# SECAT: Quantifying differential protein-protein interaction states by network-centric analysis

**DOI:** 10.1101/819755

**Authors:** George Rosenberger, Moritz Heusel, Isabell Bludau, Ben Collins, Claudia Martelli, Evan Williams, Peng Xue, Yansheng Liu, Ruedi Aebersold, Andrea Califano

## Abstract

Protein-protein interactions (PPIs) play critical functional and regulatory roles in virtually all cellular processes. They are essential for the formation of macromolecular complexes, which in turn constitute the basis for extended protein interaction networks that determine the functional state of a cell. We and others have previously shown that chromatographic fractionation of native protein complexes in combination with bottom-up mass spectrometric analysis of consecutive fractions supports the multiplexed characterization and detection of state-specific changes of protein complexes.

In this study, we describe a computational approach that extends the analysis of data from the co-fractionation / mass spectrometric analysis of native complexes to the level of PPI networks, thus enabling a qualitative and quantitative comparison of the proteome organization between samples and states. The Size-Exclusion Chromatography Algorithmic Toolkit (SECAT) implements a novel, network-centric strategy for the scalable and robust differential analysis of PPI networks. SECAT and its underlying statistical framework elucidate differential quantitative abundance and stoichiometry attributes of proteins in the context of their PPIs. We validate algorithm predictions using publicly available datasets and demonstrate that SECAT represents a more scalable and effective methodology to assess protein-network state and that our approach thus obviates the need to explicitly infer individual protein complexes. Further, by differential analysis of PPI networks of HeLa cells in interphase and mitotic state, respectively, we demonstrate the ability of the algorithm to detect PPI network differences and to thus suggest molecular mechanisms that differentiate cellular states.

## Introduction

Living cells depend on many coordinated and concurrent biochemical reactions. Most of these are catalyzed and controlled by macromolecular entities of well-defined subunit composition and 3D structure, a notion that has been captured by the term “modular cell biology” by Hartwell and colleagues [1]. Most of these modules consist of or contain protein complexes. It is thus a basic assumption of the modular cell biology model that alterations in protein complex structure, composition and abundance will alter the biochemical state of cells. Elucidating protein complexes and their organization in extended protein-protein interaction (PPI) networks is, therefore, of paramount importance for both basic and translational research.

Traditionally, composition and structure of protein complexes has been determined by two broad and complementary approaches, structural biology and interaction proteomics. Structural biology encompasses a suite of powerful techniques to characterize individual, purified or reconstituted protein complexes at high, at times, atomic resolution. High resolution structures have provided a wealth of functional and mechanistic insights into biochemical reactions [2]. However, they have been solved for only a few hundred human protein complexes and the number of cases where the structure of protein complexes is assessed across different functional states is even lower. This is contrasted by the observation that, under mild lysis conditions, approximately 60% of proteins and total protein cell mass in protein cell extracts is engaged in protein complexes [3]. As a result, methodologies for the rapid elucidation of protein complexes are still critically needed.

Interaction proteomics encompasses multiple methodologies to determine composition and, when possible, subcellular location and abundance of protein complexes, albeit at lower resolution, yet higher throughput, than structural biology techniques. Among these methods, liquid chromatography coupled to tandem mass spectrometry [4, 5] (LC-MS/MS)—and more specifically affinity purification coupled to LC-MS/MS (AP-MS [6])—has been most widely used. For AP-MS analyses, individual proteins are engineered to display an affinity tag and are expressed as “bait” proteins in cells. The bait and the corresponding “prey” proteins assembled around it are then isolated and qualitatively [7–10] or quantitatively [11–14] analyzed by MS. This method has proven robust across laboratories [15] and, through process automation and integrative data analysis, efforts to map PPIs across the entire human proteome [16, 17] are underway and have so far characterized interactions of more than half of canonical human proteins [18]. This knowledge is embedded in a variety of databases, including BioPlex [17, 18], STRING [19], IntAct [20] and hu.MAP [21] that present generic human PPI maps. The data have also been used to predict PPIs for previously uncharacterized proteins, e.g. PrePPI [22–24]. Whereas it can be expected that these systematic PPI mapping projects will reach saturation in the next few years, AP-MS and related approaches are fundamentally limited in their ability to detect compositional or abundance changes across multiple cell states and to detect concurrent changes in different complexes within the same sample.

Protein correlation profiling (PCP) [25] and related methods [3, 26–31] have been proposed as a means to concurrently analyze multiple complexes from the same sample. PCP proceeds by first separating native protein complexes, e.g. according to their hydrodynamic radius by size-exclusion chromatography (SEC), by collecting 40-80 consecutive fractions and by finally performing bottom-up mass spectrometric analysis of the proteins in each consecutive fraction. The result is a set of quantitative protein abundance profiles across the SEC separation range (Fig. 1a). Under the assumption that protein subunits of the same complex have congruent SEC profiles, they can be used to infer protein-protein interactions and protein complex composition. Conditional to availability of quantitative mass spectrometric data, the method further supports comparative analysis across multiple biological conditions, thus detecting condition-specific differences, information that is critically missing in current PPI databases. For the most part, PCP datasets have been analyzed using interaction-centric algorithms [26, 27, 29, 31–34] which essentially use chromatographic co-elution of protein profiles to identify PPIs, to infer protein complexes and to conduct qualitative and quantitative comparisons across biological conditions [30, 35]. Interaction-centric algorithms are limited by the inherently low SEC resolution and the high degree of complexity of proteomic samples. This results in the presence of hundreds to thousands of proteins per SEC fraction and lowers the confidence of inferred interactions, because the probability that non-interacting proteins may show indistinguishable elution profiles by chance is relatively high [33].

**Figure 1.**
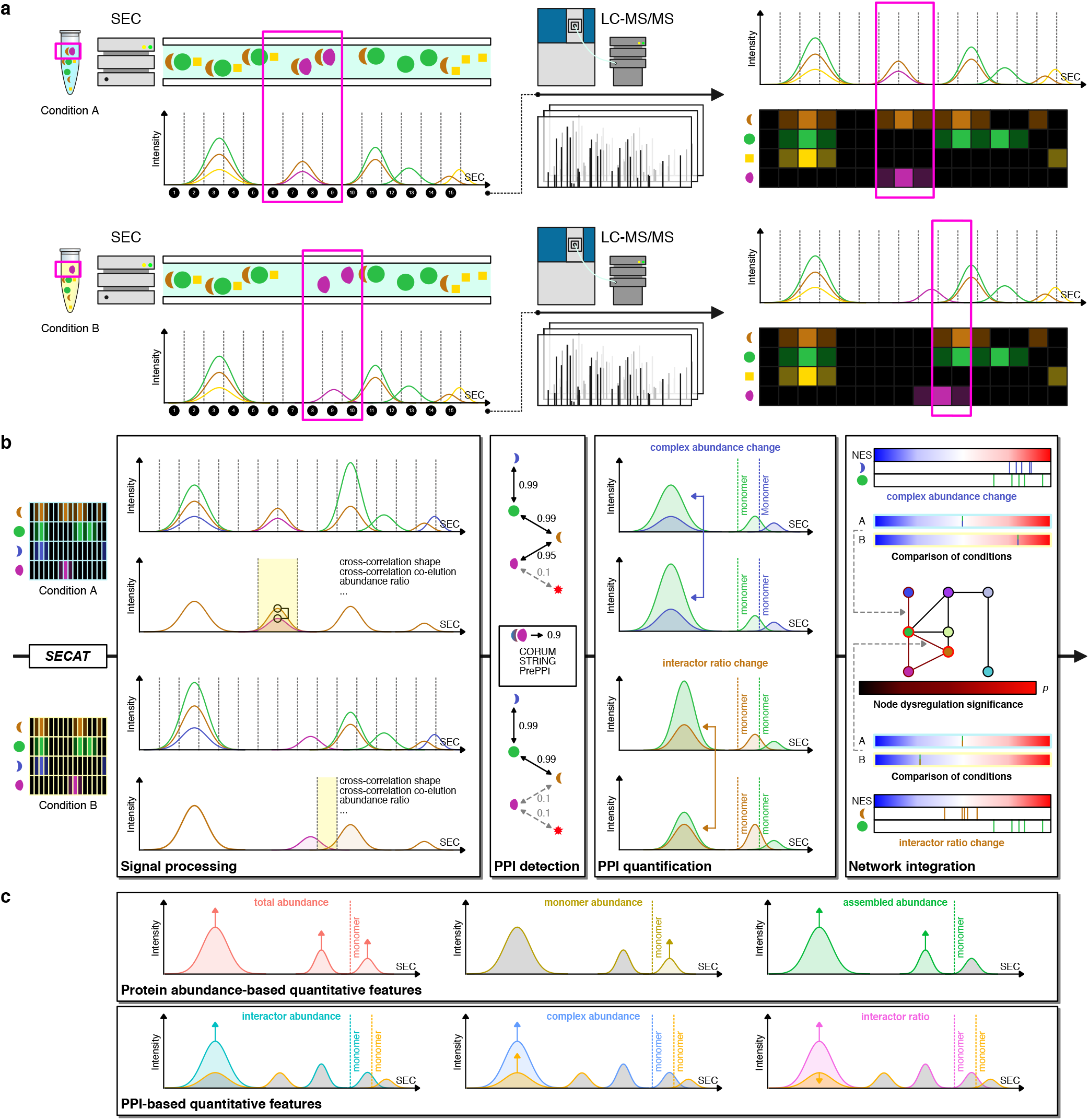
Methodological overview. **(a)** The SEC-SWATH-MS workflow is depicted comparing two different experimental conditions A and B. Native protein extracts are separated by SEC and individual fractions are sampled (1-15). Each fraction is separately measured by LC-MS/MS. Peptide or protein abundance grouped according to SEC fraction can be depicted as profile, whereby co-eluting proteins might be part of the same protein complex. Two experimental conditions can result in similar protein complex profiles but might differ for some proteins (highlighted in pink). **(b)** The SECAT algorithm uses peptide-level protein complex profiles and optional reference PPI networks as input. The signal processing module computes a set of scores for the peptide profiles of candidate interactors. Using the set of scores and ground truth positive and negative interactomes, the PPI detection module conducts semi-supervised learning and generates context-specific PPI subnetworks. Confident PPIs are quantified to assess complex abundance and interactor ratio changes using the PPI quantification module. The proteoVIPER algorithm based on VIPER [41] is used for quantitative protein inference from peptide-level abundances. Network integration: Significance of changes between the experimental conditions is integrated by a network-centric approach on all different levels. **(c)** SECAT provides quantitative insights on different levels. Protein abundance-based quantitative metrics are computed for total abundance, monomer abundance and assembled abundance. PPI-based quantitative metrics are provided separately for each interactor, integrated as positively correlated complex abundance or negatively correlated interactor ratio metric.

To address these limitations, we recently developed the algorithm CCprofiler [3], which implements a targeted, complex-centric strategy to query the protein elution profiles generated by high resolution SEC-SWATH-MS [3] to assess presence, composition and abundance of predefined protein complexes. Similar to targeted proteomic approaches [36, 37], transforming the problem from *de novo* inference of unknown protein complexes to *a posteriori* detection of established complexes significantly increases sensitivity. Using prior knowledge from reference databases like CORUM [38], BioPlex [17, 18] or STRING [19], substantially improved protein complex detection confidence, albeit at the cost of missing potential novel interactions or complexes not included in the query set. Taking advantage of the precise quantitative values generated by SWATH-MS, we have also shown that the CCprofiler strategy is well-suited to detect changes in complex-associated protein abundance across conditions [35].

However, several critical challenges remain, both for interaction- and complex-centric strategies. This is especially relevant in studies where an increasing number of experimental conditions or samples are compared, as can be soon expected. Specifically, the key assumption underlying the complex analysis methods described above, that protein complexes constitute entities of fixed subunit composition across different contexts and conditions, is both highly restrictive and biologically unrealistic. In contrast, the ability to accurately detect changes in protein complex composition between different conditions is increasingly critical to address key biological questions. For example, the spliceosome—a complex molecular machinery controlling intron removal from pre-mRNA—consists of small nuclear RNAs (snRNA) and more than 100 protein subunits, which are assembled into a variety of submodules, each with highly context-specific activity and composition [39, 40]. Since only a fraction of these protein subunits is detectable across all conditions in typical bottom-up proteomic experiments, differentiating between biological effects and technical artefacts is challenging and affecting the biological conclusions from such studies. However, if the PPIs of protein complexes are interpreted as a network rather than discrete entities, missing observations of PPIs in some conditions (e.g. missing PPI of proteins A and B) are less problematic and can be substituted by other PPIs (e.g. observed PPI of proteins A and C), due to their frequently redundant behavior within the same complexes (e.g. the PPIs of proteins A, B and C within complex A-B-C).

To address these limitations, we introduce the Size-Exclusion Chromatography Algorithmic Toolkit (SECAT). The algorithm extends the analysis of PCP data from the level of well-defined complexes to the level of PPI networks, obviating the need to explicitly infer individual protein complexes. For this purpose, SECAT generates error-rate-controlled PPI networks for each tested condition. But in contrast to existing methods, rather than using the resulting connectivity maps to directly infer protein complexes, SECAT transforms the differential protein abundances across fractions into quantitative metrics for each PPI, representing differential complex abundance and stoichiometry, which can be further integrated on network-level to derive a differential representation of the protein network state. In analogy to the extension of AP-MS from a qualitative [7–10] to a quantitative [11–14] characterization of PPIs, SECAT supports the quantitative characterization of PPI network states, while accounting for the dynamic rather than static nature of protein complexes.

We demonstrate that this novel, network-centric strategy for PPI analysis is robust against technical variability and overcomes key limitations of current protein-complex inference methods, while representing changes in network connectivity in an intuitive and unbiased way. Applying SECAT to conduct differential analysis of PPI networks of HeLa cells in interphase and mitotic state, respectively, we demonstrate the ability of the algorithm to detect PPI network differences and to thus suggest molecular mechanisms that differentiate cellular states.

## Results

### The Size-Exclusion Chromatography Algorithmic Toolkit

The primary goal of SECAT is to systematically quantify differences in abundance and interaction stoichiometry of complex-bound proteins across different conditions. SECAT explicitly is designed to follow the paradigm that complexes represent dynamic rather than static entities and that proteins can thus be constituents of several dependent or independent (sub-)complexes. Therefore, the algorithm makes two main assumptions regarding the PPIs within complexes: First, interactions of the same two proteins or proteoforms within different complexes are expected to be mediated by a similar mode of interaction (e.g. binding interface) and consequently, perturbations of these binary interactions are expected to be consistently observed across multiple protein complexes that contain the two tested proteins. Second, the quantitative measures of PPIs for any protein might be either redundant (e.g. PPIs quantified between protein in question with other subunits of the same complex) or orthogonal (e.g. PPIs quantified between protein in question with subunits in different complexes).

To implement a model that fulfills these assumptions, SECAT employs a network-centric strategy: First, similar to previous approaches [27, 33, 34], context-specific PPI networks are generated for each tested condition. Second, these networks are used to derive novel quantitative metrics representing changes in in abundance and interaction stoichiometry for proteins that are detected in complexed form based on their SEC profiles. Third, the redundant and non-redundant PPIs of the detected proteins are statistically integrated to represent the global PPI network state. In contrast to existing methods, our approach omits the explicit inference of protein complexes and provides representation of the qualitative and quantitative changes of PPI networks between experimental conditions.

The SECAT workflow consists of five consecutive steps: i) data preprocessing, ii) signal processing, iii) PPI detection, iv) PPI quantification and v) network integration. The main input data are the peptide-level intensity *vs*. SEC elution fraction profiles that are acquired by SEC-SWATH-MS [3, 35] in the conditions tested, and optional reference PPI networks (Fig. 1b, Methods). For data preprocessing and PPI detection, SECAT operates on proteotypic peptide-level profiles and uses the inferred proteins from upstream analysis pipelines to group queries and to compute meta-attributes, such as the expected monomeric weight for each subunit. For quantitative protein and PPI inference, SECAT implements a novel strategy termed proteoVIPER (Methods). The output of the system is a set of condition-specific PPI networks with protein-level metrics summarizing the associated differential protein and PPI properties (Fig. 1c, Methods). In the following we describe each step.

#### Step 1, Data preprocessing

As main input, SECAT requires quantitative peptide-level intensity *vs*. SEC elution fraction profiles acquired by SEC-SWATH-MS [3, 35] across one or several conditions. To account for varying sample amounts and batch effects between SEC runs, we first normalize peptide signal intensities within conditions and replicates and across the SEC fractions (Methods). Next, based on the molecular weight calibration curve of the SEC separation [3], for each inferred protein the partition between assembled and monomeric state is determined (Methods). This “monomer threshold” is defined by the expected molecular weight of the monomeric subunit and is multiplied by a user-defined factor to account for potential homomultimers (default: F = 2) and deviations of the protein’s chromatographic behavior from the calibration curve. Optionally, the peptide-level profiles grouped by inferred proteins can be preprocessed by detrending or local-maximum peak-picking (Methods). However, as the consistent quantification of peptides, inferred proteins and PPIs between conditions and replicates is a crucial requirement for SECAT, only minimal preprocessing is conducted by default.

#### Step 2, Signal processing

To assess PPIs, peptide-level profiles from the range of fractions representing complexed forms of each protein are queried and scored. To account for the possibility that a protein is part of multiple complexes, SECAT focuses the analysis of PPIs on those SEC fractions in which both tested interactors are detectable (Methods). To restrict the query space, candidate interactions can optionally be obtained from a comprehensive set of PPI network representations, such as CORUM [38], STRING [19] or PrePPI [22]. Alternatively, all putative PPI combinations are assessed. For each candidate PPI, peptide-level elution profiles from each protein are processed to compute chromatographic (cross-correlation shape and shift [42, 43], maximal and total information criterion [44]), interactor ratio, SEC coverage, and monomeric fraction distance metrics (Fig. 1b, Methods). The result of this step is a table in which each candidate PPI is associated with a score vector representing the different properties that can differentiate between true and false interactions.

#### Step 3, PPI detection

To generate context-specific interaction networks, partial scores from the second step are used as input to a machine learning (ML) approach based on the PyProphet algorithm [42, 45, 46]. This step is designed to discriminate true *vs*. false interactions and to estimate their confidence. For this purpose, a classifier is trained in a semi-supervised manner using a set of true negative PPIs as null model and the most confidently detected PPIs, which are evaluated and selected over multiple iterations, as a true interaction gold standard model (Fig. 1b, Methods). Since queried or tested candidate PPIs may by chance match co-eluting protein profiles that are not representative of true positive interactions detectable within the experimental context, this semi-supervised learning approach is designed to achieve high sensitivity at high confidence levels (Fig. 1b, Methods).

Classification is mediated by an XGBoost-based [47] gradient boosting approach (Methods). The first machine learning iteration is initialized by a single composite score that selects only the most confident PPIs (Methods). The full set of partial scores is then used within each subsequent iteration, thus progressively increasing the classifier sensitivity. Cross-validation and “early stopping” are employed to preclude potential overfitting of the classifier (Methods). By default, SECAT learns a classifier using high confidence PPI networks (“learning reference network”; e.g. CORUM [38]) and then applies it to integrate additional potential interactions from less stringent, optionally used networks (“query reference network”; e.g. STRING [19] or PrePPI [22]) or all potential interactions, thus maximizing sensitivity and coverage of the assessed interactions. If optionally a query reference network was used to restrict the query space, potentially available confidence metrics for individual interactions can be incorporated as priors by computing a group-based false discovery rate (FDR) metric (Methods). In summary, this step generates a set of confidence metrics for each candidate PPI (posterior error probabilities and *q*-values), that allow thresholding the list at any user-defined FDR.

#### Step 4, PPI quantification

The combined set of PPIs that are confidently detected (default: *q*-value < 0.05) across all experimental conditions and replicates is then used for quantification. Specifically, for each peptide of a protein in a candidate binary interaction, peptide-level data within the boundaries of the SEC fractions defined in Step 2 are independently summarized. In addition, for each peptide of a protein, three metrics summarizing the total, assembled and monomeric fractions are computed. This provides quantitative metrics that can be used to assess inferred protein or PPI changes across experimental conditions, as described in the following steps (Fig. 1b, Methods).

The VIPER [41] algorithm was originally developed to infer protein activity from the abundance of transcripts that are targets of specific transcription factors in gene regulatory network models (GRN). SECAT contains the proteoVIPER algorithm that extends the VIPER strategy to estimate protein- or PPI-level metrics. This is achieved by adapting VIPER to proteomic data, specifically the fractionated peptide abundance values, as computed above (Methods). To estimate protein-level metrics from the peptide-level data, proteotypic peptide-protein mappings are used from upstream analysis pipelines to compute a normalized enrichment score (NES), assessing the change of peptide-based protein abundances within individual replicates and conditions against the reference samples (Methods). When all fractions are considered, a metric for “total” protein abundance can be computed. Alternatively, considering the above-defined monomer threshold, metrics for “monomeric” (right side of threshold) or “assembled” (left side of threshold) states can be estimated (Fig. 1c, Methods), thus providing the basis to quantify differences in PPIs across samples and states. In analogy to other quantitative protein inference strategies [48], the protein-level NES provides a metric for protein abundance. However, the underlying statistical framework increases the robustness for quantification with diverse sets of peptides and enables the differential comparison to control samples (Methods).

To estimate PPI-level metrics, the peptides of two interactor proteins are quantified using only their overlapping SEC fractions instead of their full elution profiles. For each protein, a separate “interactor abundance” metric can be computed as described above to assess the quantitative changes for the individual protein within the PPI. Alternatively, complex-related metrics can be derived: First, the peptides of the two interactors are assessed in a positively correlated setting, where the resulting metric can be used to assess “complex abundance”. Second, the interactor peptides are assessed in a negatively correlated setting to derive a metric representing “interactor ratio” (Fig. 1c, Methods). This metric can represent stoichiometric changes between interacting subunits within a complex or alterations in their connectivity, if a PPI is abrogated or quantitatively changed in some conditions.

The proteoVIPER module reports six quantitative metrics. On the protein level it reports total abundance, assembled abundance, monomer abundance and at the PPI-level it reports interactor abundance, complex abundance and interactor ratio. While these values represent different properties of a protein and its interactions, in some cases they are strongly correlated. For example, proteins present in assembled state only will have strongly correlated total and assembled abundances.

#### Step 5, Network integration

The goal of this last step of the algorithm is to integrate individual quantitative protein and binary PPI metrics from the previous step into comprehensive PPI network states. In the ensuing representation, nodes are individual proteins annotated with attributes representing differential protein or complex abundance and interactor ratio across conditions and replicates, and edges indicate specific binary PPIs, representing the consensus across conditions and replicates (Fig. 1b, Methods). This transformation can be used to provide a summary overview of the different PPI network states between two or several conditions.

Using the experimental design, SECAT first statistically compares the quantitative metrics between different groups to identify PPIs that change between the conditions (Methods). However, since our approach is agnostic to protein complexes, these PPIs can be either redundant (interactions within the same complex) or orthogonal (interactions within different complexes). For this reason, differential PPI-level (edges) metrics are integrated using the Empirical Brown’s Method [49] on a protein-by-protein basis (nodes) to assess protein abundance or protein-complex-based changes for every protein of interest. This method is specifically designed to account for non-statistically-independent evidence and can integrate both redundant and orthogonal information (Methods).

In conclusion, SECAT leverages large-scale, quantitatively consistent co-fractionation mass spectrometry measurements across multiple replicates and conditions, such as those acquired by SEC-SWATH-MS, to generate context-specific PPI networks, as well as protein abundance and other PPI-related quantitative metrics. In combination, these metrics comprehensively characterize condition-specific proteome abundance and PPI network changes in a single operation.

### Parameter selection and validation of signal processing and PPI detection modules

In different fields of computational proteomics, the quantification of analytes, i.e. peptides or proteins [50], or their interactions, i.e. protein-protein interactions [11–14] or cross-linked peptides [51], have relied on their prior identification at high confidence. Therefore, by analogy, the consistent identification or detection of PPIs across multiple samples with accurate confidence estimation is also a crucial property for PCP-based PPI studies. Due to the limited resolving power of SEC and the large PPI query space, previous approaches either used ad hoc thresholds to filter out less likely interactions (e.g. by requiring a minimum correlation of 0.5 for two candidate interactors [27]) or they removed background noise by peak-picking [3, 33], assuming Gaussian elution peaks in the SEC dimension. SECAT, in contrast, maintains quantitative consistency between conditions and replicates by applying an optimized semi-supervised learning strategy that requires only minimal signal preprocessing and obviates the need for prefiltering.

To demonstrate that SECAT can retain high sensitivity for PPI detection at high confidence levels without extensive data preprocessing, we first compared the effects of different preprocessing methods. For this, we used a publicly available SEC-SWATH-MS dataset [35] of HeLa CCL2 cells that were measured in triplicate in both interphase and mitotic cell state (referred to as HeLa-CC dataset, Methods). Then, we assessed the method’s robustness based on its ability to infer bona fide PPIs using reference networks with an increasing ratio of false *vs*. true PPIs. For both assessments, the entire SECAT methodology was used, however, with different choices for the parameters at each step.

Defining reference sets of true and false PPIs is a non-trivial problem. For this benchmark, we adapted a previously applied strategy used by the PrInCE algorithm [33] (Methods). It leverages CORUM [38] PPIs as true and all other interactions of CORUM proteins that are not included in the database (CORUM-inverted) as false PPIs (Methods). For the purpose of the benchmark we further excluded any known or predicted interactions from CORUM-inverted and split the combined true/false reference set into equally sized training/validation and hold-out subsets. These were generally used for all benchmarks, but for algorithm comparisons, we generated subsets on protein complex level (see below, Methods).

#### Effect of signal preprocessing on PPI detection consistency

To better assess the effects of different preprocessing strategies on PPI detection consistency, we compared the default, minimal preprocessing approach of SECAT (none) to two different peak-picking strategies, termed “detrend”—base line removal with (detrend zero) or without (detrend drop) missing values—and “localmax”—local maximum peak-picking (Methods).

Using the above-described benchmark setup, we first processed three replicates of each HeLa-CC dataset condition with the different peak-picking options. The results (Fig. 2a) show that detrending or peak-picking based noise removal results in tighter, less variable mean peak widths across conditions and replicates. When comparing peak-width standard deviation across the three replicates, the “localmax” method introduced slightly higher variability (Fig. 2a). In contrast, the quantitative, integrated peak area-based metrics and particularly the standard deviation are more similar across all three options (Fig. 2a), suggesting that all three tested approaches perform similarly in terms of quantitative applications. However, considerable differences were found when comparing detection consistency (Fig. 2b). Indeed, decomposing the total number of detected PPIs for detectability across experimental replicates revealed that the fractions of PPIs consistently detected across replicates was substantially higher if no preprocessing or detrending was used, thus supporting the omission of peak-picking in SECAT.

**Figure 2.**
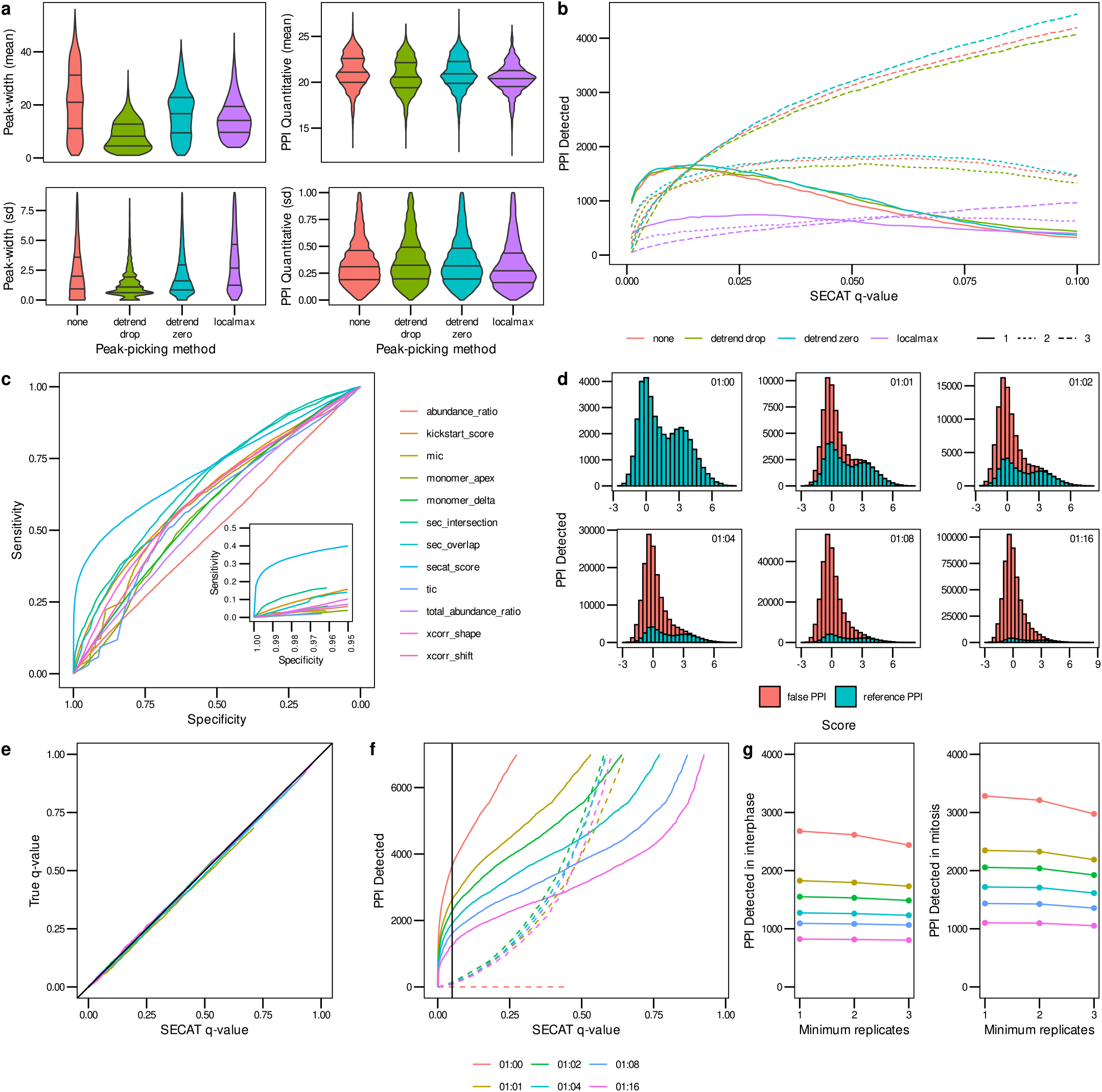
Signal processing and PPI detection performance evaluated using the HeLa-CC [35] dataset. **(a)** The effect of different peak-picking methods for signal processing on peak-width and PPI quantification within replicates of the same experimental condition is depicted in violin plots (lines representing 25, 50 and 75% quantiles respectively, Methods). **(b)** Sensitivity of PPI detection *vs*. SECAT *q*-value for peak-picking options. The PPI were filtered to a global context *q*-value < 0.05. Colors indicate different peak-picking options; line types indicate number of detections with the three replicates per condition. **(c)** The receiver-operating characteristic (ROC) illustrates the sensitivity *vs*. specificity of the different SECAT partial scores, the kickstart score and the integrated SECAT score. **(d)** The SECAT score histograms depict the dilution of the reference PPI network (true: CORUM) with false PPI (false: CORUM-inverted) for the benchmark (Methods). **(e)** The SECAT estimated *q*-value is accurate as evaluated against the ground truth for all levels of reference PPI network accuracy. For the dilution 1:0, no *a priori* false PPI are tested, thus the true FDR equals to 0. **(f)** The sensitivity of PPI detection in dependency of SECAT *q*-value for different levels of reference PPI network accuracy is depicted. Solid lines represent the true, whereas dashed lines represent the false PPI. **(g)** Consistency of PPI detection between replicates of the same conditions.

#### Error-rate estimation accuracy and PPI detection sensitivity

To optimally separate true *vs*. false candidate PPIs, SECAT integrates 11 different partial scores into a composite score (Fig. 2c), using a semi-supervised learning strategy initialized by a high confidence PPI score (kickstart score, Methods). Consistent with other evidence integration approaches, the final integrative score (SECAT score) is more discriminative than any of the initial or partial scores (Fig. 2c). To assess error-rate estimation accuracy and algorithm sensitivity, we performed cross-validation analysis by applying the classifier to the hold-out subset with increasing fractions (1:0 –1:16) of false reference interactions (Methods, Fig. 2d). The accuracy of PPI detection confidence was then assessed by comparing the *q*-value estimates with the ground truth estimated FDR (Fig. 2e). Our results show that the *q*-value estimates for the hold-out dataset were accurate within the assessed range. This shows that the SECAT PPI detection module is robust against a variable fraction of false or undetectable PPIs in reference databases, especially in the more relevant high-confidence region (*q*-value < 0.1). Moreover, assessing all pairwise PPI combinations instead of using reference PPI networks to restrict the query space substantially reduces sensitivity due to multiple hypotheses testing correction [52] (Fig. 2f). At the same high confidence level (*q*-value < 0.05), the reference-network-based approach, without additional false reference interactions, detected 3,630 PPIs, whereas a reference-network-free approach, represented by using 16 times as many false reference interactions as true interactions, only detected 1,244 bona fide PPIs, suggesting key loss of accuracy when low-quality PPIs are included and even greater loss when no prior PPI network is used and PPI interactions must be discovered *de novo*. Indeed, the substantially higher PPI recovery rate at the same confidence threshold illustrates the benefit of using a high-quality reference PPI network, rather than comparing all potential PPI interactions using the proposed scoring approach.

#### Compatibility with different data modalities

PCP datasets have been acquired by proteomic methods different from SWATH-MS, e.g. by label-based [30] or label-free [26] data-dependent acquisition methods. We tested the performance of SECAT with a publicly available SILAC-PCP dataset [30] which compared Anti-Fas IgM treated (inducing apoptosis) against control samples of Jurkat cells in triplicates per condition. We refer to this dataset as Jurkat-Fas. The assessment was conducted using the same strategy as described above.

Benchmark data shows that SECAT can be effectively applied to the Jurkat-Fas dataset, with similar performance characteristics in terms of signal processing and error-rate control robustness, resulting in 2,132 PPIs (*q*-value < 0.05) detected across both conditions and replicates (Supplementary Fig. 1). A crucial requirement for studies comparing multiple experimental conditions and replicates is high PPI detection reproducibility. For example, requiring PPIs to be detected in all three instead of just one replicate reduced recall by 43.5-44.4% (Supplementary Fig. 1g) in the Jurkat-FAS dataset. In comparison, the corresponding drop for SEC-SWATH-MS in the HeLa-CC dataset was substantially smaller (9.0-9.3%) (Fig. 2g). In conclusion, the core assumptions for SECAT are also fulfilled for datasets acquired by different methods. But because different biological systems were investigated (i.e., HeLa-CC *vs*. Jurkat-Fas dataset), no conclusions can be made in terms of performance metric differences, since they could be of technical or biological (e.g. sample complexity) origin.

#### Comparison of SECAT with established algorithms

To our knowledge, no other algorithms have been published to quantitatively assess differential PPI network states using PCP datasets and thus a direct comparison of SECAT is not possible. However, several algorithms have been developed for the interaction- and complex-centric analysis of proteomic co-fractionation profiles. They share with SECAT the requirement to accurately estimate the confidence of identified or detect PPIs. For the purpose of this benchmark, we thus compared the PPI identification or detection modules of three representative algorithms, EPIC [27] (interaction-centric), CCprofiler [3] (complex-centric) and SECAT (network-centric). As reference dataset, we used a previously published SEC-SWATH-MS profile of a HEK293 cell line [3] in exponential growth state, measured in a single replicate. In the following we refer to this dataset as HEK293-EG. We used the CORUM reference PPI network to generate a pseudo ground-truth dataset for classifier training, as described above, but restricted validation results to a previously published [3], manually curated list of detectable complexes within that dataset (Methods).

To compare binary PPI identification/detection performance, we trained the SECAT, CCprofiler and EPIC classifiers using the same input data, selecting 50% of the complexes for training and the other 50% for validation. For CCprofiler, complex hypotheses were defined as binary complexes. Since the algorithms apply very different signal processing and filtering steps, the numbers of reported PPIs with predicted scores varied substantially. SECAT reported 31,932 PPIs of which 2,506 had a *q*-value < 0.05. Similarly, CCprofiler conducts targeted analysis, but the preprocessing steps of the algorithm reduced the number of reported PPIs to 4,955 of which 2,500 had a *q*-value < 0.05. EPIC assesses all combinations of protein interactions in an unbiased manner and only reported results with a score above 0.5 (not error-rate controlled), resulting in 4,592 PPIs.

Comparison of these diverse sets of predicted PPIs is not trivial, particularly because the algorithms were designed for very different purposes. We thus only assessed the power of individual classifiers to discriminate true and false PPIs and for this purpose, we compared all combinations of the three classifiers and the full intersection of all predicted PPIs (without cutoffs, except EPIC score>0.5). The full intersection of the three algorithms resulted in a set of predictions for 2,633 ground truth “true” and 191 ground truth “false” PPIs. On this set, SECAT achieved an area under the receiver operating characteristic (AUROC) of 0.856, with CCprofiler reporting a slightly lower performance of 0.805, whereas the AUROC of EPIC was lower at 0.690 (Supplementary Fig. 2a). Further, SECAT and CCprofiler outperformed EPIC in all pairwise comparisons, especially in the high-confidence region (Supplementary Fig. 2b-d).

It should be noted though that this comparison is limited in scope and generalizability of the conclusions. First, in absence of any systematic ground truth datasets that could be used for benchmarking, only the interactions of manually curated complexes [3] and their inferred negative PPIs were used. These examples likely form characteristic elution peaks in the SEC dimension and might not be entirely representative of all PPIs detectable by PCP. Second, only a single replicate was assessed and thus no conclusions about PPI detection consistency across replicates and conditions could be drawn, which might be more variable between different preprocessing methods, as discussed above. Third, the algorithms were originally designed for different purposes, e.g. EPIC was not primarily developed for the analysis of SEC, but rather ion-exchange (IEX) profiles [27].

In summary, SECAT’s novel approach for accurate estimation of PPI detection confidence combines (optional) reference-network-based assessment of PPIs with a semi-supervised machine learning strategy. While the semi-supervised learning component provides selectivity by exclusion of spontaneously co-eluting PPIs present in the ground truth dataset during the first learning iteration, the restriction of the PPI query space using reference networks dramatically increases sensitivity, thus enabling an optimal tradeoff between the two metrics. In turn, these improvements allow SECAT to operate with minimal data preprocessing, ensuring reproducible detection of PPIs across replicates and conditions, a crucial requirement for their quantification.

### Validation of PPI quantification & network integration modules

Building on SECAT’s benchmarked ability to detect true PPIs from PCP datasets, we next validated its PPI quantification and network integration modules. For these analyses, we used the previously described HeLa-CC dataset [35]. We performed independent analyses, either assessing all potential PPI combinations or by using a restricted query space based on CORUM [38], STRING [19] or PrePPI [22] as reference PPI networks. The classifiers were trained on the full CORUM true/false reference network.

#### PPI detection with different reference networks

For the analyses involving the three reference PPI networks or all combinations, the total respective query space ranged from 17,751 (CORUM), 198,399 (PrePPI) to 528,163 (STRING) and 6,824,388 (all combinations) PPIs (Fig. 3a). Of these, between 2,891 (all combinations), 5,656 (PrePPI), 7,898 (CORUM) and 8,560 (STRING) high-confidence PPIs (*q*-value < 0.05) were detected, with a core set of 1,129 common PPIs (Fig. 3b). Notably, the full intersection of detected true PPIs might be larger, but two factors conspire to make objective comparison more complex. First, different confidence scores provided by each database are used as priors in our analyses. Second, the largely variable number of PPIs represented in each database needs to be accounted for during multiple testing correction (Methods). Assessing all pairwise protein combinations expectedly resulted in the lowest number of detected PPIs (*N* = 2, 891). While this mode discovered 1,068 unique PPIs, the much lower sensitivity represents a considerable drawback for quantitative applications. Consequently, we primarily used the STRING-based results for further analyses.

**Figure 3.**
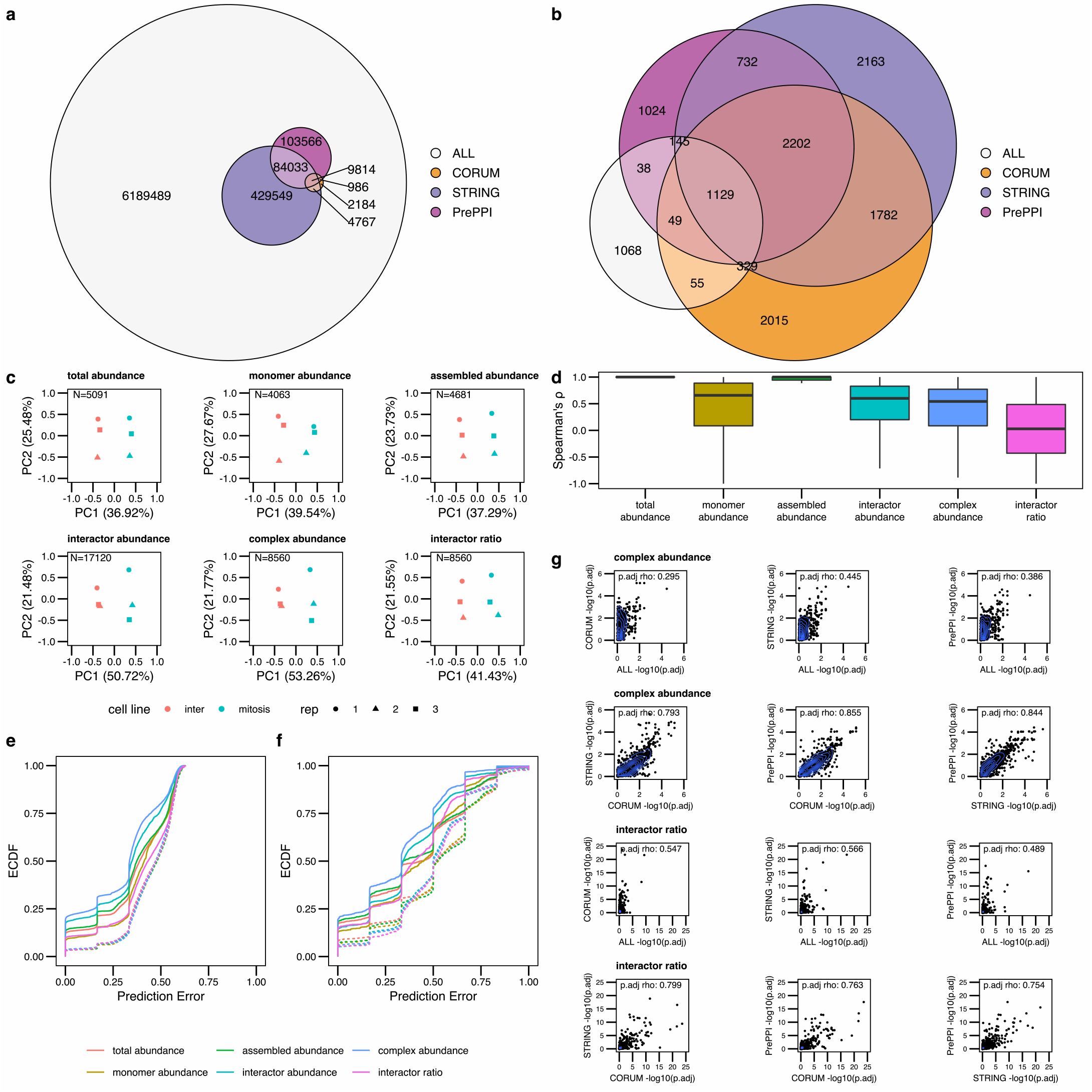
PPI quantification and Network integration assessment using the HeLa-CC [35] dataset. **(a)** The Euler diagram indicates the overlap of STRING, PrePPI and CORUM reference PPI networks resulting in at least a partial overlap of the interactors in any of the replicates. **(b)** Overlap of confident (*q*-value < 0.05) PPI detections after SECAT analysis with different reference PPI networks. **(c)** Quantitative SECAT metrics can be used to separate experimental conditions and replicates. **(d)** Spearman’s correlation of the metrics (PPI: summarized by averaging) to total-level is depicted in individual boxplots (lower and upper hinges represent the first and third quartiles; the bar represents the median; lower and upper whisker extend to 1.5 * IQR from the hinge). **(e)** Predictive error quantified by the empirical cumulative distribution function (ECDF) for leave-one-out cross-validated logistic regression classification of replicates to biological conditions. **(f)** Predictive error for group-wise leave-one-out cross-validated logistic regression classification of replicates to biological conditions. Groups represent several metrics connected to a core metric based on the PPI network. **(g)** Correlation of node-level integrated adjusted *p*-values estimated using different reference PPI networks.

#### PPI quantification and relation of metric classes

SECAT’s quantification module computes two main quantitative property categories from SEC-SWATH-MS data: one indicating protein abundance, the other representing PPIs. Some of these metrics are expected to be related. For example, if a protein is only present in an “assembled” conformation, the “total abundance” and“assembled abundance” values will be highly correlated. To assess the relation and redundancy within metric classes, we conducted dimensionality reduction using principal component analysis (PCA), separating replicates and conditions (Fig. 3c). The data show that the first two principal components of the “total abundance”, “monomer abundance” and “assembled abundance” metrics explain 62.40-67.21% of the variance between them, while the first two principal components for the “interactor abundance”, “complex abundance” and “interactor ratio” metrics explain 62.98-75.03% of the variance. The small difference in variance between protein abundance- and PPI-based metrics might potentially arise from metrics quantifying redundant interactions between same-complex proteins.

To assess the relationship between metric classes, we further computed the distribution of Spearman’s correlation of metrics aggregated over conditions, replicates and proteins to the corresponding “total” protein abundances (Fig. 3d). As expected, the “assembled abundance” metrics which cover most SEC fractions have the strongest correlation, followed by “monomer abundance”, “interactor abundance” and “complex abundance” metrics. Notably, since the “interactor ratio” metrics summarize relative changes between the interactors, they are substantially less correlated.

#### Assessment of predictive power of quantitative metrics

To assess the predictive power of the different metric classes for classifying the six samples to one of the two represented mitotic states, we estimated the prediction error using leave-one-out cross-validation using a logistic regression classifier applied to each metric separately (Fig. 3e). The empirical cumulative distribution function (ECDF) in dependency of the prediction error indicates that the “complex abundance” metrics are most informative, followed by “interactor abundance” and the other metrics, to separate the two mitotic states. While these metrics are computed for single proteins or PPIs, network-centric data integration might further boost the predictive power of the respective values.

To assess this effect, we combined individual PPI metrics based on the SECAT PPI networks by extending the score vector for the classifier from single metrics to all PPI scores connected to a specific protein (Fig. 3f). The results show that the “complex abundance”-based metrics performed best and the “assembled abundance” metrics performed second best, indicating that the fractions covering assembled protein states have a higher information content than those covering monomeric subunits and that the connectivity information calculated by SECAT has higher information content than the protein abundance.

#### Redundancy of network integration

In higher order complexes composed of three or more subunits, PPI information is expected to be partly redundant, with each subunit relating to at least two other co-complex members. Thus, a change in one subunit might be apparent in other PPIs as well. This PPI redundancy between subunits of a multiprotein complex is critically relevant to any strategy attempting to integrate individual protein or PPI properties at the network level. Consistent with this observation, SECAT assumes that changes in the individual subunits of a protein complex can be measured redundantly by assessing its interactions with all proteins in the complex. Different reference PPI networks are thus expected to provide comparable quantitative metrics if they partially overlap. To test this assumption, we compared the differential integrated metrics on “complex abundance” and “interactor ratio” levels using different reference PPI networks. The results indicate a high degree of correlation among the “complex abundance” (Spearman’s rho: 0.793-0.855) and “interactor ratio” levels (Spearman’s rho: 0.754-0.799) (Fig. 3g), confirming that different reference PPI networks provided comparable quantitative metrics from the same PCP input data. Notably, the comparison with the reference-network-free mode indicates much lower correlation. This can be explained by differences in the topologies of the generated PPI networks. Although the node degree distributions of both reference-network-free and reference-network-based PPI networks approximate a power-law [53], substantially more high degree nodes (*k* > 60) could be detected when using reference networks (Supplementary Fig. 3).

In summary, SECAT’s PPI quantification and network integration modules generate instances of PPI networks for each sample or phenotypic state. To achieve this, the signal processing and PPI detection modules first transform SEC fraction peptide abundance values to a set of protein abundance or complex-associated metrics. The presented validation results show that PPI-level quantitative metrics have higher predictive power to separate sample groups than the underlying total protein abundances. In a second step, redundant and non-redundant PPI-level metrics (edges) are integrated to protein-level (nodes). With different empirical or predicted reference PPI networks, this transformation achieves similar results attesting to the robustness of results despite different background PPI networks.

### Molecular mechanisms differentiating HeLa cell cycle states

To demonstrate SECAT’s ability to identify molecular mechanisms that differ between experimental conditions or phenotypes, we further investigated the results obtained from the HeLa-CC dataset [35]. This study was designed to compare differences at the level of protein complexes between cell cycle states of a HeLa CCL2 cell line. The HeLa cell states were induced by thymidine blocking which arrested cells in interphase and by a subsequent release in nocodazole which arrested cells in mitosis [35]. For each experimental condition, three full-process replicates were generated, and 65 SEC fractions per condition and replicates were measured by SEC-SWATH-MS. Cumulatively, 70,445 peptides associated with 5,514 proteins were quantified across the full dataset. We conducted the SECAT analysis of this quantitative matrix using STRING as the query reference PPI network with the goal to identify phenotype-associated molecular mechanisms (Methods). Since the cell cycle and its checkpoints are controlled by clearly defined events involving selected checkpoint proteins, the literature provides a reference framework for the interpretation of the results.

After network-centric data integration, SECAT identified 25 differentially abundant proteins (“total abundance”-level) across the two cell states, 44 alterations affecting “assembled abundance”, no alterations regarding “monomer abundance”, 117 alterations affecting “interactor abundance”, 129 alterations in “complex abundance”, and 141 alterations in “interactor ratio” (adj. *p*-value < 0.01; |*log*_2_(fold-change)| > 1) (Supplementary Data 2-3). To visualize the data SECAT implements a simplistic aggregation to the most significant level per protein that provides a reduced overview. This can be further augmented by integrating the context-specific PPI network with CORUM (Fig. 4, Supplementary Fig. 4) or Reactome [54] (Supplementary Fig. 5), visualizing changes affecting protein complexes or modules, respectively (Methods). Importantly, modules and complexes can represent the same entities, e.g. the anaphase-promoting complex (APC), which constitutes a well-defined complex in both complex or module representations (Supplementary Fig. 4-5), or their modularization cannot be attributed to specific complexes, e.g. the proteins part of the rRNA processing module (Supplementary Fig. 5).

**Figure 4.**
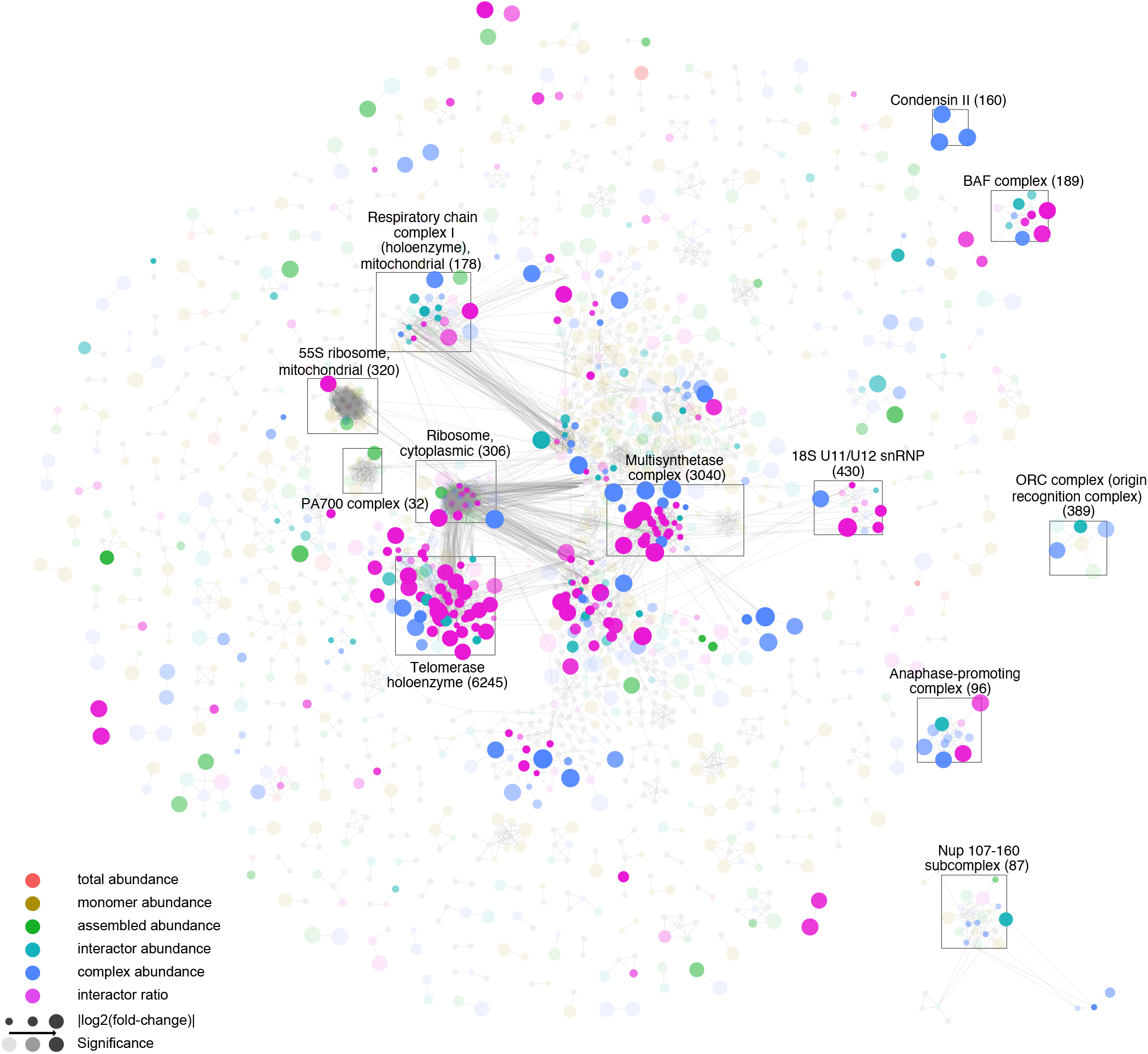
Complex-level molecular mechanisms differentiating HeLa cell cycle states. The integrated STRING-based PPI network of the HeLa cell cycle dataset [35] is depicted with proteins (nodes) and binary PPIs (edges) clustered against CORUM complexes (identifiers in brackets; Methods). Different colors indicate the most significant metric-level (legend, Methods). Node-size indicates effect-size and opacity indicates significance. The most prominent clusters highlight essential macromolecular assemblies. Interactive results (Cytoscape) are provided as Supplementary Data 1.

**Figure 5.**
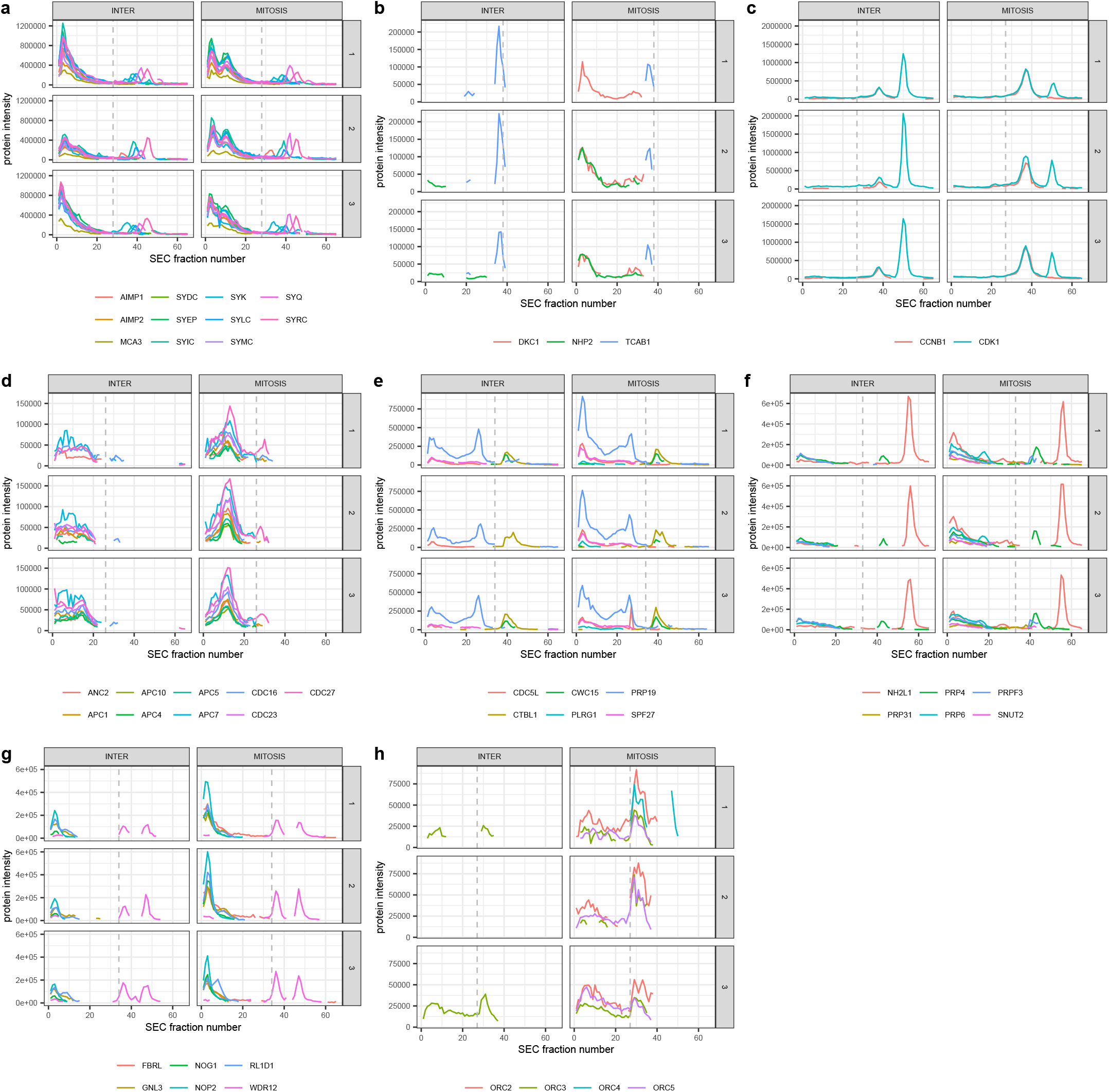
Protein-level SEC-SWATH-MS profiles of HeLa cell cycle states. Lines indicate different protein subunits, whereas the dashed line indicates the highest monomer threshold per group. **(a)** Multisynthetase complex. **(b)** Telomerase holoenzyme. **(c)** Cyclin B1-Cdk1 complex. **(d)** Subunits of the anaphase-promoting complex (APC). **(e)** Subunits of the NineTeen Complex (NTC). **(f)** Associated U4/U6.U5 tri-snRNP subcomplex proteins. **(g)** The most abundant ribosome biogenesis-associated proteins. **(h)** Subunits of the origin recognition complex (ORC).

The data show that a few primary clusters representing large protein complexes and their interactors dominate the modules covered by the network of detected PPIs. These include the cytoplasmic ribosome (80 proteins; peptide chain elongation module), the mitochondrial 55S ribosome (72 proteins; mitochondrial translation termination module), the multisynthetase complex (44 proteins; selenoamino acid metabolism module), the telomerase holoenzyme (41 proteins; rRNA modification in the nucleus and cytosol module) and the mitochondrial respiratory chain complex I (holoenzyme) (32 proteins; respiratory electron chain transport module). These structures have in common that they consist of numerous subunits with many intra-complex PPIs. For the ribosomal complexes and the respiratory chain complex, many PPIs could be detected and quantified, but most PPIs and complex subunits did not change between conditions. In contrast, for the telomerase holoenzyme and the multisynthetase complexes, a larger number of subunits were significantly altered between the conditions (Fig. 4, Supplementary Fig. 4) indicating that these structures changed composition between mitotic states. Specifically, the subunits of the multisynthetase complex form a single complex in interphase, whereas two subunits of the telomerase holoenzyme, DKC1 and NHP2, and their interaction, are only detectable in mitosis suggesting a significant quantitative difference of the two proteins between mitotic states (Fig. 5b). These results indicate SECAT’s ability to quantify different types of alterations of large molecular modules between cellular states.

Numerous changes were also detected for smaller modules and their interactions. Cyclin B1 bound to Cyclin-dependent kinase 1 (Cdk1) is a major catalytic factor promoting mitosis [55]. Counteracting Cyclin B1-Cdk1 mediated activation, sequential degradation of cell cycle regulating proteins via the ubiquitin pathway is important to progress through mitosis [56]. SECAT recalled these well-known biochemical events from the SEC-SWATH-MS data. Specifically, it detected different levels of Cdk1 “complex abundance” and “interactor ratio” as a principal factor differentiating the two cell cycle states (Fig. 5c). Further, SECAT identified several subunits of the anaphase-promoting complex (APC) to be significantly more abundant during mitosis (Fig. 5d). The network visualization further illustrates the high connectivity between the individual subunits, resulting in a distinctive complex module within the graph, primarily affected by differential protein abundance between the mitosis states (Fig. 4). This is consistent with the role of the APC as a ubiquitin ligase mediating this particular step [56]. Between transcription and translation, pre-mRNA is processed to remove introns and splice exons to produce mature mRNA molecules for translation. This process is catalyzed by the spliceosome, a multi-megadalton ribonucleoprotein complex, which highly dynamically adapts to context-dependent functions [40]. Spliceosome assembly and function typically involve several intermediate complexes, requiring the integrity of the nuclear compartment [39]. With the disassembly of the nucleus and associated nuclear pore complex (NPC) proteins during mitosis, the rate of transcription is reduced and it is currently believed that spliceosome subunits are distributed across the mitotic cytoplasm awaiting re-activation upon nuclear reassembly [39]. However, systematic screens identified spliceosome subcomplexes, including the NineTeen Complex (NTC) with five out of seven of its core proteins (PRPF19 (PRP19), CDC5L, SPF27, PLRG1 and CTNNBL1 (CTBL1)) as essential components for mitosis [57]. Correspondingly, in our analyses PRPF19 and CDC5L were identified as differentially abundant between interphase and mitosis on the “complex abundance” level. In addition, they display changes on the “interactor ratio” level. Specifically, the NTC subunits form a more distinctive SEC elution peak during mitosis (Fig. 5e). Similarly, NHP2-like protein 1 (NH2L1), a component of the U4/U6.U5 tri-snRNP subcomplex, has been found to be required during mitosis [57]. Our analysis further supports the importance of NH2L1 during mitosis, as it is among the most significantly changed proteins in terms of “interactor ratio” but not “total abundance”. SECAT further found the linked U4/U6.U5 tri-snRNP subcomplex proteins such as the tri-snRNP-associated protein 2 (SNUT2), the pre-mRNA-processing factors 3, 4, 6 and 31 (PRP3,4,6,31) to be of similar differential “interactor ratio” significance, suggesting subcomplex activity during the cell cycle (Fig. 5f).

The Ribosomal RNA Processing complex represents one of the larger and most densely connected submodules of the dataset. SECAT found several proteins known to belong to this group to be differentially abundant (Fig. 5g). Of those, the majority is associated with ribosome biogenesis, a process located in both cytoplasm and nucleolus. Similar to the origin recognition complex (ORC, Fig. 5h), which is only active in the nucleus, this difference in abundance might be explained by the experimental design, because under the conditions used for cell lysis, proteins of the nucleus are less accessible during interphase than mitosis, a cell cycle state where the NPC is disassembled. The proteins, complexes and modules involved in rRNA processing illustrate a further important asset of network-centric analysis: Reactome modularization in Supplementary Fig. 5 shows that although the proteins involved in the associated rRNA processing and modification processes cannot always be attributed to specific complexes, their functional association can provide clues about differential mechanisms, either by visualization or pathways analysis. Further examples illustrating the SEC profiles of the protein complexes covered by Fig. 4 are visualized in Supplementary Fig. 6.

In summary, SECAT provides context-specific networks and protein-level metrics that can be visualized as intuitive maps (Fig. 4), facilitating the interpretation of observed molecular differences between cell states at different levels including protein abundance, PPIs, protein complexes and PPI network modules, concurrently, from the same dataset. This representation allows to analyze changes on different levels, from larger network modules to complexes and specific PPIs, thus providing an interpretable and scalable resource from overview representations to zoomed-in molecular mechanisms for expert analysis or multi-omic data integration.

## Discussion

Protein-protein interactions are a principal characteristic of proteome organization and are significantly affected by or determine the biochemical state of the cell. Most biochemical functions are catalyzed and controlled by multiprotein complexes which, in turn, are organized in extensive interaction networks, exemplified by PPI interaction resources such as STRING [19]. The ability to accurately compare PPI networks of different cellular states and to deduce from the detected differences altered biochemical functions and mechanisms is therefore of fundamental importance for molecular biology. To address this need under the term “interaction proteomics” several powerful methods have been developed, including AP-MS that investigates the interactions of specific proteins at relatively high throughput. The cumulative results of thousands of AP-MS measurements constitute PPI network maps for investigated organisms [17, 18]. However, the extension of the AP-MS approach to compare PPI networks at different states is intrinsically limited because it would require the comparative analysis of the results of a high number of AP-MS measurements in the different states.

For this reason, PCP-based methods, such as SEC-SWATH-MS, have emerged as complementary approaches. They can measure protein profiles across the chromatographic size separation range quickly and reproducibly for thousands of proteins, thus indicating their abundance and association with complexes. They achieve, however, lower proteome coverage than typical bottom-up proteomic measurements and are limited to medium to high affinity-binding protein complexes that are soluble and remain intact under the extraction conditions used. Previous studies have already demonstrated the application of PCP to qualitatively characterize metazoan macromolecular complexes [31]. With increasing throughput and further methodological improvements, it can be expected that these developments enable the qualitative and quantitative comparison of dozen to hundreds of samples in single studies.

However, the relatively low peak capacity of SEC imposes major limitations for PPI identification, i.e. the number of proteins identified by far exceeds the number of separable peaks and thus fractions collected. In previous studies this limitation was addressed by sequentially applying orthogonal biochemical fractionation methods [26] or by focusing on protein complex detection using predetermined subunits [3]. This requires either more complex and costly experimental designs or focus on well characterized protein complexes, limiting the scalability and generic applicability of protein complex profiling studies.

With SECAT, we introduce an alternative analysis strategy which makes use of the high consistency of peptide-level quantification of SWATH-MS and prior knowledge from PPI reference databases. We demonstrate that SECAT applied to data with these qualities provides accurate estimation of PPI detection confidence while substantially increasing the coverage of binary interactions. The robustness of the scoring and semi-supervised learning strategy further permits omission of preprocessing steps such as peak-picking and obviates the need for scoring thresholds, making the algorithm robust against context-specific deviations related to SEC peak shape or different calibration of SEC fractions.

Because SEC separates stable native protein complexes, the inference of their composition from binary interactions is a key component of most previous data analysis strategies. This provides the opportunity to identify previously unknown associations in a global fashion, however, the underlying challenges of annotating and comparing context-specific related subcomplexes will become more severe with increasing number of conditions tested in a study. We show that the proposed protein abundance-level and PPI-based metrics are comparable across different PPI reference networks and thus their implementation in SECAT provides a scalable alternative to protein complex inference that makes use of redundant information to increase the consistency of quantitative comparisons. In turn, these improvements allow quantitative comparisons of the PPI network state, which can further be grouped to protein complexes or functional modules, where concurrent changes highlight differential molecular mechanisms between the investigated conditions.

Our application of SECAT to the HeLa-CC dataset [35] illustrates that different network states can efficiently be visualized in a network-centric representation to highlight the complex relations of different qualities. This provides a bird’s-eye view of alterations in PPI networks that can be used intuitively to guide follow-up investigations. SECAT is available for all platforms as open source software implemented in Python and is compatible with different LC-MS/MS profile and reference PPI database formats. We expect that our toolkit and the underlying concepts will be particularly useful for future PCP datasets, guiding the qualitative and quantitative comparison of multiple conditions, where protein complexes represent dynamic rather than static modules.

## Methods

### The Size-Exclusion Chromatography Algorithmic Toolkit

#### Input data

The primary input data for SECAT are quantitative, proteotypic/unique peptide-level profiles, e.g. acquired by SEC-SWATH-MS [3, 35]. The input can be supplied either as matrix (protein, peptide and run-wise peptide intensity columns) or as transposed long list. Protein identifiers need to be provided in UniProtKB/Swiss-Prot format. The column names can be freely specified, and example files are provided and referred to in the online documentation.

The second required input file represents the experimental design and molecular weight calibration of the experiment [3, 35]. The primary column is the run identifier (matching the quantitative profiles above), with additional columns for SEC fraction identifier (integer value), SEC molecular weight (float value, as specified previously [3, 35]), a group condition identifier (freetext value) and a replicate identifier (freetext value). The column names can be freely specified, and example files are provided and referred to in the online documentation.

The third required file covers matching UniProtKB/Swiss-Prot meta data in XML format and can be obtained from UniProt.

Optionally, reference PPI networks can be specified to support semi-supervised learning and to restrict the peptide query space. SECAT can accept three files: A positive and a negative reference network for the learning step and a separate reference network to restrict the query space. SECAT natively supports HUPO-PSI MITAB (2.5-2.7), STRING-DB, BioPlex and PrePPI formats and provides filtering options to optionally exclude lower confidence PPIs. Example files for CORUM (version 3.0), PrePPI (version 2016) and STRING (version 11.0) are provided and referred to in the online documentation.

#### Data preprocessing

By default, SECAT first normalizes the quantitative profiles on peptide-level. To reduce local fluctuations in total protein abundance between individual samples, but to still conserve the global distribution of the SEC-fraction-dependent protein abundances, SECAT implements a sliding window-based approach: Cyclic lowess [58, 59] (span=0.7, iterations=3) is applied group-wise to all conditions and replicates by a sliding window (default: *N* = 5) and a step-size (default: *N* = 1) over the SEC fraction indices and the average intensity is computed for each peptide in each SEC fraction. The sliding windows are by default “padded”, meaning that the average of the first and last window frames is computed over the restricted set of covered fractions only. Supplementary Fig. 7-10 illustrate the effects of the normalization on the total protein abundance profiles.

Using the user-provided molecular weight calibration of the experiment [3, 35], the apparent molecular weights of the SEC fractions are matched with the reference molecular weights for each monomeric subunit to identify the closest SEC fraction index, representing a protein-specific “monomer threshold”. In this process, a user-defined factor (default: *F* = 2) can be specified that is used to multiply the reference molecular weights prior to matching to account for potential homomultimers.

By default, SECAT does not conduct strict prefiltering or peak-picking of the SEC elution profiles. Optionally, peptide-level detrending including or excluding zero values can be conducted, where the quantitative matrix *I_s_*, represents sample *s*. *I_s_* consists of peptide intensities *i_s_* with rows, representing peptides of the complete set *P* with index *p* and columns, representing runs of the complete set *R* with index *r*. *I_s_* can be transformed to *I_s,detrend_zero__* or *I_s,detrend_drop__*:

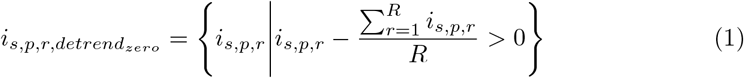

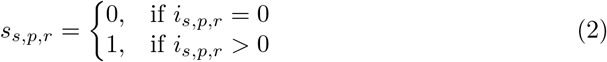

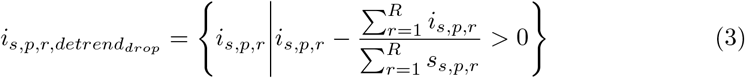

As alternative option, a local maximum (“local-max”) peak-picking algorithm can be applied on protein-level, either individually per sample or averaged over the replicates of the same conditions under the same assumptions as stated above. First, the quantitative matrix is extended to protein-level *J_s_* for sample *s* or averaged over all replicates of condition *c*, 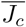, with rows, representing proteins of the complete set *O* with index *o* and *o_p_* indicating the set of peptides *p* mapping to the protein with a minimum (default: *N* = 1) and maximum (default: *N* = 3) peptides, sorted according to decreasing total intensity over the full SEC gradient.

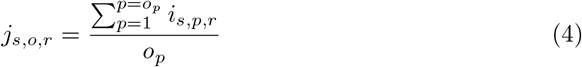

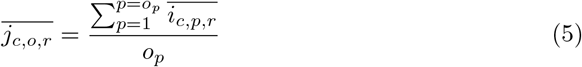

Protein level arrays are then used as input for the SciPy [60] local maxima peak-picking function “scipy.signal.find_peaks” (minimum width=3, relative height=0.9), which returns a binary vector *k_s,o_*(*r*) for each protein indicating the peak boundaries. Using this vector, the peptide-level matrix is transformed: Peptide intensities *i_s,p,r_* are set to zero if the binary vector *k_s,o_*(*r*) of mapping protein *o* at run index *r* is zero, indicating outside-peak boundaries:

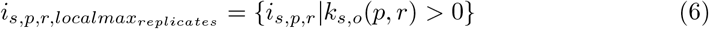

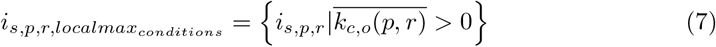

#### Signal processing

Candidate PPIs, either defined by the reference PPI networks or by computing all pairwise combinations, are evaluated in the signal processing module. First, a minimum (default: *N* = 1) and maximum (default: *N* = 3) number of peptides are selected according to total intensity computed over the full SEC profile and all samples. If there are at least 3 consecutive non-zero values on protein-level, a vector of 11 scores is computed within the assembled fractions of each candidate PPI, where the input data are the preprocessed SEC profile intensities of the peptides corresponding to the two interactor proteins. The peptide *p_A,j_* denotes the *j*-th ranking peptide with the complete set *J* of protein *A*, whereas *i_A,j,r_* denotes the peptide intensity of that peptide in run *r* of the complete set *R*.

##### Cross-correlation-based scores

Inspired by the chromatographic cross-correlation-based scores of mQuest/mProphet [42], two scores are computed by comparing the peptides of protein *A* with the peptides of protein *B*.

First, peptides intensities are normalized over all runs:

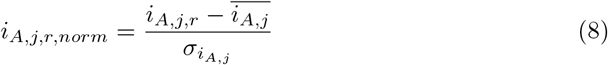

Second, the cross-correlation function between each combination of peptides of protein *A* and *B* is computed using the NumPy function “numpy.correlate”:

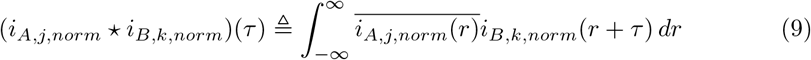

Based on this function, two scores are derived. *xcorr_shape_* describes the average of all normalized peptide pair convolution products retrieved at full signal overlap. *xcorr_shift_* describes the maximum difference between the intersection and protein *A* or *B* in *xcorr_apex_*, which represents the average delay *τ* at which the cross-correlation is maximal:

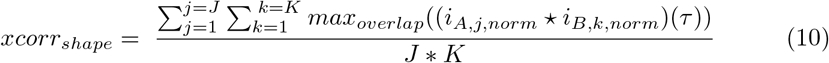

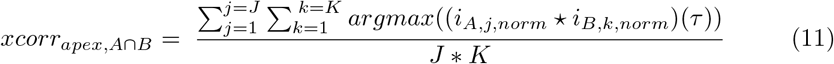

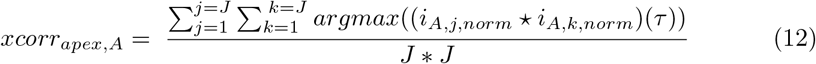

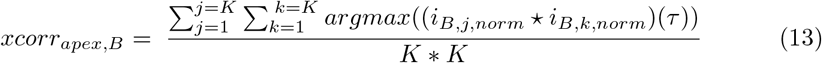

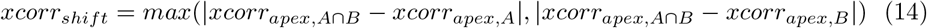

##### Monomer-based scores

Two scores are computed to measure the distance in SEC fractions between monomers of proteins *A* and *B* and their PPIs. *m_A_* denotes the monomer threshold computed for protein *A*:

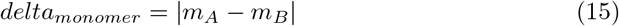

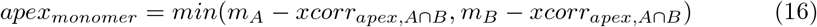

##### Maximal and total information coefficient-based scores (MIC/TIC)

Mean equicharacteristic *mic* and *tic* scores are computed for all peptide combinations between proteins *A* and *B* using the minepy package [44]:

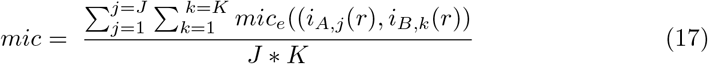

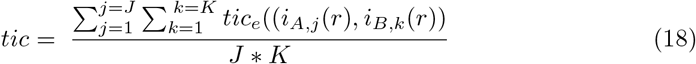

##### SEC profile intersection-based scores

Two scores are computed to describe the intersection of the protein profiles. The intersection of proteins *A* and *B* over the SEC profile is defined as true at index *r*, if any peptide of protein *A* and any peptide of protein *B* have non-zero intensities:

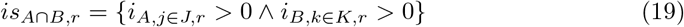

The union of proteins *A* and *B* over the SEC profile is defined as true at index *r*, if any peptide of protein *A* or *B* has non-zero intensities:

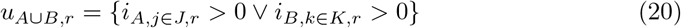

The score *sec_intersection_* describes the maximum stretch of consecutive intersecting fractions:

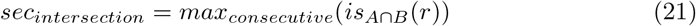

The score *sec_overlap_* represents the Jaccard Index and describes the total intersection divided by the total overlap.

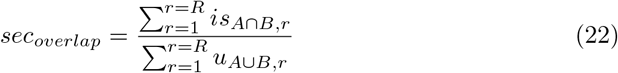

##### Protein-abundance-based scores

Two scores are derived to describe the relative ratio of proteins *A* and *B*. First, the peptide intensities are summarized over the intersection or the full SEC profile:

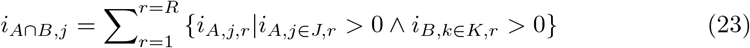

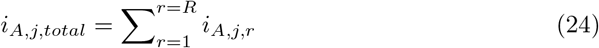

Second, for each protein an abundance metric is computed by averaging the peptide intensities:

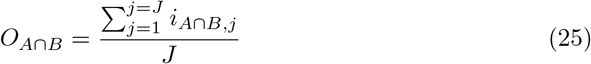

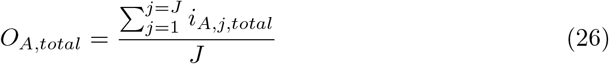

The score *abundance_ratio_* defines the relative abundance ratio between the intersection of proteins *A* and B. The score *total_abundance_ratio_* defines the abundance ratio between the full SEC profiles of proteins *A* and *B*. If the values are larger than 1, the inverse values are computed:

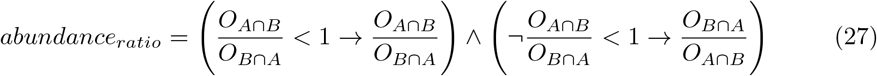

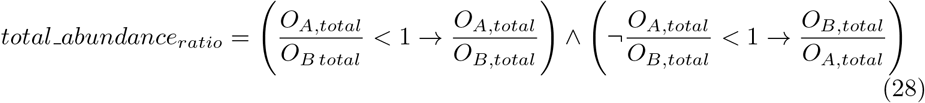

##### Kickstart score

To initialize semi-supervised learning, a *kickstart* score is computed to select PPIs that co-elute, have similar shape, and similar interactor mass, where values range between 0 and 1 with higher values indicating better signals:

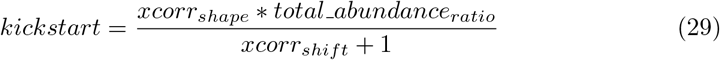

#### PPI detection

Spontaneous co-elution of protein subunits in PCP datasets represents a considerable challenge to identify or detect PPIs. While partial scores can describe properties to discriminate true from false candidate PPIs, several properties need to be combined to achieve sensitivity and selectivity. For this reason, supervised learning of classifiers for PPI identification using a ground truth dataset was a critical component of all previous approaches [3, 26, 27, 29, 31–34].

The CORUM [38] protein complex reference database has been used previously for this purpose. For the application in SECAT, the complexes were transformed to PPIs, representing the positive ground truth dataset. For the negative ground truth dataset, we adapted the approach proposed by the PrInCE algorithm [33]. It leverages CORUM PPIs as true and all other interactions of CORUM proteins that are not included in the database (CORUM-inverted) as false PPIs. Since proteins in CORUM complexes are well characterized and, for the most part, supported by 3D structure data, this strategy assumes that any true interactions within that set of proteins should be known already. As such, identified or detected interactions that are not reported are likely false positives [33]. For the purpose of the benchmark we further excluded any known or predicted interactions from CORUM-inverted and split the combined true/false reference set into equally sized training/validation and hold-out subsets.

However, since SECAT does by default not conduct strict prefiltering and uses considerably more candidate PPIs for machine learning and scoring, spontaneous co-elution events of (partially) overlapping interactors within the positive ground truth dataset would lead to an accumulation of false positives. For this reason, a semi-supervised learning approach, inspired by the solutions developed as part of Percolator [61], mProphet [42] and PyProphet [45, 46] for related challenges within computational proteomics was implemented, where the positive ground truth set is learned over several iterations, ensuring both selective and sensitive scoring of candidate signals.

The input for the semi-supervised learning step are the partial score vectors for the positive and negative ground truth dataset. Optionally, two filters can be applied to remove the most unlikely candidate PPIs. The minimum abundance ratio filter (default: *F* = 0.1) ensures that only PPIs within a maximum 10-fold difference of protein abundance ratios are considered. The maximum SEC fraction shift filter (default: *F* = 10) ensures that only PPIs with maximum elution peaks within 10 SEC fractions are considered. The PyProphet learning module is then applied to the ground truth dataset.

##### Semi-supervised learning

This step is conducted essentially as described before [42] with modifications:

1. Cross-validation is conducted with a randomly sampled fraction *f* of the data (default:*f* = 0.8) and repeated *r* (default: *r* = 10) times.

a. Initialization of semi-supervised learning:

i. All negative ground truth PPIs of the cross-validation fold are used. The *kickstart* score is used to select the initial positive ground truth PPI subset at a defined *q*-value threshold (default: *q* = 0.1).
ii. As part of the initialization step, all partial scores, except the *kickstart*score are then used to train an XGBoost [47]-based classifier. Hyperparameters can optionally be tuned as described below, but a default set (num_boost_round=100, early jstopping_rounds=10, test_sìze=0.33, eta=1.0, gamma=0, max_depth=6, min_child_weight=1, subsample=1, colsample_bytree=1, colsample_bylevel=1, colsample_bynode=1, lambda=1, alpha=0, scale_pos_weight=1, silent=1, objective=binary:logitraw, nthread=1, eval_metric=auc) is provided.
iii. Based on this classification, the positive ground truth dataset is re-scored for the next iteration.
b. Iterate *i* (default: *i* = 3) times:

i. All negative ground truth PPIs of the cross-validation fold are used. The previous classifier is applied to the ground truth dataset to select the iteration positive ground truth PPI subset at a defined *q*-value threshold (default: *q* = 0.05).
ii. Classification of the new dataset is conducted as described above in the initialization step.
iii. Based on this classification, the positive ground truth dataset is re-scored for the next iteration.
c. The classifier discriminant scores are normalized relative to the negative ground truth data points by subtracting the mean and dividing by the standard deviation of the negative ground truth data points as described previously [42].
2. Classifier discriminant scores are averaged over all cross-validation folds. In combination with the negative ground truth dataset, this score is used to select the final positive ground truth PPI subset at a defined *q*-value threshold (default: *q* = 0.05). A final classifier is trained and stored, which will later be used and applied to the full dataset for classification. Optionally, hyperparameters can be tuned at this stage using the hyperopt framework [62] in a hierarchical fashion within 10 rounds optimizing the evaluation metric “auc”: 1) Complexity hyperparameters: max_depth=(2,8) and min_child_weight=(1,5), 2) Gamma hyperparameter: gamma=(0.0,0.5), 3) Subsampling hyperparameters: subsample=(0.5,1.0), colsample_bytree=(1.0), colsample_bylevel=(1.0), colsample_bynode=(1.0), 4) Regularization hyperparameters: lambda=(0.0,1.0), alpha=(0.0,1.0), 5) Learning rate: eta=(0.5,1.0). Integer ranges are sampled by a quantized uniform distribution, floating number ranges are sampled by a uniform distribution. In our applications, we found that the default hyperparameter set described above was applicable to all tested datasets and we thus omitted autotuning in further iterations.

##### Statistical validation

The Storey-Tibshirani *q*-value framework [63] is used to assess false-discovery rates. First, empirical *p*-values are estimated for the positive ground truth dataset using the negative ground truth dataset as null model [63]. *q*-values are then estimated as described previously [63] with the following parameters in all iterations: pi0_lambda=(min=0.01,max=0.5,step=0.01), pi0_method=bootstrap.

##### Incorporation of prior information from reference PPI networks

If a reference PPI network was used to restrict the query space, the optionally provided confidence scores can be incorporated during statistical validation. Because these scores often only to some extent represent the PPIs measurable by PCP datasets and can be differently calibrated across databases, SECAT uses them to compute a grouped FDR. Queried PPIs are grouped according to their prior score in N predefined confidence bins (default: *N* = 100) and *q*-value estimation is conducted for each bin separately. This enables multiple hypothesis testing correction to account for different prior probabilities of false detection of interactions and is generically applicable to confidence scores with different statistical properties.

##### Integration across multiple replicates

If multiple replicates of the same biological condition are scored together, a global *q*-value is computed for each PPI by prior computing of the average of the classifier scores over all replicates.

#### PPI quantification

SEC peptide profiles can be partitioned into components to represent quantitative information on different levels. Based on the peptide-level SEC profiles, SECAT computes four quantitative metrics for selected peptides of a protein (default: minimum *N* = 1; maximum *N* = 3 peptides per protein, ranked according to total protein abundance):

##### Total intensity

The sum of the full elution profile corresponds to the total peptide abundance, which to some degree represents protein abundances measured by conventional, non-fractionated LC-MS/MS. Assembled intensity: Based on the protein-specific monomer threshold defined in the preprocessing step, the peptide signals of all assembled fractions (left hand-side of monomer threshold) are summarized.

##### Monomer intensity

Based on the protein-specific monomer threshold defined in the preprocessing step, the peptide signals of all monomer fractions (right hand-side of monomer threshold) are summarized.

##### Interactor intensity

To compute the PPI-level quantities, for each PPI below a specified confidence threshold (default: integrated *q*-value < 0.05 in any of the compared conditions), the intersecting fractions of the two interactor protein profiles are analogously extracted on peptide level.

These summarized peptide-level quantities are then *log*_2_-transformed and the *log*_2_-fold-change between groups is computed. The intensities are further used by the proteoVIPER module for differential quantitative protein and PPI assessment. proteoVIPER is based on the VIPER algorithm [41], which was originally developed to assess protein activity from transcriptomic profiles using gene regulatory networks. Using the peptide-protein relationships, proteoVIPER computes three differential protein-level metrics, total, assembled and monomer abundance to describe changes between the groups. In addition, proteoVIPER computes three differential PPI-level metrics based on the peptide interactor intensities of the two proteins within a PPI: interactor and complex abundance and interactor ratio.

The main component of VIPER and thus proteoVIPER is the analytic rank-based enrichment analysis (aREA) module, which tests for a global shift in the position of the peptides mapping to the same protein or PPI when projected on the rank-sorted peptide intensities of a run on separate levels [41]. The description of the algorithm below is adapted from the original publication [41]:

1. To compute total, assembled and monomer abundances, the peptides mapping to the protein-of-interest are used. To compute the interactor abundance, for each PPI, the peptides of each interactor are separately assessed with a positively correlated mode of interaction. To compute complex abundance, the peptides of proteins *A* and *B* are used with the same, positively correlated mode of interaction. To compute interactor ratio, the peptides of proteins *A* and *B* are used with negatively correlated mode of interaction.
2. The means of the quantile-transformed rank positions are used as test statistic (enrichment score), which is computed twice:

a. First, a one-tail approach is used based on the absolute value of the peptide intensities, which rank-sorts proteins from the less invariant between groups to the most differentially abundant.
b. Second, a two-tail approach is used, where the positions of the peptides of one interactor is inverted (negatively correlated mode of interaction) within the peptide intensity signature to compute the interactor ratio enrichment score.
3. The one-tail and two-tail enrichment scores are then integrated exactly as described previously [41], but with equal confidence for each peptide. The scores are normalized and calibrated against the reference exactly as described previously [41].

The quantitative metrics reported by proteoVIPER have several advantages: First, quantitative changes of proteins between different samples can also be assessed by only partially overlapping sets of peptides. Second, complex abundance changes can be estimated by contributions of the peptides of both interactor proteins. Third, by assessing the interactor proteins in a negatively correlated mode of interaction, a differential metric for changes affecting the ratio of the interactors can be derived, which to some extent represents a metric describing changes in complex stoichiometry. proteoVIPER reports six quantitative matrices, representing protein or PPI metrics on each level that can be used for differential comparisons between the samples and conditions. Experimental conditions can then be statistically compared by independent *t*-tests [41] on each level.

#### Network inference

Using the PPI-level quantitative metrics from above, SECAT conducts network-centric data integration. For each protein, the test statistics and proteoVIPER normalized enrichment scores of its PPIs are integrated using Empirical Brown’s Method [49]. The evidence of multiple measured PPI is thus summarized to protein complex metrics summarizing changes in protein complex abundance or interactor ratio for each protein. Notably, highly correlated PPI (e.g. from the same protein complex) are integrated in a dependent fashion, whereas independent PPI (e.g. from different protein complexes) combine and increase the significance of the protein complex engagement metric.

In a network context, this helps to identify the most perturbed or dysregulated proteins based on changes of their protein complexes. Instead of clustering or inferring protein complexes, which are difficult to define in presence of subcomplexes across multiple experimental conditions, SECAT’s metrics can be more robustly characterized from a partial subset of their interactions.

Integrated *p*-values are adjusted for multiple testing using the Benjamini-Hochberg [64] approach, as suggested in the original publication [49].

### Primary data analysis

Processed mass spectrometry datasets have been obtained from the repositories linked by the original publications or the authors of the corresponding publications.

### SECAT data analysis

SECAT (version 1.0.5), PyProphet (version 2.1.4) and VIPER (version 1.20) were used for all data analyses with CORUM [38] (version 3.0), PrePPI [22] (version 2016) and STRING [19] (version 11.0) and default parameters if not otherwise specified. Semi-supervised learning was conducted using CORUM as positive network and CORUM-inverted as negative network. The inverted CORUM reference PPI network was generated by using the inverted set of PPIs (i.e. all possible PPIs that are not covered by CORUM) and removing all PPI in this set covered by STRING (version 11.0), IID [65] (version 2018-11), PrePPI (version 2016) or BioPlex [18] (version 2.0).

All input and output data and used parameters are provided on the Zenodo repository to reproduce all analysis steps.

### Parameter selection and validation of signal processing and PPI detection modules

The SECAT PPI detection benchmark was conducted by using 50% of the CORUM reference PPI network for learning and the other fraction for evaluation. For all assessments, this random selection was conducted on PPI-level, except for the algorithm comparison, where the selection was conducted on complex-level. Reference false PPI from CORUM-inverted were randomly selected in predefined ratios (1:0 – 1:16) and added to the target set for evaluation but not learning.

Fig. 2a and Supplementary Fig. 1a depict violin plots with the following parameters: Lower and upper hinges represent the first and third quartiles; the bar represents the median. This represents the default parameters of the function “geom_violin” of ggplot2.

Fig. 2b and Supplementary Fig. 1b were generated by assessing the PPIs with a global-context*q*-value < 0.05 and decomposing the number of PPIs for detection amongst replicates at different confidence thresholds.

Fig. 2e and Supplementary Fig. 1e were generated by using the ground truth CORUM and CORUM-inverted reference values. Because the estimated *q*-values are dependent on the combined reference sets with unknown ratios of true and false PPIs, the “true *q*-values” were corrected by a factor, which accounts for the PyProphet estimated proportion of false targets in the 1:0 dilution step.

For Supplementary Fig. 2, the CORUM reference PPI network was similarly used as described above to generate a pseudo-ground truth dataset for classifier training. However, for the validation subset, an excess of 10 times as many false interactions as targets prior to analysis was added and subsets were generated by randomly selecting protein complexes instead of PPIs. For the downstream analysis, the CORUM targets only were reduced to the intersection with a previously published [3], manually curated annotation of the dataset.

The CCprofiler [3] analysis (git revision: 39650f2) was conducted as suggested by the software documentation. All input data and parameters are provided on the Zenodo repository.

The EPIC [27] analysis (git revision: b6432b9) was conducted as suggested by the software documentation with the provided Docker container. The input data was first aggregated from peptide-level to protein-level using the top3 method implemented in aLFQ [66] (version 1.3.5). All input data and parameters are provided on the Zenodo repository.

### Validation of PPI quantification & network integration modules

The data was analyzed as described above with the full CORUM, PrePPI and STRING reference networks and a network encompassing all potential PPIs. Fig. 3c-f were generated using the STRING-based analysis. Fig. 3d depicts boxplots with the following parameters: Lower and upper hinges represent the first and third quartiles; the bar represents the median. Lower and upper whisker extend to 1.5 * IQR from the hinge. This represents the default parameters of the function “geom_boxplot” of ggplot2.

Supplementary Fig. 3 was generated by computing the node degrees with the “NetworkAnalyzer” module of Cytoscape [67] (version 3.7.2) and visualization of the density distributions of each network using the function density.compare from the R-package “sm” (version 2.2-5.6) with default parameters.

### Molecular mechanisms differentiating HeLa cell cycle states

To annotate and visualize differential proteins in Fig. 4 and Supplementary Fig. 4-5 between the HeLa cell cycle states identified by SECAT, we used Cytoscape [67] (version 3.7.2). CORUM (version 3.0) or Reactome [54] (version 71) was used to cluster PPI using the Cytoscape App AutoAnnotate [68] (version 1.3.2) with default parameters and a maximum cluster size (“Max words per label”) of 1. Clusters were arranged according to the CoSE layout.

Visualization of protein-level SEC-SWATH-MS profiles in Fig. 5 and Supplementary Fig. 6 was conducted using the R-package “ggplot2” by averaging the three most intense peptide precursors per protein.

### Source code availability

SECAT is available as platform-independent open source software under the Modified BSD License and distributed as part of the SECAT (https://pypi.org/project/secat) and PyProphet (https://pypi.org/project/pyprophet) Python PyPI packages. SECAT further depends on the R/Bioconductor package “viper”, which is distributed under a non-commercial usage license. Further documentation and instructions for usage can be found on the SECAT source code repository (https://github.com/grosenberger/secat).

### Data availability

All analysis results are available on Zenodo with the dataset identifier 10.5281/zenodo.3631786.

## Acknowledgments

We thank E.O. Paull and A.T. Griffin for discussions regarding the methodologies and N.E. Scott and L.J. Foster for providing access to the peptide-level data of their study [30]. G.R. is supported by grants P2EZP3_175127 and P400PB_183933 from the Swiss National Science Foundation. The project was supported by the European Research Council (ERC-20I40AdG 67082I) and the Swiss National Science Foundation (grant 31003A_166435 to R.A.).

## Author contributions

Conceptualization, G.R., M.H., R.A. and A.C; Formal analysis, G.R.; Funding acquisition: G.R., R.A.; Investigation: all authors; Methodology: G.R., M.H., I.B.; Resources: R.A. and A.C.; Software: G.R.; Supervision: R.A. and A.C.; Validation: G.R.; Visualization: G.R.; Writing – original draft: G.R., R.A. and A.C.; Writing – review & editing: all authors.

---

**Supplementary Figure 1.**
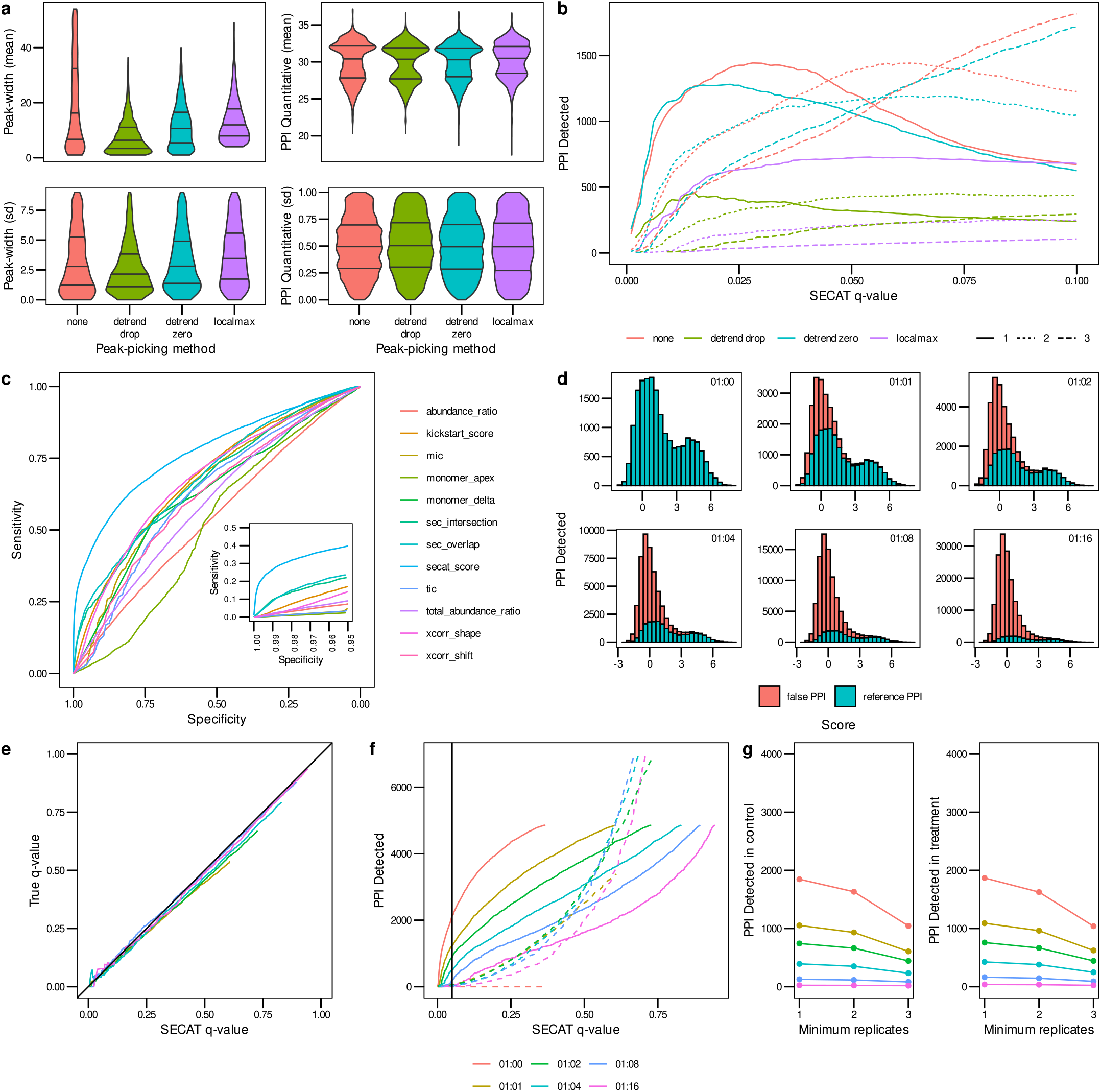
Signal processing and PPI detection performance evaluated using the Jurkat-Fas [30] dataset. **(a)** The effect of different peak-picking methods for signal processing on peak-width and PPI quantification within replicates of the same experimental condition is depicted in violin plots (lines representing 25, 50 and 75% quantiles respectively, Methods). **(b)** Sensitivity of PPI detection *vs*. SECAT *q*-value for peak-picking options. The PPI were filtered to a global context *q*-value < 0.05. Colors indicate different peak-picking options; line types indicate number of detections with the three replicates per condition. **(c)** The receiver-operating characteristic (ROC) illustrates the sensitivity *vs*. specificity of the different SECAT partial scores, the kickstart score and the integrated SECAT score. **(d)** The SECAT score histograms depict the dilution of the reference PPI network (true: CORUM) with false PPI (false: CORUM-inverted) for the benchmark (Methods). **(e)** The SECAT estimated *q*-value is accurate as evaluated against the ground truth for all levels of reference PPI network accuracy. For the dilution 1:0, no *a priori* false PPI are tested, thus the true FDR equals to 0. **(f)** The sensitivity of PPI detection in dependency of SECAT *q*-value for different levels of reference PPI network accuracy is depicted. Solid lines represent the true, whereas dashed lines represent the false PPI. **(g)** Consistency of PPI detection between replicates of the same conditions.

**Supplementary Figure 2.**
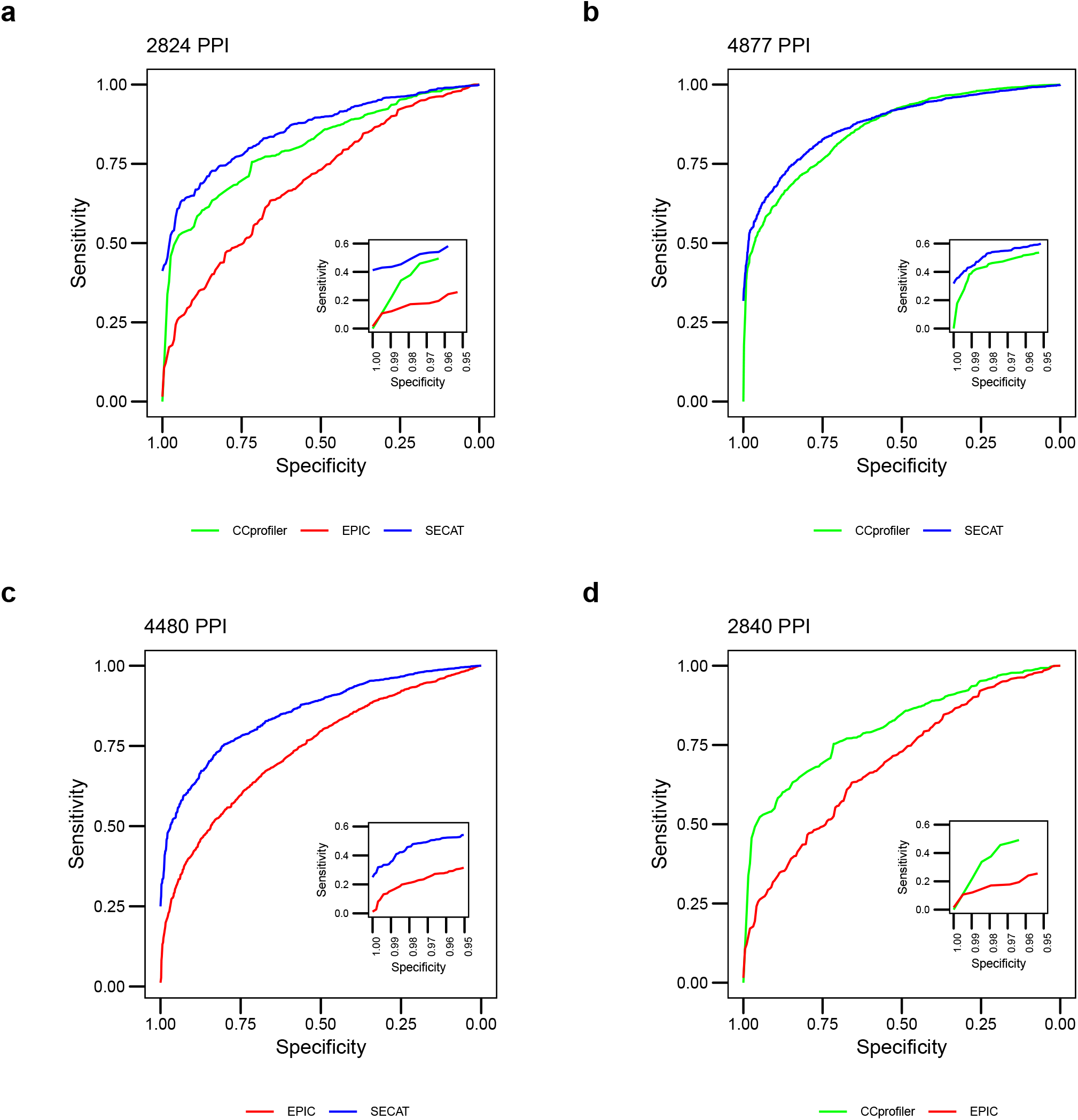
Receiver operating characteristic for the comparison of SECAT with EPIC and CCprofiler using the HEK293-EG [3] dataset.

**Supplementary Figure 3.**
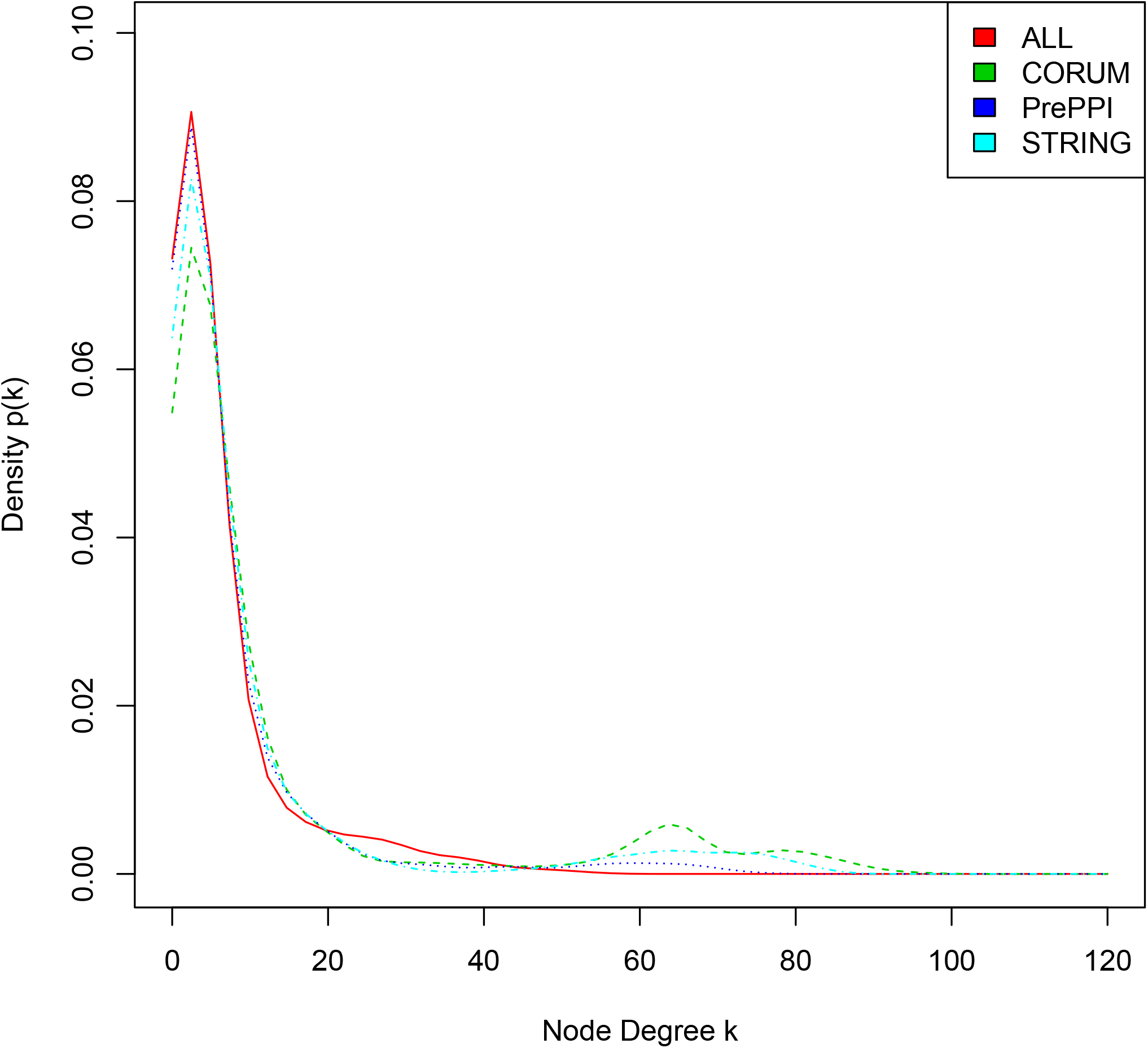
Comparison of node degree densities between different reference PPI networks and the network covering all possible PPI combinations.

**Supplementary Figure 4.**
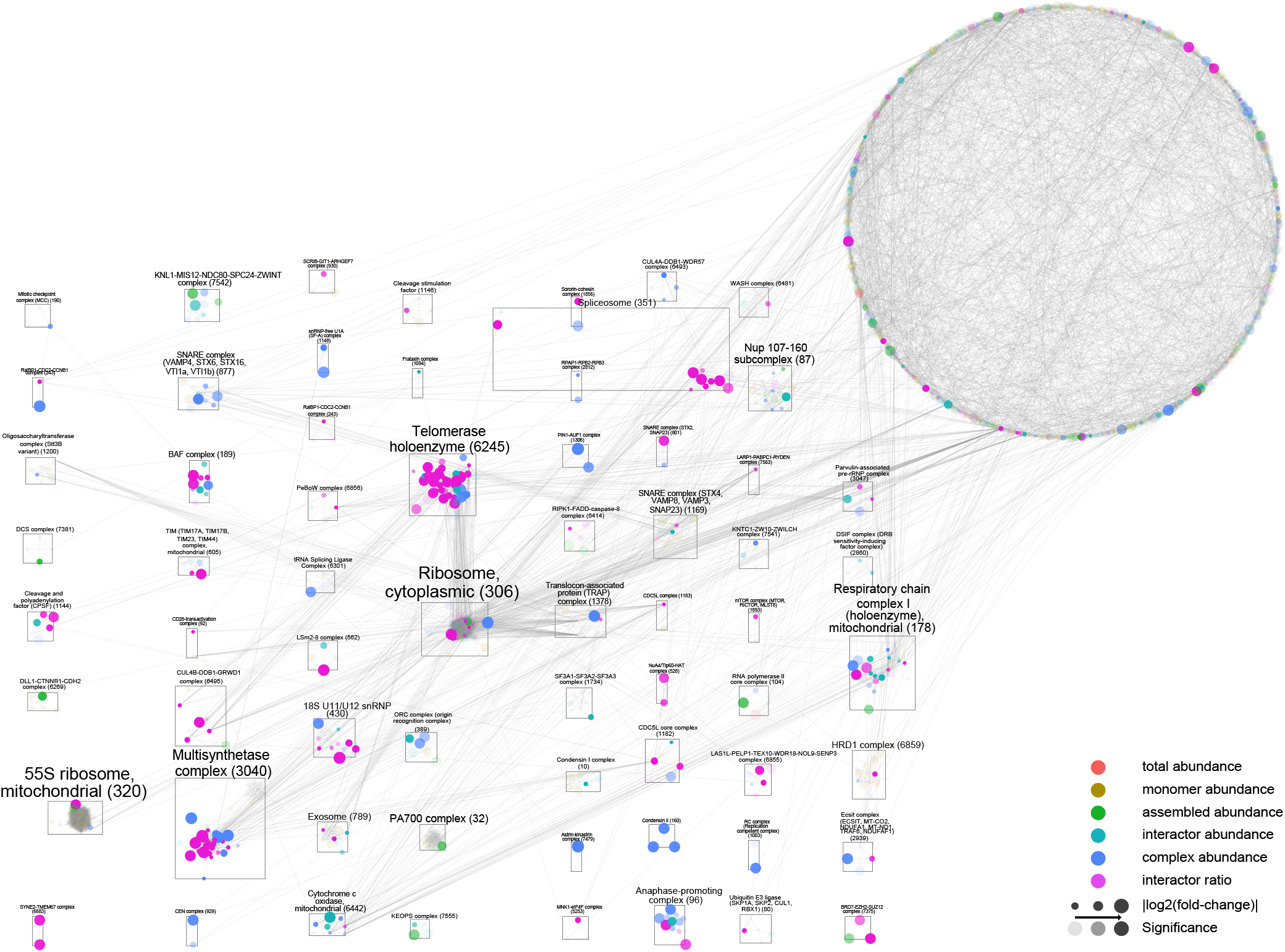
Extended complex-level molecular mechanisms differentiating HeLa cell cycle states. The integrated STRING-based PPI network of the HeLa cell cycle dataset [35] is depicted with proteins (nodes) and binary PPIs (edges) clustered against CORUM complexes (identifiers in brackets; Methods). Different colors indicate the most significant metric-level (legend, Methods). Node-size indicates effect-size and opacity indicates significance. The most prominent clusters highlight essential macromolecular assemblies. Interactive results (Cytoscape) are provided as Supplementary Data 1.

**Supplementary Figure 5.**
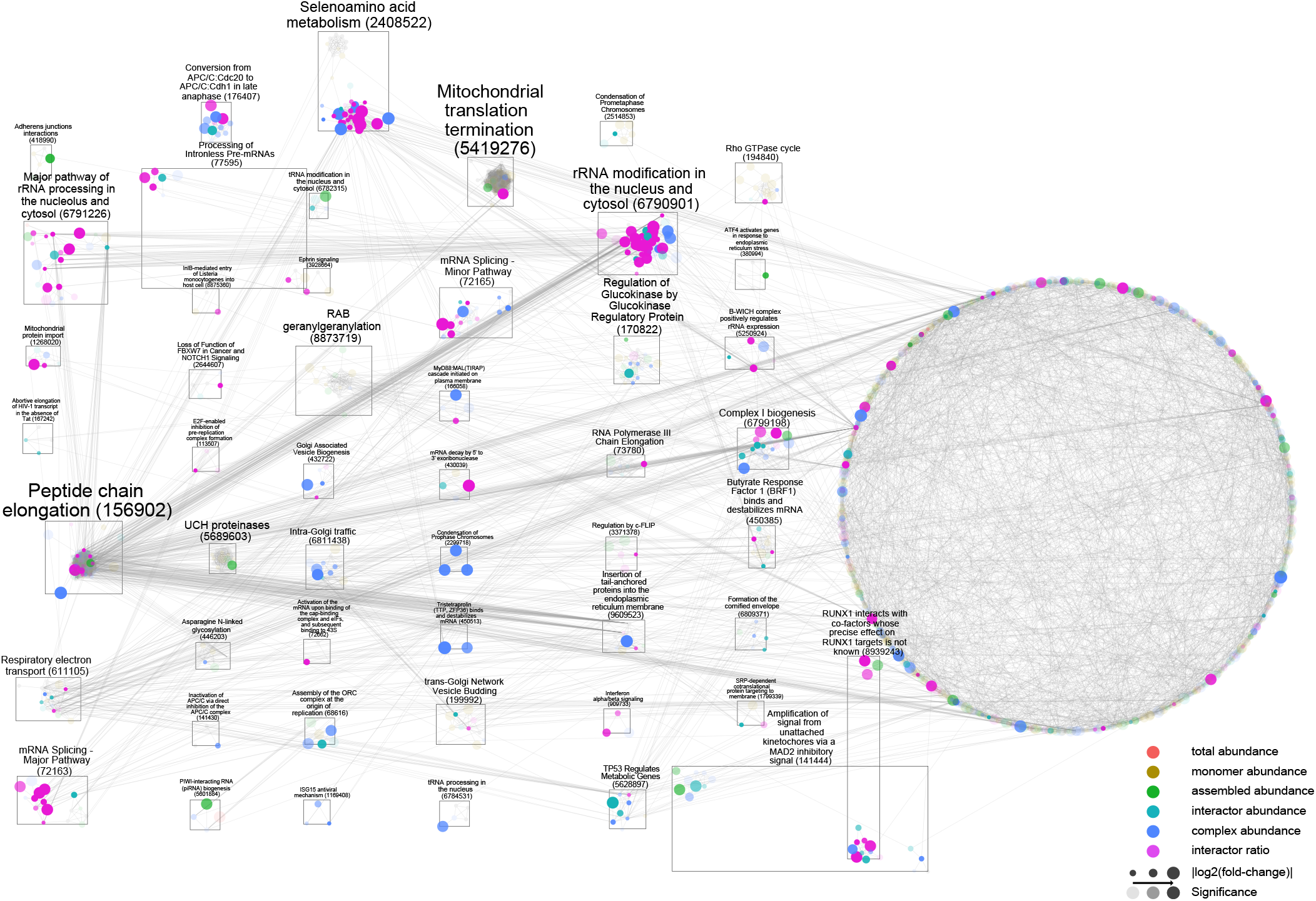
Pathway-level molecular mechanisms differentiating HeLa cell cycle states. The integrated STRING-based PPI network of the HeLa cell cycle dataset [35] is depicted with proteins (nodes) and binary PPI (edges) clustered against Reactome pathways (identifiers in brackets; Methods). Different colors indicate the most significant metric-level (legend, Methods). Node-size indicates effect-size and opacity indicates significance. The most prominent clusters highlight essential macromolecular assemblies. Interactive results (Cytoscape) are provided as Supplementary Data 1.

**Supplementary Figure 6.**
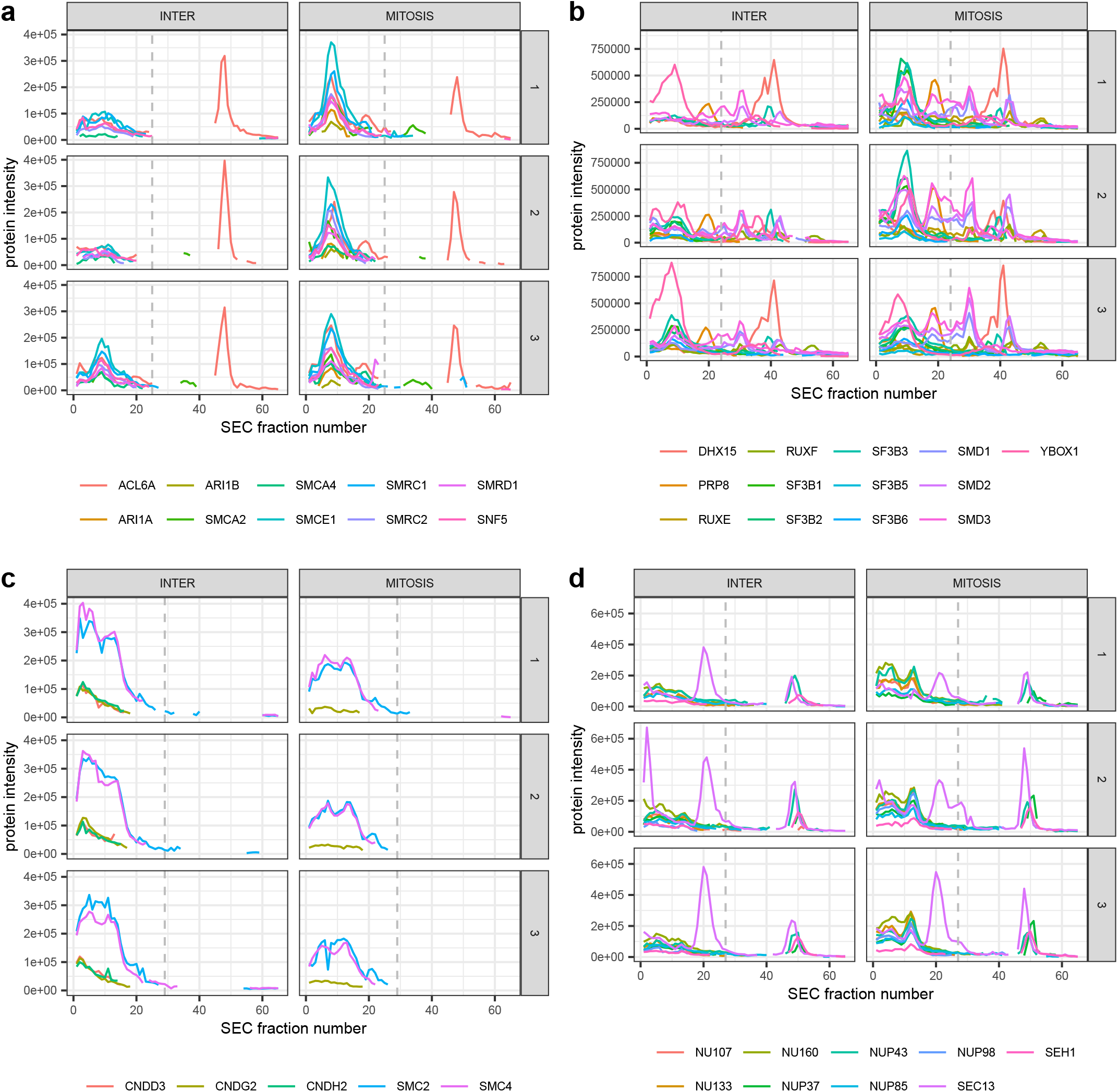
Extended protein-level SEC-SWATH-MS profiles of HeLa cell cycle states. Lines indicate different protein subunits, whereas the dashed line indicates the highest monomer threshold per group. **(a)** BAF complex. **(b)** 18S U11/U12 snRNP. **(c)** Condensin II. **(d)** Nup 107-160 subcomplex.

**Supplementary Figure 7.**
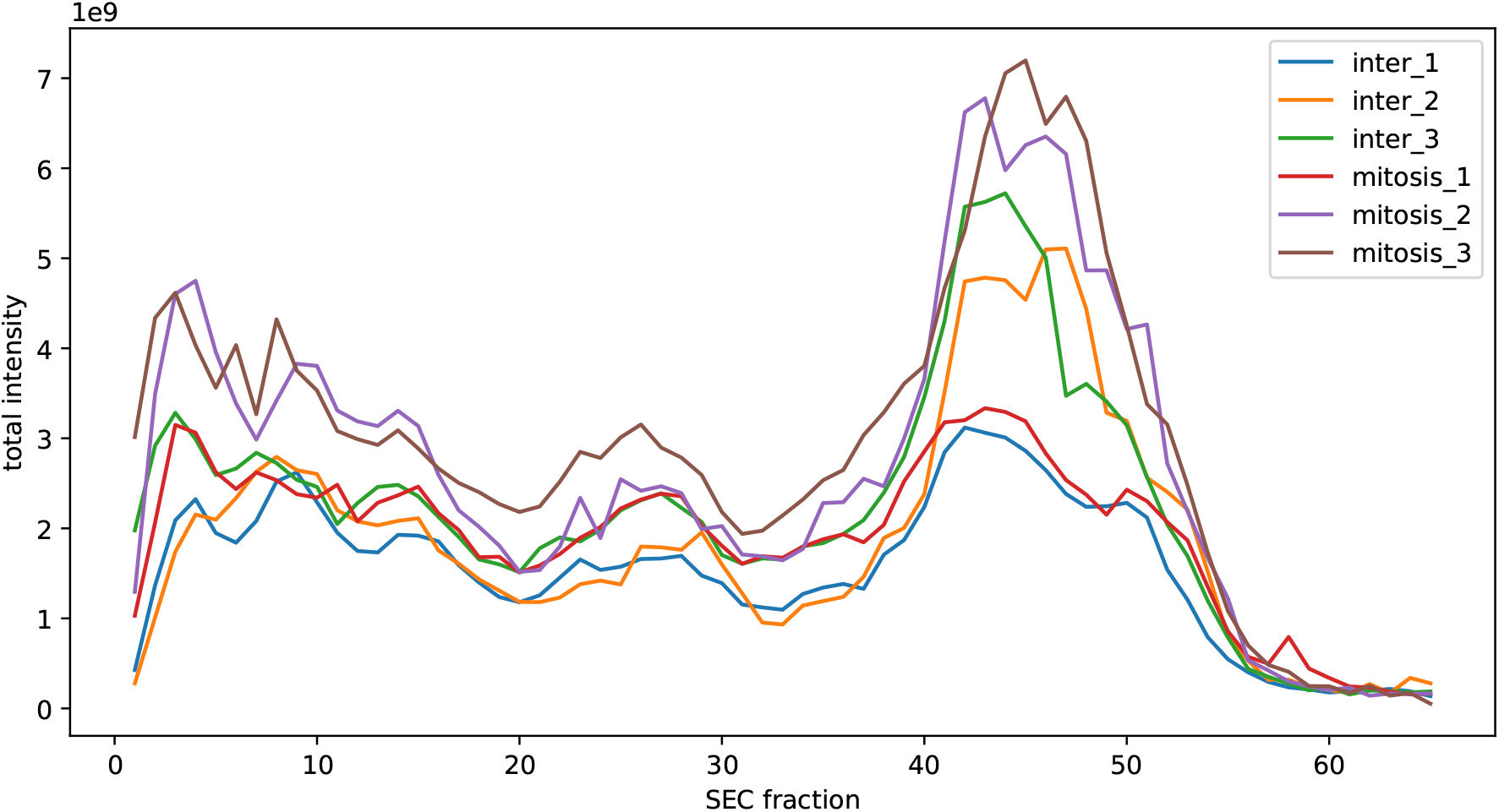
Summarized raw peptide precursor intensities over the SEC profile for the HeLa-CC [35] dataset.

**Supplementary Figure 8.**
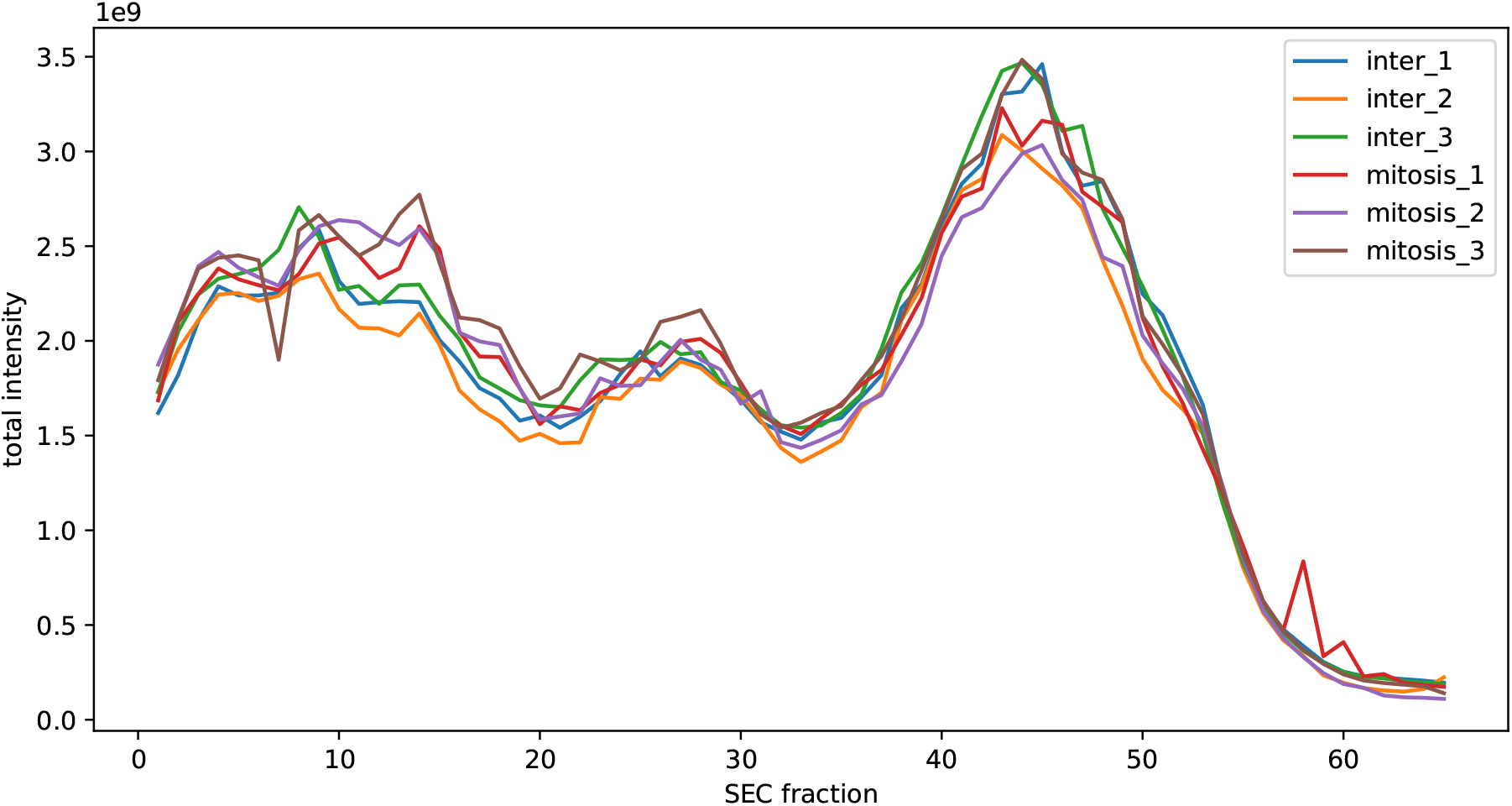
Summarized normalized peptide precursor intensities over the SEC profile for the HeLa-CC [35] dataset.

**Supplementary Figure 9.**
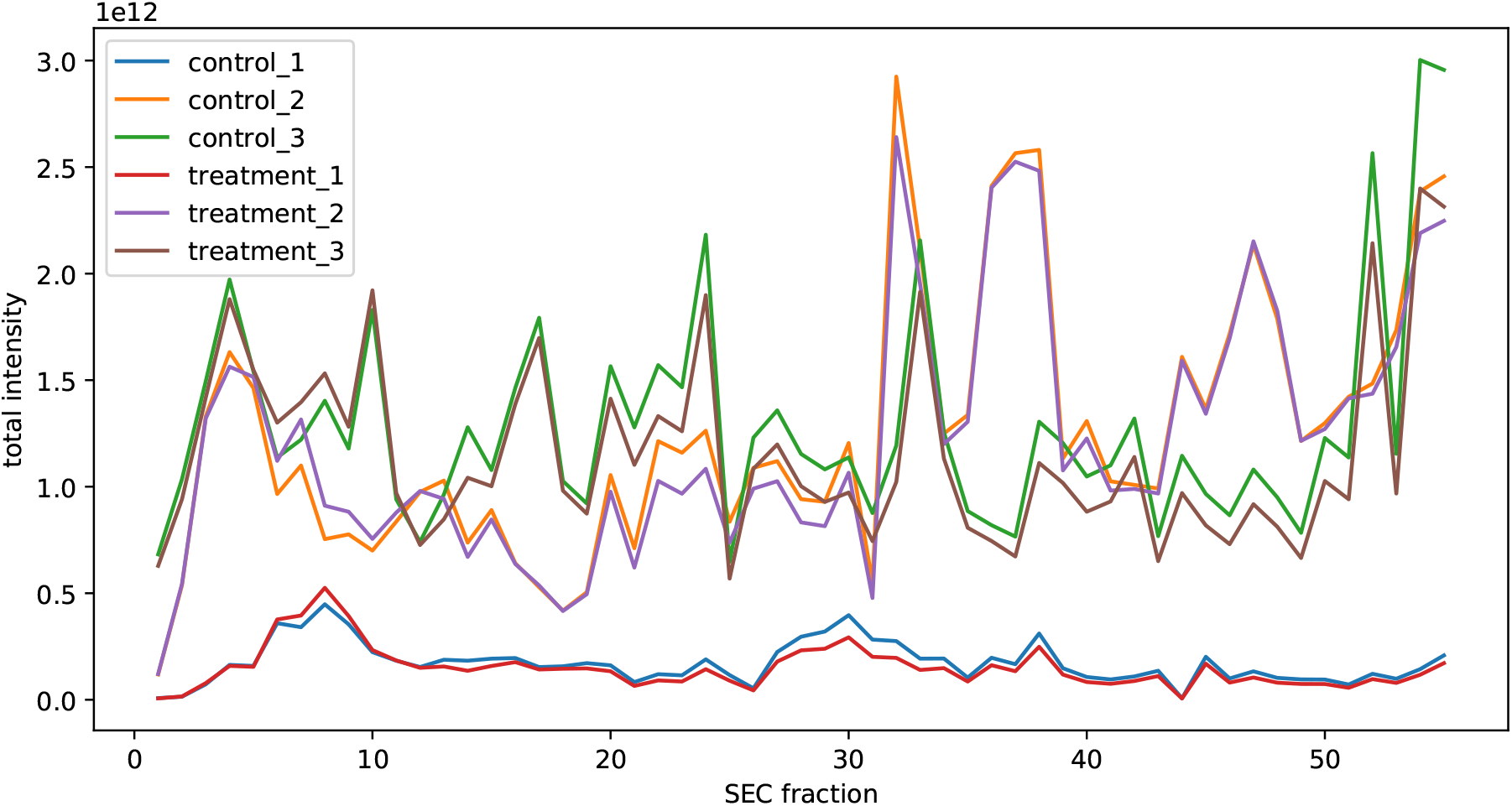
Summarized raw peptide precursor intensities over the SEC profile for the Jurkat-Fas [30] dataset.

**Supplementary Figure 10.**
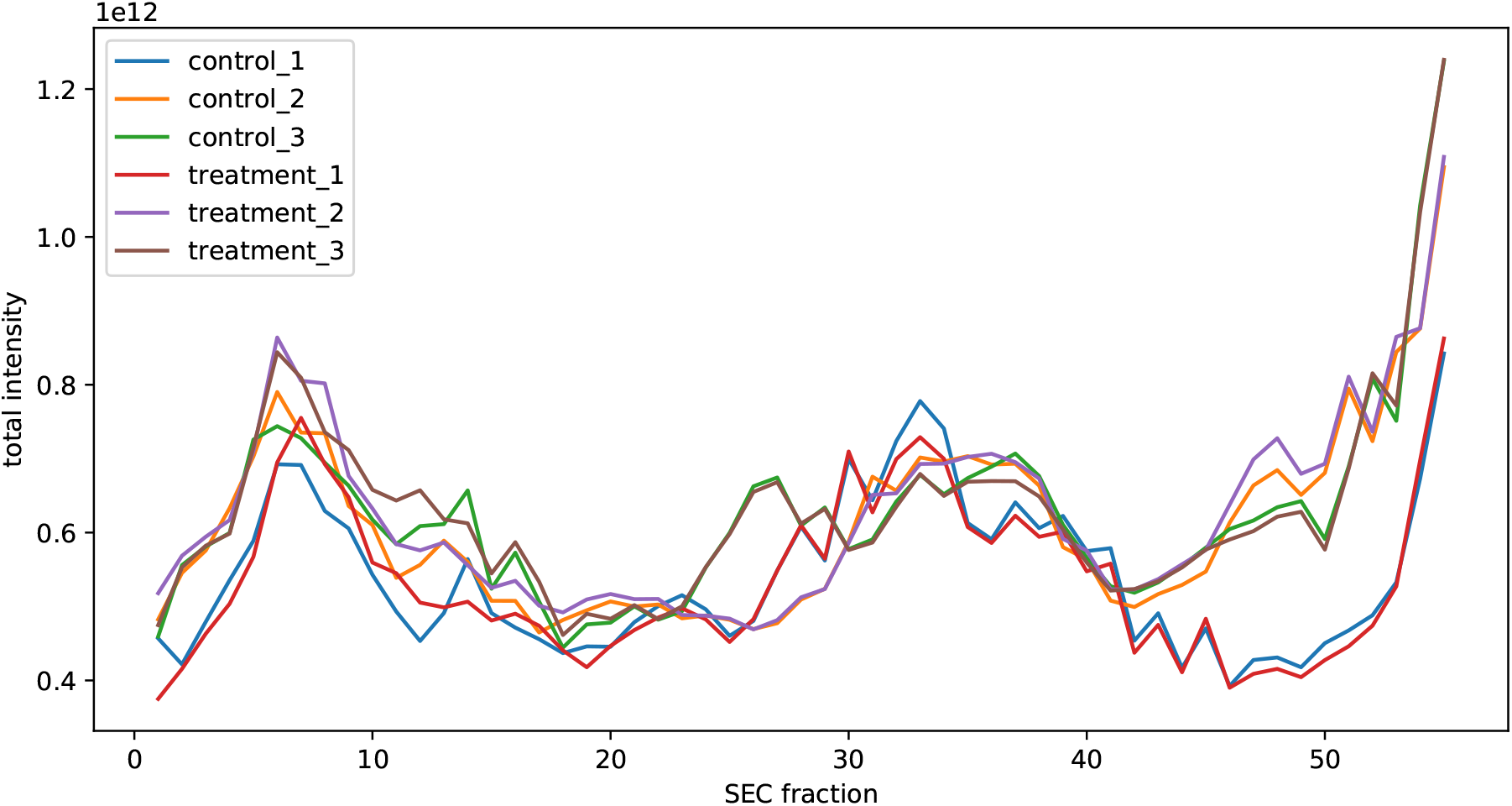
Summarized normalized peptide precursor intensities over the SEC profile for the Jurkat-Fas [30] dataset.

## References

1. Hartwell, L. H., Hopfield, J. J., Leibler, S. & Murray, A. W. From molecular to modular cell biology. Nature 402, C47–C52 (1999).

2. Campbell, I. D. The march of structural biology. Nature Reviews Molecular Cell Biology 3, 377–381 (2002).

3. Heusel, M. et al. Complex-centric proteome profiling by SEC-SWATH-MS. Molecular Systems Biology 15, e8438 (2019).

4. Aebersold, R. & Mann, M. Mass spectrometry-based proteomics. Nature 422, 198–207 (2003).

5. Aebersold, R. & Mann, M. Mass-spectrometric exploration of proteome structure and function. Nature 537, 347–355 (2016).

6. Gingras, A.-C., Gstaiger, M., Raught, B. & Aebersold, R. Analysis of protein complexes using mass spectrometry. Nature Reviews Molecular Cell Biology 8, 645–654 (2007).

7. Choi, H. et al. SAINT: probabilistic scoring of affinity purification–mass spectrometry data. Nature Methods 8, 70–73 (2011).

8. Herzog, F. et al. Structural Probing of a Protein Phosphatase 2A Network by Chemical Cross-Linking and Mass Spectrometry. Science 337, 1348–1352 (2012).

9. Krogan, N. J. et al. Global landscape of protein complexes in the yeast Saccharomyces cerevisiae. Nature 440, 637–643 (2006).

10. Sowa, M. E., Bennett, E. J., Gygi, S. P. & Harper, J. W. Defining the Human Deubiquitinating Enzyme Interaction Landscape. Cell 138, 389–403 (2009).

11. Bisson, N. et al. Selected reaction monitoring mass spectrometry reveals the dynamics of signaling through the GRB2 adaptor. Nature Biotechnology 29, 653–658 (2011).

12. Collins, B. C. et al. Quantifying protein interaction dynamics by SWATH mass spectrometry: application to the 14-3-3 system. Nature Methods 10, 1246–1253 (2013).

13. Keilhauer, E. C., Hein, M. Y. & Mann, M. Accurate Protein Complex Retrieval by Affinity Enrichment Mass Spectrometry (AE-MS) Rather than Affinity Purification Mass Spectrometry (AP-MS). Molecular & Cellular Proteomics 14, 120–135 (2015).

14. Lambert, J.-P. et al. Mapping differential interactomes by affinity purification coupled with data-independent mass spectrometry acquisition. Nature Methods 10, 1239–1245 (2013).

15. Varjosalo, M. et al. Interlaboratory reproducibility of large-scale human protein-complex analysis by standardized AP-MS. Nature Methods 10, 307–314 (2013).

16. Hein, M. Y. et al. A Human Interactome in Three Quantitative Dimensions Organized by Stoichiometries and Abundances. Cell 163, 712–723 (2015).

17. Huttlin, E. L. et al. The BioPlex Network: A Systematic Exploration of the Human Interactome. Cell 162, 425–440 (2015).

18. Huttlin, E. L. et al. Architecture of the human interactome defines protein communities and disease networks. Nature 545, 505–509 (2017).

19. Szklarczyk, D. et al. STRING v11: protein–protein association networks with increased coverage, supporting functional discovery in genome-wide experimental datasets. Nucleic Acids Research 47, D607–D613 (2019).

20. Orchard, S. et al. The MIntAct project - IntAct as a common curation platform for 11 molecular interaction databases. Nucleic Acids Research 42, D358–D363 (2014).

21. Drew, K. et al. Integration of over 9,000 mass spectrometry experiments builds a global map of human protein complexes. Molecular Systems Biology 13, 932 (2017).

22. Garzón, J. I. et al. A computational interactome and functional annotation for the human proteome. eLife 5, e18715 (2016).

23. Zhang, Q. C. et al. Structure-based prediction of protein-protein interactions on a genome-wide scale. Nature 494, 127 (2013).

24. Zhang, Q. C., Petrey, D., Garzón, J. I., Deng, L. & Honig, B. PrePPI: A structure-informed database of protein-protein interactions. Nucleic Acids Research 41, D828–33 (2013).

25. Foster, L. J. et al. A Mammalian Organelle Map by Protein Correlation Profiling. Cell 125, 187–199 (2006).

26. Havugimana, P. C. et al. A census of human soluble protein complexes. Cell 150, 1068–1081 (2012).

27. Hu, L. Z. et al. EPIC: software toolkit for elution profile-based inference of protein complexes. Nature Methods 2019 16, 737–742 (2019).

28. Kirkwood, K. J., Ahmad, Y., Larance, M. & Lamond, A. I. Characterization of Native Protein Complexes and Protein Isoform Variation Using Size-fractionation-based Quantitative Proteomics. Molecular & Cellular Proteomics 12, 3851–3873 (2013).

29. Kristensen, A. R., Gsponer, J. & Foster, L. J. A high-throughput approach for measuring temporal changes in the interactome. Nature Methods 9, 907–909 (2012).

30. Scott, N. E. et al. Interactome disassembly during apoptosis occurs independent of caspase cleavage. Molecular Systems Biology 13, 906 (2017).

31. Wan, C. et al. Panorama of ancient metazoan macromolecular complexes. Nature 525, 339–344 (2015).

32. Scott, N. E., Brown, L. M., Kristensen, A. R. & Foster, L. J. Development of a computational framework for the analysis of protein correlation profiling and spatial proteomics experiments. Journal of Proteomics 118, 112–129 (2015).

33. Stacey, R. G., Skinnider, M. A., Scott, N. E. & Foster, L. J. A rapid and accurate approach for prediction of interactomes from co-elution data (PrInCE). BMC Bioinformatics 18, 457 (2017).

34. Wan, C. et al. ComplexQuant: High-throughput computational pipeline for the global quantitative analysis of endogenous soluble protein complexes using high resolution protein HPLC and precision label-free LC/MS/MS. Journal of Proteomics 81, 102–111 (2013).

35. Heusel, M. et al. A Global Screen for Assembly State Changes of the Mitotic Proteome by SEC-SWATH-MS. Cell Systems (2020).

36. Picotti, P. & Aebersold, R. Selected reaction monitoring–based proteomics: workflows, potential, pitfalls and future directions. Nature Methods 9, 555–566 (2012).

37. Ting, Y. S. et al. Peptide-Centric Proteome Analysis: An Alternative Strategy for the Analysis of Tandem Mass Spectrometry Data. Molecular & Cellular Proteomics 14, 2301–2307 (2015).

38. Giurgiu, M. et al. CORUM: the comprehensive resource of mammalian protein complexes—2019. Nucleic Acids Research 47, D559–D563 (2019).

39. Hofmann, J. C., Husedzinovic, A. & Gruss, O. J. The function of spliceosome components in open mitosis. Nucleus 1, 447–459 (2010).

40. Will, C. L. & Lührmann, R. Spliceosome structure and function. Cold Spring Harbor Perspectives in Biology 3, 1–2 (2011).

41. Alvarez, M. J. et al. Functional characterization of somatic mutations in cancer using network-based inference of protein activity. Nature Genetics 48, 838–847 (2016).

42. Reiter, L. et al. mProphet: automated data processing and statistical validation for large-scale SRM experiments. Nature Methods 8, 430–435 (2011).

43. Röst, H. L. et al. OpenSWATH enables automated, targeted analysis of data-independent acquisition MS data. Nature Biotechnology 32, 219–223 (2014).

44. Albanese, D. et al. minerva and minepy: a C engine for the MINE suite and its R, Python and MATLAB wrappers. Bioinformatics 29, 407–408 (2013).

45. Rosenberger, G. et al. Statistical control of peptide and protein error rates in large-scale targeted data-independent acquisition analyses. Nature Methods 14, 921–927 (2017).

46. Teleman, J. et al. DIANA-algorithmic improvements for analysis of data-independent acquisition MS data. Bioinformatics 31, 555–562 (2015).

47. Chen, T. & Guestrin, C. XGBoost: A scalable tree boosting system. In Proceedings of the ACM SIGKDD International Conference on Knowledge Discovery and Data Mining, vol. 13-17-August-2016, 785–794 (Association for Computing Machinery, New York, New York, USA, 2016).

48. Ahrné, E., Molzahn, L., Glatter, T. & Schmidt, A. Critical assessment of proteome-wide label-free absolute abundance estimation strategies. Proteomics 13, 2567–2578 (2013).

49. Poole, W., Gibbs, D. L., Shmulevich, I., Bernard, B. & Knijnenburg, T. A. Combining dependent P-values with an empirical adaptation of Brown’s method. Bioinformatics 32, i430–i436 (2016).

50. Domon, B. & Aebersold, R. Options and considerations when selecting a quantitative proteomics strategy. Nature Biotechnology 28, 710–721 (2010).

51. Walzthoeni, T. et al. xTract: software for characterizing conformational changes of protein complexes by quantitative cross-linking mass spectrometry. Nature Methods 12, 1185–1190 (2015).

52. Sham, P. C. & Purcell, S. M. Statistical power and significance testing in large-scale genetic studies. Nature Reviews Genetics 15, 335–346 (2014).

53. Barabási, A. L. & Oltvai, Z. N. Network biology: Understanding the cell’s functional organization. Nature Reviews Genetics 5, 101–113 (2004).

54. Jassal, B. et al. The reactome pathway knowledgebase. Nucleic acids research 48, D498–D503 (2020).

55. Gavet, O. & Pines, J. Progressive Activation of CyclinB1-Cdk1 Coordinates Entry to Mitosis. Developmental Cell 18, 533–543 (2010).

56. Castro, A., Bernis, C., Vigneron, S., Labbée, J. C. & Lorca, T. The anaphase-promoting complex: A key factor in the regulation of cell cycle. Oncogene 24, 314–325 (2005).

57. Neumann, B. et al. Phenotypic profiling of the human genome by time-lapse microscopy reveals cell division genes. Nature 464, 721–727 (2010).

58. Ballman, K. V., Grill, D. E., Oberg, A. L. & Therneau, T. M. Faster cyclic loess: Normalizing RNA arrays via linear models. Bioinformatics 20, 2778–2786 (2004).

59. Bolstad, B. M., Irizarry, R. A., Astrand, M. & Speed, T. P. A comparison of normalization methods for high density oligonucleotide array data based on variance and bias. Bioinformatics 19, 185–193 (2003).

60. Virtanen, P. et al. SciPy 1.0: fundamental algorithms for scientific computing in Python. Nature Methods (2020).

61. Käll, L., Canterbury, J. D., Weston, J., Noble, W. S. & MacCoss, M. J. Semi-supervised learning for peptide identification from shotgun proteomics datasets. Nature Methods 4, 923–925 (2007).

62. Bergstra, J., Yamins, D. & Cox, D. Making a Science of Model Search: Hyperparameter Optimization in Hundreds of Dimensions for Vision Architectures. In Dasgupta, S. & McAllester, D. (eds.) Proceedings of the 30th International Conference on Machine Learning, vol. 28 of Proceedings of Machine Learning Research, 115–123 (PMLR, Atlanta, Georgia, USA, 2013).

63. Storey, J. D. The positive false discovery rate: A Bayesian interpretation and the q-value. Annals of Statistics 31, 2013–2035 (2003).

64. Benjamini, Y. & Hochberg, Y. Controlling the false discovery rate: a practical and powerful approach to multiple testing. Journal of the Royal Statistical Society 57, 289–300 (1995).

65. Kotlyar, M., Pastrello, C., Malik, Z. & Jurisica, I. IID 2018 update: context-specific physical protein–protein interactions in human, model organisms and domesticated species. Nucleic Acids Research (2018).

66. Rosenberger, G., Ludwig, C., Röst, H. L., Aebersold, R. & Malmström, L. ALFQ: An R-package for estimating absolute protein quantities from label-free LC-MS/MS proteomics data. Bioinformatics 30, 2511–2513 (2014).

67. Shannon, P. et al. Cytoscape: a software environment for integrated models of biomolecular interaction networks. Genome research 13, 2498–504 (2003).

68. Kucera, M., Isserlin, R., Arkhangorodsky, A. & Bader, G. D. AutoAnnotate: A Cytoscape app for summarizing networks with semantic annotations. F1000Research 5, 1717 (2016).

