## Supplementary Data 2 for "SECAT: Quantifying differential protein-protein interaction states by network-centric analysis": hela_string_000001_P61254.pdf

P61254_P61254

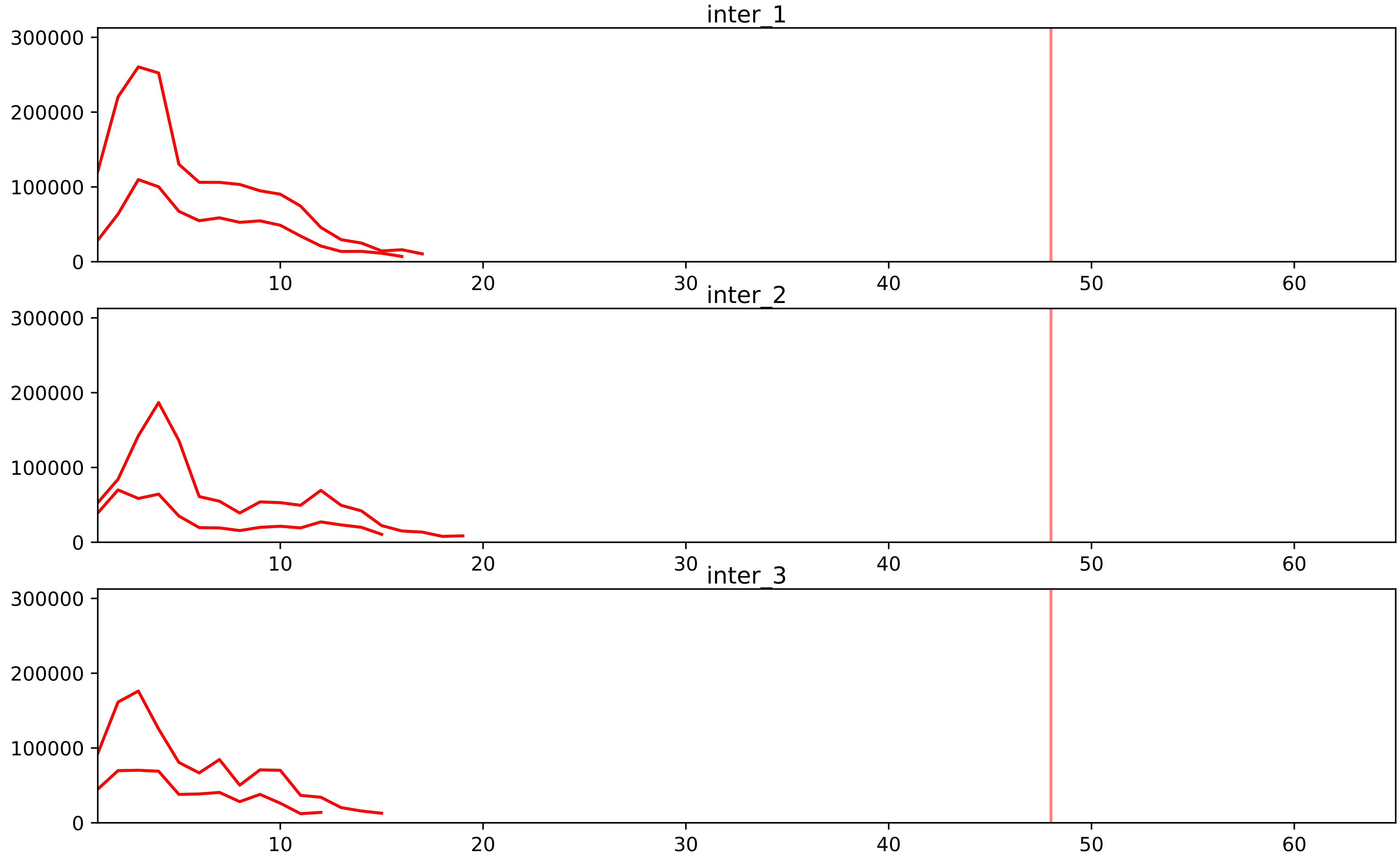


| level | condition_1 | condition_2 | pvalue | pvalue_adjusted |
|---|---|---|---|---|
| interactor_ratio | mitosis | inter | 1.2673527714633817e-106 | 2.331929099492622e-103 |
| complex_abundance | mitosis | inter | 5.689792436931684e-08 | 1.4956025834220426e-05 |
| interactor_abundance | mitosis | inter | 2.6008102022659265e-06 | 0.0002175223078258775 |
| assembled_abundance | mitosis | inter | 5.746807045066382e-05 | 0.007270487507555603 |
| total_abundance | mitosis | inter | 9.964756334129977e-05 | 0.01273261737241483 |

P05388_P61254
RLA0_HUMAN vs RL26_HUMAN
p-value: 0.05 q-value: 0.033

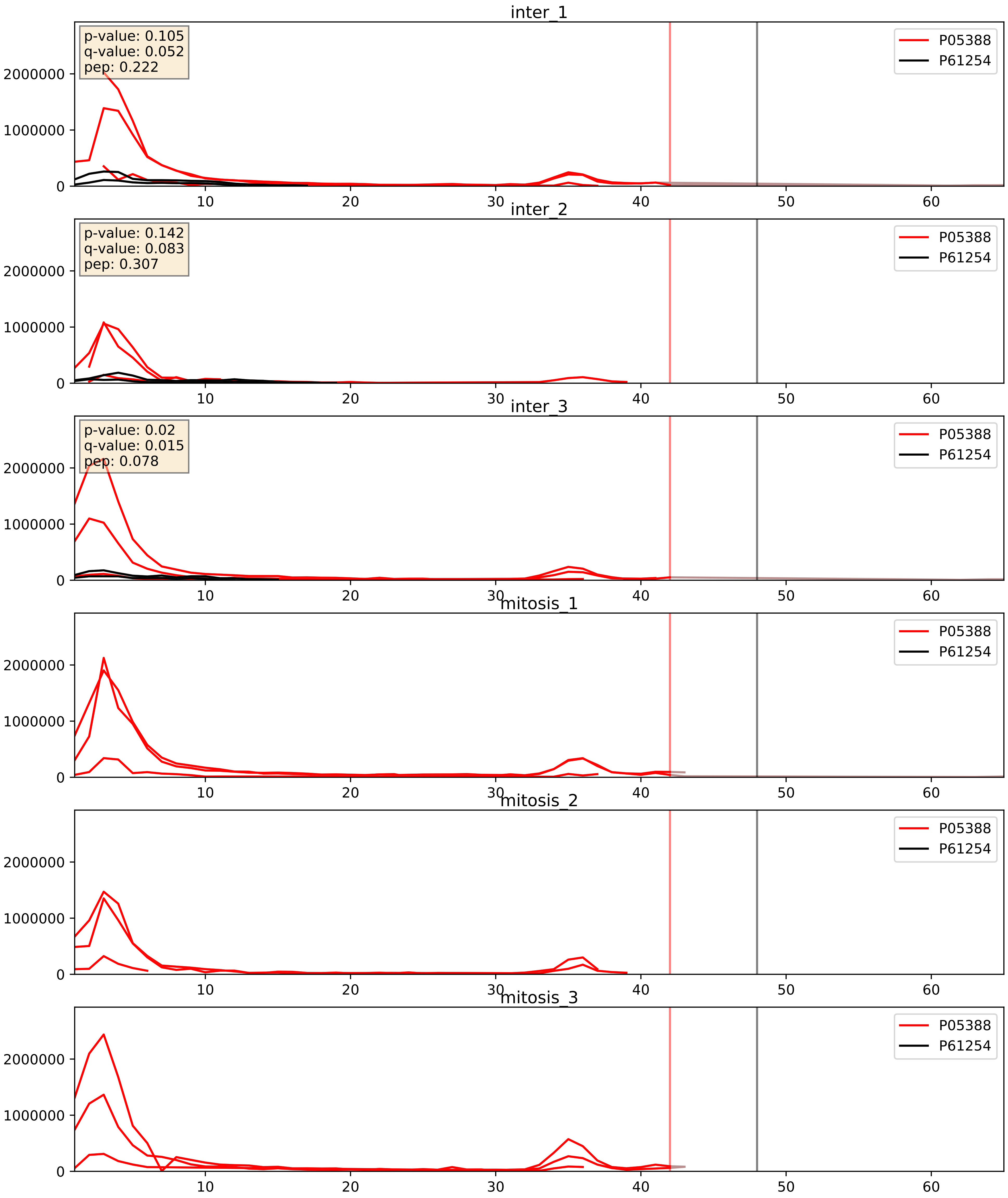


| level | condition_1 | condition_2 | pvalue | pvalue_adjusted |
|---|---|---|---|---|
| interactor_abundance | mitosis | inter | 0.0001793461765278864 | 0.010140496760639767 |
| complex_abundance | mitosis | inter | 0.0002298320204321024 | 0.010219389159334393 |
| interactor_ratio | mitosis | inter | 0.0002494714800990228 | 0.01605620954622282 |

P08708_P61254
RS17_HUMAN vs RL26_HUMAN
p-value: 0.0 q-value: 0.0

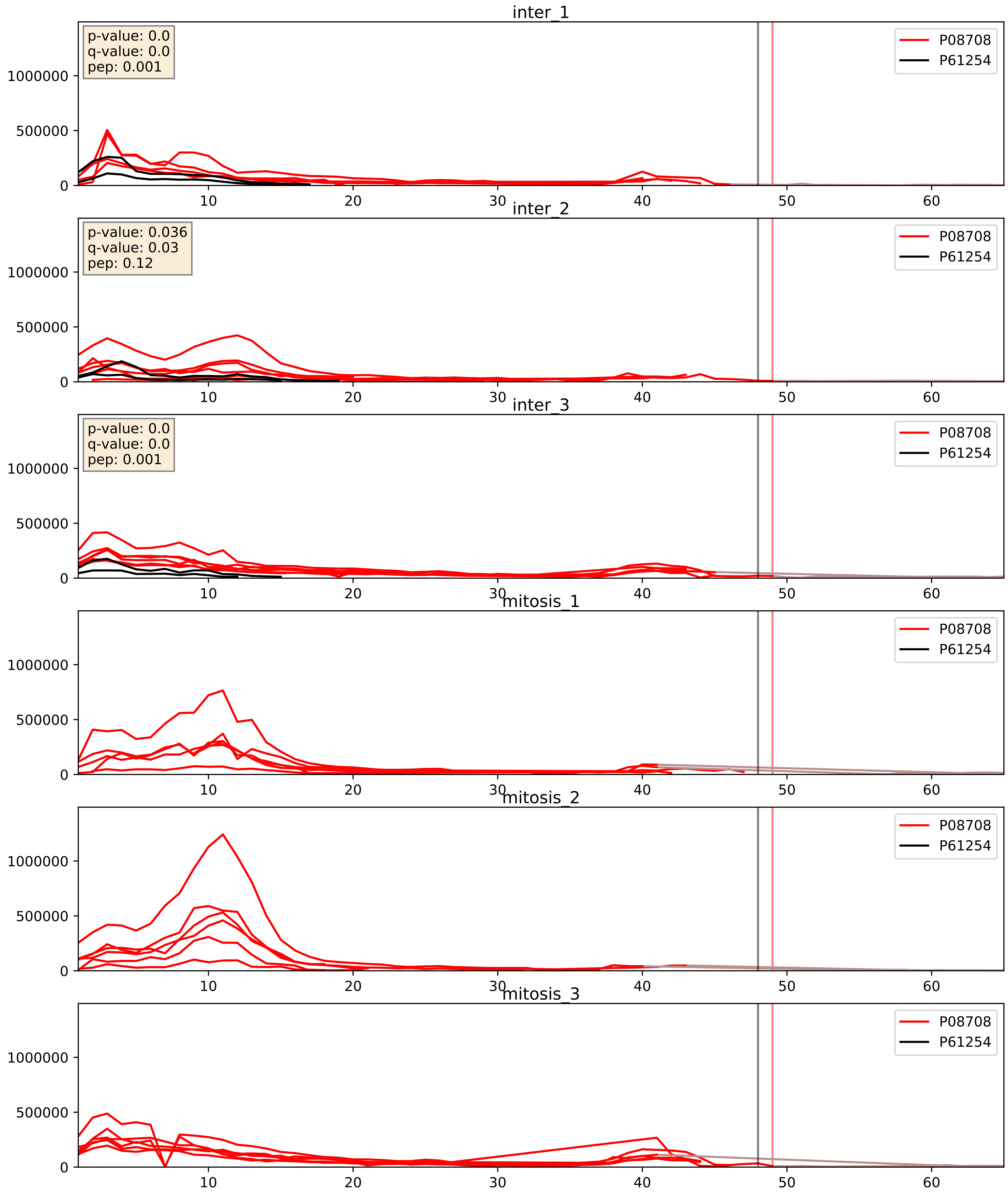


| level | condition_1 | condition_2 | pvalue | pvalue_adjusted |
|---|---|---|---|---|
| interactor_abundance | mitosis | inter | 9.170603546153091e-05 | 0.0075120681697771925 |
| complex_abundance | mitosis | inter | 0.00012786725061173694 | 0.007762721030045874 |
| interactor_ratio | mitosis | inter | 0.0003630105561275218 | 0.020578611658619777 |

P18077_P61254
RL35A_HUMAN vs RL26_HUMAN
p-value: 0.0 q-value: 0.001

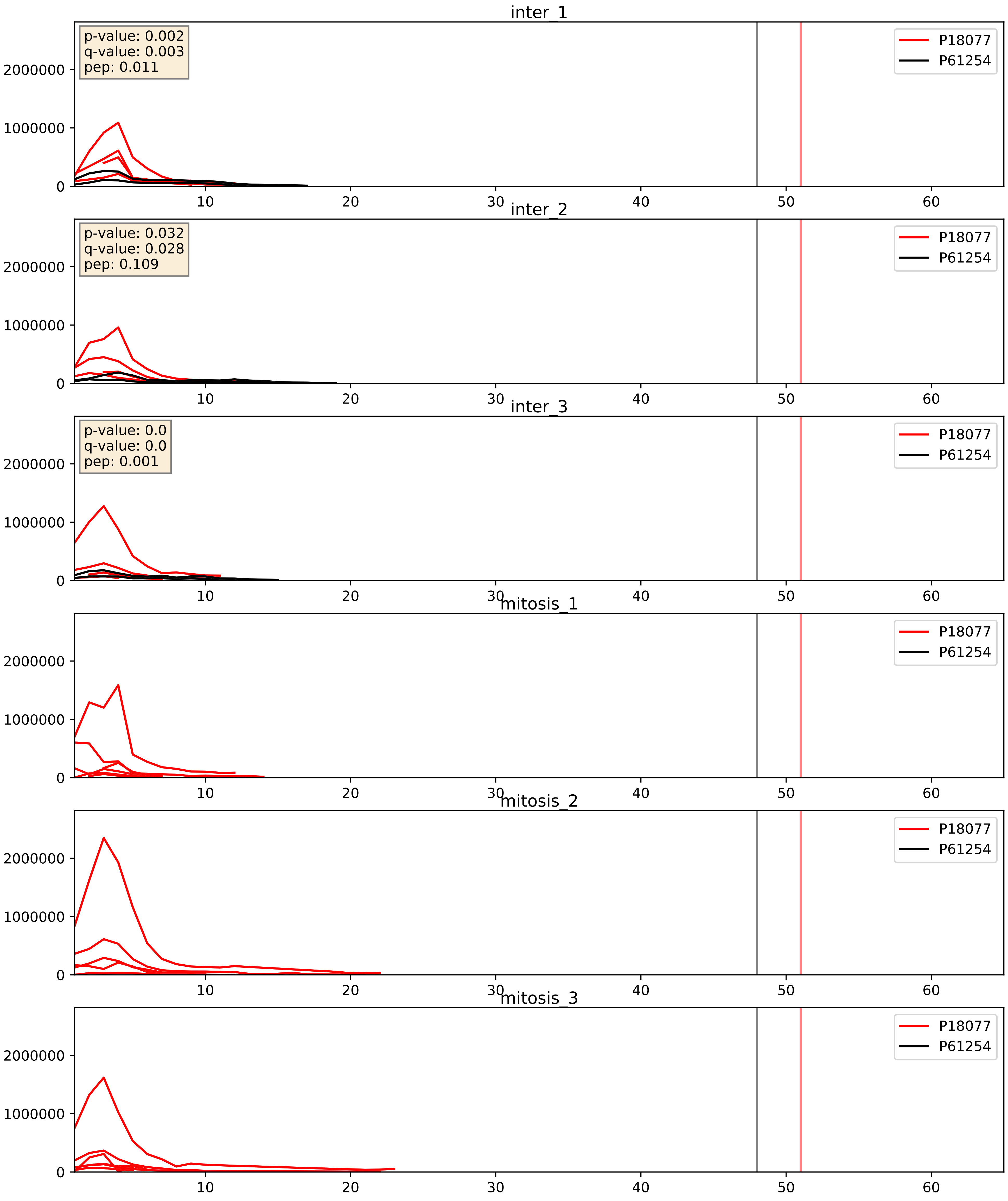


| level | condition_1 | condition_2 | pvalue | pvalue_adjusted |
|---|---|---|---|---|
| interactor_ratio | mitosis | inter | 0.00011582782362096936 | 0.011396392760867789 |
| interactor_abundance | mitosis | inter | 0.00024465216712364293 | 0.01187905696742526 |
| complex_abundance | mitosis | inter | 0.00036048069653242006 | 0.012039775462644836 |

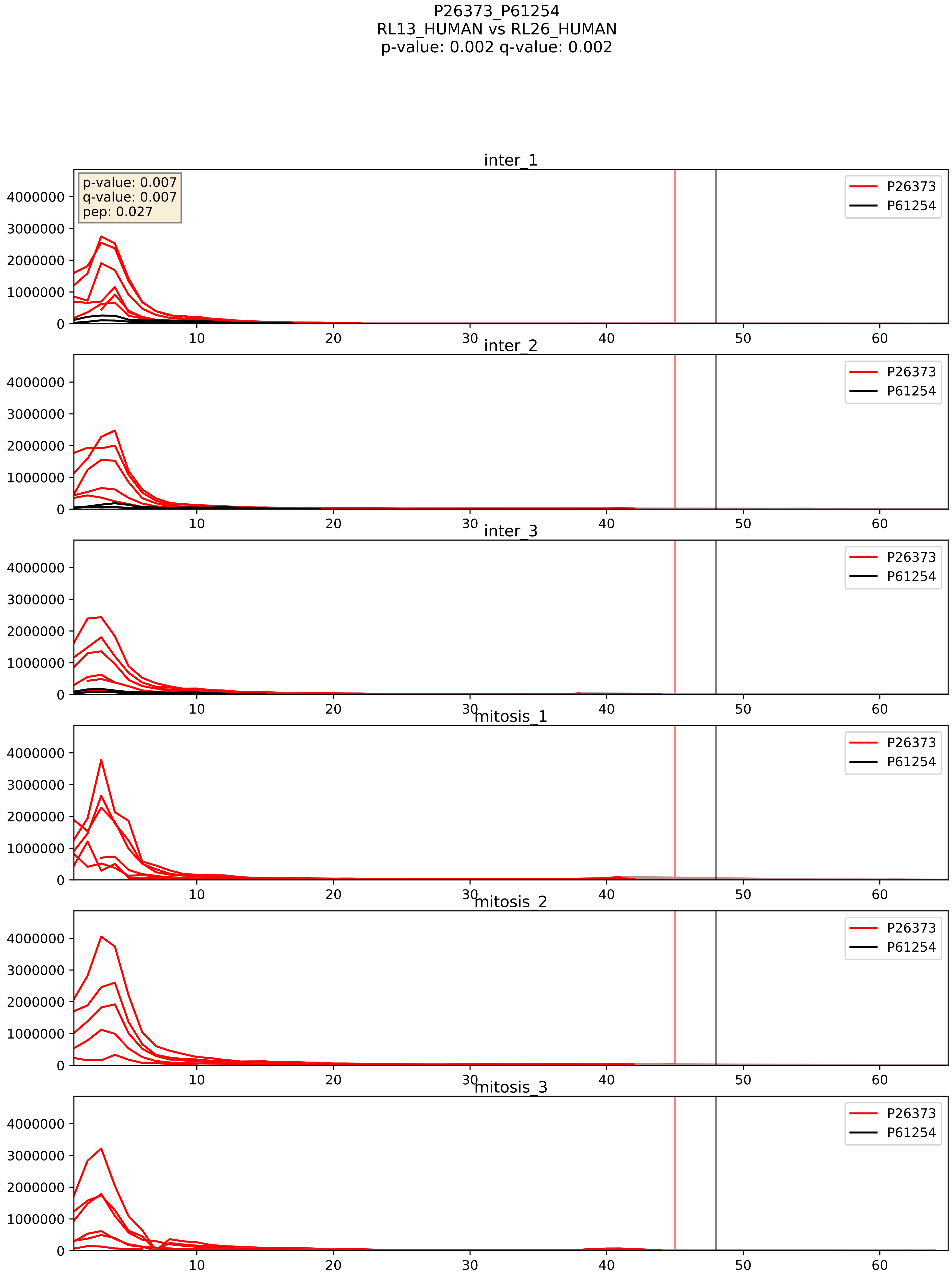


| level | condition_1 | condition_2 | pvalue | pvalue_adjusted |
|---|---|---|---|---|
| interactor_ratio | mitosis | inter | 2.041780948909418e-06 | 0.002520386835947235 |
| interactor_abundance | mitosis | inter | 1.5413956431593788e-05 | 0.0037089335794386596 |
| complex_abundance | mitosis | inter | 4.943712628168651e-05 | 0.005691857151652958 |

P27635_P61254
RL10_HUMAN vs RL26_HUMAN
p-value: 0.008 q-value: 0.007

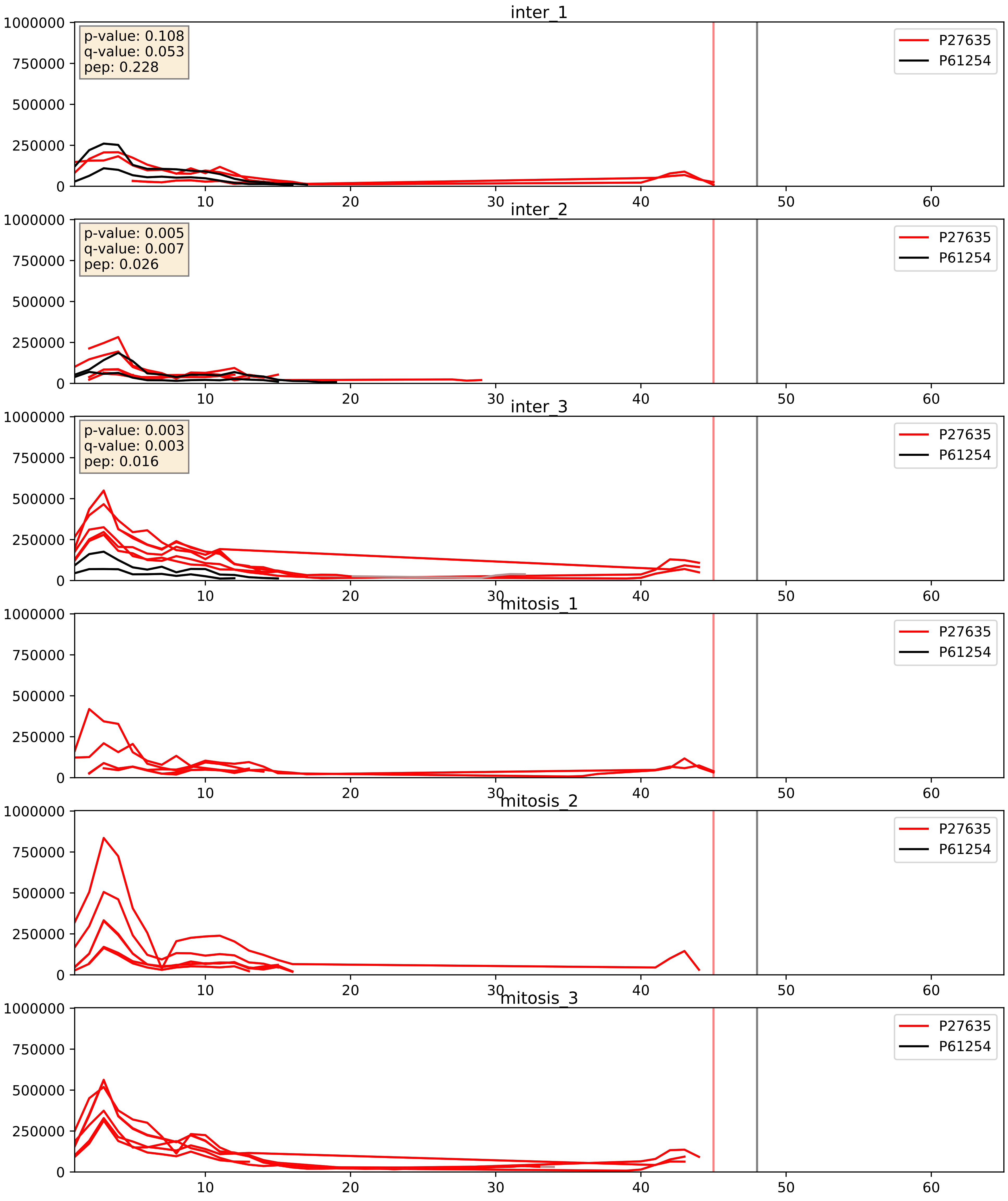


| level | condition_1 | condition_2 | pvalue | pvalue_adjusted |
|---|---|---|---|---|
| complex_abundance | mitosis | inter | 0.00037471788909165294 | 0.012384498573839959 |
| interactor_abundance | mitosis | inter | 0.0015855554986002476 | 0.029031766160815386 |
| interactor_ratio | mitosis | inter | 0.03330347482230045 | 0.29268762266826676 |

P32969_P61254
RL9_HUMAN vs RL26_HUMAN
p-value: 0.002 q-value: 0.003

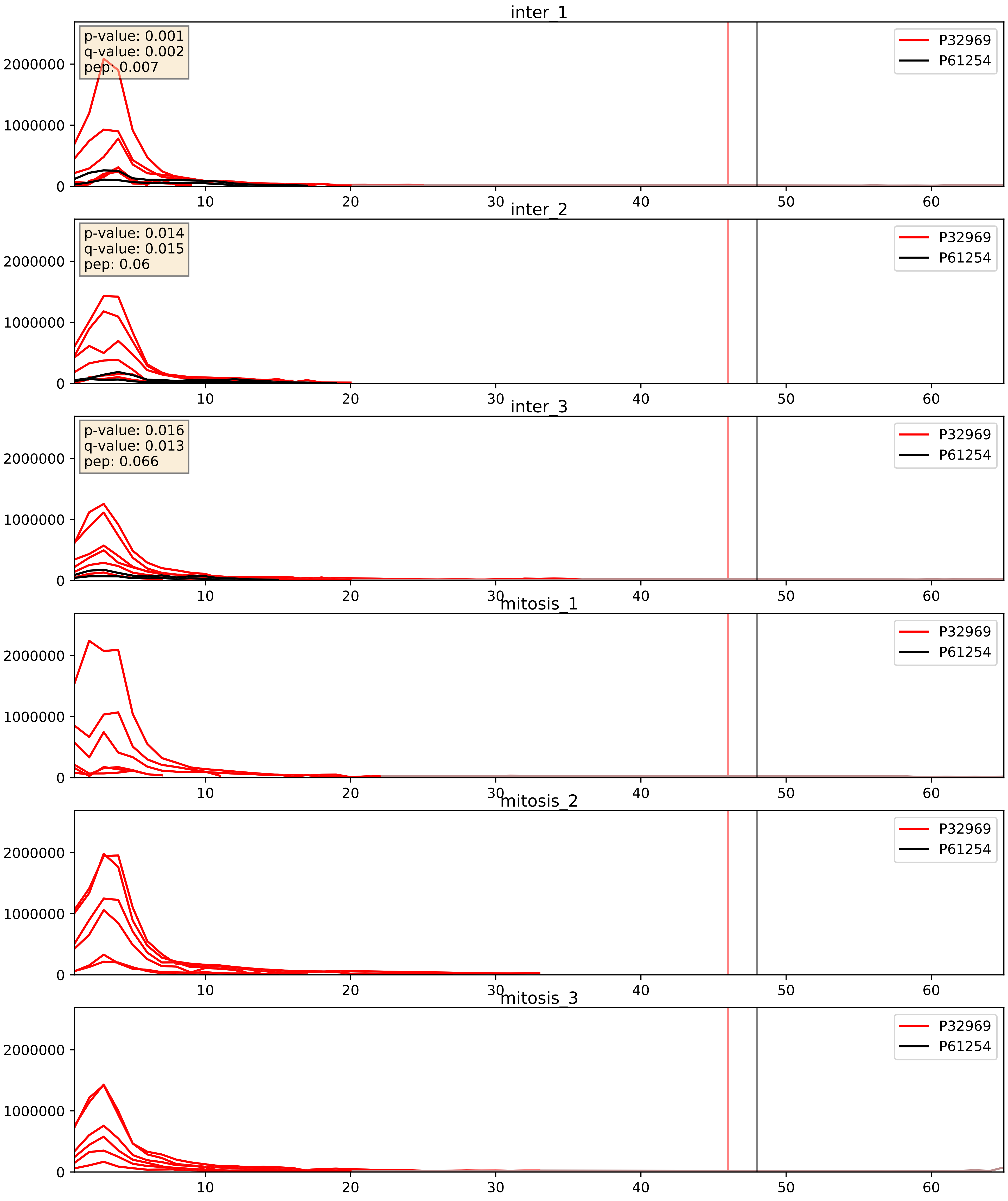


| level | condition_1 | condition_2 | pvalue | pvalue_adjusted |
|---|---|---|---|---|
| interactor_ratio | mitosis | inter | 7.860739514228402e-06 | 0.0032529045057416803 |
| interactor_abundance | mitosis | inter | 3.1428782185550784e-05 | 0.004891461372878449 |
| complex_abundance | mitosis | inter | 8.26516458512303e-05 | 0.00615215729118723 |

P35268_P61254
RL22_HUMAN vs RL26_HUMAN
p-value: 0.002 q-value: 0.003

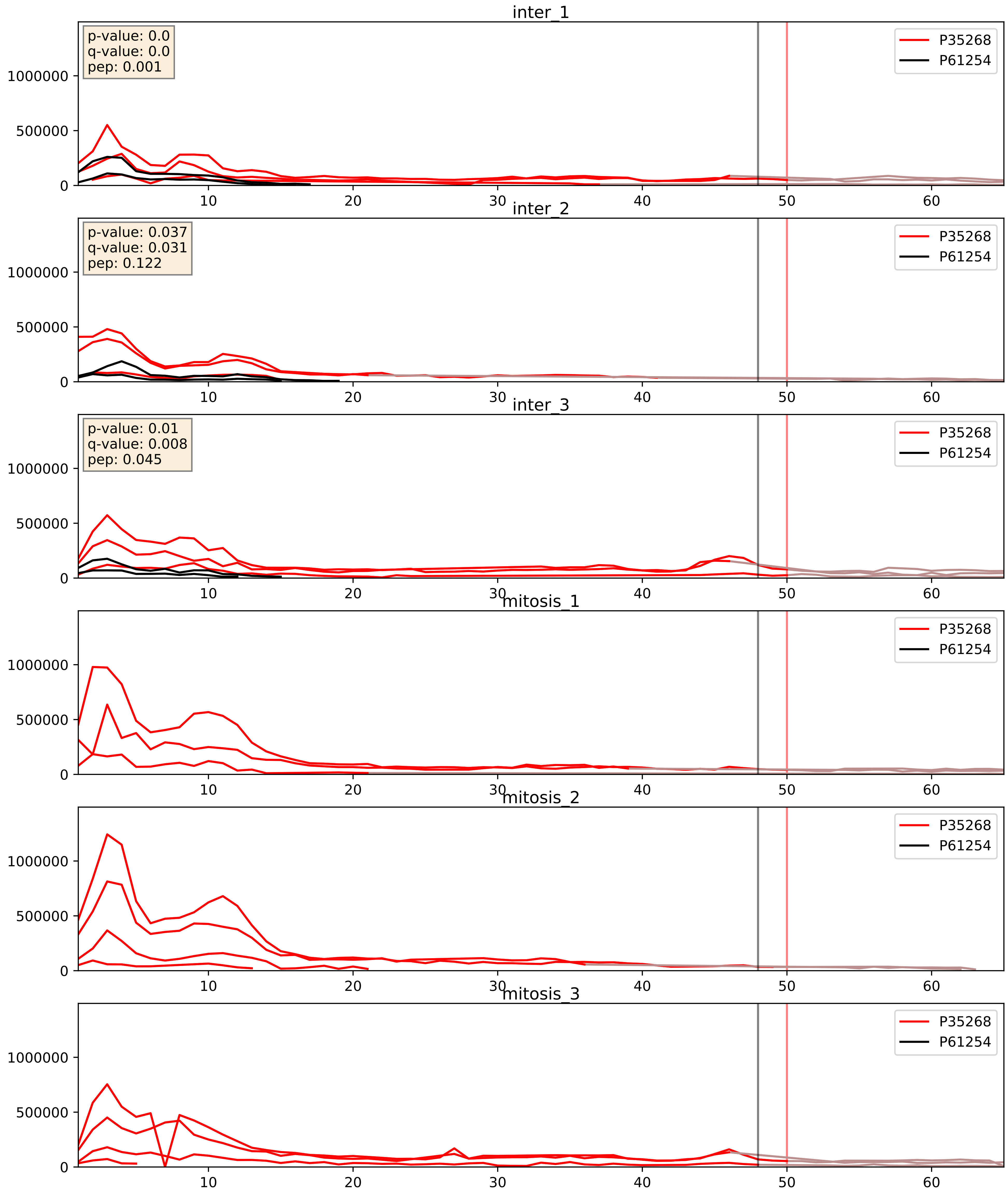


| level | condition_1 | condition_2 | pvalue | pvalue_adjusted |
|---|---|---|---|---|
| interactor_abundance | mitosis | inter | 7.421754572657264e-05 | 0.006870282762438709 |
| complex_abundance | mitosis | inter | 8.3817361471577e-05 | 0.006185143225833613 |
| interactor_ratio | mitosis | inter | 0.0008875457249814373 | 0.034218313408302205 |

P37108_P61254
SRP14_HUMAN vs RL26_HUMAN
p-value: 0.036 q-value: 0.025

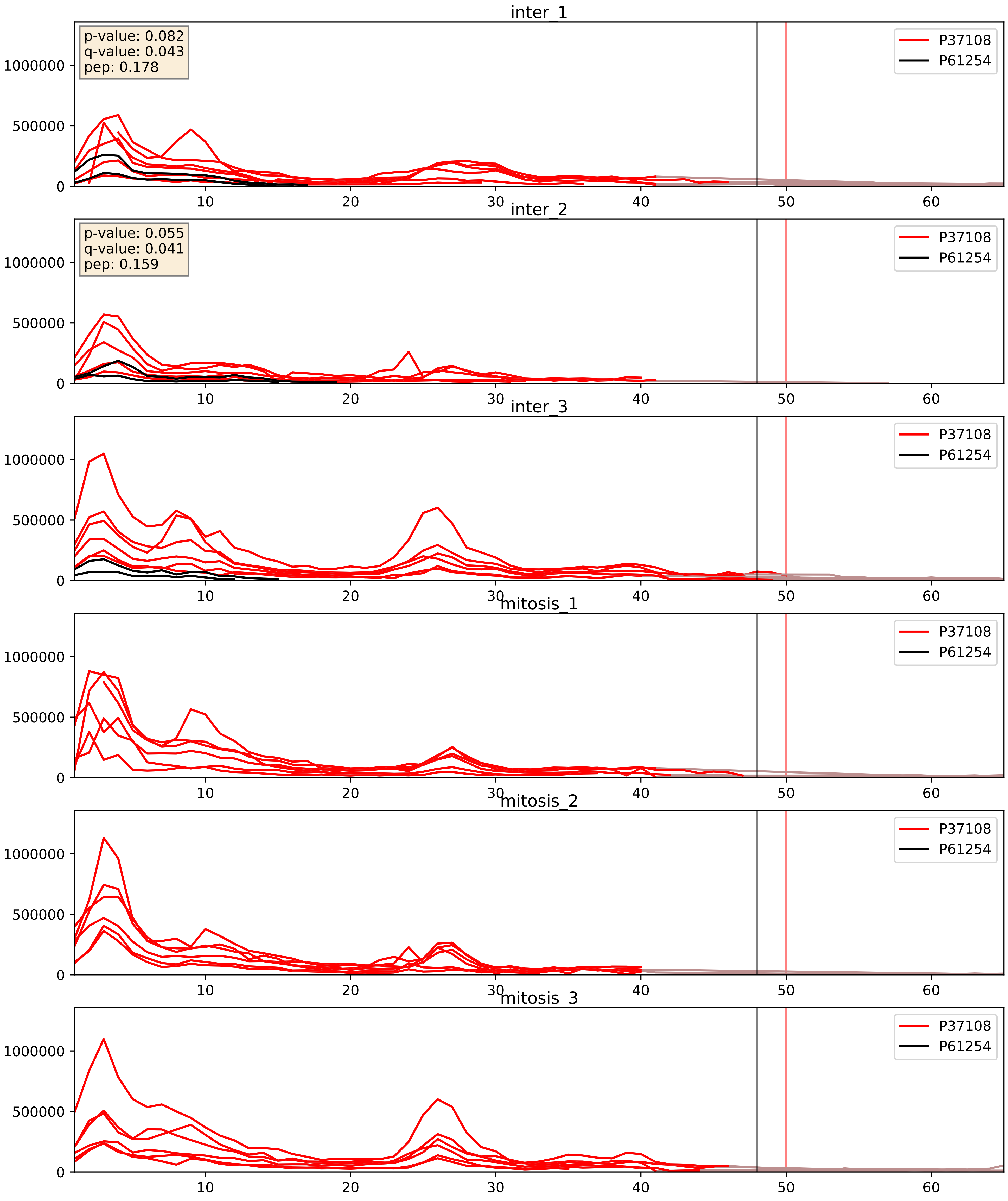


| level | condition_1 | condition_2 | pvalue | pvalue_adjusted |
|---|---|---|---|---|
| complex_abundance | mitosis | inter | 0.000612677764346236 | 0.01584447632267003 |
| interactor_abundance | mitosis | inter | 0.0015770968345815443 | 0.029025269221018554 |
| interactor_ratio | mitosis | inter | 0.011703427523279374 | 0.15628914134051708 |

P39023_P61254
RL3_HUMAN vs RL26_HUMAN
p-value: 0.039 q-value: 0.026

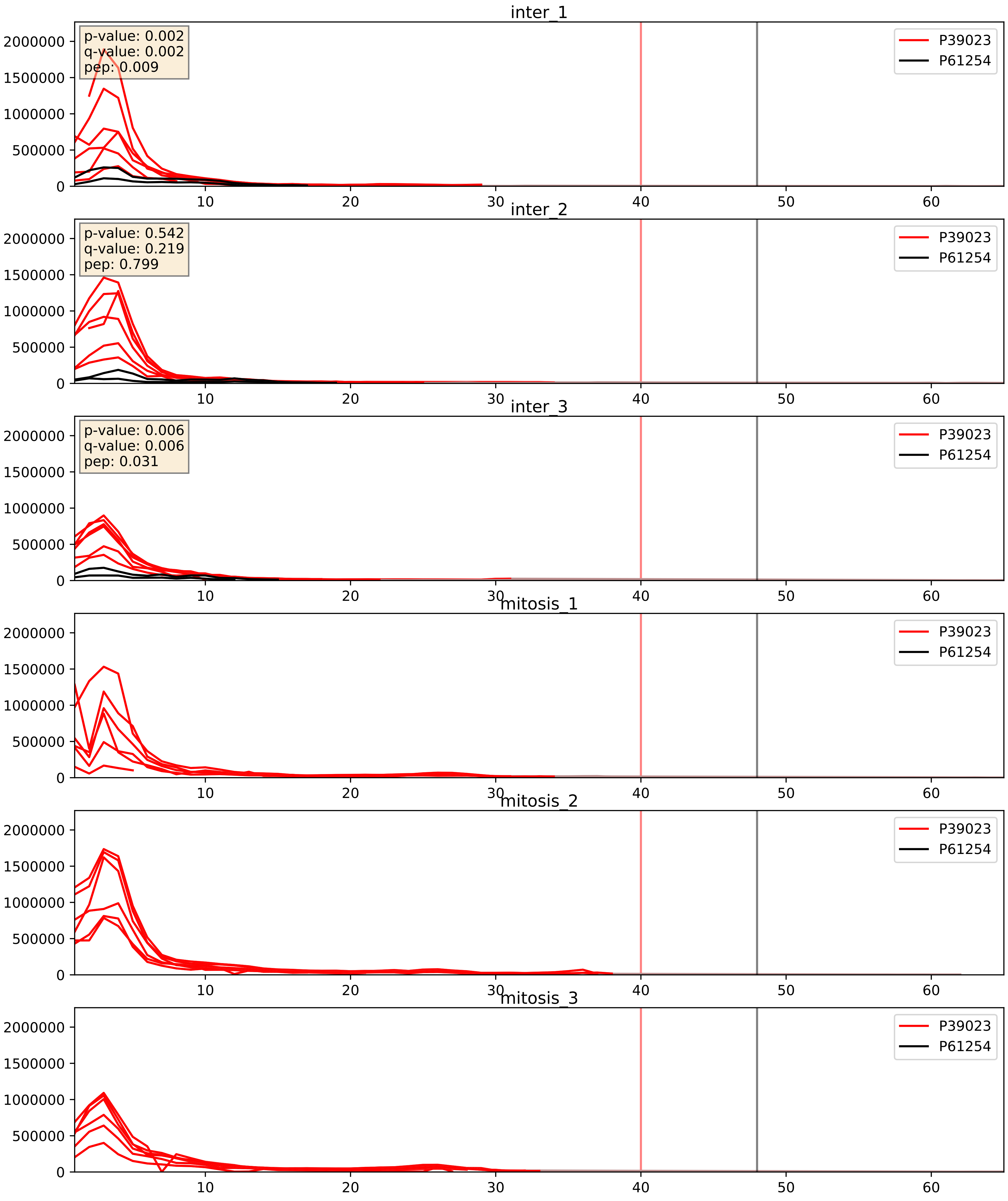


| level | condition_1 | condition_2 | pvalue | pvalue_adjusted |
|---|---|---|---|---|
| interactor_abundance | mitosis | inter | 4.2975123221088804e-05 | 0.006030607455287216 |
| interactor_ratio | mitosis | inter | 6.146887867472464e-05 | 0.008128440693030705 |
| complex_abundance | mitosis | inter | 8.212420270142806e-05 | 0.00615215729118723 |

P40429_P61254
RL13A_HUMAN vs RL26_HUMAN
p-value: 0.001 q-value: 0.001

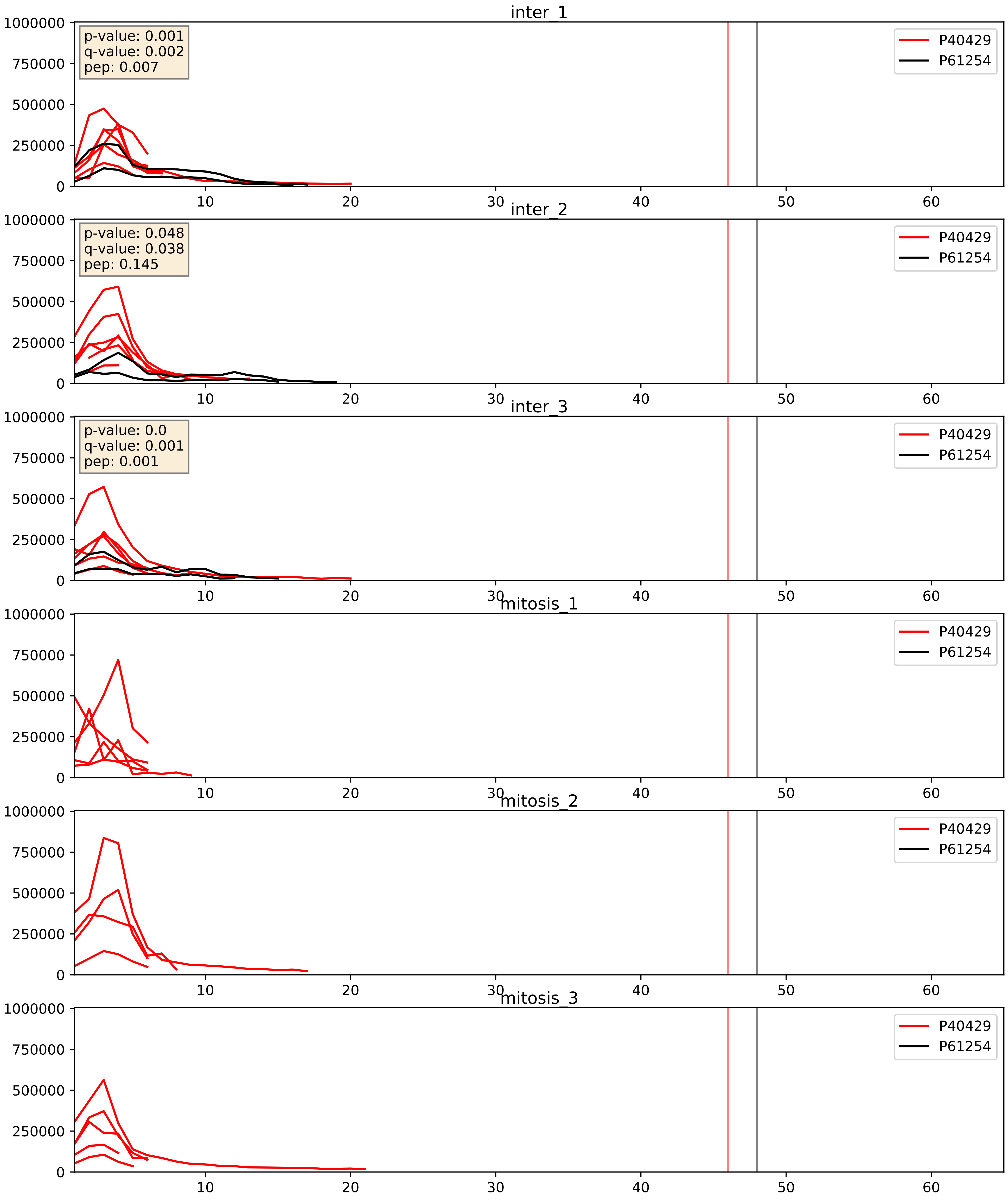


| level | condition_1 | condition_2 | pvalue | pvalue_adjusted |
|---|---|---|---|---|
| interactor_abundance | mitosis | inter | 0.00010749829679460574 | 0.007876481428251637 |
| complex_abundance | mitosis | inter | 0.000149229153604957 | 0.00845961294608233 |
| interactor_ratio | mitosis | inter | 0.000672218340536711 | 0.029848225645041686 |

P42766_P61254
RL35_HUMAN vs RL26_HUMAN
p-value: 0.002 q-value: 0.003

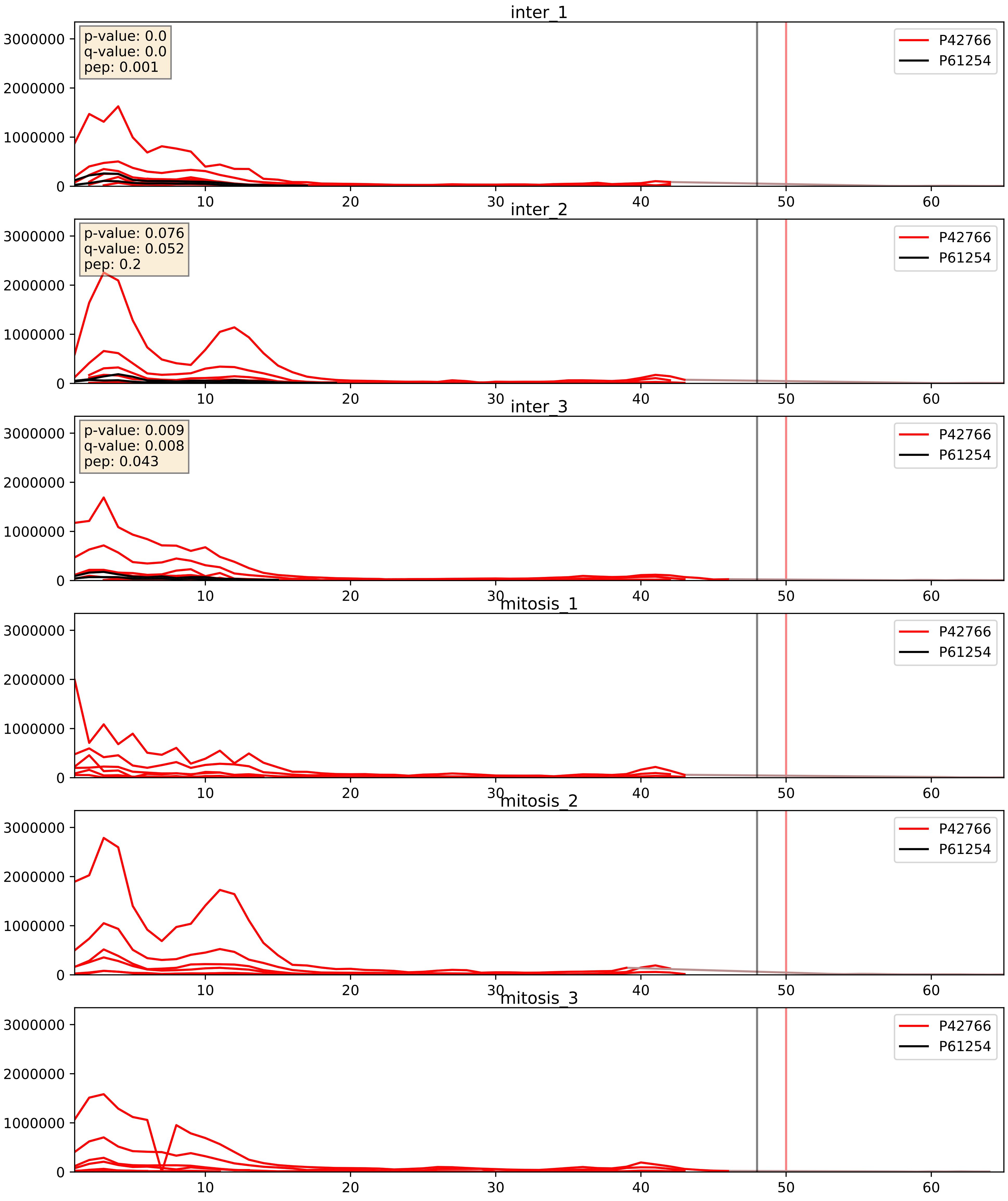


| level | condition_1 | condition_2 | pvalue | pvalue_adjusted |
|---|---|---|---|---|
| interactor_abundance | mitosis | inter | 2.4526274259084307e-05 | 0.004450836370313492 |
| interactor_ratio | mitosis | inter | 3.431898356409952e-05 | 0.006185155691845816 |
| complex_abundance | mitosis | inter | 5.896564332446068e-05 | 0.005691857151652958 |

P46776_P61254
RL27A_HUMAN vs RL26_HUMAN
p-value: 0.018 q-value: 0.014

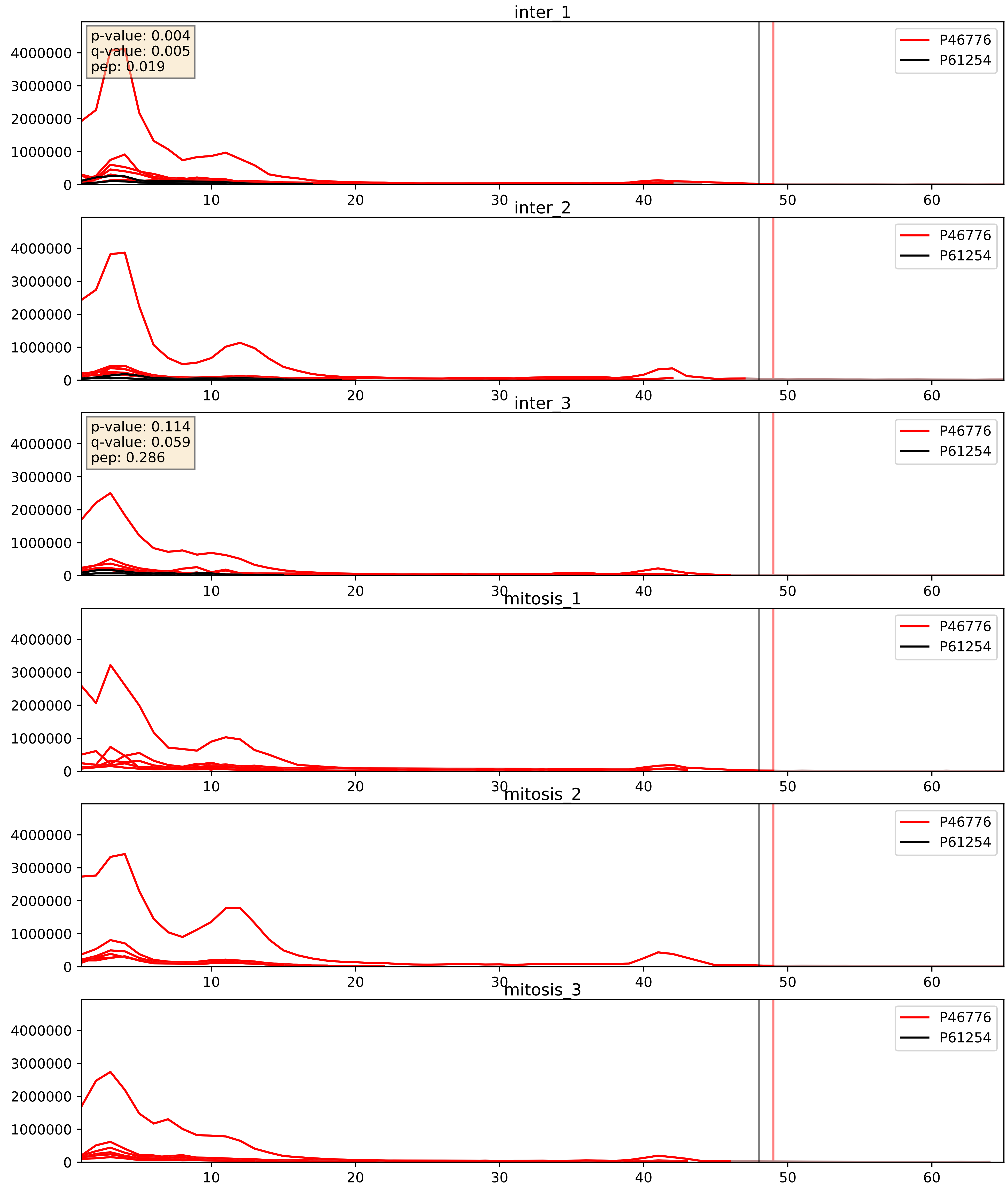


| level | condition_1 | condition_2 | pvalue | pvalue_adjusted |
|---|---|---|---|---|
| interactor_ratio | mitosis | inter | 1.5127254382454834e-06 | 0.002520386835947235 |
| interactor_abundance | mitosis | inter | 3.415709164225707e-05 | 0.005164930486161031 |
| complex_abundance | mitosis | inter | 8.961722529992237e-05 | 0.00644641553417929 |

P46777_P61254
RL5_HUMAN vs RL26_HUMAN
p-value: 0.005 q-value: 0.005

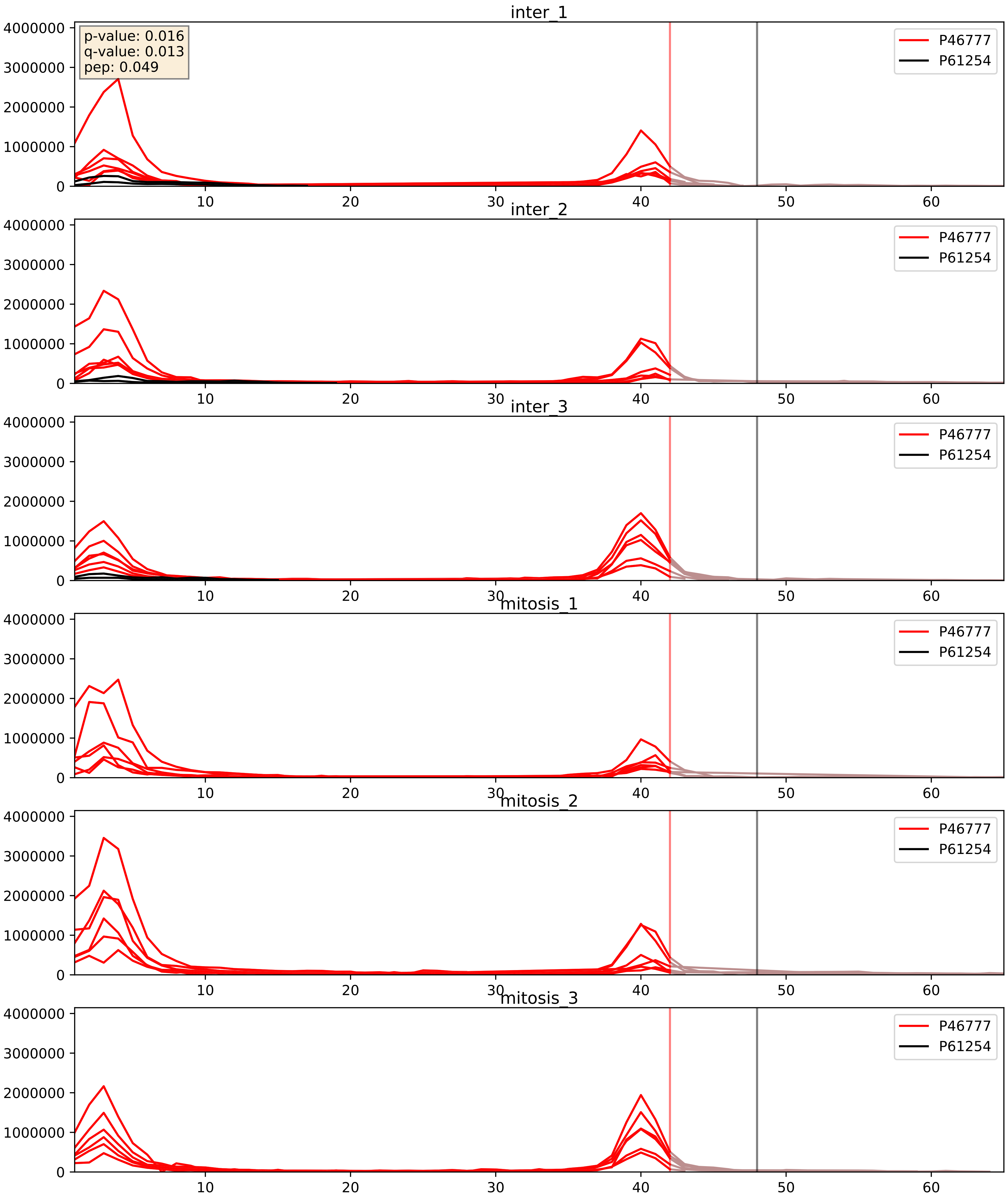


| level | condition_1 | condition_2 | pvalue | pvalue_adjusted |
|---|---|---|---|---|
| interactor_abundance | mitosis | inter | 3.4694334457273275e-05 | 0.005164930486161031 |
| complex_abundance | mitosis | inter | 6.484541610676946e-05 | 0.005691857151652958 |
| interactor_ratio | mitosis | inter | 8.478071930844215e-05 | 0.009941410373702258 |

P46778_P61254
RL21_HUMAN vs RL26_HUMAN
p-value: 0.027 q-value: 0.02

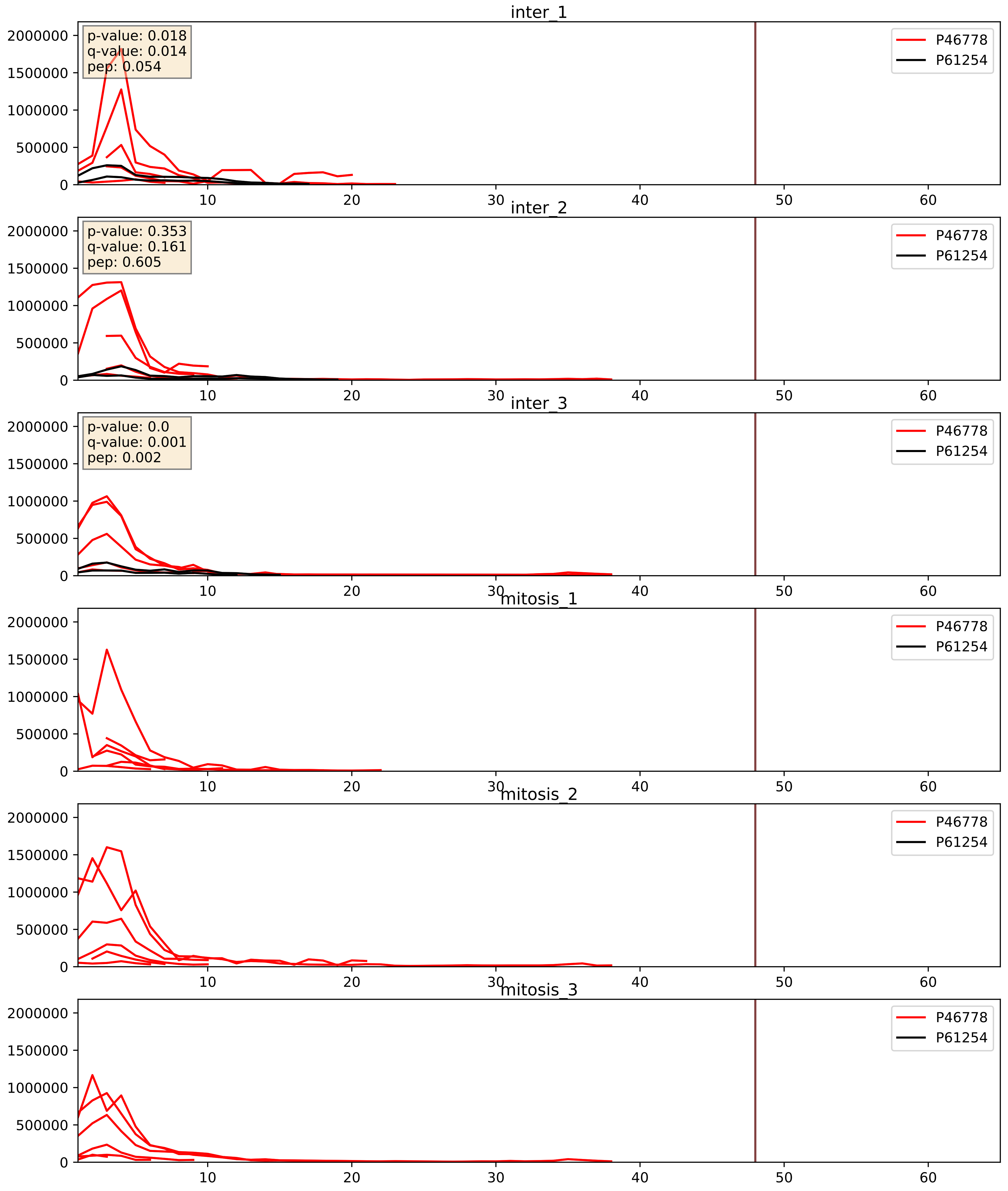


| level | condition_1 | condition_2 | pvalue | pvalue_adjusted |
|---|---|---|---|---|
| interactor_abundance | mitosis | inter | 3.297012345080543e-05 | 0.005085121743043144 |
| complex_abundance | mitosis | inter | 5.967886072755383e-05 | 0.005691857151652958 |
| interactor_ratio | mitosis | inter | 0.00015123109647191348 | 0.012836278661937291 |

P46779_P61254
RL28_HUMAN vs RL26_HUMAN
p-value: 0.003 q-value: 0.003

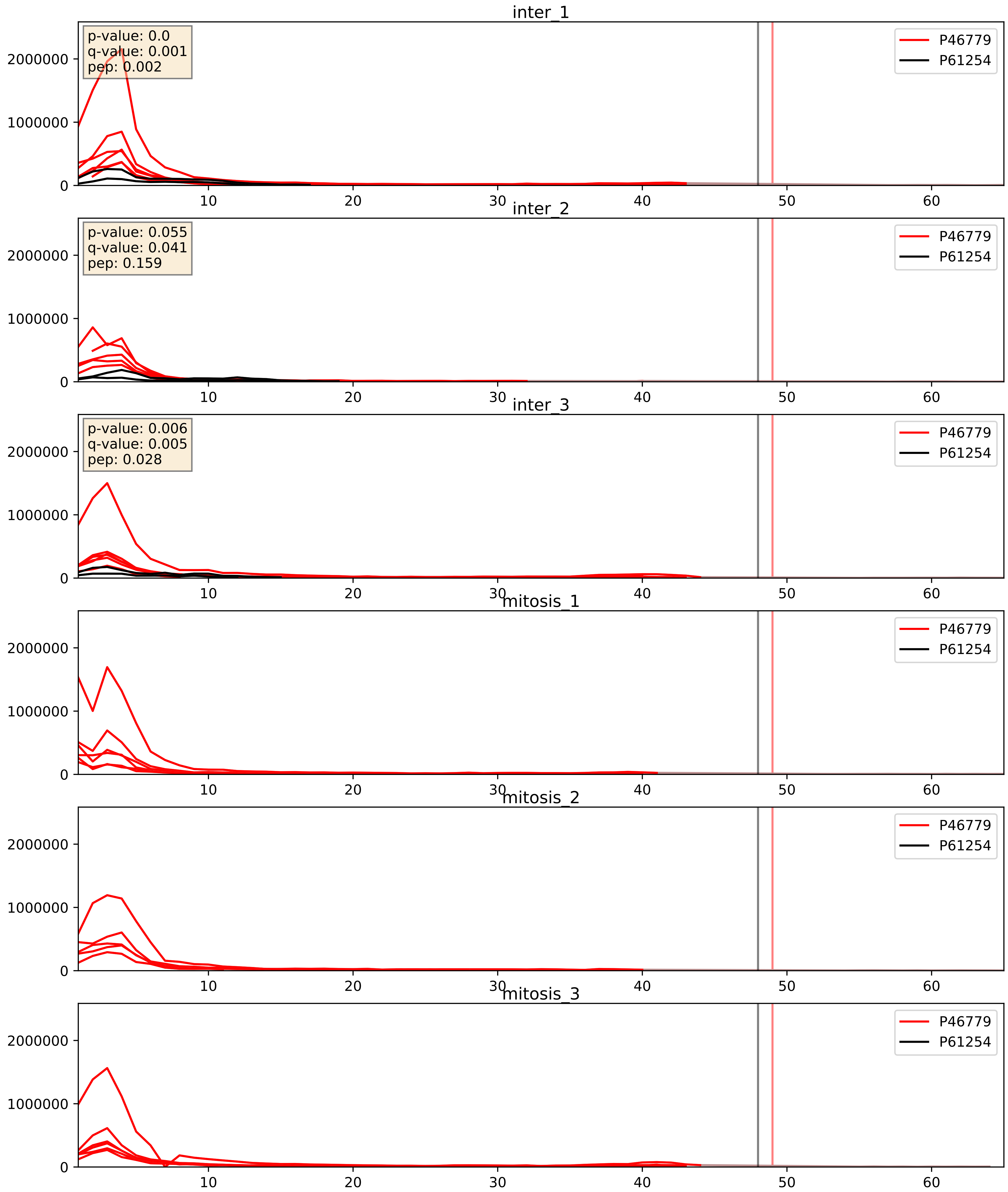


| level | condition_1 | condition_2 | pvalue | pvalue_adjusted |
|---|---|---|---|---|
| interactor_ratio | mitosis | inter | 5.727064801717087e-06 | 0.002835004461043519 |
| interactor_abundance | mitosis | inter | 5.1707692295123786e-05 | 0.006283598333582311 |
| complex_abundance | mitosis | inter | 0.00012530619449886146 | 0.0077166980209370795 |

P47914_P61254
RL29_HUMAN vs RL26_HUMAN
p-value: 0.057 q-value: 0.036

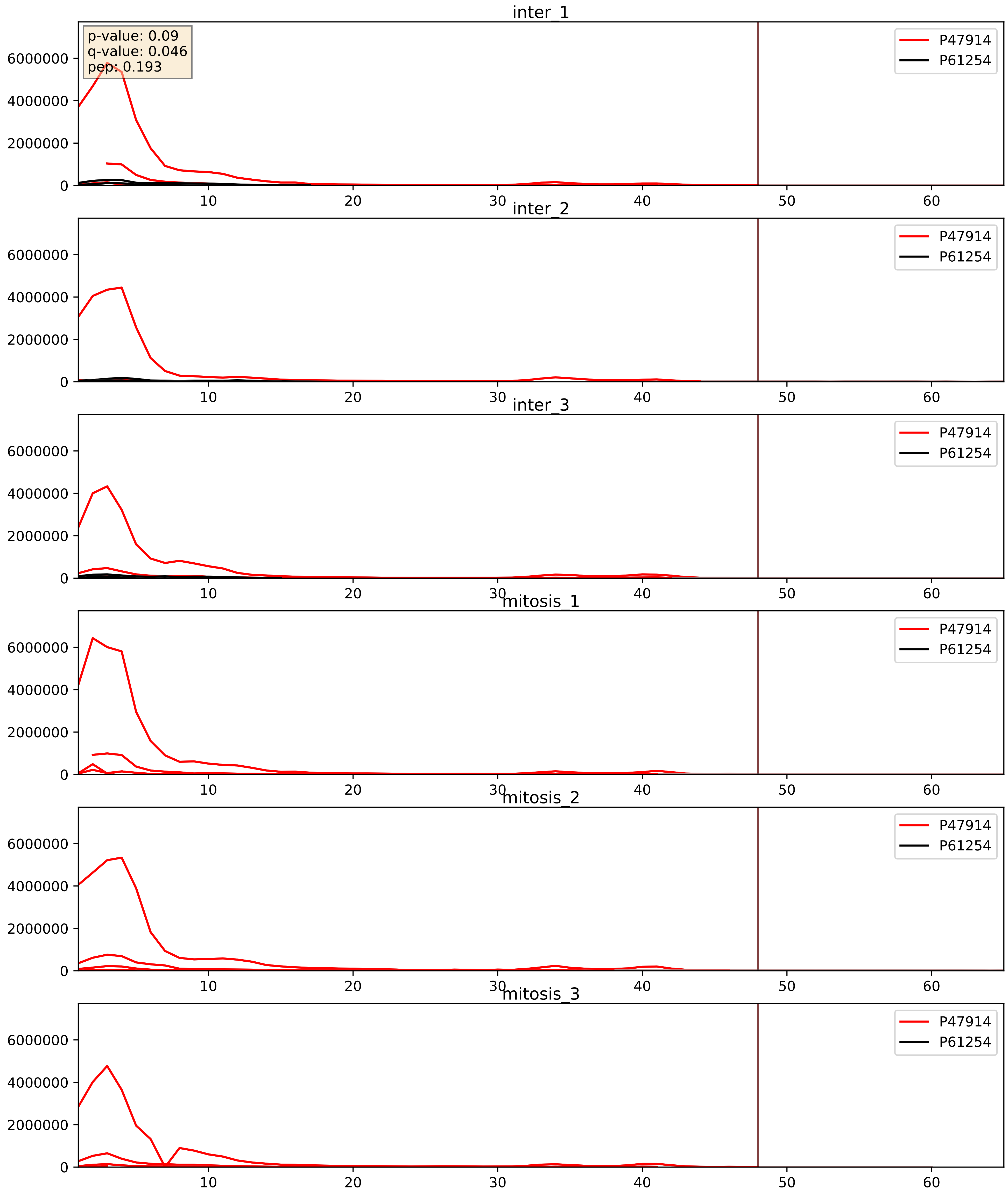


| level | condition_1 | condition_2 | pvalue | pvalue_adjusted |
|---|---|---|---|---|
| complex_abundance | mitosis | inter | 0.00048598835034400985 | 0.013857879989144268 |
| interactor_abundance | mitosis | inter | 0.0007529717761997596 | 0.020141995013343572 |
| interactor_ratio | mitosis | inter | 0.0030406750334639188 | 0.0715059843034372 |

P49207_P61254
RL34_HUMAN vs RL26_HUMAN
p-value: 0.005 q-value: 0.005

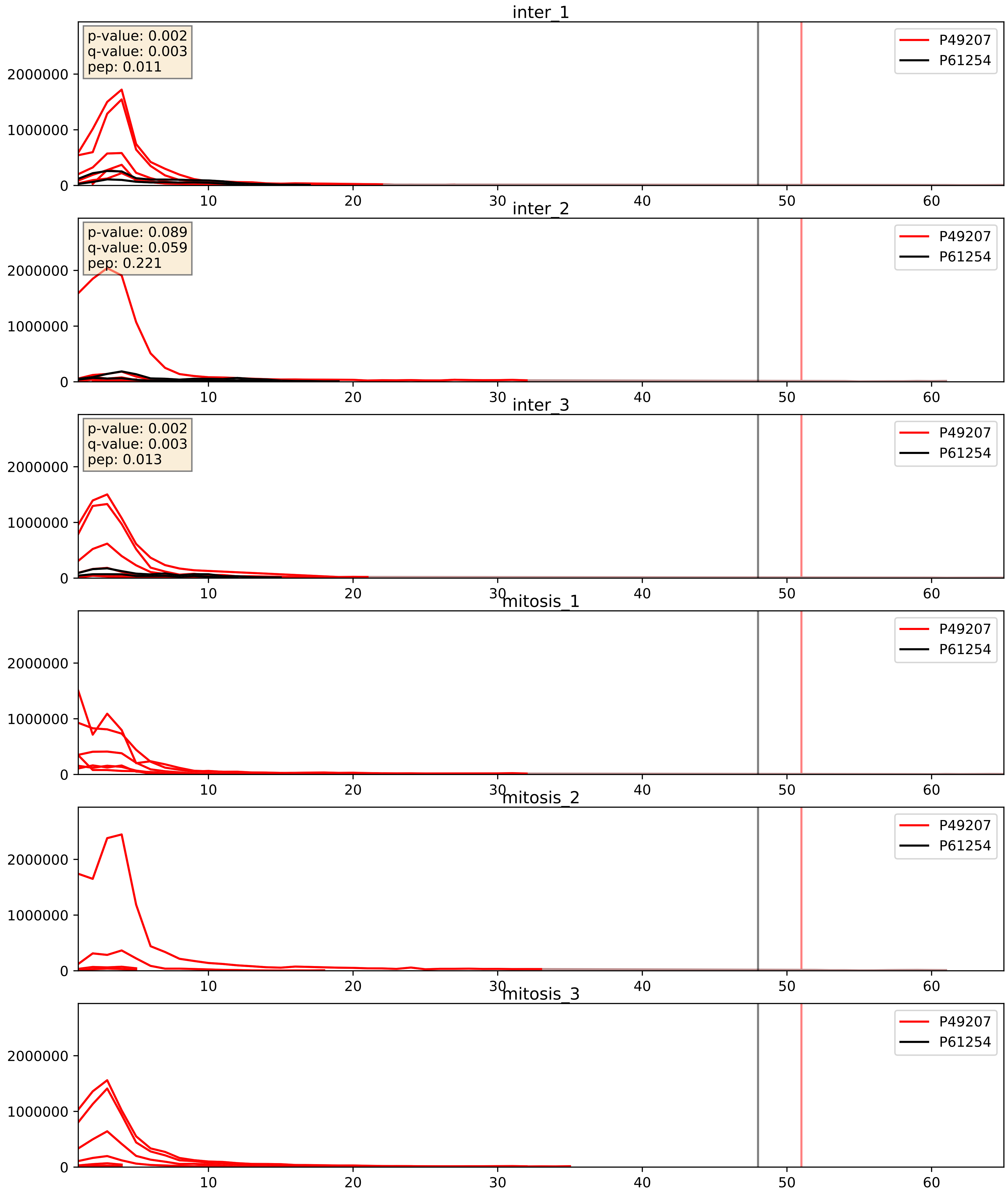


| level | condition_1 | condition_2 | pvalue | pvalue_adjusted |
|---|---|---|---|---|
| complex_abundance | mitosis | inter | 0.0005302897161240373 | 0.014642838612973419 |
| interactor_abundance | mitosis | inter | 0.0009700043449336144 | 0.02306454775731039 |
| interactor_ratio | mitosis | inter | 0.004875773163143465 | 0.09295460640647675 |

P49458_P61254
SRP09_HUMAN vs RL26_HUMAN
p-value: 0.008 q-value: 0.007

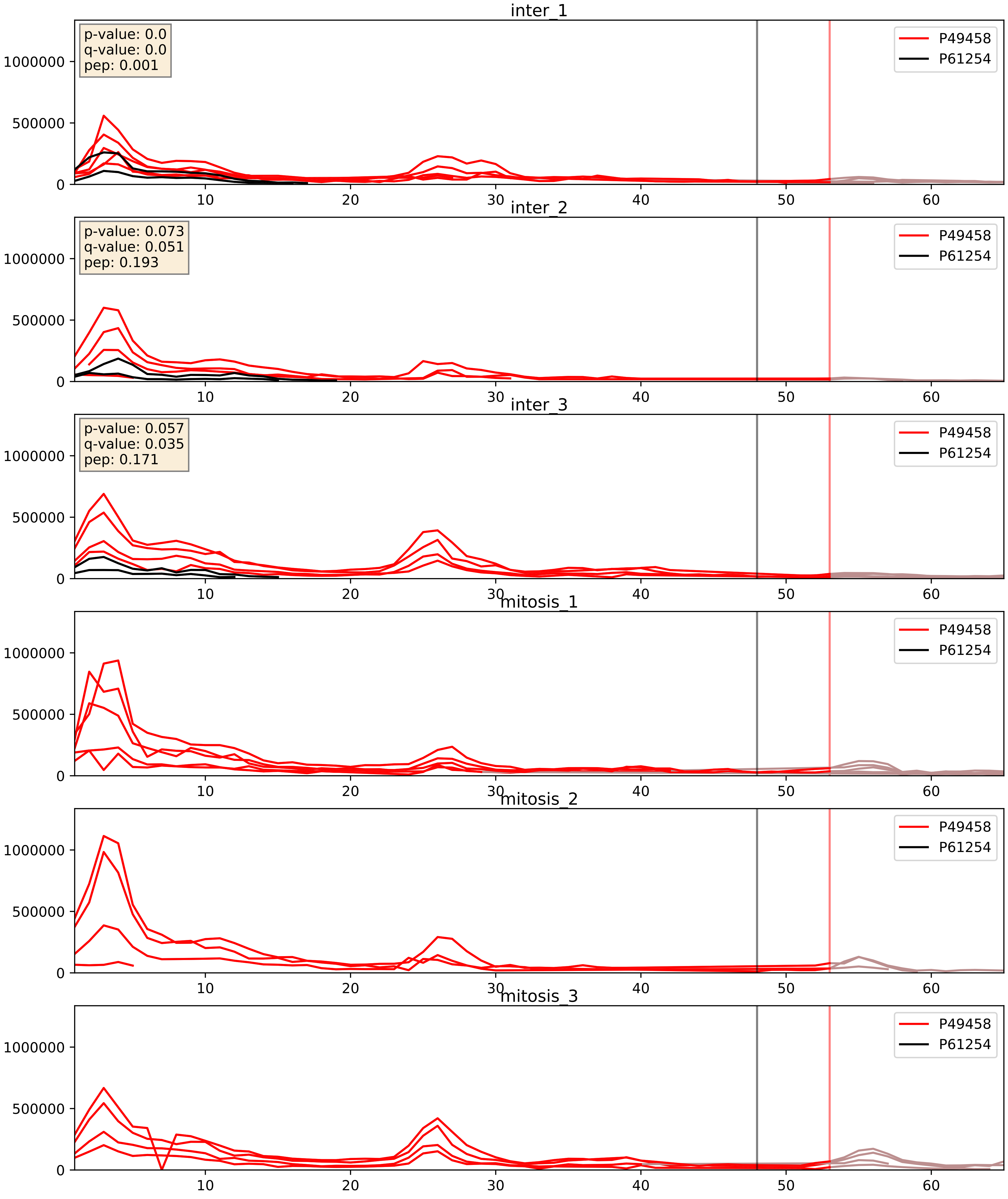


| level | condition_1 | condition_2 | pvalue | pvalue_adjusted |
|---|---|---|---|---|
| interactor_abundance | mitosis | inter | 5.518730952144667e-05 | 0.006414334379145961 |
| complex_abundance | mitosis | inter | 6.915955333606029e-05 | 0.005803978201536039 |
| interactor_ratio | mitosis | inter | 0.0004938131211380184 | 0.025464098294827937 |

P50914_P61254
RL14_HUMAN vs RL26_HUMAN
p-value: 0.001 q-value: 0.001

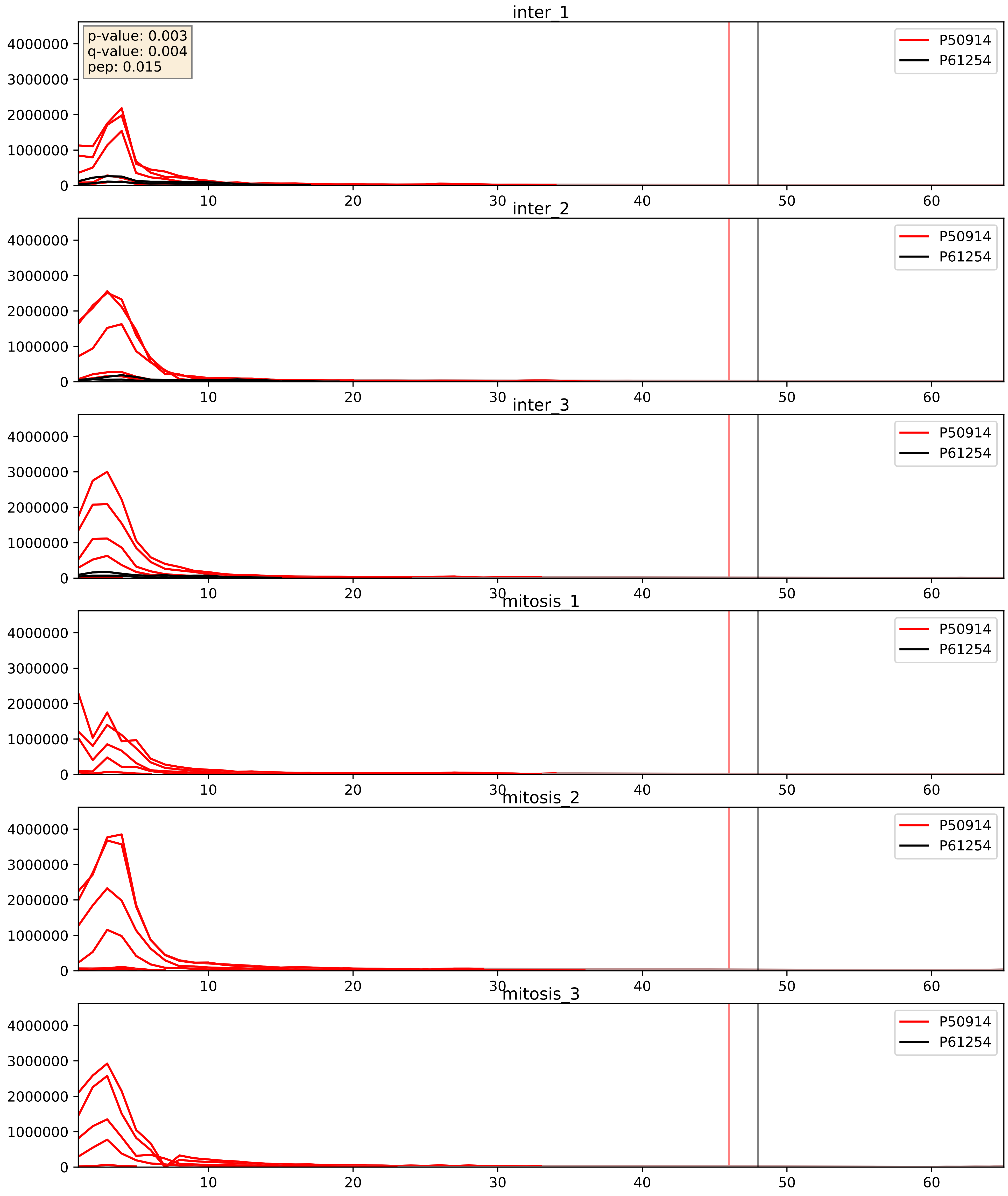


| level | condition_1 | condition_2 | pvalue | pvalue_adjusted |
| --- | --- | --- | --- | --- |
| interactor_abundance | mitosis | inter | 2.469798219508071e-05 | 0.004450836370313492 |
| complex_abundance | mitosis | inter | 2.7749091845267907e-05 | 0.005691857151652958 |
| interactor_ratio | mitosis | inter | 0.00016286192105085615 | 0.013151868341465365 |

P61247_P61254
RS3A_HUMAN vs RL26_HUMAN
p-value: 0.041 q-value: 0.028

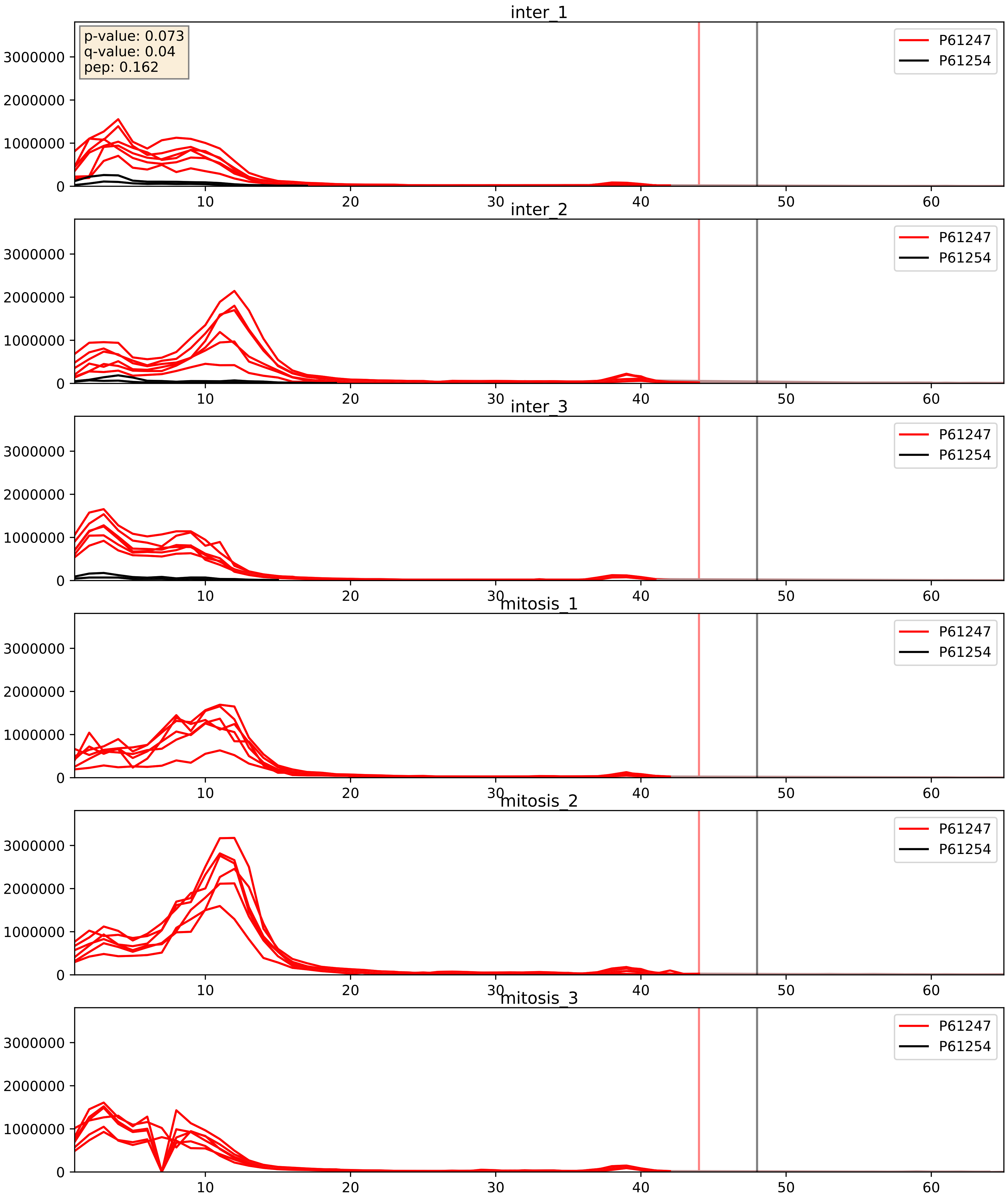


| level | condition_1 | condition_2 | pvalue | pvalue_adjusted |
|---|---|---|---|---|
| interactor_abundance | mitosis | inter | 1.3901103960192236e-05 | 0.0036058621181589558 |
| complex_abundance | mitosis | inter | 2.09741592387447e-05 | 0.005691857151652958 |
| interactor_ratio | mitosis | inter | 6.56812164772372e-05 | 0.008371379138291892 |

P61254_P61313
RL26_HUMAN vs RL15_HUMAN
p-value: 0.021 q-value: 0.016

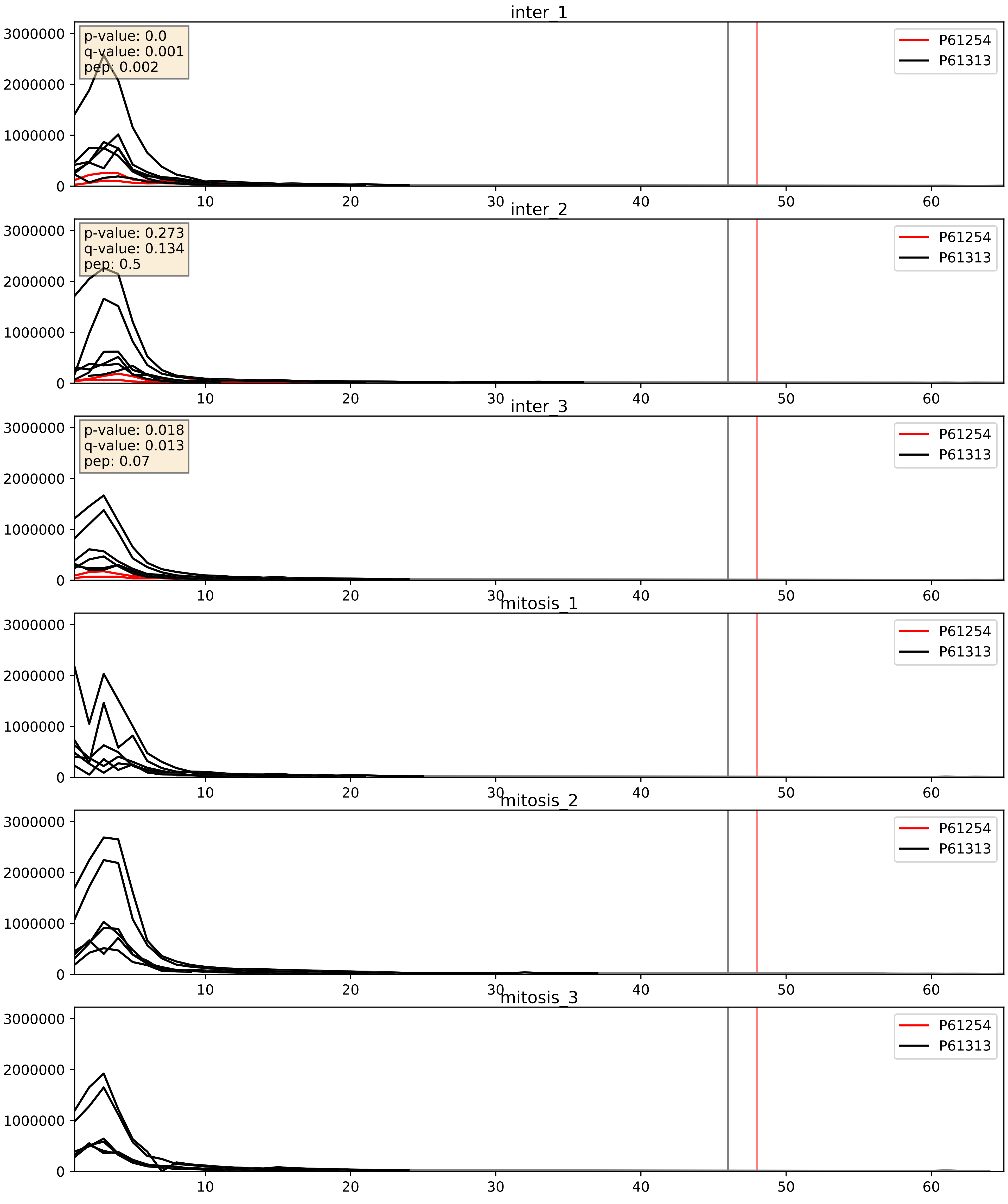


| level | condition_1 | condition_2 | pvalue | pvalue_adjusted |
|---|---|---|---|---|
| interactor_ratio | mitosis | inter | 4.416565717197257e-06 | 0.002520386835947235 |
| complex_abundance | mitosis | inter | 0.00011055789914316915 | 0.007224241348591816 |
| interactor_abundance | mitosis | inter | 0.0006144309768949645 | 0.017415659477552636 |

P61254_P61353
RL26_HUMAN vs RL27_HUMAN
p-value: 0.013 q-value: 0.011

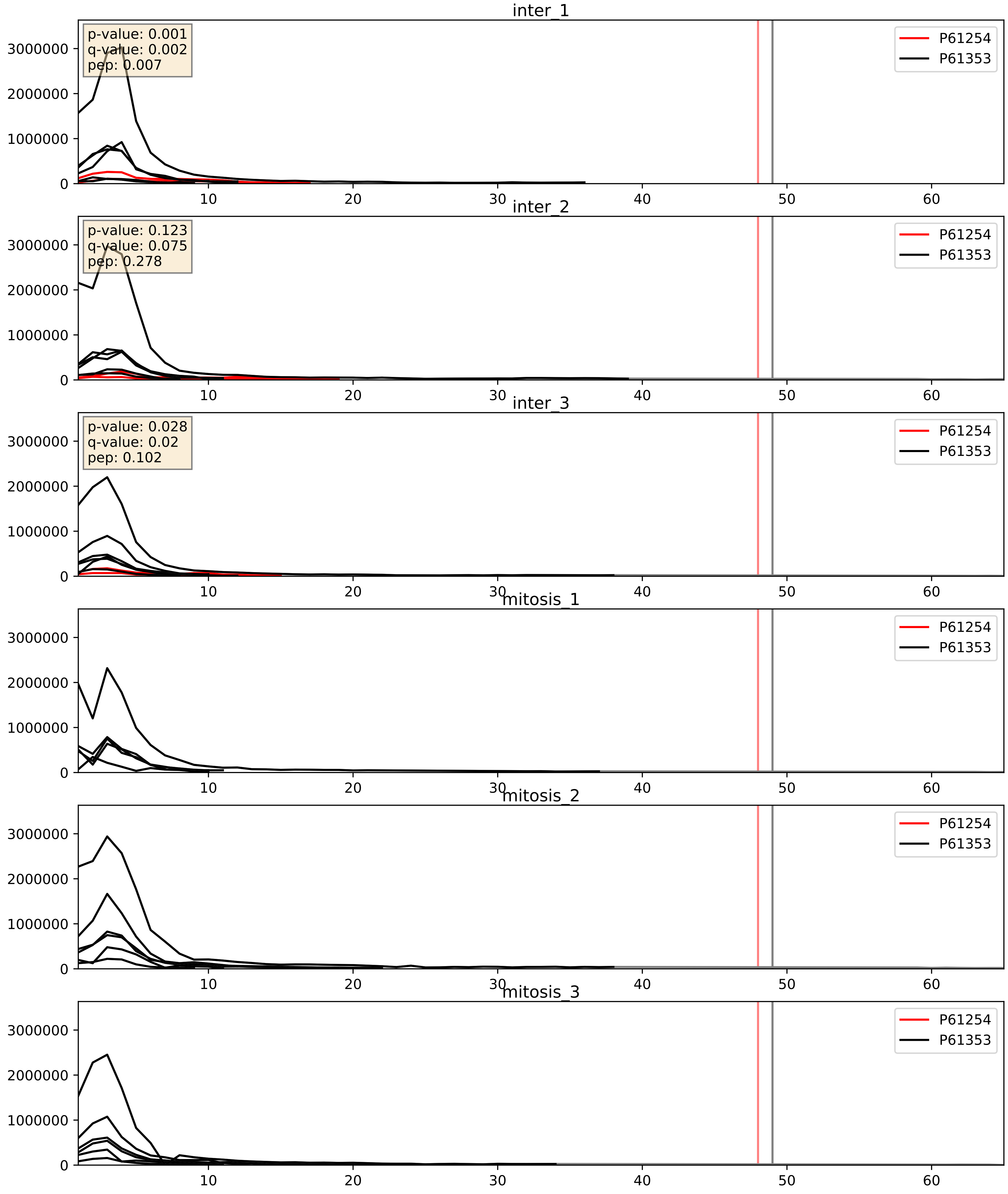


| level | condition_1 | condition_2 | pvalue | pvalue_adjusted |
|---|---|---|---|---|
| interactor_ratio | mitosis | inter | 3.454600341540311e-06 | 0.002520386835947235 |
| complex_abundance | mitosis | inter | 7.783958089694102e-05 | 0.005949167968551921 |
| interactor_abundance | mitosis | inter | 0.0006144309768949645 | 0.017415659477552636 |

P61254_P61513
RL26_HUMAN vs RL37A_HUMAN
p-value: 0.01 q-value: 0.008

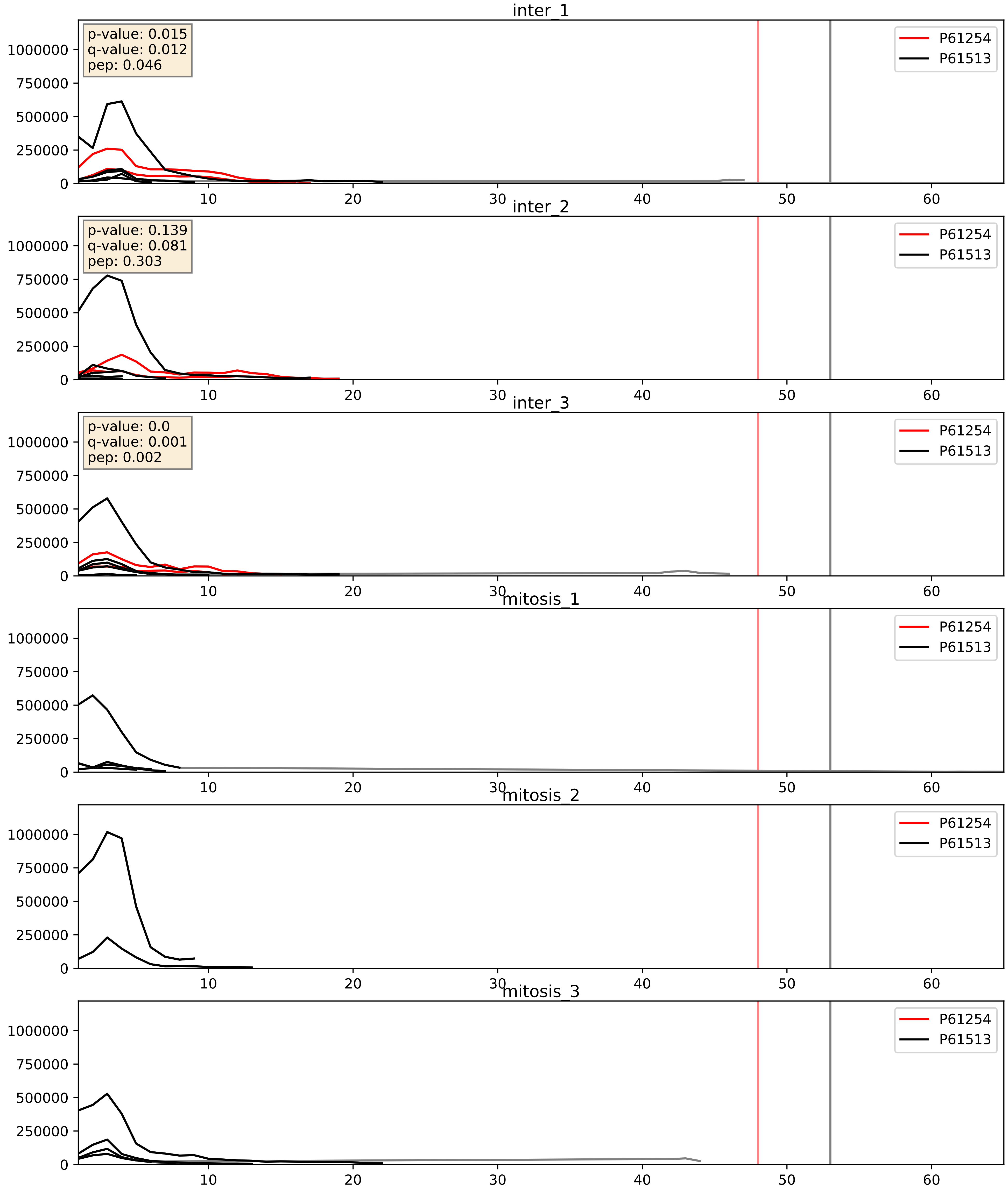


| level | condition_1 | condition_2 | pvalue | pvalue_adjusted |
|---|---|---|---|---|
| complex_abundance | mitosis | inter | 0.0002987527856444634 | 0.011624199295984575 |
| interactor_abundance | mitosis | inter | 0.0005799546565092048 | 0.017415659477552636 |
| interactor_ratio | mitosis | inter | 0.0018318325645875259 | 0.0519767486809081 |

P61254_P62081
RL26_HUMAN vs RS7_HUMAN
p-value: 0.085 q-value: 0.05

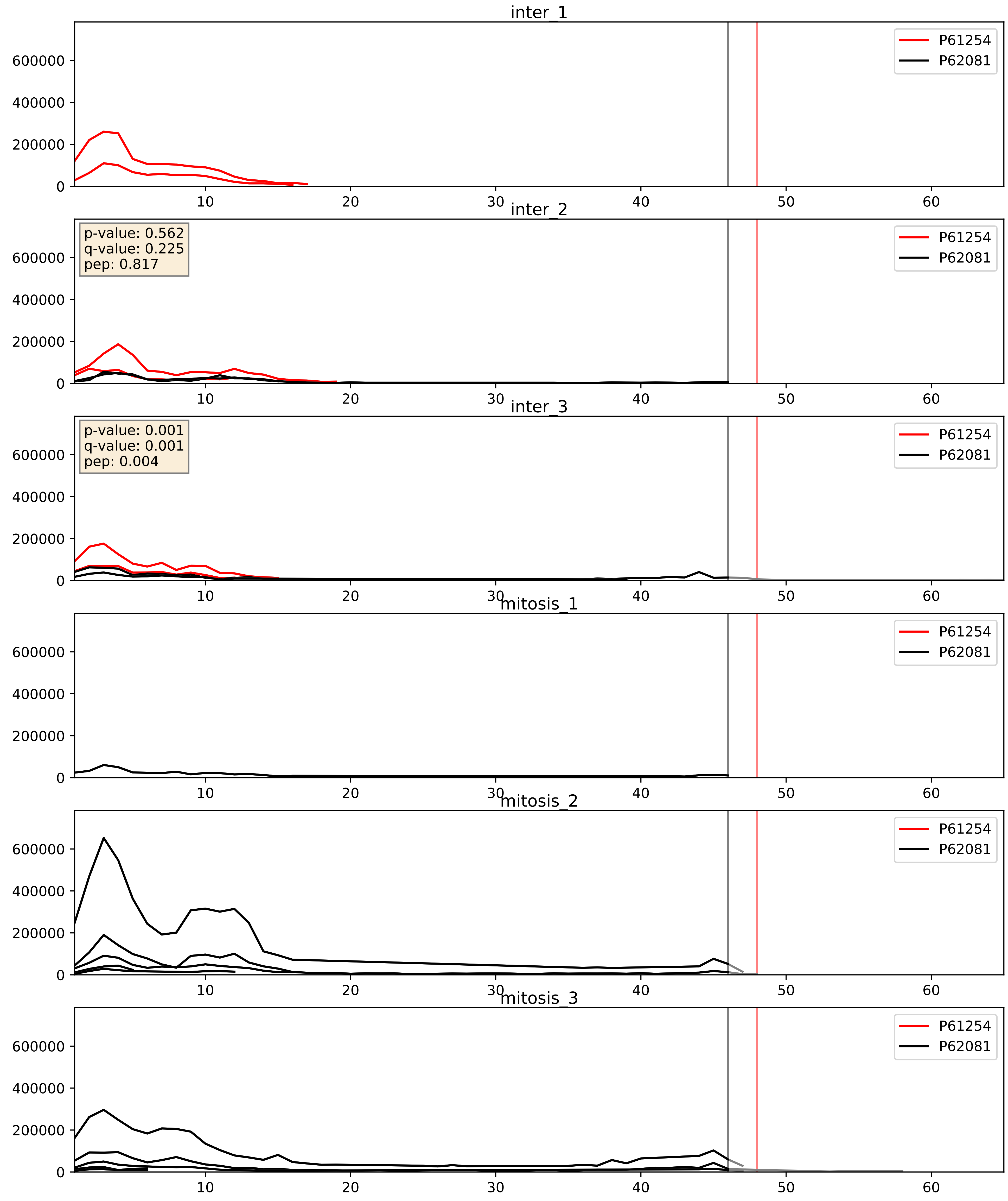


| level | condition_1 | condition_2 | pvalue | pvalue_adjusted |
|---|---|---|---|---|
| complex_abundance | mitosis | inter | 0.027086655804580792 | 0.12025018583230009 |
| interactor_abundance | mitosis | inter | 0.03198556731937464 | 0.1489898770550745 |
| interactor_ratio | mitosis | inter | 0.12596017699033923 | 0.5750062399685051 |

P61254_P62244
RL26_HUMAN vs RS15A_HUMAN
p-value: 0.015 q-value: 0.012

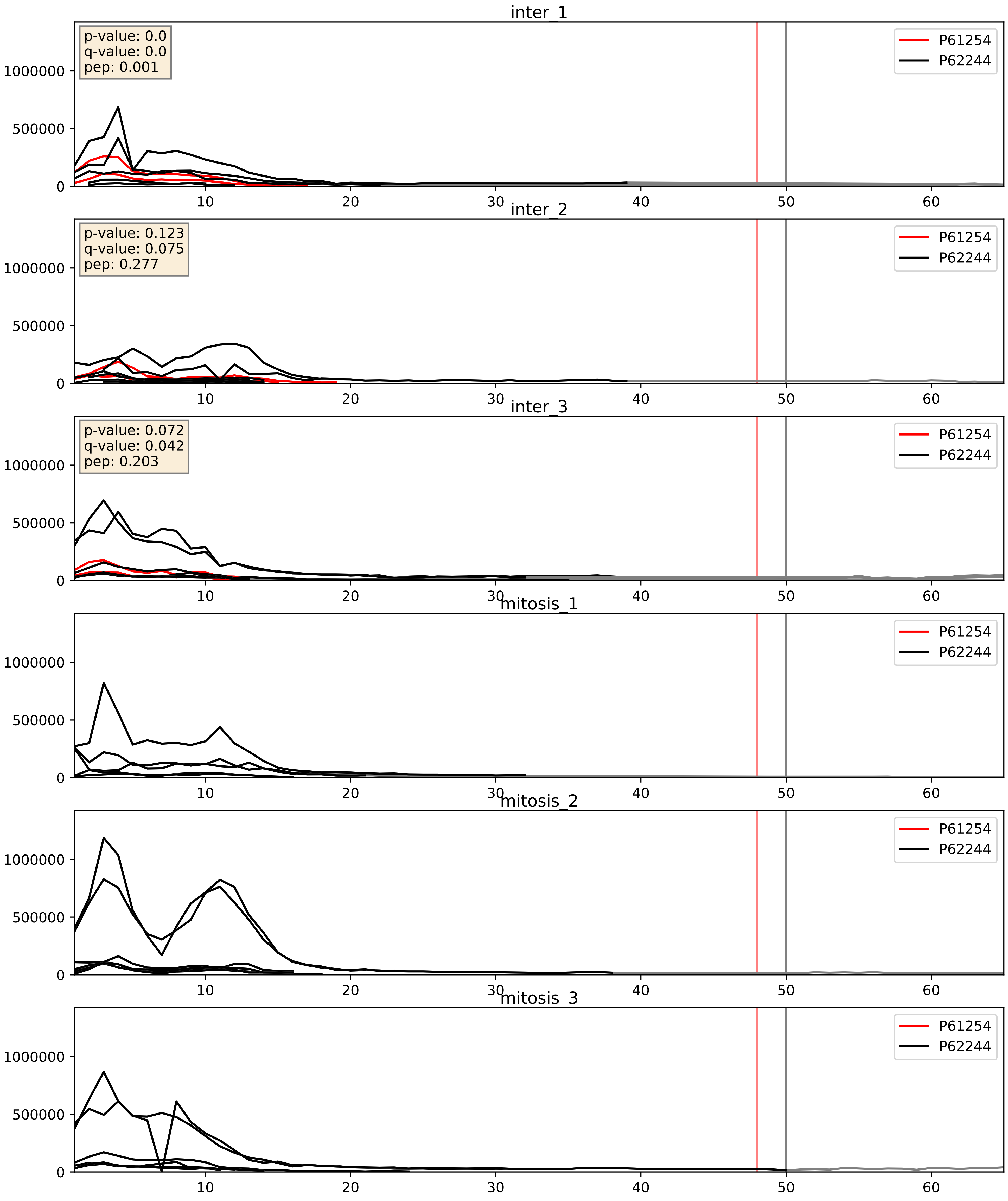


| level | condition_1 | condition_2 | pvalue | pvalue_adjusted |
|---|---|---|---|---|
| complex_abundance | mitosis | inter | 0.00015396727515901132 | 0.008563834848633118 |
| interactor_ratio | mitosis | inter | 0.00018989089546798363 | 0.014134487523529912 |
| interactor_abundance | mitosis | inter | 0.0006144309768949645 | 0.017415659477552636 |

P61254_P62249
RL26_HUMAN vs RS16_HUMAN
p-value: 0.027 q-value: 0.02

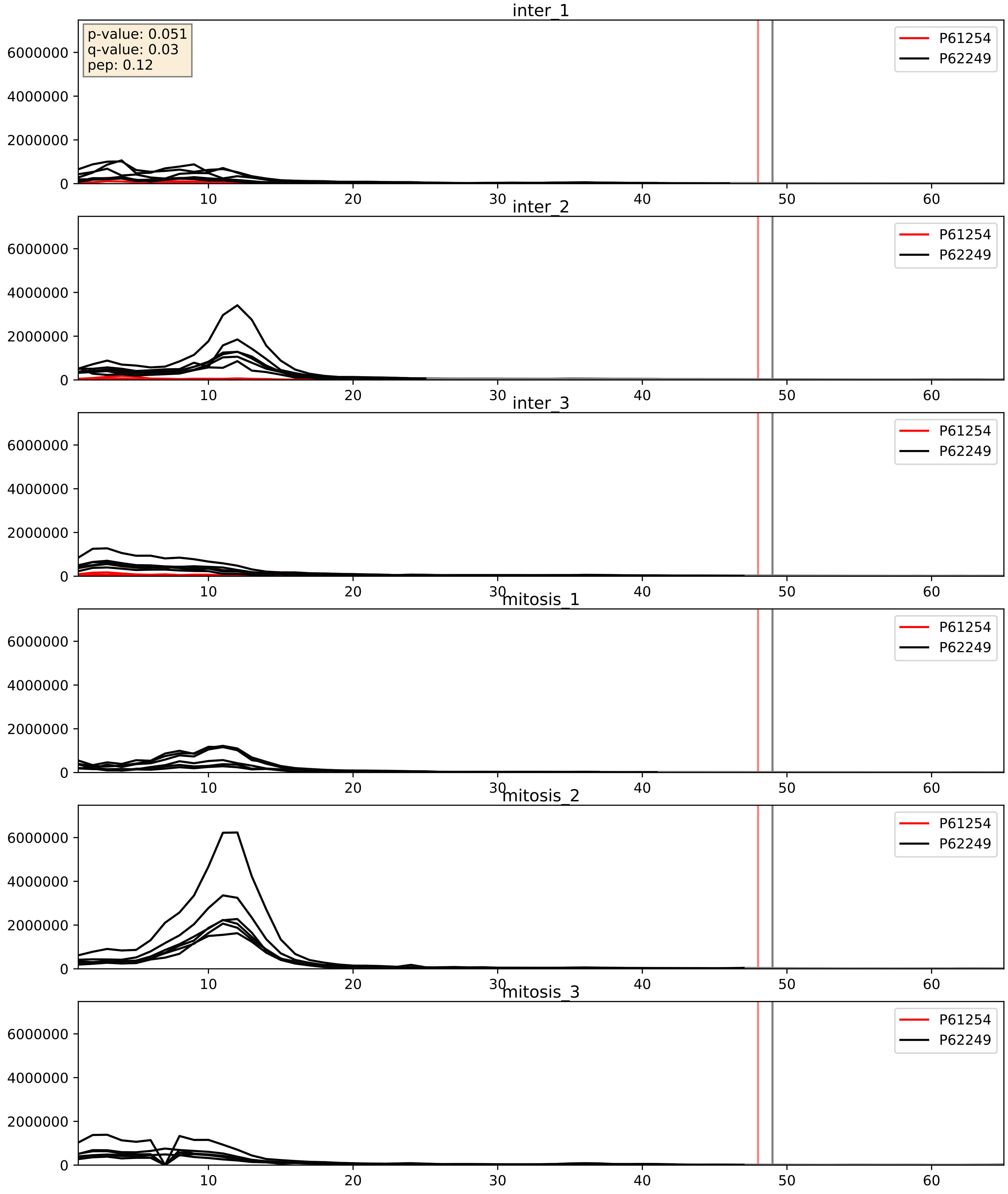


| level | condition_1 | condition_2 | pvalue | pvalue_adjusted |
|---|---|---|---|---|
| complex_abundance | mitosis | inter | 7.265303406503967e-05 | 0.005893640694944935 |
| interactor_ratio | mitosis | inter | 0.0003182223516565166 | 0.01878749154928596 |
| interactor_abundance | mitosis | inter | 0.0006144309768949645 | 0.017415659477552636 |

P61254_P62266
RL26_HUMAN vs RS23_HUMAN
p-value: 0.0 q-value: 0.0

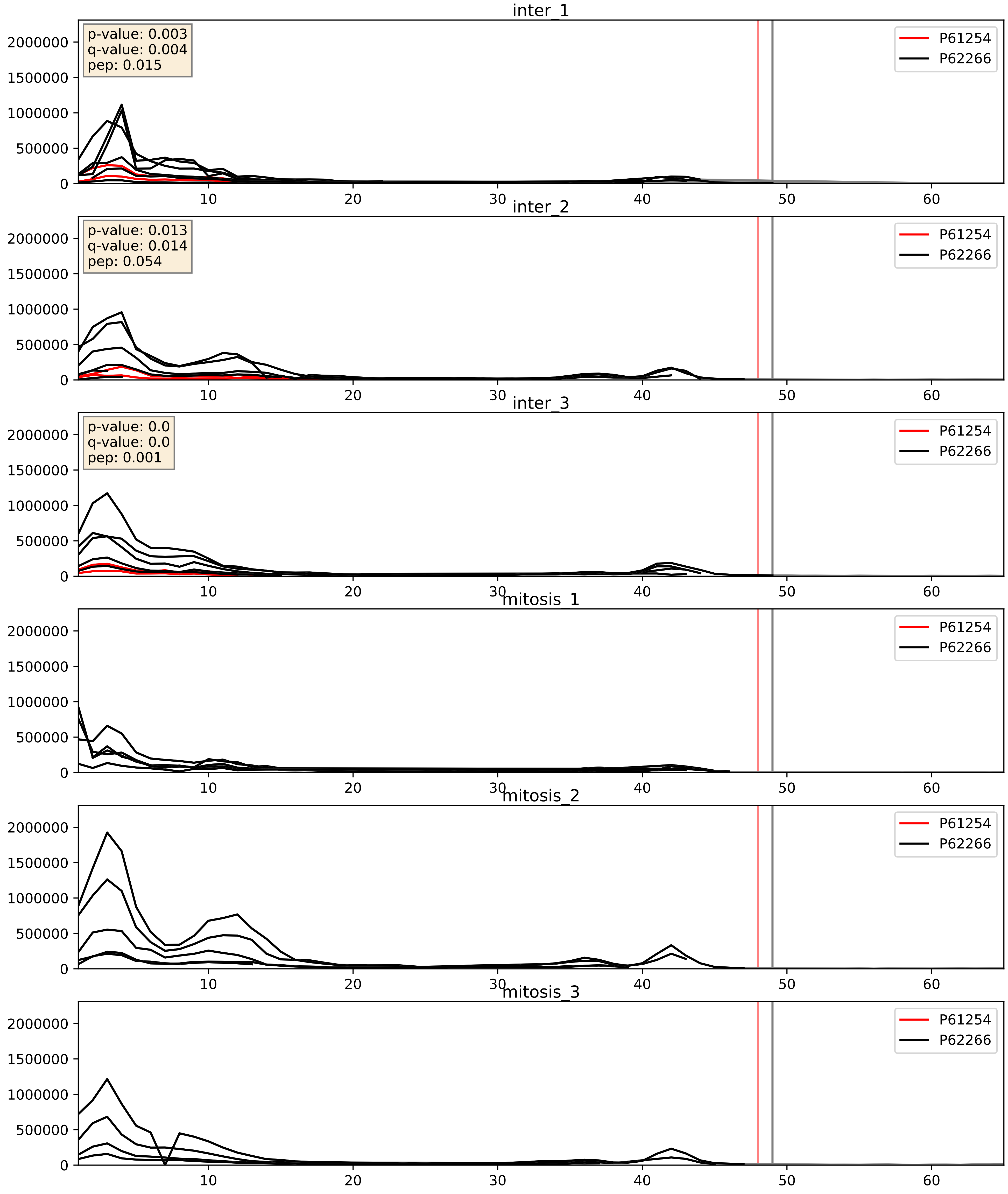


| level | condition_1 | condition_2 | pvalue | pvalue_adjusted |
| --- | --- | --- | --- | --- |
| interactor_ratio | mitosis | inter | 2.297420027729169e-05 | 0.004993481478971454 |
| complex_abundance | mitosis | inter | 5.796246078448919e-05 | 0.005691857151652958 |
| interactor_abundance | mitosis | inter | 0.0006144309768949645 | 0.017415659477552636 |

P61254_P62269
RL26_HUMAN vs RS18_HUMAN
p-value: 0.052 q-value: 0.034

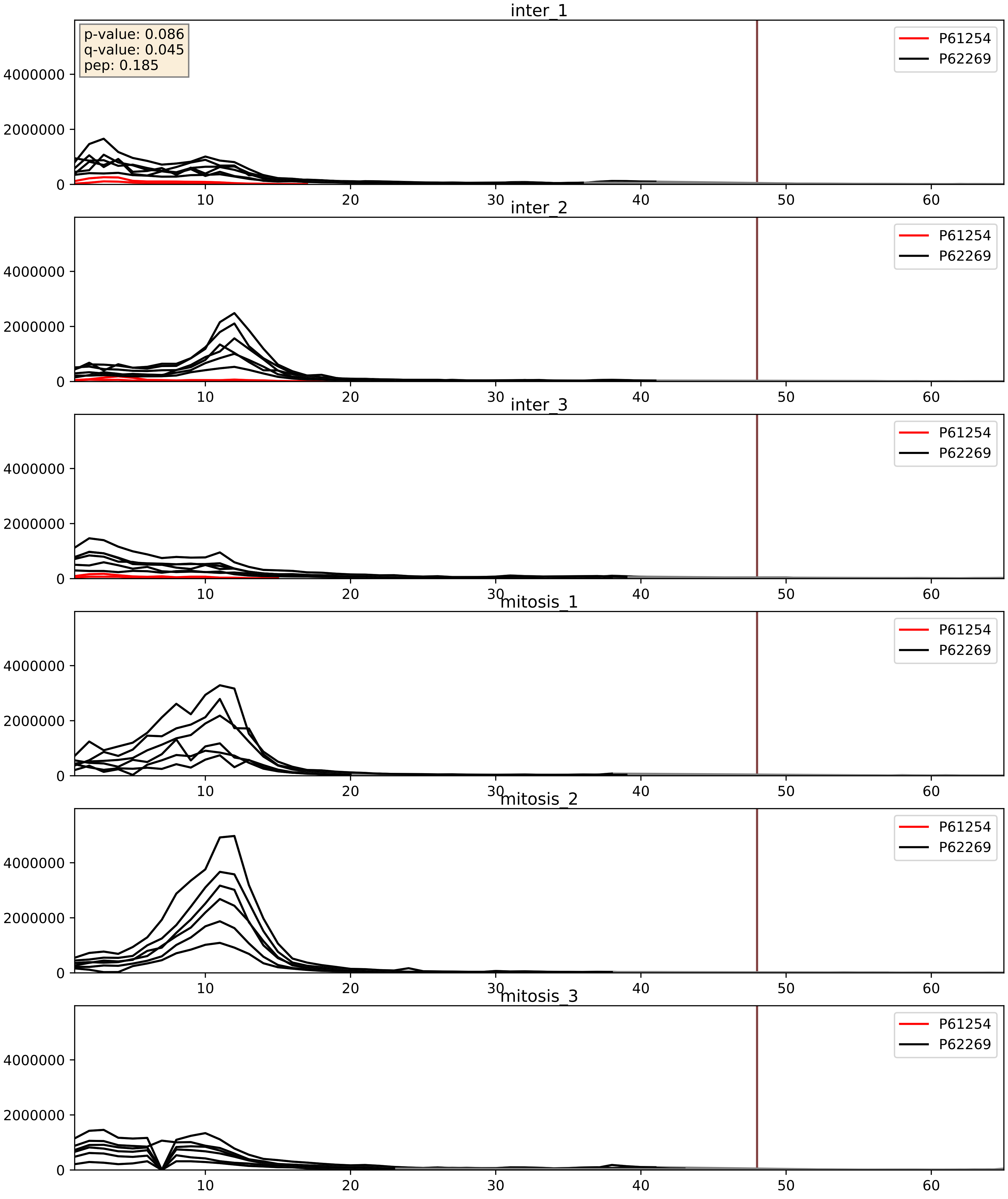


| level | condition_1 | condition_2 | pvalue | pvalue_adjusted |
|---|---|---|---|---|
| complex_abundance | mitosis | inter | 2.6885787207521043e-05 | 0.005691857151652958 |
| interactor_ratio | mitosis | inter | 2.9185735534168128e-05 | 0.005677952185738163 |
| interactor_abundance | mitosis | inter | 0.0006144309768949645 | 0.017415659477552636 |

P61254_P62277
RL26_HUMAN vs RS13_HUMAN
p-value: 0.0 q-value: 0.001

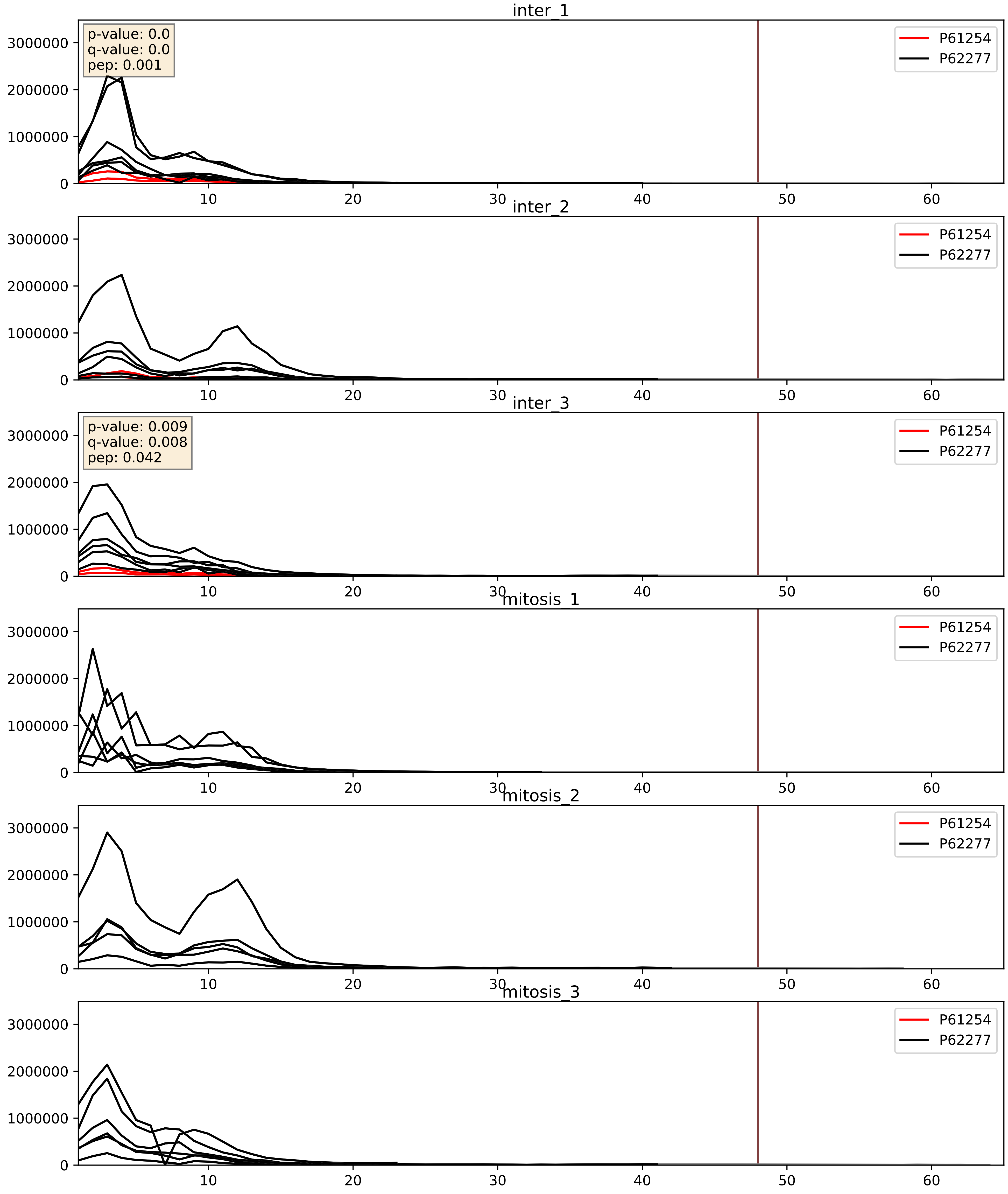


| level | condition_1 | condition_2 | pvalue | pvalue_adjusted |
|---|---|---|---|---|
| interactor_ratio | mitosis | inter | 1.3377783377575025e-05 | 0.003969306345409136 |
| complex_abundance | mitosis | inter | 3.86662153311982e-05 | 0.005691857151652958 |
| interactor_abundance | mitosis | inter | 0.0006144309768949645 | 0.017415659477552636 |

P61254_P62280
RL26_HUMAN vs RS11_HUMAN
p-value: 0.005 q-value: 0.005

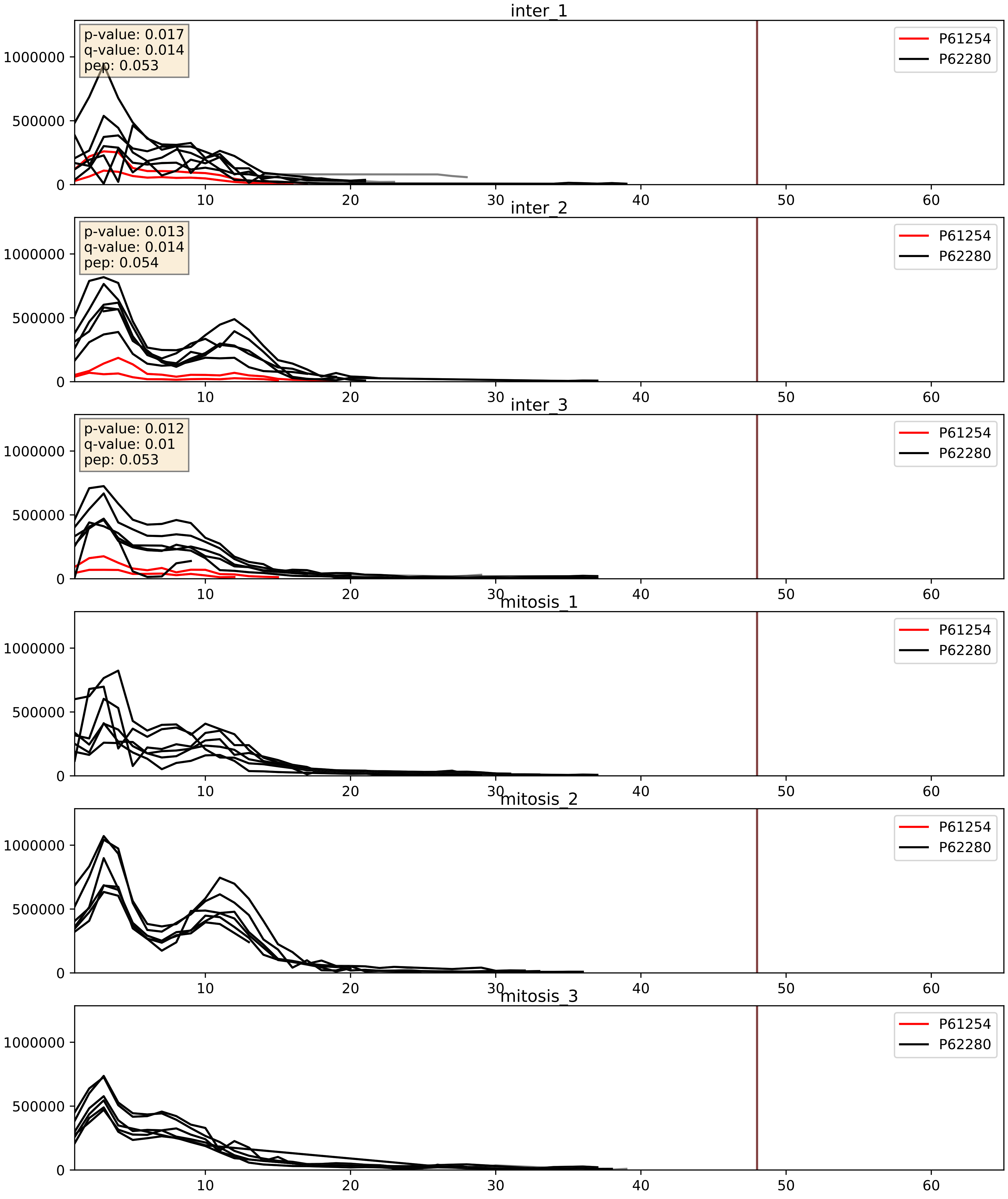


| level | condition_1 | condition_2 | pvalue | pvalue_adjusted |
|---|---|---|---|---|
| complex_abundance | mitosis | inter | 5.488019942660338e-05 | 0.005691857151652958 |
| interactor_ratio | mitosis | inter | 0.0003585396643569484 | 0.020460663512636524 |
| interactor_abundance | mitosis | inter | 0.0006144309768949645 | 0.017415659477552636 |

P61254_P62424
RL26_HUMAN vs RL7A_HUMAN
p-value: 0.034 q-value: 0.024

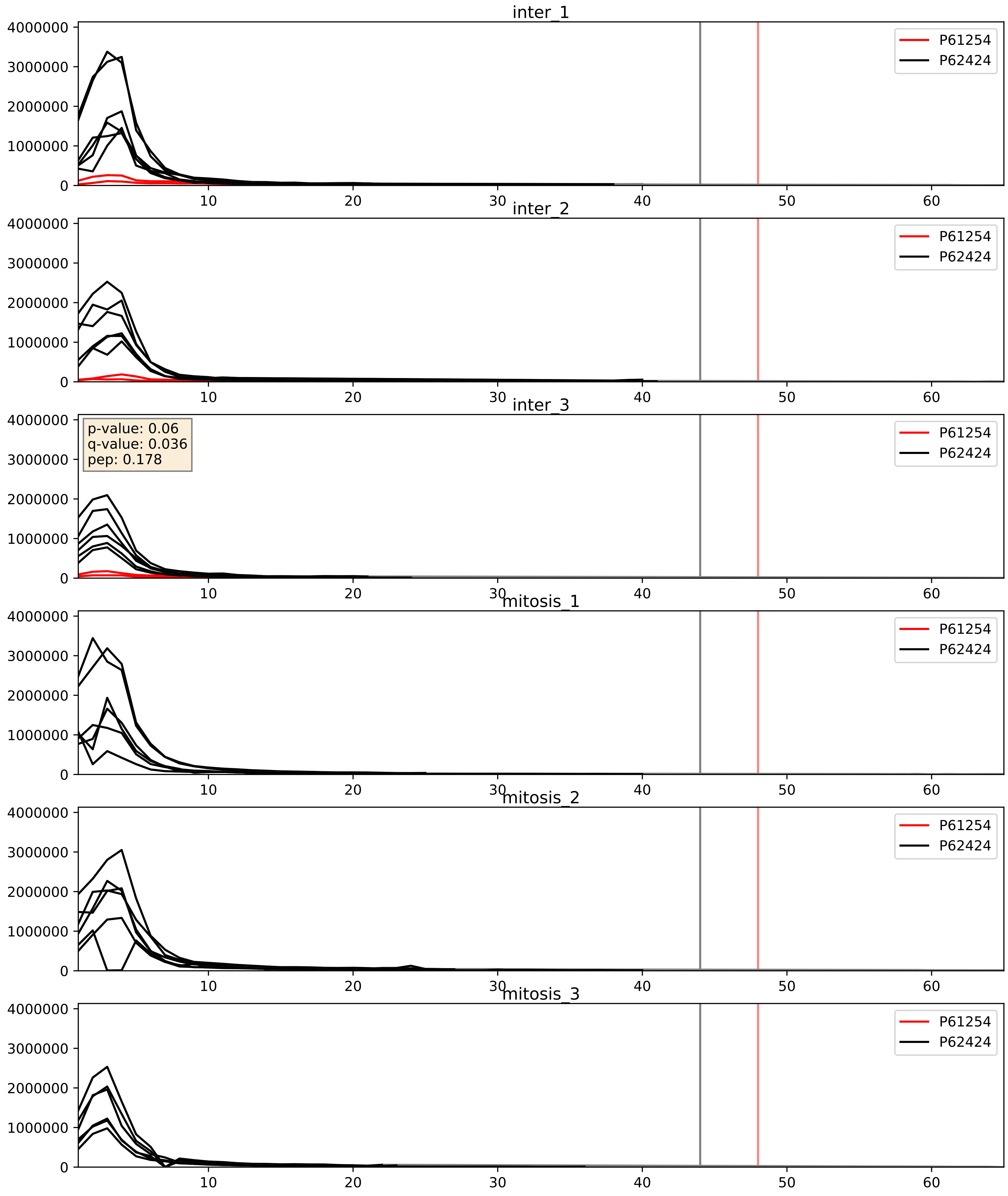


| level | condition_1 | condition_2 | pvalue | pvalue_adjusted |
|---|---|---|---|---|
| interactor_ratio | mitosis | inter | 1.2841400950072114e-05 | 0.003969306345409136 |
| complex_abundance | mitosis | inter | 0.00012096602574608935 | 0.00755816919990164 |
| interactor_abundance | mitosis | inter | 0.0006144309768949645 | 0.017415659477552636 |

P61254_P62701
RL26_HUMAN vs RS4X_HUMAN
p-value: 0.004 q-value: 0.004

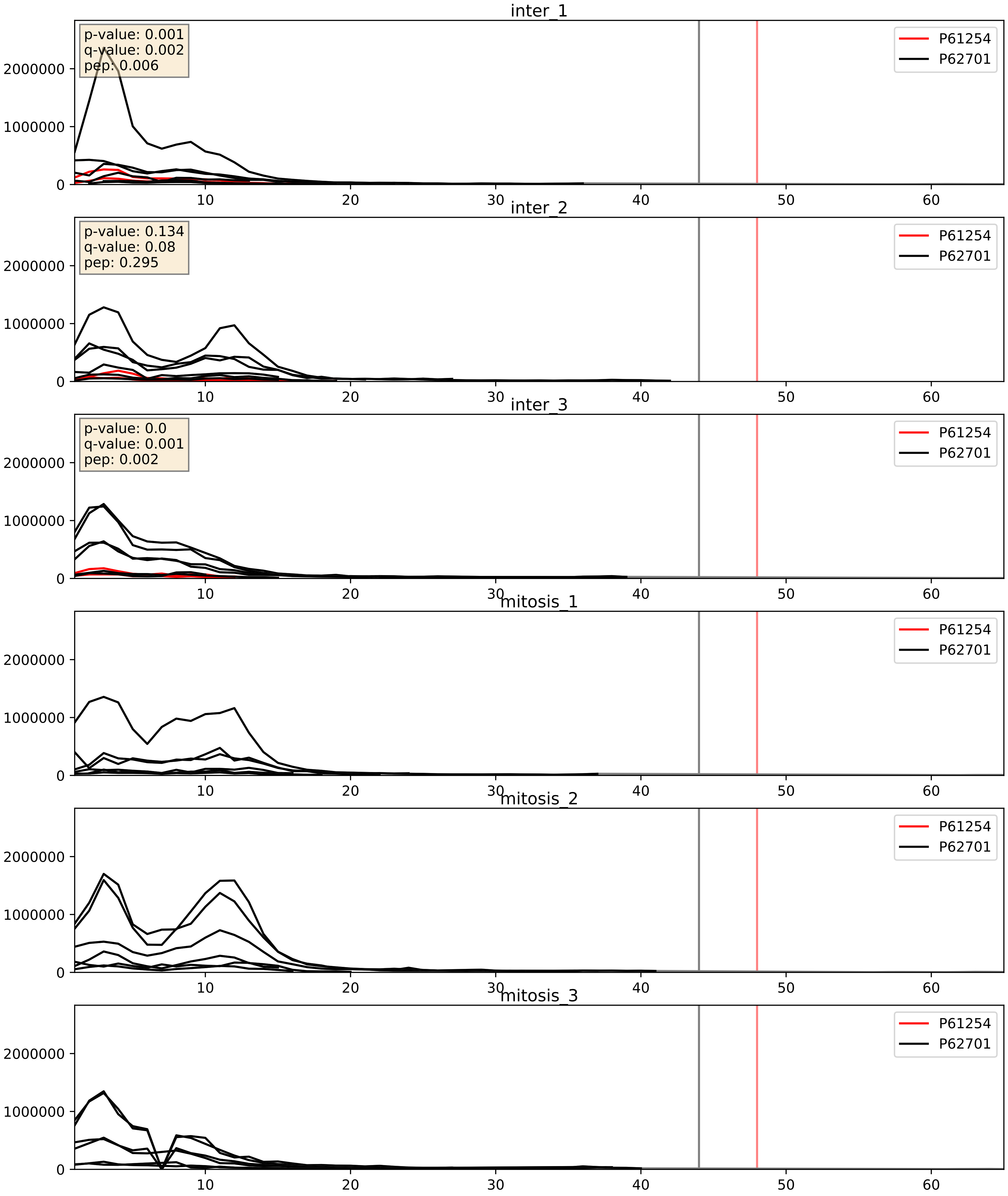


| level | condition_1 | condition_2 | pvalue | pvalue_adjusted |
|---|---|---|---|---|
| complex_abundance | mitosis | inter | 3.3573100722286054e-05 | 0.005691857151652958 |
| interactor_ratio | mitosis | inter | 0.00014876390814639185 | 0.012836278661937291 |
| interactor_abundance | mitosis | inter | 0.0006144309768949645 | 0.017415659477552636 |

P61254_P62750
RL26_HUMAN vs RL23A_HUMAN
p-value: 0.017 q-value: 0.014

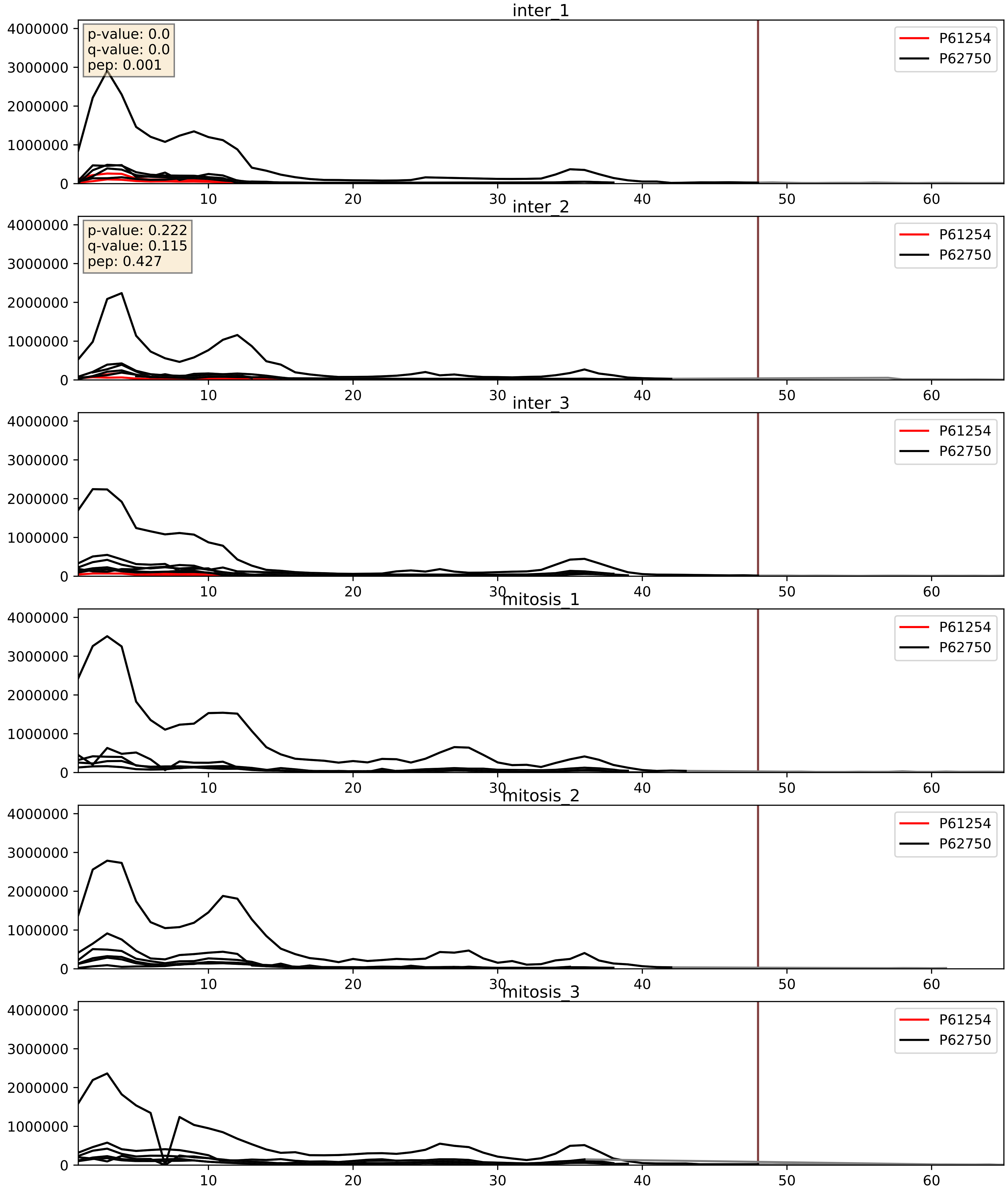


| level | condition_1 | condition_2 | pvalue | pvalue_adjusted |
|---|---|---|---|---|
| complex_abundance | mitosis | inter | 6.145318478878758e-05 | 0.005691857151652958 |
| interactor_ratio | mitosis | inter | 9.576941404330796e-05 | 0.010510079284752772 |
| interactor_abundance | mitosis | inter | 0.0006144309768949645 | 0.017415659477552636 |

P61254_P62829
RL26_HUMAN vs RL23_HUMAN
p-value: 0.048 q-value: 0.032

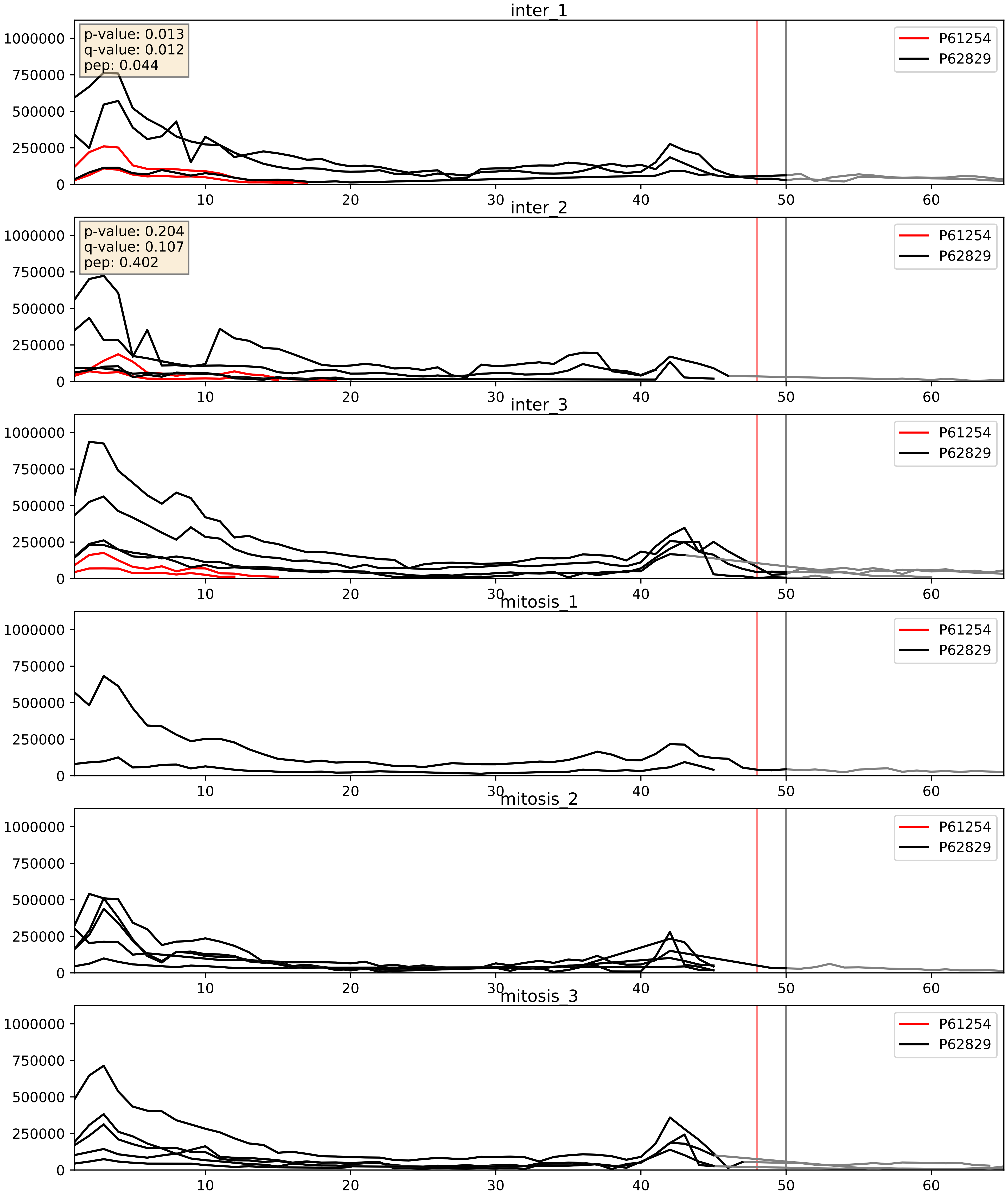


| level | condition_1 | condition_2 | pvalue | pvalue_adjusted |
|---|---|---|---|---|
| complex_abundance | mitosis | inter | 0.00010015156561264747 | 0.006746492382951911 |
| interactor_abundance | mitosis | inter | 0.0006144309768949645 | 0.017415659477552636 |
| interactor_ratio | mitosis | inter | 0.000773065761217179 | 0.03185945617443502 |

P61254_P62841
RL26_HUMAN vs RS15_HUMAN
p-value: 0.001 q-value: 0.001

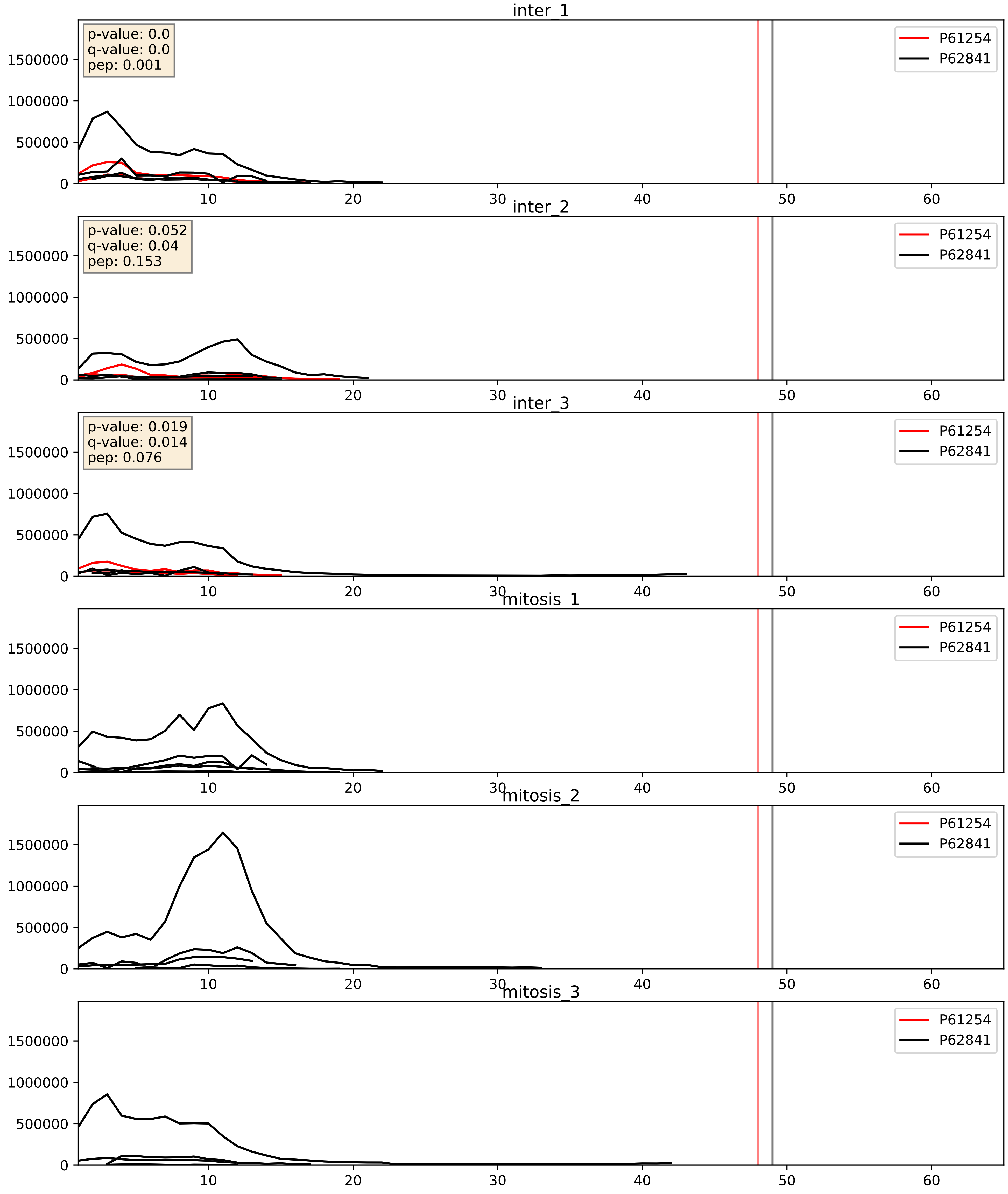


| level | condition_1 | condition_2 | pvalue | pvalue_adjusted |
|---|---|---|---|---|
| interactor_ratio | mitosis | inter | 0.00010365205704255138 | 0.010720128702277293 |
| complex_abundance | mitosis | inter | 0.00038293323433936715 | 0.01249066060683217 |
| interactor_abundance | mitosis | inter | 0.0006144309768949645 | 0.017415659477552636 |

P61254_P62847
RL26_HUMAN vs RS24_HUMAN
p-value: 0.0 q-value: 0.0

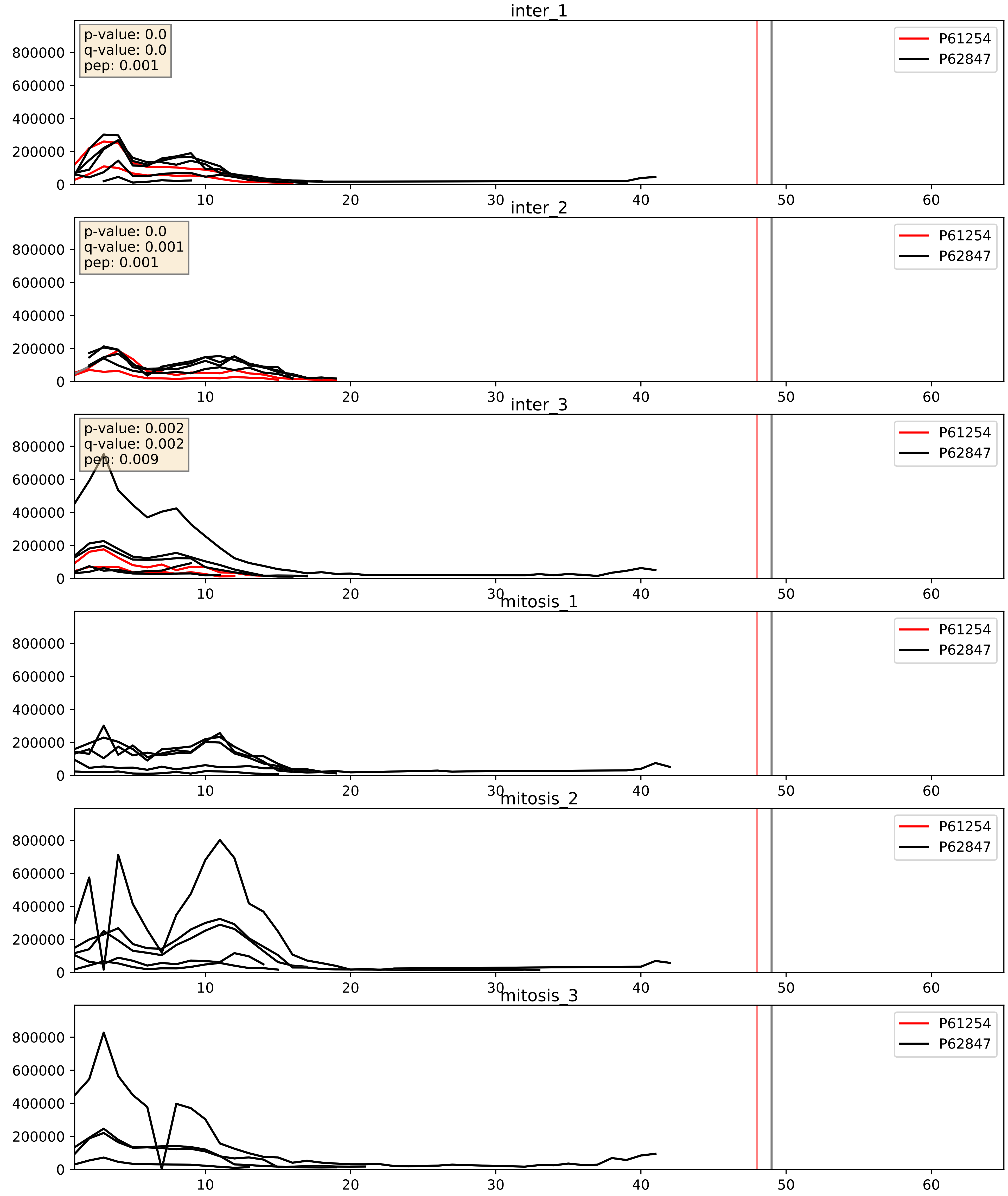


| level | condition_1 | condition_2 | pvalue | pvalue_adjusted |
|---|---|---|---|---|
| complex_abundance | mitosis | inter | 0.00012018025740401858 | 0.00755816919990164 |
| interactor_abundance | mitosis | inter | 0.0006612858544051219 | 0.018319116225591726 |
| interactor_ratio | mitosis | inter | 0.0006725566861629701 | 0.029848225645041686 |

P61254_P62861
RL26_HUMAN vs RS30_HUMAN
p-value: 0.081 q-value: 0.048

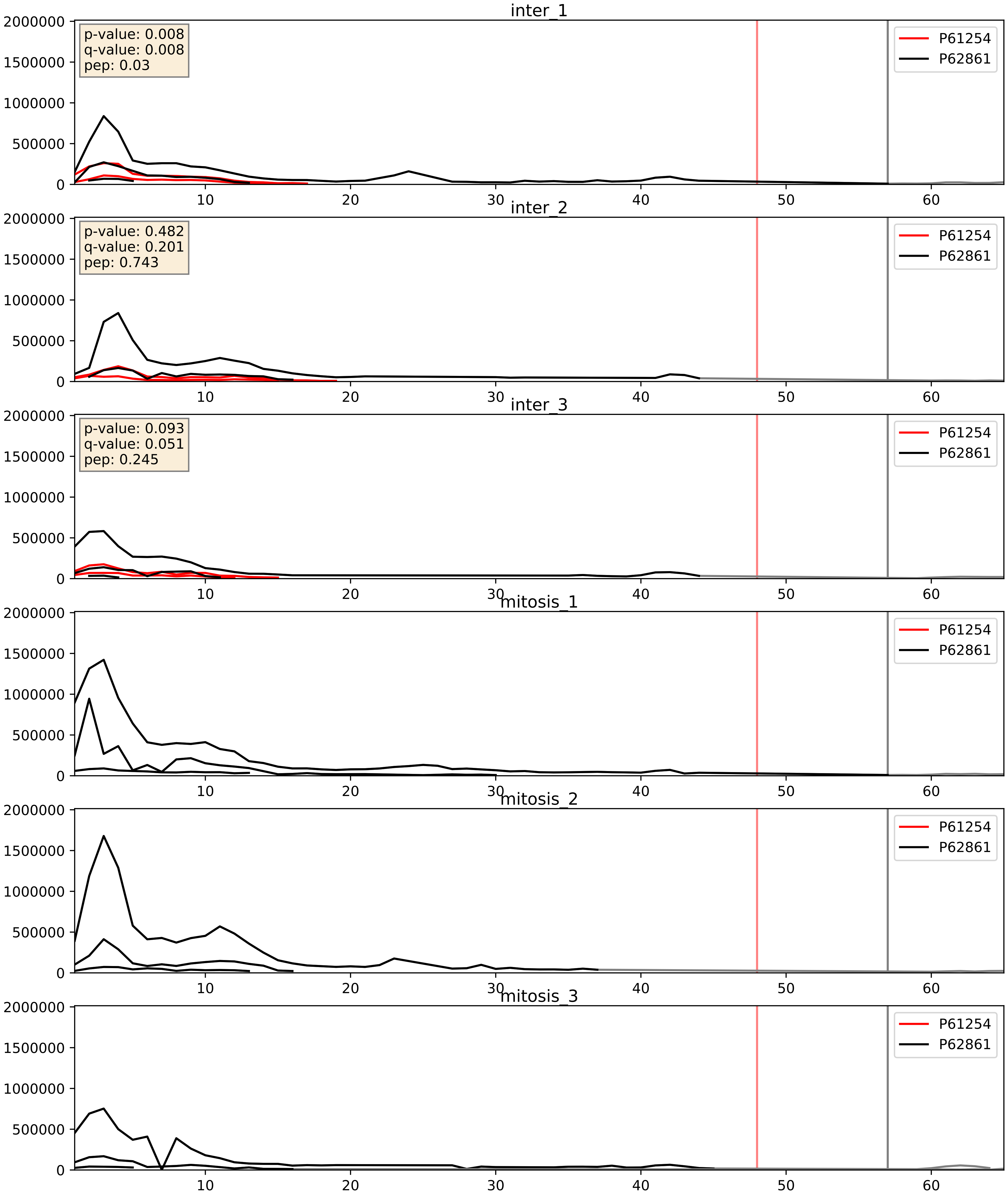


| level | condition_1 | condition_2 | pvalue | pvalue_adjusted |
|---|---|---|---|---|
| interactor_ratio | mitosis | inter | 0.00020926966620735956 | 0.014683183137172115 |
| complex_abundance | mitosis | inter | 0.0004803957564769742 | 0.013833744180327739 |
| interactor_abundance | mitosis | inter | 0.0006144309768949645 | 0.017415659477552636 |

P61254_P62888
RL26_HUMAN vs RL30_HUMAN
p-value: 0.025 q-value: 0.019

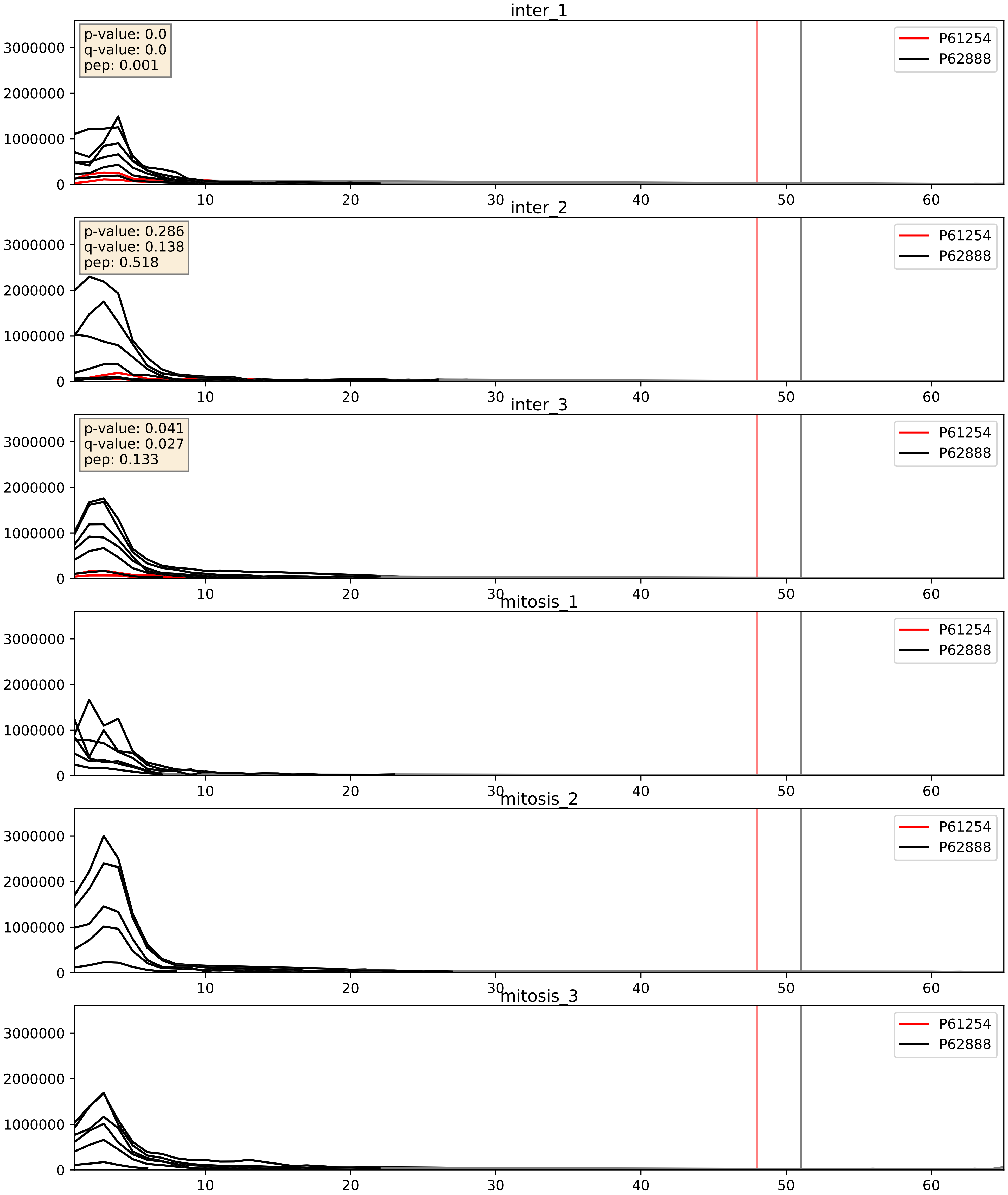


| level | condition_1 | condition_2 | pvalue | pvalue_adjusted |
|---|---|---|---|---|
| complex_abundance | mitosis | inter | 4.2546048398160366e-05 | 0.005691857151652958 |
| interactor_ratio | mitosis | inter | 0.0004573761432106749 | 0.02416752954248998 |
| interactor_abundance | mitosis | inter | 0.0006144309768949645 | 0.017415659477552636 |

P61254_P62899
RL26_HUMAN vs RL31_HUMAN
p-value: 0.005 q-value: 0.005

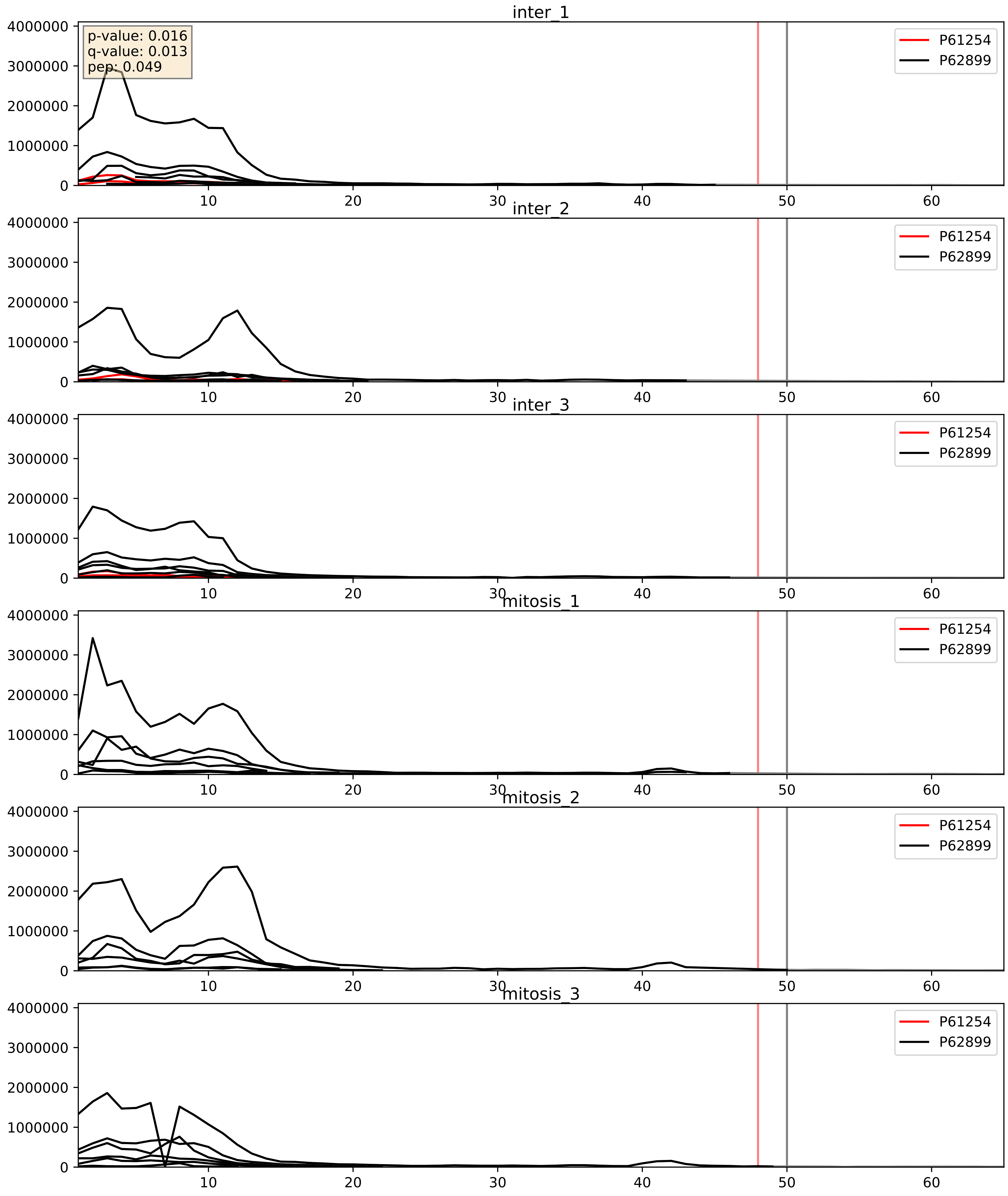


| level | condition_1 | condition_2 | pvalue | pvalue_adjusted |
|---|---|---|---|---|
| interactor_ratio | mitosis | inter | 2.278040868862456e-05 | 0.004993481478971454 |
| complex_abundance | mitosis | inter | 7.565654693739751e-05 | 0.005941468273248832 |
| interactor_abundance | mitosis | inter | 0.0006144309768949645 | 0.017415659477552636 |

P61254_P62906
RL26_HUMAN vs RL10A_HUMAN
p-value: 0.009 q-value: 0.008

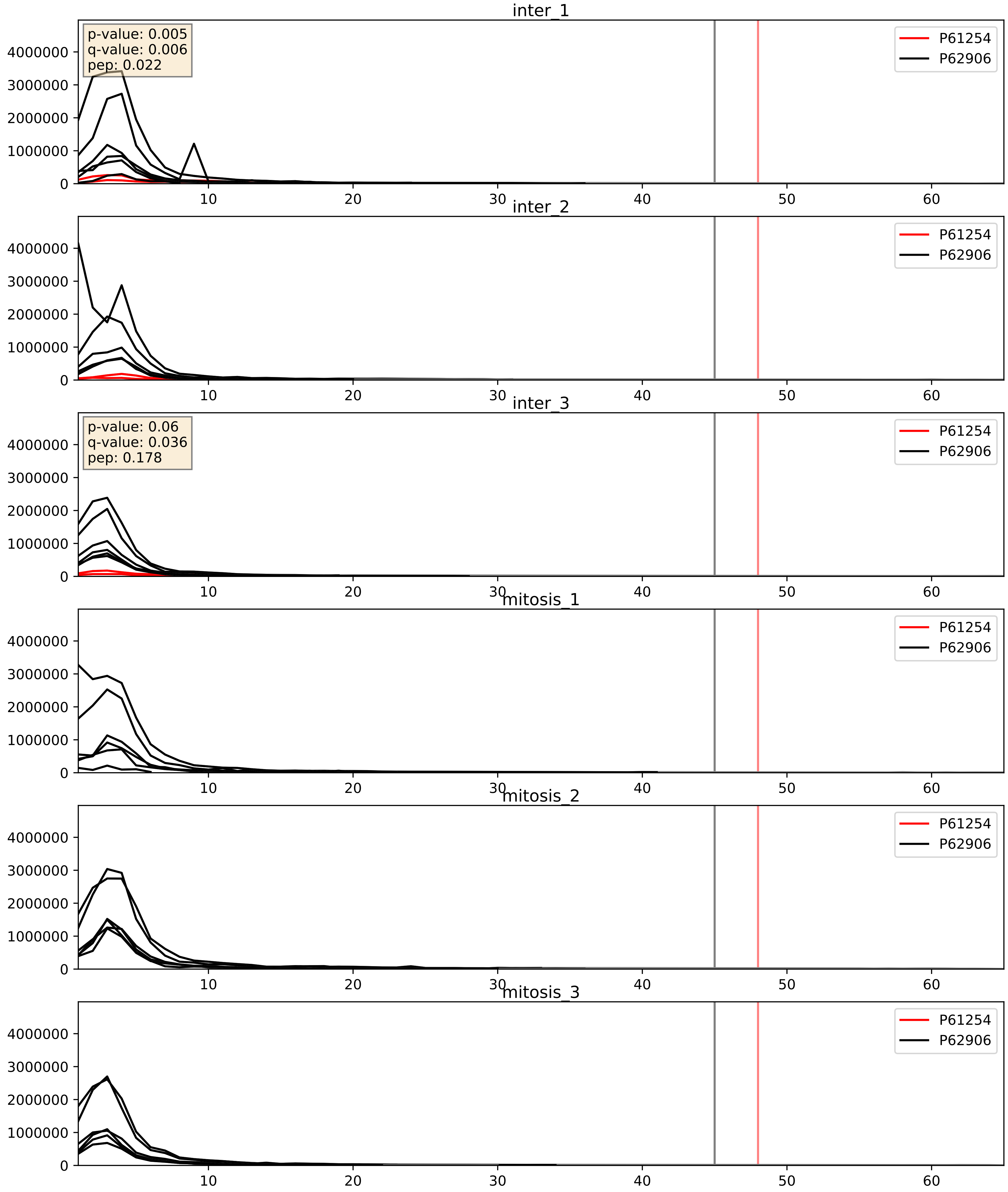


| level | condition_1 | condition_2 | pvalue | pvalue_adjusted |
|---|---|---|---|---|
| interactor_ratio | mitosis | inter | 6.337284687624959e-07 | 0.0013561789231517412 |
| complex_abundance | mitosis | inter | 6.359477139163931e-05 | 0.005691857151652958 |
| interactor_abundance | mitosis | inter | 0.0006144309768949645 | 0.017415659477552636 |

P61254_P62910
RL26_HUMAN vs RL32_HUMAN
p-value: 0.0 q-value: 0.0

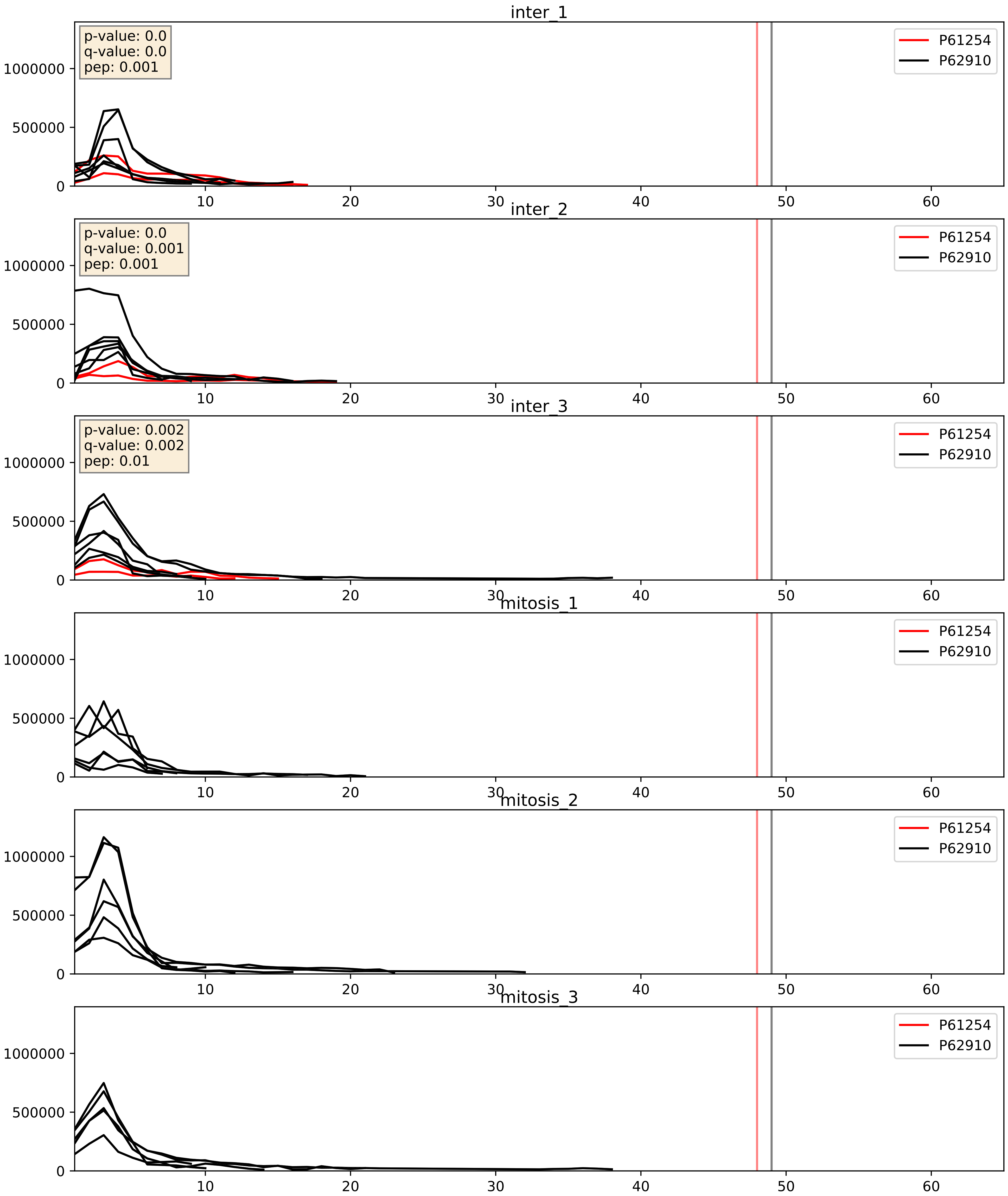


| level | condition_1 | condition_2 | pvalue | pvalue_adjusted |
|---|---|---|---|---|
| complex_abundance | mitosis | inter | 7.897500856149904e-05 | 0.005982531622003821 |
| interactor_abundance | mitosis | inter | 0.0006106728749080789 | 0.017415659477552636 |
| interactor_ratio | mitosis | inter | 0.0009113282783599036 | 0.034517566649383956 |

P61254_P62913
RL26_HUMAN vs RL11_HUMAN
p-value: 0.003 q-value: 0.004

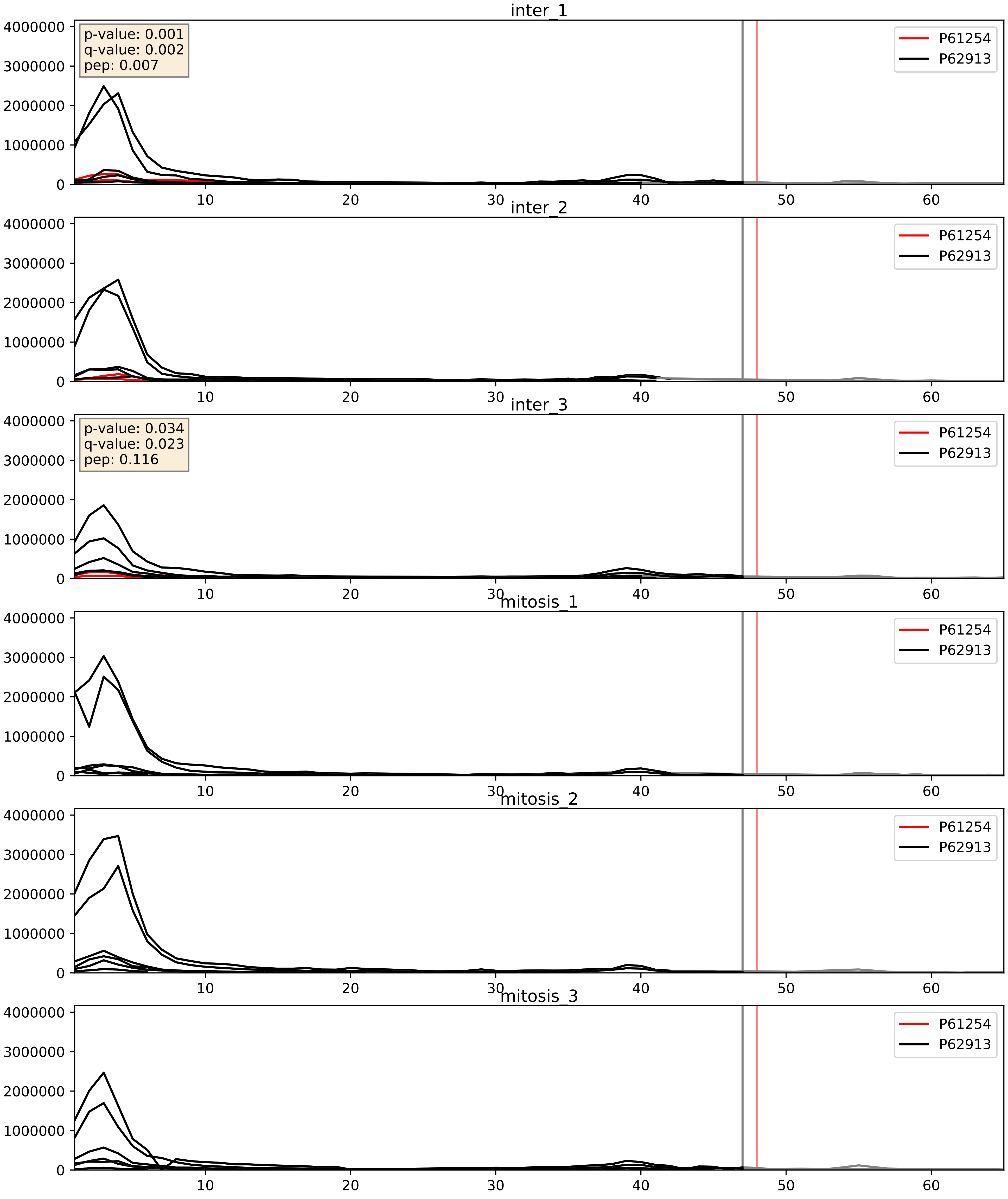


| level | condition_1 | condition_2 | pvalue | pvalue_adjusted |
|---|---|---|---|---|
| interactor_ratio | mitosis | inter | 3.68438568323015e-05 | 0.00641081637209747 |
| complex_abundance | mitosis | inter | 6.494523676077732e-05 | 0.005691857151652958 |
| interactor_abundance | mitosis | inter | 0.0006144309768949645 | 0.017415659477552636 |

P61254_P62917
RL26_HUMAN vs RL8_HUMAN
p-value: 0.05 q-value: 0.032

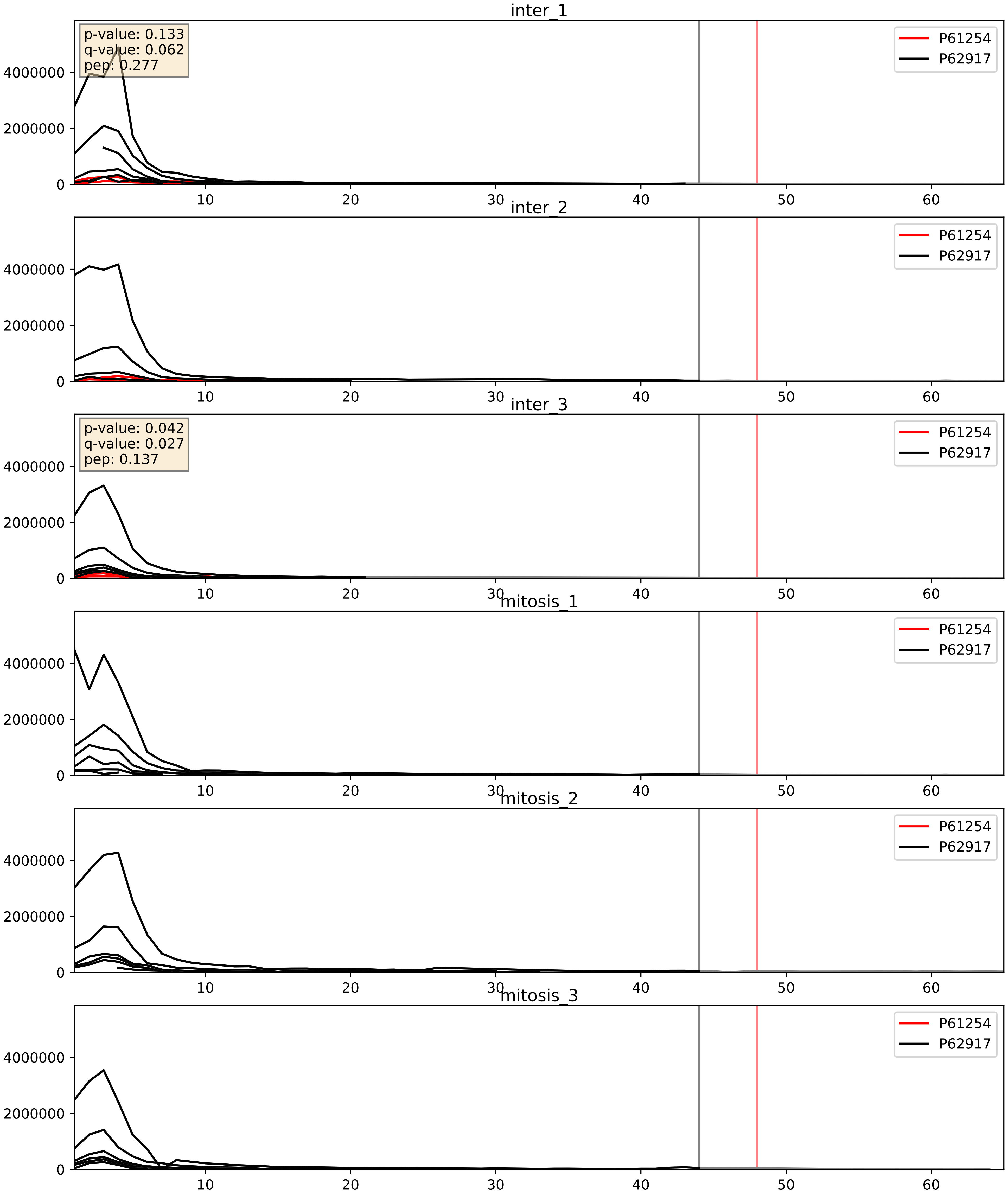


| level | condition_1 | condition_2 | pvalue | pvalue_adjusted |
|---|---|---|---|---|
| complex_abundance | mitosis | inter | 0.0003185262429502608 | 0.01165207110963347 |
| interactor_abundance | mitosis | inter | 0.0006144309768949645 | 0.017415659477552636 |
| interactor_ratio | mitosis | inter | 0.0006197193447599191 | 0.029219626944512057 |

P61254_P83731
RL26_HUMAN vs RL24_HUMAN
p-value: 0.002 q-value: 0.002

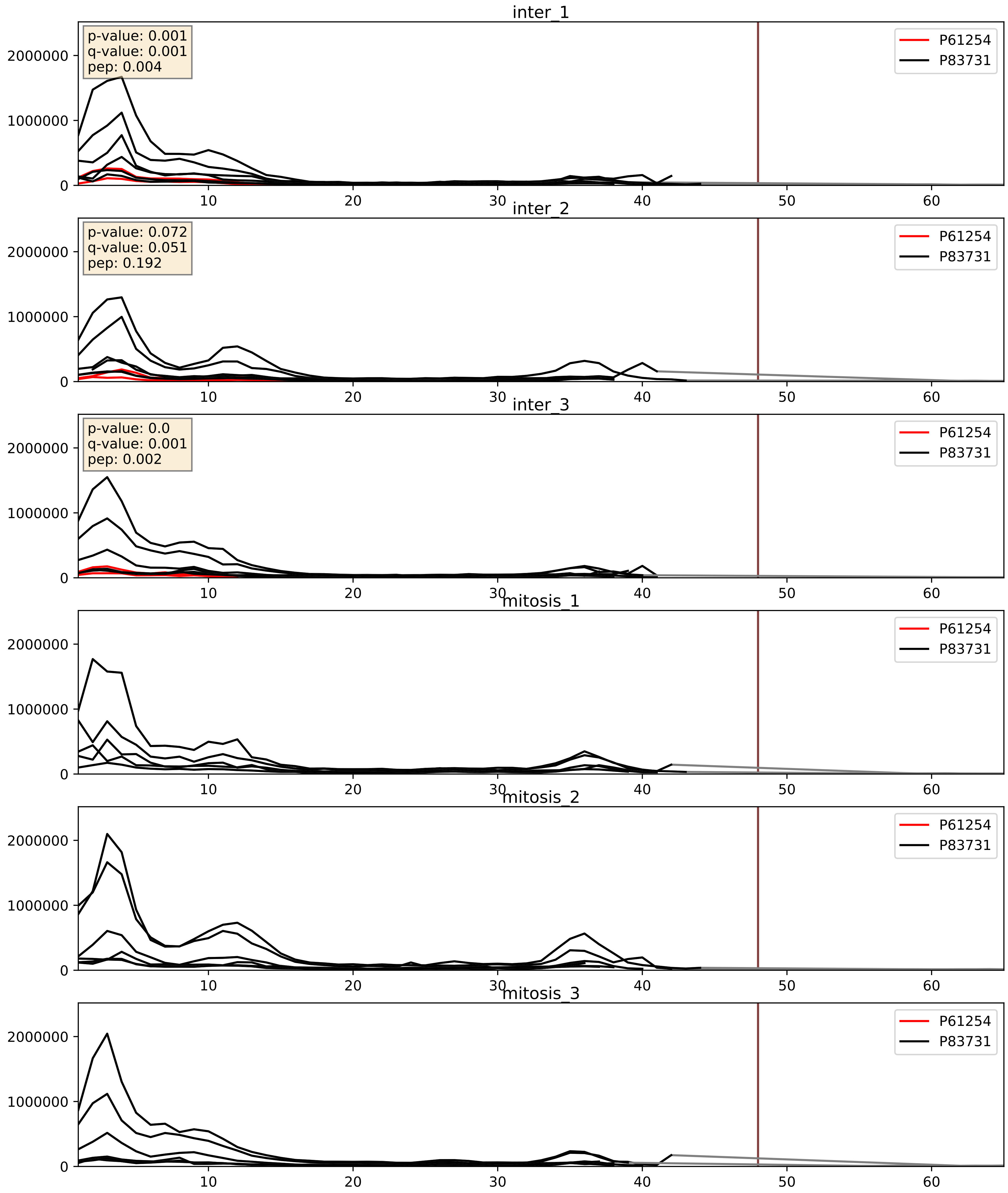


| level | condition_1 | condition_2 | pvalue | pvalue_adjusted |
|---|---|---|---|---|
| interactor_ratio | mitosis | inter | 2.189883545739685e-05 | 0.004993481478971454 |
| complex_abundance | mitosis | inter | 6.522773579750339e-05 | 0.005691857151652958 |
| interactor_abundance | mitosis | inter | 0.0006144309768949645 | 0.017415659477552636 |

P61254_P84098
RL26_HUMAN vs RL19_HUMAN
p-value: 0.003 q-value: 0.003

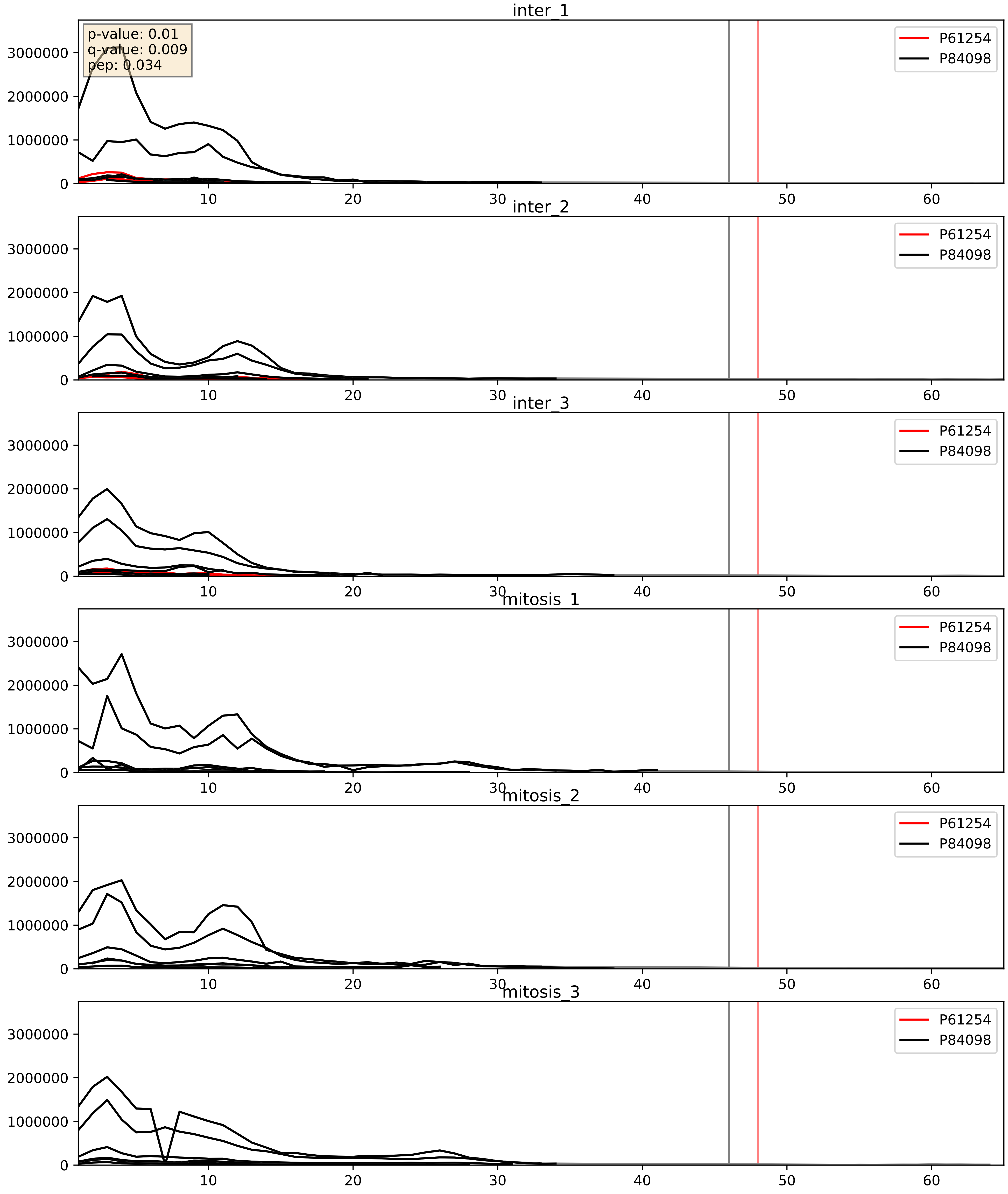


| level | condition_1 | condition_2 | pvalue | pvalue_adjusted |
|---|---|---|---|---|
| complex_abundance | mitosis | inter | 4.213236819254313e-05 | 0.005691857151652958 |
| interactor_ratio | mitosis | inter | 4.334882414149513e-05 | 0.006796360849516482 |
| interactor_abundance | mitosis | inter | 0.0006144309768949645 | 0.017415659477552636 |

P61254_Q02543
RL26_HUMAN vs RL18A_HUMAN
p-value: 0.0 q-value: 0.0

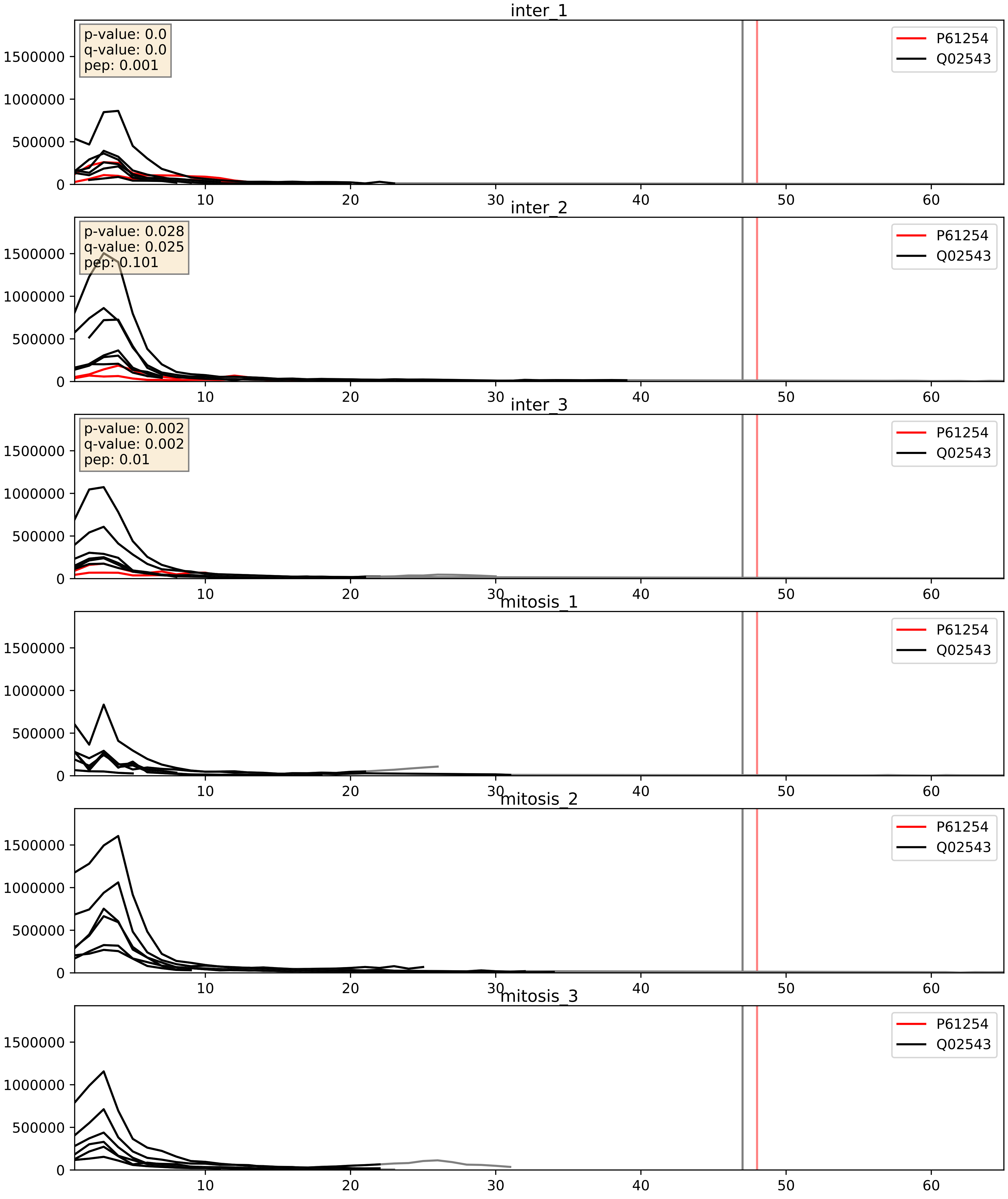


| level | condition_1 | condition_2 | pvalue | pvalue_adjusted |
|---|---|---|---|---|
| complex_abundance | mitosis | inter | 0.00017755033091332515 | 0.008807651911797798 |
| interactor_abundance | mitosis | inter | 0.0006144309768949645 | 0.017415659477552636 |
| interactor_ratio | mitosis | inter | 0.003408449719940575 | 0.07572939655107606 |

P61254_Q969Q0
RL26_HUMAN vs RL36L_HUMAN
p-value: 0.003 q-value: 0.003

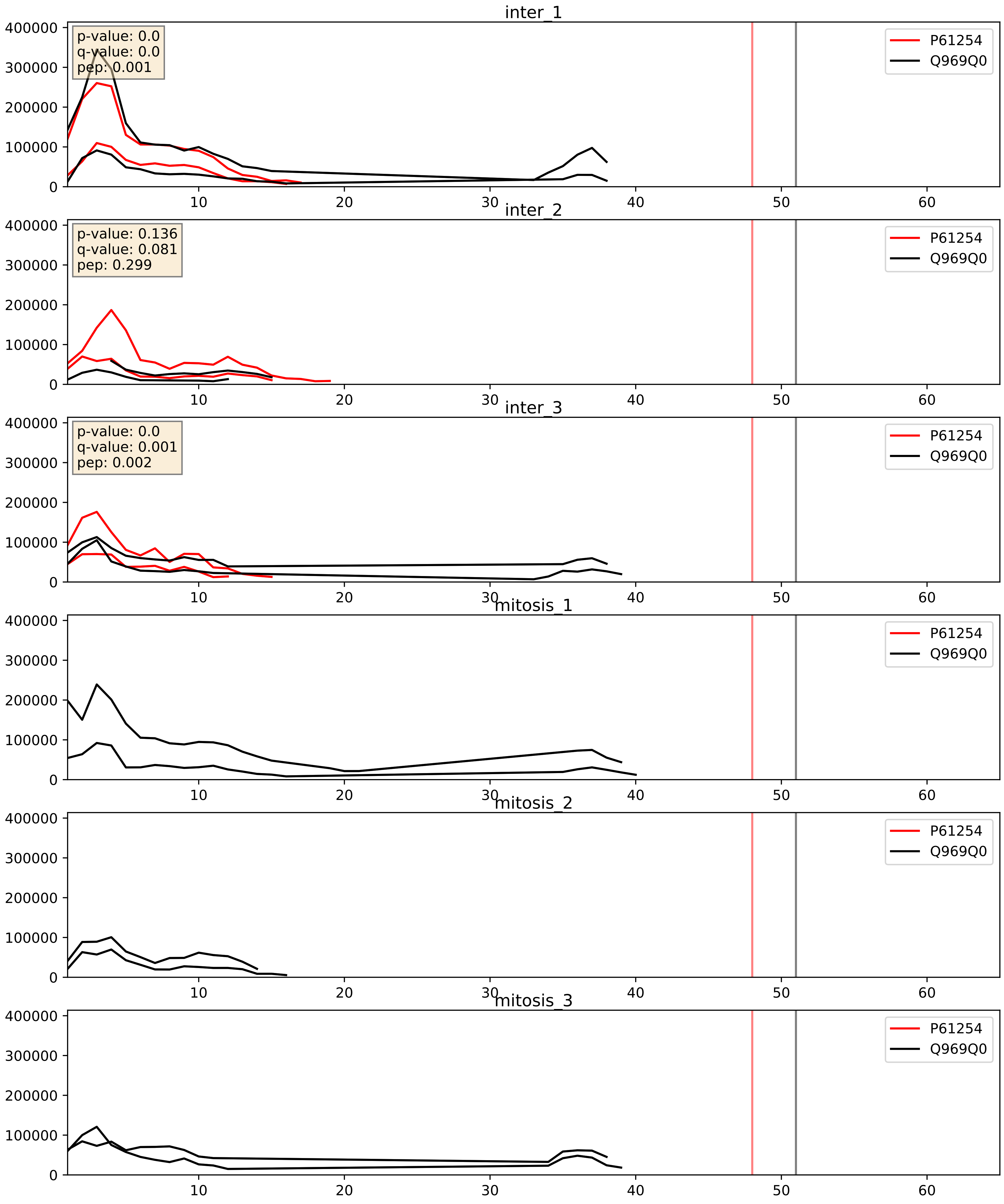


| level | condition_1 | condition_2 | pvalue | pvalue_adjusted |
|---|---|---|---|---|
| interactor_abundance | mitosis | inter | 0.000696587806406029 | 0.019073880342257484 |
| complex_abundance | mitosis | inter | 0.0015220567673542066 | 0.02714334568448335 |
| interactor_ratio | mitosis | inter | 0.2523662728653968 | 0.7251677296572153 |

P61254_Q9Y221
RL26_HUMAN vs NIP7_HUMAN
p-value: 0.0 q-value: 0.016

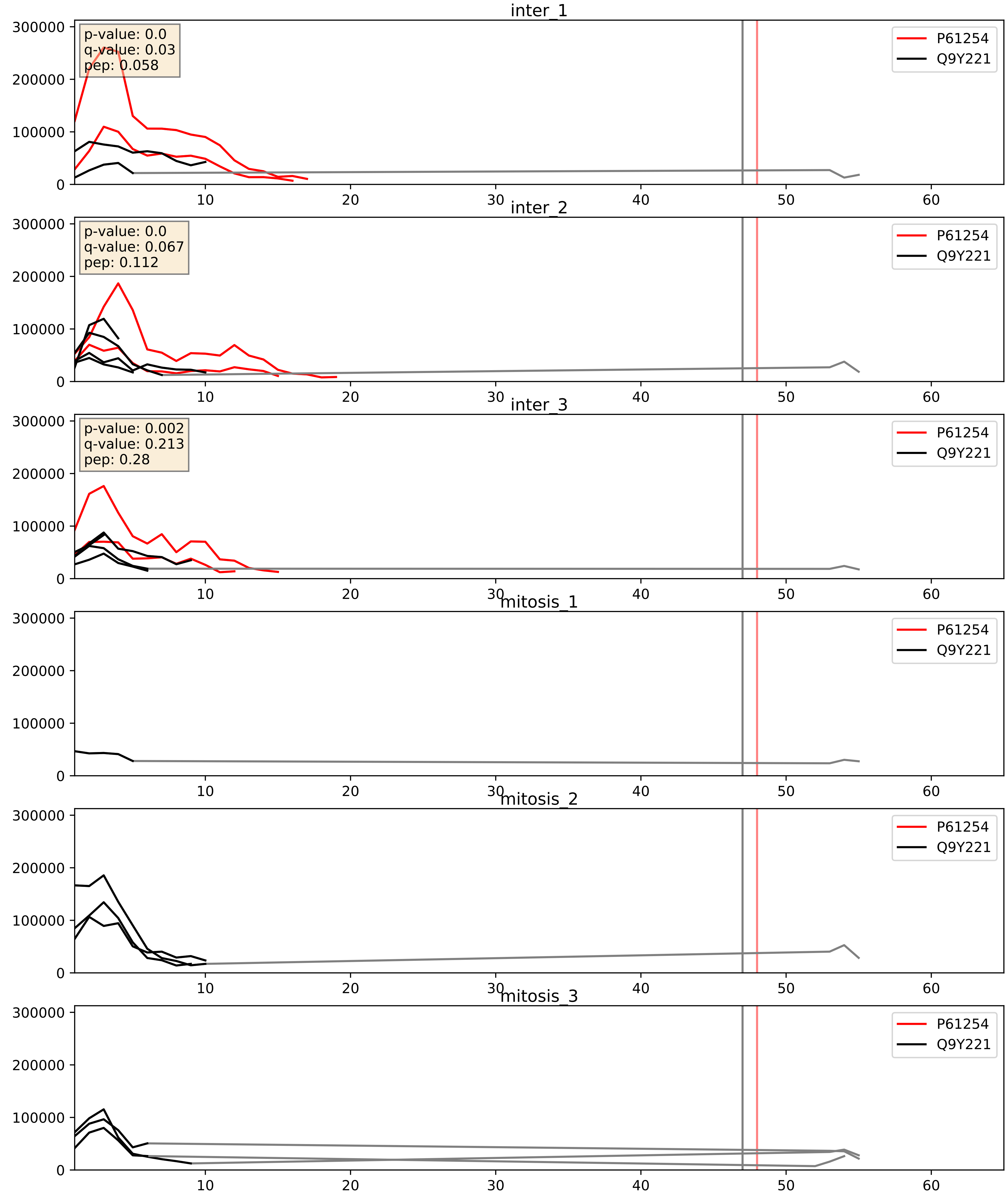


| level | condition_1 | condition_2 | pvalue | pvalue_adjusted |
|---|---|---|---|---|
| complex_abundance | mitosis | inter | 0.0008670015576876434 | 0.019513211095311817 |
| interactor_abundance | mitosis | inter | 0.0009560351713573535 | 0.022955571014919903 |
| interactor_ratio | mitosis | inter | 0.8261522456111503 | 0.969784012122695 |

P61254_Q9Y3U8
RL26_HUMAN vs RL36_HUMAN
p-value: 0.01 q-value: 0.009

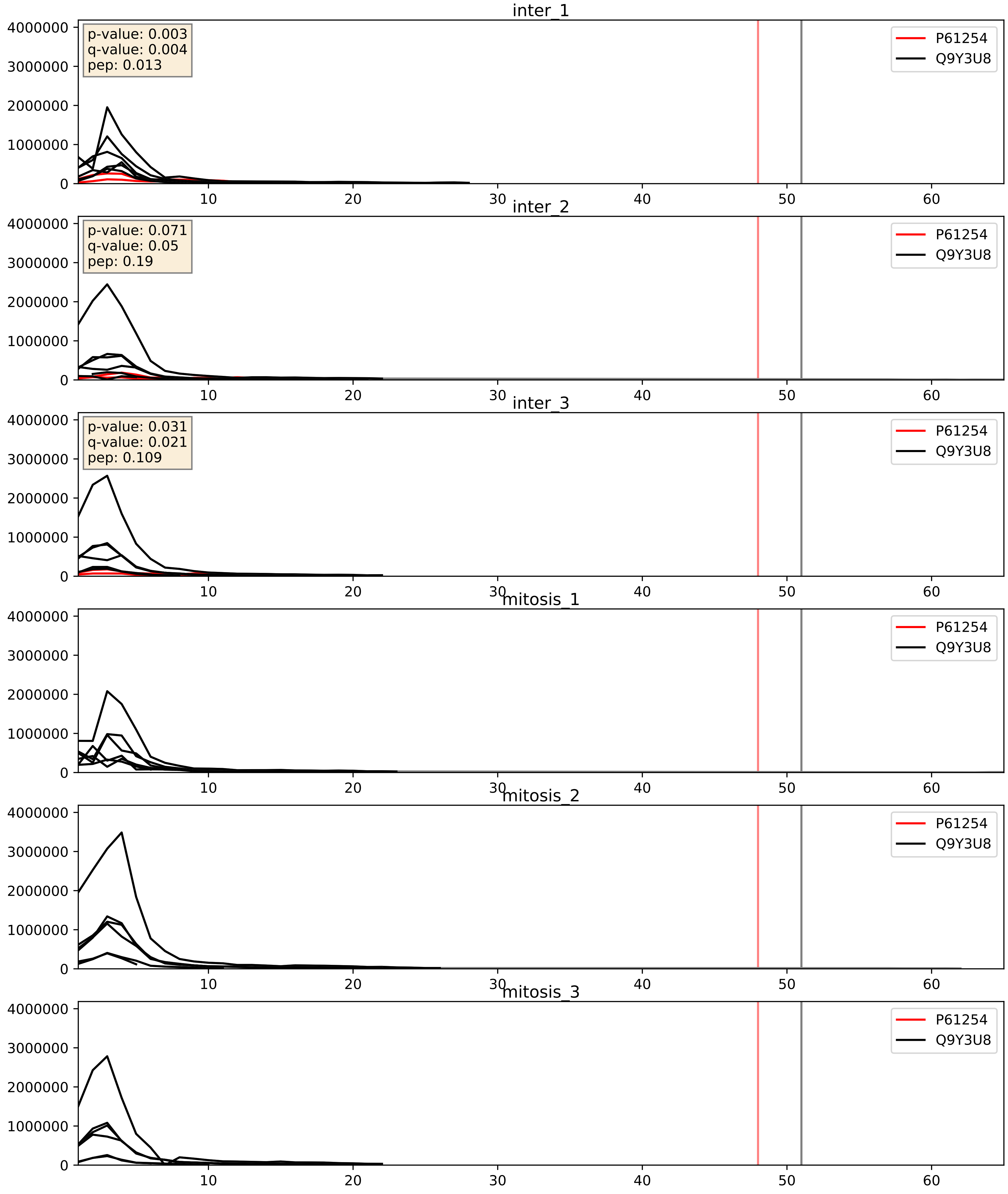


| level | condition_1 | condition_2 | pvalue | pvalue_adjusted |
|---|---|---|---|---|
| complex_abundance | mitosis | inter | 4.8380686524386975e-05 | 0.005691857151652958 |
| interactor_ratio | mitosis | inter | 6.172297255221914e-05 | 0.008128440693030705 |
| interactor_abundance | mitosis | inter | 0.0006144309768949645 | 0.017415659477552636 |
