## Supplementary Data 2 for "SECAT: Quantifying differential protein-protein interaction states by network-centric analysis": hela_string_000002_P55769.pdf

P55769_P55769

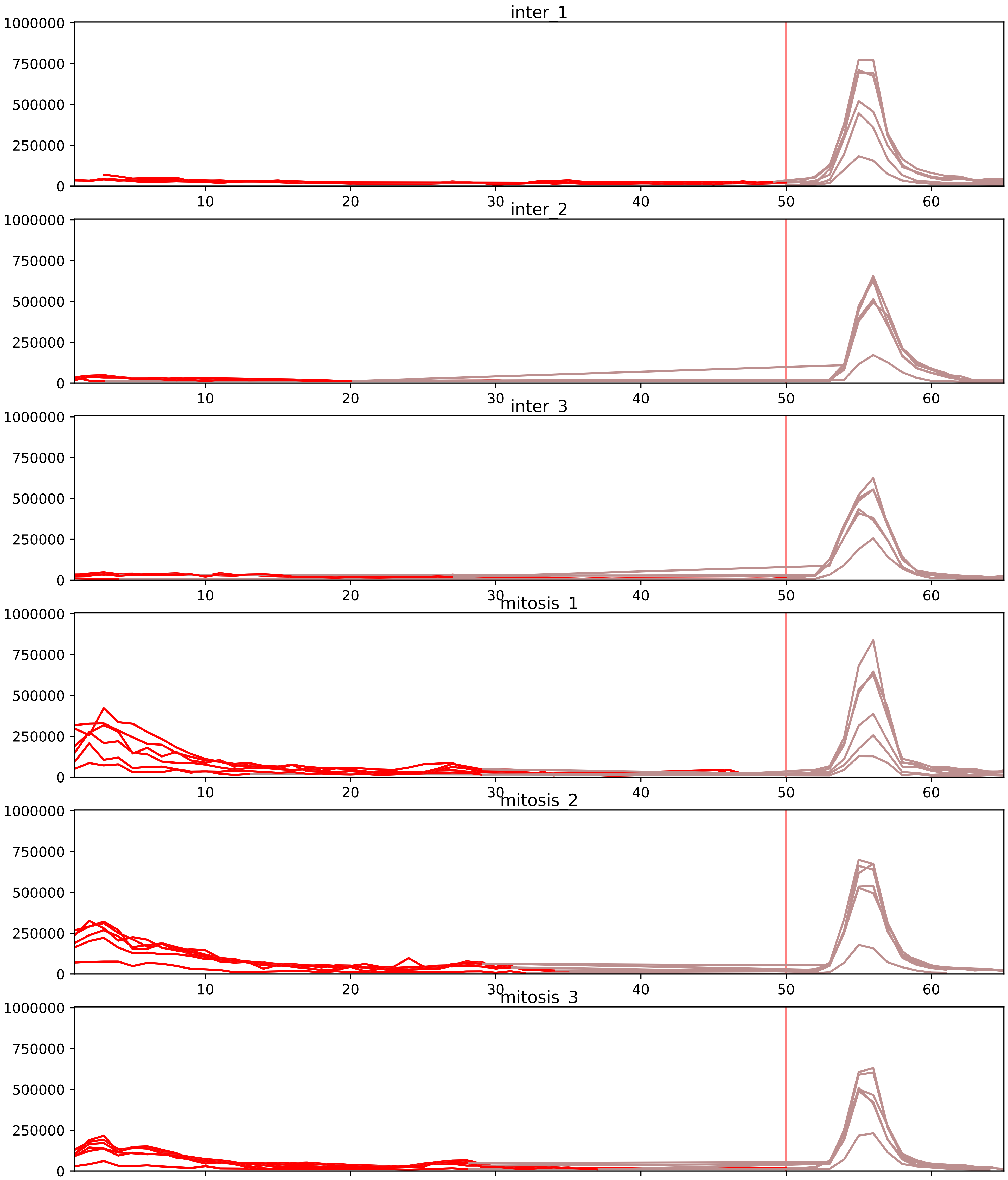


| level | condition_1 | condition_2 | pvalue | pvalue_adjusted |
|---|---|---|---|---|
| interactor_ratio | mitosis | inter | 1.9463435304507197e-25 | 1.790636048014662e-22 |
| interactor_abundance | mitosis | inter | 2.914570180647952e-06 | 0.00023316561445183617 |
| complex_abundance | mitosis | inter | 8.626760474604487e-06 | 0.00042900646684519613 |
| assembled_abundance | mitosis | inter | 0.0056771075818936485 | 0.07816041350248285 |
| total_abundance | mitosis | inter | 0.018915898353741977 | 0.17413642561621925 |
| monomer_abundance | mitosis | inter | 0.48888611234566415 | 0.7484545586191849 |

O00567_P55769
NOP56_HUMAN vs NH2L1_HUMAN
p-value: 0.0 q-value: 0.0

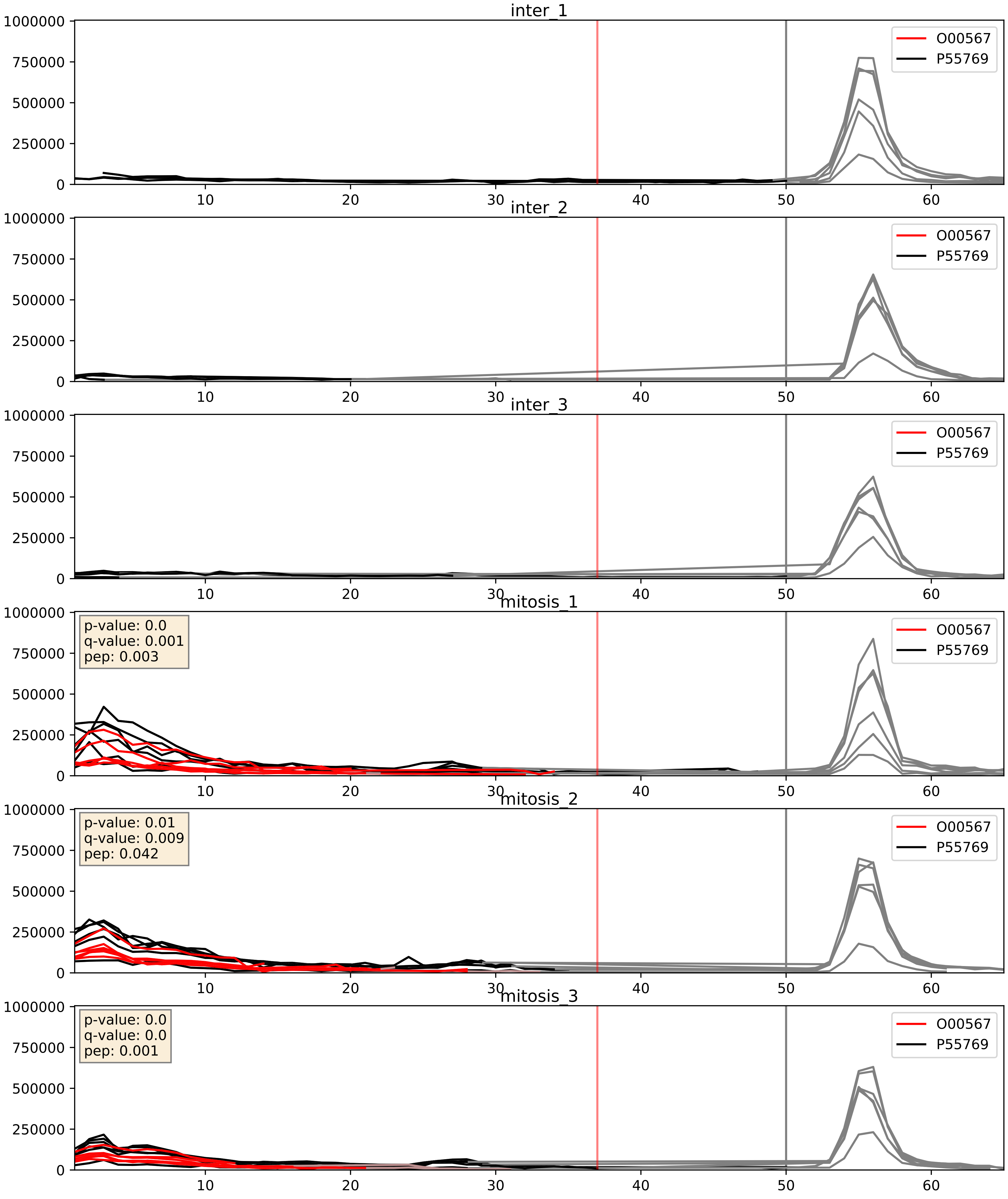


| level | condition_1 | condition_2 | pvalue | pvalue_adjusted |
|---|---|---|---|---|
| interactor_ratio | mitosis | inter | 0.000189167225482195 | 0.014134487523529912 |
| complex_abundance | mitosis | inter | 0.00048729227531920847 | 0.013857879989144268 |
| interactor_abundance | mitosis | inter | 0.0011181996263145508 | 0.024049720606162197 |

O43172_P55769
PRP4_HUMAN vs NH2L1_HUMAN
p-value: 0.009 q-value: 0.008

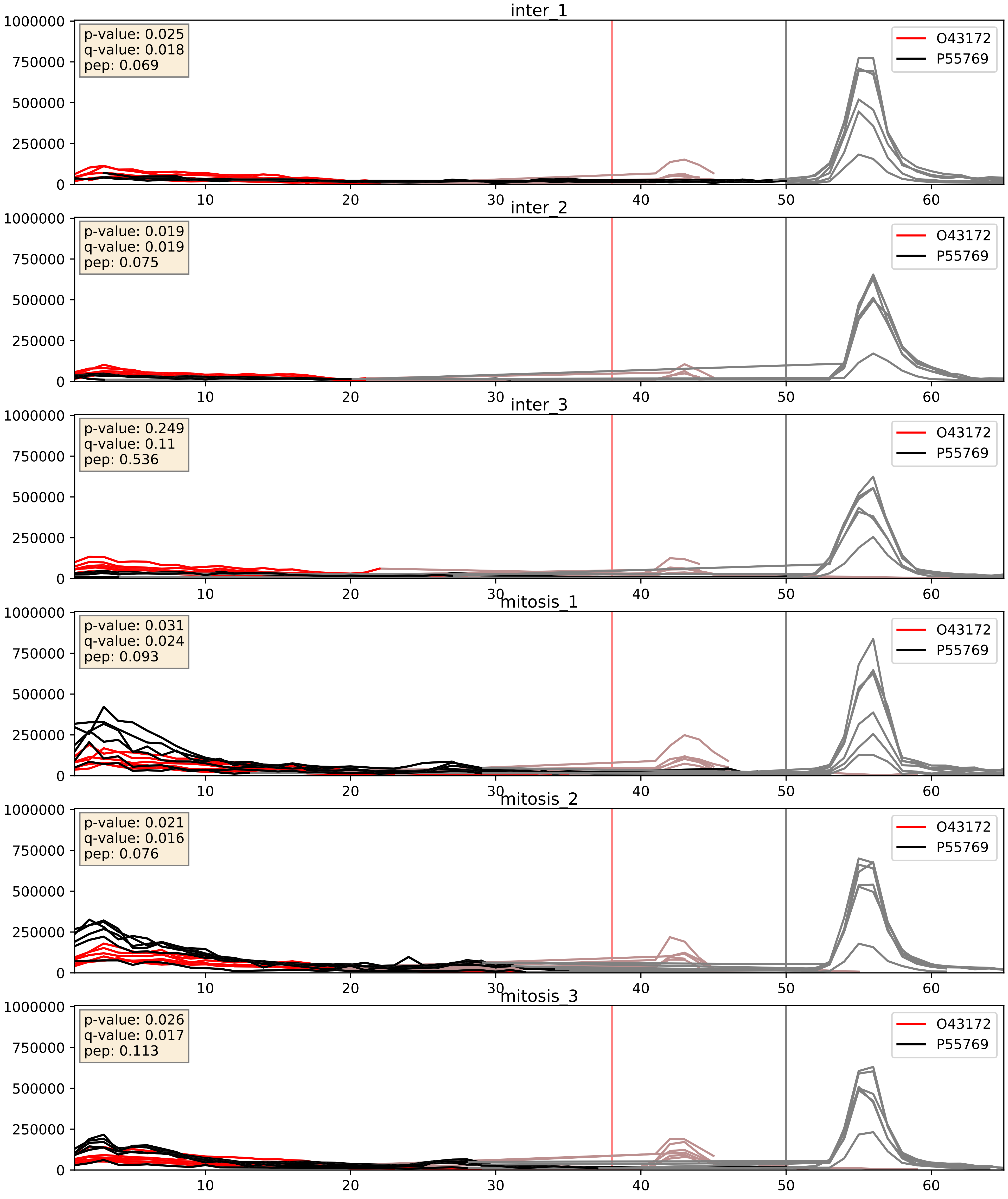


| level | condition_1 | condition_2 | pvalue | pvalue_adjusted |
|---|---|---|---|---|
| interactor_ratio | mitosis | inter | 0.0030875916908493423 | 0.07241036951690513 |
| complex_abundance | mitosis | inter | 0.008800537902912024 | 0.06968788570668541 |
| interactor_abundance | mitosis | inter | 0.20966725888123622 | 0.40652154270556234 |

O43395_P55769
PRPF3_HUMAN vs NH2L1_HUMAN
p-value: 0.066 q-value: 0.041

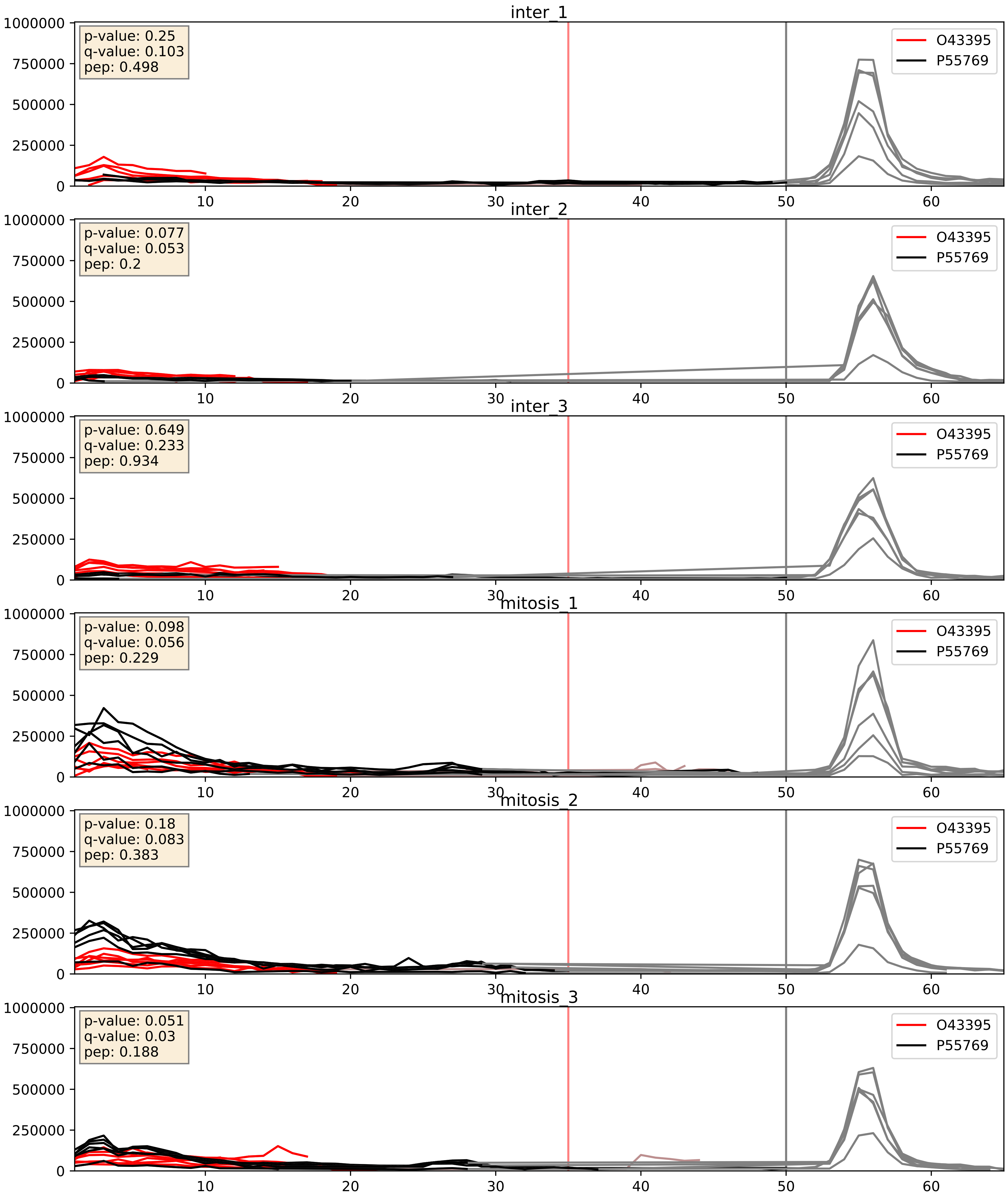


| level | condition_1 | condition_2 | pvalue | pvalue_adjusted |
|---|---|---|---|---|
| complex_abundance | mitosis | inter | 0.0049893793514525385 | 0.05279244406481302 |
| interactor_ratio | mitosis | inter | 0.016058636694013865 | 0.18702303415069207 |
| interactor_abundance | mitosis | inter | 0.3100367719395898 | 0.5063273428985764 |

O60832_P55769
DKC1_HUMAN vs NH2L1_HUMAN
p-value: 0.008 q-value: 0.007

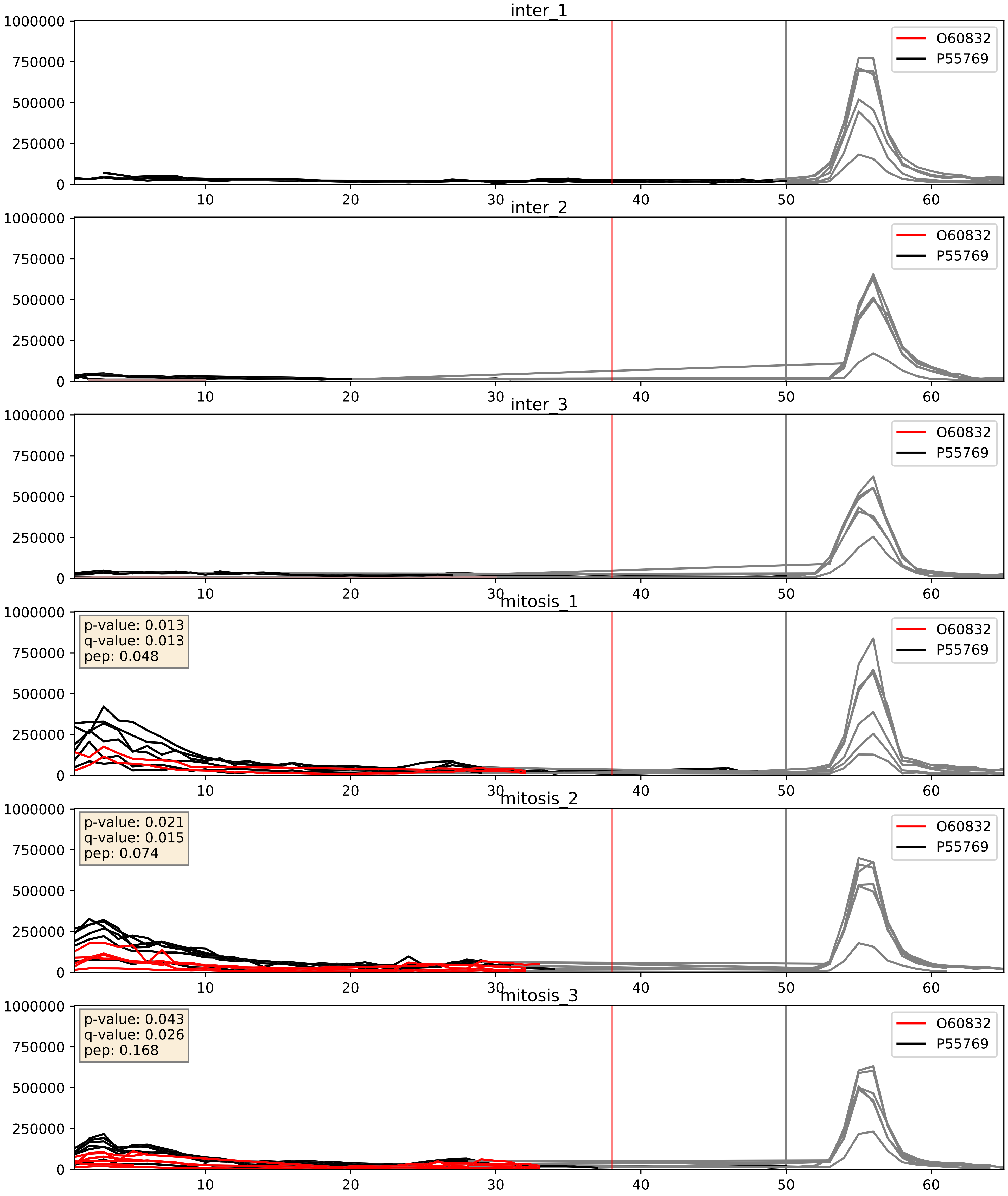


| level | condition_1 | condition_2 | pvalue | pvalue_adjusted |
|---|---|---|---|---|
| complex_abundance | mitosis | inter | 0.0010098562157940983 | 0.021187179429405588 |
| interactor_ratio | mitosis | inter | 0.043200488775377 | 0.34205635861393335 |
| interactor_abundance | mitosis | inter | 0.04824150833940766 | 0.17504902777779313 |

O76021_P55769
RL1D1_HUMAN vs NH2L1_HUMAN
p-value: 0.0 q-value: 0.013

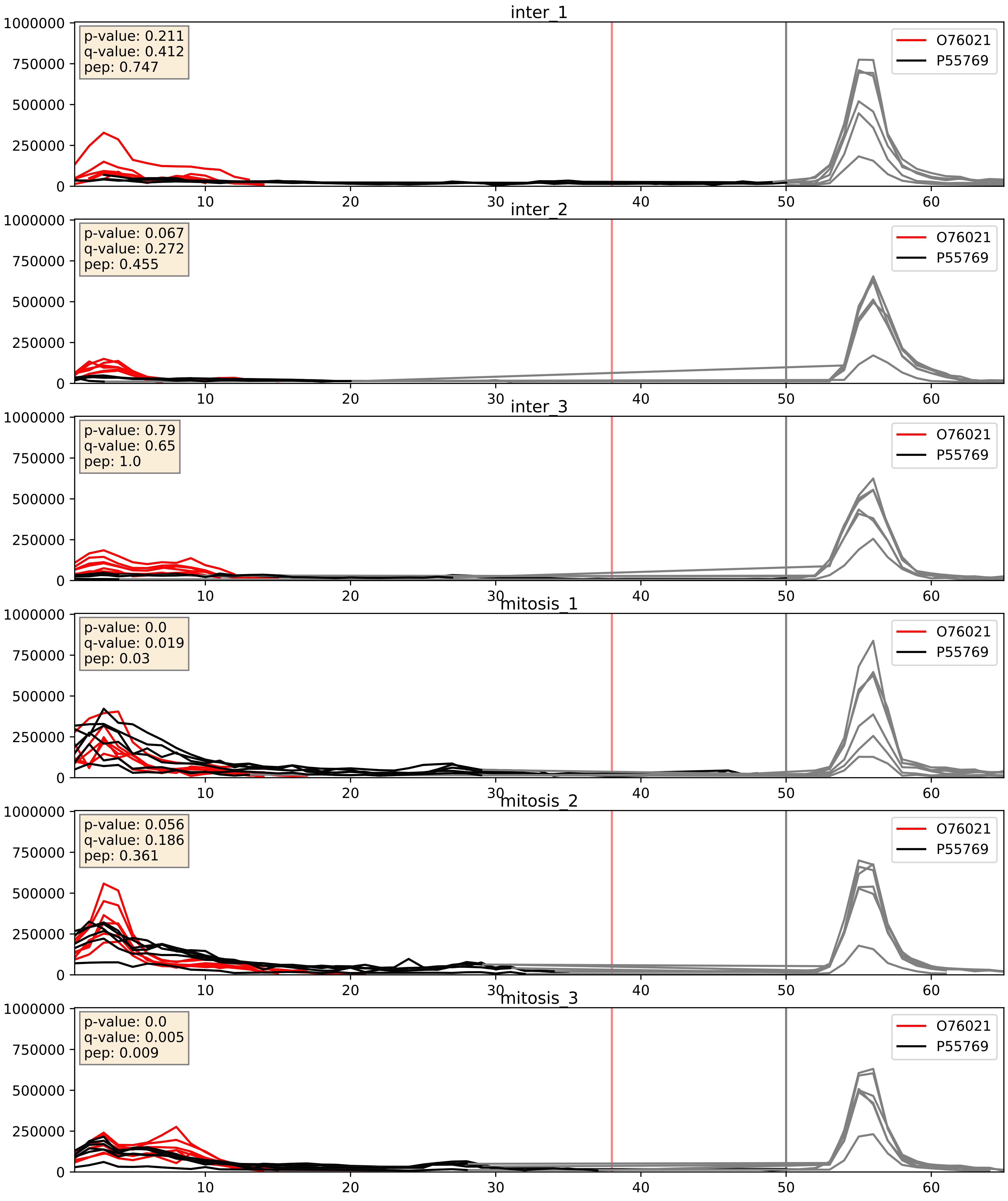


| level | condition_1 | condition_2 | pvalue | pvalue_adjusted |
|---|---|---|---|---|
| complex_abundance | mitosis | inter | 0.0010308703241052963 | 0.021560581032375372 |
| interactor_abundance | mitosis | inter | 0.007302544007558834 | 0.07362753439894419 |
| interactor_ratio | mitosis | inter | 0.018751790484042963 | 0.20631790044139817 |

O94906_P55769
PRP6_HUMAN vs NH2L1_HUMAN
p-value: 0.005 q-value: 0.005

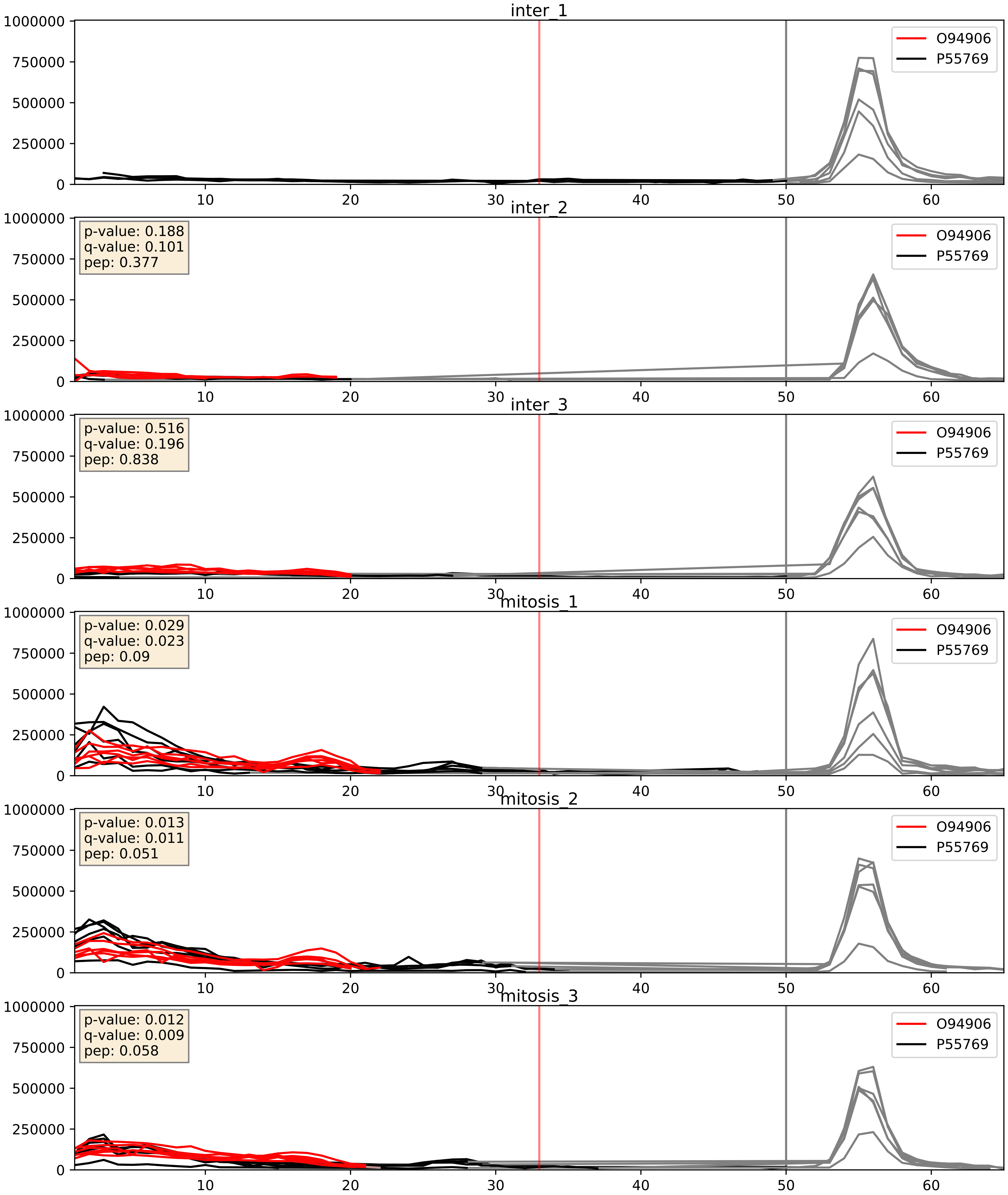


| level | condition_1 | condition_2 | pvalue | pvalue_adjusted |
|---|---|---|---|---|
| complex_abundance | mitosis | inter | 0.01249359904338843 | 0.08303199364239515 |
| interactor_abundance | mitosis | inter | 0.04665310860590202 | 0.1724311786124876 |
| interactor_ratio | mitosis | inter | 0.17781759270905903 | 0.6444111479450476 |

P09651_P55769
ROA1_HUMAN vs NH2L1_HUMAN
p-value: 0.0 q-value: 0.003

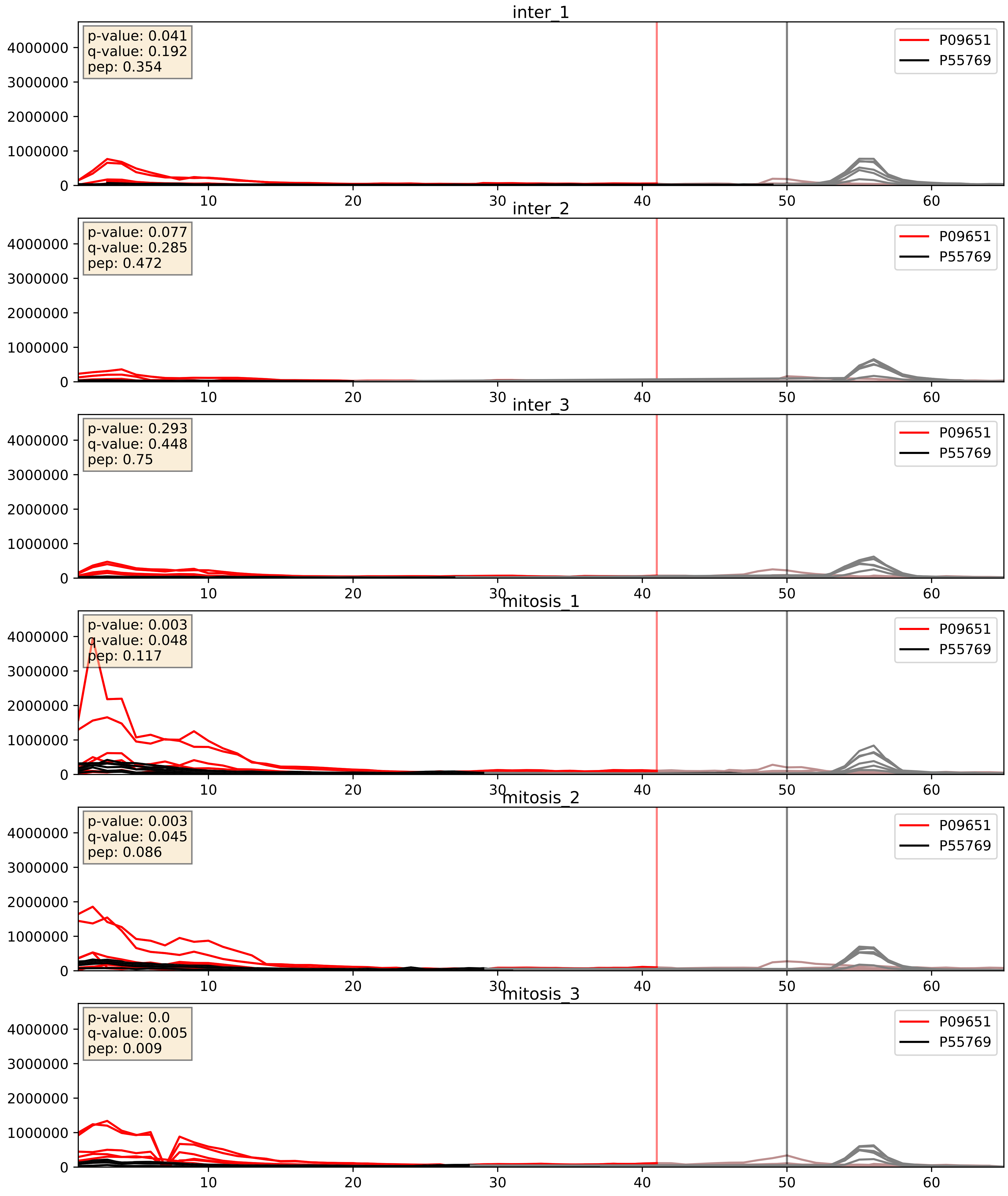


| level | condition_1 | condition_2 | pvalue | pvalue_adjusted |
|---|---|---|---|---|
| complex_abundance | mitosis | inter | 0.004746622165762423 | 0.05149694010003338 |
| interactor_abundance | mitosis | inter | 0.01925771606241603 | 0.11885079271397347 |
| interactor_ratio | mitosis | inter | 0.7812992714023176 | 0.9589625145419588 |

P10606_P55769
COX5B_HUMAN vs NH2L1_HUMAN
p-value: 0.0 q-value: 0.026

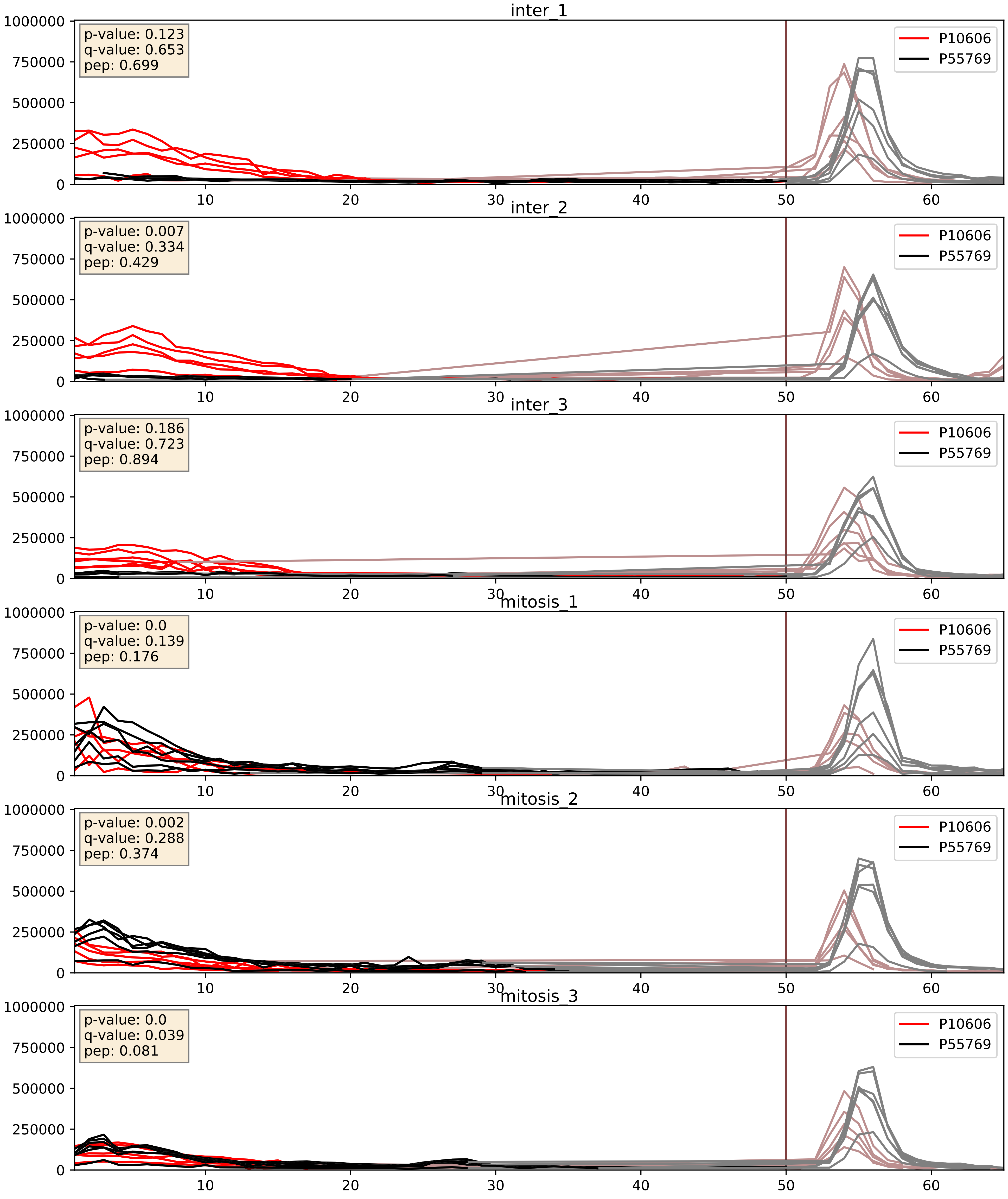


| level | condition_1 | condition_2 | pvalue | pvalue_adjusted |
|---|---|---|---|---|
| interactor_ratio | mitosis | inter | 0.0012388457487284845 | 0.041750077201243414 |
| interactor_abundance | mitosis | inter | 0.07790927811011916 | 0.23144314441180636 |
| complex_abundance | mitosis | inter | 0.1916272517241159 | 0.34283281829888396 |

P15880_P55769
RS2_HUMAN vs NH2L1_HUMAN
p-value: 0.041 q-value: 0.028

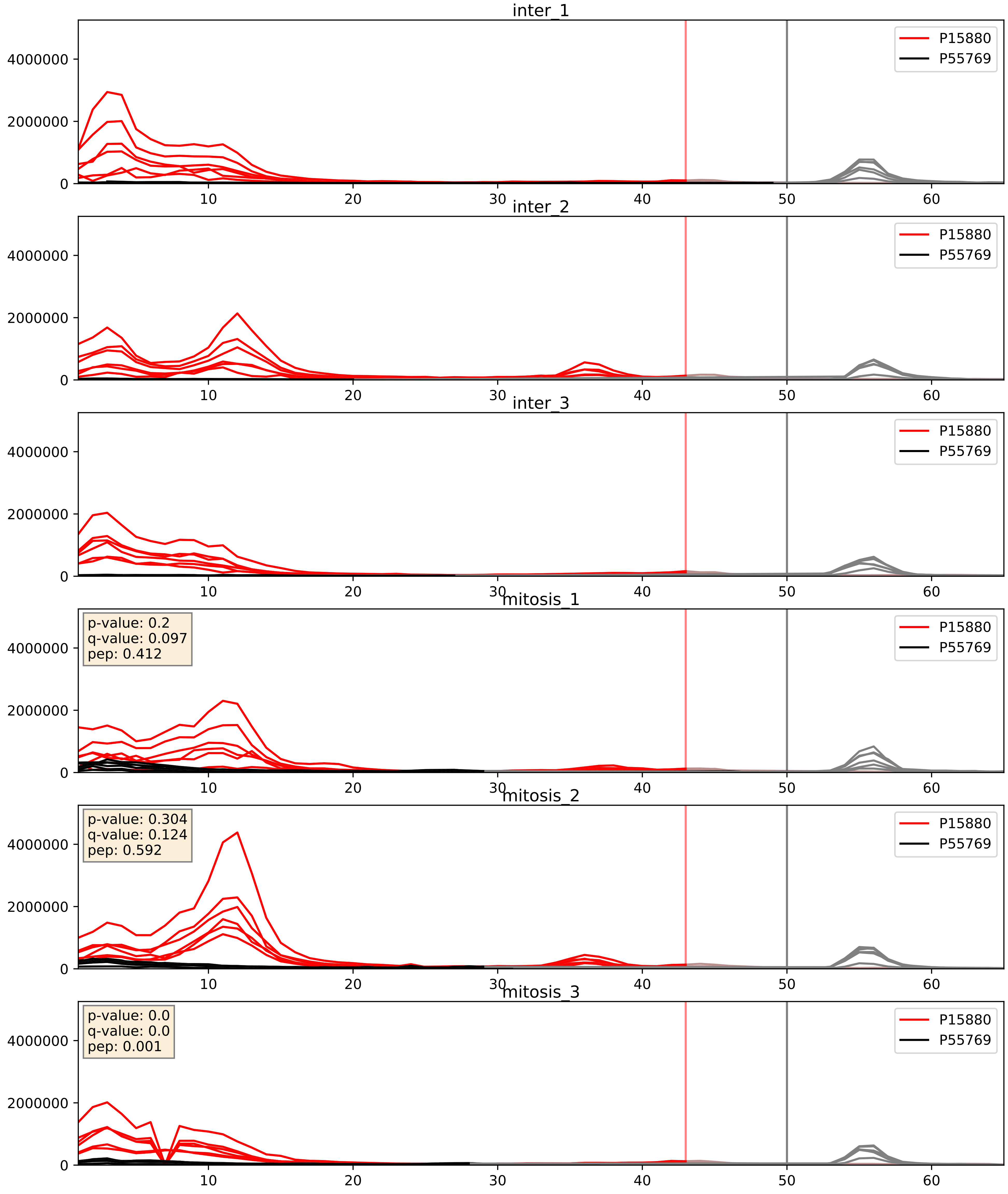


| level | condition_1 | condition_2 | pvalue | pvalue_adjusted |
|---|---|---|---|---|
| interactor_ratio | mitosis | inter | 0.0009813620140001324 | 0.03599129628116289 |
| complex_abundance | mitosis | inter | 0.00827489961282815 | 0.06742444735376033 |
| interactor_abundance | mitosis | inter | 0.6832770140112532 | 0.7940602057914786 |

P18077_P55769
RL35A_HUMAN vs NH2L1_HUMAN
p-value: 0.001 q-value: 0.039

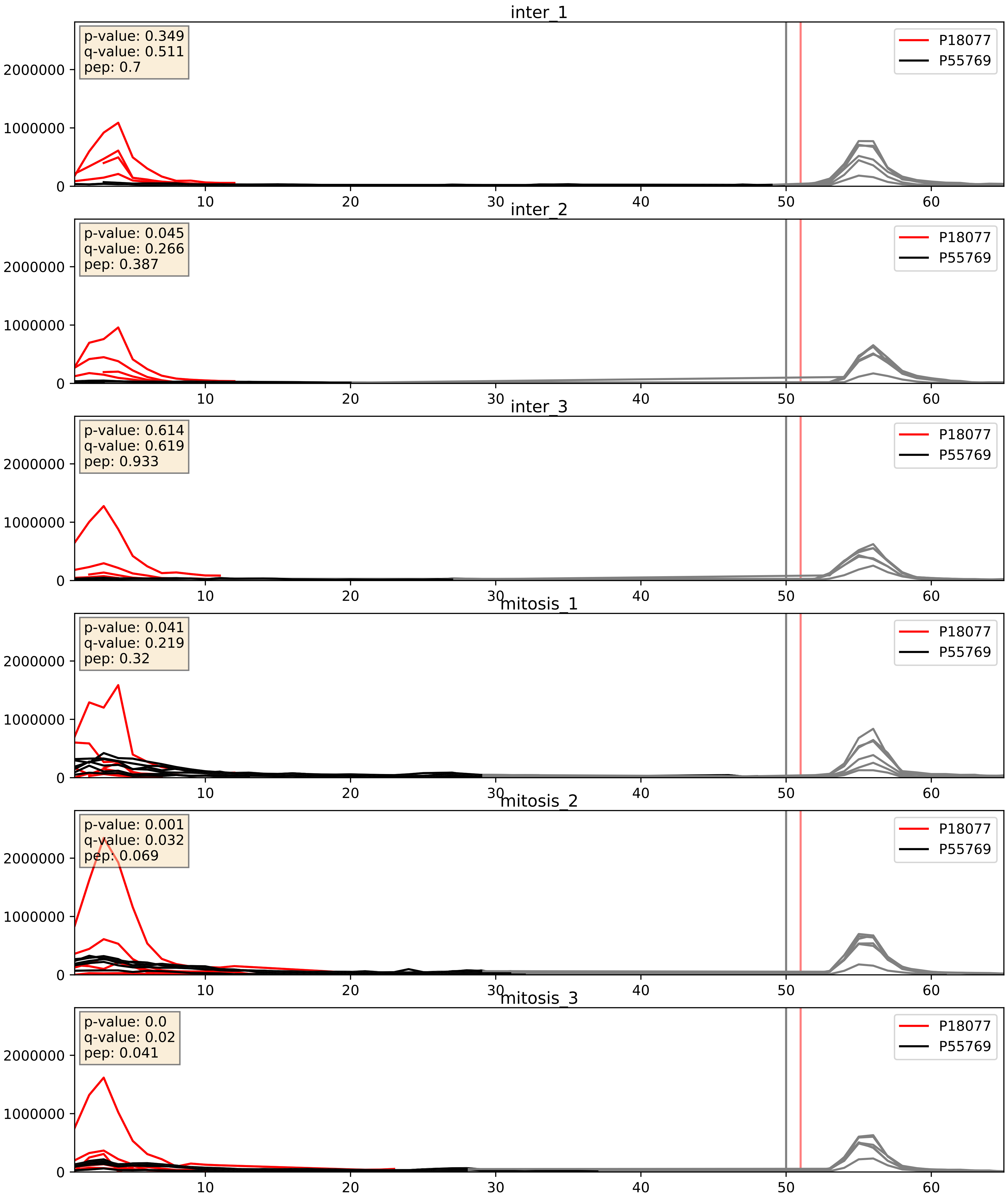


| level | condition_1 | condition_2 | pvalue | pvalue_adjusted |
|---|---|---|---|---|
| interactor_ratio | mitosis | inter | 0.00022461029121093303 | 0.01529848062979237 |
| complex_abundance | mitosis | inter | 0.027432003538751186 | 0.12057983221353287 |
| interactor_abundance | mitosis | inter | 0.8878397503920612 | 0.9361222224987428 |

P20674_P55769
COX5A_HUMAN vs NH2L1_HUMAN
p-value: 0.0 q-value: 0.037

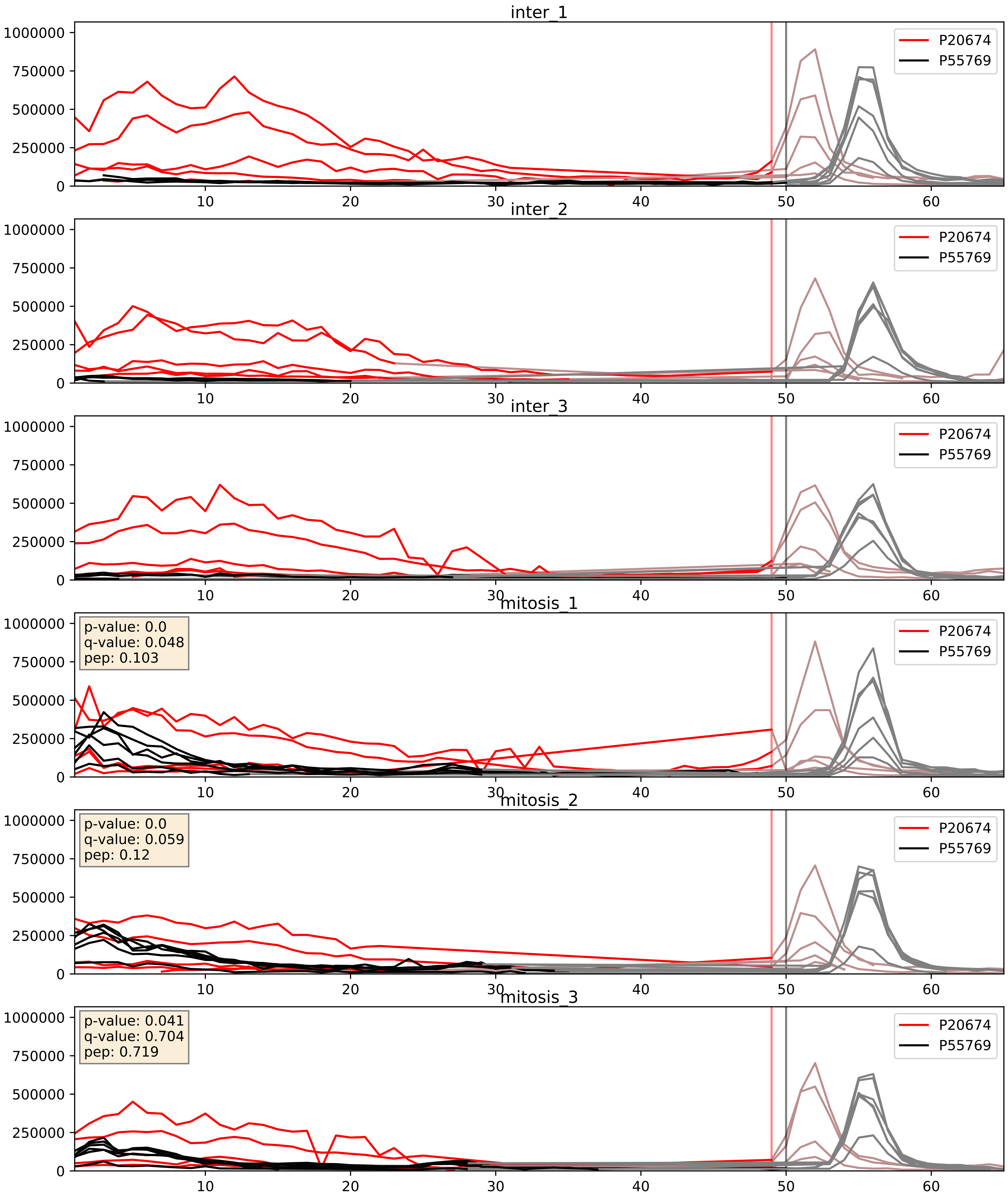


| level | condition_1 | condition_2 | pvalue | pvalue_adjusted |
|---|---|---|---|---|
| interactor_ratio | mitosis | inter | 0.00026084499122796666 | 0.016318883418974577 |
| interactor_abundance | mitosis | inter | 0.014966275857983836 | 0.10701782781851343 |
| complex_abundance | mitosis | inter | 0.10261923701165743 | 0.23799312559378397 |

P22087_P55769
FBRL_HUMAN vs NH2L1_HUMAN
p-value: 0.0 q-value: 0.0

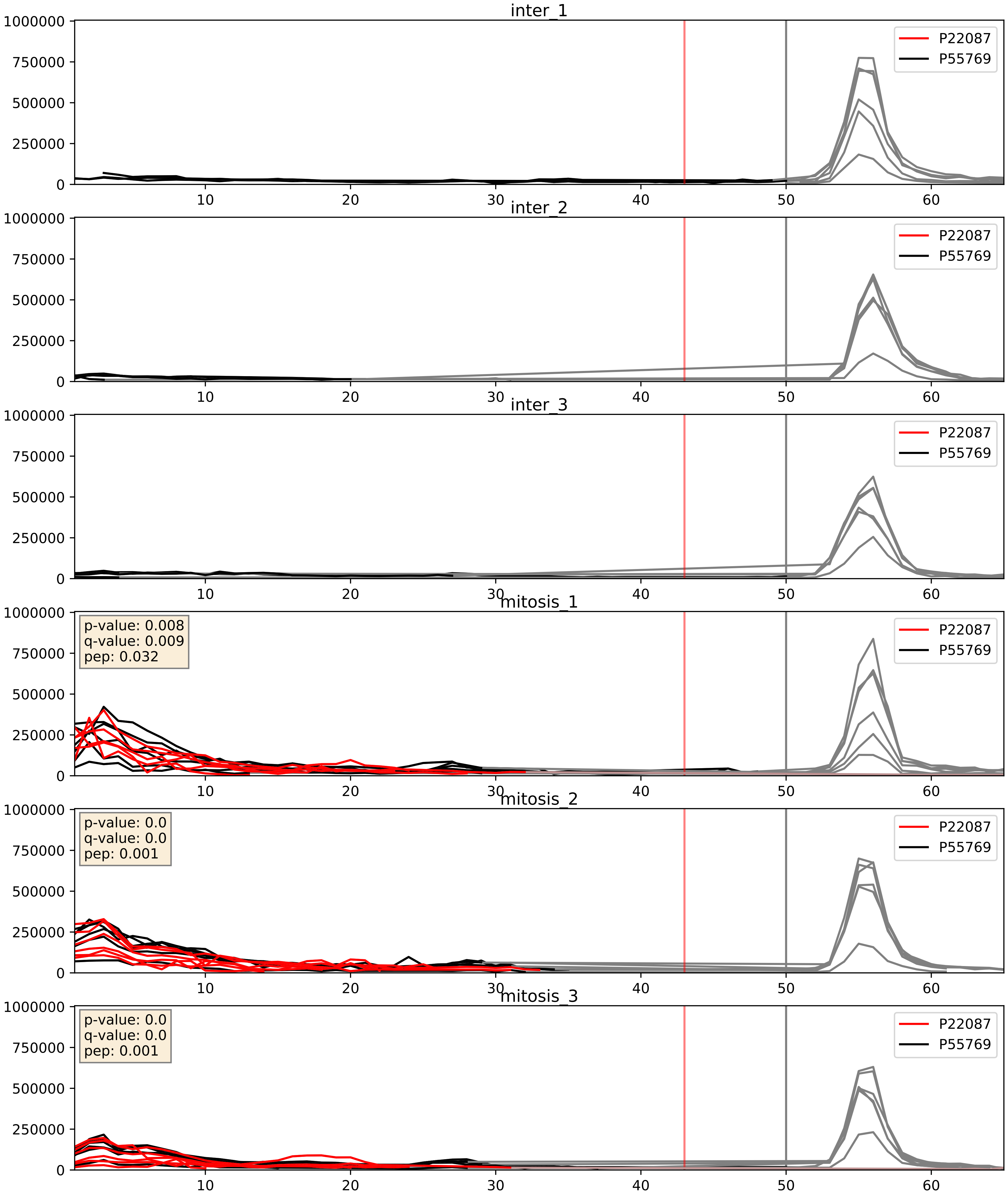


| level | condition_1 | condition_2 | pvalue | pvalue_adjusted |
|---|---|---|---|---|
| interactor_ratio | mitosis | inter | 3.3712270526144467e-05 | 0.006185155691845816 |
| complex_abundance | mitosis | inter | 0.0002827767244629666 | 0.011311068978518664 |
| interactor_abundance | mitosis | inter | 0.0003671184791785453 | 0.01356427145975096 |

P26373_P55769
RL13_HUMAN vs NH2L1_HUMAN
p-value: 0.0 q-value: 0.041

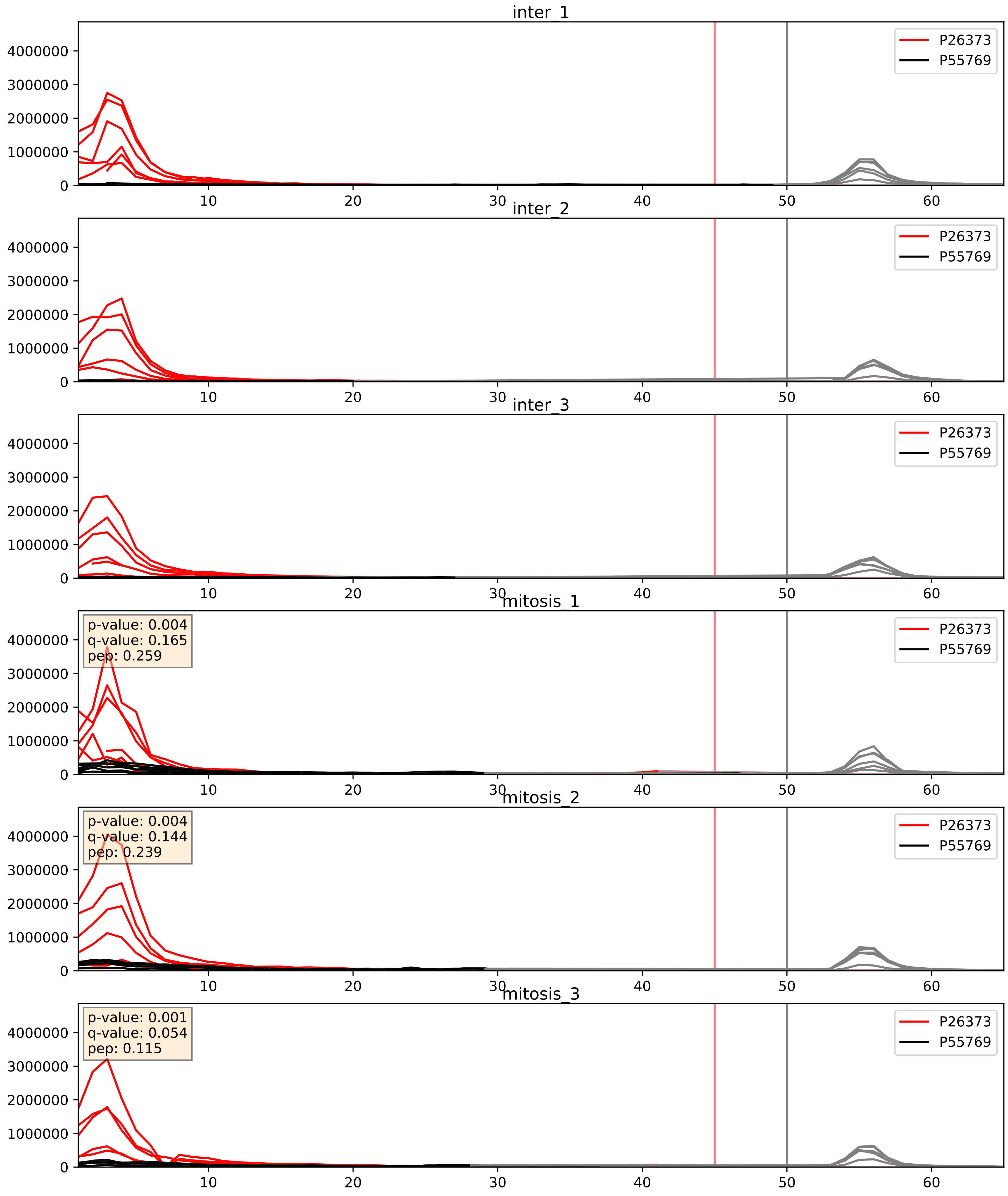


| level | condition_1 | condition_2 | pvalue | pvalue_adjusted |
|---|---|---|---|---|
| interactor_ratio | mitosis | inter | 0.0016612159709601506 | 0.04925768316782793 |
| complex_abundance | mitosis | inter | 0.006115812545326621 | 0.05820202463873419 |
| interactor_abundance | mitosis | inter | 0.6715663074712546 | 0.7848999989014116 |

P35268_P55769
RL22_HUMAN vs NH2L1_HUMAN
p-value: 0.0 q-value: 0.016

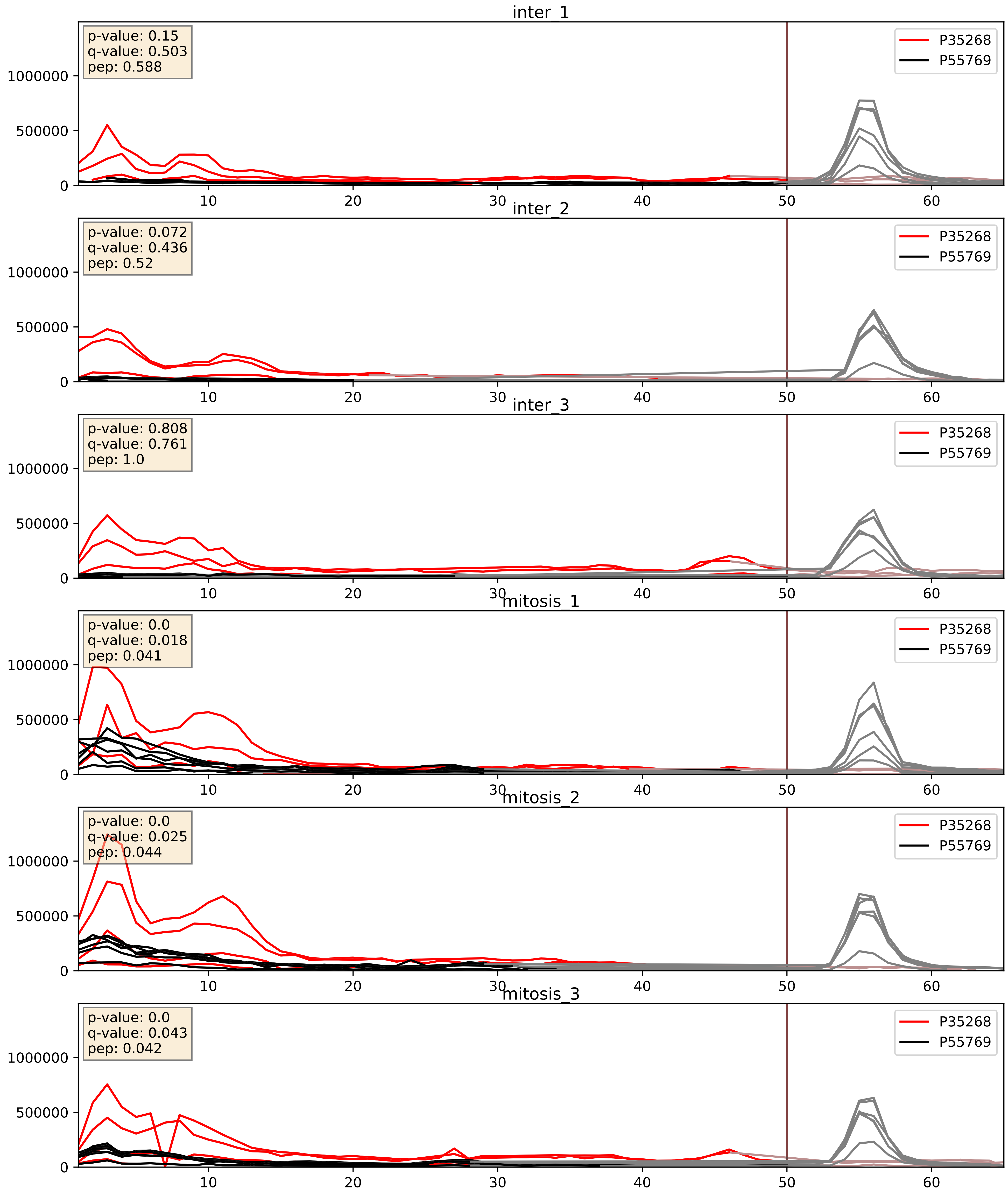


| level | condition_1 | condition_2 | pvalue | pvalue_adjusted |
|---|---|---|---|---|
| complex_abundance | mitosis | inter | 0.0010492460264906755 | 0.021612845224852777 |
| interactor_abundance | mitosis | inter | 0.014141208892344399 | 0.10541033287779904 |
| interactor_ratio | mitosis | inter | 0.024818760043489395 | 0.24732082185363125 |

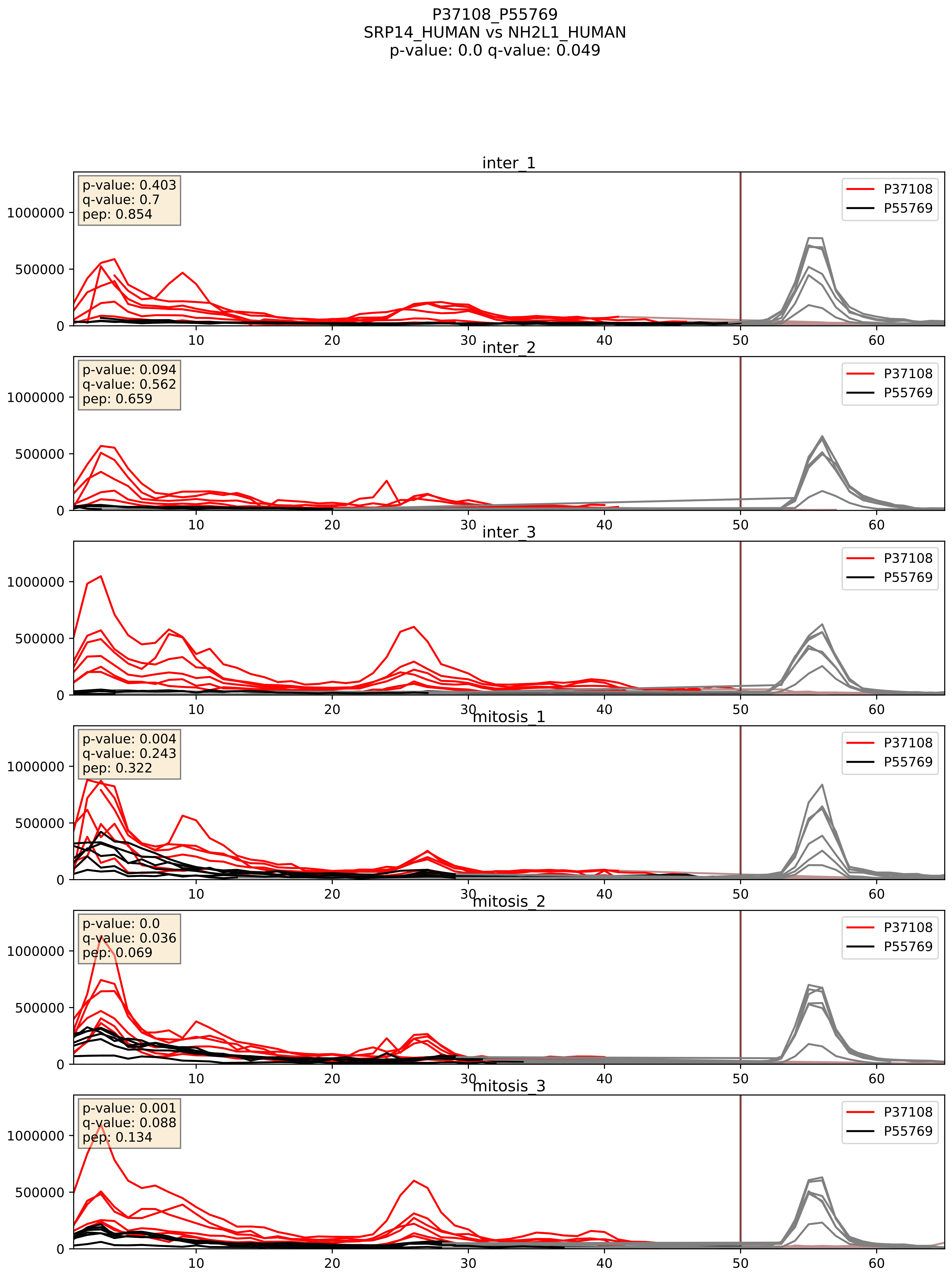


| level | condition_1 | condition_2 | pvalue | pvalue_adjusted |
|---|---|---|---|---|
| complex_abundance | mitosis | inter | 0.0536024187735455 | 0.16510856592355144 |
| interactor_ratio | mitosis | inter | 0.12403236196725721 | 0.5739294160480145 |
| interactor_abundance | mitosis | inter | 0.525406257381365 | 0.686009390357609 |

P39023_P55769
RL3_HUMAN vs NH2L1_HUMAN
p-value: 0.0 q-value: 0.041

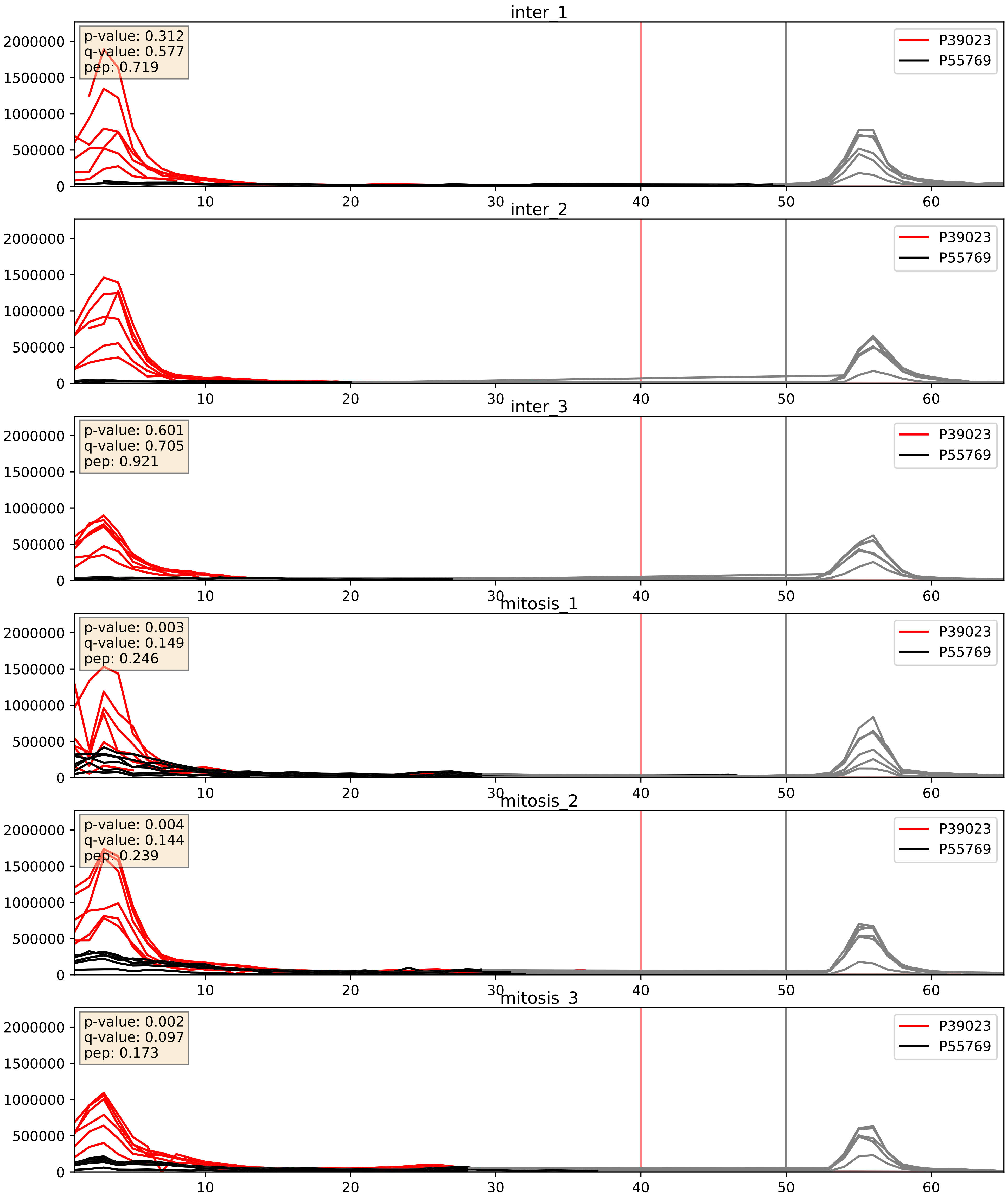


| level | condition_1 | condition_2 | pvalue | pvalue_adjusted |
|---|---|---|---|---|
| complex_abundance | mitosis | inter | 0.0016737830020827449 | 0.028828133798447276 |
| interactor_ratio | mitosis | inter | 0.004039007201836414 | 0.08271268336775048 |
| interactor_abundance | mitosis | inter | 0.4358819834363643 | 0.6141304877319197 |

P42677_P55769
RS27_HUMAN vs NH2L1_HUMAN
p-value: 0.001 q-value: 0.023

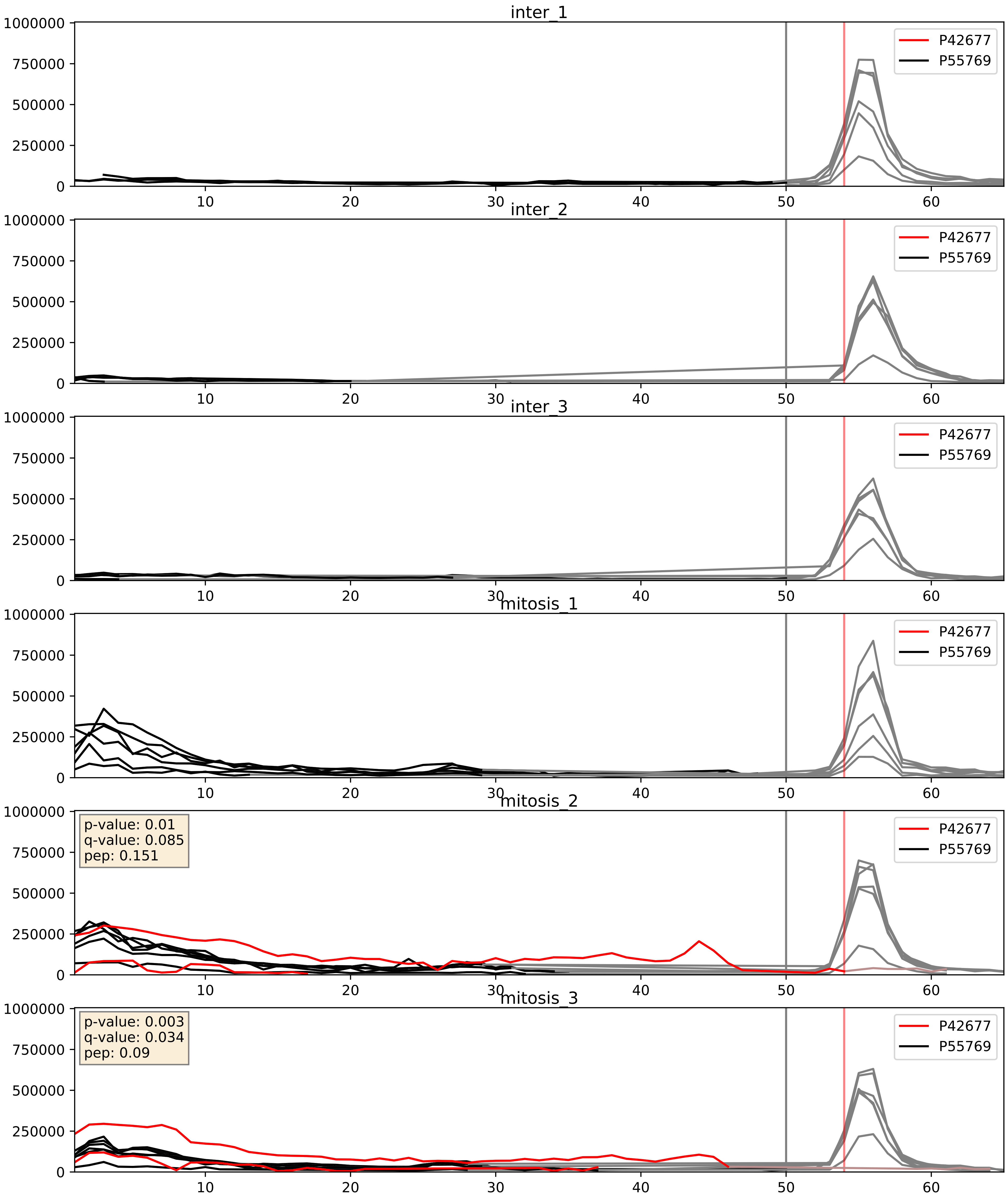


| level | condition_1 | condition_2 | pvalue | pvalue_adjusted |
|---|---|---|---|---|
| complex_abundance | mitosis | inter | 0.15162530868448854 | 0.29775467821500845 |
| interactor_abundance | mitosis | inter | 0.1580999442007717 | 0.3439663292308059 |
| interactor_ratio | mitosis | inter | 0.17202298278234604 | 0.6355272907280458 |

P42766_P55769
RL35_HUMAN vs NH2L1_HUMAN
p-value: 0.0 q-value: 0.008

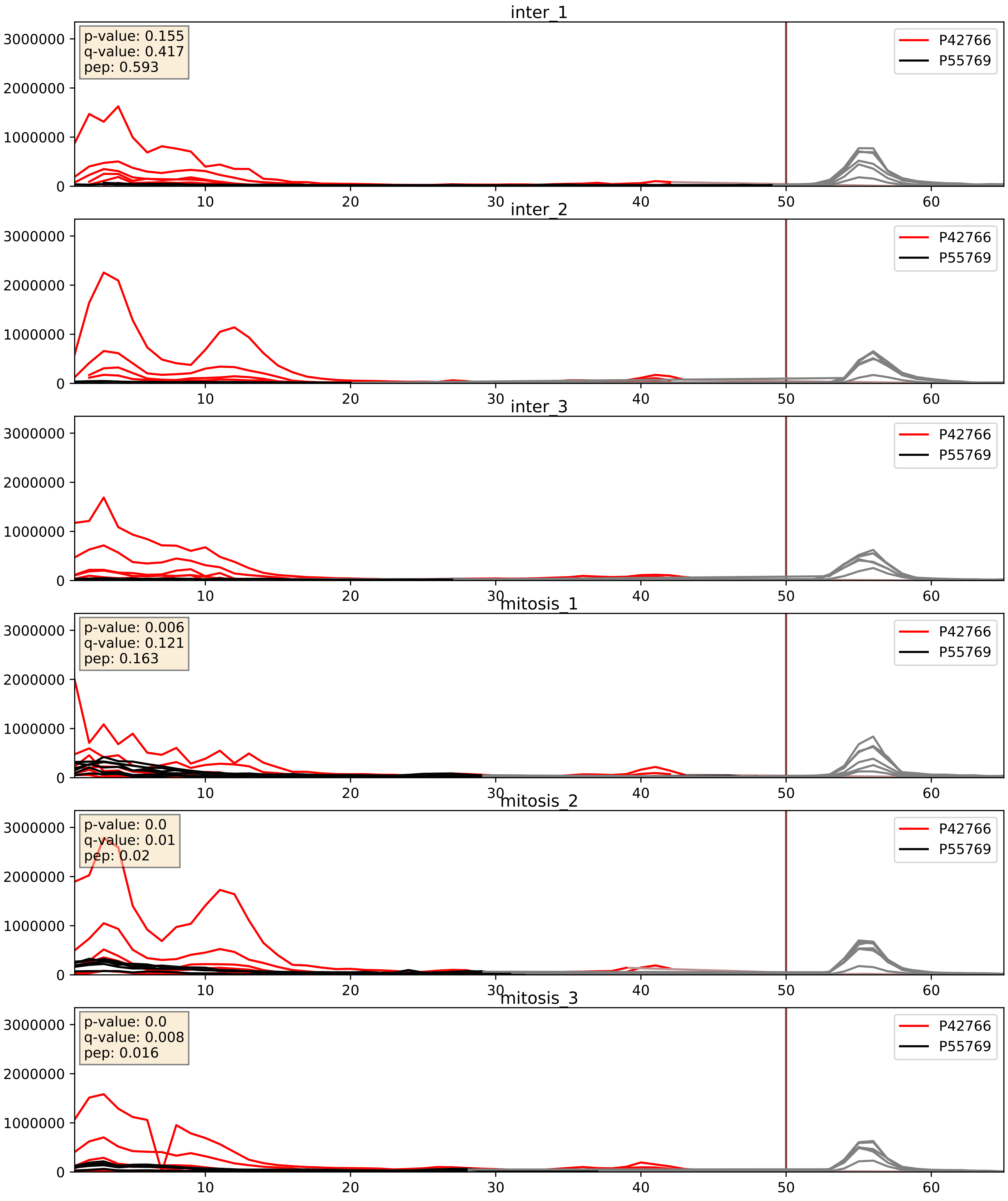


| level | condition_1 | condition_2 | pvalue | pvalue_adjusted |
|---|---|---|---|---|
| complex_abundance | mitosis | inter | 0.0017189885243836867 | 0.02936214455576784 |
| interactor_ratio | mitosis | inter | 0.007332808884641069 | 0.11910596594407505 |
| interactor_abundance | mitosis | inter | 0.8006603892902106 | 0.883601228946587 |

P46776_P55769
RL27A_HUMAN vs NH2L1_HUMAN
p-value: 0.0 q-value: 0.016

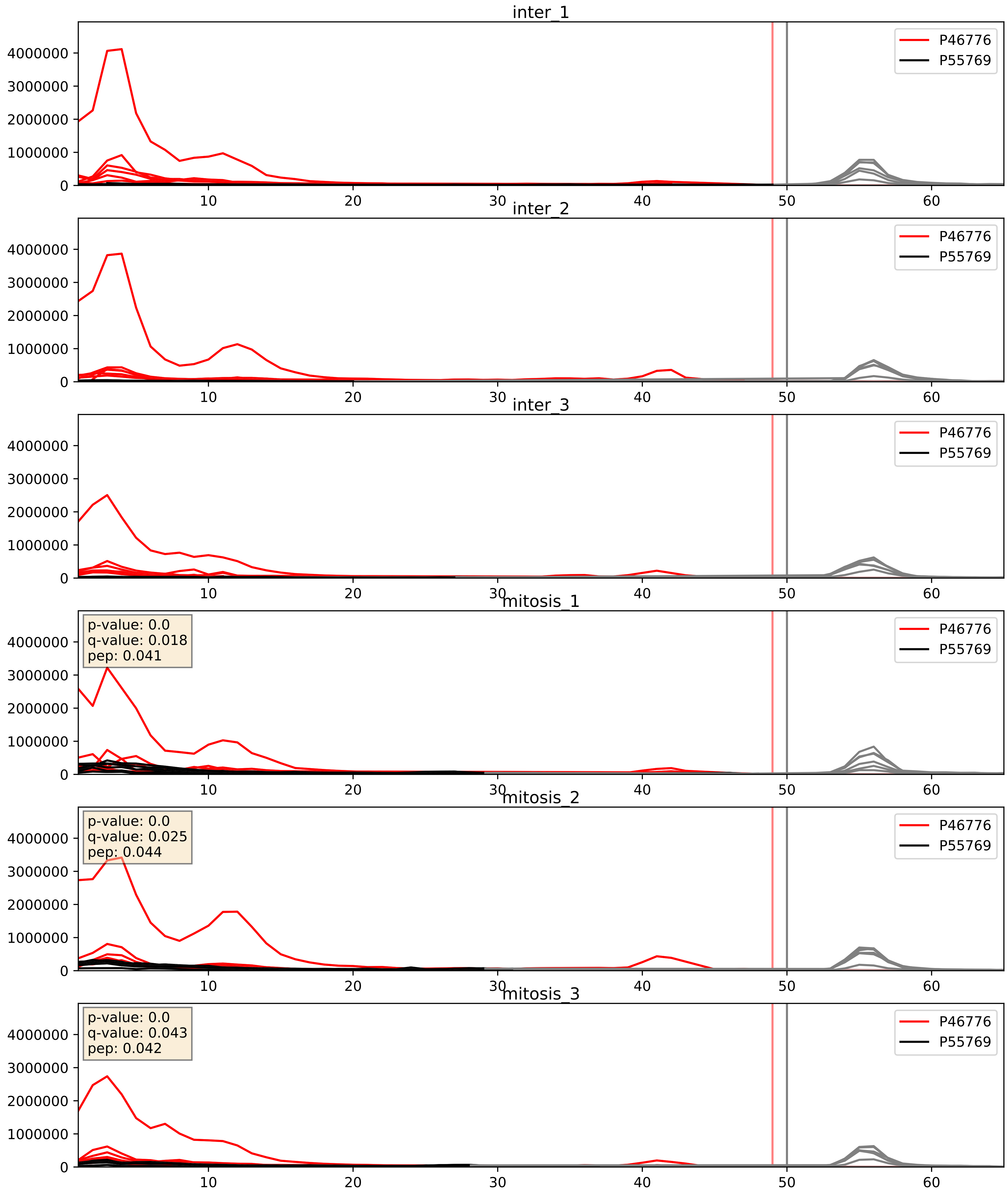


| level | condition_1 | condition_2 | pvalue | pvalue_adjusted |
|---|---|---|---|---|
| interactor_ratio | mitosis | inter | 0.0005671150149868073 | 0.02774002587592612 |
| complex_abundance | mitosis | inter | 0.010656811552043503 | 0.07661909071036627 |
| interactor_abundance | mitosis | inter | 0.8985882811758585 | 0.9441584502781086 |

P46778_P55769
RL21_HUMAN vs NH2L1_HUMAN
p-value: 0.0 q-value: 0.041

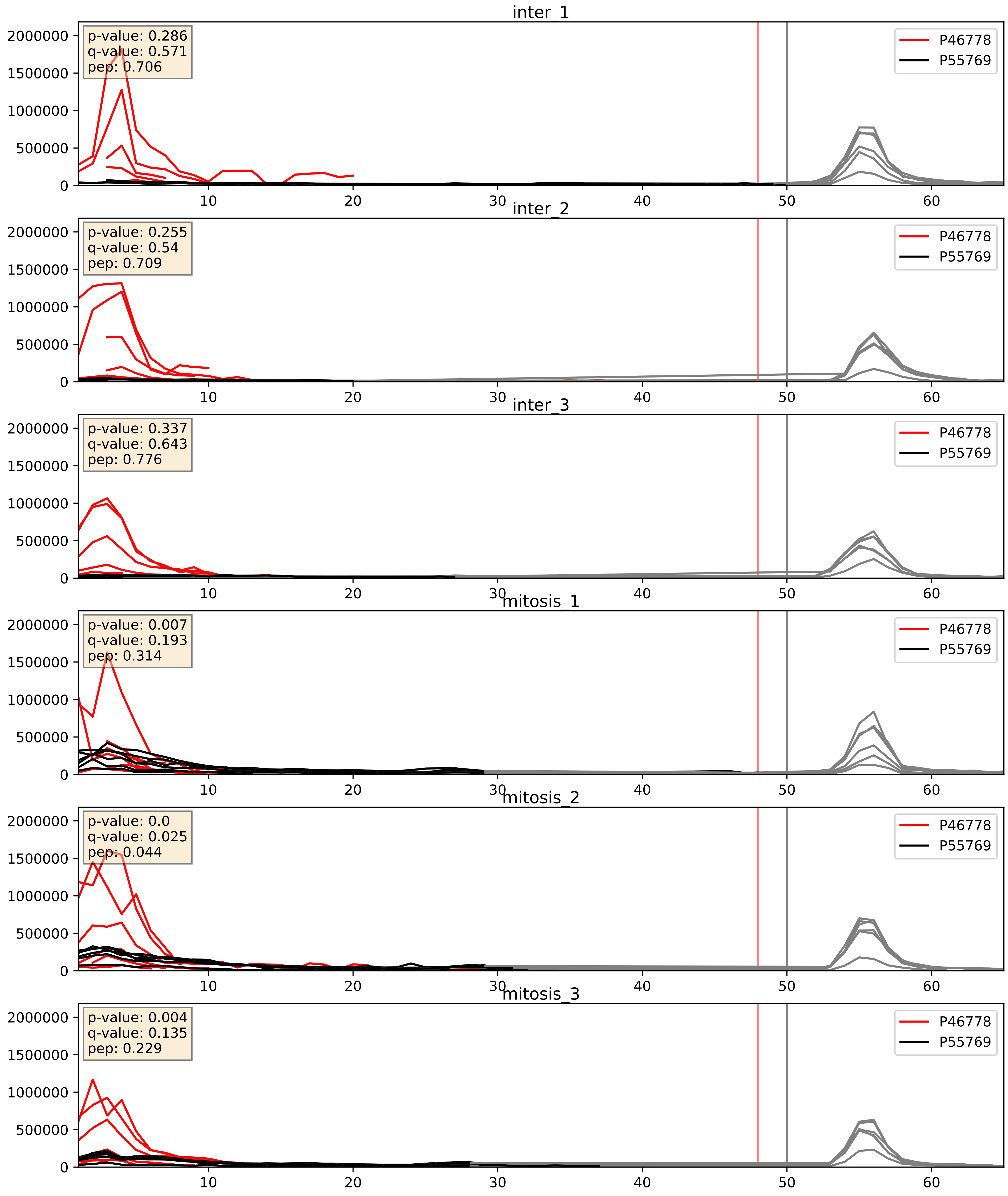


| level | condition_1 | condition_2 | pvalue | pvalue_adjusted |
|---|---|---|---|---|
| complex_abundance | mitosis | inter | 0.0030334186114187945 | 0.040382680114688776 |
| interactor_ratio | mitosis | inter | 0.003568846118141189 | 0.07714475447295095 |
| interactor_abundance | mitosis | inter | 0.48937275953919535 | 0.6579973023007024 |

P46779_P55769
RL28_HUMAN vs NH2L1_HUMAN
p-value: 0.0 q-value: 0.046

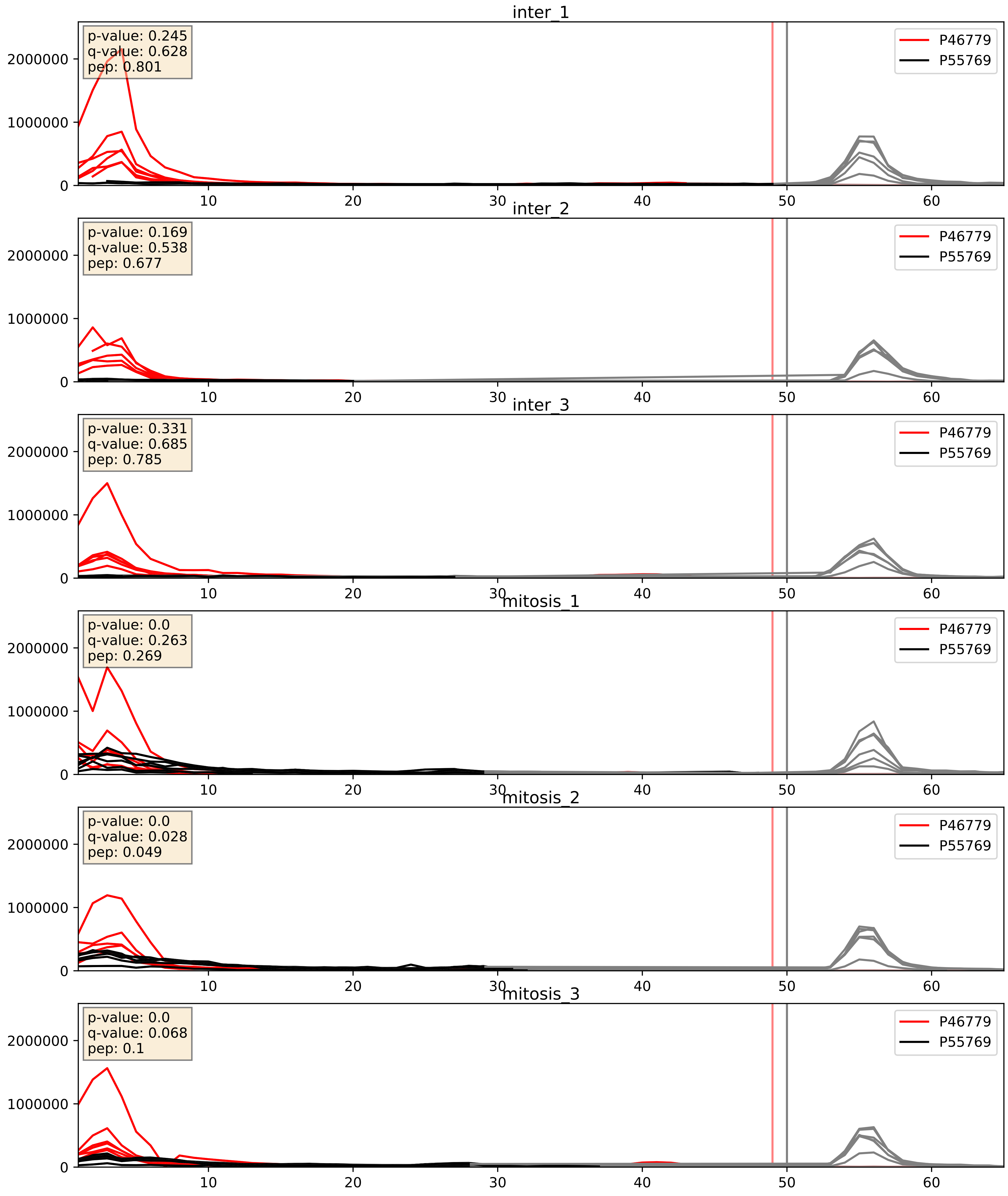


| level | condition_1 | condition_2 | pvalue | pvalue_adjusted |
|---|---|---|---|---|
| interactor_ratio | mitosis | inter | 0.0017975534568758092 | 0.051461731073100085 |
| complex_abundance | mitosis | inter | 0.0065132397257623995 | 0.05956552569714331 |
| interactor_abundance | mitosis | inter | 0.20546607270756348 | 0.40205499654286053 |

P46781_P55769
RS9_HUMAN vs NH2L1_HUMAN
p-value: 0.049 q-value: 0.032

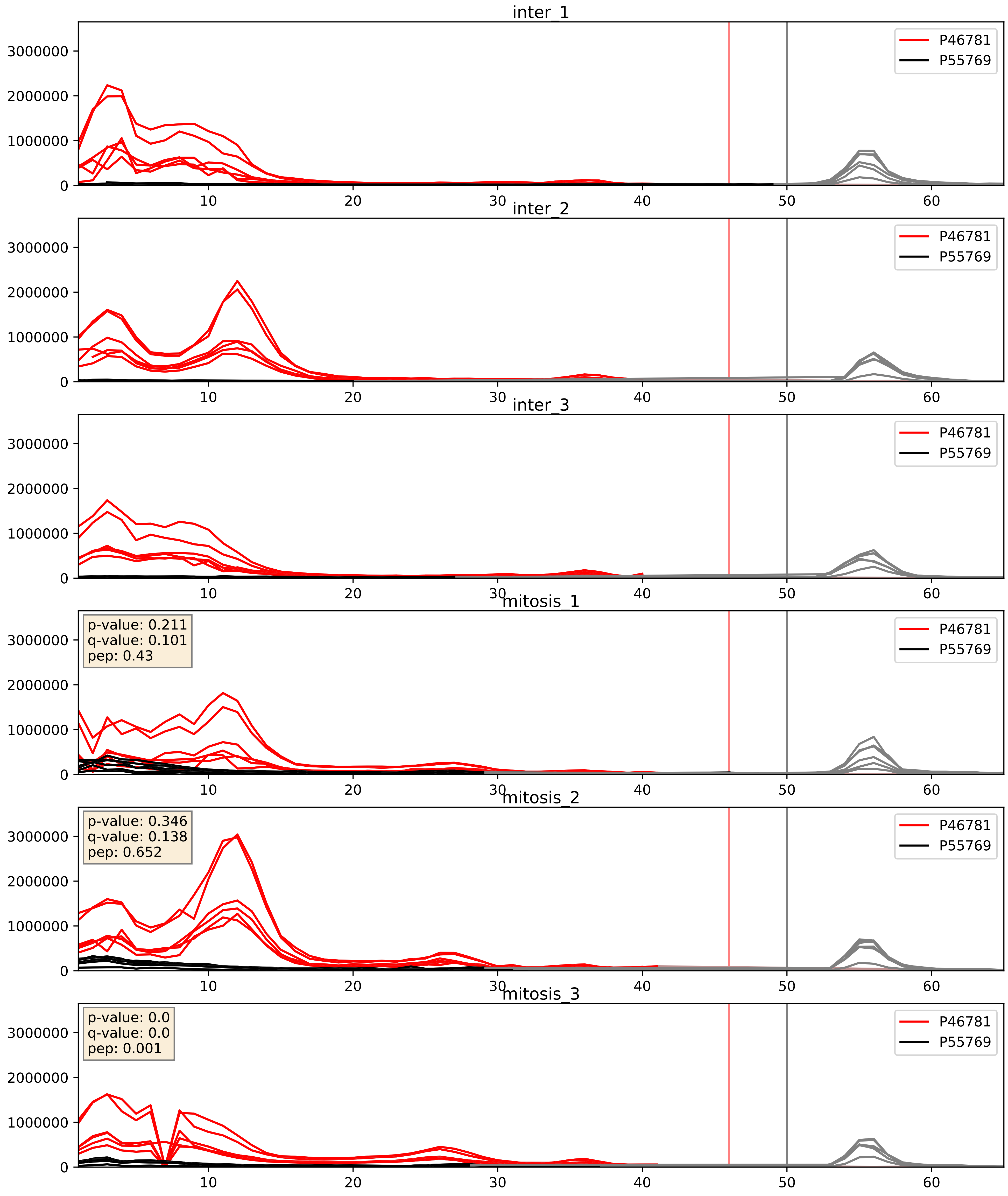


| level | condition_1 | condition_2 | pvalue | pvalue_adjusted |
|---|---|---|---|---|
| complex_abundance | mitosis | inter | 0.0008638921020909193 | 0.019513211095311817 |
| interactor_ratio | mitosis | inter | 0.010014220696020447 | 0.14263182888175546 |
| interactor_abundance | mitosis | inter | 0.5712841599732108 | 0.7174578065391263 |

P46782_P55769
RS5_HUMAN vs NH2L1_HUMAN
p-value: 0.079 q-value: 0.047

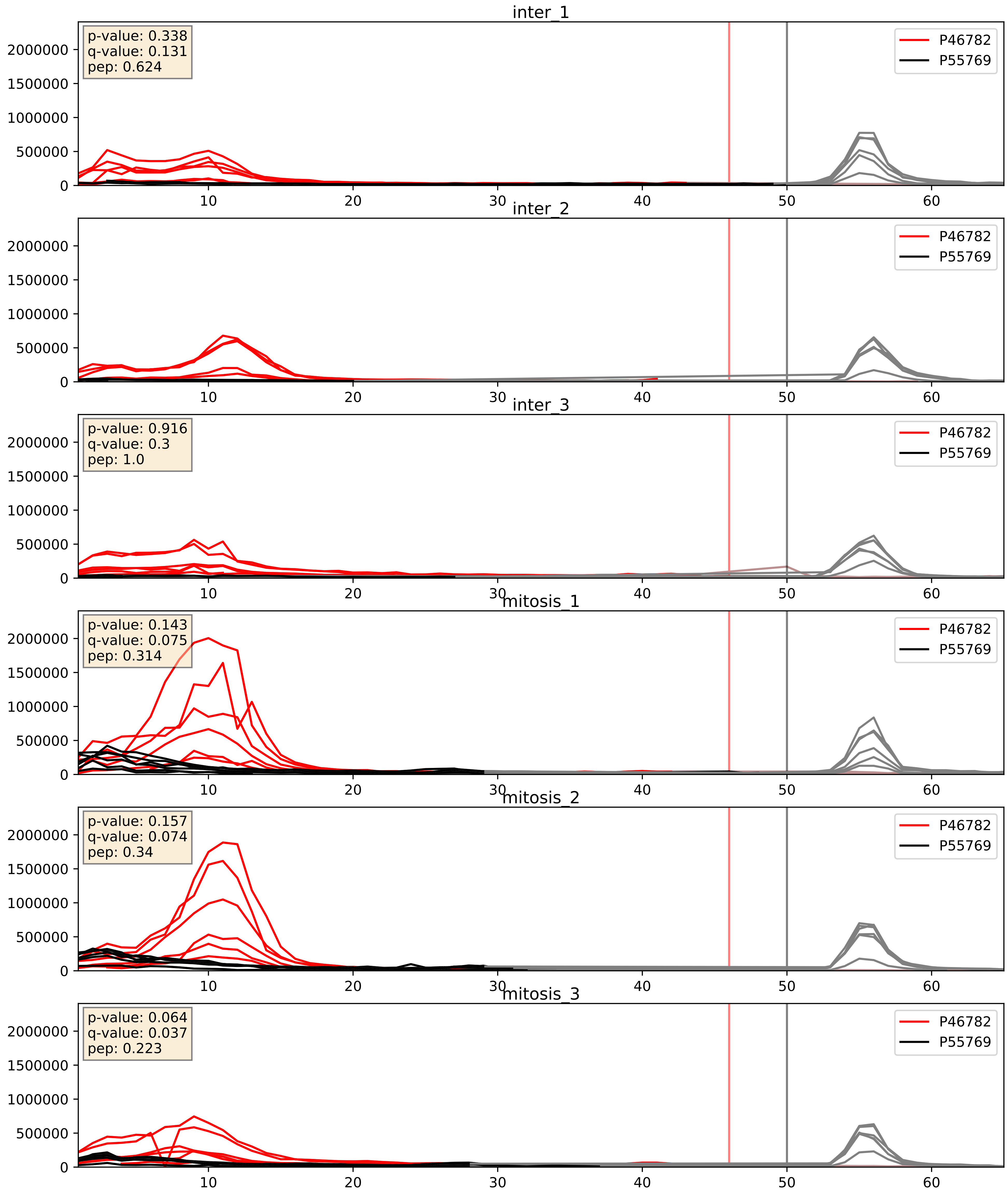


| level | condition_1 | condition_2 | pvalue | pvalue_adjusted |
|---|---|---|---|---|
| complex_abundance | mitosis | inter | 0.006070361828801624 | 0.05817678481497925 |
| interactor_ratio | mitosis | inter | 0.009513845612864247 | 0.13913572462569762 |
| interactor_abundance | mitosis | inter | 0.09473555347456417 | 0.2636762600365044 |

P49207_P55769
RL34_HUMAN vs NH2L1_HUMAN
p-value: 0.0 q-value: 0.023

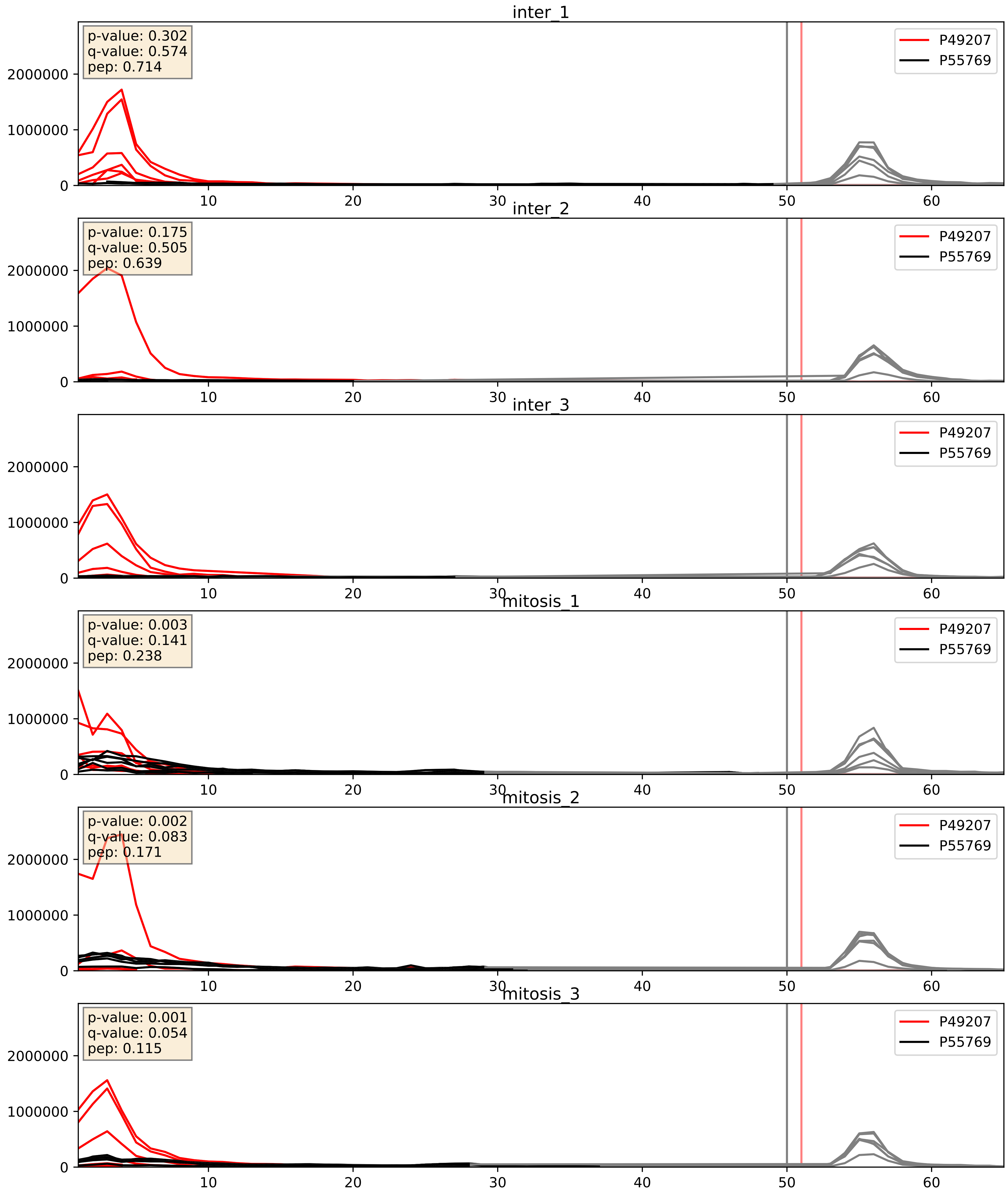


| level | condition_1 | condition_2 | pvalue | pvalue_adjusted |
|---|---|---|---|---|
| interactor_ratio | mitosis | inter | 0.0488496422782553 | 0.3645622823904668 |
| complex_abundance | mitosis | inter | 0.10606036529967314 | 0.2424236921135386 |
| interactor_abundance | mitosis | inter | 0.6942121200095892 | 0.7977893922170306 |

P51991_P55769
ROA3_HUMAN vs NH2L1_HUMAN
p-value: 0.0 q-value: 0.01

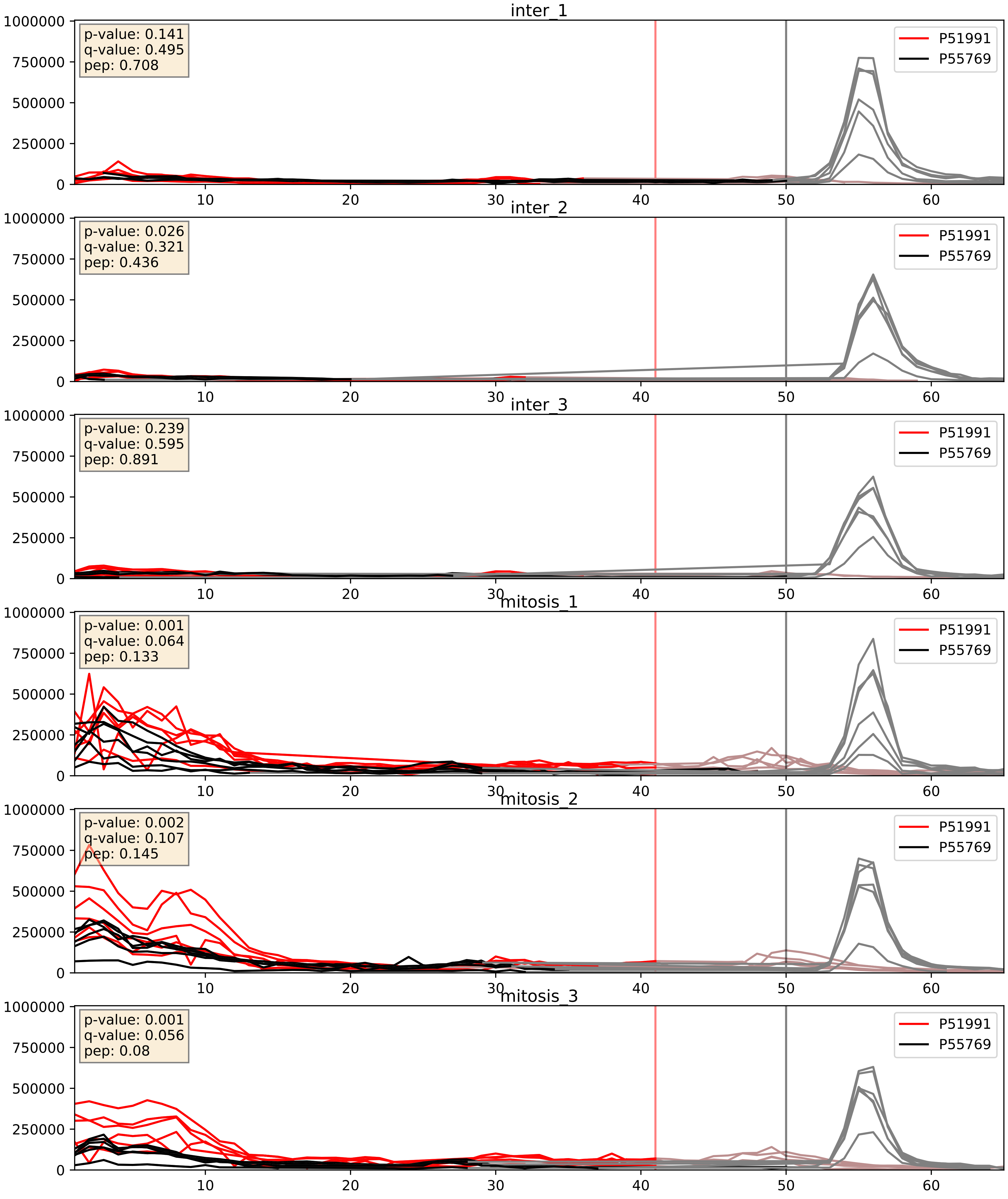


| level | condition_1 | condition_2 | pvalue | pvalue_adjusted |
|---|---|---|---|---|
| interactor_abundance | mitosis | inter | 4.454929811102081e-06 | 0.0033734327106189023 |
| complex_abundance | mitosis | inter | 9.719836869876729e-05 | 0.00670982287146329 |
| interactor_ratio | mitosis | inter | 0.09016169571238221 | 0.5008332999986967 |

P52597_P55769
HNRPF_HUMAN vs NH2L1_HUMAN
p-value: 0.0 q-value: 0.042

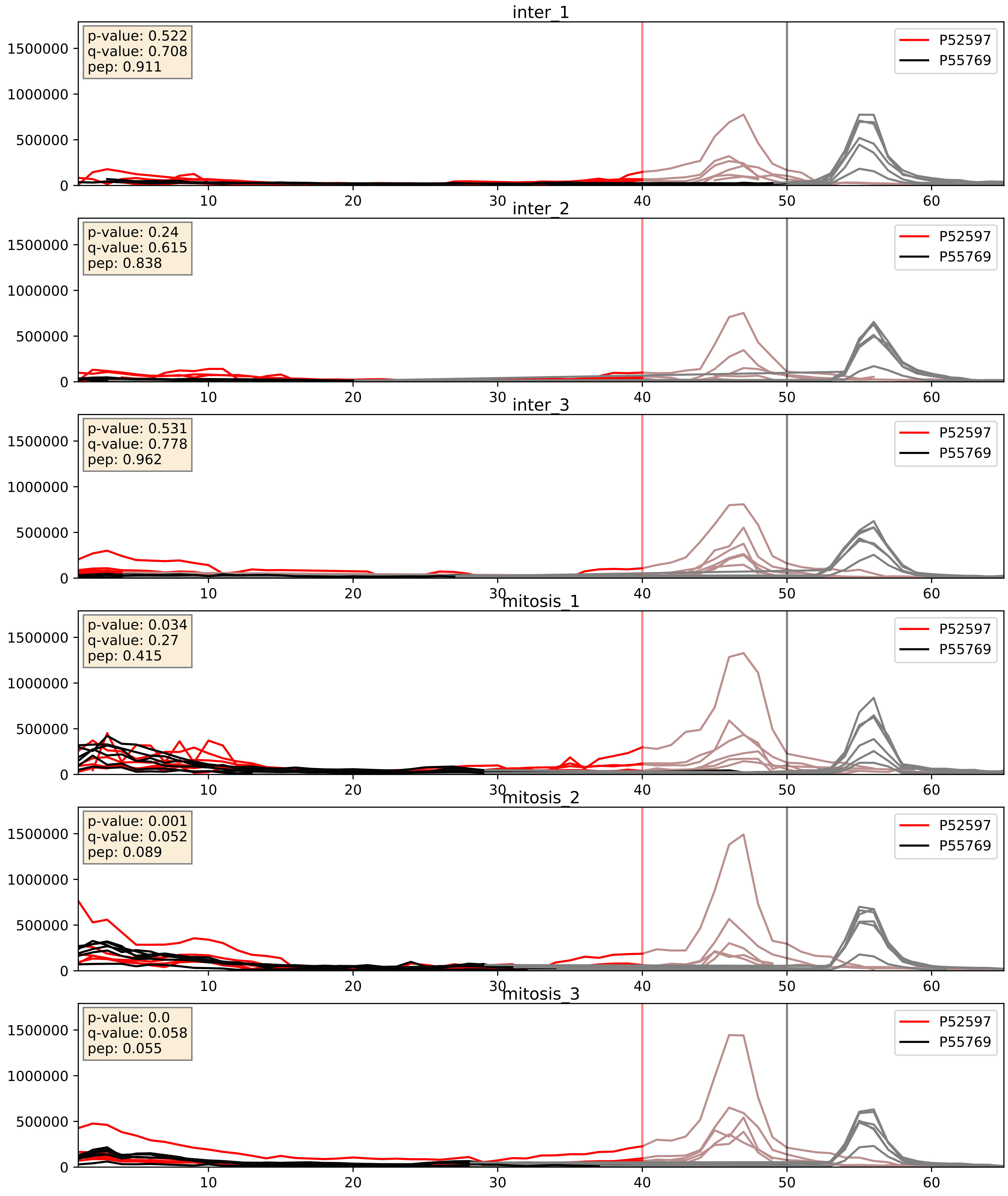


| level | condition_1 | condition_2 | pvalue | pvalue_adjusted |
|---|---|---|---|---|
| complex_abundance | mitosis | inter | 0.0012657499792246994 | 0.02413100183109895 |
| interactor_abundance | mitosis | inter | 0.006646850462384629 | 0.06976951558309309 |
| interactor_ratio | mitosis | inter | 0.03615766025694413 | 0.3092003714280137 |

P55769_P56537
NH2L1_HUMAN vs IF6_HUMAN
p-value: 0.001 q-value: 0.03

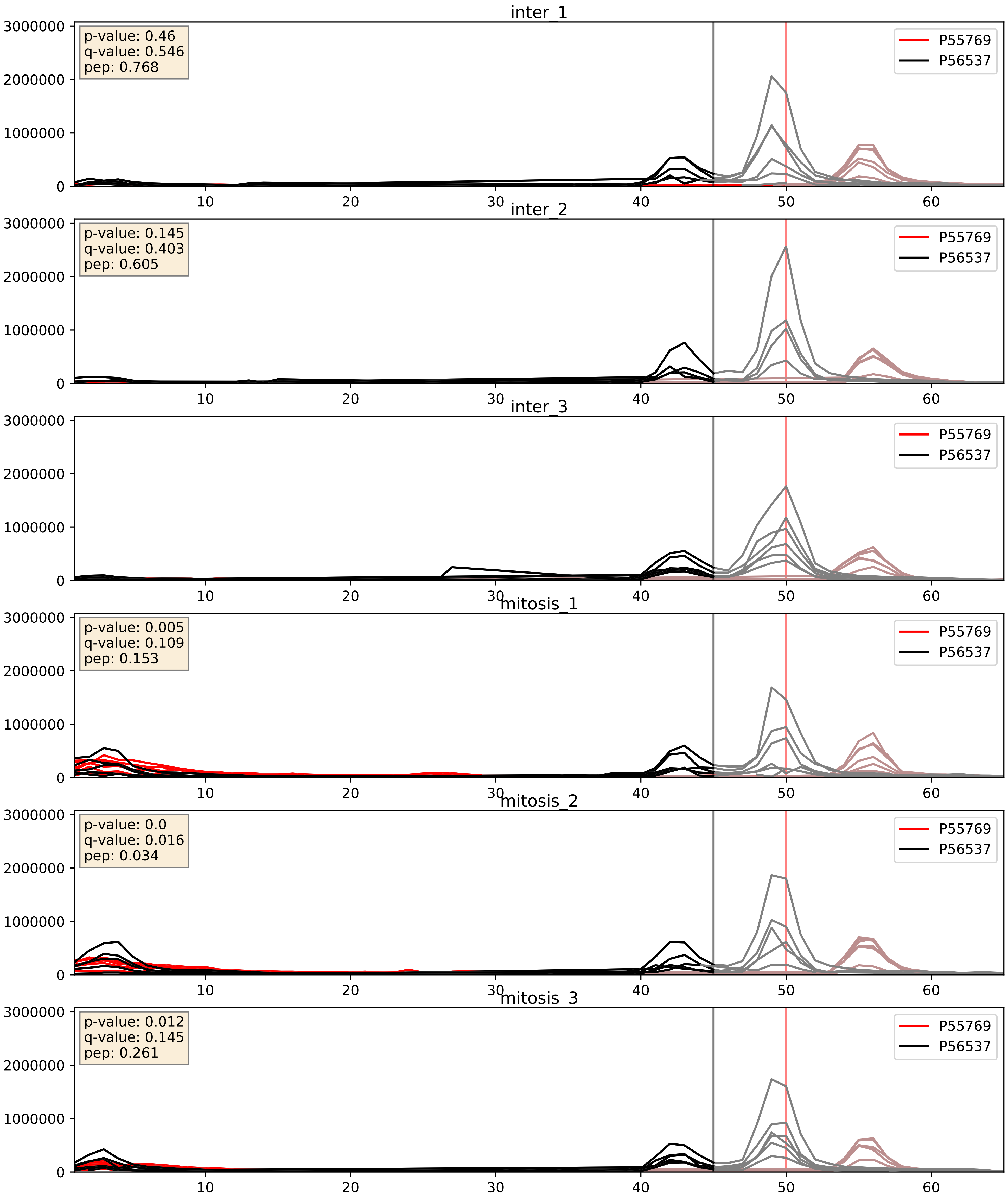


| level | condition_1 | condition_2 | pvalue | pvalue_adjusted |
|---|---|---|---|---|
| interactor_abundance | mitosis | inter | 0.001960763129046175 | 0.032370554261591626 |
| complex_abundance | mitosis | inter | 0.020954510270985917 | 0.10689403306976396 |
| interactor_ratio | mitosis | inter | 0.04772747899799155 | 0.3614455797387484 |

P55769_P61247
NH2L1_HUMAN vs RS3A_HUMAN
p-value: 0.029 q-value: 0.021

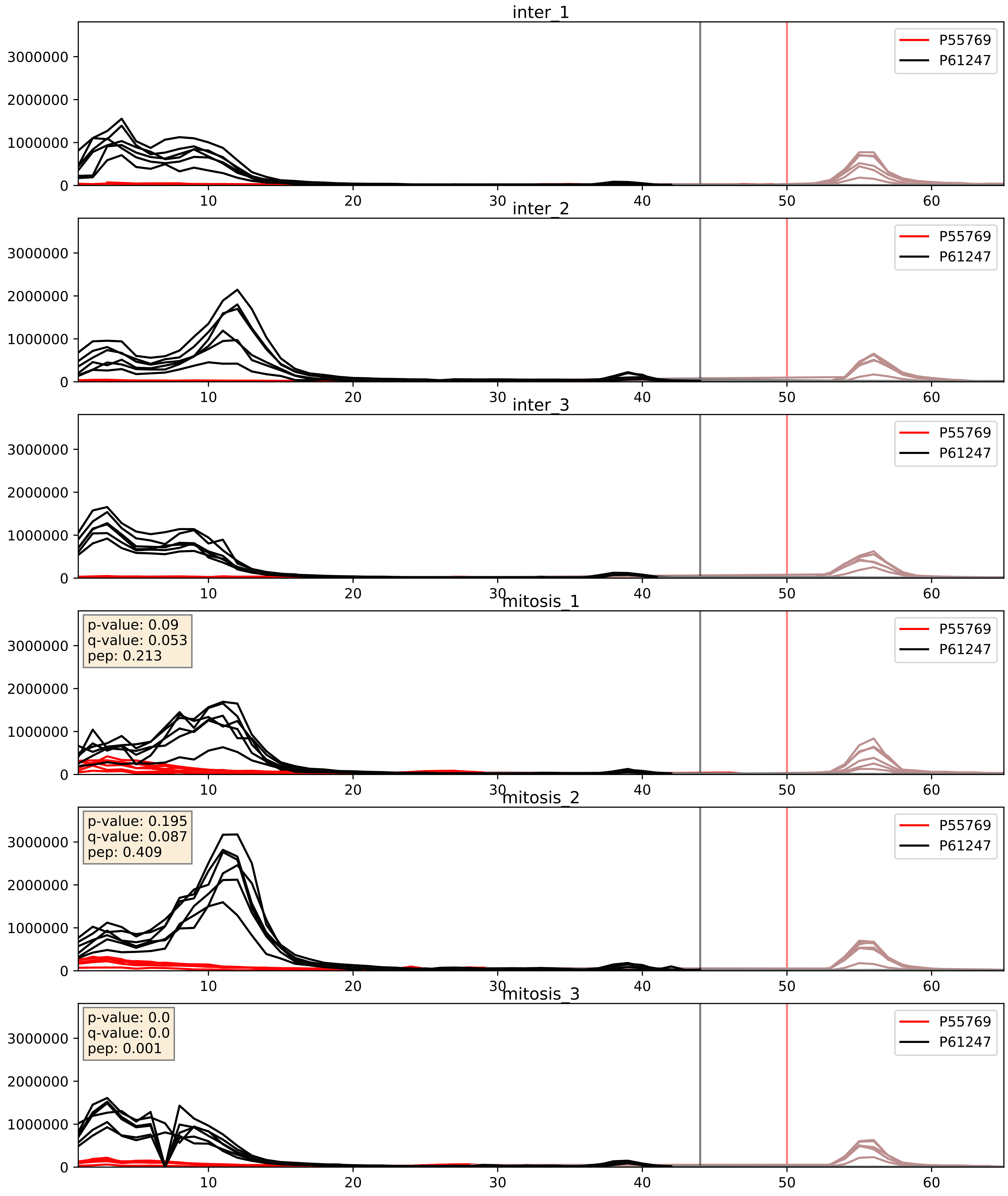


| level | condition_1 | condition_2 | pvalue | pvalue_adjusted |
|---|---|---|---|---|
| complex_abundance | mitosis | inter | 0.0015577419918366706 | 0.02760718726733313 |
| interactor_abundance | mitosis | inter | 0.0021242110805868956 | 0.03376647511573598 |
| interactor_ratio | mitosis | inter | 0.010955920585766941 | 0.15054322834266998 |

P55769_P61353
NH2L1_HUMAN vs RL27_HUMAN
p-value: 0.0 q-value: 0.008

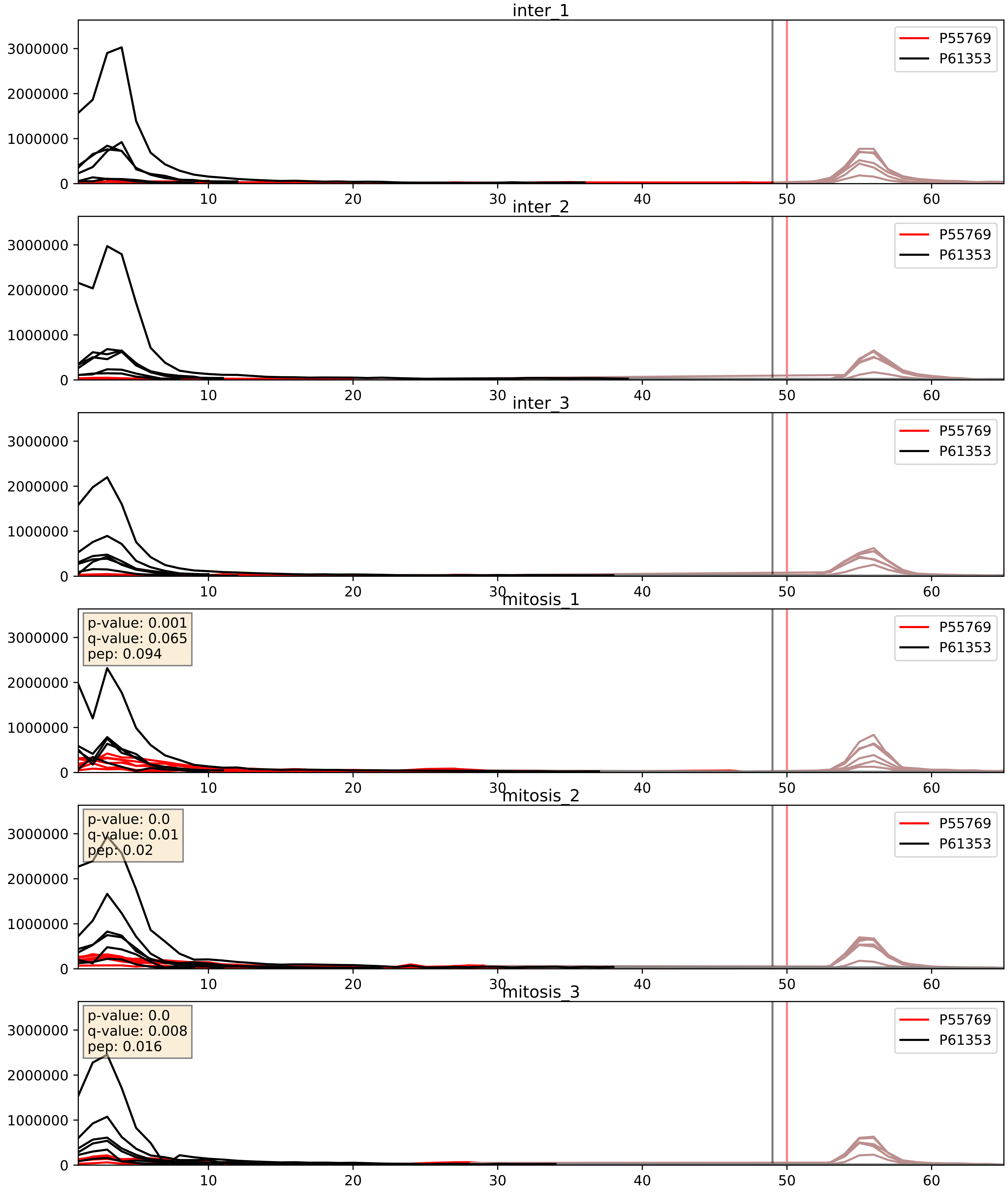


| level | condition_1 | condition_2 | pvalue | pvalue_adjusted |
|---|---|---|---|---|
| interactor_ratio | mitosis | inter | 0.0013567231783374613 | 0.04349644347029464 |
| interactor_abundance | mitosis | inter | 0.0014620432003608957 | 0.028218917238081776 |
| complex_abundance | mitosis | inter | 0.004474029937243554 | 0.049788645279664145 |

P55769_P62081
NH2L1_HUMAN vs RS7_HUMAN
p-value: 0.0 q-value: 0.001

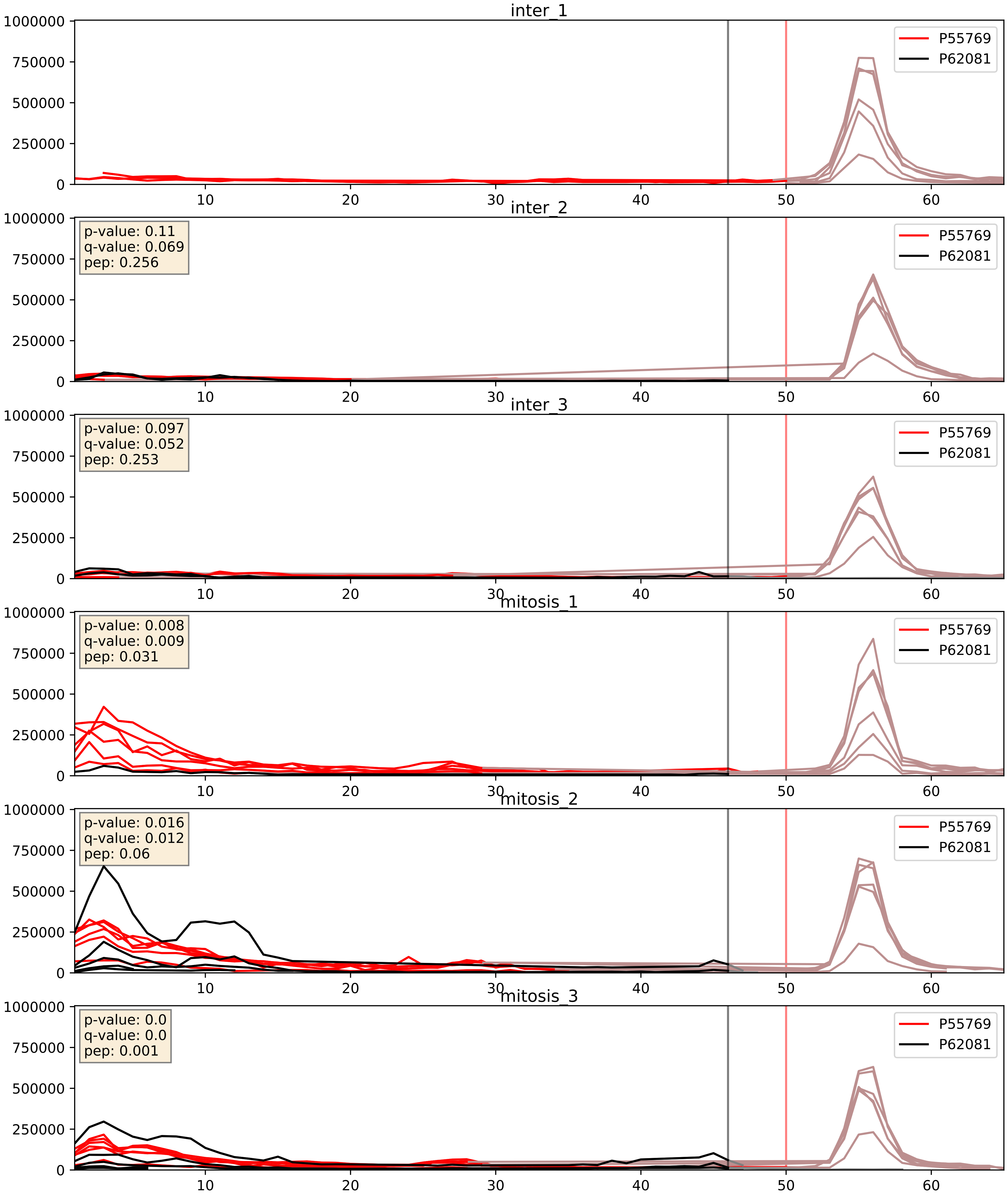


| level | condition_1 | condition_2 | pvalue | pvalue_adjusted |
|---|---|---|---|---|
| interactor_abundance | mitosis | inter | 0.002776703337451414 | 0.03993577968615009 |
| complex_abundance | mitosis | inter | 0.021496464517949856 | 0.10786033779229236 |
| interactor_ratio | mitosis | inter | 0.11064229200612882 | 0.5520385380060858 |

P55769_P62241
NH2L1_HUMAN vs RS8_HUMAN
p-value: 0.001 q-value: 0.001

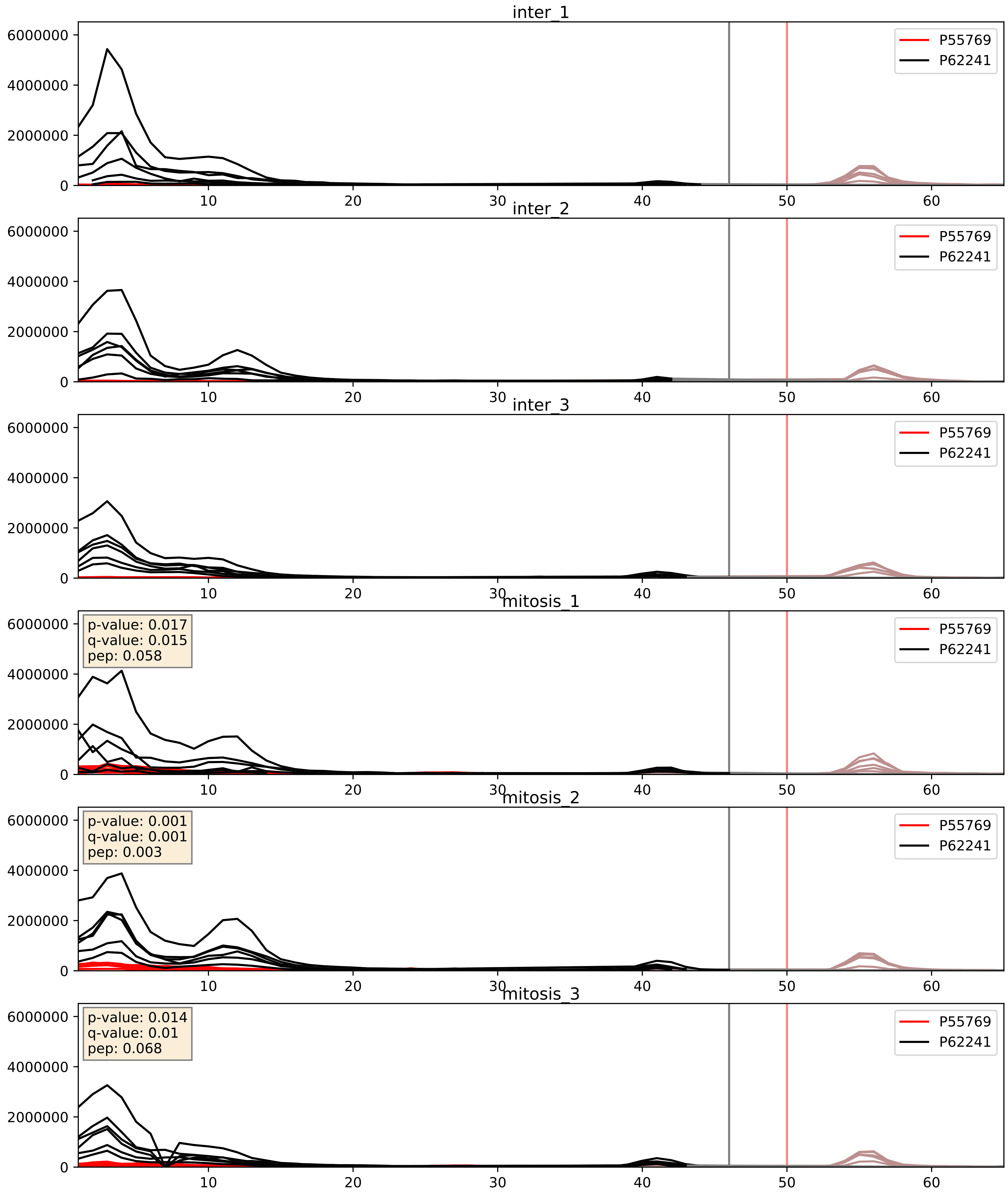


| level | condition_1 | condition_2 | pvalue | pvalue_adjusted |
|---|---|---|---|---|
| interactor_ratio | mitosis | inter | 0.0017972345362234274 | 0.051461731073100085 |
| interactor_abundance | mitosis | inter | 0.003136097951291225 | 0.043089885173439625 |
| complex_abundance | mitosis | inter | 0.008317782290370432 | 0.06742444735376033 |

P55769_P62244
NH2L1_HUMAN vs RS15A_HUMAN
p-value: 0.0 q-value: 0.0

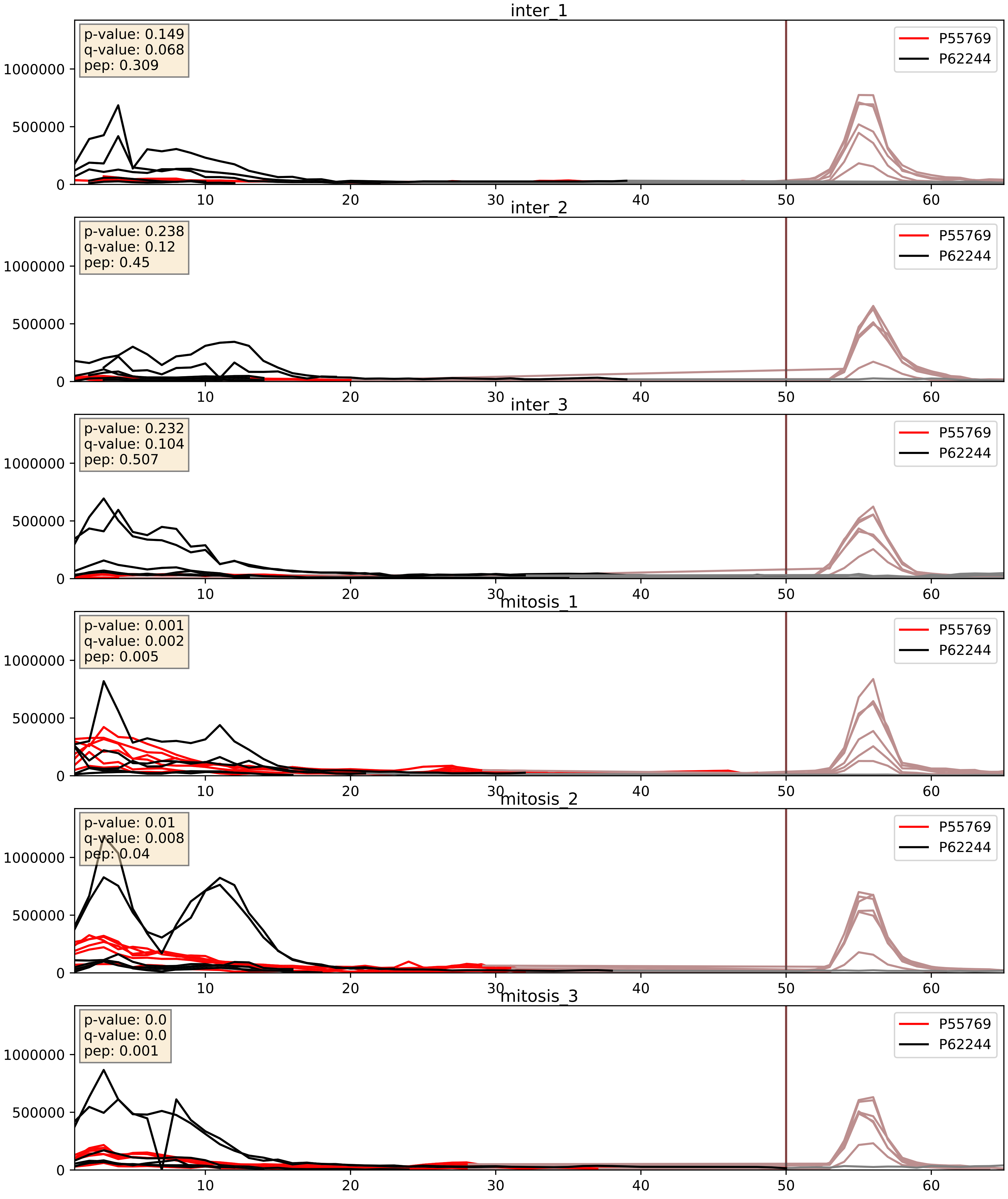


| level | condition_1 | condition_2 | pvalue | pvalue_adjusted |
|---|---|---|---|---|
| interactor_abundance | mitosis | inter | 0.0009839140650830256 | 0.023233943164443306 |
| interactor_ratio | mitosis | inter | 0.0010684903745133802 | 0.038414698994273265 |
| complex_abundance | mitosis | inter | 0.0034681439054112745 | 0.04302508960916016 |

P55769_P62249
NH2L1_HUMAN vs RS16_HUMAN
p-value: 0.05 q-value: 0.033

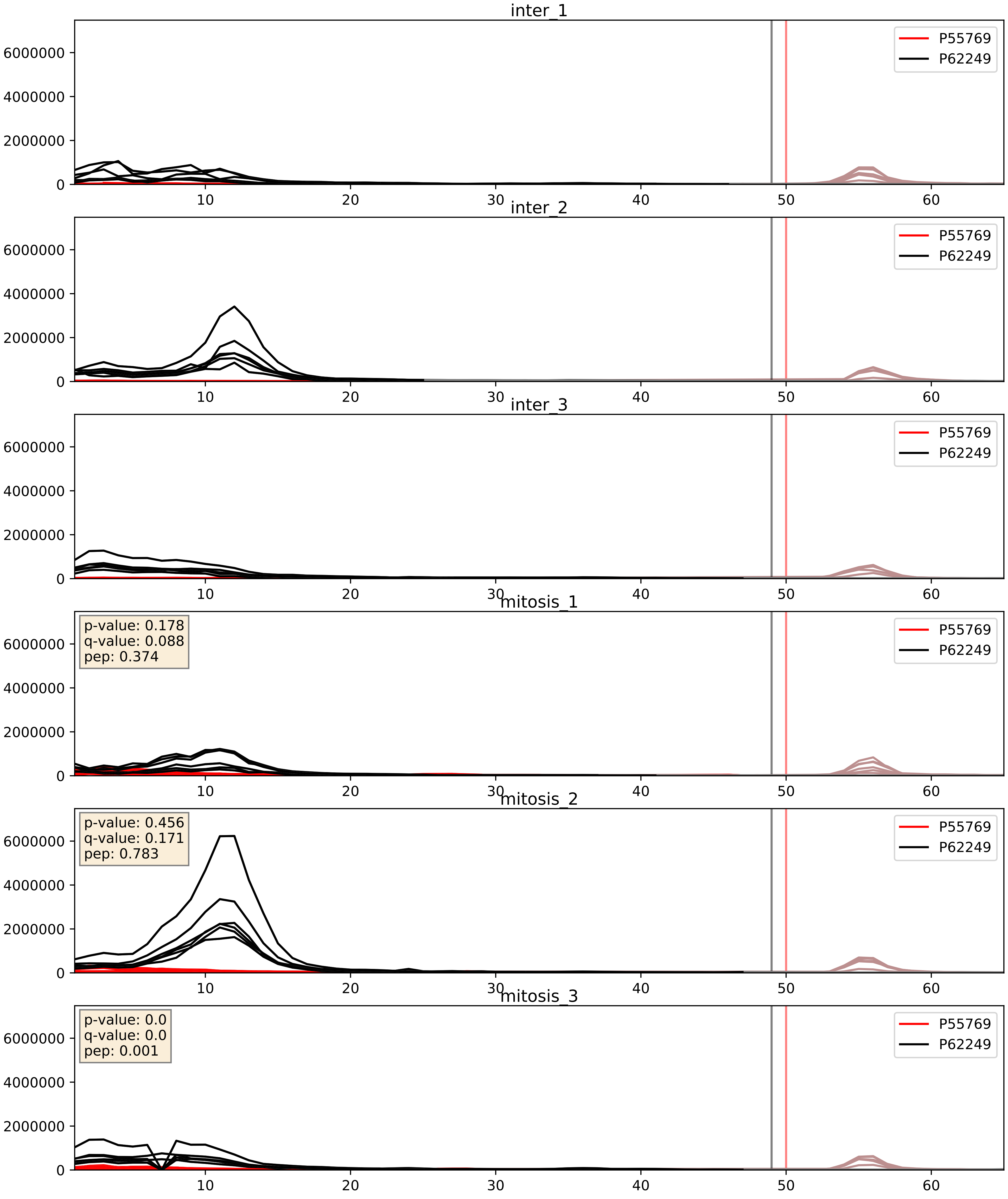


| level | condition_1 | condition_2 | pvalue | pvalue_adjusted |
|---|---|---|---|---|
| interactor_abundance | mitosis | inter | 0.0024684610377058687 | 0.037061411436434794 |
| complex_abundance | mitosis | inter | 0.007468050120599692 | 0.06431238333232732 |
| interactor_ratio | mitosis | inter | 0.05619442370902446 | 0.3926647223857623 |

P55769_P62266
NH2L1_HUMAN vs RS23_HUMAN
p-value: 0.0 q-value: 0.0

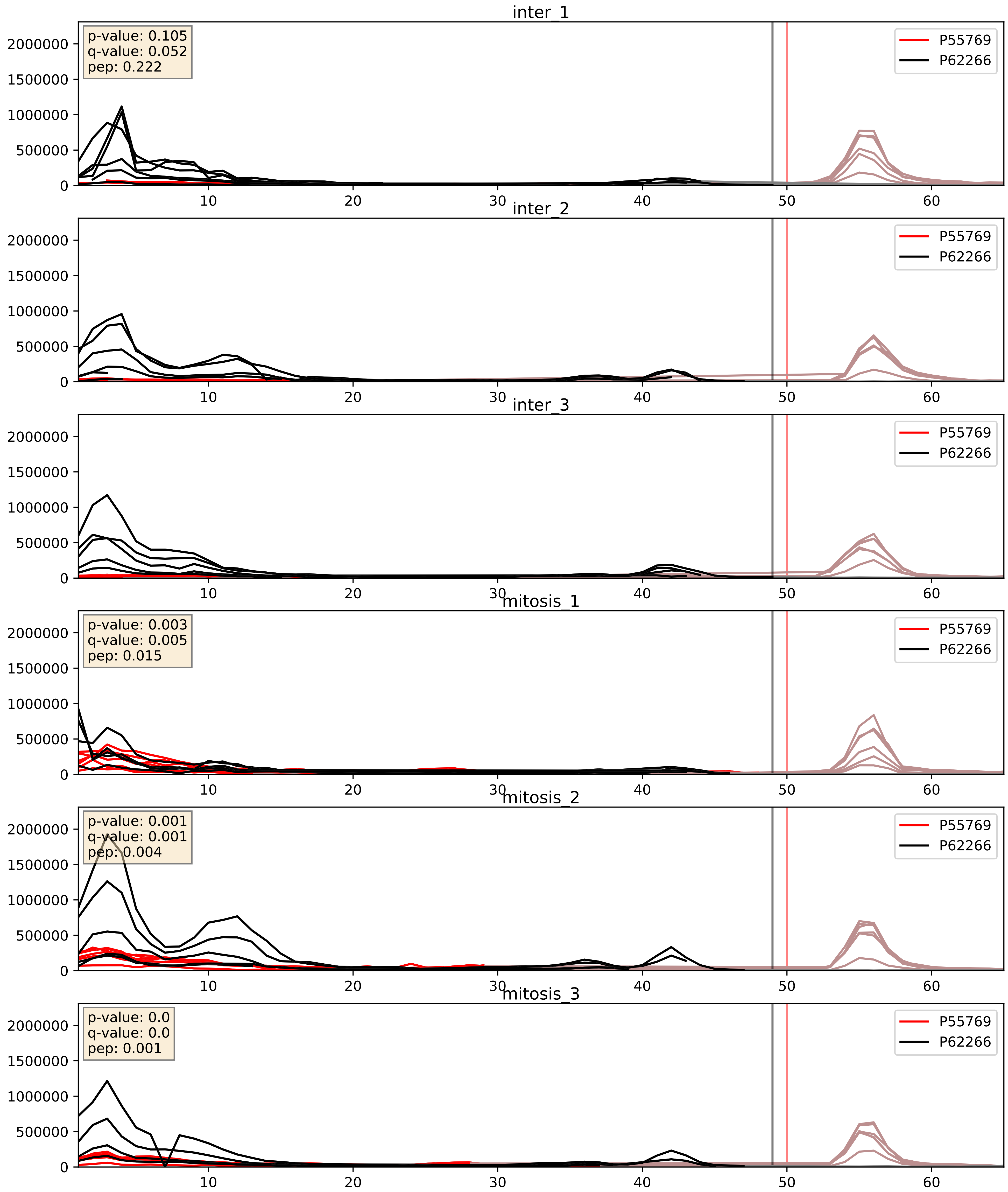


| level | condition_1 | condition_2 | pvalue | pvalue_adjusted |
|---|---|---|---|---|
| interactor_abundance | mitosis | inter | 0.0031769585353846404 | 0.043442116713885814 |
| interactor_ratio | mitosis | inter | 0.005738830207404453 | 0.10363794636156566 |
| complex_abundance | mitosis | inter | 0.15575217624509755 | 0.30259614812937696 |

P55769_P62277
NH2L1_HUMAN vs RS13_HUMAN
p-value: 0.0 q-value: 0.0

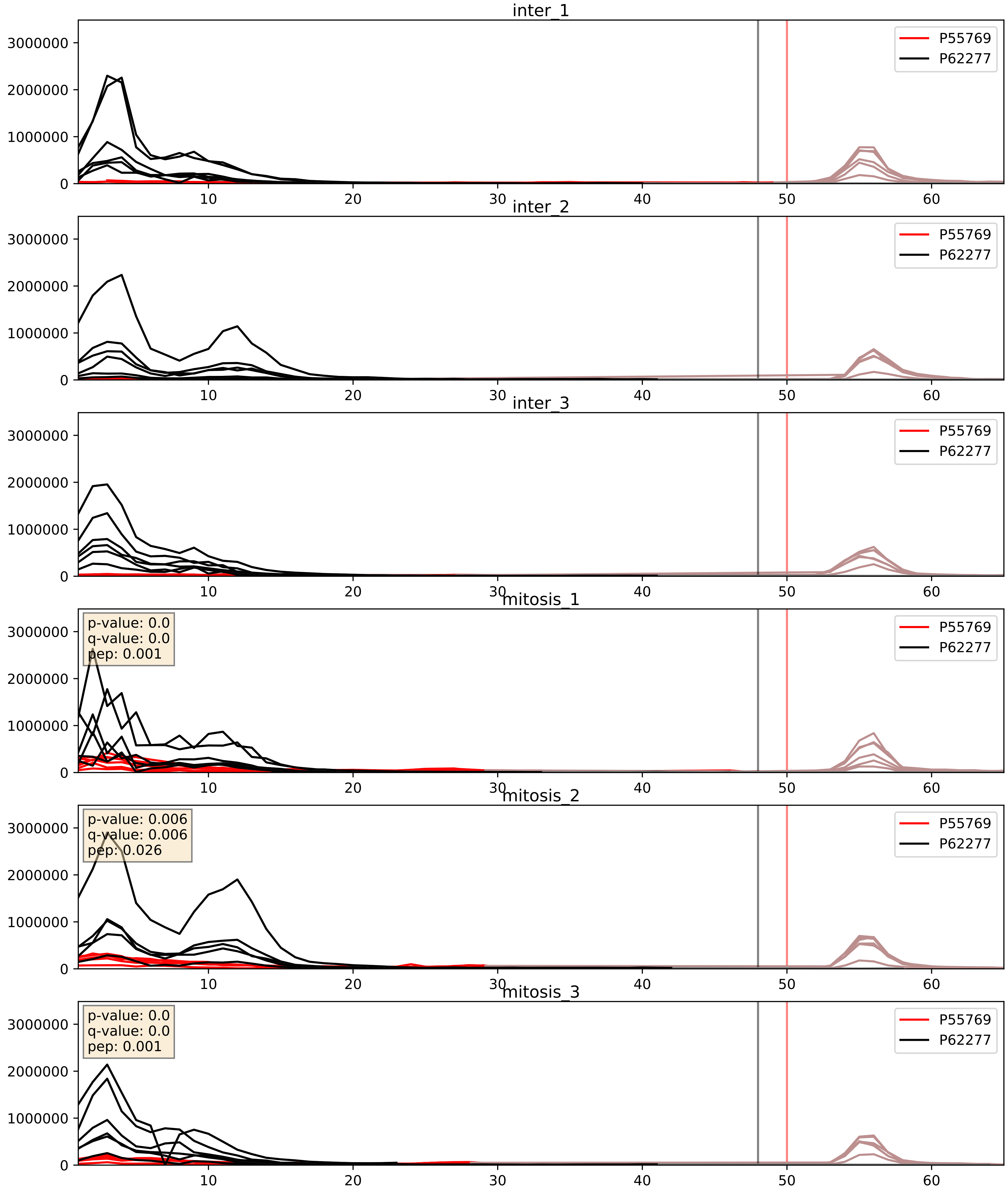


| level | condition_1 | condition_2 | pvalue | pvalue_adjusted |
|---|---|---|---|---|
| complex_abundance | mitosis | inter | 0.0016281493033817628 | 0.02826969175851499 |
| interactor_abundance | mitosis | inter | 0.002154014372967189 | 0.03396392286310966 |
| interactor_ratio | mitosis | inter | 0.004612115938377692 | 0.09096707933758767 |

P55769_P62280
NH2L1_HUMAN vs RS11_HUMAN
p-value: 0.0 q-value: 0.0

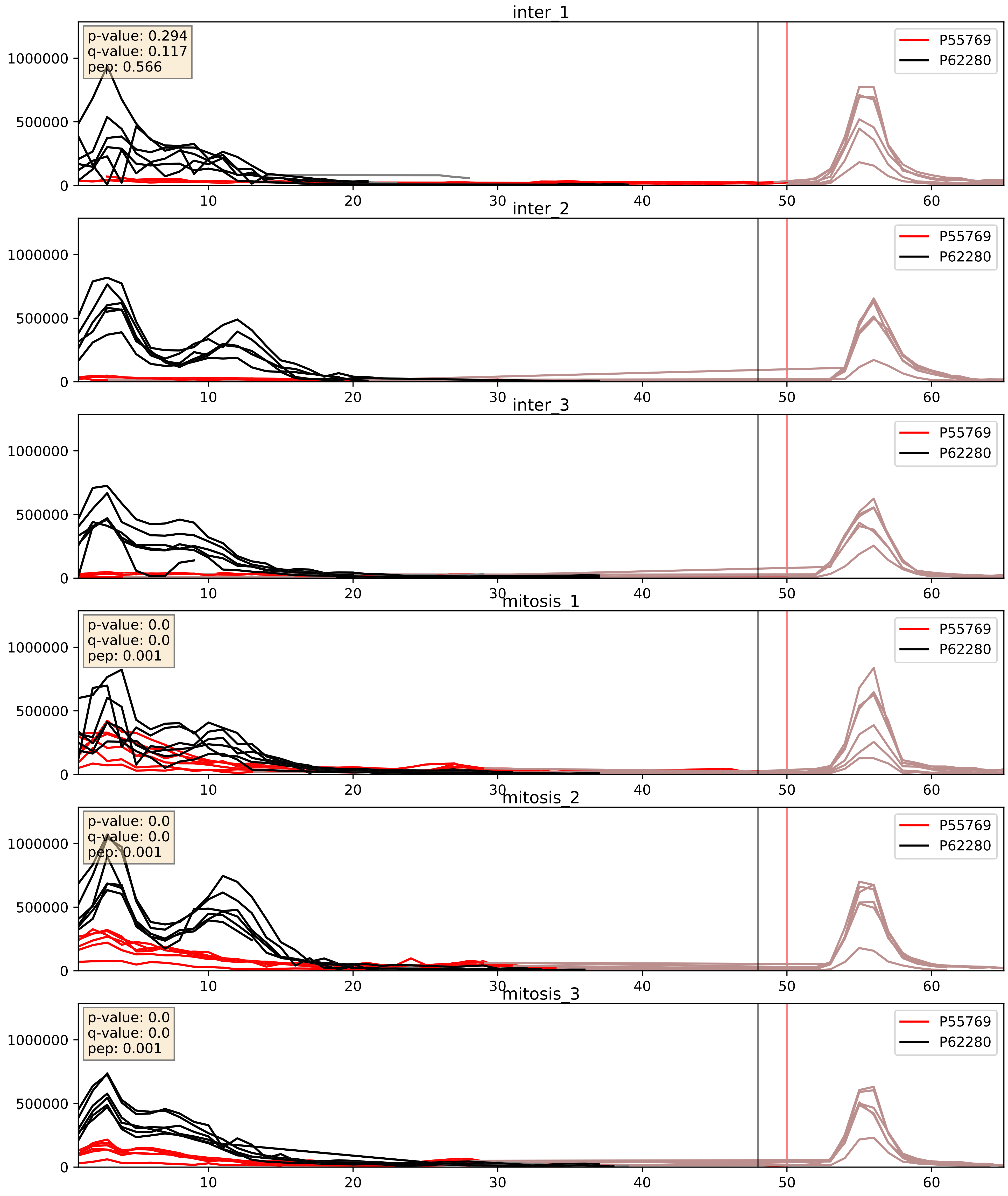


| level | condition_1 | condition_2 | pvalue | pvalue_adjusted |
|---|---|---|---|---|
| complex_abundance | mitosis | inter | 0.0003990885249934917 | 0.01272633620489383 |
| interactor_abundance | mitosis | inter | 0.0011097319494582842 | 0.024049720606162197 |
| interactor_ratio | mitosis | inter | 0.006566244961729941 | 0.1124141137448166 |

P55769_P62701
NH2L1_HUMAN vs RS4X_HUMAN
p-value: 0.0 q-value: 0.001

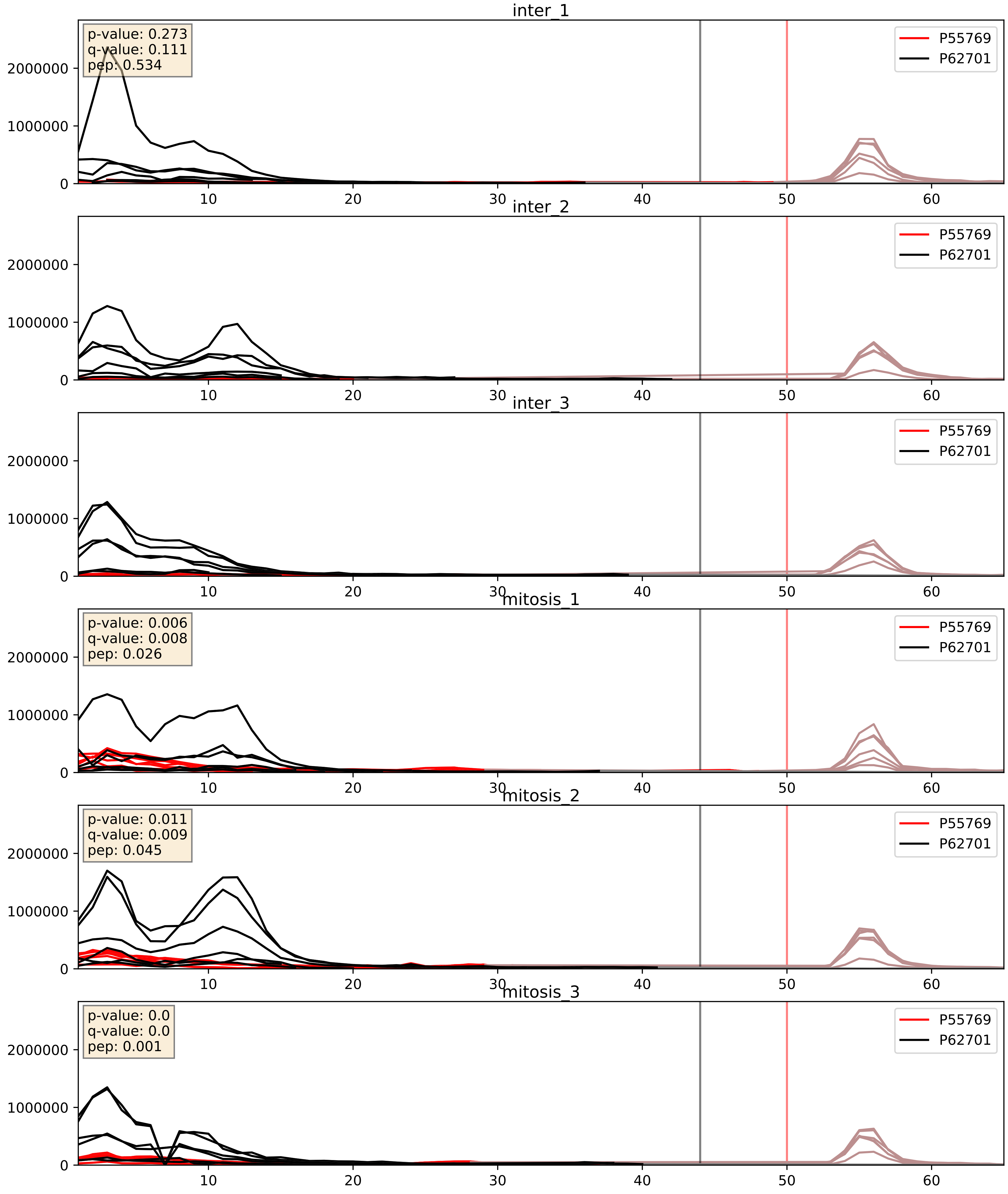


| level | condition_1 | condition_2 | pvalue | pvalue_adjusted |
|---|---|---|---|---|
| complex_abundance | mitosis | inter | 0.0011179963250585142 | 0.022133971147621622 |
| interactor_abundance | mitosis | inter | 0.0018457486603657728 | 0.03159921706546203 |
| interactor_ratio | mitosis | inter | 0.008738550111391514 | 0.13215899108394233 |

P55769_P62750
NH2L1_HUMAN vs RL23A_HUMAN
p-value: 0.0 q-value: 0.008

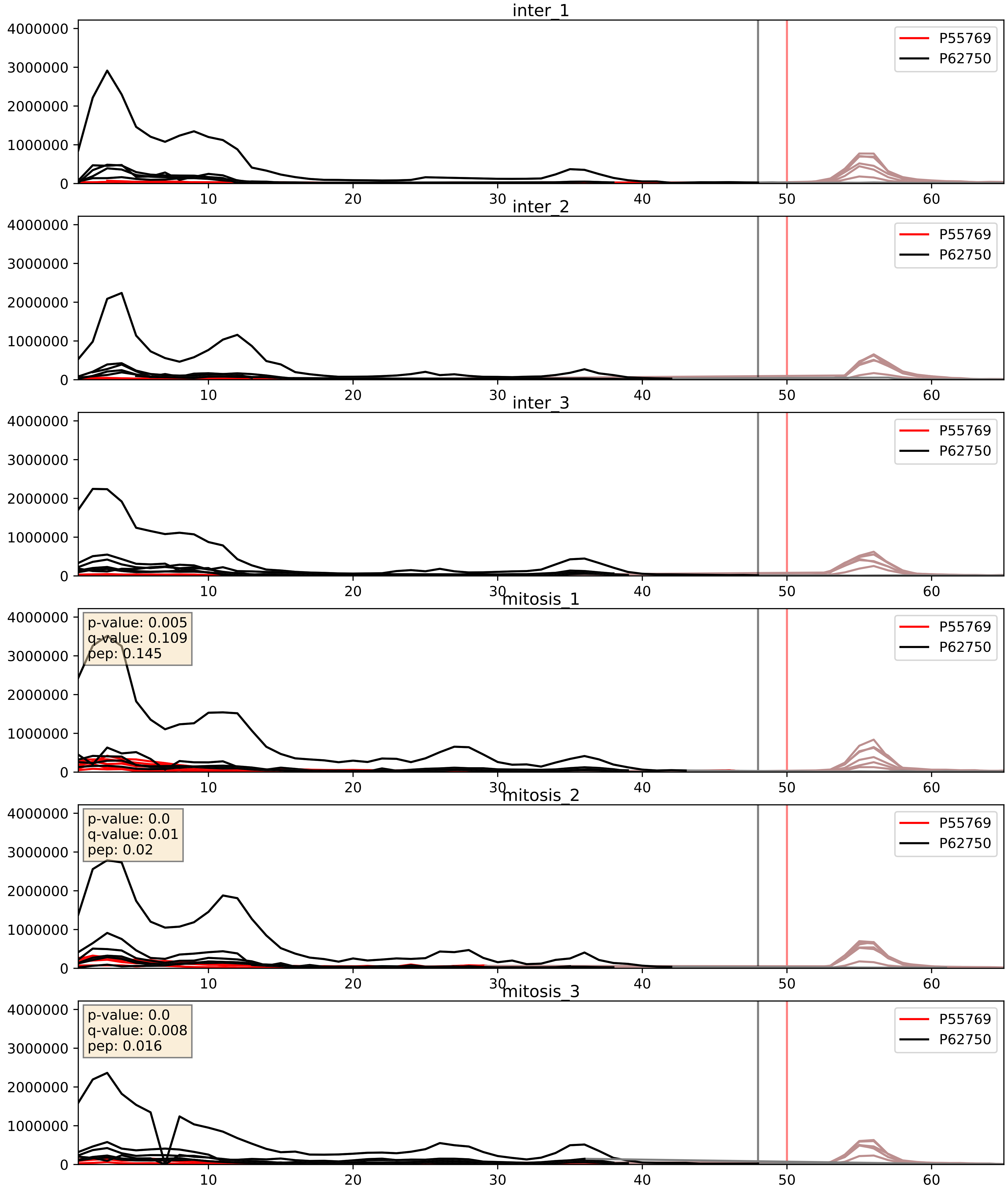


| level | condition_1 | condition_2 | pvalue | pvalue_adjusted |
|---|---|---|---|---|
| interactor_abundance | mitosis | inter | 0.0022737770306248332 | 0.03522811109891145 |
| complex_abundance | mitosis | inter | 0.005326196929449992 | 0.054883686885757 |
| interactor_ratio | mitosis | inter | 0.012218288328575378 | 0.15943376233628845 |

P55769_P62753
NH2L1_HUMAN vs RS6_HUMAN
p-value: 0.0 q-value: 0.0

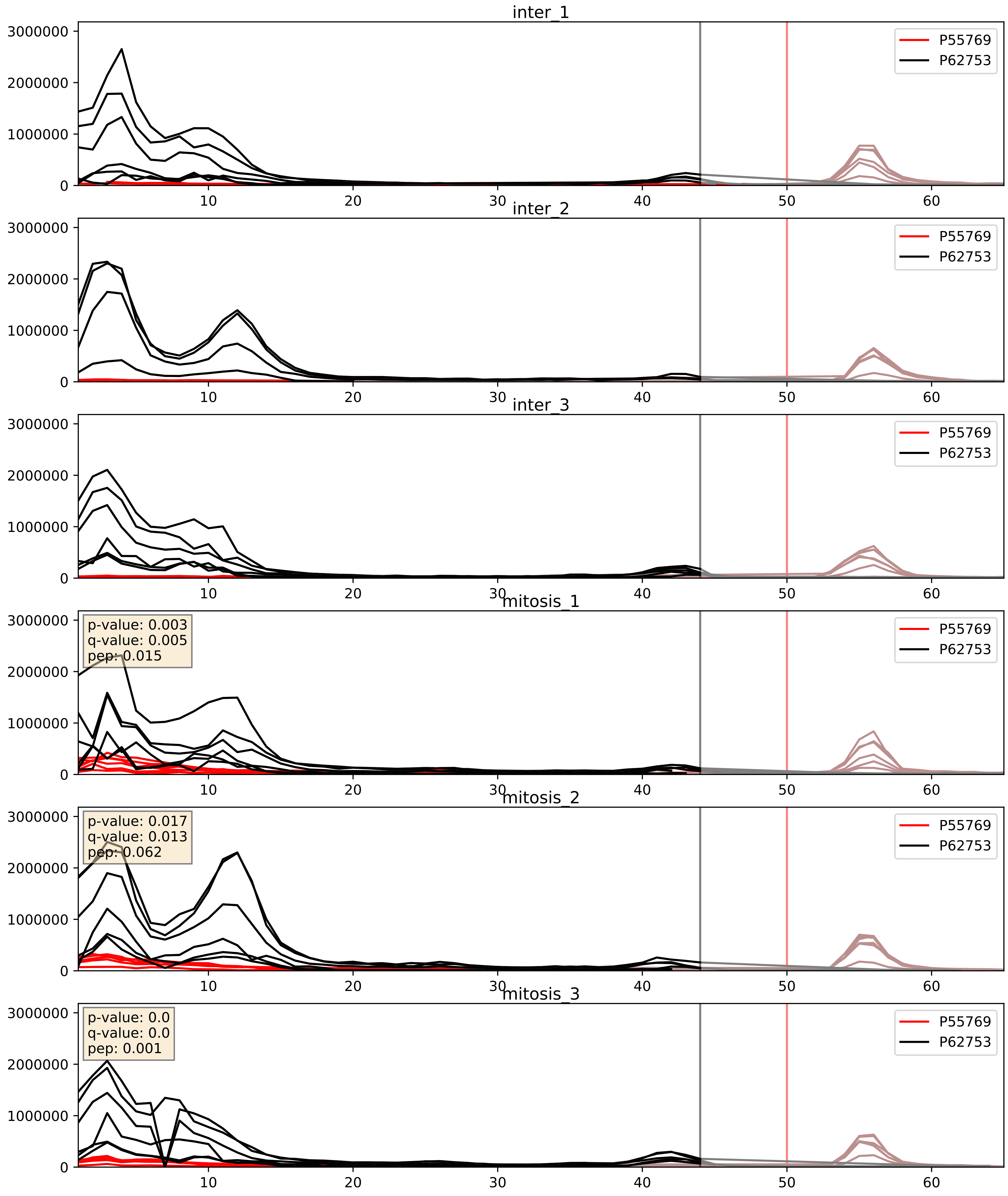


| level | condition_1 | condition_2 | pvalue | pvalue_adjusted |
|---|---|---|---|---|
| complex_abundance | mitosis | inter | 0.0006746791326246952 | 0.016643381484920432 |
| interactor_abundance | mitosis | inter | 0.0022247149364754016 | 0.03466841143289118 |
| interactor_ratio | mitosis | inter | 0.00836956527510944 | 0.129788910787929 |

P55769_P62829
NH2L1_HUMAN vs RL23_HUMAN
p-value: 0.0 q-value: 0.008

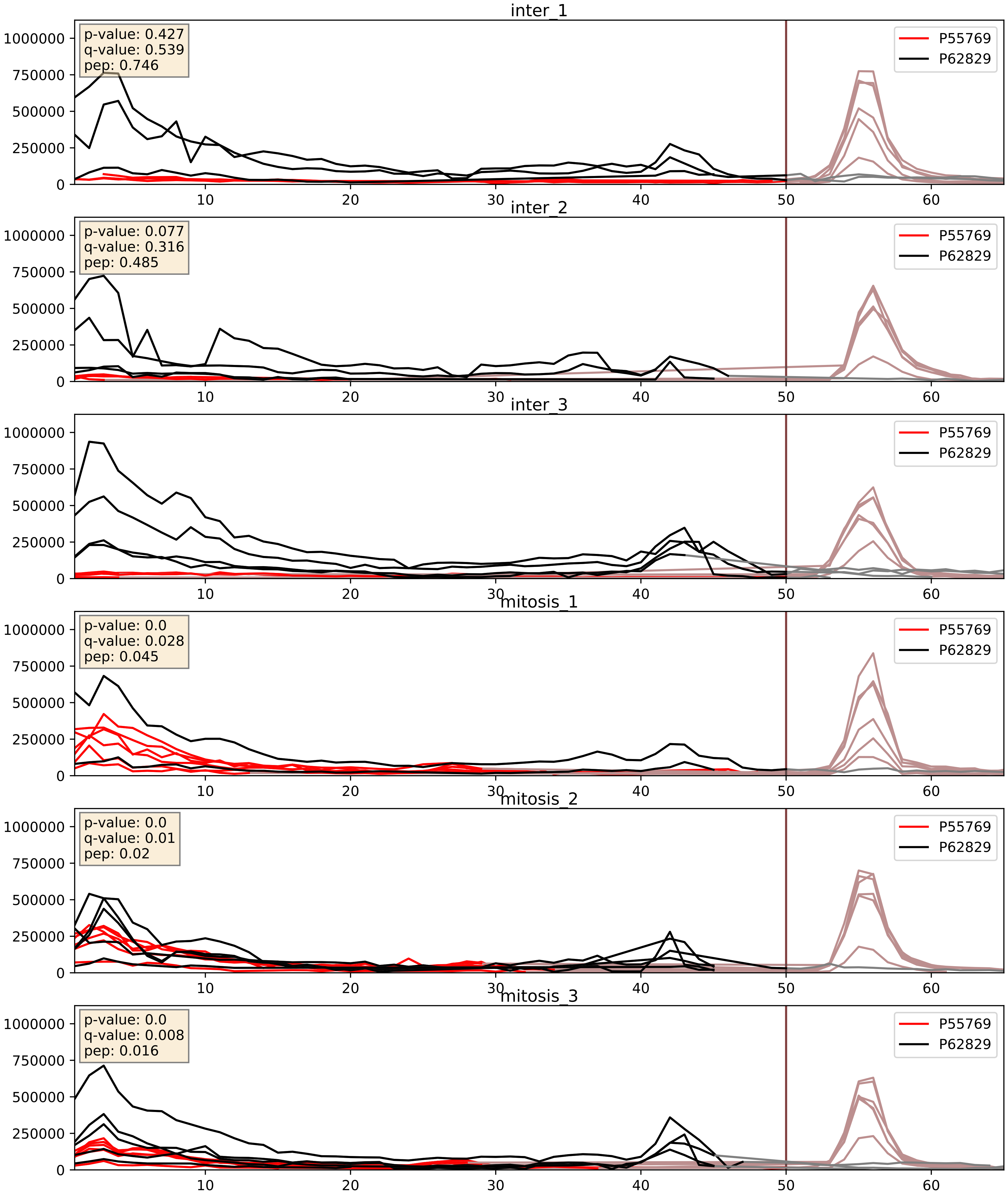


| level | condition_1 | condition_2 | pvalue | pvalue_adjusted |
|---|---|---|---|---|
| interactor_ratio | mitosis | inter | 0.0010607158442868909 | 0.038414698994273265 |
| interactor_abundance | mitosis | inter | 0.003522677249577062 | 0.04582692592154961 |
| complex_abundance | mitosis | inter | 0.8831107774455889 | 0.9275372091943853 |

P55769_P62841
NH2L1_HUMAN vs RS15_HUMAN
p-value: 0.0 q-value: 0.005

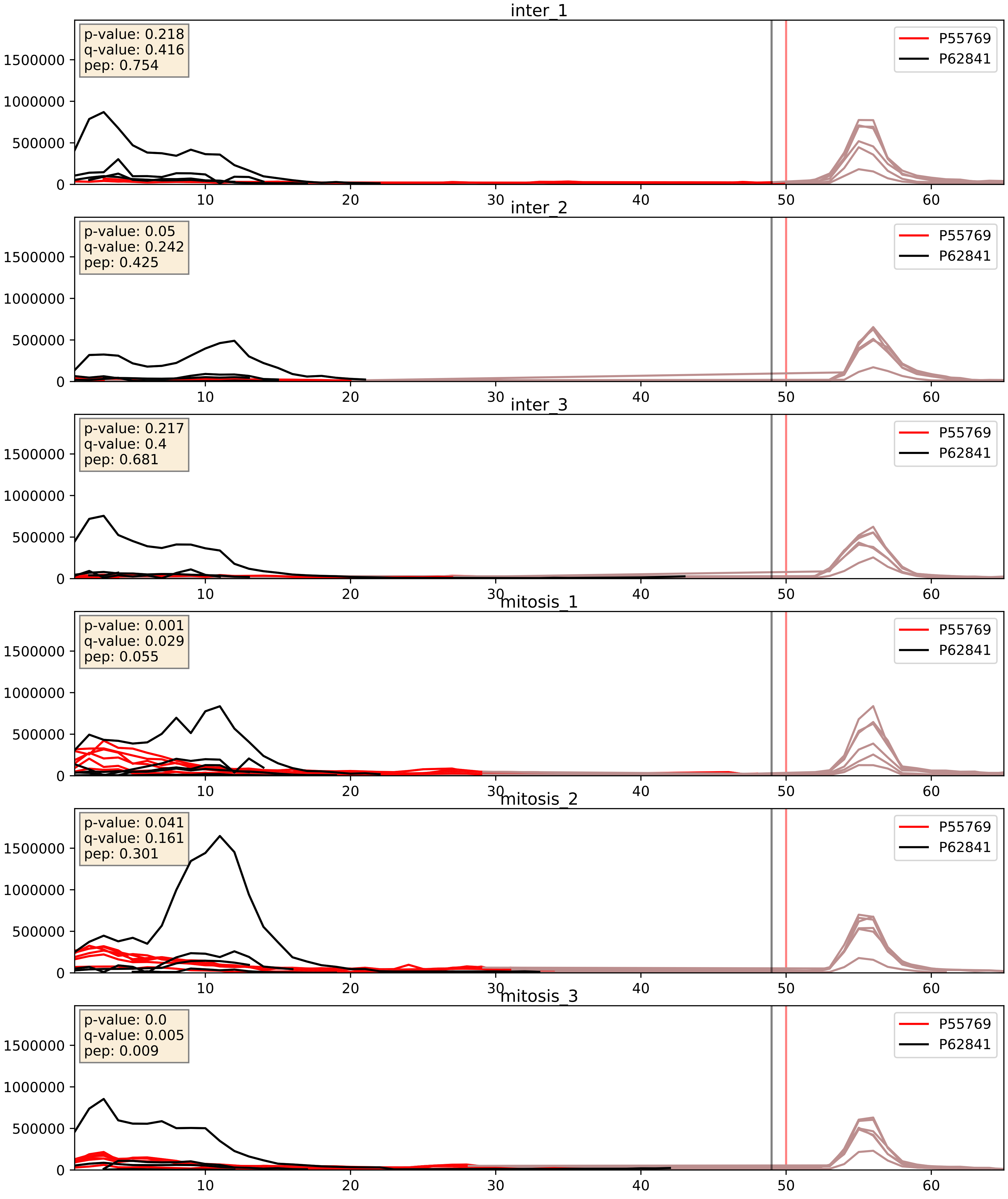


| level | condition_1 | condition_2 | pvalue | pvalue_adjusted |
|---|---|---|---|---|
| interactor_abundance | mitosis | inter | 0.0013262026640028536 | 0.02690117252100575 |
| interactor_ratio | mitosis | inter | 0.0028175245201769636 | 0.06871227889662339 |
| complex_abundance | mitosis | inter | 0.05952392987834658 | 0.1746144070454581 |

P55769_P62847
NH2L1_HUMAN vs RS24_HUMAN
p-value: 0.004 q-value: 0.004

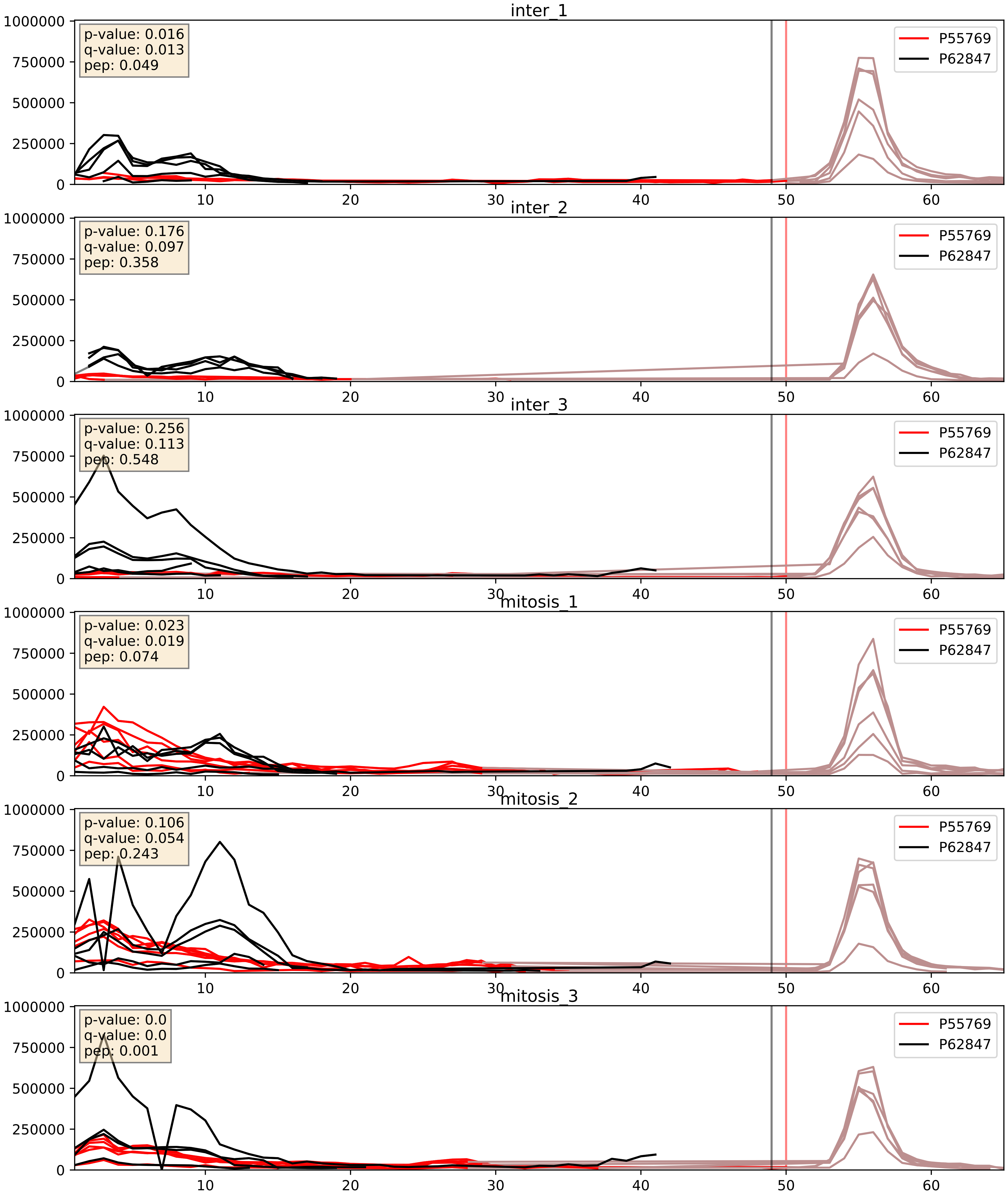


| level | condition_1 | condition_2 | pvalue | pvalue_adjusted |
|---|---|---|---|---|
| interactor_abundance | mitosis | inter | 0.0008638052713741788 | 0.02184393832485368 |
| complex_abundance | mitosis | inter | 0.0013946167086880382 | 0.02556299577381072 |
| interactor_ratio | mitosis | inter | 0.0032207078089795675 | 0.07405554735675686 |

P55769_P62861
NH2L1_HUMAN vs RS30_HUMAN
p-value: 0.0 q-value: 0.0

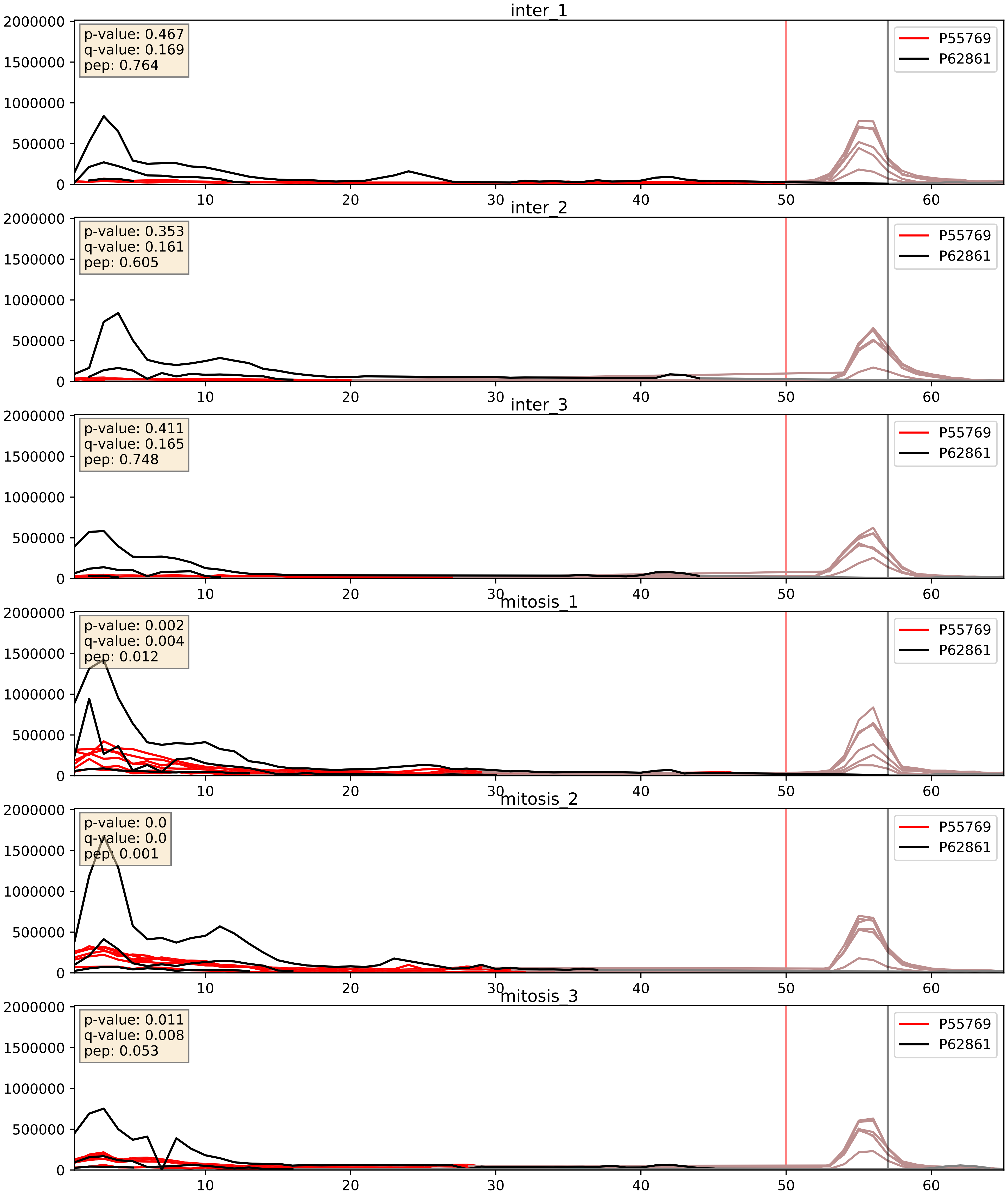


| level | condition_1 | condition_2 | pvalue | pvalue_adjusted |
|---|---|---|---|---|
| interactor_ratio | mitosis | inter | 0.0014846064640640533 | 0.04597663709000905 |
| interactor_abundance | mitosis | inter | 0.0050647865460358285 | 0.05914675693597093 |
| complex_abundance | mitosis | inter | 0.03980362482872163 | 0.1449976963395235 |

P55769_P62888
NH2L1_HUMAN vs RL30_HUMAN
p-value: 0.0 q-value: 0.005

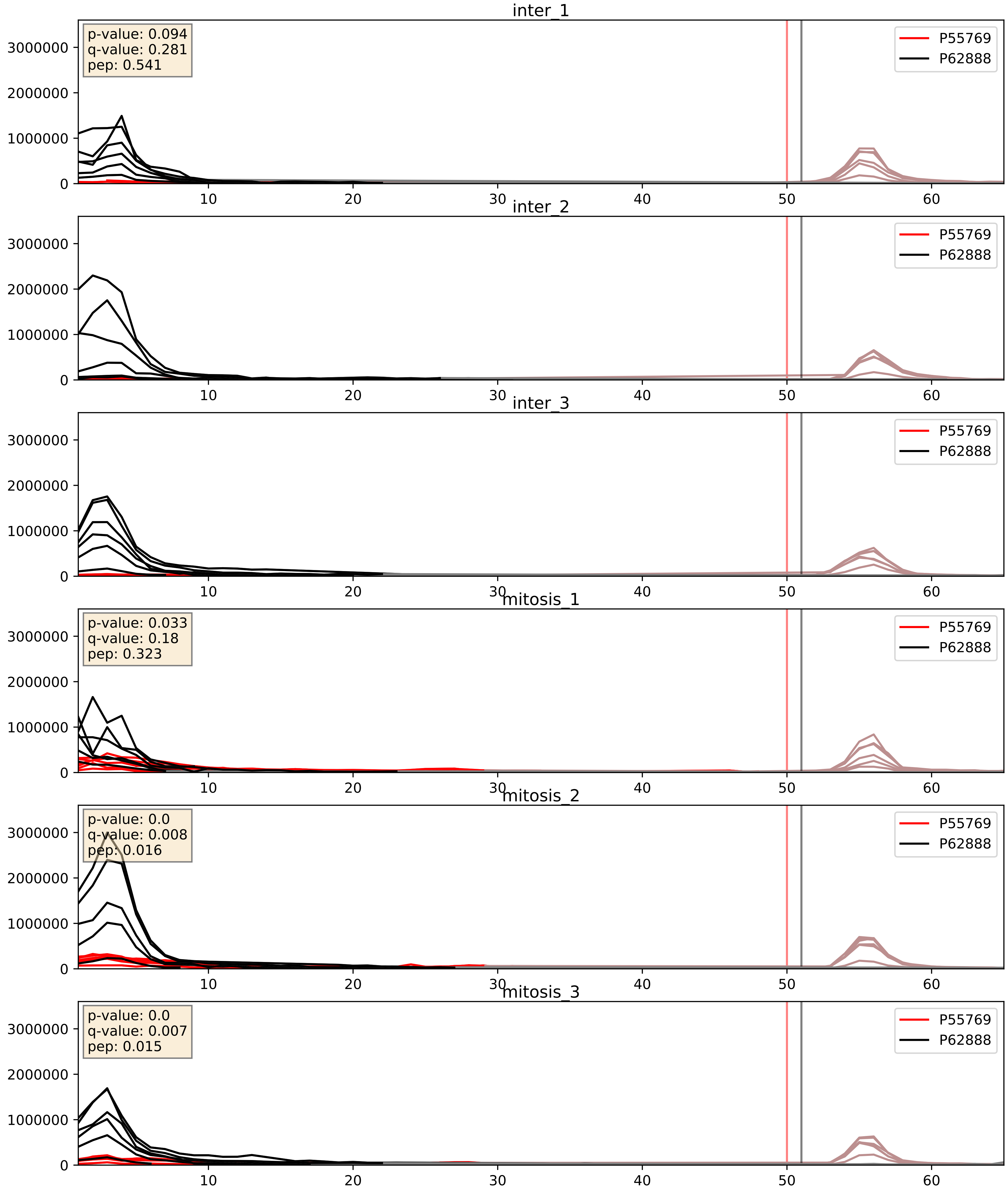


| level | condition_1 | condition_2 | pvalue | pvalue_adjusted |
|---|---|---|---|---|
| interactor_abundance | mitosis | inter | 0.0018182251455957498 | 0.03134744661893176 |
| complex_abundance | mitosis | inter | 0.007629160553136954 | 0.06504543260443459 |
| interactor_ratio | mitosis | inter | 0.01983635837482736 | 0.213852931597635 |

P55769_P62899
NH2L1_HUMAN vs RL31_HUMAN
p-value: 0.0 q-value: 0.041

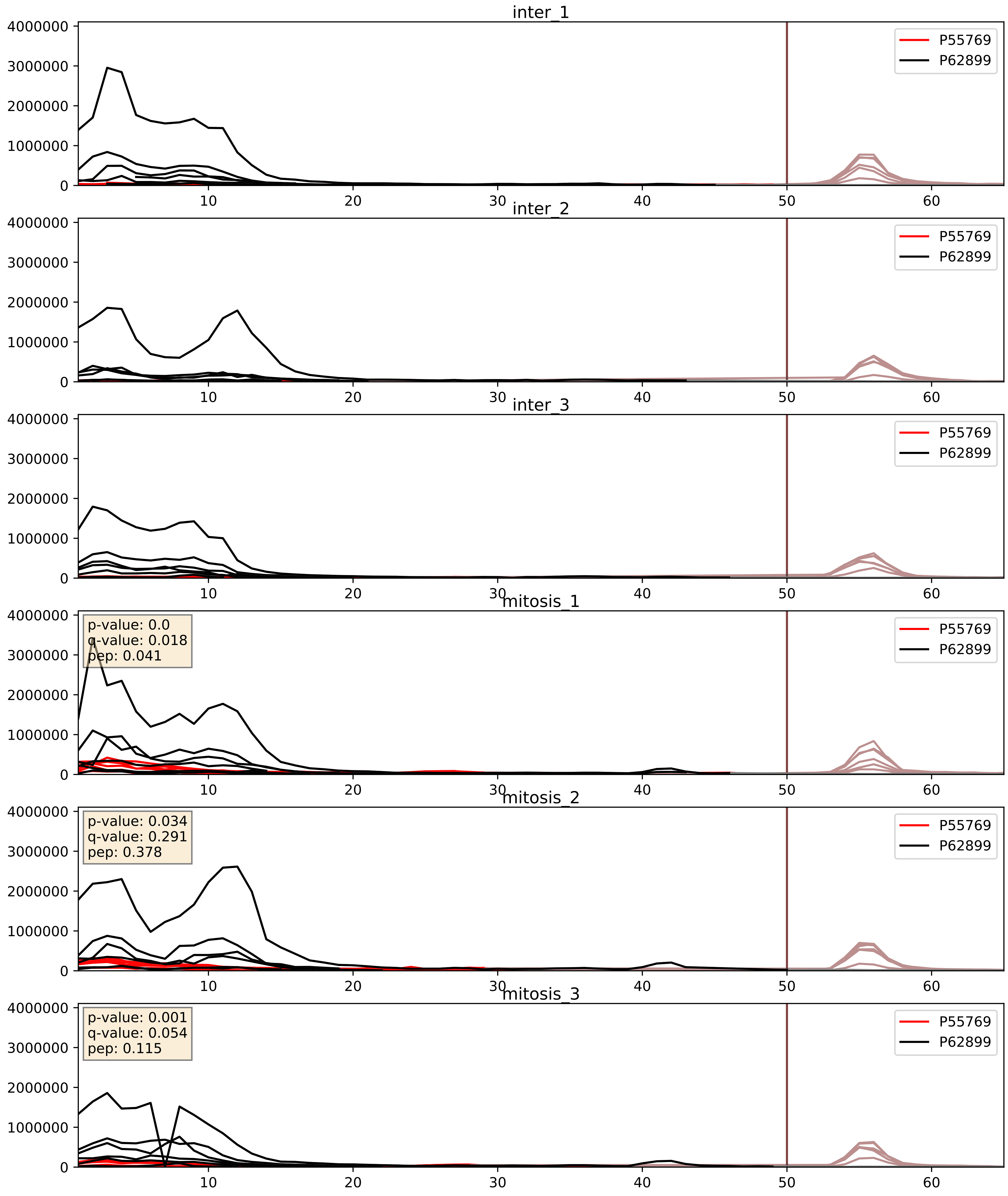


| level | condition_1 | condition_2 | pvalue | pvalue_adjusted |
|---|---|---|---|---|
| interactor_ratio | mitosis | inter | 0.0008232728353916324 | 0.03277774637652267 |
| interactor_abundance | mitosis | inter | 0.0024100075505886463 | 0.03654502149342571 |
| complex_abundance | mitosis | inter | 0.009749797605994615 | 0.07329383942917299 |

P55769_P62906
NH2L1_HUMAN vs RL10A_HUMAN
p-value: 0.0 q-value: 0.016

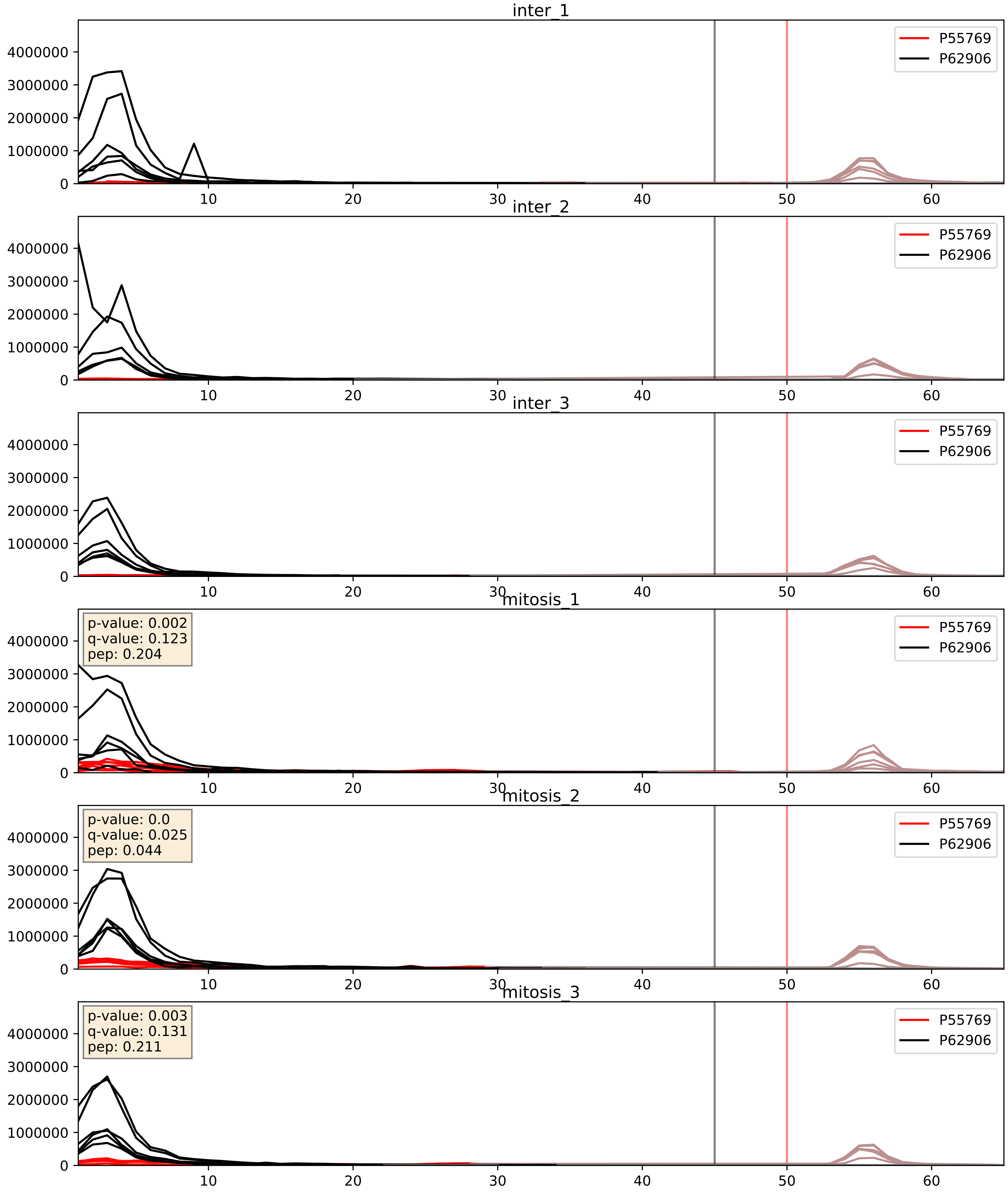


| level | condition_1 | condition_2 | pvalue | pvalue_adjusted |
|---|---|---|---|---|
| interactor_ratio | mitosis | inter | 0.002581226673103492 | 0.0651778770553566 |
| interactor_abundance | mitosis | inter | 0.002912682752214205 | 0.040906586314936176 |
| complex_abundance | mitosis | inter | 0.0072955578923150815 | 0.06327386507688666 |

P55769_P62910
NH2L1_HUMAN vs RL32_HUMAN
p-value: 0.0 q-value: 0.008

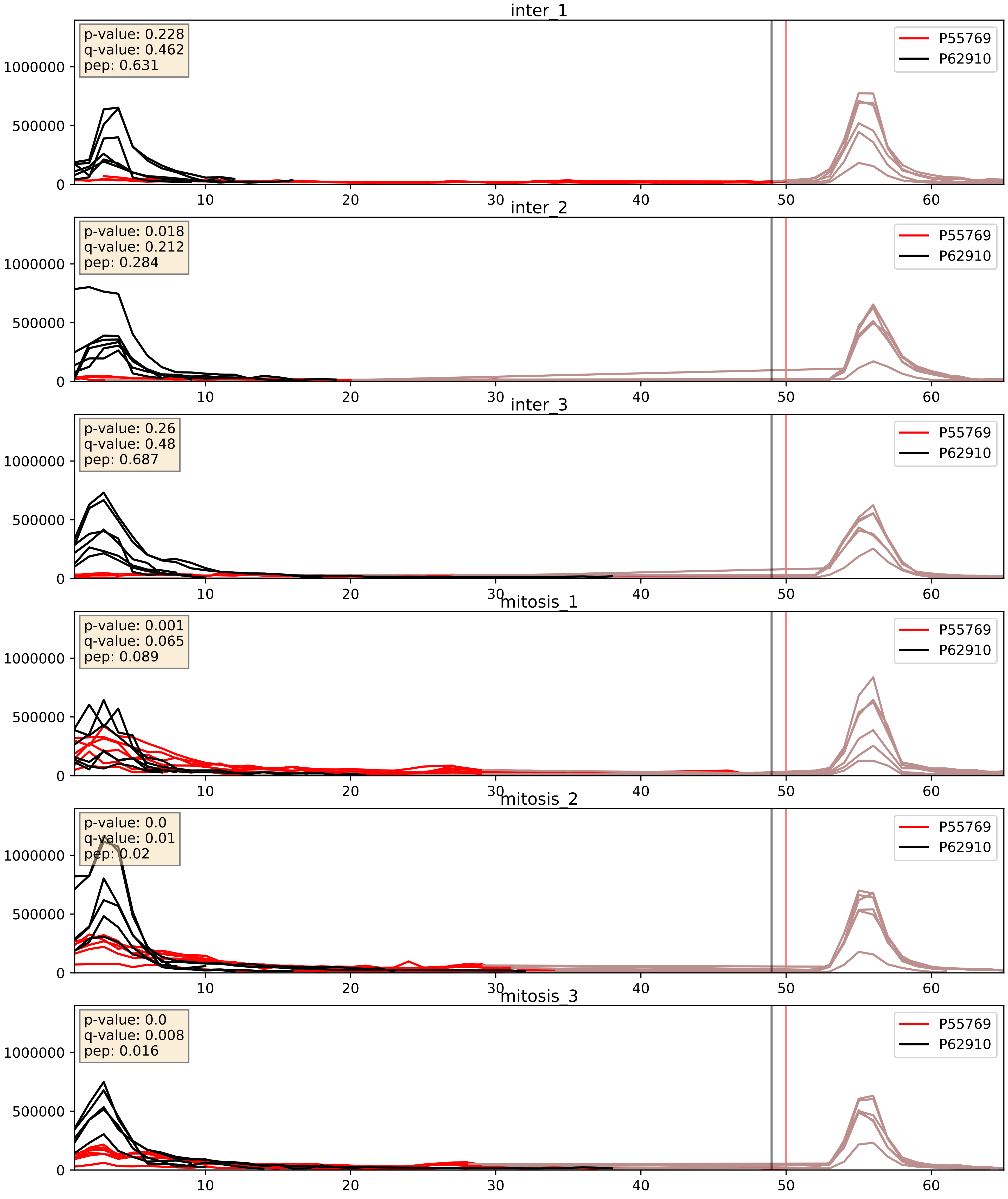


| level | condition_1 | condition_2 | pvalue | pvalue_adjusted |
|---|---|---|---|---|
| interactor_abundance | mitosis | inter | 0.0011891362735882815 | 0.025102358821000467 |
| complex_abundance | mitosis | inter | 0.0016896704090206235 | 0.028927157402433075 |
| interactor_ratio | mitosis | inter | 0.01241749524749119 | 0.16080750275117184 |

P55769_P62913
NH2L1_HUMAN vs RL11_HUMAN
p-value: 0.0 q-value: 0.008

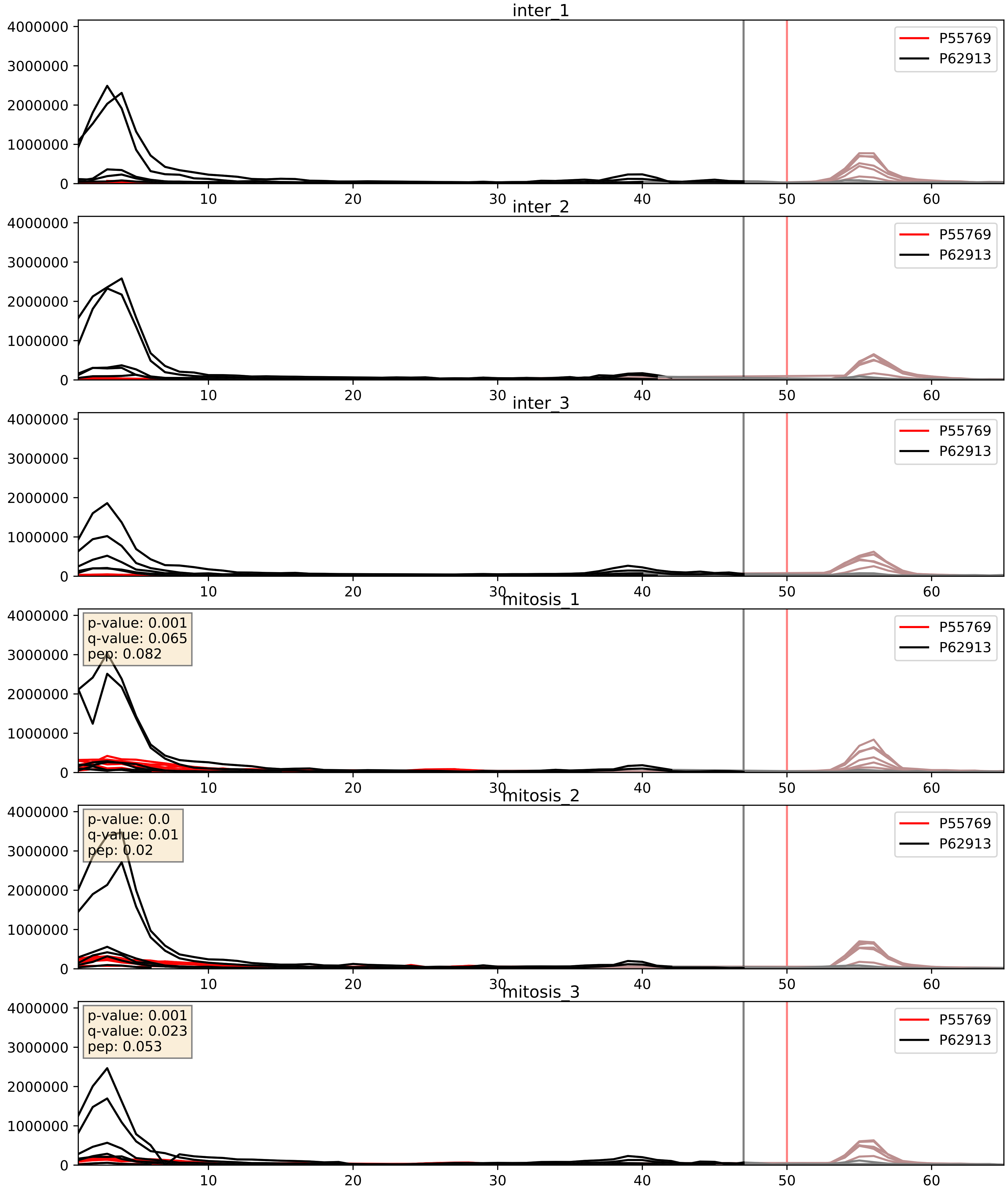


| level | condition_1 | condition_2 | pvalue | pvalue_adjusted |
|---|---|---|---|---|
| complex_abundance | mitosis | inter | 0.001422093816424617 | 0.02584527190784442 |
| interactor_abundance | mitosis | inter | 0.002500371661828678 | 0.03735284716449125 |
| interactor_ratio | mitosis | inter | 0.005129879459469436 | 0.09650938060012829 |

P55769_P62917
NH2L1_HUMAN vs RL8_HUMAN
p-value: 0.0 q-value: 0.016

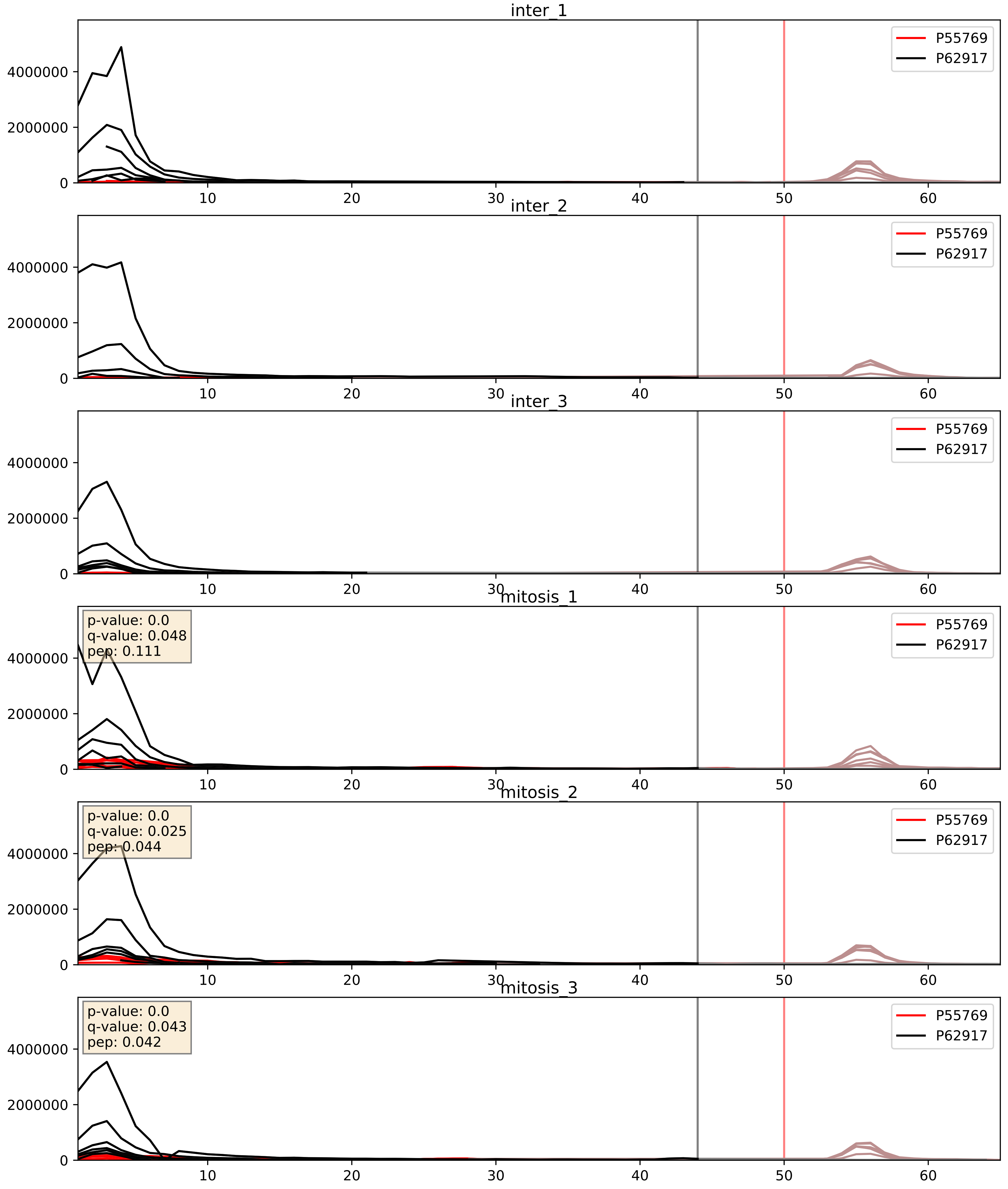


| level | condition_1 | condition_2 | pvalue | pvalue_adjusted |
|---|---|---|---|---|
| interactor_abundance | mitosis | inter | 0.0010161487469955068 | 0.023821155197003323 |
| interactor_ratio | mitosis | inter | 0.0033929072330672596 | 0.07563355707045767 |
| complex_abundance | mitosis | inter | 0.02218589020882896 | 0.10952717692621856 |

P55769_P62995
NH2L1_HUMAN vs TRA2B_HUMAN
p-value: 0.0 q-value: 0.009

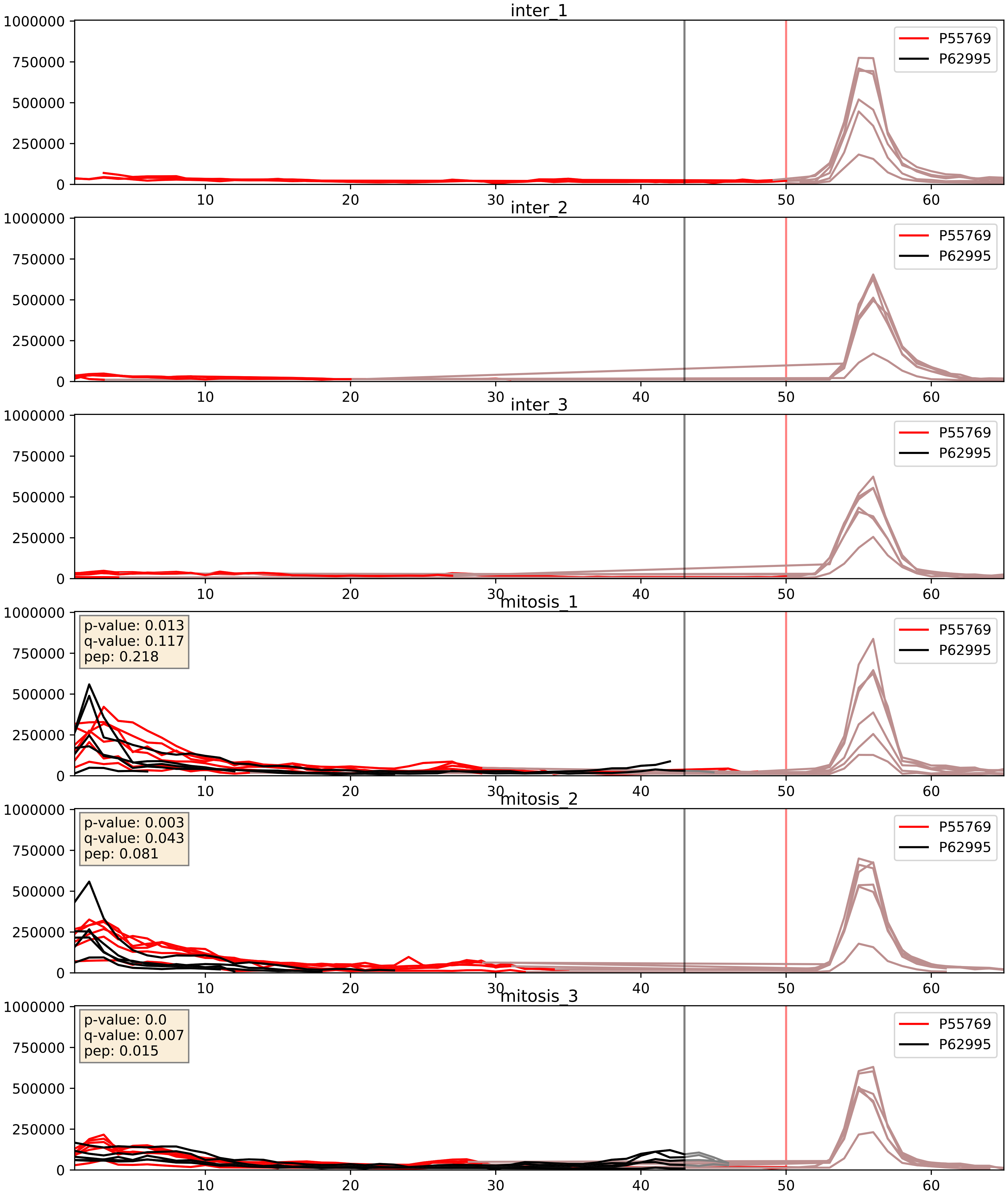


| level | condition_1 | condition_2 | pvalue | pvalue_adjusted |
|---|---|---|---|---|
| interactor_abundance | mitosis | inter | 8.443492929225096e-05 | 0.007227629947416683 |
| complex_abundance | mitosis | inter | 0.00011484000270838403 | 0.007391206189351633 |
| interactor_ratio | mitosis | inter | 0.07572511598736271 | 0.4628166812754875 |

P55769_P83731
NH2L1_HUMAN vs RL24_HUMAN
p-value: 0.0 q-value: 0.016

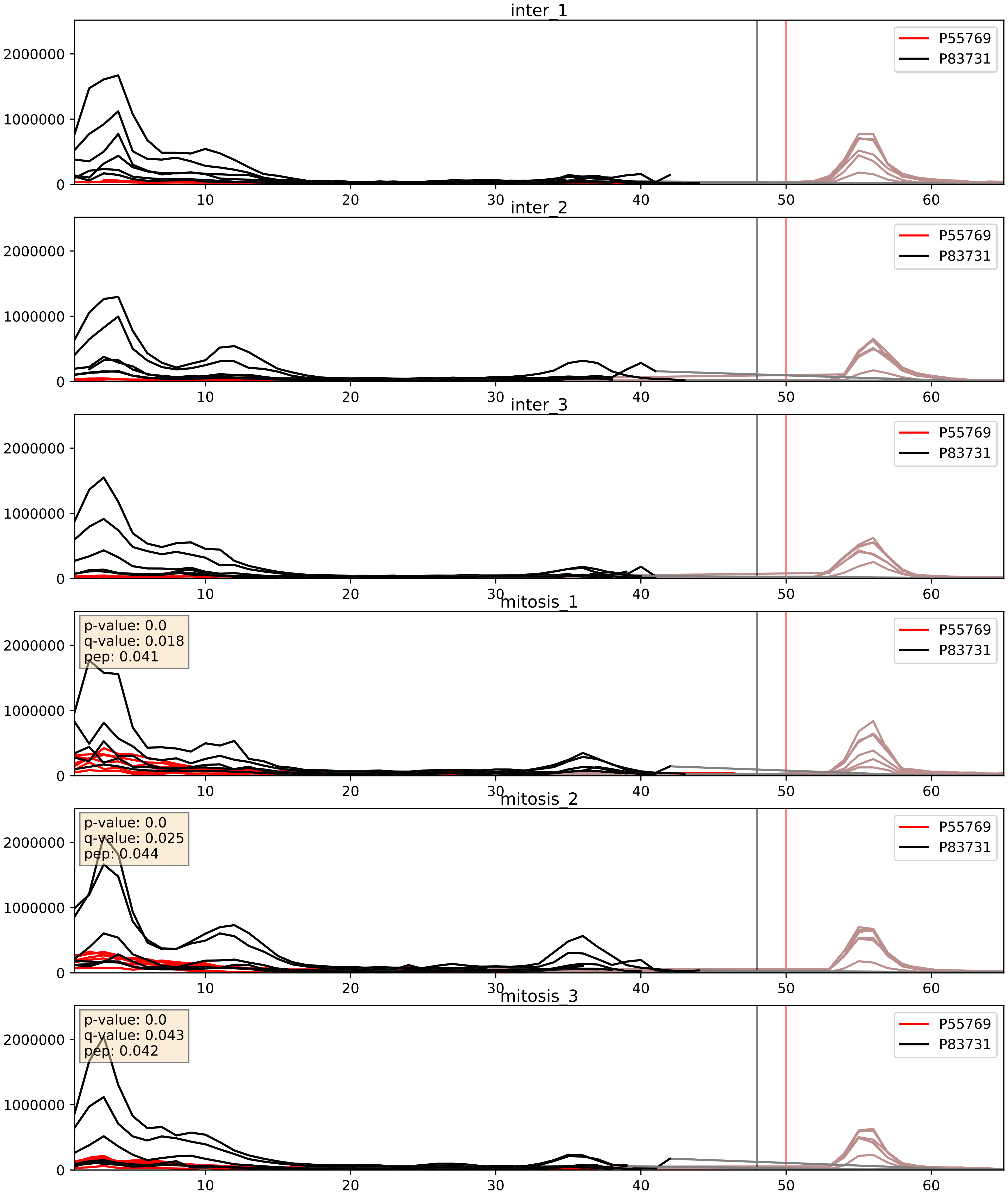


| level | condition_1 | condition_2 | pvalue | pvalue_adjusted |
|---|---|---|---|---|
| interactor_abundance | mitosis | inter | 0.0023736732188134827 | 0.036283290630434666 |
| interactor_ratio | mitosis | inter | 0.005068429617363597 | 0.09598618921378847 |
| complex_abundance | mitosis | inter | 0.0055543202669368005 | 0.05624098855659709 |

P55769_P84098
NH2L1_HUMAN vs RL19_HUMAN
p-value: 0.0 q-value: 0.008

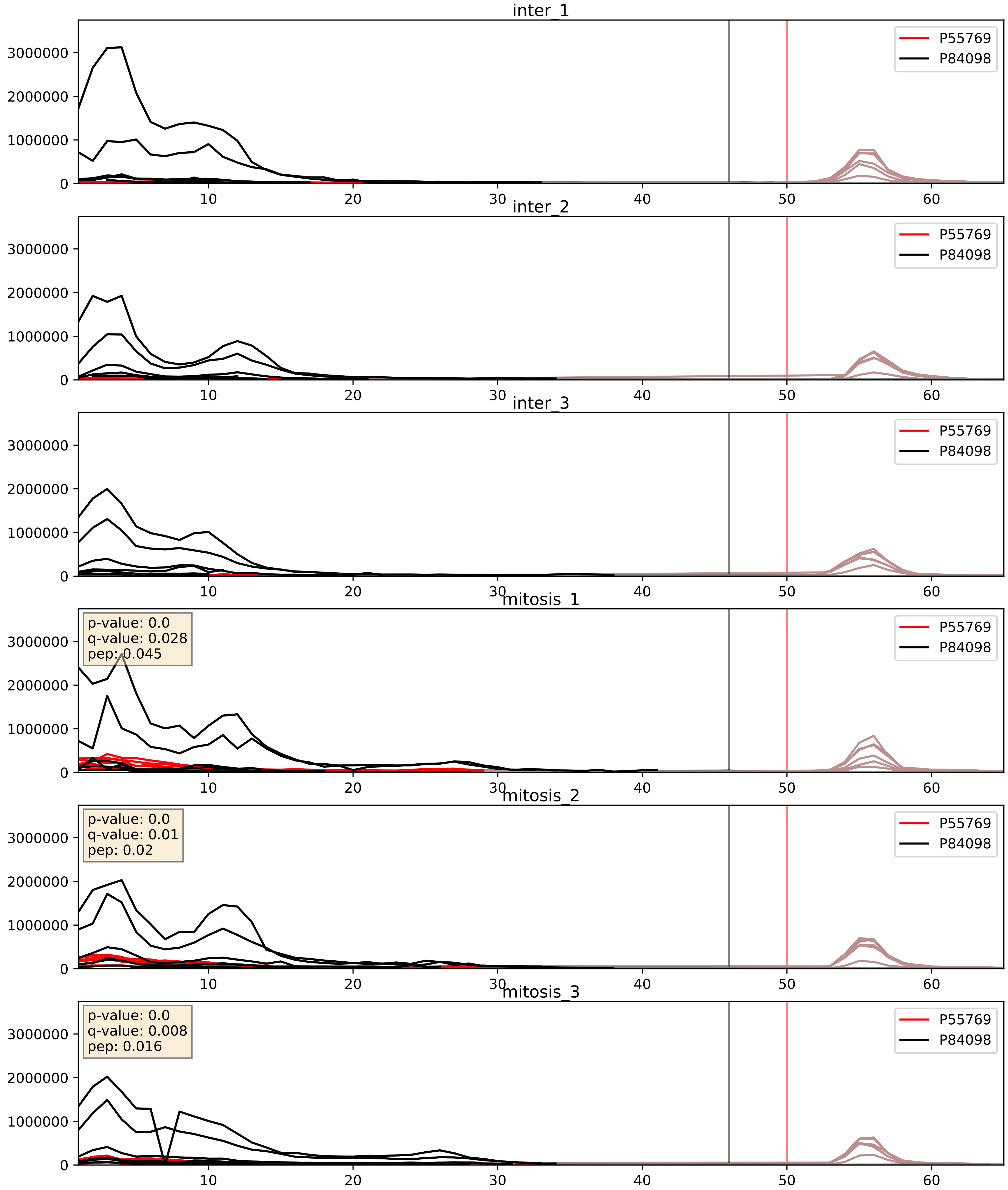


| level | condition_1 | condition_2 | pvalue | pvalue_adjusted |
|---|---|---|---|---|
| interactor_abundance | mitosis | inter | 0.0015711628412833026 | 0.029025269221018554 |
| complex_abundance | mitosis | inter | 0.0022559509527014065 | 0.034819618598671796 |
| interactor_ratio | mitosis | inter | 0.0107762695448819 | 0.14902240275313255 |

P55769_Q02543
NH2L1_HUMAN vs RL18A_HUMAN
p-value: 0.0 q-value: 0.014

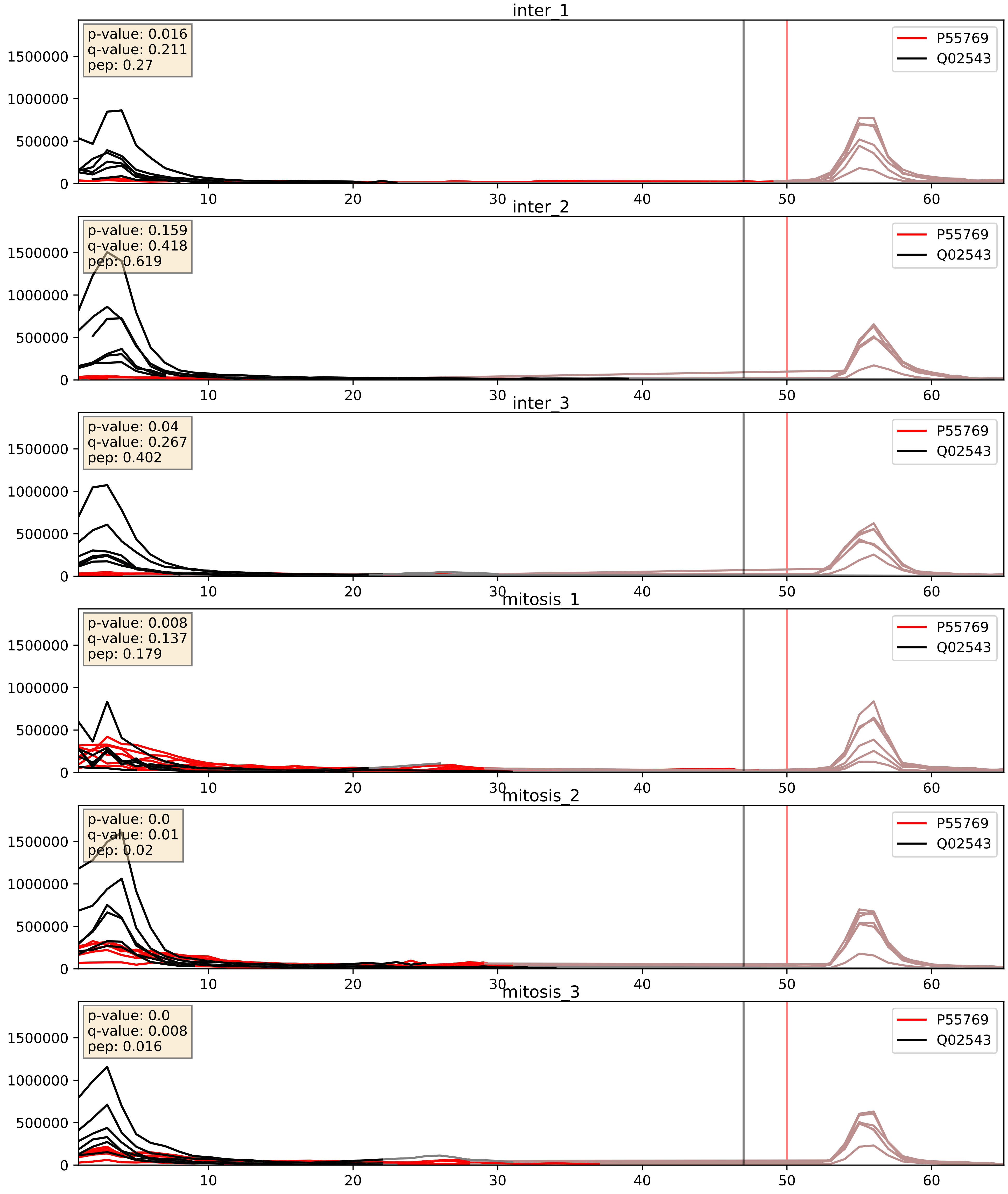


| level | condition_1 | condition_2 | pvalue | pvalue_adjusted |
|---|---|---|---|---|
| interactor_abundance | mitosis | inter | 0.0025905294417015996 | 0.03806046117740333 |
| complex_abundance | mitosis | inter | 0.024457818542900852 | 0.11409965655757742 |
| interactor_ratio | mitosis | inter | 0.02783468005572525 | 0.2618295178868221 |

P55769_Q08211
NH2L1_HUMAN vs DHX9_HUMAN
p-value: 0.001 q-value: 0.048

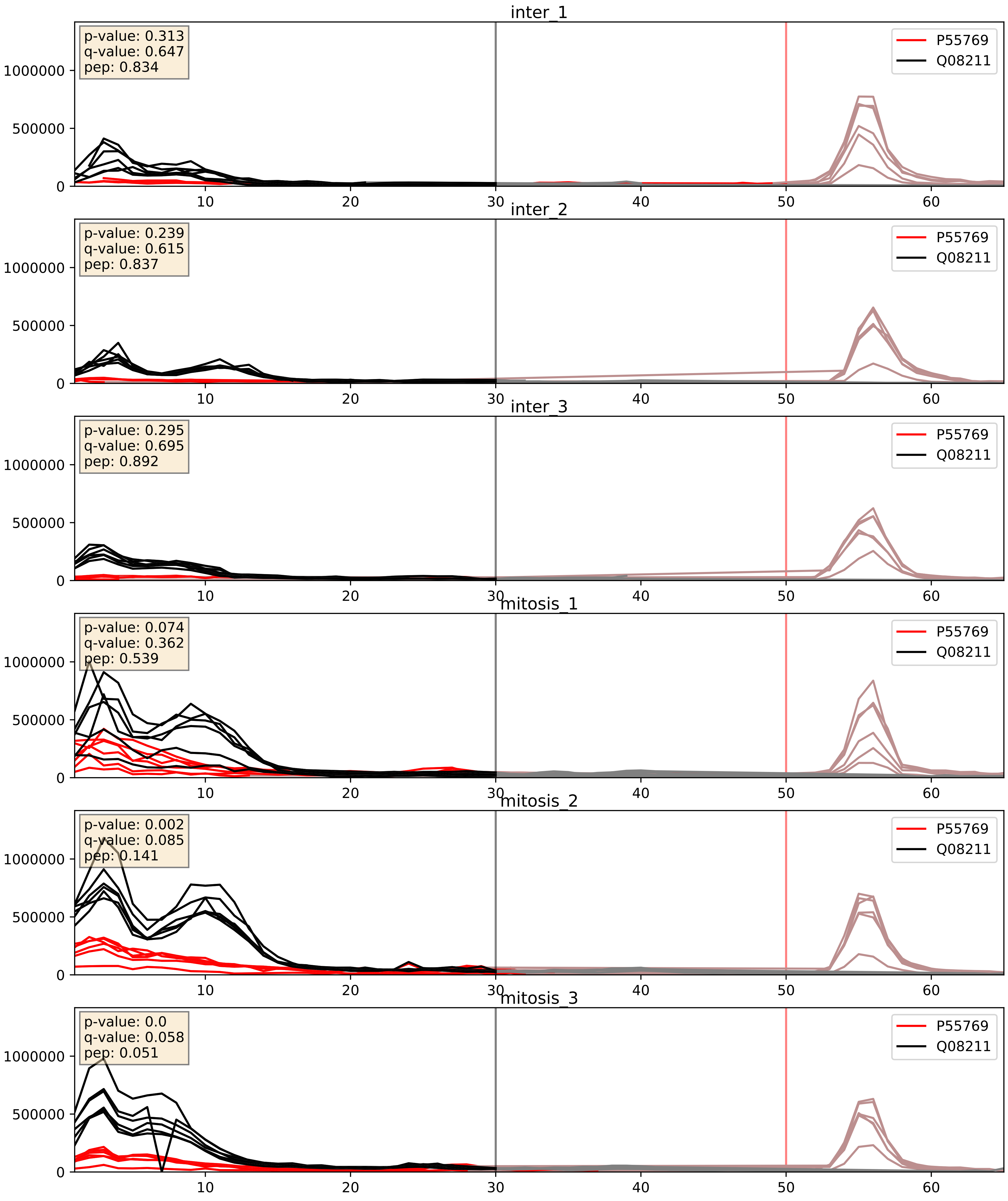


| level | condition_1 | condition_2 | pvalue | pvalue_adjusted |
|---|---|---|---|---|
| complex_abundance | mitosis | inter | 0.0007198582539638076 | 0.01750564390321078 |
| interactor_abundance | mitosis | inter | 0.0013996461911011957 | 0.02753913185123736 |
| interactor_ratio | mitosis | inter | 0.012767744743810204 | 0.16336606129598708 |

P55769_Q12788
NH2L1_HUMAN vs TBL3_HUMAN
p-value: 0.004 q-value: 0.004

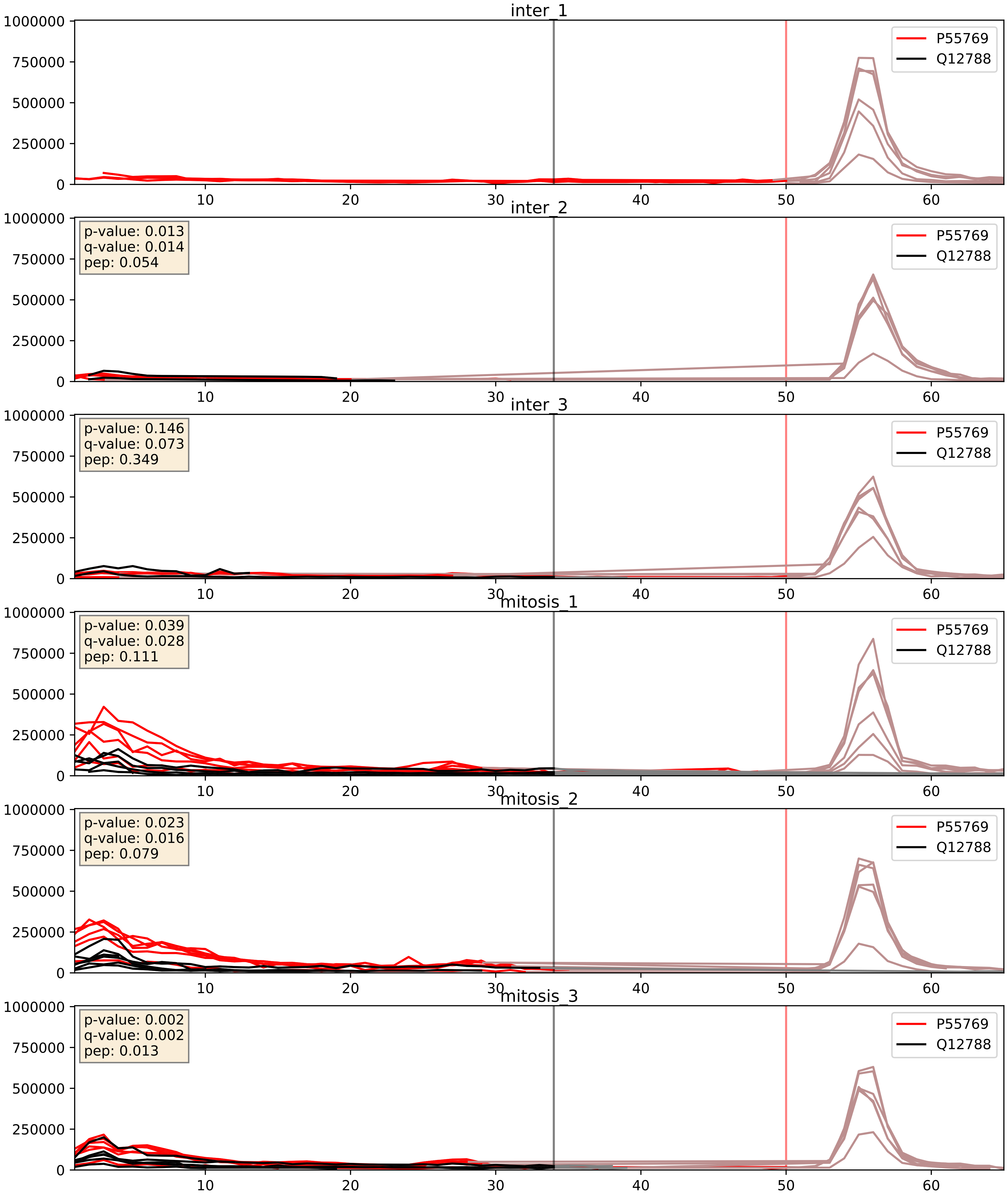


| level | condition_1 | condition_2 | pvalue | pvalue_adjusted |
|---|---|---|---|---|
| complex_abundance | mitosis | inter | 0.002265540023863082 | 0.03487953705803594 |
| interactor_abundance | mitosis | inter | 0.002465726822584136 | 0.037061411436434794 |
| interactor_ratio | mitosis | inter | 0.02210957812286063 | 0.22829672947127502 |

P55769_Q12905
NH2L1_HUMAN vs ILF2_HUMAN
p-value: 0.0 q-value: 0.047

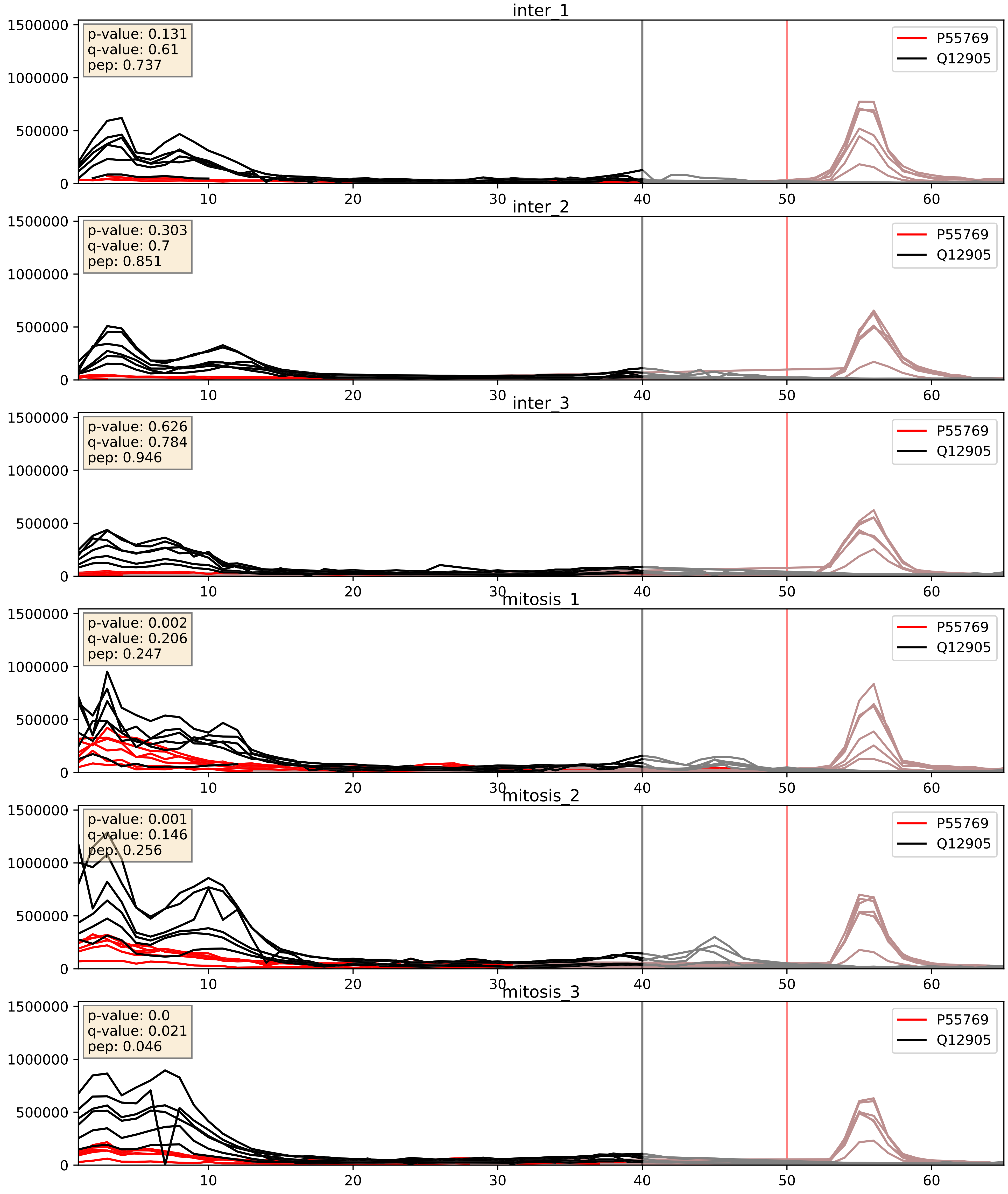


| level | condition_1 | condition_2 | pvalue | pvalue_adjusted |
|---|---|---|---|---|
| complex_abundance | mitosis | inter | 0.0011196272788458134 | 0.022133971147621622 |
| interactor_abundance | mitosis | inter | 0.0018958531698051928 | 0.03196215383917848 |
| interactor_ratio | mitosis | inter | 0.009975986588717106 | 0.14263182888175546 |

P55769_Q13151
NH2L1_HUMAN vs ROA0_HUMAN
p-value: 0.001 q-value: 0.041

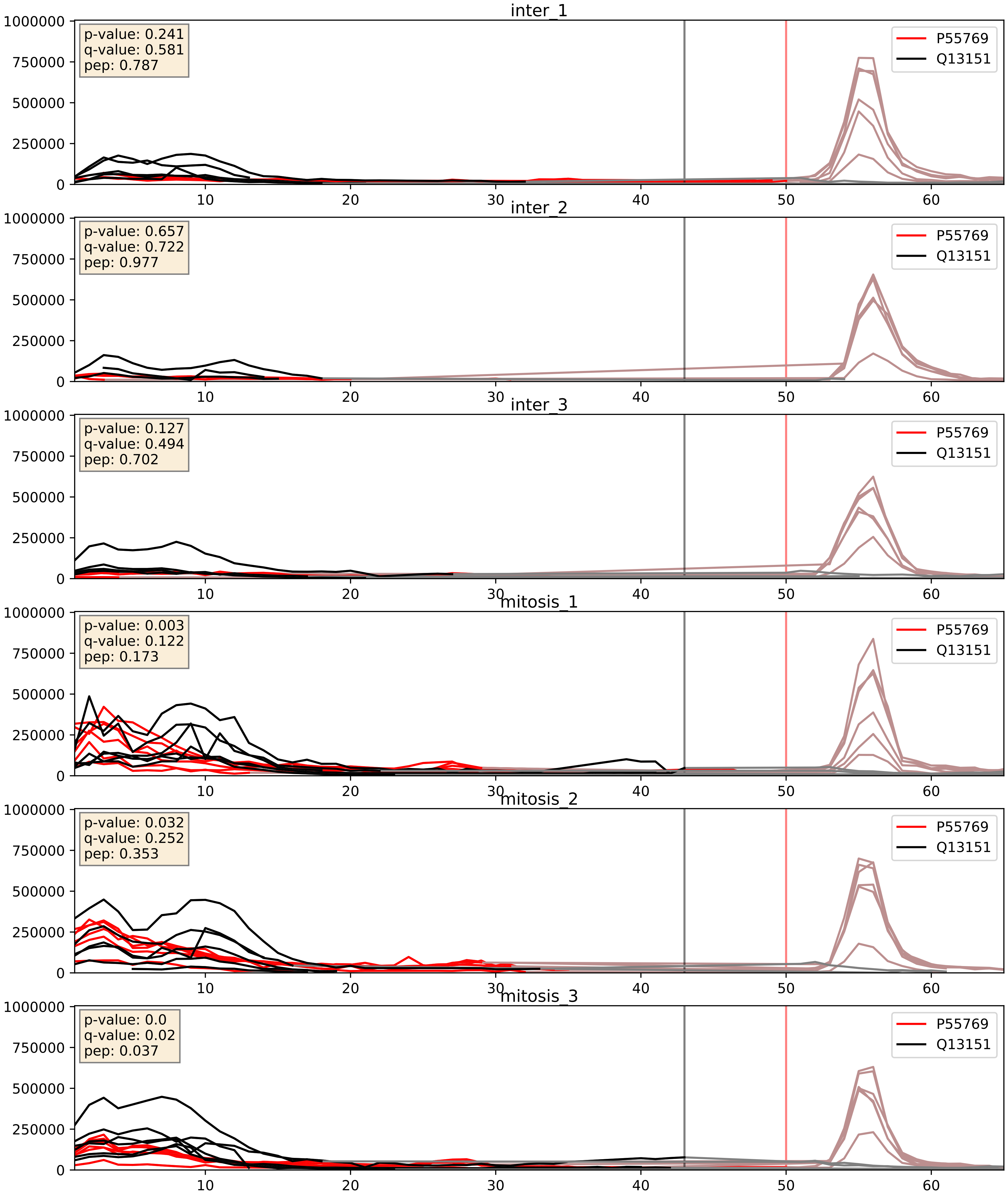


| level | condition_1 | condition_2 | pvalue | pvalue_adjusted |
|---|---|---|---|---|
| interactor_abundance | mitosis | inter | 0.0012966100224587672 | 0.026616263290760302 |
| complex_abundance | mitosis | inter | 0.002280148765337147 | 0.03502731659907978 |
| interactor_ratio | mitosis | inter | 0.06995079943241257 | 0.4380240257069873 |

P55769_Q13895
NH2L1_HUMAN vs BYST_HUMAN
p-value: 0.062 q-value: 0.039

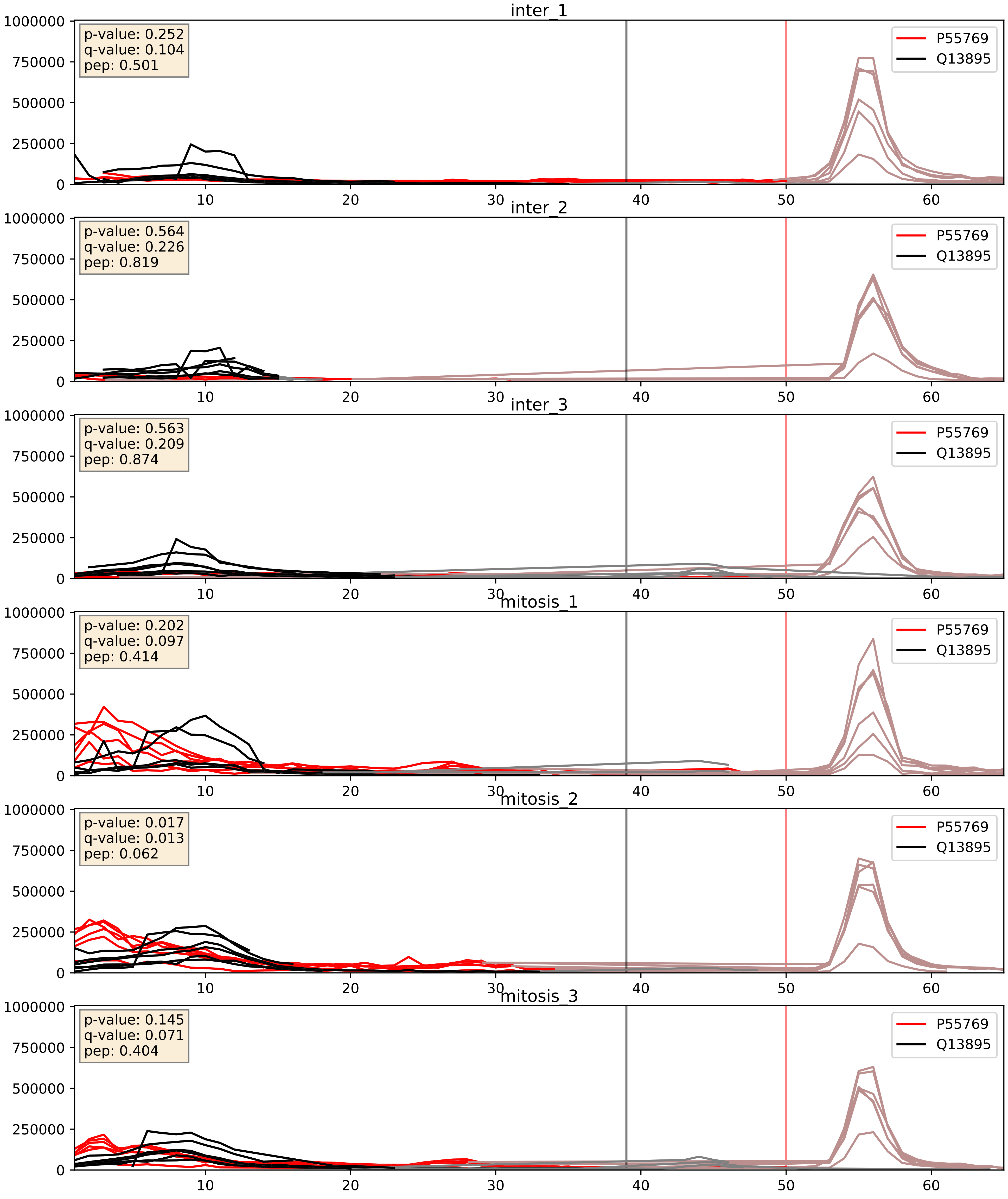


| level | condition_1 | condition_2 | pvalue | pvalue_adjusted |
|---|---|---|---|---|
| complex_abundance | mitosis | inter | 0.0017384148809047146 | 0.0295241467010272 |
| interactor_abundance | mitosis | inter | 0.00183168678013043 | 0.03145283618438612 |
| interactor_ratio | mitosis | inter | 0.0033897716918089944 | 0.07563355707045767 |

P55769_Q15061
NH2L1_HUMAN vs WDR43_HUMAN
p-value: 0.036 q-value: 0.025

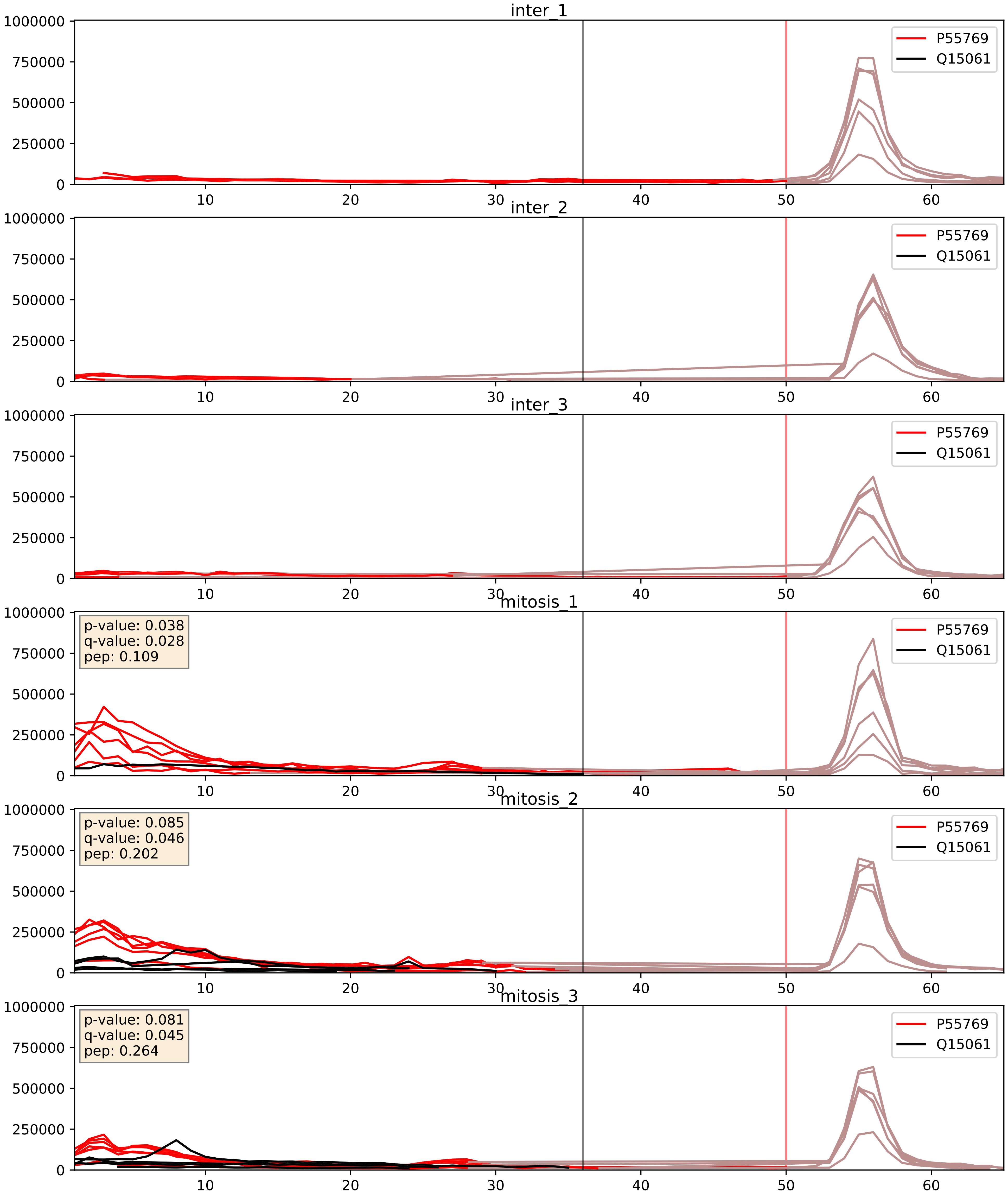


| level | condition_1 | condition_2 | pvalue | pvalue_adjusted |
|---|---|---|---|---|
| interactor_abundance | mitosis | inter | 5.175159842494777e-05 | 0.006283598333582311 |
| complex_abundance | mitosis | inter | 0.0008534383700319466 | 0.01942934161562091 |
| interactor_ratio | mitosis | inter | 0.020867400541259323 | 0.21890312332497525 |

P55769_Q15269
NH2L1_HUMAN vs PWP2_HUMAN
p-value: 0.015 q-value: 0.013

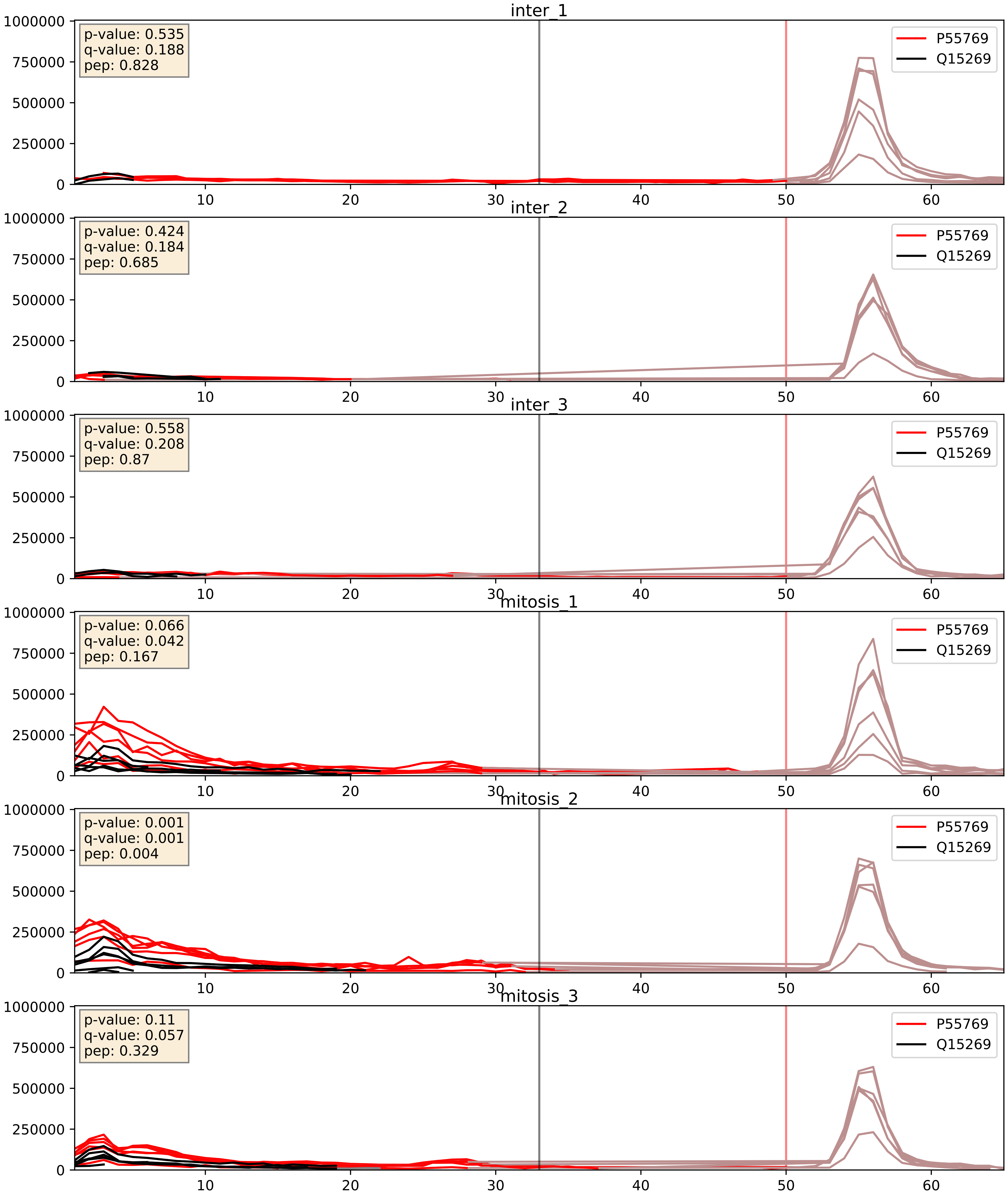


| level | condition_1 | condition_2 | pvalue | pvalue_adjusted |
|---|---|---|---|---|
| interactor_abundance | mitosis | inter | 0.0009531398130480196 | 0.022918193257559123 |
| complex_abundance | mitosis | inter | 0.0021258483587280417 | 0.033489340249041655 |
| interactor_ratio | mitosis | inter | 0.003761862384957826 | 0.07990457075741685 |

P55769_Q16629
NH2L1_HUMAN vs SRSF7_HUMAN
p-value: 0.001 q-value: 0.021

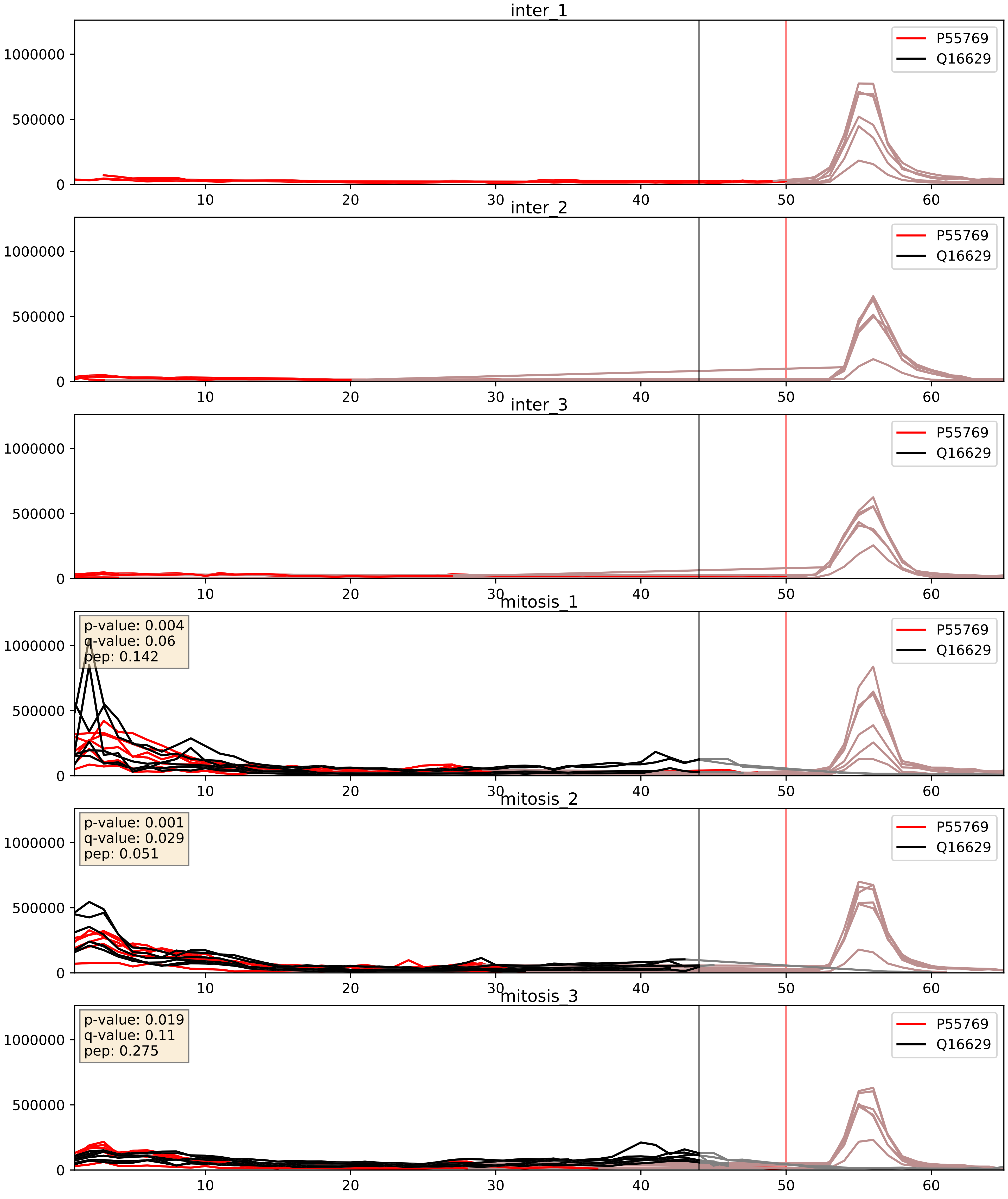


| level | condition_1 | condition_2 | pvalue | pvalue_adjusted |
|---|---|---|---|---|
| complex_abundance | mitosis | inter | 5.694619762543979e-05 | 0.005691857151652958 |
| interactor_abundance | mitosis | inter | 9.321731834197482e-05 | 0.007548664551888891 |
| interactor_ratio | mitosis | inter | 0.06548194300852817 | 0.42820888628953485 |

P55769_Q53GS9
NH2L1_HUMAN vs SNUT2_HUMAN
p-value: 0.002 q-value: 0.002

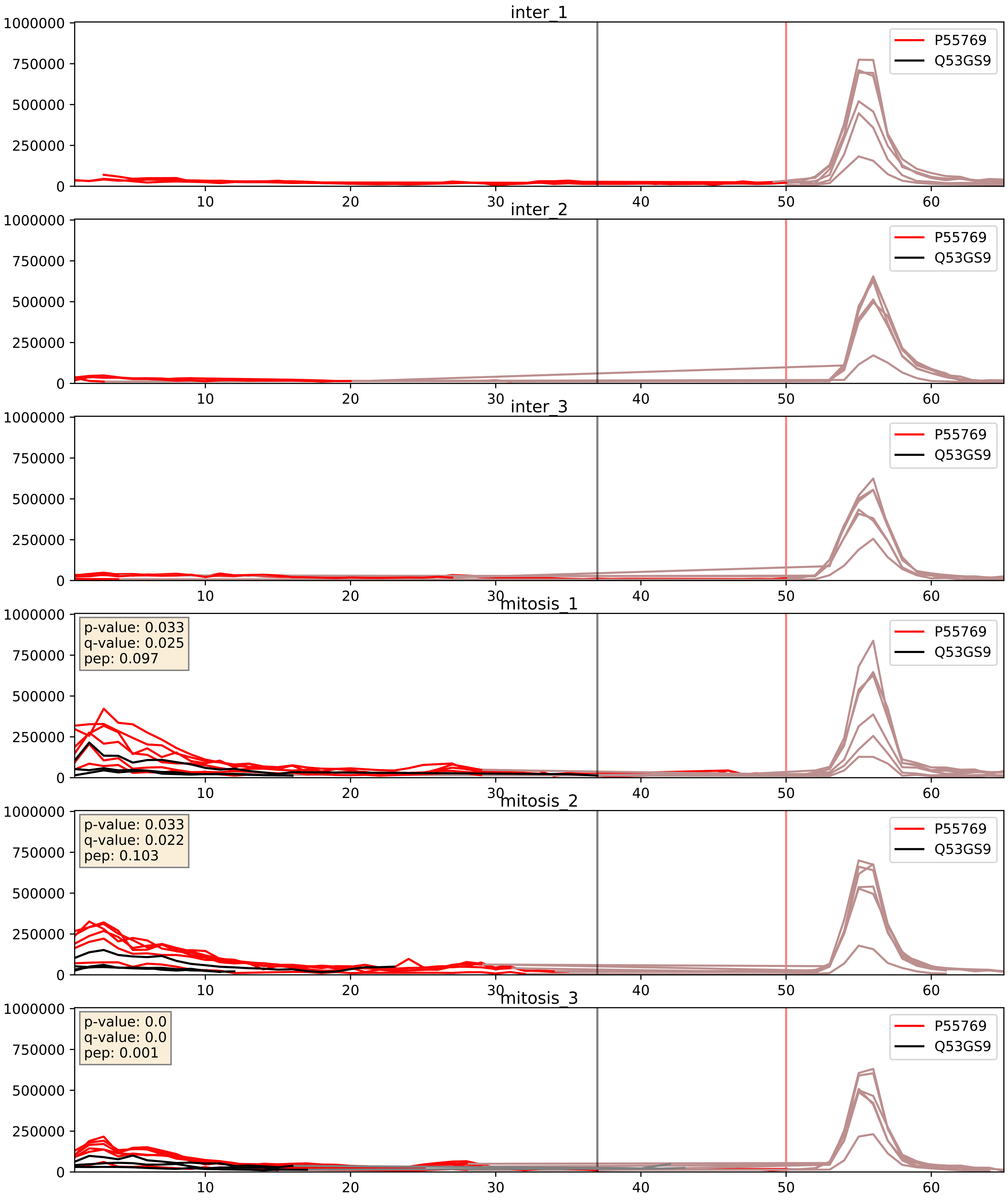


| level | condition_1 | condition_2 | pvalue | pvalue_adjusted |
|---|---|---|---|---|
| interactor_abundance | mitosis | inter | 0.00033269578556084927 | 0.013536301629984253 |
| complex_abundance | mitosis | inter | 0.0004200036215168646 | 0.013091389064164169 |
| interactor_ratio | mitosis | inter | 0.0011227623642383034 | 0.03908562924515634 |

P55769_Q7RTV0
NH2L1_HUMAN vs PHF5A_HUMAN
p-value: 0.077 q-value: 0.046

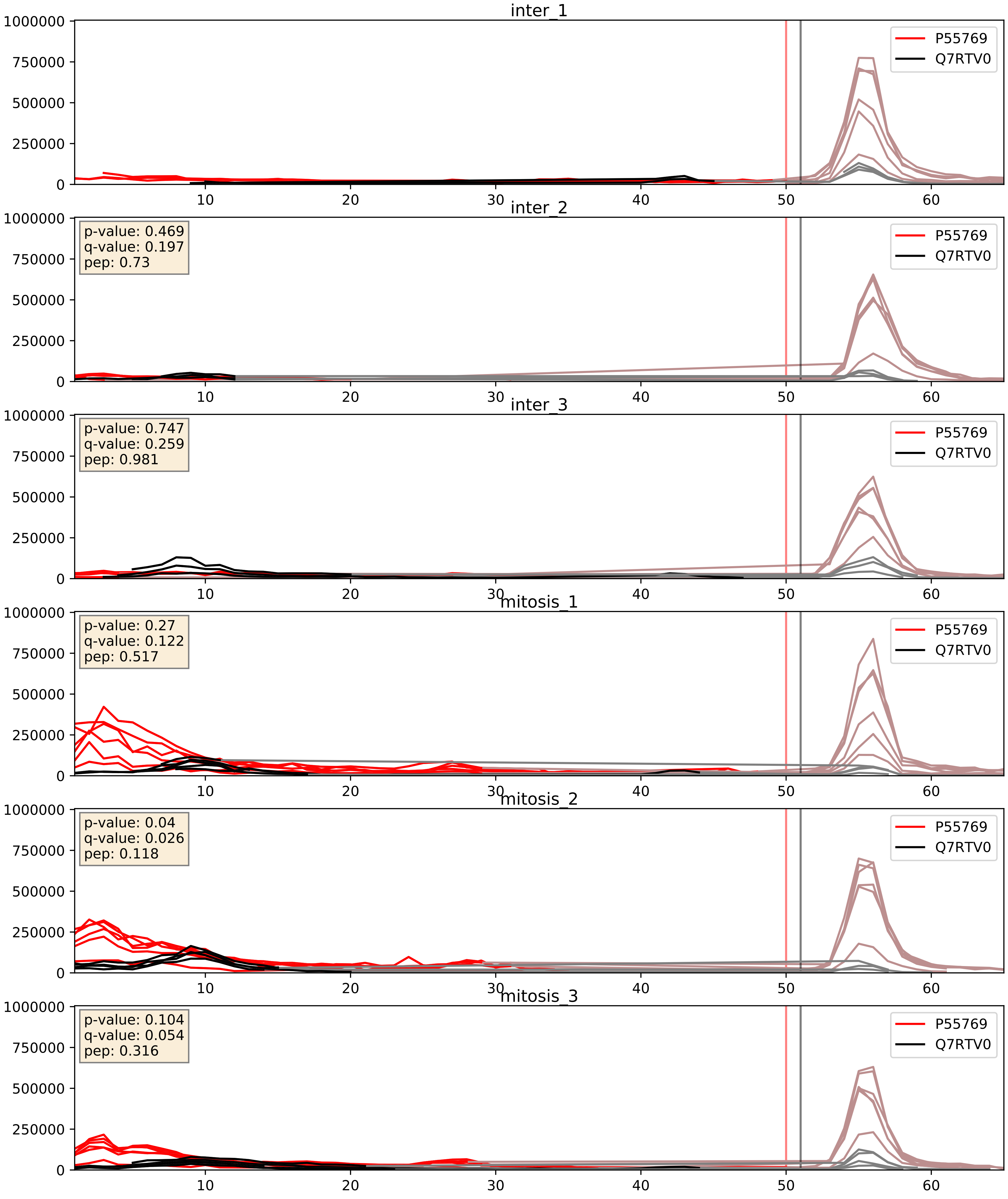


| level | condition_1 | condition_2 | pvalue | pvalue_adjusted |
|---|---|---|---|---|
| interactor_abundance | mitosis | inter | 7.426998330880908e-05 | 0.006870282762438709 |
| complex_abundance | mitosis | inter | 0.0008716073857922254 | 0.019513211095311817 |
| interactor_ratio | mitosis | inter | 0.001565331631028302 | 0.04768412370676963 |

P55769_Q8NI36
NH2L1_HUMAN vs WDR36_HUMAN
p-value: 0.017 q-value: 0.014

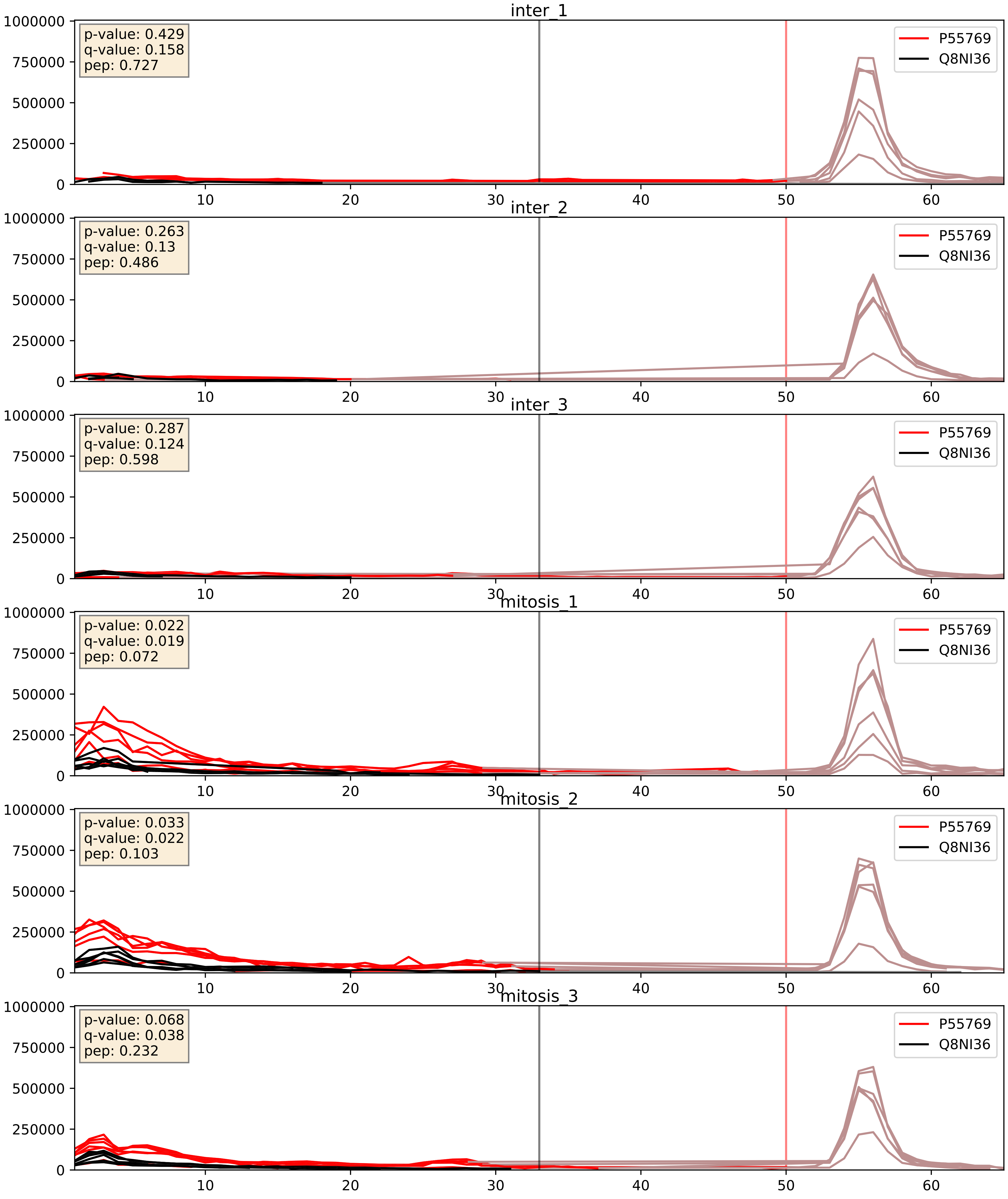


| level | condition_1 | condition_2 | pvalue | pvalue_adjusted |
|---|---|---|---|---|
| interactor_abundance | mitosis | inter | 0.0005601385574333299 | 0.017415659477552636 |
| complex_abundance | mitosis | inter | 0.0007027447438489493 | 0.01718712859242002 |
| interactor_ratio | mitosis | inter | 0.0015892366528110781 | 0.0480939461418057 |

P55769_Q8WWY3
NH2L1_HUMAN vs PRP31_HUMAN
p-value: 0.032 q-value: 0.023

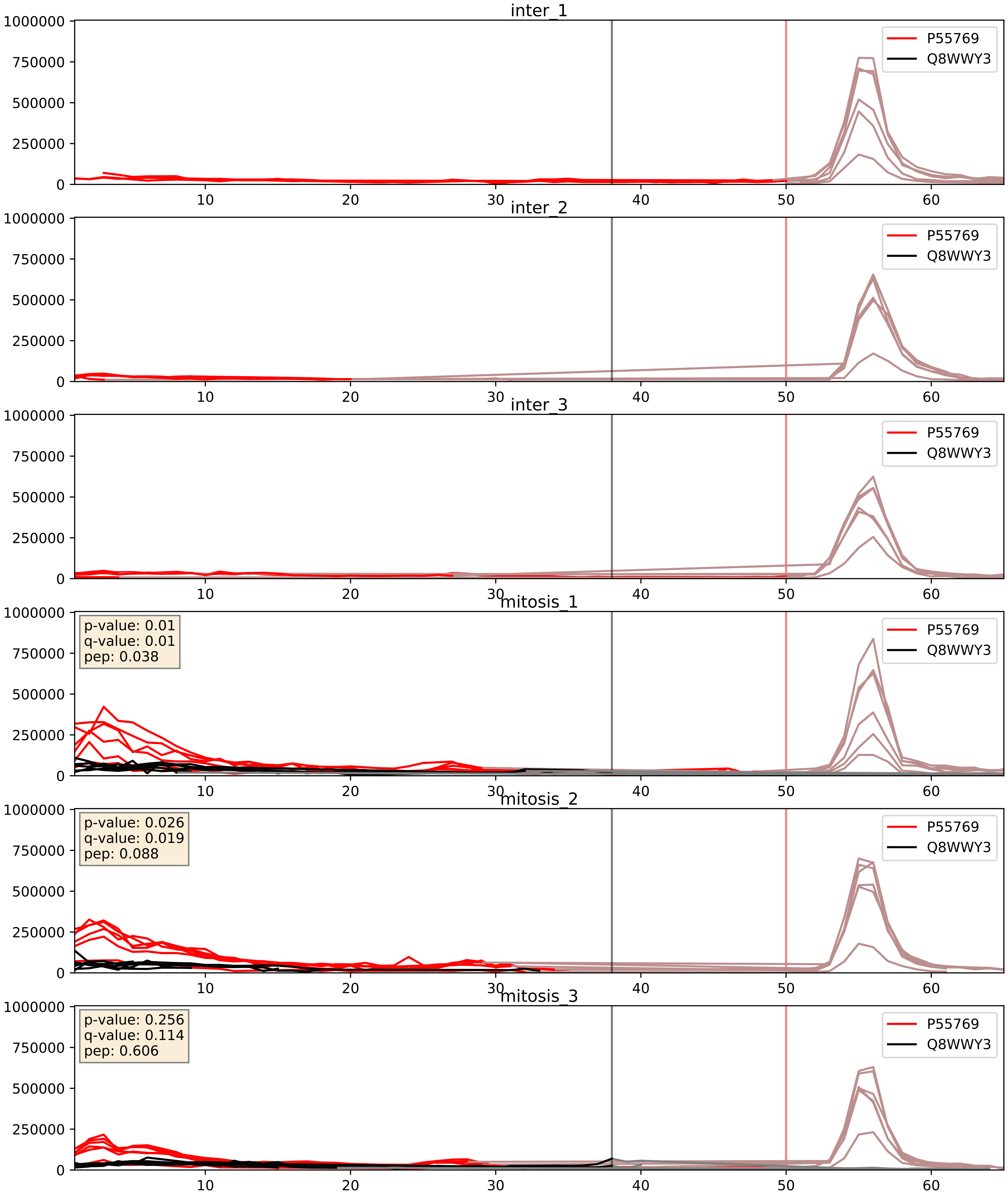


| level | condition_1 | condition_2 | pvalue | pvalue_adjusted |
|---|---|---|---|---|
| interactor_abundance | mitosis | inter | 6.0879231696132886e-05 | 0.006459477818931243 |
| interactor_ratio | mitosis | inter | 0.00020587750302125888 | 0.014683183137172115 |
| complex_abundance | mitosis | inter | 0.00030195135818513163 | 0.01165207110963347 |

P55769_Q96DI7
NH2L1_HUMAN vs SNR40_HUMAN
p-value: 0.082 q-value: 0.049

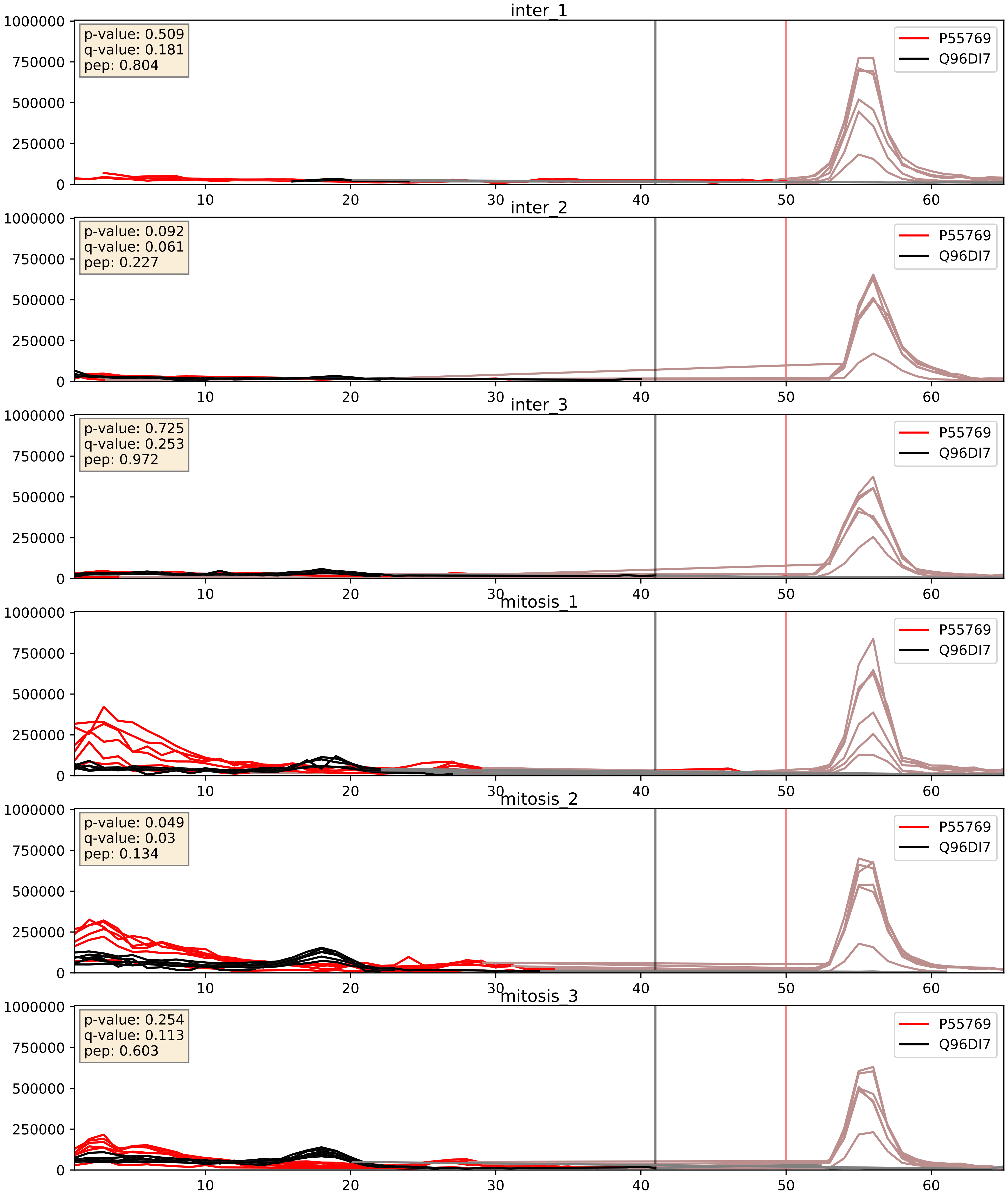


| level | condition_1 | condition_2 | pvalue | pvalue_adjusted |
|---|---|---|---|---|
| interactor_abundance | mitosis | inter | 0.00011044849743284689 | 0.008012196084959062 |
| complex_abundance | mitosis | inter | 0.00031764872867737817 | 0.011652071109633347 |
| interactor_ratio | mitosis | inter | 0.03602142916165438 | 0.30857523351954663 |

P55769_Q9BUQ8
NH2L1_HUMAN vs DDX23_HUMAN
p-value: 0.008 q-value: 0.007

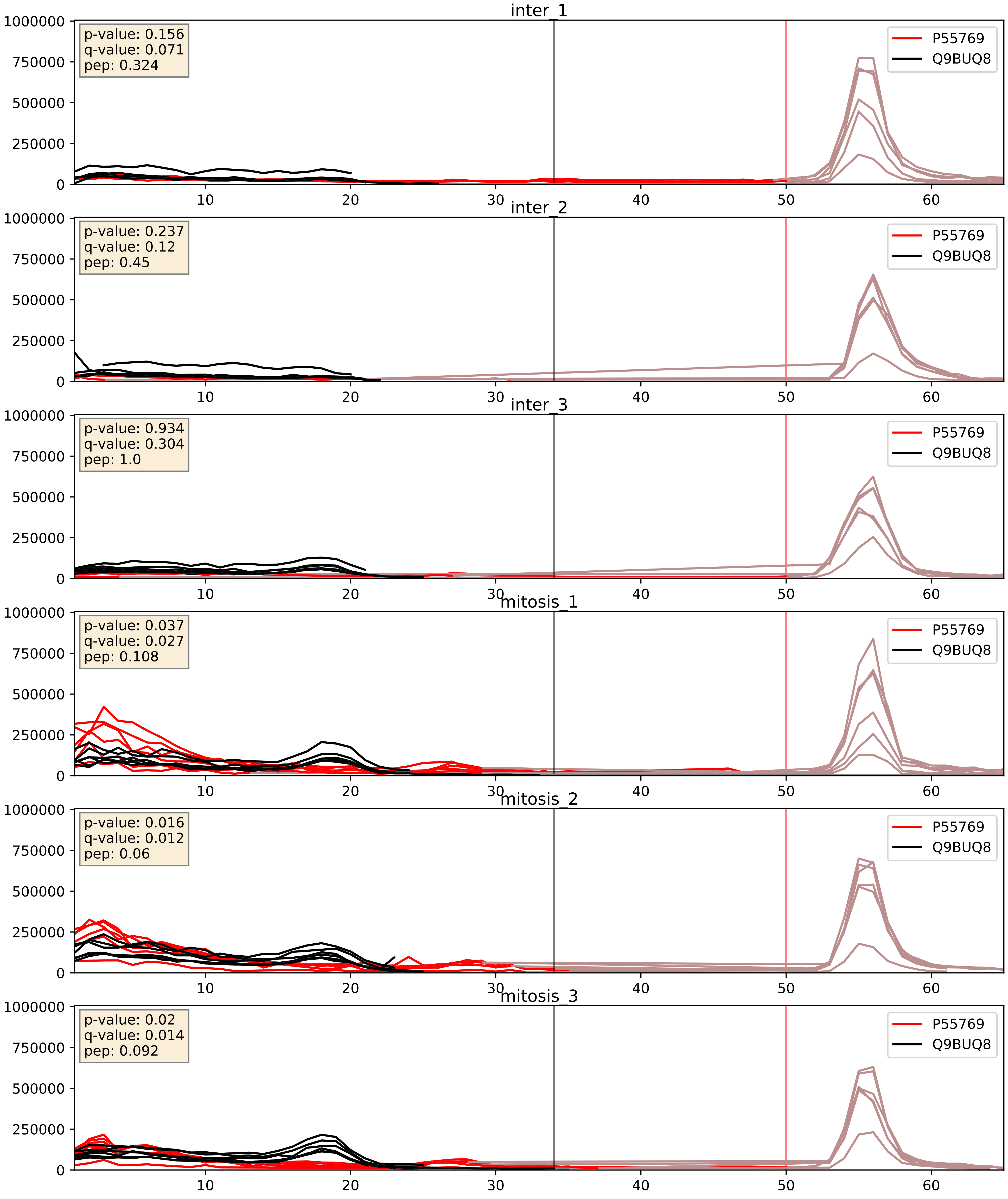


| level | condition_1 | condition_2 | pvalue | pvalue_adjusted |
|---|---|---|---|---|
| complex_abundance | mitosis | inter | 5.162211395155756e-05 | 0.005691857151652958 |
| interactor_abundance | mitosis | inter | 0.0010679120199147774 | 0.024049720606162197 |
| interactor_ratio | mitosis | inter | 0.1115601989567642 | 0.5532962393902757 |

P55769_Q9NR30
NH2L1_HUMAN vs DDX21_HUMAN
p-value: 0.0 q-value: 0.025

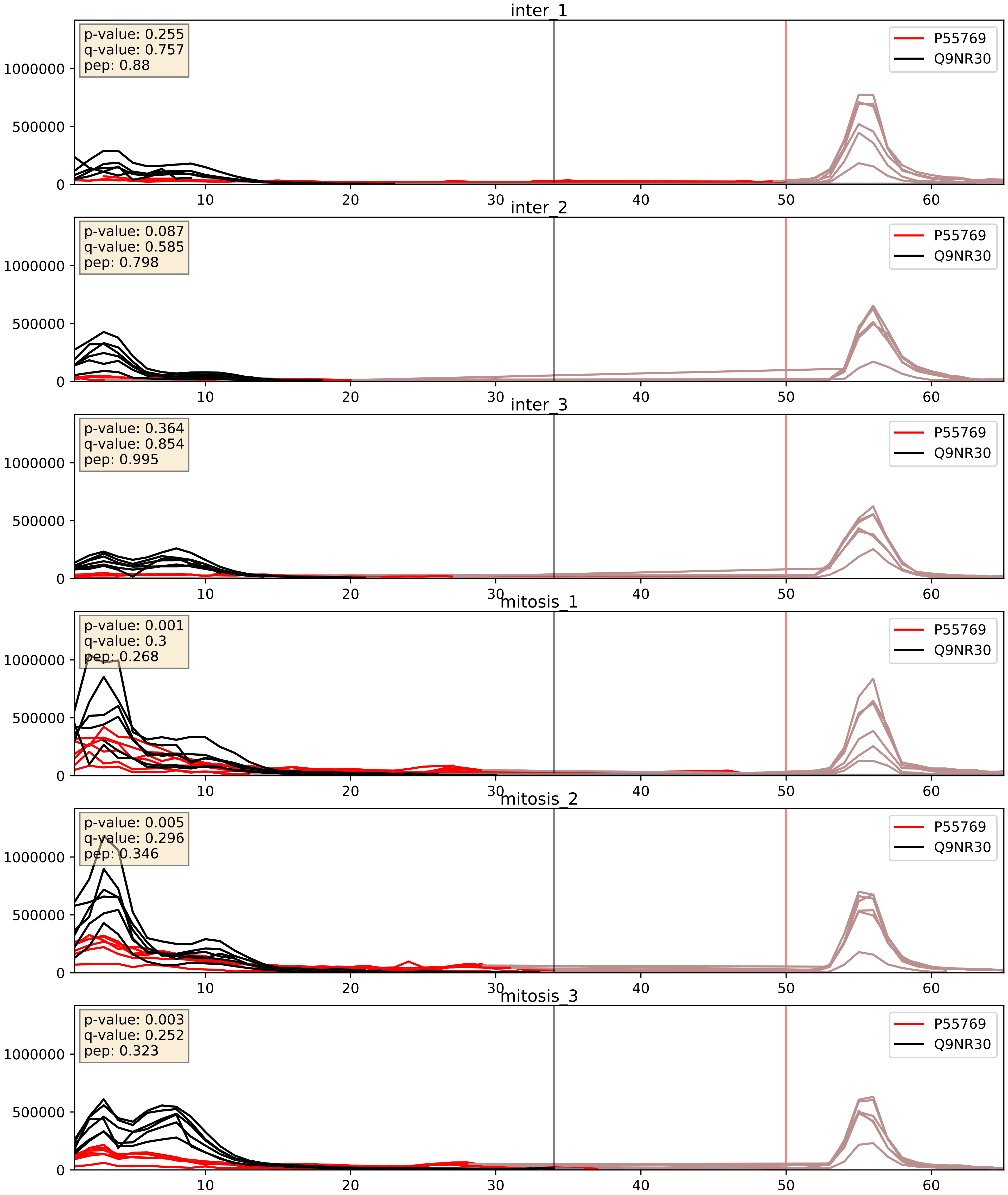


| level | condition_1 | condition_2 | pvalue | pvalue_adjusted |
|---|---|---|---|---|
| complex_abundance | mitosis | inter | 0.0002757050388916281 | 0.011185000629916286 |
| interactor_abundance | mitosis | inter | 0.0010777766016947538 | 0.024049720606162197 |
| interactor_ratio | mitosis | inter | 0.02503180125222349 | 0.24857565976685972 |

P55769_Q9NX24
NH2L1_HUMAN vs NHP2_HUMAN
p-value: 0.0 q-value: 0.0

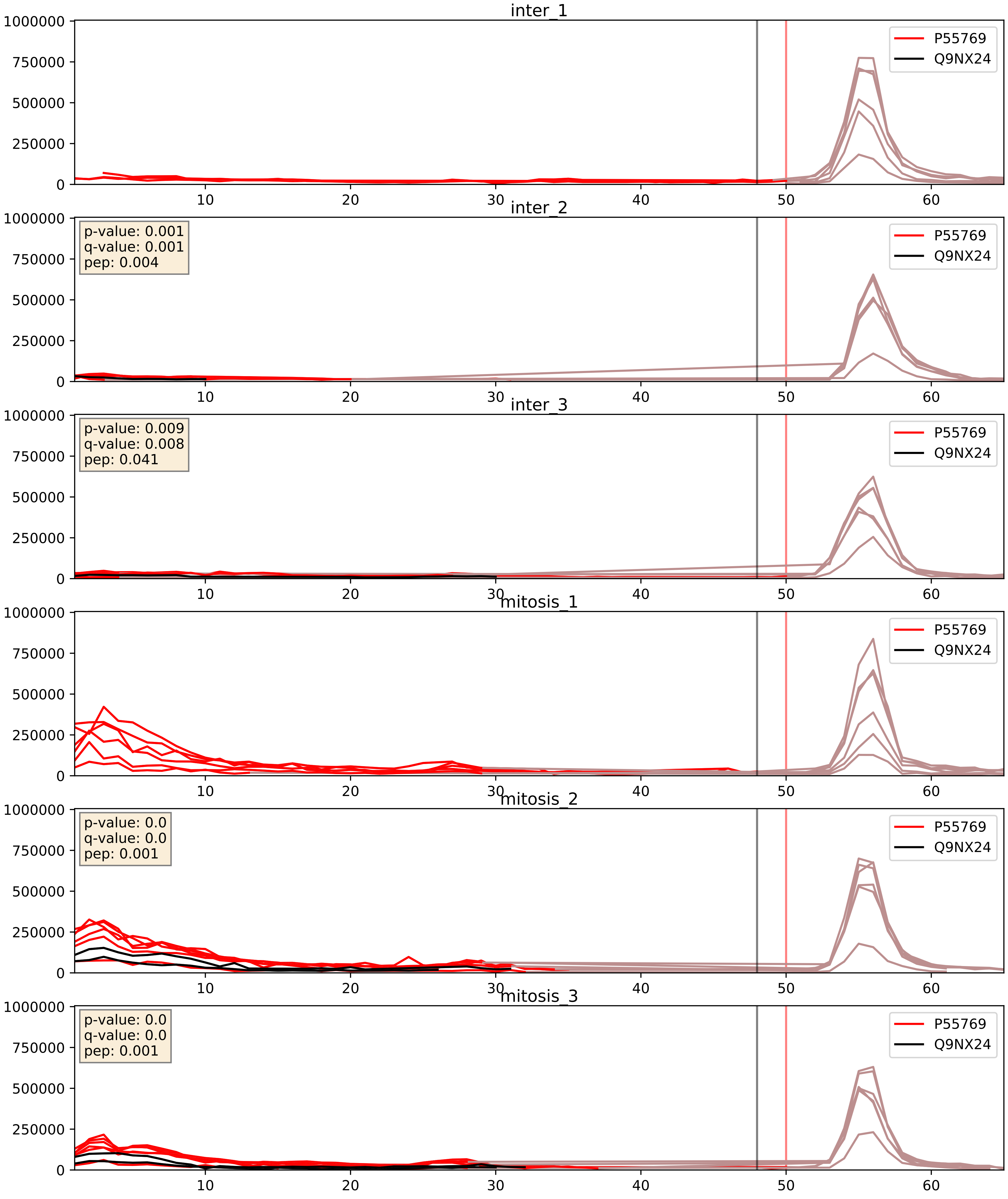


| level | condition_1 | condition_2 | pvalue | pvalue_adjusted |
|---|---|---|---|---|
| interactor_abundance | mitosis | inter | 0.325184730427145 | 0.5246595594112452 |
| complex_abundance | mitosis | inter | 0.32584222271544905 | 0.4742746856732263 |
| interactor_ratio | mitosis | inter | 0.3277826972160907 | 0.7859439462660326 |

P55769_Q9UKD2
NH2L1_HUMAN vs MRT4_HUMAN
p-value: 0.001 q-value: 0.029

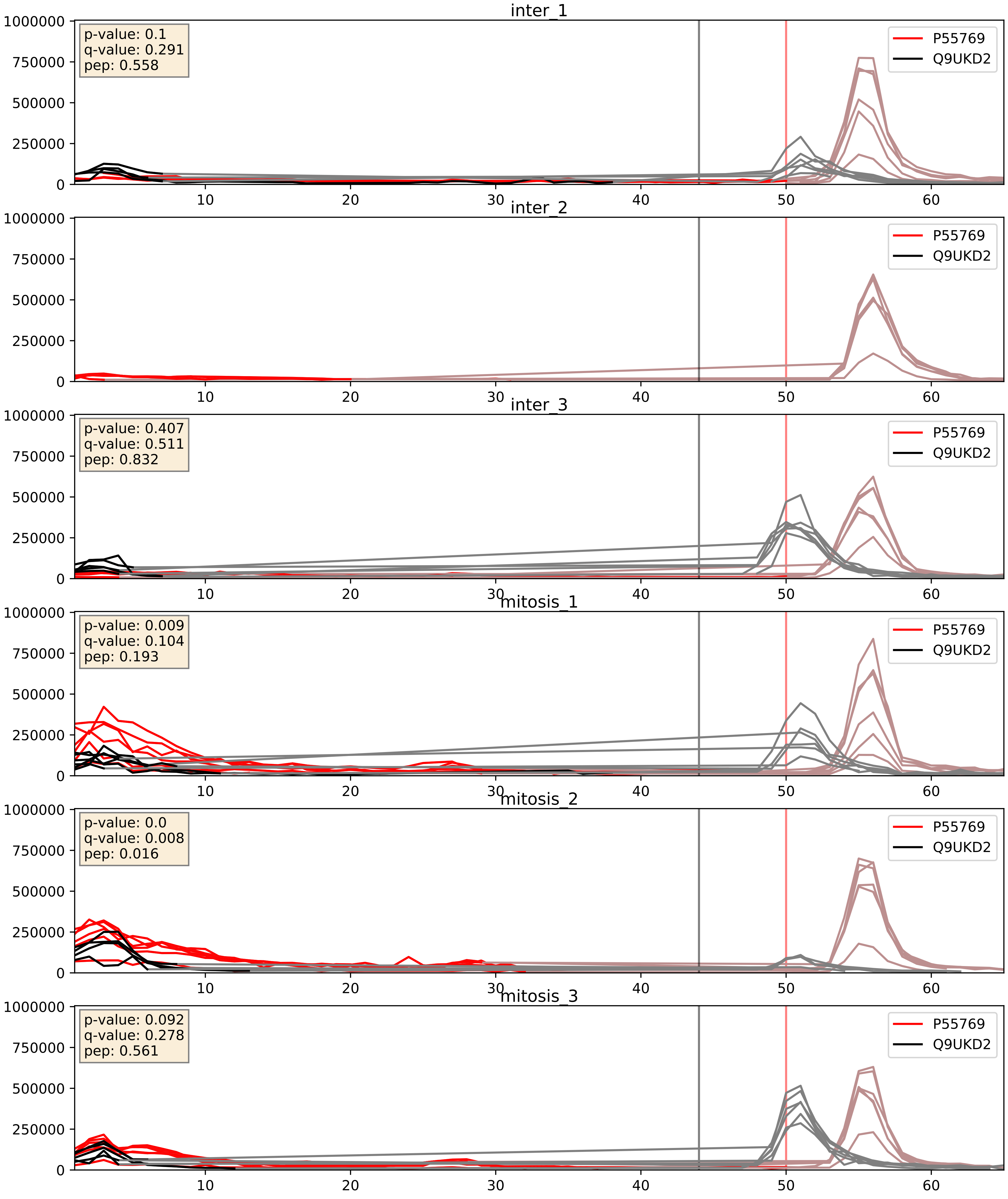


| level | condition_1 | condition_2 | pvalue | pvalue_adjusted |
|---|---|---|---|---|
| interactor_ratio | mitosis | inter | 0.0025457301912372872 | 0.06485550725294995 |
| interactor_abundance | mitosis | inter | 0.019320041195118556 | 0.1190514126751731 |
| complex_abundance | mitosis | inter | 0.051399232483414346 | 0.16145960736074377 |

P55769_Q9Y221
NH2L1_HUMAN vs NIP7_HUMAN
p-value: 0.072 q-value: 0.044

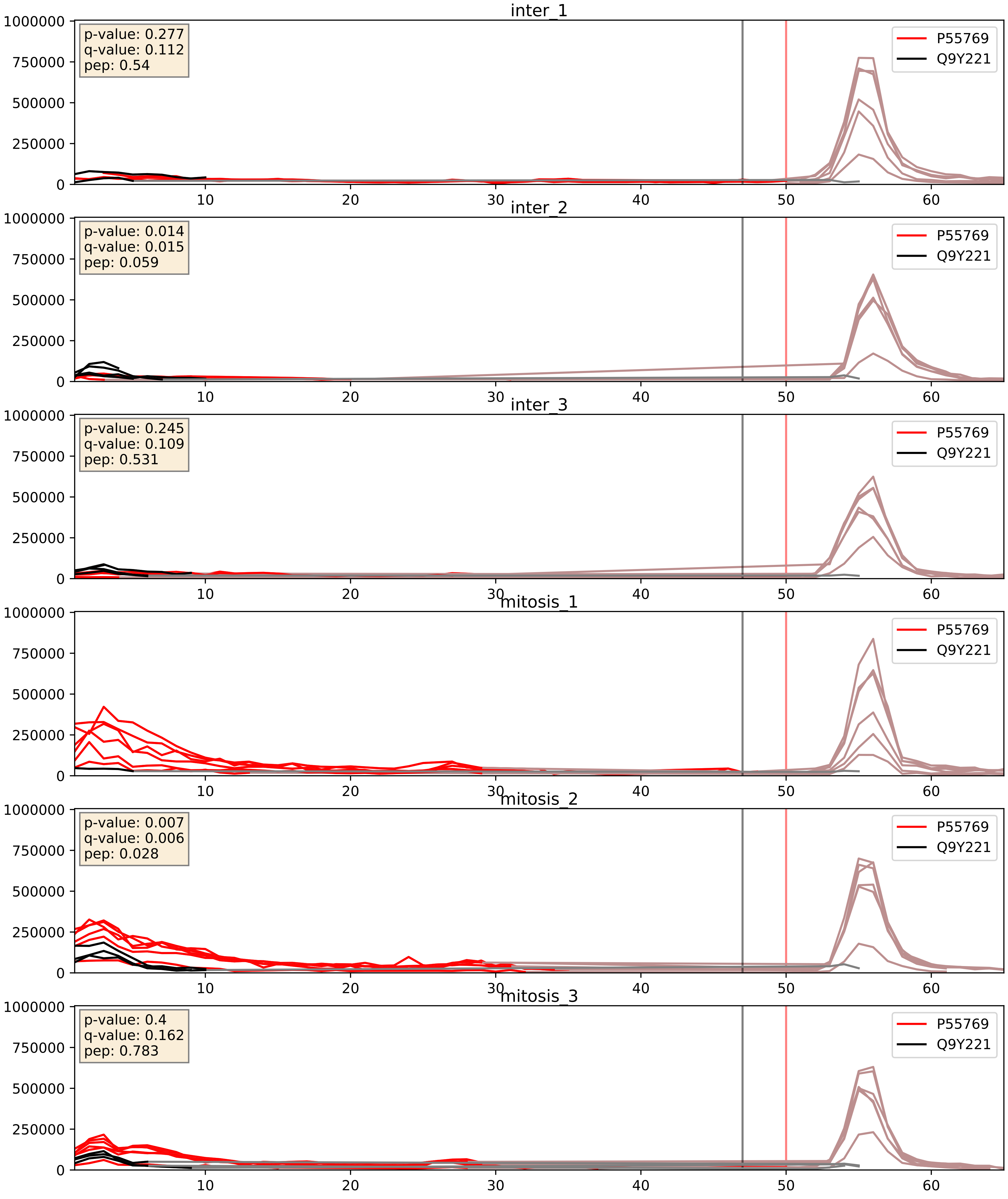


| level | condition_1 | condition_2 | pvalue | pvalue_adjusted |
|---|---|---|---|---|
| interactor_abundance | mitosis | inter | 0.006240942665075991 | 0.06728270681744393 |
| interactor_ratio | mitosis | inter | 0.009362521776968582 | 0.13838047771338102 |
| complex_abundance | mitosis | inter | 0.10568485694956464 | 0.24175612942802124 |

P55769_Q9Y2X3
NH2L1_HUMAN vs NOP58_HUMAN
p-value: 0.0 q-value: 0.0

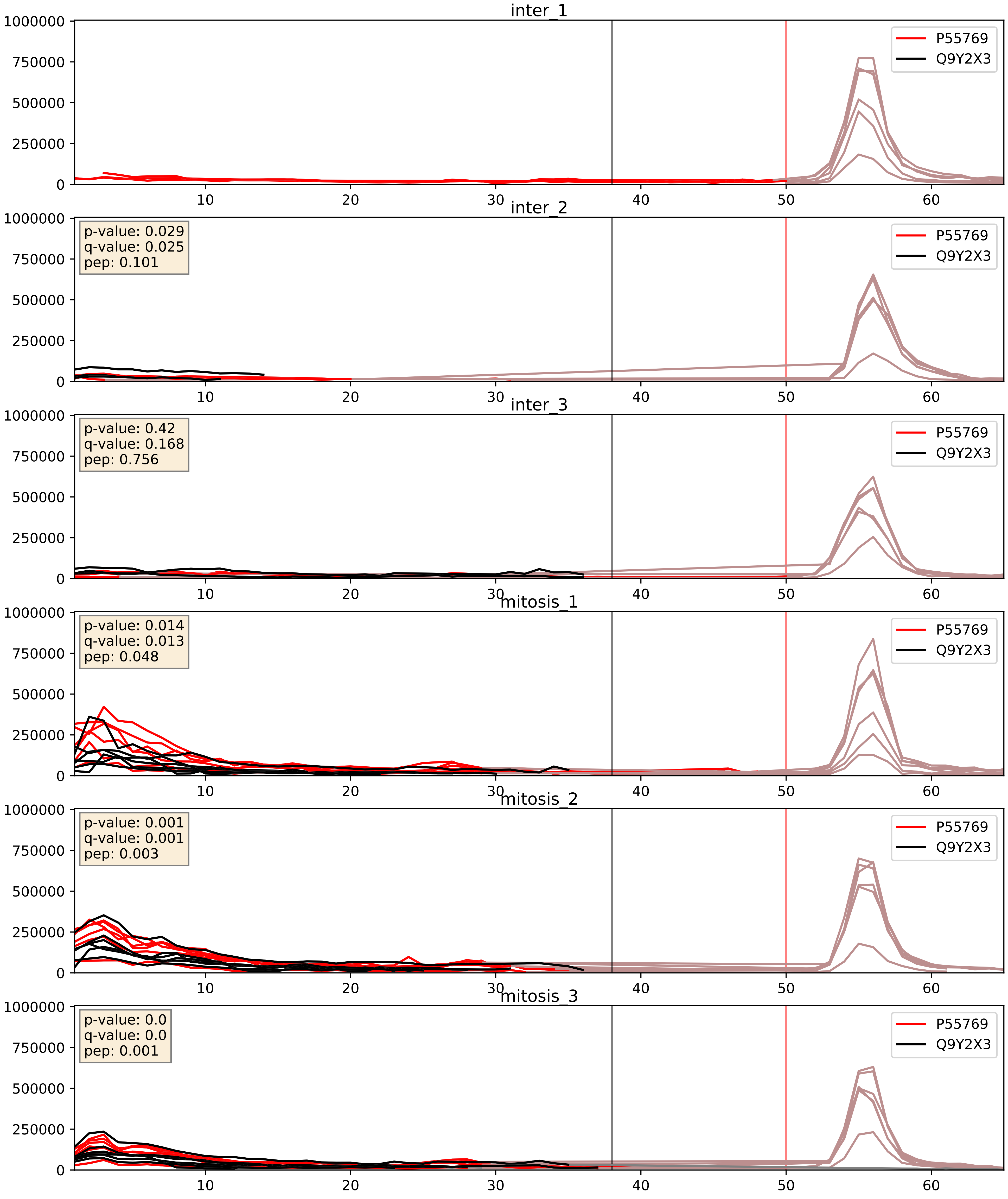


| level | condition_1 | condition_2 | pvalue | pvalue_adjusted |
|---|---|---|---|---|
| interactor_abundance | mitosis | inter | 0.0024610293995718582 | 0.037056133087660696 |
| complex_abundance | mitosis | inter | 0.0024997379458740598 | 0.03638642644133238 |
| interactor_ratio | mitosis | inter | 0.04568692907471442 | 0.35552737534505047 |

P55769_Q9Y3U8
NH2L1_HUMAN vs RL36_HUMAN
p-value: 0.0 q-value: 0.008

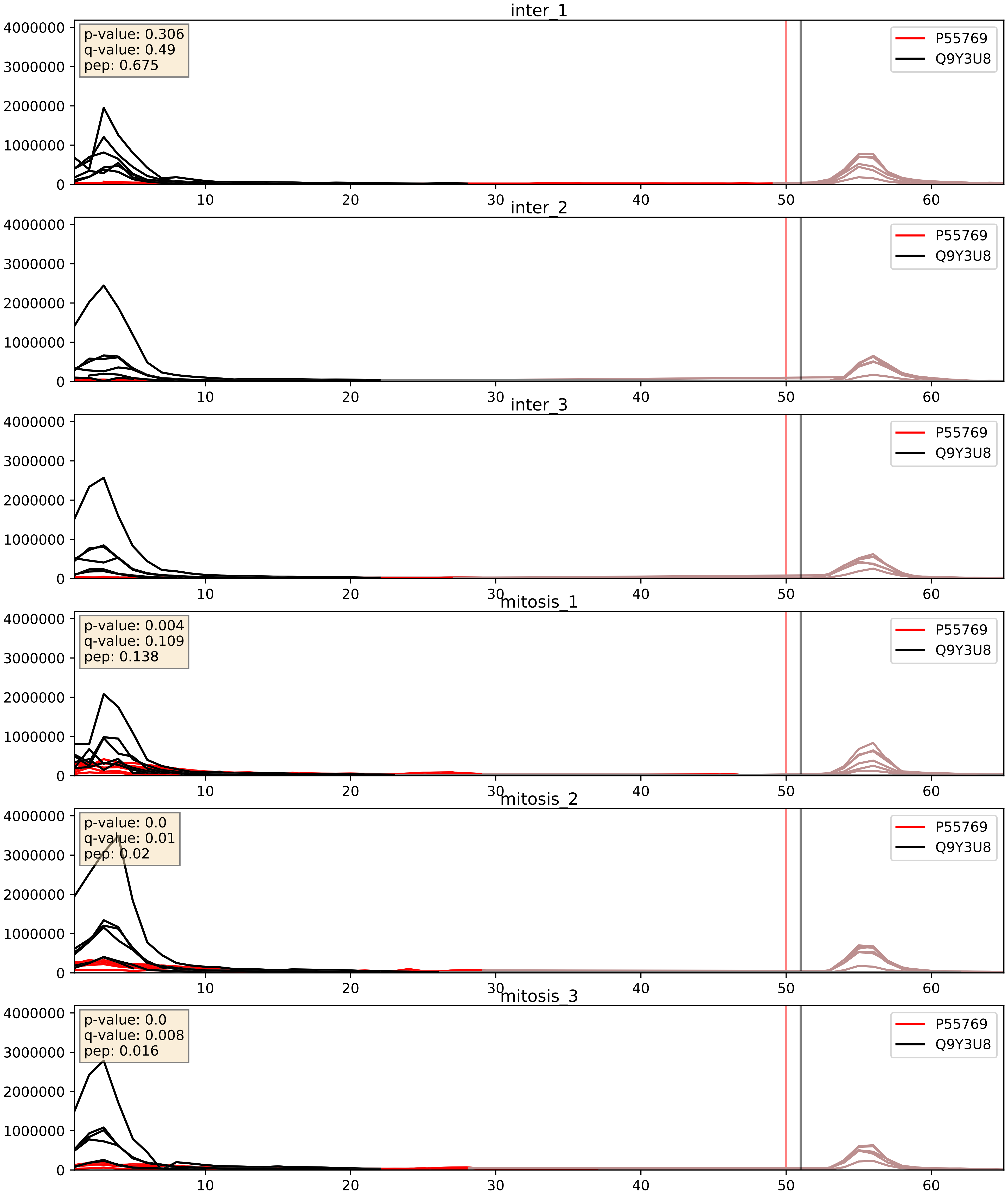


| level | condition_1 | condition_2 | pvalue | pvalue_adjusted |
|---|---|---|---|---|
| complex_abundance | mitosis | inter | 0.0013593972713581571 | 0.025187100958497455 |
| interactor_abundance | mitosis | inter | 0.002521189739124521 | 0.03753284202940157 |
| interactor_ratio | mitosis | inter | 0.006719299349918978 | 0.1136703605440839 |
