## Supplementary Data 2 for "SECAT: Quantifying differential protein-protein interaction states by network-centric analysis": hela_string_000003_P56537.pdf

P56537_P56537

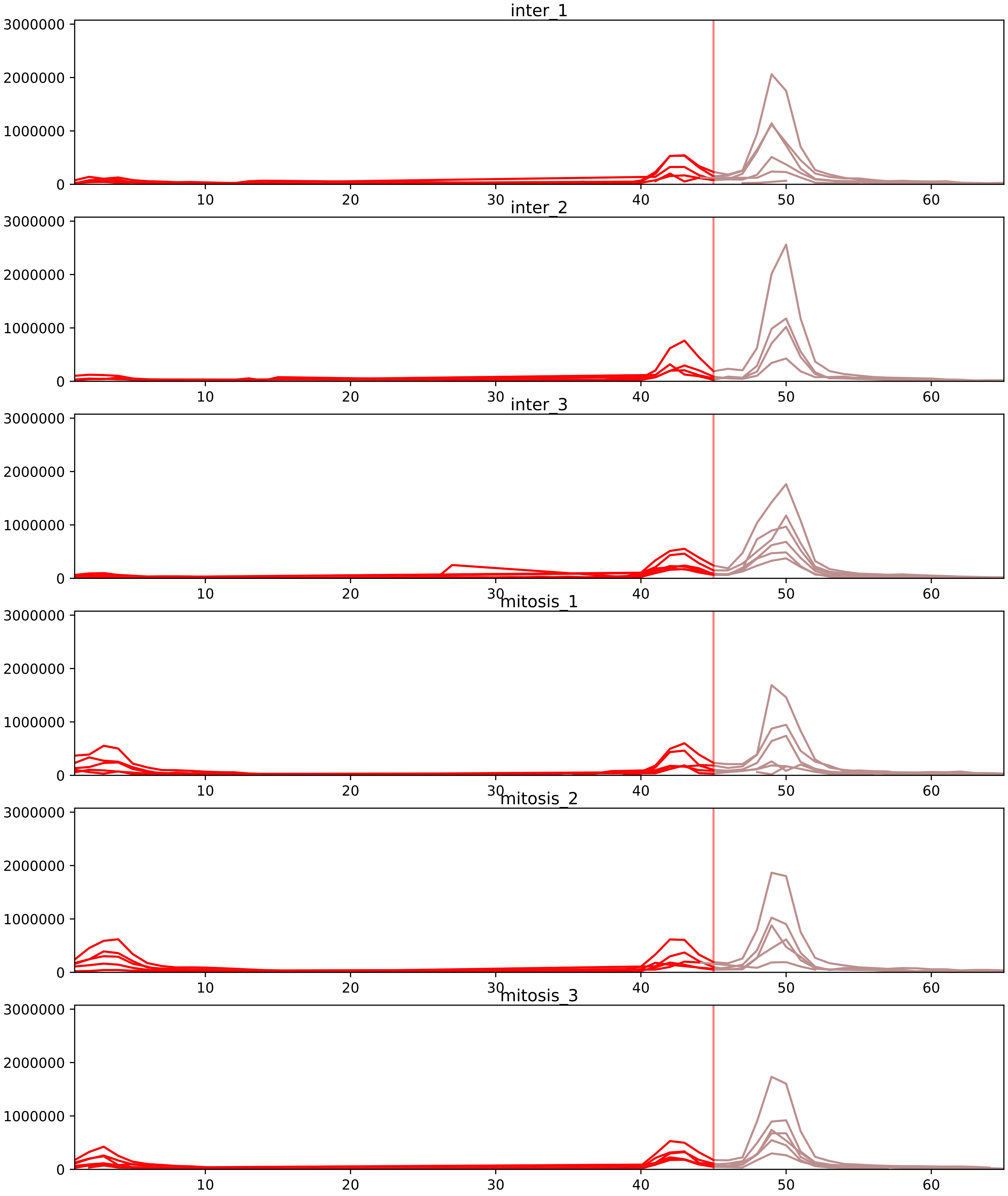


| level | condition_1 | condition_2 | pvalue | pvalue_adjusted |
|---|---|---|---|---|
| interactor_ratio | mitosis | inter | 2.30079008425788e-22 | 1.4111512516781665e-19 |
| interactor_abundance | mitosis | inter | 0.00016175083766852022 | 0.003166186609681672 |
| complex_abundance | mitosis | inter | 0.00046613853656582606 | 0.006547289368558167 |
| assembled_abundance | mitosis | inter | 0.01036040926470159 | 0.113310924691748 |
| total_abundance | mitosis | inter | 0.32634671896499784 | 0.6482200305904395 |
| monomer_abundance | mitosis | inter | 0.40058755835559556 | 0.6958474773829776 |

O76021_P56537
RL1D1_HUMAN vs IF6_HUMAN
p-value: 0.001 q-value: 0.039

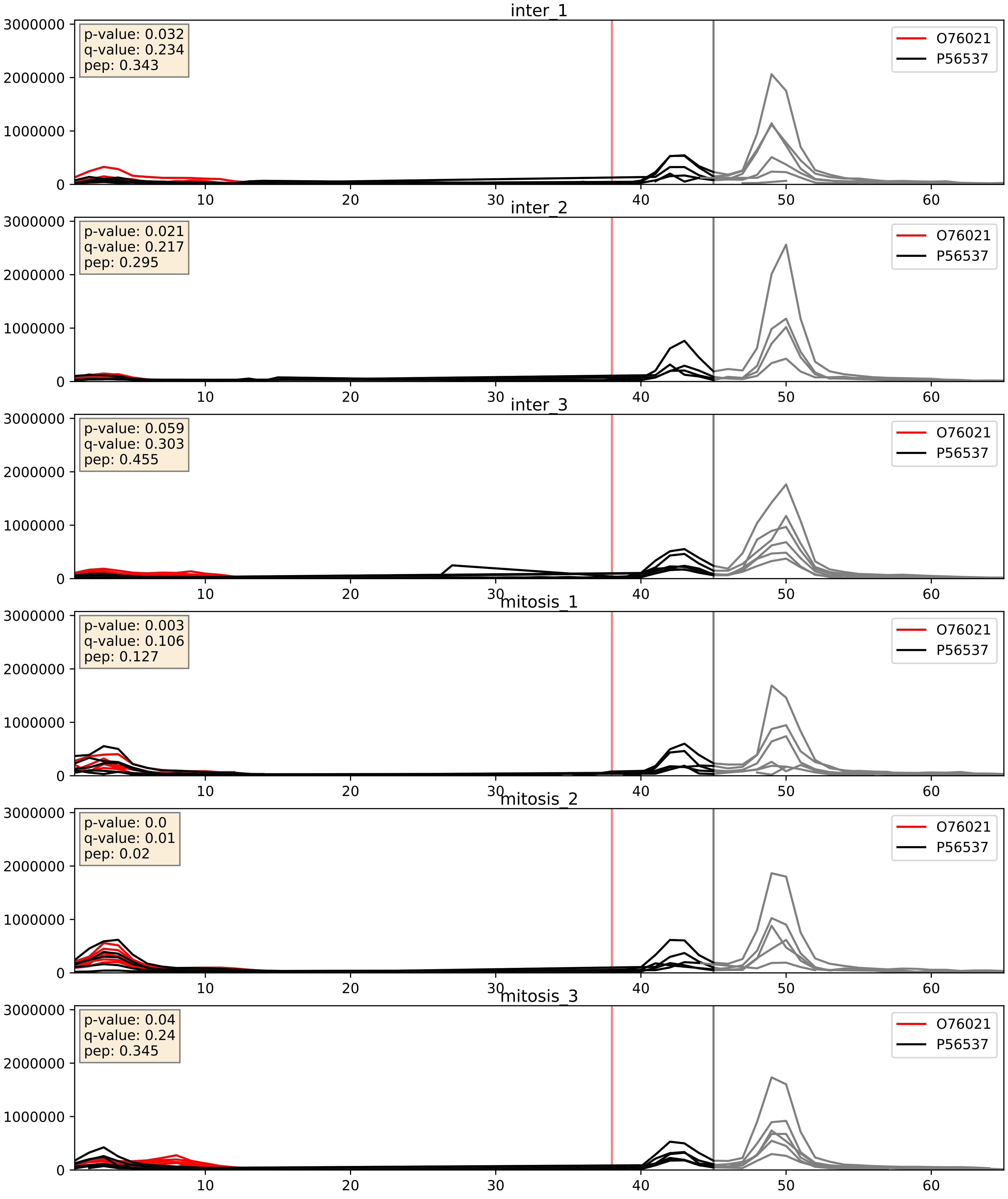


| level | condition_1 | condition_2 | pvalue | pvalue_adjusted |
|---|---|---|---|---|
| complex_abundance | mitosis | inter | 0.001587159867408426 | 0.027897512248493073 |
| interactor_abundance | mitosis | inter | 0.007302544007558834 | 0.07362753439894419 |
| interactor_ratio | mitosis | inter | 0.17049014932935702 | 0.6336933036297422 |

P18077_P56537
RL35A_HUMAN vs IF6_HUMAN
p-value: 0.001 q-value: 0.03

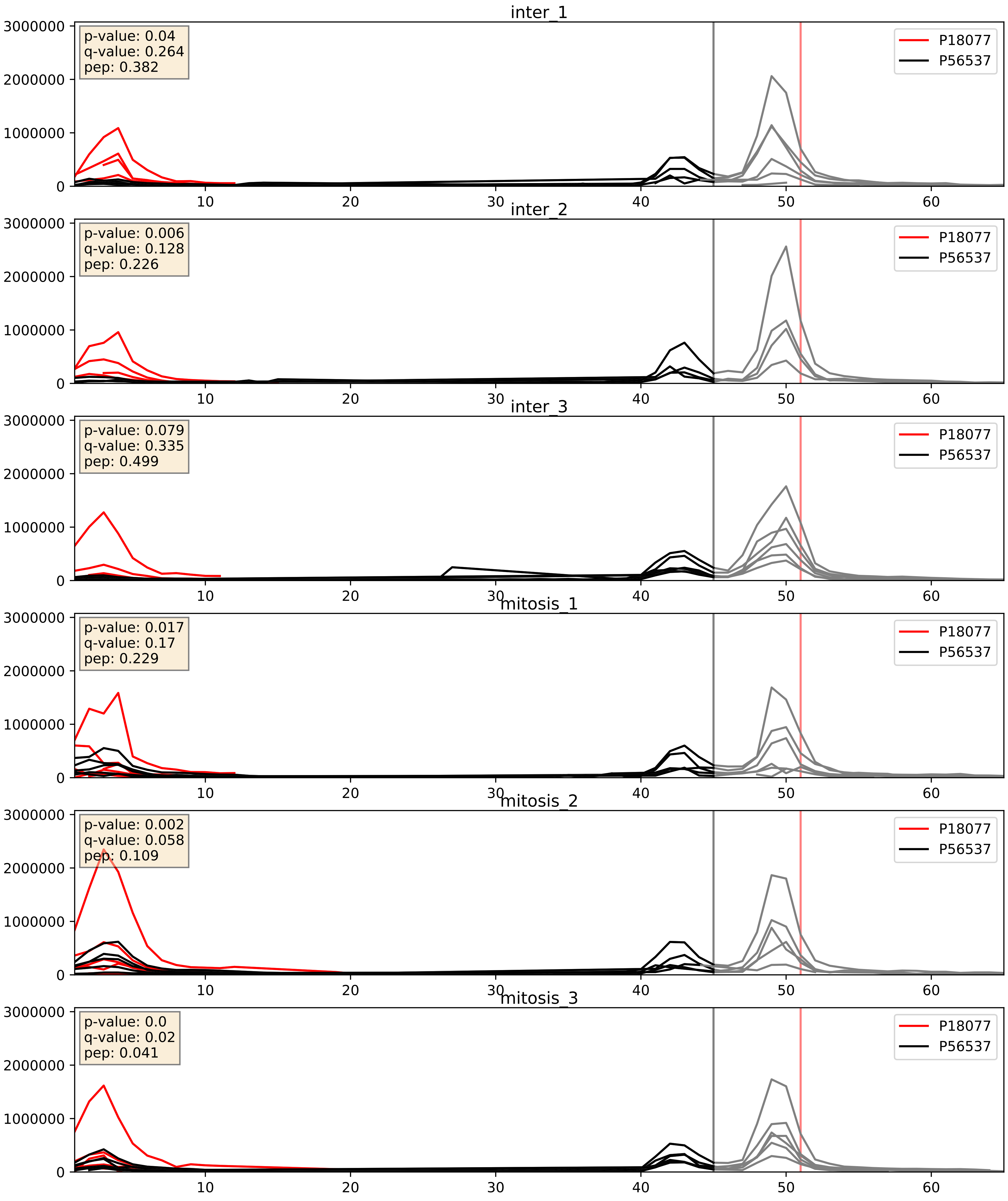


| level | condition_1 | condition_2 | pvalue | pvalue_adjusted |
|---|---|---|---|---|
| interactor_ratio | mitosis | inter | 9.619201967251694e-06 | 0.0035800160365075866 |
| complex_abundance | mitosis | inter | 0.068877227309555 | 0.1900061443022207 |
| interactor_abundance | mitosis | inter | 0.8824024184766702 | 0.9349380742864584 |

P18124_P56537
RL7_HUMAN vs IF6_HUMAN
p-value: 0.0 q-value: 0.013

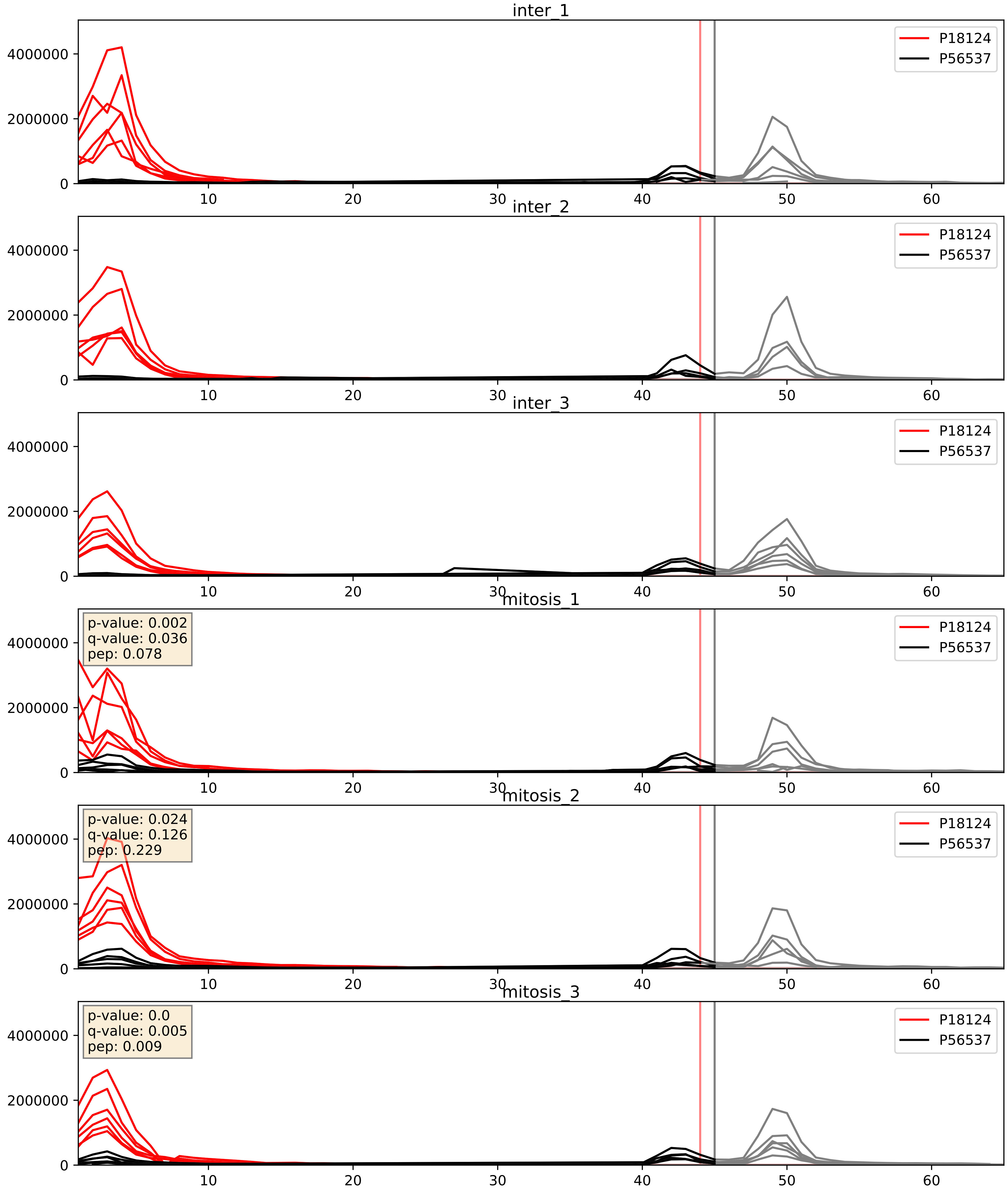


| level | condition_1 | condition_2 | pvalue | pvalue_adjusted |
|---|---|---|---|---|
| interactor_ratio | mitosis | inter | 0.006211836518656897 | 0.10851698081572049 |
| complex_abundance | mitosis | inter | 0.04274581029093276 | 0.1481949160937304 |
| interactor_abundance | mitosis | inter | 0.5623313926707146 | 0.7112621176332597 |

P22087_P56537
FBRL_HUMAN vs IF6_HUMAN
p-value: 0.001 q-value: 0.015

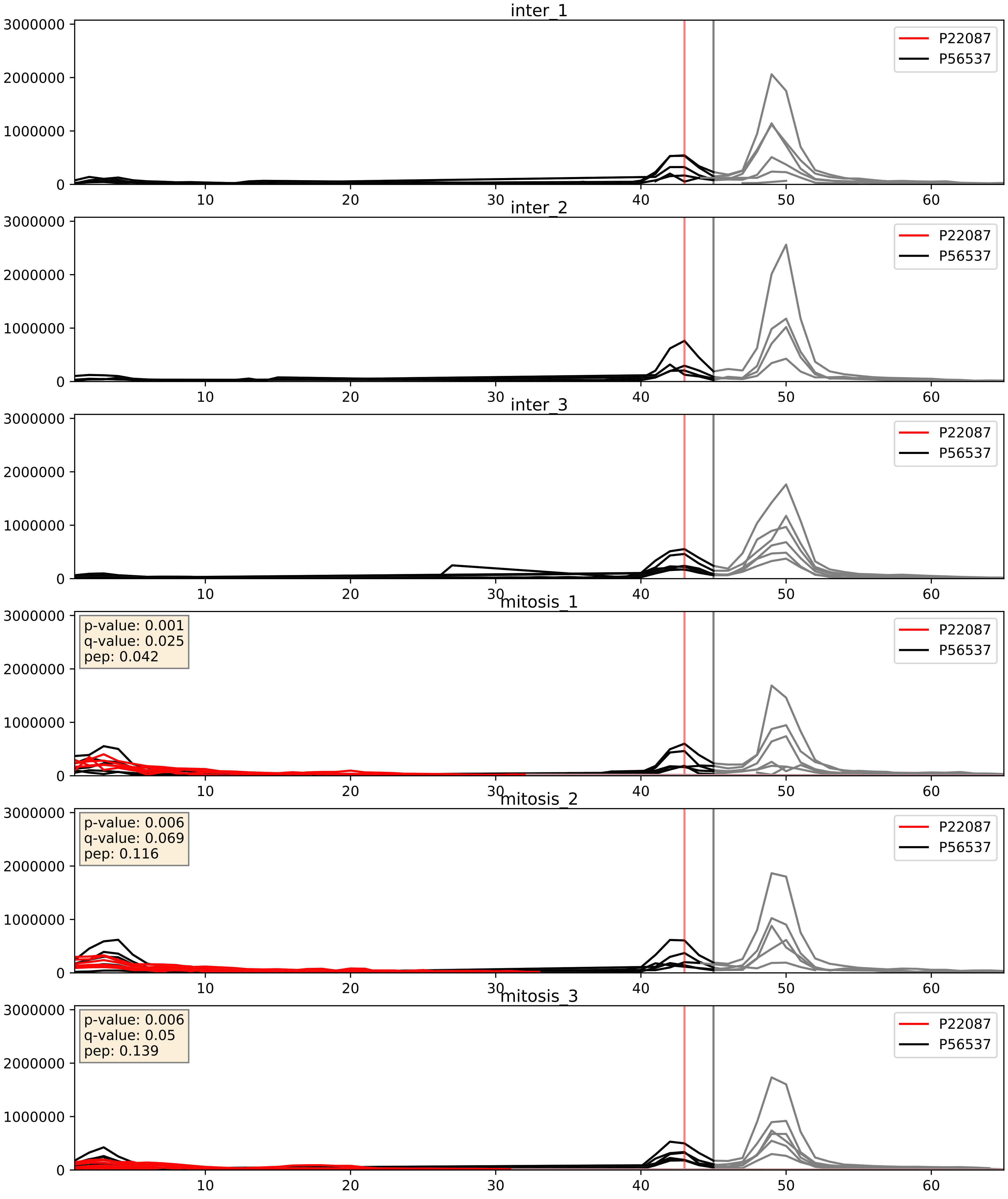


| level | condition_1 | condition_2 | pvalue | pvalue_adjusted |
|---|---|---|---|---|
| complex_abundance | mitosis | inter | 0.00029723422483891155 | 0.011617922212881656 |
| interactor_abundance | mitosis | inter | 0.00036693098007483434 | 0.01356427145975096 |
| interactor_ratio | mitosis | inter | 0.12685036942305242 | 0.5750062399685051 |

P32969_P56537
RL9_HUMAN vs IF6_HUMAN
p-value: 0.0 q-value: 0.003

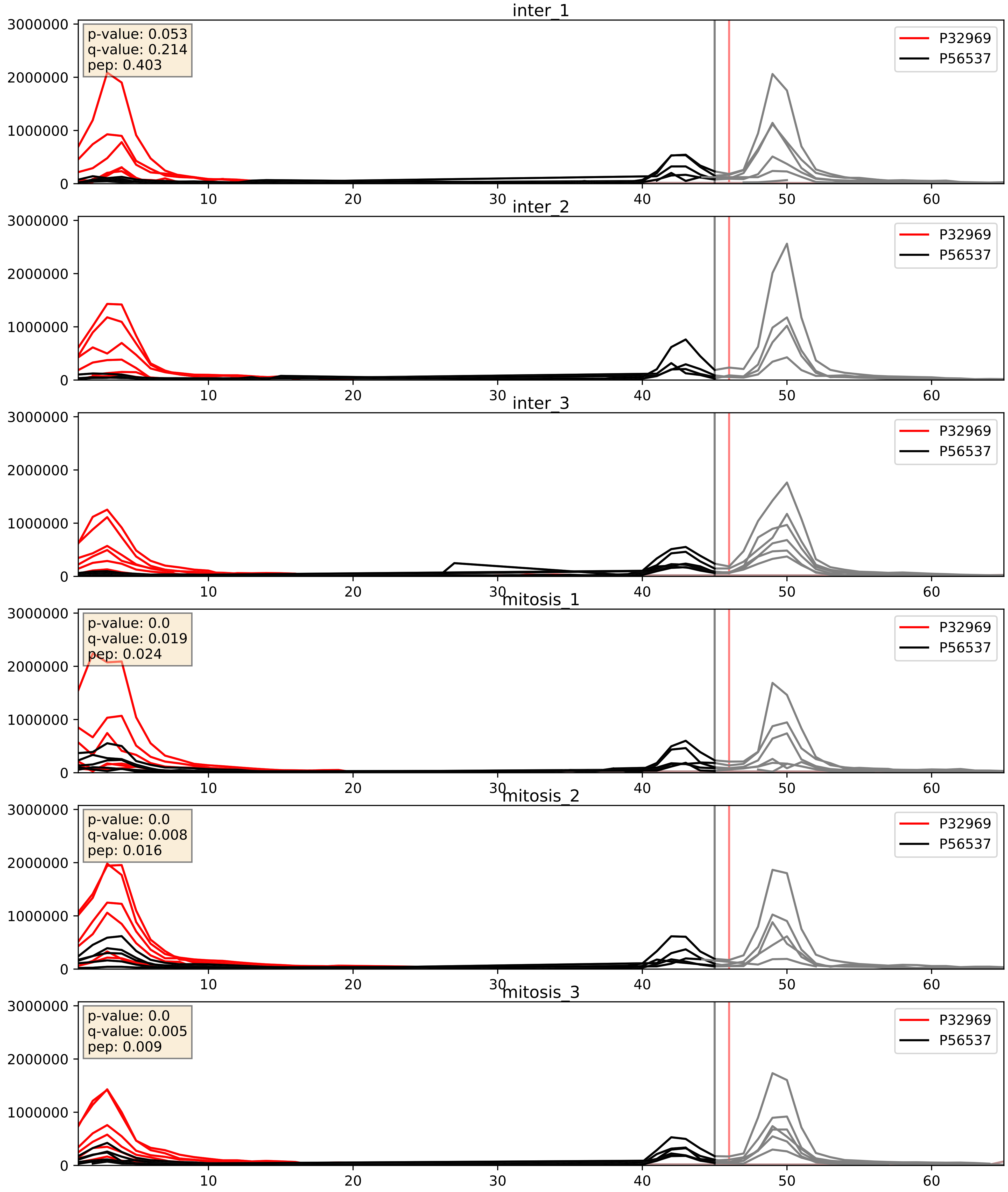


| level | condition_1 | condition_2 | pvalue | pvalue_adjusted |
|---|---|---|---|---|
| interactor_ratio | mitosis | inter | 0.0011232552329799603 | 0.03908562924515634 |
| complex_abundance | mitosis | inter | 0.008368800867374564 | 0.06770976883244449 |
| interactor_abundance | mitosis | inter | 0.18662723254004443 | 0.37950407698664385 |

P36578_P56537
RL4_HUMAN vs IF6_HUMAN
p-value: 0.004 q-value: 0.004

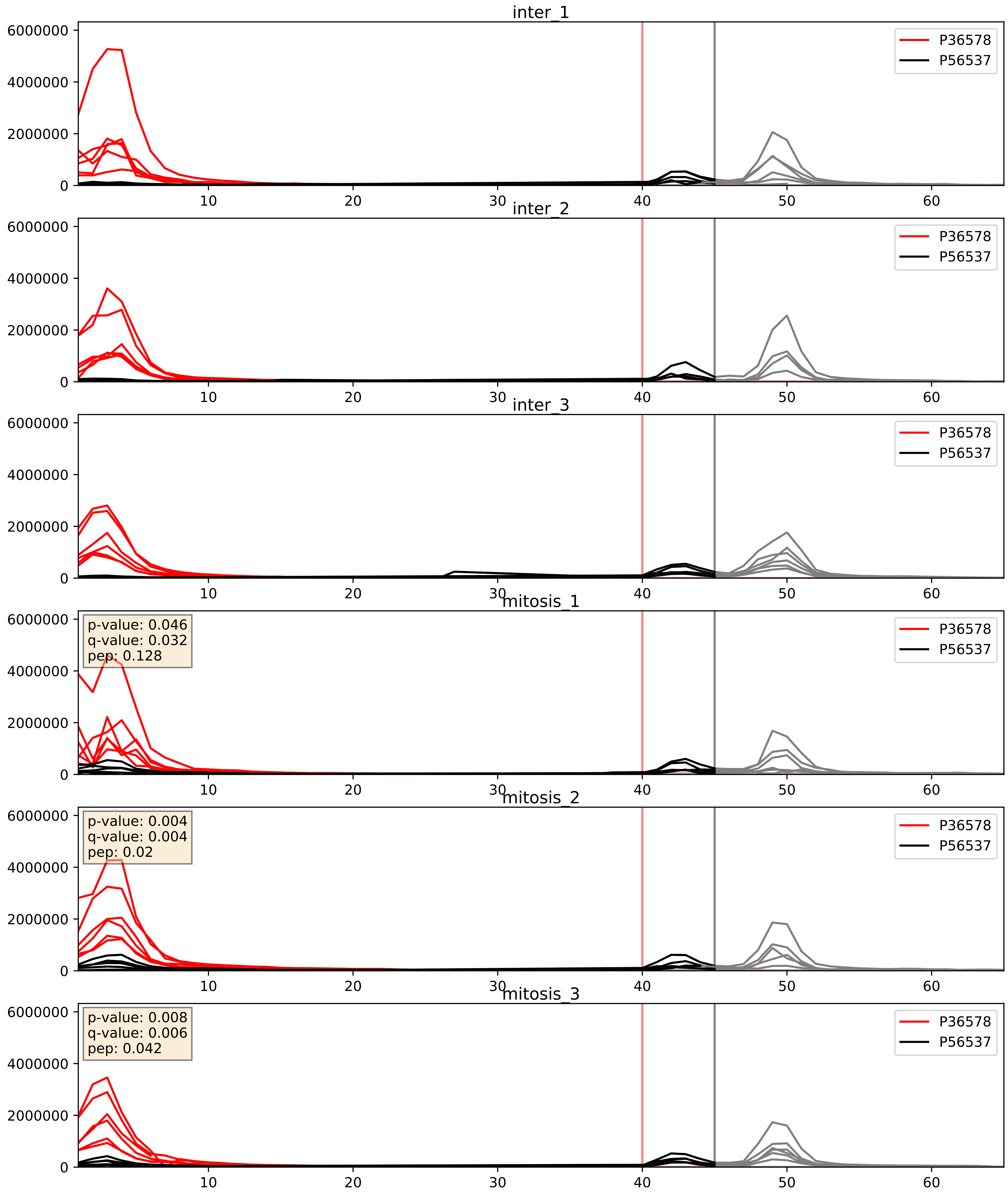


| level | condition_1 | condition_2 | pvalue | pvalue_adjusted |
|---|---|---|---|---|
| complex_abundance | mitosis | inter | 0.0013077920738187984 | 0.024459508269436846 |
| interactor_ratio | mitosis | inter | 0.018083508166905432 | 0.20237106227759052 |
| interactor_abundance | mitosis | inter | 0.827671747607865 | 0.9044322664866693 |

P39023_P56537
RL3_HUMAN vs IF6_HUMAN
p-value: 0.0 q-value: 0.015

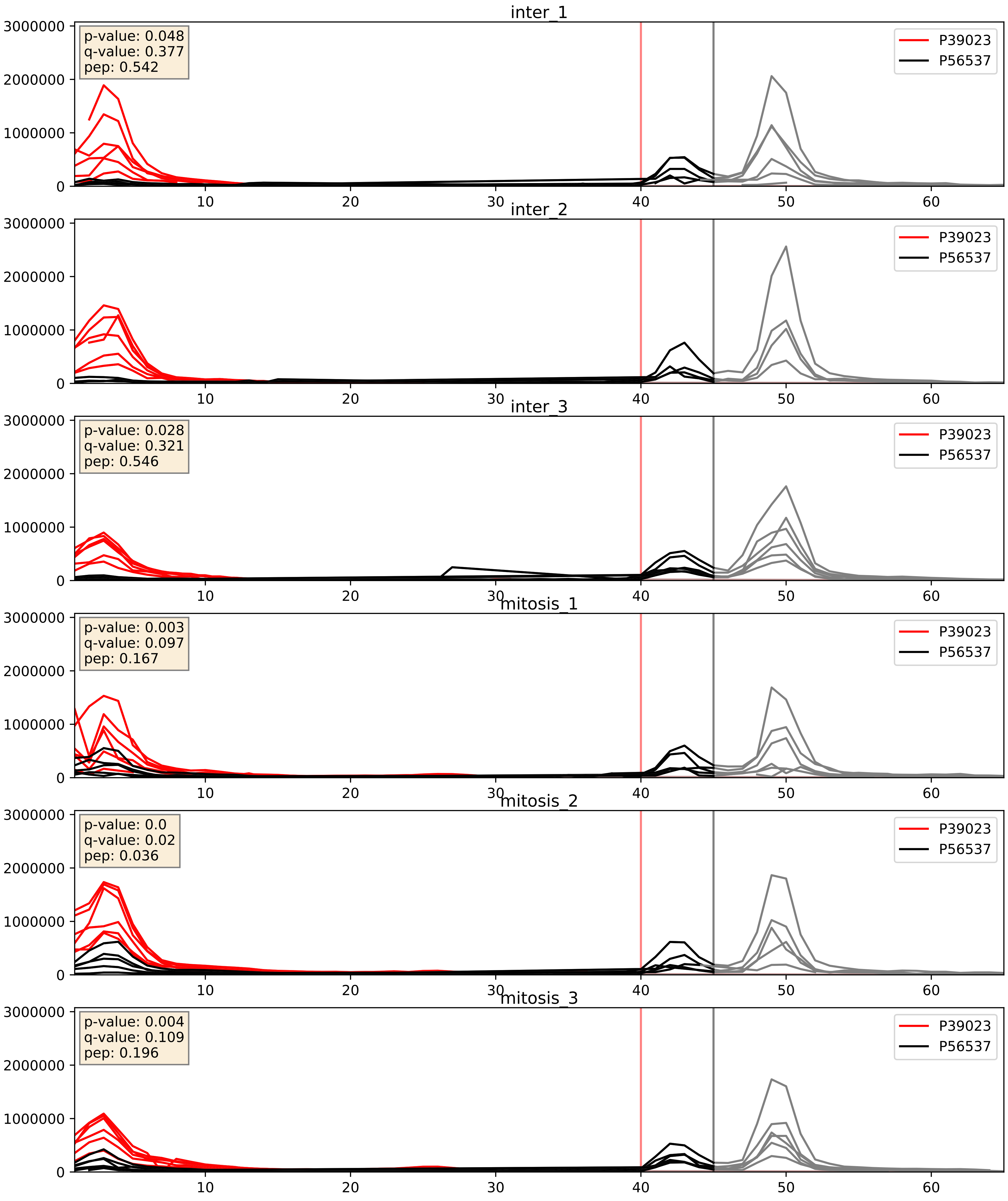


| level | condition_1 | condition_2 | pvalue | pvalue_adjusted |
|---|---|---|---|---|
| interactor_ratio | mitosis | inter | 0.004233880729778598 | 0.08567853202577967 |
| complex_abundance | mitosis | inter | 0.018043815404843450 | 0.10031583038215336 |
| interactor_abundance | mitosis | inter | 0.5383686742633752 | 0.6958755532947515 |

P40429_P56537
RL13A_HUMAN vs IF6_HUMAN
p-value: 0.0 q-value: 0.017

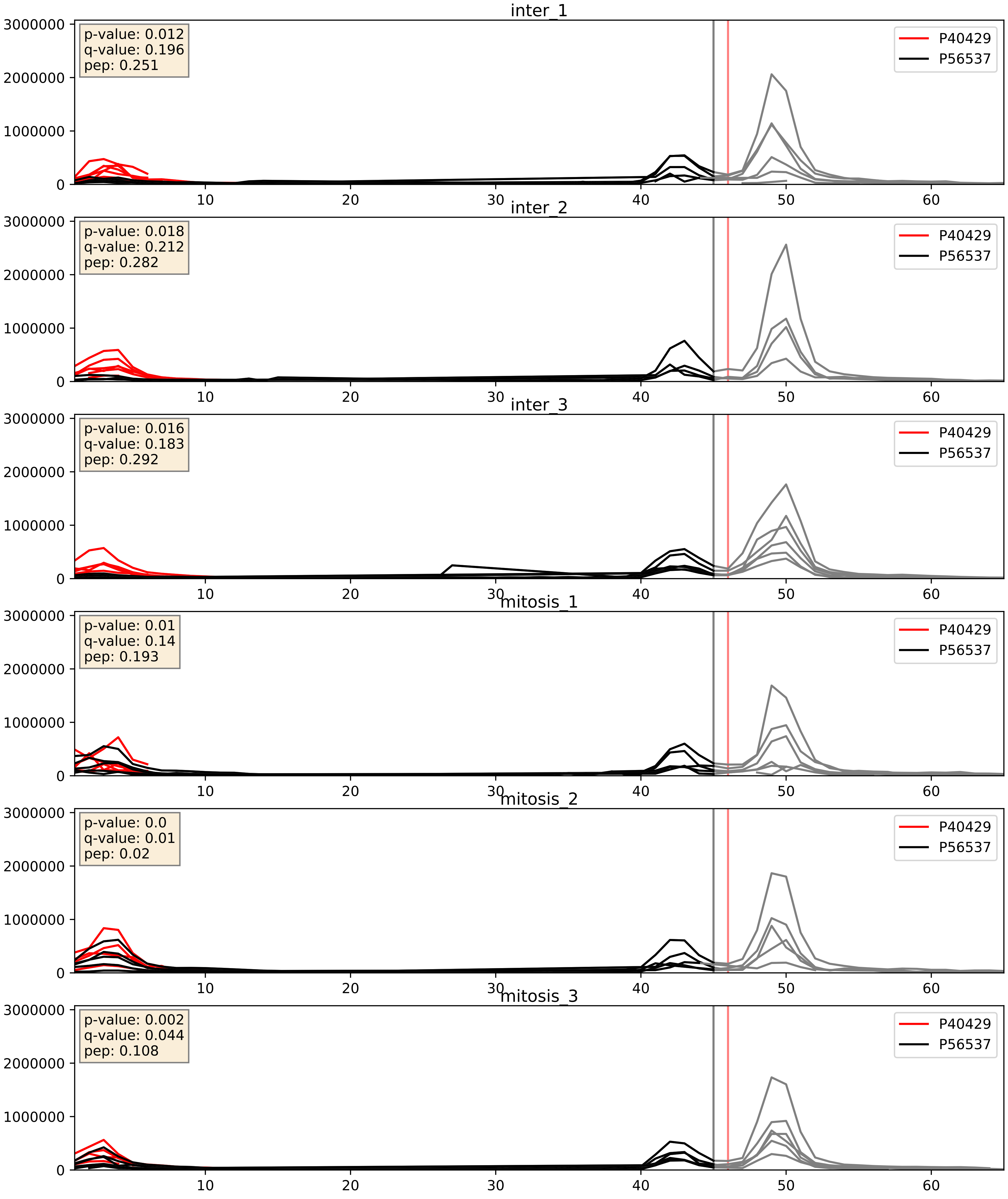


| level | condition_1 | condition_2 | pvalue | pvalue_adjusted |
|---|---|---|---|---|
| interactor_ratio | mitosis | inter | 0.0059272323470886505 | 0.1048287373782621 |
| complex_abundance | mitosis | inter | 0.023500031369345923 | 0.11152773505083037 |
| interactor_abundance | mitosis | inter | 0.9842056238999787 | 0.99062850732951 |

P46777_P56537
RL5_HUMAN vs IF6_HUMAN
p-value: 0.011 q-value: 0.01

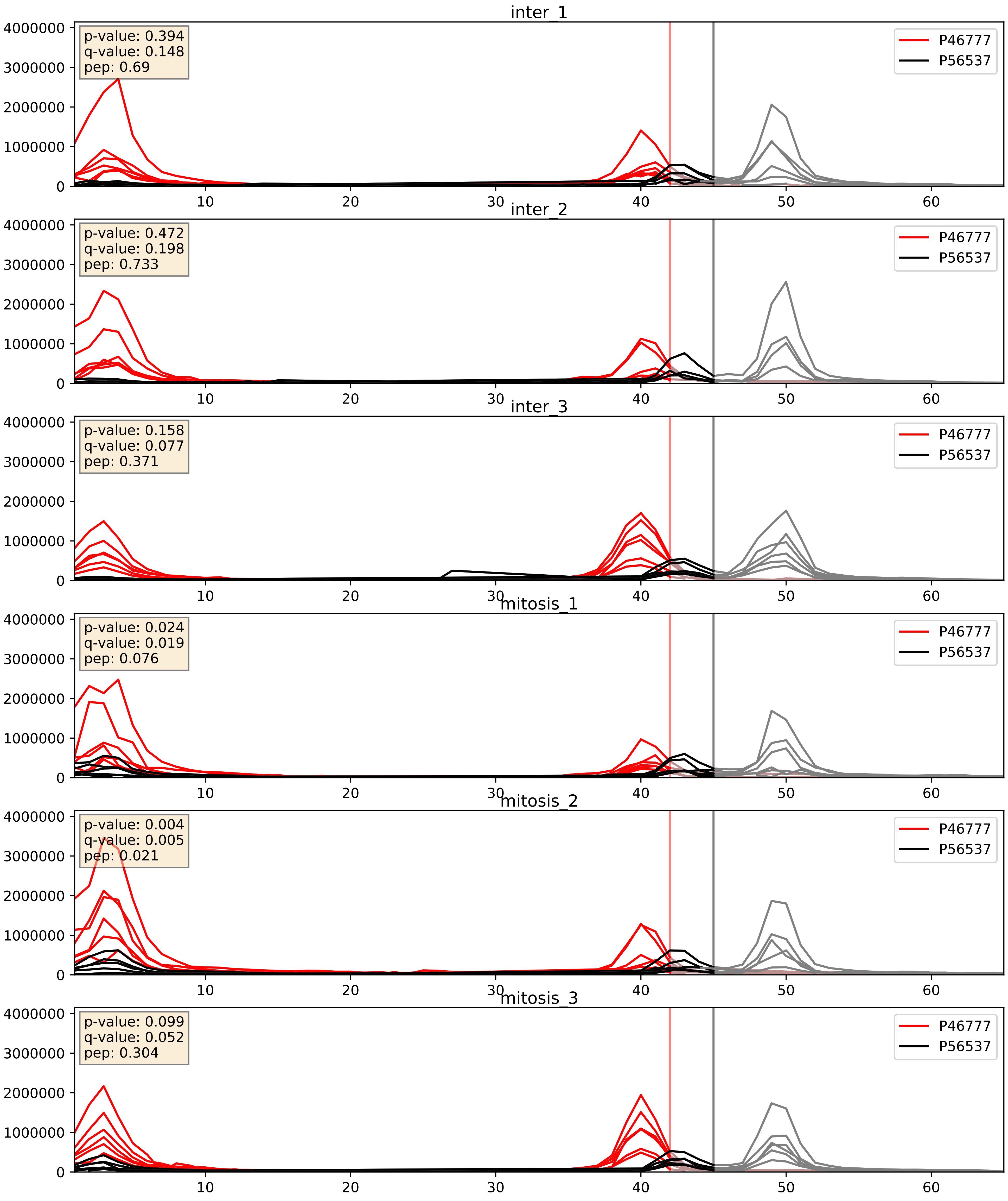


| level | condition_1 | condition_2 | pvalue | pvalue_adjusted |
|---|---|---|---|---|
| complex_abundance | mitosis | inter | 0.010357552994704032 | 0.07507252636296911 |
| interactor_ratio | mitosis | inter | 0.08217105395520377 | 0.48078210653215603 |
| interactor_abundance | mitosis | inter | 0.6738003221176949 | 0.7866655701330614 |

P46778_P56537
RL21_HUMAN vs IF6_HUMAN
p-value: 0.0 q-value: 0.008

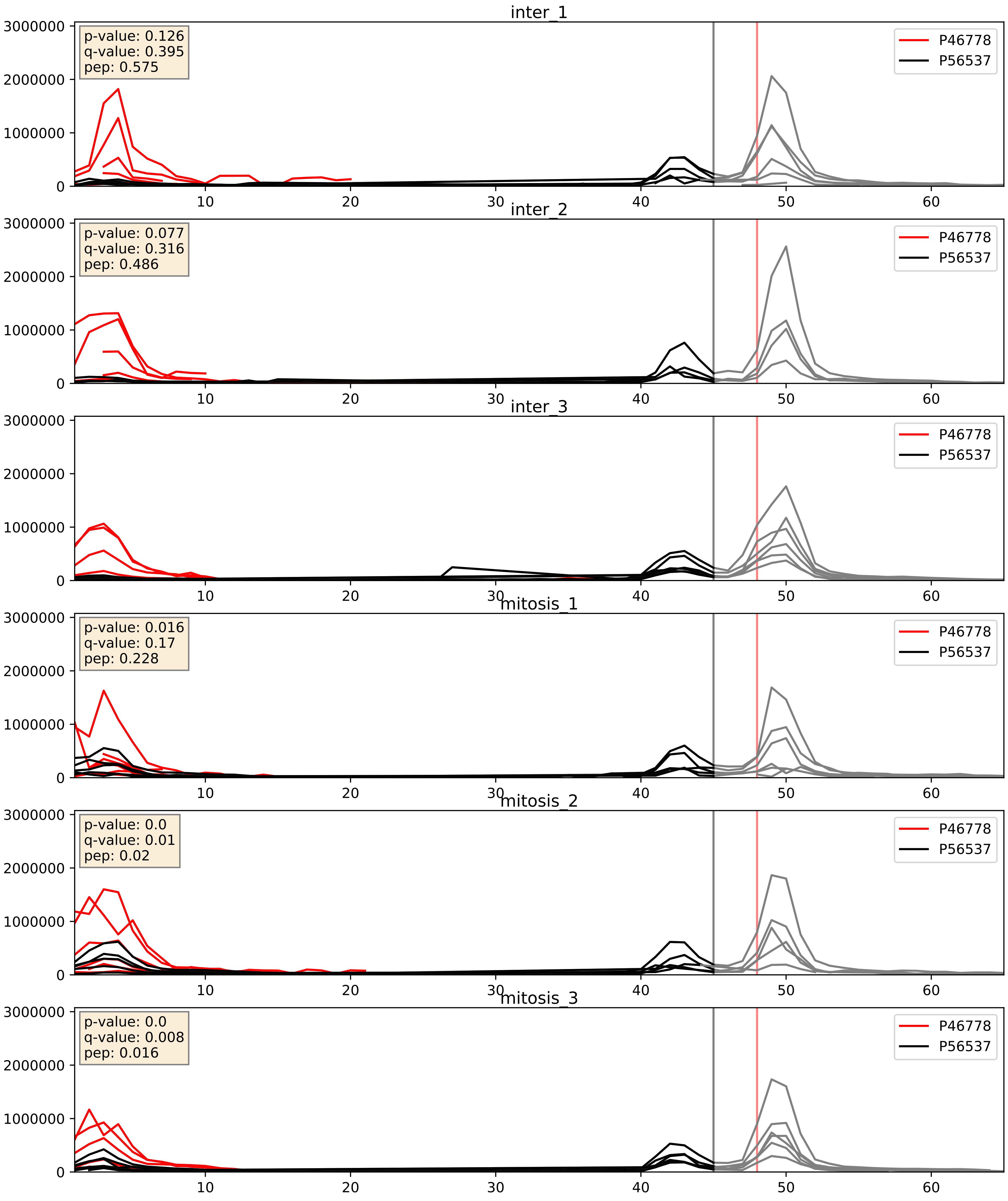


| level | condition_1 | condition_2 | pvalue | pvalue_adjusted |
|---|---|---|---|---|
| interactor_ratio | mitosis | inter | 0.010569061392807683 | 0.14758754571359506 |
| complex_abundance | mitosis | inter | 0.01957886056576645 | 0.10412204938653433 |
| interactor_abundance | mitosis | inter | 0.5575258192658037 | 0.7081794053888232 |

P46779_P56537
RL28_HUMAN vs IF6_HUMAN
p-value: 0.0 q-value: 0.018

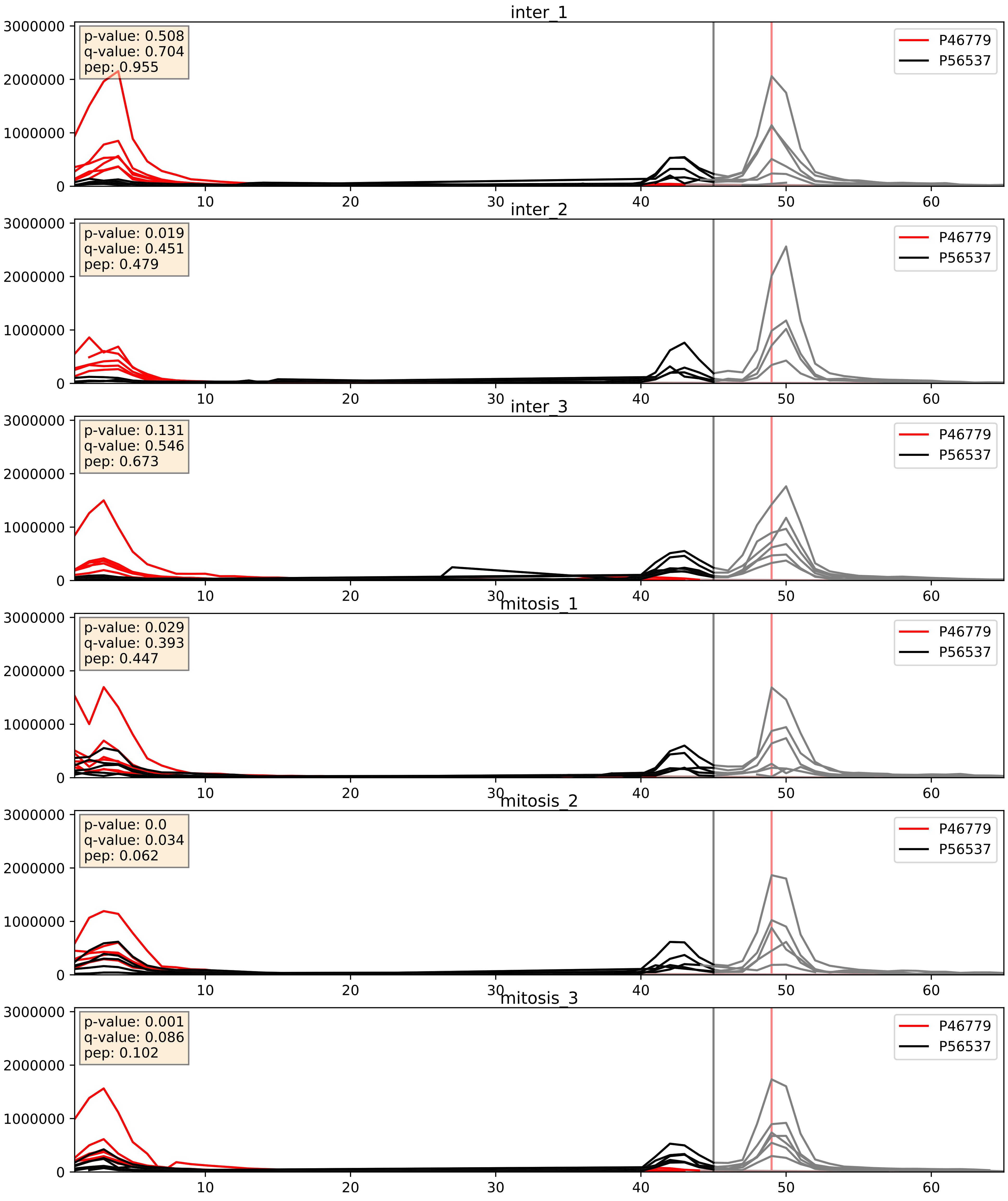


| level | condition_1 | condition_2 | pvalue | pvalue_adjusted |
|---|---|---|---|---|
| interactor_ratio | mitosis | inter | 0.13829234755191047 | 0.5876014454413842 |
| interactor_abundance | mitosis | inter | 0.25273075170977777 | 0.44596479790469956 |
| complex_abundance | mitosis | inter | 0.41675593933220534 | 0.5610932432657562 |

P50914_P56537
RL14_HUMAN vs IF6_HUMAN
p-value: 0.0 q-value: 0.008

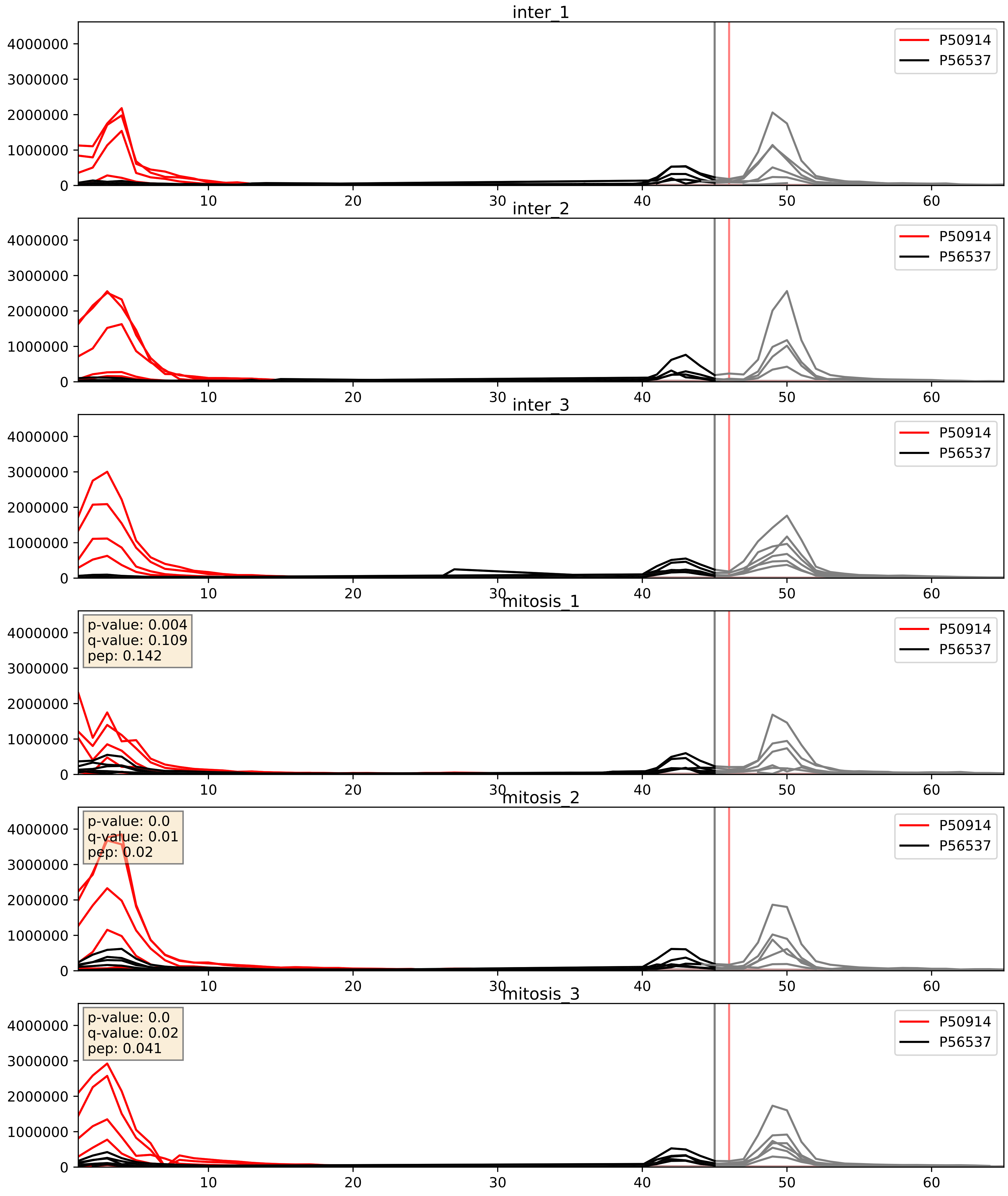


| level | condition_1 | condition_2 | pvalue | pvalue_adjusted |
|---|---|---|---|---|
| complex_abundance | mitosis | inter | 0.021659129890377753 | 0.10835894322713825 |
| interactor_ratio | mitosis | inter | 0.05032663055579263 | 0.37073662440411786 |
| interactor_abundance | mitosis | inter | 0.930853808405876 | 0.960890506798932 |

P55769_P56537
NH2L1_HUMAN vs IF6_HUMAN
p-value: 0.001 q-value: 0.03

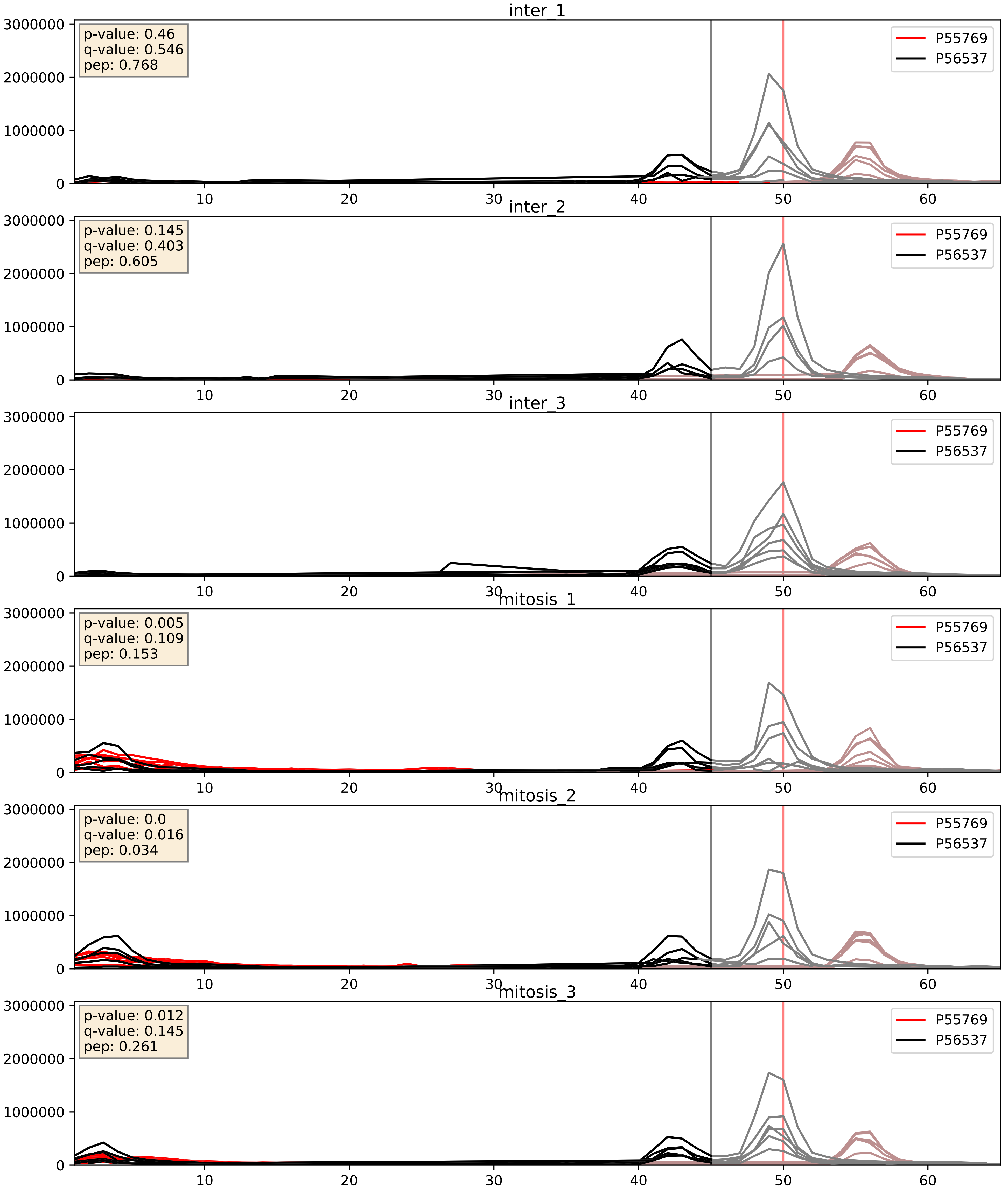


| level | condition_1 | condition_2 | pvalue | pvalue_adjusted |
|---|---|---|---|---|
| interactor_abundance | mitosis | inter | 0.001960763129046175 | 0.032370554261591626 |
| complex_abundance | mitosis | inter | 0.020954510270985917 | 0.10689403306976396 |
| interactor_ratio | mitosis | inter | 0.04772747899799155 | 0.3614455797387484 |

P56537_P61313
IF6_HUMAN vs RL15_HUMAN
p-value: 0.0 q-value: 0.017

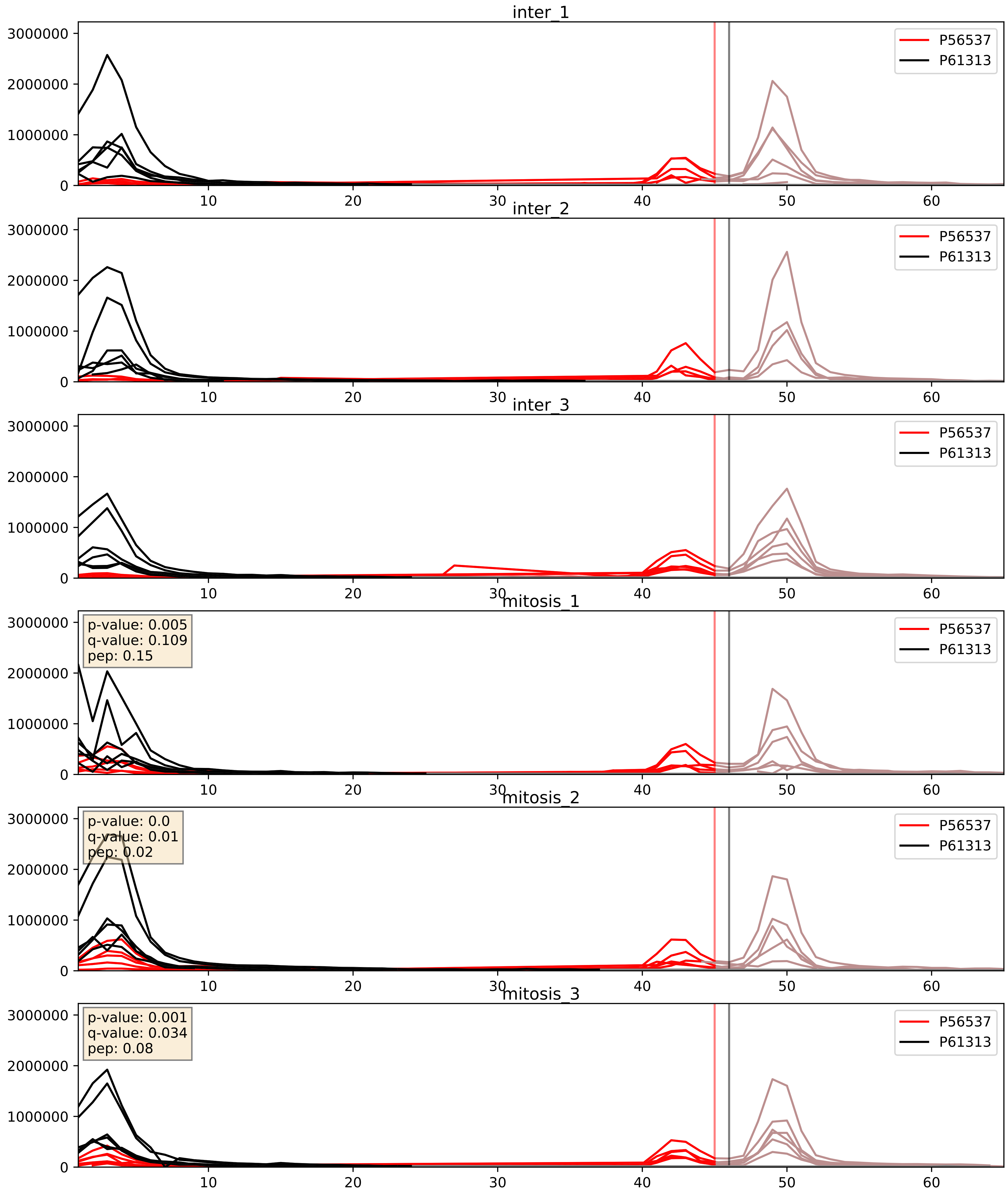


| level | condition_1 | condition_2 | pvalue | pvalue_adjusted |
|---|---|---|---|---|
| interactor_ratio | mitosis | inter | 0.001963936653798777 | 0.05388236460422286 |
| interactor_abundance | mitosis | inter | 0.00640084648112694 | 0.06847945796917453 |
| complex_abundance | mitosis | inter | 0.027013783390175718 | 0.12018606331595849 |

P56537_P61353
IF6_HUMAN vs RL27_HUMAN
p-value: 0.0 q-value: 0.016

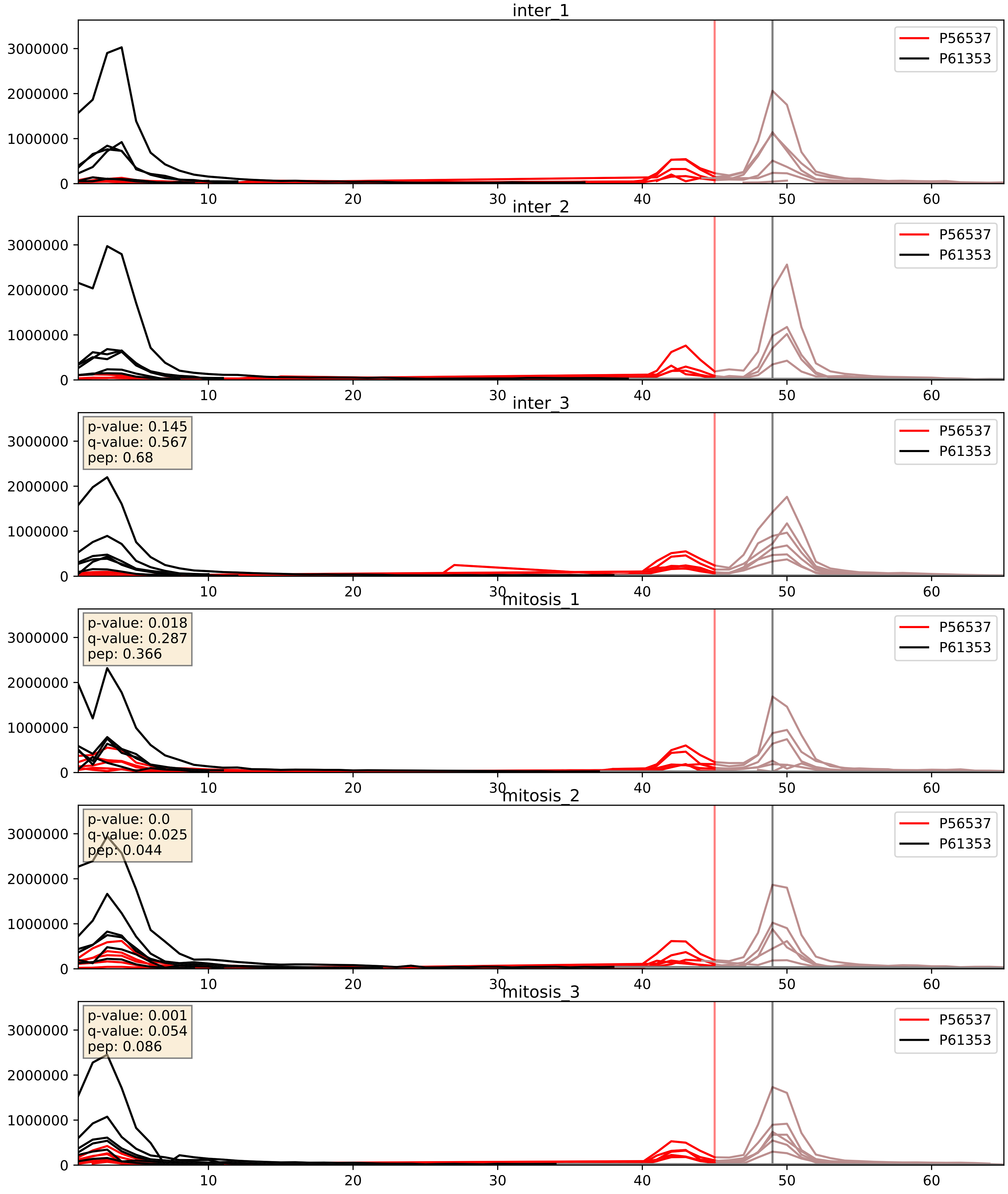


| level | condition_1 | condition_2 | pvalue | pvalue_adjusted |
|---|---|---|---|---|
| interactor_abundance | mitosis | inter | 0.004292299499120139 | 0.05222755325155421 |
| interactor_ratio | mitosis | inter | 0.005174572490593194 | 0.09671253388532258 |
| complex_abundance | mitosis | inter | 0.013303836127421224 | 0.08556035856553394 |

P56537_P62424
IF6_HUMAN vs RL7A_HUMAN
p-value: 0.0 q-value: 0.017

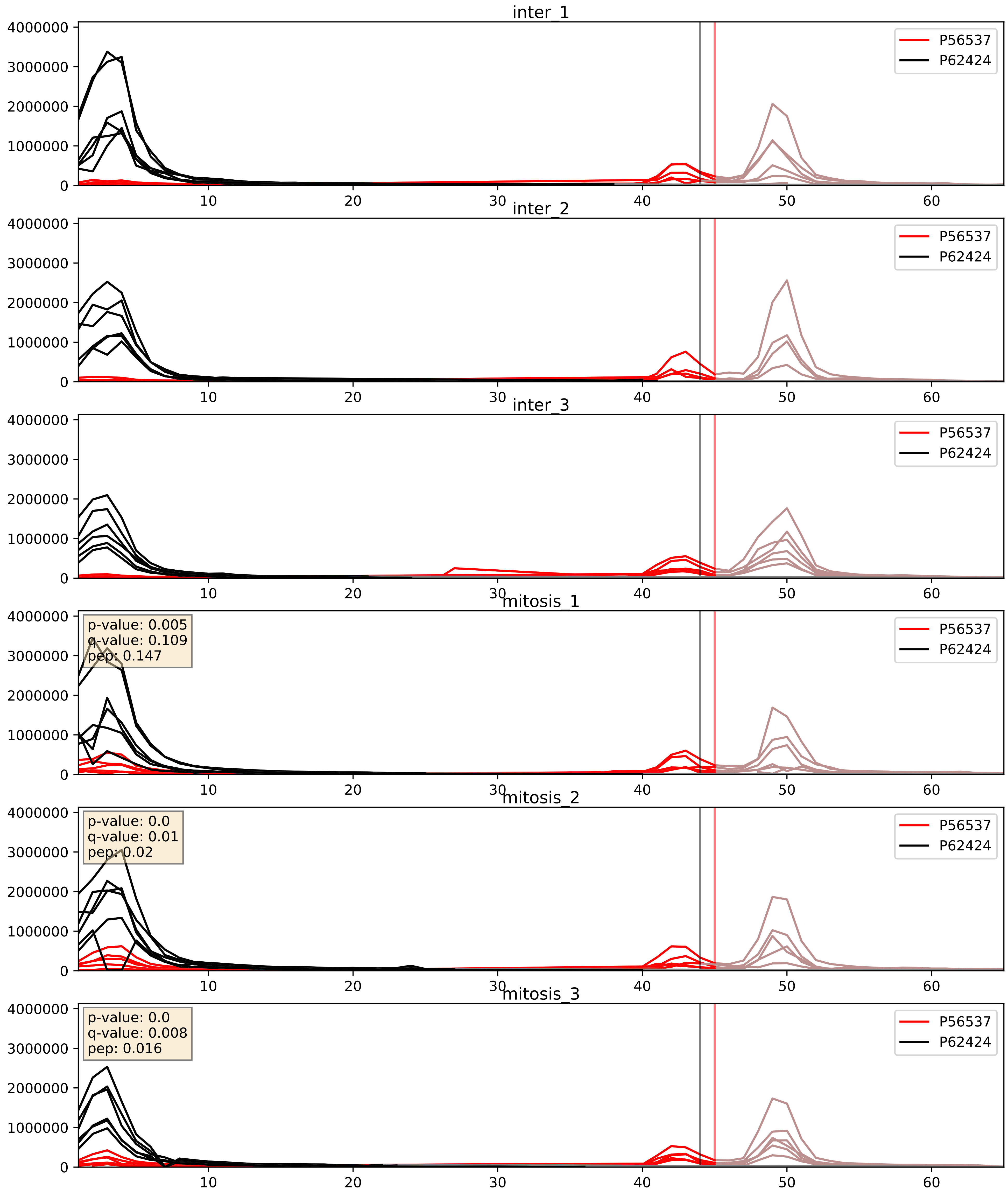


| level | condition_1 | condition_2 | pvalue | pvalue_adjusted |
|---|---|---|---|---|
| interactor_abundance | mitosis | inter | 0.011802011355852736 | 0.0959859545901182 |
| interactor_ratio | mitosis | inter | 0.01319828471635139 | 0.16610126135854406 |
| complex_abundance | mitosis | inter | 0.05340417648424683 | 0.16497650946897674 |

P56537_P62888
IF6_HUMAN vs RL30_HUMAN
p-value: 0.0 q-value: 0.008

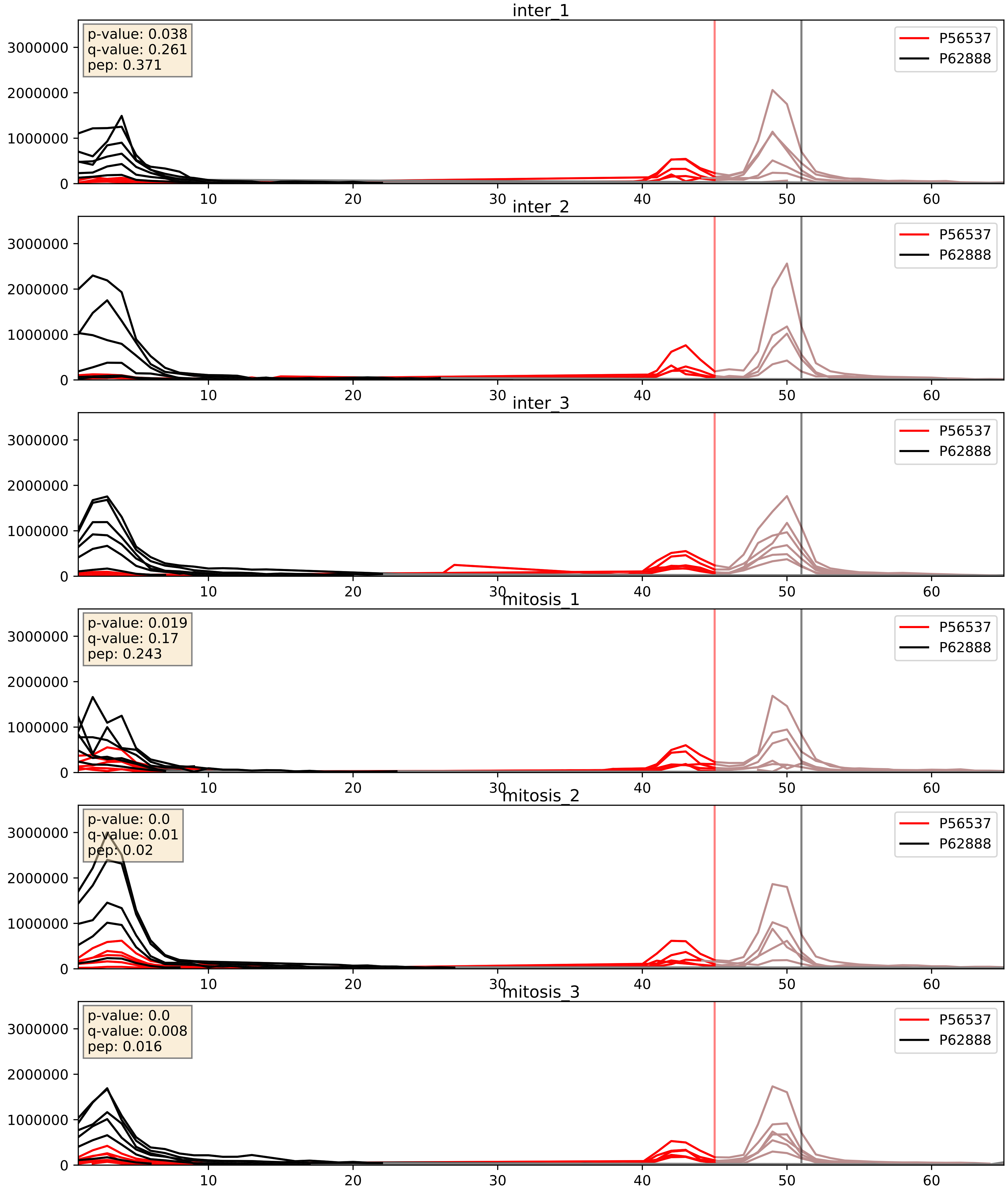


| level | condition_1 | condition_2 | pvalue | pvalue_adjusted |
|---|---|---|---|---|
| interactor_abundance | mitosis | inter | 0.0035466111482954548 | 0.04601489785035819 |
| complex_abundance | mitosis | inter | 0.044301904917346376 | 0.15060165151763358 |
| interactor_ratio | mitosis | inter | 0.046665667966653945 | 0.35838626305591875 |

P56537_P62906
IF6_HUMAN vs RL10A_HUMAN
p-value: 0.0 q-value: 0.014

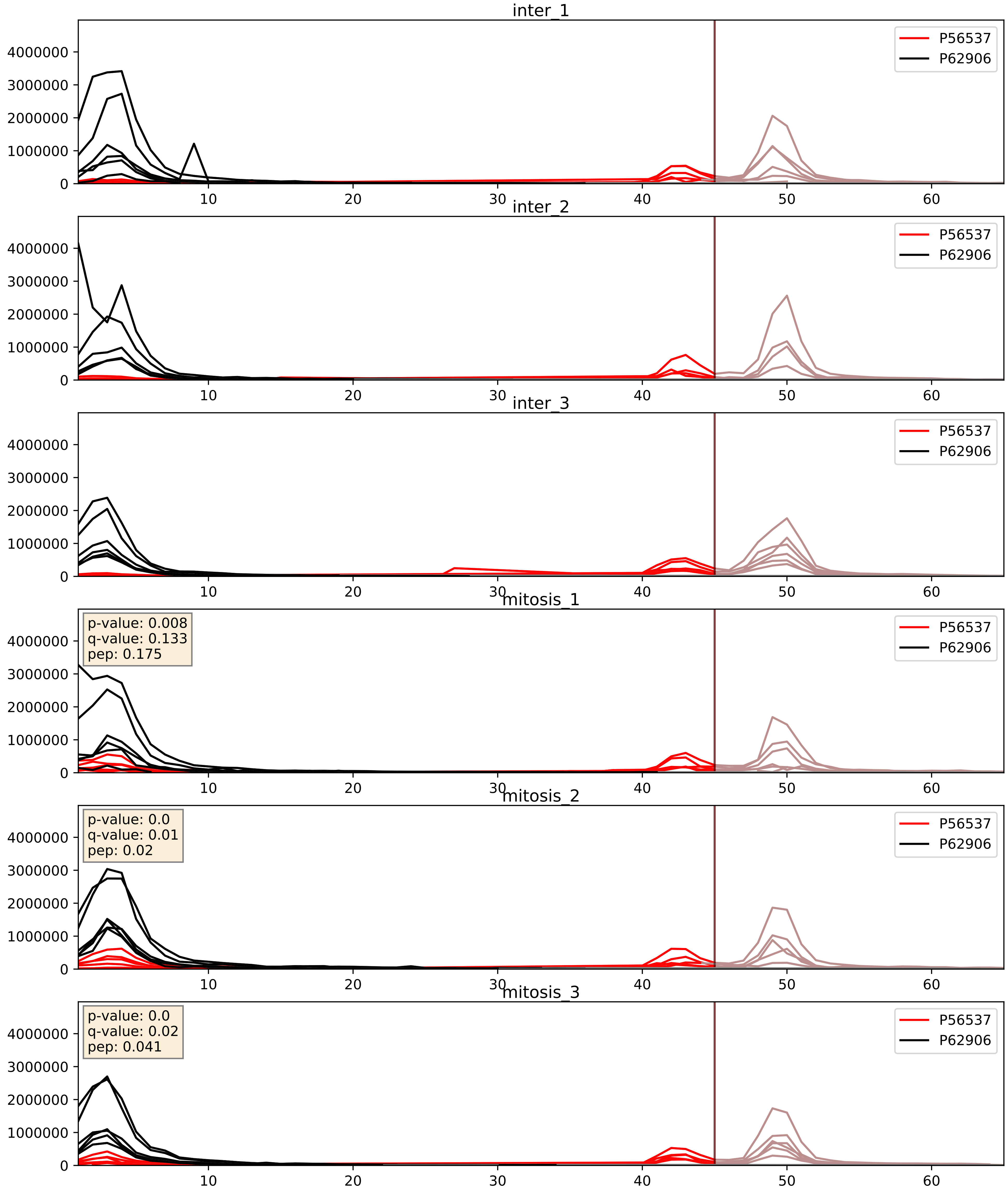


| level | condition_1 | condition_2 | pvalue | pvalue_adjusted |
|---|---|---|---|---|
| interactor_abundance | mitosis | inter | 0.004768879721580431 | 0.05653962661596744 |
| interactor_ratio | mitosis | inter | 0.007006627641535918 | 0.1166862502170184 |
| complex_abundance | mitosis | inter | 0.011403424595661052 | 0.07923158647634627 |

P56537_P62910
IF6_HUMAN vs RL32_HUMAN
p-value: 0.0 q-value: 0.008

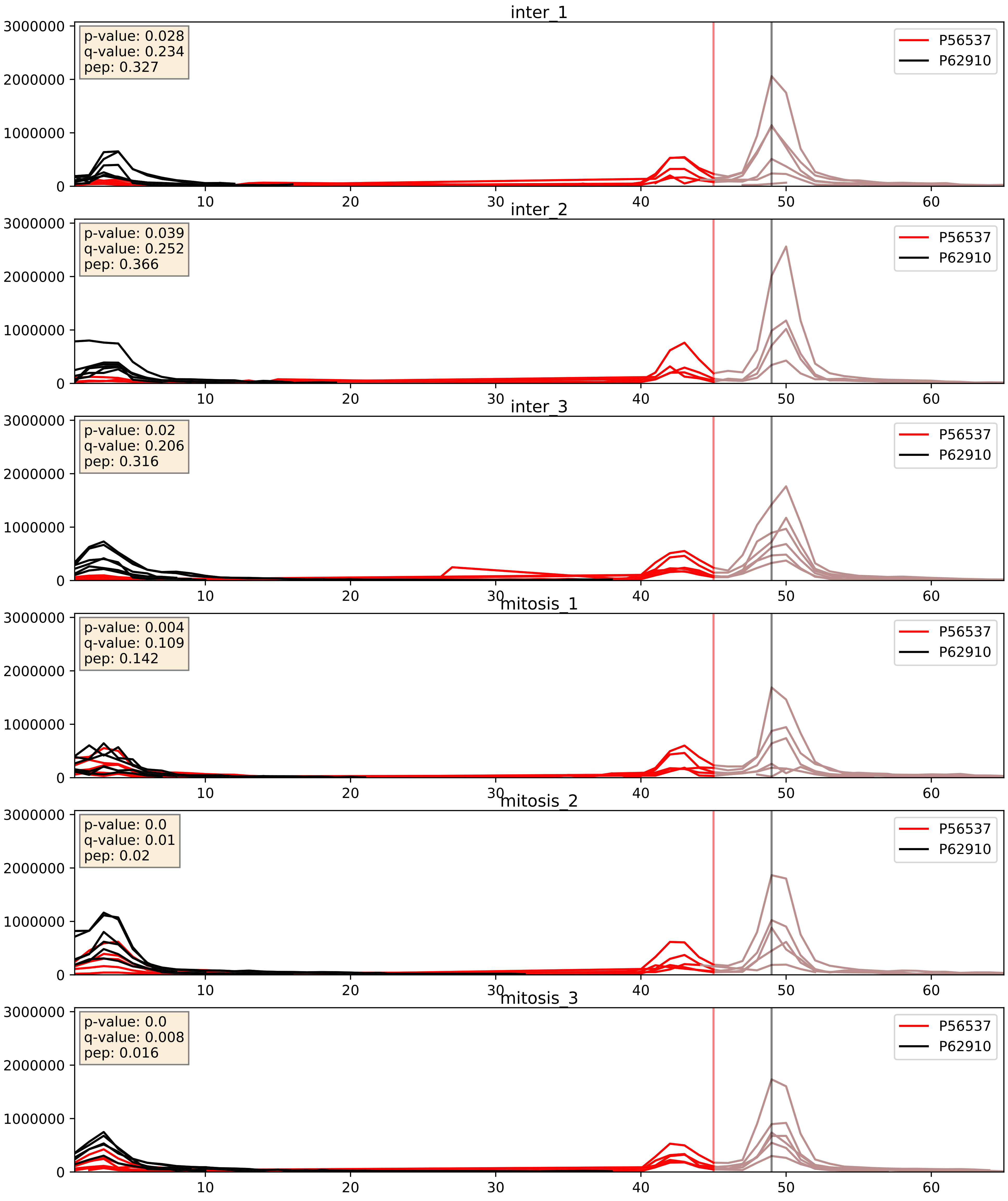


| level | condition_1 | condition_2 | pvalue | pvalue_adjusted |
|---|---|---|---|---|
| interactor_abundance | mitosis | inter | 0.004083792706615565 | 0.050736234497284816 |
| complex_abundance | mitosis | inter | 0.007170048891153688 | 0.06275154604611013 |
| interactor_ratio | mitosis | inter | 0.0436227532311622 | 0.3444748779139746 |

P56537_Q02543
IF6_HUMAN vs RL18A_HUMAN
p-value: 0.0 q-value: 0.003

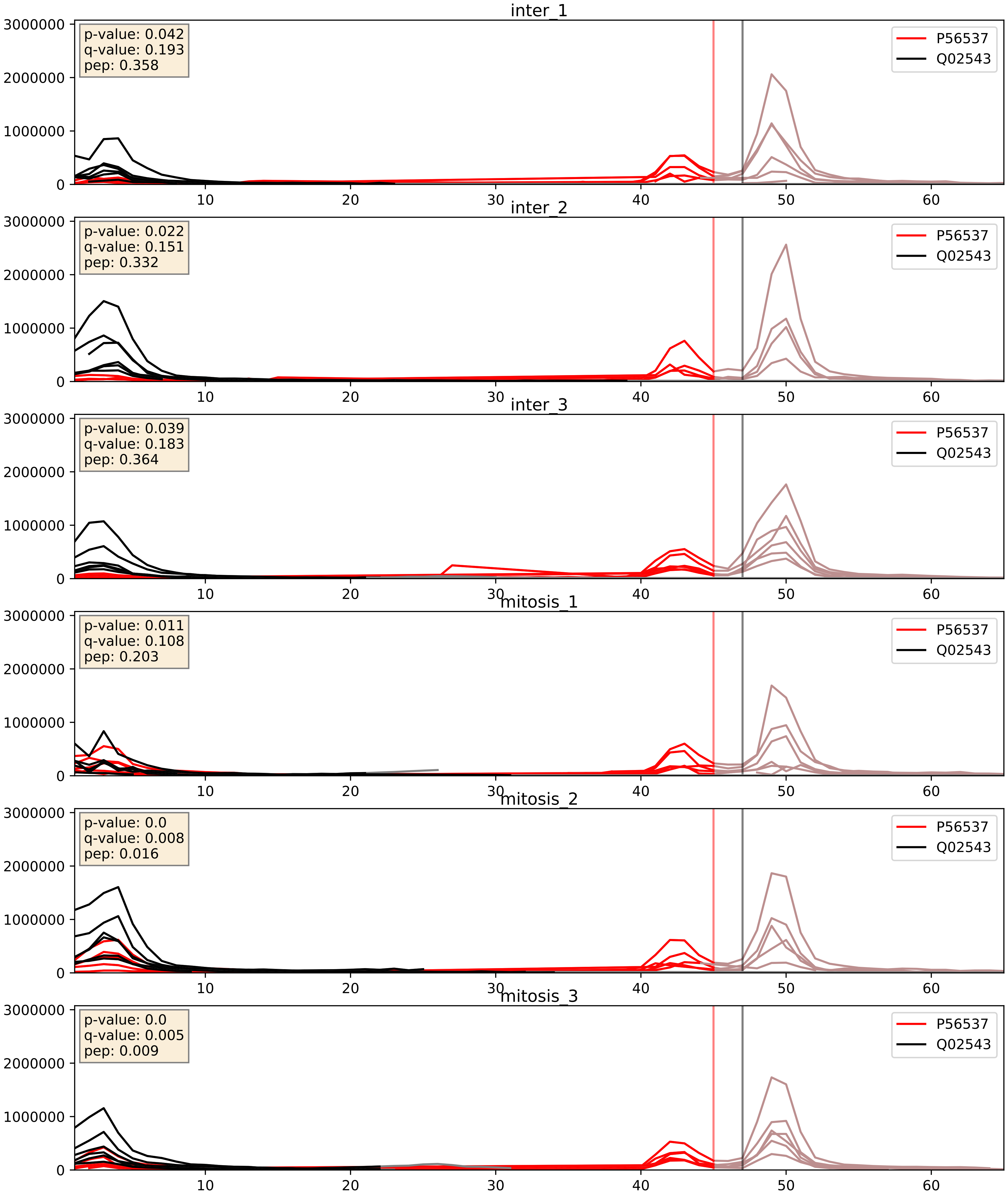


| level | condition_1 | condition_2 | pvalue | pvalue_adjusted |
|---|---|---|---|---|
| interactor_abundance | mitosis | inter | 0.006979780935512527 | 0.07224537461667138 |
| interactor_ratio | mitosis | inter | 0.05112928813464553 | 0.37414961988671497 |
| complex_abundance | mitosis | inter | 0.131756932062296 | 0.2755532221972279 |

P56537_Q07020
IF6_HUMAN vs RL18_HUMAN
p-value: 0.001 q-value: 0.039

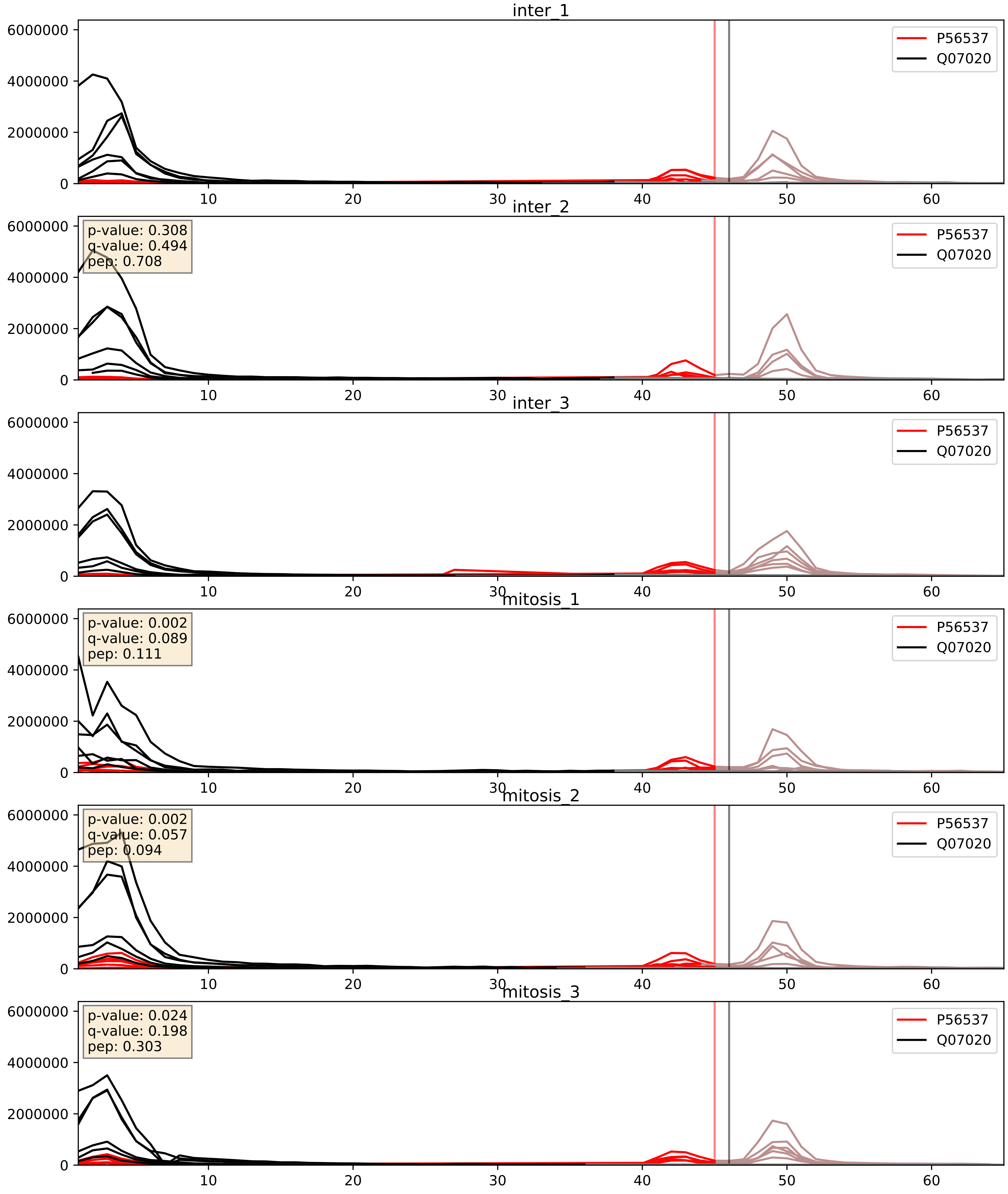


| level | condition_1 | condition_2 | pvalue | pvalue_adjusted |
|---|---|---|---|---|
| interactor_ratio | mitosis | inter | 0.09296681379441395 | 0.5070243509919888 |
| interactor_abundance | mitosis | inter | 0.11396154735529197 | 0.29128421778480124 |
| complex_abundance | mitosis | inter | 0.2392466528158904 | 0.39127843869010737 |

P56537_Q9BQ67
IF6_HUMAN vs GRWD1_HUMAN
p-value: 0.0 q-value: 0.046

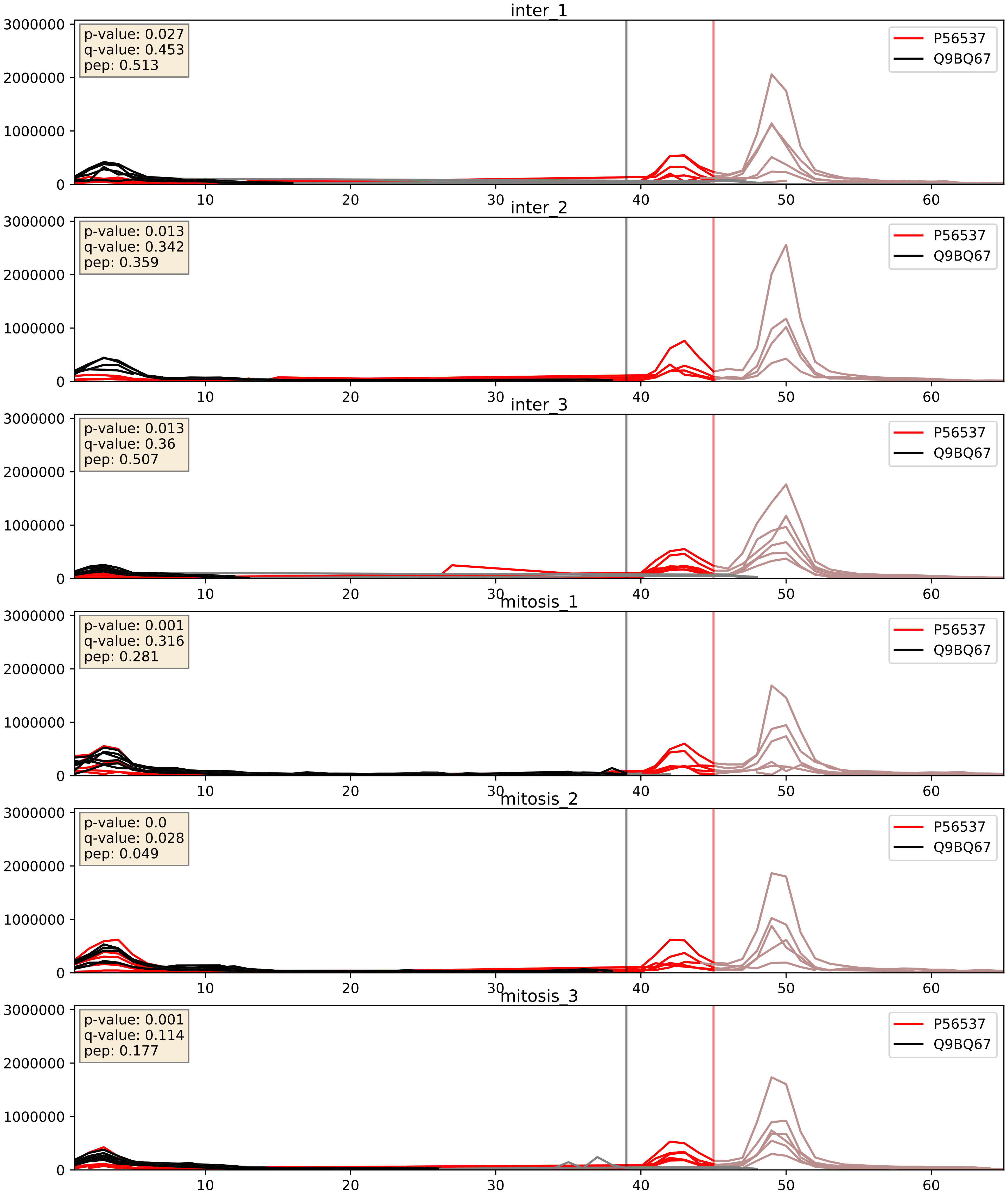


| level | condition_1 | condition_2 | pvalue | pvalue_adjusted |
|---|---|---|---|---|
| interactor_ratio | mitosis | inter | 0.0052956920235738775 | 0.0974780445871482 |
| interactor_abundance | mitosis | inter | 0.007702330616543858 | 0.07617787415091325 |
| complex_abundance | mitosis | inter | 0.03073328011425226 | 0.12751204260889337 |

P56537_Q9BZE4
IF6_HUMAN vs NOG1_HUMAN
p-value: 0.002 q-value: 0.036

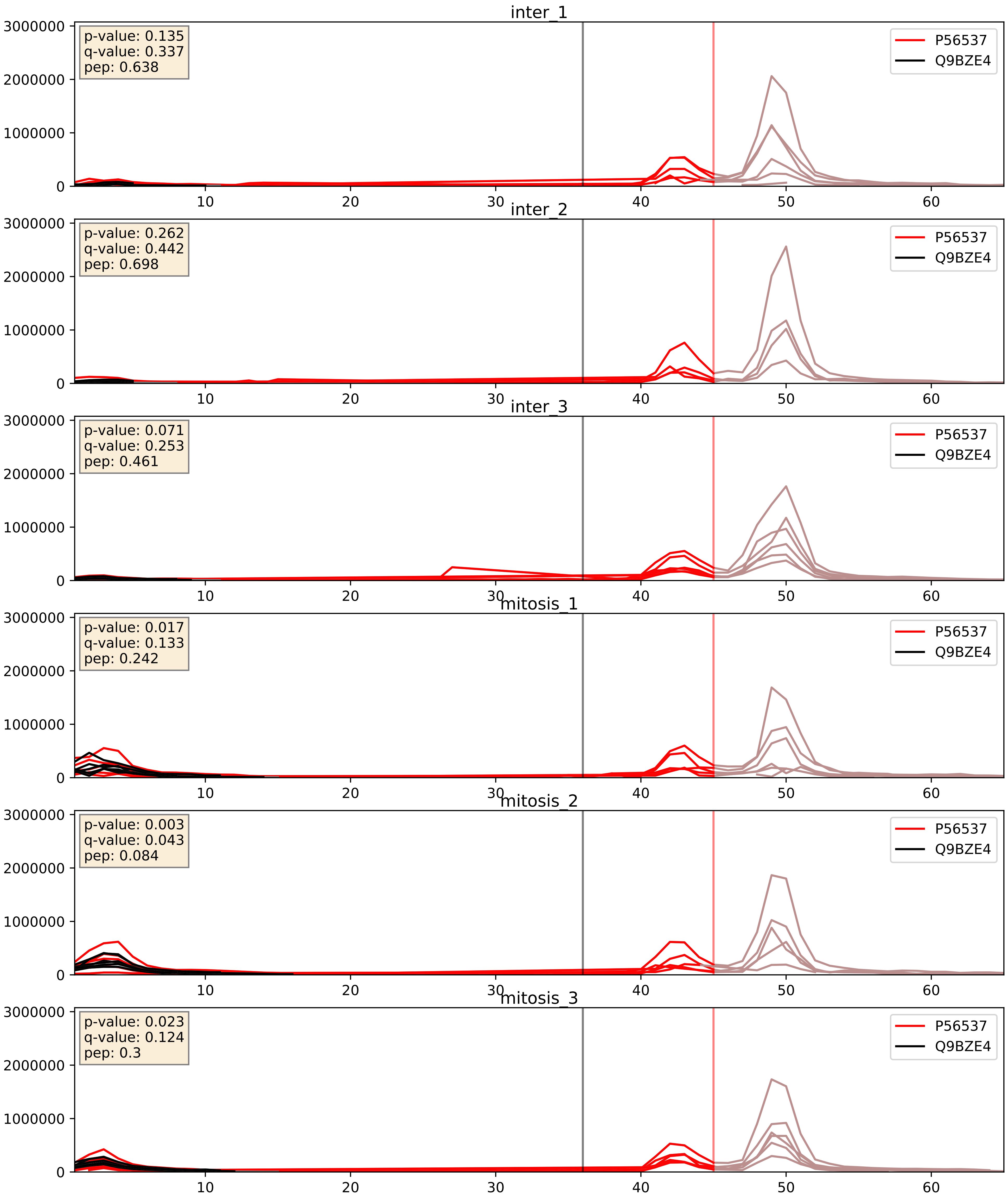


| level | condition_1 | condition_2 | pvalue | pvalue_adjusted |
|---|---|---|---|---|
| complex_abundance | mitosis | inter | 0.0021330438577376568 | 0.033489340249041655 |
| interactor_abundance | mitosis | inter | 0.004459234357215569 | 0.05368749863632952 |
| interactor_ratio | mitosis | inter | 0.17659636703522258 | 0.6420034343641515 |

P56537_Q9NX24
IF6_HUMAN vs NHP2_HUMAN
p-value: 0.0 q-value: 0.041

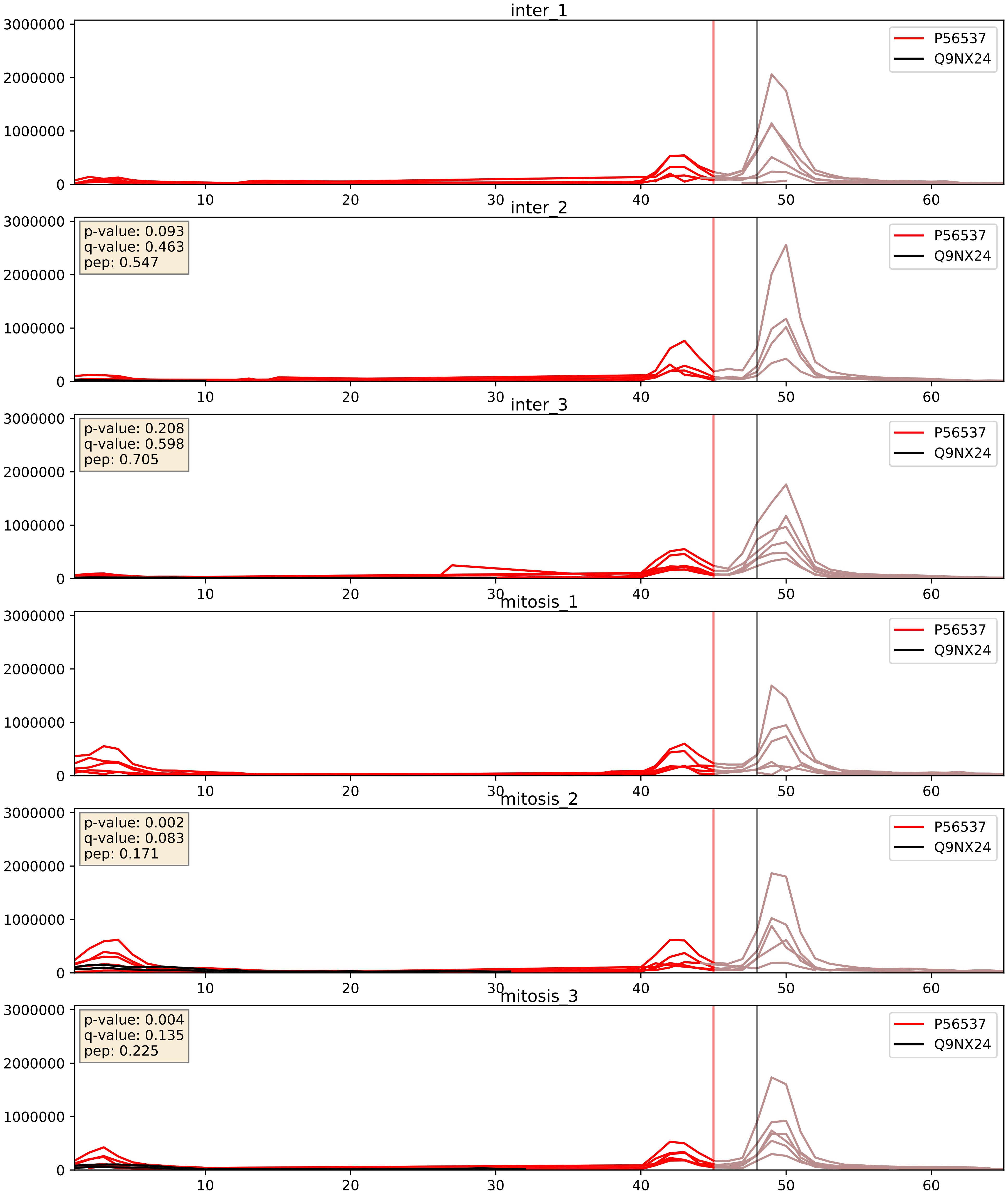


| level | condition_1 | condition_2 | pvalue | pvalue_adjusted |
|---|---|---|---|---|
| complex_abundance | mitosis | inter | 0.4559467124206309 | 0.5978713018260724 |
| interactor_abundance | mitosis | inter | 0.5163408382462942 | 0.6792496658042536 |
| interactor_ratio | mitosis | inter | 0.6981465253616923 | 0.9359699707113329 |

P56537_Q9UKD2
IF6_HUMAN vs MRT4_HUMAN
p-value: 0.002 q-value: 0.035

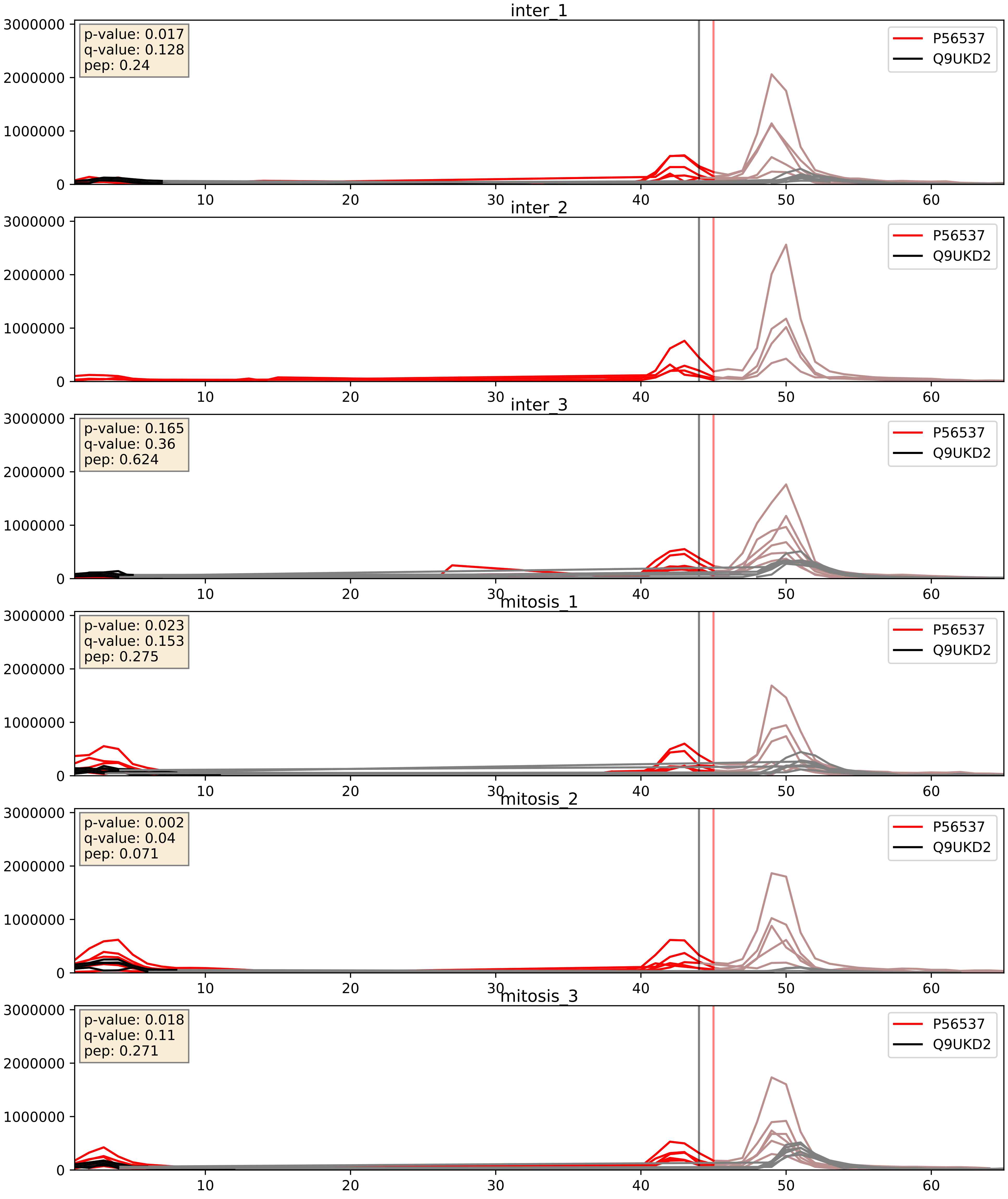


| level | condition_1 | condition_2 | pvalue | pvalue_adjusted |
|---|---|---|---|---|
| interactor_ratio | mitosis | inter | 0.0007143459035063286 | 0.03088283300007158 |
| interactor_abundance | mitosis | inter | 0.025852706349435345 | 0.13331275081395577 |
| complex_abundance | mitosis | inter | 0.060938893075678645 | 0.17730690847308267 |

P56537_Q9Y221
IF6_HUMAN vs NIP7_HUMAN
p-value: 0.004 q-value: 0.004

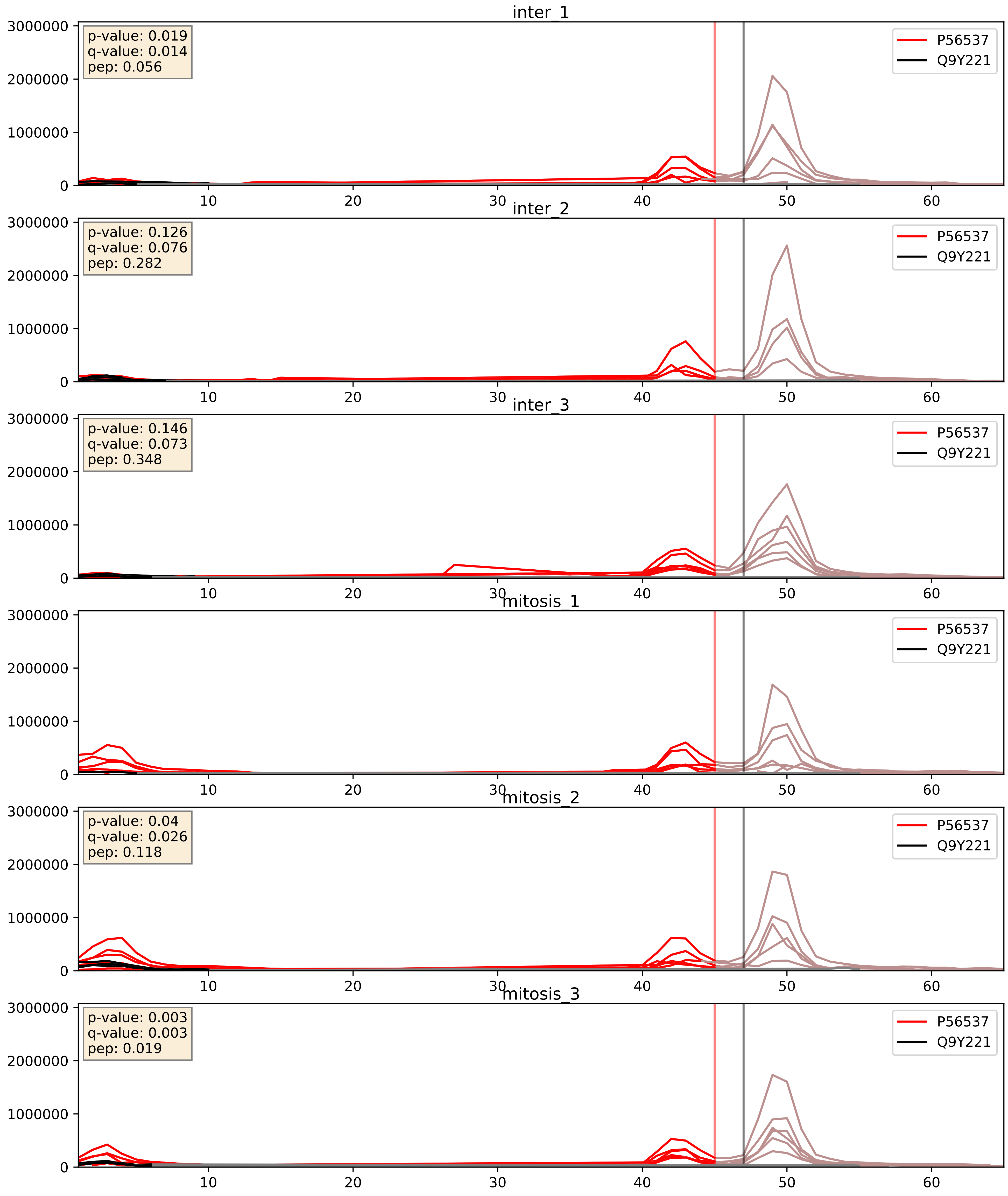


| level | condition_1 | condition_2 | pvalue | pvalue_adjusted |
|---|---|---|---|---|
| interactor_abundance | mitosis | inter | 0.009435635913037536 | 0.08417826306993362 |
| interactor_ratio | mitosis | inter | 0.017423488439053336 | 0.19780512074044637 |
| complex_abundance | mitosis | inter | 0.19616618706993139 | 0.3486286077123025 |

P56537_Q9Y2X3
IF6_HUMAN vs NOP58_HUMAN
p-value: 0.002 q-value: 0.03

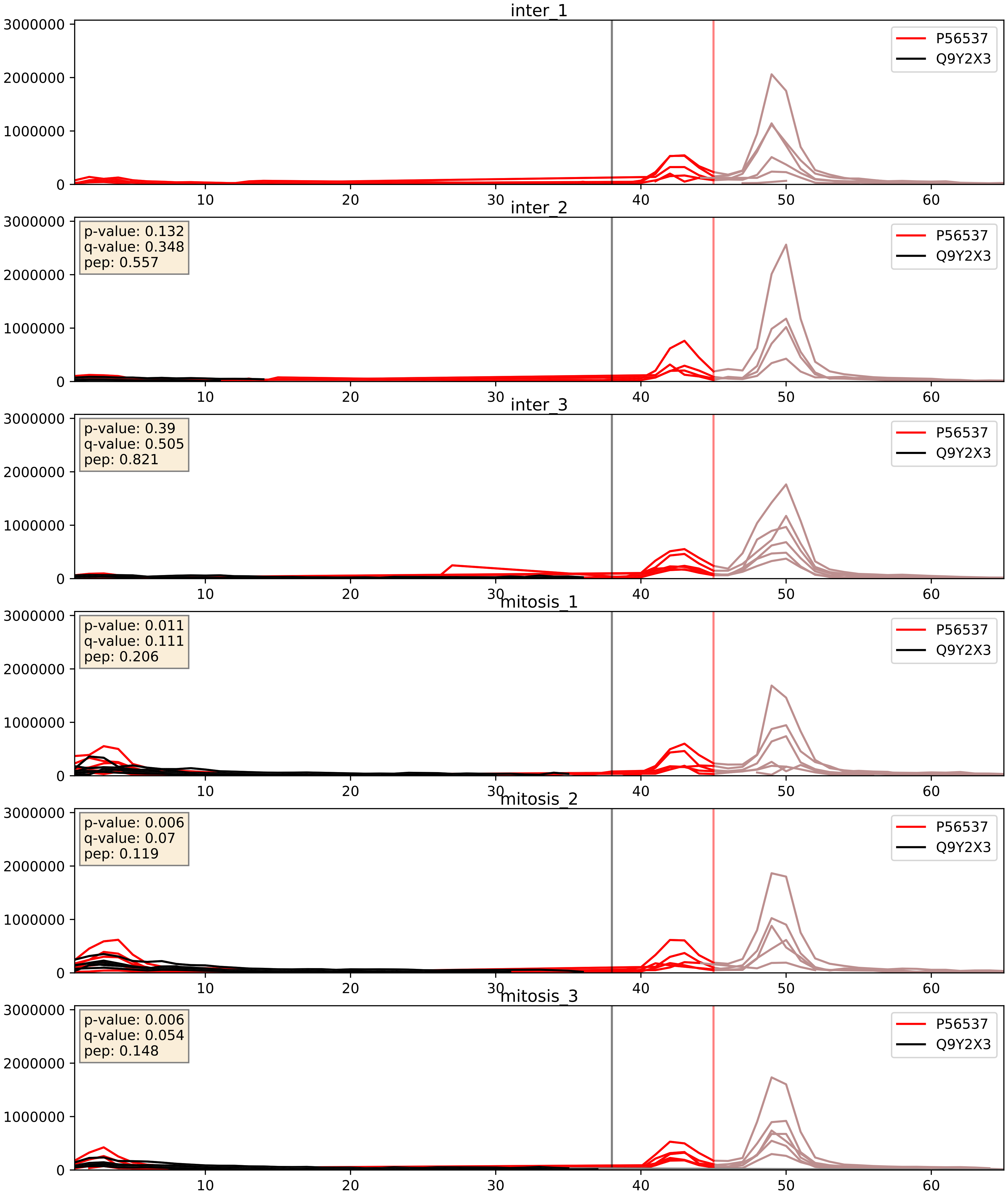


| level | condition_1 | condition_2 | pvalue | pvalue_adjusted |
|---|---|---|---|---|
| complex_abundance | mitosis | inter | 0.008291925197678492 | 0.06742444735376033 |
| interactor_abundance | mitosis | inter | 0.018773691505101508 | 0.11803363884220999 |
| interactor_ratio | mitosis | inter | 0.2376377875942226 | 0.7165126670681738 |

P56537_Q9Y3U8
IF6_HUMAN vs RL36_HUMAN
p-value: 0.0 q-value: 0.008

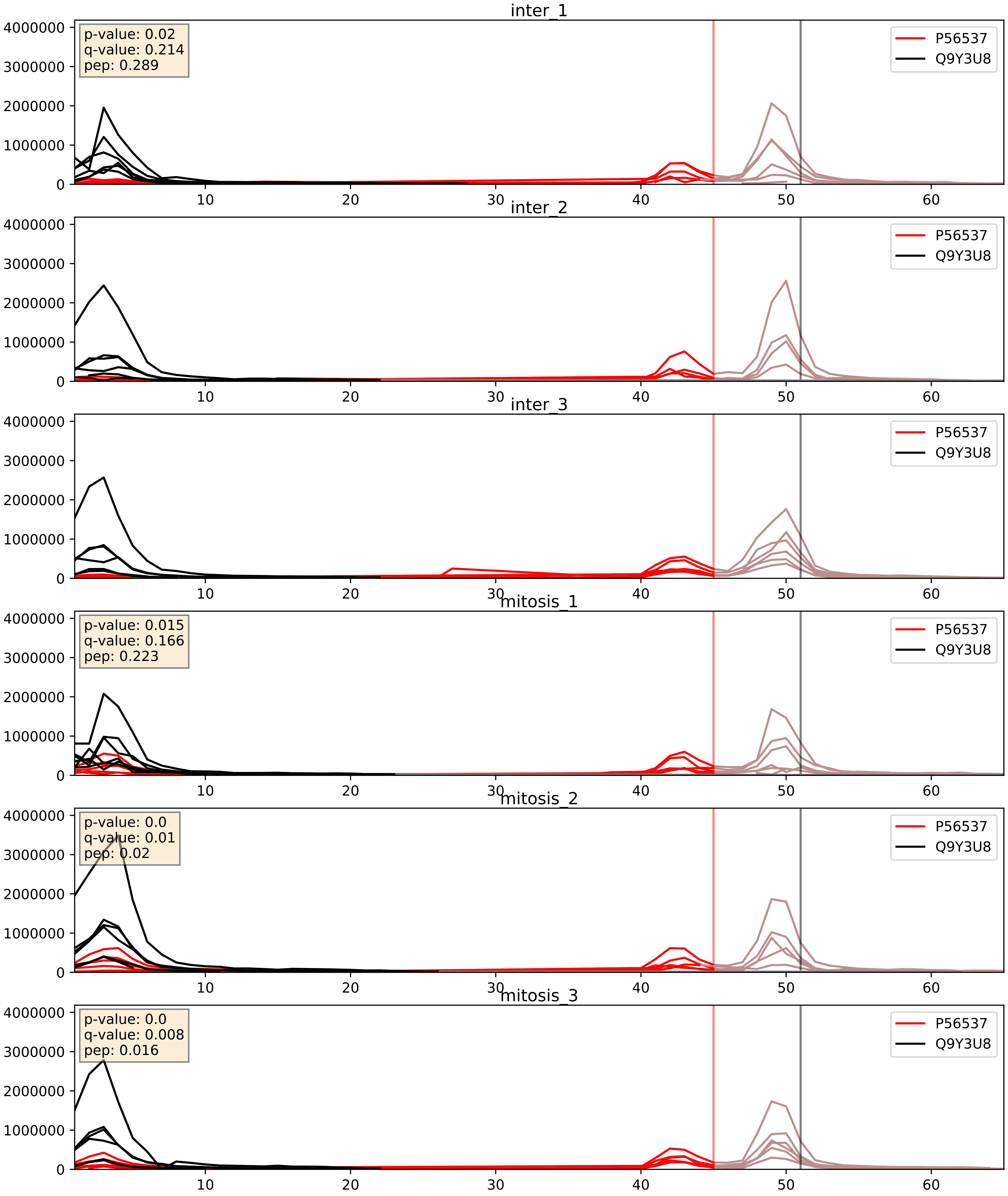


| level | condition_1 | condition_2 | pvalue | pvalue_adjusted |
|---|---|---|---|---|
| complex_abundance | mitosis | inter | 0.0025874317613308133 | 0.03664546283589145 |
| interactor_abundance | mitosis | inter | 0.005818635892822735 | 0.06462369741607146 |
| interactor_ratio | mitosis | inter | 0.02003757512968488 | 0.21413438590524667 |
