## Supplementary Data 2 for "SECAT: Quantifying differential protein-protein interaction states by network-centric analysis": hela_string_000004_Q9BZE4.pdf

Q9BZE4_Q9BZE4

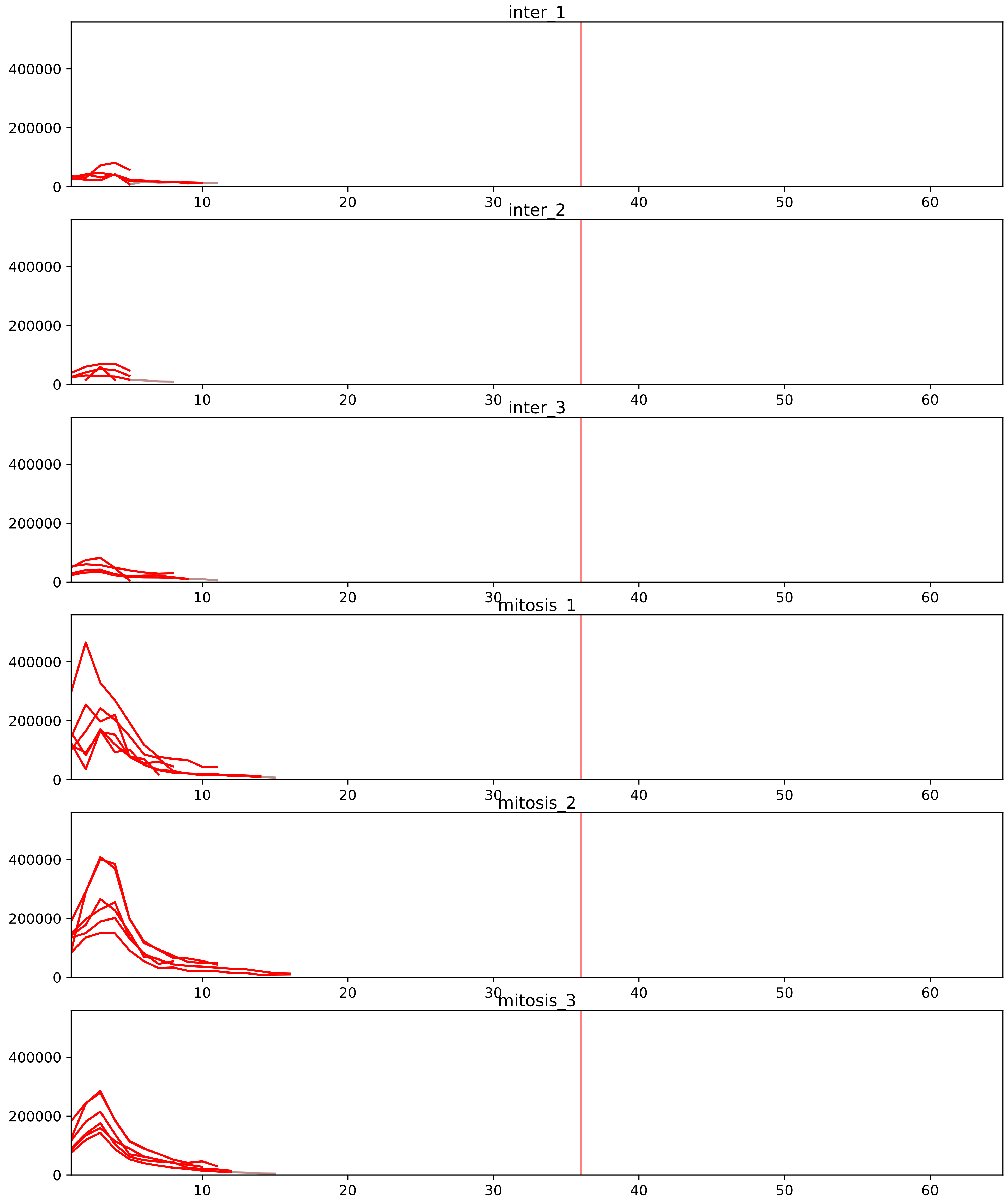


| level | condition_1 | condition_2 | pvalue | pvalue_adjusted |
|---|---|---|---|---|
| interactor_ratio | mitosis | inter | 1.9340128453493062e-20 | 8.89645908860681e-18 |
| interactor_abundance | mitosis | inter | 4.741416738217906e-06 | 0.000300834717183481 |
| complex_abundance | mitosis | inter | 0.00010643045779761653 | 0.002323510983773901 |
| assembled_abundance | mitosis | inter | 0.00039517783776387815 | 0.018876015153929147 |
| total_abundance | mitosis | inter | 0.0006086975537824142 | 0.028961488283236175 |

P46087_Q9BZE4
NOP2_HUMAN vs NOG1_HUMAN
p-value: 0.004 q-value: 0.004

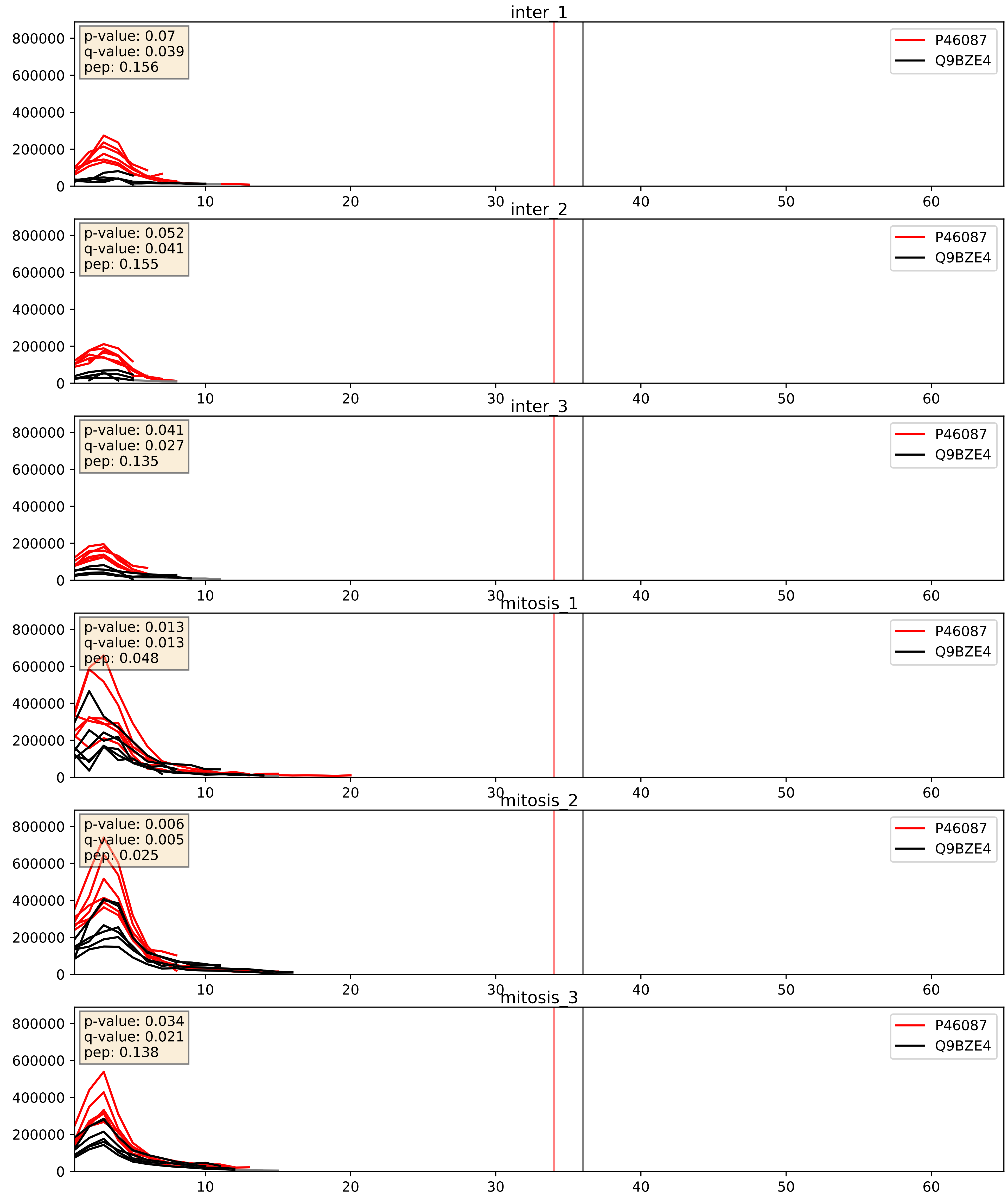


| level | condition_1 | condition_2 | pvalue | pvalue_adjusted |
|---|---|---|---|---|
| complex_abundance | mitosis | inter | 0.00198928380982054 | 0.03207280622328027 |
| interactor_ratio | mitosis | inter | 0.004188742047813209 | 0.08517183734867877 |
| interactor_abundance | mitosis | inter | 0.004516185423873405 | 0.05406789822147741 |

P56537_Q9BZE4
IF6_HUMAN vs NOG1_HUMAN
p-value: 0.002 q-value: 0.036

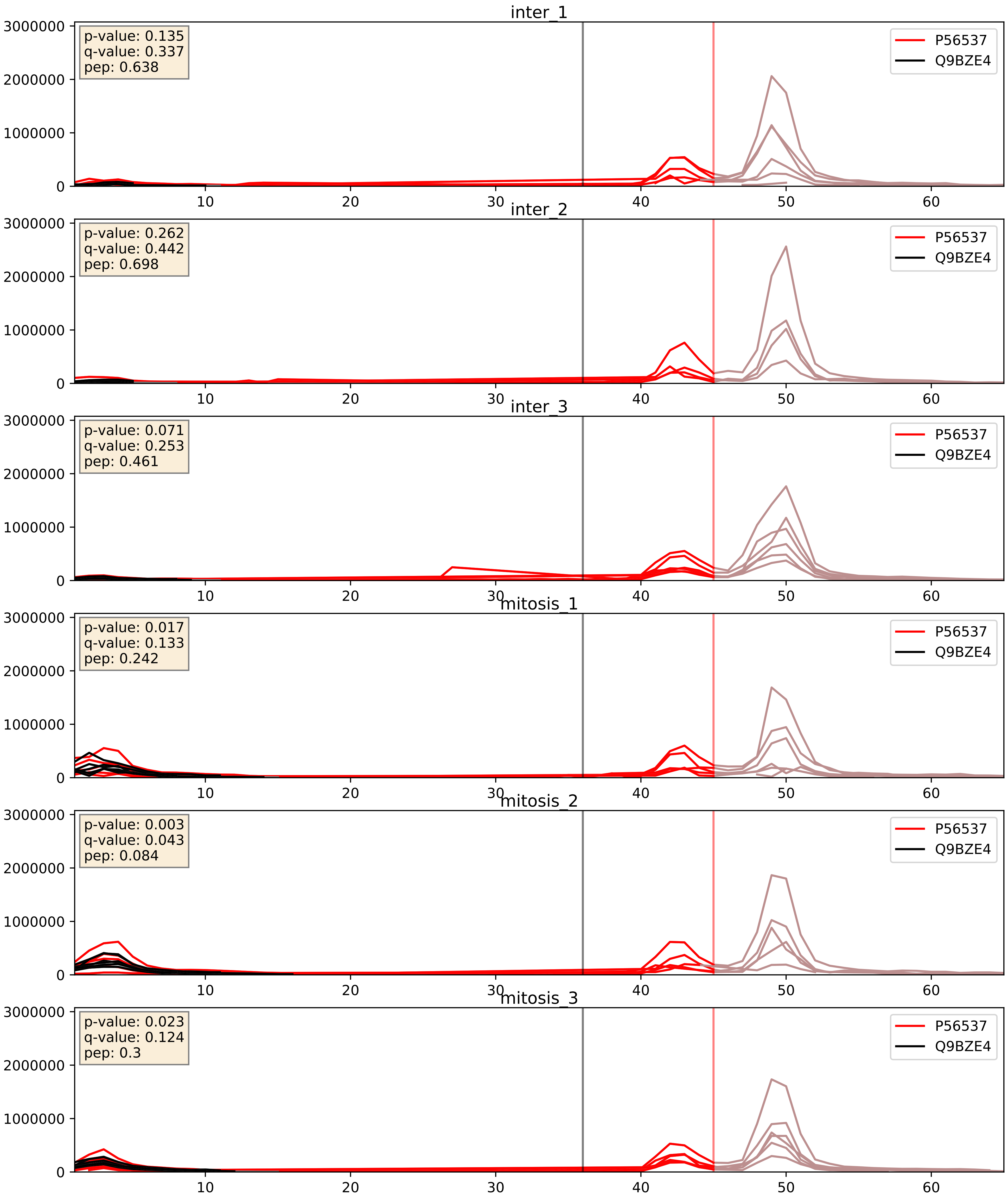


| level | condition_1 | condition_2 | pvalue | pvalue_adjusted |
|---|---|---|---|---|
| complex_abundance | mitosis | inter | 0.0021330438577376568 | 0.033489340249041655 |
| interactor_abundance | mitosis | inter | 0.004459234357215569 | 0.05368749863632952 |
| interactor_ratio | mitosis | inter | 0.17659636703522258 | 0.6420034343641515 |

P62906_Q9BZE4
RL10A_HUMAN vs NOG1_HUMAN
p-value: 0.001 q-value: 0.041

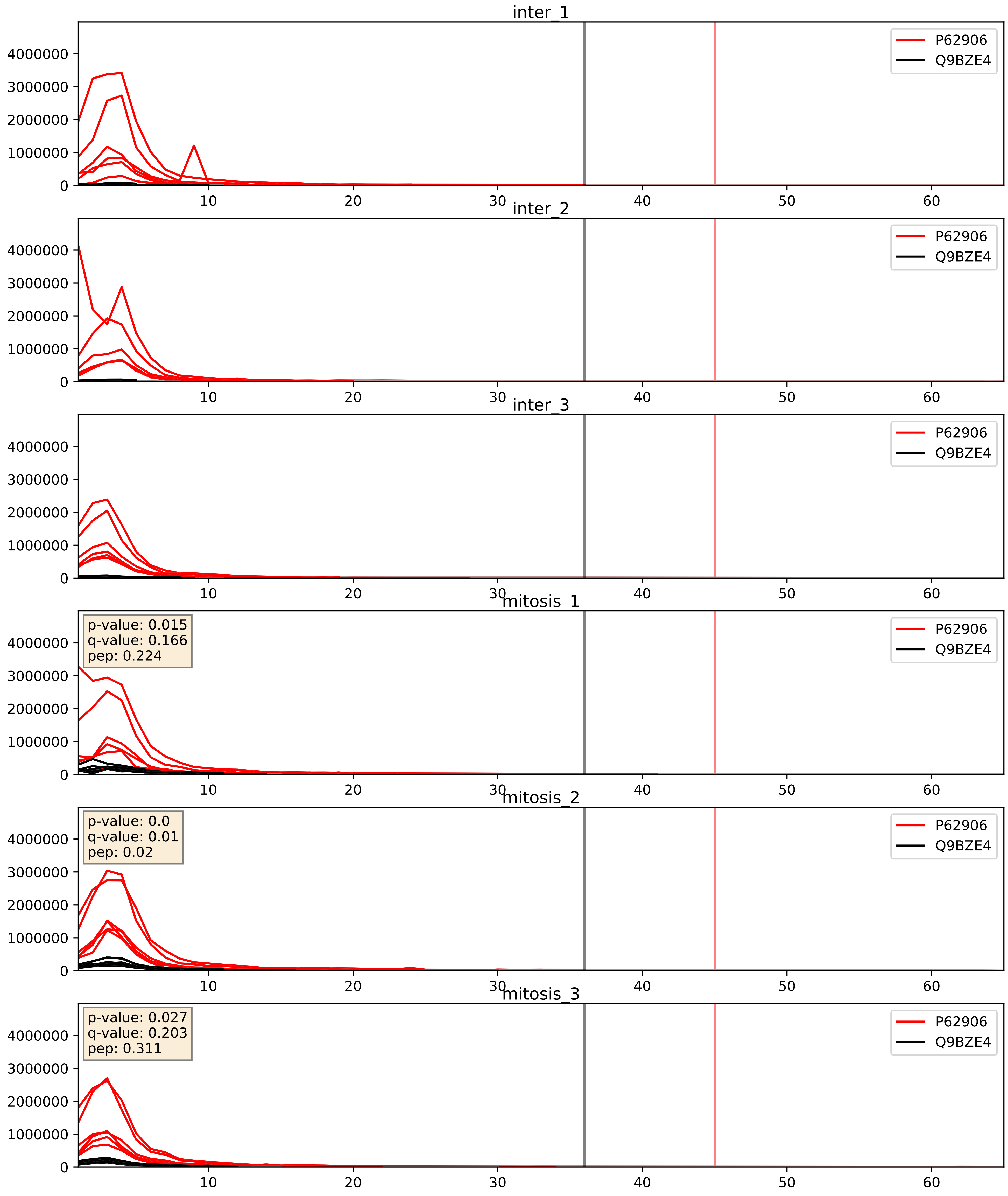


| level | condition_1 | condition_2 | pvalue | pvalue_adjusted |
|---|---|---|---|---|
| complex_abundance | mitosis | inter | 0.0017417866920582635 | 0.0295241467010272 |
| interactor_ratio | mitosis | inter | 0.002923484829602876 | 0.07007095913365735 |
| interactor_abundance | mitosis | inter | 0.40955067394198835 | 0.594379996833825 |

Q03701_Q9BZE4
CEBPZ_HUMAN vs NOG1_HUMAN
p-value: 0.034 q-value: 0.024

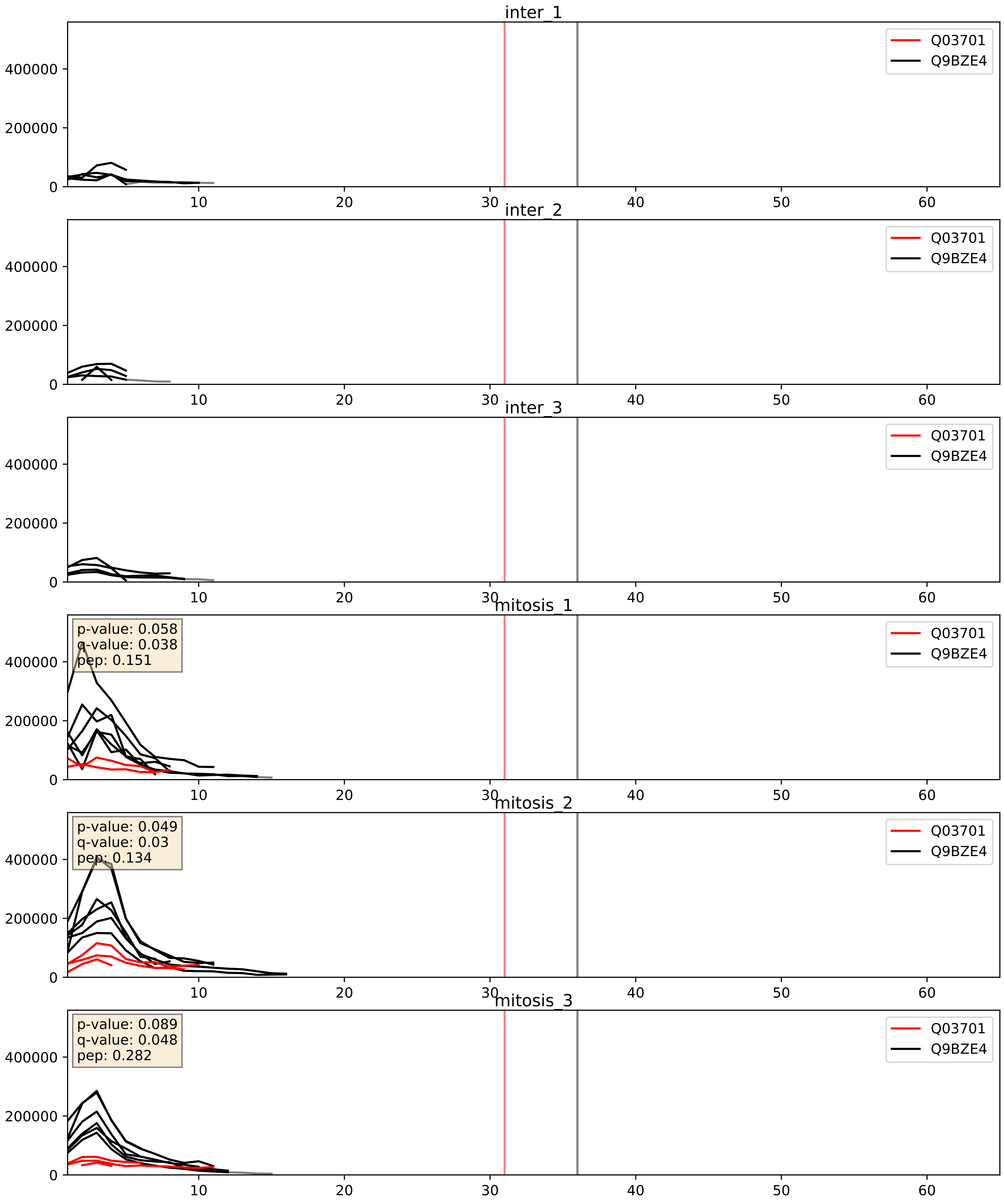


| level | condition_1 | condition_2 | pvalue | pvalue_adjusted |
|---|---|---|---|---|
| interactor_ratio | mitosis | inter | 2.782884935953381e-06 | 0.002520386835947235 |
| complex_abundance | mitosis | inter | 0.0011733846300989105 | 0.02303709273772173 |
| interactor_abundance | mitosis | inter | 0.019842839707373484 | 0.1190514126751731 |

Q13823_Q9BZE4
NOG2_HUMAN vs NOG1_HUMAN
p-value: 0.007 q-value: 0.006

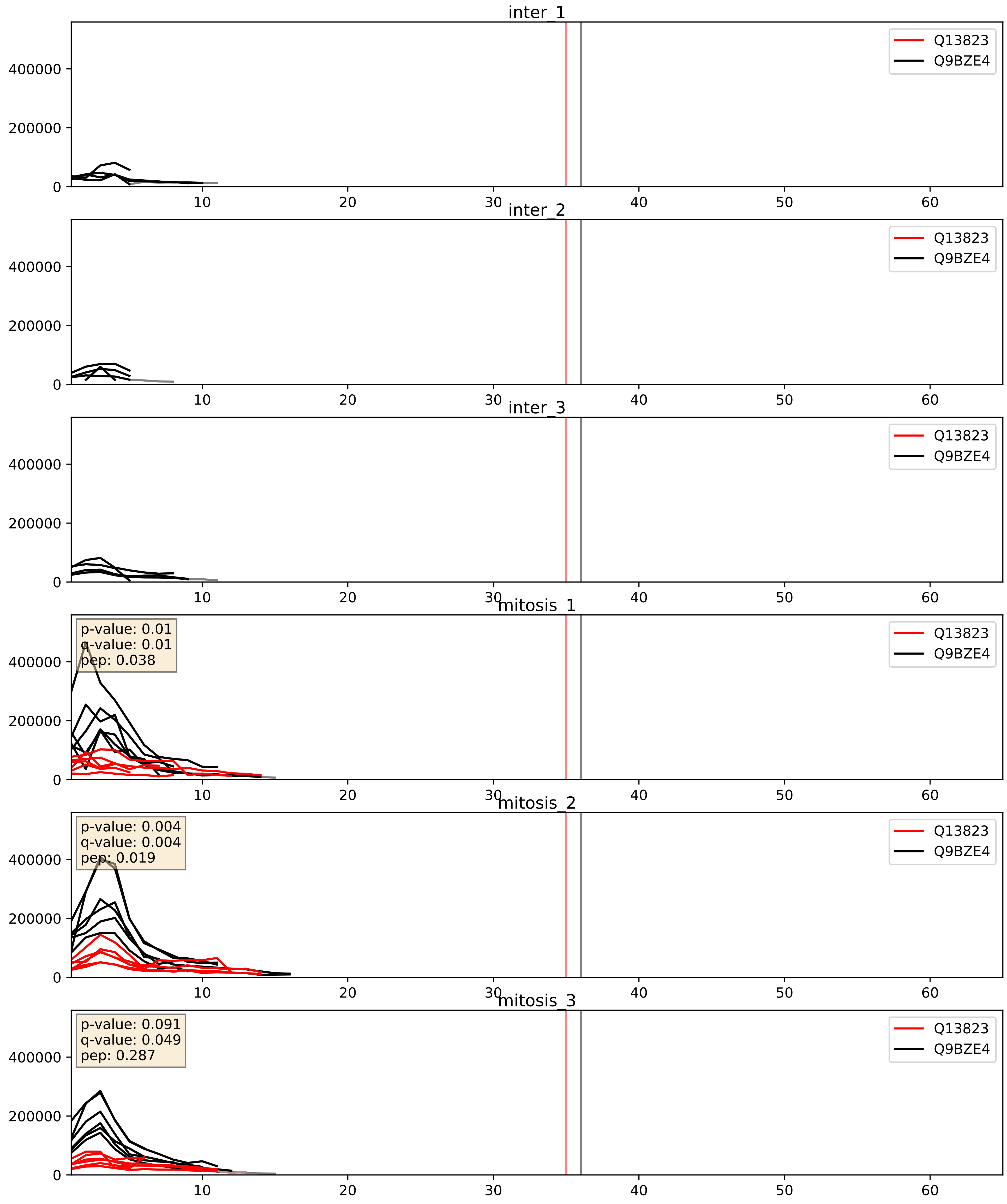


| level | condition_1 | condition_2 | pvalue | pvalue_adjusted |
|---|---|---|---|---|
| interactor_ratio | mitosis | inter | 0.0009758498273769684 | 0.03599129628116289 |
| complex_abundance | mitosis | inter | 0.0012618827871512452 | 0.024110974683068435 |
| interactor_abundance | mitosis | inter | 0.009576535749111607 | 0.08503645851908231 |

Q14137_Q9BZE4
BOP1_HUMAN vs NOG1_HUMAN
p-value: 0.053 q-value: 0.034

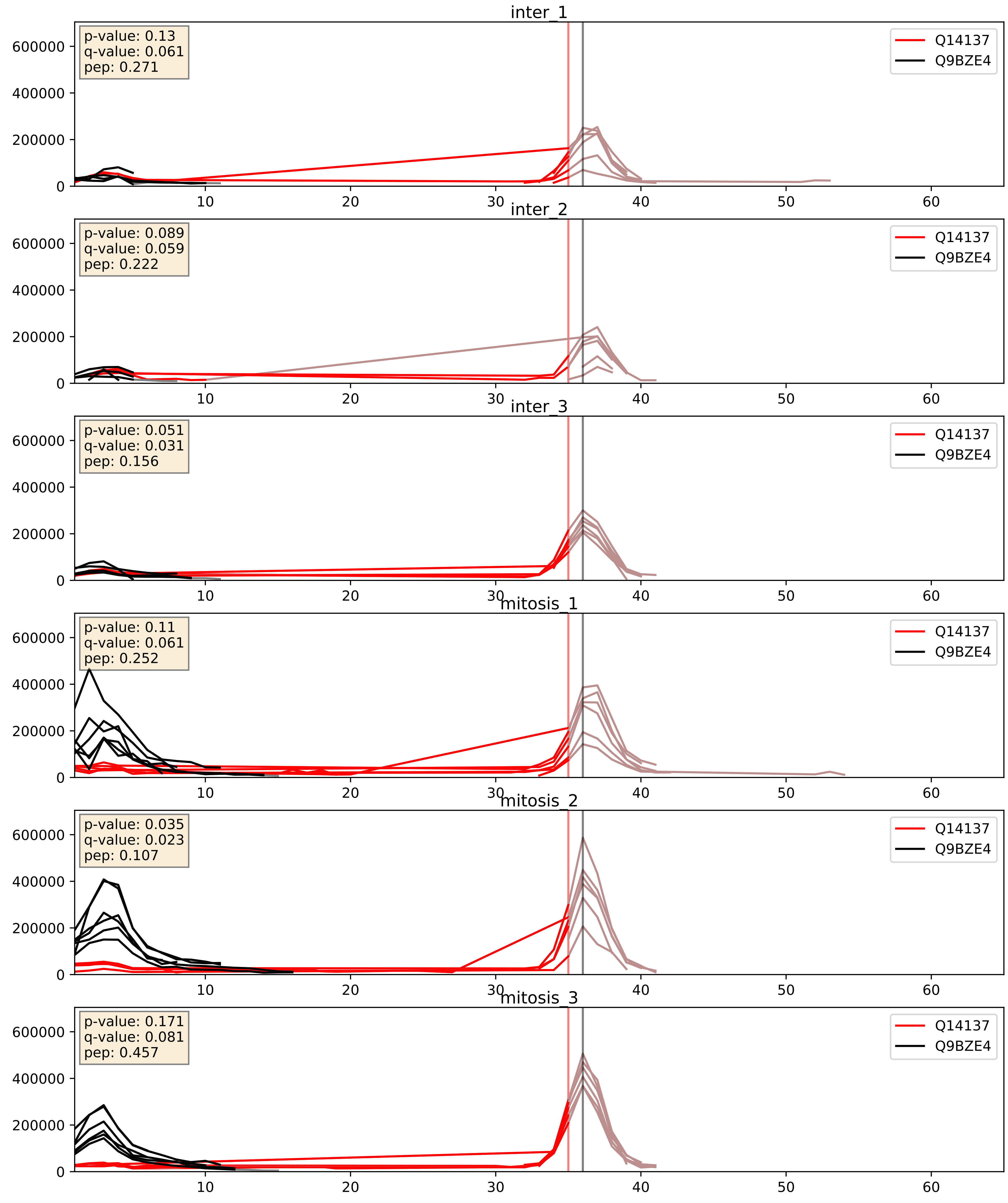


| level | condition_1 | condition_2 | pvalue | pvalue_adjusted |
|---|---|---|---|---|
| interactor_ratio | mitosis | inter | 0.0007612082988845985 | 0.03178508799244958 |
| complex_abundance | mitosis | inter | 0.0967292144756393 | 0.23055517679920082 |
| interactor_abundance | mitosis | inter | 0.09980342138358718 | 0.2711705935253959 |

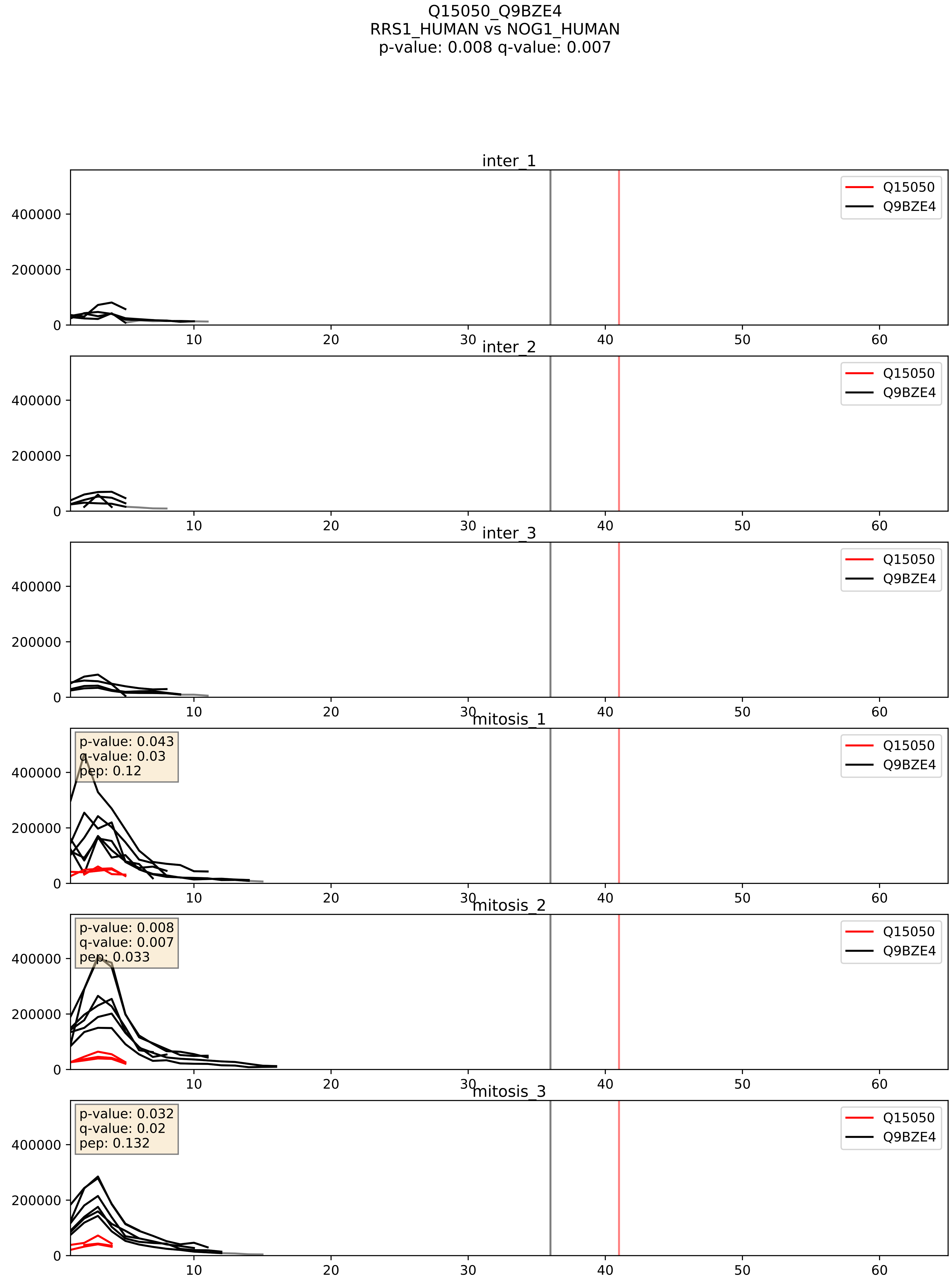


| level | condition_1 | condition_2 | pvalue | pvalue_adjusted |
|---|---|---|---|---|
| interactor_ratio | mitosis | inter | 0.0022417858763910544 | 0.05896690574521502 |
| complex_abundance | mitosis | inter | 0.005630838424790495 | 0.05650642076929266 |
| interactor_abundance | mitosis | inter | 0.2162538244564617 | 0.4119120465837366 |

Q15397_Q9BZE4
PUM3_HUMAN vs NOG1_HUMAN
p-value: 0.006 q-value: 0.006

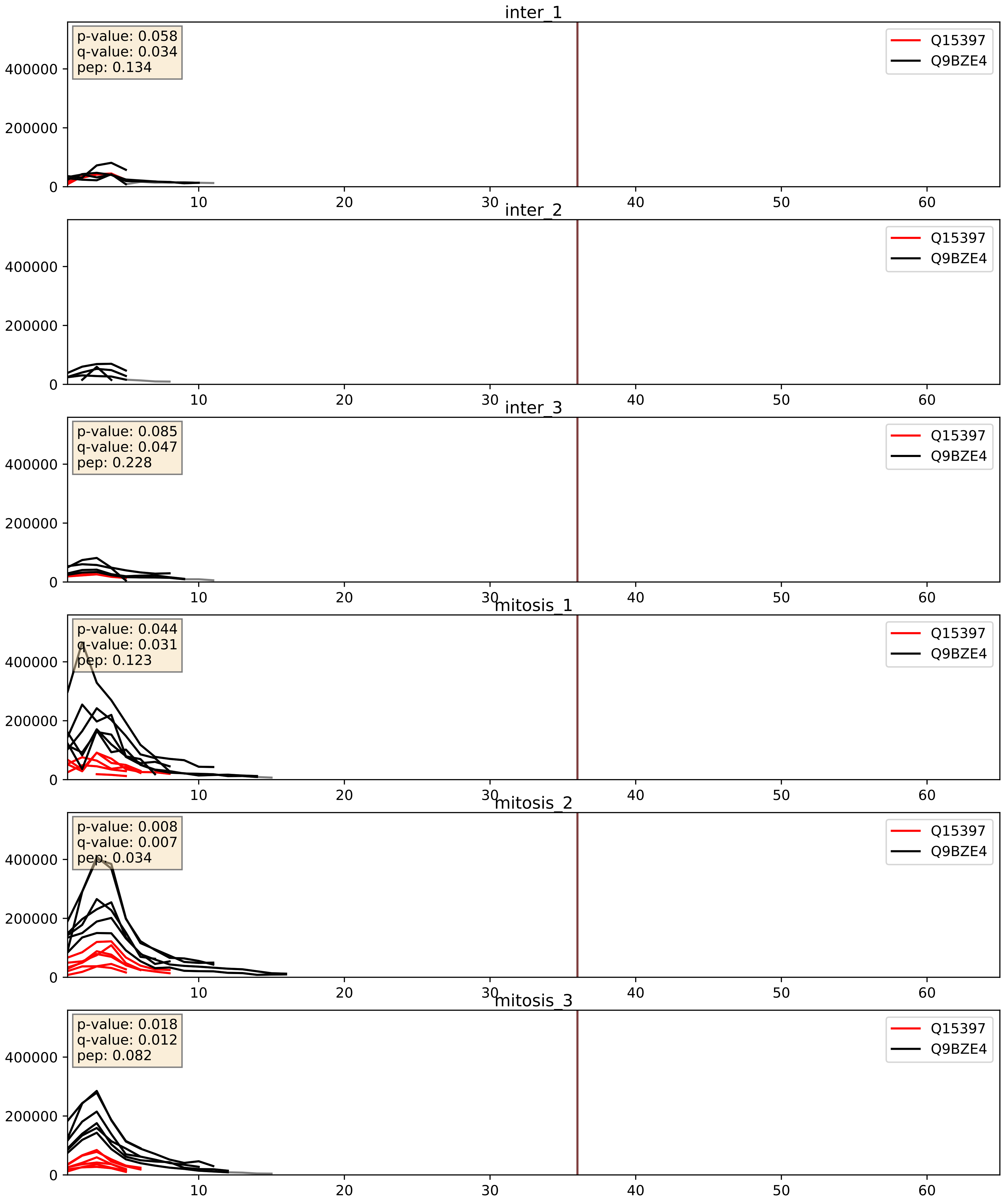


| level | condition_1 | condition_2 | pvalue | pvalue_adjusted |
|---|---|---|---|---|
| interactor_ratio | mitosis | inter | 0.00047537542148074825 | 0.02481227809680003 |
| complex_abundance | mitosis | inter | 0.022095800715059954 | 0.10952717692621856 |
| interactor_abundance | mitosis | inter | 0.2744724807239444 | 0.46751127239370466 |

Q8IY81_Q9BZE4
SPB1_HUMAN vs NOG1_HUMAN
p-value: 0.005 q-value: 0.005

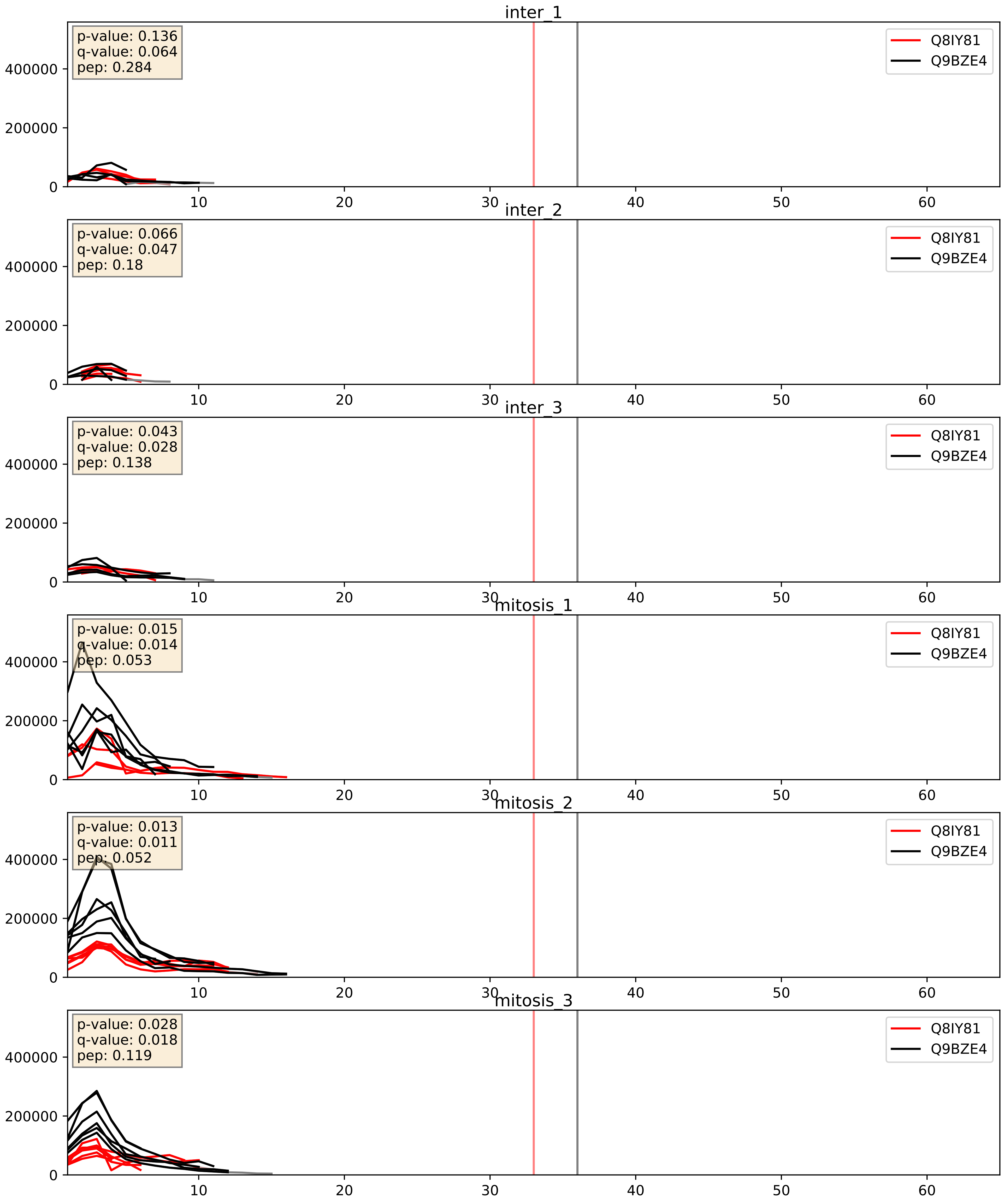


| level | condition_1 | condition_2 | pvalue | pvalue_adjusted |
|---|---|---|---|---|
| interactor_ratio | mitosis | inter | 0.0007974295630027685 | 0.03250474790144618 |
| complex_abundance | mitosis | inter | 0.012254193945439154 | 0.0822068183173661 |
| interactor_abundance | mitosis | inter | 0.33114478037662615 | 0.5307908853629082 |

Q8TDN6_Q9BZE4
BRX1_HUMAN vs NOG1_HUMAN
p-value: 0.017 q-value: 0.014

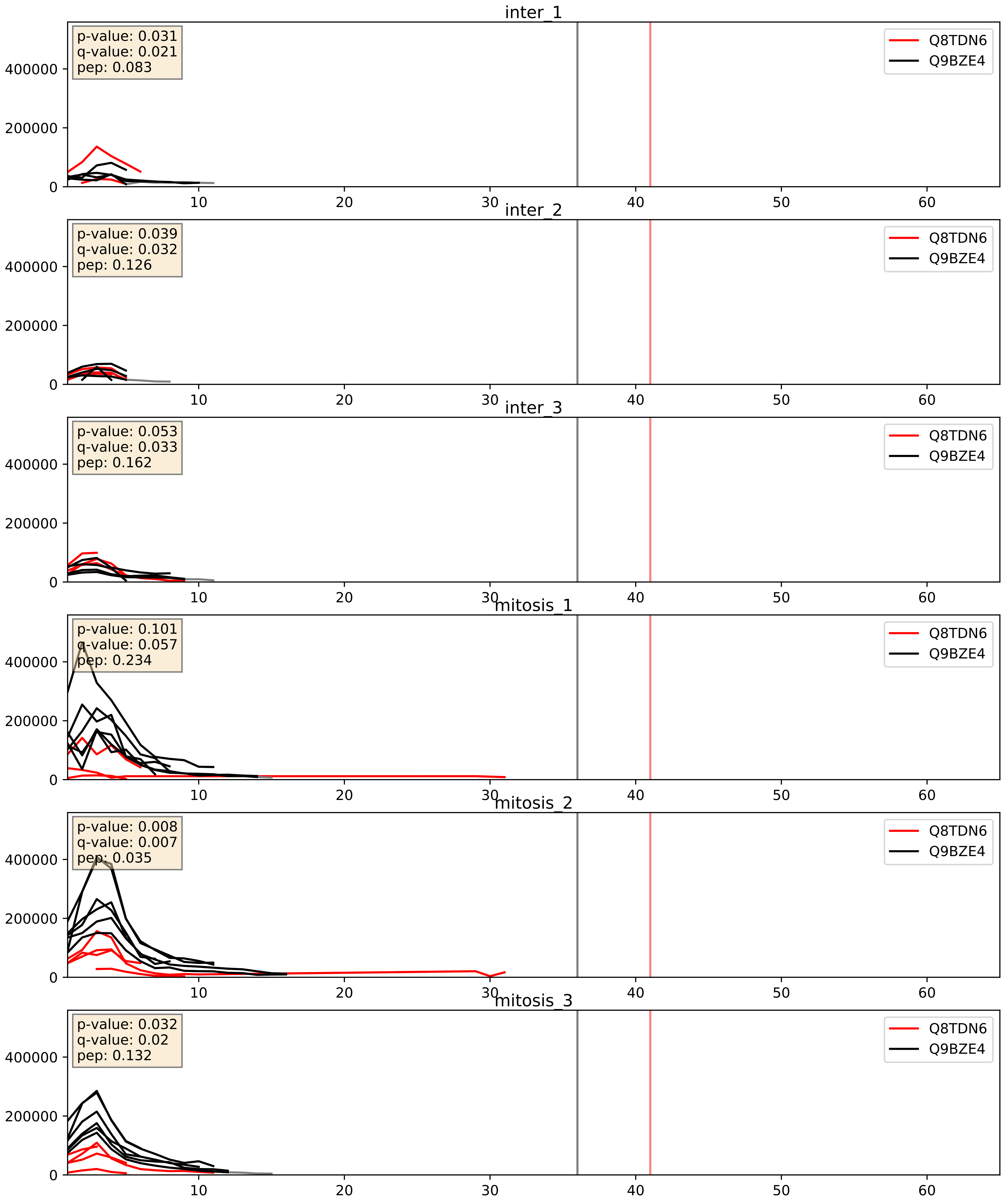


| level | condition_1 | condition_2 | pvalue | pvalue_adjusted |
|---|---|---|---|---|
| interactor_ratio | mitosis | inter | 0.00011836259628161376 | 0.011513452547393338 |
| complex_abundance | mitosis | inter | 0.01080016423974069 | 0.07693511812192051 |
| interactor_abundance | mitosis | inter | 0.0462072460121471 | 0.17159827586289772 |

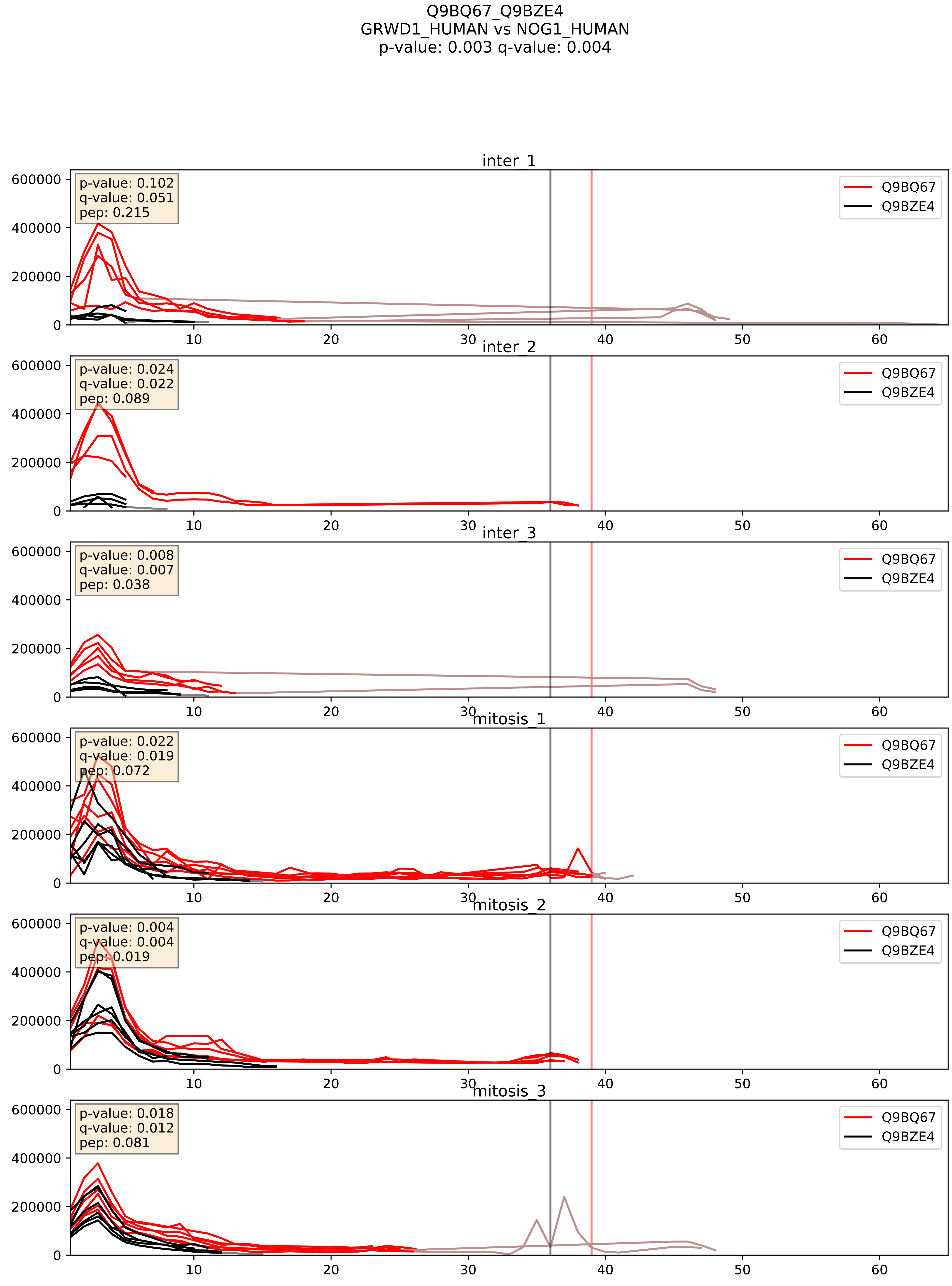


| level | condition_1 | condition_2 | pvalue | pvalue_adjusted |
|---|---|---|---|---|
| interactor_ratio | mitosis | inter | 0.0016120197660940117 | 0.048587638020298377 |
| complex_abundance | mitosis | inter | 0.007234422190094093 | 0.06298977015239762 |
| interactor_abundance | mitosis | inter | 0.08268426471817841 | 0.24090473795481276 |

Q9BVP2_Q9BZE4
GNL3_HUMAN vs NOG1_HUMAN
p-value: 0.004 q-value: 0.004

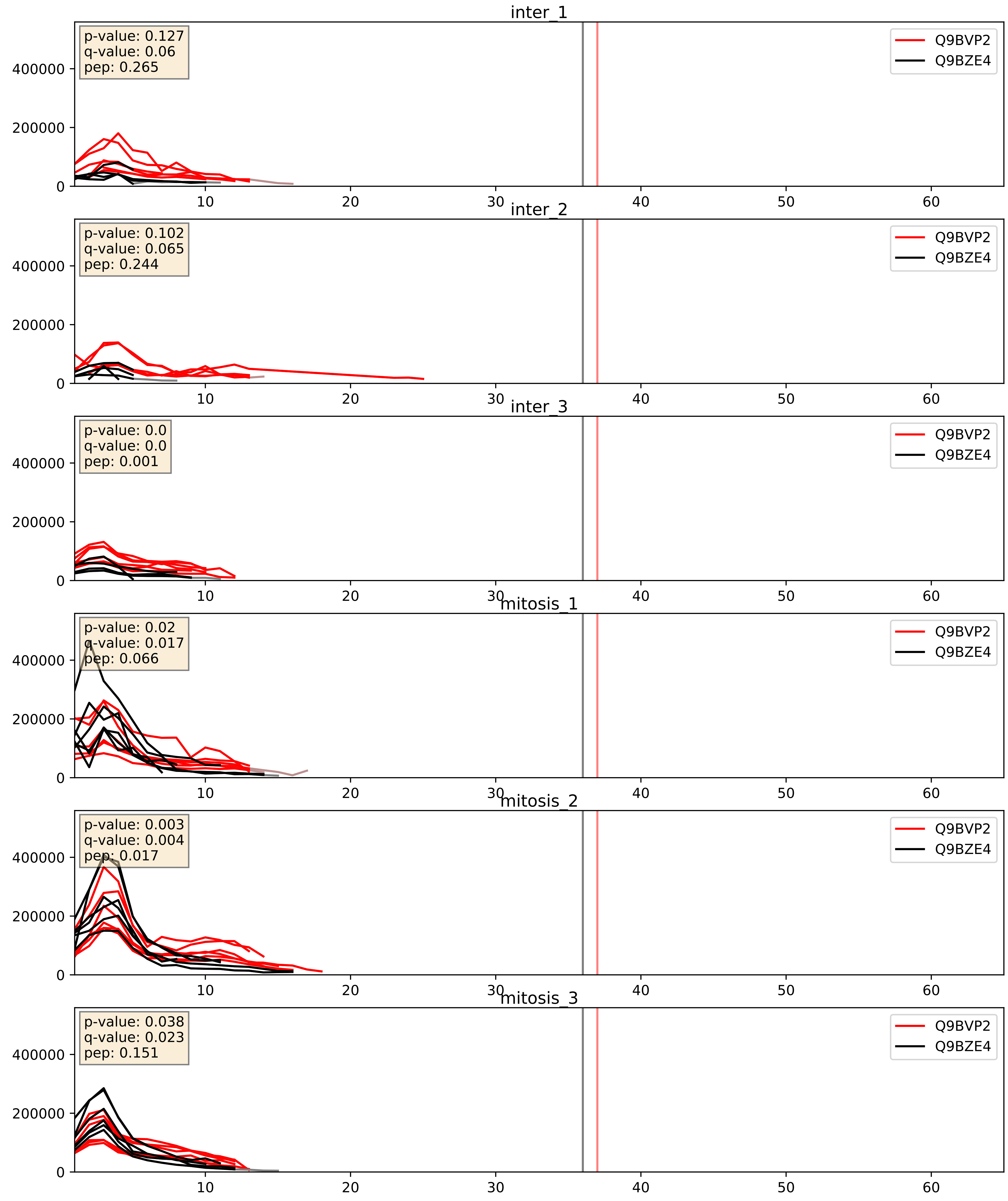


| level | condition_1 | condition_2 | pvalue | pvalue_adjusted |
|---|---|---|---|---|
| complex_abundance | mitosis | inter | 0.0013895549742411744 | 0.02552487248820698 |
| interactor_abundance | mitosis | inter | 0.006897488603137285 | 0.07148798621825644 |
| interactor_ratio | mitosis | inter | 0.011563542728058424 | 0.15494522140242886 |

Q9BYG3_Q9BZE4
MK67I_HUMAN vs NOG1_HUMAN
p-value: 0.068 q-value: 0.041

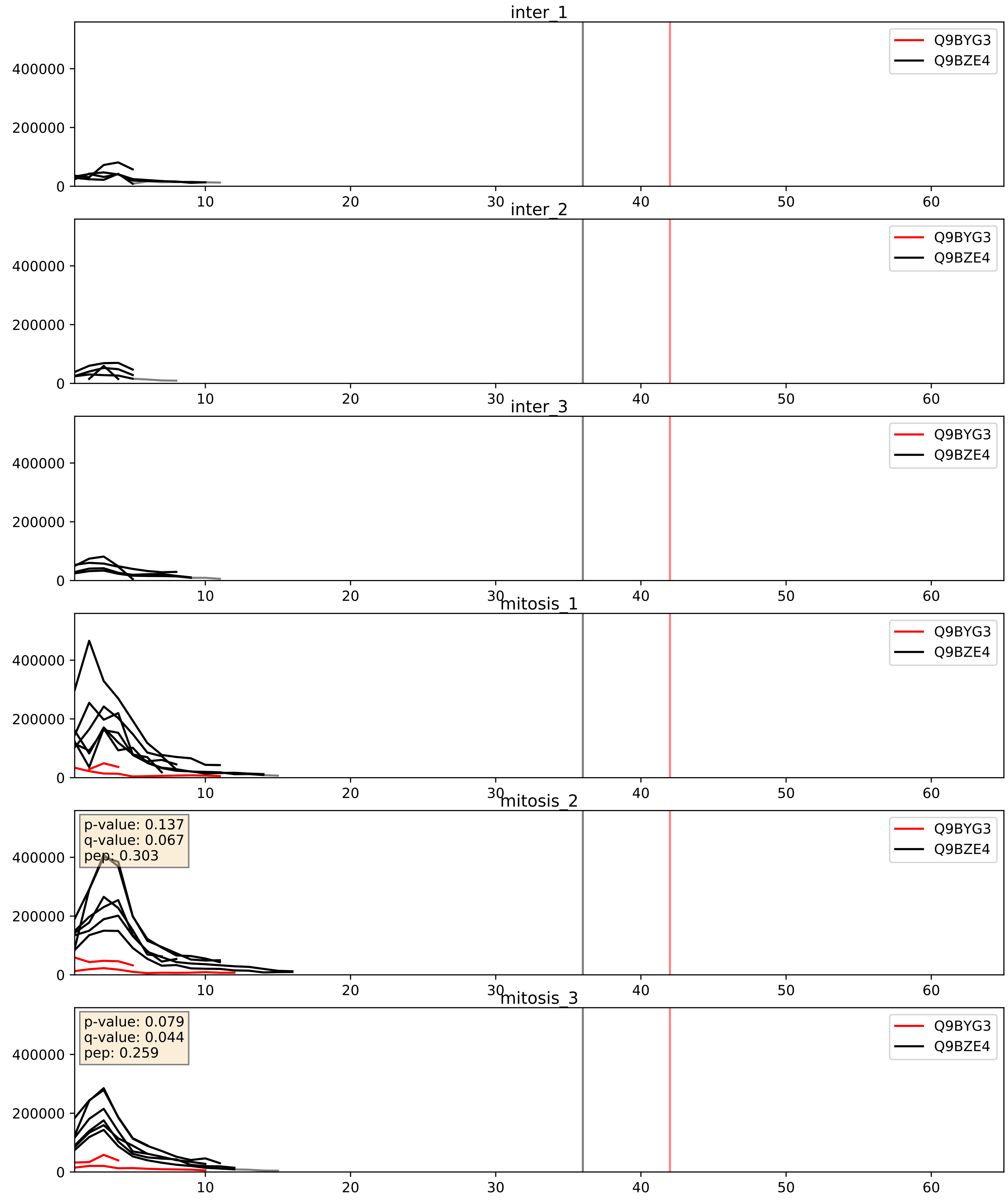


| level | condition_1 | condition_2 | pvalue | pvalue_adjusted |
|---|---|---|---|---|
| interactor_ratio | mitosis | inter | 0.0006141803584999456 | 0.029219626944512057 |
| complex_abundance | mitosis | inter | 0.0011171790006910644 | 0.022133971147621622 |
| interactor_abundance | mitosis | inter | 0.21337749899904426 | 0.40861552381025035 |

Q9BZE4_Q9GZR7
NOG1_HUMAN vs DDX24_HUMAN
p-value: 0.012 q-value: 0.01

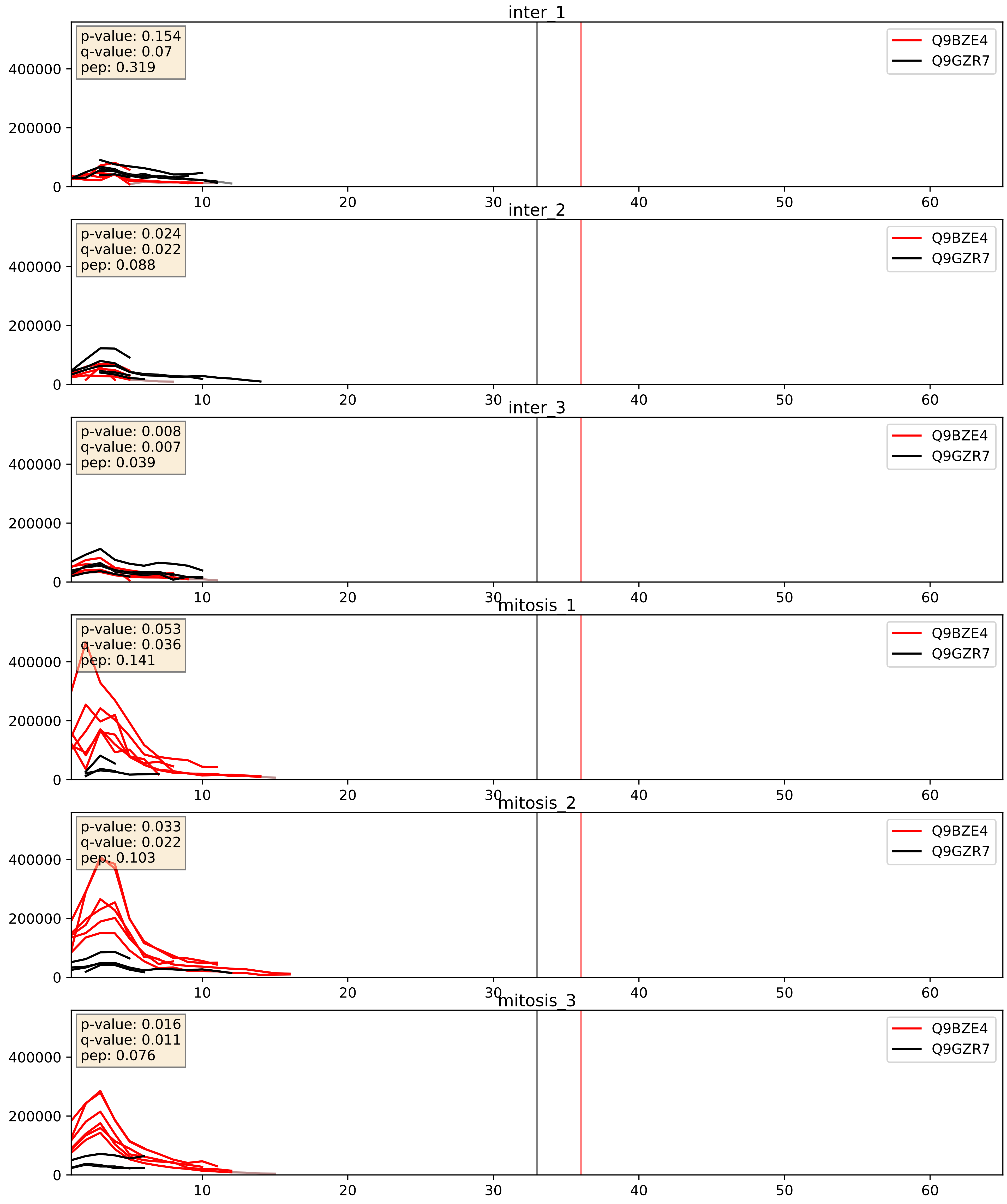


| level | condition_1 | condition_2 | pvalue | pvalue_adjusted |
|---|---|---|---|---|
| interactor_ratio | mitosis | inter | 0.00020599297265075348 | 0.014683183137172115 |
| interactor_abundance | mitosis | inter | 0.004638788434277854 | 0.05526517605764569 |
| complex_abundance | mitosis | inter | 0.2407443405492743 | 0.39200524160201405 |

Q9BZE4_Q9NVP1
NOG1_HUMAN vs DDX18_HUMAN
p-value: 0.008 q-value: 0.007

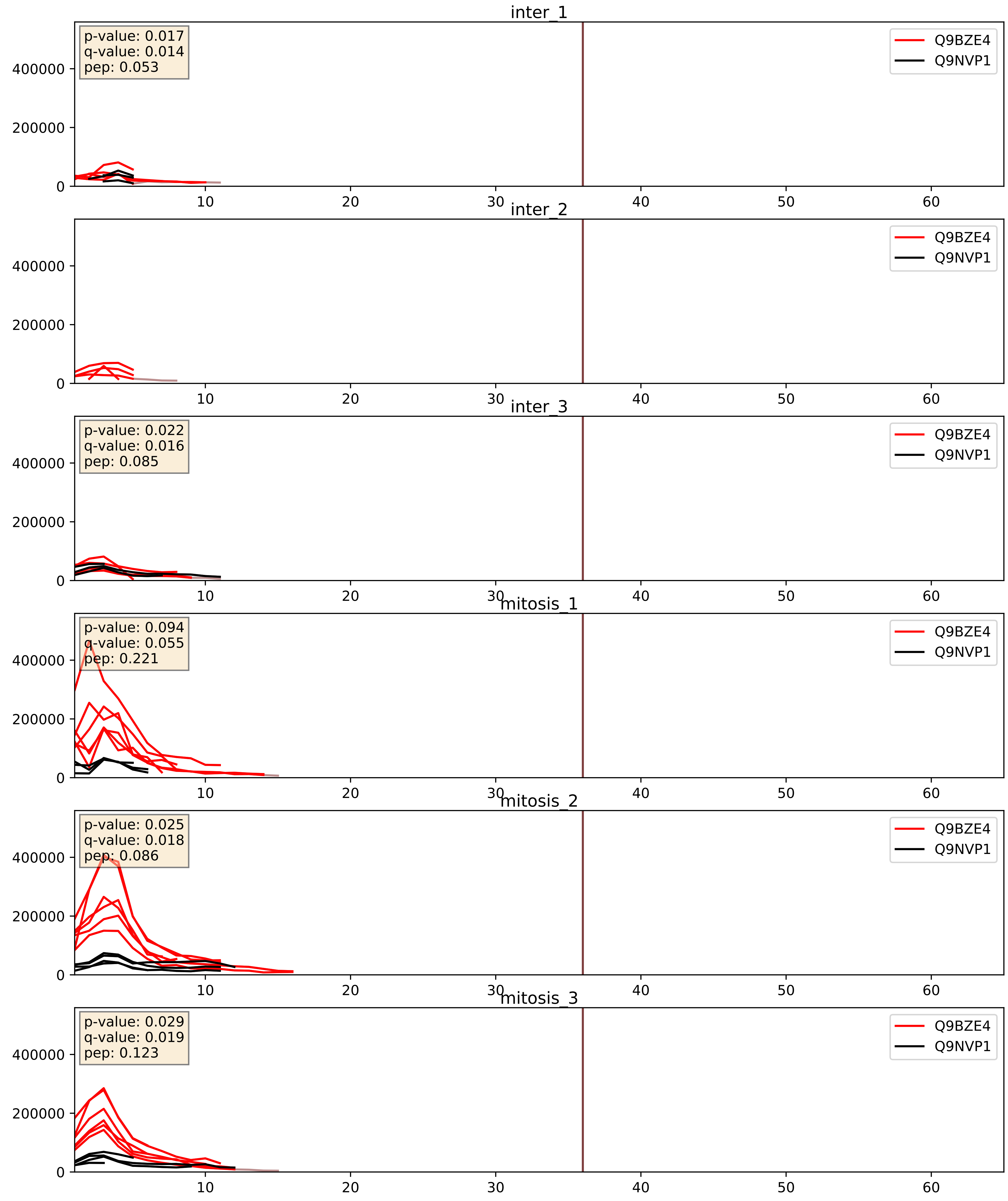


| level | condition_1 | condition_2 | pvalue | pvalue_adjusted |
|---|---|---|---|---|
| interactor_ratio | mitosis | inter | 0.0007256664780815352 | 0.031214598253155482 |
| interactor_abundance | mitosis | inter | 0.0031709934969099437 | 0.043395210765066534 |
| complex_abundance | mitosis | inter | 0.028801127868799545 | 0.1234203919435165 |

Q9BZE4_Q9NY93
NOG1_HUMAN vs DDX56_HUMAN
p-value: 0.051 q-value: 0.033

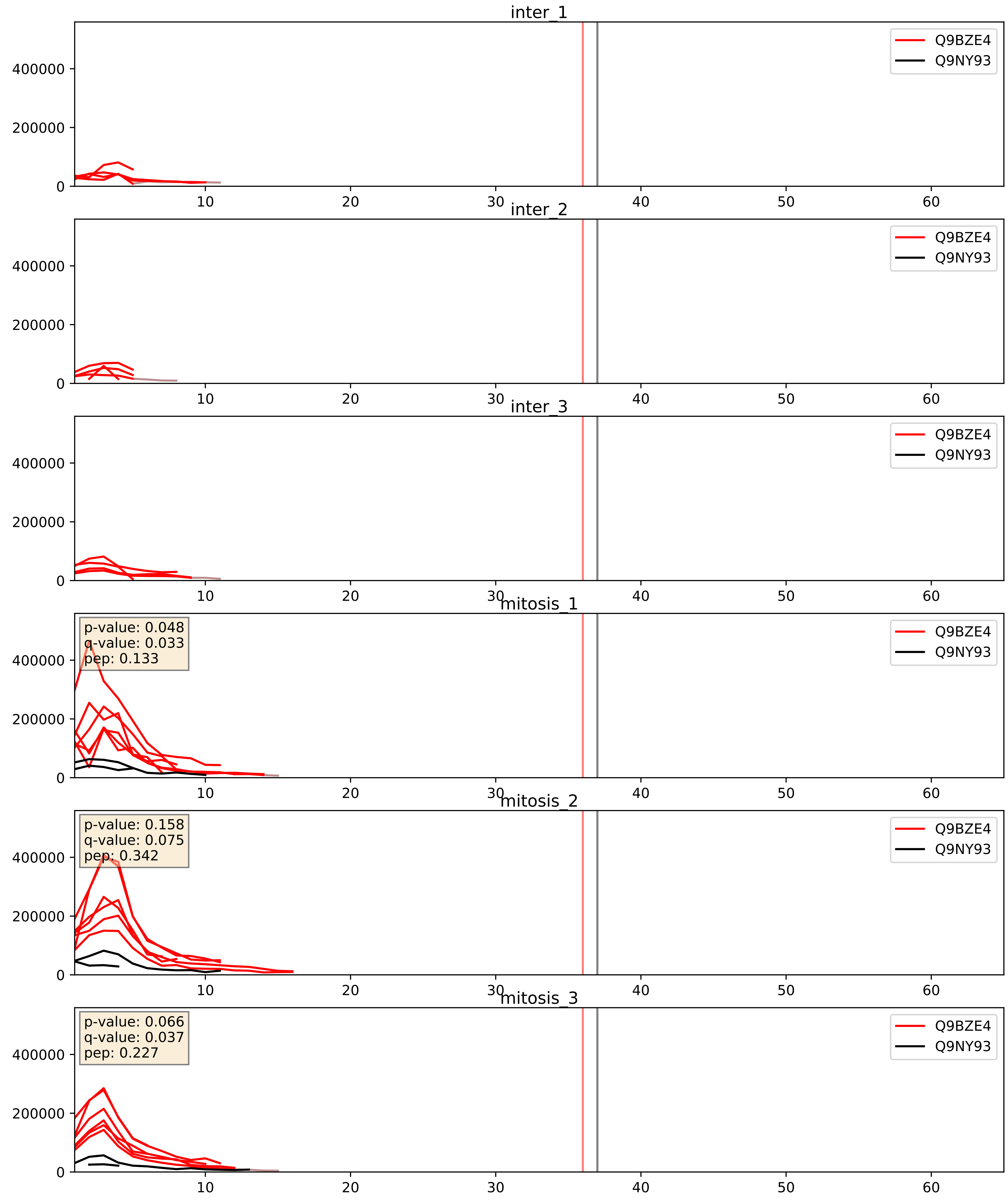


| level | condition_1 | condition_2 | pvalue | pvalue_adjusted |
|---|---|---|---|---|
| interactor_abundance | mitosis | inter | 0.0002829560238761811 | 0.012681170494136702 |
| interactor_ratio | mitosis | inter | 0.0002899670013636358 | 0.017729410940519448 |
| complex_abundance | mitosis | inter | 0.0011975320997738852 | 0.02329744266832831 |

Q9BZE4_Q9UKD2
NOG1_HUMAN vs MRT4_HUMAN
p-value: 0.008 q-value: 0.008

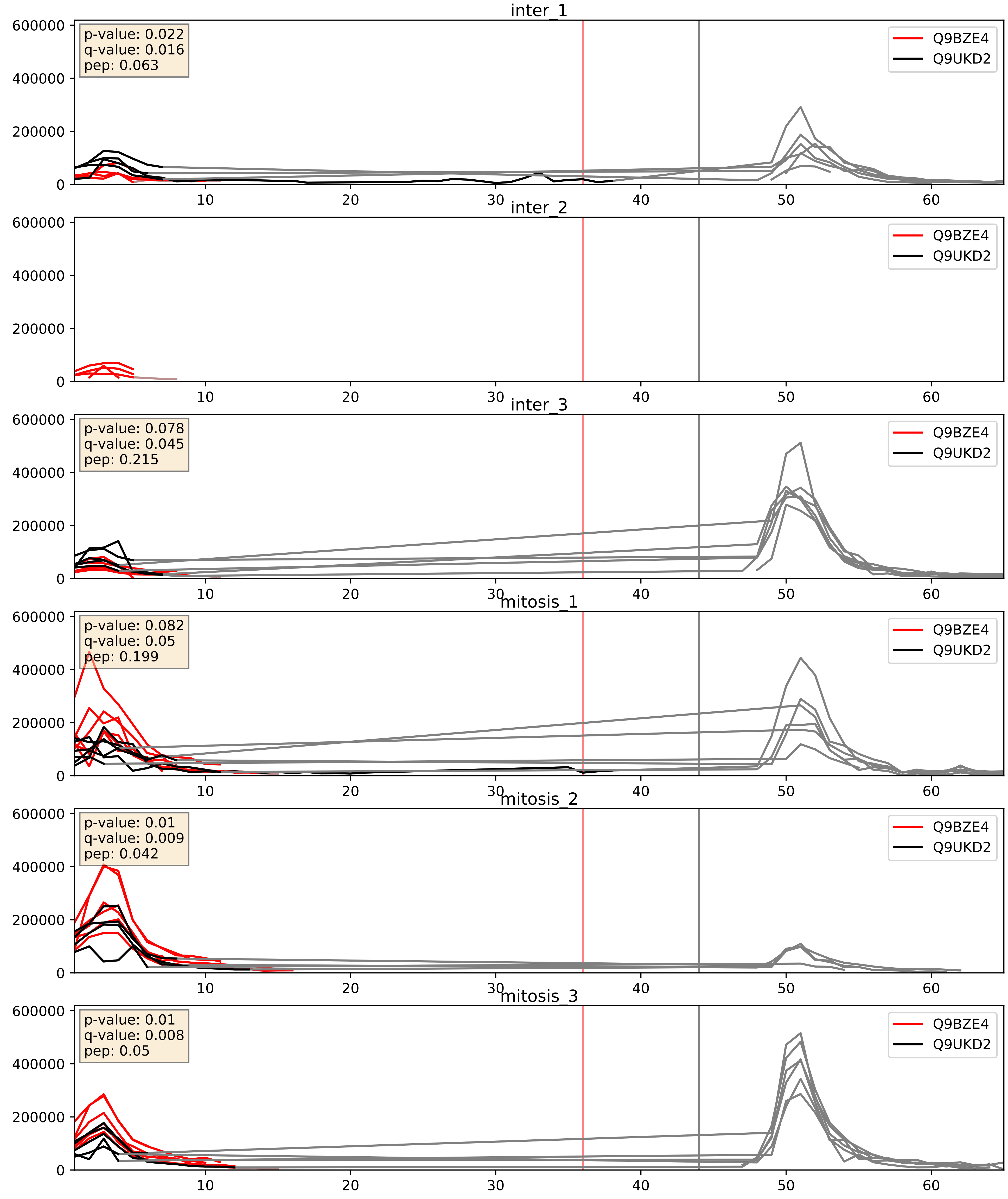


| level | condition_1 | condition_2 | pvalue | pvalue_adjusted |
|---|---|---|---|---|
| interactor_abundance | mitosis | inter | 0.0030352039949890198 | 0.0421728093362545 |
| interactor_ratio | mitosis | inter | 0.0047183833657339245 | 0.09159088101537749 |
| complex_abundance | mitosis | inter | 0.02221883361752925 | 0.10952717692621856 |

Q9BZE4_Q9Y221
NOG1_HUMAN vs NIP7_HUMAN
p-value: 0.011 q-value: 0.009

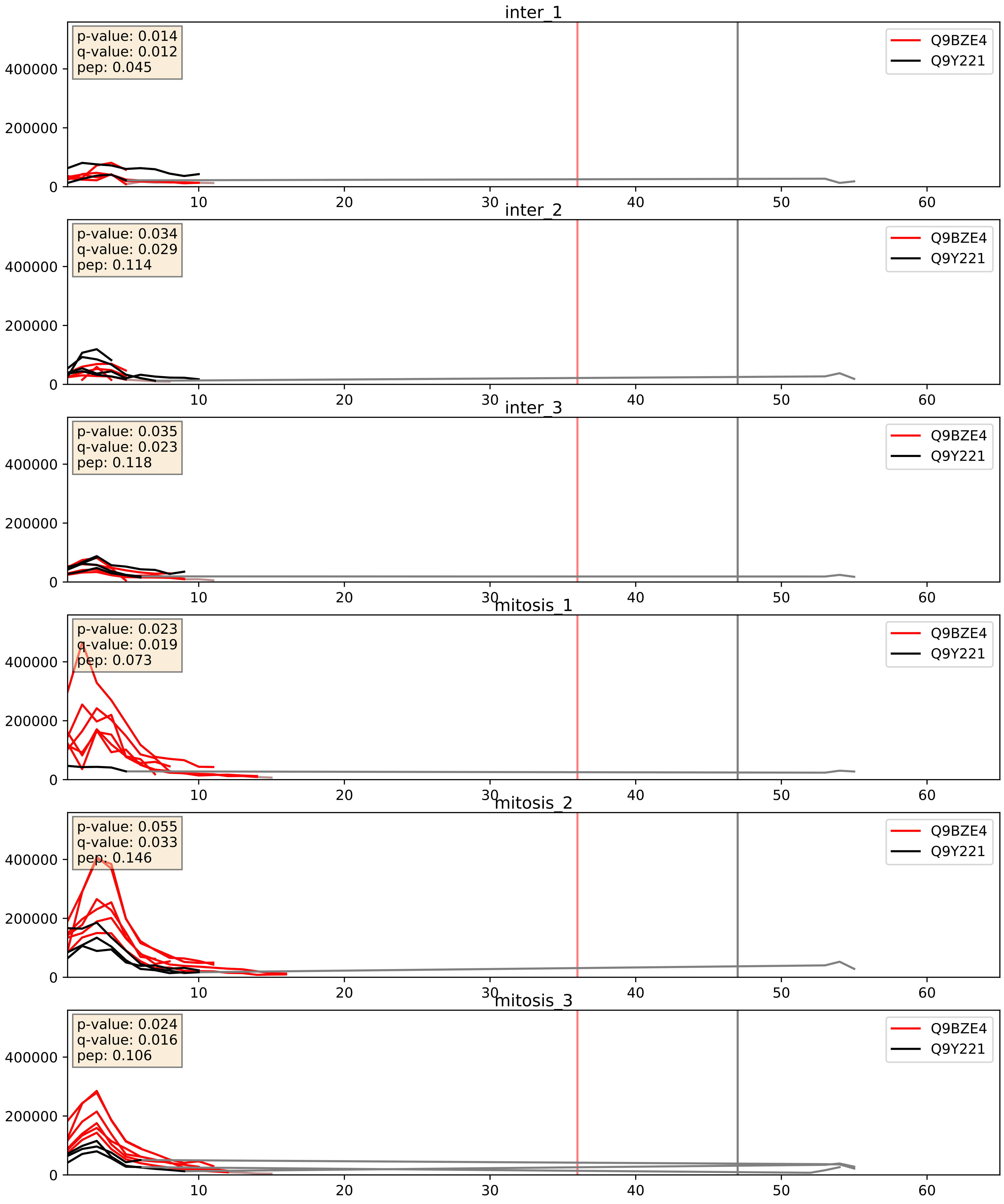


| level | condition_1 | condition_2 | pvalue | pvalue_adjusted |
|---|---|---|---|---|
| interactor_abundance | mitosis | inter | 0.001937848367081554 | 0.032272338564626656 |
| interactor_ratio | mitosis | inter | 0.004133771637519246 | 0.08445127736793497 |
| complex_abundance | mitosis | inter | 0.09716538449166384 | 0.23097353269887322 |

Q9BZE4_Q9Y2X3
NOG1_HUMAN vs NOP58_HUMAN
p-value: 0.034 q-value: 0.024

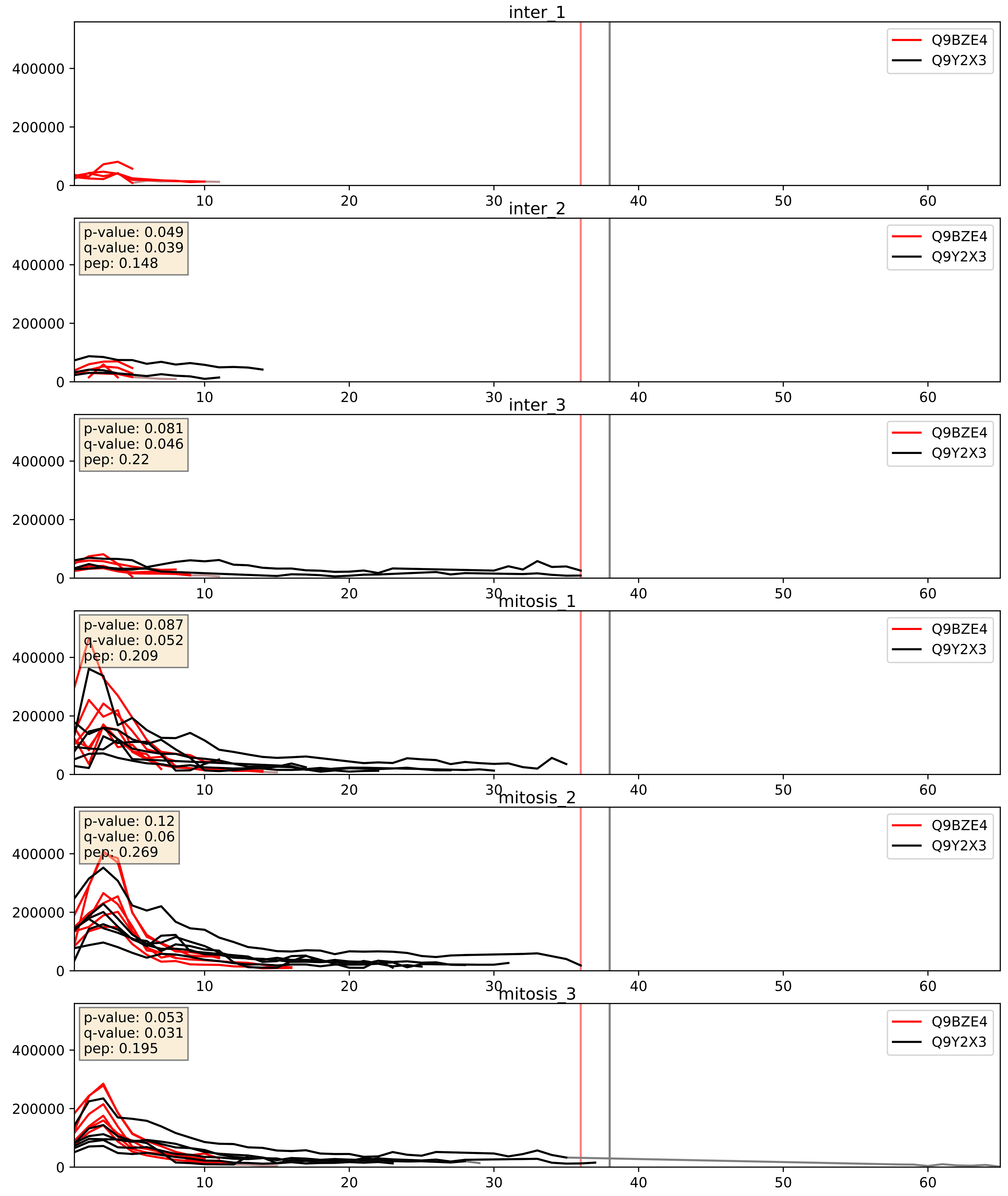


| level | condition_1 | condition_2 | pvalue | pvalue_adjusted |
|---|---|---|---|---|
| complex_abundance | mitosis | inter | 0.0015925256220138068 | 0.027934465828766776 |
| interactor_abundance | mitosis | inter | 0.001698666342849207 | 0.030042528708242176 |
| interactor_ratio | mitosis | inter | 0.04024854859360579 | 0.33223488520854927 |

Q9BZE4_Q9Y3T9
NOG1_HUMAN vs NOC2L_HUMAN
p-value: 0.015 q-value: 0.012

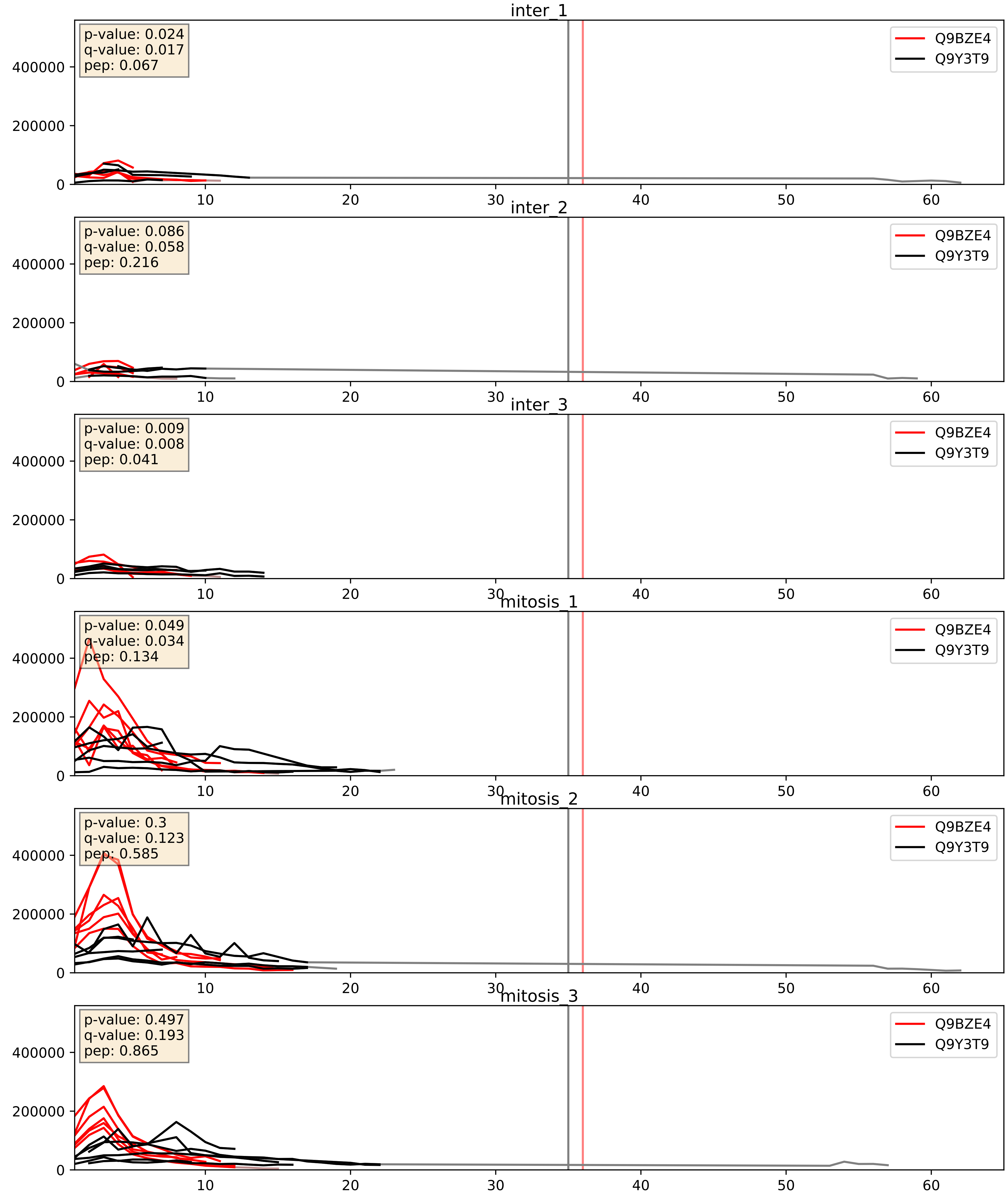


| level | condition_1 | condition_2 | pvalue | pvalue_adjusted |
|---|---|---|---|---|
| complex_abundance | mitosis | inter | 0.0007505306717870162 | 0.017895661700548352 |
| interactor_abundance | mitosis | inter | 0.0009449316519668341 | 0.022846219799594687 |
| interactor_ratio | mitosis | inter | 0.13624795726384295 | 0.5866546850118911 |
