## Supplementary Data 2 for "SECAT: Quantifying differential protein-protein interaction states by network-centric analysis": hela_string_000005_P62269.pdf

P62269_P62269

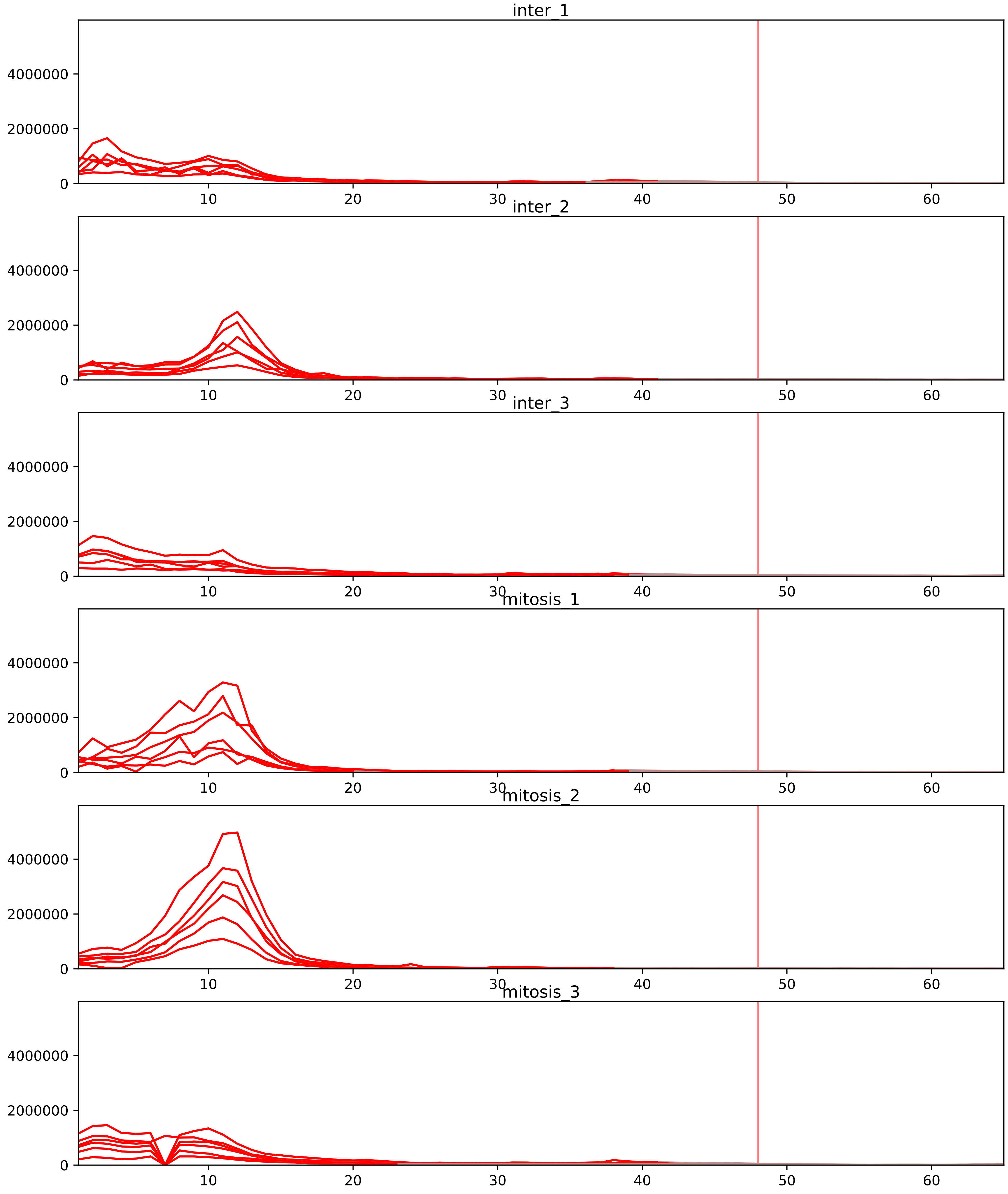


| level | condition_1 | condition_2 | pvalue | pvalue_adjusted |
| --- | --- | --- | --- | --- |
| interactor_ratio | mitosis | inter | 8.72197458617678e-20 | 3.2096866477130555e-17 |
| interactor_abundance | mitosis | inter | 0.000740443991247939 | 0.009461228777056997 |
| complex_abundance | mitosis | inter | 0.0018121806467932913 | 0.015802902322747184 |
| total_abundance | mitosis | inter | 0.7675684840055483 | 0.9041054674160921 |
| assembled_abundance | mitosis | inter | 0.7854887946572418 | 0.9106320699503349 |
| monomer_abundance | mitosis | inter | 0.9886927089370946 | 0.9981662859499865 |

O00303_P62269
EIF3F_HUMAN vs RS18_HUMAN
p-value: 0.082 q-value: 0.049

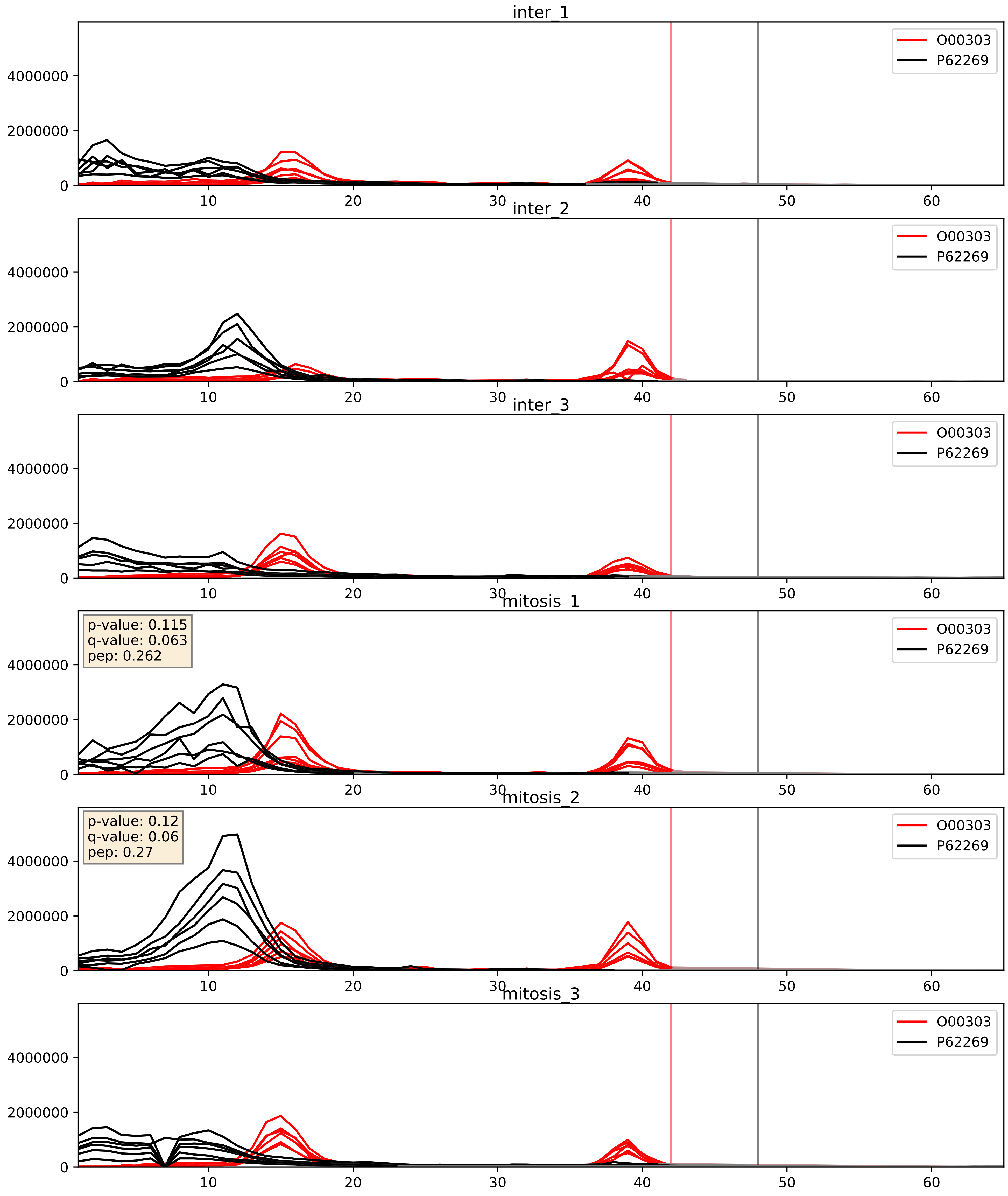


| level | condition_1 | condition_2 | pvalue | pvalue_adjusted |
|---|---|---|---|---|
| complex_abundance | mitosis | inter | 0.005055645522624262 | 0.05313654189341448 |
| interactor_abundance | mitosis | inter | 0.27816544418952294 | 0.47145751950545817 |
| interactor_ratio | mitosis | inter | 0.34490127371089785 | 0.7969645820582403 |

P05387_P62269
RLA2_HUMAN vs RS18_HUMAN
p-value: 0.01 q-value: 0.009

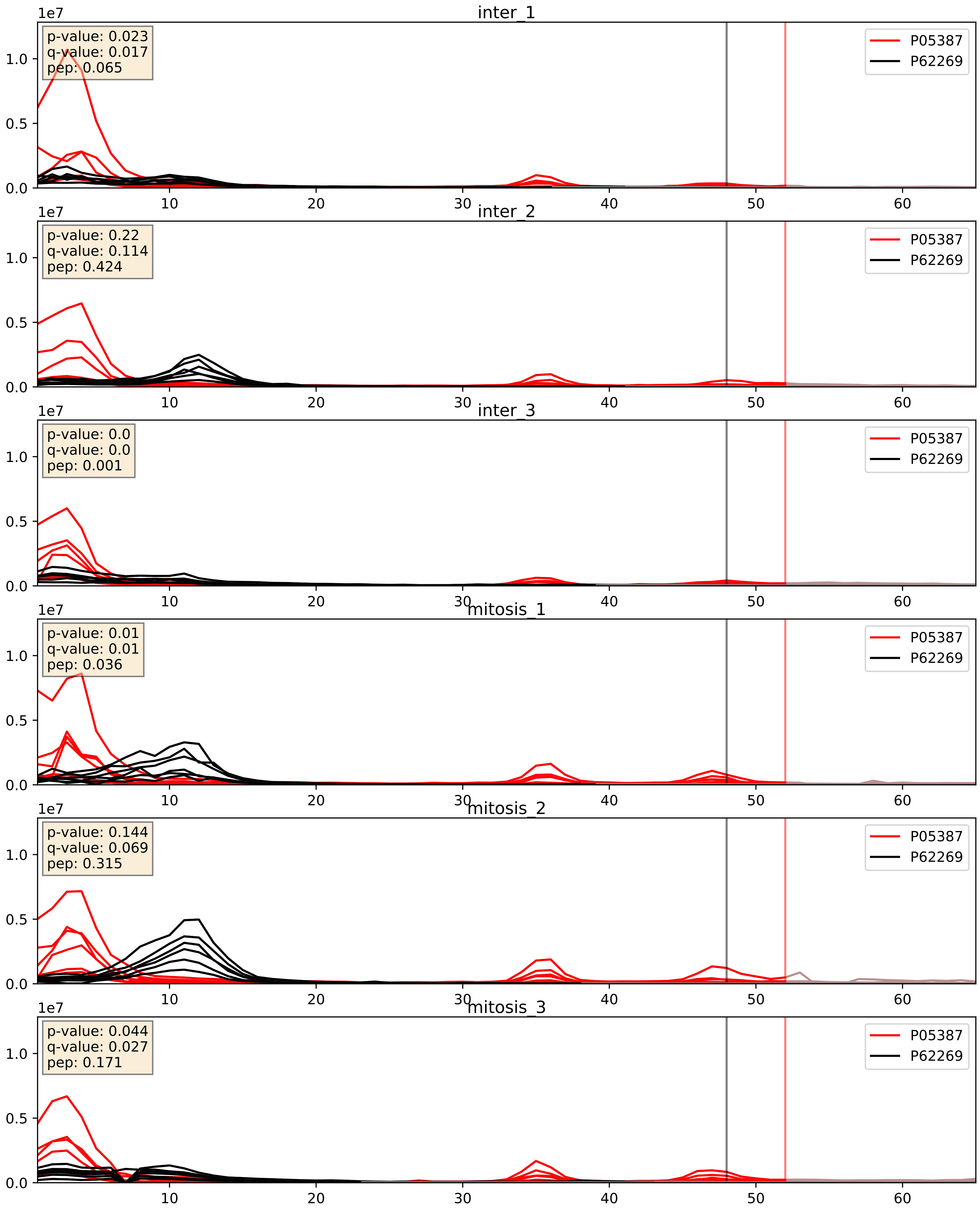


| level | condition_1 | condition_2 | pvalue | pvalue_adjusted |
|---|---|---|---|---|
| complex_abundance | mitosis | inter | 0.02787747575457837 | 0.12144081041180196 |
| interactor_ratio | mitosis | inter | 0.13439924587159455 | 0.5847231583055633 |
| interactor_abundance | mitosis | inter | 0.8248718694852493 | 0.9021789053591943 |

P08708_P62269
RS17_HUMAN vs RS18_HUMAN
p-value: 0.0 q-value: 0.0

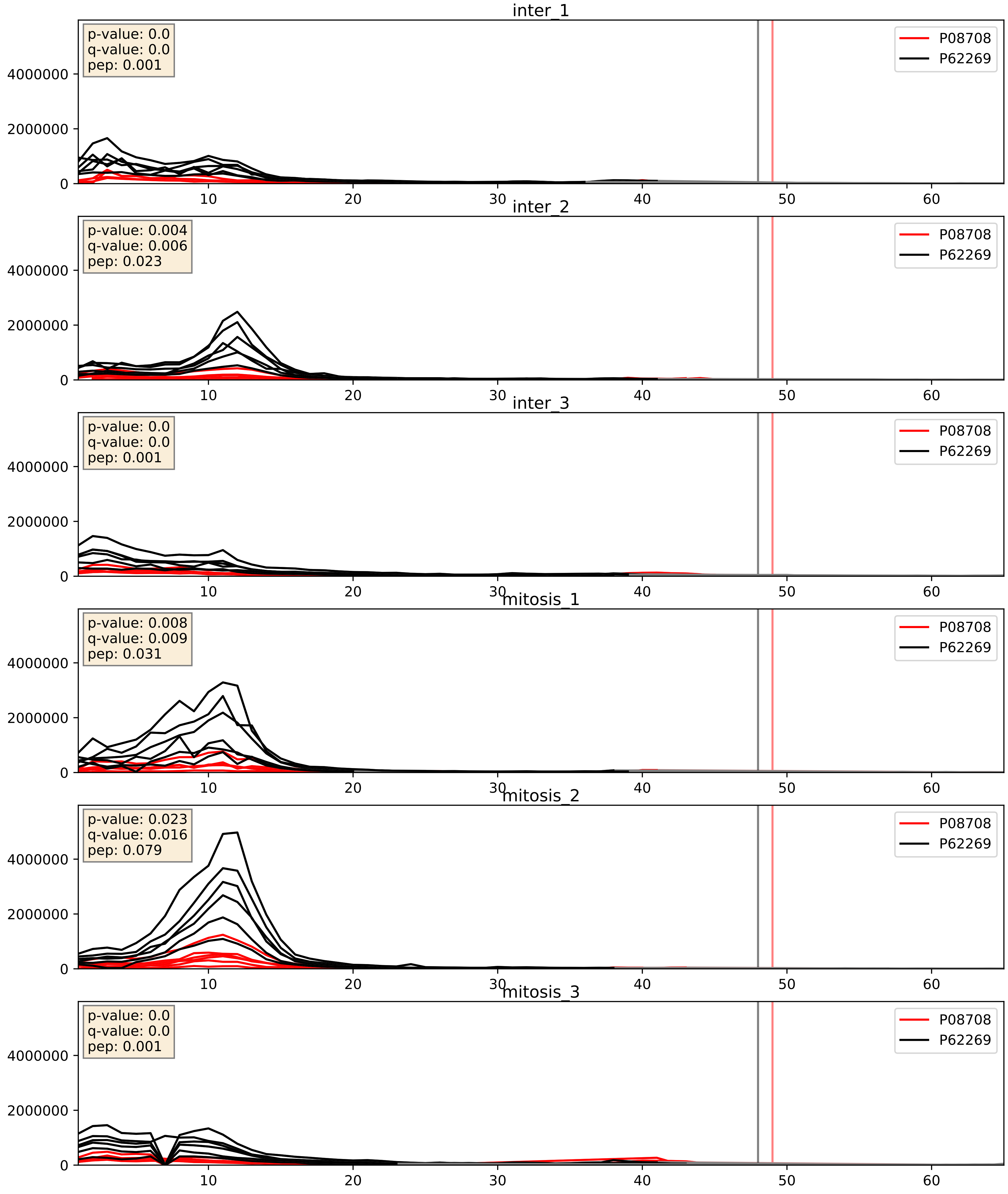


| level | condition_1 | condition_2 | pvalue | pvalue_adjusted |
|---|---|---|---|---|
| interactor_ratio | mitosis | inter | 0.009180527608022524 | 0.13667011534725704 |
| complex_abundance | mitosis | inter | 0.016795488374930955 | 0.09665323655973425 |
| interactor_abundance | mitosis | inter | 0.04090370098055168 | 0.16258912486348848 |

P08865_P62269
RSSA_HUMAN vs RS18_HUMAN
p-value: 0.0 q-value: 0.0

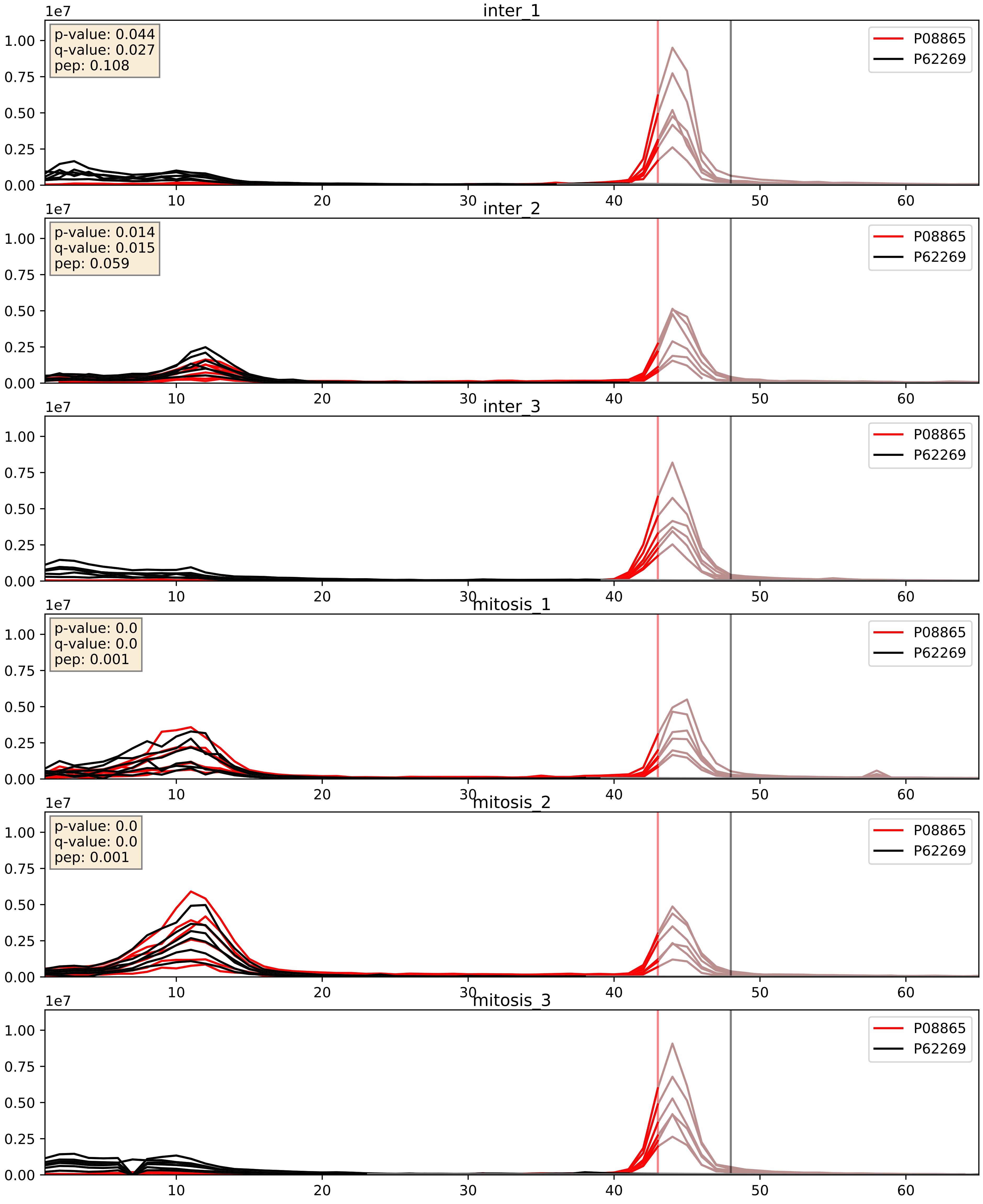


| level | condition_1 | condition_2 | pvalue | pvalue_adjusted |
|---|---|---|---|---|
| complex_abundance | mitosis | inter | 0.14693643767978928 | 0.29271024122387623 |
| interactor_abundance | mitosis | inter | 0.25504636911208617 | 0.4481110261903648 |
| interactor_ratio | mitosis | inter | 0.4922891558694187 | 0.86984863222988 |

P15880_P62269
RS2_HUMAN vs RS18_HUMAN
p-value: 0.0 q-value: 0.0

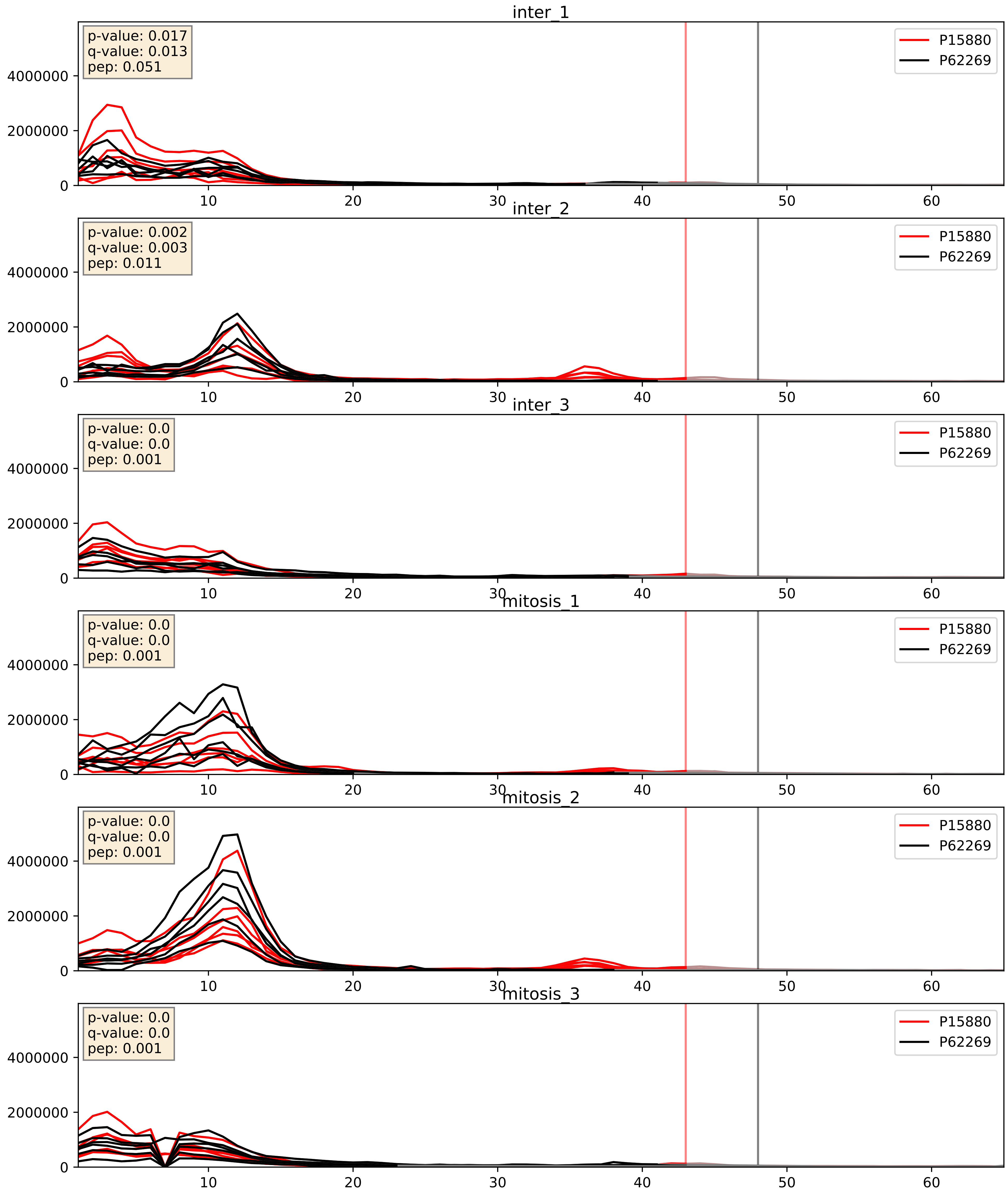


| level | condition_1 | condition_2 | pvalue | pvalue_adjusted |
|---|---|---|---|---|
| interactor_ratio | mitosis | inter | 0.00041647645495156035 | 0.022852810605034336 |
| complex_abundance | mitosis | inter | 0.12185934348542515 | 0.2641110761636074 |
| interactor_abundance | mitosis | inter | 0.29737981089028054 | 0.49163725584376783 |

P18124_P62269
RL7_HUMAN vs RS18_HUMAN
p-value: 0.002 q-value: 0.002

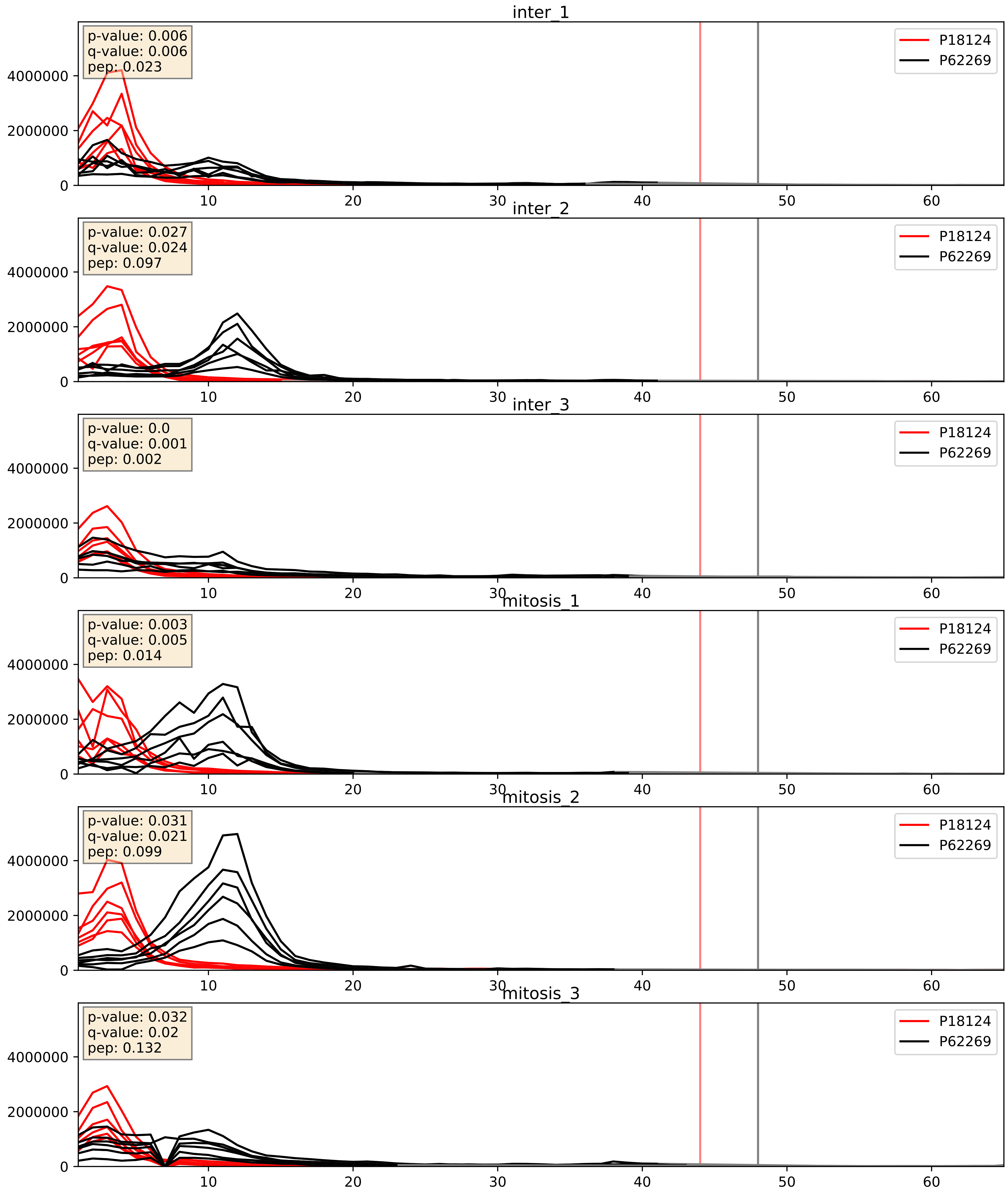


| level | condition_1 | condition_2 | pvalue | pvalue_adjusted |
|---|---|---|---|---|
| complex_abundance | mitosis | inter | 0.04469308772666204 | 0.15121455768388423 |
| interactor_ratio | mitosis | inter | 0.06545770872892186 | 0.42820888628953485 |
| interactor_abundance | mitosis | inter | 0.49648279161018033 | 0.6622168268689187 |

P23396_P62269
RS3_HUMAN vs RS18_HUMAN
p-value: 0.0 q-value: 0.0

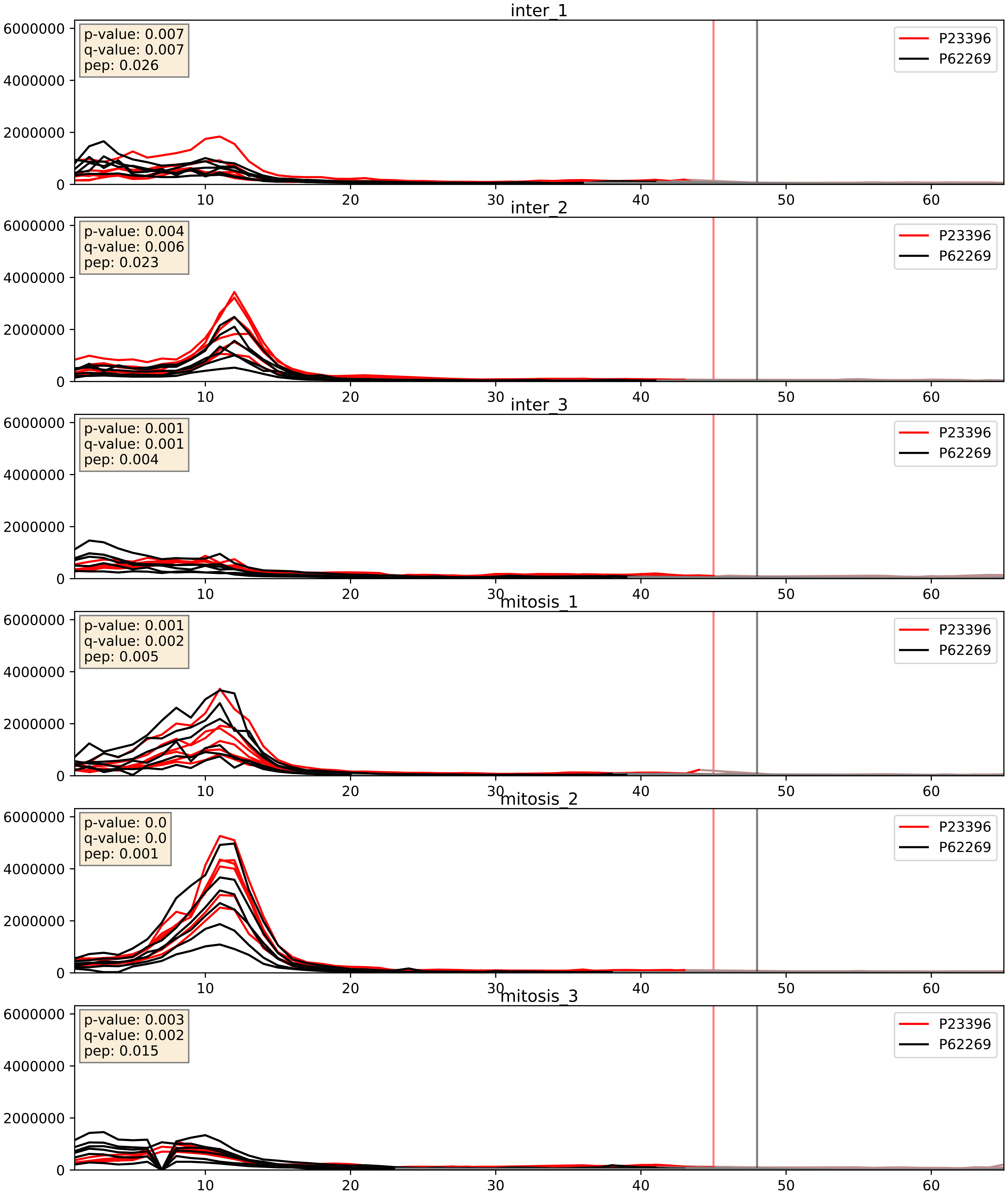


| level | condition_1 | condition_2 | pvalue | pvalue_adjusted |
|---|---|---|---|---|
| interactor_ratio | mitosis | inter | 0.19910927275719648 | 0.6736811410050544 |
| complex_abundance | mitosis | inter | 0.2554300728887485 | 0.40603183359845635 |
| interactor_abundance | mitosis | inter | 0.8016282914082349 | 0.8843273631618649 |

P25398_P62269
RS12_HUMAN vs RS18_HUMAN
p-value: 0.0 q-value: 0.0

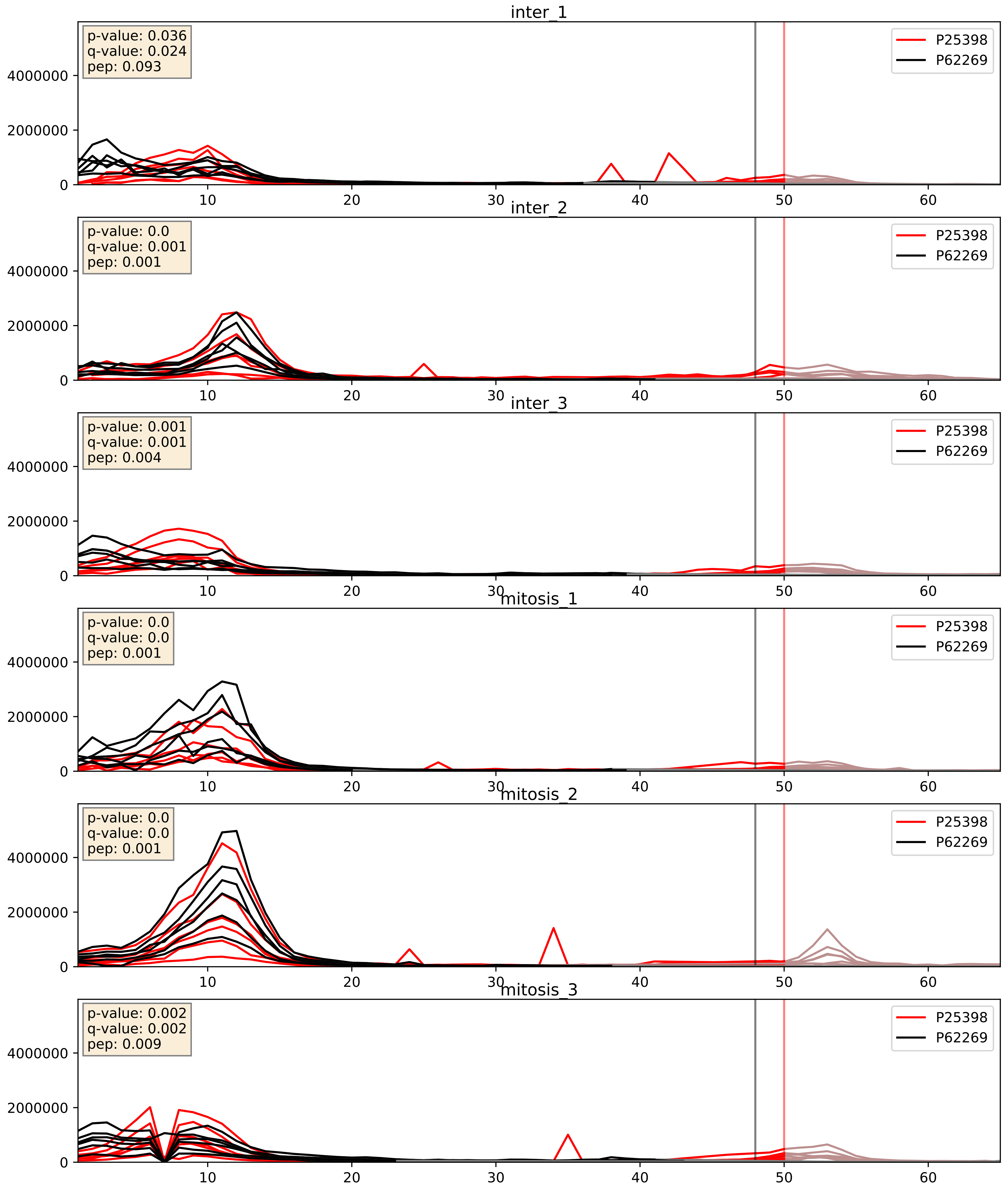


| level | condition_1 | condition_2 | pvalue | pvalue_adjusted |
|---|---|---|---|---|
| interactor_ratio | mitosis | inter | 0.044519419165674576 | 0.3496203927139214 |
| complex_abundance | mitosis | inter | 0.13129815867057482 | 0.2750641796916594 |
| interactor_abundance | mitosis | inter | 0.716021616373337 | 0.8171648604967355 |

P26373_P62269
RL13_HUMAN vs RS18_HUMAN
p-value: 0.002 q-value: 0.003

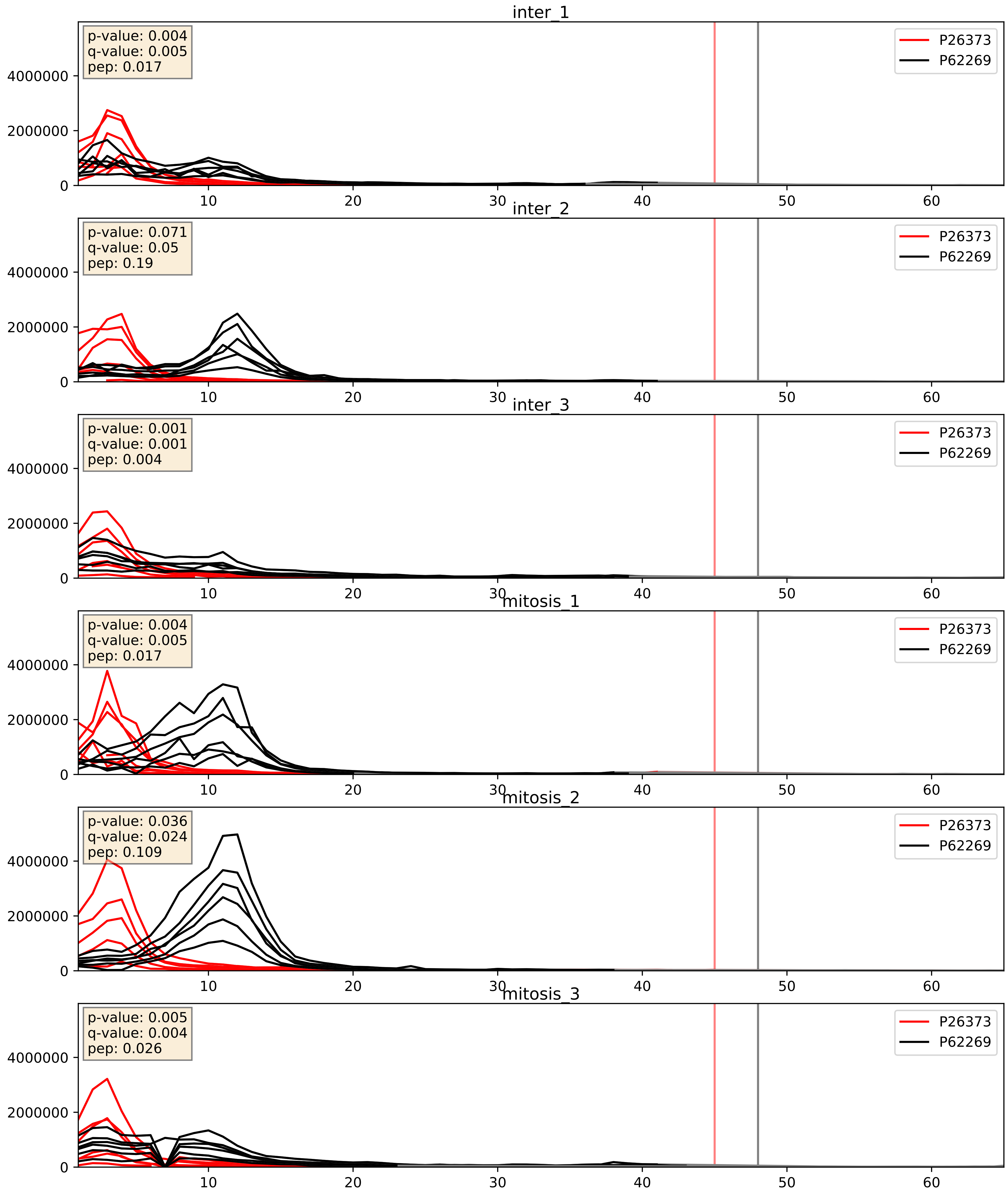


| level | condition_1 | condition_2 | pvalue | pvalue_adjusted |
|---|---|---|---|---|
| complex_abundance | mitosis | inter | 0.002475493221641855 | 0.03616078835708921 |
| interactor_ratio | mitosis | inter | 0.08068962102934402 | 0.47583520079221503 |
| interactor_abundance | mitosis | inter | 0.6092878016969288 | 0.7434257832692909 |

P27635_P62269
RL10_HUMAN vs RS18_HUMAN
p-value: 0.02 q-value: 0.016

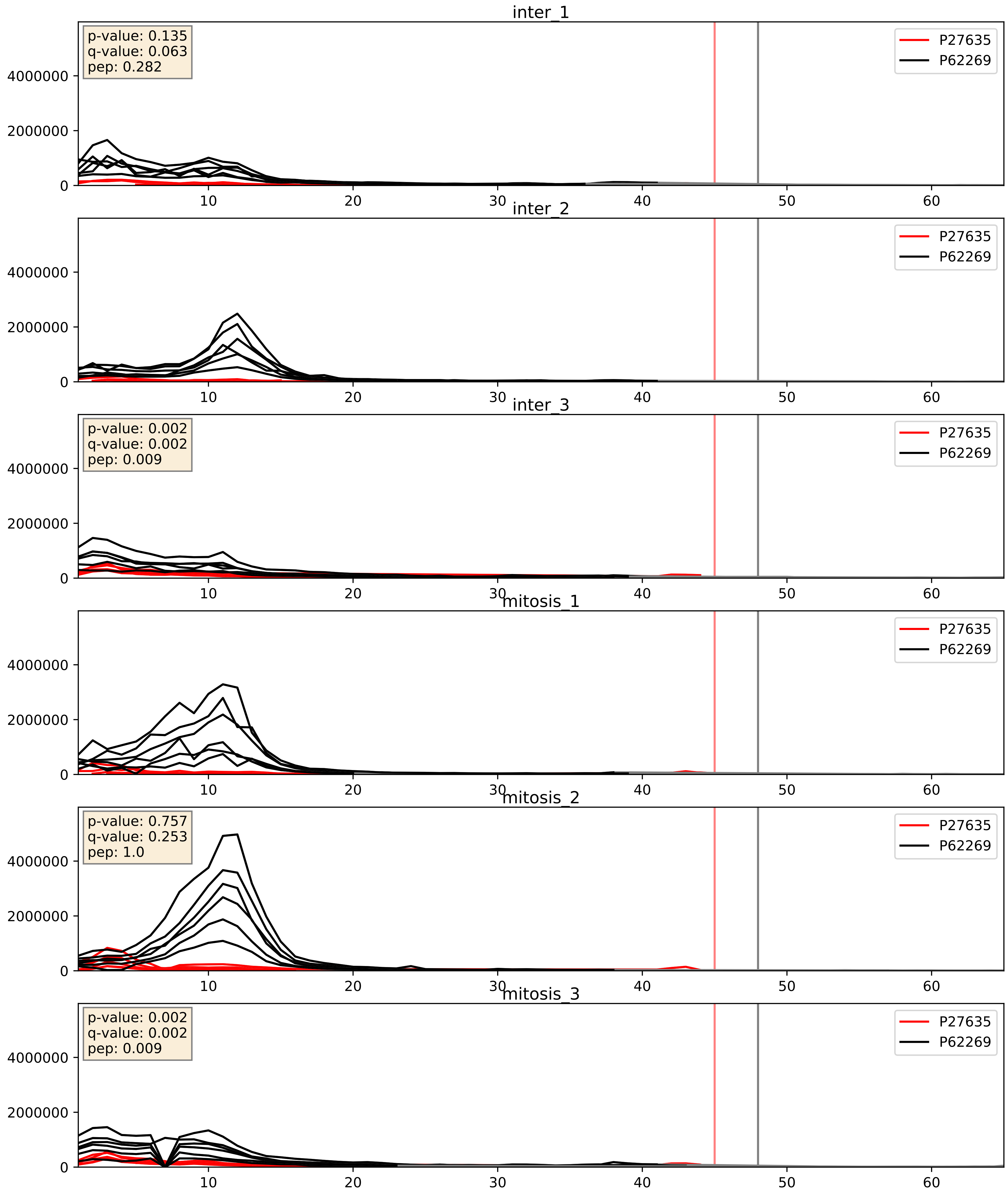


| level | condition_1 | condition_2 | pvalue | pvalue_adjusted |
|---|---|---|---|---|
| complex_abundance | mitosis | inter | 0.16402193607763574 | 0.3129101343491335 |
| interactor_ratio | mitosis | inter | 0.3982186192306028 | 0.82998572695738 |
| interactor_abundance | mitosis | inter | 0.6562920514903565 | 0.7718430941481695 |

P30050_P62269
RL12_HUMAN vs RS18_HUMAN
p-value: 0.0 q-value: 0.0

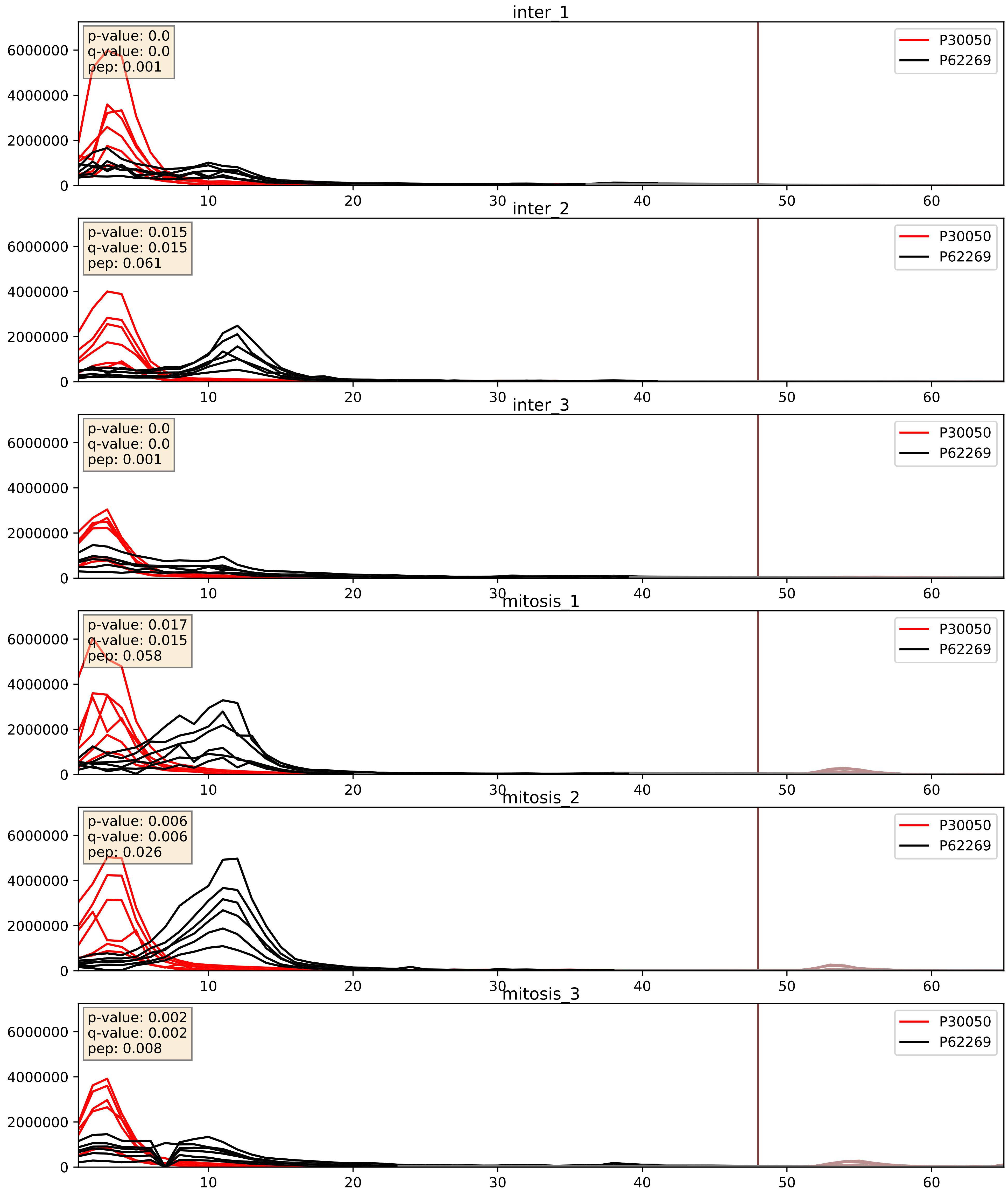


| level | condition_1 | condition_2 | pvalue | pvalue_adjusted |
|---|---|---|---|---|
| complex_abundance | mitosis | inter | 0.007747198218692334 | 0.06546497211451764 |
| interactor_ratio | mitosis | inter | 0.14450137340224645 | 0.5972630402333315 |
| interactor_abundance | mitosis | inter | 0.8816242532233419 | 0.9344027248921942 |

P32969_P62269
RL9_HUMAN vs RS18_HUMAN
p-value: 0.01 q-value: 0.009

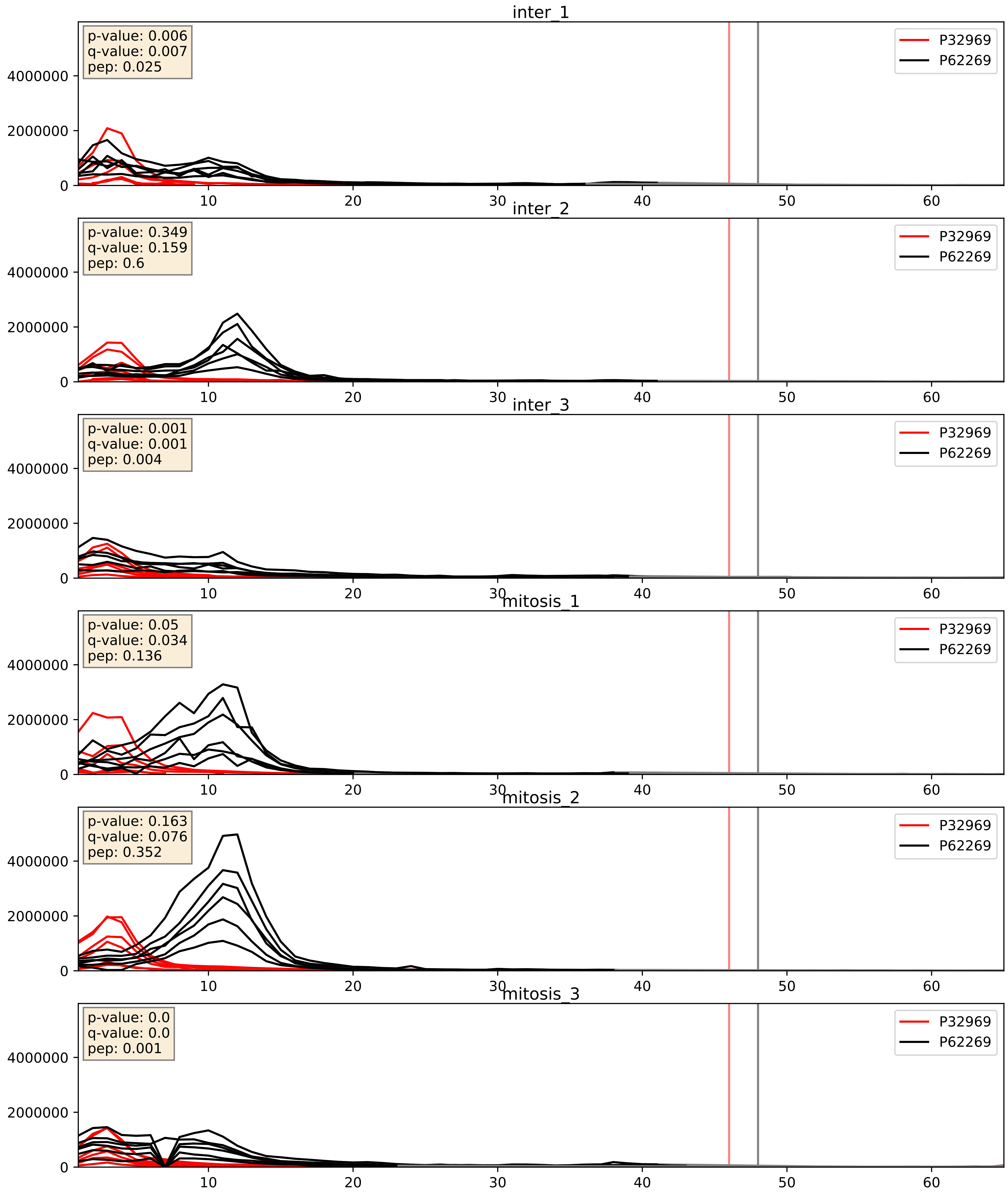


| level | condition_1 | condition_2 | pvalue | pvalue_adjusted |
|---|---|---|---|---|
| complex_abundance | mitosis | inter | 0.01461098170913859 | 0.0898251565234827 |
| interactor_ratio | mitosis | inter | 0.04929111276857474 | 0.3665785623796696 |
| interactor_abundance | mitosis | inter | 0.2103491843605969 | 0.40652154270556234 |

P35268_P62269
RL22_HUMAN vs RS18_HUMAN
p-value: 0.0 q-value: 0.0

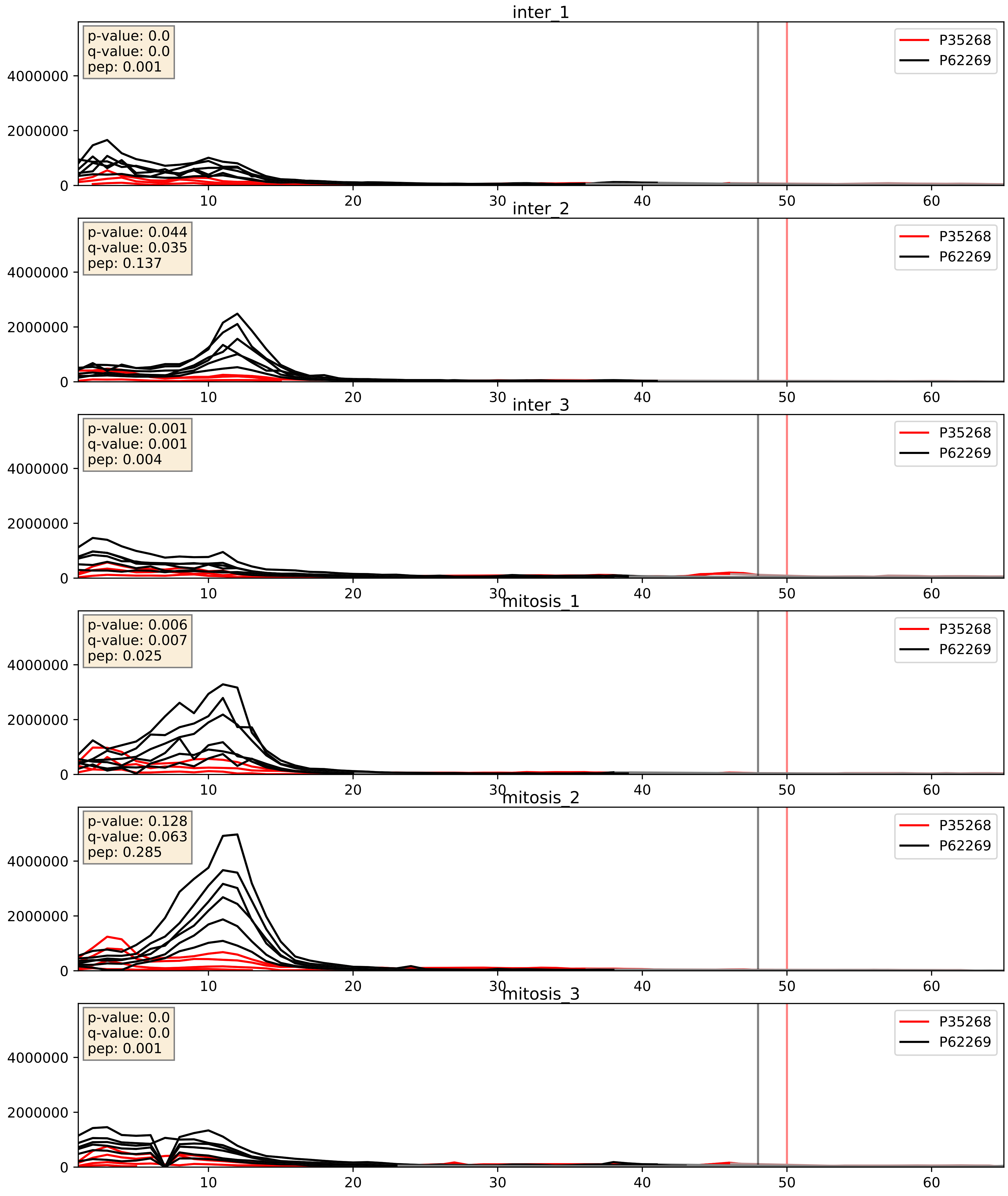


| level | condition_1 | condition_2 | pvalue | pvalue_adjusted |
|---|---|---|---|---|
| complex_abundance | mitosis | inter | 0.012824079295520931 | 0.08405635484106207 |
| interactor_abundance | mitosis | inter | 0.03731151707436548 | 0.15802829520754763 |
| interactor_ratio | mitosis | inter | 0.4899321816095601 | 0.8692763386835093 |

P36578_P62269
RL4_HUMAN vs RS18_HUMAN
p-value: 0.009 q-value: 0.008

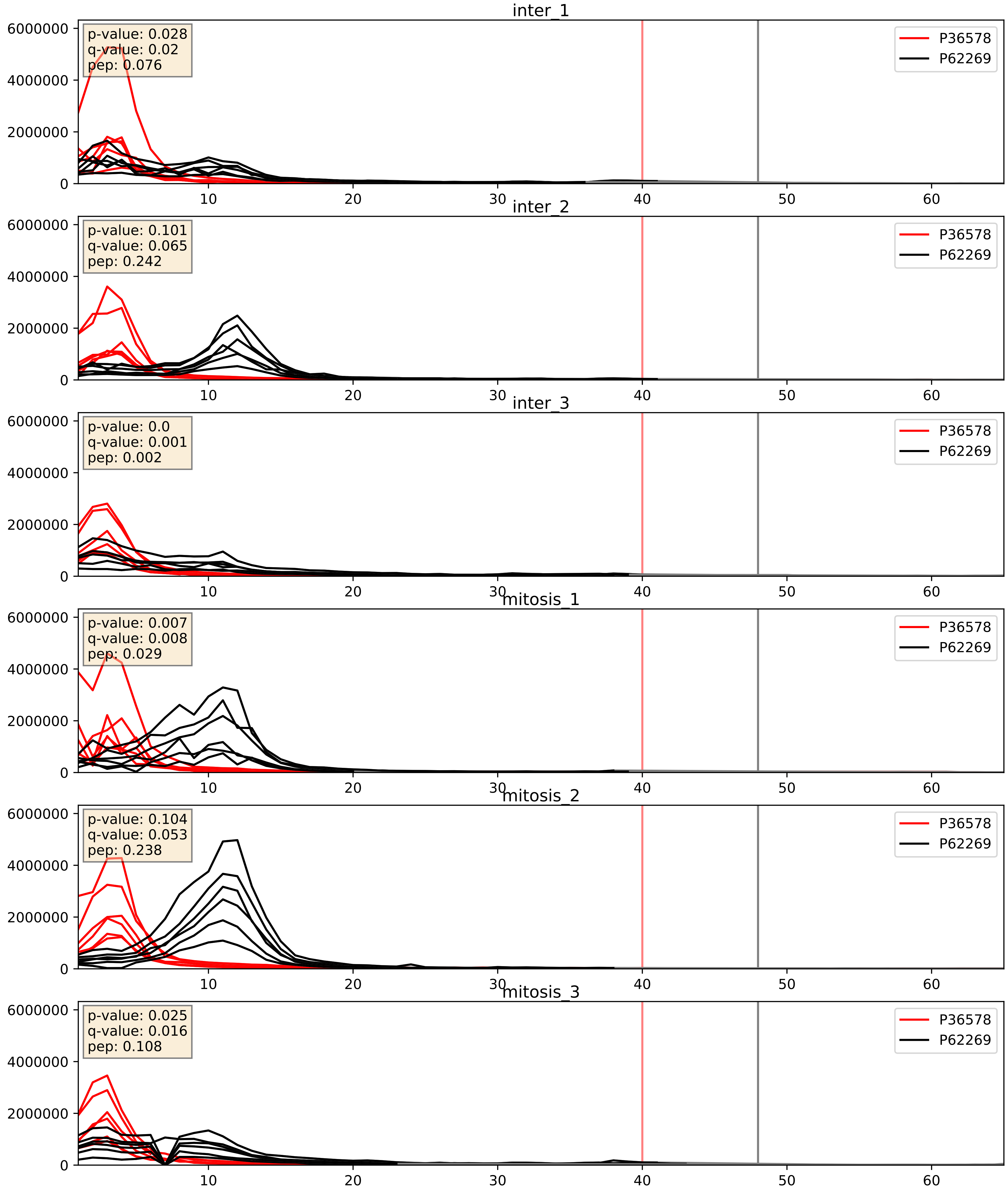


| level | condition_1 | condition_2 | pvalue | pvalue_adjusted |
|---|---|---|---|---|
| complex_abundance | mitosis | inter | 0.015814427027425828 | 0.09374757296036364 |
| interactor_ratio | mitosis | inter | 0.04334152288613971 | 0.3425701162560997 |
| interactor_abundance | mitosis | inter | 0.9858211760603667 | 0.9908564864764562 |

P37108_P62269
SRP14_HUMAN vs RS18_HUMAN
p-value: 0.007 q-value: 0.006

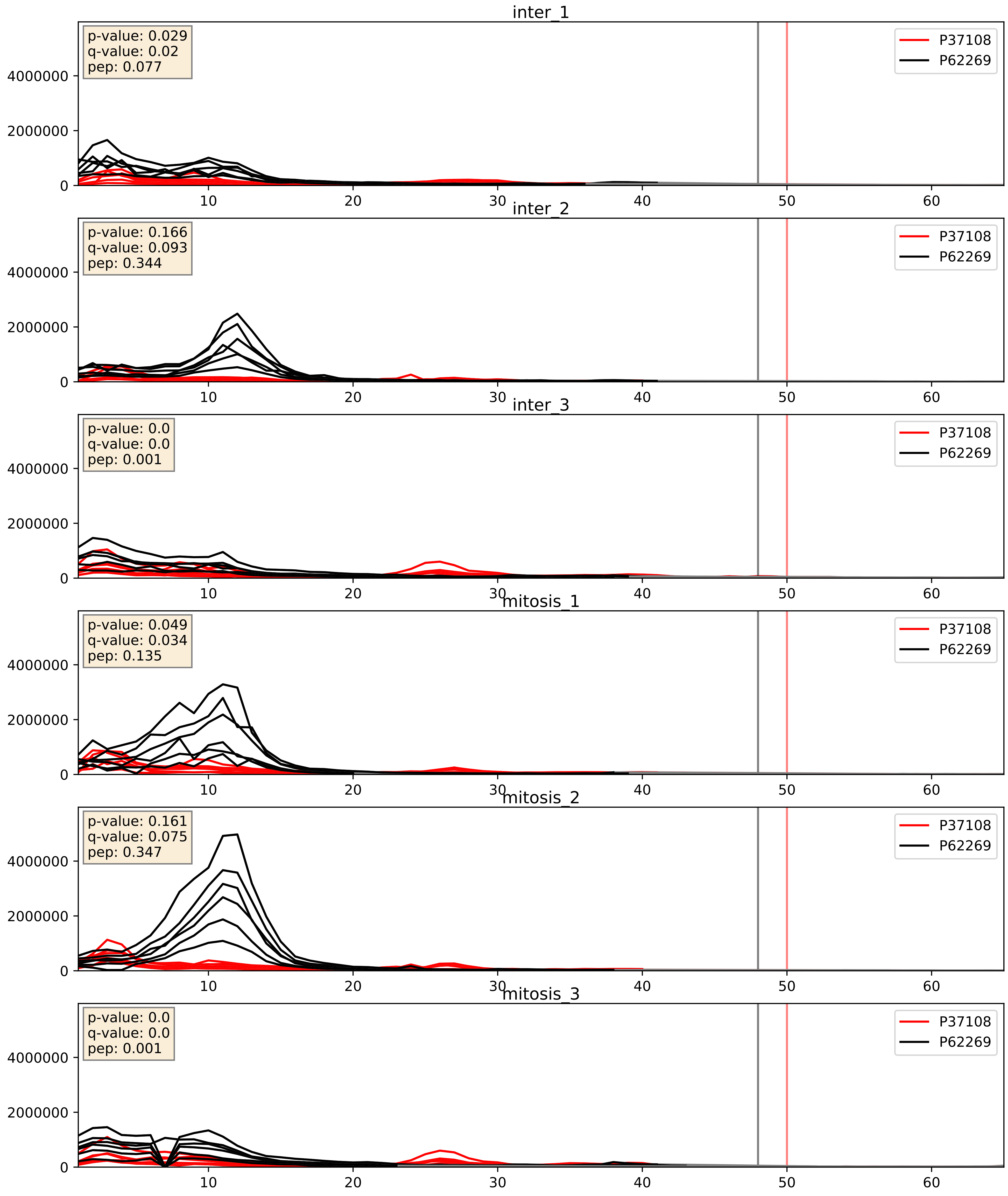


| level | condition_1 | condition_2 | pvalue | pvalue_adjusted |
|---|---|---|---|---|
| complex_abundance | mitosis | inter | 0.18976272331156624 | 0.3413517188001596 |
| interactor_abundance | mitosis | inter | 0.5558107473188234 | 0.707733729572202 |
| interactor_ratio | mitosis | inter | 0.8043147253663596 | 0.9634160836222222 |

P39019_P62269
RS19_HUMAN vs RS18_HUMAN
p-value: 0.0 q-value: 0.0

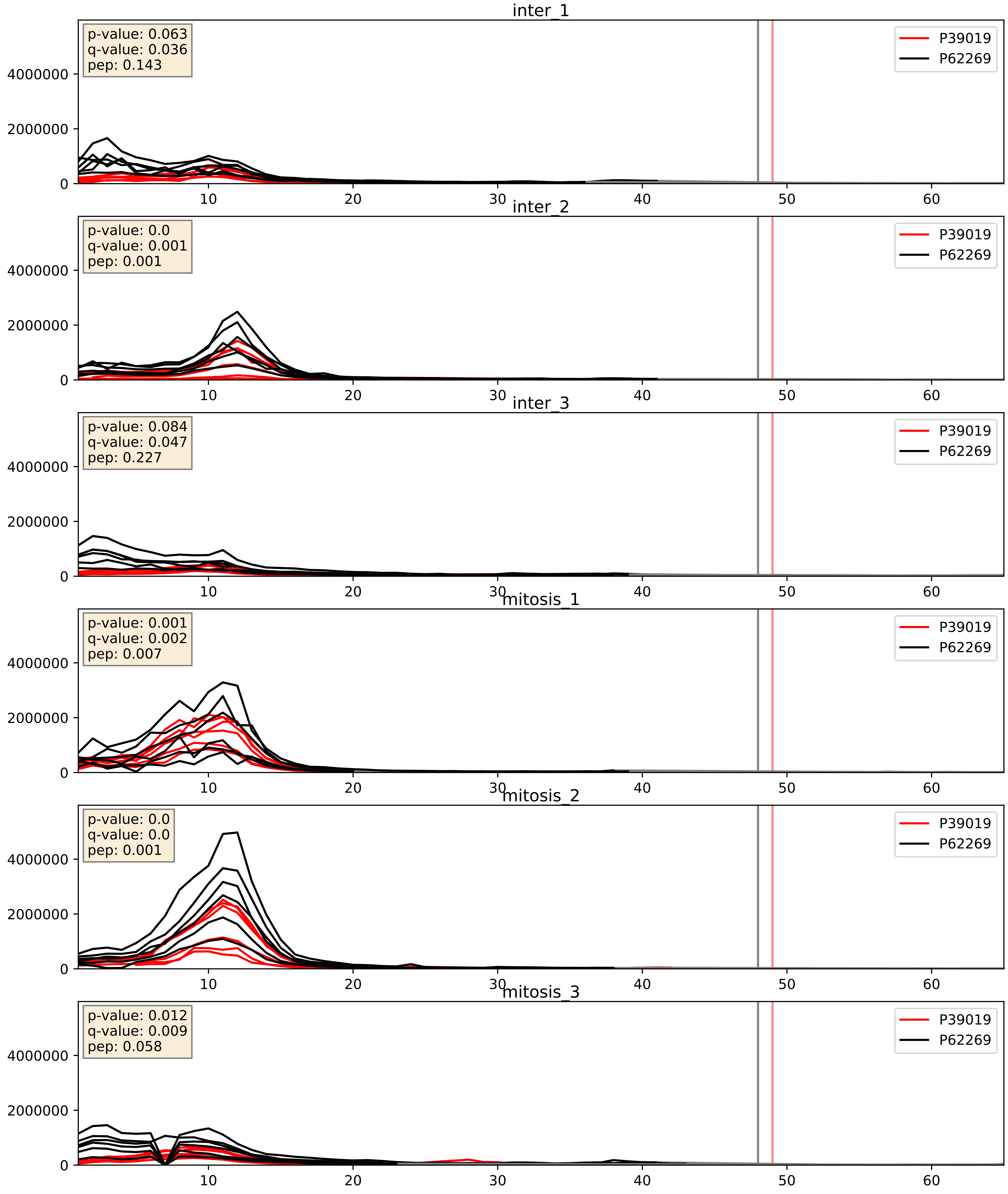


| level | condition_1 | condition_2 | pvalue | pvalue_adjusted |
|---|---|---|---|---|
| complex_abundance | mitosis | inter | 0.07091664589726682 | 0.19345012392626001 |
| interactor_abundance | mitosis | inter | 0.2118751697034527 | 0.40715039907095185 |
| interactor_ratio | mitosis | inter | 0.8392488947963327 | 0.9737835196474741 |

P39023_P62269
RL3_HUMAN vs RS18_HUMAN
p-value: 0.036 q-value: 0.025

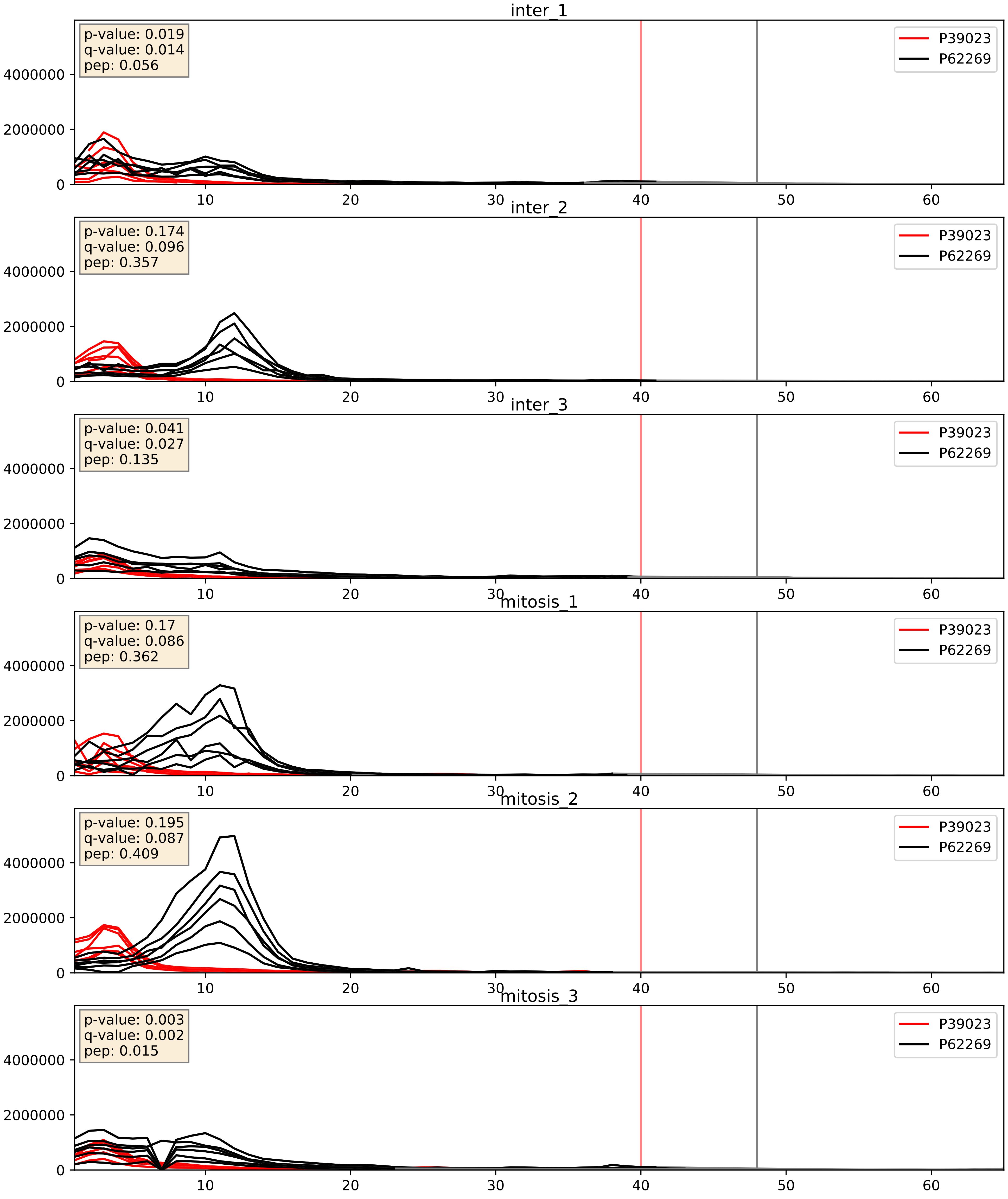


| level | condition_1 | condition_2 | pvalue | pvalue_adjusted |
|---|---|---|---|---|
| interactor_ratio | mitosis | inter | 0.030996755775698798 | 0.2804780438054775 |
| complex_abundance | mitosis | inter | 0.04362861134872484 | 0.1496214737595309 |
| interactor_abundance | mitosis | inter | 0.4577908838109454 | 0.629609570279835 |

P42677_P62269
RS27_HUMAN vs RS18_HUMAN
p-value: 0.011 q-value: 0.009

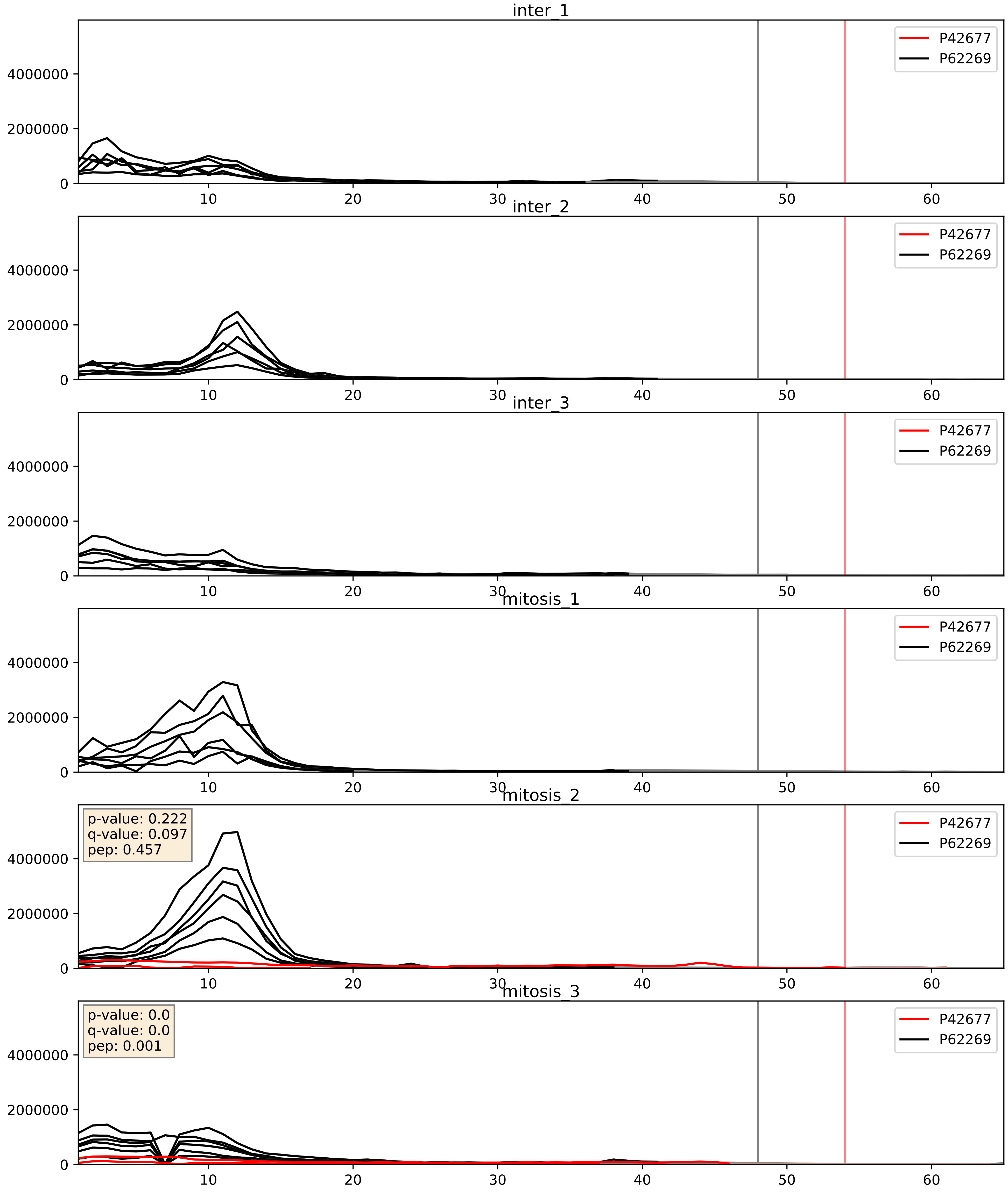


| level | condition_1 | condition_2 | pvalue | pvalue_adjusted |
|---|---|---|---|---|
| interactor_ratio | mitosis | inter | 0.1261182666820657 | 0.5750062399685051 |
| complex_abundance | mitosis | inter | 0.13705024440173494 | 0.2820982332168934 |
| interactor_abundance | mitosis | inter | 0.15575859694663505 | 0.34249153565282997 |

P42766_P62269
RL35_HUMAN vs RS18_HUMAN
p-value: 0.0 q-value: 0.0

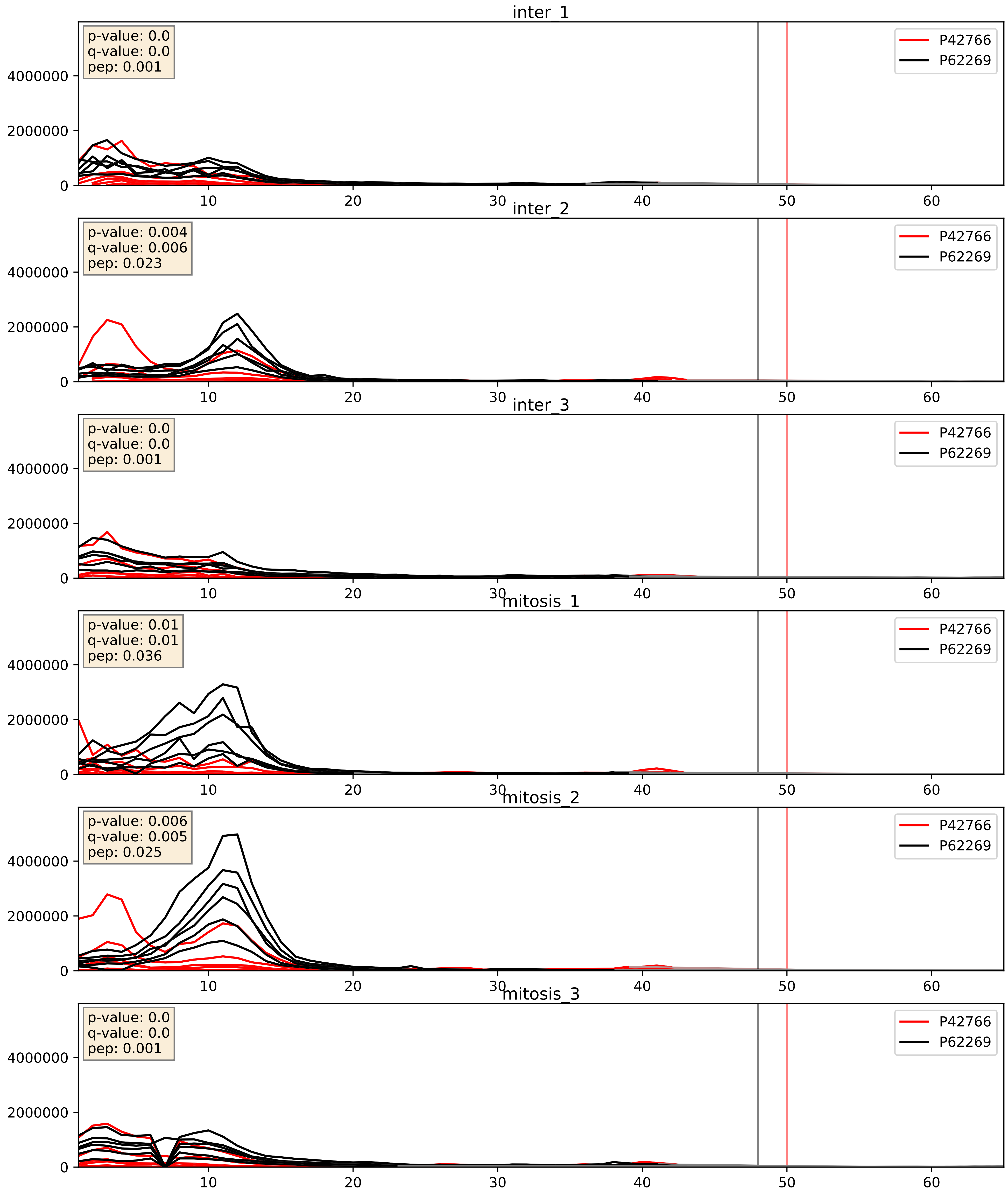


| level | condition_1 | condition_2 | pvalue | pvalue_adjusted |
|---|---|---|---|---|
| interactor_ratio | mitosis | inter | 0.009905798910117317 | 0.14263182888175546 |
| complex_abundance | mitosis | inter | 0.06345210036503628 | 0.18165551141294664 |
| interactor_abundance | mitosis | inter | 0.8084321695920049 | 0.8894496230032257 |

P46776_P62269
RL27A_HUMAN vs RS18_HUMAN
p-value: 0.0 q-value: 0.0

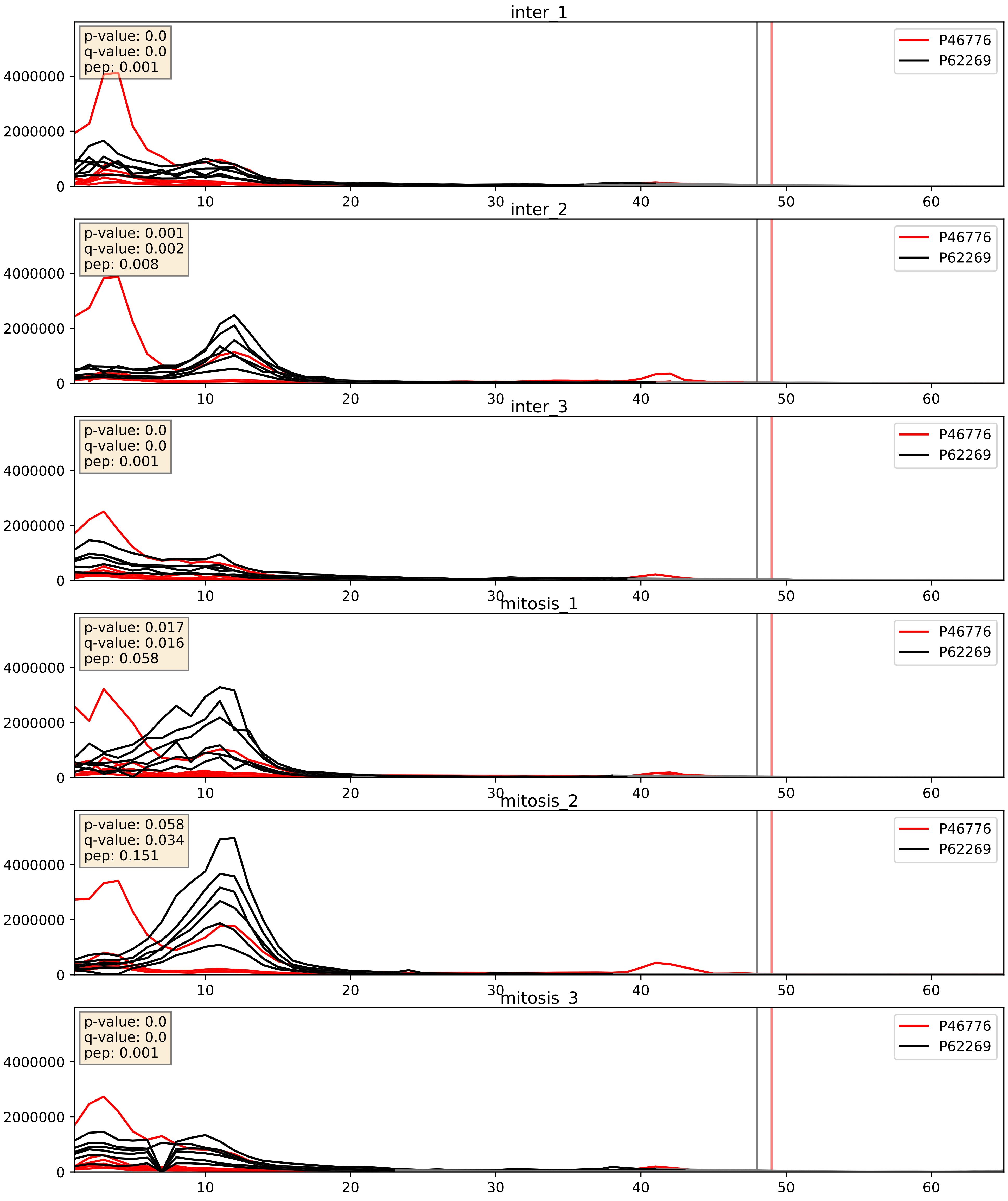


| level | condition_1 | condition_2 | pvalue | pvalue_adjusted |
|---|---|---|---|---|
| complex_abundance | mitosis | inter | 0.005805741536474478 | 0.057519846703960105 |
| interactor_ratio | mitosis | inter | 0.050499148773522248 | 0.37168762983779235 |
| interactor_abundance | mitosis | inter | 0.692224768235187 | 0.7964306473243549 |

P46778_P62269
RL21_HUMAN vs RS18_HUMAN
p-value: 0.006 q-value: 0.006

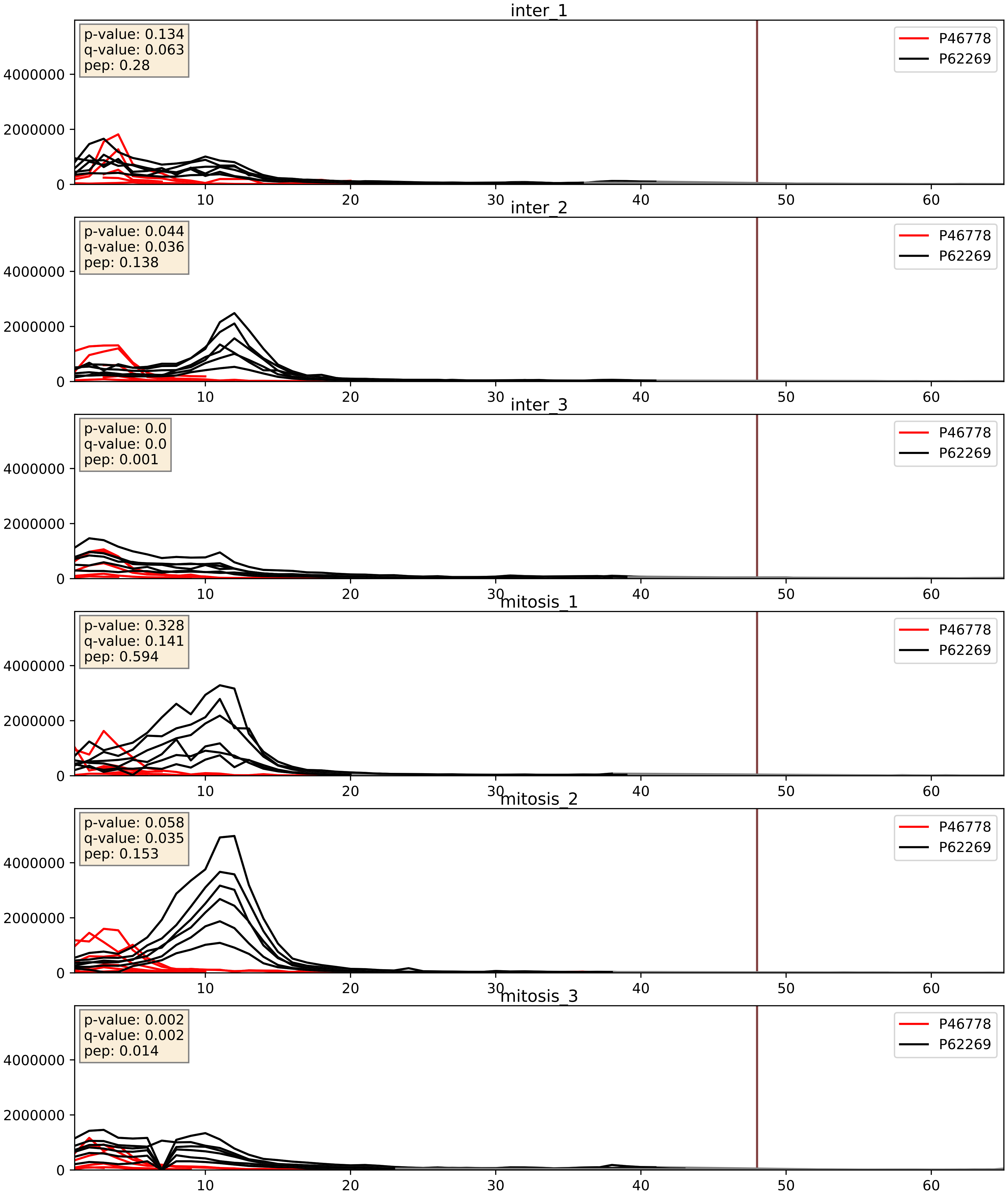


| level | condition_1 | condition_2 | pvalue | pvalue_adjusted |
|---|---|---|---|---|
| interactor_ratio | mitosis | inter | 0.0059474269897681195 | 0.10496902068539196 |
| complex_abundance | mitosis | inter | 0.23555744923466201 | 0.3879129983548878 |
| interactor_abundance | mitosis | inter | 0.4845822245322886 | 0.6544702818064209 |

P46779_P62269
RL28_HUMAN vs RS18_HUMAN
p-value: 0.0 q-value: 0.0

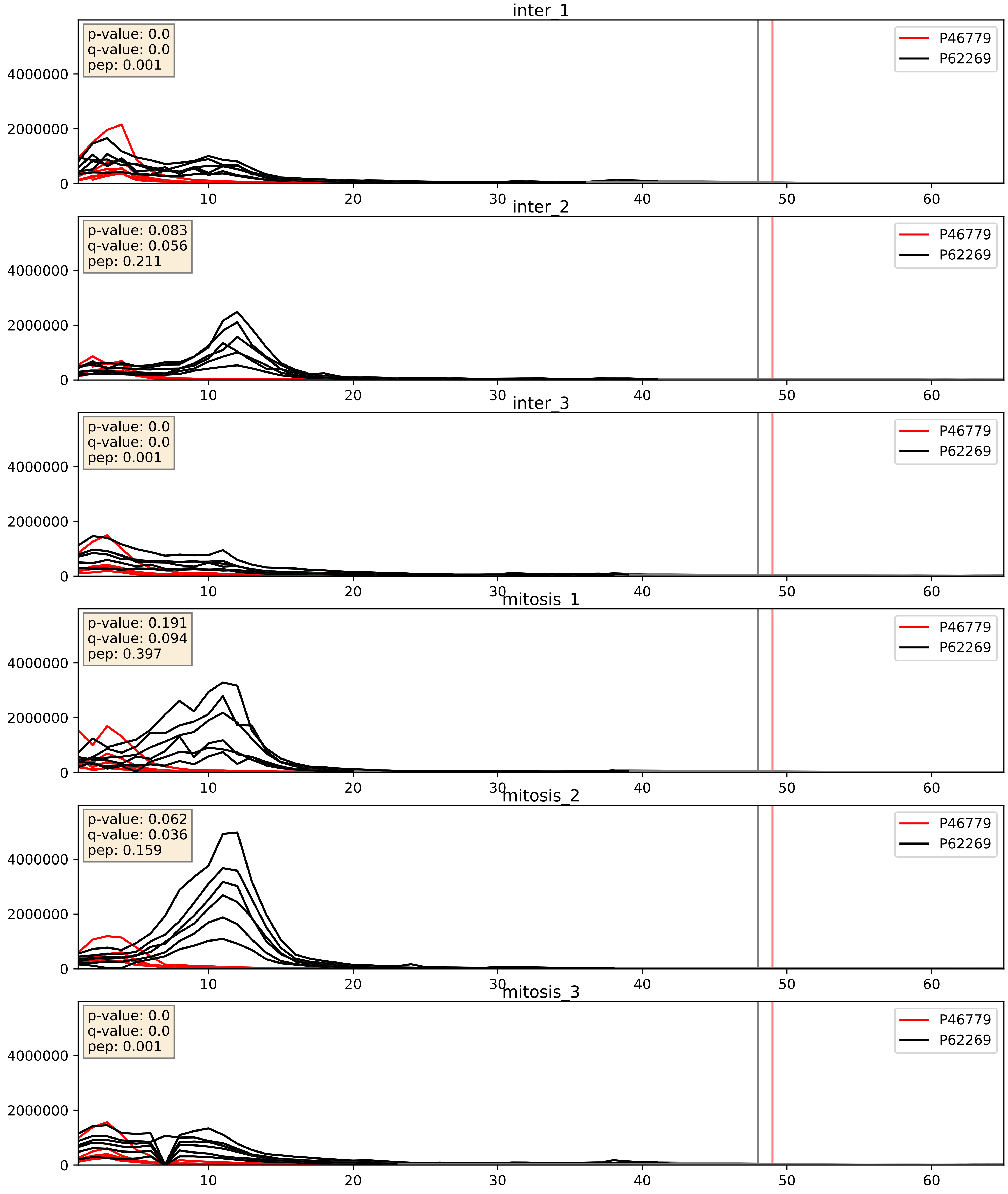


| level | condition_1 | condition_2 | pvalue | pvalue_adjusted |
|---|---|---|---|---|
| interactor_ratio | mitosis | inter | 0.014059519207909826 | 0.1741671265118786 |
| complex_abundance | mitosis | inter | 0.06934300042322808 | 0.19079912684758352 |
| interactor_abundance | mitosis | inter | 0.20770102547817784 | 0.4048550103821479 |

P46781_P62269
RS9_HUMAN vs RS18_HUMAN
p-value: 0.0 q-value: 0.0

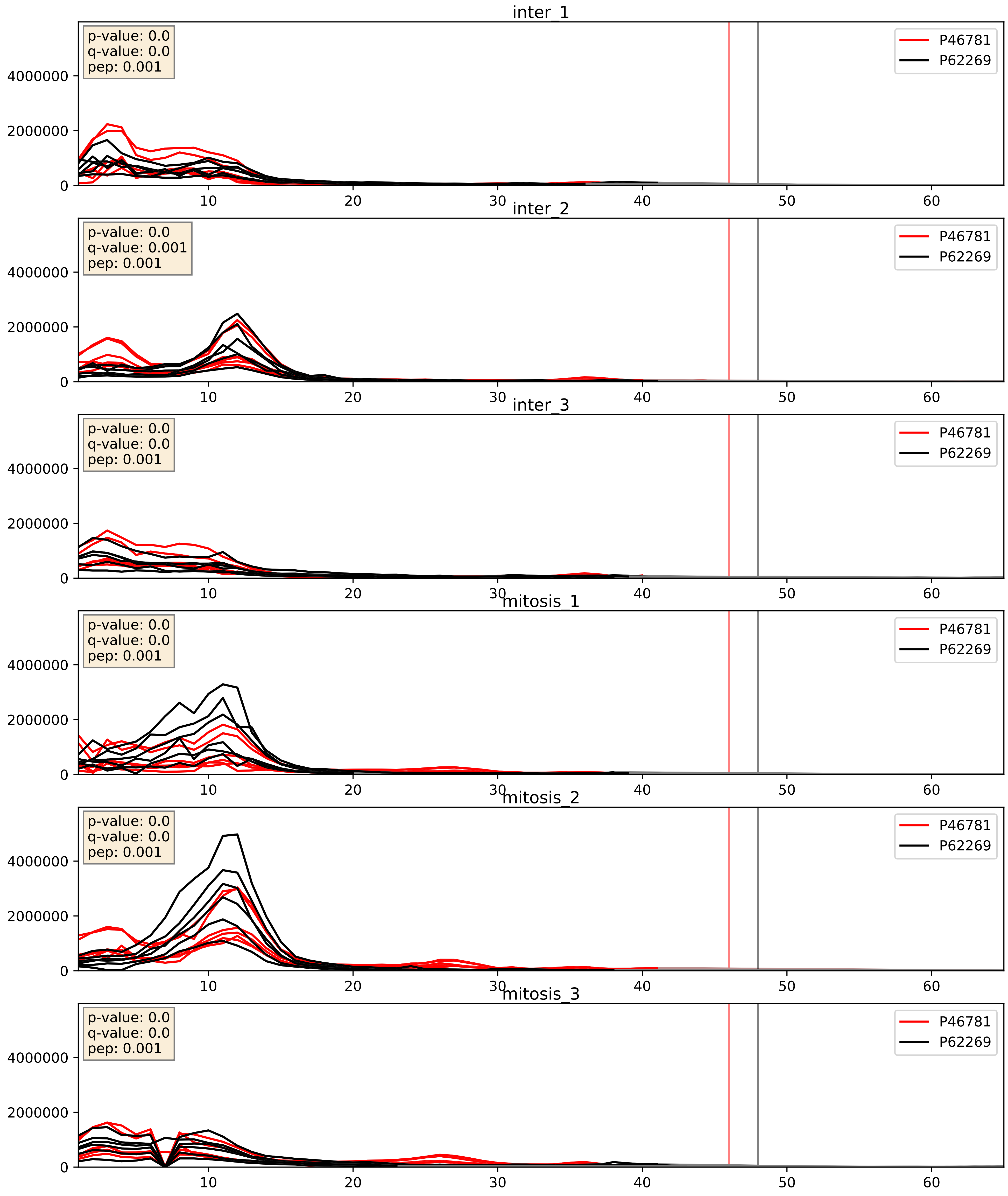


| level | condition_1 | condition_2 | pvalue | pvalue_adjusted |
|---|---|---|---|---|
| interactor_ratio | mitosis | inter | 0.0260912184418928 | 0.25543205403291264 |
| complex_abundance | mitosis | inter | 0.053133538181358894 | 0.16455249161810137 |
| interactor_abundance | mitosis | inter | 0.7657979703540582 | 0.8606053073691399 |

P46782_P62269
RS5_HUMAN vs RS18_HUMAN
p-value: 0.0 q-value: 0.001

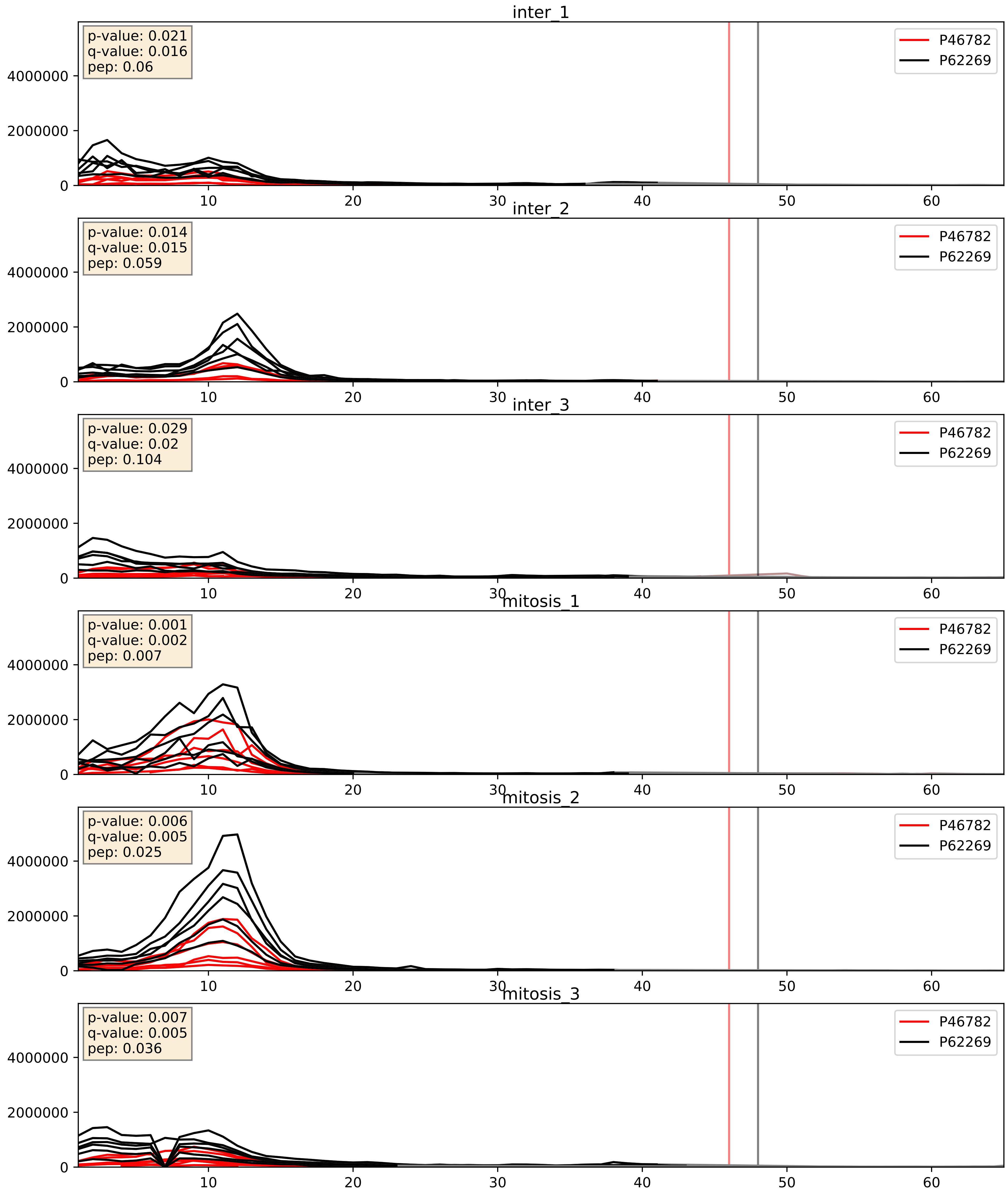


| level | condition_1 | condition_2 | pvalue | pvalue_adjusted |
|---|---|---|---|---|
| complex_abundance | mitosis | inter | 0.020352272483554926 | 0.10515559050023798 |
| interactor_abundance | mitosis | inter | 0.11668647902250837 | 0.29516437956048214 |
| interactor_ratio | mitosis | inter | 0.7442831453801275 | 0.9491588884573259 |

P46783_P62269
RS10_HUMAN vs RS18_HUMAN
p-value: 0.004 q-value: 0.004

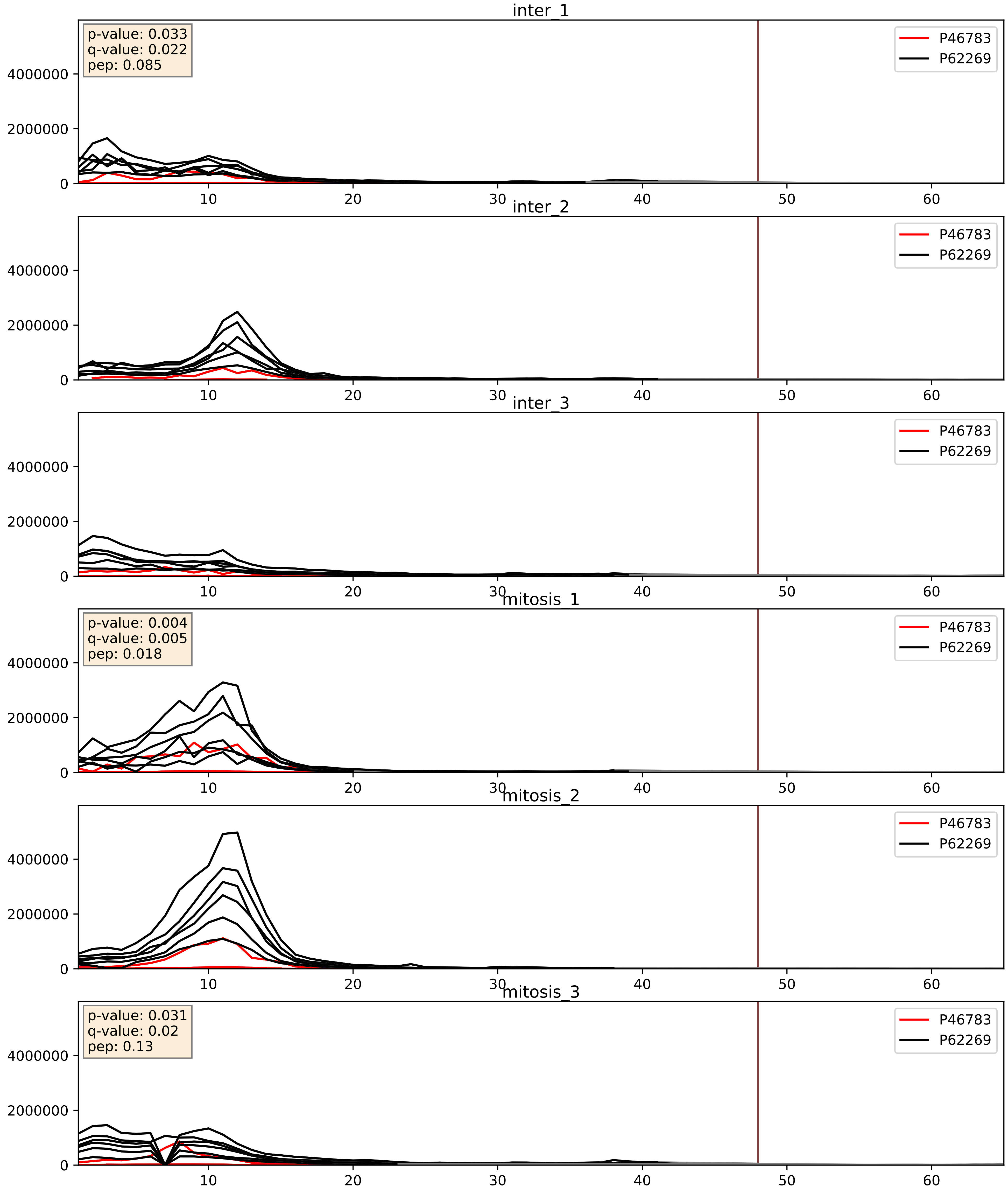


| level | condition_1 | condition_2 | pvalue | pvalue_adjusted |
|---|---|---|---|---|
| complex_abundance | mitosis | inter | 0.08100911760185286 | 0.20854635595369386 |
| interactor_ratio | mitosis | inter | 0.15362109570191307 | 0.6107740730182889 |
| interactor_abundance | mitosis | inter | 0.9635437341838033 | 0.9767804789925814 |

P47914_P62269
RL29_HUMAN vs RS18_HUMAN
p-value: 0.0 q-value: 0.0

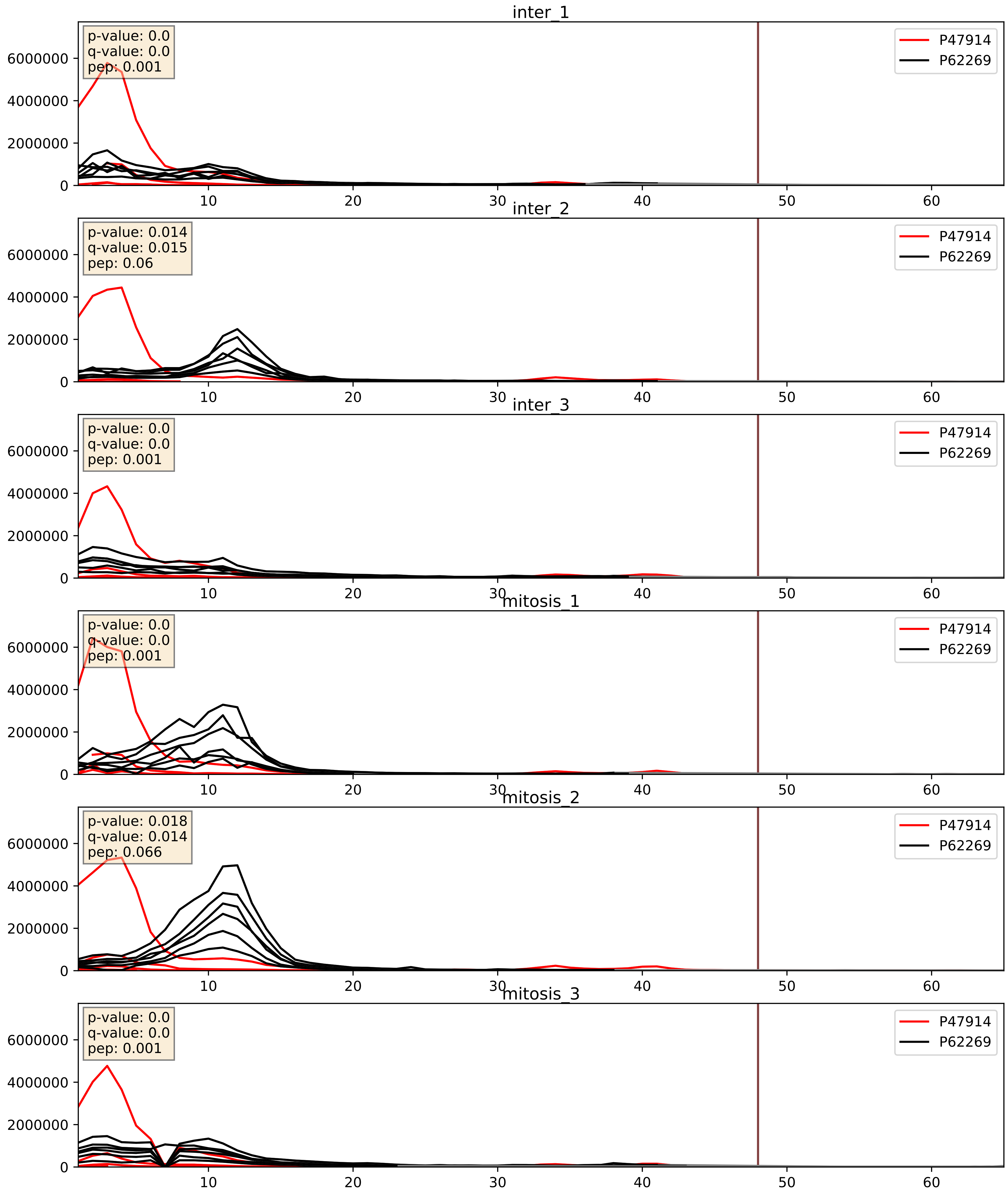


| level | condition_1 | condition_2 | pvalue | pvalue_adjusted |
|---|---|---|---|---|
| complex_abundance | mitosis | inter | 0.06509373018616871 | 0.18450408291178944 |
| interactor_abundance | mitosis | inter | 0.41490076134959875 | 0.597652590181332 |
| interactor_ratio | mitosis | inter | 0.6186608912950338 | 0.9118004871703665 |

P49207_P62269
RL34_HUMAN vs RS18_HUMAN
p-value: 0.024 q-value: 0.019

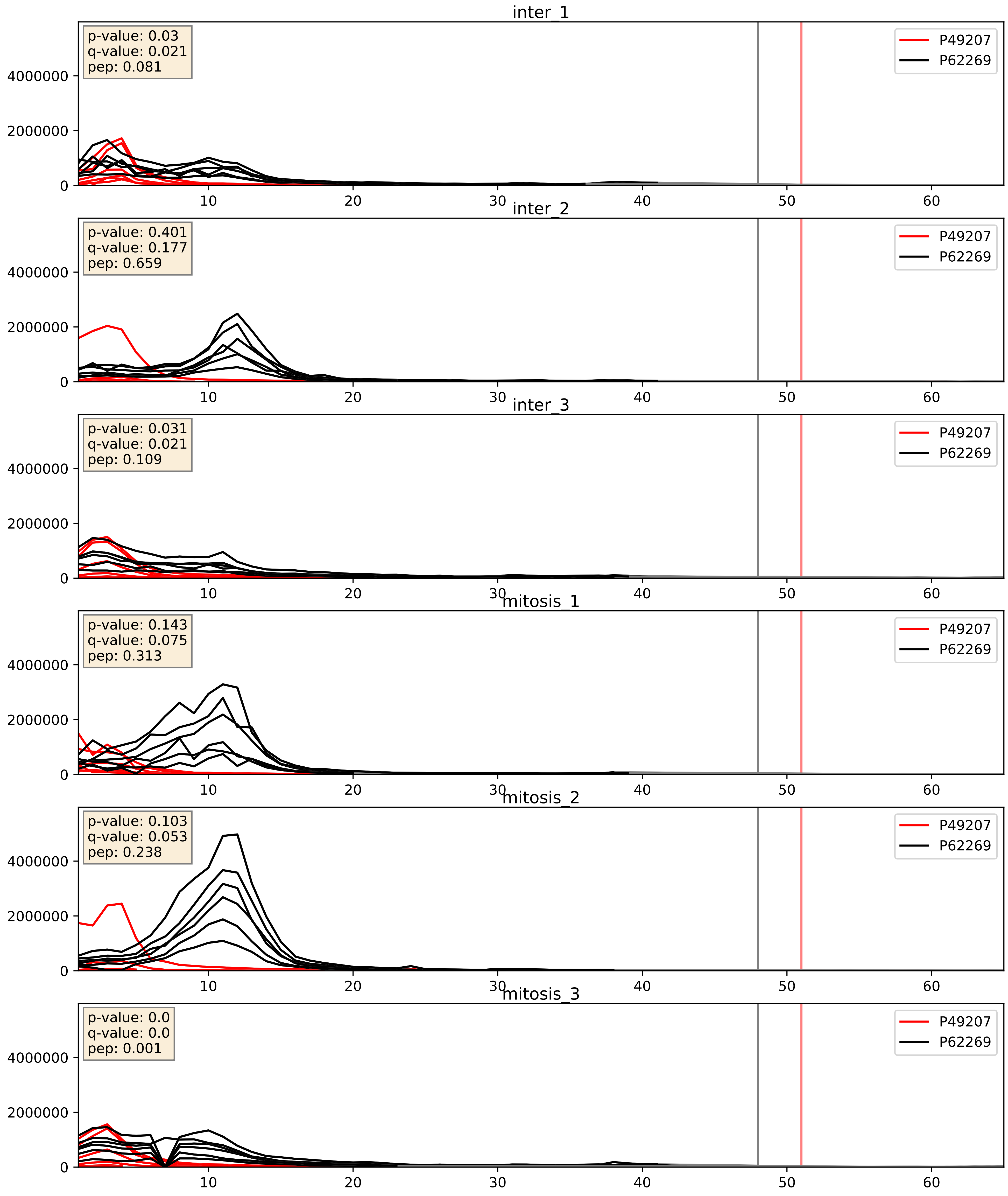


| level | condition_1 | condition_2 | pvalue | pvalue_adjusted |
|---|---|---|---|---|
| interactor_ratio | mitosis | inter | 0.3148293516058483 | 0.7783421708721073 |
| complex_abundance | mitosis | inter | 0.5513752263473786 | 0.6848846768360524 |
| interactor_abundance | mitosis | inter | 0.6896689856436142 | 0.7949860647871448 |

P49458_P62269
SRP09_HUMAN vs RS18_HUMAN
p-value: 0.009 q-value: 0.008

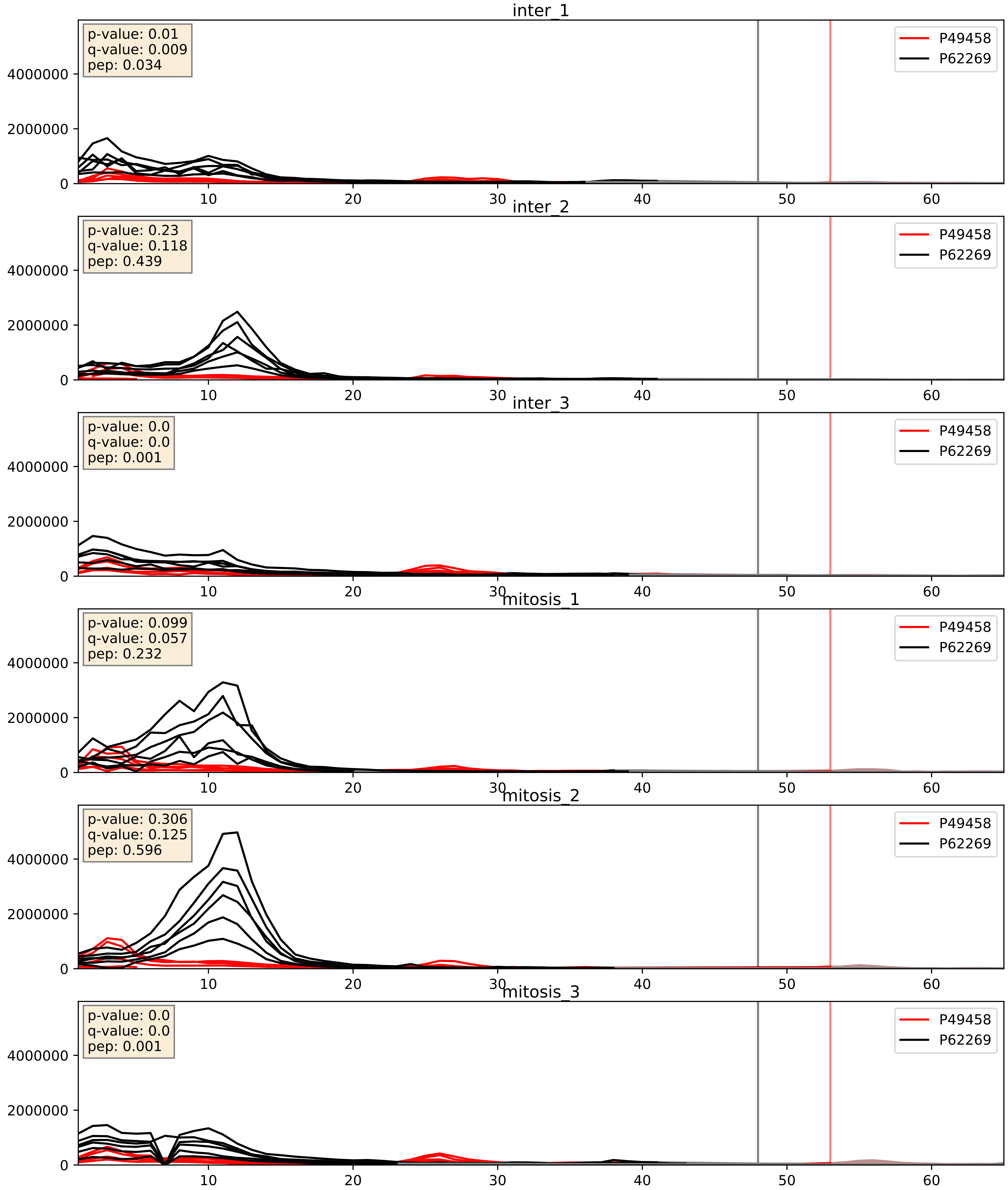


| level | condition_1 | condition_2 | pvalue | pvalue_adjusted |
|---|---|---|---|---|
| complex_abundance | mitosis | inter | 0.03035345538192613 | 0.12643580441327867 |
| interactor_ratio | mitosis | inter | 0.2024027702385481 | 0.6778433932871564 |
| interactor_abundance | mitosis | inter | 0.43867321308572516 | 0.6167941366645544 |

P50914_P62269
RL14_HUMAN vs RS18_HUMAN
p-value: 0.006 q-value: 0.006

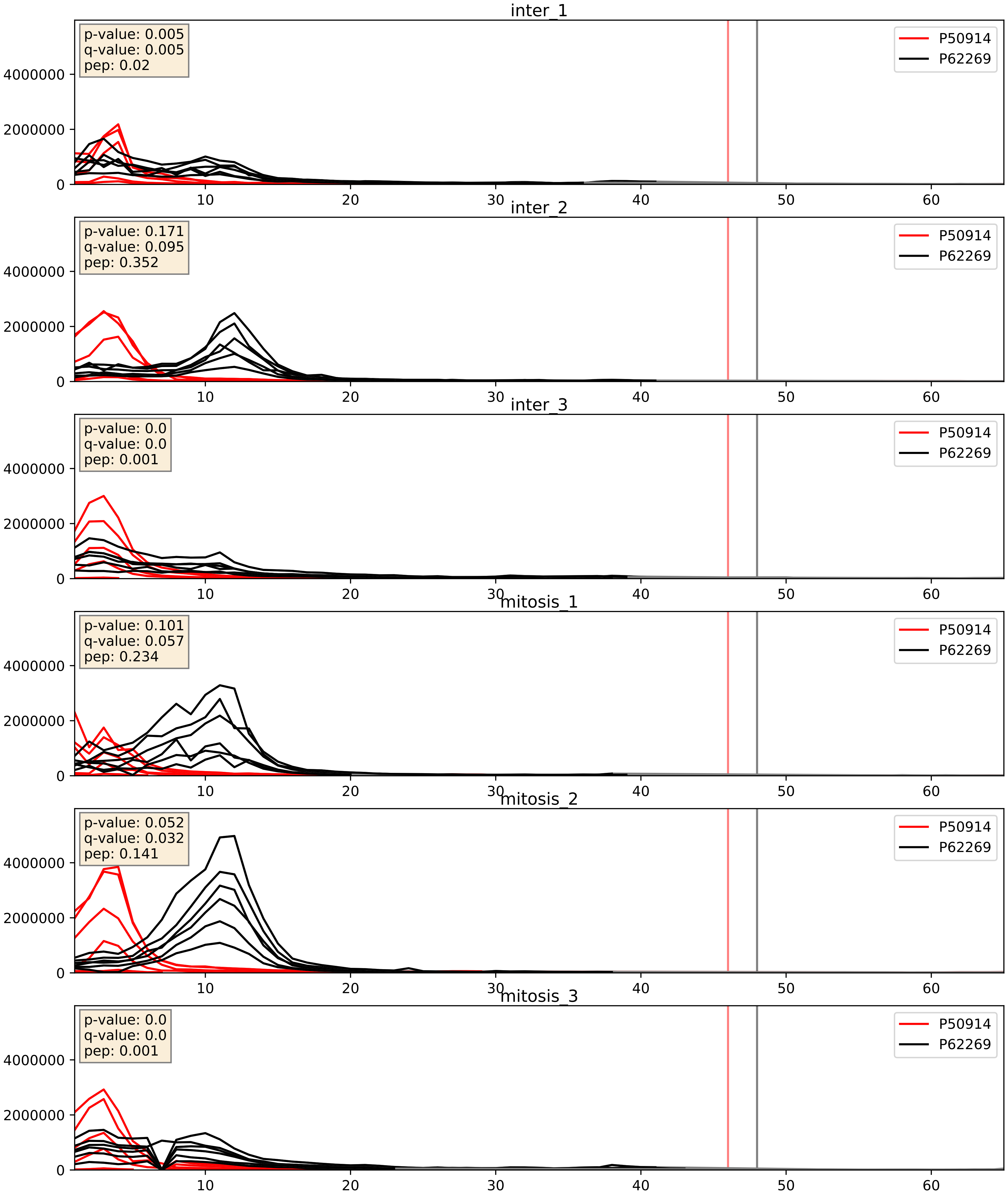


| level | condition_1 | condition_2 | pvalue | pvalue_adjusted |
|---|---|---|---|---|
| interactor_ratio | mitosis | inter | 0.05338320847374129 | 0.38327943971738976 |
| complex_abundance | mitosis | inter | 0.24158985751800655 | 0.39293353227325406 |
| interactor_abundance | mitosis | inter | 0.8451161396574962 | 0.9151994630233623 |

P51571_P62269
SSRD_HUMAN vs RS18_HUMAN
p-value: 0.001 q-value: 0.042

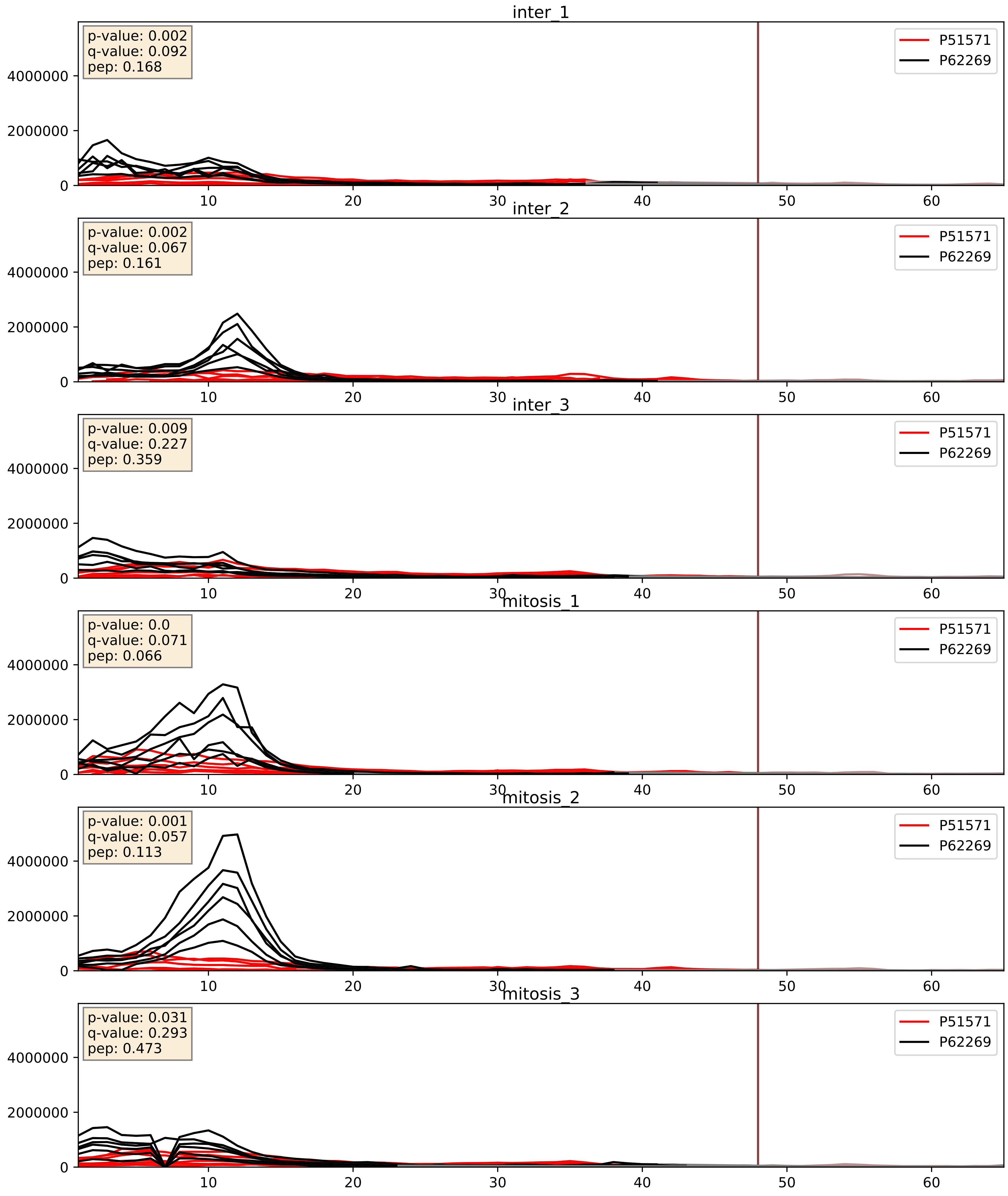


| level | condition_1 | condition_2 | pvalue | pvalue_adjusted |
|---|---|---|---|---|
| complex_abundance | mitosis | inter | 0.05979943868484623 | 0.17506265223744313 |
| interactor_ratio | mitosis | inter | 0.06860291211775972 | 0.43402877141760765 |
| interactor_abundance | mitosis | inter | 0.488070337426603 | 0.6572613998854279 |

P52815_P62269
RM12_HUMAN vs RS18_HUMAN
p-value: 0.0 q-value: 0.015

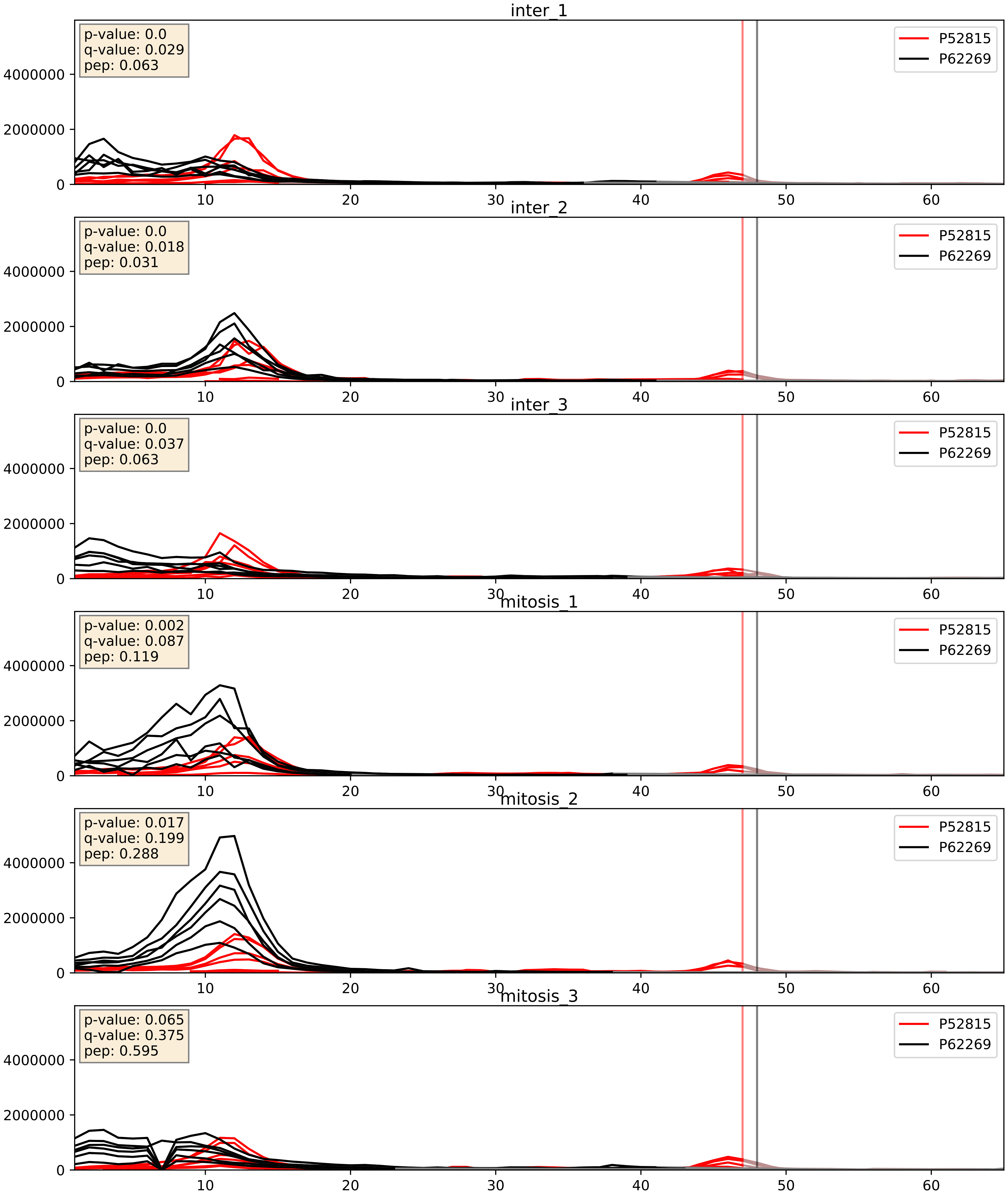


| level | condition_1 | condition_2 | pvalue | pvalue_adjusted |
|---|---|---|---|---|
| complex_abundance | mitosis | inter | 0.00925955438140137 | 0.07106059027160733 |
| interactor_ratio | mitosis | inter | 0.014758850915369316 | 0.17767618346953945 |
| interactor_abundance | mitosis | inter | 0.027918492319173822 | 0.1388624603440603 8 |

P60468_P62269
SC61B_HUMAN vs RS18_HUMAN
p-value: 0.003 q-value: 0.003

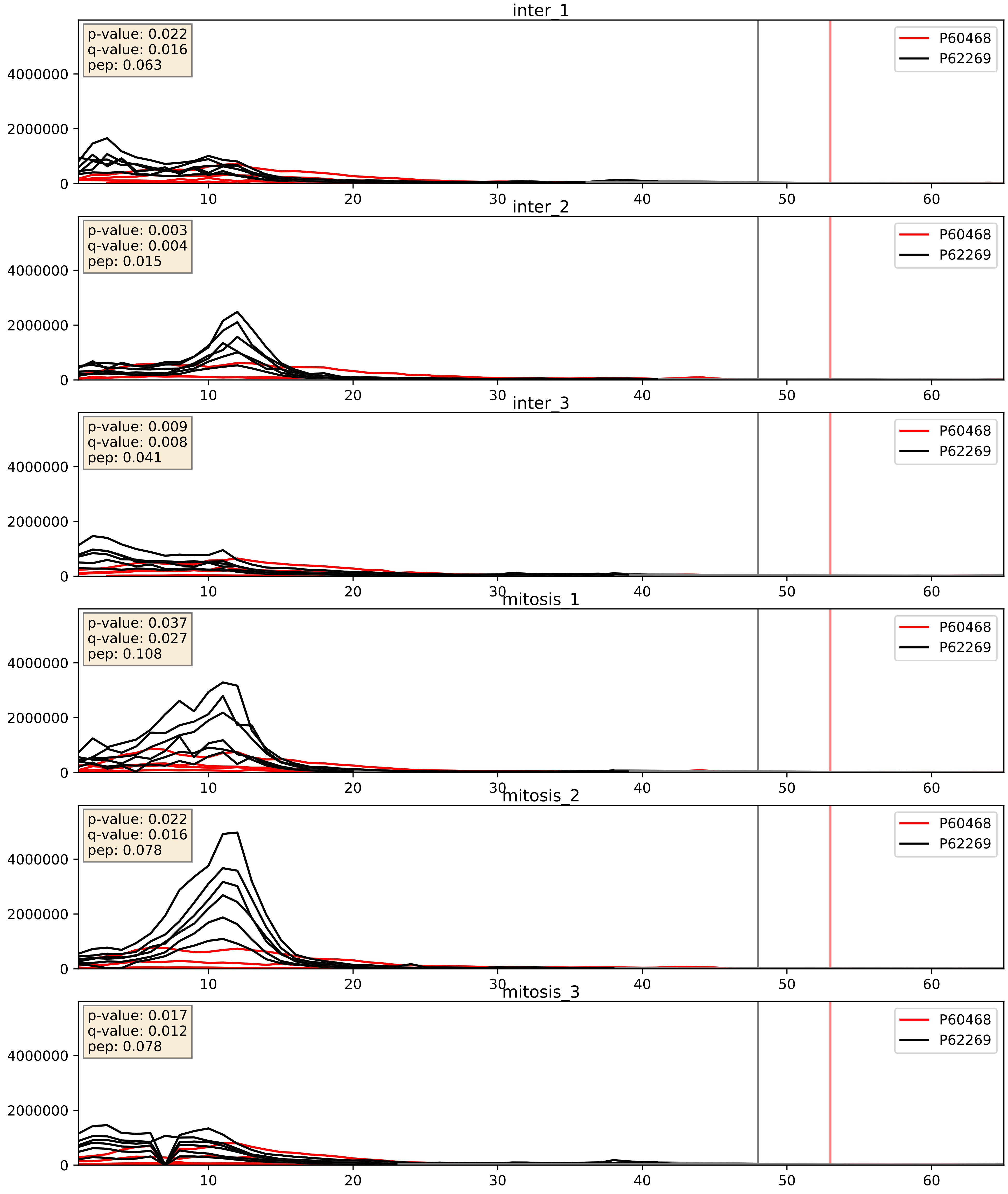


| level | condition_1 | condition_2 | pvalue | pvalue_adjusted |
|---|---|---|---|---|
| interactor_ratio | mitosis | inter | 0.06806144227815981 | 0.4323044818991264 |
| interactor_abundance | mitosis | inter | 0.2647477545276368 | 0.45687793447647373 |
| complex_abundance | mitosis | inter | 0.30513507089515246 | 0.4552196842234692 |

P60866_P62269
RS20_HUMAN vs RS18_HUMAN
p-value: 0.0 q-value: 0.0

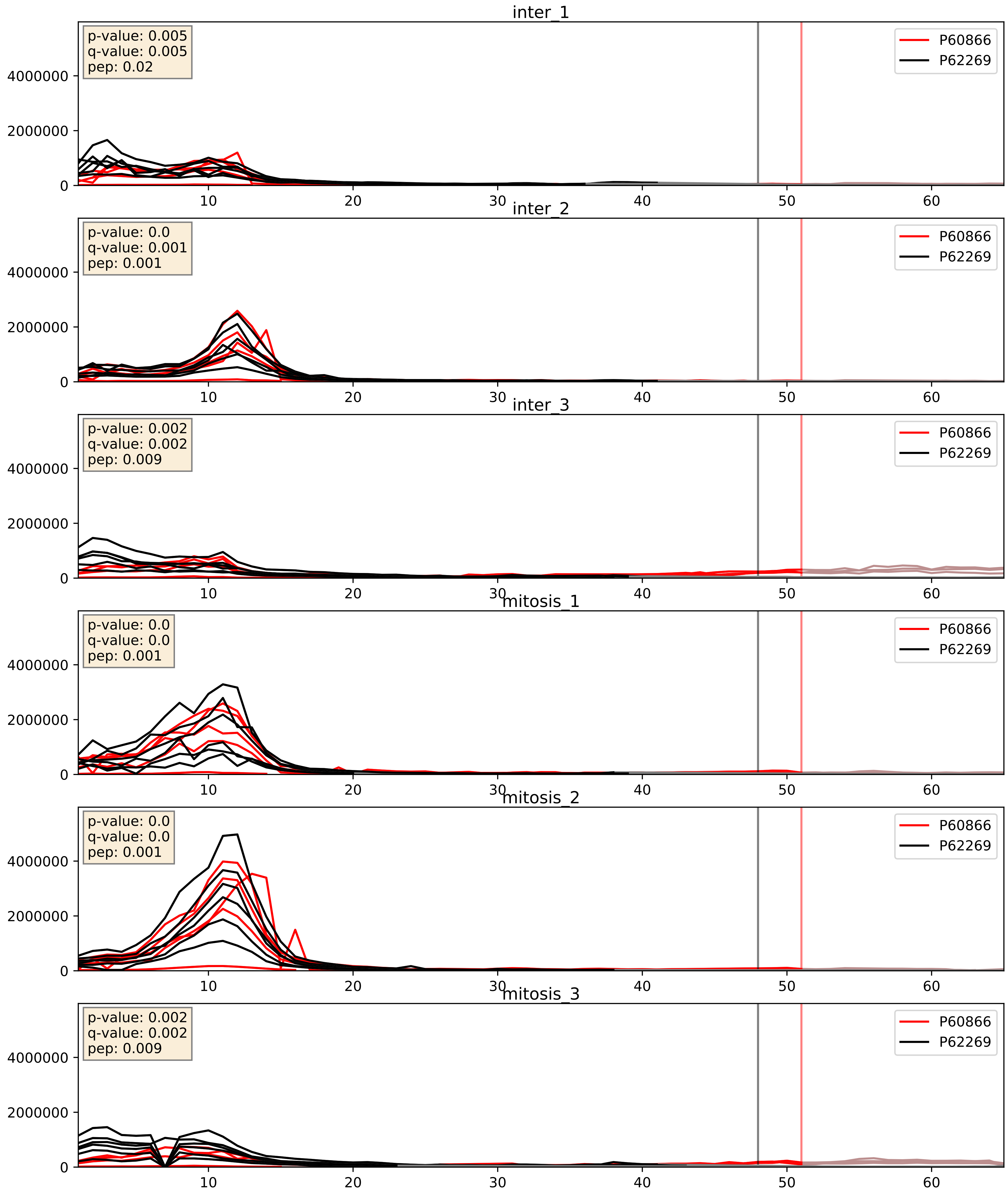


| level | condition_1 | condition_2 | pvalue | pvalue_adjusted |
|---|---|---|---|---|
| complex_abundance | mitosis | inter | 0.10737258421972205 | 0.2443542206845082 |
| interactor_abundance | mitosis | inter | 0.3905076992182803 | 0.5825871649997609 |
| interactor_ratio | mitosis | inter | 0.47827247030619513 | 0.8660416446969278 |

P61247_P62269
RS3A_HUMAN vs RS18_HUMAN
p-value: 0.0 q-value: 0.0

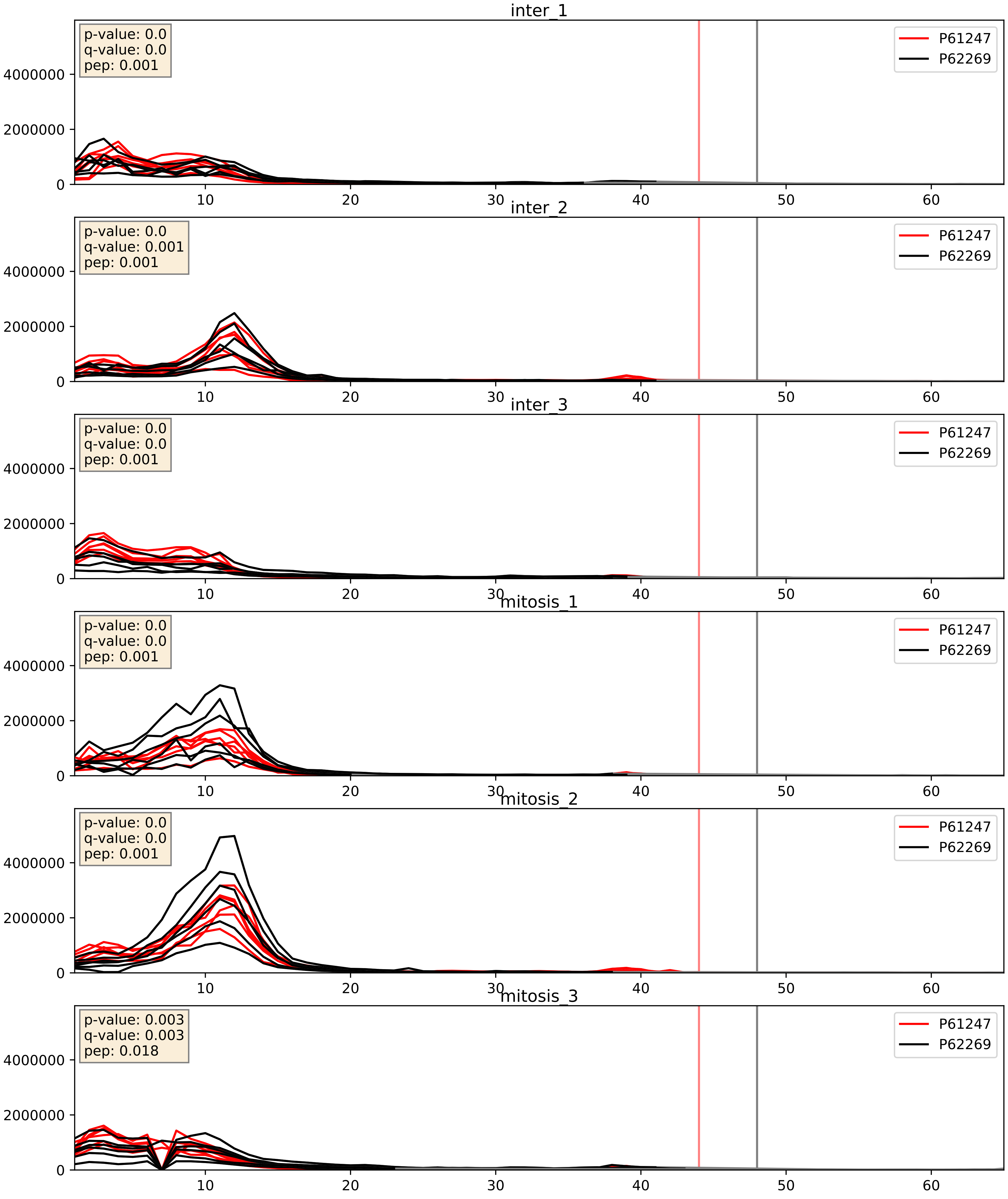


| level | condition_1 | condition_2 | pvalue | pvalue_adjusted |
|---|---|---|---|---|
| interactor_ratio | mitosis | inter | 0.010631950211521158 | 0.14788976221250855 |
| complex_abundance | mitosis | inter | 0.07571191507518986 | 0.1998767548072416 |
| interactor_abundance | mitosis | inter | 0.5746620223826564 | 0.720300118516995 |

P61254_P62269
RL26_HUMAN vs RS18_HUMAN
p-value: 0.052 q-value: 0.034

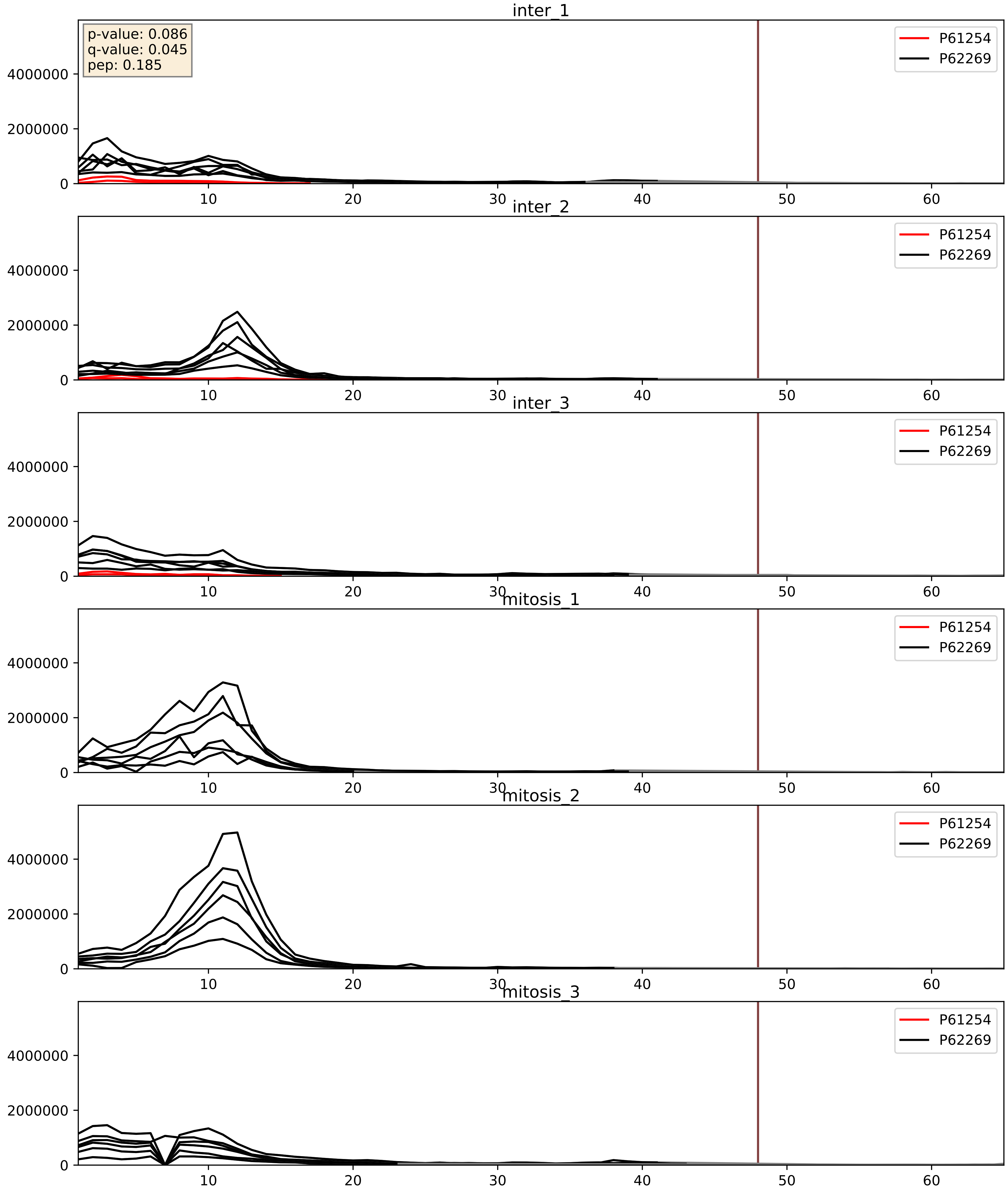


| level | condition_1 | condition_2 | pvalue | pvalue_adjusted |
|---|---|---|---|---|
| complex_abundance | mitosis | inter | 2.6885787207521043e-05 | 0.005691857151652958 |
| interactor_ratio | mitosis | inter | 2.9185735534168128e-05 | 0.005677952185738163 |
| interactor_abundance | mitosis | inter | 0.0006144309768949645 | 0.017415659477552636 |

P61313_P62269
RL15_HUMAN vs RS18_HUMAN
p-value: 0.0 q-value: 0.0

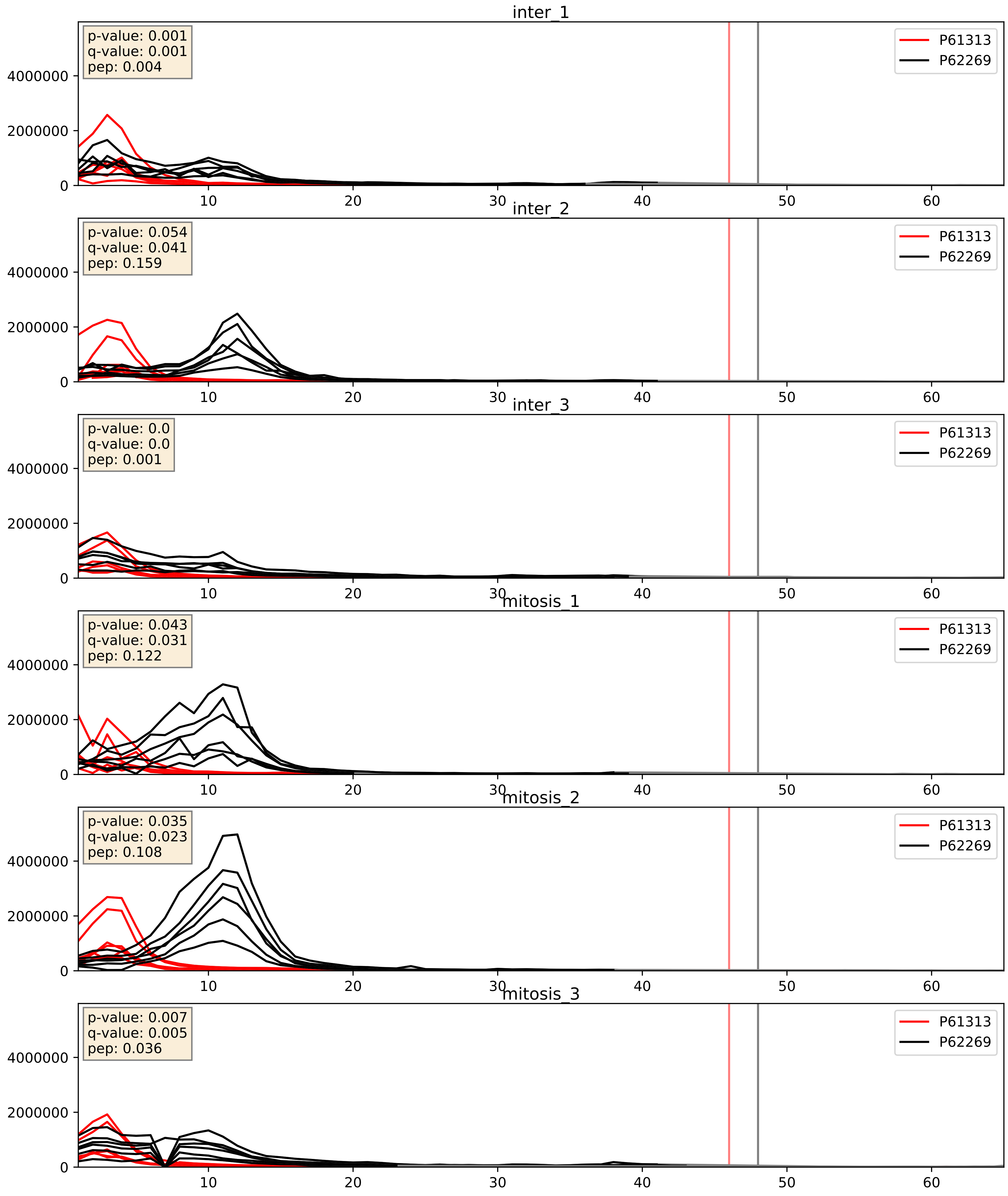


| level | condition_1 | condition_2 | pvalue | pvalue_adjusted |
|---|---|---|---|---|
| interactor_ratio | mitosis | inter | 0.056732177139738084 | 0.39385842361407786 |
| complex_abundance | mitosis | inter | 0.0702658124841788 | 0.19241213284599734 |
| interactor_abundance | mitosis | inter | 0.9175497077049597 | 0.951681267170054 |

P61353_P62269
RL27_HUMAN vs RS18_HUMAN
p-value: 0.0 q-value: 0.001

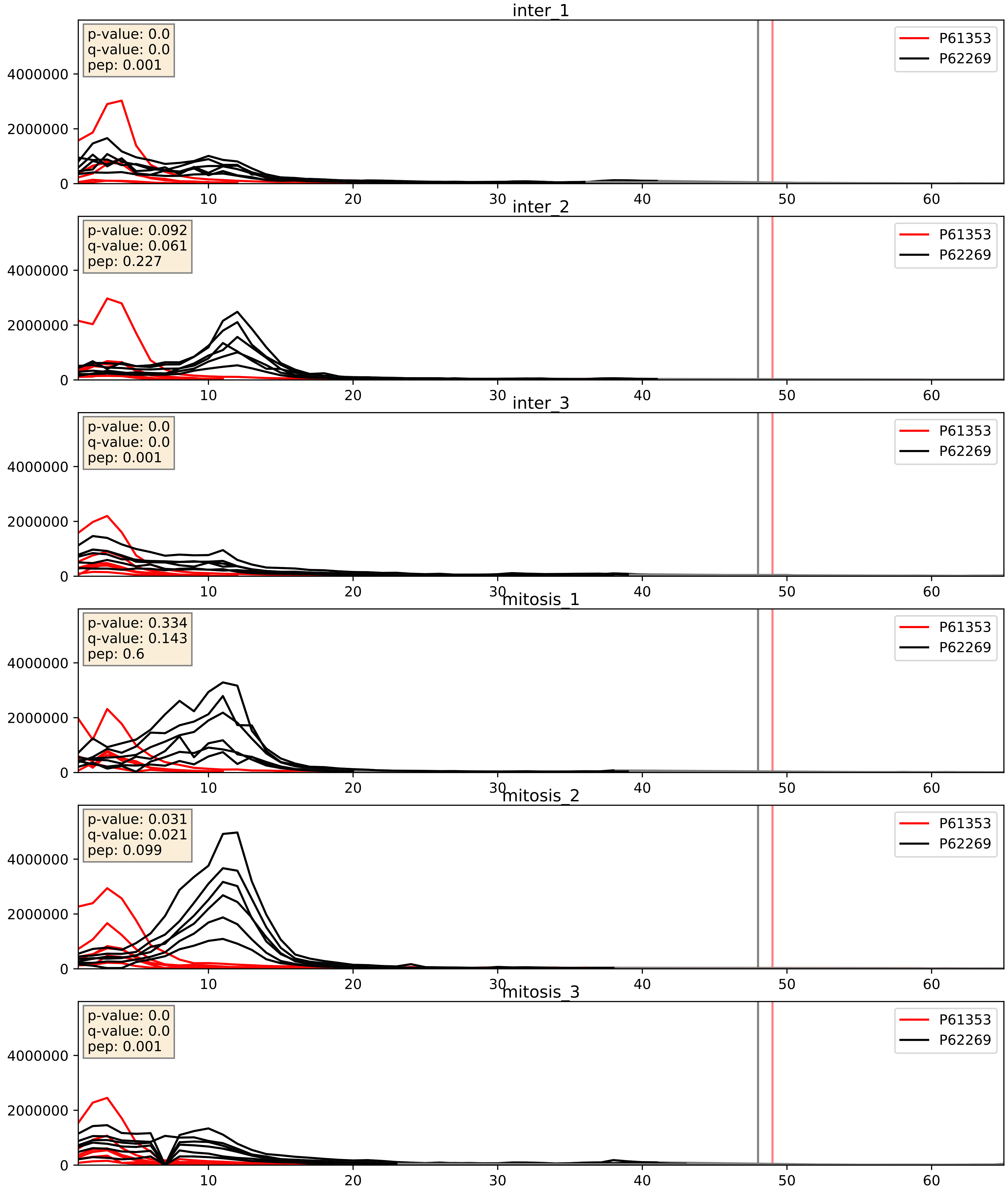


| level | condition_1 | condition_2 | pvalue | pvalue_adjusted |
|---|---|---|---|---|
| interactor_ratio | mitosis | inter | 0.02262796585455358 | 0.2314162338291262 |
| complex_abundance | mitosis | inter | 0.08513350504315893 | 0.2139201058931811 |
| interactor_abundance | mitosis | inter | 0.5867616308489645 | 0.7283996171513504 |

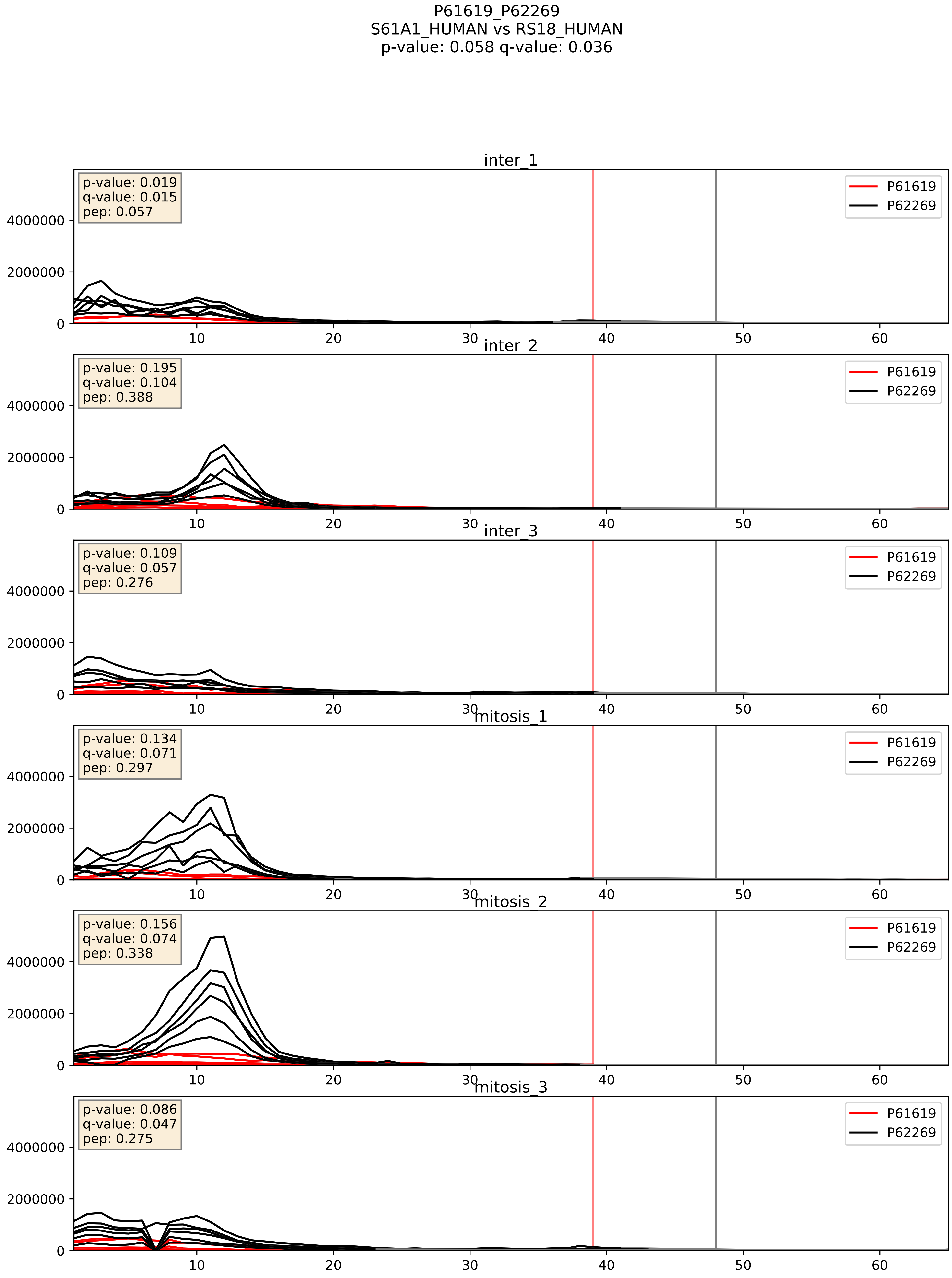


| level | condition_1 | condition_2 | pvalue | pvalue_adjusted |
|---|---|---|---|---|
| interactor_ratio | mitosis | inter | 0.025778133197931933 | 0.2542175347630154 |
| complex_abundance | mitosis | inter | 0.3067017251231653 | 0.4570624594453856 |
| interactor_abundance | mitosis | inter | 0.6440730353911753 | 0.7618163856499186 |

P62081_P62269
RS7_HUMAN vs RS18_HUMAN
p-value: 0.02 q-value: 0.016

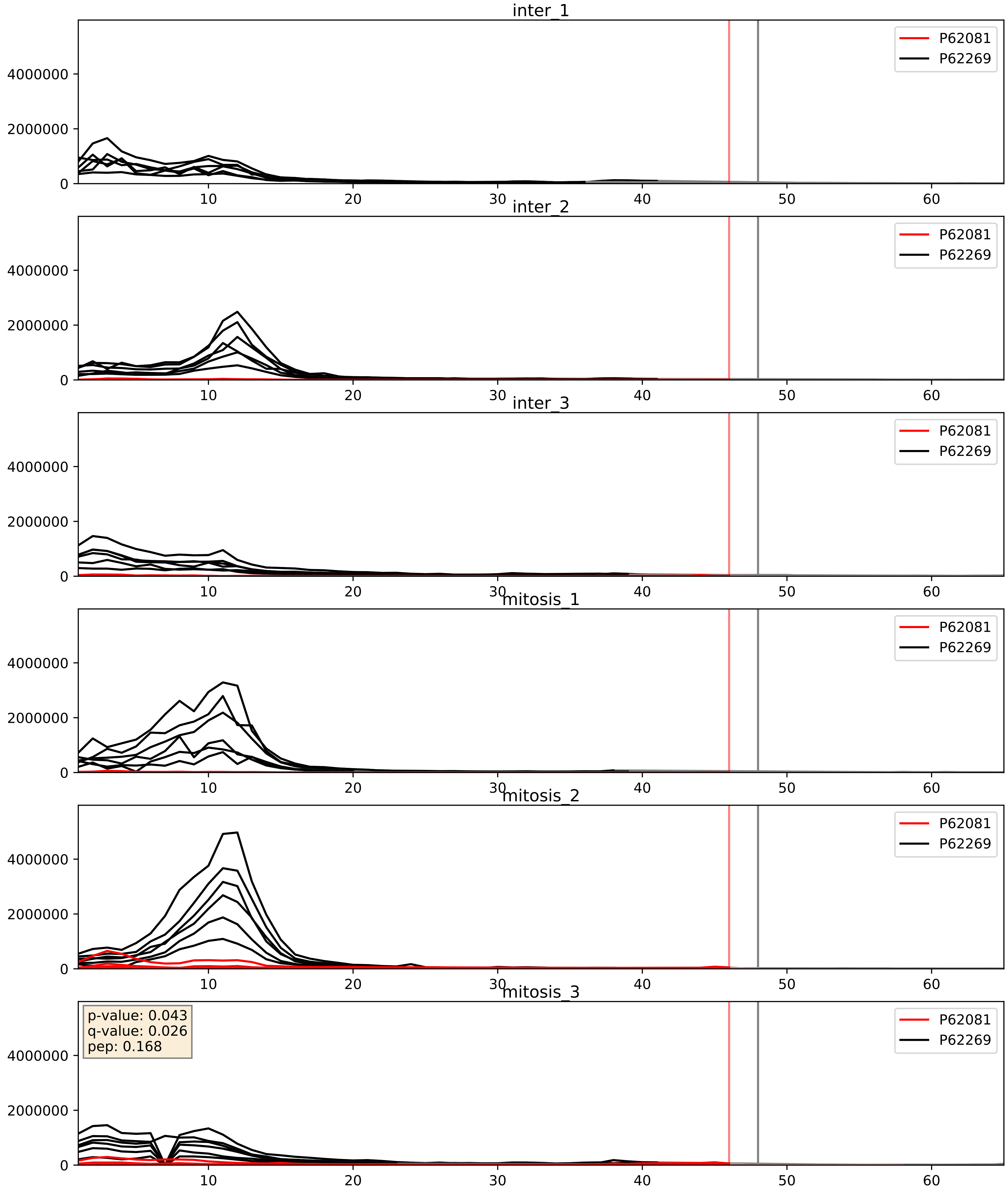


| level | condition_1 | condition_2 | pvalue | pvalue_adjusted |
|---|---|---|---|---|
| complex_abundance | mitosis | inter | 0.19633457081326972 | 0.34874951777580177 |
| interactor_abundance | mitosis | inter | 0.2581997179165959 | 0.4510560417467788 |
| interactor_ratio | mitosis | inter | 0.4357889724765848 | 0.8435878041032946 |

P62241_P62269
RS8_HUMAN vs RS18_HUMAN
p-value: 0.0 q-value: 0.0

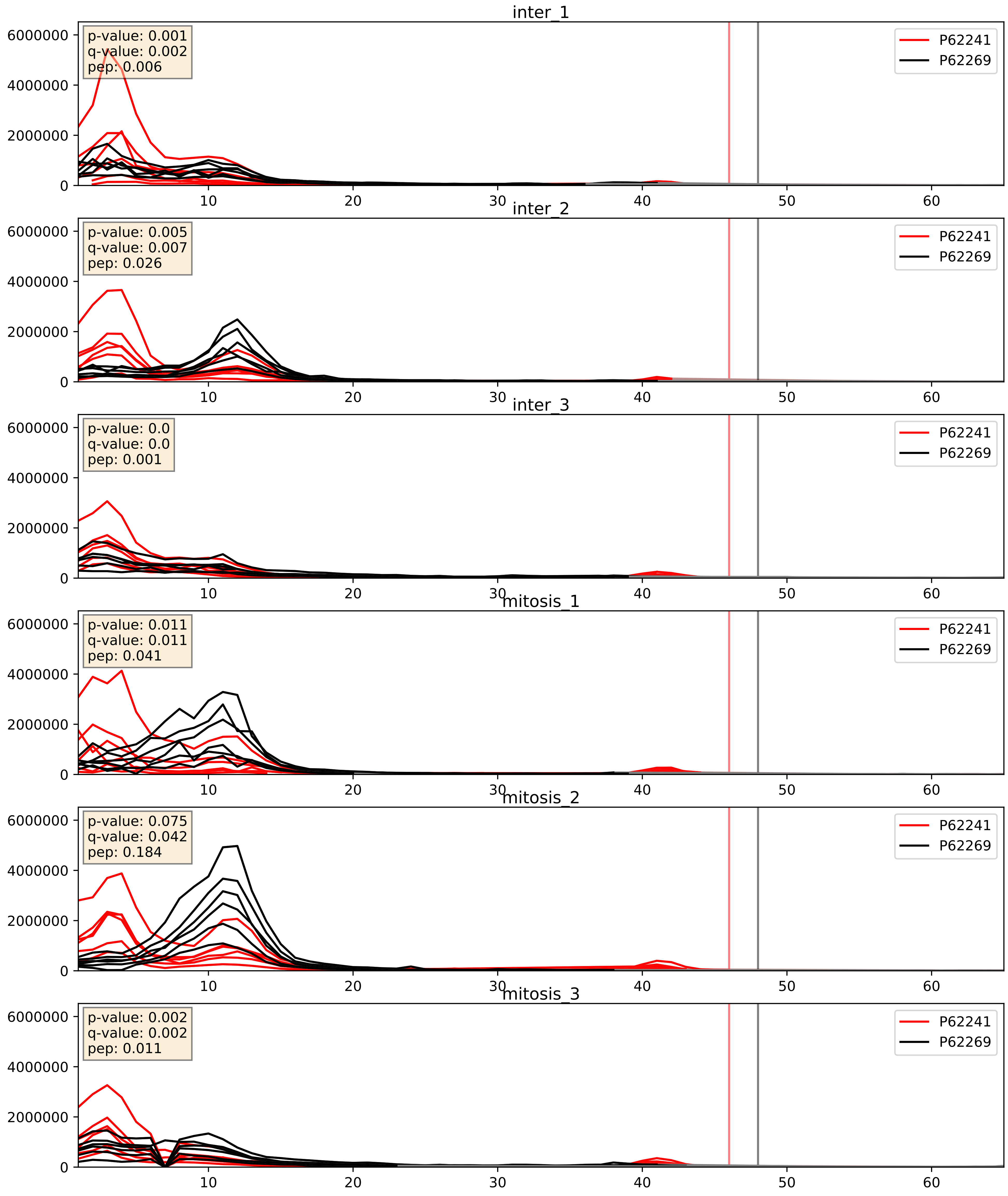


| level | condition_1 | condition_2 | pvalue | pvalue_adjusted |
|---|---|---|---|---|
| complex_abundance | mitosis | inter | 0.014550147396708365 | 0.08966829497179525 |
| interactor_ratio | mitosis | inter | 0.0199796376451993 | 0.21404307217424748 |
| interactor_abundance | mitosis | inter | 0.1923539599836578 | 0.38466298270298116 |

P62244_P62269
RS15A_HUMAN vs RS18_HUMAN
p-value: 0.017 q-value: 0.014

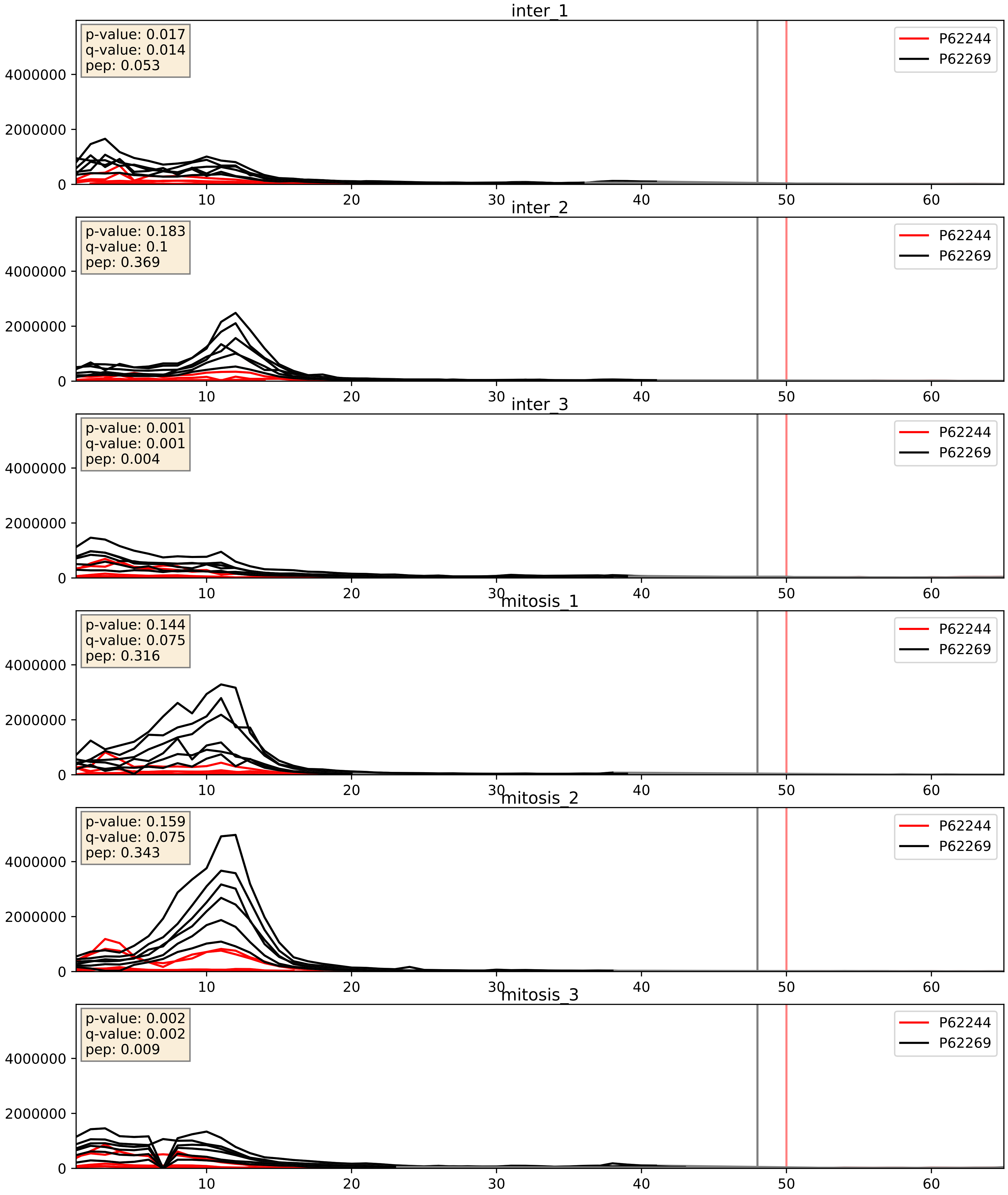


| level | condition_1 | condition_2 | pvalue | pvalue_adjusted |
|---|---|---|---|---|
| complex_abundance | mitosis | inter | 0.02445944740457413 | 0.11409965655757742 |
| interactor_ratio | mitosis | inter | 0.1554762875705694 | 0.6121789427801628 |
| interactor_abundance | mitosis | inter | 0.16118758951289522 | 0.3478547248784528 |

P62249_P62269
RS16_HUMAN vs RS18_HUMAN
p-value: 0.0 q-value: 0.0

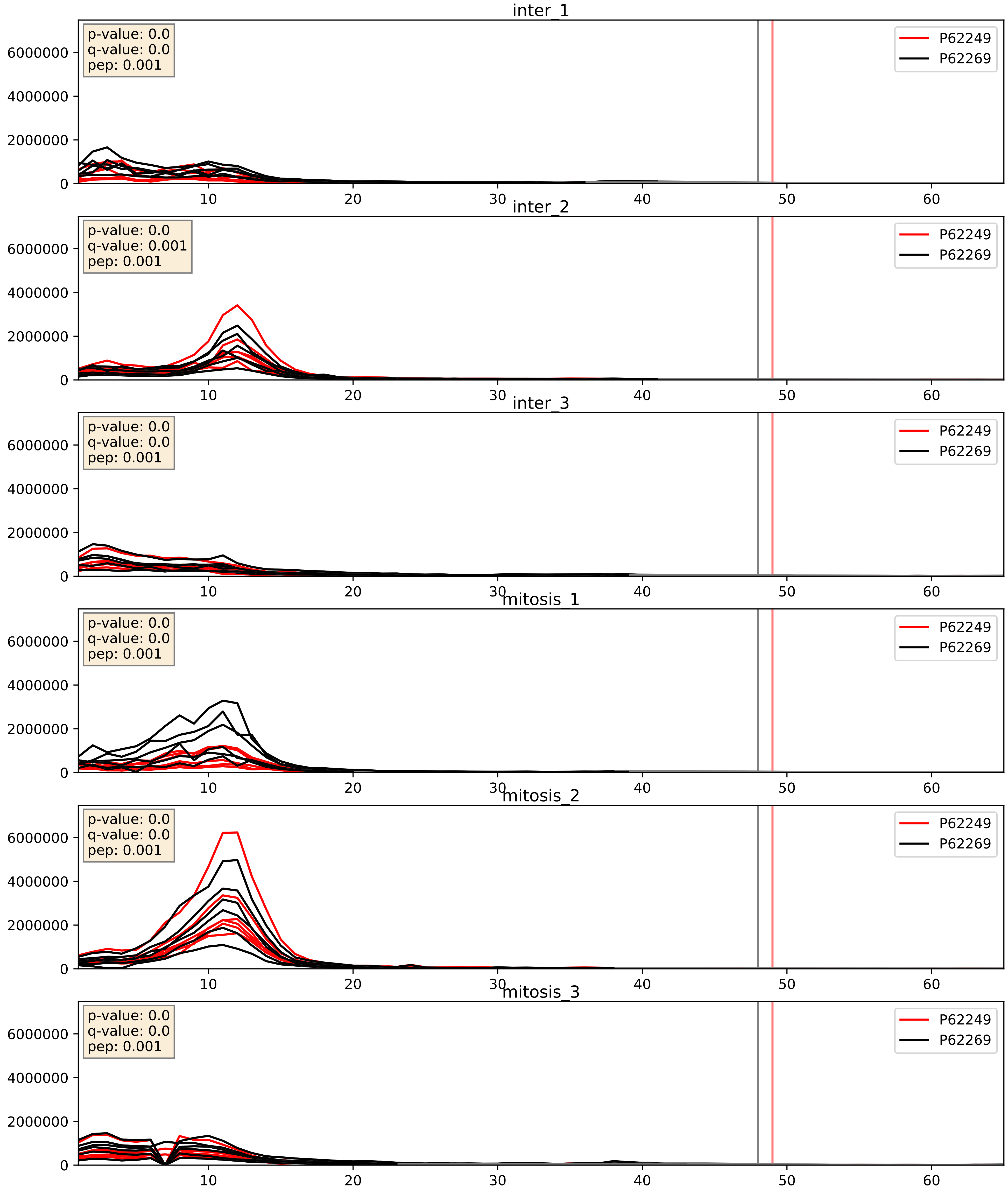


| level | condition_1 | condition_2 | pvalue | pvalue_adjusted |
|---|---|---|---|---|
| complex_abundance | mitosis | inter | 0.1855415390962496 | 0.3370221298338614 |
| interactor_ratio | mitosis | inter | 0.3209339586813907 | 0.7815632108997735 |
| interactor_abundance | mitosis | inter | 0.5894138718393125 | 0.7305267129435337 |

P62263_P62269
RS14_HUMAN vs RS18_HUMAN
p-value: 0.0 q-value: 0.0

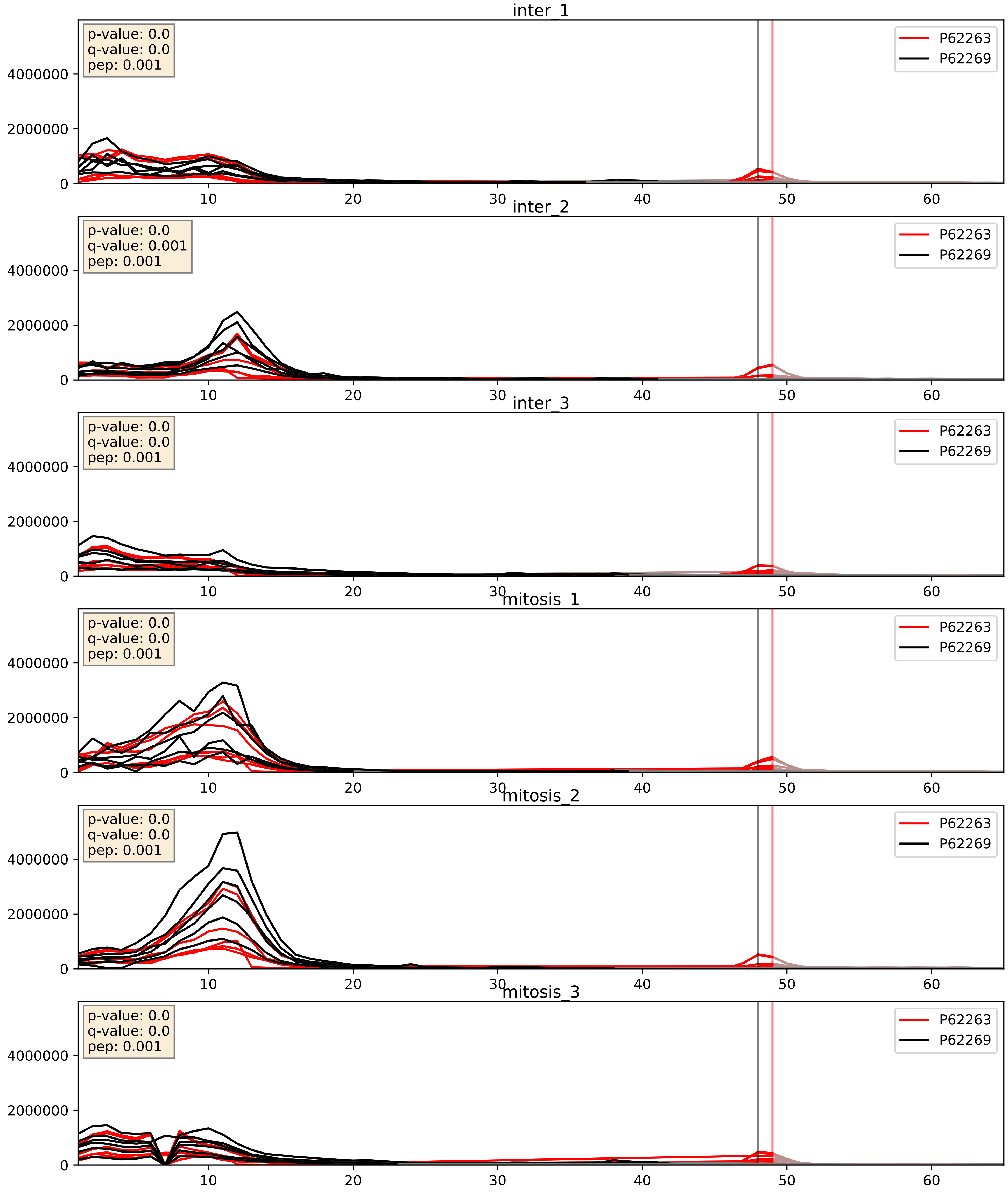


| level | condition_1 | condition_2 | pvalue | pvalue_adjusted |
|---|---|---|---|---|
| complex_abundance | mitosis | inter | 0.04091910649389891 | 0.1455676624066993 |
| interactor_abundance | mitosis | inter | 0.3792505275137932 | 0.5721983160982499 |
| interactor_ratio | mitosis | inter | 0.522186792432599 | 0.8786895401167326 |

P62266_P62269
RS23_HUMAN vs RS18_HUMAN
p-value: 0.0 q-value: 0.0

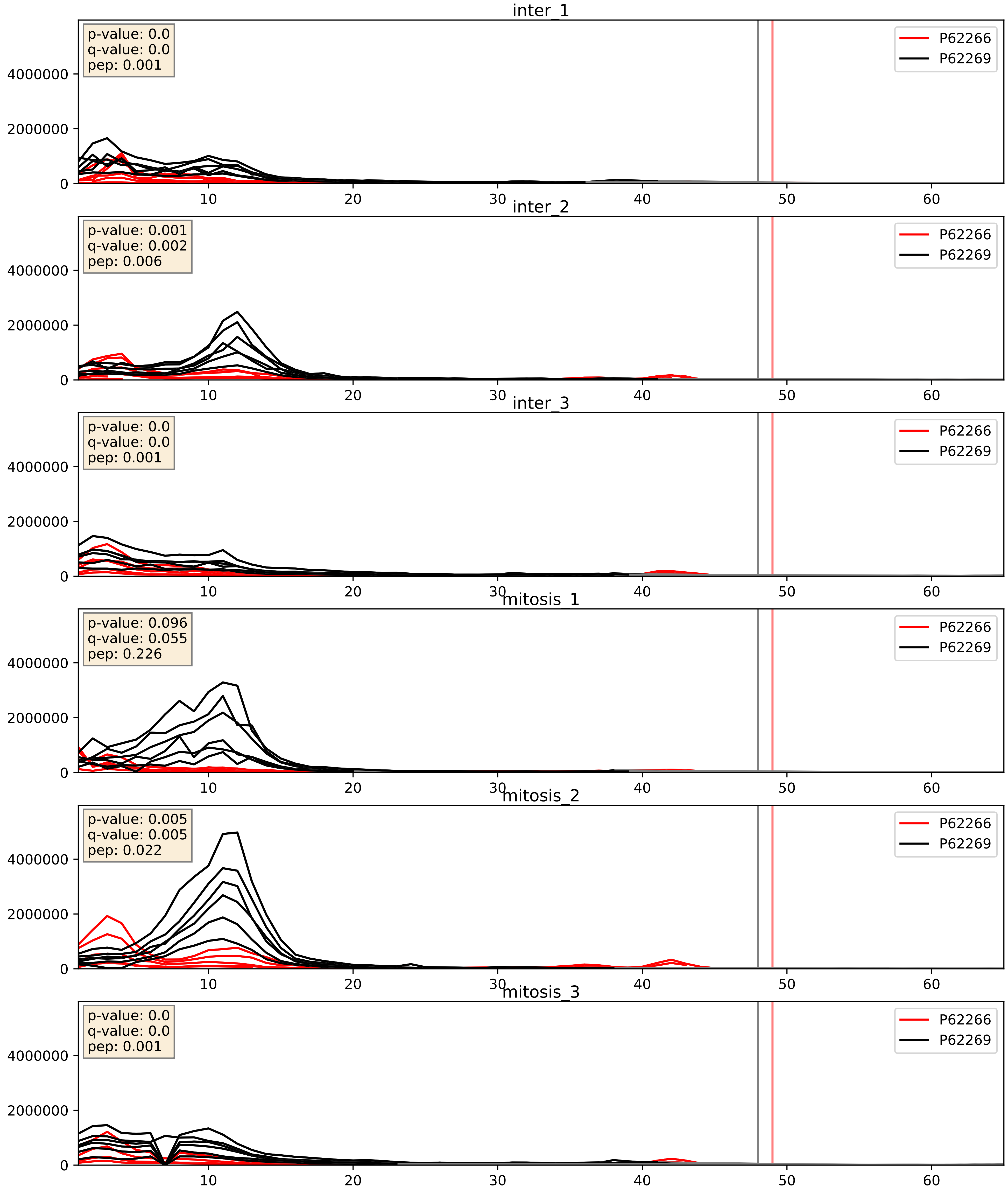


| level | condition_1 | condition_2 | pvalue | pvalue_adjusted |
|---|---|---|---|---|
| interactor_ratio | mitosis | inter | 0.03726128020612114 | 0.3165269945684102 |
| interactor_abundance | mitosis | inter | 0.2799218243492273 | 0.4731086541433709 |
| complex_abundance | mitosis | inter | 0.9940529907575837 | 0.9966143828630729 |

P62269_P62273
RS18_HUMAN vs RS29_HUMAN
p-value: 0.004 q-value: 0.005

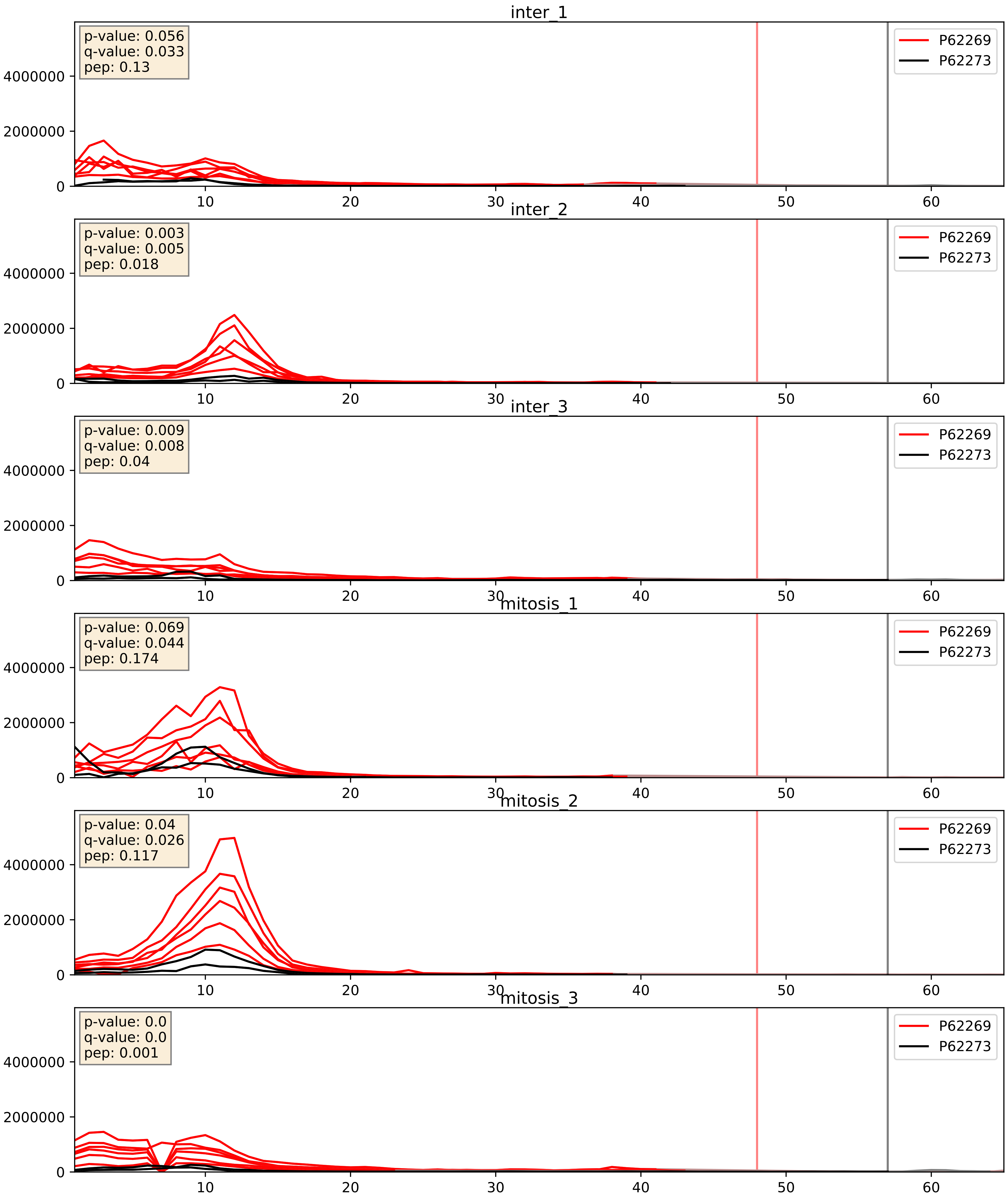


| level | condition_1 | condition_2 | pvalue | pvalue_adjusted |
|---|---|---|---|---|
| interactor_abundance | mitosis | inter | 0.01041014945690444 | 0.0889330133244531 |
| complex_abundance | mitosis | inter | 0.022361198436343783 | 0.10987189501272765 |
| interactor_ratio | mitosis | inter | 0.46452371700824635 | 0.8591803704157076 |

P62269_P62277
RS18_HUMAN vs RS13_HUMAN
p-value: 0.0 q-value: 0.001

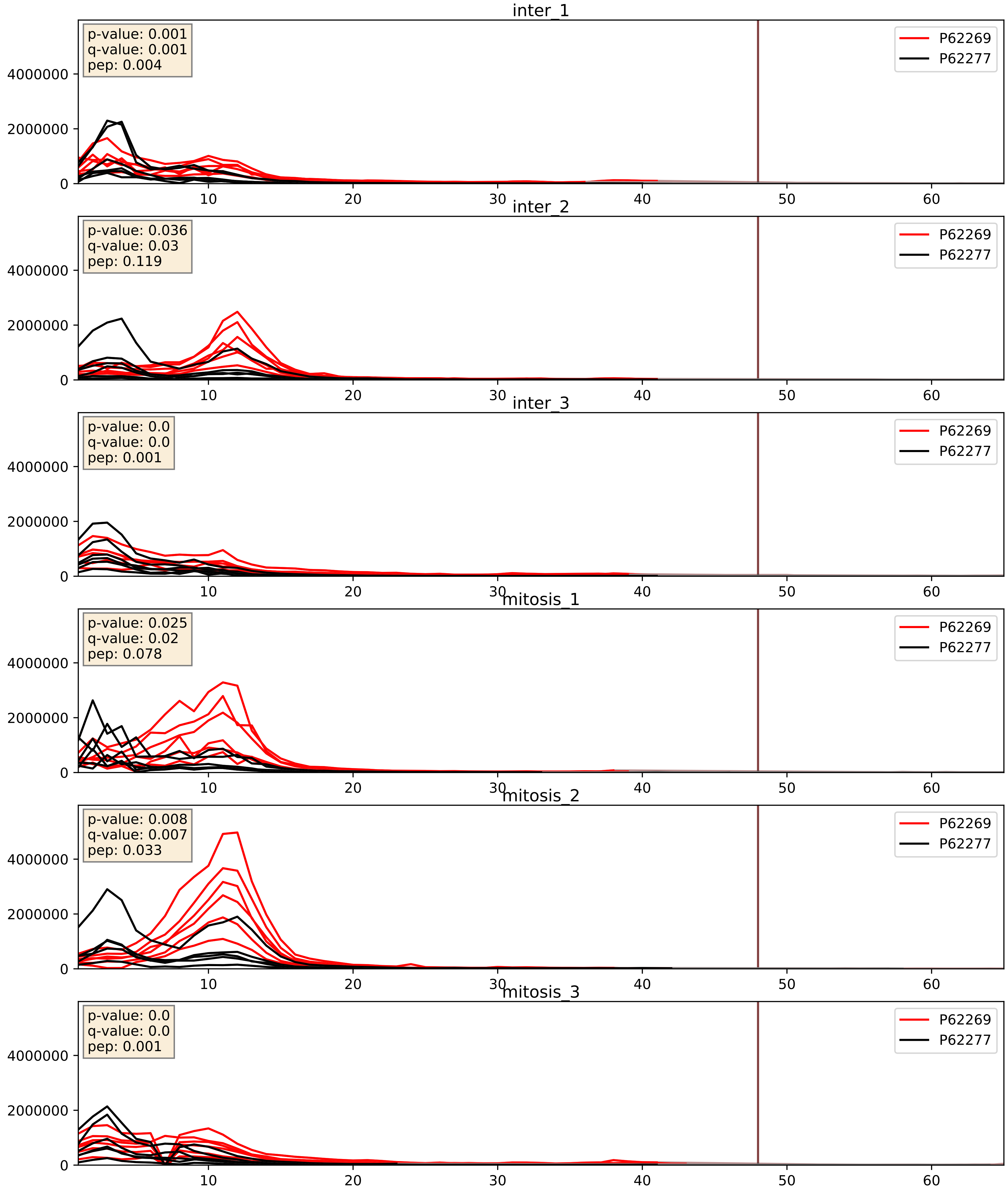


| level | condition_1 | condition_2 | pvalue | pvalue_adjusted |
|---|---|---|---|---|
| complex_abundance | mitosis | inter | 0.008943530751085981 | 0.07017105703876811 |
| interactor_abundance | mitosis | inter | 0.011627613253151009 | 0.09510976535783339 |
| interactor_ratio | mitosis | inter | 0.046921710294060934 | 0.35838626305591875 |

P62269_P62280
RS18_HUMAN vs RS11_HUMAN
p-value: 0.0 q-value: 0.0

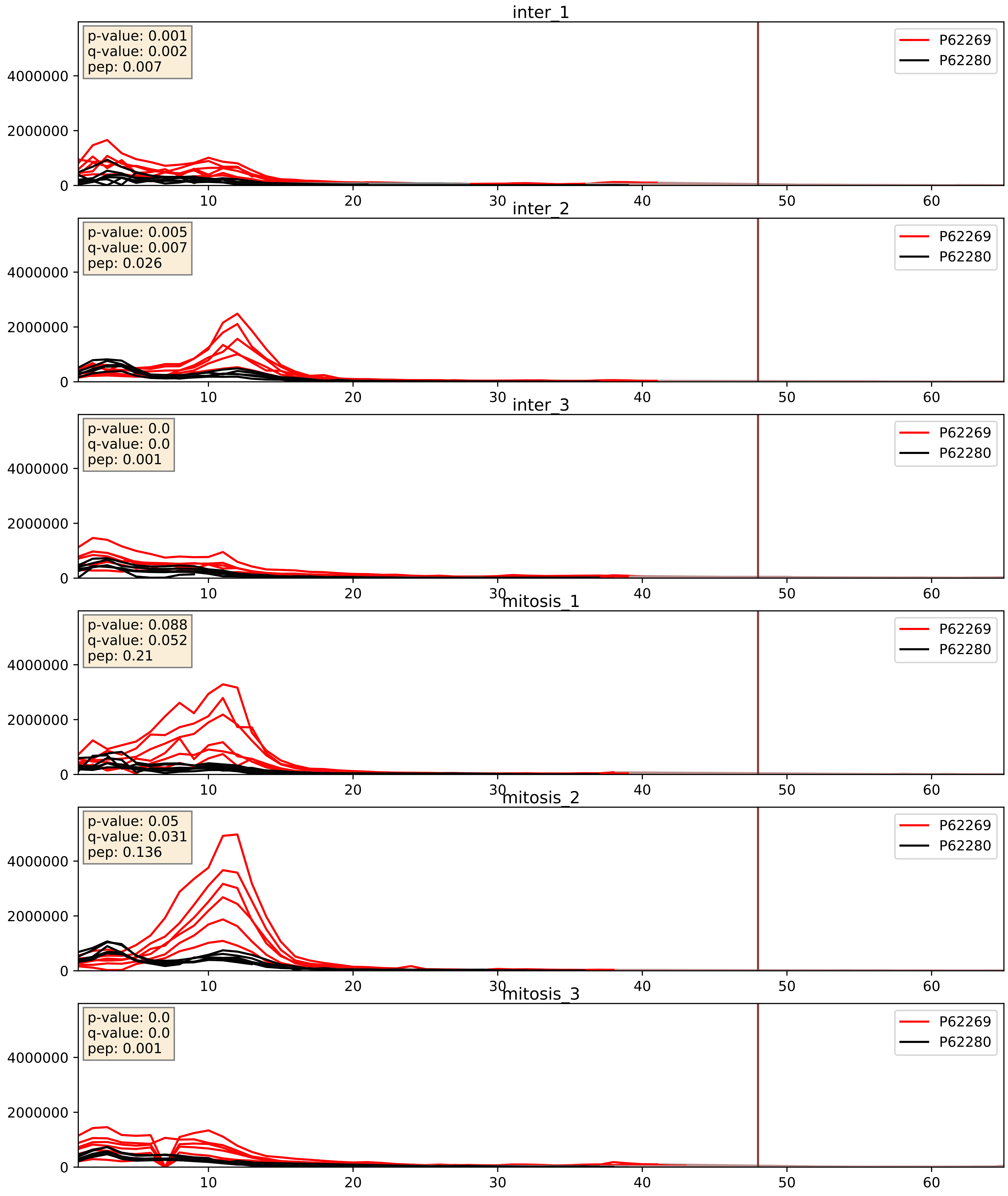


| level | condition_1 | condition_2 | pvalue | pvalue_adjusted |
|---|---|---|---|---|
| interactor_ratio | mitosis | inter | 0.0034151958076039566 | 0.07572939655107606 |
| interactor_abundance | mitosis | inter | 0.010842572684605211 | 0.09135080923250061 |
| complex_abundance | mitosis | inter | 0.07280693201883015 | 0.19598343964817172 |

P62269_P62424
RS18_HUMAN vs RL7A_HUMAN
p-value: 0.0 q-value: 0.001

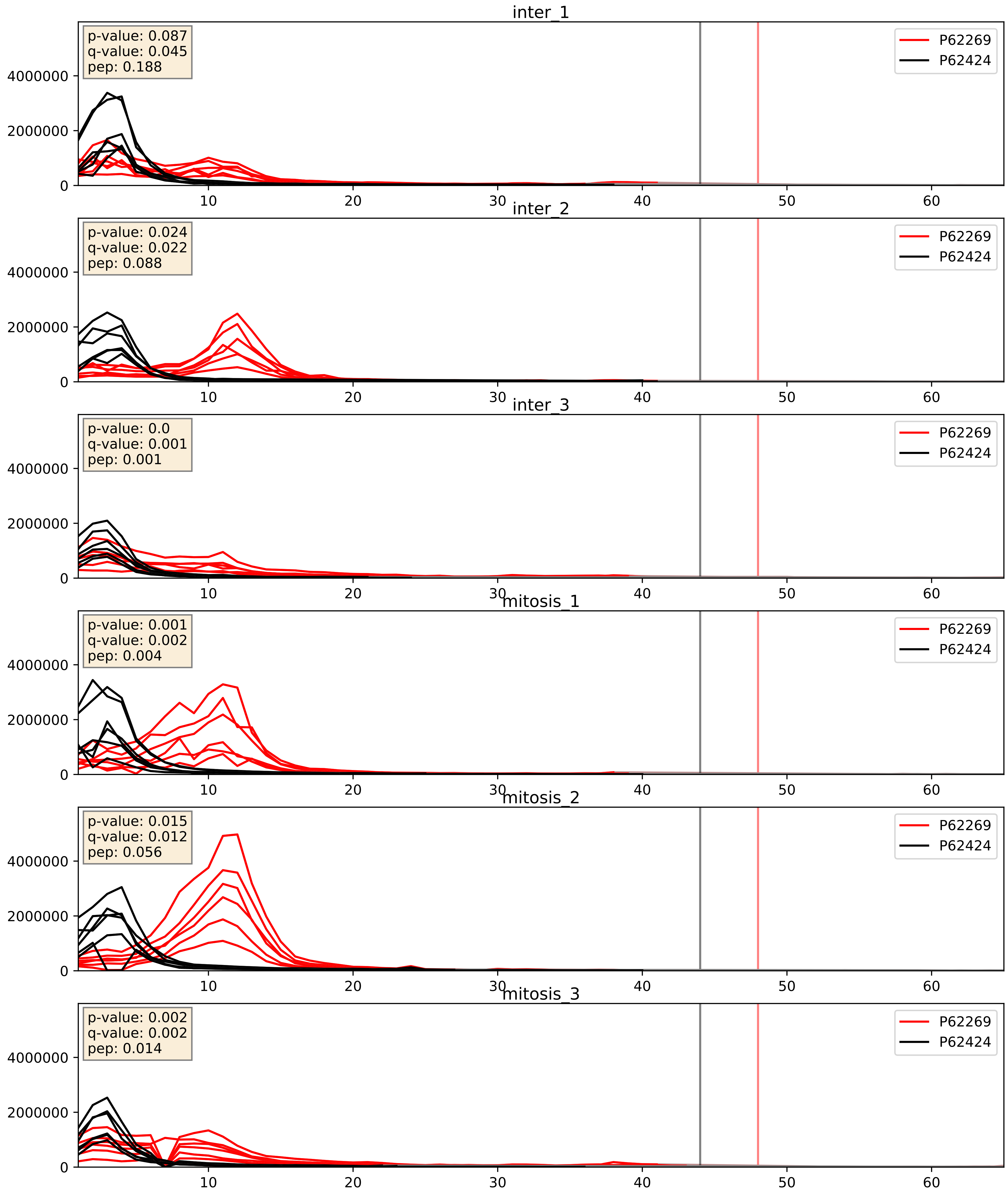


| level | condition_1 | condition_2 | pvalue | pvalue_adjusted |
|---|---|---|---|---|
| interactor_abundance | mitosis | inter | 0.013163539444812453 | 0.10304923015527873 |
| complex_abundance | mitosis | inter | 0.04281211183435931 | 0.1481949160937304 |
| interactor_ratio | mitosis | inter | 0.12924410892295735 | 0.5779454961622694 |

P62269_P62701
RS18_HUMAN vs RS4X_HUMAN
p-value: 0.001 q-value: 0.001

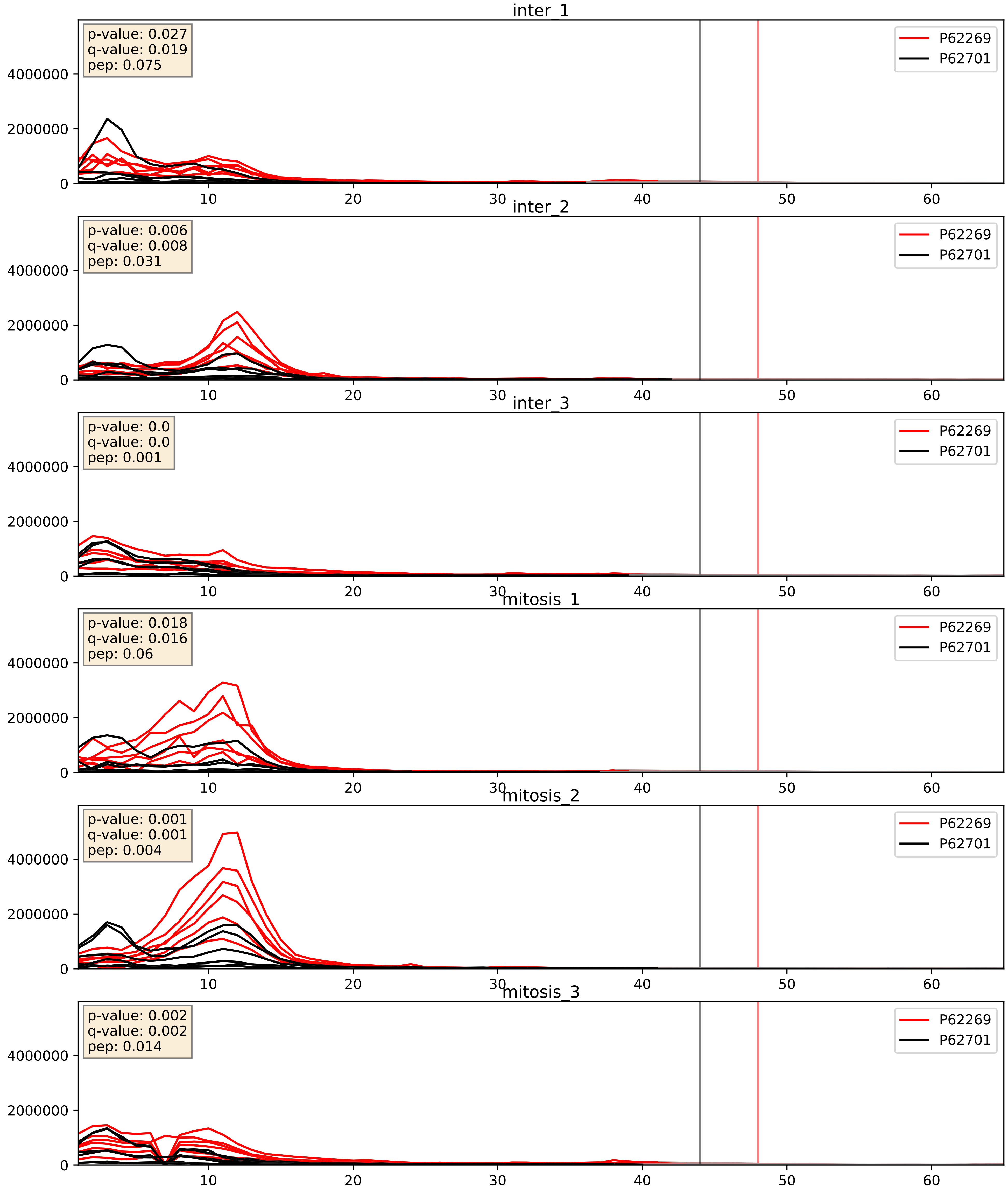


| level | condition_1 | condition_2 | pvalue | pvalue_adjusted |
|---|---|---|---|---|
| interactor_ratio | mitosis | inter | 0.0010811141035510382 | 0.038414698994273265 |
| interactor_abundance | mitosis | inter | 0.01463403608248235 | 0.10605571190893752 |
| complex_abundance | mitosis | inter | 0.09243572012492991 | 0.22415007486385272 |

P62269_P62750
RS18_HUMAN vs RL23A_HUMAN
p-value: 0.0 q-value: 0.0

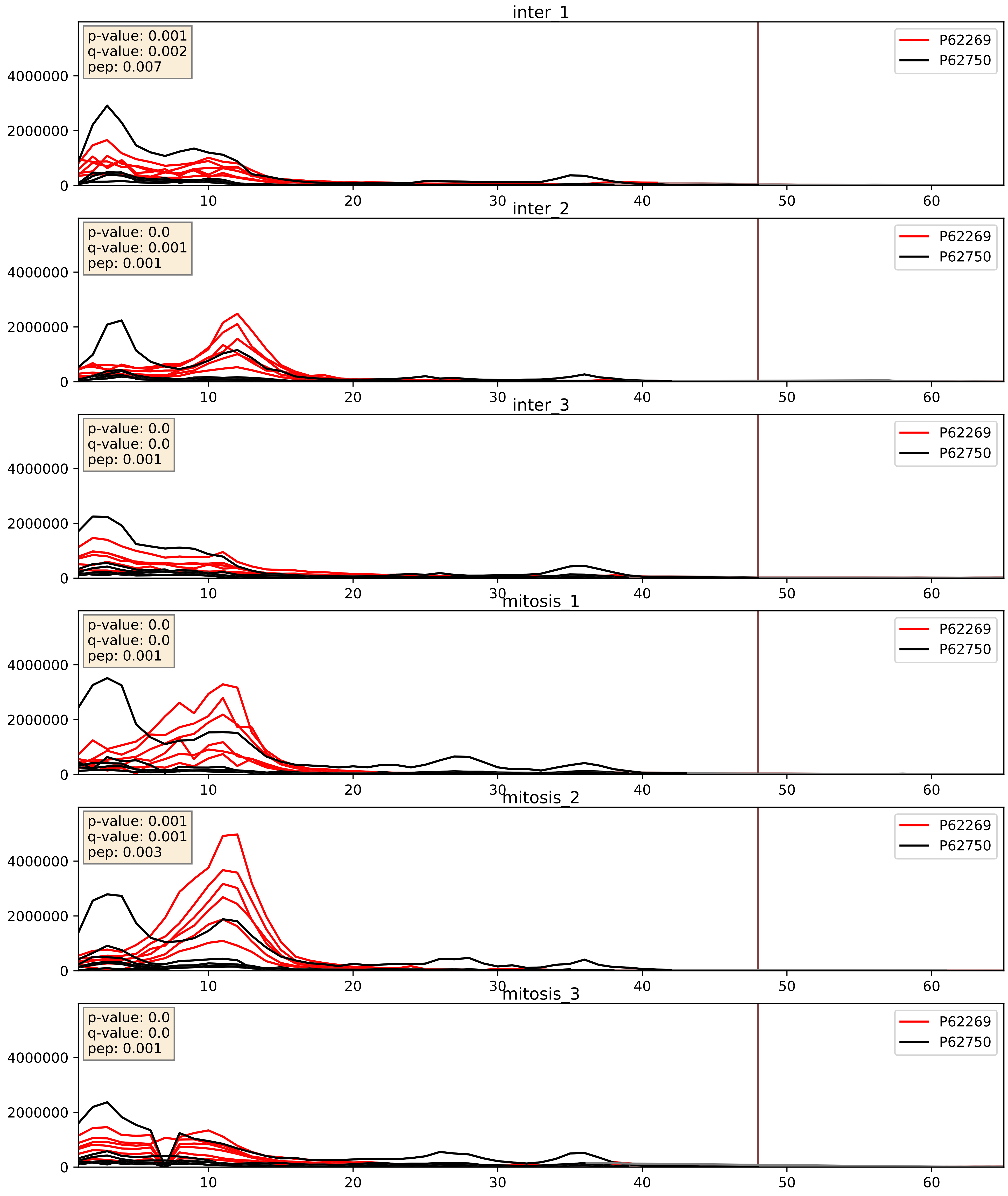


| level | condition_1 | condition_2 | pvalue | pvalue_adjusted |
|---|---|---|---|---|
| complex_abundance | mitosis | inter | 0.009183845920069116 | 0.07082317214035282 |
| interactor_abundance | mitosis | inter | 0.012169990301213955 | 0.0981395355425261 |
| interactor_ratio | mitosis | inter | 0.49164995762918695 | 0.86984863222988 |

P62269_P62753
RS18_HUMAN vs RS6_HUMAN
p-value: 0.0 q-value: 0.0

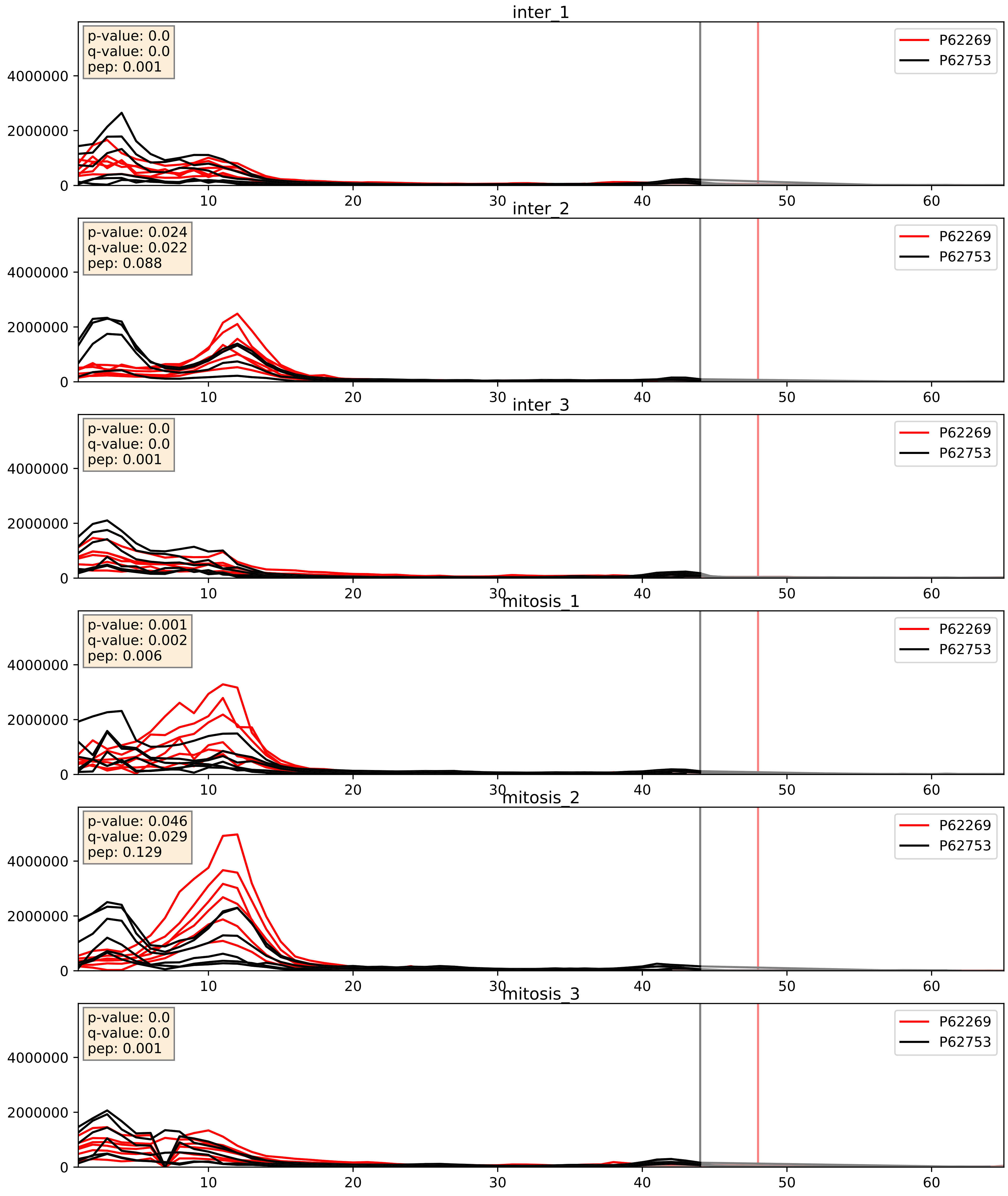


| level | condition_1 | condition_2 | pvalue | pvalue_adjusted |
|---|---|---|---|---|
| interactor_abundance | mitosis | inter | 0.01170822175968245 | 0.095549545119116 |
| interactor_ratio | mitosis | inter | 0.022317791267322973 | 0.2287907703572271 |
| complex_abundance | mitosis | inter | 0.07196154056847812 | 0.19493379343866224 |

P62269_P62829
RS18_HUMAN vs RL23_HUMAN
p-value: 0.001 q-value: 0.001

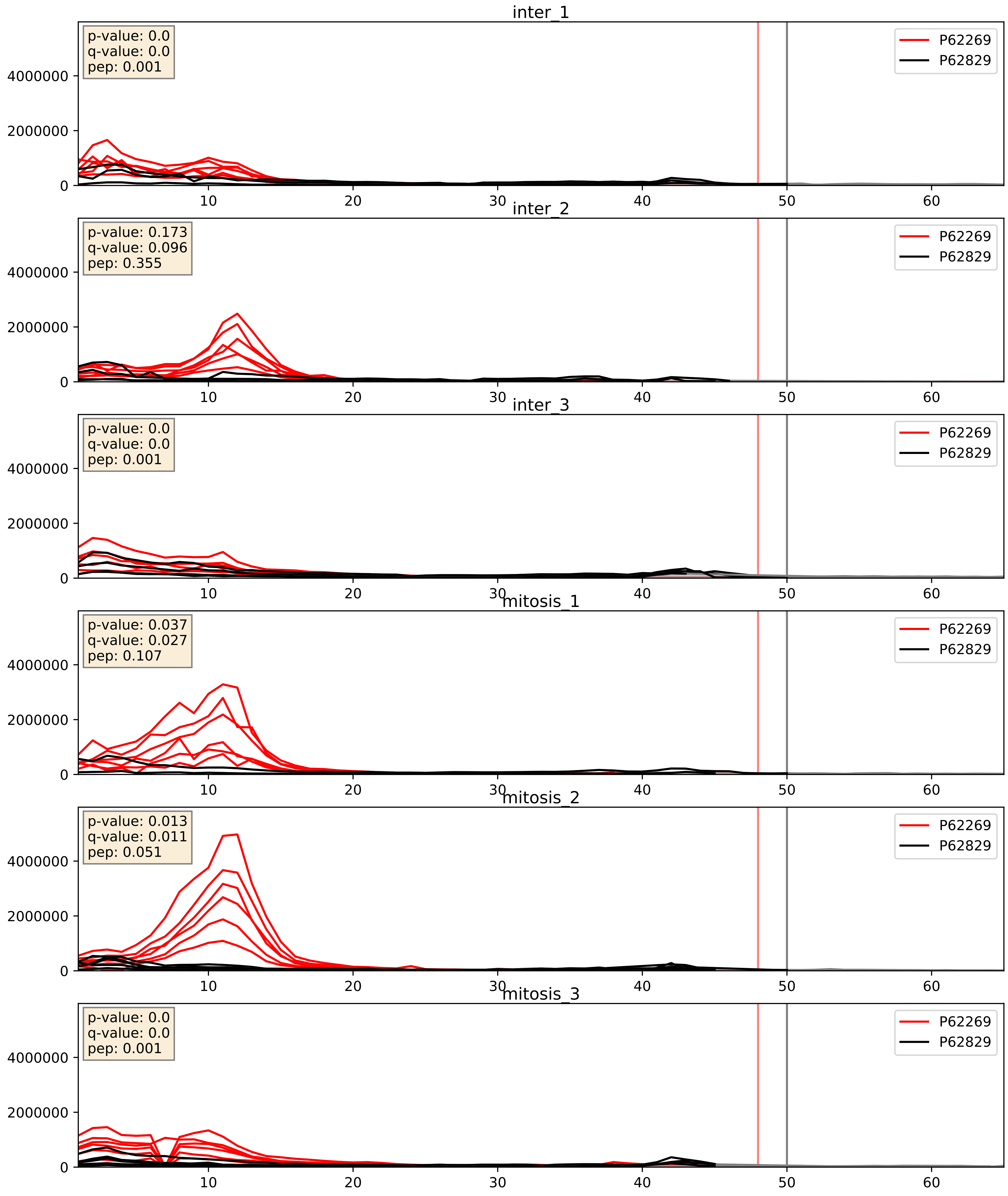


| level | condition_1 | condition_2 | pvalue | pvalue_adjusted |
|---|---|---|---|---|
| interactor_ratio | mitosis | inter | 0.003926685926982767 | 0.08138603277233046 |
| interactor_abundance | mitosis | inter | 0.01140063841385368 | 0.09438052690772485 |
| complex_abundance | mitosis | inter | 0.046698979628882156 | 0.15446030356384516 |

P62269_P62841
RS18_HUMAN vs RS15_HUMAN
p-value: 0.0 q-value: 0.0

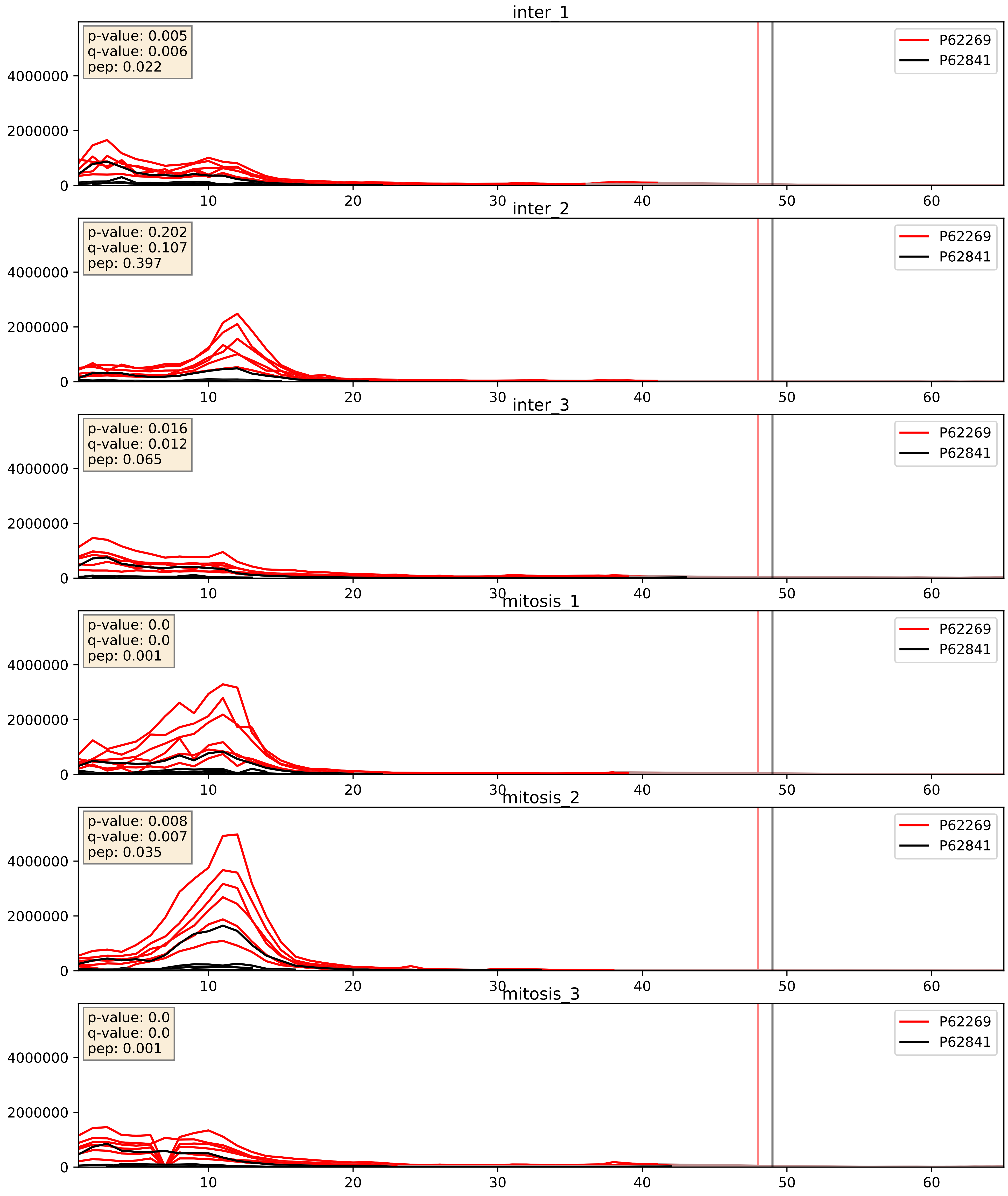


| level | condition_1 | condition_2 | pvalue | pvalue_adjusted |
|---|---|---|---|---|
| interactor_abundance | mitosis | inter | 0.013420049270359122 | 0.10304923015527873 |
| interactor_ratio | mitosis | inter | 0.15103768003384063 | 0.606604569117897 |
| complex_abundance | mitosis | inter | 0.3050644474334144 | 0.4552196842234692 |

P62269_P62847
RS18_HUMAN vs RS24_HUMAN
p-value: 0.0 q-value: 0.0

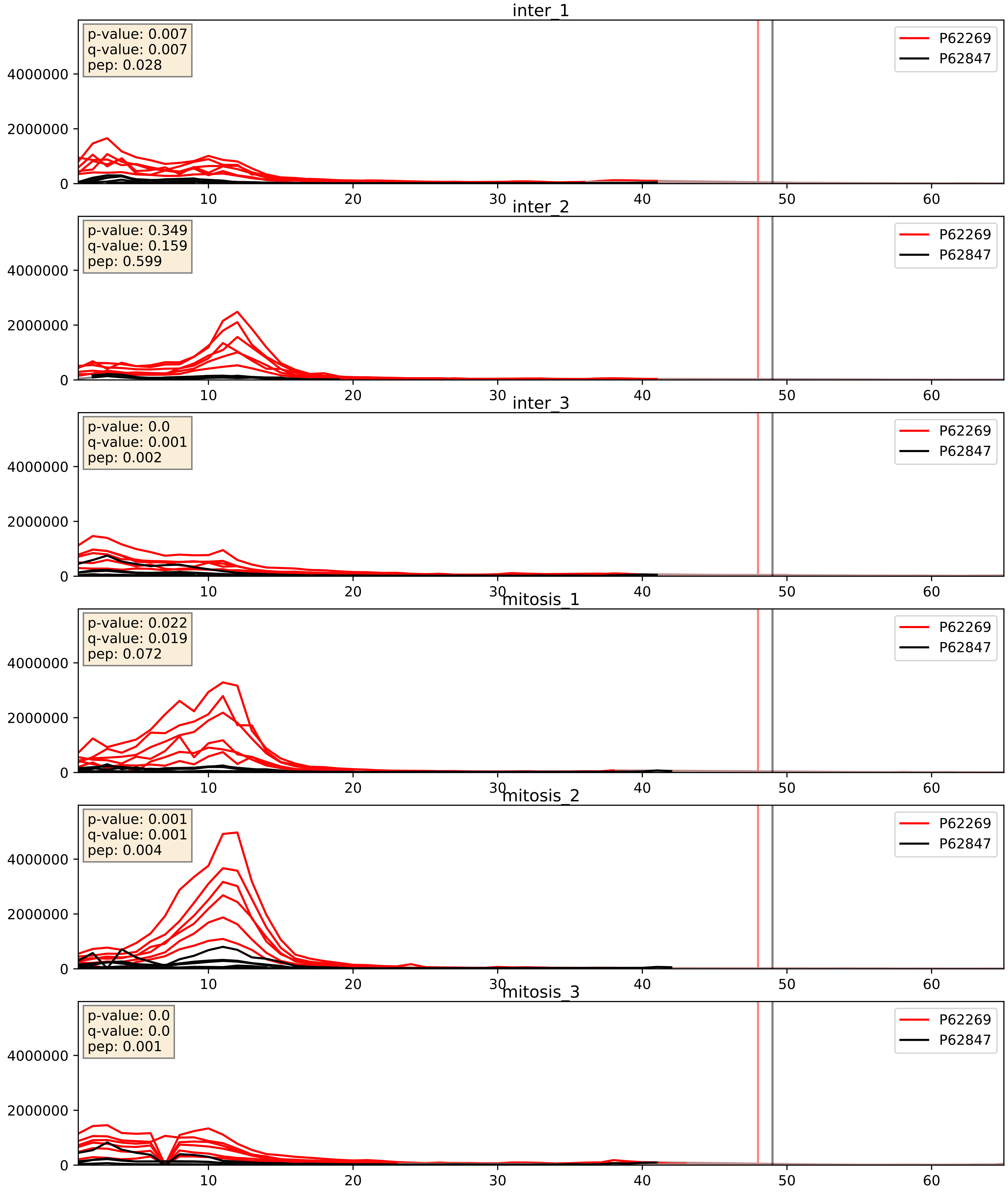


| level | condition_1 | condition_2 | pvalue | pvalue_adjusted |
|---|---|---|---|---|
| interactor_abundance | mitosis | inter | 0.008851192463549136 | 0.08170219532413543 |
| complex_abundance | mitosis | inter | 0.02144671259324164 | 0.10779265935309998 |
| interactor_ratio | mitosis | inter | 0.023564796737102236 | 0.23825177699606717 |

P62269_P62851
RS18_HUMAN vs RS25_HUMAN
p-value: 0.0 q-value: 0.0

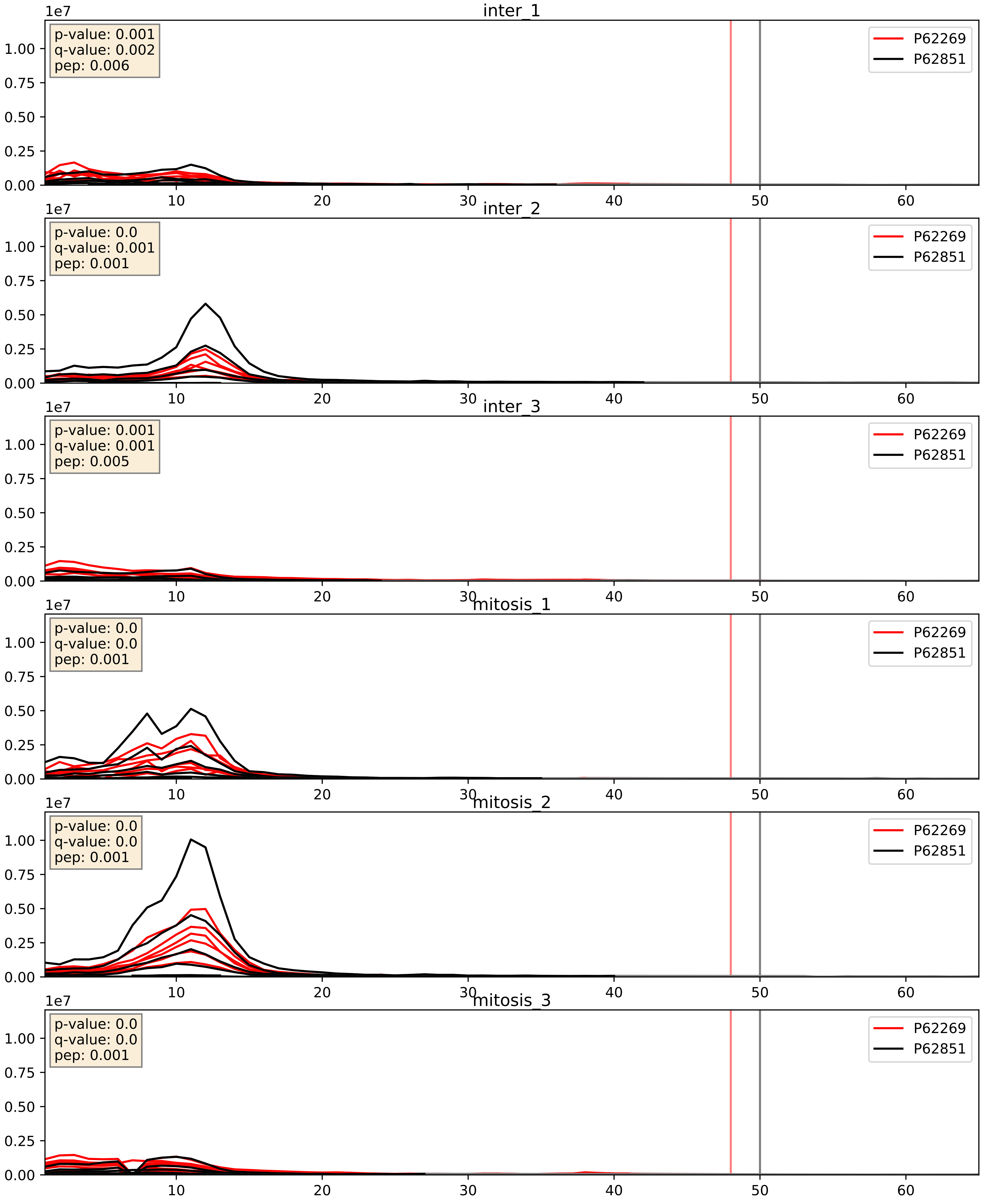


| level | condition_1 | condition_2 | pvalue | pvalue_adjusted |
|---|---|---|---|---|
| interactor_abundance | mitosis | inter | 0.013976128429178225 | 0.10480565865419676 |
| complex_abundance | mitosis | inter | 0.20774905780824526 | 0.36188177097973334 |
| interactor_ratio | mitosis | inter | 0.8491904994995086 | 0.9755634896635457 |

P62269_P62857
RS18_HUMAN vs RS28_HUMAN
p-value: 0.0 q-value: 0.0

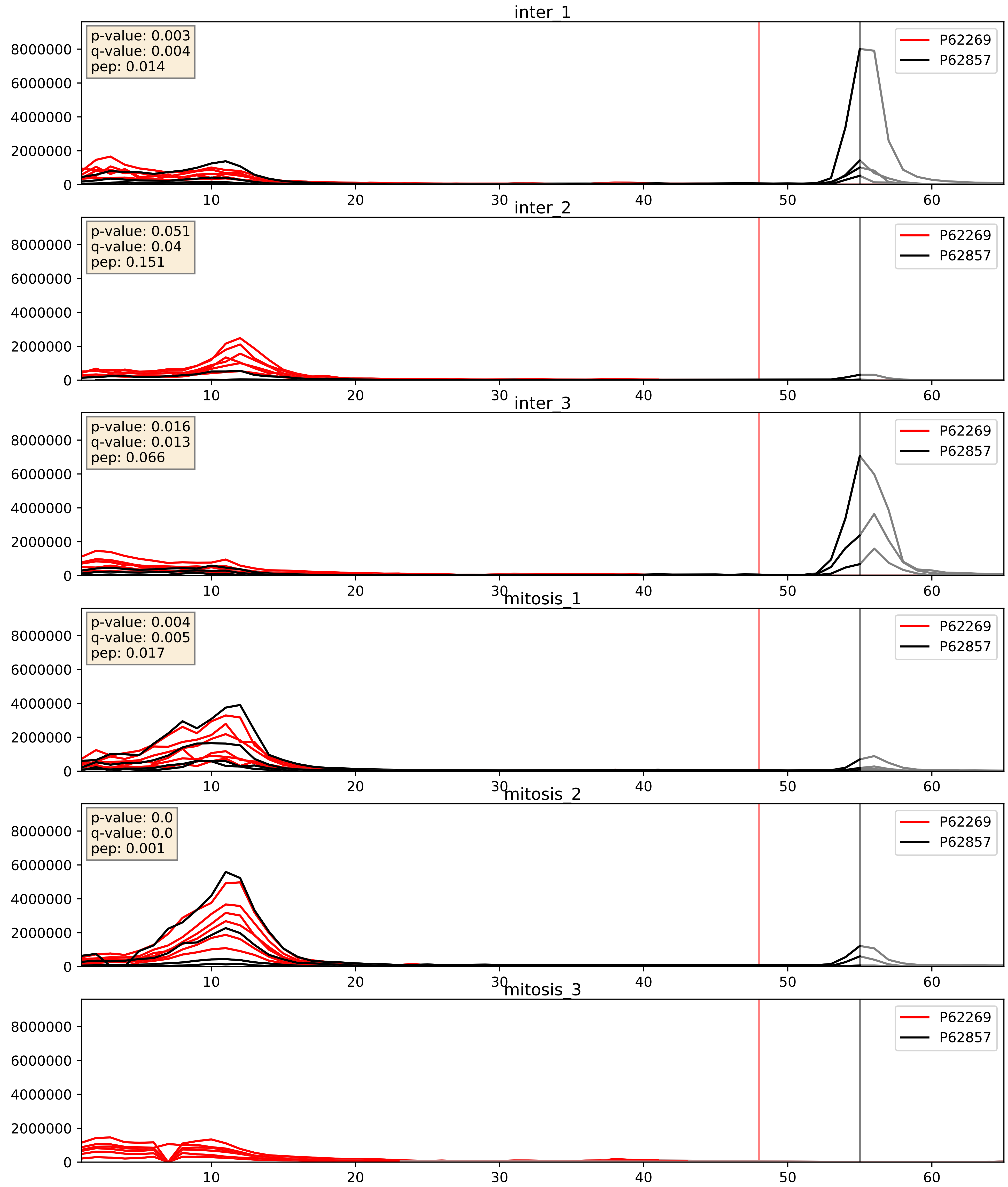


| level | condition_1 | condition_2 | pvalue | pvalue_adjusted |
|---|---|---|---|---|
| interactor_ratio | mitosis | inter | 0.0735062628030063 | 0.4539780732999524 |
| interactor_abundance | mitosis | inter | 0.5314249146780222 | 0.6900760554415658 |
| complex_abundance | mitosis | inter | 0.7942825341989727 | 0.8664532296091763 |

P62269_P62861
RS18_HUMAN vs RS30_HUMAN
p-value: 0.016 q-value: 0.013

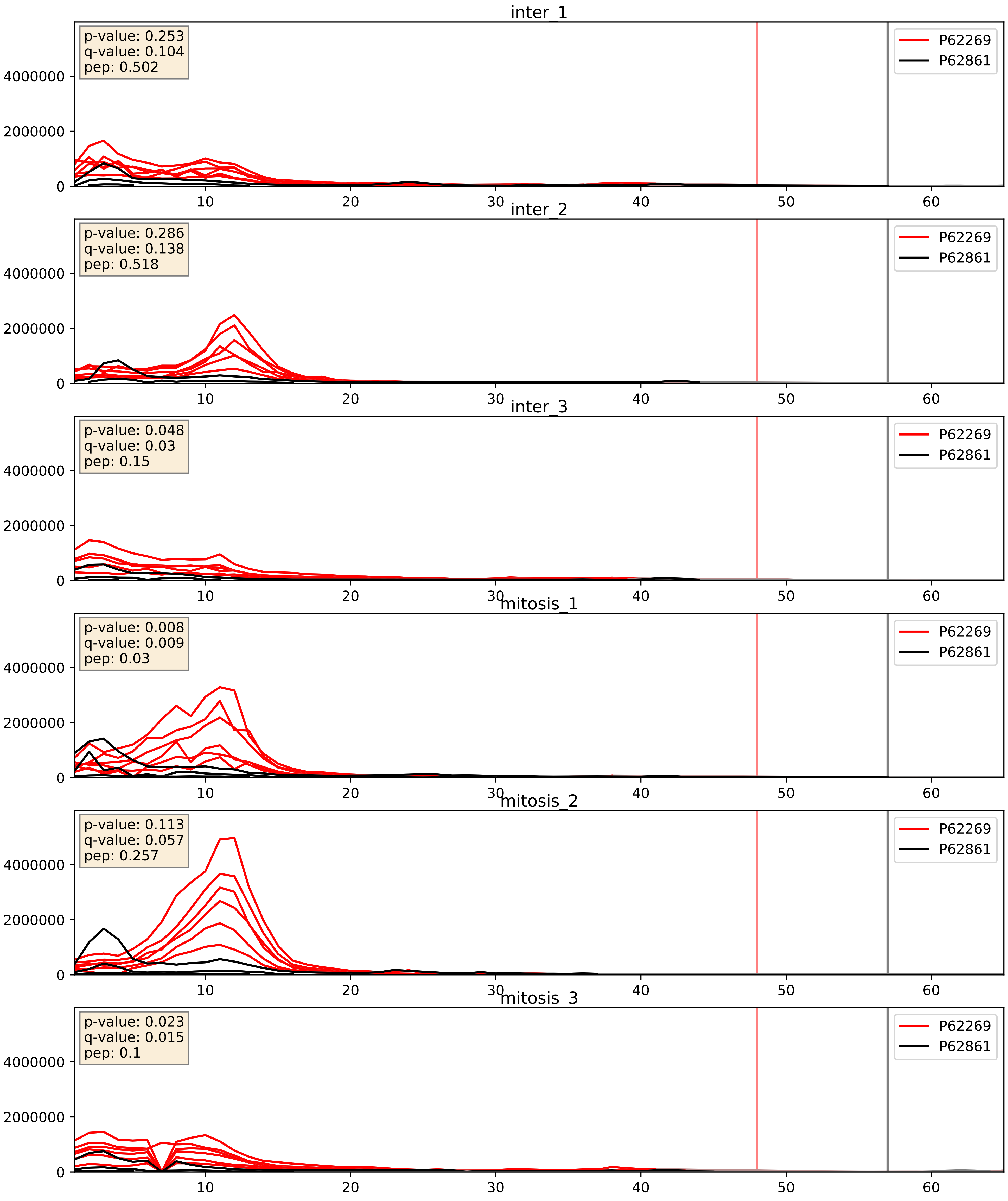


| level | condition_1 | condition_2 | pvalue | pvalue_adjusted |
|---|---|---|---|---|
| interactor_abundance | mitosis | inter | 0.010345526662824988 | 0.08869074435030737 |
| complex_abundance | mitosis | inter | 0.09439111511289694 | 0.22721820735837958 |
| interactor_ratio | mitosis | inter | 0.49754009251343034 | 0.8702488381316982 |

P62269_P62888
RS18_HUMAN vs RL30_HUMAN
p-value: 0.044 q-value: 0.029

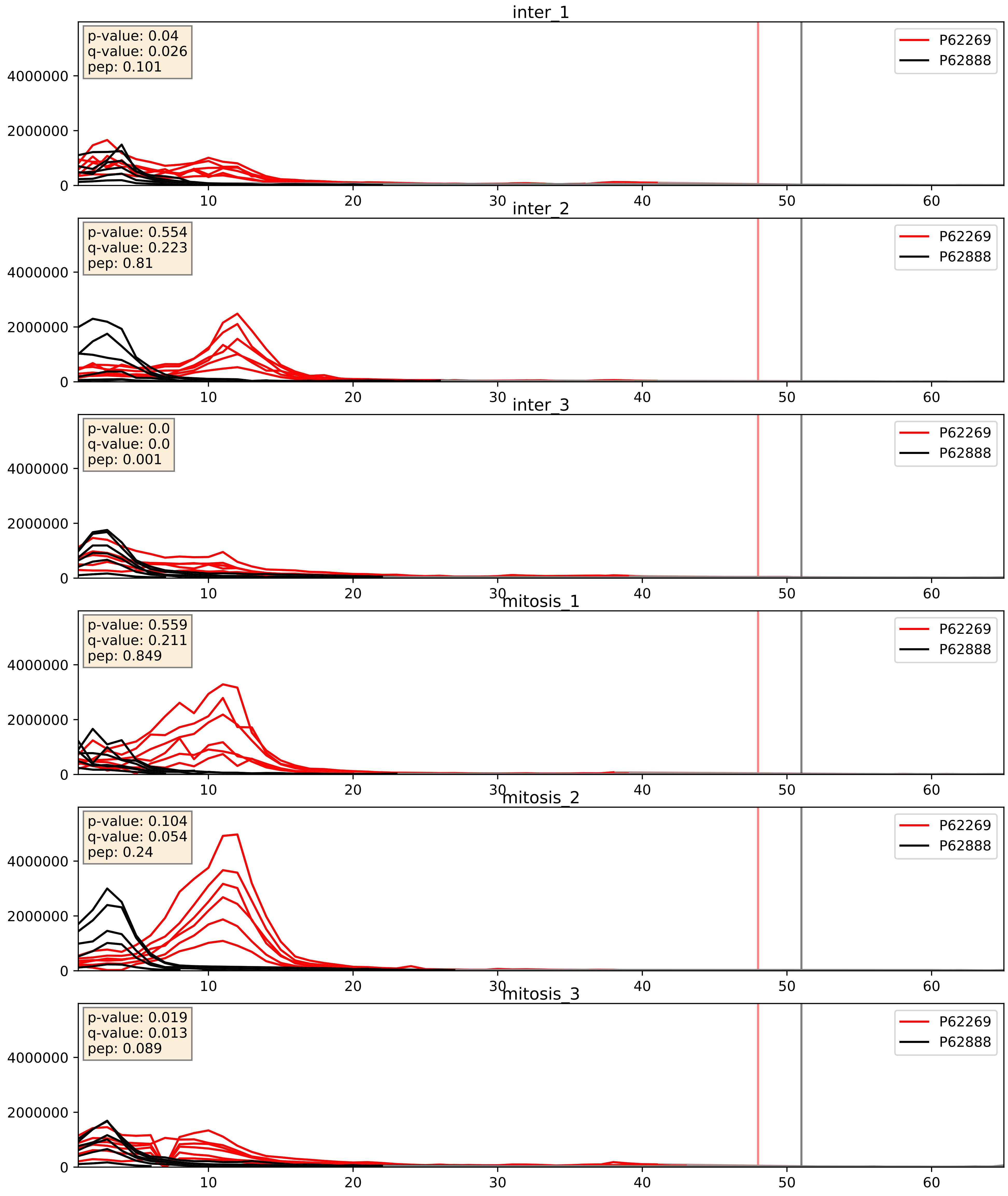


| level | condition_1 | condition_2 | pvalue | pvalue_adjusted |
|---|---|---|---|---|
| interactor_abundance | mitosis | inter | 0.021602061675789177 | 0.12219131964586558 |
| interactor_ratio | mitosis | inter | 0.047757703089379856 | 0.3614455797387484 |
| complex_abundance | mitosis | inter | 0.3887739945367265 | 0.5349271429503147 |

P62269_P62899
RS18_HUMAN vs RL31_HUMAN
p-value: 0.0 q-value: 0.0

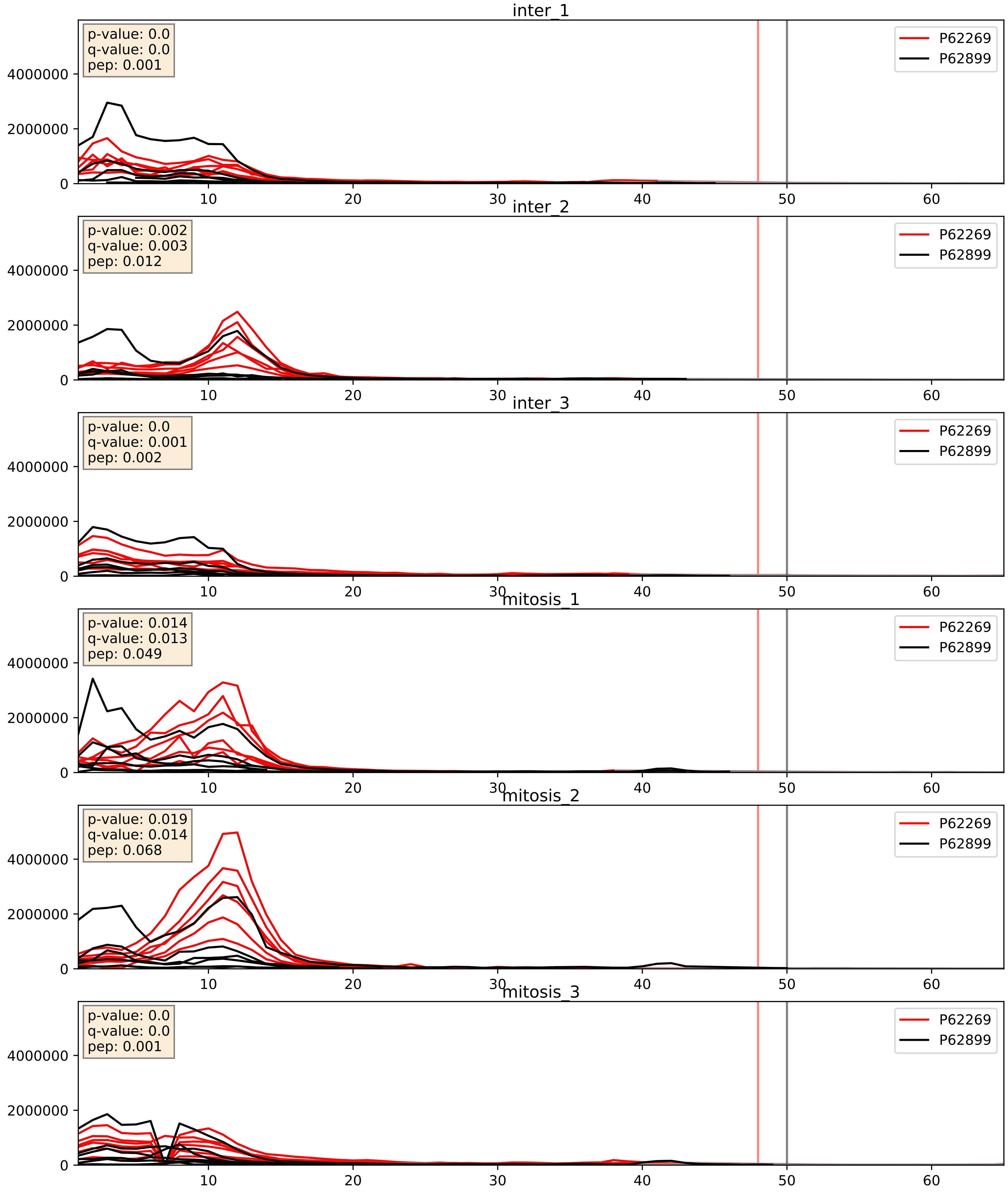


| level | condition_1 | condition_2 | pvalue | pvalue_adjusted |
|---|---|---|---|---|
| complex_abundance | mitosis | inter | 0.003997864986950128 | 0.04687907436752479 |
| interactor_abundance | mitosis | inter | 0.009876120958878595 | 0.08628103889292982 |
| interactor_ratio | mitosis | inter | 0.26184665990223577 | 0.7356112270308953 |

P62269_P62906
RS18_HUMAN vs RL10A_HUMAN
p-value: 0.006 q-value: 0.006

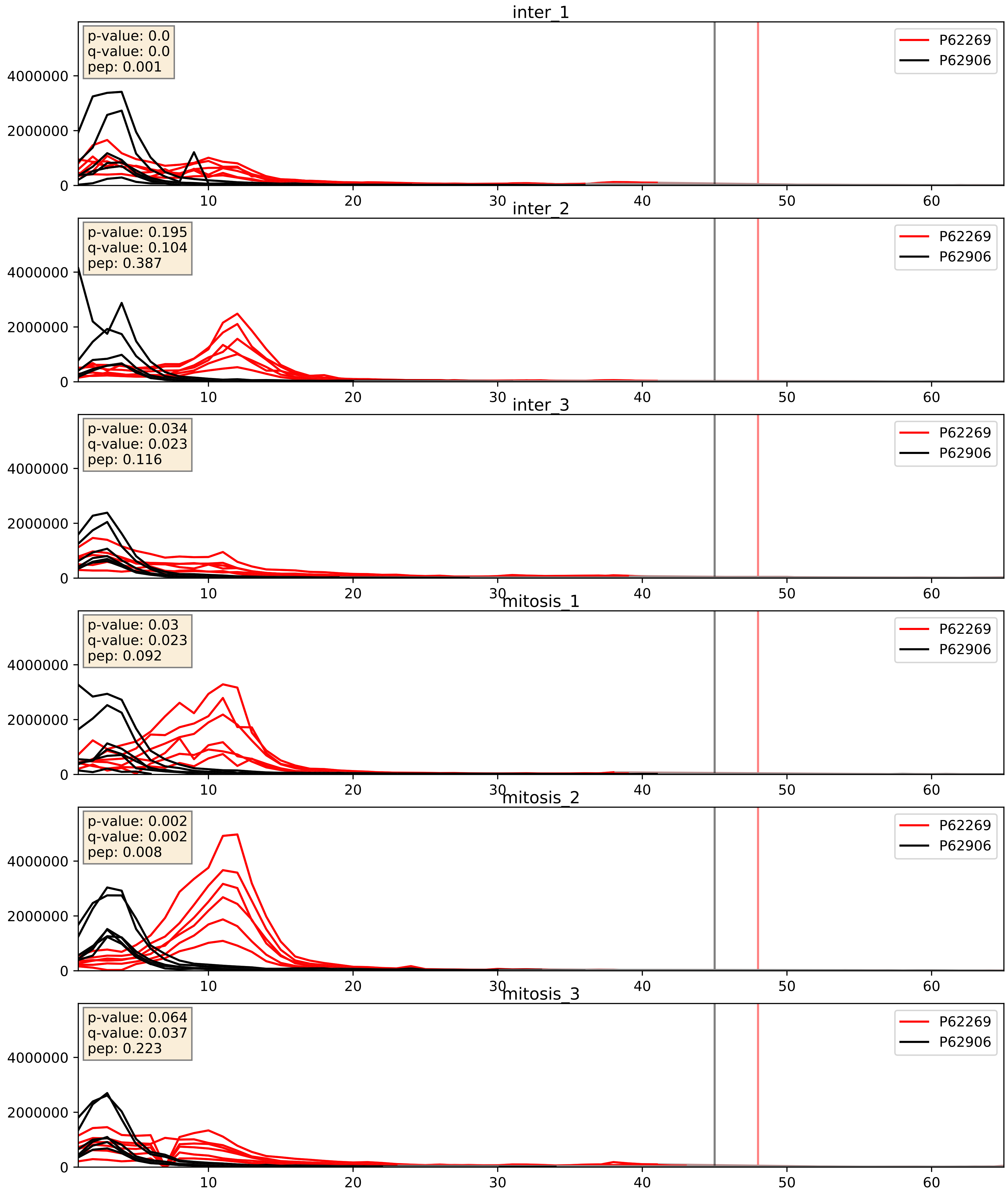


| level | condition_1 | condition_2 | pvalue | pvalue_adjusted |
|---|---|---|---|---|
| complex_abundance | mitosis | inter | 0.006023236527422844 | 0.05806974312565789 |
| interactor_abundance | mitosis | inter | 0.01739271702981435 | 0.11421684524373674 |
| interactor_ratio | mitosis | inter | 0.0760206439167499 | 0.46316861816154964 |

P62269_P62910
RS18_HUMAN vs RL32_HUMAN
p-value: 0.048 q-value: 0.031

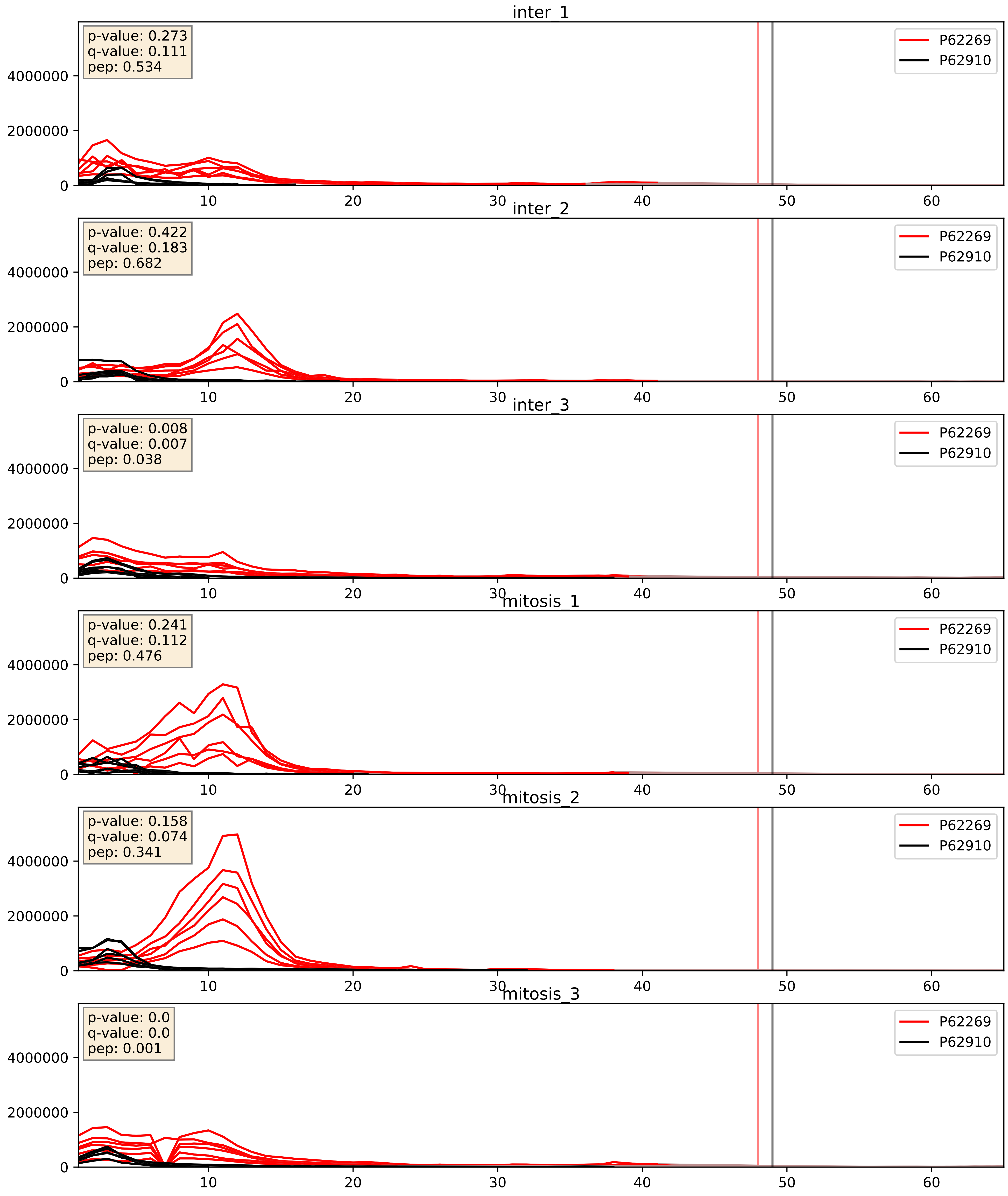


| level | condition_1 | condition_2 | pvalue | pvalue_adjusted |
|---|---|---|---|---|
| interactor_abundance | mitosis | inter | 0.020820214267006282 | 0.12061307420112273 |
| interactor_ratio | mitosis | inter | 0.04034051759816009 | 0.33235306125144404 |
| complex_abundance | mitosis | inter | 0.10260332196110886 | 0.23799312559378397 |

P62269_P62913
RS18_HUMAN vs RL11_HUMAN
p-value: 0.0 q-value: 0.0

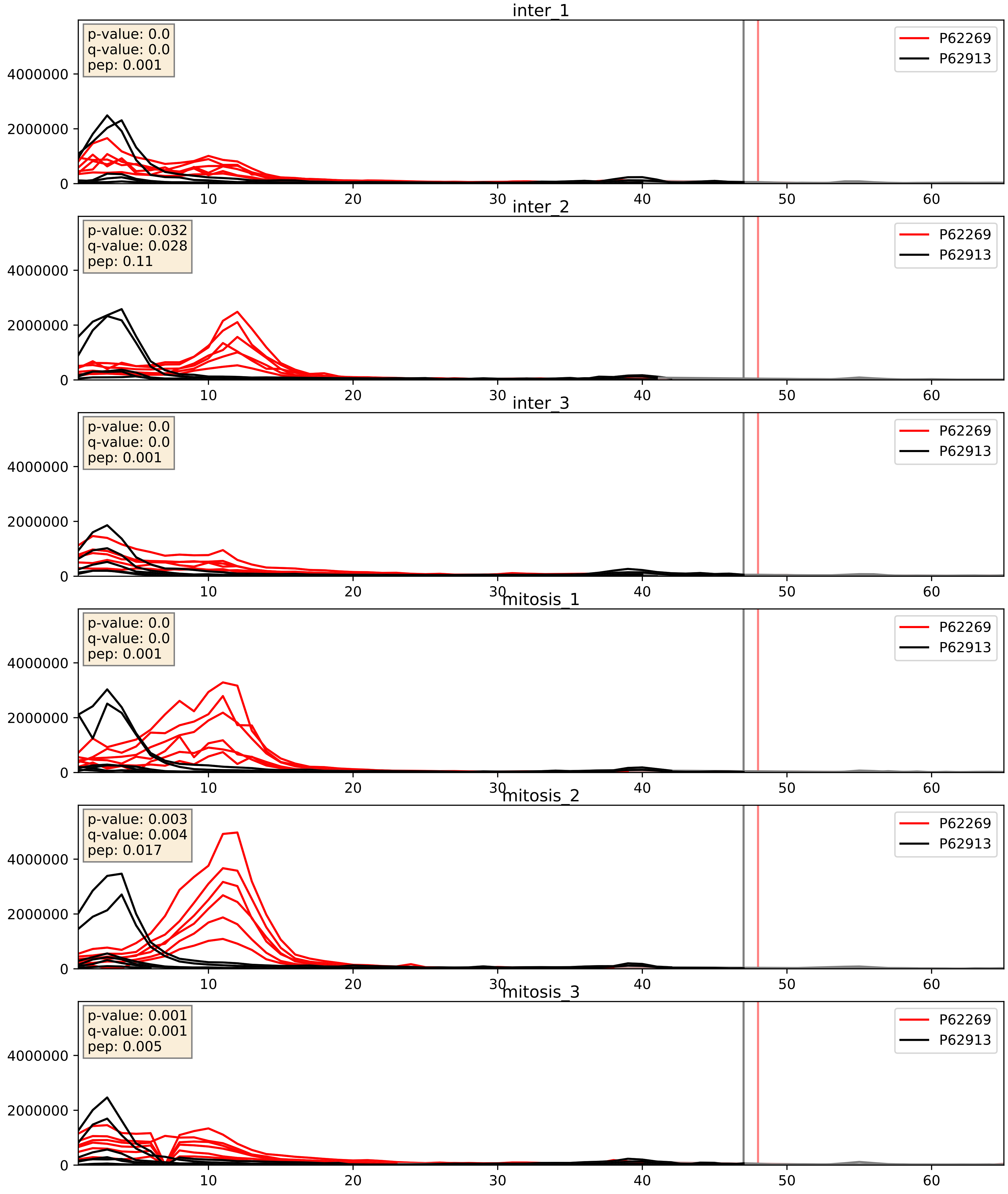


| level | condition_1 | condition_2 | pvalue | pvalue_adjusted |
|---|---|---|---|---|
| complex_abundance | mitosis | inter | 0.006833585943598483 | 0.061187756984521974 |
| interactor_abundance | mitosis | inter | 0.011243111228547091 | 0.09348327548942506 |
| interactor_ratio | mitosis | inter | 0.03479843403474809 | 0.3021223786073155 |

P62269_P62917
RS18_HUMAN vs RL8_HUMAN
p-value: 0.003 q-value: 0.003

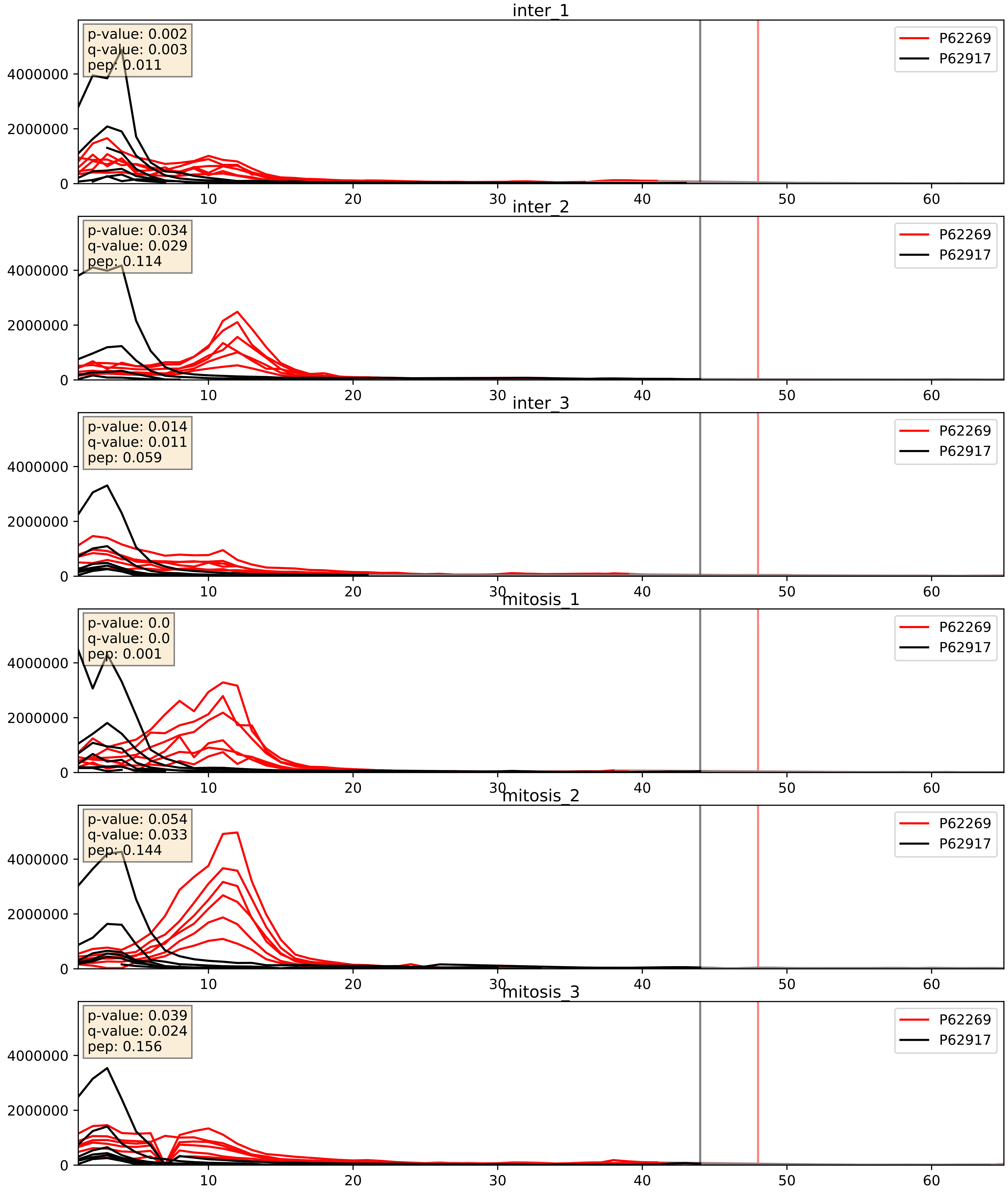


| level | condition_1 | condition_2 | pvalue | pvalue_adjusted |
|---|---|---|---|---|
| interactor_abundance | mitosis | inter | 0.005719315236138822 | 0.06408028589181716 |
| complex_abundance | mitosis | inter | 0.03239025221310241 | 0.13109246285775727 |
| interactor_ratio | mitosis | inter | 0.40486229727328615 | 0.8313271231500527 |

P62269_P63173
RS18_HUMAN vs RL38_HUMAN
p-value: 0.0 q-value: 0.0

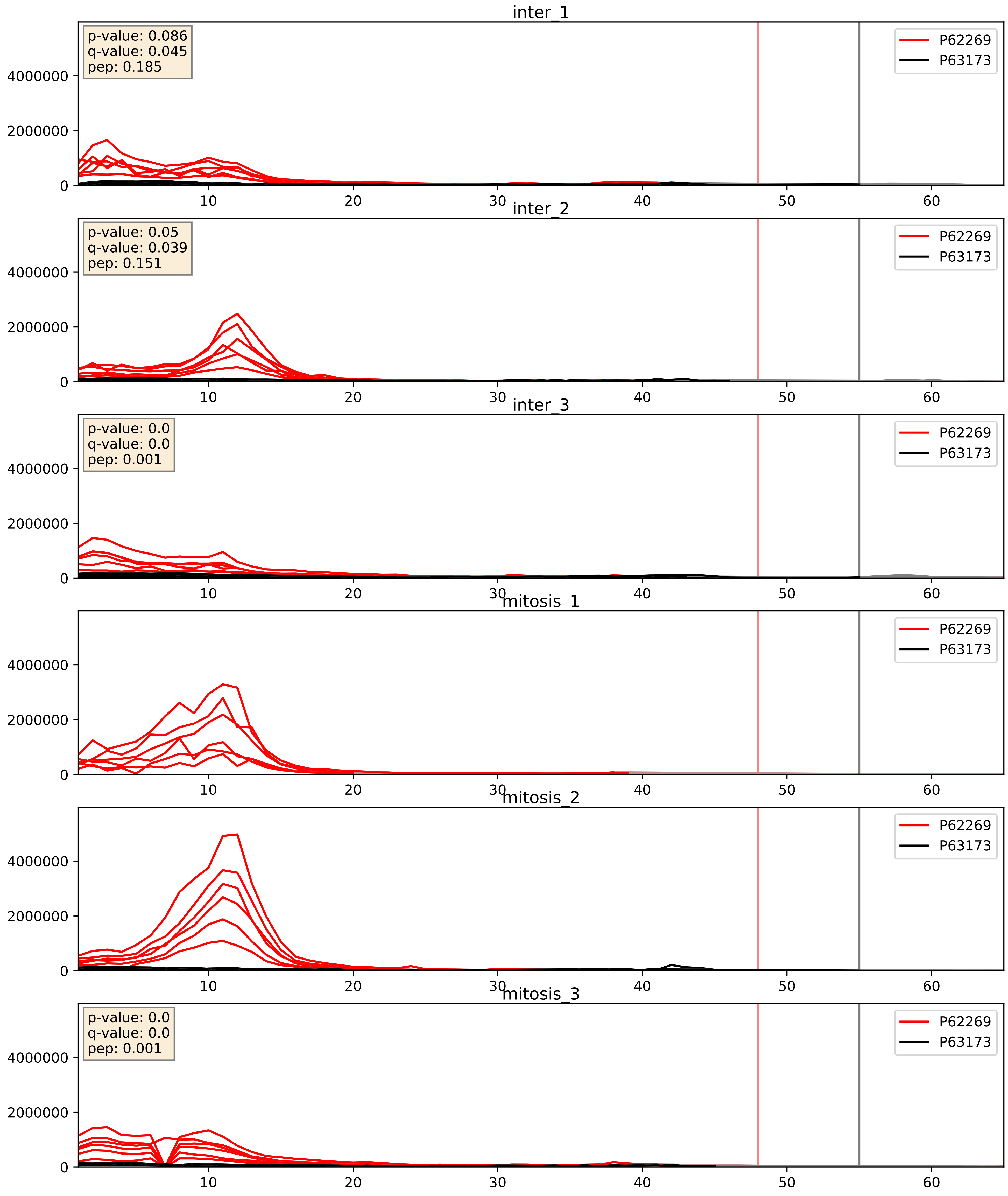


| level | condition_1 | condition_2 | pvalue | pvalue_adjusted |
|---|---|---|---|---|
| complex_abundance | mitosis | inter | 0.4085180707644842 | 0.5535084016101673 |
| interactor_abundance | mitosis | inter | 0.5413241733716816 | 0.6985881085574542 |
| interactor_ratio | mitosis | inter | 0.9511693208167908 | 0.9919597934691097 |

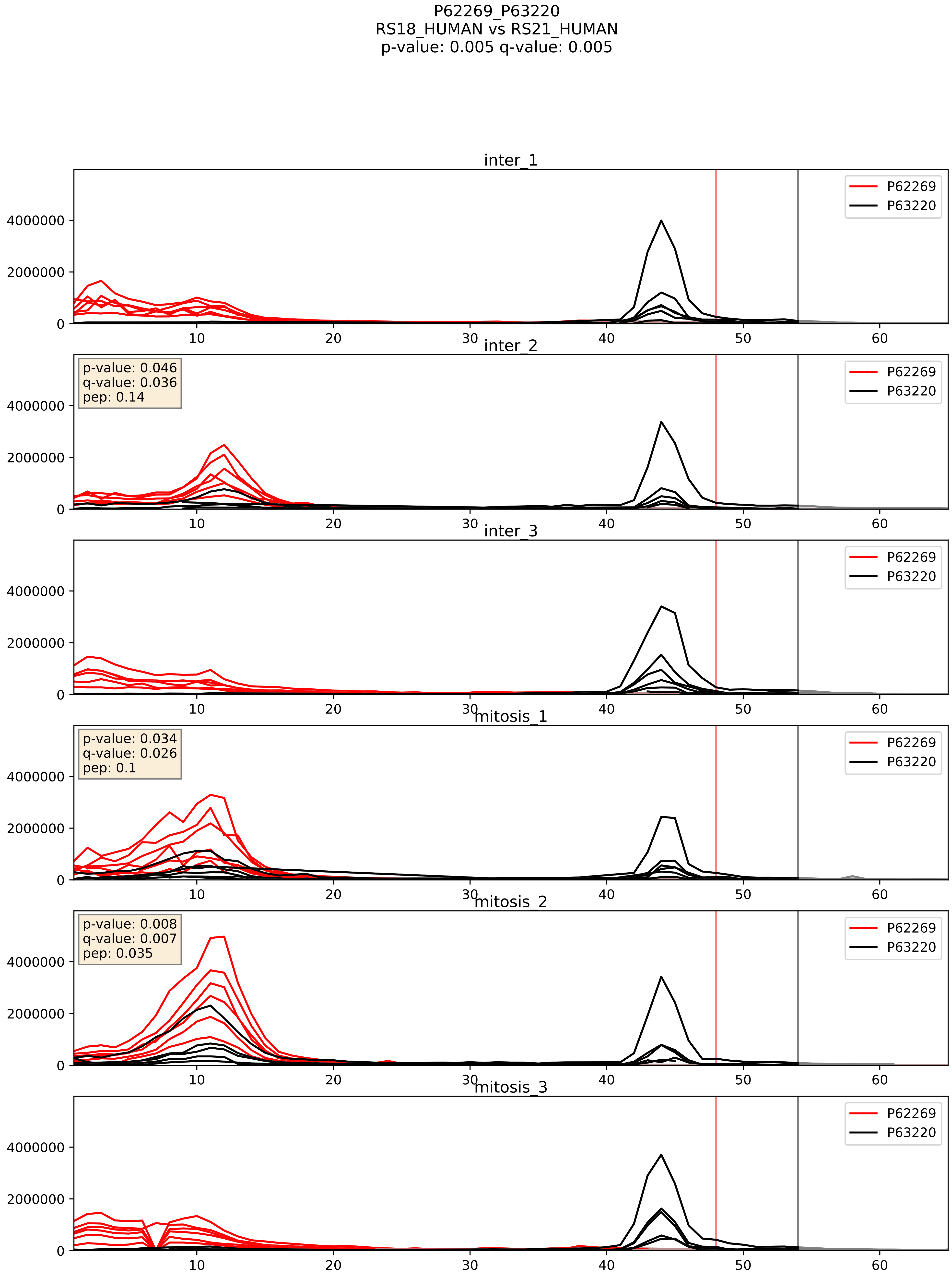


| level | condition_1 | condition_2 | pvalue | pvalue_adjusted |
|---|---|---|---|---|
| interactor_ratio | mitosis | inter | 0.052031111294175036 | 0.37764936302518703 |
| interactor_abundance | mitosis | inter | 0.06231095264275949 | 0.2044301789122679 |
| complex_abundance | mitosis | inter | 0.11915464425668097 | 0.26092702861017886 |

P62269_P63244
RS18_HUMAN vs RACK1_HUMAN
p-value: 0.042 q-value: 0.028

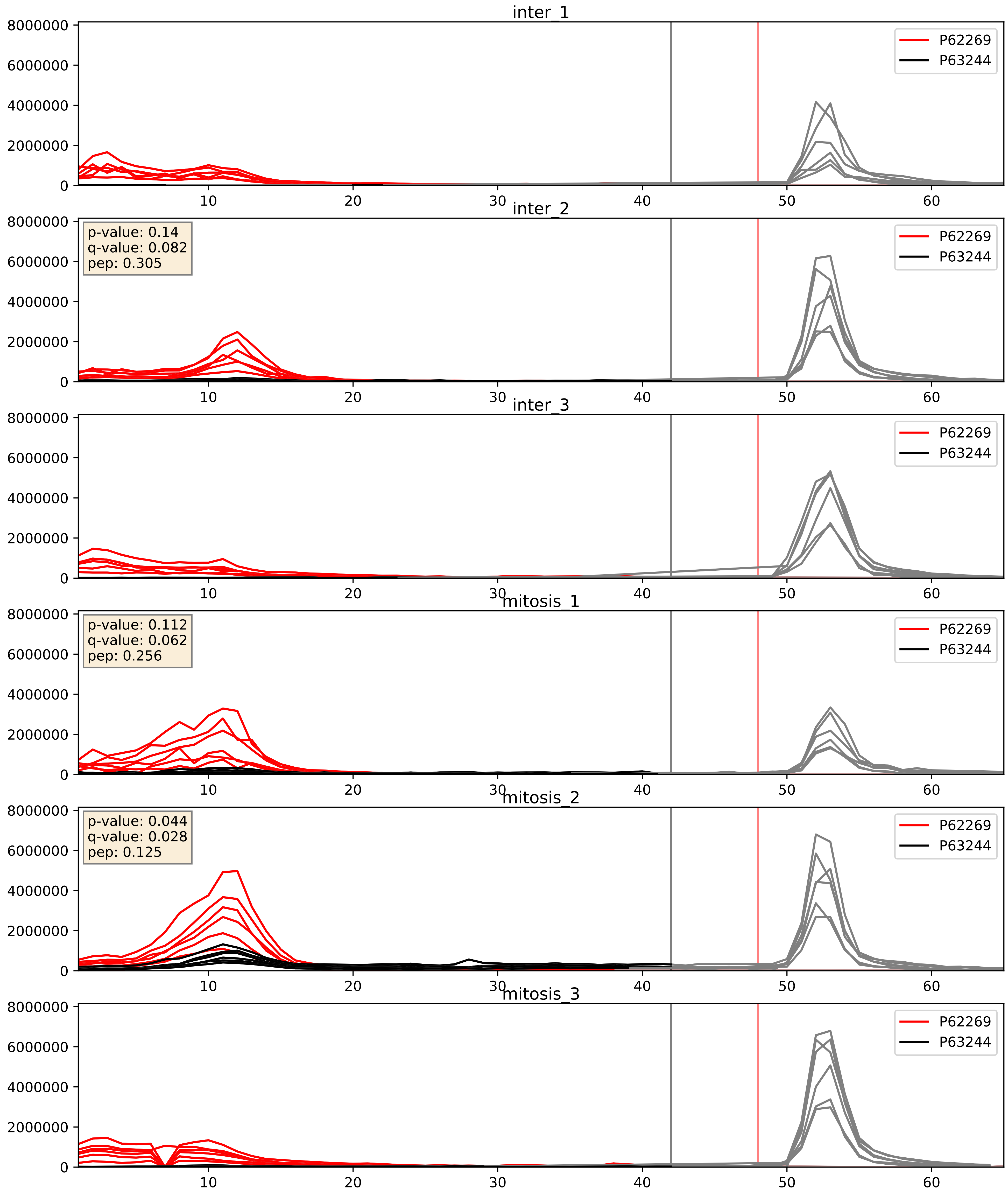


| level | condition_1 | condition_2 | pvalue | pvalue_adjusted |
|---|---|---|---|---|
| interactor_abundance | mitosis | inter | 0.048861486004349274 | 0.1764041839718388 |
| complex_abundance | mitosis | inter | 0.1080175857714414 | 0.24506507665081856 |
| interactor_ratio | mitosis | inter | 0.45836324508957677 | 0.8567124985945661 |

P62269_P67809
RS18_HUMAN vs YBOX1_HUMAN
p-value: 0.0 q-value: 0.018

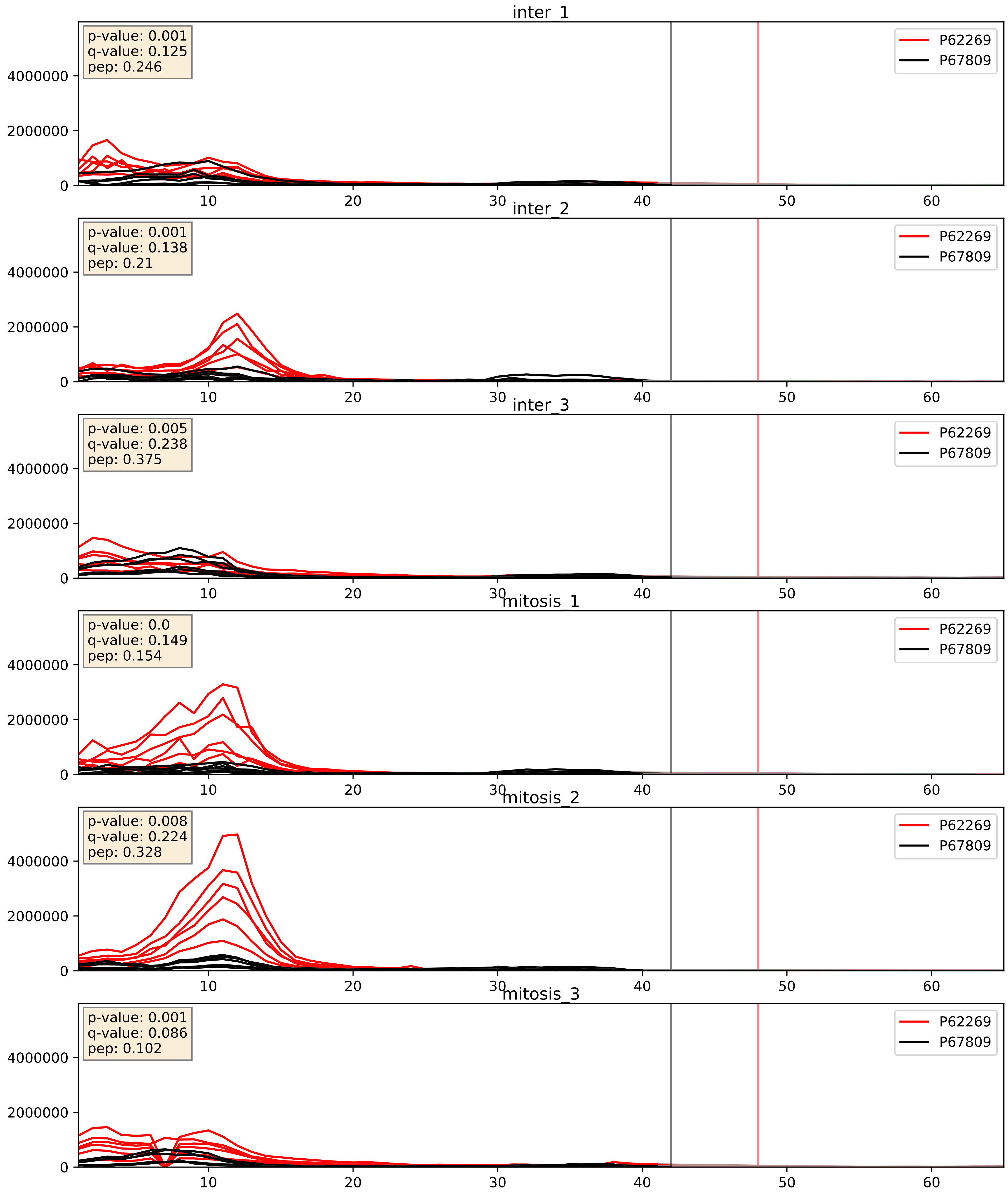


| level | condition_1 | condition_2 | pvalue | pvalue_adjusted |
|---|---|---|---|---|
| interactor_abundance | mitosis | inter | 0.012121353991992215 | 0.09779339318704369 |
| interactor_ratio | mitosis | inter | 0.023011904626773733 | 0.23394525368786598 |
| complex_abundance | mitosis | inter | 0.8324050809378118 | 0.8915650016050638 |

P62269_P83731
RS18_HUMAN vs RL24_HUMAN
p-value: 0.0 q-value: 0.0

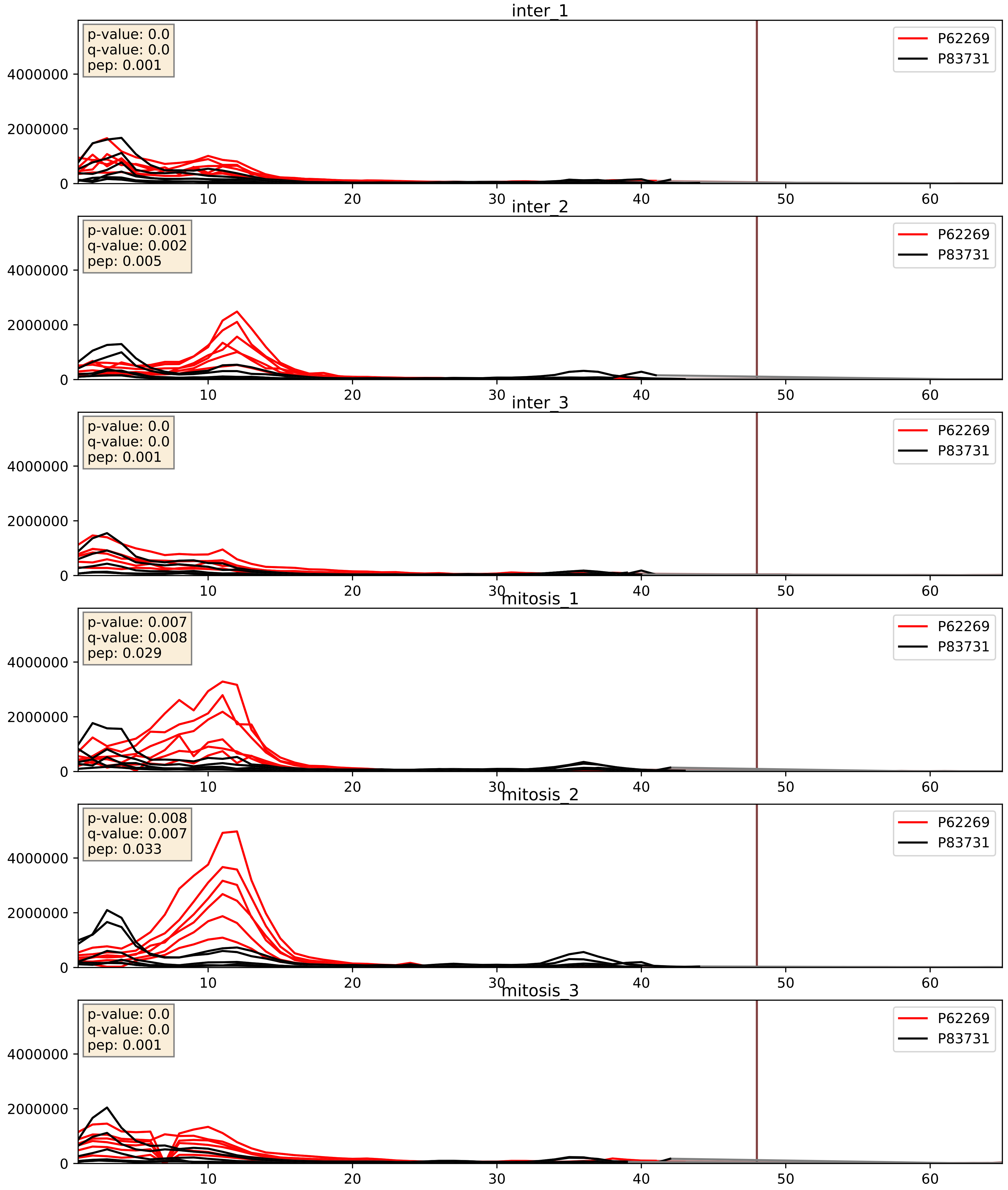


| level | condition_1 | condition_2 | pvalue | pvalue_adjusted |
|---|---|---|---|---|
| complex_abundance | mitosis | inter | 0.008859979272681418 | 0.06989992863977229 |
| interactor_abundance | mitosis | inter | 0.011776276231423785 | 0.09594604221213761 |
| interactor_ratio | mitosis | inter | 0.10892471031819799 | 0.5478234549493389 |

P62269_P84098
RS18_HUMAN vs RL19_HUMAN
p-value: 0.0 q-value: 0.0

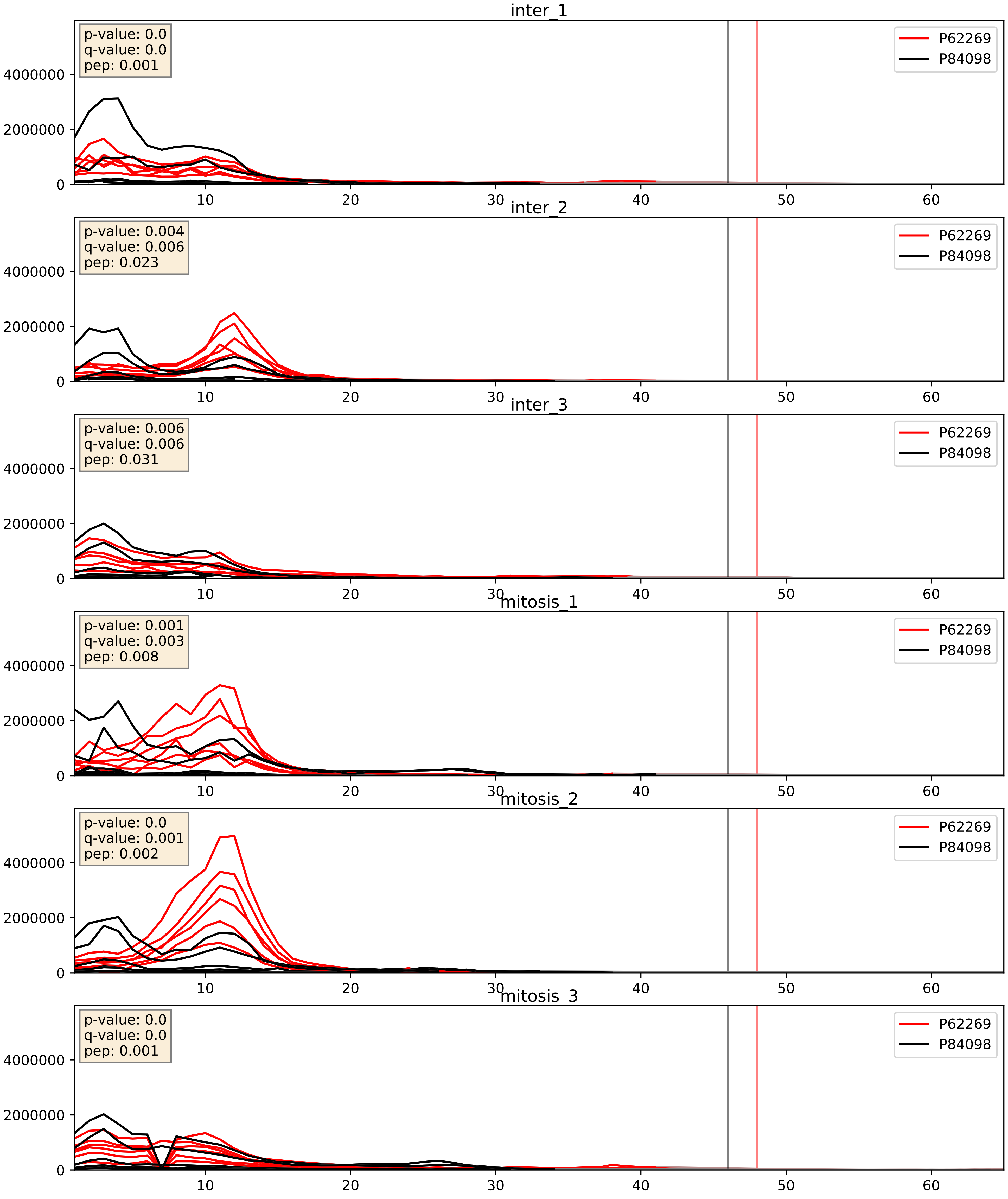


| level | condition_1 | condition_2 | pvalue | pvalue_adjusted |
|---|---|---|---|---|
| complex_abundance | mitosis | inter | 0.011061334382965202 | 0.07799425232140209 |
| interactor_abundance | mitosis | inter | 0.01770297816139897 | 0.11493173535197207 |
| interactor_ratio | mitosis | inter | 0.2637278489220485 | 0.7382309963285595 |

P62269_Q02543
RS18_HUMAN vs RL18A_HUMAN
p-value: 0.0 q-value: 0.0

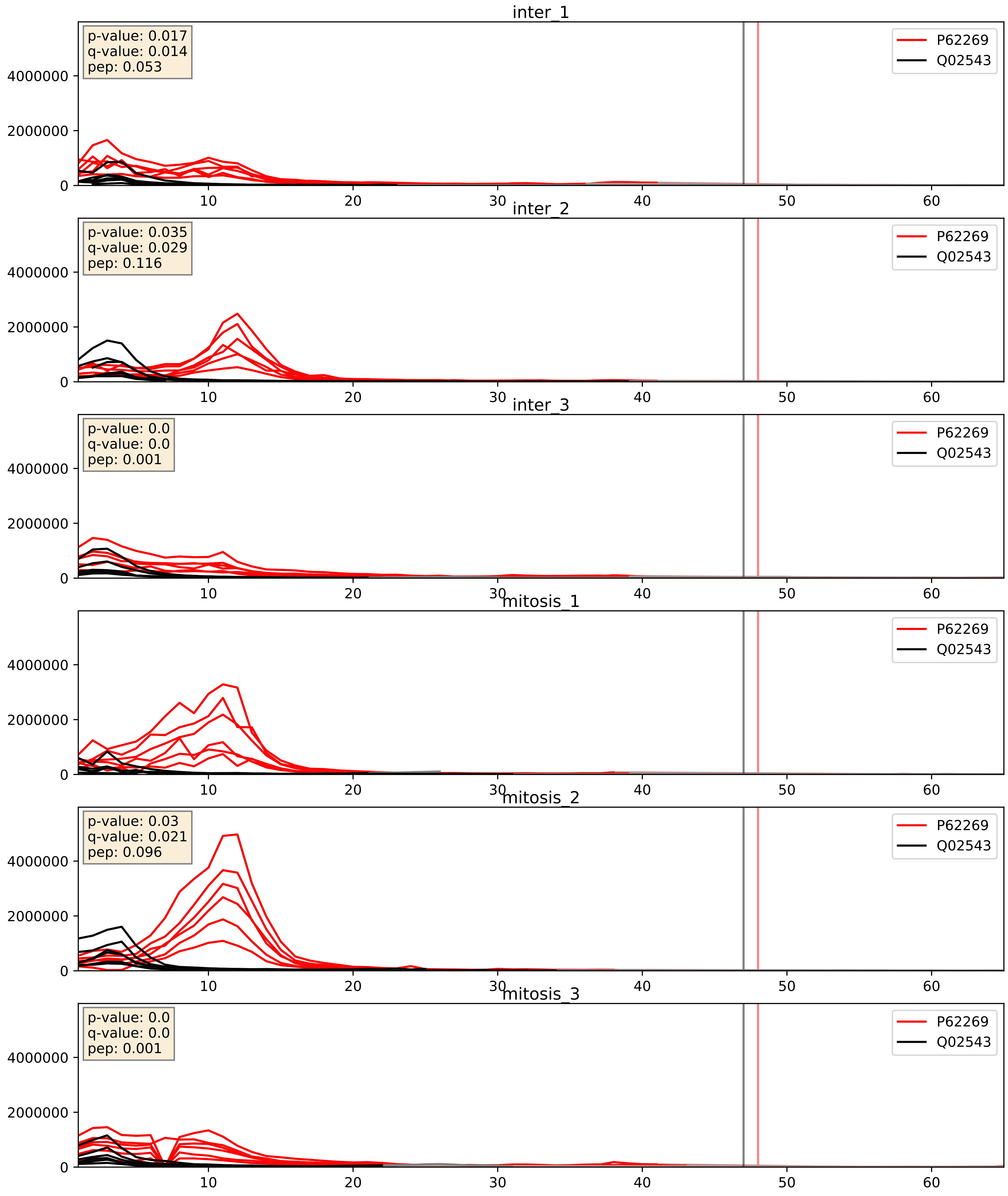


| level | condition_1 | condition_2 | pvalue | pvalue_adjusted |
|---|---|---|---|---|
| interactor_abundance | mitosis | inter | 0.029845563529237444 | 0.14365318846106628 |
| interactor_ratio | mitosis | inter | 0.07745430765925769 | 0.46684312337322714 |
| complex_abundance | mitosis | inter | 0.5396916050089915 | 0.6743190977779838 |

P62269_Q02878
RS18_HUMAN vs RL6_HUMAN
p-value: 0.0 q-value: 0.0

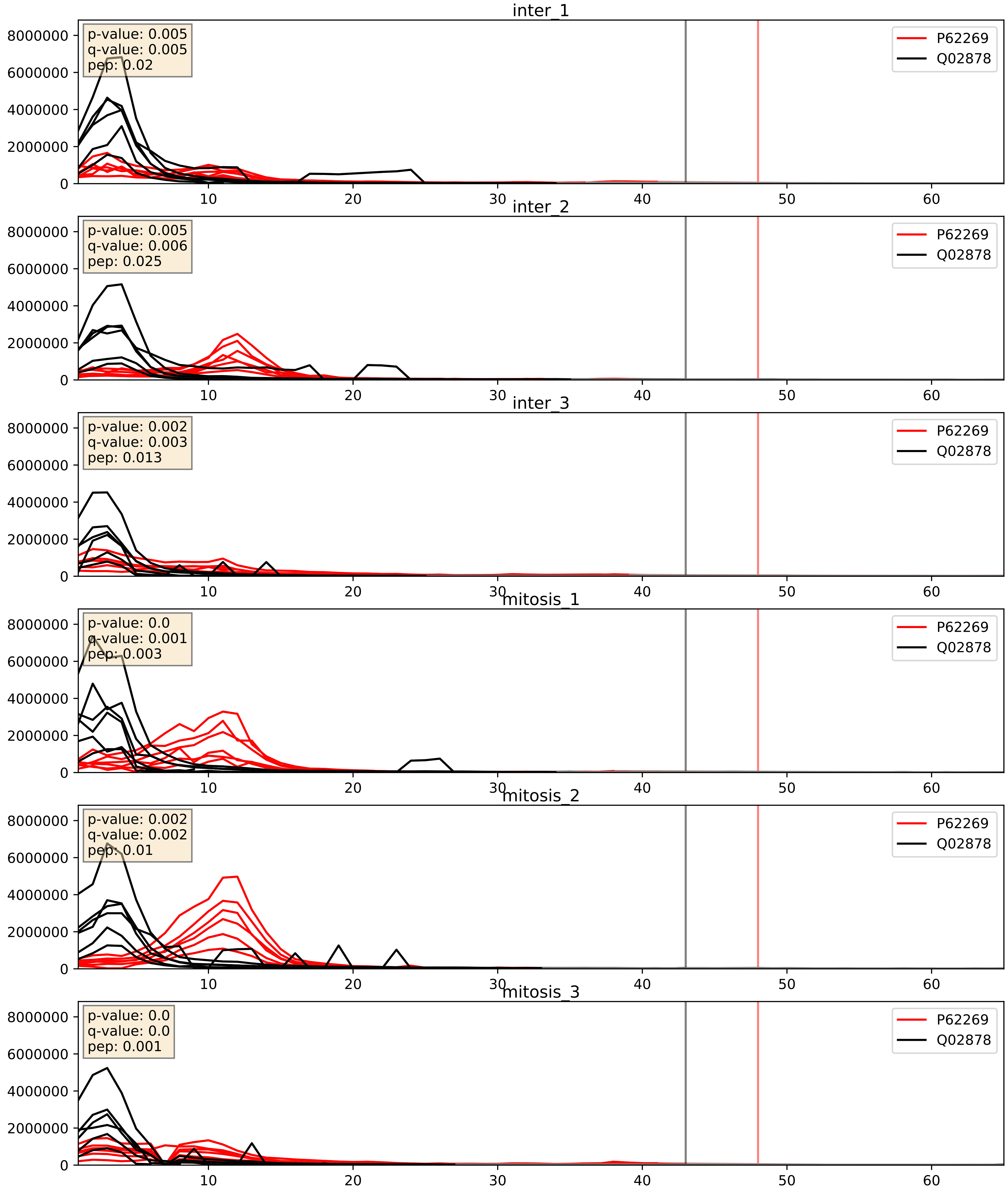


| level | condition_1 | condition_2 | pvalue | pvalue_adjusted |
|---|---|---|---|---|
| interactor_abundance | mitosis | inter | 0.015416986876980718 | 0.10875105699790272 |
| interactor_ratio | mitosis | inter | 0.09429955073244997 | 0.5094488449611217 |
| complex_abundance | mitosis | inter | 0.2144424969573108 | 0.36911920488133304 |

P62269_Q07020
RS18_HUMAN vs RL18_HUMAN
p-value: 0.003 q-value: 0.003

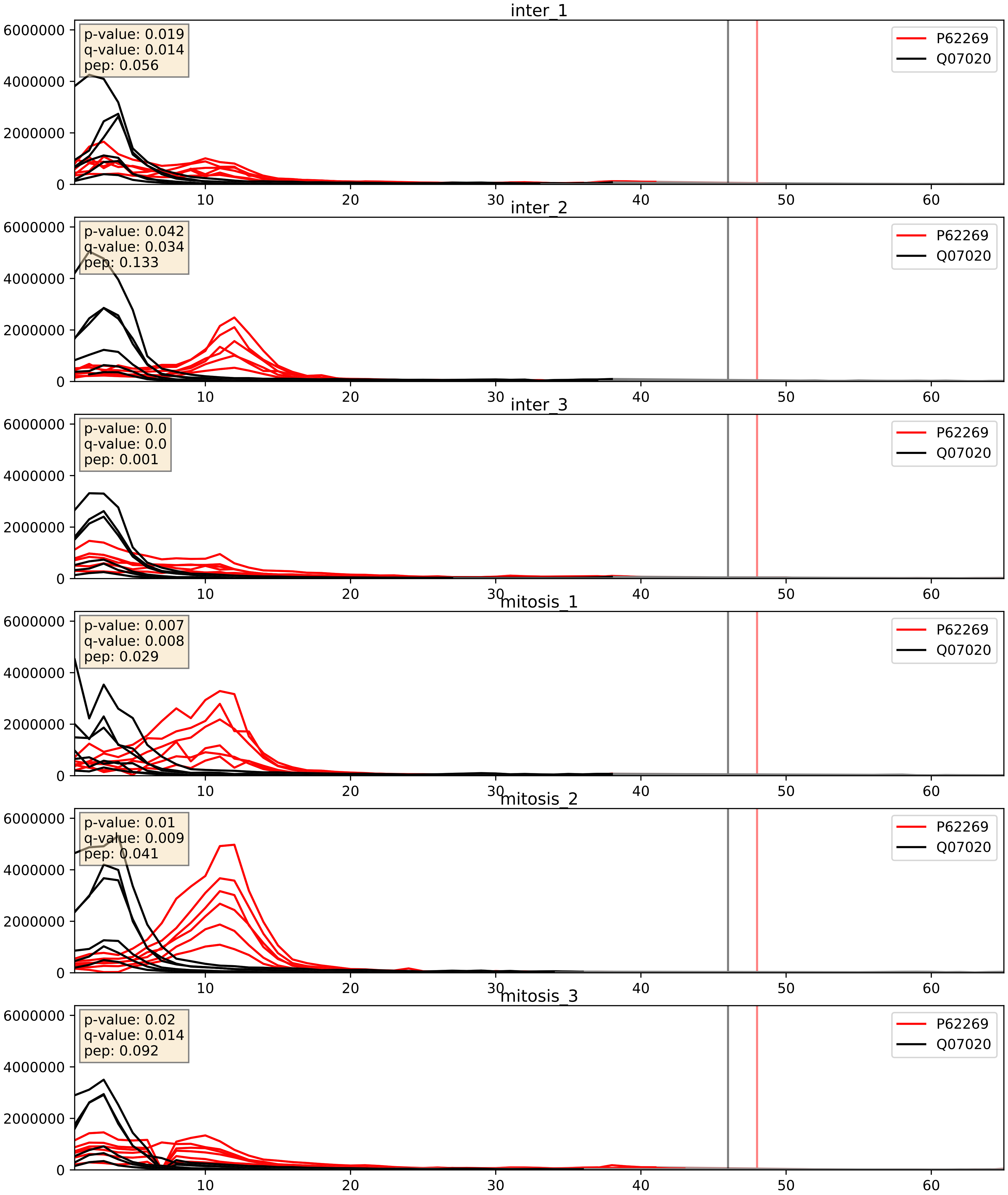


| level | condition_1 | condition_2 | pvalue | pvalue_adjusted |
|---|---|---|---|---|
| interactor_abundance | mitosis | inter | 0.01663738694456197 | 0.11277132084292767 |
| interactor_ratio | mitosis | inter | 0.021123001578685775 | 0.2202349494683925 |
| complex_abundance | mitosis | inter | 0.2558632975721986 | 0.40649402880809576 |

P62269_Q969Q0
RS18_HUMAN vs RL36L_HUMAN
p-value: 0.036 q-value: 0.025

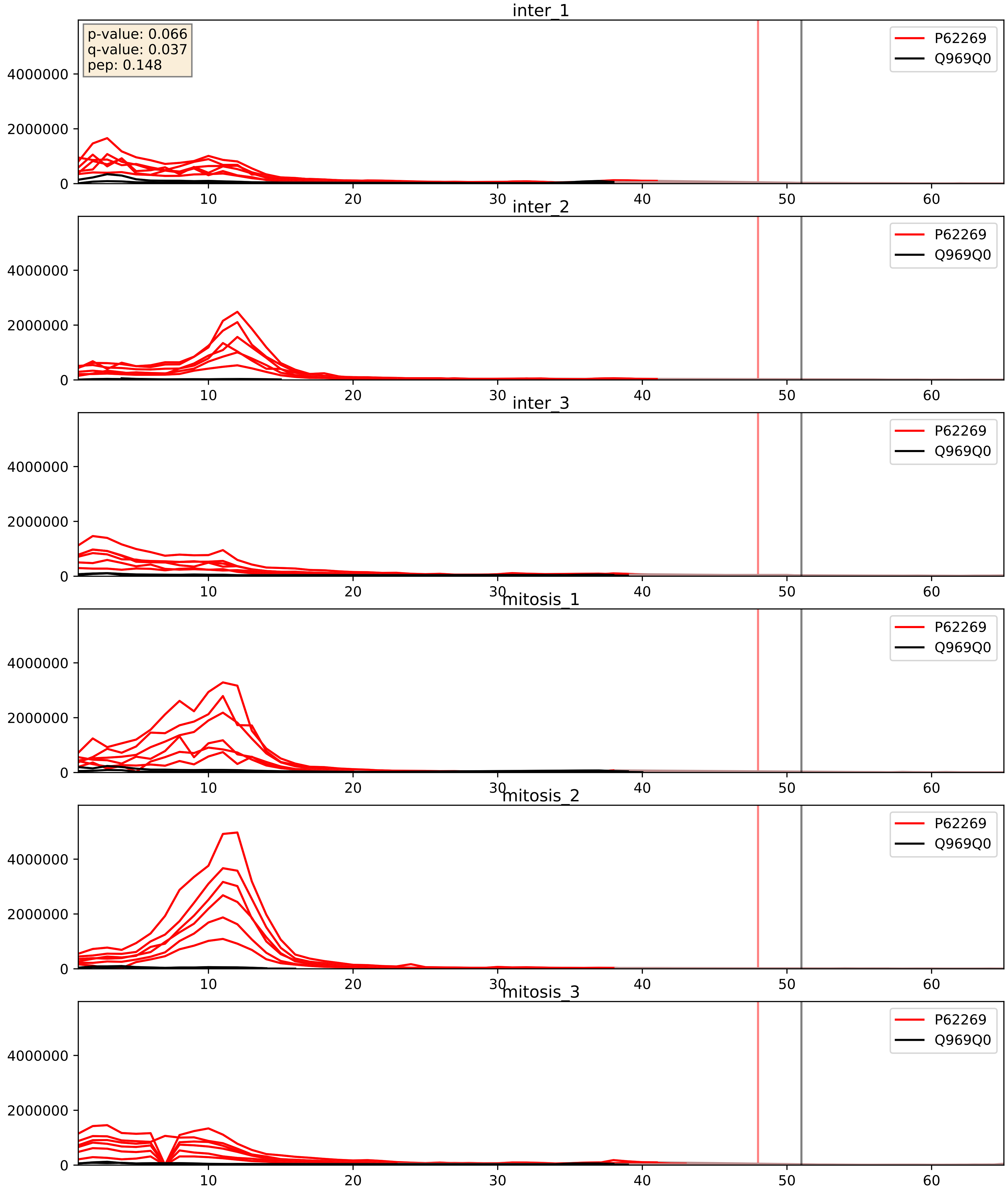


| level | condition_1 | condition_2 | pvalue | pvalue_adjusted |
|---|---|---|---|---|
| interactor_abundance | mitosis | inter | 0.04474339125385491 | 0.16824222672216033 |
| complex_abundance | mitosis | inter | 0.12571192979521037 | 0.2678183471993531 |
| interactor_ratio | mitosis | inter | 0.2076987810064808 | 0.6851258440907421 |

P62269_Q9Y2R9
RS18_HUMAN vs RT07_HUMAN
p-value: 0.004 q-value: 0.047

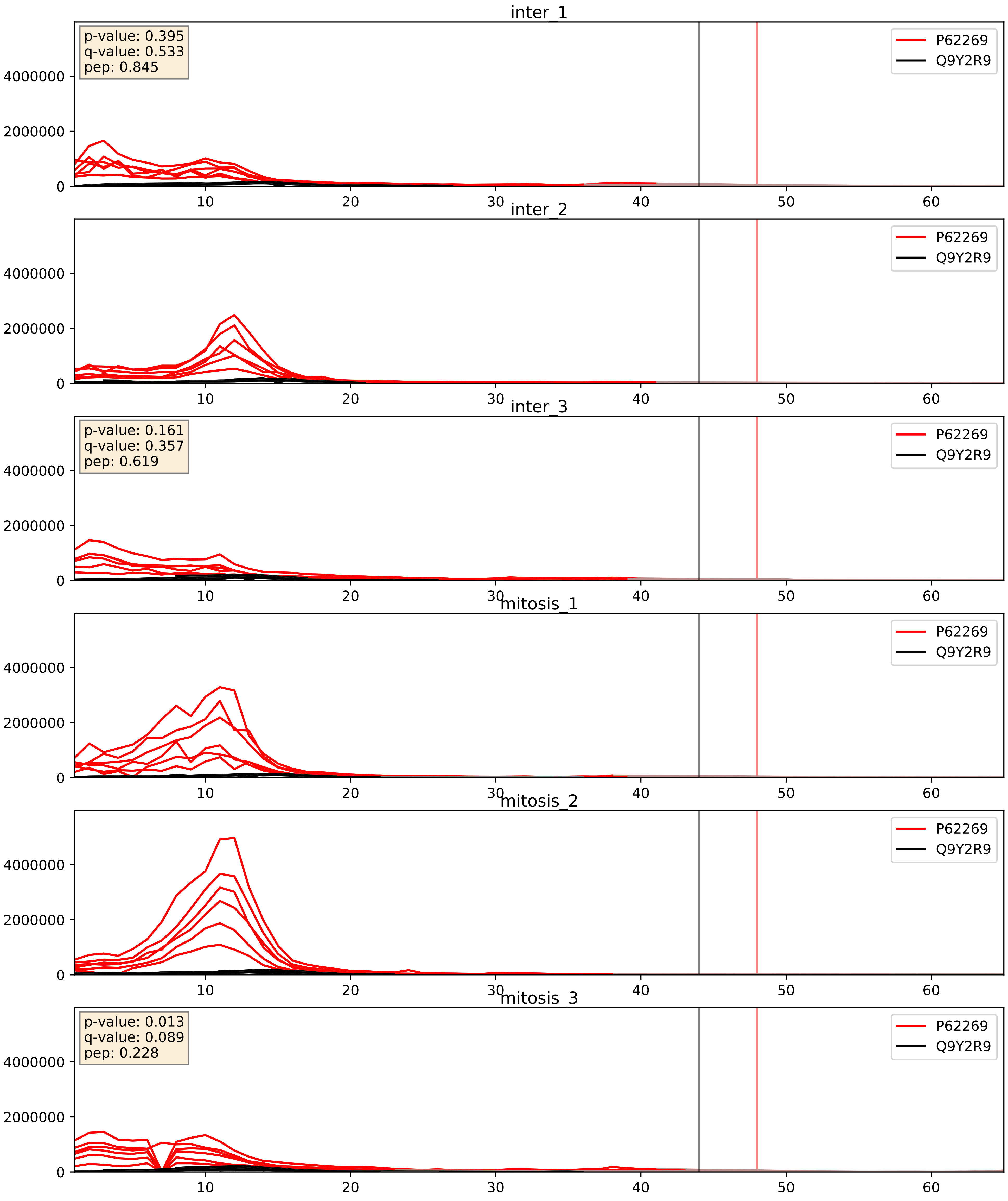


| level | condition_1 | condition_2 | pvalue | pvalue_adjusted |
|---|---|---|---|---|
| interactor_abundance | mitosis | inter | 0.020333471193953786 | 0.12002651953515421 |
| interactor_ratio | mitosis | inter | 0.048939097505324713 | 0.3649117374961494 |
| complex_abundance | mitosis | inter | 0.42079892649253725 | 0.5644059559348353 |

P62269_Q9Y2X3
RS18_HUMAN vs NOP58_HUMAN
p-value: 0.0 q-value: 0.024

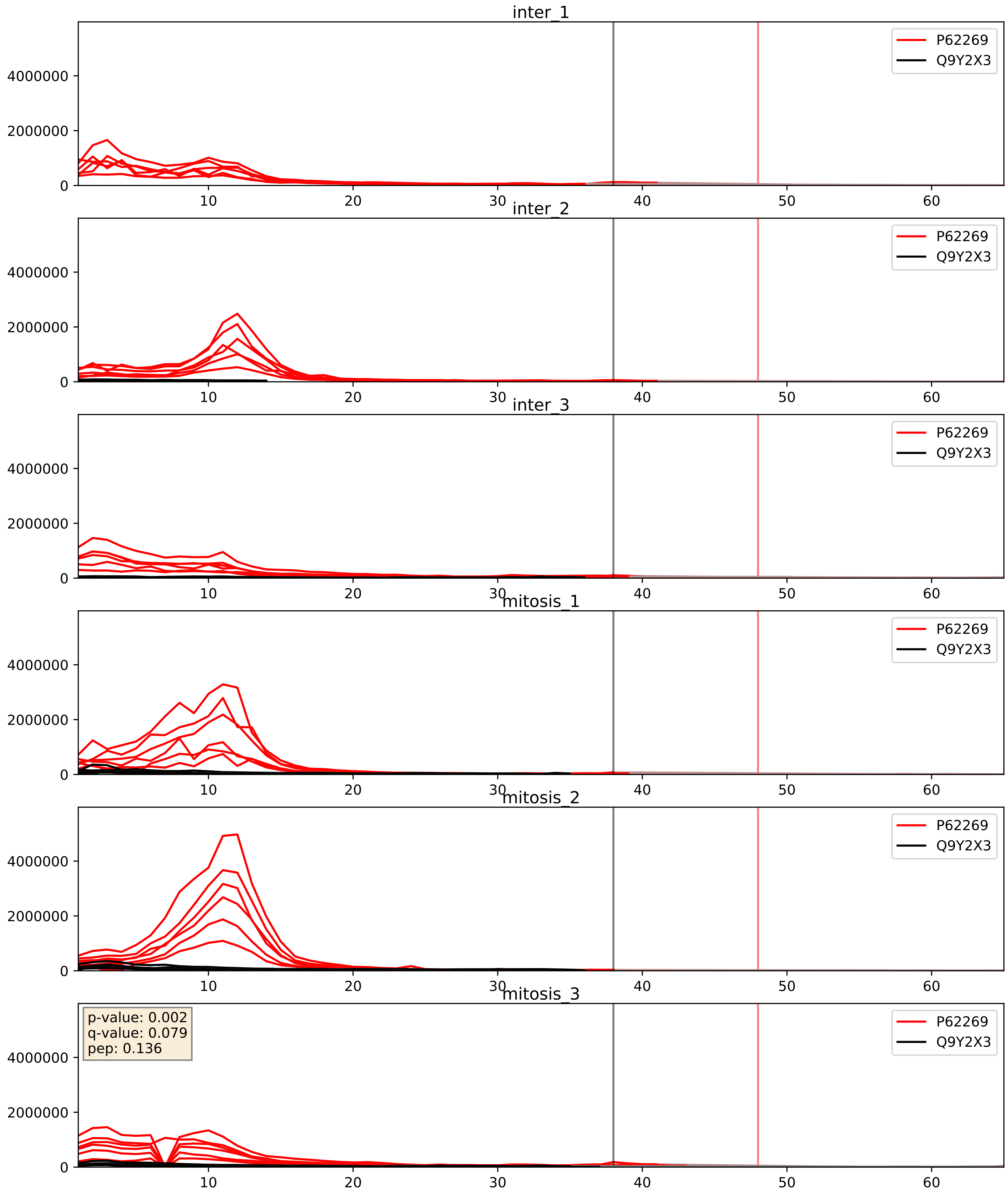


| level | condition_1 | condition_2 | pvalue | pvalue_adjusted |
|---|---|---|---|---|
| complex_abundance | mitosis | inter | 0.09773888757235956 | 0.23169340283007417 |
| interactor_abundance | mitosis | inter | 0.22976059832030665 | 0.42246864822439595 |
| interactor_ratio | mitosis | inter | 0.7394020622773316 | 0.9491588884573259 |

P62269_Q9Y3U8
RS18_HUMAN vs RL36_HUMAN
p-value: 0.001 q-value: 0.001

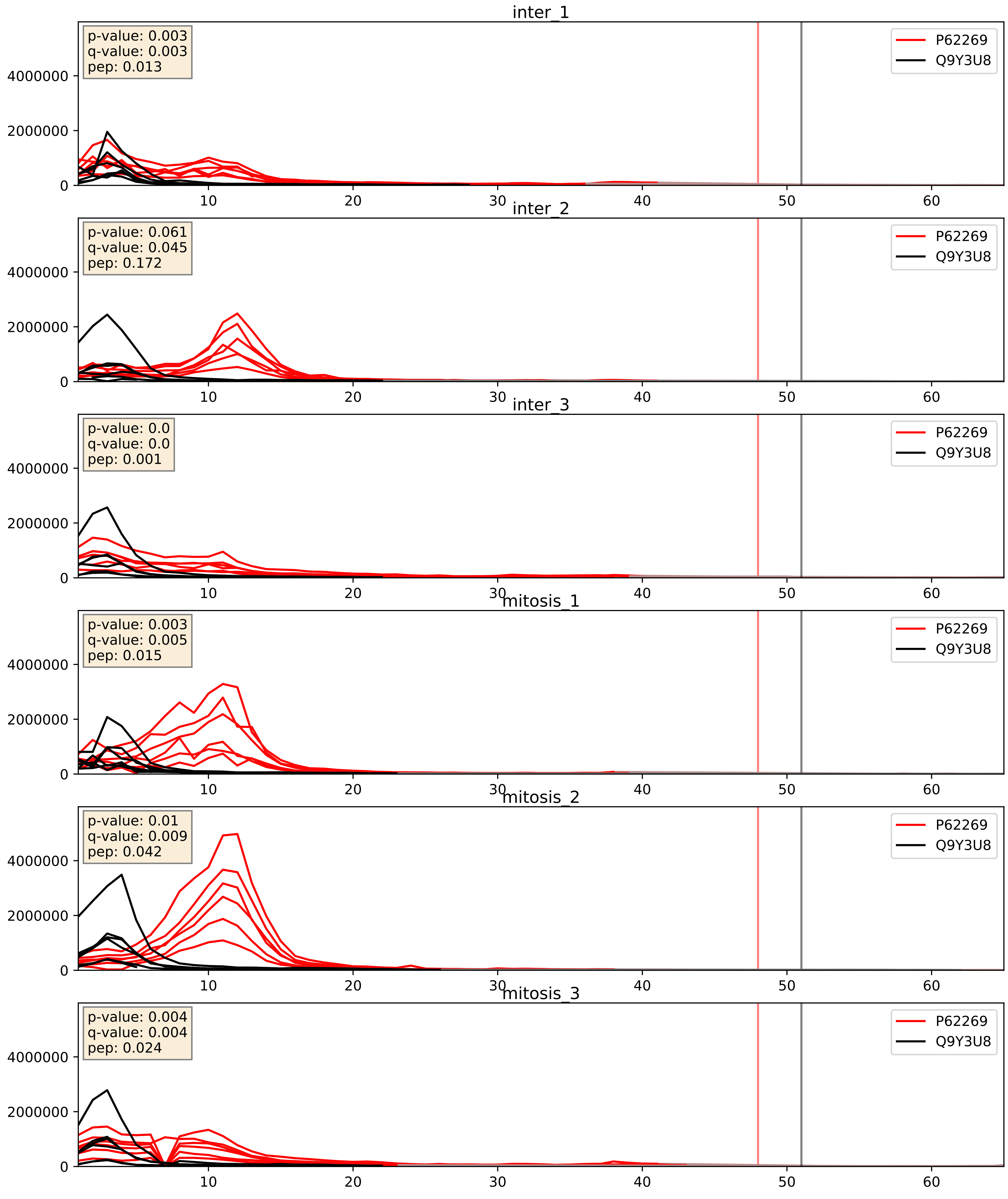


| level | condition_1 | condition_2 | pvalue | pvalue_adjusted |
|---|---|---|---|---|
| interactor_abundance | mitosis | inter | 0.021824689875546744 | 0.12294790742657462 |
| complex_abundance | mitosis | inter | 0.03173805408262735 | 0.12967911357865877 |
| interactor_ratio | mitosis | inter | 0.03758459947362565 | 0.3178686235304411 |
