## Supplementary Data 2 for "SECAT: Quantifying differential protein-protein interaction states by network-centric analysis": hela_string_000006_P46087.pdf

P46087_P46087

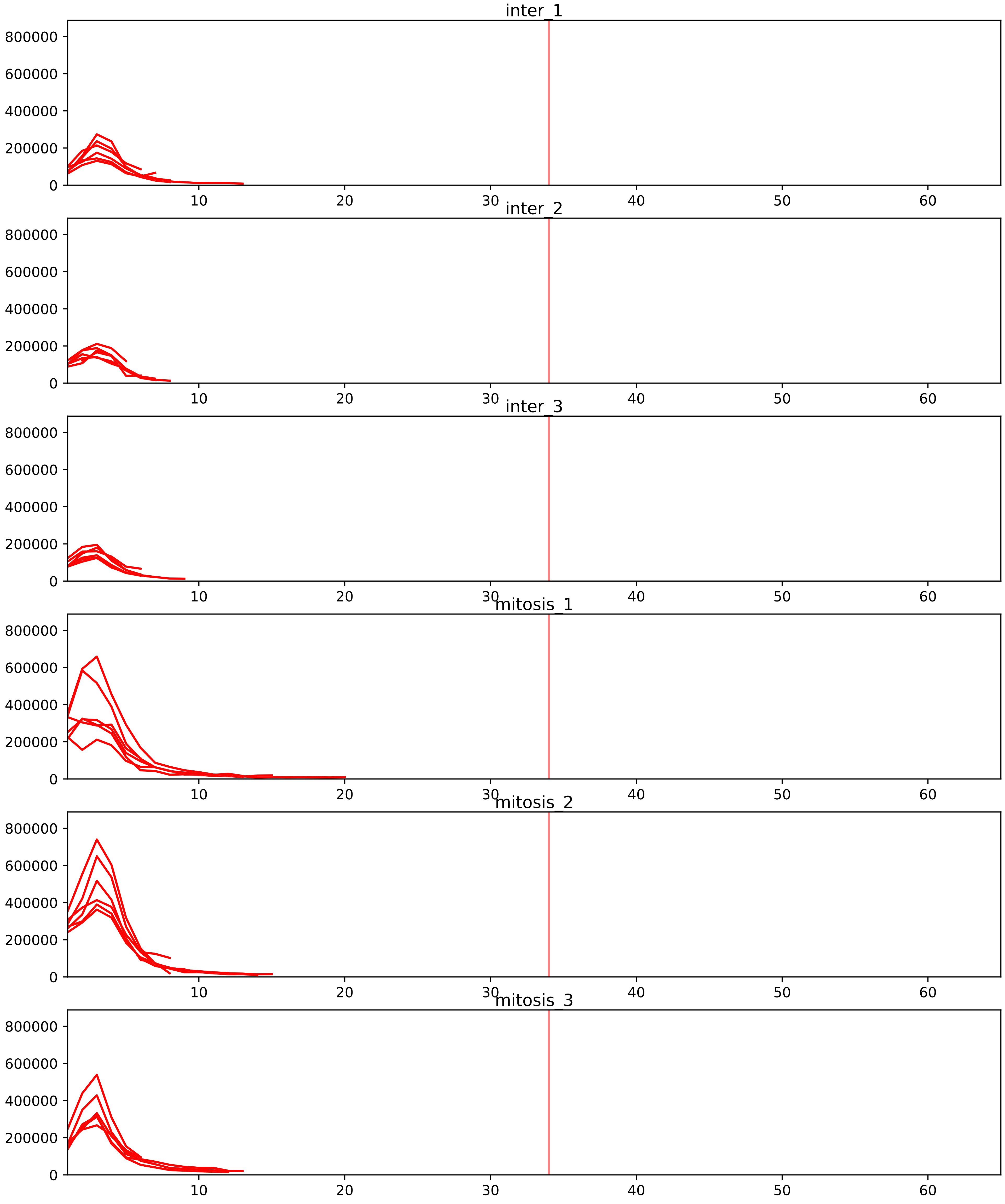


| level | condition_1 | condition_2 | pvalue | pvalue_adjusted |
|---|---|---|---|---|
| interactor_ratio | mitosis | inter | 1.871092711642598e-17 | 5.738017649037301e-15 |
| interactor_abundance | mitosis | inter | 0.00013355125004408088 | 0.0028245321848403312 |
| complex_abundance | mitosis | inter | 0.00039045830344758655 | 0.005888879330684911 |
| assembled_abundance | mitosis | inter | 0.007519948729417653 | 0.09331793469809328 |
| total_abundance | mitosis | inter | 0.008888099848097101 | 0.11468713451692311 |

O00567_P46087
NOP56_HUMAN vs NOP2_HUMAN
p-value: 0.01 q-value: 0.009

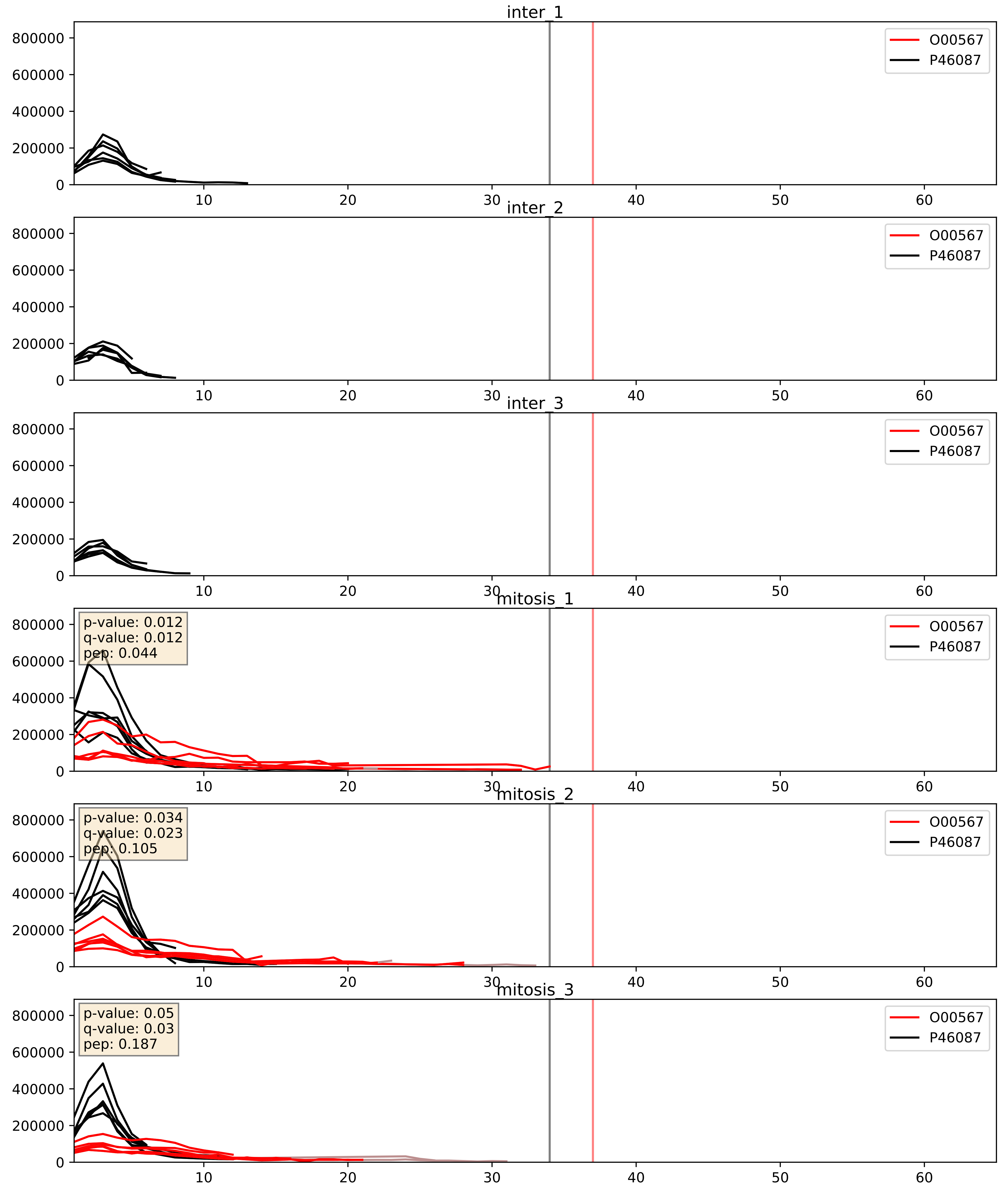


| level | condition_1 | condition_2 | pvalue | pvalue_adjusted |
|---|---|---|---|---|
| complex_abundance | mitosis | inter | 0.0004100837102177417 | 0.012905575586264225 |
| interactor_abundance | mitosis | inter | 0.001371443612175878 | 0.02742887224351756 |
| interactor_ratio | mitosis | inter | 0.0037884888716758825 | 0.08027095233055831 |

O43159_P46087
RRP8_HUMAN vs NOP2_HUMAN
p-value: 0.0 q-value: 0.009

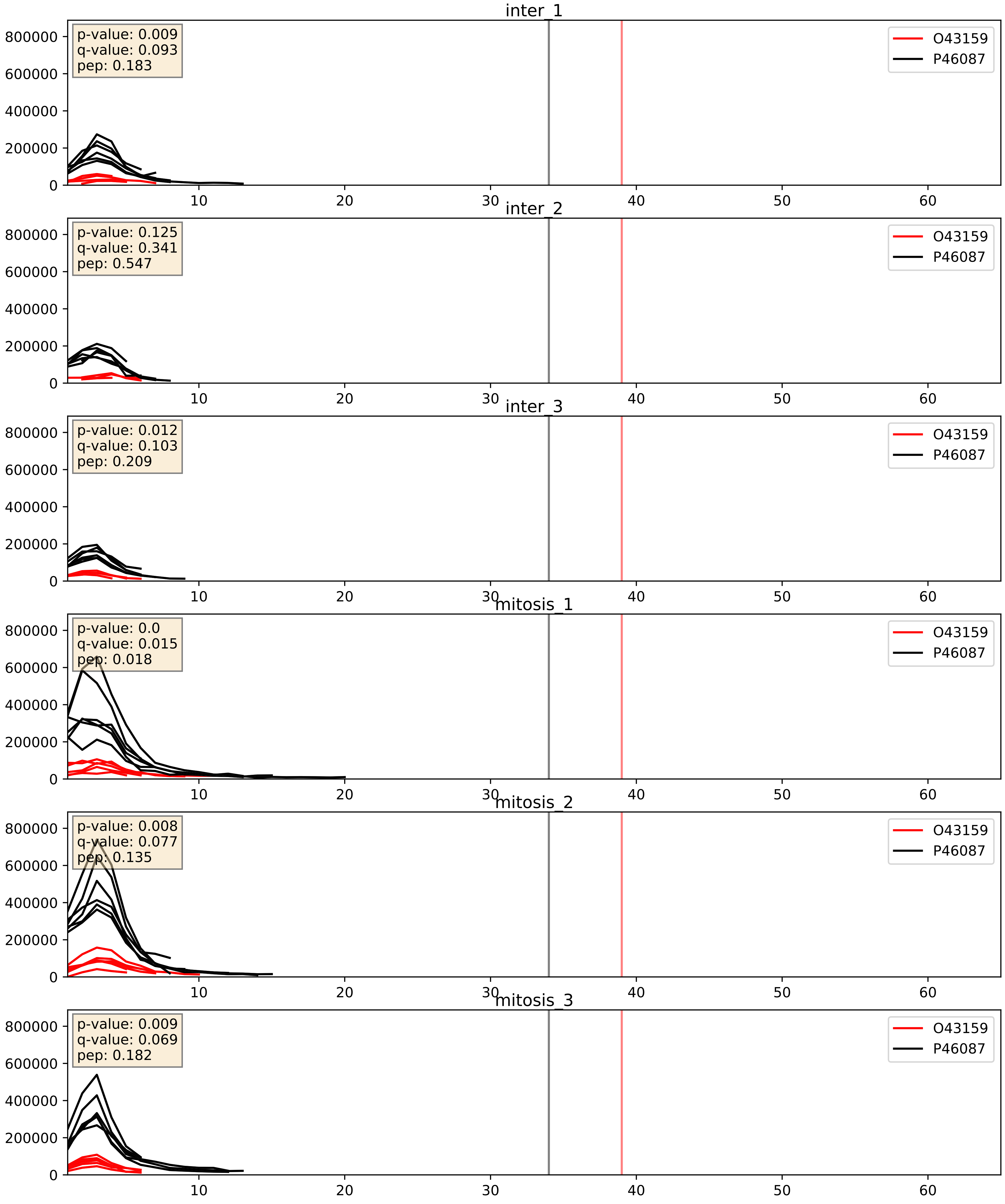


| level | condition_1 | condition_2 | pvalue | pvalue_adjusted |
|---|---|---|---|---|
| interactor_ratio | mitosis | inter | 0.002493678187167931 | 0.06390983617412421 |
| complex_abundance | mitosis | inter | 0.07519460130734286 | 0.1994814550301895 |
| interactor_abundance | mitosis | inter | 0.7068938551500539 | 0.8089046721588746 |

O76021_P46087
RL1D1_HUMAN vs NOP2_HUMAN
p-value: 0.018 q-value: 0.014

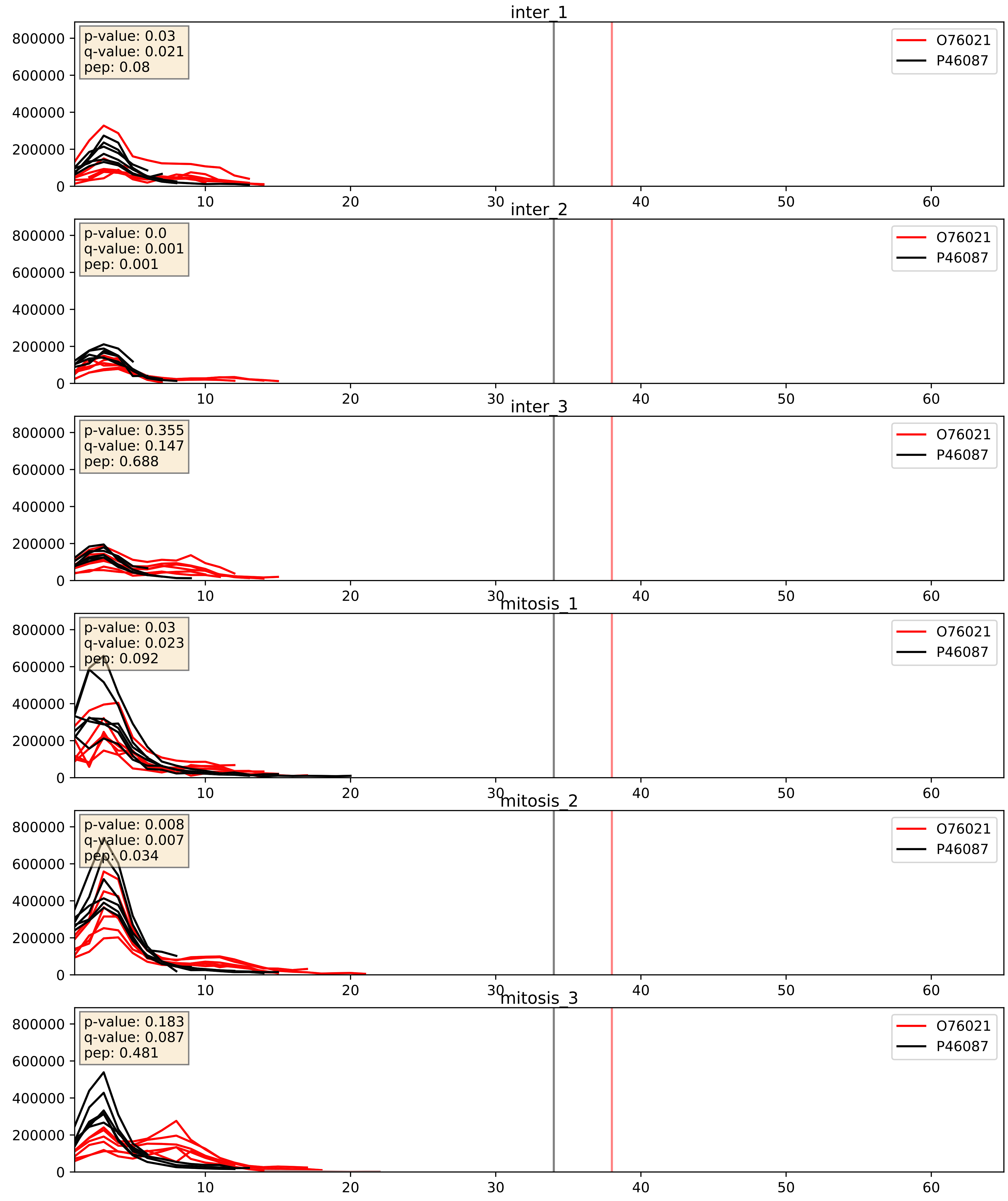


| level | condition_1 | condition_2 | pvalue | pvalue_adjusted |
|---|---|---|---|---|
| complex_abundance | mitosis | inter | 0.0026377819363957414 | 0.03679993741510981 |
| interactor_abundance | mitosis | inter | 0.007992044606047556 | 0.07749253525281795 |
| interactor_ratio | mitosis | inter | 0.4410474716757986 | 0.8466845385837265 |

P22087_P46087
FBRL_HUMAN vs NOP2_HUMAN
p-value: 0.004 q-value: 0.048

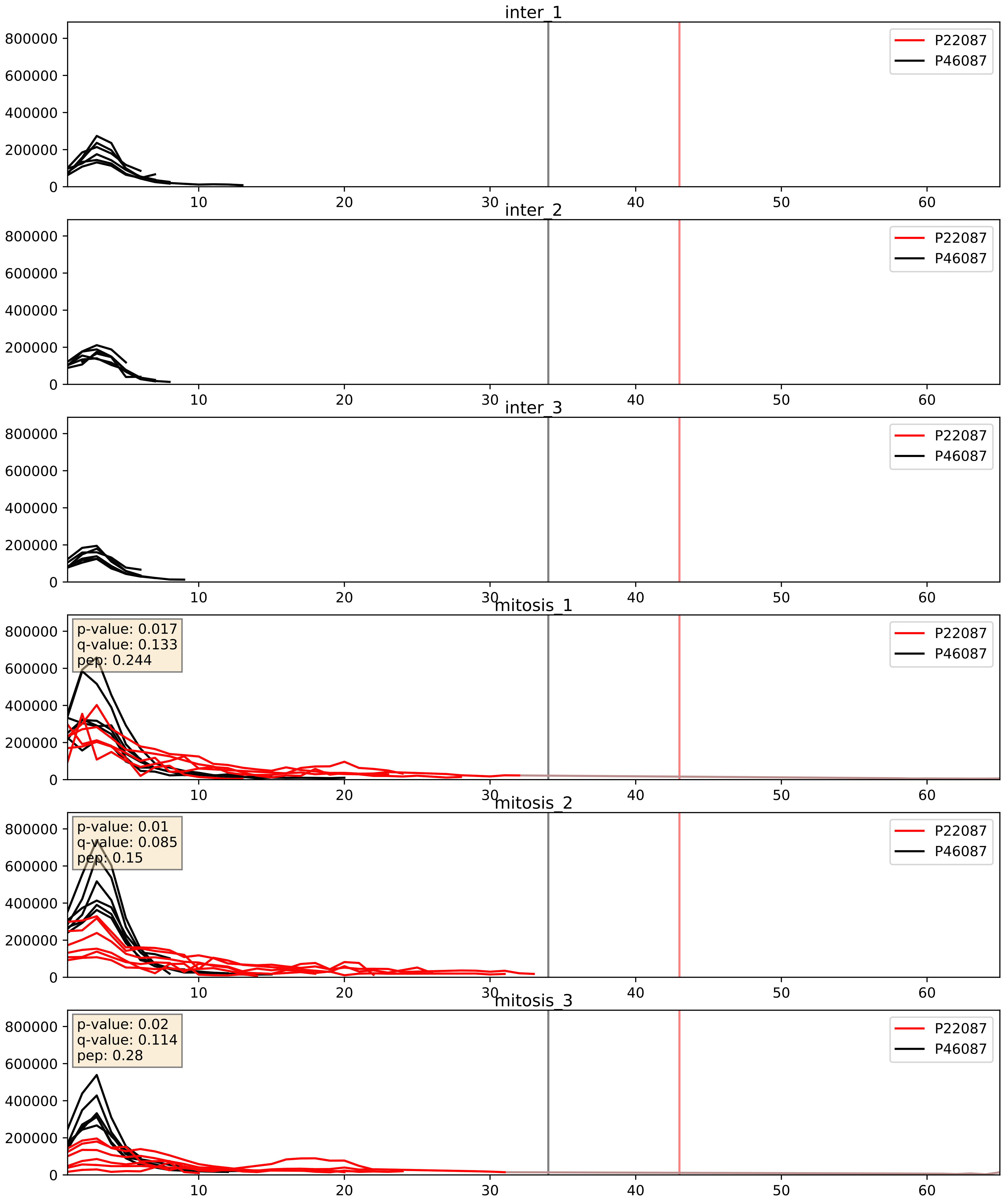


| level | condition_1 | condition_2 | pvalue | pvalue_adjusted |
|---|---|---|---|---|
| complex_abundance | mitosis | inter | 0.00034747852944409986 | 0.012015488415248196 |
| interactor_abundance | mitosis | inter | 0.0008503677168821298 | 0.021631939543866364 |
| interactor_ratio | mitosis | inter | 0.13219010083240598 | 0.58057838025931 |

P36578_P46087
RL4_HUMAN vs NOP2_HUMAN
p-value: 0.001 q-value: 0.047

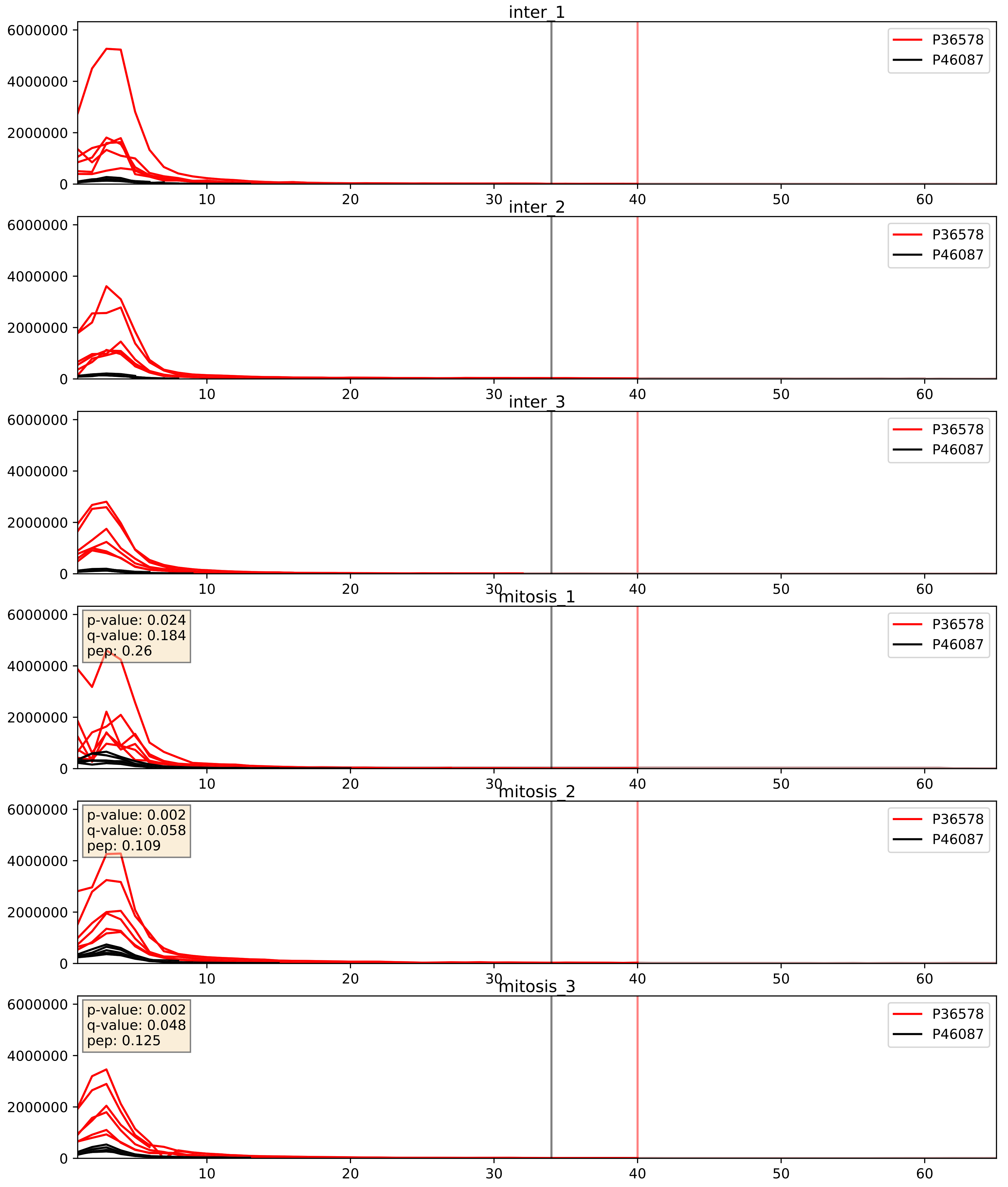


| level | condition_1 | condition_2 | pvalue | pvalue_adjusted |
|---|---|---|---|---|
| complex_abundance | mitosis | inter | 0.005861170144914151 | 0.05756820427608883 |
| interactor_ratio | mitosis | inter | 0.025025449747705465 | 0.24857565976685972 |
| interactor_abundance | mitosis | inter | 0.6670566057629634 | 0.7813930741742554 |

P46087_P62917
NOP2_HUMAN vs RL8_HUMAN
p-value: 0.0 q-value: 0.026

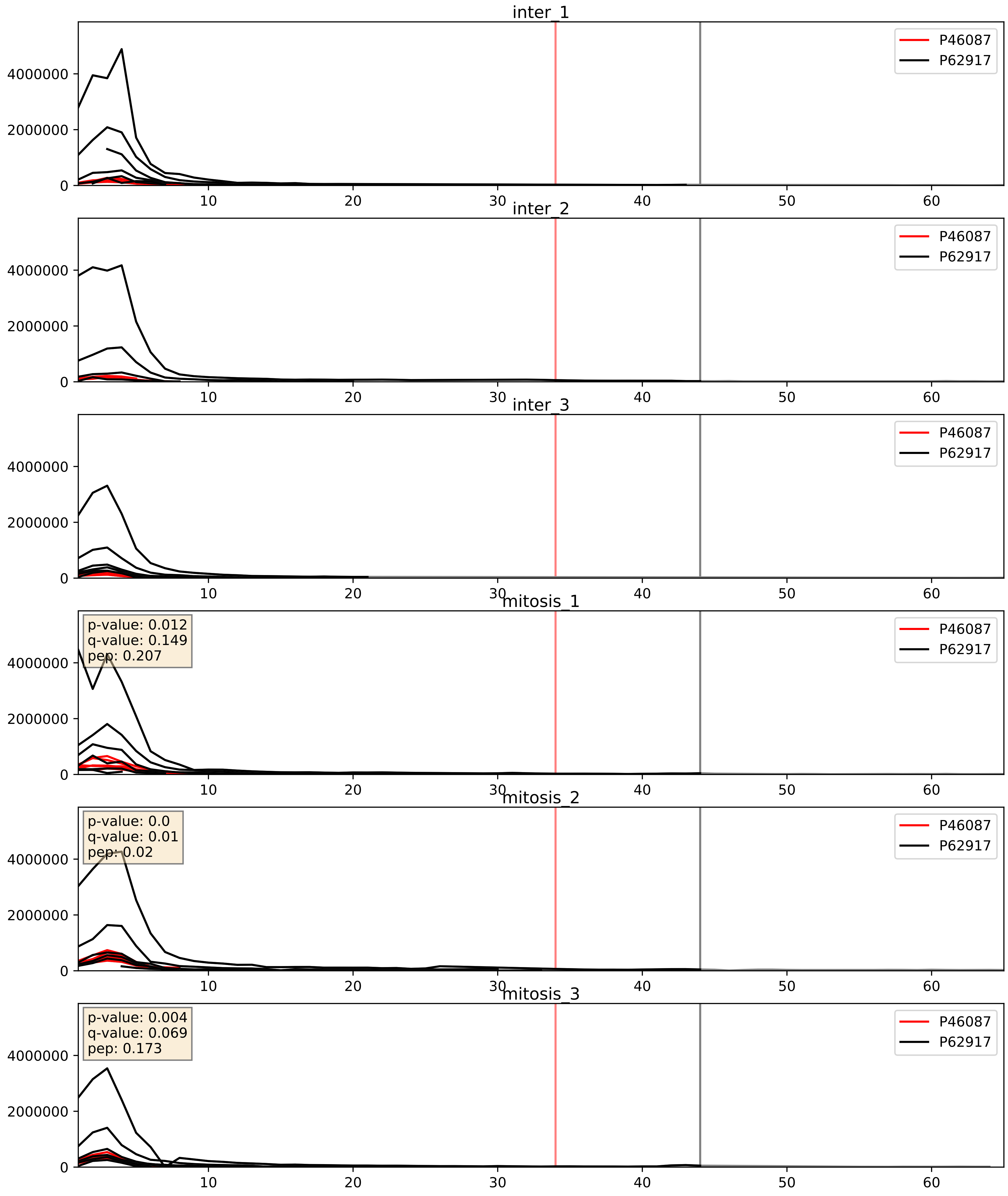


| level | condition_1 | condition_2 | pvalue | pvalue_adjusted |
|---|---|---|---|---|
| interactor_abundance | mitosis | inter | 0.008044903744847214 | 0.07776688517202765 |
| complex_abundance | mitosis | inter | 0.0701361532034238 | 0.19218777394728123 |
| interactor_ratio | mitosis | inter | 0.14495121437875413 | 0.5979674193166917 |

P46087_Q03701
NOP2_HUMAN vs CEBPZ_HUMAN
p-value: 0.043 q-value: 0.029

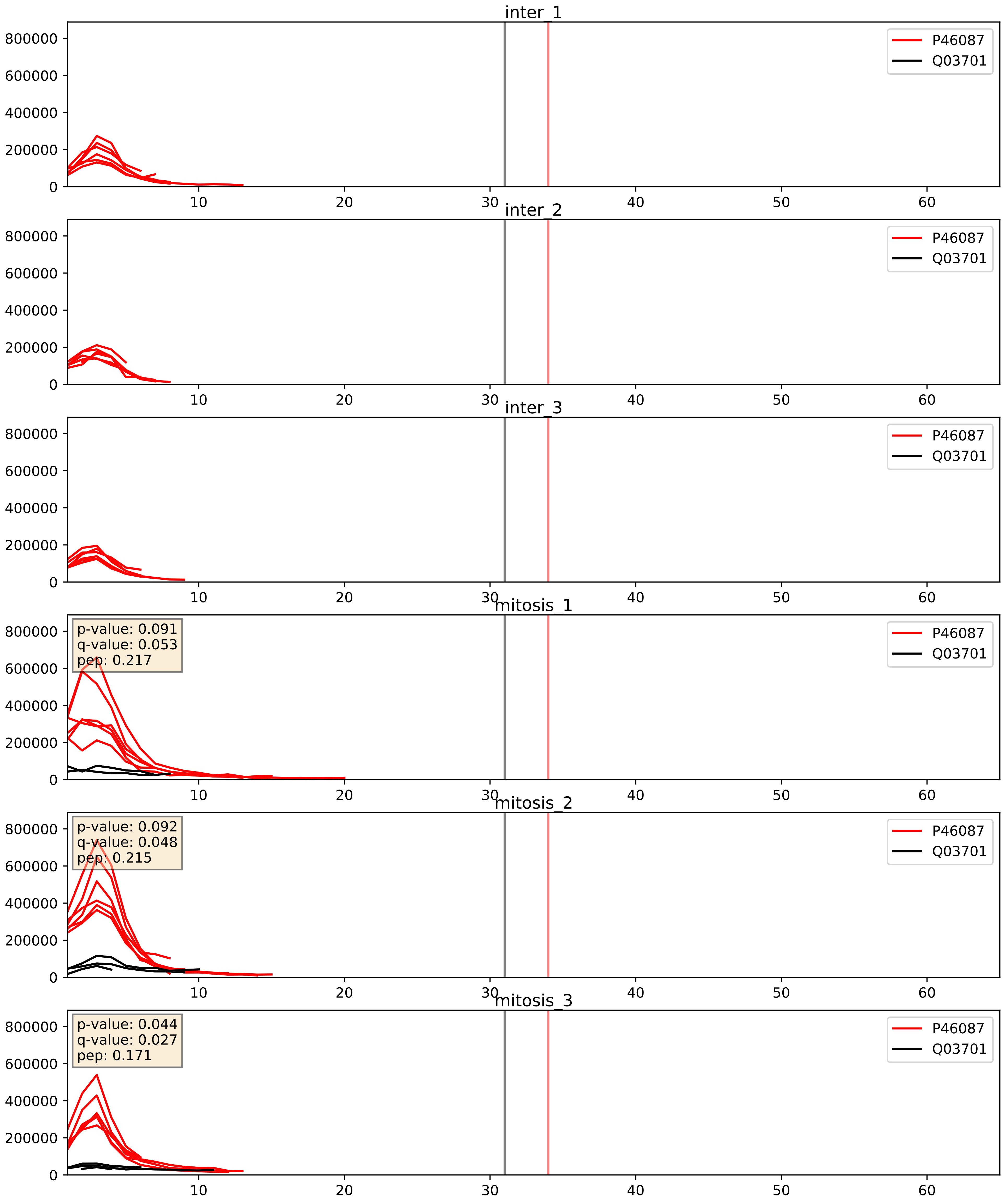


| level | condition_1 | condition_2 | pvalue | pvalue_adjusted |
|---|---|---|---|---|
| interactor_ratio | mitosis | inter | 7.980256380908328e-06 | 0.0032529045057416803 |
| interactor_abundance | mitosis | inter | 0.00013476672746345162 | 0.00890813271881966 |
| complex_abundance | mitosis | inter | 0.000587508565109972 | 0.015666895069599254 |

P46087_Q14137
NOP2_HUMAN vs BOP1_HUMAN
p-value: 0.044 q-value: 0.029

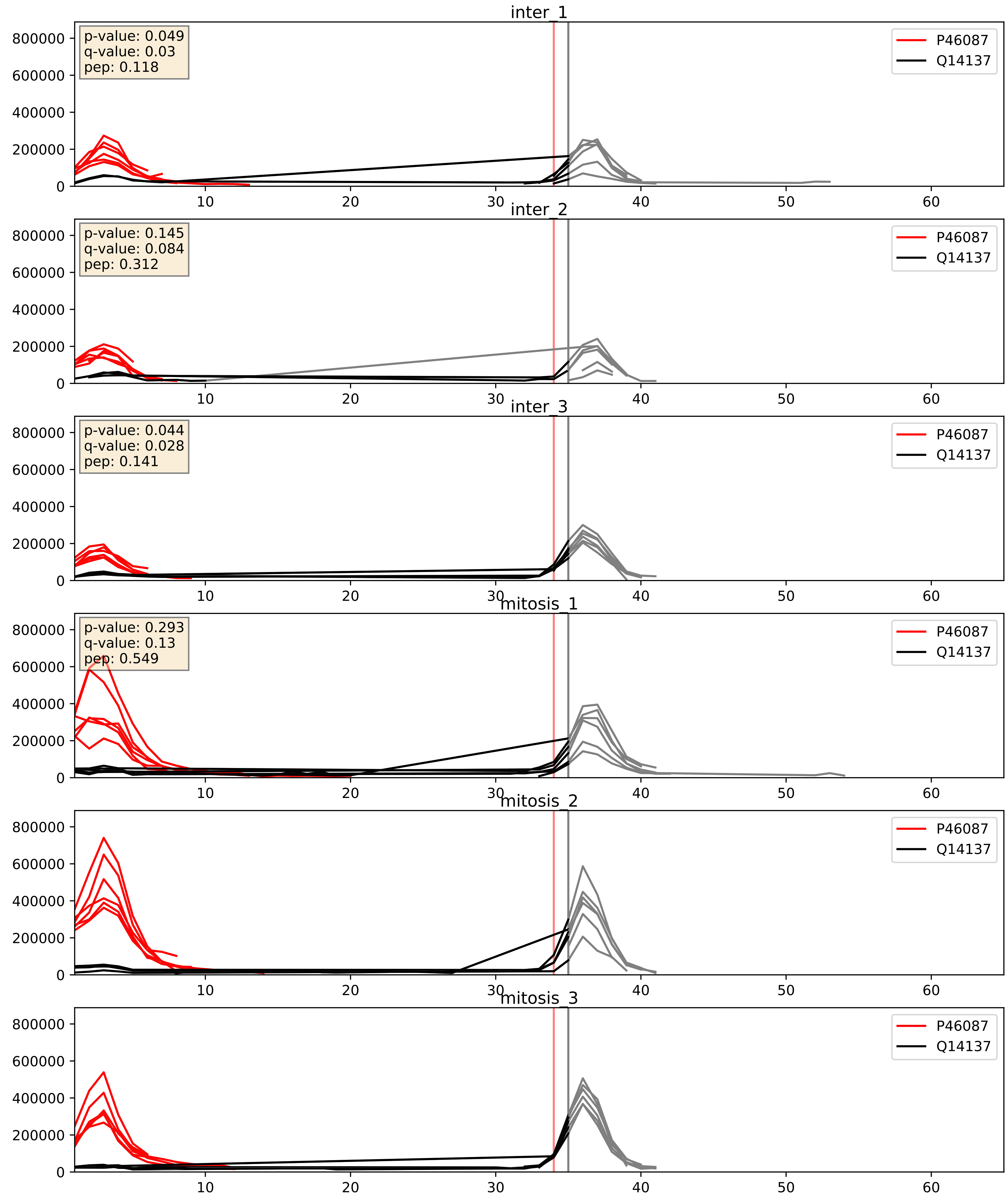


| level | condition_1 | condition_2 | pvalue | pvalue_adjusted |
|---|---|---|---|---|
| interactor_ratio | mitosis | inter | 0.00214770110748558 | 0.05734759220344108 |
| interactor_abundance | mitosis | inter | 0.017814919958046103 | 0.11522154502521695 |
| complex_abundance | mitosis | inter | 0.3752348769755155 | 0.5222928817512863 |

P46087_Q15397
NOP2_HUMAN vs PUM3_HUMAN
p-value: 0.032 q-value: 0.023

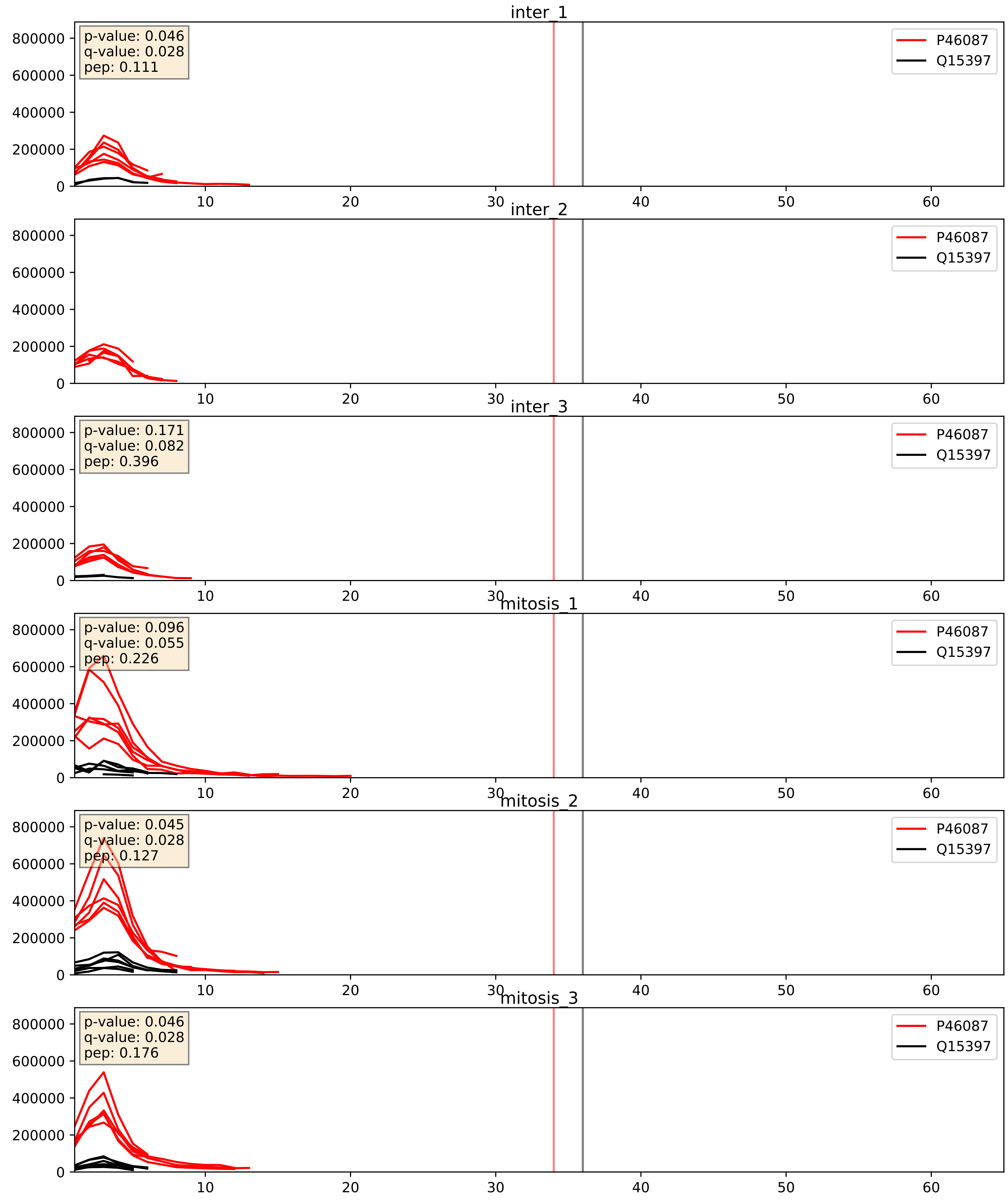


| level | condition_1 | condition_2 | pvalue | pvalue_adjusted |
|---|---|---|---|---|
| interactor_ratio | mitosis | inter | 0.0396609606963561 | 0.32897075926434904 |
| interactor_abundance | mitosis | inter | 0.05779753858331698 | 0.19593937832601716 |
| complex_abundance | mitosis | inter | 0.08310913278336679 | 0.21135299364991675 |

P46087_Q8IY81
NOP2_HUMAN vs SPB1_HUMAN
p-value: 0.026 q-value: 0.019

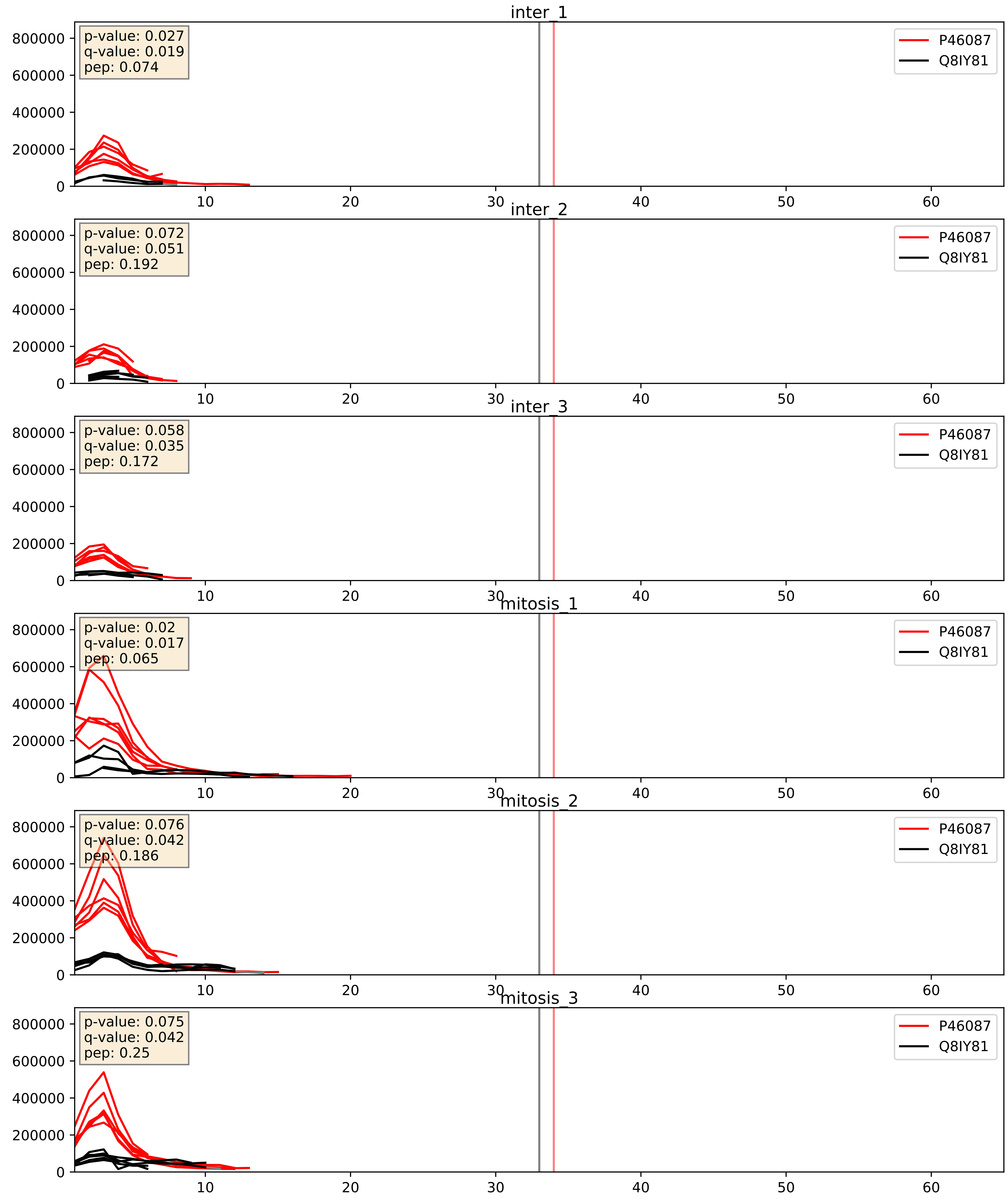


| level | condition_1 | condition_2 | pvalue | pvalue_adjusted |
|---|---|---|---|---|
| interactor_abundance | mitosis | inter | 0.007209065250286371 | 0.07343152394042615 |
| interactor_ratio | mitosis | inter | 0.013389338509009651 | 0.1678078149884665 |
| complex_abundance | mitosis | inter | 0.03232225811702476 | 0.13090015450091652 |

P46087_Q8TDD1
NOP2_HUMAN vs DDX54_HUMAN
p-value: 0.049 q-value: 0.032

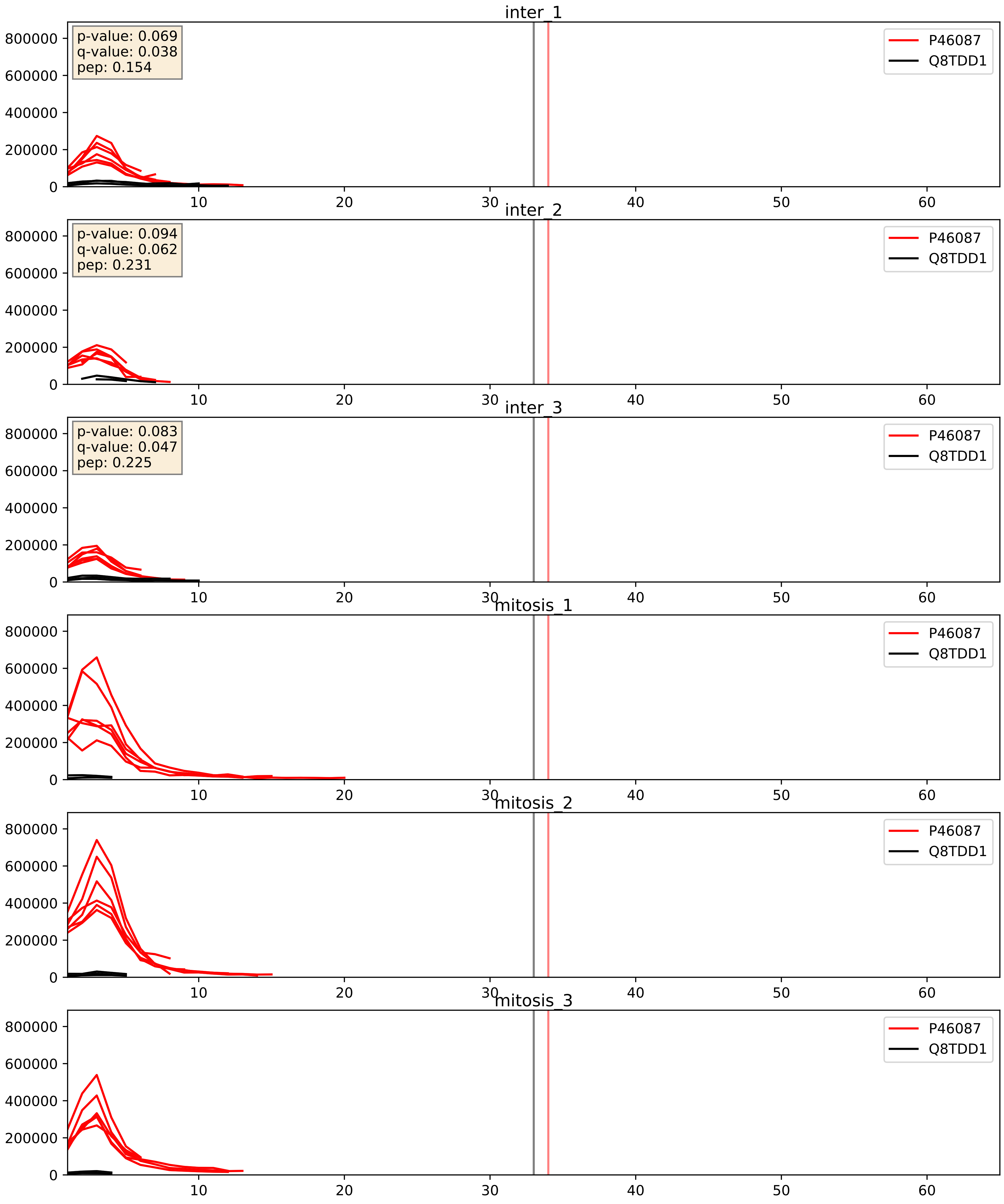


| level | condition_1 | condition_2 | pvalue | pvalue_adjusted |
|---|---|---|---|---|
| interactor_ratio | mitosis | inter | 0.0014545102410756308 | 0.045440173954771526 |
| interactor_abundance | mitosis | inter | 0.04147498932271661 | 0.1637021393260352 |
| complex_abundance | mitosis | inter | 0.8141791930681086 | 0.8791943853491875 |

P46087_Q8TDN6
NOP2_HUMAN vs BRX1_HUMAN
p-value: 0.026 q-value: 0.02

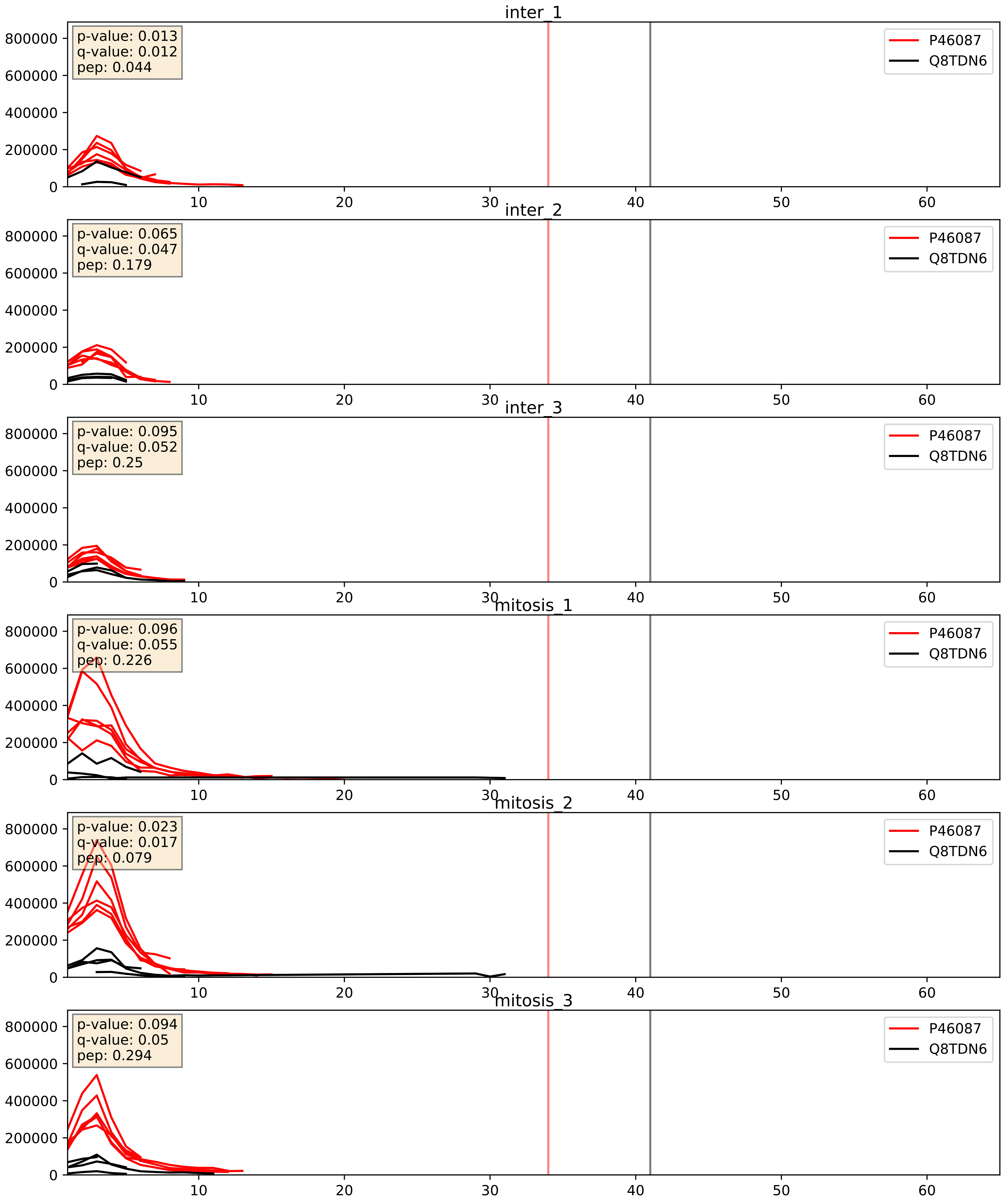


| level | condition_1 | condition_2 | pvalue | pvalue_adjusted |
|---|---|---|---|---|
| interactor_ratio | mitosis | inter | 0.001498537587396323 | 0.04597663709000905 |
| interactor_abundance | mitosis | inter | 0.005651157289129827 | 0.0634828167912747 |
| complex_abundance | mitosis | inter | 0.04071320618289467 | 0.1453315450064942 |

P46087_Q99459
NOP2_HUMAN vs CDC5L_HUMAN
p-value: 0.017 q-value: 0.014

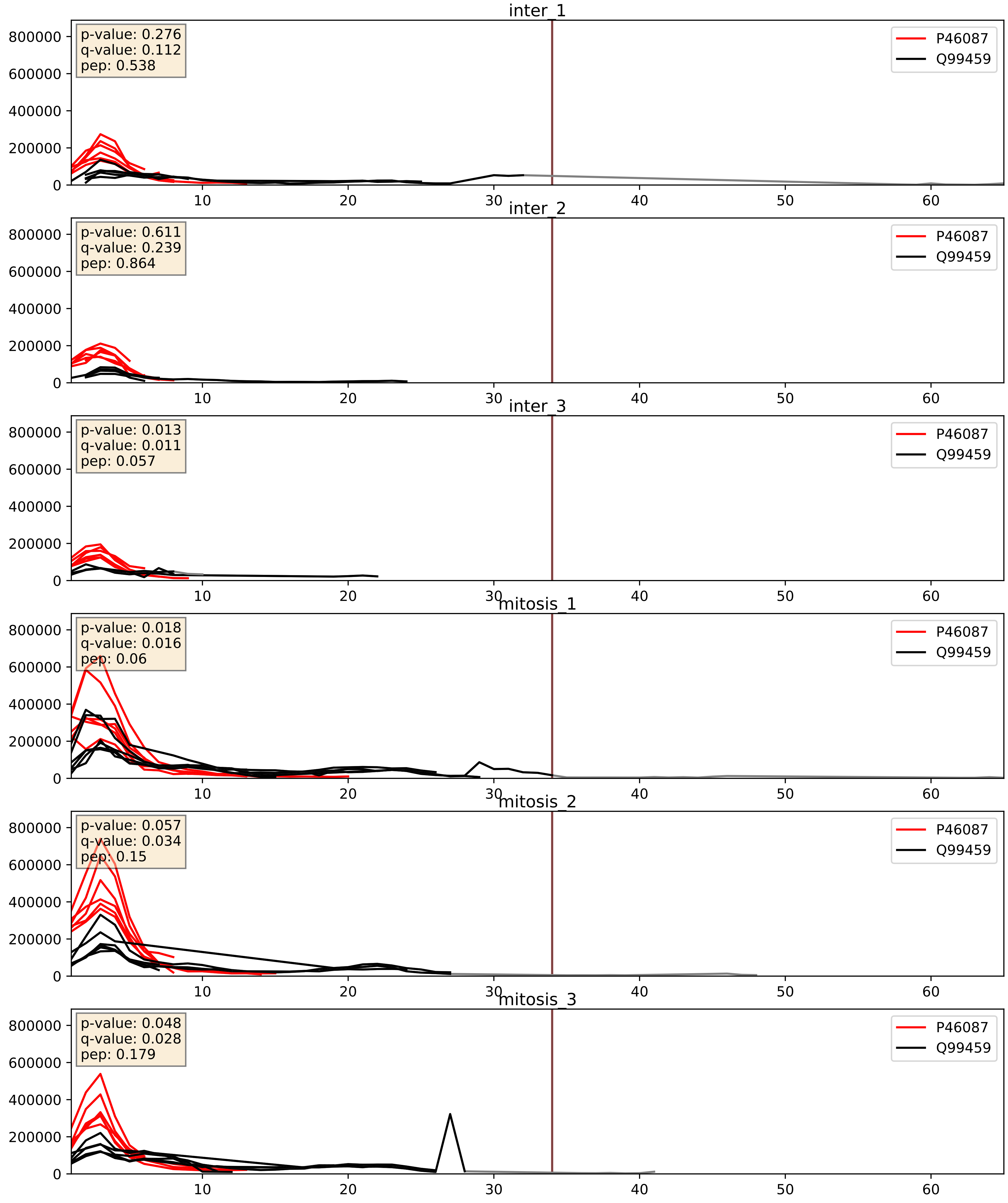


| level | condition_1 | condition_2 | pvalue | pvalue_adjusted |
|---|---|---|---|---|
| interactor_abundance | mitosis | inter | 0.008163501708093478 | 0.07851637597896649 |
| complex_abundance | mitosis | inter | 0.017128092485186742 | 0.09767919498547536 |
| interactor_ratio | mitosis | inter | 0.3721881012830839 | 0.8138388063636502 |

P46087_Q9BVP2
NOP2_HUMAN vs GNL3_HUMAN
p-value: 0.026 q-value: 0.019

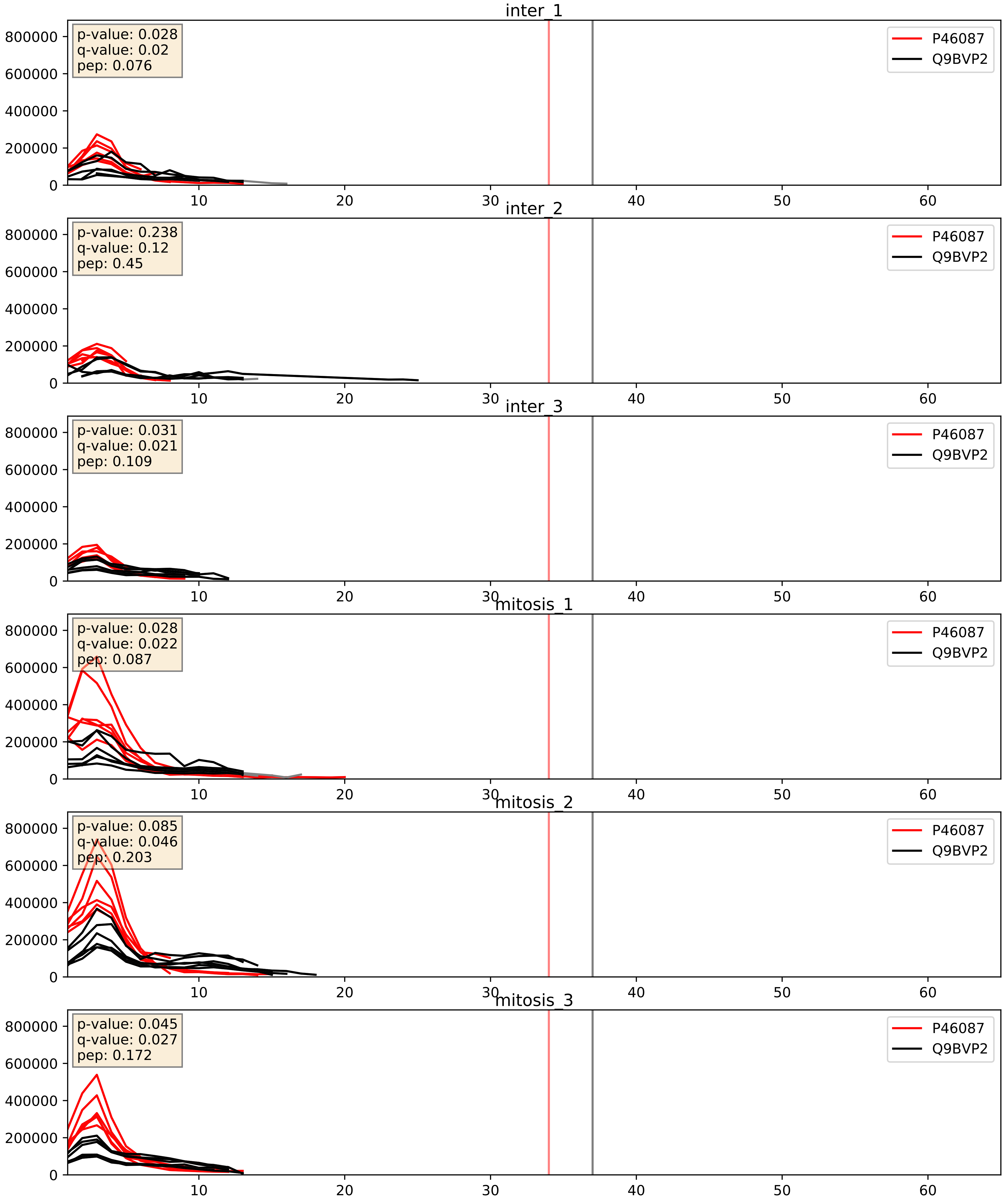


| level | condition_1 | condition_2 | pvalue | pvalue_adjusted |
|---|---|---|---|---|
| complex_abundance | mitosis | inter | 0.00781659329937401 | 0.0656624520536227 |
| interactor_abundance | mitosis | inter | 0.007951593035973885 | 0.07739128639901814 |
| interactor_ratio | mitosis | inter | 0.05096474016989619 | 0.373742437899162 |

P46087_Q9BZE4
NOP2_HUMAN vs NOG1_HUMAN
p-value: 0.004 q-value: 0.004

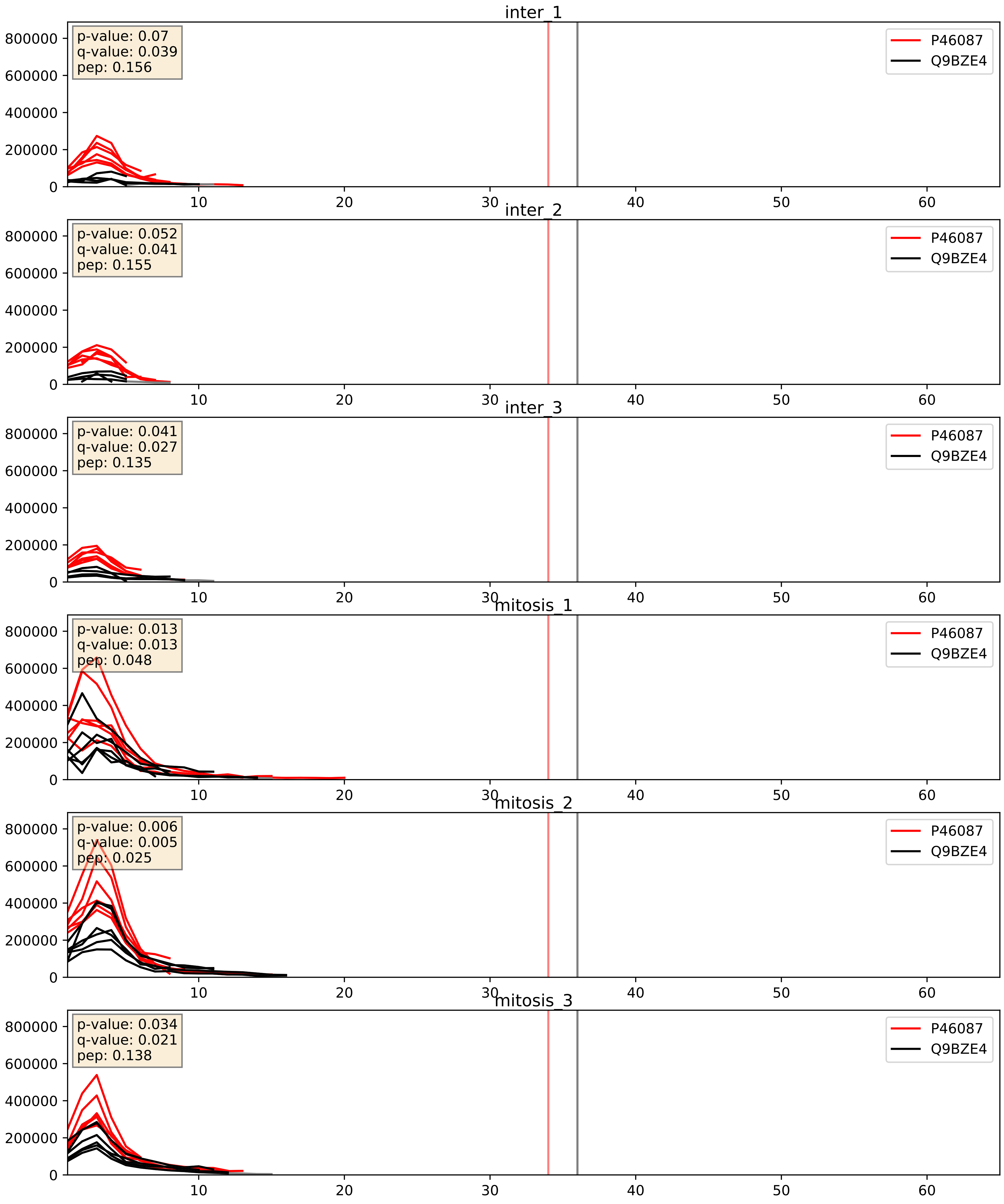


| level | condition_1 | condition_2 | pvalue | pvalue_adjusted |
|---|---|---|---|---|
| complex_abundance | mitosis | inter | 0.00198928380982054 | 0.03207280622328027 |
| interactor_ratio | mitosis | inter | 0.004188742047813209 | 0.08517183734867877 |
| interactor_abundance | mitosis | inter | 0.004516185423873405 | 0.05406789822147741 |

P46087_Q9GZL7
NOP2_HUMAN vs WDR12_HUMAN
p-value: 0.045 q-value: 0.03

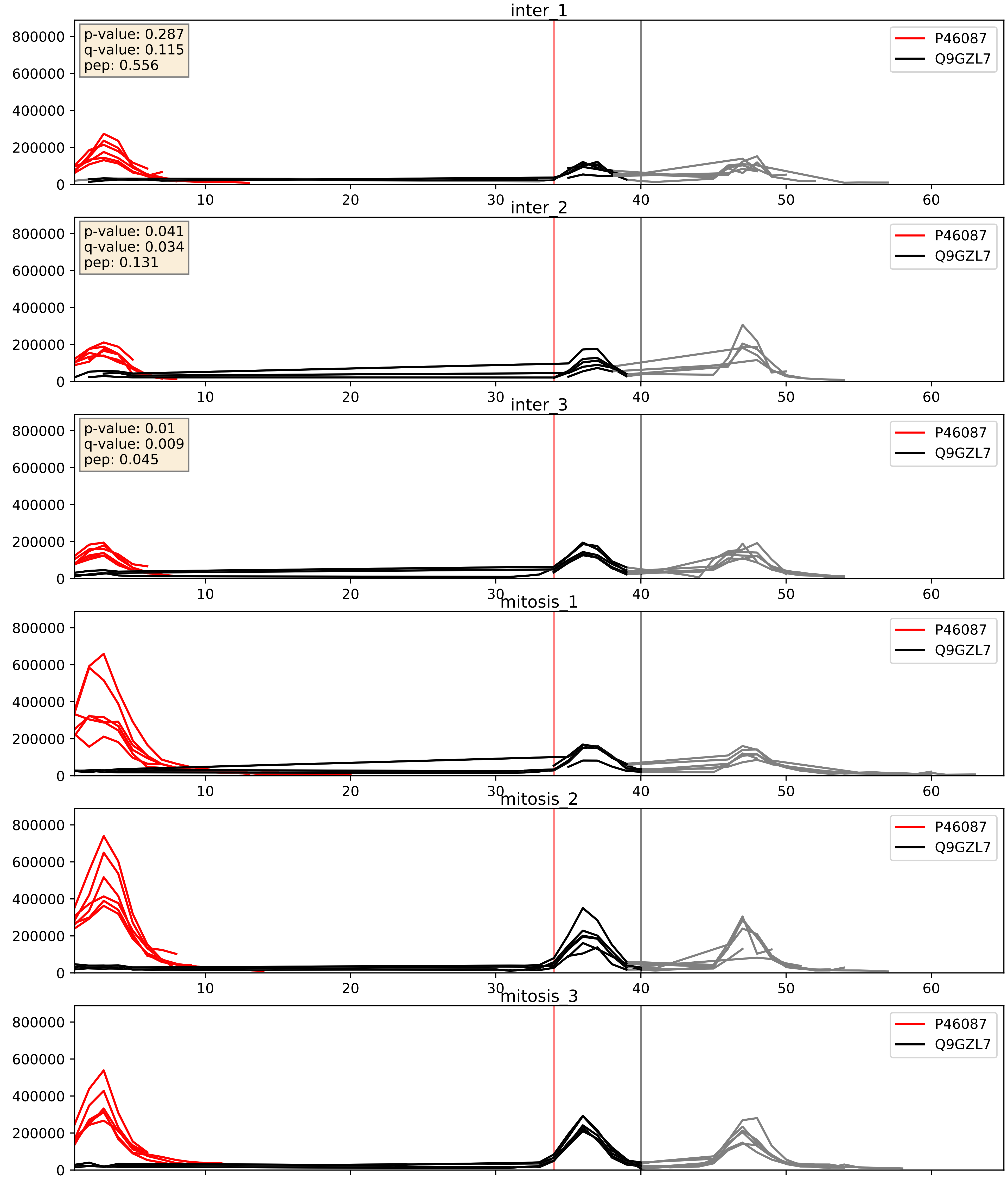


| level | condition_1 | condition_2 | pvalue | pvalue_adjusted |
|---|---|---|---|---|
| interactor_ratio | mitosis | inter | 0.0017565937997646257 | 0.051144363693827195 |
| interactor_abundance | mitosis | inter | 0.03342104099149199 | 0.15249686081405728 |
| complex_abundance | mitosis | inter | 0.5242060841311404 | 0.6597859256230794 |

P46087_Q9GZR7
NOP2_HUMAN vs DDX24_HUMAN
p-value: 0.039 q-value: 0.027

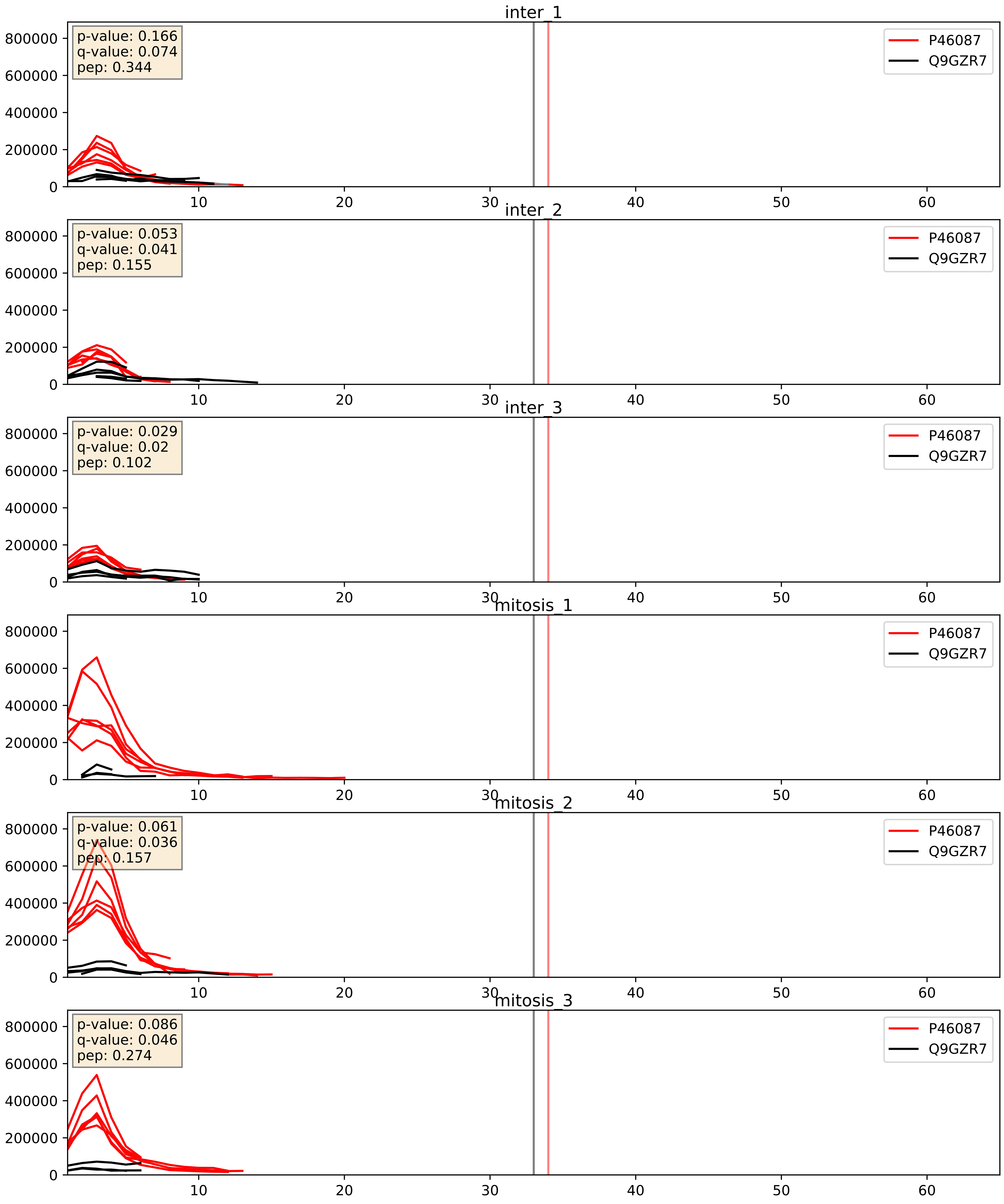


| level | condition_1 | condition_2 | pvalue | pvalue_adjusted |
|---|---|---|---|---|
| interactor_ratio | mitosis | inter | 0.0003942176979097222 | 0.021937350419330353 |
| interactor_abundance | mitosis | inter | 0.021324997778811212 | 0.1216554664472706 |
| complex_abundance | mitosis | inter | 0.8062023626534441 | 0.8745098220636244 |

P46087_Q9H6R4
NOP2_HUMAN vs NOL6_HUMAN
p-value: 0.06 q-value: 0.038

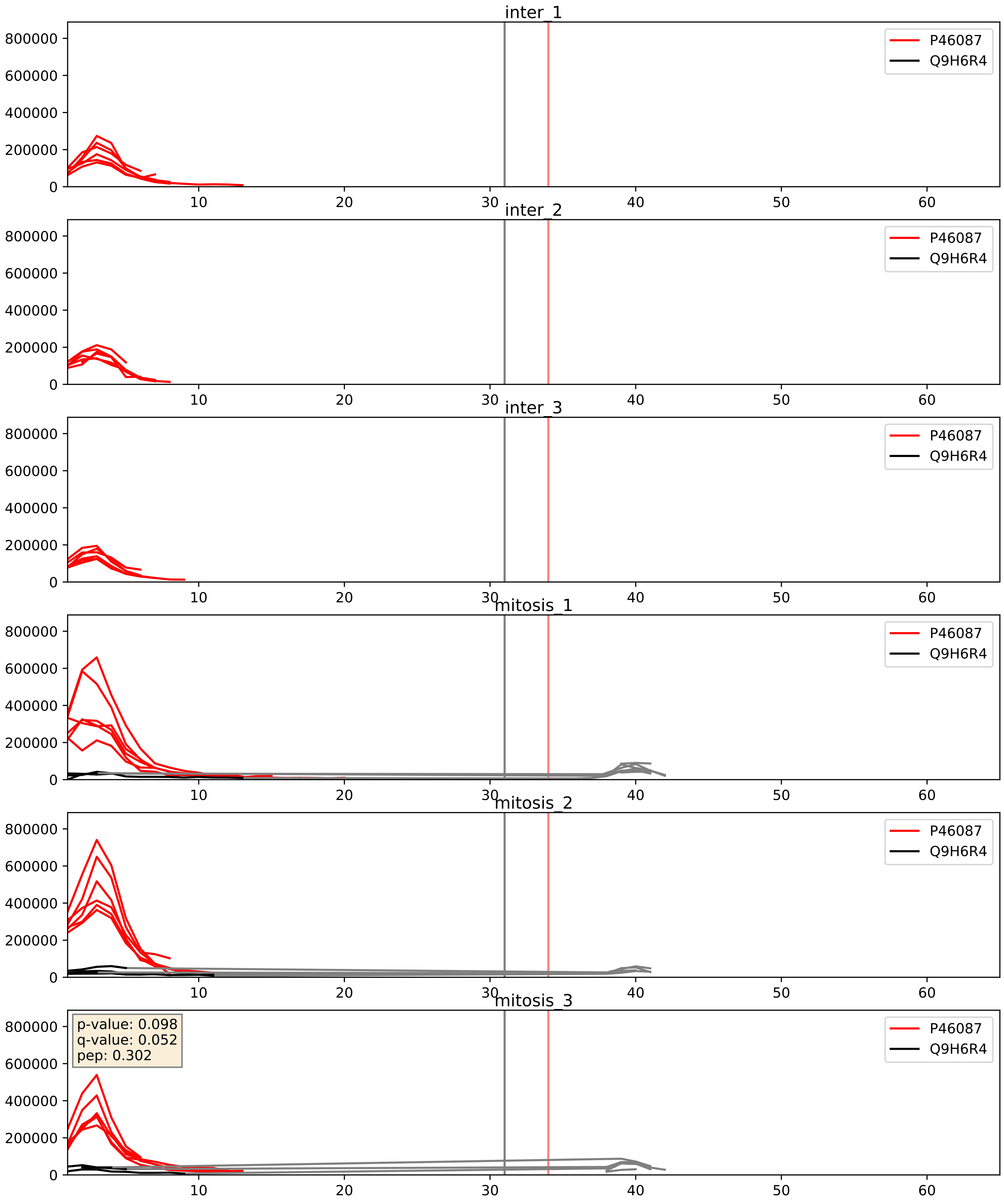


| level | condition_1 | condition_2 | pvalue | pvalue_adjusted |
|---|---|---|---|---|
| interactor_abundance | mitosis | inter | 0.00019761530192014963 | 0.01052608162799415 |
| interactor_ratio | mitosis | inter | 0.00027831638304536276 | 0.017163005662466667 |
| complex_abundance | mitosis | inter | 0.000983162917257506 | 0.02083137270228775 |

P46087_Q9NVP1
NOP2_HUMAN vs DDX18_HUMAN
p-value: 0.046 q-value: 0.031

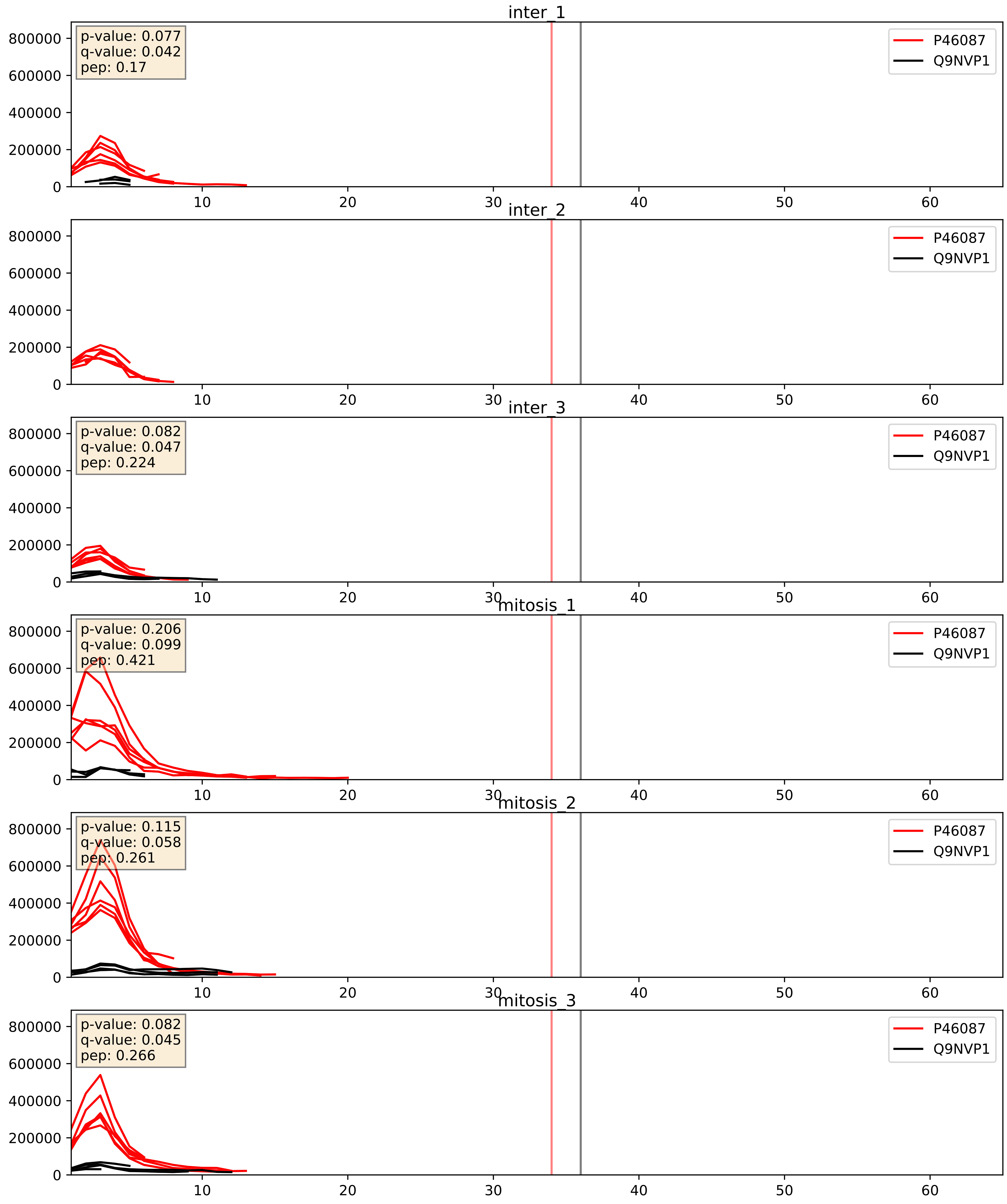


| level | condition_1 | condition_2 | pvalue | pvalue_adjusted |
|---|---|---|---|---|
| interactor_ratio | mitosis | inter | 0.005387411984646833 | 0.0985389884371301 |
| interactor_abundance | mitosis | inter | 0.03915622151066956 | 0.16083596262333216 |
| complex_abundance | mitosis | inter | 0.1003973827814877 | 0.2350018038308818 |

P46087_Q9NW13
NOP2_HUMAN vs RBM28_HUMAN
p-value: 0.083 q-value: 0.049

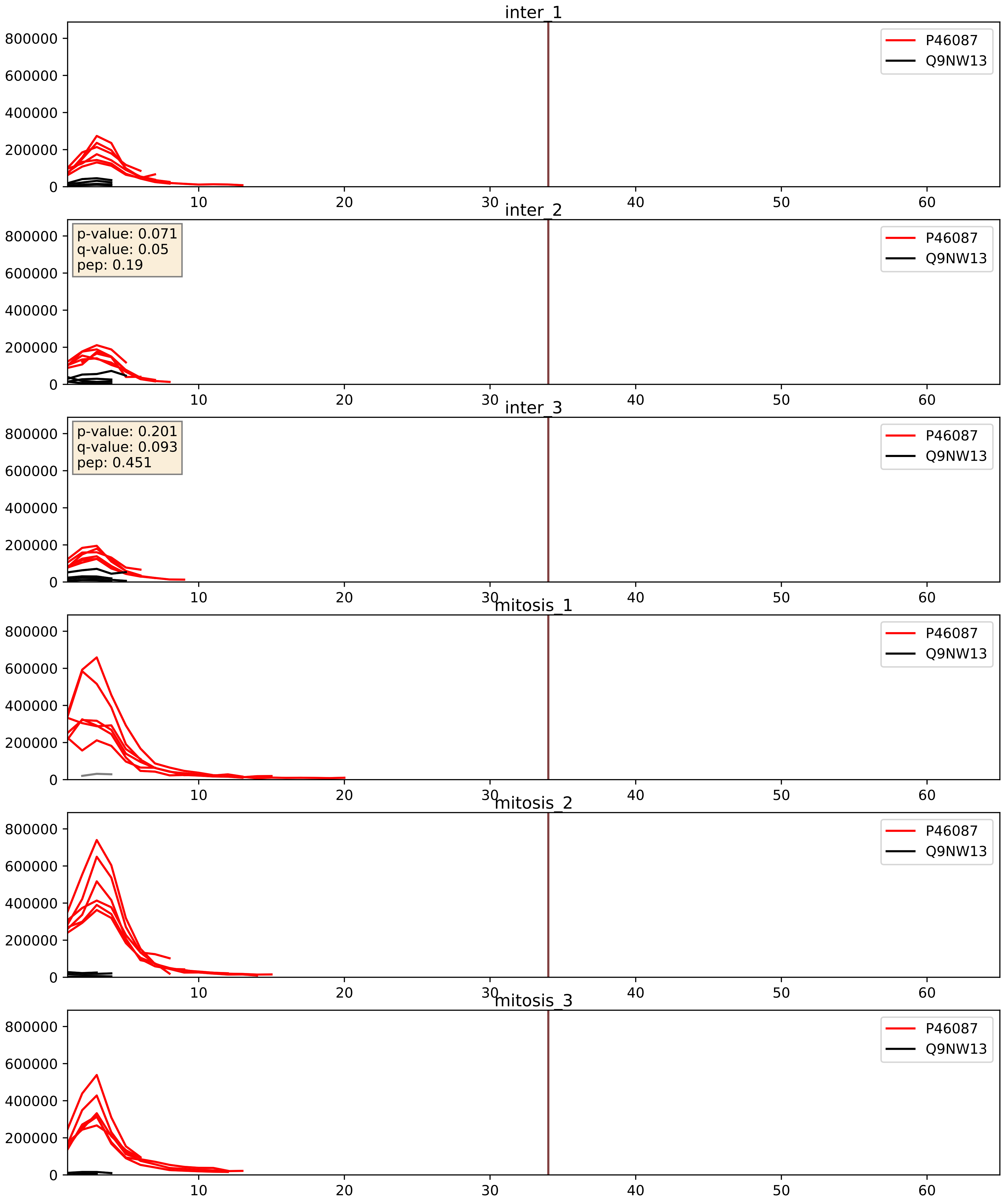


| level | condition_1 | condition_2 | pvalue | pvalue_adjusted |
|---|---|---|---|---|
| complex_abundance | mitosis | inter | 0.13763659808947898 | 0.2828737766256759 |
| interactor_ratio | mitosis | inter | 0.34802209469128087 | 0.7998854020815096 |
| interactor_abundance | mitosis | inter | 0.8352324841573822 | 0.9101381279851304 |

P46087_Q9NY93
NOP2_HUMAN vs DDX56_HUMAN
p-value: 0.046 q-value: 0.031

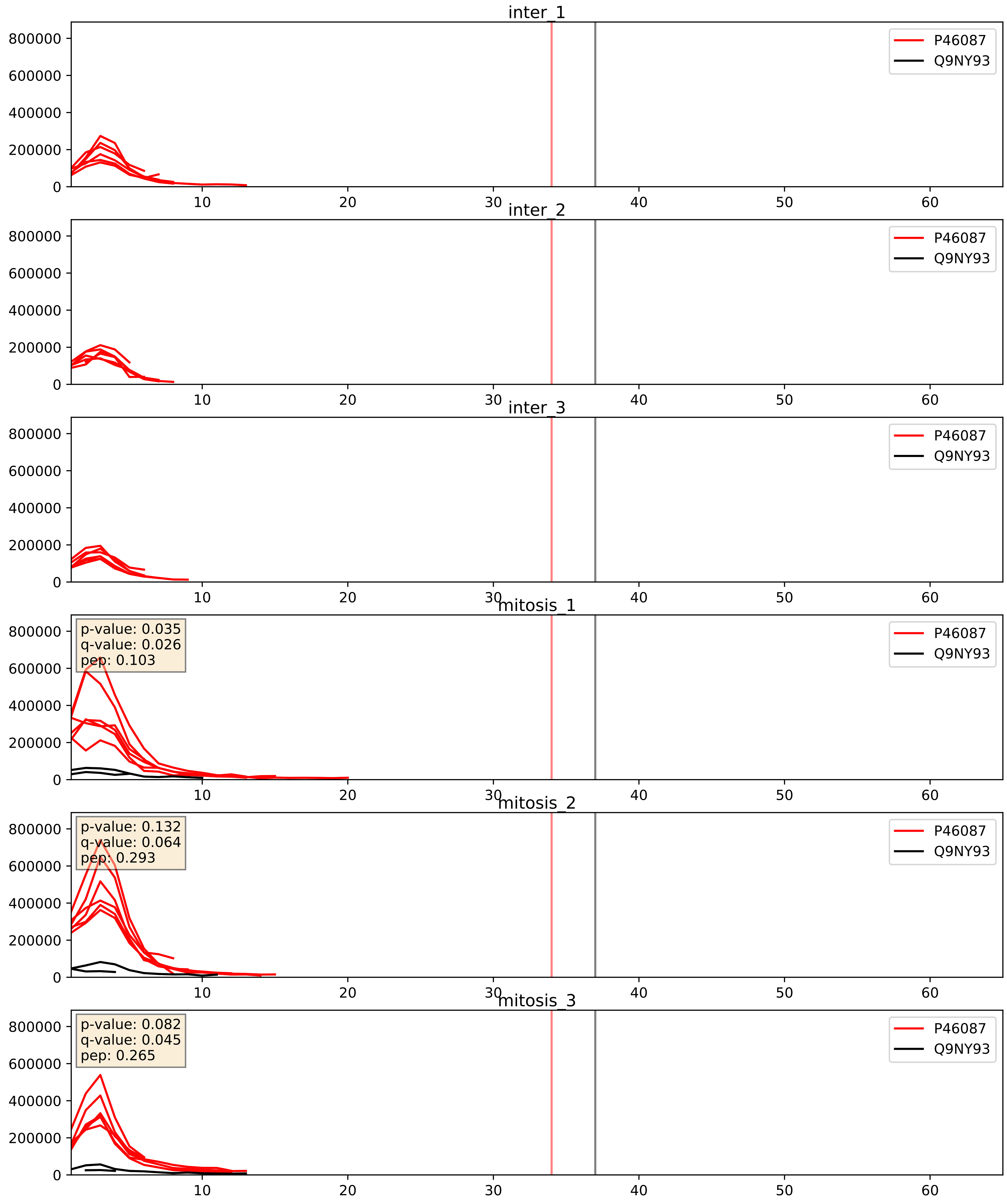


| level | condition_1 | condition_2 | pvalue | pvalue_adjusted |
|---|---|---|---|---|
| interactor_ratio | mitosis | inter | 2.6497983805864818e-05 | 0.005274947473911693 |
| interactor_abundance | mitosis | inter | 0.00012951041339078335 | 0.008661008895508637 |
| complex_abundance | mitosis | inter | 0.0006897093281200277 | 0.016916652861625893 |

P46087_Q9UKD2
NOP2_HUMAN vs MRT4_HUMAN
p-value: 0.002 q-value: 0.003

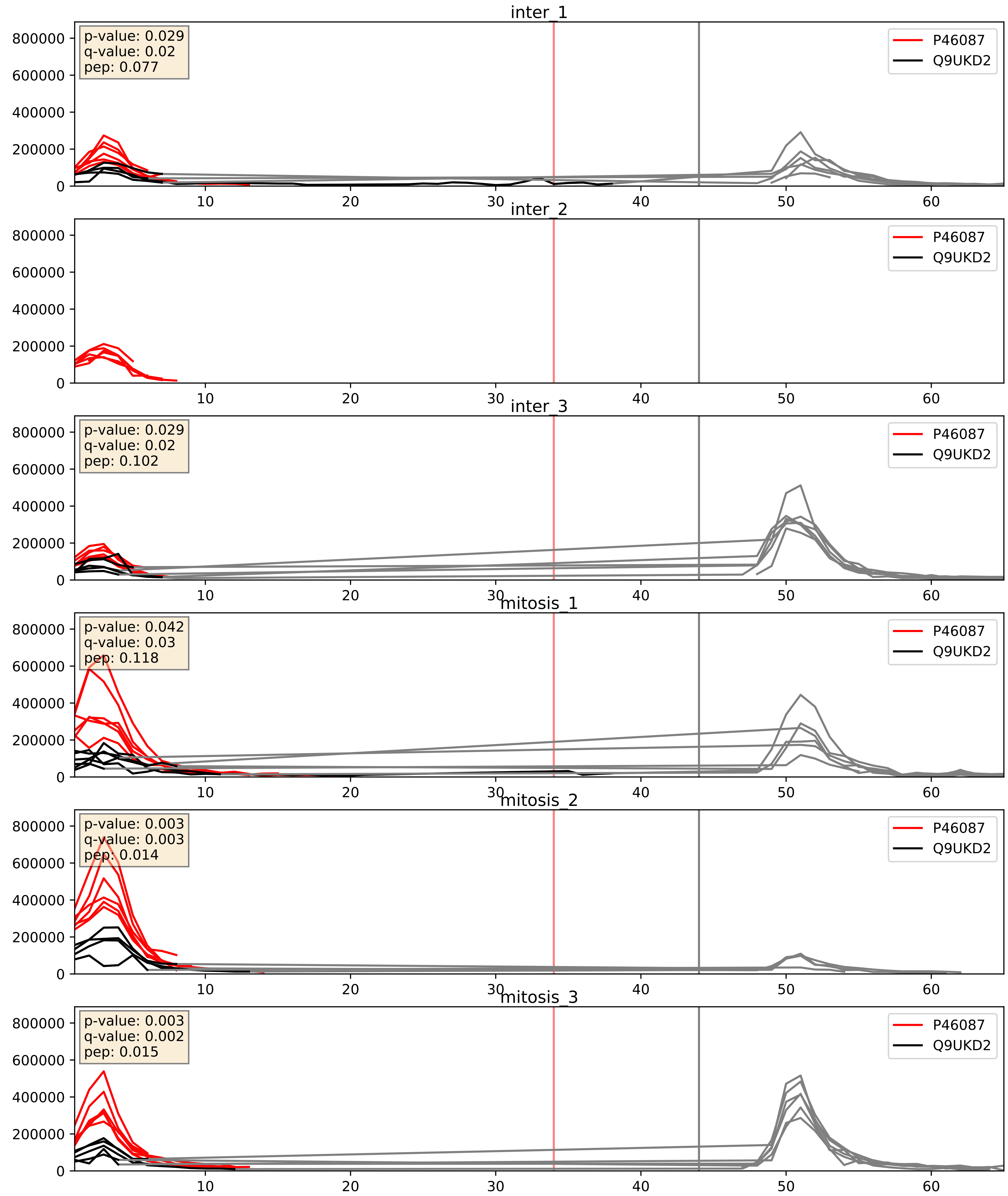


| level | condition_1 | condition_2 | pvalue | pvalue_adjusted |
|---|---|---|---|---|
| interactor_ratio | mitosis | inter | 0.008169453022261623 | 0.12756688776860511 |
| interactor_abundance | mitosis | inter | 0.05561217096474677 | 0.1913344788819262 |
| complex_abundance | mitosis | inter | 0.08848513121730248 | 0.21834324682044082 |

P46087_Q9Y221
NOP2_HUMAN vs NIP7_HUMAN
p-value: 0.015 q-value: 0.012

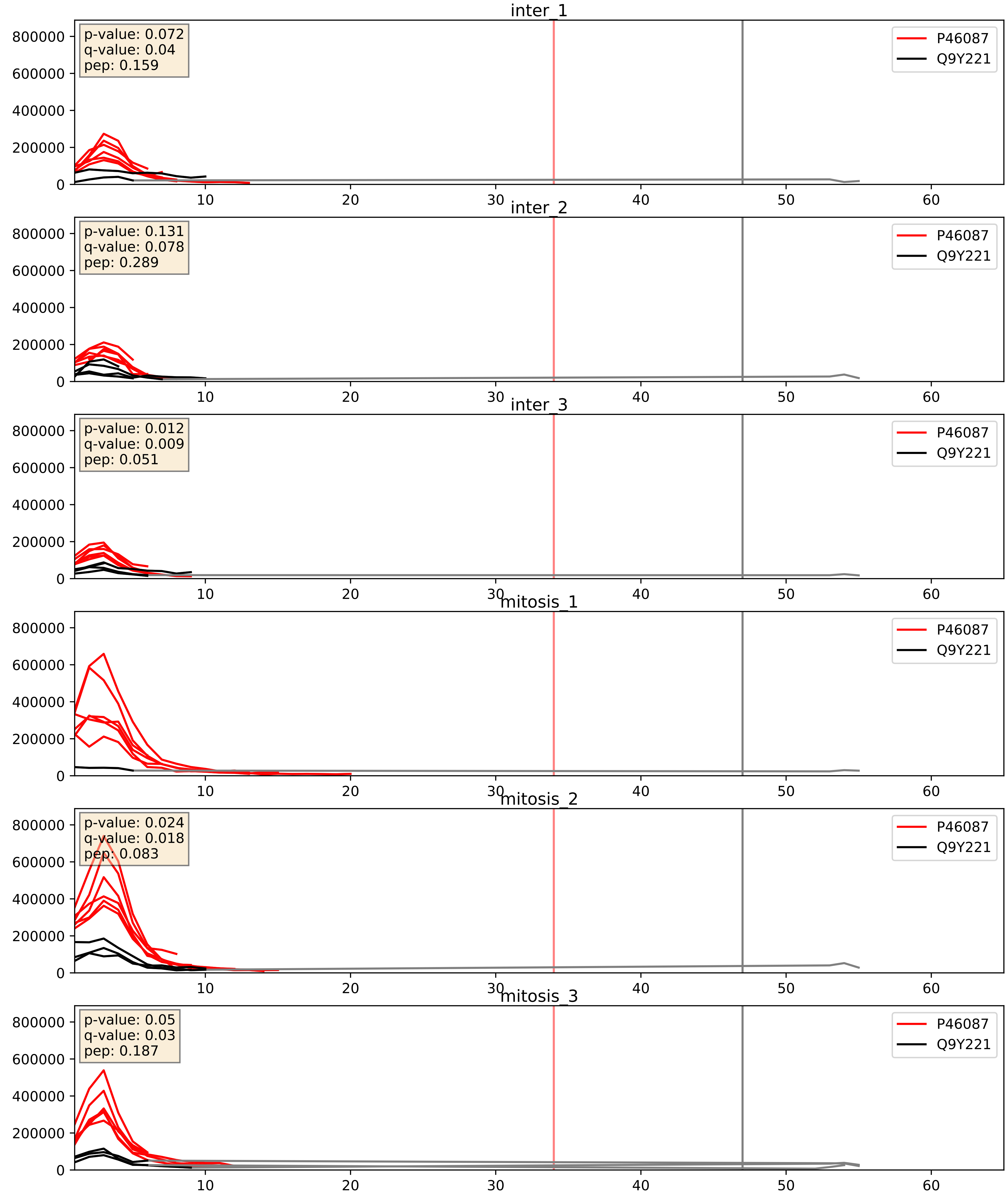


| level | condition_1 | condition_2 | pvalue | pvalue_adjusted |
|---|---|---|---|---|
| interactor_abundance | mitosis | inter | 0.010363624052286788 | 0.08871262188757491 |
| interactor_ratio | mitosis | inter | 0.0262708202591697 | 0.25543205403291264 |
| complex_abundance | mitosis | inter | 0.22060645660497263 | 0.3746808072497155 |

P46087_Q9Y2X3
NOP2_HUMAN vs NOP58_HUMAN
p-value: 0.014 q-value: 0.011

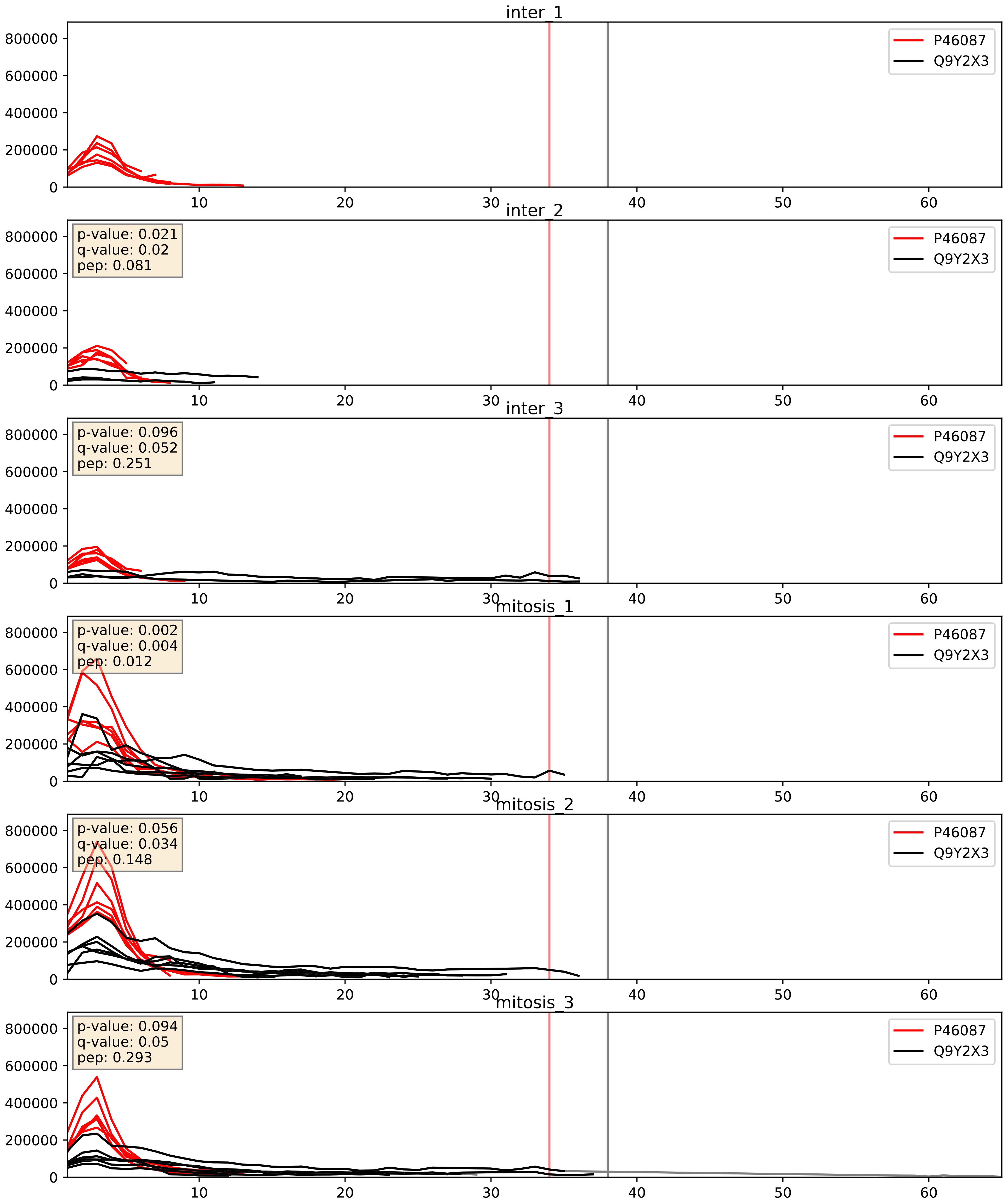


| level | condition_1 | condition_2 | pvalue | pvalue_adjusted |
|---|---|---|---|---|
| complex_abundance | mitosis | inter | 0.012113264791281528 | 0.0818150085787963 |
| interactor_abundance | mitosis | inter | 0.03839338005658009 | 0.15936046986653518 |
| interactor_ratio | mitosis | inter | 0.6680032234742932 | 0.9302273617927362 |
