## Supplementary Data 2 for "SECAT: Quantifying differential protein-protein interaction states by network-centric analysis": hela_string_000007_O00567.pdf

O00567_O00567

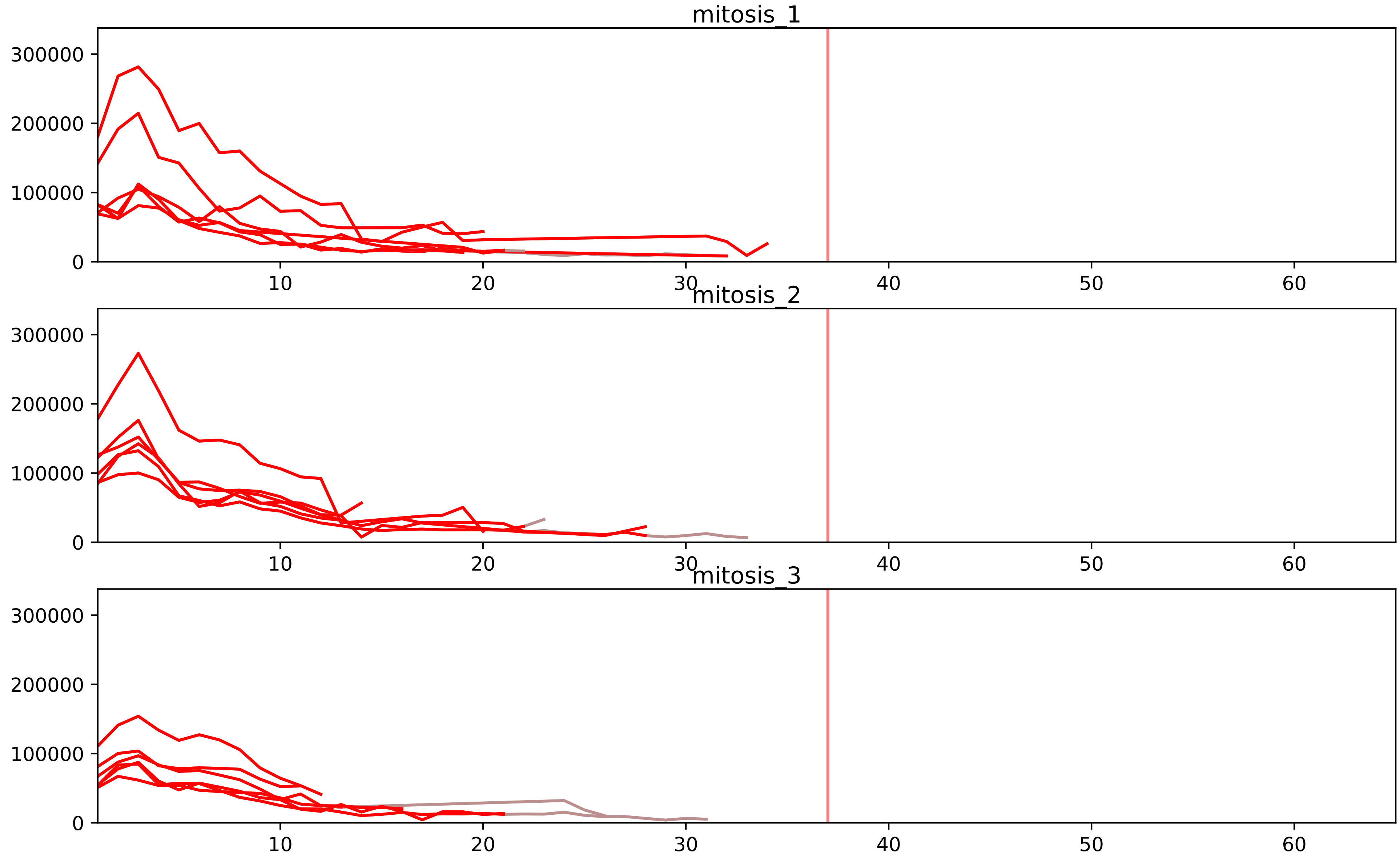


| level | condition_1 | condition_2 | pvalue | pvalue_adjusted |
|---|---|---|---|---|
| interactor_ratio | mitosis | inter | 5.466319871370034e-16 | 1.4368612233315517e-13 |
| complex_abundance | mitosis | inter | 3.336275691799981e-06 | 0.00021168094044524016 |
| interactor_abundance | mitosis | inter | 1.1656810902371506e-05 | 0.000579690055685502 |
| assembled_abundance | mitosis | inter | 0.00018514547961713212 | 0.014378367623948875 |
| total_abundance | mitosis | inter | 0.0003119854042913784 | 0.023706234227573245 |

O00567_O60832
NOP56_HUMAN vs DKC1_HUMAN
p-value: 0.008 q-value: 0.007

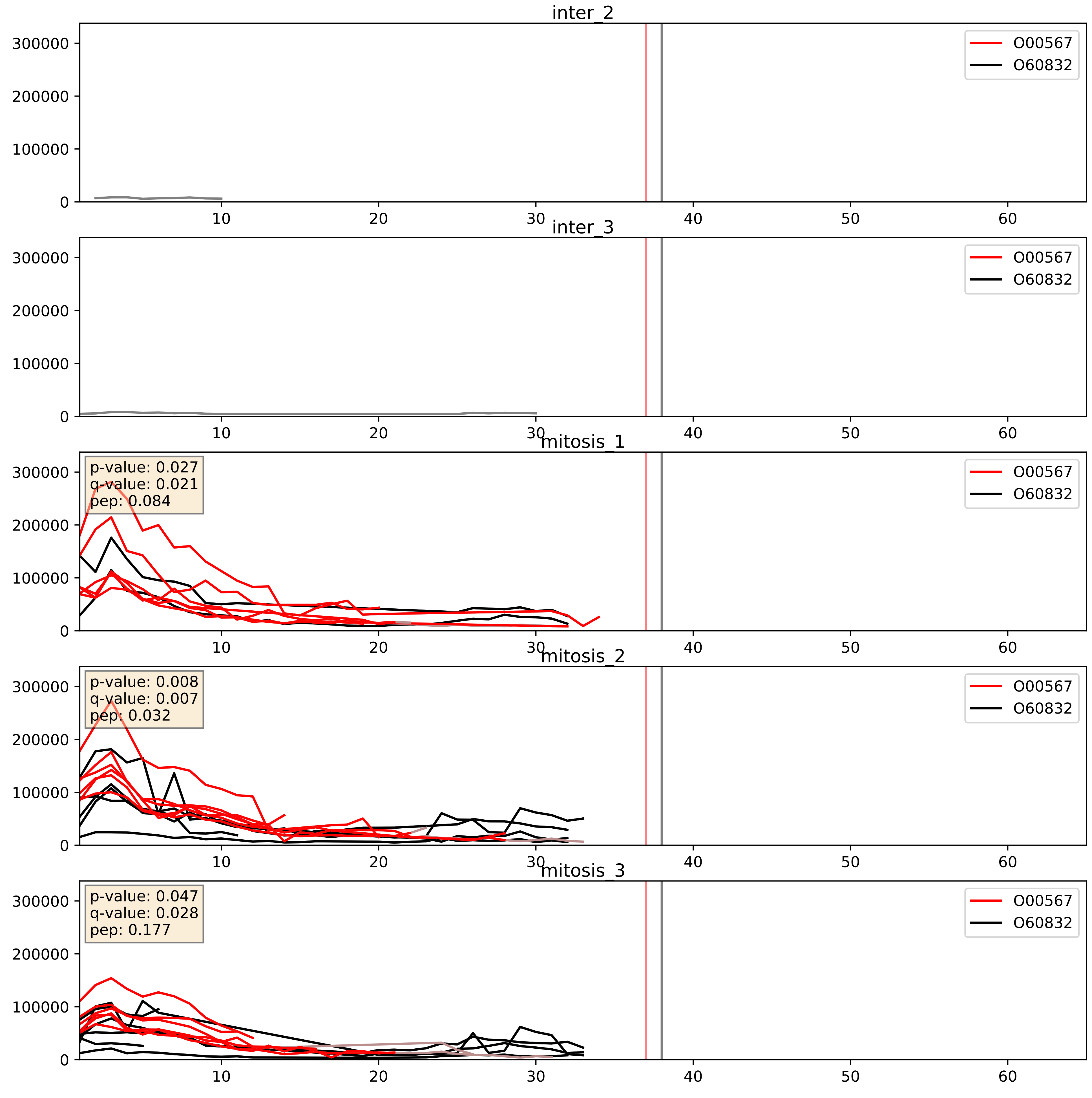


| level | condition_1 | condition_2 | pvalue | pvalue_adjusted |
|---|---|---|---|---|
| interactor_abundance | mitosis | inter | 0.0013960261808965133 | 0.027539131851237736 |
| complex_abundance | mitosis | inter | 0.0023625900336761763 | 0.035168510428347066 |
| interactor_ratio | mitosis | inter | 0.1363687124743647 | 0.5866546850118911 |

O00567_O76021
NOP56_HUMAN vs RL1D1_HUMAN
p-value: 0.001 q-value: 0.001

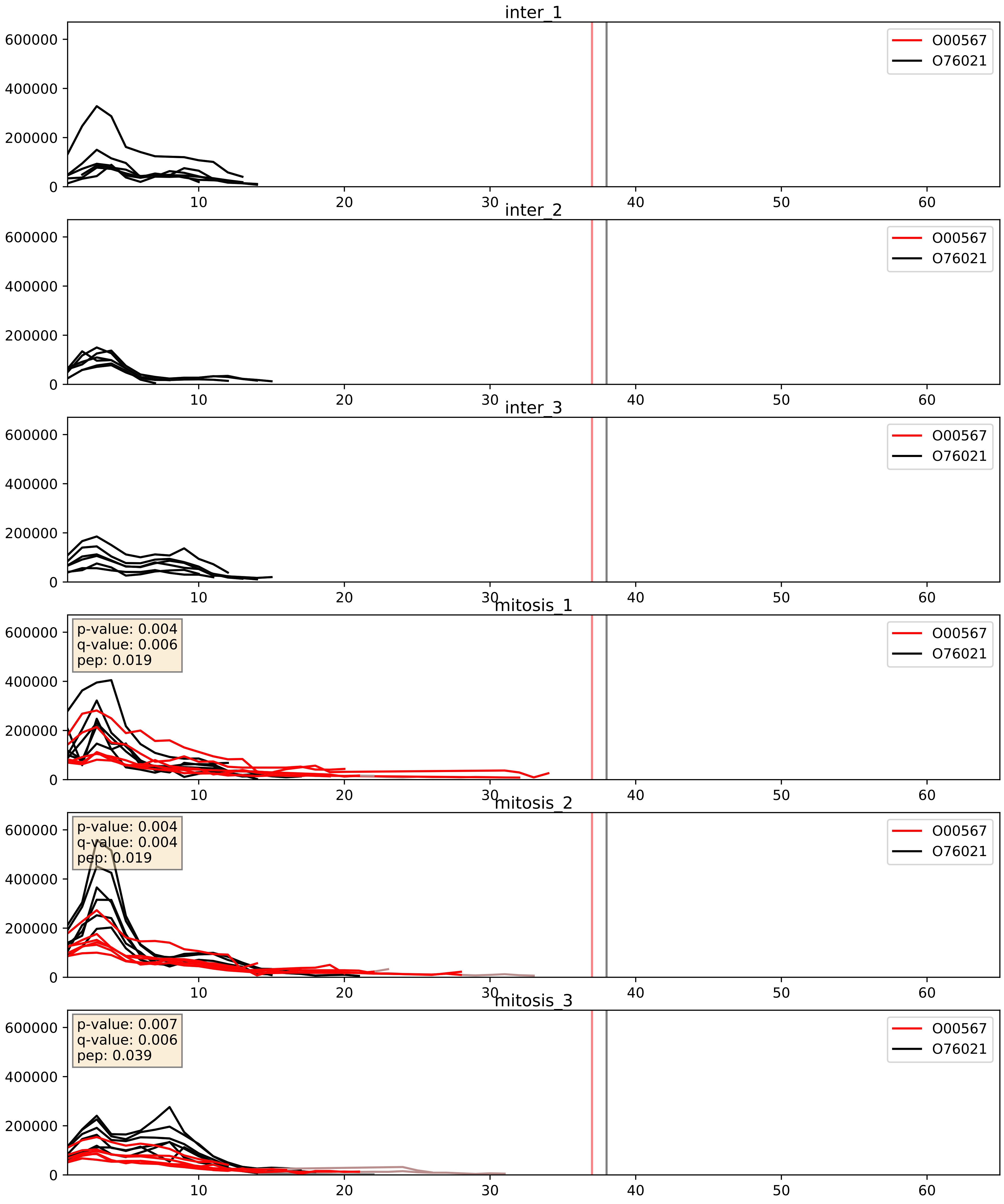


| level | condition_1 | condition_2 | pvalue | pvalue_adjusted |
|---|---|---|---|---|
| complex_abundance | mitosis | inter | 0.00010088212909086971 | 0.006746492382951911 |
| interactor_abundance | mitosis | inter | 0.0009352754344708583 | 0.022711936791689494 |
| interactor_ratio | mitosis | inter | 0.13519290058957723 | 0.5858060166978465 |

O00567_P22087
NOP56_HUMAN vs FBRL_HUMAN
p-value: 0.001 q-value: 0.002

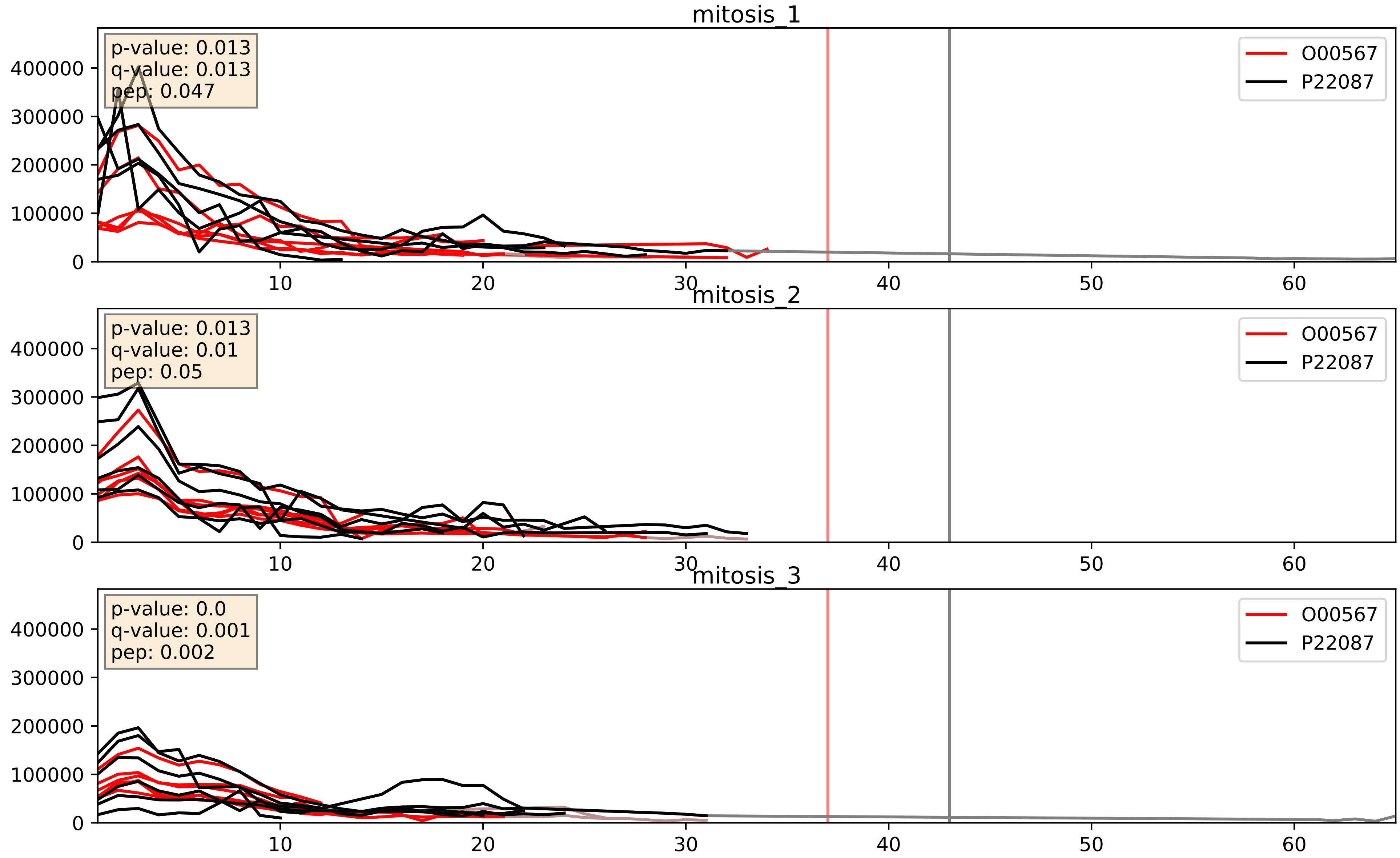


| level | condition_1 | condition_2 | pvalue | pvalue_adjusted |
|---|---|---|---|---|
| interactor_ratio | mitosis | inter | 0.00011298744803945258 | 0.011246192502531558 |
| complex_abundance | mitosis | inter | 0.0006517472043178059 | 0.01642175463112936 |
| interactor_abundance | mitosis | inter | 0.0011056112671619452 | 0.024049720606162197 |

O00567_P42285
NOP56_HUMAN vs MTREX_HUMAN
p-value: 0.031 q-value: 0.023

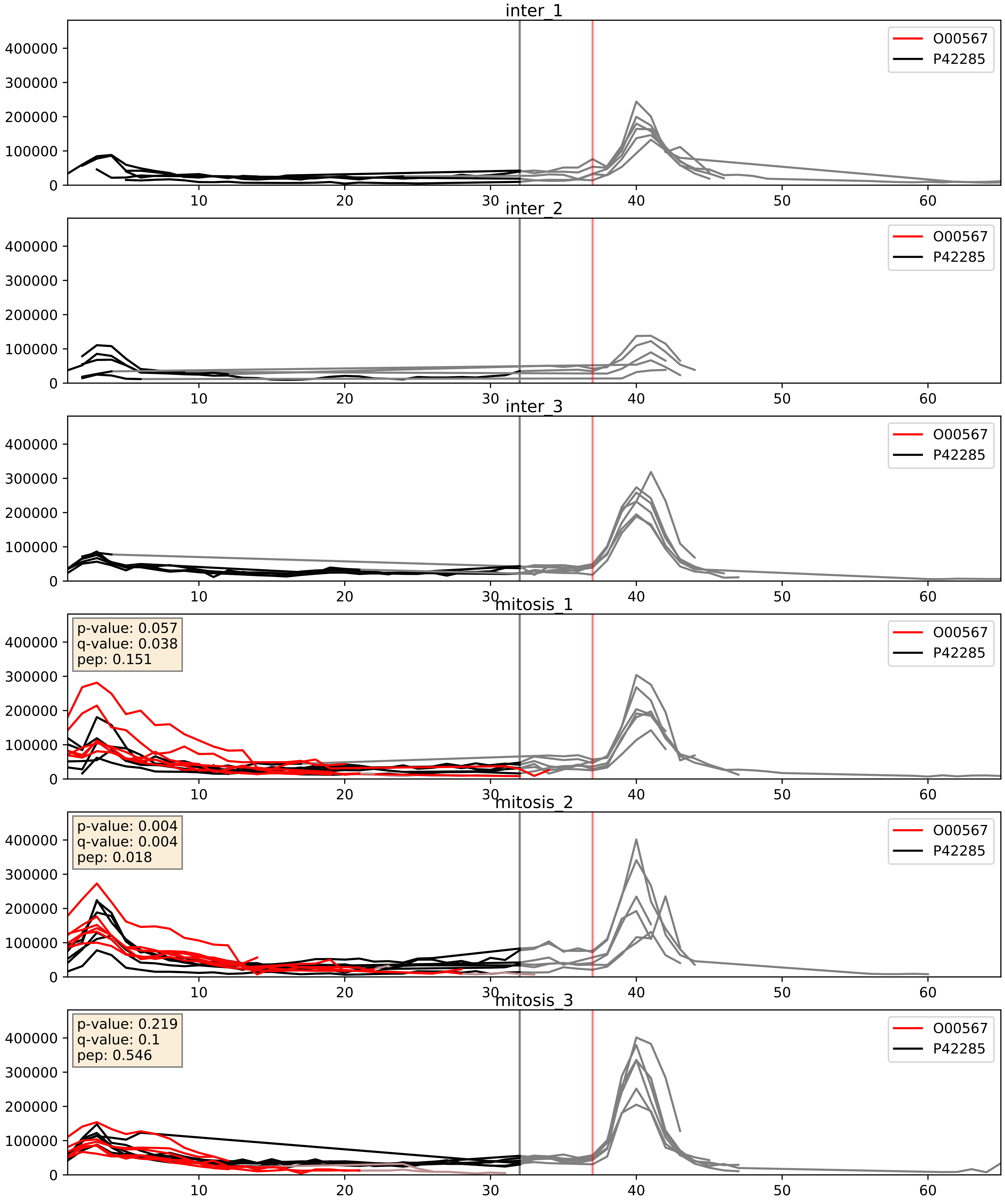


| level | condition_1 | condition_2 | pvalue | pvalue_adjusted |
|---|---|---|---|---|
| complex_abundance | mitosis | inter | 0.0005777242975888954 | 0.015600378509025063 |
| interactor_abundance | mitosis | inter | 0.0010892229270805026 | 0.024049720606162197 |
| interactor_ratio | mitosis | inter | 0.2336592312420024 | 0.7130563349132052 |

O00567_P46087
NOP56_HUMAN vs NOP2_HUMAN
p-value: 0.01 q-value: 0.009

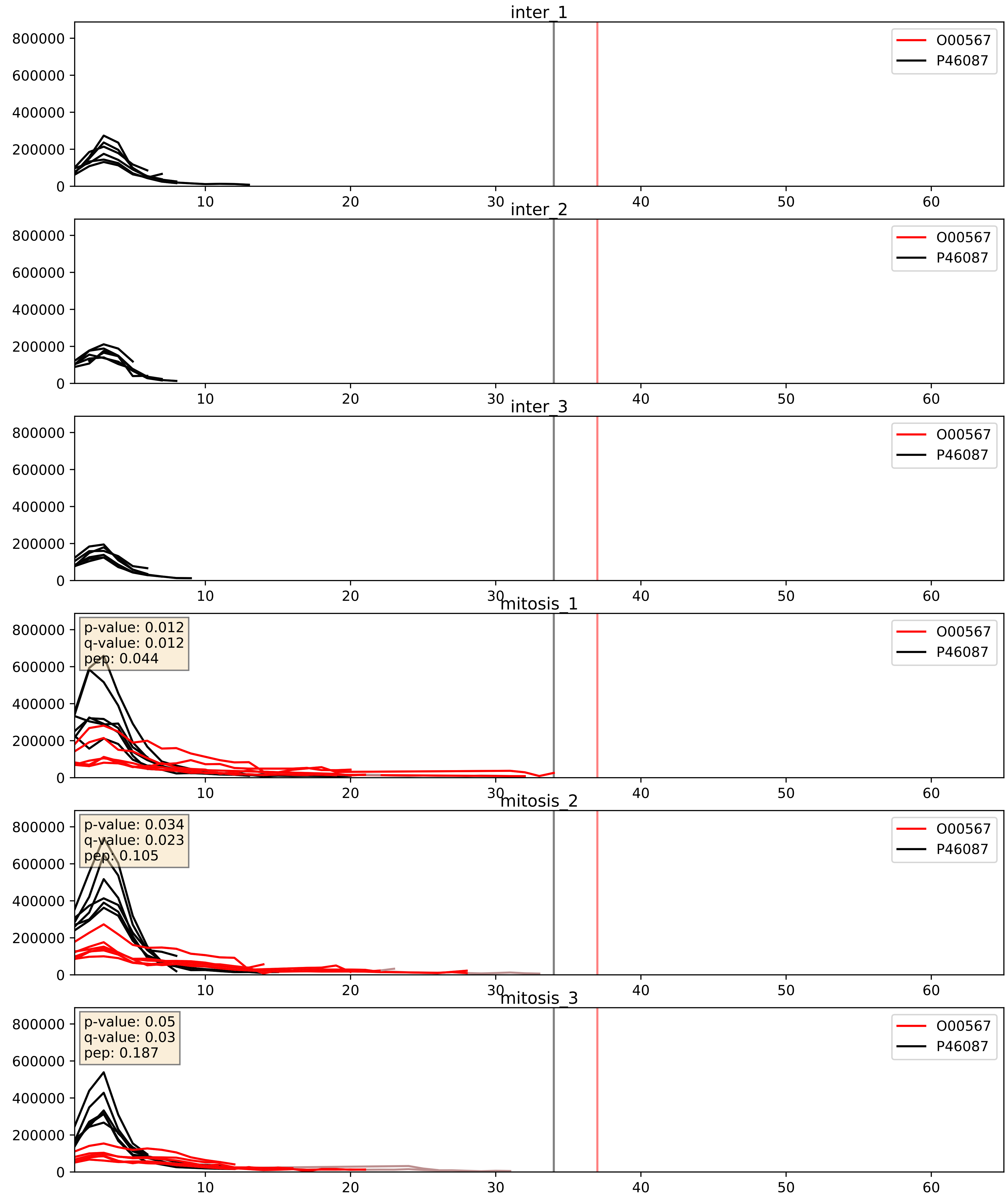


| level | condition_1 | condition_2 | pvalue | pvalue_adjusted |
|---|---|---|---|---|
| complex_abundance | mitosis | inter | 0.0004100837102177417 | 0.012905575586264225 |
| interactor_abundance | mitosis | inter | 0.001371443612175878 | 0.02742887224351756 |
| interactor_ratio | mitosis | inter | 0.0037884888716758825 | 0.08027095233055831 |

O00567_P55769
NOP56_HUMAN vs NH2L1_HUMAN
p-value: 0.0 q-value: 0.0

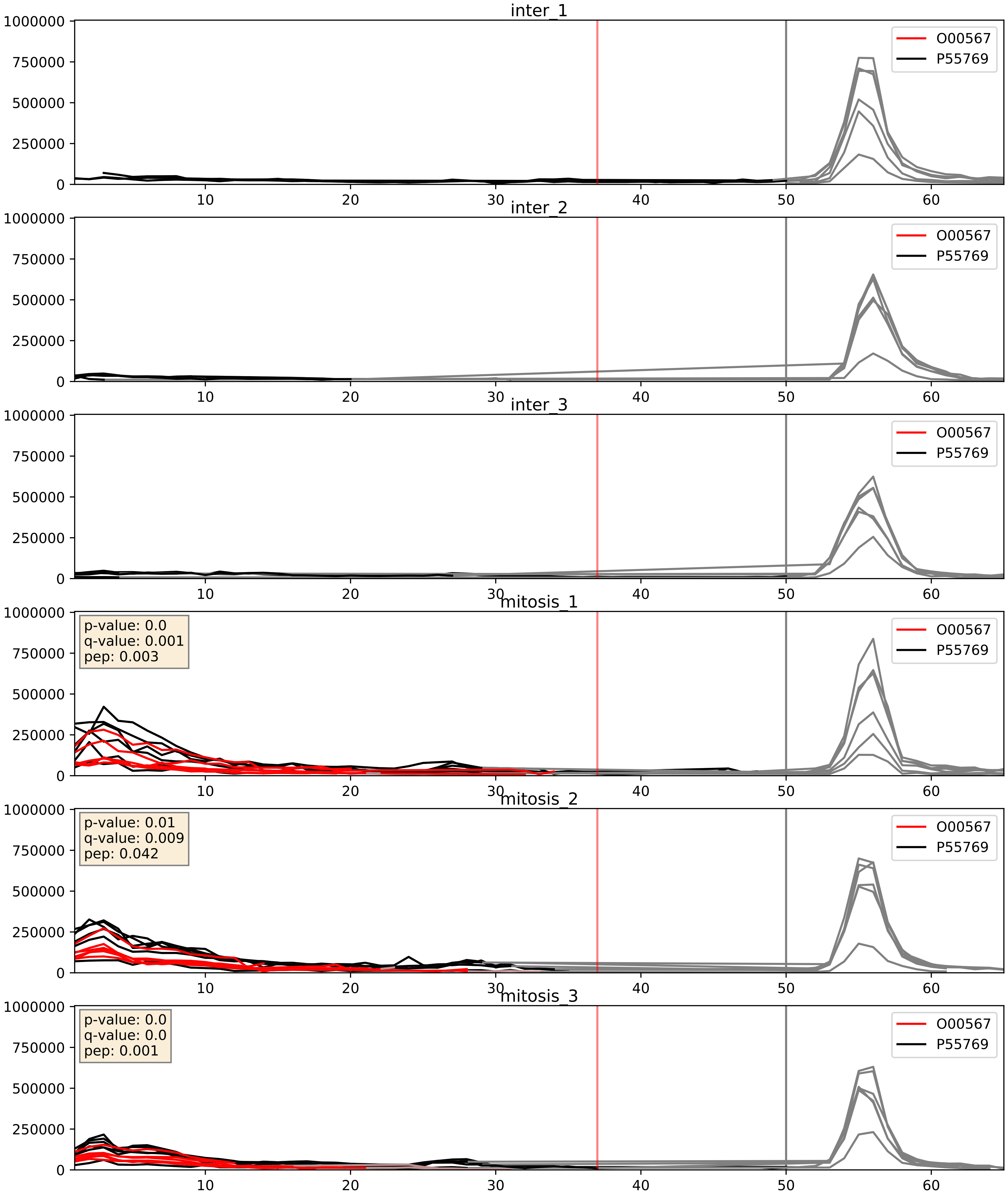


| level | condition_1 | condition_2 | pvalue | pvalue_adjusted |
|---|---|---|---|---|
| interactor_ratio | mitosis | inter | 0.000189167225482195 | 0.014134487523529912 |
| complex_abundance | mitosis | inter | 0.00048729227531920847 | 0.013857879989144268 |
| interactor_abundance | mitosis | inter | 0.0011181996263145508 | 0.024049720606162197 |

O00567_P56182
NOP56_HUMAN vs RRP1_HUMAN
p-value: 0.011 q-value: 0.01

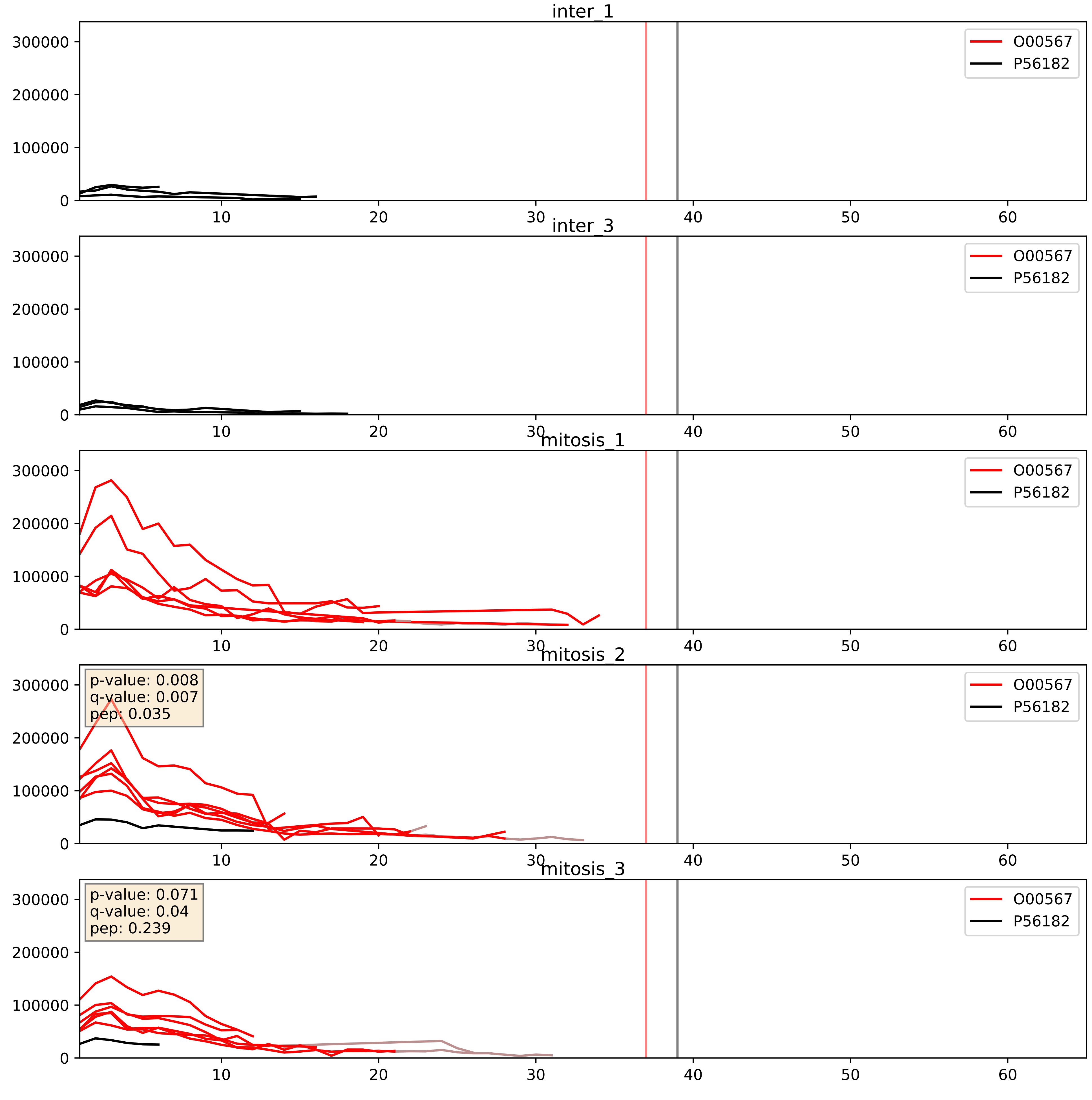


| level | condition_1 | condition_2 | pvalue | pvalue_adjusted |
|---|---|---|---|---|
| interactor_ratio | mitosis | inter | 0.1856723899874445 | 0.6559196457151685 |
| interactor_abundance | mitosis | inter | 0.21154728512611398 | 0.40715039907095185 |
| complex_abundance | mitosis | inter | 0.23486897194862114 | 0.3870770889257214 |

O00567_P61313
NOP56_HUMAN vs RL15_HUMAN
p-value: 0.004 q-value: 0.047

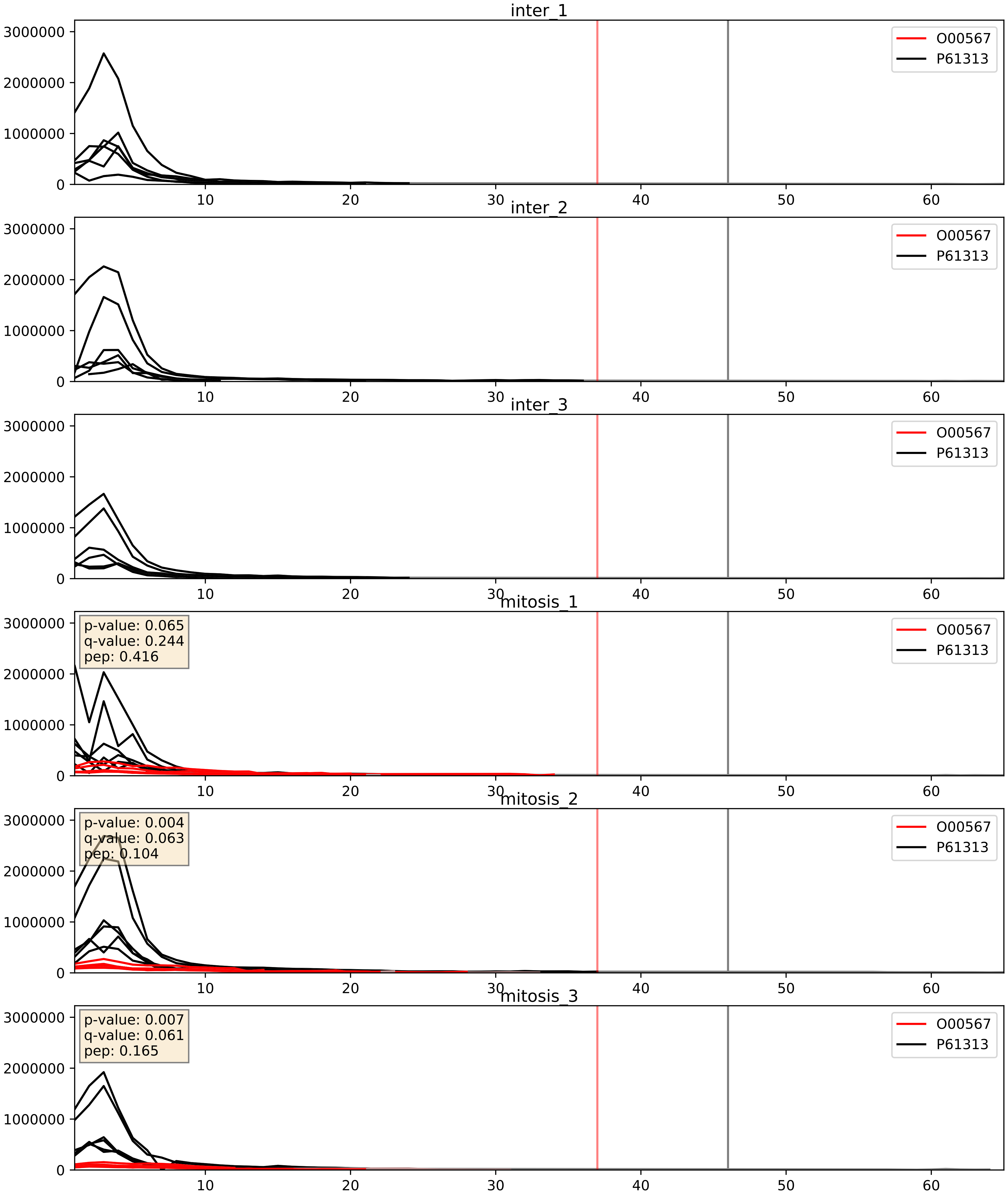


| level | condition_1 | condition_2 | pvalue | pvalue_adjusted |
|---|---|---|---|---|
| complex_abundance | mitosis | inter | 5.906513236339796e-05 | 0.005691857151652958 |
| interactor_abundance | mitosis | inter | 0.0010562059318354998 | 0.024049720606162197 |
| interactor_ratio | mitosis | inter | 0.0014917736926577606 | 0.04597663709000905 |

O00567_P62081
NOP56_HUMAN vs RS7_HUMAN
p-value: 0.002 q-value: 0.002

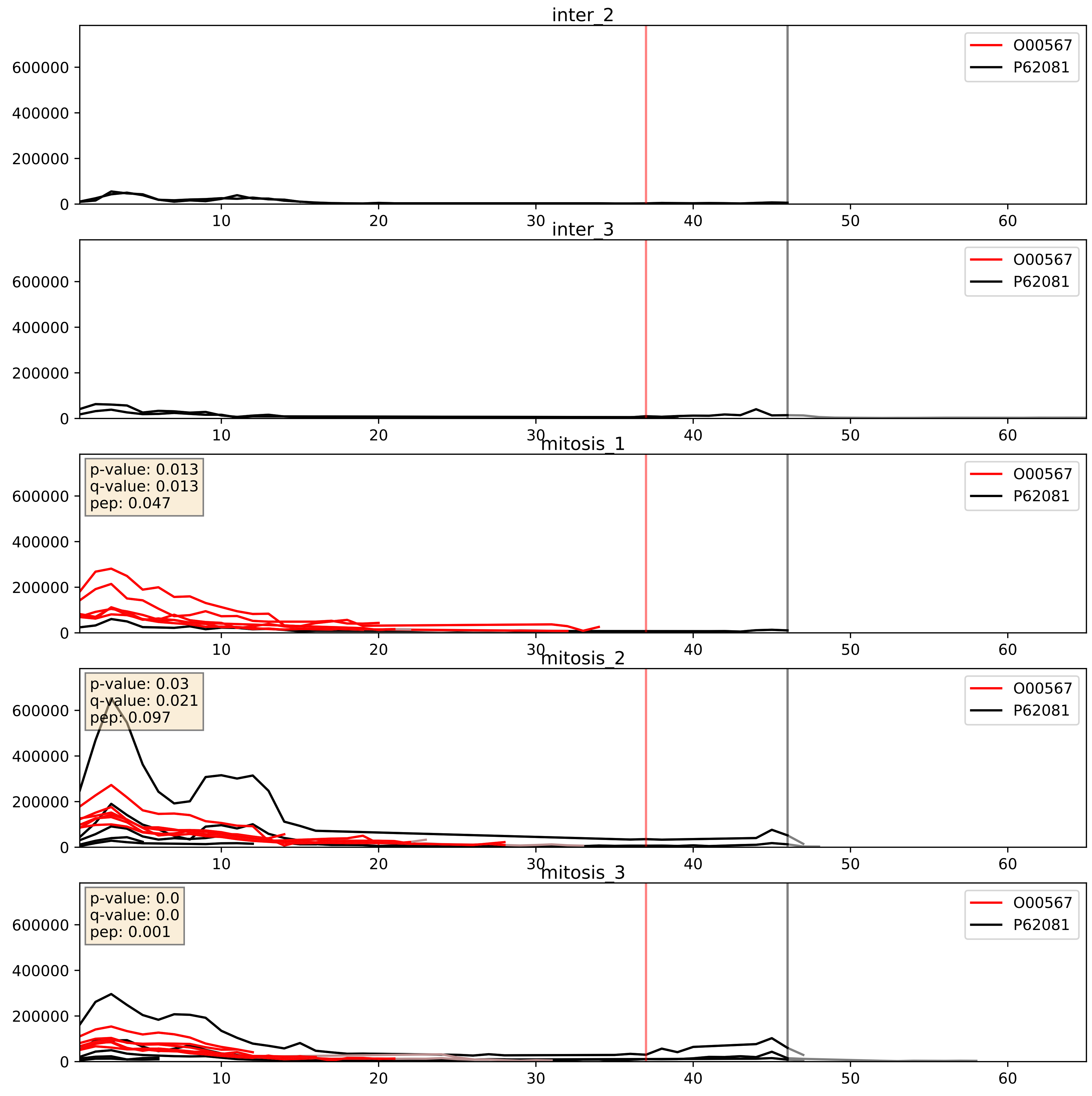


| level | condition_1 | condition_2 | pvalue | pvalue_adjusted |
|---|---|---|---|---|
| interactor_abundance | mitosis | inter | 0.0007896418237053181 | 0.020607725643041228 |
| complex_abundance | mitosis | inter | 0.004425707233152511 | 0.049683977369634376 |
| interactor_ratio | mitosis | inter | 0.41099655815660535 | 0.8340755186867097 |

O00567_P62244
NOP56_HUMAN vs RS15A_HUMAN
p-value: 0.006 q-value: 0.006

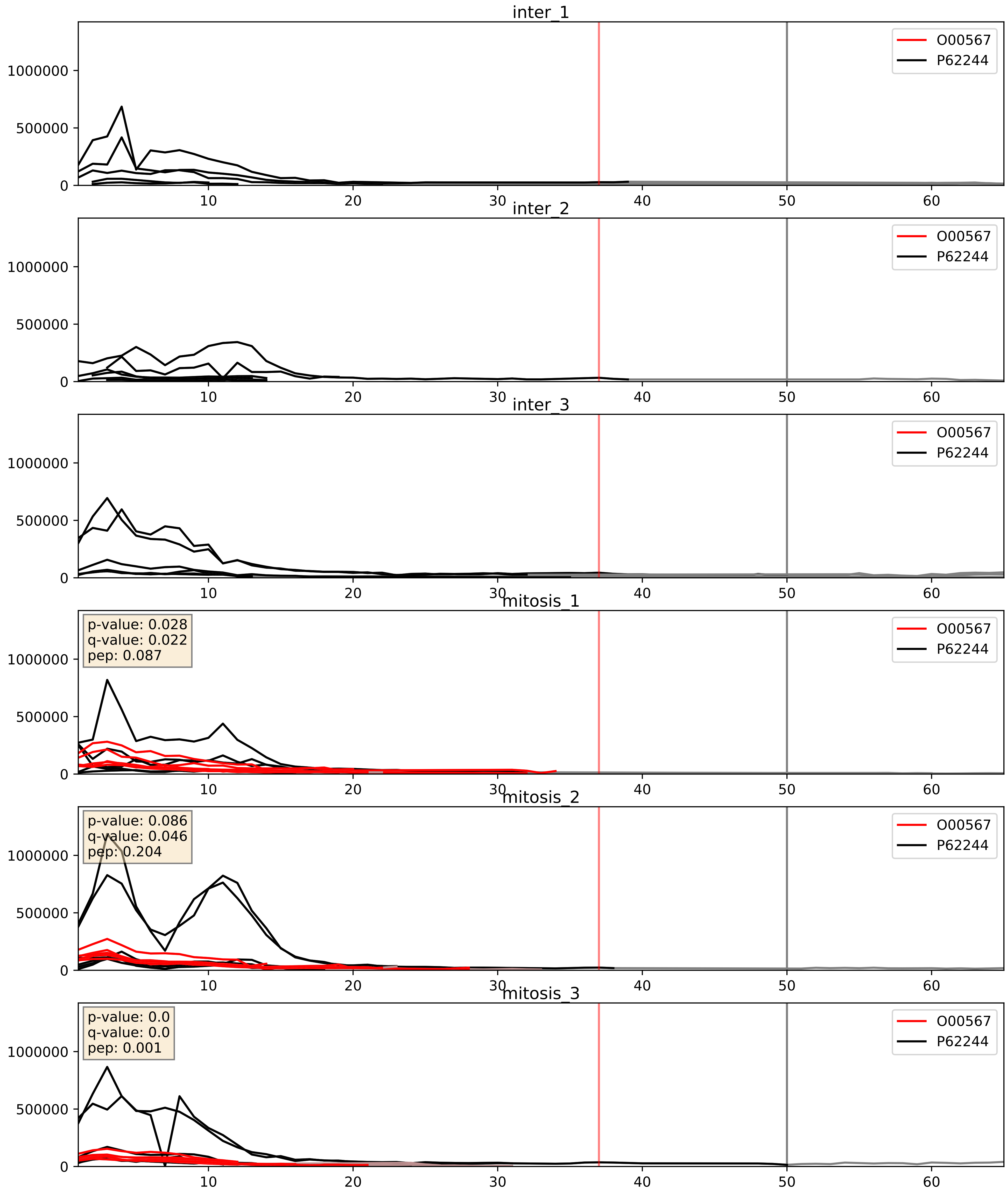


| level | condition_1 | condition_2 | pvalue | pvalue_adjusted |
|---|---|---|---|---|
| complex_abundance | mitosis | inter | 0.00014575867540400682 | 0.008317961743055323 |
| interactor_abundance | mitosis | inter | 0.0011052779688176132 | 0.024049720606162197 |
| interactor_ratio | mitosis | inter | 0.007167137700217959 | 0.11775565971951195 |

O00567_P62277
NOP56_HUMAN vs RS13_HUMAN
p-value: 0.078 q-value: 0.047

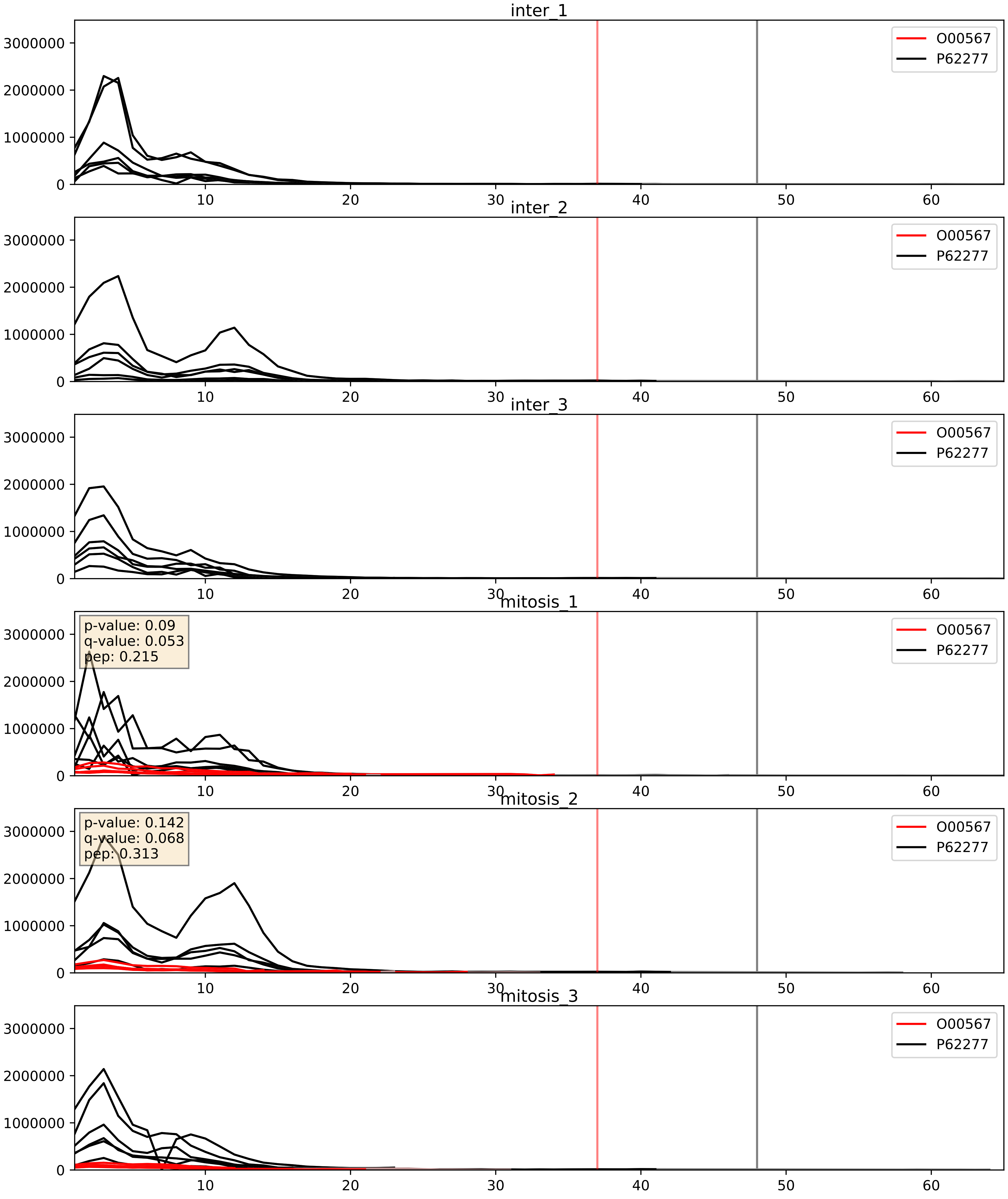


| level | condition_1 | condition_2 | pvalue | pvalue_adjusted |
|---|---|---|---|---|
| complex_abundance | mitosis | inter | 2.346006912375981e-05 | 0.005691857151652958 |
| interactor_abundance | mitosis | inter | 0.0011181996263145508 | 0.024049720606162197 |
| interactor_ratio | mitosis | inter | 0.0011704914723791608 | 0.04023858234363702 |

O00567_P62280
NOP56_HUMAN vs RS11_HUMAN
p-value: 0.005 q-value: 0.005

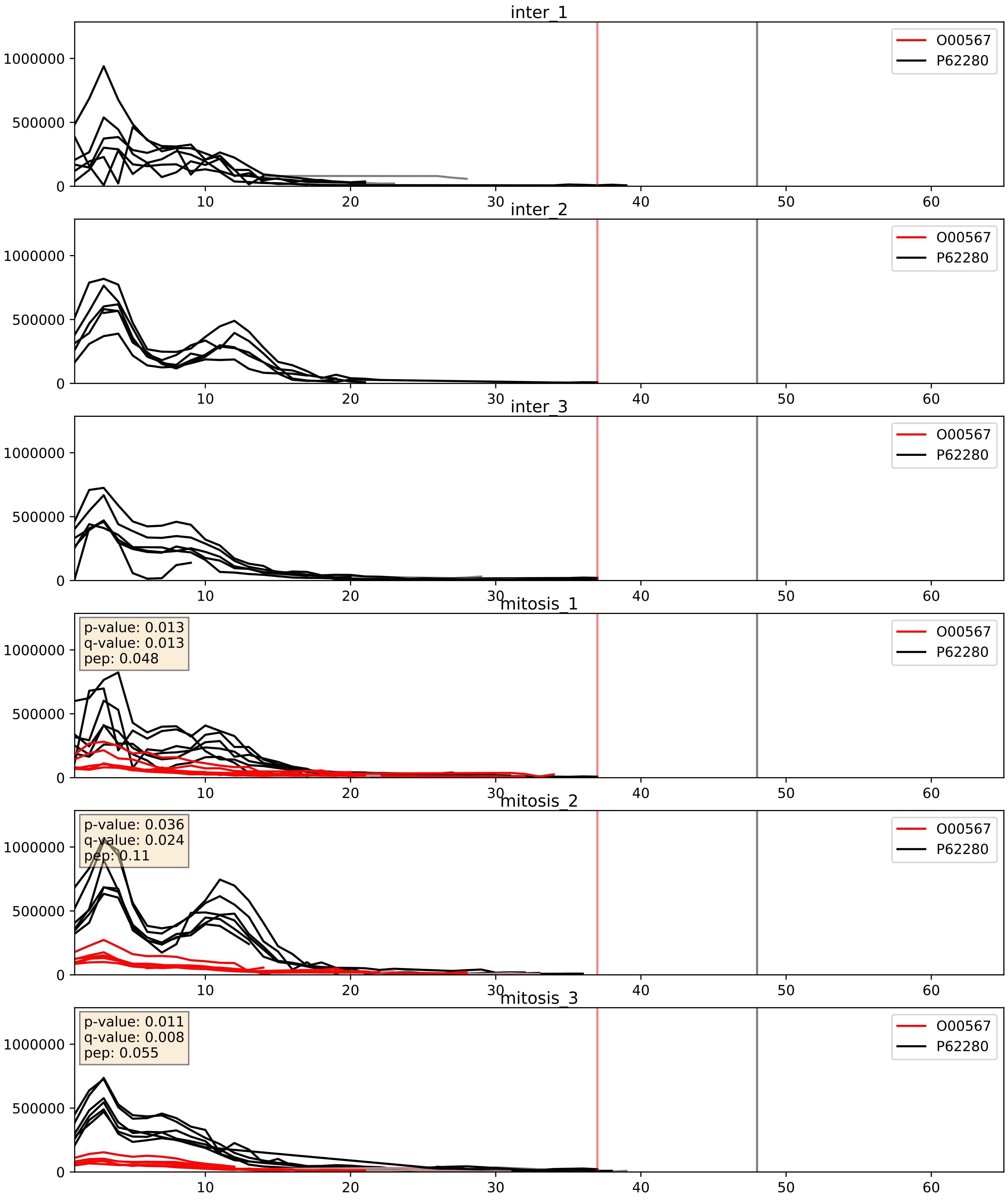


| level | condition_1 | condition_2 | pvalue | pvalue_adjusted |
|---|---|---|---|---|
| complex_abundance | mitosis | inter | 7.338827789298022e-05 | 0.005893640694944935 |
| interactor_abundance | mitosis | inter | 0.0010892646924931246 | 0.024049720606162197 |
| interactor_ratio | mitosis | inter | 0.0015900218175386696 | 0.0480939461418057 |

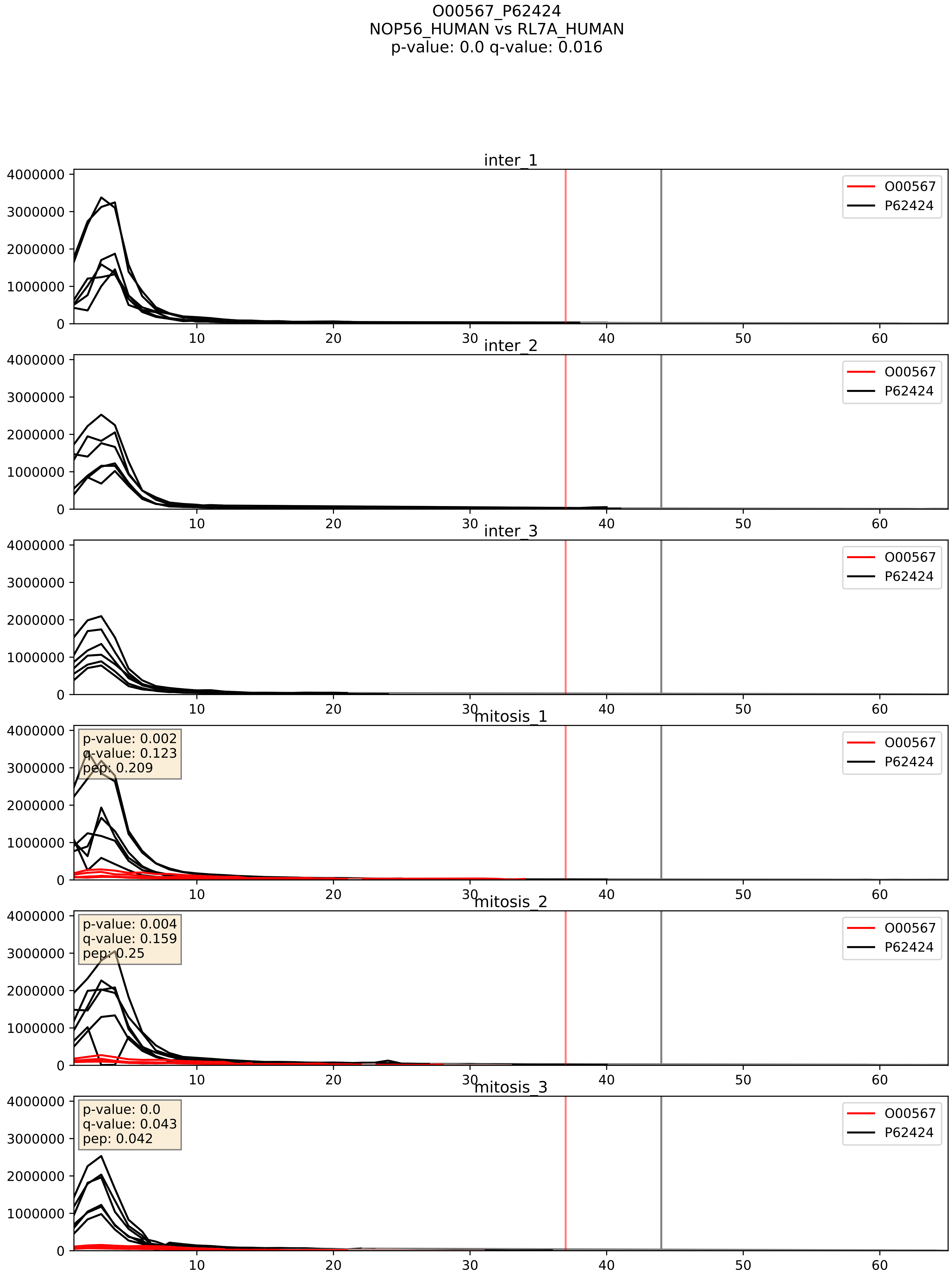


| level | condition_1 | condition_2 | pvalue | pvalue_adjusted |
|---|---|---|---|---|
| complex_abundance | mitosis | inter | 5.760963949085291e-05 | 0.005691857151652958 |
| interactor_ratio | mitosis | inter | 0.00014691320261744684 | 0.012836278661937291 |
| interactor_abundance | mitosis | inter | 0.0011181996263145508 | 0.024049720606162197 |

O00567_P62701
NOP56_HUMAN vs RS4X_HUMAN
p-value: 0.062 q-value: 0.039

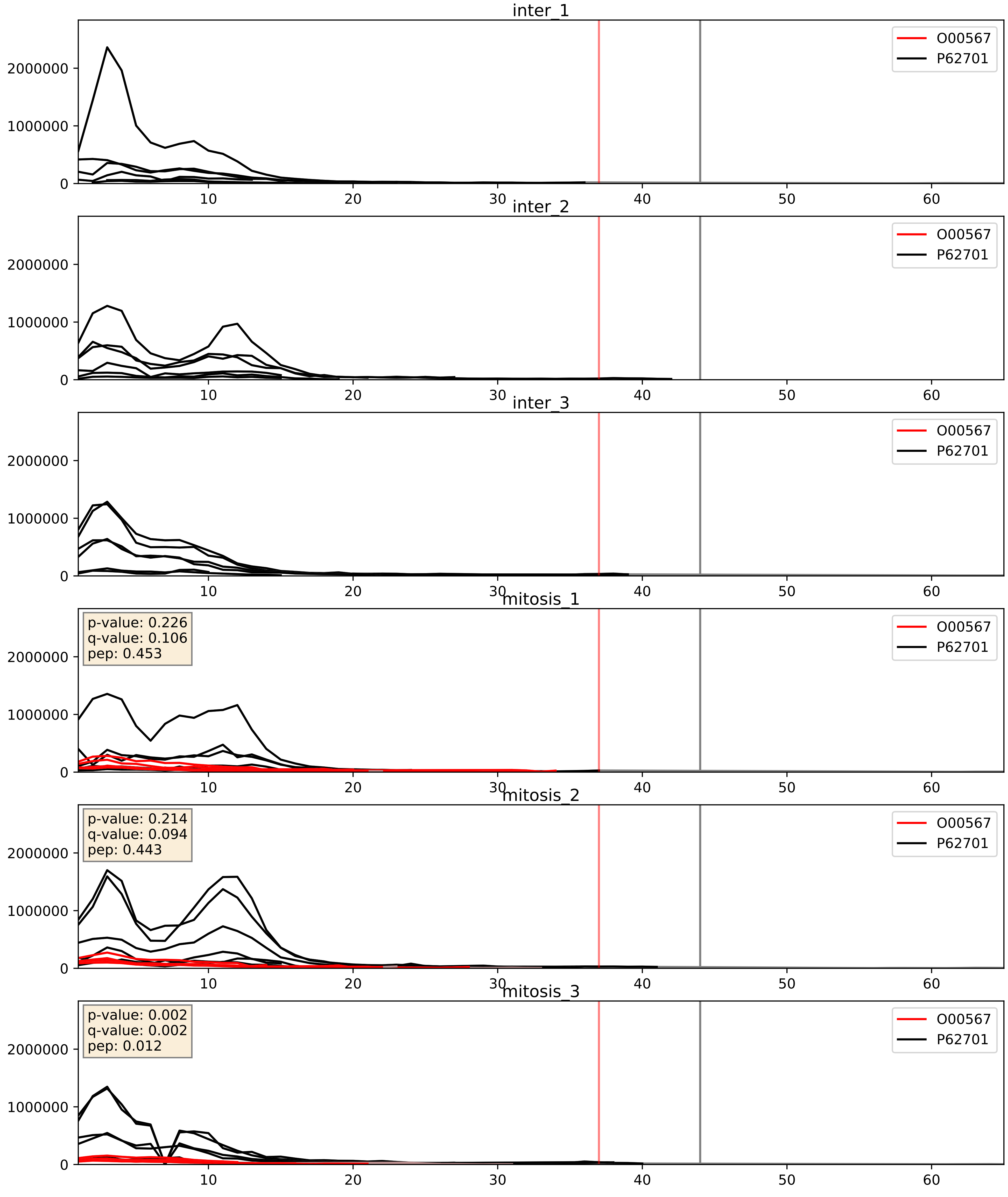


| level | condition_1 | condition_2 | pvalue | pvalue_adjusted |
|---|---|---|---|---|
| complex_abundance | mitosis | inter | 4.9797560413752016e-05 | 0.005691857151652958 |
| interactor_abundance | mitosis | inter | 0.0011181996263145508 | 0.024049720606162197 |
| interactor_ratio | mitosis | inter | 0.0012256721316524868 | 0.041469381213222474 |

O00567_P62753
NOP56_HUMAN vs RS6_HUMAN
p-value: 0.0 q-value: 0.0

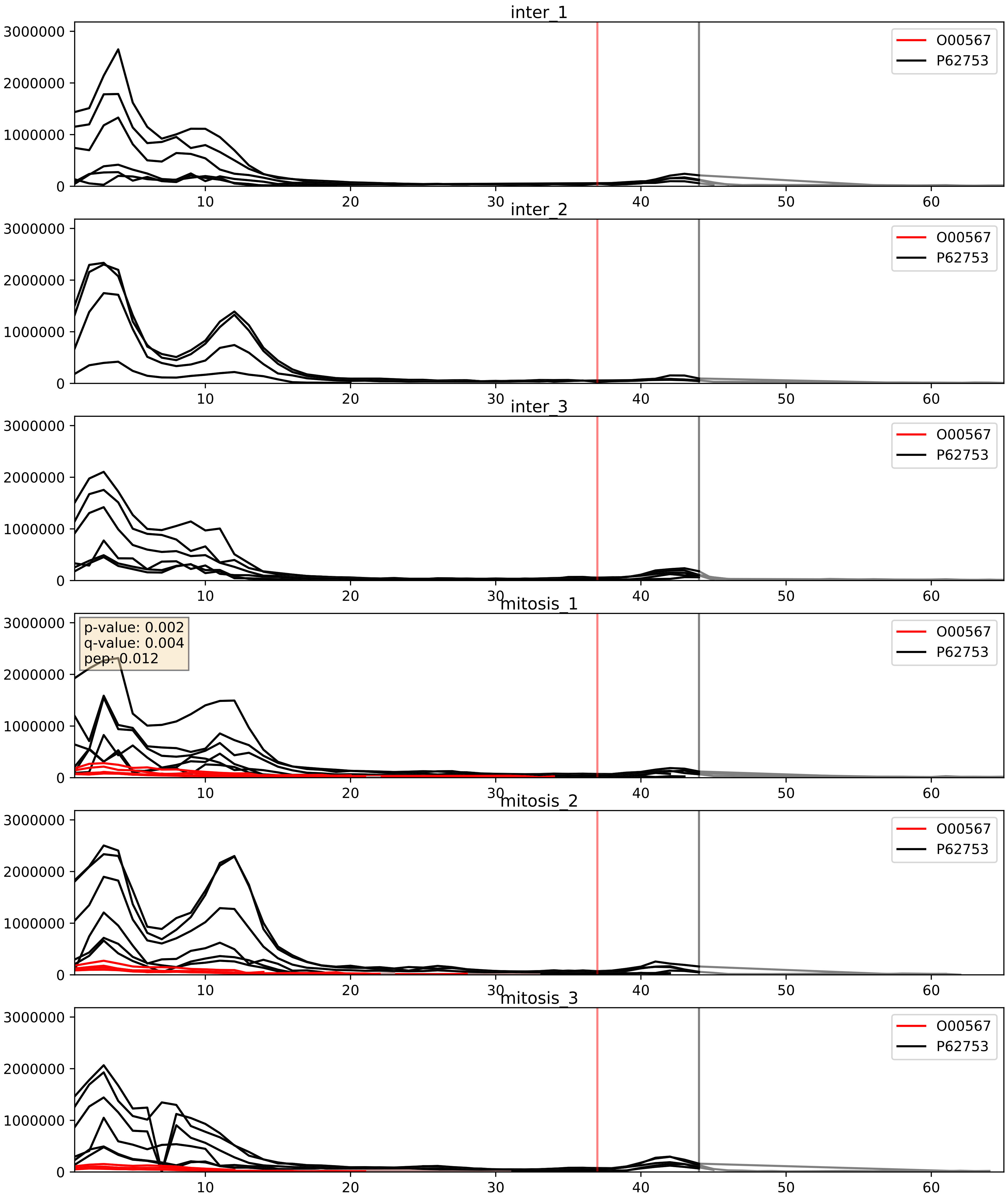


| level | condition_1 | condition_2 | pvalue | pvalue_adjusted |
|---|---|---|---|---|
| complex_abundance | mitosis | inter | 1.5692732711379486e-05 | 0.005691857151652958 |
| interactor_ratio | mitosis | inter | 0.0004213753580807645 | 0.02297435073357544 |
| interactor_abundance | mitosis | inter | 0.0011181996263145508 | 0.024049720606162197 |

O00567_P62847
NOP56_HUMAN vs RS24_HUMAN
p-value: 0.035 q-value: 0.025

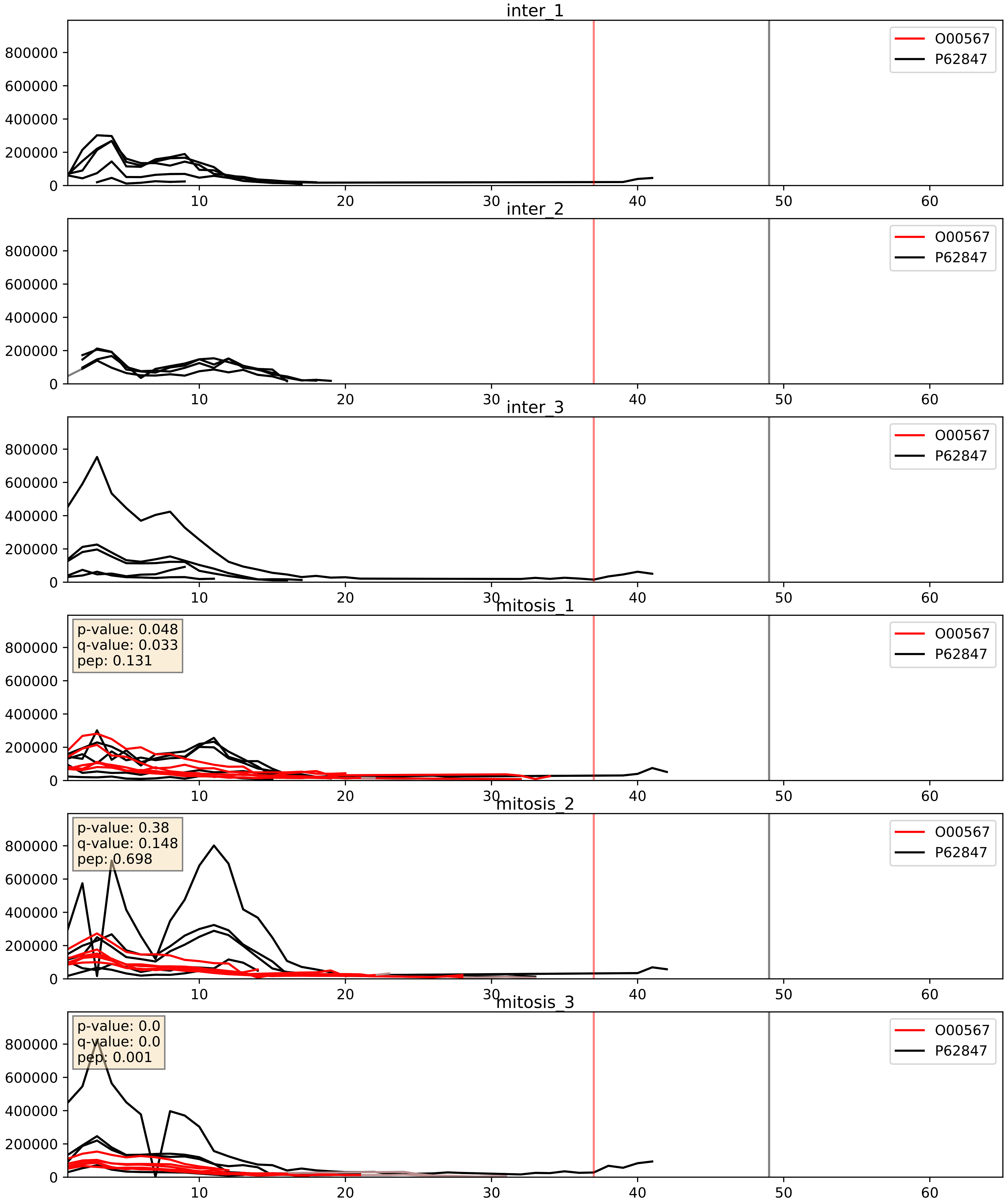


| level | condition_1 | condition_2 | pvalue | pvalue_adjusted |
|---|---|---|---|---|
| complex_abundance | mitosis | inter | 0.0001307170847499598 | 0.00787984679901166 |
| interactor_abundance | mitosis | inter | 0.001056251194238277 | 0.024049720606162197 |
| interactor_ratio | mitosis | inter | 0.012080002499307237 | 0.15864619391321497 |

O00567_P62913
NOP56_HUMAN vs RL11_HUMAN
p-value: 0.001 q-value: 0.046

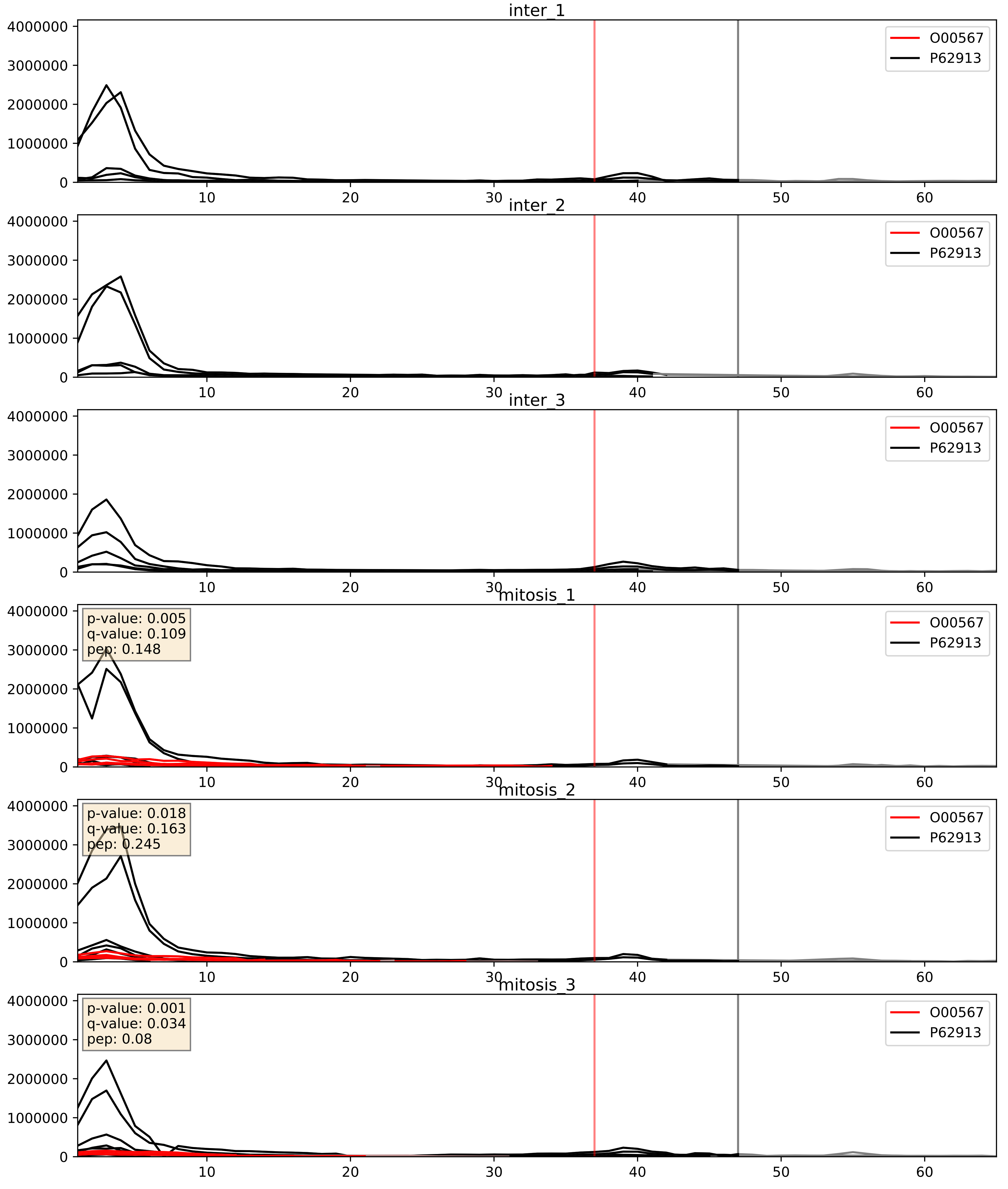


| level | condition_1 | condition_2 | pvalue | pvalue_adjusted |
|---|---|---|---|---|
| complex_abundance | mitosis | inter | 5.641121545173787e-05 | 0.005691857151652958 |
| interactor_ratio | mitosis | inter | 0.0005452171487126521 | 0.026977218456533537 |
| interactor_abundance | mitosis | inter | 0.001118199626314508 | 0.024049720606162197 |

O00567_Q02543
NOP56_HUMAN vs RL18A_HUMAN
p-value: 0.001 q-value: 0.039

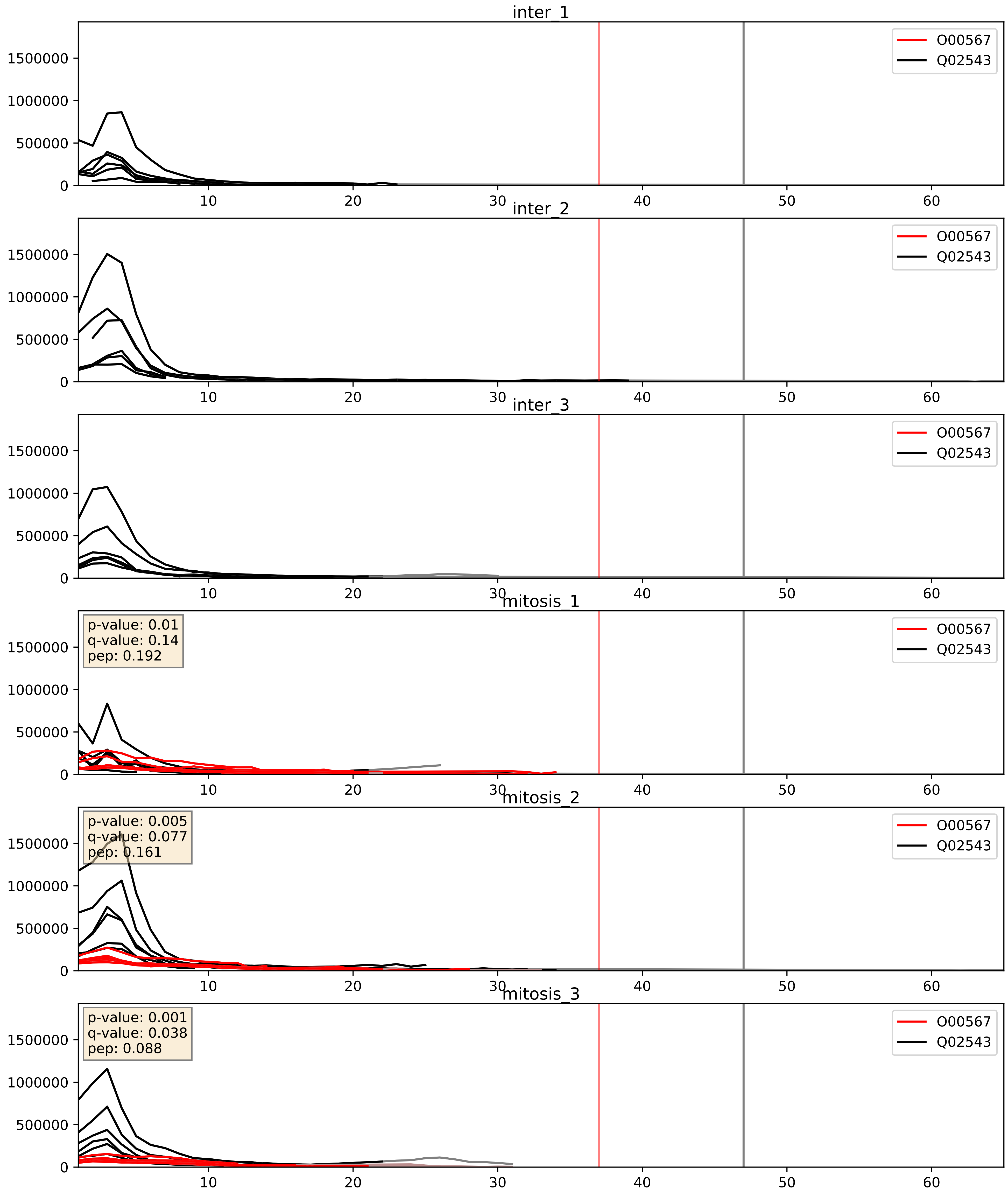


| level | condition_1 | condition_2 | pvalue | pvalue_adjusted |
|---|---|---|---|---|
| complex_abundance | mitosis | inter | 0.00020225689468314427 | 0.009565298444683509 |
| interactor_abundance | mitosis | inter | 0.0010892229270805026 | 0.024049720606162197 |
| interactor_ratio | mitosis | inter | 0.12192059394147745 | 0.5706647308029218 |

O00567_Q12788
NOP56_HUMAN vs TBL3_HUMAN
p-value: 0.007 q-value: 0.007

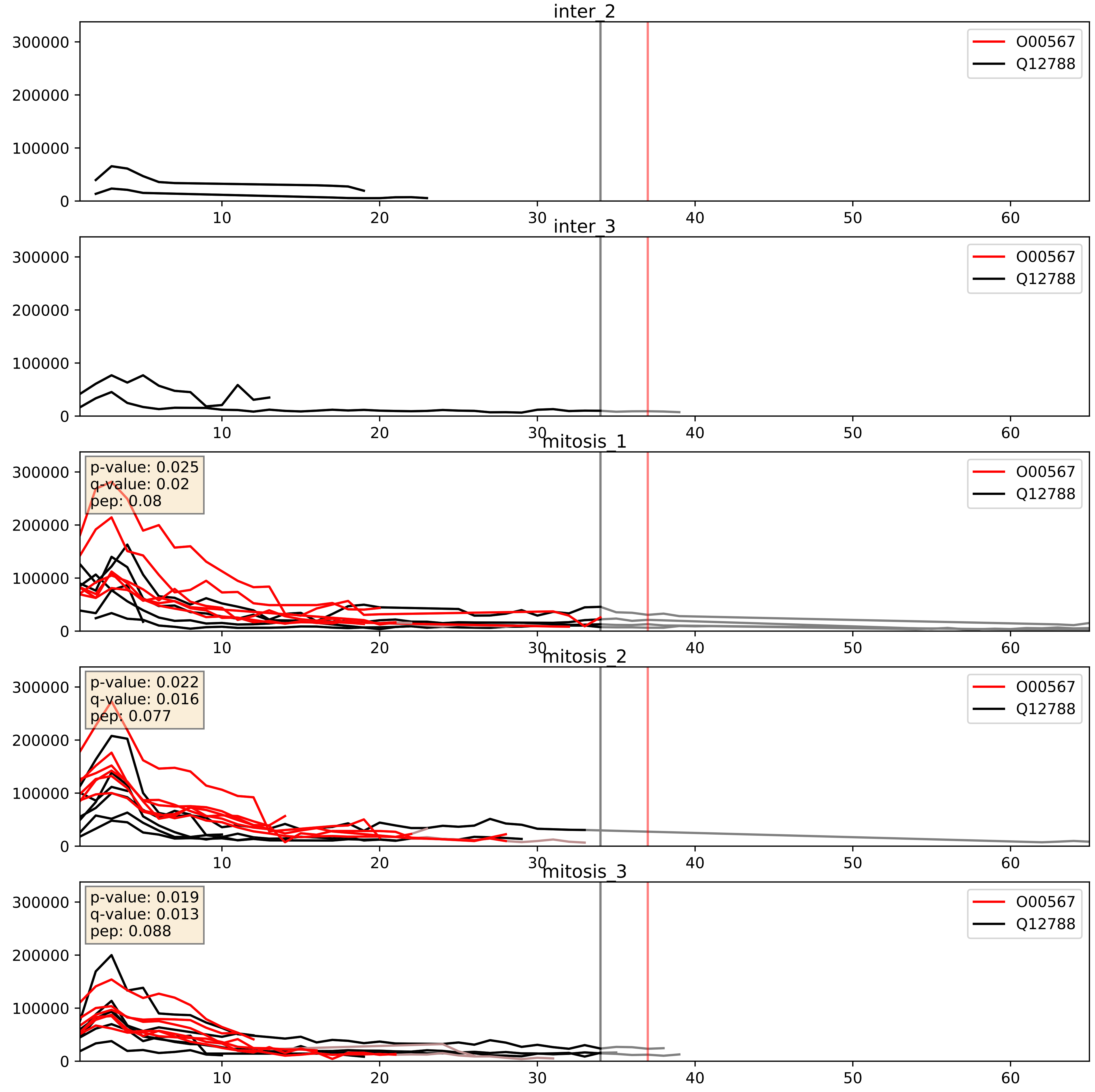


| level | condition_1 | condition_2 | pvalue | pvalue_adjusted |
|---|---|---|---|---|
| complex_abundance | mitosis | inter | 0.00030967416140721313 | 0.011652071109633347 |
| interactor_abundance | mitosis | inter | 0.0011111765577599547 | 0.024049720606162197 |
| interactor_ratio | mitosis | inter | 0.1581429879180654 | 0.6144687163742579 |

O00567_Q13895
NOP56_HUMAN vs BYST_HUMAN
p-value: 0.067 q-value: 0.041

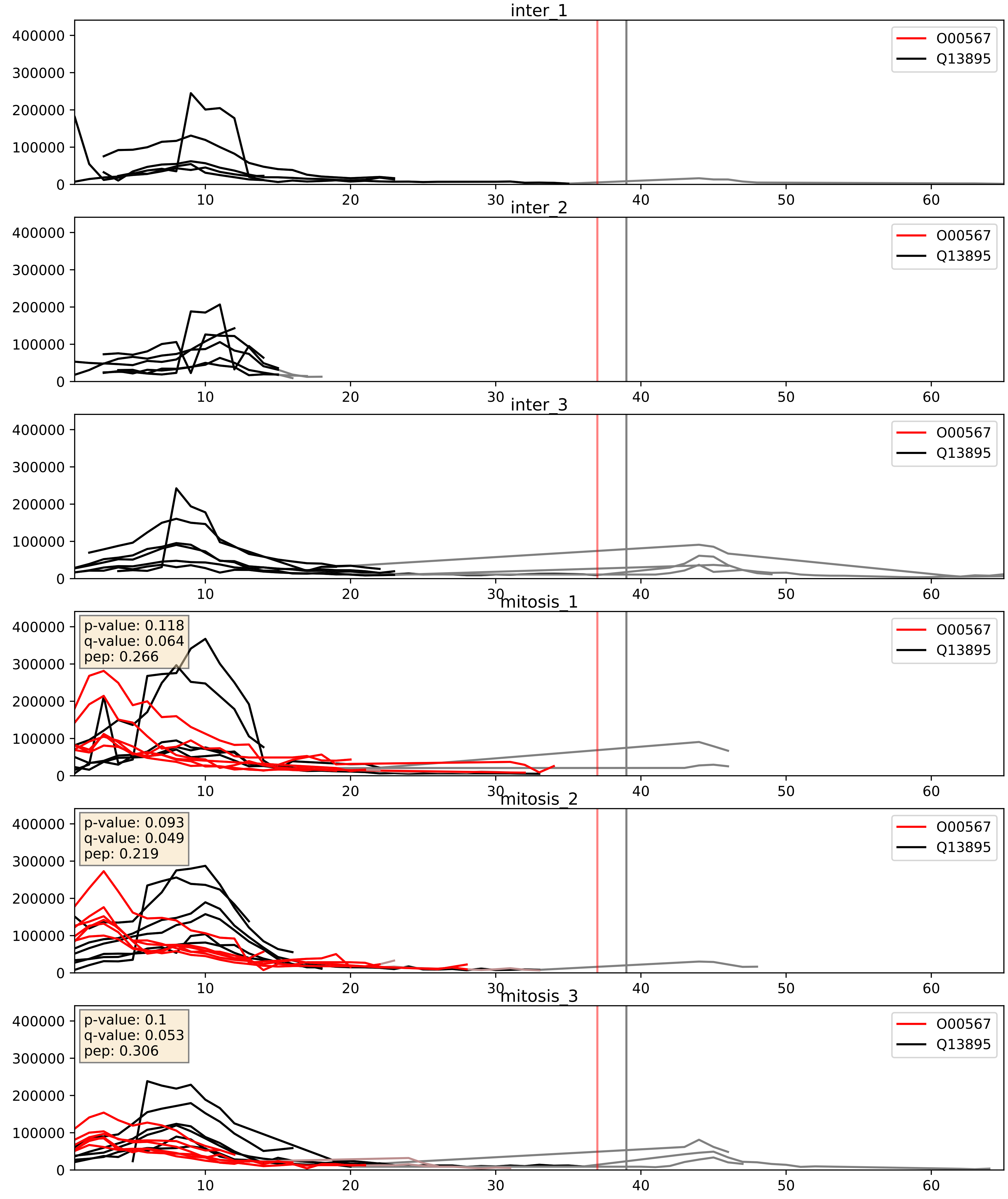


| level | condition_1 | condition_2 | pvalue | pvalue_adjusted |
|---|---|---|---|---|
| complex_abundance | mitosis | inter | 0.0003517935453936726 | 0.012039775462644836 |
| interactor_abundance | mitosis | inter | 0.0011111765577599547 | 0.024049720606162197 |
| interactor_ratio | mitosis | inter | 0.04276010684486895 | 0.3405601671696144 |

O00567_Q15061
NOP56_HUMAN vs WDR43_HUMAN
p-value: 0.021 q-value: 0.017

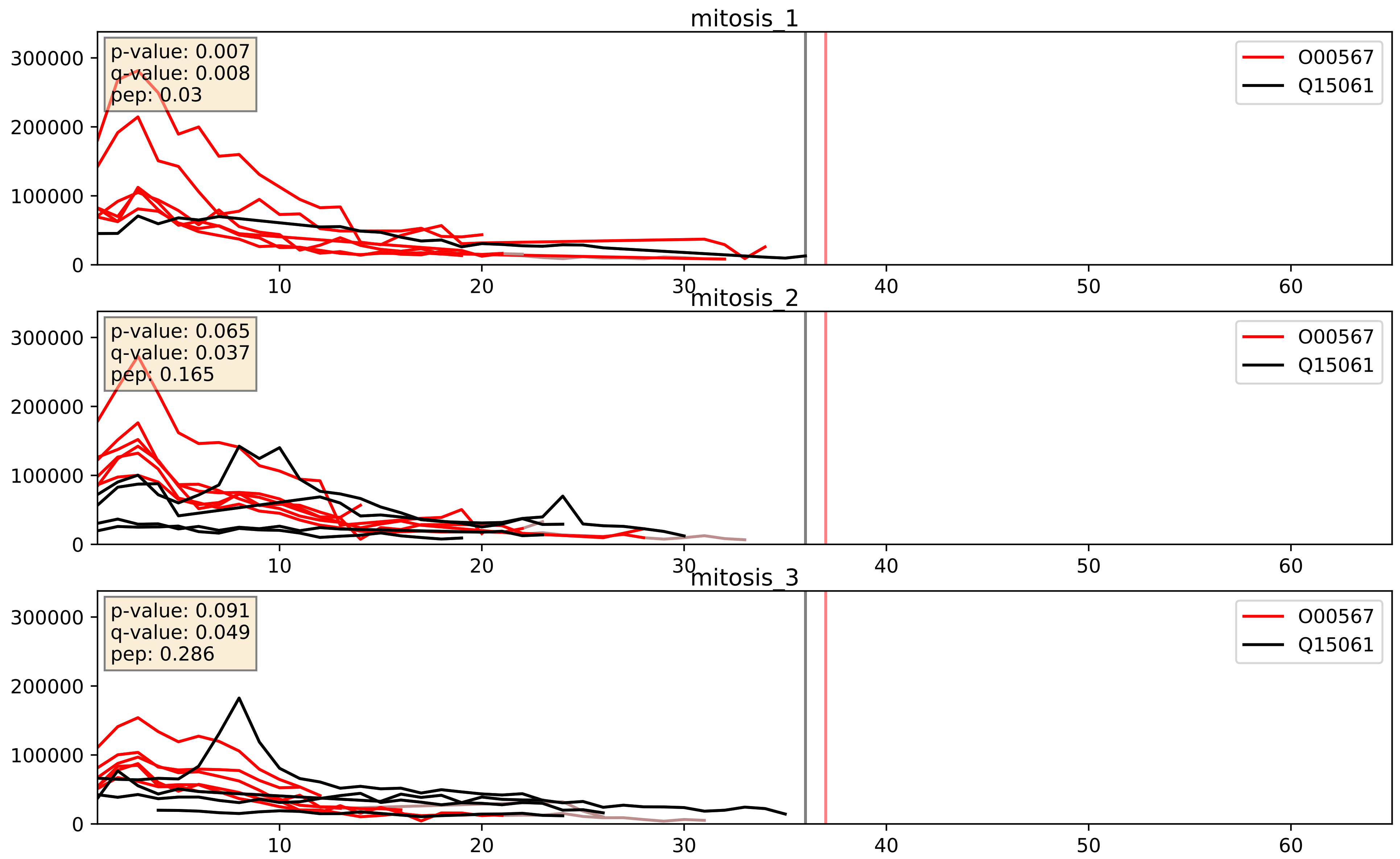


| level | condition_1 | condition_2 | pvalue | pvalue_adjusted |
|---|---|---|---|---|
| interactor_abundance | mitosis | inter | 0.0005738807154731015 | 0.017415659477552636 |
| complex_abundance | mitosis | inter | 0.0012557787786977876 | 0.02410194247904274 |
| interactor_ratio | mitosis | inter | 0.15614177644181174 | 0.6132288760320385 |

O00567_Q15269
NOP56_HUMAN vs PWP2_HUMAN
p-value: 0.001 q-value: 0.001

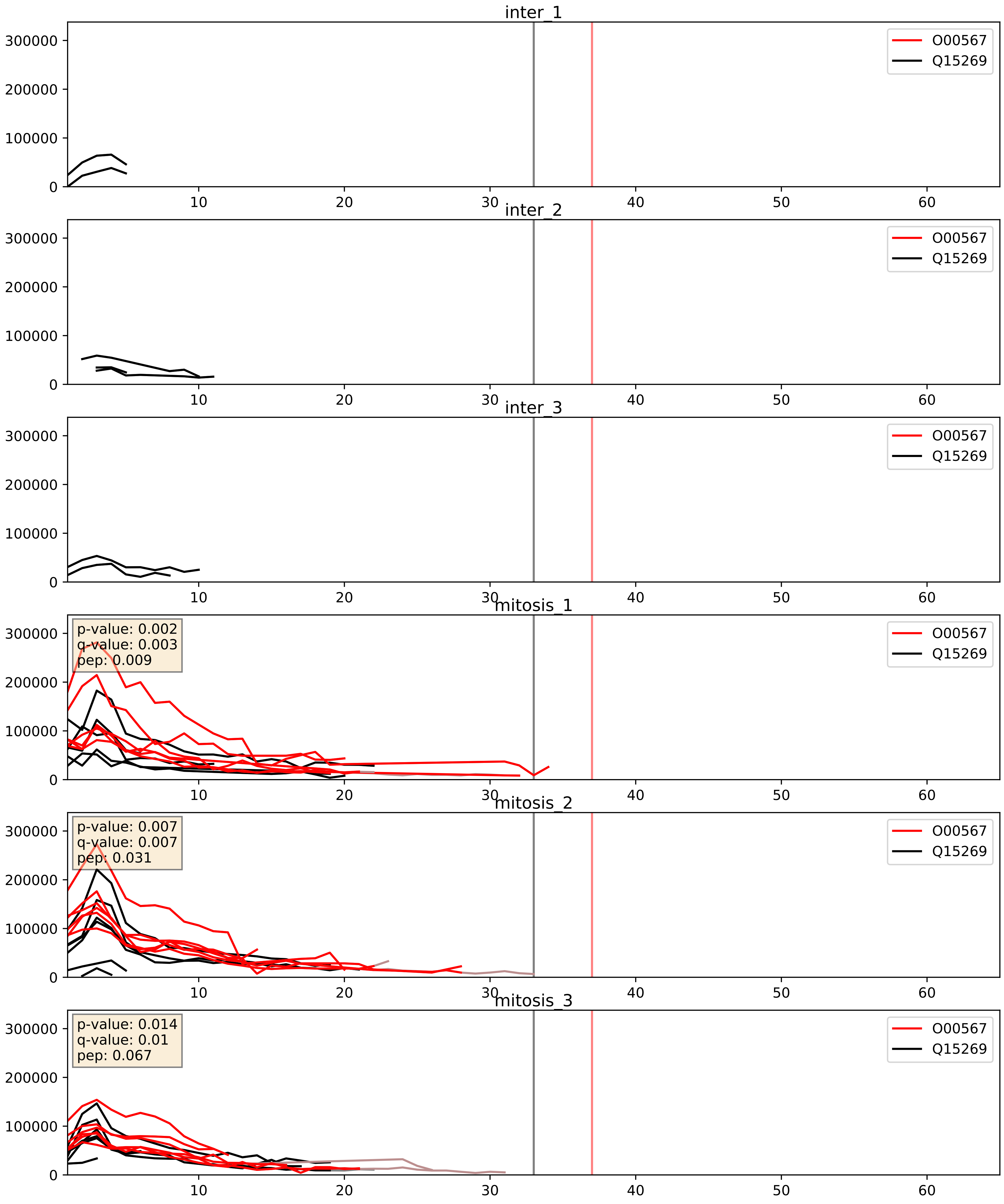


| level | condition_1 | condition_2 | pvalue | pvalue_adjusted |
|---|---|---|---|---|
| complex_abundance | mitosis | inter | 0.0010352101406899857 | 0.021560581032375372 |
| interactor_abundance | mitosis | inter | 0.0011045983974246115 | 0.024049720606162197 |
| interactor_ratio | mitosis | inter | 0.008196470592608973 | 0.12756688776860511 |

O00567_Q15397
NOP56_HUMAN vs PUM3_HUMAN
p-value: 0.045 q-value: 0.03

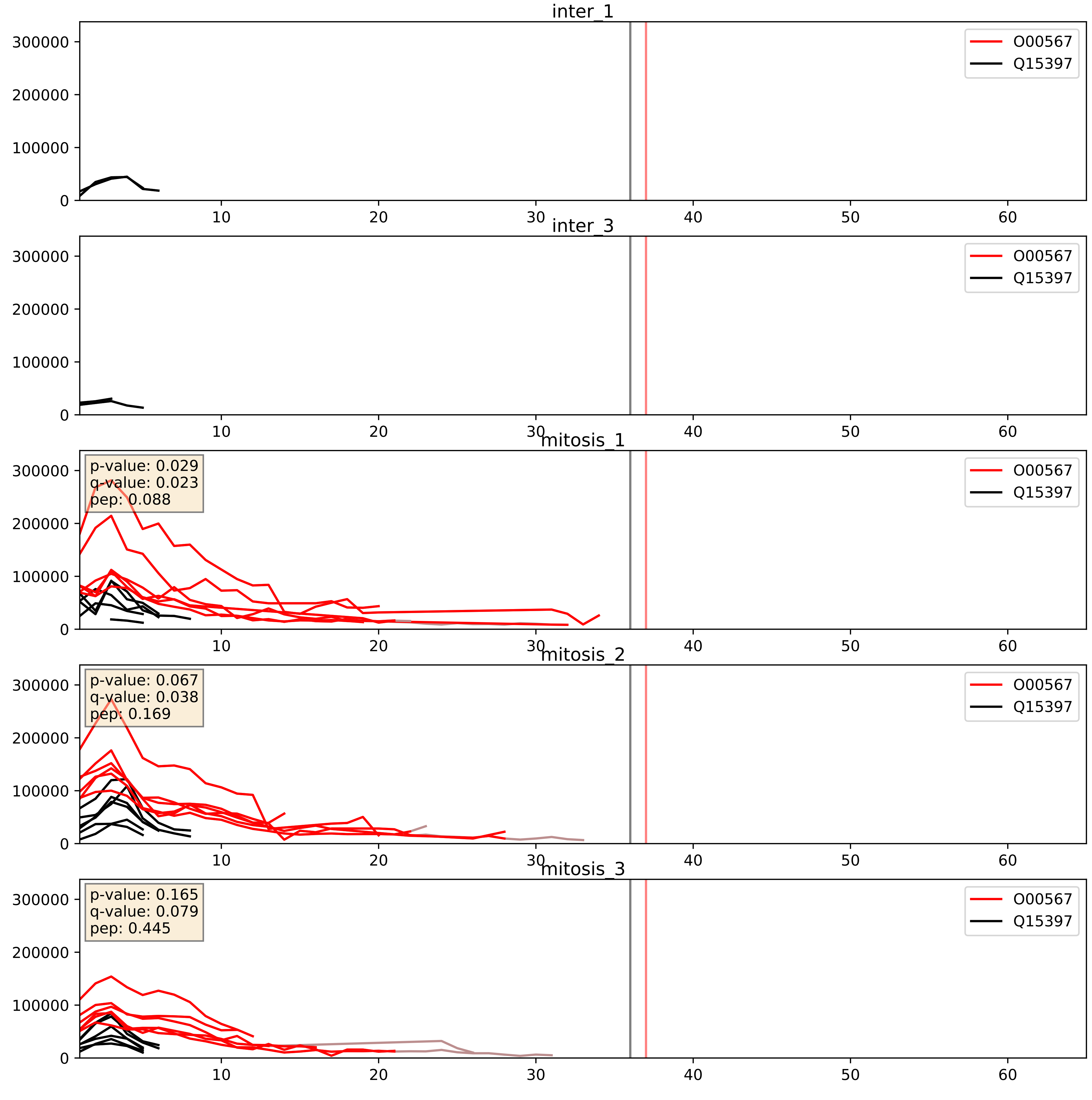


| level | condition_1 | condition_2 | pvalue | pvalue_adjusted |
|---|---|---|---|---|
| interactor_ratio | mitosis | inter | 0.0006892251594627307 | 0.030255217256415255 |
| interactor_abundance | mitosis | inter | 0.00467469140185995 | 0.05557688666655718 |
| complex_abundance | mitosis | inter | 0.008906695963925746 | 0.07001039251717574 |

O00567_Q5JTH9
NOP56_HUMAN vs RRP12_HUMAN
p-value: 0.002 q-value: 0.03

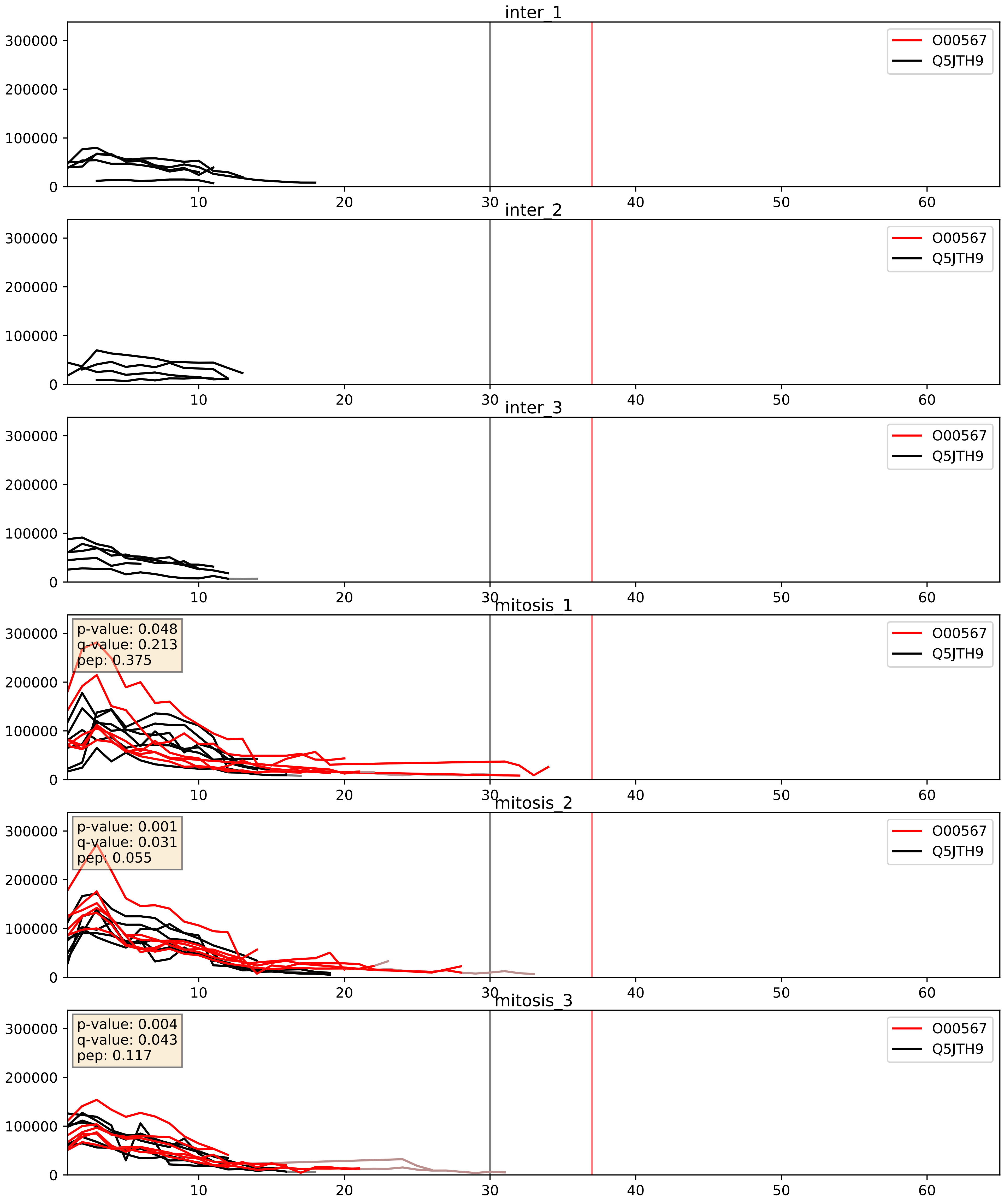


| level | condition_1 | condition_2 | pvalue | pvalue_adjusted |
|---|---|---|---|---|
| complex_abundance | mitosis | inter | 0.0007100535285470436 | 0.017316405140634452 |
| interactor_abundance | mitosis | inter | 0.0008734408149987851 | 0.021954049575556266 |
| interactor_ratio | mitosis | inter | 0.11524458091398708 | 0.5598715168125593 |

O00567_Q8IY81
NOP56_HUMAN vs SPB1_HUMAN
p-value: 0.01 q-value: 0.009

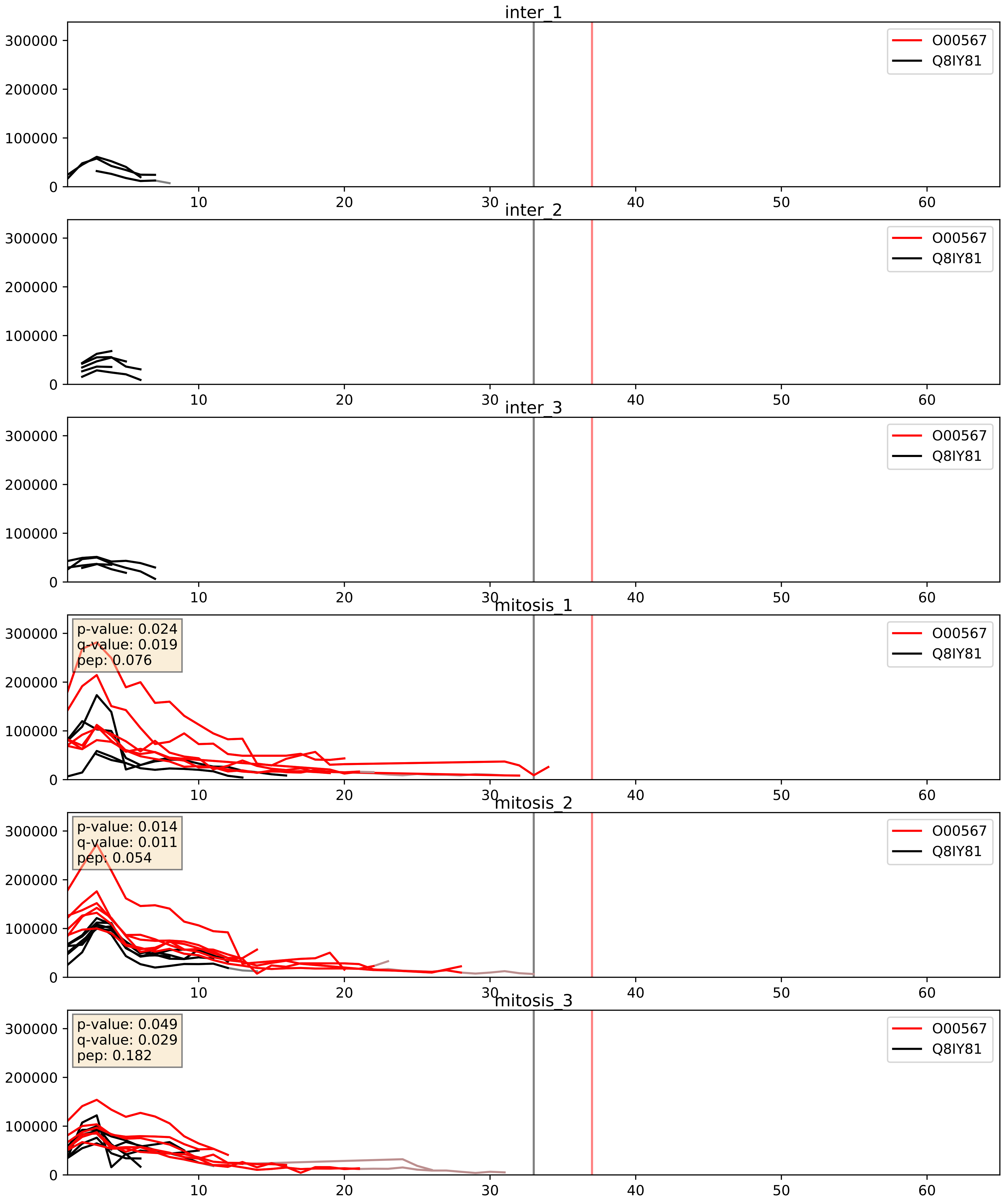


| level | condition_1 | condition_2 | pvalue | pvalue_adjusted |
|---|---|---|---|---|
| interactor_abundance | mitosis | inter | 0.0015635866039636899 | 0.029025269221018554 |
| complex_abundance | mitosis | inter | 0.001688865399875853 | 0.028927157402433075 |
| interactor_ratio | mitosis | inter | 0.010319605669621772 | 0.146251365119143 |

O00567_Q8NI36
NOP56_HUMAN vs WDR36_HUMAN
p-value: 0.005 q-value: 0.005

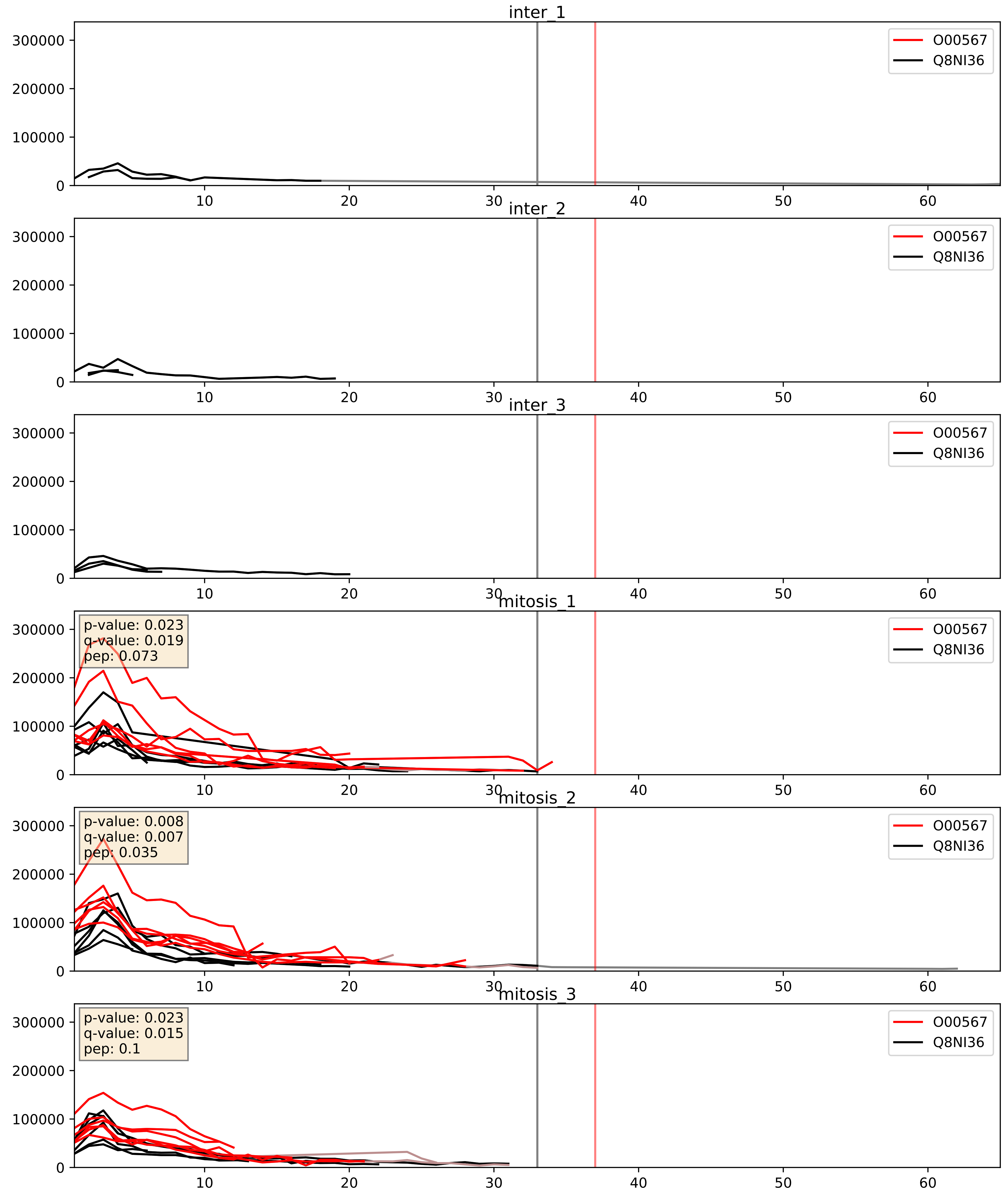


| level | condition_1 | condition_2 | pvalue | pvalue_adjusted |
|---|---|---|---|---|
| complex_abundance | mitosis | inter | 0.0007499876579213406 | 0.017895661700548352 |
| interactor_abundance | mitosis | inter | 0.0011052779688176132 | 0.024049720606162197 |
| interactor_ratio | mitosis | inter | 0.010958138448517993 | 0.15054322834266998 |

O00567_Q8TED0
NOP56_HUMAN vs UTP15_HUMAN
p-value: 0.05 q-value: 0.033

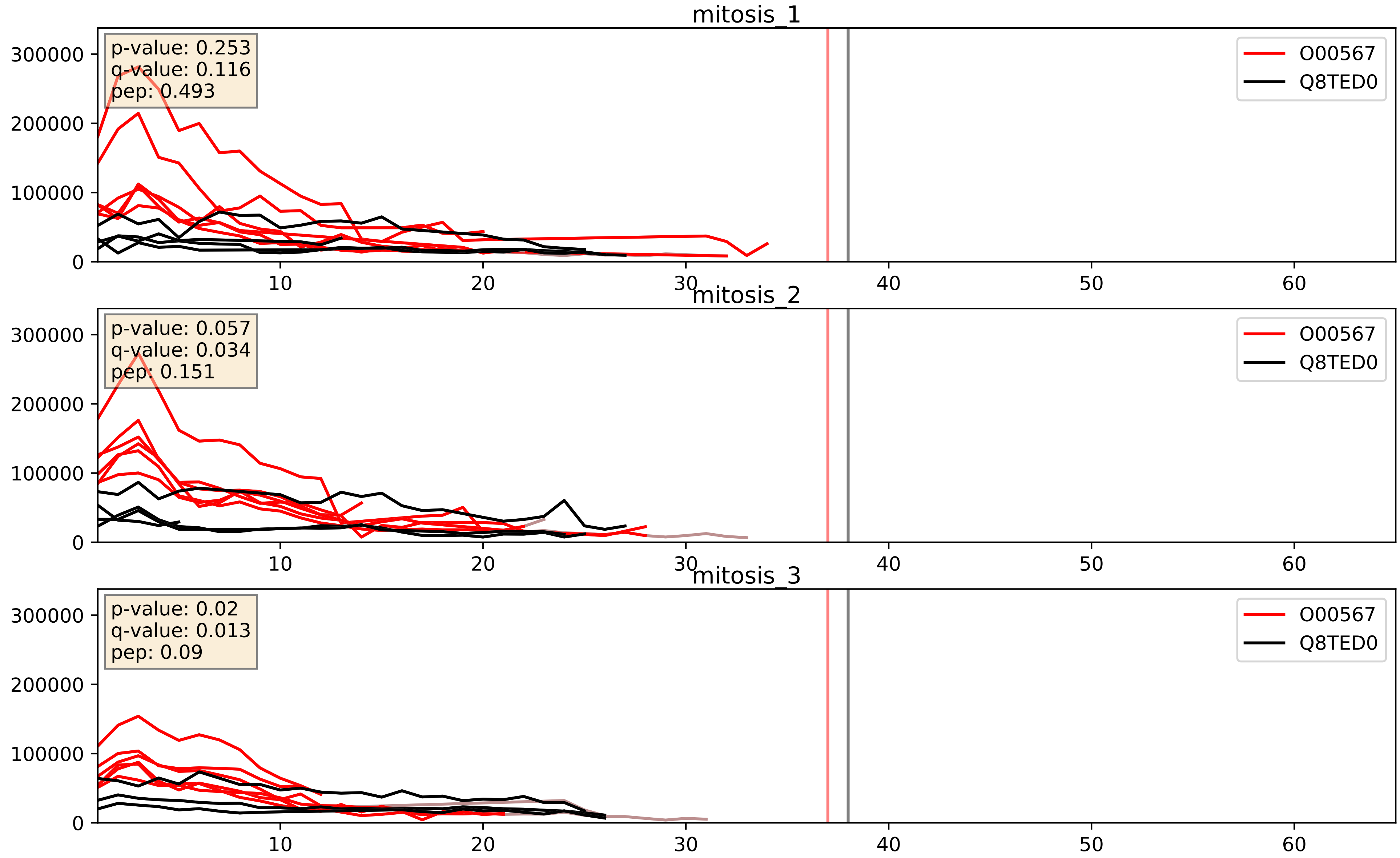


| level | condition_1 | condition_2 | pvalue | pvalue_adjusted |
|---|---|---|---|---|
| complex_abundance | mitosis | inter | 0.0008768577861510543 | 0.019513211095311817 |
| interactor_abundance | mitosis | inter | 0.0010560524122632741 | 0.024049720606162197 |
| interactor_ratio | mitosis | inter | 0.0075917466514874325 | 0.12238295920288592 |

O00567_Q92979
NOP56_HUMAN vs NEP1_HUMAN
p-value: 0.068 q-value: 0.042

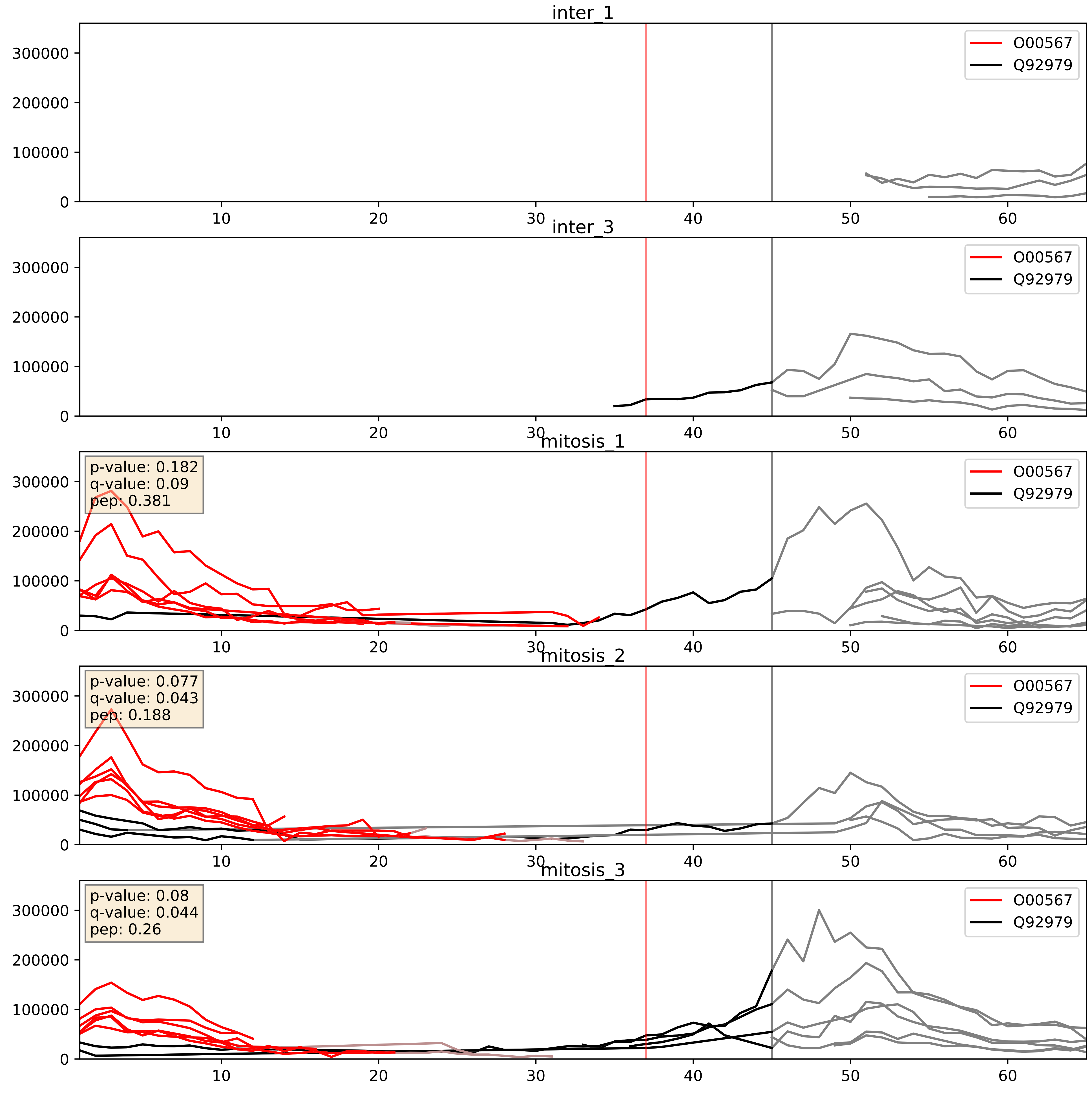


| level | condition_1 | condition_2 | pvalue | pvalue_adjusted |
|---|---|---|---|---|
| interactor_ratio | mitosis | inter | 0.00021846409665752416 | 0.015168670877391028 |
| interactor_abundance | mitosis | inter | 0.0013322765484747097 | 0.026992395869688792 |
| complex_abundance | mitosis | inter | 0.0060296068177722595 | 0.05806974312565789 |

O00567_Q969X6
NOP56_HUMAN vs UTP4_HUMAN
p-value: 0.006 q-value: 0.006

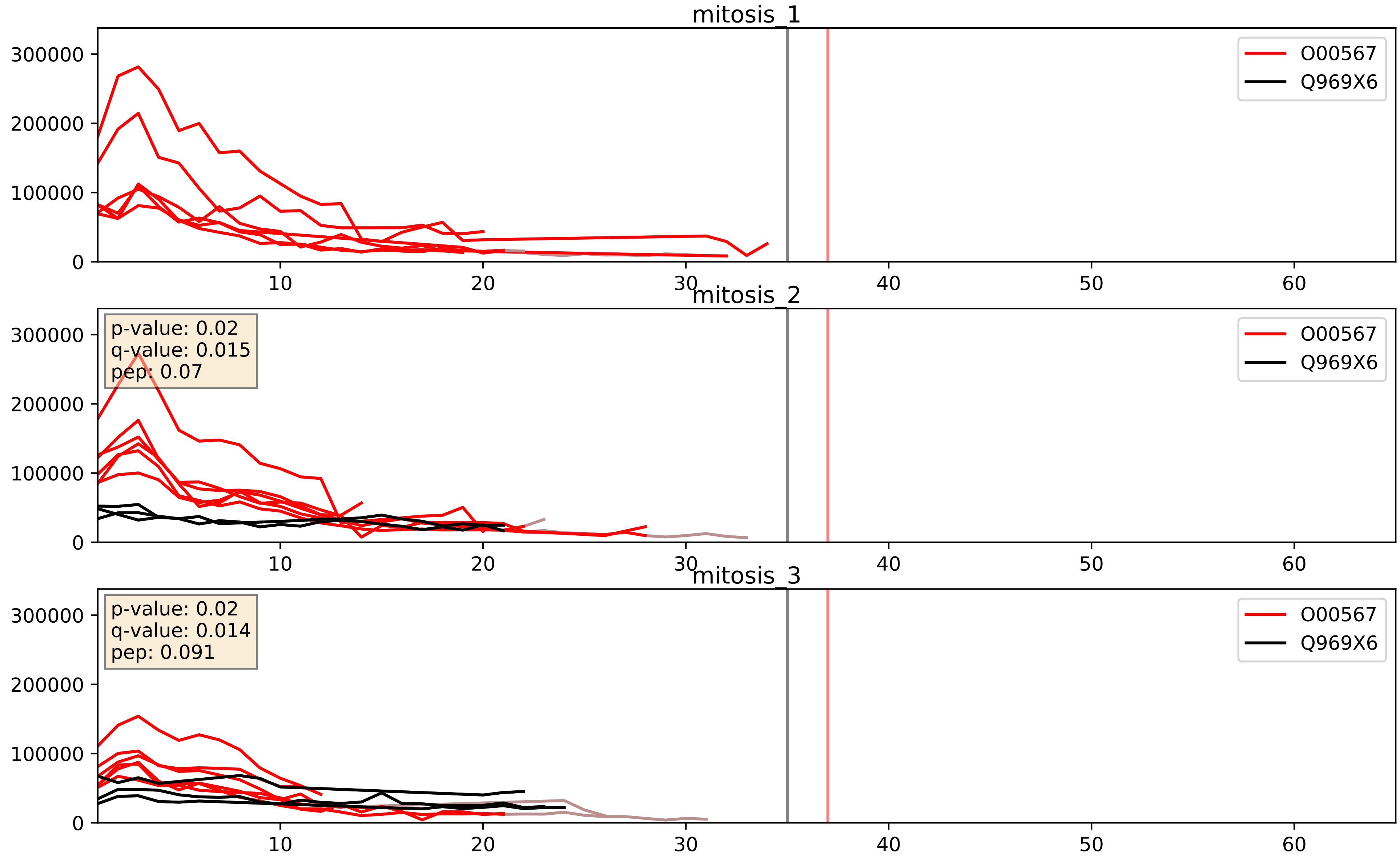


| level | condition_1 | condition_2 | pvalue | pvalue_adjusted |
|---|---|---|---|---|
| interactor_ratio | mitosis | inter | 0.1635008352446056 | 0.6234152114449104 |
| interactor_abundance | mitosis | inter | 0.17640681945440545 | 0.367125475823002 |
| complex_abundance | mitosis | inter | 0.1962735824510143 | 0.348713546239245 |

O00567_Q96GQ7
NOP56_HUMAN vs DDX27_HUMAN
p-value: 0.036 q-value: 0.025

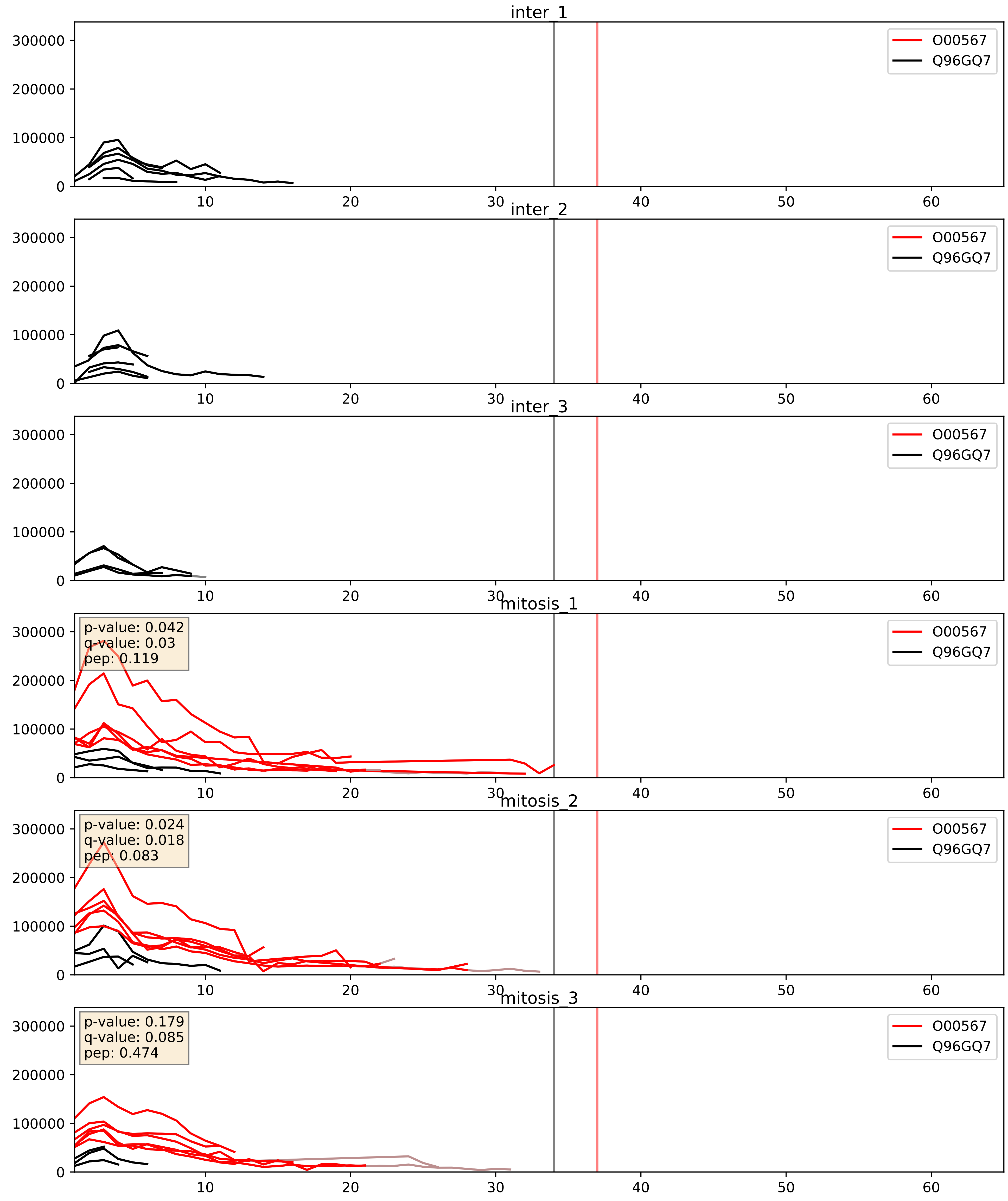


| level | condition_1 | condition_2 | pvalue | pvalue_adjusted |
|---|---|---|---|---|
| interactor_ratio | mitosis | inter | 0.0007576191550616581 | 0.03178508799244958 |
| interactor_abundance | mitosis | inter | 0.007061028371252754 | 0.07263199229395266 |
| complex_abundance | mitosis | inter | 0.02076315597200204 | 0.1061724104661514 |

O00567_Q9BVP2
NOP56_HUMAN vs GNL3_HUMAN
p-value: 0.0 q-value: 0.013

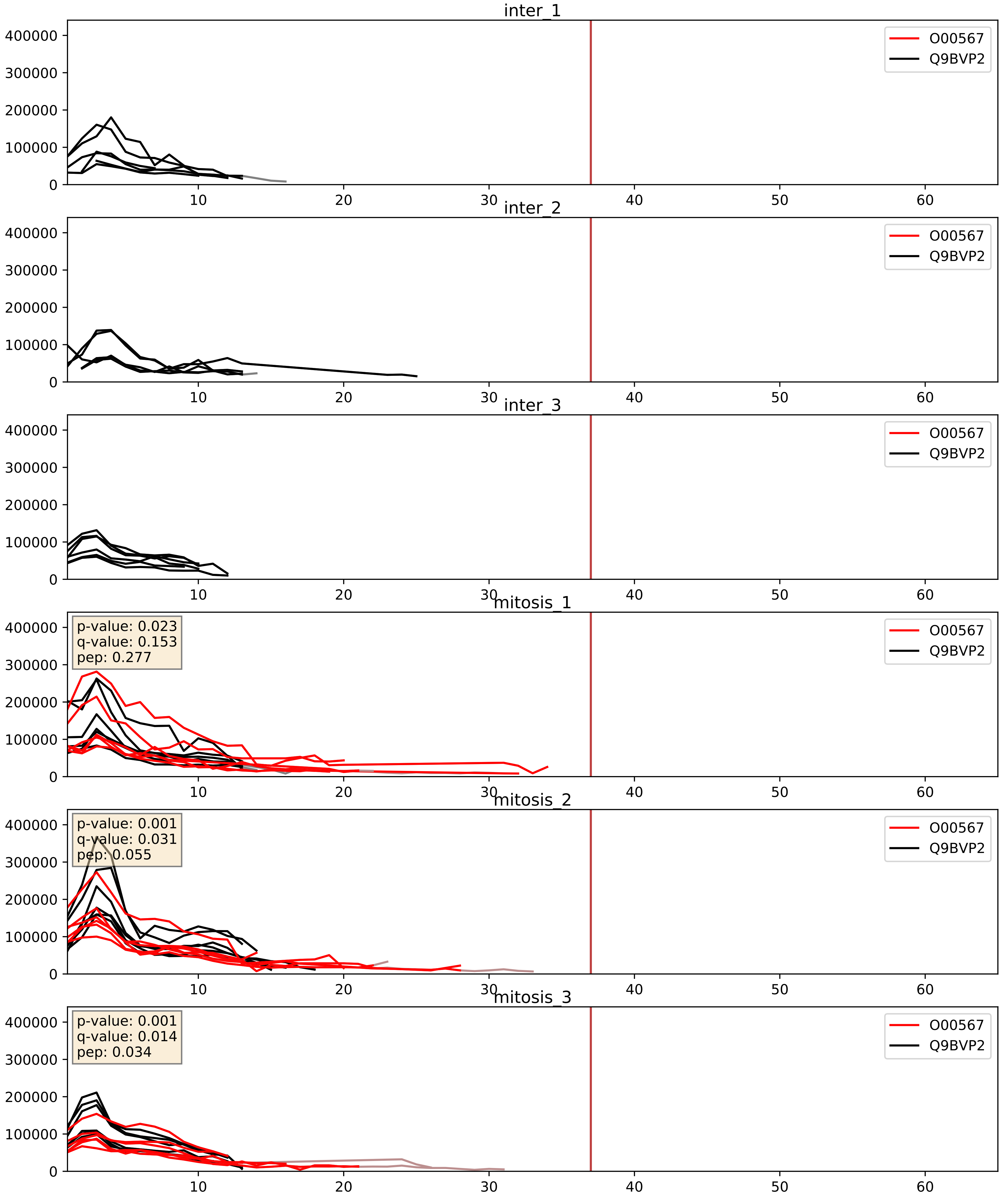


| level | condition_1 | condition_2 | pvalue | pvalue_adjusted |
|---|---|---|---|---|
| complex_abundance | mitosis | inter | 0.0003457116354080251 | 0.012015488415248196 |
| interactor_abundance | mitosis | inter | 0.0008877885552668727 | 0.02197564842852207 |
| interactor_ratio | mitosis | inter | 0.22699272553855118 | 0.7059145275954049 |

O00567_Q9BYG3
NOP56_HUMAN vs MK67I_HUMAN
p-value: 0.03 q-value: 0.022

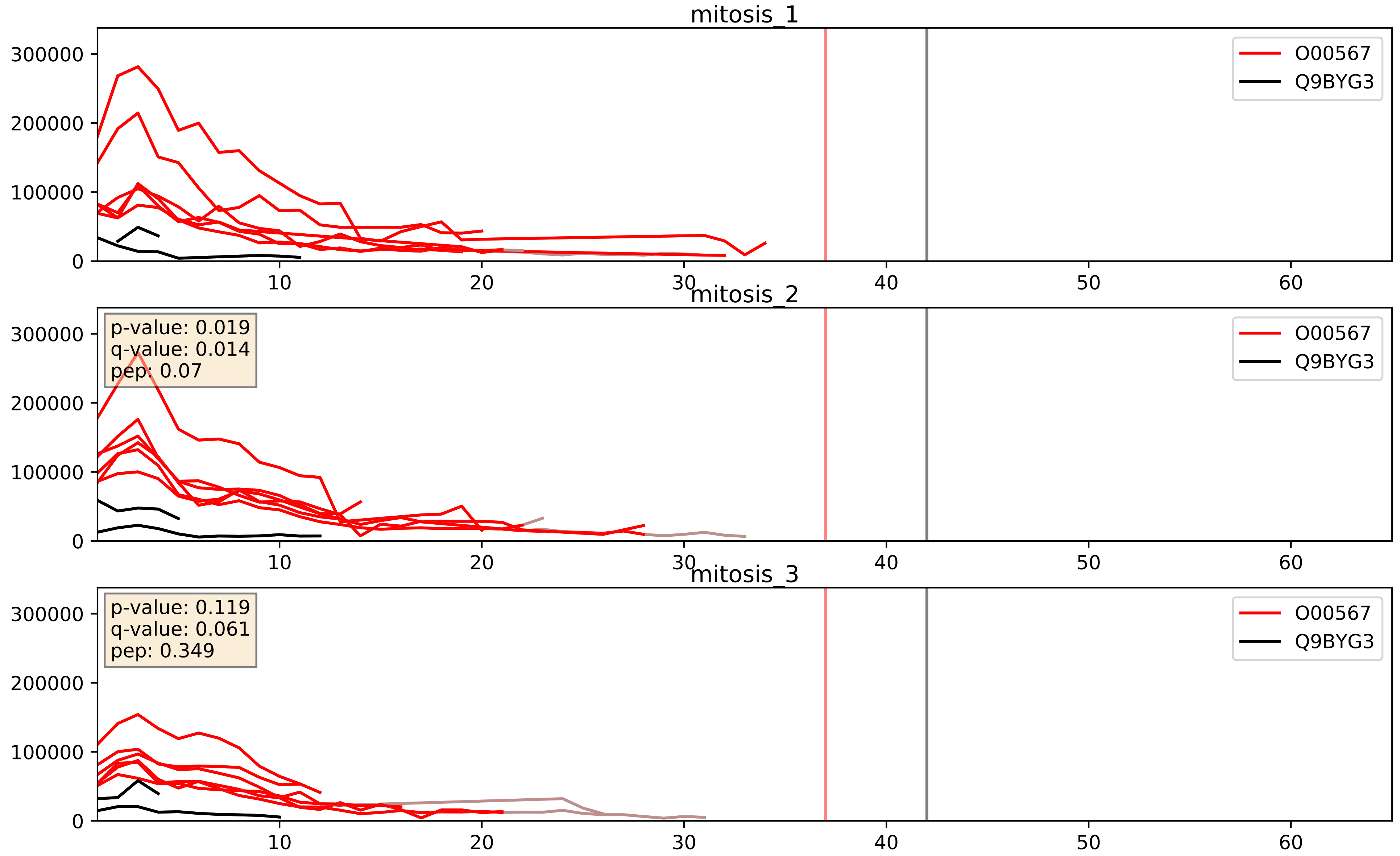


| level | condition_1 | condition_2 | pvalue | pvalue_adjusted |
|---|---|---|---|---|
| interactor_ratio | mitosis | inter | 0.00025624310259385303 | 0.016247710801506533 |
| interactor_abundance | mitosis | inter | 0.000695734930284271 | 0.019073880342257484 |
| complex_abundance | mitosis | inter | 0.002507942943970339 | 0.03638642644133238 |

O00567_Q9H0A0
NOP56_HUMAN vs NAT10_HUMAN
p-value: 0.005 q-value: 0.005

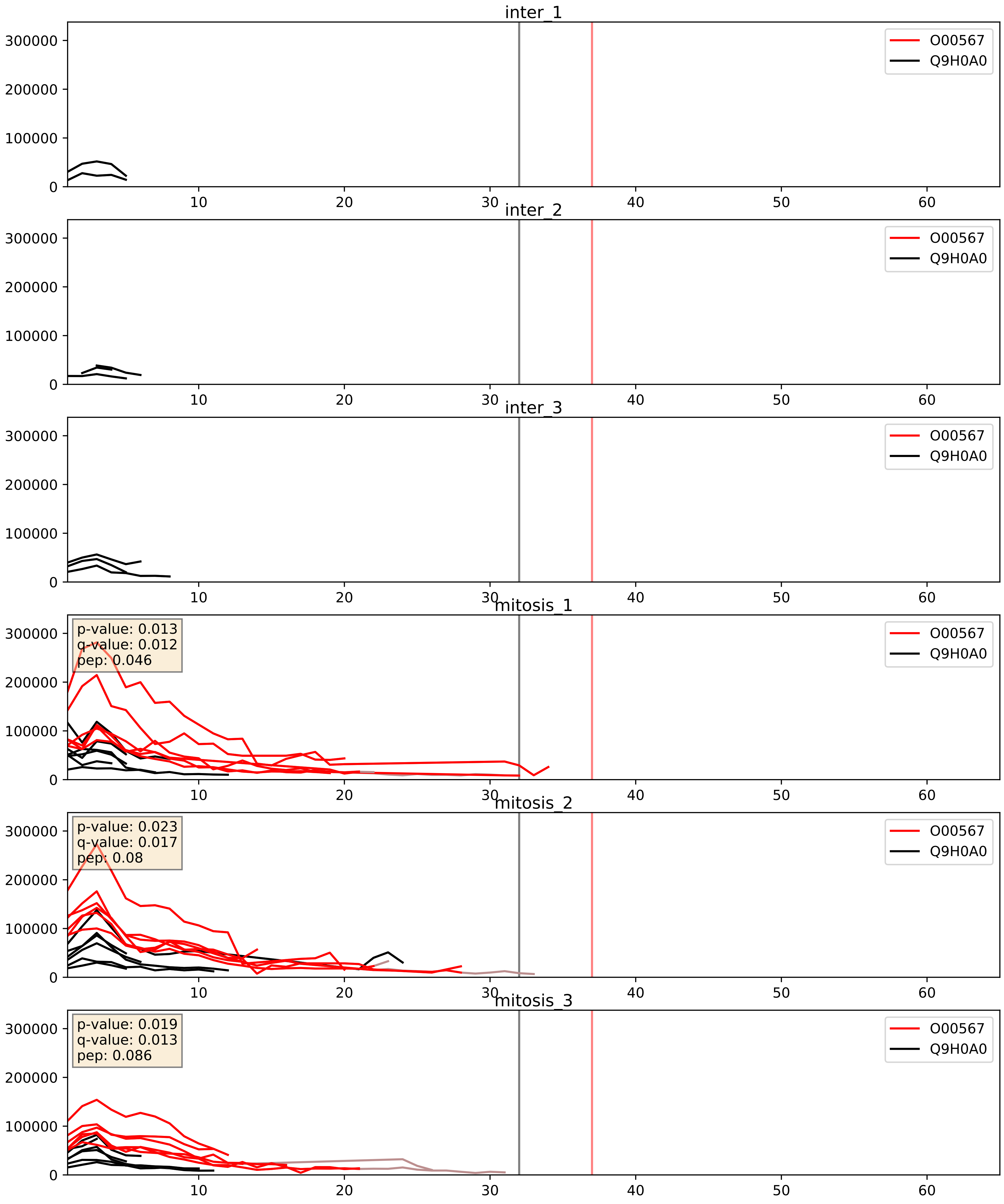


| level | condition_1 | condition_2 | pvalue | pvalue_adjusted |
|---|---|---|---|---|
| interactor_ratio | mitosis | inter | 2.3043765380856052e-08 | 0.0001972546316601278 |
| interactor_abundance | mitosis | inter | 0.0009620251184093979 | 0.02306454775731039 |
| complex_abundance | mitosis | inter | 0.0035552641482872074 | 0.043600374082146835 |

O00567_Q9H0S4
NOP56_HUMAN vs DDX47_HUMAN
p-value: 0.01 q-value: 0.009

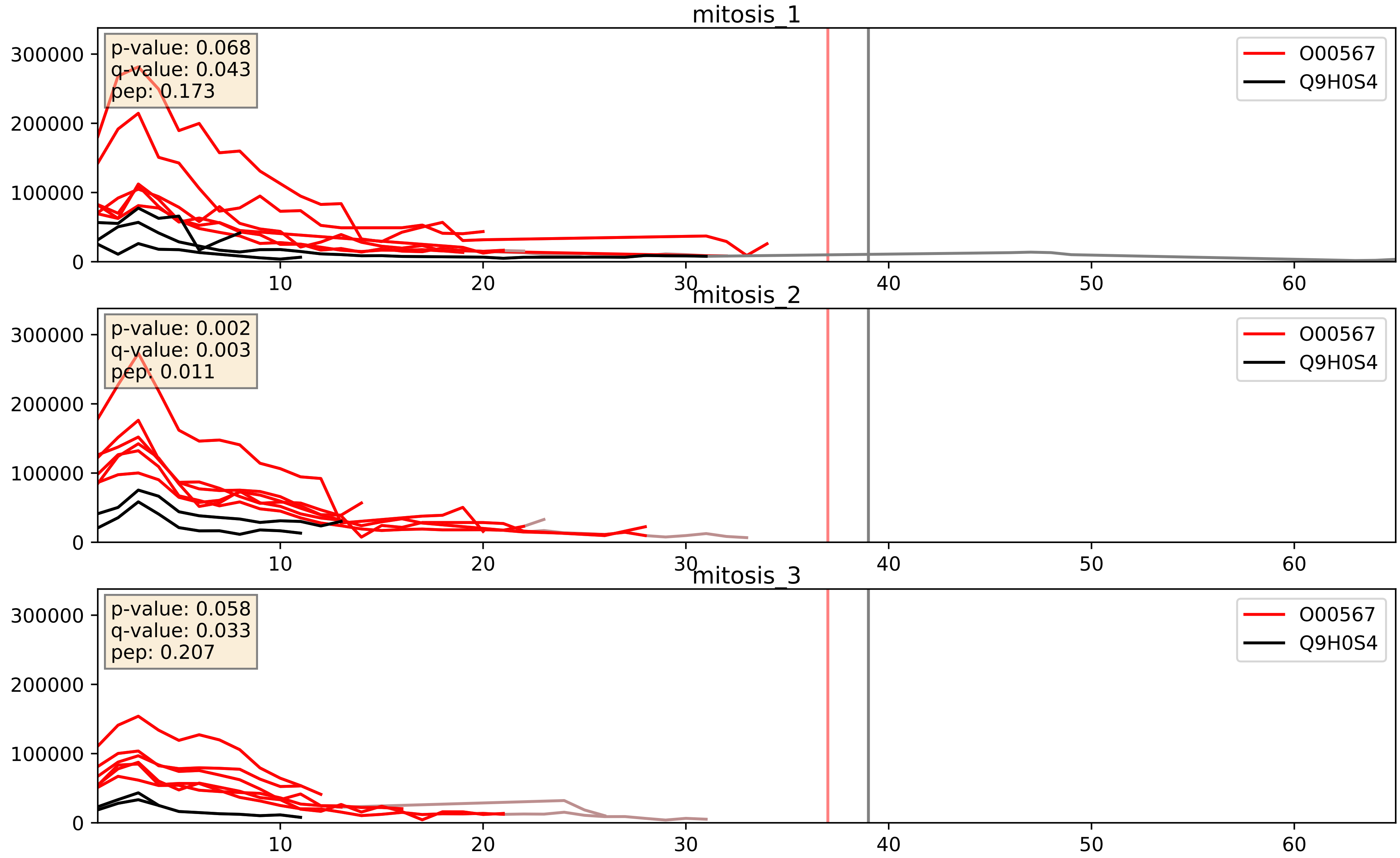


| level | condition_1 | condition_2 | pvalue | pvalue_adjusted |
|---|---|---|---|---|
| interactor_ratio | mitosis | inter | 0.0001712185653981663 | 0.013577643810159747 |
| interactor_abundance | mitosis | inter | 0.0014024757199137346 | 0.02753913185123736 |
| complex_abundance | mitosis | inter | 0.01641041278178478 | 0.09575537383236381 |

O00567_Q9H583
NOP56_HUMAN vs HEAT1_HUMAN
p-value: 0.012 q-value: 0.01

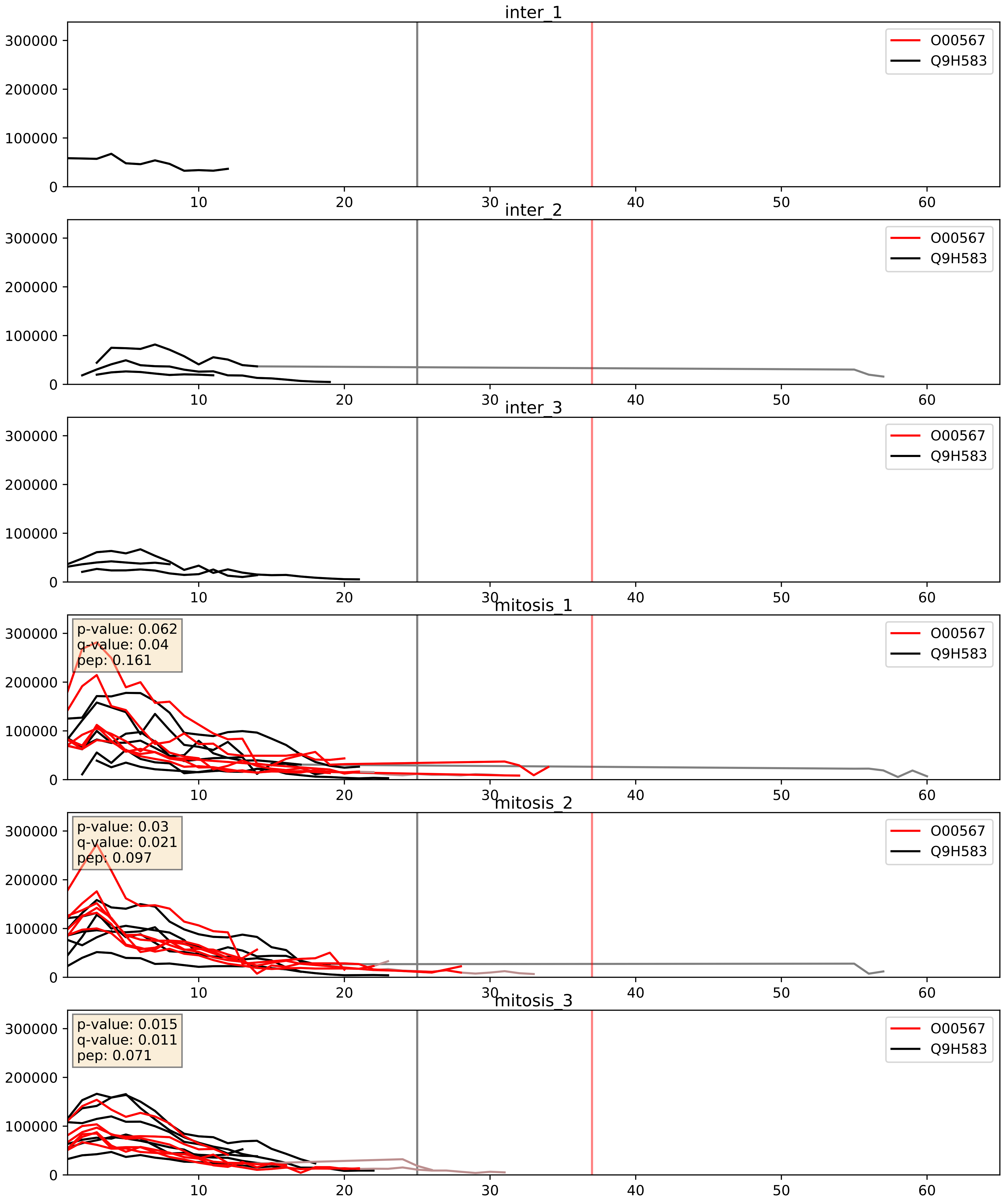


| level | condition_1 | condition_2 | pvalue | pvalue_adjusted |
|---|---|---|---|---|
| complex_abundance | mitosis | inter | 0.00016025150516764934 | 0.008563834848633118 |
| interactor_abundance | mitosis | inter | 0.001056251194238277 | 0.024049720606162197 |
| interactor_ratio | mitosis | inter | 0.8485851427407834 | 0.9755634896635457 |

O00567_Q9H6R4
NOP56_HUMAN vs NOL6_HUMAN
p-value: 0.045 q-value: 0.03

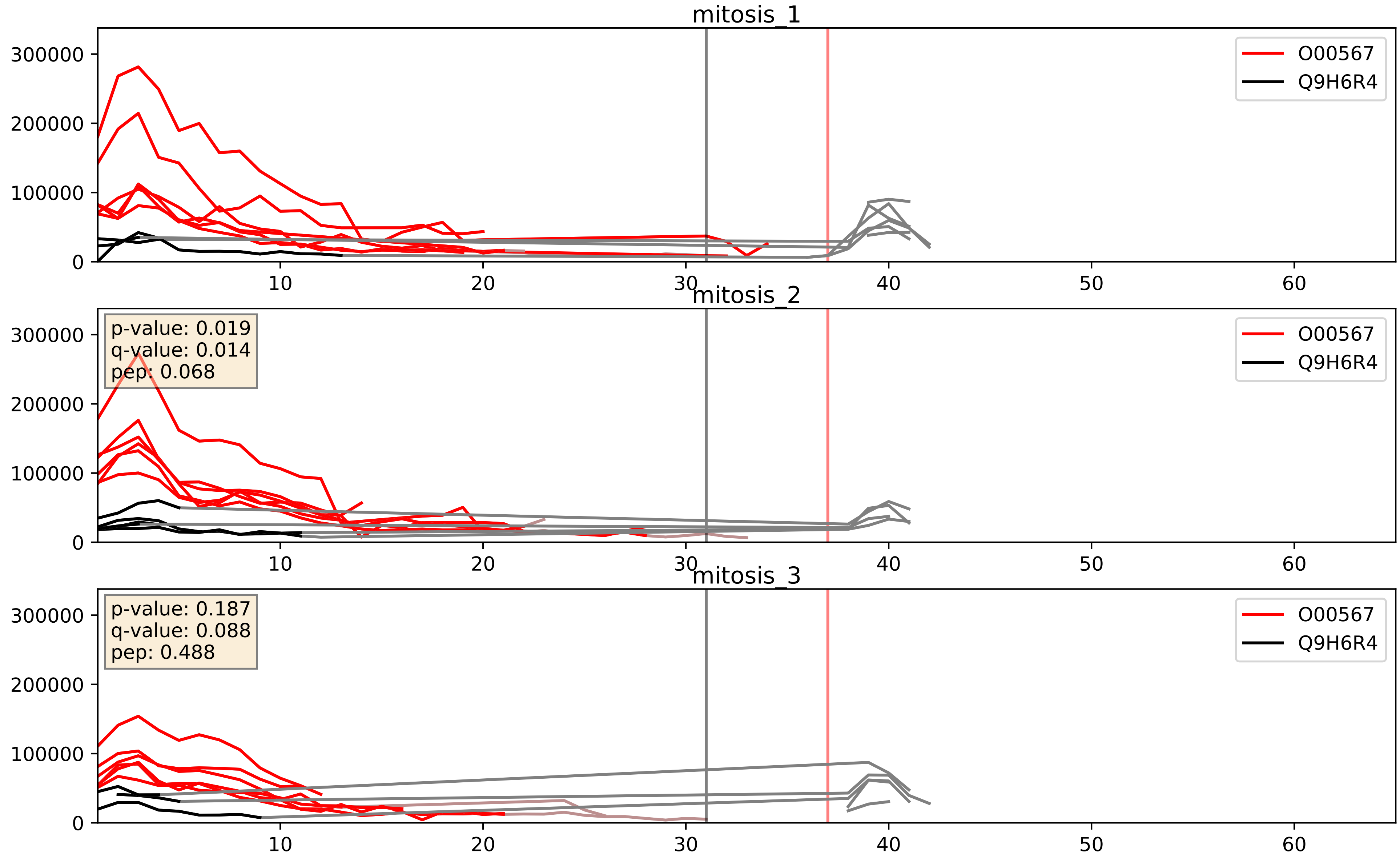


| level | condition_1 | condition_2 | pvalue | pvalue_adjusted |
|---|---|---|---|---|
| interactor_ratio | mitosis | inter | 0.0013886285934761222 | 0.044212523606194314 |
| interactor_abundance | mitosis | inter | 0.0019094235057377626 | 0.031986606092438766 |
| complex_abundance | mitosis | inter | 0.006619064498898424 | 0.06002696841014766 |

O00567_Q9NX24
NOP56_HUMAN vs NHP2_HUMAN
p-value: 0.008 q-value: 0.008

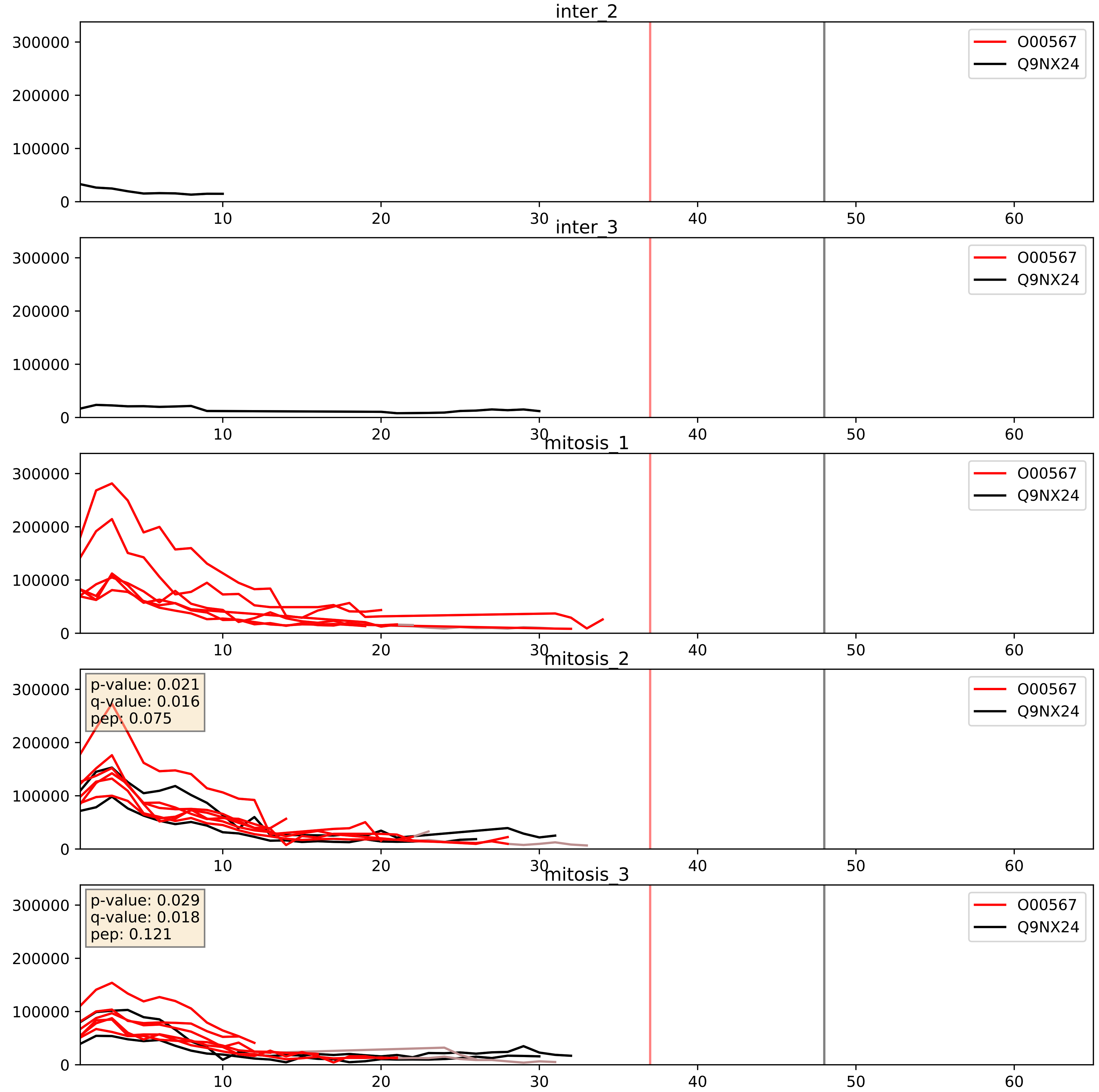


| level | condition_1 | condition_2 | pvalue | pvalue_adjusted |
|---|---|---|---|---|
| interactor_ratio | mitosis | inter | 0.15819140191817735 | 0.6144687163742579 |
| interactor_abundance | mitosis | inter | 0.1764432485439054 | 0.367125475823002 |
| complex_abundance | mitosis | inter | 0.18218672033295358 | 0.3330880662217178 |

O00567_Q9NY93
NOP56_HUMAN vs DDX56_HUMAN
p-value: 0.032 q-value: 0.023

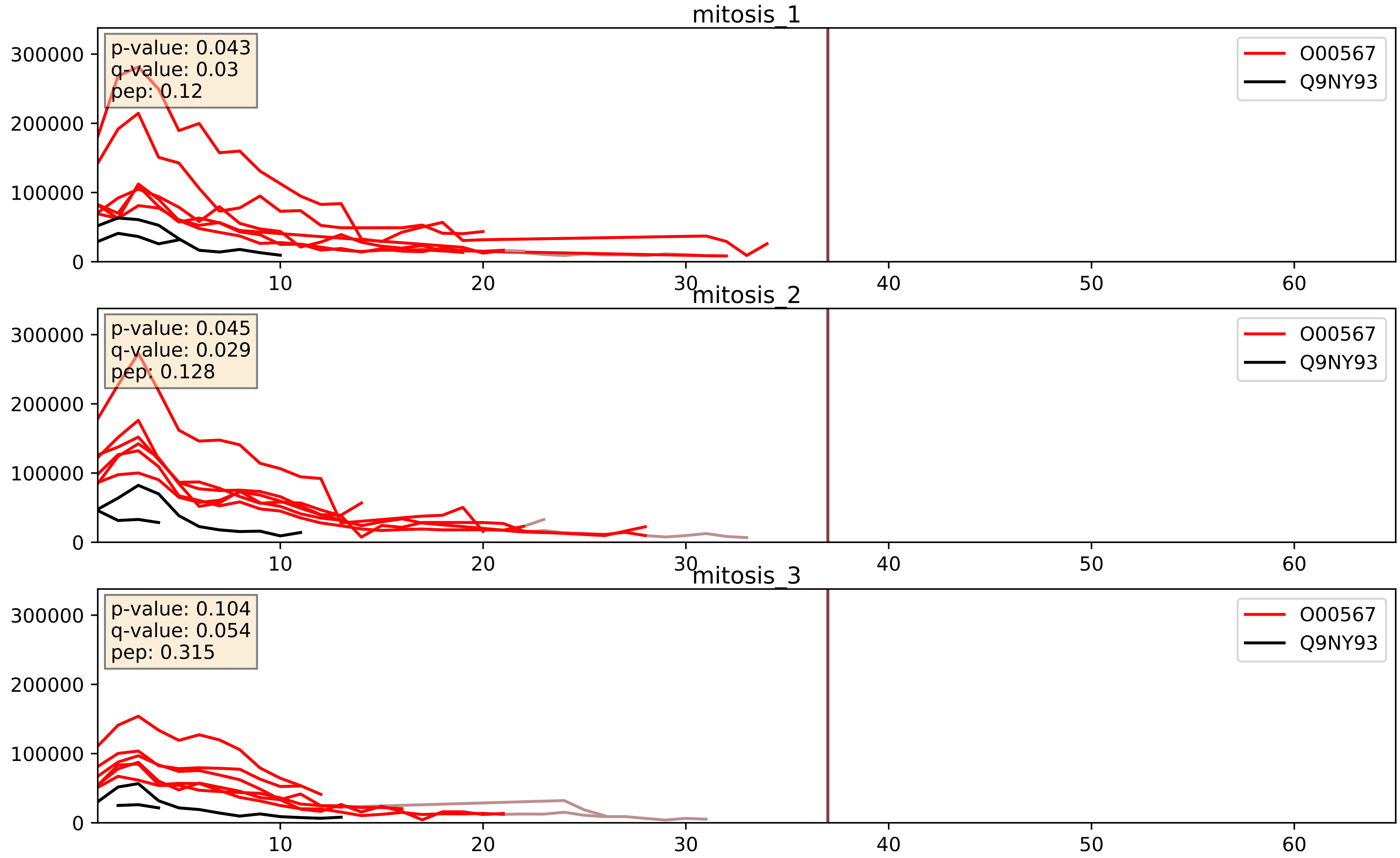


| level | condition_1 | condition_2 | pvalue | pvalue_adjusted |
|---|---|---|---|---|
| interactor_ratio | mitosis | inter | 2.3383393070343584e-05 | 0.004993481478971454 |
| interactor_abundance | mitosis | inter | 0.0005414629468052343 | 0.017071538948997444 |
| complex_abundance | mitosis | inter | 0.0026325520572434646 | 0.03679993741510981 |

O00567_Q9UNX4
NOP56_HUMAN vs WDR3_HUMAN
p-value: 0.017 q-value: 0.014

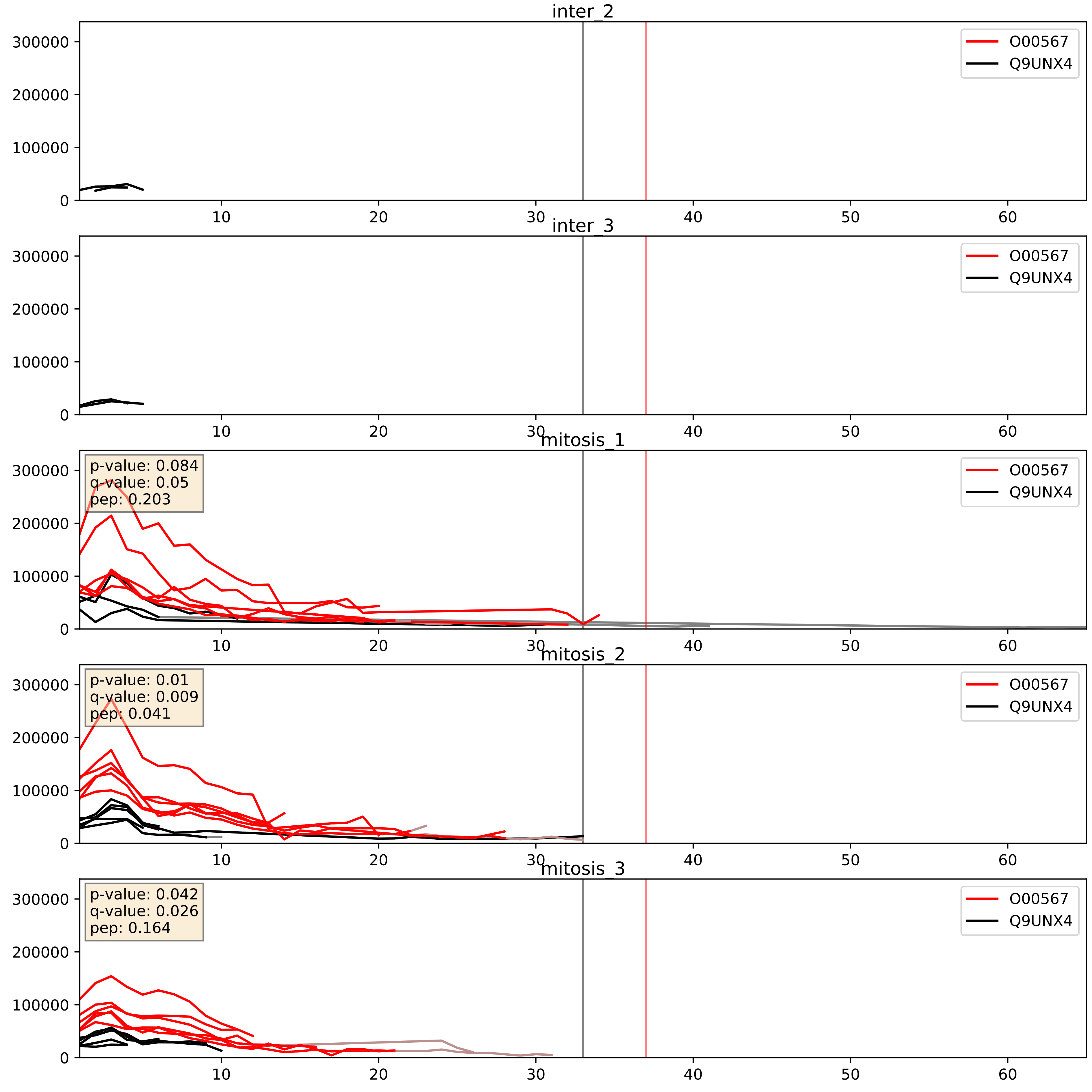


| level | condition_1 | condition_2 | pvalue | pvalue_adjusted |
|---|---|---|---|---|
| interactor_ratio | mitosis | inter | 0.00014655098812599722 | 0.012836278661937291 |
| interactor_abundance | mitosis | inter | 0.001146348816560698 | 0.024325226673177953 |
| complex_abundance | mitosis | inter | 0.003340135899144592 | 0.04242071705738532 |

O00567_Q9Y221
NOP56_HUMAN vs NIP7_HUMAN
p-value: 0.065 q-value: 0.04

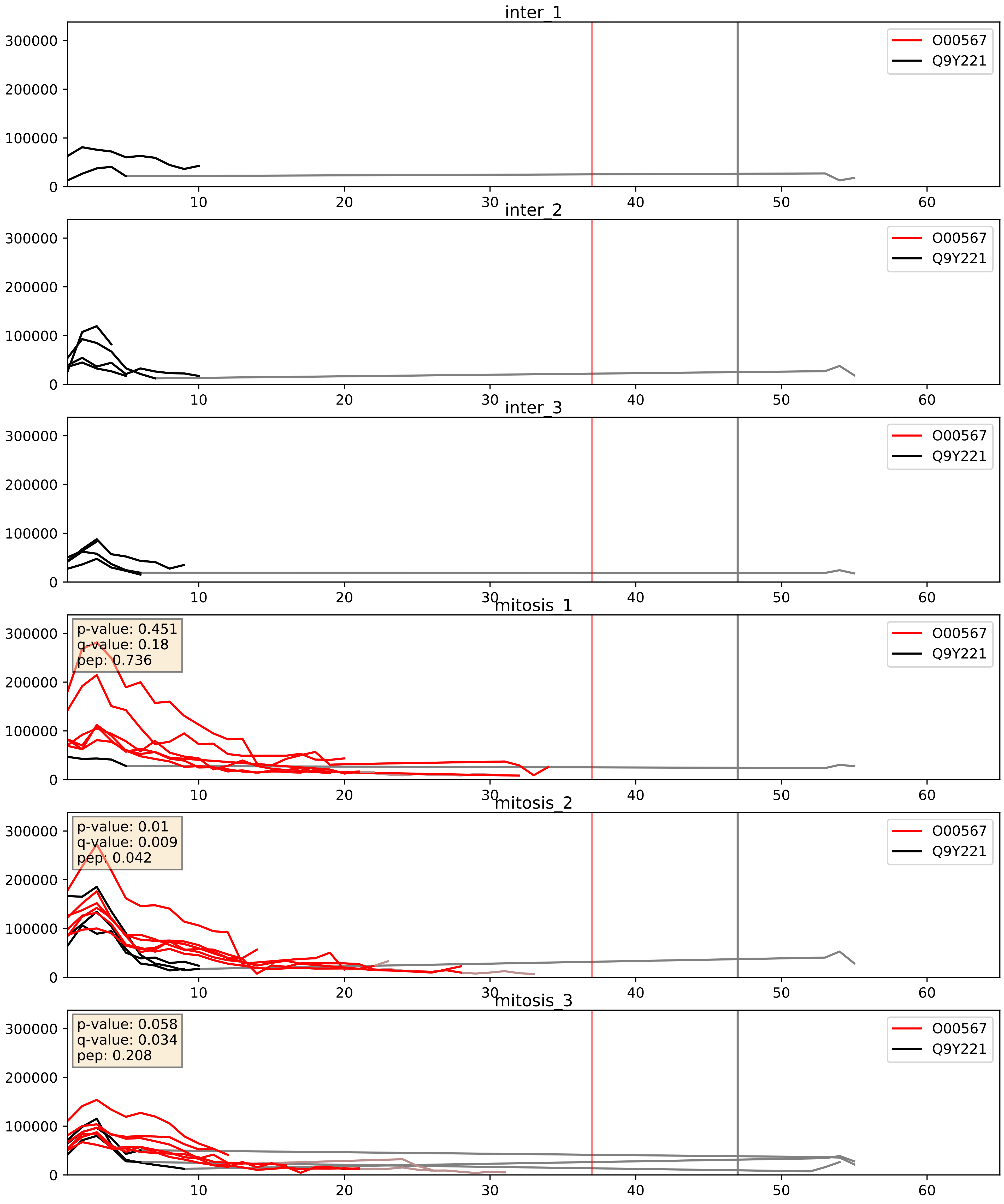


| level | condition_1 | condition_2 | pvalue | pvalue_adjusted |
|---|---|---|---|---|
| interactor_abundance | mitosis | inter | 0.0006422914560265031 | 0.017938058282502015 |
| complex_abundance | mitosis | inter | 0.006912891195033164 | 0.0616199142025868 |
| interactor_ratio | mitosis | inter | 0.026081672786619357 | 0.25543205403291264 |

O00567_Q9Y2X3
NOP56_HUMAN vs NOP58_HUMAN
p-value: 0.001 q-value: 0.001

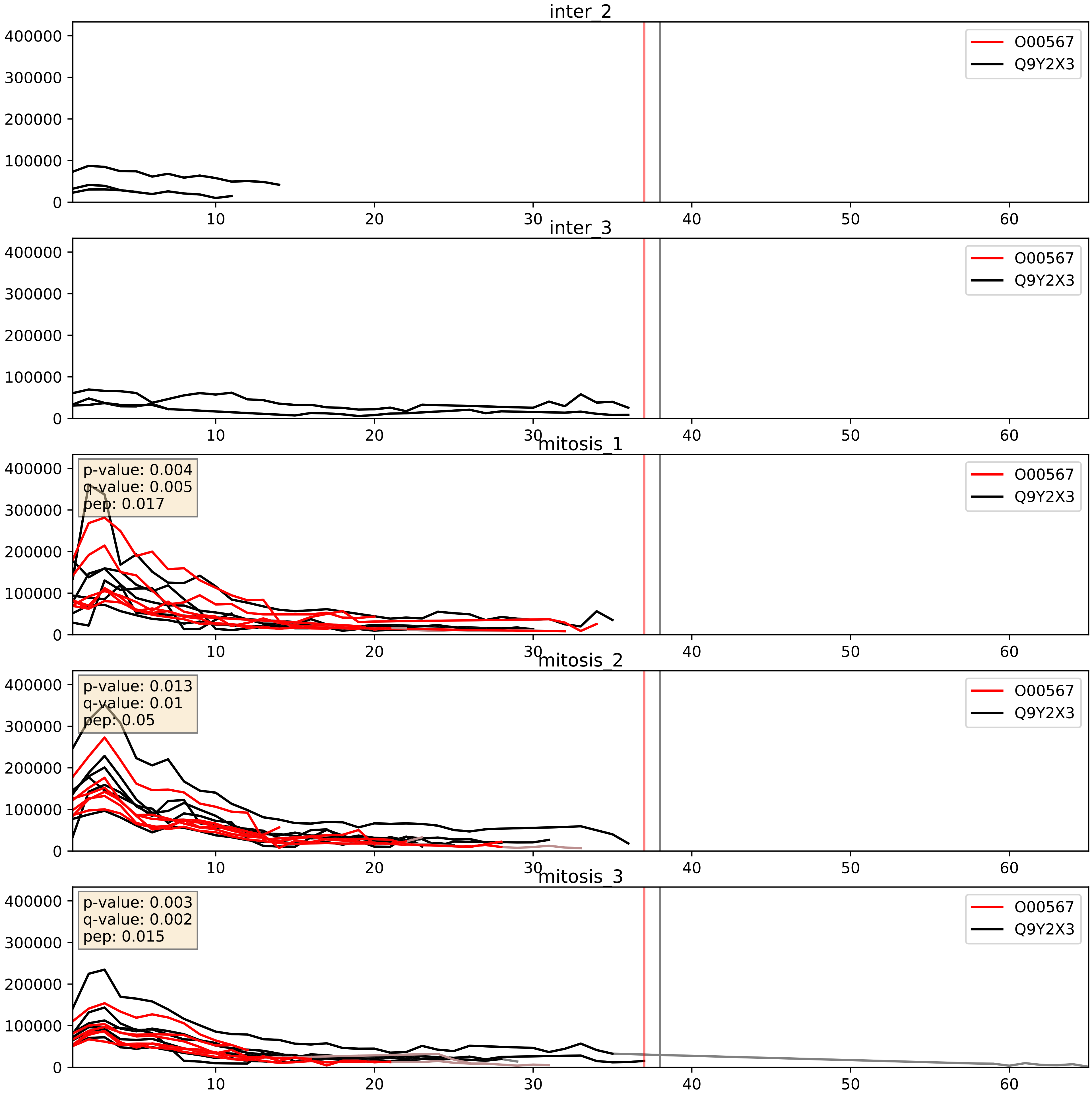


| level | condition_1 | condition_2 | pvalue | pvalue_adjusted |
|---|---|---|---|---|
| complex_abundance | mitosis | inter | 0.00028226689255234446 | 0.011311068978518664 |
| interactor_abundance | mitosis | inter | 0.0011181996263145508 | 0.024049720606162197 |
| interactor_ratio | mitosis | inter | 0.12583682042012595 | 0.5750062399685051 |
