## Supplementary Data 2 for "SECAT: Quantifying differential protein-protein interaction states by network-centric analysis": hela_string_000008_P22087.pdf

P22087_P22087

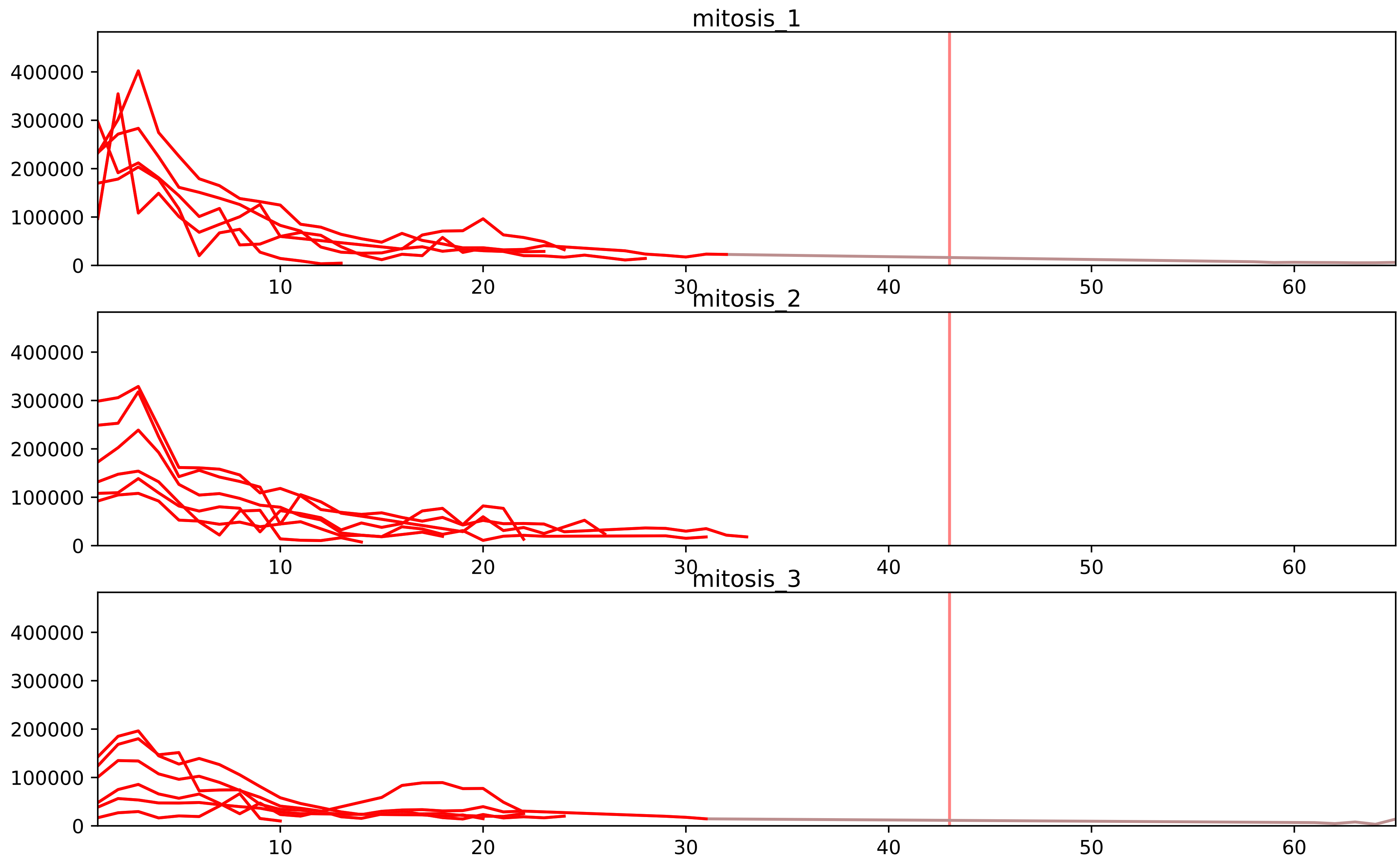


| level | condition_1 | condition_2 | pvalue | pvalue_adjusted |
|---|---|---|---|---|
| interactor_ratio | mitosis | inter | 3.649419048223071e-15 | 8.393663810913063e-13 |
| complex_abundance | mitosis | inter | 3.230032160661462e-07 | 4.245185125440779e-05 |
| interactor_abundance | mitosis | inter | 1.3510226340135832e-06 | 0.00013353840650924488 |
| monomer_abundance | mitosis | inter | 0.07323385102926859 | 0.37224850260048076 |
| assembled_abundance | mitosis | inter | 0.10789205200512232 | 0.38523470284971595 |
| total_abundance | mitosis | inter | 0.1150754859414043 | 0.4209856739593246 |

O00541_P22087
PESC_HUMAN vs FBRL_HUMAN
p-value: 0.073 q-value: 0.044

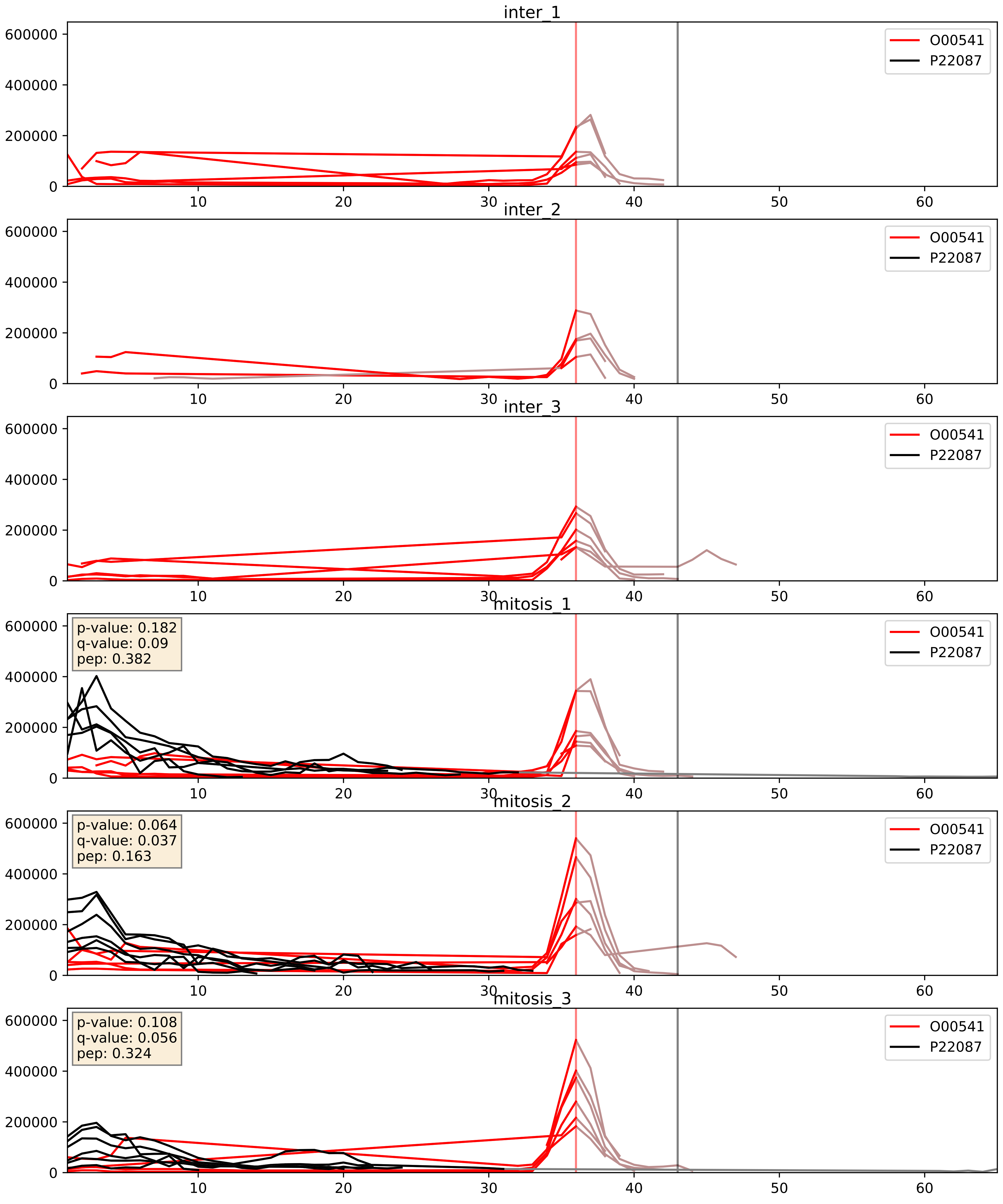


| level | condition_1 | condition_2 | pvalue | pvalue_adjusted |
|---|---|---|---|---|
| interactor_ratio | mitosis | inter | 0.0006227650099035394 | 0.029219626944512057 |
| complex_abundance | mitosis | inter | 0.006526491567282415 | 0.05962301794657148 |
| interactor_abundance | mitosis | inter | 0.1085544053990082 | 0.2833005214071677 |

O00567_P22087
NOP56_HUMAN vs FBRL_HUMAN
p-value: 0.001 q-value: 0.002

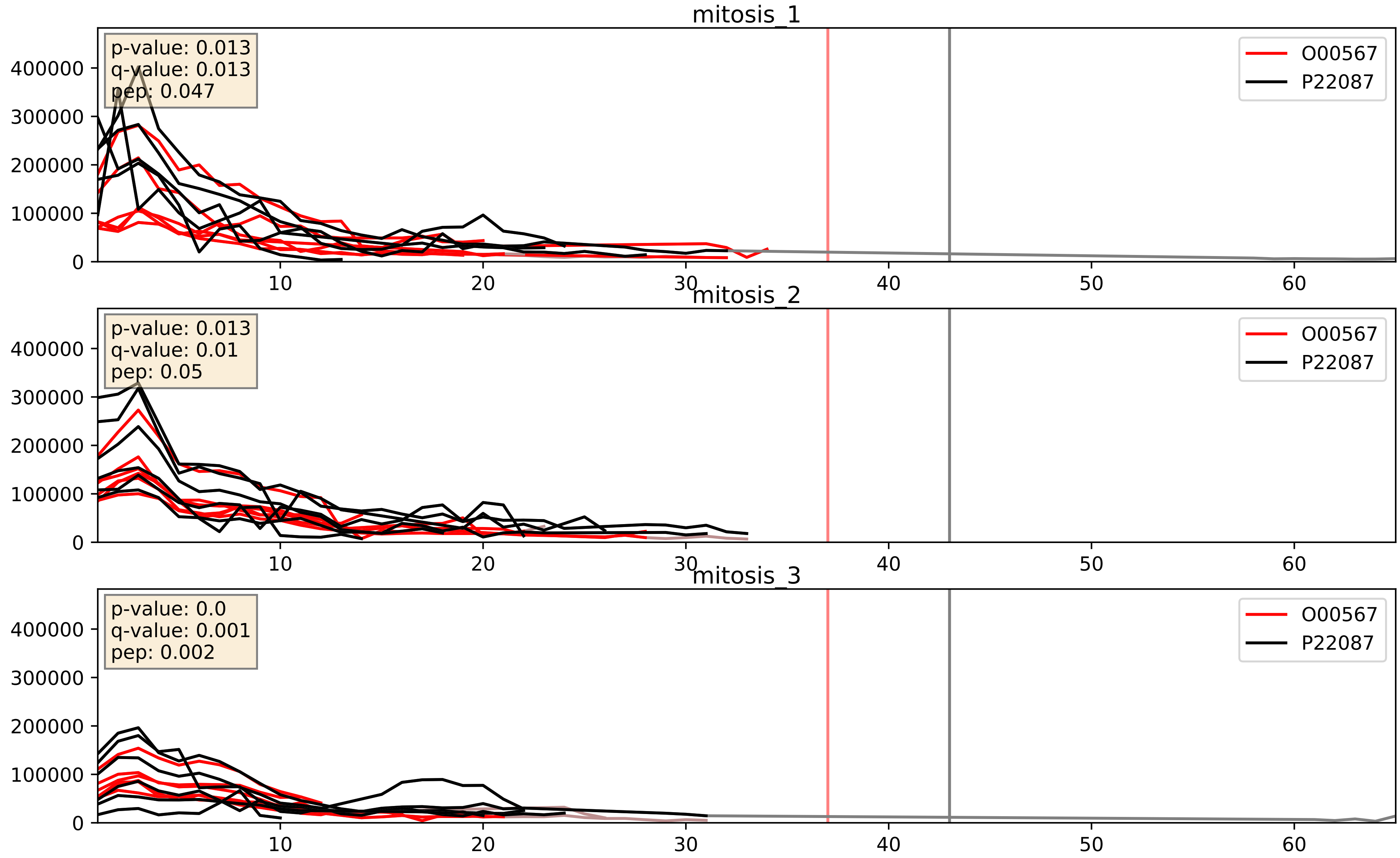


| level | condition_1 | condition_2 | pvalue | pvalue_adjusted |
|---|---|---|---|---|
| interactor_ratio | mitosis | inter | 0.00011298744803945258 | 0.011246192502531558 |
| complex_abundance | mitosis | inter | 0.0006517472043178059 | 0.01642175463112936 |
| interactor_abundance | mitosis | inter | 0.0011056112671619452 | 0.024049720606162197 |

O60832_P22087
DKC1_HUMAN vs FBRL_HUMAN
p-value: 0.005 q-value: 0.005

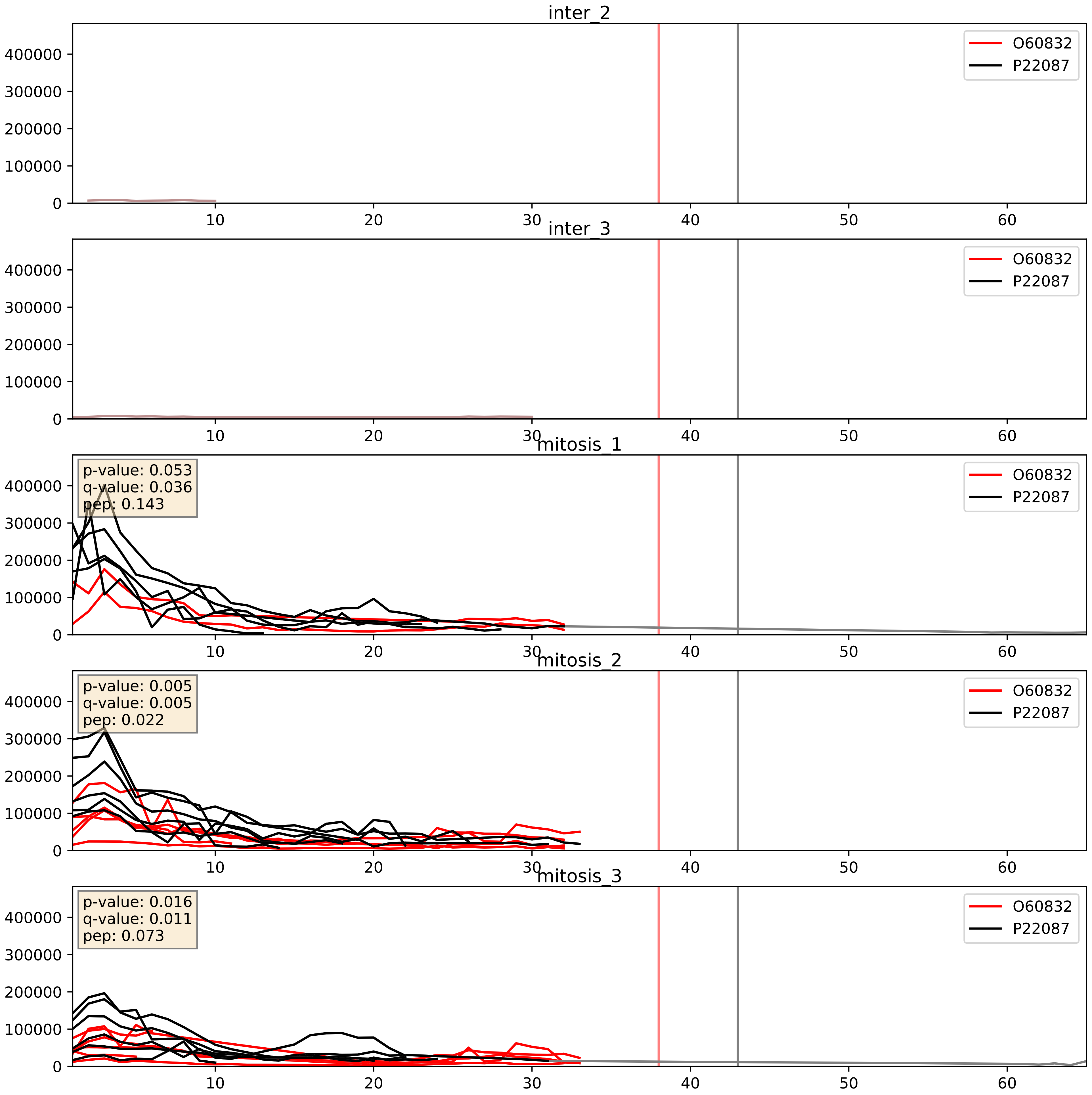


| level | condition_1 | condition_2 | pvalue | pvalue_adjusted |
|---|---|---|---|---|
| complex_abundance | mitosis | inter | 0.0015667645418376422 | 0.027691822035811595 |
| interactor_abundance | mitosis | inter | 0.050819620539780355 | 0.18065446504174412 |
| interactor_ratio | mitosis | inter | 0.07249205172524499 | 0.44970456769966505 |

O76021_P22087
RL1D1_HUMAN vs FBRL_HUMAN
p-value: 0.003 q-value: 0.004

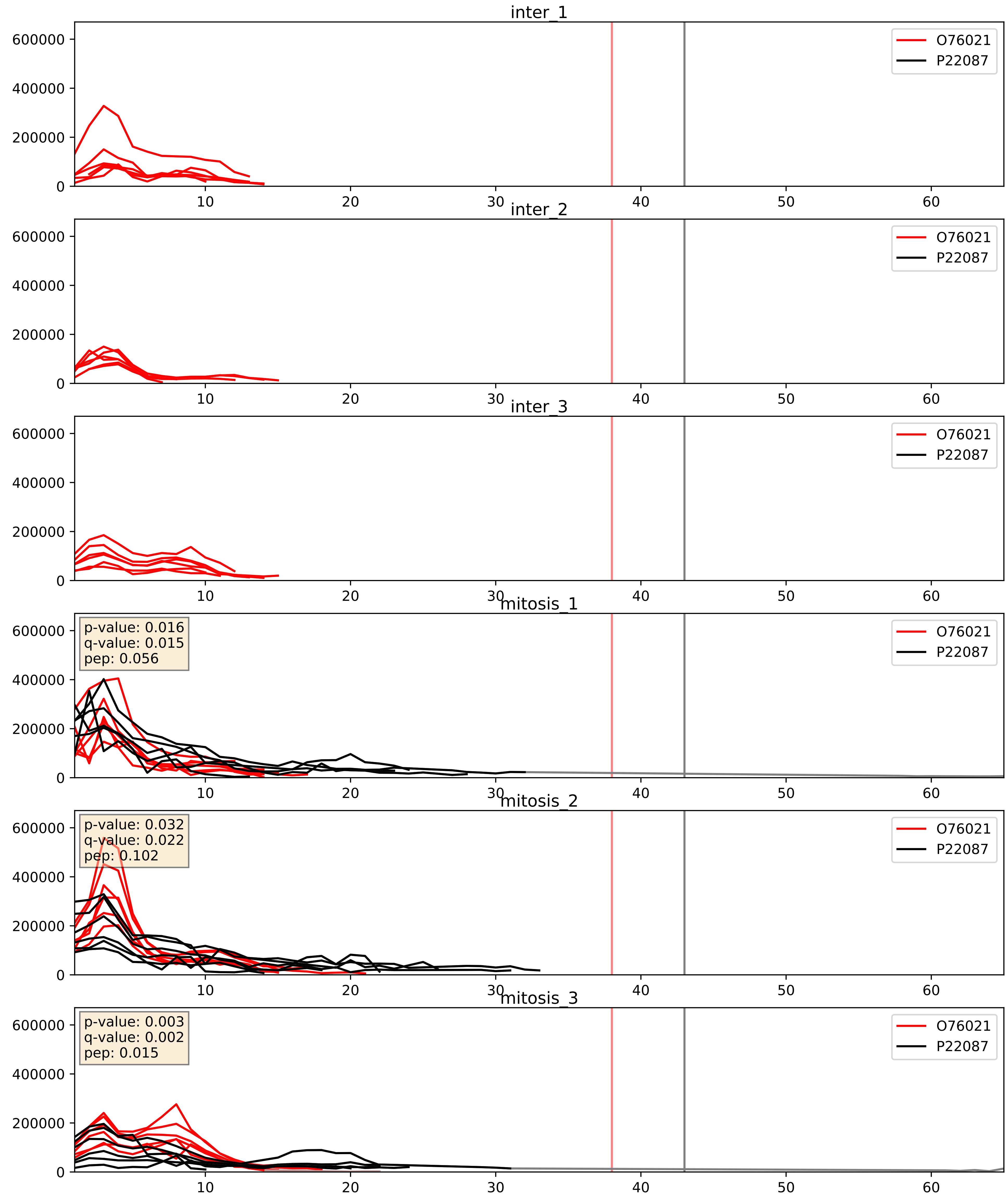


| level | condition_1 | condition_2 | pvalue | pvalue_adjusted |
|---|---|---|---|---|
| complex_abundance | mitosis | inter | 7.351496278048426e-05 | 0.005893640694944935 |
| interactor_abundance | mitosis | inter | 8.109392026765088e-05 | 0.0071196303332419645 |
| interactor_ratio | mitosis | inter | 0.9974833049507463 | 0.9996442035996809 |

P18124_P22087
RL7_HUMAN vs FBRL_HUMAN
p-value: 0.001 q-value: 0.037

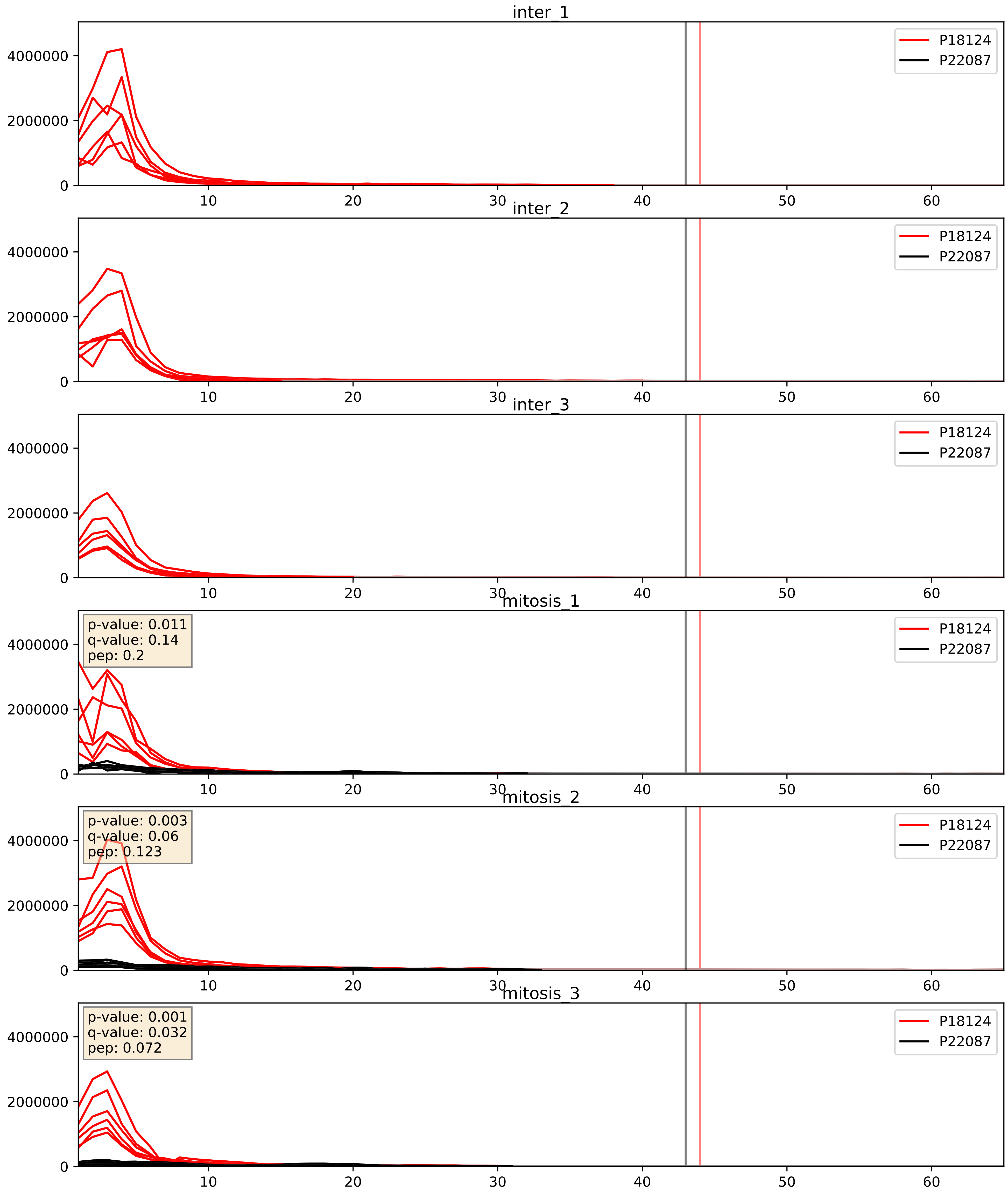


| level | condition_1 | condition_2 | pvalue | pvalue_adjusted |
|---|---|---|---|---|
| interactor_abundance | mitosis | inter | 4.739527379151582e-06 | 0.0033734327106189023 |
| complex_abundance | mitosis | inter | 3.150426976970383e-05 | 0.005691857151652958 |
| interactor_ratio | mitosis | inter | 0.0001558387989749646 | 0.012836278661937291 |

P22087_P26373
FBRL_HUMAN vs RL13_HUMAN
p-value: 0.0 q-value: 0.016

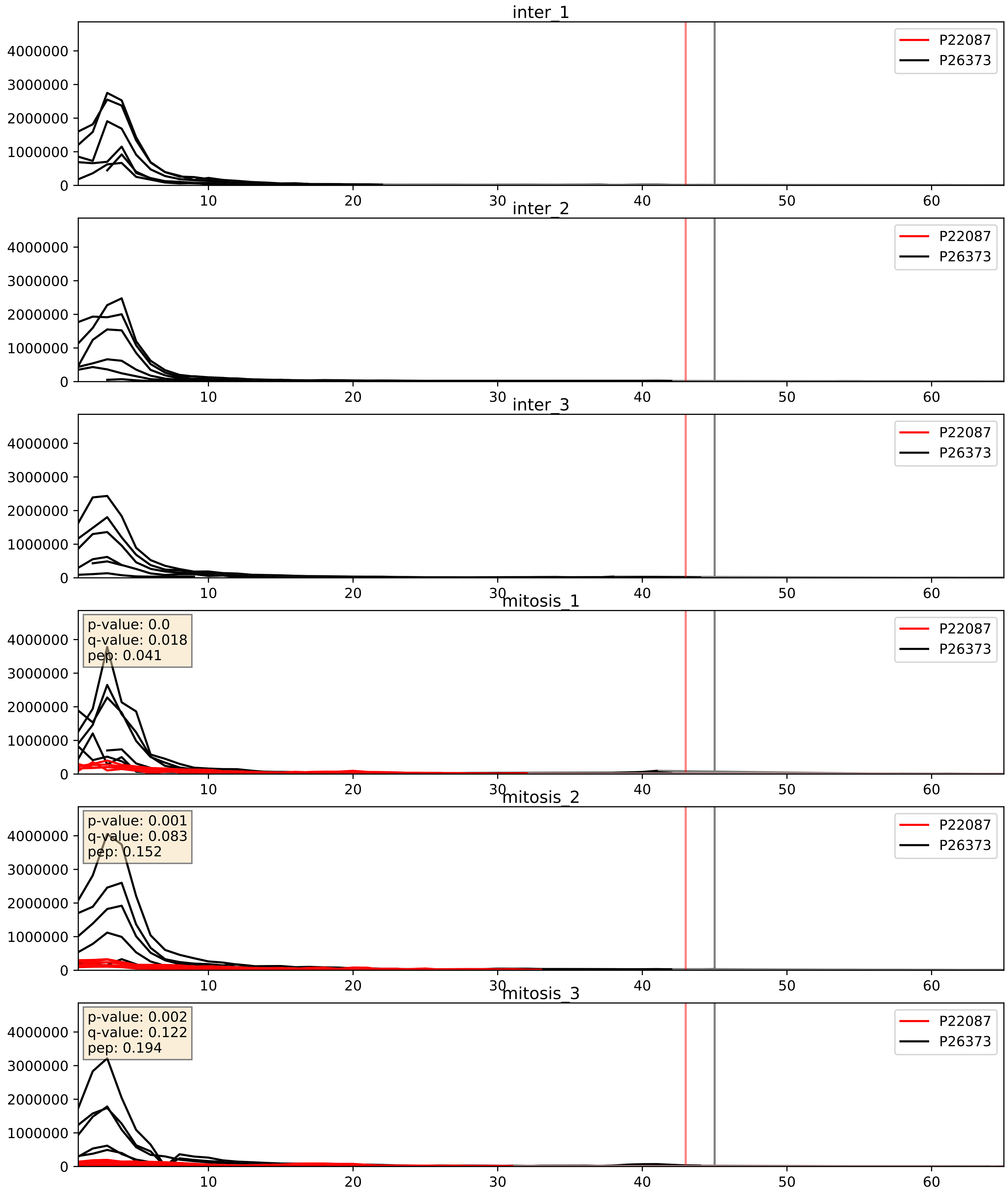


| level | condition_1 | condition_2 | pvalue | pvalue_adjusted |
|---|---|---|---|---|
| complex_abundance | mitosis | inter | 1.8594605808112858e-05 | 0.005691857151652958 |
| interactor_abundance | mitosis | inter | 0.0003671184791785453 | 0.01356427145975096 |
| interactor_ratio | mitosis | inter | 0.0008838330286563458 | 0.034218313408302205 |

P22087_P36578
FBRL_HUMAN vs RL4_HUMAN
p-value: 0.0 q-value: 0.039

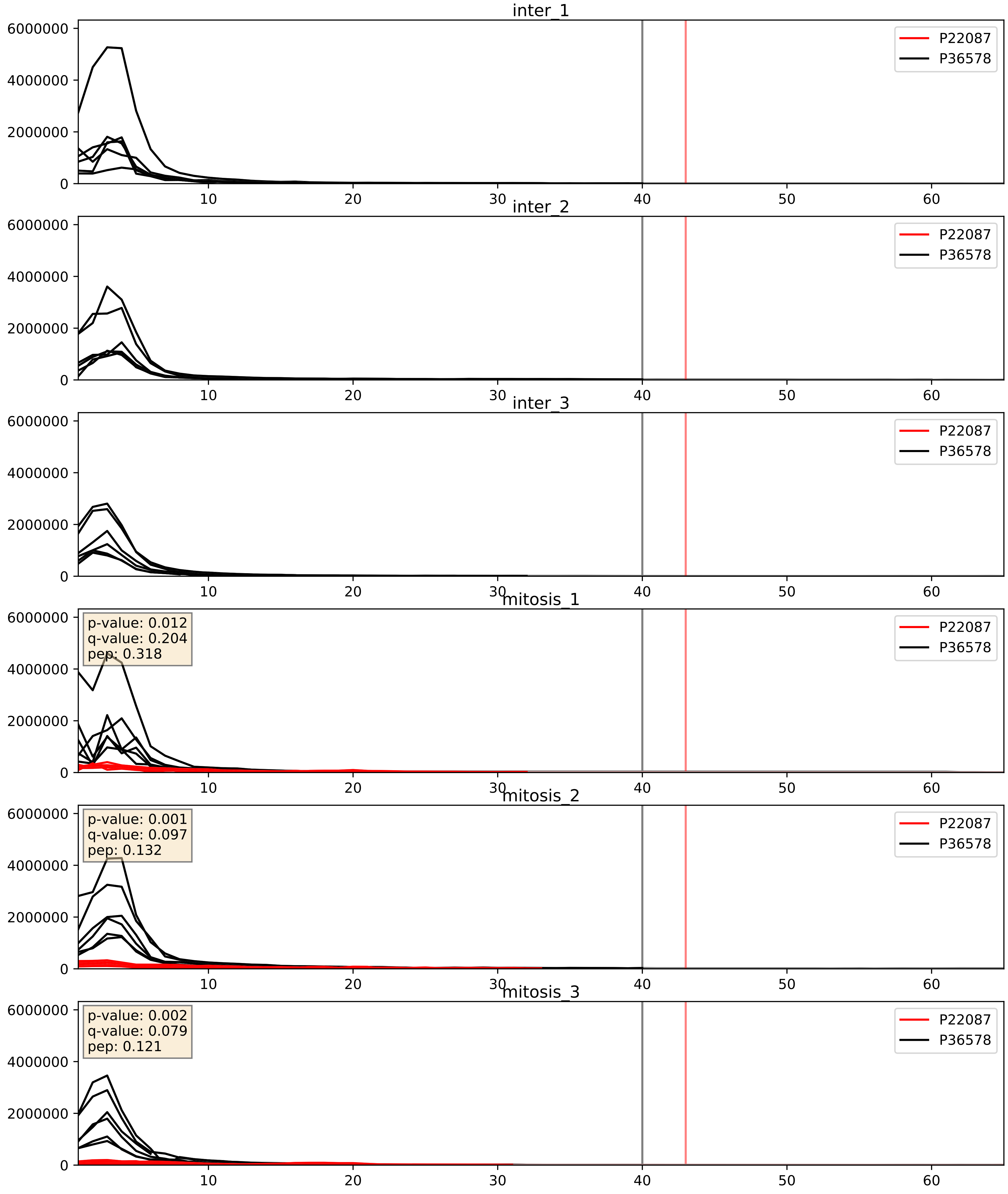


| level | condition_1 | condition_2 | pvalue | pvalue_adjusted |
|---|---|---|---|---|
| complex_abundance | mitosis | inter | 1.1986800633791924e-05 | 0.005691857151652958 |
| interactor_abundance | mitosis | inter | 0.0003671184791785453 | 0.01356427145975096 |
| interactor_ratio | mitosis | inter | 0.001313510109738704 | 0.04307910551480194 |

P22087_P39023
FBRL_HUMAN vs RL3_HUMAN
p-value: 0.0 q-value: 0.023

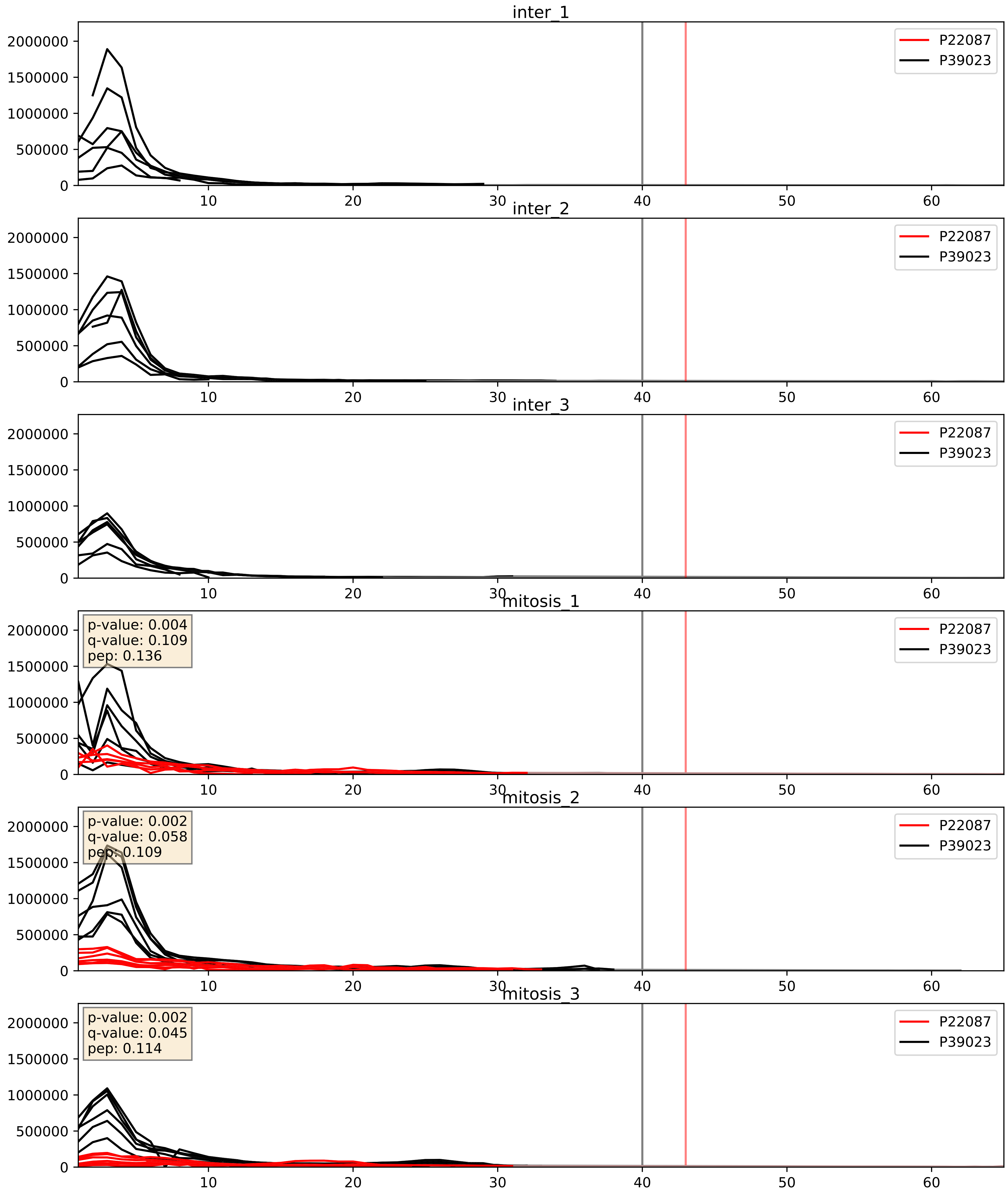


| level | condition_1 | condition_2 | pvalue | pvalue_adjusted |
|---|---|---|---|---|
| complex_abundance | mitosis | inter | 4.93530119560175e-05 | 0.005691857151652958 |
| interactor_abundance | mitosis | inter | 0.0003671184791785453 | 0.01356427145975096 |
| interactor_ratio | mitosis | inter | 0.00360829452514137 | 0.07764180691118368 |

P22087_P46087
FBRL_HUMAN vs NOP2_HUMAN
p-value: 0.004 q-value: 0.048

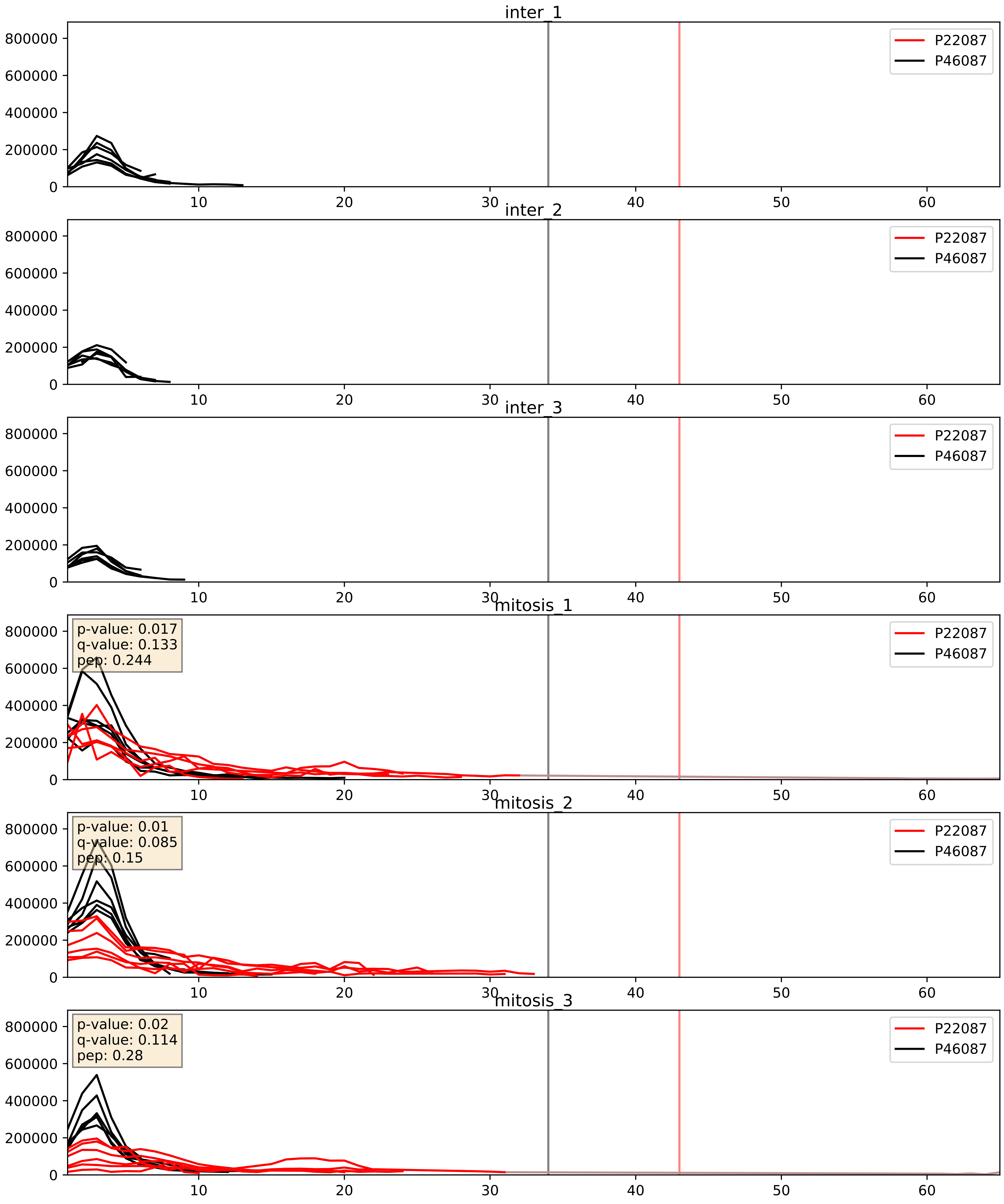


| level | condition_1 | condition_2 | pvalue | pvalue_adjusted |
|---|---|---|---|---|
| complex_abundance | mitosis | inter | 0.00034747852944409986 | 0.012015488415248196 |
| interactor_abundance | mitosis | inter | 0.0008503677168821298 | 0.021631939543866364 |
| interactor_ratio | mitosis | inter | 0.13219010083240598 | 0.58057838025931 |

P22087_P55769
FBRL_HUMAN vs NH2L1_HUMAN
p-value: 0.0 q-value: 0.0

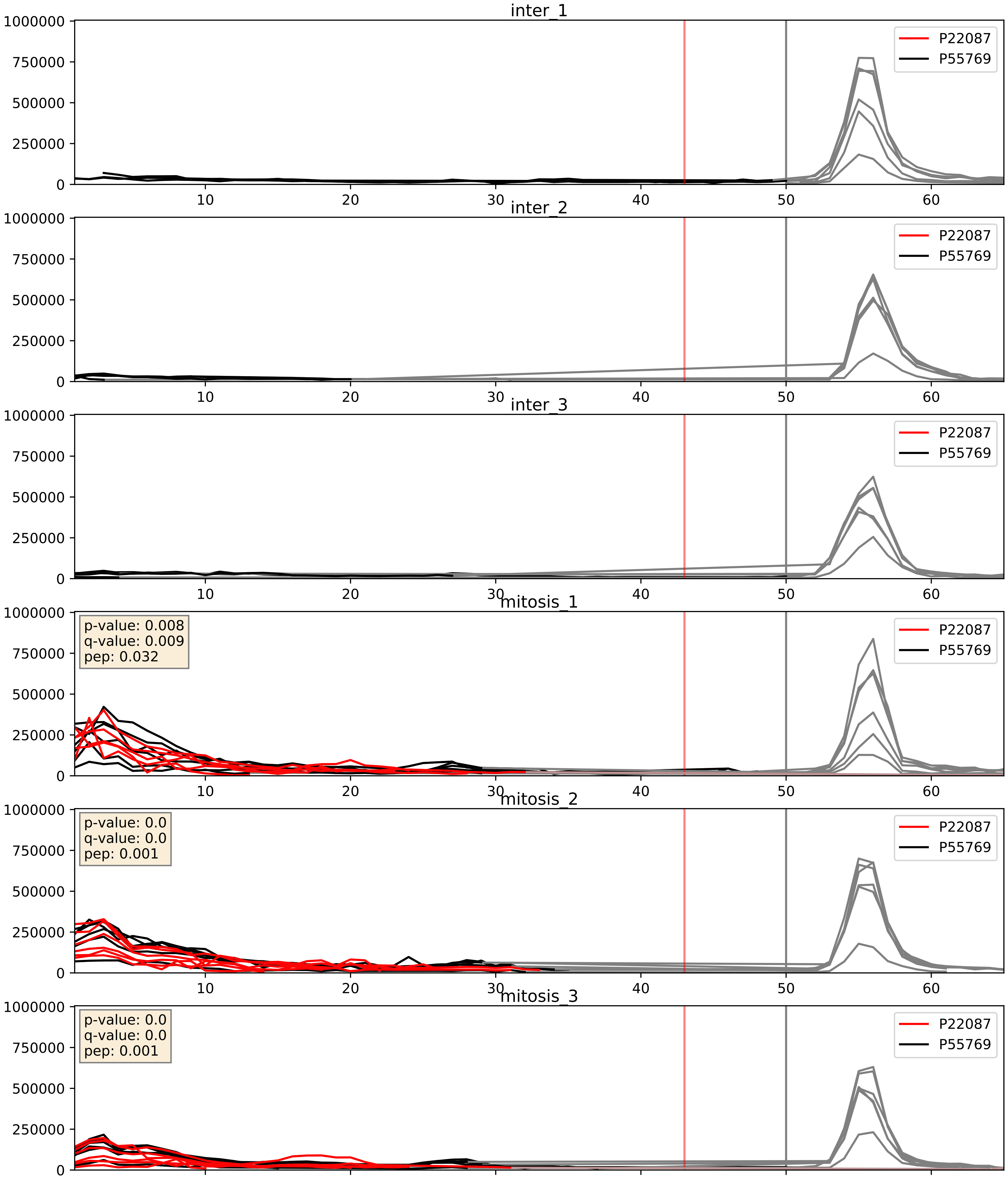


| level | condition_1 | condition_2 | pvalue | pvalue_adjusted |
|---|---|---|---|---|
| interactor_ratio | mitosis | inter | 3.3712270526144467e-05 | 0.006185155691845816 |
| complex_abundance | mitosis | inter | 0.0002827767244629666 | 0.011311068978518664 |
| interactor_abundance | mitosis | inter | 0.0003671184791785453 | 0.01356427145975096 |

P22087_P56537
FBRL_HUMAN vs IF6_HUMAN
p-value: 0.001 q-value: 0.015

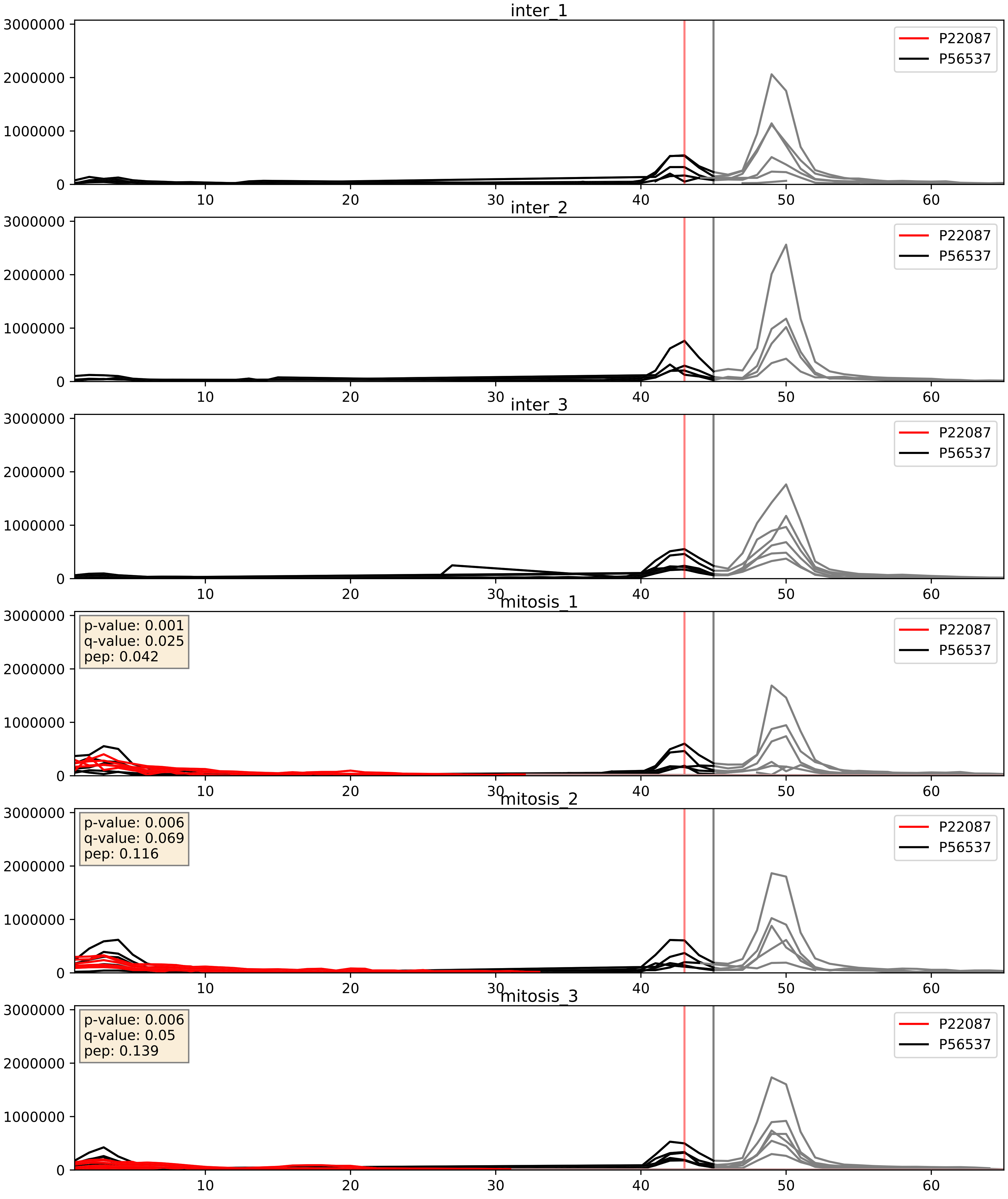


| level | condition_1 | condition_2 | pvalue | pvalue_adjusted |
|---|---|---|---|---|
| complex_abundance | mitosis | inter | 0.00029723422483891155 | 0.011617922212881656 |
| interactor_abundance | mitosis | inter | 0.00036693098007483434 | 0.01356427145975096 |
| interactor_ratio | mitosis | inter | 0.12685036942305242 | 0.5750062399685051 |

P22087_P61313
FBRL_HUMAN vs RL15_HUMAN
p-value: 0.001 q-value: 0.016

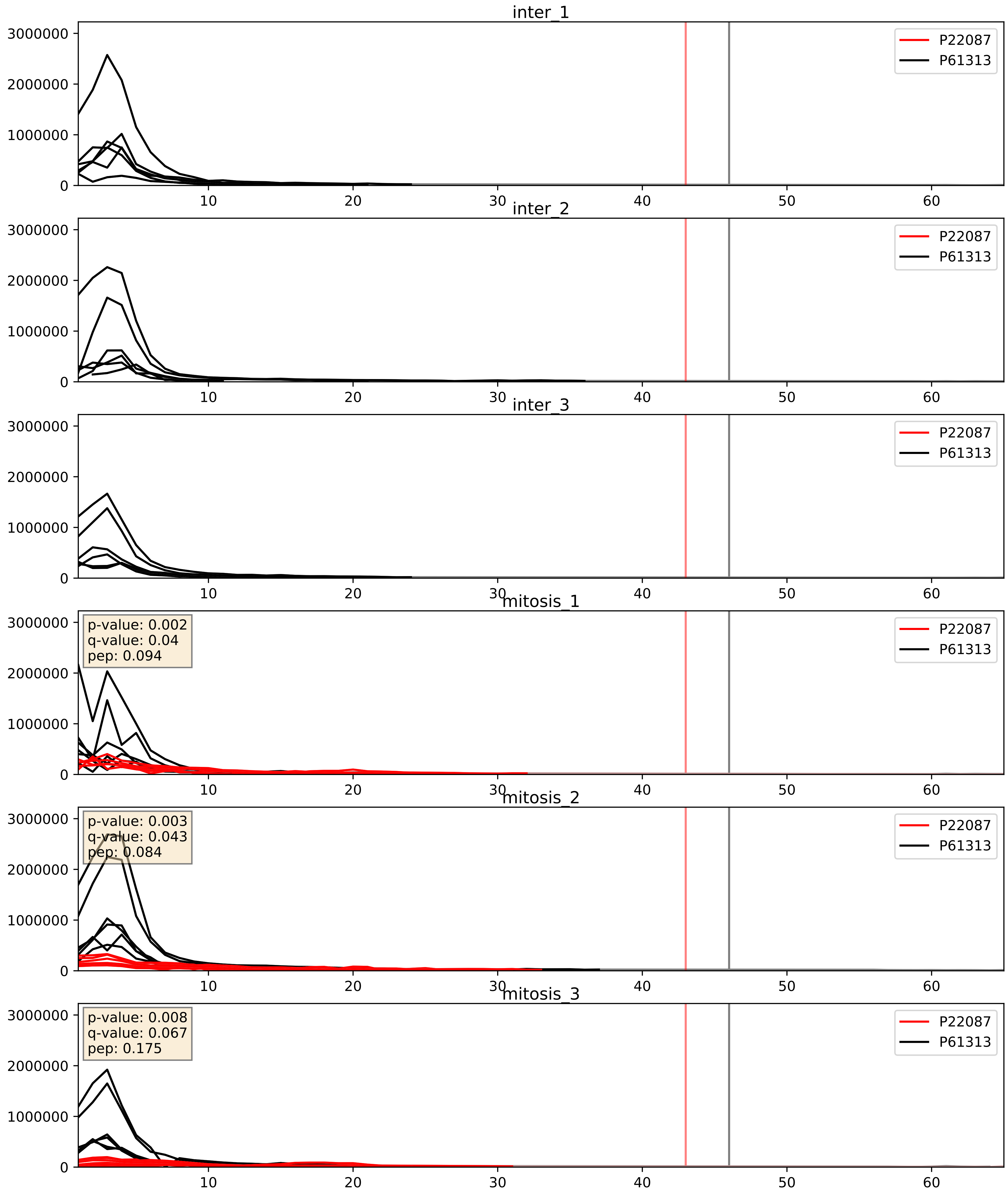


| level | condition_1 | condition_2 | pvalue | pvalue_adjusted |
|---|---|---|---|---|
| complex_abundance | mitosis | inter | 4.346136759748988e-05 | 0.005691857151652958 |
| interactor_abundance | mitosis | inter | 0.0003812827100365066 | 0.013573913378163039 |
| interactor_ratio | mitosis | inter | 0.004250536428321106 | 0.08581271657176573 |

P22087_P61353
FBRL_HUMAN vs RL27_HUMAN
p-value: 0.0 q-value: 0.041

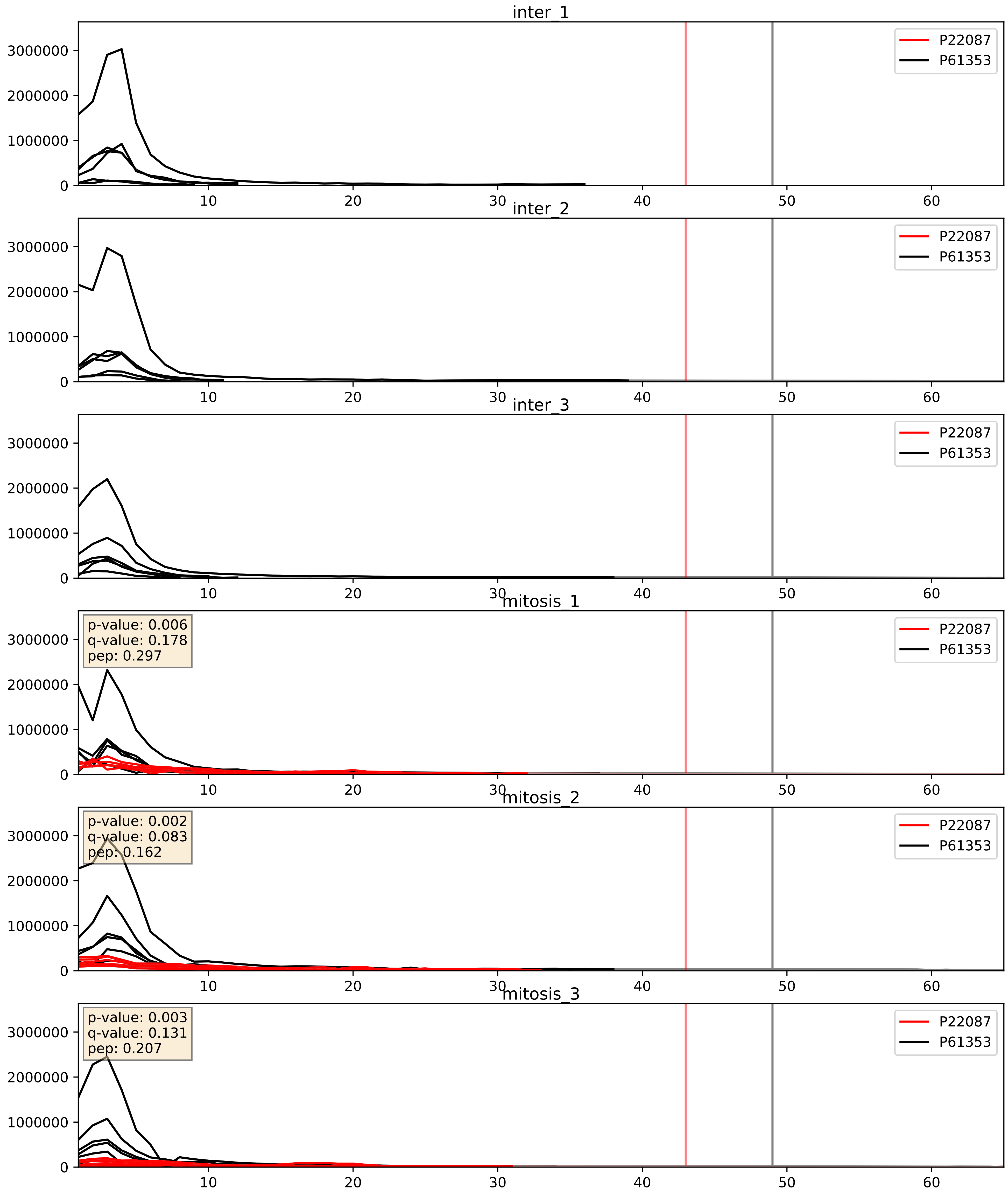


| level | condition_1 | condition_2 | pvalue | pvalue_adjusted |
|---|---|---|---|---|
| complex_abundance | mitosis | inter | 3.316358693033294e-05 | 0.005691857151652958 |
| interactor_abundance | mitosis | inter | 0.0003158592306741209 | 0.013227619301987383 |
| interactor_ratio | mitosis | inter | 0.007119864116795093 | 0.11742974342922158 |

P22087_P62081
FBRL_HUMAN vs RS7_HUMAN
p-value: 0.002 q-value: 0.002

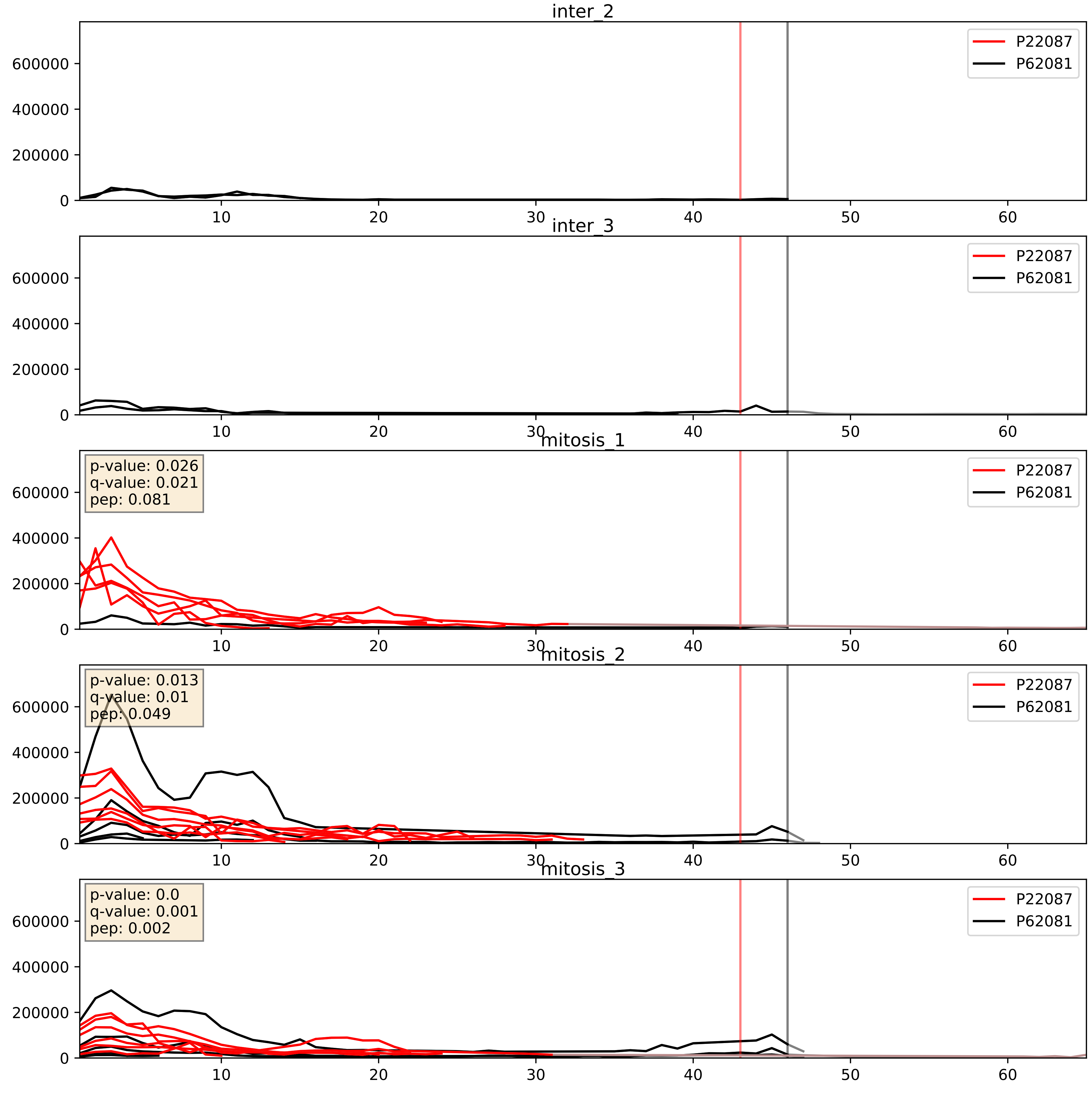


| level | condition_1 | condition_2 | pvalue | pvalue_adjusted |
|---|---|---|---|---|
| interactor_abundance | mitosis | inter | 0.00024745763614672354 | 0.011921759831644983 |
| complex_abundance | mitosis | inter | 0.0033855088768150795 | 0.042806434247469834 |
| interactor_ratio | mitosis | inter | 0.14943468907022306 | 0.6039771666075939 |

P22087_P62241
FBRL_HUMAN vs RS8_HUMAN
p-value: 0.012 q-value: 0.01

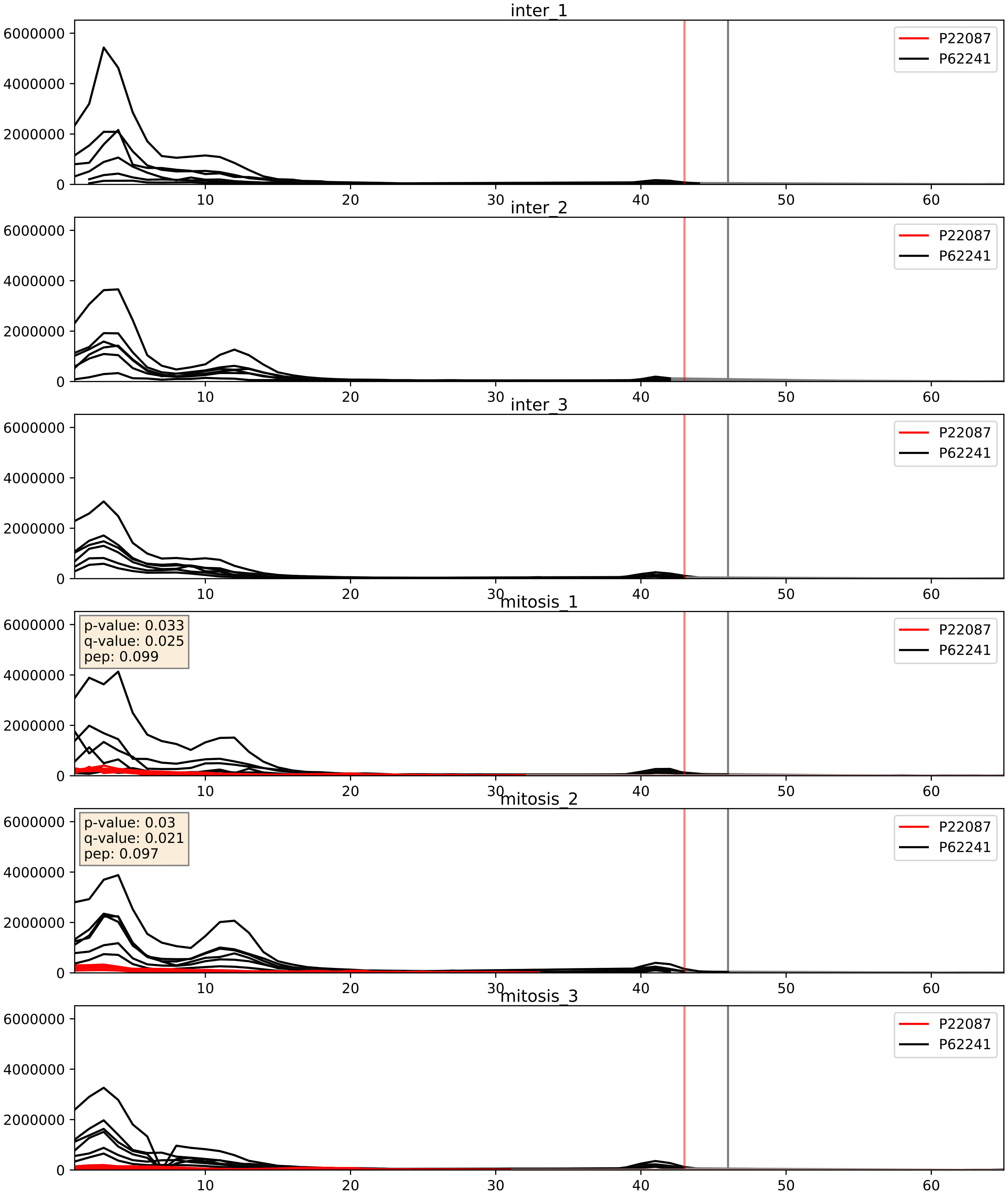


| level | condition_1 | condition_2 | pvalue | pvalue_adjusted |
|---|---|---|---|---|
| complex_abundance | mitosis | inter | 1.4441646991084911e-05 | 0.005691857151652958 |
| interactor_ratio | mitosis | inter | 0.0002611783911681679 | 0.016318883418974577 |
| interactor_abundance | mitosis | inter | 0.00037582732769035214 | 0.013573913378163039 |

P22087_P62244
FBRL_HUMAN vs RS15A_HUMAN
p-value: 0.004 q-value: 0.004

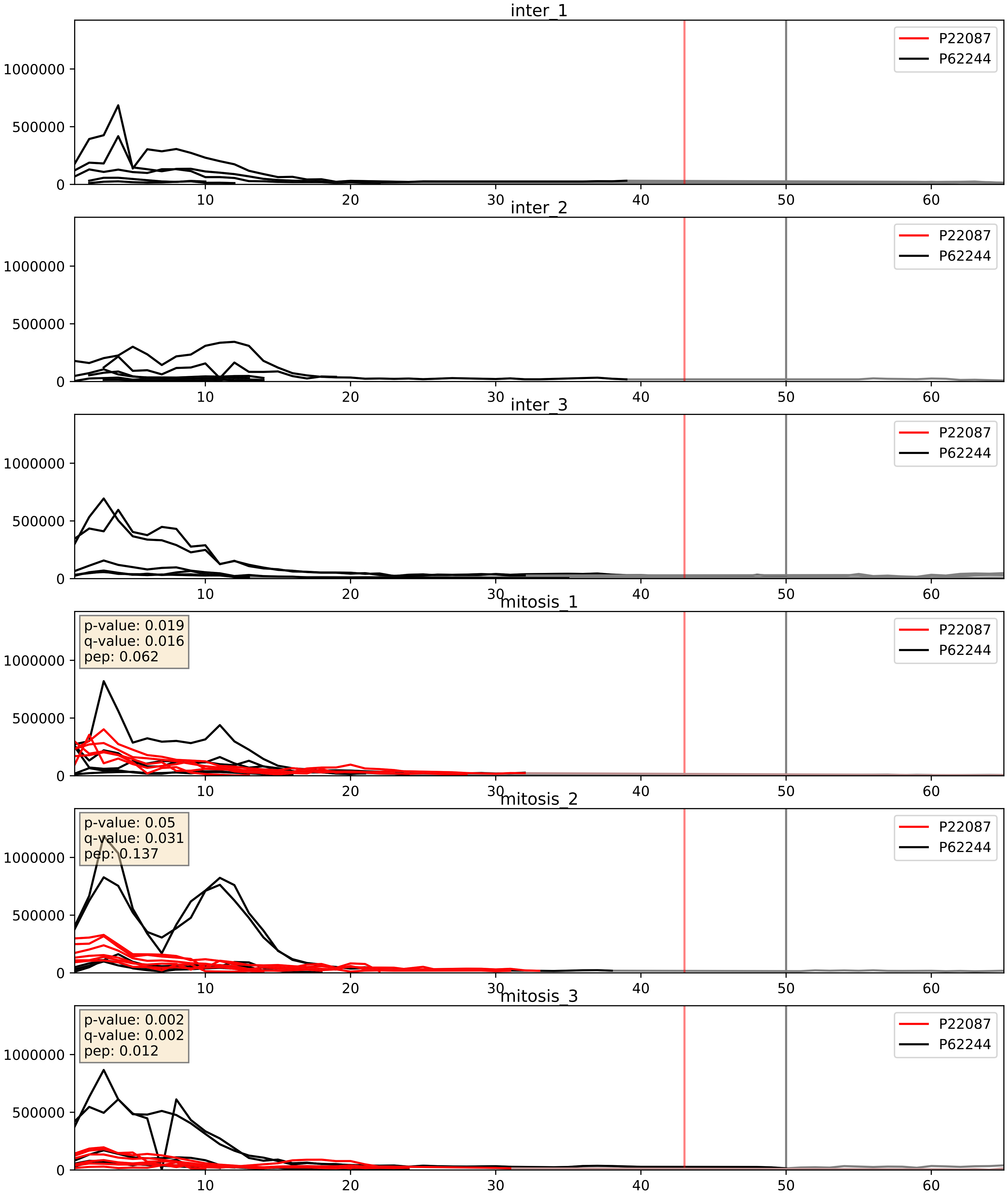


| level | condition_1 | condition_2 | pvalue | pvalue_adjusted |
|---|---|---|---|---|
| complex_abundance | mitosis | inter | 9.047875791541205e-05 | 0.00645415139796606 |
| interactor_abundance | mitosis | inter | 0.0003623659339208455 | 0.01356427145975096 |
| interactor_ratio | mitosis | inter | 0.04499390548439066 | 0.35269947888863007 |

P22087_P62249
FBRL_HUMAN vs RS16_HUMAN
p-value: 0.005 q-value: 0.005

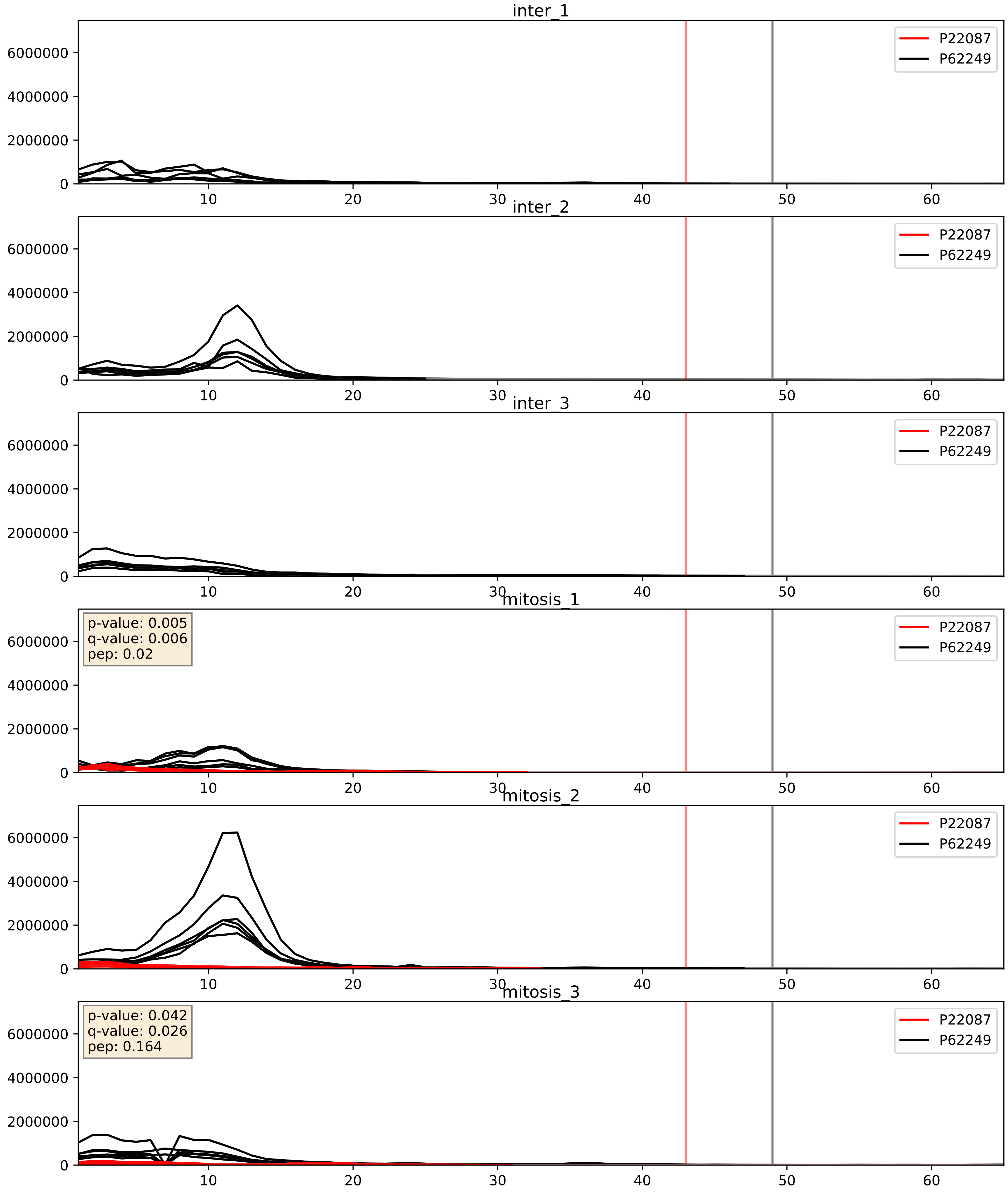


| level | condition_1 | condition_2 | pvalue | pvalue_adjusted |
|---|---|---|---|---|
| complex_abundance | mitosis | inter | 0.00010224102514160368 | 0.006784365699318818 |
| interactor_abundance | mitosis | inter | 0.0003671184791785453 | 0.01356427145975096 |
| interactor_ratio | mitosis | inter | 0.0029287716764384536 | 0.07007095913365735 |

P22087_P62266
FBRL_HUMAN vs RS23_HUMAN
p-value: 0.003 q-value: 0.004

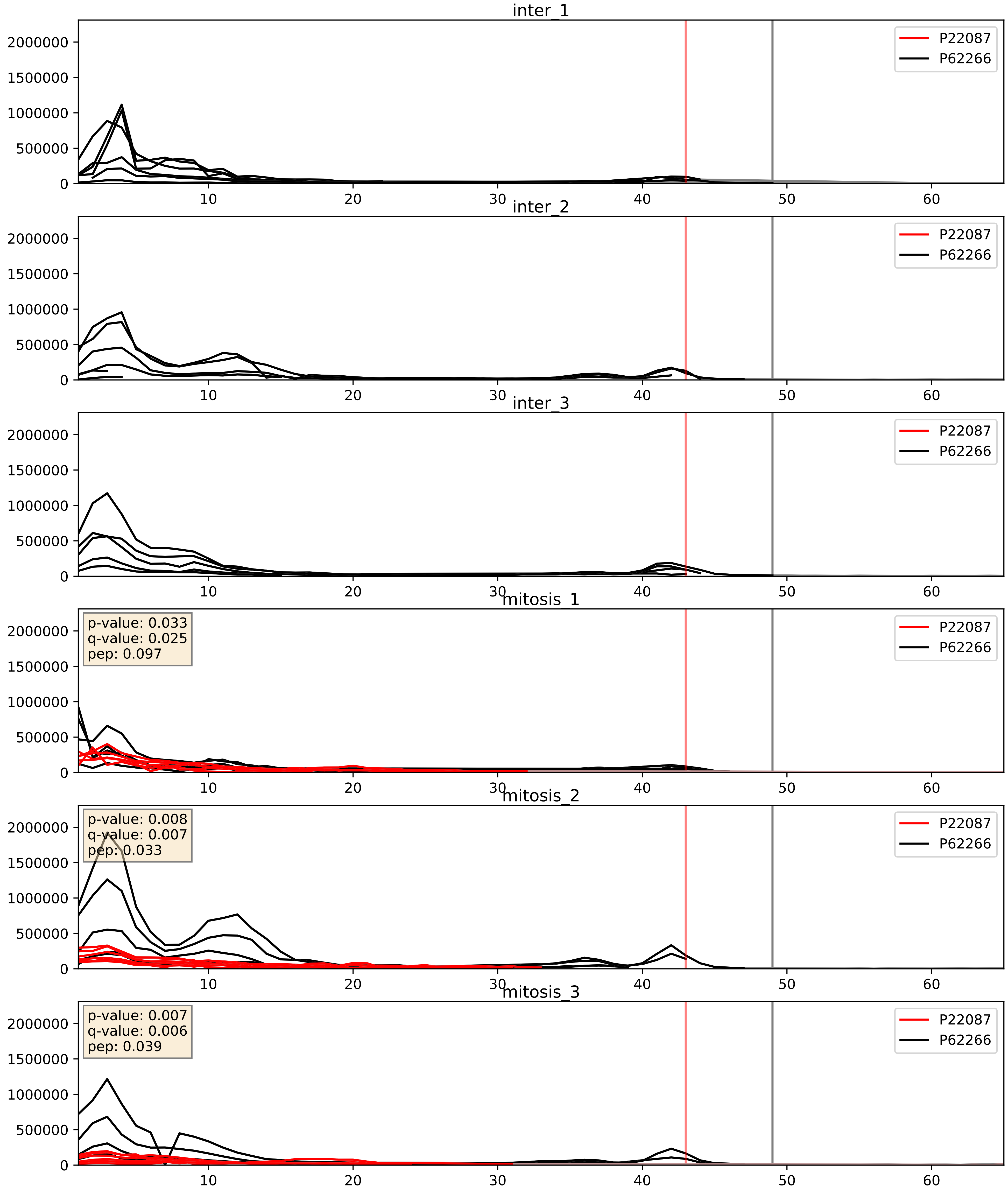


| level | condition_1 | condition_2 | pvalue | pvalue_adjusted |
|---|---|---|---|---|
| interactor_abundance | mitosis | inter | 0.0003671184791785453 | 0.01356427145975096 |
| complex_abundance | mitosis | inter | 0.0011910106738603468 | 0.02322335163609241 |
| interactor_ratio | mitosis | inter | 0.18291817842614824 | 0.6518805262931596 |

P22087_P62277
FBRL_HUMAN vs RS13_HUMAN
p-value: 0.012 q-value: 0.01

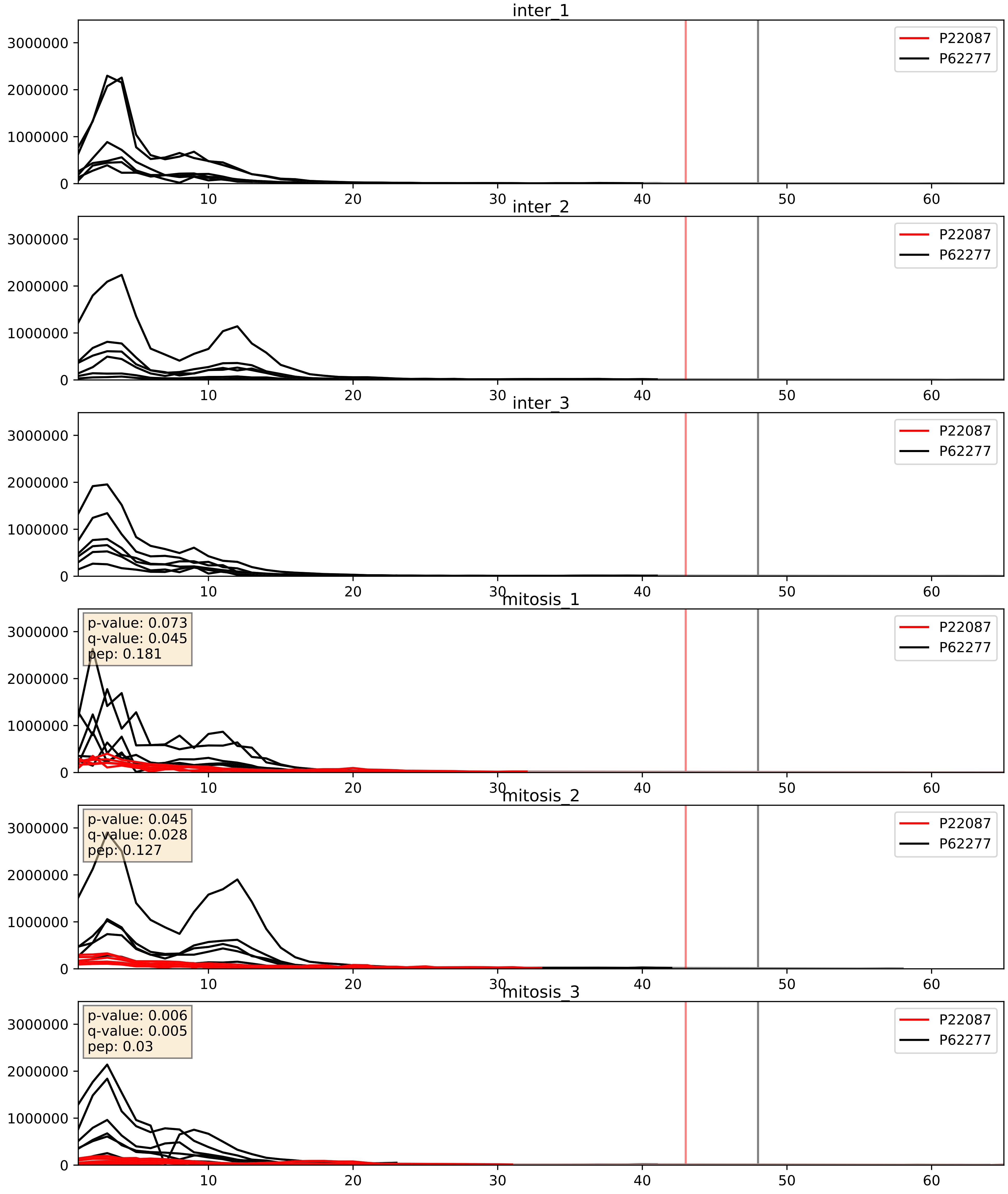


| level | condition_1 | condition_2 | pvalue | pvalue_adjusted |
|---|---|---|---|---|
| complex_abundance | mitosis | inter | 1.536099136567195e-05 | 0.005691857151652958 |
| interactor_abundance | mitosis | inter | 0.0003671184791785453 | 0.01356427145975096 |
| interactor_ratio | mitosis | inter | 0.002614307383523888 | 0.06562601525796036 |

P22087_P62280
FBRL_HUMAN vs RS11_HUMAN
p-value: 0.0 q-value: 0.001

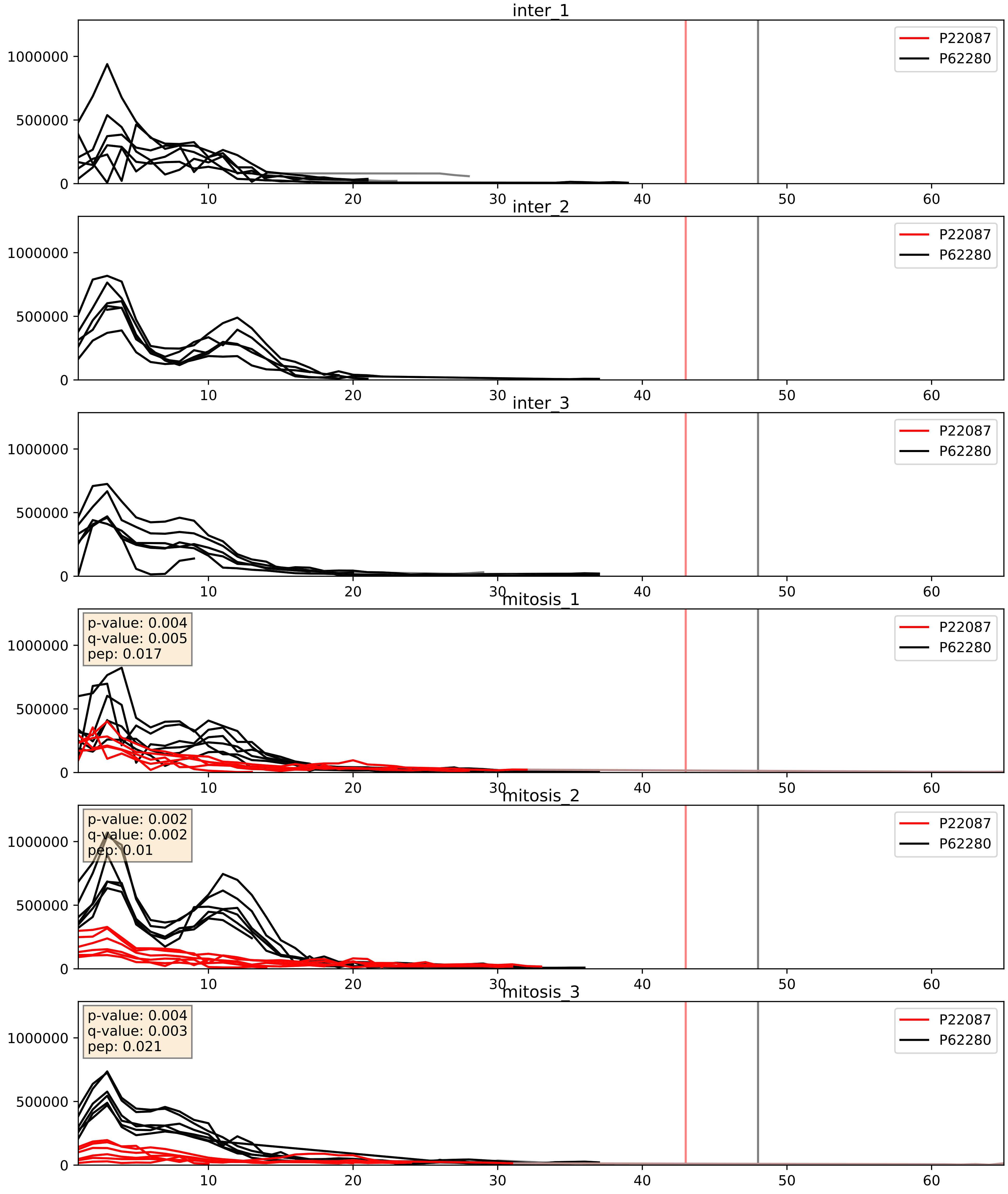


| level | condition_1 | condition_2 | pvalue | pvalue_adjusted |
| --- | --- | --- | --- | --- |
| complex_abundance | mitosis | inter | 5.170860890879182e-05 | 0.005691857151652958 |
| interactor_abundance | mitosis | inter | 0.00038131627813842286 | 0.013573913378163039 |
| interactor_ratio | mitosis | inter | 0.003925103685787187 | 0.08138603277233046 |

P22087_P62424
FBRL_HUMAN vs RL7A_HUMAN
p-value: 0.0 q-value: 0.023

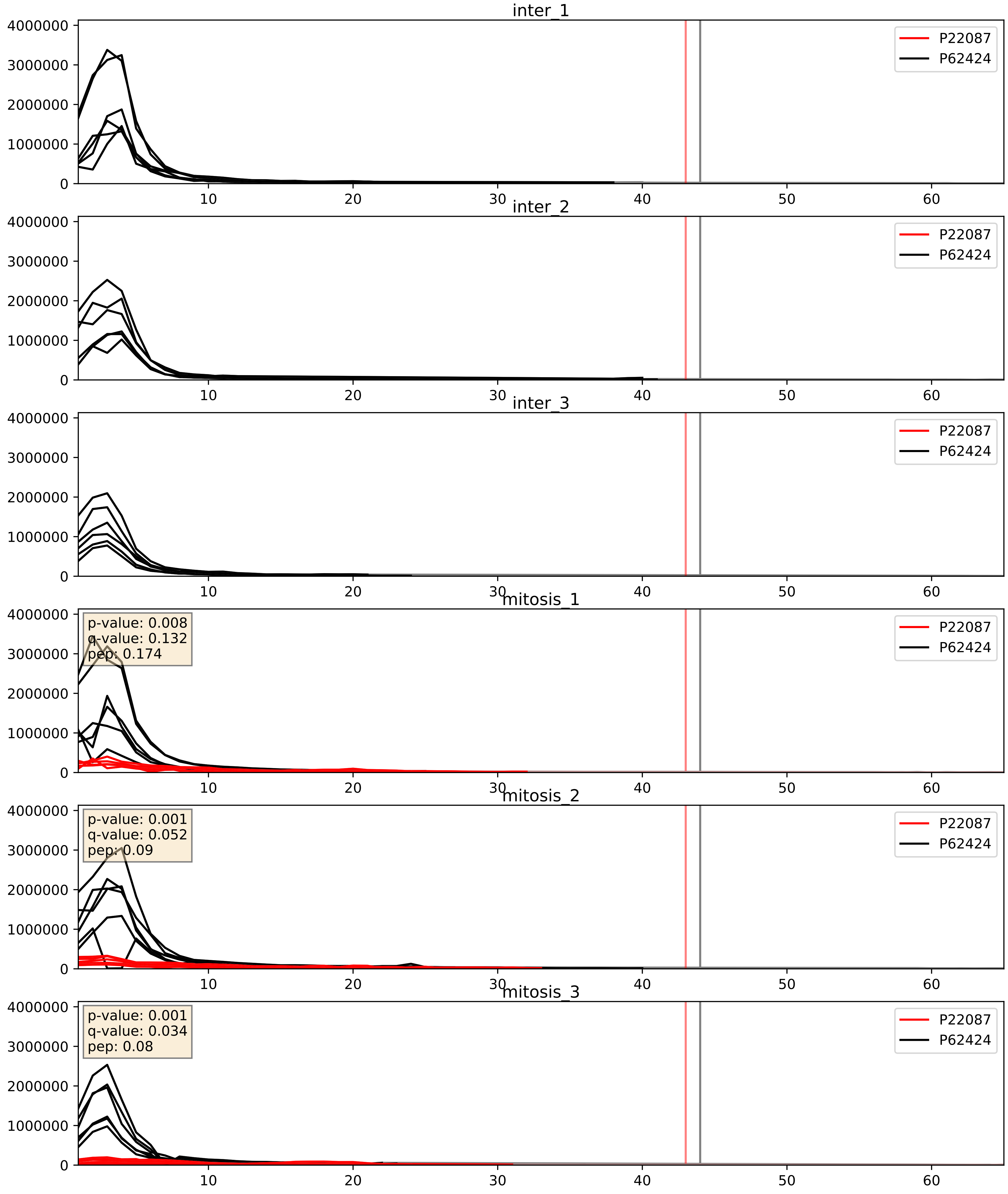


| level | condition_1 | condition_2 | pvalue | pvalue_adjusted |
|---|---|---|---|---|
| complex_abundance | mitosis | inter | 3.630302465341418e-05 | 0.005691857151652958 |
| interactor_abundance | mitosis | inter | 0.00038188998169085025 | 0.013573913378163039 |
| interactor_ratio | mitosis | inter | 0.0003946672855814106 | 0.021937350419330353 |

P22087_P62701
FBRL_HUMAN vs RS4X_HUMAN
p-value: 0.019 q-value: 0.015

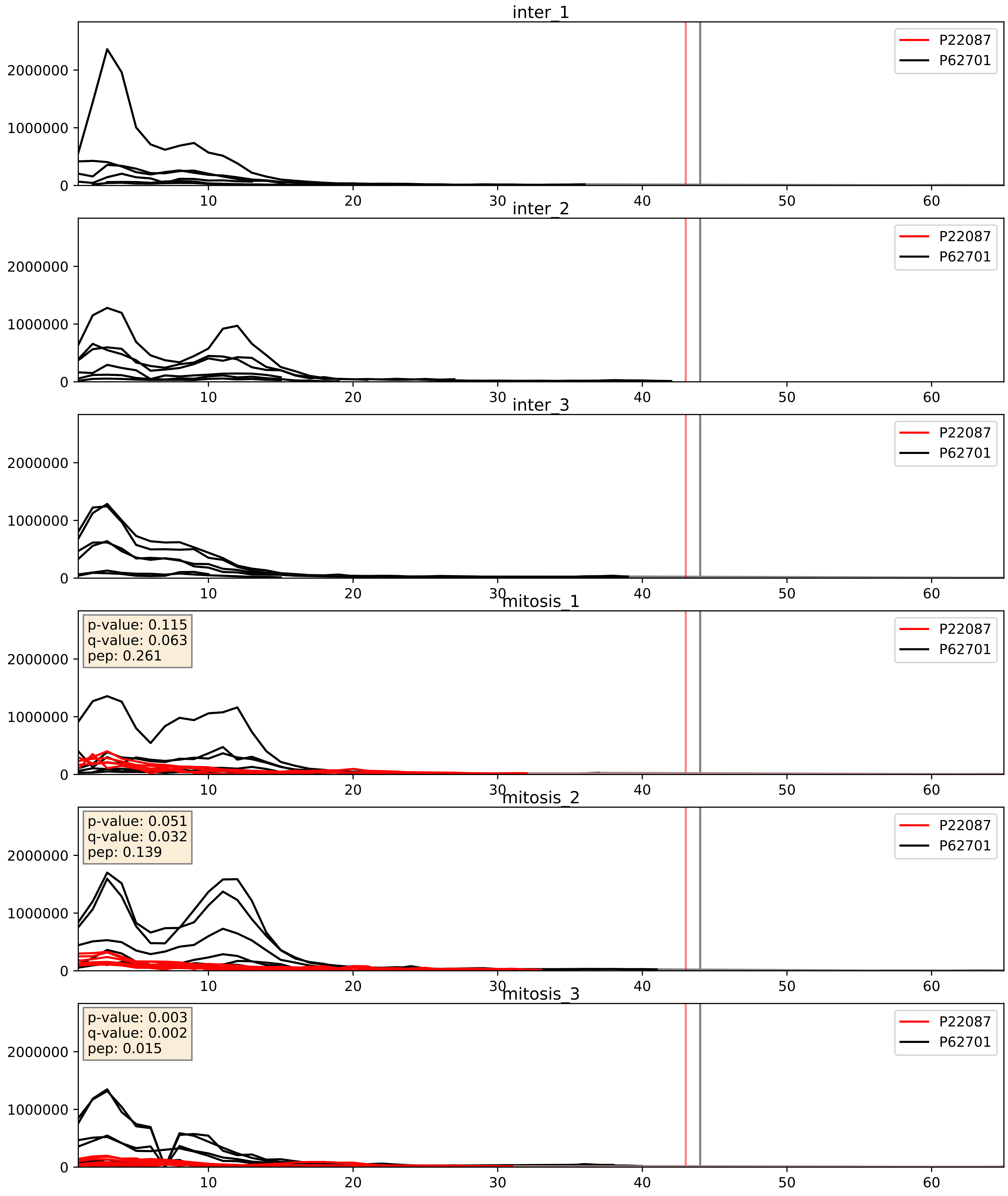


| level | condition_1 | condition_2 | pvalue | pvalue_adjusted |
|---|---|---|---|---|
| complex_abundance | mitosis | inter | 3.7130534683429326e-05 | 0.005691857151652958 |
| interactor_abundance | mitosis | inter | 0.0003671184791785453 | 0.013564271459750960 |
| interactor_ratio | mitosis | inter | 0.0023173846963125285 | 0.06011155454677347 |

P22087_P62753
FBRL_HUMAN vs RS6_HUMAN
p-value: 0.026 q-value: 0.019

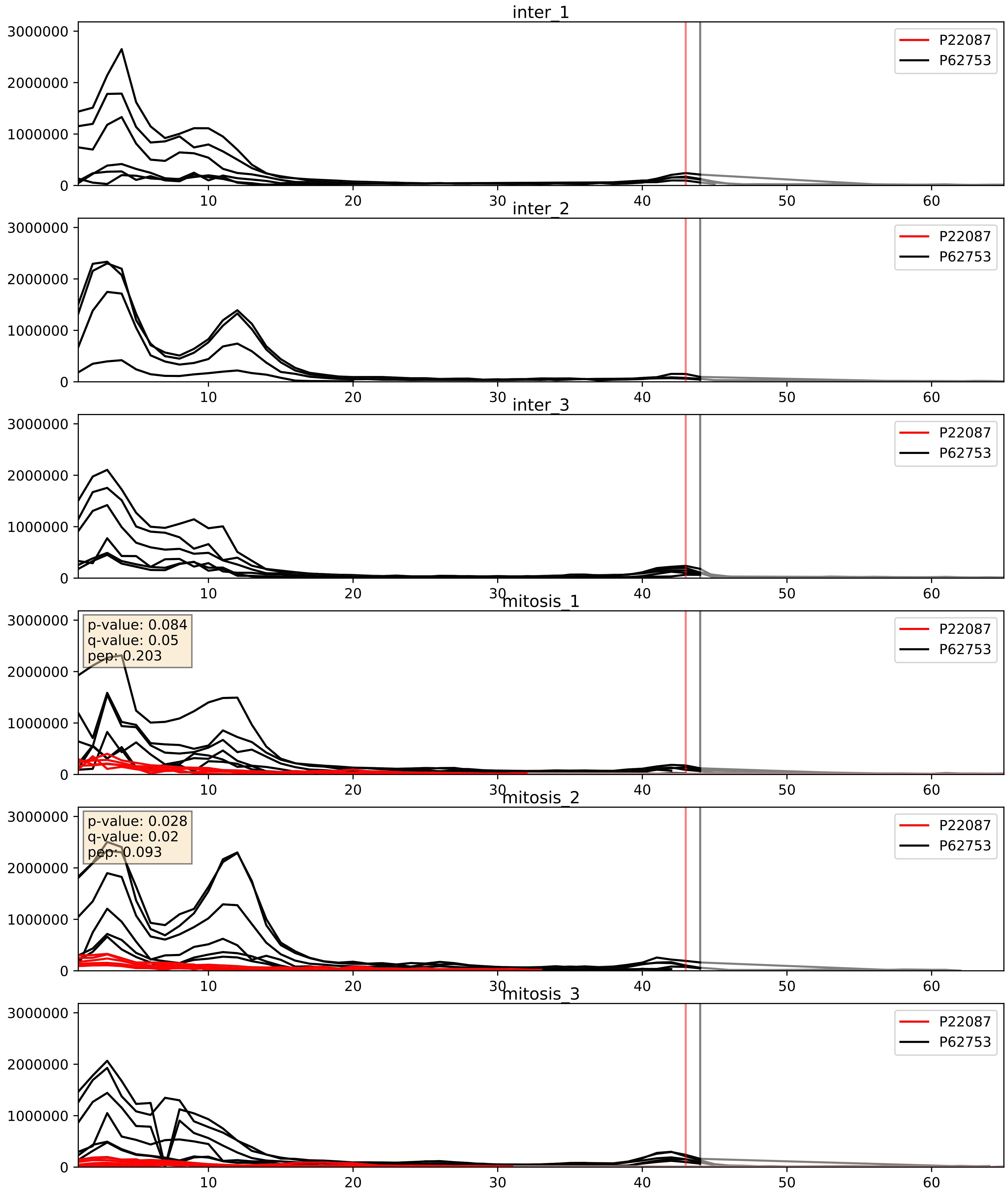


| level | condition_1 | condition_2 | pvalue | pvalue_adjusted |
|---|---|---|---|---|
| complex_abundance | mitosis | inter | 1.1949045645942093e-05 | 0.005691857151652958 |
| interactor_abundance | mitosis | inter | 0.0003671184791785453 | 0.01356427145975096 |
| interactor_ratio | mitosis | inter | 0.0005429526486218265 | 0.026977218456533537 |

P22087_P62847
FBRL_HUMAN vs RS24_HUMAN
p-value: 0.023 q-value: 0.018

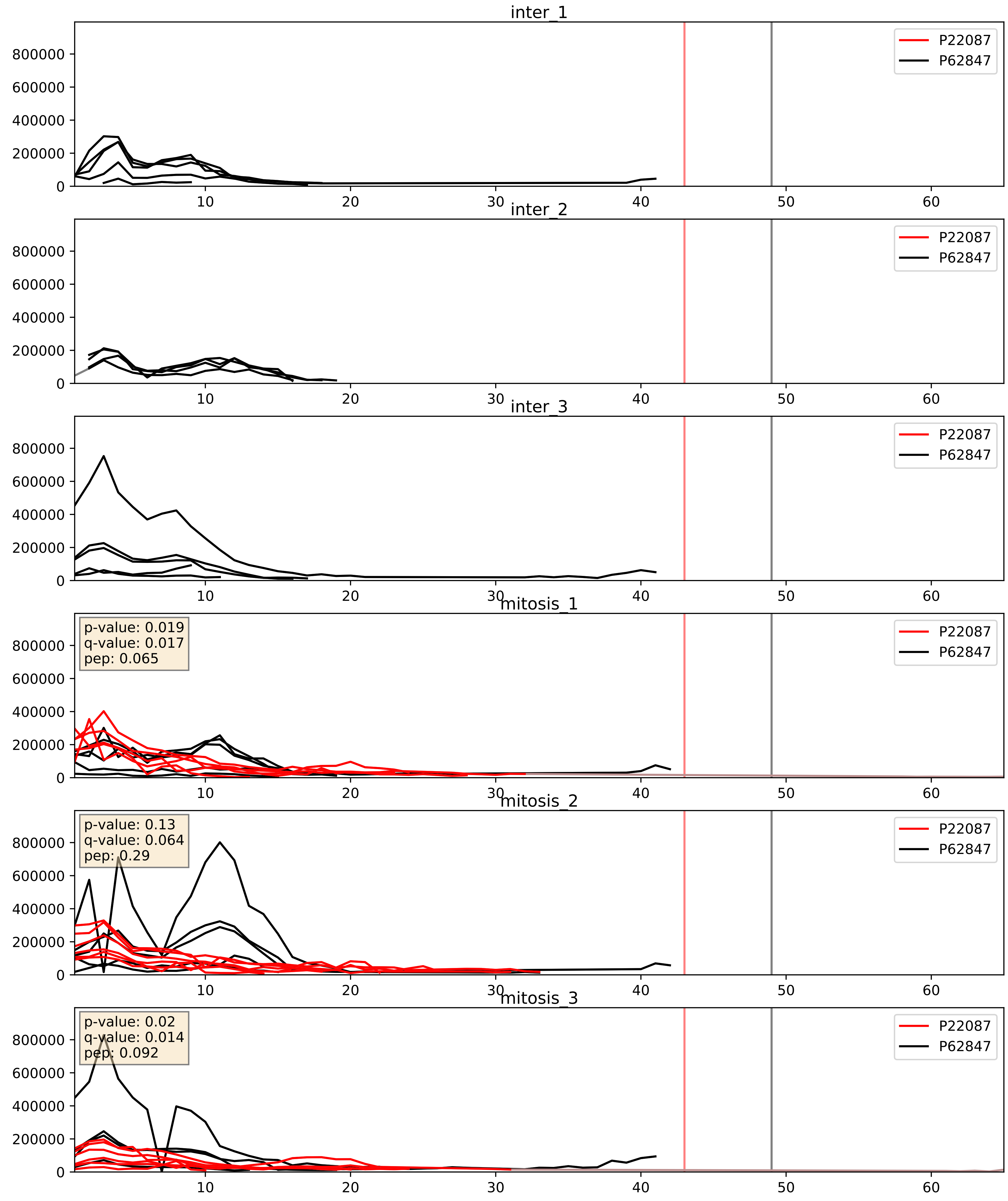


| level | condition_1 | condition_2 | pvalue | pvalue_adjusted |
|---|---|---|---|---|
| complex_abundance | mitosis | inter | 8.832364348528197e-05 | 0.006407206679949268 |
| interactor_abundance | mitosis | inter | 0.0003433386879125071 | 0.01356427145975096 |
| interactor_ratio | mitosis | inter | 0.07801173968815125 | 0.46705246033499004 |

P22087_P62861
FBRL_HUMAN vs RS30_HUMAN
p-value: 0.001 q-value: 0.028

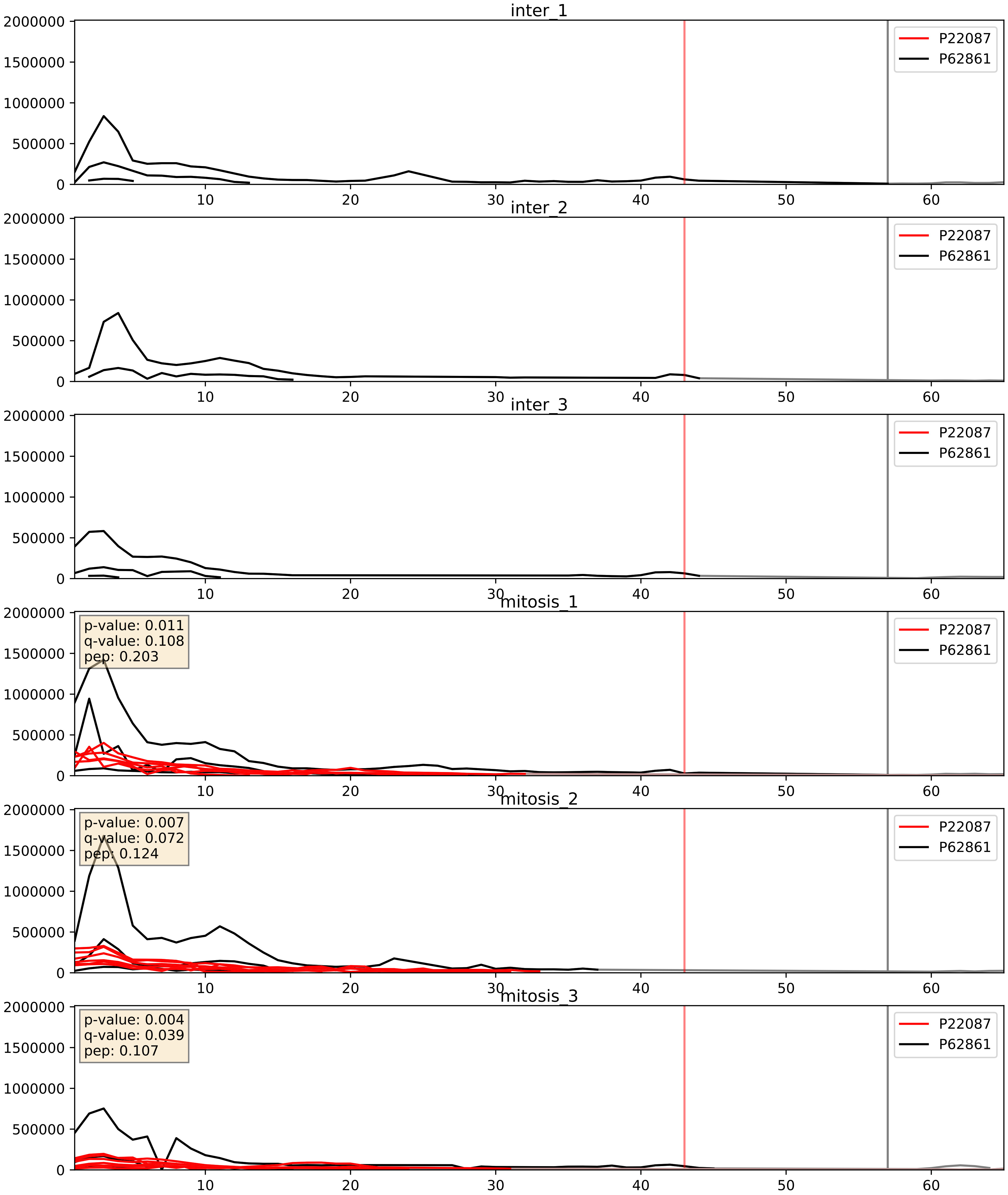


| level | condition_1 | condition_2 | pvalue | pvalue_adjusted |
|---|---|---|---|---|
| interactor_abundance | mitosis | inter | 0.0005590298359306732 | 0.017415659477552636 |
| complex_abundance | mitosis | inter | 0.0012837664514777853 | 0.024365944178824483 |
| interactor_ratio | mitosis | inter | 0.731875723500164 | 0.948492556694308 |

P22087_P62906
FBRL_HUMAN vs RL10A_HUMAN
p-value: 0.0 q-value: 0.017

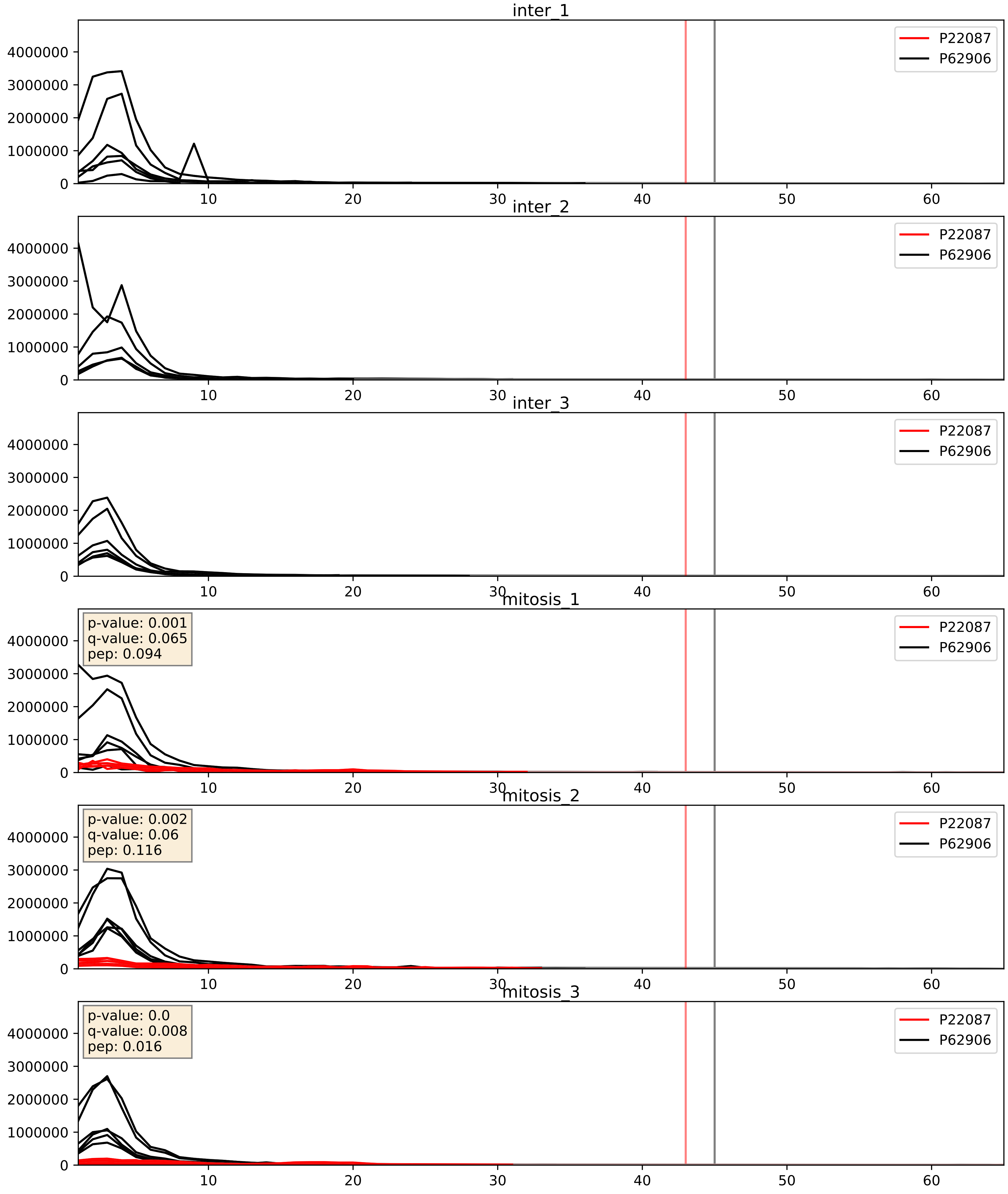


| level | condition_1 | condition_2 | pvalue | pvalue_adjusted |
|---|---|---|---|---|
| complex_abundance | mitosis | inter | 2.264226142142095e-05 | 0.005691857151652958 |
| interactor_abundance | mitosis | inter | 0.0003841155059274424 | 0.013586895581565731 |
| interactor_ratio | mitosis | inter | 0.0014007263931960893 | 0.04424434658951485 |

P22087_P62913
FBRL_HUMAN vs RL11_HUMAN
p-value: 0.0 q-value: 0.015

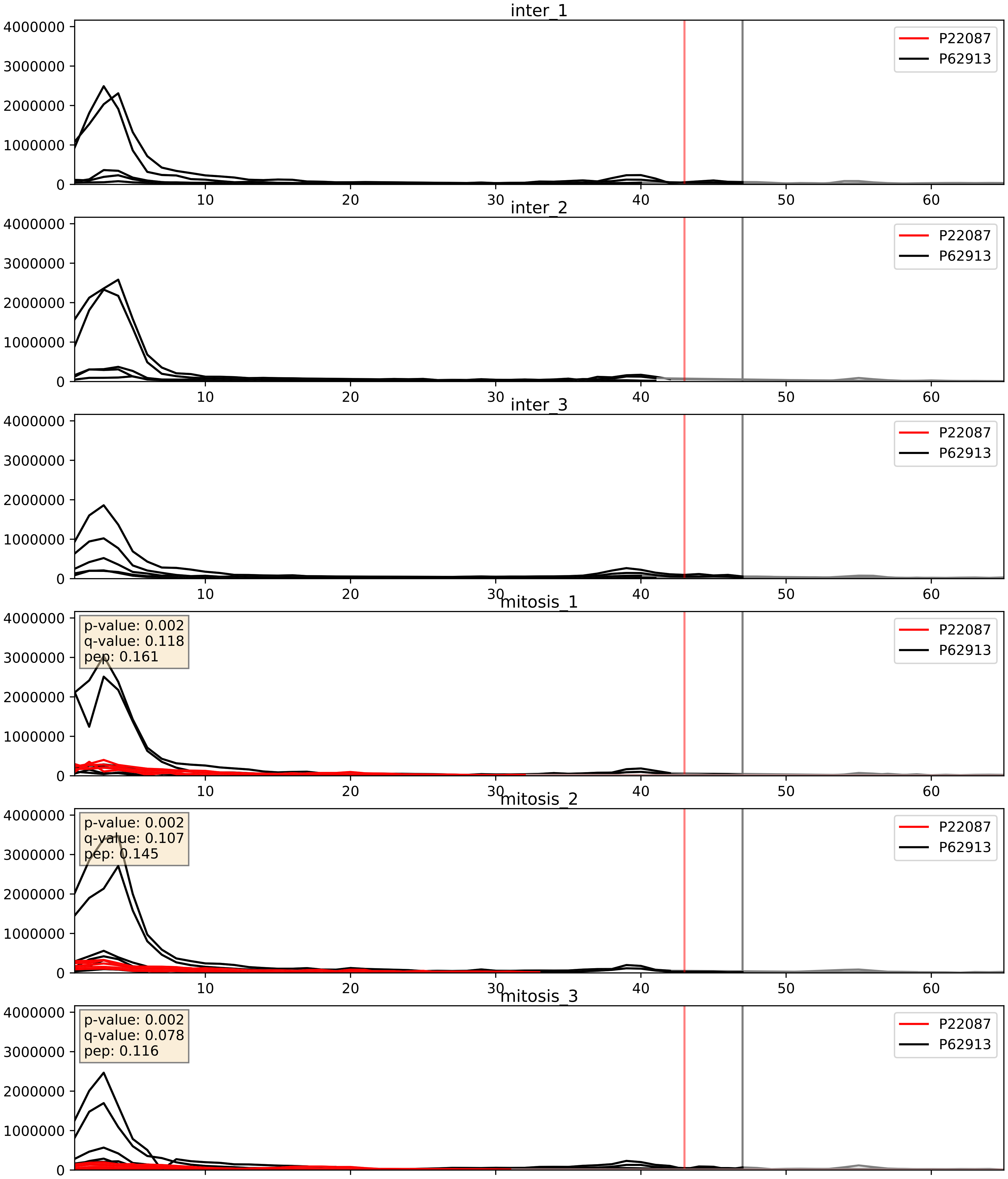


| level | condition_1 | condition_2 | pvalue | pvalue_adjusted |
|---|---|---|---|---|
| complex_abundance | mitosis | inter | 3.642014135100391e-05 | 0.005691857151652958 |
| interactor_abundance | mitosis | inter | 0.0003671184791785453 | 0.013564271459750096 |
| interactor_ratio | mitosis | inter | 0.0013469600971013035 | 0.04334578357589157 |

P22087_Q01780
FBRL_HUMAN vs EXOSX_HUMAN
p-value: 0.002 q-value: 0.034

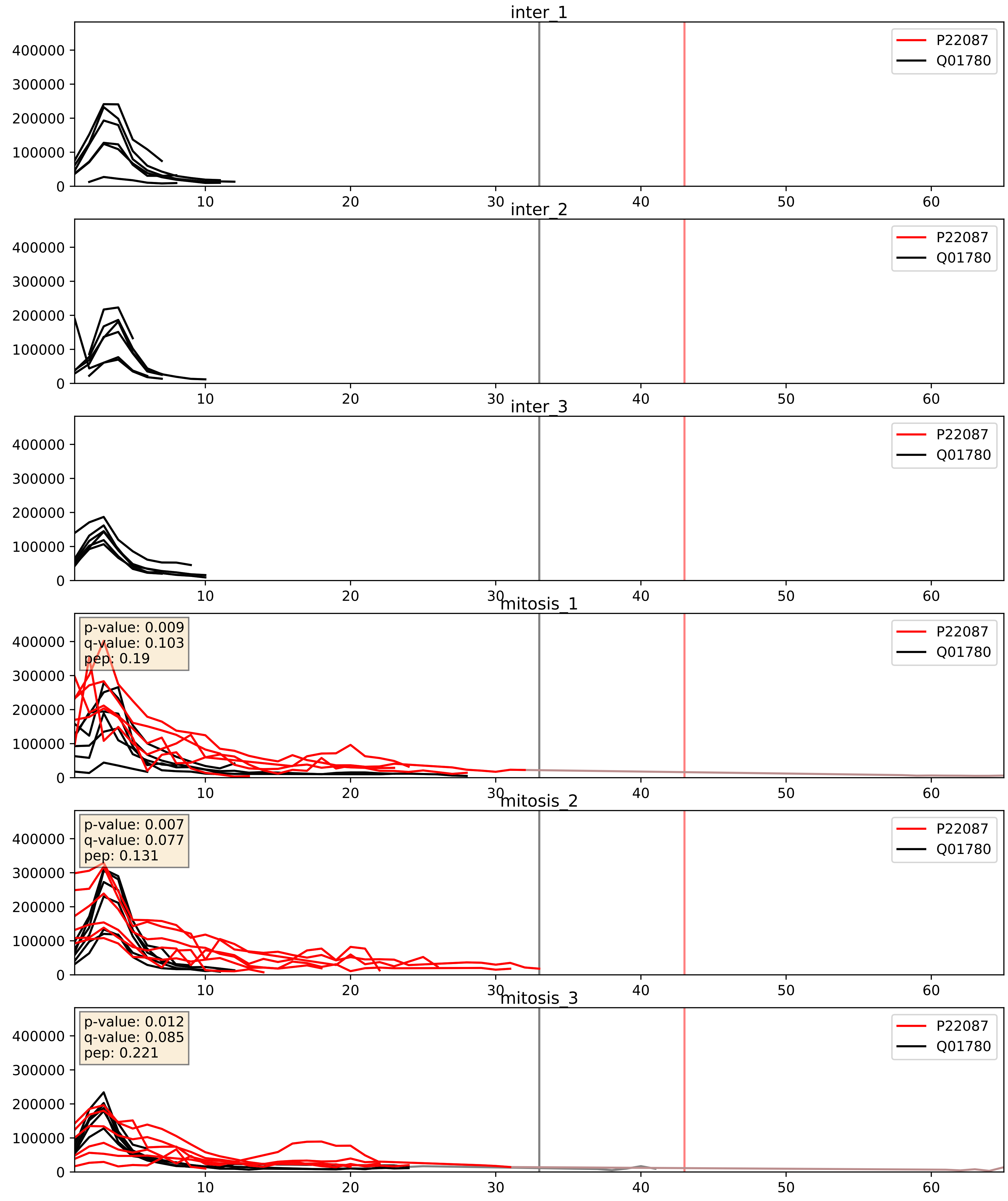


| level | condition_1 | condition_2 | pvalue | pvalue_adjusted |
|---|---|---|---|---|
| interactor_ratio | mitosis | inter | 3.1797532631946304e-06 | 0.002520386835947235 |
| interactor_abundance | mitosis | inter | 0.0004135262530475819 | 0.014019253191624928 |
| complex_abundance | mitosis | inter | 0.0006473146188650986 | 0.016393529992559892 |

P22087_Q02543
FBRL_HUMAN vs RL18A_HUMAN
p-value: 0.0 q-value: 0.01

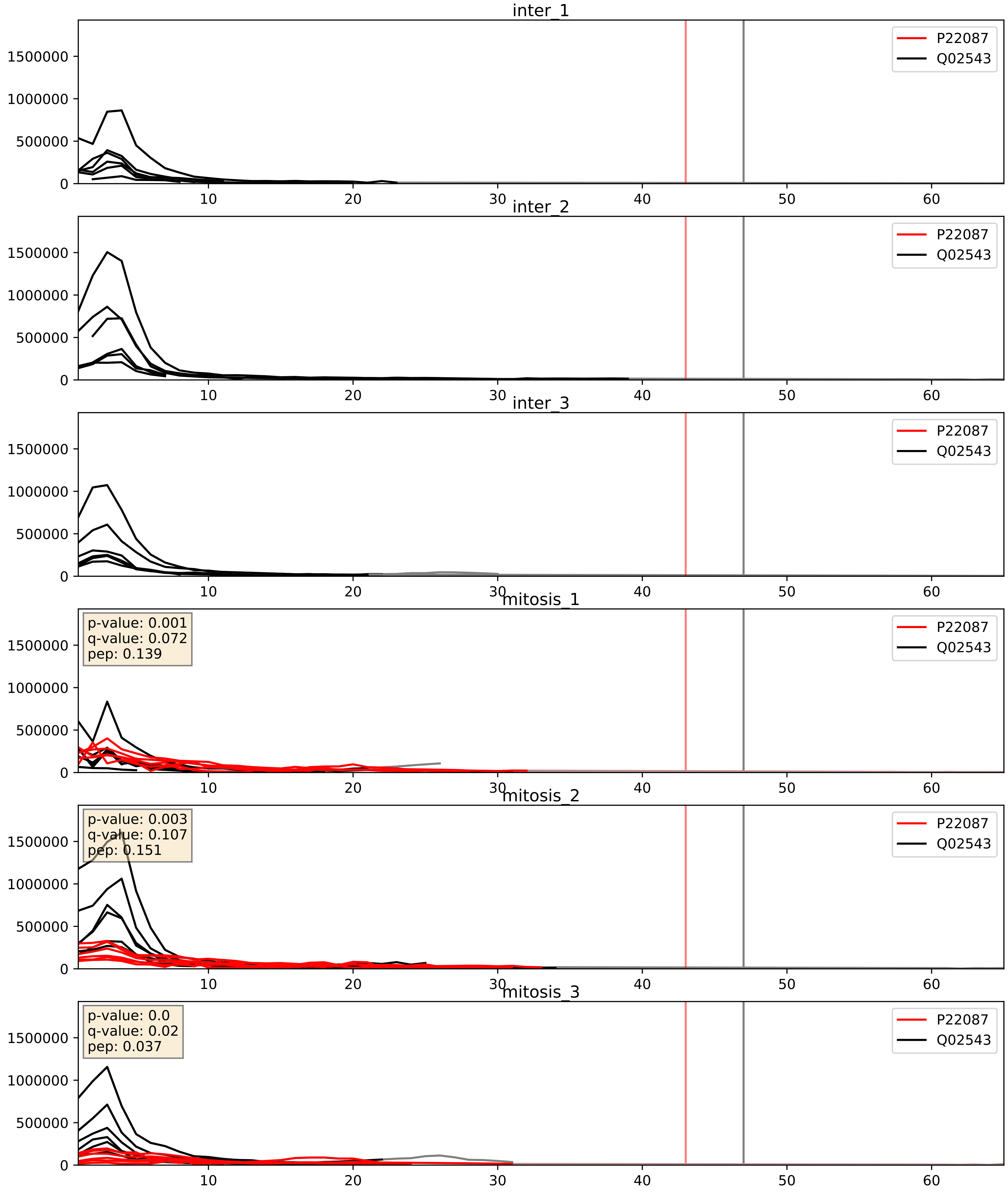


| level | condition_1 | condition_2 | pvalue | pvalue_adjusted |
|---|---|---|---|---|
| complex_abundance | mitosis | inter | 0.00018840423370753738 | 0.009111526782692203 |
| interactor_abundance | mitosis | inter | 0.00037774436755851884 | 0.013573913378163039 |
| interactor_ratio | mitosis | inter | 0.4211998682973811 | 0.8388785723072948 |

P22087_Q12788
FBRL_HUMAN vs TBL3_HUMAN
p-value: 0.009 q-value: 0.008

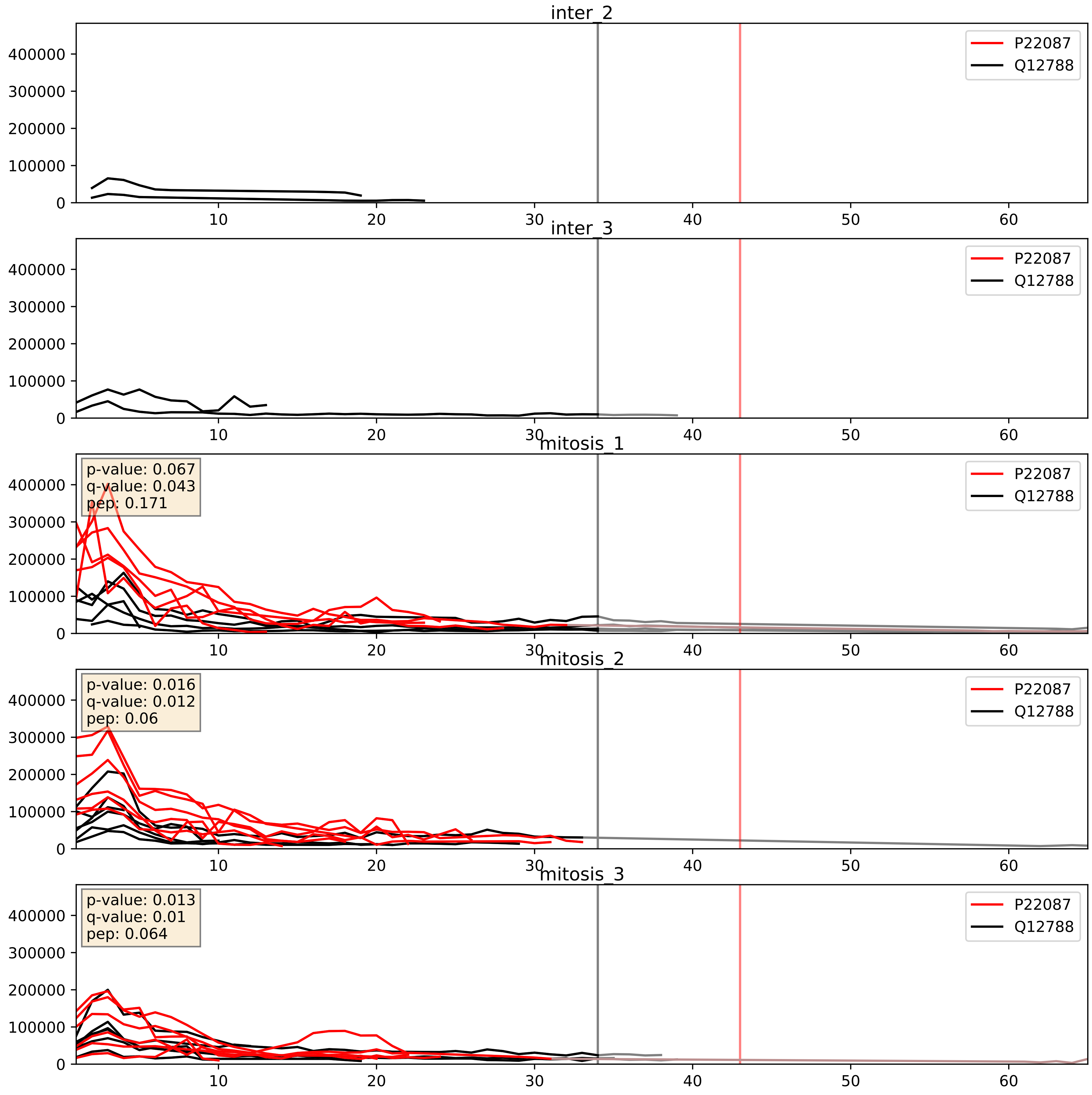


| level | condition_1 | condition_2 | pvalue | pvalue_adjusted |
|---|---|---|---|---|
| complex_abundance | mitosis | inter | 0.0001327659247700999 | 0.007892196639111494 |
| interactor_abundance | mitosis | inter | 0.0003671184791785453 | 0.01356427145975096 |
| interactor_ratio | mitosis | inter | 0.02640428302391608 | 0.25543205403291264 |

P22087_Q13895
FBRL_HUMAN vs BYST_HUMAN
p-value: 0.062 q-value: 0.039

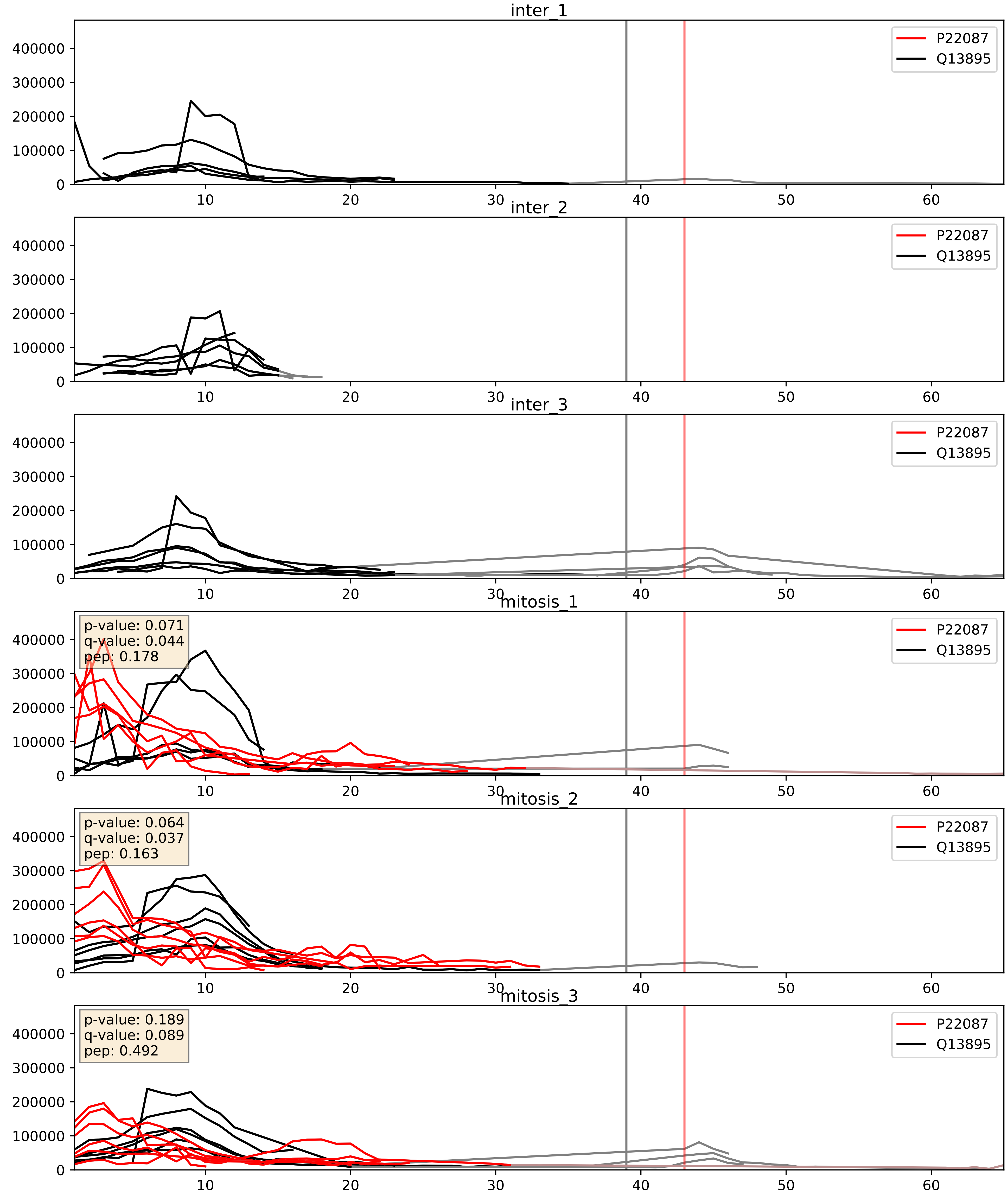


| level | condition_1 | condition_2 | pvalue | pvalue_adjusted |
|---|---|---|---|---|
| complex_abundance | mitosis | inter | 0.0001916121831474646 | 0.009214608358102793 |
| interactor_abundance | mitosis | inter | 0.00035333092874419484 | 0.01356427145975096 |
| interactor_ratio | mitosis | inter | 0.26434880442933373 | 0.7392439614227693 |

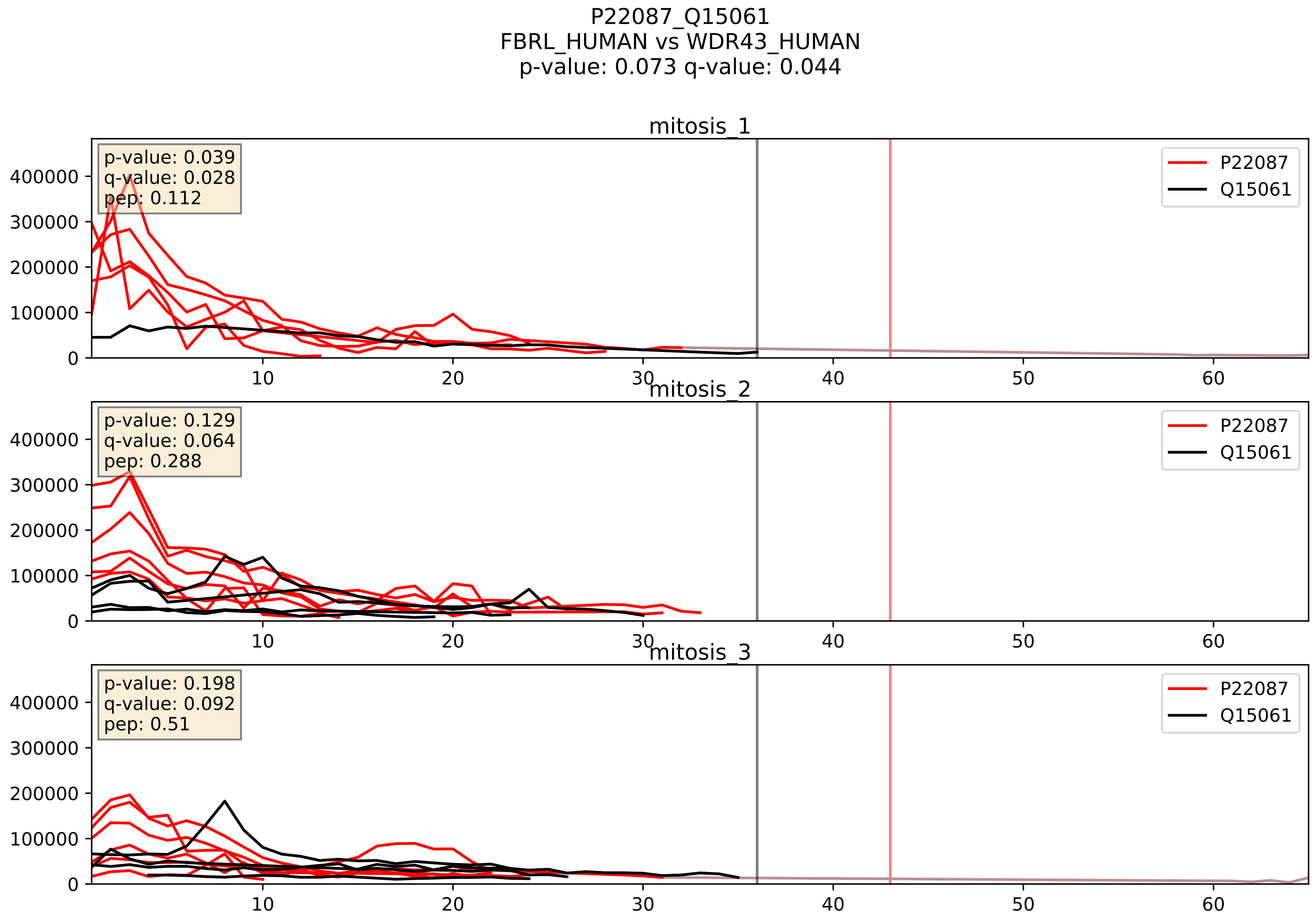


| level | condition_1 | condition_2 | pvalue | pvalue_adjusted |
|---|---|---|---|---|
| interactor_abundance | mitosis | inter | 0.00022145143300215012 | 0.011215116209912415 |
| complex_abundance | mitosis | inter | 0.000804026877839027 | 0.018753324453139156 |
| interactor_ratio | mitosis | inter | 0.05603485152930944 | 0.3926647223857623 |

P22087_Q15269
FBRL_HUMAN vs PWP2_HUMAN
p-value: 0.001 q-value: 0.002

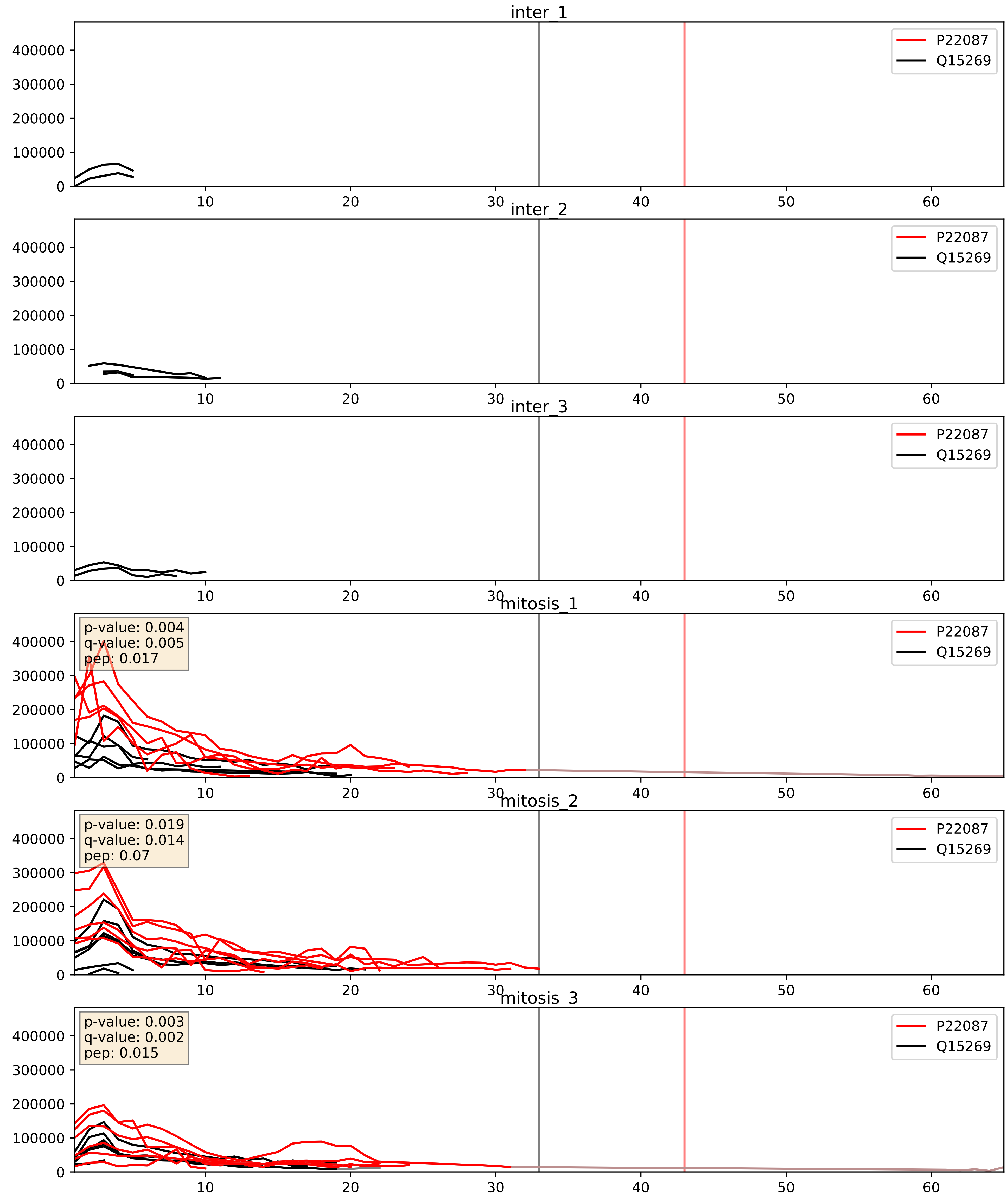


| level | condition_1 | condition_2 | pvalue | pvalue_adjusted |
|---|---|---|---|---|
| interactor_abundance | mitosis | inter | 0.0005008244807677675 | 0.015907449185054136 |
| interactor_ratio | mitosis | inter | 0.0005887273659099557 | 0.028471786735532318 |
| complex_abundance | mitosis | inter | 0.000661858101371452 | 0.01642175463112936 |

P22087_Q8IY81
FBRL_HUMAN vs SPB1_HUMAN
p-value: 0.02 q-value: 0.016

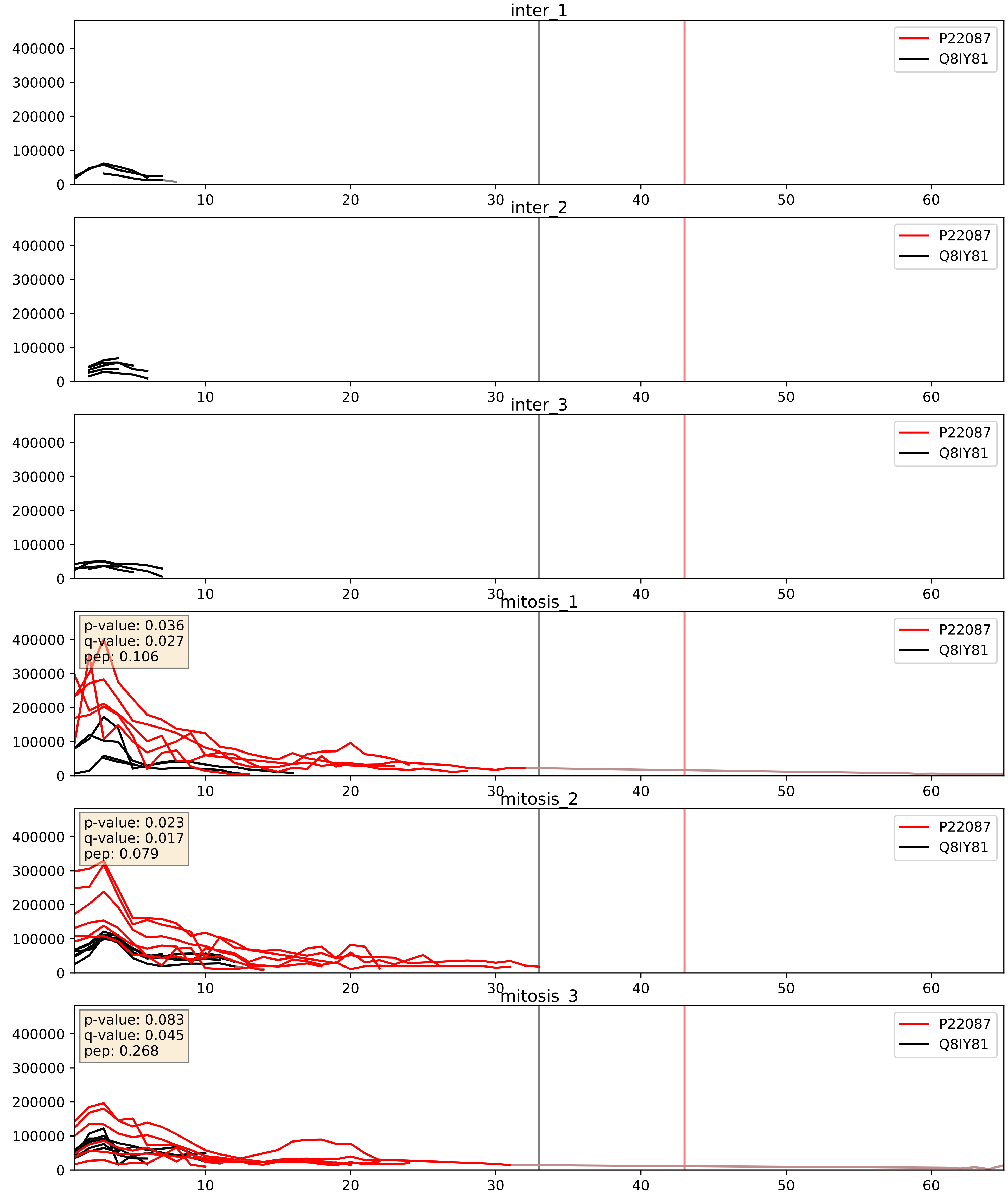


| level | condition_1 | condition_2 | pvalue | pvalue_adjusted |
|---|---|---|---|---|
| interactor_abundance | mitosis | inter | 0.0010710204172406286 | 0.024049720606162197 |
| complex_abundance | mitosis | inter | 0.0012994811847285065 | 0.024459508269436846 |
| interactor_ratio | mitosis | inter | 0.0034502879269765964 | 0.07572939655107606 |

P22087_Q8NI36
FBRL_HUMAN vs WDR36_HUMAN
p-value: 0.005 q-value: 0.005

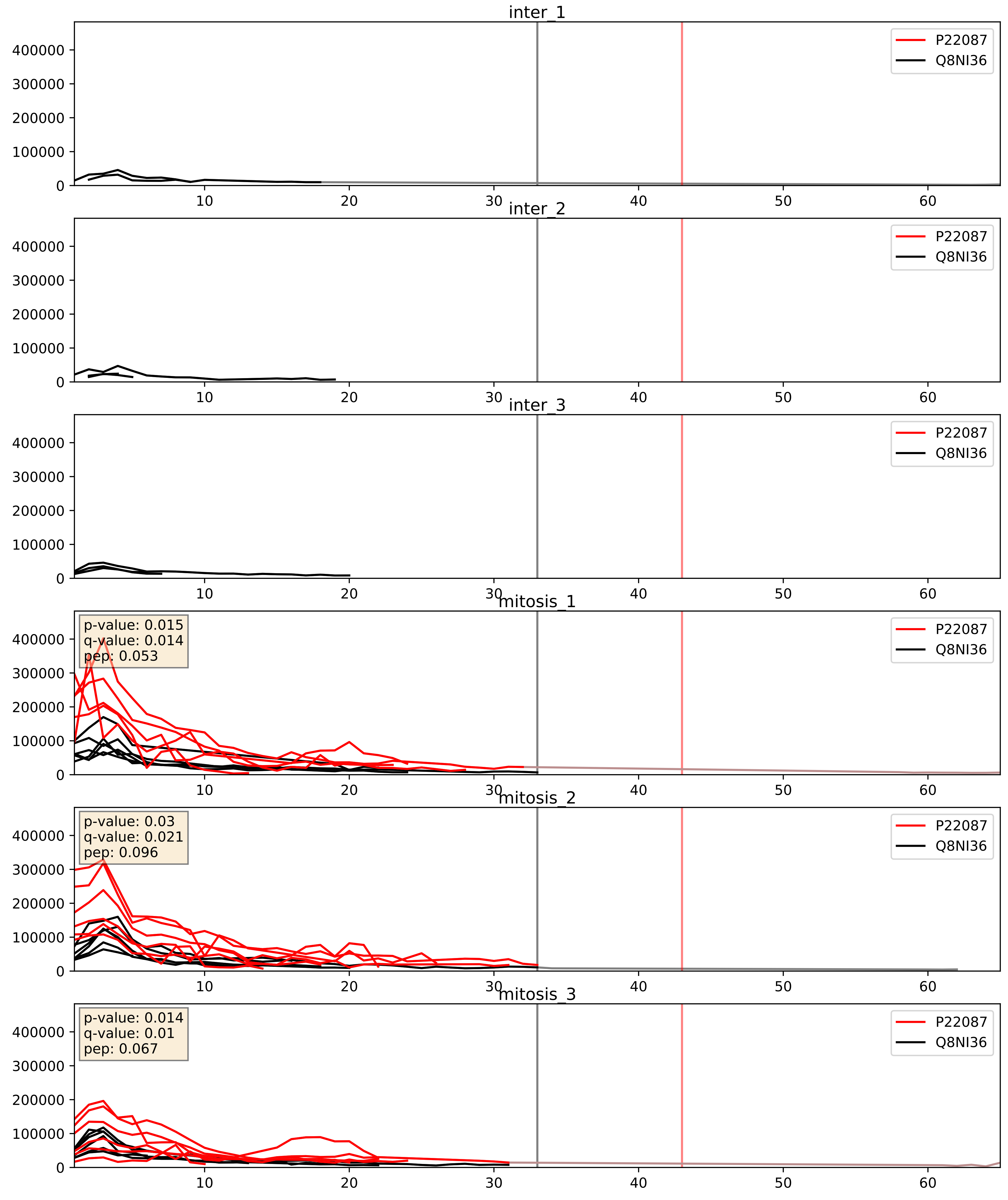


| level | condition_1 | condition_2 | pvalue | pvalue_adjusted |
|---|---|---|---|---|
| complex_abundance | mitosis | inter | 0.00034440768556571094 | 0.012015488415248196 |
| interactor_abundance | mitosis | inter | 0.0003496674327824524 | 0.01356427145975096 |
| interactor_ratio | mitosis | inter | 0.0010740795022735833 | 0.038414698994273265 |

P22087_Q92979
FBRL_HUMAN vs NEP1_HUMAN
p-value: 0.035 q-value: 0.024

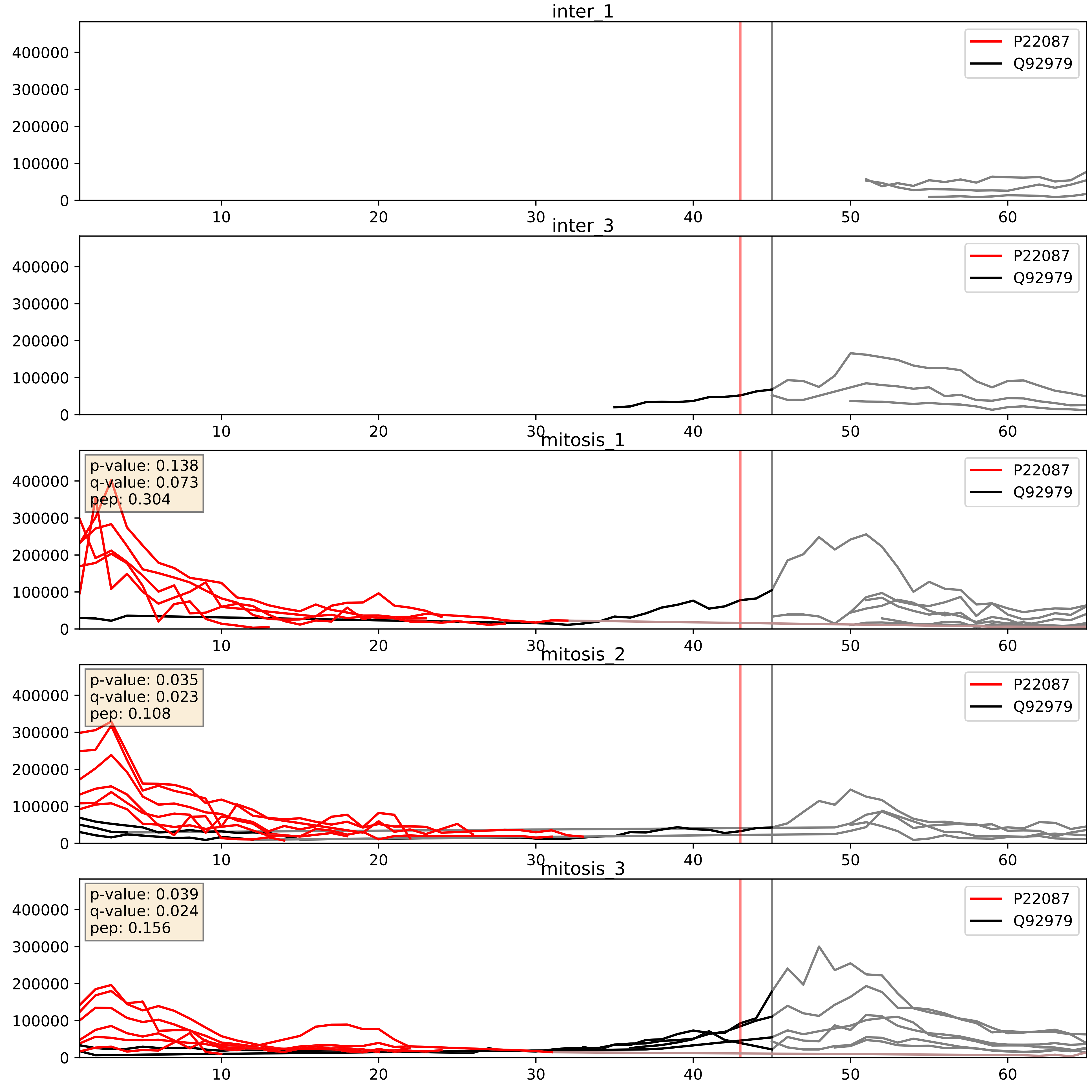


| level | condition_1 | condition_2 | pvalue | pvalue_adjusted |
|---|---|---|---|---|
| interactor_ratio | mitosis | inter | 9.565040877399547e-05 | 0.010510079284752772 |
| interactor_abundance | mitosis | inter | 0.0010078483561365378 | 0.023668537526827883 |
| complex_abundance | mitosis | inter | 0.0040748418684974745 | 0.04732787841836958 |

P22087_Q969X6
FBRL_HUMAN vs UTP4_HUMAN
p-value: 0.081 q-value: 0.048

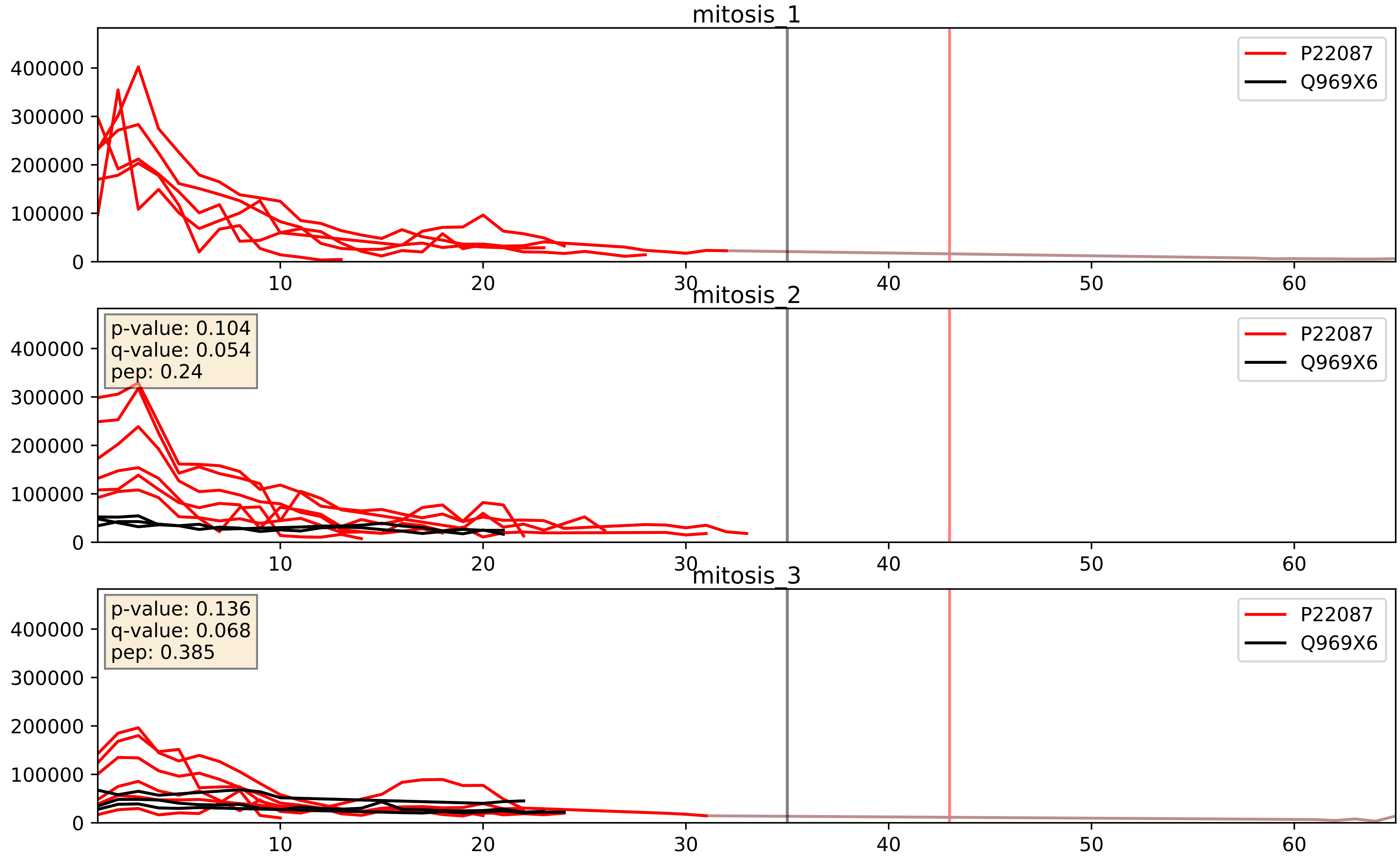


| level | condition_1 | condition_2 | pvalue | pvalue_adjusted |
|---|---|---|---|---|
| interactor_ratio | mitosis | inter | 0.14264333123055517 | 0.5930193857860866 |
| interactor_abundance | mitosis | inter | 0.16030116872089478 | 0.34690380590339004 |
| complex_abundance | mitosis | inter | 0.1796374809803189 | 0.3300486879569714 |

P22087_Q9BQ67
FBRL_HUMAN vs GRWD1_HUMAN
p-value: 0.001 q-value: 0.029

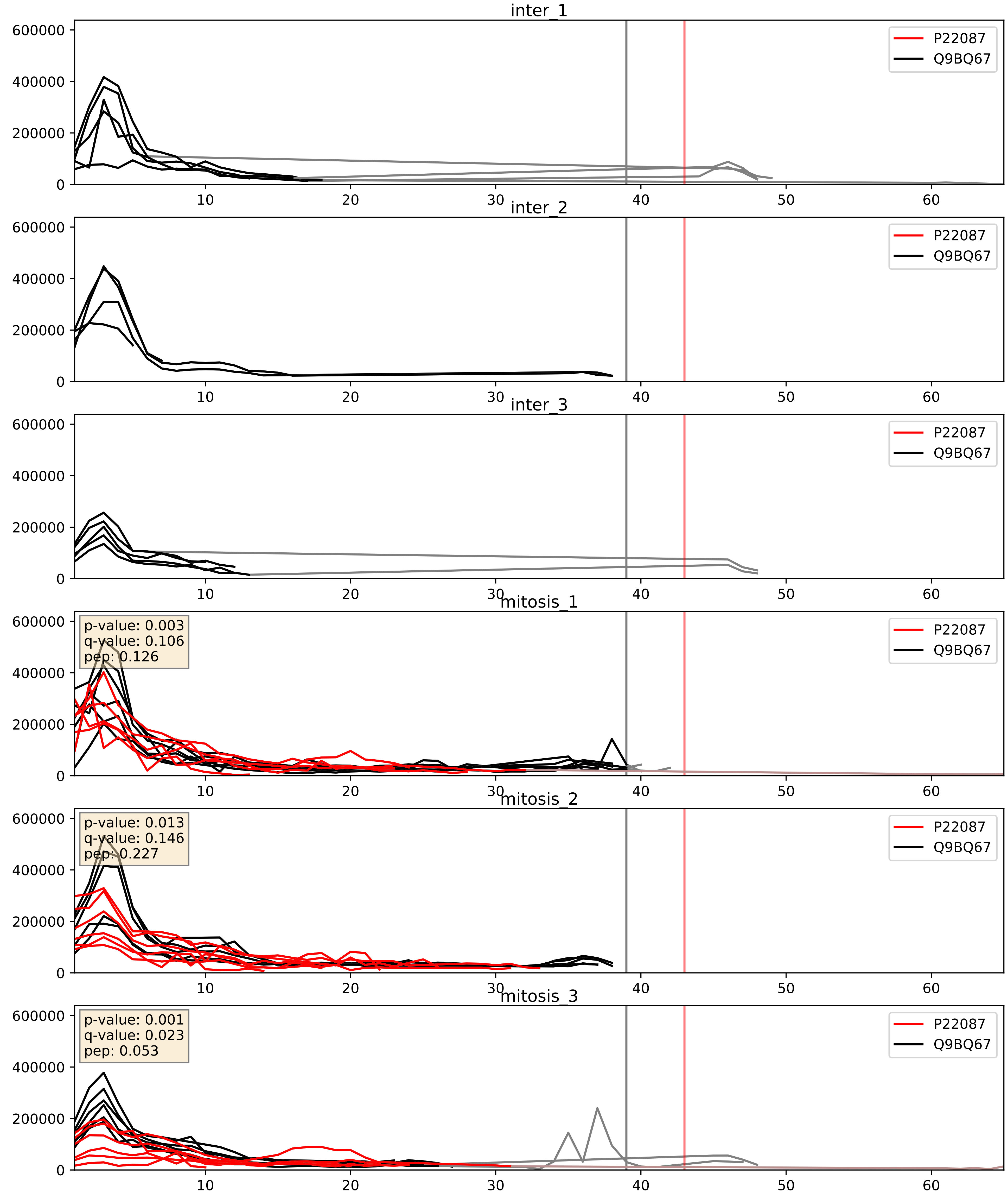


| level | condition_1 | condition_2 | pvalue | pvalue_adjusted |
|---|---|---|---|---|
| complex_abundance | mitosis | inter | 0.00022948761081750734 | 0.010219389159334393 |
| interactor_abundance | mitosis | inter | 0.0003900245617774174 | 0.01362698060740691 |
| interactor_ratio | mitosis | inter | 0.011537958352666234 | 0.15494522140242886 |

P22087_Q9BVP2
FBRL_HUMAN vs GNL3_HUMAN
p-value: 0.001 q-value: 0.027

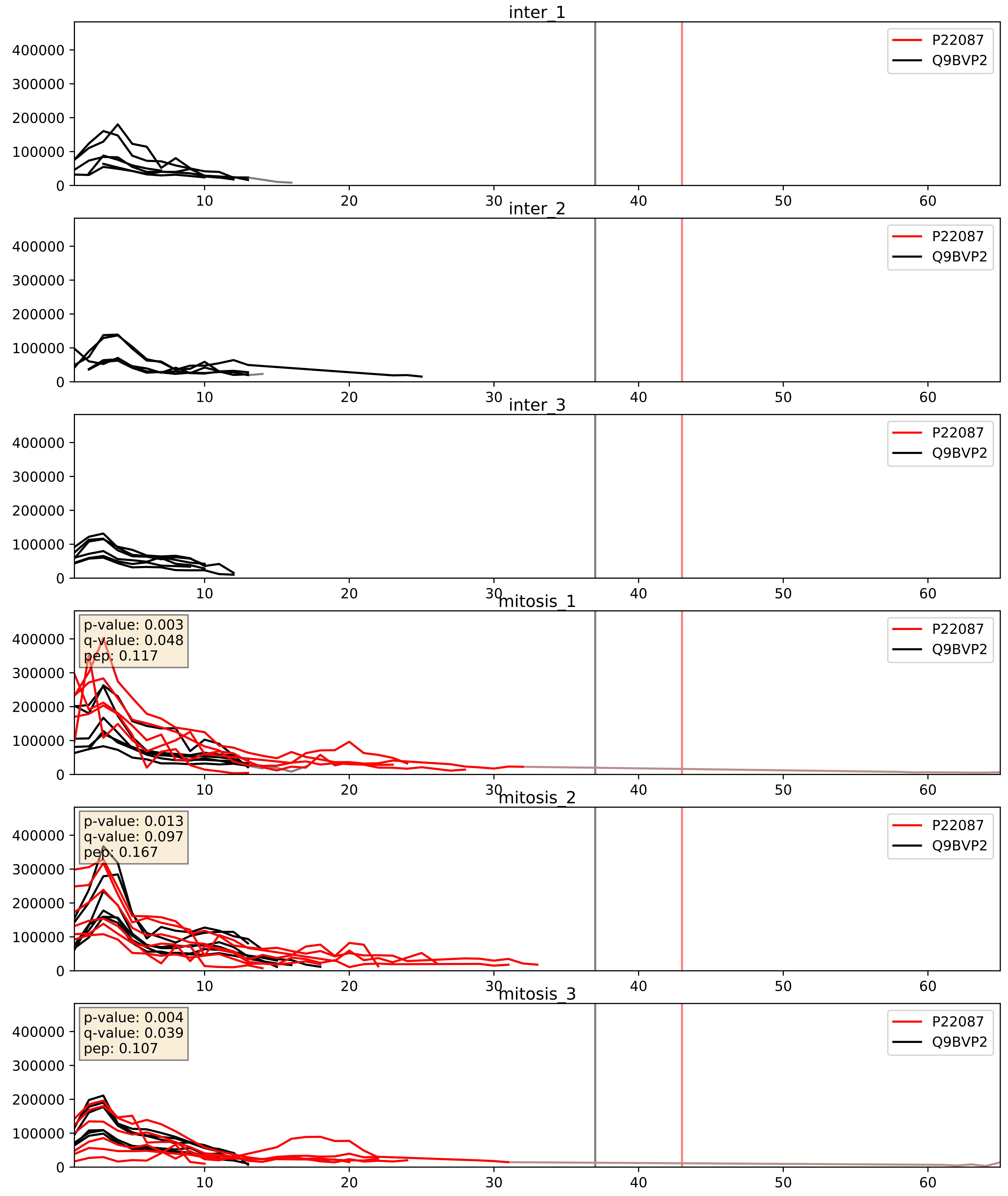


| level | condition_1 | condition_2 | pvalue | pvalue_adjusted |
|---|---|---|---|---|
| complex_abundance | mitosis | inter | 0.00030575611516796747 | 0.01165207110963347 |
| interactor_abundance | mitosis | inter | 0.0006136131553603794 | 0.017415659477552636 |
| interactor_ratio | mitosis | inter | 0.22854497040560845 | 0.7070495646135039 |

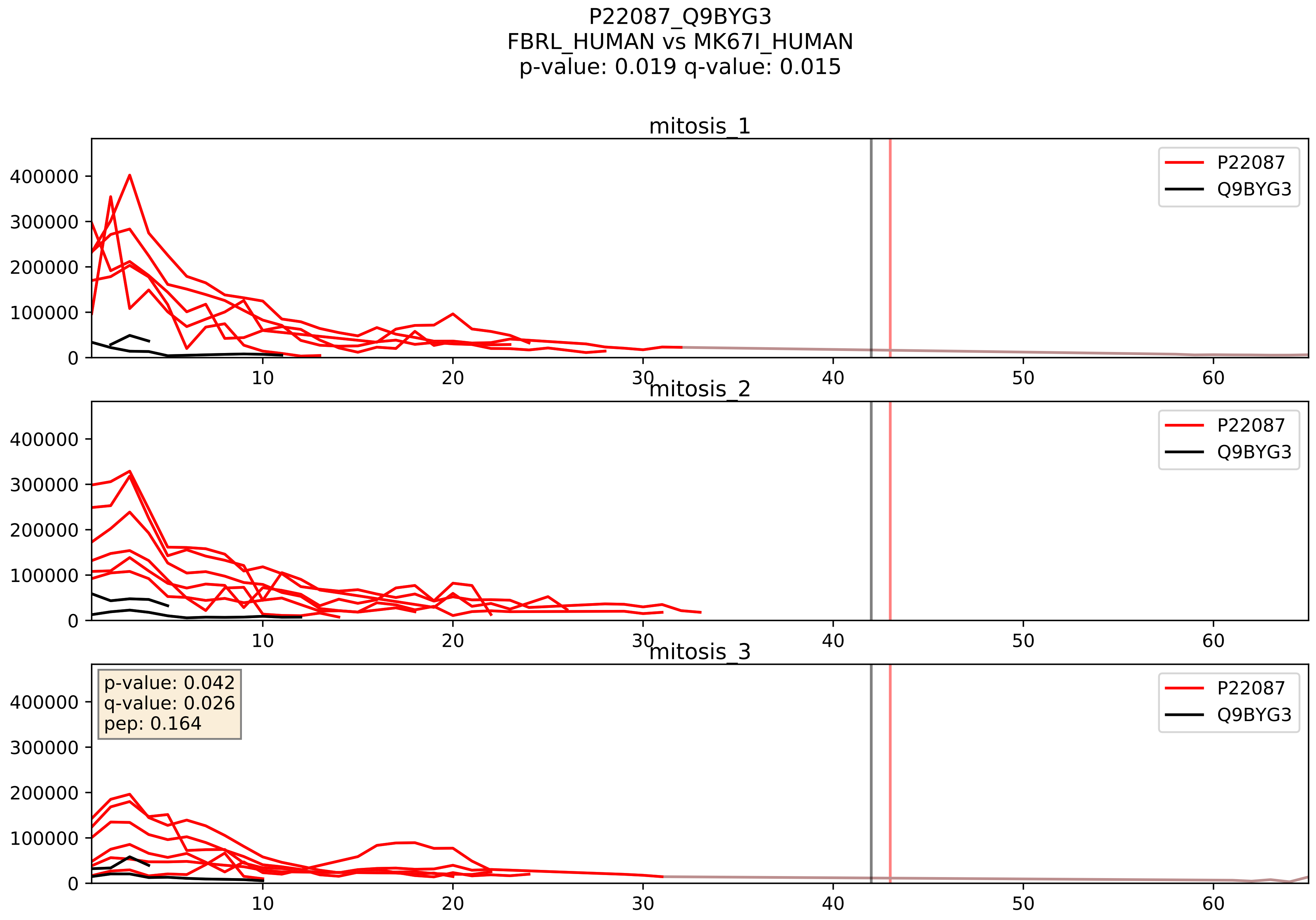


| level | condition_1 | condition_2 | pvalue | pvalue_adjusted |
|---|---|---|---|---|
| interactor_ratio | mitosis | inter | 0.00022528201190776103 | 0.01529848062979237 |
| interactor_abundance | mitosis | inter | 0.0005233714416599235 | 0.016562142479145826 |
| complex_abundance | mitosis | inter | 0.0016144918603450086 | 0.02814408910325431 |

P22087_Q9H0S4
FBRL_HUMAN vs DDX47_HUMAN
p-value: 0.003 q-value: 0.042

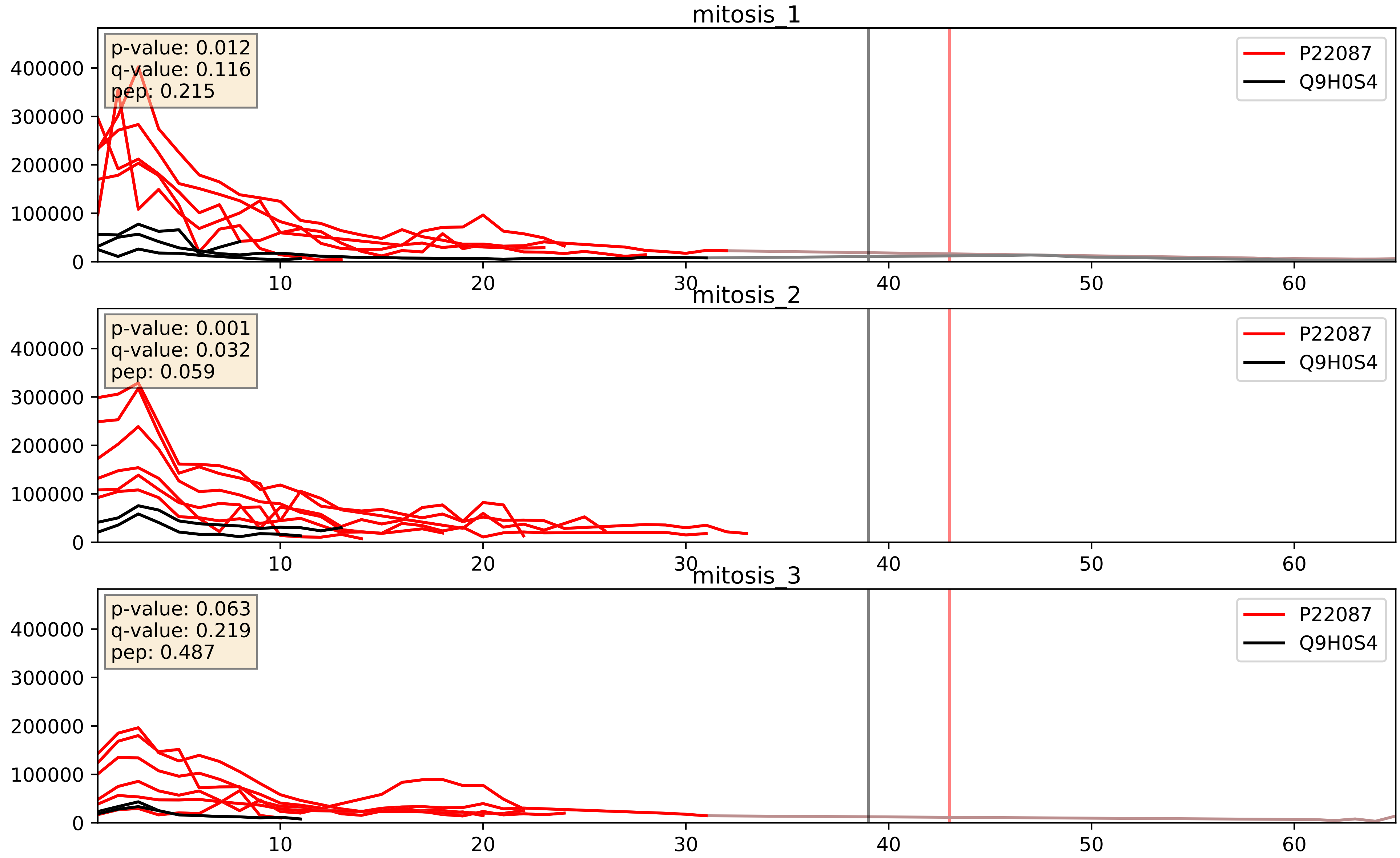


| level | condition_1 | condition_2 | pvalue | pvalue_adjusted |
|---|---|---|---|---|
| interactor_ratio | mitosis | inter | 6.650160997708512e-05 | 0.008371379138291892 |
| interactor_abundance | mitosis | inter | 0.0010898797318194015 | 0.024049720606162197 |
| complex_abundance | mitosis | inter | 0.011229383006423614 | 0.0785322863847926 |

P22087_Q9H583
FBRL_HUMAN vs HEAT1_HUMAN
p-value: 0.069 q-value: 0.042

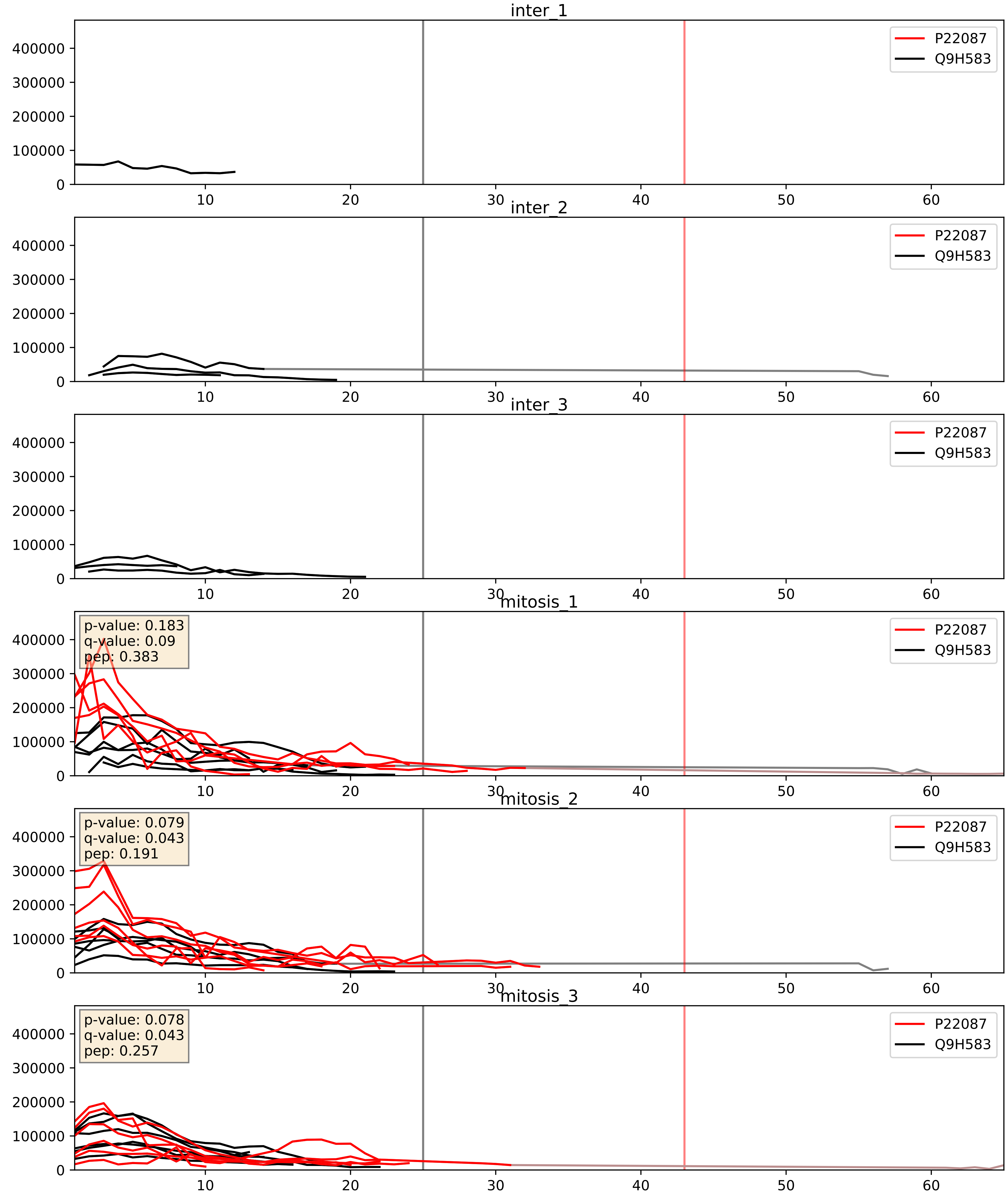


| level | condition_1 | condition_2 | pvalue | pvalue_adjusted |
|---|---|---|---|---|
| complex_abundance | mitosis | inter | 9.233901065022553e-05 | 0.006532412654263889 |
| interactor_abundance | mitosis | inter | 0.00038515934981581823 | 0.013595727977003728 |
| interactor_ratio | mitosis | inter | 0.08842363453252496 | 0.4953575337685953 |

P22087_Q9NQT4
FBRL_HUMAN vs EXOS5_HUMAN
p-value: 0.003 q-value: 0.044

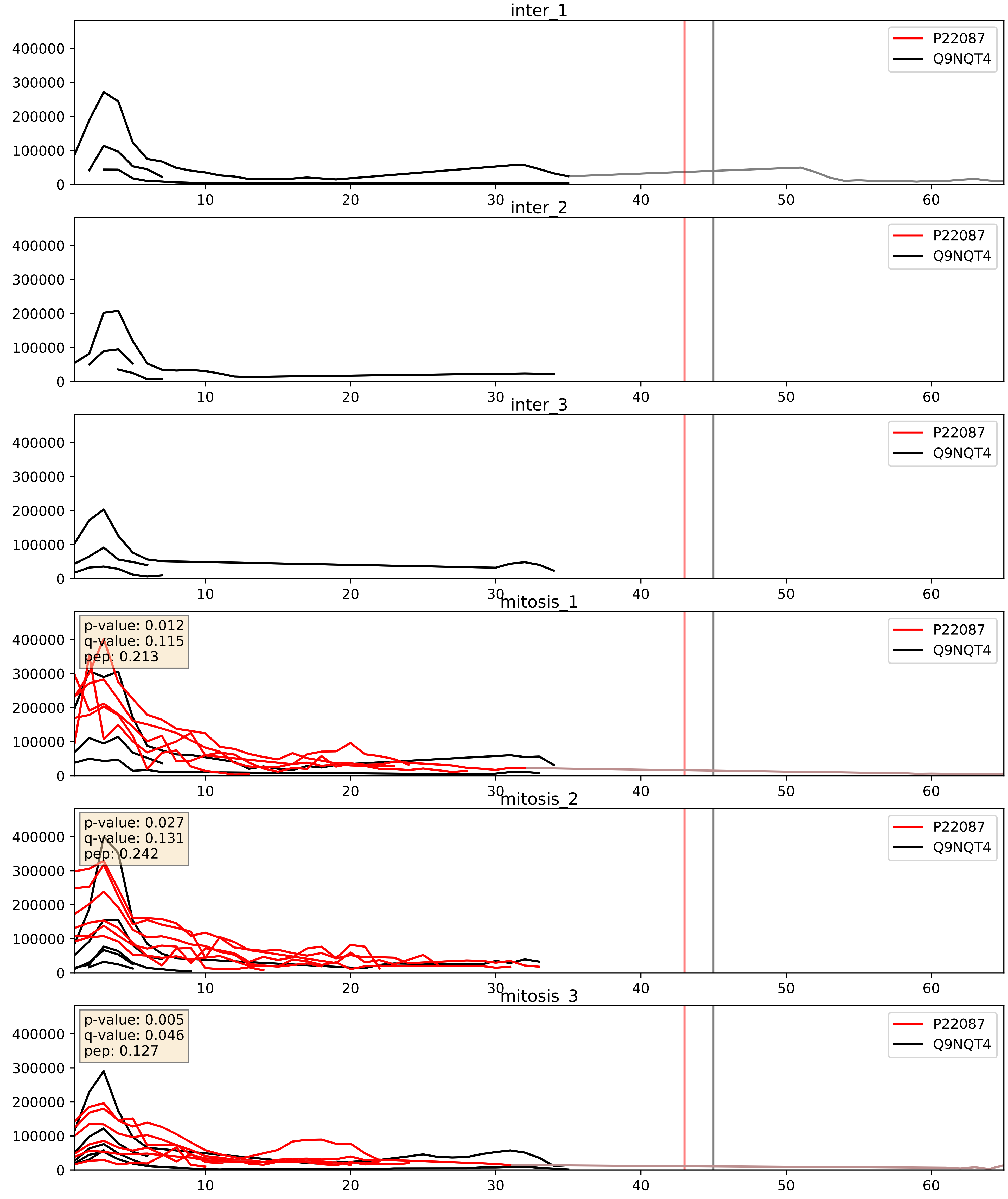


| level | condition_1 | condition_2 | pvalue | pvalue_adjusted |
|---|---|---|---|---|
| interactor_ratio | mitosis | inter | 6.0081390880697794e-05 | 0.008128440693030705 |
| interactor_abundance | mitosis | inter | 0.0003046041652790056 | 0.013073369371061003 |
| complex_abundance | mitosis | inter | 0.0006369301444058181 | 0.01632371867099941 |

P22087_Q9NX24
FBRL_HUMAN vs NHP2_HUMAN
p-value: 0.0 q-value: 0.0

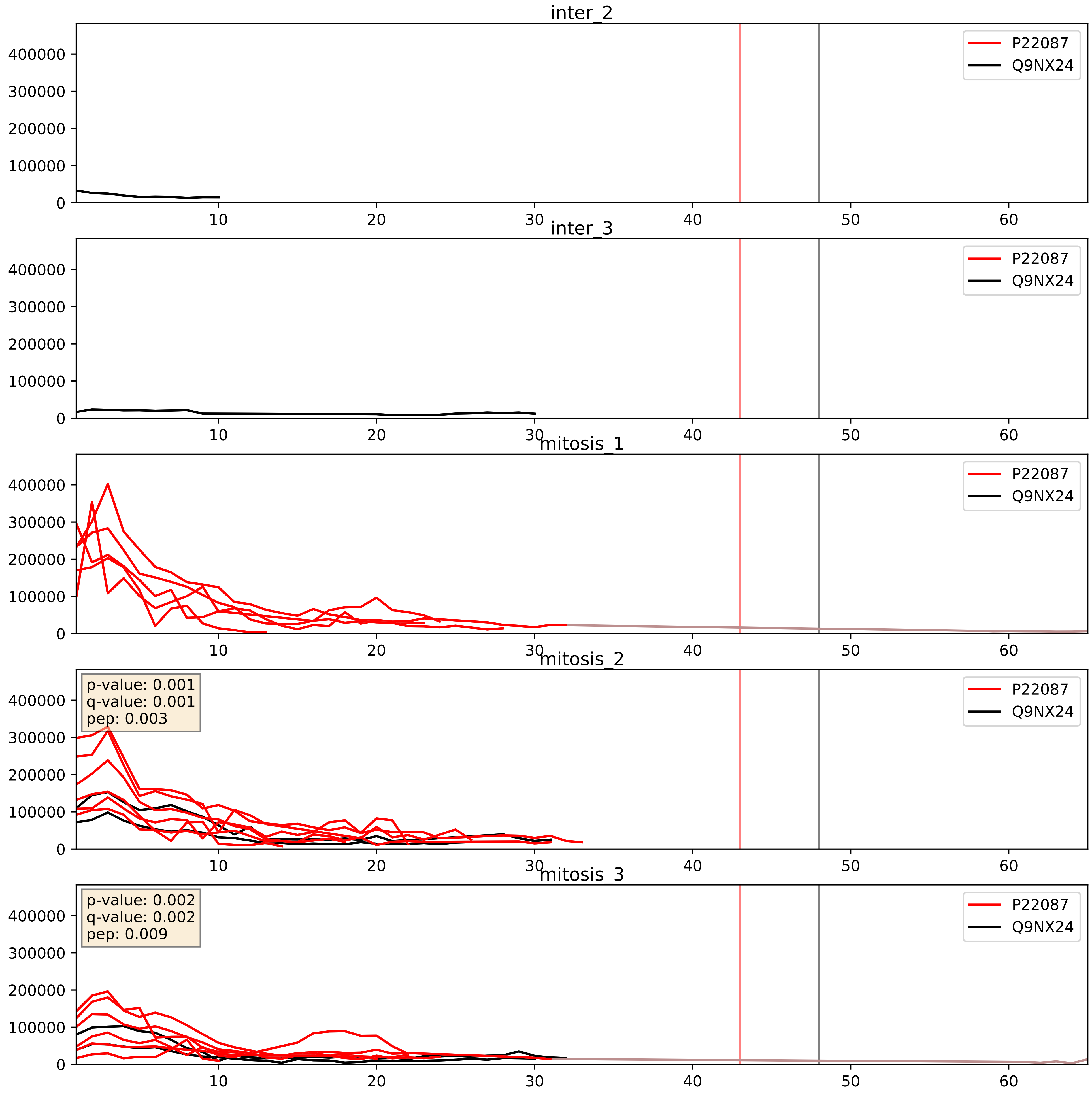


| level | condition_1 | condition_2 | pvalue | pvalue_adjusted |
|---|---|---|---|---|
| interactor_ratio | mitosis | inter | 0.14261129595203648 | 0.5930193857860866 |
| interactor_abundance | mitosis | inter | 0.16018705696216212 | 0.3467445208233931 |
| complex_abundance | mitosis | inter | 0.16754336790949512 | 0.31624503402541965 |

P22087_Q9UKD2
FBRL_HUMAN vs MRT4_HUMAN
p-value: 0.002 q-value: 0.034

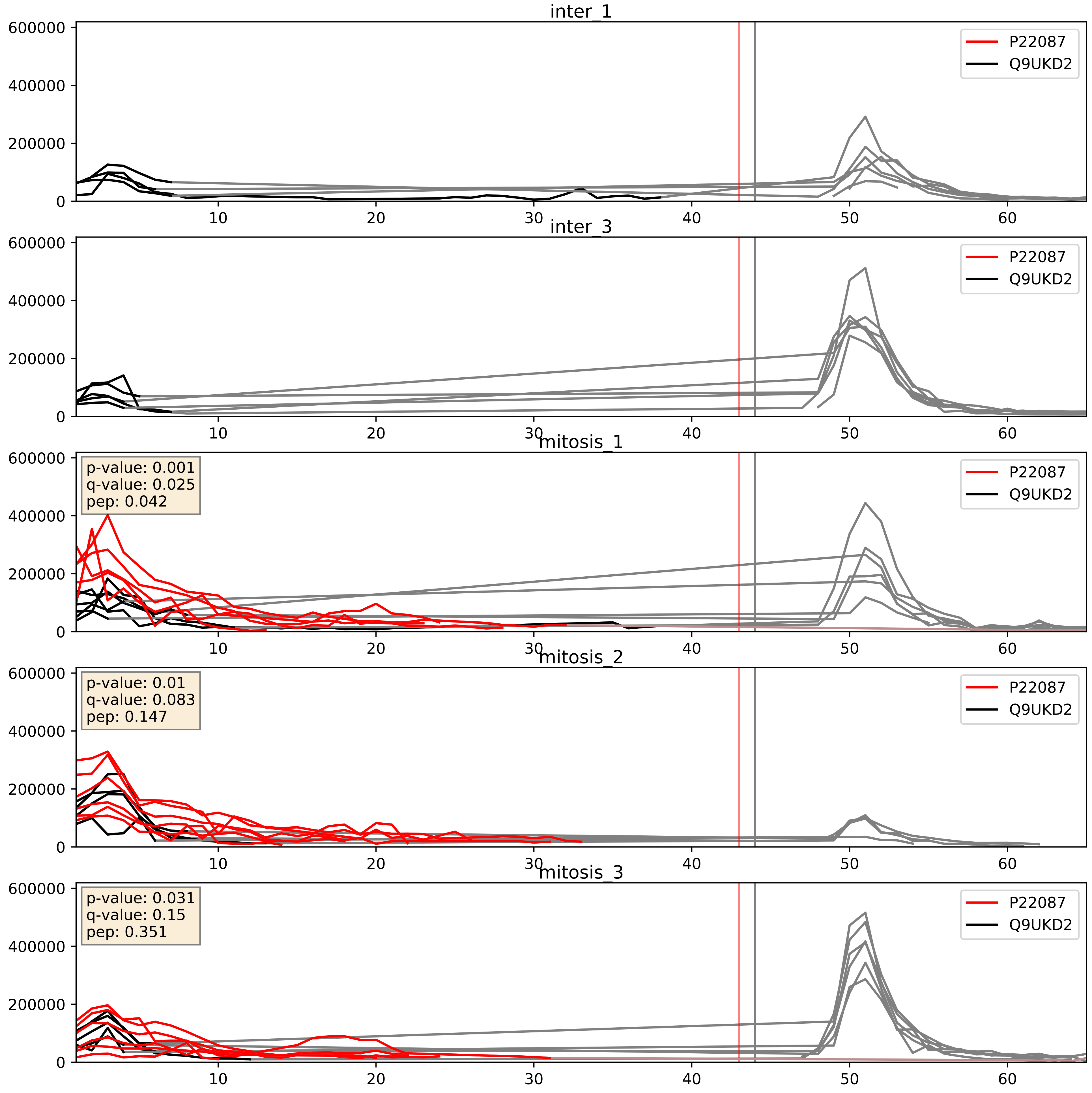


| level | condition_1 | condition_2 | pvalue | pvalue_adjusted |
|---|---|---|---|---|
| complex_abundance | mitosis | inter | 0.0008743237108210411 | 0.019513211095311817 |
| interactor_abundance | mitosis | inter | 0.0009724779092061511 | 0.023091292379485866 |
| interactor_ratio | mitosis | inter | 0.004228737105466719 | 0.08567853202577967 |

P22087_Q9UNX4
FBRL_HUMAN vs WDR3_HUMAN
p-value: 0.025 q-value: 0.019

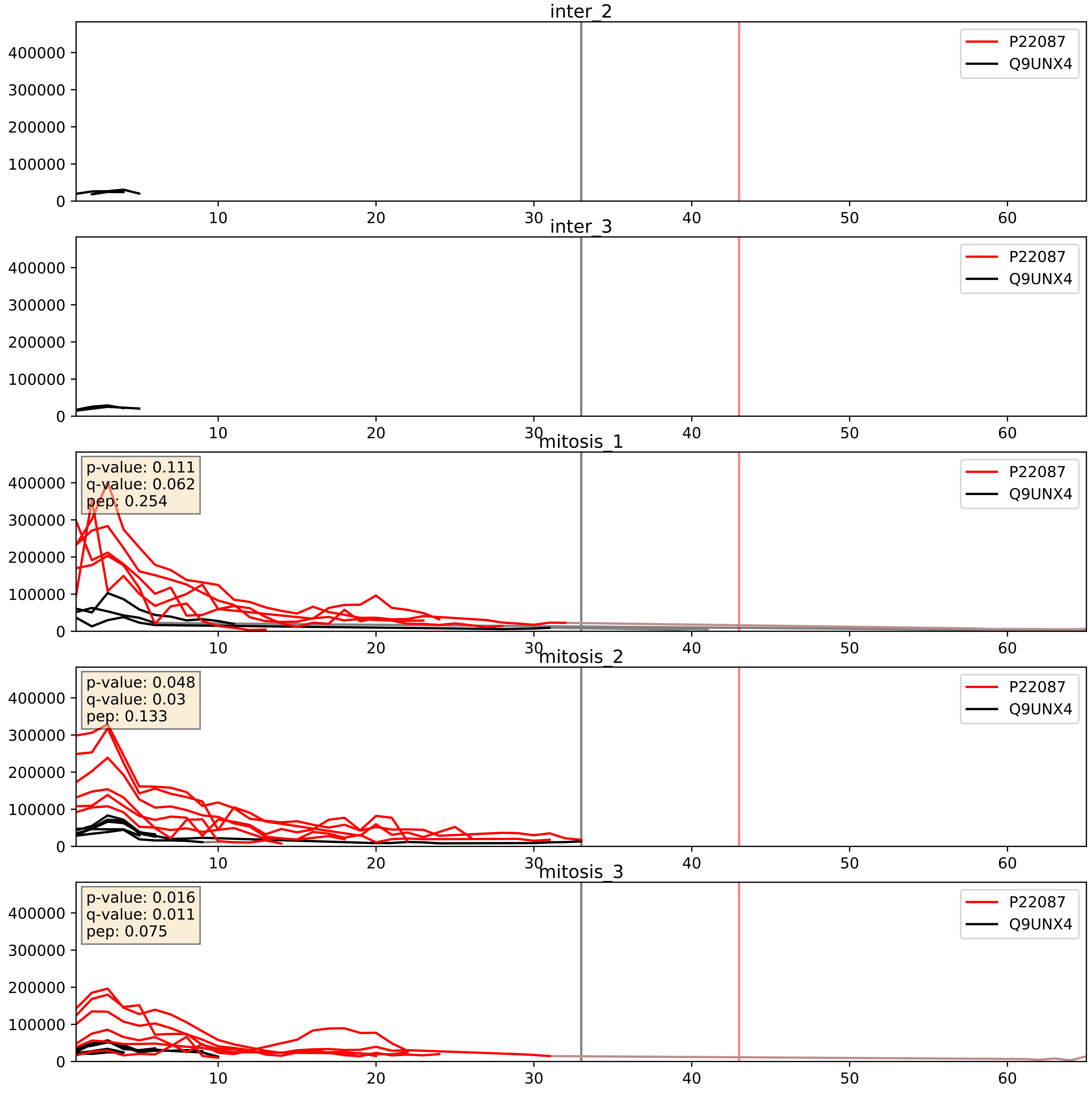


| level | condition_1 | condition_2 | pvalue | pvalue_adjusted |
|---|---|---|---|---|
| interactor_ratio | mitosis | inter | 0.00040930987289006183 | 0.022604467818960836 |
| interactor_abundance | mitosis | inter | 0.0008065885048889307 | 0.020834500976147856 |
| complex_abundance | mitosis | inter | 0.002005177899684022 | 0.032203232310122384 |

P22087_Q9Y221
FBRL_HUMAN vs NIP7_HUMAN
p-value: 0.01 q-value: 0.008

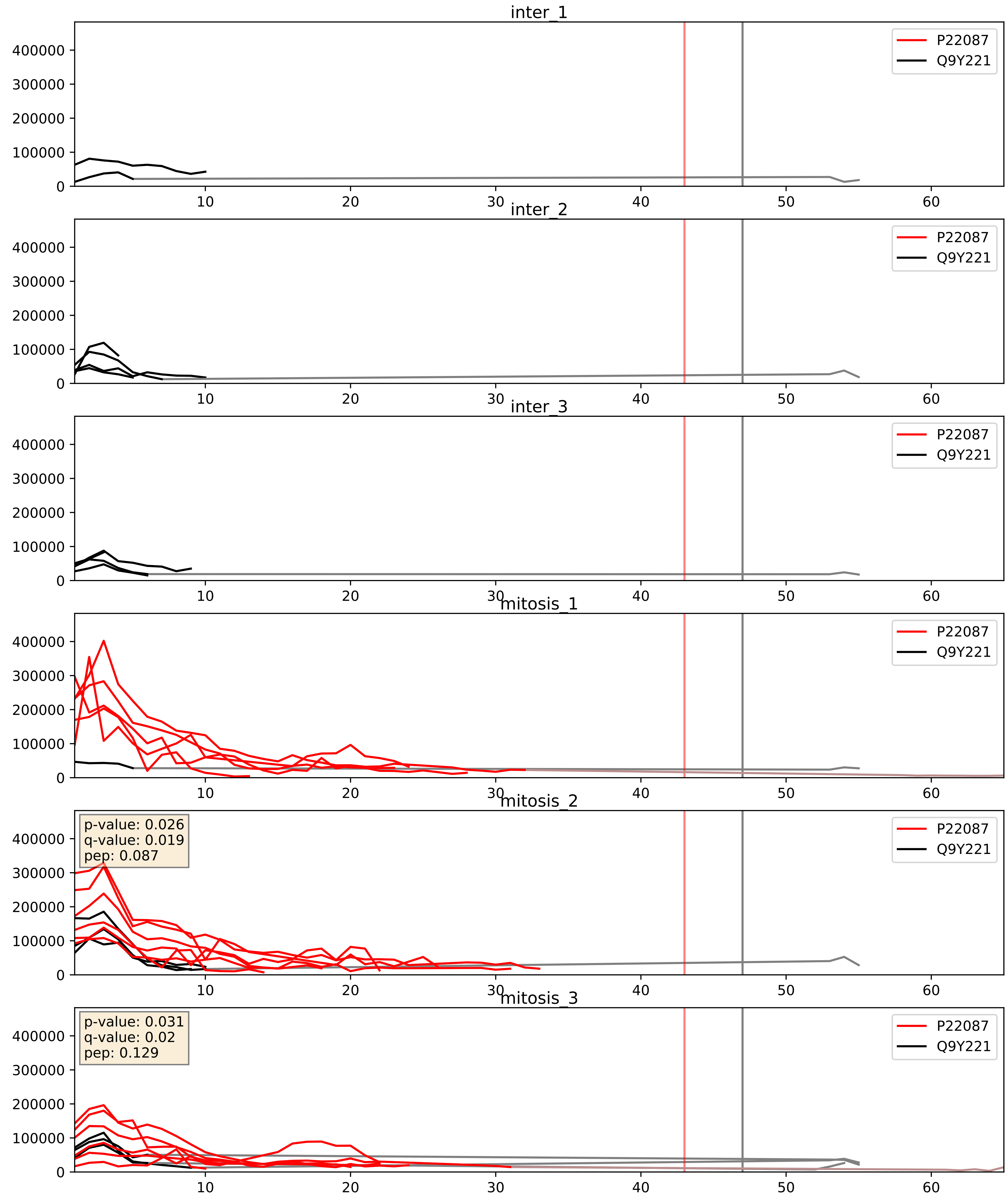


| level | condition_1 | condition_2 | pvalue | pvalue_adjusted |
|---|---|---|---|---|
| interactor_abundance | mitosis | inter | 0.00042214500914833364 | 0.014167094422781207 |
| complex_abundance | mitosis | inter | 0.003888247479744257 | 0.046162827221374256 |
| interactor_ratio | mitosis | inter | 0.008710686291152805 | 0.13211821468820312 |

P22087_Q9Y2X3
FBRL_HUMAN vs NOP58_HUMAN
p-value: 0.0 q-value: 0.0

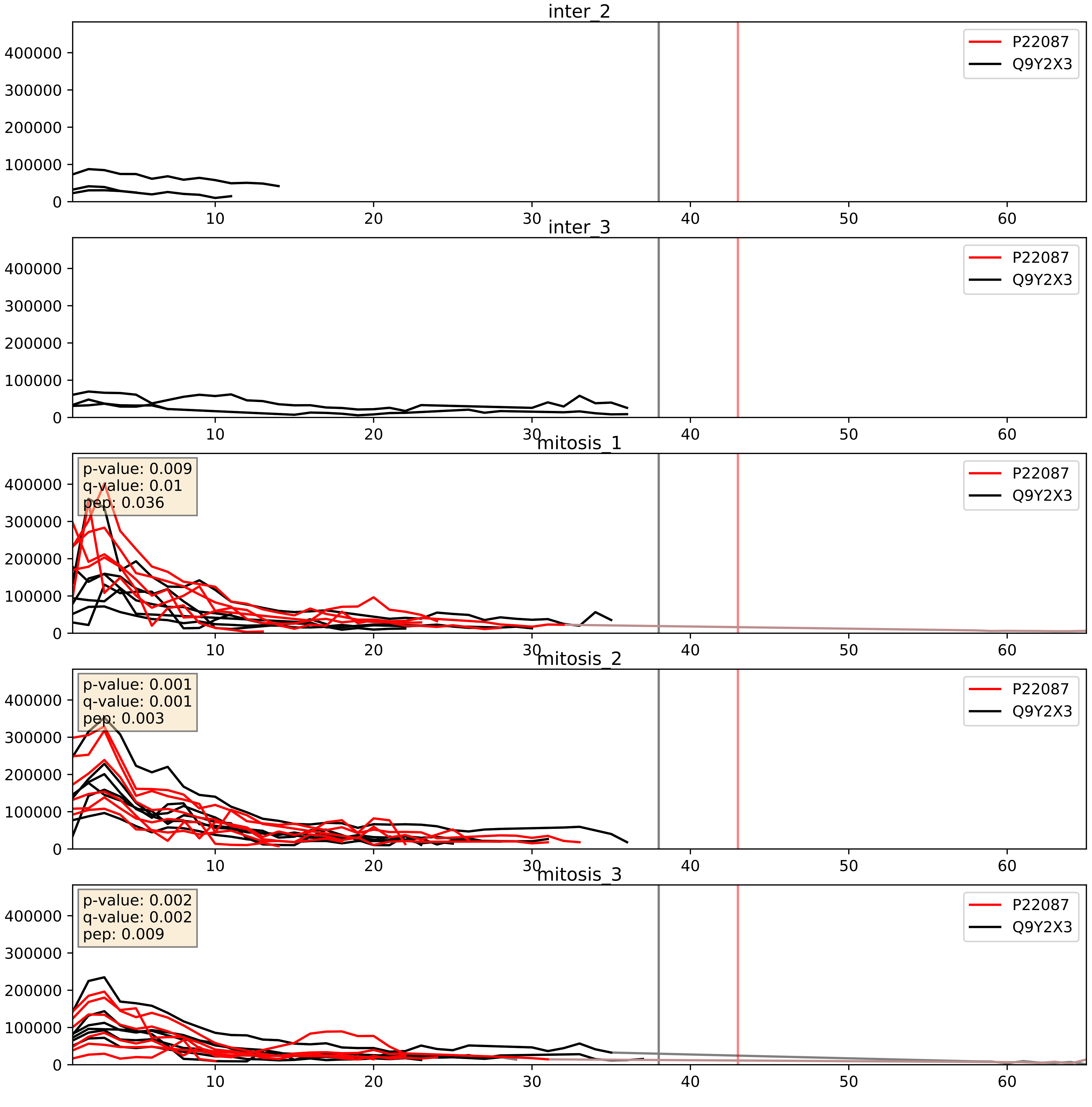


| level | condition_1 | condition_2 | pvalue | pvalue_adjusted |
|---|---|---|---|---|
| complex_abundance | mitosis | inter | 0.00015183324738868412 | 0.008550609195046946 |
| interactor_abundance | mitosis | inter | 0.00038188998169085025 | 0.013573913378163039 |
| interactor_ratio | mitosis | inter | 0.42563529182084064 | 0.841246386050888 |

P22087_Q9Y3T9
FBRL_HUMAN vs NOC2L_HUMAN
p-value: 0.029 q-value: 0.021

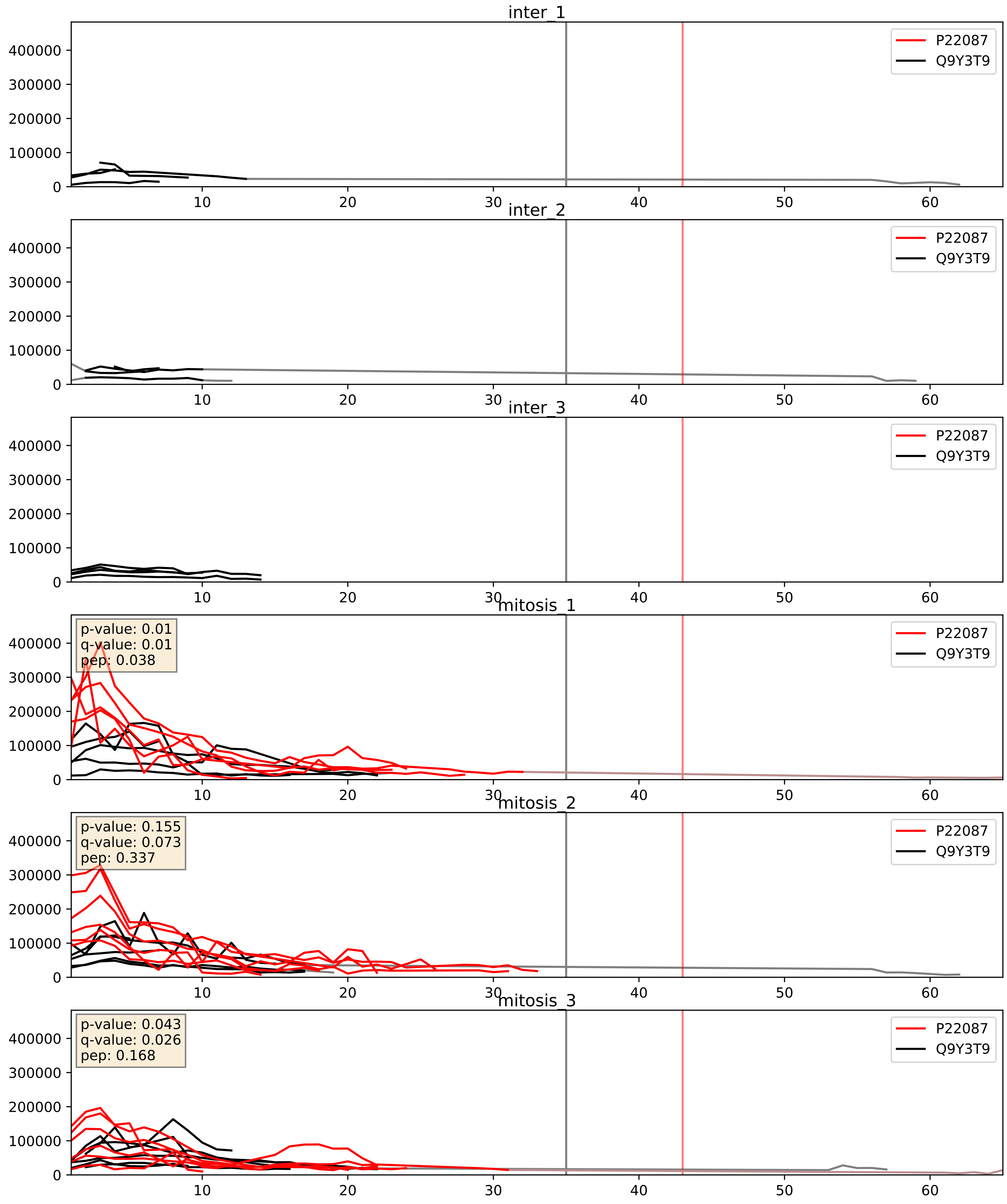


| level | condition_1 | condition_2 | pvalue | pvalue_adjusted |
|---|---|---|---|---|
| complex_abundance | mitosis | inter | 0.00035590071739036747 | 0.012039775462644836 |
| interactor_abundance | mitosis | inter | 0.00035742084283765416 | 0.01356427145975096 |
| interactor_ratio | mitosis | inter | 0.009198067791017207 | 0.13668839544866102 |

P22087_Q9Y3U8
FBRL_HUMAN vs RL36_HUMAN
p-value: 0.0 q-value: 0.018

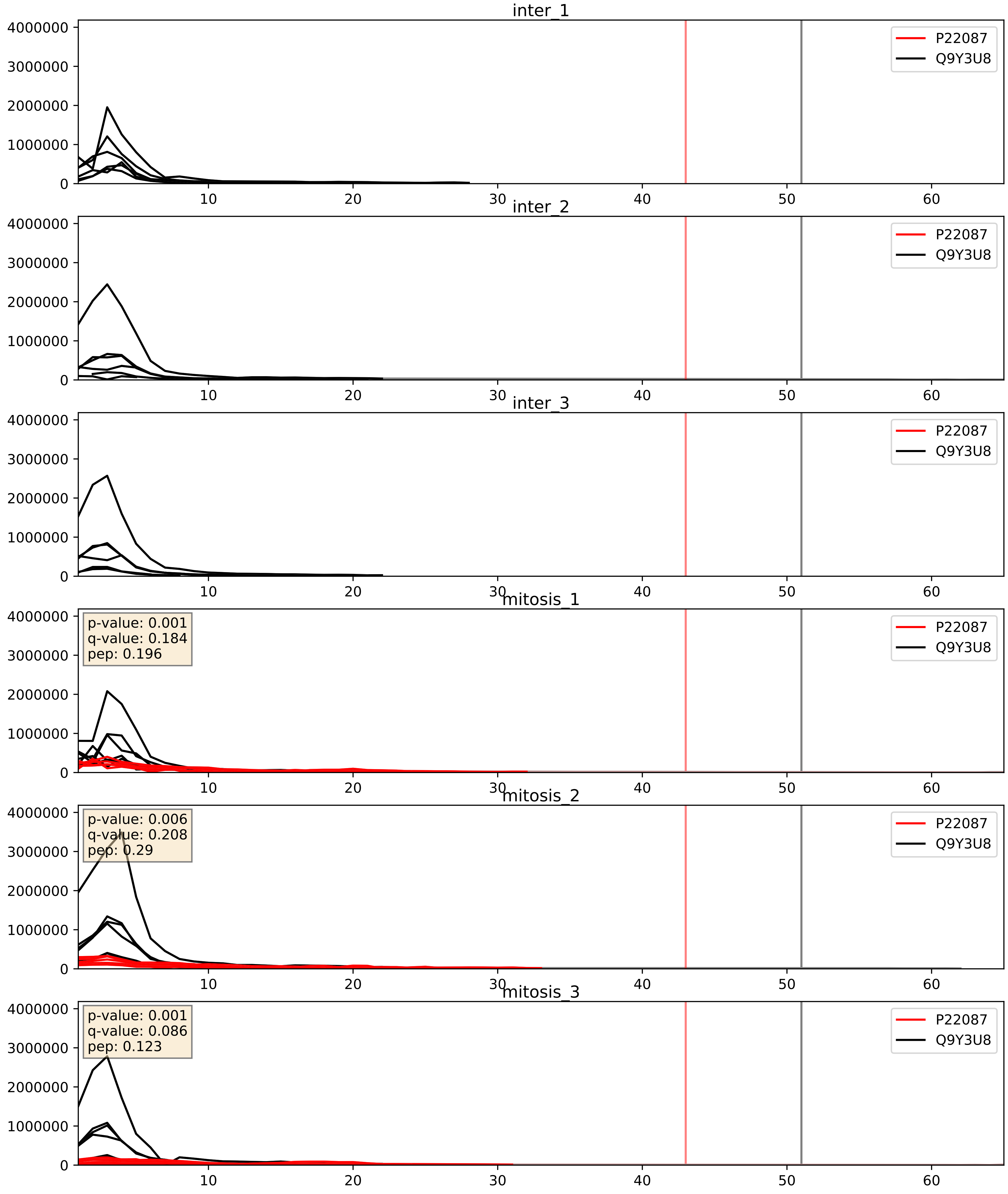


| level | condition_1 | condition_2 | pvalue | pvalue_adjusted |
|---|---|---|---|---|
| complex_abundance | mitosis | inter | 2.8538907879179594e-05 | 0.005691857151652958 |
| interactor_abundance | mitosis | inter | 0.00039298031109972966 | 0.013702287018385686 |
| interactor_ratio | mitosis | inter | 0.007968812527067133 | 0.1259990886723217 |

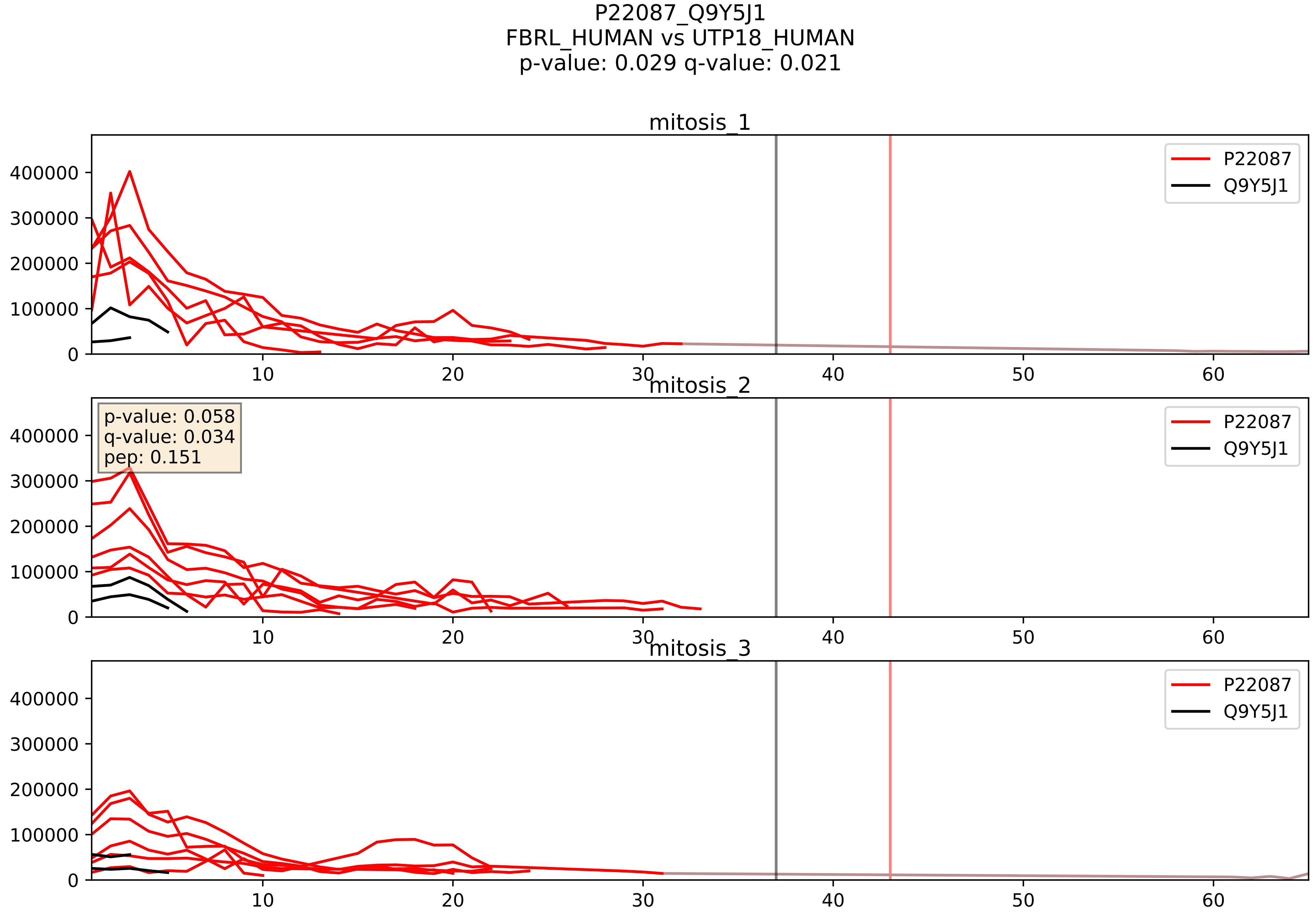


| level | condition_1 | condition_2 | pvalue | pvalue_adjusted |
|---|---|---|---|---|
| interactor_ratio | mitosis | inter | 0.00023104780740332214 | 0.015451322120097167 |
| interactor_abundance | mitosis | inter | 0.0020671551707800175 | 0.03307448273248028 |
| complex_abundance | mitosis | inter | 0.0066943970868762175 | 0.06051112889510077 |
