## Supplementary Data 2 for "SECAT: Quantifying differential protein-protein interaction states by network-centric analysis": hela_string_000009_Q03701.pdf

Q03701_Q03701

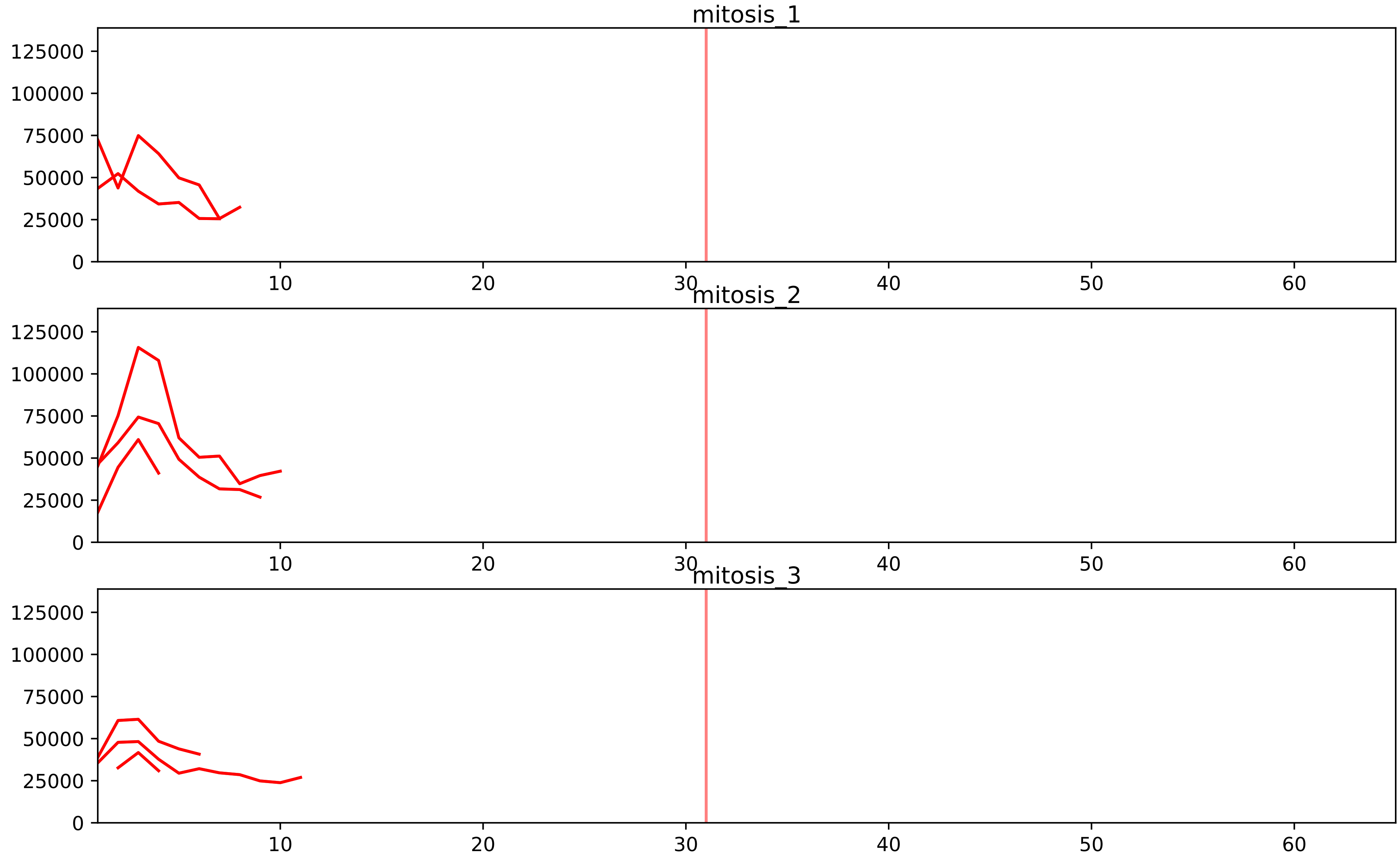


| level | condition_1 | condition_2 | pvalue | pvalue_adjusted |
|---|---|---|---|---|
| interactor_ratio | mitosis | inter | 4.517381837178695e-15 | 9.235536200454219e-13 |
| complex_abundance | mitosis | inter | 0.0002700437359397445 | 0.004476400667829999 |
| assembled_abundance | mitosis | inter | 0.0037666130624475205 | 0.06079833015626498 |
| interactor_abundance | mitosis | inter | 0.005749321087905118 | 0.03764680000621145 |
| total_abundance | mitosis | inter | 0.009154065417066065 | 0.1154817130772922 |

P46087_Q03701
NOP2_HUMAN vs CEBPZ_HUMAN
p-value: 0.043 q-value: 0.029

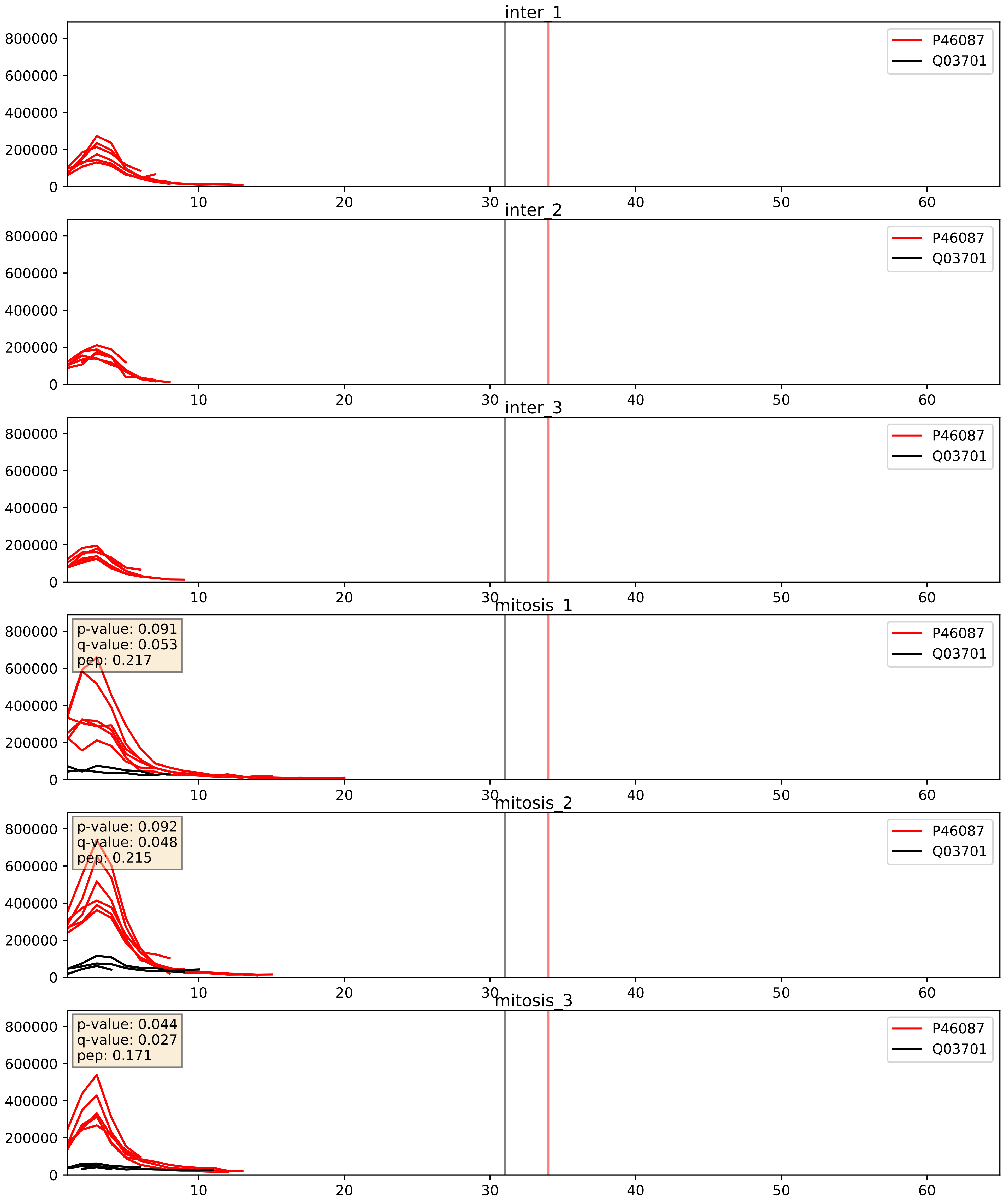


| level | condition_1 | condition_2 | pvalue | pvalue_adjusted |
|---|---|---|---|---|
| interactor_ratio | mitosis | inter | 7.980256380908328e-06 | 0.0032529045057416803 |
| interactor_abundance | mitosis | inter | 0.00013476672746345162 | 0.00890813271881966 |
| complex_abundance | mitosis | inter | 0.000587508565109972 | 0.015666895069599254 |

Q03701_Q15397
CEBPZ_HUMAN vs PUM3_HUMAN
p-value: 0.031 q-value: 0.022

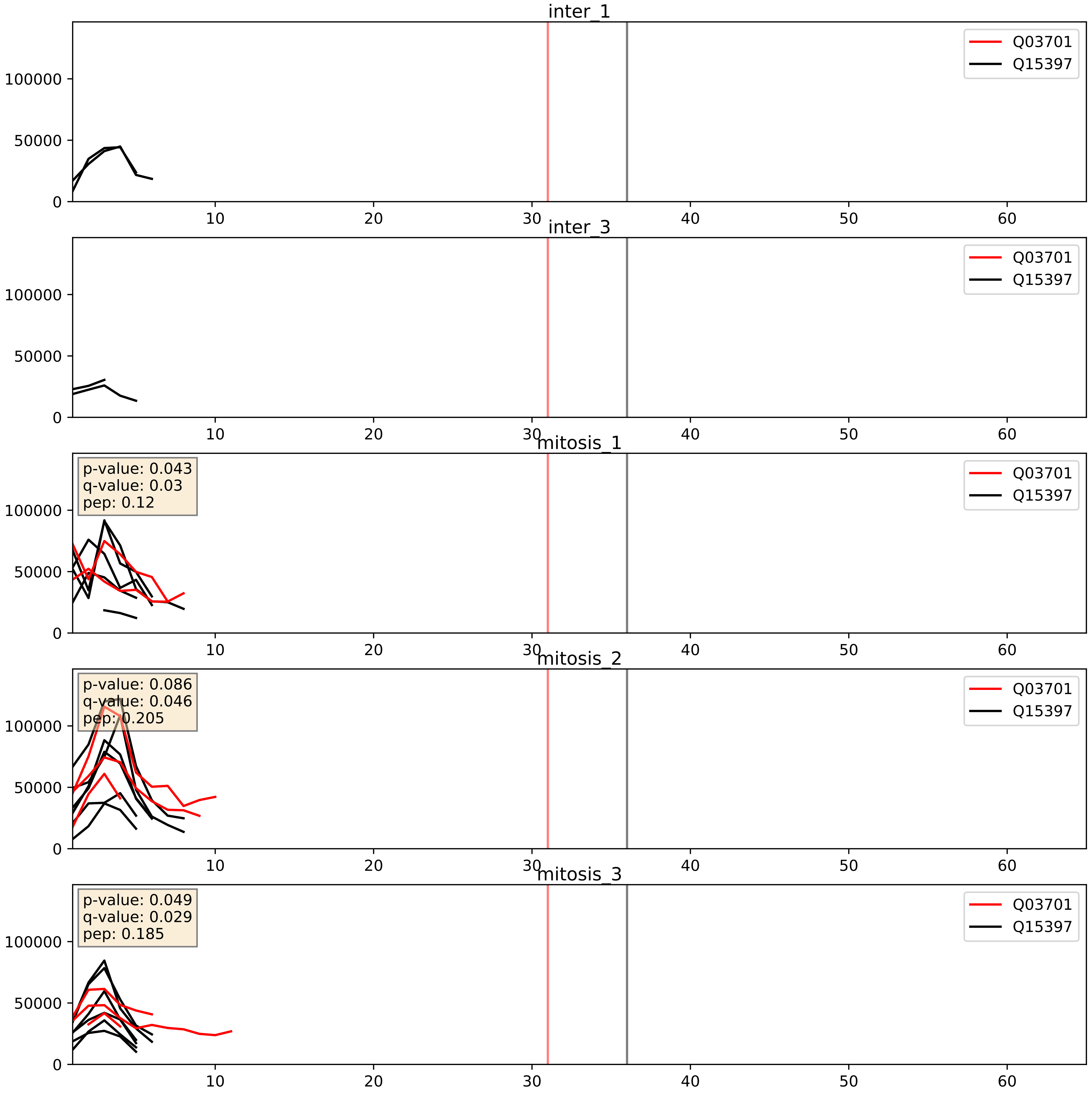


| level | condition_1 | condition_2 | pvalue | pvalue_adjusted |
|---|---|---|---|---|
| complex_abundance | mitosis | inter | 0.028263516439963447 | 0.1224370955091534 |
| interactor_abundance | mitosis | inter | 0.03470855862624382 | 0.1553074576333373 |
| interactor_ratio | mitosis | inter | 0.0981551795291016 | 0.5212210525862964 |

Q03701_Q9BZE4
CEBPZ_HUMAN vs NOG1_HUMAN
p-value: 0.034 q-value: 0.024

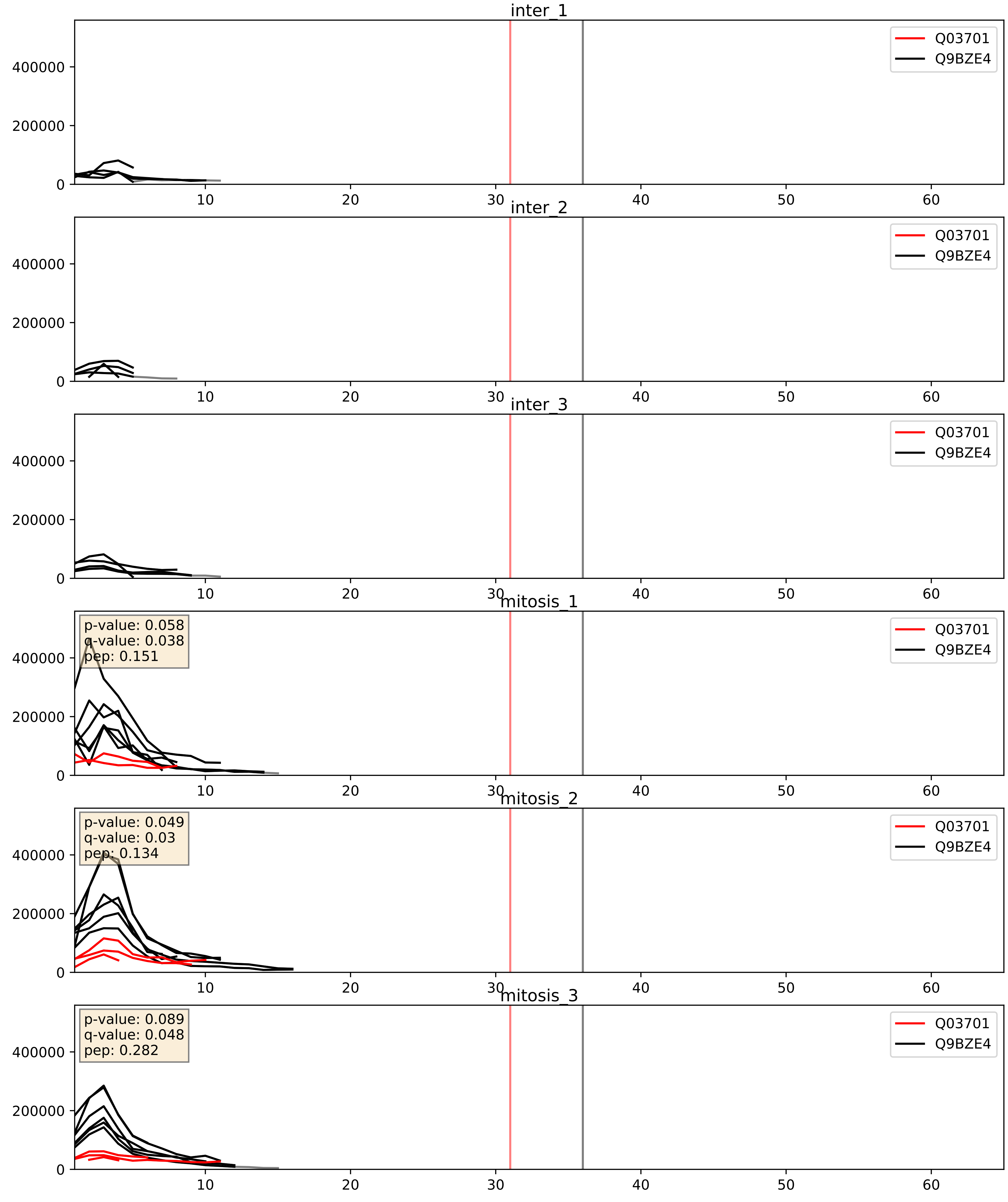


| level | condition_1 | condition_2 | pvalue | pvalue_adjusted |
|---|---|---|---|---|
| interactor_ratio | mitosis | inter | 2.782884935953381e-06 | 0.002520386835947235 |
| complex_abundance | mitosis | inter | 0.0011733846300989105 | 0.02303709273772173 |
| interactor_abundance | mitosis | inter | 0.019842839707373484 | 0.1190514126751731 |

Q03701_Q9NY93
CEBPZ_HUMAN vs DDX56_HUMAN
p-value: 0.011 q-value: 0.01

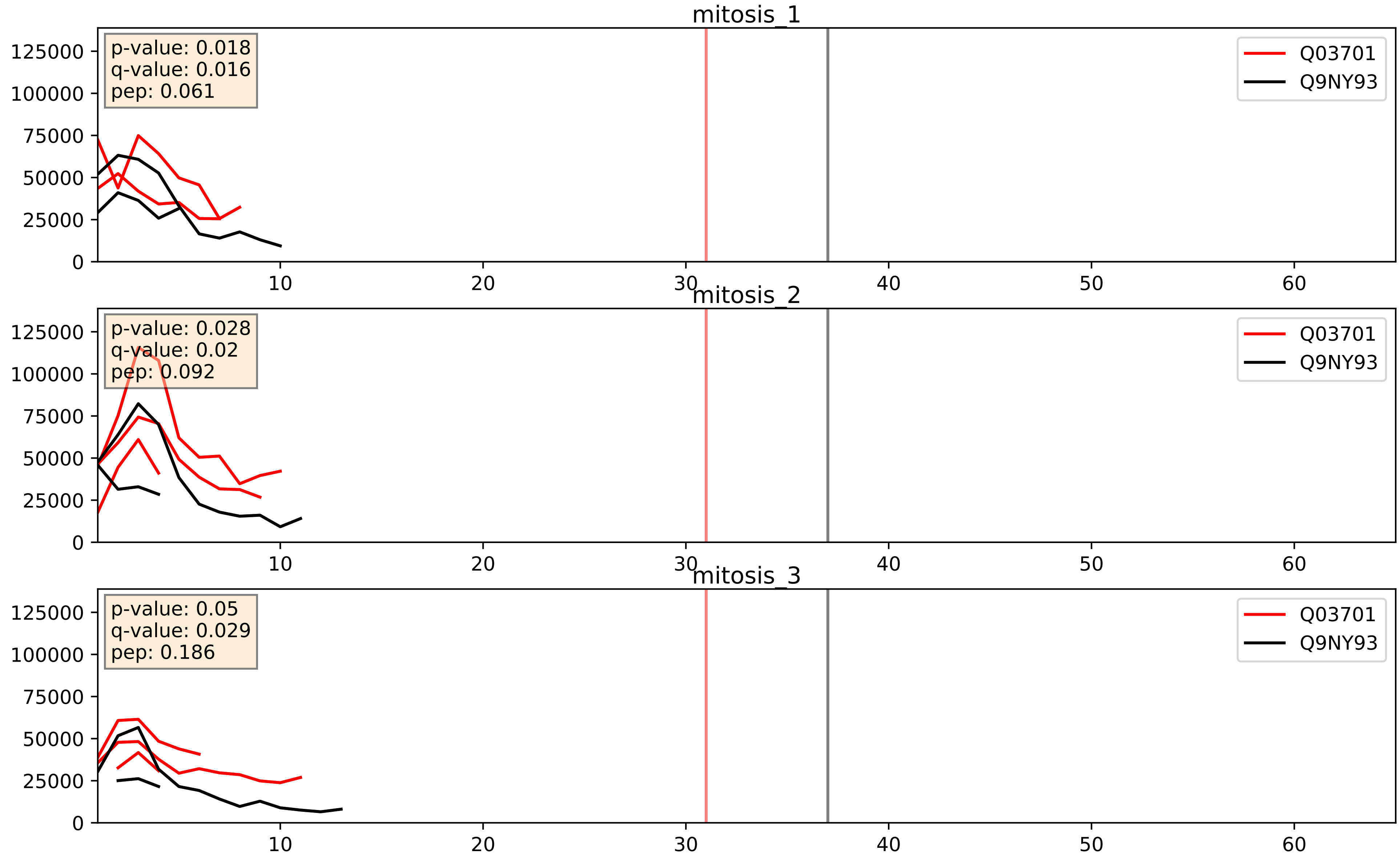


| level | condition_1 | condition_2 | pvalue | pvalue_adjusted |
|---|---|---|---|---|
| interactor_abundance | mitosis | inter | 0.019842839707373484 | 0.1190514126751731 |
| complex_abundance | mitosis | inter | 0.03402980185974759 | 0.1342992641398983 |
| interactor_ratio | mitosis | inter | 0.10349888889113636 | 0.5362896421962029 |

Q03701_Q9Y2X3
CEBPZ_HUMAN vs NOP58_HUMAN
p-value: 0.076 q-value: 0.046

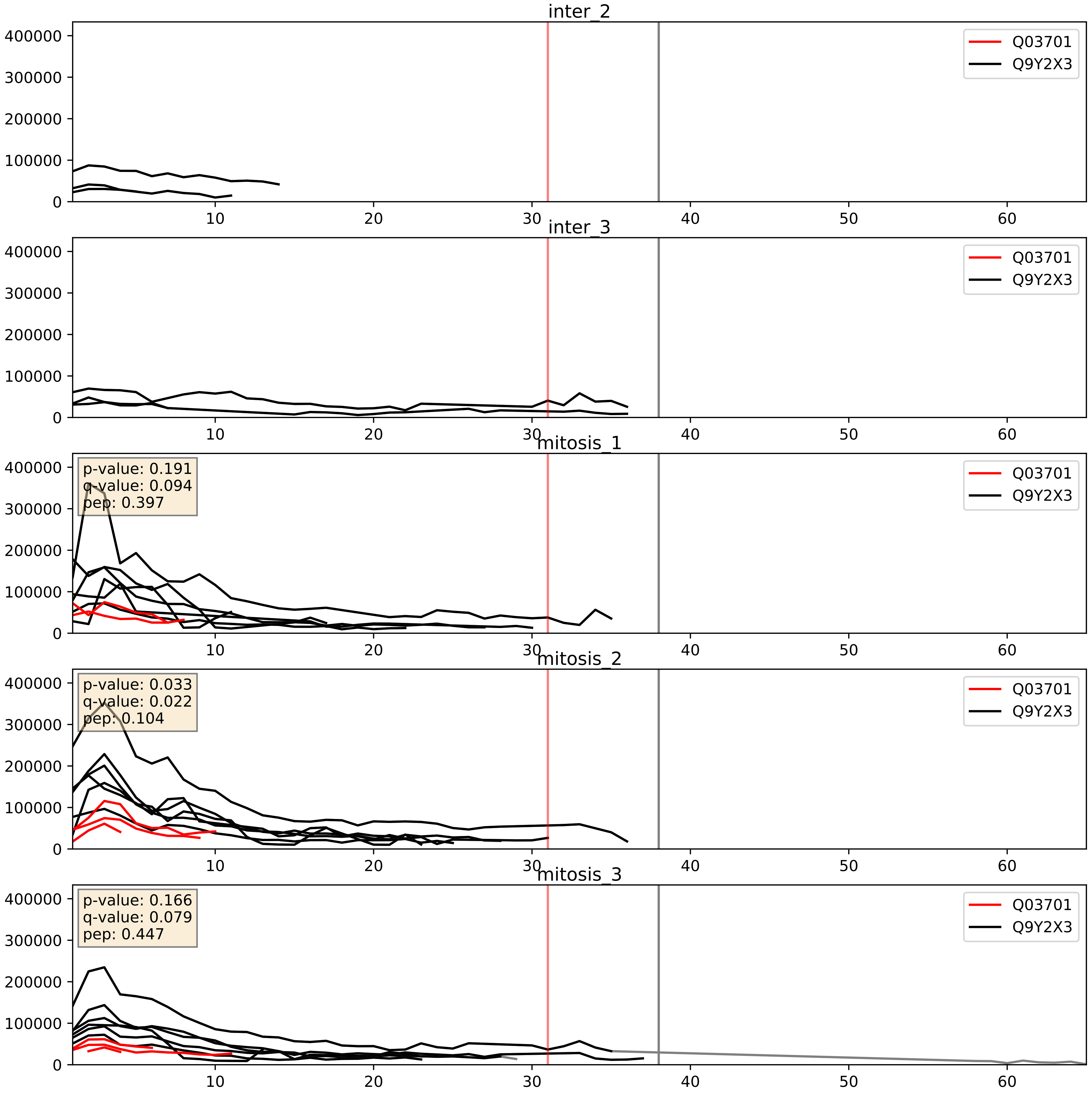


| level | condition_1 | condition_2 | pvalue | pvalue_adjusted |
|---|---|---|---|---|
| interactor_ratio | mitosis | inter | 1.052988896845605e-07 | 0.0004506792478499189 |
| complex_abundance | mitosis | inter | 0.000883253249181854 | 0.019570164268135633 |
| interactor_abundance | mitosis | inter | 0.019842839707373484 | 0.1190514126751731 |
