## Supplementary Data 2 for "SECAT: Quantifying differential protein-protein interaction states by network-centric analysis": hela_string_000010_P52597.pdf

P52597_P52597

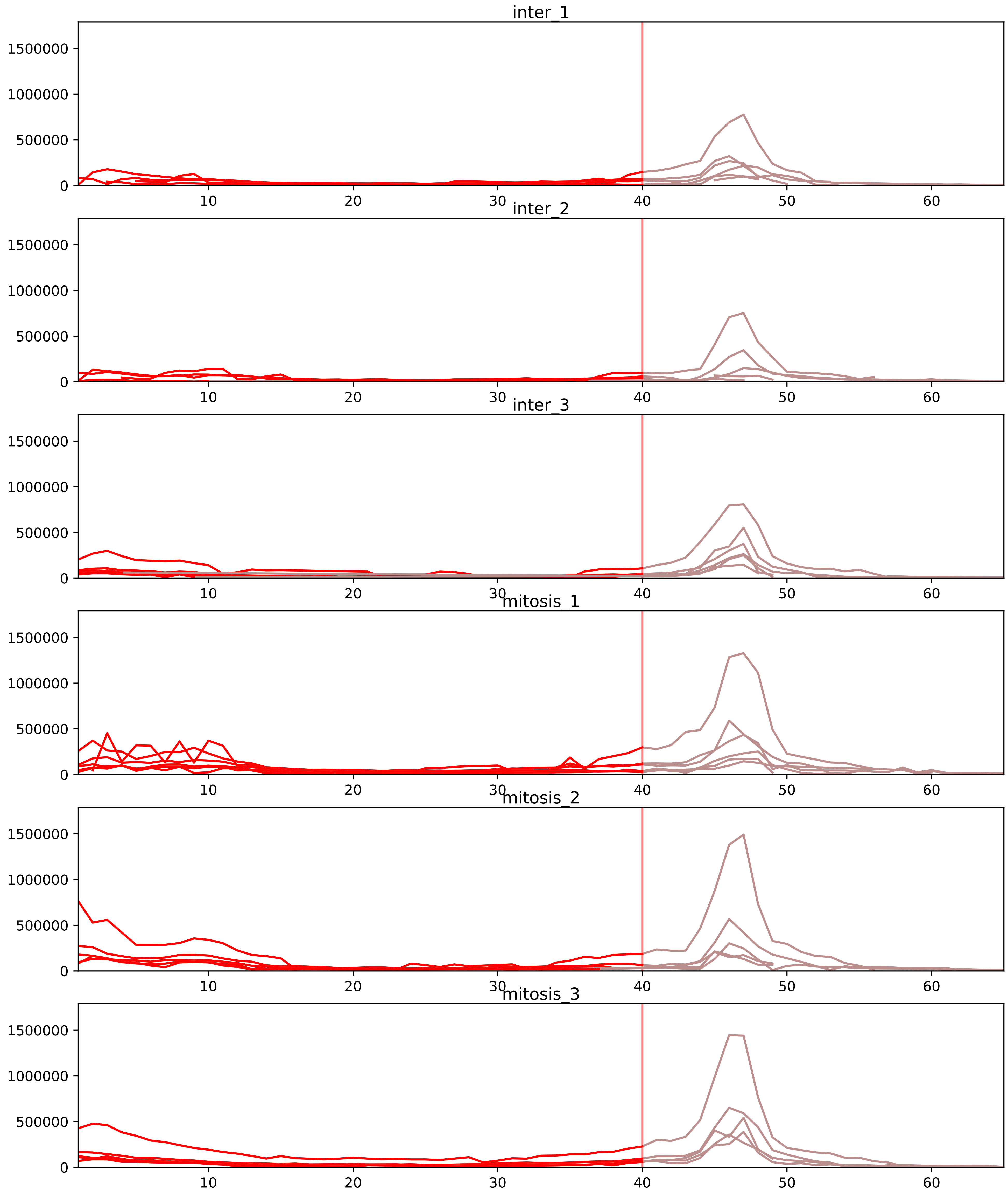


| level | condition_1 | condition_2 | pvalue | pvalue_adjusted |
|---|---|---|---|---|
| interactor_ratio | mitosis | inter | 7.773056638889959e-15 | 1.4302424215557526e-12 |
| complex_abundance | mitosis | inter | 2.832630674361764e-07 | 4.009261877558189e-05 |
| interactor_abundance | mitosis | inter | 4.669397727902558e-06 | 0.000300834717183481 |
| total_abundance | mitosis | inter | 0.0019655939371504834 | 0.04766014179201654 |
| assembled_abundance | mitosis | inter | 0.00296509605400293 | 0.05379695592553378 |
| monomer_abundance | mitosis | inter | 0.016050973601114356 | 0.21518226668971946 |

O14979_P52597
HNRDL_HUMAN vs HNRPF_HUMAN
p-value: 0.001 q-value: 0.024

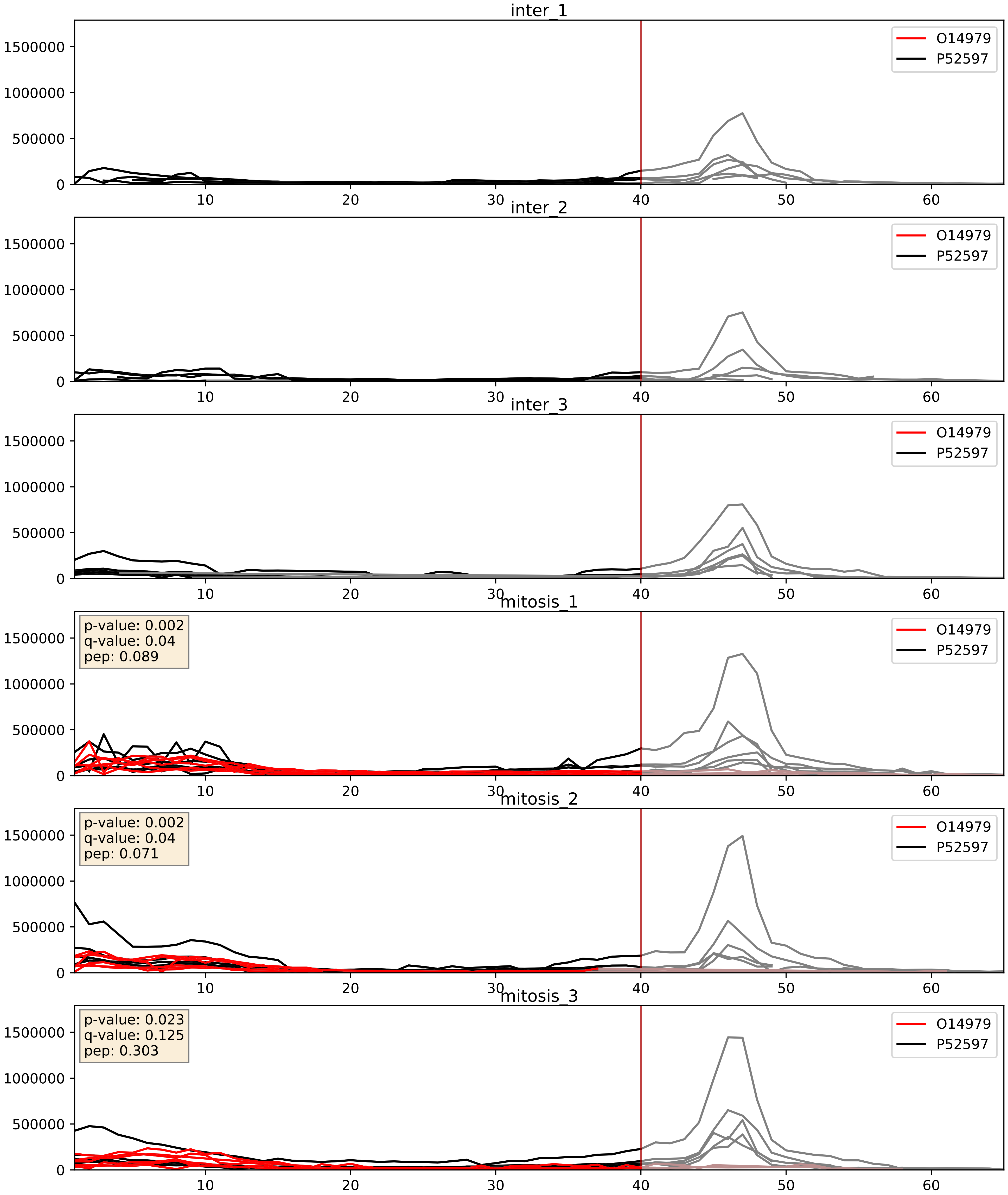


| level | condition_1 | condition_2 | pvalue | pvalue_adjusted |
|---|---|---|---|---|
| complex_abundance | mitosis | inter | 0.00023969958657414883 | 0.010310695784295044 |
| interactor_abundance | mitosis | inter | 0.0008164873709066391 | 0.021004694279574423 |
| interactor_ratio | mitosis | inter | 0.009098125486910772 | 0.13591615038037733 |

O43390_P52597
HNRPR_HUMAN vs HNRPF_HUMAN
p-value: 0.003 q-value: 0.003

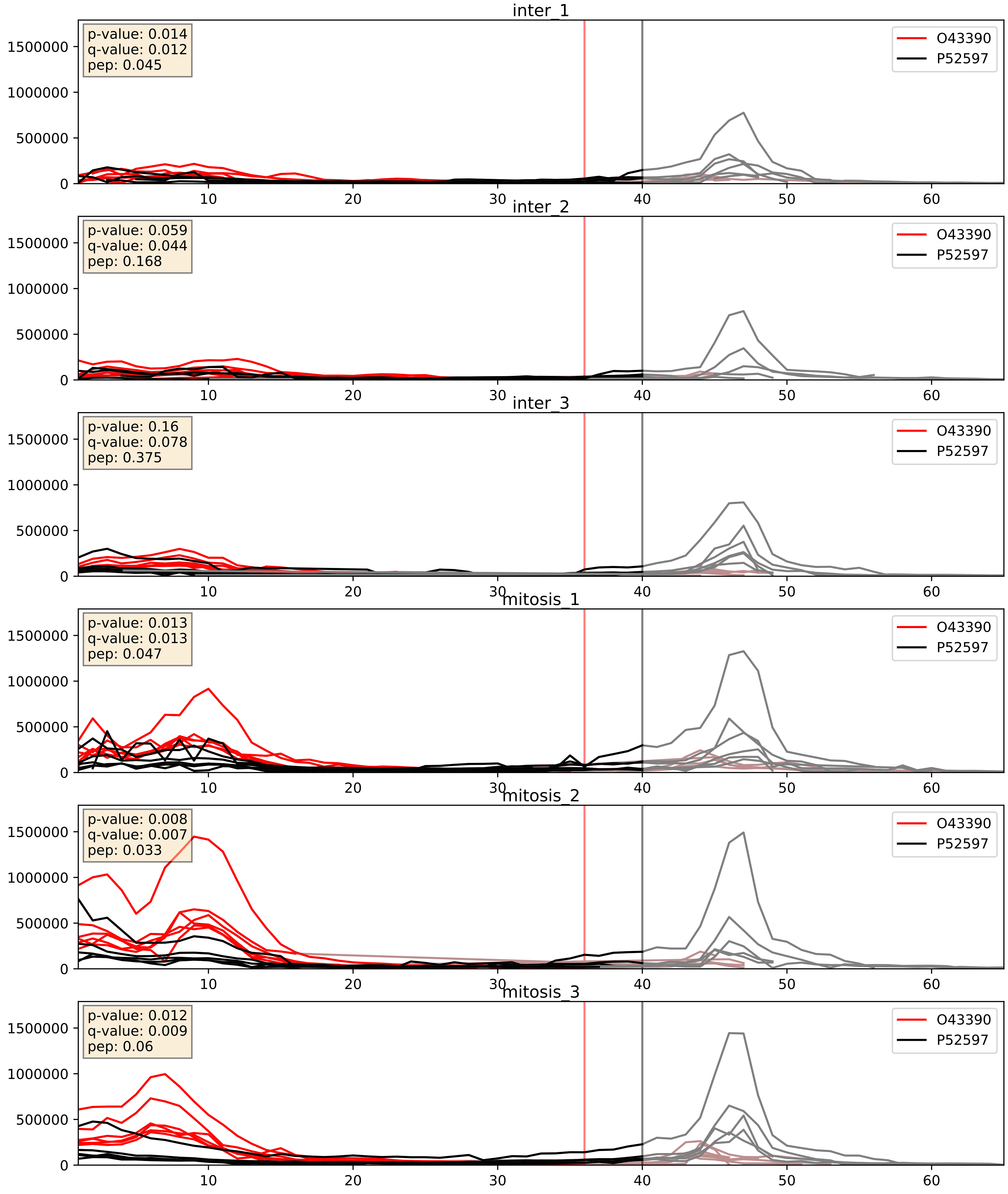


| level | condition_1 | condition_2 | pvalue | pvalue_adjusted |
|---|---|---|---|---|
| complex_abundance | mitosis | inter | 7.40732189271377e-05 | 0.005893640694944935 |
| interactor_abundance | mitosis | inter | 0.00010796061997593195 | 0.007876481428251637 |
| interactor_ratio | mitosis | inter | 0.29205419947420935 | 0.7616429249926009 |

P07910_P52597
HNRPC_HUMAN vs HNRPF_HUMAN
p-value: 0.007 q-value: 0.007

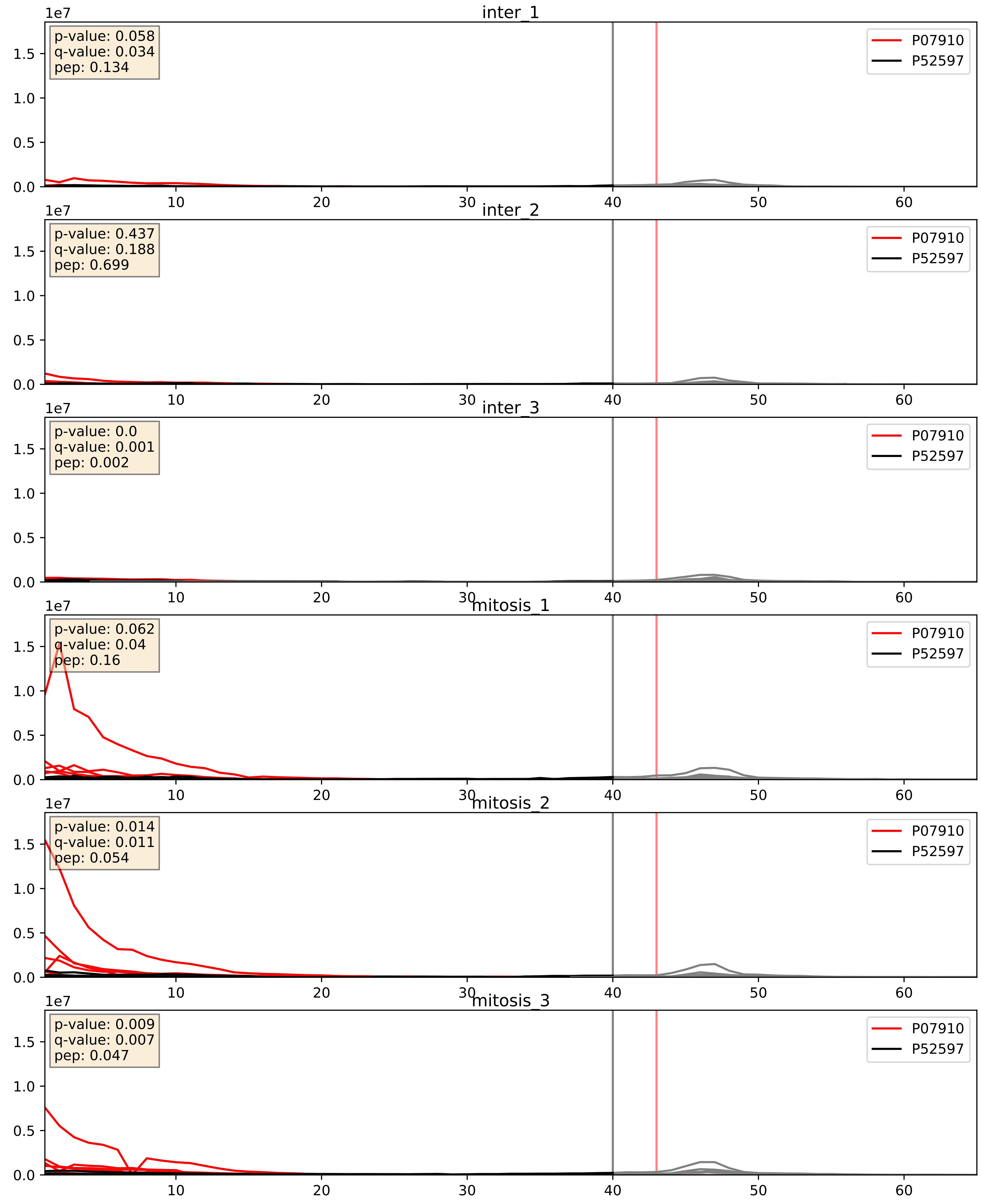


| level | condition_1 | condition_2 | pvalue | pvalue_adjusted |
|---|---|---|---|---|
| complex_abundance | mitosis | inter | 2.867619584776087e-06 | 0.004091137274280551 |
| interactor_abundance | mitosis | inter | 9.49140558984193e-06 | 0.0033734327106189023 |
| interactor_ratio | mitosis | inter | 0.0007484433338408752 | 0.03166103300210851 |

P09651_P52597
ROA1_HUMAN vs HNRPF_HUMAN
p-value: 0.001 q-value: 0.001

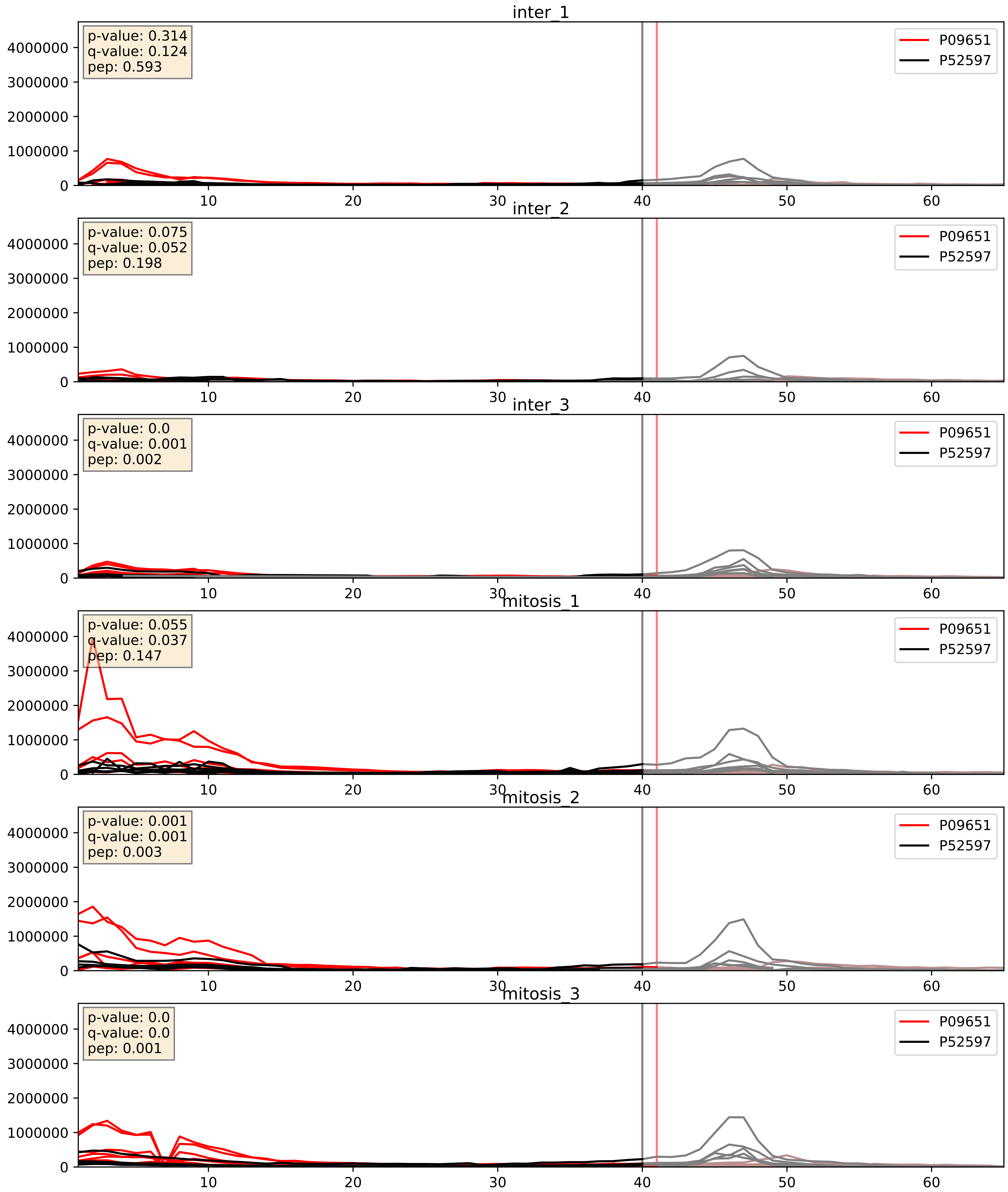


| level | condition_1 | condition_2 | pvalue | pvalue_adjusted |
|---|---|---|---|---|
| complex_abundance | mitosis | inter | 0.006743082334176311 | 0.06075872082163076 |
| interactor_abundance | mitosis | inter | 0.017050532289726136 | 0.11334462047152971 |
| interactor_ratio | mitosis | inter | 0.11810419448591807 | 0.566001132973789 |

P11940_P52597
PABP1_HUMAN vs HNRPF_HUMAN
p-value: 0.0 q-value: 0.018

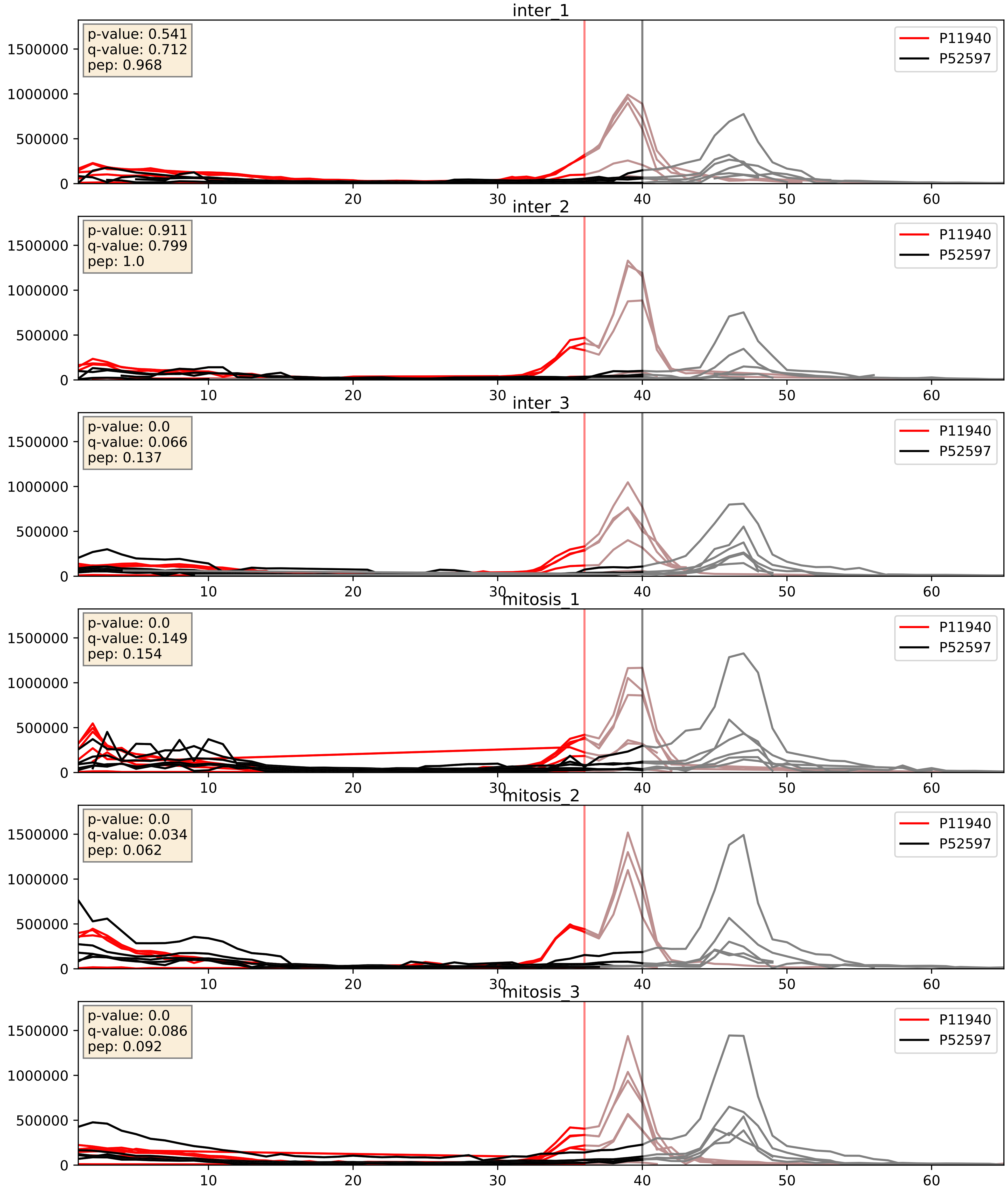


| level | condition_1 | condition_2 | pvalue | pvalue_adjusted |
|---|---|---|---|---|
| complex_abundance | mitosis | inter | 0.0061419595529173445 | 0.05828645200067901 |
| interactor_ratio | mitosis | inter | 0.016353146604940094 | 0.1886562465475569 |
| interactor_abundance | mitosis | inter | 0.15090604372175434 | 0.33732834796845007 |

P14866_P52597
HNRPL_HUMAN vs HNRPF_HUMAN
p-value: 0.001 q-value: 0.002

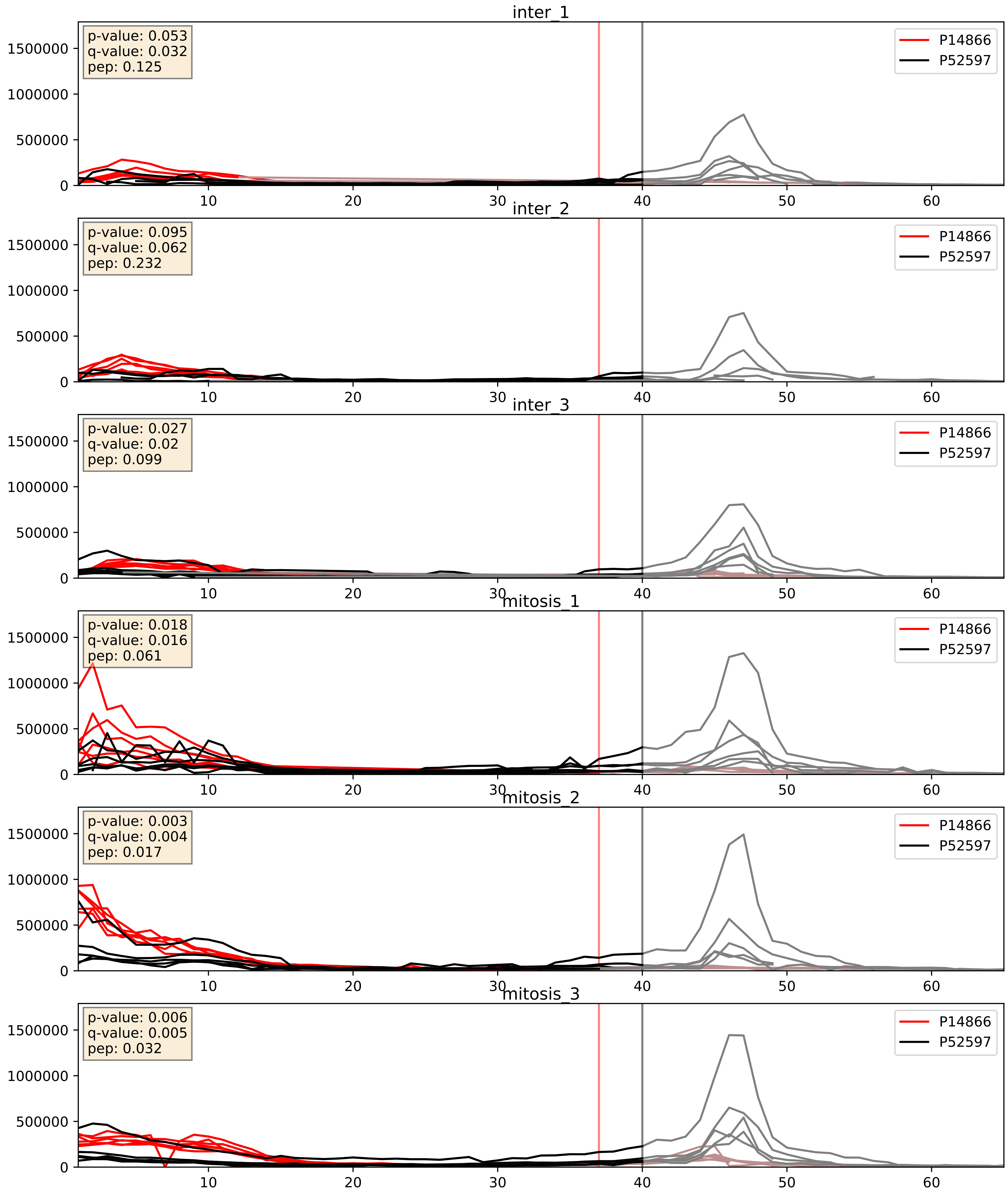


| level | condition_1 | condition_2 | pvalue | pvalue_adjusted |
|---|---|---|---|---|
| complex_abundance | mitosis | inter | 0.005276617137903093 | 0.054740033961712564 |
| interactor_abundance | mitosis | inter | 0.009697640385696541 | 0.08525122834595471 |
| interactor_ratio | mitosis | inter | 0.48618414815202177 | 0.8688384776996464 |

P17844_P52597
DDX5_HUMAN vs HNRPF_HUMAN
p-value: 0.0 q-value: 0.005

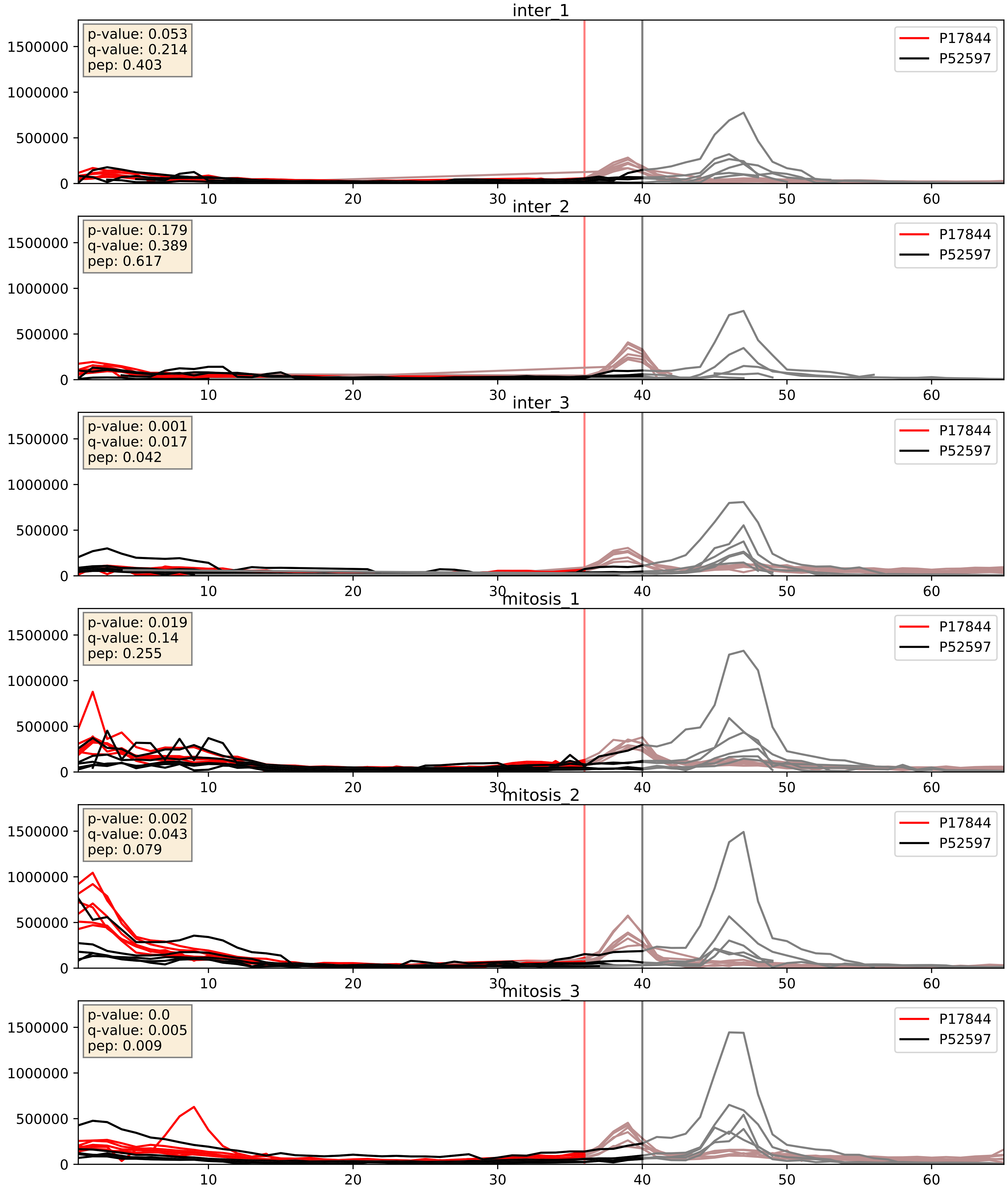


| level | condition_1 | condition_2 | pvalue | pvalue_adjusted |
|---|---|---|---|---|
| interactor_abundance | mitosis | inter | 0.0021426459467421676 | 0.033870820506210444 |
| complex_abundance | mitosis | inter | 0.002394620564026592 | 0.035463584823646414 |
| interactor_ratio | mitosis | inter | 0.413906743817922 | 0.8357726189348997 |

P22626_P52597
ROA2_HUMAN vs HNRPF_HUMAN
p-value: 0.006 q-value: 0.006

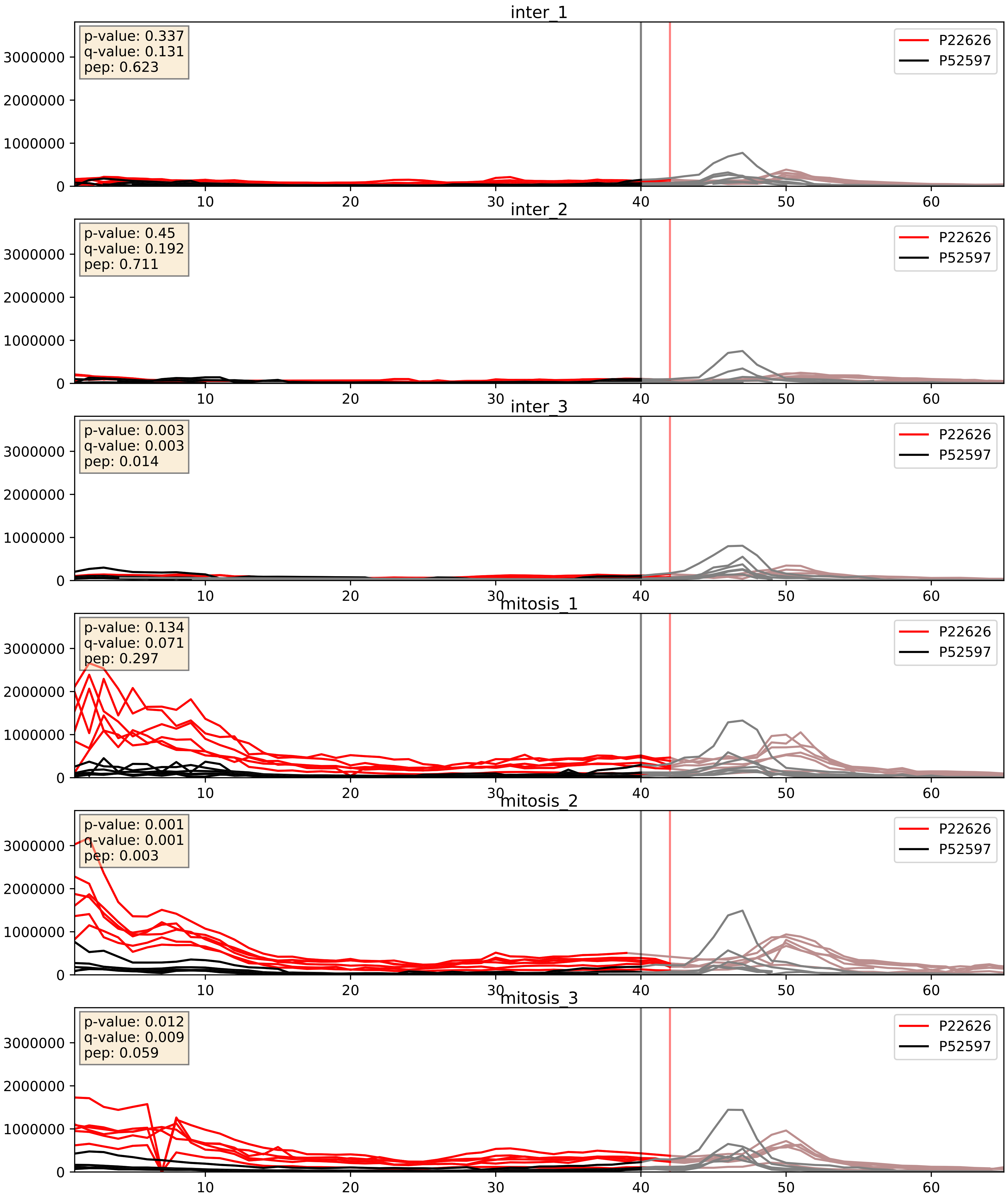


| level | condition_1 | condition_2 | pvalue | pvalue_adjusted |
|---|---|---|---|---|
| complex_abundance | mitosis | inter | 0.0010673061249328343 | 0.021804631096479864 |
| interactor_abundance | mitosis | inter | 0.003361167395046095 | 0.04478068934100323 |
| interactor_ratio | mitosis | inter | 0.01572743448355028 | 0.1848097318528163 |

P31943_P52597
HNRH1_HUMAN vs HNRPF_HUMAN
p-value: 0.001 q-value: 0.002

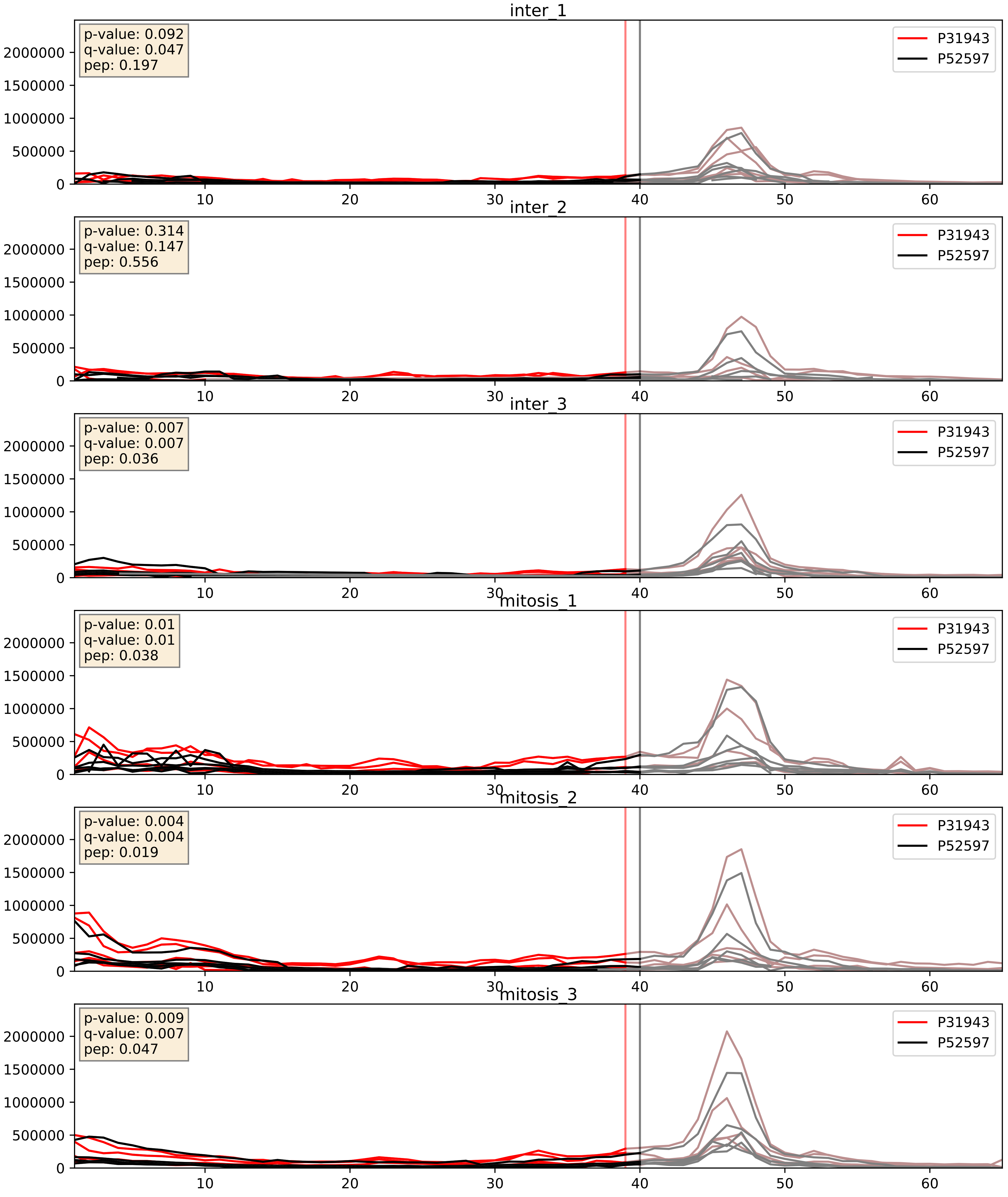


| level | condition_1 | condition_2 | pvalue | pvalue_adjusted |
|---|---|---|---|---|
| complex_abundance | mitosis | inter | 0.0023078745983505836 | 0.03502731659907978 |
| interactor_abundance | mitosis | inter | 0.0035613165684418575 | 0.04601489785035819 |
| interactor_ratio | mitosis | inter | 0.11518859280013378 | 0.5598715168125593 |

P38159_P52597
RBMX_HUMAN vs HNRPF_HUMAN
p-value: 0.005 q-value: 0.005

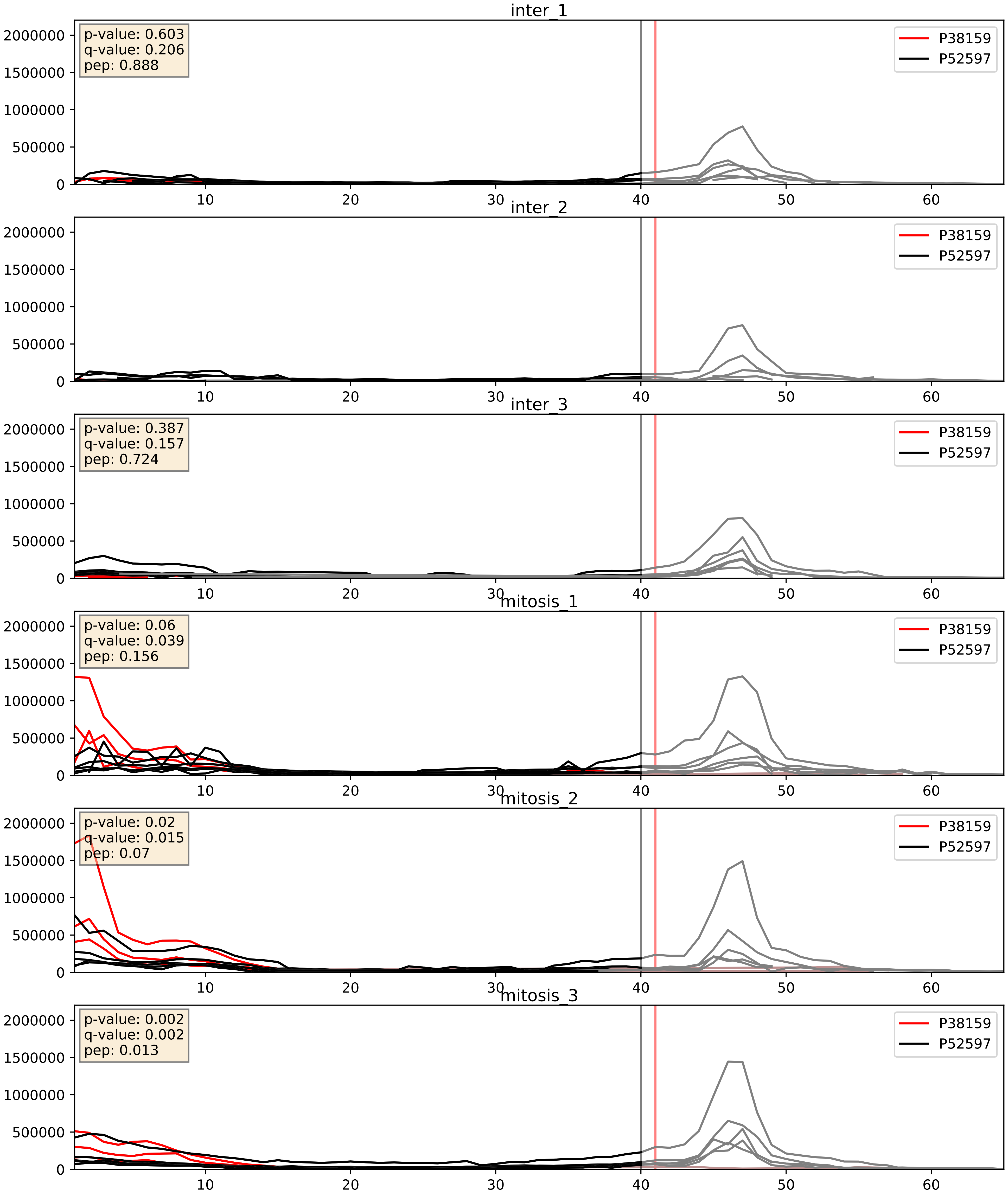


| level | condition_1 | condition_2 | pvalue | pvalue_adjusted |
|---|---|---|---|---|
| interactor_abundance | mitosis | inter | 0.0010973699494020942 | 0.024049720606162197 |
| complex_abundance | mitosis | inter | 0.0032164973285077132 | 0.0416428285245195 |
| interactor_ratio | mitosis | inter | 0.00676965061563321 | 0.11425781573990595 |

P51991_P52597
ROA3_HUMAN vs HNRPF_HUMAN
p-value: 0.0 q-value: 0.0

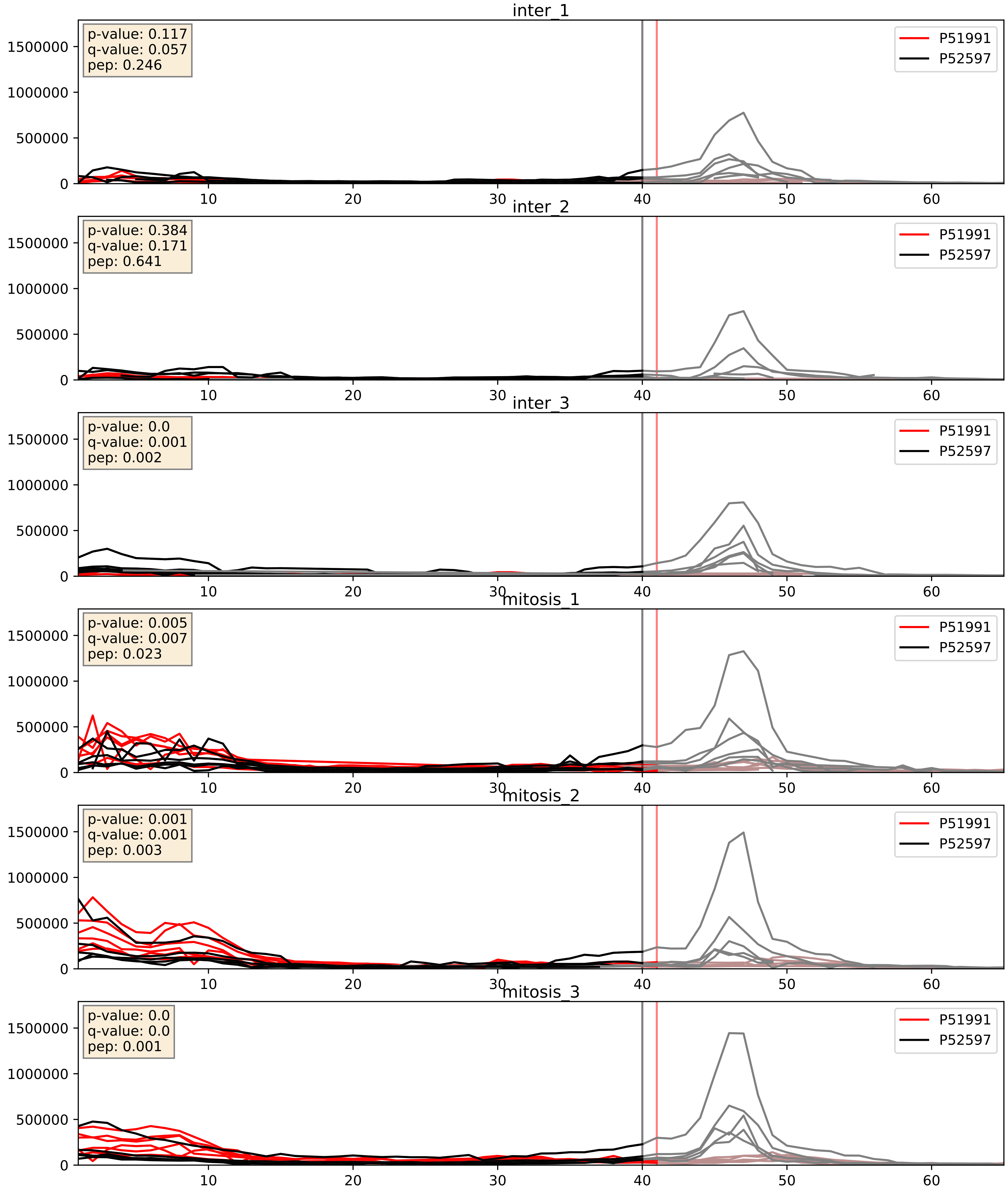


| level | condition_1 | condition_2 | pvalue | pvalue_adjusted |
|---|---|---|---|---|
| interactor_abundance | mitosis | inter | 1.0516040165513909e-05 | 0.0033734327106189023 |
| interactor_ratio | mitosis | inter | 0.00014720601620827773 | 0.012836278661937291 |
| complex_abundance | mitosis | inter | 0.0001689637015938096 | 0.008722442186052247 |

P52272_P52597
HNRPM_HUMAN vs HNRPF_HUMAN
p-value: 0.012 q-value: 0.01

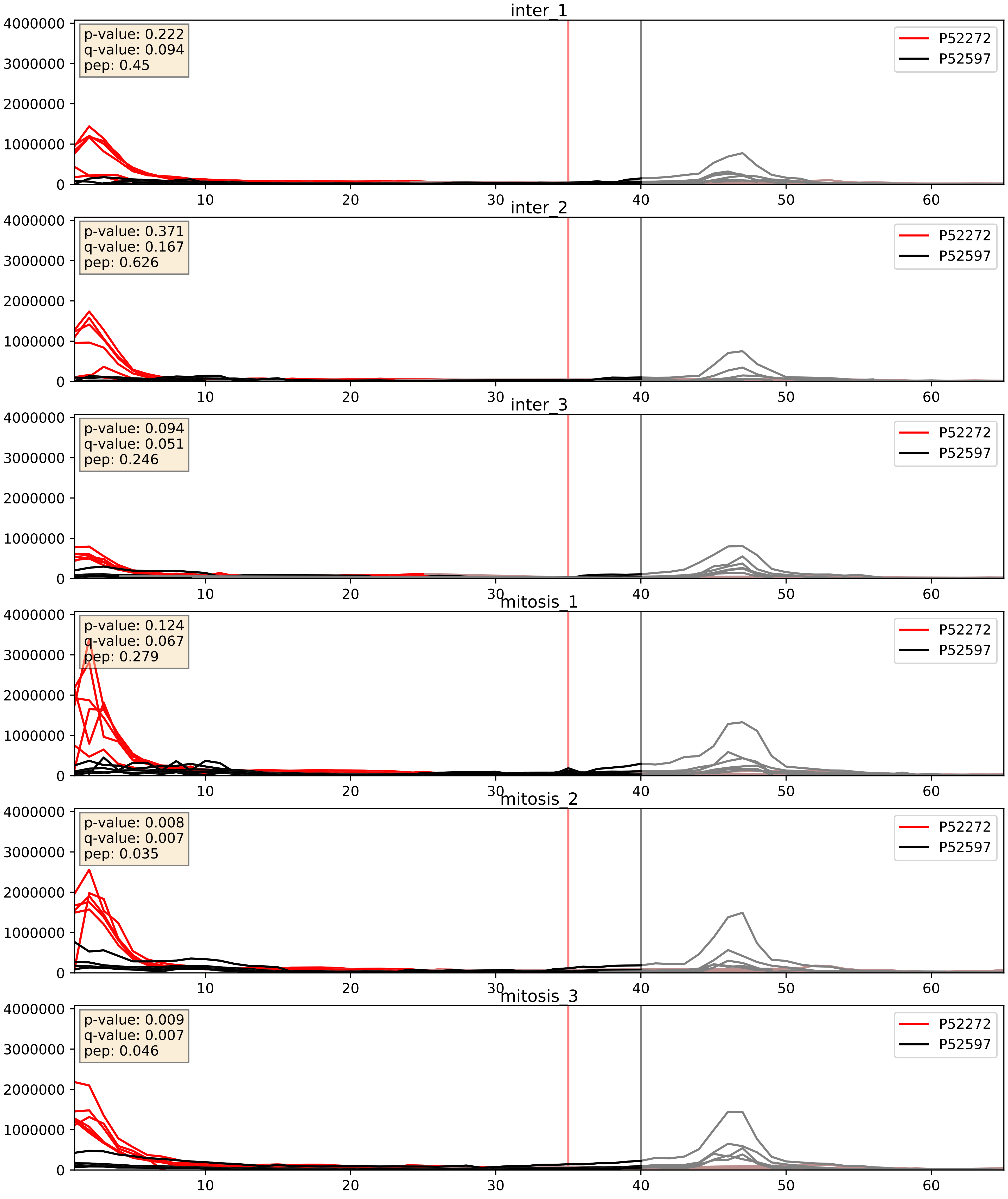


| level | condition_1 | condition_2 | pvalue | pvalue_adjusted |
|---|---|---|---|---|
| complex_abundance | mitosis | inter | 0.009939955657301148 | 0.07371585975104354 |
| interactor_ratio | mitosis | inter | 0.15013875473174929 | 0.6053639851642835 |
| interactor_abundance | mitosis | inter | 0.19757579480384646 | 0.391223701458744 |

P52597_P55769
HNRPF_HUMAN vs NH2L1_HUMAN
p-value: 0.0 q-value: 0.042

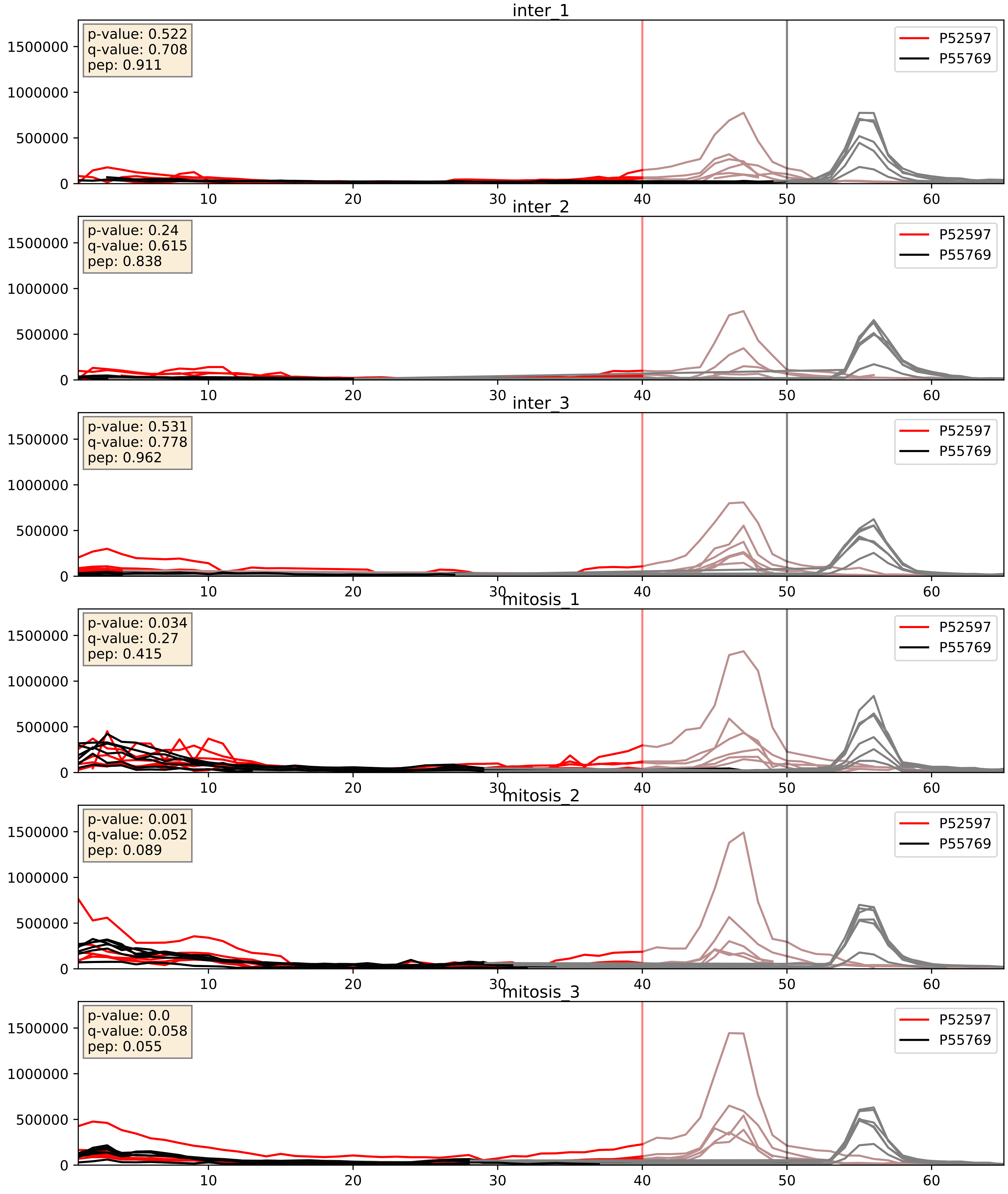


| level | condition_1 | condition_2 | pvalue | pvalue_adjusted |
|---|---|---|---|---|
| complex_abundance | mitosis | inter | 0.0012657499792246994 | 0.02413100183109895 |
| interactor_abundance | mitosis | inter | 0.006646850462384629 | 0.06976951558309309 |
| interactor_ratio | mitosis | inter | 0.03615766025694413 | 0.3092003714280137 |

P52597_P61978
HNRPF_HUMAN vs HNRPK_HUMAN
p-value: 0.028 q-value: 0.02

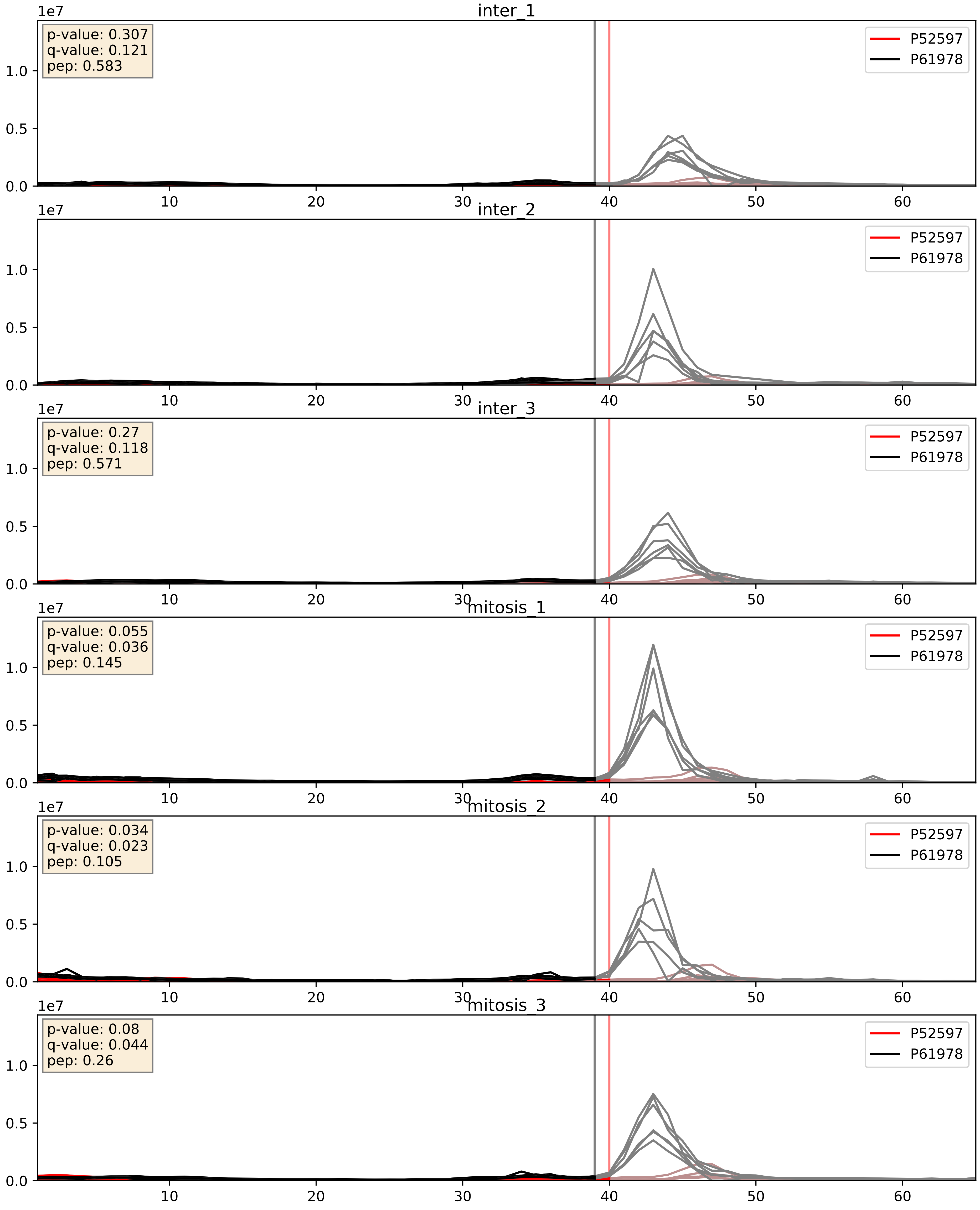


| level | condition_1 | condition_2 | pvalue | pvalue_adjusted |
|---|---|---|---|---|
| interactor_abundance | mitosis | inter | 0.004157412060800134 | 0.05116814844061703 |
| interactor_ratio | mitosis | inter | 0.2162728164558602 | 0.6924580509898395 |
| complex_abundance | mitosis | inter | 0.4194249258357087 | 0.5632691191016107 |

P52597_P62995
HNRPF_HUMAN vs TRA2B_HUMAN
p-value: 0.003 q-value: 0.003

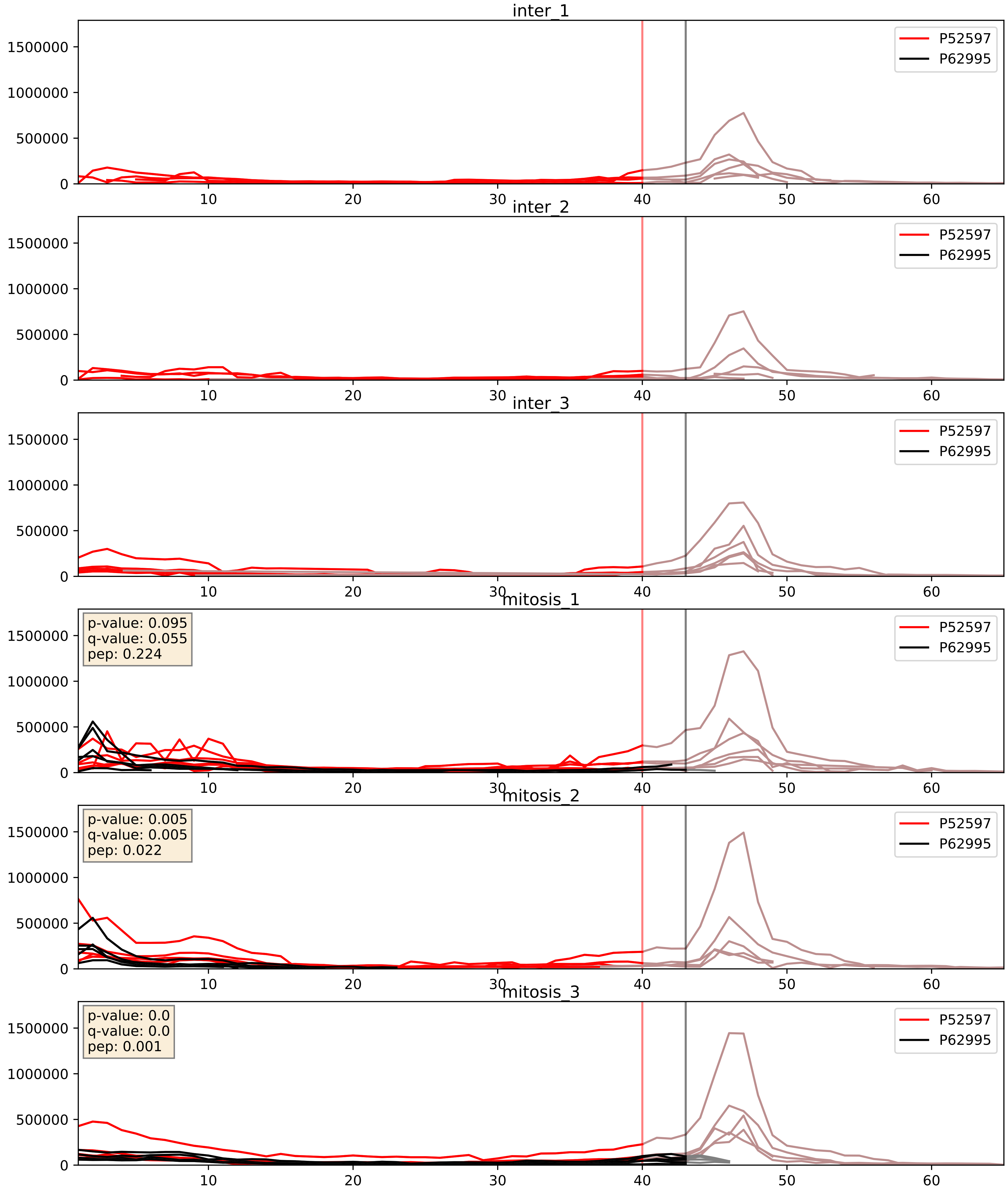


| level | condition_1 | condition_2 | pvalue | pvalue_adjusted |
|---|---|---|---|---|
| interactor_ratio | mitosis | inter | 4.6063738991628886e-05 | 0.0067983725132472975 |
| interactor_abundance | mitosis | inter | 0.00010371553266387402 | 0.007876481428251637 |
| complex_abundance | mitosis | inter | 0.0001881994972916382 | 0.009111526782692203 |

P52597_P84103
HNRPF_HUMAN vs SRSF3_HUMAN
p-value: 0.001 q-value: 0.002

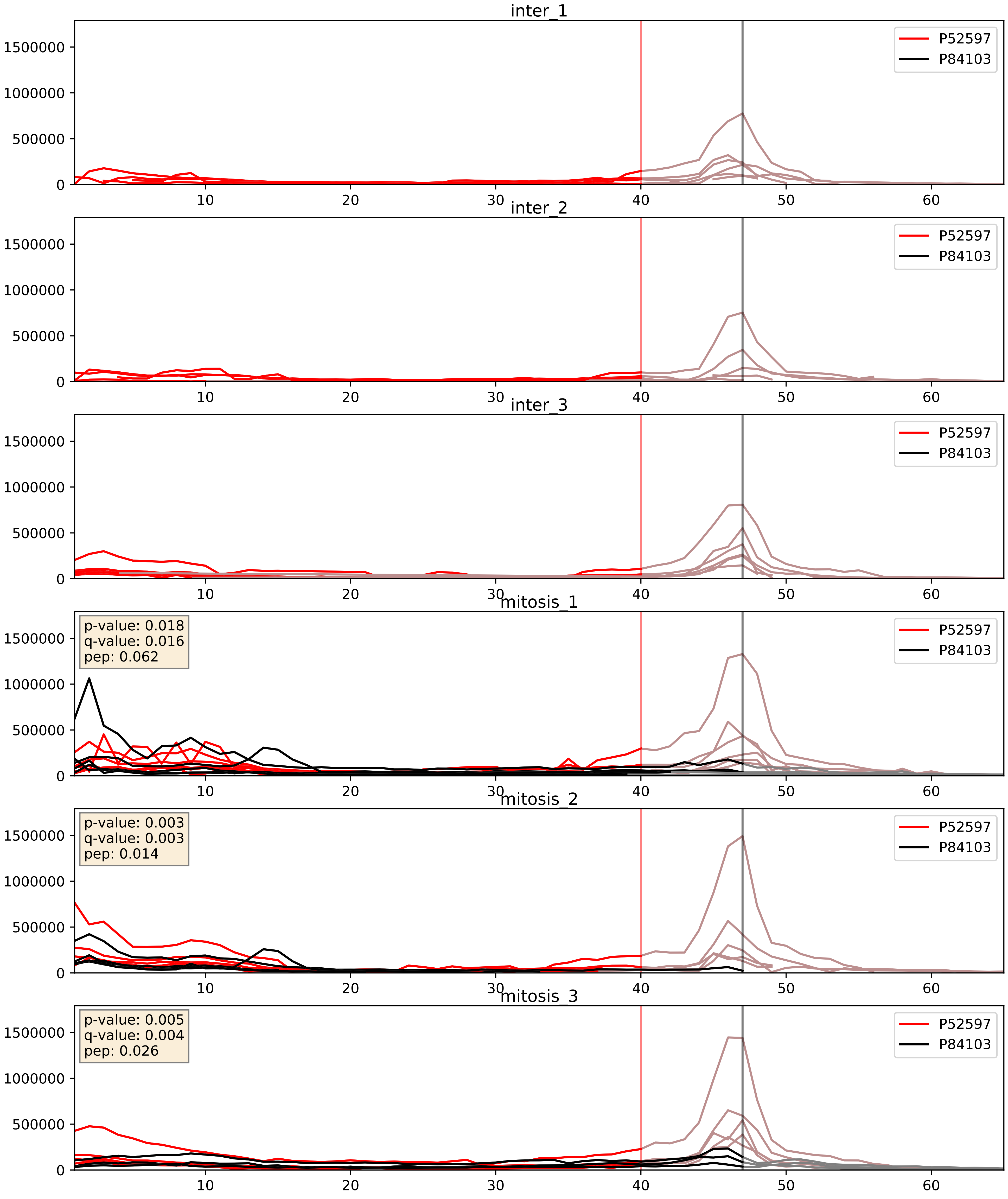


| level | condition_1 | condition_2 | pvalue | pvalue_adjusted |
|---|---|---|---|---|
| interactor_abundance | mitosis | inter | 4.920255045596975e-05 | 0.006283598333582311 |
| complex_abundance | mitosis | inter | 0.00044192382616051125 | 0.013273220883978866 |
| interactor_ratio | mitosis | inter | 0.0357560041616608 | 0.30712516692473574 |

P52597_Q07955
HNRPF_HUMAN vs SRSF1_HUMAN
p-value: 0.0 q-value: 0.0

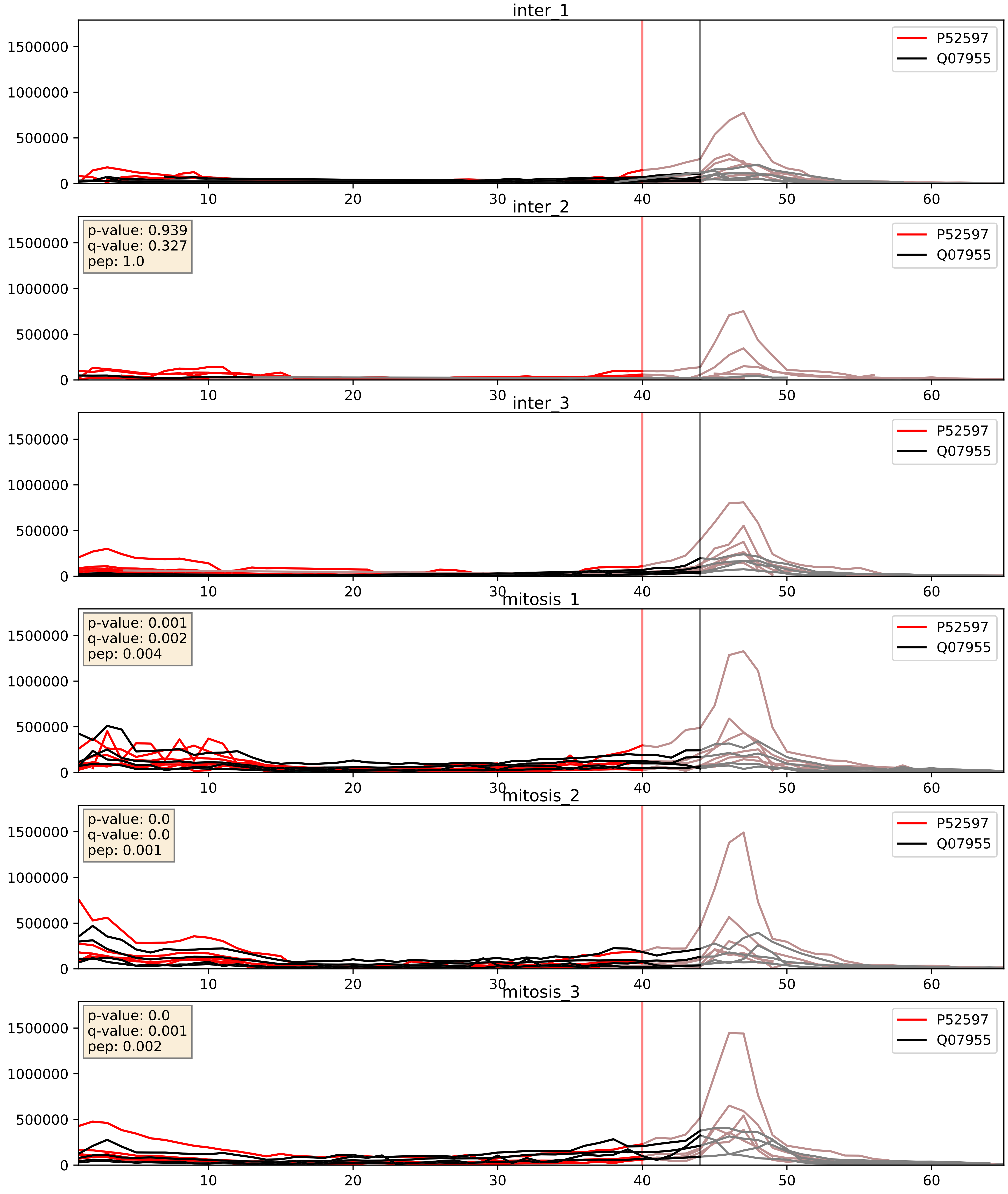


| level | condition_1 | condition_2 | pvalue | pvalue_adjusted |
|---|---|---|---|---|
| interactor_abundance | mitosis | inter | 0.0030437040272975205 | 0.042192884977598016 |
| complex_abundance | mitosis | inter | 0.003478199315974476 | 0.043087389500349516 |
| interactor_ratio | mitosis | inter | 0.17421734291907873 | 0.6394941918470471 |

P52597_Q08211
HNRPF_HUMAN vs DHX9_HUMAN
p-value: 0.002 q-value: 0.03

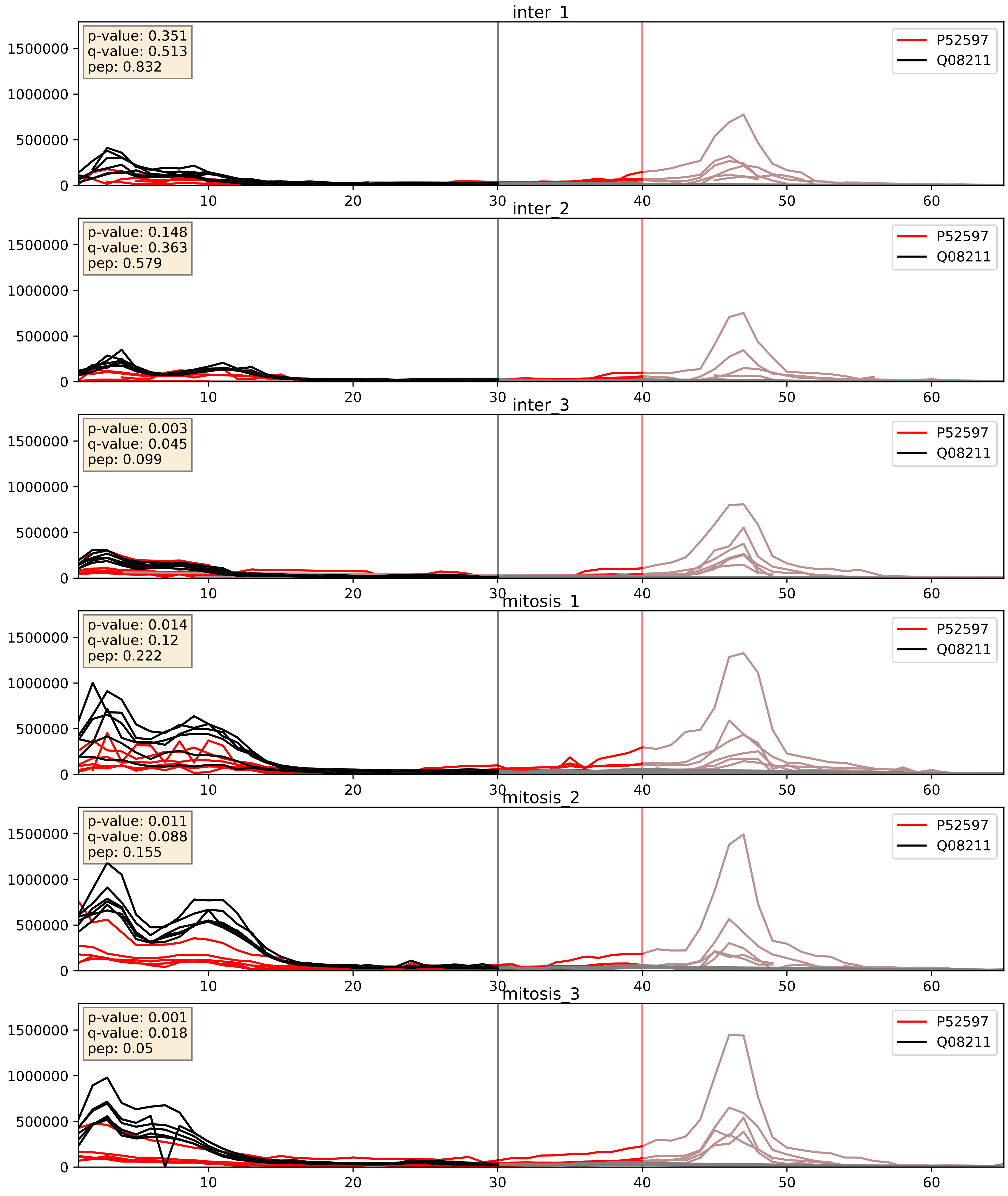


| level | condition_1 | condition_2 | pvalue | pvalue_adjusted |
|---|---|---|---|---|
| complex_abundance | mitosis | inter | 0.0005487925892557542 | 0.015047650716485379 |
| interactor_abundance | mitosis | inter | 0.007718766641450282 | 0.0762963538693007 |
| interactor_ratio | mitosis | inter | 0.1472602964791206 | 0.6016936218908221 |

P52597_Q13151
HNRPF_HUMAN vs ROA0_HUMAN
p-value: 0.005 q-value: 0.005

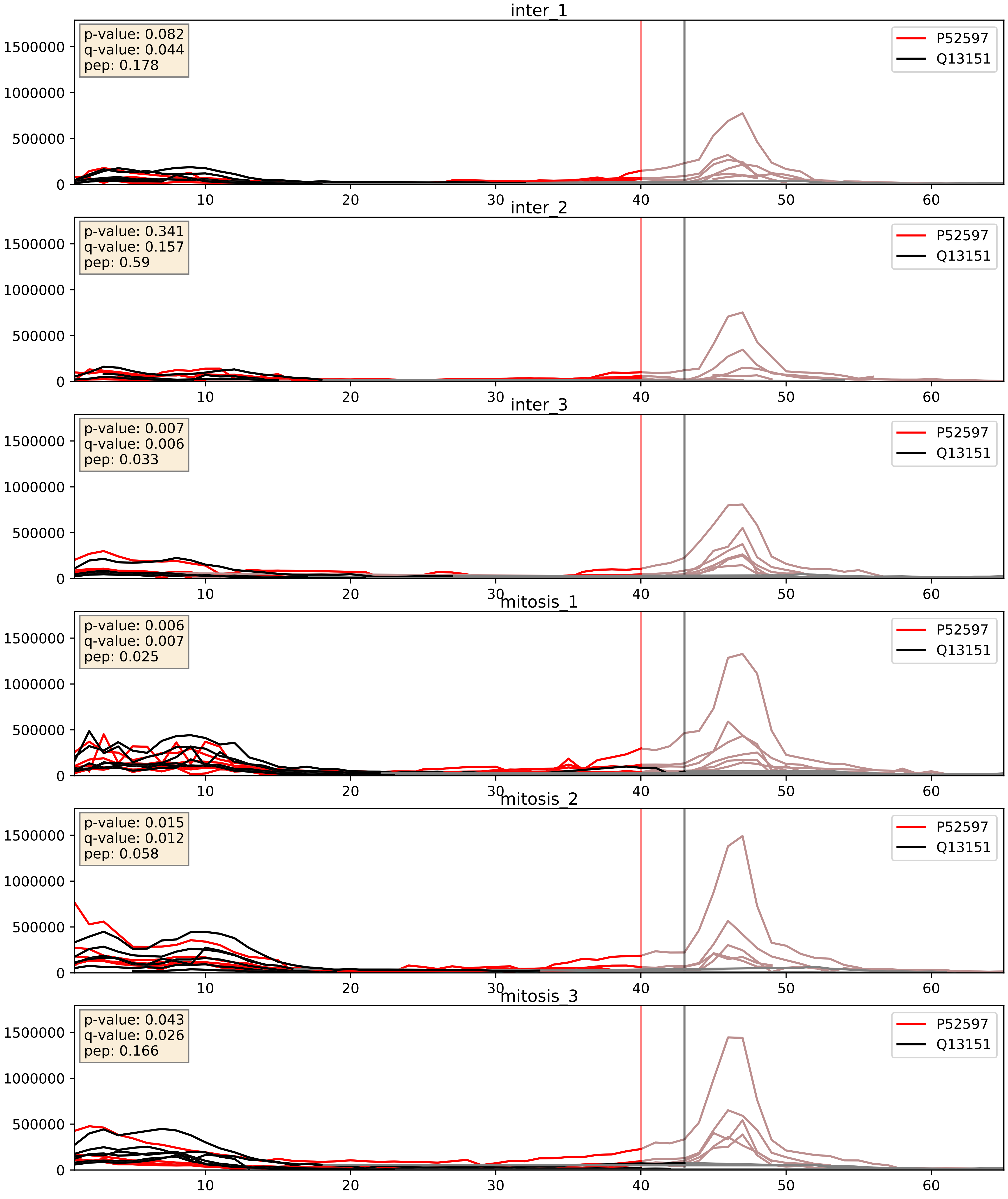


| level | condition_1 | condition_2 | pvalue | pvalue_adjusted |
|---|---|---|---|---|
| complex_abundance | mitosis | inter | 0.001076328295497409 | 0.021833569470040306 |
| interactor_abundance | mitosis | inter | 0.003070826023648689 | 0.042465703978082026 |
| interactor_ratio | mitosis | inter | 0.5732357411651294 | 0.8946257776299471 |

P52597_Q13247
HNRPF_HUMAN vs SRSF6_HUMAN
p-value: 0.002 q-value: 0.03

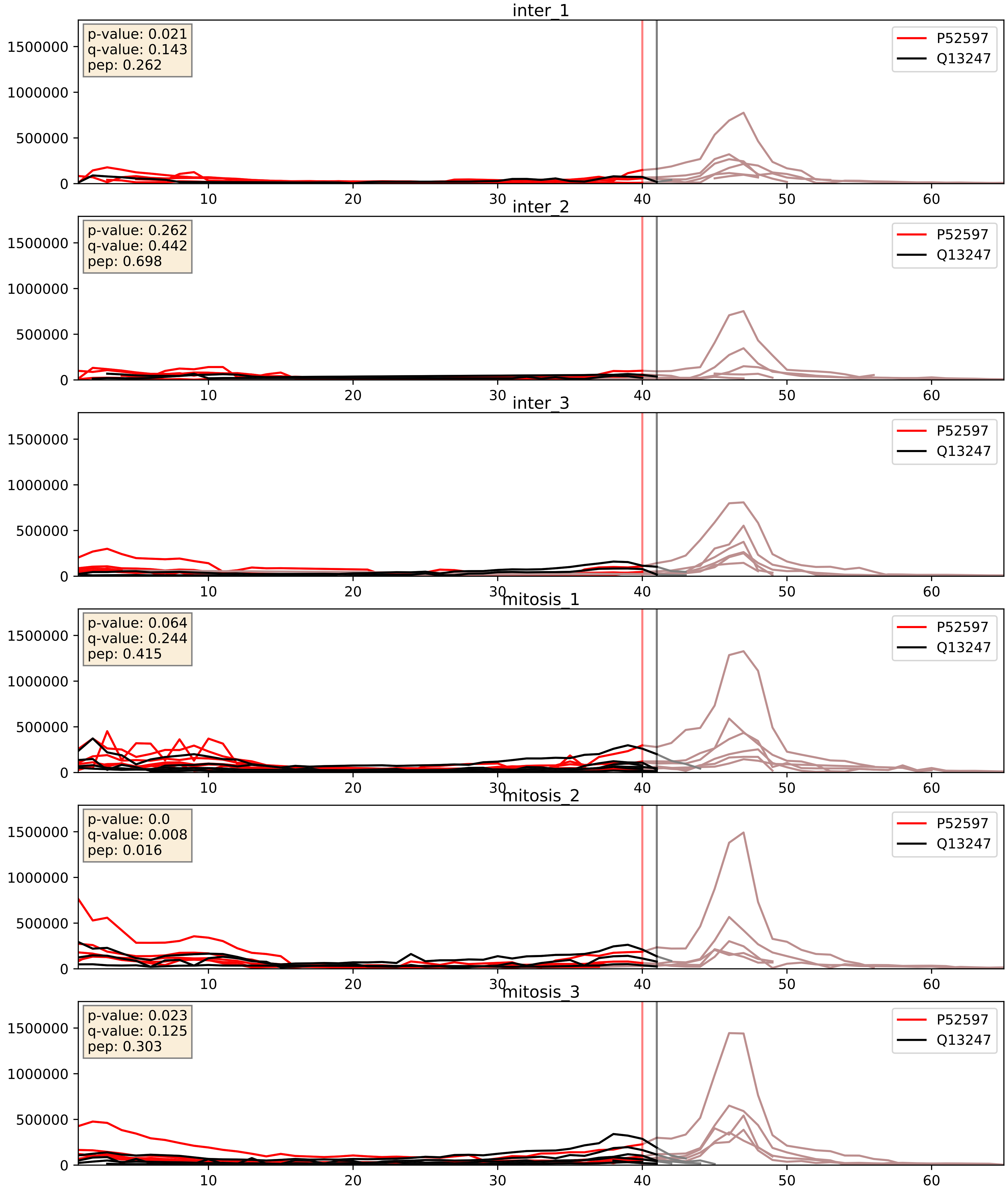


| level | condition_1 | condition_2 | pvalue | pvalue_adjusted |
|---|---|---|---|---|
| interactor_abundance | mitosis | inter | 0.005847440986472472 | 0.06479494478214157 |
| complex_abundance | mitosis | inter | 0.010125200125526758 | 0.07439632023563009 |
| interactor_ratio | mitosis | inter | 0.2895904474858344 | 0.7585355662419652 |

P52597_Q15365
HNRPF_HUMAN vs PCBP1_HUMAN
p-value: 0.002 q-value: 0.035

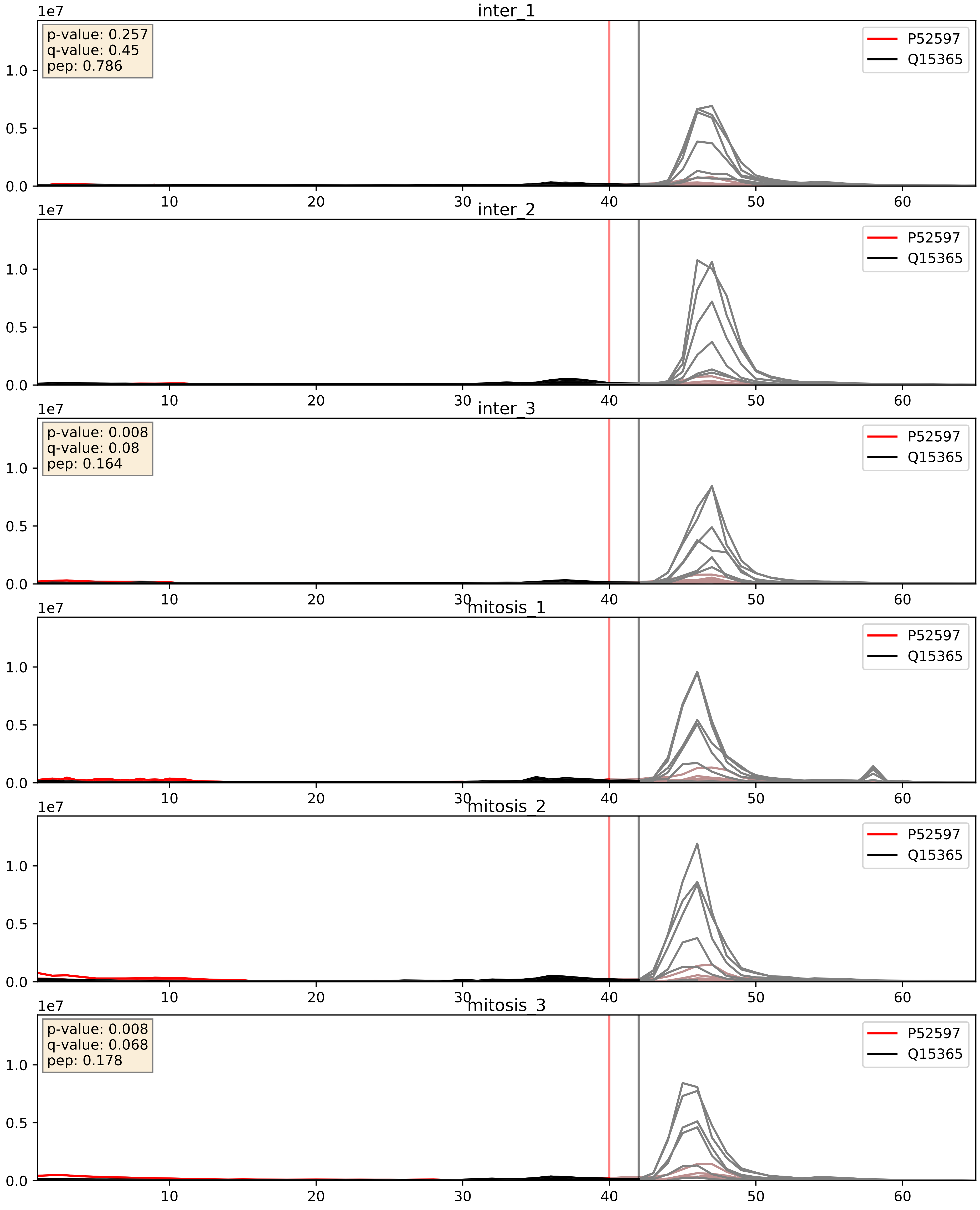


| level | condition_1 | condition_2 | pvalue | pvalue_adjusted |
|---|---|---|---|---|
| interactor_ratio | mitosis | inter | 0.0031499845140135186 | 0.07327137891292315 |
| interactor_abundance | mitosis | inter | 0.003829085421829145 | 0.04830798999389459 |
| complex_abundance | mitosis | inter | 0.01784052834109063 | 0.1000532792033756 |

P52597_Q15717
HNRPF_HUMAN vs ELAV1_HUMAN
p-value: 0.005 q-value: 0.005

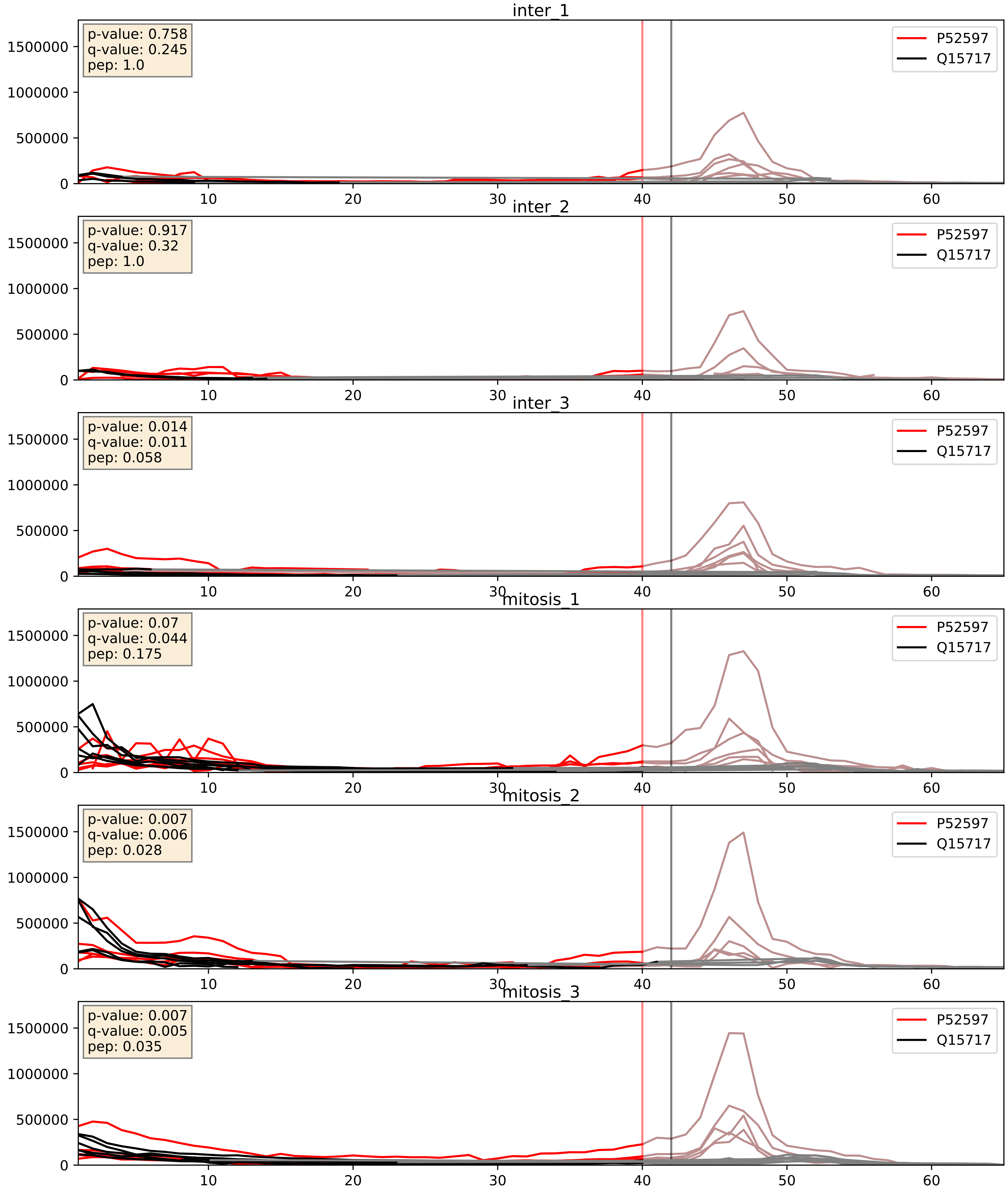


| level | condition_1 | condition_2 | pvalue | pvalue_adjusted |
|---|---|---|---|---|
| complex_abundance | mitosis | inter | 0.00016915016388839636 | 0.008722442186052247 |
| interactor_abundance | mitosis | inter | 0.002413019482392022 | 0.036558312865974704 |
| interactor_ratio | mitosis | inter | 0.014023604107891958 | 0.1741671265118786 |

P52597_Q16629
HNRPF_HUMAN vs SRSF7_HUMAN
p-value: 0.0 q-value: 0.003

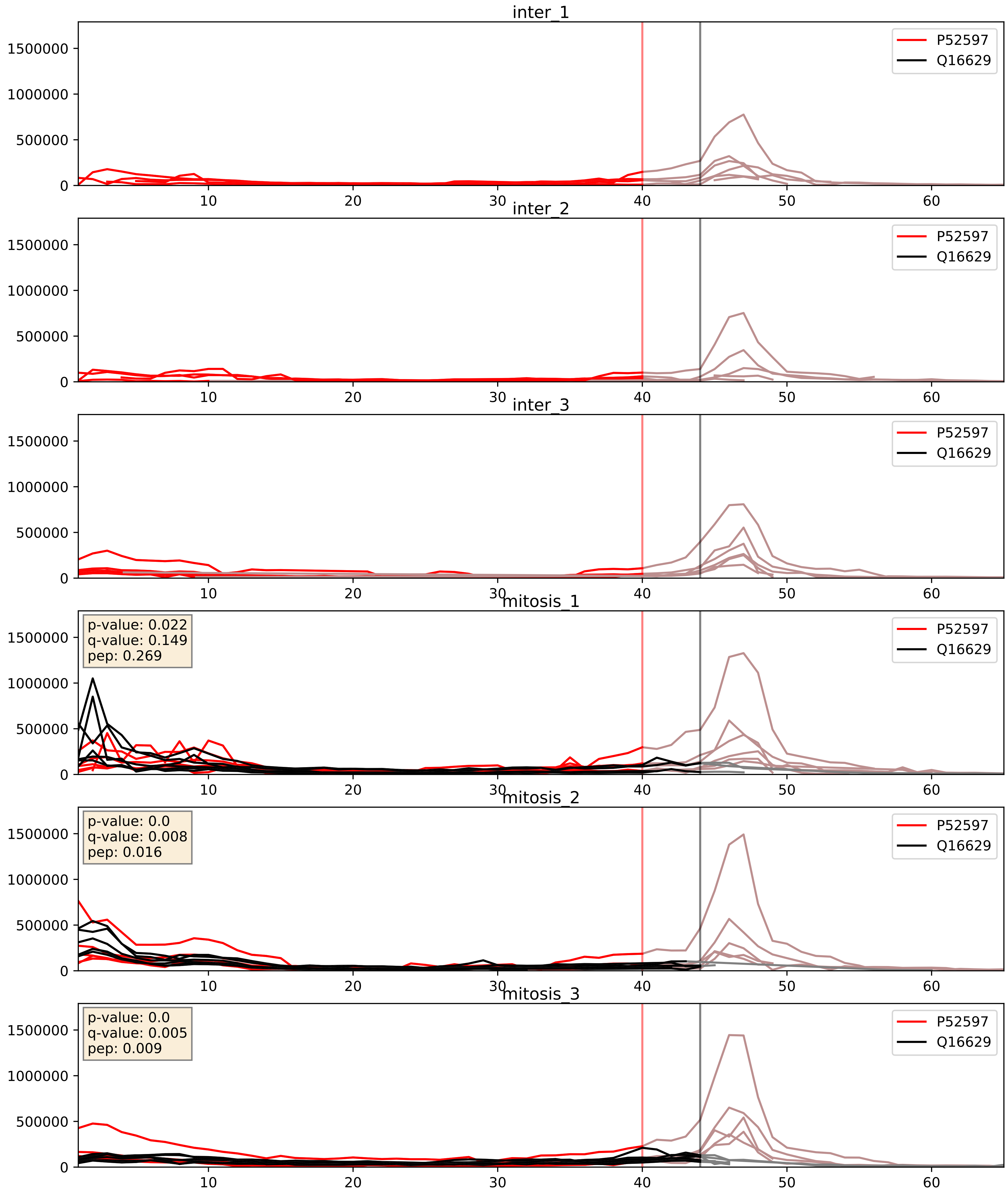


| level | condition_1 | condition_2 | pvalue | pvalue_adjusted |
|---|---|---|---|---|
| interactor_abundance | mitosis | inter | 4.447475947098241e-05 | 0.006058954526139308 |
| complex_abundance | mitosis | inter | 5.25369688885978365e-05 | 0.005691857151652958 |
| interactor_ratio | mitosis | inter | 0.4299776830859134 | 0.8432094269126428 |

P52597_Q99459
HNRPF_HUMAN vs CDC5L_HUMAN
p-value: 0.016 q-value: 0.013

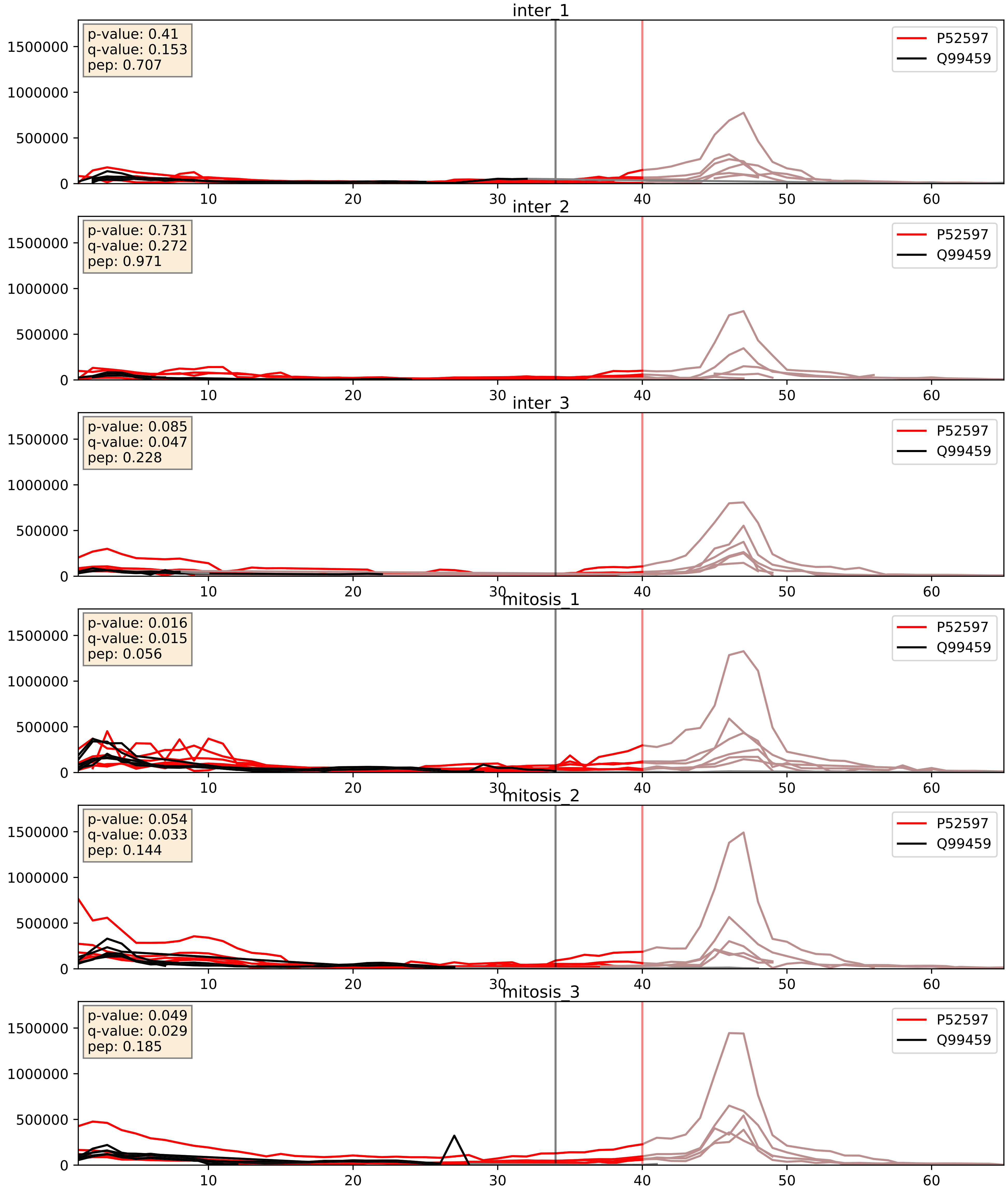


| level | condition_1 | condition_2 | pvalue | pvalue_adjusted |
|---|---|---|---|---|
| interactor_abundance | mitosis | inter | 0.005872037501971724 | 0.06494139666263302 |
| complex_abundance | mitosis | inter | 0.008301617280340168 | 0.06742444735376033 |
| interactor_ratio | mitosis | inter | 0.13750648150689043 | 0.5873558393508659 |

P52597_Q9BUJ2
HNRPF_HUMAN vs HNRL1_HUMAN
p-value: 0.0 q-value: 0.0

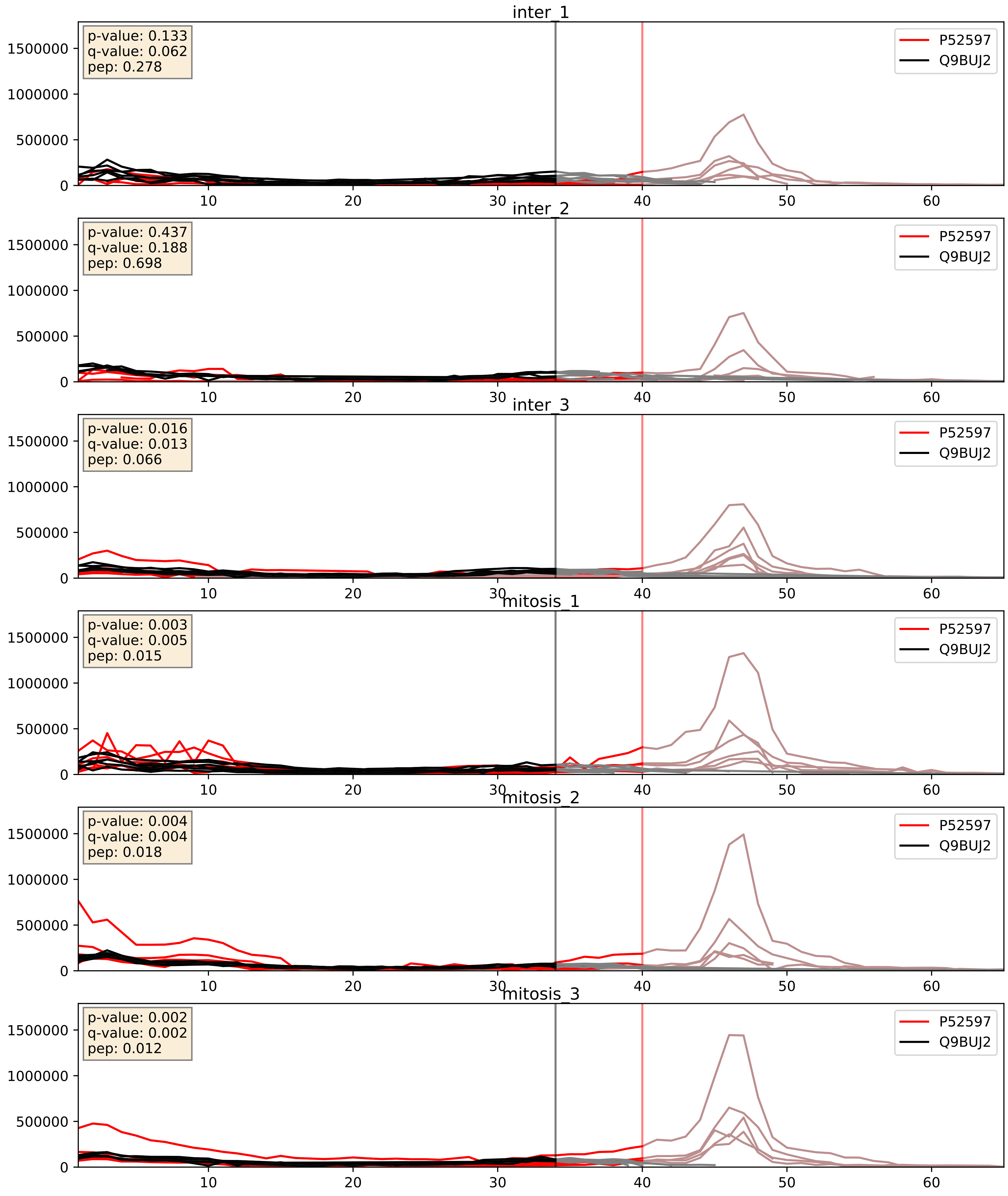


| level | condition_1 | condition_2 | pvalue | pvalue_adjusted |
|---|---|---|---|---|
| interactor_abundance | mitosis | inter | 0.005867622300035323 | 0.06493451439987377 |
| interactor_ratio | mitosis | inter | 0.00655810316750251 | 0.1124141137448166 |
| complex_abundance | mitosis | inter | 0.00962378637334349 | 0.07290231093435423 |
