## Supplementary Data 2 for "SECAT: Quantifying differential protein-protein interaction states by network-centric analysis": hela_string_000011_P24941.pdf

P24941_P24941

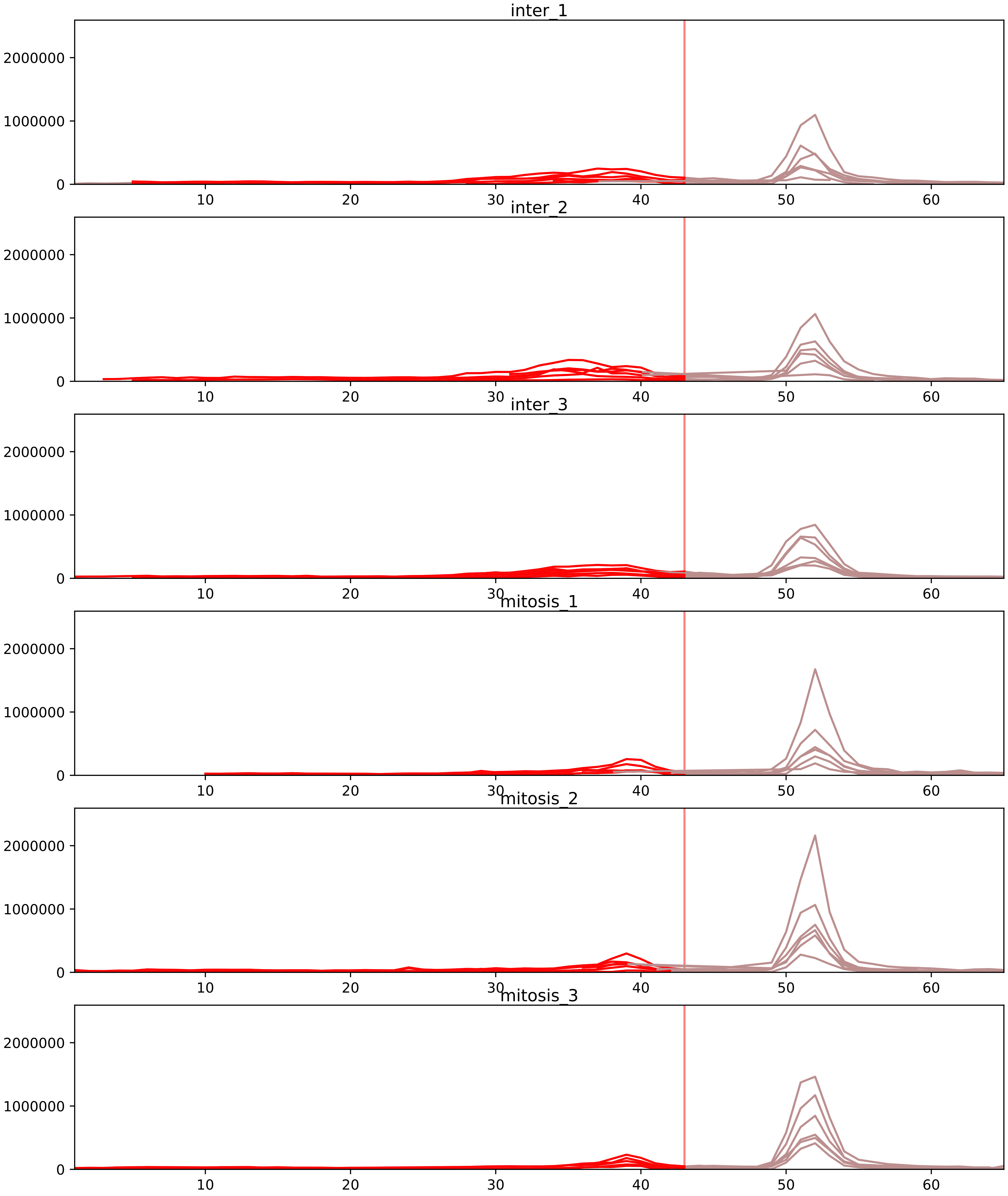


| level | condition_1 | condition_2 | pvalue | pvalue_adjusted |
|---|---|---|---|---|
| interactor_ratio | mitosis | inter | 1.471400026943556e-14 | 2.46125095416013e-12 |
| complex_abundance | mitosis | inter | 3.893004405086517e-09 | 2.3877093684530637e-06 |
| interactor_abundance | mitosis | inter | 3.7481673747059376e-06 | 0.0002733228684700698 |
| assembled_abundance | mitosis | inter | 0.004386881880806217 | 0.06581728873094198 |
| monomer_abundance | mitosis | inter | 0.08862614520131269 | 0.38770285874899957 |
| total_abundance | mitosis | inter | 0.6057532772278731 | 0.8204426728836646 |

O95067_P24941
CCNB2_HUMAN vs CDK2_HUMAN
p-value: 0.004 q-value: 0.004

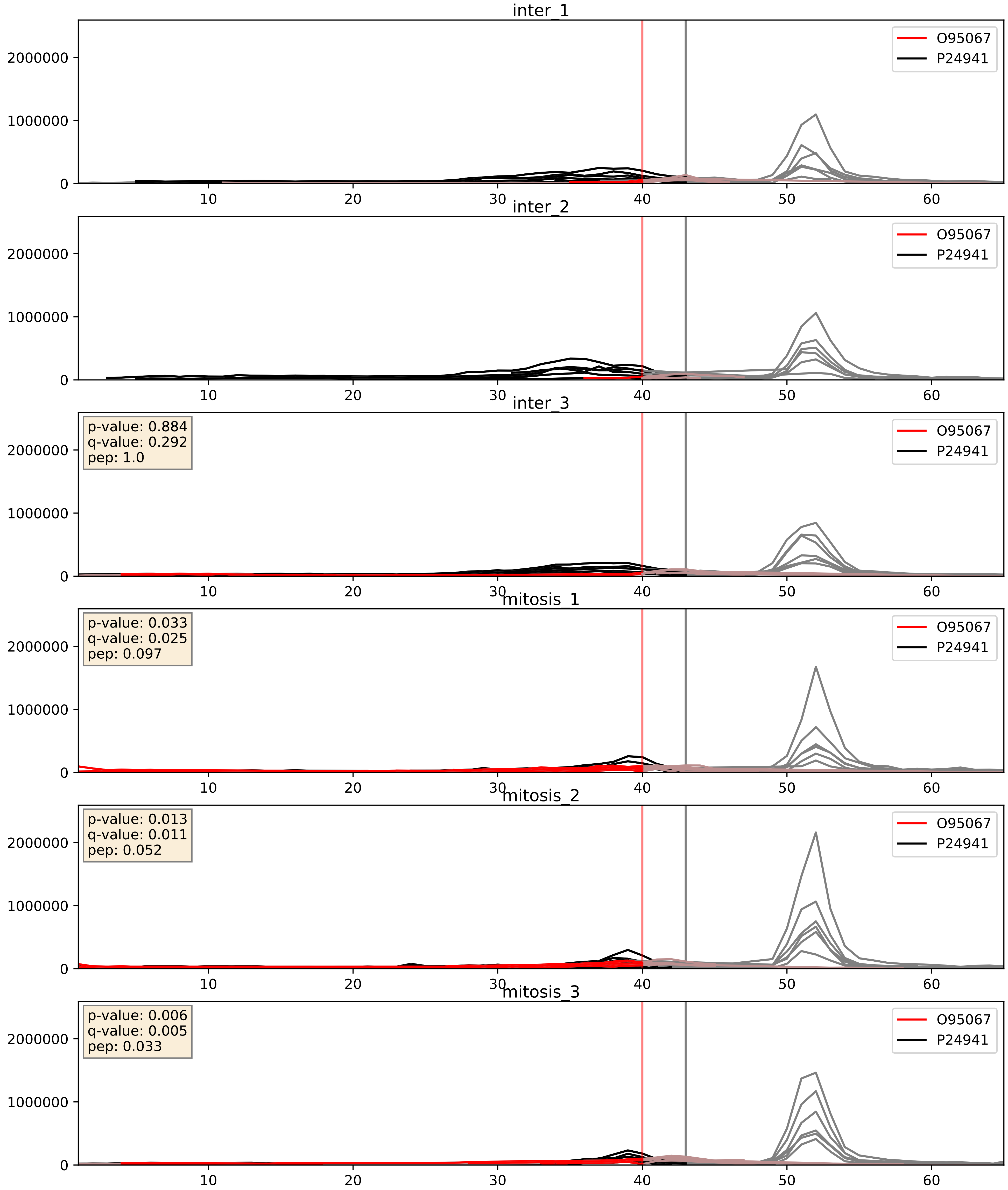


| level | condition_1 | condition_2 | pvalue | pvalue_adjusted |
|---|---|---|---|---|
| interactor_abundance | mitosis | inter | 0.001443970468219757 | 0.027996347016899476 |
| interactor_ratio | mitosis | inter | 0.011385932666237232 | 0.15421453104903593 |
| complex_abundance | mitosis | inter | 0.04027580454763698 | 0.14508340529050803 |

P06493_P24941
CDK1_HUMAN vs CDK2_HUMAN
p-value: 0.001 q-value: 0.001

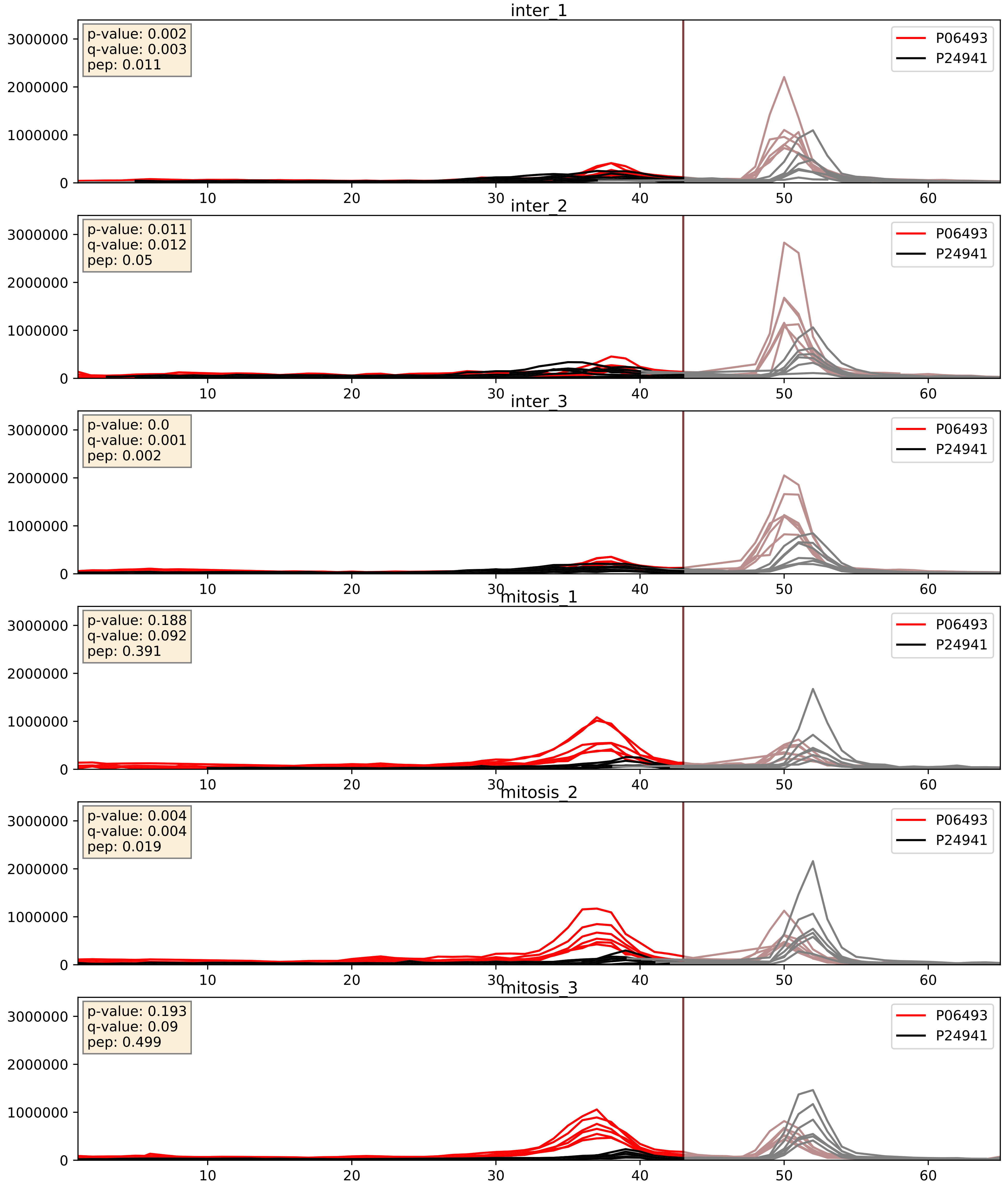


| level | condition_1 | condition_2 | pvalue | pvalue_adjusted |
|---|---|---|---|---|
| interactor_ratio | mitosis | inter | 0.00019753625235696597 | 0.014576813104962318 |
| interactor_abundance | mitosis | inter | 0.017640941221411546 | 0.11492120004207217 |
| complex_abundance | mitosis | inter | 0.34351583213978315 | 0.4915572589629796 |

P14635_P24941
CCNB1_HUMAN vs CDK2_HUMAN
p-value: 0.023 q-value: 0.018

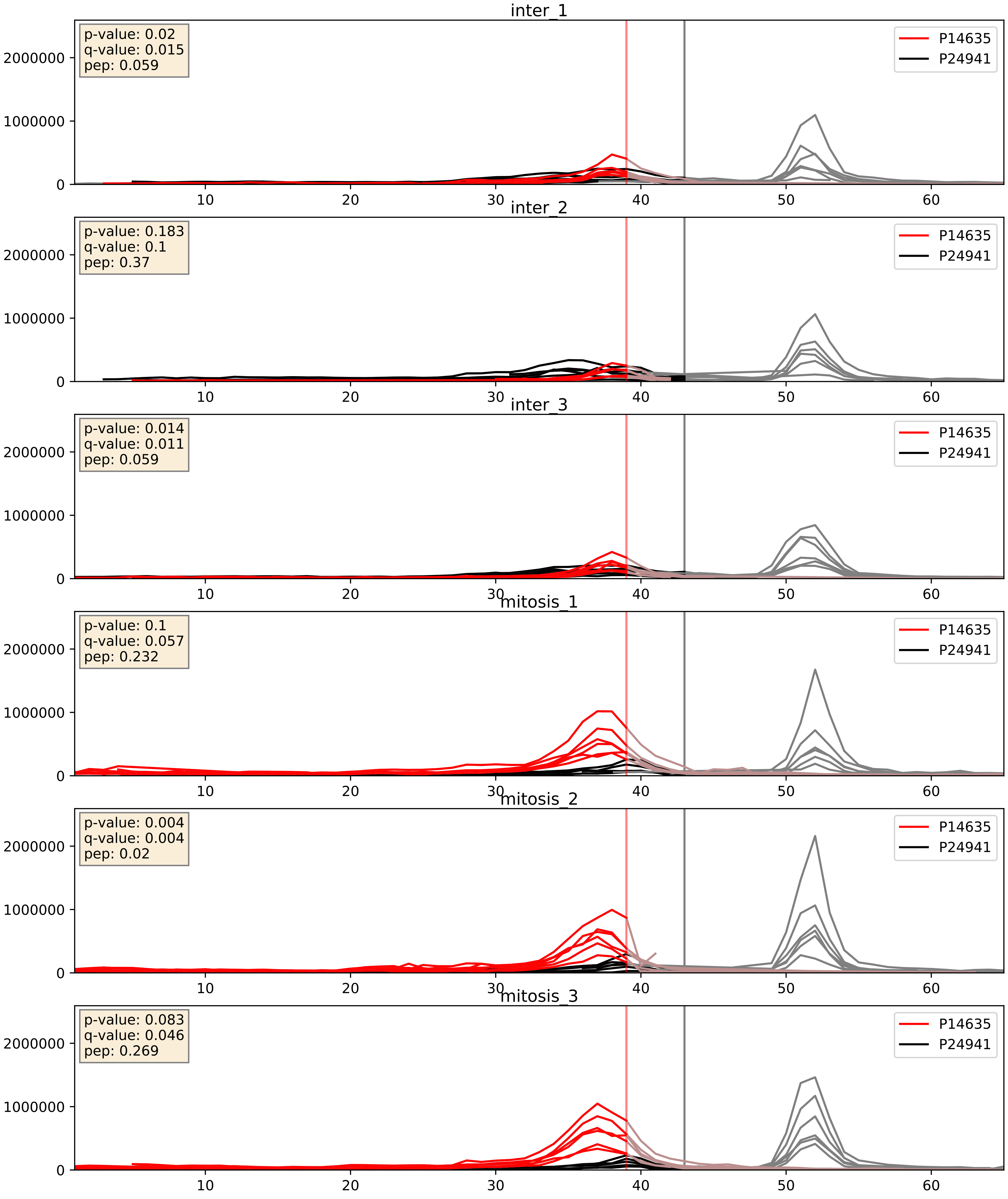


| level | condition_1 | condition_2 | pvalue | pvalue_adjusted |
|---|---|---|---|---|
| interactor_ratio | mitosis | inter | 0.0008074983906906686 | 0.032604652001472285 |
| interactor_abundance | mitosis | inter | 0.002332060110087462 | 0.03586830170101165 |
| complex_abundance | mitosis | inter | 0.0425001476113568 | 0.1476929246638287 |

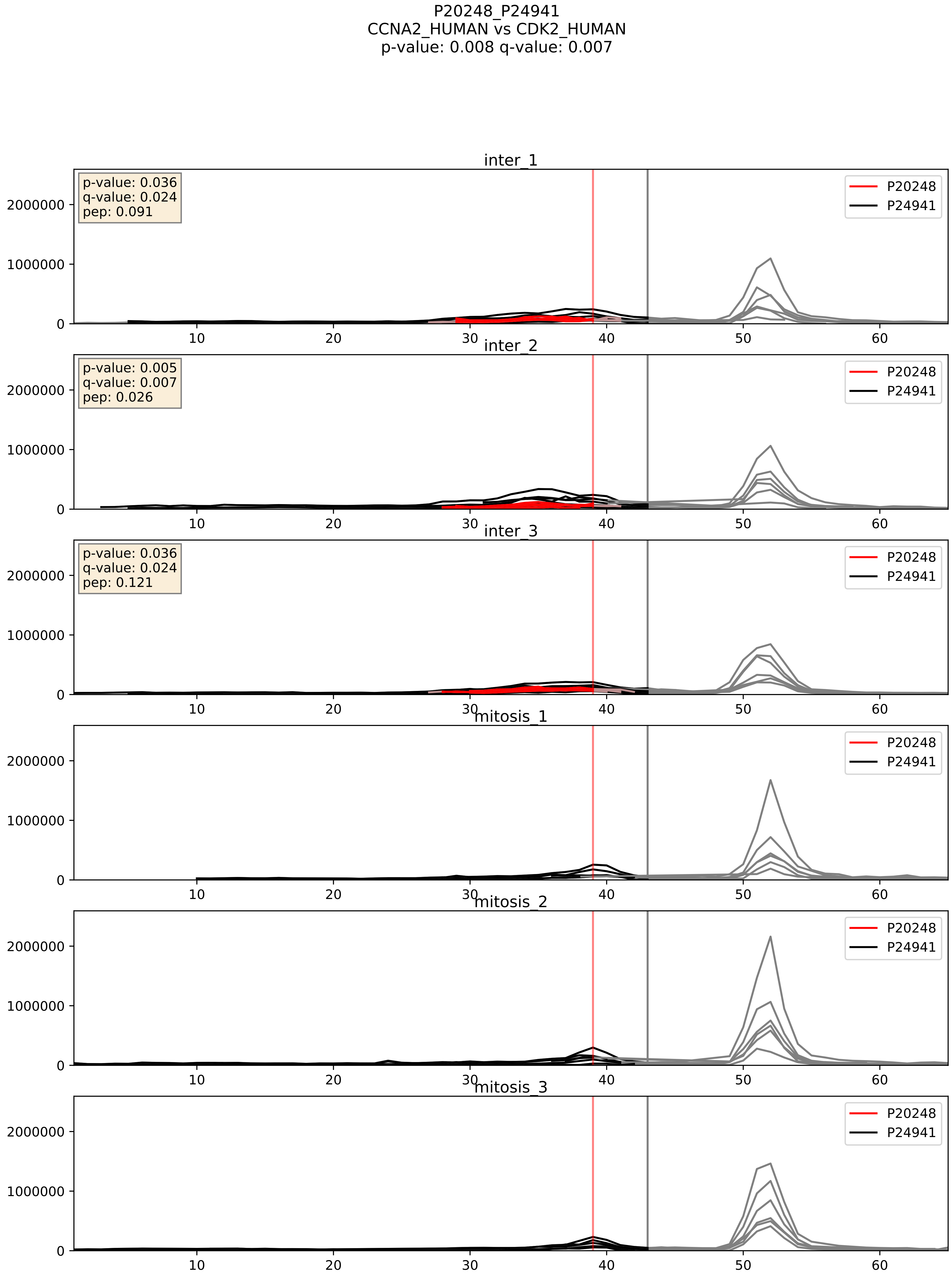


| level | condition_1 | condition_2 | pvalue | pvalue_adjusted |
|---|---|---|---|---|
| complex_abundance | mitosis | inter | 0.00017174227842104968 | 0.008803077265174762 |
| interactor_abundance | mitosis | inter | 0.0004058346268662555 | 0.013951583959739548 |
| interactor_ratio | mitosis | inter | 0.01412332765988096 | 0.17445264757370998 |

P24941_P25205
CDK2_HUMAN vs MCM3_HUMAN
p-value: 0.051 q-value: 0.033

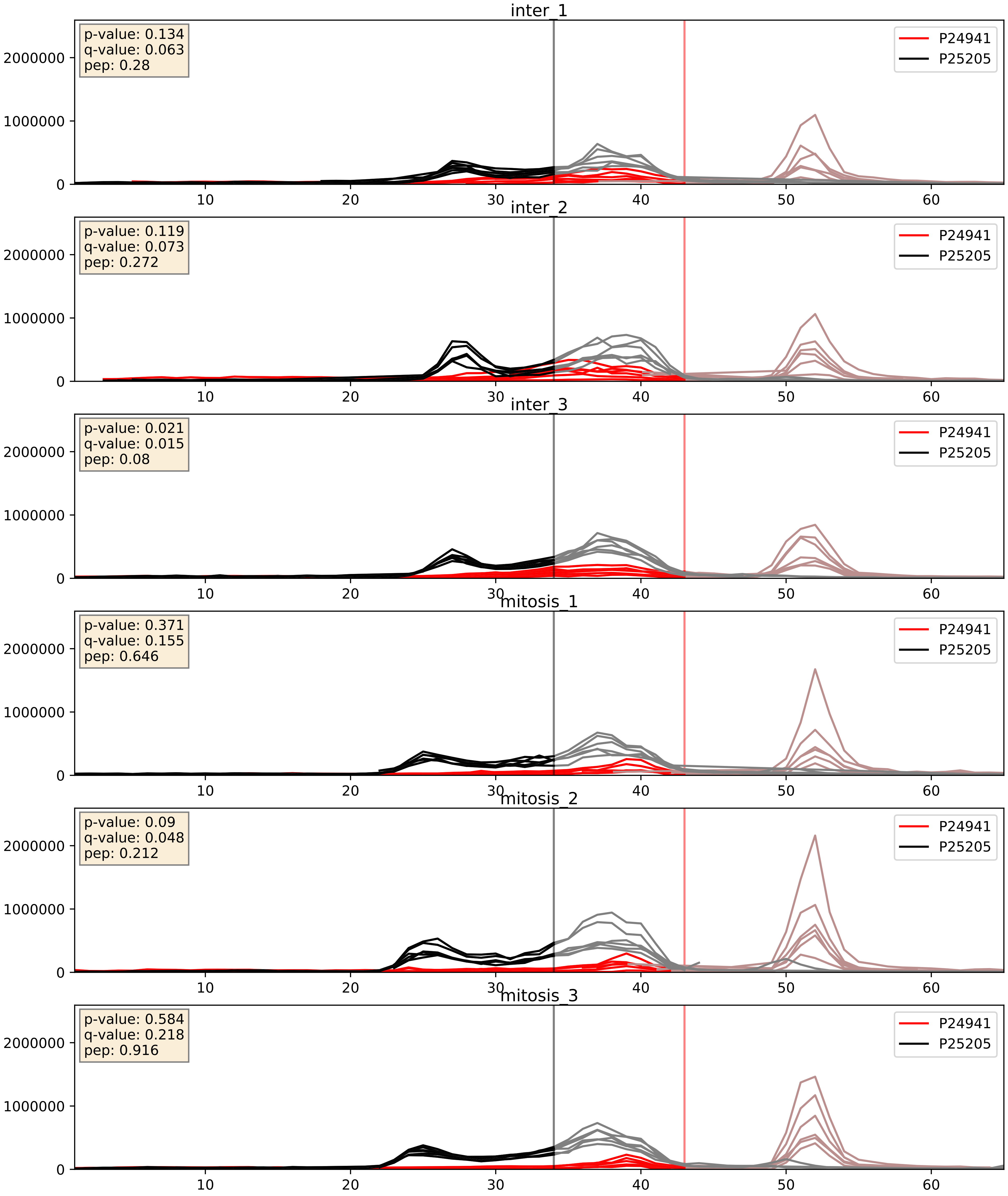


| level | condition_1 | condition_2 | pvalue | pvalue_adjusted |
|---|---|---|---|---|
| complex_abundance | mitosis | inter | 0.000877638583141945 | 0.019513211095311817 |
| interactor_abundance | mitosis | inter | 0.0012740450977966962 | 0.026310798642074112 |
| interactor_ratio | mitosis | inter | 0.004021962211288759 | 0.08271268336775048 |

P24941_P33992
CDK2_HUMAN vs MCM5_HUMAN
p-value: 0.067 q-value: 0.041

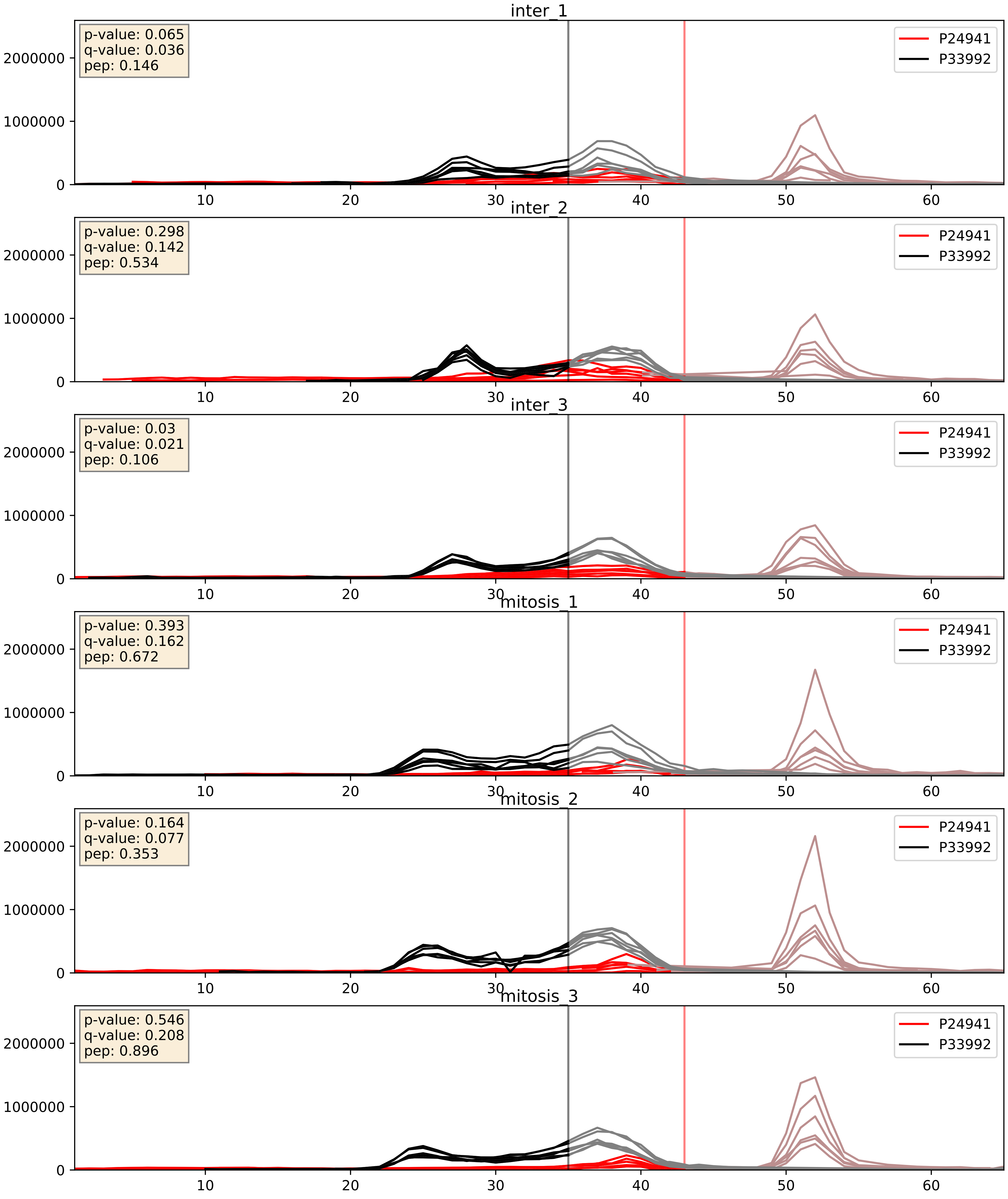


| level | condition_1 | condition_2 | pvalue | pvalue_adjusted |
|---|---|---|---|---|
| interactor_ratio | mitosis | inter | 0.00015356953979651935 | 0.012836278661937291 |
| interactor_abundance | mitosis | inter | 0.0009207487723061253 | 0.022454727894417187 |
| complex_abundance | mitosis | inter | 0.003410888998198883 | 0.04302508960916016 |

P24941_Q13309
CDK2_HUMAN vs SKP2_HUMAN
p-value: 0.0 q-value: 0.0

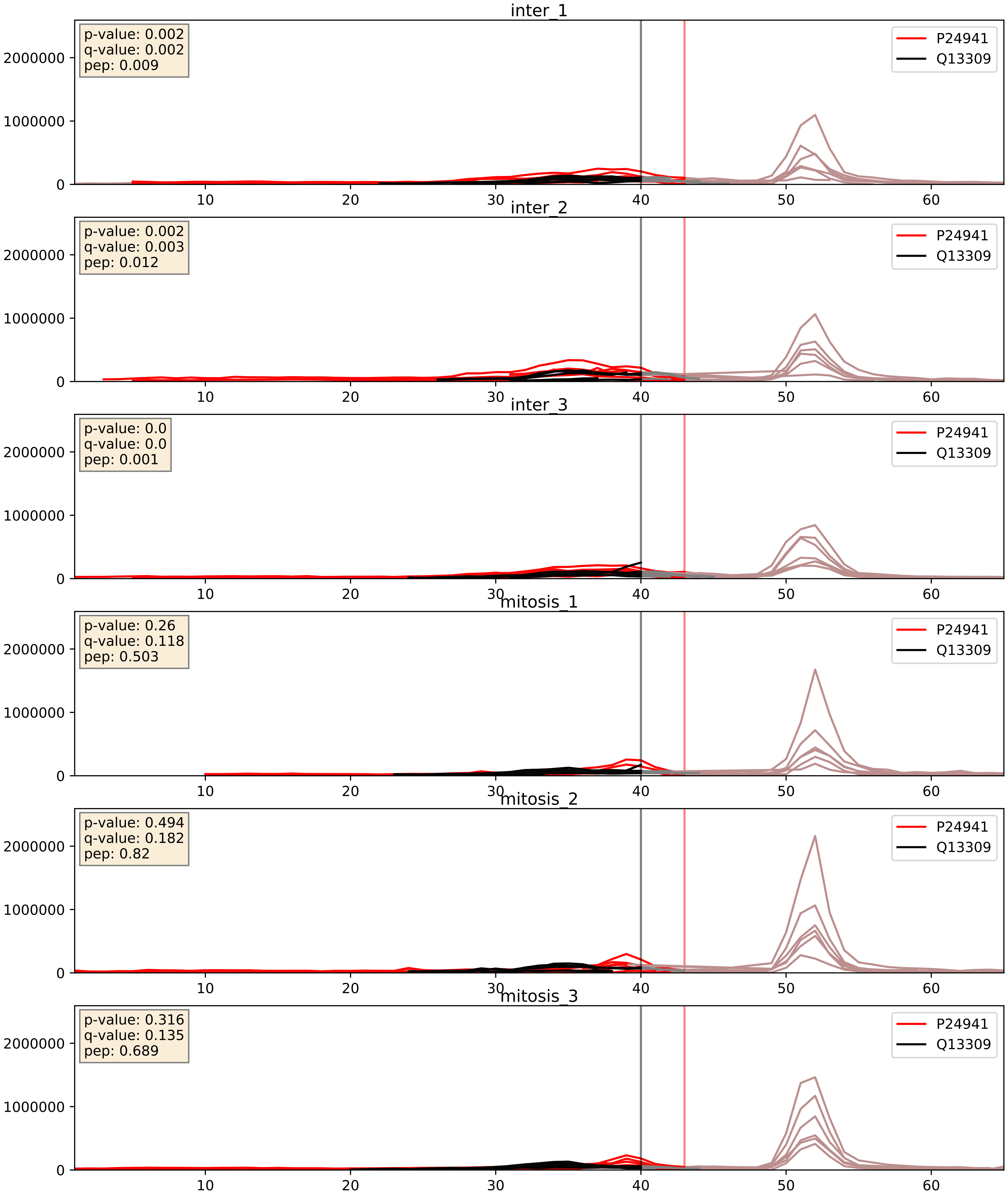


| level | condition_1 | condition_2 | pvalue | pvalue_adjusted |
|---|---|---|---|---|
| interactor_ratio | mitosis | inter | 4.400737078739184e-05 | 0.006796360849516482 |
| interactor_abundance | mitosis | inter | 0.0009100691980148597 | 0.022289534577989125 |
| complex_abundance | mitosis | inter | 0.004846689670913454 | 0.05231735634680853 |
