## Supplementary Data 2 for "SECAT: Quantifying differential protein-protein interaction states by network-centric analysis": hela_string_000012_P62995.pdf

# P62995_P62995

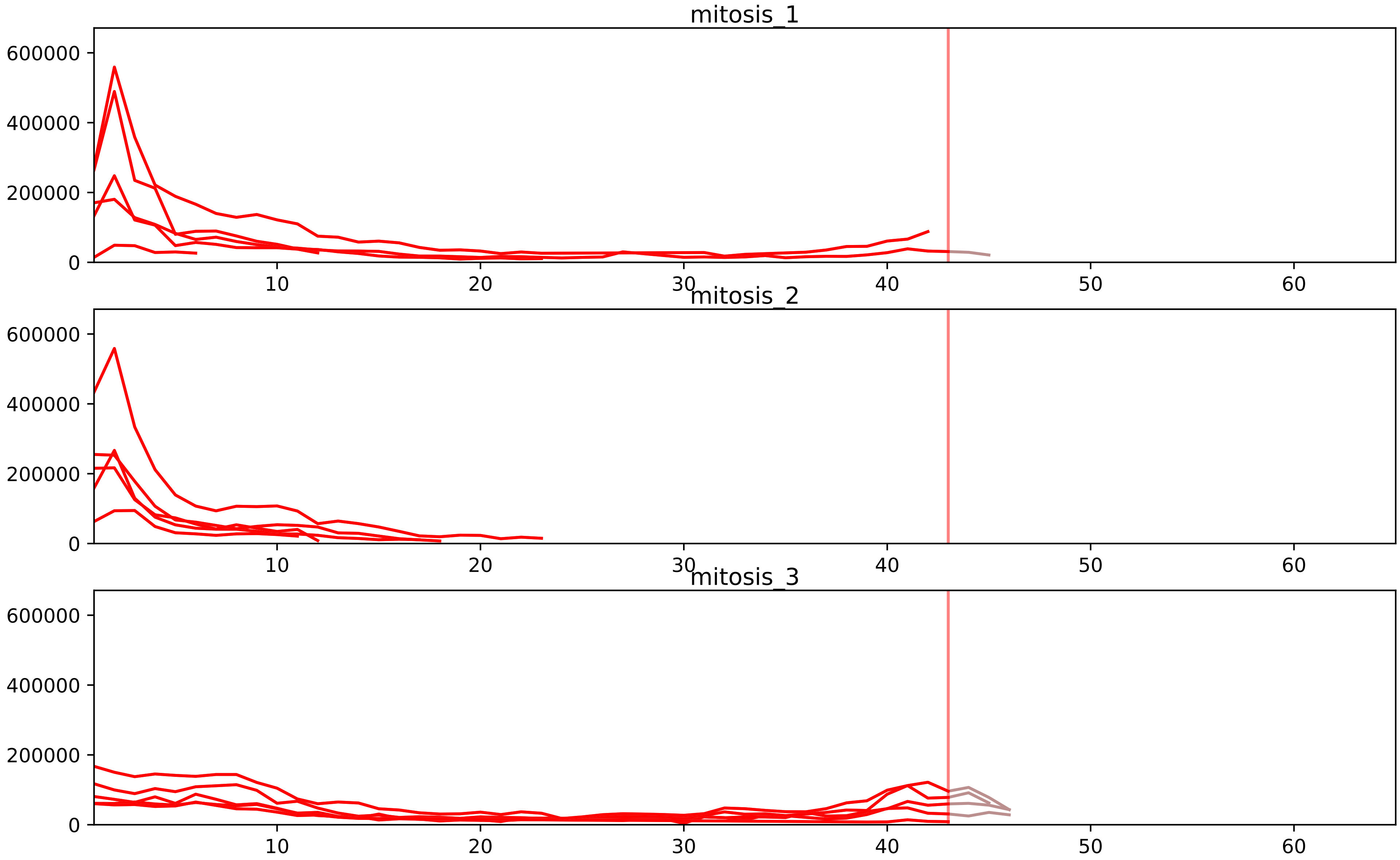


| level | condition_1 | condition_2 | pvalue | pvalue_adjusted |
|---|---|---|---|---|
| interactor_ratio | mitosis | inter | 1.904197674536582e-14 | 2.919769767622759e-12 |
| complex_abundance | mitosis | inter | 4.940989694069434e-07 | 5.6821381481798486e-05 |
| interactor_abundance | mitosis | inter | 2.0155346074324335e-06 | 0.00017659922274646084 |
| assembled_abundance | mitosis | inter | 0.03434953900947724 | 0.22171564497781057 |
| total_abundance | mitosis | inter | 0.0359075753692699 | 0.23802795078769934 |
| monomer_abundance | mitosis | inter | 0.21698986178111387 | 0.5950843593520406 |

O75934_P62995
SPF27_HUMAN vs TRA2B_HUMAN
p-value: 0.004 q-value: 0.047

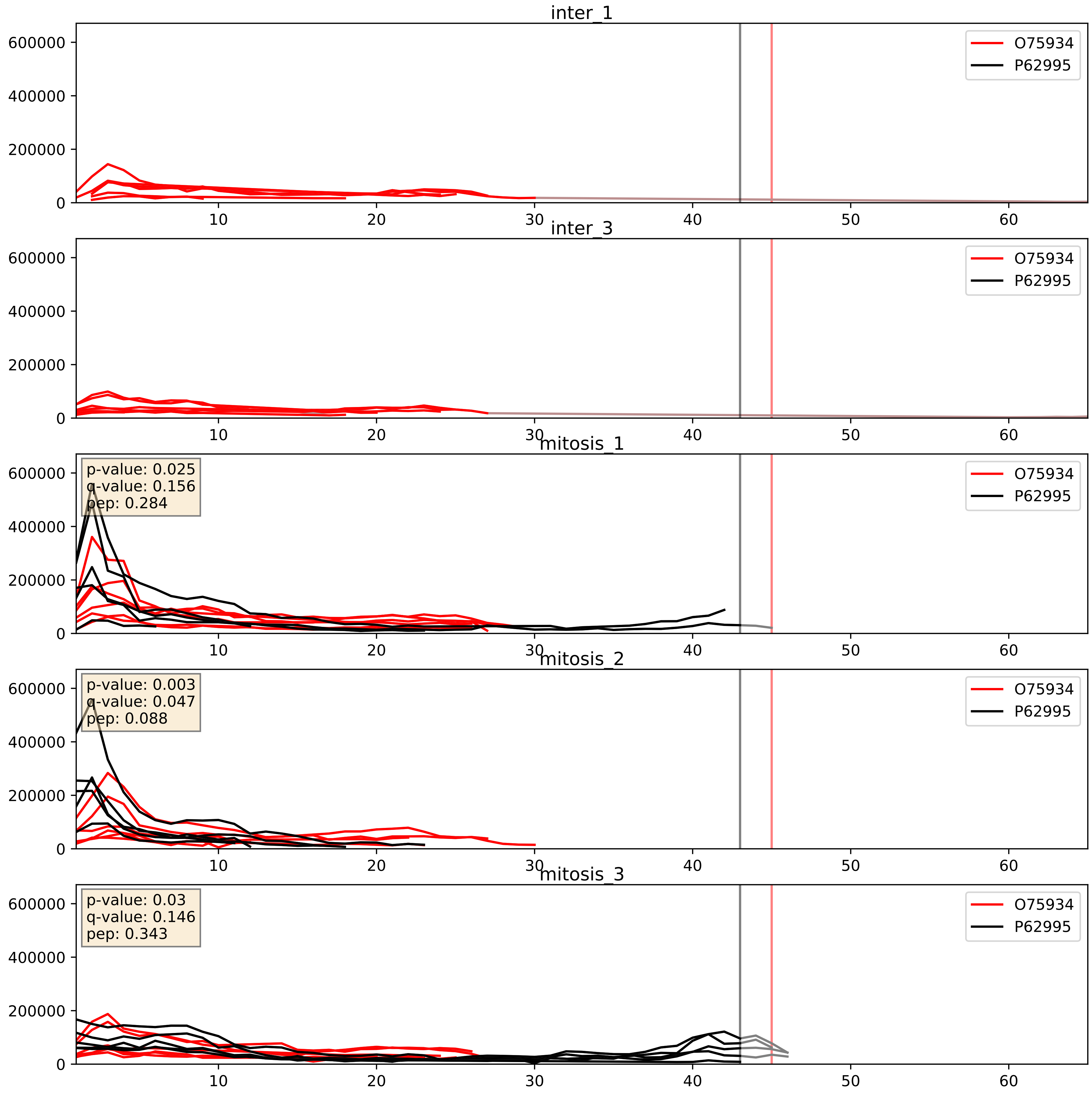


| level | condition_1 | condition_2 | pvalue | pvalue_adjusted |
|---|---|---|---|---|
| complex_abundance | mitosis | inter | 0.000556330786787743 | 0.01507022637627557 |
| interactor_abundance | mitosis | inter | 0.0007609849117950717 | 0.020324589219862133 |
| interactor_ratio | mitosis | inter | 0.23601861432310012 | 0.7155827200888129 |

P09651_P62995
ROA1_HUMAN vs TRA2B_HUMAN
p-value: 0.006 q-value: 0.005

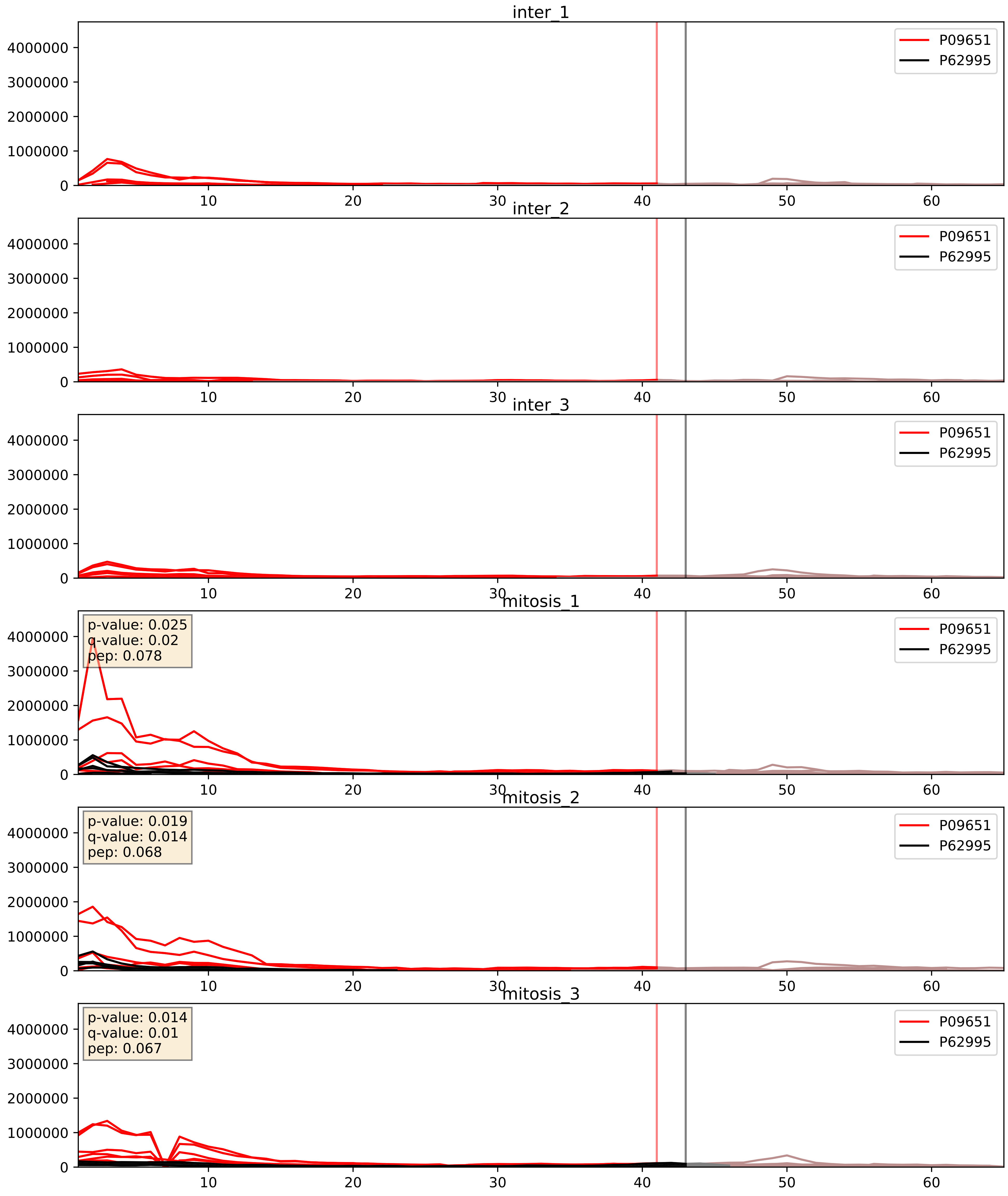


| level | condition_1 | condition_2 | pvalue | pvalue_adjusted |
|---|---|---|---|---|
| interactor_ratio | mitosis | inter | 9.380515584716844e-06 | 0.0035800160365075866 |
| interactor_abundance | mitosis | inter | 9.497380795475017e-05 | 0.007548664551888891 |
| complex_abundance | mitosis | inter | 0.0001646369536294834 | 0.008645965172198638 |

P11940_P62995
PABP1_HUMAN vs TRA2B_HUMAN
p-value: 0.0 q-value: 0.047

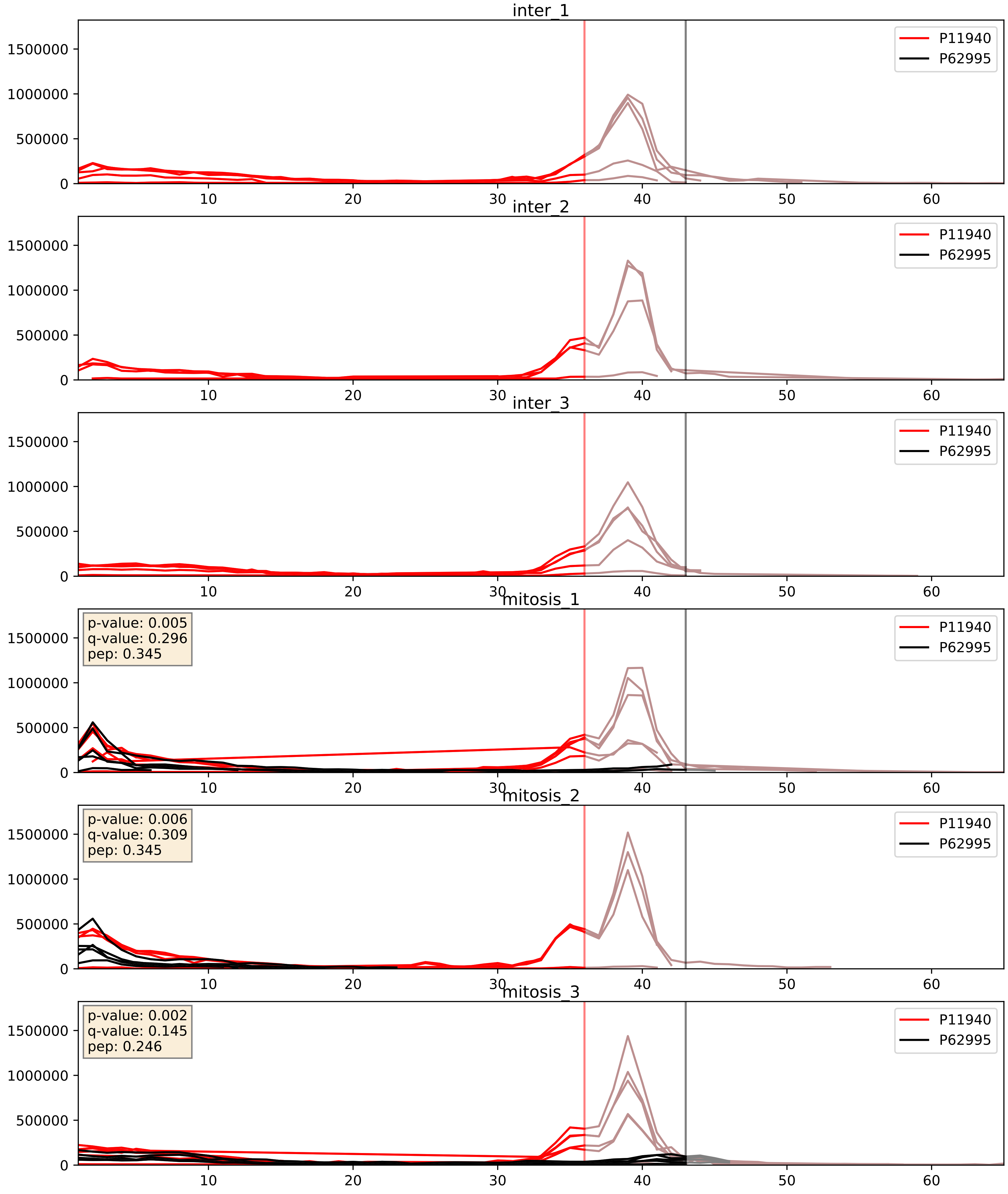


| level | condition_1 | condition_2 | pvalue | pvalue_adjusted |
|---|---|---|---|---|
| interactor_ratio | mitosis | inter | 2.219737766599098e-05 | 0.004993481478971454 |
| interactor_abundance | mitosis | inter | 7.82818796061288e-05 | 0.006980134264879818 |
| complex_abundance | mitosis | inter | 0.00015804606083051257 | 0.008563834848633118 |

P14866_P62995
HNRPL_HUMAN vs TRA2B_HUMAN
p-value: 0.007 q-value: 0.007

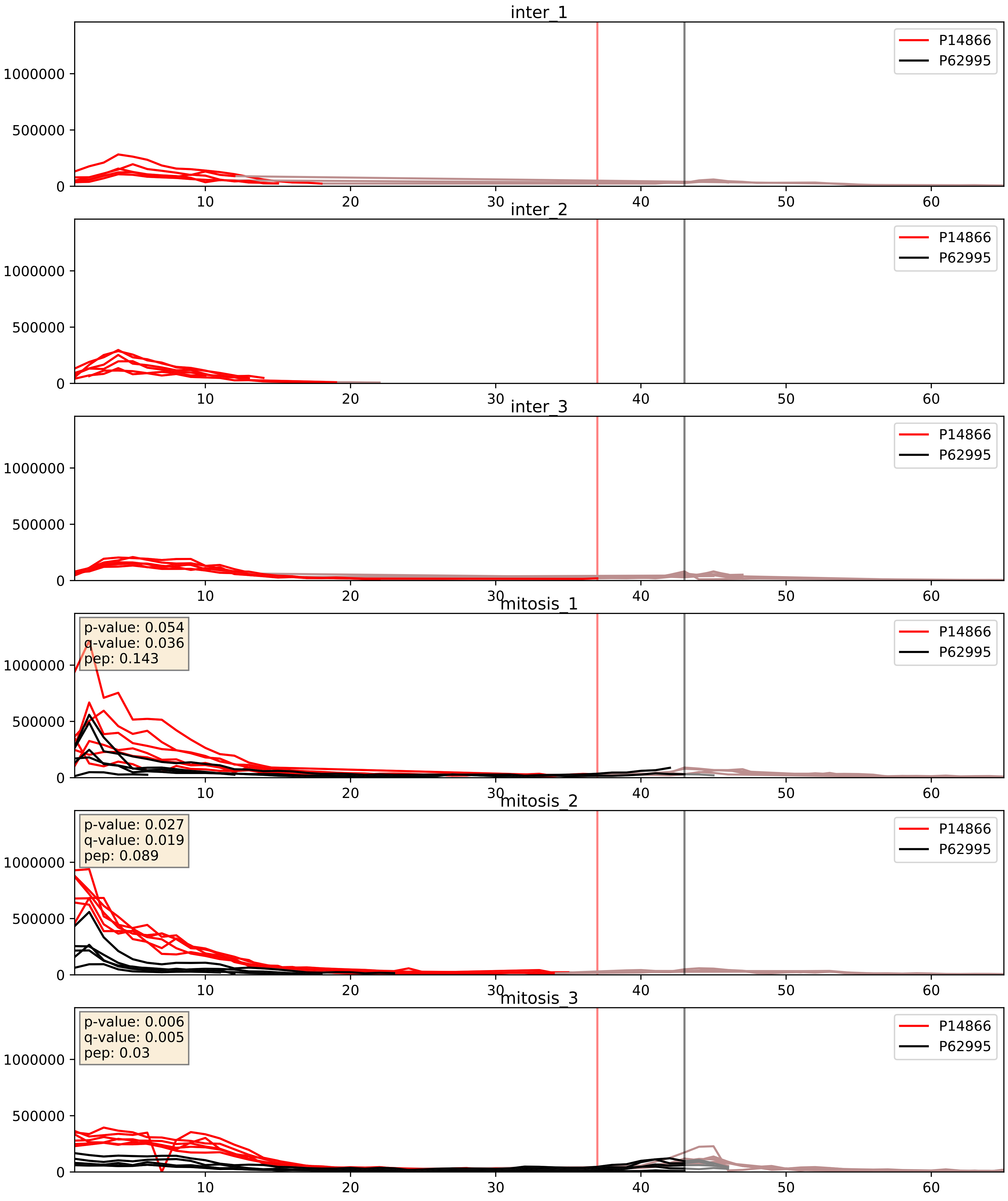


| level | condition_1 | condition_2 | pvalue | pvalue_adjusted |
|---|---|---|---|---|
| interactor_abundance | mitosis | inter | 1.2674740761041976e-05 | 0.003391165020149013 |
| complex_abundance | mitosis | inter | 5.6324767565087076e-05 | 0.005691857151652958 |
| interactor_ratio | mitosis | inter | 0.000967869032068439 | 0.03586562300651878 |

P31942_P62995
HNRH3_HUMAN vs TRA2B_HUMAN
p-value: 0.0 q-value: 0.023

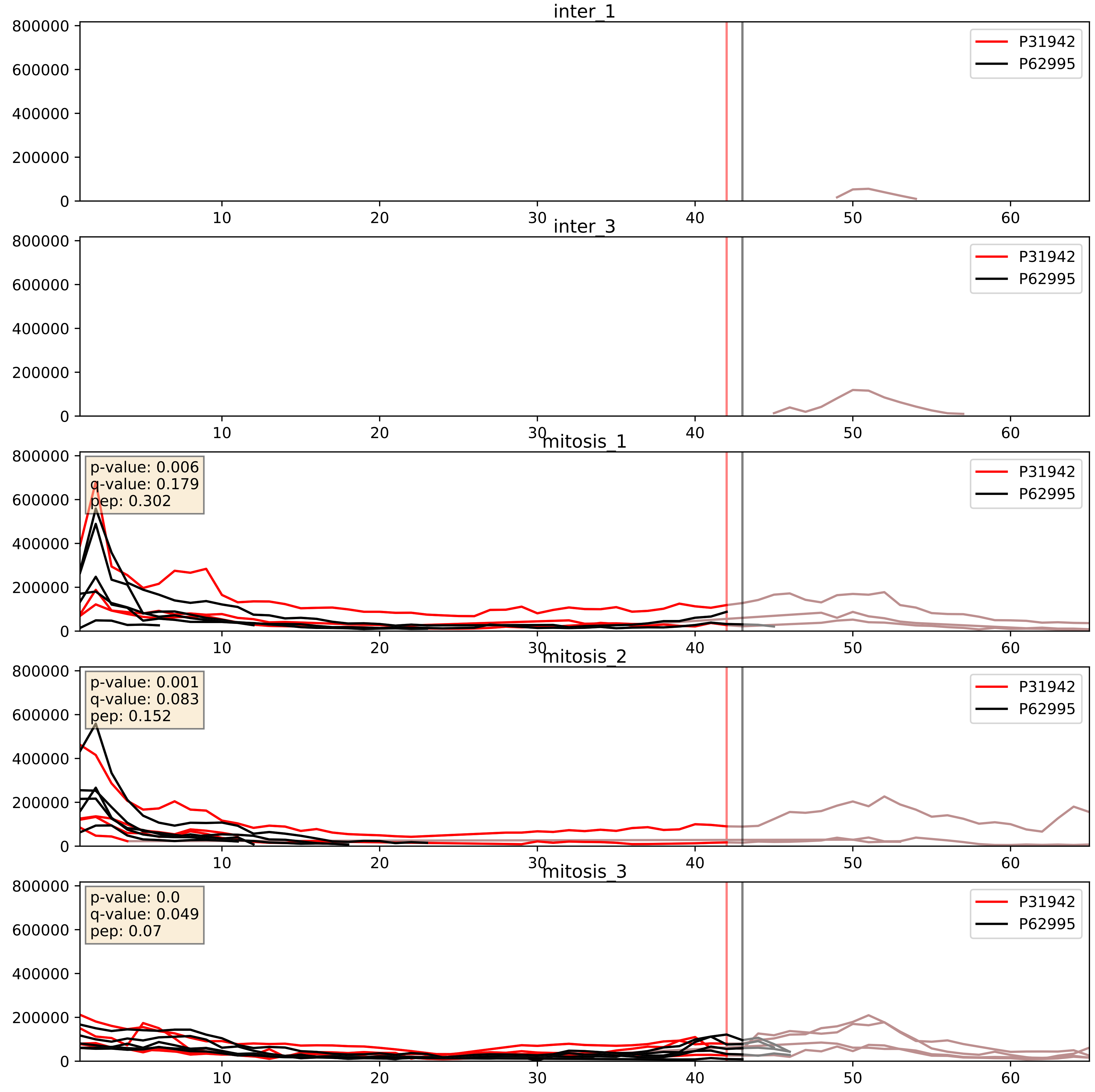


| level | condition_1 | condition_2 | pvalue | pvalue_adjusted |
|---|---|---|---|---|
| complex_abundance | mitosis | inter | 0.0010827209575059045 | 0.021833569470040306 |
| interactor_abundance | mitosis | inter | 0.004530454063144402 | 0.05416297036384927 |
| interactor_ratio | mitosis | inter | 0.20927687488332142 | 0.686697008467737 |

P31943_P62995
HNRH1_HUMAN vs TRA2B_HUMAN
p-value: 0.007 q-value: 0.006

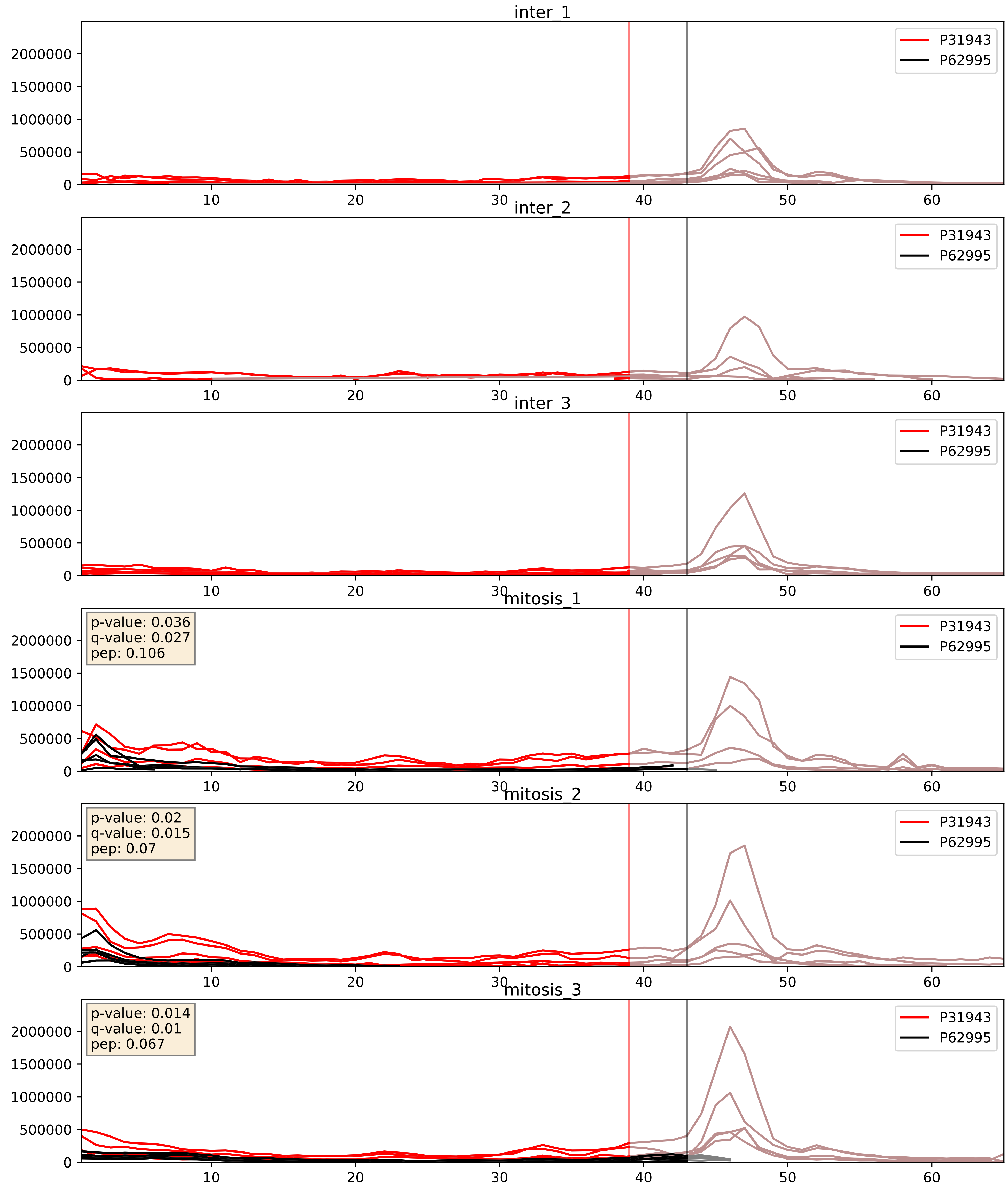


| level | condition_1 | condition_2 | pvalue | pvalue_adjusted |
|---|---|---|---|---|
| interactor_ratio | mitosis | inter | 2.5032546650867087e-05 | 0.005101871412652911 |
| interactor_abundance | mitosis | inter | 4.9215985987051895e-05 | 0.006283598333582311 |
| complex_abundance | mitosis | inter | 0.00011780432600576765 | 0.007493687219947184 |

P38159_P62995
RBMX_HUMAN vs TRA2B_HUMAN
p-value: 0.0 q-value: 0.001

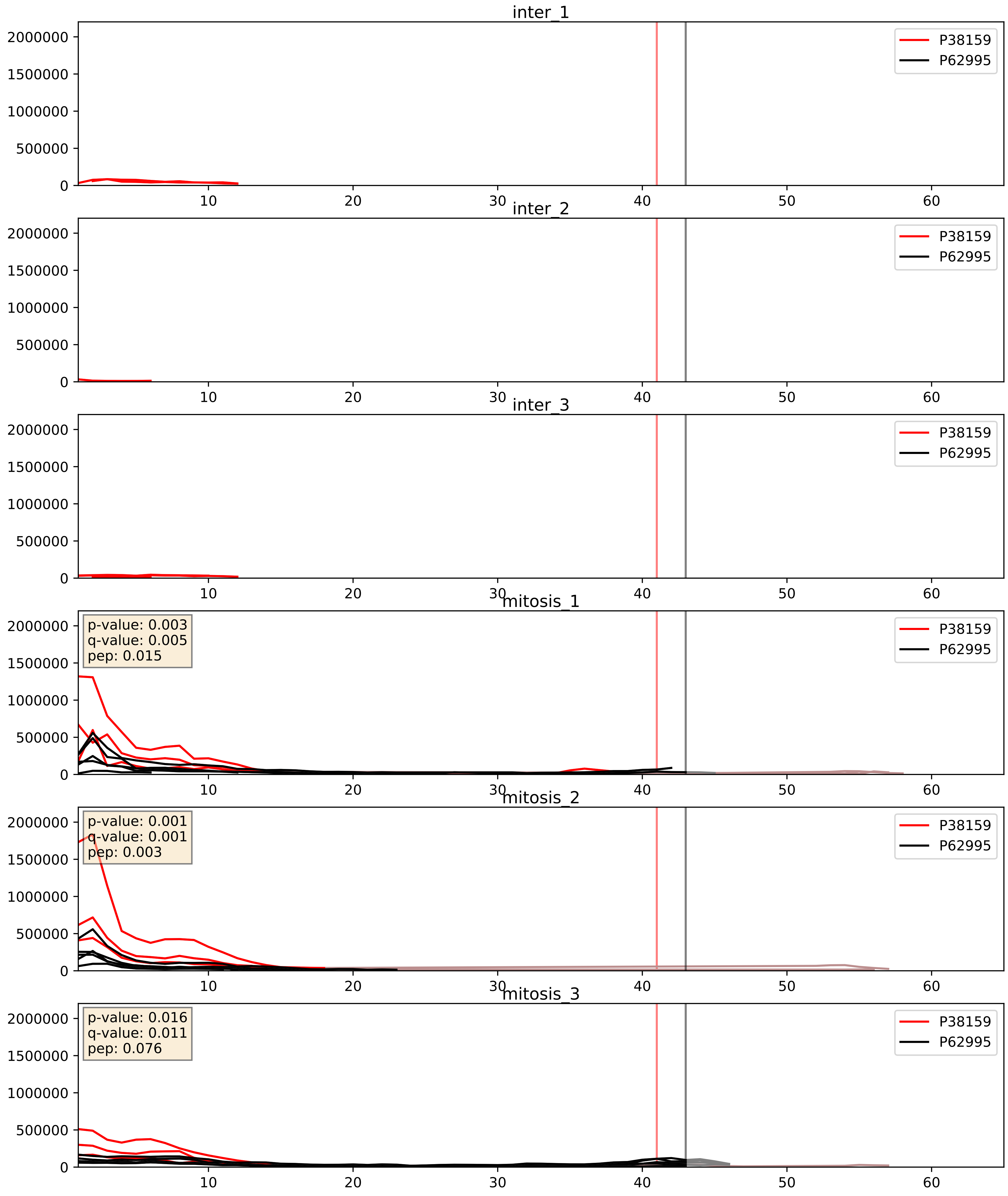


| level | condition_1 | condition_2 | pvalue | pvalue_adjusted |
|---|---|---|---|---|
| interactor_abundance | mitosis | inter | 0.0001647383149740902 | 0.009910743628815524 |
| complex_abundance | mitosis | inter | 0.00022597518718696217 | 0.010219389159334393 |
| interactor_ratio | mitosis | inter | 0.011485414155784251 | 0.15494522140242886 |

P51991_P62995
ROA3_HUMAN vs TRA2B_HUMAN
p-value: 0.002 q-value: 0.002

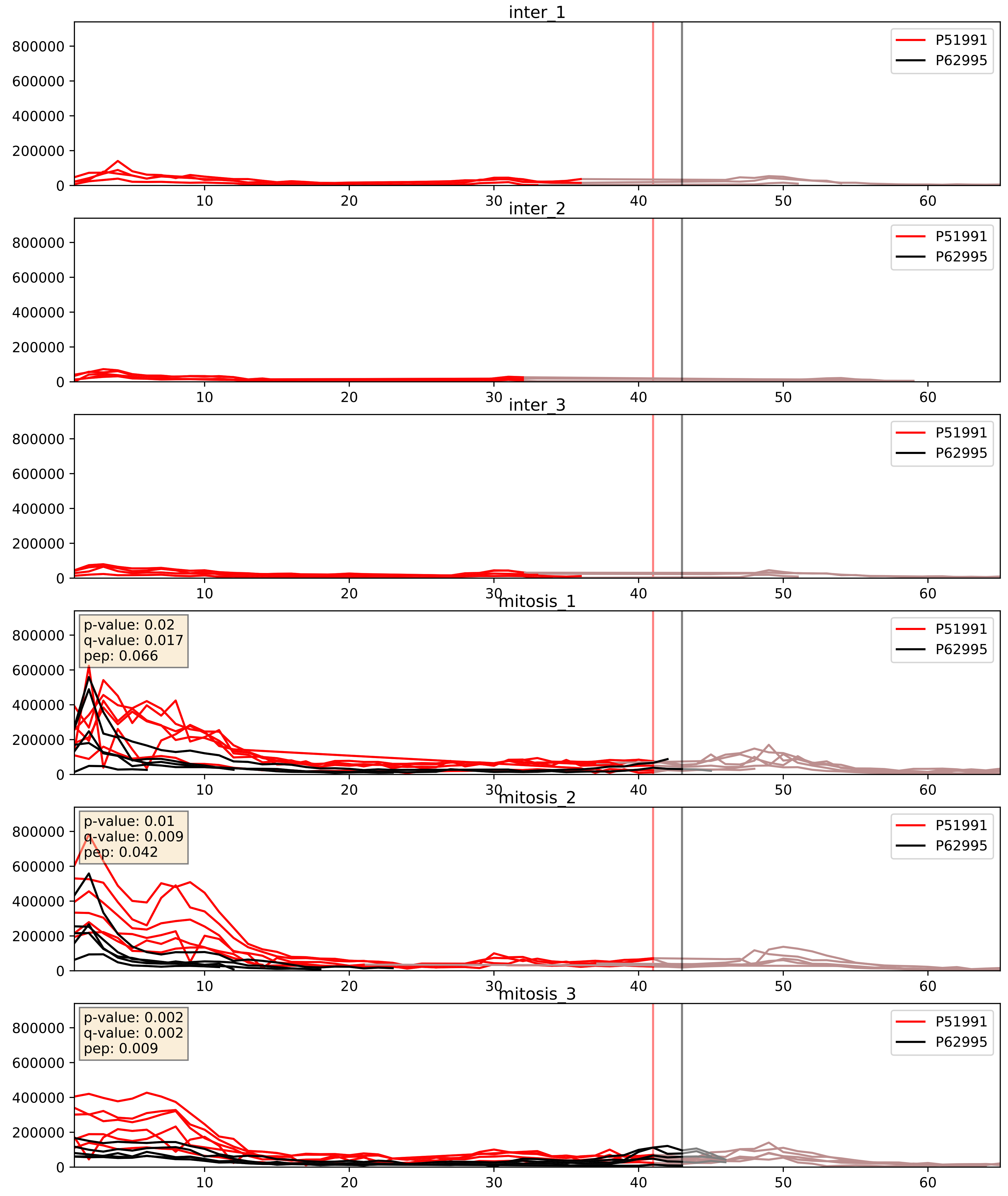


| level | condition_1 | condition_2 | pvalue | pvalue_adjusted |
|---|---|---|---|---|
| interactor_abundance | mitosis | inter | 1.7334412988378383e-05 | 0.003956868671480506 |
| complex_abundance | mitosis | inter | 7.644256938603826e-05 | 0.005948621763131705 |
| interactor_ratio | mitosis | inter | 0.0003545280013586819 | 0.0203675147089283 |

P52272_P62995
HNRPM_HUMAN vs TRA2B_HUMAN
p-value: 0.03 q-value: 0.022

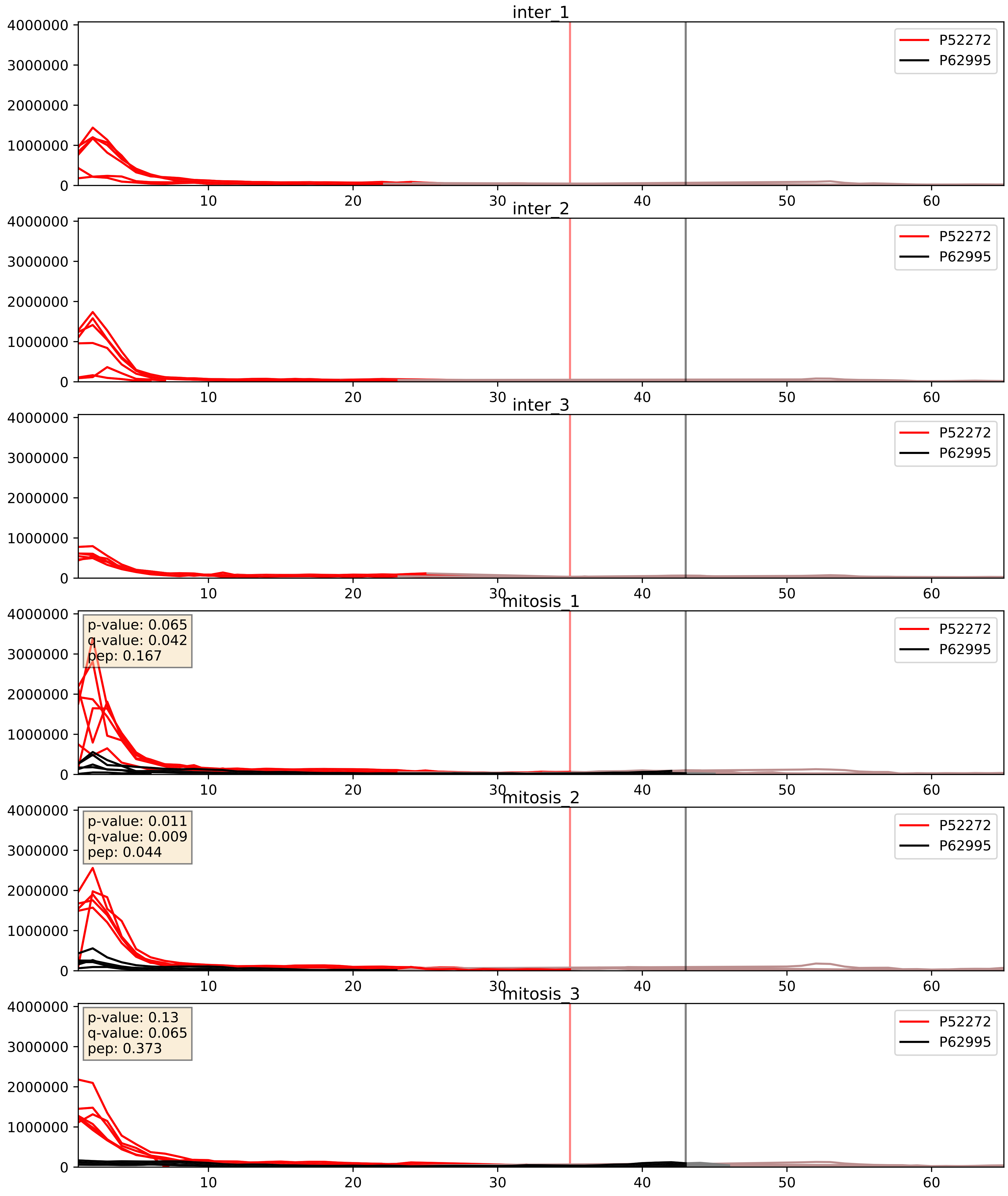


| level | condition_1 | condition_2 | pvalue | pvalue_adjusted |
|---|---|---|---|---|
| interactor_ratio | mitosis | inter | 3.4737747771941747e-07 | 0.0009911837364260712 |
| interactor_abundance | mitosis | inter | 2.4013033340954548e-05 | 0.004450836370313492 |
| complex_abundance | mitosis | inter | 7.744600555564312e-05 | 0.005949167968551921 |

P52597_P62995
HNRPF_HUMAN vs TRA2B_HUMAN
p-value: 0.003 q-value: 0.003

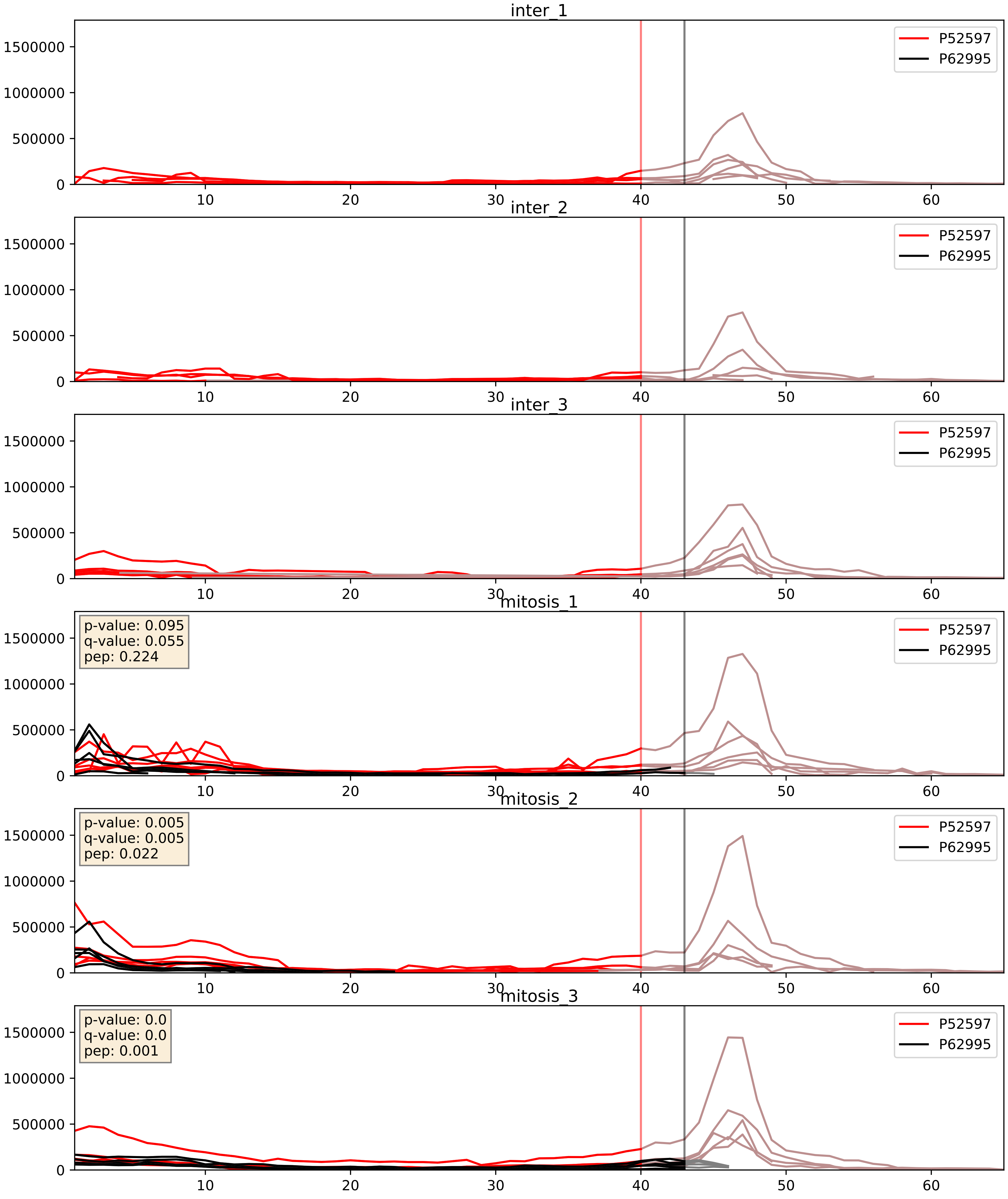


| level | condition_1 | condition_2 | pvalue | pvalue_adjusted |
|---|---|---|---|---|
| interactor_ratio | mitosis | inter | 4.6063738991628886e-05 | 0.0067983725132472975 |
| interactor_abundance | mitosis | inter | 0.00010371553266387402 | 0.007876481428251637 |
| complex_abundance | mitosis | inter | 0.0001881994972916382 | 0.009111526782692203 |

P55769_P62995
NH2L1_HUMAN vs TRA2B_HUMAN
p-value: 0.0 q-value: 0.009

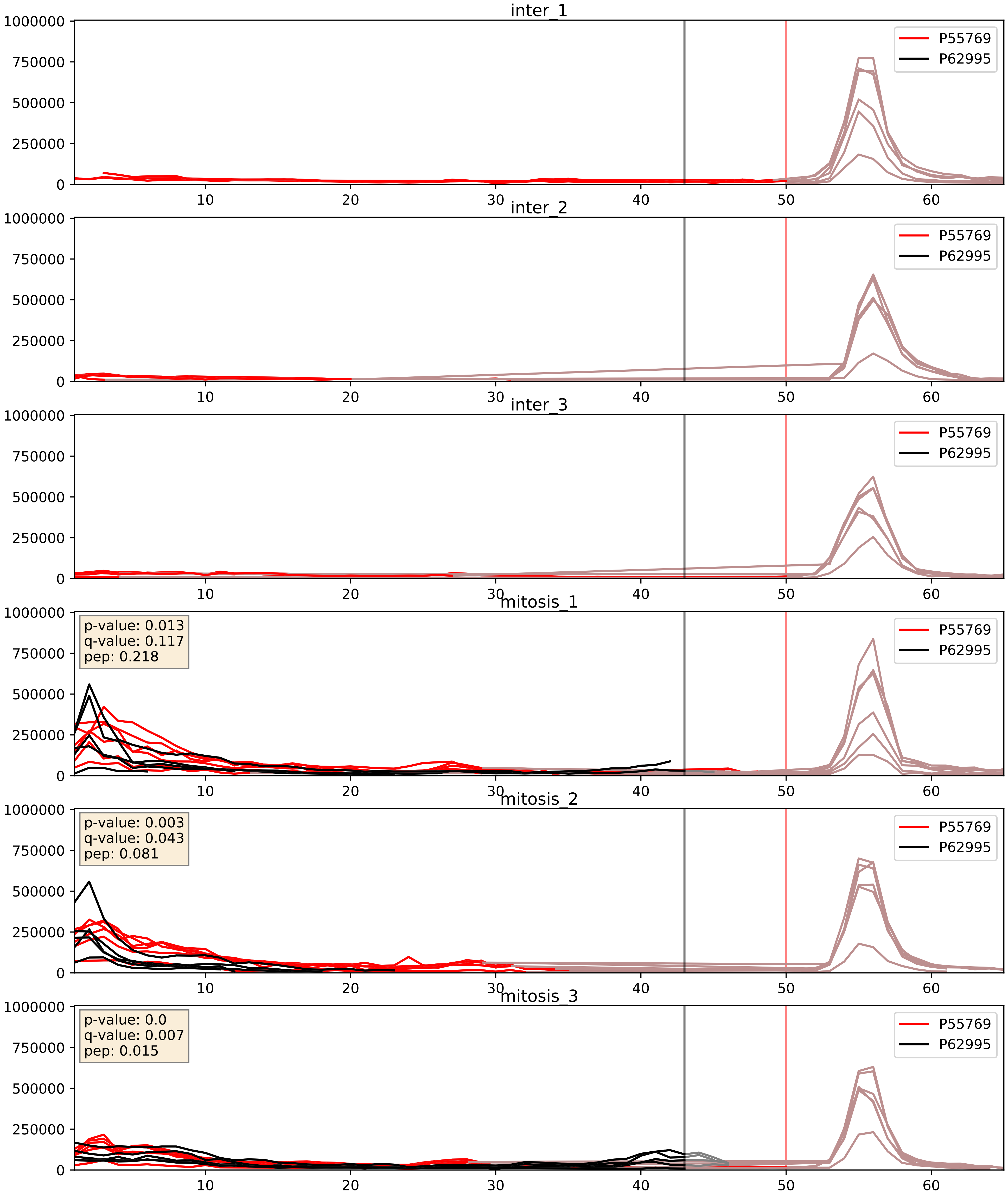


| level | condition_1 | condition_2 | pvalue | pvalue_adjusted |
|---|---|---|---|---|
| interactor_abundance | mitosis | inter | 8.443492929225096e-05 | 0.007227629947416683 |
| complex_abundance | mitosis | inter | 0.00011484000270838403 | 0.007391206189351633 |
| interactor_ratio | mitosis | inter | 0.07572511598736271 | 0.4628166812754875 |

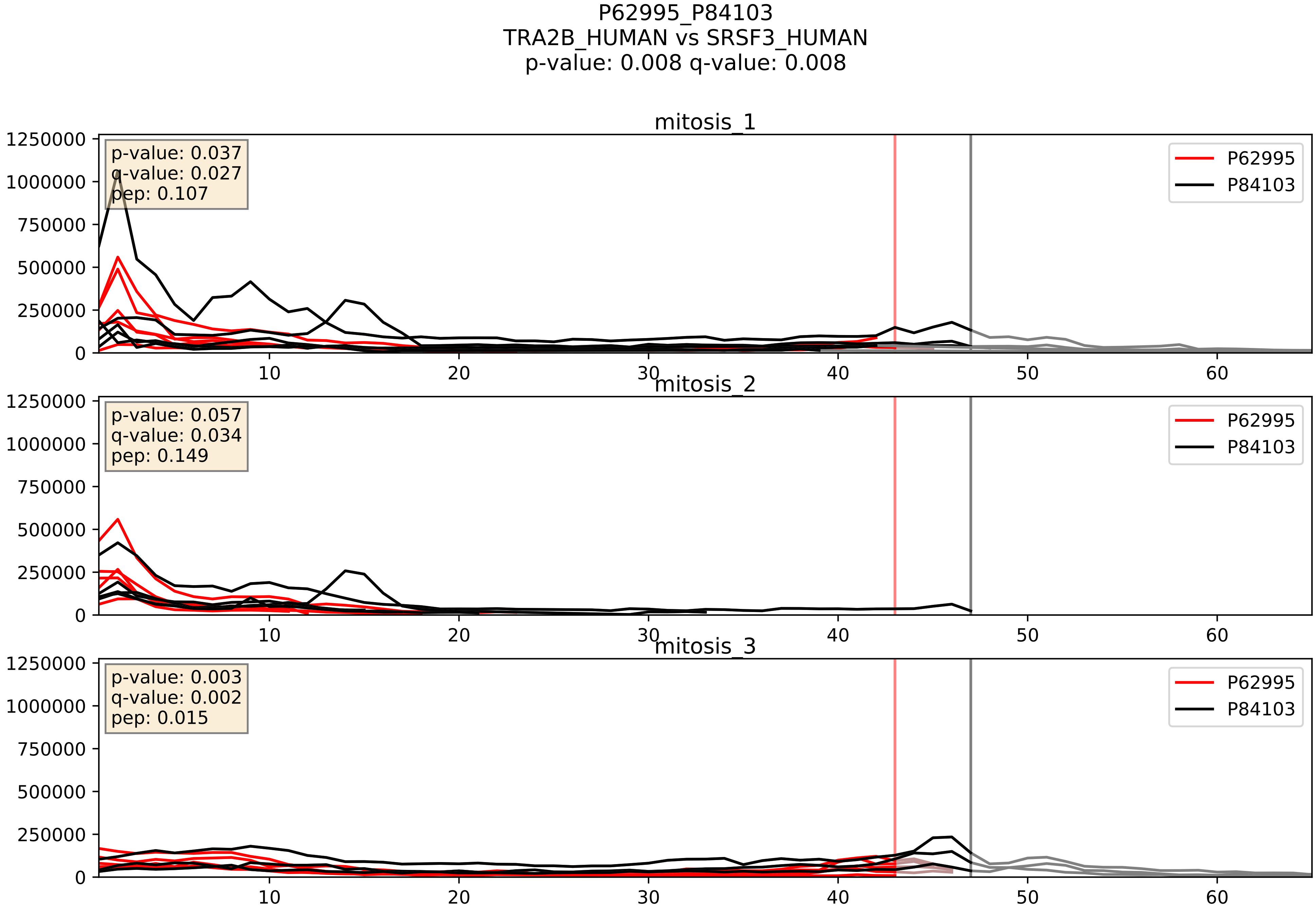


| level | condition_1 | condition_2 | pvalue | pvalue_adjusted |
|---|---|---|---|---|
| interactor_abundance | mitosis | inter | 0.0004584932841066445 | 0.014951247664582388 |
| complex_abundance | mitosis | inter | 0.000642767496886288 | 0.016393529992559892 |
| interactor_ratio | mitosis | inter | 0.8736089821848678 | 0.980069417404396 |

P62995_Q07666
TRA2B_HUMAN vs KHDR1_HUMAN
p-value: 0.0 q-value: 0.023

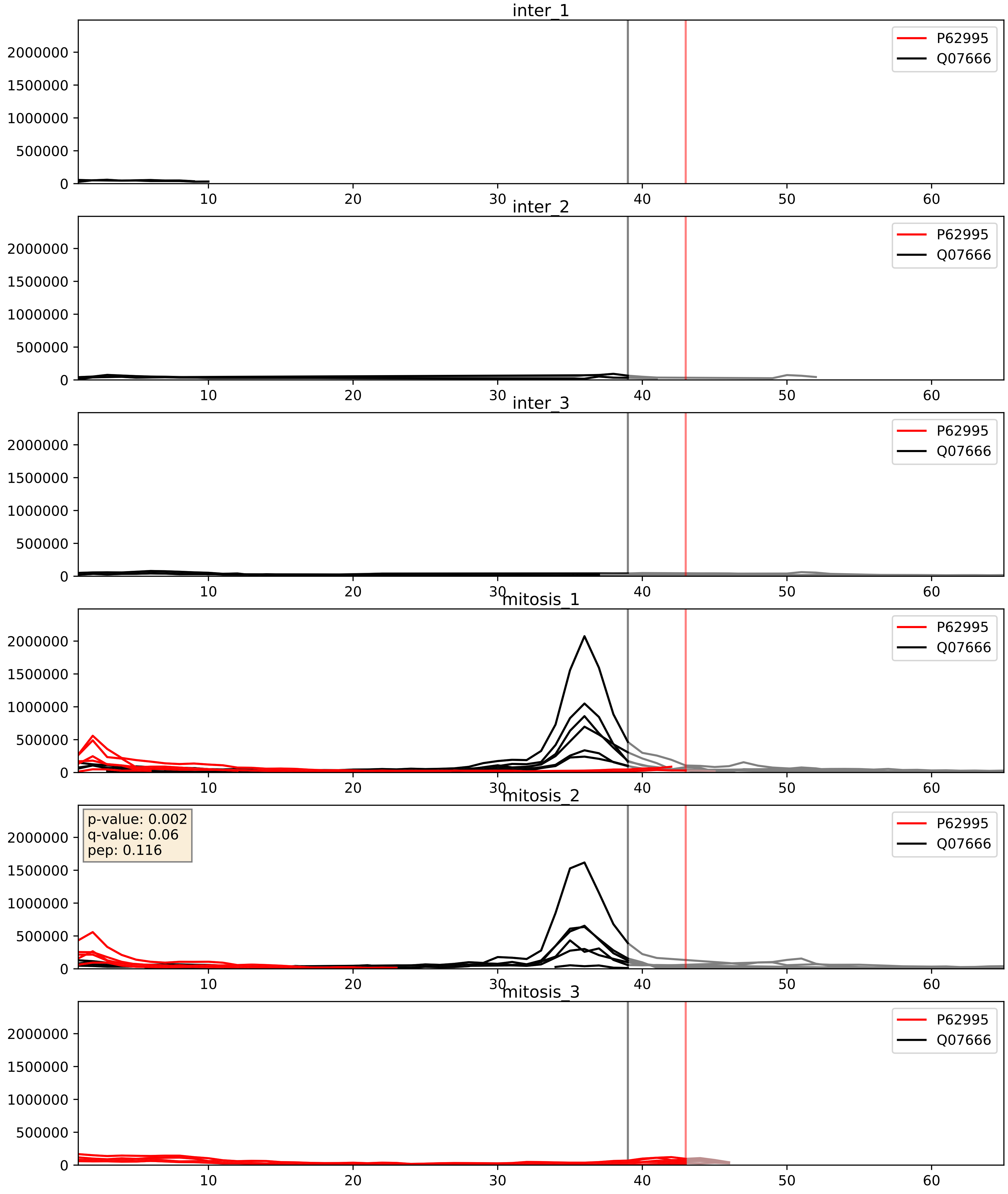


| level | condition_1 | condition_2 | pvalue | pvalue_adjusted |
|---|---|---|---|---|
| interactor_abundance | mitosis | inter | 0.270194466891906 | 0.46243419705982514 |
| complex_abundance | mitosis | inter | 0.30363085198893075 | 0.4536708139335394 |
| interactor_ratio | mitosis | inter | 0.9886088667776481 | 0.998078263757106 |

P62995_Q07955
TRA2B_HUMAN vs SRSF1_HUMAN
p-value: 0.019 q-value: 0.015

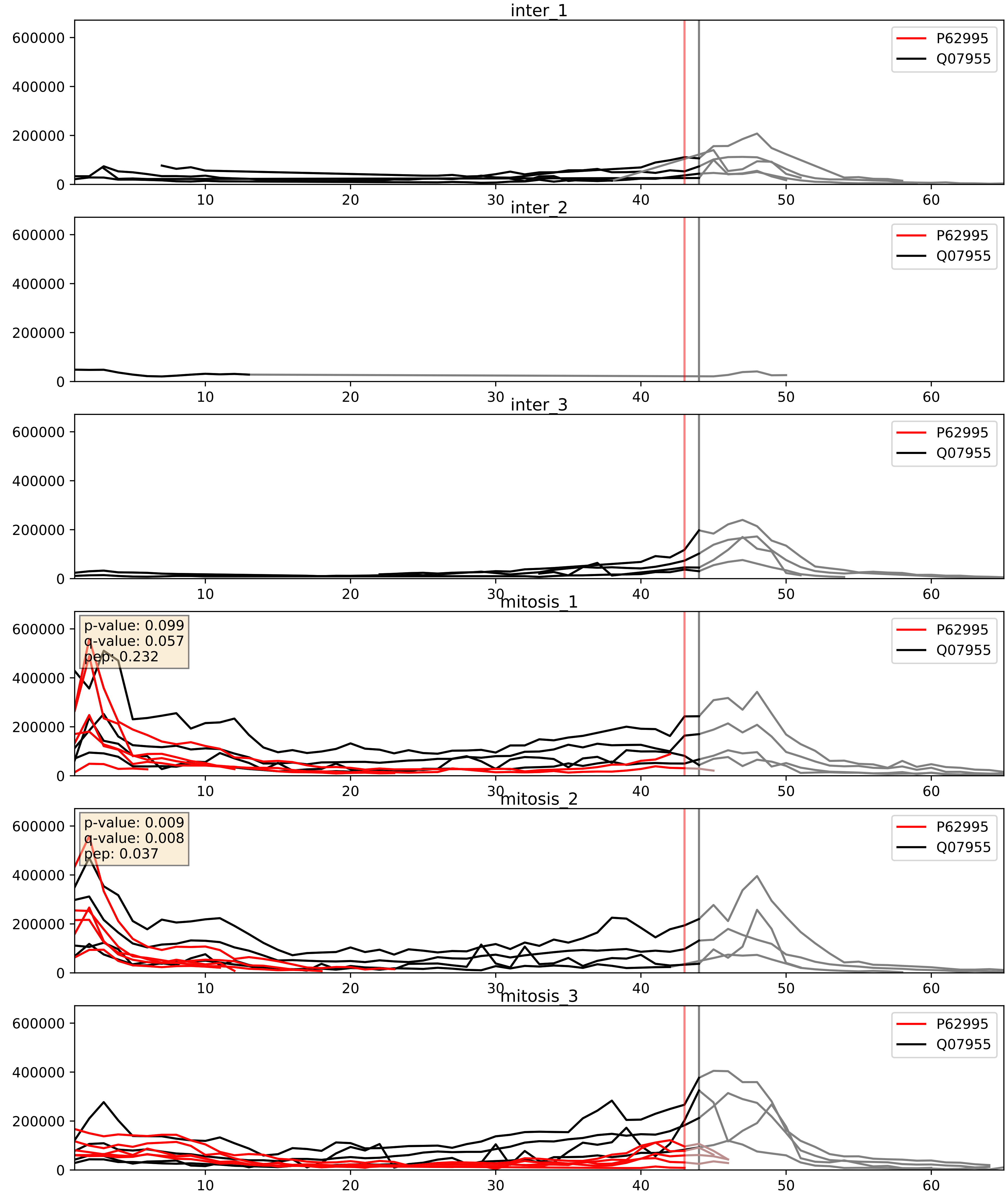


| level | condition_1 | condition_2 | pvalue | pvalue_adjusted |
|---|---|---|---|---|
| complex_abundance | mitosis | inter | 0.0002950478593069016 | 0.011617922212881656 |
| interactor_abundance | mitosis | inter | 0.0004584932841066445 | 0.014951247664582388 |
| interactor_ratio | mitosis | inter | 0.4511140392976068 | 0.8525838327540917 |

P62995_Q12905
TRA2B_HUMAN vs ILF2_HUMAN
p-value: 0.001 q-value: 0.045

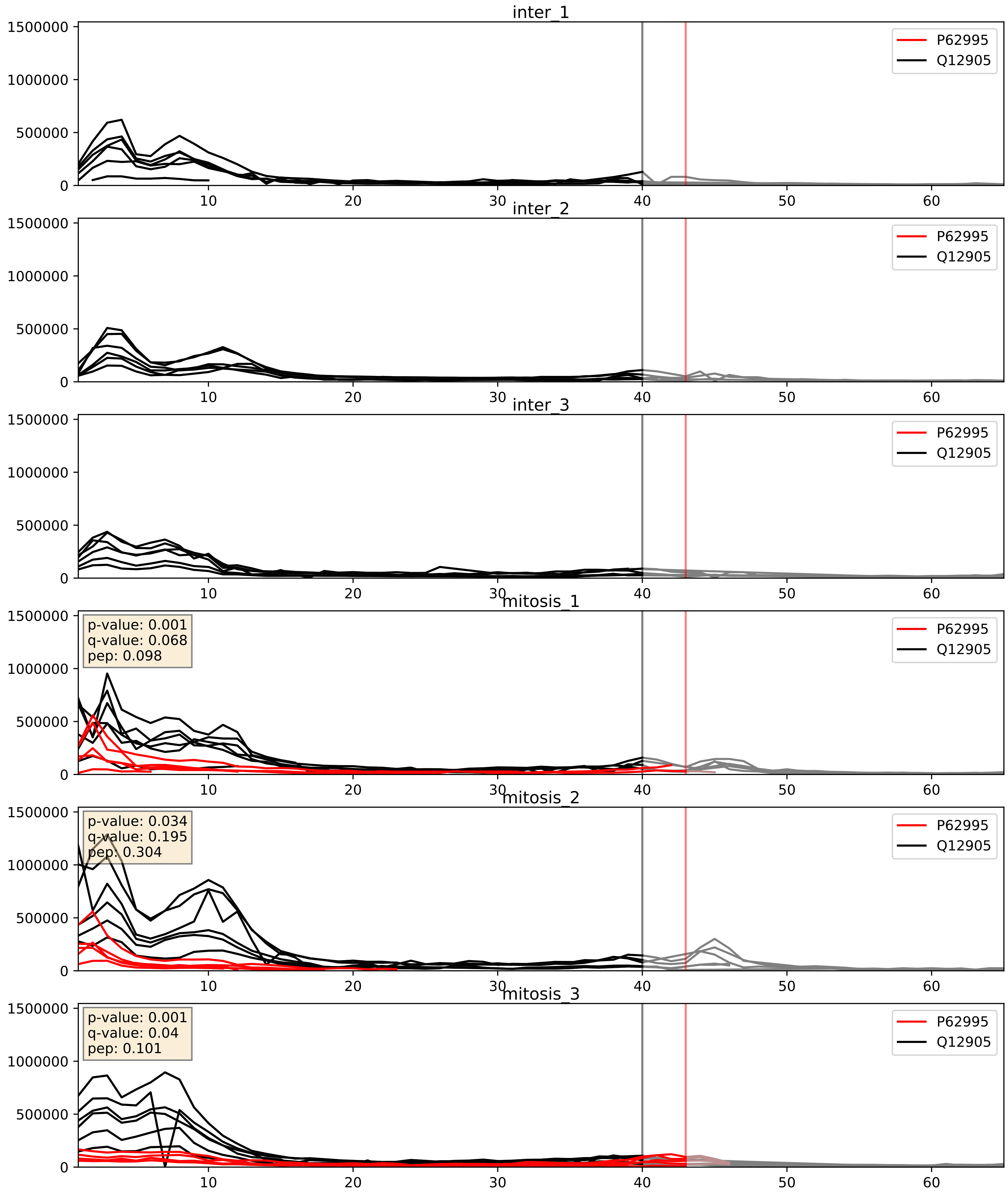


| level | condition_1 | condition_2 | pvalue | pvalue_adjusted |
|---|---|---|---|---|
| complex_abundance | mitosis | inter | 2.505847793058248e-05 | 0.005691857151652958 |
| interactor_abundance | mitosis | inter | 0.00037334314755474554 | 0.013573913378163039 |
| interactor_ratio | mitosis | inter | 0.0014969399724690845 | 0.04597663709000905 |

P62995_Q13151
TRA2B_HUMAN vs ROA0_HUMAN
p-value: 0.005 q-value: 0.005

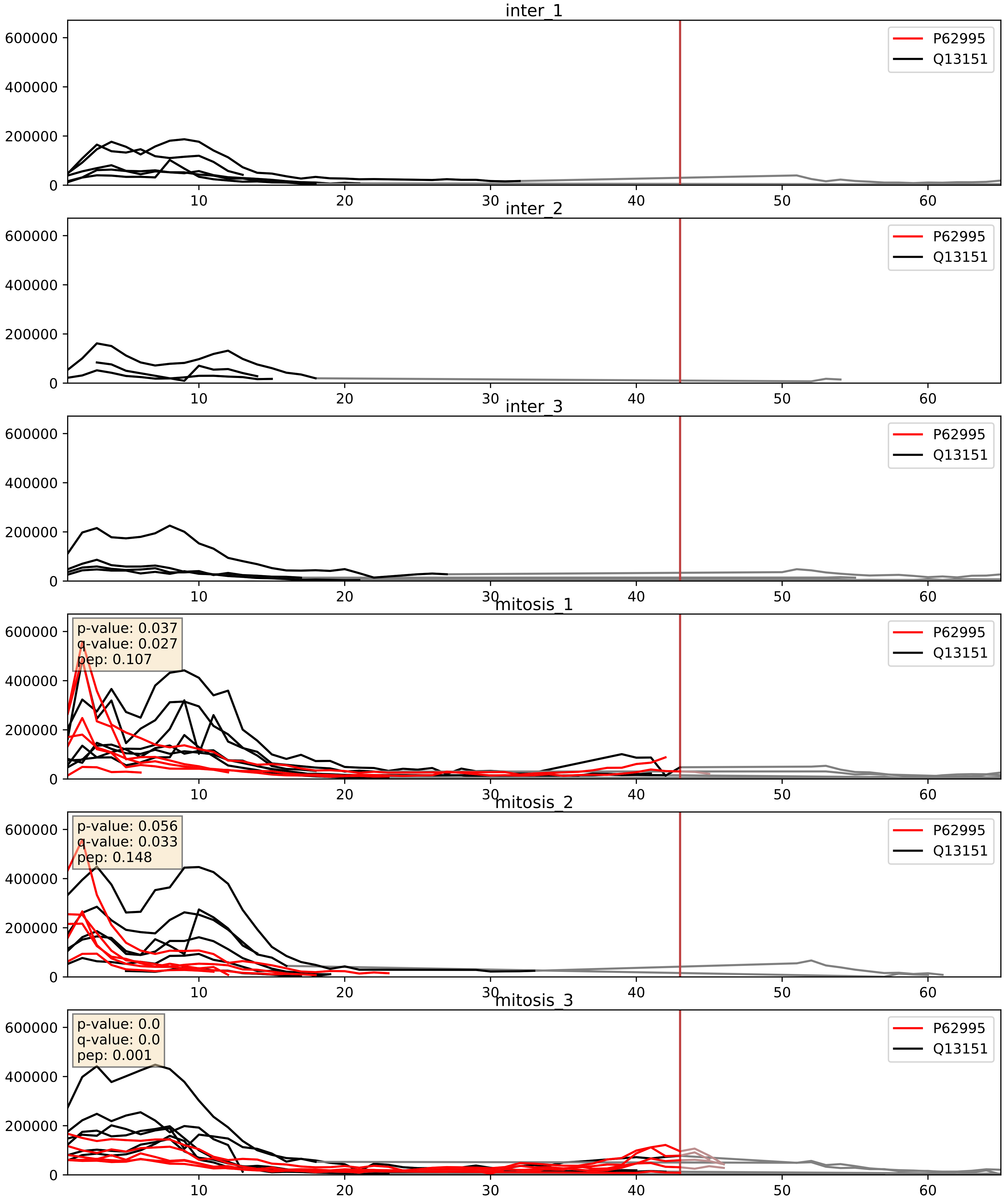


| level | condition_1 | condition_2 | pvalue | pvalue_adjusted |
|---|---|---|---|---|
| complex_abundance | mitosis | inter | 0.00011818315124916703 | 0.007493687219947184 |
| interactor_abundance | mitosis | inter | 0.0004573130702955831 | 0.014951247664582388 |
| interactor_ratio | mitosis | inter | 0.008403991269352195 | 0.13008709812957467 |

P62995_Q13242
TRA2B_HUMAN vs SRSF9_HUMAN
p-value: 0.001 q-value: 0.002

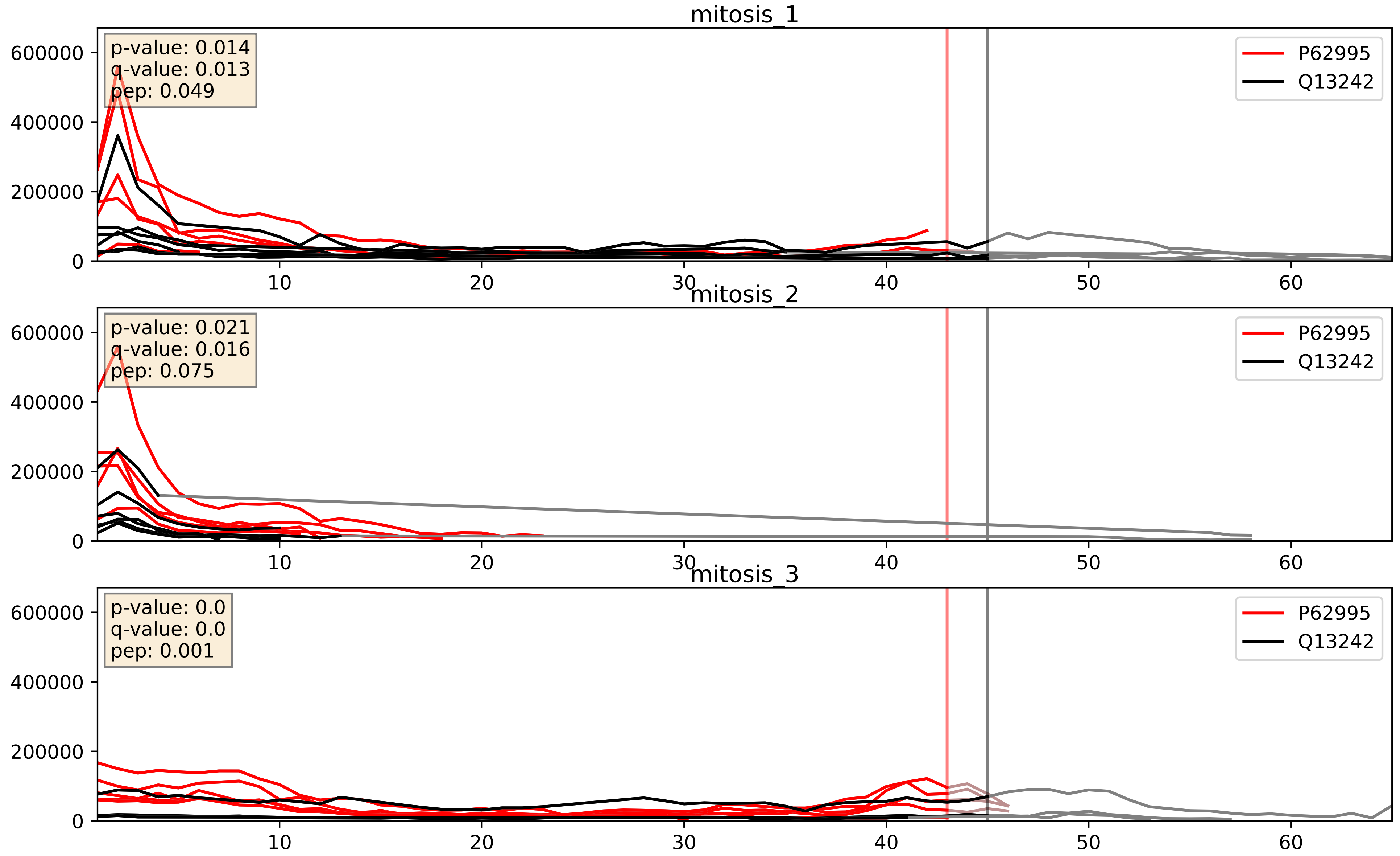


| level | condition_1 | condition_2 | pvalue | pvalue_adjusted |
|---|---|---|---|---|
| interactor_abundance | mitosis | inter | 0.0006196284287297792 | 0.01748938971196646 |
| complex_abundance | mitosis | inter | 0.004301196506507519 | 0.04889540782962067 |
| interactor_ratio | mitosis | inter | 0.008613258867848741 | 0.13203195690147984 |

P62995_Q13247
TRA2B_HUMAN vs SRSF6_HUMAN
p-value: 0.007 q-value: 0.006

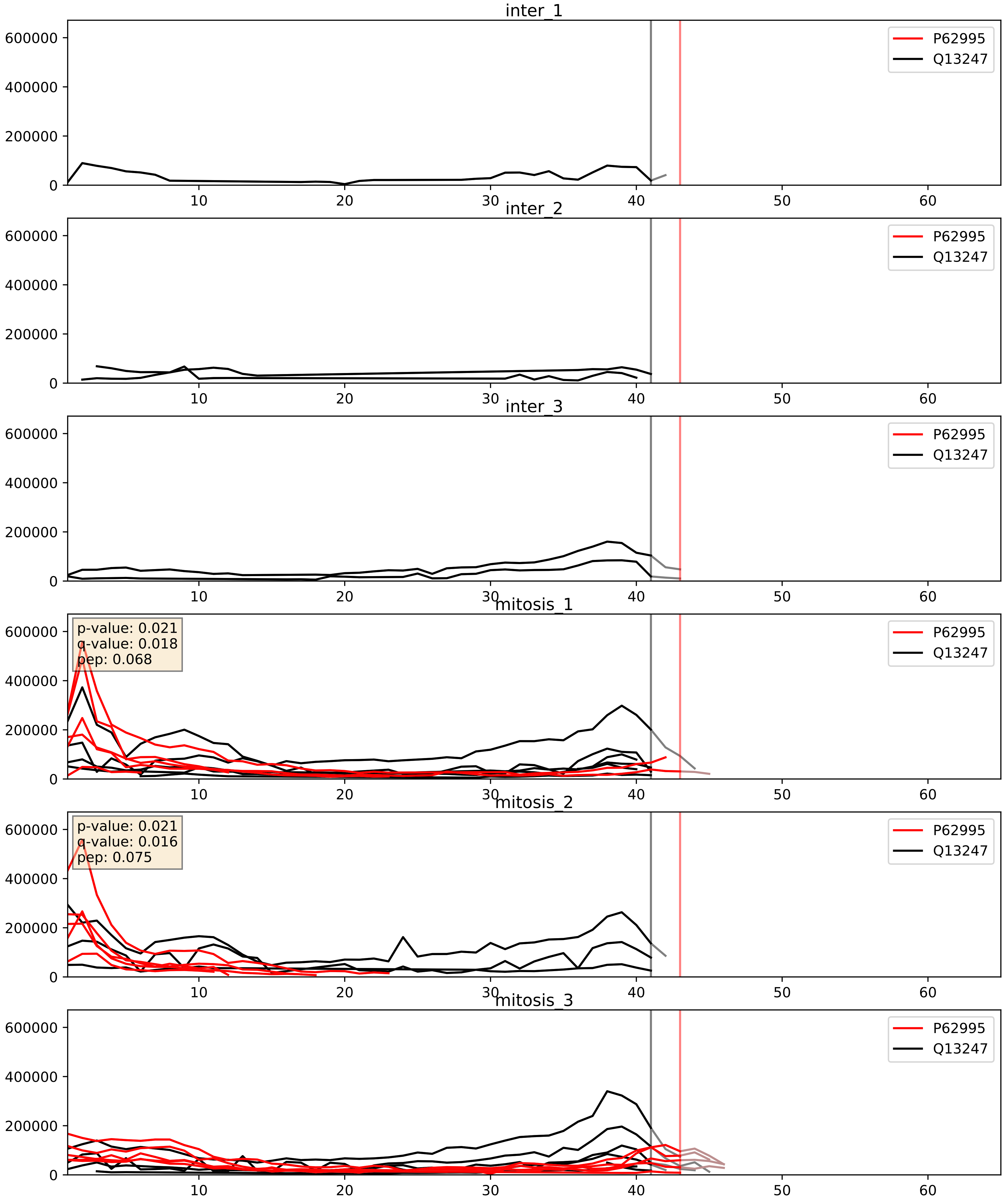


| level | condition_1 | condition_2 | pvalue | pvalue_adjusted |
|---|---|---|---|---|
| interactor_abundance | mitosis | inter | 0.00038990687589259026 | 0.01362698060740691 |
| complex_abundance | mitosis | inter | 0.002674131184473713 | 0.037220427543243874 |
| interactor_ratio | mitosis | inter | 0.9430800845236742 | 0.9912117205440141 |

P62995_Q13838
TRA2B_HUMAN vs DX39B_HUMAN
p-value: 0.003 q-value: 0.047

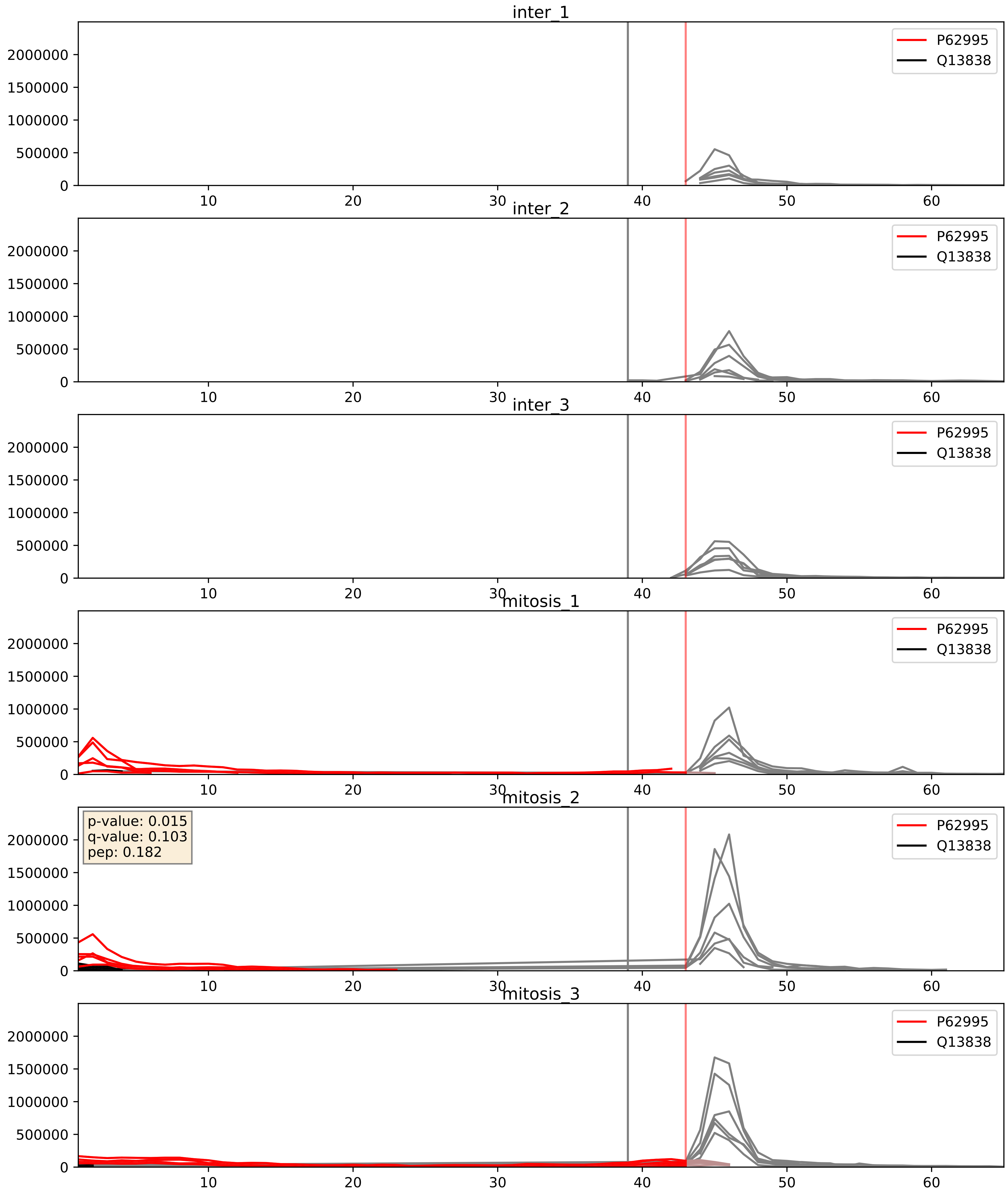


| level | condition_1 | condition_2 | pvalue | pvalue_adjusted |
|---|---|---|---|---|
| interactor_ratio | mitosis | inter | 0.014722011230281114 | 0.1776545333938431 |
| interactor_abundance | mitosis | inter | 0.06364300330806487 | 0.2075368031683944 |
| complex_abundance | mitosis | inter | 0.1430857576735257 | 0.2885309978057432 |

P62995_Q15717
TRA2B_HUMAN vs ELAV1_HUMAN
p-value: 0.0 q-value: 0.0

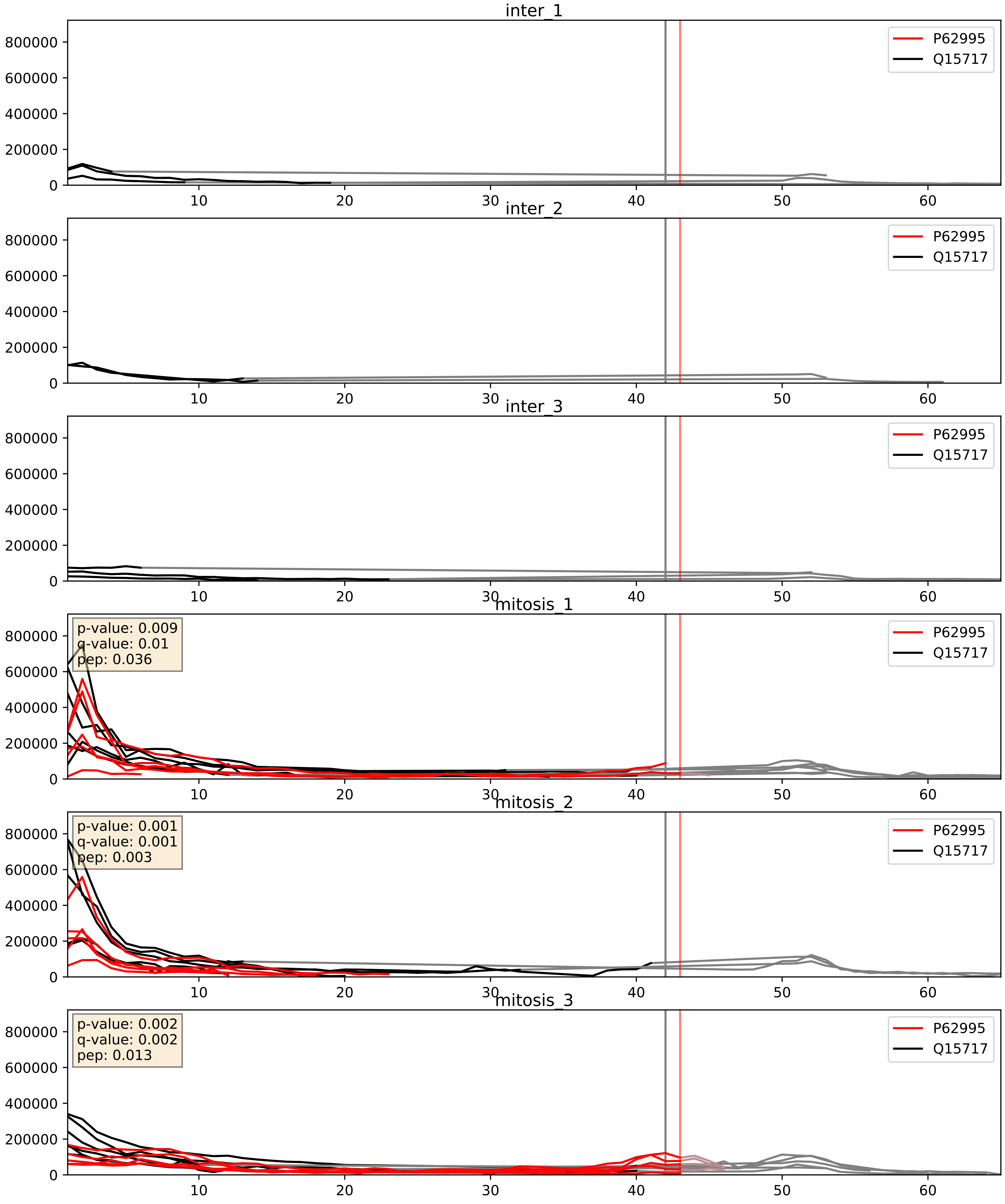


| level | condition_1 | condition_2 | pvalue | pvalue_adjusted |
|---|---|---|---|---|
| complex_abundance | mitosis | inter | 0.0001242130151056852 | 0.007704807313801923 |
| interactor_abundance | mitosis | inter | 0.0004121602900123102 | 0.014019253191624928 |
| interactor_ratio | mitosis | inter | 0.01618022122043033 | 0.18789564671895648 |

P62995_Q16629
TRA2B_HUMAN vs SRSF7_HUMAN
p-value: 0.0 q-value: 0.001

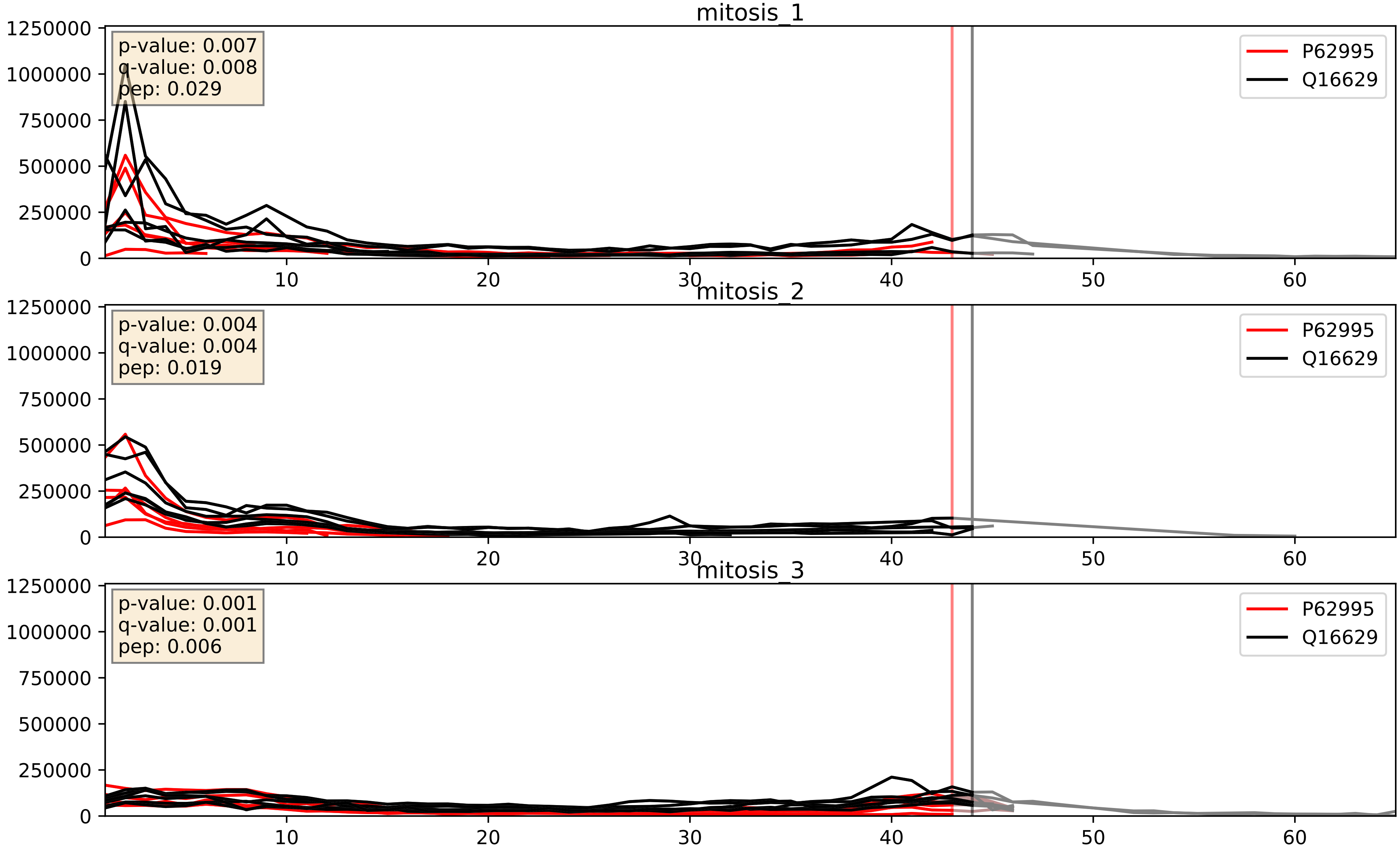


| level | condition_1 | condition_2 | pvalue | pvalue_adjusted |
|---|---|---|---|---|
| complex_abundance | mitosis | inter | 0.00017630276153620522 | 0.008807651911797798 |
| interactor_abundance | mitosis | inter | 0.0004584932841066445 | 0.014951247664582388 |
| interactor_ratio | mitosis | inter | 0.002039014647753622 | 0.05540941391990795 |
