## Supplementary Data 2 for "SECAT: Quantifying differential protein-protein interaction states by network-centric analysis": hela_string_000013_Q12905.pdf

Q12905_Q12905

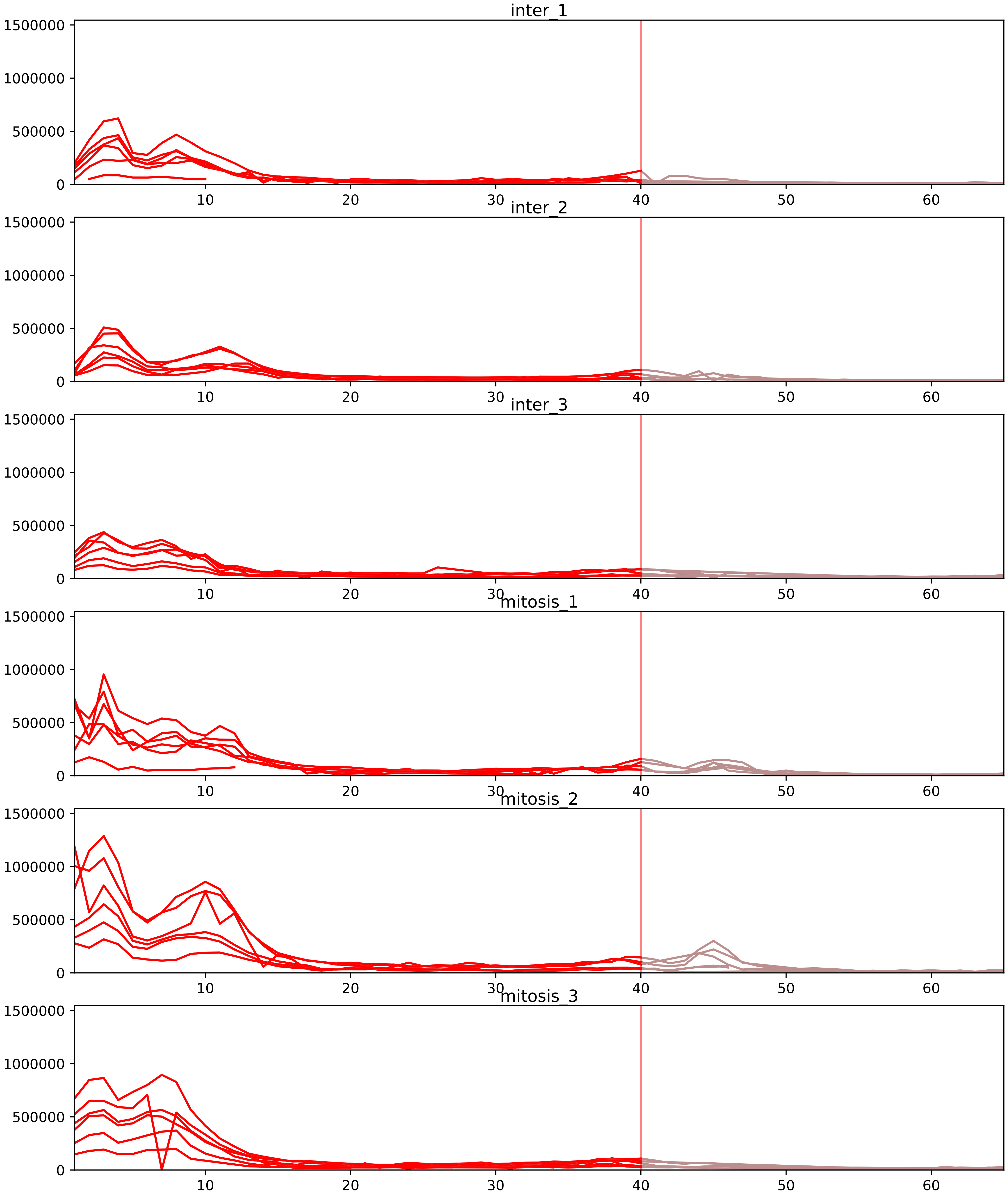


| level | condition_1 | condition_2 | pvalue | pvalue_adjusted |
|---|---|---|---|---|
| interactor_ratio | mitosis | inter | 2.1932399310796645e-14 | 3.1042780562973715e-12 |
| complex_abundance | mitosis | inter | 8.912389736371743e-06 | 0.0004315472924980002 |
| interactor_abundance | mitosis | inter | 7.231128098793699e-05 | 0.0018479549585806118 |
| assembled_abundance | mitosis | inter | 0.00514197226894421 | 0.0729380975482662 |
| total_abundance | mitosis | inter | 0.00515604220760259 | 0.08280571255175012 |
| monomer_abundance | mitosis | inter | 0.08337771936980803 | 0.37967498034042735 |

O43390_Q12905
HNRPR_HUMAN vs ILF2_HUMAN
p-value: 0.003 q-value: 0.041

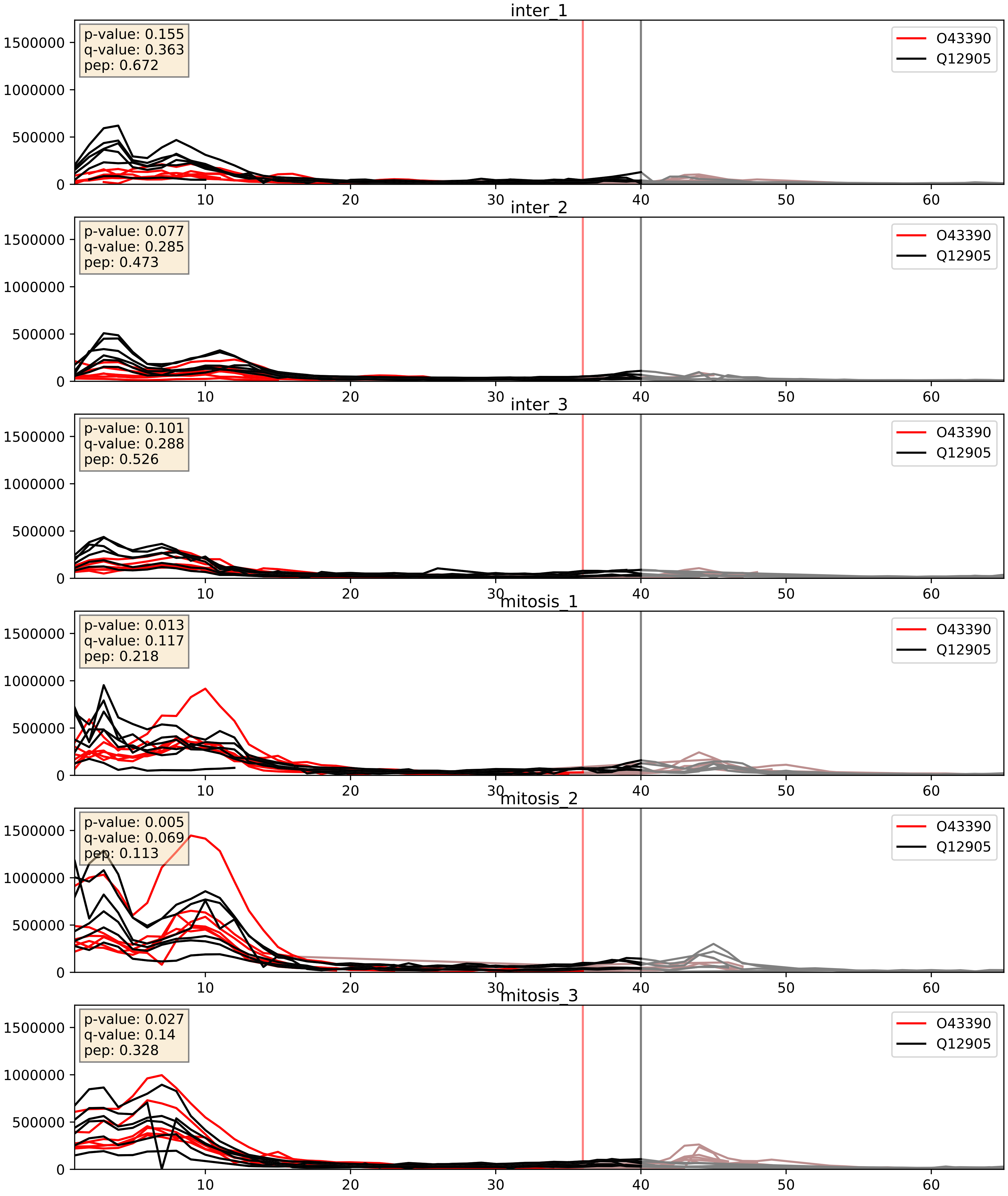


| level | condition_1 | condition_2 | pvalue | pvalue_adjusted |
|---|---|---|---|---|
| interactor_abundance | mitosis | inter | 0.00010796061997593195 | 0.007876481428251637 |
| complex_abundance | mitosis | inter | 0.0003909354311201745 | 0.012627952039202618 |
| interactor_ratio | mitosis | inter | 0.004324784485630598 | 0.08690177276290592 |

P09651_Q12905
ROA1_HUMAN vs ILF2_HUMAN
p-value: 0.0 q-value: 0.0

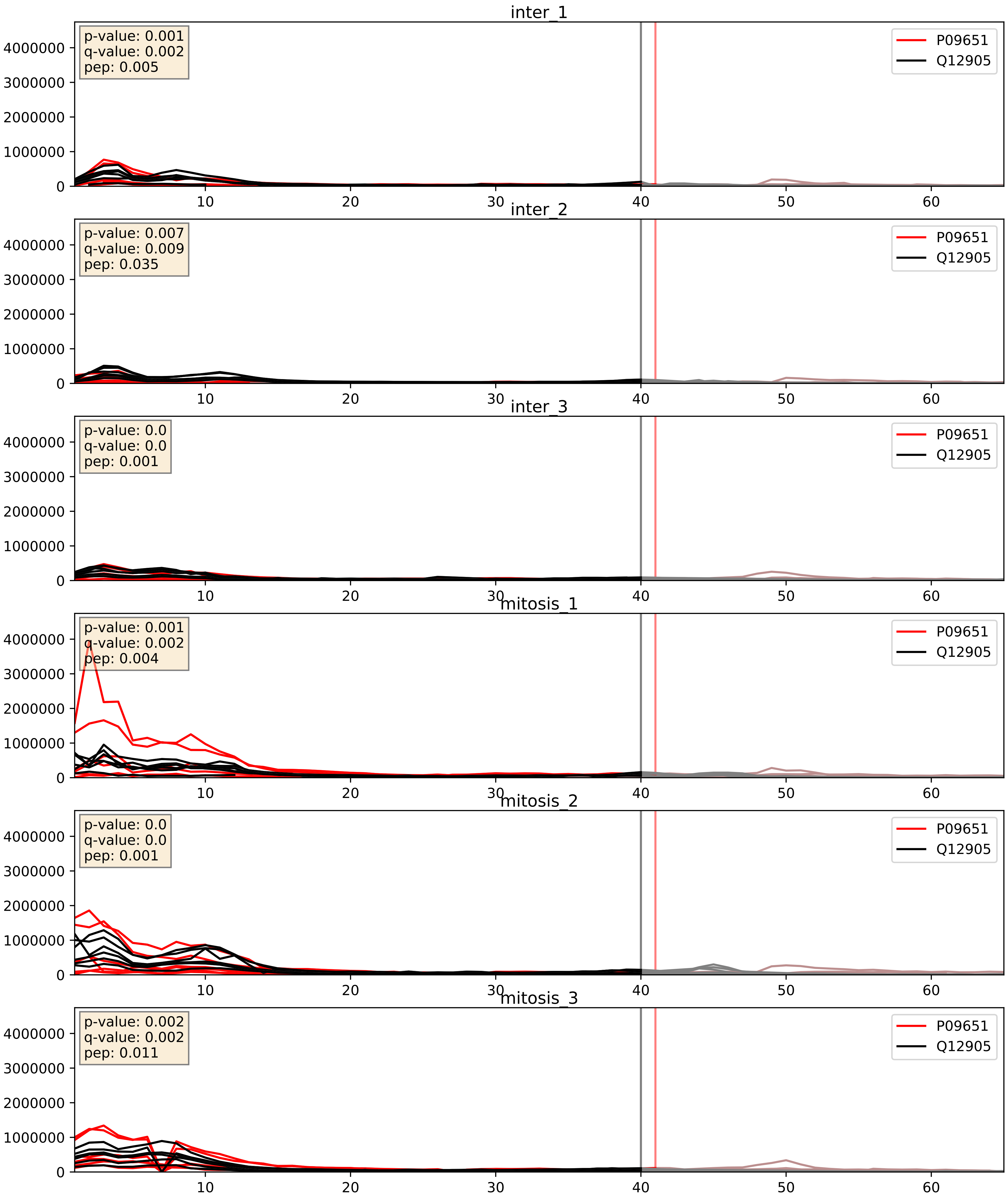


| level | condition_1 | condition_2 | pvalue | pvalue_adjusted |
|---|---|---|---|---|
| complex_abundance | mitosis | inter | 0.005618403472428057 | 0.05646492674842808 |
| interactor_abundance | mitosis | inter | 0.017050532289726136 | 0.11334462047152971 |
| interactor_ratio | mitosis | inter | 0.07777248885697595 | 0.4669171100029366 |

P22626_Q12905
ROA2_HUMAN vs ILF2_HUMAN
p-value: 0.002 q-value: 0.035

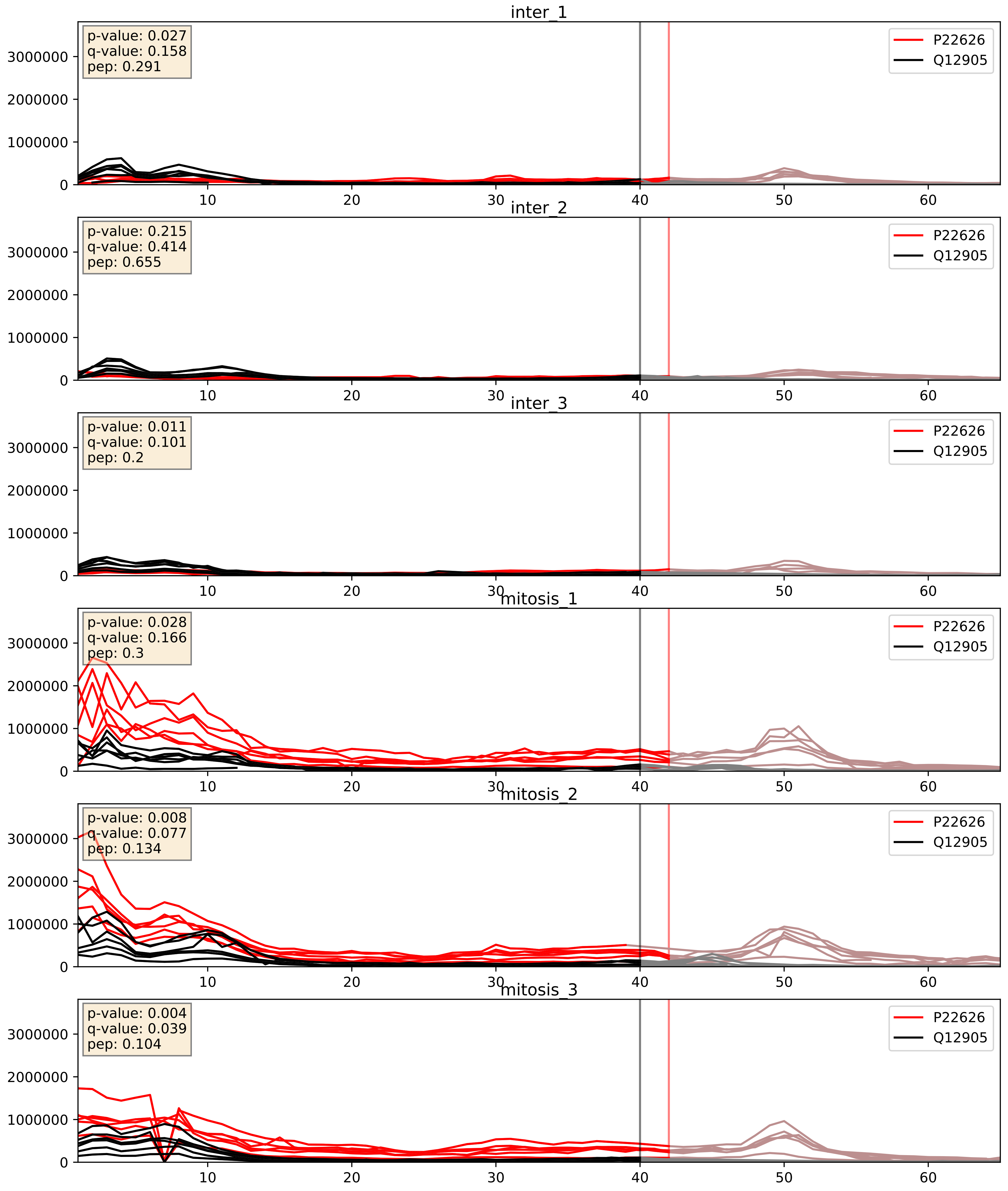


| level | condition_1 | condition_2 | pvalue | pvalue_adjusted |
|---|---|---|---|---|
| complex_abundance | mitosis | inter | 0.0022575804114793045 | 0.034819618598671796 |
| interactor_abundance | mitosis | inter | 0.003361167395046095 | 0.04478068934100323 |
| interactor_ratio | mitosis | inter | 0.006434293228359545 | 0.11104344765072117 |

P32969_Q12905
RL9_HUMAN vs ILF2_HUMAN
p-value: 0.0 q-value: 0.049

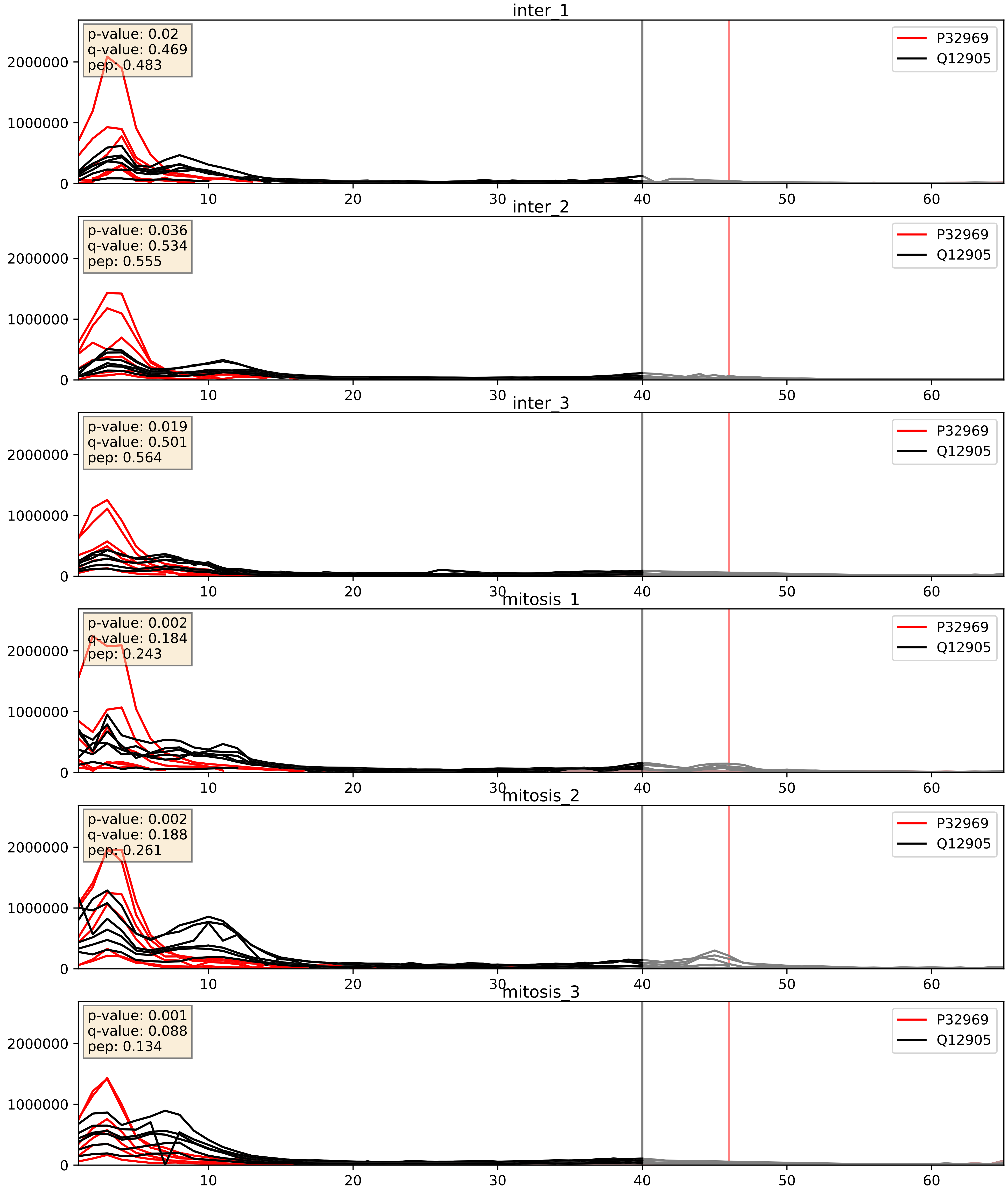


| level | condition_1 | condition_2 | pvalue | pvalue_adjusted |
|---|---|---|---|---|
| complex_abundance | mitosis | inter | 0.005256311849077548 | 0.05467075264654169 |
| interactor_ratio | mitosis | inter | 0.02202480886494059 | 0.2276960916472119 |
| interactor_abundance | mitosis | inter | 0.2103491843605969 | 0.40652154270556234 |

P39023_Q12905
RL3_HUMAN vs ILF2_HUMAN
p-value: 0.0 q-value: 0.047

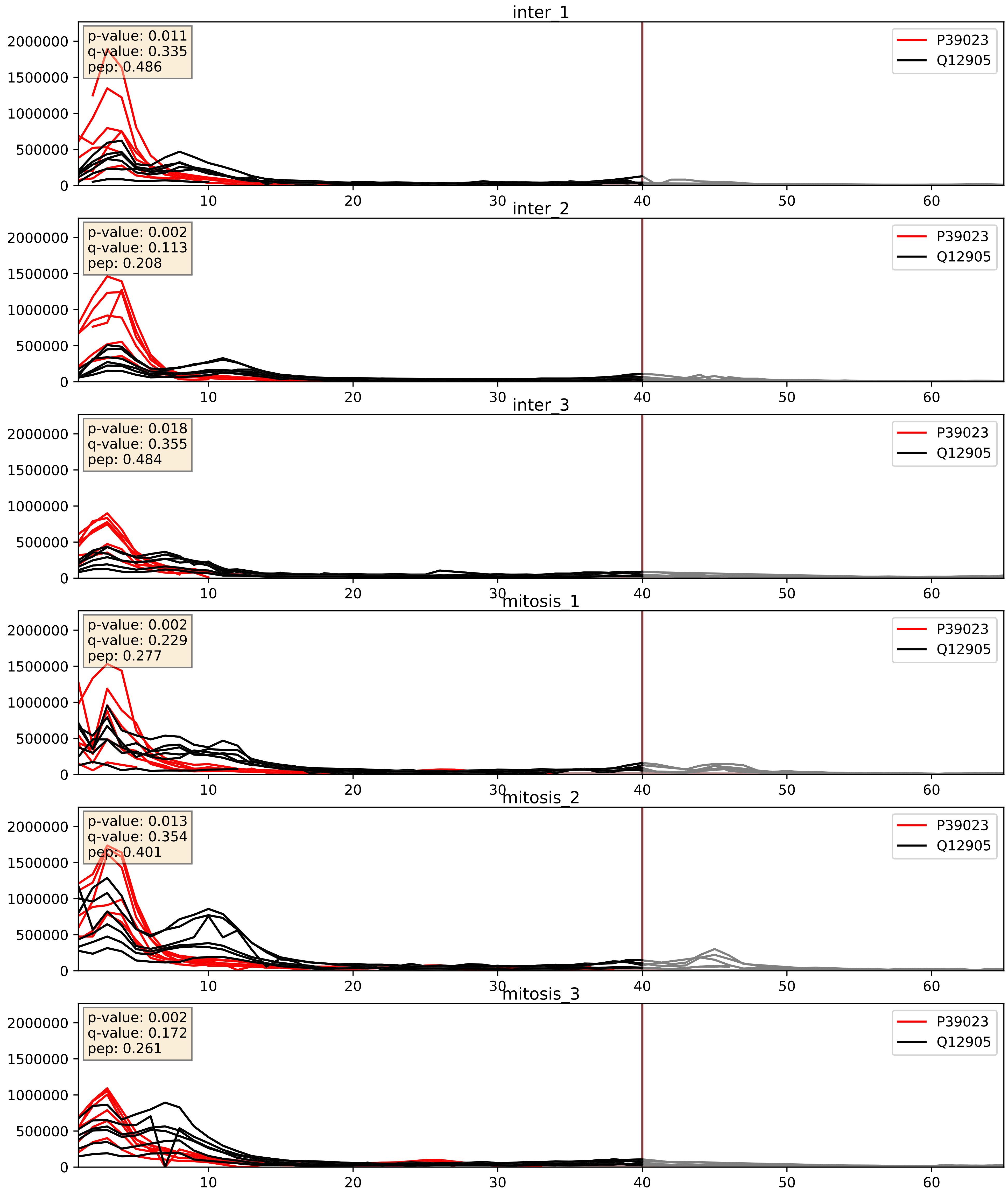


| level | condition_1 | condition_2 | pvalue | pvalue_adjusted |
|---|---|---|---|---|
| interactor_ratio | mitosis | inter | 0.009975649083884552 | 0.14263182888175546 |
| complex_abundance | mitosis | inter | 0.03702231914352979 | 0.14016674328897277 |
| interactor_abundance | mitosis | inter | 0.4577908838109454 | 0.629609570279835 |

P42766_Q12905
RL35_HUMAN vs ILF2_HUMAN
p-value: 0.0 q-value: 0.047

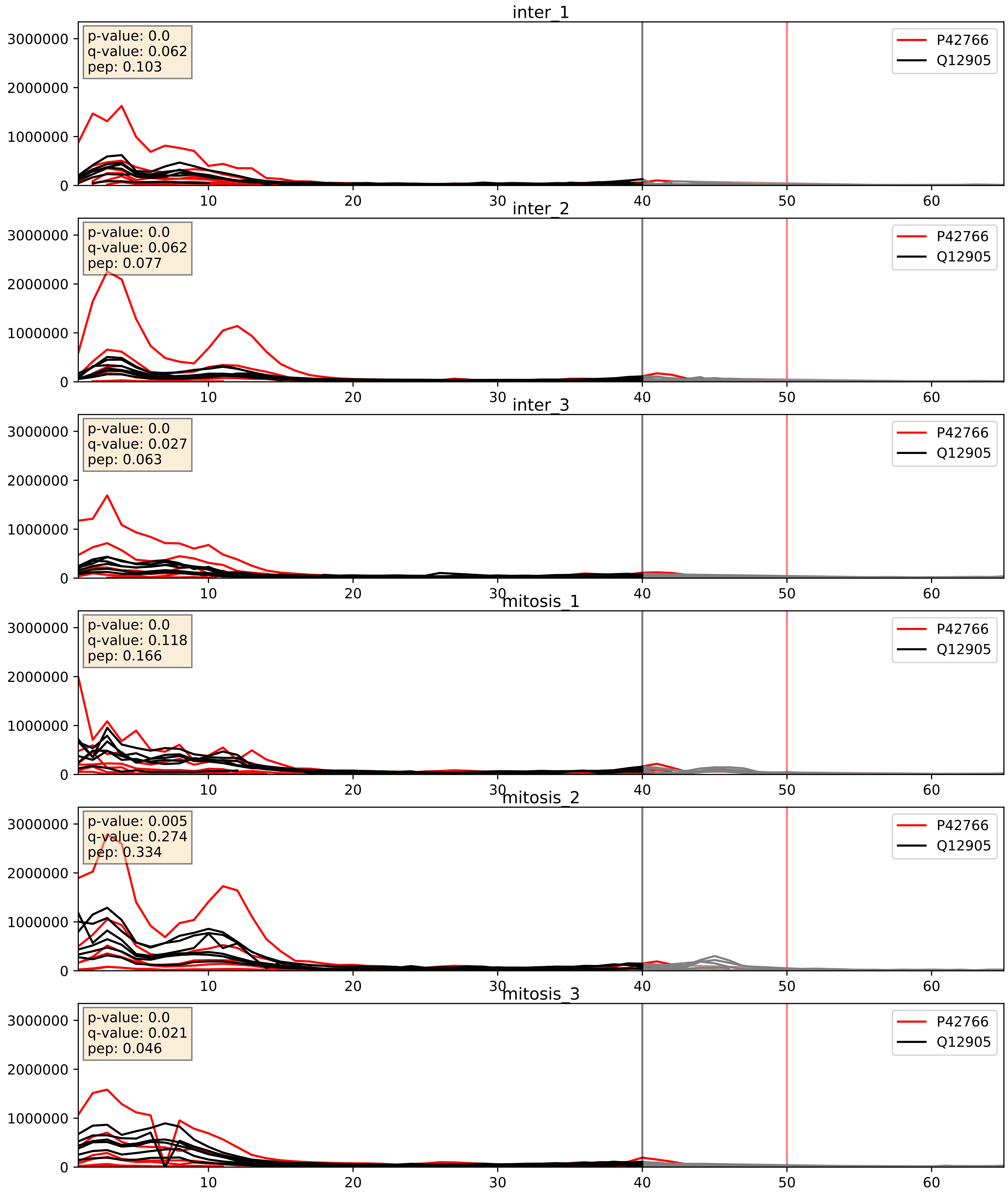


| level | condition_1 | condition_2 | pvalue | pvalue_adjusted |
|---|---|---|---|---|
| interactor_ratio | mitosis | inter | 0.008599101484341015 | 0.13203195690147984 |
| complex_abundance | mitosis | inter | 0.07138512115416014 | 0.19410947810661078 |
| interactor_abundance | mitosis | inter | 0.8496713088875427 | 0.9182736448554214 |

P46781_Q12905
RS9_HUMAN vs ILF2_HUMAN
p-value: 0.0 q-value: 0.042

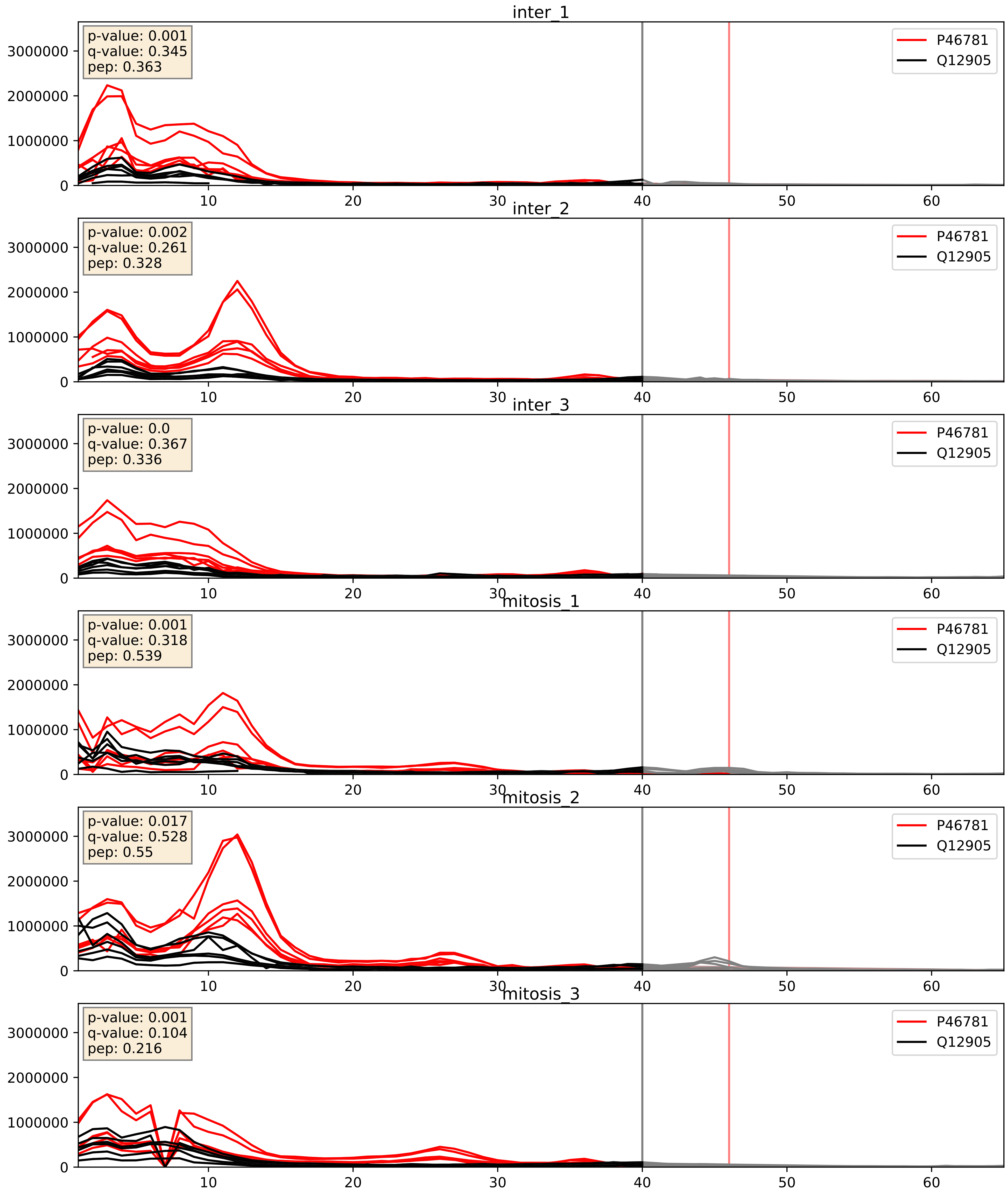


| level | condition_1 | condition_2 | pvalue | pvalue_adjusted |
|---|---|---|---|---|
| interactor_ratio | mitosis | inter | 0.011510917397957572 | 0.15494522140242886 |
| complex_abundance | mitosis | inter | 0.05934455932812005 | 0.17426738519681223 |
| interactor_abundance | mitosis | inter | 0.8270269397762977 | 0.9040528689253451 |

P50914_Q12905
RL14_HUMAN vs ILF2_HUMAN
p-value: 0.0 q-value: 0.018

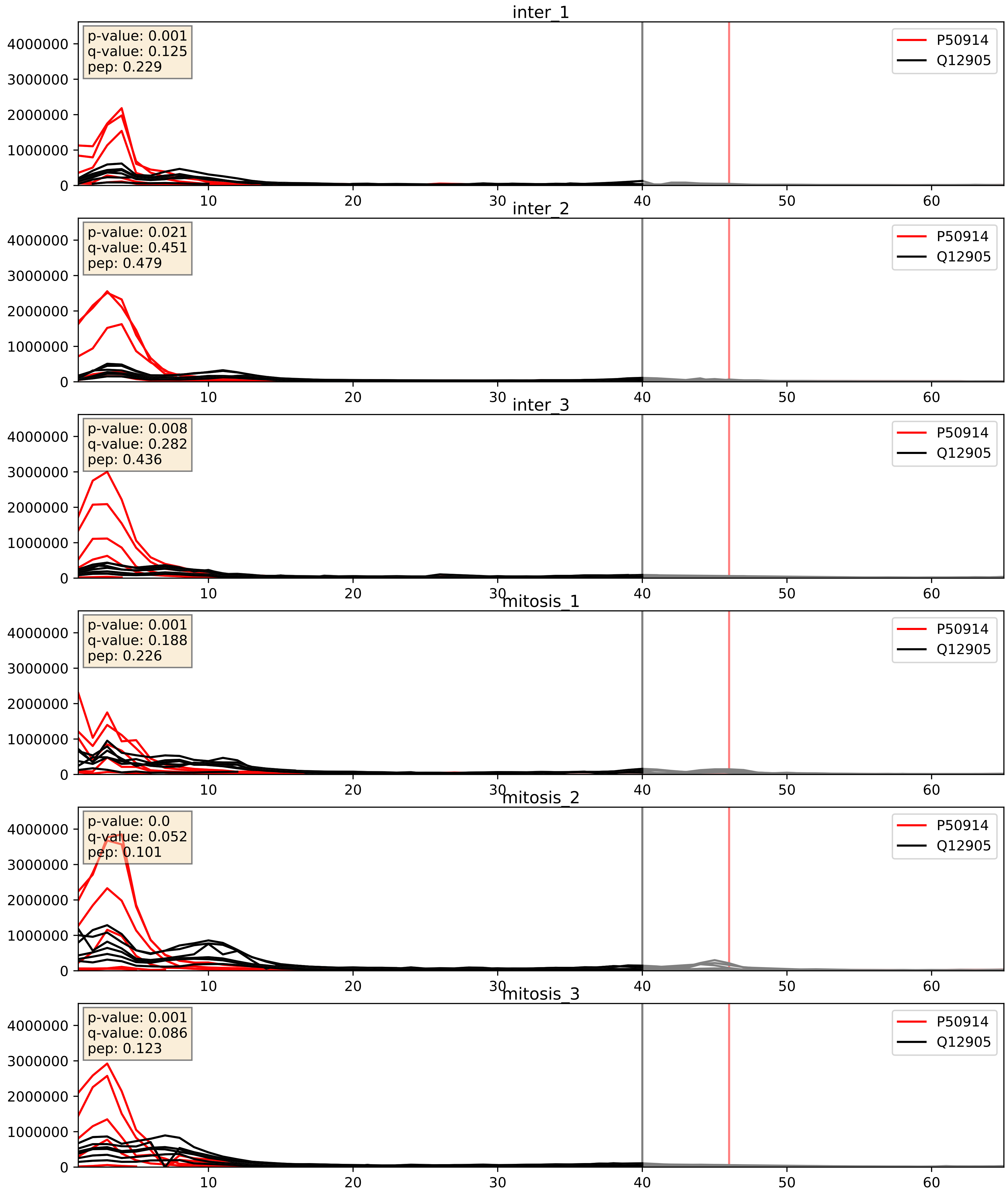


| level | condition_1 | condition_2 | pvalue | pvalue_adjusted |
|---|---|---|---|---|
| interactor_ratio | mitosis | inter | 0.06791275554537803 | 0.4323044818991264 |
| complex_abundance | mitosis | inter | 0.2388434594574438 | 0.3909925440726179 |
| interactor_abundance | mitosis | inter | 0.8453806192959343 | 0.9152981529527123 |

P55769_Q12905
NH2L1_HUMAN vs ILF2_HUMAN
p-value: 0.0 q-value: 0.047

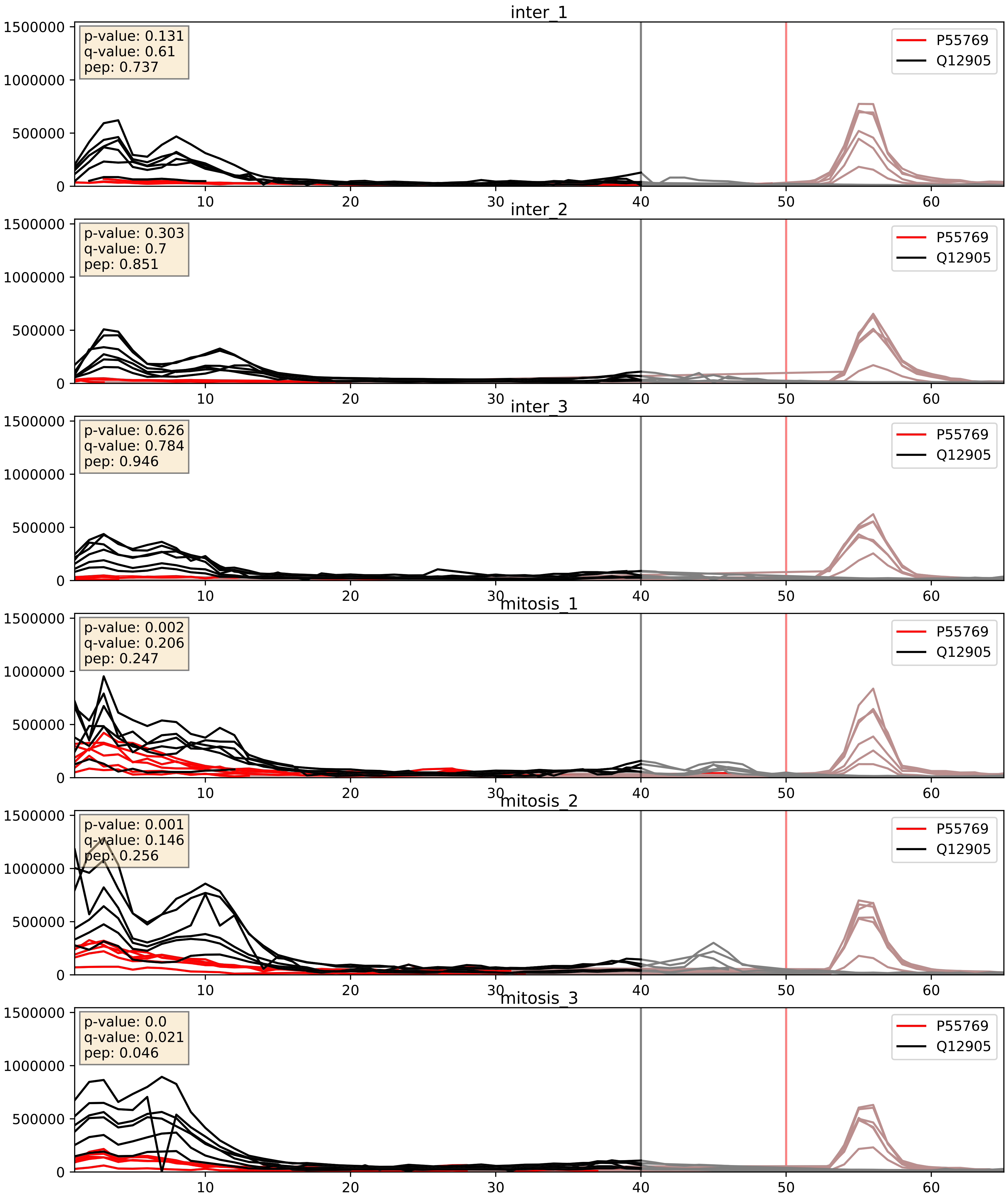


| level | condition_1 | condition_2 | pvalue | pvalue_adjusted |
|---|---|---|---|---|
| complex_abundance | mitosis | inter | 0.0011196272788458134 | 0.022133971147621622 |
| interactor_abundance | mitosis | inter | 0.0018958531698051928 | 0.03196215383917848 |
| interactor_ratio | mitosis | inter | 0.009975986588717106 | 0.14263182888175546 |

P62241_Q12905
RS8_HUMAN vs ILF2_HUMAN
p-value: 0.0 q-value: 0.049

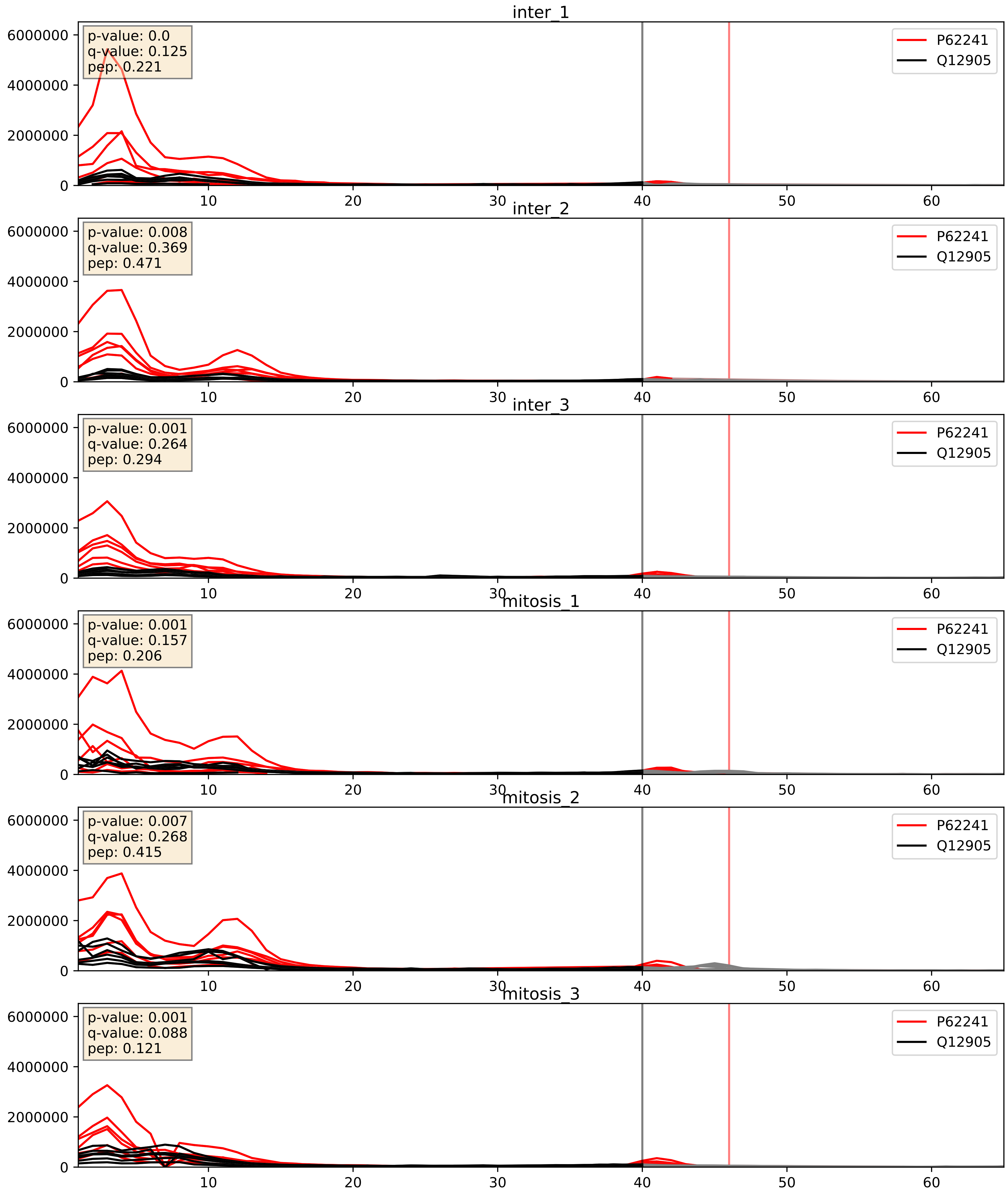


| level | condition_1 | condition_2 | pvalue | pvalue_adjusted |
|---|---|---|---|---|
| interactor_ratio | mitosis | inter | 0.0027642173648887137 | 0.06779856917893234 |
| complex_abundance | mitosis | inter | 0.04197968837428533 | 0.14726323635770058 |
| interactor_abundance | mitosis | inter | 0.13238802431073732 | 0.3178814938442557 |

P62266_Q12905
RS23_HUMAN vs ILF2_HUMAN
p-value: 0.0 q-value: 0.047

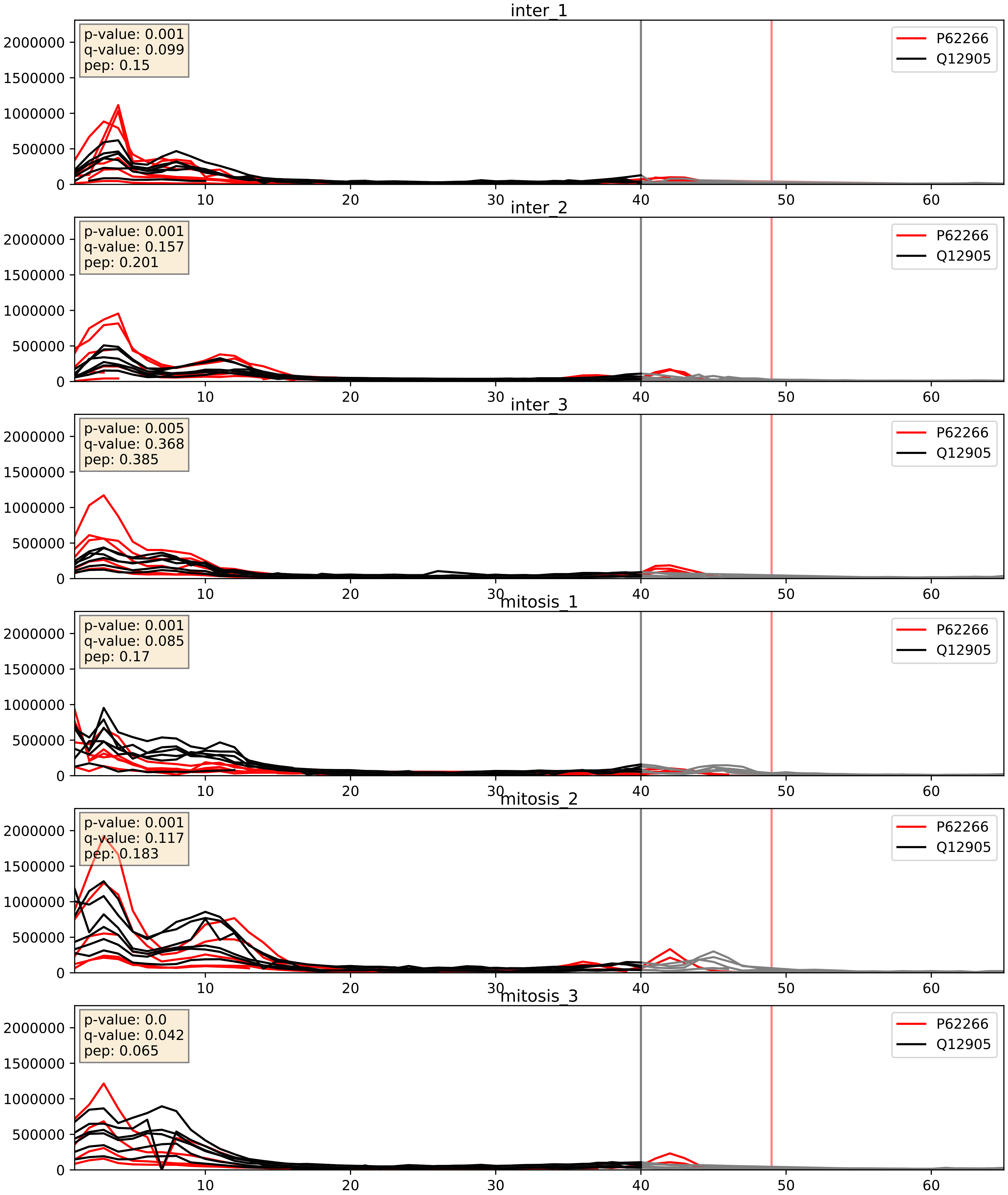


| level | condition_1 | condition_2 | pvalue | pvalue_adjusted |
|---|---|---|---|---|
| interactor_ratio | mitosis | inter | 0.08832457928501604 | 0.4953575337685953 |
| interactor_abundance | mitosis | inter | 0.301469321193808 | 0.49640807721823527 |
| complex_abundance | mitosis | inter | 0.9673493135661784 | 0.9807544858612445 |

P62277_Q12905
RS13_HUMAN vs ILF2_HUMAN
p-value: 0.0 q-value: 0.049

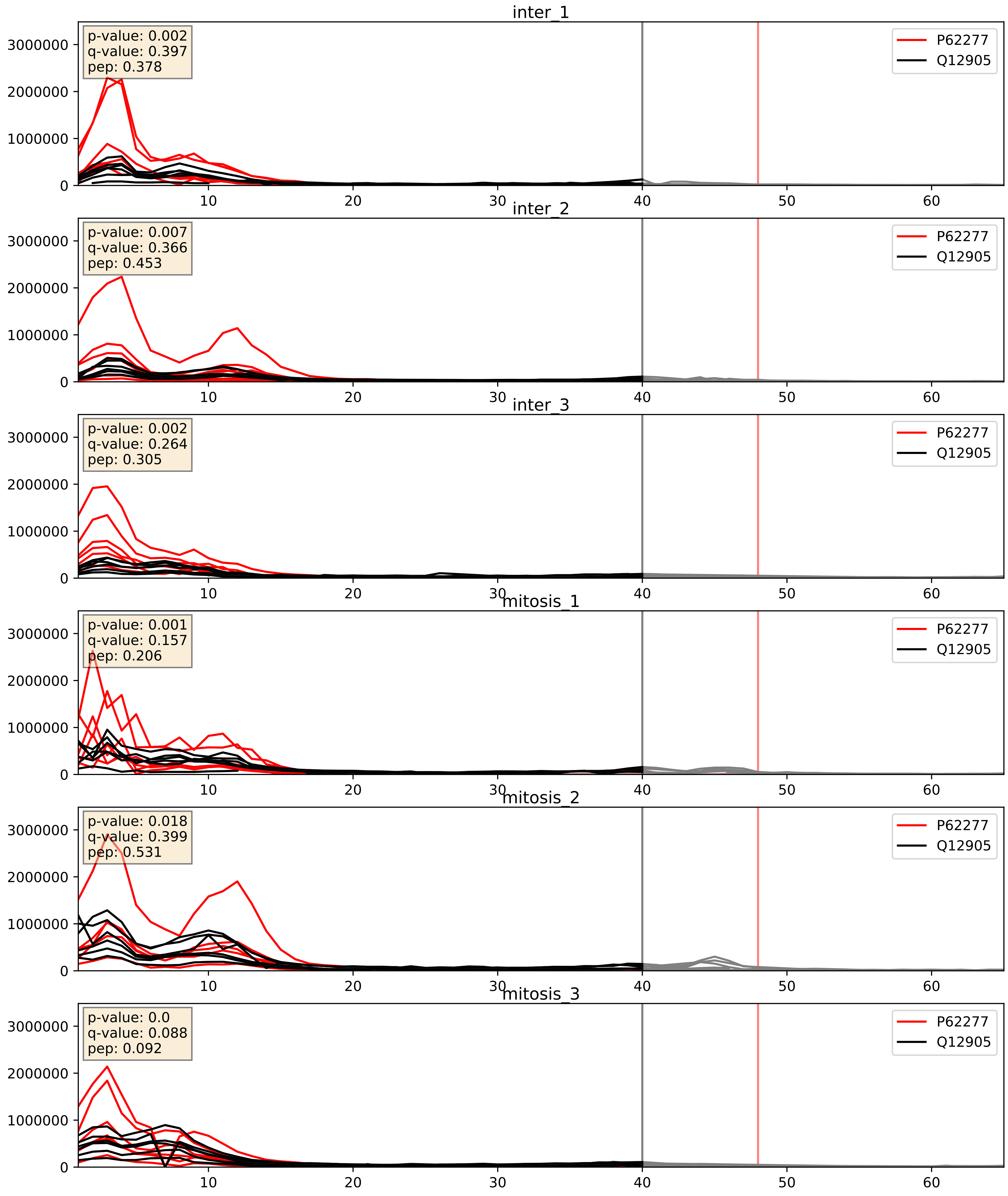


| level | condition_1 | condition_2 | pvalue | pvalue_adjusted |
|---|---|---|---|---|
| interactor_ratio | mitosis | inter | 0.005251363139102532 | 0.0974780445871482 |
| complex_abundance | mitosis | inter | 0.020352789722355272 | 0.10515559050023798 |
| interactor_abundance | mitosis | inter | 0.3699370722830677 | 0.5640650763703348 |

P62280_Q12905
RS11_HUMAN vs ILF2_HUMAN
p-value: 0.0 q-value: 0.018

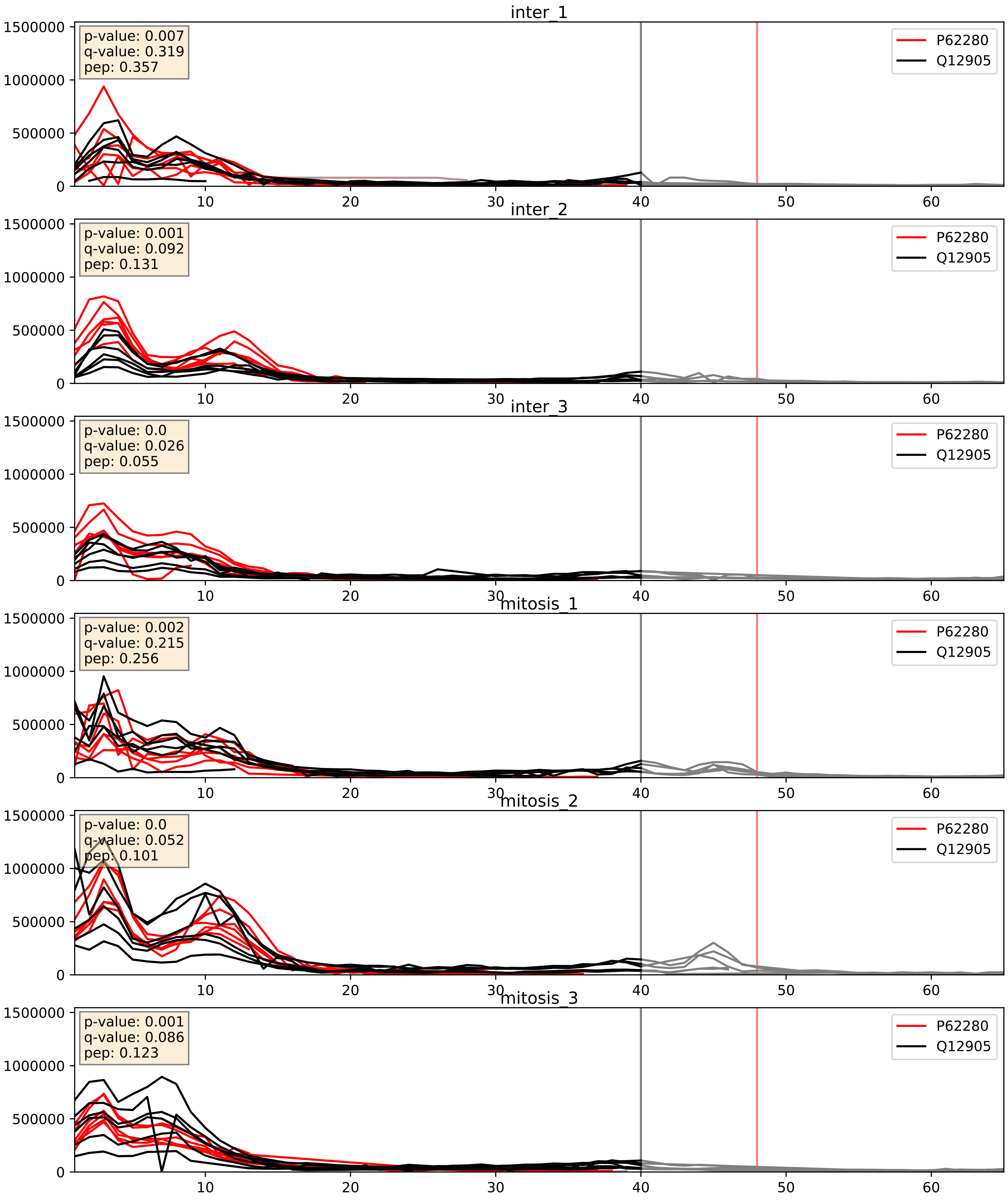


| level | condition_1 | condition_2 | pvalue | pvalue_adjusted |
|---|---|---|---|---|
| complex_abundance | mitosis | inter | 0.02324464678263421 | 0.11100971021145899 |
| interactor_ratio | mitosis | inter | 0.042242787641378815 | 0.3399226239910554 |
| interactor_abundance | mitosis | inter | 0.7956222020674533 | 0.8808985116259767 |

P62753_Q12905
RS6_HUMAN vs ILF2_HUMAN
p-value: 0.0 q-value: 0.049

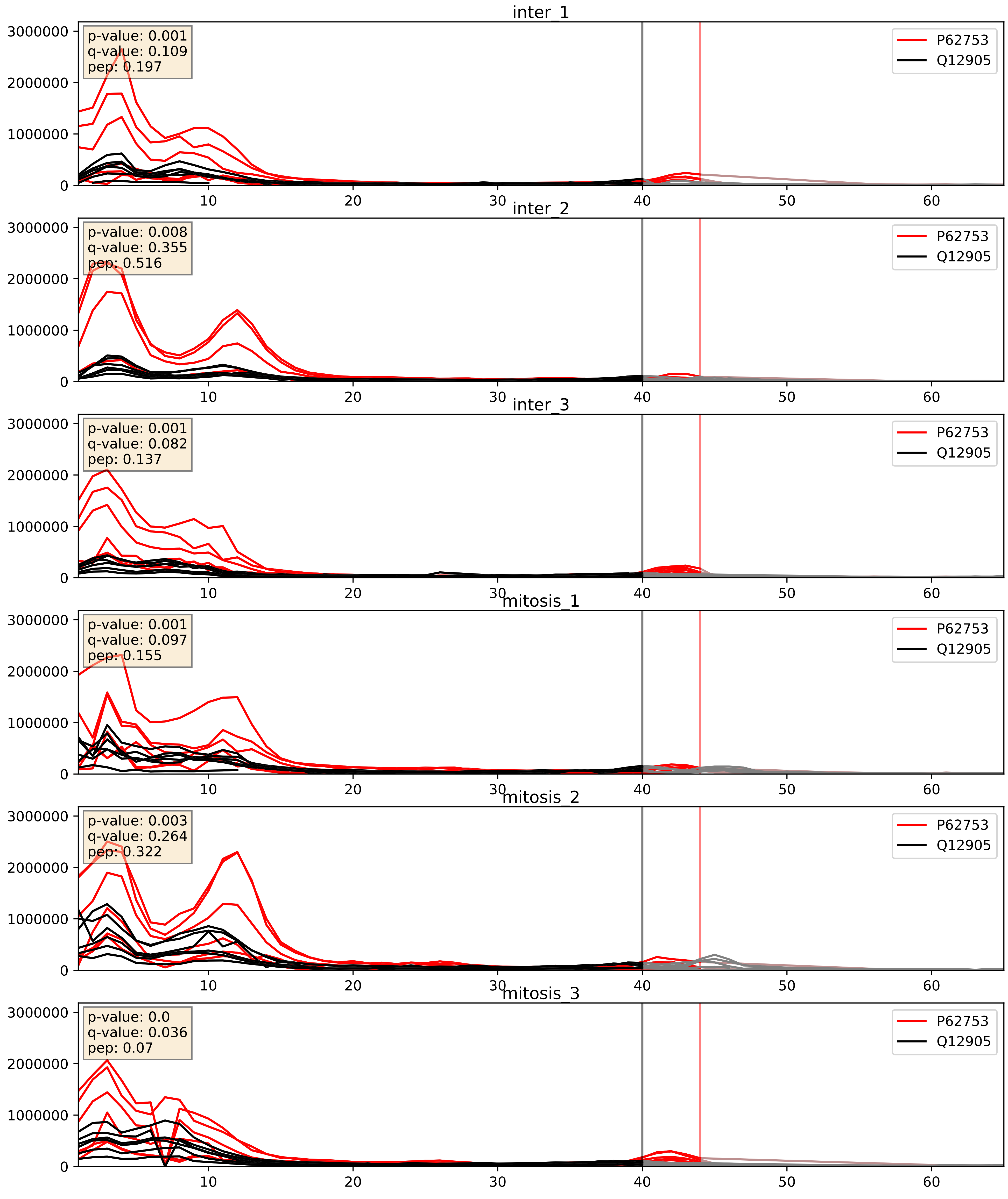


| level | condition_1 | condition_2 | pvalue | pvalue_adjusted |
|---|---|---|---|---|
| interactor_ratio | mitosis | inter | 0.015583696141321714 | 0.1843379164928052 |
| complex_abundance | mitosis | inter | 0.04990125352069983 | 0.15897087091075196 |
| interactor_abundance | mitosis | inter | 0.8343197771242803 | 0.9094329927650375 |

P62829_Q12905
RL23_HUMAN vs ILF2_HUMAN
p-value: 0.0 q-value: 0.047

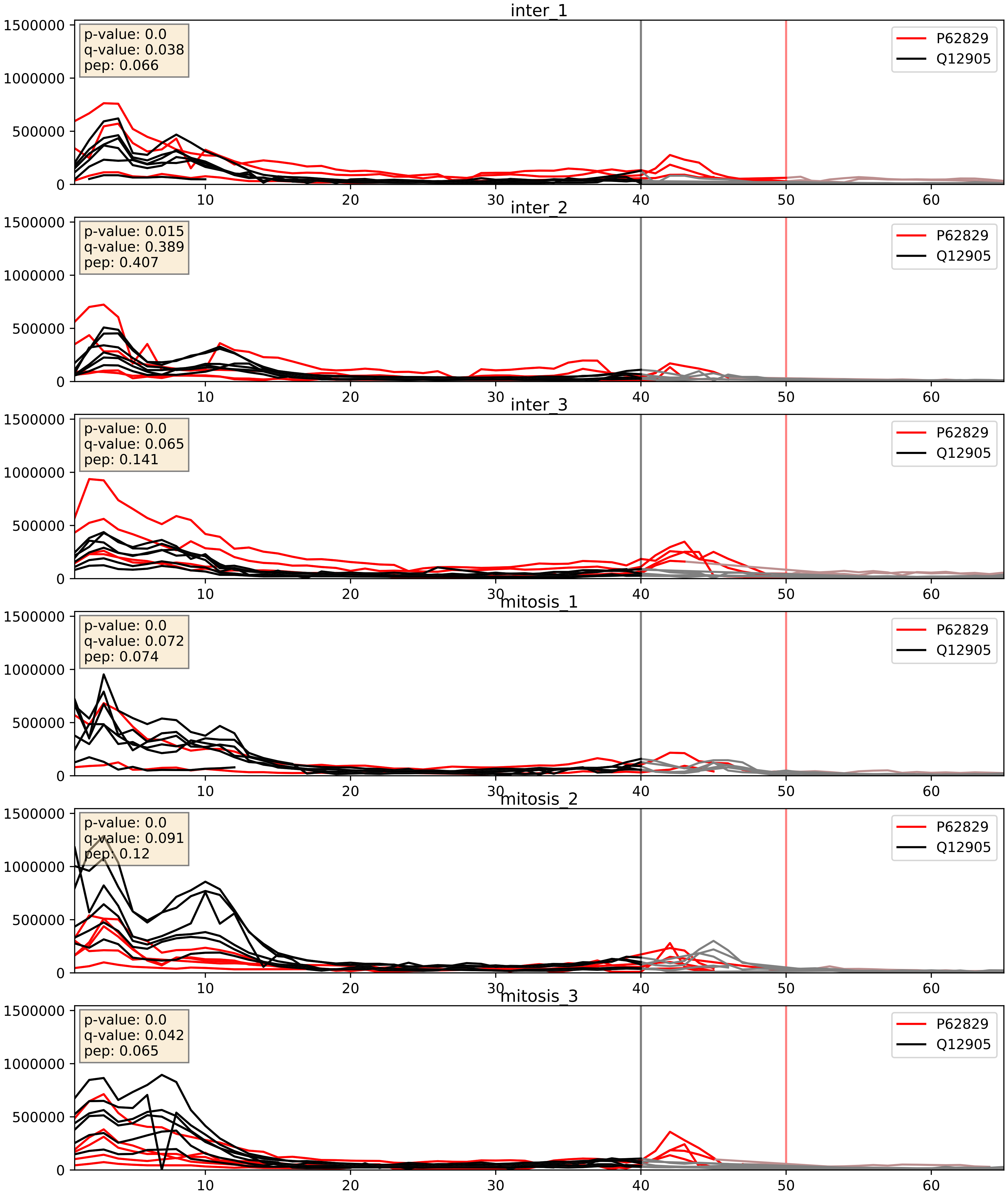


| level | condition_1 | condition_2 | pvalue | pvalue_adjusted |
|---|---|---|---|---|
| interactor_ratio | mitosis | inter | 0.0022056667910768353 | 0.05863511717893699 |
| interactor_abundance | mitosis | inter | 0.004946130873219955 | 0.05819777357355713 |
| complex_abundance | mitosis | inter | 0.023997797201938377 | 0.11305511505150936 |

P62899_Q12905
RL31_HUMAN vs ILF2_HUMAN
p-value: 0.0 q-value: 0.047

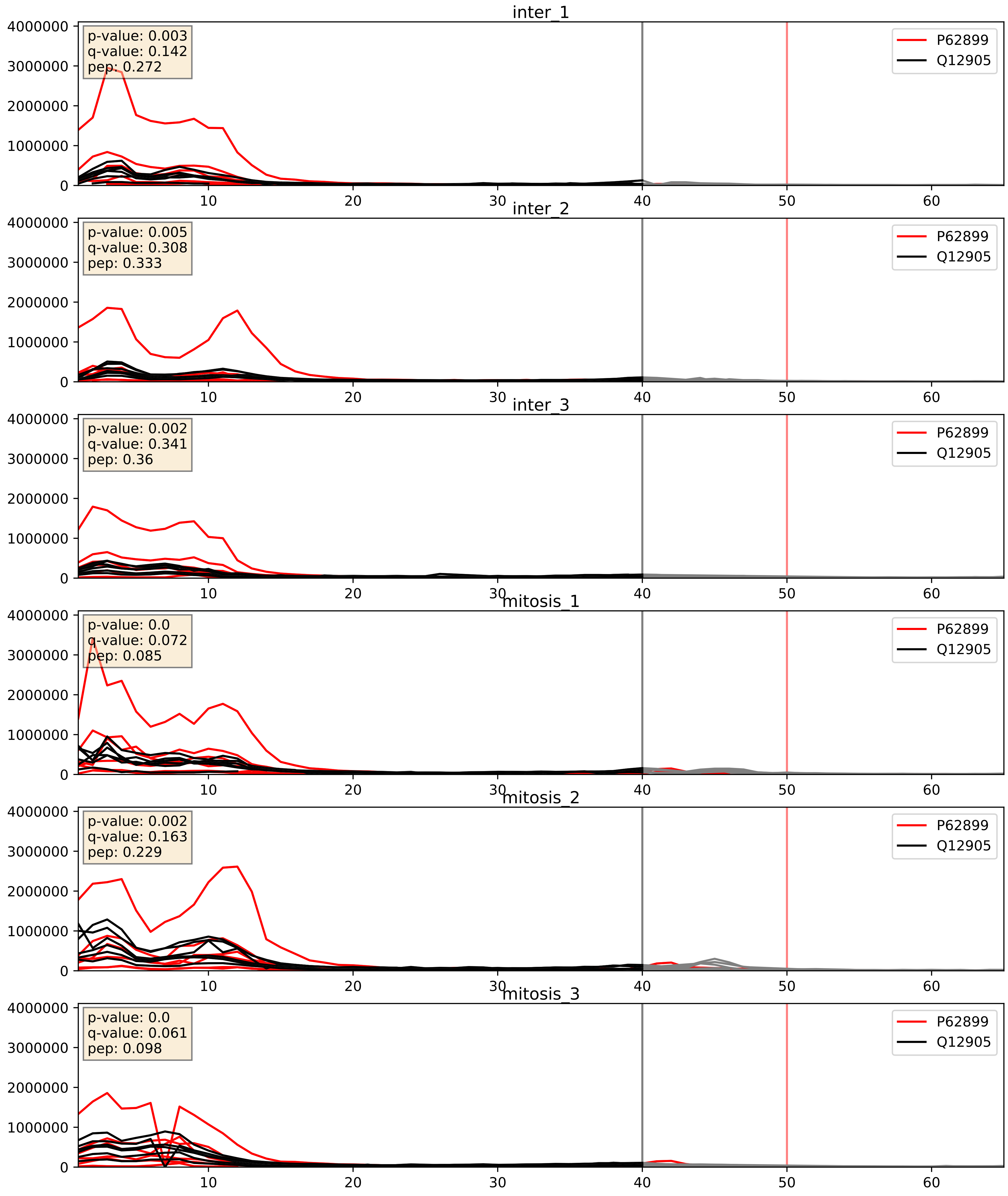


| level | condition_1 | condition_2 | pvalue | pvalue_adjusted |
|---|---|---|---|---|
| complex_abundance | mitosis | inter | 0.01816970602023441 | 0.10074914023469604 |
| interactor_ratio | mitosis | inter | 0.21552192622893387 | 0.6924580509898395 |
| interactor_abundance | mitosis | inter | 0.2904832686279918 | 0.4848940677565542 |

P62906_Q12905
RL10A_HUMAN vs ILF2_HUMAN
p-value: 0.0 q-value: 0.041

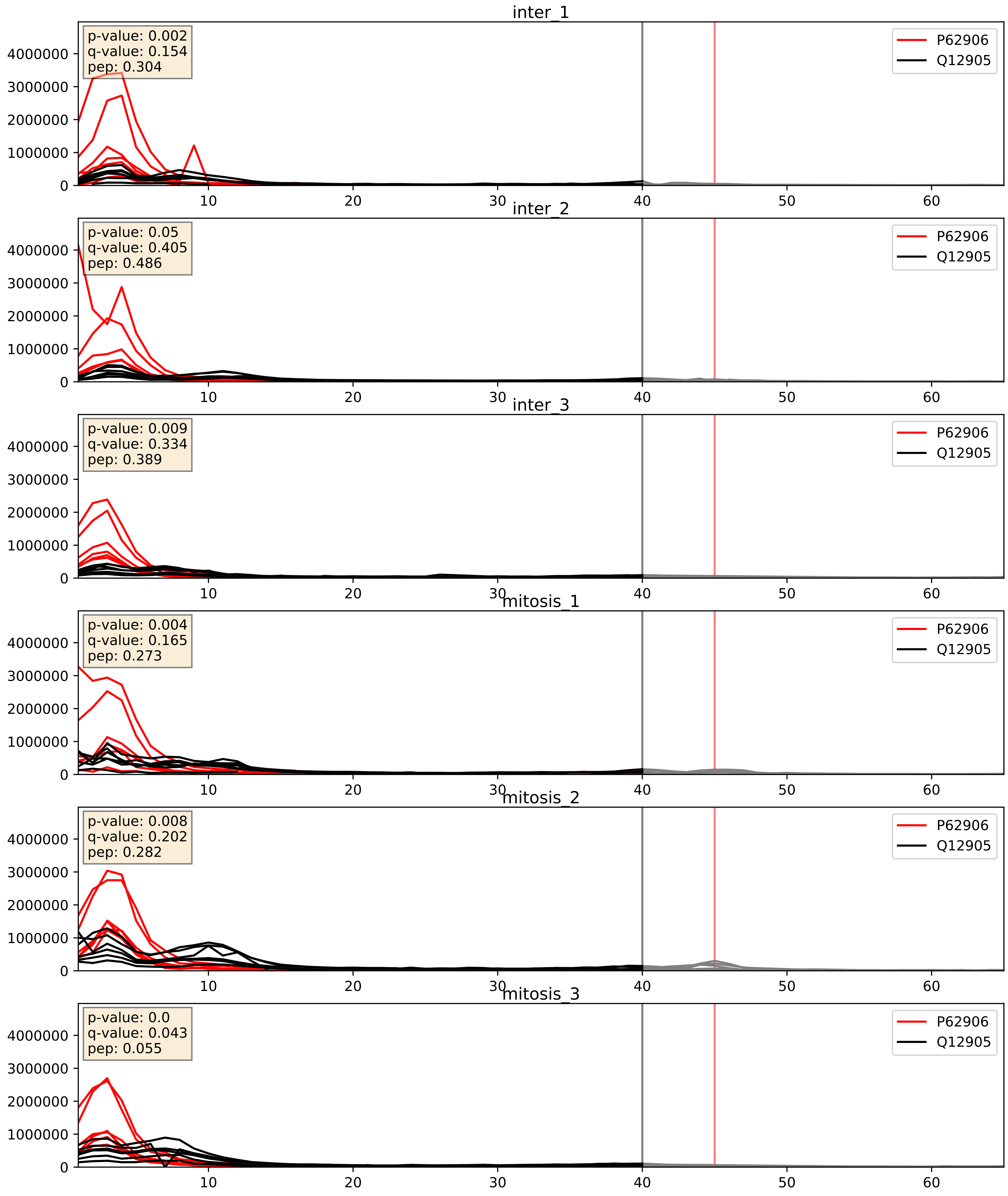


| level | condition_1 | condition_2 | pvalue | pvalue_adjusted |
|---|---|---|---|---|
| interactor_ratio | mitosis | inter | 0.019821751658541914 | 0.213852931597635 |
| complex_abundance | mitosis | inter | 0.05391602986386887 | 0.16559785275734393 |
| interactor_abundance | mitosis | inter | 0.8597142149458314 | 0.9247491429927516 |

P62995_Q12905
TRA2B_HUMAN vs ILF2_HUMAN
p-value: 0.001 q-value: 0.045

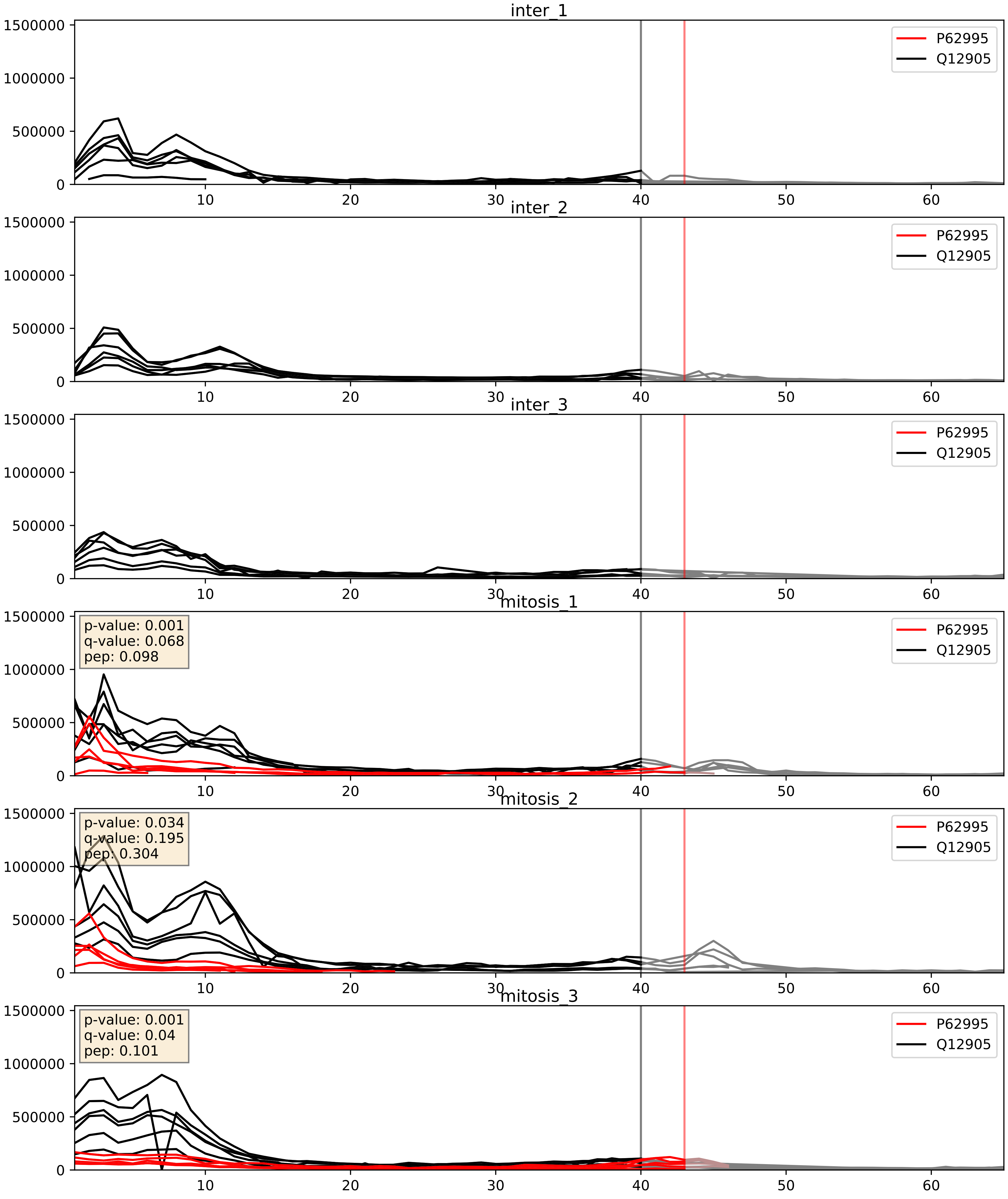


| level | condition_1 | condition_2 | pvalue | pvalue_adjusted |
|---|---|---|---|---|
| complex_abundance | mitosis | inter | 2.505847793058248e-05 | 0.005691857151652958 |
| interactor_abundance | mitosis | inter | 0.00037334314755474554 | 0.013573913378163039 |
| interactor_ratio | mitosis | inter | 0.0014969399724690845 | 0.04597663709000905 |

P83731_Q12905
RL24_HUMAN vs ILF2_HUMAN
p-value: 0.0 q-value: 0.044

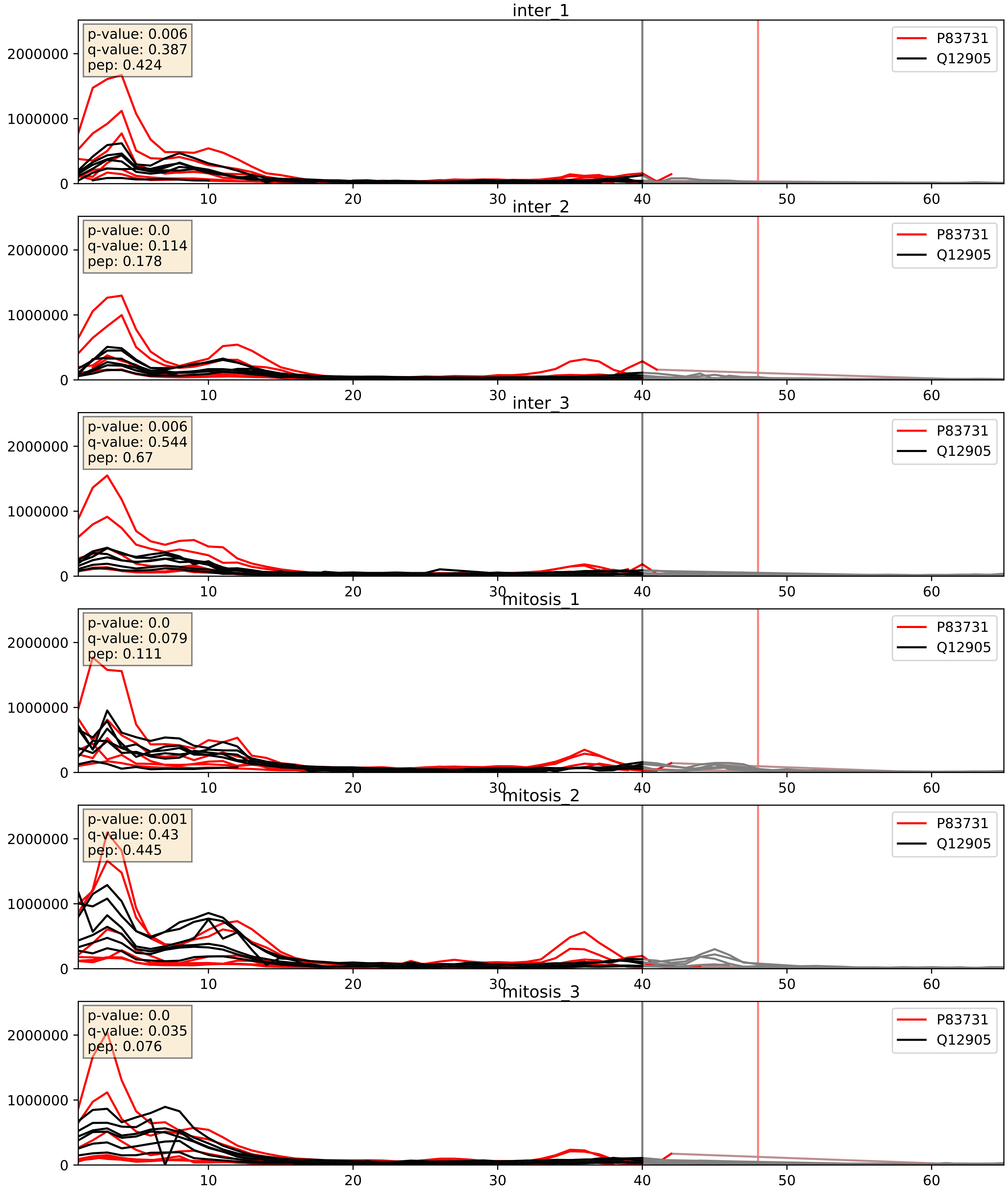


| level | condition_1 | condition_2 | pvalue | pvalue_adjusted |
|---|---|---|---|---|
| complex_abundance | mitosis | inter | 0.02533428749824106 | 0.11659220483061478 |
| interactor_ratio | mitosis | inter | 0.03516369047609029 | 0.30434902980316775 |
| interactor_abundance | mitosis | inter | 0.2783968434326379 | 0.47161626356290925 |

Q08211_Q12905
DHX9_HUMAN vs ILF2_HUMAN
p-value: 0.001 q-value: 0.039

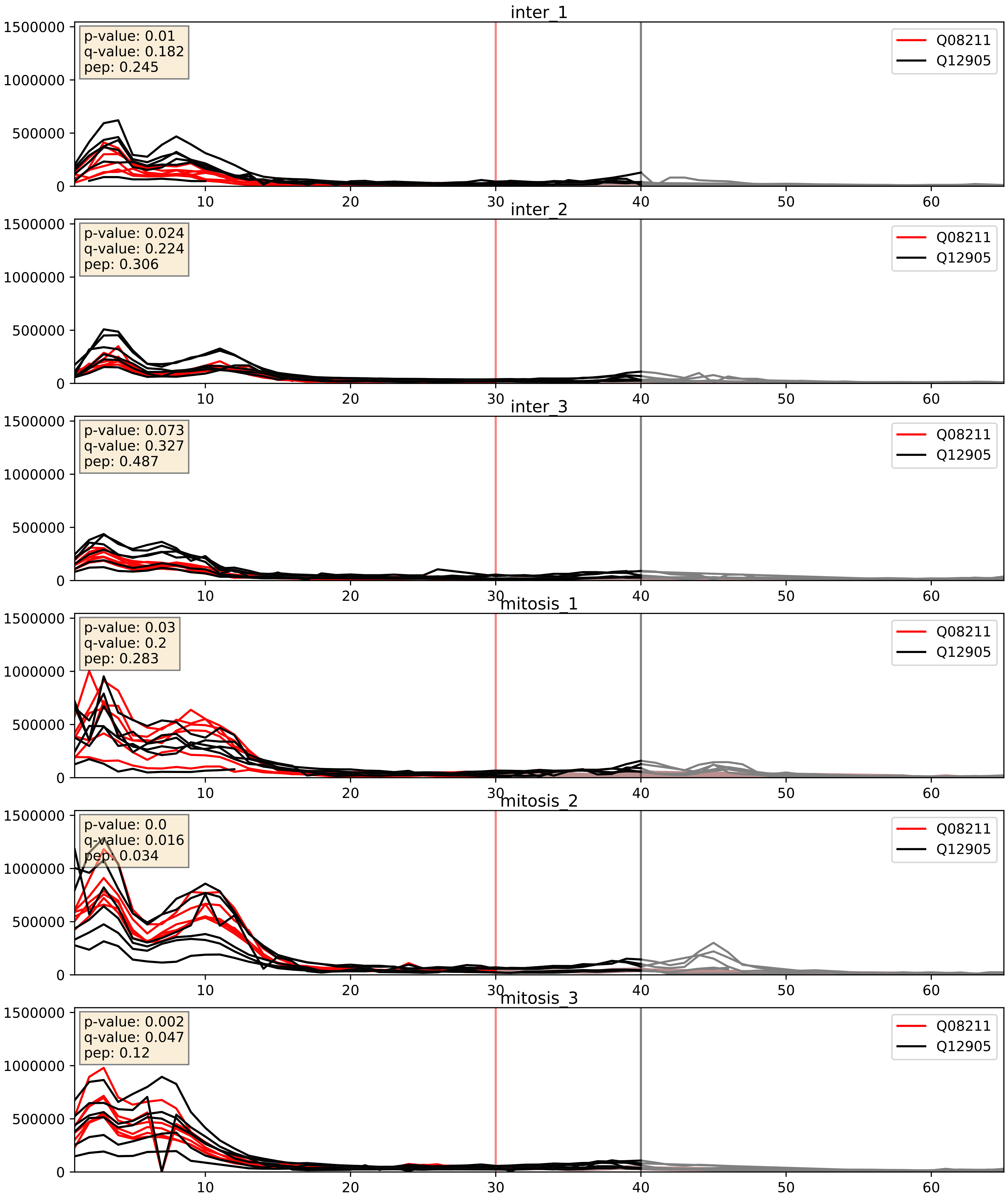


| level | condition_1 | condition_2 | pvalue | pvalue_adjusted |
|---|---|---|---|---|
| interactor_abundance | mitosis | inter | 0.0001655881985577161 | 0.009910743628815524 |
| complex_abundance | mitosis | inter | 0.00023397135977256306 | 0.010221981956574496 |
| interactor_ratio | mitosis | inter | 0.007958384649893444 | 0.1259990886723217 |

Q12905_Q12906
ILF2_HUMAN vs ILF3_HUMAN
p-value: 0.0 q-value: 0.0

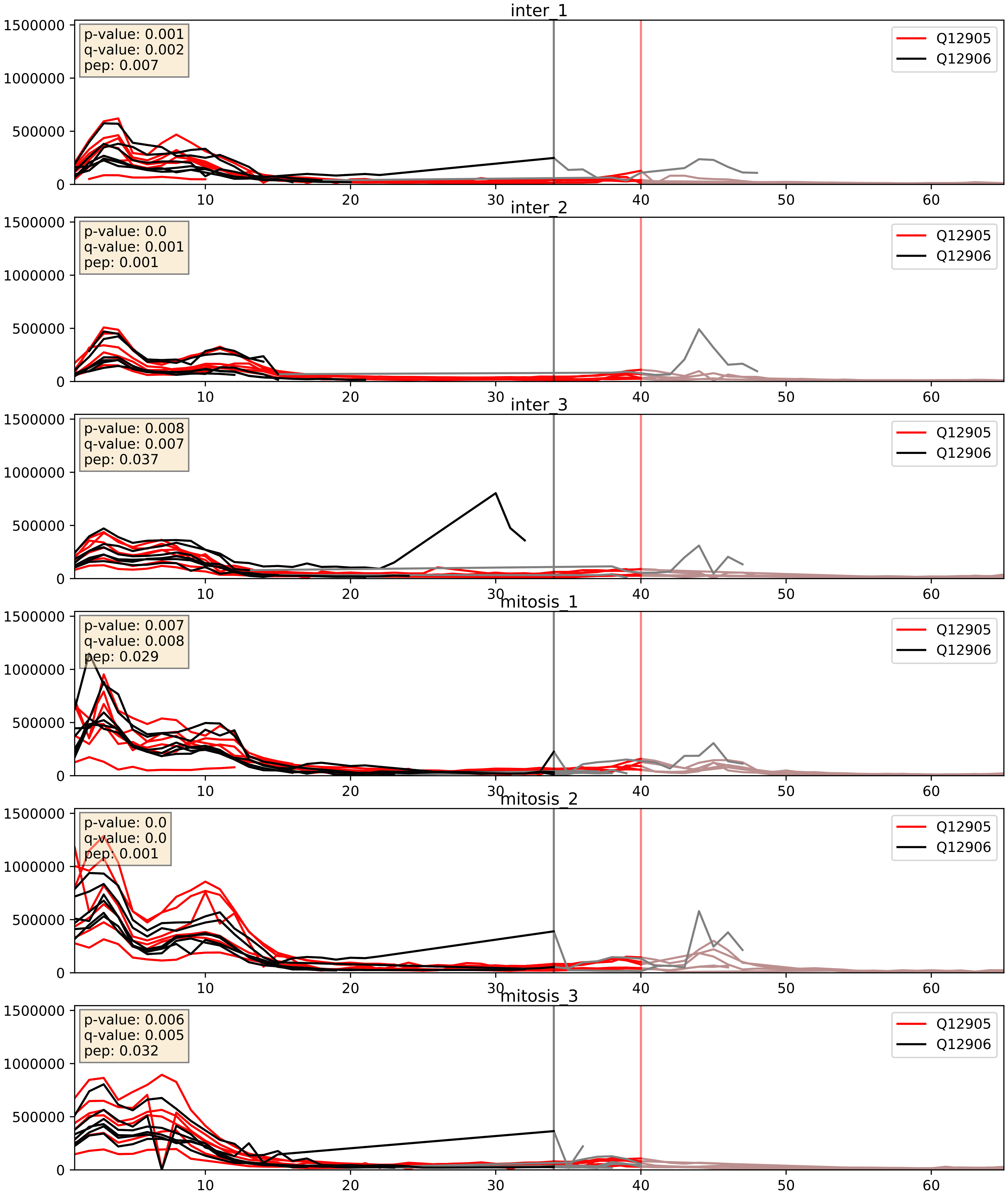


| level | condition_1 | condition_2 | pvalue | pvalue_adjusted |
|---|---|---|---|---|
| complex_abundance | mitosis | inter | 0.0014631415558530808 | 0.026478840841654062 |
| interactor_abundance | mitosis | inter | 0.0016872493479083844 | 0.030026724361945468 |
| interactor_ratio | mitosis | inter | 0.3605319558504416 | 0.805243537848486 |

Q12905_Q14257
ILF2_HUMAN vs RCN2_HUMAN
p-value: 0.0 q-value: 0.025

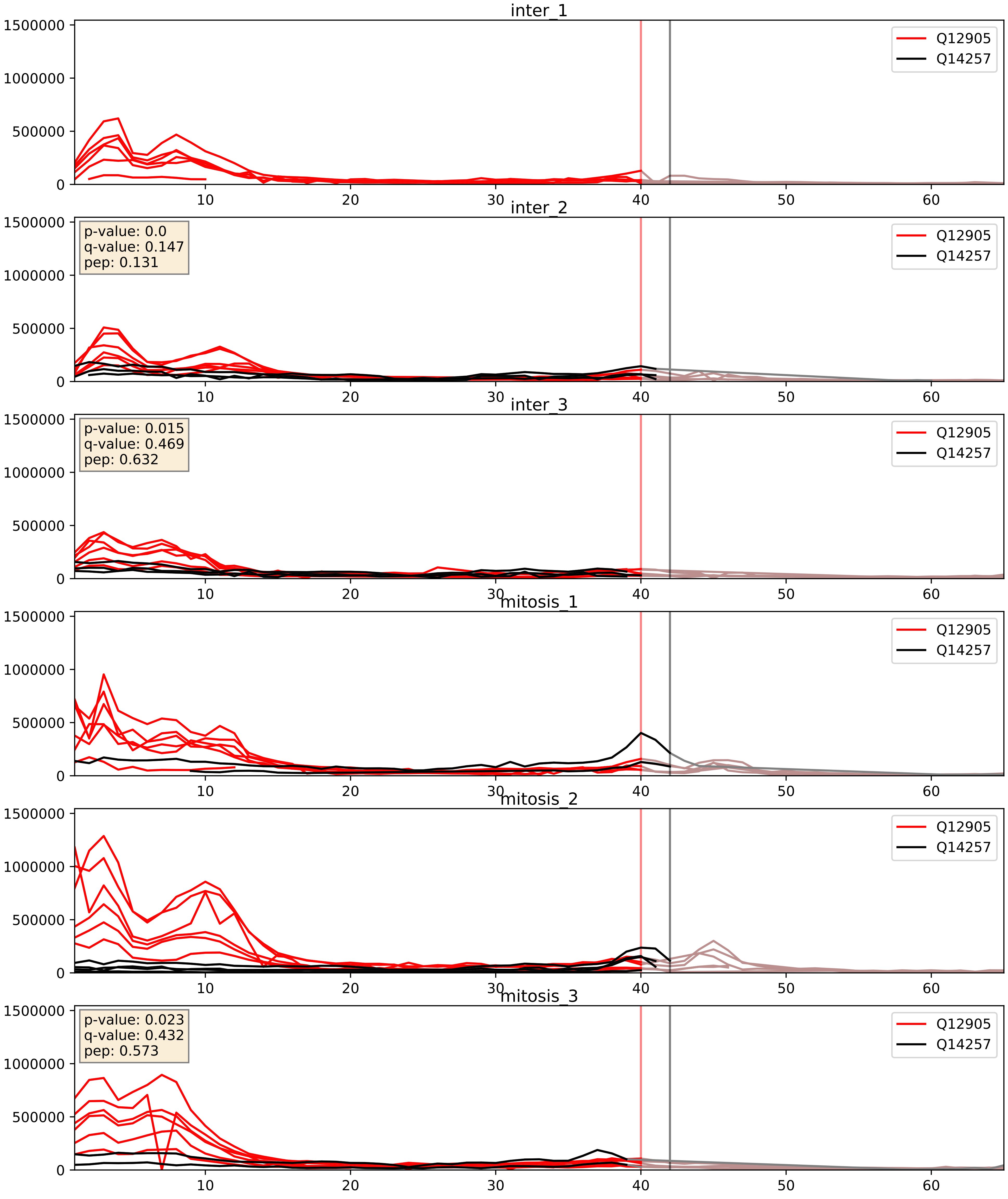


| level | condition_1 | condition_2 | pvalue | pvalue_adjusted |
|---|---|---|---|---|
| interactor_ratio | mitosis | inter | 0.014566409246335154 | 0.17686306829592754 |
| interactor_abundance | mitosis | inter | 0.20459910743912274 | 0.4009542947982808 |
| complex_abundance | mitosis | inter | 0.5608712321999432 | 0.6926512096954721 |

Q12905_Q15717
ILF2_HUMAN vs ELAV1_HUMAN
p-value: 0.004 q-value: 0.047

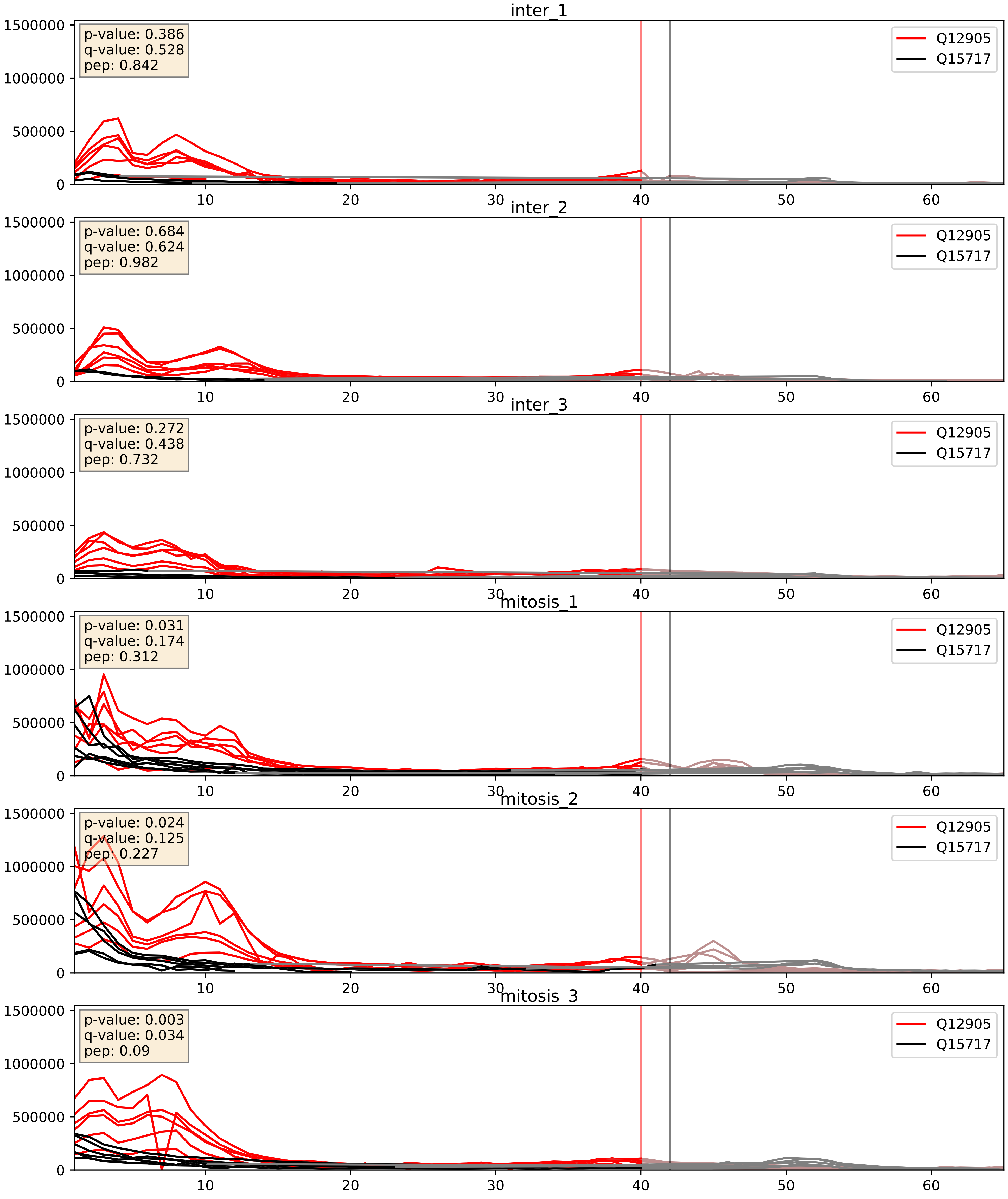


| level | condition_1 | condition_2 | pvalue | pvalue_adjusted |
|---|---|---|---|---|
| complex_abundance | mitosis | inter | 4.5501477611336626e-05 | 0.005691857151652958 |
| interactor_abundance | mitosis | inter | 0.0015109761494022987 | 0.028710223837699618 |
| interactor_ratio | mitosis | inter | 0.003190844185944654 | 0.07382061143698984 |

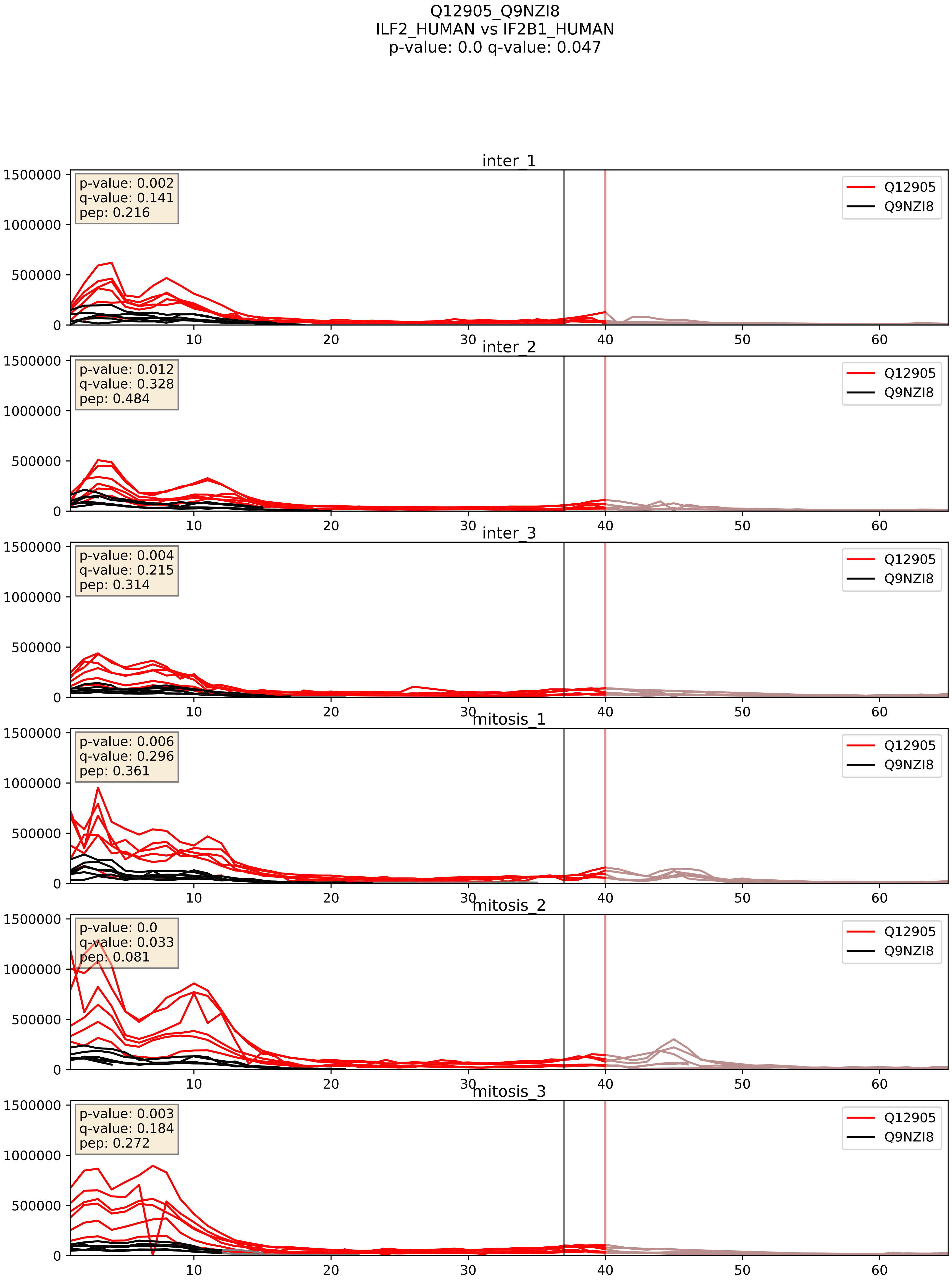


| level | condition_1 | condition_2 | pvalue | pvalue_adjusted |
|---|---|---|---|---|
| interactor_abundance | mitosis | inter | 0.007285978219052328 | 0.07362753439894419 |
| interactor_ratio | mitosis | inter | 0.046008117716284255 | 0.35622692584467375 |
| complex_abundance | mitosis | inter | 0.06607339195523967 | 0.1855604445987046 |
