## Supplementary Data 2 for "SECAT: Quantifying differential protein-protein interaction states by network-centric analysis": hela_string_000014_P62304.pdf

P62304_P62304

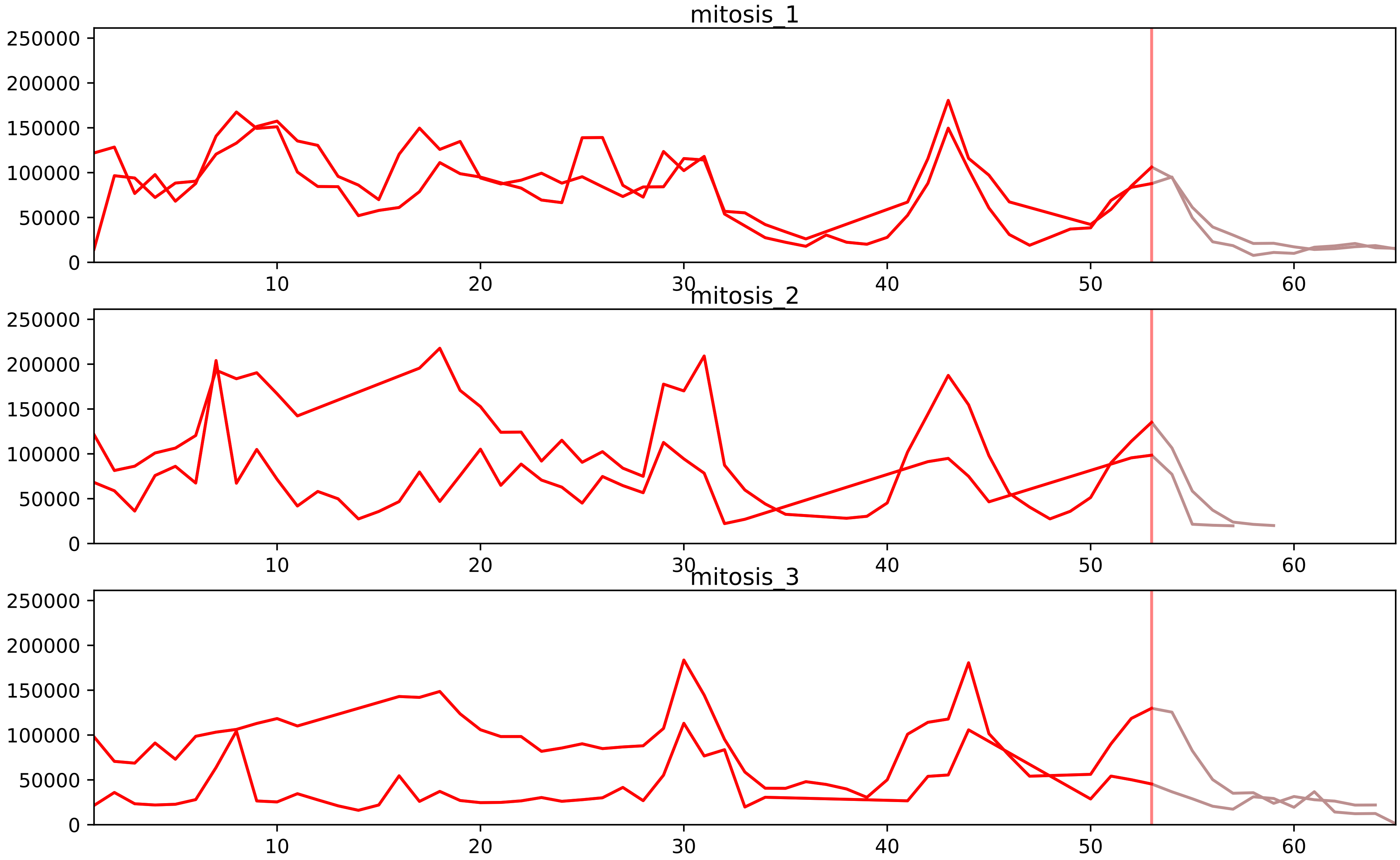


| level | condition_1 | condition_2 | pvalue | pvalue_adjusted |
|---|---|---|---|---|
| interactor_ratio | mitosis | inter | 1.1607704891948622e-13 | 1.5255840715132475e-11 |
| complex_abundance | mitosis | inter | 1.533987000220793e-07 | 2.3521134003385493e-05 |
| interactor_abundance | mitosis | inter | 4.7399140413755866e-07 | 7.2678681967759e-05 |
| total_abundance | mitosis | inter | 1.863768989903585e-05 | 0.0058684532992606024 |
| assembled_abundance | mitosis | inter | 2.0852663284143003e-05 | 0.005741842166651376 |
| monomer_abundance | mitosis | inter | 0.0003072853809853298 | 0.05240402371705197 |

P08579_P62304
RU2B_HUMAN vs RUXE_HUMAN
p-value: 0.009 q-value: 0.008

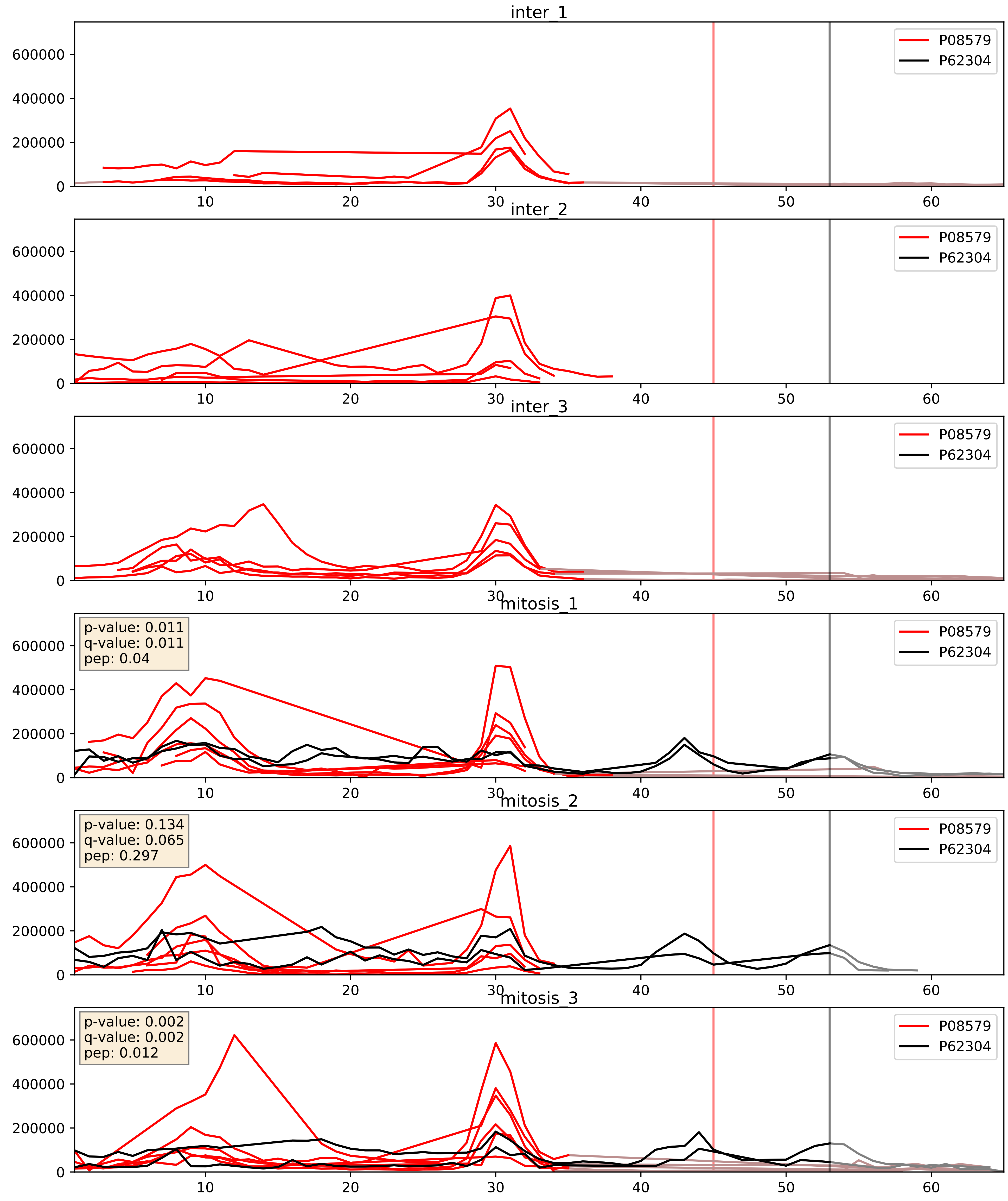


| level | condition_1 | condition_2 | pvalue | pvalue_adjusted |
|---|---|---|---|---|
| complex_abundance | mitosis | inter | 6.639607125748741e-05 | 0.005691857151652958 |
| interactor_abundance | mitosis | inter | 8.77072104426966e-05 | 0.007396785432408699 |
| interactor_ratio | mitosis | inter | 0.0022594795659381457 | 0.05896690574521502 |

P09661_P62304
RU2A_HUMAN vs RUXE_HUMAN
p-value: 0.072 q-value: 0.044

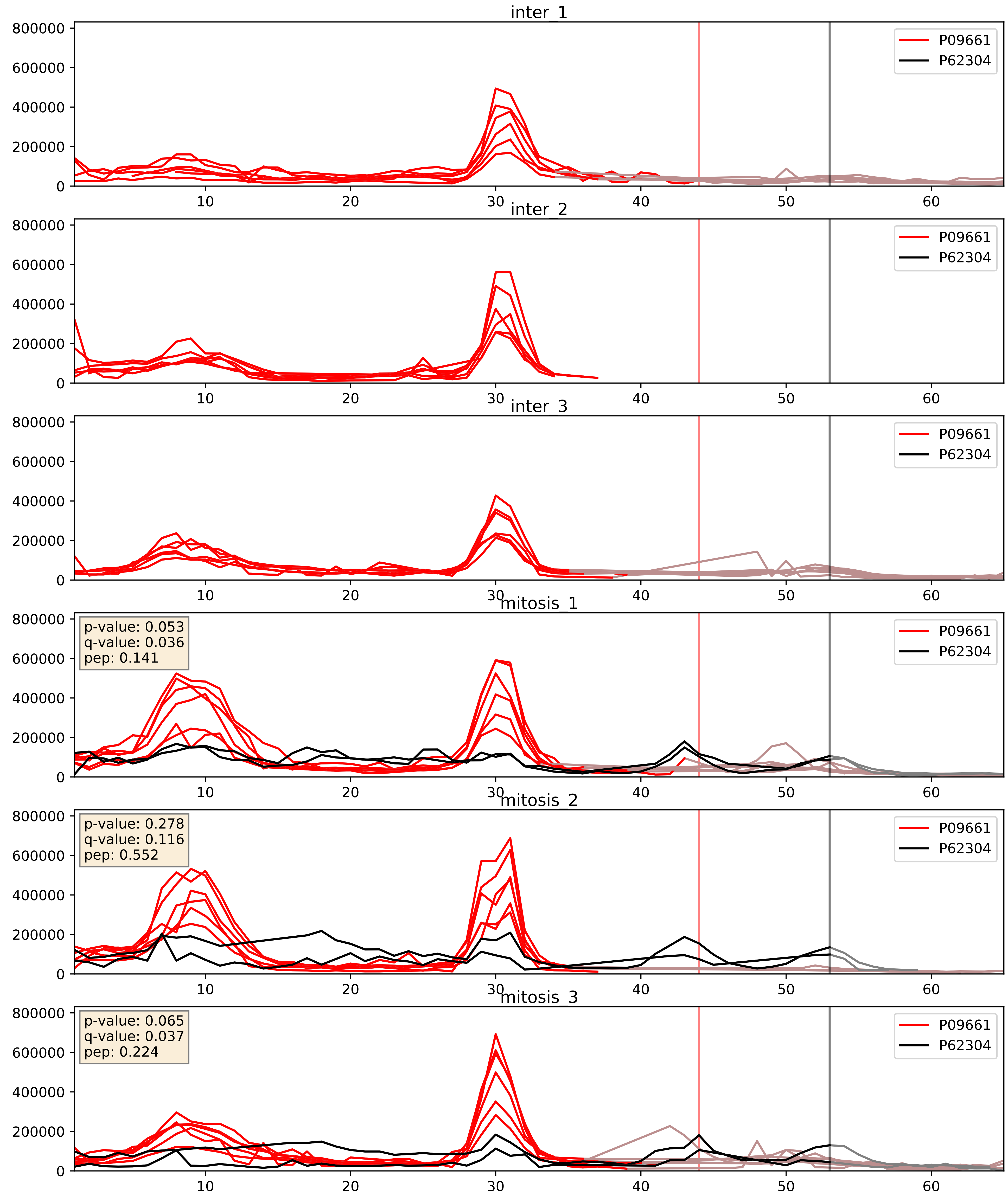


| level | condition_1 | condition_2 | pvalue | pvalue_adjusted |
|---|---|---|---|---|
| interactor_ratio | mitosis | inter | 1.9350871757215154e-05 | 0.004993481478971454 |
| interactor_abundance | mitosis | inter | 5.2162128658301765e-05 | 0.0062888425537332825 |
| complex_abundance | mitosis | inter | 8.64707192201536e-05 | 0.006326404756619784 |

P62304_P62306
RUXE_HUMAN vs RUXF_HUMAN
p-value: 0.0 q-value: 0.0

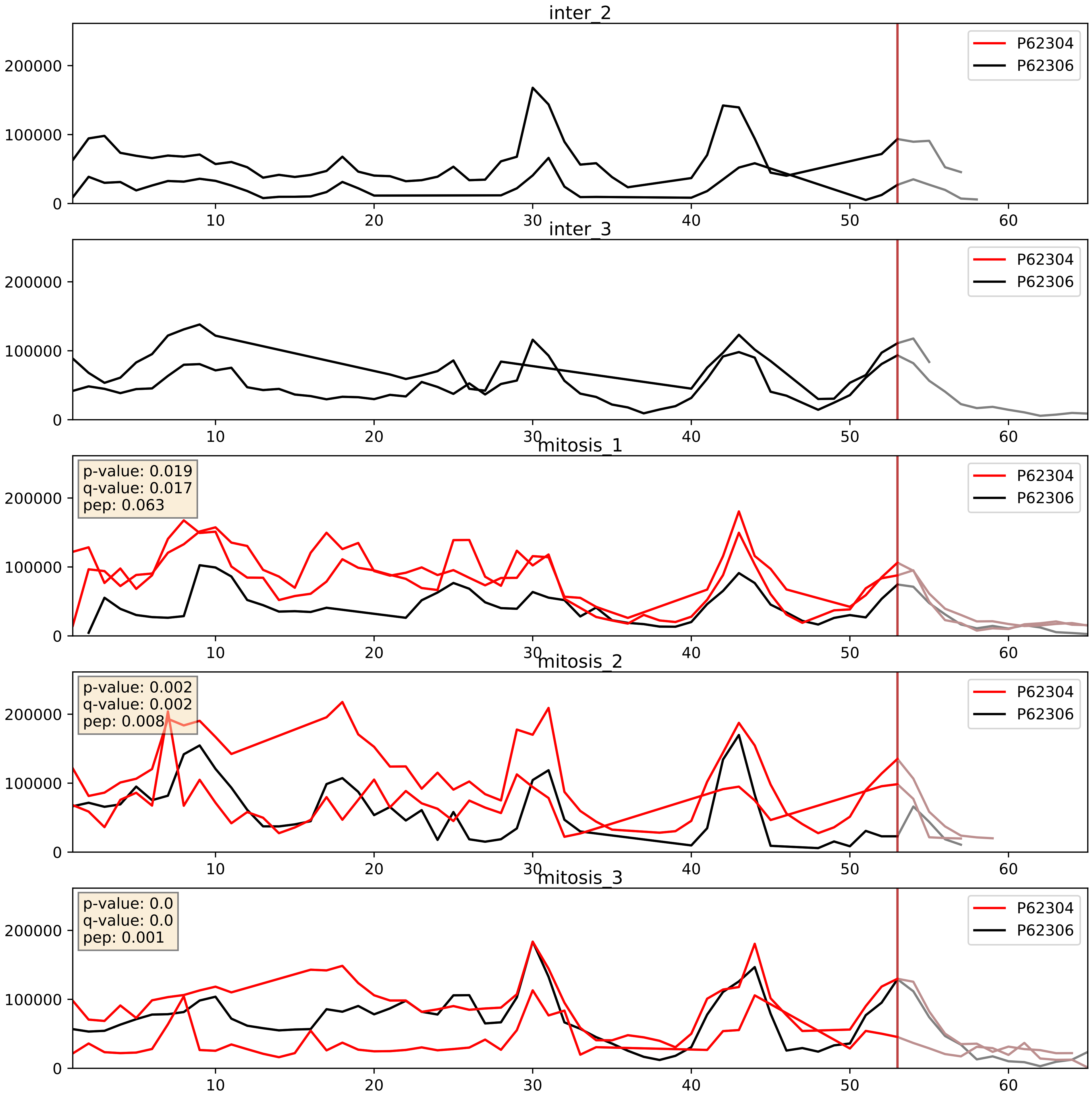


| level | condition_1 | condition_2 | pvalue | pvalue_adjusted |
|---|---|---|---|---|
| complex_abundance | mitosis | inter | 1.9058781942551378e-05 | 0.005691857151652958 |
| interactor_abundance | mitosis | inter | 5.613578522364216e-05 | 0.006414334379145961 |
| interactor_ratio | mitosis | inter | 0.0016271261094656015 | 0.048854197059354385 |

P62304_P62314
RUXE_HUMAN vs SMD1_HUMAN
p-value: 0.0 q-value: 0.0

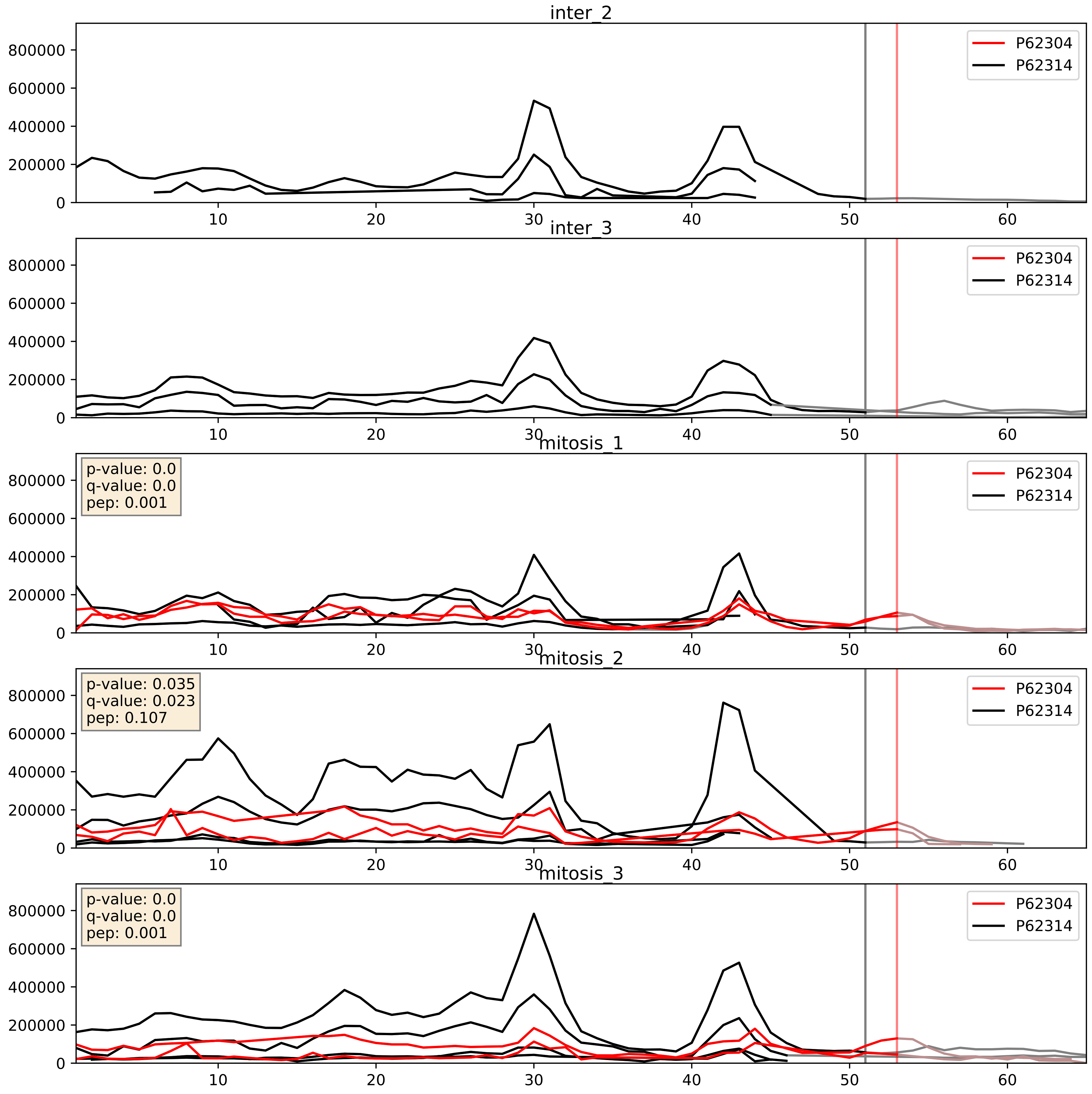


| level | condition_1 | condition_2 | pvalue | pvalue_adjusted |
|---|---|---|---|---|
| interactor_abundance | mitosis | inter | 0.00010811758969854758 | 0.007876481428251637 |
| complex_abundance | mitosis | inter | 0.00020038690159054093 | 0.009529510431194613 |
| interactor_ratio | mitosis | inter | 0.11847273284144728 | 0.567184895482544 |

P62304_P62316
RUXE_HUMAN vs SMD2_HUMAN
p-value: 0.001 q-value: 0.002

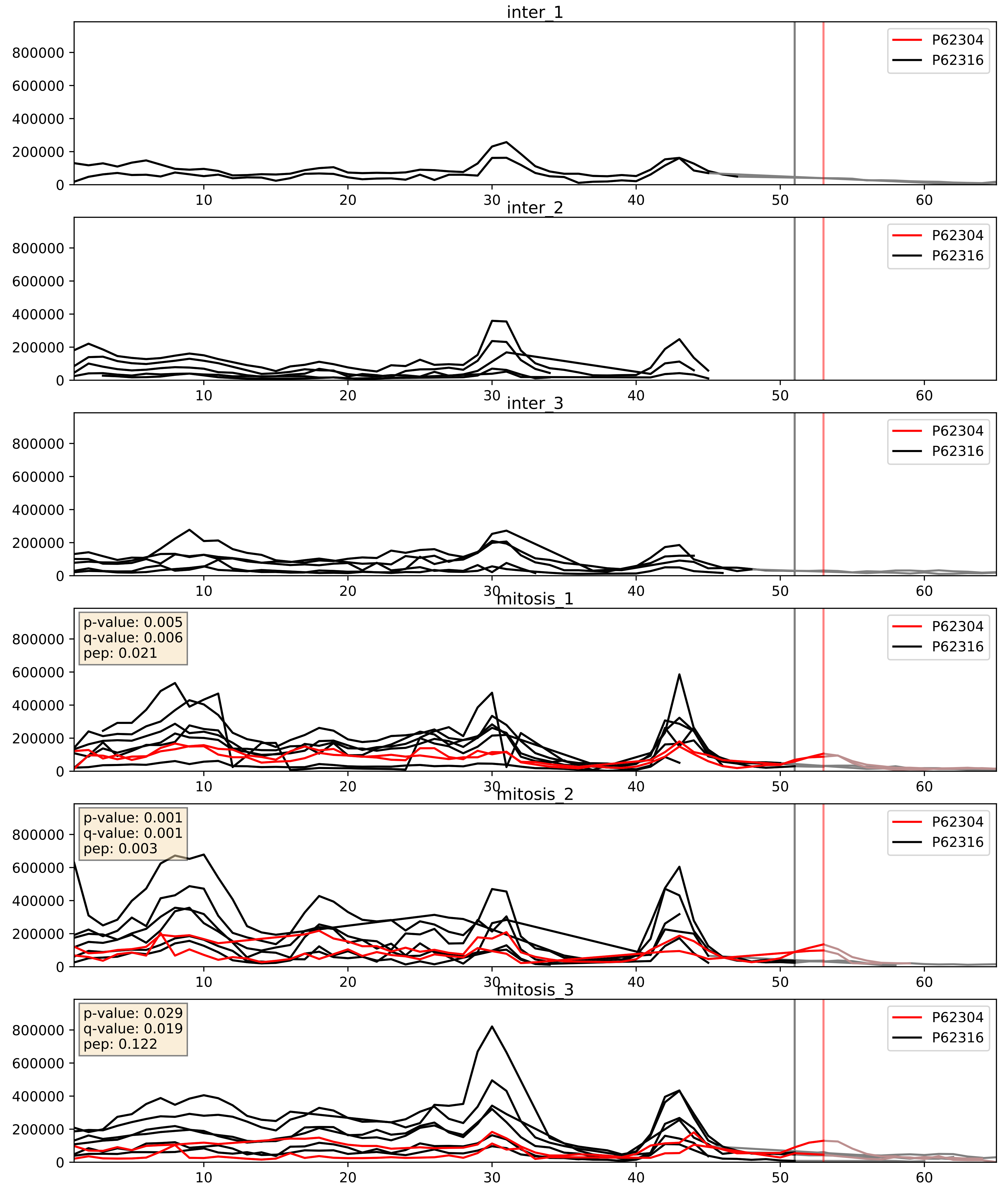


| level | condition_1 | condition_2 | pvalue | pvalue_adjusted |
|---|---|---|---|---|
| complex_abundance | mitosis | inter | 1.149781427818535e-05 | 0.005691857151652958 |
| interactor_abundance | mitosis | inter | 0.00011451049632638166 | 0.008168415404615225 |
| interactor_ratio | mitosis | inter | 0.001270295386270424 | 0.0423373831474418 |

P62304_P62318
RUXE_HUMAN vs SMD3_HUMAN
p-value: 0.002 q-value: 0.002

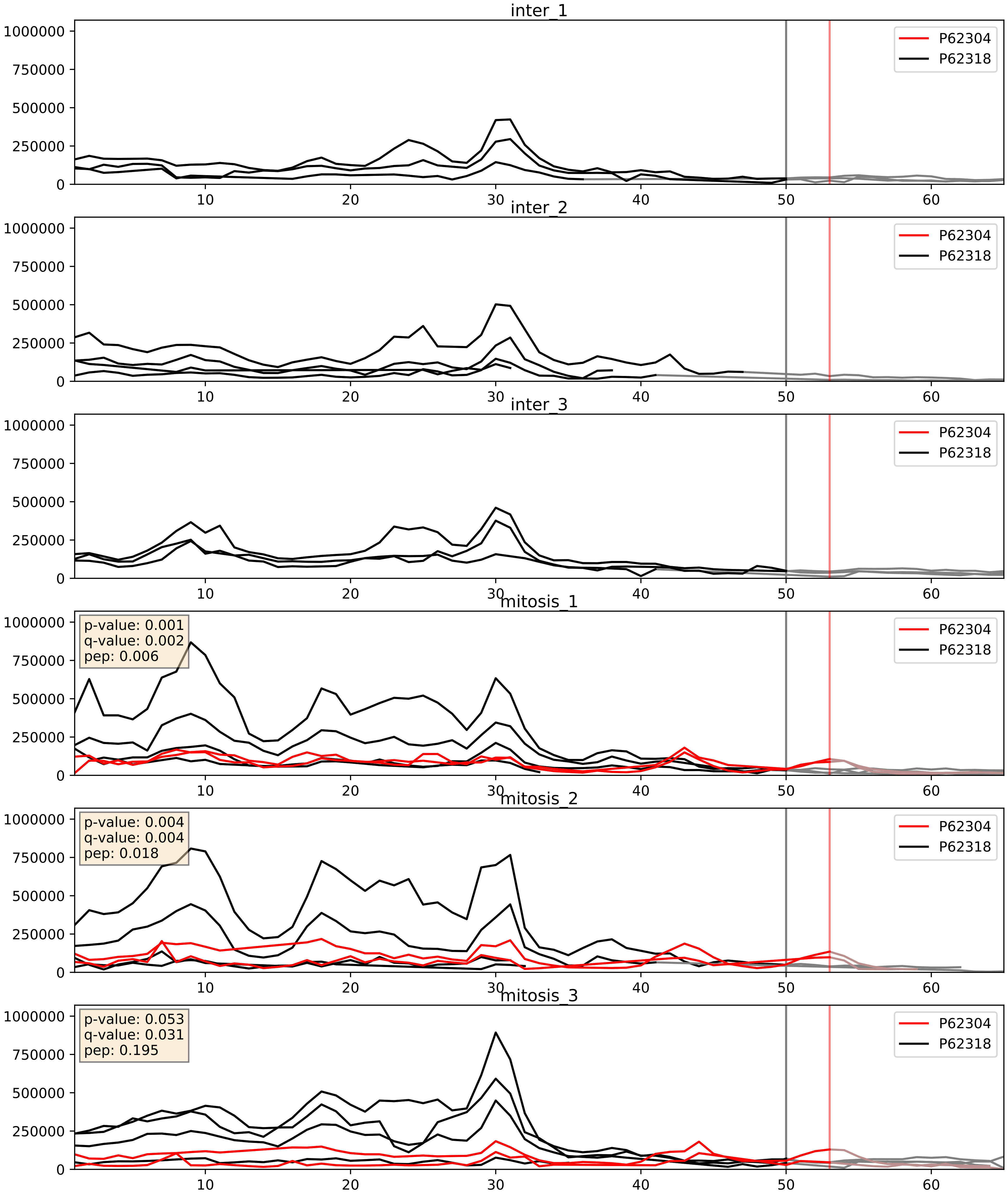


| level | condition_1 | condition_2 | pvalue | pvalue_adjusted |
|---|---|---|---|---|
| interactor_abundance | mitosis | inter | 0.00012182372888288902 | 0.008484493377800183 |
| complex_abundance | mitosis | inter | 0.0008863328576890988 | 0.019570164268135633 |
| interactor_ratio | mitosis | inter | 0.049675343900825265 | 0.3678381866704708 |

P62304_Q15717
RUXE_HUMAN vs ELAV1_HUMAN
p-value: 0.082 q-value: 0.048

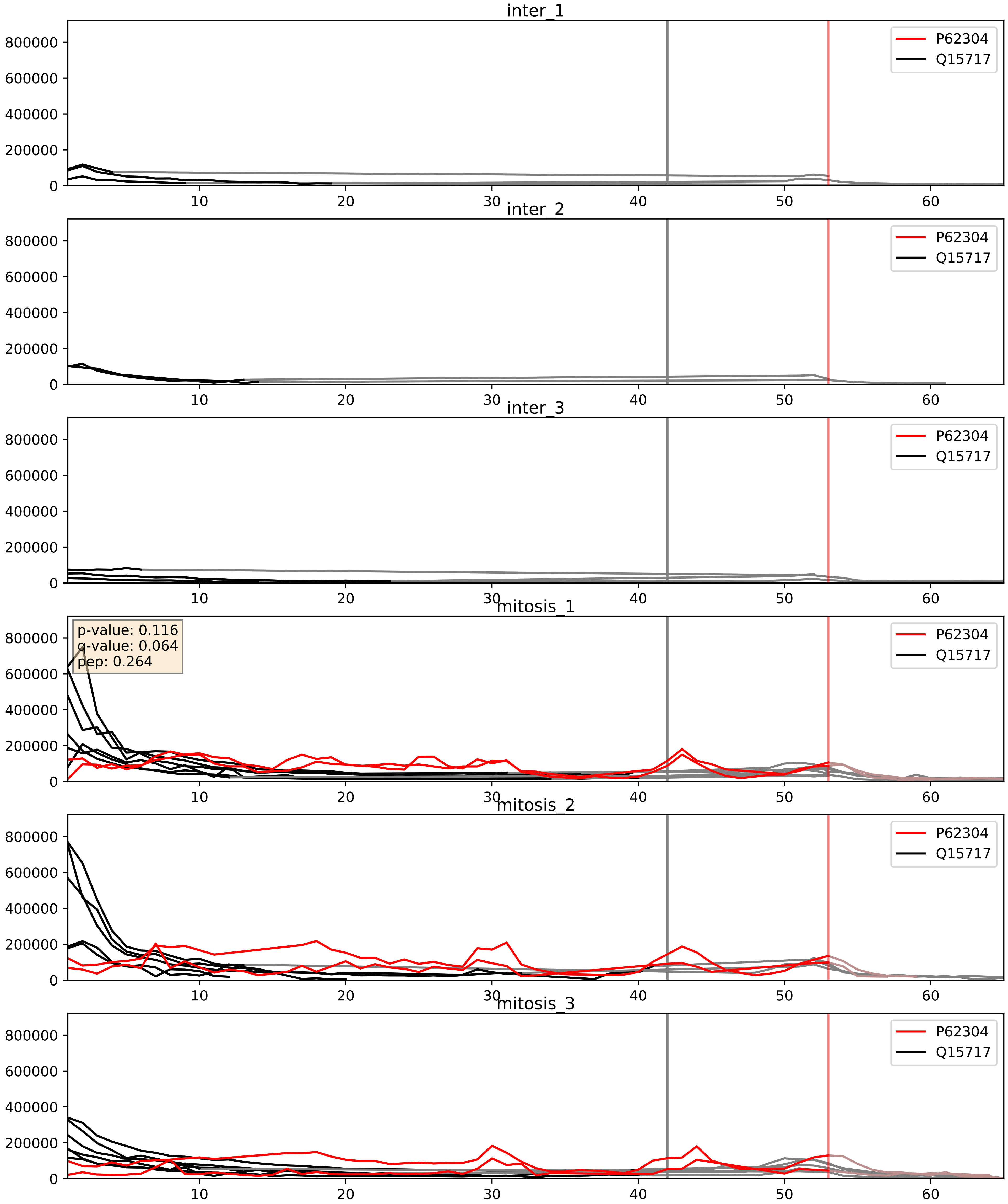


| level | condition_1 | condition_2 | pvalue | pvalue_adjusted |
|---|---|---|---|---|
| interactor_ratio | mitosis | inter | 1.3829541673086077e-05 | 0.003969306345409136 |
| complex_abundance | mitosis | inter | 9.823710518782713e-05 | 0.006719978526560179 |
| interactor_abundance | mitosis | inter | 0.00019797887174147875 | 0.01052608162799415 |
