## Supplementary Data 2 for "SECAT: Quantifying differential protein-protein interaction states by network-centric analysis": hela_string_000015_Q9H6R4.pdf

Q9H6R4_Q9H6R4

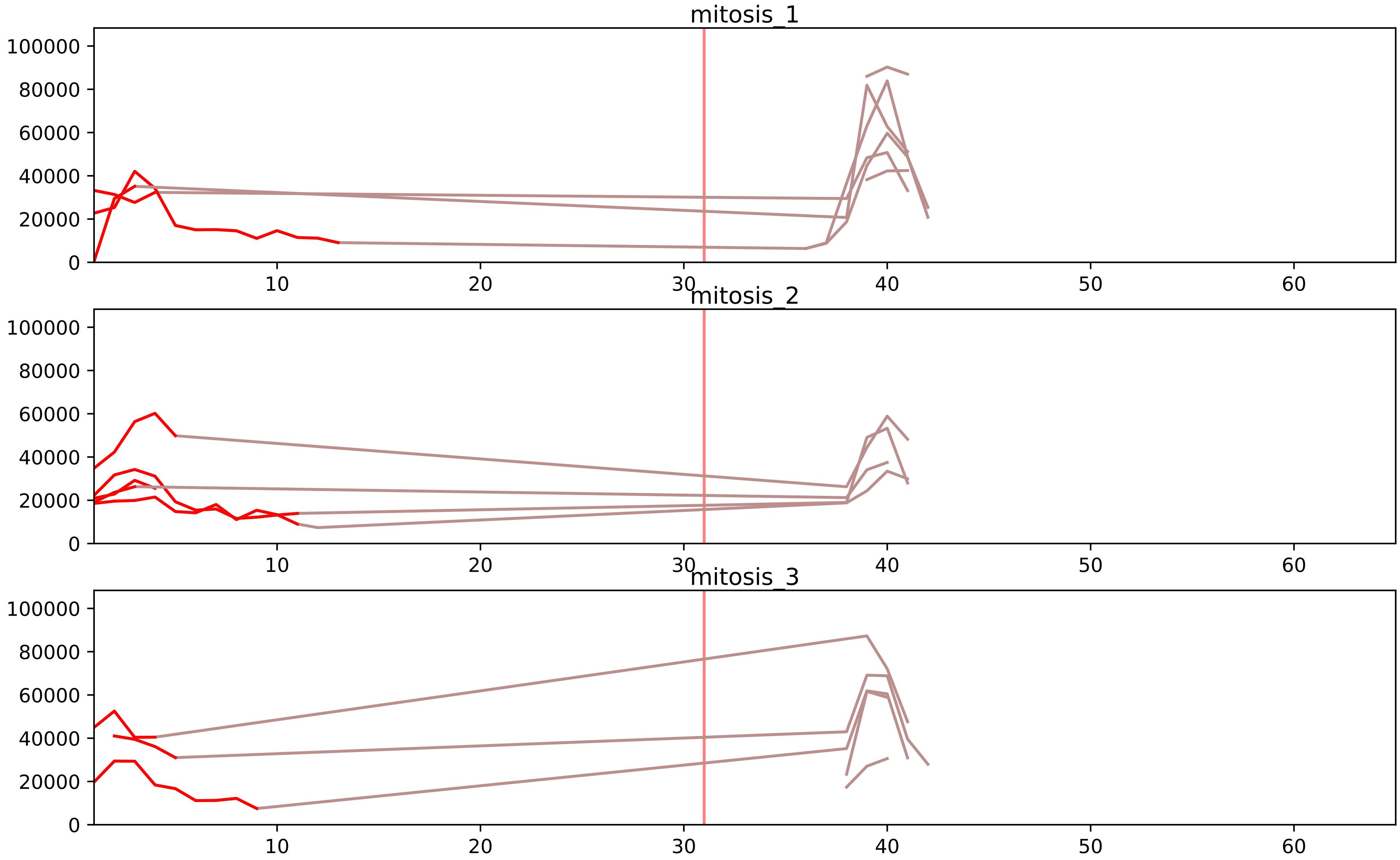


| level | condition_1 | condition_2 | pvalue | pvalue_adjusted |
|---|---|---|---|---|
| interactor_ratio | mitosis | inter | 2.384147980600694e-13 | 2.924554856203518e-11 |
| total_abundance | mitosis | inter | 0.002620454154985446 | 0.055355734867348155 |
| complex_abundance | mitosis | inter | 0.0029996901663464645 | 0.021730039000305096 |
| monomer_abundance | mitosis | inter | 0.003239869742306546 | 0.11999603554765999 |
| assembled_abundance | mitosis | inter | 0.0036925879753224 | 0.06015016024434401 |
| interactor_abundance | mitosis | inter | 0.6044237779369844 | 0.7419211150127094 |

O00567_Q9H6R4
NOP56_HUMAN vs NOL6_HUMAN
p-value: 0.045 q-value: 0.03

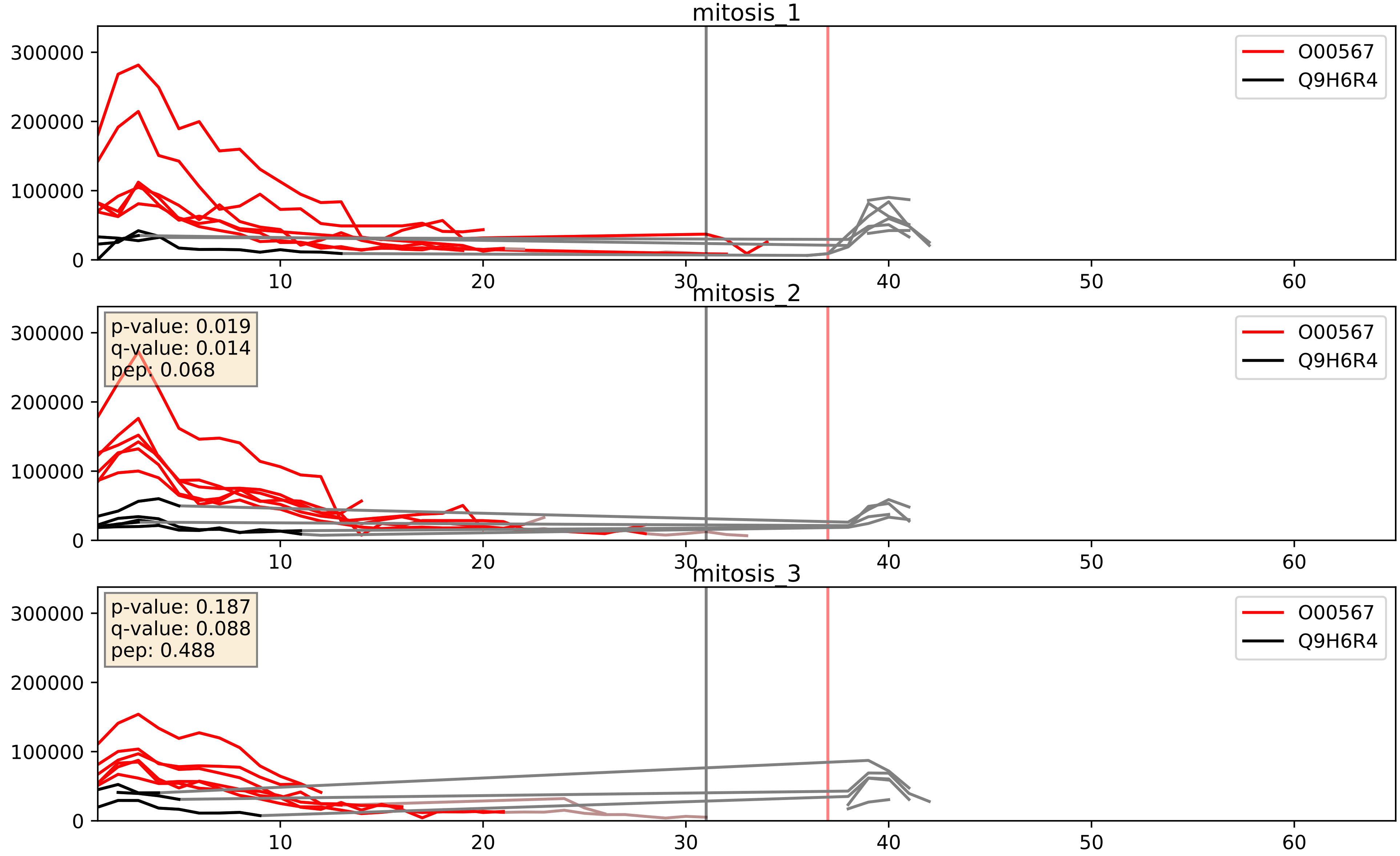


| level | condition_1 | condition_2 | pvalue | pvalue_adjusted |
|---|---|---|---|---|
| interactor_ratio | mitosis | inter | 0.0013886285934761222 | 0.044212523606194314 |
| interactor_abundance | mitosis | inter | 0.0019094235057377626 | 0.031986606092438766 |
| complex_abundance | mitosis | inter | 0.006619064498898424 | 0.06002696841014766 |

O76021_Q9H6R4
RL1D1_HUMAN vs NOL6_HUMAN
p-value: 0.02 q-value: 0.016

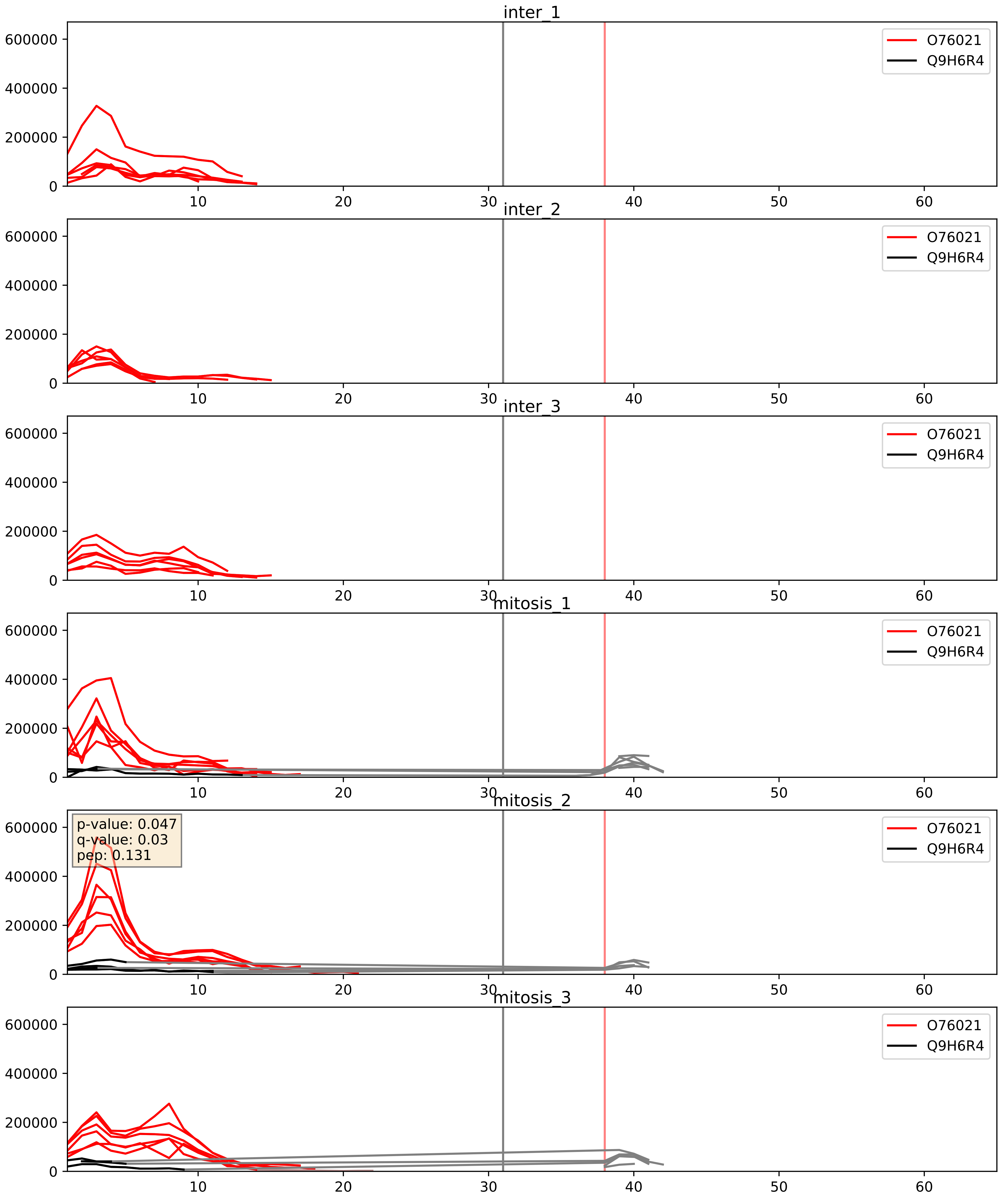


| level | condition_1 | condition_2 | pvalue | pvalue_adjusted |
|---|---|---|---|---|
| interactor_abundance | mitosis | inter | 6.462019261636246e-05 | 0.0065503332782105754 |
| interactor_ratio | mitosis | inter | 0.0002037188630223042 | 0.014683183137172115 |
| complex_abundance | mitosis | inter | 0.0007402222580168641 | 0.017798602608495383 |

P46087_Q9H6R4
NOP2_HUMAN vs NOL6_HUMAN
p-value: 0.06 q-value: 0.038

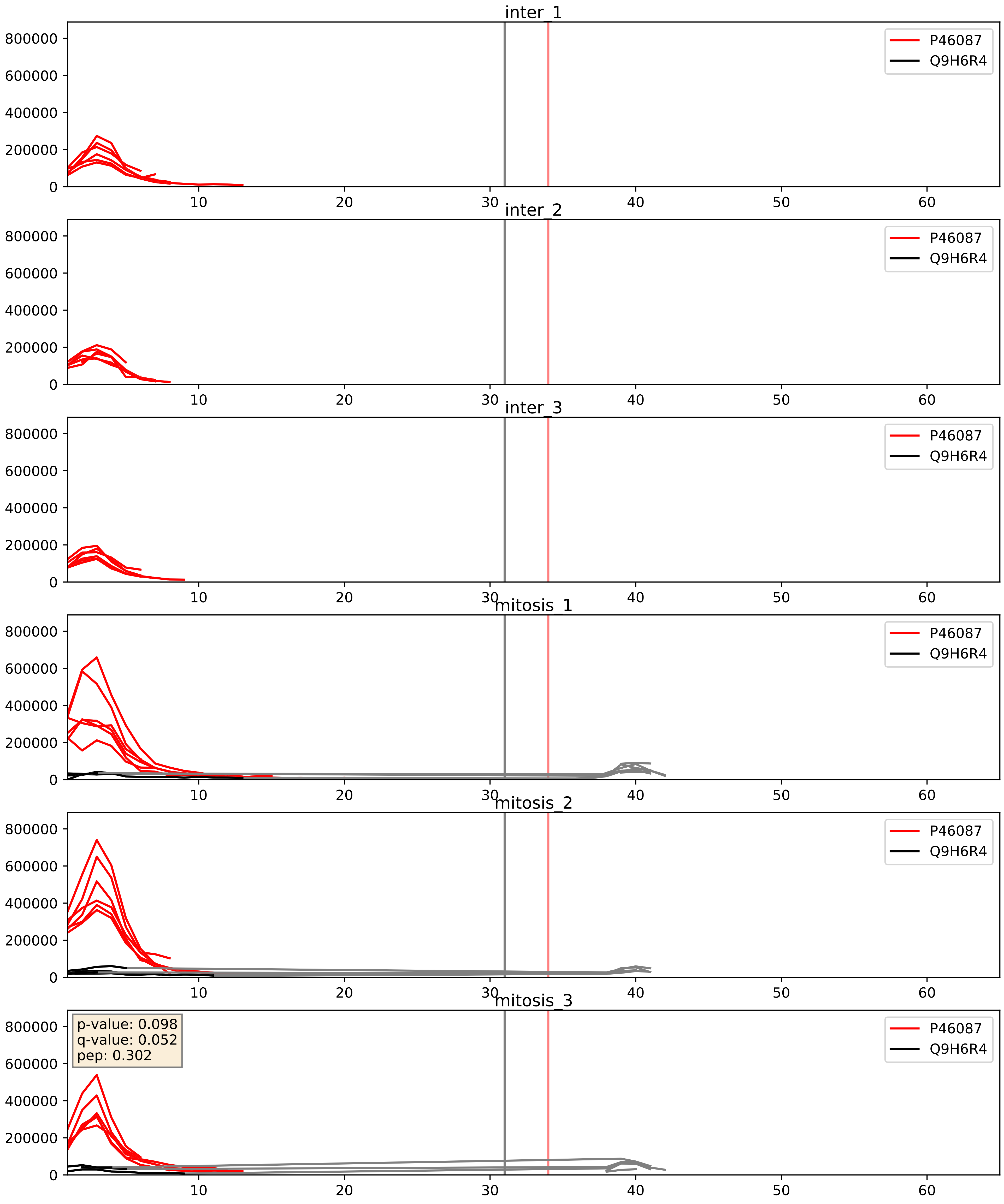


| level | condition_1 | condition_2 | pvalue | pvalue_adjusted |
|---|---|---|---|---|
| interactor_abundance | mitosis | inter | 0.00019761530192014963 | 0.01052608162799415 |
| interactor_ratio | mitosis | inter | 0.00027831638304536276 | 0.017163005662466667 |
| complex_abundance | mitosis | inter | 0.000983162917257506 | 0.02083137270228775 |

Q15269_Q9H6R4
PWP2_HUMAN vs NOL6_HUMAN
p-value: 0.028 q-value: 0.021

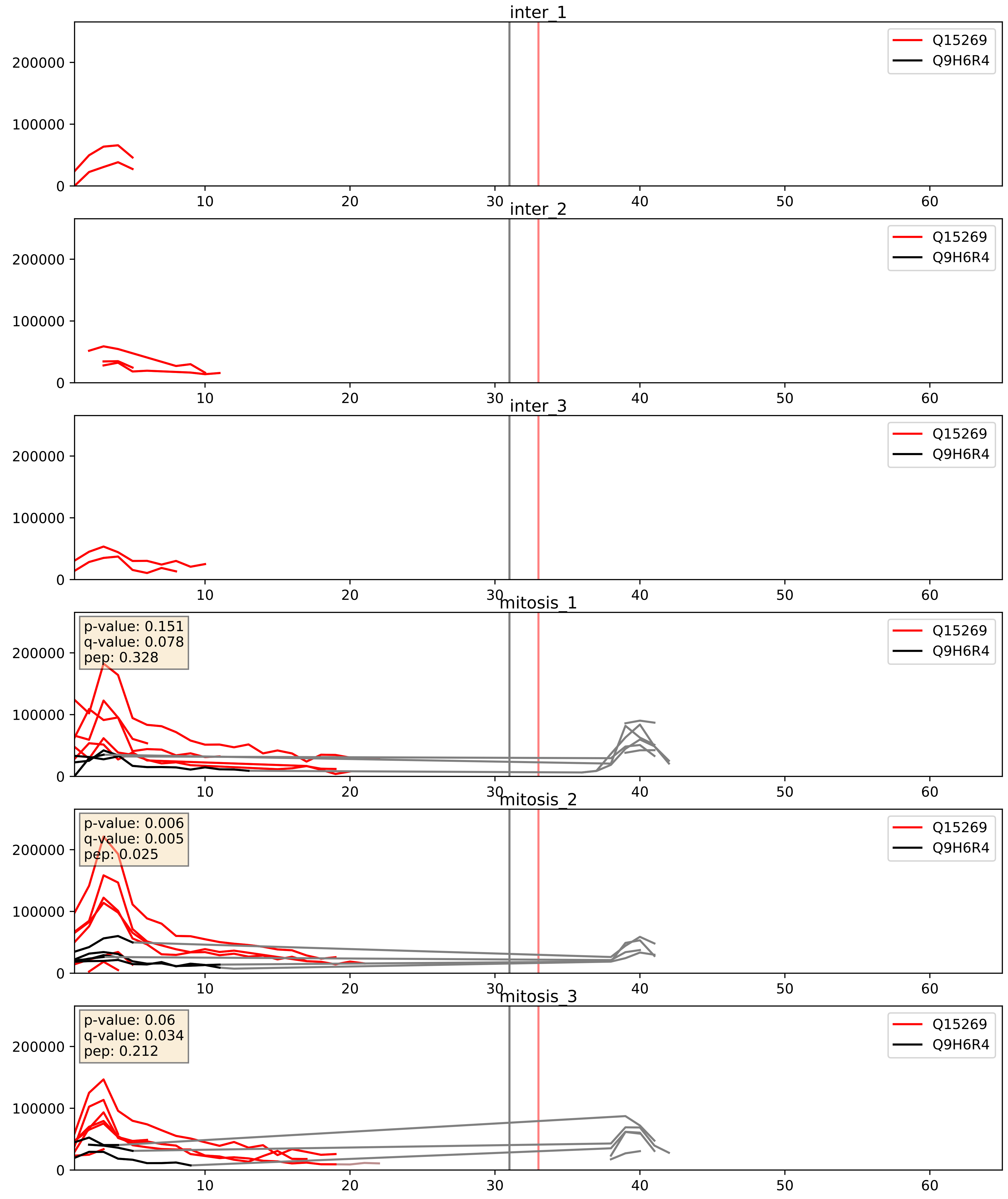


| level | condition_1 | condition_2 | pvalue | pvalue_adjusted |
|---|---|---|---|---|
| interactor_abundance | mitosis | inter | 0.003100387467477749 | 0.04271207144514066 |
| interactor_ratio | mitosis | inter | 0.004656516393809501 | 0.09159088101537749 |
| complex_abundance | mitosis | inter | 0.012416401996518596 | 0.08277281176263881 |

Q8IY81_Q9H6R4
SPB1_HUMAN vs NOL6_HUMAN
p-value: 0.063 q-value: 0.039

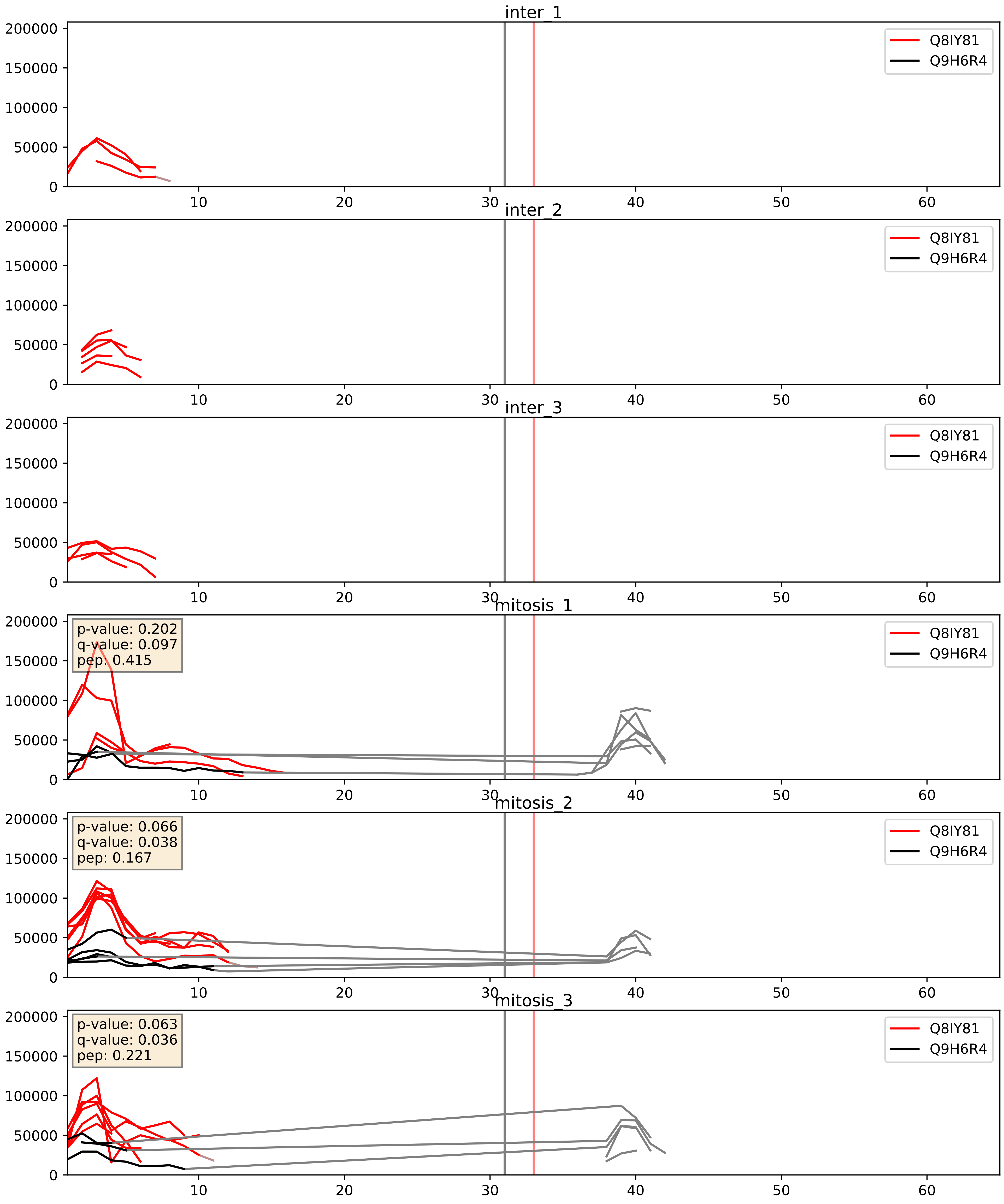


| level | condition_1 | condition_2 | pvalue | pvalue_adjusted |
|---|---|---|---|---|
| interactor_abundance | mitosis | inter | 0.006105255729208133 | 0.0666372269553884 |
| complex_abundance | mitosis | inter | 0.015707262428466266 | 0.09352166546417016 |
| interactor_ratio | mitosis | inter | 0.026384193828163788 | 0.25543205403291264 |

Q8NI36_Q9H6R4
WDR36_HUMAN vs NOL6_HUMAN
p-value: 0.032 q-value: 0.023

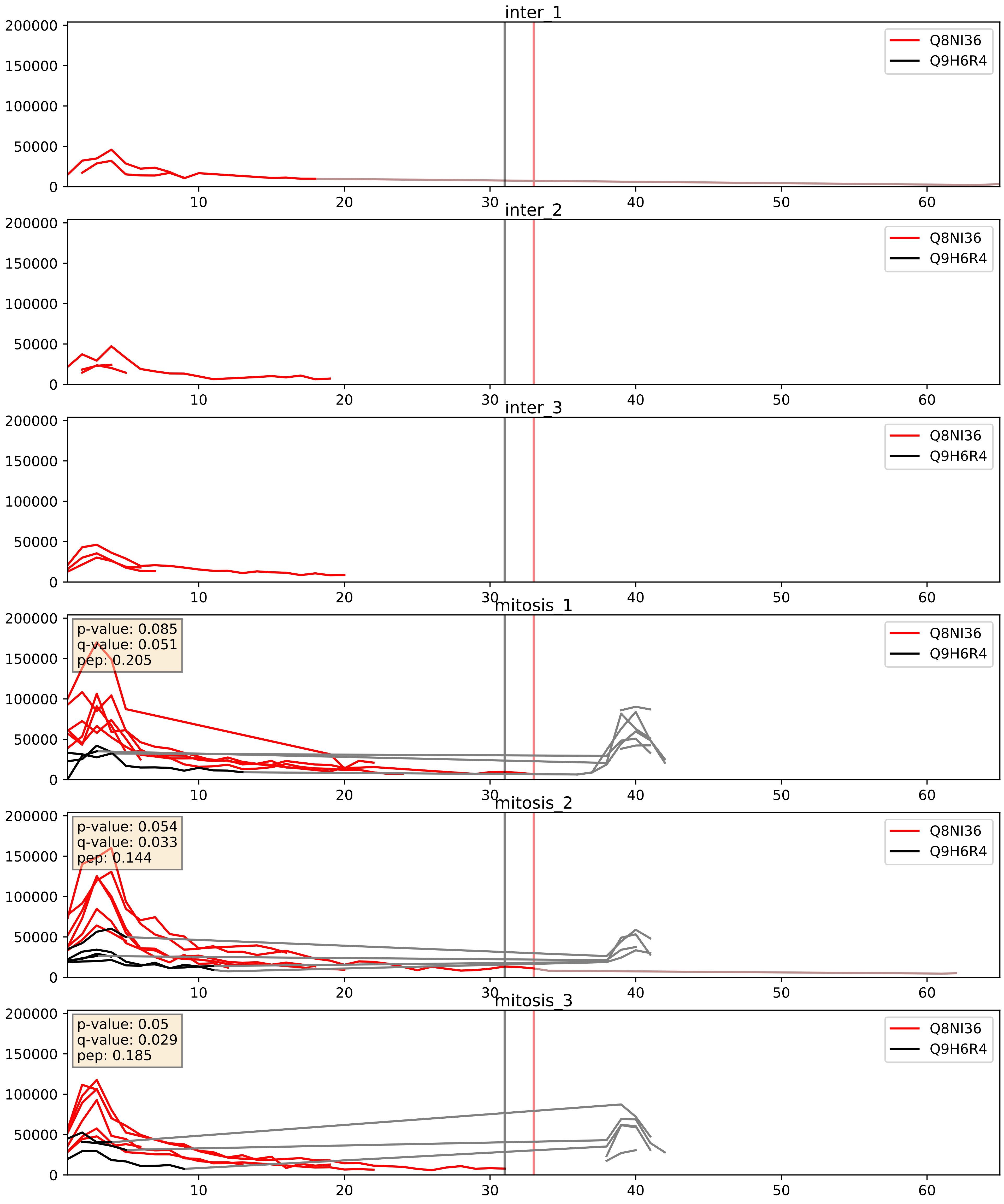


| level | condition_1 | condition_2 | pvalue | pvalue_adjusted |
|---|---|---|---|---|
| interactor_ratio | mitosis | inter | 0.0015565326714841756 | 0.04758542738537337 |
| interactor_abundance | mitosis | inter | 0.0022775002900272527 | 0.03523946087610491 |
| complex_abundance | mitosis | inter | 0.01595418682162696 | 0.09419630360707916 |

Q9H0A0_Q9H6R4
NAT10_HUMAN vs NOL6_HUMAN
p-value: 0.029 q-value: 0.021

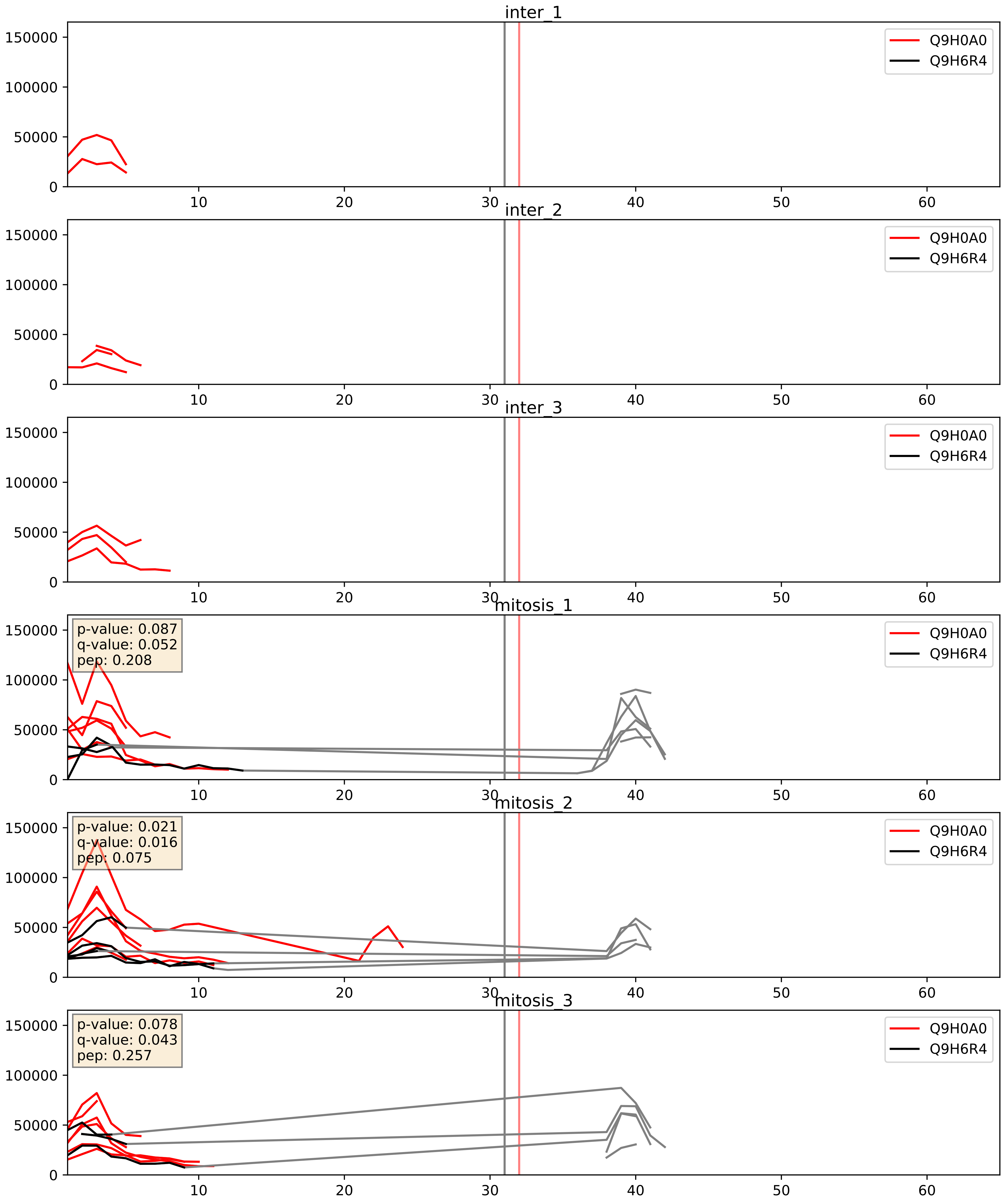


| level | condition_1 | condition_2 | pvalue | pvalue_adjusted |
|---|---|---|---|---|
| interactor_abundance | mitosis | inter | 0.025428031496039612 | 0.13211772358488563 |
| interactor_ratio | mitosis | inter | 0.0271500034485924 | 0.2577730310940899 |
| complex_abundance | mitosis | inter | 0.06262730312099983 | 0.18019822343386843 |

Q9H6R4_Q9NVP1
NOL6_HUMAN vs DDX18_HUMAN
p-value: 0.074 q-value: 0.045

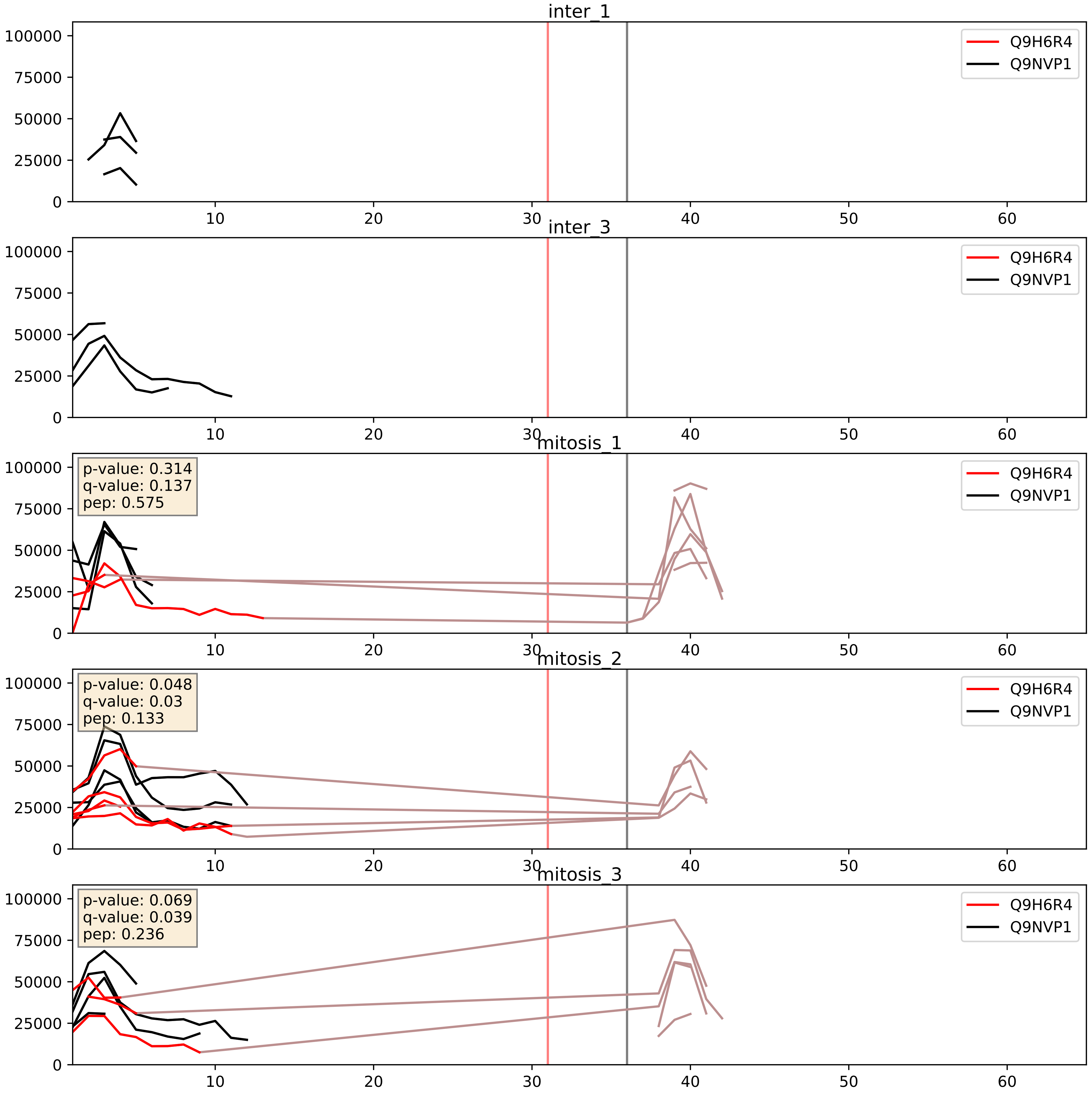


| level | condition_1 | condition_2 | pvalue | pvalue_adjusted |
|---|---|---|---|---|
| interactor_ratio | mitosis | inter | 0.009838253155822414 | 0.14263182888175546 |
| complex_abundance | mitosis | inter | 0.06470209978002295 | 0.1836981672029839 |
| interactor_abundance | mitosis | inter | 0.5002996189915532 | 0.6656579265768231 |

Q9H6R4_Q9NW13
NOL6_HUMAN vs RBM28_HUMAN
p-value: 0.055 q-value: 0.035

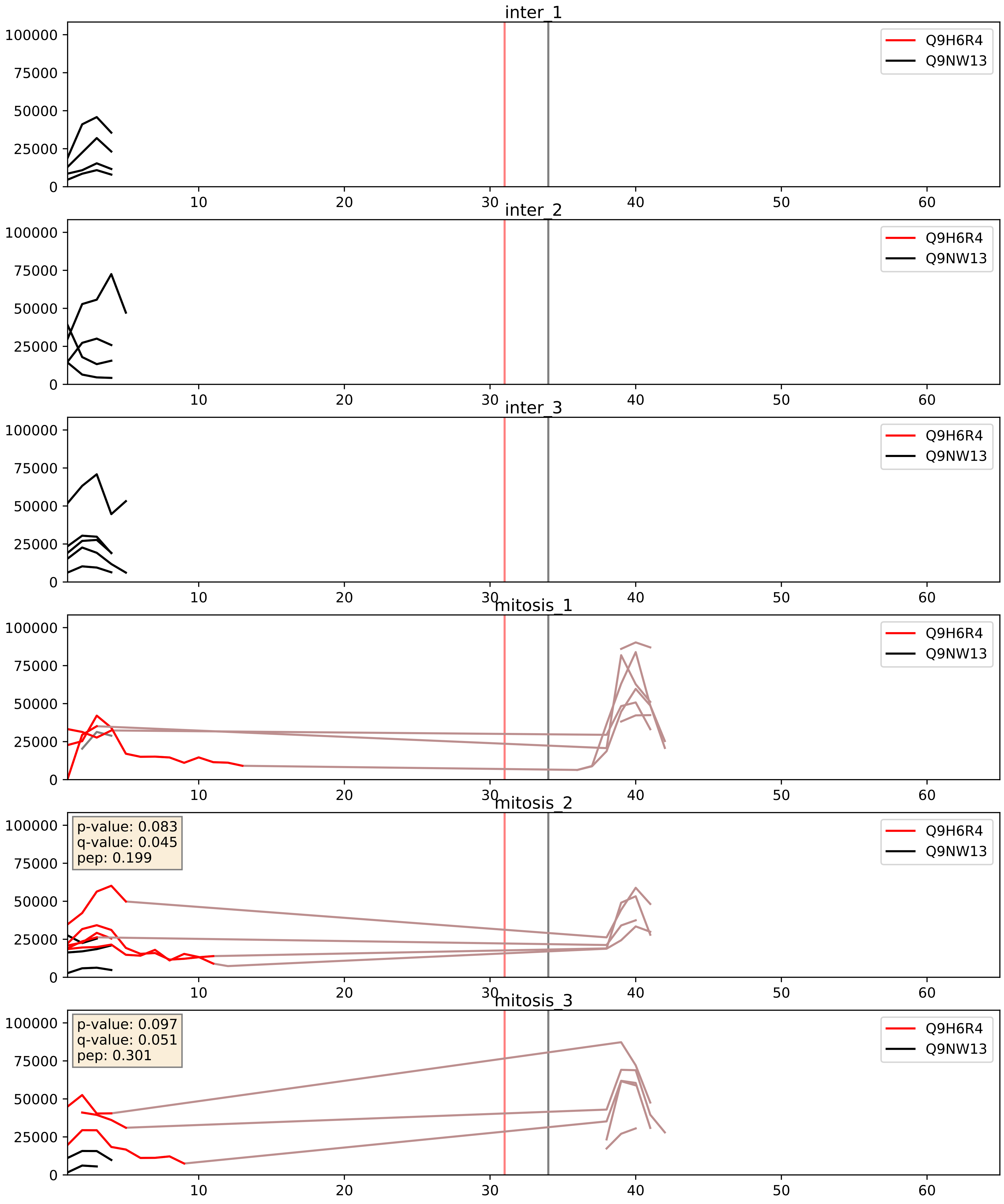


| level | condition_1 | condition_2 | pvalue | pvalue_adjusted |
|---|---|---|---|---|
| complex_abundance | mitosis | inter | 0.4550360875637394 | 0.5969913112594304 |
| interactor_abundance | mitosis | inter | 0.5274626077760354 | 0.6872987496111436 |
| interactor_ratio | mitosis | inter | 0.6699802415148992 | 0.9304647856701197 |

Q9H6R4_Q9UNX4
NOL6_HUMAN vs WDR3_HUMAN
p-value: 0.053 q-value: 0.034

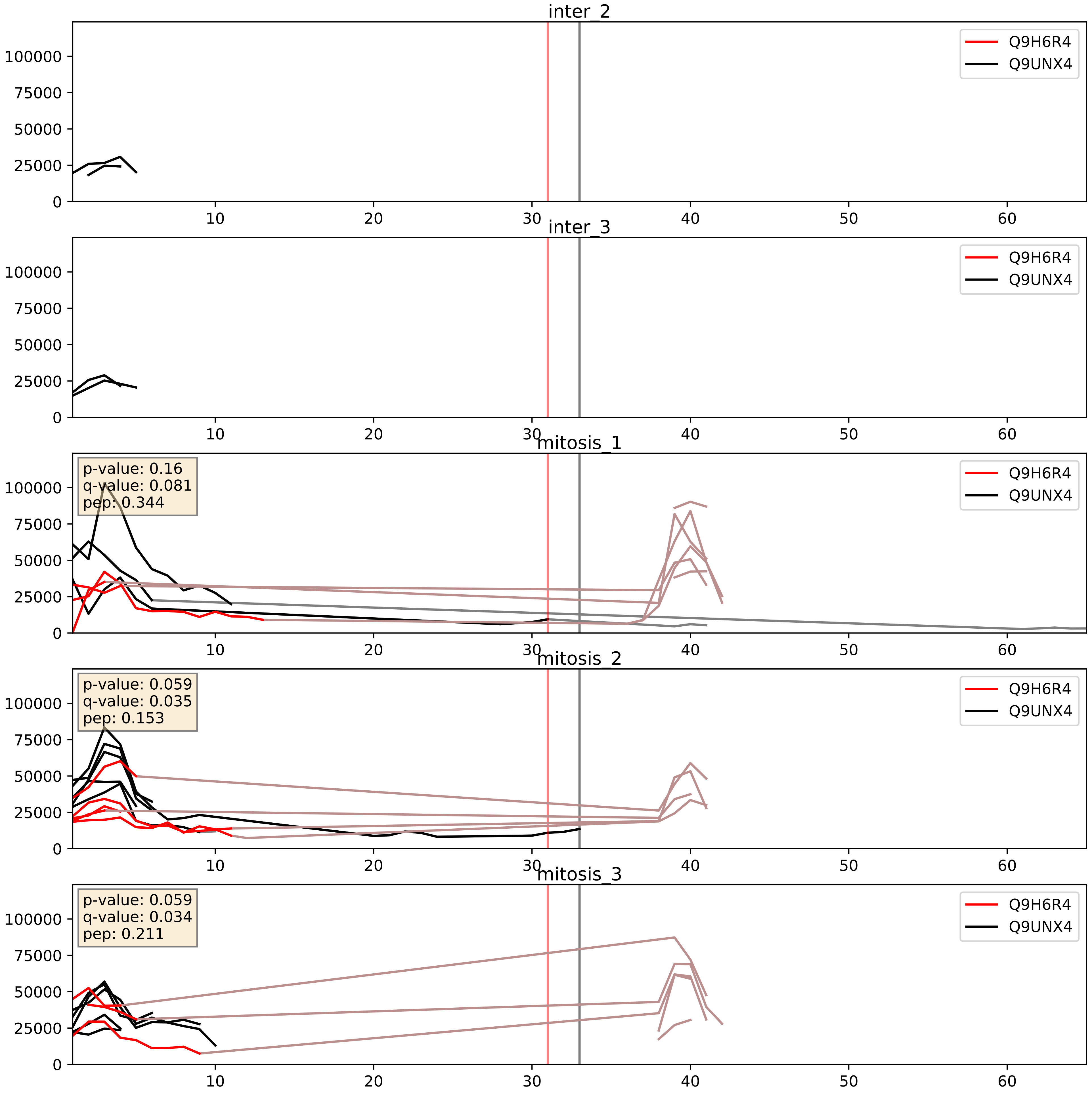


| level | condition_1 | condition_2 | pvalue | pvalue_adjusted |
|---|---|---|---|---|
| interactor_ratio | mitosis | inter | 0.002692662930541276 | 0.06718455808461757 |
| complex_abundance | mitosis | inter | 0.09645403910829943 | 0.23018381938539362 |
| interactor_abundance | mitosis | inter | 0.5045521248563168 | 0.6689848495616593 |

Q9H6R4_Q9Y221
NOL6_HUMAN vs NIP7_HUMAN
p-value: 0.071 q-value: 0.043

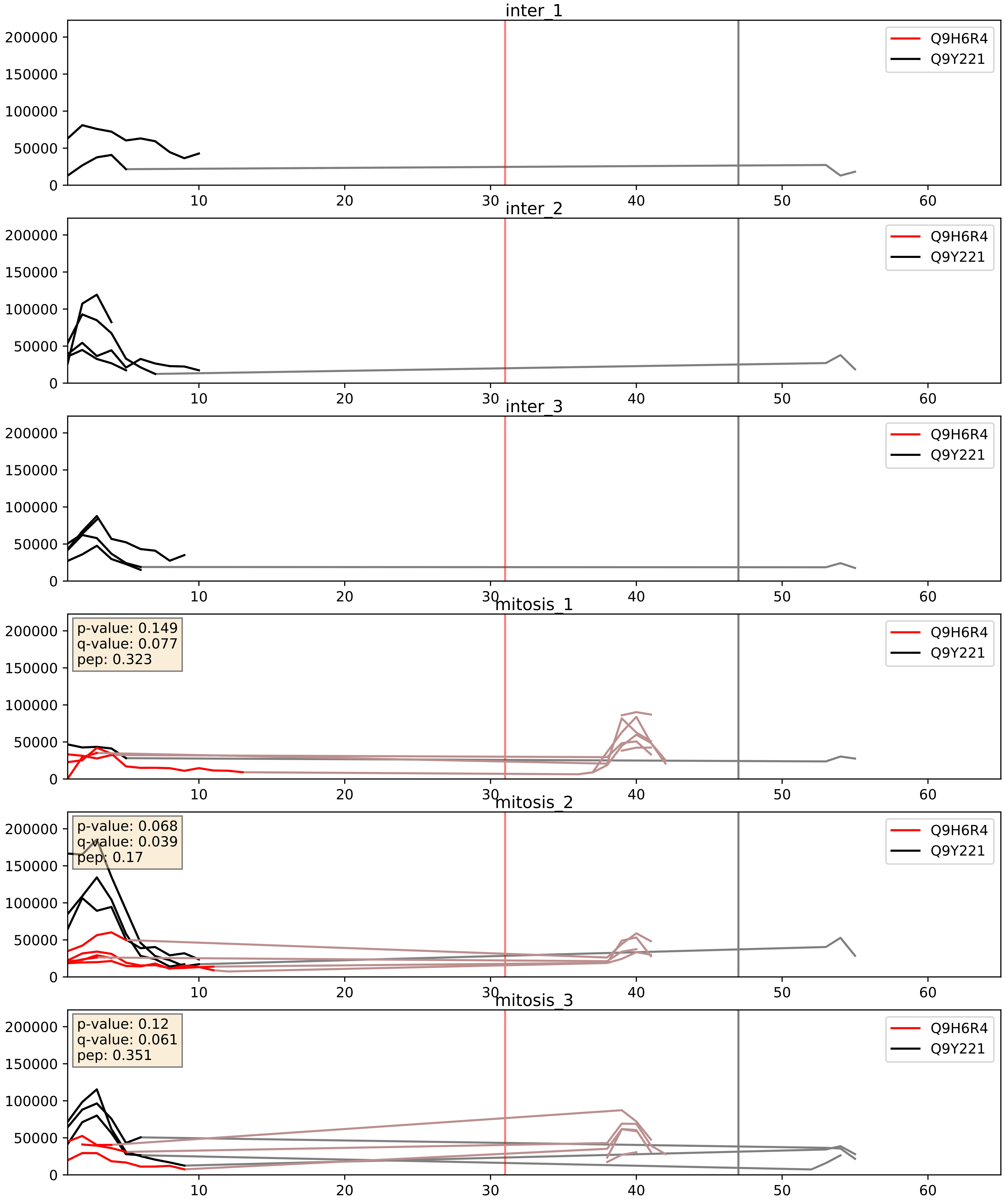


| level | condition_1 | condition_2 | pvalue | pvalue_adjusted |
|---|---|---|---|---|
| complex_abundance | mitosis | inter | 0.07398502436357436 | 0.19784675062063986 |
| interactor_ratio | mitosis | inter | 0.24988597224484135 | 0.7241110096194455 |
| interactor_abundance | mitosis | inter | 0.5249344519496493 | 0.6856499350875437 |

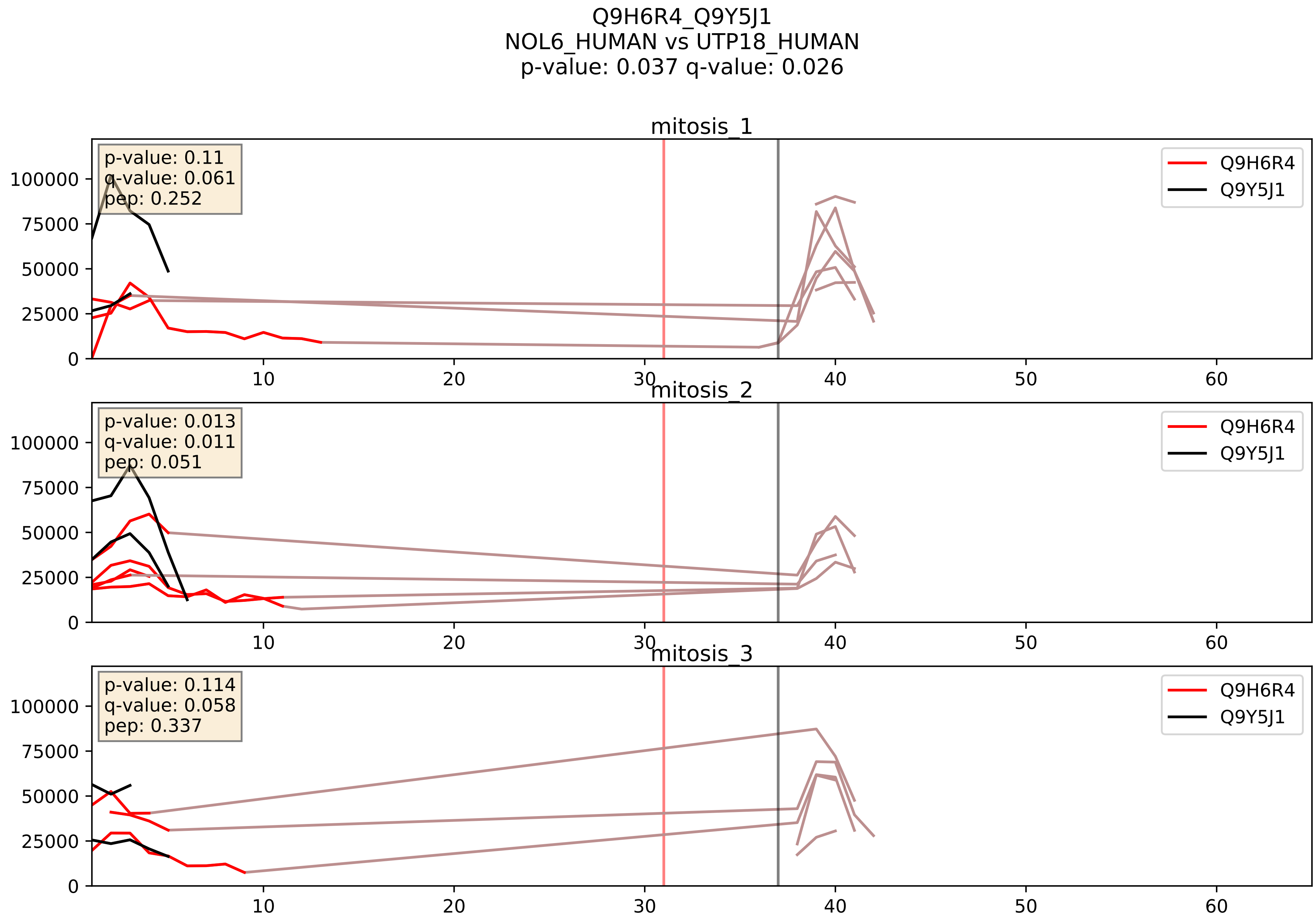


| level | condition_1 | condition_2 | pvalue | pvalue_adjusted |
|---|---|---|---|---|
| interactor_ratio | mitosis | inter | 0.2463865098672941 | 0.7216070053502578 |
| complex_abundance | mitosis | inter | 0.30028671272589114 | 0.4500094994631702 |
| interactor_abundance | mitosis | inter | 0.6347233985504436 | 0.7573495751476084 |
