## Supplementary Data 2 for "SECAT: Quantifying differential protein-protein interaction states by network-centric analysis": hela_string_000016_P52272.pdf

P52272_P52272

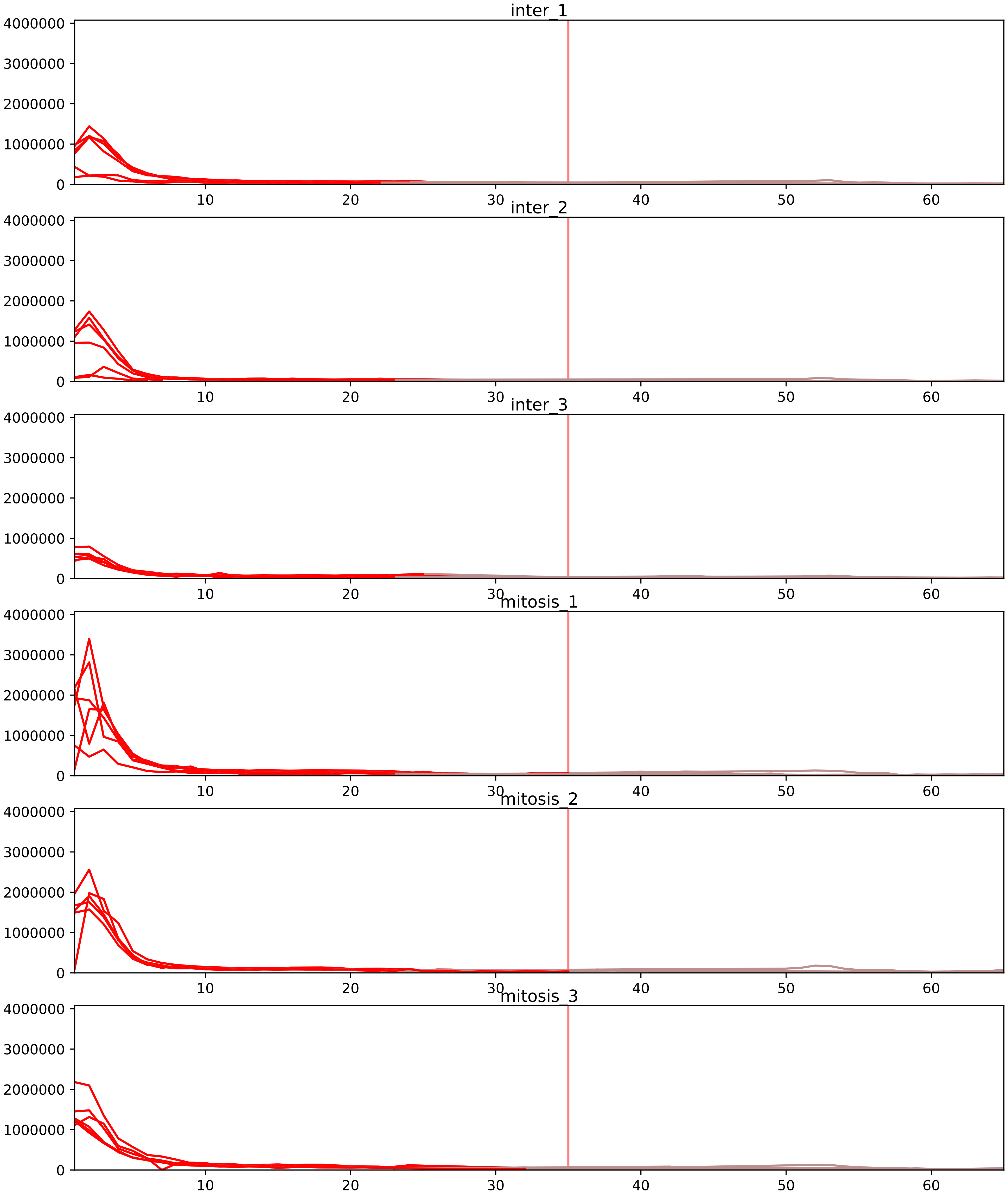


| level | condition_1 | condition_2 | pvalue | pvalue_adjusted |
|---|---|---|---|---|
| interactor_ratio | mitosis | inter | 1.126611005203759e-12 | 1.295602655984323e-10 |
| complex_abundance | mitosis | inter | 2.8786254593558792e-05 | 0.0010185905471566958 |
| monomer_abundance | mitosis | inter | 0.003257404115552365 | 0.11999603554765999 |
| interactor_abundance | mitosis | inter | 0.0222012463123562 | 0.0841051214605524 |
| total_abundance | mitosis | inter | 0.03643158582431479 | 0.23901186009225078 |
| assembled_abundance | mitosis | inter | 0.04885427995723825 | 0.2625593533374663 |

O43390_P52272
HNRPR_HUMAN vs HNRPM_HUMAN
p-value: 0.058 q-value: 0.037

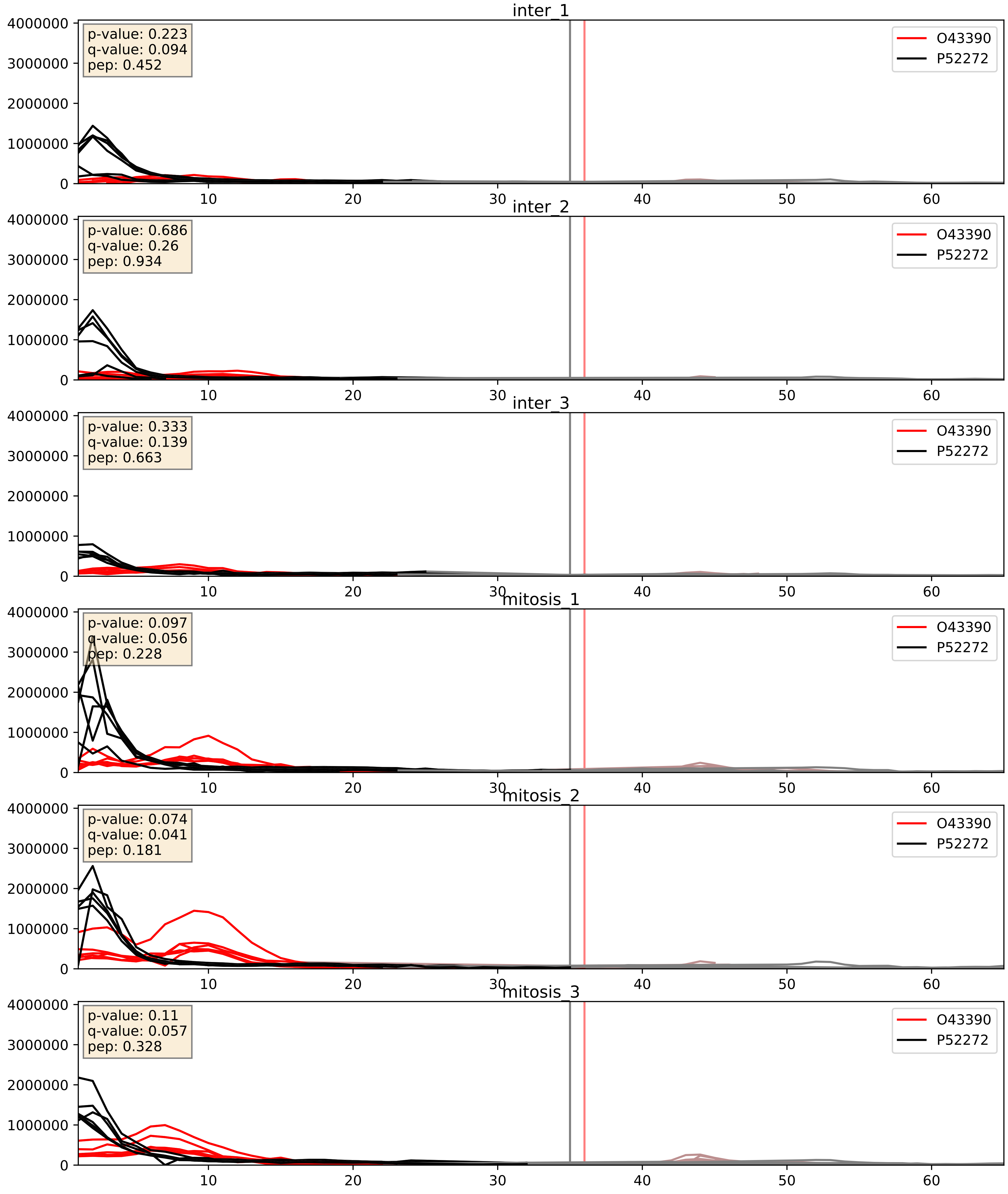


| level | condition_1 | condition_2 | pvalue | pvalue_adjusted |
| --- | --- | --- | --- | --- |
| interactor_abundance | mitosis | inter | 6.140823405978327e-05 | 0.006459477818931243 |
| complex_abundance | mitosis | inter | 0.0025869915277018023 | 0.03664546283589145 |
| interactor_ratio | mitosis | inter | 0.04696217208361891 | 0.35838626305591875 |

P07910_P52272
HNRPC_HUMAN vs HNRPM_HUMAN
p-value: 0.001 q-value: 0.002

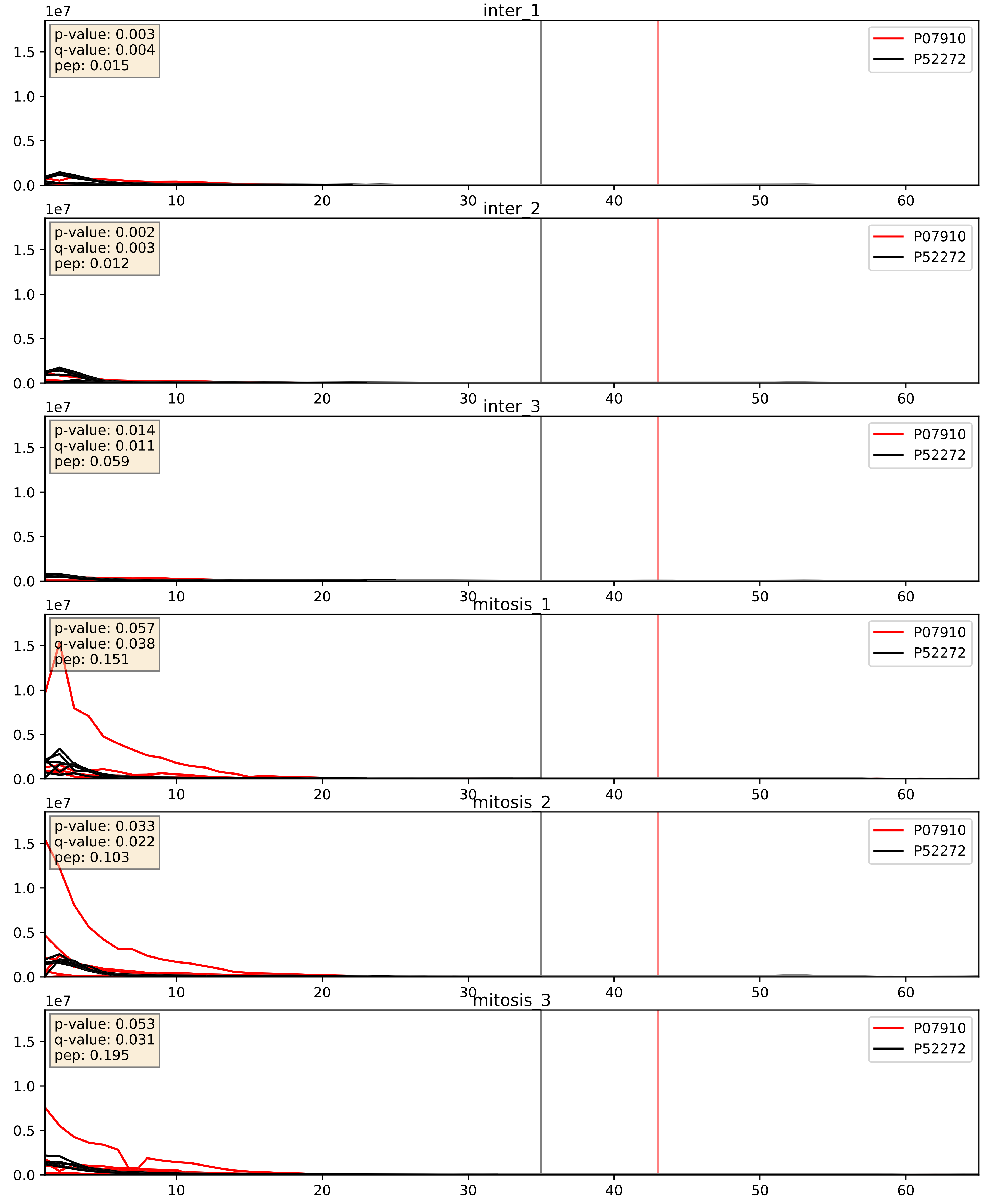


| level | condition_1 | condition_2 | pvalue | pvalue_adjusted |
|---|---|---|---|---|
| interactor_abundance | mitosis | inter | 9.77406216392995e-06 | 0.0033734327106189023 |
| interactor_ratio | mitosis | inter | 0.00022697512149341483 | 0.01529848062979237 |
| complex_abundance | mitosis | inter | 0.000330922405409354 | 0.011886635542316795 |

P09651_P52272
ROA1_HUMAN vs HNRPM_HUMAN
p-value: 0.0 q-value: 0.001

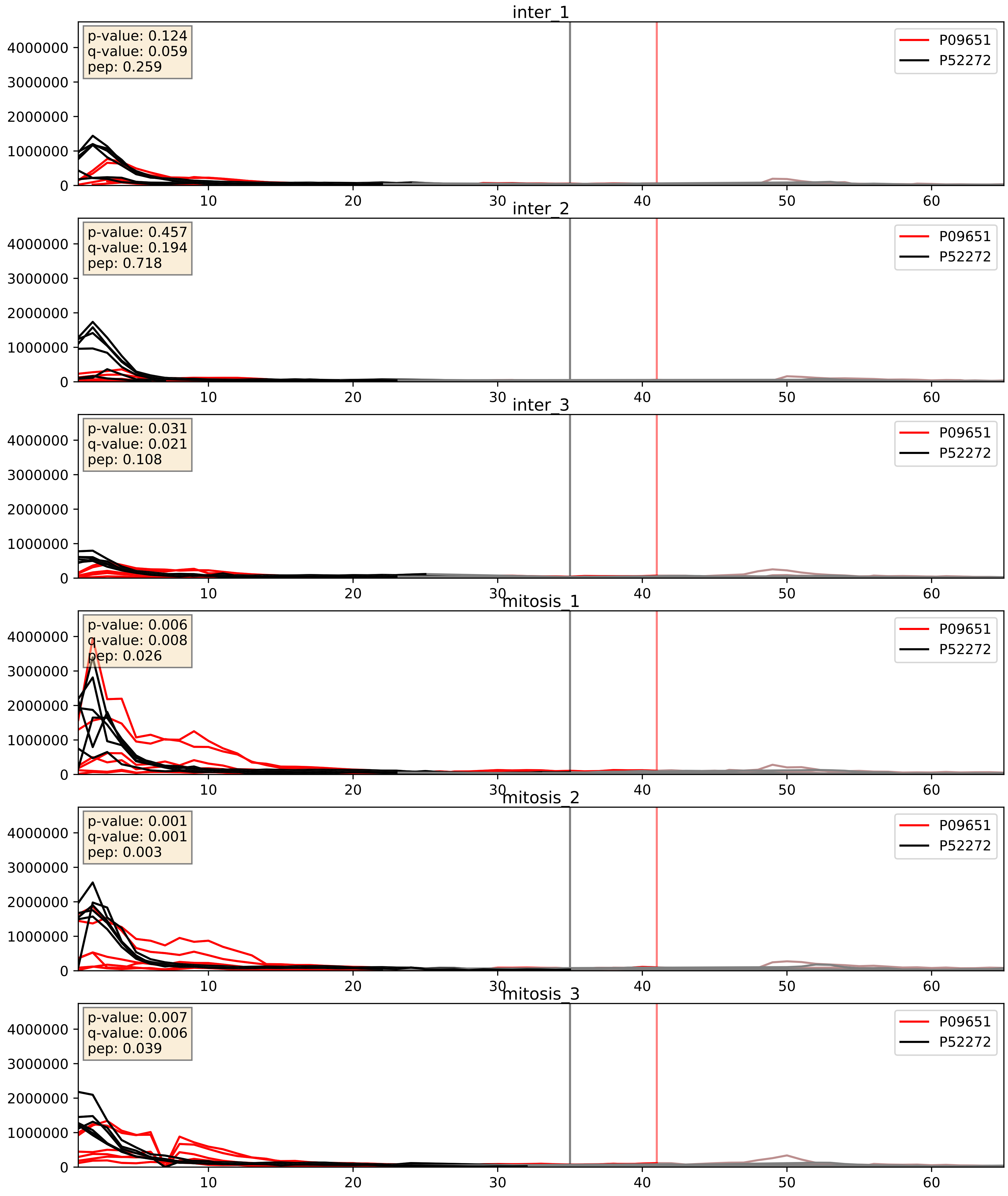


| level | condition_1 | condition_2 | pvalue | pvalue_adjusted |
|---|---|---|---|---|
| interactor_abundance | mitosis | inter | 0.01476418936640577 | 0.1063369465514795 |
| complex_abundance | mitosis | inter | 0.016581499584325202 | 0.09584596180627945 |
| interactor_ratio | mitosis | inter | 0.04323656308648082 | 0.34205635861393335 |

P14866_P52272
HNRPL_HUMAN vs HNRPM_HUMAN
p-value: 0.01 q-value: 0.009

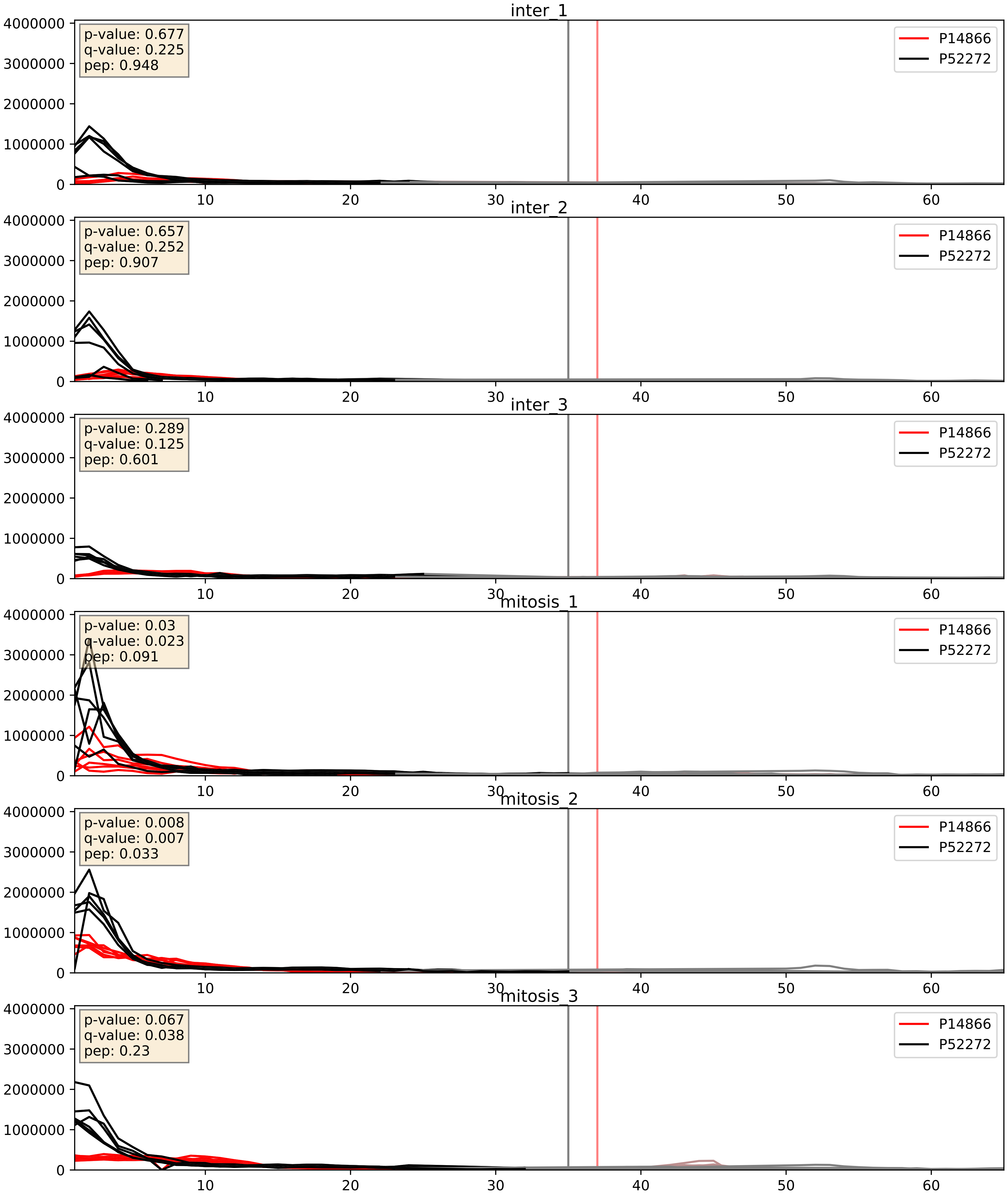


| level | condition_1 | condition_2 | pvalue | pvalue_adjusted |
|---|---|---|---|---|
| complex_abundance | mitosis | inter | 0.00290128716918596 | 0.03923383596877065 |
| interactor_abundance | mitosis | inter | 0.00986072689417613 | 0.08626246521629807 |
| interactor_ratio | mitosis | inter | 0.16723764333693136 | 0.6293444626080091 |

P17844_P52272
DDX5_HUMAN vs HNRPM_HUMAN
p-value: 0.0 q-value: 0.005

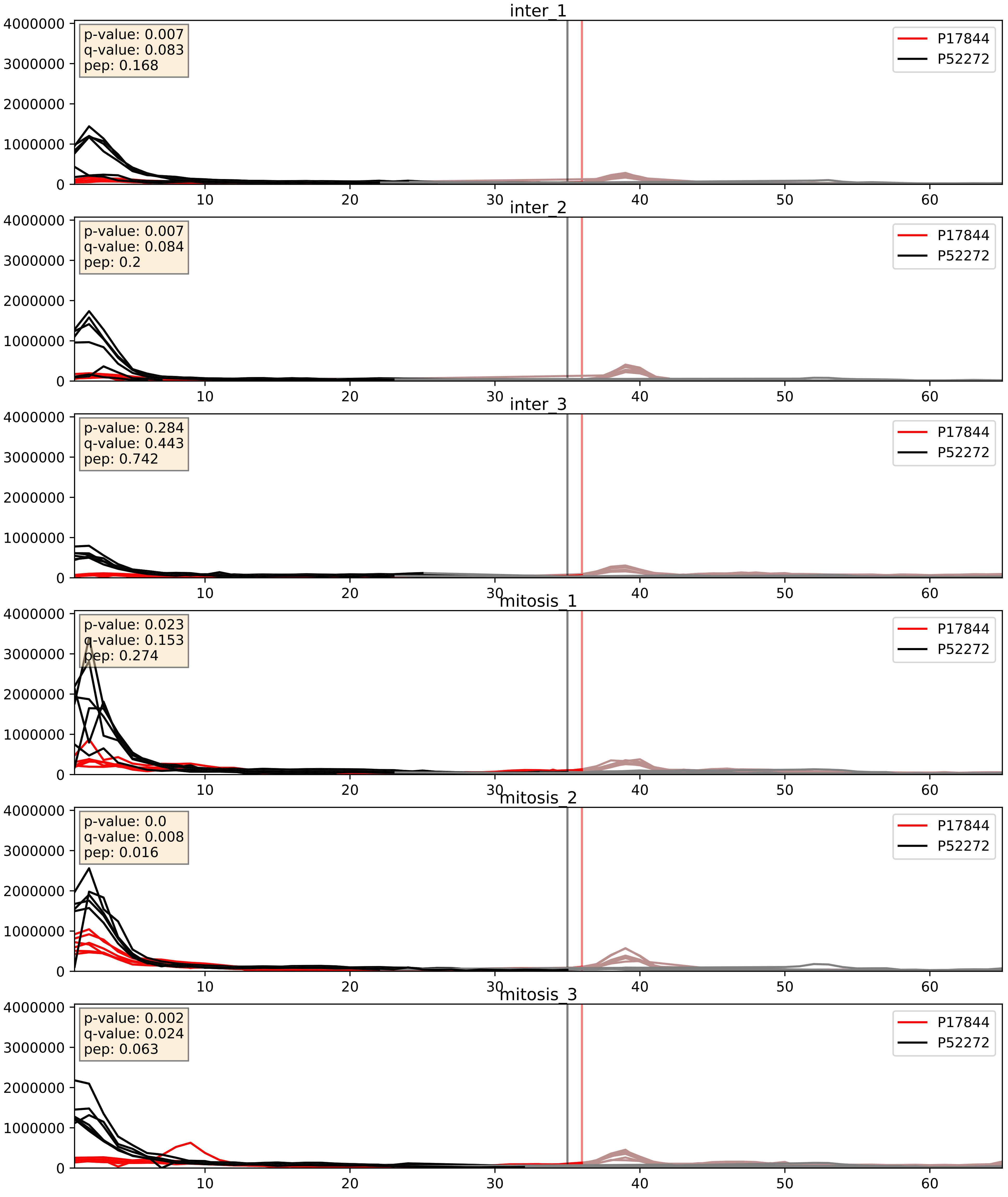


| level | condition_1 | condition_2 | pvalue | pvalue_adjusted |
|---|---|---|---|---|
| interactor_abundance | mitosis | inter | 0.0008880993781604356 | 0.02197564842852207 |
| complex_abundance | mitosis | inter | 0.003908349803170464 | 0.0462092186673193 |
| interactor_ratio | mitosis | inter | 0.08332897328355368 | 0.4841037832929192 |

P22626_P52272
ROA2_HUMAN vs HNRPM_HUMAN
p-value: 0.003 q-value: 0.003

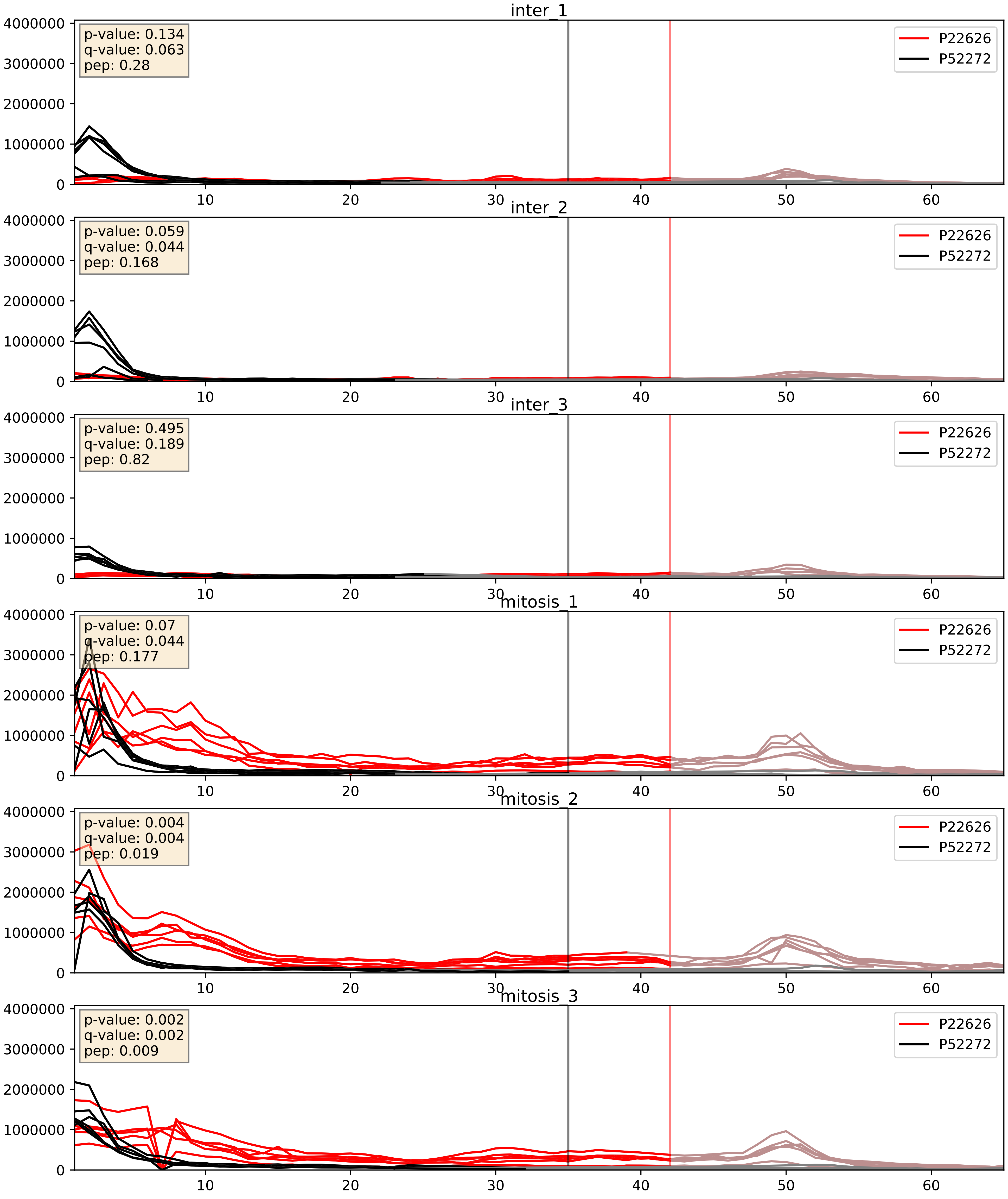


| level | condition_1 | condition_2 | pvalue | pvalue_adjusted |
|---|---|---|---|---|
| interactor_ratio | mitosis | inter | 0.0006314989468148049 | 0.029219626944512057 |
| interactor_abundance | mitosis | inter | 0.0029874185243059705 | 0.04171664366730687 |
| complex_abundance | mitosis | inter | 0.007981194454066766 | 0.06600871934957633 |

P31943_P52272
HNRH1_HUMAN vs HNRPM_HUMAN
p-value: 0.006 q-value: 0.006

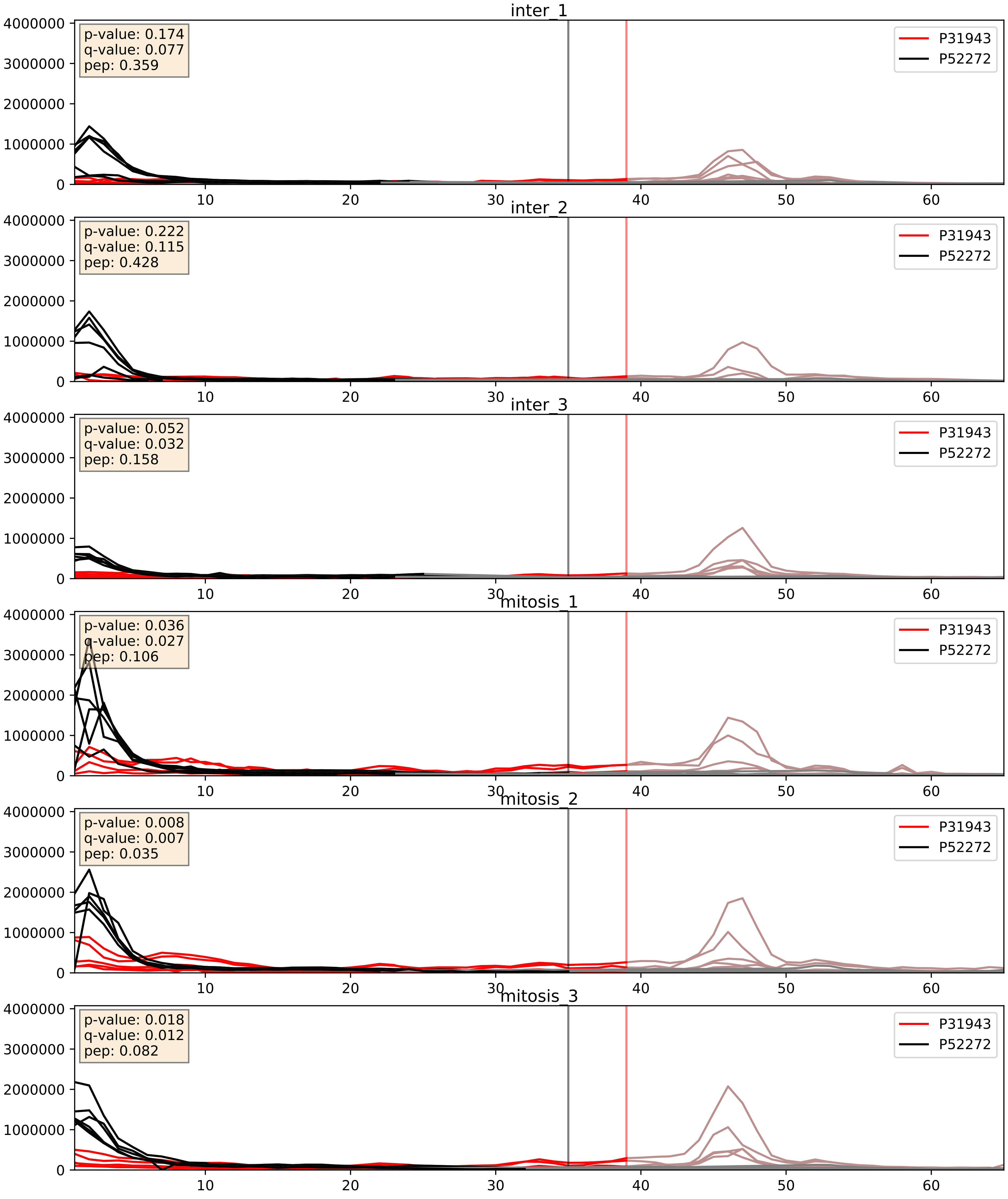


| level | condition_1 | condition_2 | pvalue | pvalue_adjusted |
|---|---|---|---|---|
| interactor_abundance | mitosis | inter | 0.002632509495742343 | 0.03838967005823521 |
| complex_abundance | mitosis | inter | 0.006583262263537986 | 0.06001355162501082 |
| interactor_ratio | mitosis | inter | 0.04049740663601096 | 0.33268502956262364 |

P38159_P52272
RBMX_HUMAN vs HNRPM_HUMAN
p-value: 0.001 q-value: 0.002

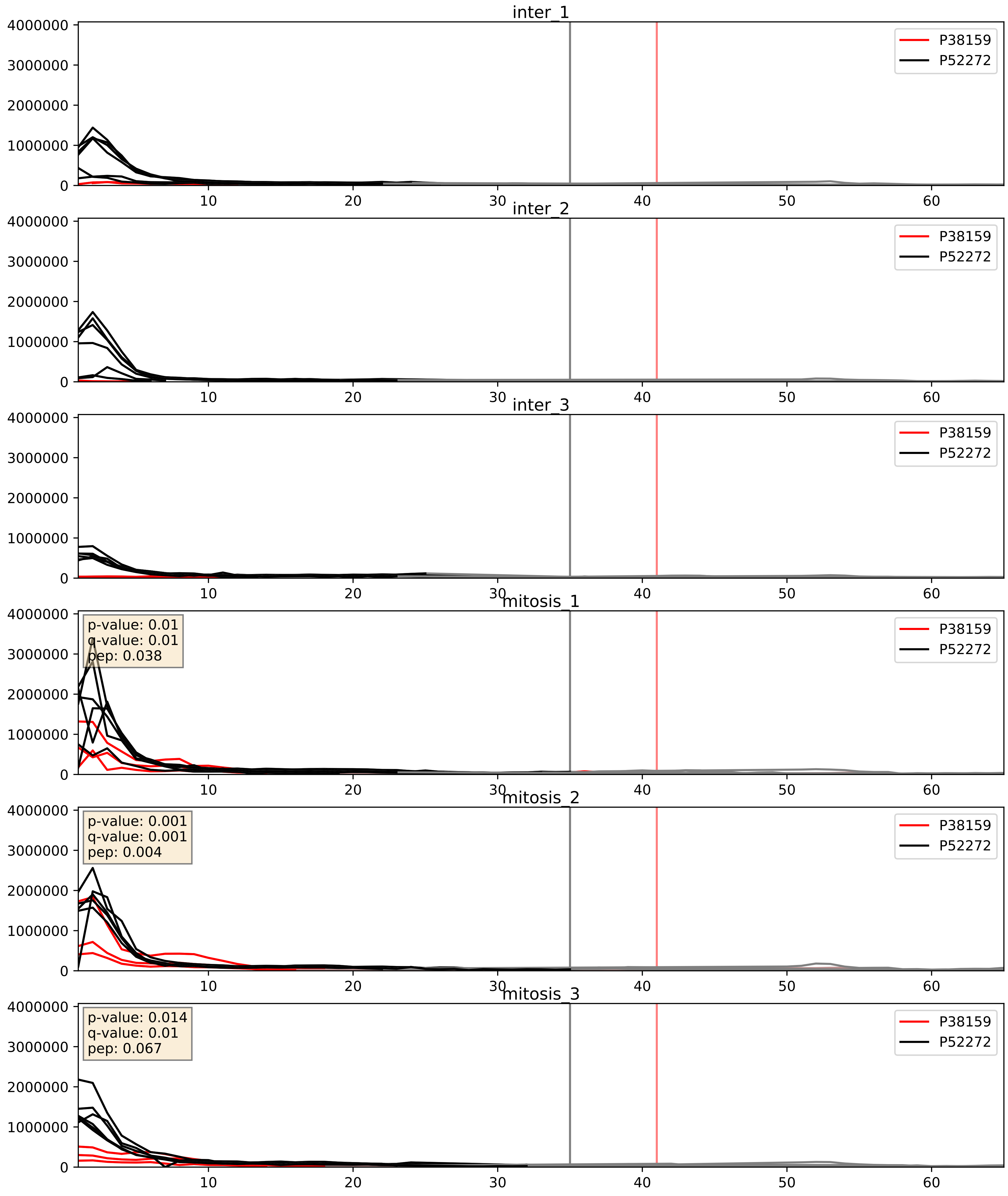


| level | condition_1 | condition_2 | pvalue | pvalue_adjusted |
|---|---|---|---|---|
| interactor_abundance | mitosis | inter | 0.0010812776417605033 | 0.024049720606162197 |
| complex_abundance | mitosis | inter | 0.002146905767739123 | 0.033596916584729235 |
| interactor_ratio | mitosis | inter | 0.00772024973379348 | 0.12375531408477938 |

P51991_P52272
ROA3_HUMAN vs HNRPM_HUMAN
p-value: 0.006 q-value: 0.006

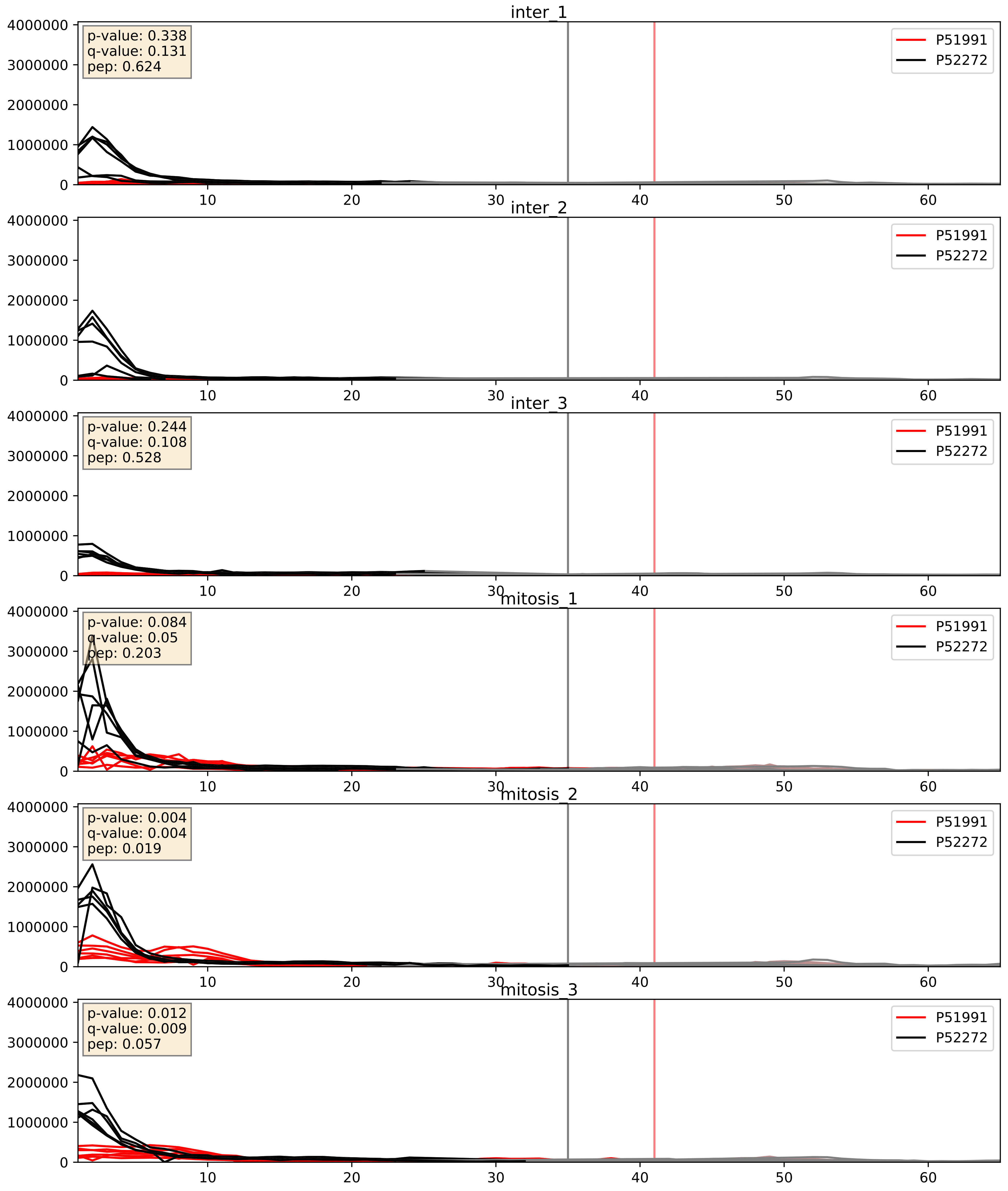


| level | condition_1 | condition_2 | pvalue | pvalue_adjusted |
|---|---|---|---|---|
| interactor_abundance | mitosis | inter | 2.6272351615725858e-06 | 0.0033734327106189023 |
| complex_abundance | mitosis | inter | 0.00045014421143136155 | 0.013379286284209911 |
| interactor_ratio | mitosis | inter | 0.0018482357365531494 | 0.05221418450460382 |

P52272_P52597
HNRPM_HUMAN vs HNRPF_HUMAN
p-value: 0.012 q-value: 0.01

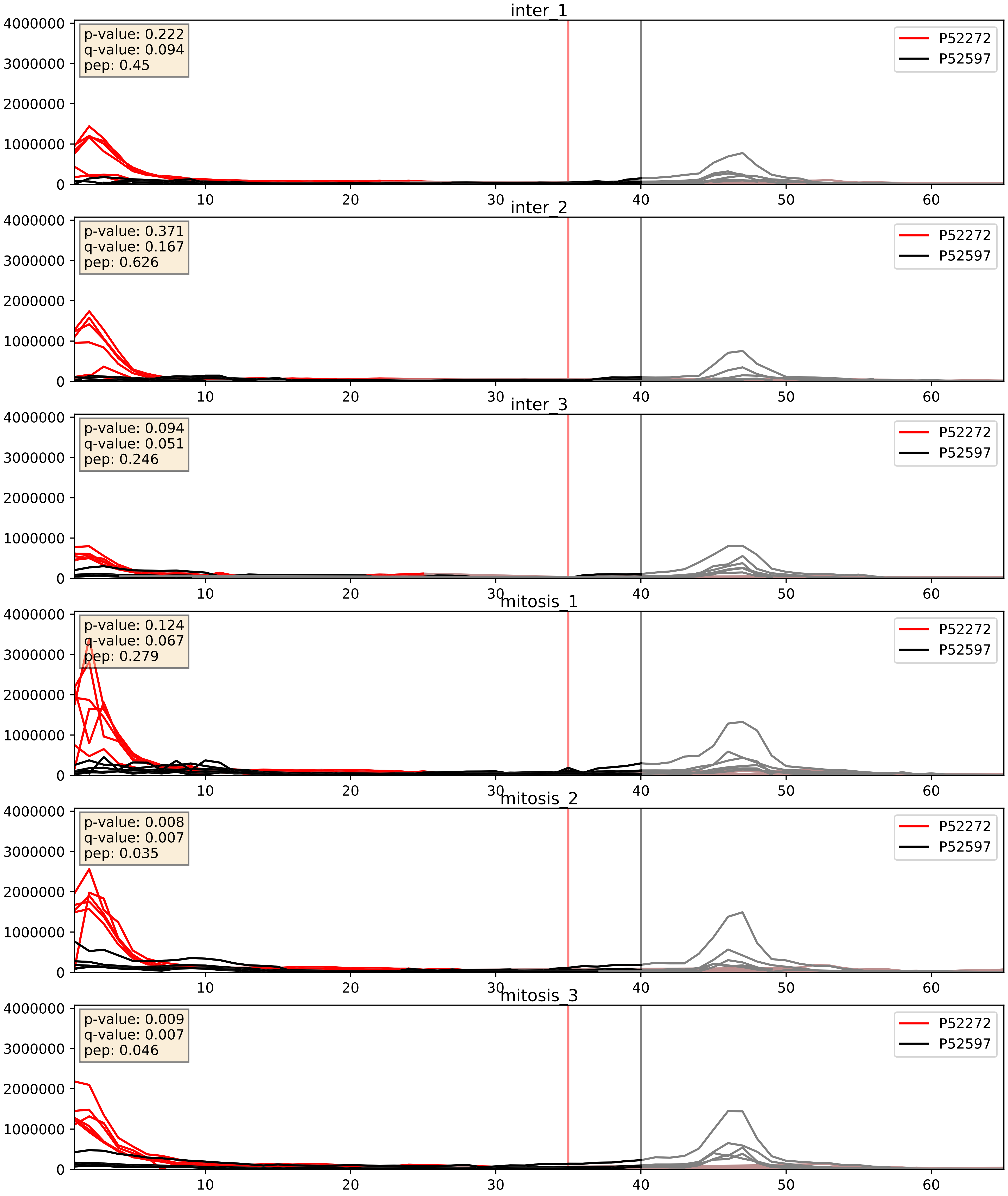


| level | condition_1 | condition_2 | pvalue | pvalue_adjusted |
|---|---|---|---|---|
| complex_abundance | mitosis | inter | 0.009939955657301148 | 0.07371585975104354 |
| interactor_ratio | mitosis | inter | 0.15013875473174929 | 0.6053639851642835 |
| interactor_abundance | mitosis | inter | 0.19757579480384646 | 0.391223701458744 |

P52272_P61978
HNRPM_HUMAN vs HNRPK_HUMAN
p-value: 0.027 q-value: 0.02

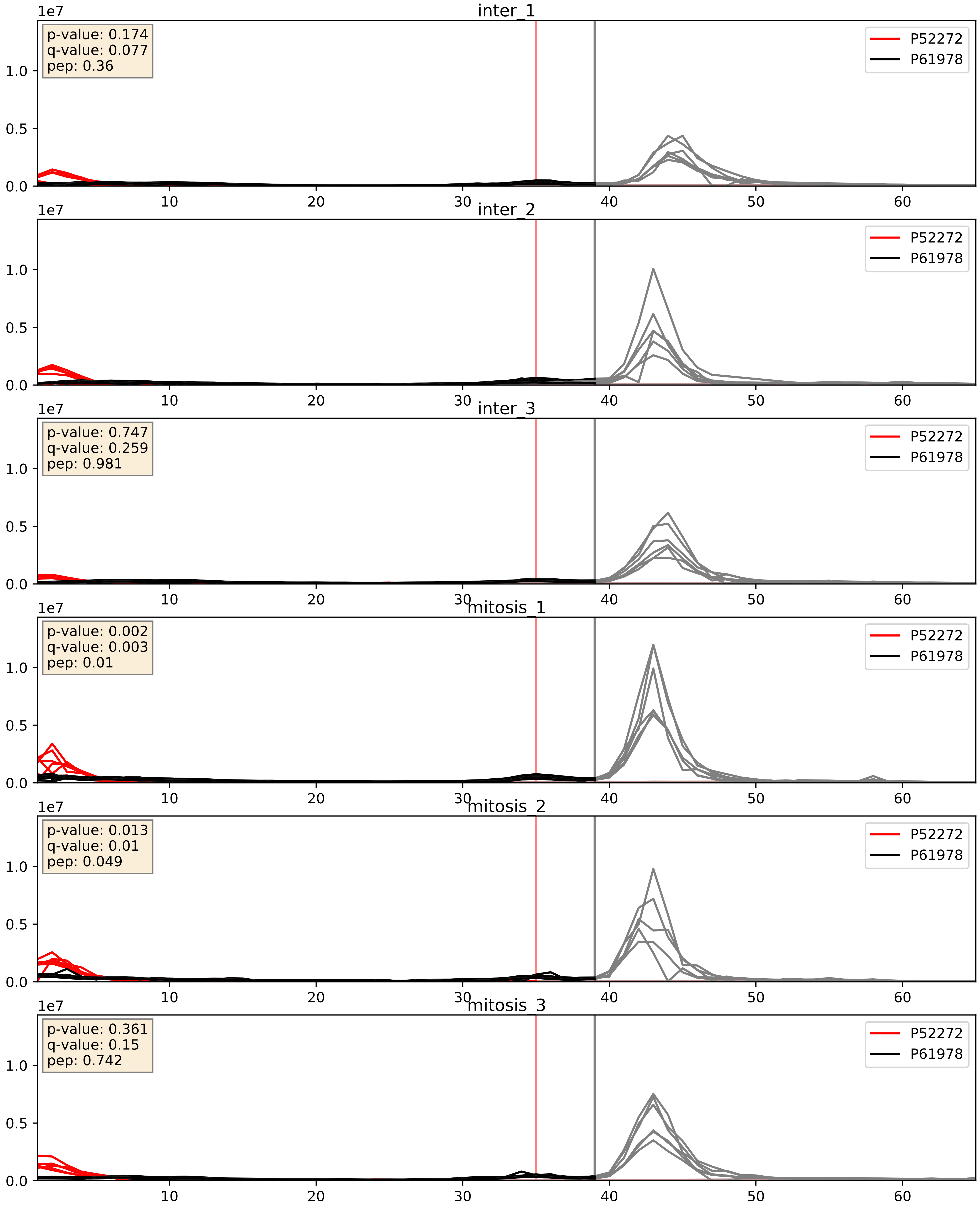


| level | condition_1 | condition_2 | pvalue | pvalue_adjusted |
|---|---|---|---|---|
| interactor_abundance | mitosis | inter | 0.19757579480384646 | 0.391223701458744 |
| interactor_ratio | mitosis | inter | 0.5278843604875636 | 0.8813144502088409 |
| complex_abundance | mitosis | inter | 0.692895826460435 | 0.7936588942398386 |

P52272_P62995
HNRPM_HUMAN vs TRA2B_HUMAN
p-value: 0.03 q-value: 0.022

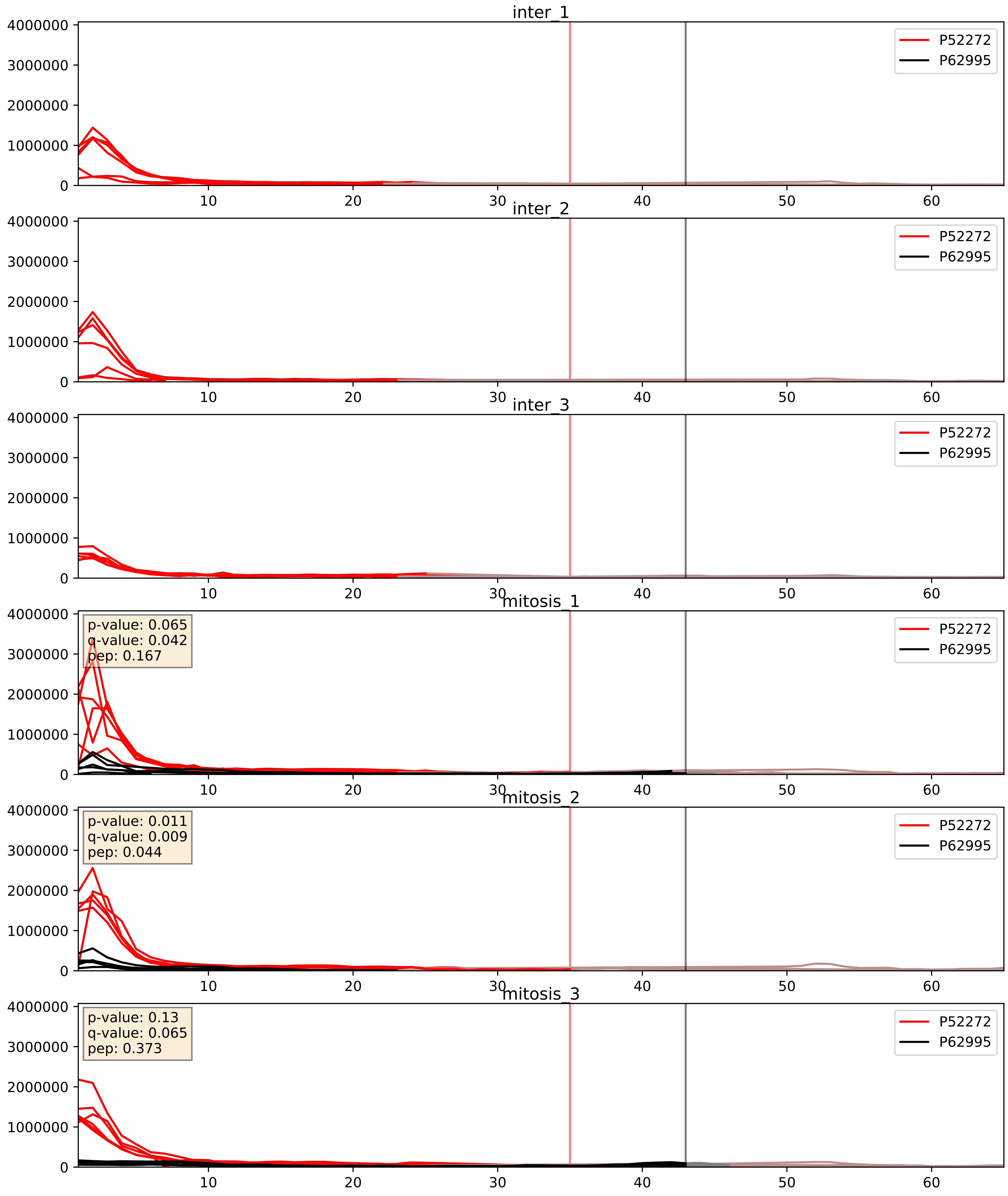


| level | condition_1 | condition_2 | pvalue | pvalue_adjusted |
|---|---|---|---|---|
| interactor_ratio | mitosis | inter | 3.4737747771941747e-07 | 0.0009911837364260712 |
| interactor_abundance | mitosis | inter | 2.4013033340954548e-05 | 0.004450836370313492 |
| complex_abundance | mitosis | inter | 7.744600555564312e-05 | 0.005949167968551921 |

P52272_Q07955
HNRPM_HUMAN vs SRSF1_HUMAN
p-value: 0.003 q-value: 0.004

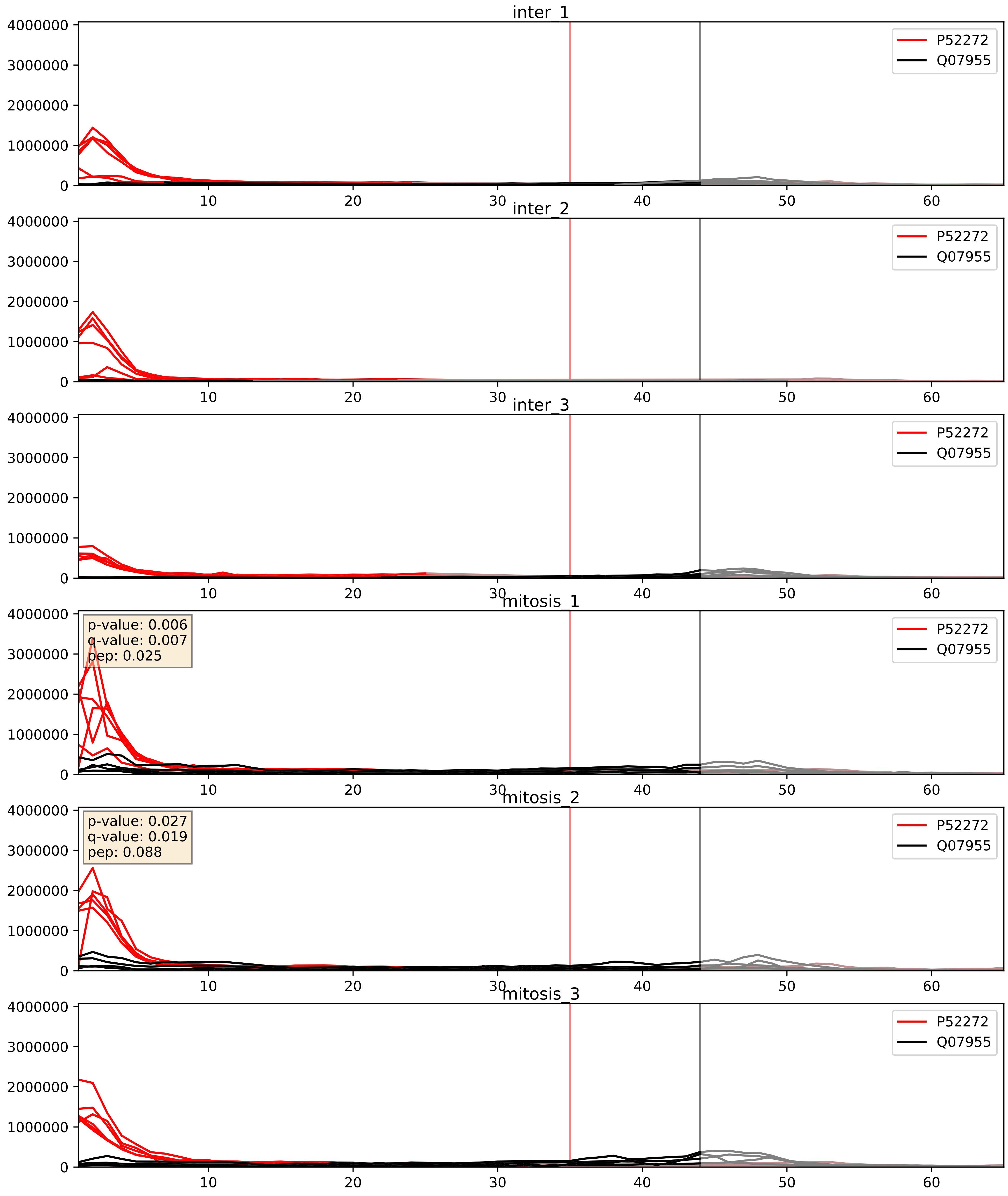


| level | condition_1 | condition_2 | pvalue | pvalue_adjusted |
|---|---|---|---|---|
| complex_abundance | mitosis | inter | 0.015294076257498249 | 0.09174685245214603 |
| interactor_ratio | mitosis | inter | 0.026601540420708367 | 0.25614196664208316 |
| interactor_abundance | mitosis | inter | 0.18545035871799714 | 0.37784844239126014 |

P52272_Q08211
HNRPM_HUMAN vs DHX9_HUMAN
p-value: 0.001 q-value: 0.002

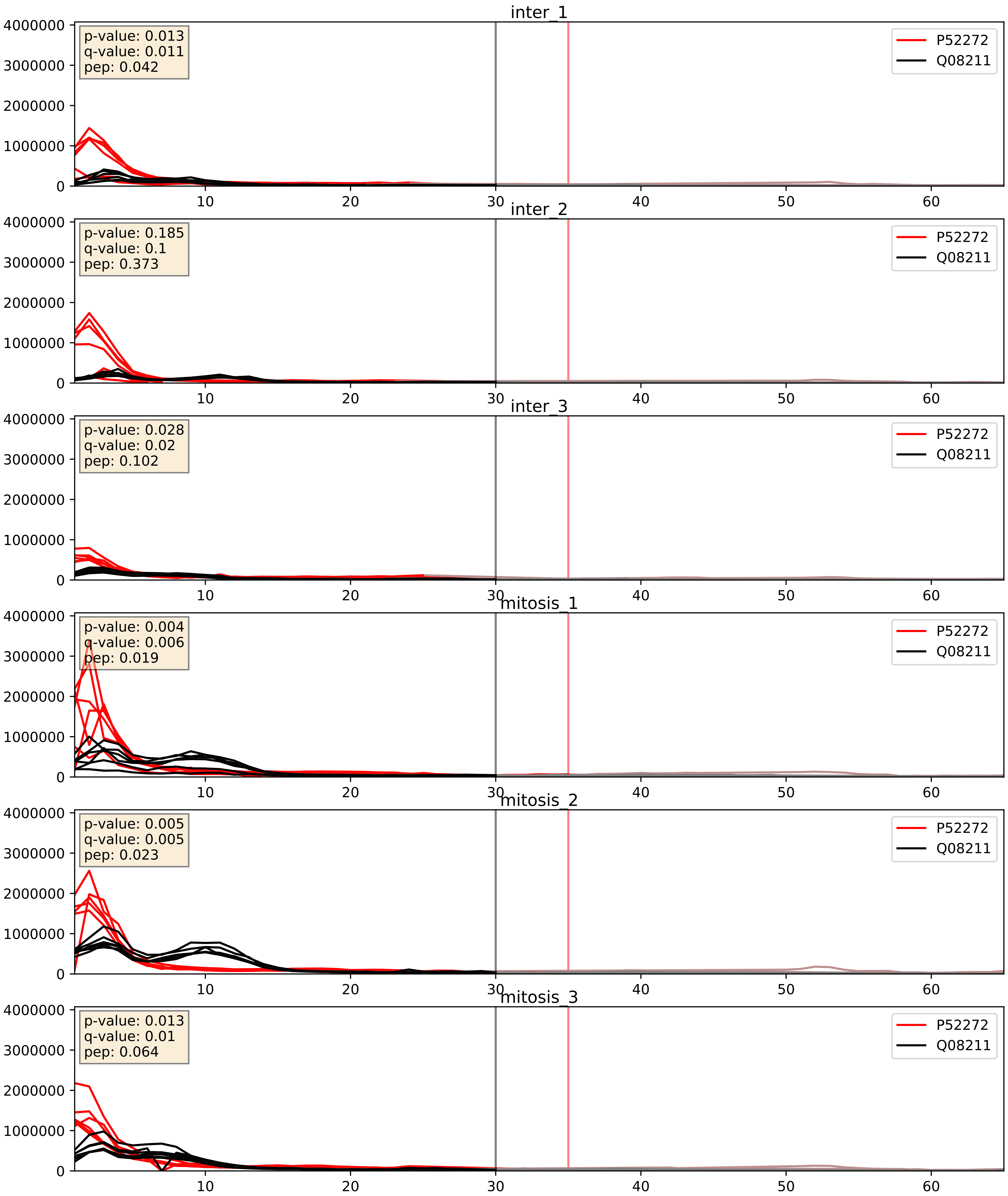


| level | condition_1 | condition_2 | pvalue | pvalue_adjusted |
|---|---|---|---|---|
| complex_abundance | mitosis | inter | 0.006357588474845068 | 0.05948545934570088 |
| interactor_ratio | mitosis | inter | 0.024442223536284675 | 0.2447639398105627 |
| interactor_abundance | mitosis | inter | 0.19956587550211832 | 0.39378164892295875 |

P52272_Q12906
HNRPM_HUMAN vs ILF3_HUMAN
p-value: 0.004 q-value: 0.047

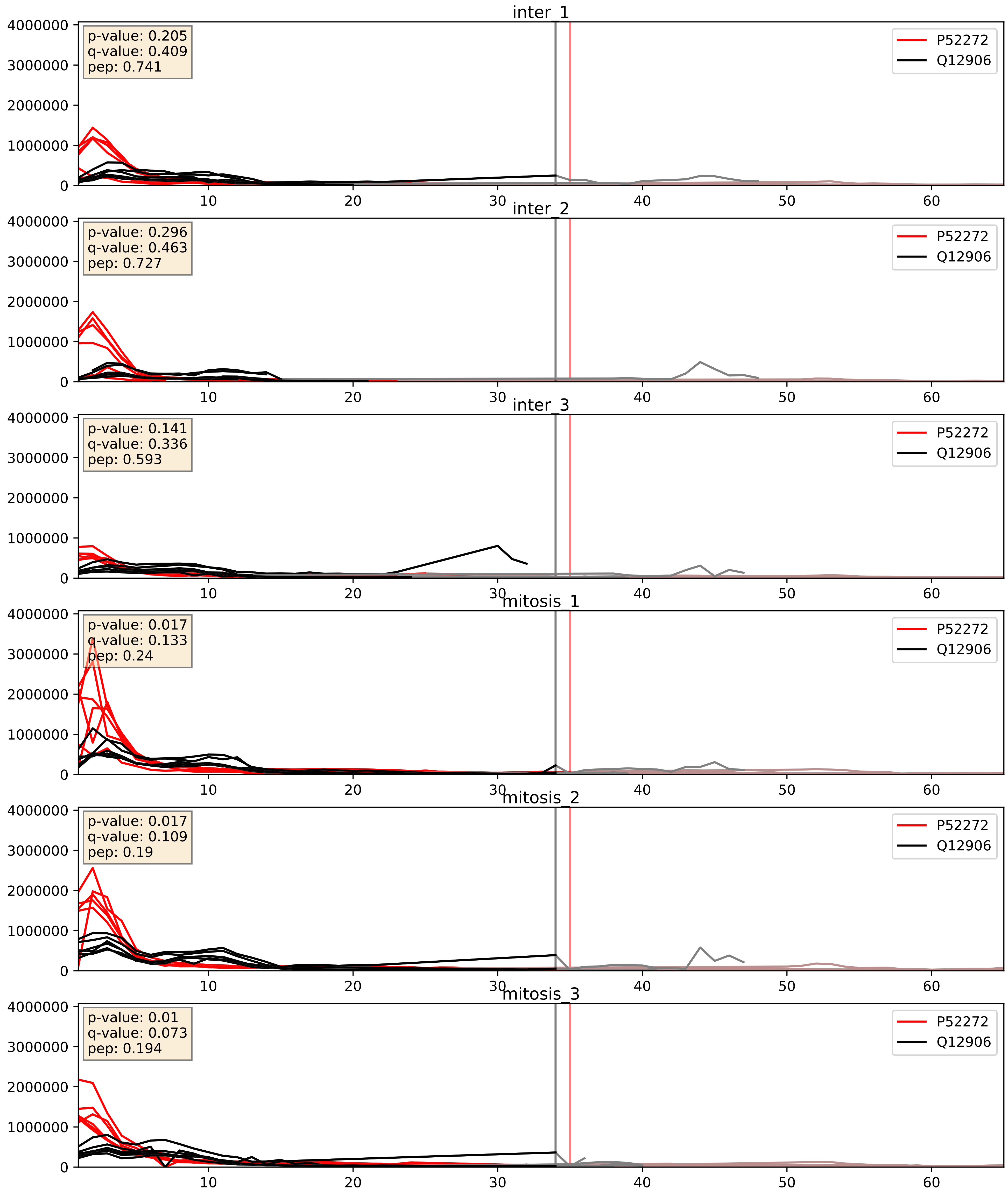


| level | condition_1 | condition_2 | pvalue | pvalue_adjusted |
|---|---|---|---|---|
| complex_abundance | mitosis | inter | 0.02102241322044666 | 0.10710544759151497 |
| interactor_abundance | mitosis | inter | 0.17718829693708443 | 0.3683175866394956 |
| interactor_ratio | mitosis | inter | 0.8491323203026978 | 0.9755634896635457 |

P52272_Q15717
HNRPM_HUMAN vs ELAV1_HUMAN
p-value: 0.001 q-value: 0.002

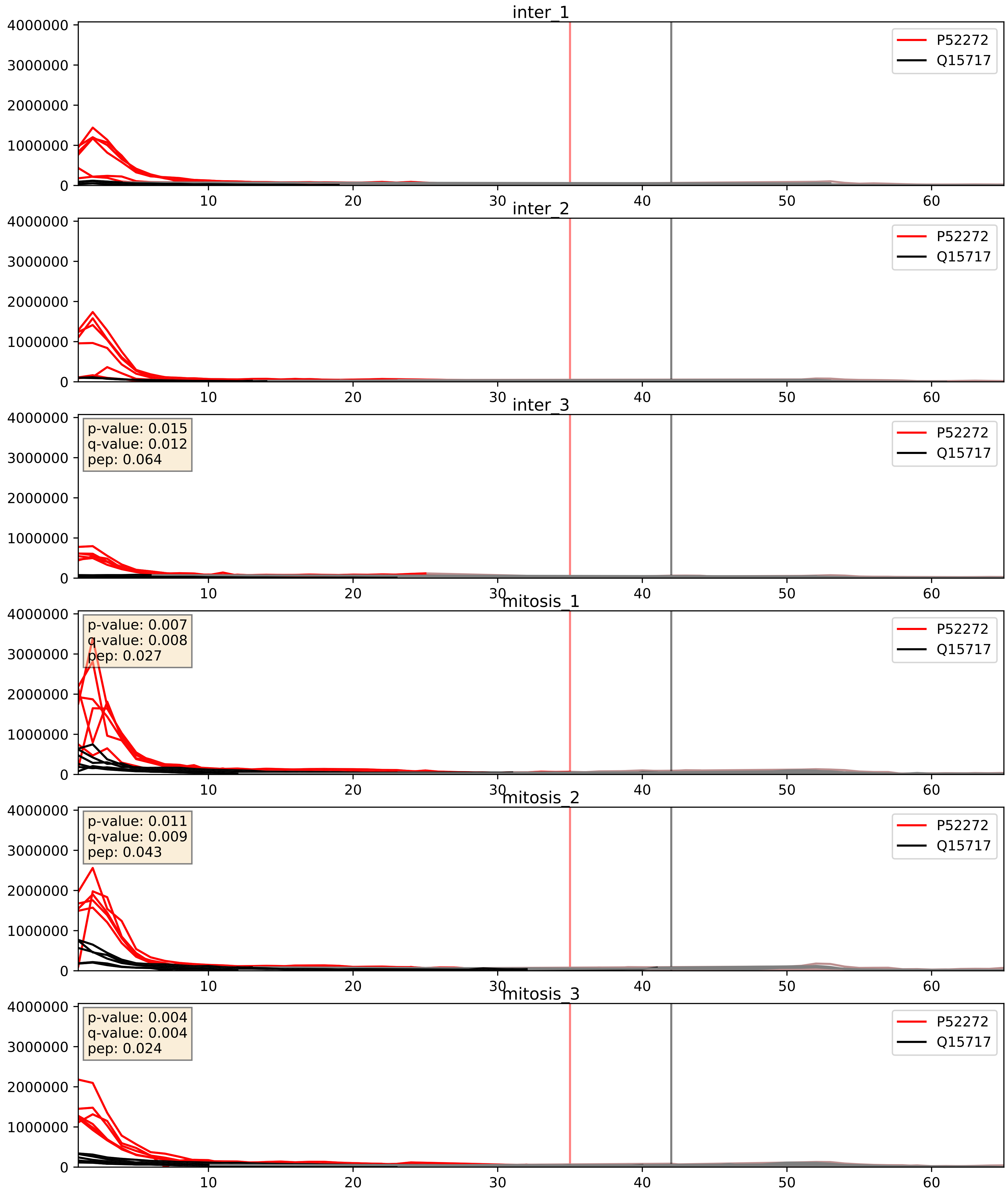


| level | condition_1 | condition_2 | pvalue | pvalue_adjusted |
|---|---|---|---|---|
| interactor_ratio | mitosis | inter | 0.0003243129433665509 | 0.019014512296011477 |
| complex_abundance | mitosis | inter | 0.002303087840743302 | 0.03502731659907978 |
| interactor_abundance | mitosis | inter | 0.1314788701529947 | 0.31658484627556527 |

P52272_Q16629
HNRPM_HUMAN vs SRSF7_HUMAN
p-value: 0.025 q-value: 0.019

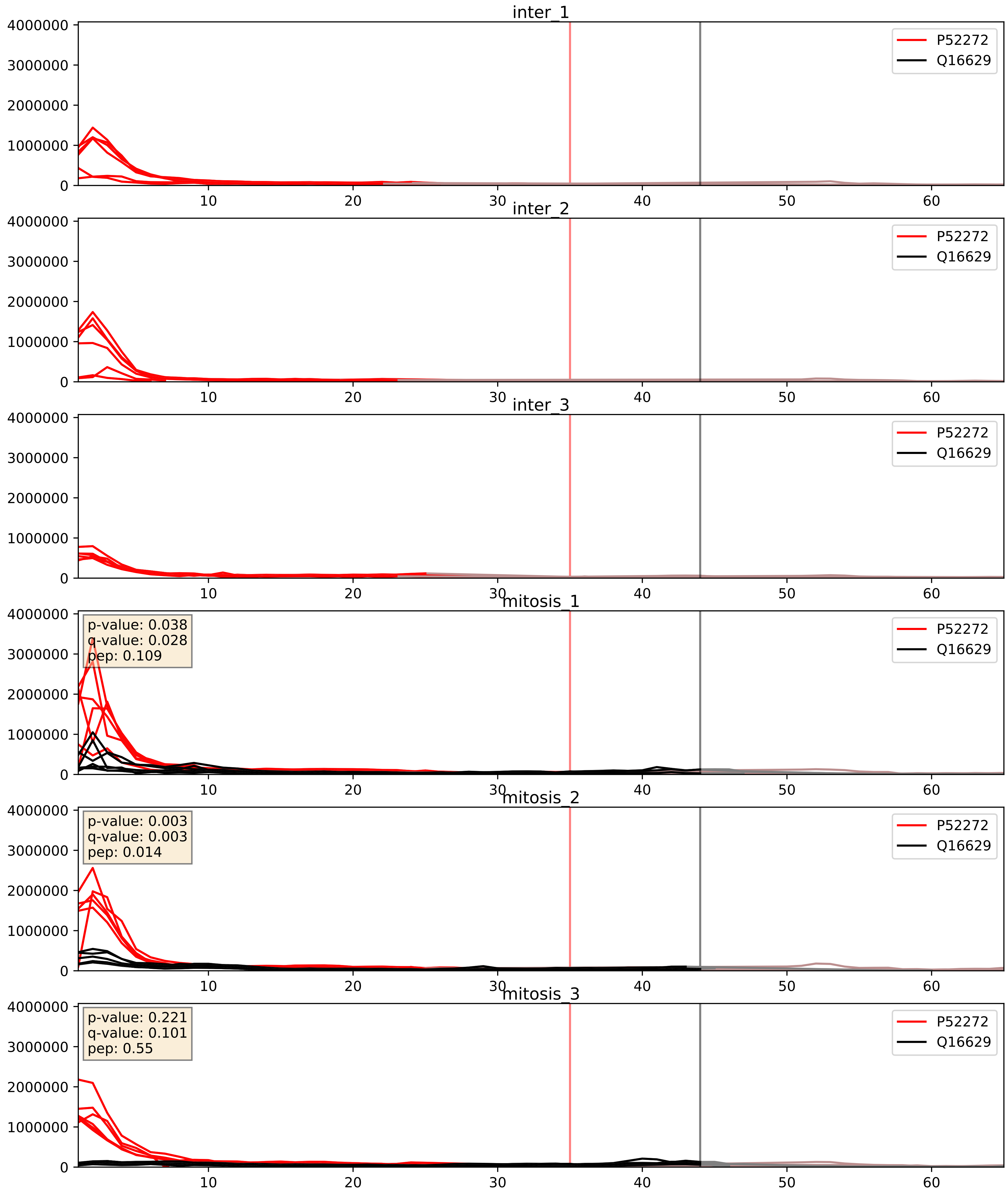


| level | condition_1 | condition_2 | pvalue | pvalue_adjusted |
|---|---|---|---|---|
| interactor_abundance | mitosis | inter | 2.2309750162464603e-05 | 0.0042438102531266 |
| complex_abundance | mitosis | inter | 3.735702009528526e-05 | 0.005691857151652958 |
| interactor_ratio | mitosis | inter | 0.0008463813255265196 | 0.033234055717922054 |
