## Supplementary Data 2 for "SECAT: Quantifying differential protein-protein interaction states by network-centric analysis": hela_string_000017_Q92925.pdf

# Q92925_Q92925

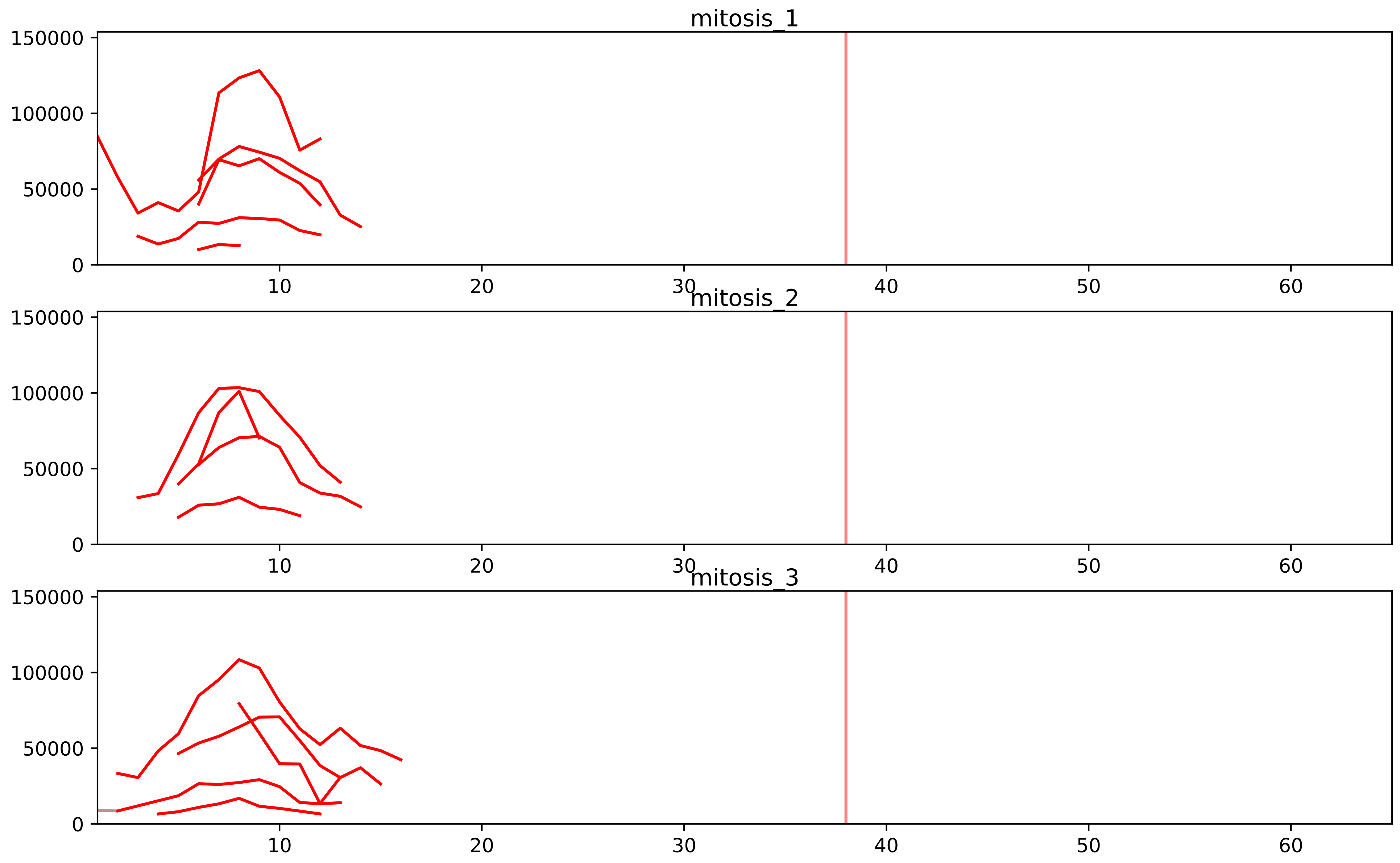


| level | condition_1 | condition_2 | pvalue | pvalue_adjusted |
|---|---|---|---|---|
| interactor_ratio | mitosis | inter | 1.2729044708927083e-12 | 1.3777318979074018e-10 |
| complex_abundance | mitosis | inter | 5.5040005819661445e-05 | 0.0015824001673152665 |
| assembled_abundance | mitosis | inter | 7.53181951614952e-05 | 0.008577738896244046 |
| total_abundance | mitosis | inter | 0.00013638168216436778 | 0.014169778446914211 |
| interactor_abundance | mitosis | inter | 0.00035646423641045916 | 0.005753457850835481 |

O14497_Q92925
ARI1A_HUMAN vs SMRD2_HUMAN
p-value: 0.035 q-value: 0.025

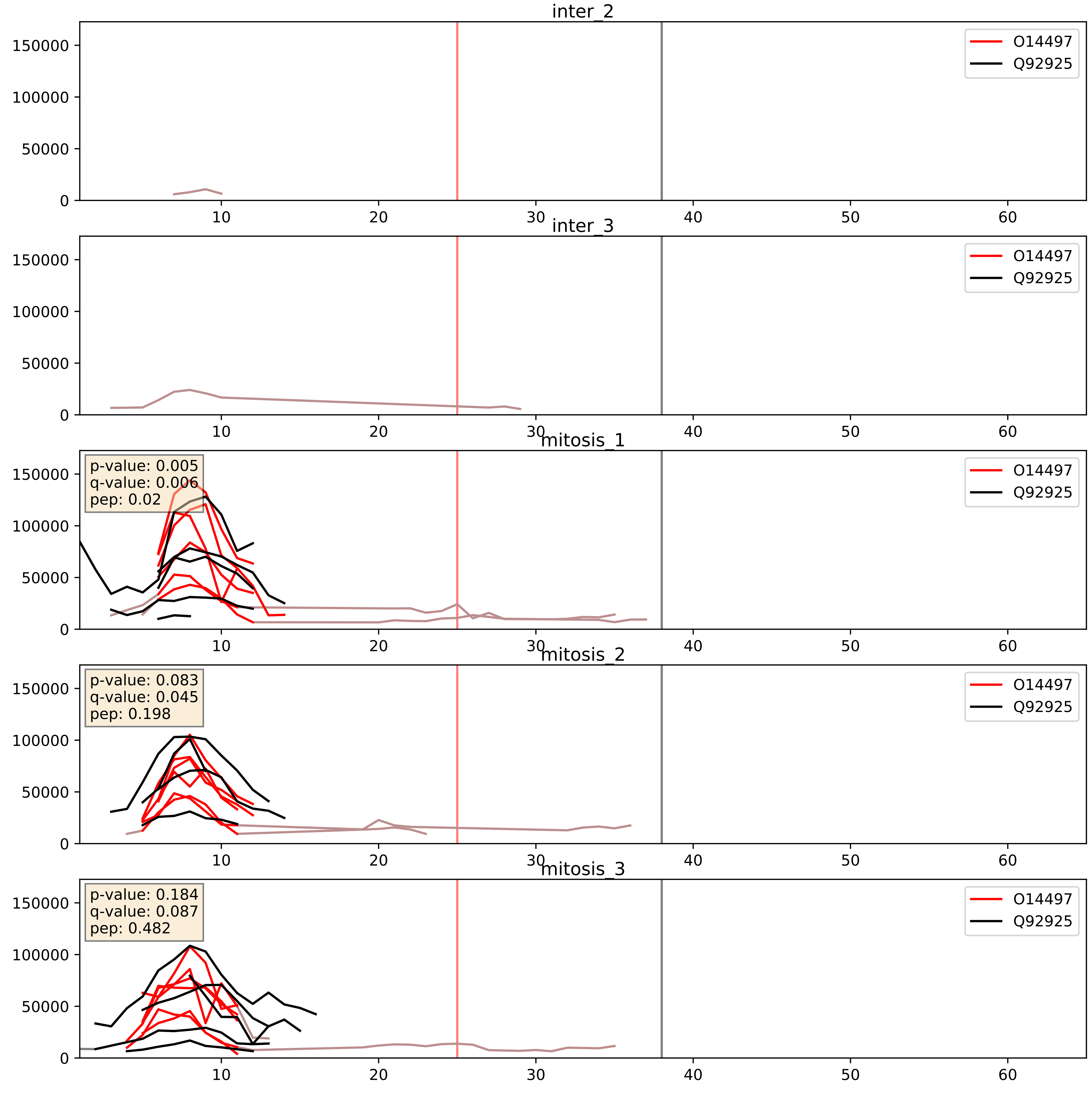


| level | condition_1 | condition_2 | pvalue | pvalue_adjusted |
|---|---|---|---|---|
| complex_abundance | mitosis | inter | 0.009992203841417977 | 0.07371585975104354 |
| interactor_abundance | mitosis | inter | 0.012640483206095423 | 0.10027749130318384 |
| interactor_ratio | mitosis | inter | 0.8861547475939666 | 0.9802747470617126 |

O96019_Q92925
ACL6A_HUMAN vs SMRD2_HUMAN
p-value: 0.036 q-value: 0.025

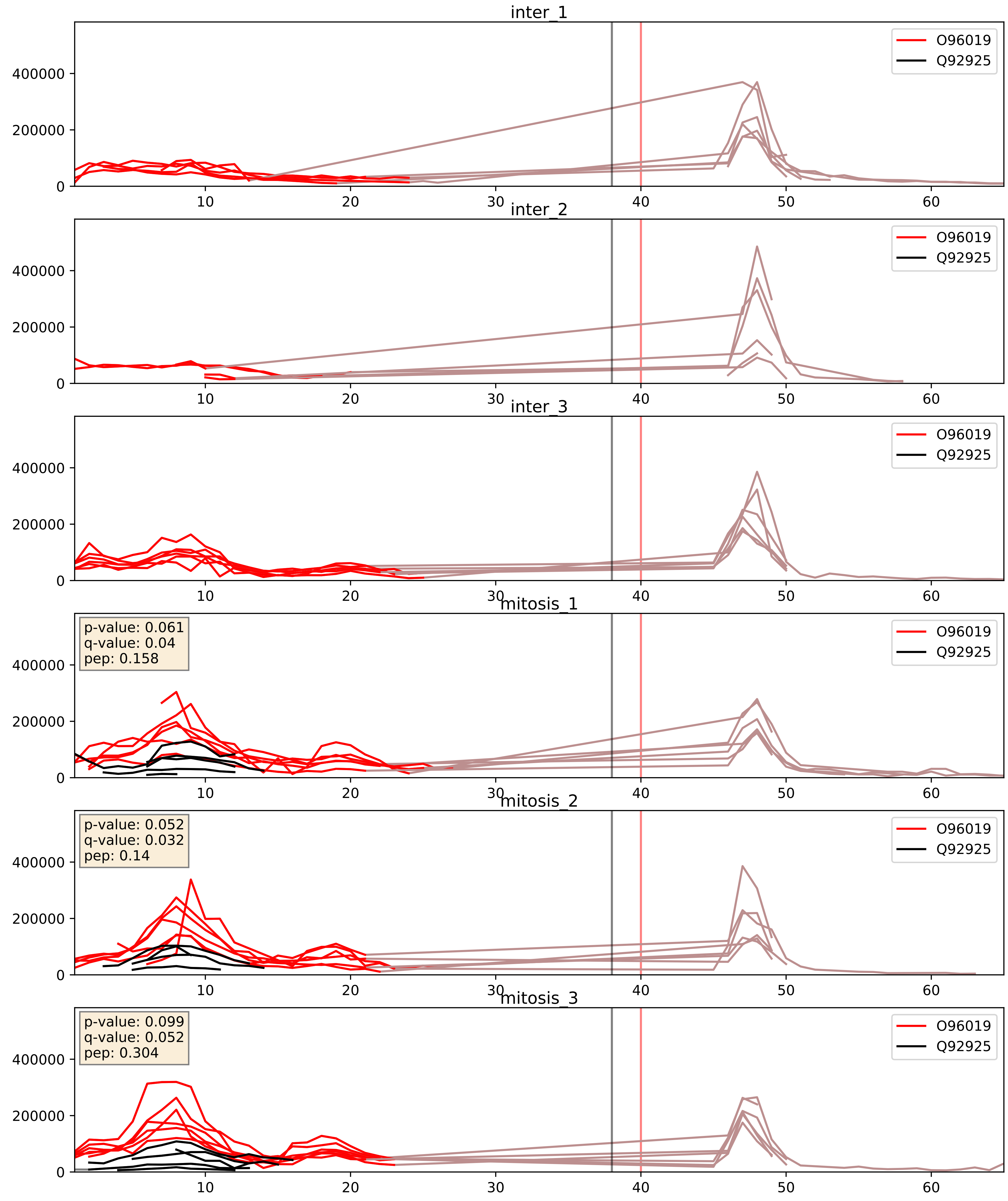


| level | condition_1 | condition_2 | pvalue | pvalue_adjusted |
|---|---|---|---|---|
| interactor_abundance | mitosis | inter | 0.0006118287183261804 | 0.017415659477552636 |
| complex_abundance | mitosis | inter | 0.0009958216364759 89 | 0.02099825493417971 |
| interactor_ratio | mitosis | inter | 0.001661415325362099 | 0.04925768316782793 |

P51531_Q92925
SMCA2_HUMAN vs SMRD2_HUMAN
p-value: 0.085 q-value: 0.05

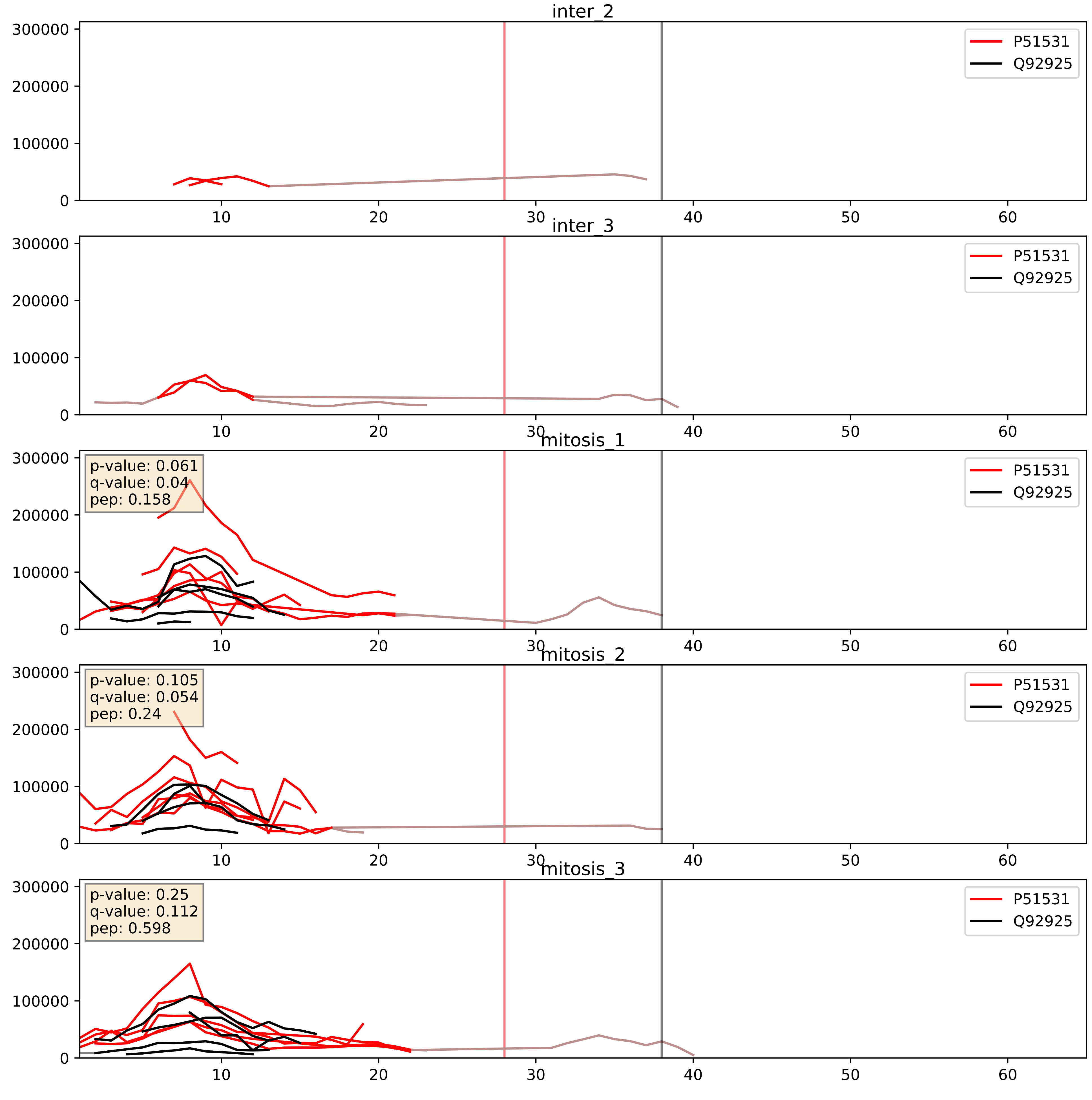


| level | condition_1 | condition_2 | pvalue | pvalue_adjusted |
|---|---|---|---|---|
| complex_abundance | mitosis | inter | 0.004621740992278264 | 0.050629452773887605 |
| interactor_abundance | mitosis | inter | 0.01019836828153345 | 0.0879625269700301 |
| interactor_ratio | mitosis | inter | 0.28817519340286885 | 0.7581846425018174 |

Q92922_Q92925
SMRC1_HUMAN vs SMRD2_HUMAN
p-value: 0.043 q-value: 0.029

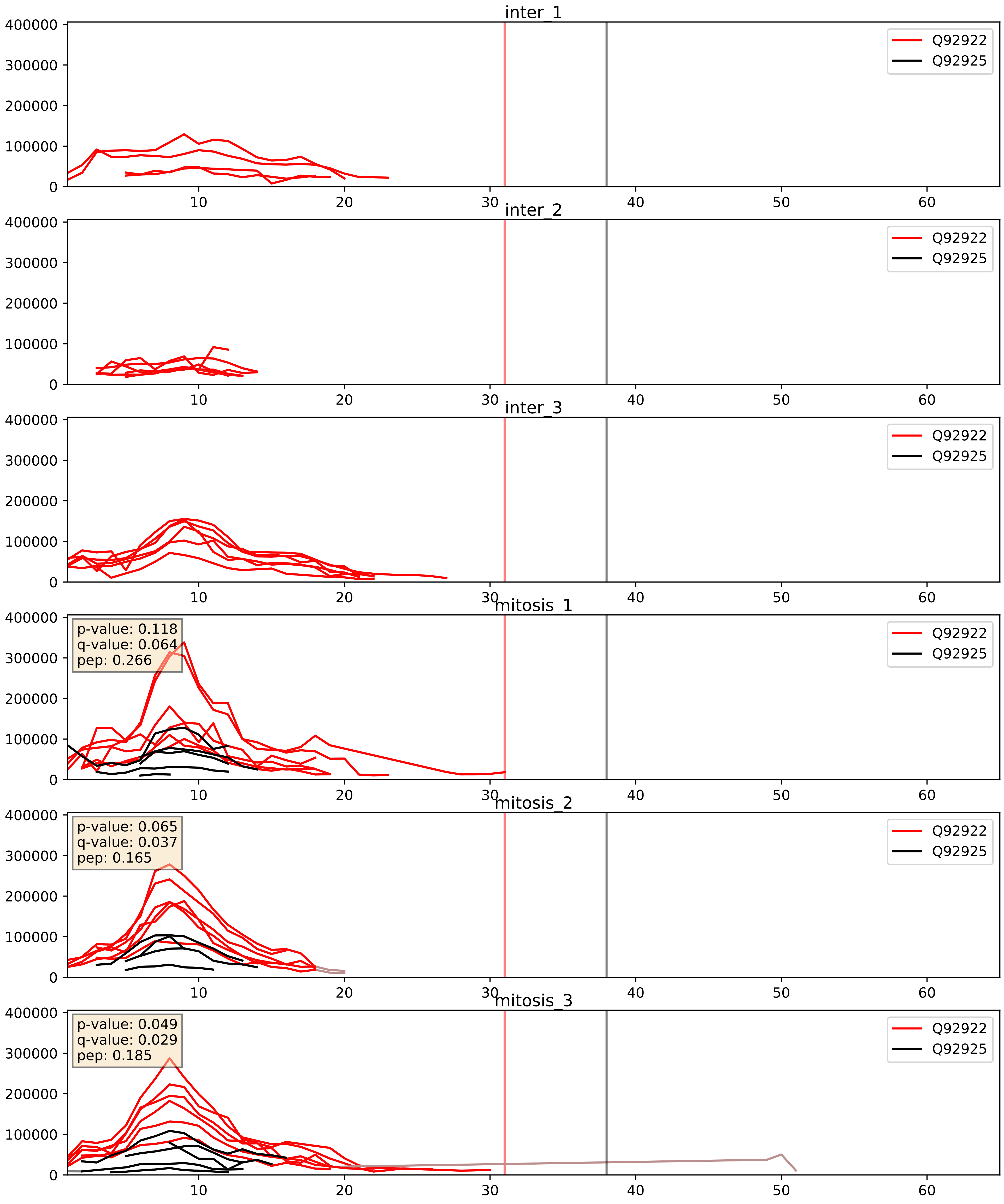


| level | condition_1 | condition_2 | pvalue | pvalue_adjusted |
|---|---|---|---|---|
| interactor_ratio | mitosis | inter | 0.00010380497123051964 | 0.010720128702277293 |
| interactor_abundance | mitosis | inter | 0.00018540588357067092 | 0.01016145604161317 |
| complex_abundance | mitosis | inter | 0.0005971387895009652 | 0.01581504091436693 |

Q92925_Q969G3
SMRD2_HUMAN vs SMCE1_HUMAN
p-value: 0.017 q-value: 0.014

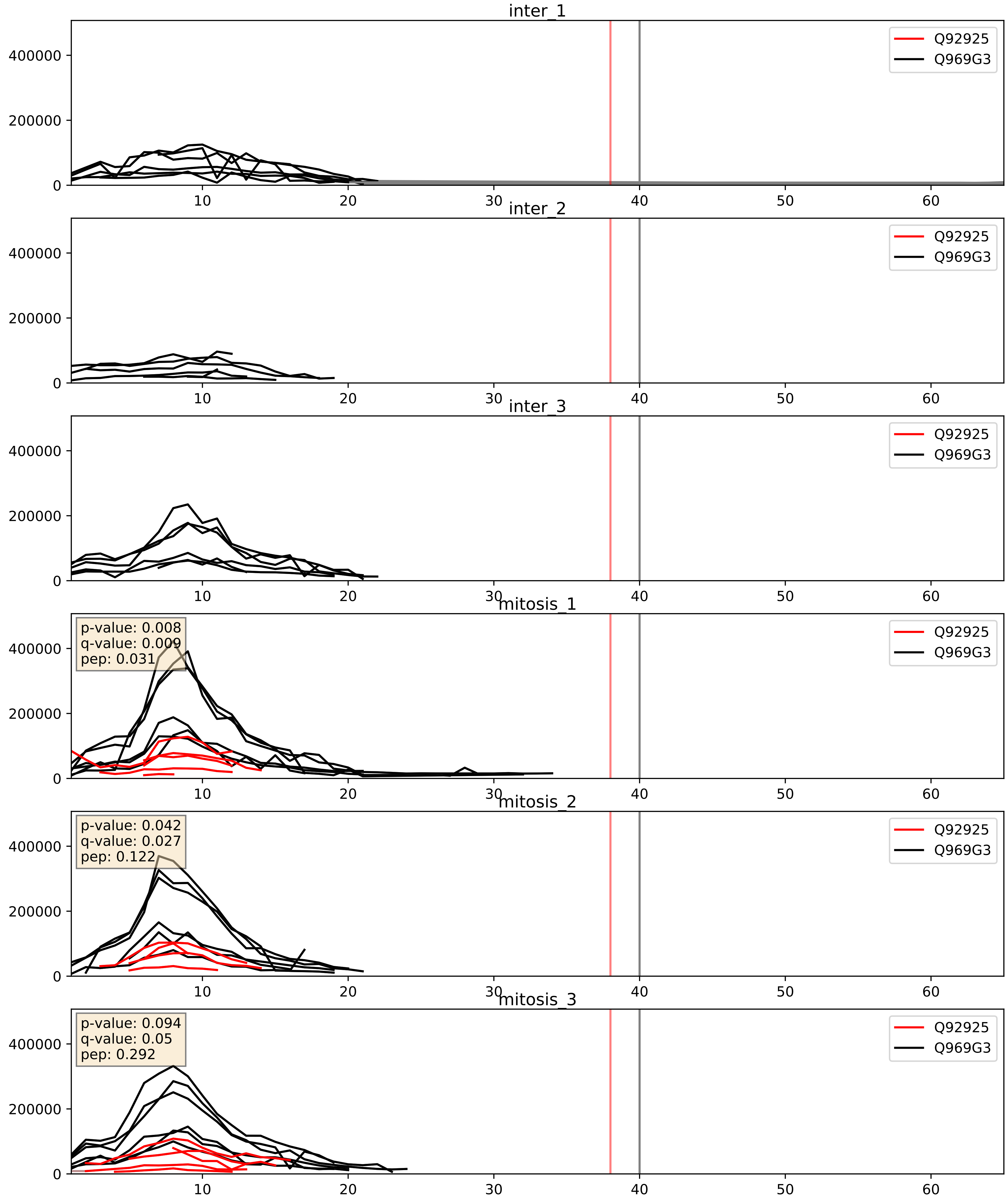


| level | condition_1 | condition_2 | pvalue | pvalue_adjusted |
|---|---|---|---|---|
| interactor_ratio | mitosis | inter | 3.4231436616441946e-06 | 0.002520386835947235 |
| complex_abundance | mitosis | inter | 0.0003362431142253742 | 0.011893558089955385 |
| interactor_abundance | mitosis | inter | 0.004123135752406863 | 0.050966125690401085 |

Q92925_Q96GM5
SMRD2_HUMAN vs SMRD1_HUMAN
p-value: 0.021 q-value: 0.017

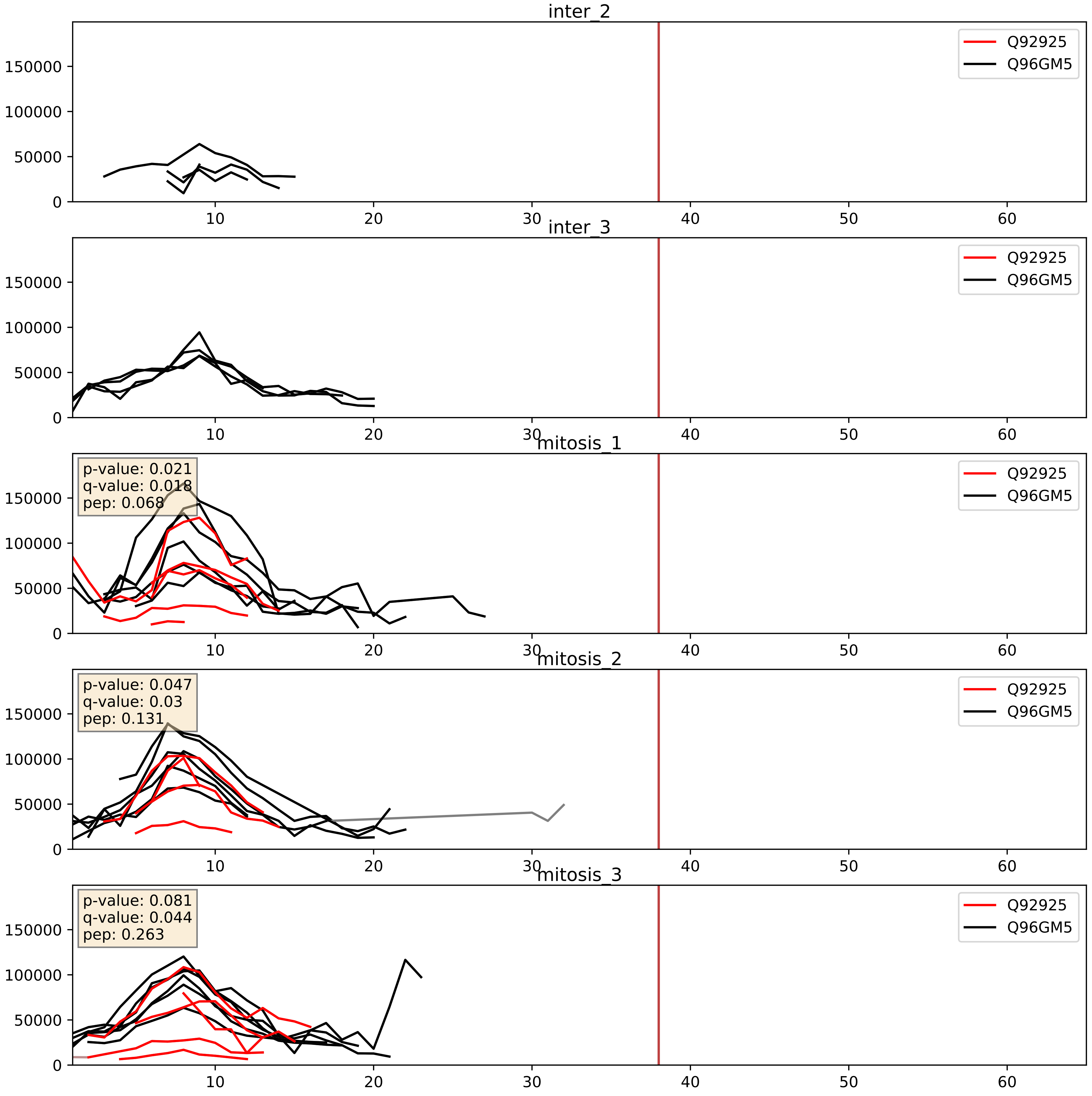


| level | condition_1 | condition_2 | pvalue | pvalue_adjusted |
|---|---|---|---|---|
| interactor_ratio | mitosis | inter | 5.661017680550811e-06 | 0.002835004461043519 |
| complex_abundance | mitosis | inter | 0.0019049156108668468 | 0.031118468757672154 |
| interactor_abundance | mitosis | inter | 0.004123135752406863 | 0.050966125690401085 |
