## Supplementary Data 2 for "SECAT: Quantifying differential protein-protein interaction states by network-centric analysis": hela_string_000018_P51991.pdf

P51991_P51991

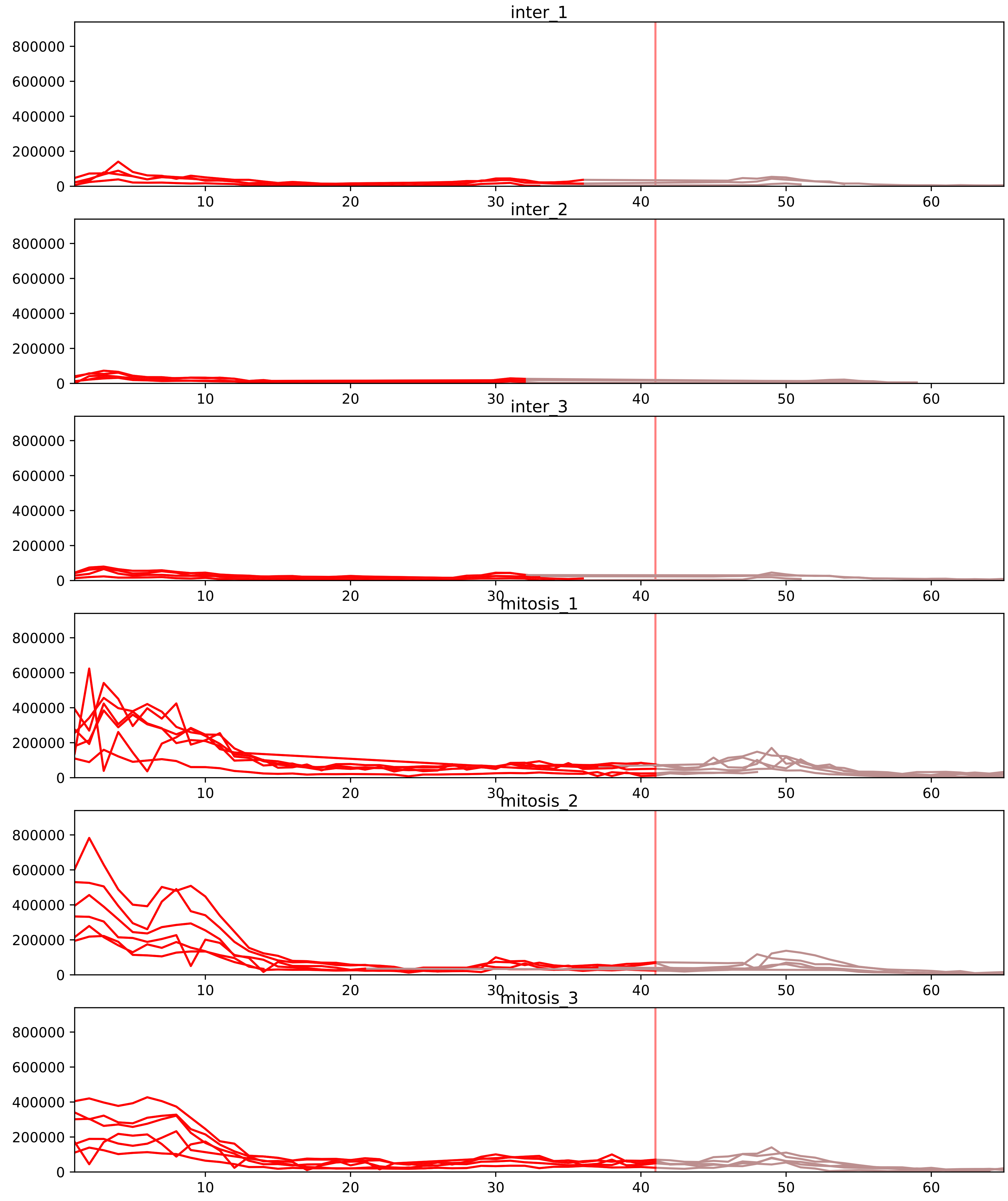


| level | condition_1 | condition_2 | pvalue | pvalue_adjusted |
|---|---|---|---|---|
| interactor_abundance | mitosis | inter | 1.4728131751340888e-12 | 2.7099762422467234e-09 |
| interactor_ratio | mitosis | inter | 3.904695906804043e-12 | 3.7813897202733894e-10 |
| complex_abundance | mitosis | inter | 5.699721939310722e-10 | 5.243744184165864e-07 |
| assembled_abundance | mitosis | inter | 7.674753266623976e-06 | 0.0032983295440323994 |
| total_abundance | mitosis | inter | 2.0748803650891936e-05 | 0.0058684532992606024 |
| monomer_abundance | mitosis | inter | 0.00010900477042974257 | 0.03406818325046493 |

O43390_P51991
HNRPR_HUMAN vs ROA3_HUMAN
p-value: 0.003 q-value: 0.041

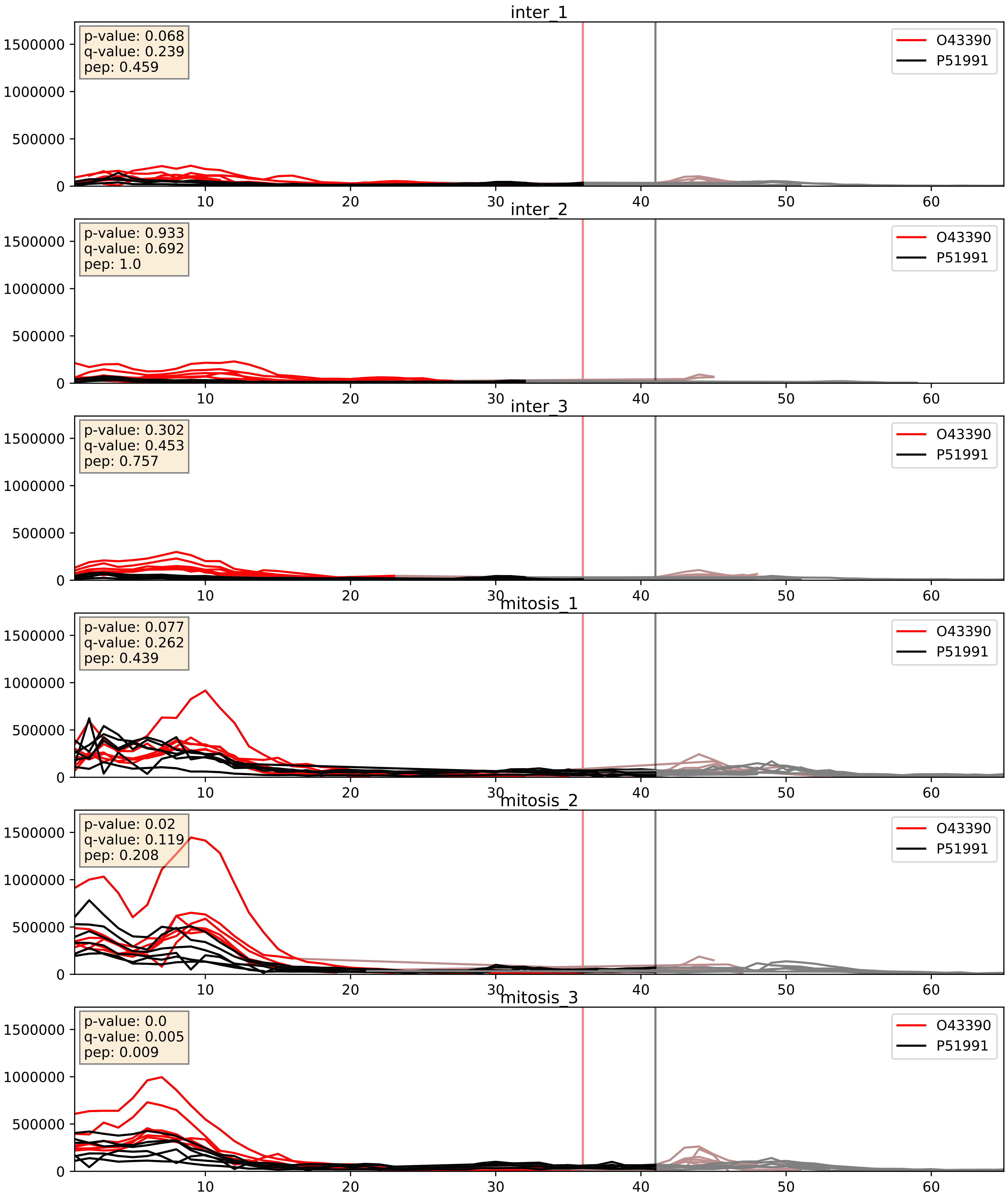


| level | condition_1 | condition_2 | pvalue | pvalue_adjusted |
|---|---|---|---|---|
| complex_abundance | mitosis | inter | 1.1164489343079466e-07 | 0.0009556802877676023 |
| interactor_abundance | mitosis | inter | 3.72135292356026e-05 | 0.005353744710197617 |
| interactor_ratio | mitosis | inter | 0.0004914306397162247 | 0.025464098294827937 |

P07910_P51991
HNRPC_HUMAN vs ROA3_HUMAN
p-value: 0.002 q-value: 0.002

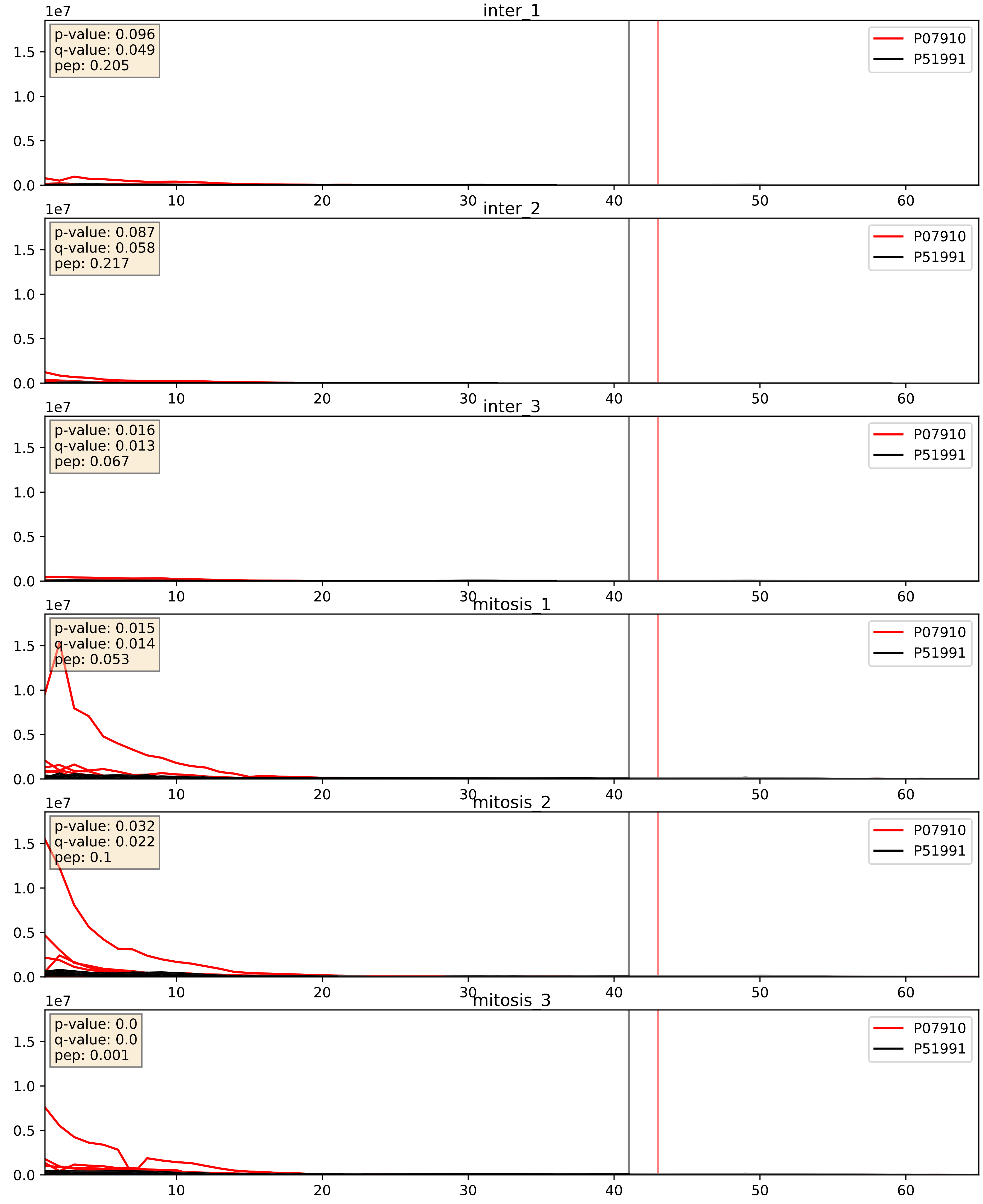


| level | condition_1 | condition_2 | pvalue | pvalue_adjusted |
|---|---|---|---|---|
| complex_abundance | mitosis | inter | 7.94637328064476e-07 | 0.0024732922429087437 |
| interactor_abundance | mitosis | inter | 9.1373717647951e-06 | 0.0033734327106189023 |
| interactor_ratio | mitosis | inter | 0.0027887838155807804 | 0.06820568417534709 |

P09651_P51991
ROA1_HUMAN vs ROA3_HUMAN
p-value: 0.0 q-value: 0.0

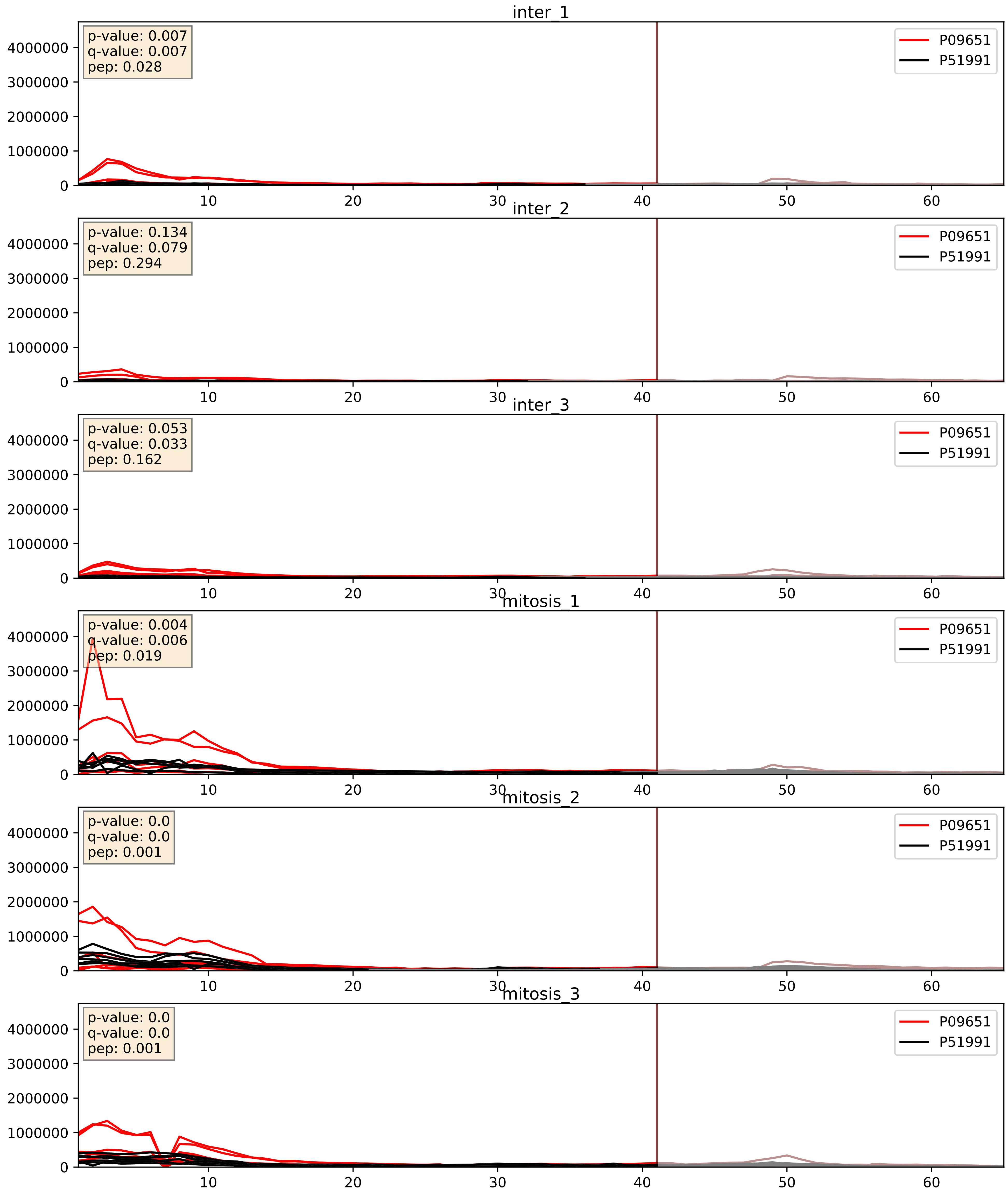


| level | condition_1 | condition_2 | pvalue | pvalue_adjusted |
|---|---|---|---|---|
| complex_abundance | mitosis | inter | 0.0011795042359659484 | 0.023096652182529428 |
| interactor_abundance | mitosis | inter | 0.016104784849936434 | 0.11126469597696198 |
| interactor_ratio | mitosis | inter | 0.19038795235618836 | 0.6627604950548505 |

P14866_P51991
HNRPL_HUMAN vs ROA3_HUMAN
p-value: 0.001 q-value: 0.002

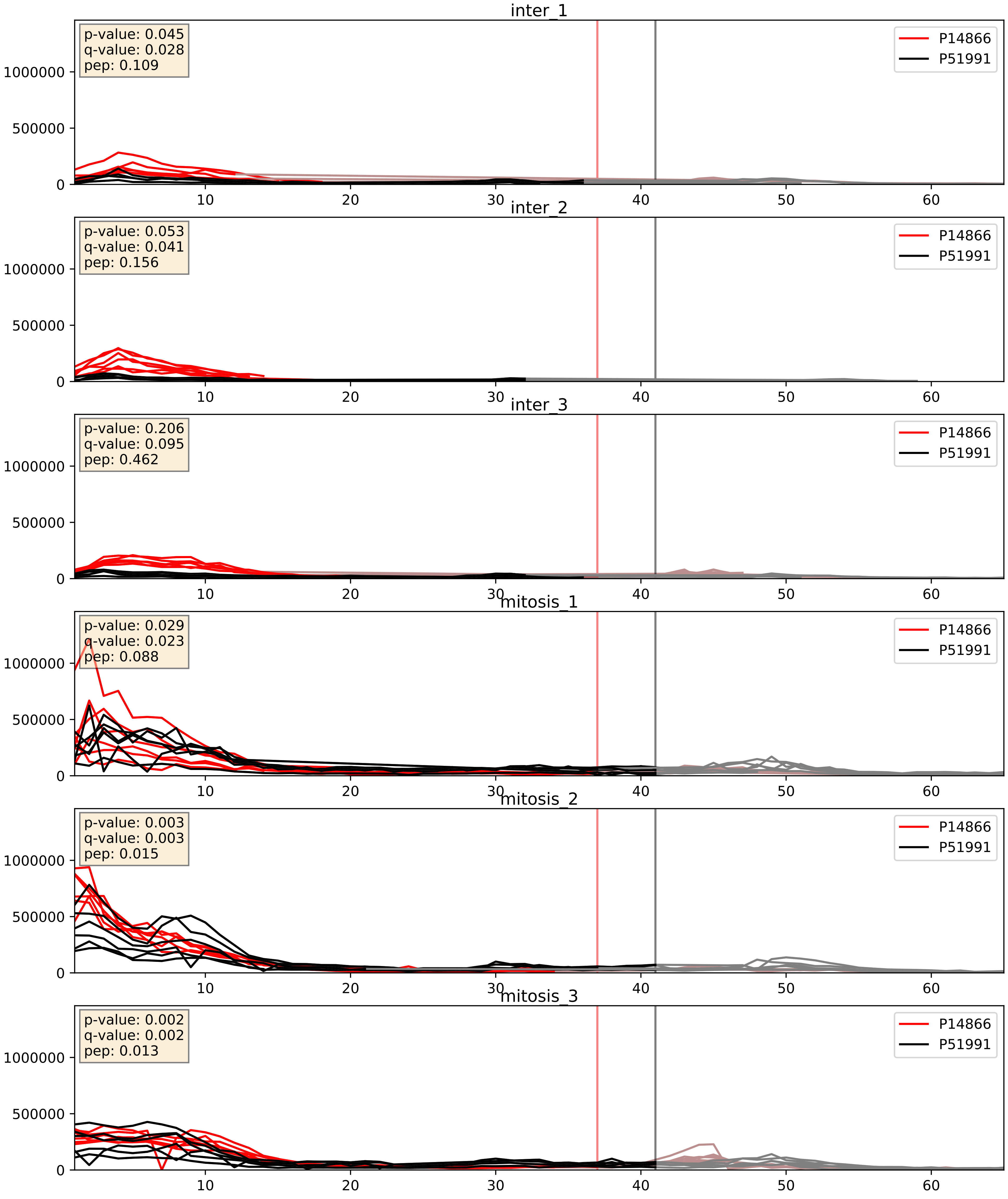


| level | condition_1 | condition_2 | pvalue | pvalue_adjusted |
|---|---|---|---|---|
| complex_abundance | mitosis | inter | 0.000306389187647456 | 0.01165207110963347 |
| interactor_ratio | mitosis | inter | 0.005592202135892223 | 0.10141790314245218 |
| interactor_abundance | mitosis | inter | 0.009697640385696541 | 0.08525122834595471 |

P22626_P51991
ROA2_HUMAN vs ROA3_HUMAN
p-value: 0.0 q-value: 0.001

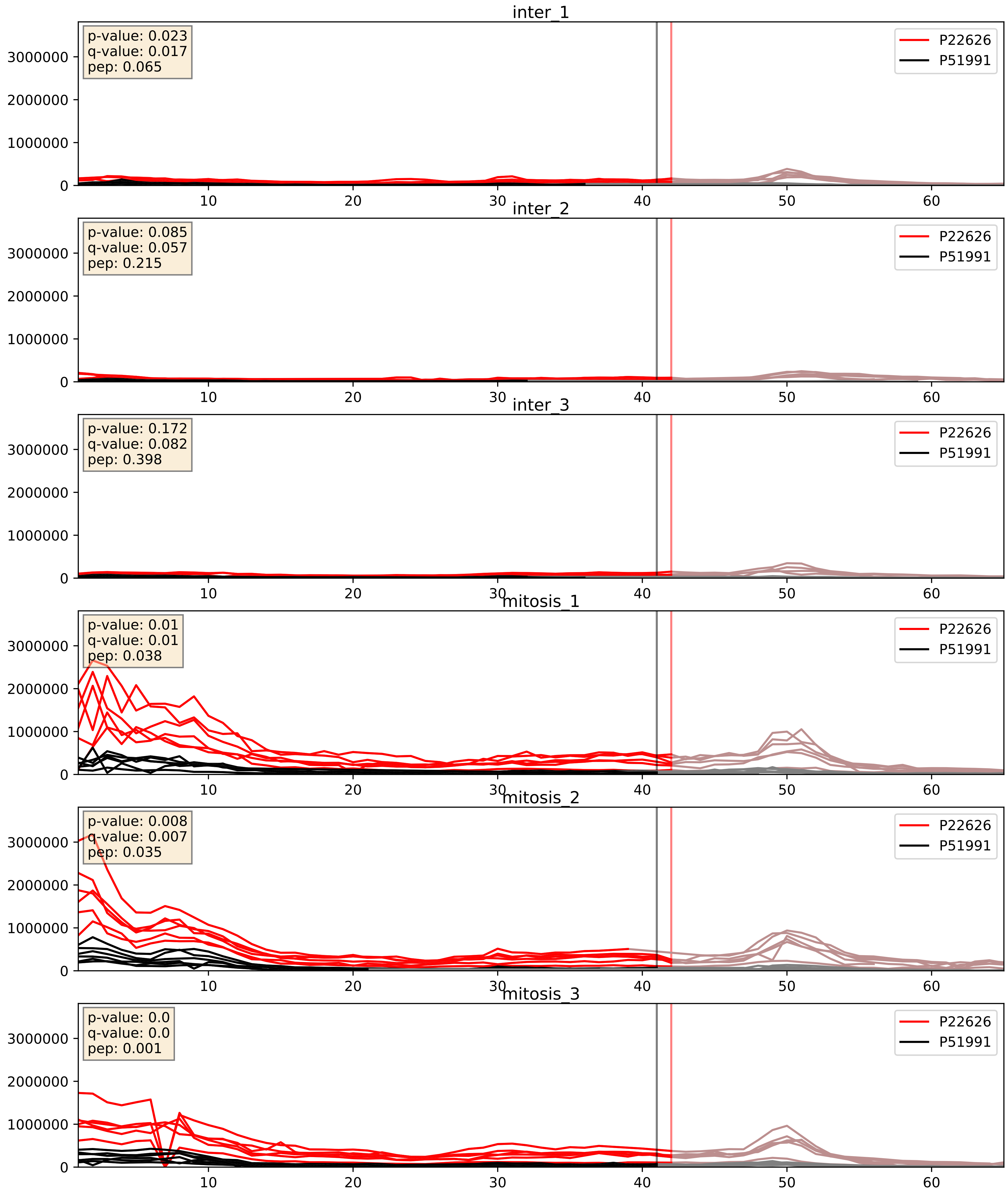


| level | condition_1 | condition_2 | pvalue | pvalue_adjusted |
|---|---|---|---|---|
| complex_abundance | mitosis | inter | 0.00023249210353232296 | 0.010221981956574496 |
| interactor_abundance | mitosis | inter | 0.0019123553121689623 | 0.031986606092438766 |
| interactor_ratio | mitosis | inter | 0.08015668685278068 | 0.47352742543809706 |

P31943_P51991
HNRH1_HUMAN vs ROA3_HUMAN
p-value: 0.0 q-value: 0.0

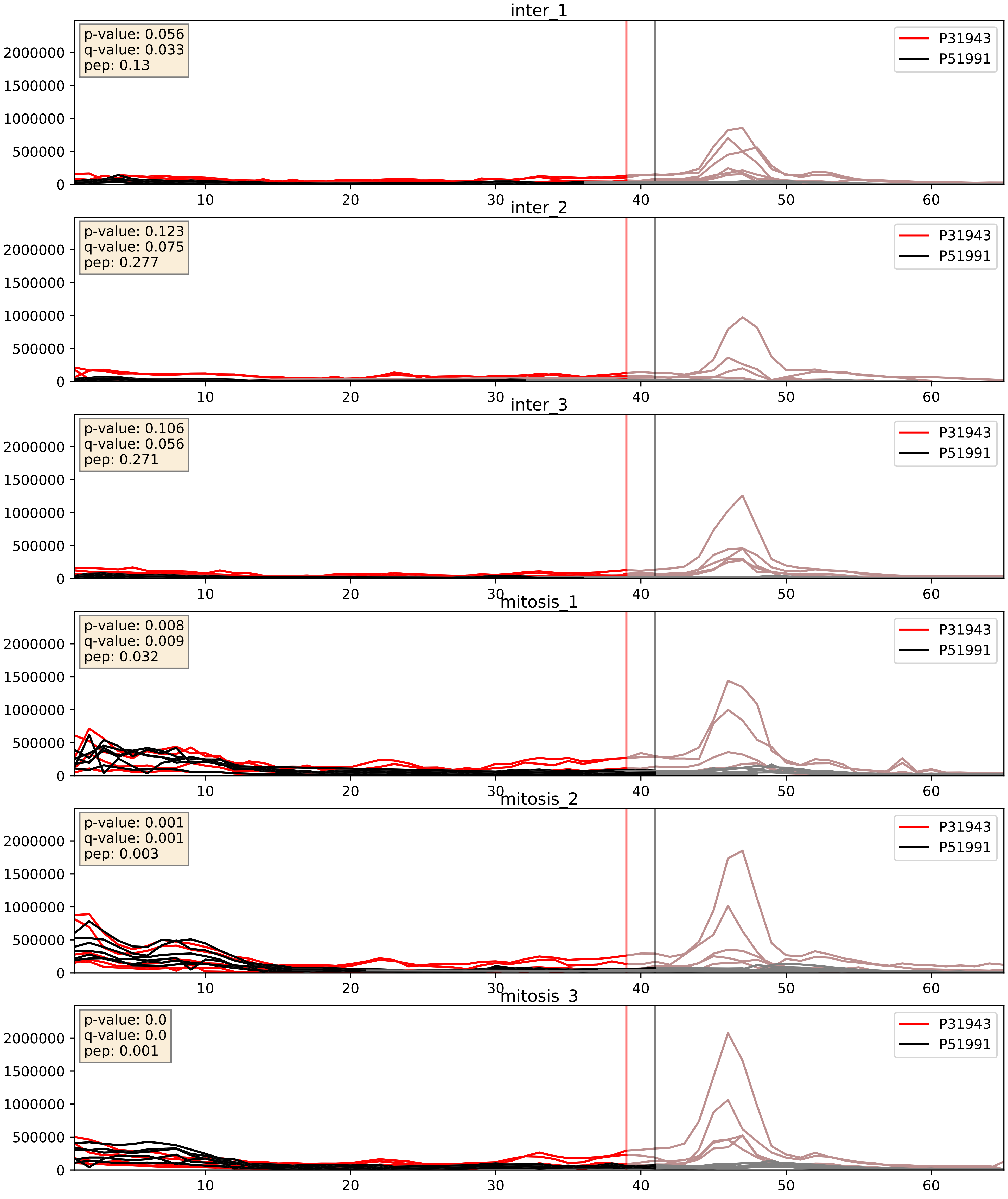


| level | condition_1 | condition_2 | pvalue | pvalue_adjusted |
|---|---|---|---|---|
| complex_abundance | mitosis | inter | 0.0003987949630443645 | 0.01272633620489383 |
| interactor_abundance | mitosis | inter | 0.006828293176471166 | 0.07097776513733234 |
| interactor_ratio | mitosis | inter | 0.01669269551176852 | 0.19102870799563973 |

P38159_P51991
RBMX_HUMAN vs ROA3_HUMAN
p-value: 0.003 q-value: 0.039

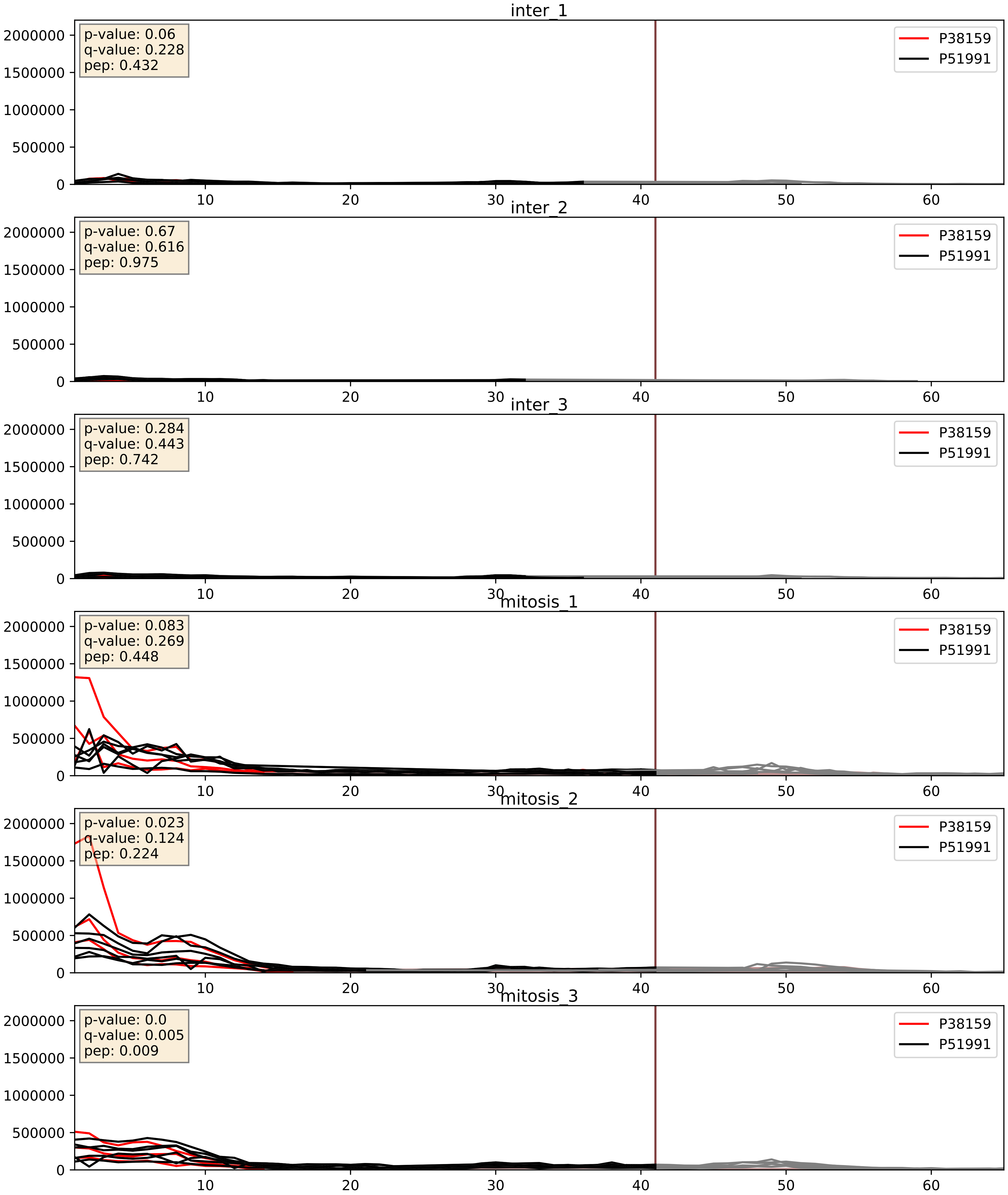


| level | condition_1 | condition_2 | pvalue | pvalue_adjusted |
|---|---|---|---|---|
| complex_abundance | mitosis | inter | 0.000160188868584408 | 0.008563834848633118 |
| interactor_abundance | mitosis | inter | 0.0010979128258399222 | 0.024049720606162197 |
| interactor_ratio | mitosis | inter | 0.9349452147444461 | 0.9881103513214468 |

P51991_P52272
ROA3_HUMAN vs HNRPM_HUMAN
p-value: 0.006 q-value: 0.006

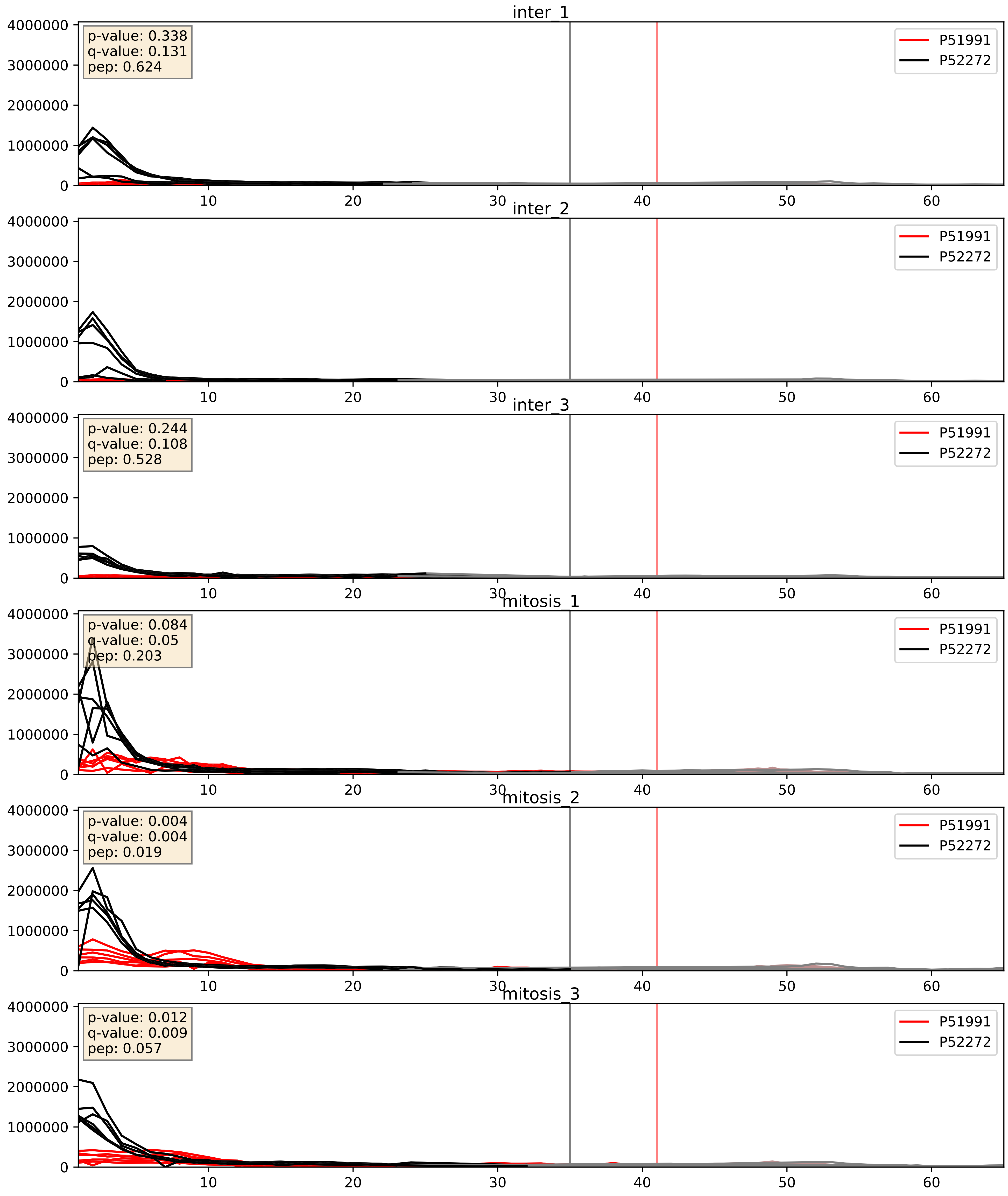


| level | condition_1 | condition_2 | pvalue | pvalue_adjusted |
|---|---|---|---|---|
| interactor_abundance | mitosis | inter | 2.6272351615725858e-06 | 0.0033734327106189023 |
| complex_abundance | mitosis | inter | 0.00045014421143136155 | 0.013379286284209911 |
| interactor_ratio | mitosis | inter | 0.0018482357365531494 | 0.05221418450460382 |

P51991_P52597
ROA3_HUMAN vs HNRPF_HUMAN
p-value: 0.0 q-value: 0.0

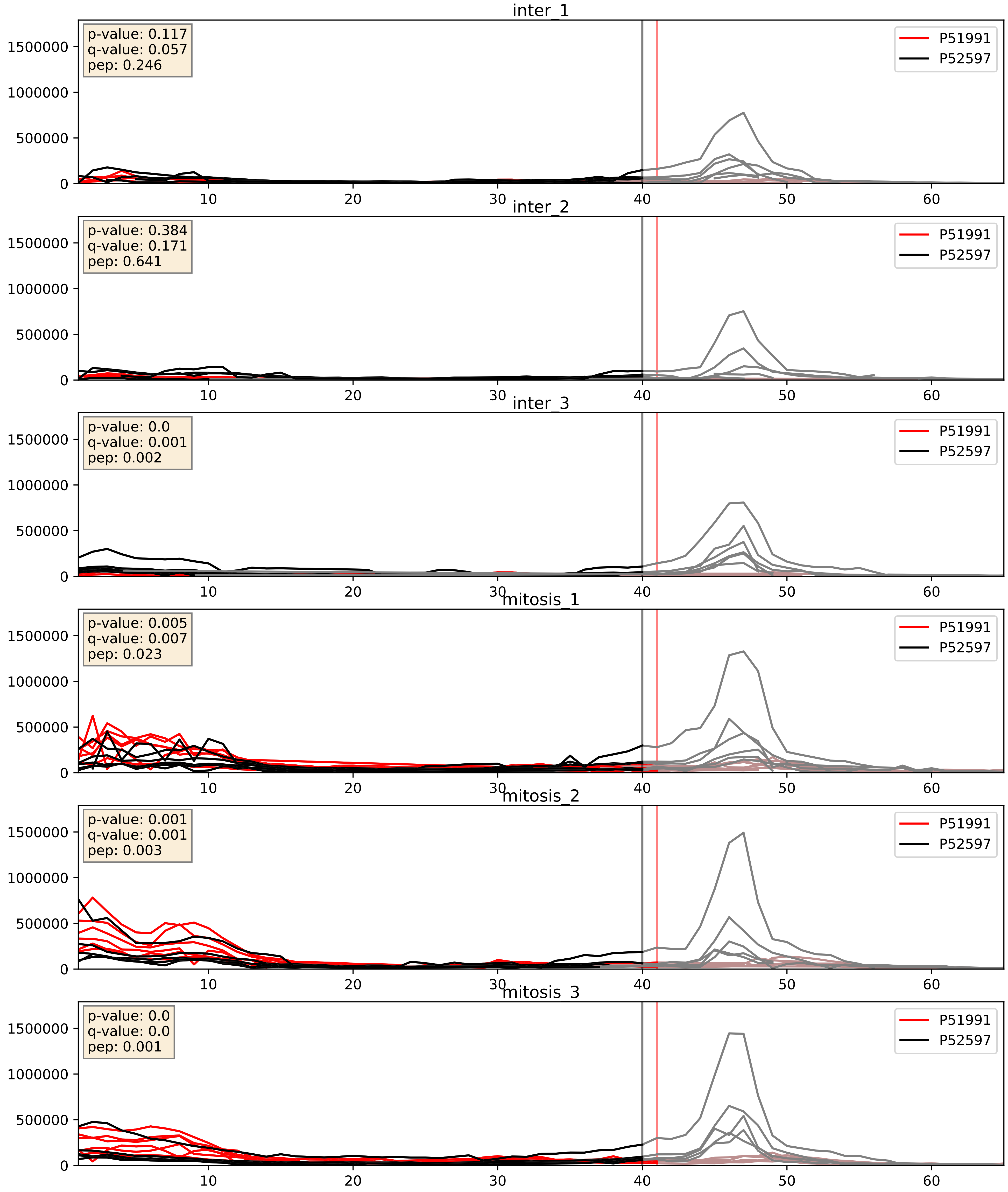


| level | condition_1 | condition_2 | pvalue | pvalue_adjusted |
|---|---|---|---|---|
| interactor_abundance | mitosis | inter | 1.0516040165513909e-05 | 0.0033734327106189023 |
| interactor_ratio | mitosis | inter | 0.00014720601620827773 | 0.012836278661937291 |
| complex_abundance | mitosis | inter | 0.0001689637015938096 | 0.008722442186052247 |

P51991_P55769
ROA3_HUMAN vs NH2L1_HUMAN
p-value: 0.0 q-value: 0.01

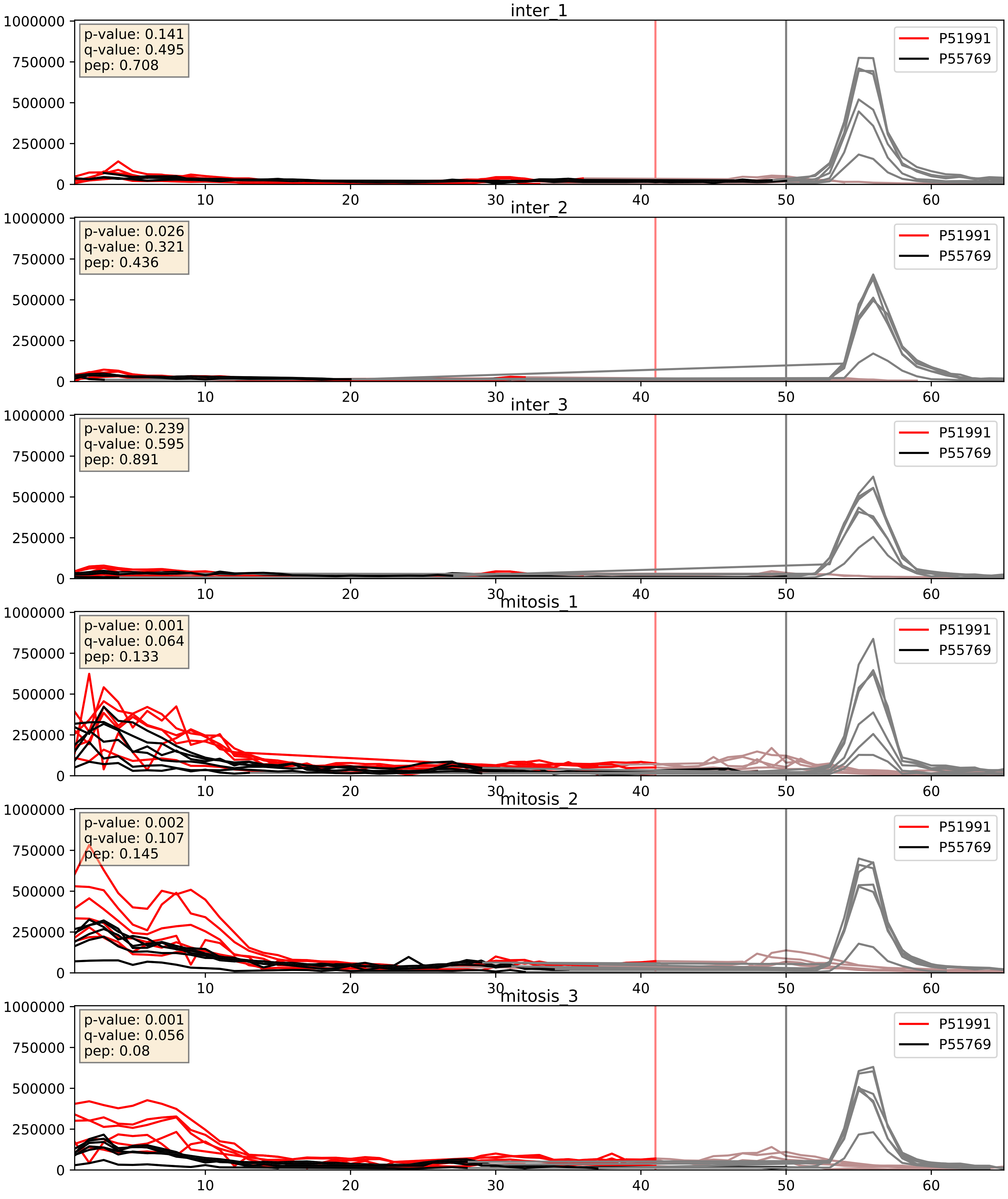


| level | condition_1 | condition_2 | pvalue | pvalue_adjusted |
|---|---|---|---|---|
| interactor_abundance | mitosis | inter | 4.454929811102081e-06 | 0.0033734327106189023 |
| complex_abundance | mitosis | inter | 9.719836869876729e-05 | 0.00670982287146329 |
| interactor_ratio | mitosis | inter | 0.09016169571238221 | 0.5008332999986967 |

P51991_P61978
ROA3_HUMAN vs HNRPK_HUMAN
p-value: 0.031 q-value: 0.022

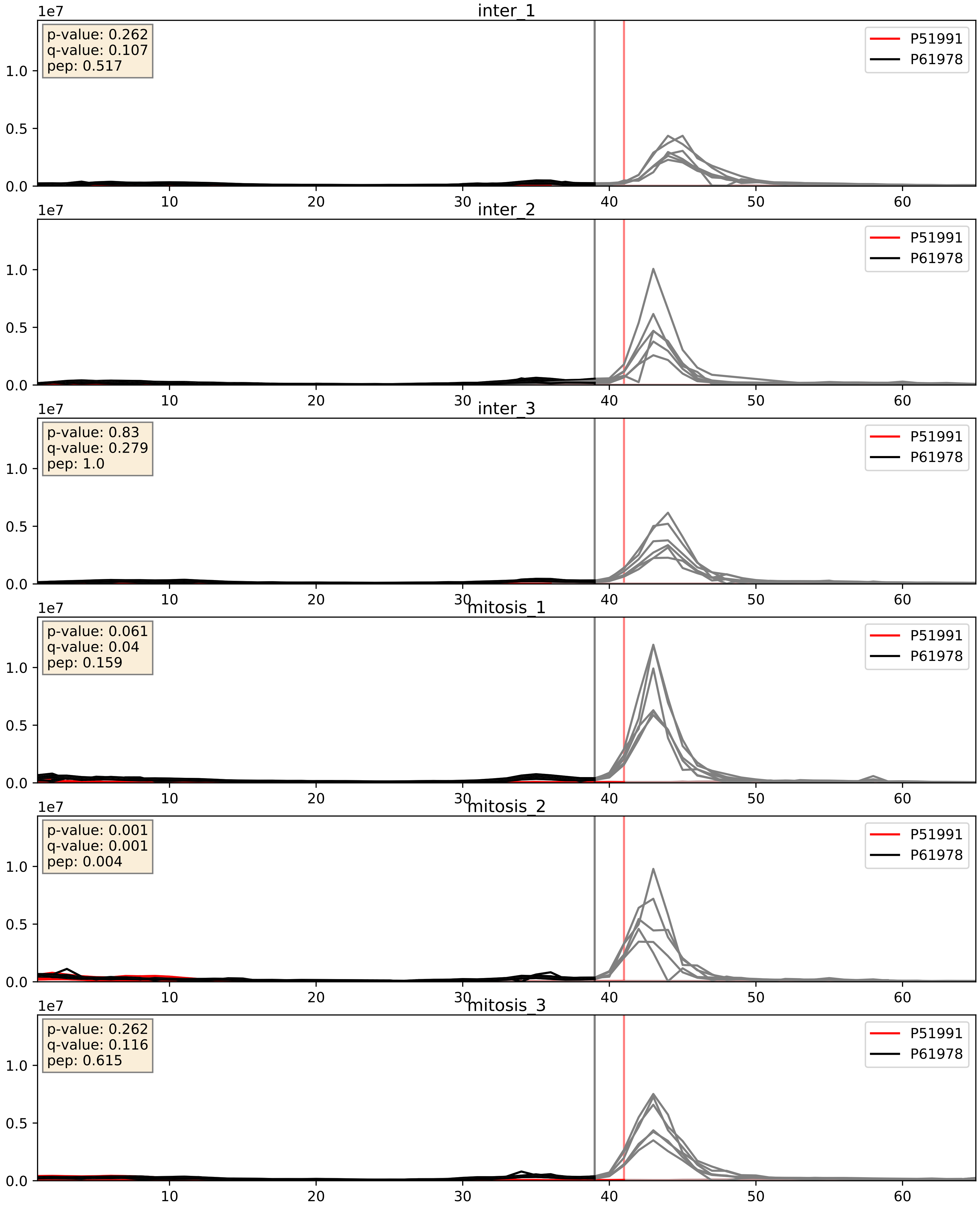


| level | condition_1 | condition_2 | pvalue | pvalue_adjusted |
|---|---|---|---|---|
| interactor_abundance | mitosis | inter | 1.0640500372279248e-05 | 0.0033734327106189023 |
| interactor_ratio | mitosis | inter | 0.04261489076837818 | 0.3405601671696144 |
| complex_abundance | mitosis | inter | 0.0461066276451223 | 0.153569156670135 |

P51991_P62995
ROA3_HUMAN vs TRA2B_HUMAN
p-value: 0.002 q-value: 0.002

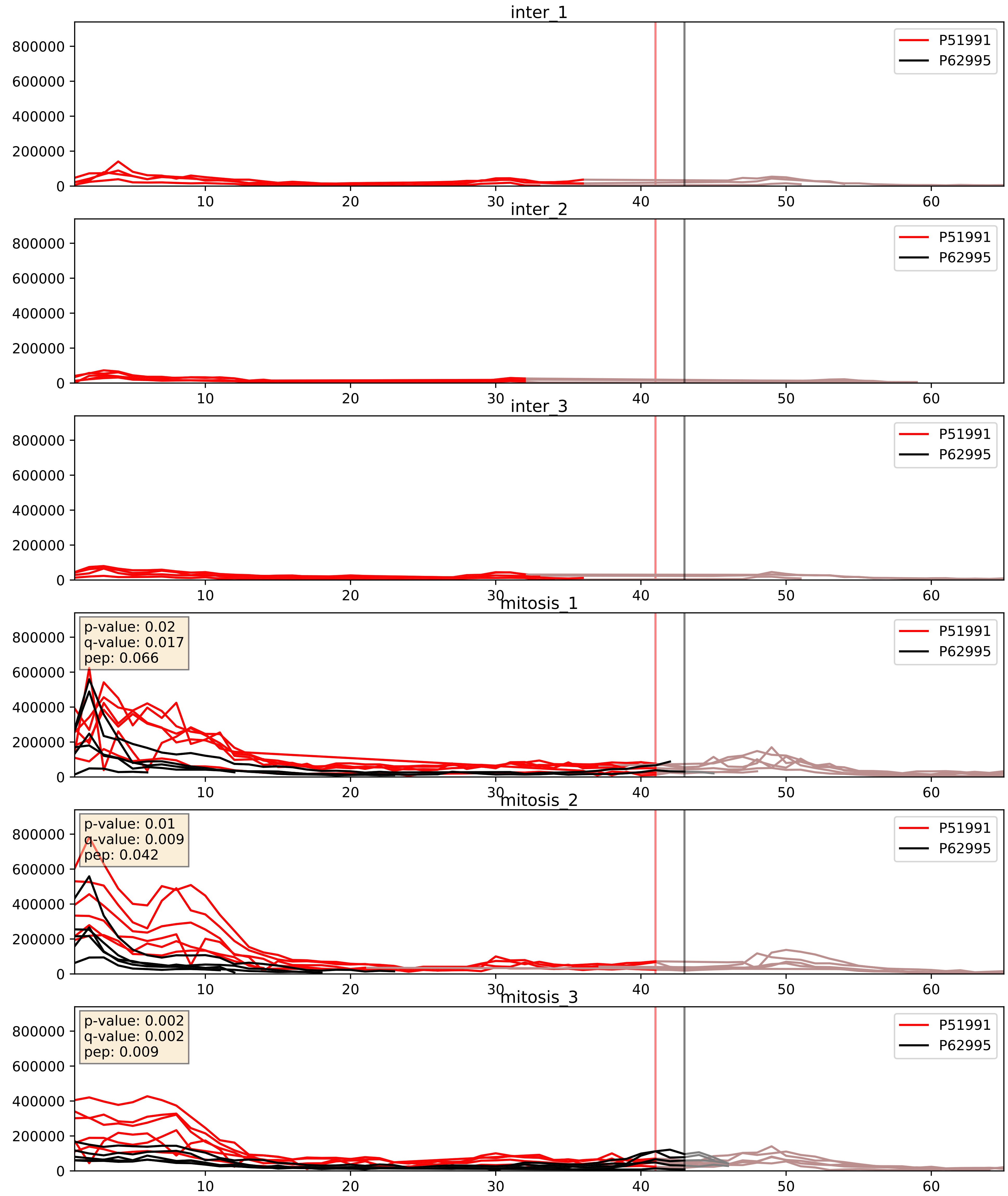


| level | condition_1 | condition_2 | pvalue | pvalue_adjusted |
|---|---|---|---|---|
| interactor_abundance | mitosis | inter | 1.7334412988378383e-05 | 0.003956868671480506 |
| complex_abundance | mitosis | inter | 7.644256938603826e-05 | 0.005948621763131705 |
| interactor_ratio | mitosis | inter | 0.0003545280013586819 | 0.0203675147089283 |

P51991_Q07955
ROA3_HUMAN vs SRSF1_HUMAN
p-value: 0.0 q-value: 0.0

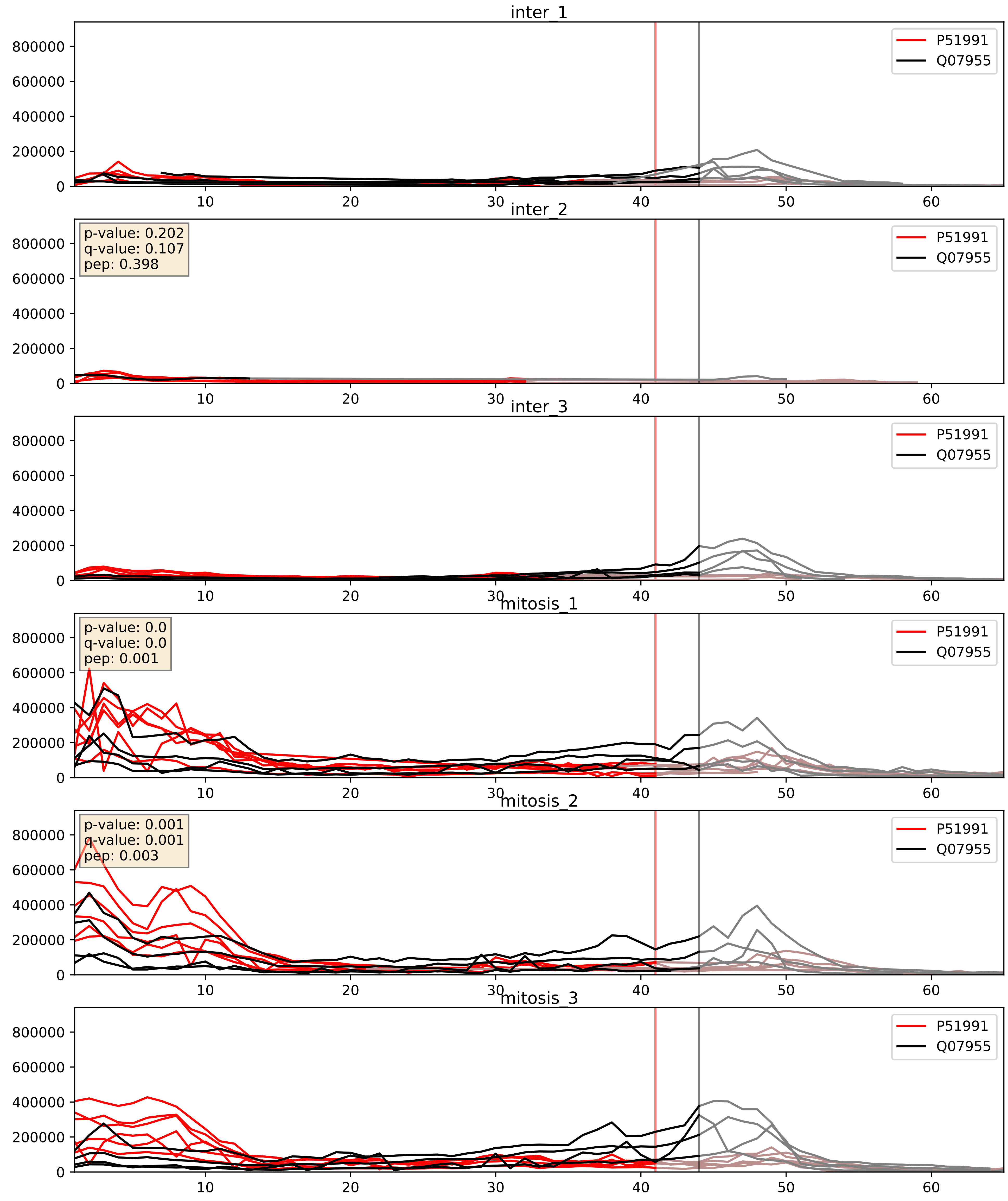


| level | condition_1 | condition_2 | pvalue | pvalue_adjusted |
|---|---|---|---|---|
| interactor_abundance | mitosis | inter | 4.395510848628261e-06 | 0.0033734327106189023 |
| complex_abundance | mitosis | inter | 0.00047430318734800036 | 0.013816623868928207 |
| interactor_ratio | mitosis | inter | 0.39080113396936994 | 0.825249705827432 |

P51991_Q08211
ROA3_HUMAN vs DHX9_HUMAN
p-value: 0.003 q-value: 0.045

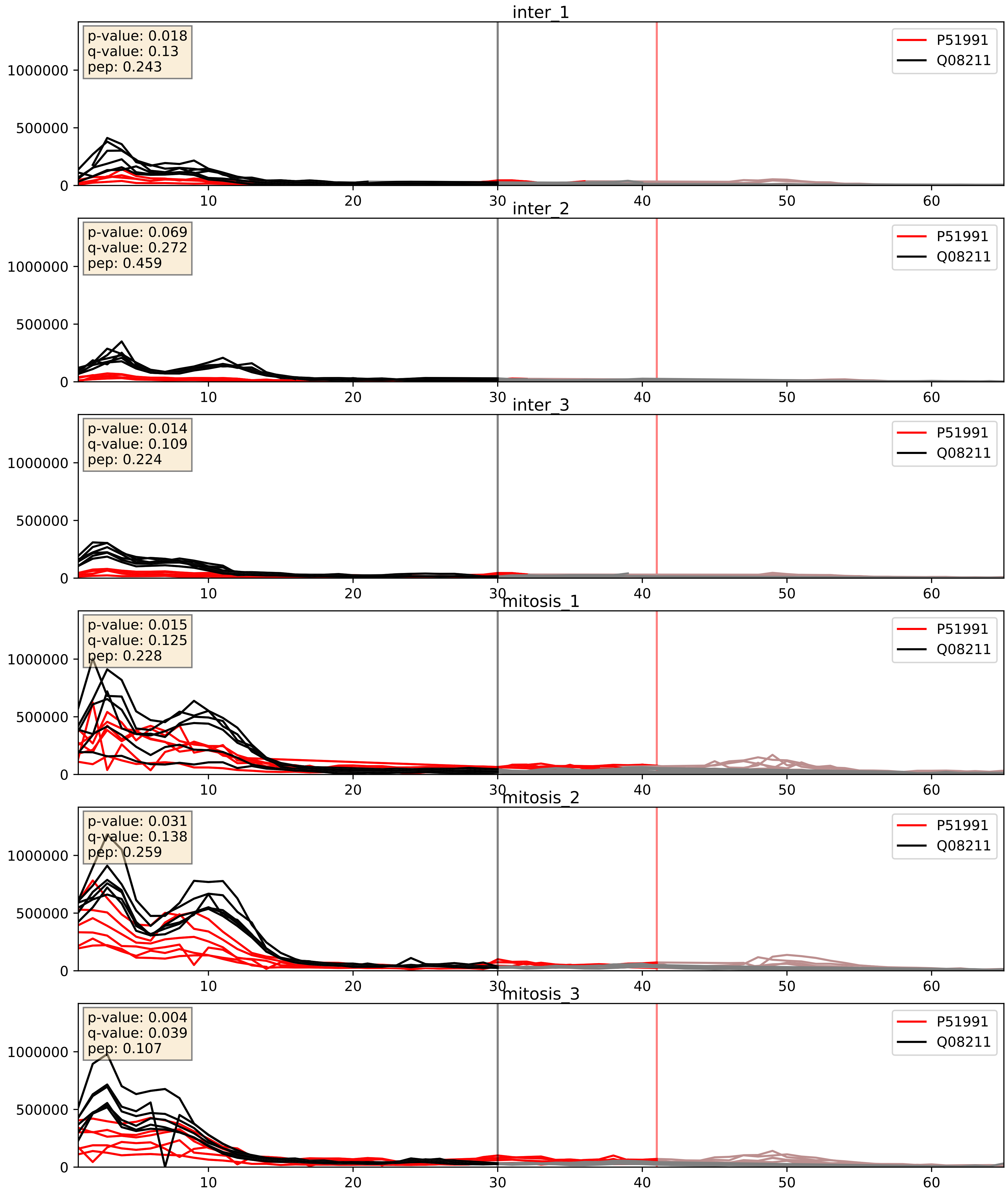


| level | condition_1 | condition_2 | pvalue | pvalue_adjusted |
|---|---|---|---|---|
| interactor_abundance | mitosis | inter | 4.591141207513425e-06 | 0.0033734327106189023 |
| complex_abundance | mitosis | inter | 7.431718998456032e-06 | 0.005691857151652958 |
| interactor_ratio | mitosis | inter | 0.0005736461993080531 | 0.0279000651481644 |

P51991_Q13151
ROA3_HUMAN vs ROA0_HUMAN
p-value: 0.0 q-value: 0.005

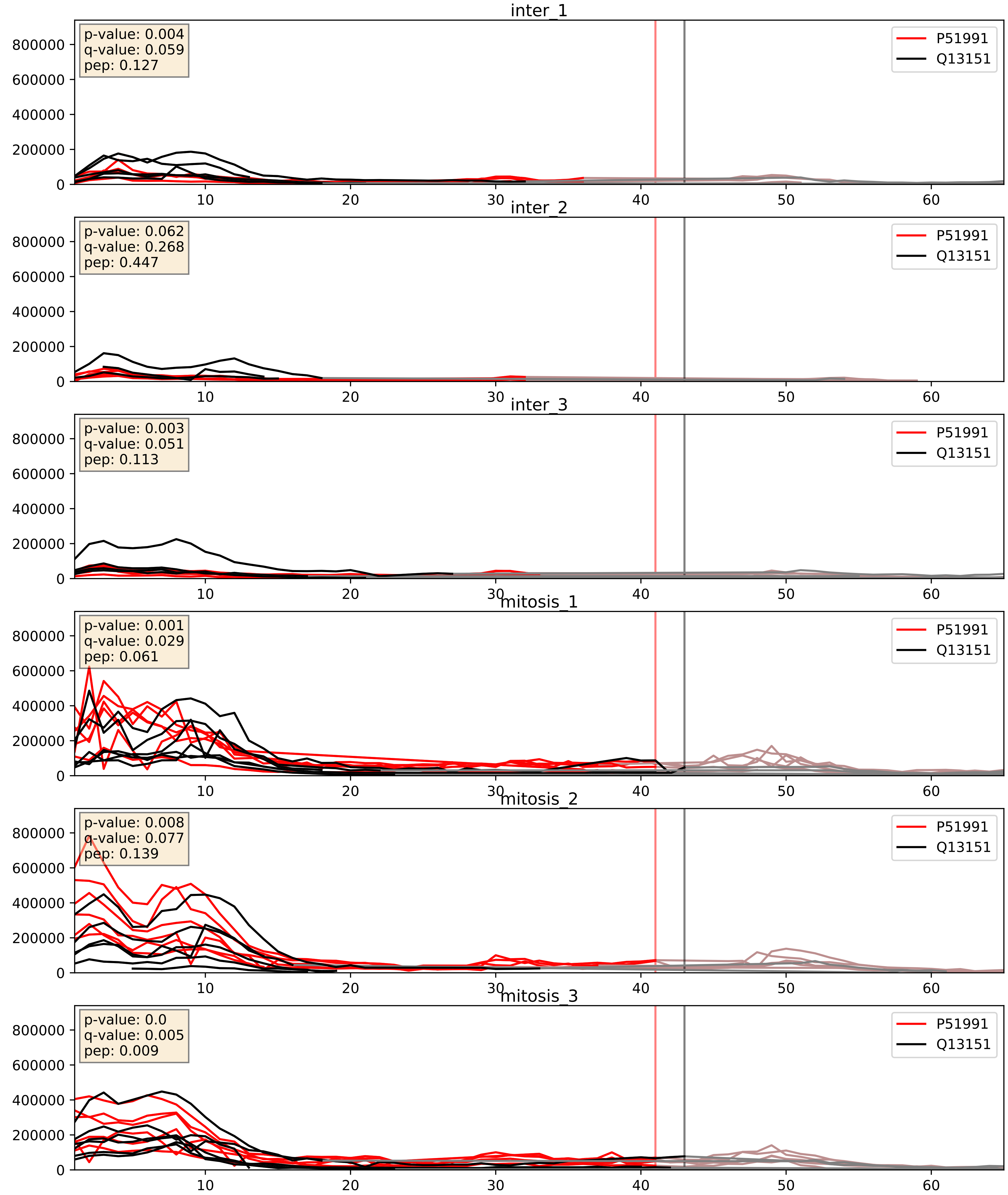


| level | condition_1 | condition_2 | pvalue | pvalue_adjusted |
|---|---|---|---|---|
| interactor_abundance | mitosis | inter | 4.098374668489626e-06 | 0.0033734327106189023 |
| complex_abundance | mitosis | inter | 0.00021831661319998973 | 0.010087086090925927 |
| interactor_ratio | mitosis | inter | 0.014115225863805373 | 0.17445264757370998 |

P51991_Q14103
ROA3_HUMAN vs HNRPD_HUMAN
p-value: 0.037 q-value: 0.026

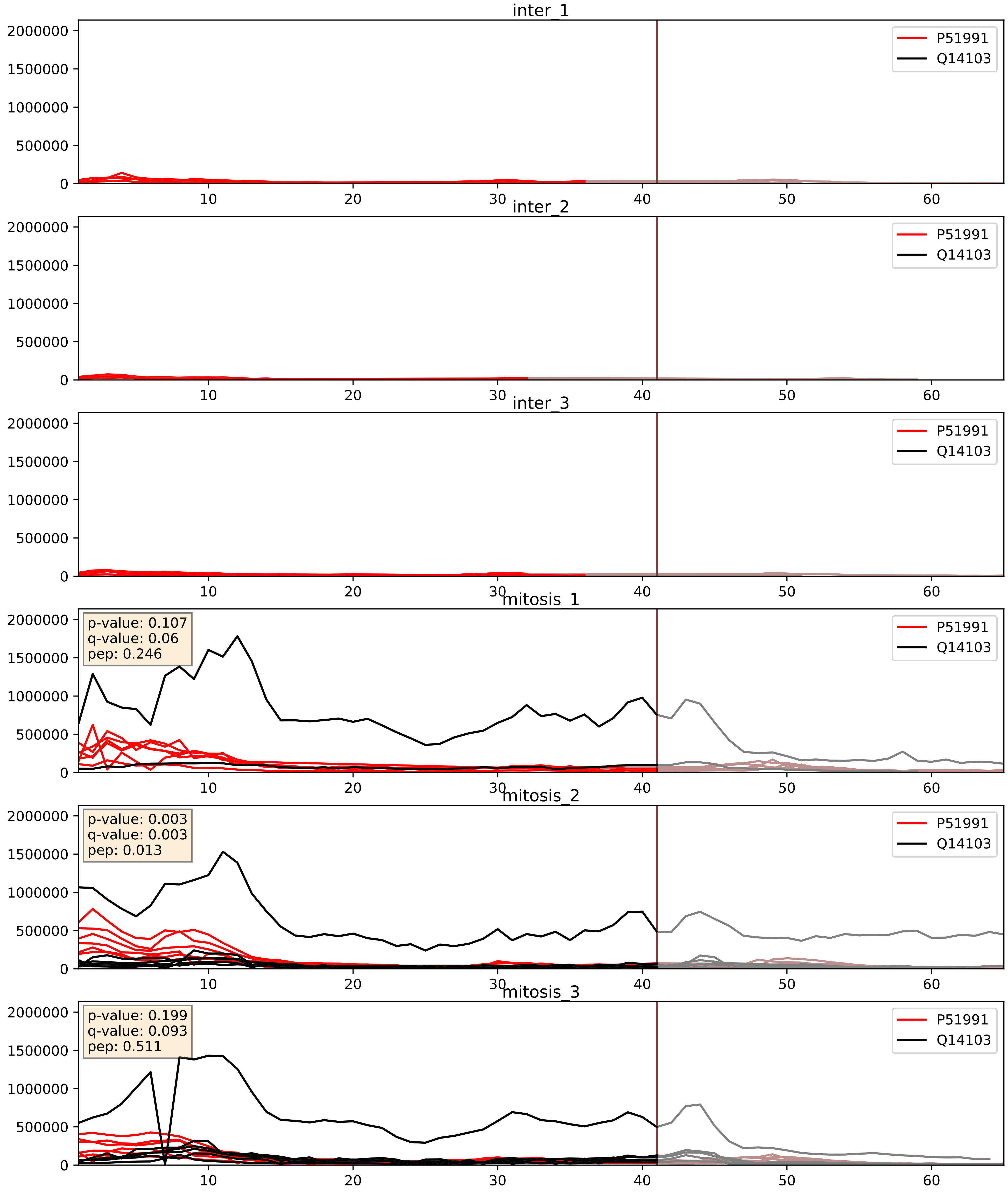


| level | condition_1 | condition_2 | pvalue | pvalue_adjusted |
|---|---|---|---|---|
| interactor_abundance | mitosis | inter | 1.150670630946923e-05 | 0.0033803141163598503 |
| complex_abundance | mitosis | inter | 0.0005845801120302033 | 0.015650423933737376 |
| interactor_ratio | mitosis | inter | 0.748127922946446 | 0.9491588884573259 |

P51991_Q15717
ROA3_HUMAN vs ELAV1_HUMAN
p-value: 0.001 q-value: 0.024

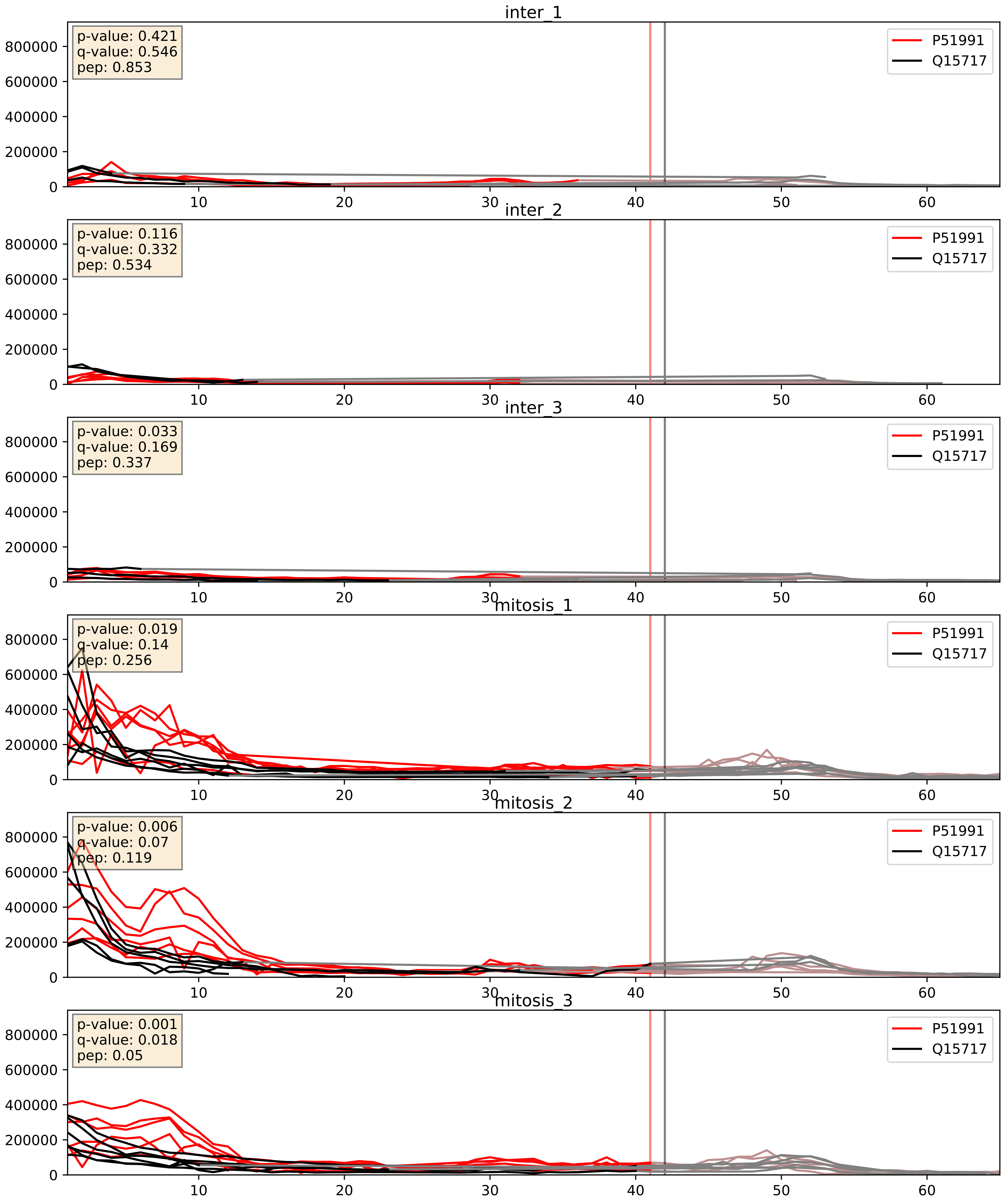


| level | condition_1 | condition_2 | pvalue | pvalue_adjusted |
|---|---|---|---|---|
| interactor_abundance | mitosis | inter | 3.736100323192025e-06 | 0.0033734327106189023 |
| complex_abundance | mitosis | inter | 1.684569173590749e-05 | 0.005691857151652958 |
| interactor_ratio | mitosis | inter | 0.24176845376887507 | 0.7200661897289671 |

P51991_Q16629
ROA3_HUMAN vs SRSF7_HUMAN
p-value: 0.0 q-value: 0.007

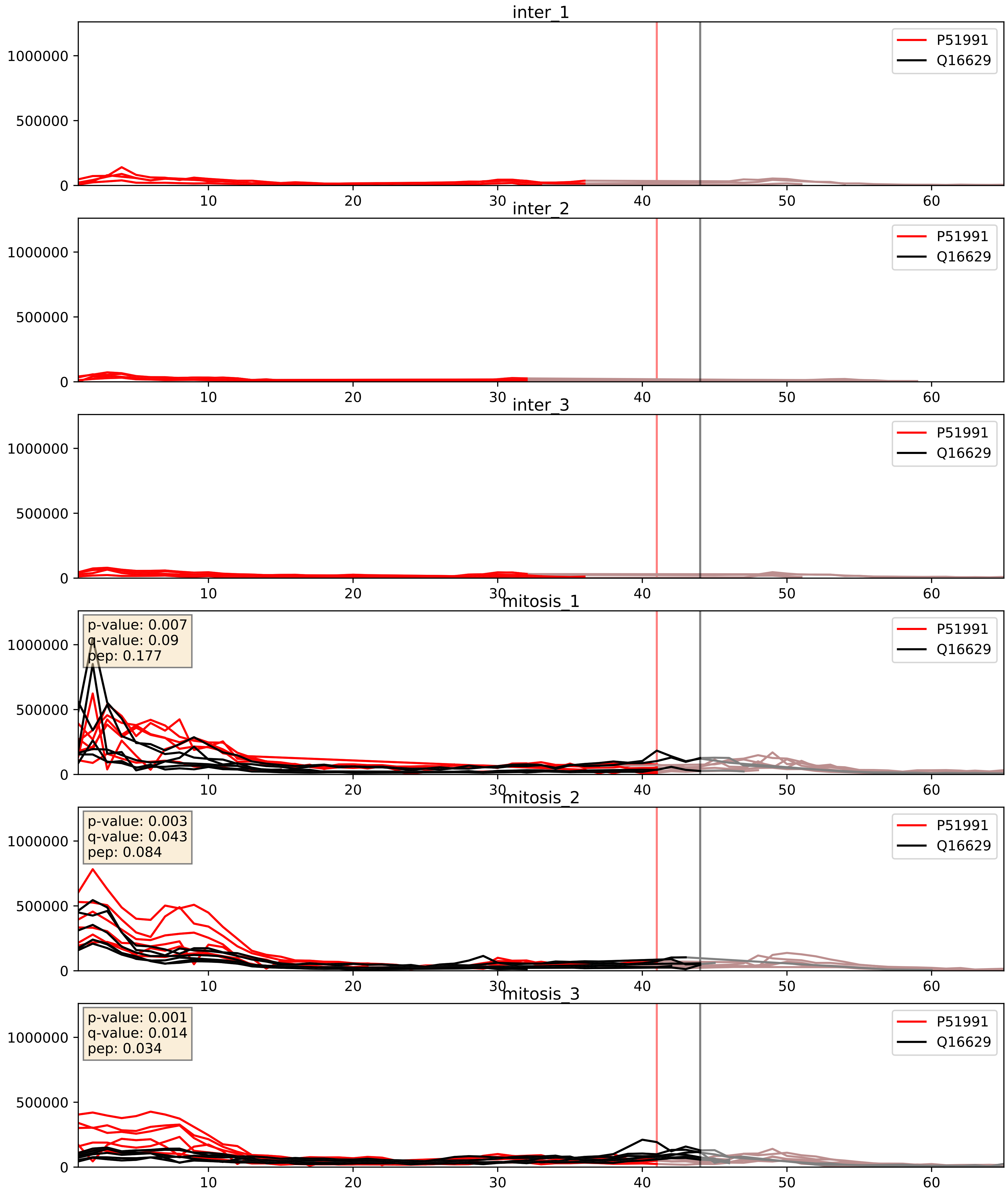


| level | condition_1 | condition_2 | pvalue | pvalue_adjusted |
|---|---|---|---|---|
| interactor_abundance | mitosis | inter | 1.150670630946923e-05 | 0.0033803141163598503 |
| complex_abundance | mitosis | inter | 2.3929388745179548e-05 | 0.005691857151652958 |
| interactor_ratio | mitosis | inter | 0.007352680833101884 | 0.11920255290028813 |

P51991_Q1KMD3
ROA3_HUMAN vs HNRL2_HUMAN
p-value: 0.0 q-value: 0.047

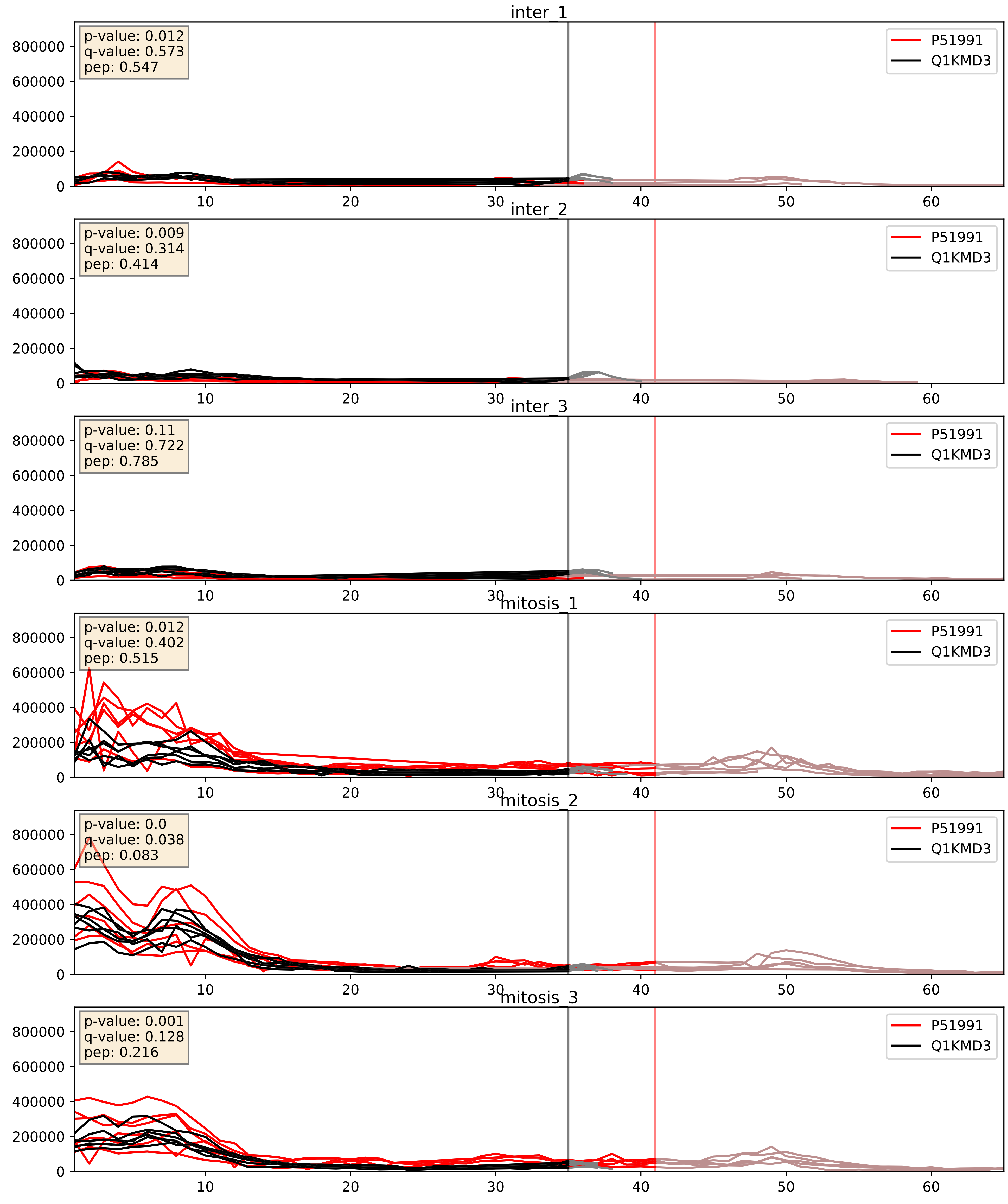


| level | condition_1 | condition_2 | pvalue | pvalue_adjusted |
| --- | --- | --- | --- | --- |
| complex_abundance | mitosis | inter | 8.668080290568026e-07 | 0.0024732922429087437 |
| interactor_abundance | mitosis | inter | 3.1946371197007753e-06 | 0.0033734327106189023 |
| interactor_ratio | mitosis | inter | 0.005567869221764732 | 0.10119099901975818 |

P51991_Q9BUJ2
ROA3_HUMAN vs HNRL1_HUMAN
p-value: 0.003 q-value: 0.039

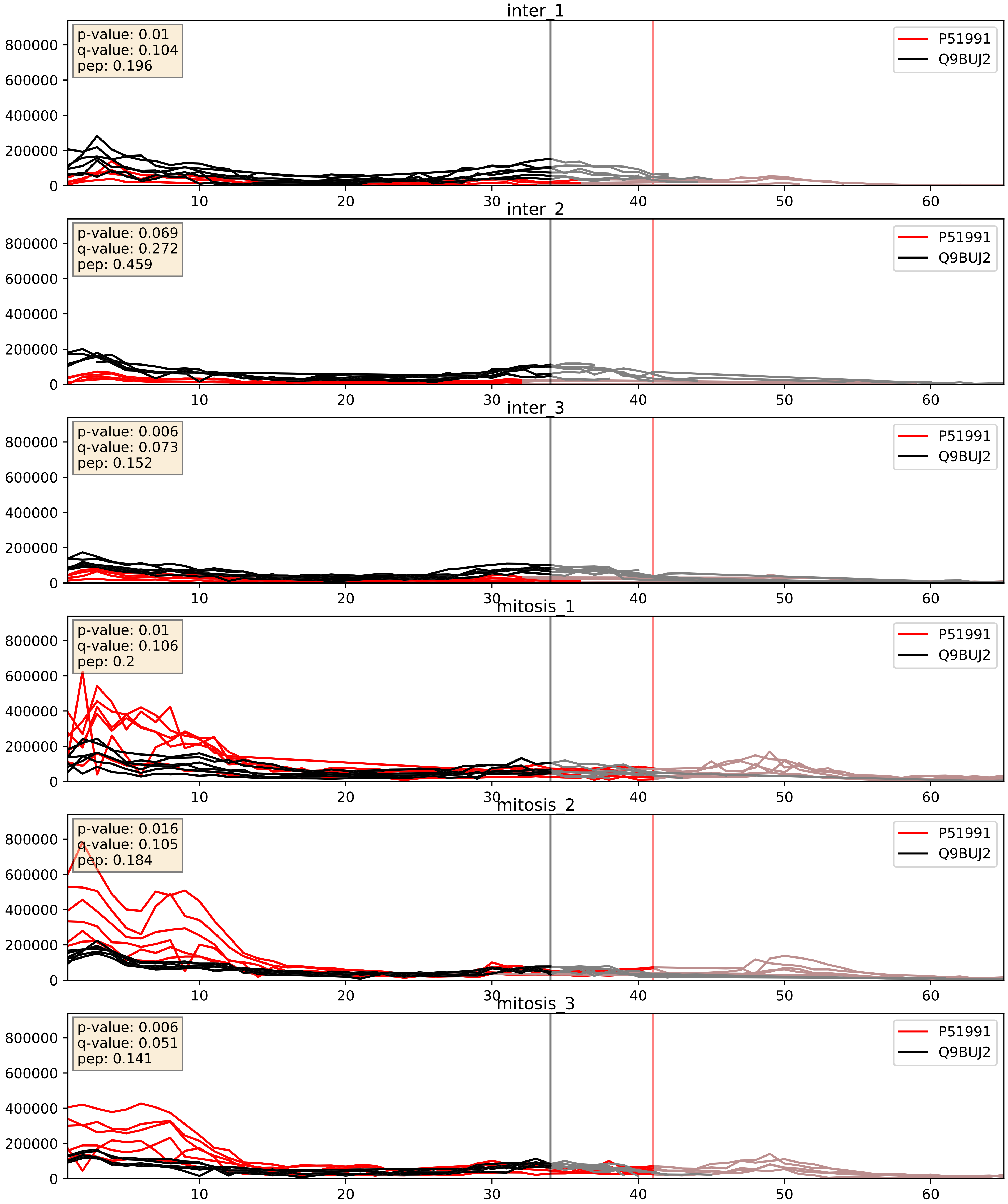


| level | condition_1 | condition_2 | pvalue | pvalue_adjusted |
|---|---|---|---|---|
| interactor_abundance | mitosis | inter | 1.1649447013156027e-05 | 0.0033803141163598503 |
| interactor_ratio | mitosis | inter | 7.24788381117057e-05 | 0.008863126489088583 |
| complex_abundance | mitosis | inter | 0.0001877343549764503 | 0.009111526782692203 |
