## Supplementary Data 2 for "SECAT: Quantifying differential protein-protein interaction states by network-centric analysis": hela_string_000019_Q9UNX4.pdf

Q9UNX4_Q9UNX4

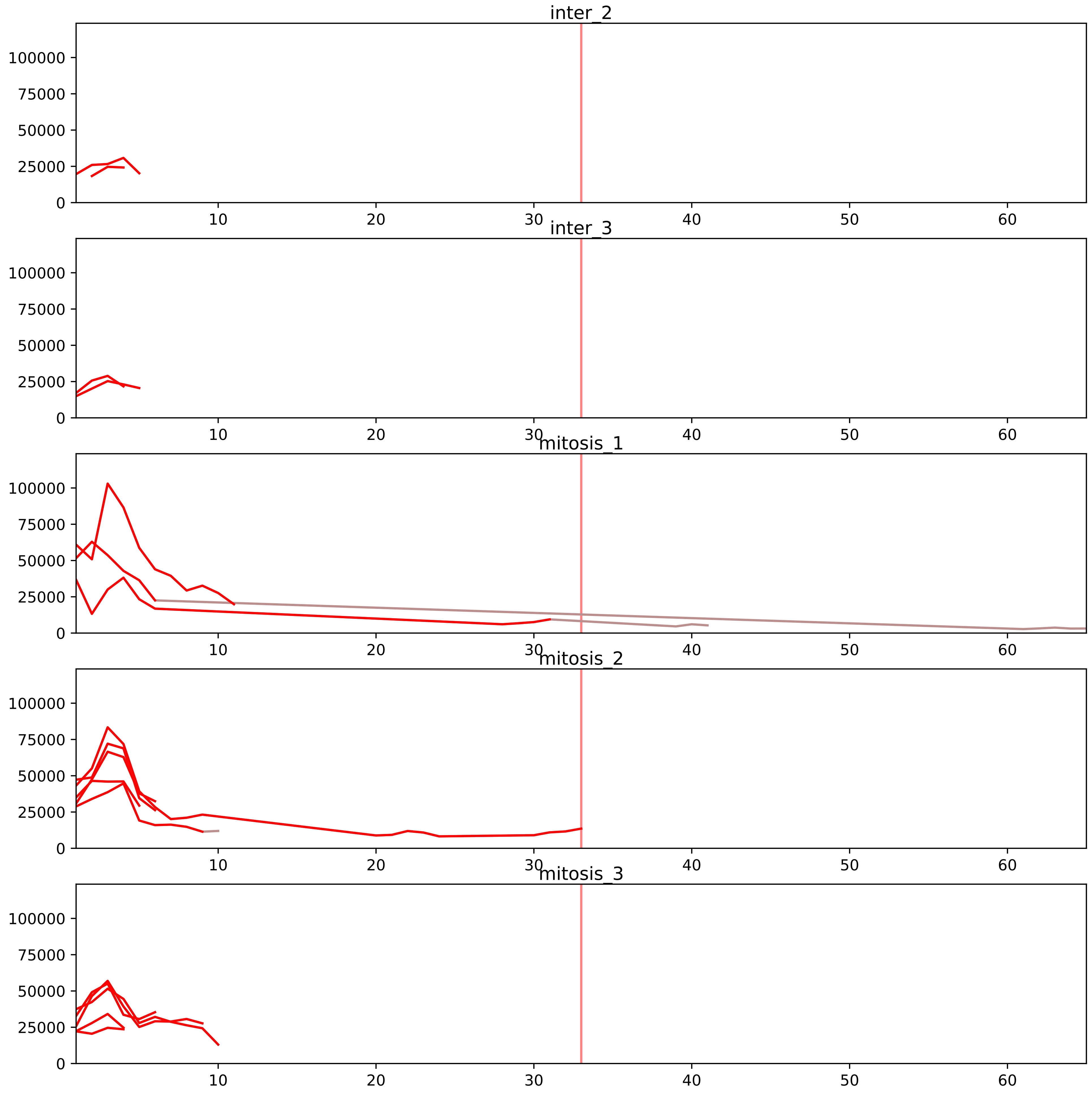


| level | condition_1 | condition_2 | pvalue | pvalue_adjusted |
|---|---|---|---|---|
| interactor_ratio | mitosis | inter | 2.8568744684025442e-12 | 2.9203605677003784e-10 |
| complex_abundance | mitosis | inter | 0.0015494707625799665 | 0.014044446405491201 |
| interactor_abundance | mitosis | inter | 0.06223371756249363 | 0.16692425701893335 |
| monomer_abundance | mitosis | inter | 0.30787750232205874 | 0.6718670942432774 |
| total_abundance | mitosis | inter | 0.36705873281484935 | 0.6763286314731807 |
| assembled_abundance | mitosis | inter | 0.37504378248513 | 0.6549319440249798 |

O00567_Q9UNX4
NOP56_HUMAN vs WDR3_HUMAN
p-value: 0.017 q-value: 0.014

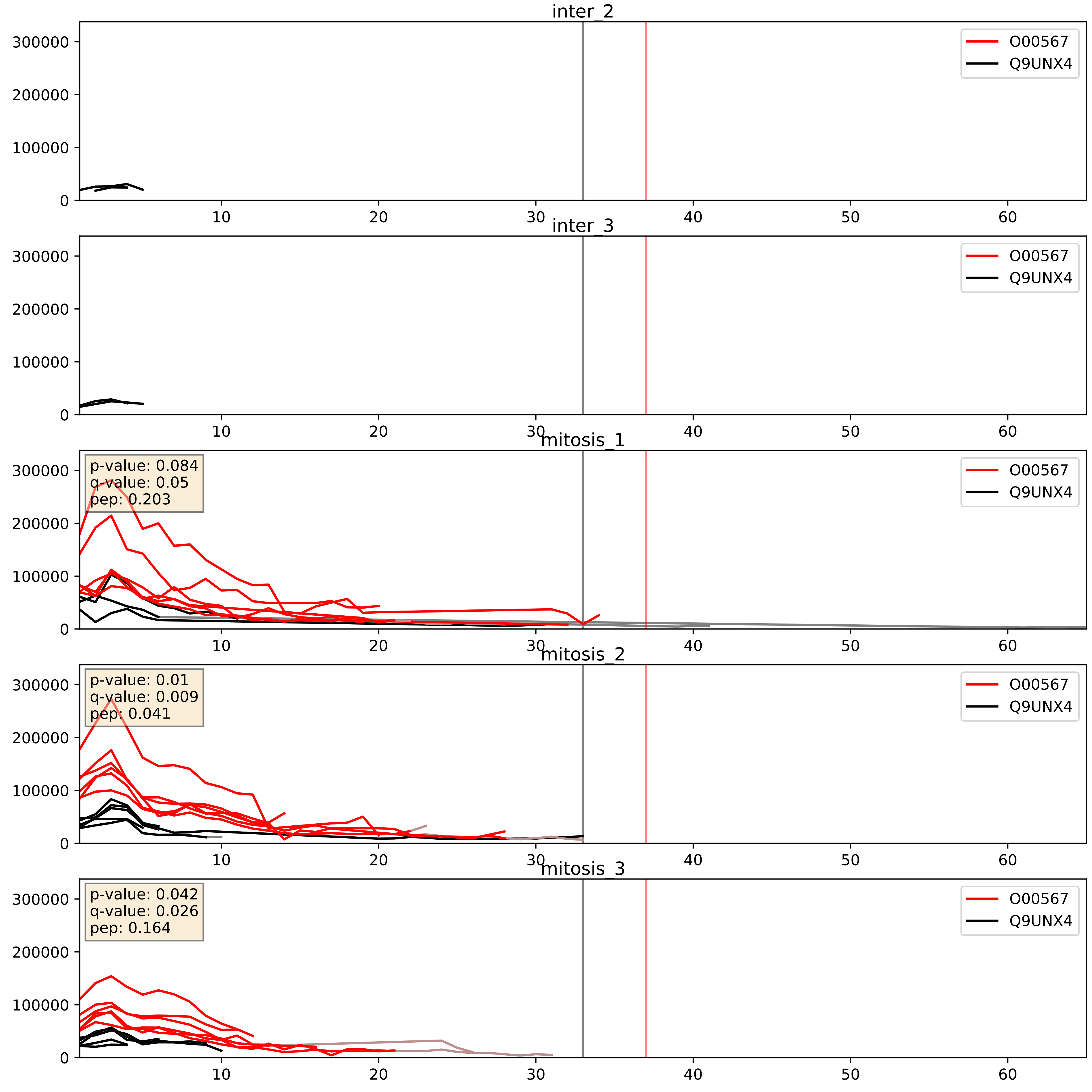


| level | condition_1 | condition_2 | pvalue | pvalue_adjusted |
|---|---|---|---|---|
| interactor_ratio | mitosis | inter | 0.00014655098812599722 | 0.012836278661937291 |
| interactor_abundance | mitosis | inter | 0.001146348816560698 | 0.024325226673177953 |
| complex_abundance | mitosis | inter | 0.003340135899144592 | 0.04242071705738532 |

O76021_Q9UNX4
RL1D1_HUMAN vs WDR3_HUMAN
p-value: 0.014 q-value: 0.011

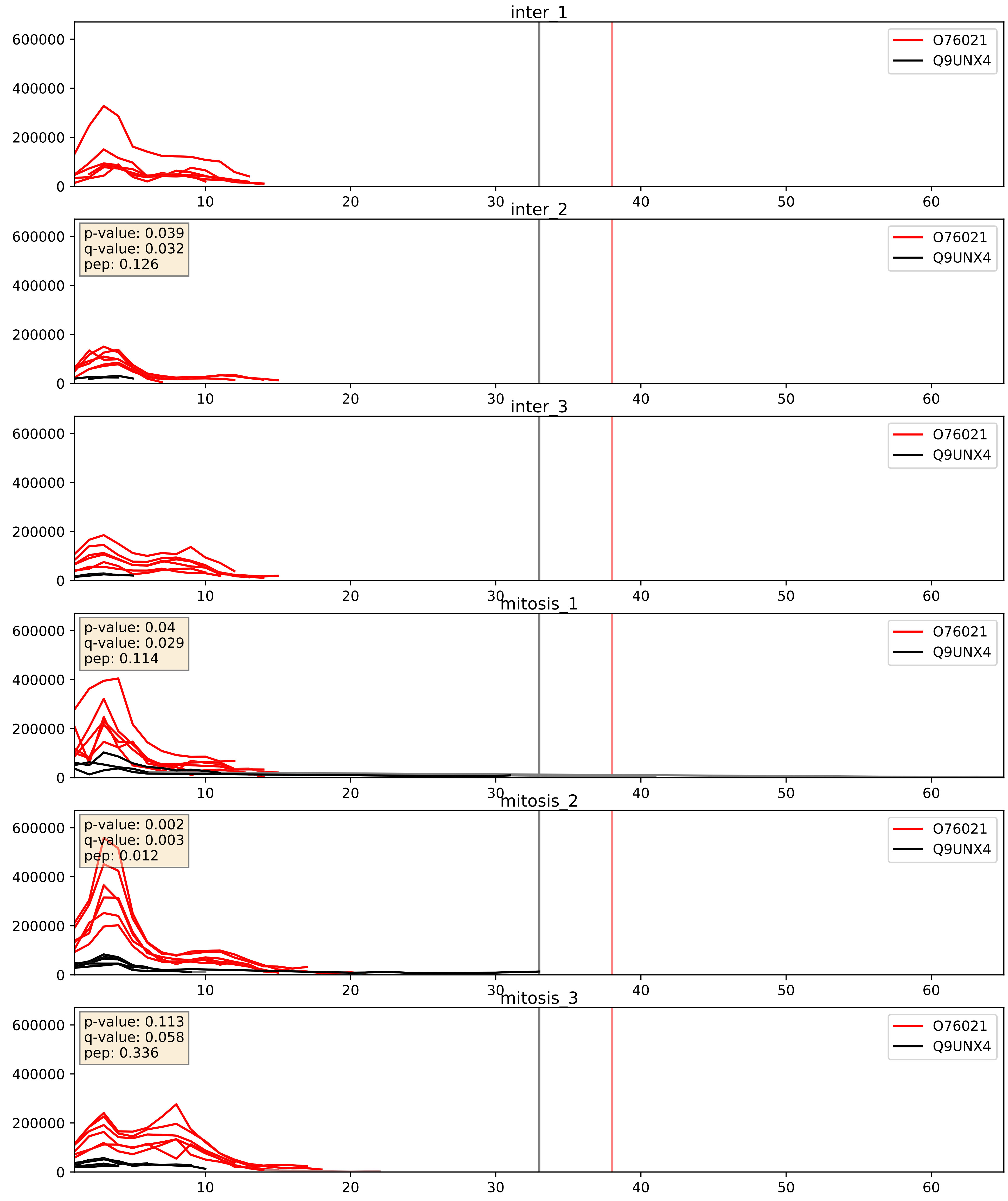


| level | condition_1 | condition_2 | pvalue | pvalue_adjusted |
|---|---|---|---|---|
| interactor_abundance | mitosis | inter | 0.022557396454085495 | 0.12329937058940194 |
| complex_abundance | mitosis | inter | 0.02766391673322131 | 0.12094133158139654 |
| interactor_ratio | mitosis | inter | 0.033938259800389954 | 0.296589834089654 |

P22087_Q9UNX4
FBRL_HUMAN vs WDR3_HUMAN
p-value: 0.025 q-value: 0.019

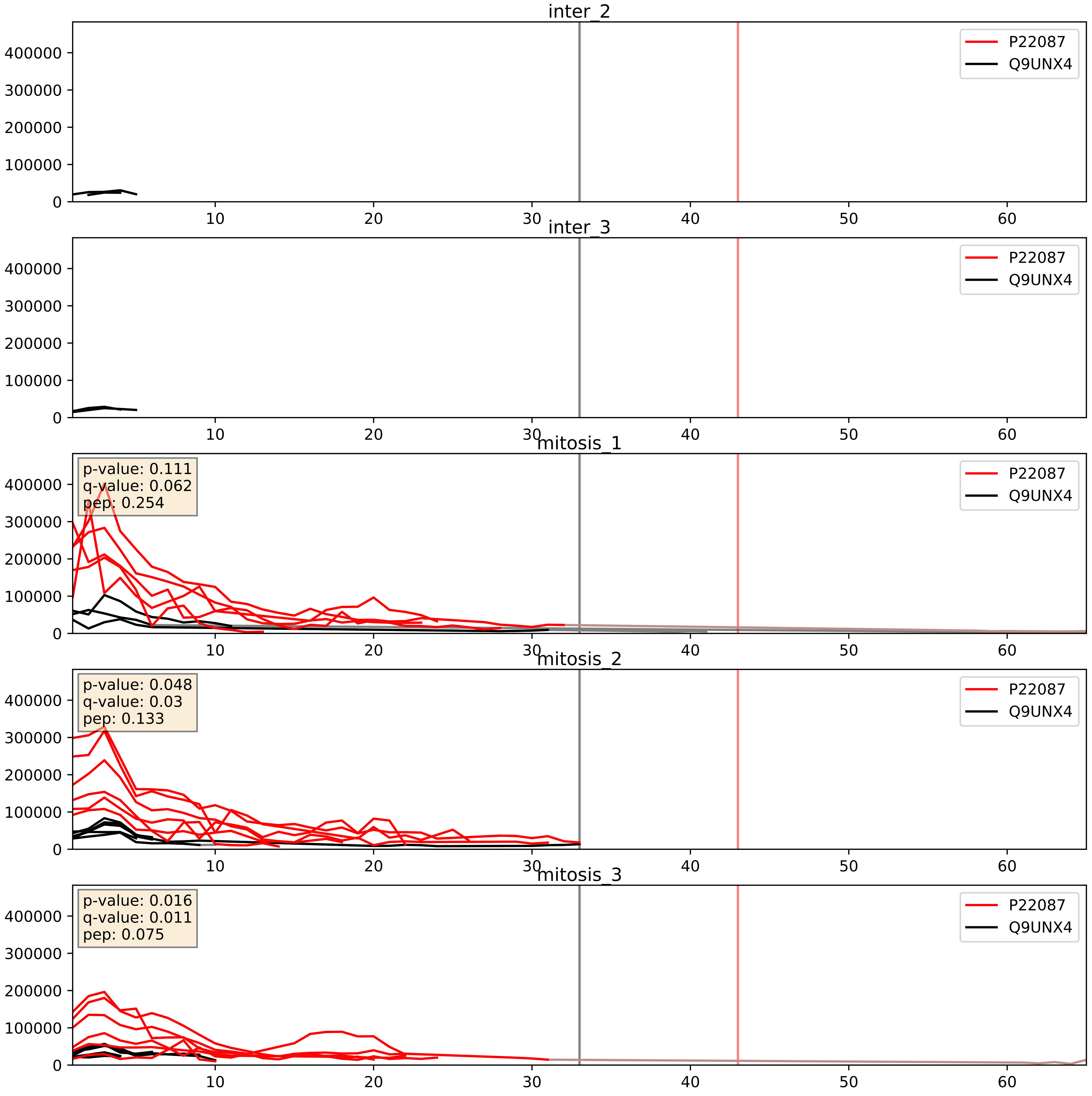


| level | condition_1 | condition_2 | pvalue | pvalue_adjusted |
|---|---|---|---|---|
| interactor_ratio | mitosis | inter | 0.00040930987289006183 | 0.022604467818960836 |
| interactor_abundance | mitosis | inter | 0.0008065885048889307 | 0.020834500976147856 |
| complex_abundance | mitosis | inter | 0.002005177899684022 | 0.032203232310122384 |

P56182_Q9UNX4
RRP1_HUMAN vs WDR3_HUMAN
p-value: 0.003 q-value: 0.042

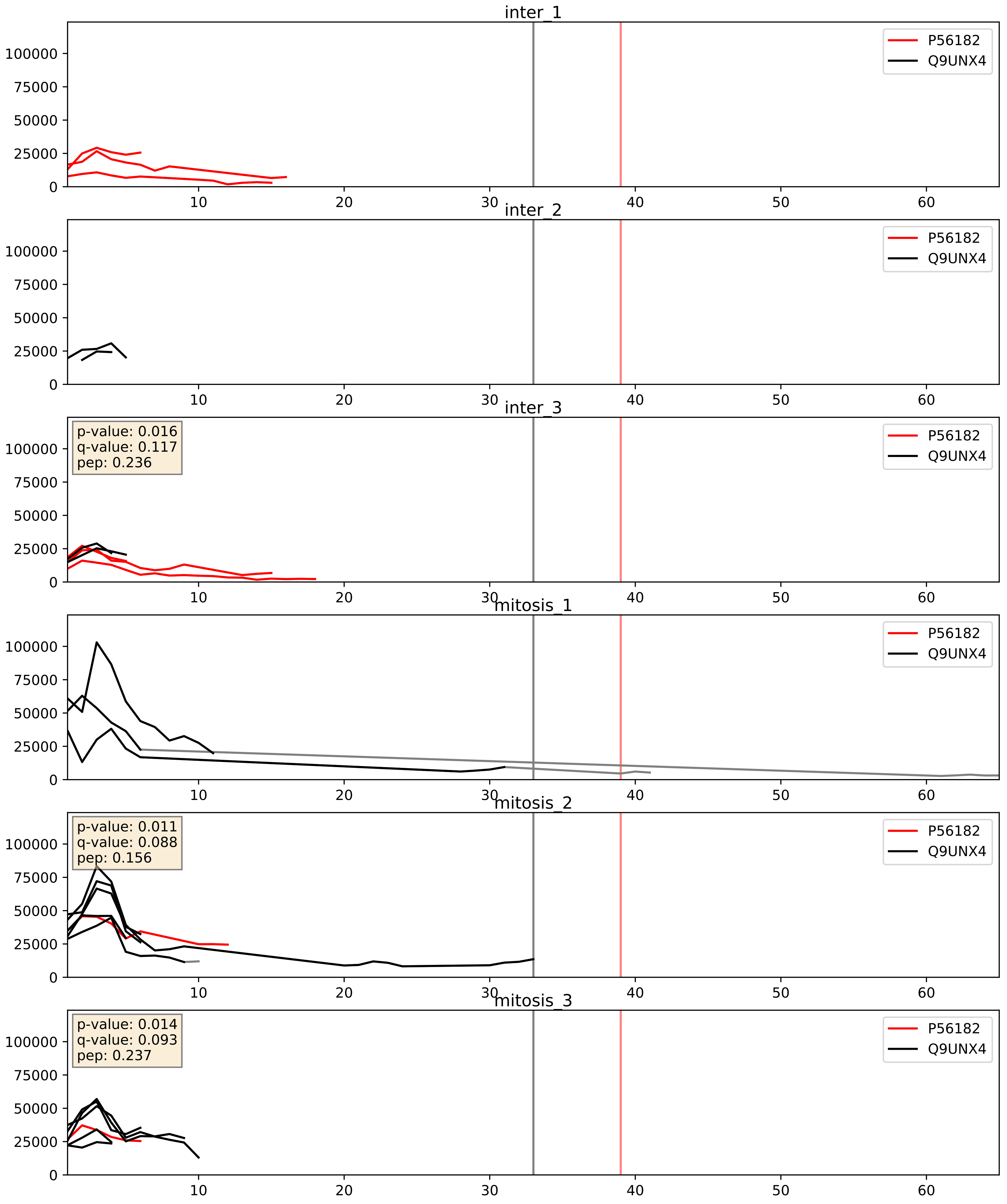


| level | condition_1 | condition_2 | pvalue | pvalue_adjusted |
|---|---|---|---|---|
| interactor_abundance | mitosis | inter | 0.011165509238416844 | 0.0931547359462458 |
| complex_abundance | mitosis | inter | 0.058870177199176074 | 0.17347193590338766 |
| interactor_ratio | mitosis | inter | 0.09493145750992908 | 0.511721206728585 |

Q12788_Q9UNX4
TBL3_HUMAN vs WDR3_HUMAN
p-value: 0.052 q-value: 0.034

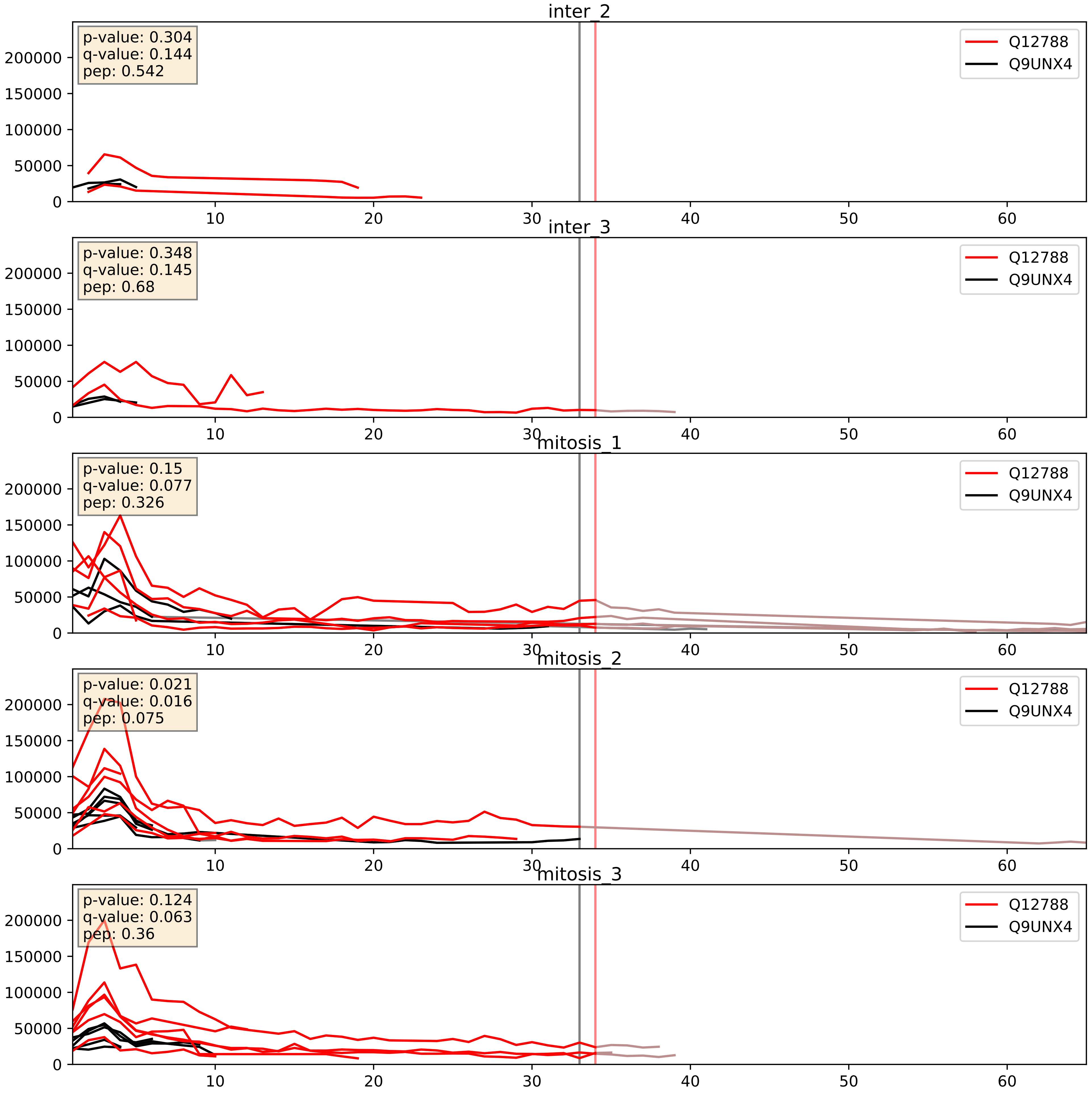


| level | condition_1 | condition_2 | pvalue | pvalue_adjusted |
|---|---|---|---|---|
| interactor_abundance | mitosis | inter | 0.005053479358839491 | 0.05906252380726447 |
| interactor_ratio | mitosis | inter | 0.006624735222091087 | 0.11281850825965009 |
| complex_abundance | mitosis | inter | 0.02123880532057976 | 0.10740063977781633 |

Q13601_Q9UNX4
KRR1_HUMAN vs WDR3_HUMAN
p-value: 0.054 q-value: 0.035

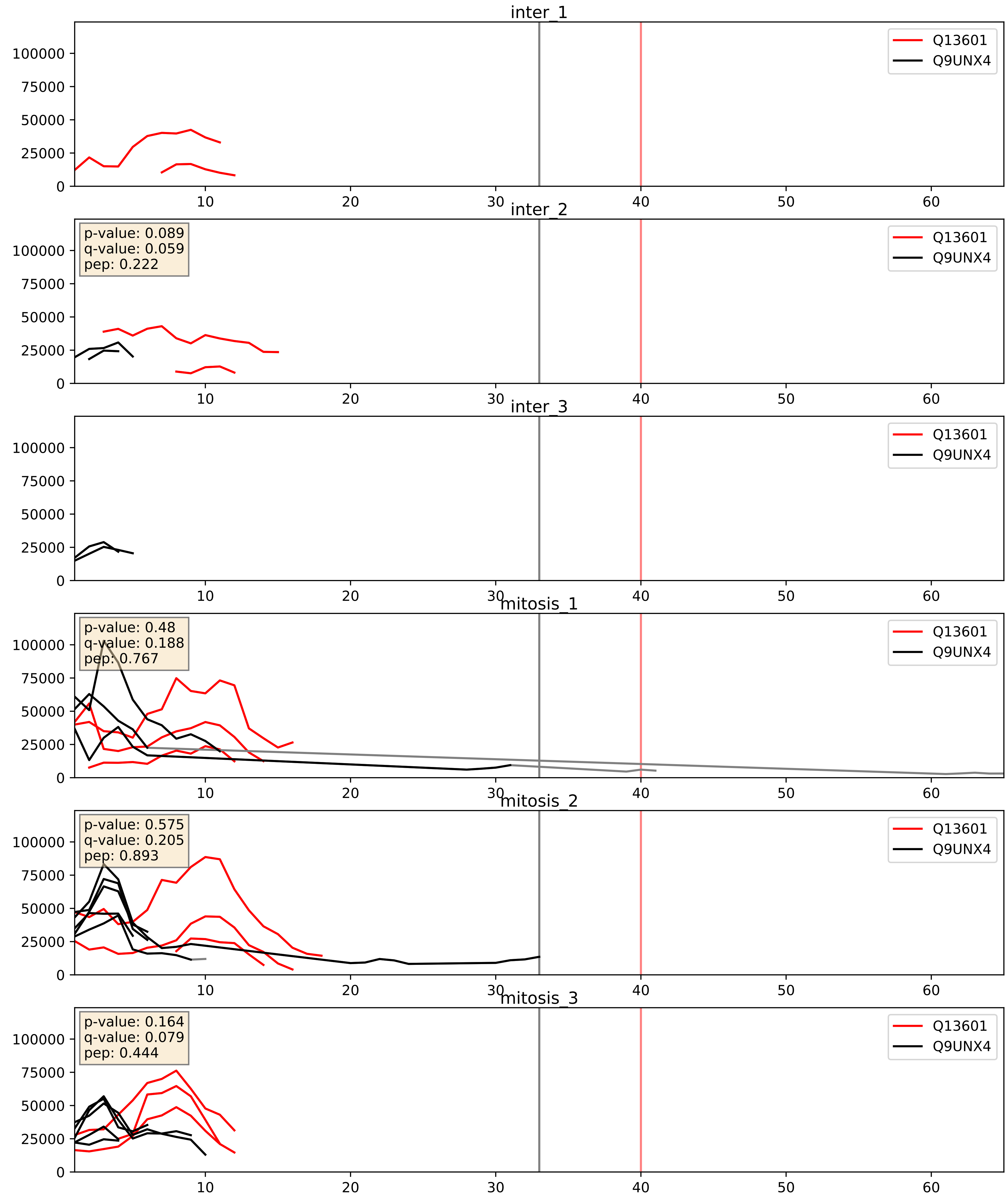


| level | condition_1 | condition_2 | pvalue | pvalue_adjusted |
|---|---|---|---|---|
| interactor_abundance | mitosis | inter | 0.10673447658103845 | 0.2798735241334627 |
| complex_abundance | mitosis | inter | 0.11913963786757466 | 0.26092702861017886 |
| interactor_ratio | mitosis | inter | 0.579276707995459 | 0.8969051975628951 |

Q14137_Q9UNX4
BOP1_HUMAN vs WDR3_HUMAN
p-value: 0.058 q-value: 0.037

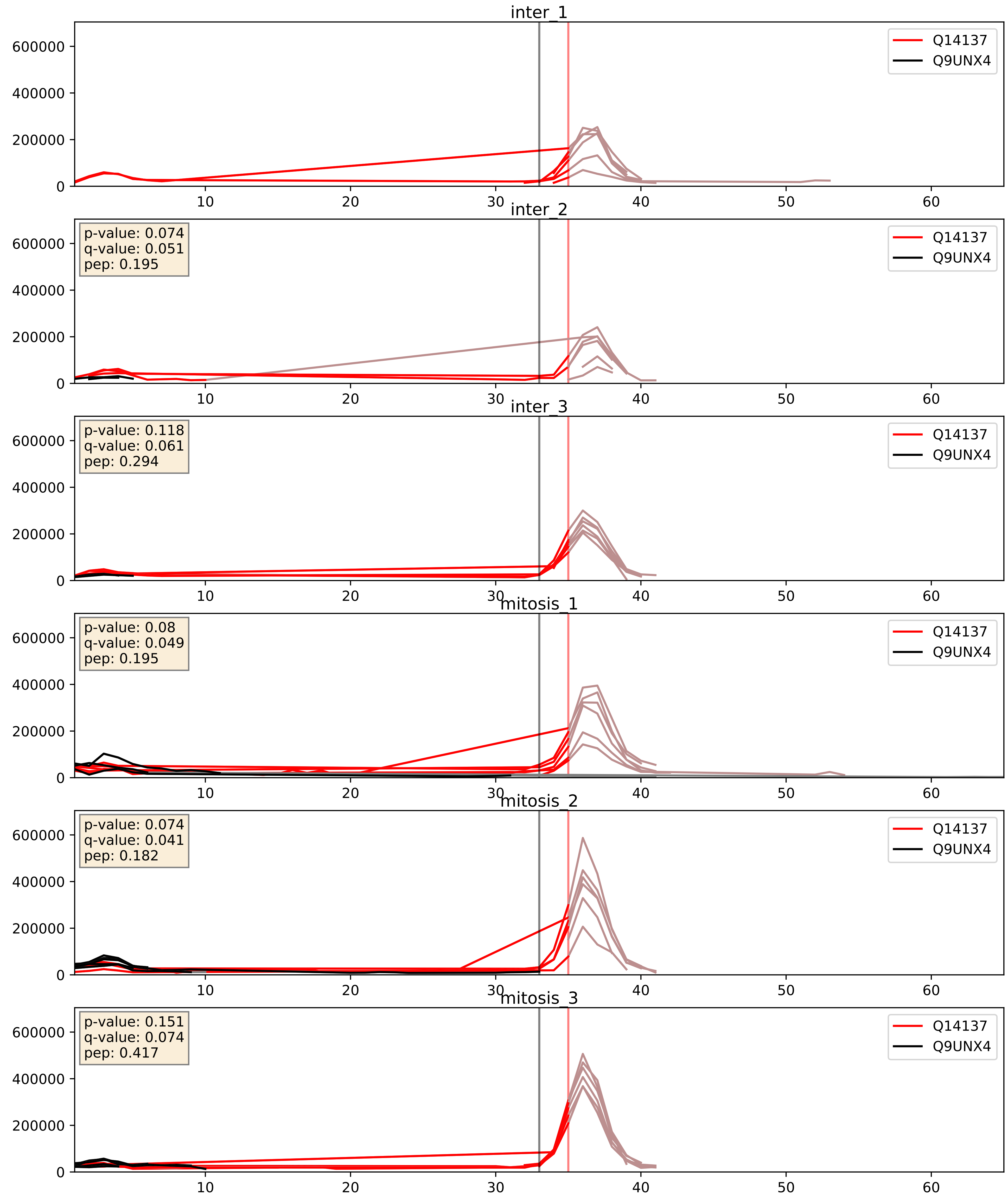


| level | condition_1 | condition_2 | pvalue | pvalue_adjusted |
|---|---|---|---|---|
| interactor_ratio | mitosis | inter | 0.025700362102627074 | 0.25374290611128925 |
| interactor_abundance | mitosis | inter | 0.5664835021564785 | 0.7138375943558745 |
| complex_abundance | mitosis | inter | 0.8968520804744315 | 0.9361137352430374 |

Q15269_Q9UNX4
PWP2_HUMAN vs WDR3_HUMAN
p-value: 0.009 q-value: 0.008

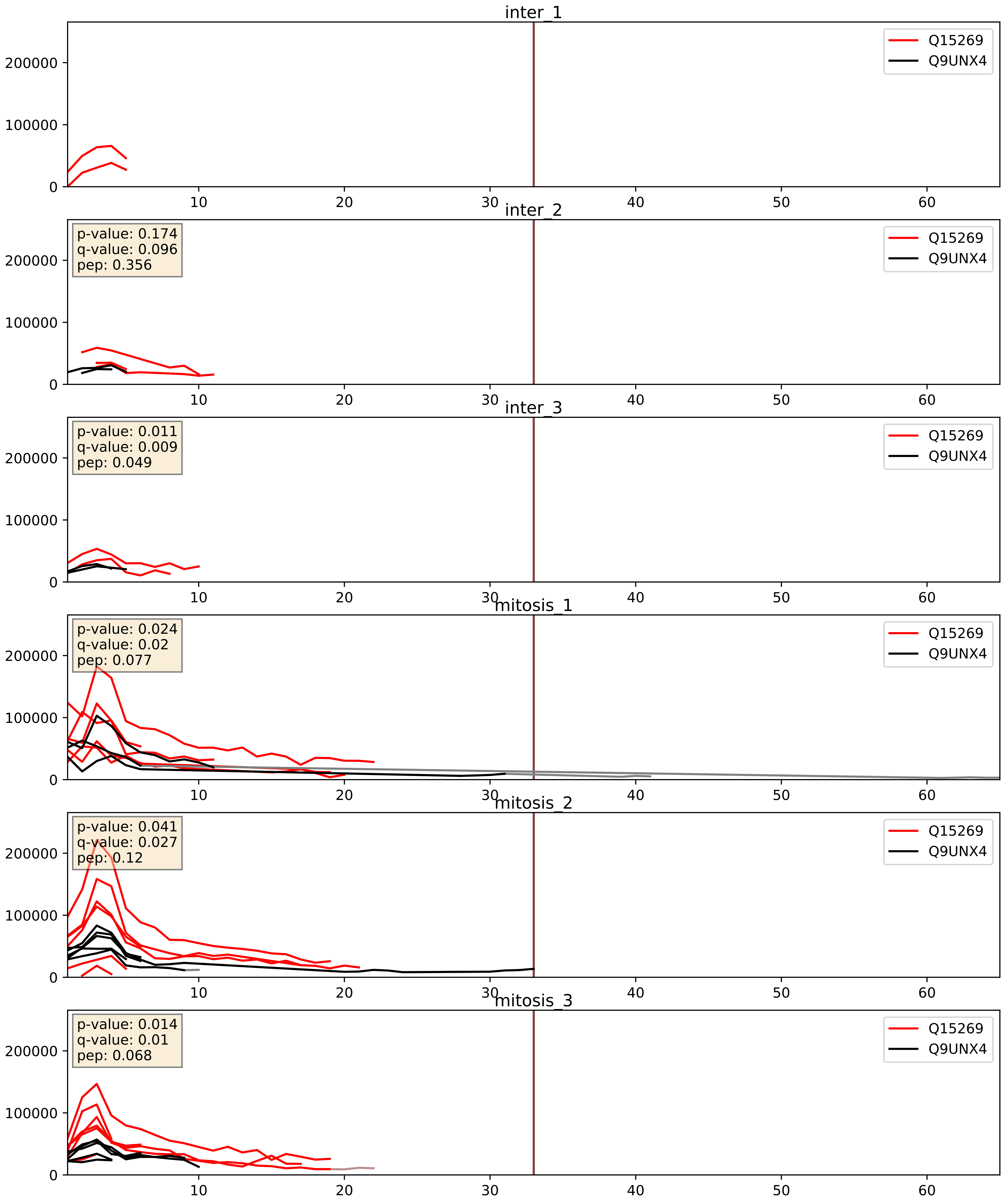


| level | condition_1 | condition_2 | pvalue | pvalue_adjusted |
|---|---|---|---|---|
| interactor_ratio | mitosis | inter | 0.027928400658634143 | 0.26184787474031573 |
| interactor_abundance | mitosis | inter | 0.041099264451958 | 0.16296731794217187 |
| complex_abundance | mitosis | inter | 0.0673384386695605 | 0.18739919037777122 |

Q8IY81_Q9UNX4
SPB1_HUMAN vs WDR3_HUMAN
p-value: 0.007 q-value: 0.006

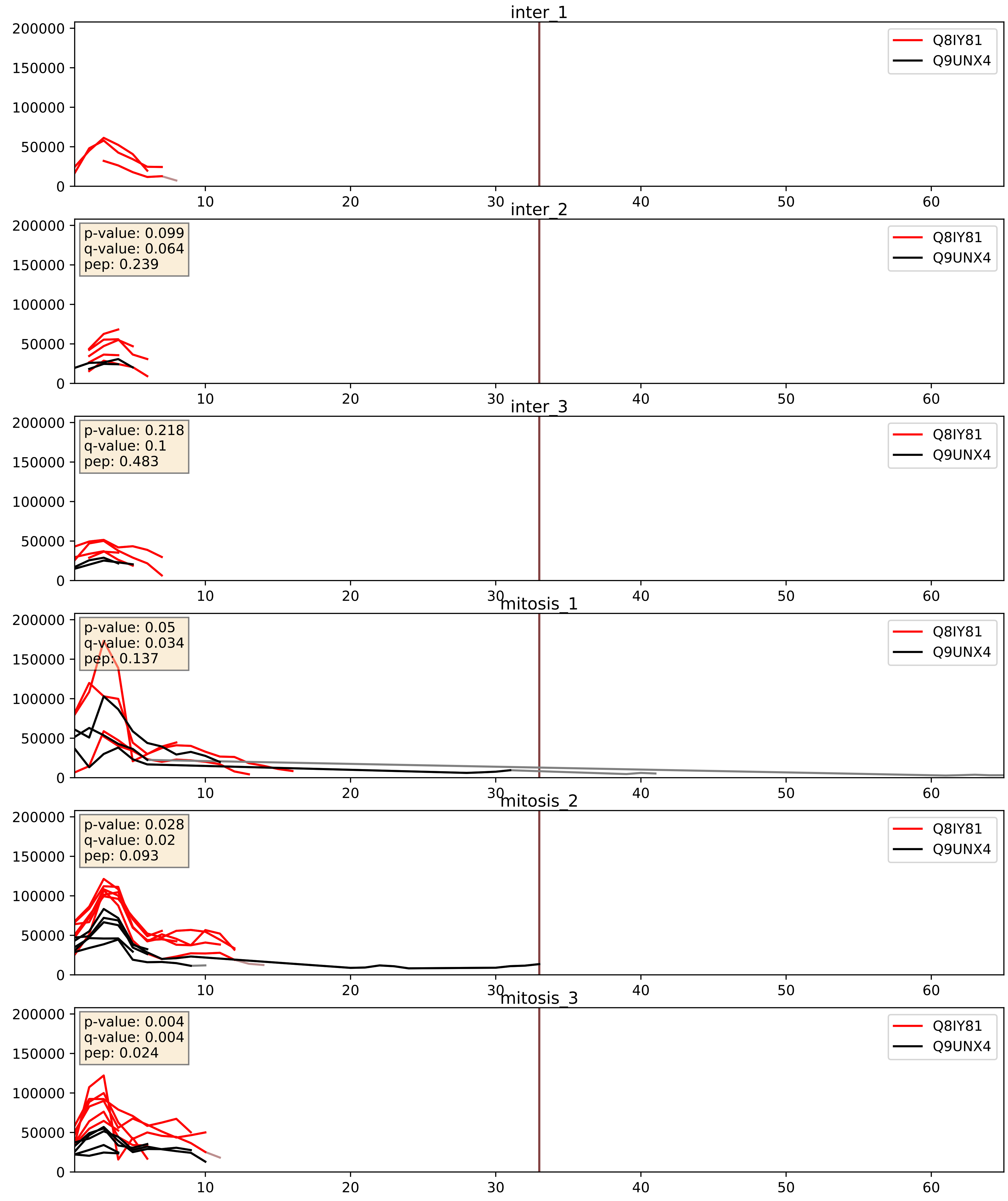


| level | condition_1 | condition_2 | pvalue | pvalue_adjusted |
|---|---|---|---|---|
| complex_abundance | mitosis | inter | 0.1603807662778196 | 0.30804508691617344 |
| interactor_abundance | mitosis | inter | 0.20267537112676712 | 0.39818709590202583 |
| interactor_ratio | mitosis | inter | 0.6731573851839978 | 0.9319765113016878 |

Q8NI36_Q9UNX4
WDR36_HUMAN vs WDR3_HUMAN
p-value: 0.005 q-value: 0.005

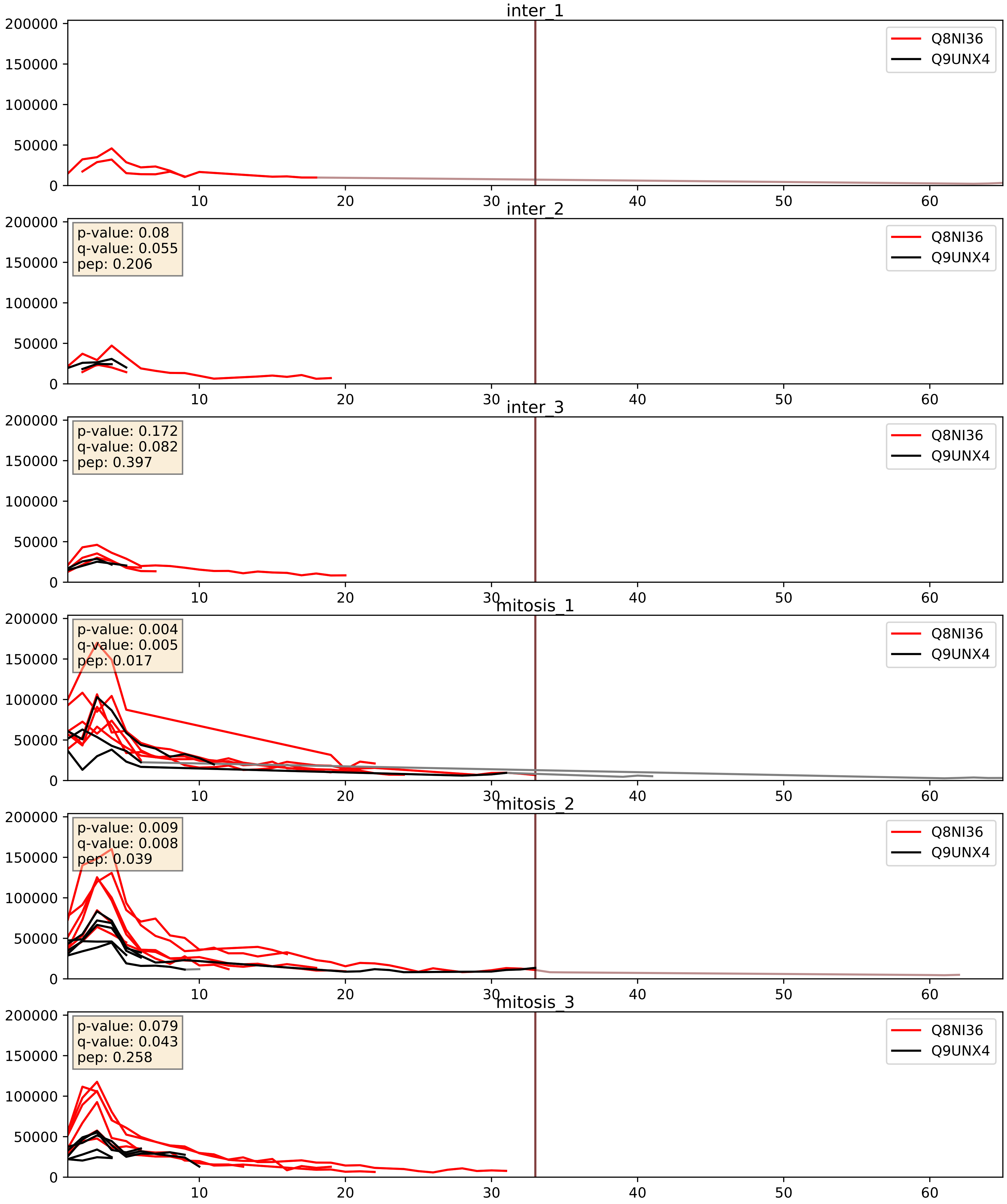


| level | condition_1 | condition_2 | pvalue | pvalue_adjusted |
|---|---|---|---|---|
| interactor_ratio | mitosis | inter | 0.0034412210964951434 | 0.07572939655107606 |
| interactor_abundance | mitosis | inter | 0.00922348089309837 | 0.08324021891678159 |
| complex_abundance | mitosis | inter | 0.03510041369517333 | 0.13586662078716946 |

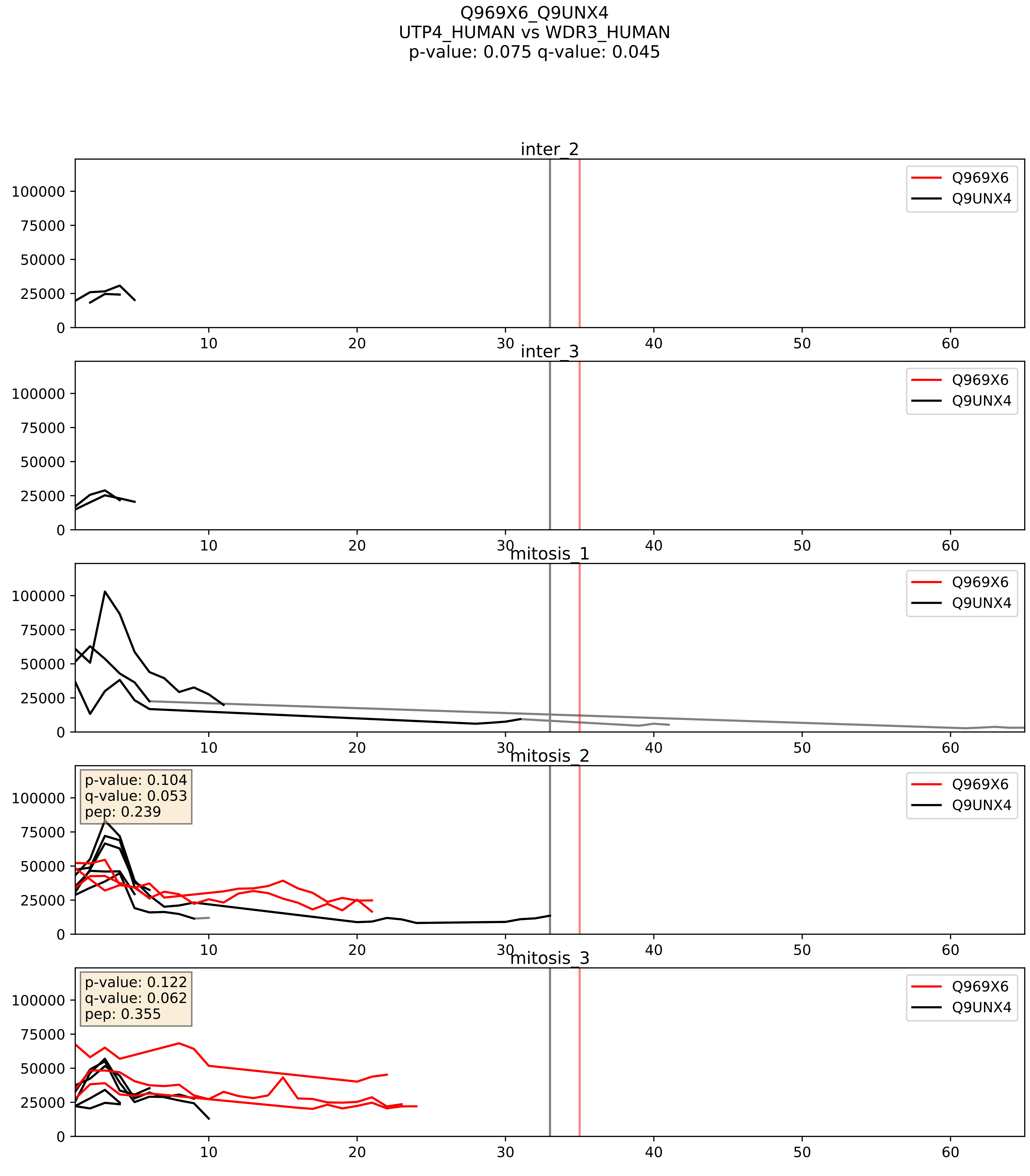


| level | condition_1 | condition_2 | pvalue | pvalue_adjusted |
|---|---|---|---|---|
| complex_abundance | mitosis | inter | 0.3813462081111618 | 0.5285497962162476 |
| interactor_abundance | mitosis | inter | 0.39515313323189816 | 0.5863253285604175 |
| interactor_ratio | mitosis | inter | 0.9109315500612556 | 0.9852158746906624 |

Q9GZL7_Q9UNX4
WDR12_HUMAN vs WDR3_HUMAN
p-value: 0.057 q-value: 0.036

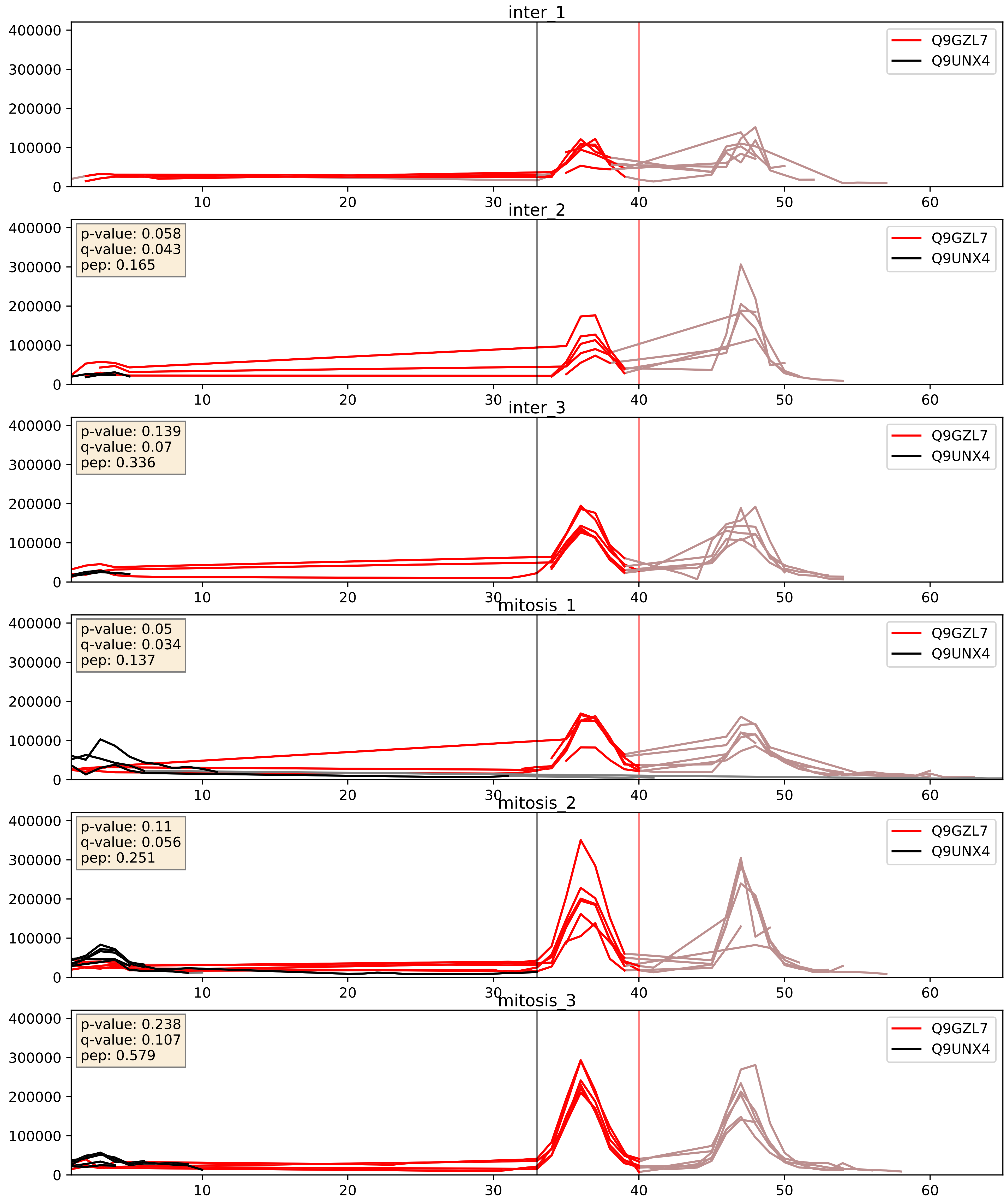


| level | condition_1 | condition_2 | pvalue | pvalue_adjusted |
|---|---|---|---|---|
| interactor_ratio | mitosis | inter | 0.015813892154450518 | 0.18518046079630154 |
| interactor_abundance | mitosis | inter | 0.18824399672868047 | 0.3800752453282745 |
| complex_abundance | mitosis | inter | 0.6867853693406996 | 0.7905974665890786 |

Q9H0A0_Q9UNX4
NAT10_HUMAN vs WDR3_HUMAN
p-value: 0.006 q-value: 0.005

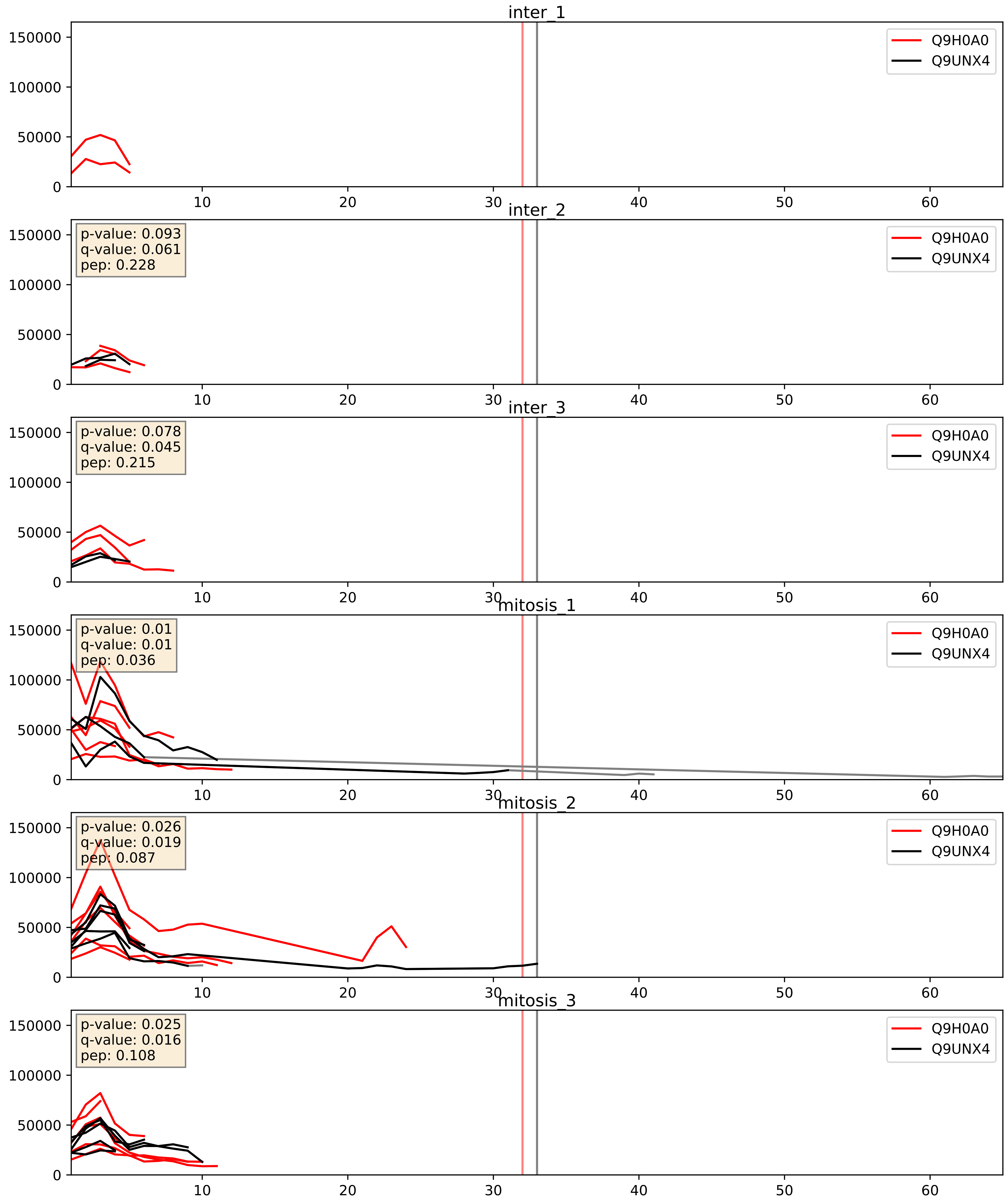


| level | condition_1 | condition_2 | pvalue | pvalue_adjusted |
|---|---|---|---|---|
| complex_abundance | mitosis | inter | 0.2858862643454428 | 0.4346099345487244 |
| interactor_abundance | mitosis | inter | 0.45197017672950507 | 0.625321595733726 |
| interactor_ratio | mitosis | inter | 0.6060595505852381 | 0.9052533265322077 |

Q9H6R4_Q9UNX4
NOL6_HUMAN vs WDR3_HUMAN
p-value: 0.053 q-value: 0.034

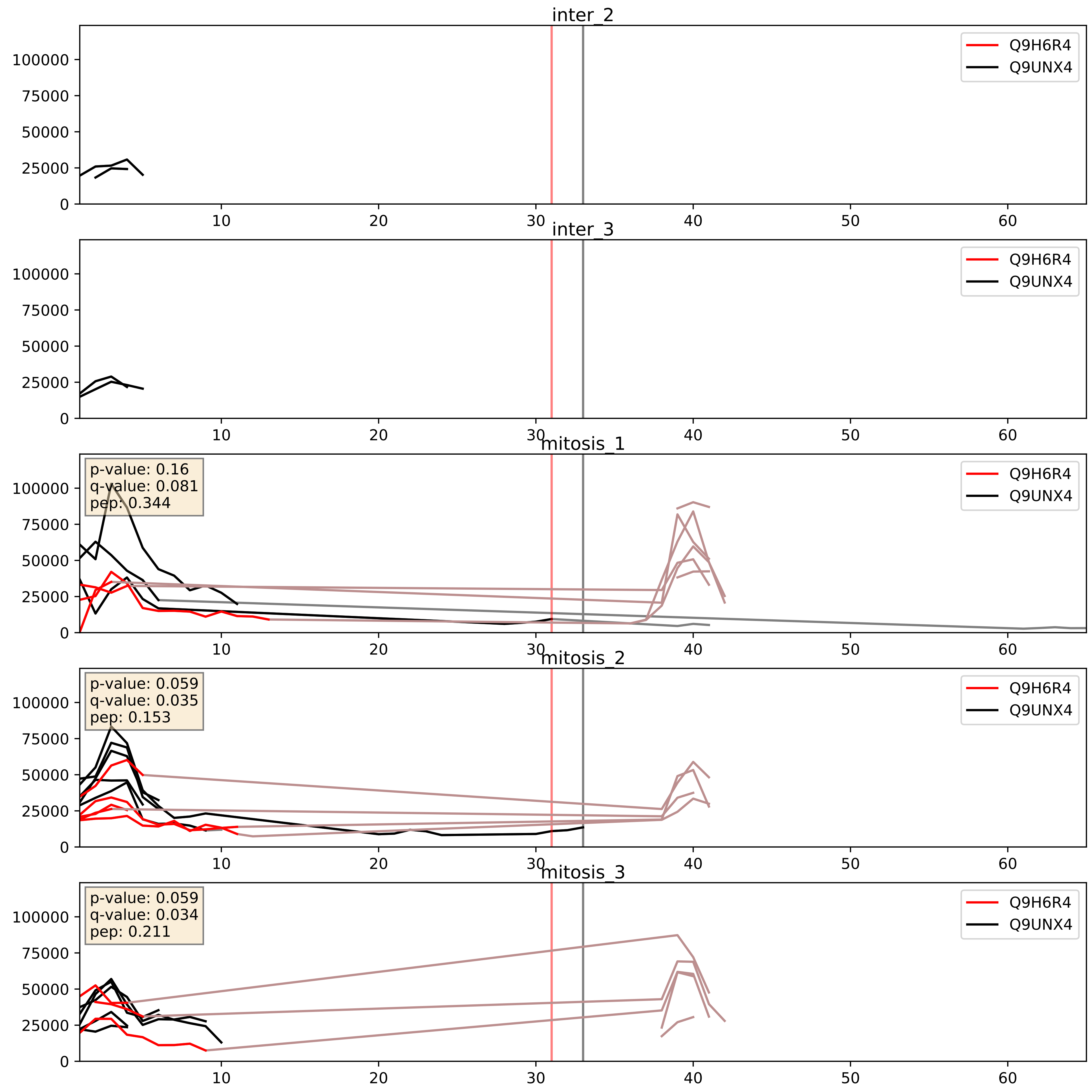


| level | condition_1 | condition_2 | pvalue | pvalue_adjusted |
|---|---|---|---|---|
| interactor_ratio | mitosis | inter | 0.002692662930541276 | 0.06718455808461757 |
| complex_abundance | mitosis | inter | 0.09645403910829943 | 0.23018381938539362 |
| interactor_abundance | mitosis | inter | 0.5045521248563168 | 0.6689848495616593 |

Q9NW13_Q9UNX4
RBM28_HUMAN vs WDR3_HUMAN
p-value: 0.028 q-value: 0.021

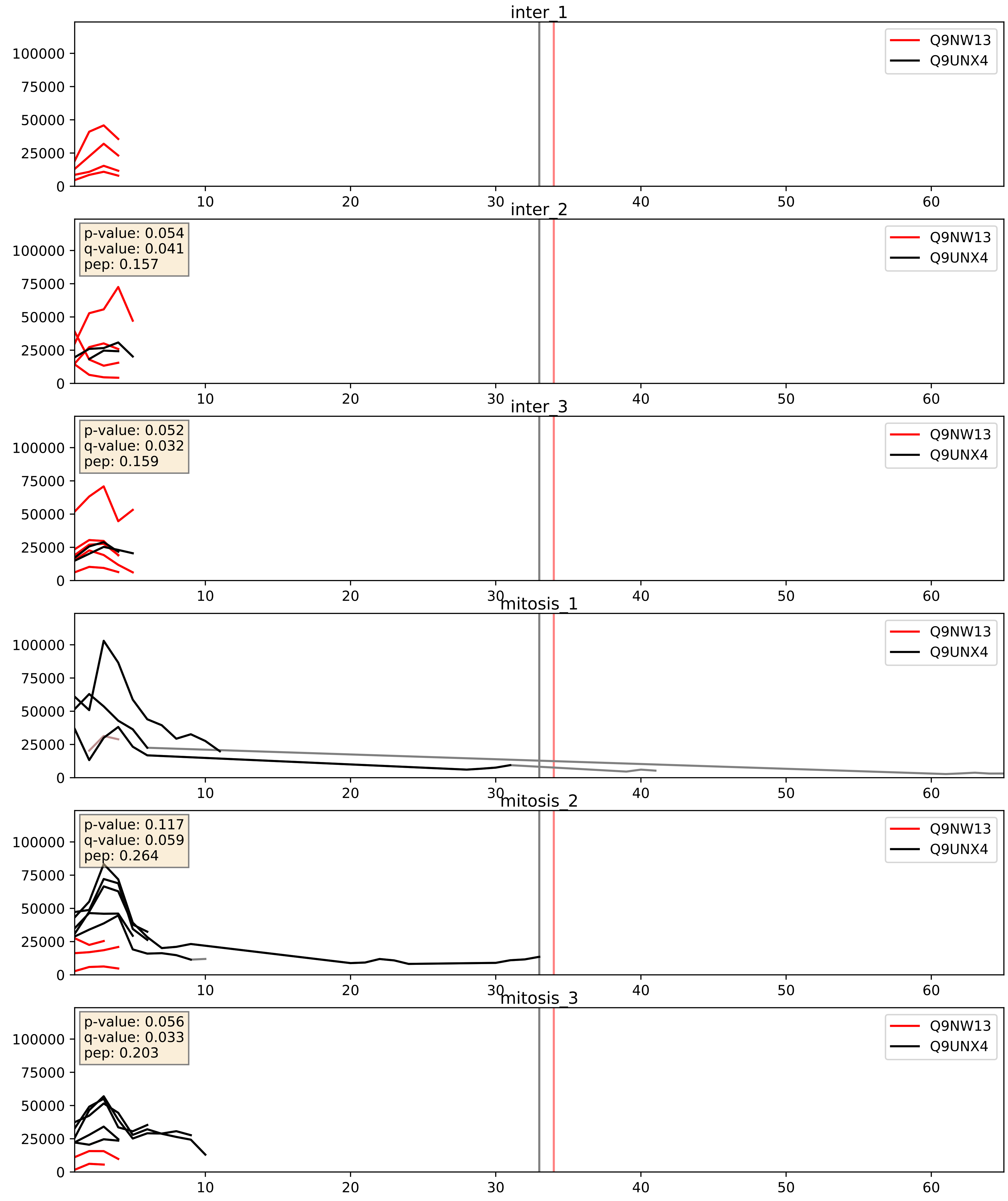


| level | condition_1 | condition_2 | pvalue | pvalue_adjusted |
|---|---|---|---|---|
| interactor_abundance | mitosis | inter | 0.017414770002192395 | 0.11427399863454725 |
| complex_abundance | mitosis | inter | 0.027075430836367088 | 0.12025018583230009 |
| interactor_ratio | mitosis | inter | 0.0596208358066013 | 0.4034421774739187 |

Q9UNX4_Q9Y221
WDR3_HUMAN vs NIP7_HUMAN
p-value: 0.013 q-value: 0.011

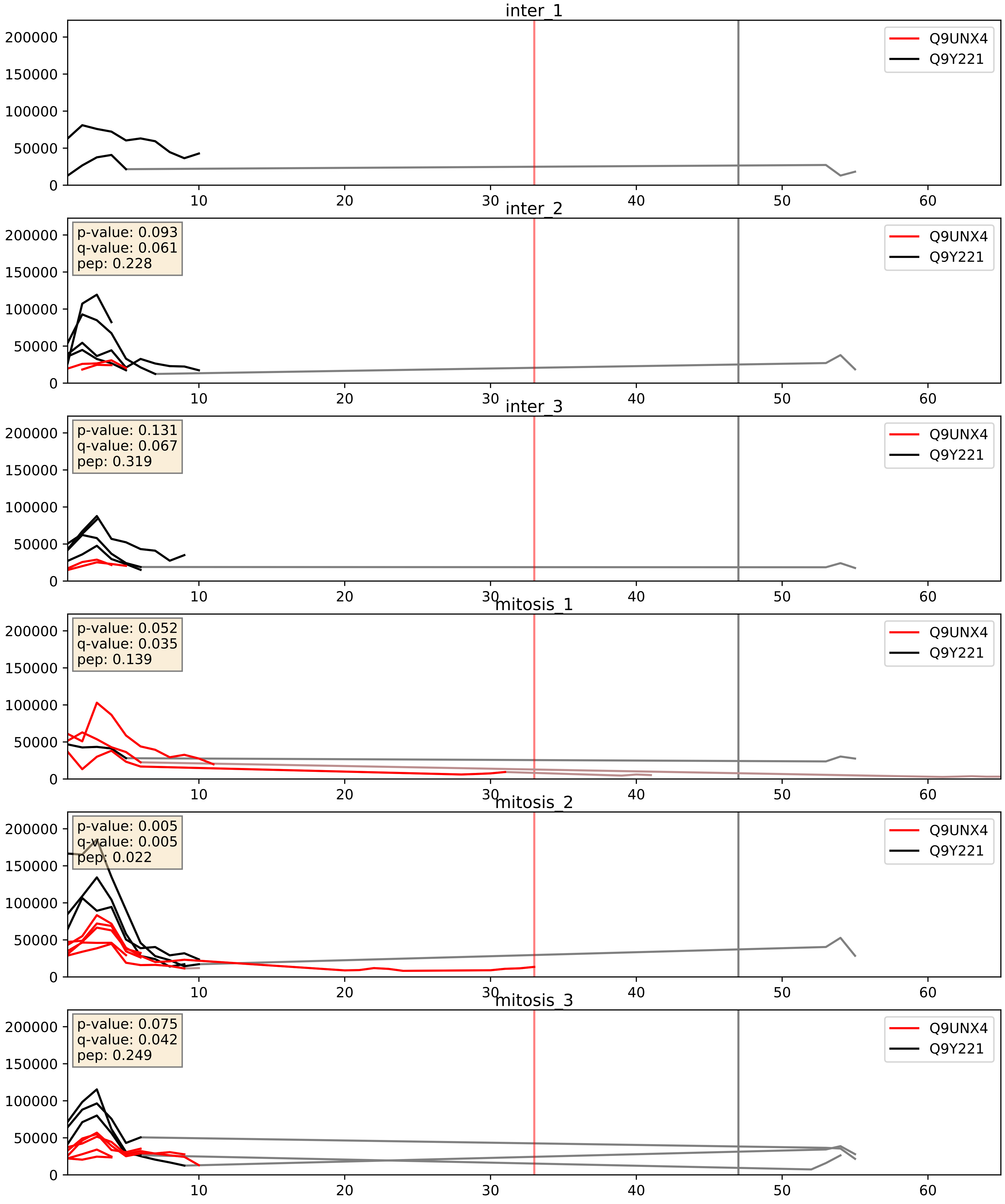


| level | condition_1 | condition_2 | pvalue | pvalue_adjusted |
|---|---|---|---|---|
| interactor_abundance | mitosis | inter | 0.19689113756785914 | 0.39076933400901326 |
| complex_abundance | mitosis | inter | 0.5704524399458758 | 0.6996806392404022 |
| interactor_ratio | mitosis | inter | 0.5801719302609729 | 0.8972209608226245 |

Q9UNX4_Q9Y2X3
WDR3_HUMAN vs NOP58_HUMAN
p-value: 0.052 q-value: 0.034

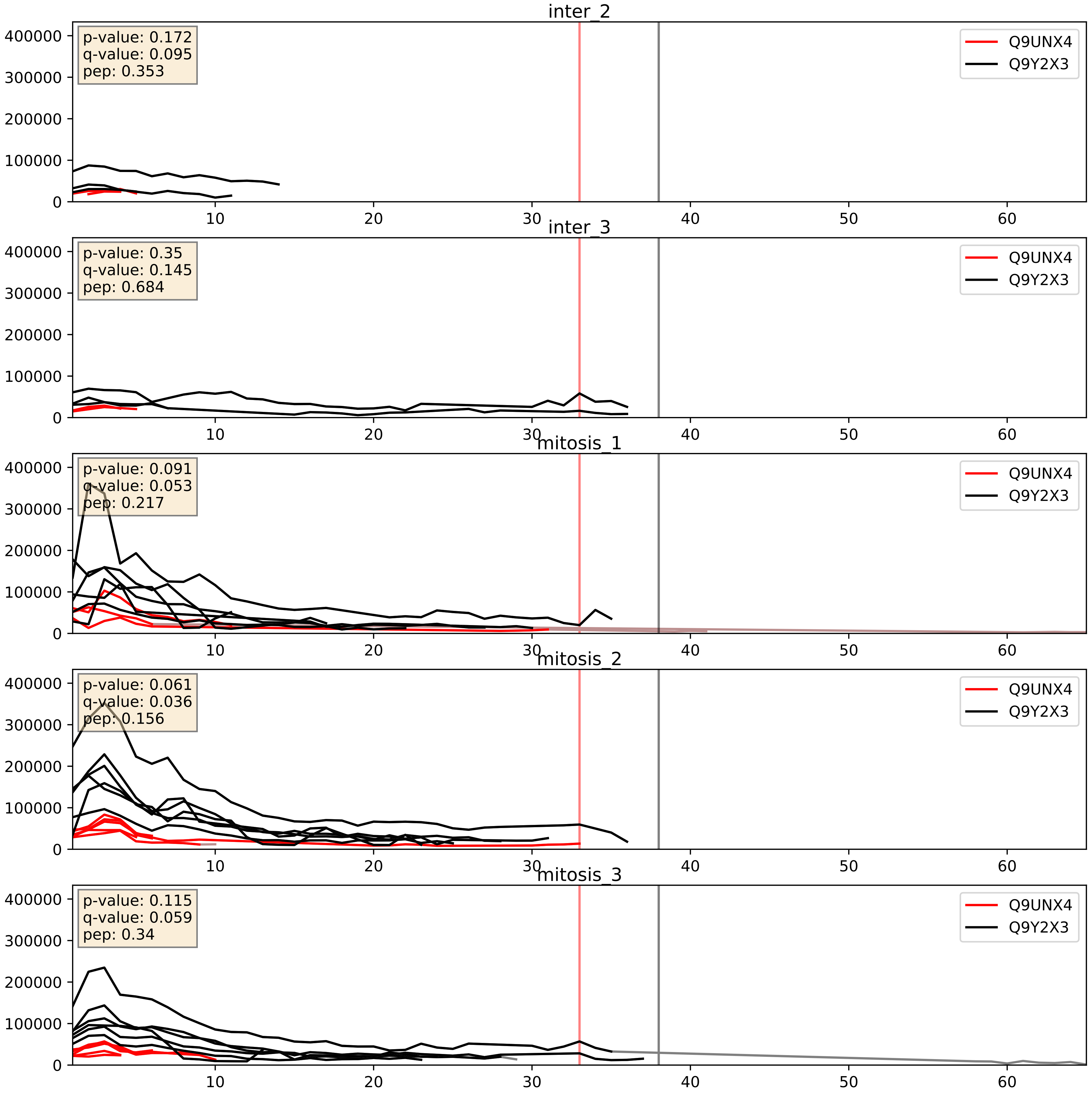


| level | condition_1 | condition_2 | pvalue | pvalue_adjusted |
|---|---|---|---|---|
| interactor_ratio | mitosis | inter | 0.0008632207210684006 | 0.033740499417102776 |
| complex_abundance | mitosis | inter | 0.00703579685561239 | 0.06202515044700521 |
| interactor_abundance | mitosis | inter | 0.15972059028570332 | 0.34604106627325243 |
