## Supplementary Data 2 for "SECAT: Quantifying differential protein-protein interaction states by network-centric analysis": hela_string_000020_P31943.pdf

P31943_P31943

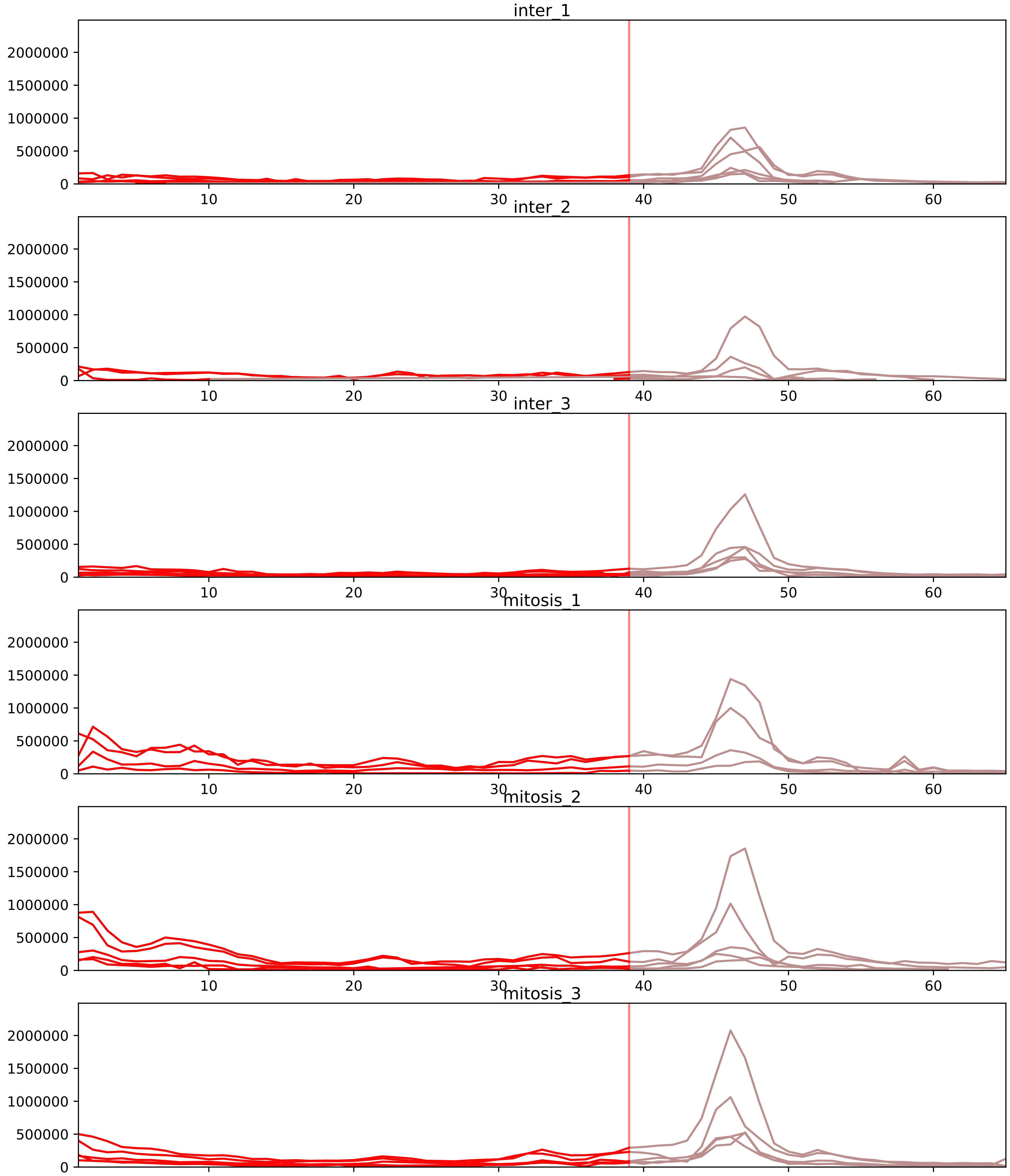


| level | condition_1 | condition_2 | pvalue | pvalue_adjusted |
|---|---|---|---|---|
| interactor_ratio | mitosis | inter | 8.658412907790795e-12 | 7.965739875167531e-10 |
| complex_abundance | mitosis | inter | 1.2429633256376748e-06 | 0.00010890726281777722 |
| interactor_abundance | mitosis | inter | 3.8621709675118555e-06 | 0.0002733228684700698 |
| assembled_abundance | mitosis | inter | 0.007849939387193614 | 0.09499304500648614 |
| total_abundance | mitosis | inter | 0.043808997115043206 | 0.26055093961762266 |
| monomer_abundance | mitosis | inter | 0.11487168136563952 | 0.4453469860578181 |

O43390_P31943
HNRPR_HUMAN vs HNRH1_HUMAN
p-value: 0.005 q-value: 0.005

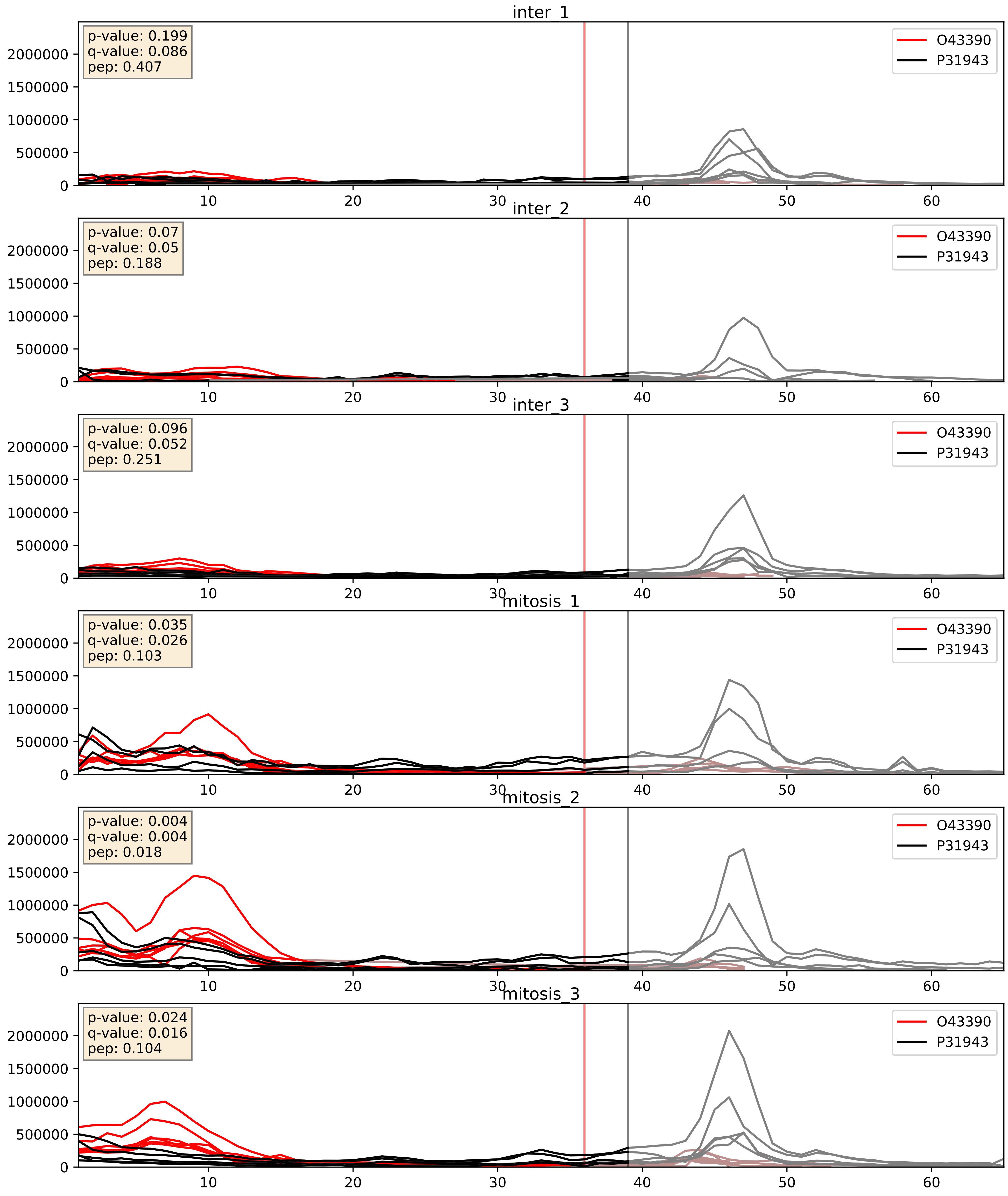


| level | condition_1 | condition_2 | pvalue | pvalue_adjusted |
|---|---|---|---|---|
| complex_abundance | mitosis | inter | 9.891557176946058e-05 | 0.006719978526560179 |
| interactor_abundance | mitosis | inter | 0.00010796061997593195 | 0.007876481428251637 |
| interactor_ratio | mitosis | inter | 0.4332742845030043 | 0.8435878041032946 |

P07910_P31943
HNRPC_HUMAN vs HNRH1_HUMAN
p-value: 0.012 q-value: 0.01

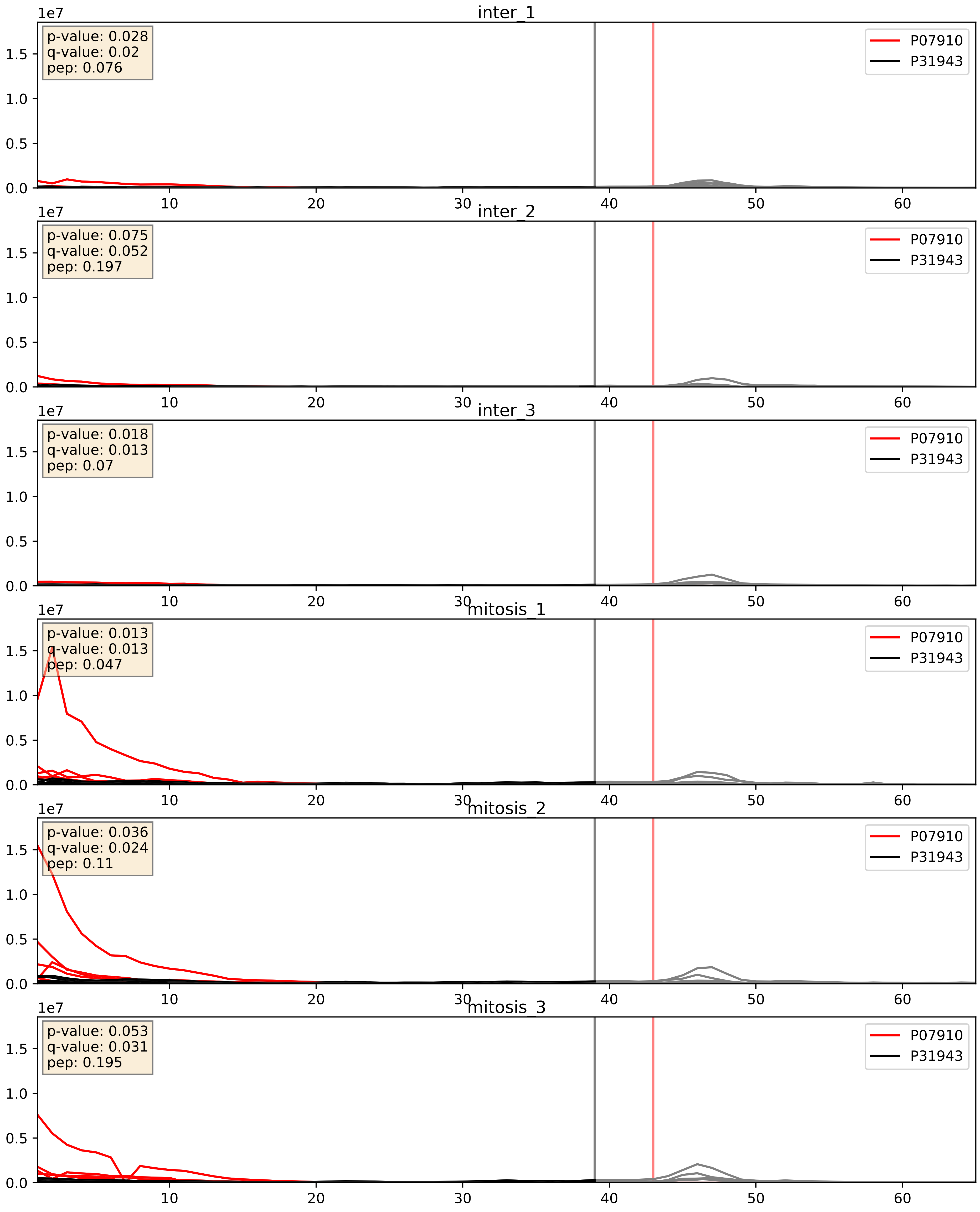


| level | condition_1 | condition_2 | pvalue | pvalue_adjusted |
|---|---|---|---|---|
| complex_abundance | mitosis | inter | 4.189010509383275e-06 | 0.005122561422902976 |
| interactor_abundance | mitosis | inter | 9.711637813774278e-06 | 0.0033734327106189023 |
| interactor_ratio | mitosis | inter | 0.0025928684235758614 | 0.0652792756053217 |

P09651_P31943
ROA1_HUMAN vs HNRH1_HUMAN
p-value: 0.0 q-value: 0.001

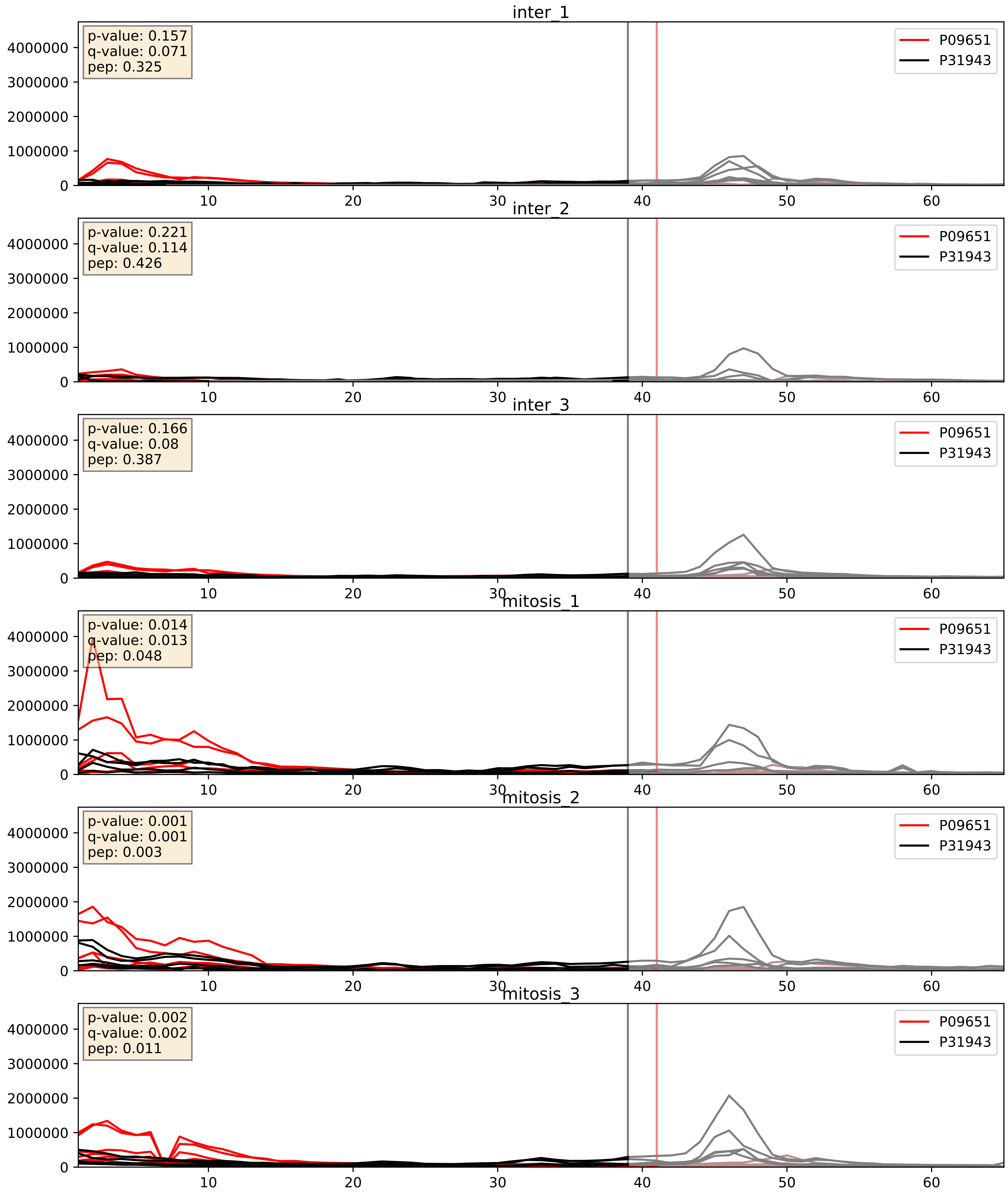


| level | condition_1 | condition_2 | pvalue | pvalue_adjusted |
|---|---|---|---|---|
| complex_abundance | mitosis | inter | 0.007967826015523555 | 0.06596188655017565 |
| interactor_abundance | mitosis | inter | 0.016854435383293762 | 0.11277132084292767 |
| interactor_ratio | mitosis | inter | 0.2314283726697915 | 0.7093426594678715 |

P14866_P31943
HNRPL_HUMAN vs HNRH1_HUMAN
p-value: 0.001 q-value: 0.002

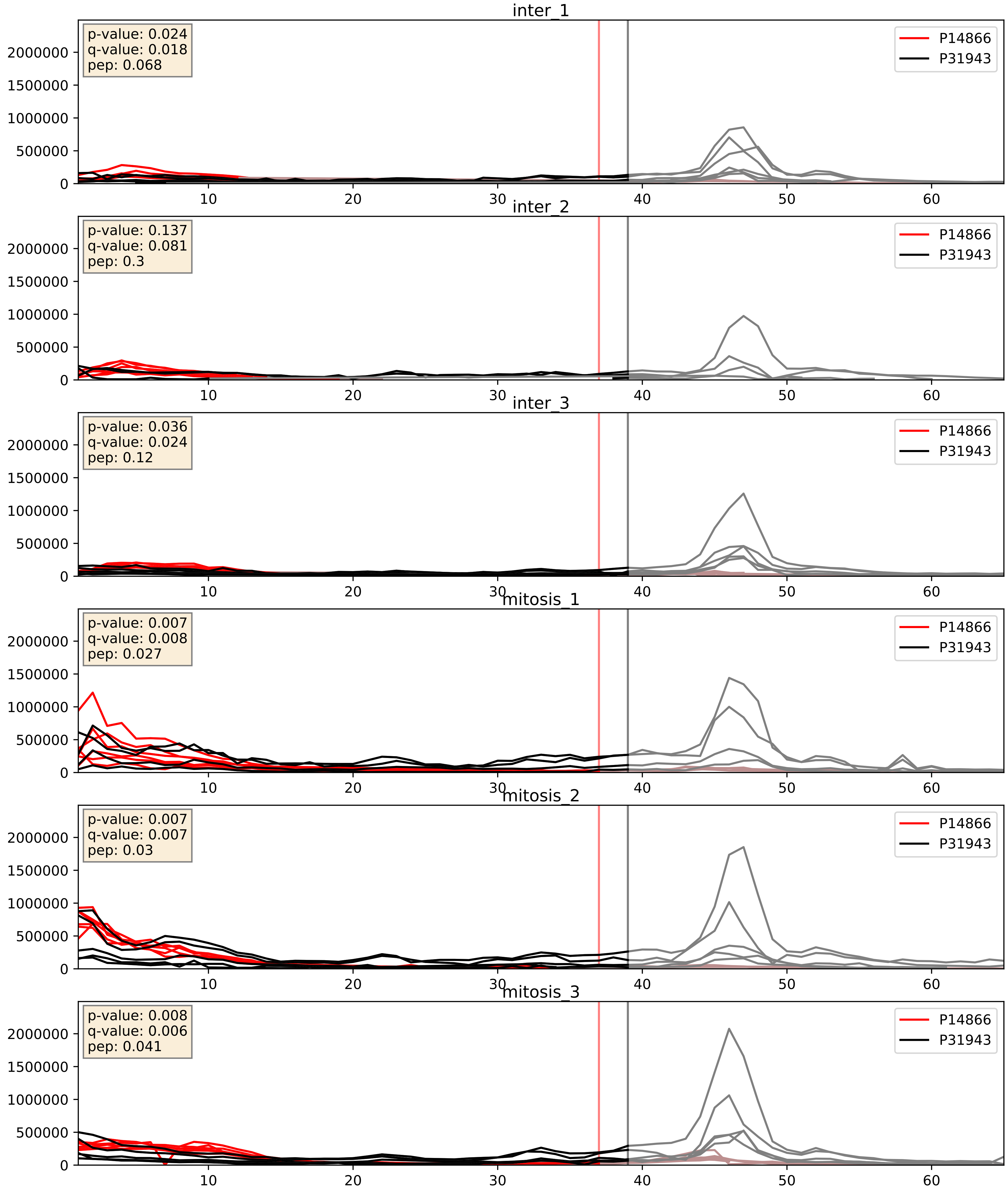


| level | condition_1 | condition_2 | pvalue | pvalue_adjusted |
|---|---|---|---|---|
| complex_abundance | mitosis | inter | 0.0010805657853290274 | 0.021833569470040306 |
| interactor_abundance | mitosis | inter | 0.009697640385696541 | 0.08525122834595471 |
| interactor_ratio | mitosis | inter | 0.20279183069839515 | 0.678106014366403 |

P17844_P31943
DDX5_HUMAN vs HNRH1_HUMAN
p-value: 0.0 q-value: 0.0

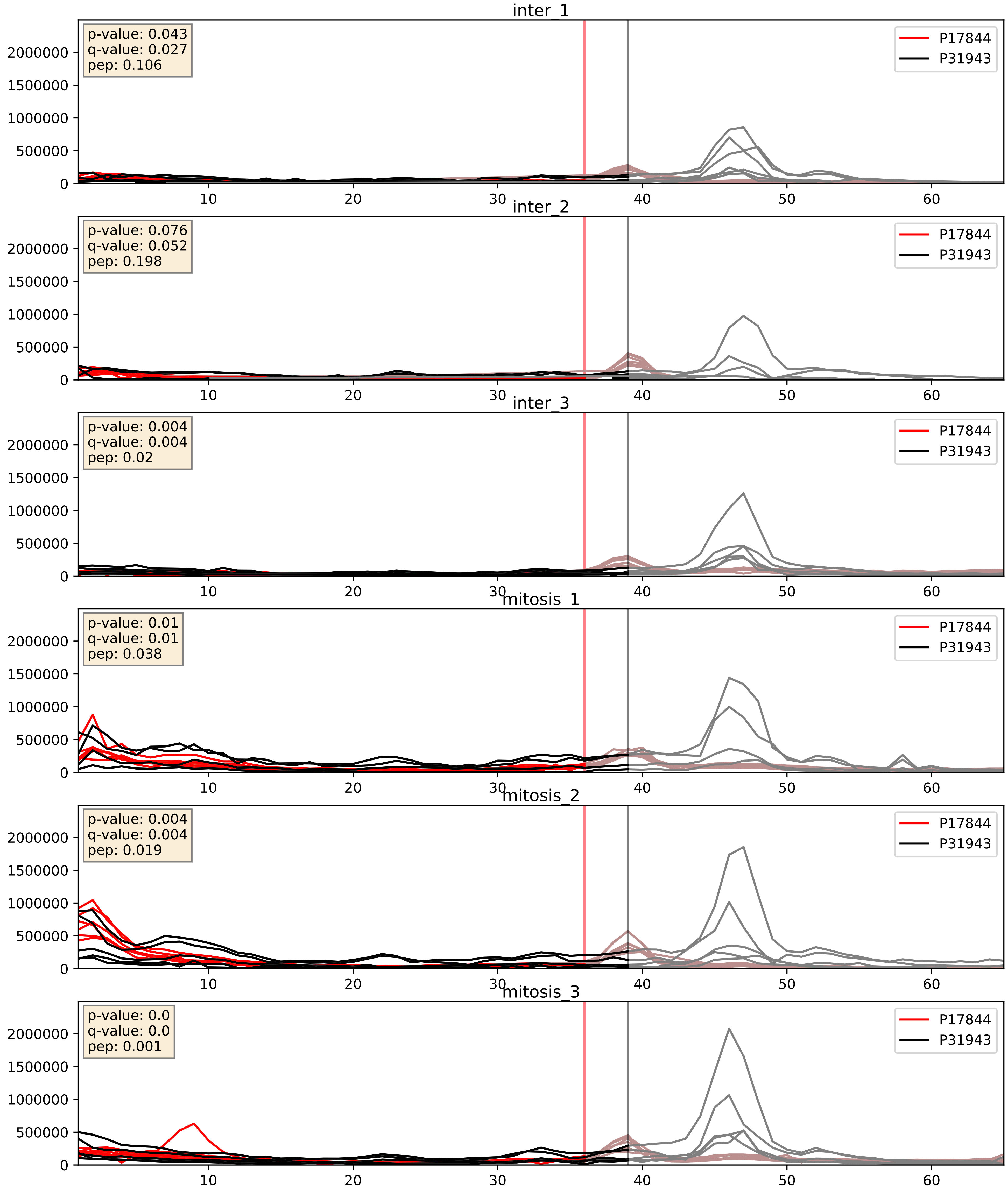


| level | condition_1 | condition_2 | pvalue | pvalue_adjusted |
|---|---|---|---|---|
| interactor_abundance | mitosis | inter | 0.0021426459467421676 | 0.033870820506210444 |
| complex_abundance | mitosis | inter | 0.0032360173229818456 | 0.04165459892439789 |
| interactor_ratio | mitosis | inter | 0.12617781079044613 | 0.5750062399685051 |

P22626_P31943
ROA2_HUMAN vs HNRH1_HUMAN
p-value: 0.003 q-value: 0.003

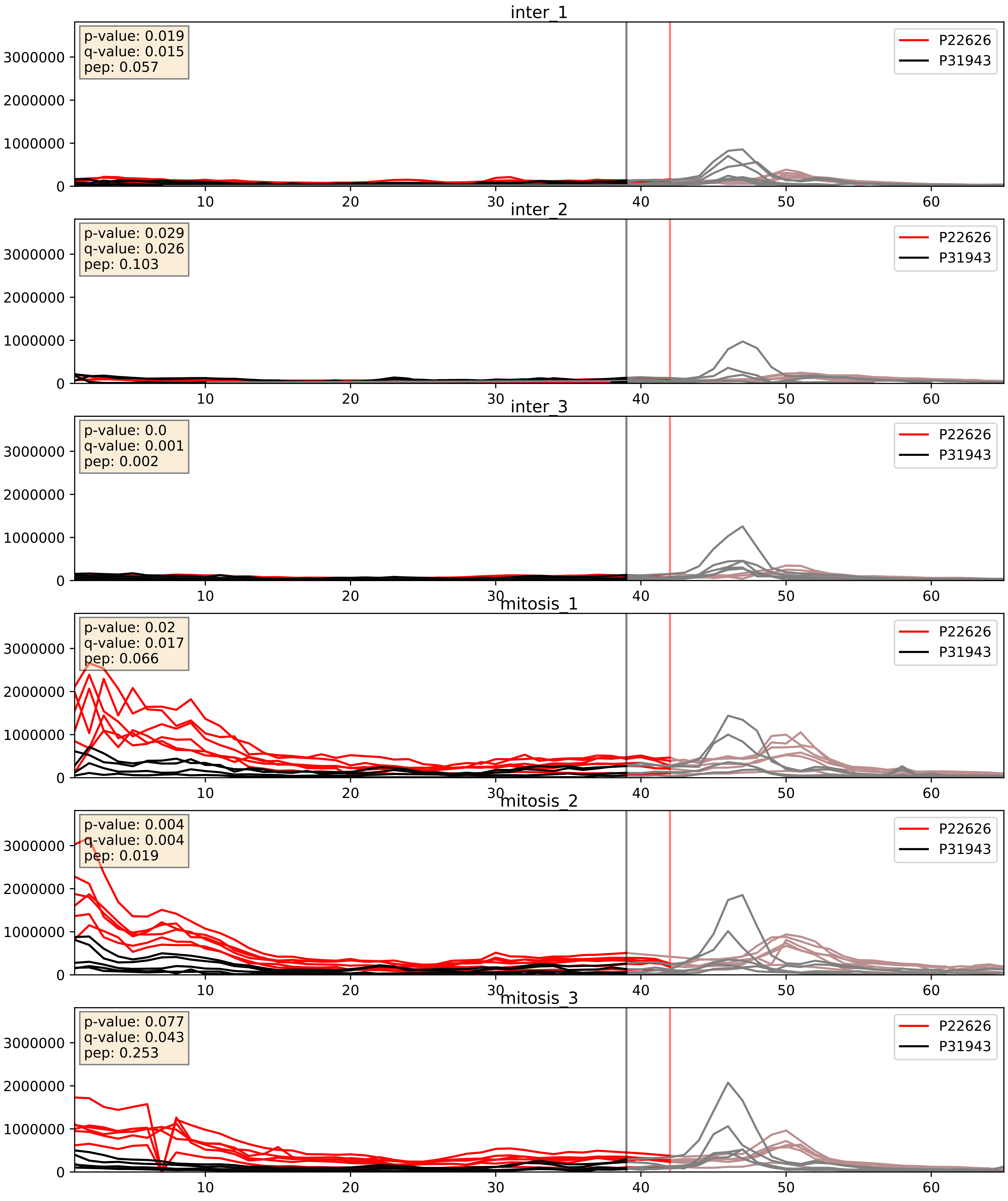


| level | condition_1 | condition_2 | pvalue | pvalue_adjusted |
|---|---|---|---|---|
| complex_abundance | mitosis | inter | 0.0008688599728642934 | 0.019513211095311817 |
| interactor_abundance | mitosis | inter | 0.003279687783791118 | 0.0442112242980346 |
| interactor_ratio | mitosis | inter | 0.02932575485422149 | 0.27108905135219863 |

P26599_P31943
PTBP1_HUMAN vs HNRH1_HUMAN
p-value: 0.042 q-value: 0.028

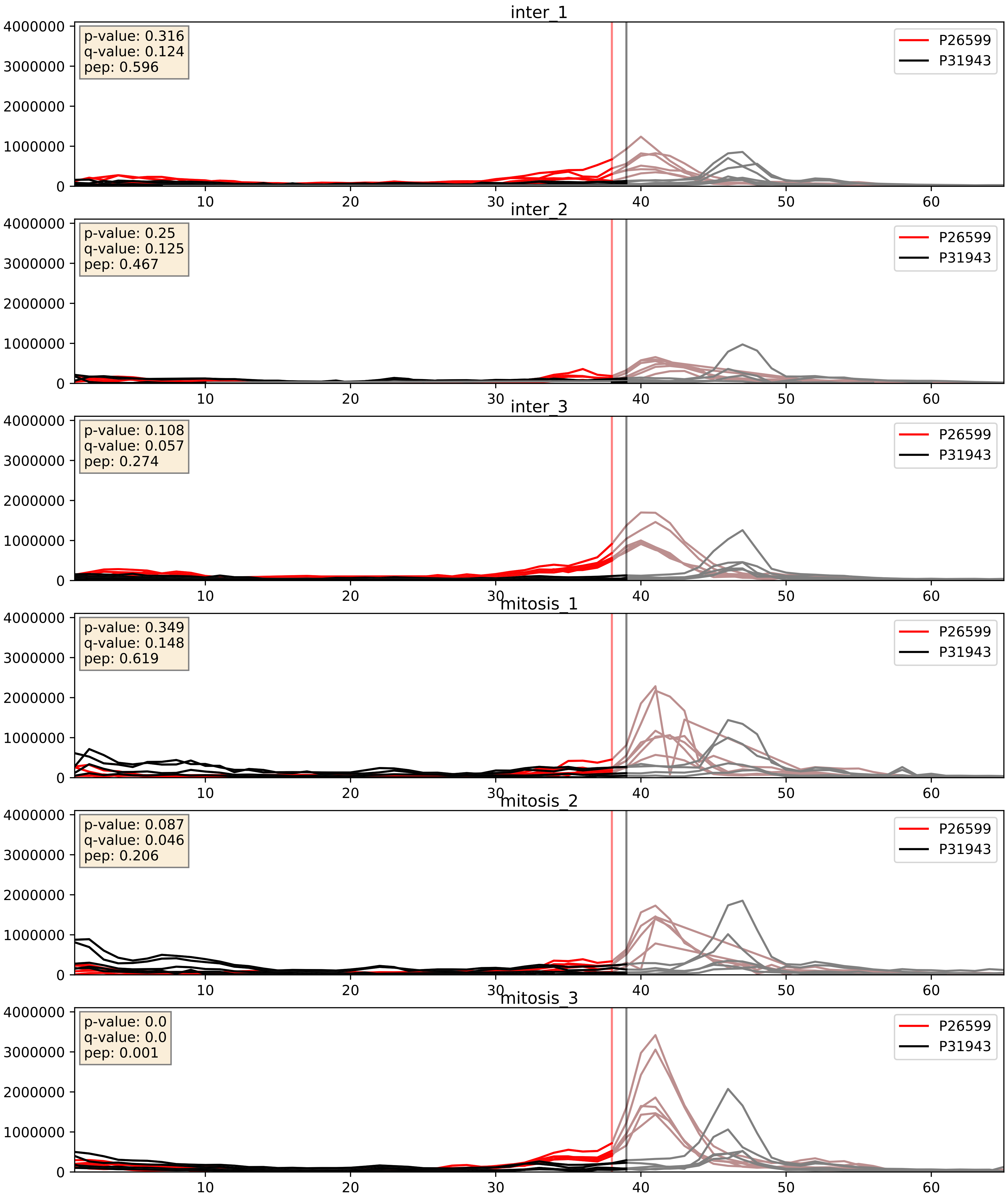


| level | condition_1 | condition_2 | pvalue | pvalue_adjusted |
|---|---|---|---|---|
| interactor_ratio | mitosis | inter | 0.047474514457461155 | 0.3605872615402551 |
| interactor_abundance | mitosis | inter | 0.3985986073620091 | 0.5886697292275717 |
| complex_abundance | mitosis | inter | 0.47360467855733784 | 0.6143439988560103 |

P31942_P31943
HNRH3_HUMAN vs HNRH1_HUMAN
p-value: 0.002 q-value: 0.03

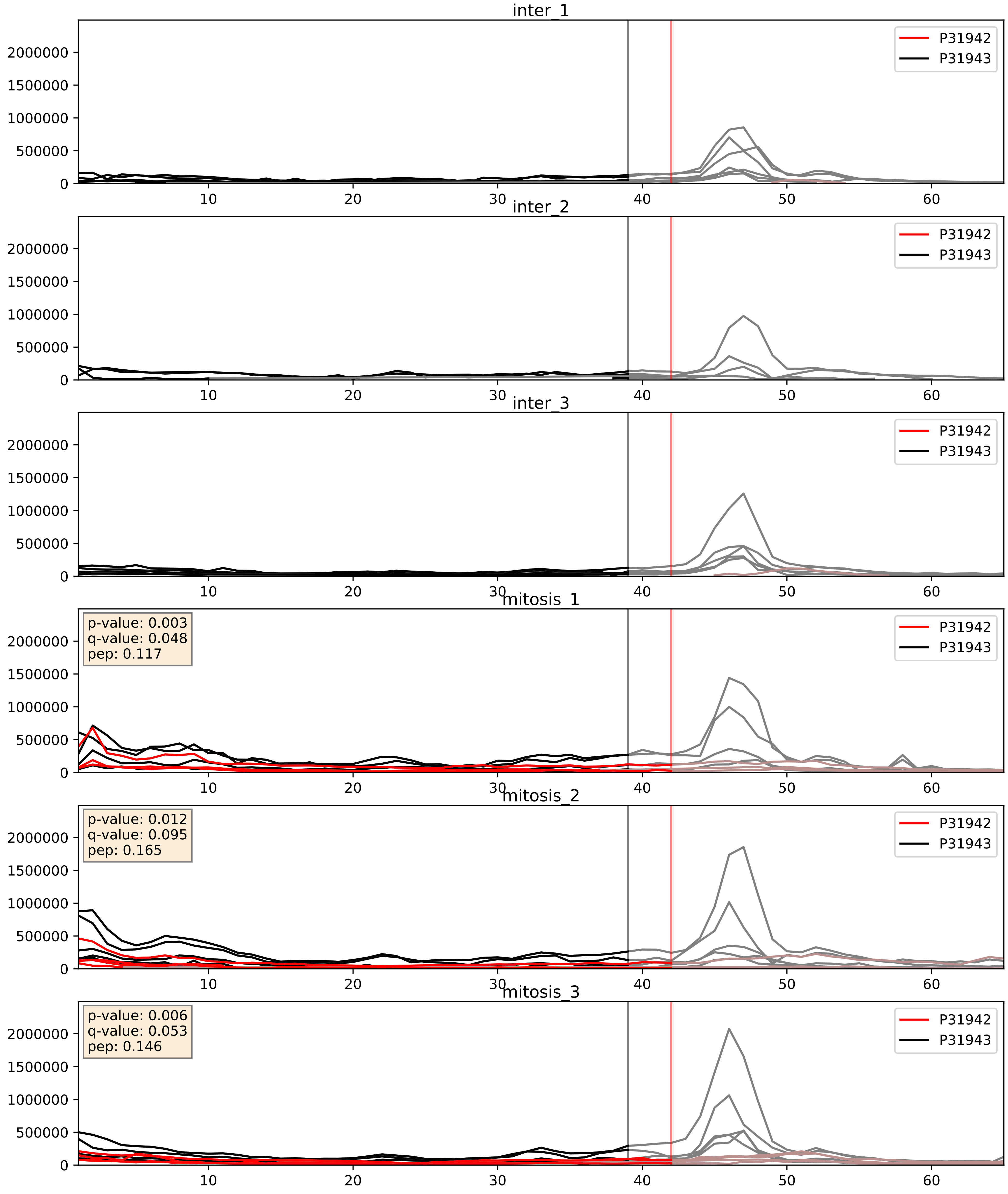


| level | condition_1 | condition_2 | pvalue | pvalue_adjusted |
|---|---|---|---|---|
| complex_abundance | mitosis | inter | 7.183772469446166e-05 | 0.005893640694944935 |
| interactor_abundance | mitosis | inter | 0.002362609793349813 | 0.03614645188753243 |
| interactor_ratio | mitosis | inter | 0.008559803010908383 | 0.13203195690147984 |

P31943_P38159
HNRH1_HUMAN vs RBMX_HUMAN
p-value: 0.004 q-value: 0.004

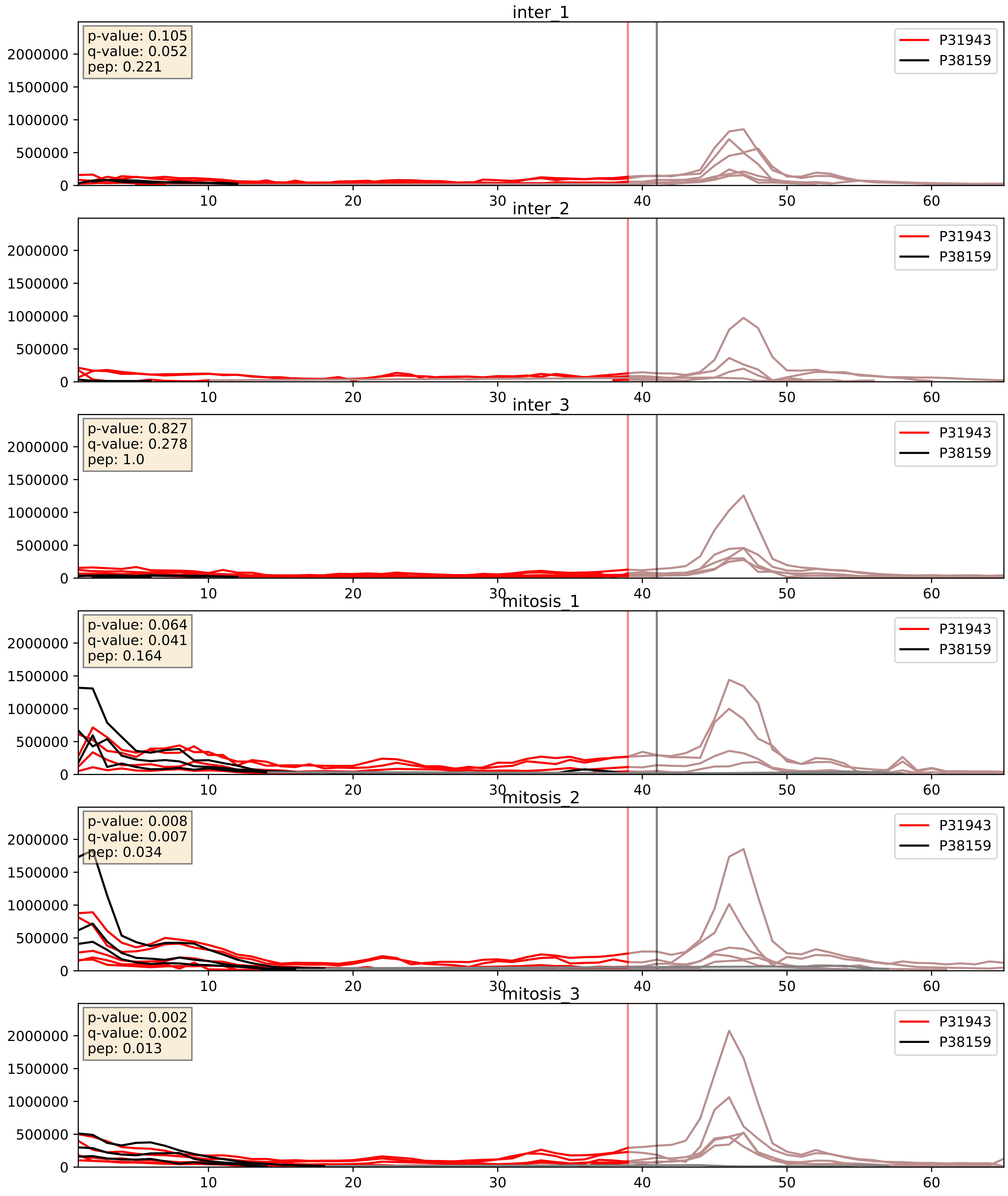


| level | condition_1 | condition_2 | pvalue | pvalue_adjusted |
|---|---|---|---|---|
| complex_abundance | mitosis | inter | 0.002318702437818041 | 0.03512936790747333 |
| interactor_ratio | mitosis | inter | 0.006225715441810617 | 0.10853793112403032 |
| interactor_abundance | mitosis | inter | 0.0062904982907834375 | 0.06768908280214485 |

P31943_P51991
HNRH1_HUMAN vs ROA3_HUMAN
p-value: 0.0 q-value: 0.0

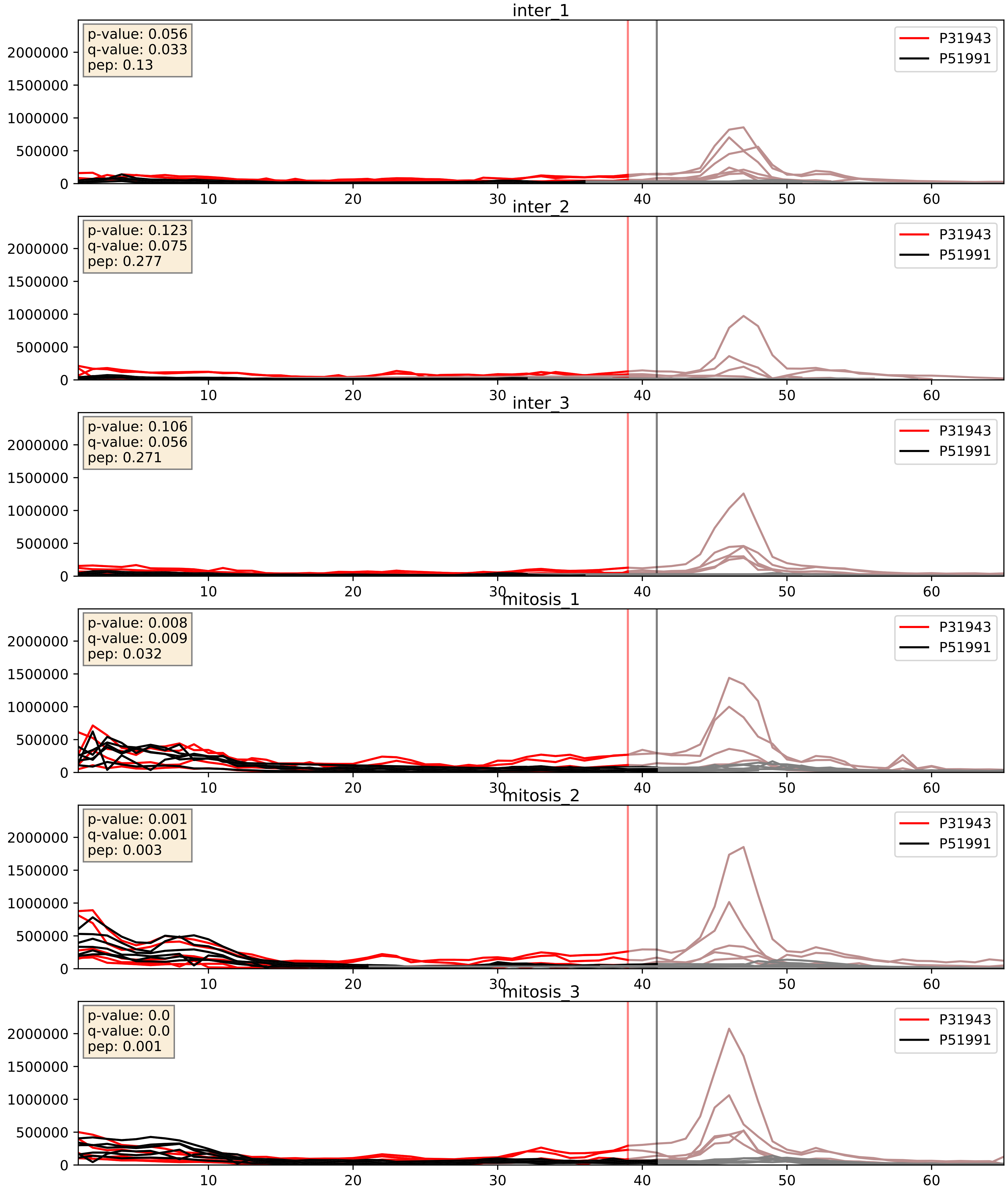


| level | condition_1 | condition_2 | pvalue | pvalue_adjusted |
|---|---|---|---|---|
| complex_abundance | mitosis | inter | 0.0003987949630443645 | 0.01272633620489383 |
| interactor_abundance | mitosis | inter | 0.006828293176471166 | 0.07097776513733234 |
| interactor_ratio | mitosis | inter | 0.01669269551176852 | 0.19102870799563973 |

P31943_P52272
HNRH1_HUMAN vs HNRPM_HUMAN
p-value: 0.006 q-value: 0.006

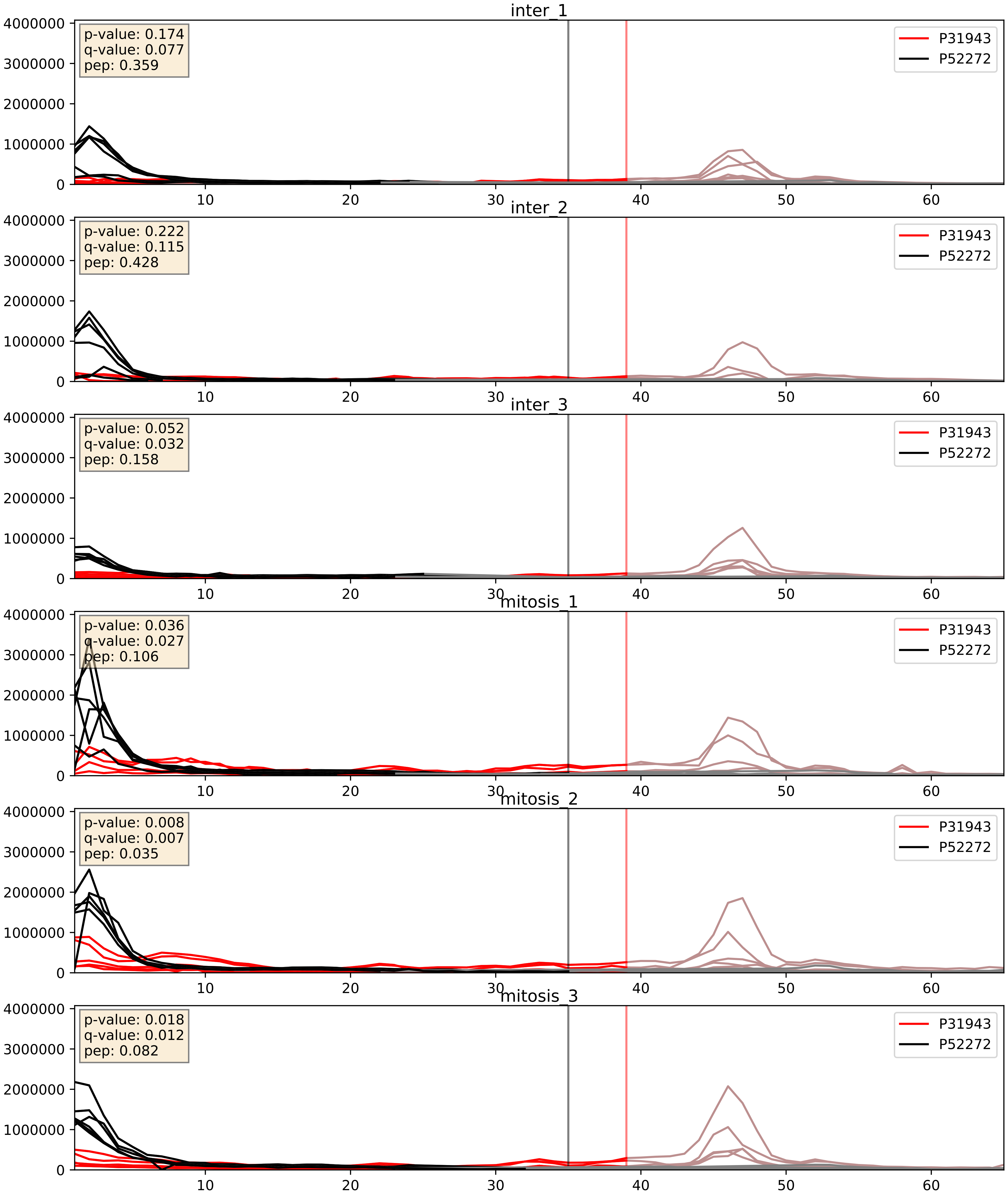


| level | condition_1 | condition_2 | pvalue | pvalue_adjusted |
|---|---|---|---|---|
| interactor_abundance | mitosis | inter | 0.002632509495742343 | 0.03838967005823521 |
| complex_abundance | mitosis | inter | 0.006583262263537986 | 0.06001355162501082 |
| interactor_ratio | mitosis | inter | 0.04049740663601096 | 0.33268502956262364 |

P31943_P52597
HNRH1_HUMAN vs HNRPF_HUMAN
p-value: 0.001 q-value: 0.002

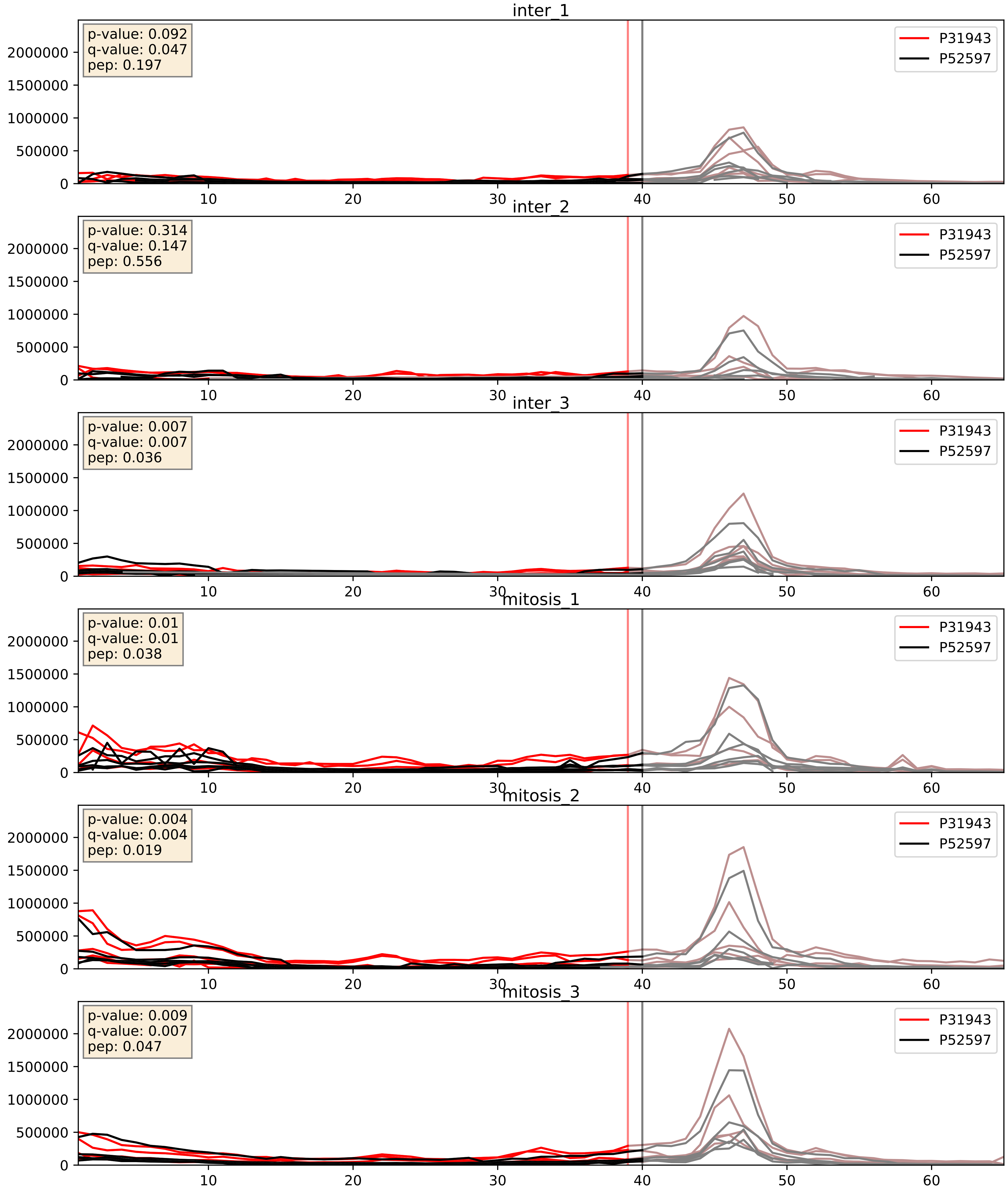


| level | condition_1 | condition_2 | pvalue | pvalue_adjusted |
|---|---|---|---|---|
| complex_abundance | mitosis | inter | 0.0023078745983505836 | 0.03502731659907978 |
| interactor_abundance | mitosis | inter | 0.0035613165684418575 | 0.04601489785035819 |
| interactor_ratio | mitosis | inter | 0.11518859280013378 | 0.5598715168125593 |

P31943_P55795
HNRH1_HUMAN vs HNRH2_HUMAN
p-value: 0.017 q-value: 0.014

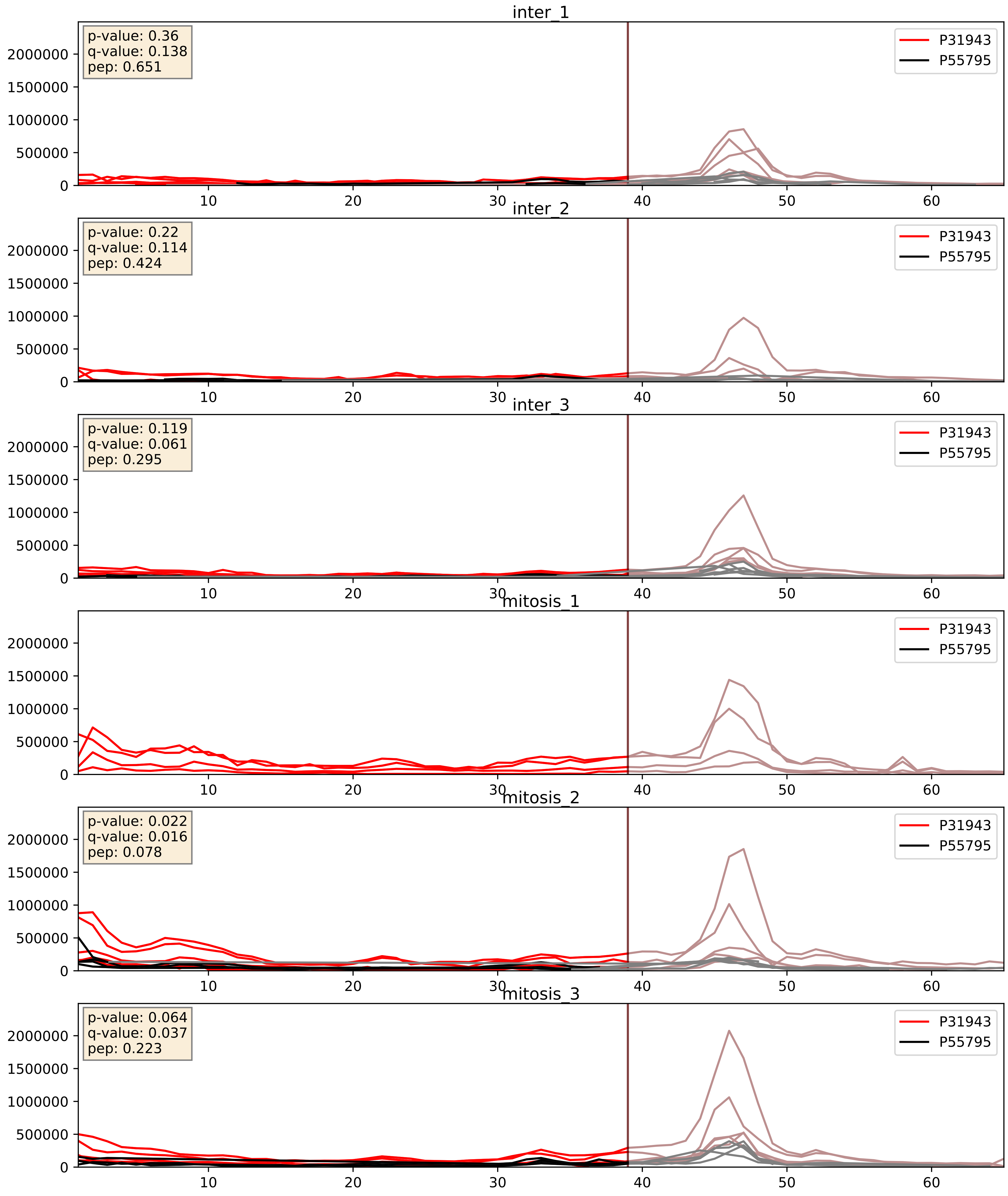


| level | condition_1 | condition_2 | pvalue | pvalue_adjusted |
|---|---|---|---|---|
| interactor_ratio | mitosis | inter | 0.6646036331503247 | 0.9283996733722243 |
| complex_abundance | mitosis | inter | 0.7506364016463556 | 0.835375407525308 |
| interactor_abundance | mitosis | inter | 0.87899693080352 | 0.9328308613535993 |

P31943_P61978
HNRH1_HUMAN vs HNRPK_HUMAN
p-value: 0.001 q-value: 0.002

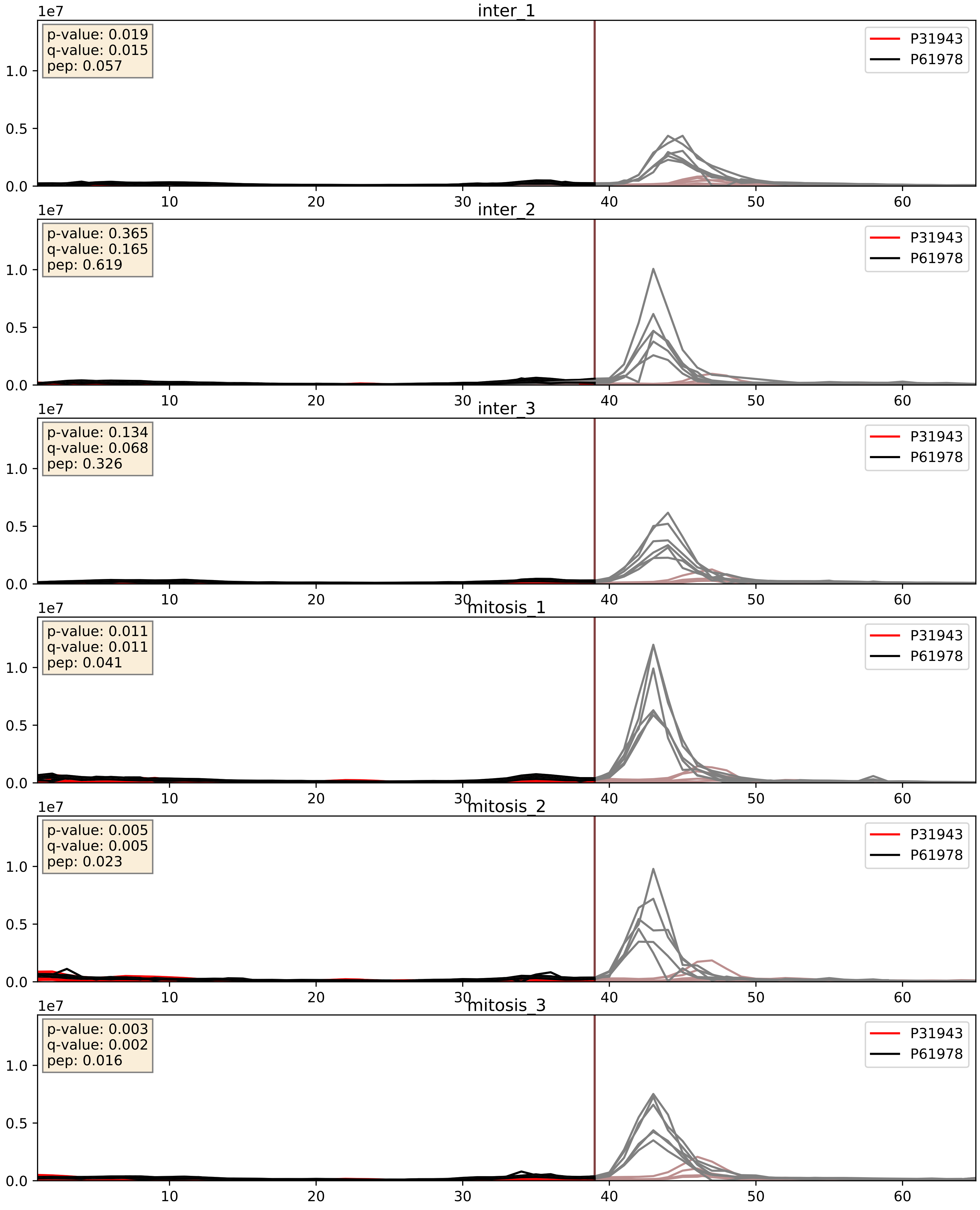


| level | condition_1 | condition_2 | pvalue | pvalue_adjusted |
|---|---|---|---|---|
| interactor_abundance | mitosis | inter | 0.0035613165684418575 | 0.04601489785035819 |
| interactor_ratio | mitosis | inter | 0.12016334003914726 | 0.5692297679773661 |
| complex_abundance | mitosis | inter | 0.26011154828329874 | 0.41017652655535747 |

P31943_P62995
HNRH1_HUMAN vs TRA2B_HUMAN
p-value: 0.007 q-value: 0.006

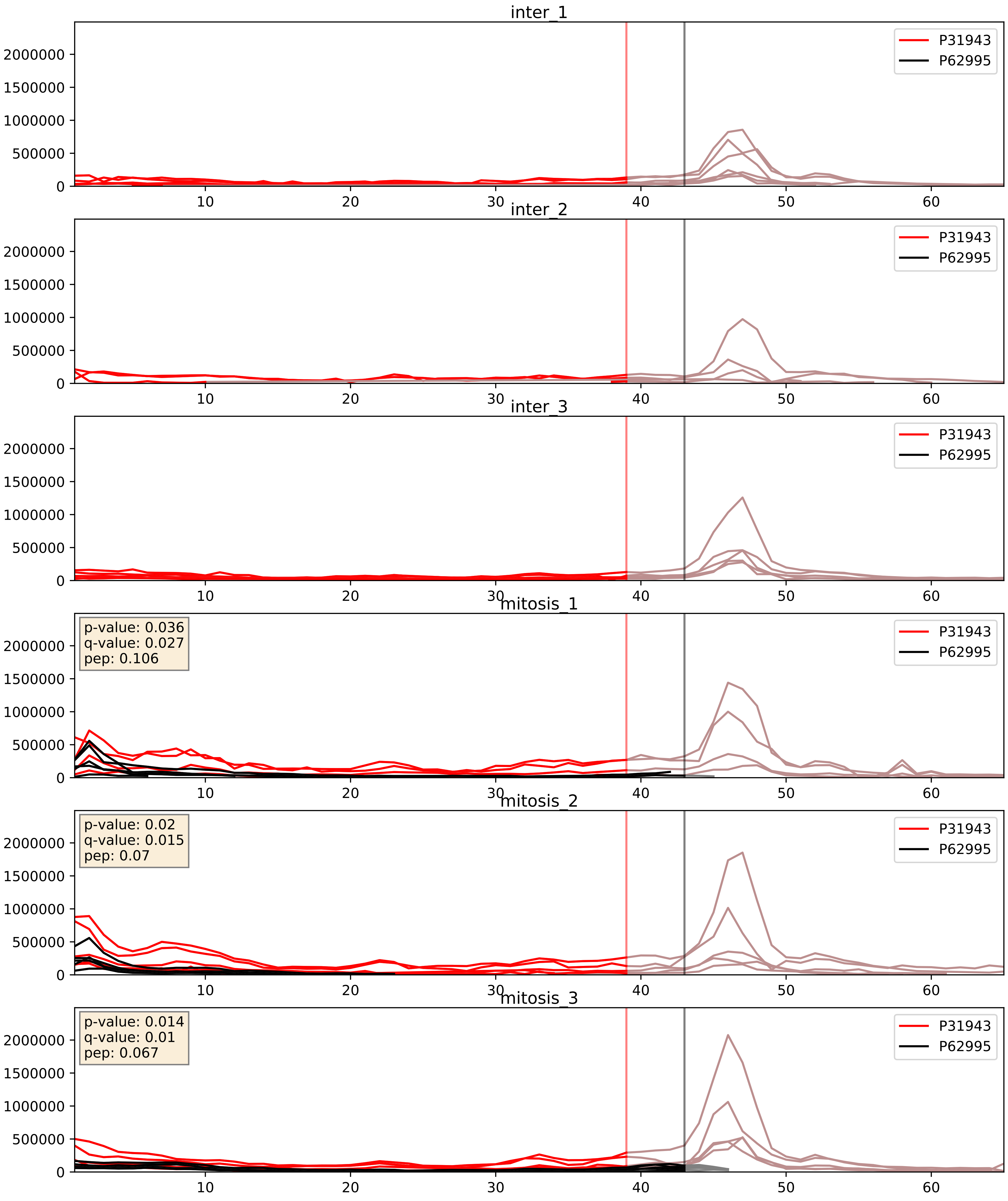


| level | condition_1 | condition_2 | pvalue | pvalue_adjusted |
|---|---|---|---|---|
| interactor_ratio | mitosis | inter | 2.5032546650867087e-05 | 0.005101871412652911 |
| interactor_abundance | mitosis | inter | 4.9215985987051895e-05 | 0.006283598333582311 |
| complex_abundance | mitosis | inter | 0.00011780432600576765 | 0.007493687219947184 |

P31943_P84103
HNRH1_HUMAN vs SRSF3_HUMAN
p-value: 0.005 q-value: 0.005

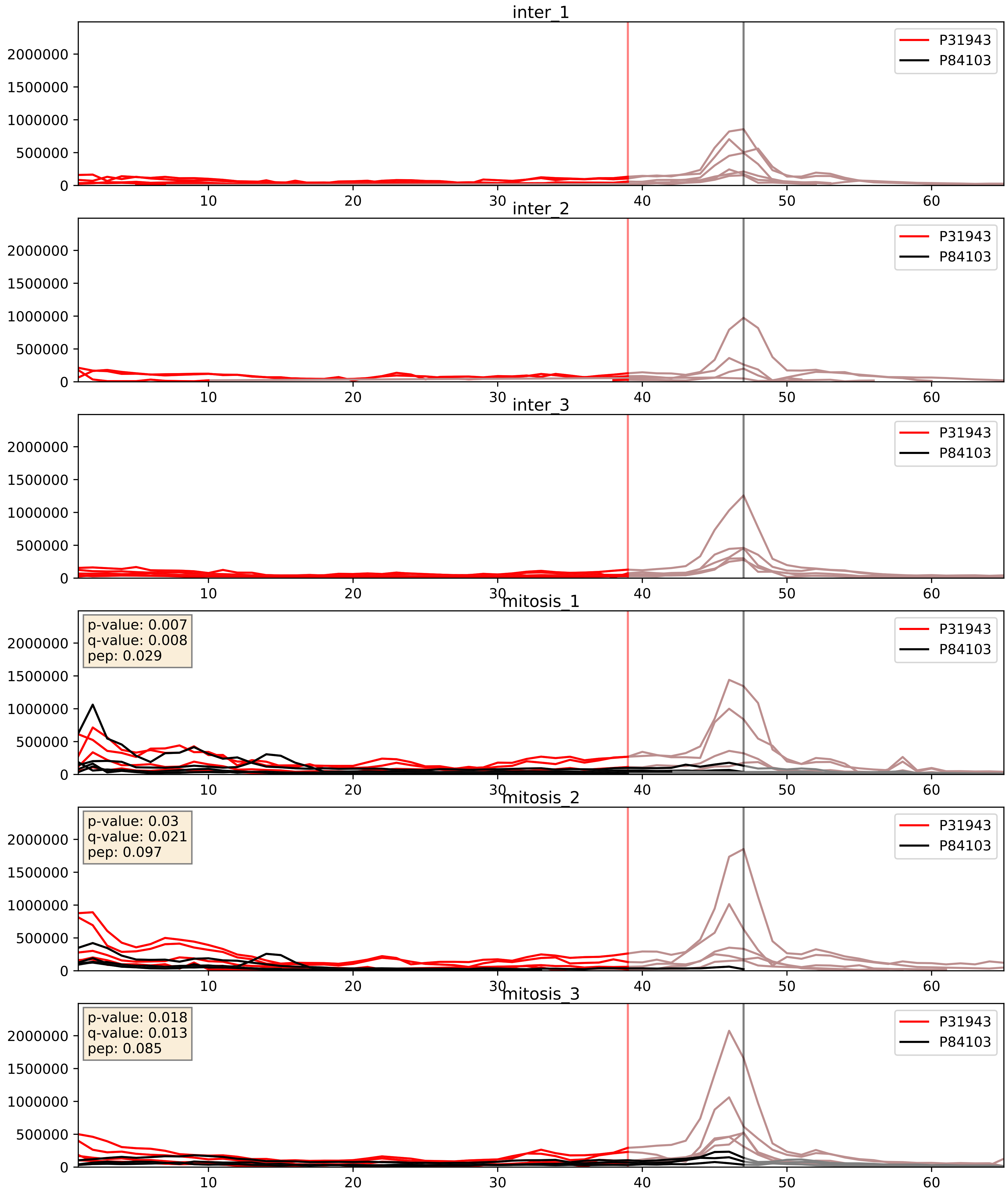


| level | condition_1 | condition_2 | pvalue | pvalue_adjusted |
|---|---|---|---|---|
| interactor_abundance | mitosis | inter | 2.5609271594898103e-05 | 0.004519904429944902 |
| complex_abundance | mitosis | inter | 0.0003358253138333869 | 0.011893558089955385 |
| interactor_ratio | mitosis | inter | 0.005707332319897169 | 0.10328702887594031 |

P31943_Q07955
HNRH1_HUMAN vs SRSF1_HUMAN
p-value: 0.001 q-value: 0.002

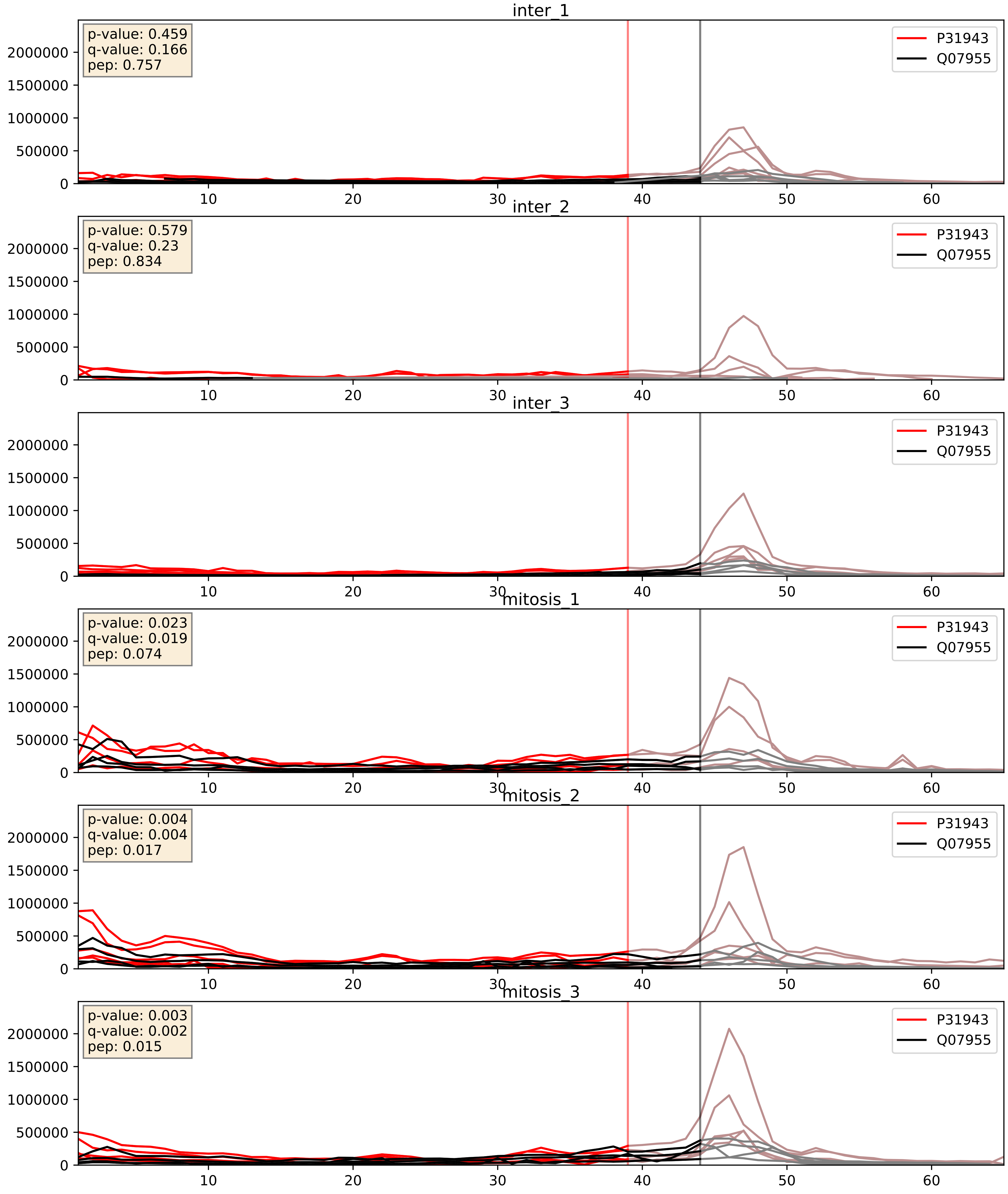


| level | condition_1 | condition_2 | pvalue | pvalue_adjusted |
|---|---|---|---|---|
| complex_abundance | mitosis | inter | 0.007690789144795531 | 0.06519315066658214 |
| interactor_abundance | mitosis | inter | 0.013391224463716843 | 0.10304923015527873 |
| interactor_ratio | mitosis | inter | 0.43412575717983676 | 0.8435878041032946 |

P31943_Q12906
HNRH1_HUMAN vs ILF3_HUMAN
p-value: 0.004 q-value: 0.048

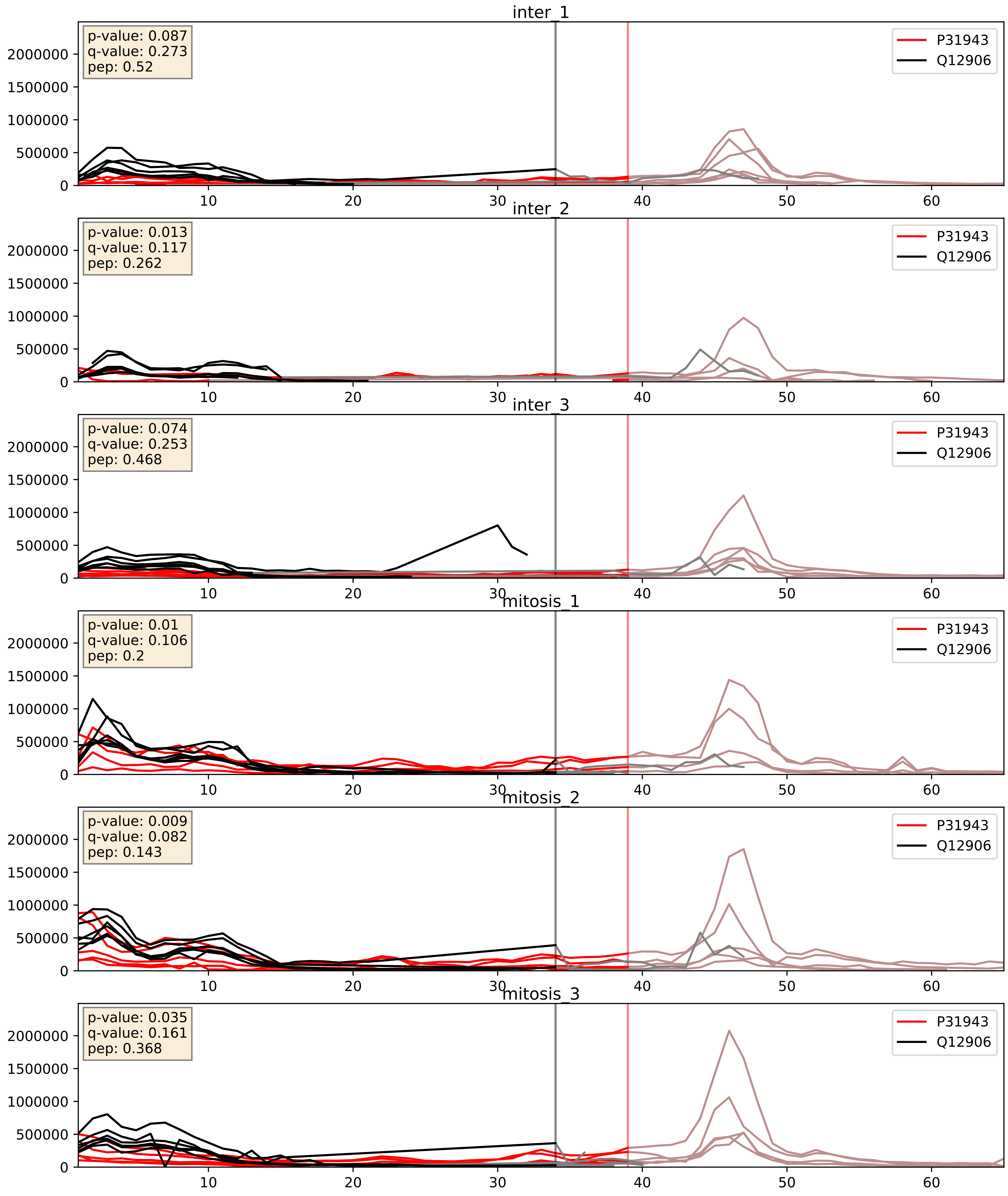


| level | condition_1 | condition_2 | pvalue | pvalue_adjusted |
|---|---|---|---|---|
| interactor_ratio | mitosis | inter | 0.004516521131172633 | 0.08949402982138366 |
| interactor_abundance | mitosis | inter | 0.005235127518247087 | 0.060728341499721825 |
| complex_abundance | mitosis | inter | 0.007172904954993601 | 0.06275154604611013 |

P31943_Q13151
HNRH1_HUMAN vs ROA0_HUMAN
p-value: 0.021 q-value: 0.016

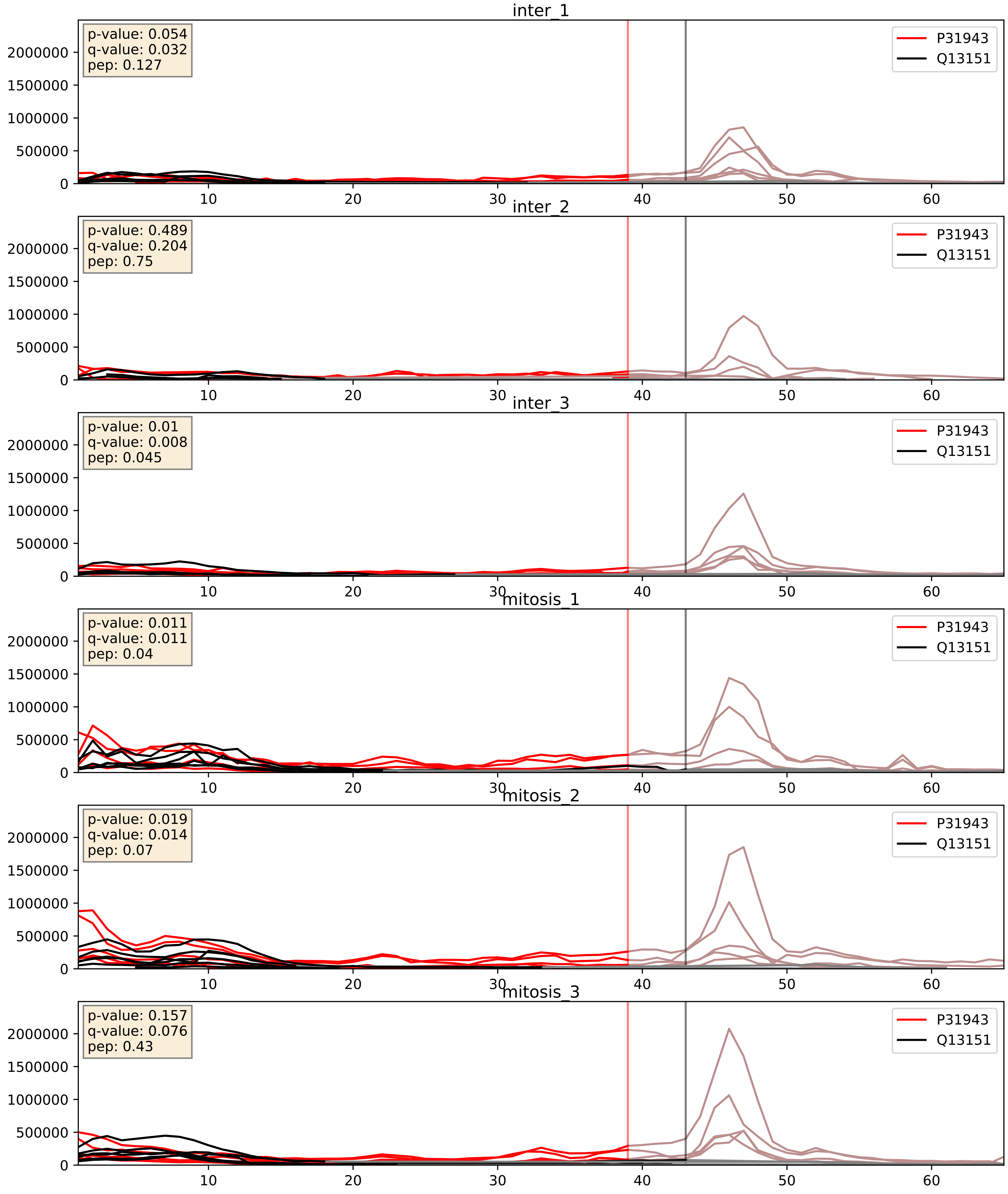


| level | condition_1 | condition_2 | pvalue | pvalue_adjusted |
|---|---|---|---|---|
| complex_abundance | mitosis | inter | 0.002219411769080441 | 0.03446495495307955 |
| interactor_abundance | mitosis | inter | 0.0024193513526238088 | 0.0366218347983374 |
| interactor_ratio | mitosis | inter | 0.413220111473216 | 0.8352217601442099 |

P31943_Q13247
HNRH1_HUMAN vs SRSF6_HUMAN
p-value: 0.005 q-value: 0.005

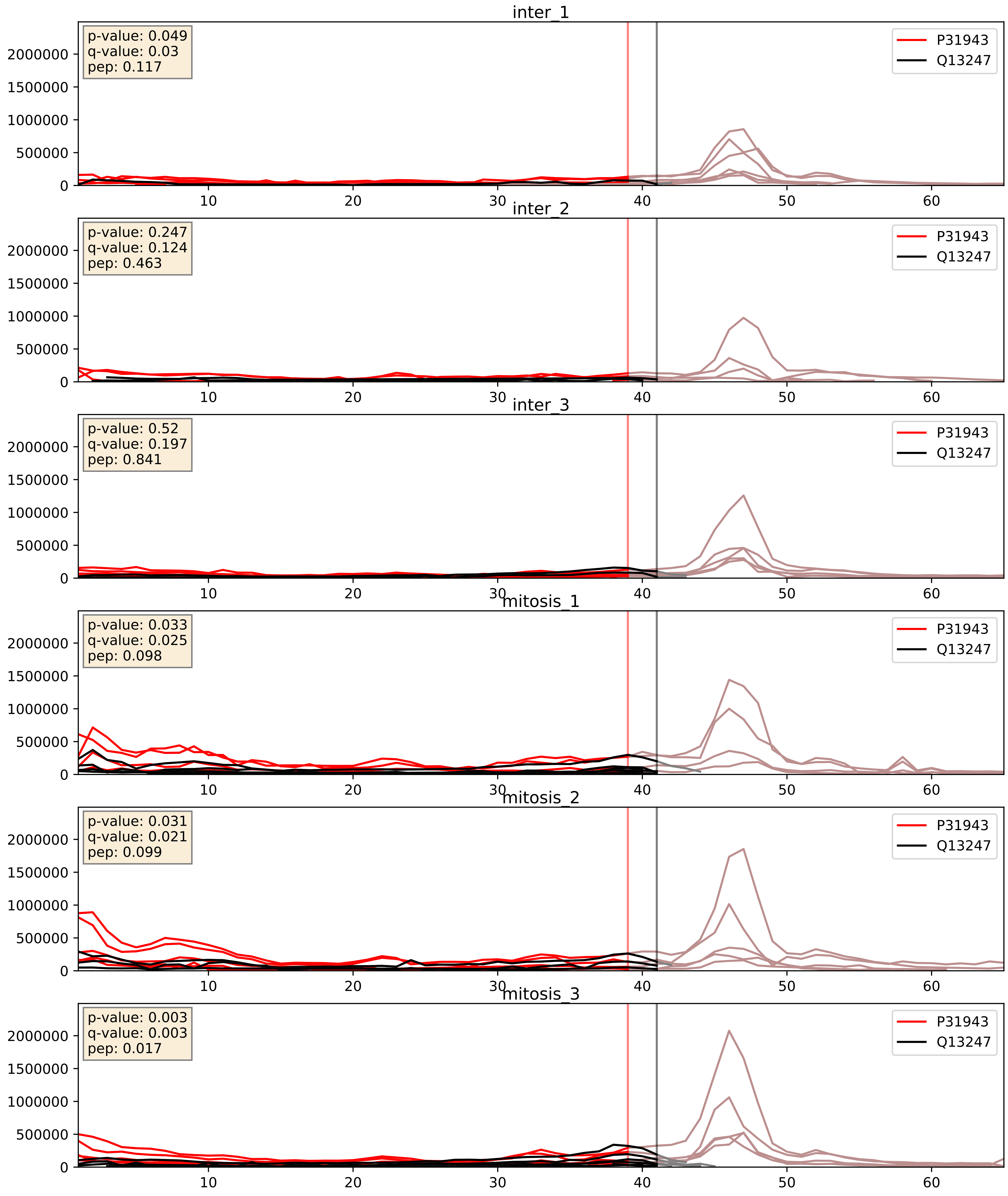


| level | condition_1 | condition_2 | pvalue | pvalue_adjusted |
|---|---|---|---|---|
| complex_abundance | mitosis | inter | 0.004753245691634782 | 0.05150352293720726 |
| interactor_abundance | mitosis | inter | 0.0056571357548407175 | 0.06348577161904892 |
| interactor_ratio | mitosis | inter | 0.9922029355017021 | 0.9981863229331772 |

P31943_Q15365
HNRH1_HUMAN vs PCBP1_HUMAN
p-value: 0.028 q-value: 0.021

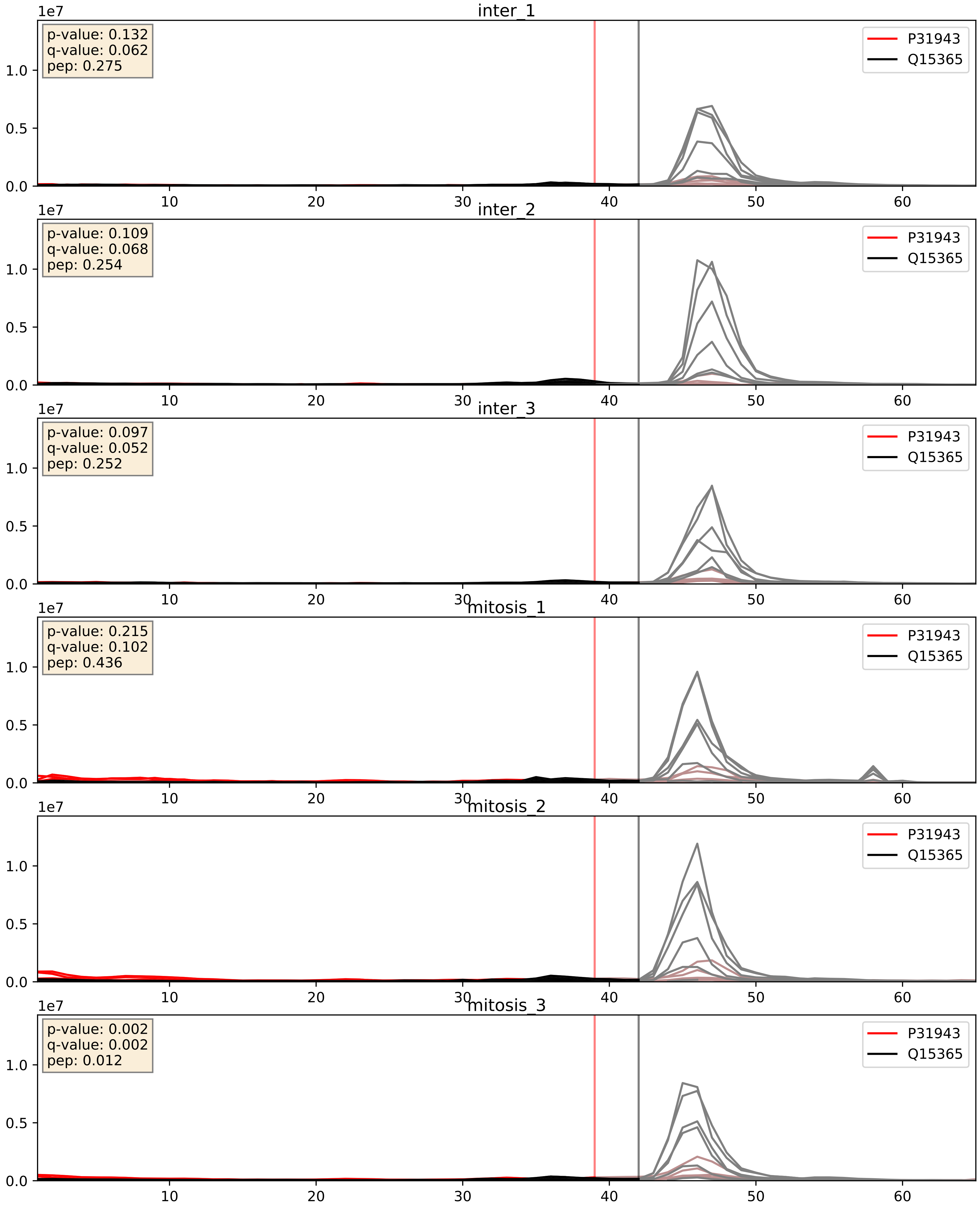


| level | condition_1 | condition_2 | pvalue | pvalue_adjusted |
|---|---|---|---|---|
| interactor_abundance | mitosis | inter | 0.0035613165684418575 | 0.04601489785035819 |
| complex_abundance | mitosis | inter | 0.004390077064485453 | 0.04944613114736247 |
| interactor_ratio | mitosis | inter | 0.005505267863595088 | 0.1004799422438677 |

P31943_Q15717
HNRH1_HUMAN vs ELAV1_HUMAN
p-value: 0.006 q-value: 0.006

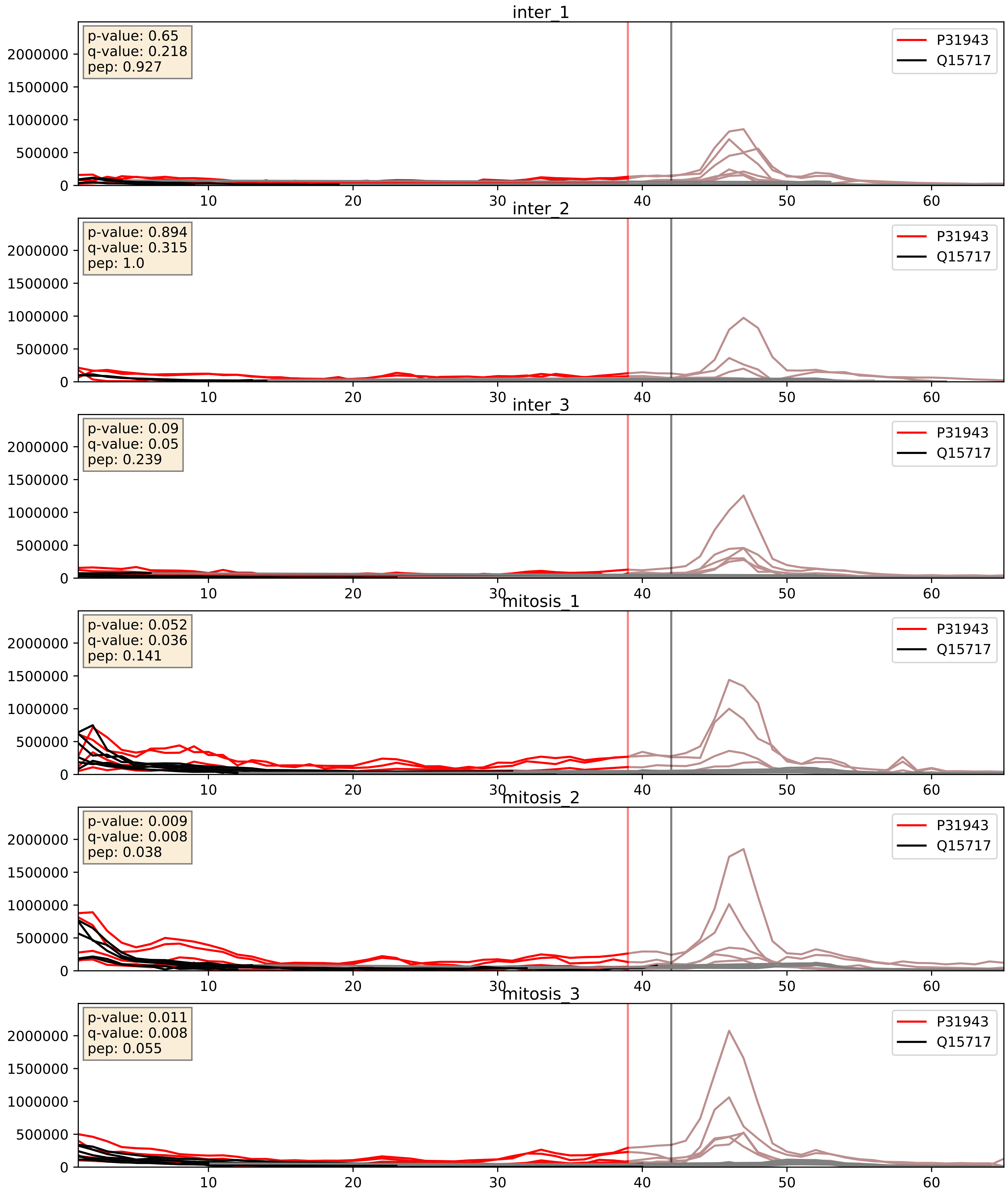


| level | condition_1 | condition_2 | pvalue | pvalue_adjusted |
|---|---|---|---|---|
| complex_abundance | mitosis | inter | 0.00017736744901018607 | 0.008807651911797798 |
| interactor_abundance | mitosis | inter | 0.0014904490749389305 | 0.0285056509549173 |
| interactor_ratio | mitosis | inter | 0.12665350951524584 | 0.5750062399685051 |

P31943_Q16629
HNRH1_HUMAN vs SRSF7_HUMAN
p-value: 0.002 q-value: 0.003

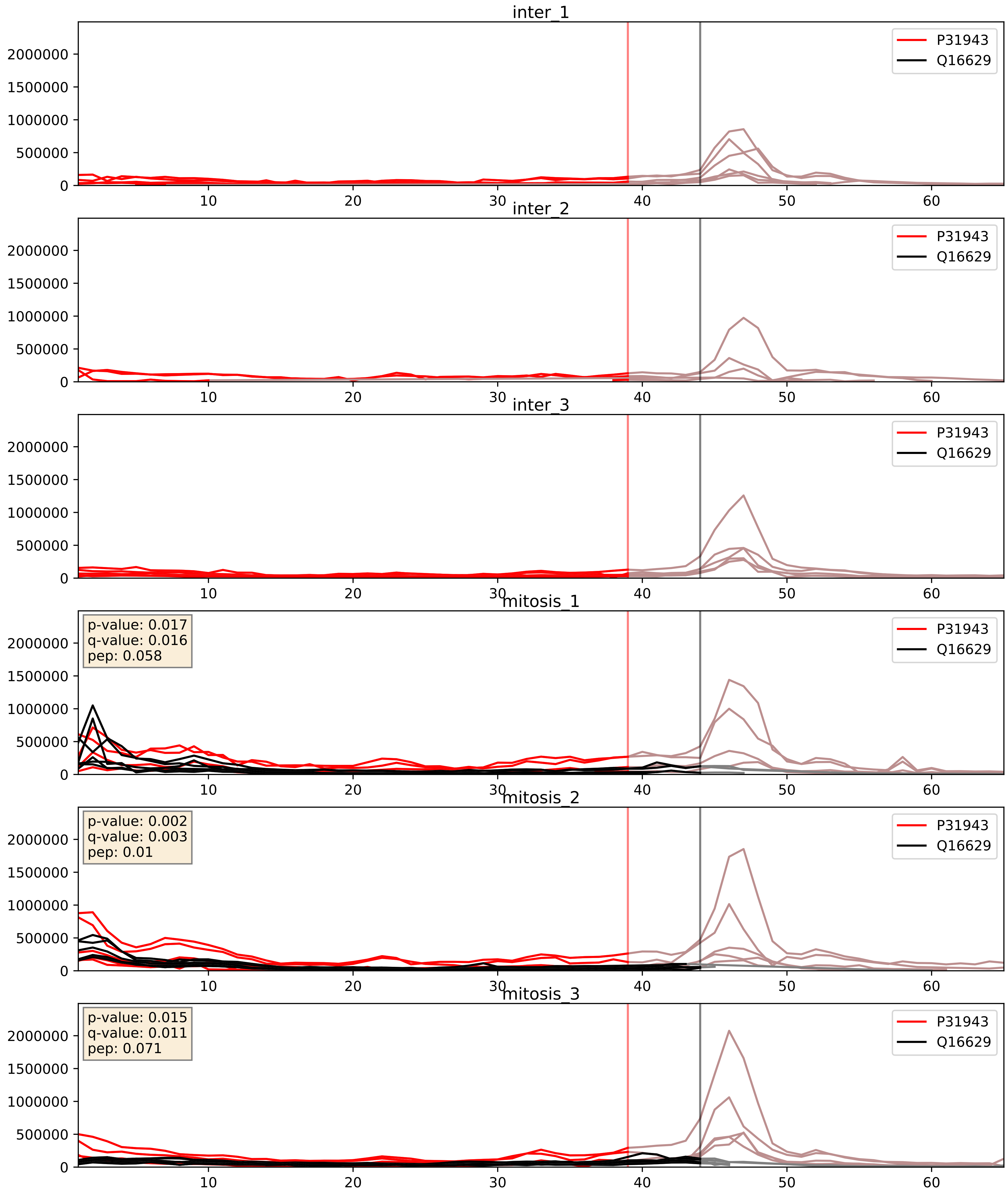


| level | condition_1 | condition_2 | pvalue | pvalue_adjusted |
|---|---|---|---|---|
| interactor_abundance | mitosis | inter | 2.510205397937702e-05 | 0.004476532959655569 |
| complex_abundance | mitosis | inter | 4.065873462401448e-05 | 0.005691857151652958 |
| interactor_ratio | mitosis | inter | 0.00017130672096930523 | 0.013577643810159747 |

P31943_Q99459
HNRH1_HUMAN vs CDC5L_HUMAN
p-value: 0.066 q-value: 0.041

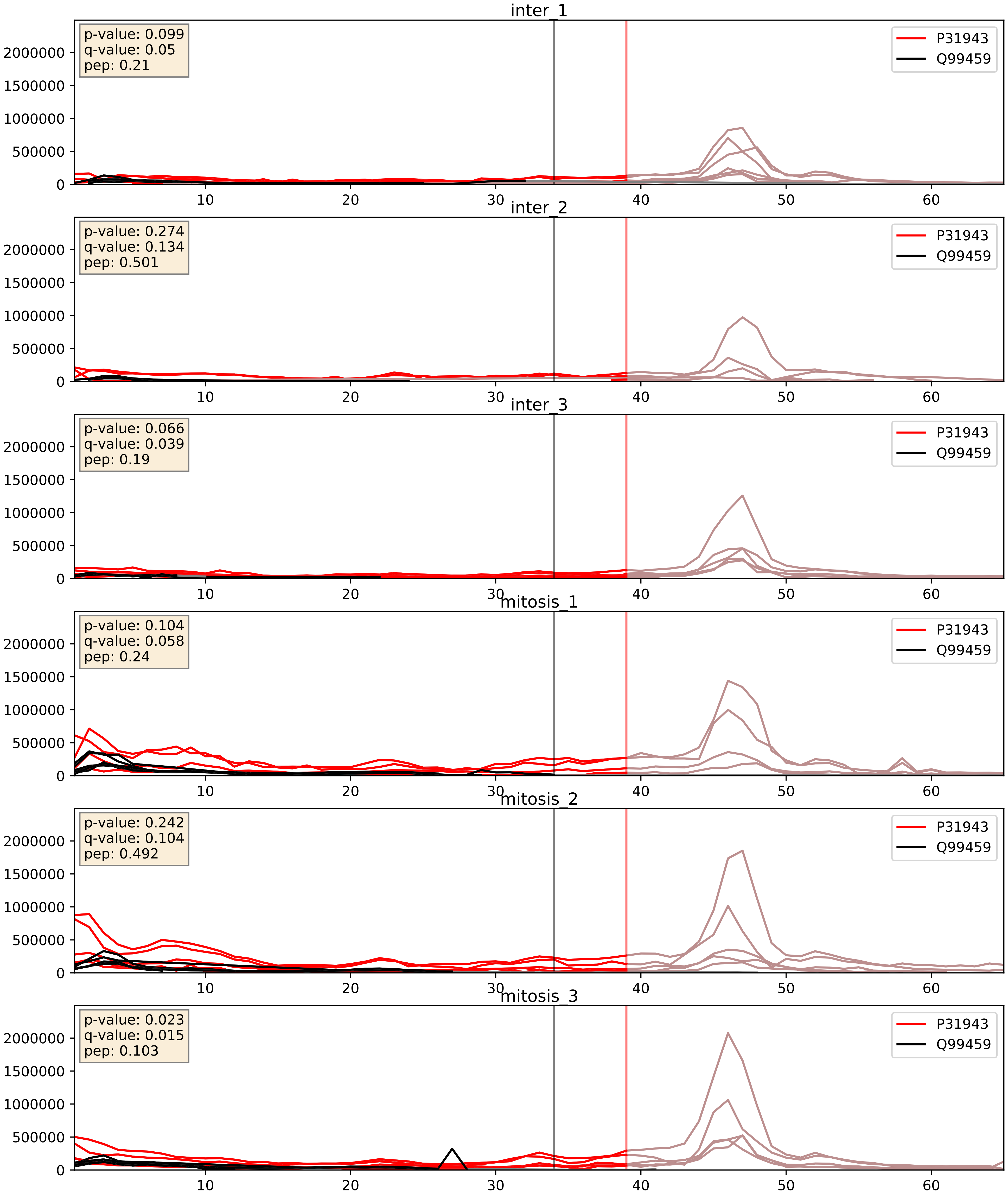


| level | condition_1 | condition_2 | pvalue | pvalue_adjusted |
|---|---|---|---|---|
| interactor_abundance | mitosis | inter | 0.0050208300212190625 | 0.05879385086407001 |
| complex_abundance | mitosis | inter | 0.008080613653928214 | 0.06653038413698939 |
| interactor_ratio | mitosis | inter | 0.6358488676126866 | 0.9197052662231202 |

P31943_Q9BUJ2
HNRH1_HUMAN vs HNRL1_HUMAN
p-value: 0.001 q-value: 0.002

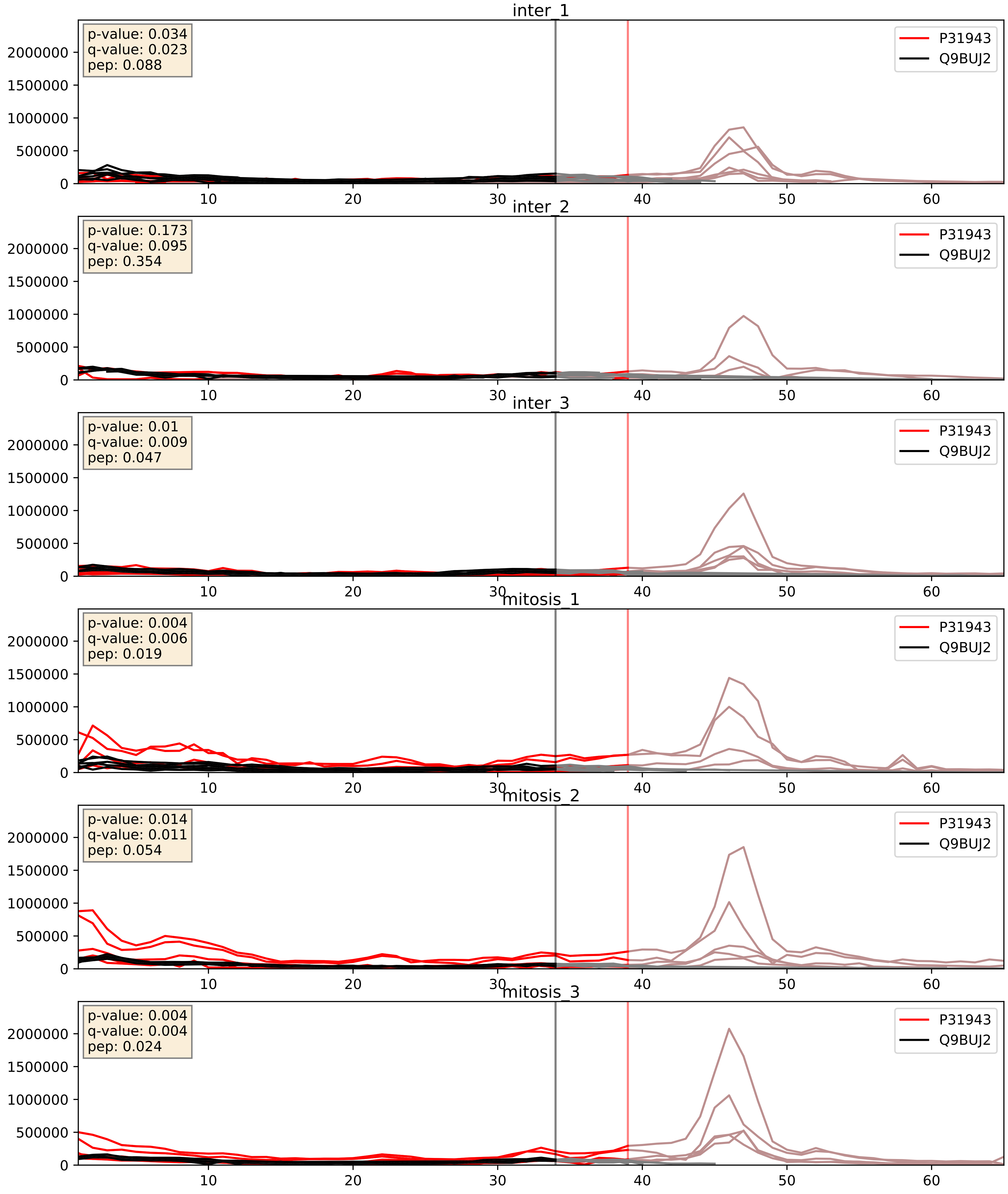


| level | condition_1 | condition_2 | pvalue | pvalue_adjusted |
|---|---|---|---|---|
| interactor_ratio | mitosis | inter | 0.001394553898793512 | 0.044212523606194314 |
| interactor_abundance | mitosis | inter | 0.0027782426171848577 | 0.03993577968615009 |
| complex_abundance | mitosis | inter | 0.007692182496874762 | 0.06519315066658214 |
