## Supplementary Data 2 for "SECAT: Quantifying differential protein-protein interaction states by network-centric analysis": hela_string_000021_Q9NVP1.pdf

Q9NVP1_Q9NVP1

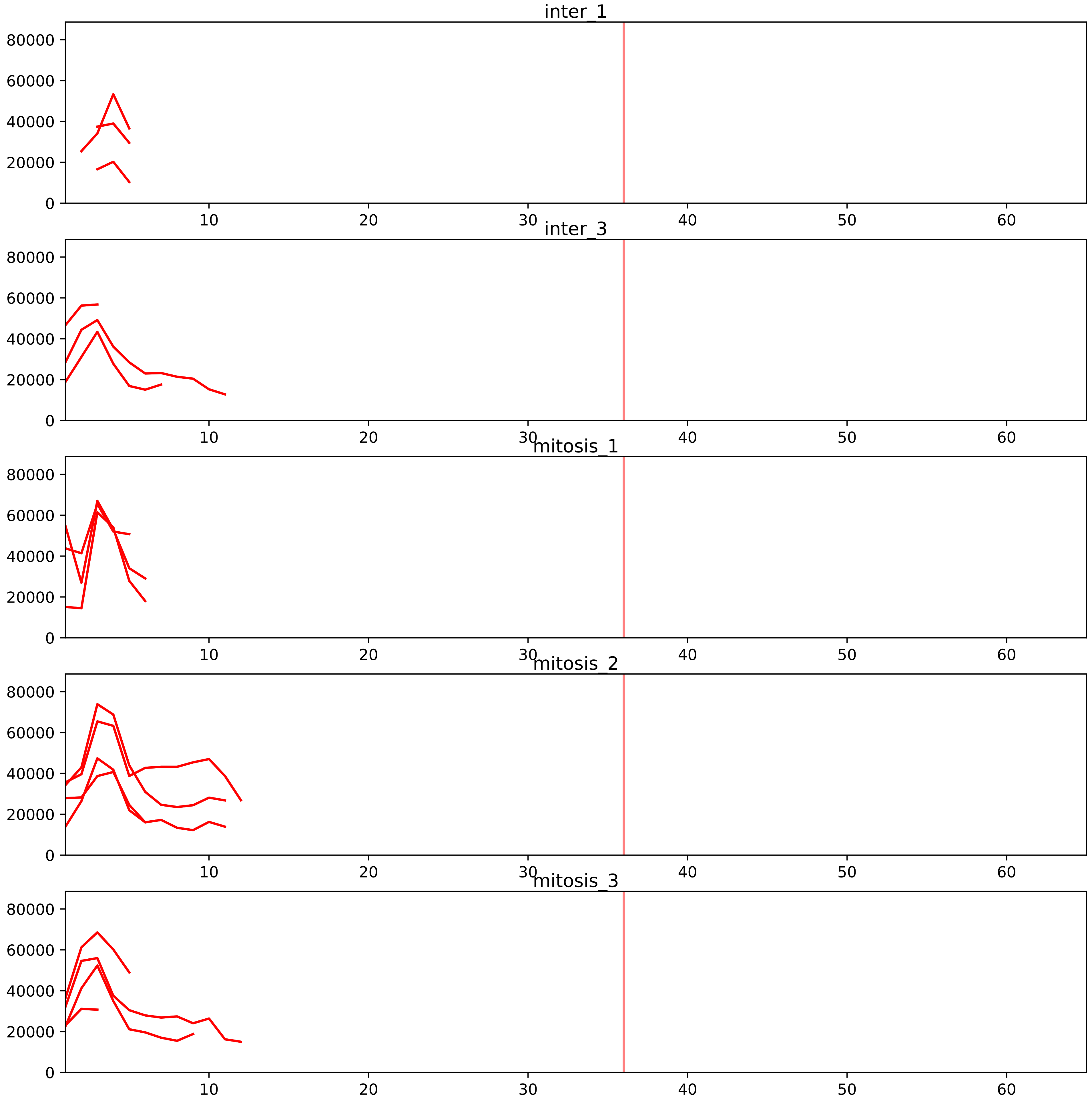


| level | condition_1 | condition_2 | pvalue | pvalue_adjusted |
|---|---|---|---|---|
| interactor_ratio | mitosis | inter | 1.0085305374926425e-11 | 8.836648518983154e-10 |
| complex_abundance | mitosis | inter | 0.025147141399425914 | 0.08131940276791508 |
| interactor_abundance | mitosis | inter | 0.16519544448921122 | 0.31207352963054275 |
| total_abundance | mitosis | inter | 0.20115180766142166 | 0.5381355939626228 |
| assembled_abundance | mitosis | inter | 0.21424393572021597 | 0.5217876498992357 |

O60832_Q9NVP1
DKC1_HUMAN vs DDX18_HUMAN
p-value: 0.076 q-value: 0.046

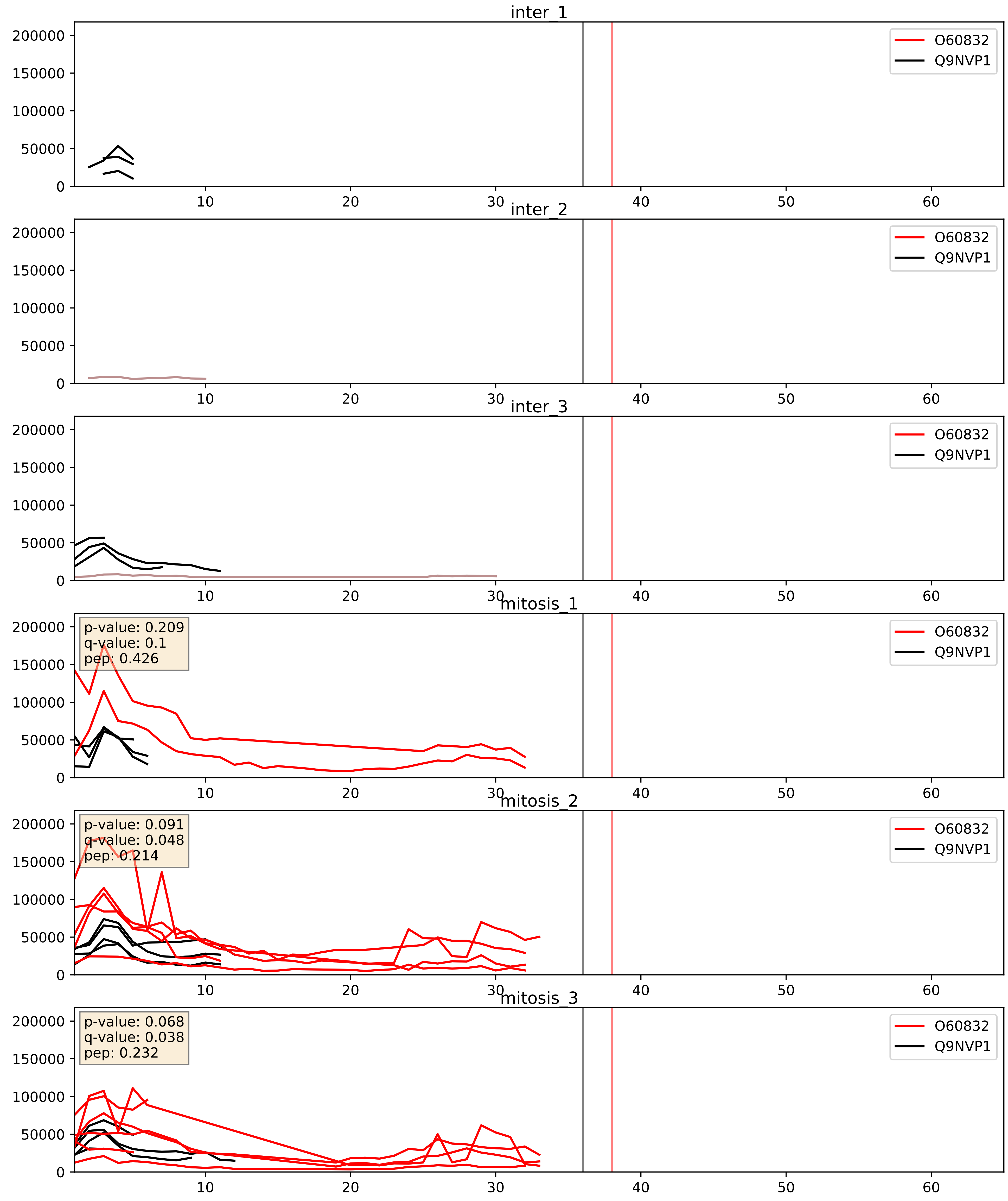


| level | condition_1 | condition_2 | pvalue | pvalue_adjusted |
|---|---|---|---|---|
| complex_abundance | mitosis | inter | 0.019061609538449552 | 0.10333589464795959 |
| interactor_abundance | mitosis | inter | 0.07954363368256376 | 0.23458863198027416 |
| interactor_ratio | mitosis | inter | 0.5961816441313526 | 0.9019028605875125 |

P46087_Q9NVP1
NOP2_HUMAN vs DDX18_HUMAN
p-value: 0.046 q-value: 0.031

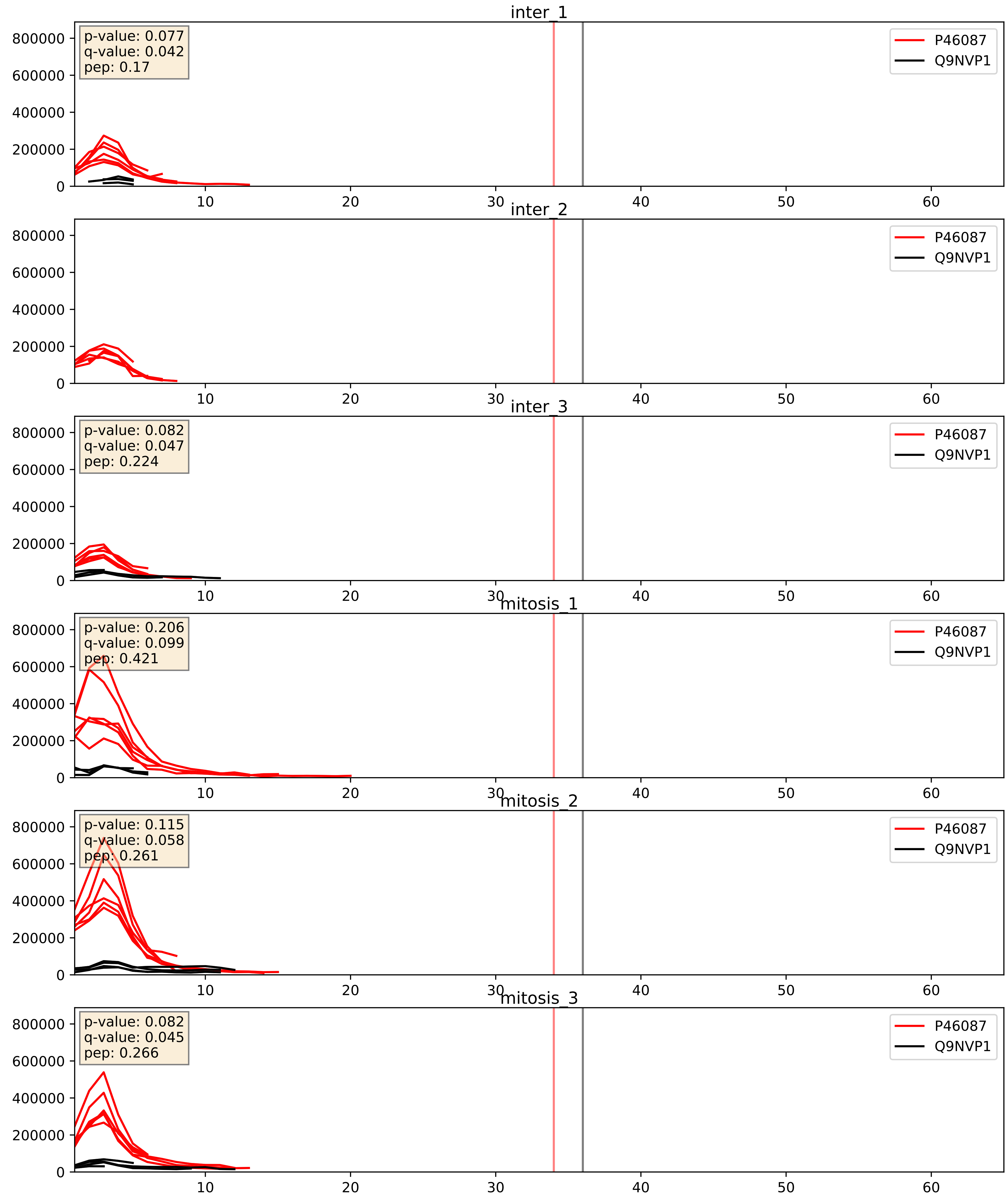


| level | condition_1 | condition_2 | pvalue | pvalue_adjusted |
|---|---|---|---|---|
| interactor_ratio | mitosis | inter | 0.005387411984646833 | 0.0985389884371301 |
| interactor_abundance | mitosis | inter | 0.03915622151066956 | 0.16083596262333216 |
| complex_abundance | mitosis | inter | 0.1003973827814877 | 0.2350018038308818 |

Q12788_Q9NVP1
TBL3_HUMAN vs DDX18_HUMAN
p-value: 0.08 q-value: 0.047

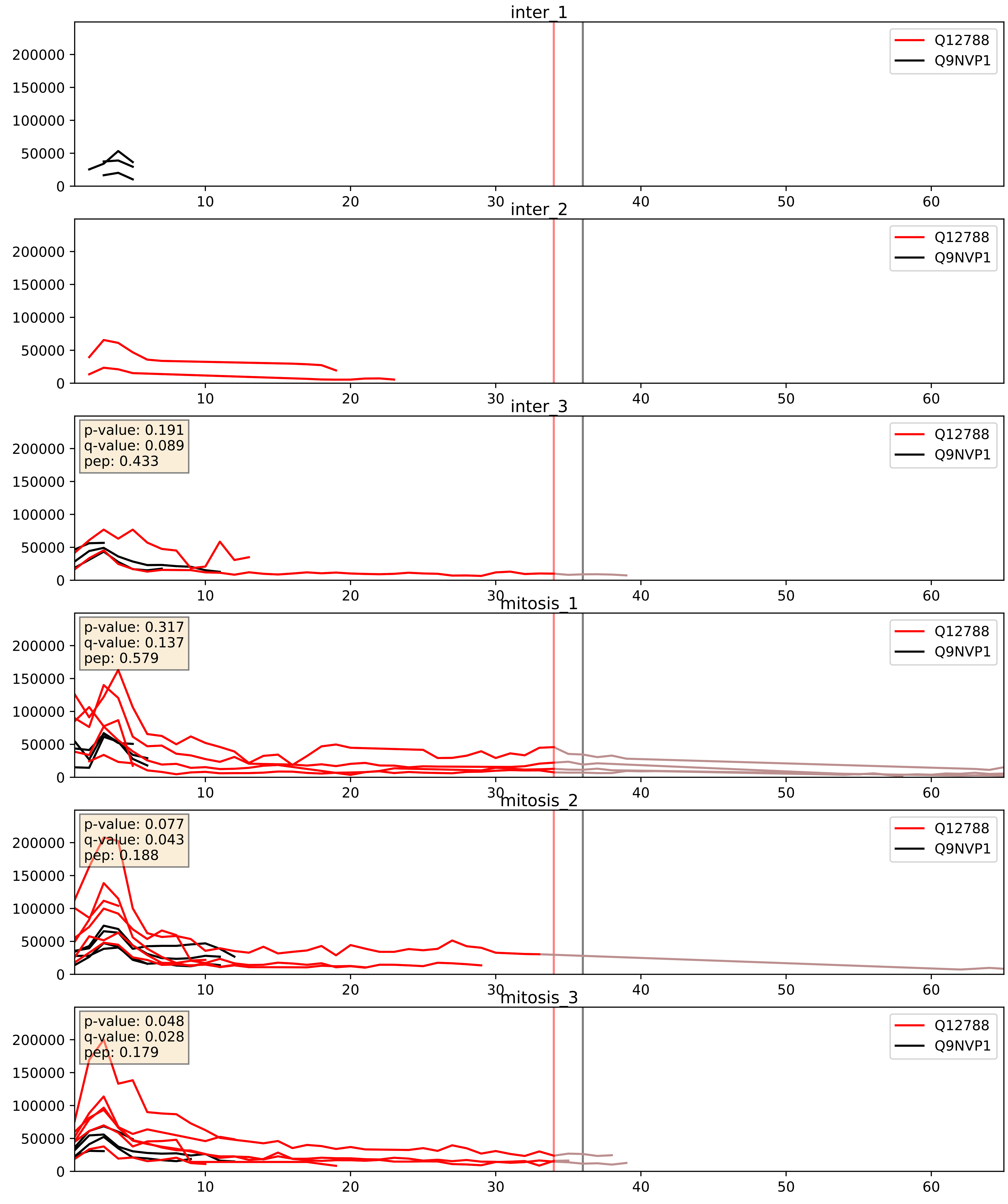


| level | condition_1 | condition_2 | pvalue | pvalue_adjusted |
|---|---|---|---|---|
| interactor_abundance | mitosis | inter | 0.007893035270042996 | 0.07711604472197223 |
| complex_abundance | mitosis | inter | 0.022357984284711807 | 0.10987189501272765 |
| interactor_ratio | mitosis | inter | 0.02782927283201316 | 0.2618295178868221 |

Q15397_Q9NVP1
PUM3_HUMAN vs DDX18_HUMAN
p-value: 0.002 q-value: 0.035

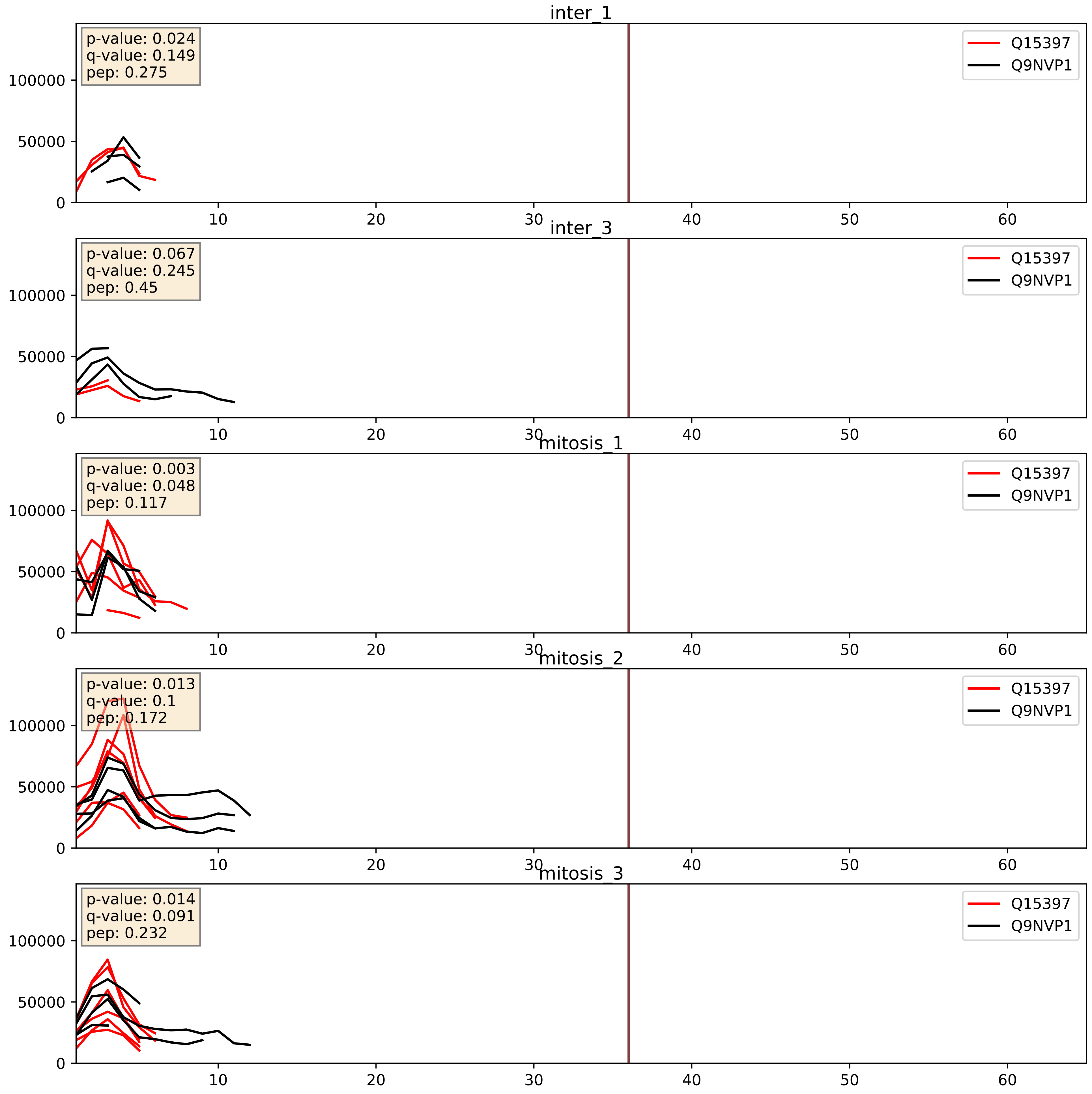


| level | condition_1 | condition_2 | pvalue | pvalue_adjusted |
|---|---|---|---|---|
| interactor_ratio | mitosis | inter | 0.07027102052804113 | 0.4393863664865099 |
| interactor_abundance | mitosis | inter | 0.26031152529654955 | 0.4522065263326465 |
| complex_abundance | mitosis | inter | 0.5653035680779996 | 0.6961586164217634 |

Q8NI36_Q9NVP1
WDR36_HUMAN vs DDX18_HUMAN
p-value: 0.034 q-value: 0.024

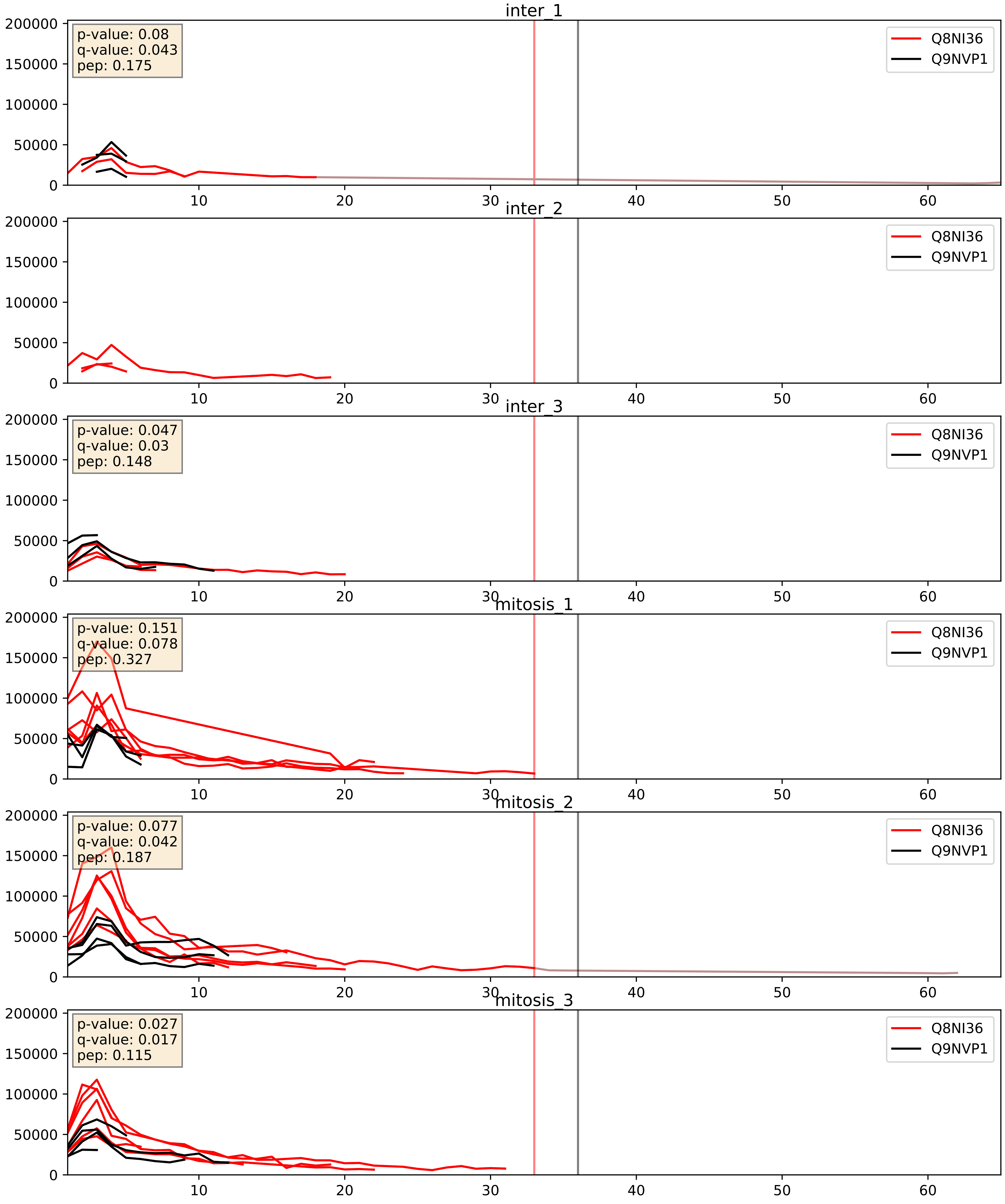


| level | condition_1 | condition_2 | pvalue | pvalue_adjusted |
|---|---|---|---|---|
| interactor_ratio | mitosis | inter | 0.009055793579369052 | 0.135757606023466 |
| interactor_abundance | mitosis | inter | 0.016895114881607966 | 0.11285383018850112 |
| complex_abundance | mitosis | inter | 0.13961312159519626 | 0.28468378070900036 |

Q8TDN6_Q9NVP1
BRX1_HUMAN vs DDX18_HUMAN
p-value: 0.02 q-value: 0.016

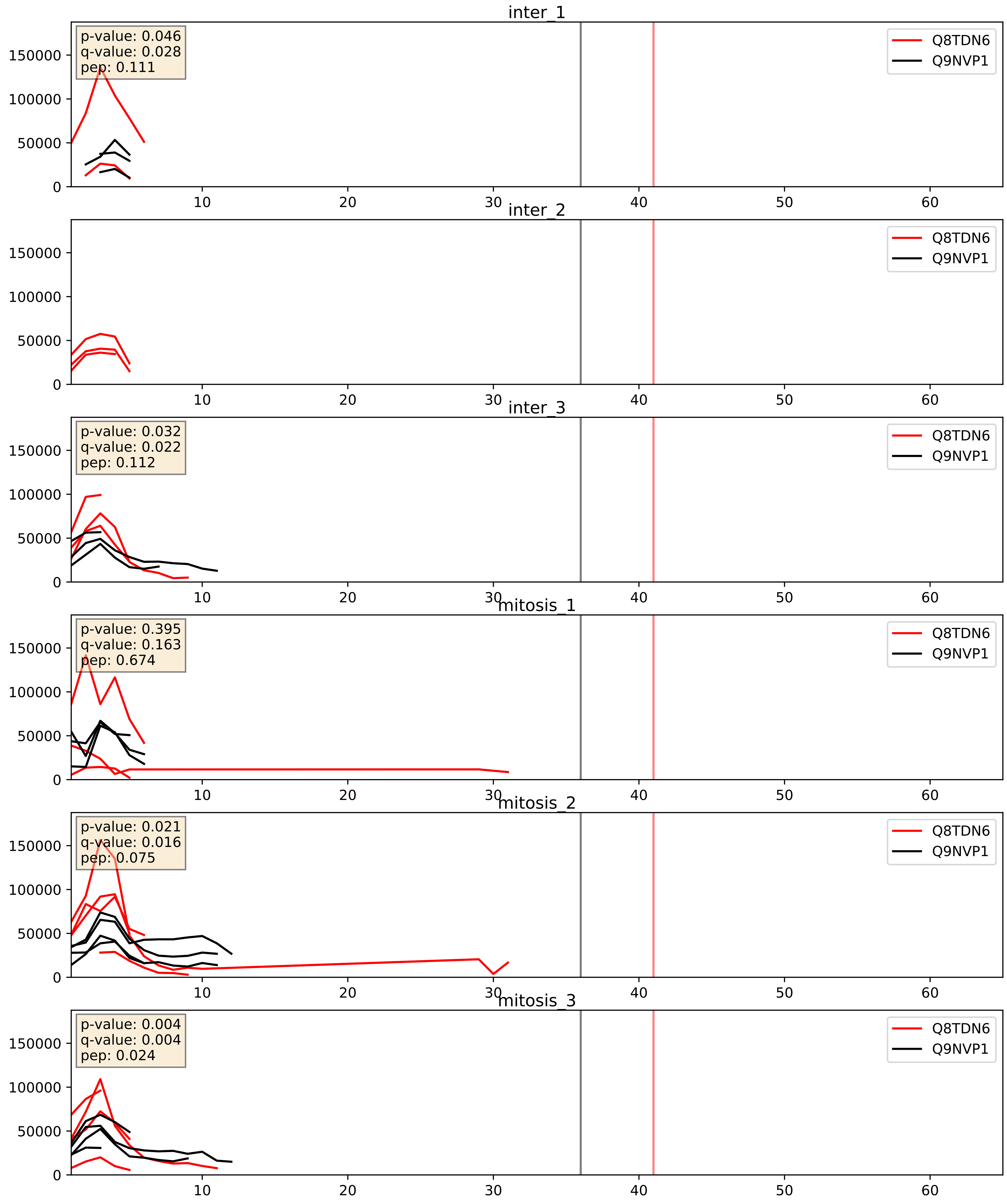


| level | condition_1 | condition_2 | pvalue | pvalue_adjusted |
|---|---|---|---|---|
| complex_abundance | mitosis | inter | 0.8425323439690111 | 0.8997101876714989 |
| interactor_abundance | mitosis | inter | 0.8683139378488682 | 0.9286903237679228 |
| interactor_ratio | mitosis | inter | 0.9136271445401767 | 0.9852905071335931 |

Q9BQ67_Q9NVP1
GRWD1_HUMAN vs DDX18_HUMAN
p-value: 0.019 q-value: 0.015

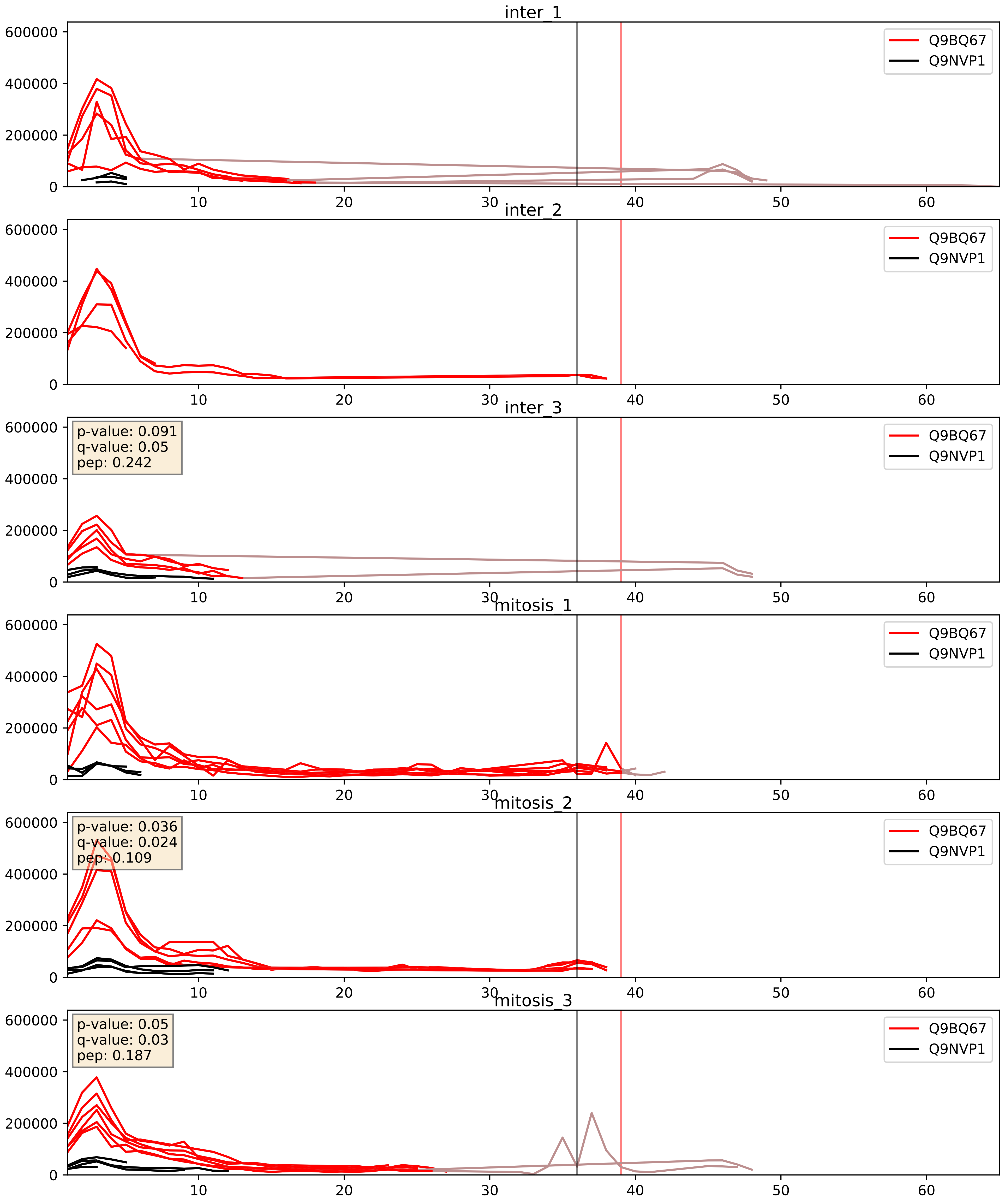


| level | condition_1 | condition_2 | pvalue | pvalue_adjusted |
|---|---|---|---|---|
| interactor_ratio | mitosis | inter | 0.05392358354238998 | 0.38487308410443505 |
| interactor_abundance | mitosis | inter | 0.14019369867573772 | 0.3269656225748469 |
| complex_abundance | mitosis | inter | 0.22993119905718284 | 0.3831280826025919 |

Q9BZE4_Q9NVP1
NOG1_HUMAN vs DDX18_HUMAN
p-value: 0.008 q-value: 0.007

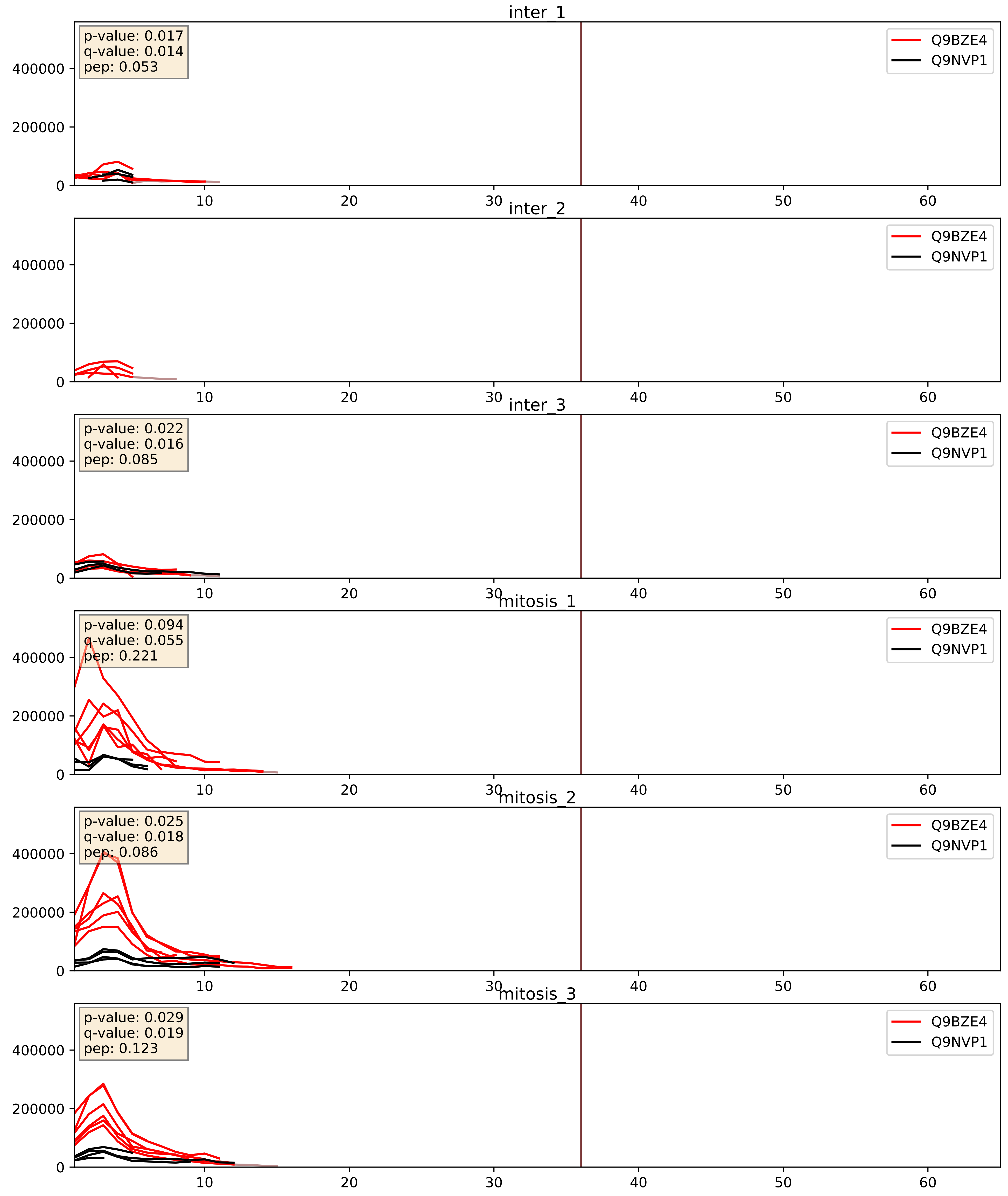


| level | condition_1 | condition_2 | pvalue | pvalue_adjusted |
|---|---|---|---|---|
| interactor_ratio | mitosis | inter | 0.0007256664780815352 | 0.031214598253155482 |
| interactor_abundance | mitosis | inter | 0.0031709934969099437 | 0.043395210765066534 |
| complex_abundance | mitosis | inter | 0.028801127868799545 | 0.1234203919435165 |

Q9H6R4_Q9NVP1
NOL6_HUMAN vs DDX18_HUMAN
p-value: 0.074 q-value: 0.045

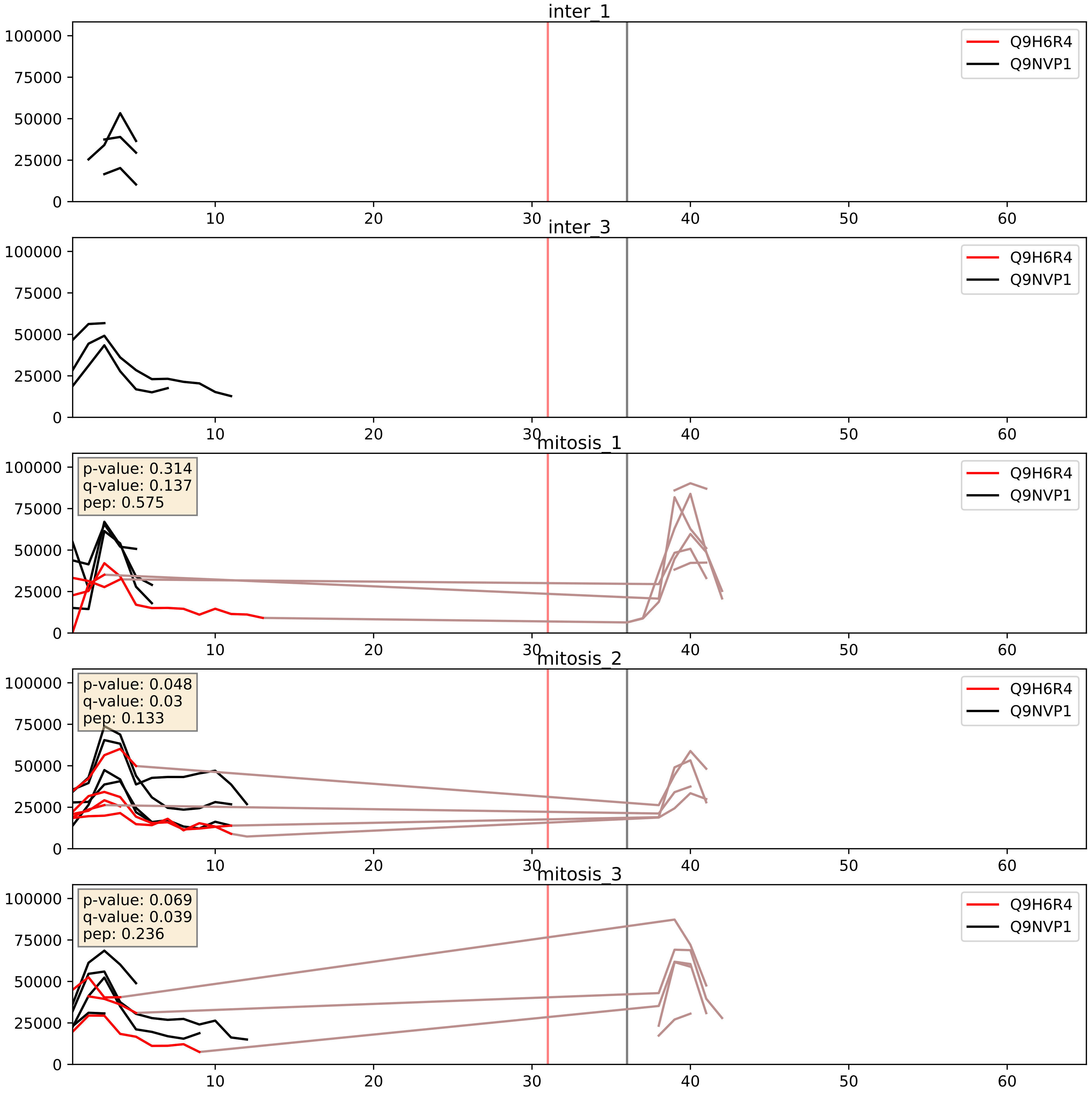


| level | condition_1 | condition_2 | pvalue | pvalue_adjusted |
|---|---|---|---|---|
| interactor_ratio | mitosis | inter | 0.009838253155822414 | 0.14263182888175546 |
| complex_abundance | mitosis | inter | 0.06470209978002295 | 0.1836981672029839 |
| interactor_abundance | mitosis | inter | 0.5002996189915532 | 0.6656579265768231 |

Q9H7B2_Q9NVP1
RPF2_HUMAN vs DDX18_HUMAN
p-value: 0.038 q-value: 0.026

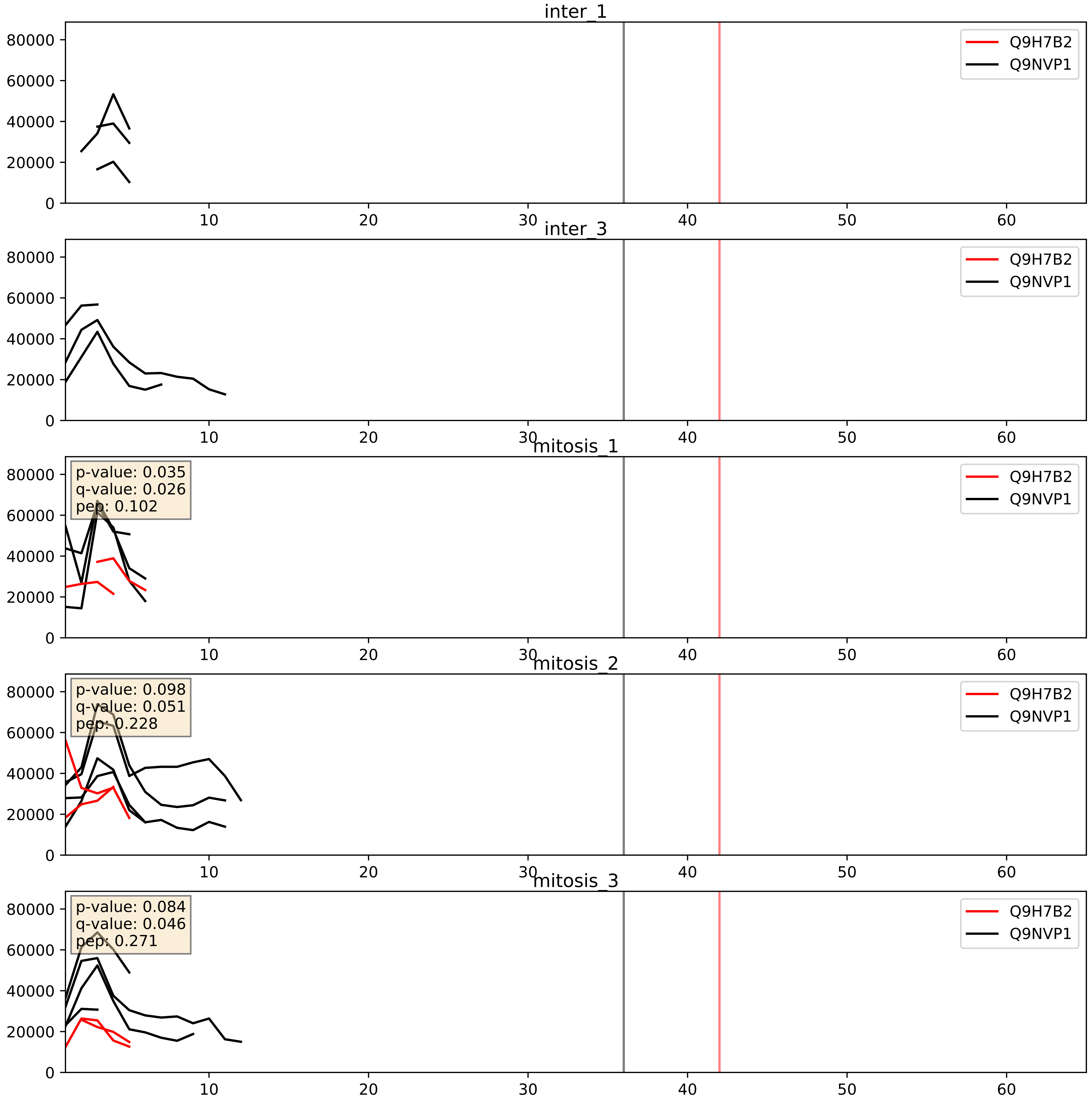


| level | condition_1 | condition_2 | pvalue | pvalue_adjusted |
|---|---|---|---|---|
| interactor_ratio | mitosis | inter | 0.009946672546552762 | 0.14263182888175546 |
| complex_abundance | mitosis | inter | 0.17688335514007095 | 0.3273476593888062 |
| interactor_abundance | mitosis | inter | 0.4740076787521983 | 0.6453802656463842 |

Q9NVP1_Q9NY93
DDX18_HUMAN vs DDX56_HUMAN
p-value: 0.085 q-value: 0.05

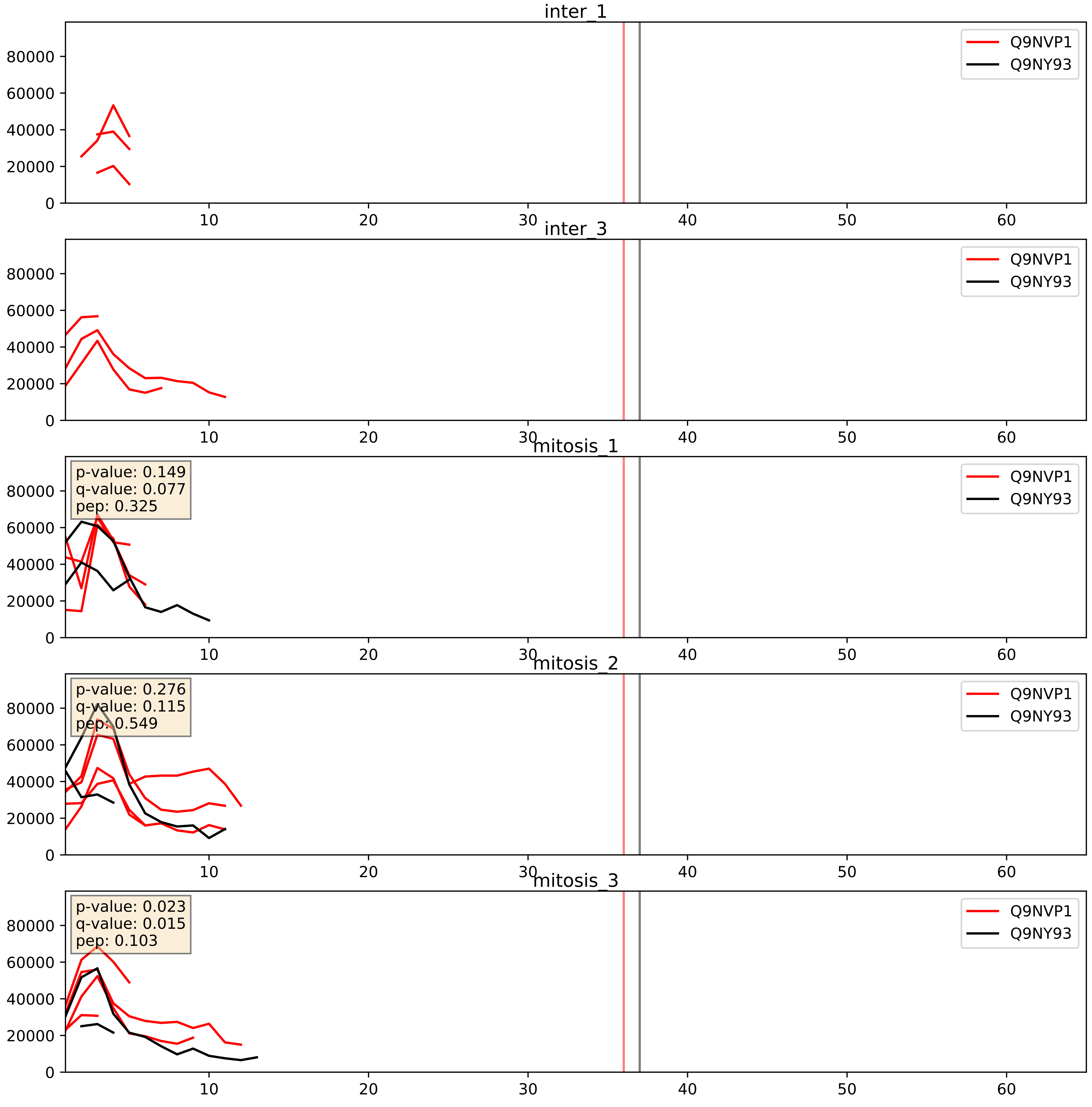


| level | condition_1 | condition_2 | pvalue | pvalue_adjusted |
|---|---|---|---|---|
| interactor_ratio | mitosis | inter | 0.003673014720945405 | 0.0787300451287945 |
| interactor_abundance | mitosis | inter | 0.010894066262585354 | 0.09160432928067842 |
| complex_abundance | mitosis | inter | 0.02682633767318529 | 0.11972879191111971 |

Q9NVP1_Q9Y221
DDX18_HUMAN vs NIP7_HUMAN
p-value: 0.022 q-value: 0.017

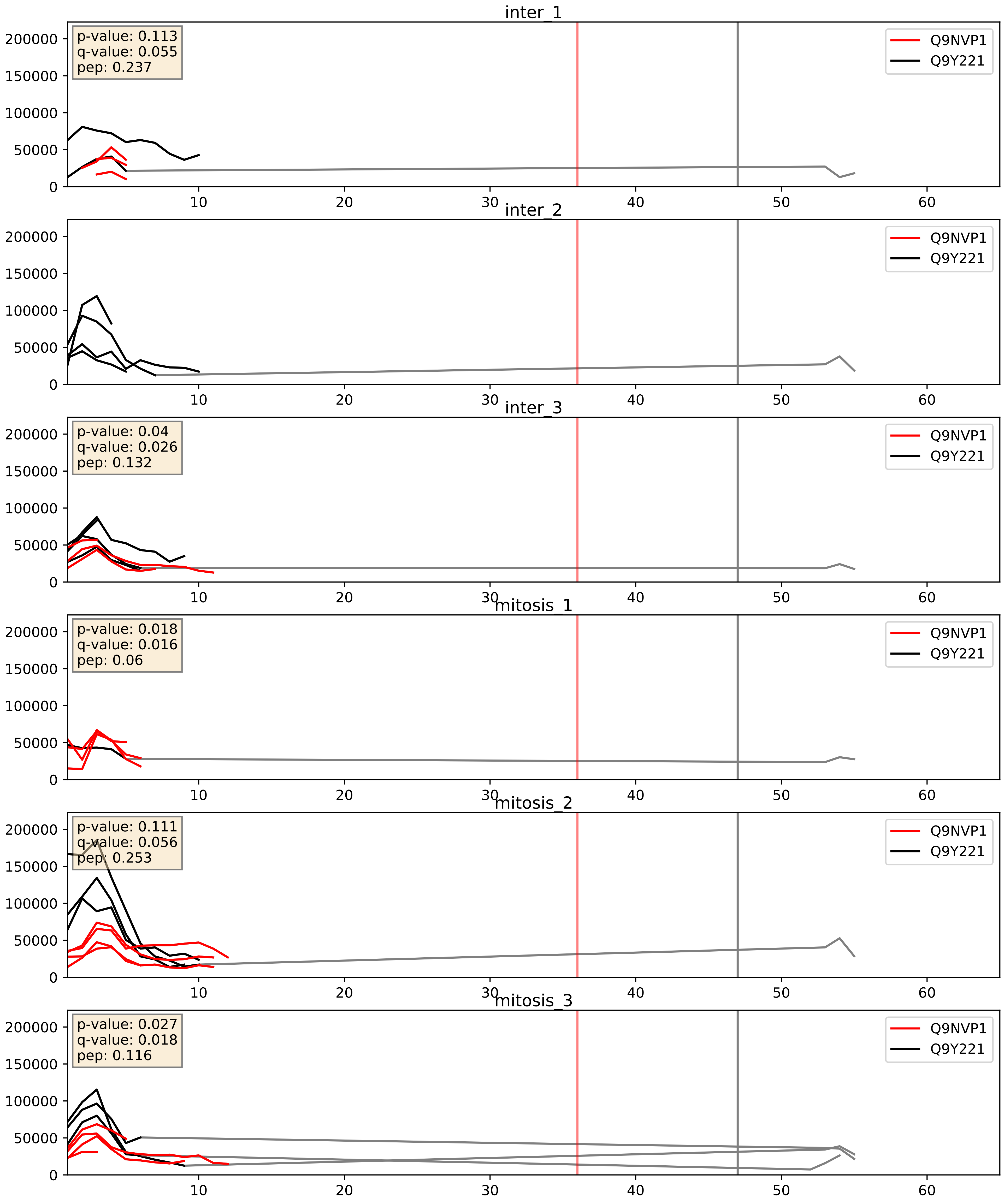


| level | condition_1 | condition_2 | pvalue | pvalue_adjusted |
|---|---|---|---|---|
| complex_abundance | mitosis | inter | 0.8108975430394811 | 0.8772319064307784 |
| interactor_ratio | mitosis | inter | 0.8821925279316942 | 0.9802747470617126 |
| interactor_abundance | mitosis | inter | 0.8871771073970744 | 0.9361222224987428 |

Q9NVP1_Q9Y2X3
DDX18_HUMAN vs NOP58_HUMAN
p-value: 0.049 q-value: 0.032

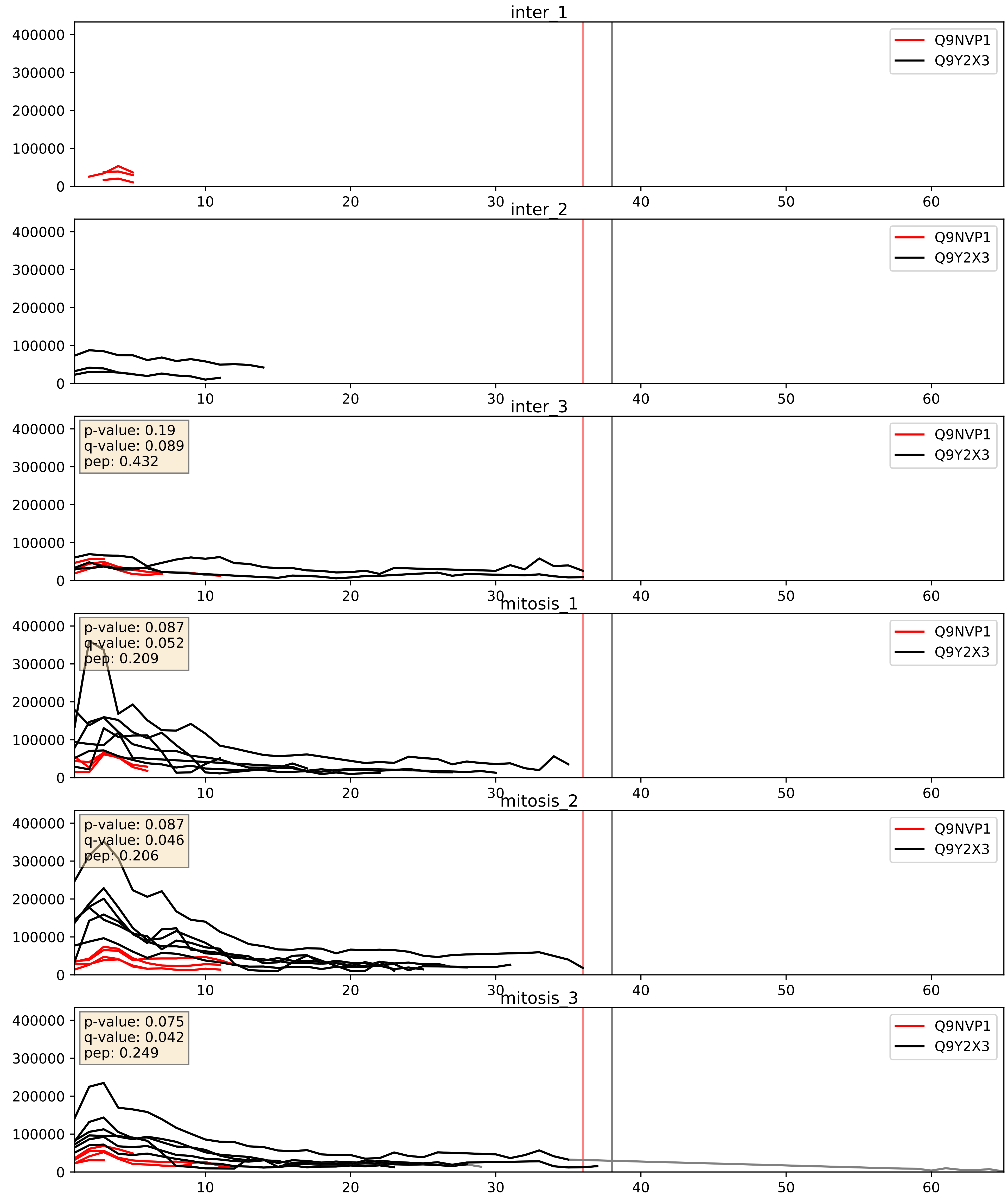


| level | condition_1 | condition_2 | pvalue | pvalue_adjusted |
|---|---|---|---|---|
| interactor_ratio | mitosis | inter | 0.0012725532055782418 | 0.0423373831474418 |
| complex_abundance | mitosis | inter | 0.006119371749399622 | 0.05820202463873419 |
| interactor_abundance | mitosis | inter | 0.16570946498784456 | 0.3537339202733041 |

Q9NVP1_Q9Y3T9
DDX18_HUMAN vs NOC2L_HUMAN
p-value: 0.037 q-value: 0.026

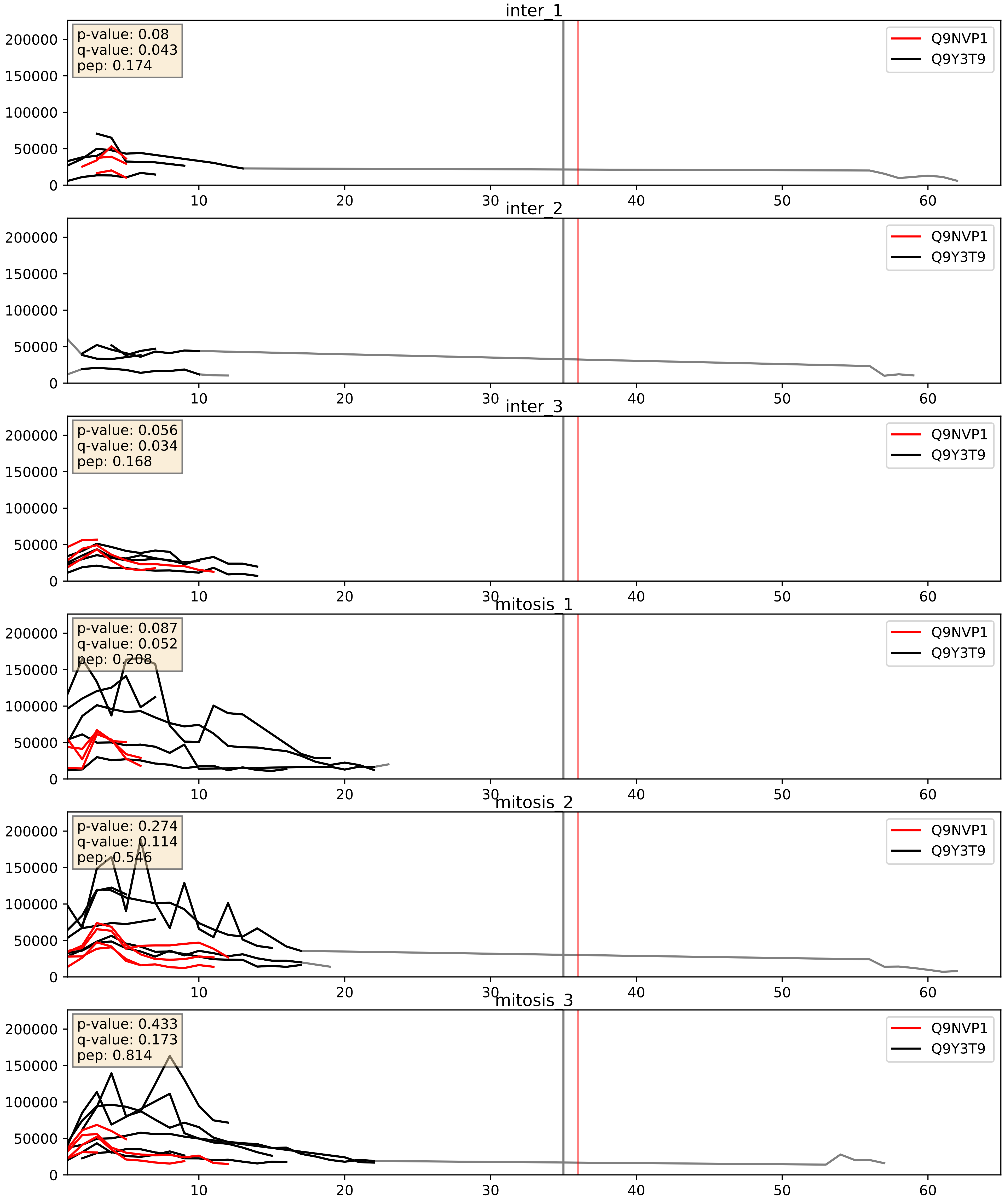


| level | condition_1 | condition_2 | pvalue | pvalue_adjusted |
|---|---|---|---|---|
| interactor_ratio | mitosis | inter | 0.0013447851463711195 | 0.04334578357589157 |
| complex_abundance | mitosis | inter | 0.07126313143207737 | 0.19408603406254607 |
| interactor_abundance | mitosis | inter | 0.7680100411481787 | 0.8624119050542318 |
