## Supplementary Data 2 for "SECAT: Quantifying differential protein-protein interaction states by network-centric analysis": hela_string_000022_Q8TDN6.pdf

Q8TDN6_Q8TDN6

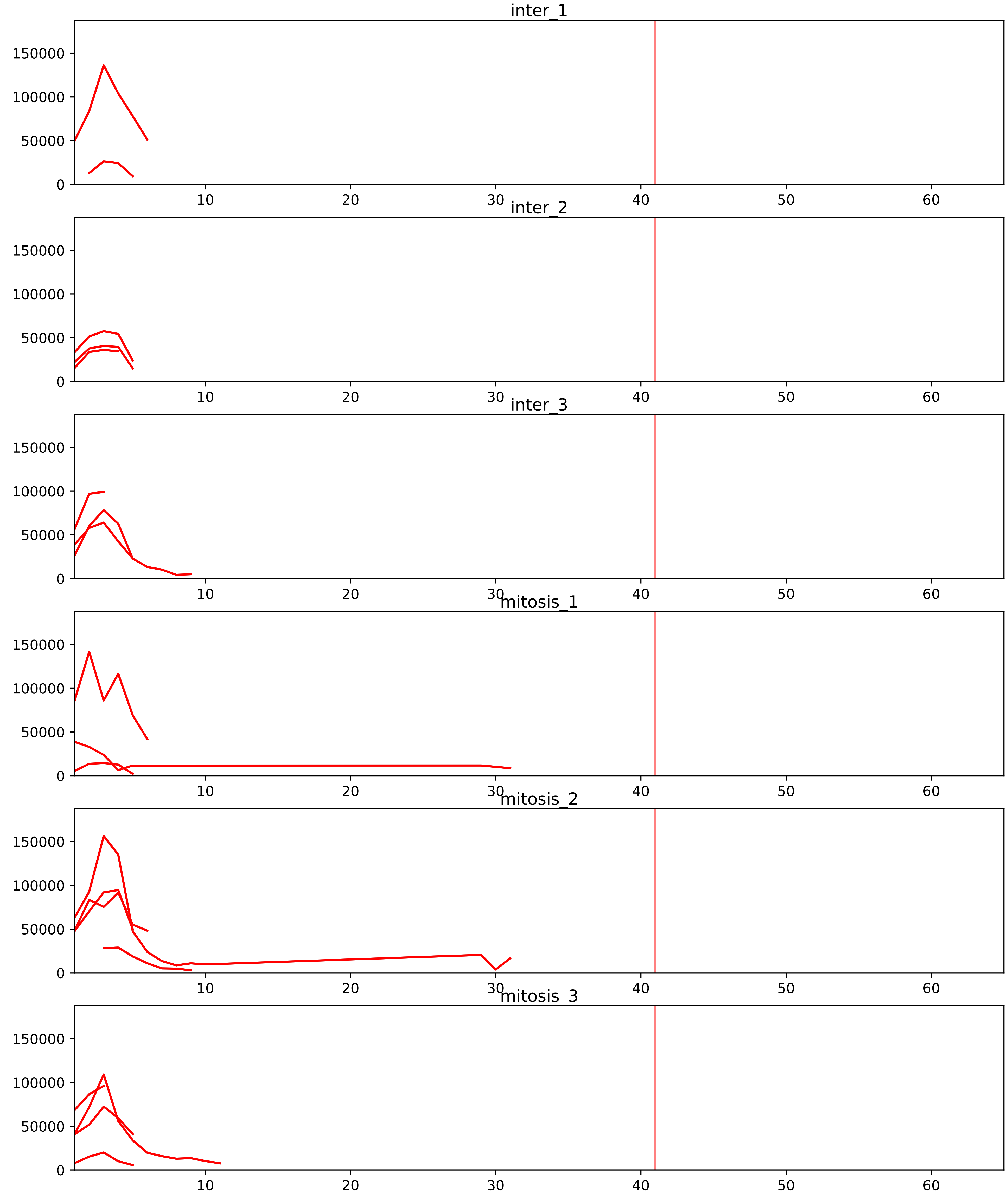


| level | condition_1 | condition_2 | pvalue | pvalue_adjusted |
|---|---|---|---|---|
| interactor_ratio | mitosis | inter | 2.7874730866231318e-11 | 2.331341126993892e-09 |
| complex_abundance | mitosis | inter | 3.6760973768188805e-05 | 0.0011866700304117087 |
| interactor_abundance | mitosis | inter | 8.86265427873271e-05 | 0.0020906774195984857 |
| assembled_abundance | mitosis | inter | 0.8508773429433129 | 0.9431394253985627 |
| total_abundance | mitosis | inter | 0.8645704643501073 | 0.9514067073643321 |

P42696_Q8TDN6
RBM34_HUMAN vs BRX1_HUMAN
p-value: 0.04 q-value: 0.028

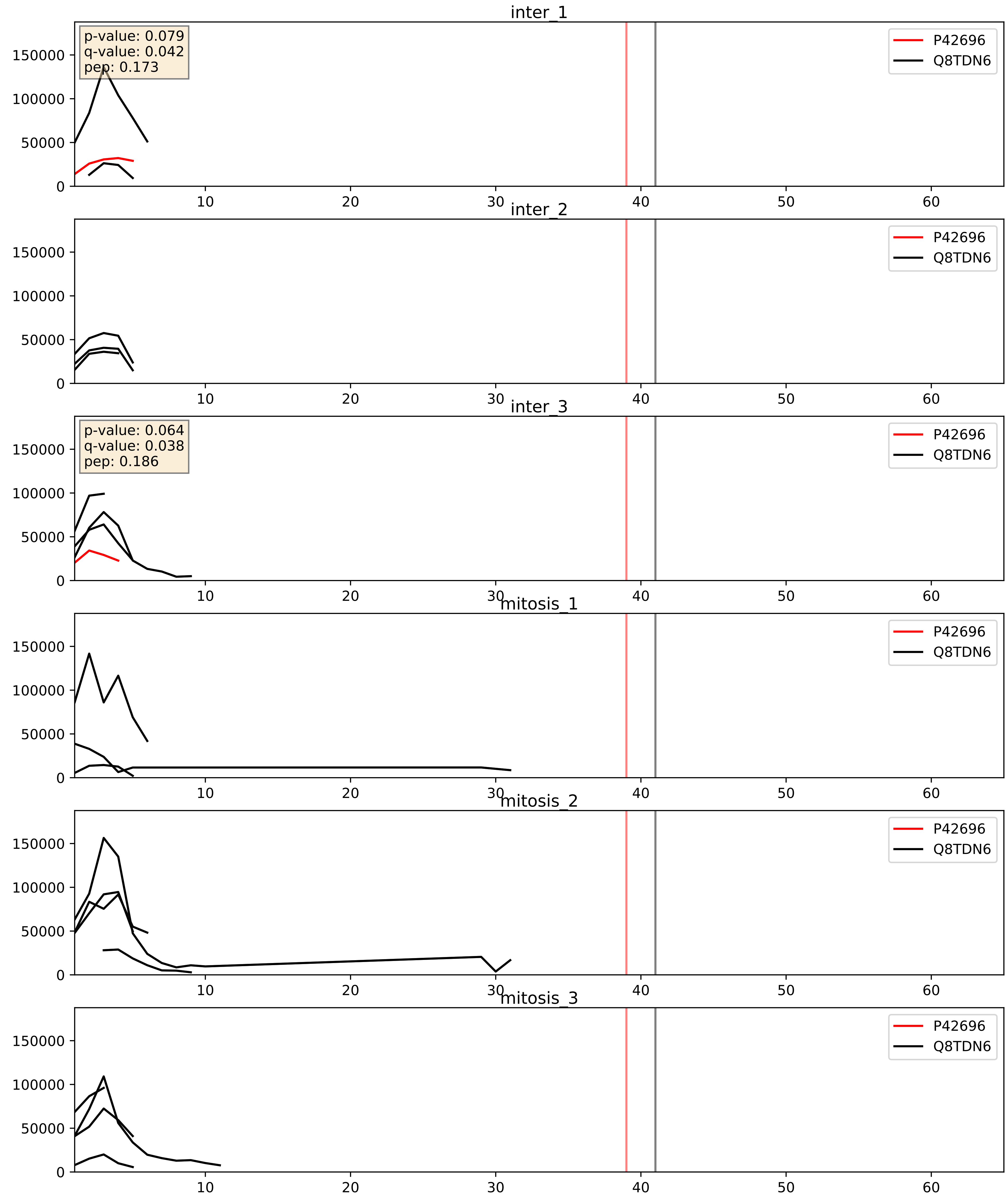


| level | condition_1 | condition_2 | pvalue | pvalue_adjusted |
|---|---|---|---|---|
| complex_abundance | mitosis | inter | 0.01884207196420192 | 0.10234018782586832 |
| interactor_ratio | mitosis | inter | 0.018860194337726403 | 0.20671352564780796 |
| interactor_abundance | mitosis | inter | 0.024446118744046258 | 0.12905259108790376 |

P46087_Q8TDN6
NOP2_HUMAN vs BRX1_HUMAN
p-value: 0.026 q-value: 0.02

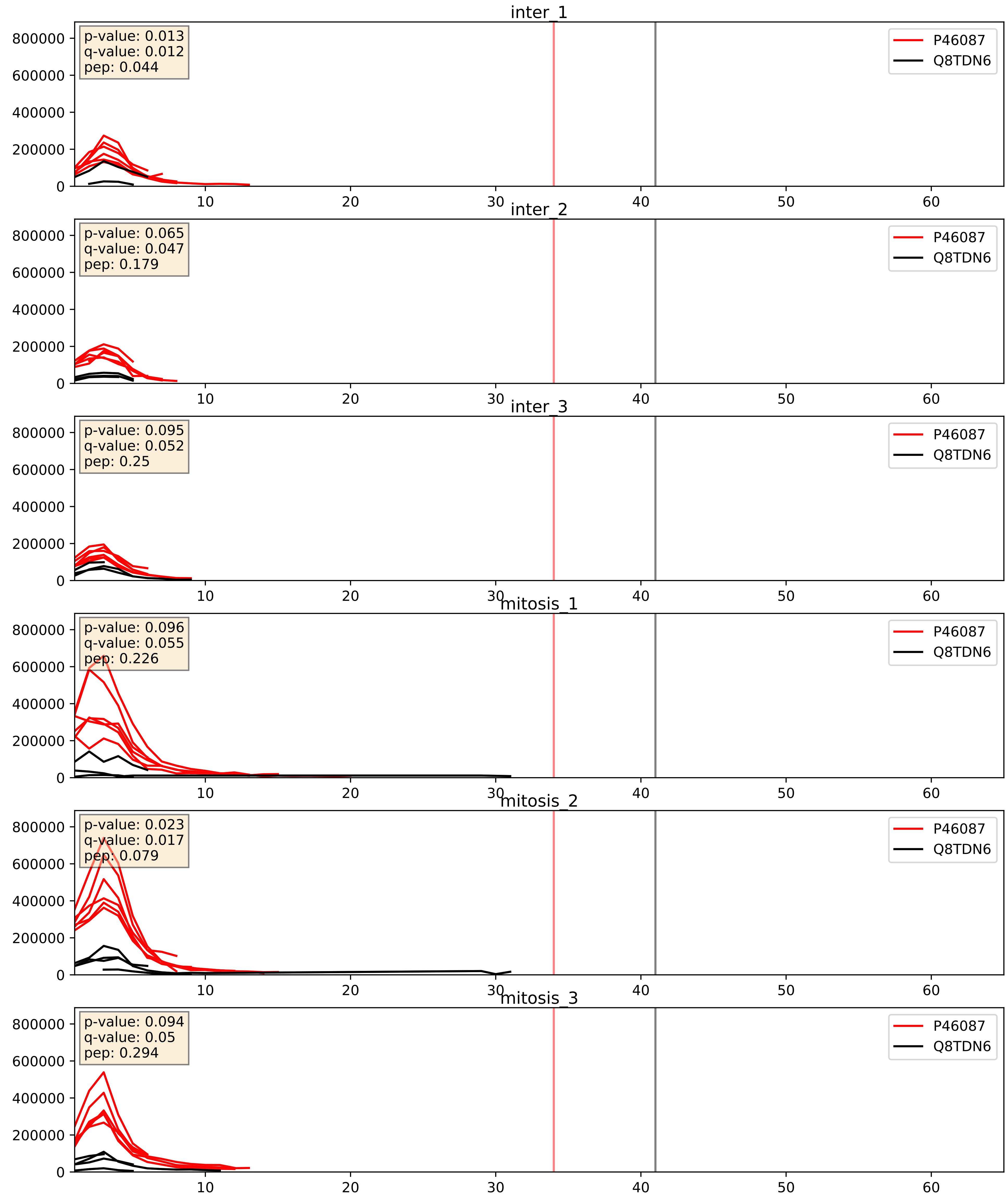


| level | condition_1 | condition_2 | pvalue | pvalue_adjusted |
|---|---|---|---|---|
| interactor_ratio | mitosis | inter | 0.001498537587396323 | 0.04597663709000905 |
| interactor_abundance | mitosis | inter | 0.005651157289129827 | 0.0634828167912747 |
| complex_abundance | mitosis | inter | 0.04071320618289467 | 0.14533154500649642 |

Q14137_Q8TDN6
BOP1_HUMAN vs BRX1_HUMAN
p-value: 0.017 q-value: 0.014

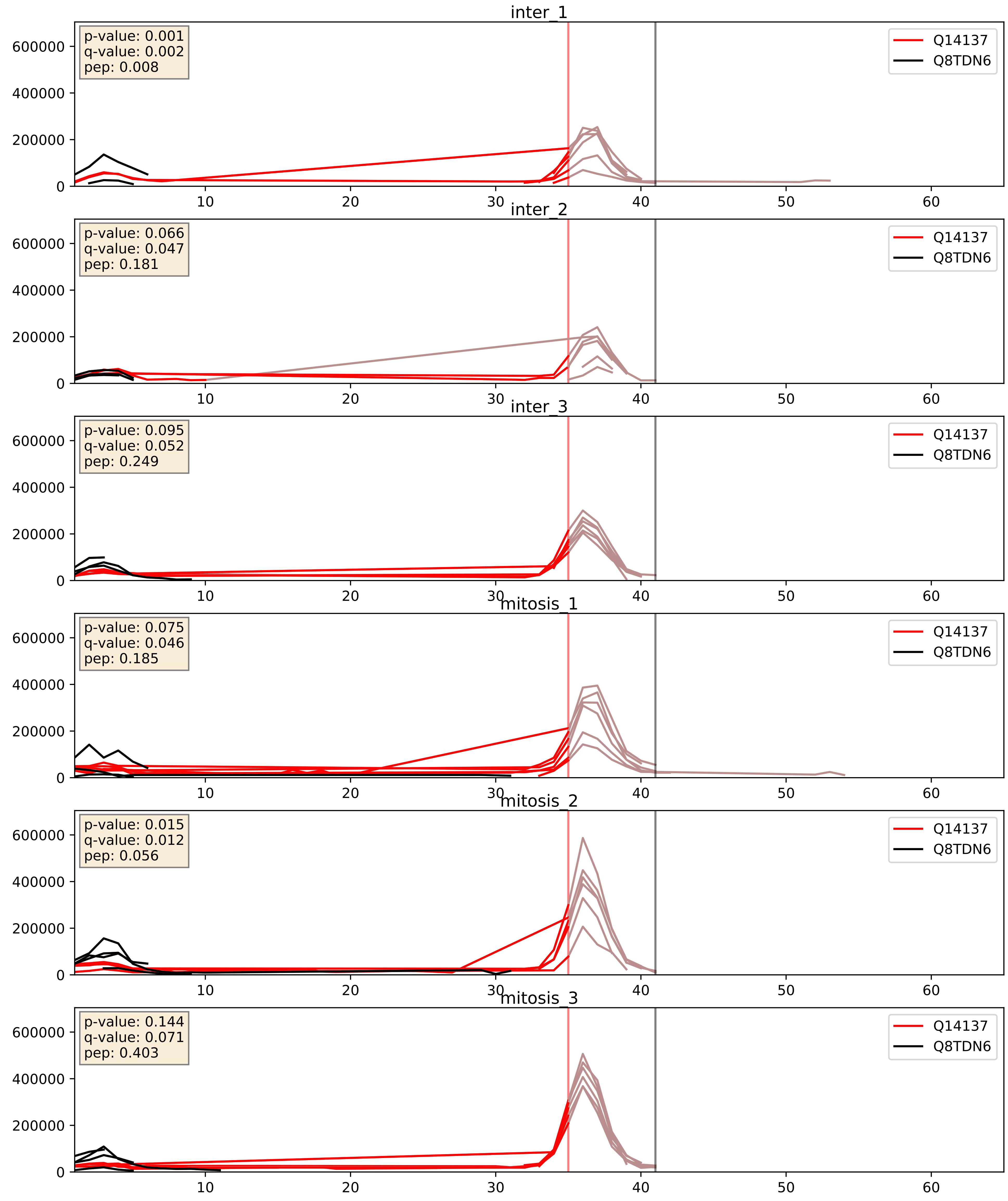


| level | condition_1 | condition_2 | pvalue | pvalue_adjusted |
|---|---|---|---|---|
| complex_abundance | mitosis | inter | 0.04192376036340907 | 0.14726323635770058 |
| interactor_abundance | mitosis | inter | 0.09173442235831389 | 0.2580077724288375 |
| interactor_ratio | mitosis | inter | 0.4069359185939671 | 0.8325457607945408 |

Q8IY81_Q8TDN6
SPB1_HUMAN vs BRX1_HUMAN
p-value: 0.013 q-value: 0.011

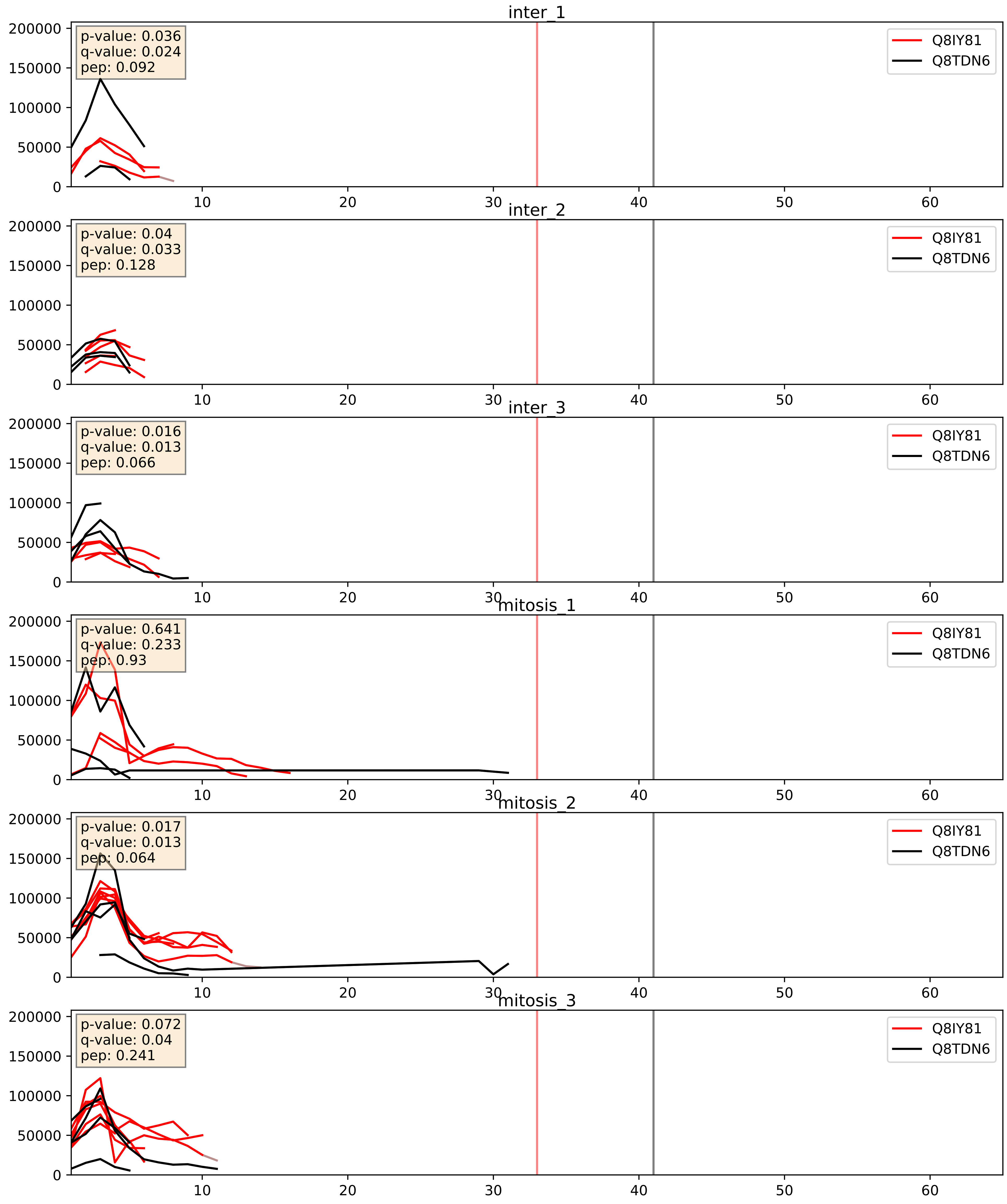


| level | condition_1 | condition_2 | pvalue | pvalue_adjusted |
|---|---|---|---|---|
| interactor_ratio | mitosis | inter | 0.05495294519905986 | 0.3887580255404565 |
| interactor_abundance | mitosis | inter | 0.5434694040854053 | 0.7001953791347185 |
| complex_abundance | mitosis | inter | 0.7217391892829187 | 0.814621236848864 |

Q8TDN6_Q9BYG3
BRX1_HUMAN vs MK67I_HUMAN
p-value: 0.044 q-value: 0.029

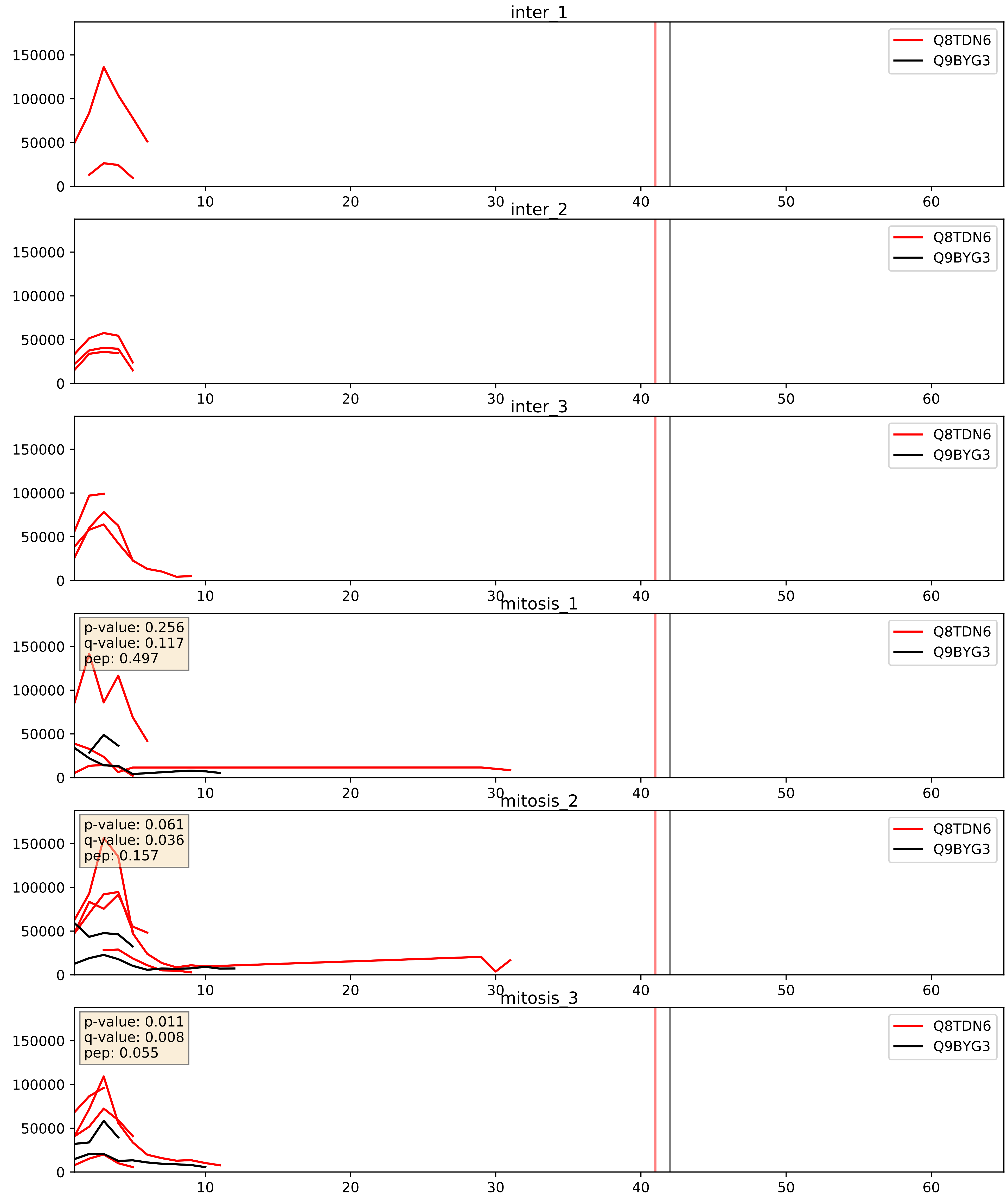


| level | condition_1 | condition_2 | pvalue | pvalue_adjusted |
|---|---|---|---|---|
| interactor_abundance | mitosis | inter | 0.015356739953974529 | 0.10851551243015083 |
| complex_abundance | mitosis | inter | 0.02656820842384979 | 0.11938260583105208 |
| interactor_ratio | mitosis | inter | 0.045559994696426284 | 0.3548621970895441 |

Q8TDN6_Q9BZE4
BRX1_HUMAN vs NOG1_HUMAN
p-value: 0.017 q-value: 0.014

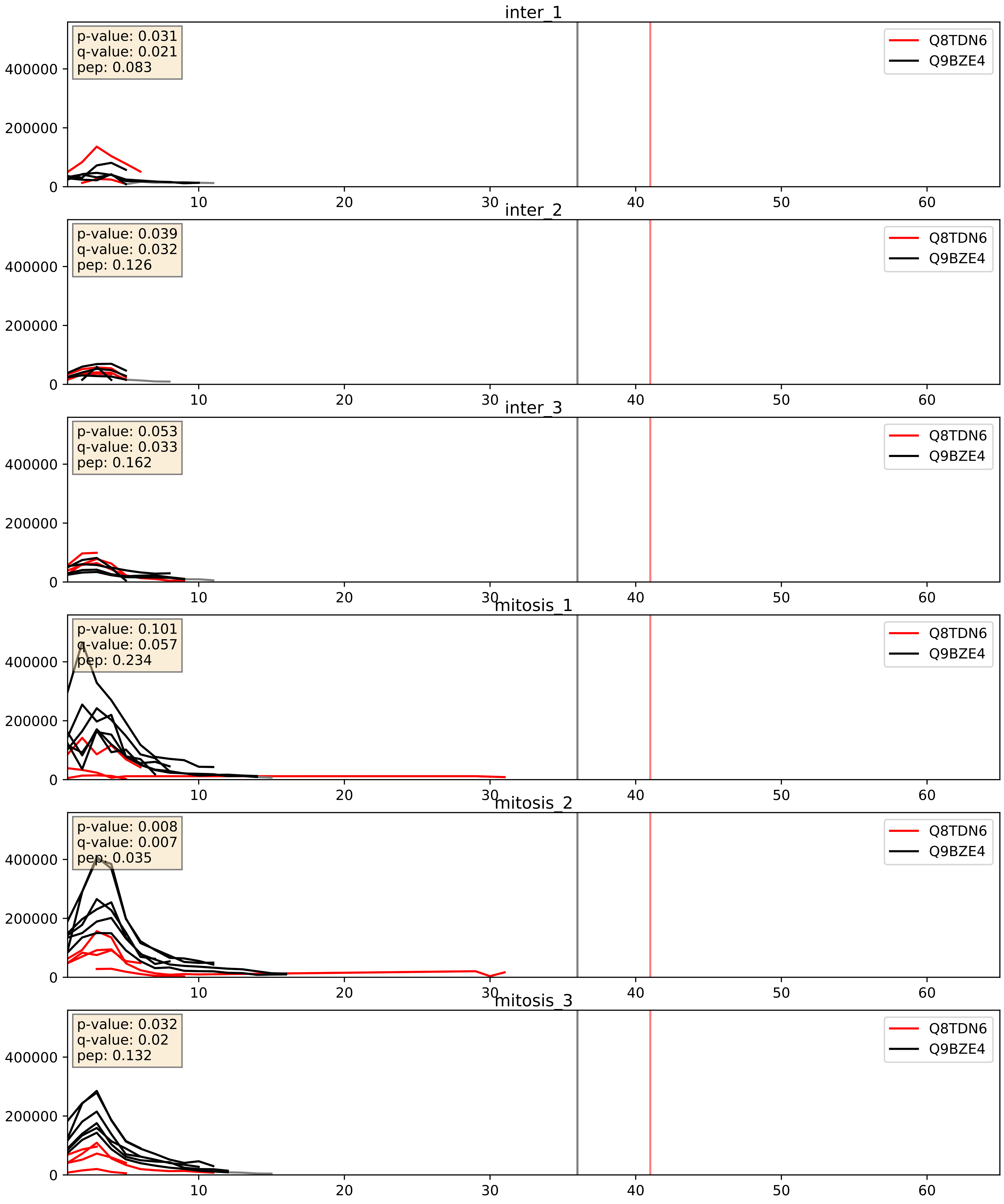


| level | condition_1 | condition_2 | pvalue | pvalue_adjusted |
|---|---|---|---|---|
| interactor_ratio | mitosis | inter | 0.00011836259628161376 | 0.011513452547393338 |
| complex_abundance | mitosis | inter | 0.01080016423974069 | 0.07693511812192051 |
| interactor_abundance | mitosis | inter | 0.0462072460121471 | 0.17159827586289772 |

Q8TDN6_Q9GZR7
BRX1_HUMAN vs DDX24_HUMAN
p-value: 0.01 q-value: 0.009

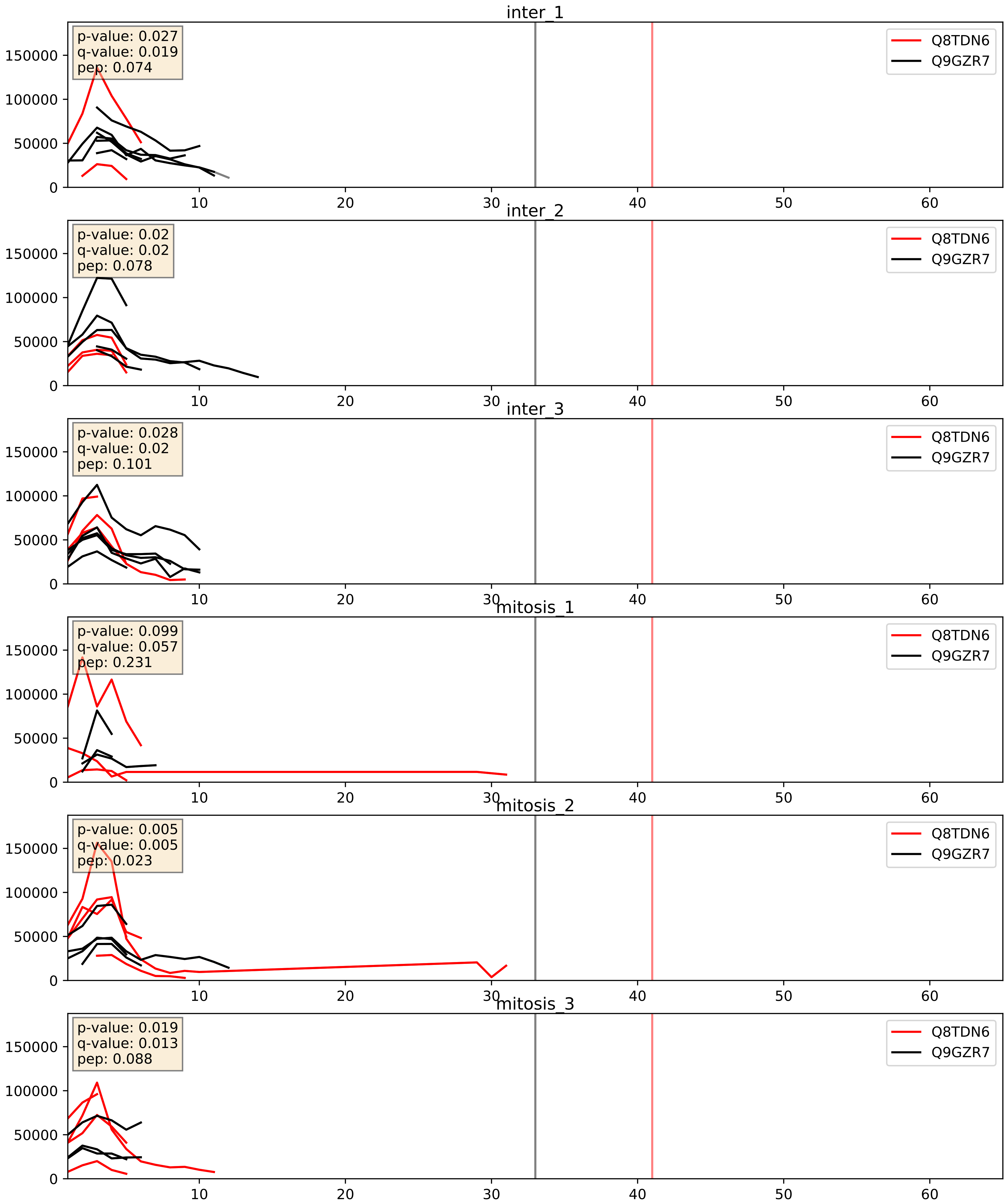


| level | condition_1 | condition_2 | pvalue | pvalue_adjusted |
|---|---|---|---|---|
| complex_abundance | mitosis | inter | 0.001617627551261813 | 0.02814408910325431 |
| interactor_ratio | mitosis | inter | 0.0017028547996455828 | 0.049919305085500645 |
| interactor_abundance | mitosis | inter | 0.04383239640025894 | 0.16685321787696922 |

Q8TDN6_Q9H7B2
BRX1_HUMAN vs RPF2_HUMAN
p-value: 0.075 q-value: 0.045

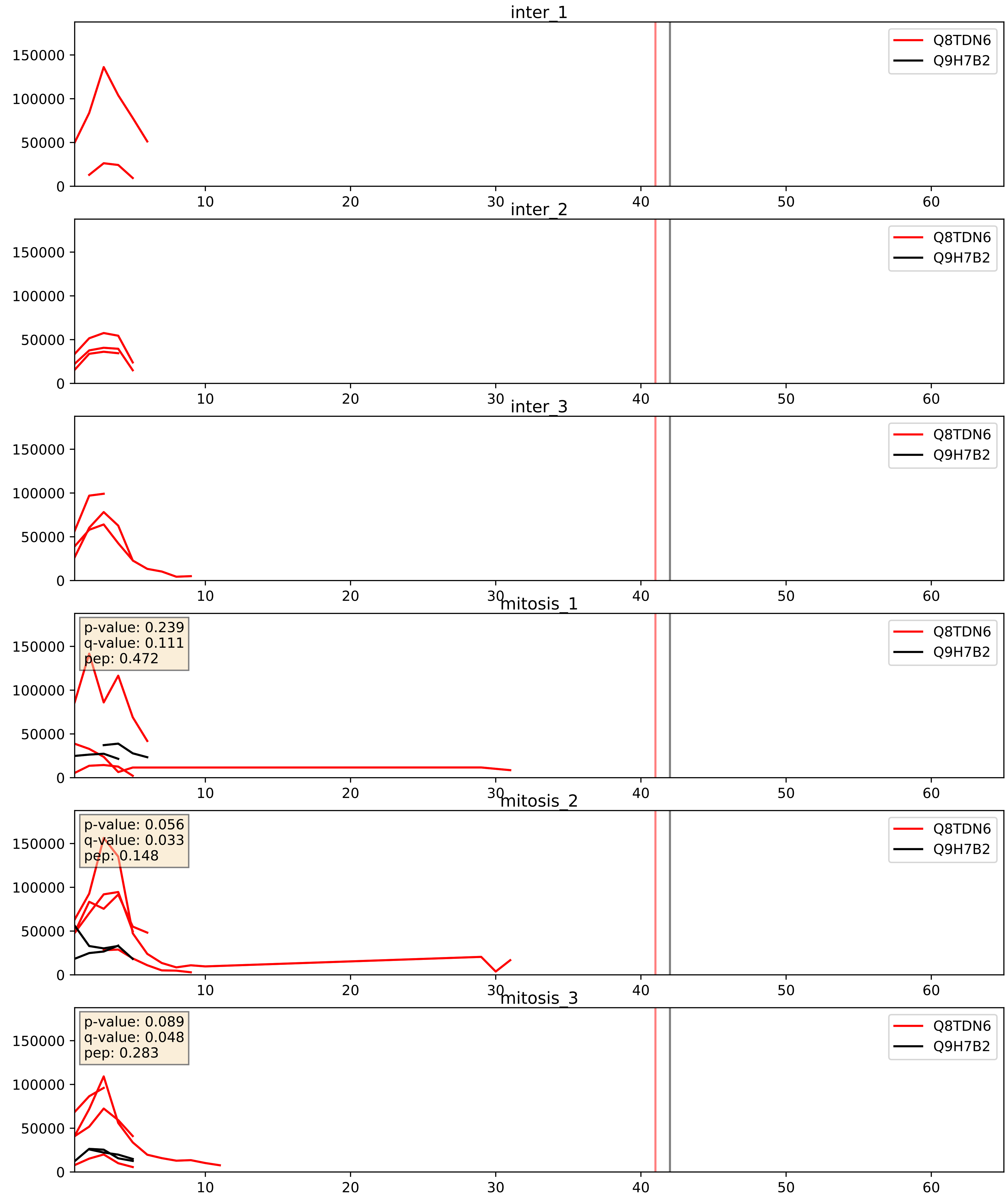


| level | condition_1 | condition_2 | pvalue | pvalue_adjusted |
|---|---|---|---|---|
| interactor_abundance | mitosis | inter | 0.015110876907849336 | 0.10761156932711341 |
| interactor_ratio | mitosis | inter | 0.039906808552959576 | 0.3303697110380406 |
| complex_abundance | mitosis | inter | 0.05121810455375765 | 0.161227792796871 |

Q8TDN6_Q9NVP1
BRX1_HUMAN vs DDX18_HUMAN
p-value: 0.02 q-value: 0.016

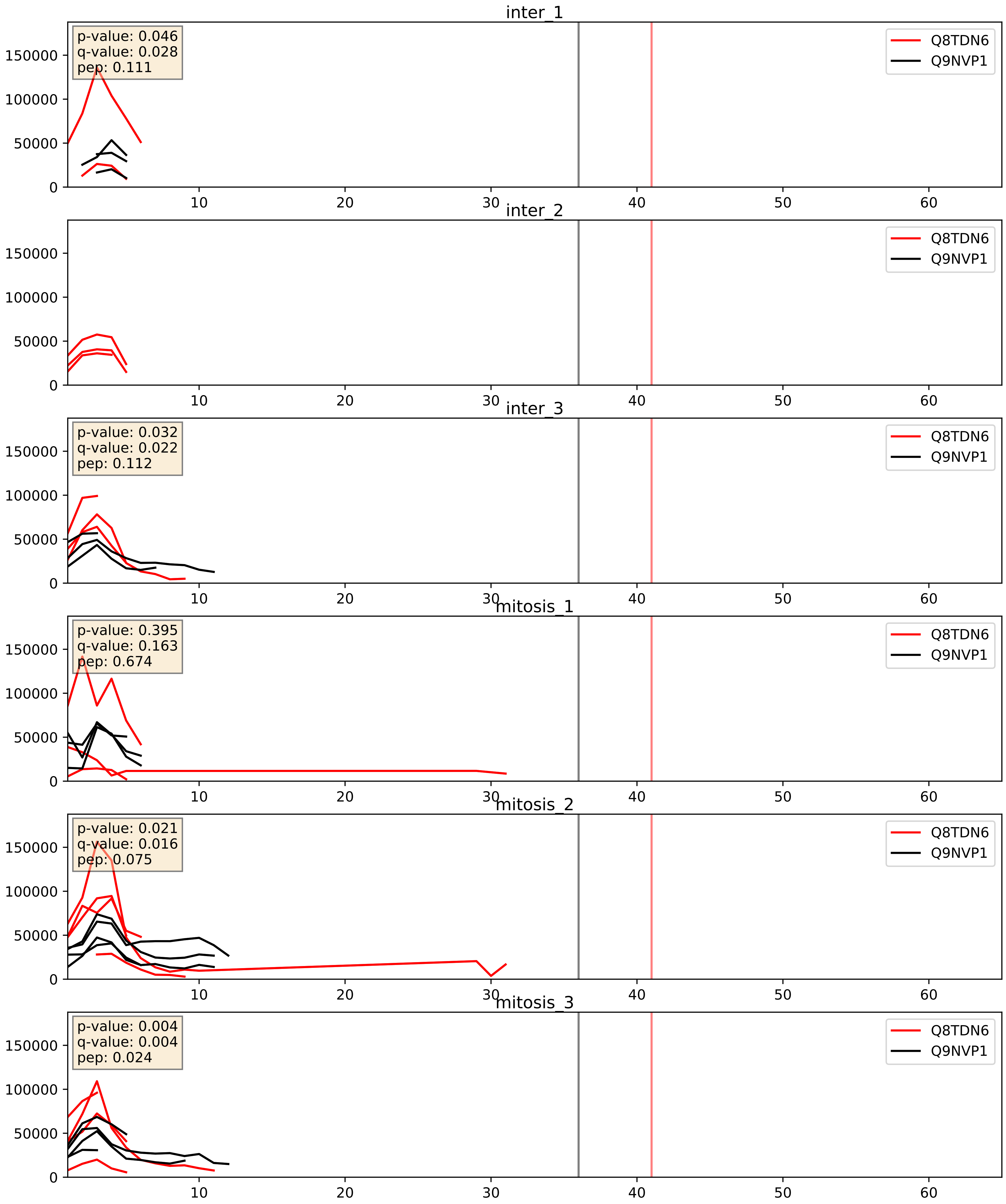


| level | condition_1 | condition_2 | pvalue | pvalue_adjusted |
|---|---|---|---|---|
| complex_abundance | mitosis | inter | 0.8425323439690111 | 0.8997101876714989 |
| interactor_abundance | mitosis | inter | 0.8683139378488682 | 0.9286903237679228 |
| interactor_ratio | mitosis | inter | 0.9136271445401767 | 0.9852905071335931 |

Q8TDN6_Q9NY93
BRX1_HUMAN vs DDX56_HUMAN
p-value: 0.019 q-value: 0.015

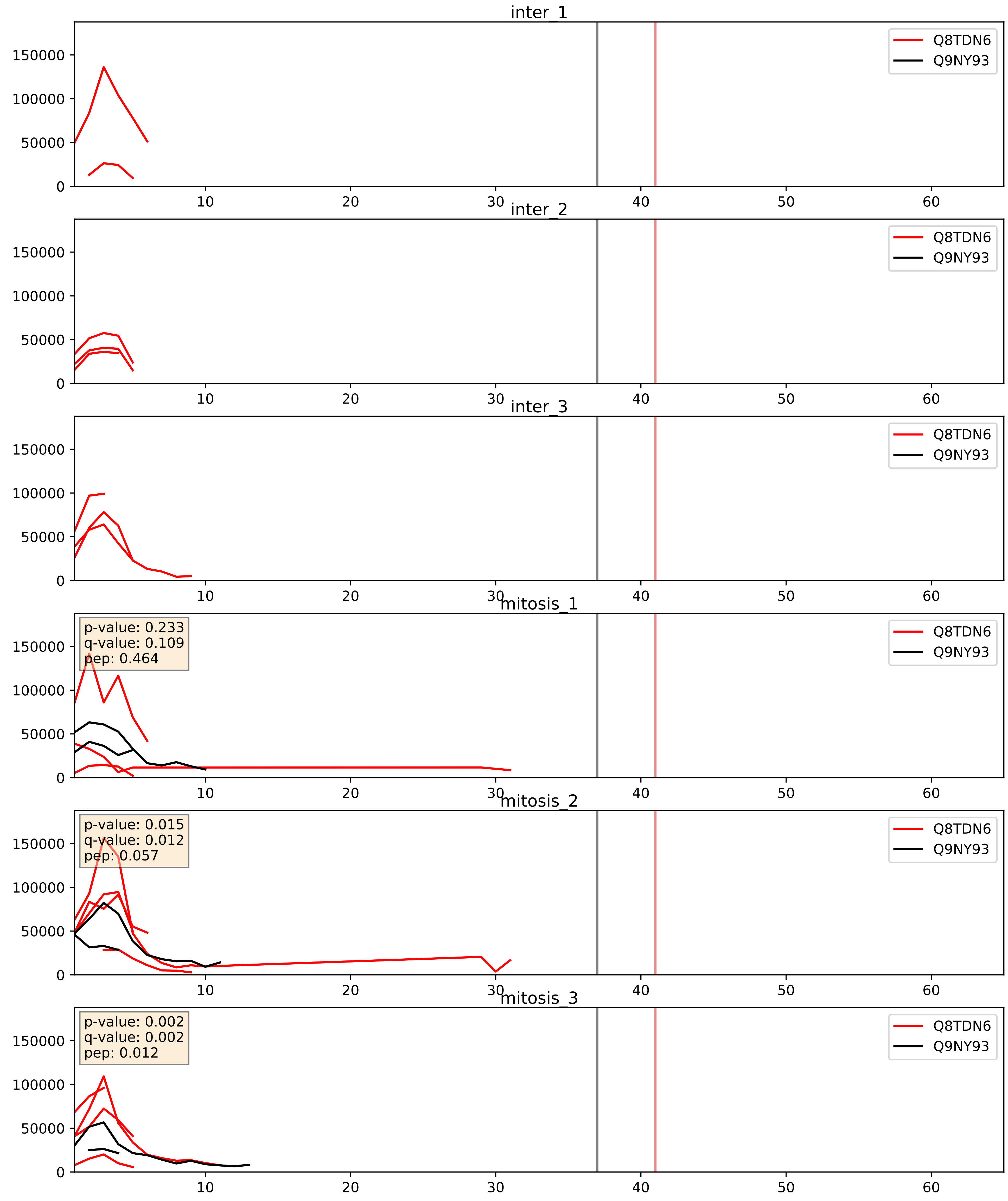


| level | condition_1 | condition_2 | pvalue | pvalue_adjusted |
|---|---|---|---|---|
| interactor_abundance | mitosis | inter | 0.013920743574672224 | 0.10466540623556807 |
| complex_abundance | mitosis | inter | 0.019947043532560785 | 0.10475257217099405 |
| interactor_ratio | mitosis | inter | 0.10974221992695789 | 0.5496158365704675 |

Q8TDN6_Q9UKD2
BRX1_HUMAN vs MRT4_HUMAN
p-value: 0.026 q-value: 0.019

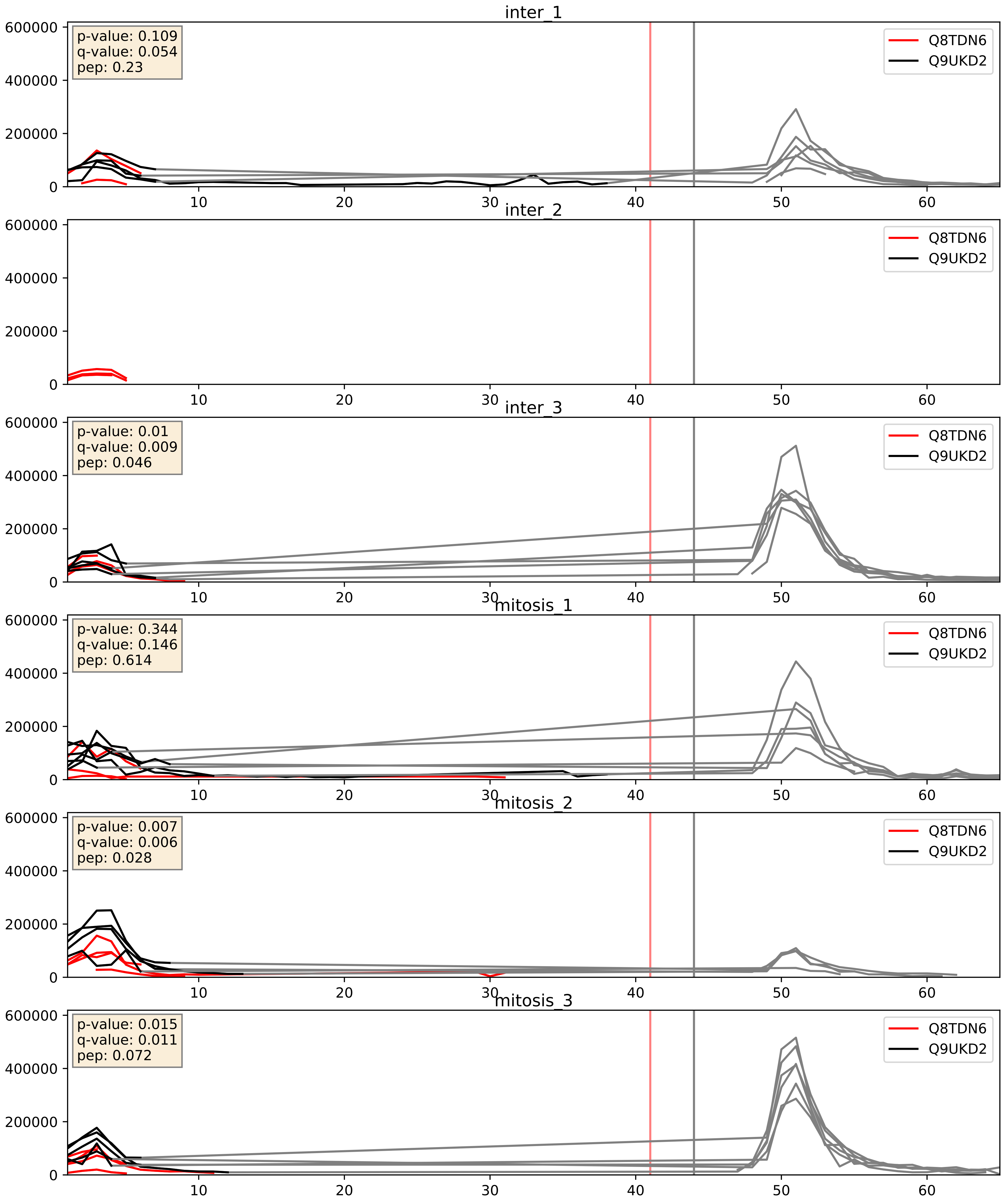


| level | condition_1 | condition_2 | pvalue | pvalue_adjusted |
|---|---|---|---|---|
| interactor_ratio | mitosis | inter | 0.026427216938707084 | 0.25543205403291264 |
| complex_abundance | mitosis | inter | 0.3724276336005471 | 0.5196382304190192 |
| interactor_abundance | mitosis | inter | 0.9044560948065035 | 0.9461832168094921 |

Q8TDN6_Q9Y2X3
BRX1_HUMAN vs NOP58_HUMAN
p-value: 0.043 q-value: 0.029

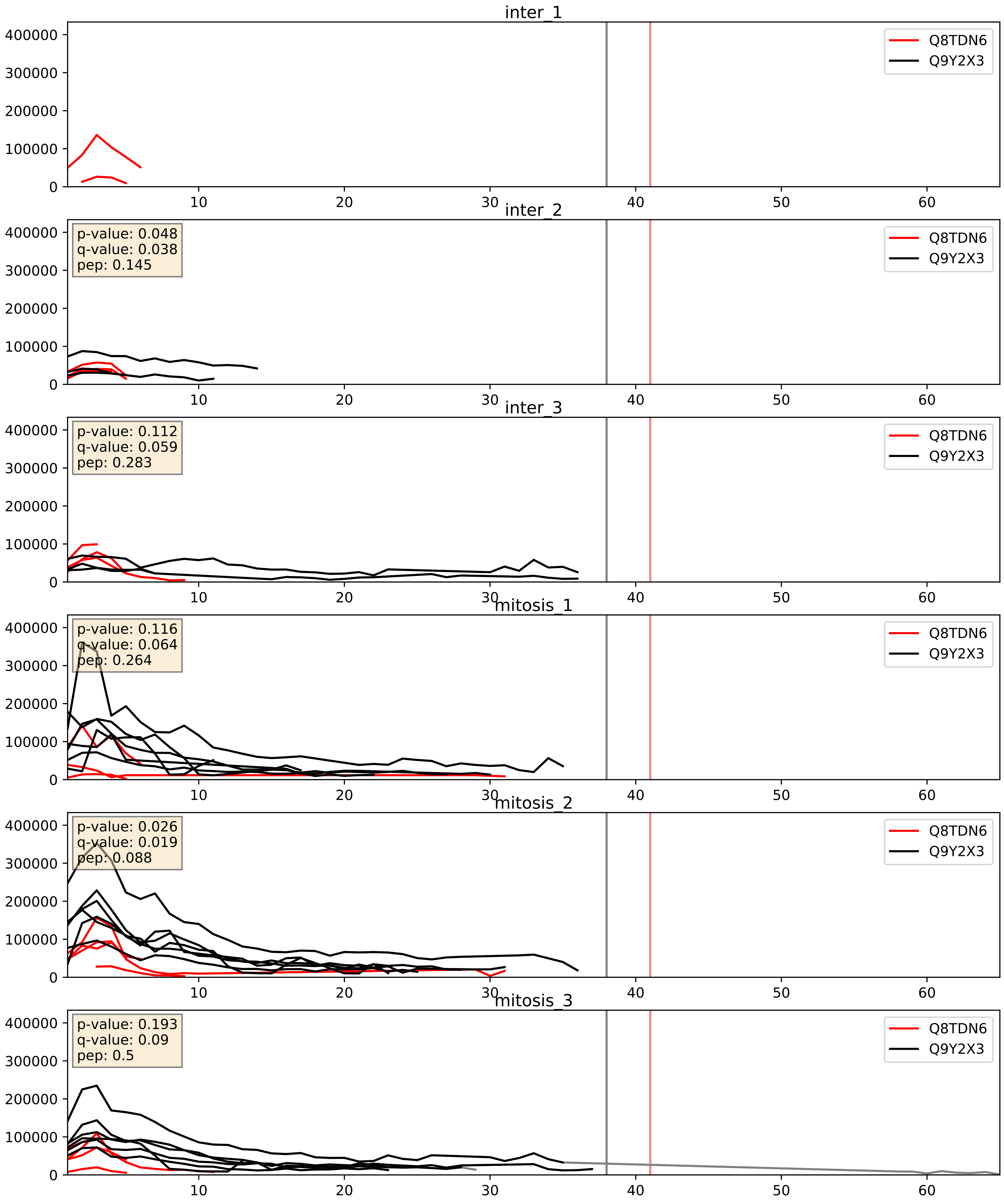


| level | condition_1 | condition_2 | pvalue | pvalue_adjusted |
|---|---|---|---|---|
| interactor_ratio | mitosis | inter | 4.525614117084573e-05 | 0.006796360849516482 |
| complex_abundance | mitosis | inter | 0.03832260841992085 | 0.14299979427834458 |
| interactor_abundance | mitosis | inter | 0.9343585822556804 | 0.962096690999235 |
