## Supplementary Data 2 for "SECAT: Quantifying differential protein-protein interaction states by network-centric analysis": hela_string_000023_Q15397.pdf

Q15397_Q15397

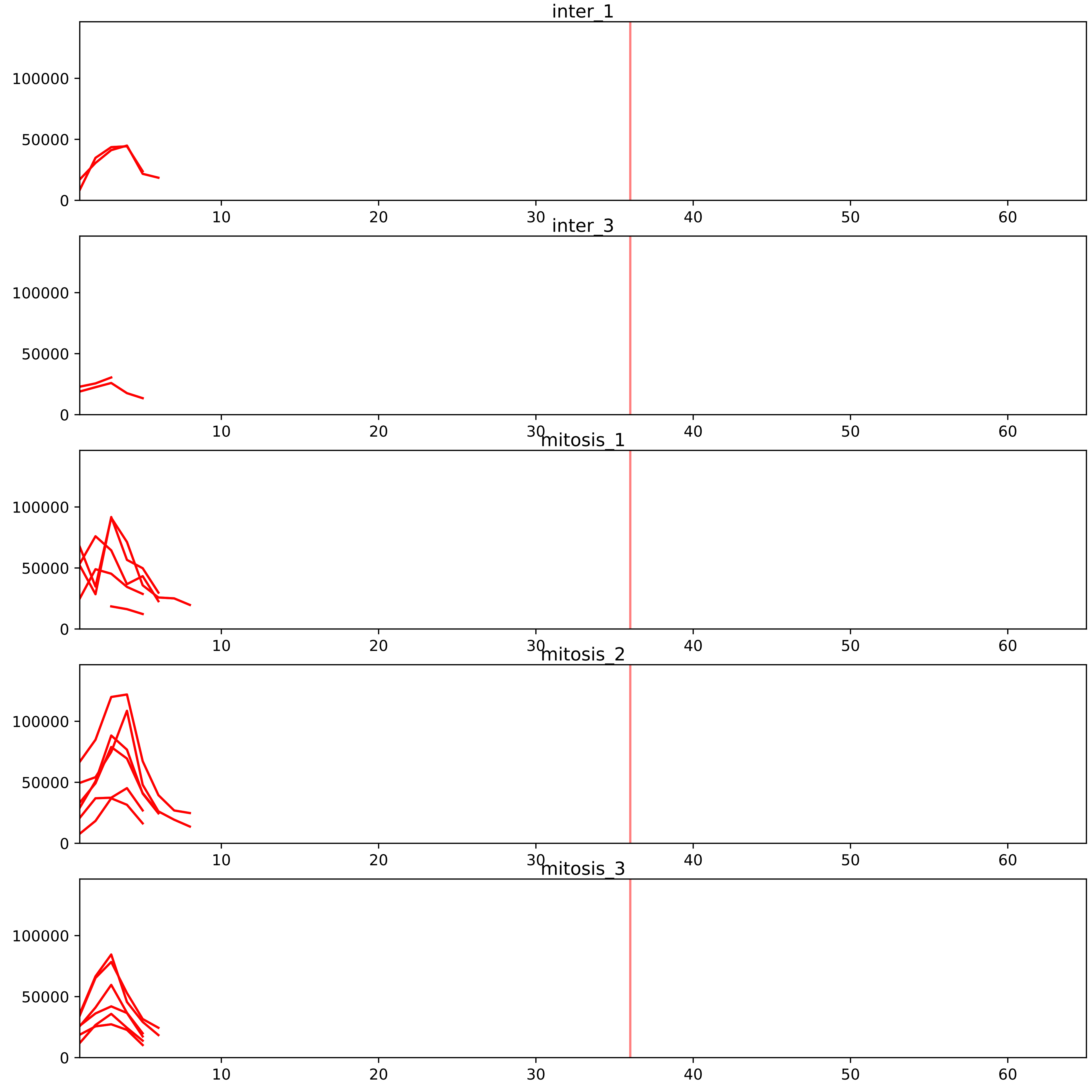


| level | condition_1 | condition_2 | pvalue | pvalue_adjusted |
|---|---|---|---|---|
| interactor_ratio | mitosis | inter | 4.530601594574162e-11 | 3.6244812756593294e-09 |
| complex_abundance | mitosis | inter | 0.0182721807640577 | 0.0659231619722866 |
| assembled_abundance | mitosis | inter | 0.02779143573240384 | 0.20263506333860185 |
| total_abundance | mitosis | inter | 0.03752636294043602 | 0.24337160984682776 |
| interactor_abundance | mitosis | inter | 0.04536441148899132 | 0.13312682159448808 |

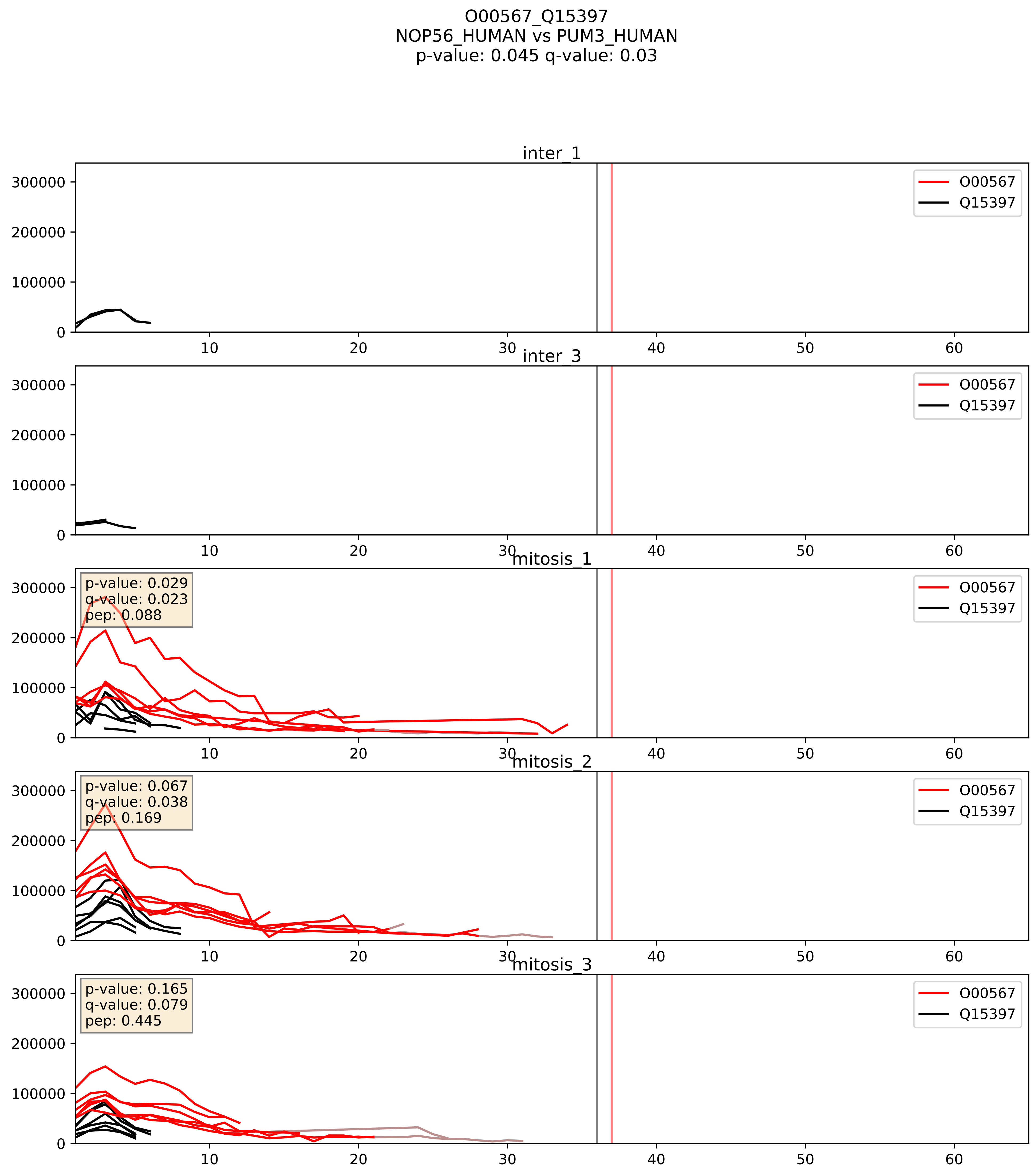


| level | condition_1 | condition_2 | pvalue | pvalue_adjusted |
|---|---|---|---|---|
| interactor_ratio | mitosis | inter | 0.0006892251594627307 | 0.030255217256415255 |
| interactor_abundance | mitosis | inter | 0.00467469140185995 | 0.05557688666655718 |
| complex_abundance | mitosis | inter | 0.008906695963925746 | 0.07001039251717574 |

O76021_Q15397
RL1D1_HUMAN vs PUM3_HUMAN
p-value: 0.003 q-value: 0.039

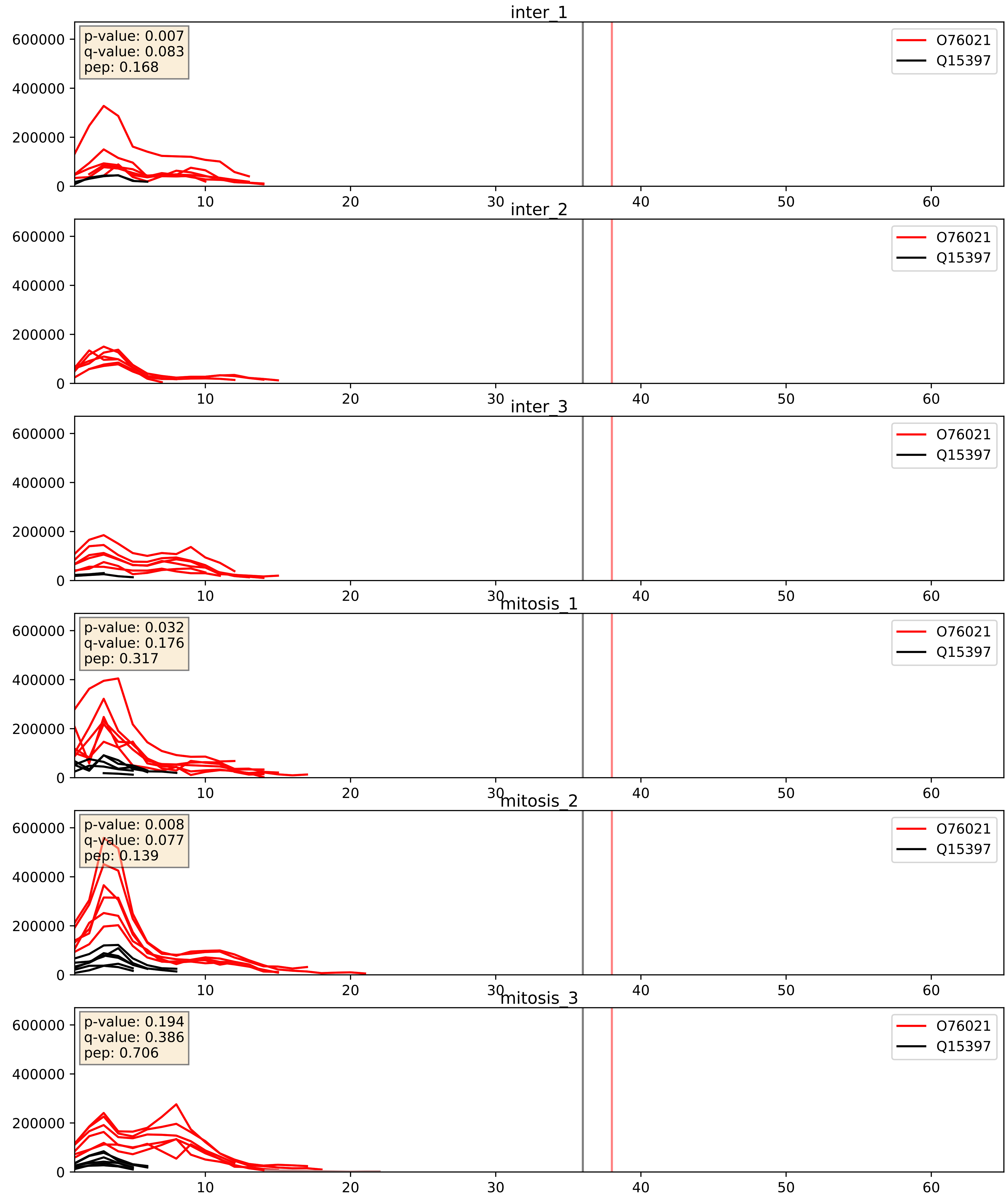


| level | condition_1 | condition_2 | pvalue | pvalue_adjusted |
|---|---|---|---|---|
| interactor_ratio | mitosis | inter | 0.09439087244256297 | 0.5094488449611217 |
| interactor_abundance | mitosis | inter | 0.09527655722192445 | 0.264708643239102 |
| complex_abundance | mitosis | inter | 0.1206192827943469 | 0.2626560826048358 |

P46087_Q15397
NOP2_HUMAN vs PUM3_HUMAN
p-value: 0.032 q-value: 0.023

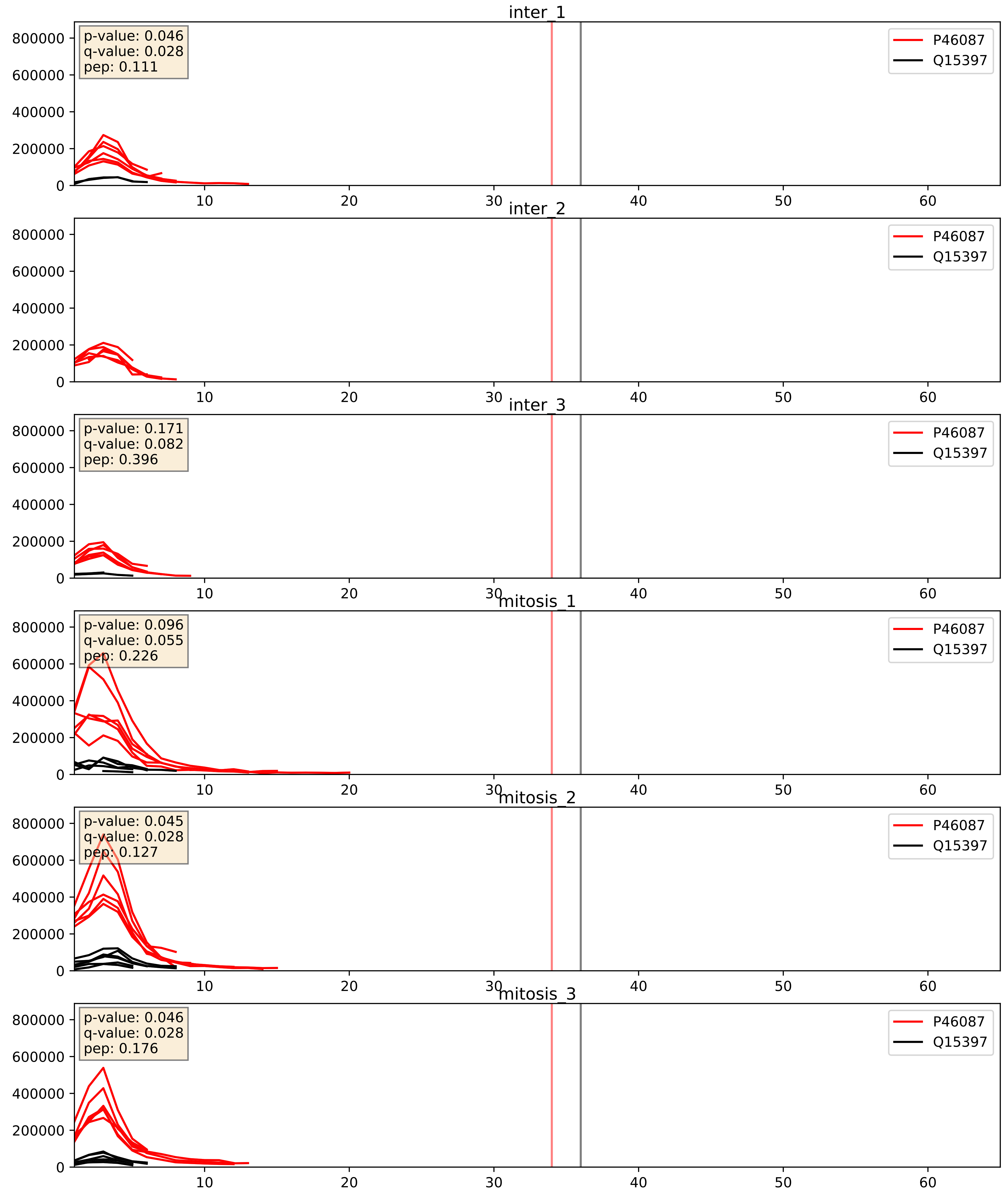


| level | condition_1 | condition_2 | pvalue | pvalue_adjusted |
|---|---|---|---|---|
| interactor_ratio | mitosis | inter | 0.0396609606963561 | 0.32897075926434904 |
| interactor_abundance | mitosis | inter | 0.05779753858331698 | 0.19593937832601716 |
| complex_abundance | mitosis | inter | 0.08310913278336679 | 0.21135299364991675 |

Q03701_Q15397
CEBPZ_HUMAN vs PUM3_HUMAN
p-value: 0.031 q-value: 0.022

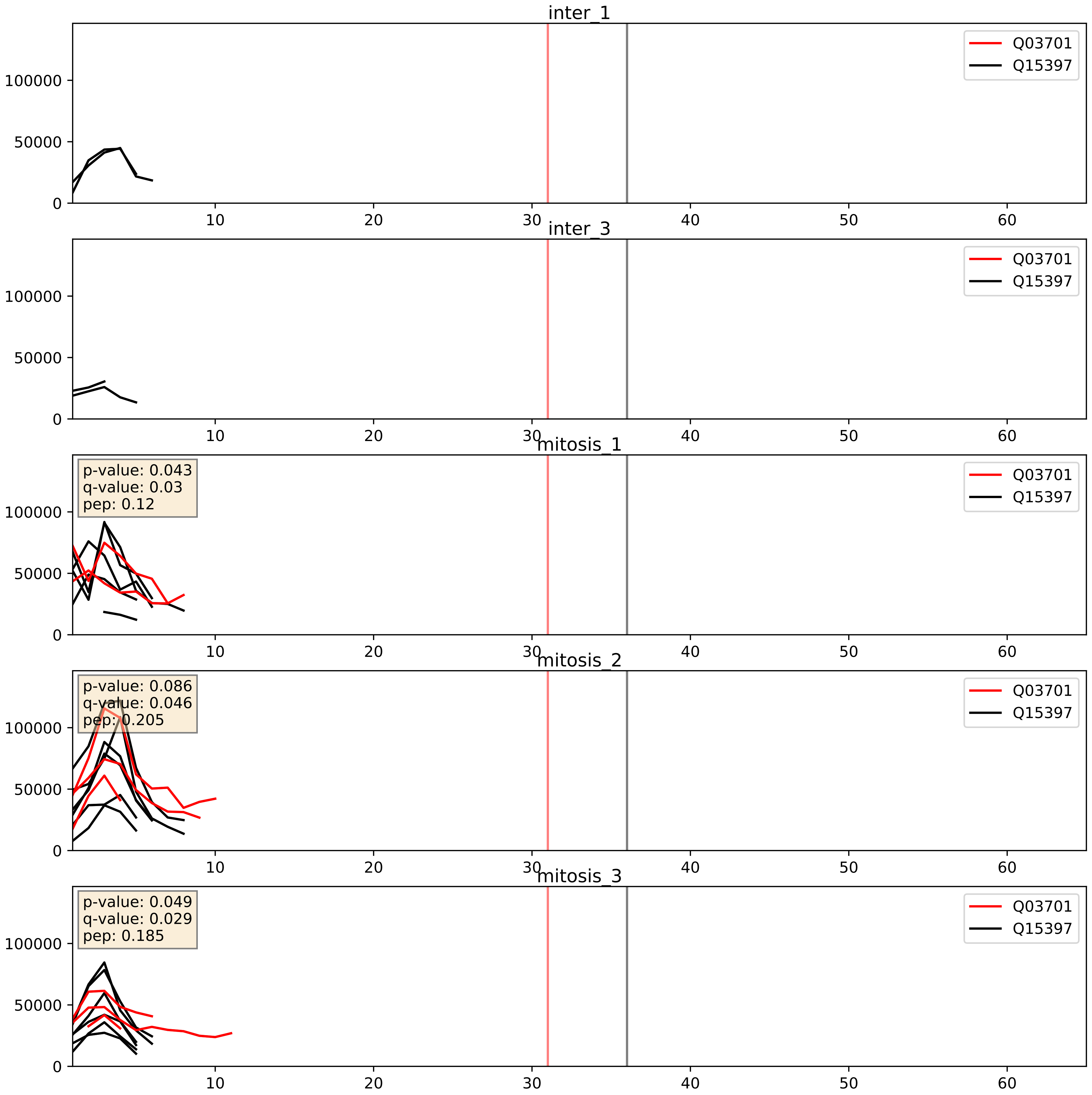


| level | condition_1 | condition_2 | pvalue | pvalue_adjusted |
|---|---|---|---|---|
| complex_abundance | mitosis | inter | 0.028263516439963447 | 0.1224370955091534 |
| interactor_abundance | mitosis | inter | 0.03470855862624382 | 0.1553074576333373 |
| interactor_ratio | mitosis | inter | 0.0981551795291016 | 0.5212210525862964 |

Q15050_Q15397
RRS1_HUMAN vs PUM3_HUMAN
p-value: 0.003 q-value: 0.044

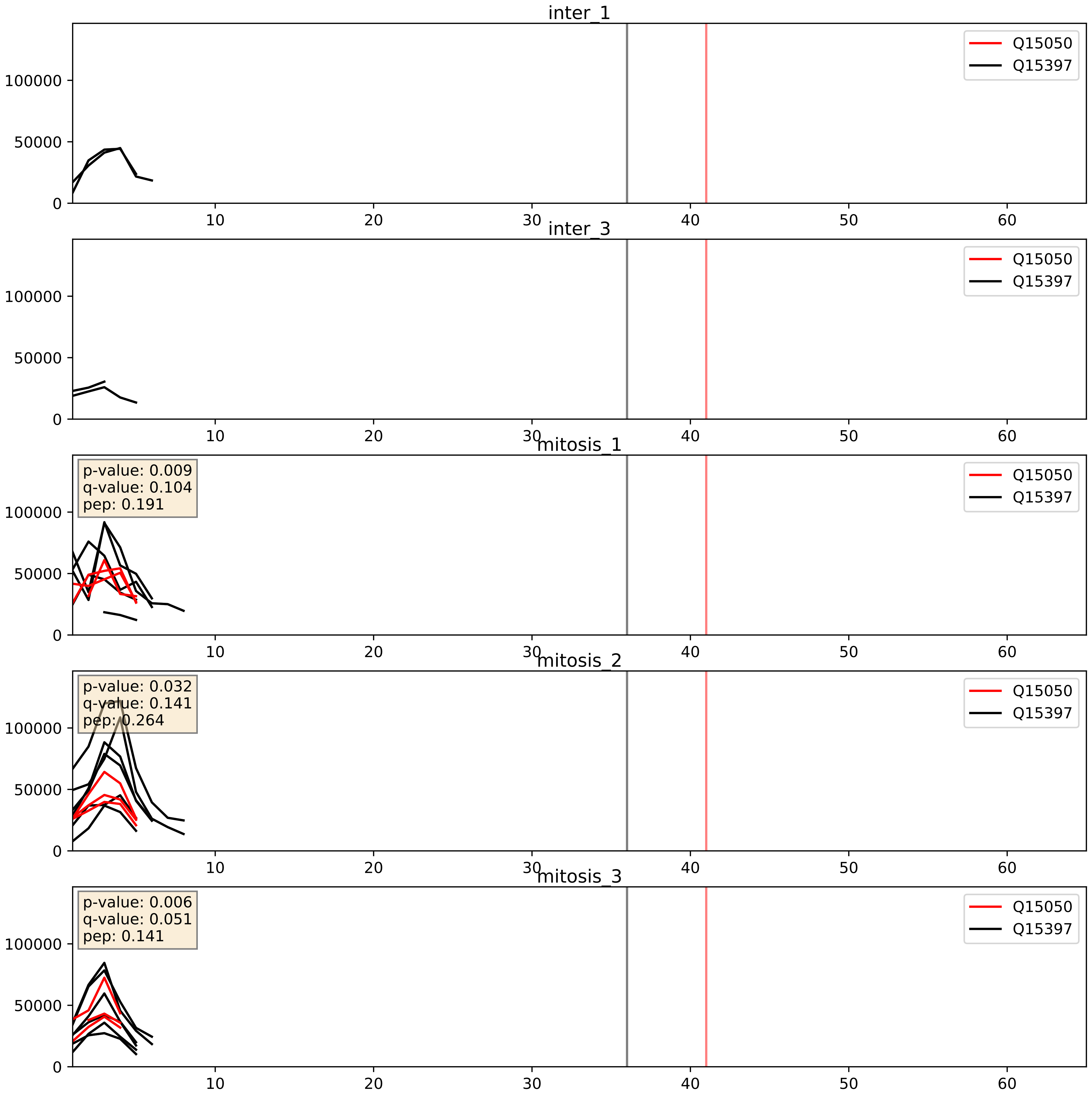


| level | condition_1 | condition_2 | pvalue | pvalue_adjusted |
|---|---|---|---|---|
| interactor_ratio | mitosis | inter | 0.05255410305415452 | 0.3793112328360562 |
| complex_abundance | mitosis | inter | 0.09861422506736761 | 0.23280136971226328 |
| interactor_abundance | mitosis | inter | 0.2162538244564617 | 0.4119120465837366 |

Q15397_Q8IY81
PUM3_HUMAN vs SPB1_HUMAN
p-value: 0.001 q-value: 0.024

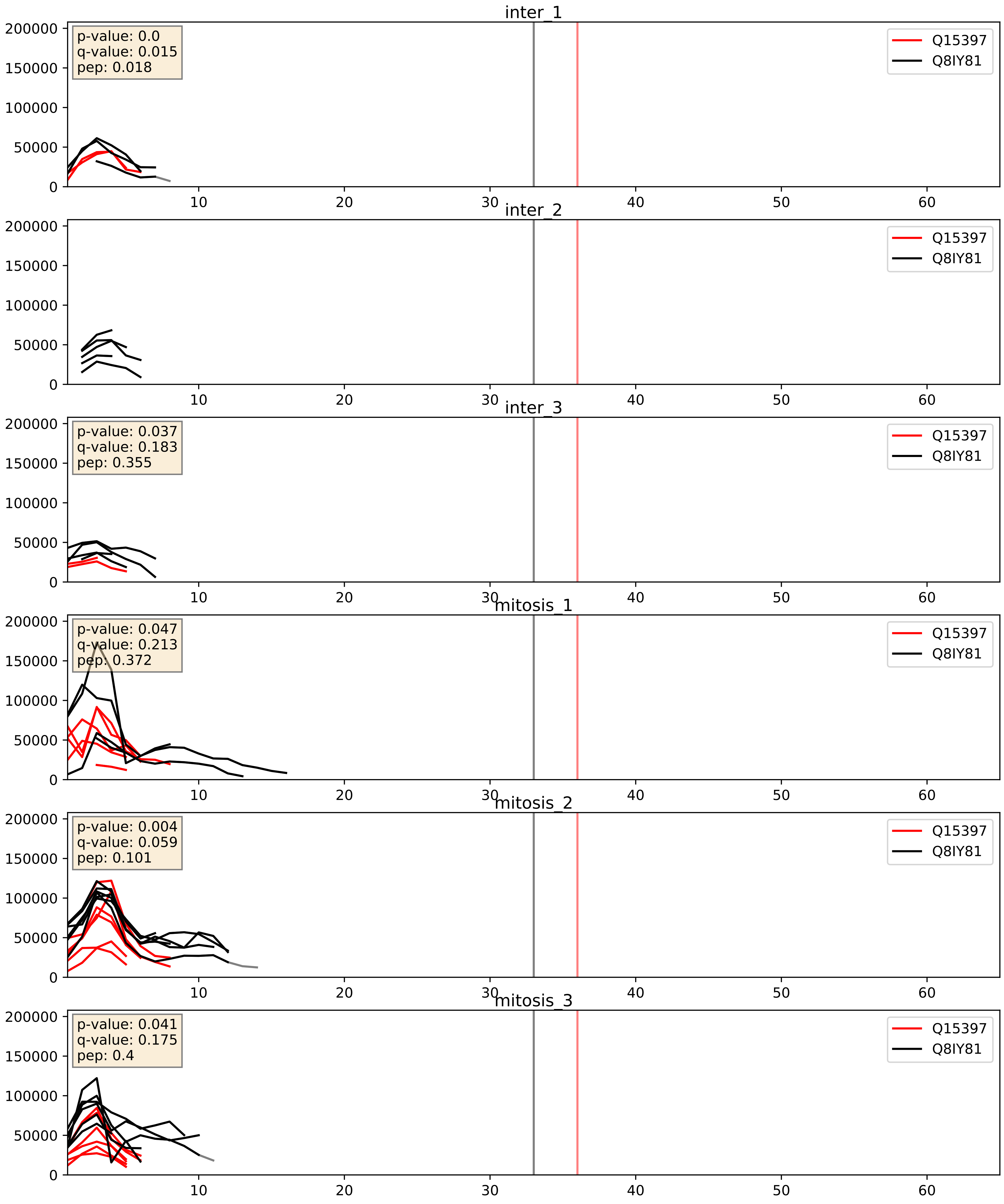


| level | condition_1 | condition_2 | pvalue | pvalue_adjusted |
|---|---|---|---|---|
| complex_abundance | mitosis | inter | 0.20380638457269512 | 0.35720365518883507 |
| interactor_abundance | mitosis | inter | 0.28137845341557466 | 0.4746944346151595 |
| interactor_ratio | mitosis | inter | 0.582051239770375 | 0.8988000350294031 |

Q15397_Q96GQ7
PUM3_HUMAN vs DDX27_HUMAN
p-value: 0.005 q-value: 0.005

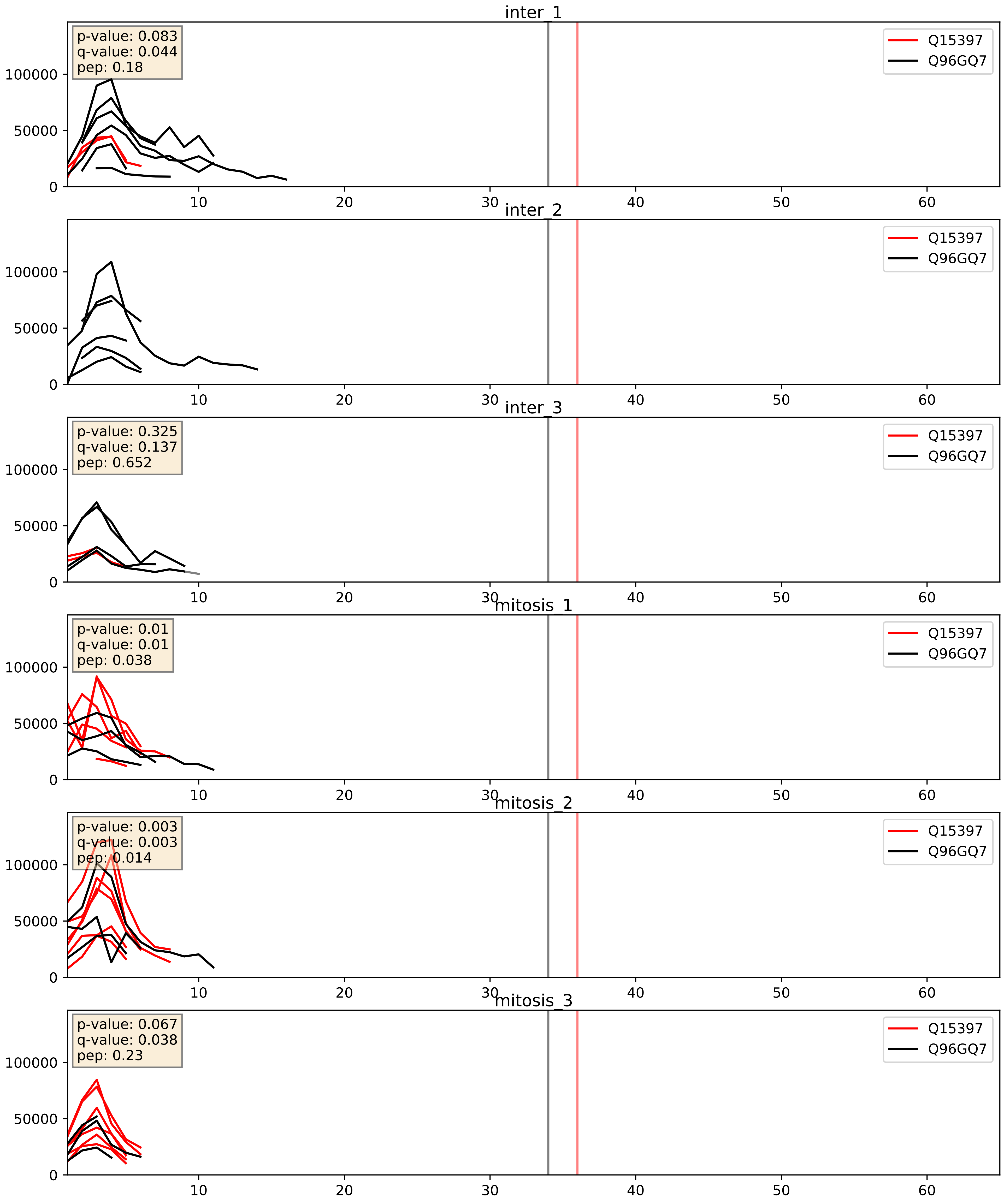


| level | condition_1 | condition_2 | pvalue | pvalue_adjusted |
|---|---|---|---|---|
| interactor_ratio | mitosis | inter | 0.020131089045294728 | 0.21442368996464944 |
| interactor_abundance | mitosis | inter | 0.28137845341557466 | 0.4746944346151595 |
| complex_abundance | mitosis | inter | 0.9286061950427655 | 0.955852456657777 |

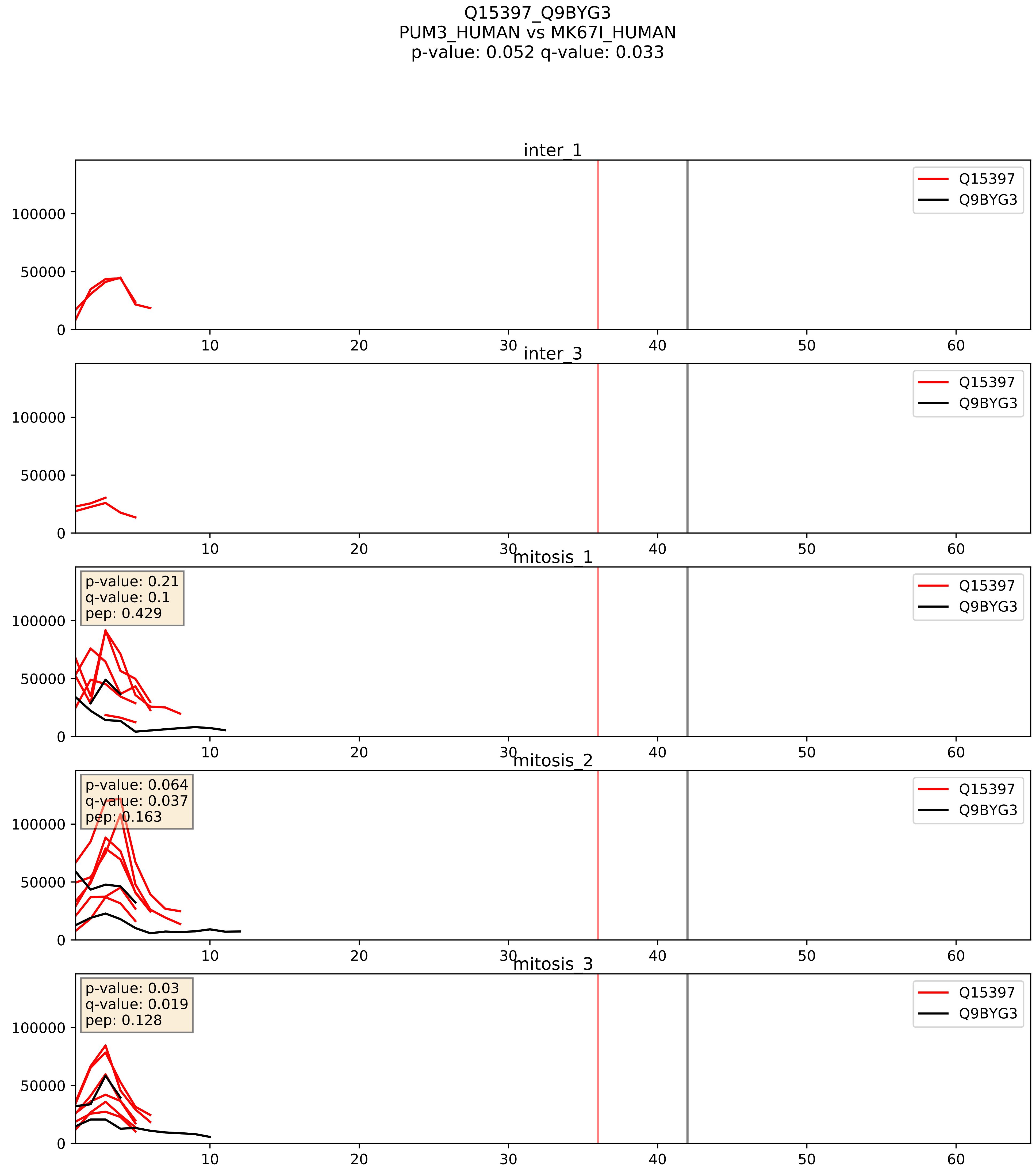


| level | condition_1 | condition_2 | pvalue | pvalue_adjusted |
|---|---|---|---|---|
| interactor_ratio | mitosis | inter | 0.004748651113544903 | 0.09196482699534925 |
| interactor_abundance | mitosis | inter | 0.024994758678114595 | 0.1308194034146505 |
| complex_abundance | mitosis | inter | 0.05783440640842851 | 0.1717178351911717 |

Q15397_Q9BZE4
PUM3_HUMAN vs NOG1_HUMAN
p-value: 0.006 q-value: 0.006

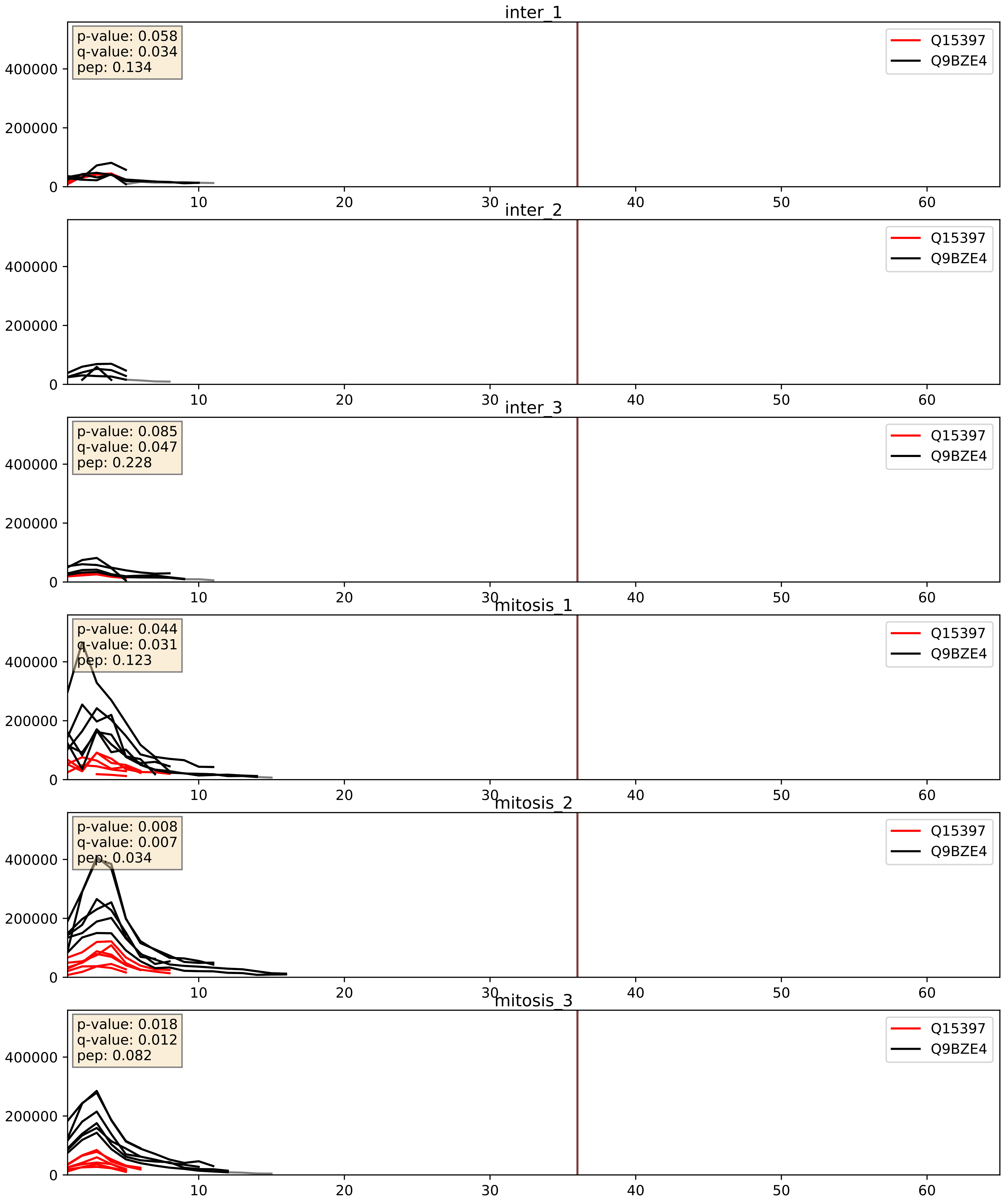


| level | condition_1 | condition_2 | pvalue | pvalue_adjusted |
|---|---|---|---|---|
| interactor_ratio | mitosis | inter | 0.00047537542148074825 | 0.02481227809680003 |
| complex_abundance | mitosis | inter | 0.022095800715059954 | 0.10952717692621856 |
| interactor_abundance | mitosis | inter | 0.2744724807239444 | 0.46751127239370466 |

Q15397_Q9GZR7
PUM3_HUMAN vs DDX24_HUMAN
p-value: 0.024 q-value: 0.018

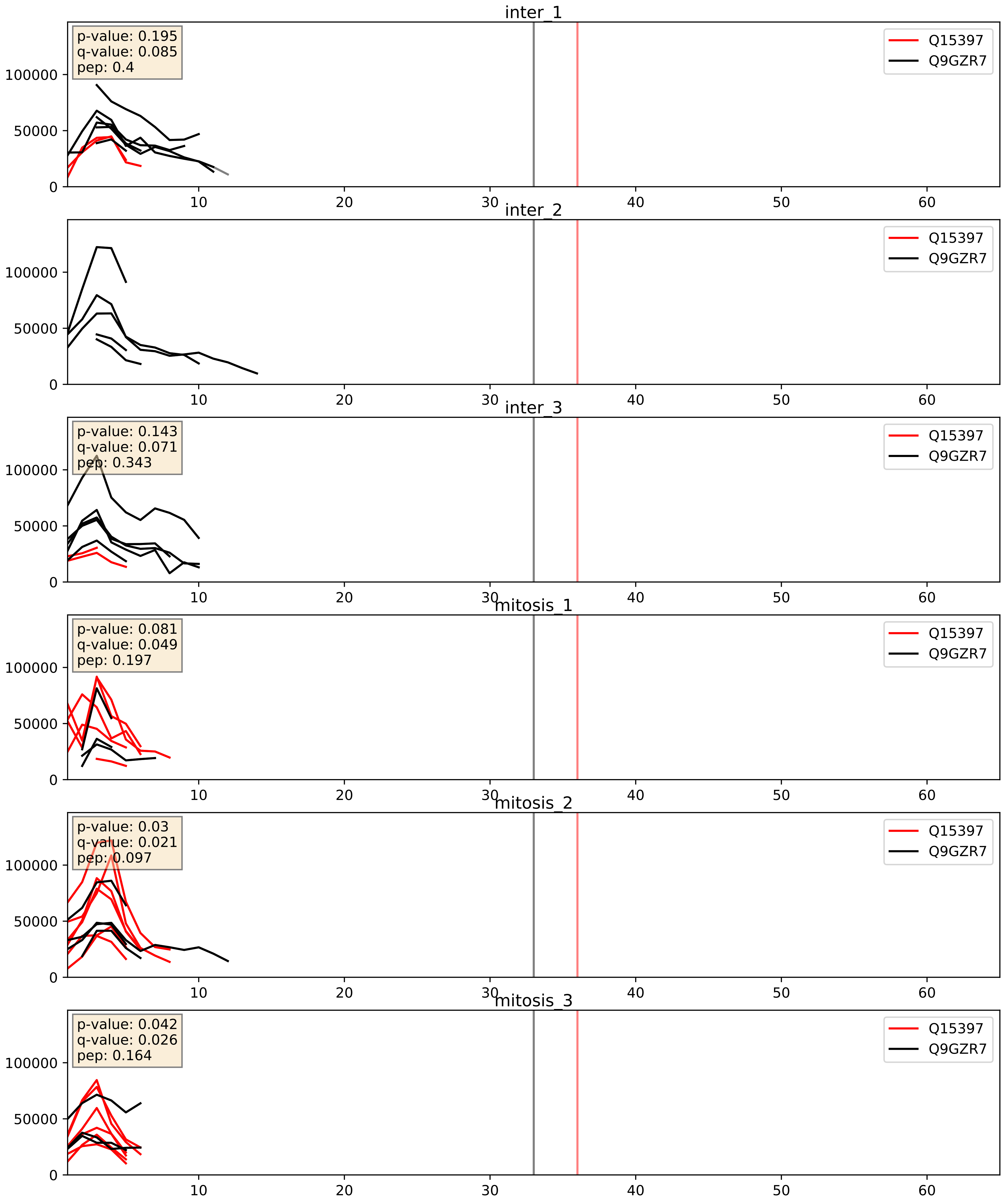


| level | condition_1 | condition_2 | pvalue | pvalue_adjusted |
|---|---|---|---|---|
| interactor_ratio | mitosis | inter | 0.04271903862311729 | 0.3405601671696144 |
| interactor_abundance | mitosis | inter | 0.30313654120020245 | 0.4984342667448584 |
| complex_abundance | mitosis | inter | 0.9077937729670189 | 0.9424158106875262 |

Q15397_Q9H7B2
PUM3_HUMAN vs RPF2_HUMAN
p-value: 0.066 q-value: 0.041

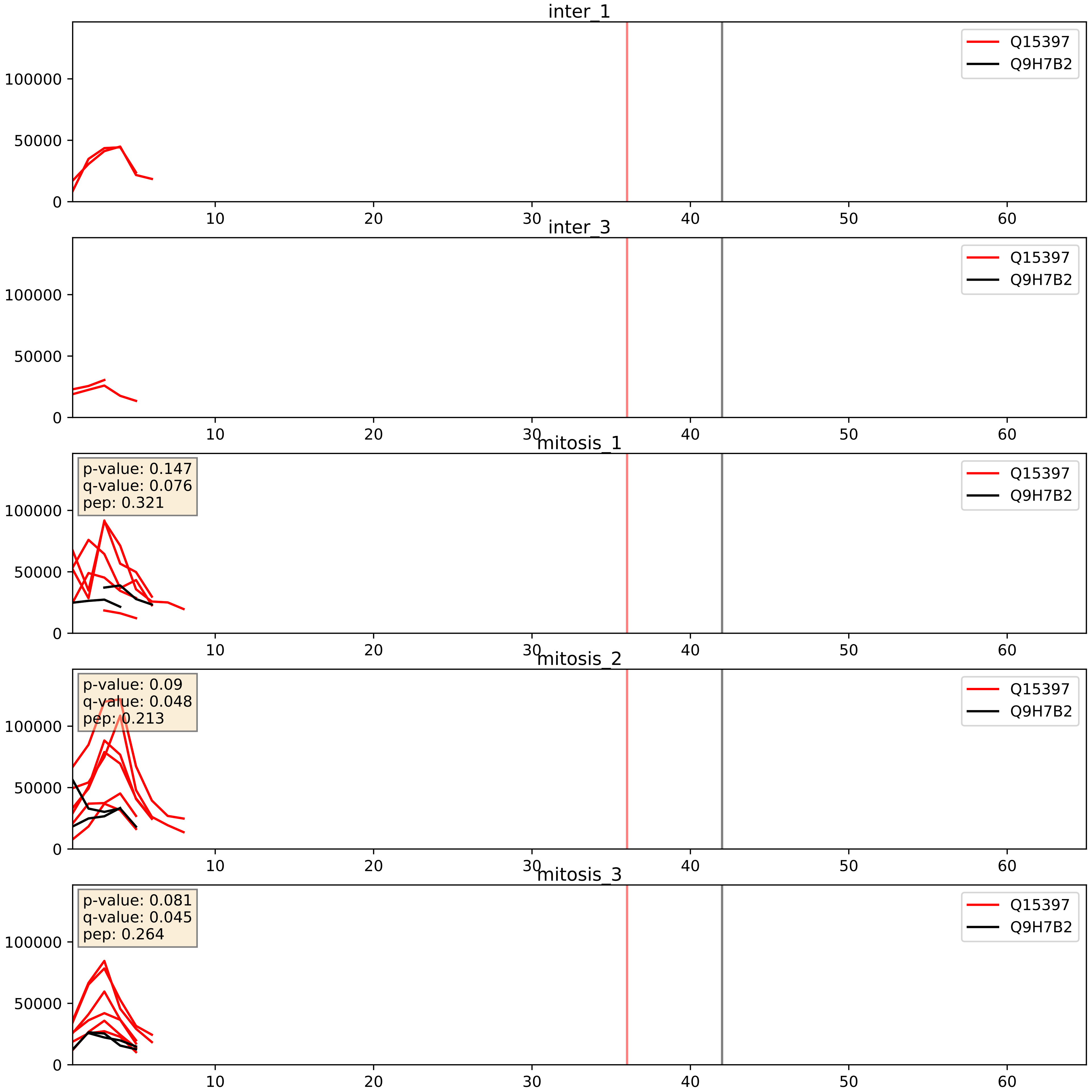


| level | condition_1 | condition_2 | pvalue | pvalue_adjusted |
|---|---|---|---|---|
| interactor_ratio | mitosis | inter | 0.00133235664863674 | 0.04334578357589157 |
| interactor_abundance | mitosis | inter | 0.037615662043017056 | 0.15827677864782586 |
| complex_abundance | mitosis | inter | 0.10346993290830749 | 0.23908191575103507 |

Q15397_Q9NVP1
PUM3_HUMAN vs DDX18_HUMAN
p-value: 0.002 q-value: 0.035

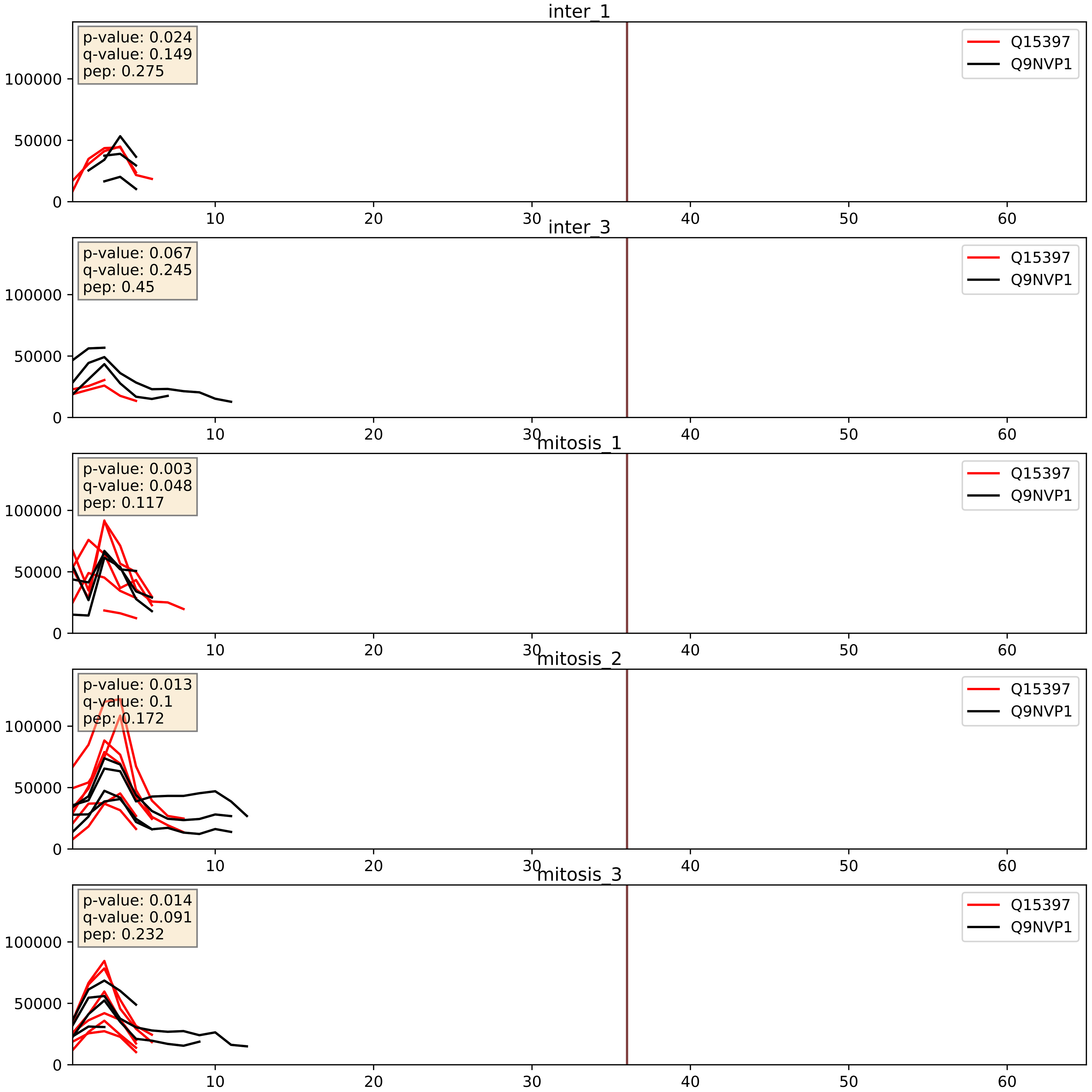


| level | condition_1 | condition_2 | pvalue | pvalue_adjusted |
|---|---|---|---|---|
| interactor_ratio | mitosis | inter | 0.07027102052804113 | 0.4393863664865099 |
| interactor_abundance | mitosis | inter | 0.26031152529654955 | 0.4522065263326465 |
| complex_abundance | mitosis | inter | 0.5653035680779996 | 0.6961586164217634 |

Q15397_Q9NW13
PUM3_HUMAN vs RBM28_HUMAN
p-value: 0.019 q-value: 0.015

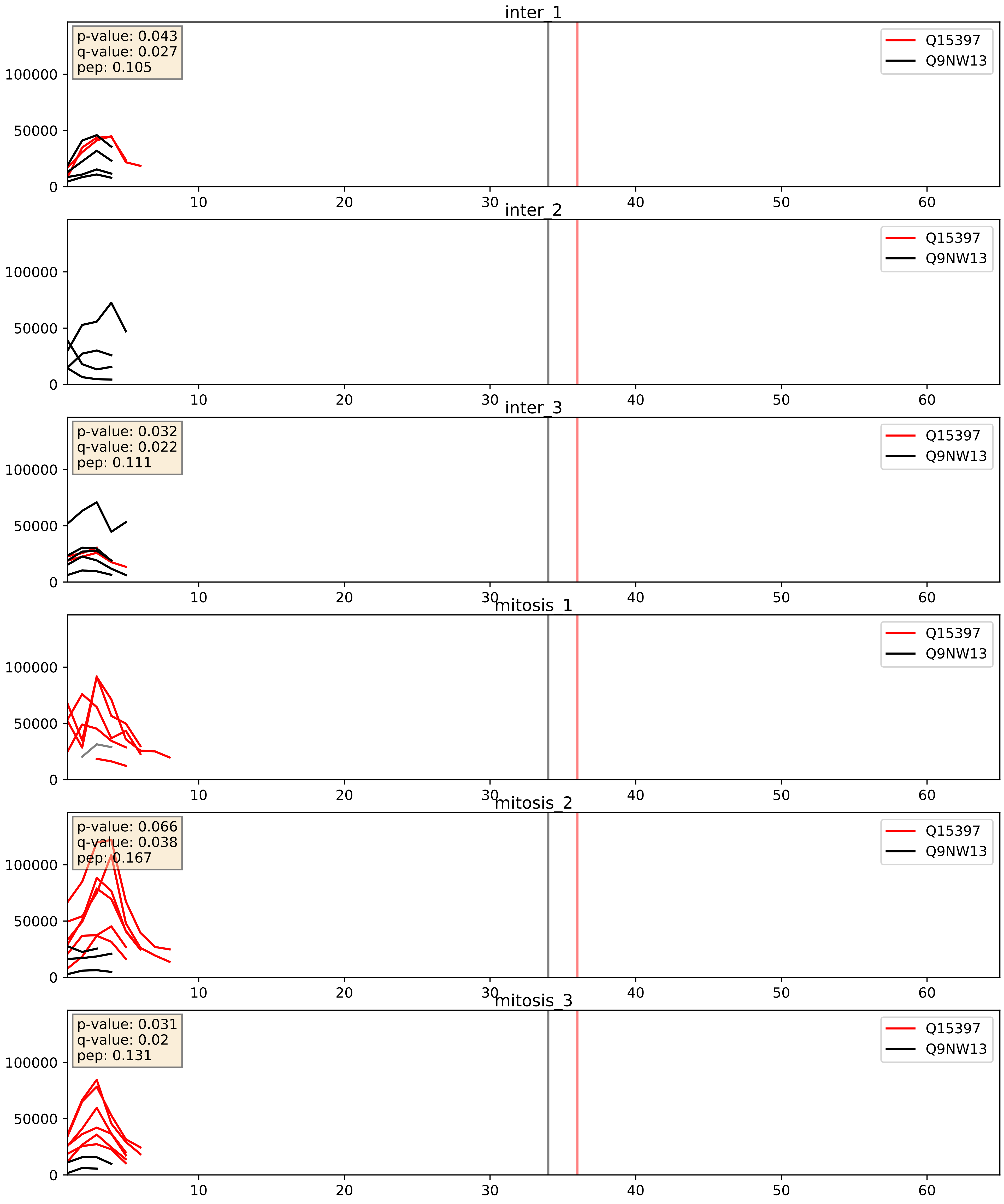


| level | condition_1 | condition_2 | pvalue | pvalue_adjusted |
|---|---|---|---|---|
| complex_abundance | mitosis | inter | 0.04097648996993001 | 0.1455676624066993 |
| interactor_ratio | mitosis | inter | 0.10396082227601368 | 0.5370731830913723 |
| interactor_abundance | mitosis | inter | 0.6013176984583364 | 0.7380141227046182 |

Q15397_Q9NY93
PUM3_HUMAN vs DDX56_HUMAN
p-value: 0.02 q-value: 0.016

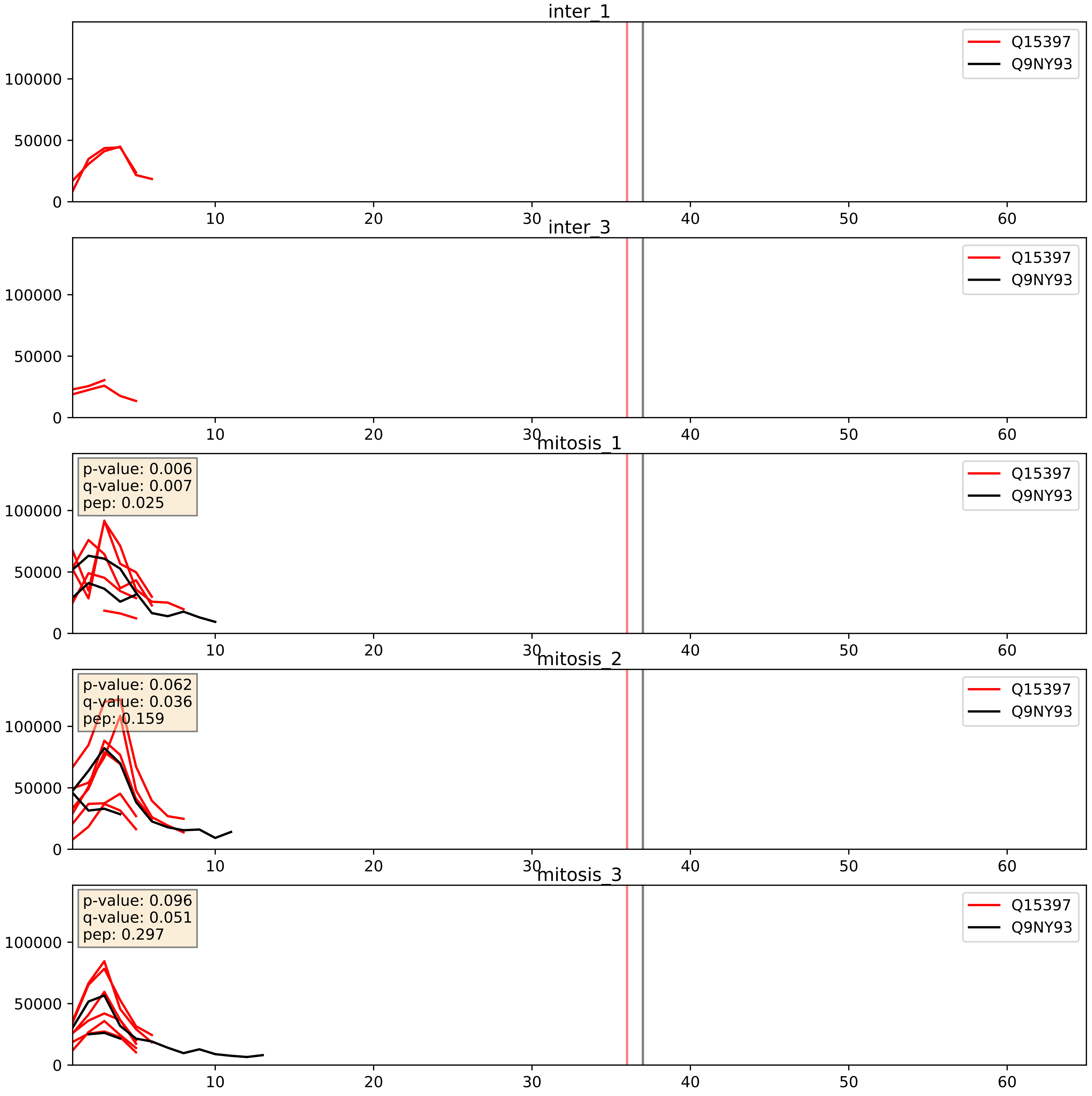


| level | condition_1 | condition_2 | pvalue | pvalue_adjusted |
|---|---|---|---|---|
| interactor_ratio | mitosis | inter | 0.0035194169325132555 | 0.07646419083591573 |
| interactor_abundance | mitosis | inter | 0.027567912416214223 | 0.13802761546789935 |
| complex_abundance | mitosis | inter | 0.05413600038038046 | 0.16599932138049858 |

Q15397_Q9UKD2
PUM3_HUMAN vs MRT4_HUMAN
p-value: 0.003 q-value: 0.041

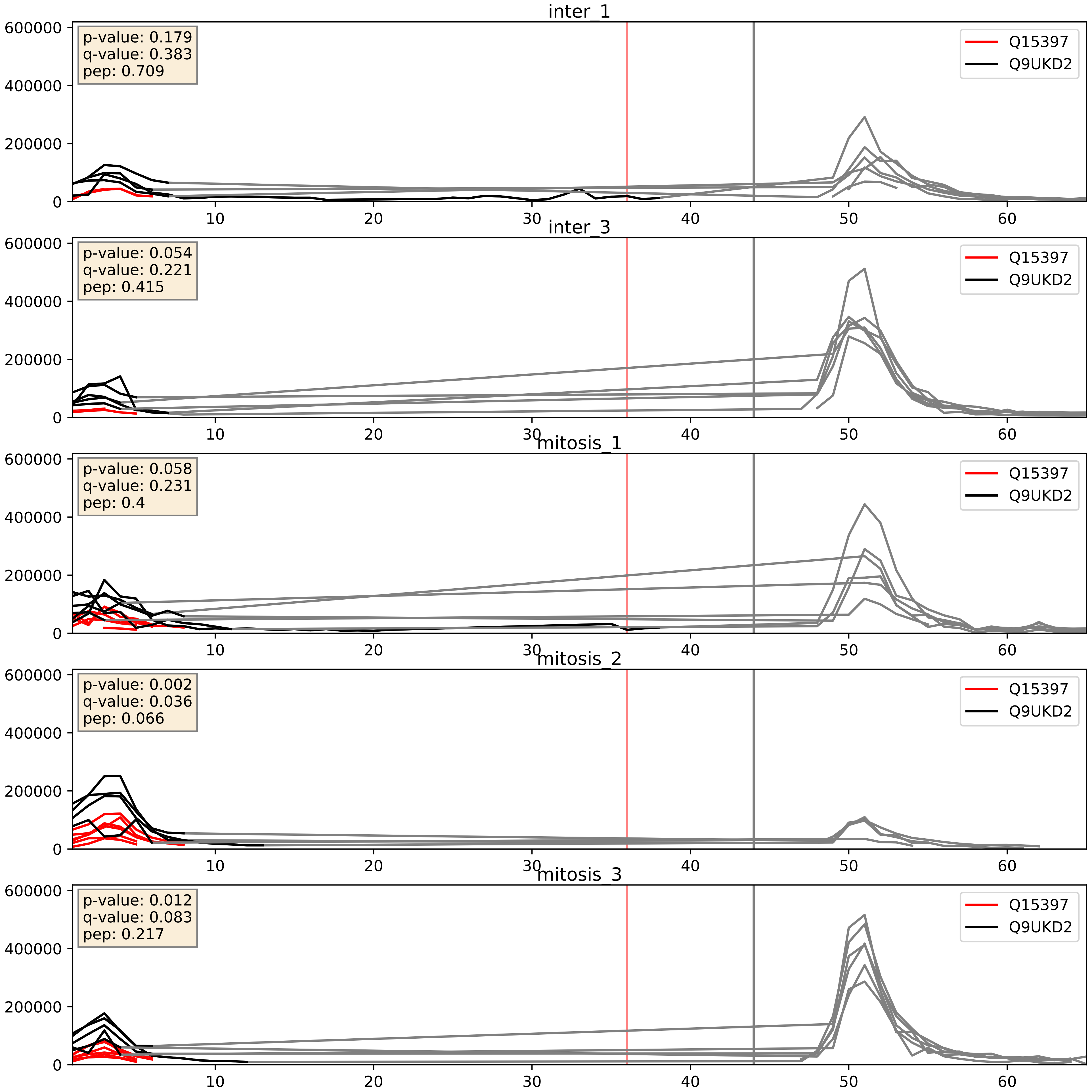


| level | condition_1 | condition_2 | pvalue | pvalue_adjusted |
|---|---|---|---|---|
| complex_abundance | mitosis | inter | 0.19909423997113293 | 0.3518989663747466 |
| interactor_ratio | mitosis | inter | 0.27205097674397627 | 0.7468898440591442 |
| interactor_abundance | mitosis | inter | 0.28137845341557466 | 0.4746944346151595 |

Q15397_Q9Y2X3
PUM3_HUMAN vs NOP58_HUMAN
p-value: 0.067 q-value: 0.041

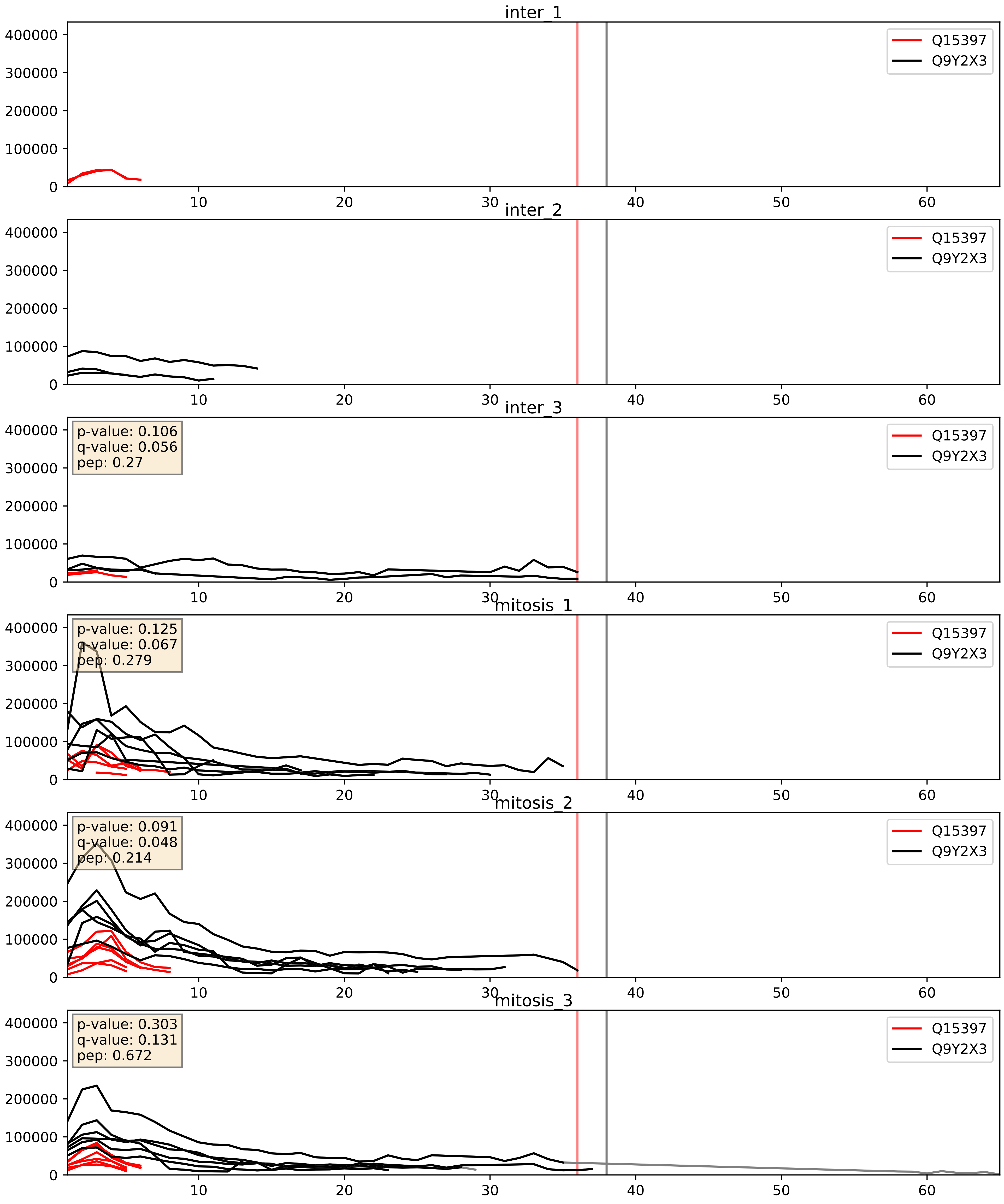


| level | condition_1 | condition_2 | pvalue | pvalue_adjusted |
|---|---|---|---|---|
| interactor_ratio | mitosis | inter | 0.0005521647453949105 | 0.02716396678494502 |
| complex_abundance | mitosis | inter | 0.004349412054734241 | 0.049247311095932673 |
| interactor_abundance | mitosis | inter | 0.02975596763925335 | 0.1434587907586644 |
