## Supplementary Data 2 for "SECAT: Quantifying differential protein-protein interaction states by network-centric analysis": hela_string_000024_Q8IY81.pdf

Q8IY81_Q8IY81

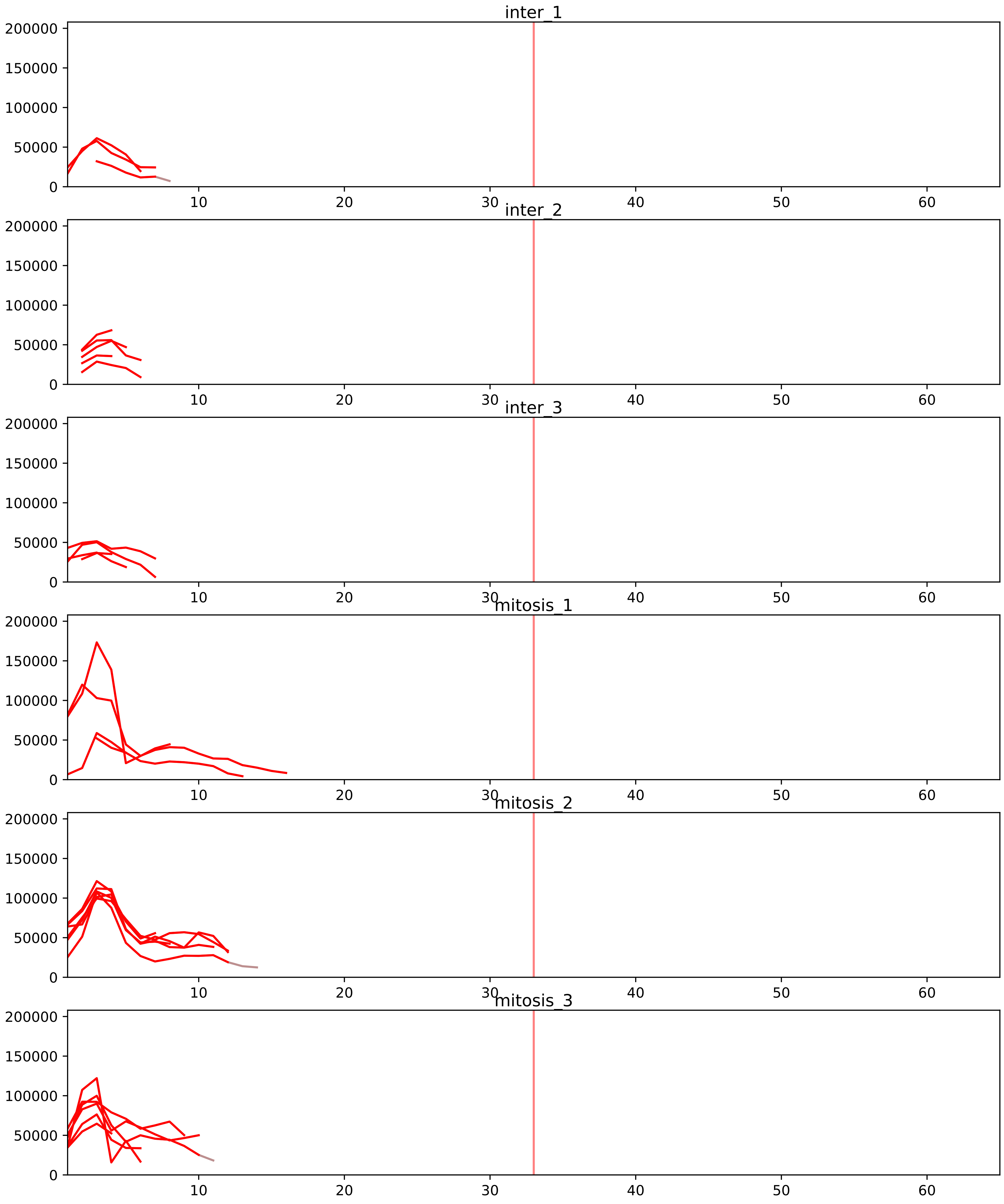


| level | condition_1 | condition_2 | pvalue | pvalue_adjusted |
|---|---|---|---|---|
| interactor_ratio | mitosis | inter | 4.807817823333567e-11 | 3.6859936645557346e-09 |
| complex_abundance | mitosis | inter | 0.0064730212326331305 | 0.03424828163185712 |
| interactor_abundance | mitosis | inter | 0.032245159926812796 | 0.10777824245034859 |
| total_abundance | mitosis | inter | 0.12473054051828428 | 0.4379332288128174 |
| assembled_abundance | mitosis | inter | 0.13081724536484782 | 0.4284507279651614 |

O00567_Q8IY81
NOP56_HUMAN vs SPB1_HUMAN
p-value: 0.01 q-value: 0.009

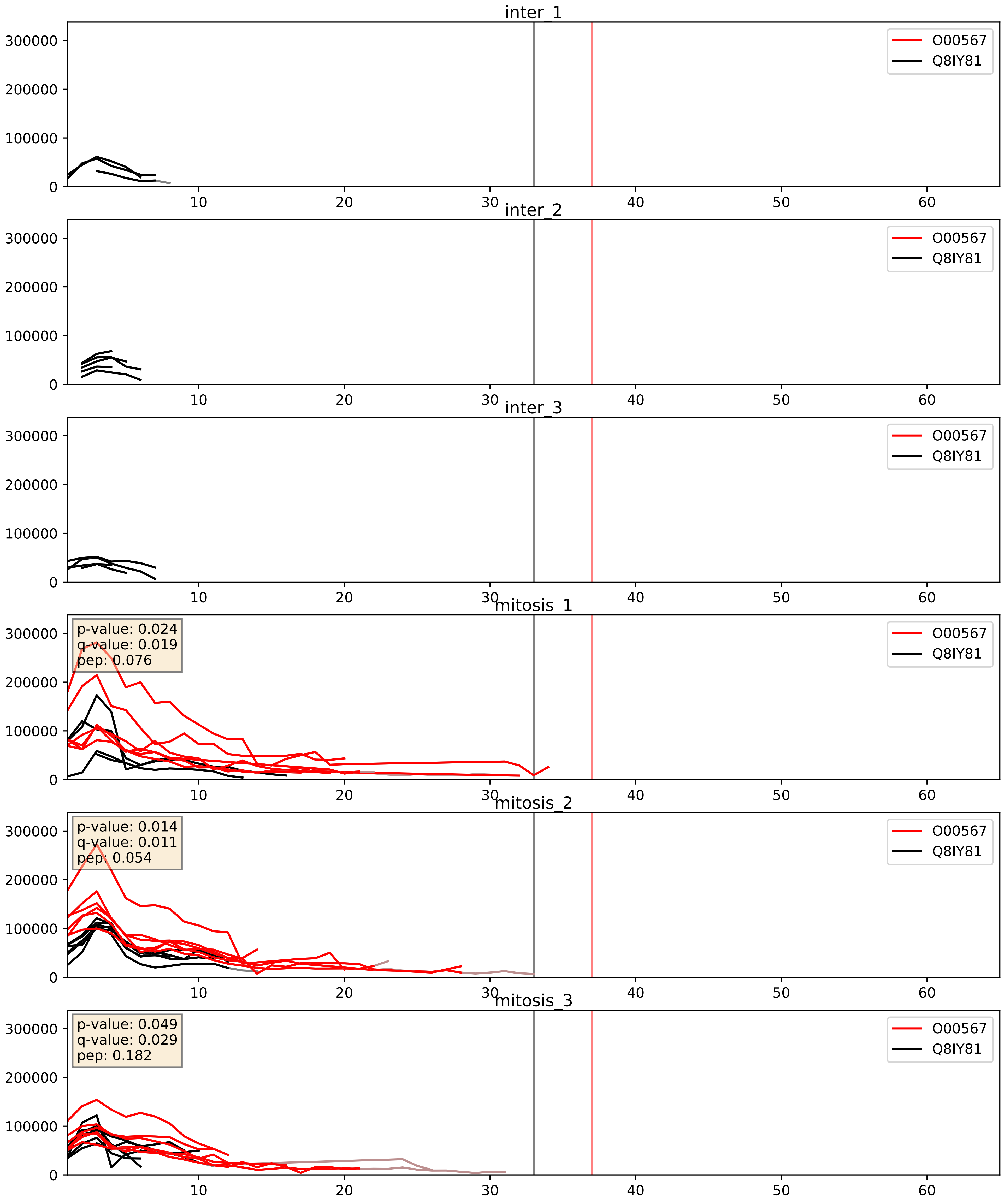


| level | condition_1 | condition_2 | pvalue | pvalue_adjusted |
|---|---|---|---|---|
| interactor_abundance | mitosis | inter | 0.0015635866039636899 | 0.029025269221018554 |
| complex_abundance | mitosis | inter | 0.001688865399875853 | 0.028927157402433075 |
| interactor_ratio | mitosis | inter | 0.010319605669621772 | 0.146251365119143 |

P22087_Q8IY81
FBRL_HUMAN vs SPB1_HUMAN
p-value: 0.02 q-value: 0.016

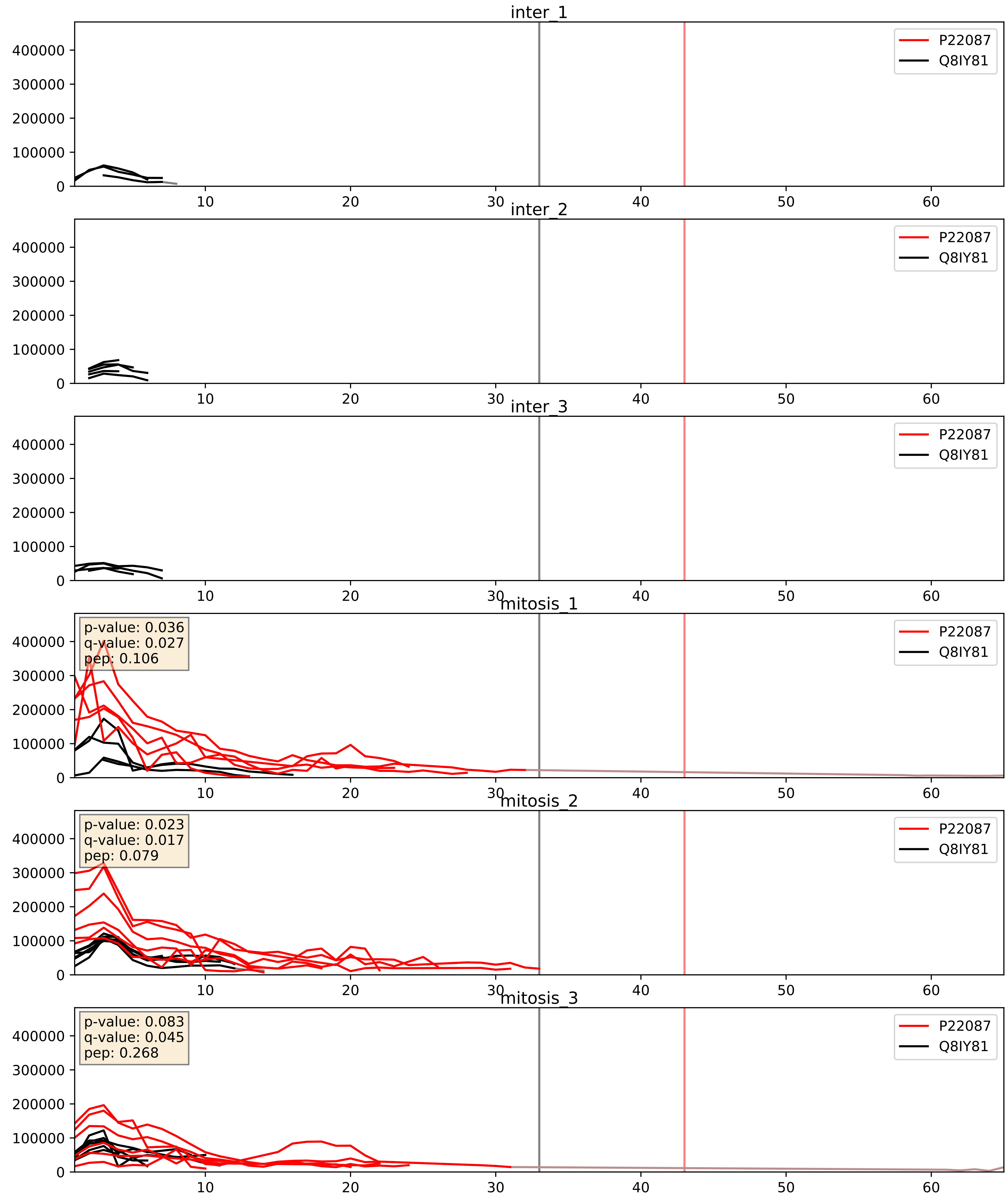


| level | condition_1 | condition_2 | pvalue | pvalue_adjusted |
|---|---|---|---|---|
| interactor_abundance | mitosis | inter | 0.0010710204172406286 | 0.024049720606162197 |
| complex_abundance | mitosis | inter | 0.0012994811847285065 | 0.024459508269436846 |
| interactor_ratio | mitosis | inter | 0.0034502879269765964 | 0.07572939655107606 |

P46087_Q8IY81
NOP2_HUMAN vs SPB1_HUMAN
p-value: 0.026 q-value: 0.019

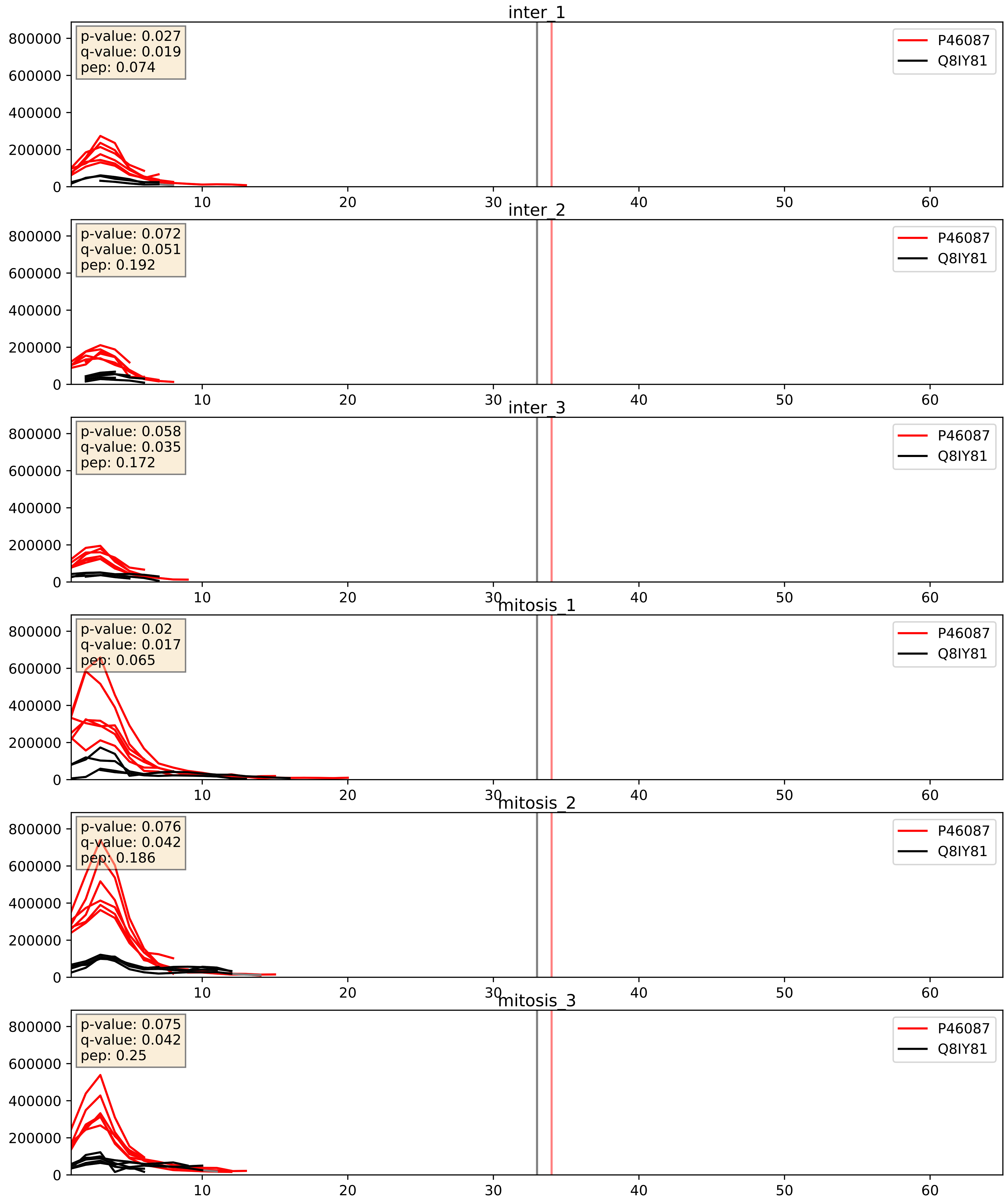


| level | condition_1 | condition_2 | pvalue | pvalue_adjusted |
|---|---|---|---|---|
| interactor_abundance | mitosis | inter | 0.007209065250286371 | 0.07343152394042615 |
| interactor_ratio | mitosis | inter | 0.013389338509009651 | 0.1678078149884665 |
| complex_abundance | mitosis | inter | 0.03232225811702476 | 0.13090015450091652 |

Q12788_Q8IY81
TBL3_HUMAN vs SPB1_HUMAN
p-value: 0.049 q-value: 0.032

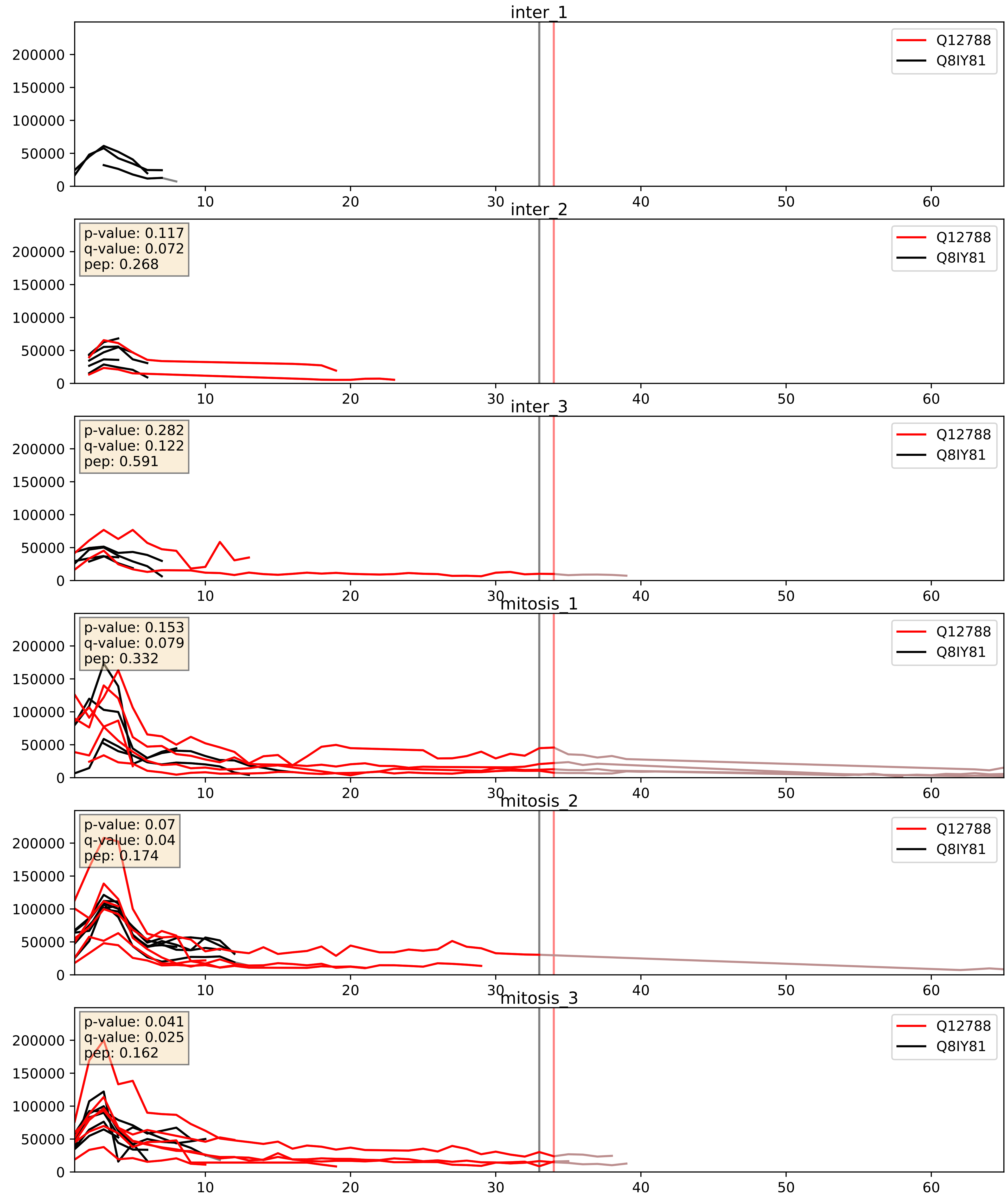


| level | condition_1 | condition_2 | pvalue | pvalue_adjusted |
|---|---|---|---|---|
| interactor_abundance | mitosis | inter | 0.007901002763394643 | 0.07711604472197223 |
| complex_abundance | mitosis | inter | 0.030978129908028652 | 0.127942946637983 |
| interactor_ratio | mitosis | inter | 0.1691647109442906 | 0.6320565072989578 |

Q13610_Q8IY81
PWP1_HUMAN vs SPB1_HUMAN
p-value: 0.008 q-value: 0.008

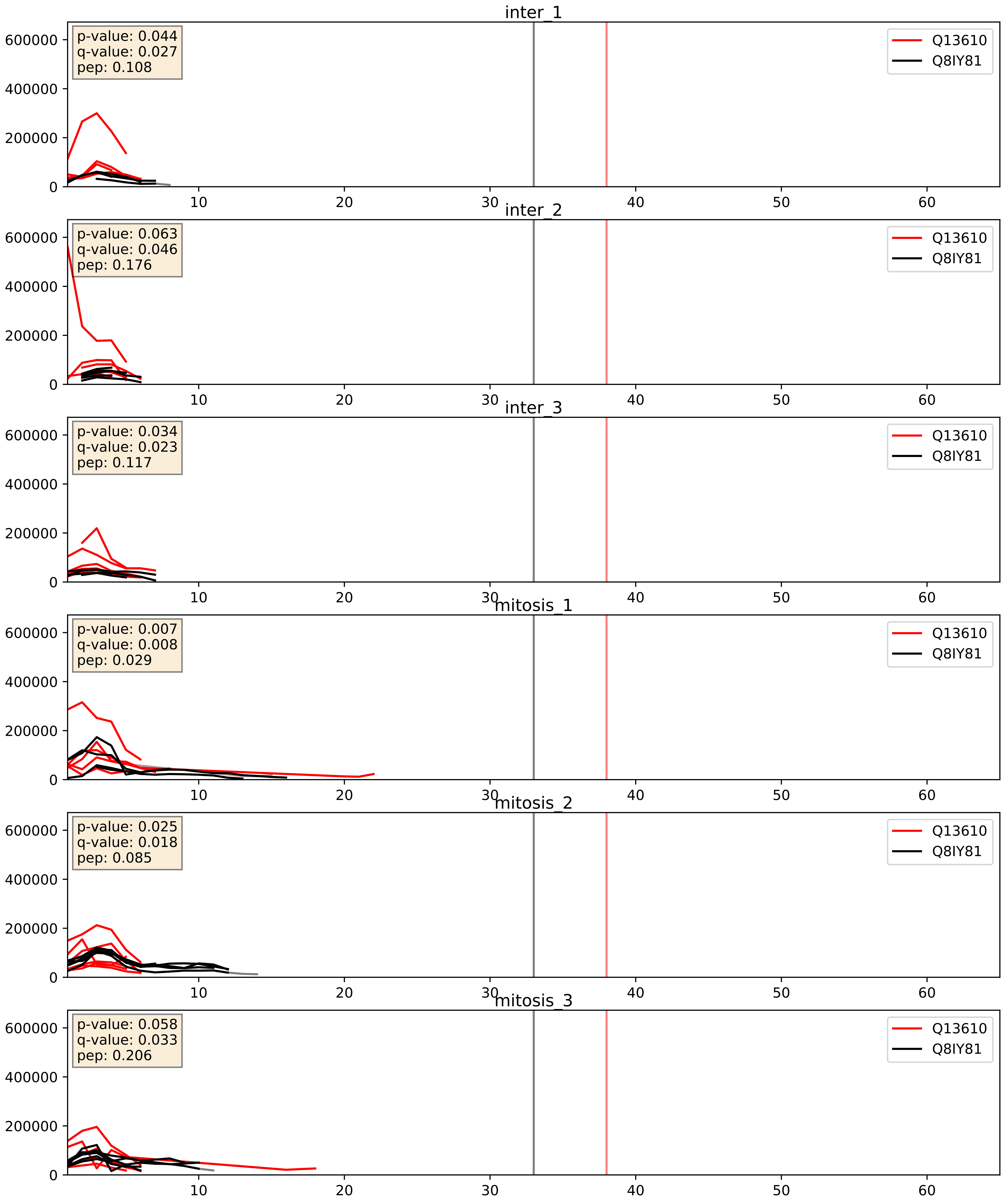


| level | condition_1 | condition_2 | pvalue | pvalue_adjusted |
|---|---|---|---|---|
| interactor_abundance | mitosis | inter | 0.4965684600005845 | 0.6622168268689187 |
| interactor_ratio | mitosis | inter | 0.49939290201744846 | 0.8703055576894411 |
| complex_abundance | mitosis | inter | 0.6902208774206902 | 0.7920085929588117 |

Q14137_Q8IY81
BOP1_HUMAN vs SPB1_HUMAN
p-value: 0.041 q-value: 0.028

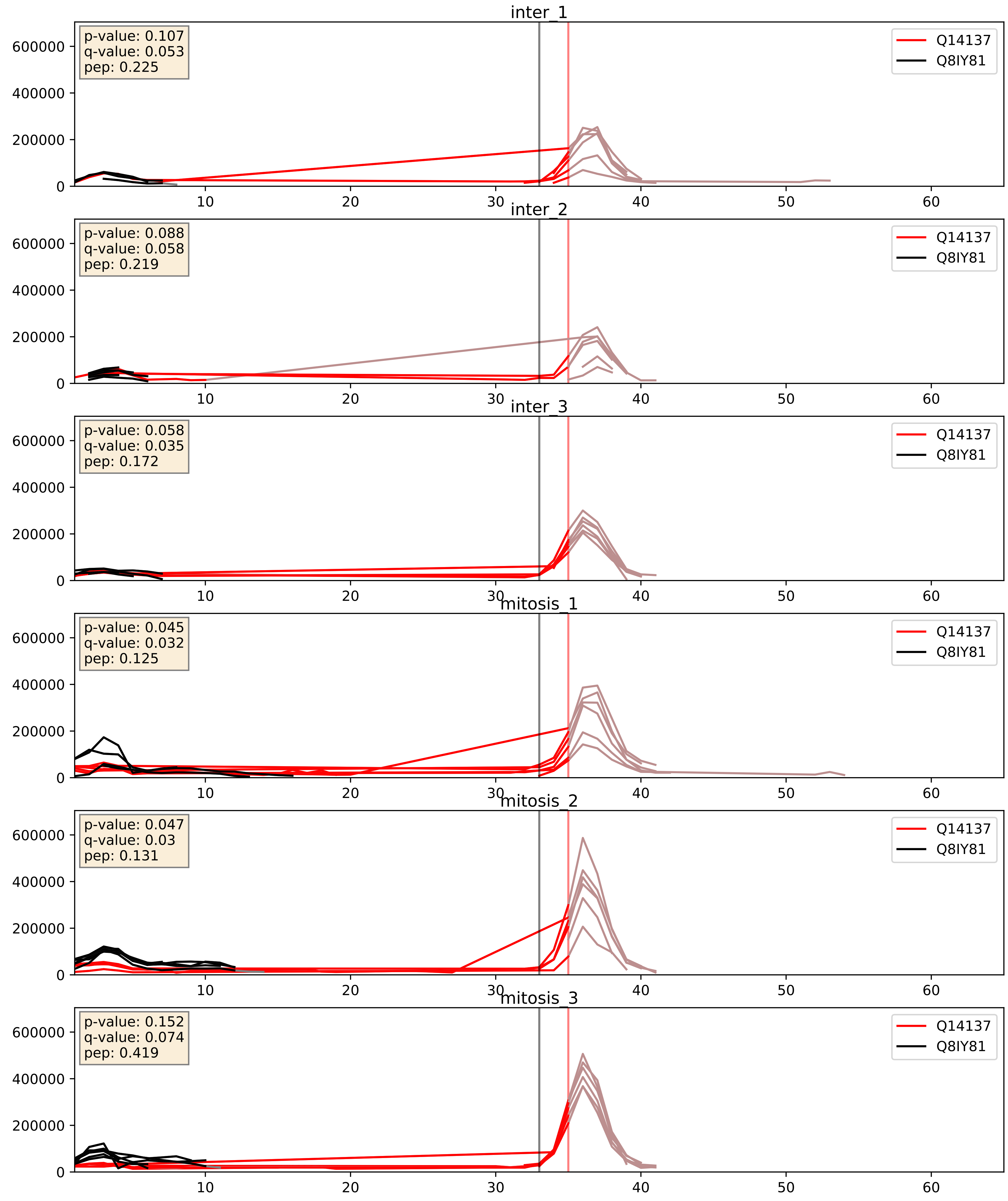


| level | condition_1 | condition_2 | pvalue | pvalue_adjusted |
|---|---|---|---|---|
| interactor_ratio | mitosis | inter | 0.06357934216723345 | 0.4189677974992443 |
| interactor_abundance | mitosis | inter | 0.11208372992542083 | 0.2886587657103679 |
| complex_abundance | mitosis | inter | 0.4002338471426604 | 0.5465861090525165 |

Q15050_Q8IY81
RRS1_HUMAN vs SPB1_HUMAN
p-value: 0.017 q-value: 0.014

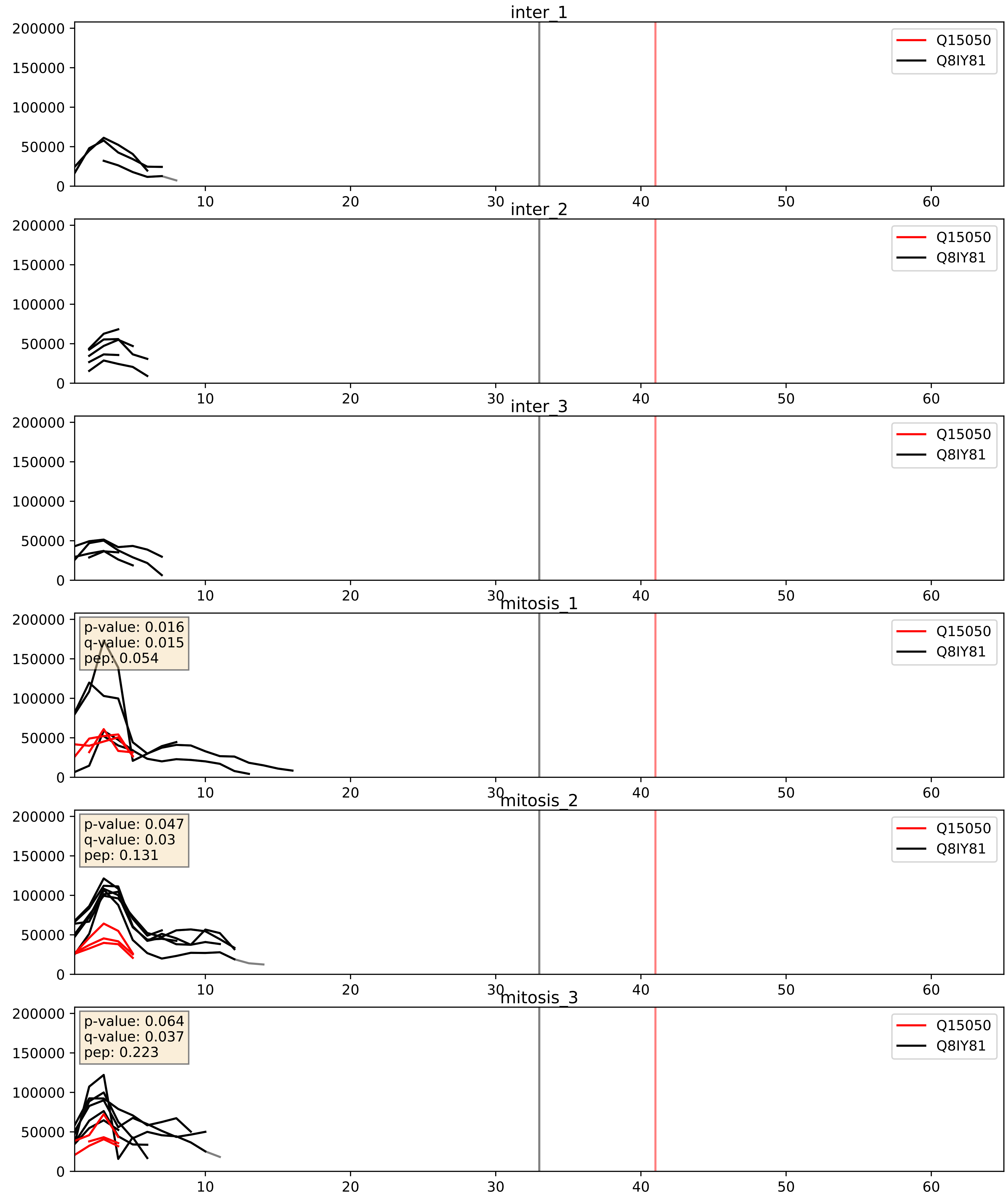


| level | condition_1 | condition_2 | pvalue | pvalue_adjusted |
|---|---|---|---|---|
| complex_abundance | mitosis | inter | 0.05036965452854774 | 0.15980547590484867 |
| interactor_ratio | mitosis | inter | 0.13938380188657049 | 0.5877140456747689 |
| interactor_abundance | mitosis | inter | 0.2162538244564617 | 0.4119120465837366 |

Q15269_Q8IY81
PWP2_HUMAN vs SPB1_HUMAN
p-value: 0.001 q-value: 0.001

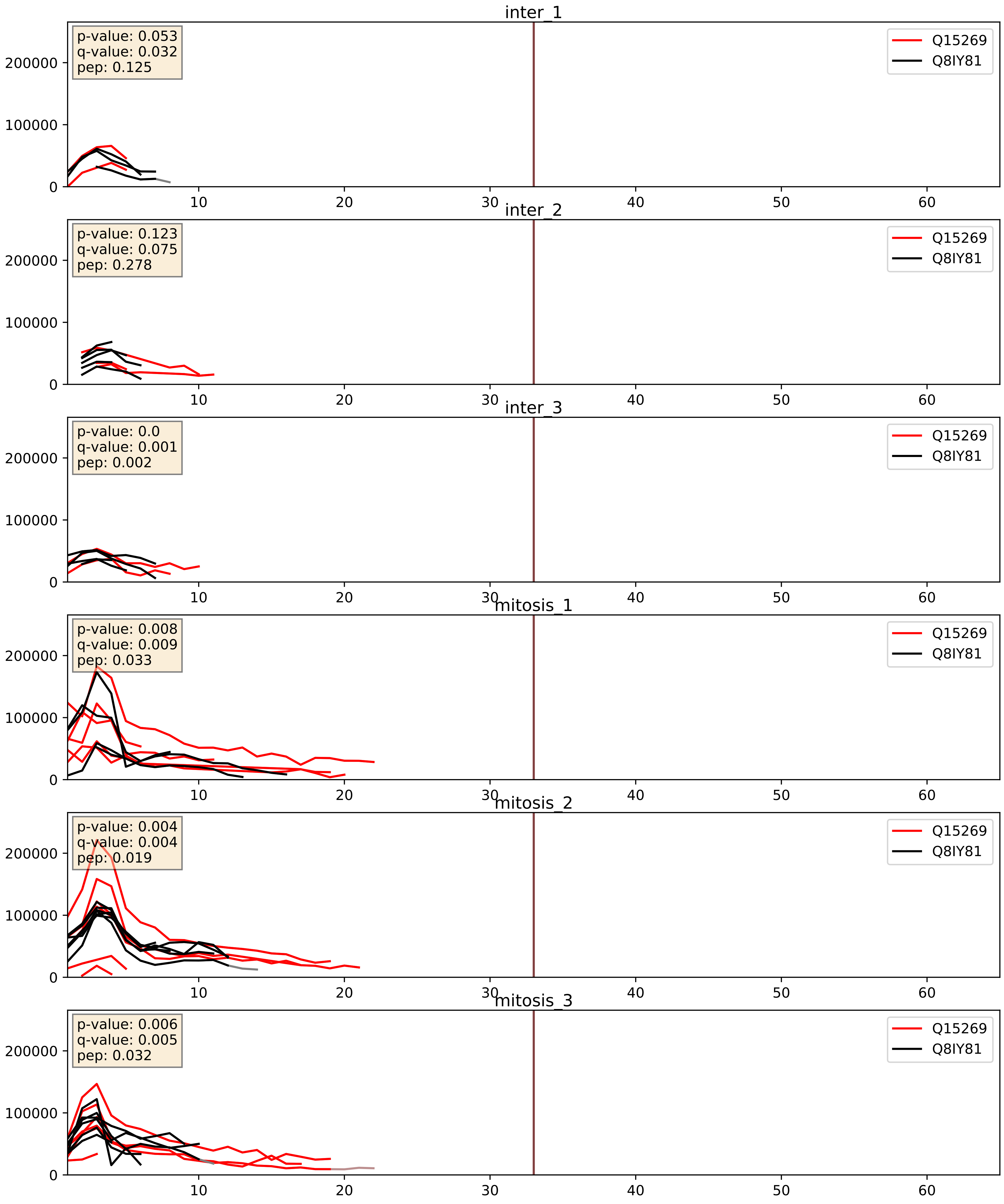


| level | condition_1 | condition_2 | pvalue | pvalue_adjusted |
|---|---|---|---|---|
| interactor_ratio | mitosis | inter | 0.04107050221712384 | 0.3361027714900383 |
| interactor_abundance | mitosis | inter | 0.05103025519733094 | 0.181177513268002 |
| complex_abundance | mitosis | inter | 0.11609552713244926 | 0.2570557972720553 |

Q15397_Q8IY81
PUM3_HUMAN vs SPB1_HUMAN
p-value: 0.001 q-value: 0.024

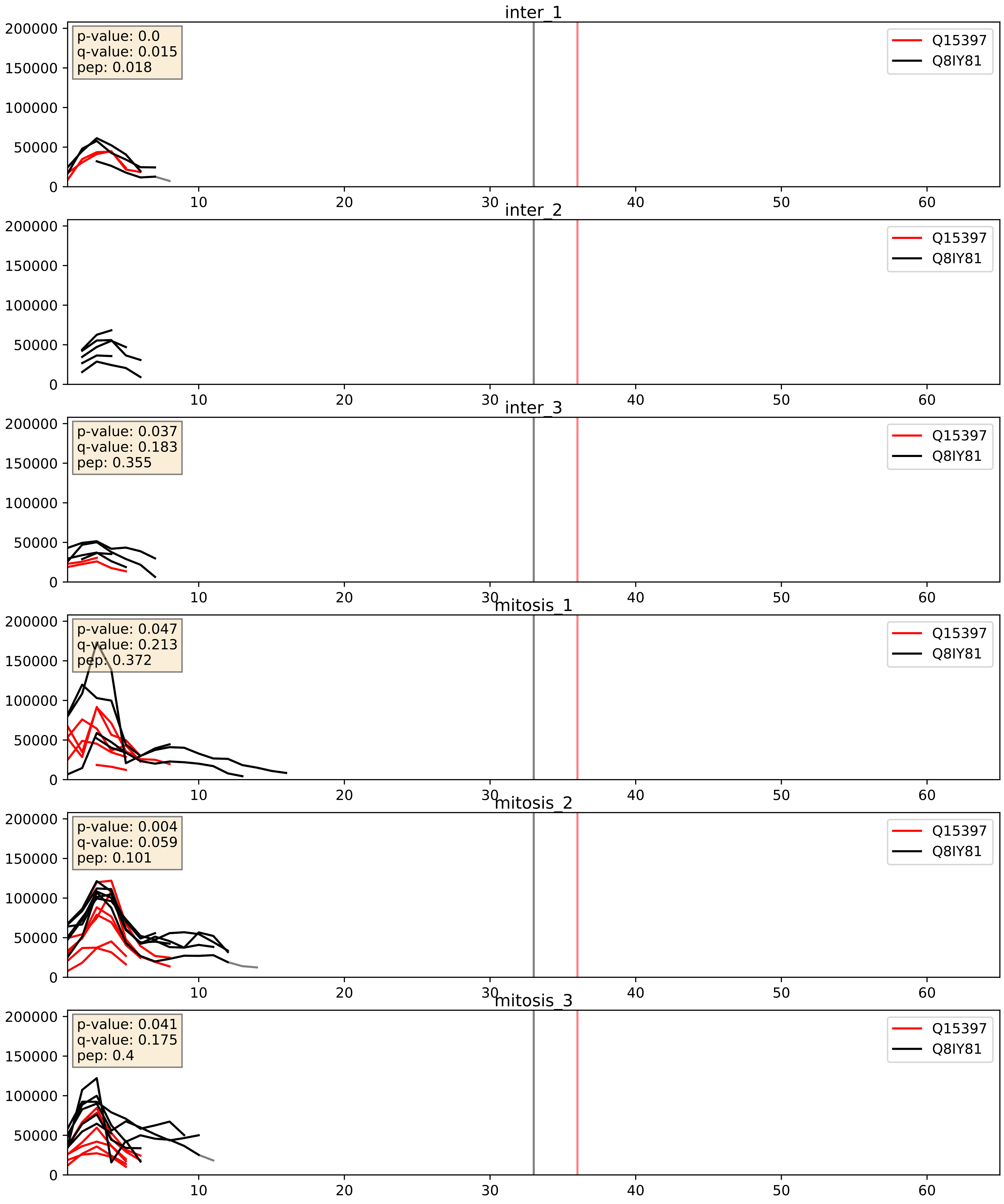


| level | condition_1 | condition_2 | pvalue | pvalue_adjusted |
|---|---|---|---|---|
| complex_abundance | mitosis | inter | 0.20380638457269512 | 0.35720365518883507 |
| interactor_abundance | mitosis | inter | 0.28137845341557466 | 0.4746944346151595 |
| interactor_ratio | mitosis | inter | 0.582051239770375 | 0.8988000350294031 |

Q8IWA0_Q8IY81
WDR75_HUMAN vs SPB1_HUMAN
p-value: 0.038 q-value: 0.026

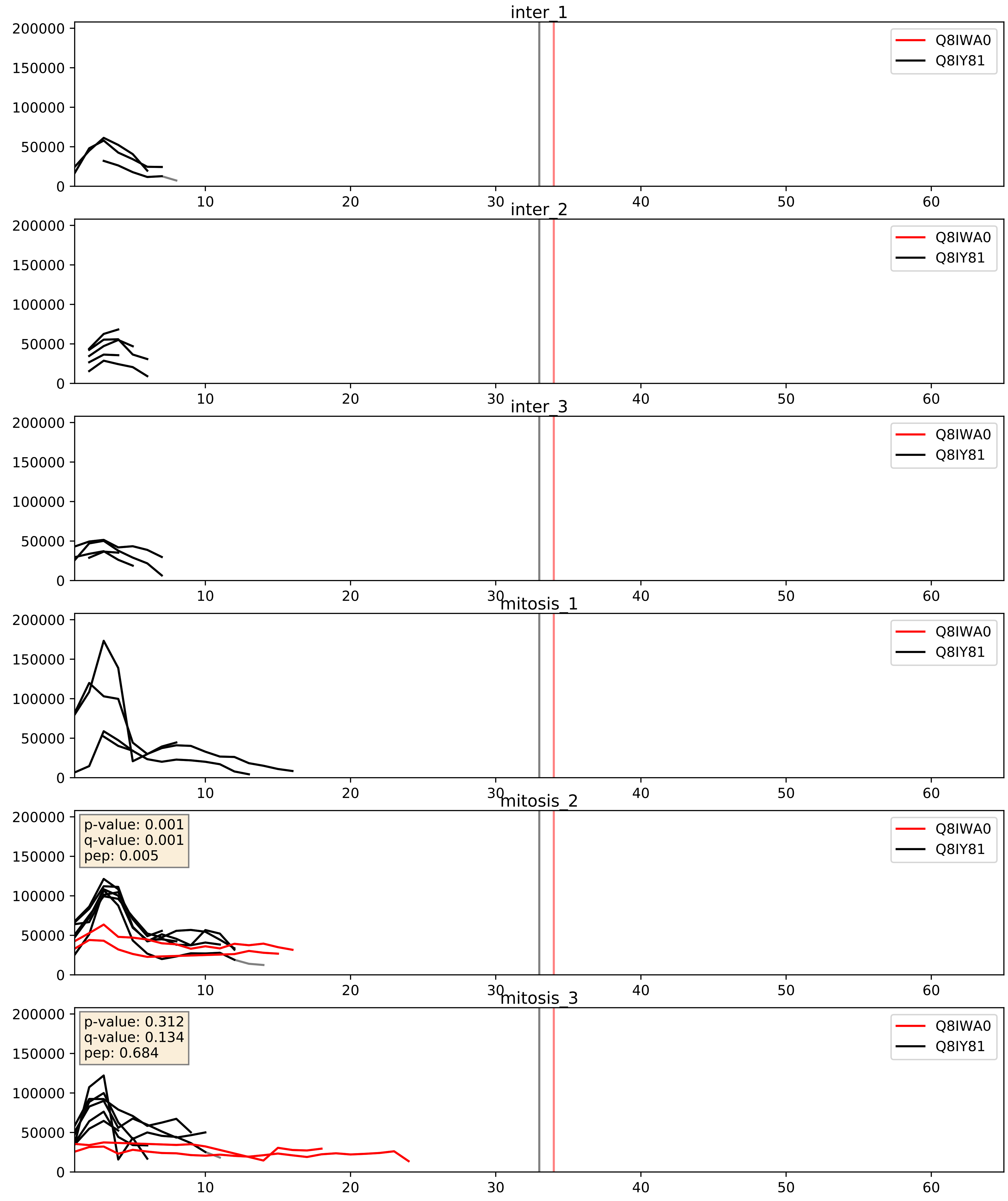


| level | condition_1 | condition_2 | pvalue | pvalue_adjusted |
|---|---|---|---|---|
| interactor_ratio | mitosis | inter | 0.18459715636898918 | 0.6543071049766241 |
| complex_abundance | mitosis | inter | 0.28354960017013753 | 0.43203712663872856 |
| interactor_abundance | mitosis | inter | 0.40120020807242107 | 0.5898786982308355 |

Q8IY81_Q8NI36
SPB1_HUMAN vs WDR36_HUMAN
p-value: 0.008 q-value: 0.008

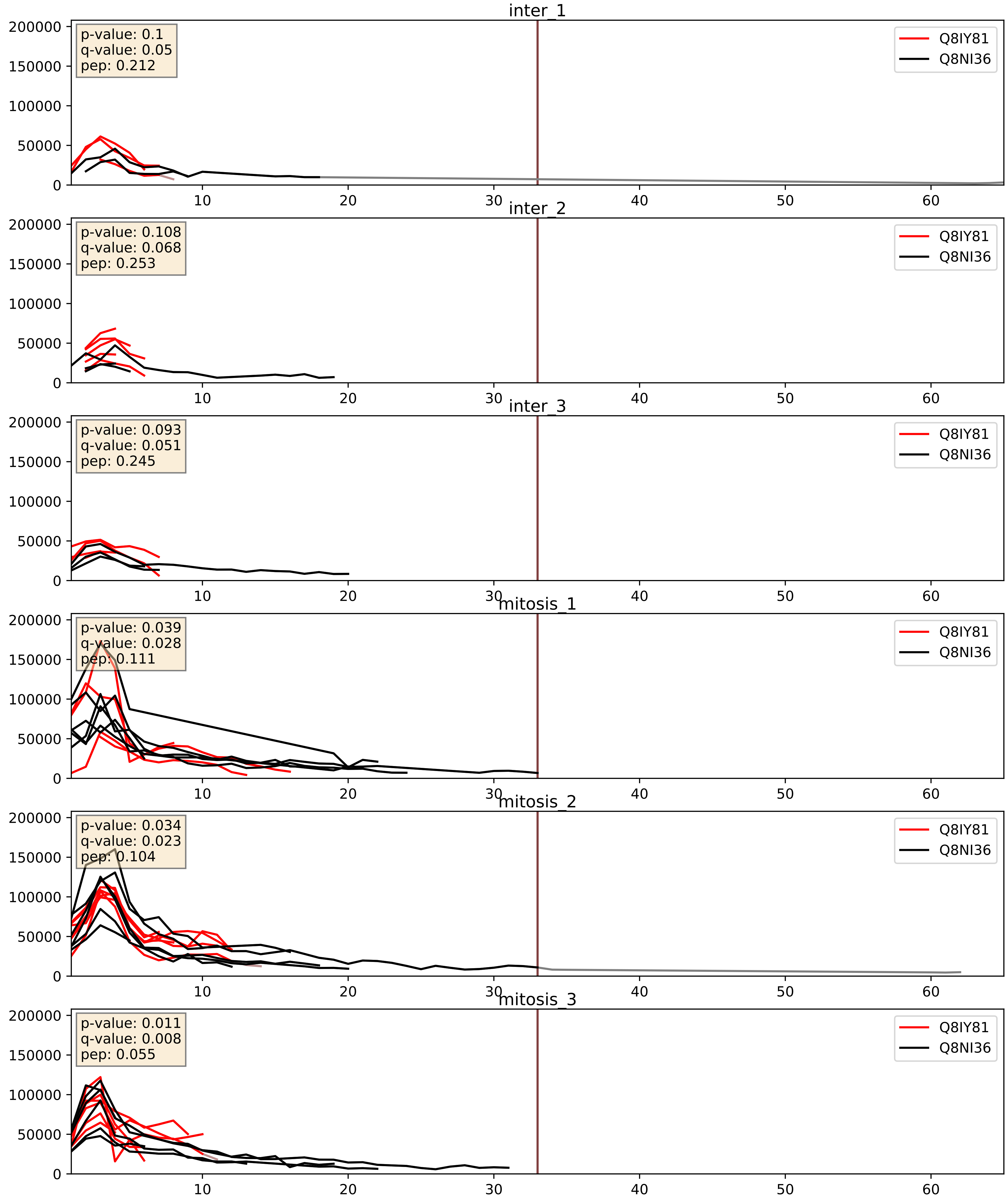


| level | condition_1 | condition_2 | pvalue | pvalue_adjusted |
|---|---|---|---|---|
| complex_abundance | mitosis | inter | 0.07129833141638876 | 0.1941068522539015 |
| interactor_ratio | mitosis | inter | 0.09852732059893228 | 0.5220660934699379 |
| interactor_abundance | mitosis | inter | 0.36653218238088725 | 0.5624025973314919 |

Q8IY81_Q8TDN6
SPB1_HUMAN vs BRX1_HUMAN
p-value: 0.013 q-value: 0.011

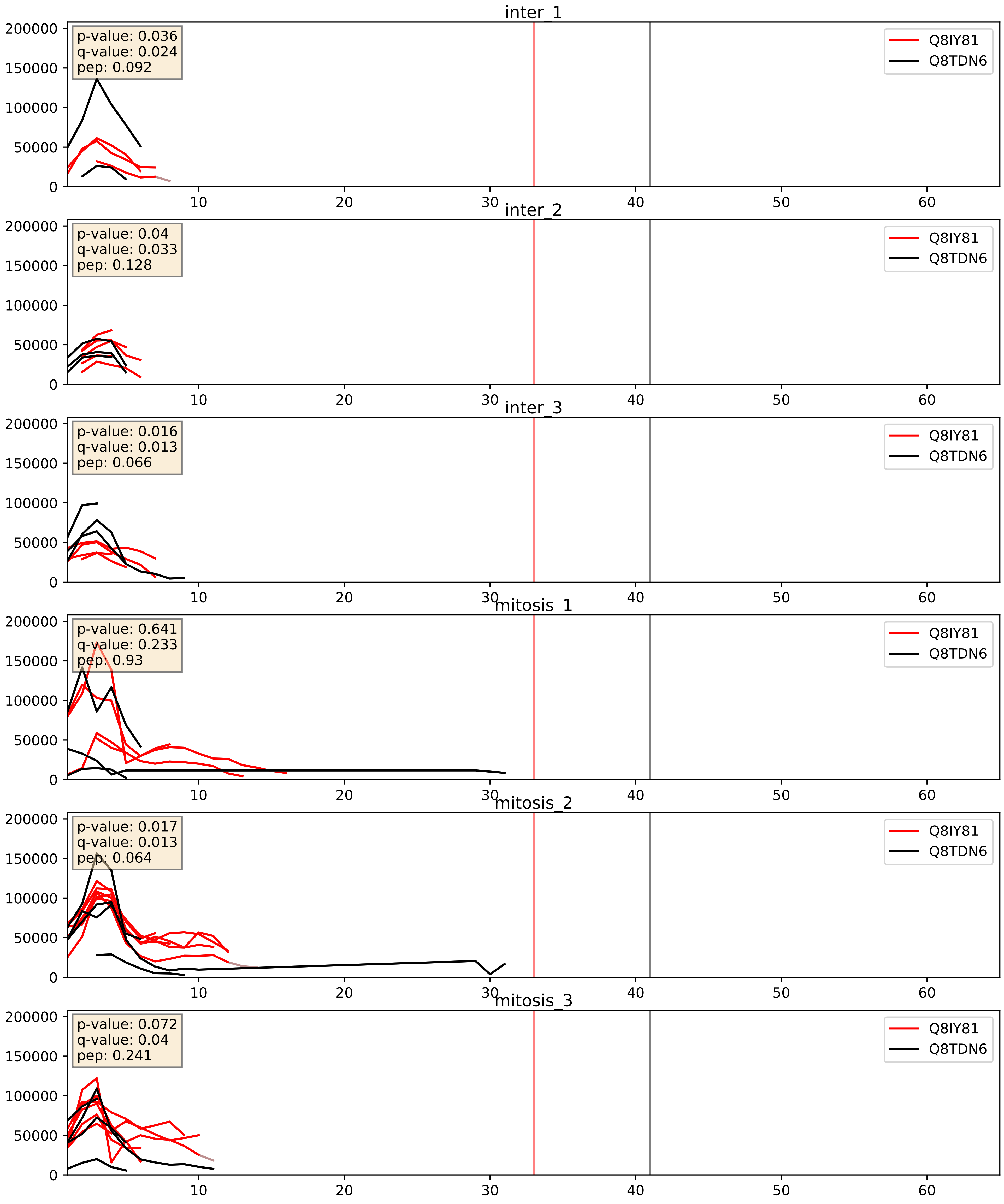


| level | condition_1 | condition_2 | pvalue | pvalue_adjusted |
|---|---|---|---|---|
| interactor_ratio | mitosis | inter | 0.05495294519905986 | 0.3887580255404565 |
| interactor_abundance | mitosis | inter | 0.5434694040854053 | 0.7001953791347185 |
| complex_abundance | mitosis | inter | 0.7217391892829187 | 0.814621236848864 |

Q8IY81_Q969X6
SPB1_HUMAN vs UTP4_HUMAN
p-value: 0.054 q-value: 0.035

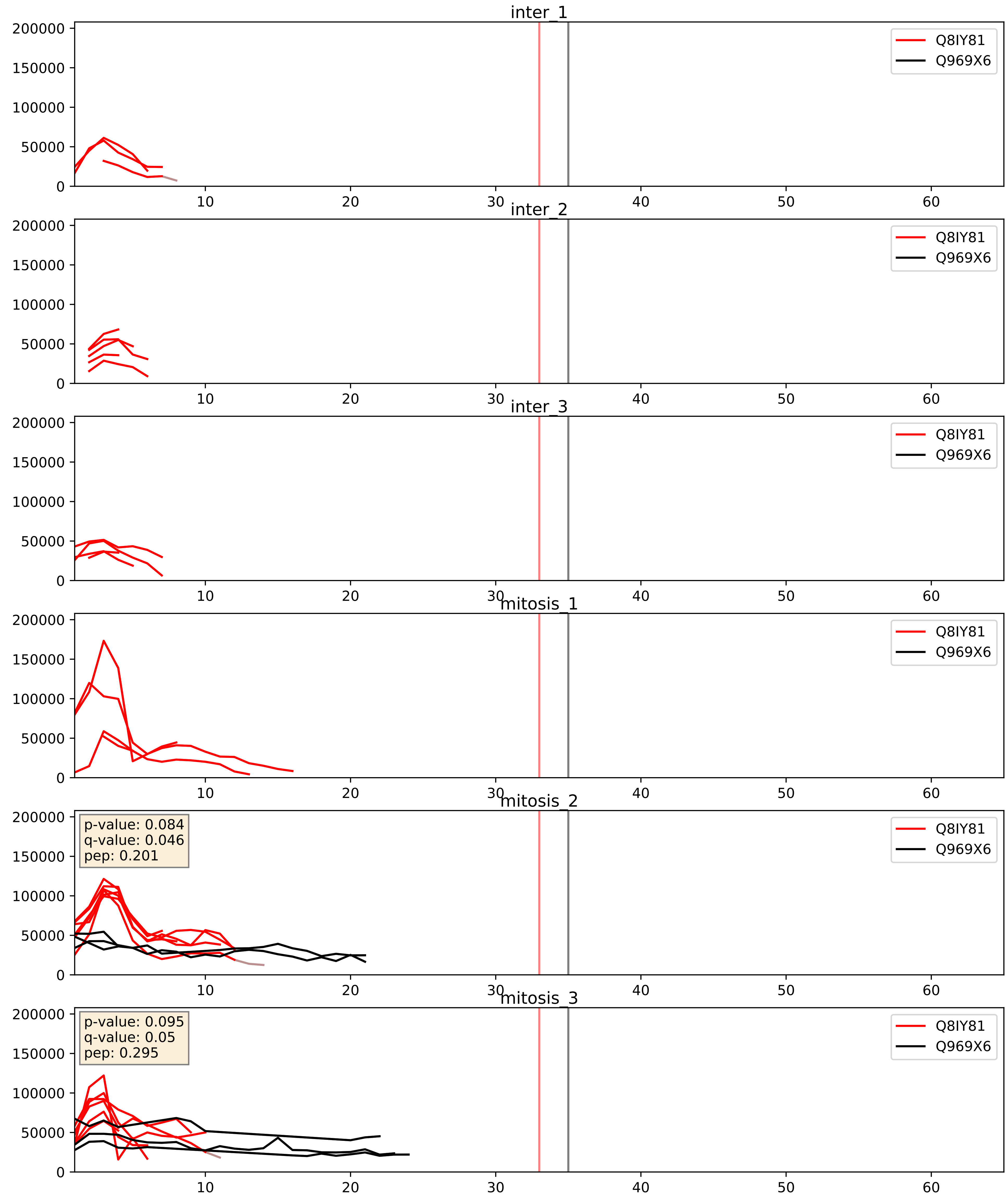


| level | condition_1 | condition_2 | pvalue | pvalue_adjusted |
|---|---|---|---|---|
| interactor_abundance | mitosis | inter | 0.24763076185556032 | 0.4410109895940074 |
| interactor_ratio | mitosis | inter | 0.26046486758792725 | 0.7338970594314211 |
| complex_abundance | mitosis | inter | 0.27894394715853627 | 0.4274224075967518 |

Q8IY81_Q96GQ7
SPB1_HUMAN vs DDX27_HUMAN
p-value: 0.012 q-value: 0.01

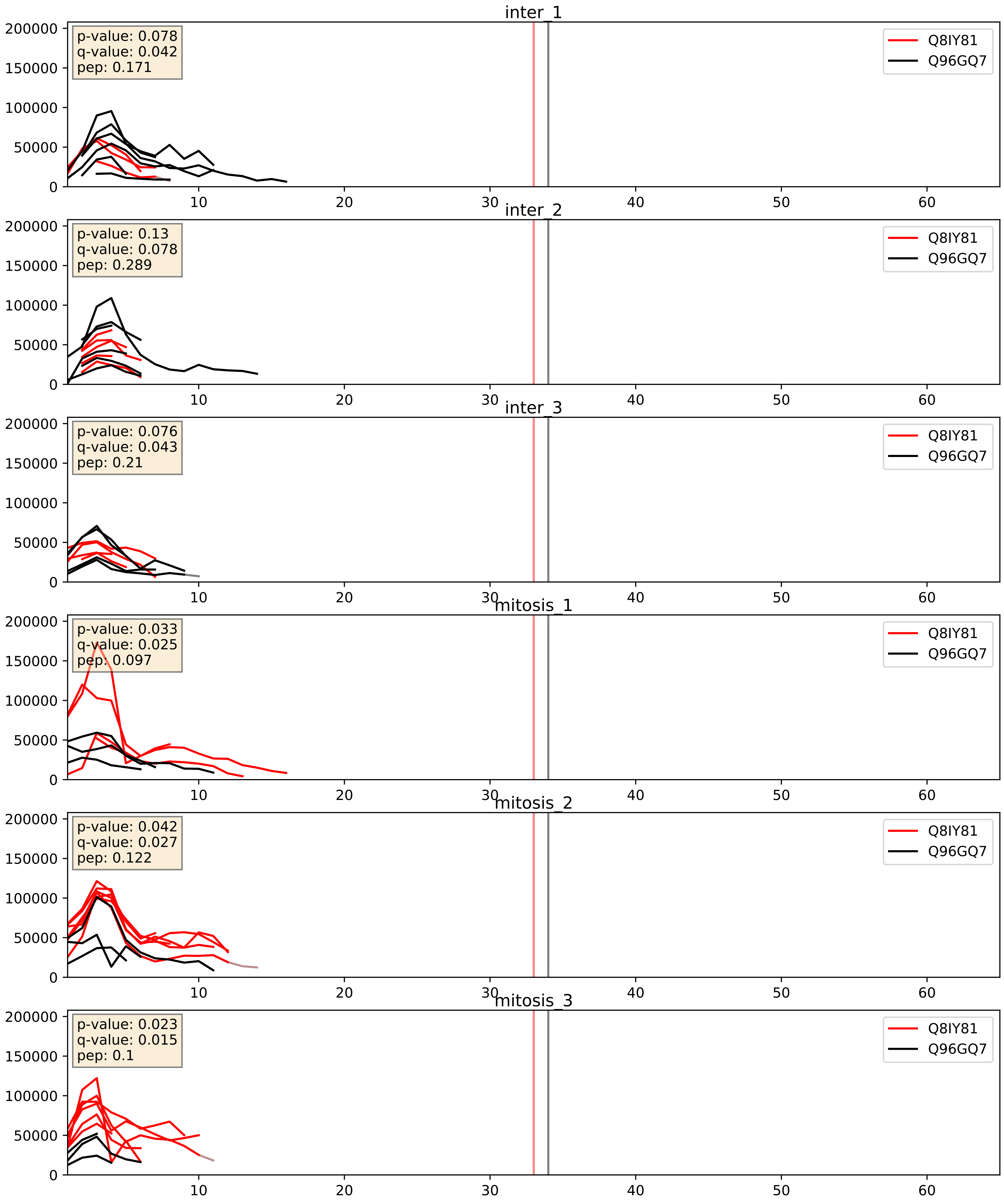


| level | condition_1 | condition_2 | pvalue | pvalue_adjusted |
|---|---|---|---|---|
| interactor_ratio | mitosis | inter | 0.0313990091864875 | 0.28316826136323564 |
| complex_abundance | mitosis | inter | 0.3682569777080367 | 0.515921395937937 |
| interactor_abundance | mitosis | inter | 0.5392572340101863 | 0.6967084632295215 |

Q8IY81_Q9BVP2
SPB1_HUMAN vs GNL3_HUMAN
p-value: 0.027 q-value: 0.02

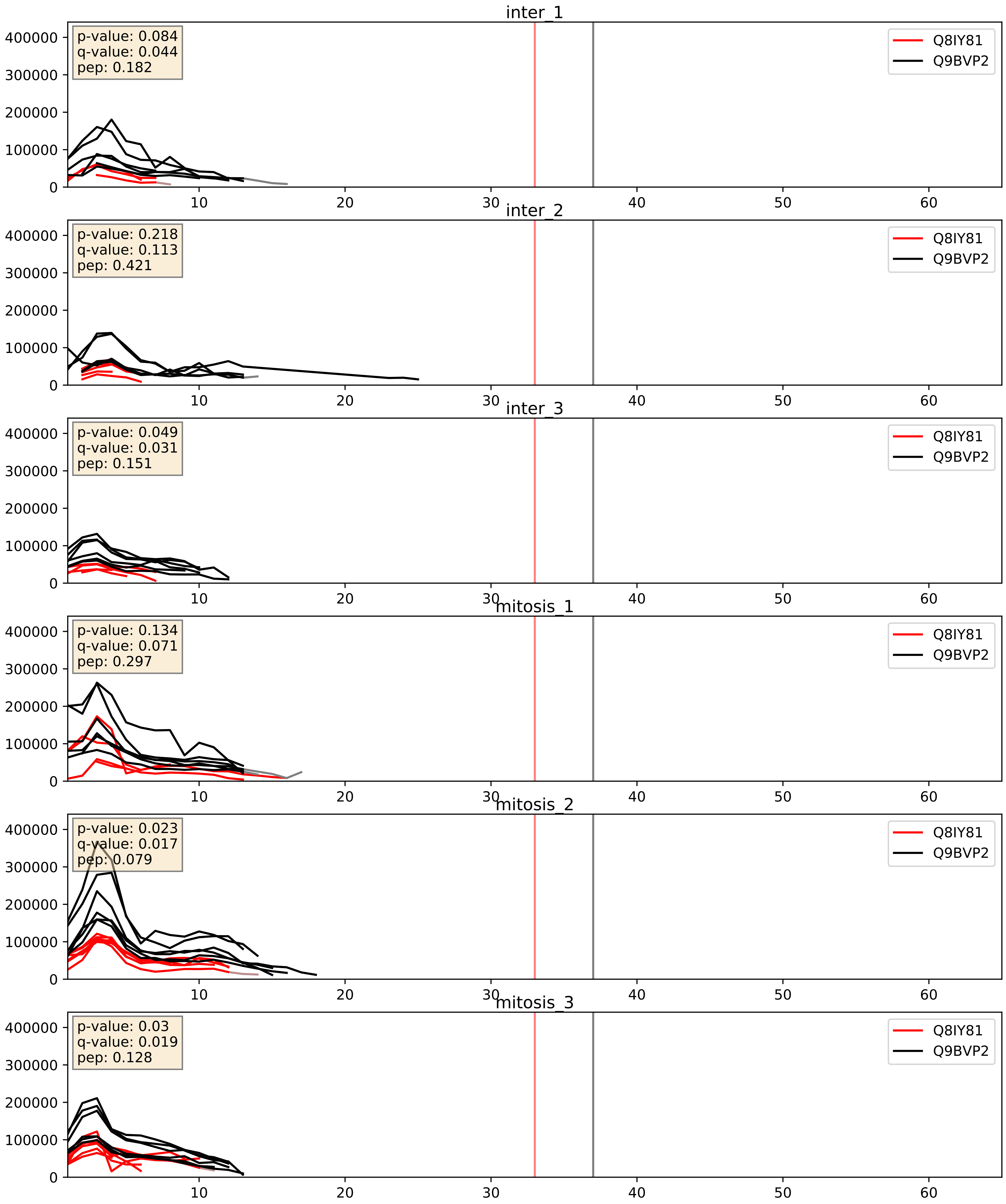


| level | condition_1 | condition_2 | pvalue | pvalue_adjusted |
|---|---|---|---|---|
| complex_abundance | mitosis | inter | 0.04433993120036613 | 0.15060165151763358 |
| interactor_ratio | mitosis | inter | 0.07249910086746936 | 0.44970456769966505 |
| interactor_abundance | mitosis | inter | 0.3789887745019212 | 0.5719072560134765 |

Q8IY81_Q9BZE4
SPB1_HUMAN vs NOG1_HUMAN
p-value: 0.005 q-value: 0.005

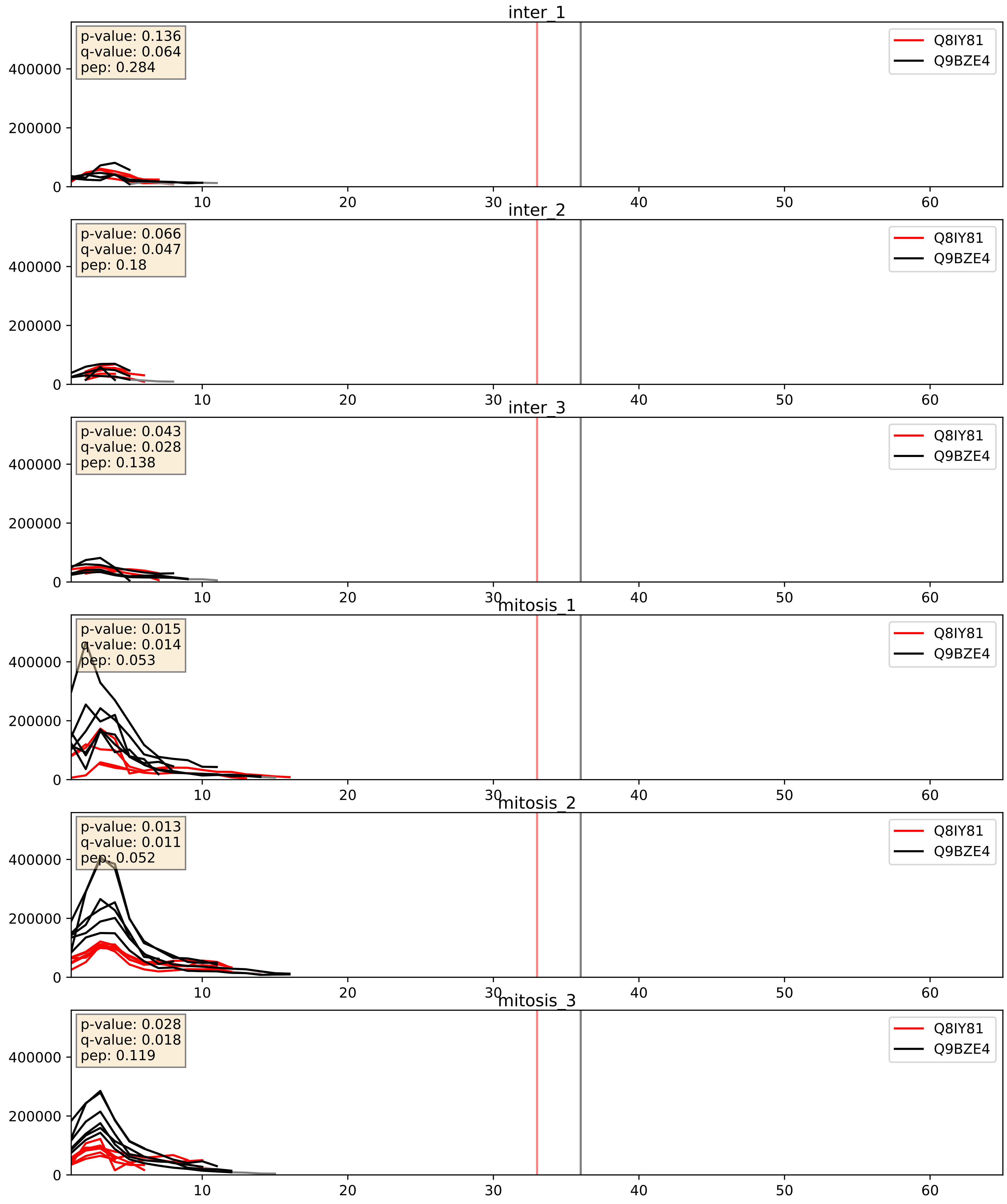


| level | condition_1 | condition_2 | pvalue | pvalue_adjusted |
|---|---|---|---|---|
| interactor_ratio | mitosis | inter | 0.0007974295630027685 | 0.03250474790144618 |
| complex_abundance | mitosis | inter | 0.012254193945439154 | 0.0822068183173661 |
| interactor_abundance | mitosis | inter | 0.33114478037662615 | 0.5307908853629082 |

Q8IY81_Q9H0S4
SPB1_HUMAN vs DDX47_HUMAN
p-value: 0.065 q-value: 0.04

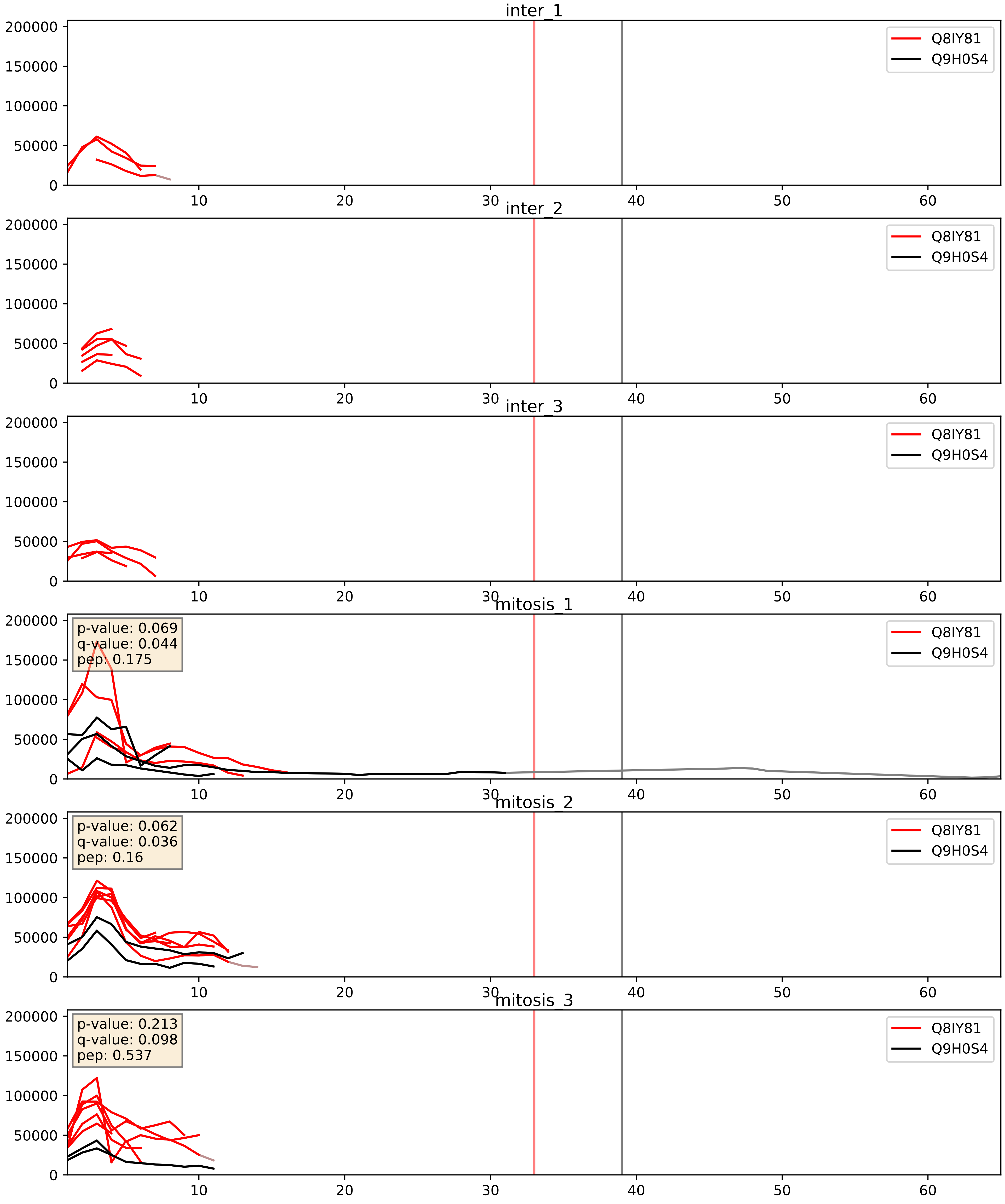


| level | condition_1 | condition_2 | pvalue | pvalue_adjusted |
|---|---|---|---|---|
| interactor_abundance | mitosis | inter | 0.0056588368861371875 | 0.06348577161904892 |
| complex_abundance | mitosis | inter | 0.03774864580132628 | 0.14178517247009784 |
| interactor_ratio | mitosis | inter | 0.10205027280289247 | 0.5320038582172714 |

Q8IY81_Q9H6R4
SPB1_HUMAN vs NOL6_HUMAN
p-value: 0.063 q-value: 0.039

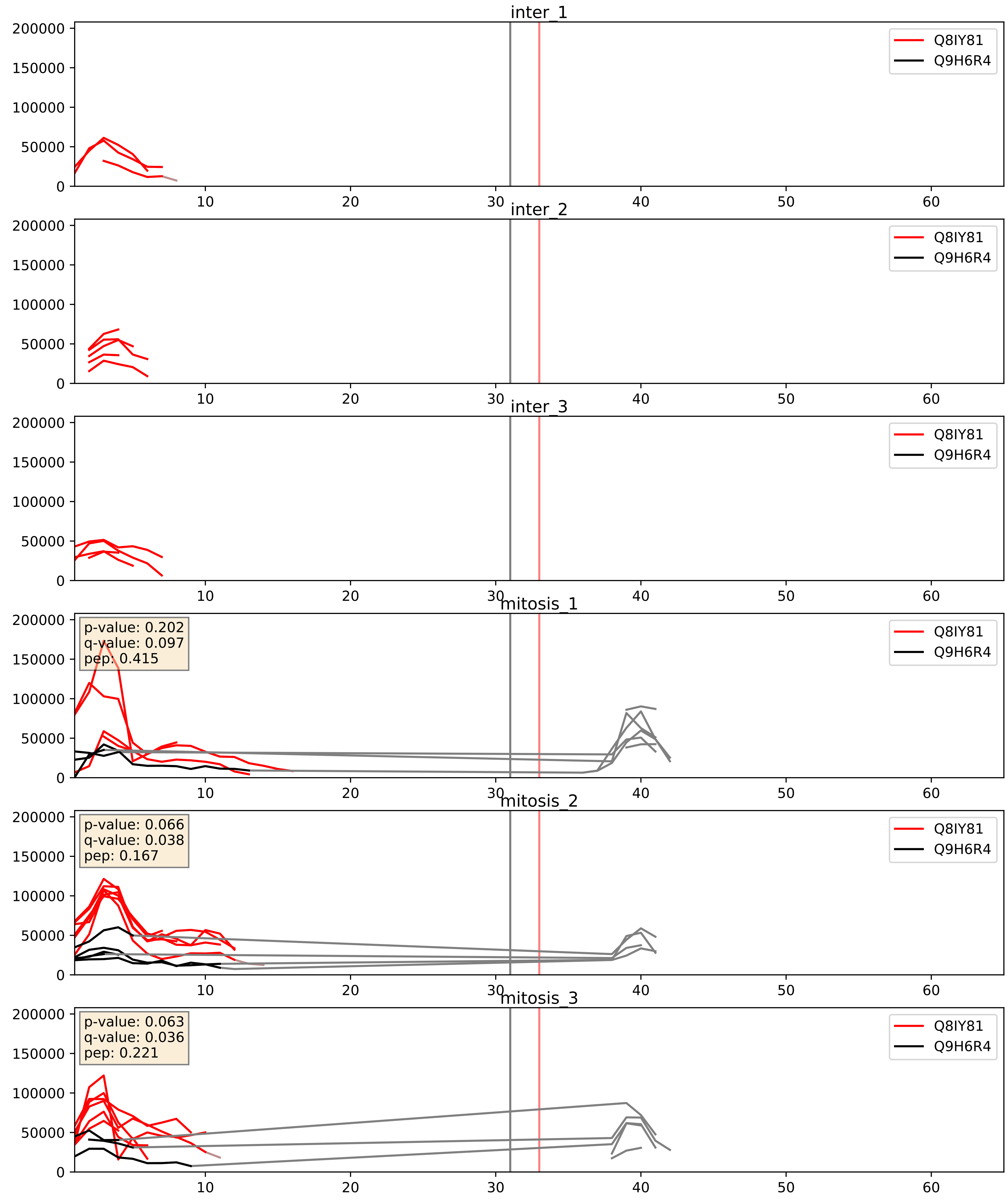


| level | condition_1 | condition_2 | pvalue | pvalue_adjusted |
|---|---|---|---|---|
| interactor_abundance | mitosis | inter | 0.006105255729208133 | 0.0666372269553884 |
| complex_abundance | mitosis | inter | 0.015707262428466266 | 0.09352166546417016 |
| interactor_ratio | mitosis | inter | 0.026384193828163788 | 0.25543205403291264 |

Q8IY81_Q9H7B2
SPB1_HUMAN vs RPF2_HUMAN
p-value: 0.059 q-value: 0.037

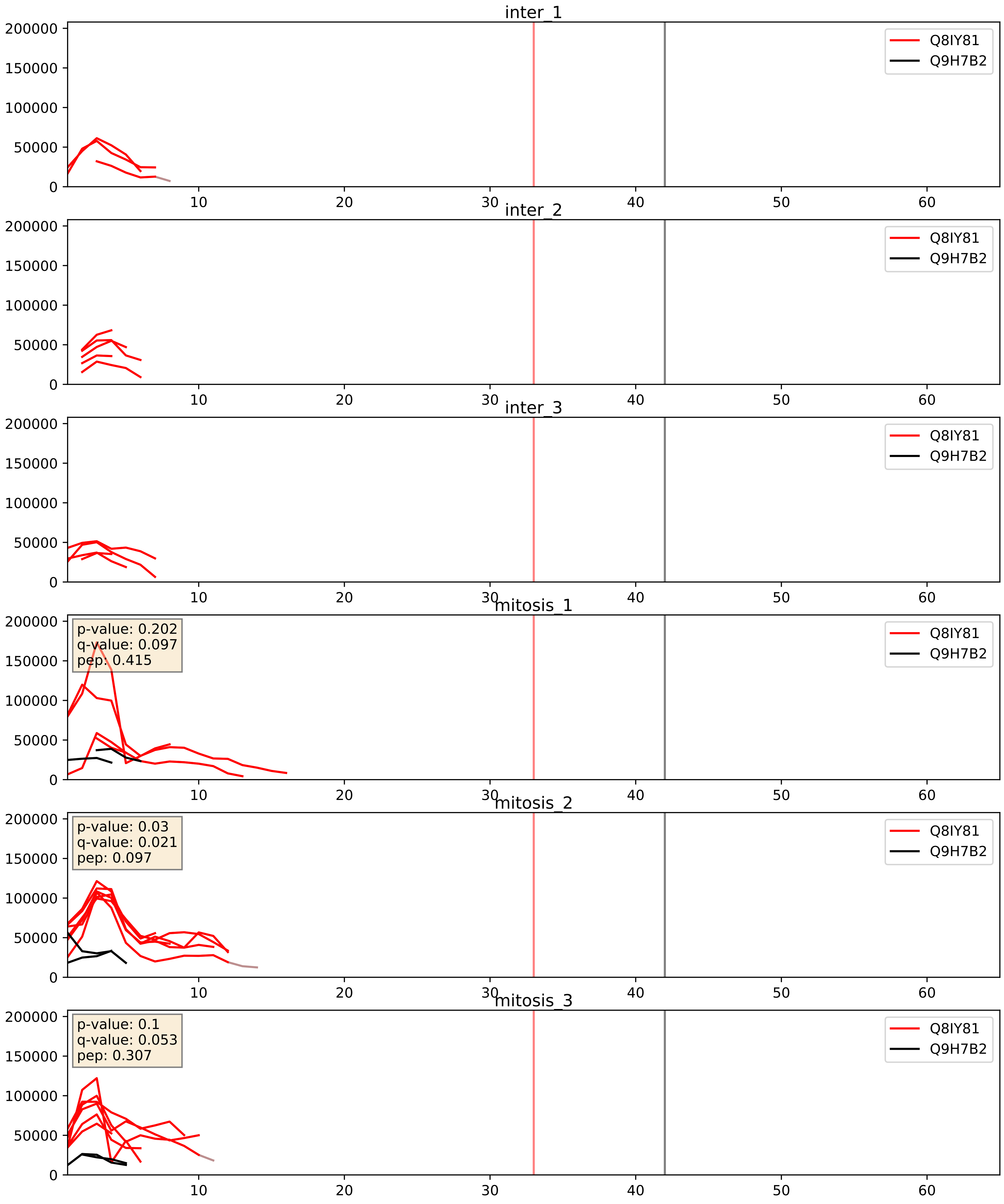


| level | condition_1 | condition_2 | pvalue | pvalue_adjusted |
|---|---|---|---|---|
| interactor_ratio | mitosis | inter | 0.006296065976282134 | 0.10931911715410765 |
| interactor_abundance | mitosis | inter | 0.00774173526098488 | 0.07639107070205253 |
| complex_abundance | mitosis | inter | 0.03990807958361974 | 0.1449976963395235 |

Q8IY81_Q9UKD2
SPB1_HUMAN vs MRT4_HUMAN
p-value: 0.024 q-value: 0.018

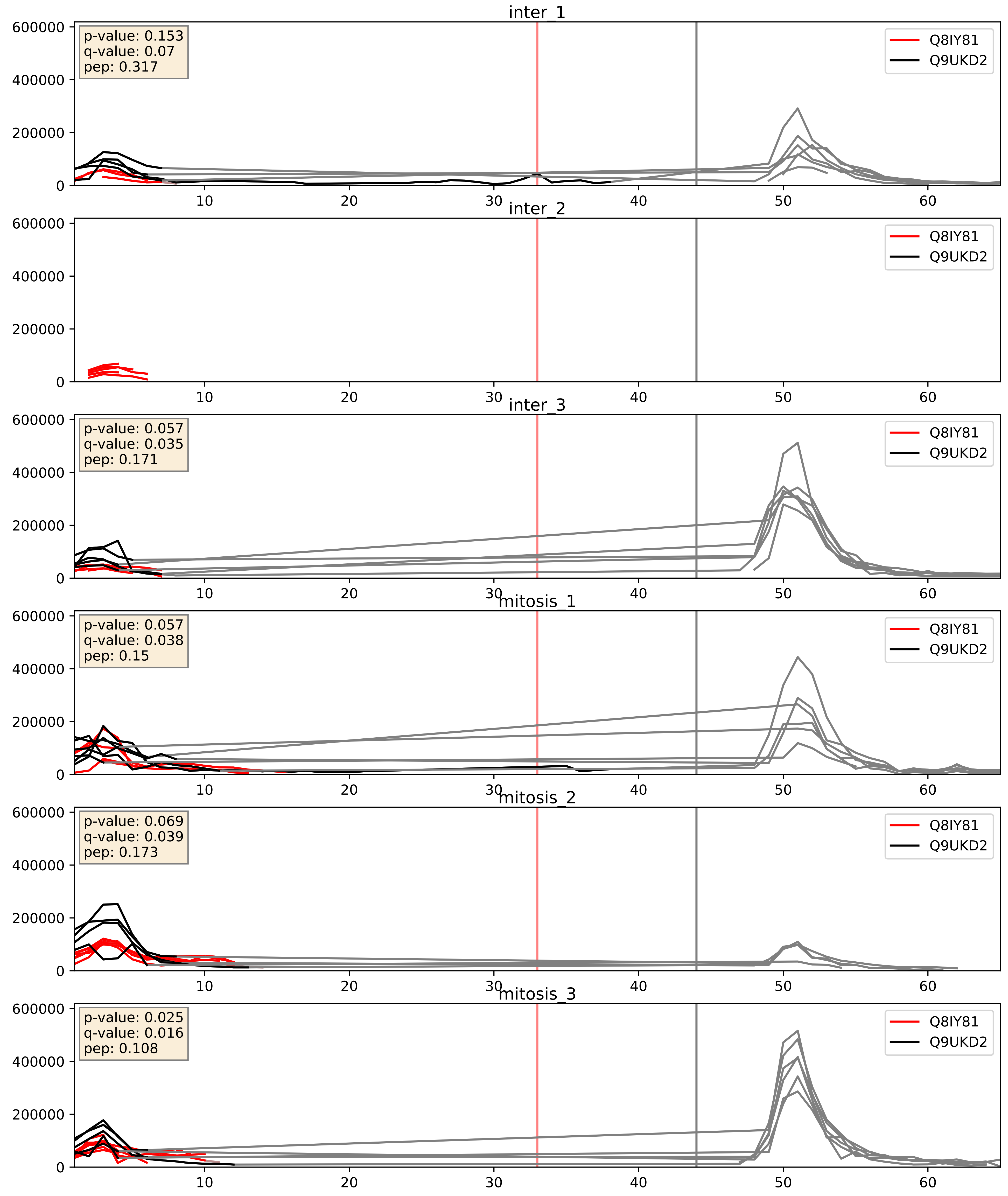


| level | condition_1 | condition_2 | pvalue | pvalue_adjusted |
|---|---|---|---|---|
| interactor_abundance | mitosis | inter | 0.10319084594938516 | 0.2756657235420093 |
| complex_abundance | mitosis | inter | 0.1365404073747575 | 0.28185013765792694 |
| interactor_ratio | mitosis | inter | 0.5766472200424113 | 0.8958439570894811 |

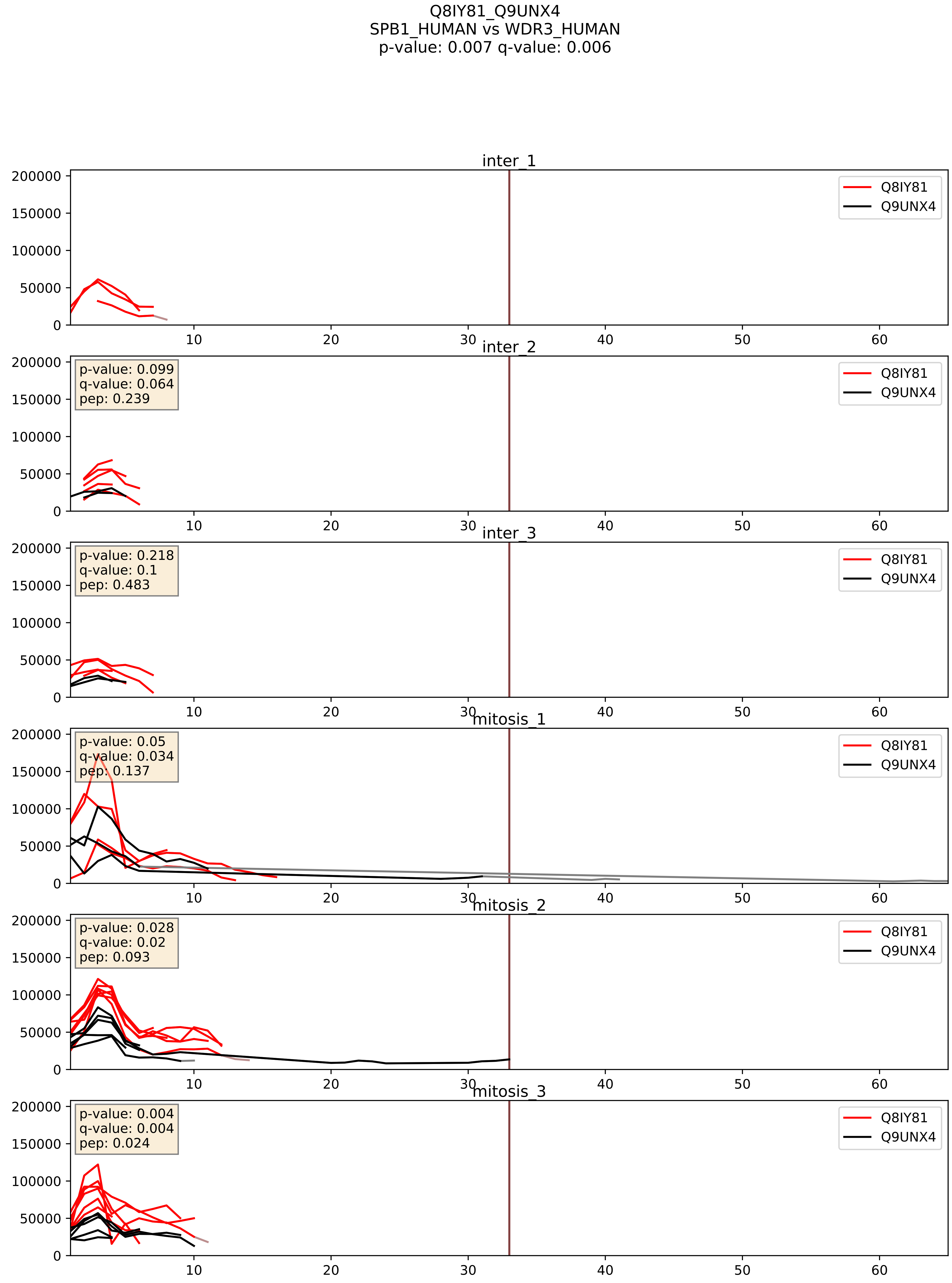


| level | condition_1 | condition_2 | pvalue | pvalue_adjusted |
|---|---|---|---|---|
| complex_abundance | mitosis | inter | 0.1603807662778196 | 0.30804508691617344 |
| interactor_abundance | mitosis | inter | 0.20267537112676712 | 0.39818709590202583 |
| interactor_ratio | mitosis | inter | 0.6731573851839978 | 0.9319765113016878 |

Q8IY81_Q9Y221
SPB1_HUMAN vs NIP7_HUMAN
p-value: 0.012 q-value: 0.01

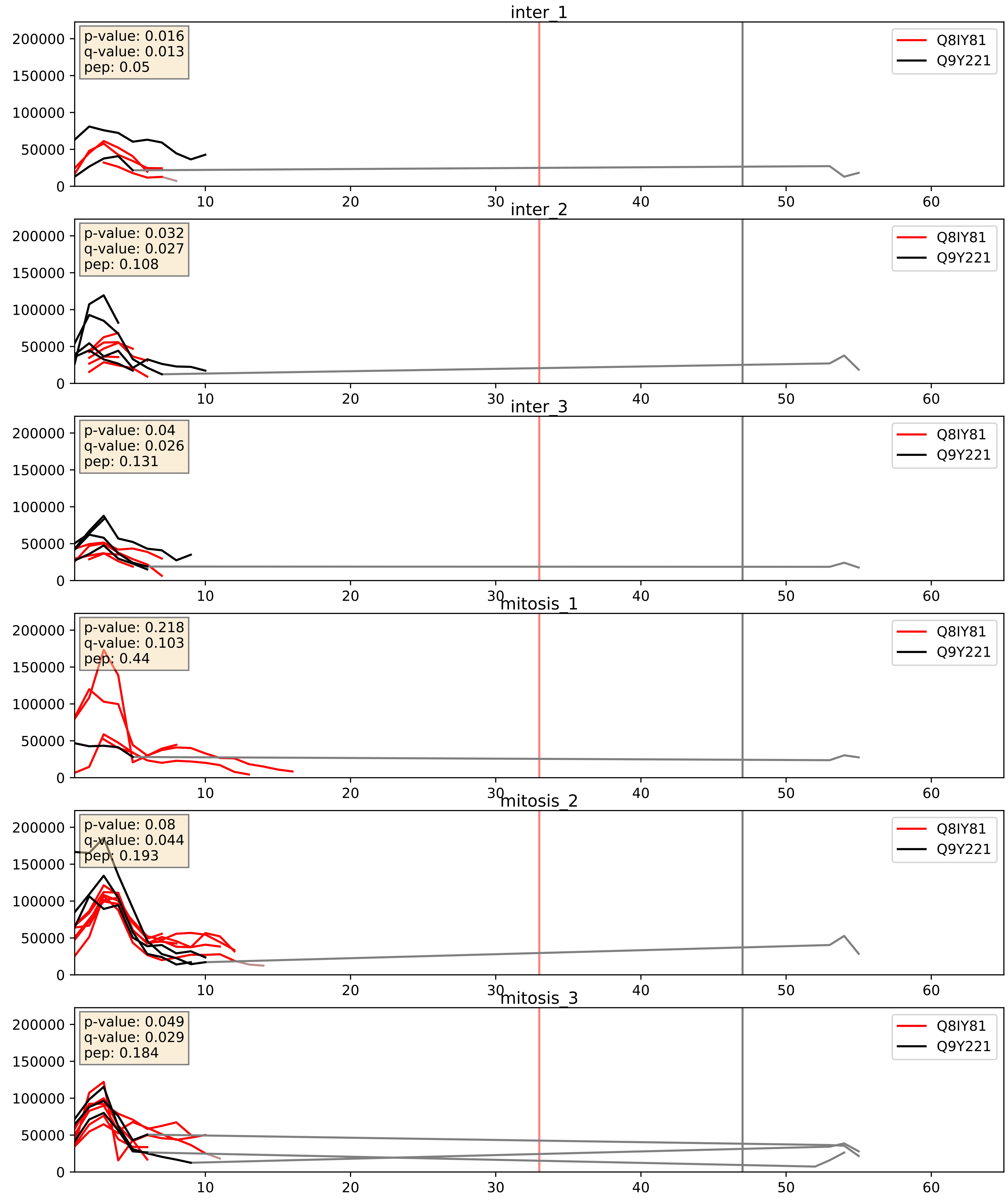


| level | condition_1 | condition_2 | pvalue | pvalue_adjusted |
|---|---|---|---|---|
| interactor_ratio | mitosis | inter | 0.22386253191740801 | 0.7026891317077497 |
| interactor_abundance | mitosis | inter | 0.6286537721490107 | 0.7560627031395197 |
| complex_abundance | mitosis | inter | 0.8269896457348944 | 0.8884326515425071 |

Q8IY81_Q9Y2X3
SPB1_HUMAN vs NOP58_HUMAN
p-value: 0.063 q-value: 0.039

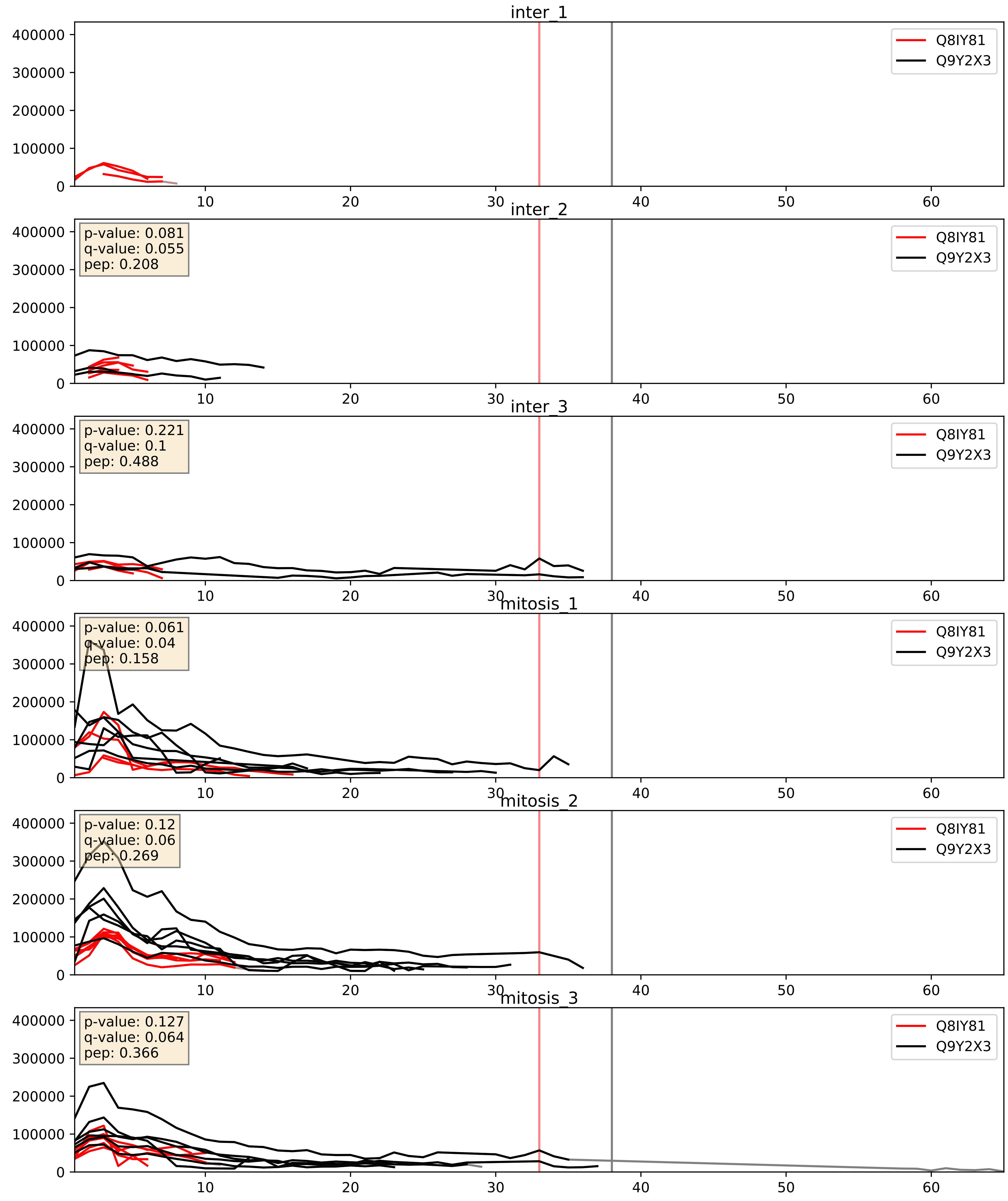


| level | condition_1 | condition_2 | pvalue | pvalue_adjusted |
|---|---|---|---|---|
| interactor_ratio | mitosis | inter | 0.0019003411607799942 | 0.052986711193083874 |
| complex_abundance | mitosis | inter | 0.01284951872274085 | 0.08405635484106207 |
| interactor_abundance | mitosis | inter | 0.18344633002469837 | 0.3757436970203683 |

Q8IY81_Q9Y5J1
SPB1_HUMAN vs UTP18_HUMAN
p-value: 0.076 q-value: 0.046

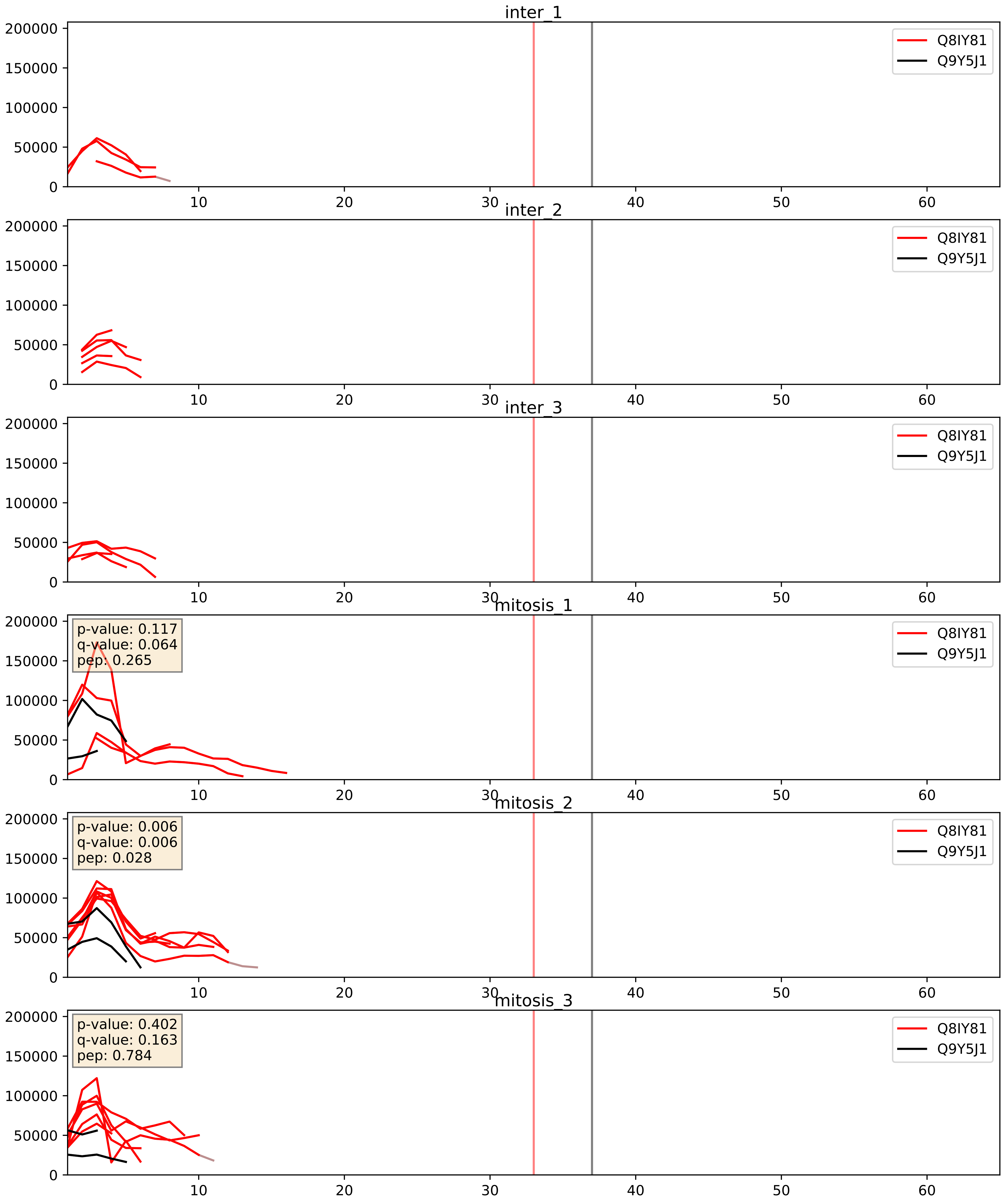


| level | condition_1 | condition_2 | pvalue | pvalue_adjusted |
|---|---|---|---|---|
| interactor_abundance | mitosis | inter | 0.008355526262714321 | 0.07951451340615297 |
| complex_abundance | mitosis | inter | 0.023105806231831845 | 0.11091884921279084 |
| interactor_ratio | mitosis | inter | 0.0241747810665698 | 0.24345426579980883 |
