## Supplementary Data 2 for "SECAT: Quantifying differential protein-protein interaction states by network-centric analysis": hela_string_000025_P62829.pdf

P62829_P62829

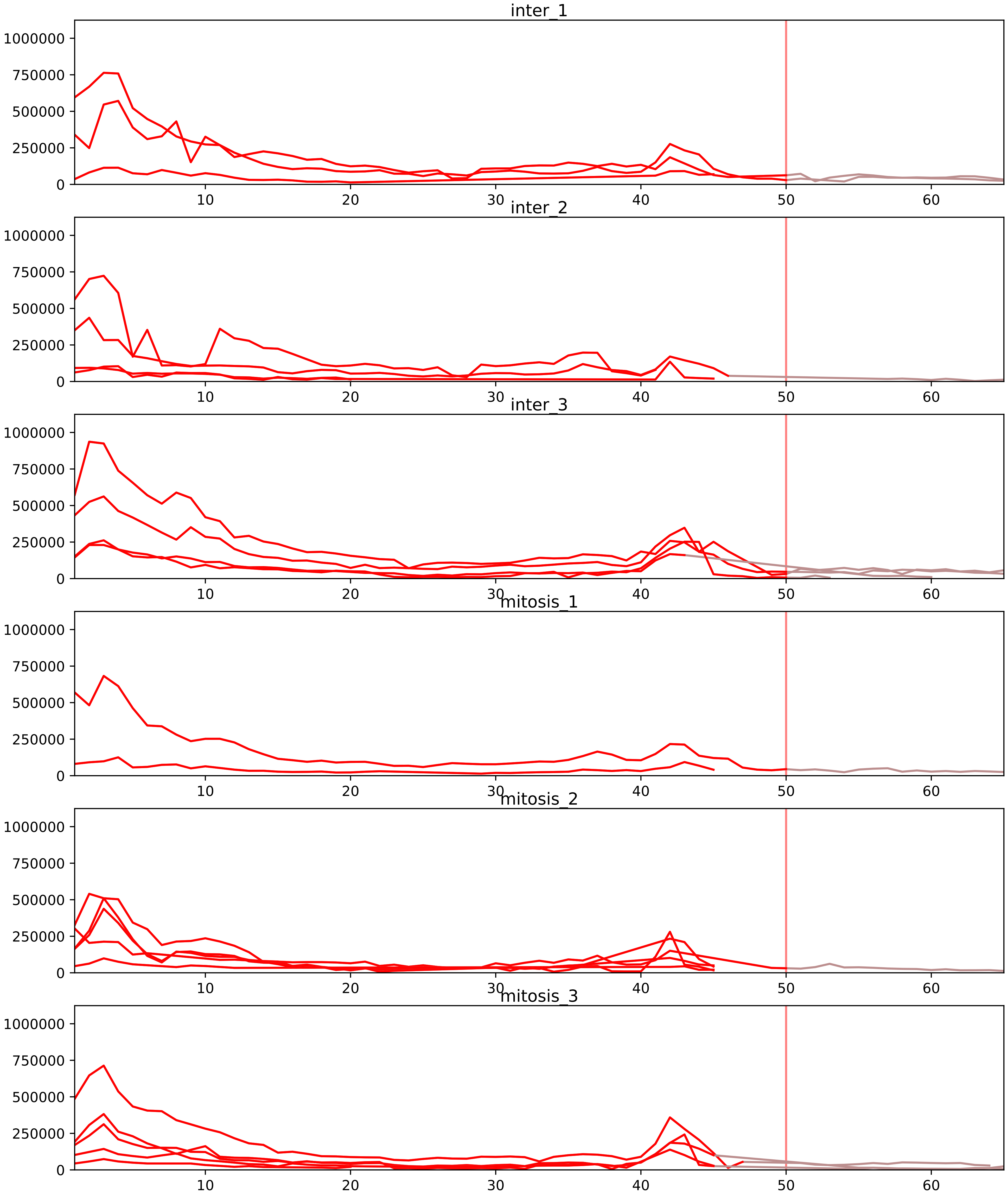


| level | condition_1 | condition_2 | pvalue | pvalue_adjusted |
|---|---|---|---|---|
| interactor_ratio | mitosis | inter | 5.250348574488934e-11 | 3.864256550823855e-09 |
| interactor_abundance | mitosis | inter | 0.0001418965315140261 | 0.002948200227185908 |
| complex_abundance | mitosis | inter | 0.0003664298431206481 | 0.005665805977663803 |
| monomer_abundance | mitosis | inter | 0.2289641714309316 | 0.6048643878568759 |
| total_abundance | mitosis | inter | 0.2859655754694843 | 0.617934662839628 |
| assembled_abundance | mitosis | inter | 0.2925918092240996 | 0.5895920185010807 |

O76021_P62829
RL1D1_HUMAN vs RL23_HUMAN
p-value: 0.0 q-value: 0.041

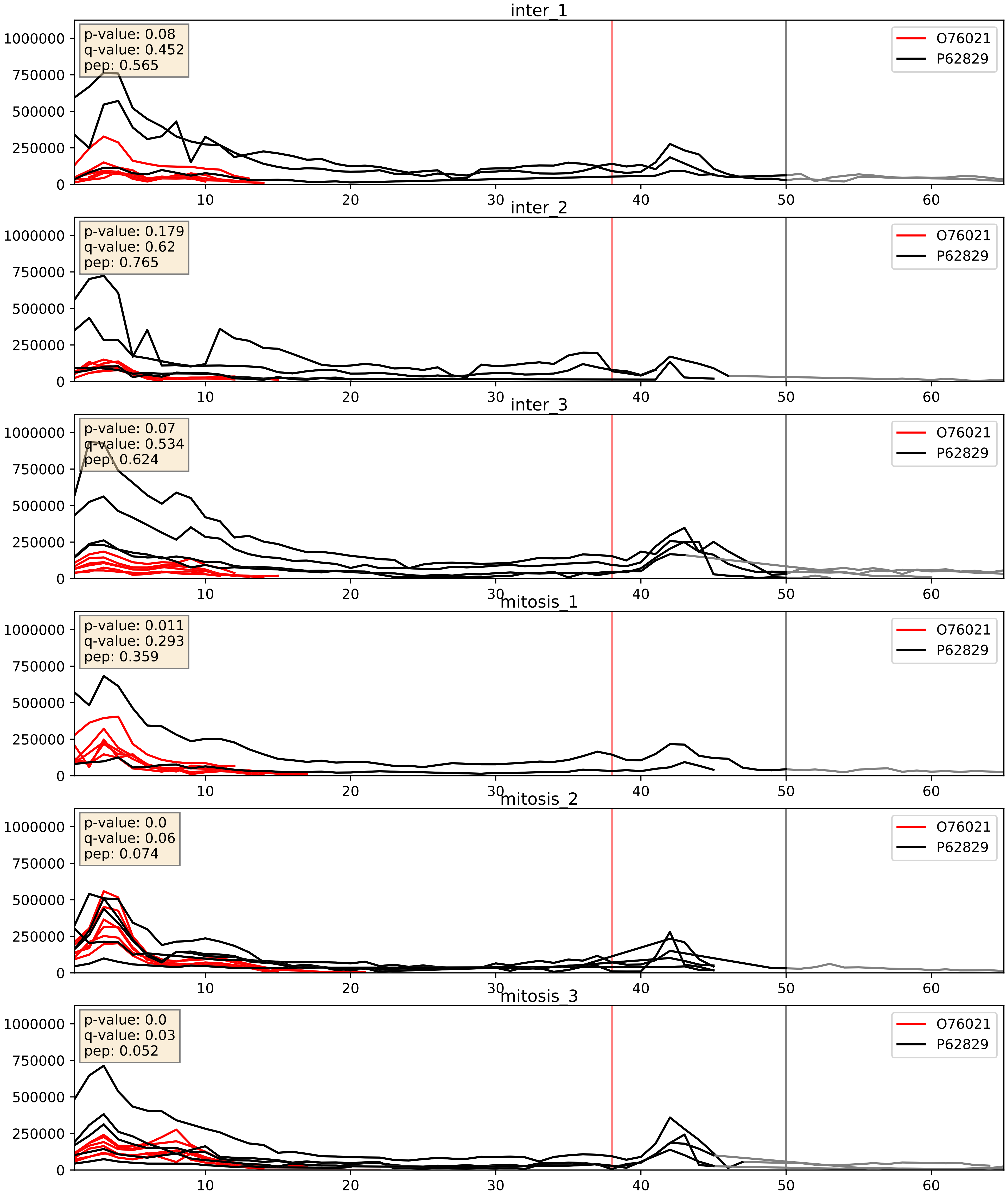


| level | condition_1 | condition_2 | pvalue | pvalue_adjusted |
|---|---|---|---|---|
| interactor_ratio | mitosis | inter | 0.0003092702819777167 | 0.018775557544179113 |
| interactor_abundance | mitosis | inter | 0.007302544007558834 | 0.07362753439894419 |
| complex_abundance | mitosis | inter | 0.086333506104879 | 0.2153229384262426 |

P04843_P62829
RPN1_HUMAN vs RL23_HUMAN
p-value: 0.03 q-value: 0.022

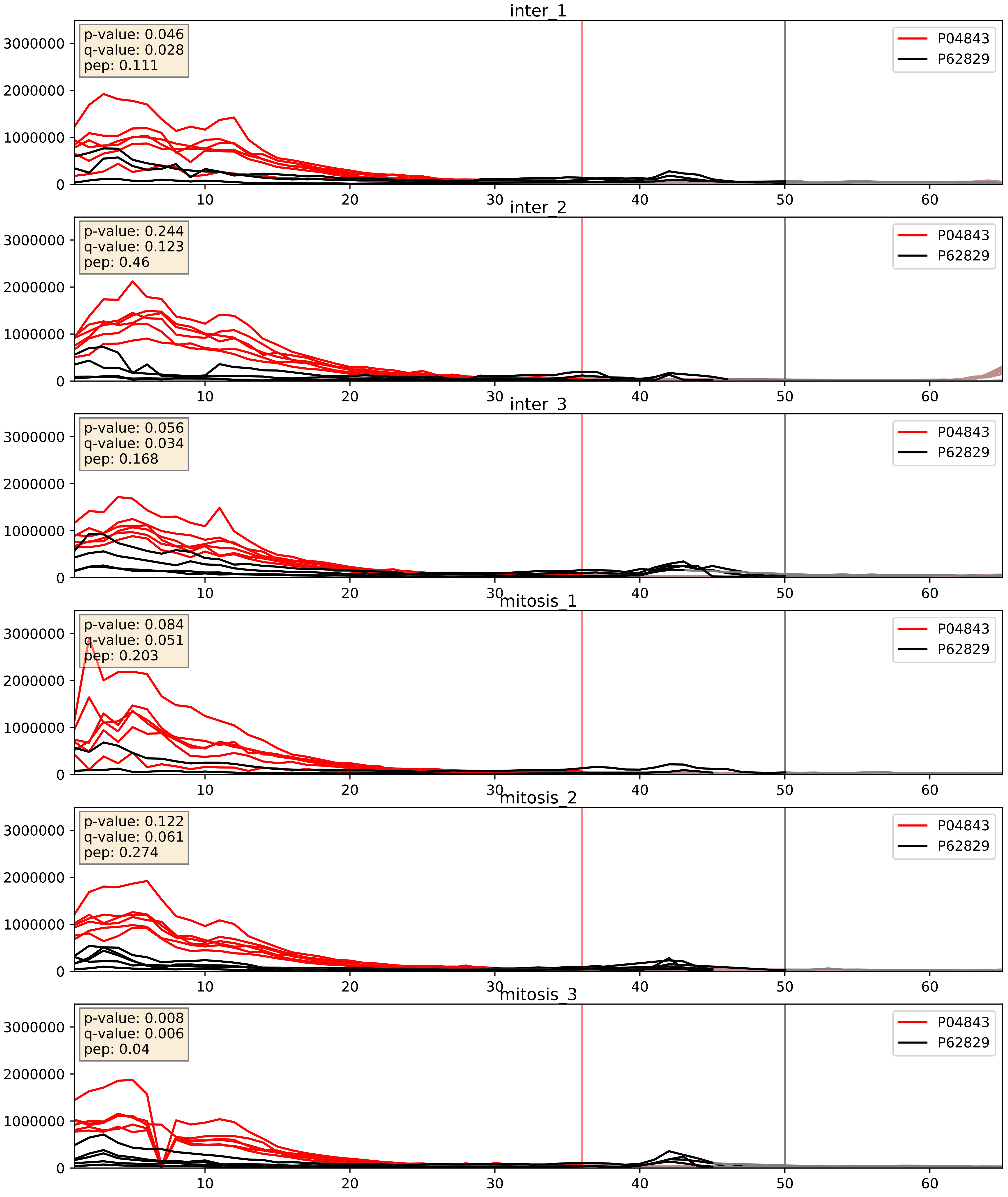


| level | condition_1 | condition_2 | pvalue | pvalue_adjusted |
|---|---|---|---|---|
| complex_abundance | mitosis | inter | 0.006841995247159383 | 0.061198487674337915 |
| interactor_ratio | mitosis | inter | 0.027492846179584727 | 0.2594694192913399 |
| interactor_abundance | mitosis | inter | 0.055512062499012145 | 0.19106684961461357 |

P05387_P62829
RLA2_HUMAN vs RL23_HUMAN
p-value: 0.001 q-value: 0.002

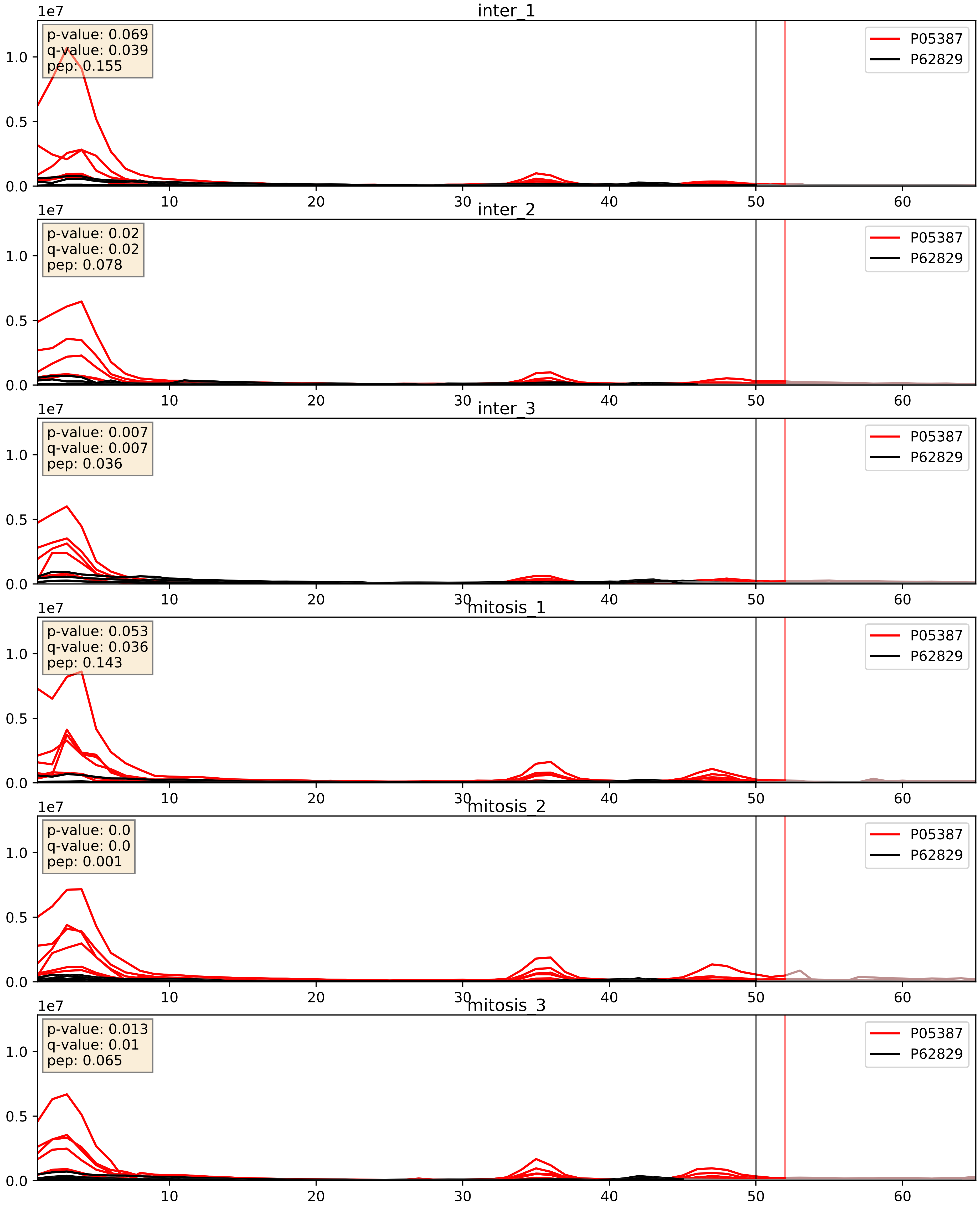


| level | condition_1 | condition_2 | pvalue | pvalue_adjusted |
|---|---|---|---|---|
| interactor_ratio | mitosis | inter | 0.001422033883744837 | 0.04475224281196987 |
| complex_abundance | mitosis | inter | 0.0409331697420589 | 0.1455676624066993 |
| interactor_abundance | mitosis | inter | 0.8921348191716616 | 0.9395514335764547 |

P05388_P62829
RLA0_HUMAN vs RL23_HUMAN
p-value: 0.0 q-value: 0.0

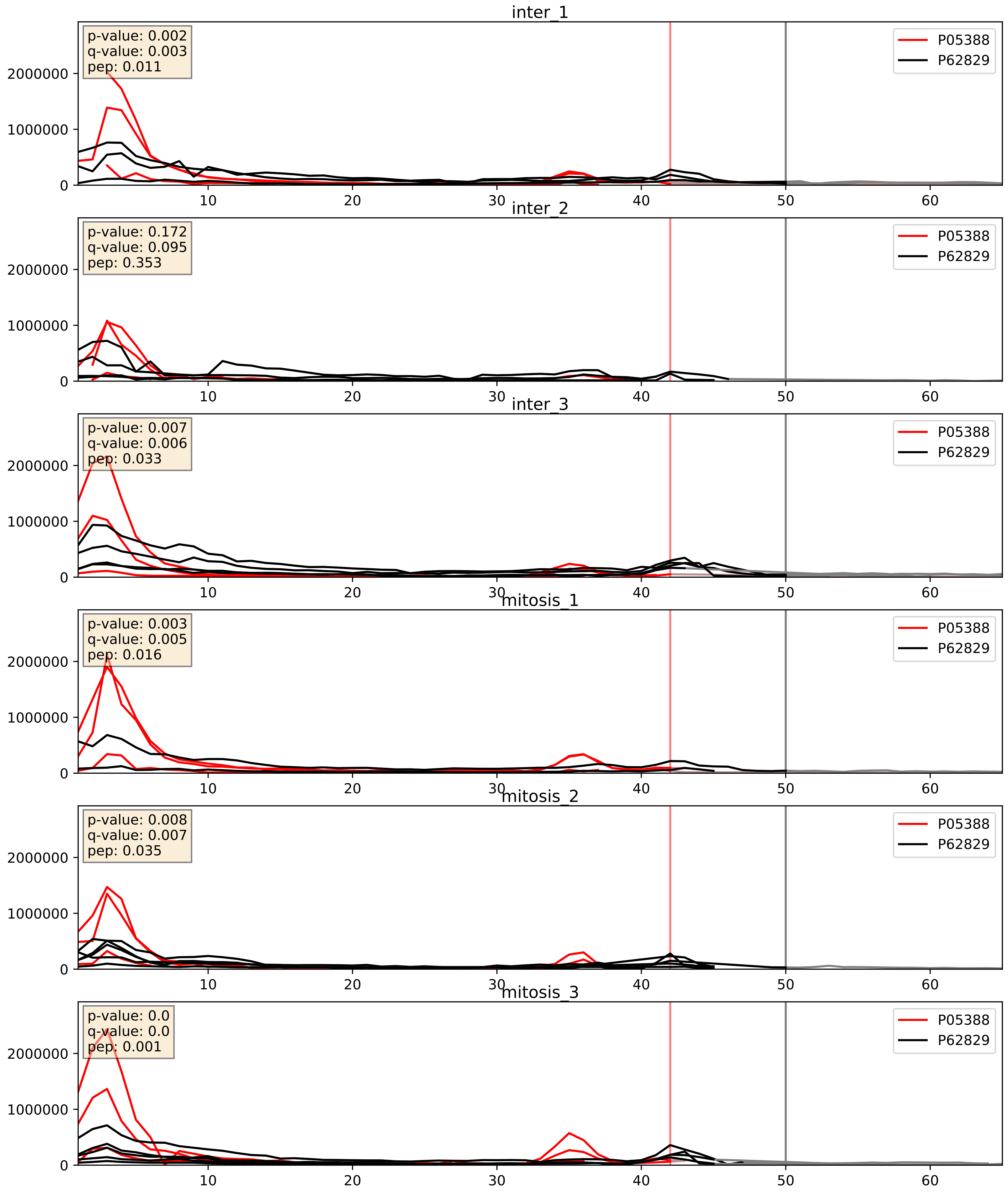


| level | condition_1 | condition_2 | pvalue | pvalue_adjusted |
|---|---|---|---|---|
| interactor_ratio | mitosis | inter | 0.004713933671340788 | 0.09159088101537749 |
| complex_abundance | mitosis | inter | 0.0993722696094373 | 0.23389445840390383 |
| interactor_abundance | mitosis | inter | 0.5648057672917774 | 0.712615132731611 |

P08708_P62829
RS17_HUMAN vs RL23_HUMAN
p-value: 0.0 q-value: 0.0

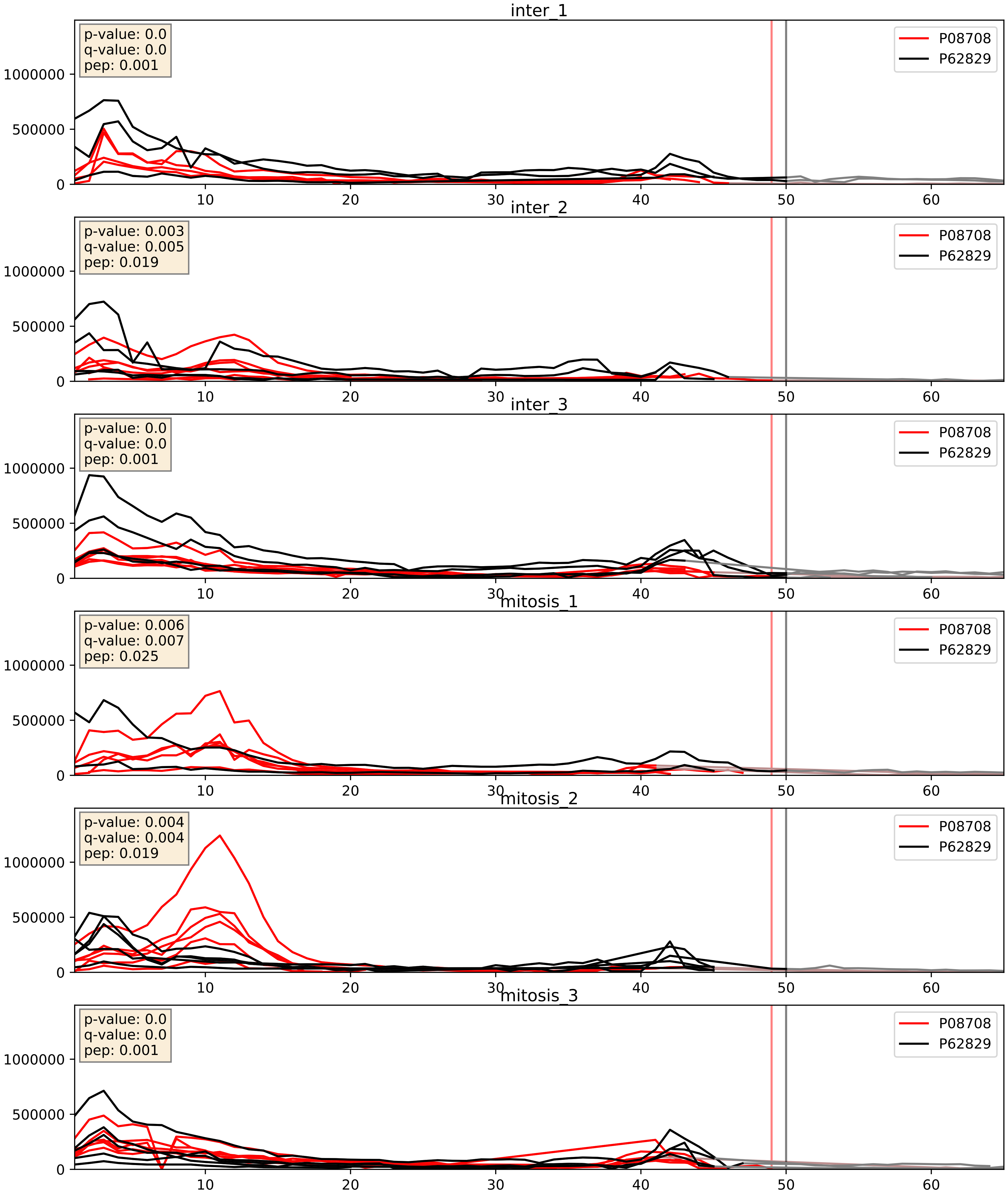


| level | condition_1 | condition_2 | pvalue | pvalue_adjusted |
|---|---|---|---|---|
| interactor_ratio | mitosis | inter | 0.008564159722841082 | 0.13203195690147984 |
| complex_abundance | mitosis | inter | 0.011934790502275522 | 0.08114978643416496 |
| interactor_abundance | mitosis | inter | 0.03809673850378837 | 0.15907711297191632 |

P11940_P62829
PABP1_HUMAN vs RL23_HUMAN
p-value: 0.003 q-value: 0.039

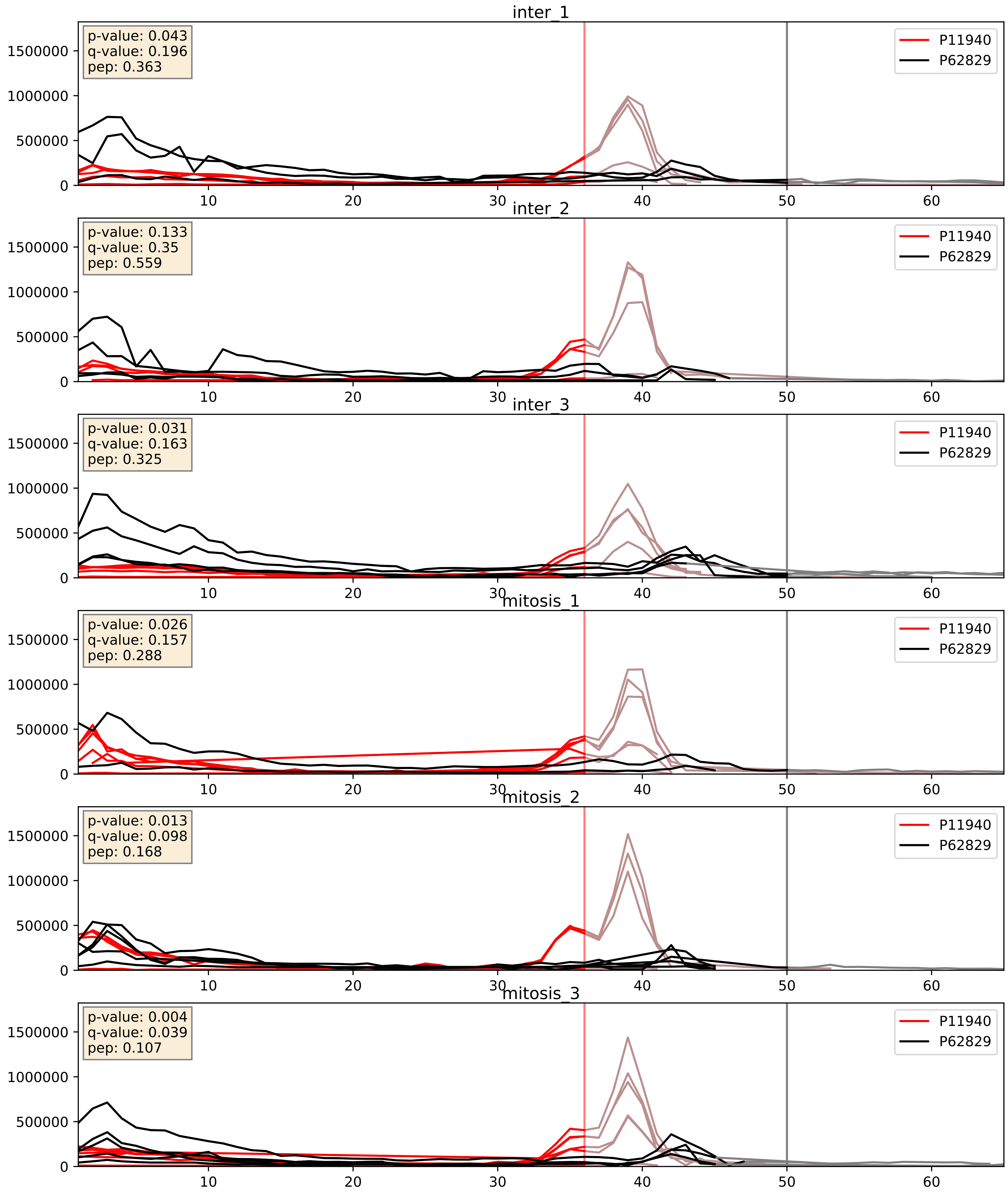


| level | condition_1 | condition_2 | pvalue | pvalue_adjusted |
|---|---|---|---|---|
| complex_abundance | mitosis | inter | 0.00659781688147954 | 0.06001839798667892 |
| interactor_ratio | mitosis | inter | 0.007455063952191203 | 0.12063392709027732 |
| interactor_abundance | mitosis | inter | 0.15090604372175434 | 0.33732834796845007 |

P15880_P62829
RS2_HUMAN vs RL23_HUMAN
p-value: 0.0 q-value: 0.0

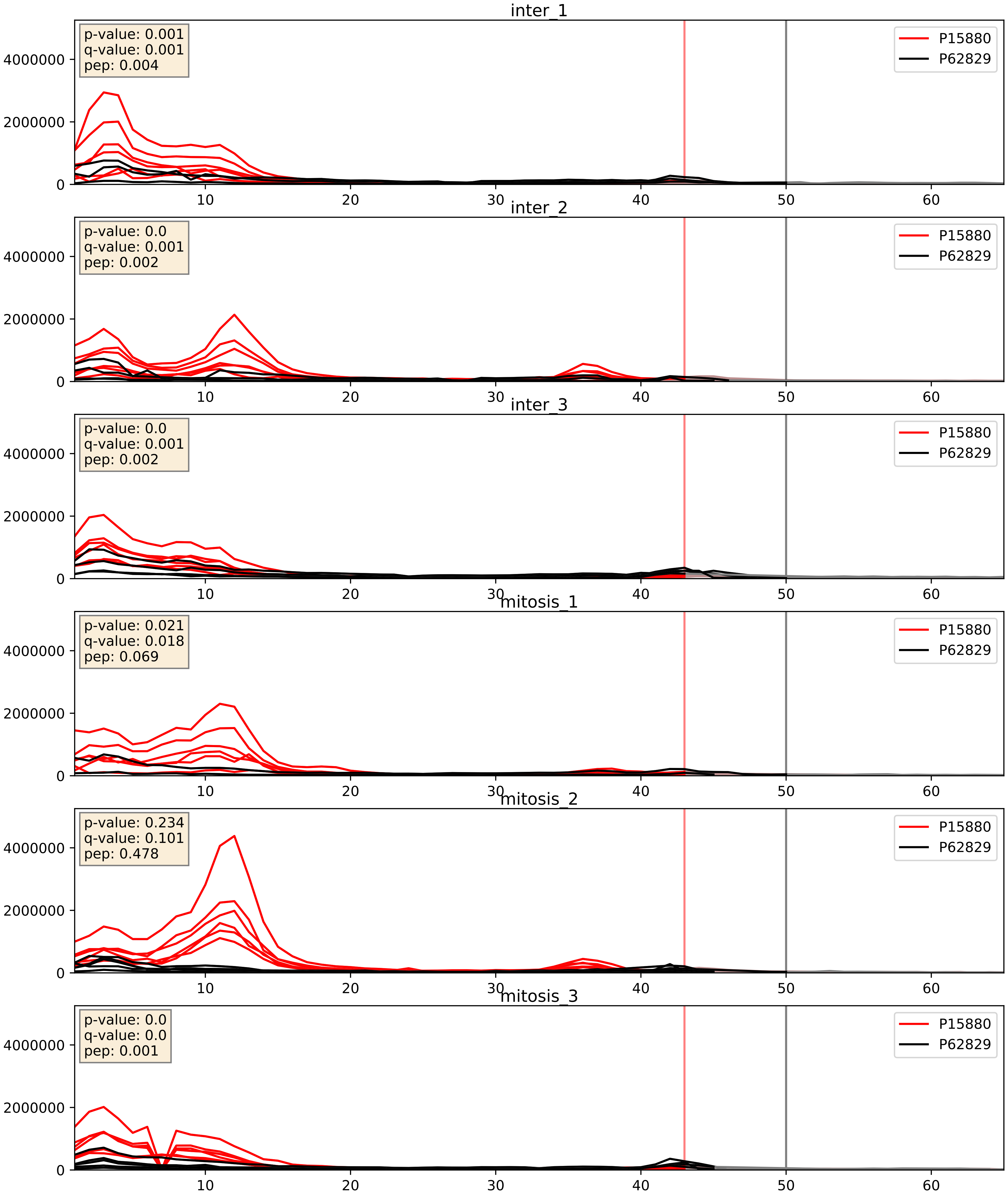


| level | condition_1 | condition_2 | pvalue | pvalue_adjusted |
|---|---|---|---|---|
| complex_abundance | mitosis | inter | 0.0058990235747750155 | 0.05764342671241339 |
| interactor_ratio | mitosis | inter | 0.011651148831008065 | 0.15583411561473287 |
| interactor_abundance | mitosis | inter | 0.29737981089028054 | 0.49163725584376783 |

P18077_P62829
RL35A_HUMAN vs RL23_HUMAN
p-value: 0.002 q-value: 0.003

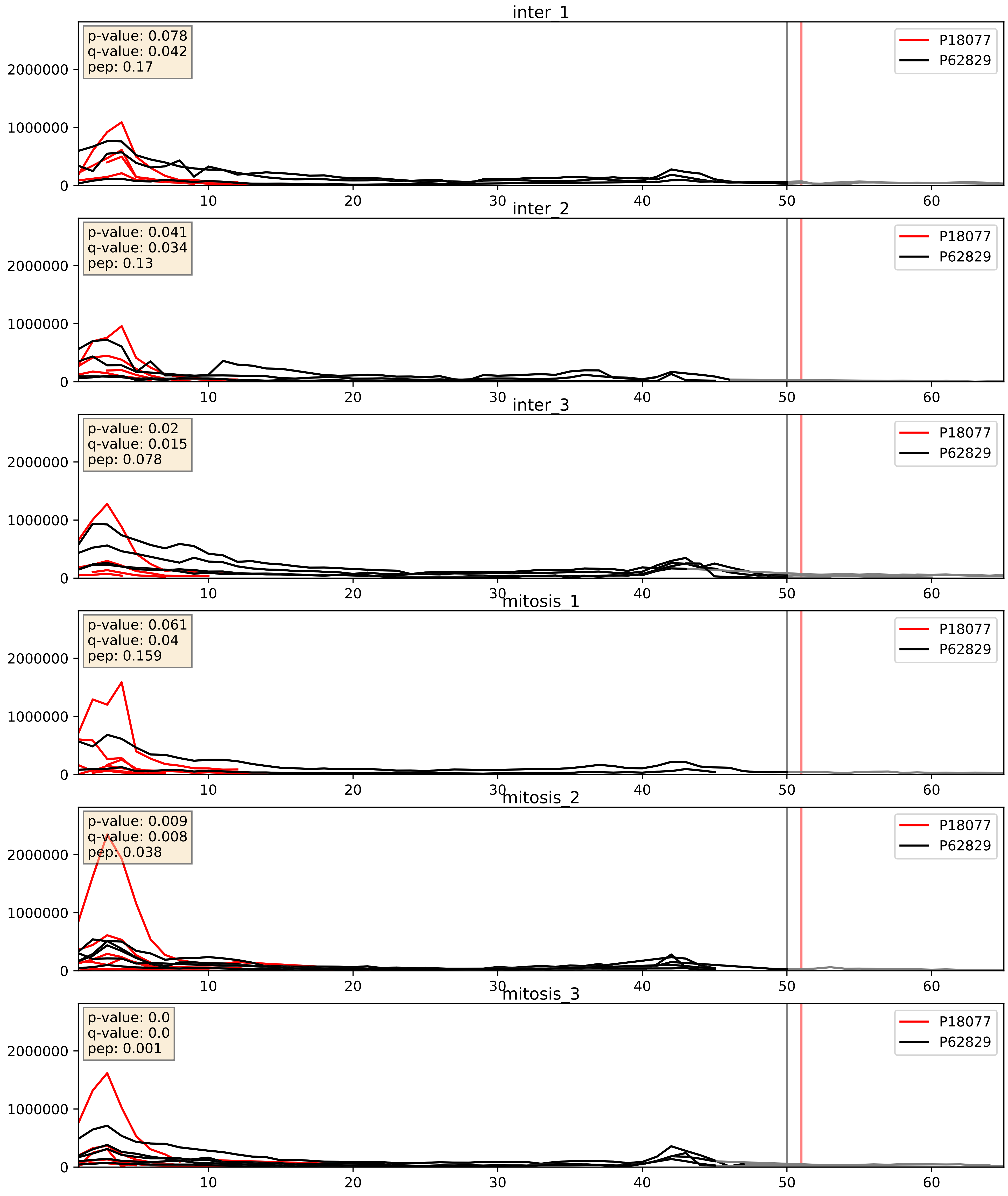


| level | condition_1 | condition_2 | pvalue | pvalue_adjusted |
|---|---|---|---|---|
| complex_abundance | mitosis | inter | 0.002794052435496297 | 0.03839019076701172 |
| interactor_ratio | mitosis | inter | 0.018588188814186153 | 0.2047810688472776 |
| interactor_abundance | mitosis | inter | 0.8878397503920612 | 0.9361222224987428 |

P18124_P62829
RL7_HUMAN vs RL23_HUMAN
p-value: 0.0 q-value: 0.0

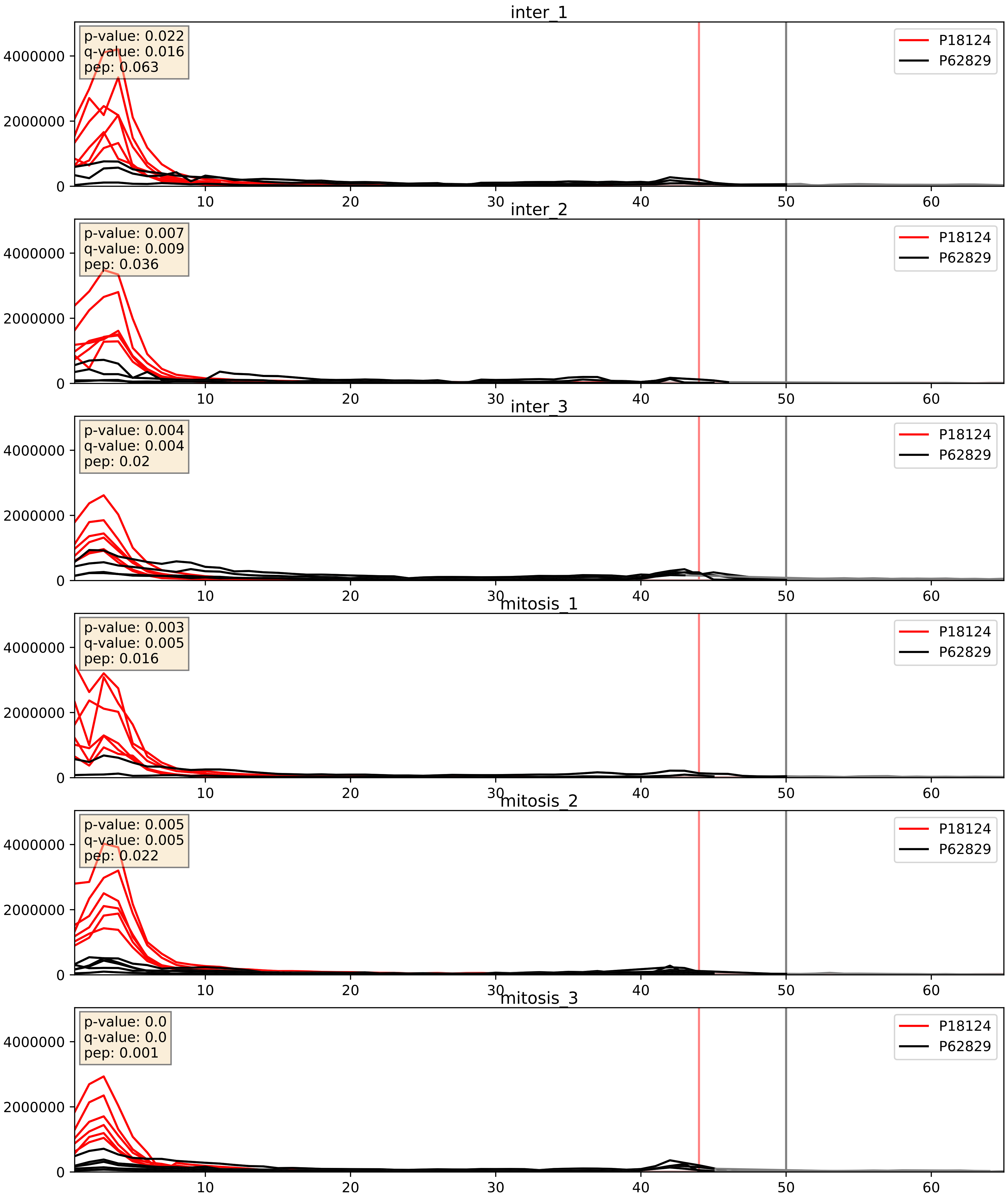


| level | condition_1 | condition_2 | pvalue | pvalue_adjusted |
|---|---|---|---|---|
| complex_abundance | mitosis | inter | 0.0017367077652753612 | 0.0295241467010272 |
| interactor_ratio | mitosis | inter | 0.024412817803191836 | 0.2447639398105627 |
| interactor_abundance | mitosis | inter | 0.49648279161018033 | 0.6622168268689187 |

P23396_P62829
RS3_HUMAN vs RL23_HUMAN
p-value: 0.023 q-value: 0.018

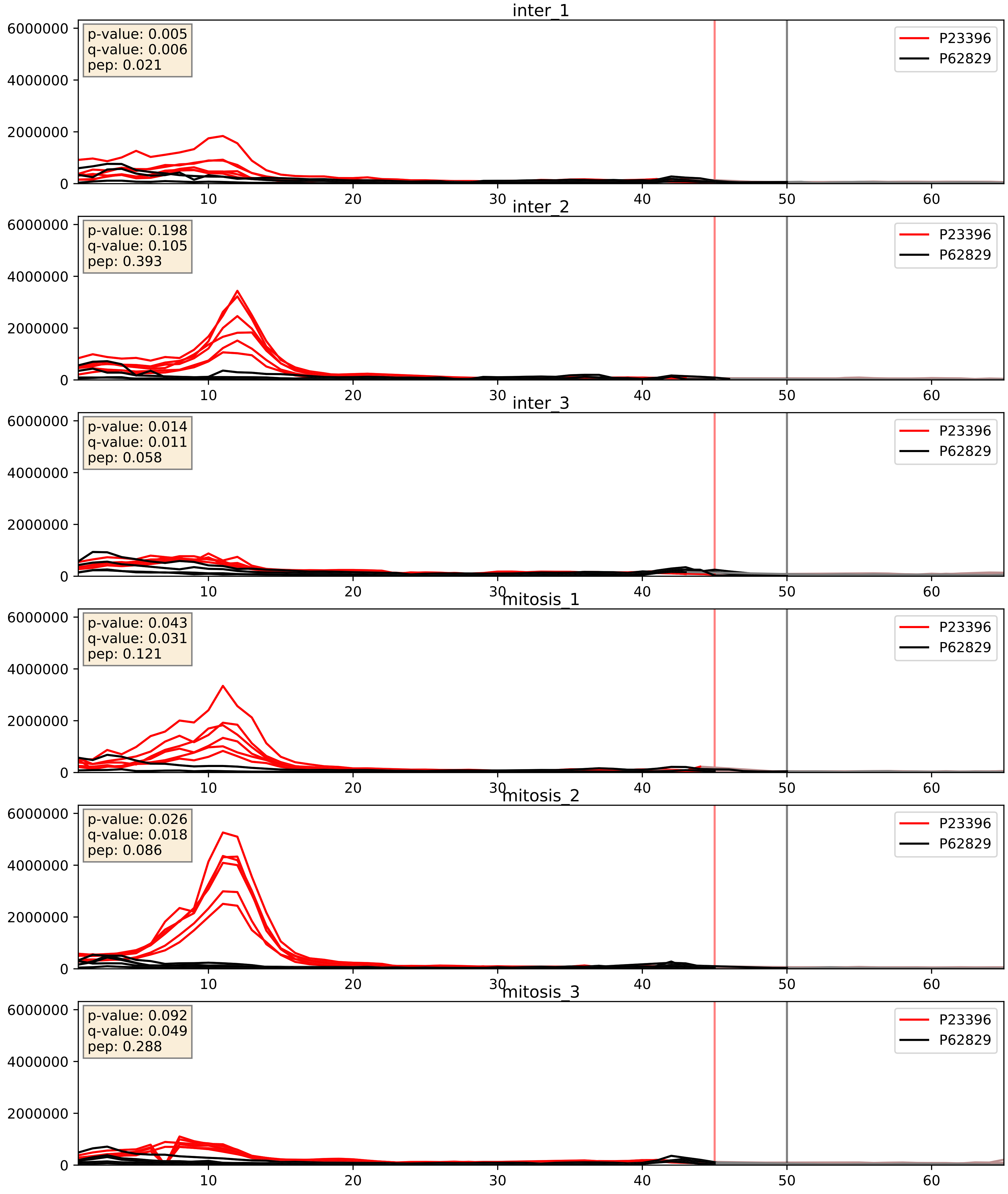


| level | condition_1 | condition_2 | pvalue | pvalue_adjusted |
|---|---|---|---|---|
| complex_abundance | mitosis | inter | 0.012811309746031889 | 0.08405635484106207 |
| interactor_ratio | mitosis | inter | 0.0599954289102443 | 0.405234575955053 |
| interactor_abundance | mitosis | inter | 0.795159623367833 | 0.8808985116259767 |

P25398_P62829
RS12_HUMAN vs RL23_HUMAN
p-value: 0.01 q-value: 0.008

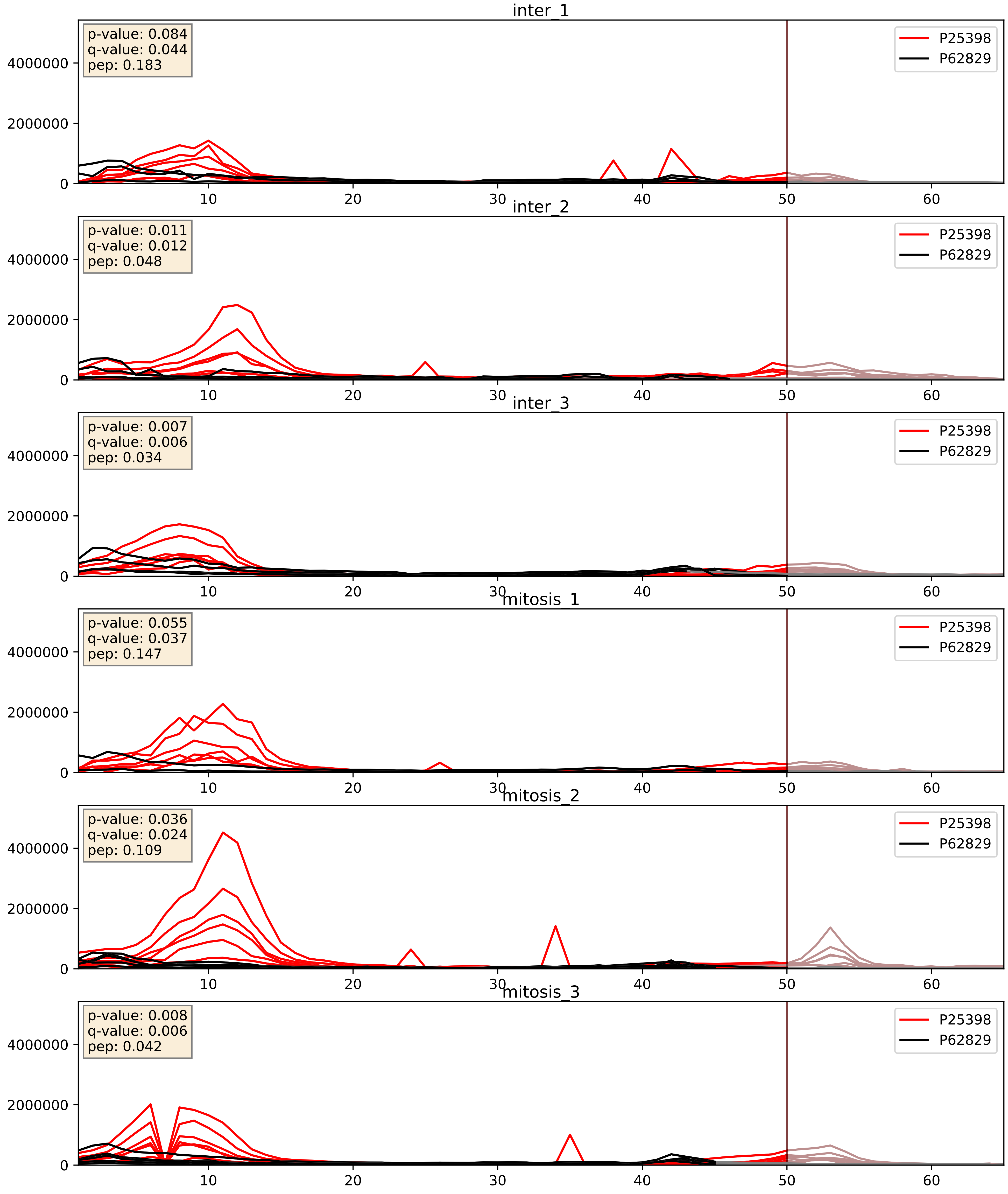


| level | condition_1 | condition_2 | pvalue | pvalue_adjusted |
|---|---|---|---|---|
| interactor_ratio | mitosis | inter | 0.012137837285786688 | 0.1588683290005108 |
| complex_abundance | mitosis | inter | 0.023377633186083567 | 0.11117363337381964 |
| interactor_abundance | mitosis | inter | 0.8077983580243697 | 0.8889001085857571 |

P26373_P62829
RL13_HUMAN vs RL23_HUMAN
p-value: 0.0 q-value: 0.0

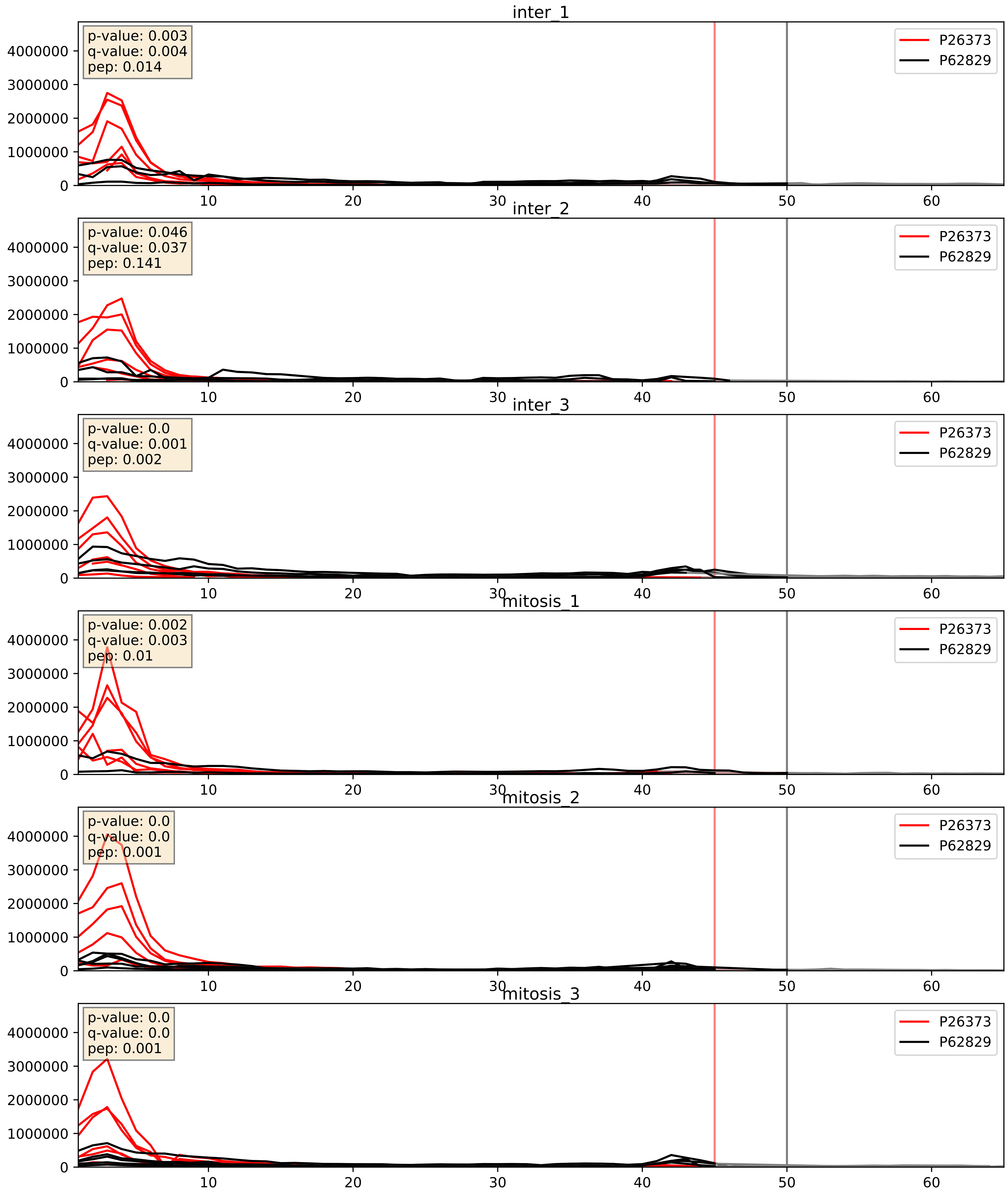


| level | condition_1 | condition_2 | pvalue | pvalue_adjusted |
|---|---|---|---|---|
| interactor_ratio | mitosis | inter | 0.0081880791659375 | 0.12756688776860511 |
| complex_abundance | mitosis | inter | 0.026610701264047975 | 0.11941204595039318 |
| interactor_abundance | mitosis | inter | 0.6092878016969288 | 0.7434257832692909 |

P27635_P62829
RL10_HUMAN vs RL23_HUMAN
p-value: 0.005 q-value: 0.005

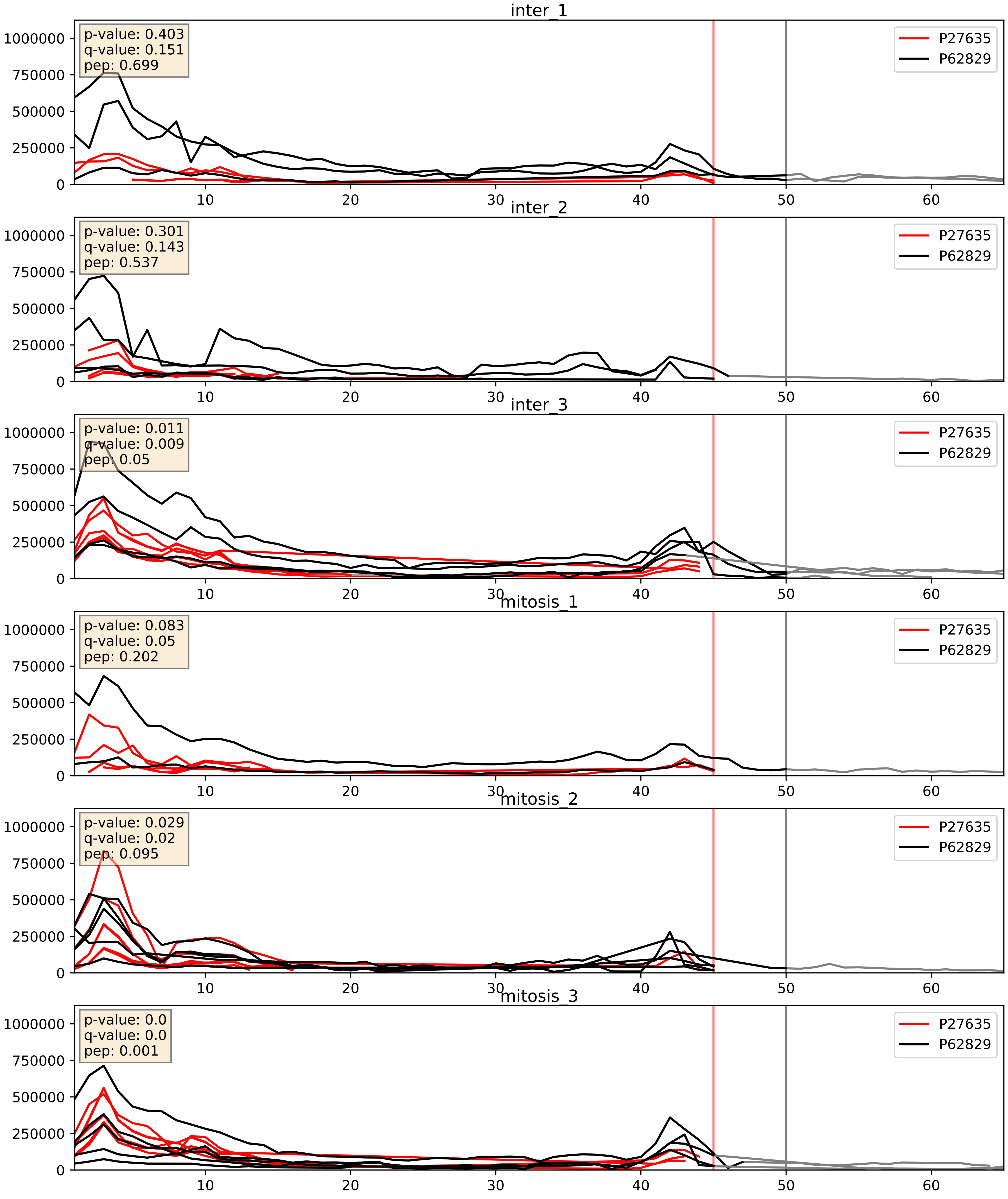


| level | condition_1 | condition_2 | pvalue | pvalue_adjusted |
|---|---|---|---|---|
| interactor_ratio | mitosis | inter | 0.0032302118195719543 | 0.07405554735675686 |
| complex_abundance | mitosis | inter | 0.1857047322297019 | 0.3370221298338614 |
| interactor_abundance | mitosis | inter | 0.6203093299259387 | 0.7499500470712932 |

P30050_P62829
RL12_HUMAN vs RL23_HUMAN
p-value: 0.0 q-value: 0.0

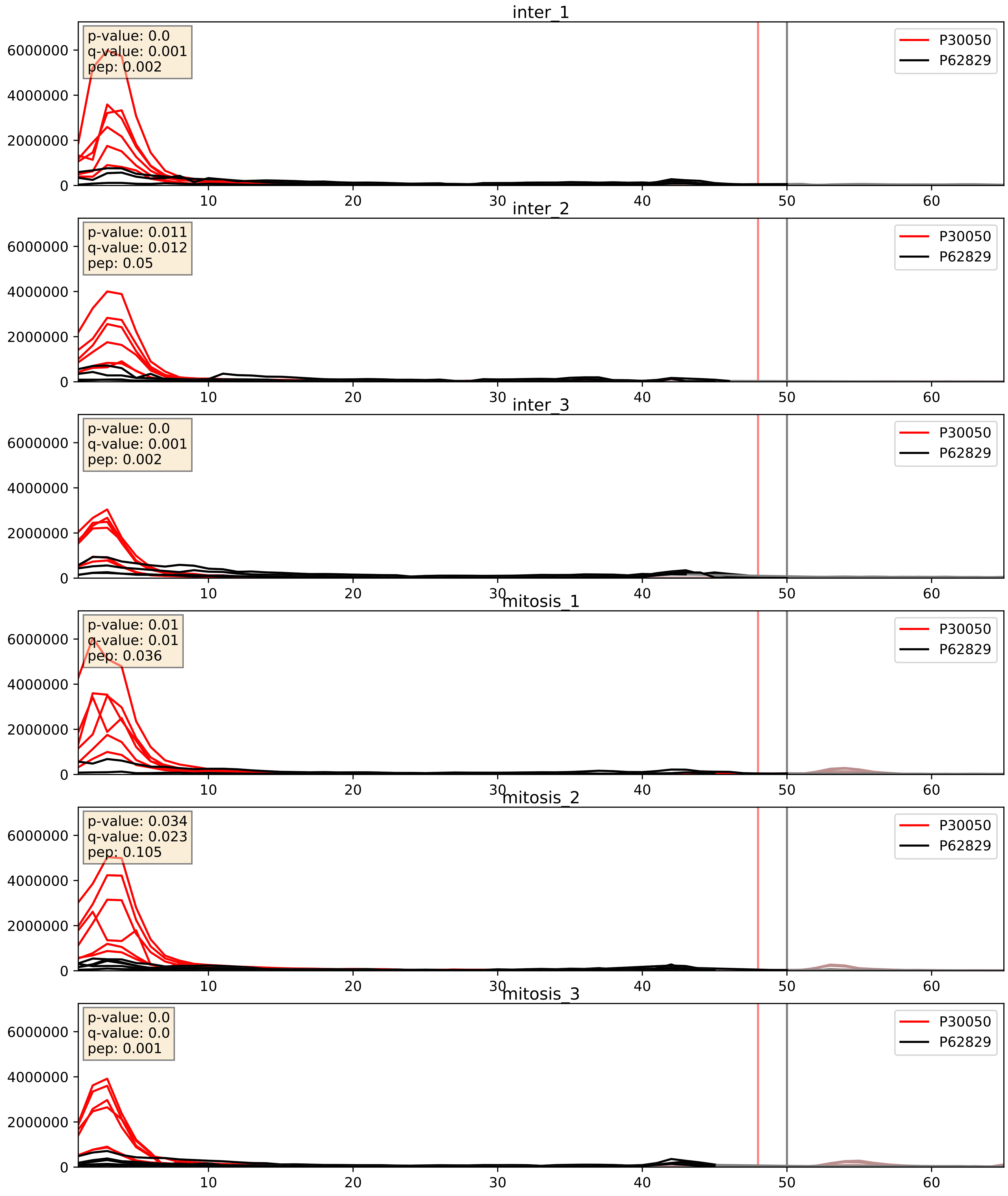


| level | condition_1 | condition_2 | pvalue | pvalue_adjusted |
|---|---|---|---|---|
| interactor_ratio | mitosis | inter | 0.01514649833891717 | 0.1807308493118951 |
| complex_abundance | mitosis | inter | 0.022128429361632312 | 0.10952717692621856 |
| interactor_abundance | mitosis | inter | 0.8832806096565684 | 0.9352897103736053 |

P32969_P62829
RL9_HUMAN vs RL23_HUMAN
p-value: 0.0 q-value: 0.0

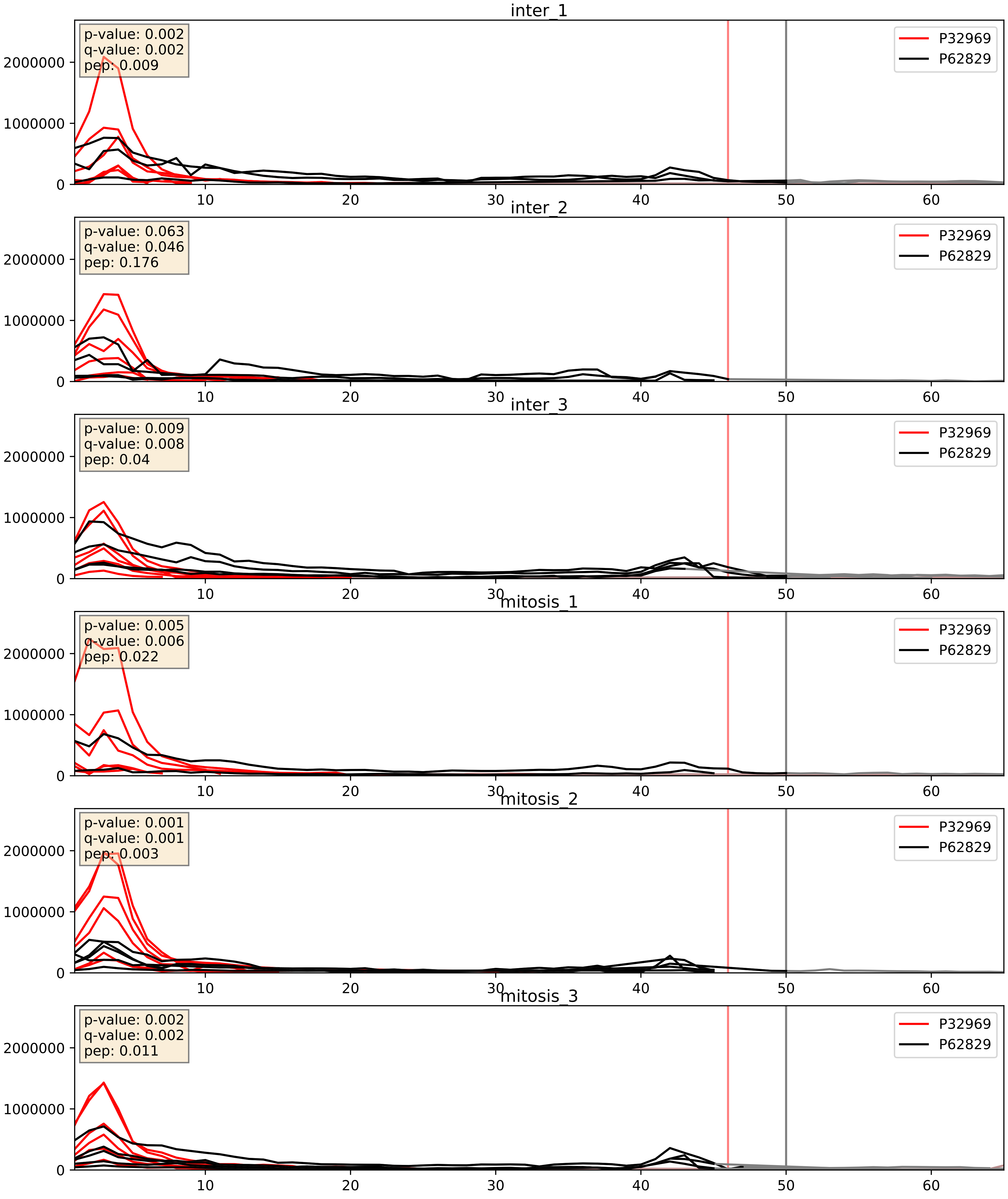


| level | condition_1 | condition_2 | pvalue | pvalue_adjusted |
|---|---|---|---|---|
| complex_abundance | mitosis | inter | 0.0054462970174589094 | 0.055632819175952586 |
| interactor_ratio | mitosis | inter | 0.015247830918452124 | 0.1810283393369628 |
| interactor_abundance | mitosis | inter | 0.2103491843605969 | 0.40652154270556234 |

P35268_P62829
RL22_HUMAN vs RL23_HUMAN
p-value: 0.0 q-value: 0.0

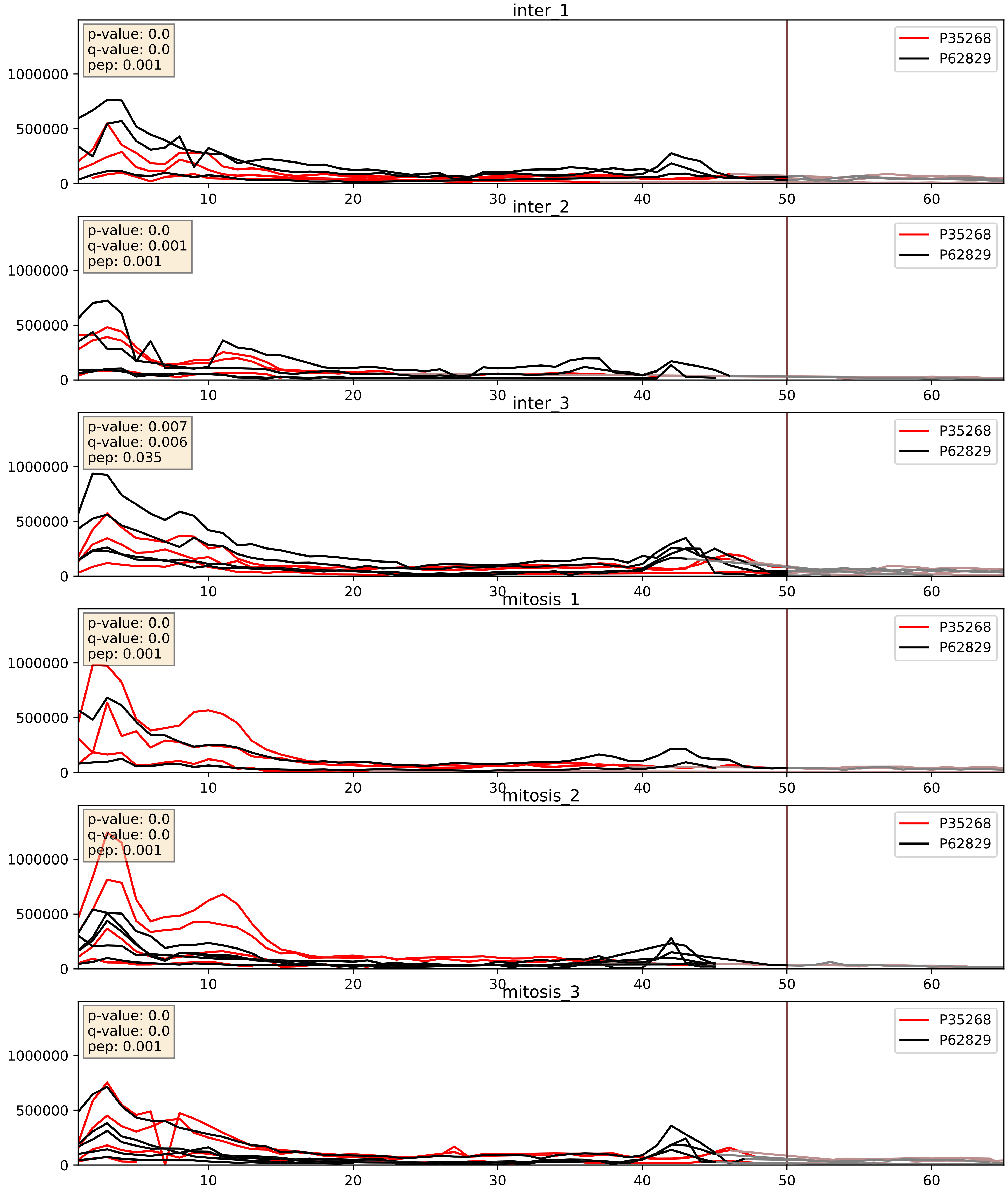


| level | condition_1 | condition_2 | pvalue | pvalue_adjusted |
|---|---|---|---|---|
| interactor_ratio | mitosis | inter | 0.0012700844385174963 | 0.0423373831474418 |
| interactor_abundance | mitosis | inter | 0.04055078168160862 | 0.1623548602406781 |
| complex_abundance | mitosis | inter | 0.07592611735990688 | 0.20016247754875358 |

P36578_P62829
RL4_HUMAN vs RL23_HUMAN
p-value: 0.001 q-value: 0.001

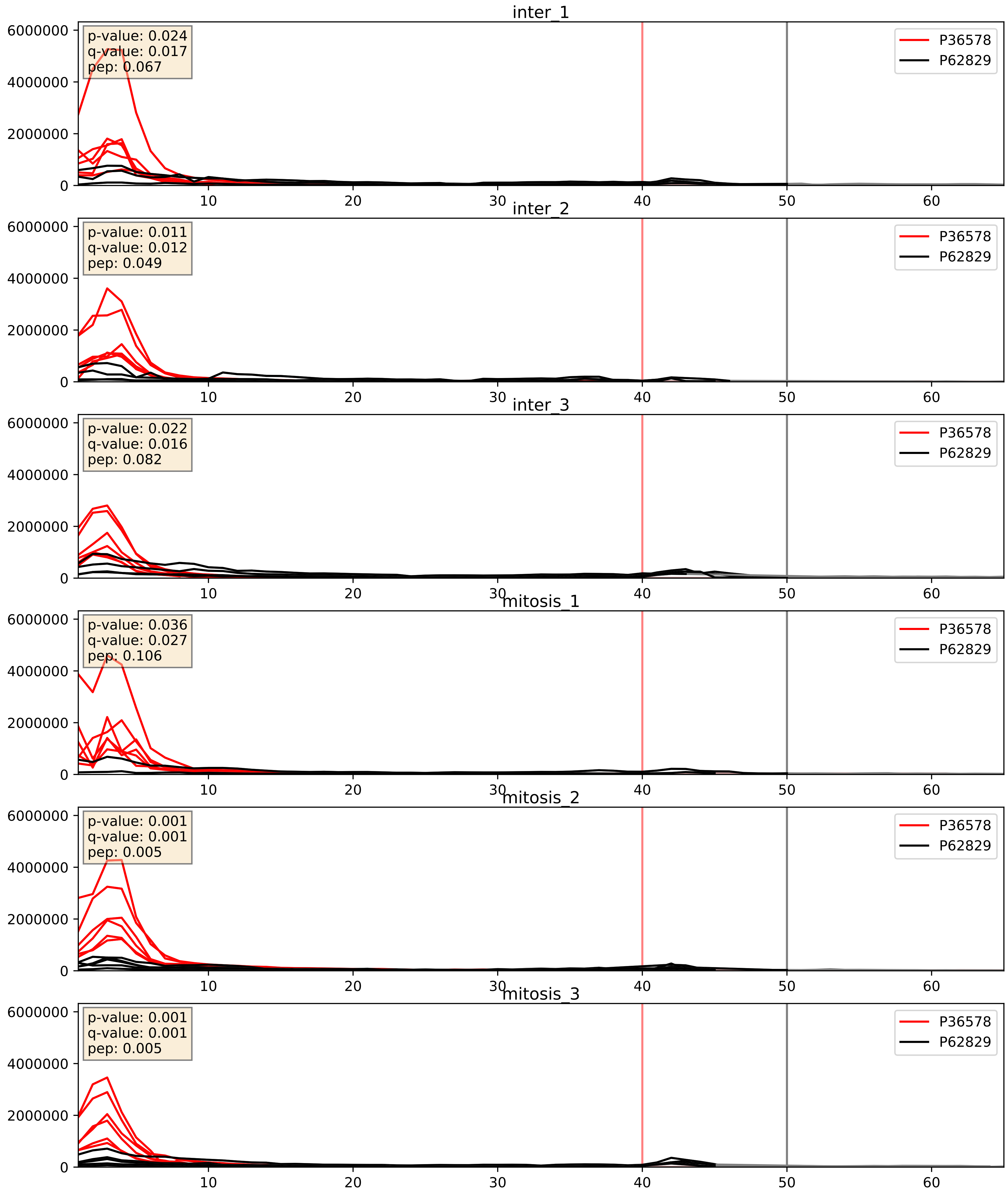


| level | condition_1 | condition_2 | pvalue | pvalue_adjusted |
|---|---|---|---|---|
| interactor_ratio | mitosis | inter | 0.0018337591240226922 | 0.0519767486809081 |
| complex_abundance | mitosis | inter | 0.012890916700927684 | 0.08411248670667204 |
| interactor_abundance | mitosis | inter | 0.9858211760603667 | 0.9908564864764562 |

P37108_P62829
SRP14_HUMAN vs RL23_HUMAN
p-value: 0.0 q-value: 0.0

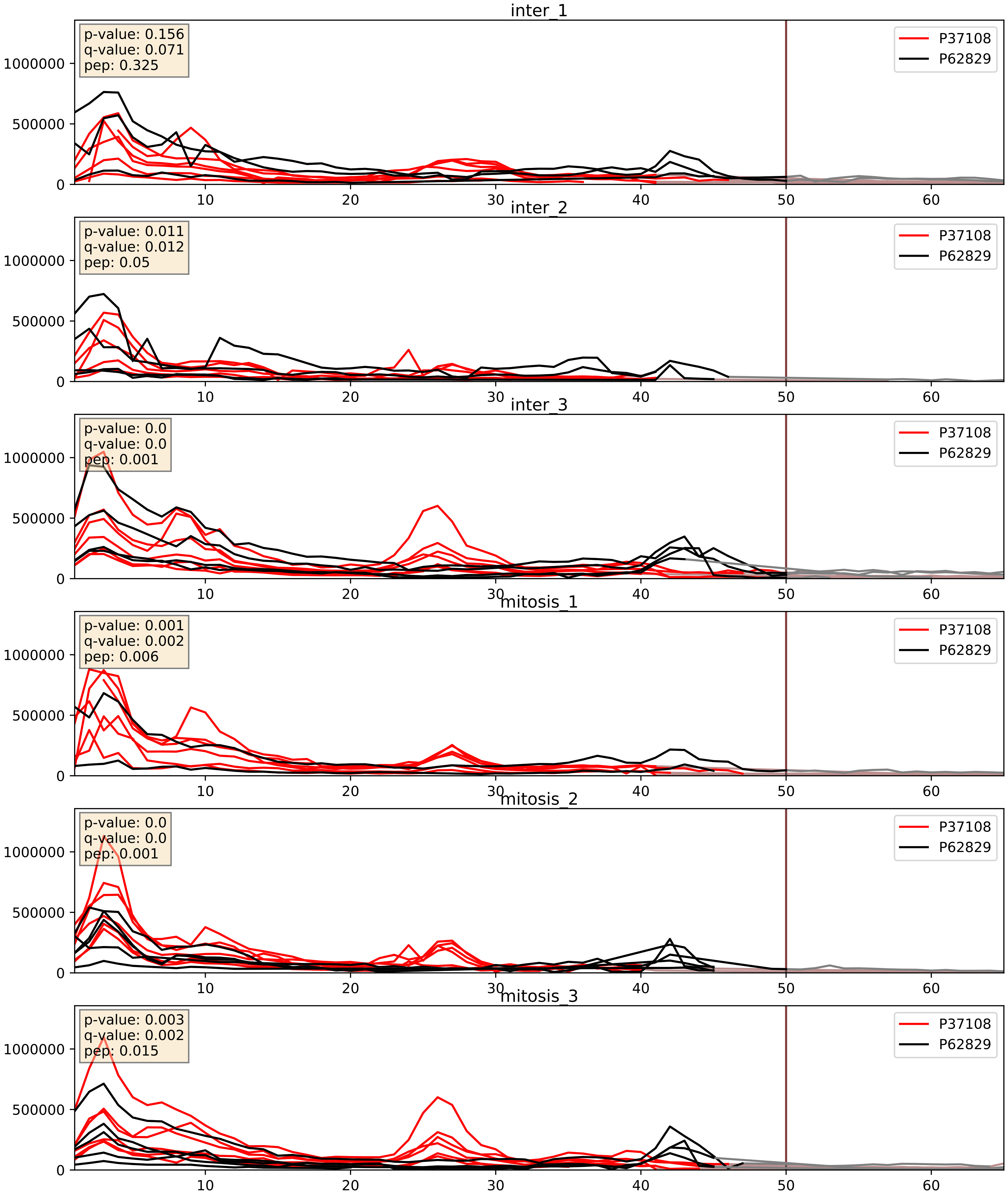


| level | condition_1 | condition_2 | pvalue | pvalue_adjusted |
|---|---|---|---|---|
| interactor_ratio | mitosis | inter | 0.0029899561580882966 | 0.07070172572717077 |
| complex_abundance | mitosis | inter | 0.2834223013021206 | 0.4319200461360428 |
| interactor_abundance | mitosis | inter | 0.5588616209015151 | 0.7088243406307556 |

P39019_P62829
RS19_HUMAN vs RL23_HUMAN
p-value: 0.003 q-value: 0.003

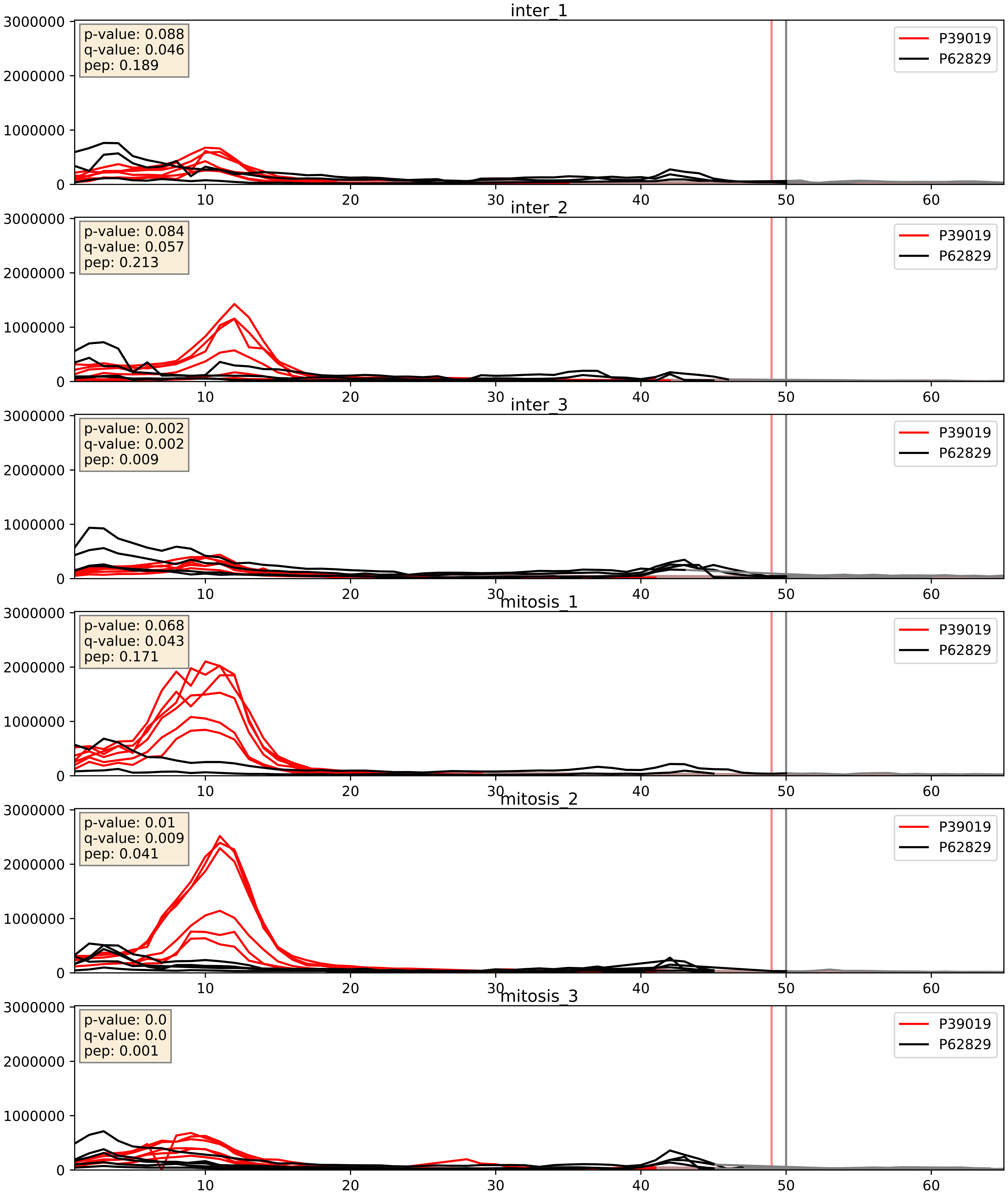


| level | condition_1 | condition_2 | pvalue | pvalue_adjusted |
|---|---|---|---|---|
| interactor_ratio | mitosis | inter | 0.035672312484526854 | 0.30712516692473574 |
| complex_abundance | mitosis | inter | 0.04688035280058762 | 0.15478990092568878 |
| interactor_abundance | mitosis | inter | 0.2118751697034527 | 0.40715039907095185 |

P39023_P62829
RL3_HUMAN vs RL23_HUMAN
p-value: 0.0 q-value: 0.0

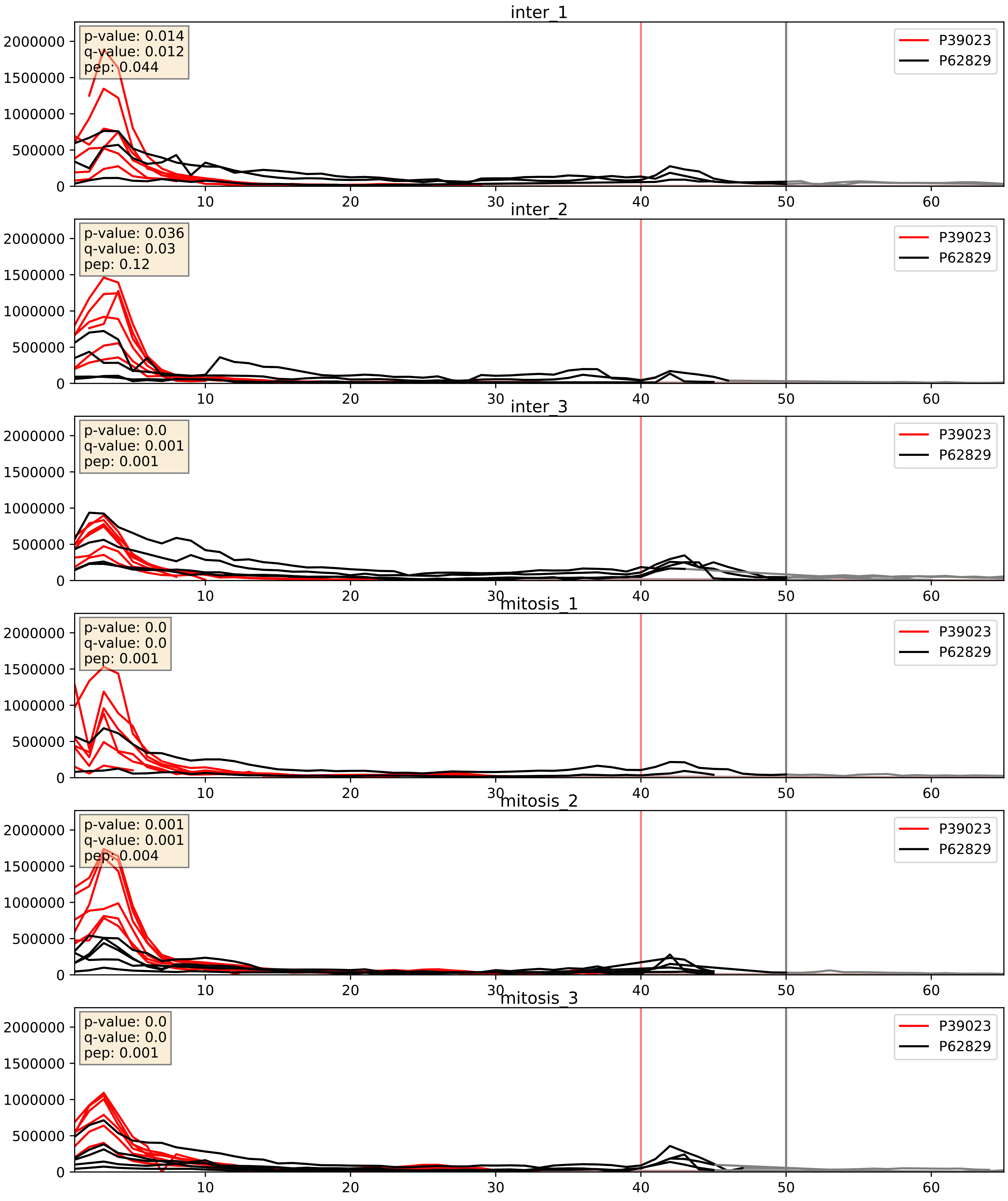


| level | condition_1 | condition_2 | pvalue | pvalue_adjusted |
|---|---|---|---|---|
| complex_abundance | mitosis | inter | 0.0007459442474047649 | 0.0178859461002375 |
| interactor_ratio | mitosis | inter | 0.012561102096494269 | 0.1616887728511142 |
| interactor_abundance | mitosis | inter | 0.4577908838109454 | 0.629609570279835 |

P40429_P62829
RL13A_HUMAN vs RL23_HUMAN
p-value: 0.024 q-value: 0.018

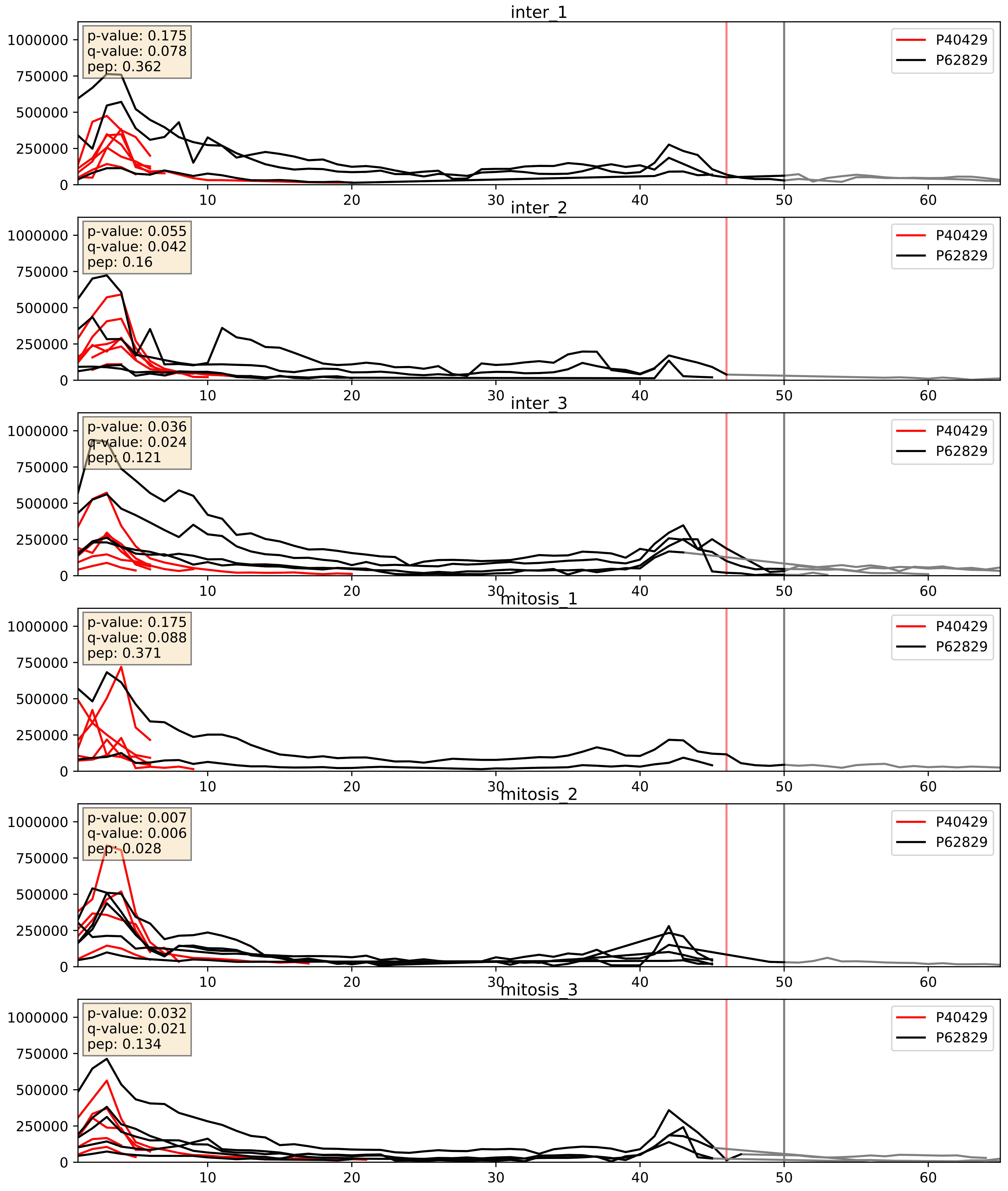


| level | condition_1 | condition_2 | pvalue | pvalue_adjusted |
|---|---|---|---|---|
| complex_abundance | mitosis | inter | 0.010243479704215621 | 0.07485435966639949 |
| interactor_ratio | mitosis | inter | 0.016036761390099 | 0.18702272138862047 |
| interactor_abundance | mitosis | inter | 0.9364707510987881 | 0.962096690999235 |

P42677_P62829
RS27_HUMAN vs RL23_HUMAN
p-value: 0.001 q-value: 0.001

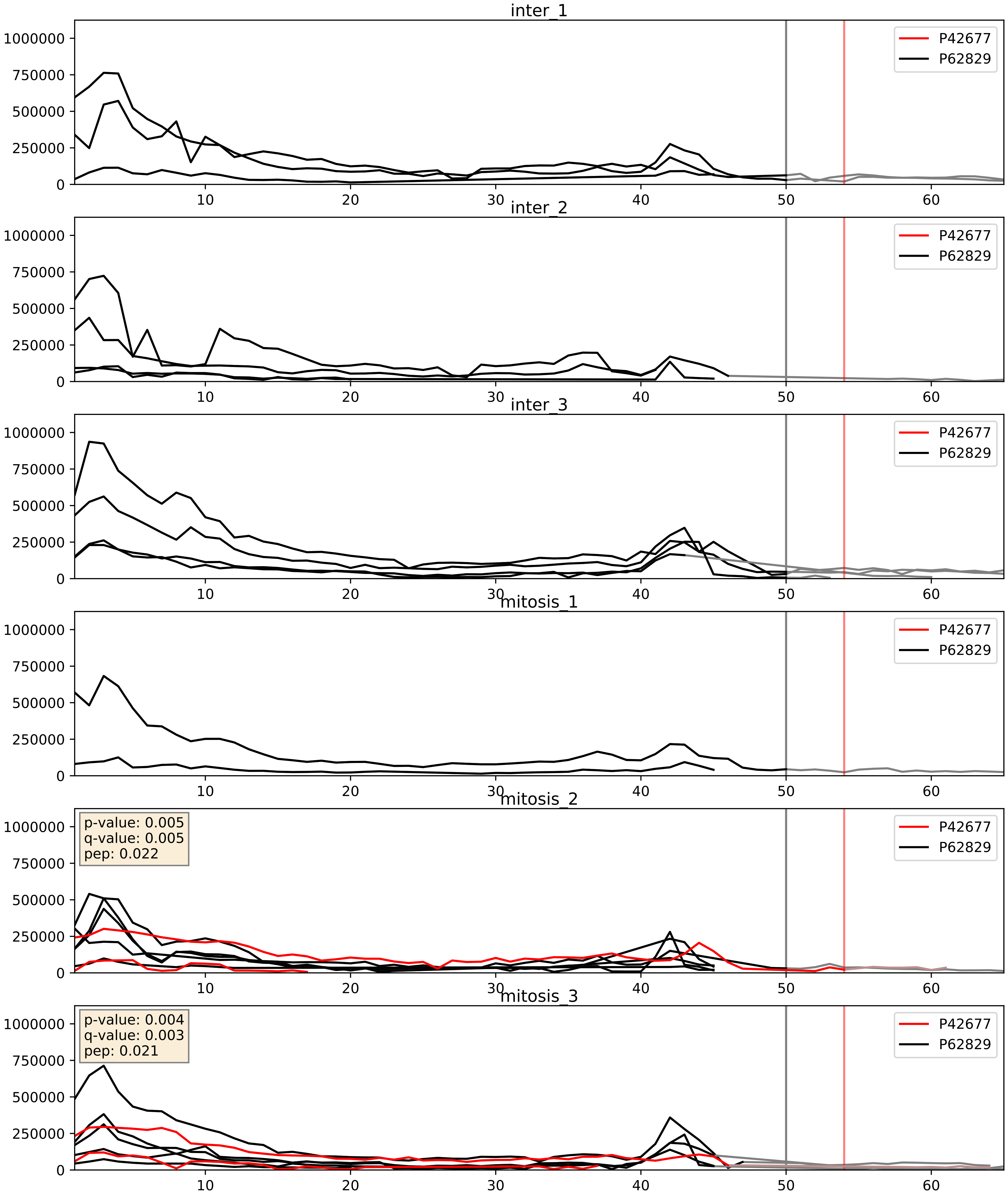


| level | condition_1 | condition_2 | pvalue | pvalue_adjusted |
|---|---|---|---|---|
| interactor_ratio | mitosis | inter | 0.11915358099553813 | 0.5681399906918084 |
| complex_abundance | mitosis | inter | 0.14898450276421568 | 0.29488459277225637 |
| interactor_abundance | mitosis | inter | 0.15591746956724173 | 0.34249153565282997 |

P42766_P62829
RL35_HUMAN vs RL23_HUMAN
p-value: 0.0 q-value: 0.0

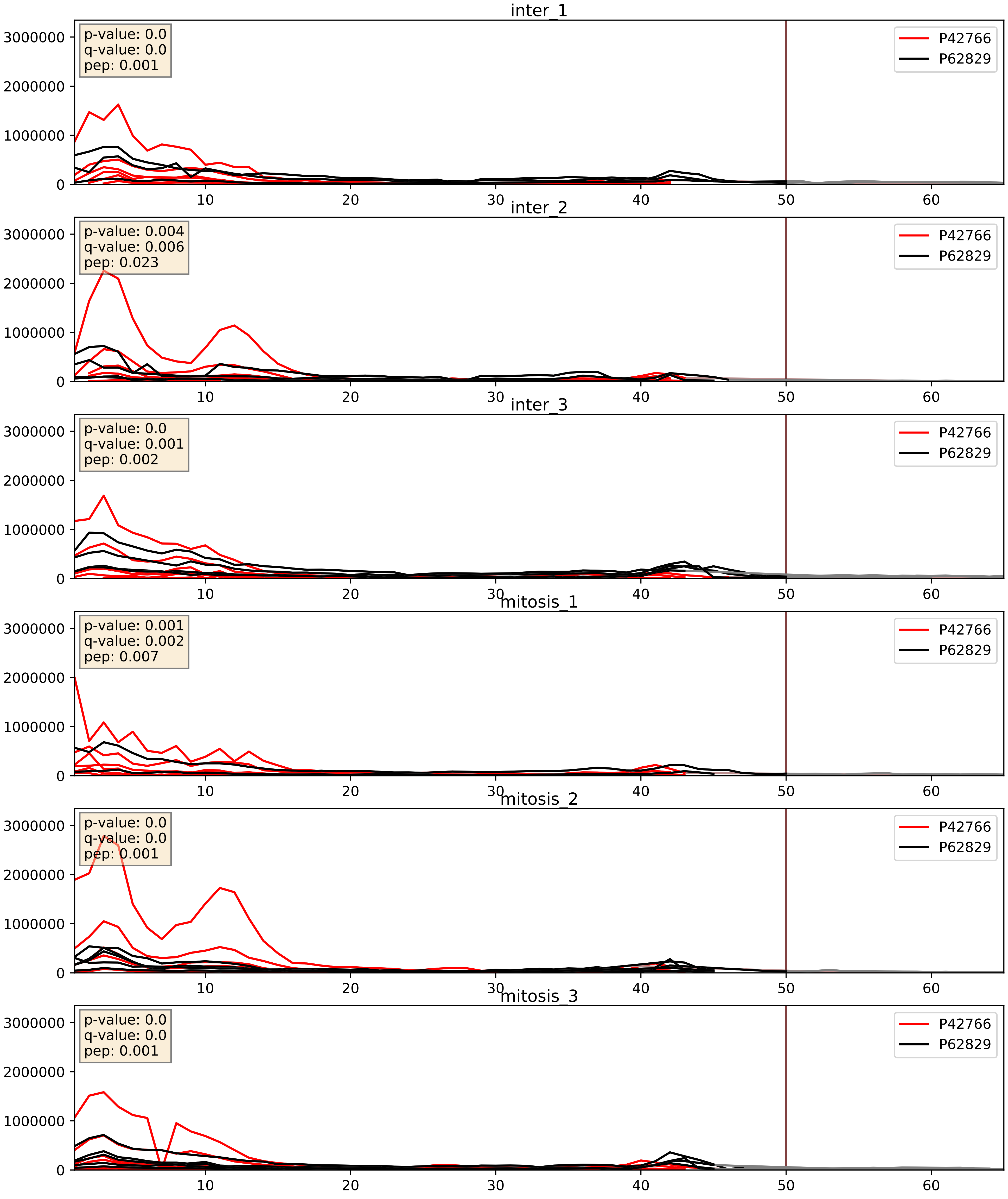


| level | condition_1 | condition_2 | pvalue | pvalue_adjusted |
|---|---|---|---|---|
| complex_abundance | mitosis | inter | 0.009790433606854661 | 0.07332118256752047 |
| interactor_ratio | mitosis | inter | 0.014244423284353047 | 0.1749386848121407 |
| interactor_abundance | mitosis | inter | 0.781558646296928 | 0.8726814827254089 |

P46776_P62829
RL27A_HUMAN vs RL23_HUMAN
p-value: 0.0 q-value: 0.0

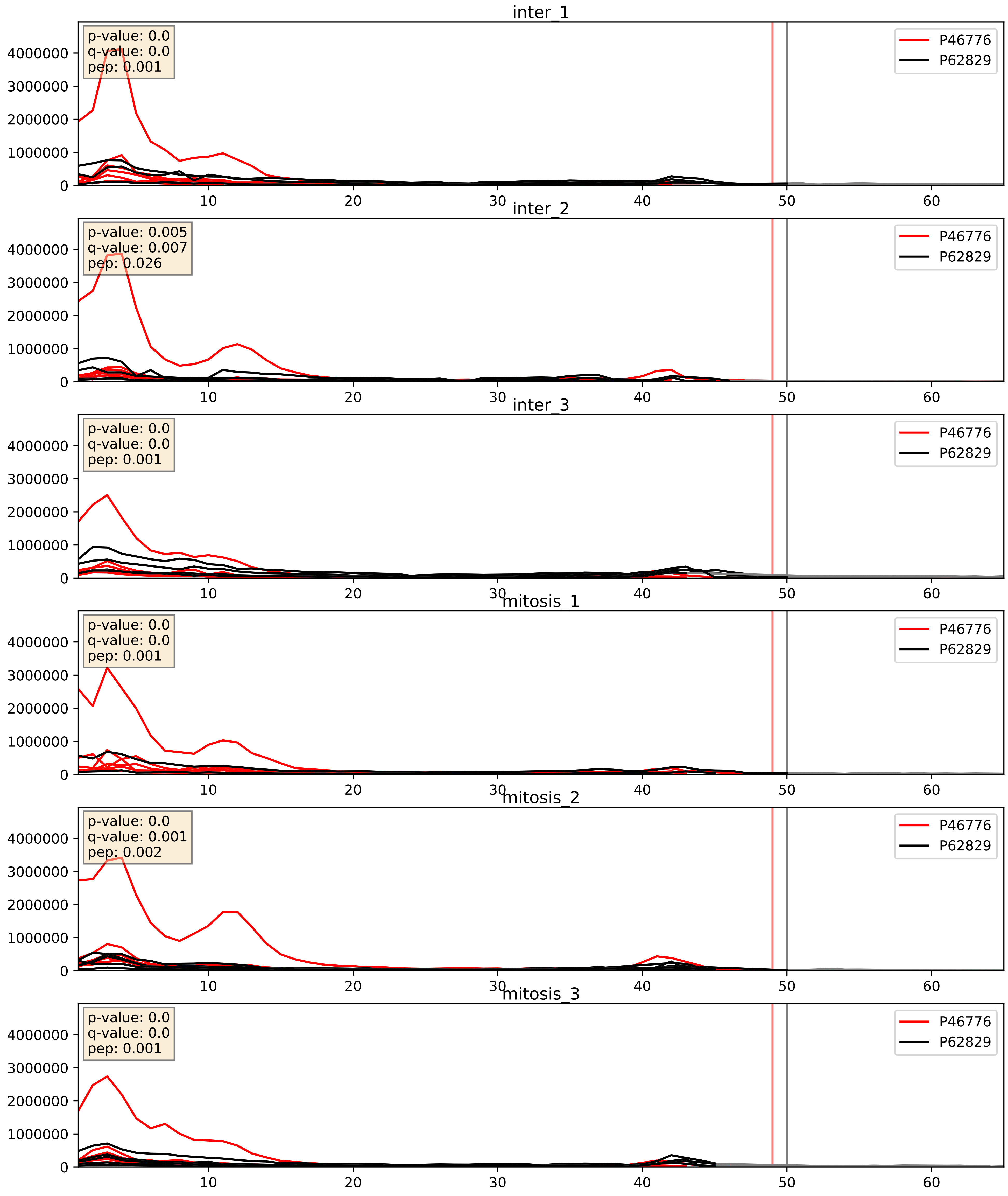


| level | condition_1 | condition_2 | pvalue | pvalue_adjusted |
|---|---|---|---|---|
| complex_abundance | mitosis | inter | 0.007141567620195767 | 0.06265939198178253 |
| interactor_ratio | mitosis | inter | 0.018562733522100039 | 0.2047641739035816 |
| interactor_abundance | mitosis | inter | 0.6997236190644978 | 0.8022547788899144 |

P46777_P62829
RL5_HUMAN vs RL23_HUMAN
p-value: 0.007 q-value: 0.007

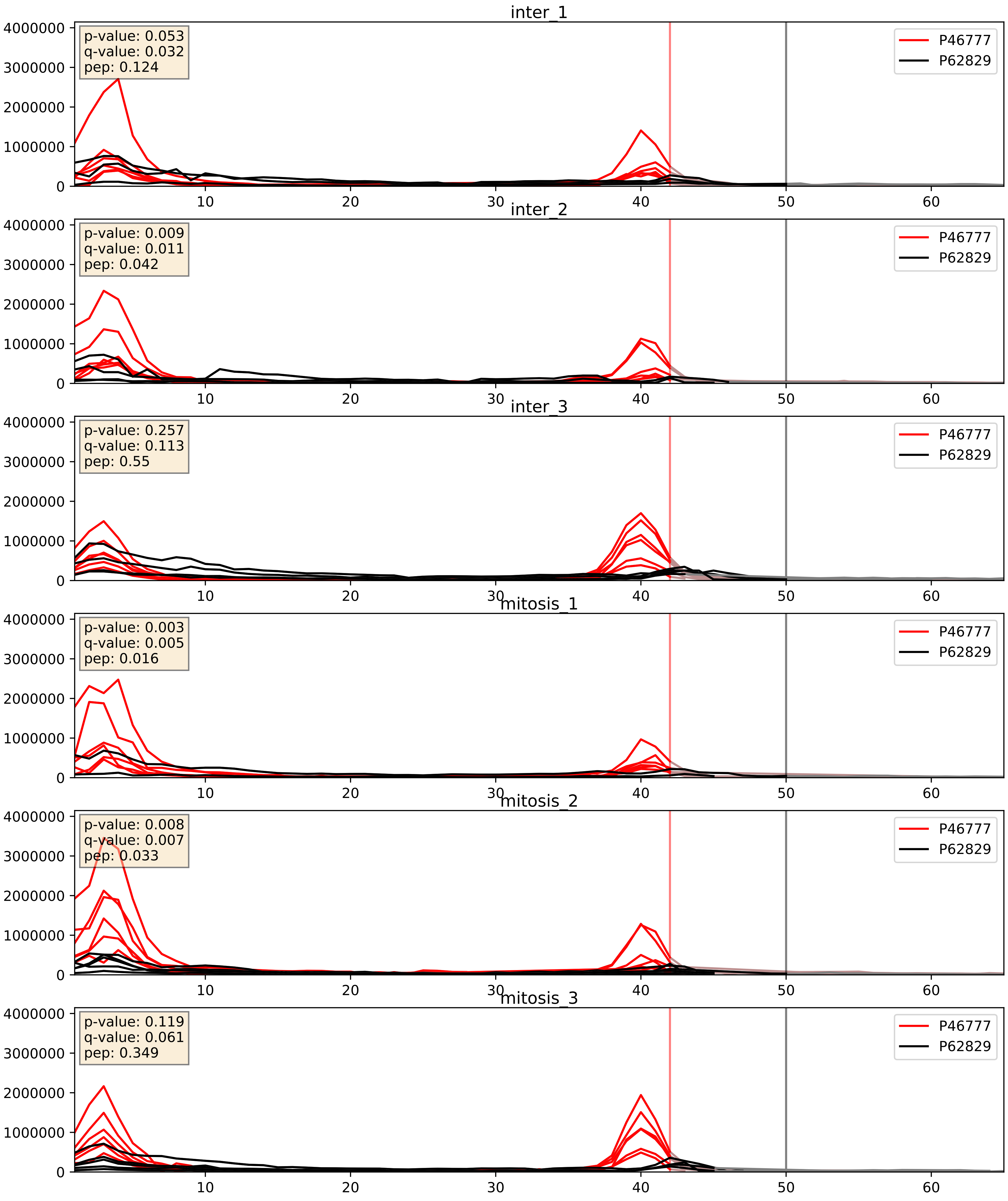


| level | condition_1 | condition_2 | pvalue | pvalue_adjusted |
|---|---|---|---|---|
| interactor_ratio | mitosis | inter | 0.0027480618992069233 | 0.06759600533681397 |
| complex_abundance | mitosis | inter | 0.04098341897197959 | 0.1455676624066993 |
| interactor_abundance | mitosis | inter | 0.6841811971746554 | 0.7940602057914786 |

P46778_P62829
RL21_HUMAN vs RL23_HUMAN
p-value: 0.0 q-value: 0.0

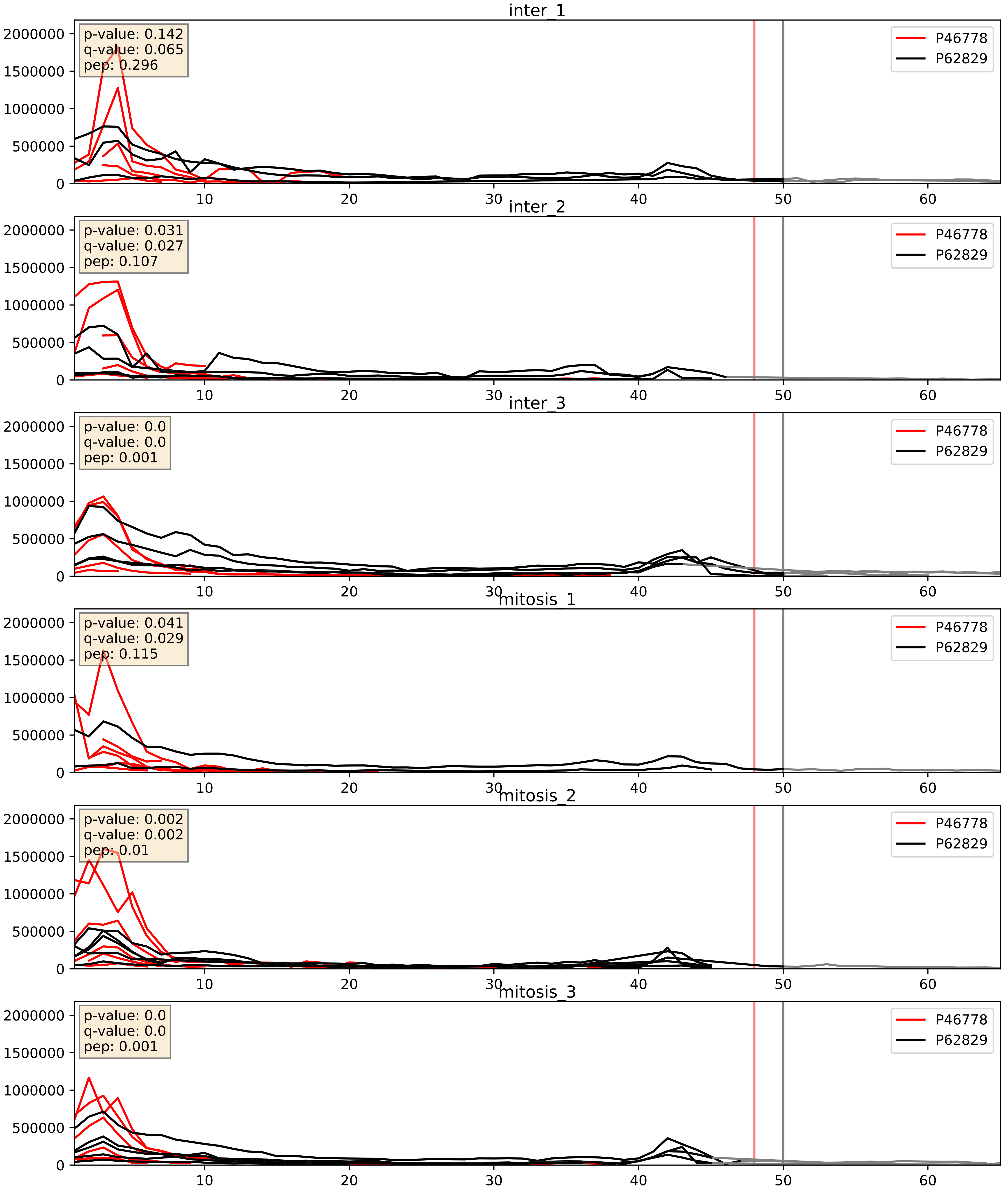


| level | condition_1 | condition_2 | pvalue | pvalue_adjusted |
|---|---|---|---|---|
| complex_abundance | mitosis | inter | 0.007925503391106073 | 0.0659424994487145 |
| interactor_ratio | mitosis | inter | 0.012807565728687513 | 0.16355562016423936 |
| interactor_abundance | mitosis | inter | 0.4845822245322886 | 0.6544702818064209 |

P46779_P62829
RL28_HUMAN vs RL23_HUMAN
p-value: 0.0 q-value: 0.0

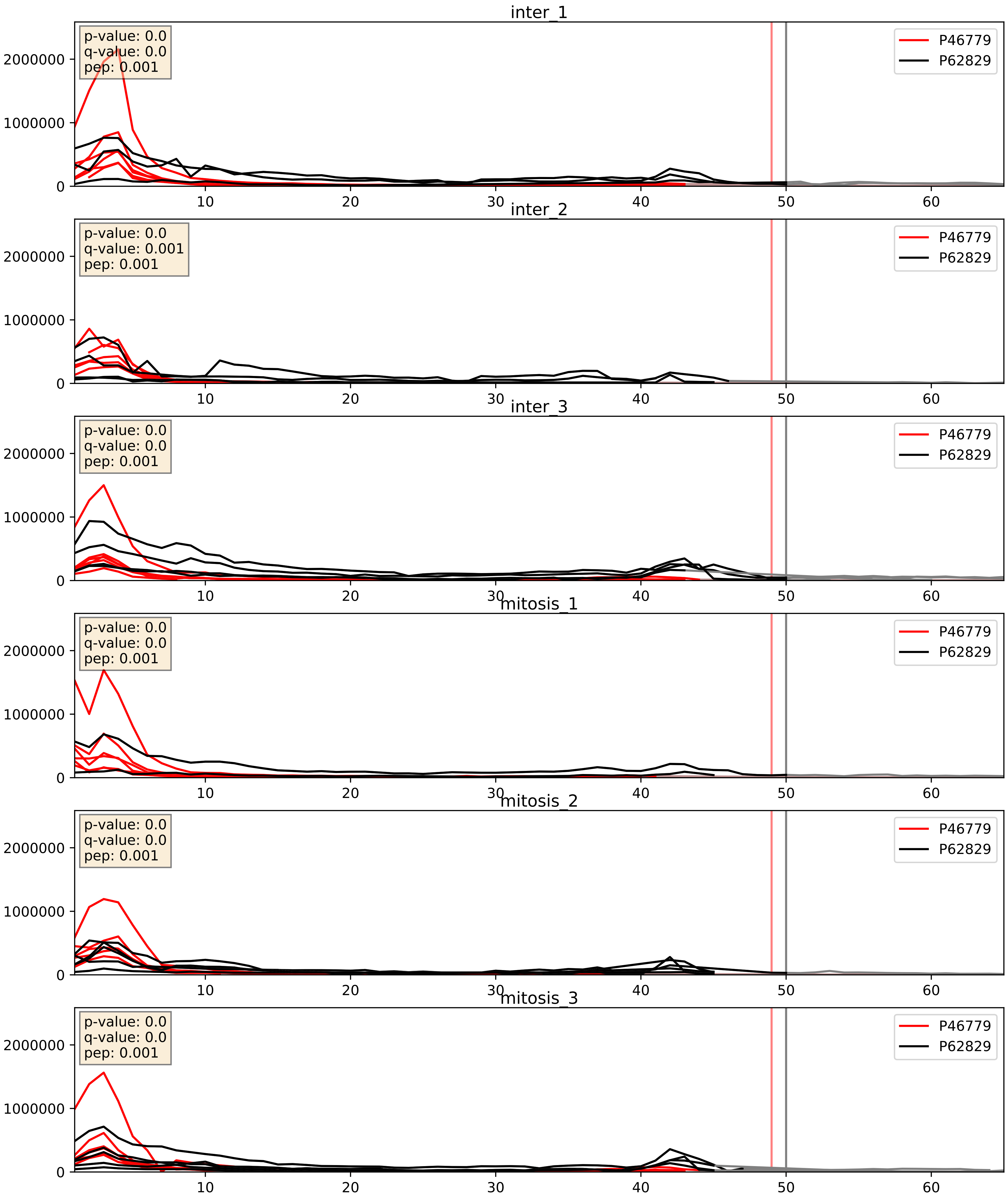


| level | condition_1 | condition_2 | pvalue | pvalue_adjusted |
|---|---|---|---|---|
| complex_abundance | mitosis | inter | 0.010372260056551404 | 0.0751155212217259 |
| interactor_ratio | mitosis | inter | 0.021478590302180325 | 0.2228566460444407 |
| interactor_abundance | mitosis | inter | 0.20770102547817784 | 0.4048550103821479 |

P46781_P62829
RS9_HUMAN vs RL23_HUMAN
p-value: 0.0 q-value: 0.0

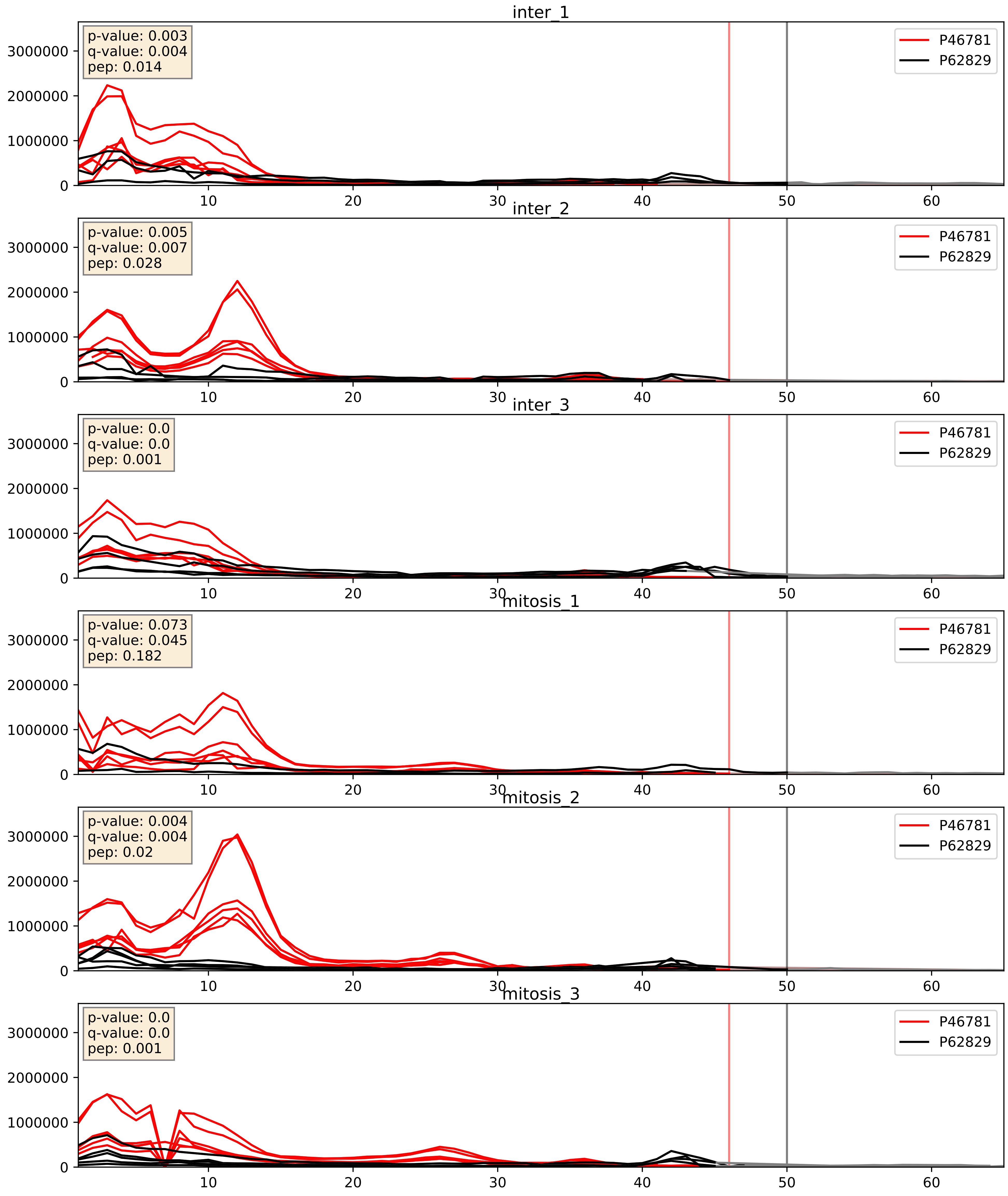


| level | condition_1 | condition_2 | pvalue | pvalue_adjusted |
|---|---|---|---|---|
| interactor_ratio | mitosis | inter | 0.010540908816021011 | 0.14745206851952483 |
| complex_abundance | mitosis | inter | 0.012067062177543546 | 0.08173689978386886 |
| interactor_abundance | mitosis | inter | 0.7628511259158188 | 0.8582513817229952 |

P46782_P62829
RS5_HUMAN vs RL23_HUMAN
p-value: 0.007 q-value: 0.006

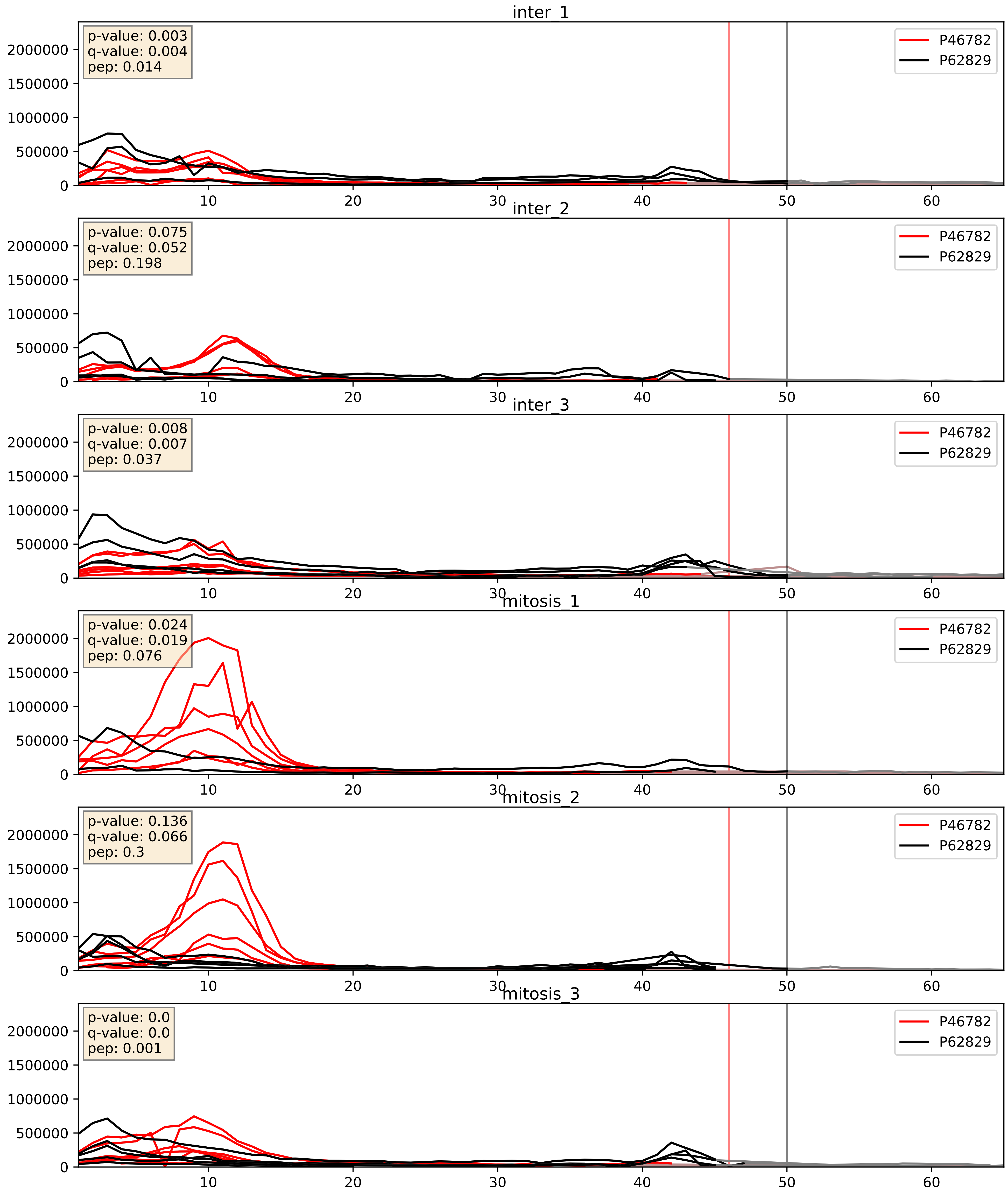


| level | condition_1 | condition_2 | pvalue | pvalue_adjusted |
|---|---|---|---|---|
| interactor_ratio | mitosis | inter | 0.009371256194171899 | 0.13838047771338102 |
| complex_abundance | mitosis | inter | 0.03900875660870847 | 0.14411521647412365 |
| interactor_abundance | mitosis | inter | 0.11668647902250837 | 0.29516437956048214 |

P46783_P62829
RS10_HUMAN vs RL23_HUMAN
p-value: 0.005 q-value: 0.005

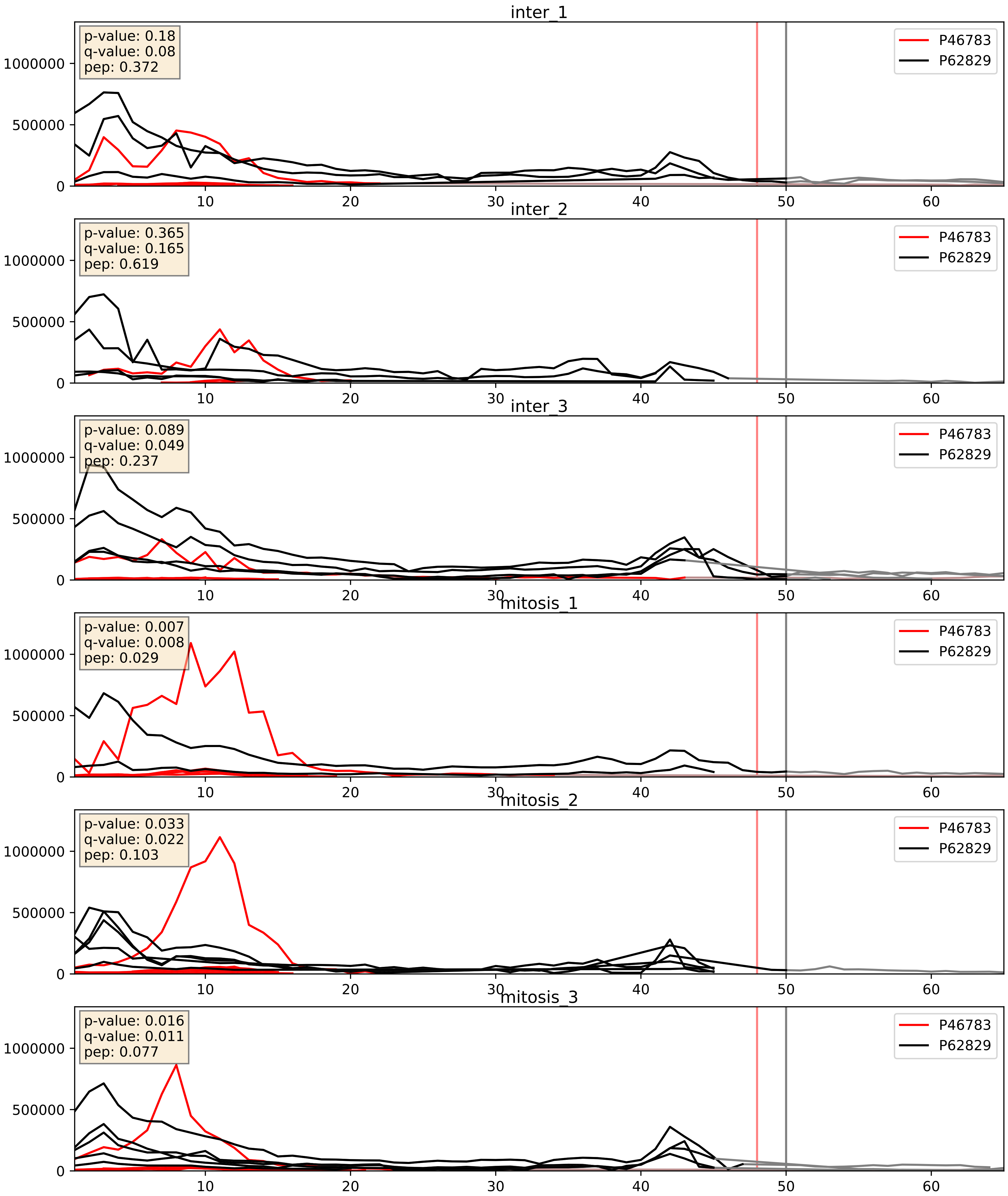


| level | condition_1 | condition_2 | pvalue | pvalue_adjusted |
|---|---|---|---|---|
| complex_abundance | mitosis | inter | 0.02219381103934062 | 0.10952717692621856 |
| interactor_ratio | mitosis | inter | 0.04422280807049488 | 0.3479294458487465 |
| interactor_abundance | mitosis | inter | 0.9635437341838033 | 0.9767804789925814 |

P47914_P62829
RL29_HUMAN vs RL23_HUMAN
p-value: 0.0 q-value: 0.0

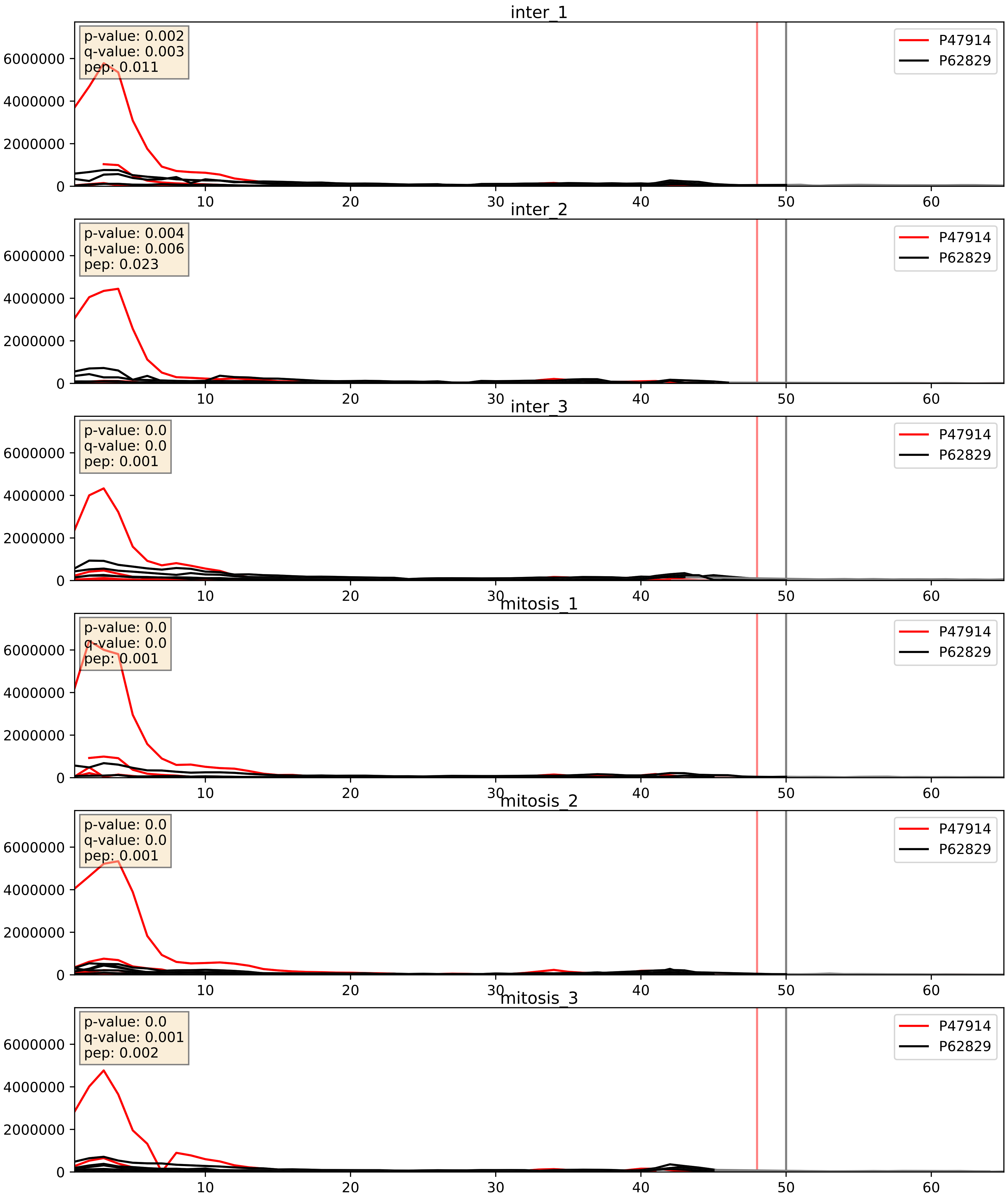


| level | condition_1 | condition_2 | pvalue | pvalue_adjusted |
|---|---|---|---|---|
| interactor_ratio | mitosis | inter | 0.003861407671655312 | 0.08075090146433855 |
| complex_abundance | mitosis | inter | 0.12584202559194957 | 0.2679166881368564 |
| interactor_abundance | mitosis | inter | 0.422038001585224 | 0.6031263351271641 |

P48047_P62829
ATPO_HUMAN vs RL23_HUMAN
p-value: 0.0 q-value: 0.049

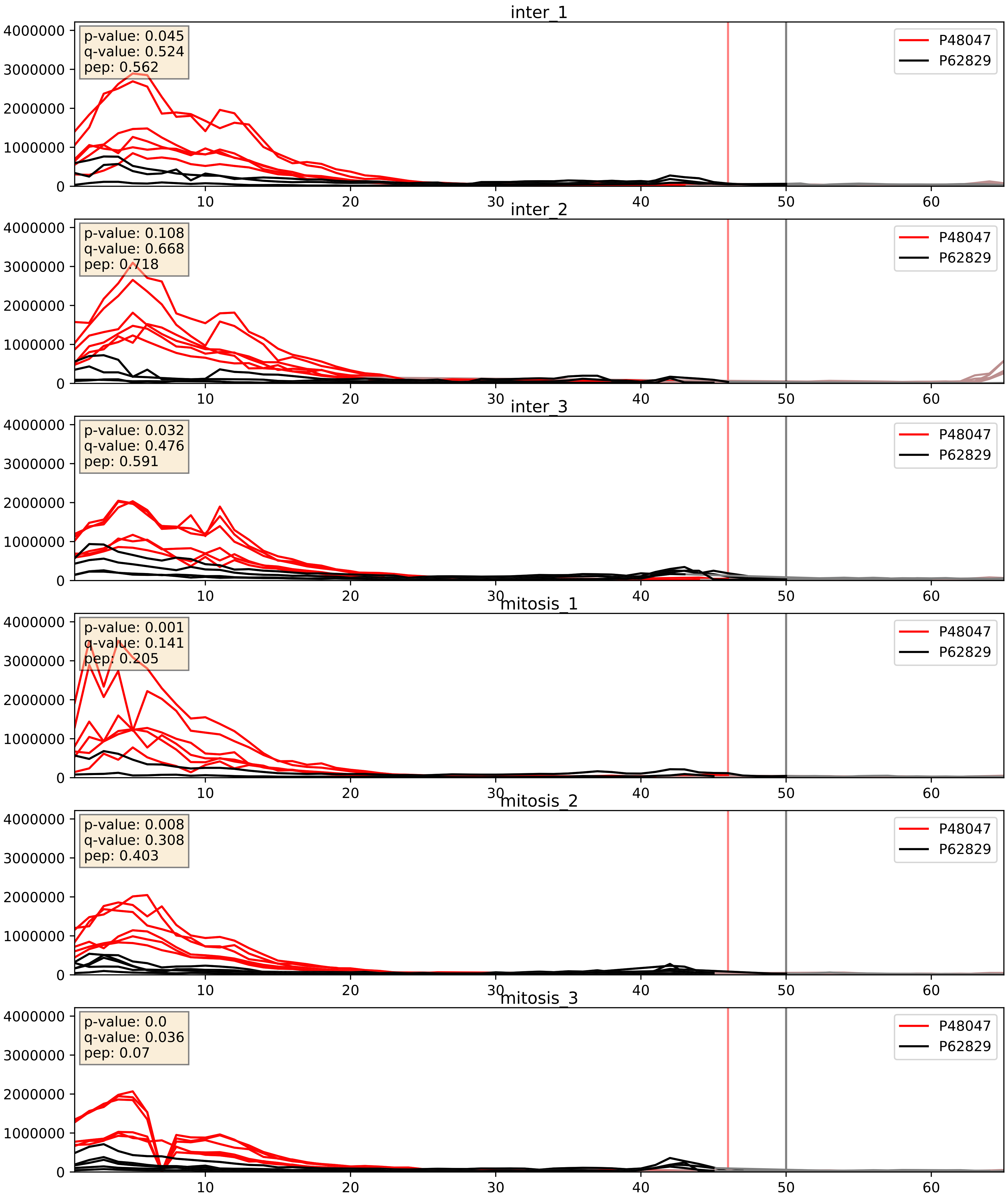


| level | condition_1 | condition_2 | pvalue | pvalue_adjusted |
|---|---|---|---|---|
| complex_abundance | mitosis | inter | 0.014561276663149561 | 0.0896723224723455 |
| interactor_ratio | mitosis | inter | 0.03802866468283547 | 0.3198713130230032 |
| interactor_abundance | mitosis | inter | 0.06144715264869247 | 0.20277086610362666 |

P49207_P62829
RL34_HUMAN vs RL23_HUMAN
p-value: 0.0 q-value: 0.0

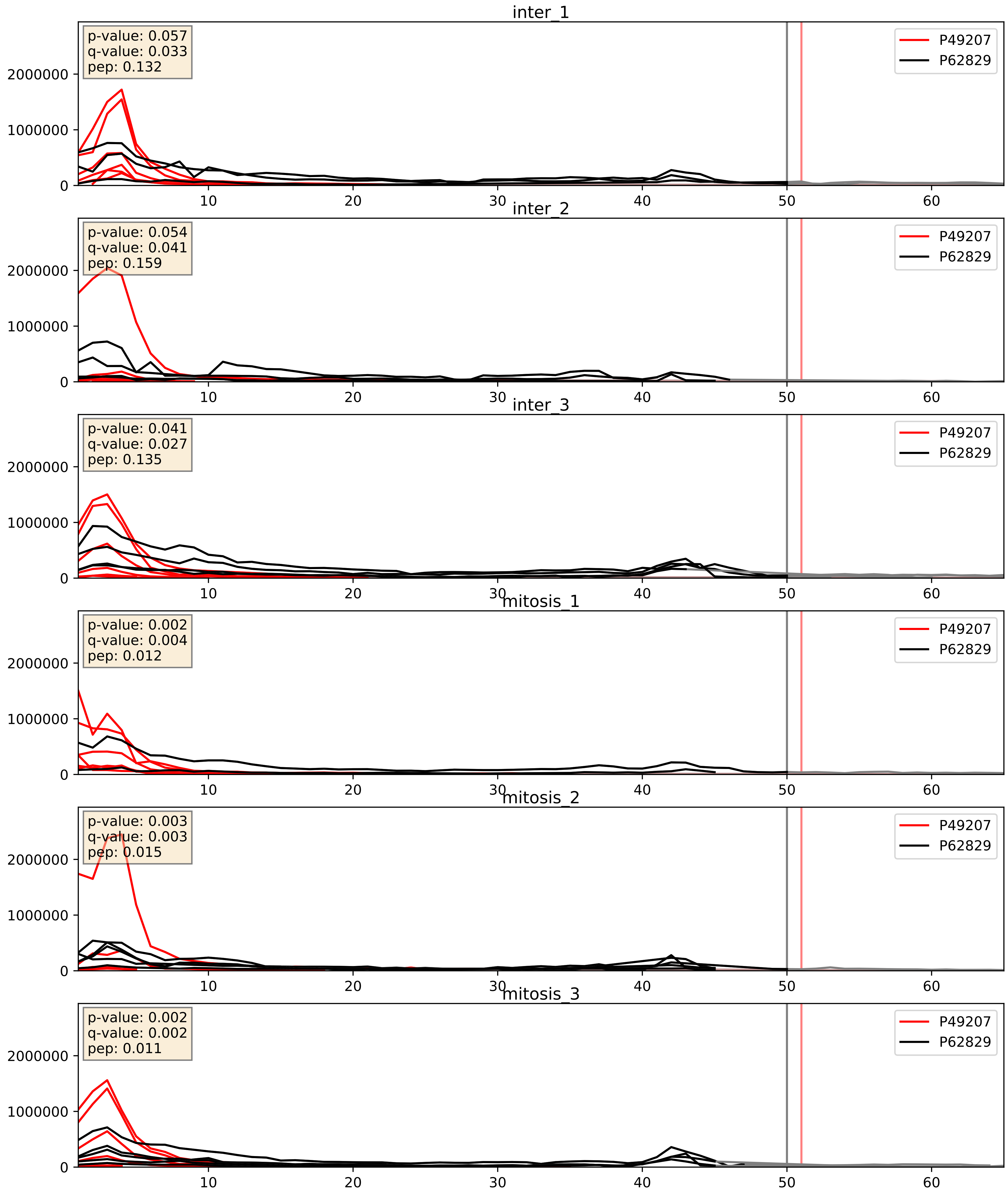


| level | condition_1 | condition_2 | pvalue | pvalue_adjusted |
|---|---|---|---|---|
| interactor_ratio | mitosis | inter | 0.09316781323348611 | 0.5070243509919888 |
| complex_abundance | mitosis | inter | 0.10447071787227112 | 0.2402721956553067 |
| interactor_abundance | mitosis | inter | 0.6896689856436142 | 0.7949860647871448 |

P49458_P62829
SRP09_HUMAN vs RL23_HUMAN
p-value: 0.001 q-value: 0.001

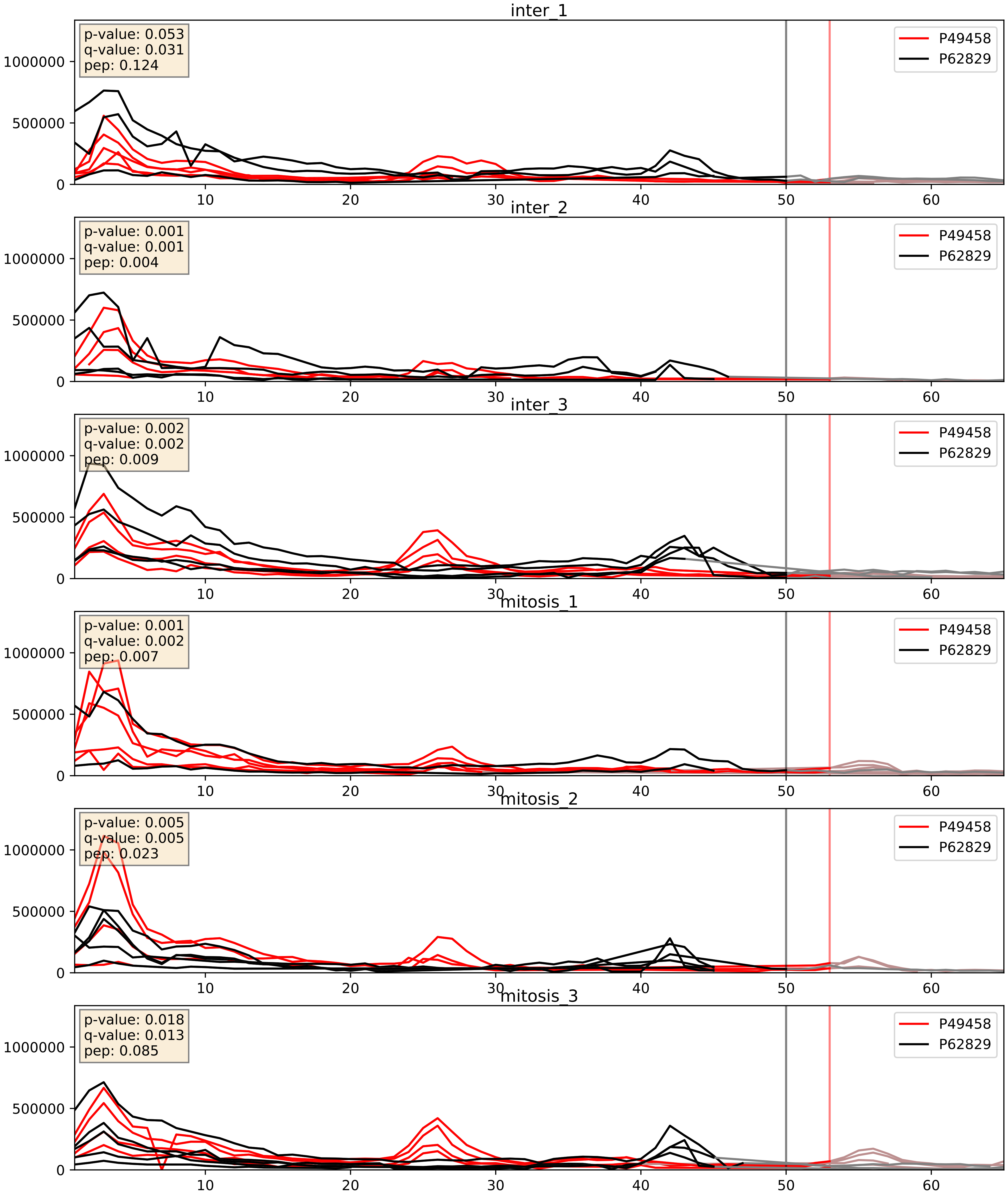


| level | condition_1 | condition_2 | pvalue | pvalue_adjusted |
|---|---|---|---|---|
| interactor_ratio | mitosis | inter | 0.00043361263857275473 | 0.02319827616364238 |
| complex_abundance | mitosis | inter | 0.06396586560369387 | 0.18245511814982324 |
| interactor_abundance | mitosis | inter | 0.45169823405767384 | 0.6250463762582749 |

P50914_P62829
RL14_HUMAN vs RL23_HUMAN
p-value: 0.0 q-value: 0.0

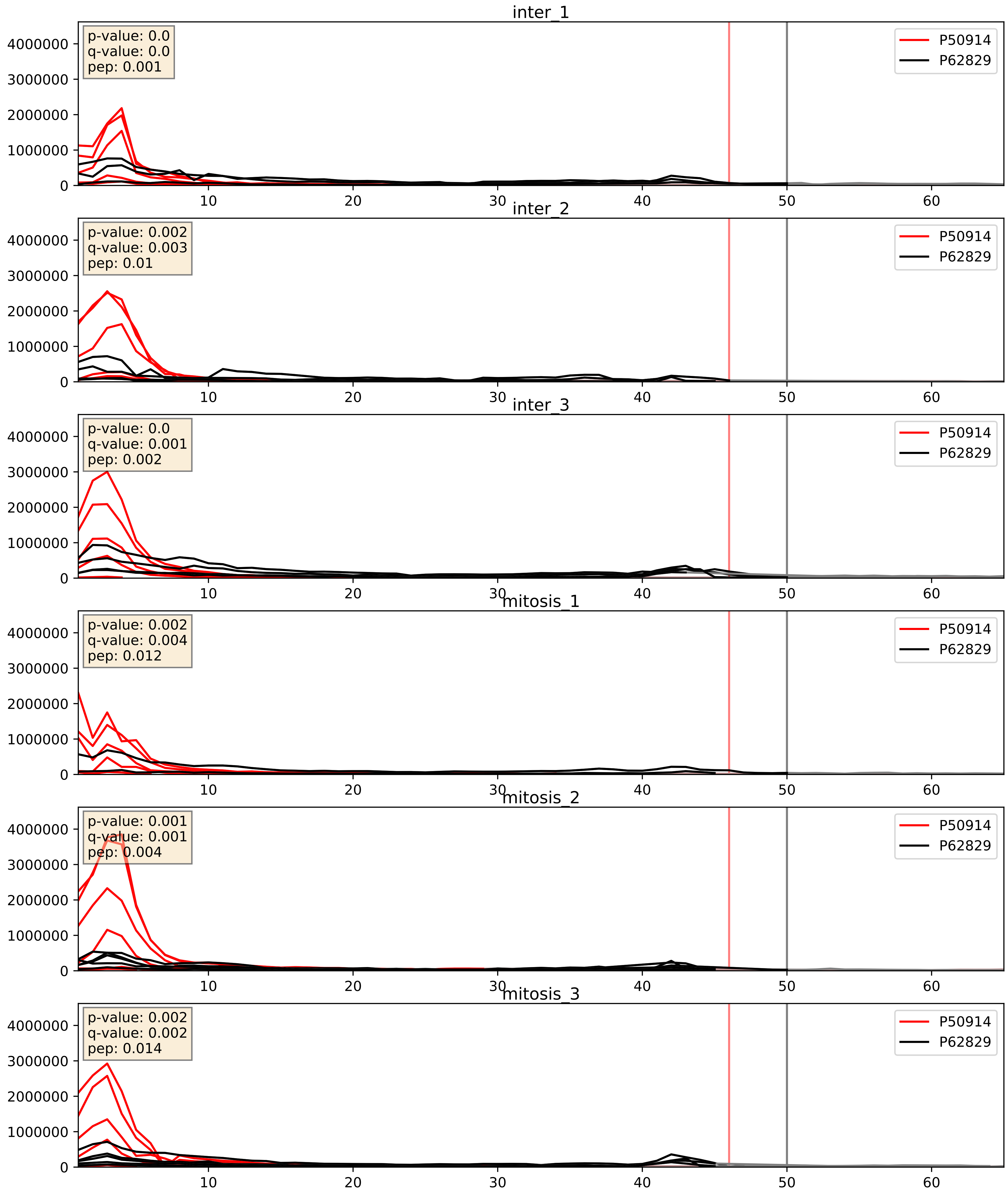


| level | condition_1 | condition_2 | pvalue | pvalue_adjusted |
|---|---|---|---|---|
| interactor_ratio | mitosis | inter | 0.010462332127968803 | 0.14681567707444748 |
| complex_abundance | mitosis | inter | 0.01609353608411727 | 0.0946158440110191 |
| interactor_abundance | mitosis | inter | 0.8451161396574962 | 0.9151994630233623 |

P55769_P62829
NH2L1_HUMAN vs RL23_HUMAN
p-value: 0.0 q-value: 0.008

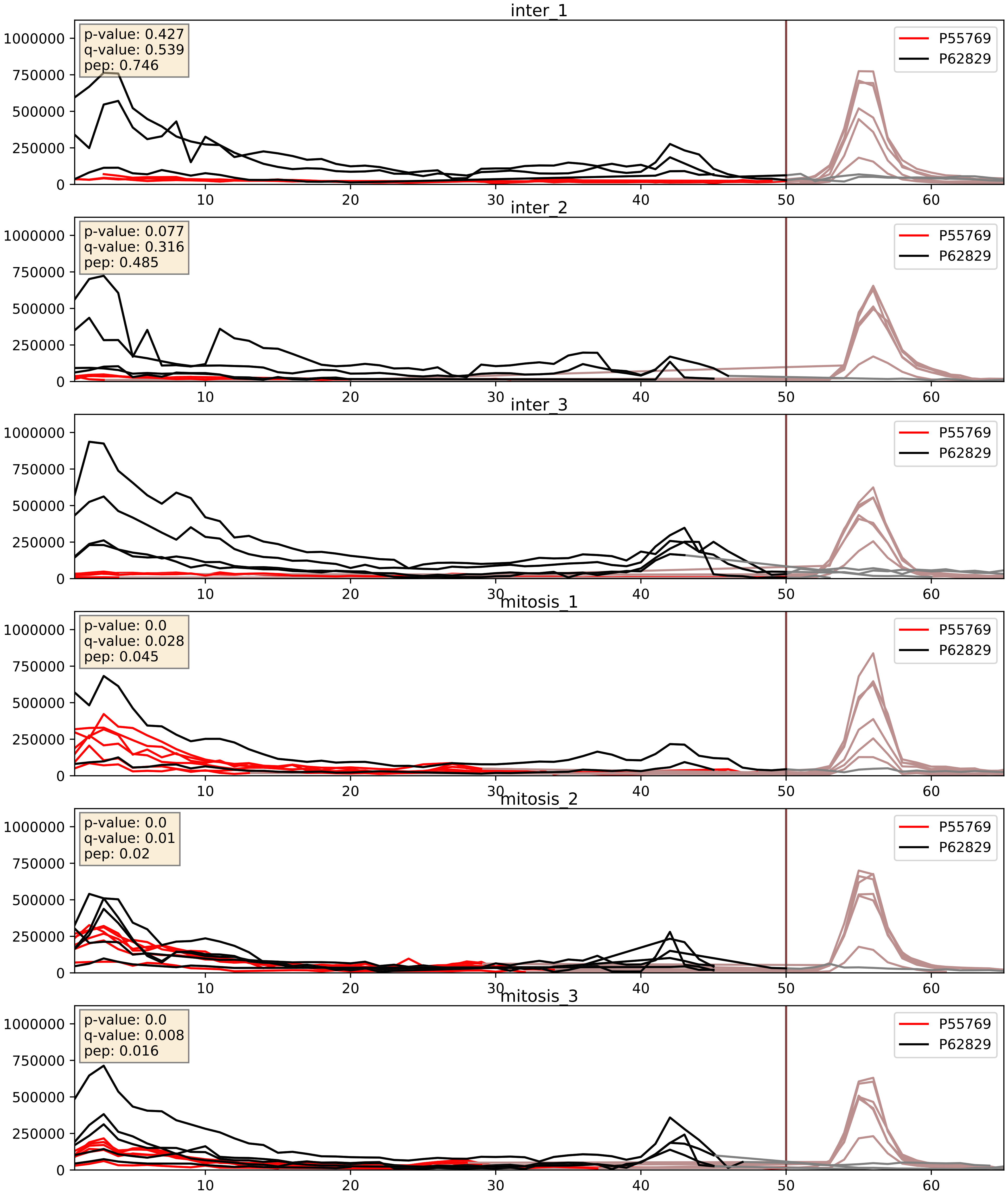


| level | condition_1 | condition_2 | pvalue | pvalue_adjusted |
|---|---|---|---|---|
| interactor_ratio | mitosis | inter | 0.0010607158442868909 | 0.038414698994273265 |
| interactor_abundance | mitosis | inter | 0.003522677249577062 | 0.04582692592154961 |
| complex_abundance | mitosis | inter | 0.8831107774455889 | 0.9275372091943853 |

P60468_P62829
SC61B_HUMAN vs RL23_HUMAN
p-value: 0.024 q-value: 0.019

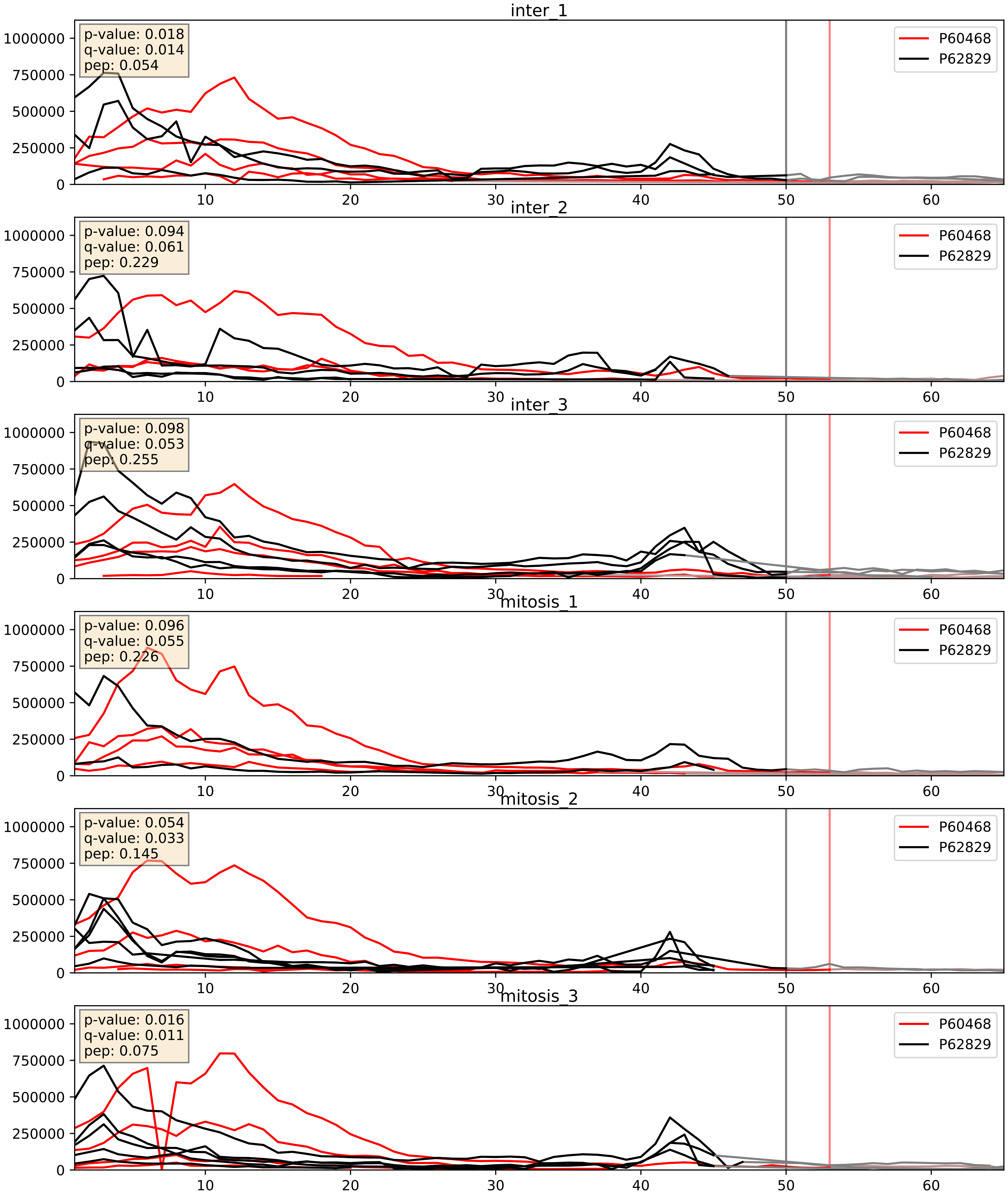


| level | condition_1 | condition_2 | pvalue | pvalue_adjusted |
|---|---|---|---|---|
| interactor_ratio | mitosis | inter | 0.01366265226048229 | 0.17048440721534752 |
| complex_abundance | mitosis | inter | 0.022476339242519364 | 0.10987189501272765 |
| interactor_abundance | mitosis | inter | 0.26587079075564074 | 0.45810265073838263 |

P60866_P62829
RS20_HUMAN vs RL23_HUMAN
p-value: 0.003 q-value: 0.003

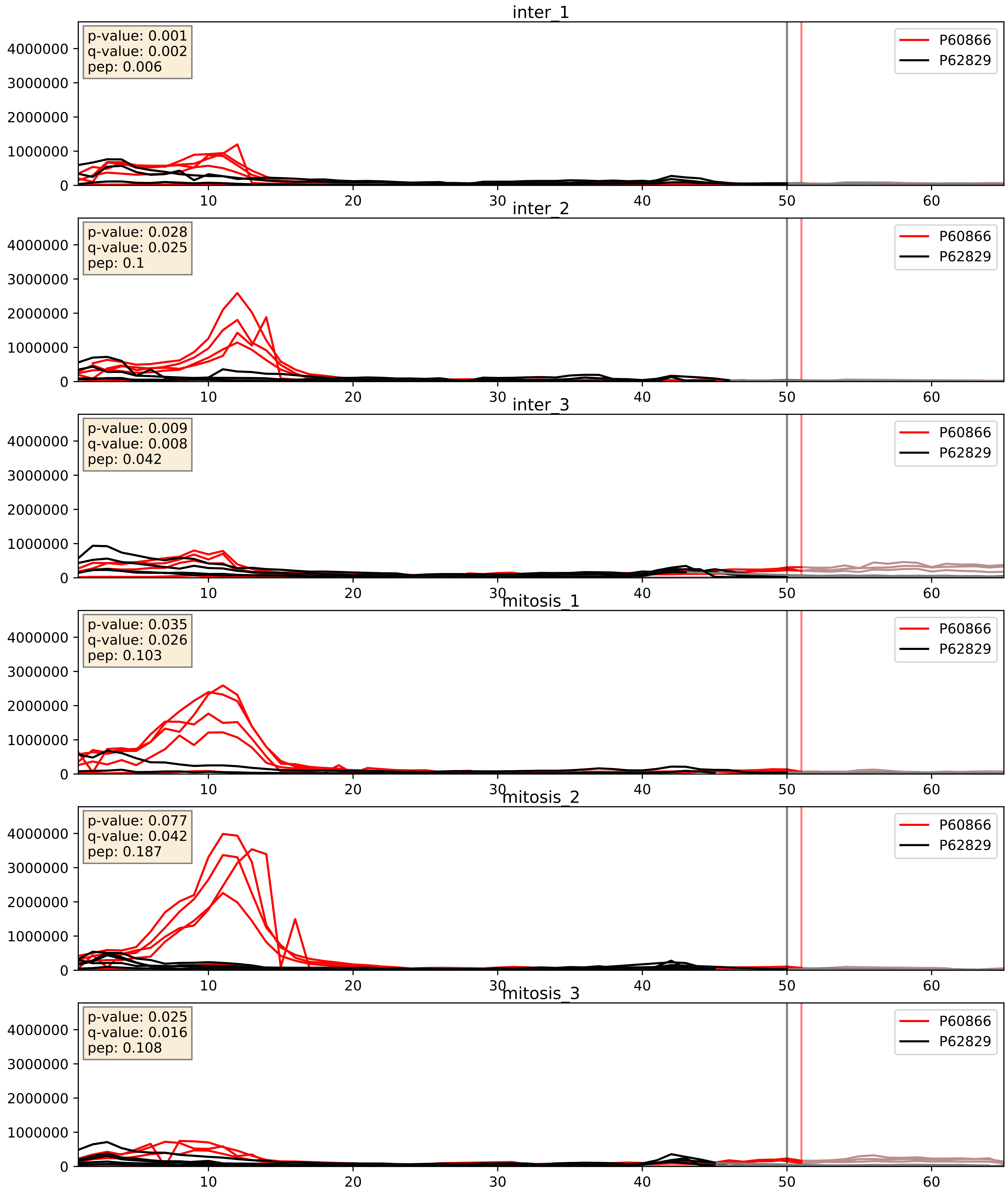


| level | condition_1 | condition_2 | pvalue | pvalue_adjusted |
|---|---|---|---|---|
| complex_abundance | mitosis | inter | 0.030201064844060637 | 0.1259849488621633 |
| interactor_ratio | mitosis | inter | 0.0302839077871649 | 0.27665981927228556 |
| interactor_abundance | mitosis | inter | 0.4180557405404574 | 0.5994735135315044 |

P61247_P62829
RS3A_HUMAN vs RL23_HUMAN
p-value: 0.0 q-value: 0.0

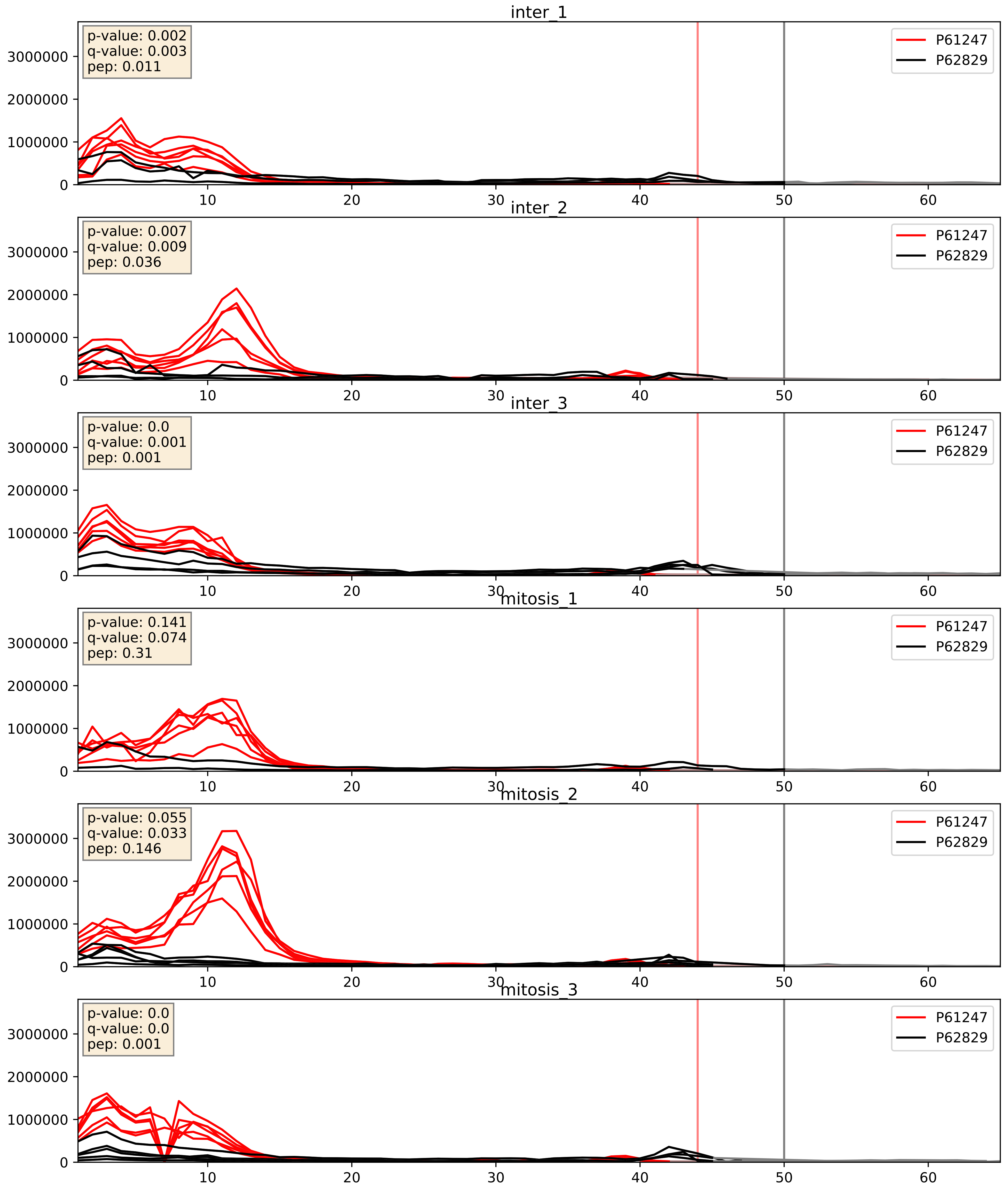


| level | condition_1 | condition_2 | pvalue | pvalue_adjusted |
|---|---|---|---|---|
| interactor_ratio | mitosis | inter | 0.0035067635692578036 | 0.07646419083591573 |
| complex_abundance | mitosis | inter | 0.025678451766092972 | 0.11728295414418871 |
| interactor_abundance | mitosis | inter | 0.5746620223826564 | 0.720300118516995 |

P61254_P62829
RL26_HUMAN vs RL23_HUMAN
p-value: 0.048 q-value: 0.032

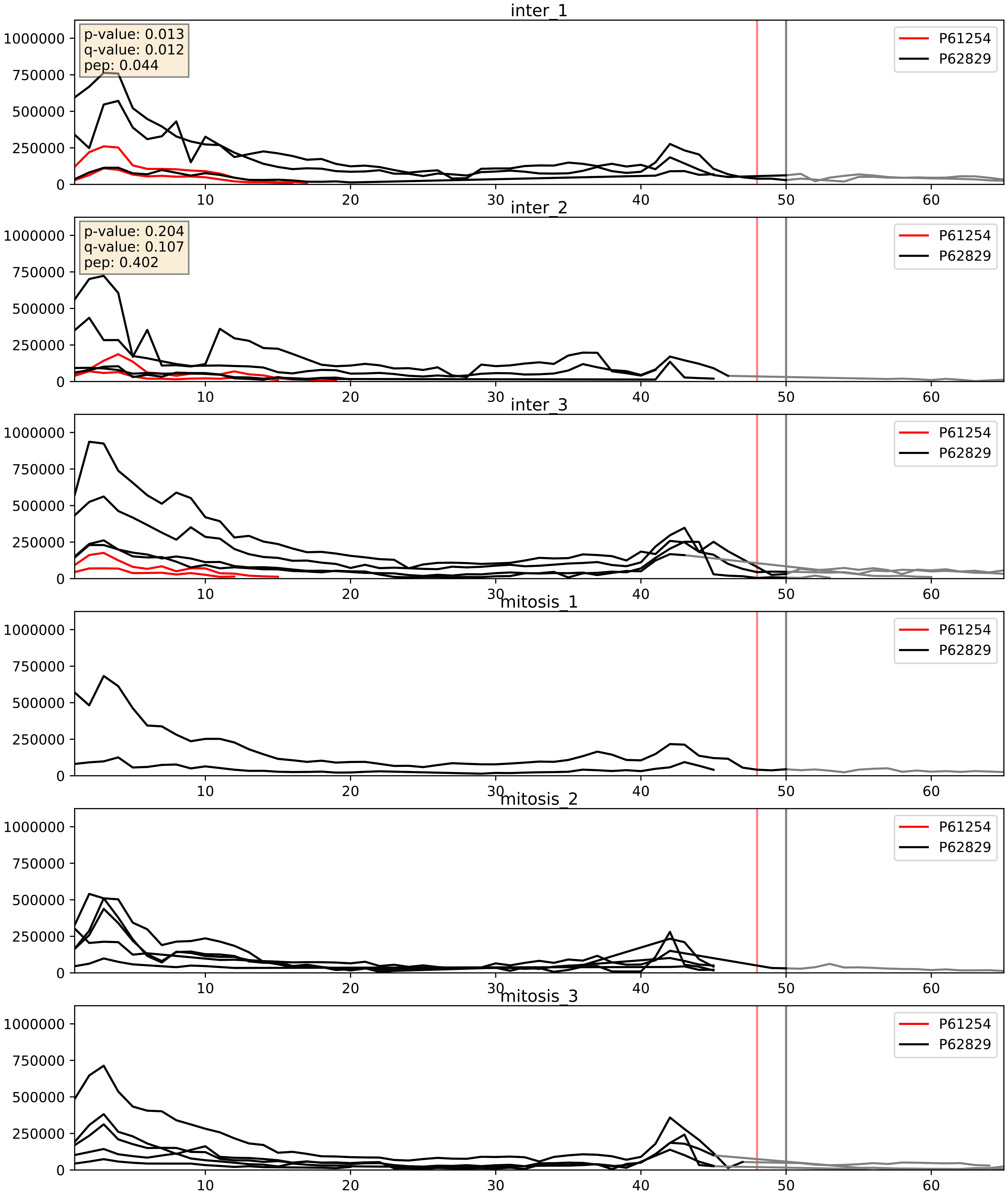


| level | condition_1 | condition_2 | pvalue | pvalue_adjusted |
|---|---|---|---|---|
| complex_abundance | mitosis | inter | 0.00010015156561264747 | 0.006746492382951911 |
| interactor_abundance | mitosis | inter | 0.0006144309768949645 | 0.017415659477552636 |
| interactor_ratio | mitosis | inter | 0.000773065761217179 | 0.03185945617443502 |

P61313_P62829
RL15_HUMAN vs RL23_HUMAN
p-value: 0.001 q-value: 0.002

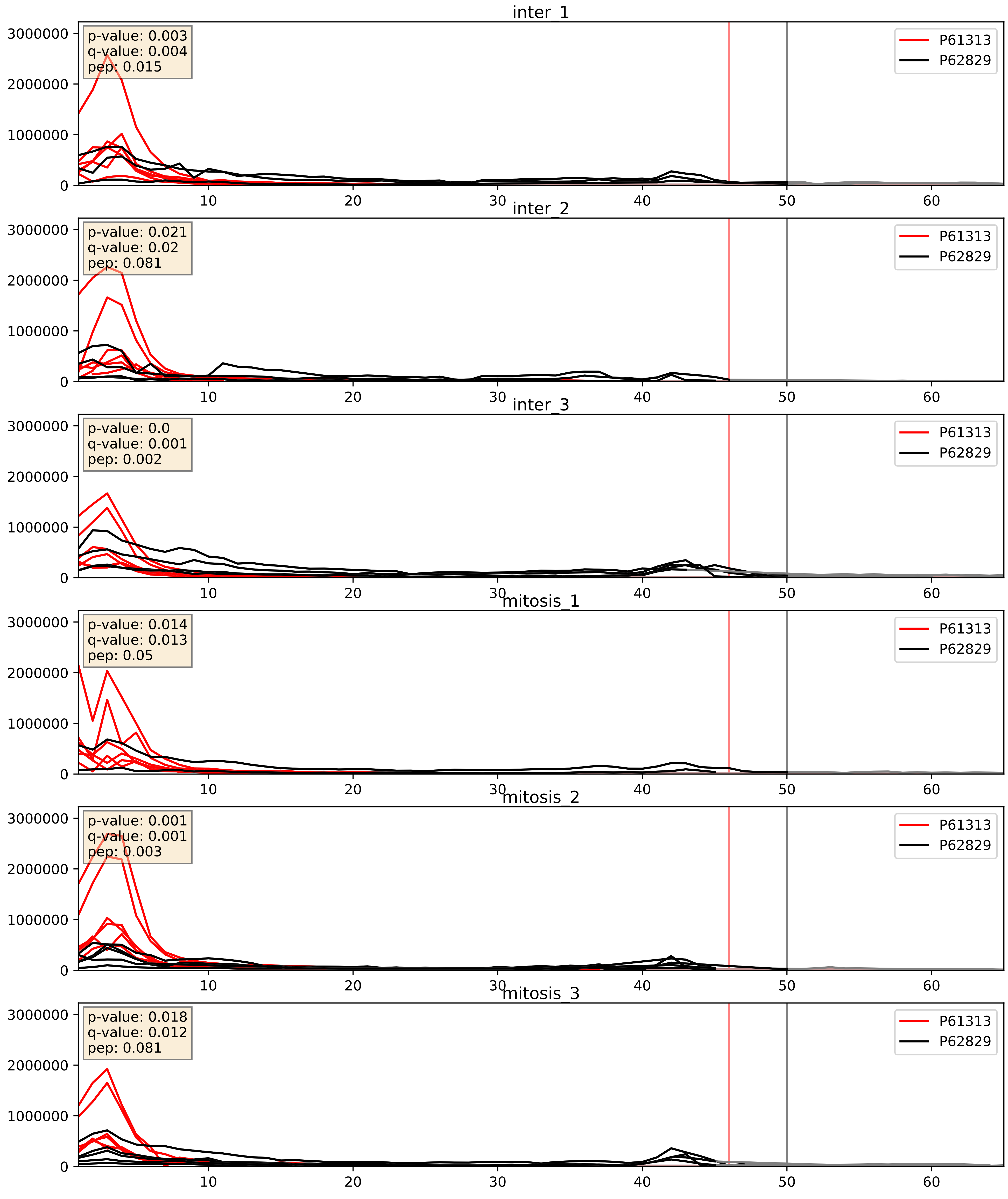


| level | condition_1 | condition_2 | pvalue | pvalue_adjusted |
|---|---|---|---|---|
| complex_abundance | mitosis | inter | 0.0007897053739137978 | 0.018520213700553727 |
| interactor_ratio | mitosis | inter | 0.0088341713603176 | 0.13290071501637724 |
| interactor_abundance | mitosis | inter | 0.9175497077049597 | 0.951681267170054 |

P61353_P62829
RL27_HUMAN vs RL23_HUMAN
p-value: 0.0 q-value: 0.0

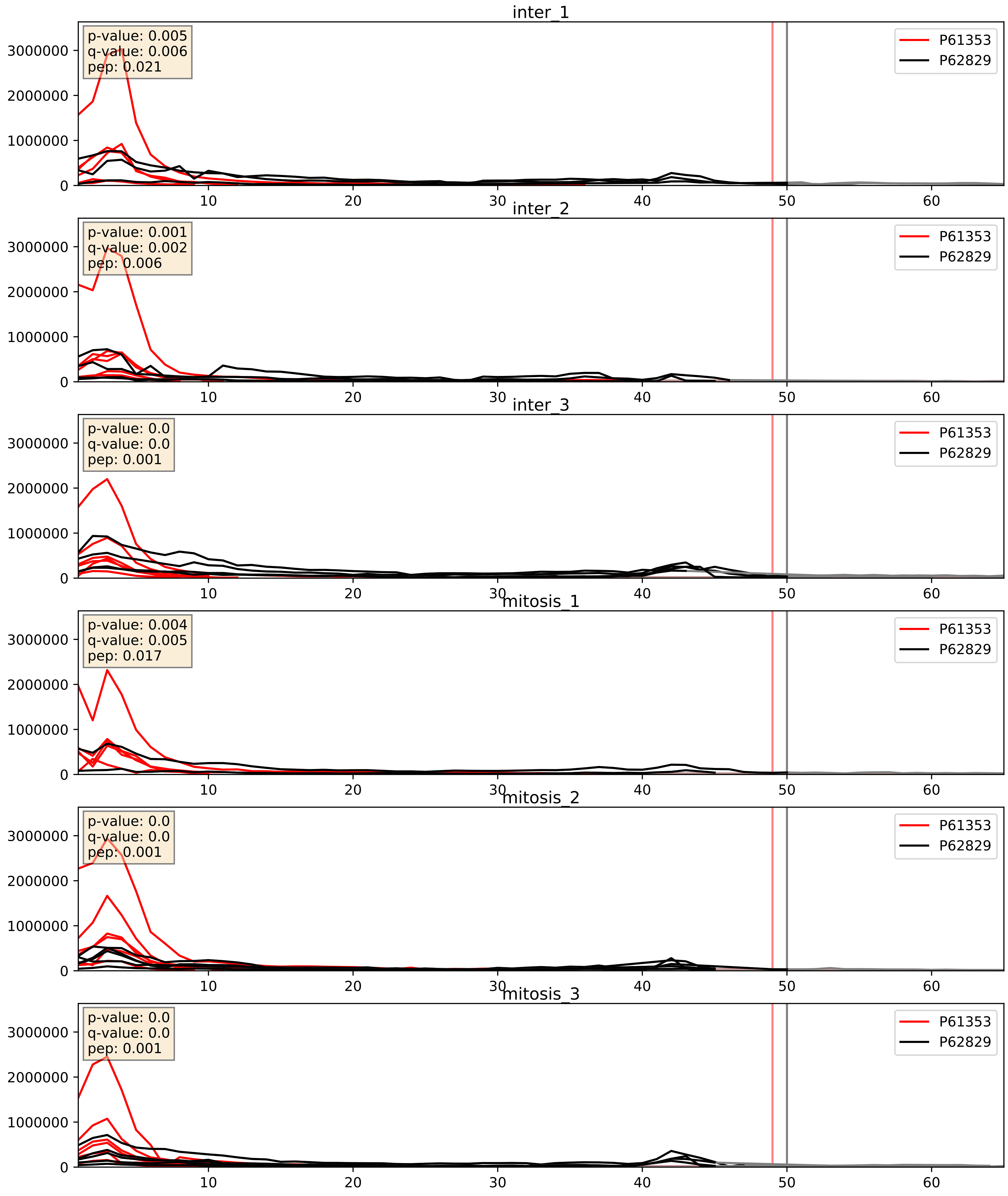


| level | condition_1 | condition_2 | pvalue | pvalue_adjusted |
|---|---|---|---|---|
| complex_abundance | mitosis | inter | 0.001721938851284516 | 0.02936214455576784 |
| interactor_ratio | mitosis | inter | 0.010417023732674625 | 0.14681567707444748 |
| interactor_abundance | mitosis | inter | 0.5867616308489645 | 0.7283996171513504 |

## P61513_P62829
### RL37A_HUMAN vs RL23_HUMAN
p-value: 0.008 q-value: 0.008

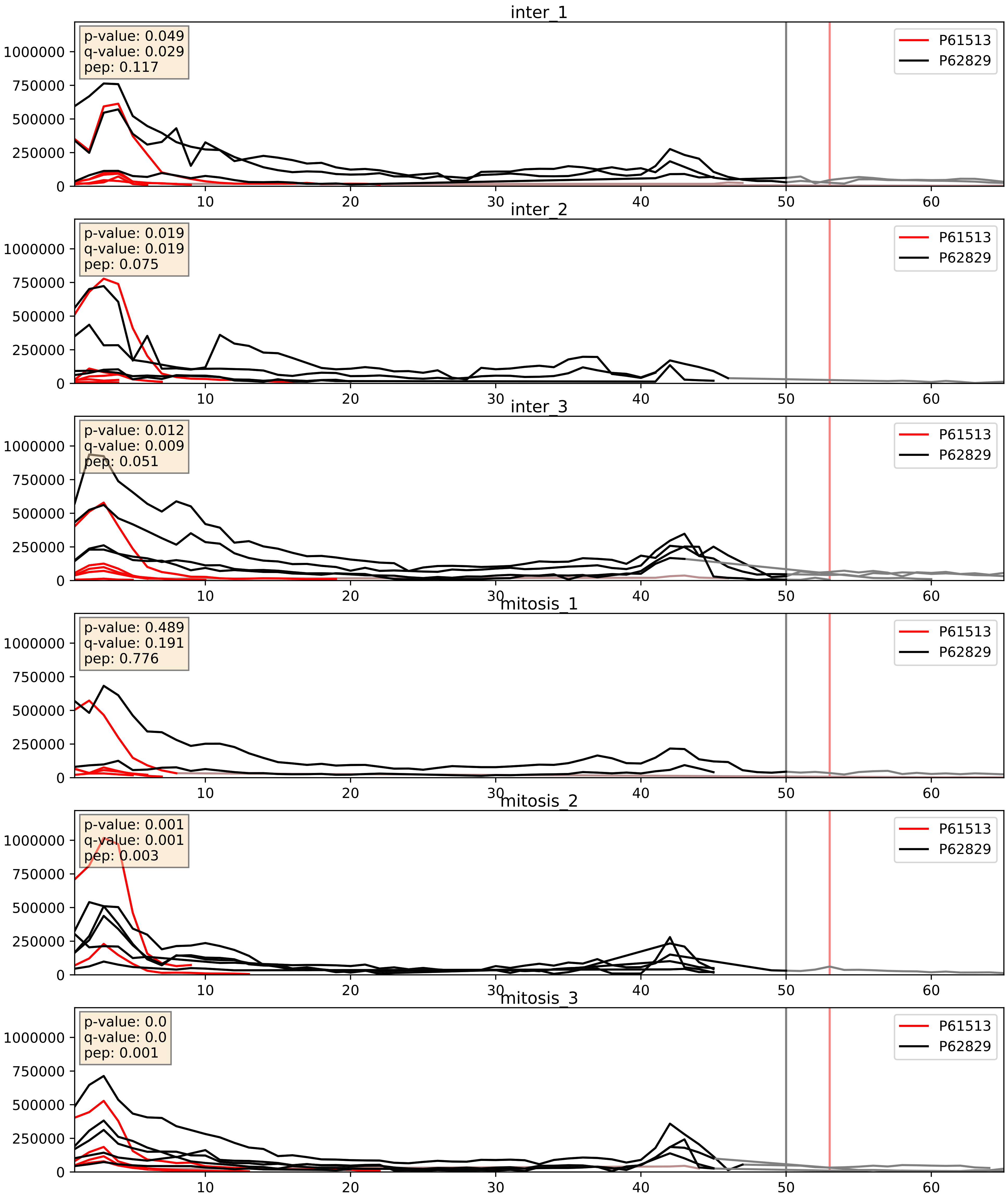


| level | condition_1 | condition_2 | pvalue | pvalue_adjusted |
|---|---|---|---|---|
| complex_abundance | mitosis | inter | 0.0034424480084898072 | 0.04302508960916016 |
| interactor_ratio | mitosis | inter | 0.01779698740545046 | 0.20091601471792916 |
| interactor_abundance | mitosis | inter | 0.04383858707081264 | 0.16685321787696922 |

P61619_P62829
S61A1_HUMAN vs RL23_HUMAN
p-value: 0.023 q-value: 0.018

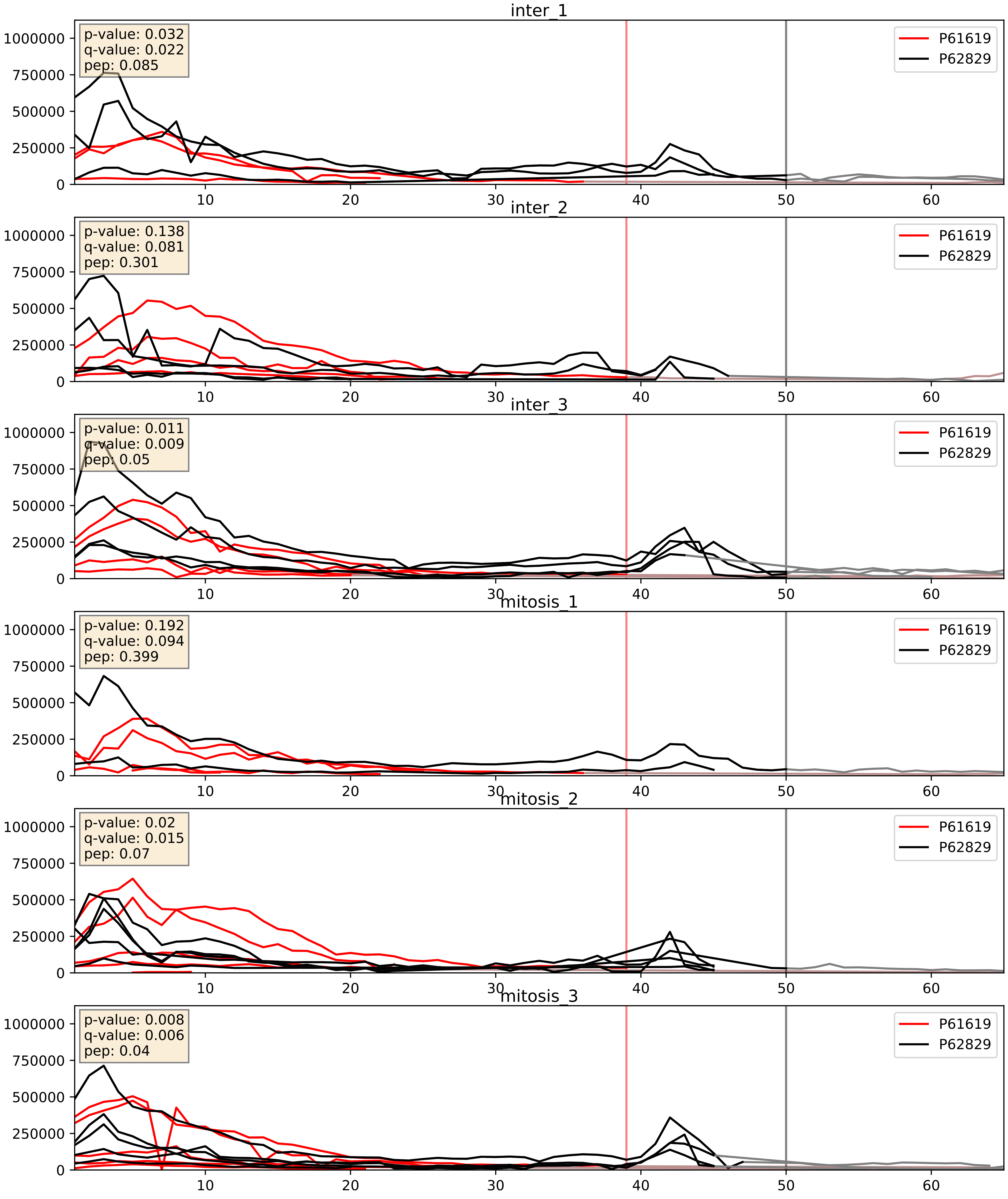


| level | condition_1 | condition_2 | pvalue | pvalue_adjusted |
|---|---|---|---|---|
| complex_abundance | mitosis | inter | 0.01644181332149885 | 0.09579267087919495 |
| interactor_ratio | mitosis | inter | 0.01992001460681836 | 0.21404307217424748 |
| interactor_abundance | mitosis | inter | 0.6440730353911753 | 0.7618163856499186 |

P62081_P62829
RS7_HUMAN vs RL23_HUMAN
p-value: 0.001 q-value: 0.001

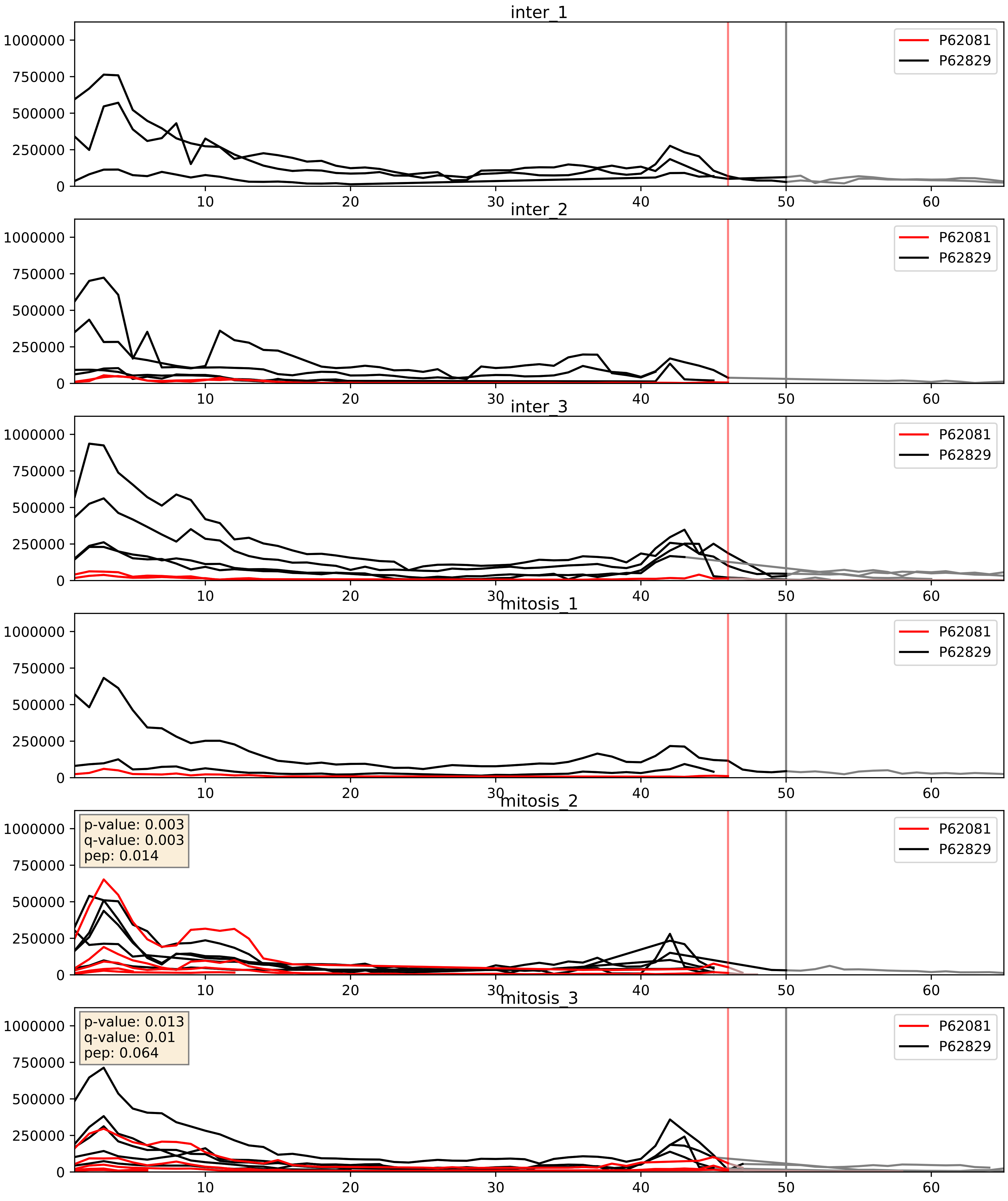


| level | condition_1 | condition_2 | pvalue | pvalue_adjusted |
|---|---|---|---|---|
| interactor_abundance | mitosis | inter | 0.25264231175823587 | 0.44596479790469956 |
| interactor_ratio | mitosis | inter | 0.33532092856761814 | 0.7914334552464135 |
| complex_abundance | mitosis | inter | 0.7957719715558504 | 0.8674141043952796 |

P62241_P62829
RS8_HUMAN vs RL23_HUMAN
p-value: 0.0 q-value: 0.0

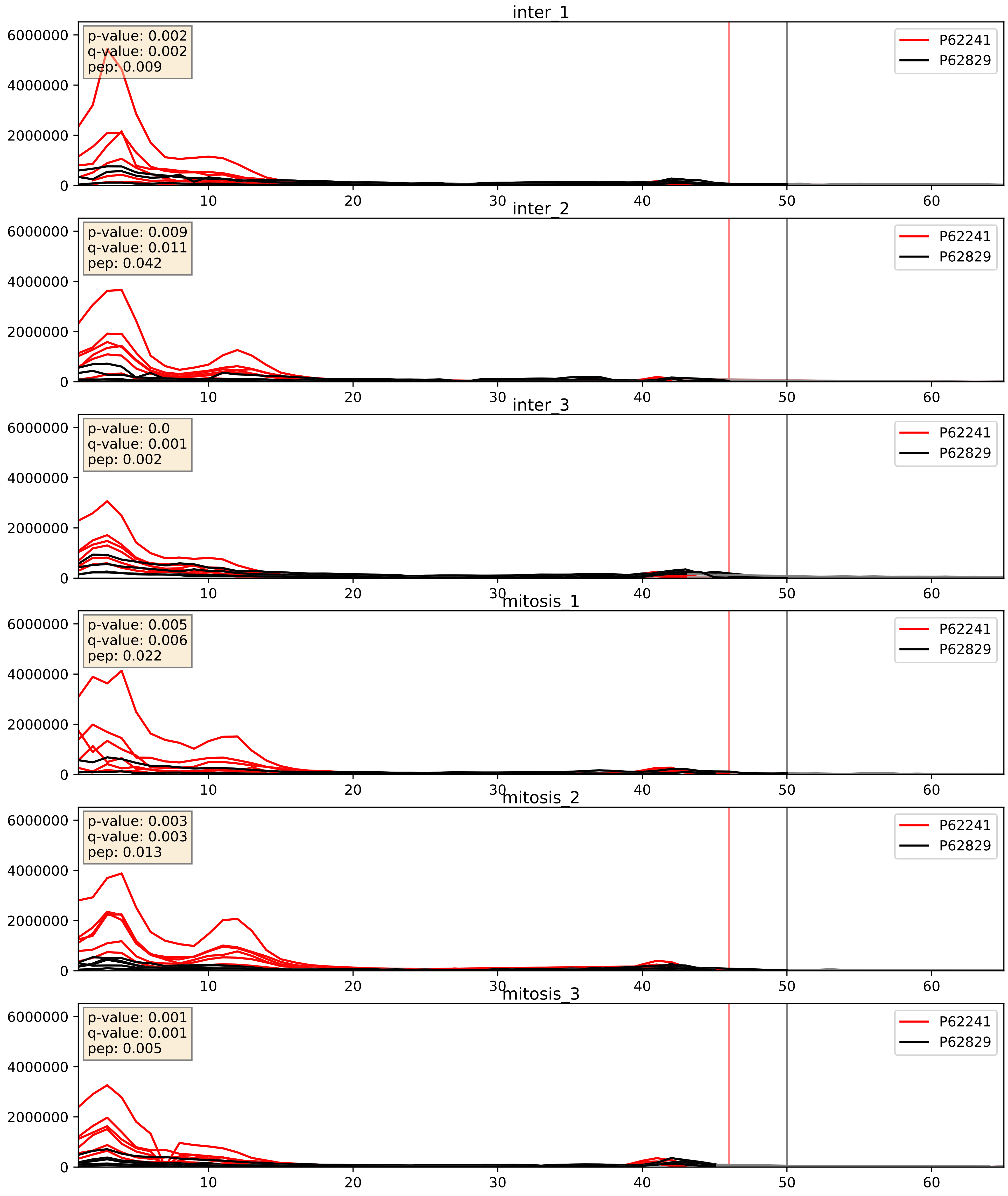


| level | condition_1 | condition_2 | pvalue | pvalue_adjusted |
|---|---|---|---|---|
| interactor_ratio | mitosis | inter | 0.006780720840639278 | 0.11425781573990595 |
| complex_abundance | mitosis | inter | 0.009059243667869806 | 0.07056153393718428 |
| interactor_abundance | mitosis | inter | 0.21849374089031023 | 0.4138761721666421 |

P62244_P62829
RS15A_HUMAN vs RL23_HUMAN
p-value: 0.0 q-value: 0.0

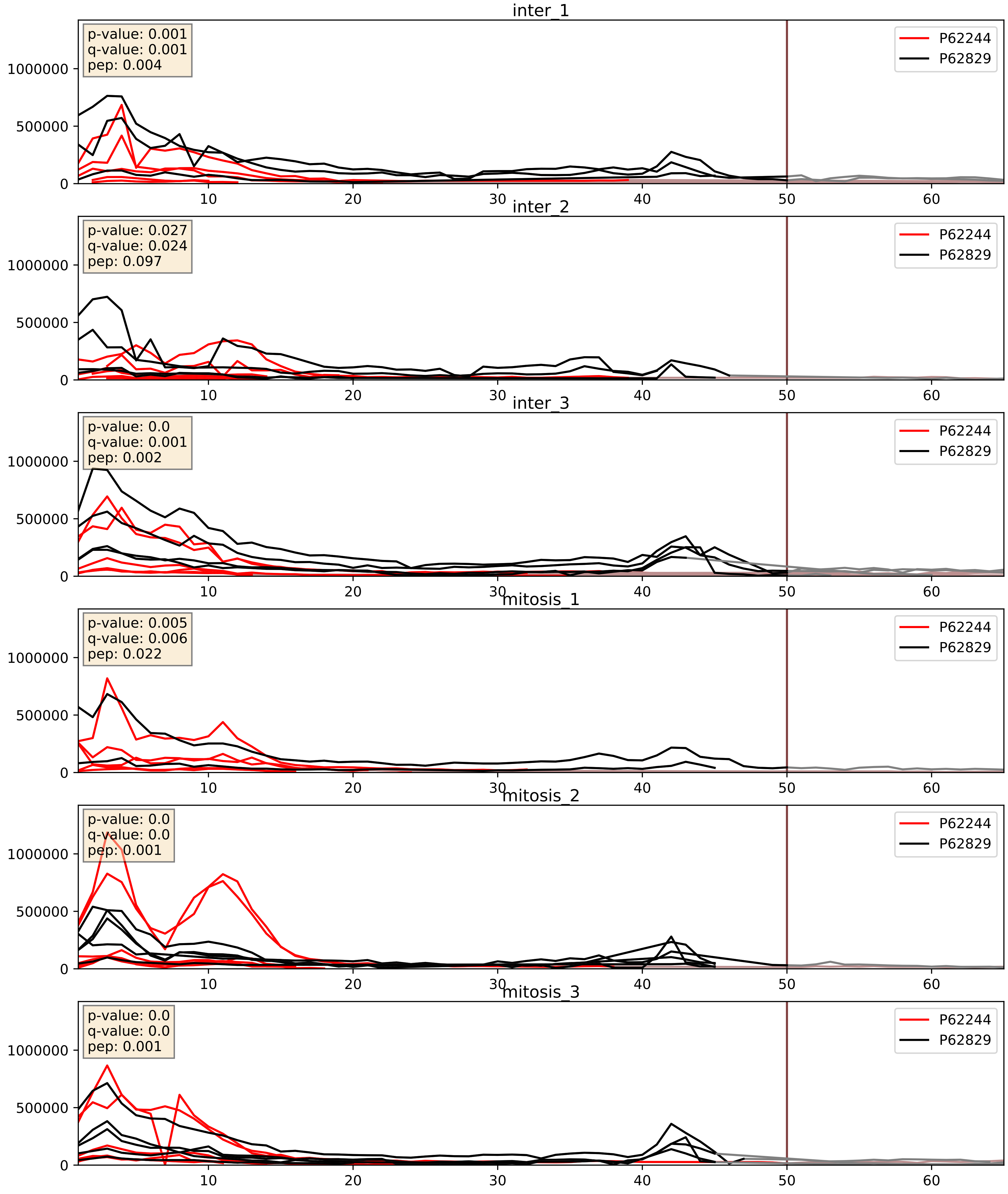


| level | condition_1 | condition_2 | pvalue | pvalue_adjusted |
|---|---|---|---|---|
| interactor_ratio | mitosis | inter | 0.004494945292431036 | 0.08927315940419876 |
| complex_abundance | mitosis | inter | 0.039470628274736276 | 0.14457363202042897 |
| interactor_abundance | mitosis | inter | 0.1664344375016143 | 0.35470653180973943 |

P62249_P62829
RS16_HUMAN vs RL23_HUMAN
p-value: 0.0 q-value: 0.0

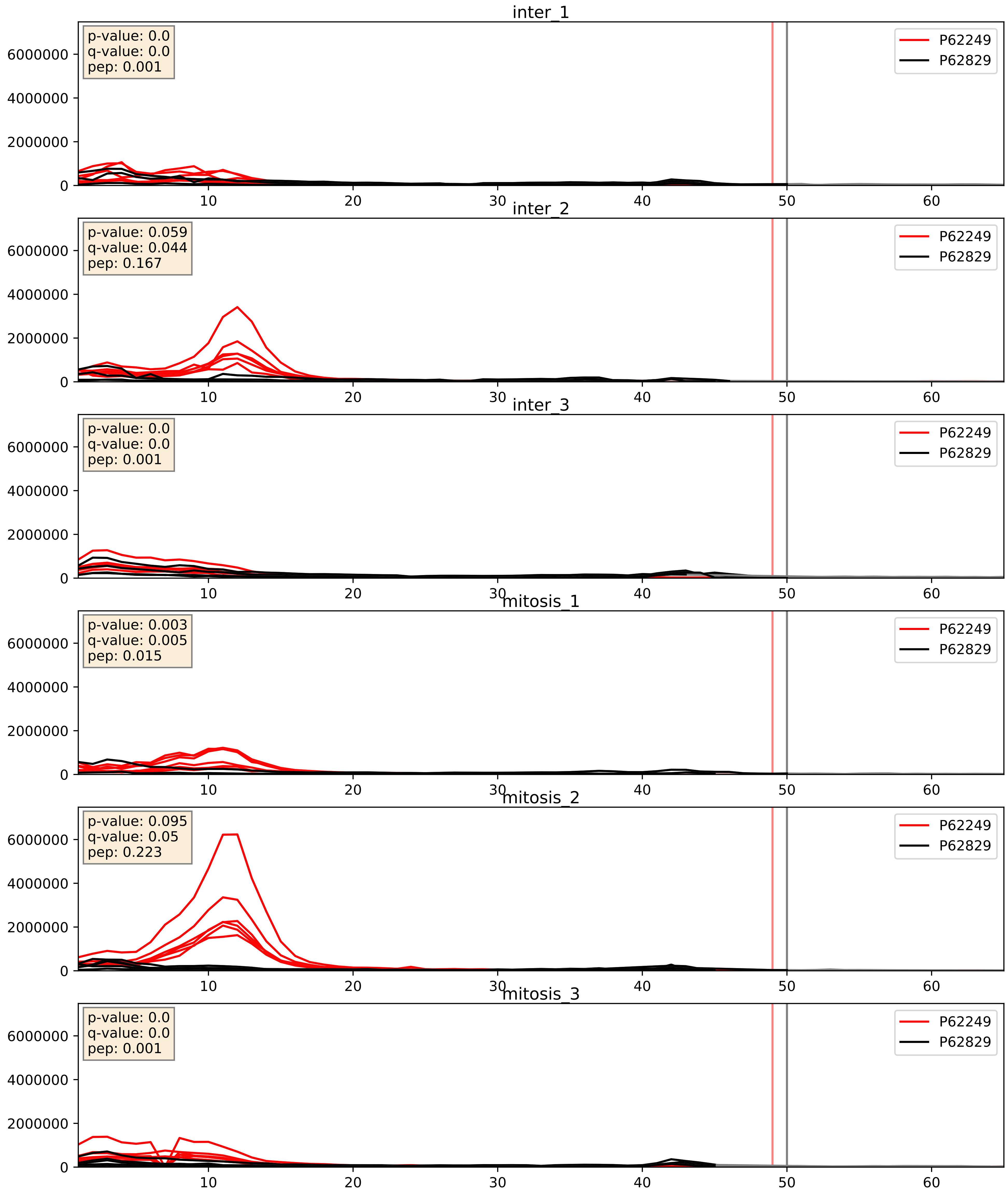


| level | condition_1 | condition_2 | pvalue | pvalue_adjusted |
|---|---|---|---|---|
| complex_abundance | mitosis | inter | 0.01753410318600555 | 0.09900522643285456 |
| interactor_ratio | mitosis | inter | 0.04672934863460885 | 0.35838626305591875 |
| interactor_abundance | mitosis | inter | 0.5933935659124083 | 0.7327008906181343 |

P62263_P62829
RS14_HUMAN vs RL23_HUMAN
p-value: 0.0 q-value: 0.0

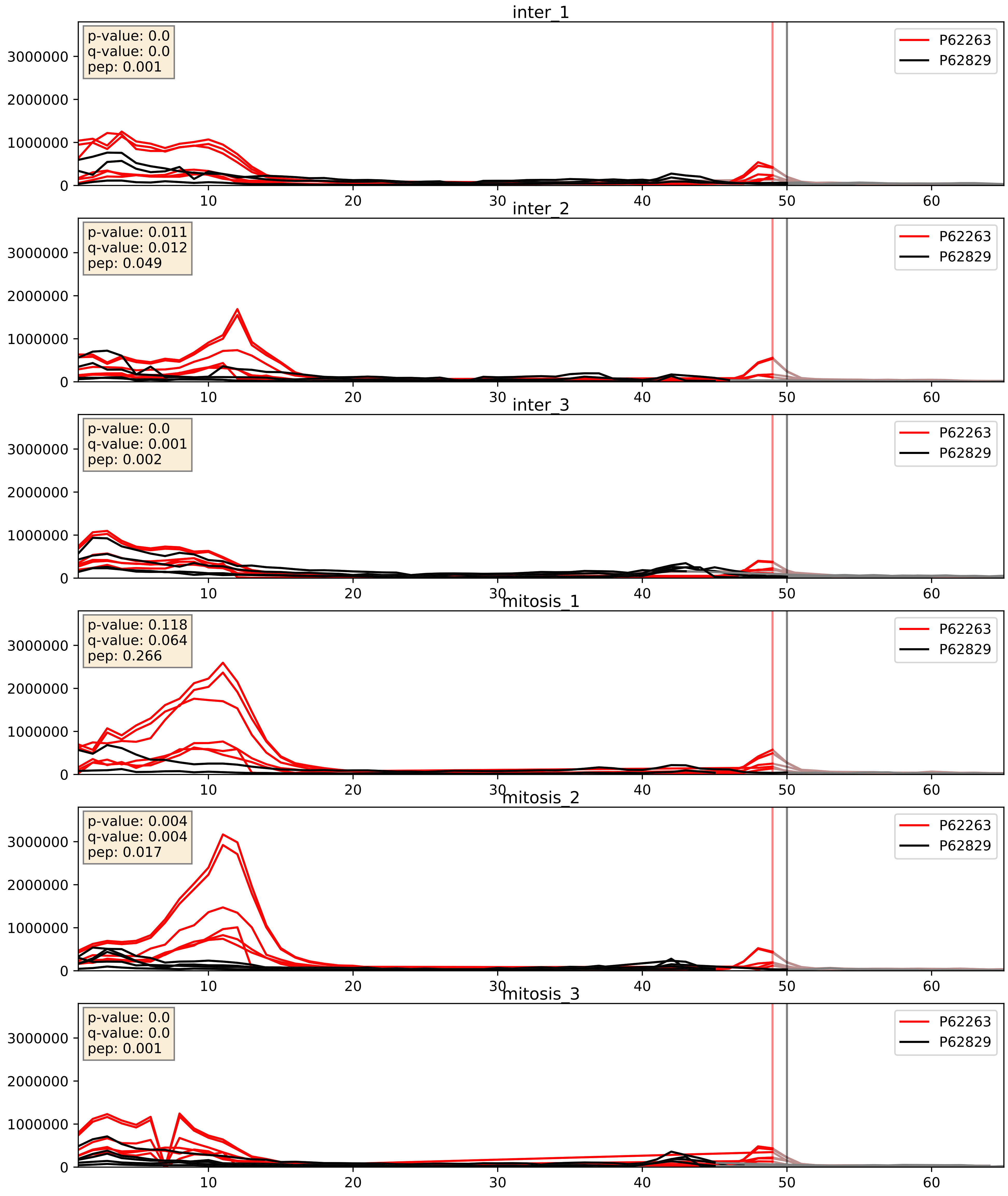


| level | condition_1 | condition_2 | pvalue | pvalue_adjusted |
|---|---|---|---|---|
| interactor_ratio | mitosis | inter | 0.020565732012535844 | 0.216534644560033 |
| complex_abundance | mitosis | inter | 0.034599625184919554 | 0.13505371253210735 |
| interactor_abundance | mitosis | inter | 0.469004134505803 | 0.6410147519351227 |

P62266_P62829
RS23_HUMAN vs RL23_HUMAN
p-value: 0.0 q-value: 0.0

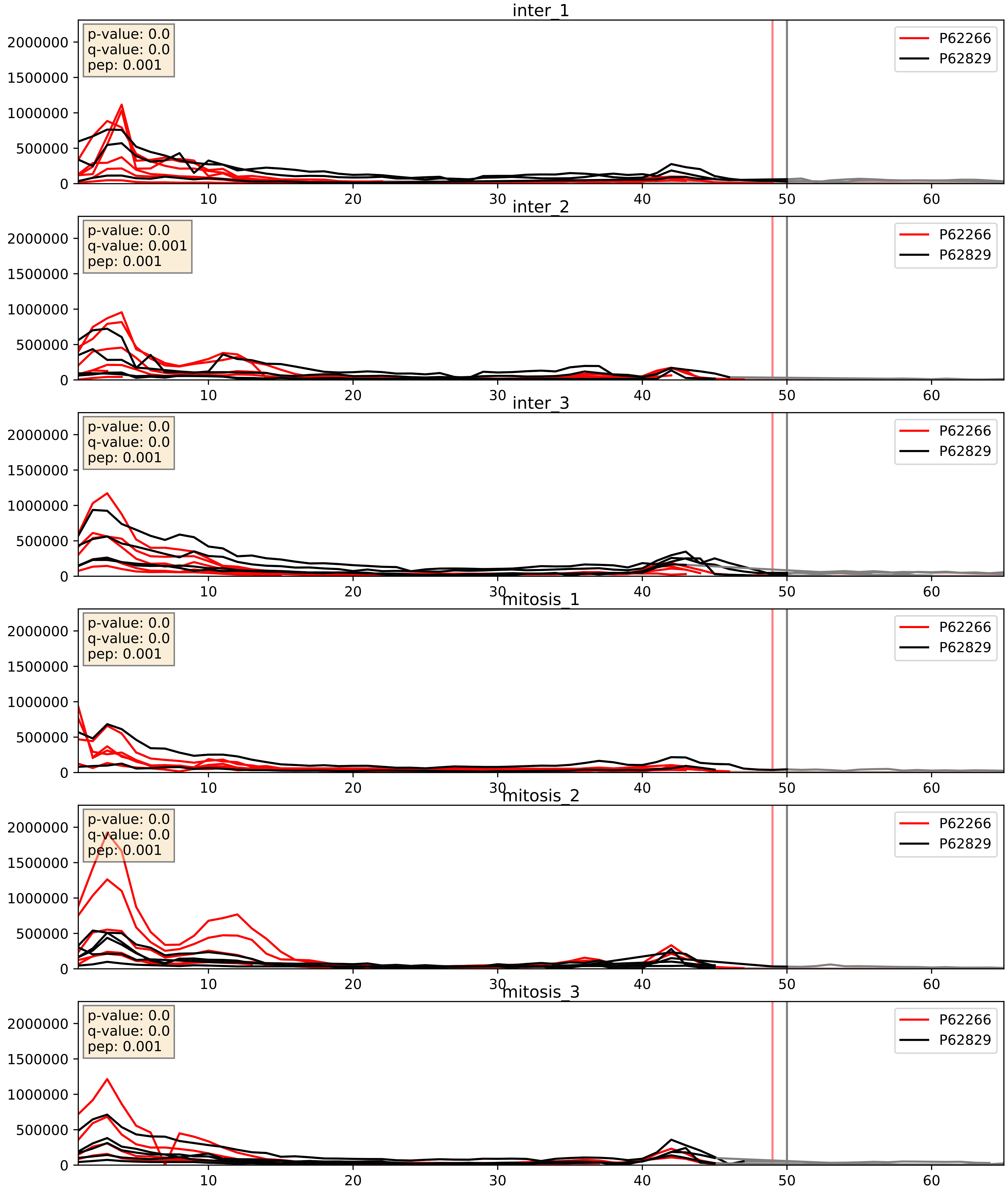


| level | condition_1 | condition_2 | pvalue | pvalue_adjusted |
|---|---|---|---|---|
| complex_abundance | mitosis | inter | 0.00997525469052954 | 0.07371585975104354 |
| interactor_ratio | mitosis | inter | 0.19323639900602177 | 0.6660896702330326 |
| interactor_abundance | mitosis | inter | 0.2858619069867244 | 0.479799592903208 |

P62269_P62829
RS18_HUMAN vs RL23_HUMAN
p-value: 0.001 q-value: 0.001

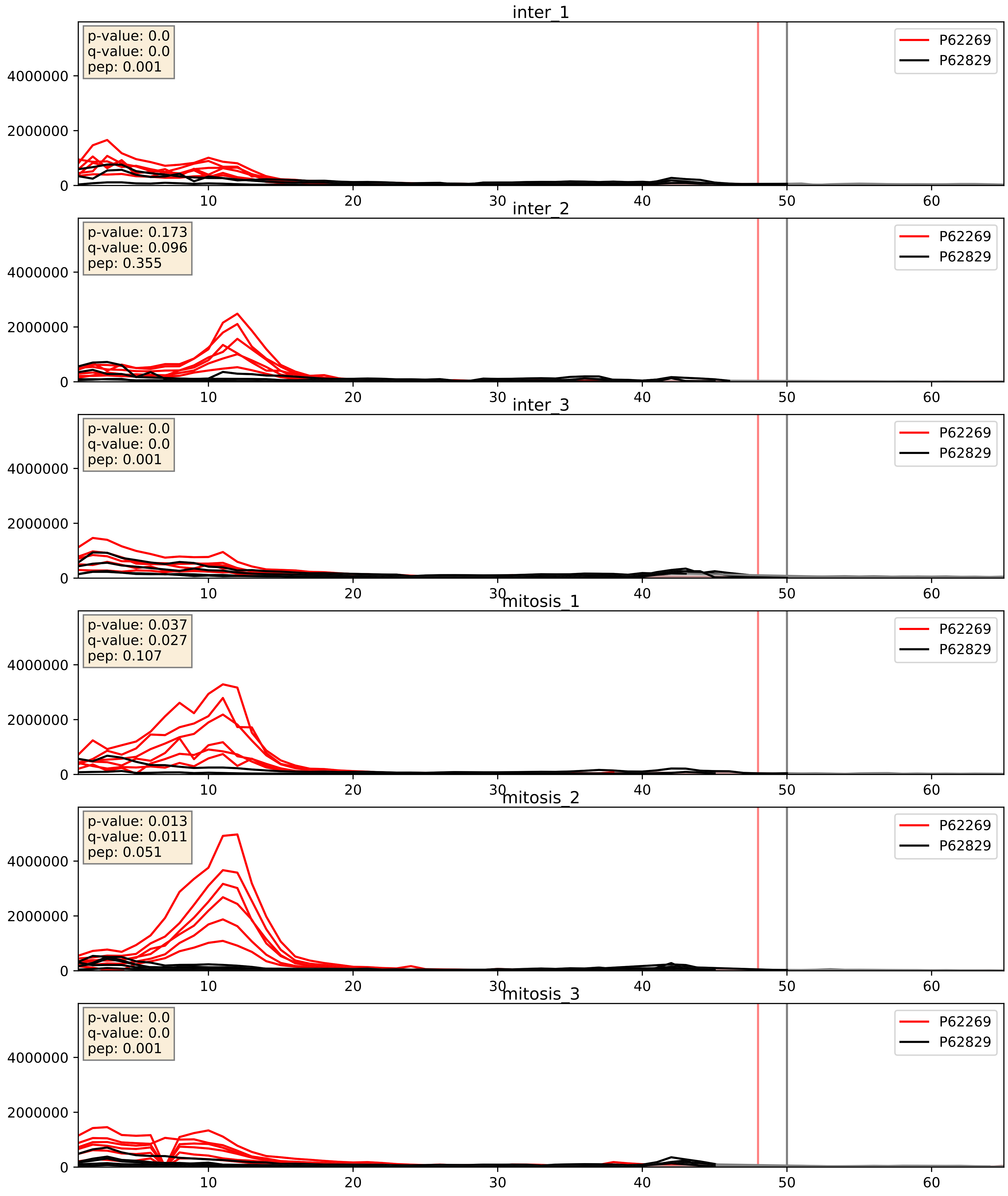


| level | condition_1 | condition_2 | pvalue | pvalue_adjusted |
|---|---|---|---|---|
| interactor_ratio | mitosis | inter | 0.003926685926982767 | 0.08138603277233046 |
| interactor_abundance | mitosis | inter | 0.01140063841385368 | 0.09438052690772485 |
| complex_abundance | mitosis | inter | 0.046698979628882156 | 0.15446030356384516 |

P62273_P62829
RS29_HUMAN vs RL23_HUMAN
p-value: 0.024 q-value: 0.019

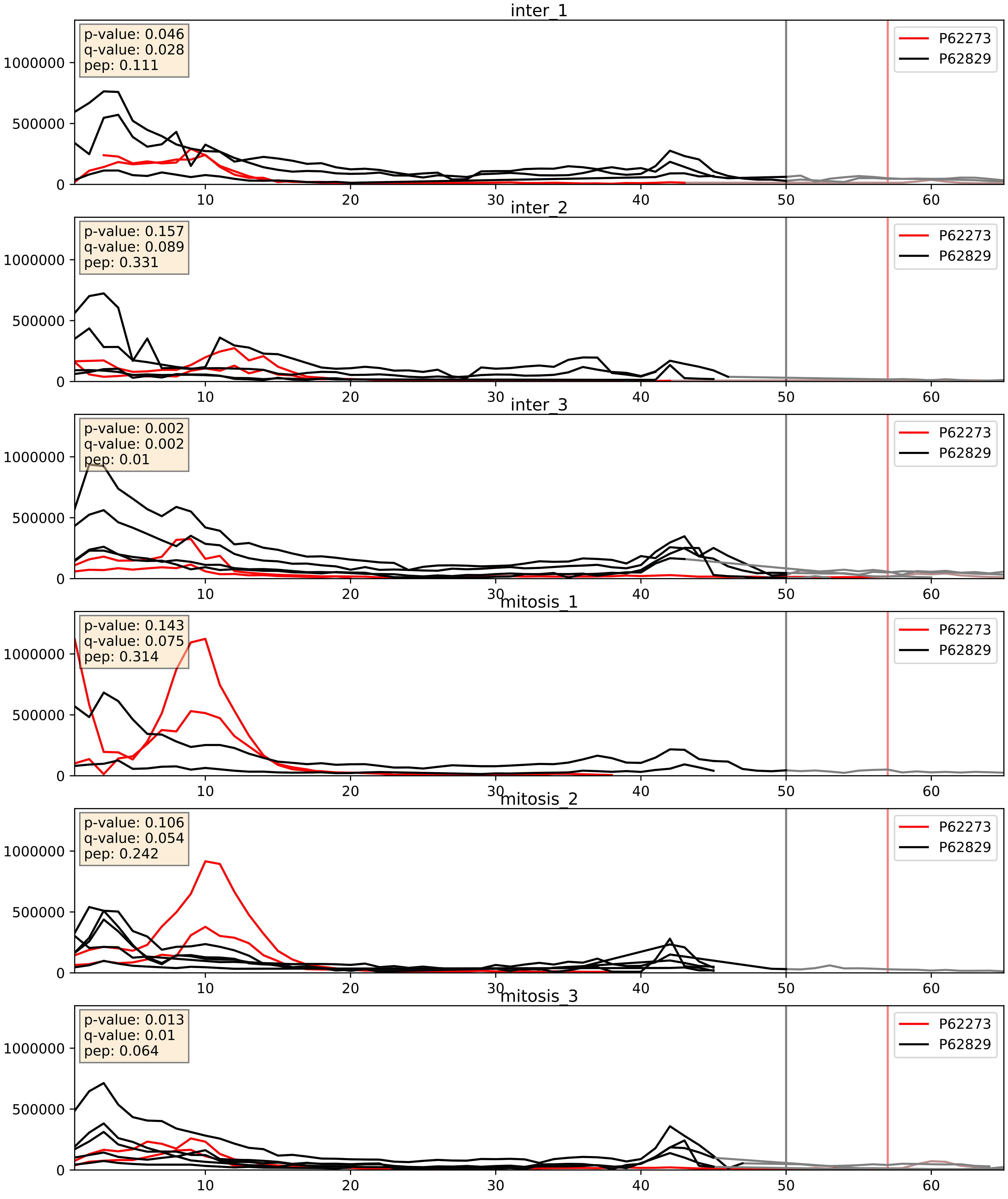


| level | condition_1 | condition_2 | pvalue | pvalue_adjusted |
|---|---|---|---|---|
| interactor_ratio | mitosis | inter | 0.023376627081019234 | 0.23708996186436568 |
| complex_abundance | mitosis | inter | 0.026923501795645594 | 0.11998266880940797 |
| interactor_abundance | mitosis | inter | 0.23108155713227638 | 0.42318727300216413 |

P62277_P62829
RS13_HUMAN vs RL23_HUMAN
p-value: 0.0 q-value: 0.0

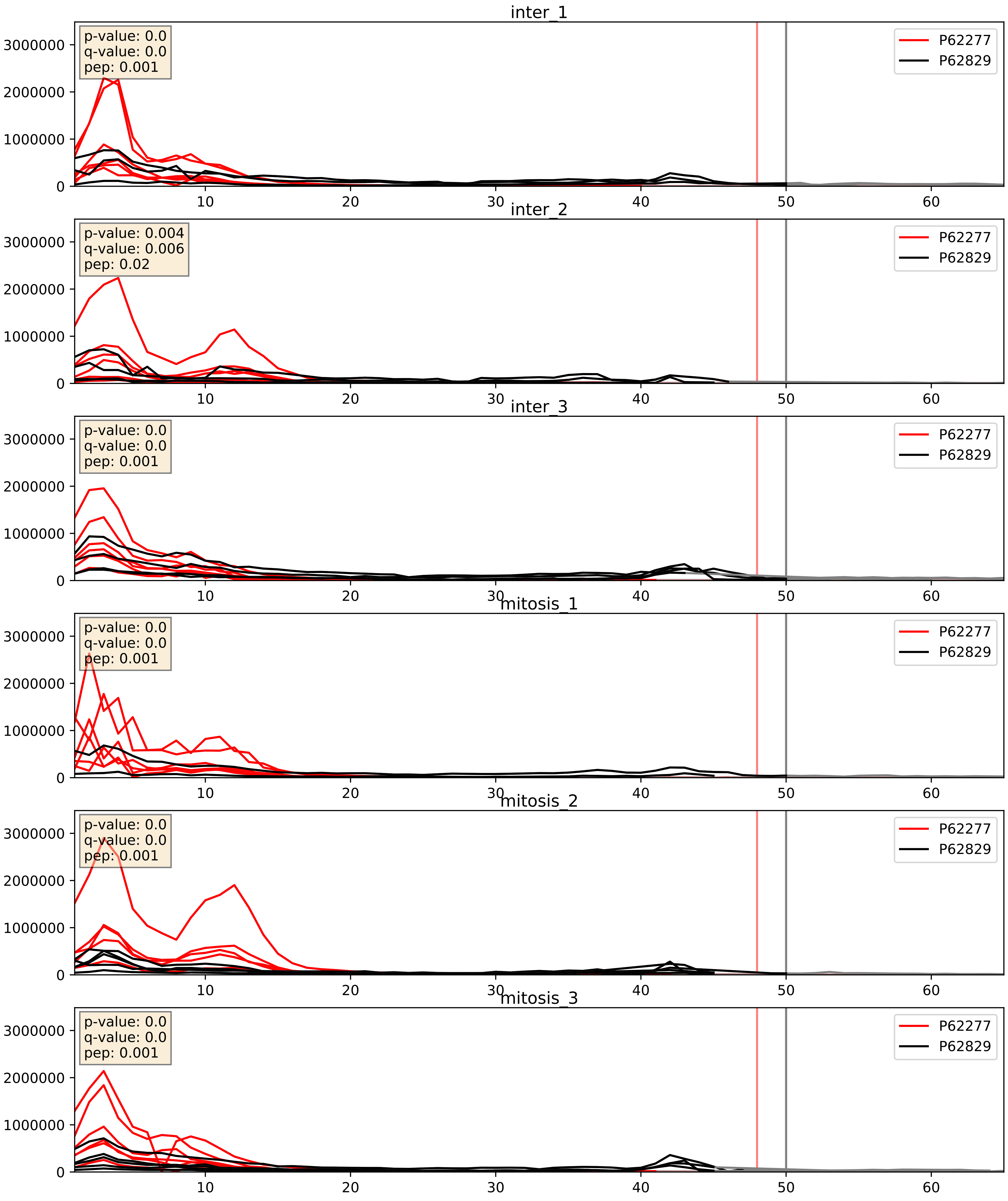


| level | condition_1 | condition_2 | pvalue | pvalue_adjusted |
|---|---|---|---|---|
| interactor_ratio | mitosis | inter | 0.0030062448102562067 | 0.07089106219226757 |
| complex_abundance | mitosis | inter | 0.014628068714221367 | 0.0898251565234827 |
| interactor_abundance | mitosis | inter | 0.354976116715731 | 0.5526728917945902 |

P62280_P62829
RS11_HUMAN vs RL23_HUMAN
p-value: 0.0 q-value: 0.0

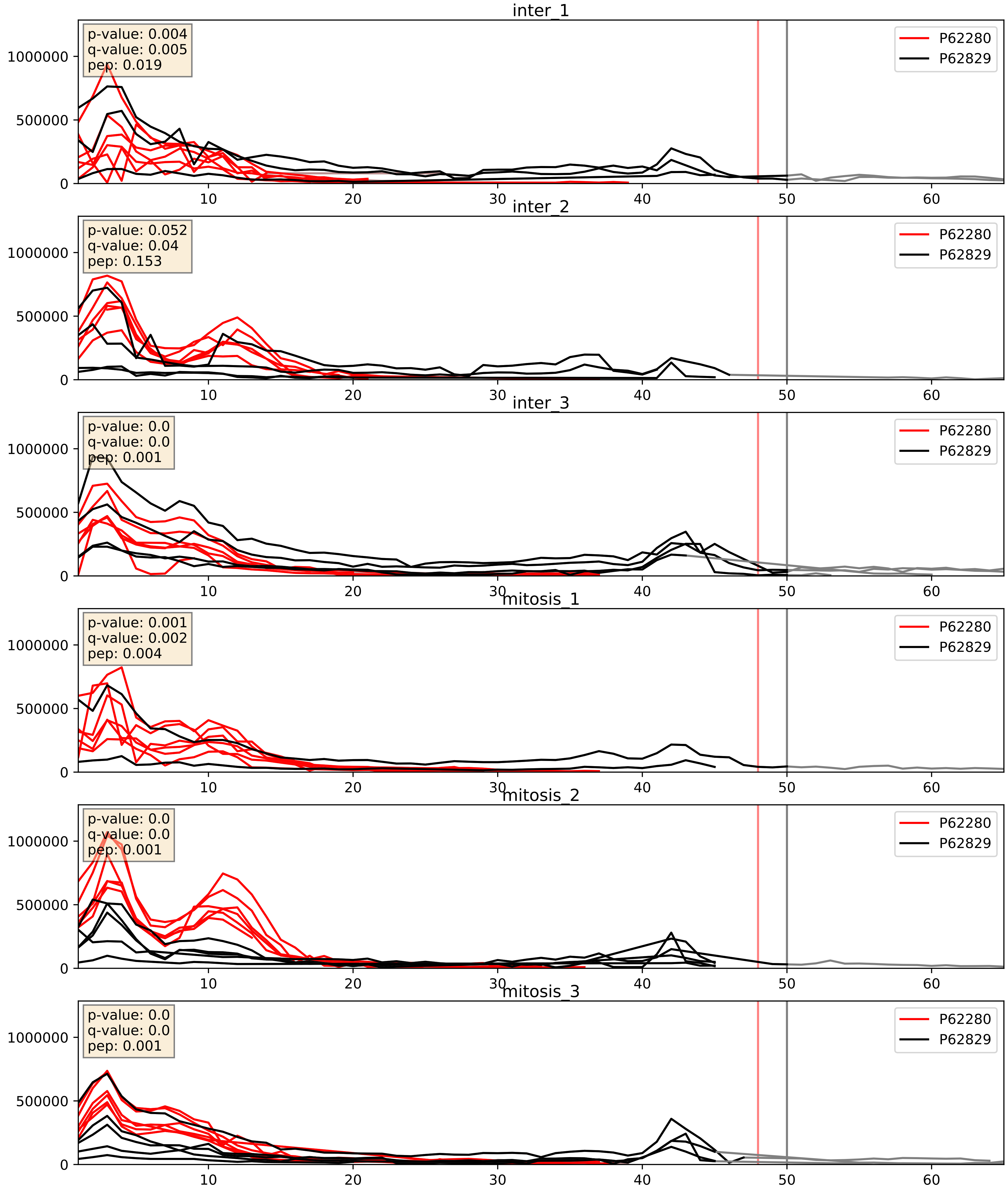


| level | condition_1 | condition_2 | pvalue | pvalue_adjusted |
|---|---|---|---|---|
| complex_abundance | mitosis | inter | 0.006264424751260746 | 0.0589917226301342 |
| interactor_ratio | mitosis | inter | 0.01227349419189634 | 0.15991036572699036 |
| interactor_abundance | mitosis | inter | 0.7956222020674533 | 0.8808985116259767 |

P62424_P62829
RL7A_HUMAN vs RL23_HUMAN
p-value: 0.0 q-value: 0.0

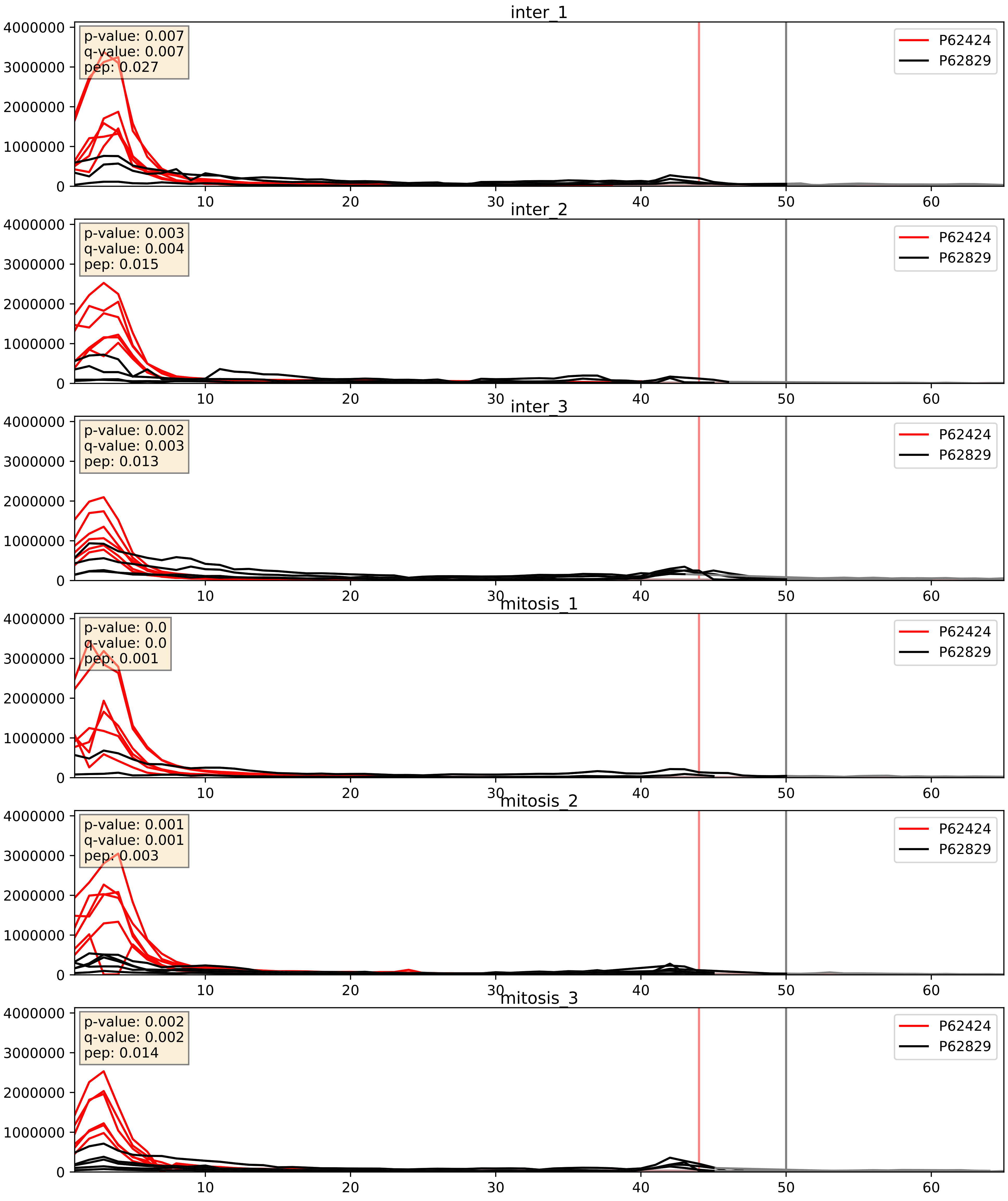


| level | condition_1 | condition_2 | pvalue | pvalue_adjusted |
|---|---|---|---|---|
| complex_abundance | mitosis | inter | 0.006497159100805557 | 0.05955783692156859 |
| interactor_ratio | mitosis | inter | 0.009086789286877207 | 0.13591615038037733 |
| interactor_abundance | mitosis | inter | 0.9391367776796664 | 0.9623524052119405 |

P62701_P62829
RS4X_HUMAN vs RL23_HUMAN
p-value: 0.0 q-value: 0.001

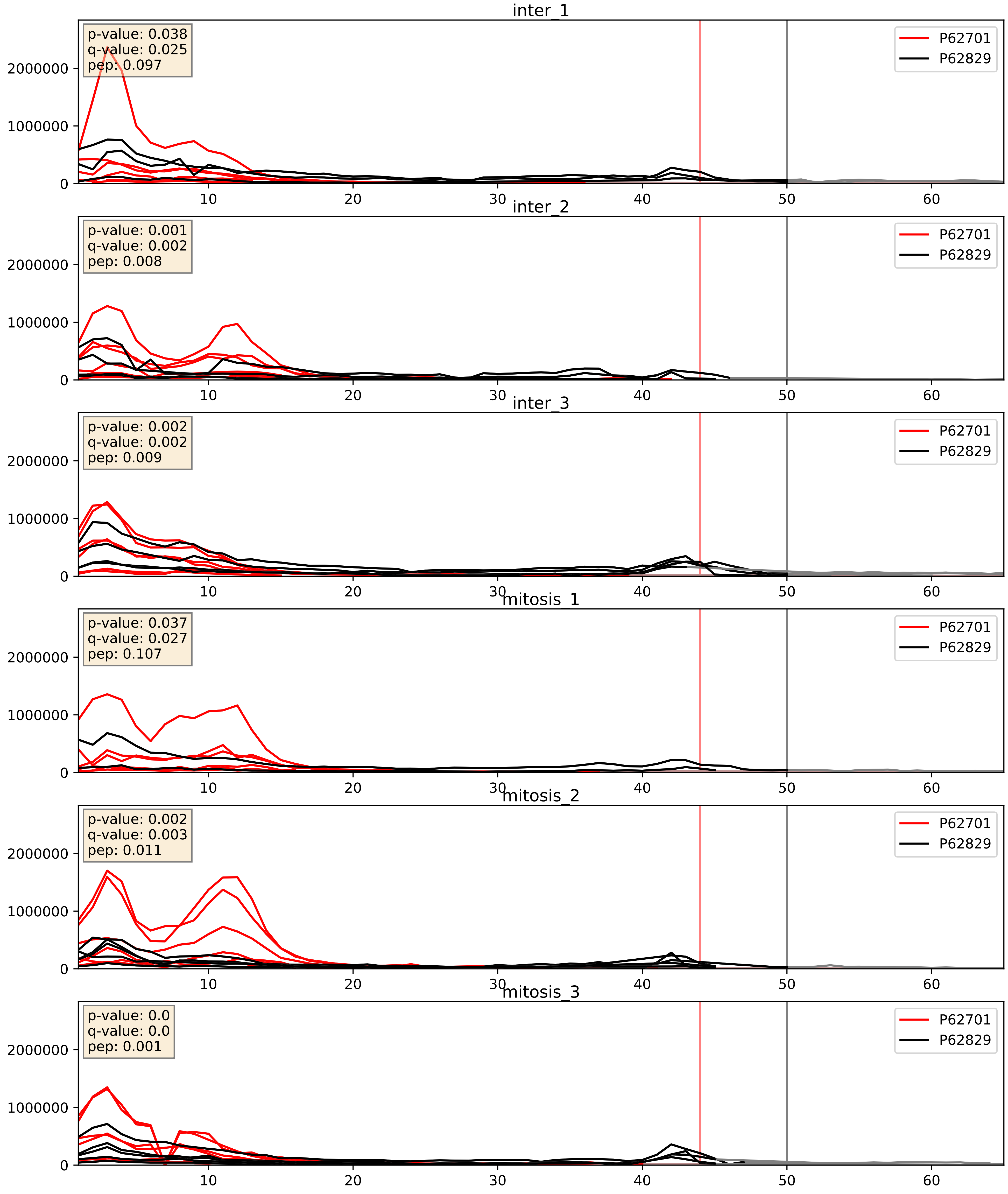


| level | condition_1 | condition_2 | pvalue | pvalue_adjusted |
|---|---|---|---|---|
| interactor_ratio | mitosis | inter | 0.005553910183506508 | 0.10115206632088448 |
| complex_abundance | mitosis | inter | 0.012791309949244463 | 0.08405635484106207 |
| interactor_abundance | mitosis | inter | 0.6893650719187185 | 0.7949860647871448 |

P62750_P62829
RL23A_HUMAN vs RL23_HUMAN
p-value: 0.0 q-value: 0.0

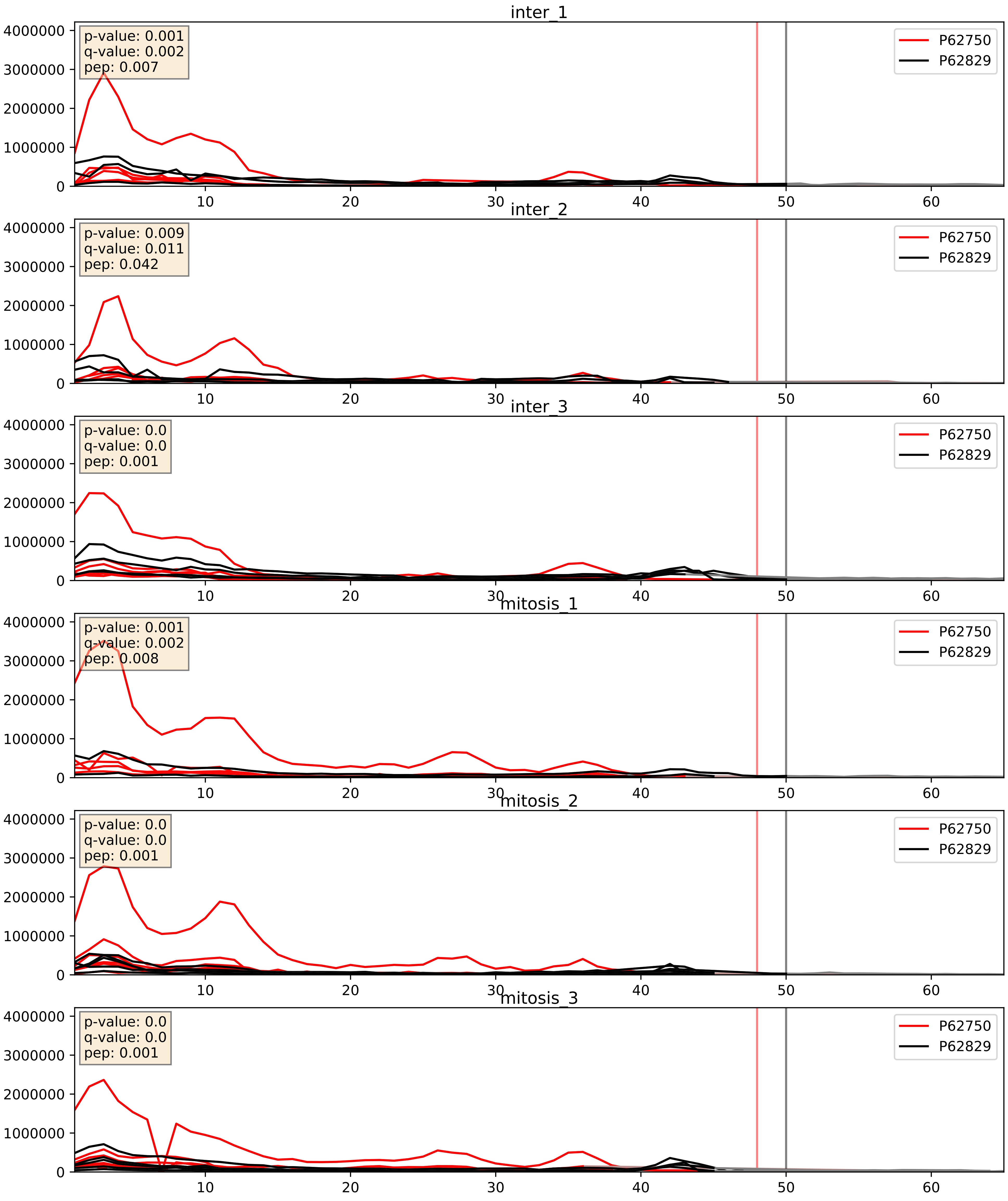


| level | condition_1 | condition_2 | pvalue | pvalue_adjusted |
|---|---|---|---|---|
| interactor_ratio | mitosis | inter | 0.00018711717641028034 | 0.014134487523529912 |
| complex_abundance | mitosis | inter | 0.07885414827655653 | 0.20485326532543974 |
| interactor_abundance | mitosis | inter | 0.15063142067463092 | 0.33732834796845007 |

P62753_P62829
RS6_HUMAN vs RL23_HUMAN
p-value: 0.0 q-value: 0.0

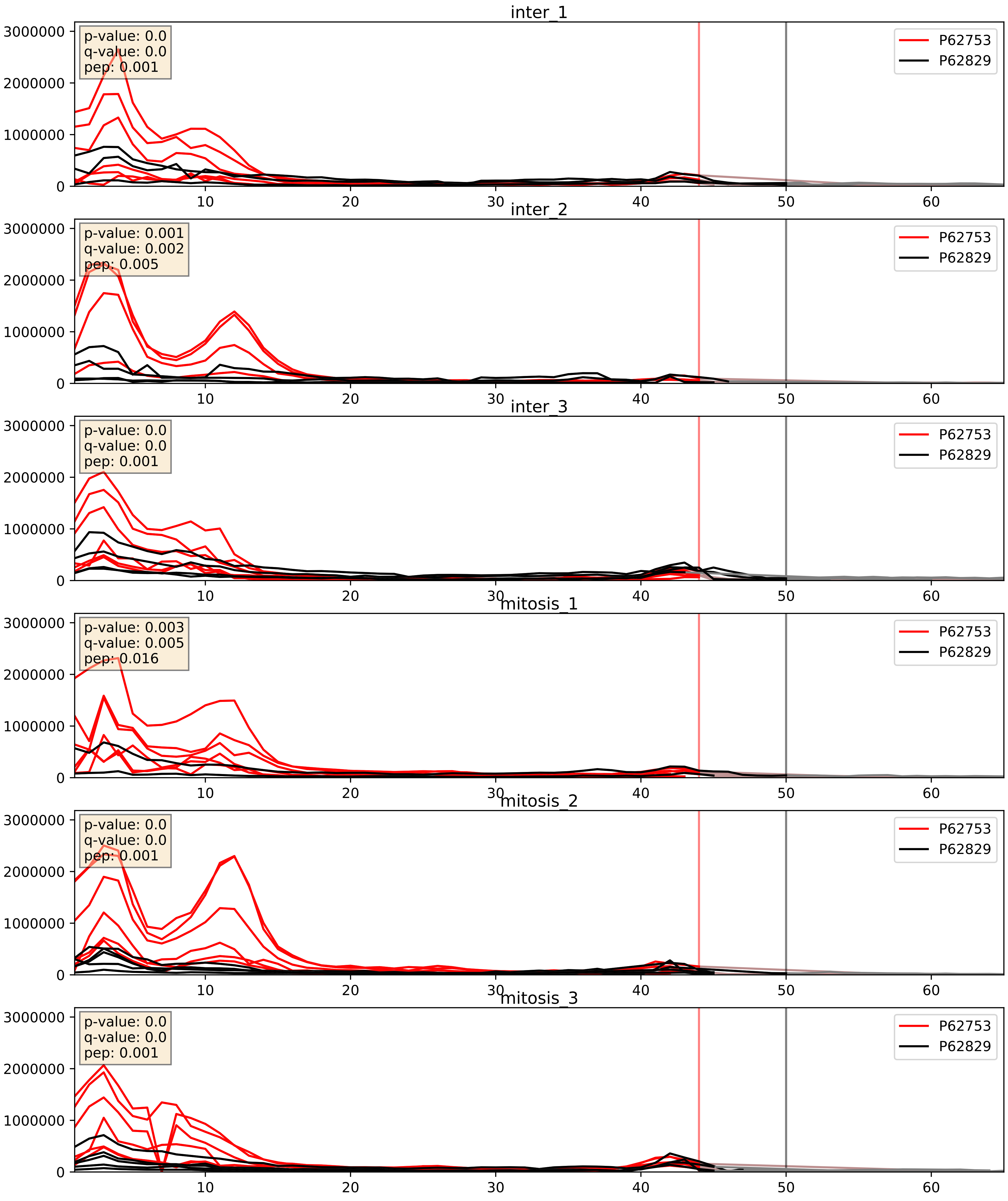


| level | condition_1 | condition_2 | pvalue | pvalue_adjusted |
|---|---|---|---|---|
| interactor_ratio | mitosis | inter | 0.006178300037298539 | 0.1083734596706465 |
| complex_abundance | mitosis | inter | 0.016541519325991585 | 0.09584596180627945 |
| interactor_abundance | mitosis | inter | 0.998848456004532 | 0.9991986424446411 |

P62829_P62841
RL23_HUMAN vs RS15_HUMAN
p-value: 0.005 q-value: 0.005

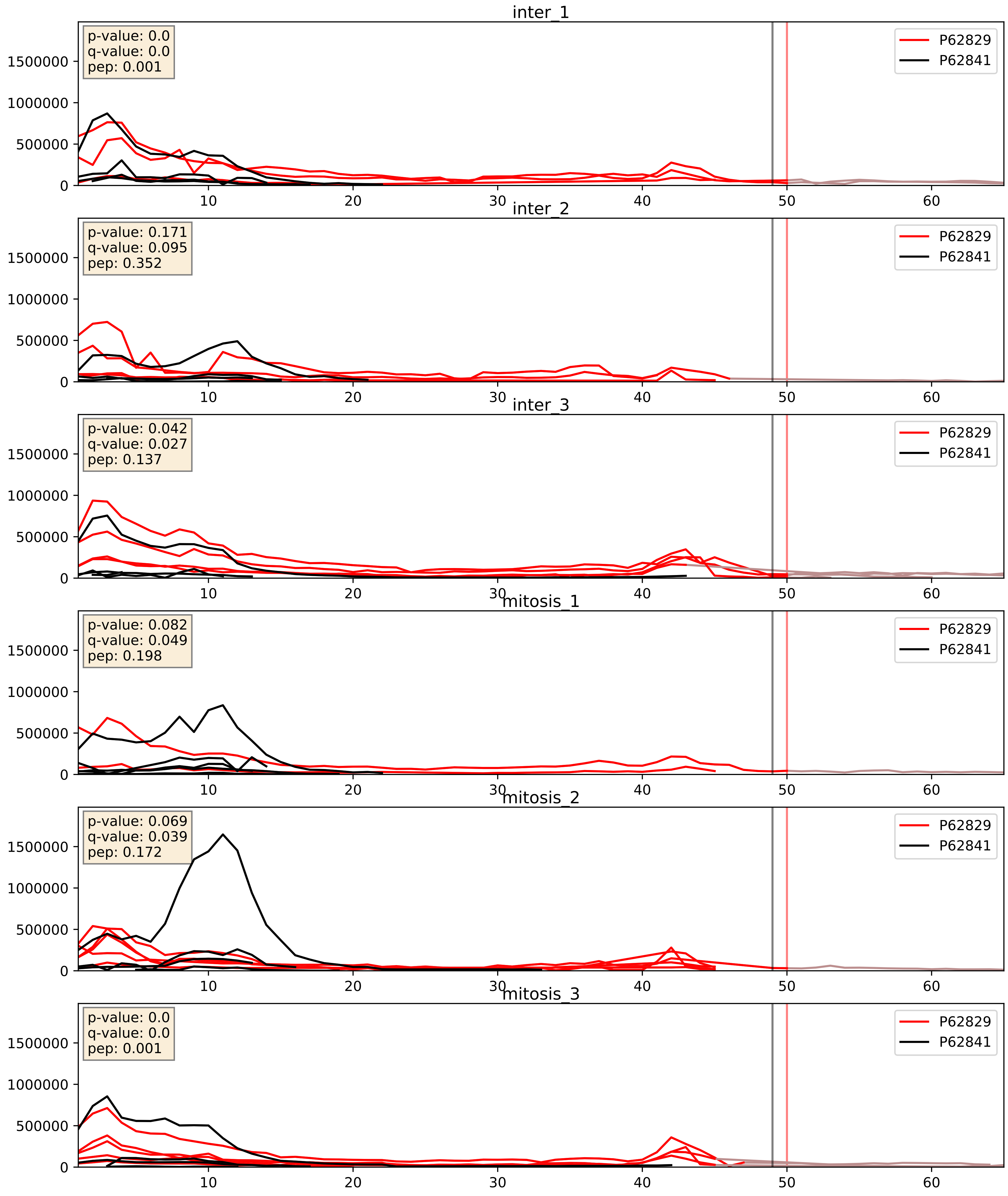


| level | condition_1 | condition_2 | pvalue | pvalue_adjusted |
|---|---|---|---|---|
| interactor_abundance | mitosis | inter | 0.010945048037109413 | 0.09180755629363702 |
| complex_abundance | mitosis | inter | 0.01562779222070651 | 0.09341575776239408 |
| interactor_ratio | mitosis | inter | 0.08789547287309328 | 0.4946978435260244 |

P62829_P62847
RL23_HUMAN vs RS24_HUMAN
p-value: 0.001 q-value: 0.002

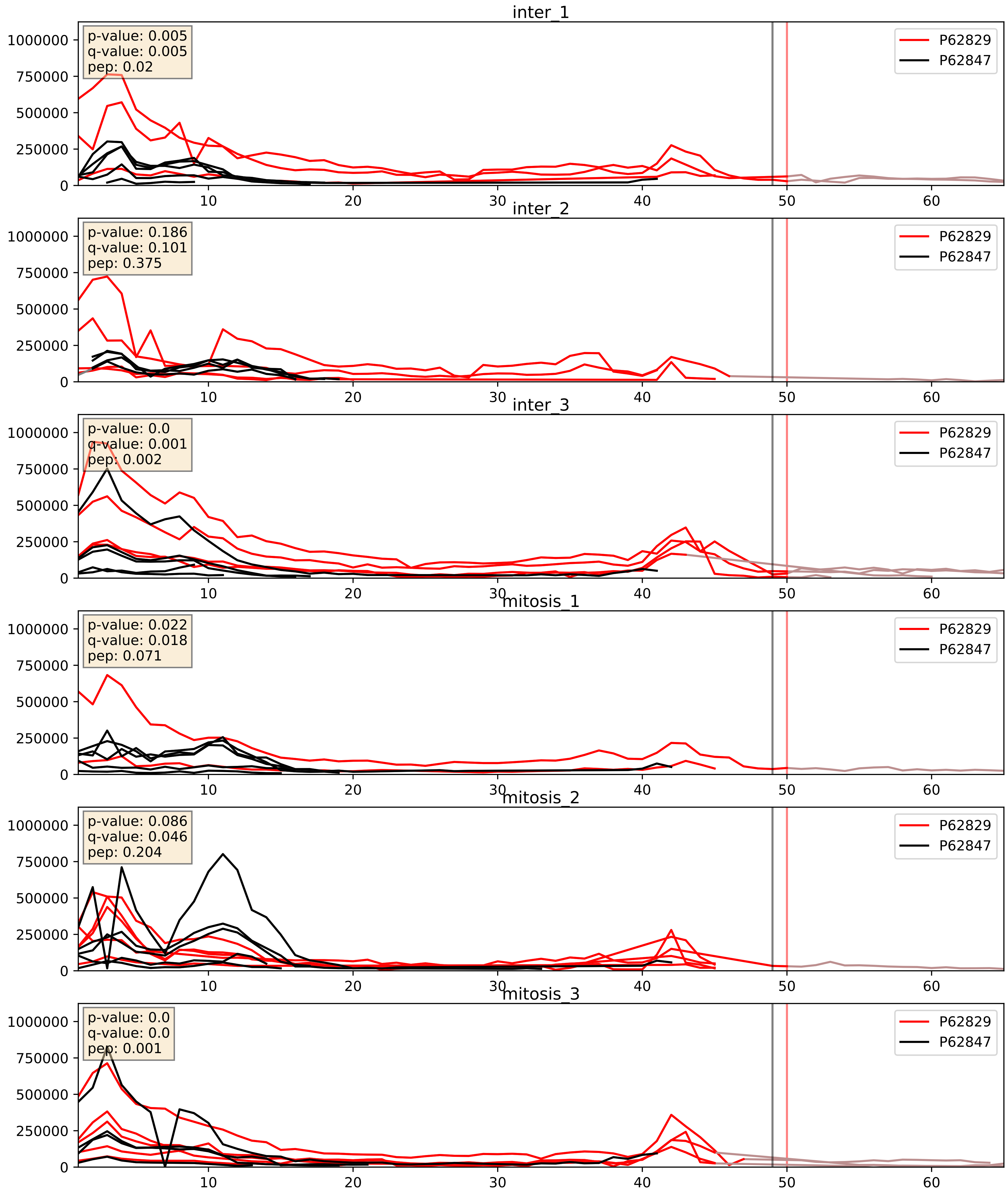


| level | condition_1 | condition_2 | pvalue | pvalue_adjusted |
|---|---|---|---|---|
| interactor_ratio | mitosis | inter | 0.0045350850798949694 | 0.0896543378381084 |
| interactor_abundance | mitosis | inter | 0.016646236821465304 | 0.11277132084292767 |
| complex_abundance | mitosis | inter | 0.07085702177127214 | 0.19345012392626001 |

P62829_P62851
RL23_HUMAN vs RS25_HUMAN
p-value: 0.005 q-value: 0.005

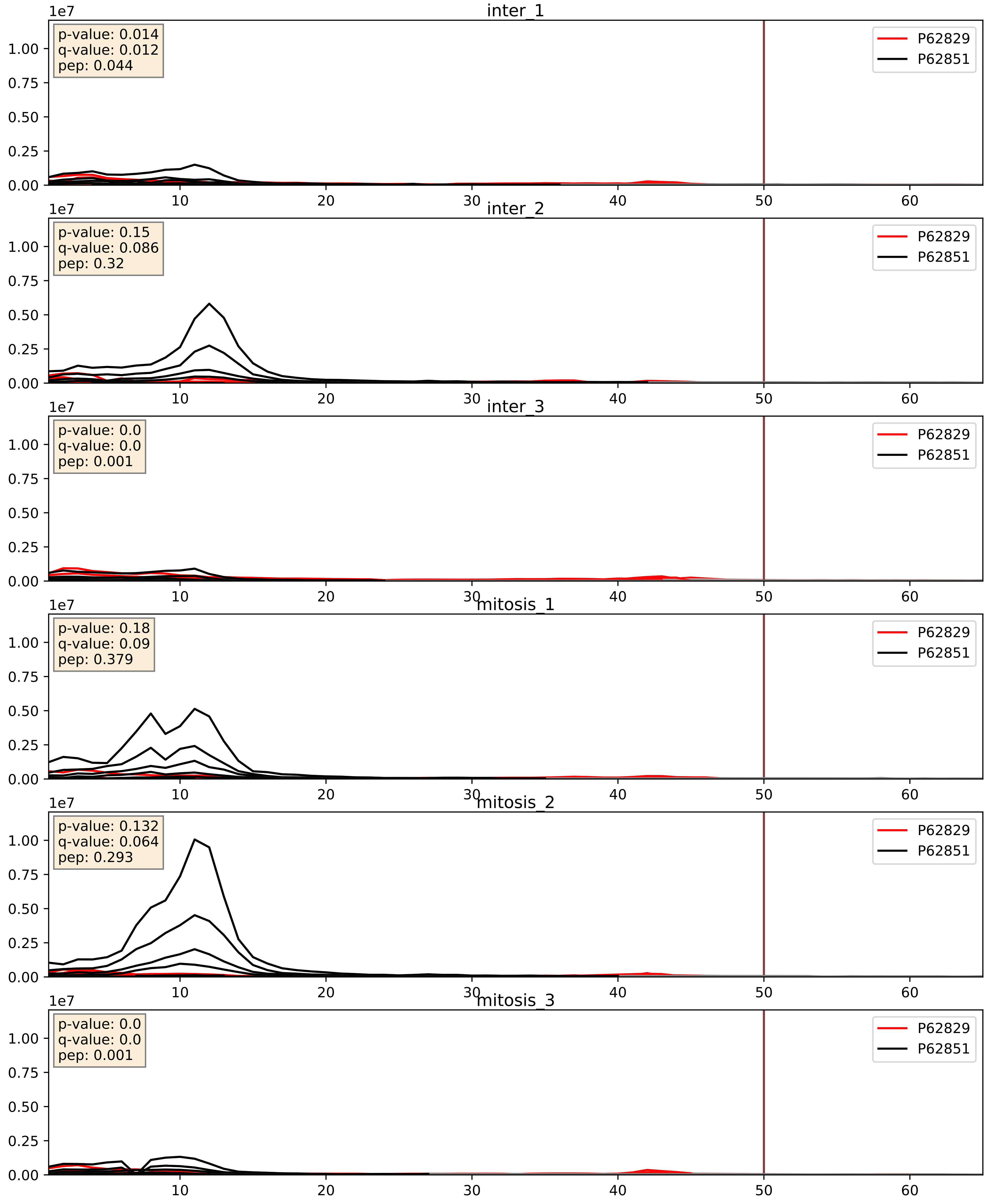


| level | condition_1 | condition_2 | pvalue | pvalue_adjusted |
|---|---|---|---|---|
| interactor_abundance | mitosis | inter | 0.007202039658835953 | 0.07343152394042615 |
| interactor_ratio | mitosis | inter | 0.10091915107347 | 0.5286829456480435 |
| complex_abundance | mitosis | inter | 0.2027828592934269 | 0.3558469199573051 |

P62829_P62857
RL23_HUMAN vs RS28_HUMAN
p-value: 0.023 q-value: 0.018

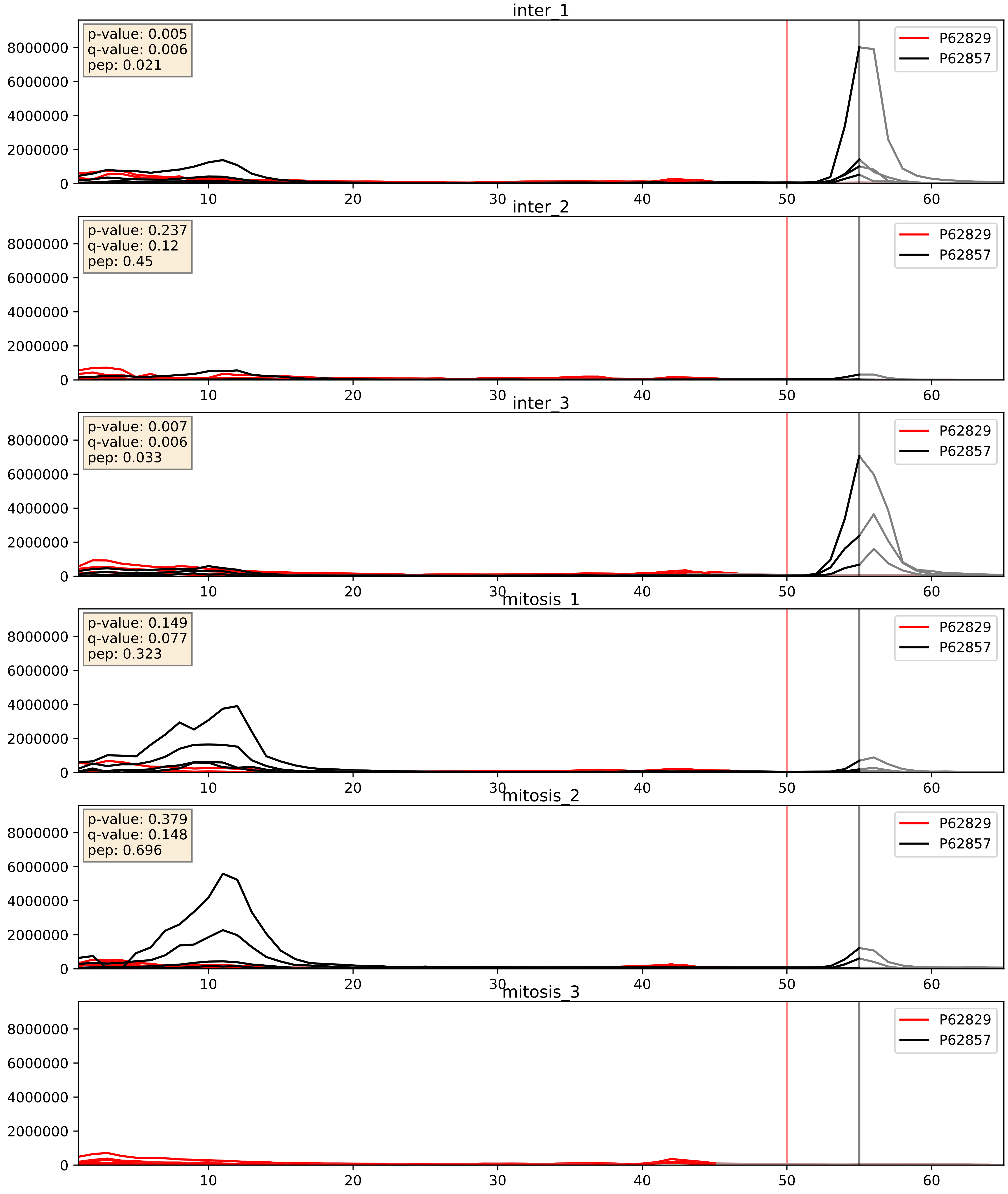


| level | condition_1 | condition_2 | pvalue | pvalue_adjusted |
|---|---|---|---|---|
| interactor_ratio | mitosis | inter | 0.047999143153140654 | 0.361488803041275 |
| interactor_abundance | mitosis | inter | 0.05463187832433133 | 0.18883459659046078 |
| complex_abundance | mitosis | inter | 0.4720766228631067 | 0.612826189218713 |

P62829_P62861
RL23_HUMAN vs RS30_HUMAN
p-value: 0.0 q-value: 0.0

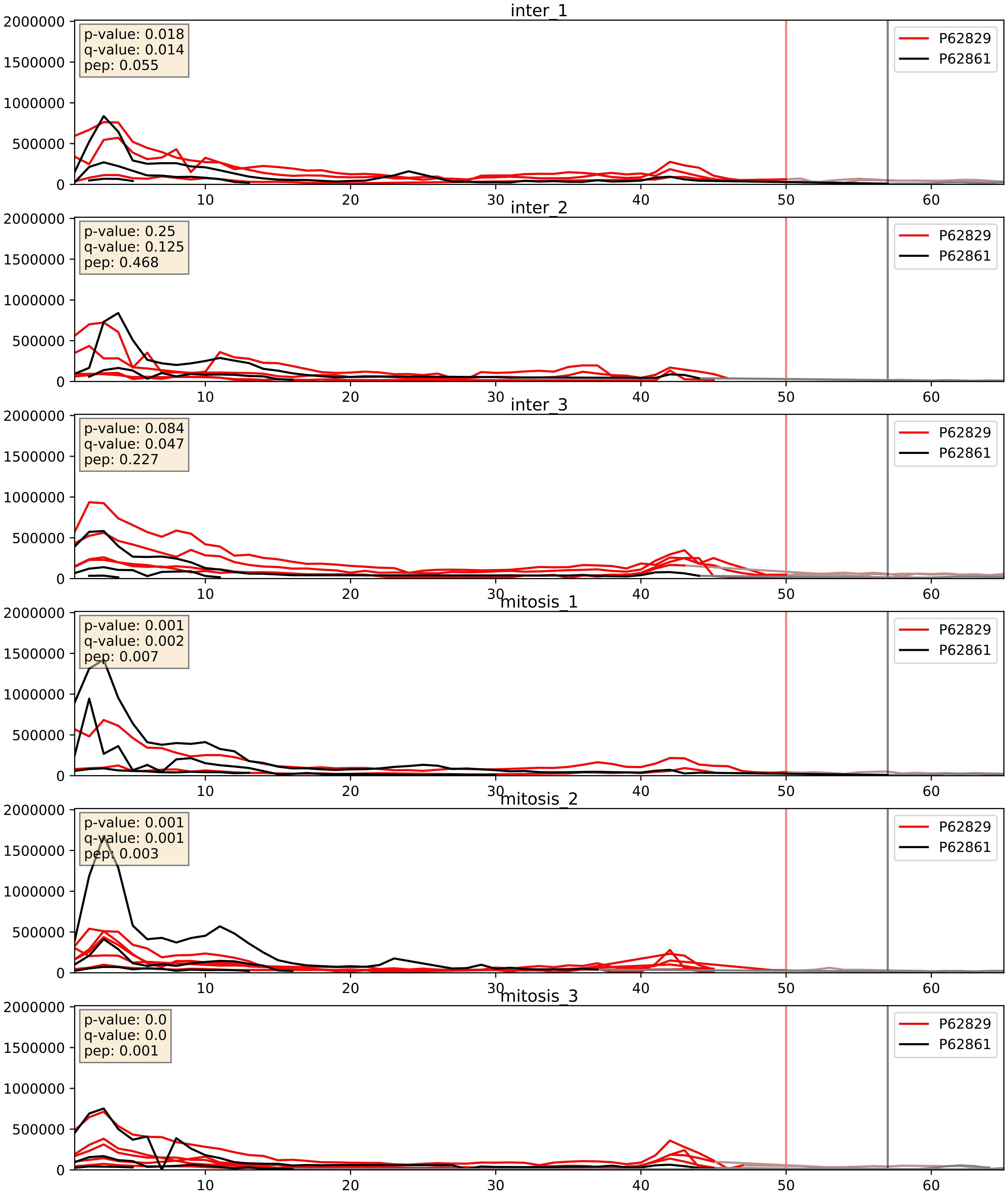


| level | condition_1 | condition_2 | pvalue | pvalue_adjusted |
|---|---|---|---|---|
| interactor_abundance | mitosis | inter | 0.006338045608738357 | 0.06815787740050293 |
| interactor_ratio | mitosis | inter | 0.024872287022184865 | 0.24756601966267727 |
| complex_abundance | mitosis | inter | 0.058570857278846304 | 0.17298981144500772 |

P62829_P62888
RL23_HUMAN vs RL30_HUMAN
p-value: 0.0 q-value: 0.0

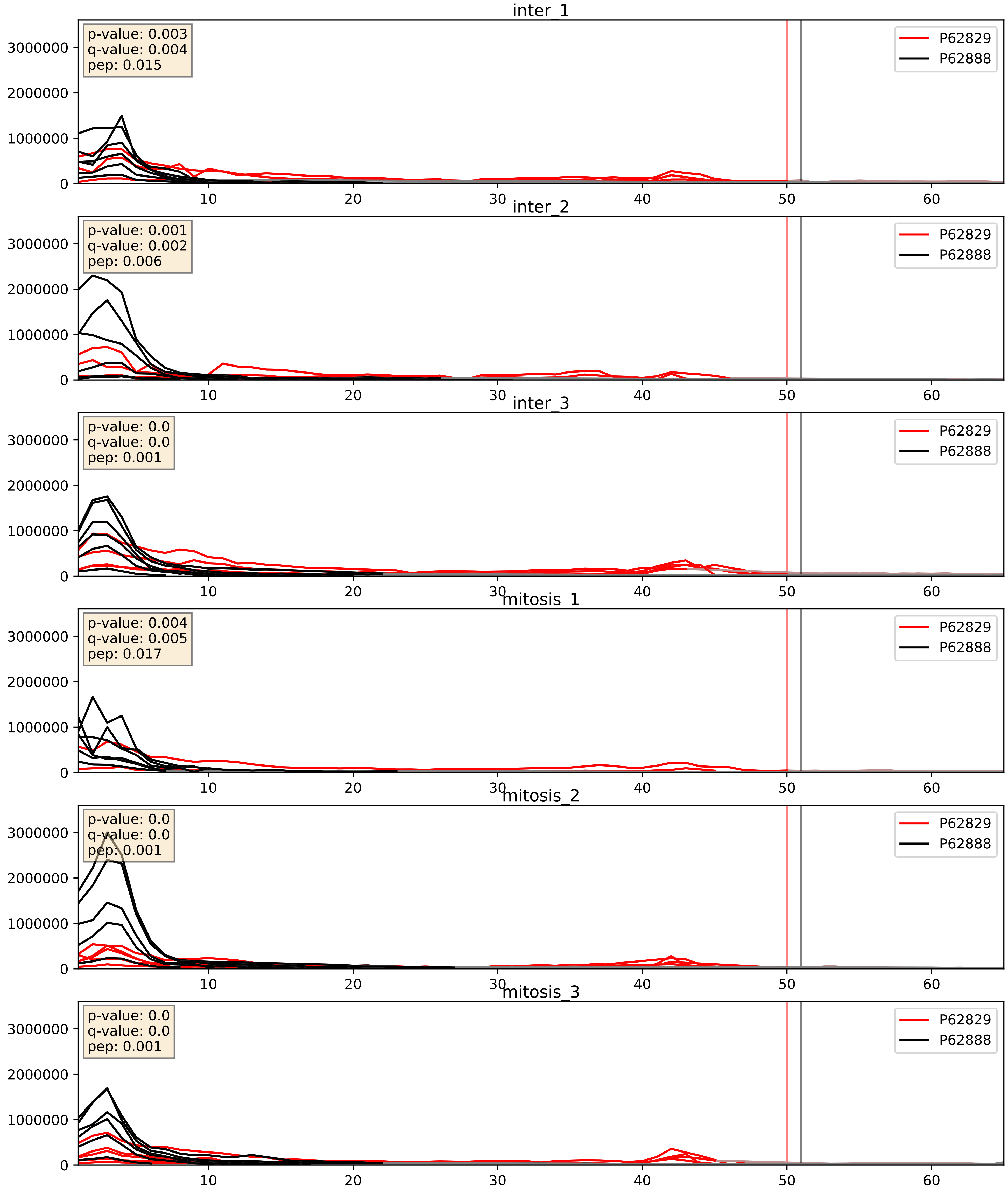


| level | condition_1 | condition_2 | pvalue | pvalue_adjusted |
|---|---|---|---|---|
| interactor_abundance | mitosis | inter | 0.003241446913957478 | 0.04397271883276706 |
| complex_abundance | mitosis | inter | 0.018334502135924035 | 0.10112328497648823 |
| interactor_ratio | mitosis | inter | 0.026438411200135587 | 0.25543205403291264 |

P62829_P62899
RL23_HUMAN vs RL31_HUMAN
p-value: 0.0 q-value: 0.0

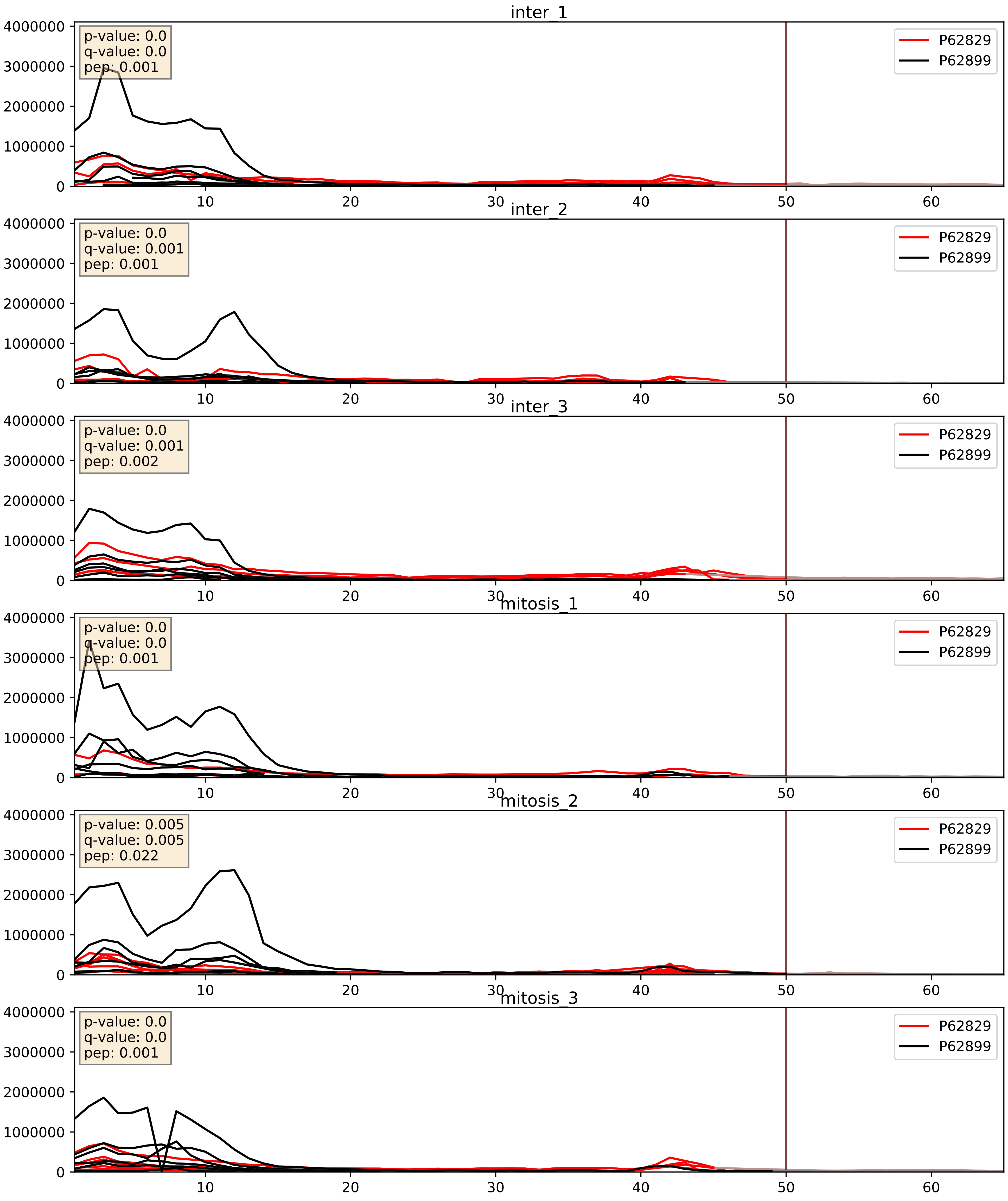


| level | condition_1 | condition_2 | pvalue | pvalue_adjusted |
|---|---|---|---|---|
| interactor_ratio | mitosis | inter | 0.003373183082024111 | 0.07563355707045767 |
| interactor_abundance | mitosis | inter | 0.009001428066350159 | 0.082762861705647 |
| complex_abundance | mitosis | inter | 0.03964276099289684 | 0.1448340829507414 |

P62829_P62906
RL23_HUMAN vs RL10A_HUMAN
p-value: 0.0 q-value: 0.0

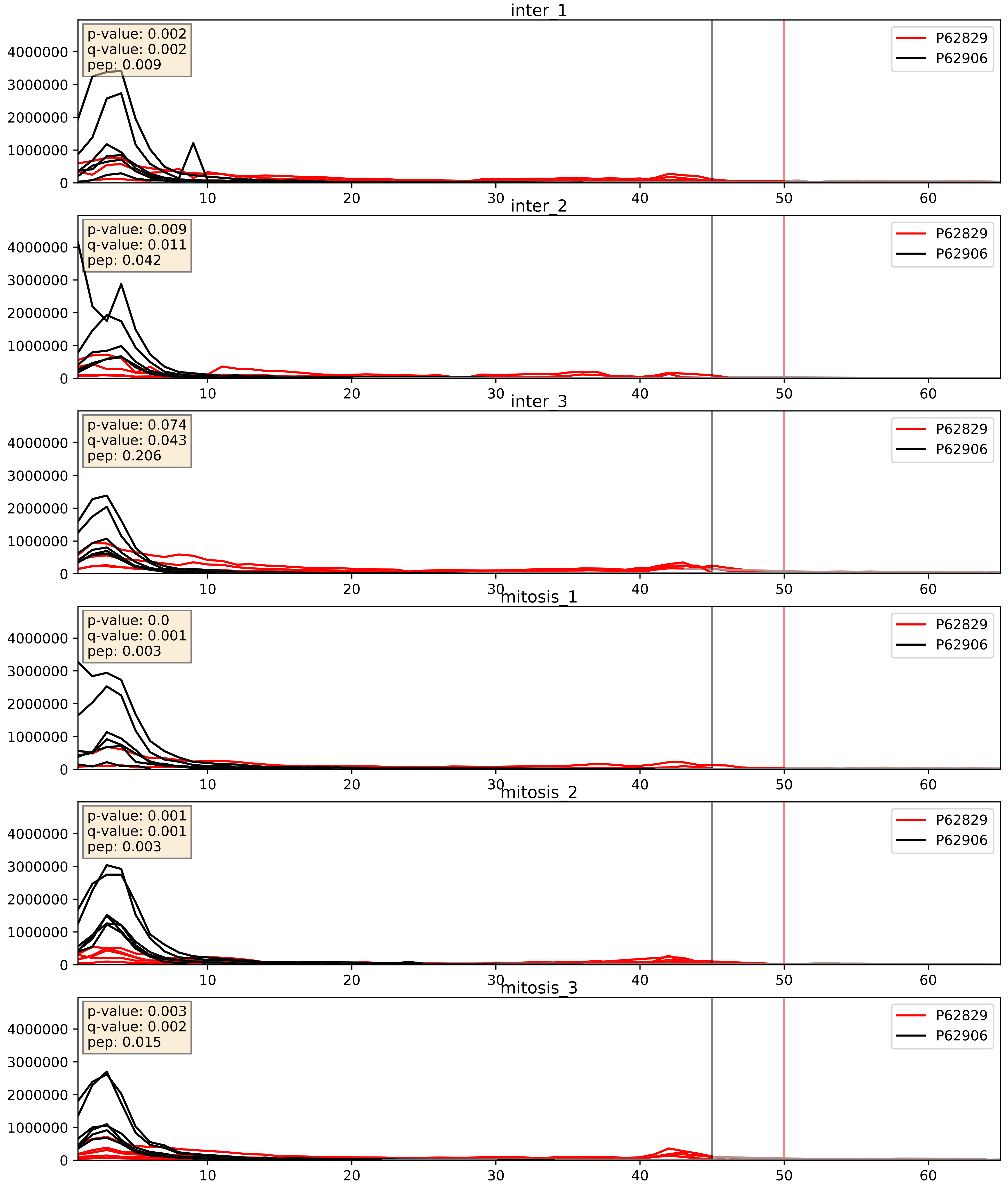


| level | condition_1 | condition_2 | pvalue | pvalue_adjusted |
|---|---|---|---|---|
| interactor_abundance | mitosis | inter | 0.002632665877349012 | 0.03838967005823521 |
| complex_abundance | mitosis | inter | 0.003632835025813253 | 0.04428054295684183 |
| interactor_ratio | mitosis | inter | 0.005822434720010065 | 0.10426786862612165 |

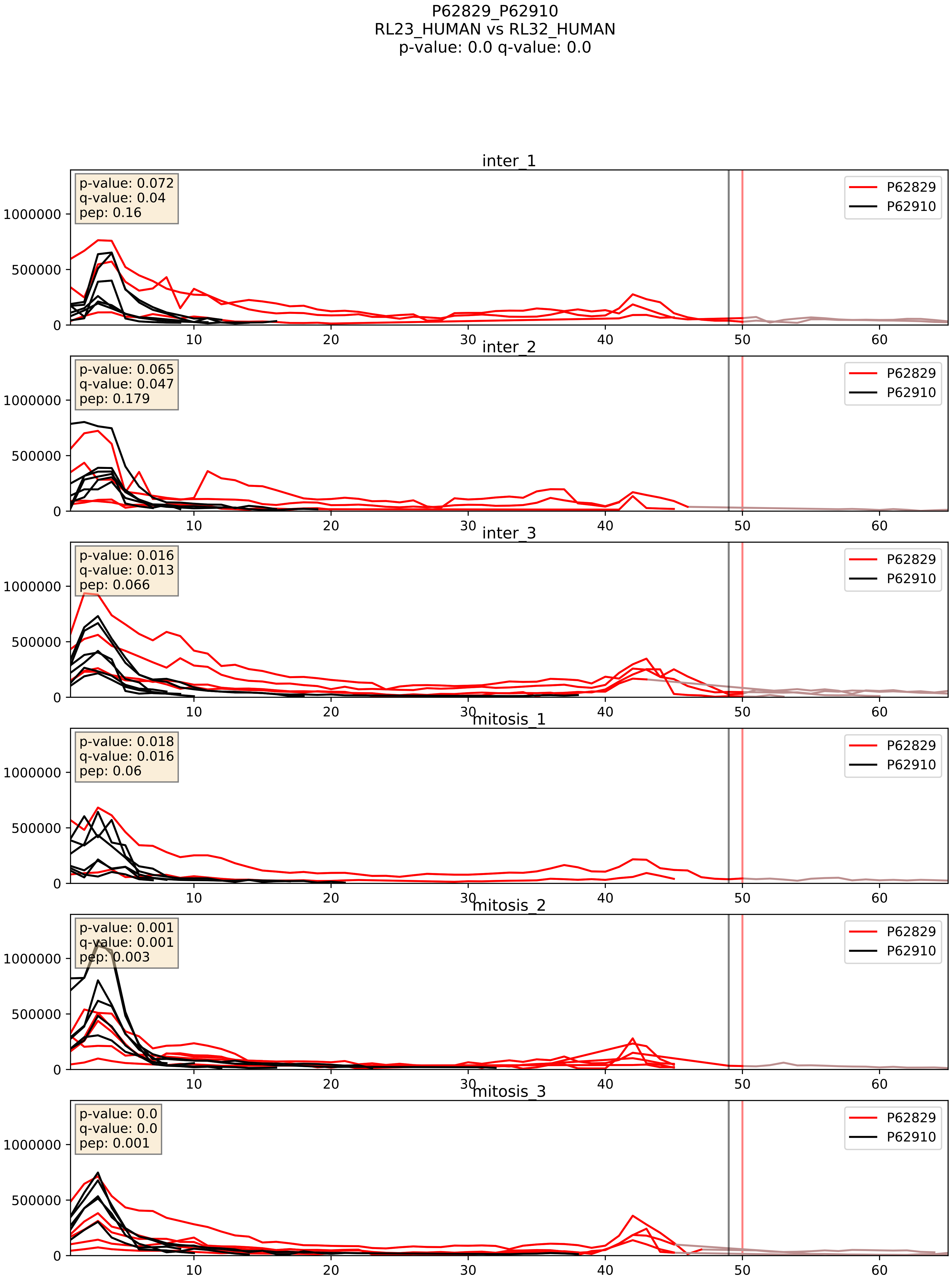


| level | condition_1 | condition_2 | pvalue | pvalue_adjusted |
|---|---|---|---|---|
| interactor_ratio | mitosis | inter | 0.006107894693682131 | 0.10745632562987323 |
| interactor_abundance | mitosis | inter | 0.007803776035269969 | 0.07671437955535183 |
| complex_abundance | mitosis | inter | 0.035995297098866307 | 0.13781615458280247 |

P62829_P62913
RL23_HUMAN vs RL11_HUMAN
p-value: 0.0 q-value: 0.0

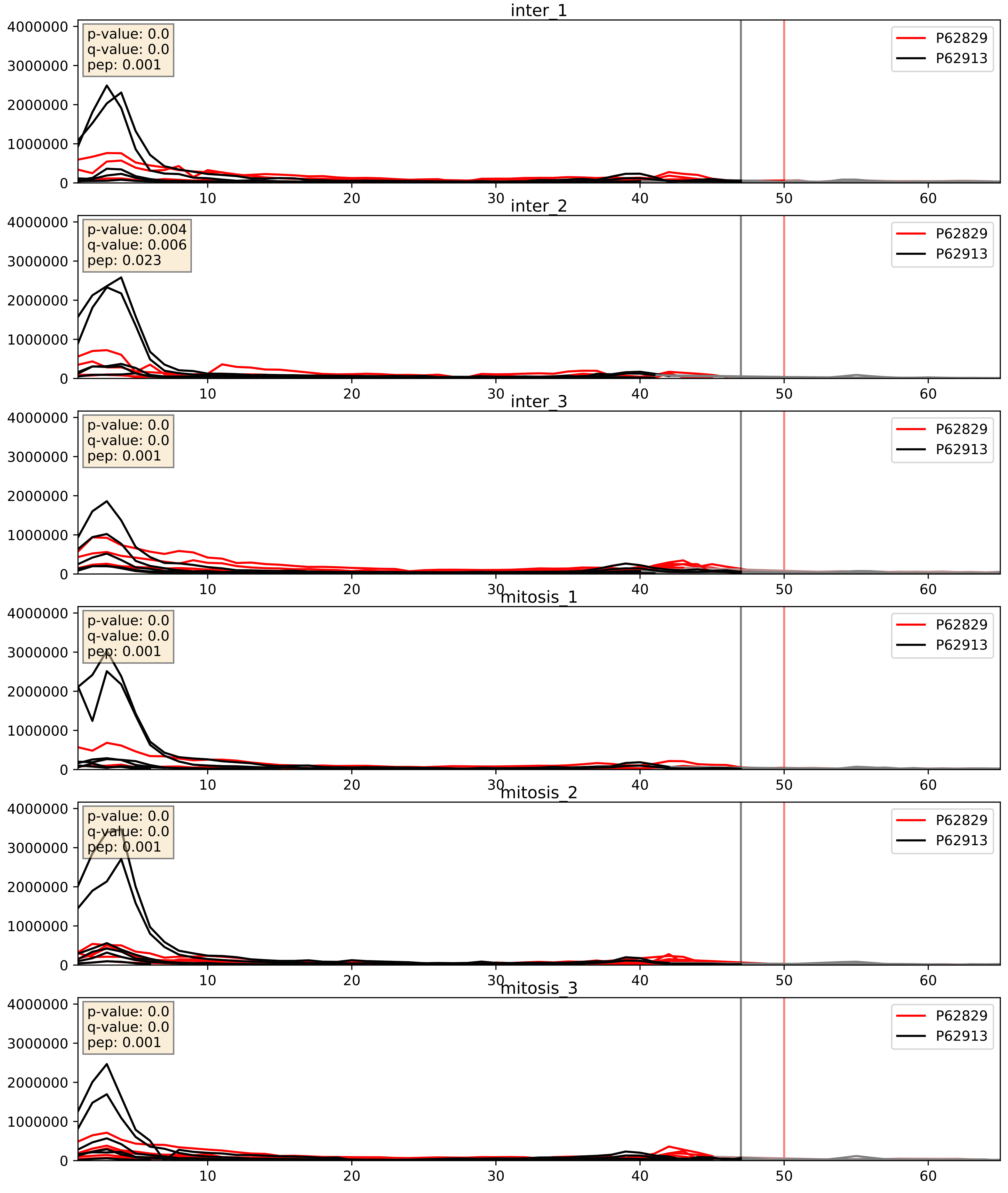


| level | condition_1 | condition_2 | pvalue | pvalue_adjusted |
|---|---|---|---|---|
| interactor_ratio | mitosis | inter | 0.00865411090032815 | 0.13211821468820312 |
| interactor_abundance | mitosis | inter | 0.009700315001046716 | 0.08525122834595471 |
| complex_abundance | mitosis | inter | 0.011919195887493676 | 0.08114978643416496 |

P62829_P62917
RL23_HUMAN vs RL8_HUMAN
p-value: 0.0 q-value: 0.0

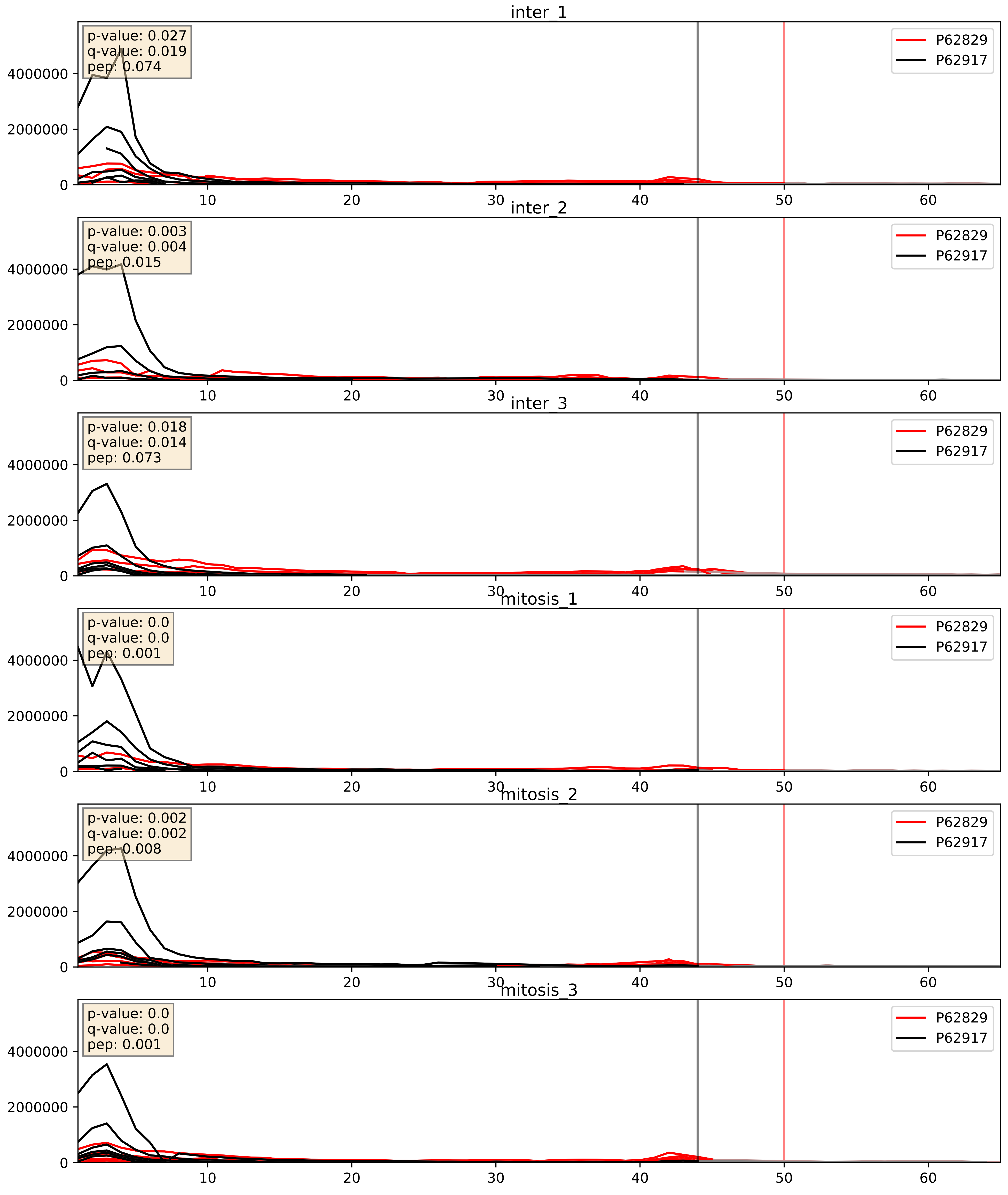


| level | condition_1 | condition_2 | pvalue | pvalue_adjusted |
|---|---|---|---|---|
| interactor_abundance | mitosis | inter | 0.0023827280082054686 | 0.03637808073281721 |
| interactor_ratio | mitosis | inter | 0.008930961999905052 | 0.13412111354243378 |
| complex_abundance | mitosis | inter | 0.08810475199817998 | 0.21753004819856378 |

P62829_P63173
RL23_HUMAN vs RL38_HUMAN
p-value: 0.0 q-value: 0.0

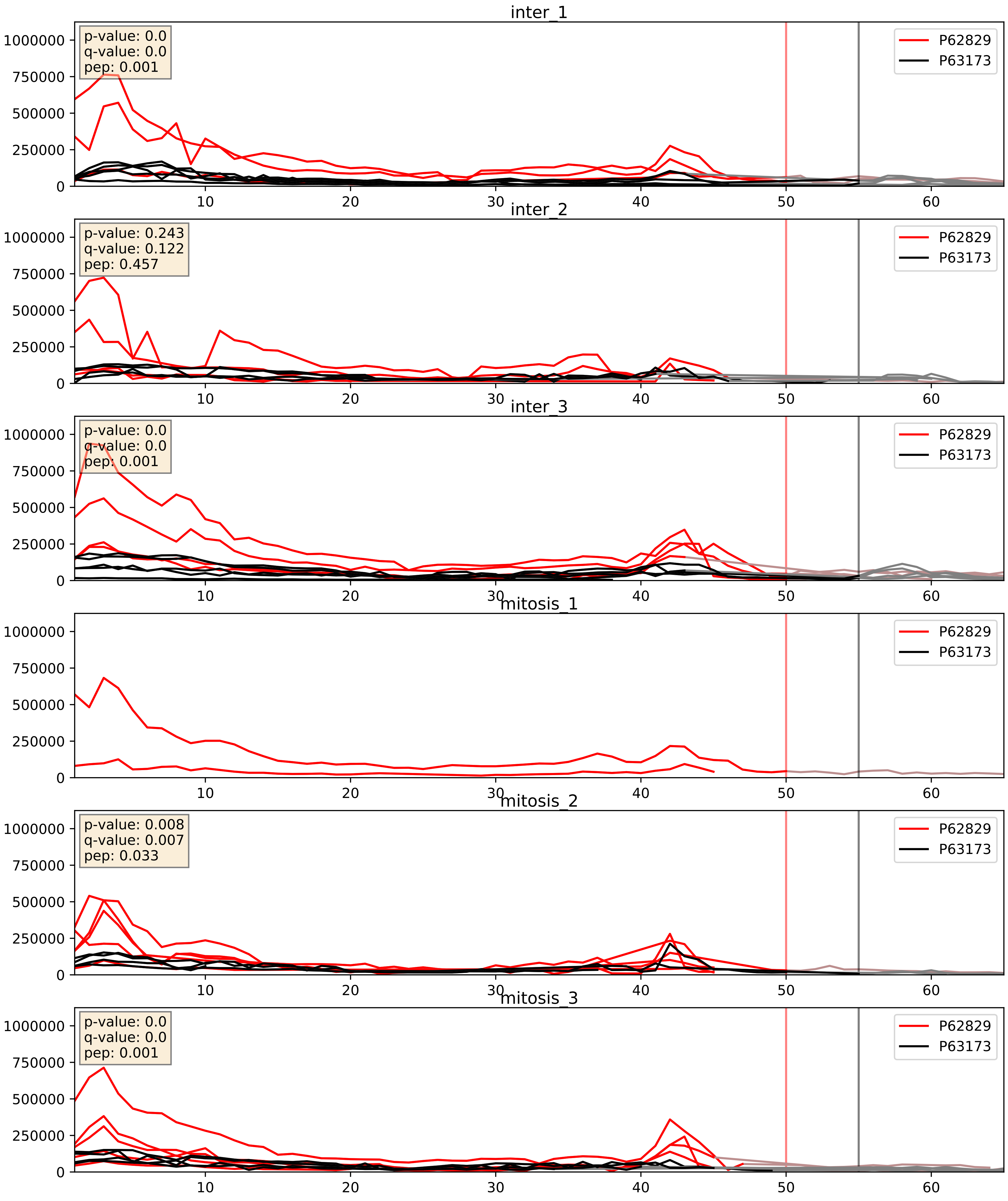


| level | condition_1 | condition_2 | pvalue | pvalue_adjusted |
|---|---|---|---|---|
| interactor_ratio | mitosis | inter | 0.018438893184676397 | 0.20476127879411543 |
| interactor_abundance | mitosis | inter | 0.04929798222404982 | 0.17730702850330524 |
| complex_abundance | mitosis | inter | 0.10207001591294664 | 0.23720313860614606 |

P62829_P83731
RL23_HUMAN vs RL24_HUMAN
p-value: 0.0 q-value: 0.0

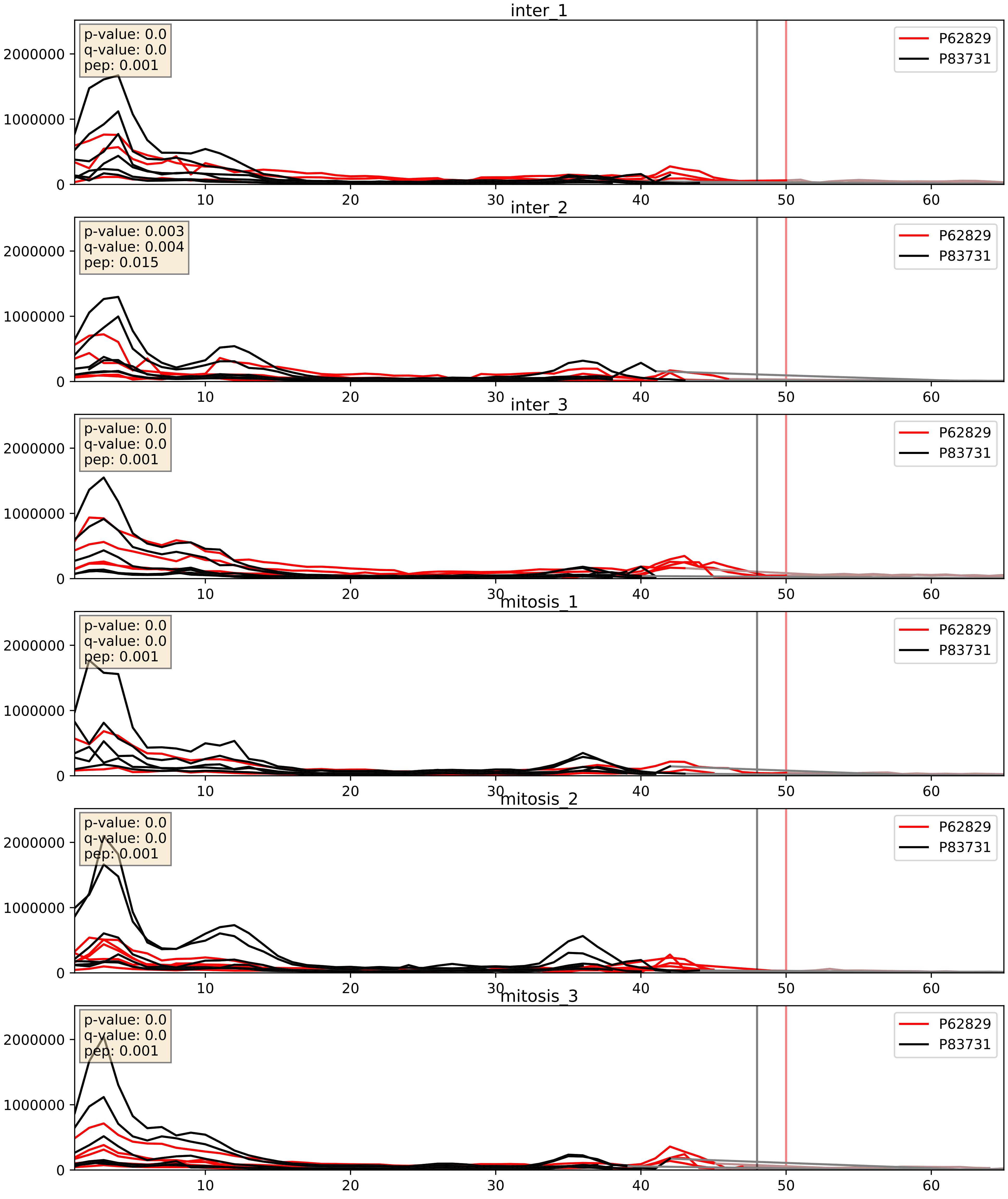


| level | condition_1 | condition_2 | pvalue | pvalue_adjusted |
|---|---|---|---|---|
| interactor_ratio | mitosis | inter | 0.0009102410313168072 | 0.034517566649383956 |
| interactor_abundance | mitosis | inter | 0.005256329787123167 | 0.06086766944637465 |
| complex_abundance | mitosis | inter | 0.022973119616356036 | 0.11091884921279084 |

P62829_P84098
RL23_HUMAN vs RL19_HUMAN
p-value: 0.0 q-value: 0.0

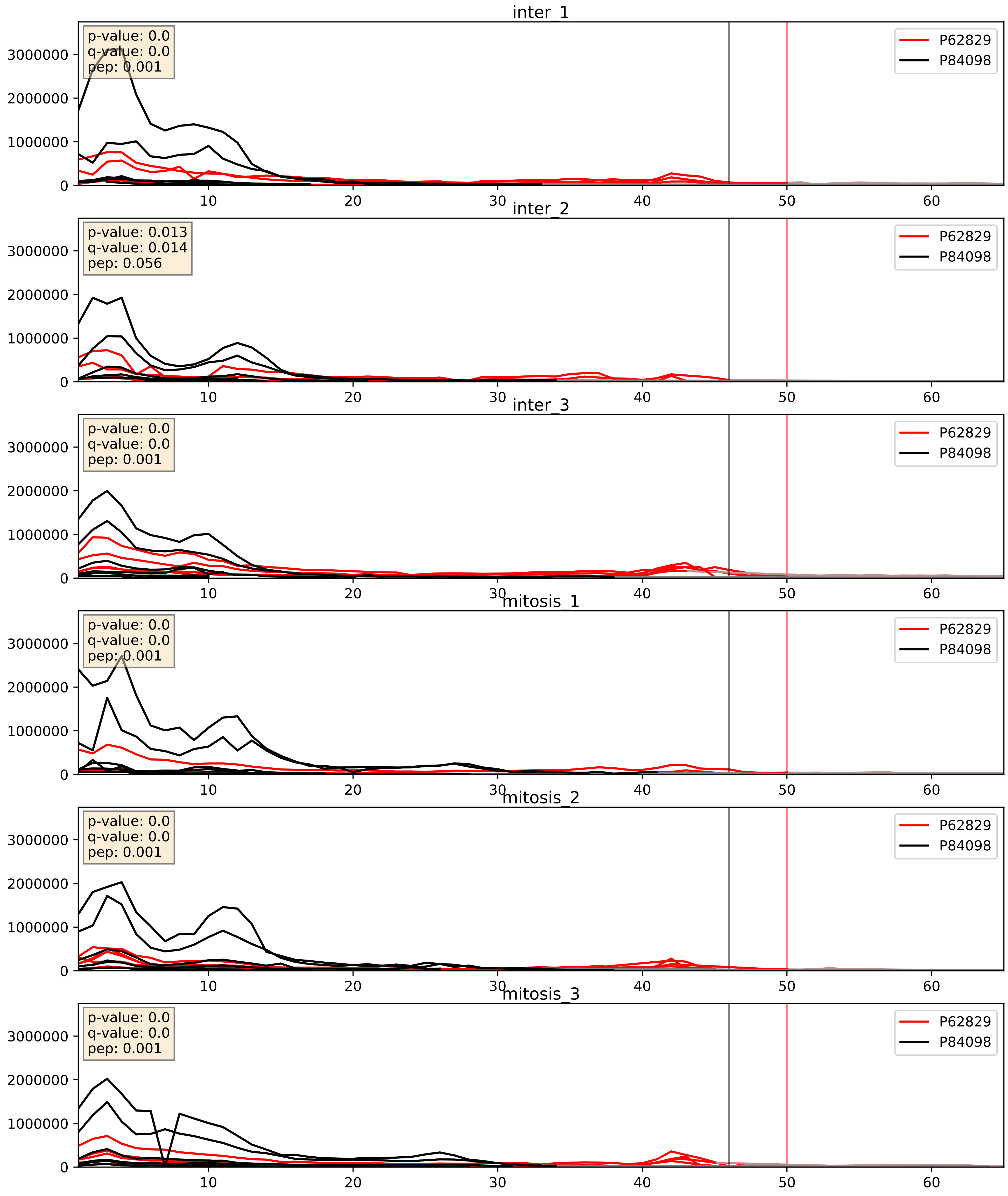


| level | condition_1 | condition_2 | pvalue | pvalue_adjusted |
|---|---|---|---|---|
| interactor_ratio | mitosis | inter | 0.0008902784219558255 | 0.034218313408302205 |
| interactor_abundance | mitosis | inter | 0.0064639807270440225 | 0.0687067394609051 |
| complex_abundance | mitosis | inter | 0.044058649064271906 | 0.15013616082411127 |

P62829_Q02543
RL23_HUMAN vs RL18A_HUMAN
p-value: 0.001 q-value: 0.001

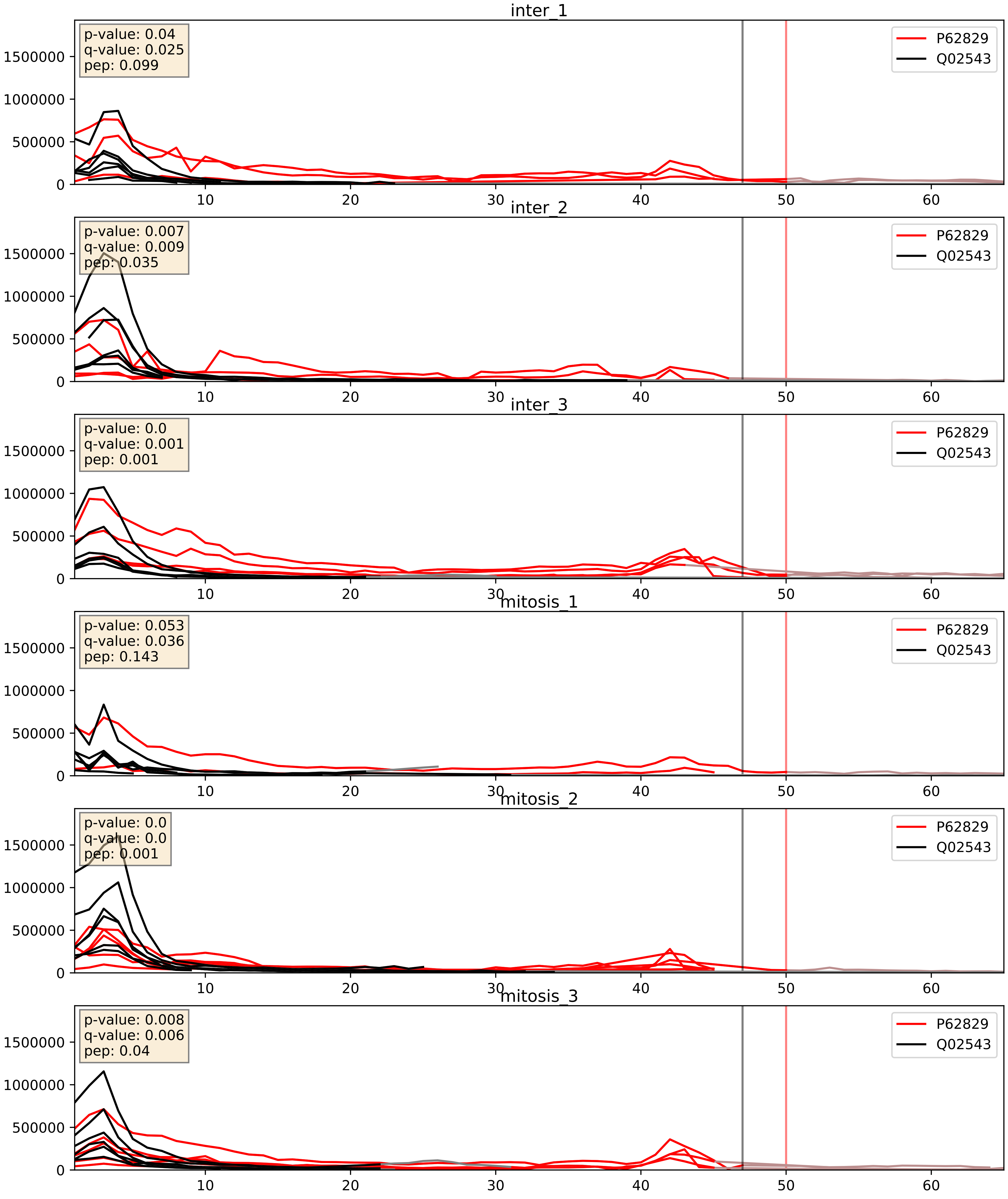


| level | condition_1 | condition_2 | pvalue | pvalue_adjusted |
|---|---|---|---|---|
| interactor_abundance | mitosis | inter | 0.001144573901567356 | 0.024325226673177953 |
| complex_abundance | mitosis | inter | 0.012282098923945444 | 0.08232949631086374 |
| interactor_ratio | mitosis | inter | 0.06310422845948546 | 0.41679953365215705 |

P62829_Q02878
RL23_HUMAN vs RL6_HUMAN
p-value: 0.0 q-value: 0.0

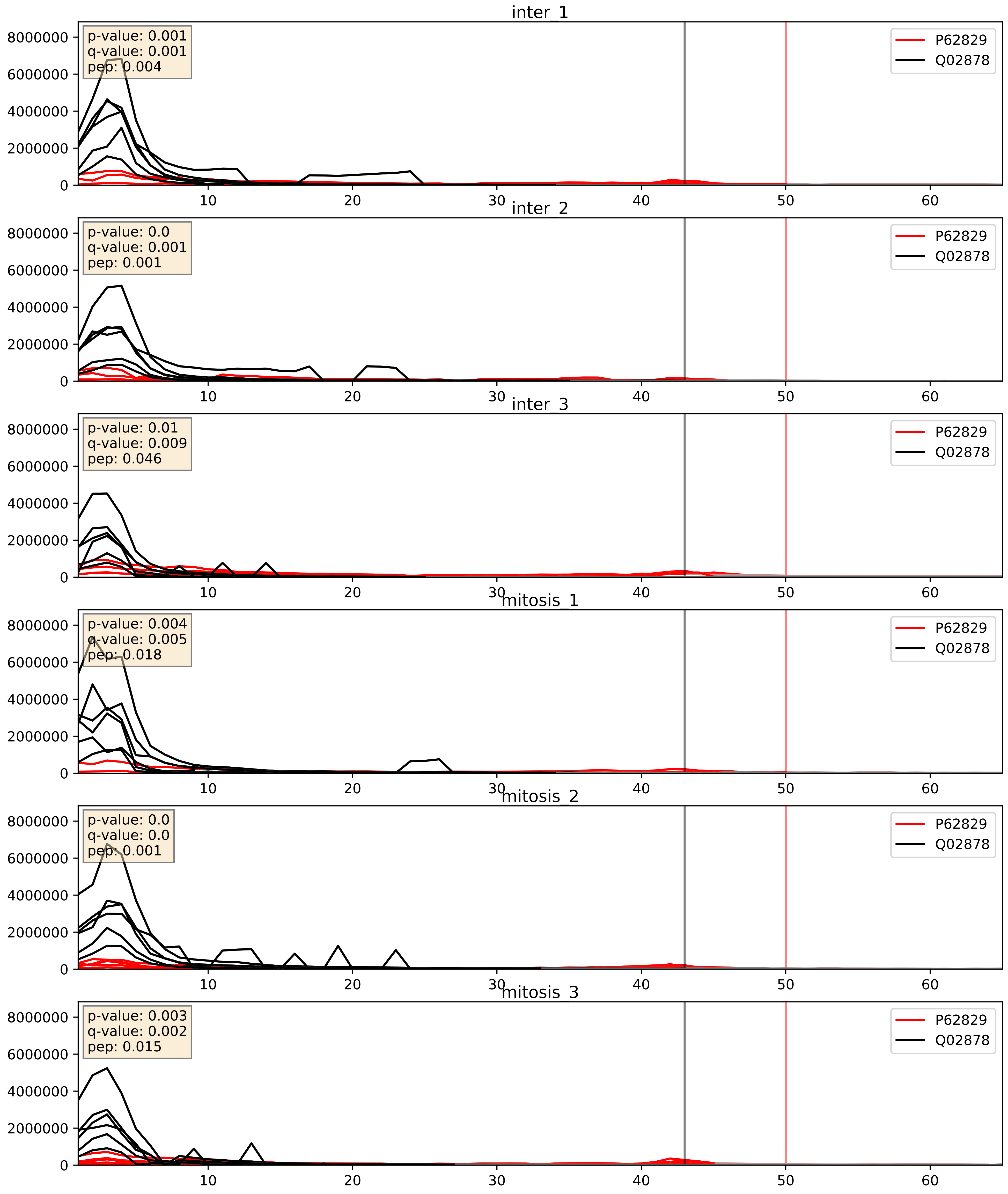


| level | condition_1 | condition_2 | pvalue | pvalue_adjusted |
|---|---|---|---|---|
| interactor_abundance | mitosis | inter | 0.00172439891736332 | 0.030275142475756547 |
| complex_abundance | mitosis | inter | 0.002950950622090906 | 0.0397179444531655 |
| interactor_ratio | mitosis | inter | 0.030976644353692084 | 0.2804780438054775 |

P62829_Q07020
RL23_HUMAN vs RL18_HUMAN
p-value: 0.0 q-value: 0.0

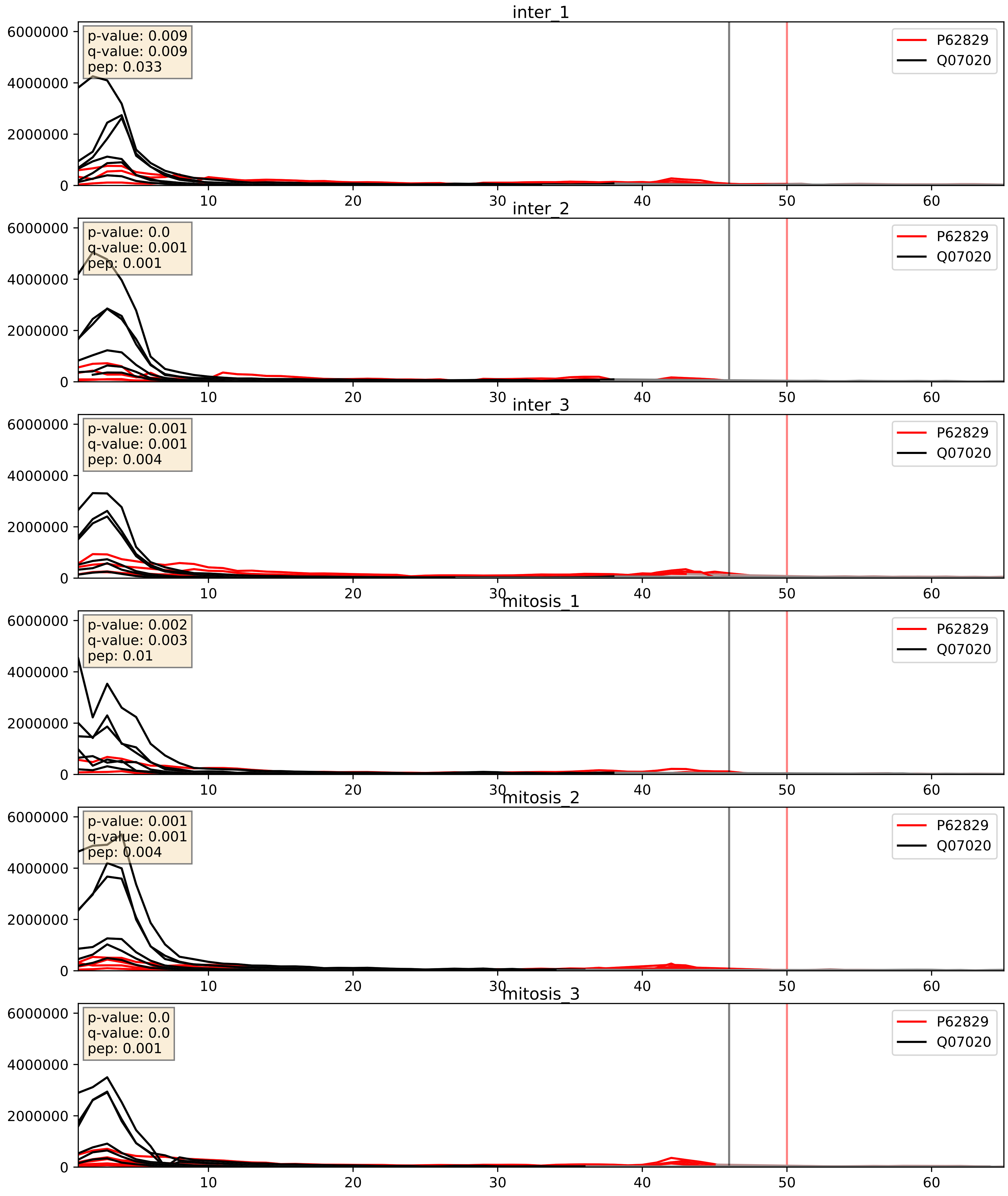


| level | condition_1 | condition_2 | pvalue | pvalue_adjusted |
|---|---|---|---|---|
| interactor_abundance | mitosis | inter | 0.0037036092869359924 | 0.047282469047236535 |
| complex_abundance | mitosis | inter | 0.005843255454504769 | 0.05756820427608883 |
| interactor_ratio | mitosis | inter | 0.01599245985203092 | 0.18676051341525876 |

P62829_Q12905
RL23_HUMAN vs ILF2_HUMAN
p-value: 0.0 q-value: 0.047

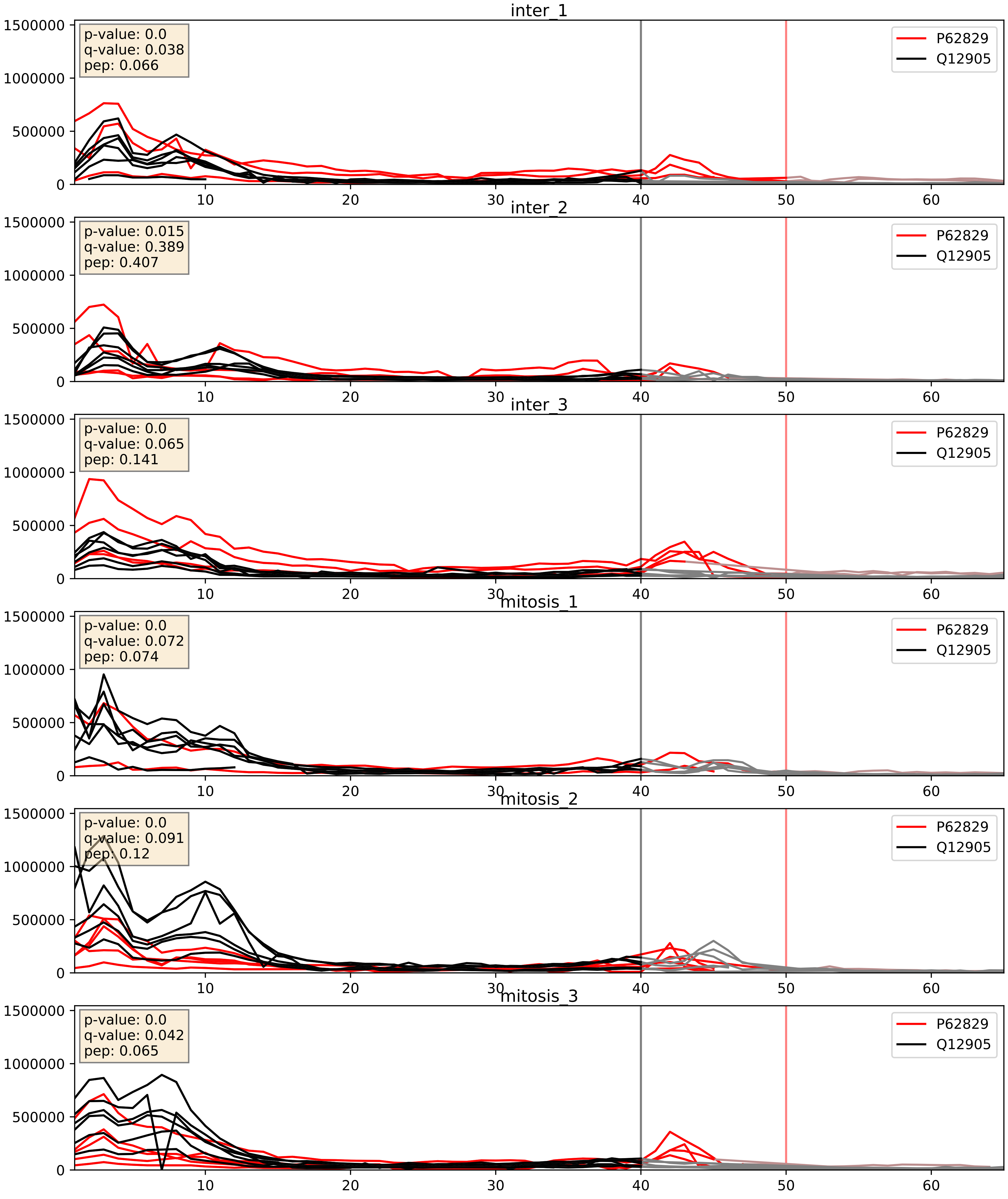


| level | condition_1 | condition_2 | pvalue | pvalue_adjusted |
|---|---|---|---|---|
| interactor_ratio | mitosis | inter | 0.0022056667910768353 | 0.05863511717893699 |
| interactor_abundance | mitosis | inter | 0.004946130873219955 | 0.05819777357355713 |
| complex_abundance | mitosis | inter | 0.023997797201938377 | 0.11305511505150936 |

P62829_Q969Q0
RL23_HUMAN vs RL36L_HUMAN
p-value: 0.0 q-value: 0.0

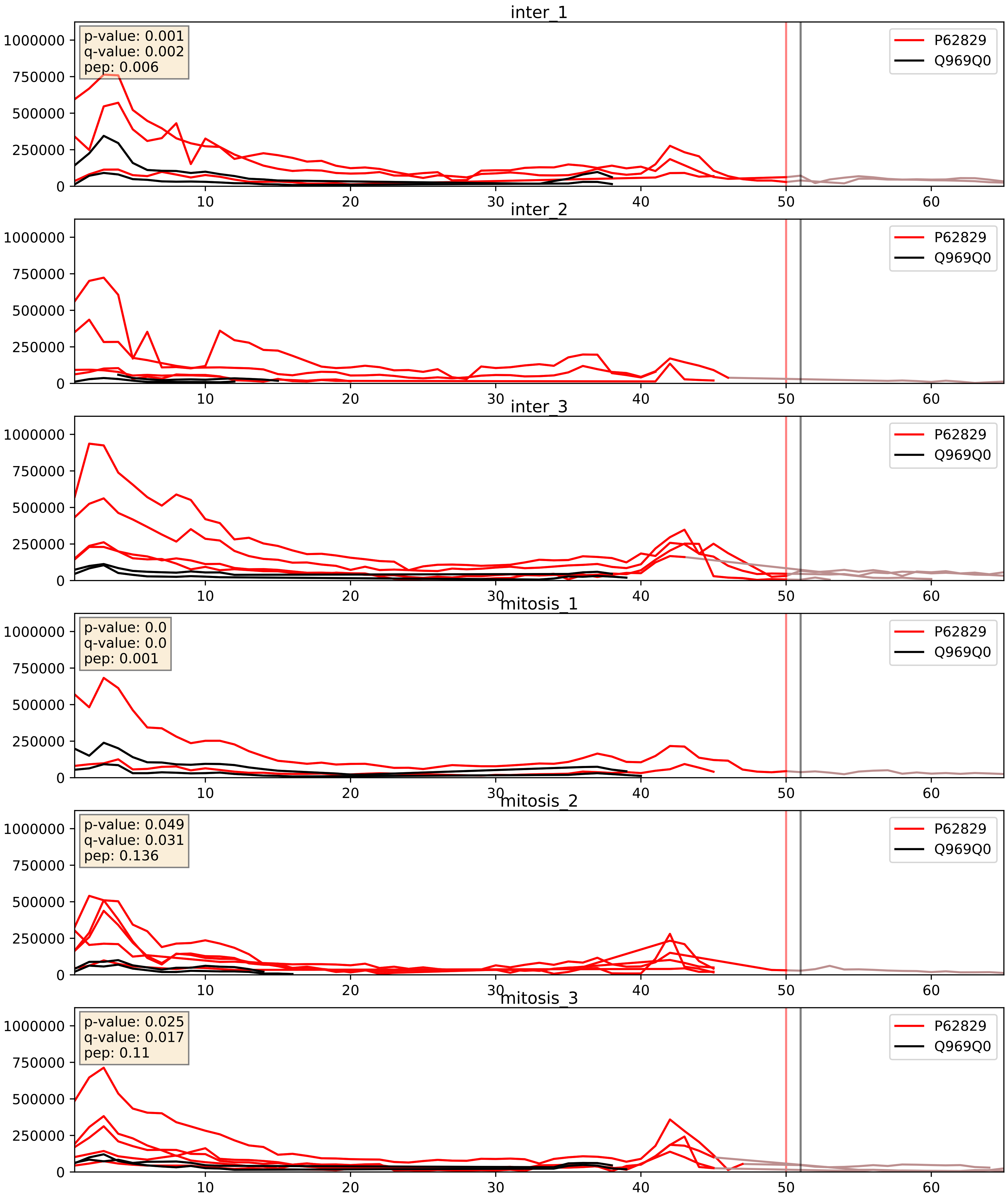


| level | condition_1 | condition_2 | pvalue | pvalue_adjusted |
|---|---|---|---|---|
| interactor_abundance | mitosis | inter | 0.0036918670855603177 | 0.047220768188103 |
| interactor_ratio | mitosis | inter | 0.006629405333481775 | 0.11281850825965009 |
| complex_abundance | mitosis | inter | 0.0358986842228082 | 0.1376167508298101 |

P62829_Q9Y2X3
RL23_HUMAN vs NOP58_HUMAN
p-value: 0.0 q-value: 0.047

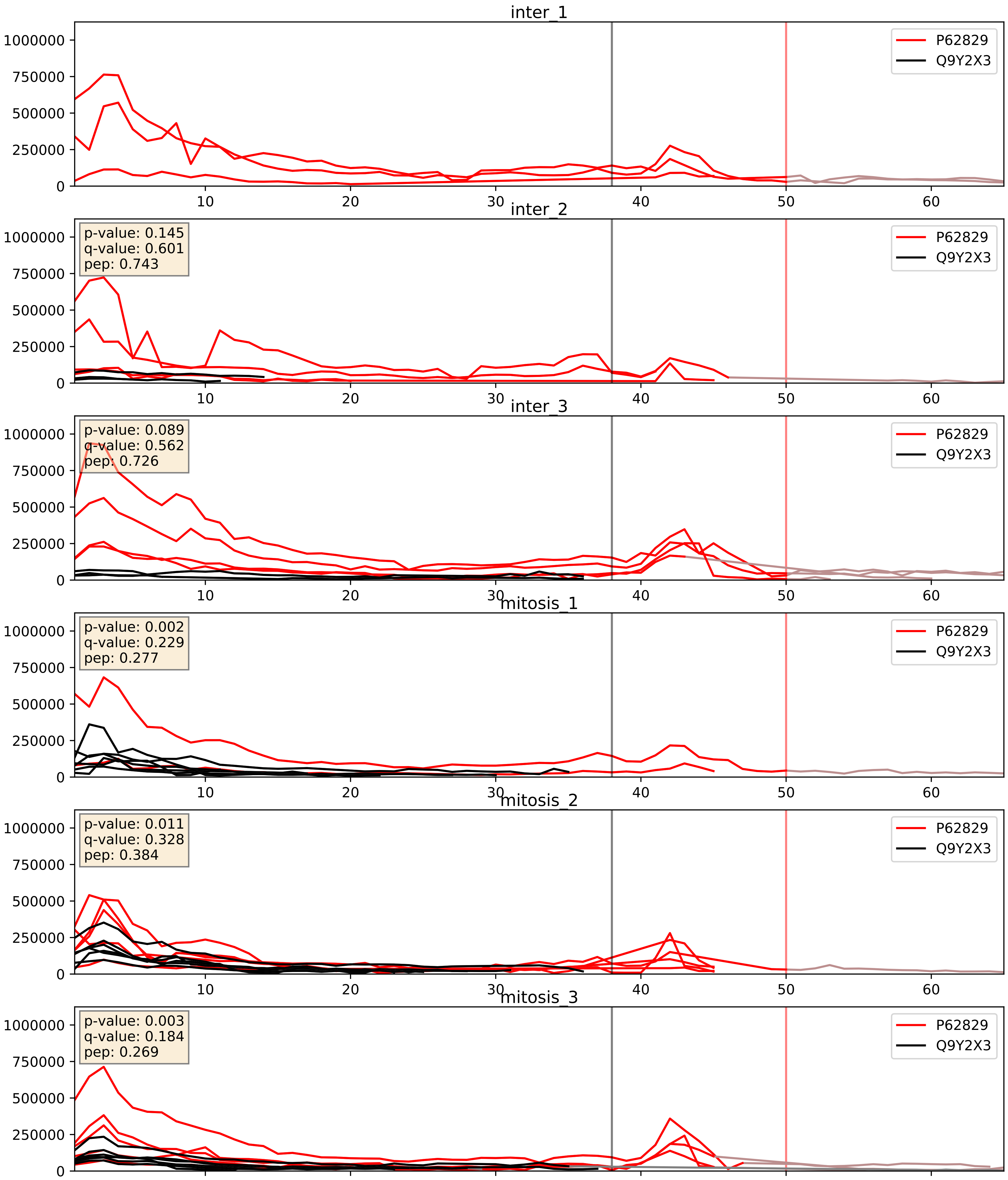


| level | condition_1 | condition_2 | pvalue | pvalue_adjusted |
|---|---|---|---|---|
| interactor_ratio | mitosis | inter | 0.08336086174927138 | 0.4841037832929192 |
| complex_abundance | mitosis | inter | 0.36594266613487514 | 0.5137722194709745 |
| interactor_abundance | mitosis | inter | 0.7978512935014155 | 0.8820363001901222 |

P62829_Q9Y3U8
RL23_HUMAN vs RL36_HUMAN
p-value: 0.0 q-value: 0.0

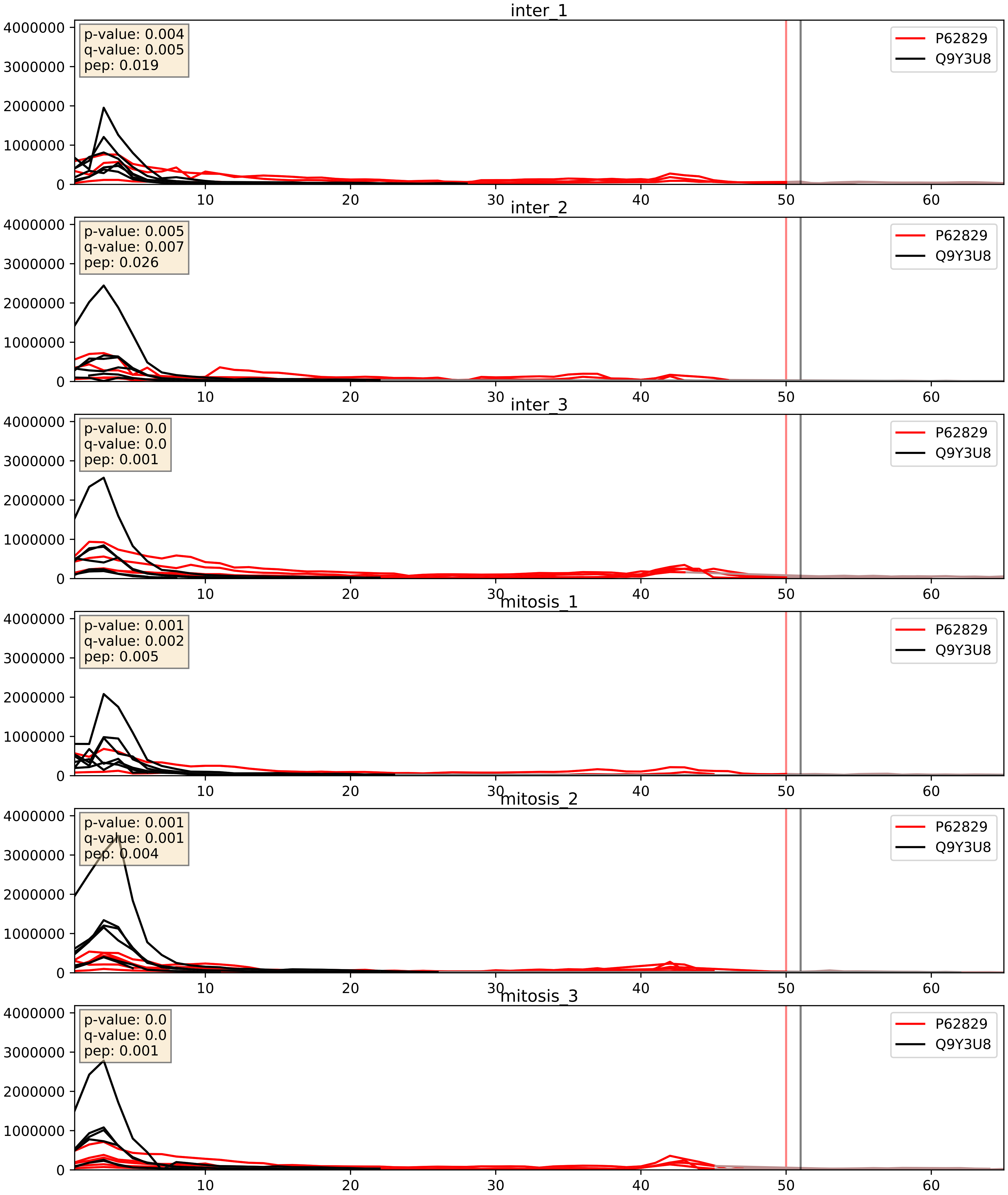


| level | condition_1 | condition_2 | pvalue | pvalue_adjusted |
|---|---|---|---|---|
| interactor_ratio | mitosis | inter | 0.0016322780793195505 | 0.048854197059354385 |
| interactor_abundance | mitosis | inter | 0.0030079256162717613 | 0.04193459816821869 |
| complex_abundance | mitosis | inter | 0.008112602748244093 | 0.06664479800860791 |
