## Supplementary Data 2 for "SECAT: Quantifying differential protein-protein interaction states by network-centric analysis": hela_string_000026_Q15269.pdf

Q15269_Q15269

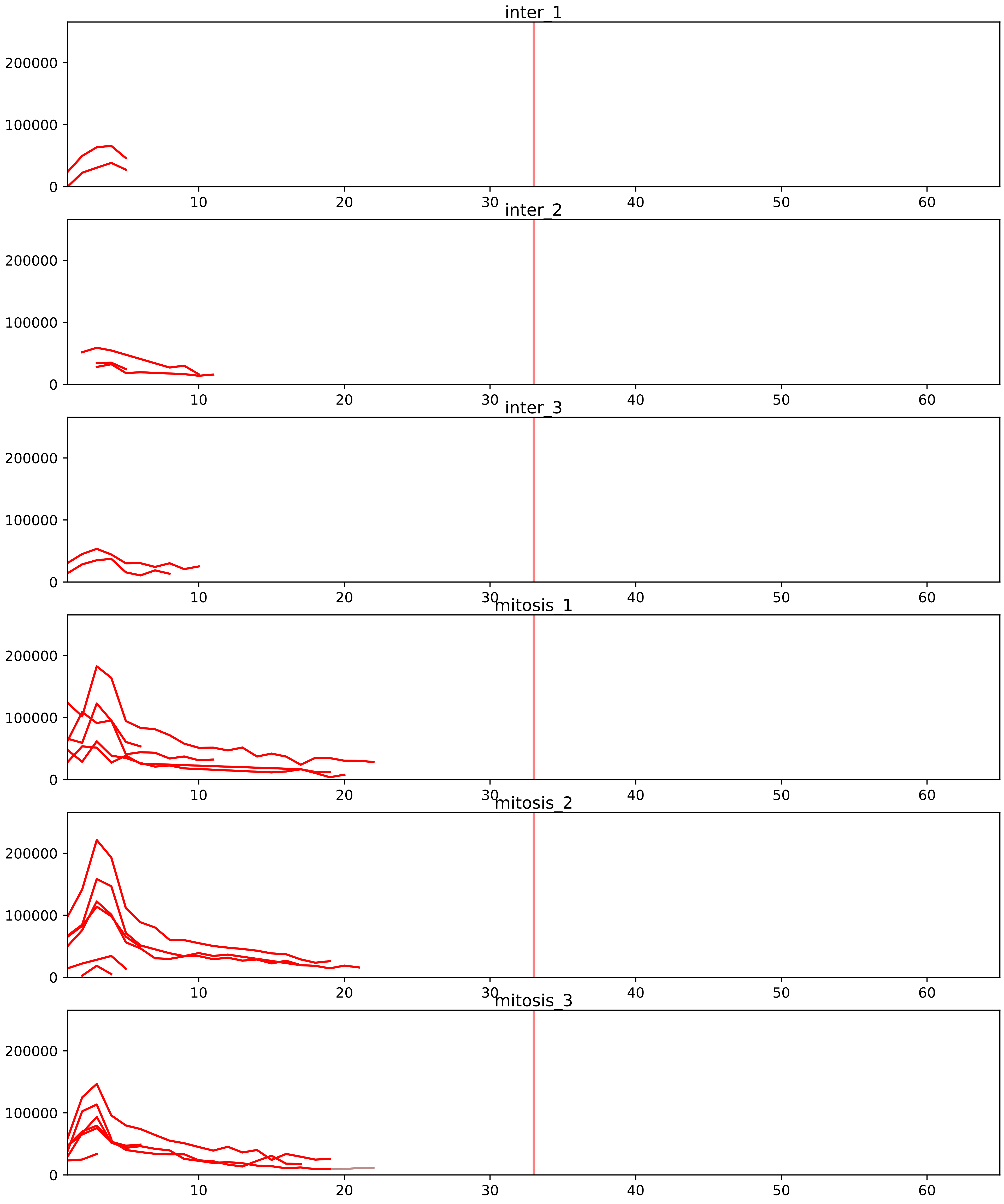


| level | condition_1 | condition_2 | pvalue | pvalue_adjusted |
|---|---|---|---|---|
| interactor_ratio | mitosis | inter | 6.422016174765646e-11 | 4.544811446757226e-09 |
| complex_abundance | mitosis | inter | 0.0004402145894501205 | 0.006328084723345483 |
| interactor_abundance | mitosis | inter | 0.002008993211699035 | 0.019153095904280958 |
| assembled_abundance | mitosis | inter | 0.007710590014530972 | 0.09448500486392535 |
| total_abundance | mitosis | inter | 0.009369278965516448 | 0.11580419412688246 |

O00567_Q15269
NOP56_HUMAN vs PWP2_HUMAN
p-value: 0.001 q-value: 0.001

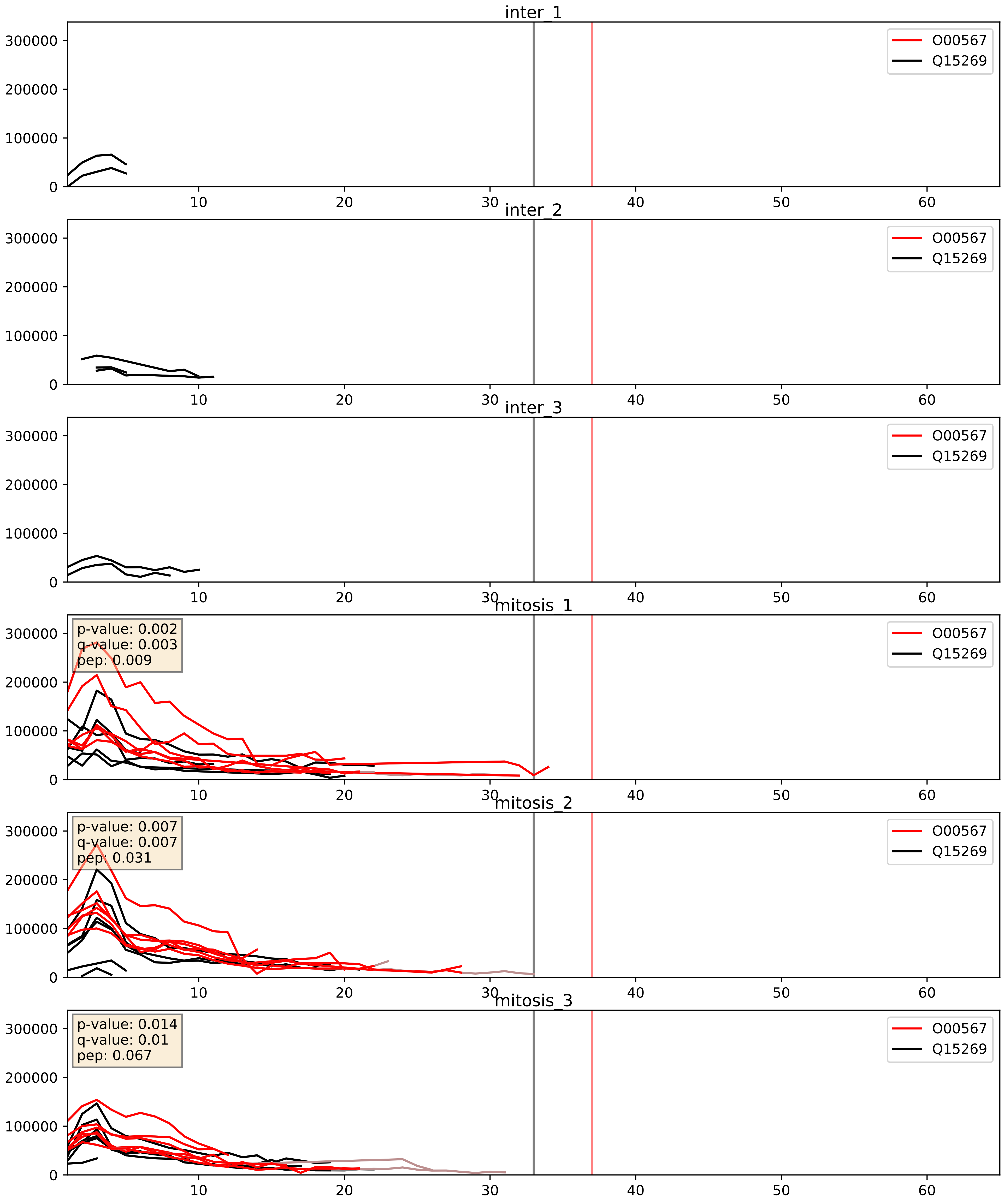


| level | condition_1 | condition_2 | pvalue | pvalue_adjusted |
|---|---|---|---|---|
| complex_abundance | mitosis | inter | 0.0010352101406899857 | 0.021560581032375372 |
| interactor_abundance | mitosis | inter | 0.0011045983974246115 | 0.024049720606162197 |
| interactor_ratio | mitosis | inter | 0.008196470592608973 | 0.12756688776860511 |

P22087_Q15269
FBRL_HUMAN vs PWP2_HUMAN
p-value: 0.001 q-value: 0.002

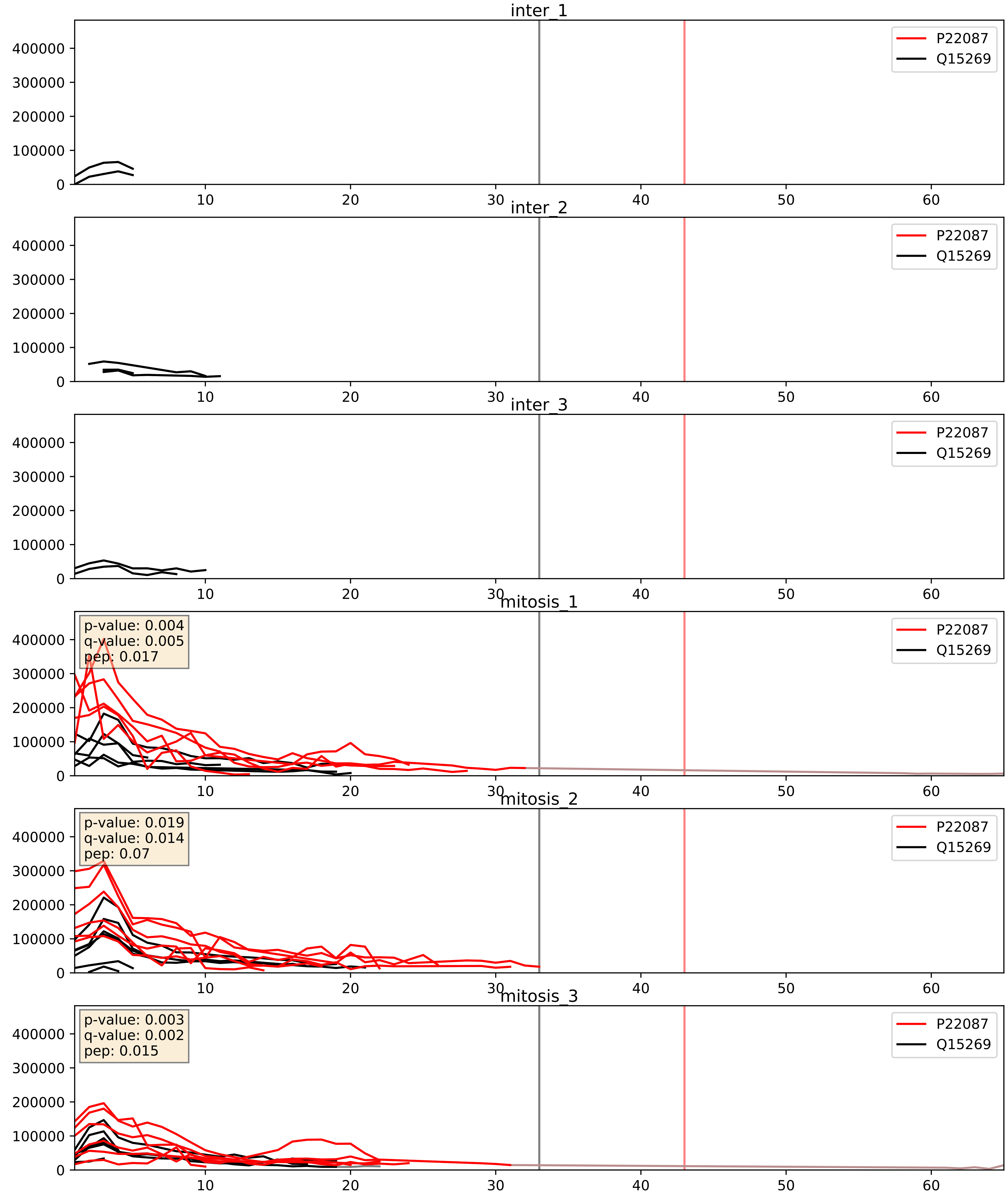


| level | condition_1 | condition_2 | pvalue | pvalue_adjusted |
|---|---|---|---|---|
| interactor_abundance | mitosis | inter | 0.0005008244807677675 | 0.015907449185054136 |
| interactor_ratio | mitosis | inter | 0.0005887273659099557 | 0.028471786735532318 |
| complex_abundance | mitosis | inter | 0.000661858101371452 | 0.01642175463112936 |

P42285_Q15269
MTREX_HUMAN vs PWP2_HUMAN
p-value: 0.08 q-value: 0.048

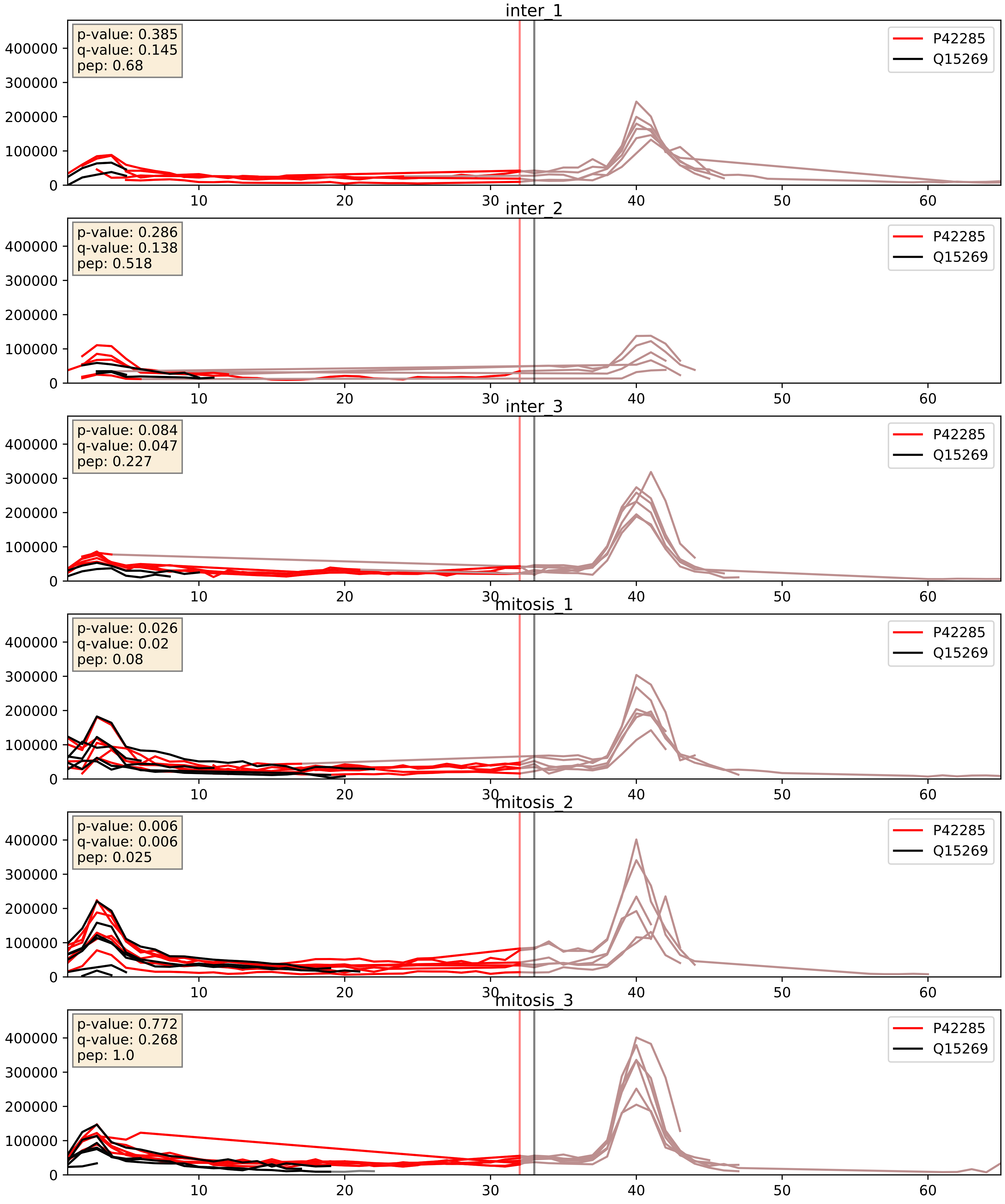


| level | condition_1 | condition_2 | pvalue | pvalue_adjusted |
|---|---|---|---|---|
| interactor_abundance | mitosis | inter | 0.0023540448297948925 | 0.036079899271341594 |
| complex_abundance | mitosis | inter | 0.005088813183140754 | 0.053382648097653 |
| interactor_ratio | mitosis | inter | 0.5603810643672665 | 0.8918094730984788 |

P55769_Q15269
NH2L1_HUMAN vs PWP2_HUMAN
p-value: 0.015 q-value: 0.013

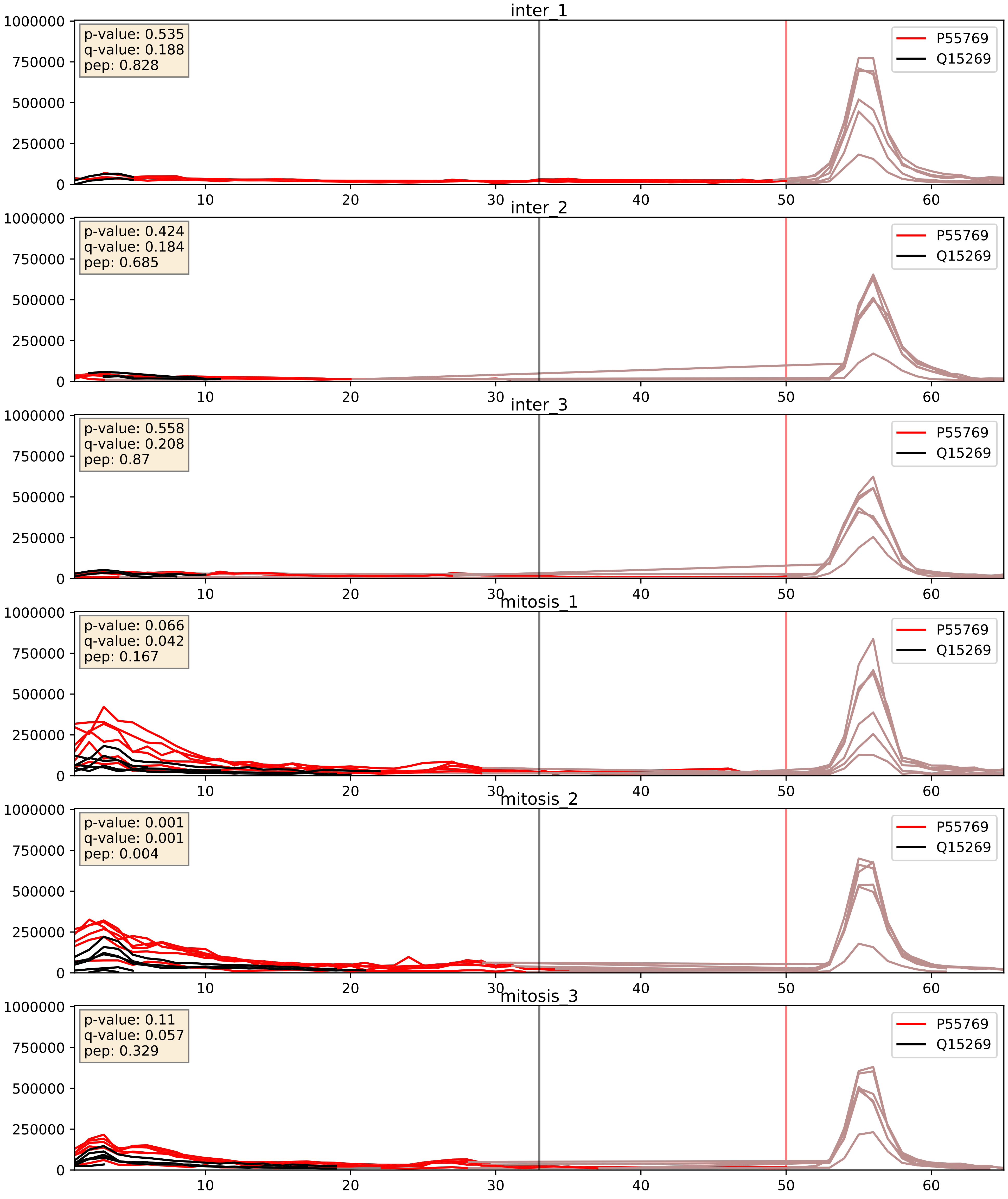


| level | condition_1 | condition_2 | pvalue | pvalue_adjusted |
|---|---|---|---|---|
| interactor_abundance | mitosis | inter | 0.0009531398130480196 | 0.022918193257559123 |
| complex_abundance | mitosis | inter | 0.0021258483587280417 | 0.033489340249041655 |
| interactor_ratio | mitosis | inter | 0.003761862384957826 | 0.07990457075741685 |

Q12788_Q15269
TBL3_HUMAN vs PWP2_HUMAN
p-value: 0.015 q-value: 0.012

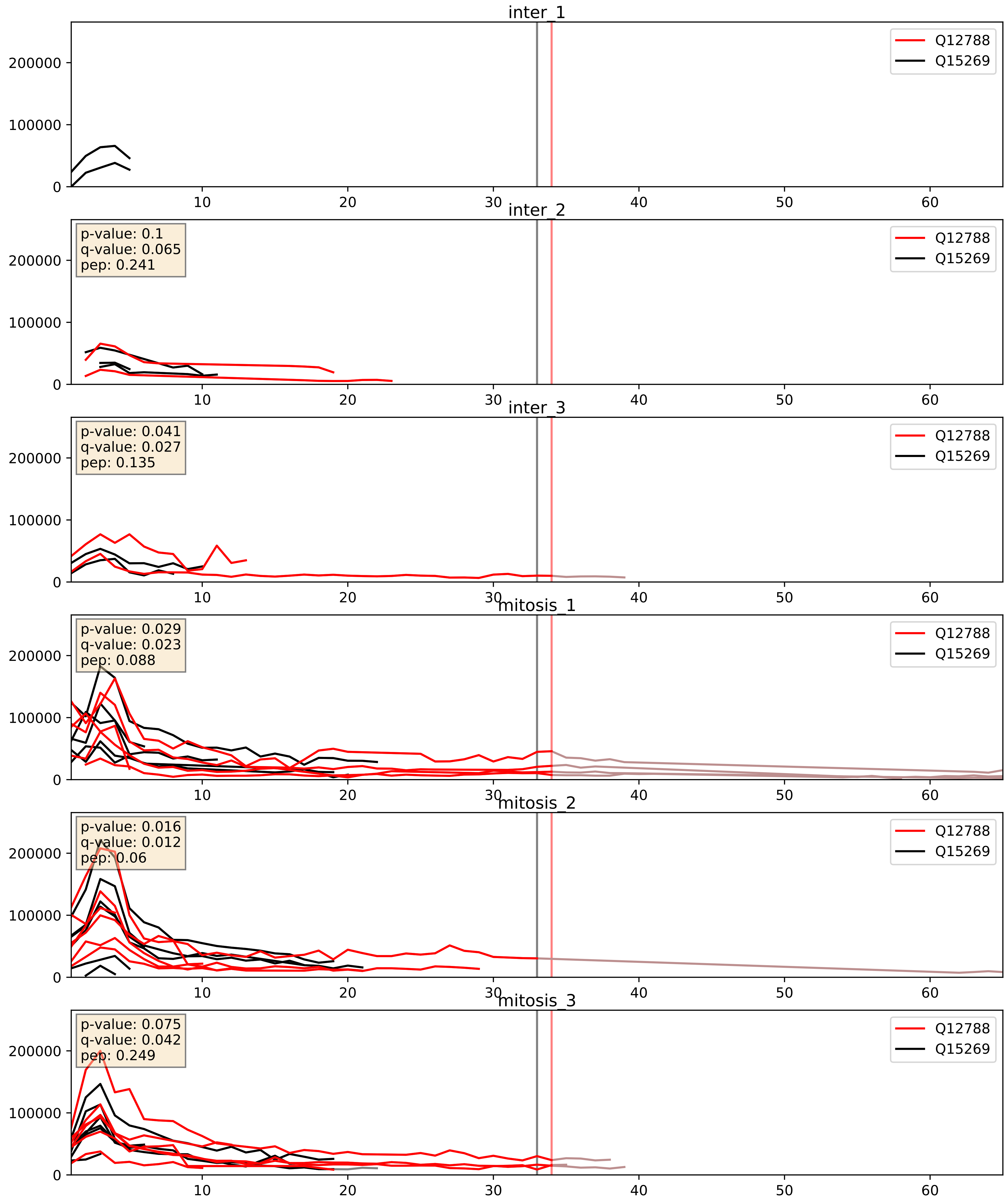


| level | condition_1 | condition_2 | pvalue | pvalue_adjusted |
|---|---|---|---|---|
| interactor_abundance | mitosis | inter | 0.0045441861105829885 | 0.05425137113889872 |
| complex_abundance | mitosis | inter | 0.005891801045557714 | 0.05764342671241339 |
| interactor_ratio | mitosis | inter | 0.6966472160731734 | 0.9356388300120577 |

Q14137_Q15269
BOP1_HUMAN vs PWP2_HUMAN
p-value: 0.058 q-value: 0.037

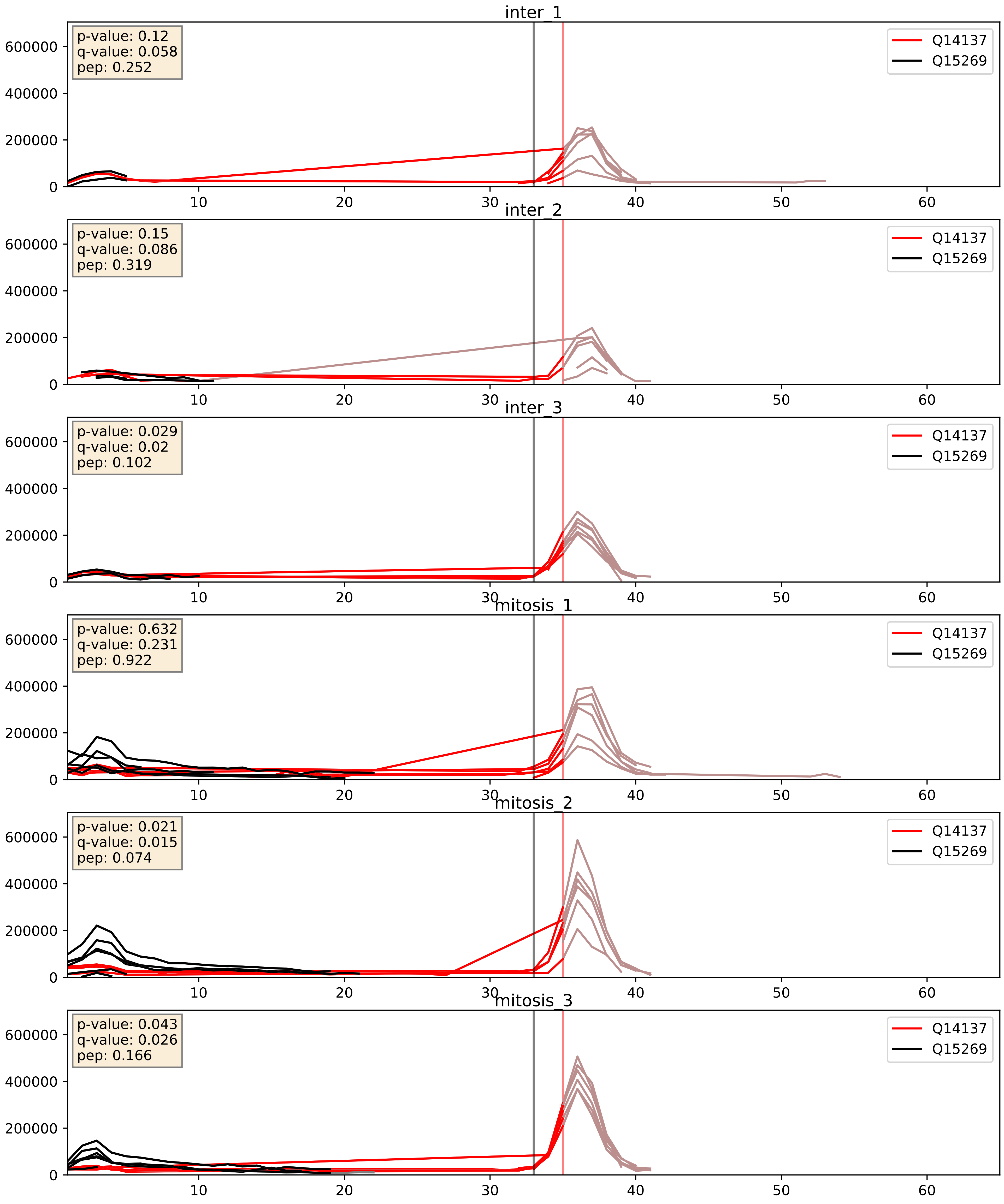


| level | condition_1 | condition_2 | pvalue | pvalue_adjusted |
|---|---|---|---|---|
| interactor_ratio | mitosis | inter | 0.0035194966342699525 | 0.07646419083591573 |
| interactor_abundance | mitosis | inter | 0.16567605704869157 | 0.3537339202733041 |
| complex_abundance | mitosis | inter | 0.7966276306979535 | 0.8676845042339333 |

Q15269_Q8IY81
PWP2_HUMAN vs SPB1_HUMAN
p-value: 0.001 q-value: 0.001

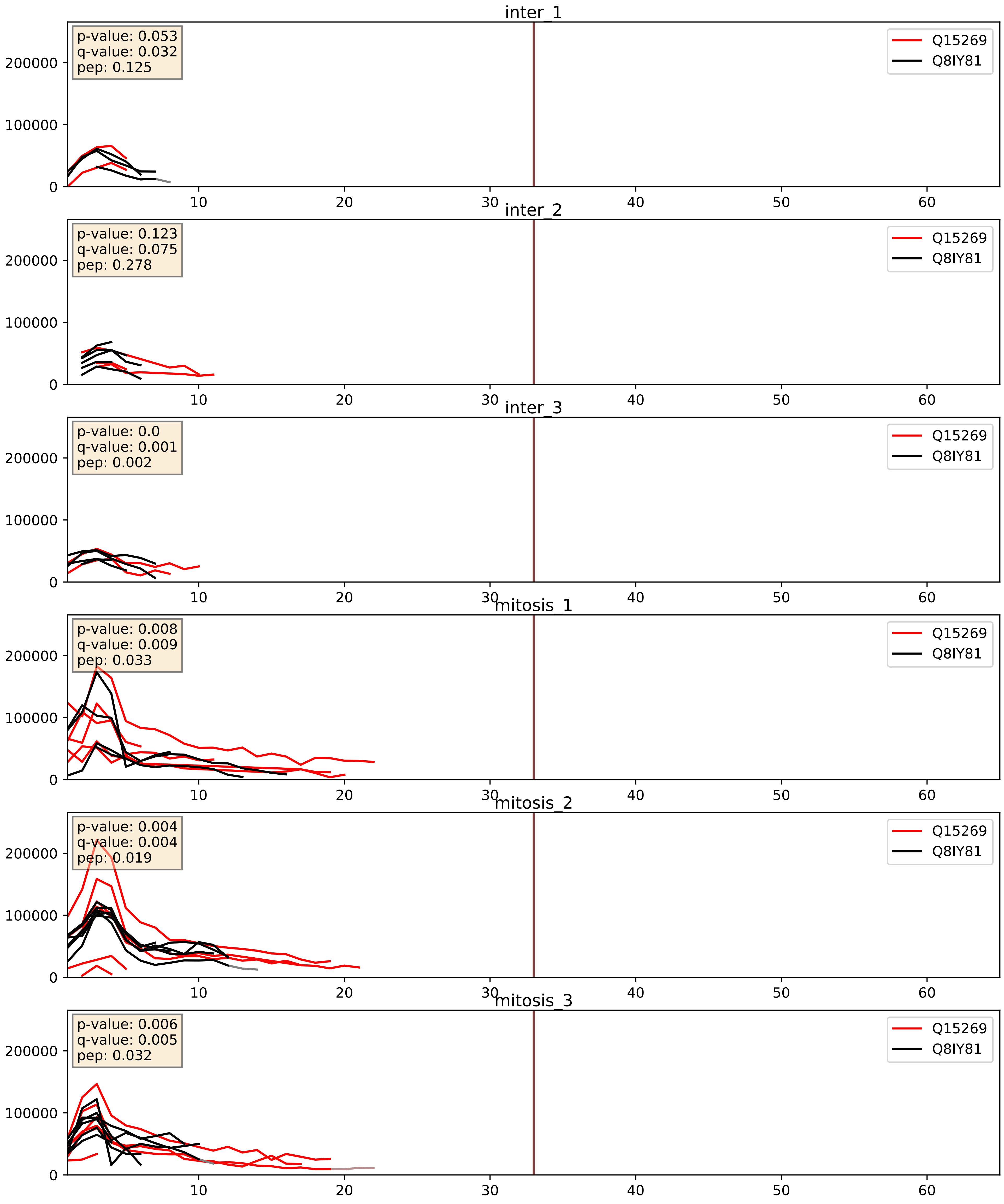


| level | condition_1 | condition_2 | pvalue | pvalue_adjusted |
|---|---|---|---|---|
| interactor_ratio | mitosis | inter | 0.04107050221712384 | 0.3361027714900383 |
| interactor_abundance | mitosis | inter | 0.05103025519733094 | 0.181177513268002 |
| complex_abundance | mitosis | inter | 0.11609552713244926 | 0.2570557972720553 |

Q15269_Q8NI36
PWP2_HUMAN vs WDR36_HUMAN
p-value: 0.001 q-value: 0.001

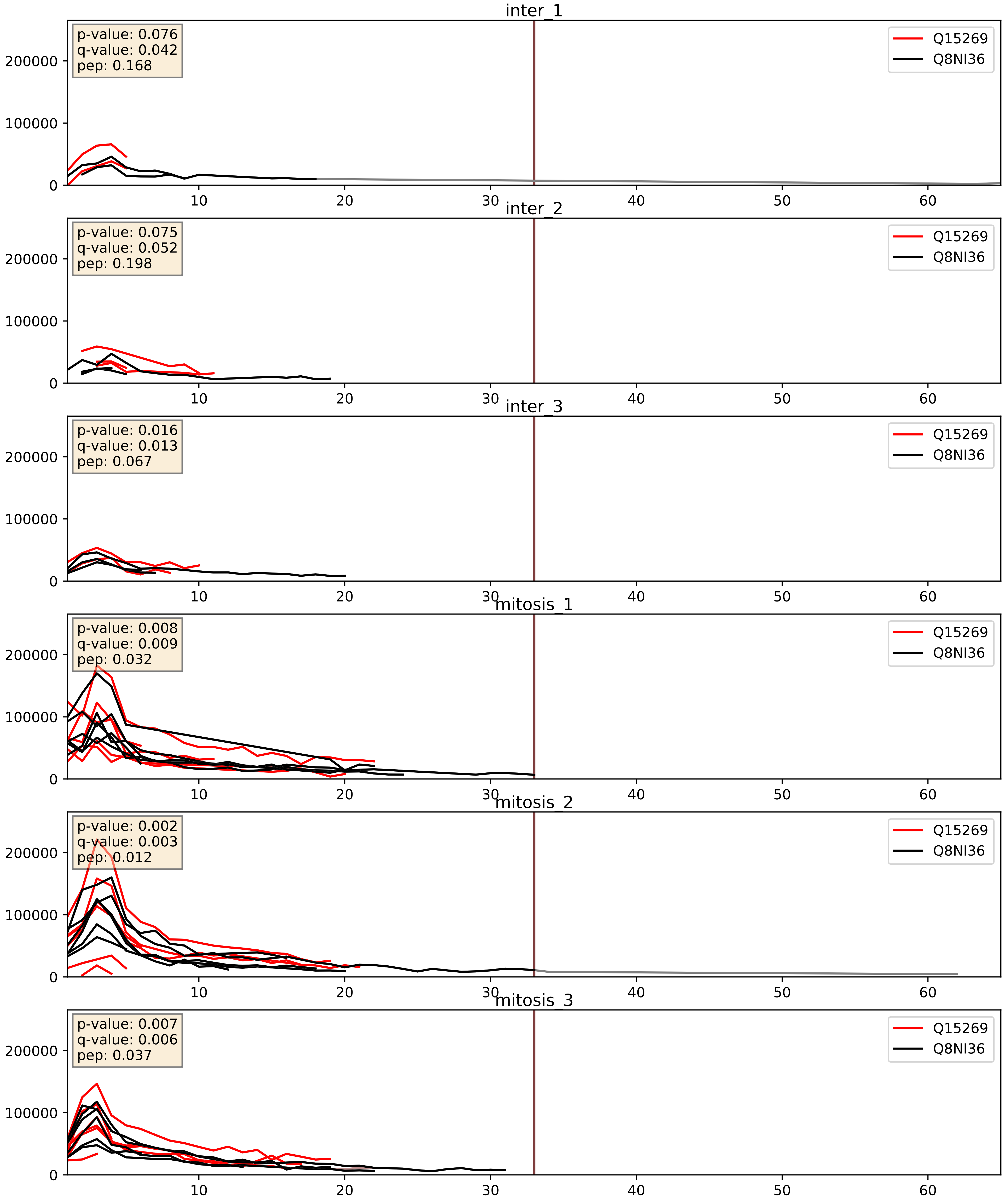


| level | condition_1 | condition_2 | pvalue | pvalue_adjusted |
|---|---|---|---|---|
| complex_abundance | mitosis | inter | 0.008082465810691268 | 0.06653038413698939 |
| interactor_abundance | mitosis | inter | 0.02134655497707949 | 0.1216554664472706 |
| interactor_ratio | mitosis | inter | 0.7936552315502841 | 0.9608508478447312 |

Q15269_Q969X6
PWP2_HUMAN vs UTP4_HUMAN
p-value: 0.073 q-value: 0.044

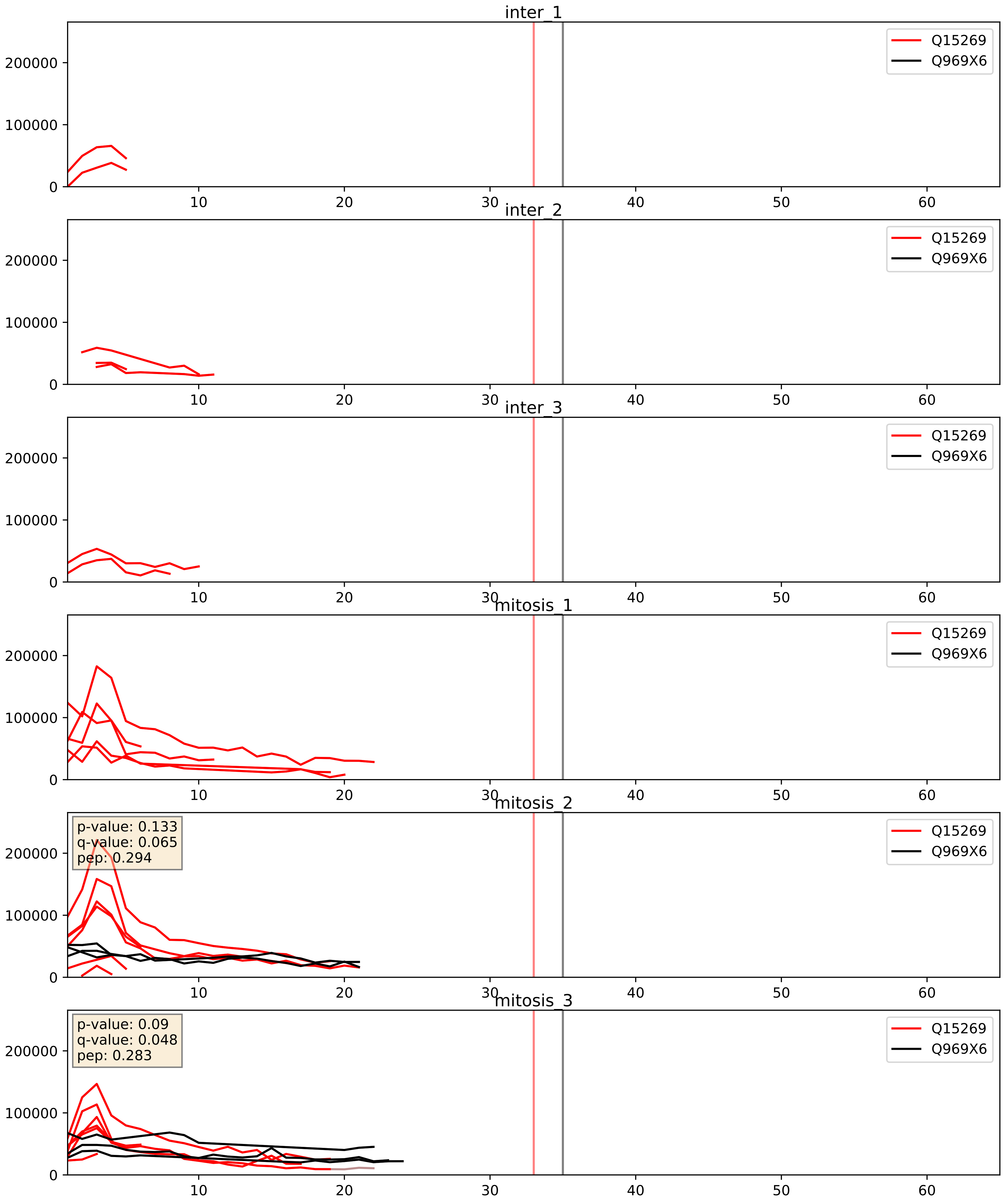


| level | condition_1 | condition_2 | pvalue | pvalue_adjusted |
|---|---|---|---|---|
| interactor_ratio | mitosis | inter | 0.1971867894585755 | 0.6719422443333624 |
| interactor_abundance | mitosis | inter | 0.20124937063532916 | 0.39608168202589294 |
| complex_abundance | mitosis | inter | 0.22045672824532372 | 0.3746808072497155 |

Q15269_Q9GZL7
PWP2_HUMAN vs WDR12_HUMAN
p-value: 0.074 q-value: 0.045

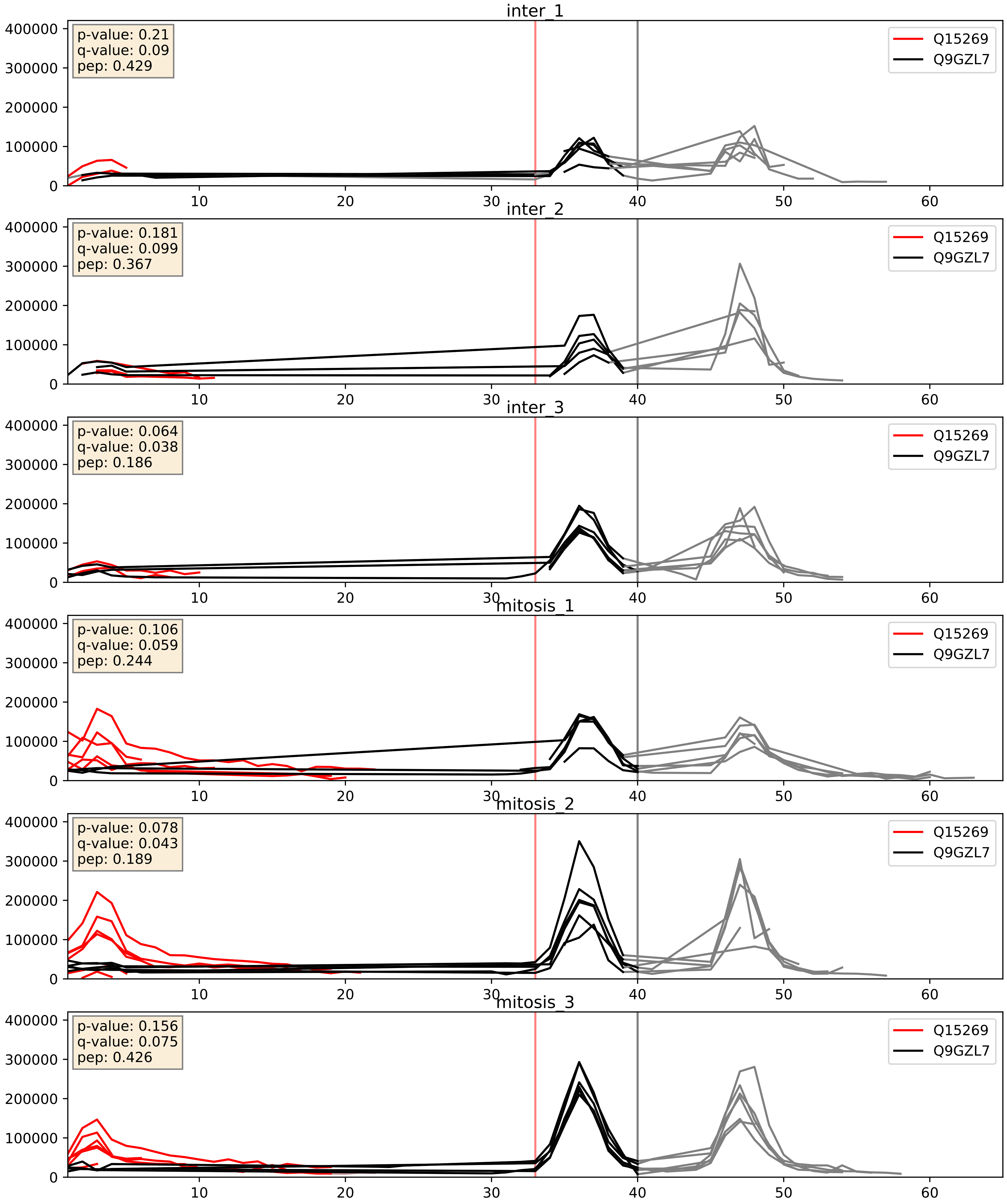


| level | condition_1 | condition_2 | pvalue | pvalue_adjusted |
|---|---|---|---|---|
| interactor_ratio | mitosis | inter | 0.007695757326345688 | 0.1235941514324936 |
| complex_abundance | mitosis | inter | 0.3683484939112922 | 0.515965162474335 |
| interactor_abundance | mitosis | inter | 0.3710710153088357 | 0.5652137254615444 |

Q15269_Q9H0A0
PWP2_HUMAN vs NAT10_HUMAN
p-value: 0.002 q-value: 0.002

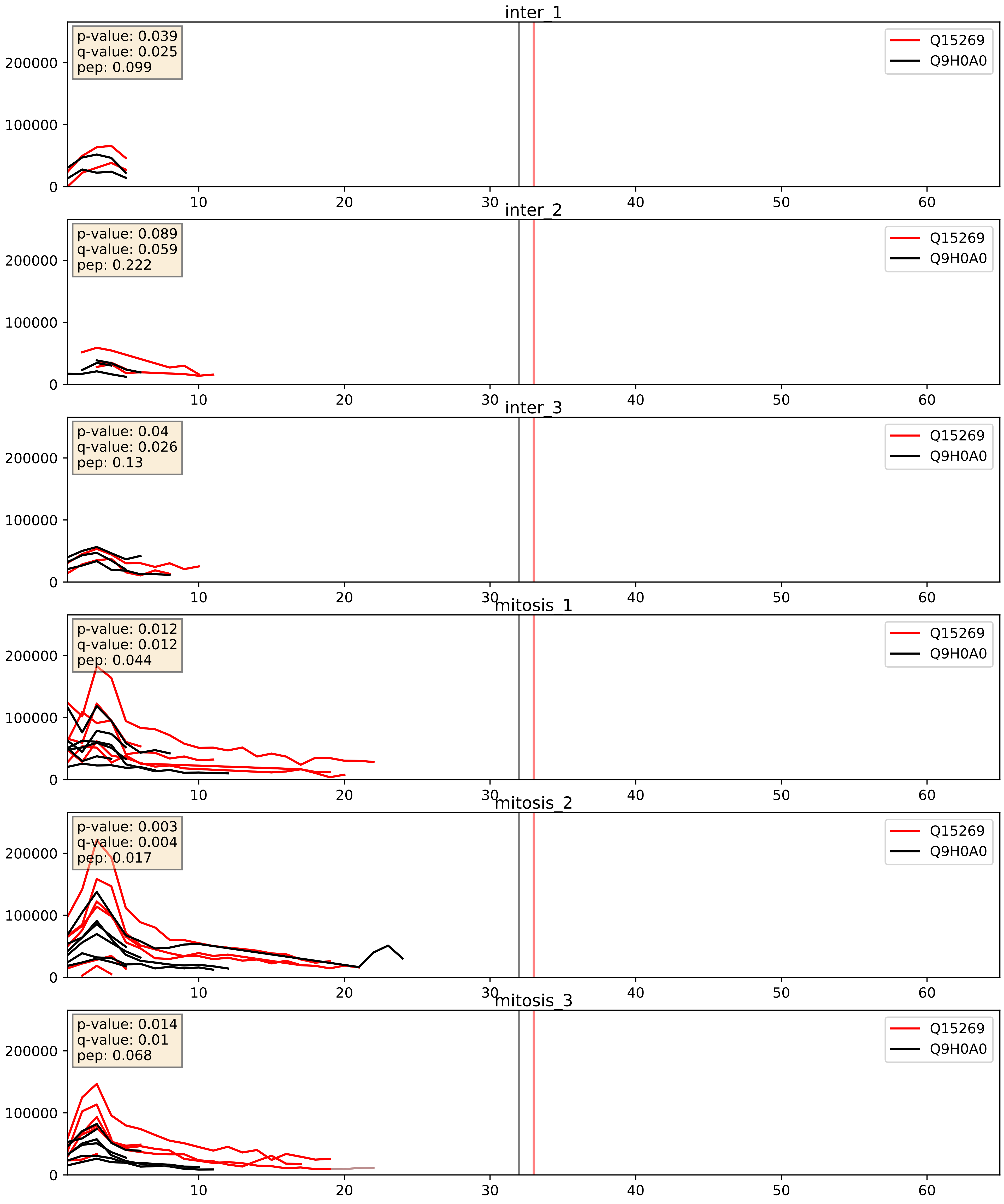


| level | condition_1 | condition_2 | pvalue | pvalue_adjusted |
|---|---|---|---|---|
| interactor_ratio | mitosis | inter | 0.013552472765333098 | 0.1693564479872282 |
| interactor_abundance | mitosis | inter | 0.0423912444936641 | 0.16595886250435155 |
| complex_abundance | mitosis | inter | 0.16592341030748023 | 0.3147952291083829 |

Q15269_Q9H583
PWP2_HUMAN vs HEAT1_HUMAN
p-value: 0.029 q-value: 0.021

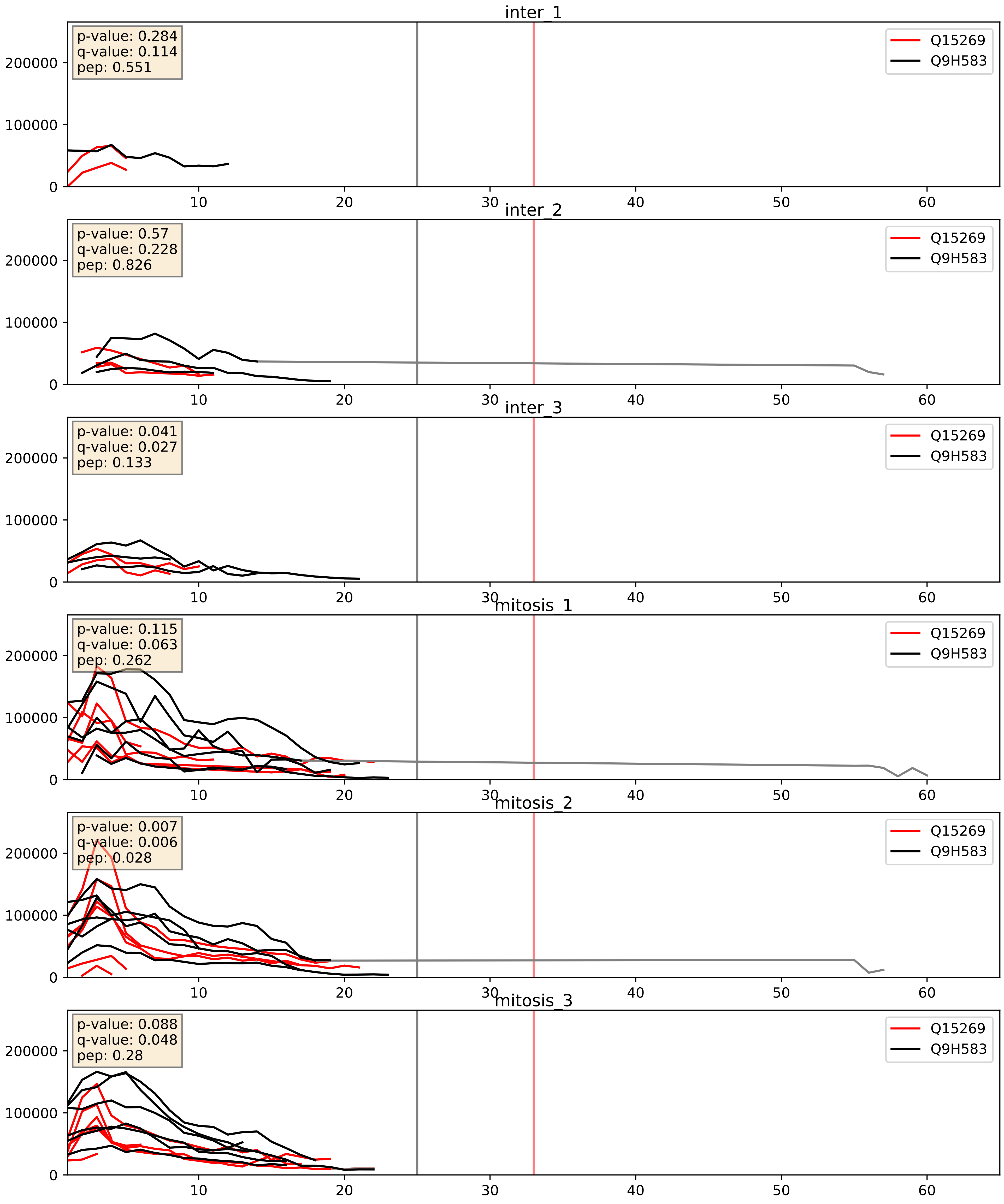


| level | condition_1 | condition_2 | pvalue | pvalue_adjusted |
|---|---|---|---|---|
| complex_abundance | mitosis | inter | 0.0013443121273097713 | 0.025015895238634005 |
| interactor_abundance | mitosis | inter | 0.02134655497707949 | 0.1216554664472706 |
| interactor_ratio | mitosis | inter | 0.06289218988803522 | 0.4166850970910073 |

Q15269_Q9H6R4
PWP2_HUMAN vs NOL6_HUMAN
p-value: 0.028 q-value: 0.021

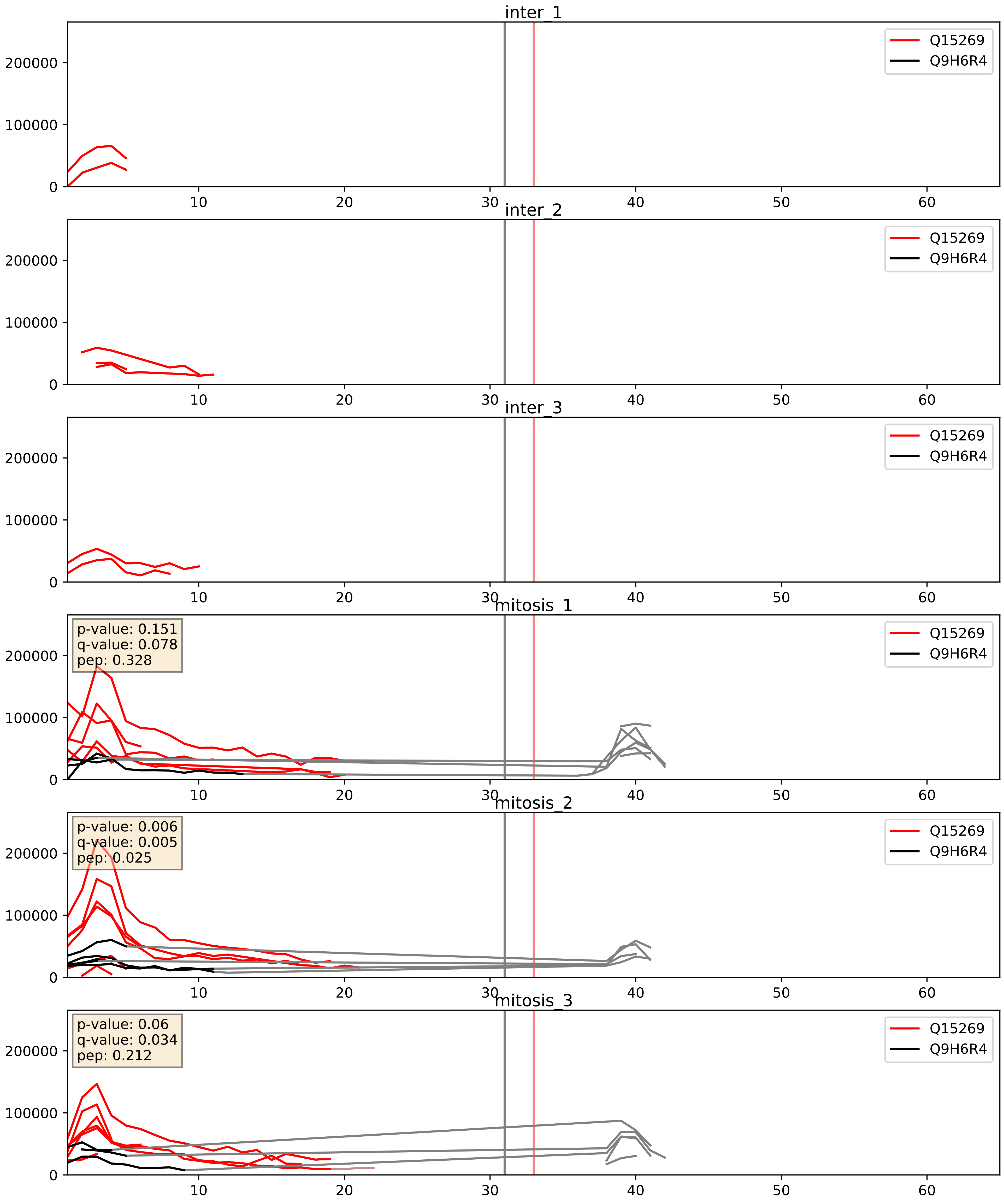


| level | condition_1 | condition_2 | pvalue | pvalue_adjusted |
|---|---|---|---|---|
| interactor_abundance | mitosis | inter | 0.003100387467477749 | 0.04271207144514066 |
| interactor_ratio | mitosis | inter | 0.004656516393809501 | 0.09159088101537749 |
| complex_abundance | mitosis | inter | 0.012416401996518596 | 0.08277281176263881 |

Q15269_Q9UNX4
PWP2_HUMAN vs WDR3_HUMAN
p-value: 0.009 q-value: 0.008

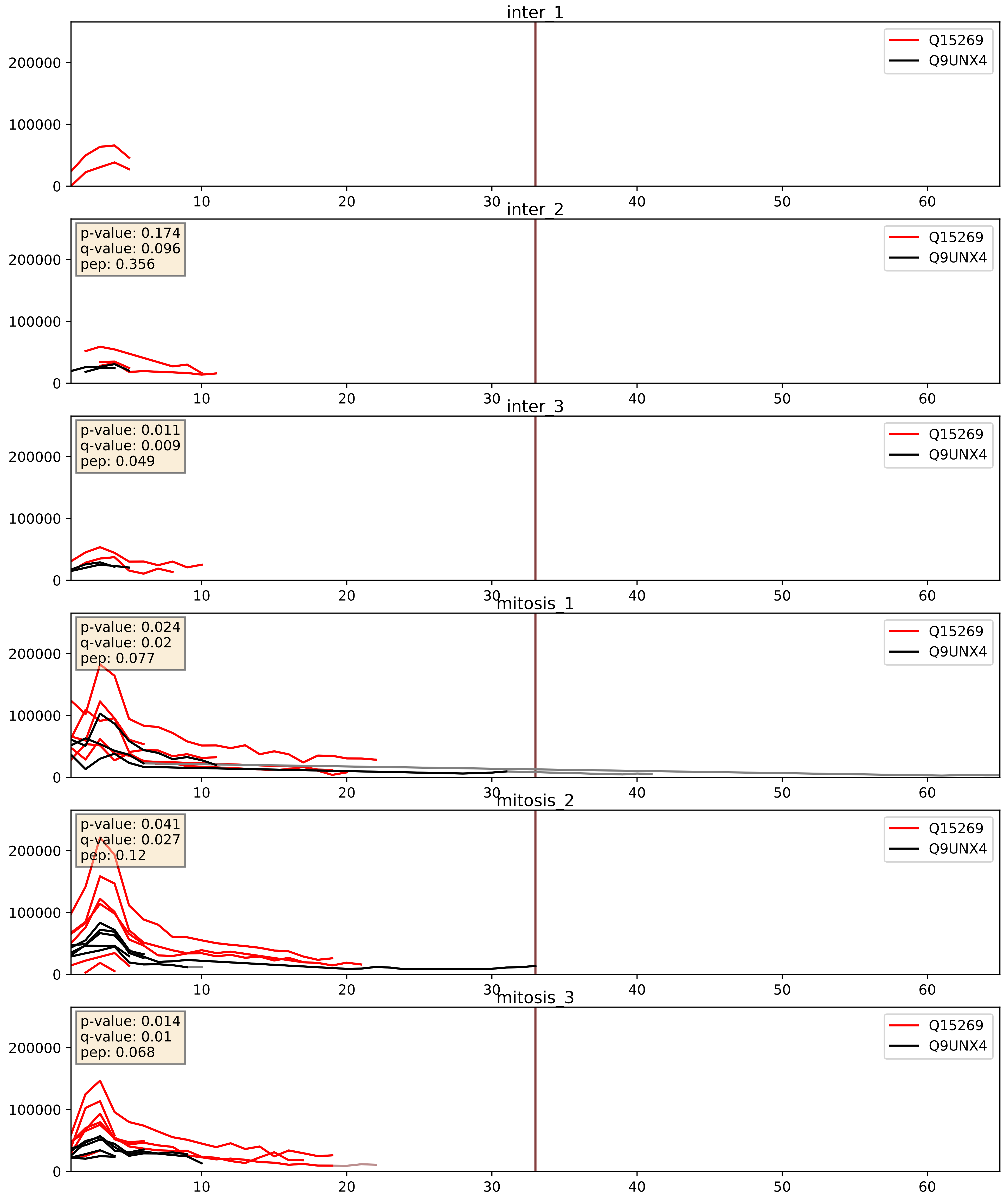


| level | condition_1 | condition_2 | pvalue | pvalue_adjusted |
|---|---|---|---|---|
| interactor_ratio | mitosis | inter | 0.027928400658634143 | 0.26184787474031573 |
| interactor_abundance | mitosis | inter | 0.041099264451958 | 0.16296731794217187 |
| complex_abundance | mitosis | inter | 0.0673384386695605 | 0.18739919037777122 |

Q15269_Q9Y2X3
PWP2_HUMAN vs NOP58_HUMAN
p-value: 0.017 q-value: 0.014

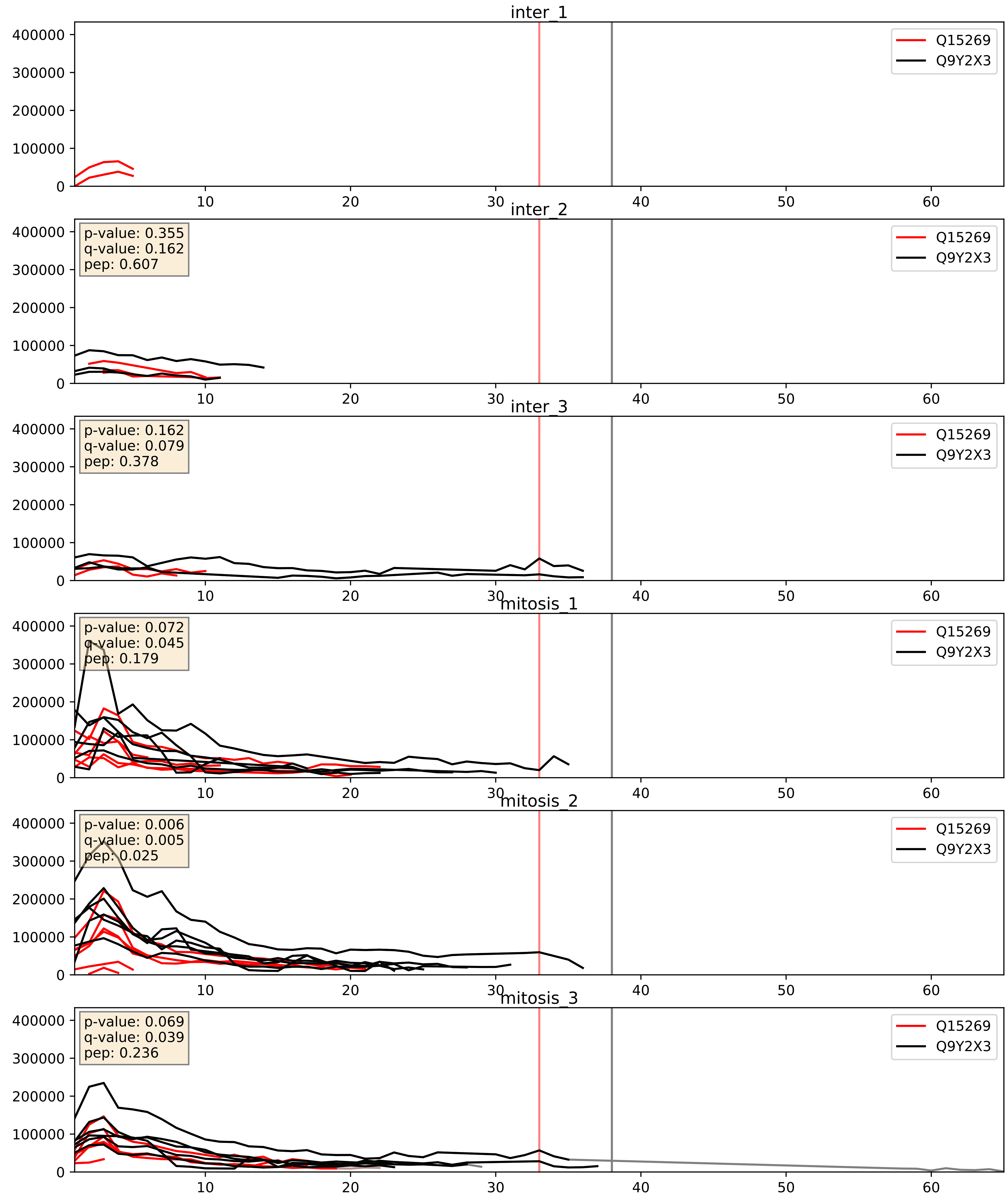


| level | condition_1 | condition_2 | pvalue | pvalue_adjusted |
|---|---|---|---|---|
| interactor_ratio | mitosis | inter | 0.0006251497771619007 | 0.029219626944512057 |
| complex_abundance | mitosis | inter | 0.0053638205370739305 | 0.055047043516043566 |
| interactor_abundance | mitosis | inter | 0.024889437139602743 | 0.13047955794942243 |
