## Supplementary Data 2 for "SECAT: Quantifying differential protein-protein interaction states by network-centric analysis": hela_string_000027_P07910.pdf

P07910_P07910

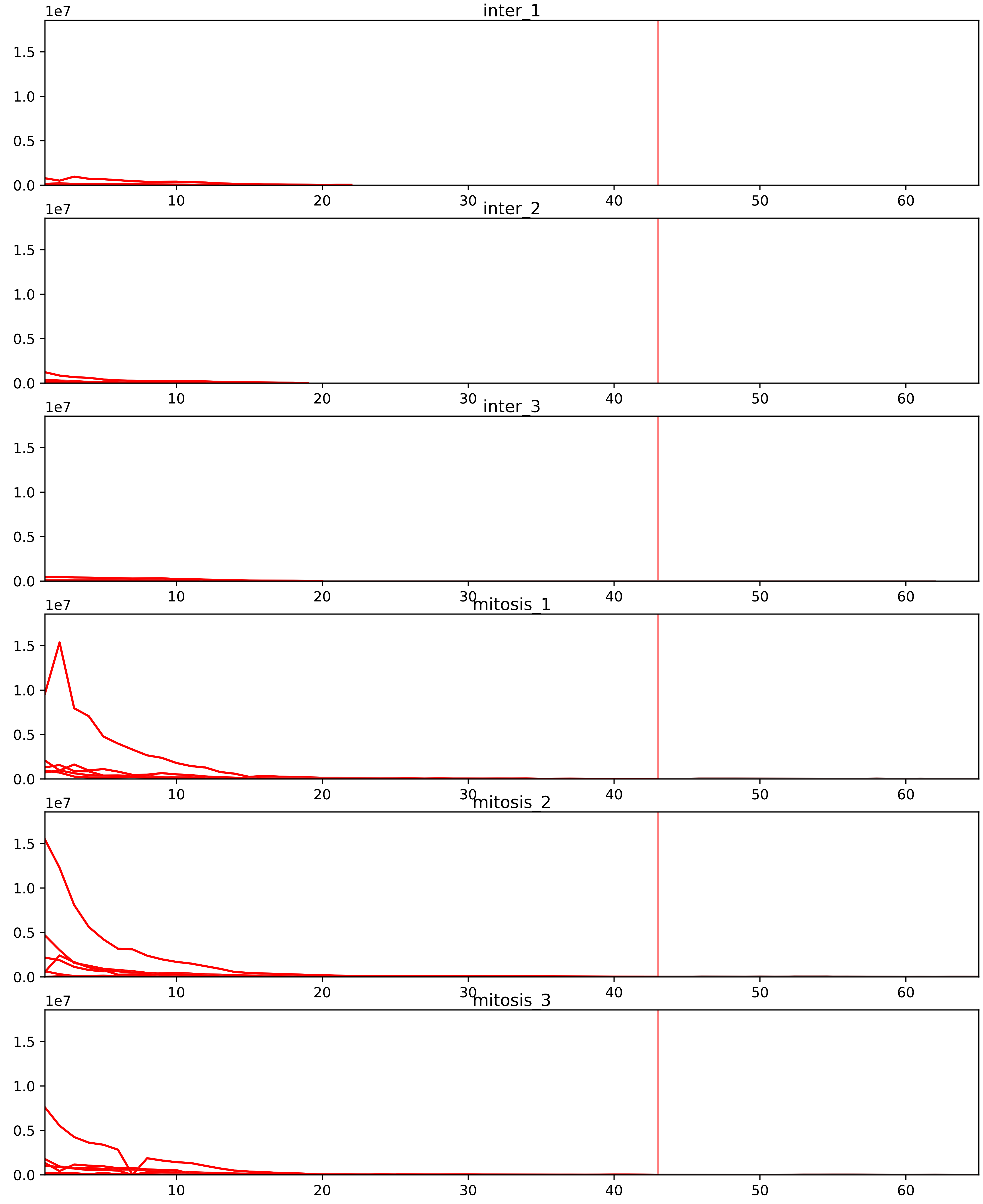


| level | condition_1 | condition_2 | pvalue | pvalue_adjusted |
|---|---|---|---|---|
| complex_abundance | mitosis | inter | 7.348485306509792e-11 | 1.3521212963978016e-07 |
| interactor_abundance | mitosis | inter | 3.6084711028307863e-10 | 3.319793414604323e-07 |
| interactor_ratio | mitosis | inter | 2.4999003706557918e-08 | 1.1794401748735018e-06 |
| monomer_abundance | mitosis | inter | 0.003979121482814966 | 0.1321415642137402 |
| total_abundance | mitosis | inter | 0.021495677763582754 | 0.18835452556753943 |
| assembled_abundance | mitosis | inter | 0.025022640455948313 | 0.19044654828687715 |

P05387_P07910
RLA2_HUMAN vs HNRPC_HUMAN
p-value: 0.0 q-value: 0.047

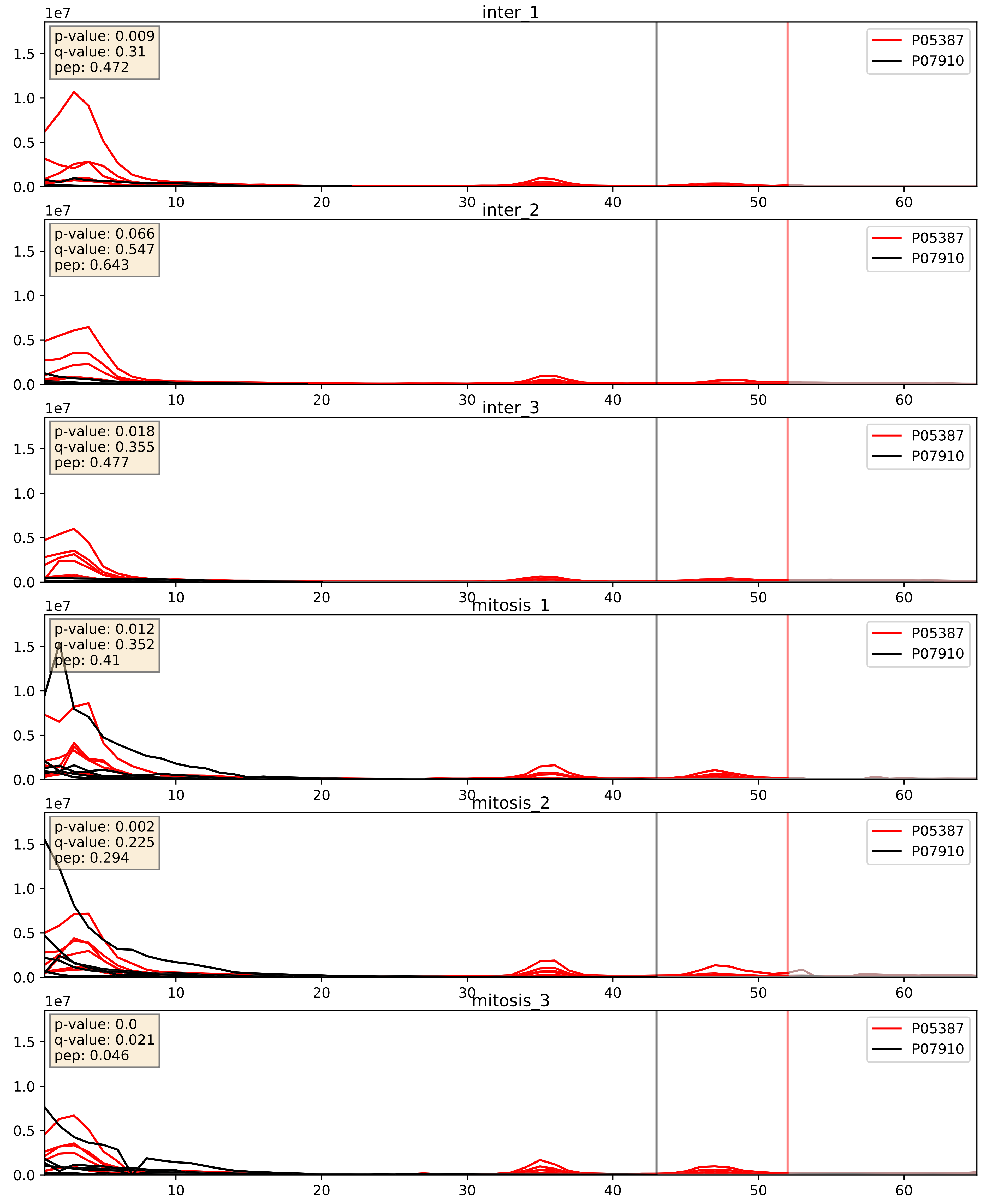


| level | condition_1 | condition_2 | pvalue | pvalue_adjusted |
|---|---|---|---|---|
| complex_abundance | mitosis | inter | 2.5859446416347255e-05 | 0.005691857151652958 |
| interactor_ratio | mitosis | inter | 0.0004311319774378024 | 0.02319827616364238 |
| interactor_abundance | mitosis | inter | 0.24758986860935167 | 0.44098403564212446 |

P07910_P08621
HNRPC_HUMAN vs RU17_HUMAN
p-value: 0.031 q-value: 0.022

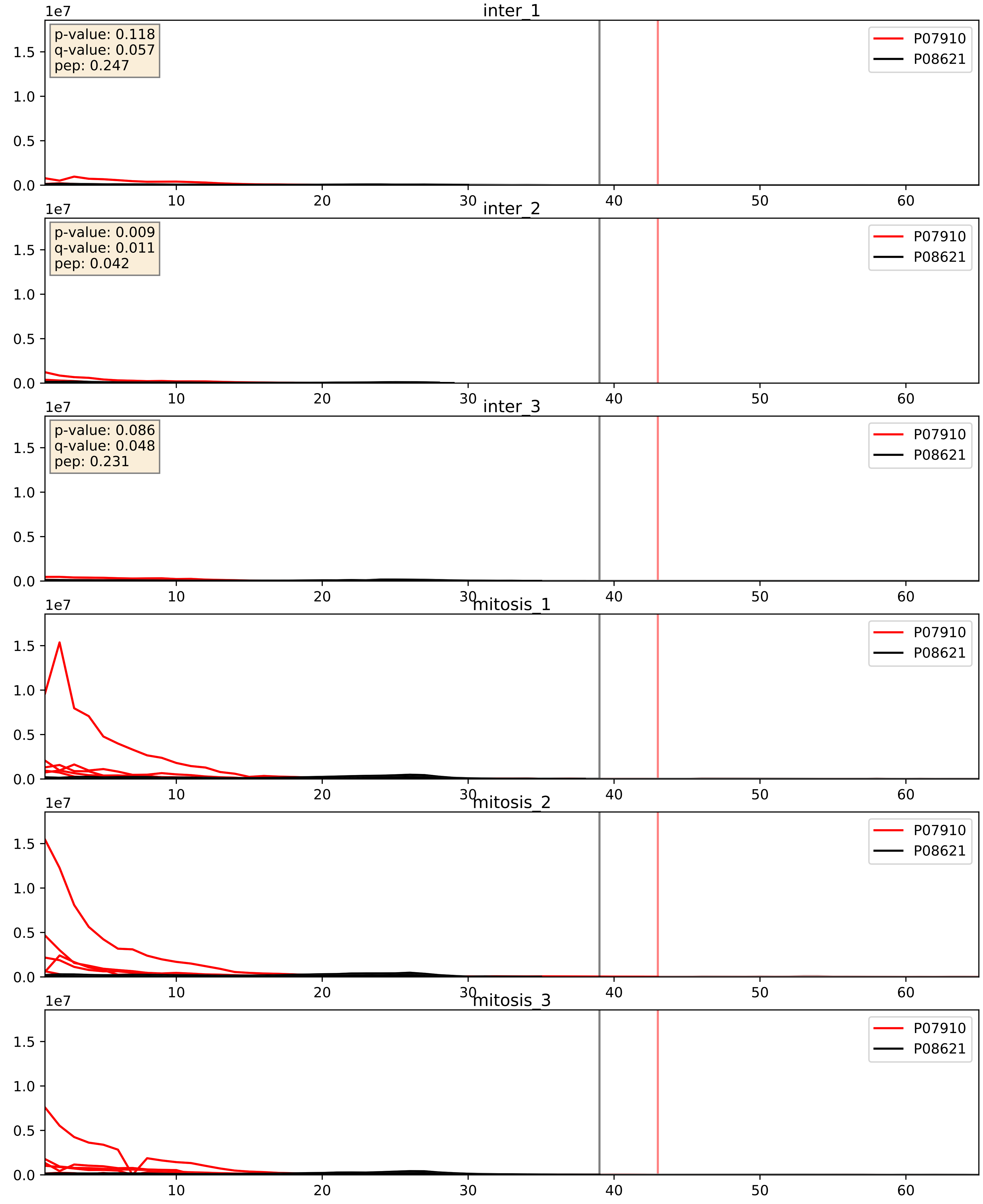


| level | condition_1 | condition_2 | pvalue | pvalue_adjusted |
|---|---|---|---|---|
| interactor_abundance | mitosis | inter | 9.496688410058796e-06 | 0.0033734327106189023 |
| complex_abundance | mitosis | inter | 1.5065308283938132e-05 | 0.005691857151652958 |
| interactor_ratio | mitosis | inter | 0.0001285712217163946 | 0.01222855175435931 |

P07910_P09651
HNRPC_HUMAN vs ROA1_HUMAN
p-value: 0.004 q-value: 0.004

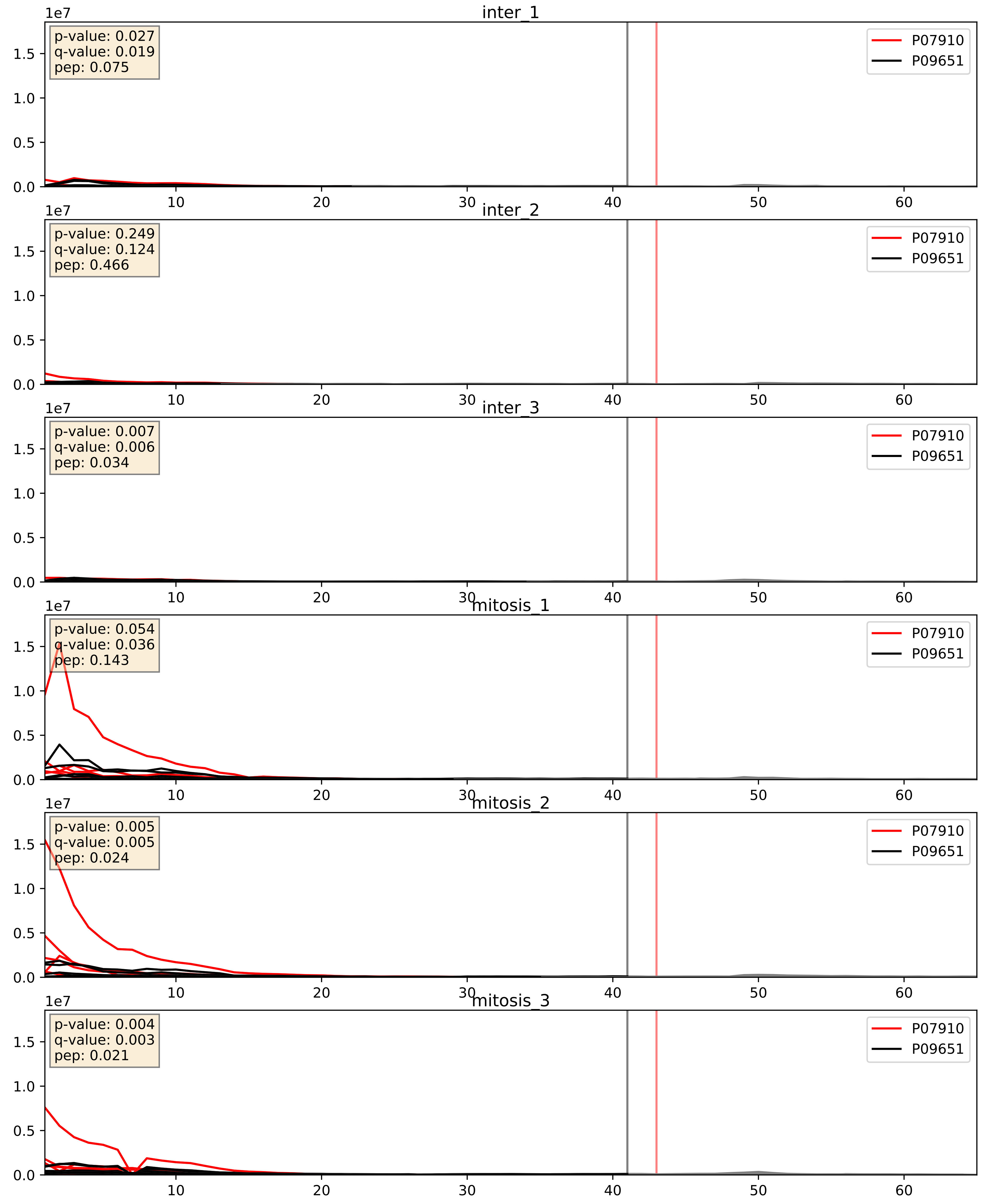


| level | condition_1 | condition_2 | pvalue | pvalue_adjusted |
|---|---|---|---|---|
| interactor_abundance | mitosis | inter | 9.357211976477008e-06 | 0.0033734327106189023 |
| complex_abundance | mitosis | inter | 7.4359018113791 24e-05 | 0.005893640694944935 |
| interactor_ratio | mitosis | inter | 0.02654415193625456 | 0.25614196664208316 |

P07910_P14866
HNRPC_HUMAN vs HNRPL_HUMAN
p-value: 0.011 q-value: 0.01

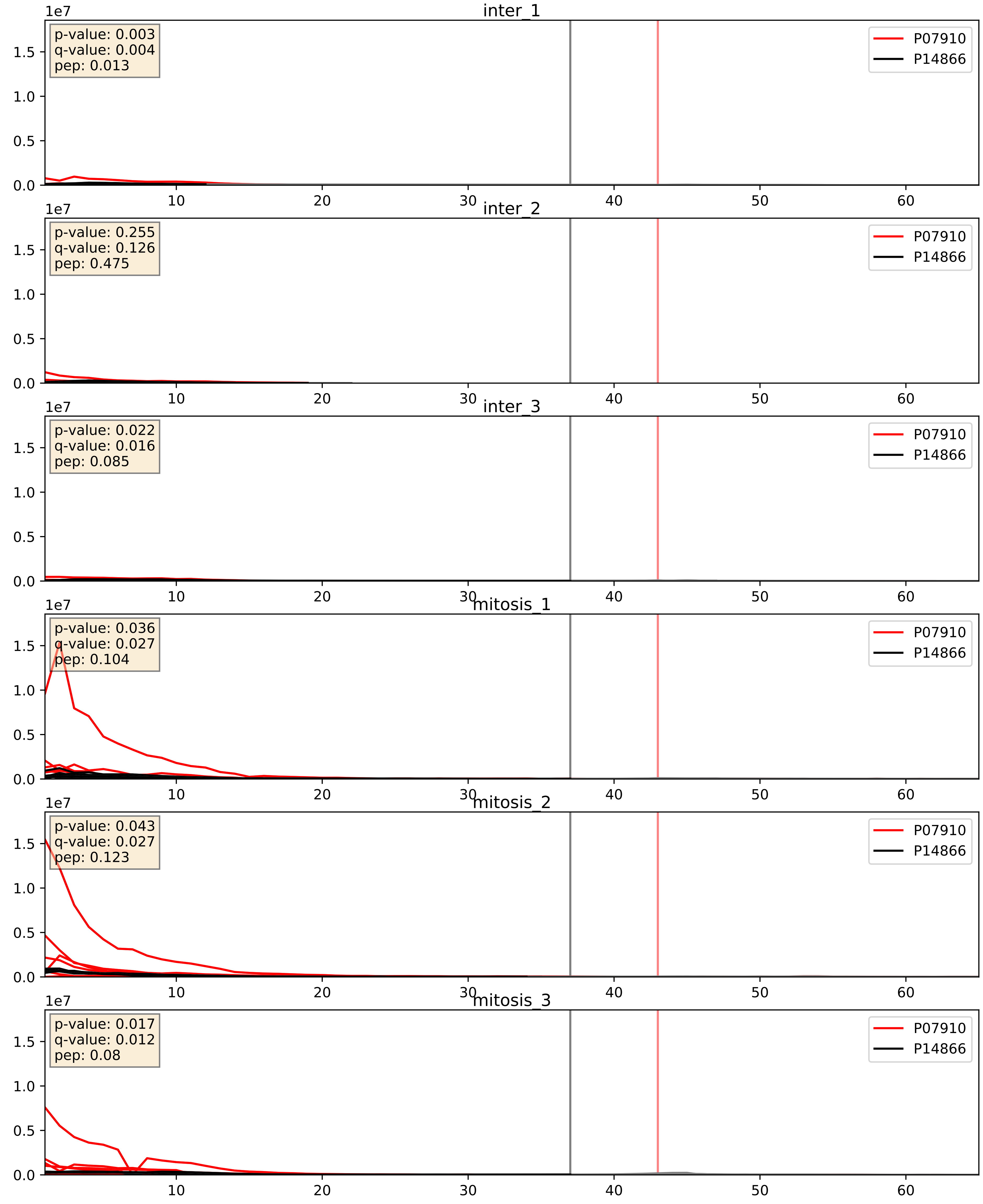


| level | condition_1 | condition_2 | pvalue | pvalue_adjusted |
|---|---|---|---|---|
| interactor_abundance | mitosis | inter | 8.645329666757008e-06 | 0.0033734327106189023 |
| complex_abundance | mitosis | inter | 3.986857495835857e-05 | 0.005691857151652958 |
| interactor_ratio | mitosis | inter | 0.0022228260684612118 | 0.05890833172144883 |

P07910_P17844
HNRPC_HUMAN vs DDX5_HUMAN
p-value: 0.002 q-value: 0.034

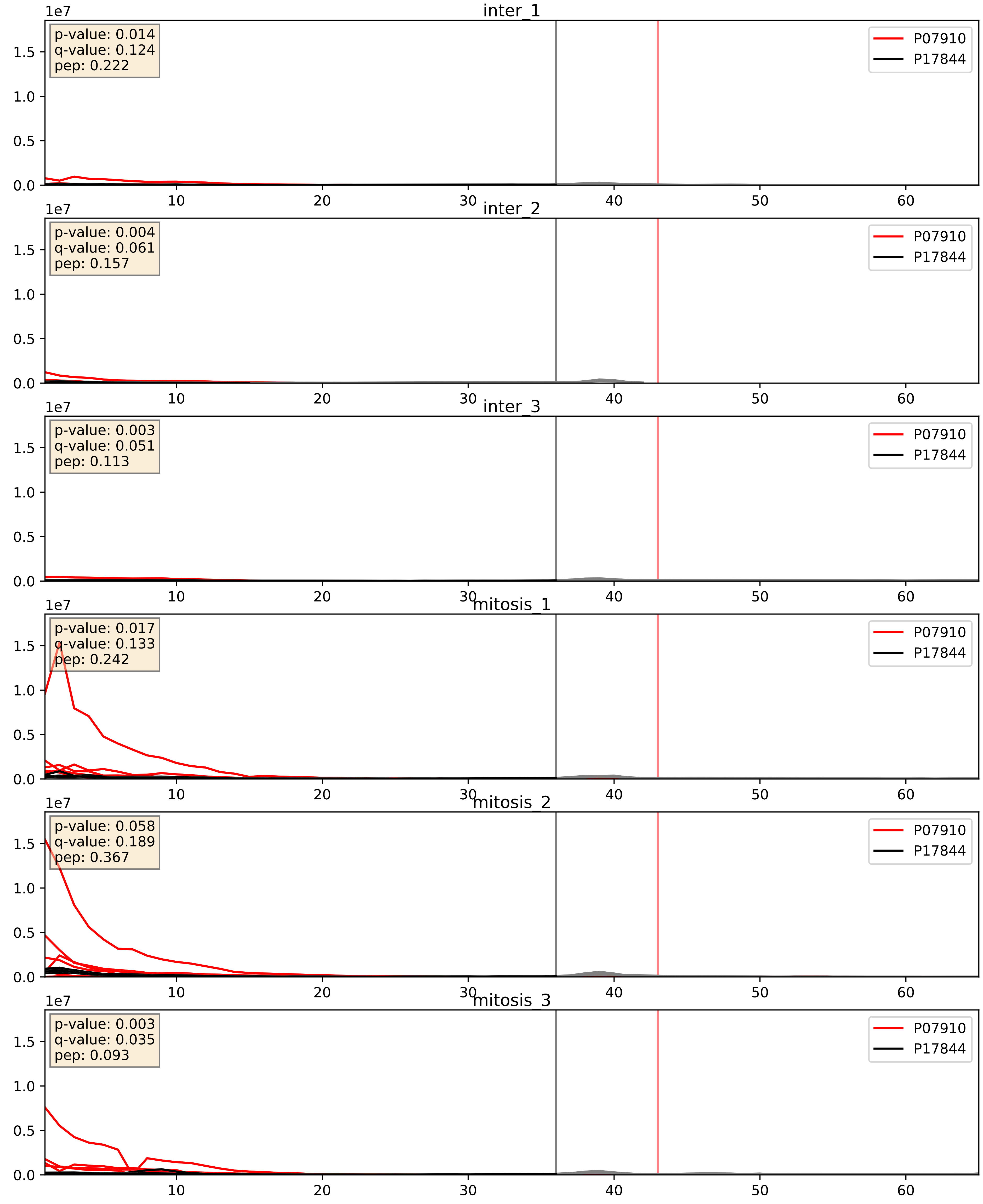


| level | condition_1 | condition_2 | pvalue | pvalue_adjusted |
|---|---|---|---|---|
| complex_abundance | mitosis | inter | 8.901605986839528e-06 | 0.005691857151652958 |
| interactor_abundance | mitosis | inter | 1.005842017873748e-05 | 0.0033734327106189023 |
| interactor_ratio | mitosis | inter | 0.00013562784083812567 | 0.012616771133069759 |

P07910_P22626
HNRPC_HUMAN vs ROA2_HUMAN
p-value: 0.0 q-value: 0.0

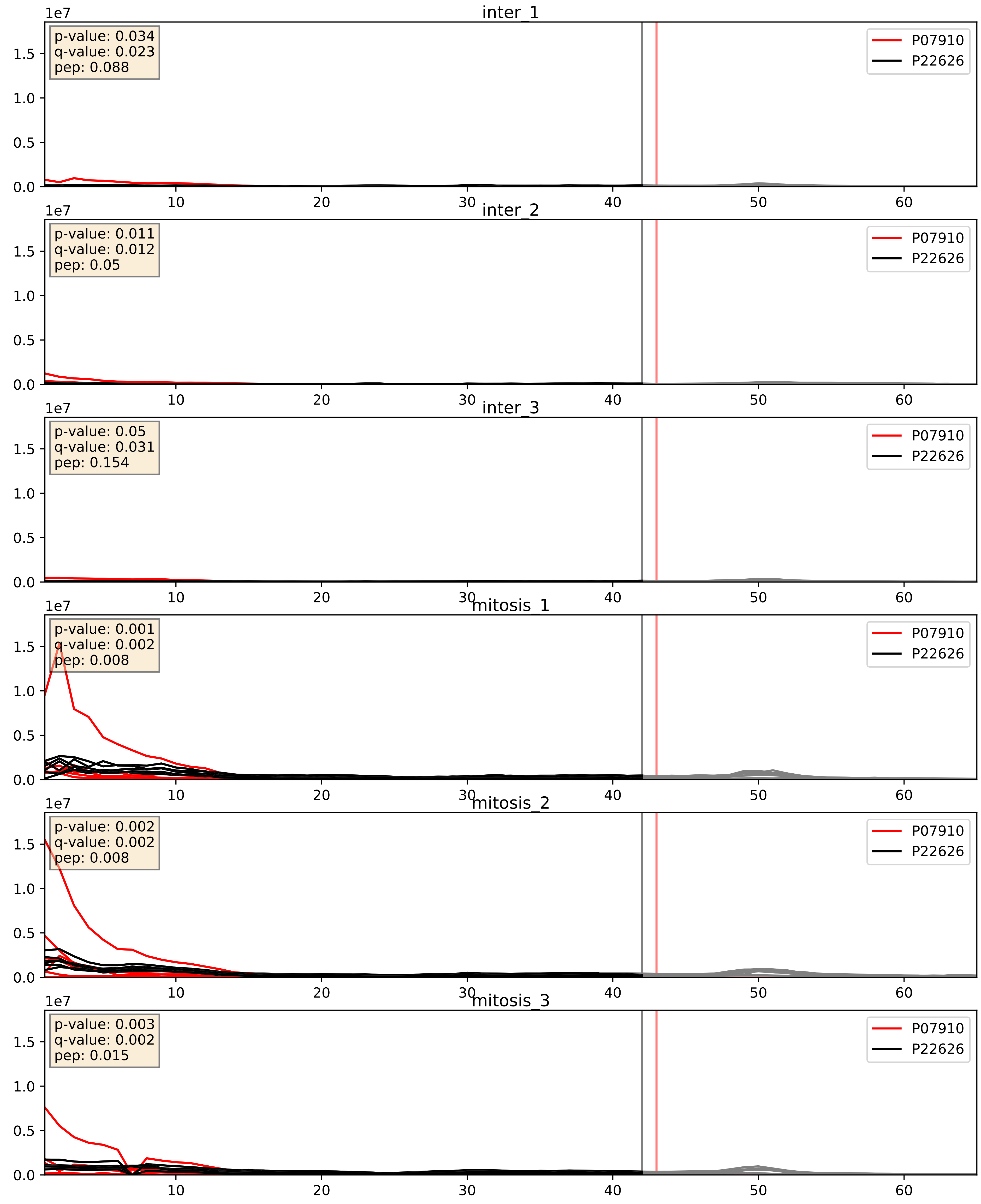


| level | condition_1 | condition_2 | pvalue | pvalue_adjusted |
|---|---|---|---|---|
| interactor_abundance | mitosis | inter | 9.148564965046394e-06 | 0.0033734327106189023 |
| complex_abundance | mitosis | inter | 0.00014044884889 71927 | 0.00812325774702682 |
| interactor_ratio | mitosis | inter | 0.417882469042839 | 0.8387043223931305 |

P07910_P26599
HNRPC_HUMAN vs PTBP1_HUMAN
p-value: 0.025 q-value: 0.019

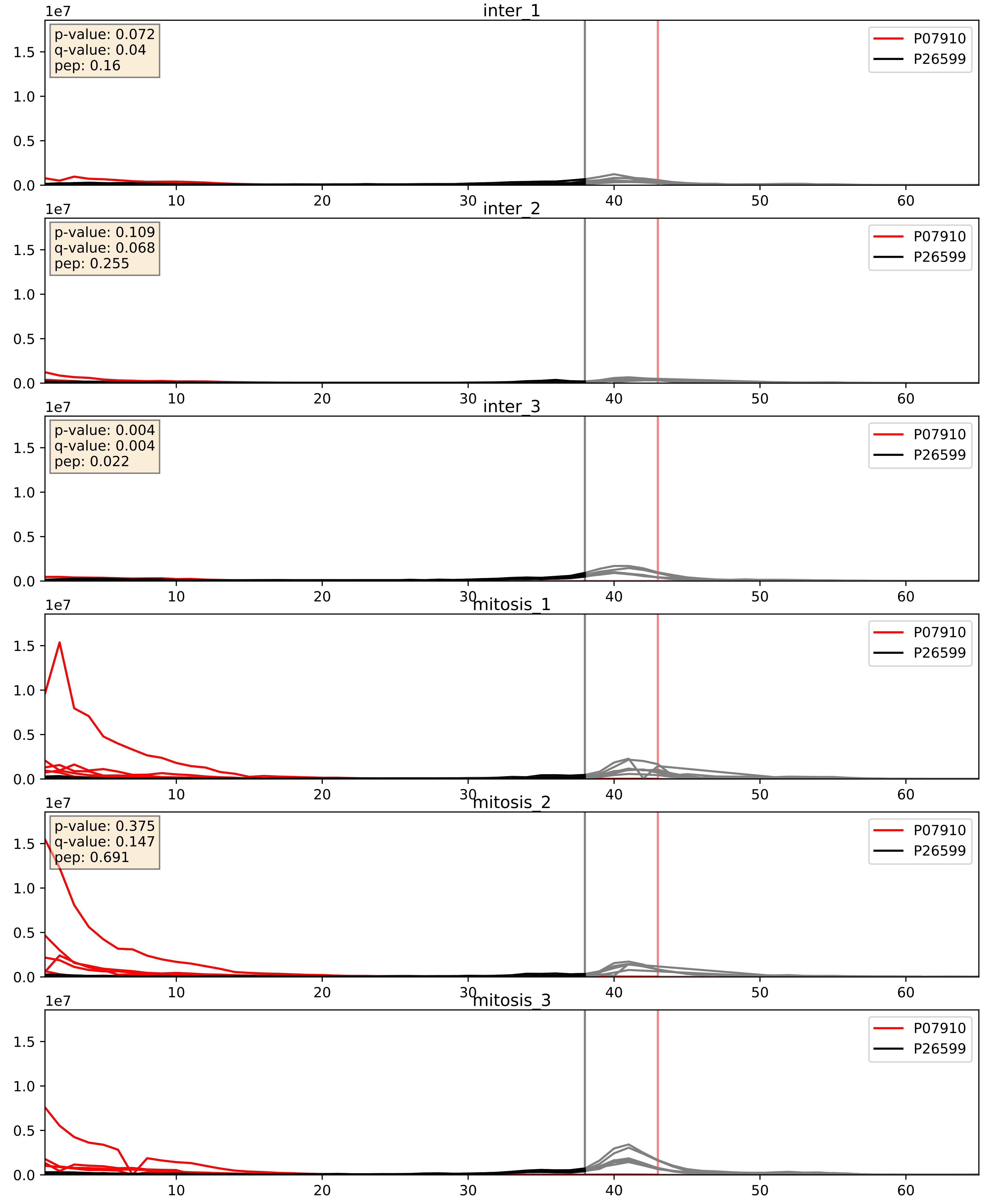


| level | condition_1 | condition_2 | pvalue | pvalue_adjusted |
|---|---|---|---|---|
| interactor_abundance | mitosis | inter | 8.777325786432812e-06 | 0.0033734327106189023 |
| complex_abundance | mitosis | inter | 0.0030170274690254867 | 0.04033974863104599 |
| interactor_ratio | mitosis | inter | 0.009939805764469163 | 0.14263182888175546 |

P07910_P31943
HNRPC_HUMAN vs HNRH1_HUMAN
p-value: 0.012 q-value: 0.01

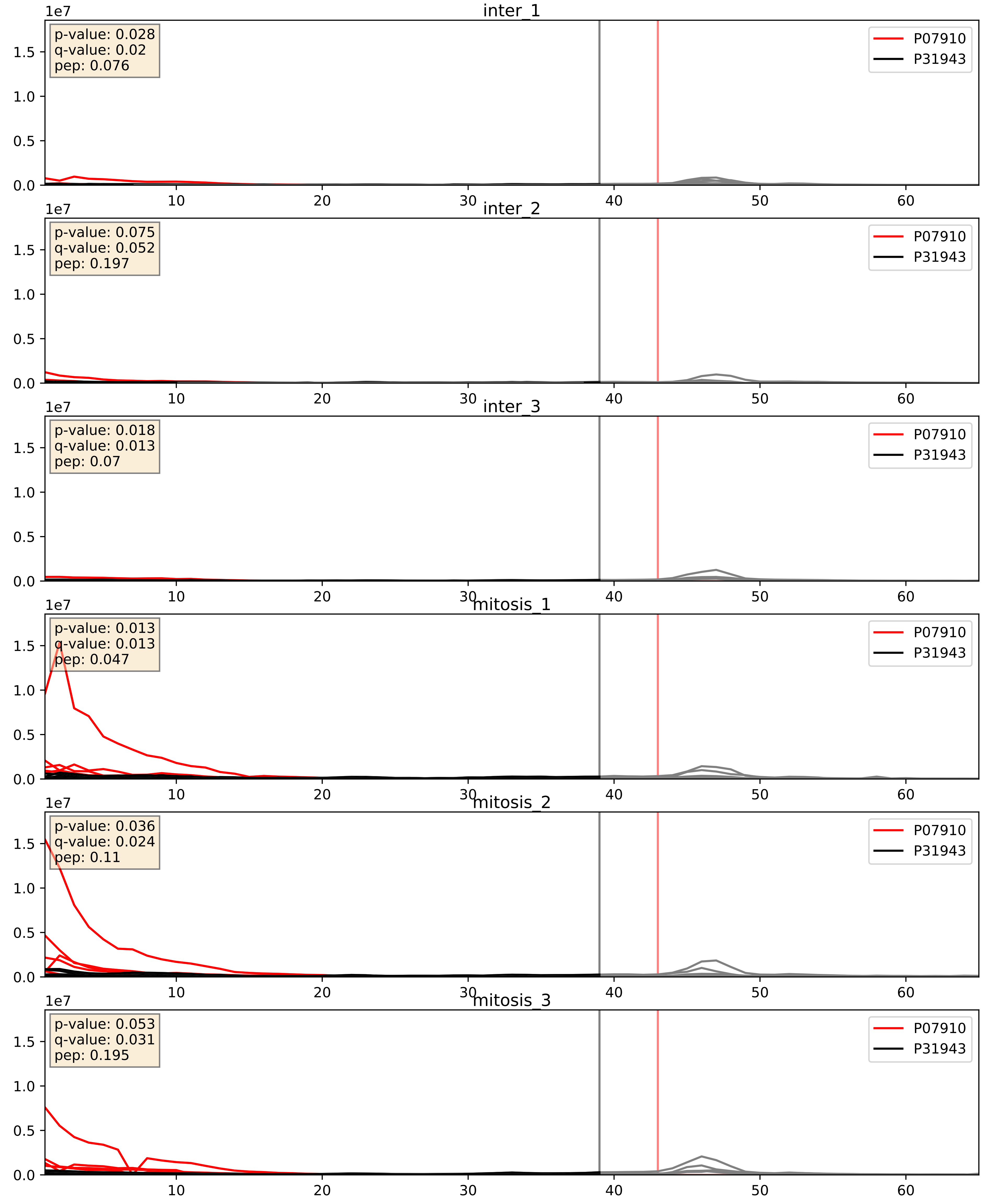


| level | condition_1 | condition_2 | pvalue | pvalue_adjusted |
|---|---|---|---|---|
| complex_abundance | mitosis | inter | 4.189010509383275e-06 | 0.005122561422902976 |
| interactor_abundance | mitosis | inter | 9.711637813774278e-06 | 0.0033734327106189023 |
| interactor_ratio | mitosis | inter | 0.0025928684235758614 | 0.0652792756053217 |

P07910_P38159
HNRPC_HUMAN vs RBMX_HUMAN
p-value: 0.022 q-value: 0.017

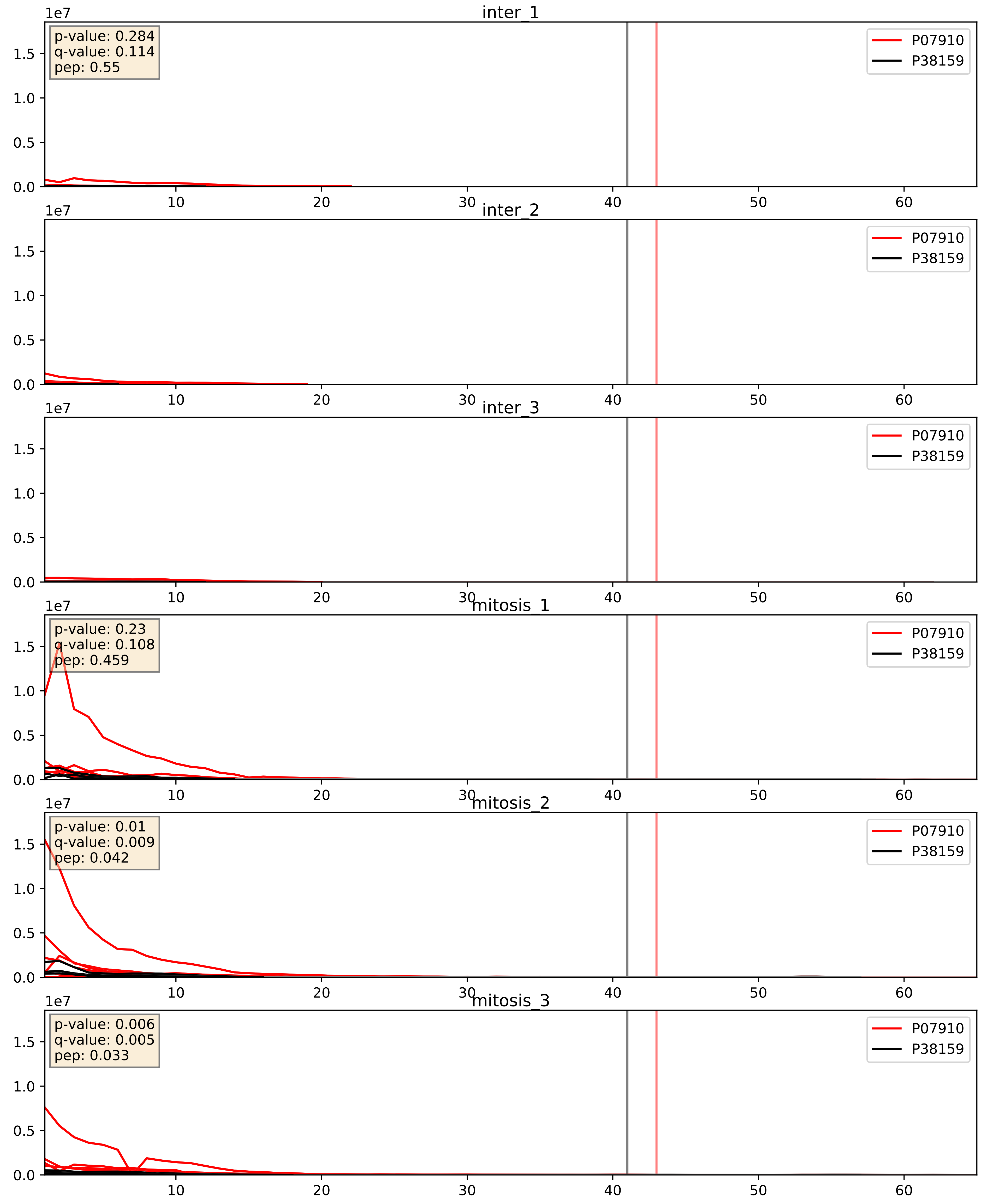


| level | condition_1 | condition_2 | pvalue | pvalue_adjusted |
|---|---|---|---|---|
| interactor_abundance | mitosis | inter | 2.354035126084445e-06 | 0.0033734327106189023 |
| complex_abundance | mitosis | inter | 4.105555000846496e-05 | 0.005691857151652958 |
| interactor_ratio | mitosis | inter | 0.018442867517554088 | 0.20476127879411543 |

P07910_P51991
HNRPC_HUMAN vs ROA3_HUMAN
p-value: 0.002 q-value: 0.002

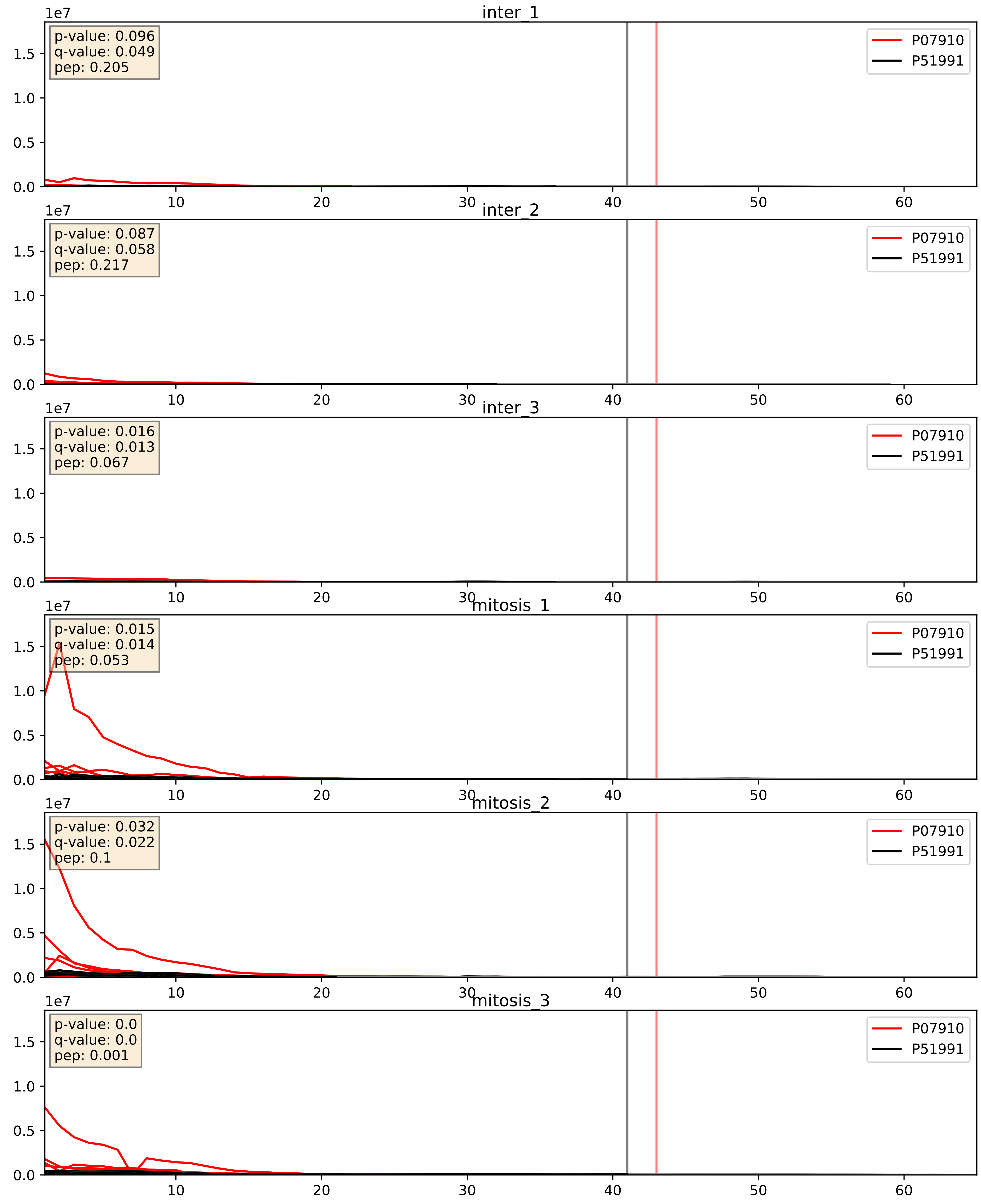


| level | condition_1 | condition_2 | pvalue | pvalue_adjusted |
|---|---|---|---|---|
| complex_abundance | mitosis | inter | 7.94637328064476e-07 | 0.0024732922429087437 |
| interactor_abundance | mitosis | inter | 9.1373717647951e-06 | 0.0033734327106189023 |
| interactor_ratio | mitosis | inter | 0.0027887838155807804 | 0.06820568417534709 |

P07910_P52272
HNRPC_HUMAN vs HNRPM_HUMAN
p-value: 0.001 q-value: 0.002

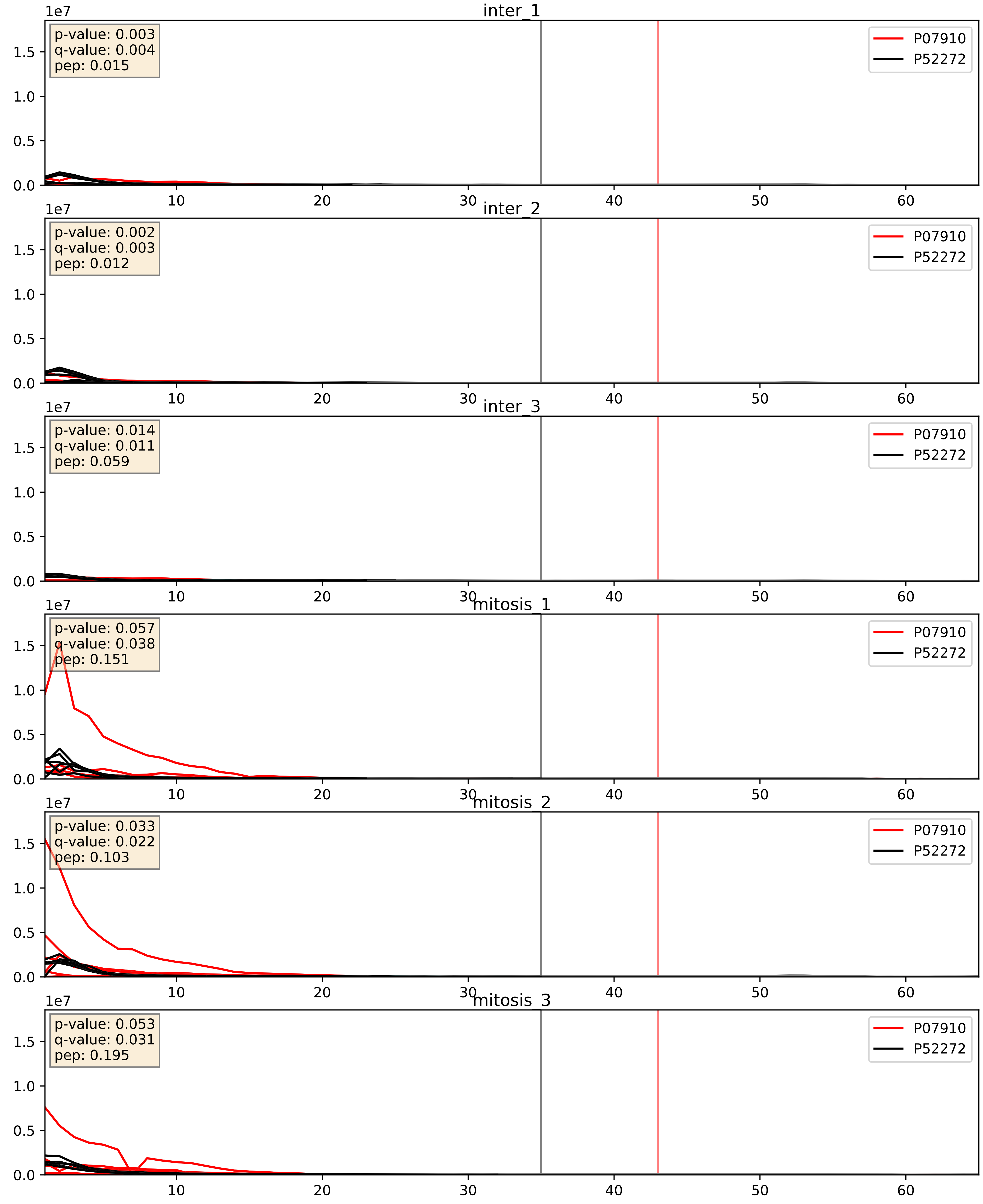


| level | condition_1 | condition_2 | pvalue | pvalue_adjusted |
|---|---|---|---|---|
| interactor_abundance | mitosis | inter | 9.77406216392995e-06 | 0.0033734327106189023 |
| interactor_ratio | mitosis | inter | 0.00022697512149341483 | 0.01529848062979237 |
| complex_abundance | mitosis | inter | 0.000330922405409354 | 0.011886635542316795 |

P07910_P52597
HNRPC_HUMAN vs HNRPF_HUMAN
p-value: 0.007 q-value: 0.007

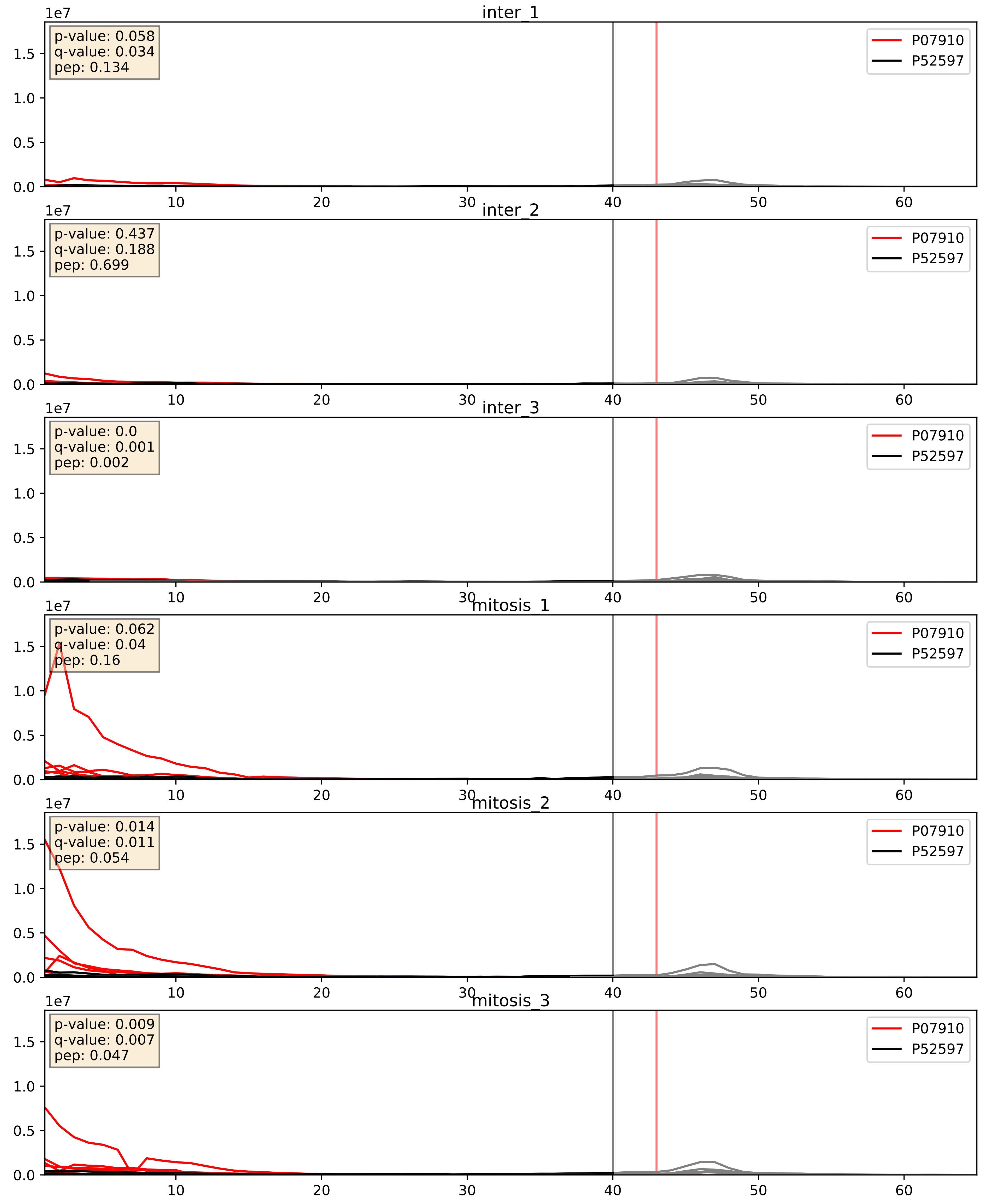


| level | condition_1 | condition_2 | pvalue | pvalue_adjusted |
|---|---|---|---|---|
| complex_abundance | mitosis | inter | 2.867619584776087e-06 | 0.004091137274280551 |
| interactor_abundance | mitosis | inter | 9.49140558984193e-06 | 0.0033734327106189023 |
| interactor_ratio | mitosis | inter | 0.0007484433338408752 | 0.03166103300210851 |

P07910_P62750
HNRPC_HUMAN vs RL23A_HUMAN
p-value: 0.0 q-value: 0.018

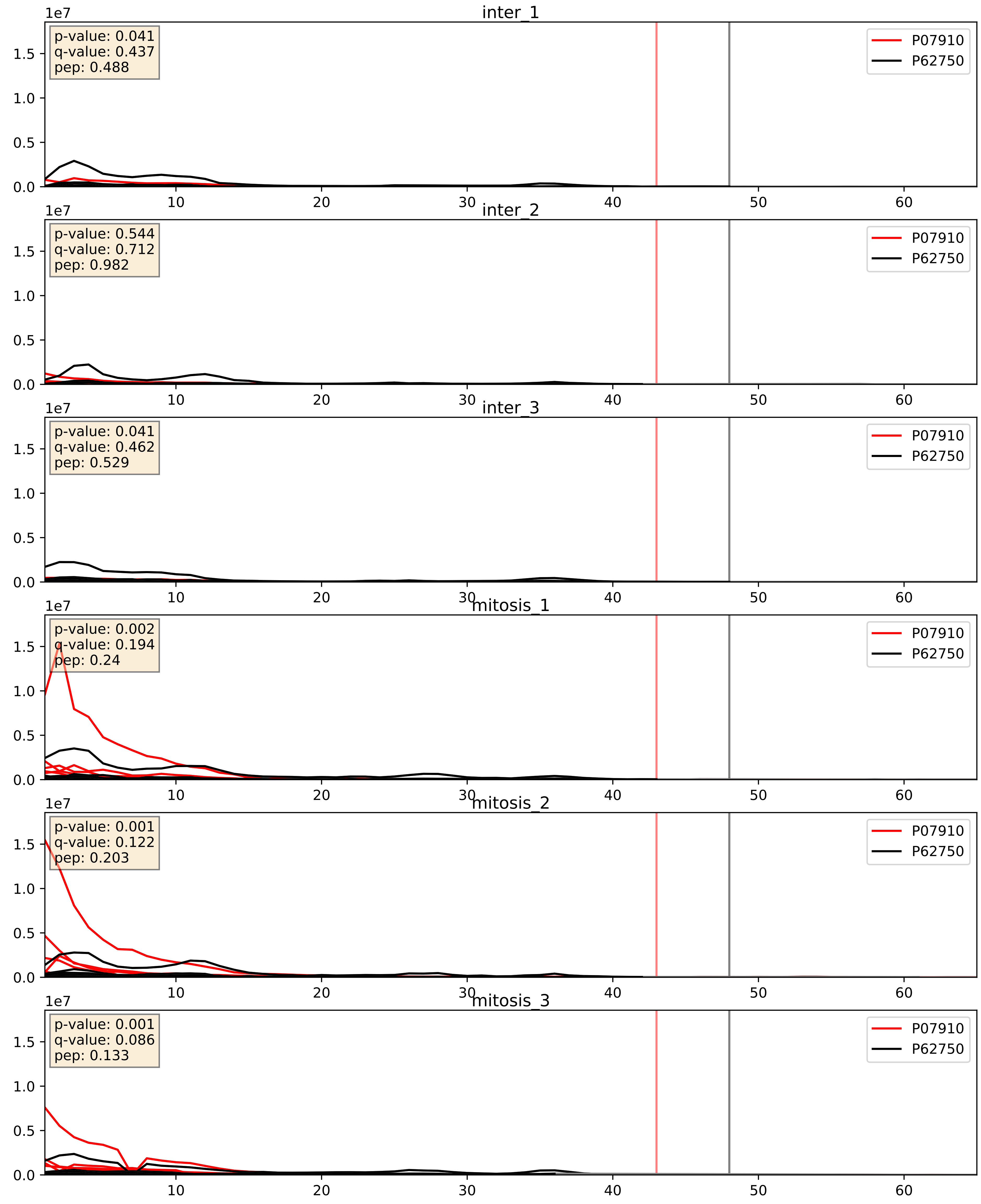


| level | condition_1 | condition_2 | pvalue | pvalue_adjusted |
|---|---|---|---|---|
| complex_abundance | mitosis | inter | 2.7095771167571986e-06 | 0.004091137274280551 |
| interactor_abundance | mitosis | inter | 9.512264939762444e-06 | 0.0033734327106189023 |
| interactor_ratio | mitosis | inter | 0.0006225976094504257 | 0.029219626944512057 |

P07910_P84103
HNRPC_HUMAN vs SRSF3_HUMAN
p-value: 0.043 q-value: 0.029

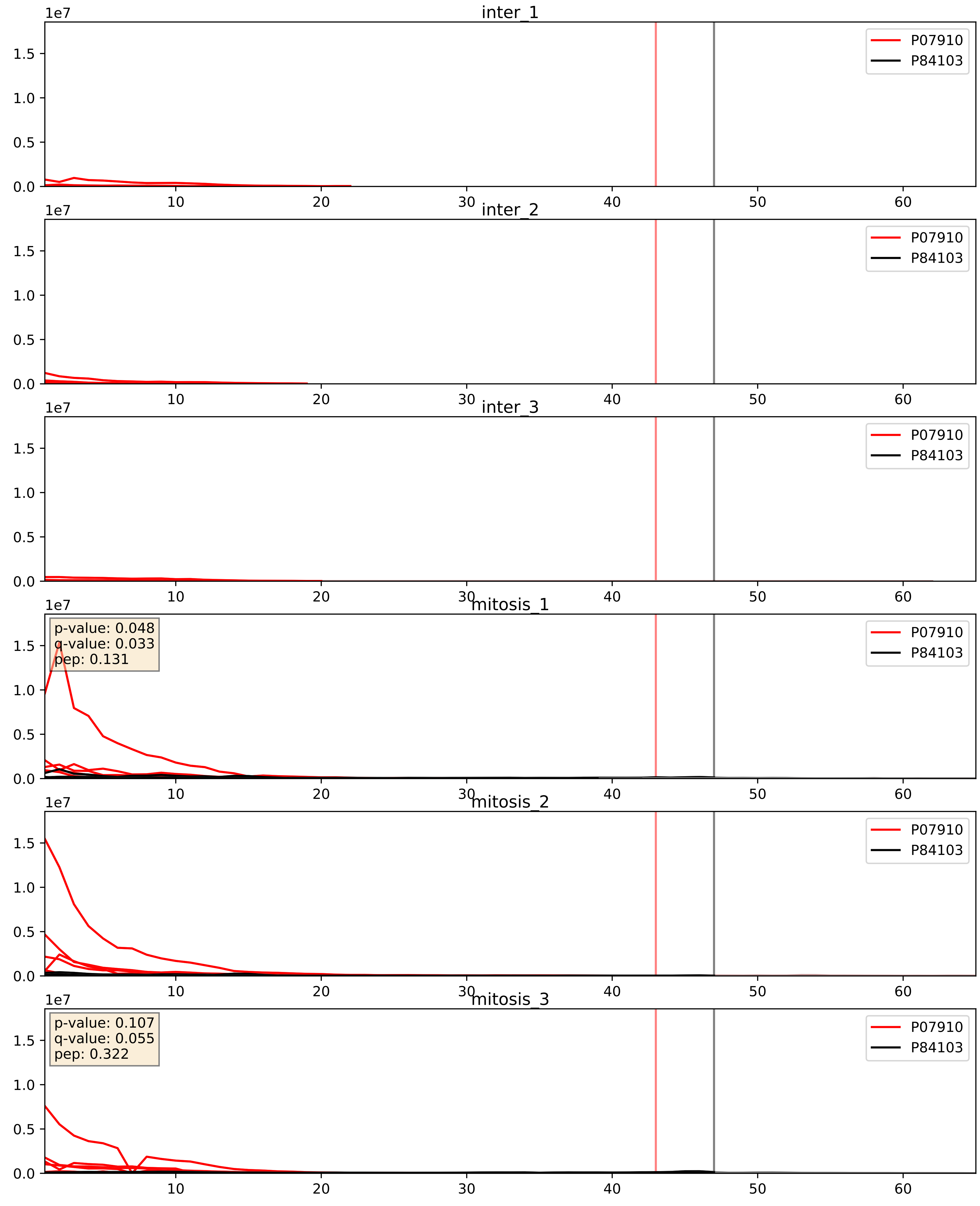


| level | condition_1 | condition_2 | pvalue | pvalue_adjusted |
|---|---|---|---|---|
| interactor_abundance | mitosis | inter | 2.2213691150803744e-06 | 0.0033734327106189023 |
| complex_abundance | mitosis | inter | 4.449045517390577e-05 | 0.005691857151652958 |
| interactor_ratio | mitosis | inter | 0.0013366807319858036 | 0.04334578357589157 |

P07910_Q07955
HNRPC_HUMAN vs SRSF1_HUMAN
p-value: 0.038 q-value: 0.026

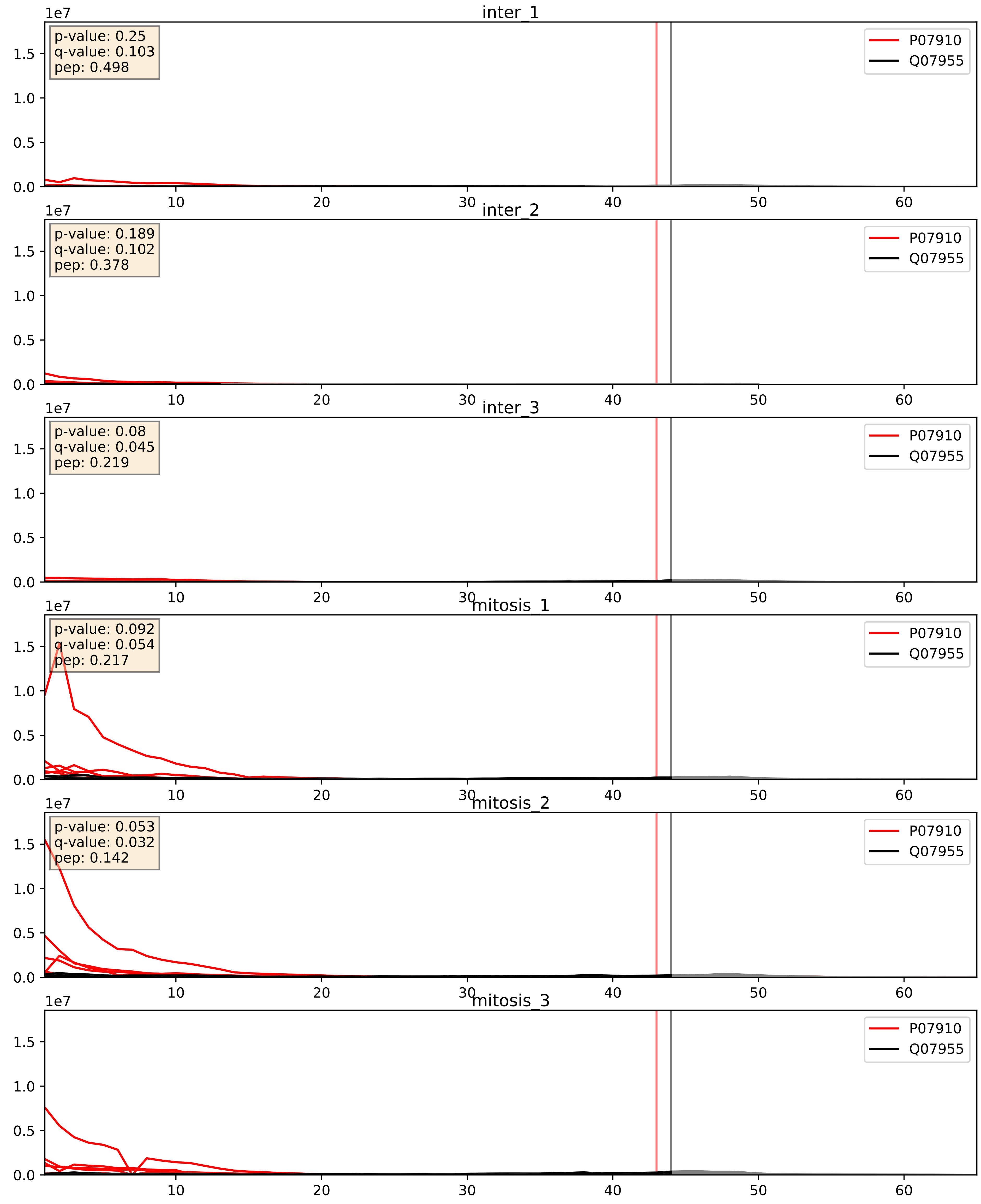


| level | condition_1 | condition_2 | pvalue | pvalue_adjusted |
|---|---|---|---|---|
| interactor_abundance | mitosis | inter | 4.923890450505053e-05 | 0.006283598333582311 |
| complex_abundance | mitosis | inter | 0.00034811228118943375 | 0.012015488415248196 |
| interactor_ratio | mitosis | inter | 0.01880601904879865 | 0.20643721891135677 |

P07910_Q08211
HNRPC_HUMAN vs DHX9_HUMAN
p-value: 0.015 q-value: 0.012

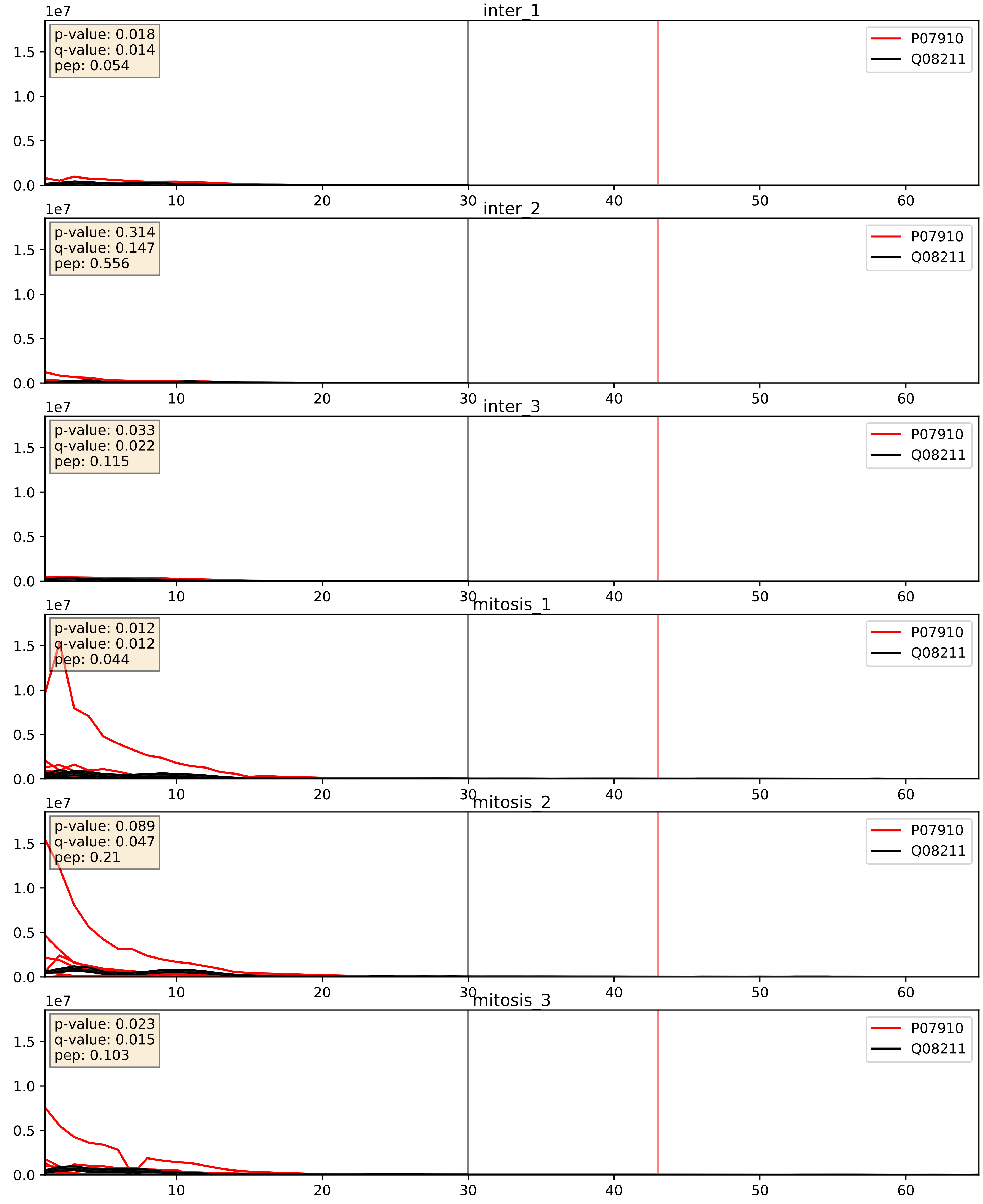


| level | condition_1 | condition_2 | pvalue | pvalue_adjusted |
|---|---|---|---|---|
| complex_abundance | mitosis | inter | 9.436874100373223e-06 | 0.005691857151652958 |
| interactor_abundance | mitosis | inter | 9.783708847801534e-06 | 0.0033734327106189023 |
| interactor_ratio | mitosis | inter | 8.825216175232504e-05 | 0.010208628440539222 |

P07910_Q12906
HNRPC_HUMAN vs ILF3_HUMAN
p-value: 0.002 q-value: 0.03

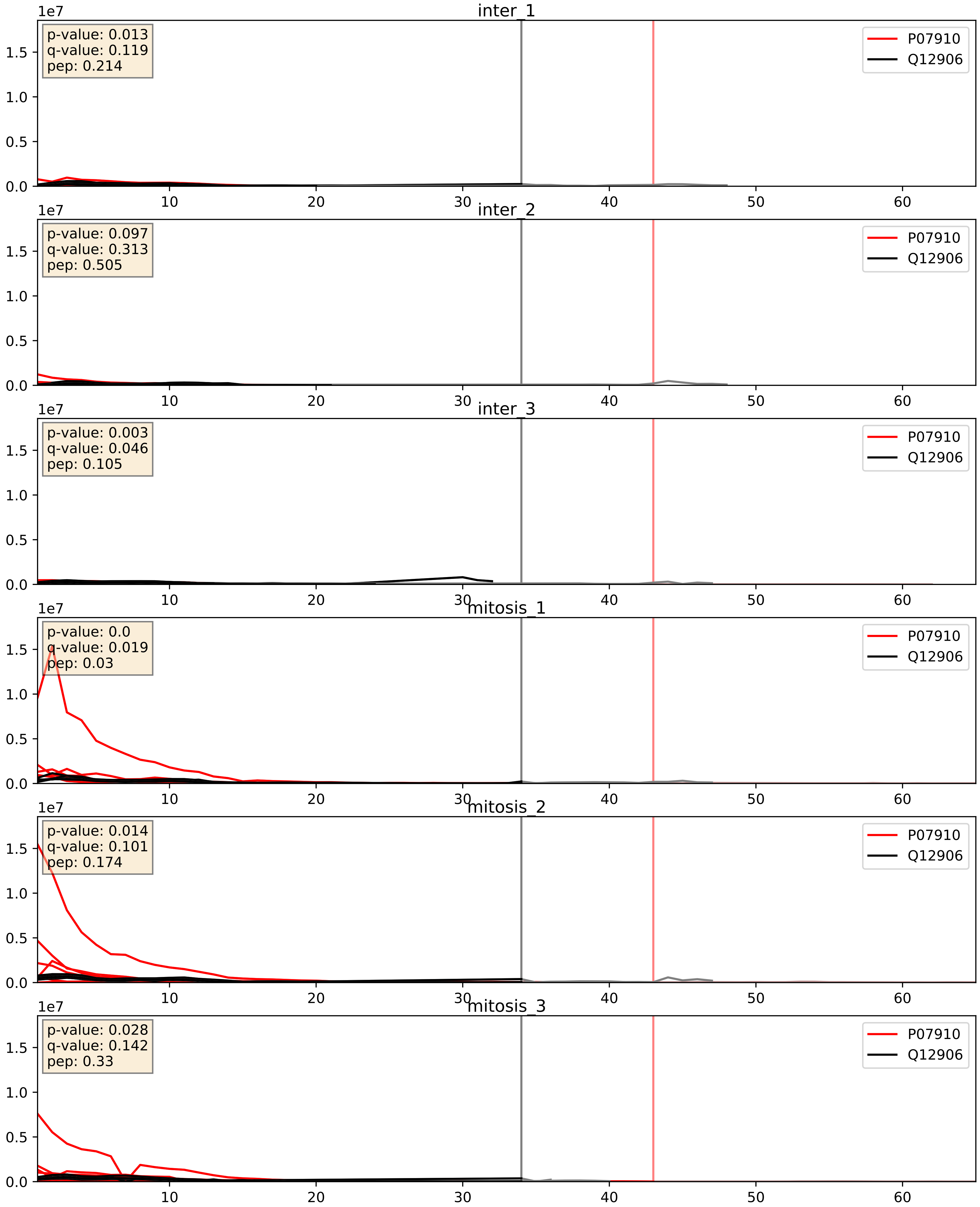


| level | condition_1 | condition_2 | pvalue | pvalue_adjusted |
|---|---|---|---|---|
| complex_abundance | mitosis | inter | 1.8856863202429171e-06 | 0.0040353687253198425 |
| interactor_abundance | mitosis | inter | 9.155180531930296e-06 | 0.0033734327106189023 |
| interactor_ratio | mitosis | inter | 0.00010644547717431898 | 0.010720128702277293 |

P07910_Q13151
HNRPC_HUMAN vs ROA0_HUMAN
p-value: 0.048 q-value: 0.032

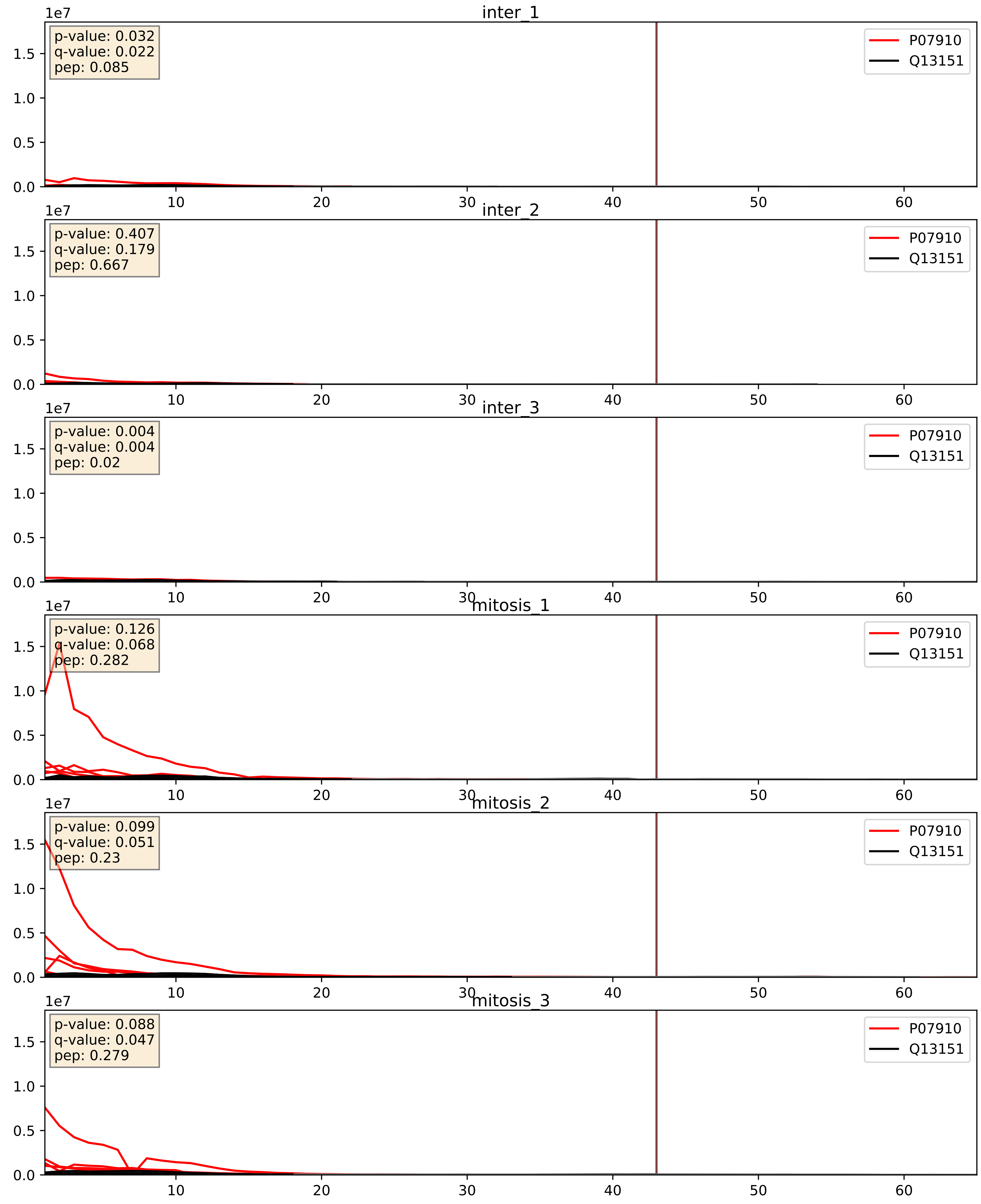


| level | condition_1 | condition_2 | pvalue | pvalue_adjusted |
|---|---|---|---|---|
| interactor_abundance | mitosis | inter | 6.849684494635132e-06 | 0.0033734327106189023 |
| complex_abundance | mitosis | inter | 0.00014564121967907843 | 0.008317961743055323 |
| interactor_ratio | mitosis | inter | 0.0007352419354070836 | 0.03146835483542318 |

P07910_Q13435
HNRPC_HUMAN vs SF3B2_HUMAN
p-value: 0.002 q-value: 0.031

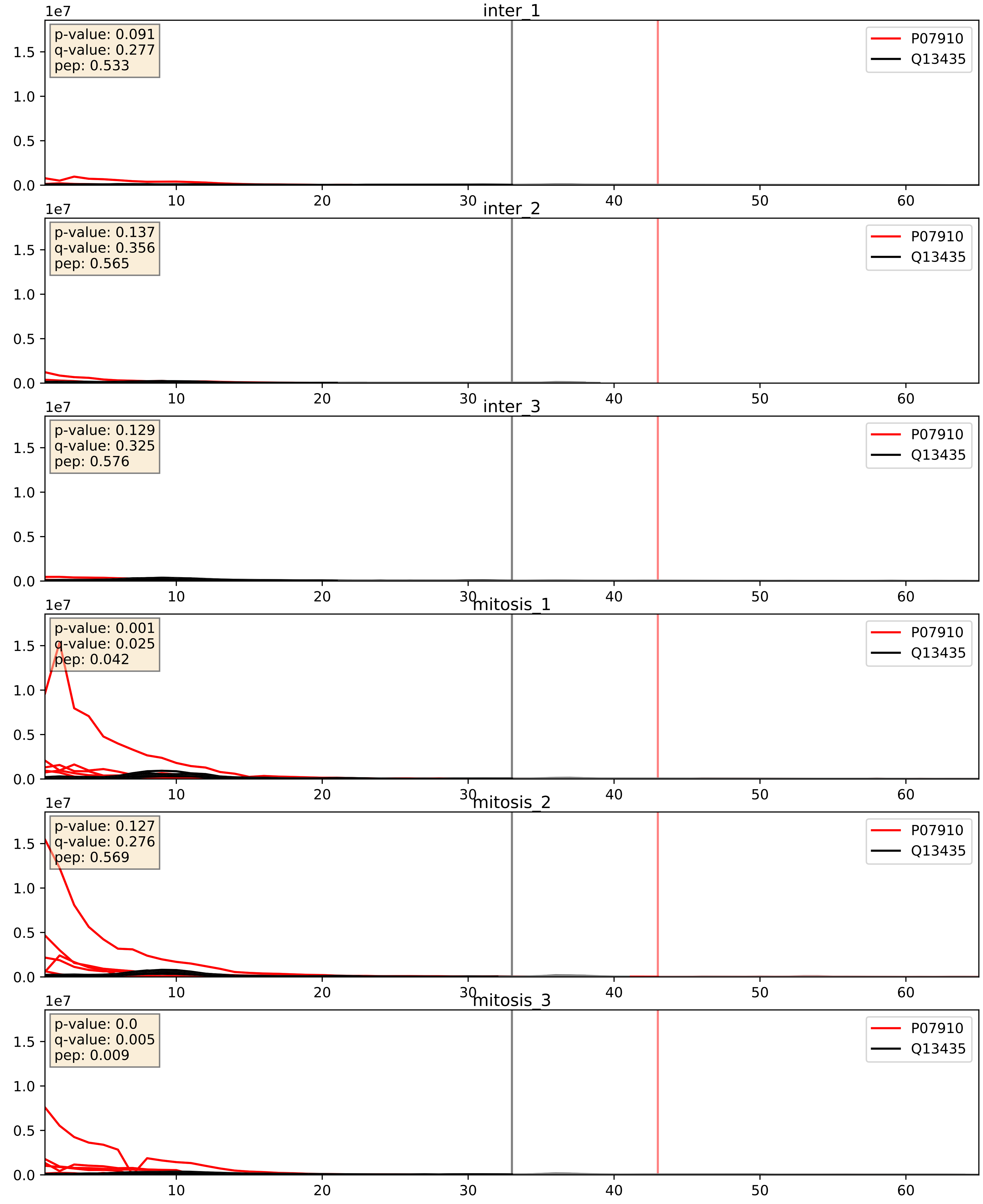


| level | condition_1 | condition_2 | pvalue | pvalue_adjusted |
|---|---|---|---|---|
| interactor_abundance | mitosis | inter | 9.628495089762232e-06 | 0.0033734327106189023 |
| complex_abundance | mitosis | inter | 3.2188906277444467e-05 | 0.005691857151652958 |
| interactor_ratio | mitosis | inter | 0.0016860726879665898 | 0.049768214513772446 |

P07910_Q15365
HNRPC_HUMAN vs PCBP1_HUMAN
p-value: 0.026 q-value: 0.019

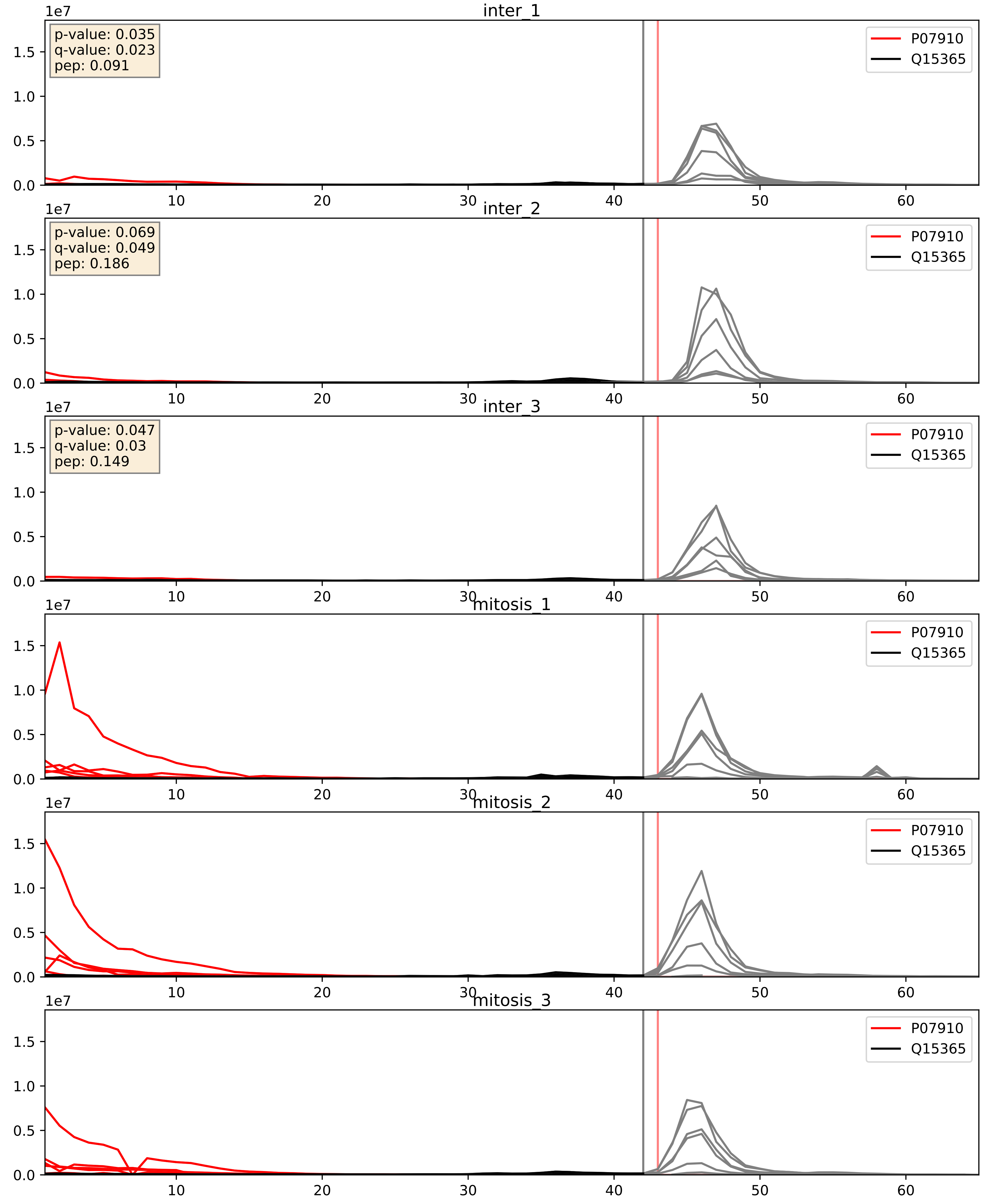


| level | condition_1 | condition_2 | pvalue | pvalue_adjusted |
|---|---|---|---|---|
| interactor_abundance | mitosis | inter | 9.148564965046394e-06 | 0.0033734327106189023 |
| complex_abundance | mitosis | inter | 1.030468151281165e-05 | 0.005691857151652958 |
| interactor_ratio | mitosis | inter | 4.476005324973264e-05 | 0.006796360849516482 |

P07910_Q15717
HNRPC_HUMAN vs ELAV1_HUMAN
p-value: 0.026 q-value: 0.02

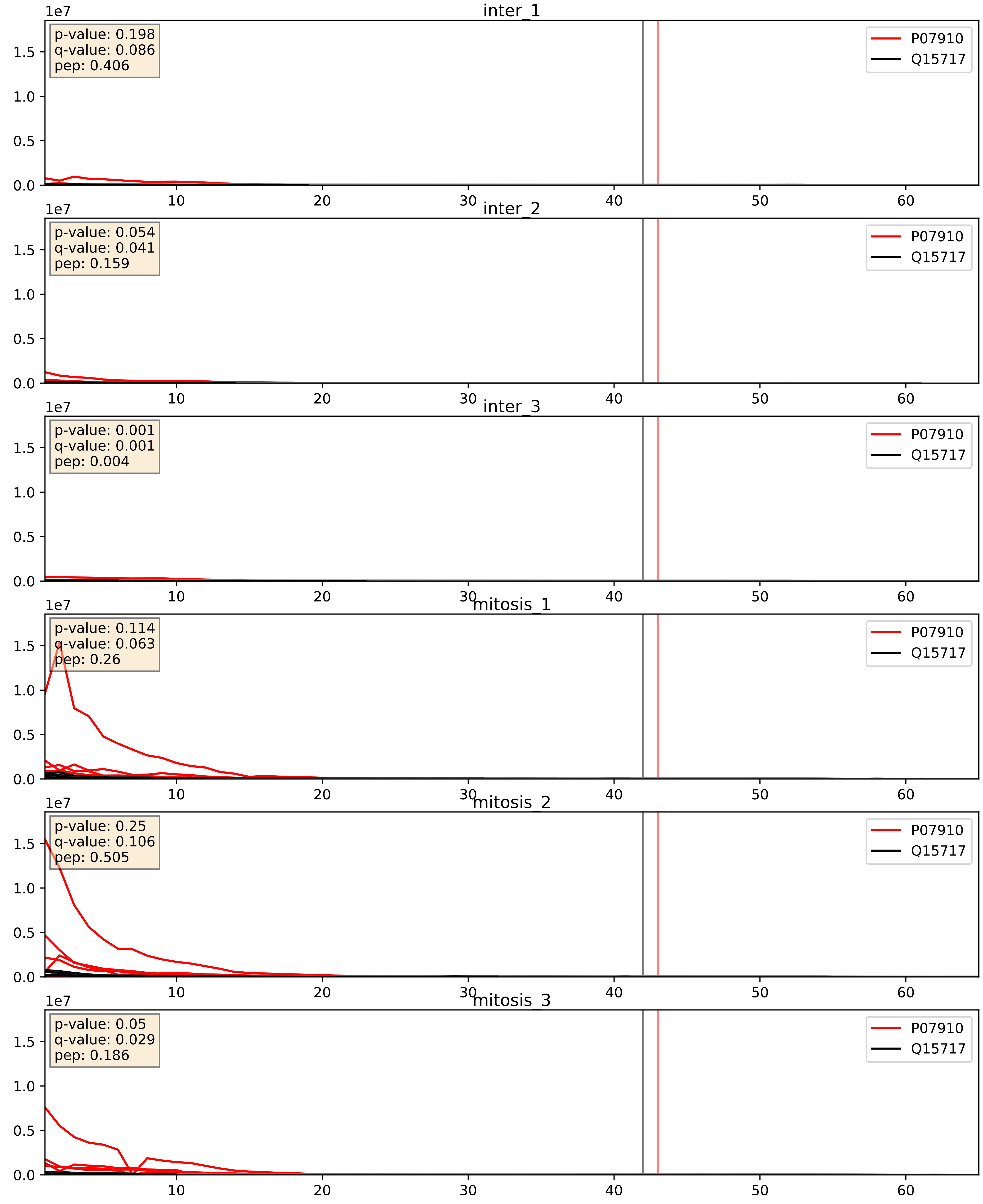


| level | condition_1 | condition_2 | pvalue | pvalue_adjusted |
|---|---|---|---|---|
| interactor_abundance | mitosis | inter | 5.855680651902104e-06 | 0.0033734327106189023 |
| complex_abundance | mitosis | inter | 1.6805290568155707e-05 | 0.005691857151652958 |
| interactor_ratio | mitosis | inter | 0.0014640241868224817 | 0.04557108014254707 |

P07910_Q16629
HNRPC_HUMAN vs SRSF7_HUMAN
p-value: 0.046 q-value: 0.031

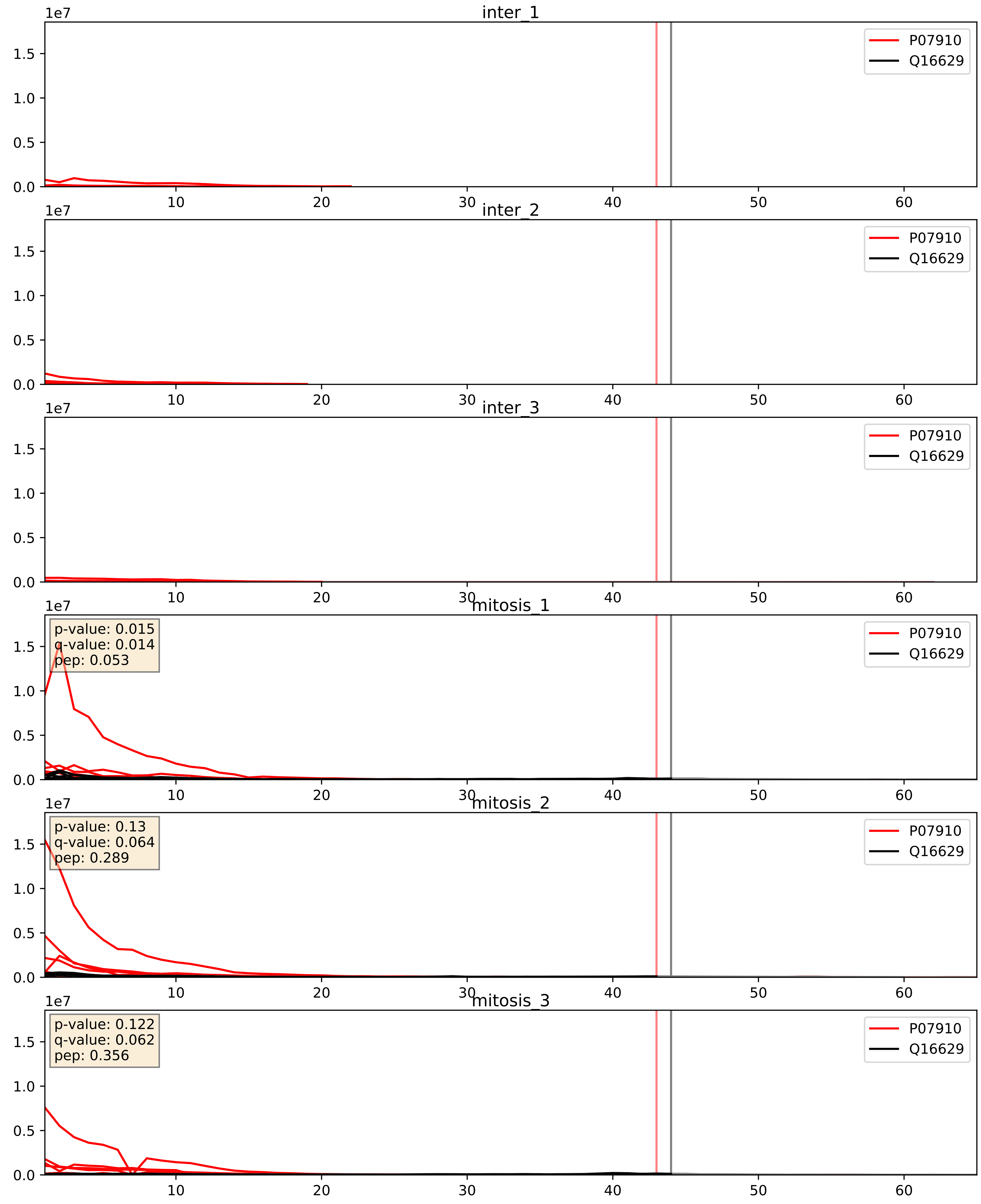


| level | condition_1 | condition_2 | pvalue | pvalue_adjusted |
|---|---|---|---|---|
| interactor_abundance | mitosis | inter | 2.525501836279769e-06 | 0.0033734327106189023 |
| complex_abundance | mitosis | inter | 5.288643111257365e-06 | 0.0056588481290453805 |
| interactor_ratio | mitosis | inter | 0.00024711905536094626 | 0.016038997600655195 |

P07910_Q99459
HNRPC_HUMAN vs CDC5L_HUMAN
p-value: 0.036 q-value: 0.025

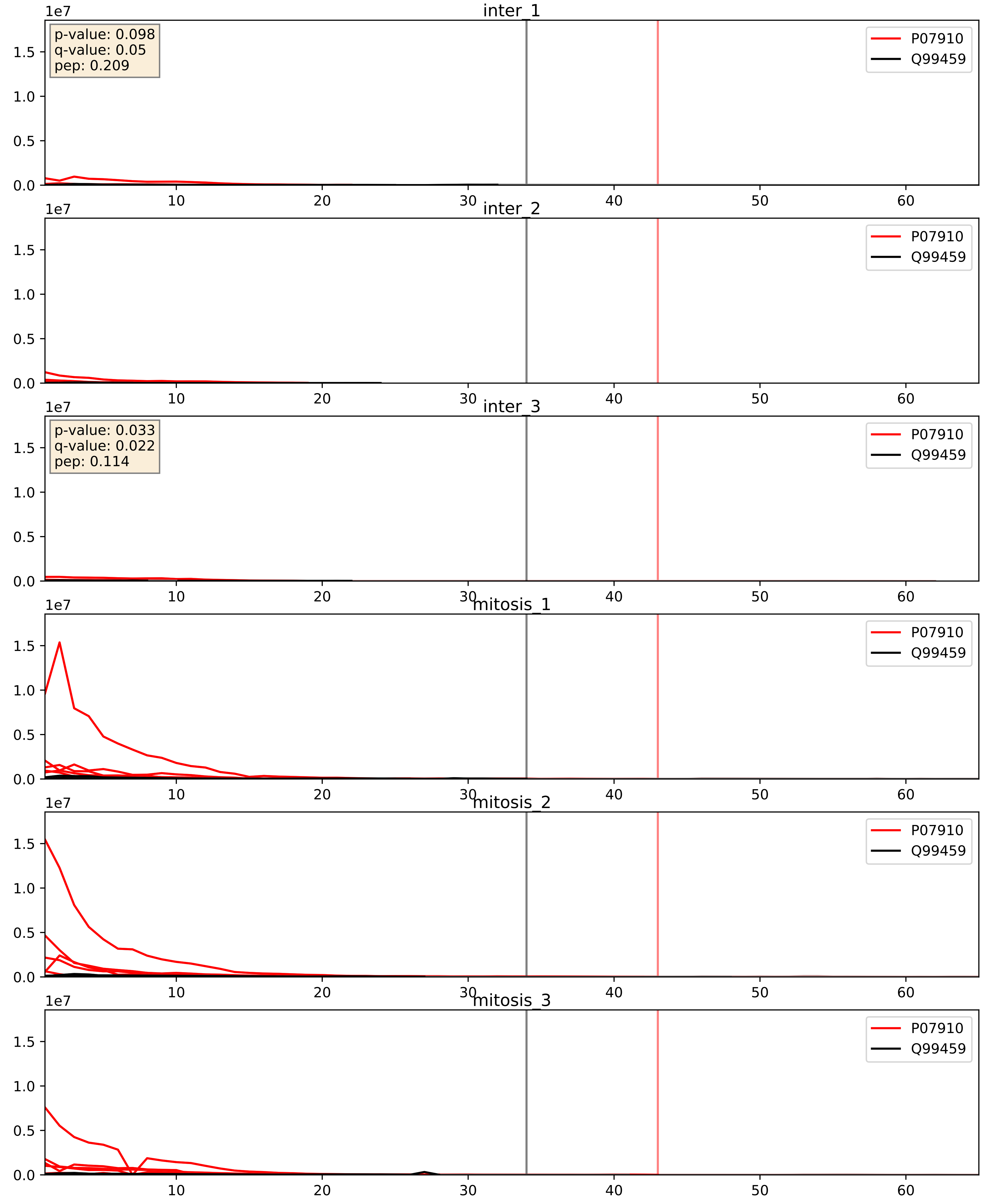


| level | condition_1 | condition_2 | pvalue | pvalue_adjusted |
|---|---|---|---|---|
| interactor_abundance | mitosis | inter | 7.673678712415949e-05 | 0.006950972463310108 |
| complex_abundance | mitosis | inter | 0.0006052291130691676 | 0.015834093001149168 |
| interactor_ratio | mitosis | inter | 0.003252907220343525 | 0.07405554735675686 |

P07910_Q9UMS4
HNRPC_HUMAN vs PRP19_HUMAN
p-value: 0.059 q-value: 0.037

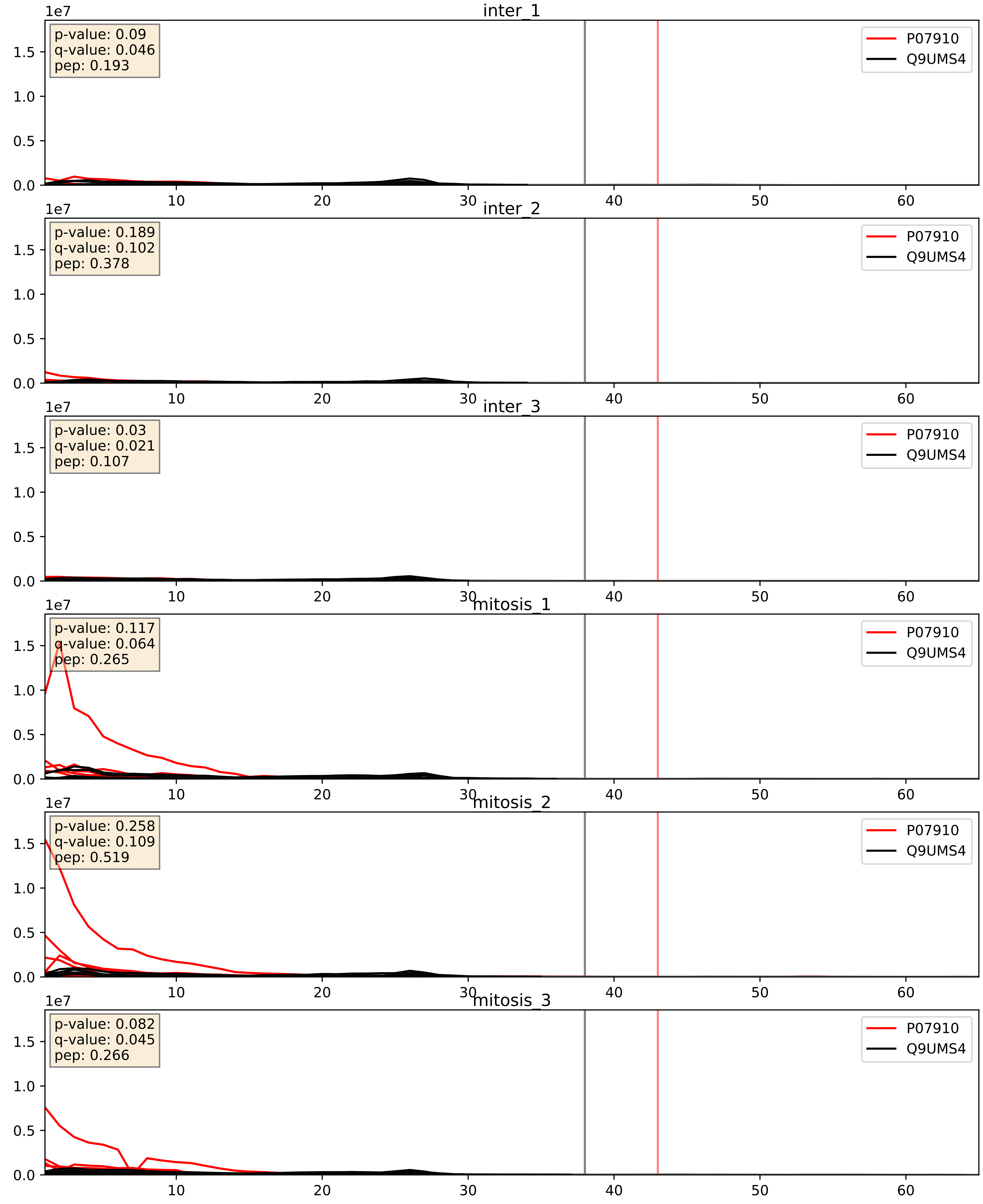


| level | condition_1 | condition_2 | pvalue | pvalue_adjusted |
|---|---|---|---|---|
| interactor_abundance | mitosis | inter | 9.831572417299737e-06 | 0.0033734327106189023 |
| complex_abundance | mitosis | inter | 1.561120506983598e-05 | 0.005691857151652958 |
| interactor_ratio | mitosis | inter | 0.002019920931775477 | 0.05524128810223029 |
