## Supplementary Data 2 for "SECAT: Quantifying differential protein-protein interaction states by network-centric analysis": hela_string_000028_Q9H0A0.pdf

Q9H0A0_Q9H0A0

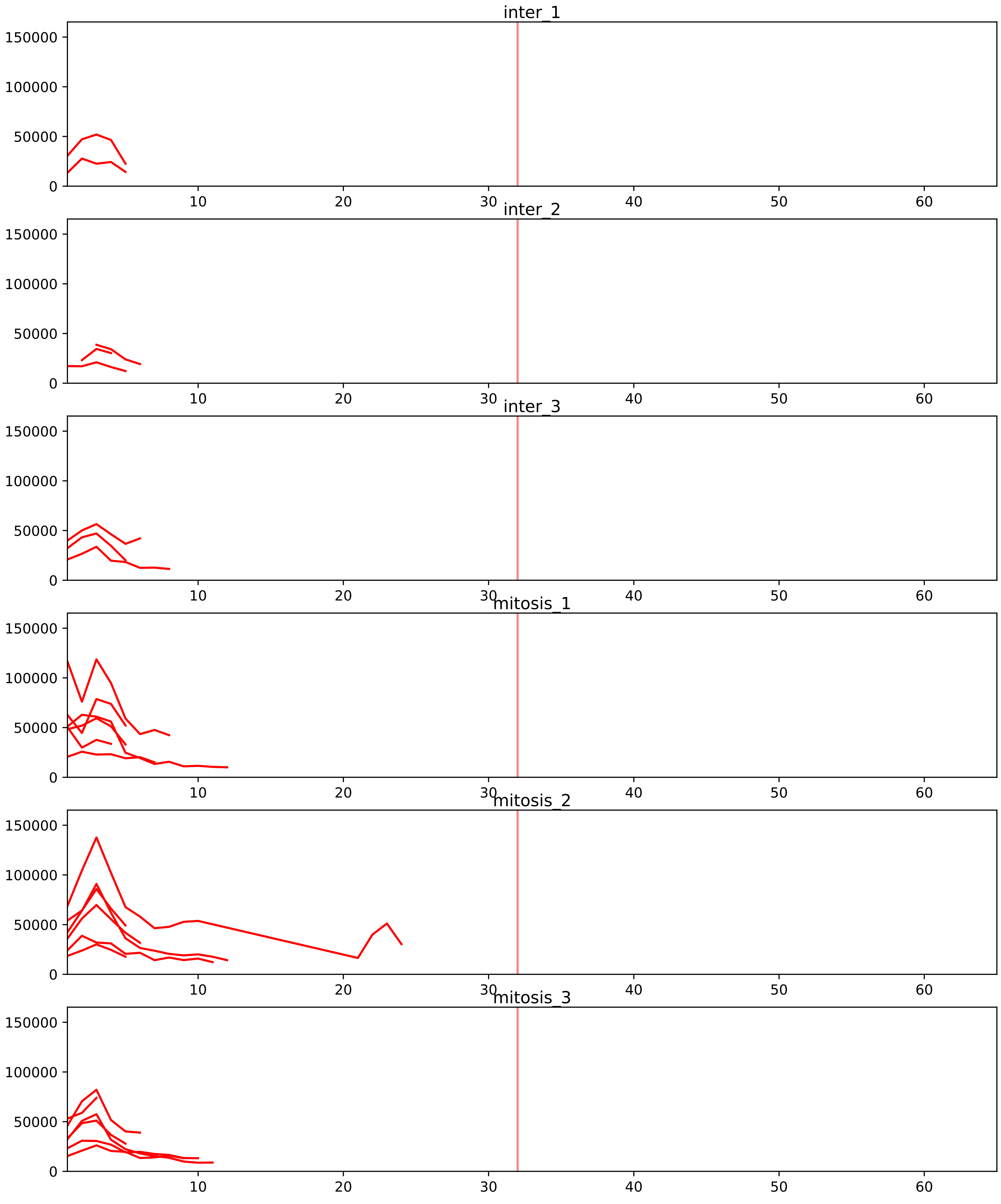


| level | condition_1 | condition_2 | pvalue | pvalue_adjusted |
|---|---|---|---|---|
| interactor_ratio | mitosis | inter | 5.873506549184944e-10 | 4.00268594462974e-08 |
| complex_abundance | mitosis | inter | 0.00509660530035969 | 0.028854626931267165 |
| assembled_abundance | mitosis | inter | 0.10246946381222684 | 0.3750270211923642 |
| interactor_abundance | mitosis | inter | 0.1048285869426164 | 0.2340832523961337 |
| total_abundance | mitosis | inter | 0.15015852973649838 | 0.47897645623548735 |

O00567_Q9H0A0
NOP56_HUMAN vs NAT10_HUMAN
p-value: 0.005 q-value: 0.005

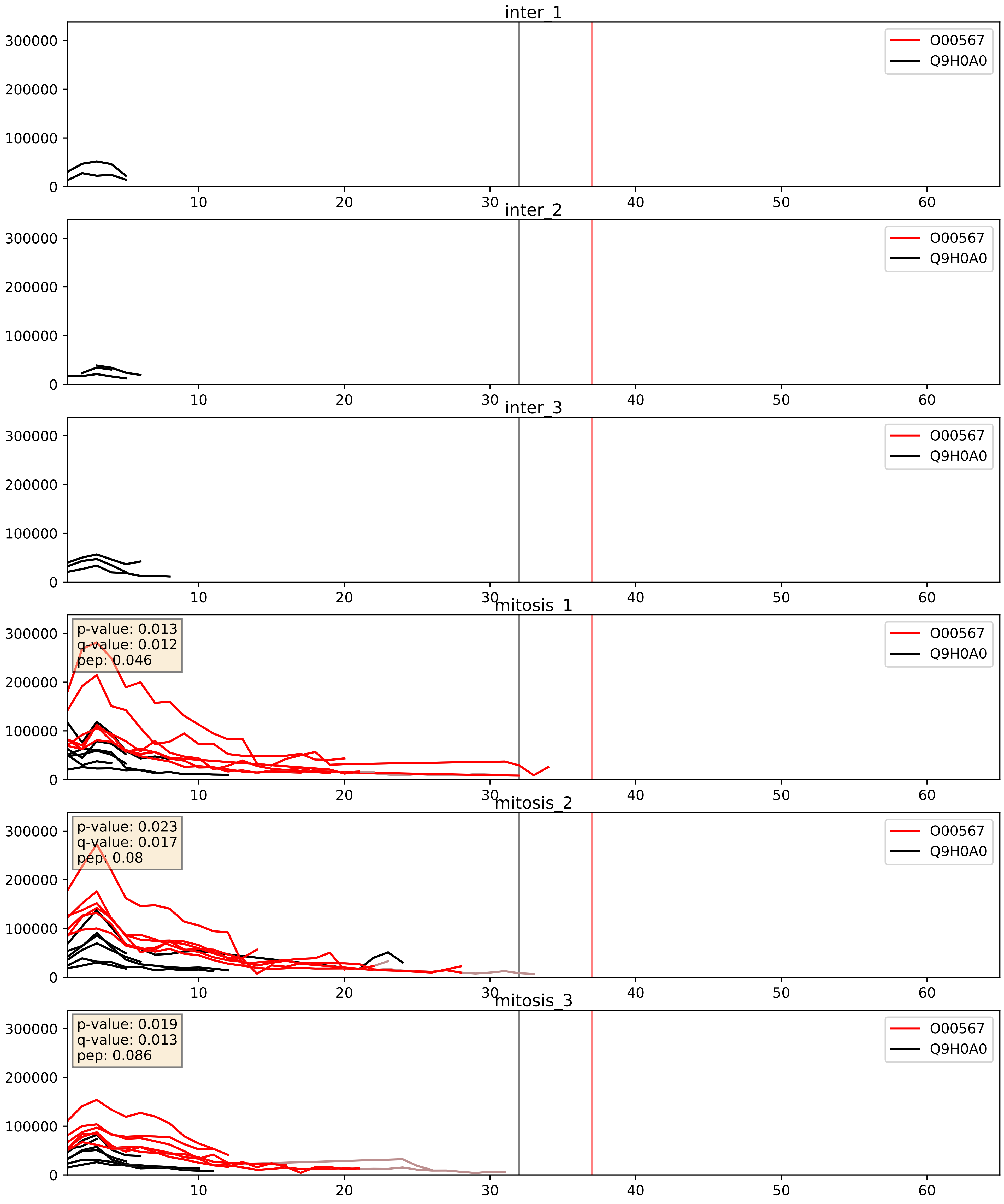


| level | condition_1 | condition_2 | pvalue | pvalue_adjusted |
|---|---|---|---|---|
| interactor_ratio | mitosis | inter | 2.3043765380856052e-08 | 0.0001972546316601278 |
| interactor_abundance | mitosis | inter | 0.0009620251184093979 | 0.02306454775731039 |
| complex_abundance | mitosis | inter | 0.0035552641482872074 | 0.043600374082146835 |

O60832_Q9H0A0
DKC1_HUMAN vs NAT10_HUMAN
p-value: 0.076 q-value: 0.046

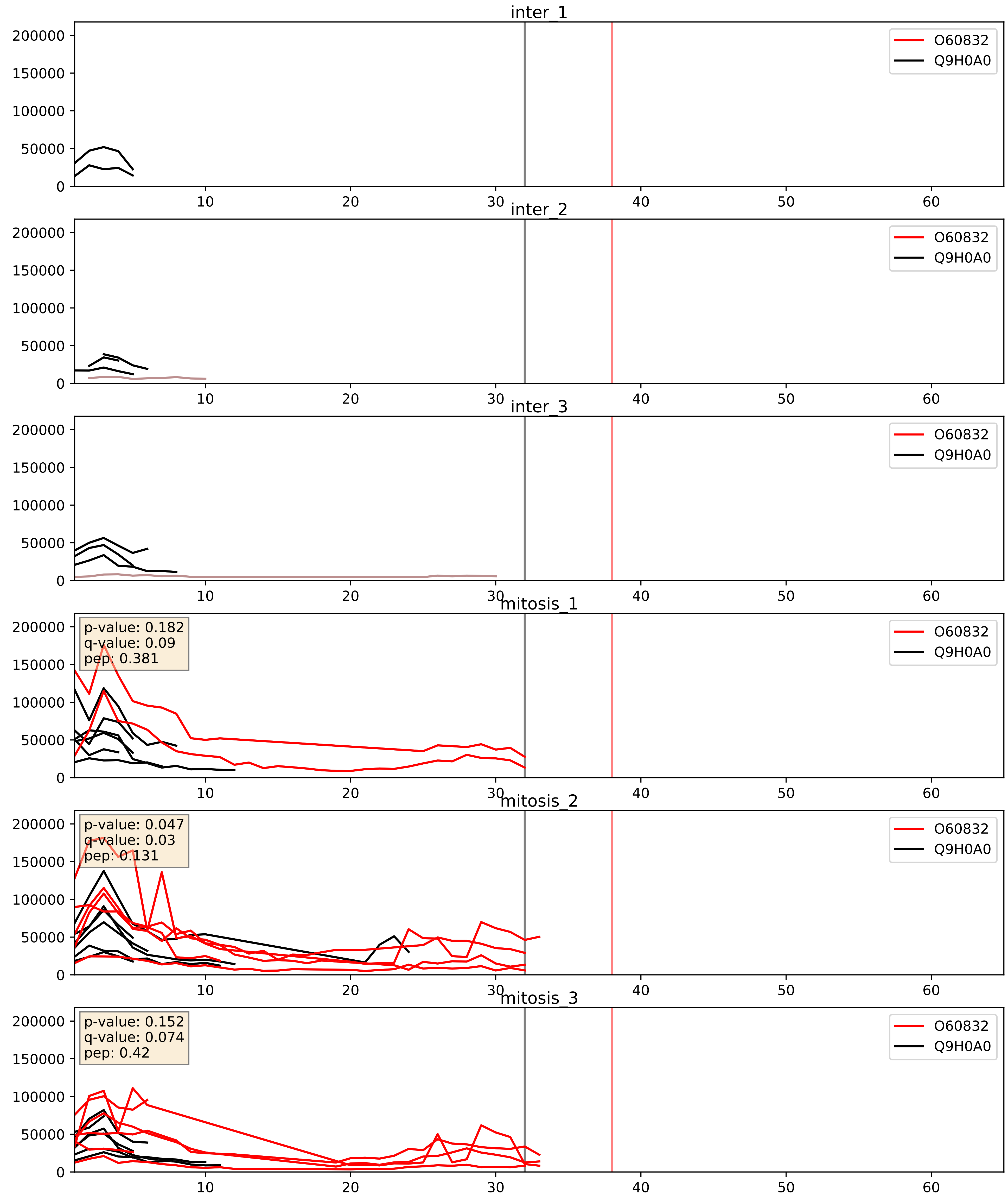


| level | condition_1 | condition_2 | pvalue | pvalue_adjusted |
|---|---|---|---|---|
| complex_abundance | mitosis | inter | 0.014138361720307944 | 0.08846811134929532 |
| interactor_abundance | mitosis | inter | 0.06548599163945651 | 0.21201213632138718 |
| interactor_ratio | mitosis | inter | 0.7825031437437657 | 0.95900240448312 |

O76021_Q9H0A0
RL1D1_HUMAN vs NAT10_HUMAN
p-value: 0.078 q-value: 0.047

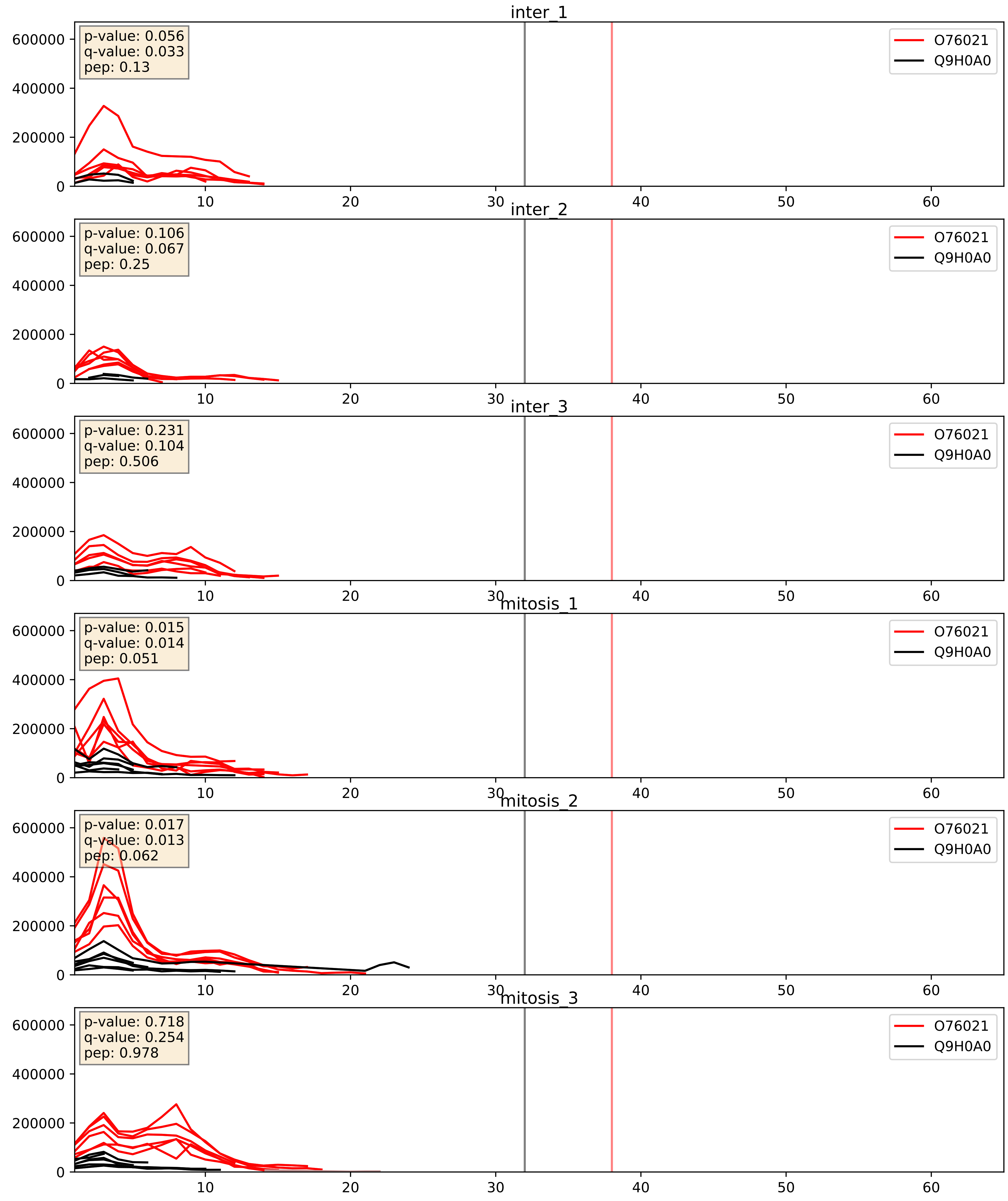


| level | condition_1 | condition_2 | pvalue | pvalue_adjusted |
|---|---|---|---|---|
| interactor_abundance | mitosis | inter | 0.0004232024324935983 | 0.014167094422781207 |
| complex_abundance | mitosis | inter | 0.002537322887836132 | 0.036409709098919725 |
| interactor_ratio | mitosis | inter | 0.02791082622771571 | 0.26184787474031573 |

Q12788_Q9H0A0
TBL3_HUMAN vs NAT10_HUMAN
p-value: 0.028 q-value: 0.02

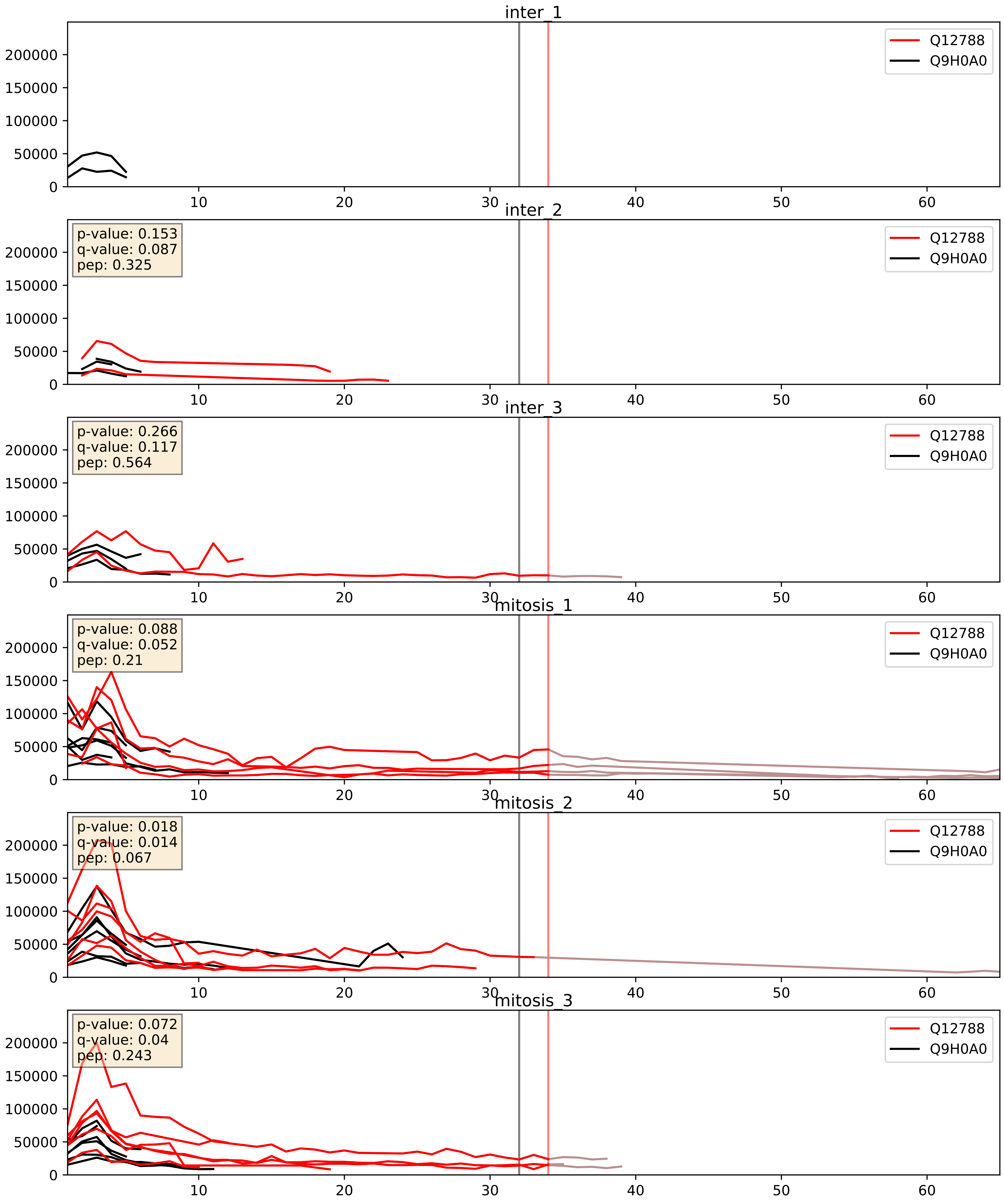


| level | condition_1 | condition_2 | pvalue | pvalue_adjusted |
|---|---|---|---|---|
| interactor_abundance | mitosis | inter | 0.005345390931153837 | 0.061666504542691174 |
| interactor_ratio | mitosis | inter | 0.04444676005266599 | 0.3493703085866124 |
| complex_abundance | mitosis | inter | 0.05040593282045461 | 0.15980547590484867 |

Q15269_Q9H0A0
PWP2_HUMAN vs NAT10_HUMAN
p-value: 0.002 q-value: 0.002

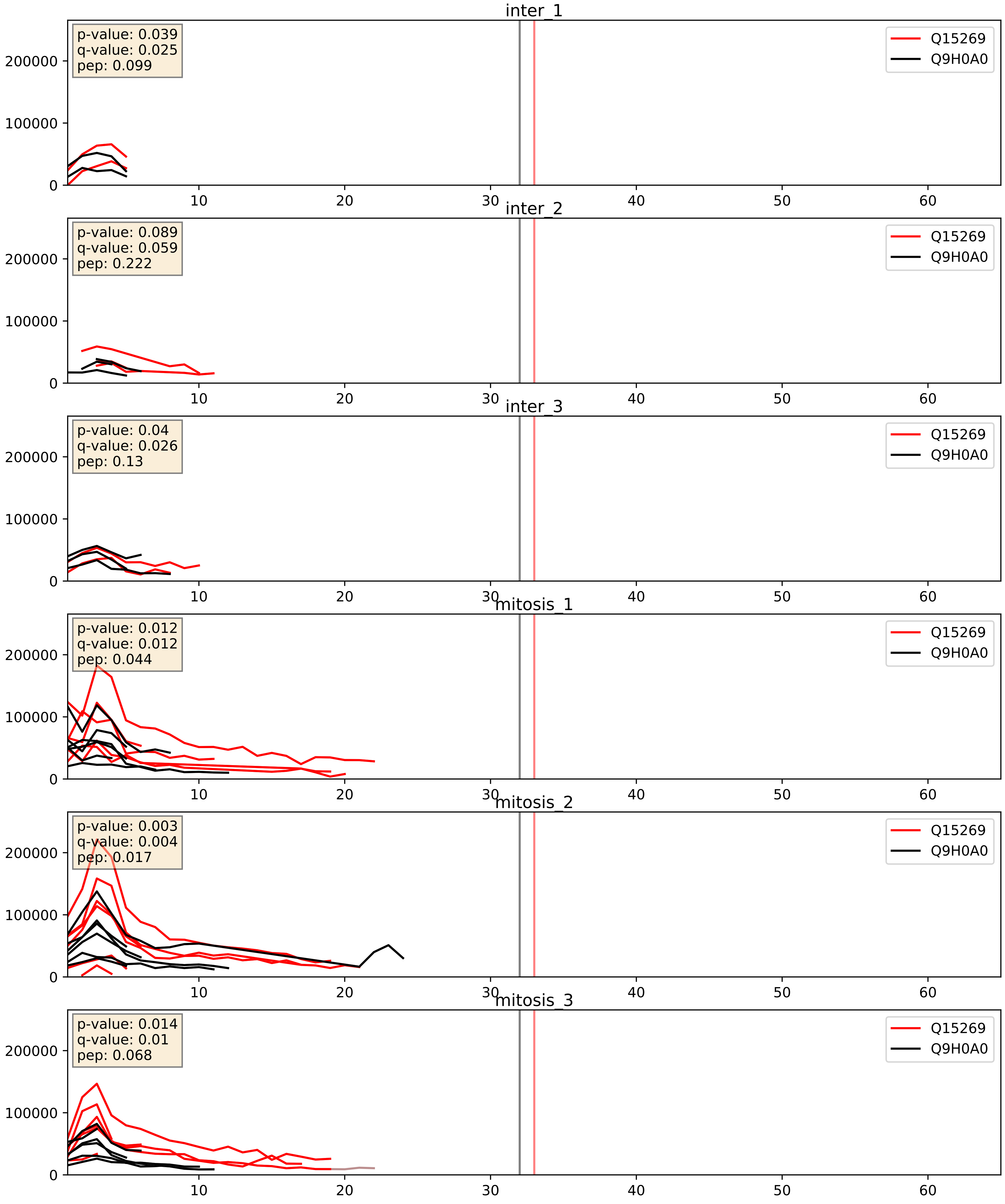


| level | condition_1 | condition_2 | pvalue | pvalue_adjusted |
|---|---|---|---|---|
| interactor_ratio | mitosis | inter | 0.013552472765333098 | 0.1693564479872282 |
| interactor_abundance | mitosis | inter | 0.0423912444936641 | 0.16595886250435155 |
| complex_abundance | mitosis | inter | 0.16592341030748023 | 0.3147952291083829 |

Q8NI36_Q9H0A0
WDR36_HUMAN vs NAT10_HUMAN
p-value: 0.008 q-value: 0.007

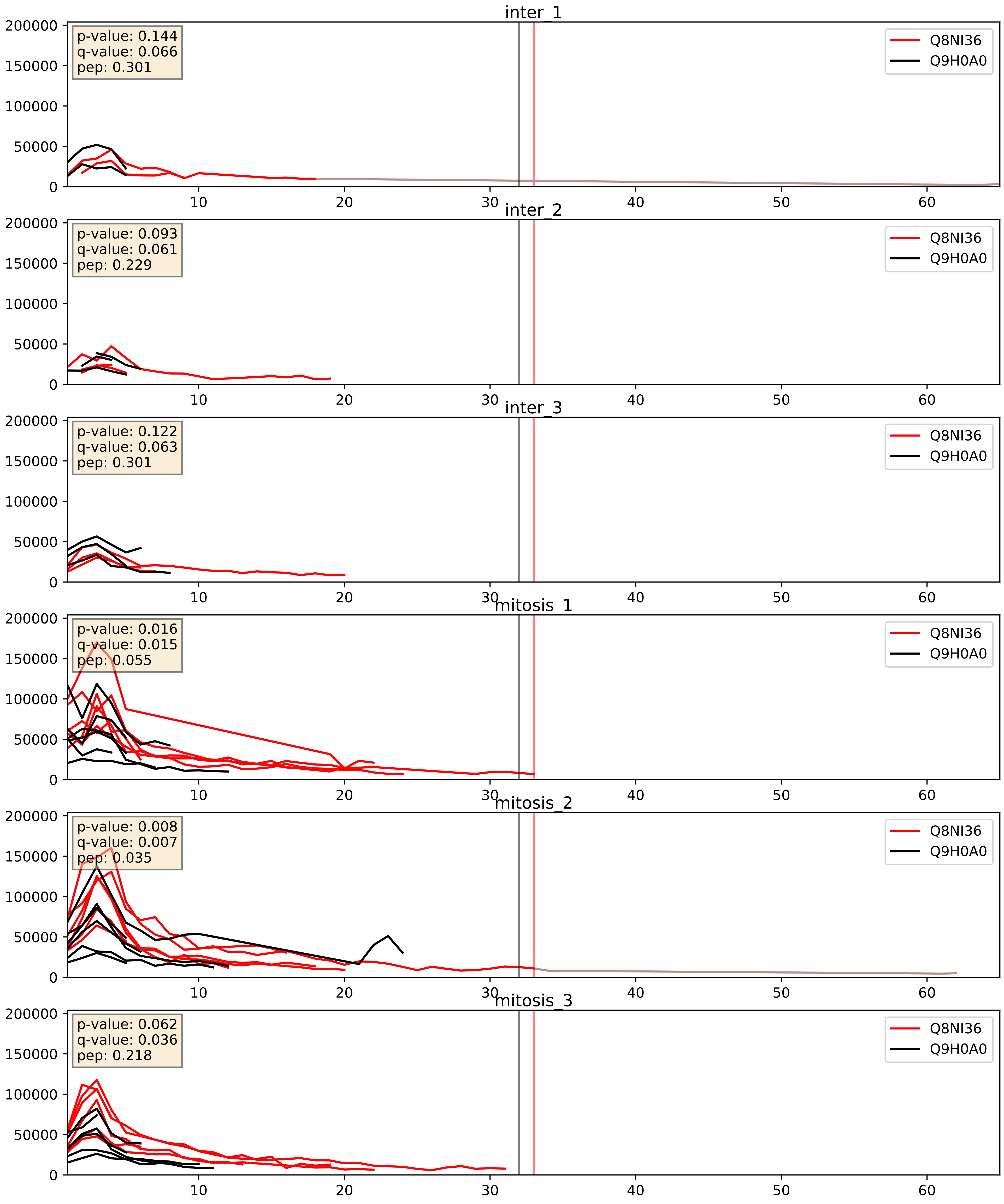


| level | condition_1 | condition_2 | pvalue | pvalue_adjusted |
|---|---|---|---|---|
| interactor_abundance | mitosis | inter | 0.00851704615394727 | 0.0803238047623781 |
| interactor_ratio | mitosis | inter | 0.018810868078371294 | 0.20643721891135677 |
| complex_abundance | mitosis | inter | 0.08899095423186006 | 0.2191545360681031 |

Q969X6_Q9H0A0
UTP4_HUMAN vs NAT10_HUMAN
p-value: 0.08 q-value: 0.048

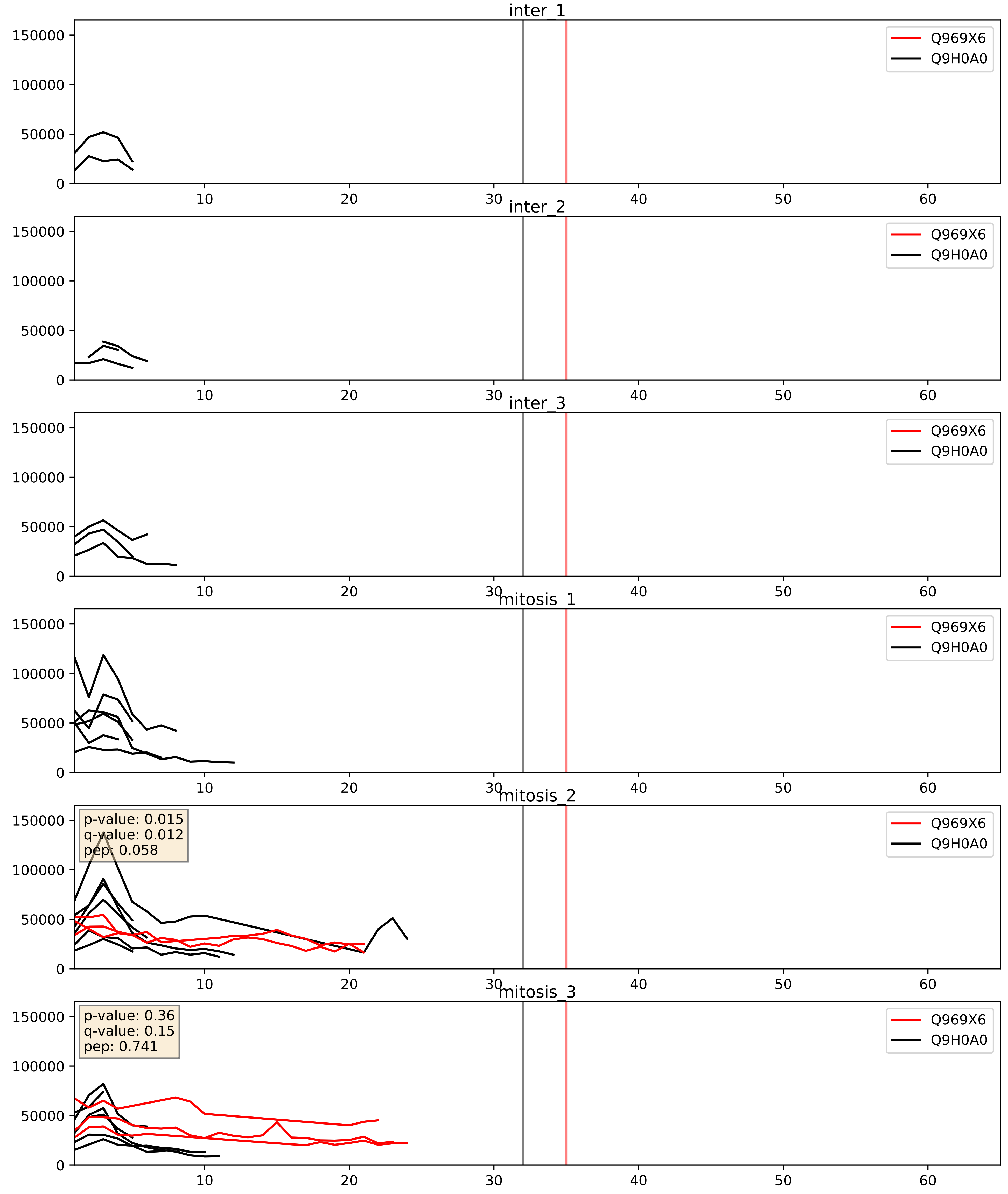


| level | condition_1 | condition_2 | pvalue | pvalue_adjusted |
|---|---|---|---|---|
| complex_abundance | mitosis | inter | 0.3288133465323389 | 0.4772279452235212 |
| interactor_abundance | mitosis | inter | 0.3460140955778534 | 0.544589568782782 |
| interactor_ratio | mitosis | inter | 0.8016188227647435 | 0.9629325179436155 |

Q9H0A0_Q9H6R4
NAT10_HUMAN vs NOL6_HUMAN
p-value: 0.029 q-value: 0.021

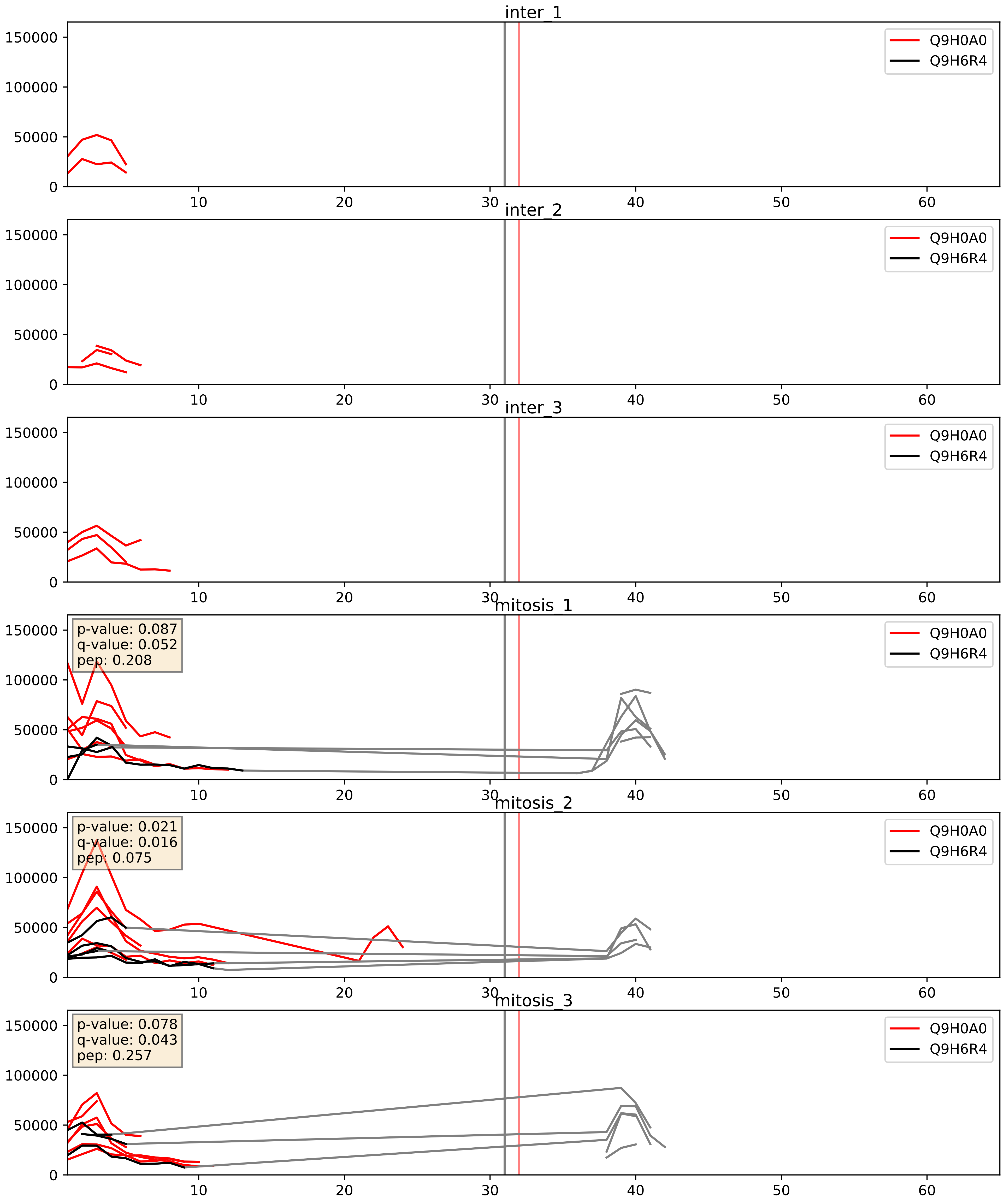


| level | condition_1 | condition_2 | pvalue | pvalue_adjusted |
|---|---|---|---|---|
| interactor_abundance | mitosis | inter | 0.025428031496039612 | 0.13211772358488563 |
| interactor_ratio | mitosis | inter | 0.0271500034485924 | 0.2577730310940899 |
| complex_abundance | mitosis | inter | 0.06262730312099983 | 0.18019822343386843 |

Q9H0A0_Q9UNX4
NAT10_HUMAN vs WDR3_HUMAN
p-value: 0.006 q-value: 0.005

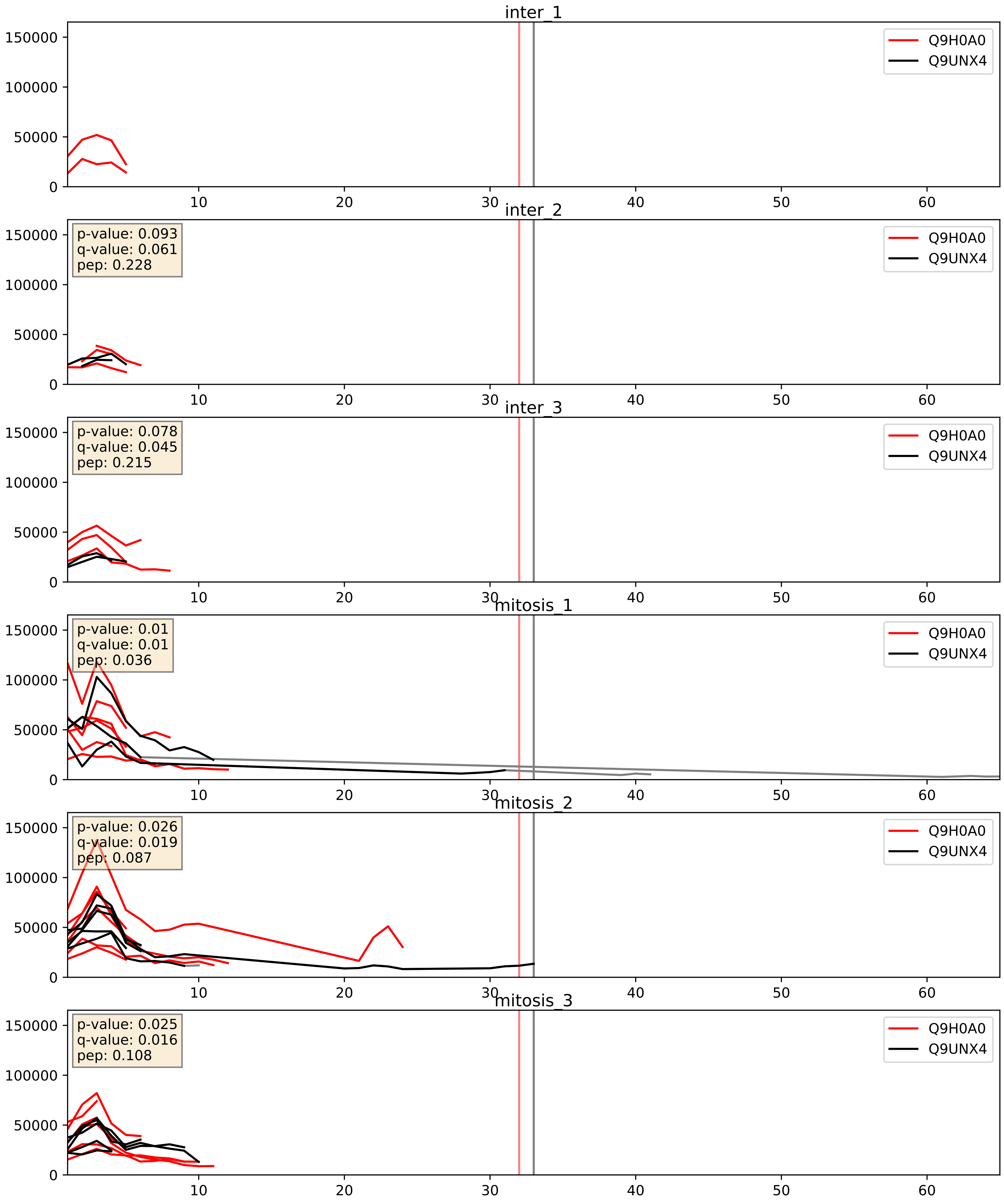


| level | condition_1 | condition_2 | pvalue | pvalue_adjusted |
|---|---|---|---|---|
| complex_abundance | mitosis | inter | 0.2858862643454428 | 0.4346099345487244 |
| interactor_abundance | mitosis | inter | 0.45197017672950507 | 0.625321595733726 |
| interactor_ratio | mitosis | inter | 0.6060595505852381 | 0.9052533265322077 |

Q9H0A0_Q9Y2X3
NAT10_HUMAN vs NOP58_HUMAN
p-value: 0.055 q-value: 0.035

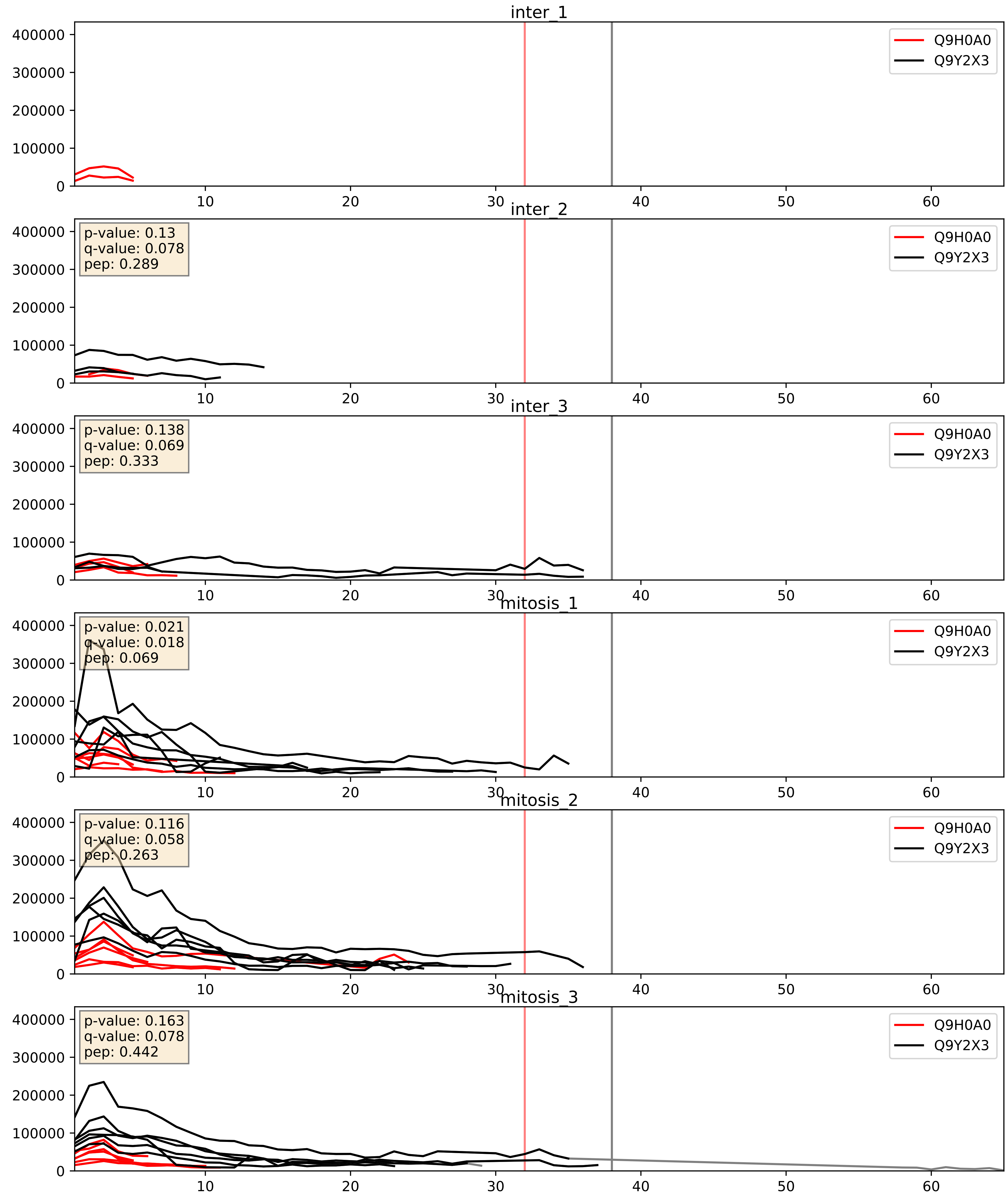


| level | condition_1 | condition_2 | pvalue | pvalue_adjusted |
|---|---|---|---|---|
| interactor_ratio | mitosis | inter | 0.0002532636495883736 | 0.016178633137884164 |
| complex_abundance | mitosis | inter | 0.02044007375838056 | 0.10540182612755276 |
| interactor_abundance | mitosis | inter | 0.45474744236861303 | 0.6283515910694637 |

Q9H0A0_Q9Y5J1
NAT10_HUMAN vs UTP18_HUMAN
p-value: 0.064 q-value: 0.04

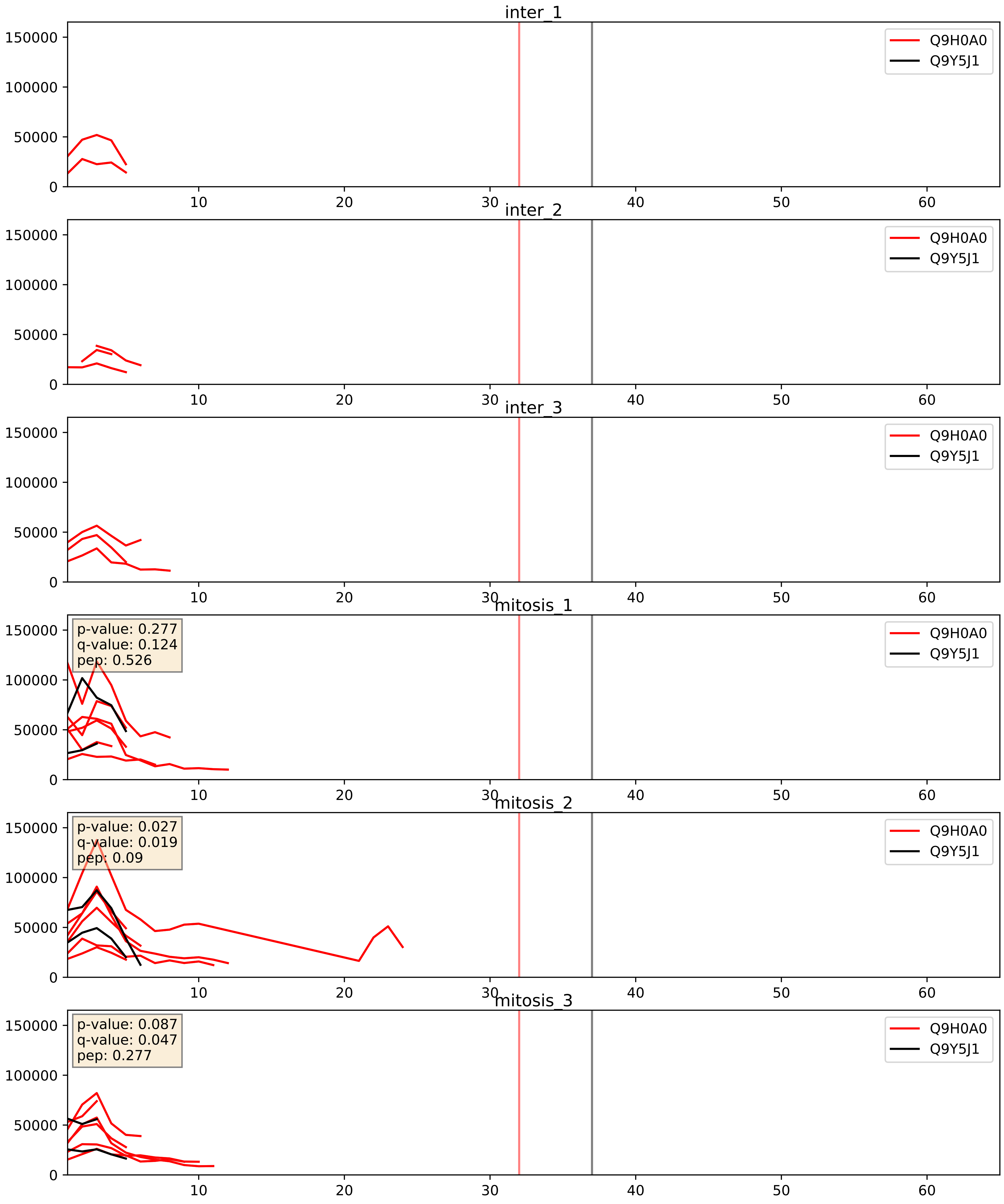


| level | condition_1 | condition_2 | pvalue | pvalue_adjusted |
|---|---|---|---|---|
| interactor_ratio | mitosis | inter | 0.0038335460165161953 | 0.08075090146433855 |
| interactor_abundance | mitosis | inter | 0.03832306629411007 | 0.15936046986653518 |
| complex_abundance | mitosis | inter | 0.06582643627069286 | 0.18541437791284332 |
