## Supplementary Data 2 for "SECAT: Quantifying differential protein-protein interaction states by network-centric analysis": hela_string_000029_Q99459.pdf

Q99459_Q99459

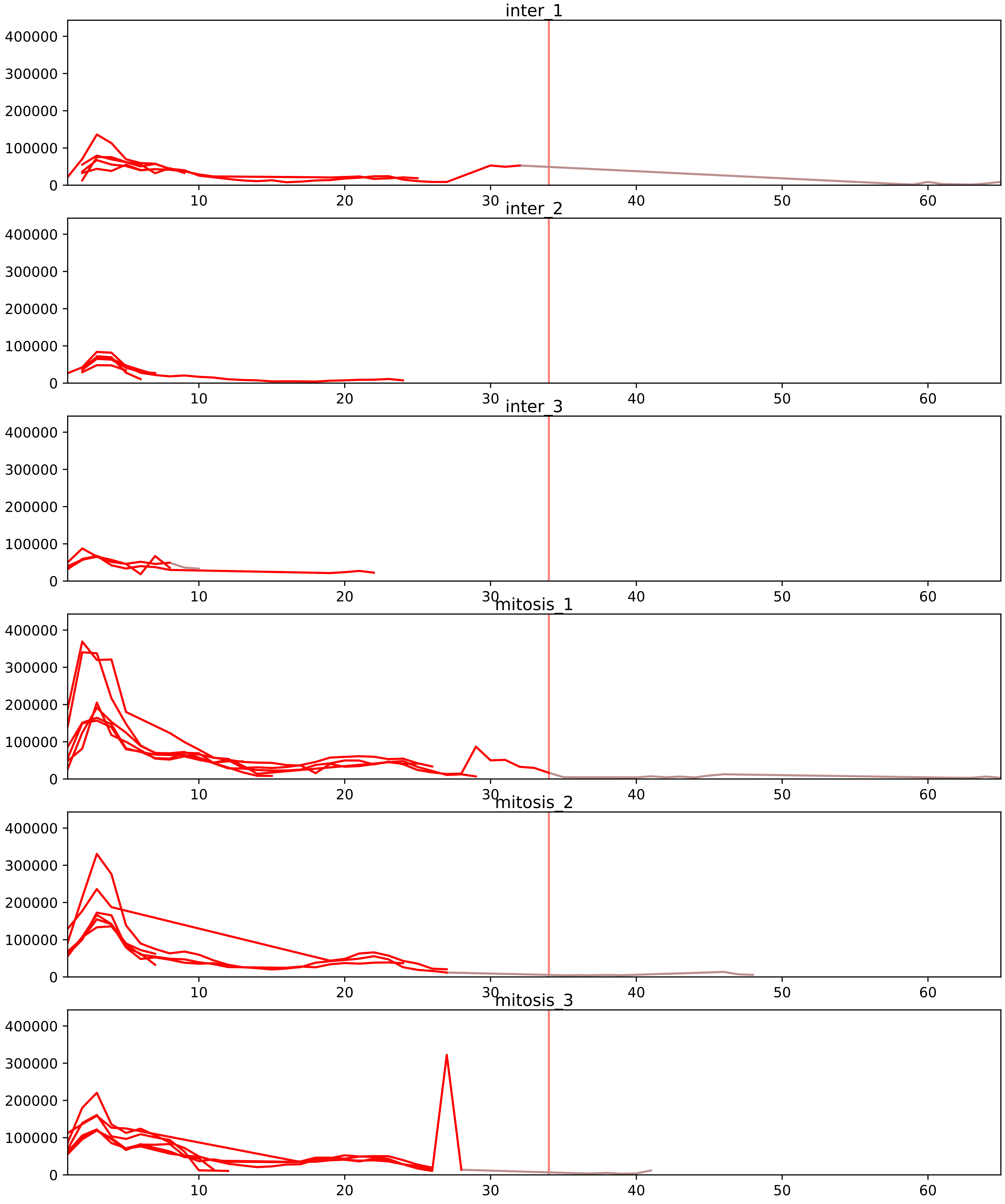


| level | condition_1 | condition_2 | pvalue | pvalue_adjusted |
| --- | --- | --- | --- | --- |
| interactor_ratio | mitosis | inter | 6.41161045517092e-10 | 4.213344013398033e-08 |
| complex_abundance | mitosis | inter | 0.0003800947150668352 | 0.005779952691925428 |
| interactor_abundance | mitosis | inter | 0.000561894262193722 | 0.007990604304242772 |
| monomer_abundance | mitosis | inter | 0.08934627042196161 | 0.3886658423173769 |
| total_abundance | mitosis | inter | 0.4408864097272551 | 0.7267771818469224 |
| assembled_abundance | mitosis | inter | 0.5815749281111882 | 0.8019526839648053 |

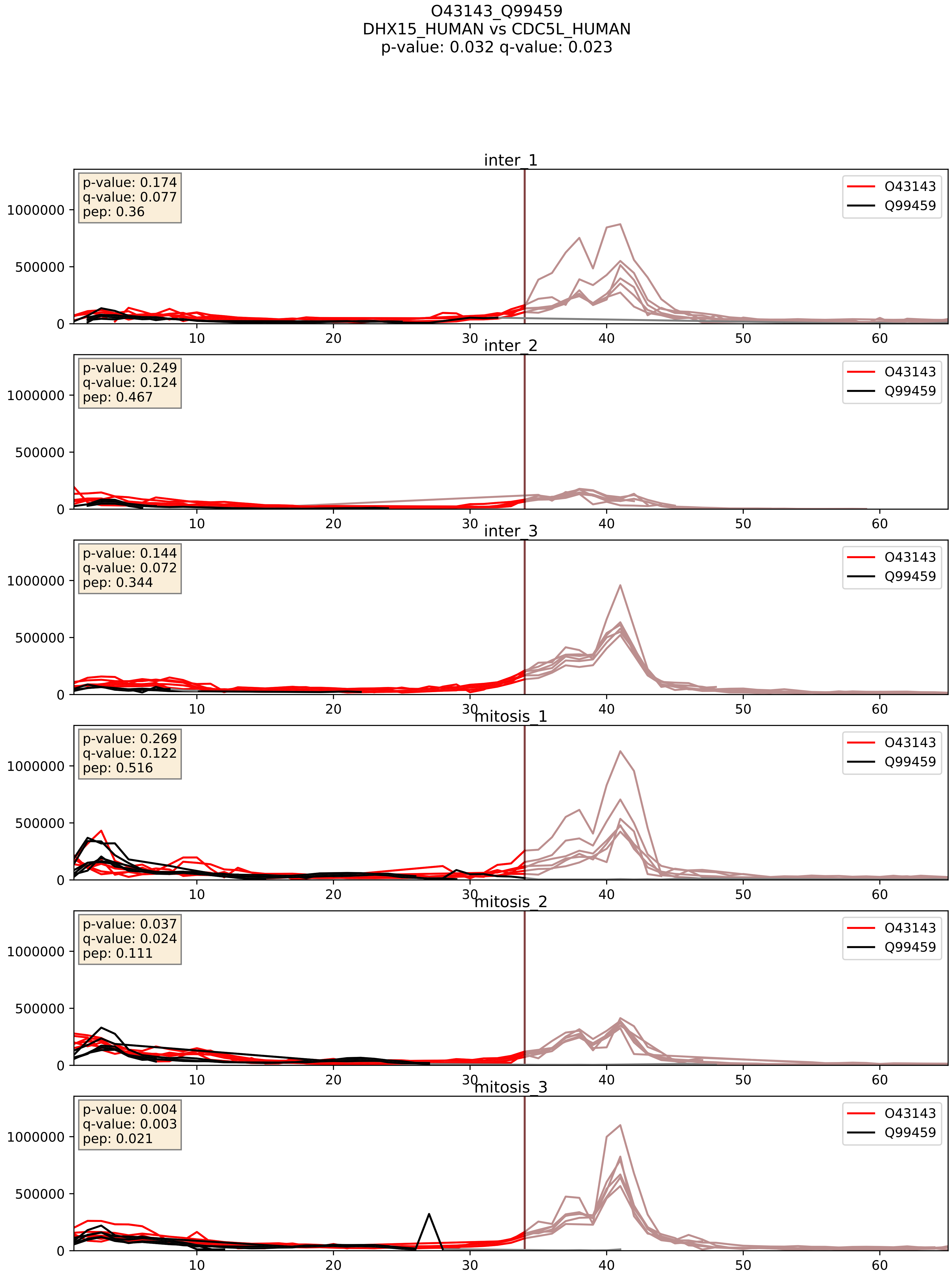


| level | condition_1 | condition_2 | pvalue | pvalue_adjusted |
|---|---|---|---|---|
| complex_abundance | mitosis | inter | 0.009703232551220584 | 0.07321717819287338 |
| interactor_abundance | mitosis | inter | 0.012897735497701254 | 0.10167976967449956 |
| interactor_ratio | mitosis | inter | 0.05512686863757614 | 0.3890548630198597 |

O43395_Q99459
PRPF3_HUMAN vs CDC5L_HUMAN
p-value: 0.003 q-value: 0.004

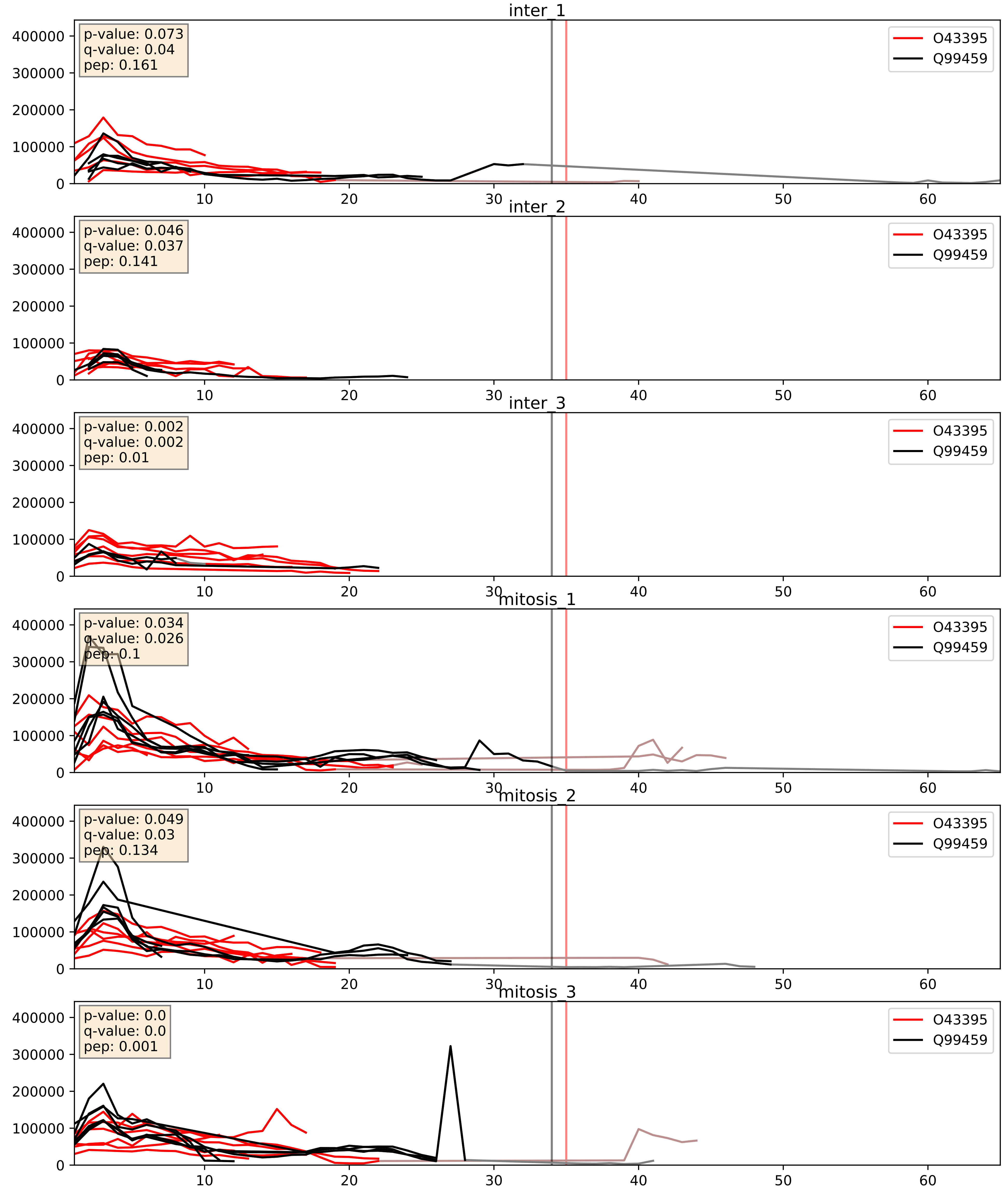


| level | condition_1 | condition_2 | pvalue | pvalue_adjusted |
|---|---|---|---|---|
| complex_abundance | mitosis | inter | 0.020735988987782956 | 0.1061724104661514 |
| interactor_ratio | mitosis | inter | 0.05638110929547731 | 0.3926647223857623 |
| interactor_abundance | mitosis | inter | 0.06577503851832661 | 0.21269195540886726 |

O43660_Q99459
PLRG1_HUMAN vs CDC5L_HUMAN
p-value: 0.075 q-value: 0.045

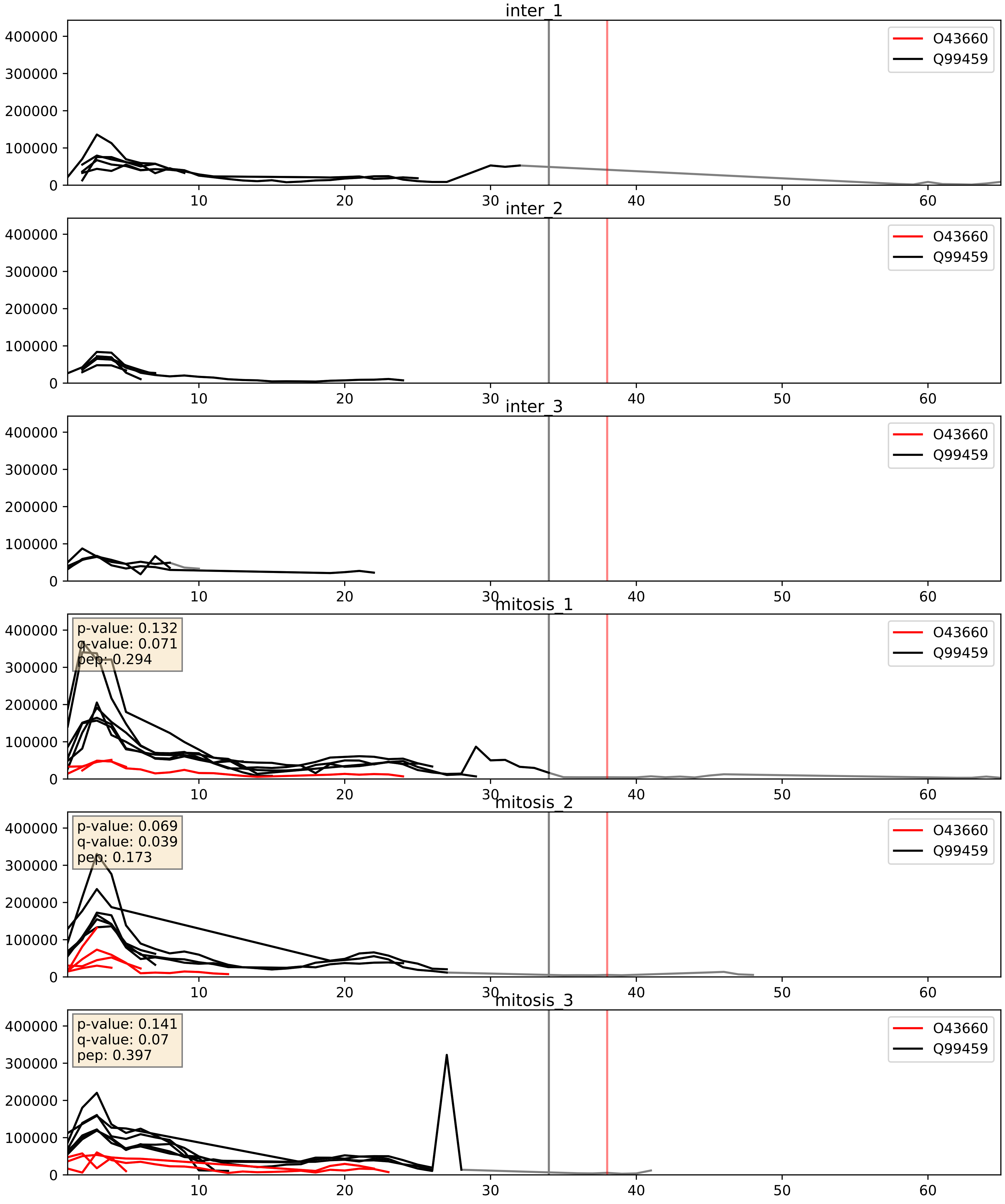


| level | condition_1 | condition_2 | pvalue | pvalue_adjusted |
|---|---|---|---|---|
| interactor_ratio | mitosis | inter | 0.0028873138380850756 | 0.0700153157337344 |
| complex_abundance | mitosis | inter | 0.0038779448692945182 | 0.046157376235893825 |
| interactor_abundance | mitosis | inter | 0.03668661671701946 | 0.15802829520754763 |

O75934_Q99459
SPF27_HUMAN vs CDC5L_HUMAN
p-value: 0.004 q-value: 0.004

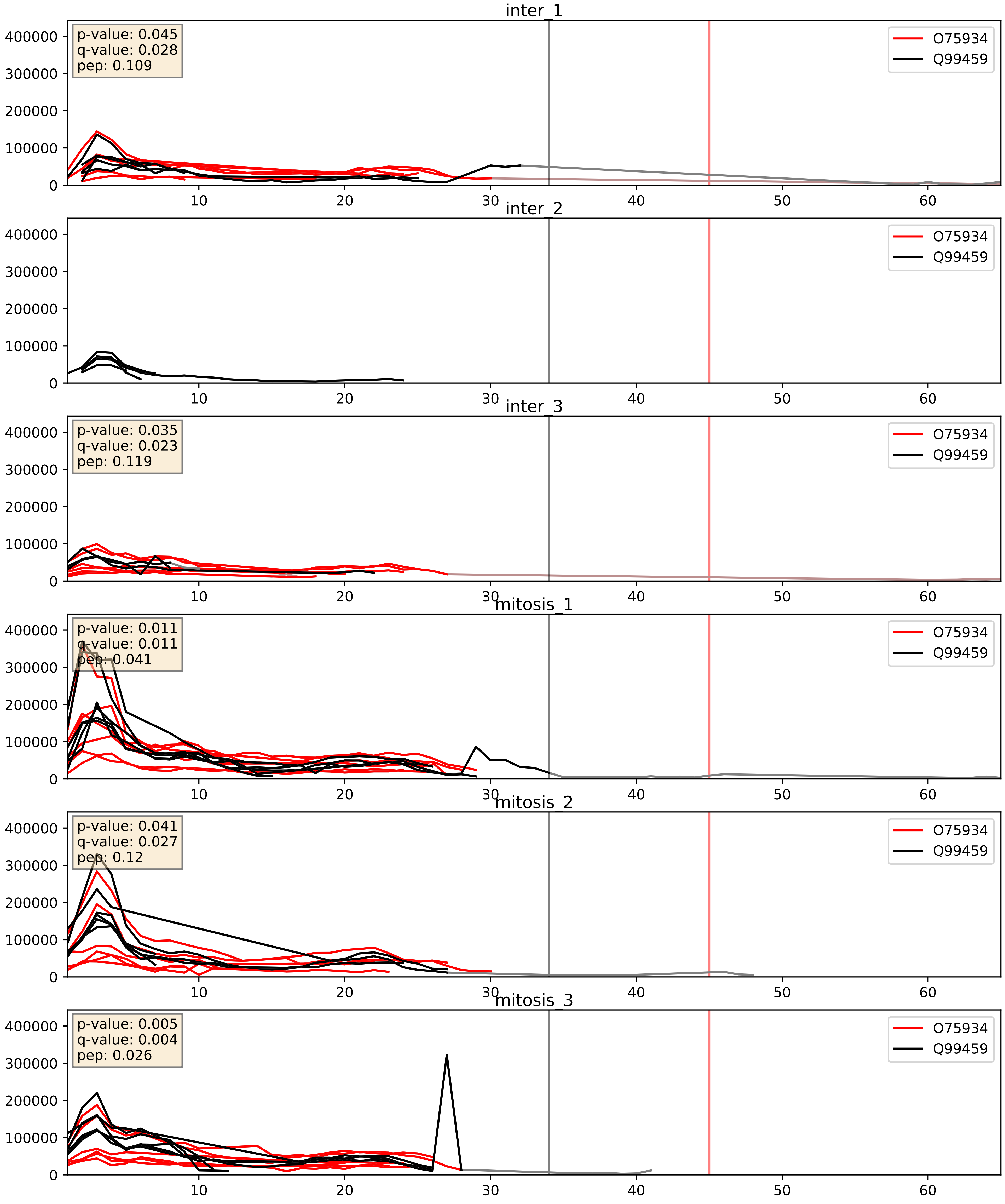


| level | condition_1 | condition_2 | pvalue | pvalue_adjusted |
|---|---|---|---|---|
| complex_abundance | mitosis | inter | 0.03282547885145826 | 0.13204233974082835 |
| interactor_abundance | mitosis | inter | 0.10154606889418748 | 0.27370850762629345 |
| interactor_ratio | mitosis | inter | 0.14845822739515527 | 0.60240057967345 |

O94906_Q99459
PRP6_HUMAN vs CDC5L_HUMAN
p-value: 0.019 q-value: 0.015

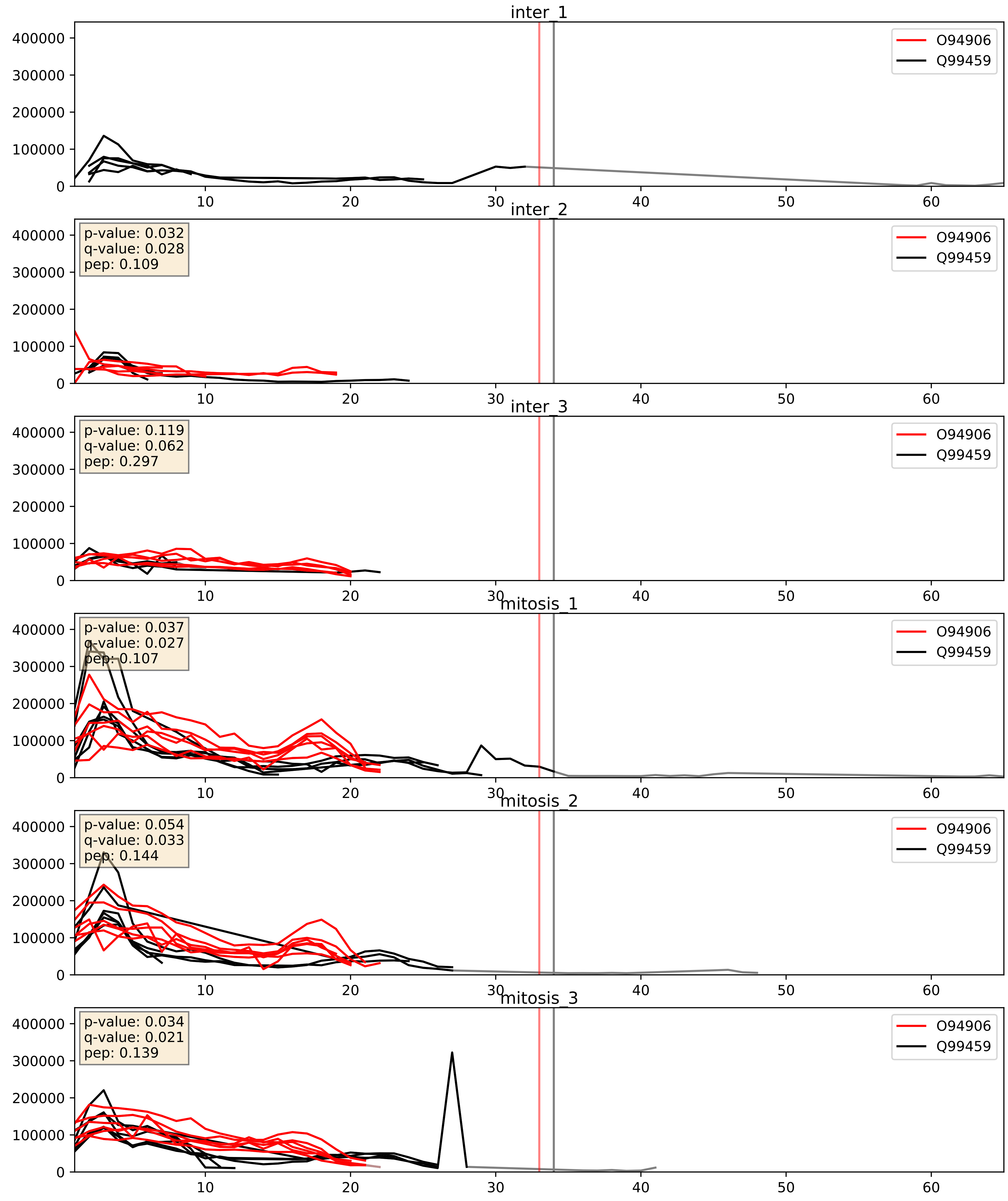


| level | condition_1 | condition_2 | pvalue | pvalue_adjusted |
|---|---|---|---|---|
| complex_abundance | mitosis | inter | 0.0063618354150012085 | 0.05948545934570088 |
| interactor_abundance | mitosis | inter | 0.014678960452509026 | 0.10605571190893752 |
| interactor_ratio | mitosis | inter | 0.9303107107087359 | 0.9878882449502933 |

P07910_Q99459
HNRPC_HUMAN vs CDC5L_HUMAN
p-value: 0.036 q-value: 0.025

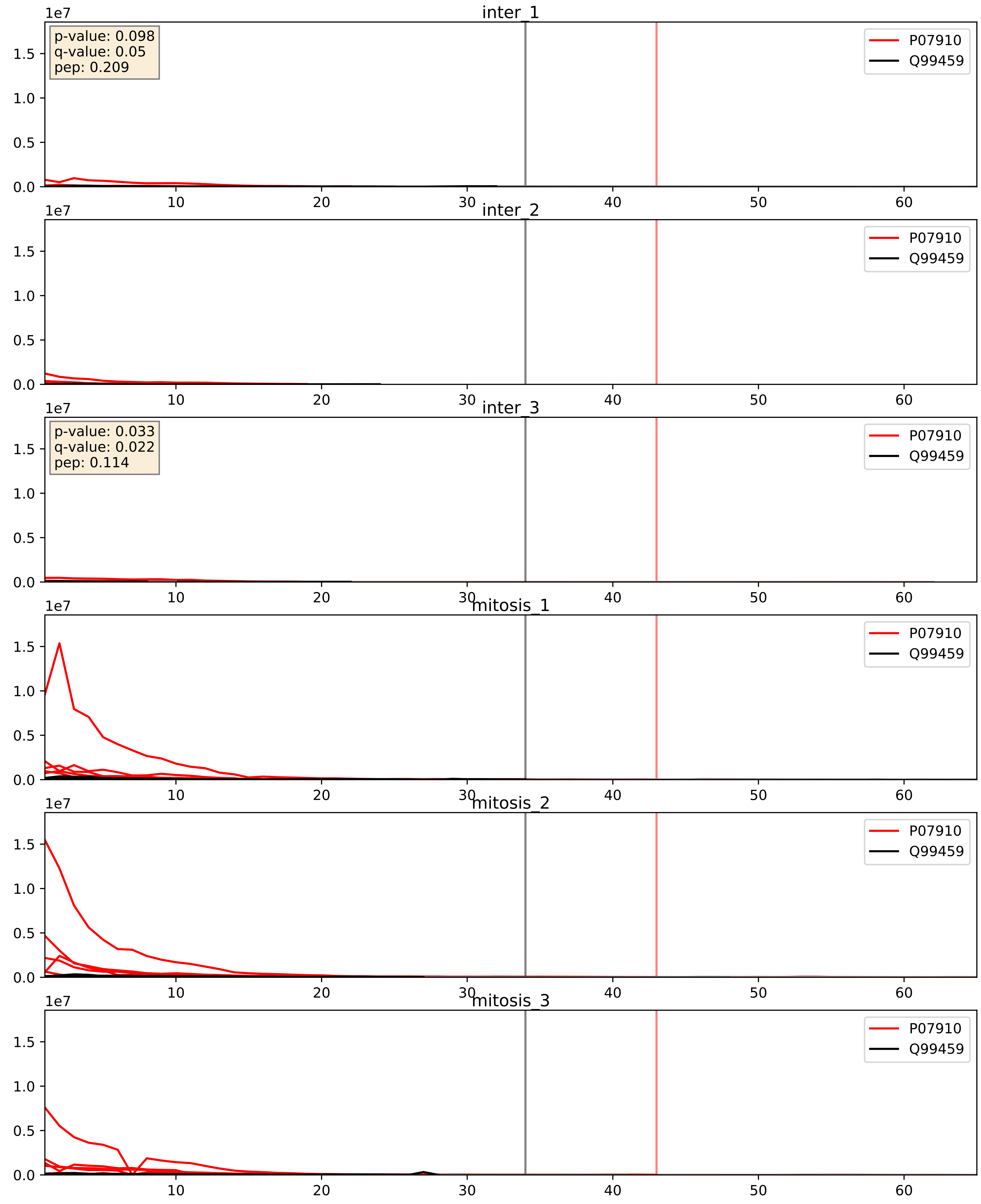


| level | condition_1 | condition_2 | pvalue | pvalue_adjusted |
|---|---|---|---|---|
| interactor_abundance | mitosis | inter | 7.673678712415949e-05 | 0.006950972463310108 |
| complex_abundance | mitosis | inter | 0.0006052291130691676 | 0.015834093001149168 |
| interactor_ratio | mitosis | inter | 0.0032529072203435253 | 0.07405554735675686 |

P17844_Q99459
DDX5_HUMAN vs CDC5L_HUMAN
p-value: 0.007 q-value: 0.006

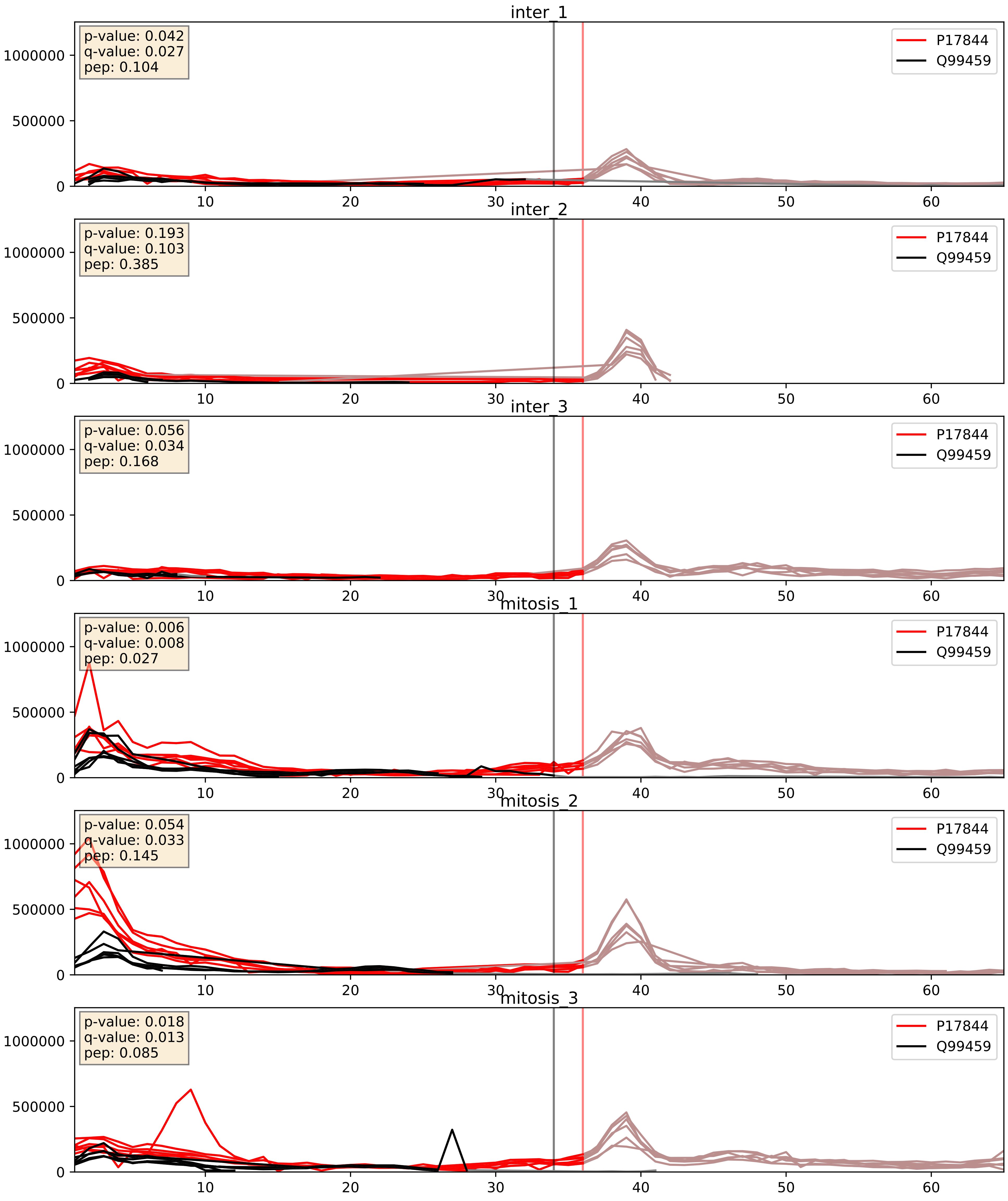


| level | condition_1 | condition_2 | pvalue | pvalue_adjusted |
|---|---|---|---|---|
| interactor_abundance | mitosis | inter | 0.006359400241119042 | 0.06830171400750187 |
| complex_abundance | mitosis | inter | 0.008211982545060673 | 0.06720322235728429 |
| interactor_ratio | mitosis | inter | 0.4083540240816134 | 0.8335535006745441 |

P31943_Q99459
HNRH1_HUMAN vs CDC5L_HUMAN
p-value: 0.066 q-value: 0.041

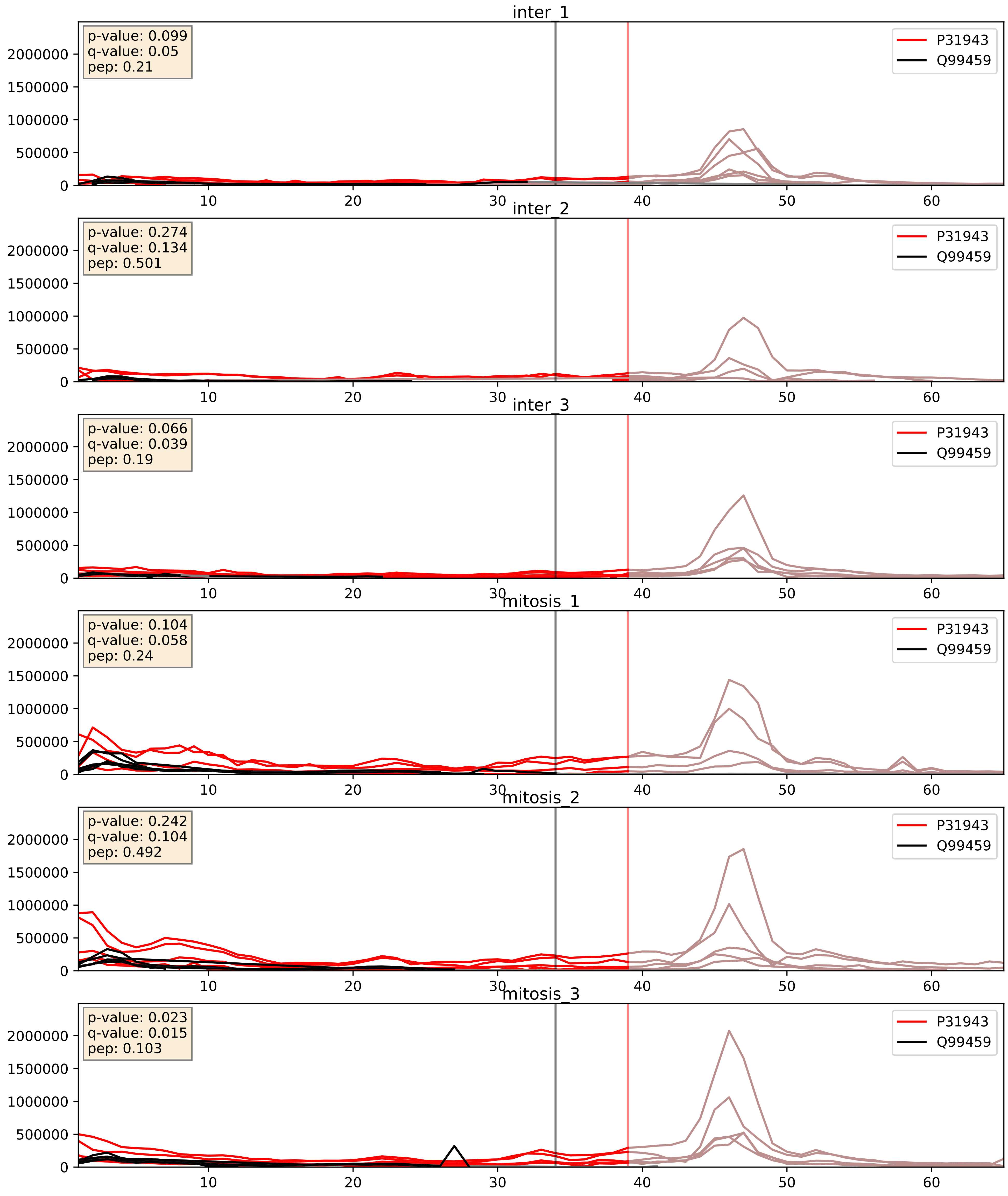


| level | condition_1 | condition_2 | pvalue | pvalue_adjusted |
|---|---|---|---|---|
| interactor_abundance | mitosis | inter | 0.0050208300212190625 | 0.05879385086407001 |
| complex_abundance | mitosis | inter | 0.008080613653928214 | 0.06653038413698939 |
| interactor_ratio | mitosis | inter | 0.6358488676126866 | 0.9197052662231202 |

P38159_Q99459
RBMX_HUMAN vs CDC5L_HUMAN
p-value: 0.036 q-value: 0.025

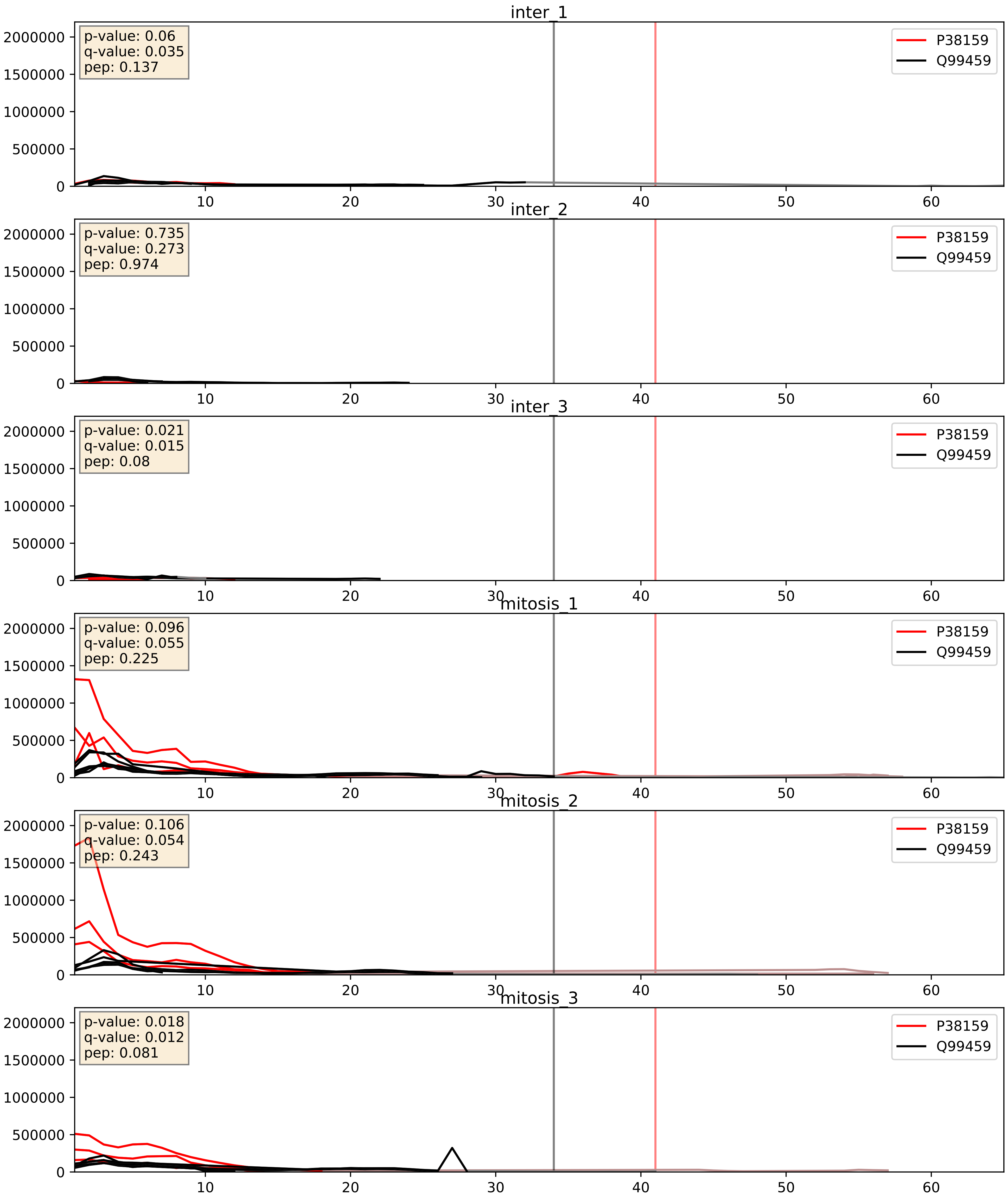


| level | condition_1 | condition_2 | pvalue | pvalue_adjusted |
|---|---|---|---|---|
| interactor_abundance | mitosis | inter | 0.0010224373165272904 | 0.0238801184978816 |
| complex_abundance | mitosis | inter | 0.0023534385428367085 | 0.035168510428347066 |
| interactor_ratio | mitosis | inter | 0.05399913247773674 | 0.38487308410443505 |

P40938_Q99459
RFC3_HUMAN vs CDC5L_HUMAN
p-value: 0.066 q-value: 0.041

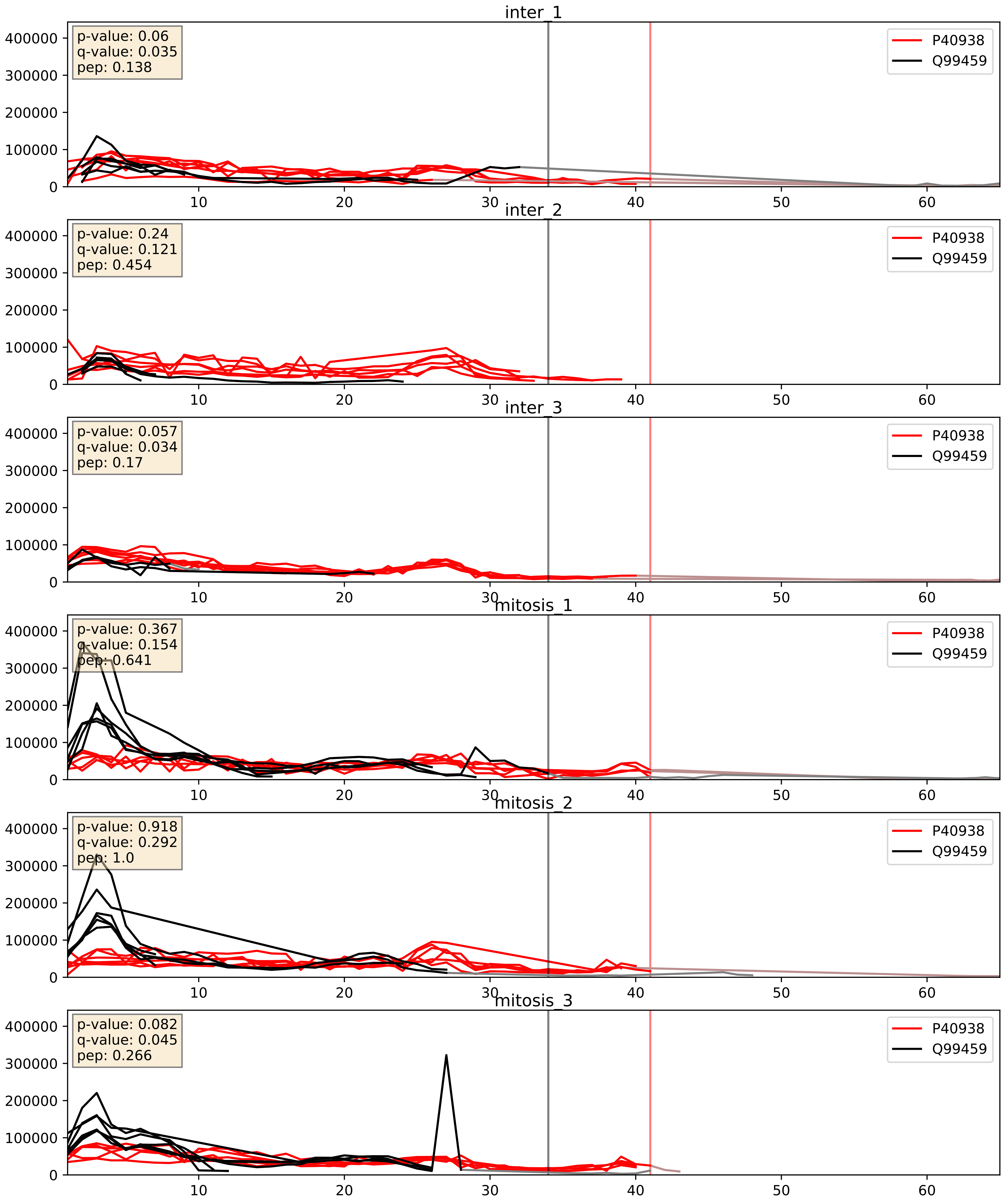


| level | condition_1 | condition_2 | pvalue | pvalue_adjusted |
|---|---|---|---|---|
| interactor_ratio | mitosis | inter | 0.0019257585573200047 | 0.05317578467954594 |
| complex_abundance | mitosis | inter | 0.05631926691748161 | 0.16922093580820607 |
| interactor_abundance | mitosis | inter | 0.7376704752982358 | 0.8375725253419417 |

P46087_Q99459
NOP2_HUMAN vs CDC5L_HUMAN
p-value: 0.017 q-value: 0.014

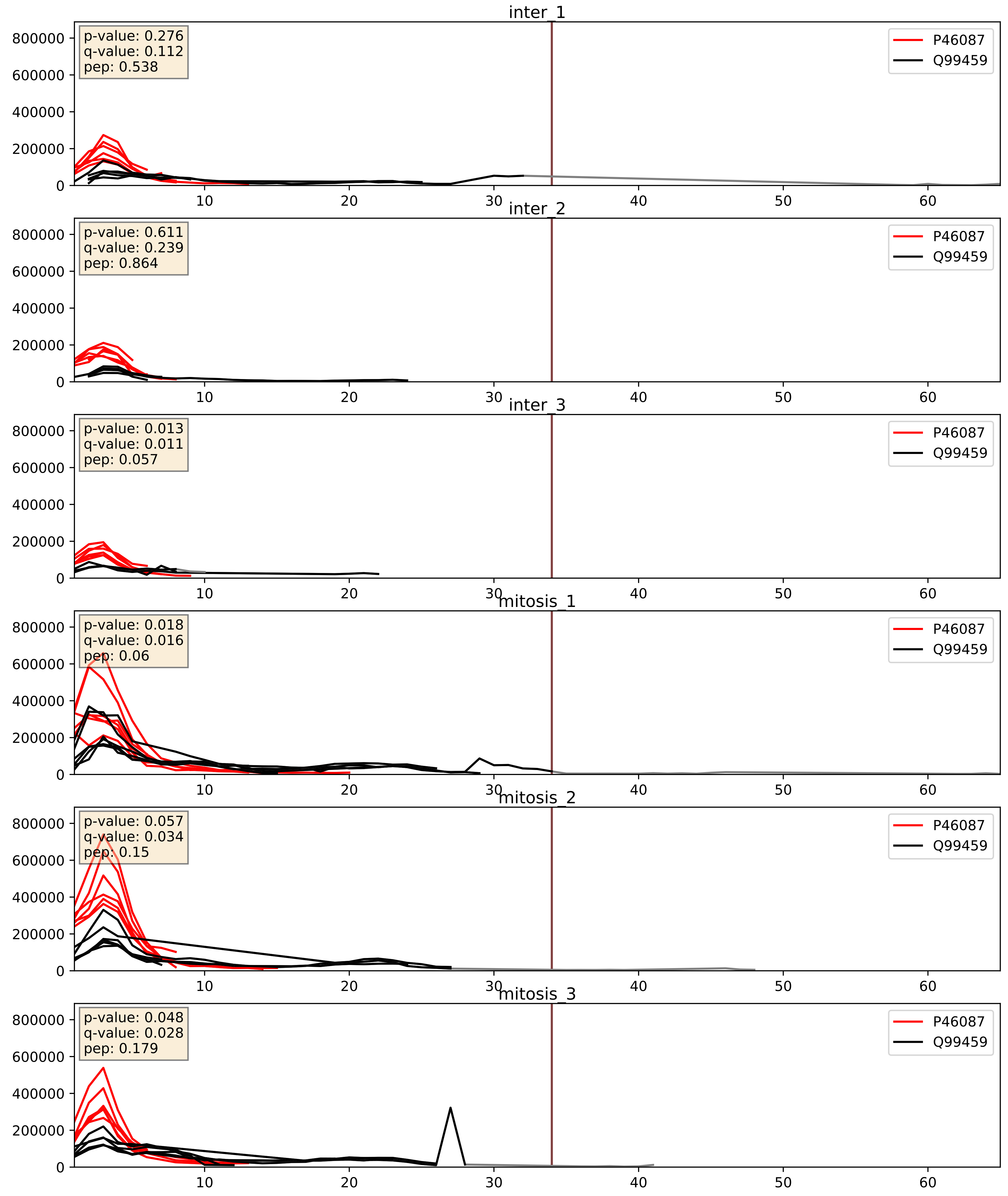


| level | condition_1 | condition_2 | pvalue | pvalue_adjusted |
|---|---|---|---|---|
| interactor_abundance | mitosis | inter | 0.008163501708093478 | 0.07851637597896649 |
| complex_abundance | mitosis | inter | 0.017128092485186742 | 0.09767919498547536 |
| interactor_ratio | mitosis | inter | 0.3721881012830839 | 0.8138388063636502 |

P52597_Q99459
HNRPF_HUMAN vs CDC5L_HUMAN
p-value: 0.016 q-value: 0.013

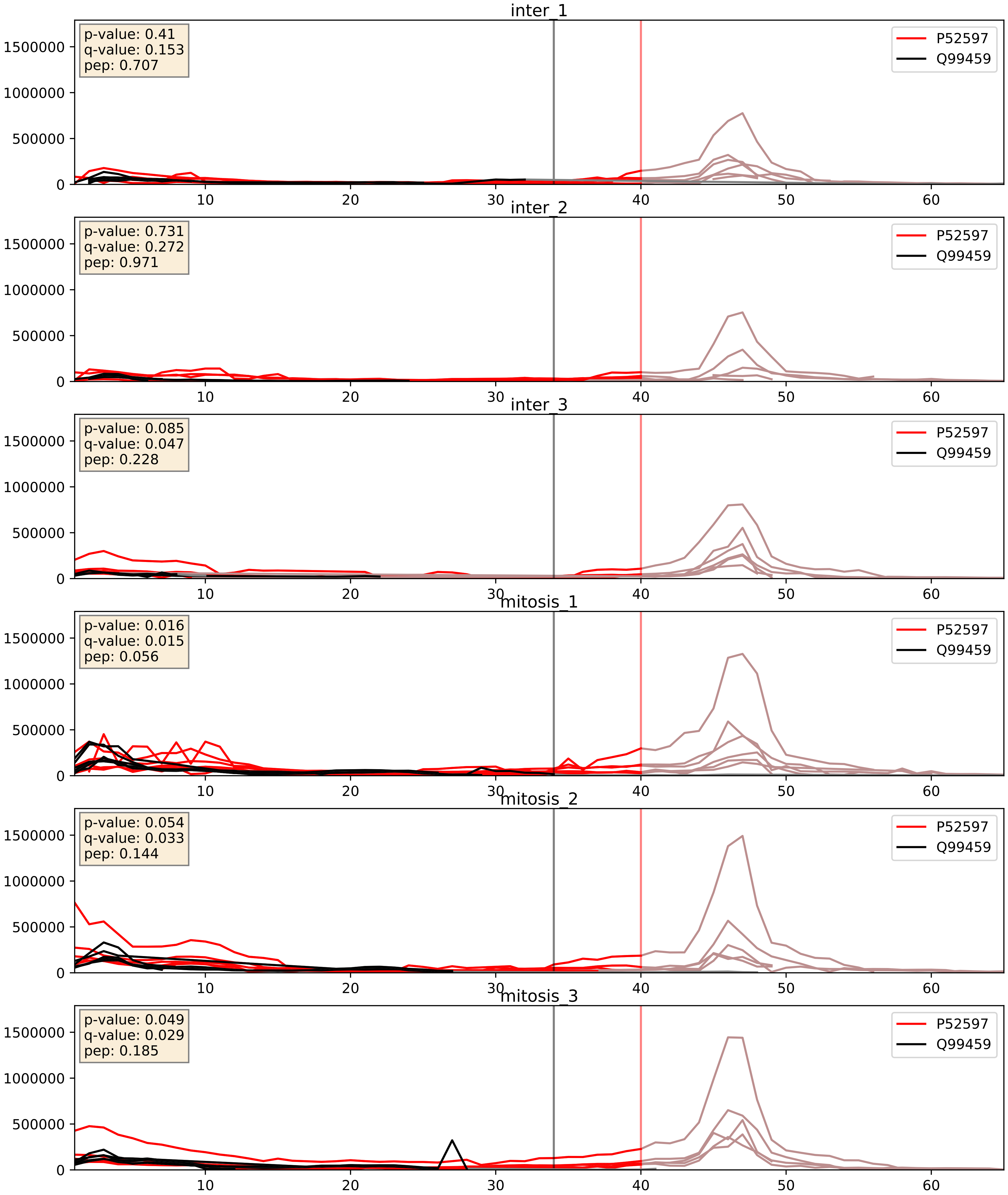


| level | condition_1 | condition_2 | pvalue | pvalue_adjusted |
|---|---|---|---|---|
| interactor_abundance | mitosis | inter | 0.005872037501971724 | 0.06494139666263302 |
| complex_abundance | mitosis | inter | 0.008301617280340168 | 0.06742444735376033 |
| interactor_ratio | mitosis | inter | 0.13750648150689043 | 0.5873558393508659 |

Q01780_Q99459
EXOSX_HUMAN vs CDC5L_HUMAN
p-value: 0.007 q-value: 0.006

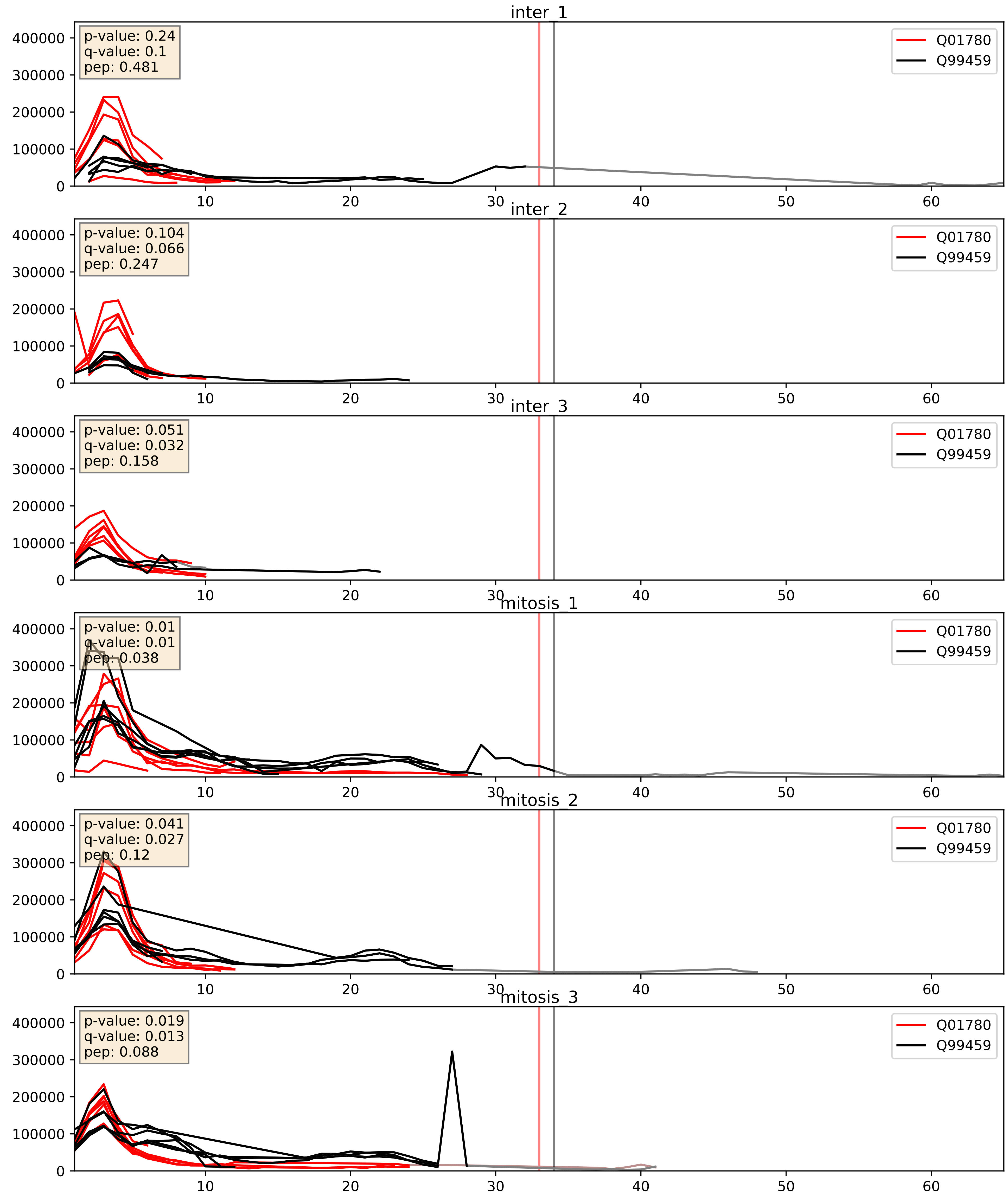


| level | condition_1 | condition_2 | pvalue | pvalue_adjusted |
|---|---|---|---|---|
| interactor_ratio | mitosis | inter | 0.010011115757435327 | 0.14263182888175546 |
| complex_abundance | mitosis | inter | 0.06311567682197815 | 0.1809950397306978 |
| interactor_abundance | mitosis | inter | 0.5783388797850517 | 0.722976387142759 |

Q07666_Q99459
KHDR1_HUMAN vs CDC5L_HUMAN
p-value: 0.054 q-value: 0.035

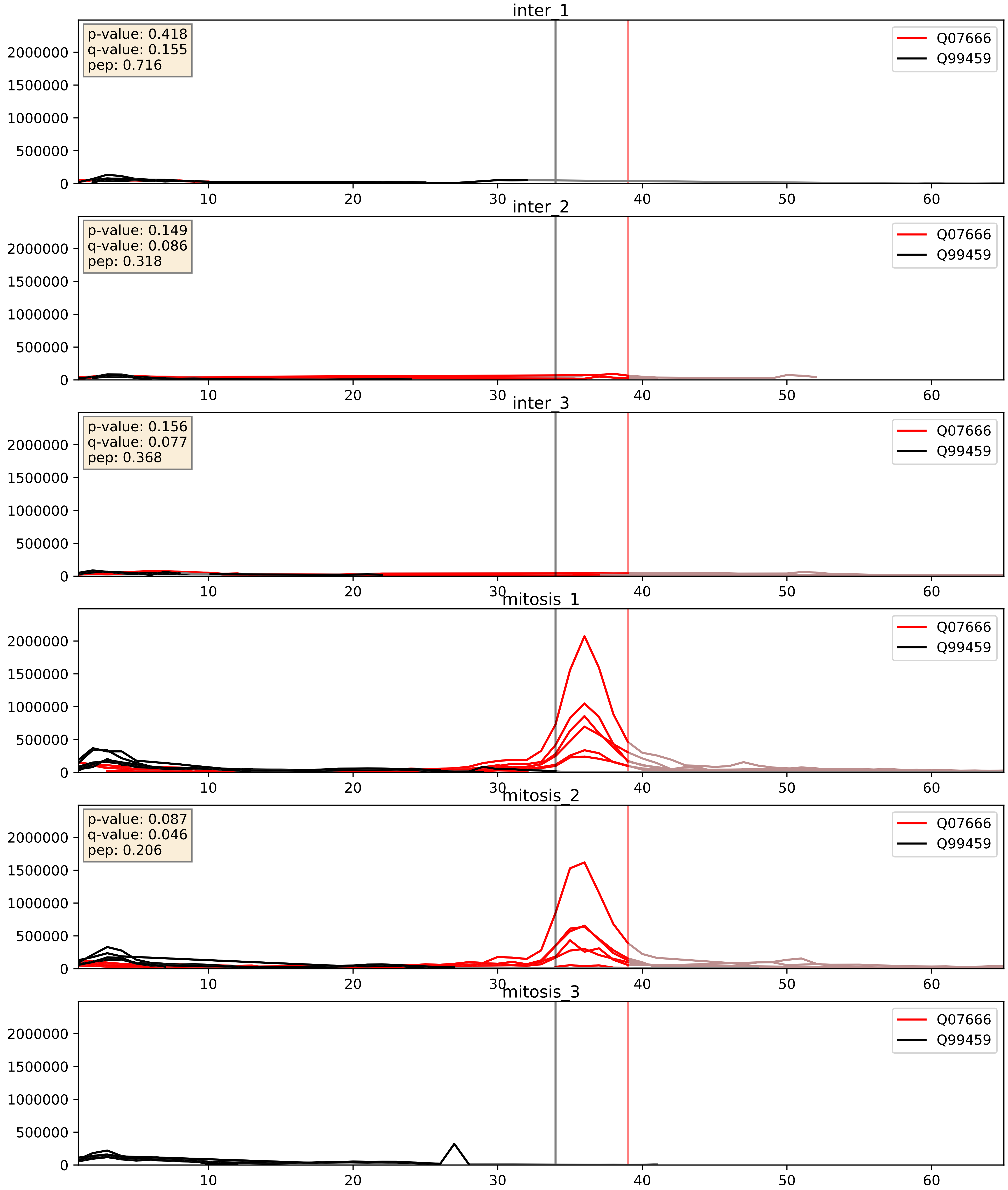


| level | condition_1 | condition_2 | pvalue | pvalue_adjusted |
|---|---|---|---|---|
| interactor_ratio | mitosis | inter | 0.4251517333812488 | 0.841246386050888 |
| complex_abundance | mitosis | inter | 0.6794414806022271 | 0.7860711585523025 |
| interactor_abundance | mitosis | inter | 0.832403034837705 | 0.9080953263506983 |

Q12906_Q99459
ILF3_HUMAN vs CDC5L_HUMAN
p-value: 0.033 q-value: 0.023

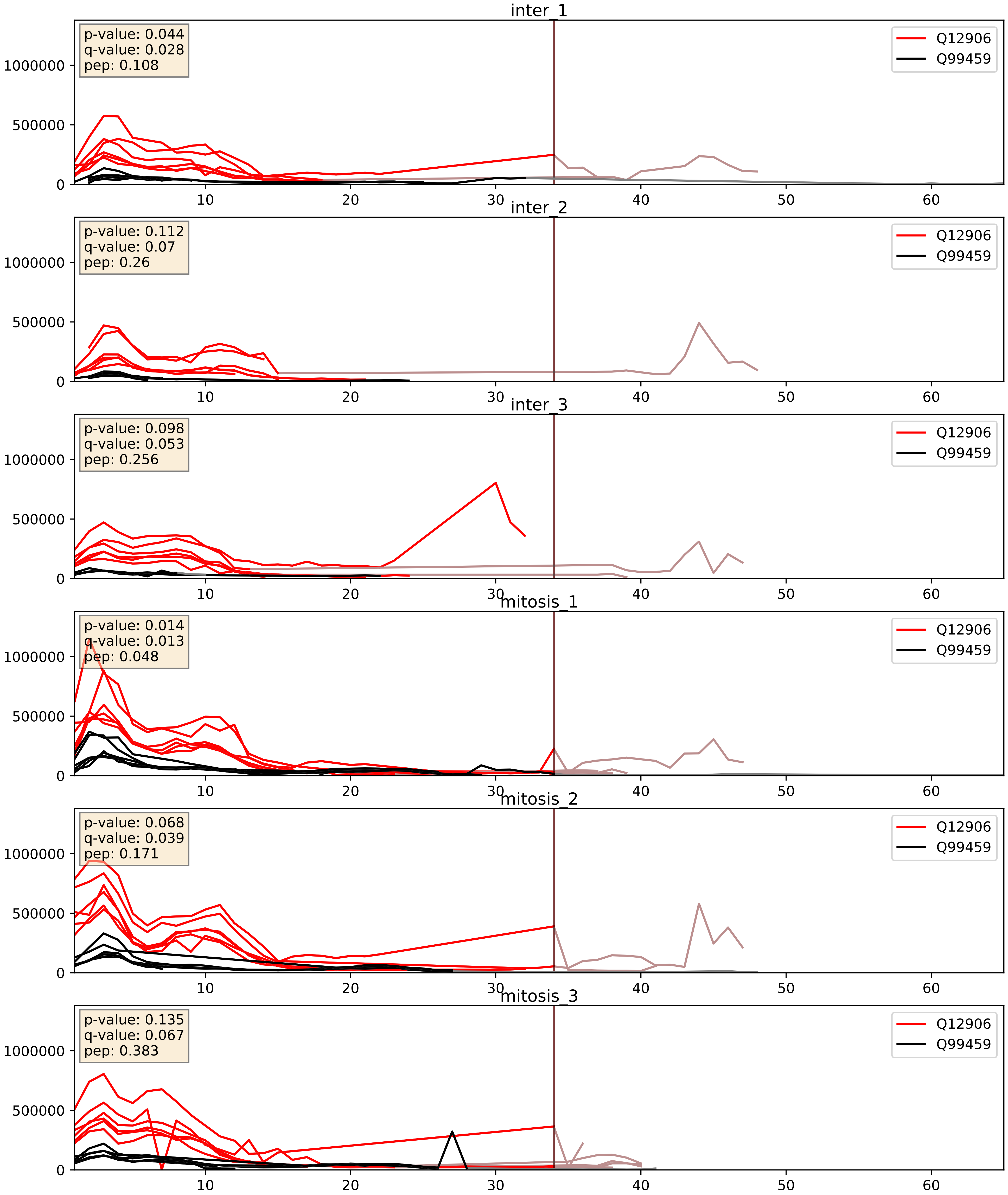


| level | condition_1 | condition_2 | pvalue | pvalue_adjusted |
|---|---|---|---|---|
| interactor_abundance | mitosis | inter | 0.008474381547771154 | 0.08017189393261118 |
| complex_abundance | mitosis | inter | 0.012844464879273177 | 0.08405635484106207 |
| interactor_ratio | mitosis | inter | 0.038268821630896746 | 0.3212185412334598 |

Q14498_Q99459
RBM39_HUMAN vs CDC5L_HUMAN
p-value: 0.013 q-value: 0.011

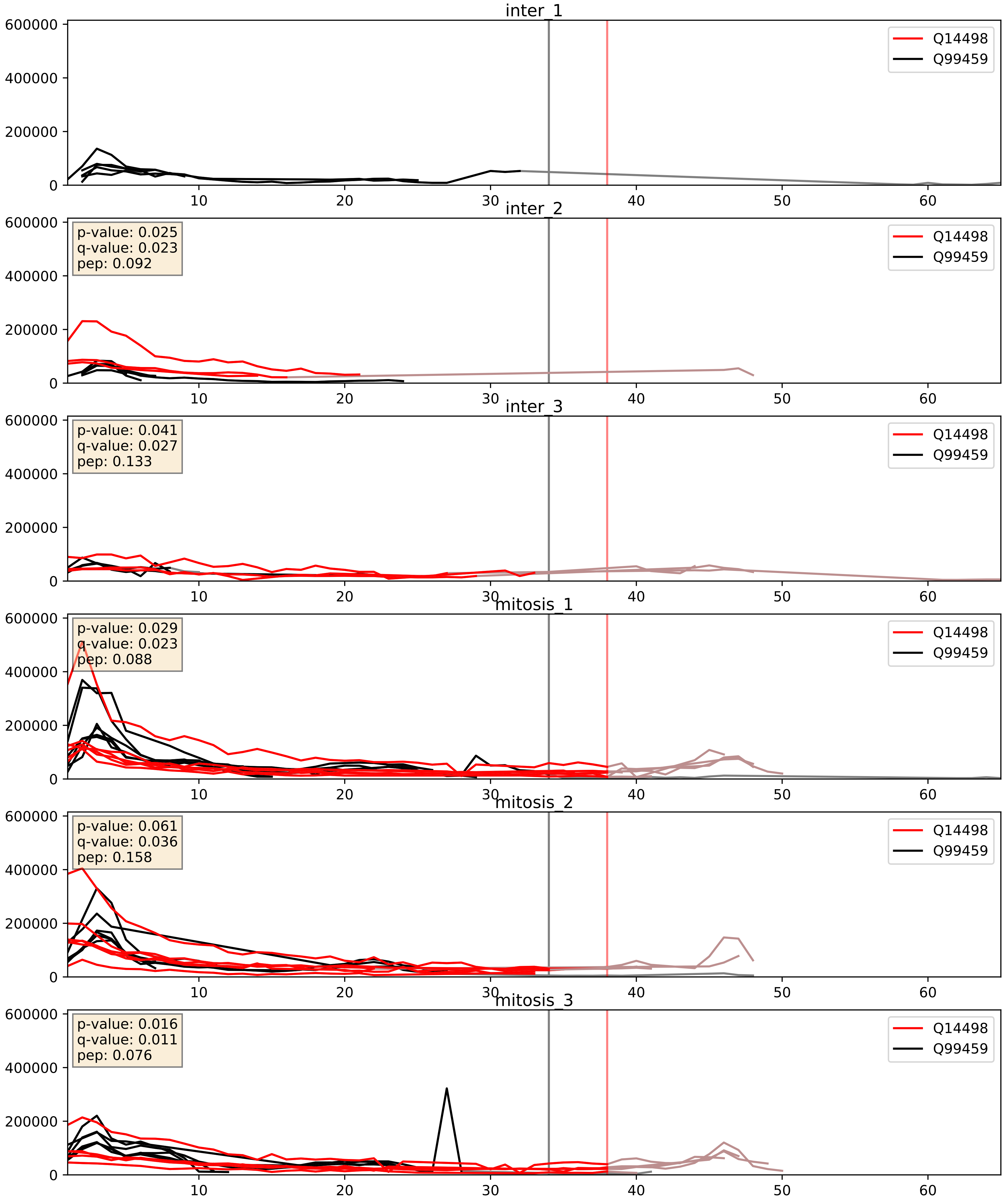


| level | condition_1 | condition_2 | pvalue | pvalue_adjusted |
|---|---|---|---|---|
| complex_abundance | mitosis | inter | 0.018459782234222916 | 0.10148730631017866 |
| interactor_abundance | mitosis | inter | 0.09910206679589971 | 0.2706939073957183 |
| interactor_ratio | mitosis | inter | 0.17627200921309574 | 0.6420034343641515 |

Q15717_Q99459
ELAV1_HUMAN vs CDC5L_HUMAN
p-value: 0.029 q-value: 0.021

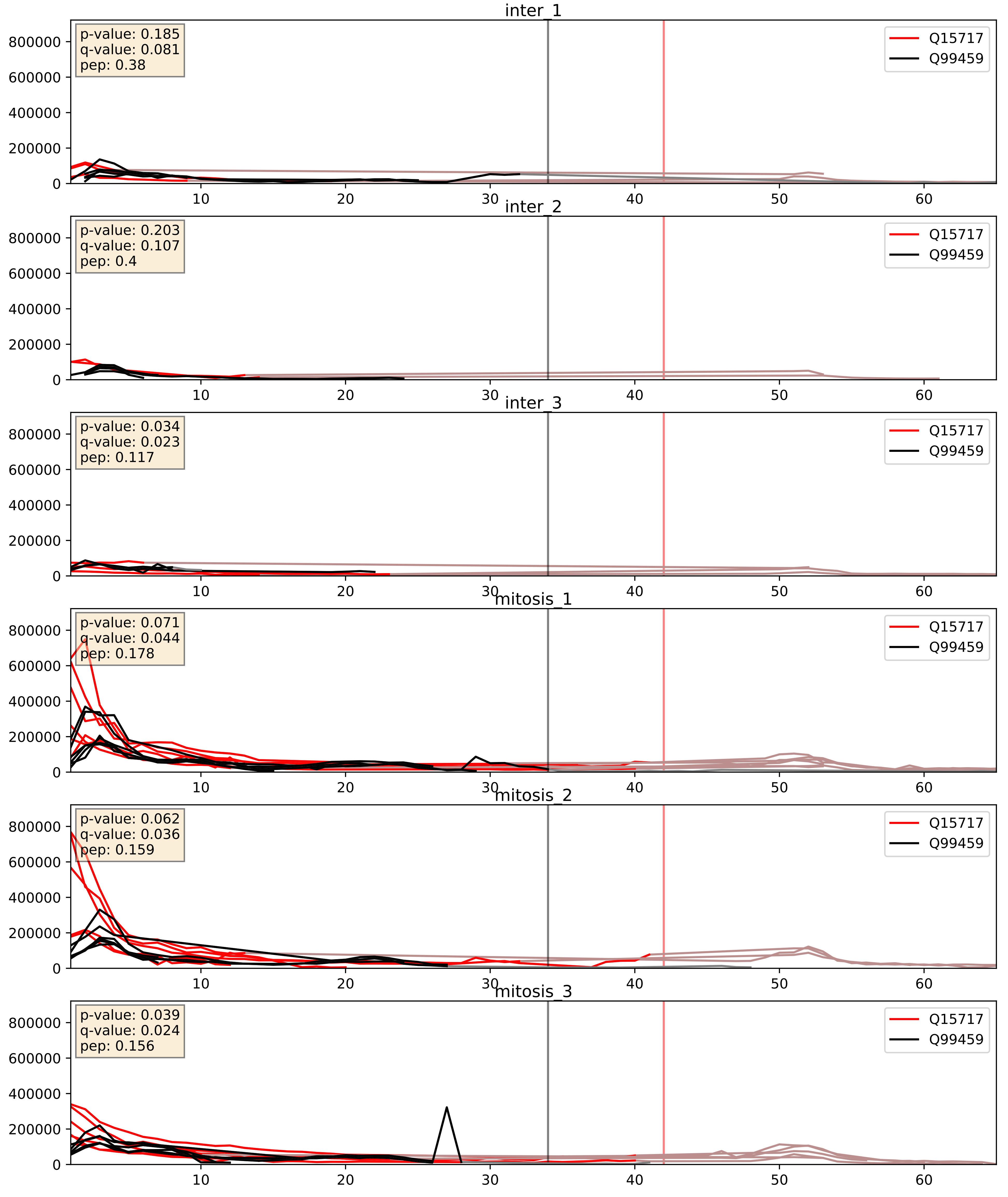


| level | condition_1 | condition_2 | pvalue | pvalue_adjusted |
|---|---|---|---|---|
| interactor_abundance | mitosis | inter | 0.00036534728597032297 | 0.01356427145975096 |
| complex_abundance | mitosis | inter | 0.002092462180363407 | 0.03329270681024306 |
| interactor_ratio | mitosis | inter | 0.046977446550768735 | 0.35838626305591875 |

Q16629_Q99459
SRSF7_HUMAN vs CDC5L_HUMAN
p-value: 0.073 q-value: 0.044

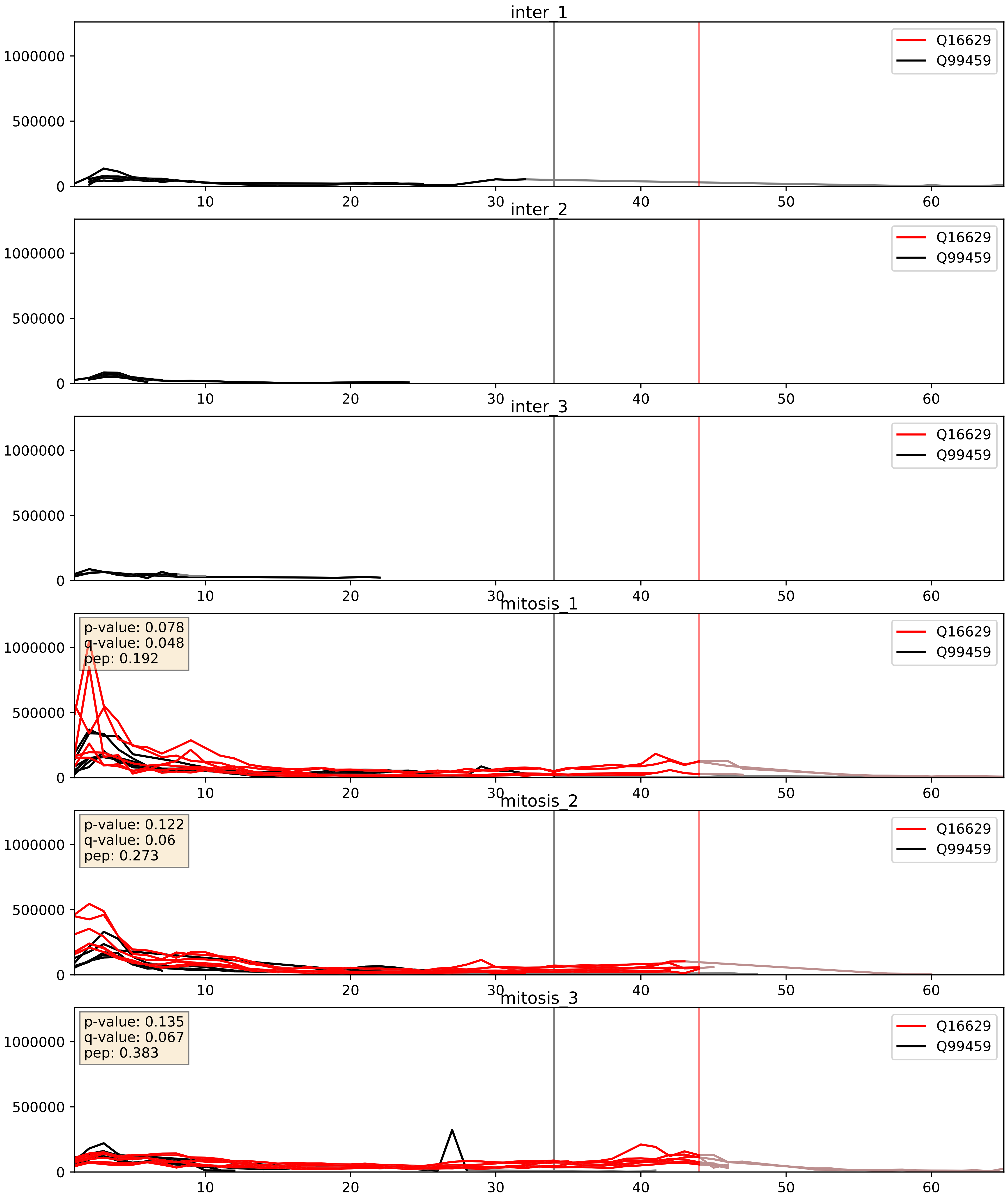


| level | condition_1 | condition_2 | pvalue | pvalue_adjusted |
|---|---|---|---|---|
| interactor_abundance | mitosis | inter | 0.00026344179809256095 | 0.01230990937725258 |
| complex_abundance | mitosis | inter | 0.0003628810828694355 | 0.012039775462644836 |
| interactor_ratio | mitosis | inter | 0.008617104652240045 | 0.13203195690147984 |

Q16637_Q99459
SMN_HUMAN vs CDC5L_HUMAN
p-value: 0.017 q-value: 0.014

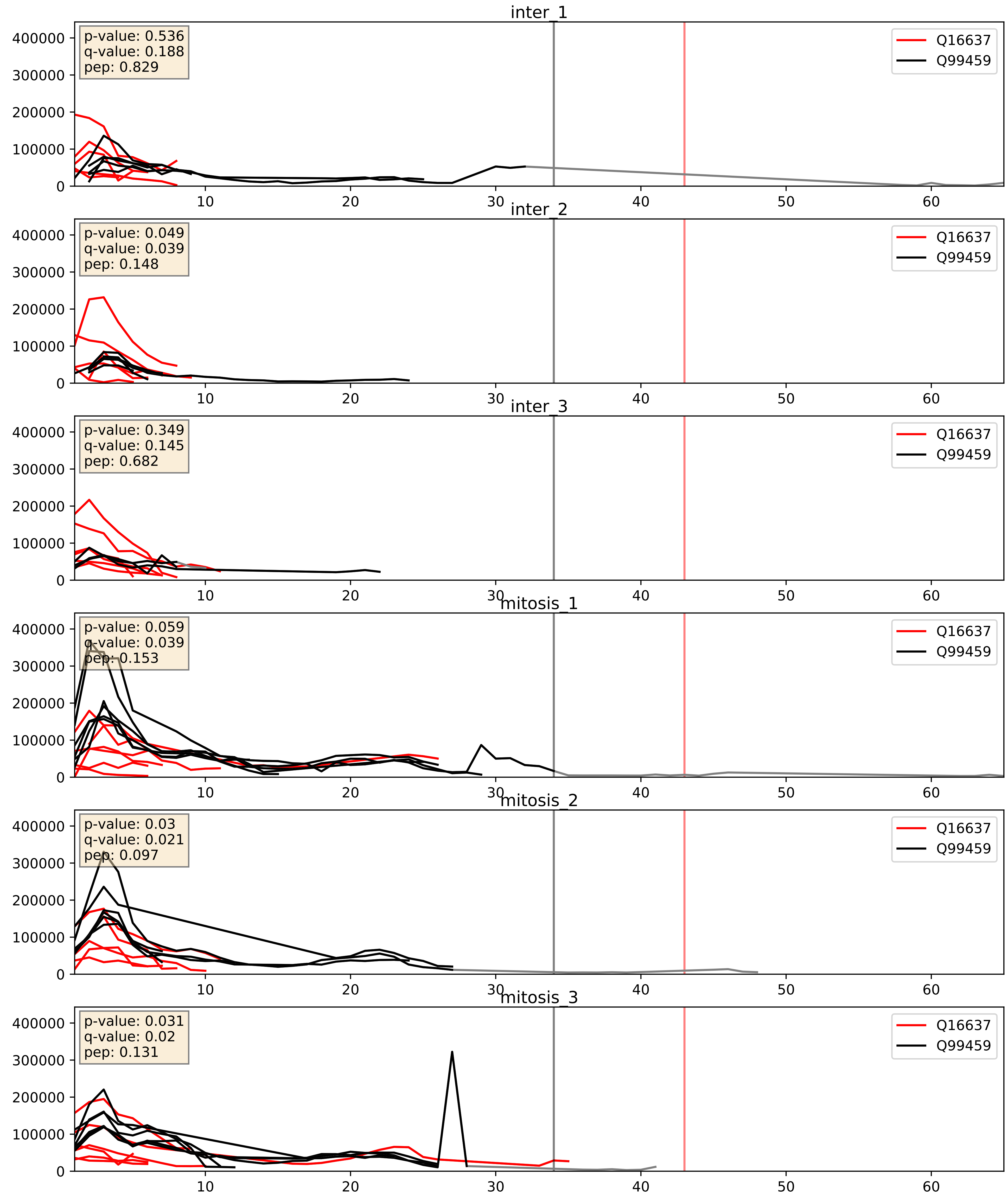


| level | condition_1 | condition_2 | pvalue | pvalue_adjusted |
|---|---|---|---|---|
| interactor_ratio | mitosis | inter | 0.020290232452811738 | 0.21527813309296567 |
| complex_abundance | mitosis | inter | 0.029789059513610345 | 0.12561298001798255 |
| interactor_abundance | mitosis | inter | 0.9572358081832746 | 0.9737300675043173 |

Q53GS9_Q99459
SNUT2_HUMAN vs CDC5L_HUMAN
p-value: 0.037 q-value: 0.026

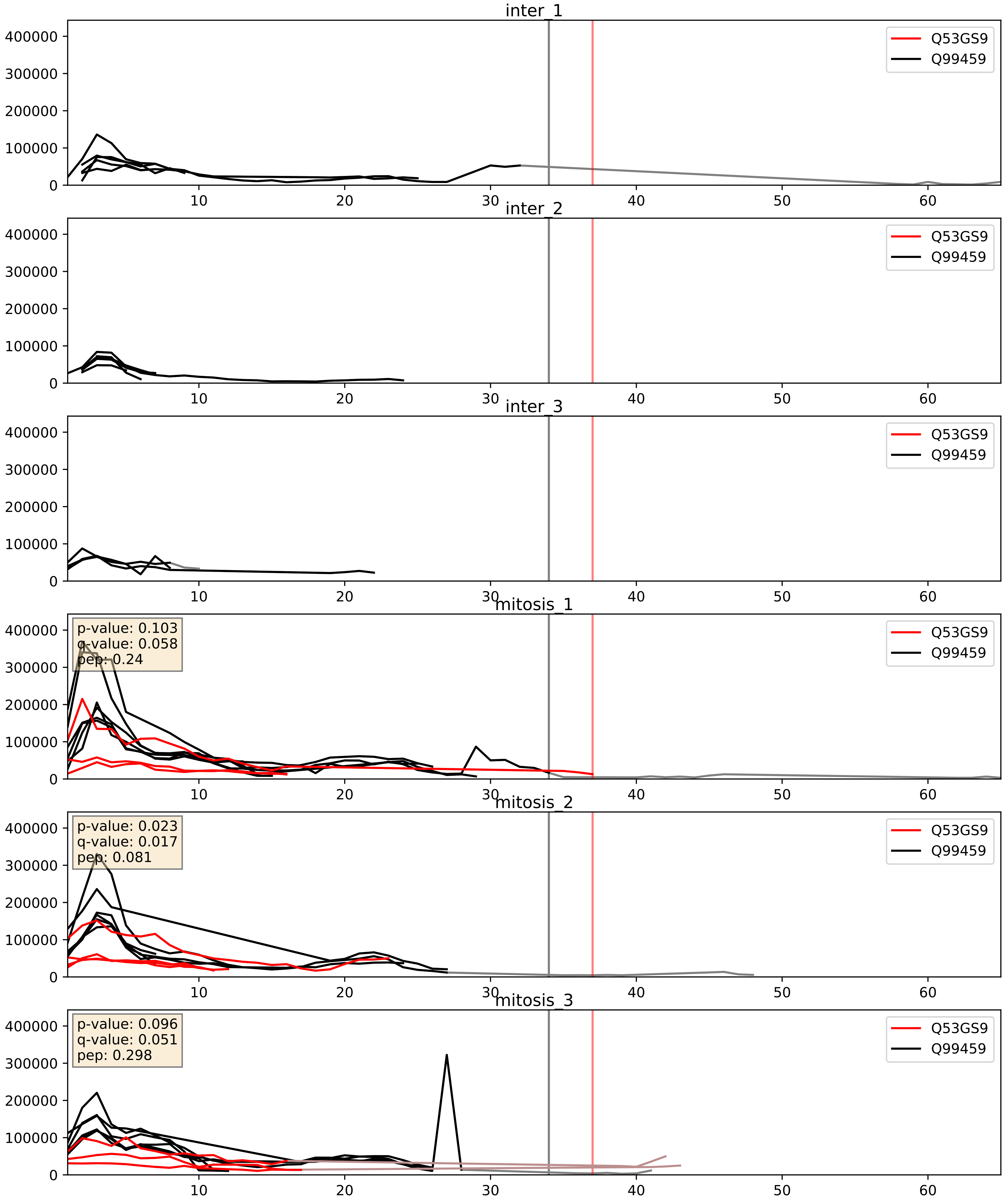


| level | condition_1 | condition_2 | pvalue | pvalue_adjusted |
|---|---|---|---|---|
| complex_abundance | mitosis | inter | 0.0010493796329760585 | 0.021612845224852777 |
| interactor_abundance | mitosis | inter | 0.0013451098335009928 | 0.027124005123129562 |
| interactor_ratio | mitosis | inter | 0.0020791470592695536 | 0.05614352942380876 |

Q8IWX8_Q99459
CHERP_HUMAN vs CDC5L_HUMAN
p-value: 0.082 q-value: 0.049

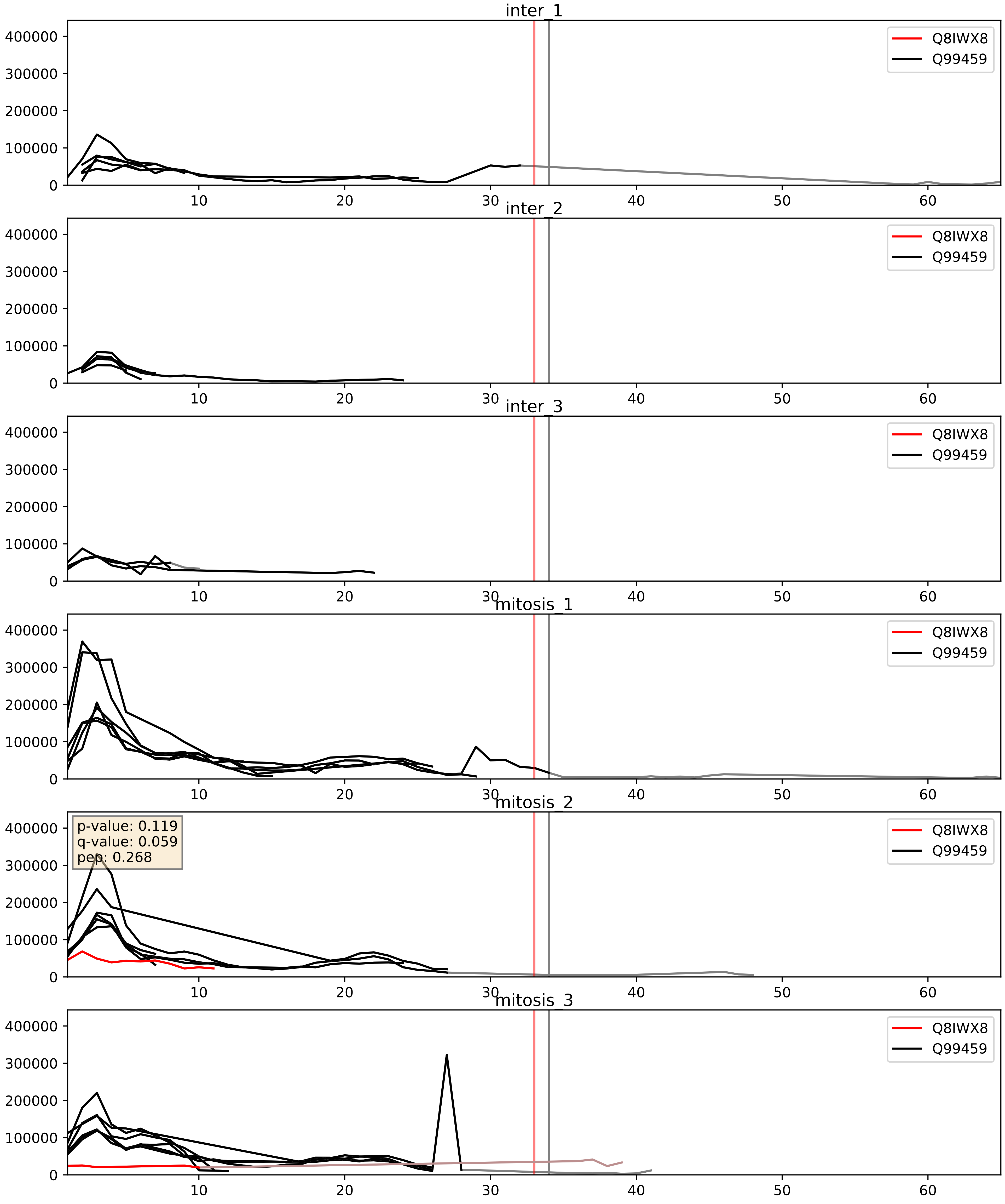


| level | condition_1 | condition_2 | pvalue | pvalue_adjusted |
|---|---|---|---|---|
| interactor_ratio | mitosis | inter | 0.1844272894939733 | 0.654246828872114 |
| complex_abundance | mitosis | inter | 0.24822890504445727 | 0.3985817721216572 |
| interactor_abundance | mitosis | inter | 0.5338195881679695 | 0.6924002840696748 |

Q99459_Q9BTC0
CDC5L_HUMAN vs DIDO1_HUMAN
p-value: 0.074 q-value: 0.045

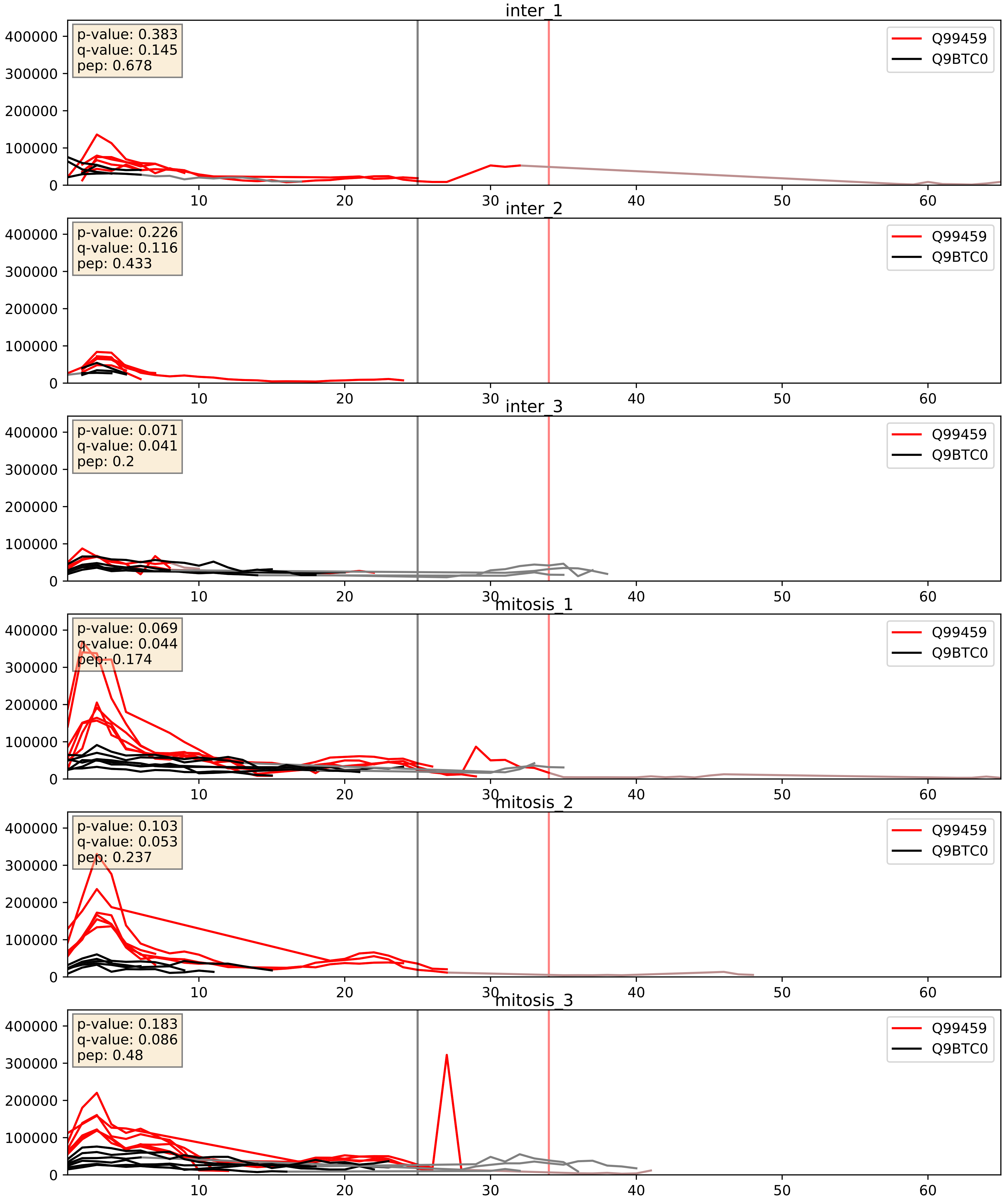


| level | condition_1 | condition_2 | pvalue | pvalue_adjusted |
|---|---|---|---|---|
| interactor_ratio | mitosis | inter | 0.011777070205603453 | 0.15667724901772678 |
| interactor_abundance | mitosis | inter | 0.016259697584858145 | 0.11171268728092203 |
| complex_abundance | mitosis | inter | 0.052794919303294864 | 0.1641728469817564 |

Q99459_Q9BZJ0
CDC5L_HUMAN vs CRNL1_HUMAN
p-value: 0.019 q-value: 0.015

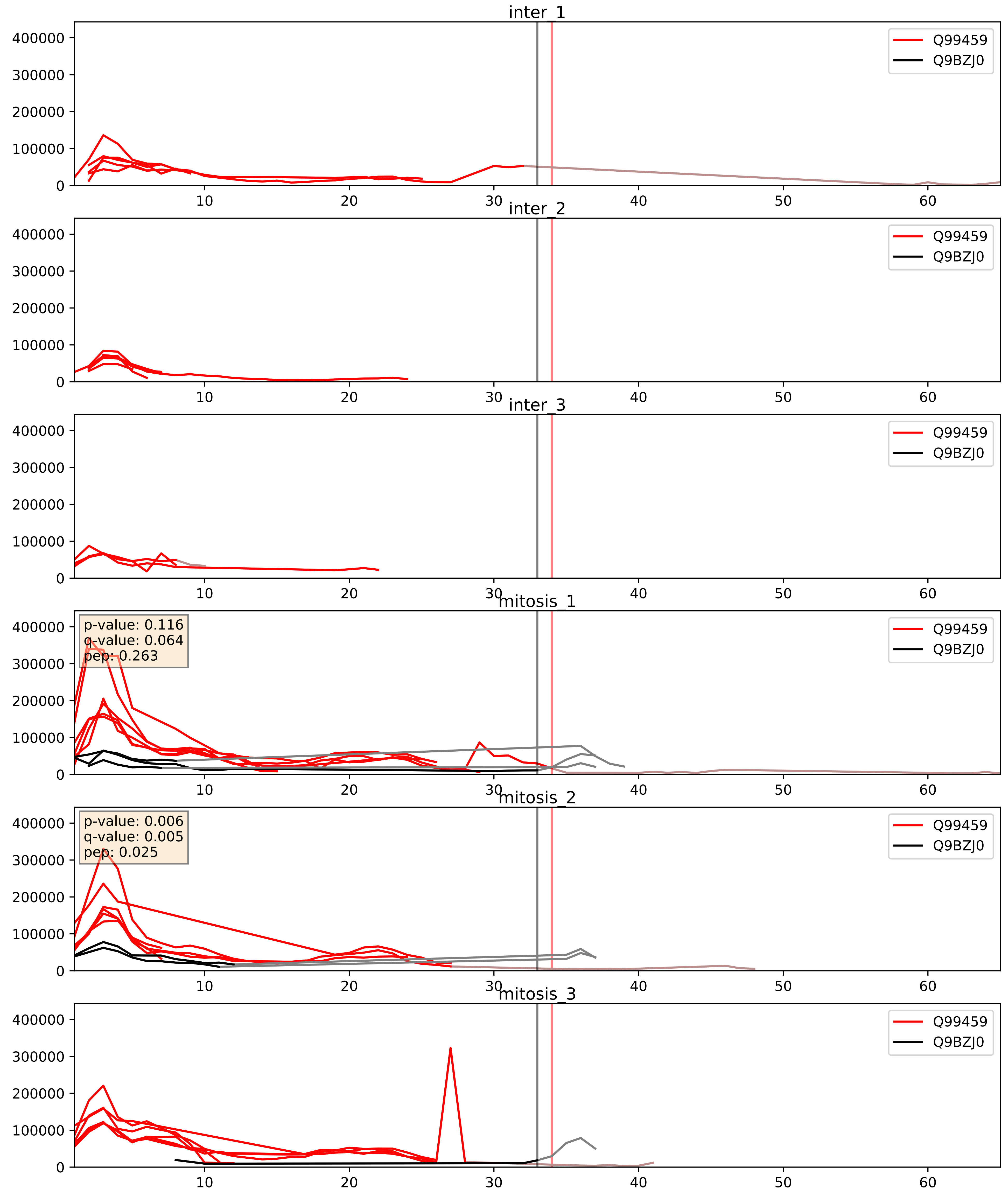


| level | condition_1 | condition_2 | pvalue | pvalue_adjusted |
|---|---|---|---|---|
| interactor_ratio | mitosis | inter | 0.022207834723738464 | 0.22875940461516397 |
| interactor_abundance | mitosis | inter | 0.14787363404624873 | 0.33422995072666123 |
| complex_abundance | mitosis | inter | 0.26937271491850834 | 0.41886111529562786 |

Q99459_Q9NR30
CDC5L_HUMAN vs DDX21_HUMAN
p-value: 0.016 q-value: 0.013

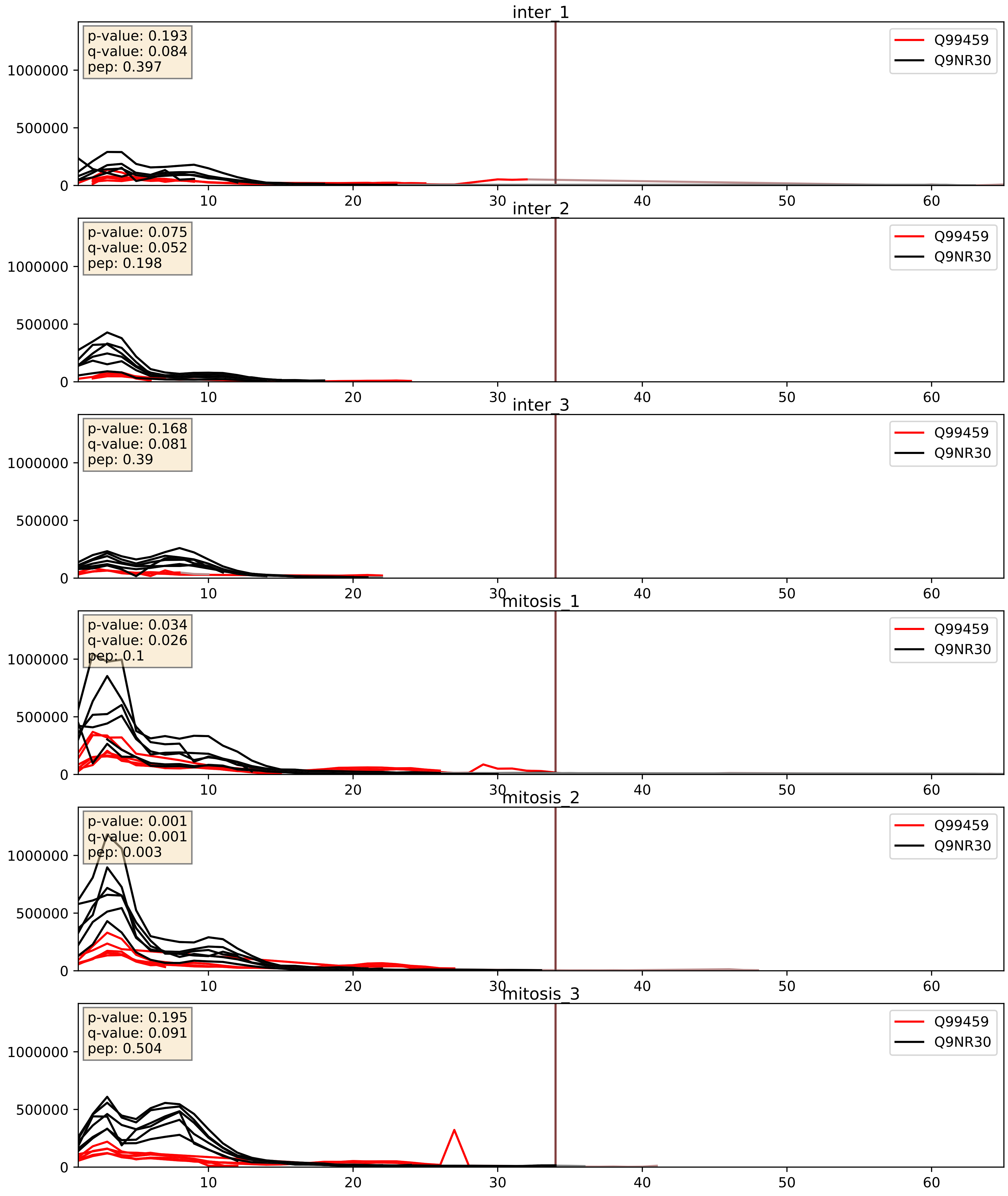


| level | condition_1 | condition_2 | pvalue | pvalue_adjusted |
|---|---|---|---|---|
| complex_abundance | mitosis | inter | 0.006365618502101007 | 0.05948545934570088 |
| interactor_abundance | mitosis | inter | 0.013030345214924086 | 0.10247106572324315 |
| interactor_ratio | mitosis | inter | 0.17752272019023063 | 0.6441689210802773 |

Q99459_Q9NTJ3
CDC5L_HUMAN vs SMC4_HUMAN
p-value: 0.077 q-value: 0.046

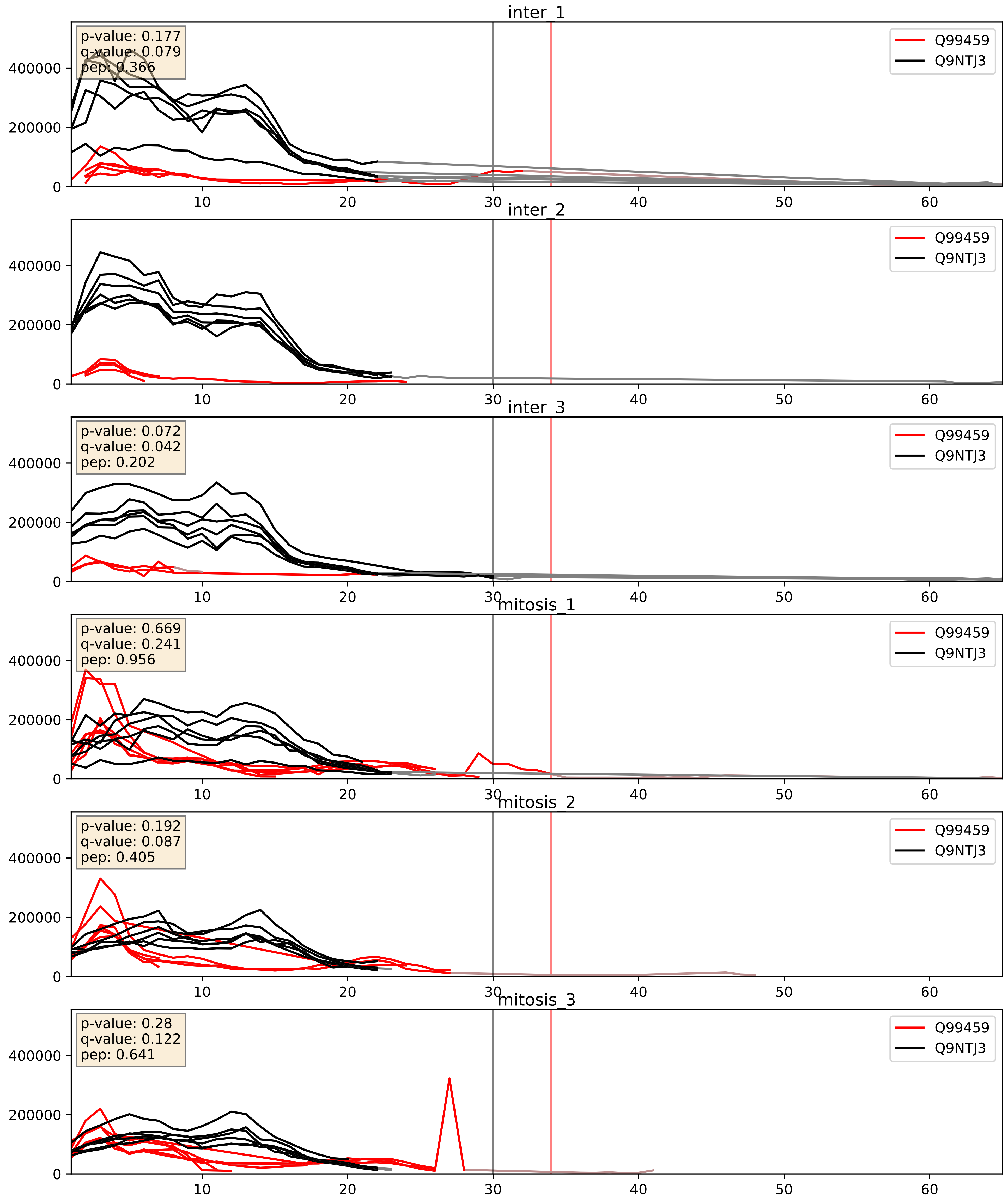


| level | condition_1 | condition_2 | pvalue | pvalue_adjusted |
|---|---|---|---|---|
| interactor_ratio | mitosis | inter | 7.860354077288164e-05 | 0.009476708577688264 |
| interactor_abundance | mitosis | inter | 0.01804490281291386 | 0.11569520685146502 |
| complex_abundance | mitosis | inter | 0.28283495543741954 | 0.4313400092130783 |

Q99459_Q9P2I0
CDC5L_HUMAN vs CPSF2_HUMAN
p-value: 0.017 q-value: 0.014

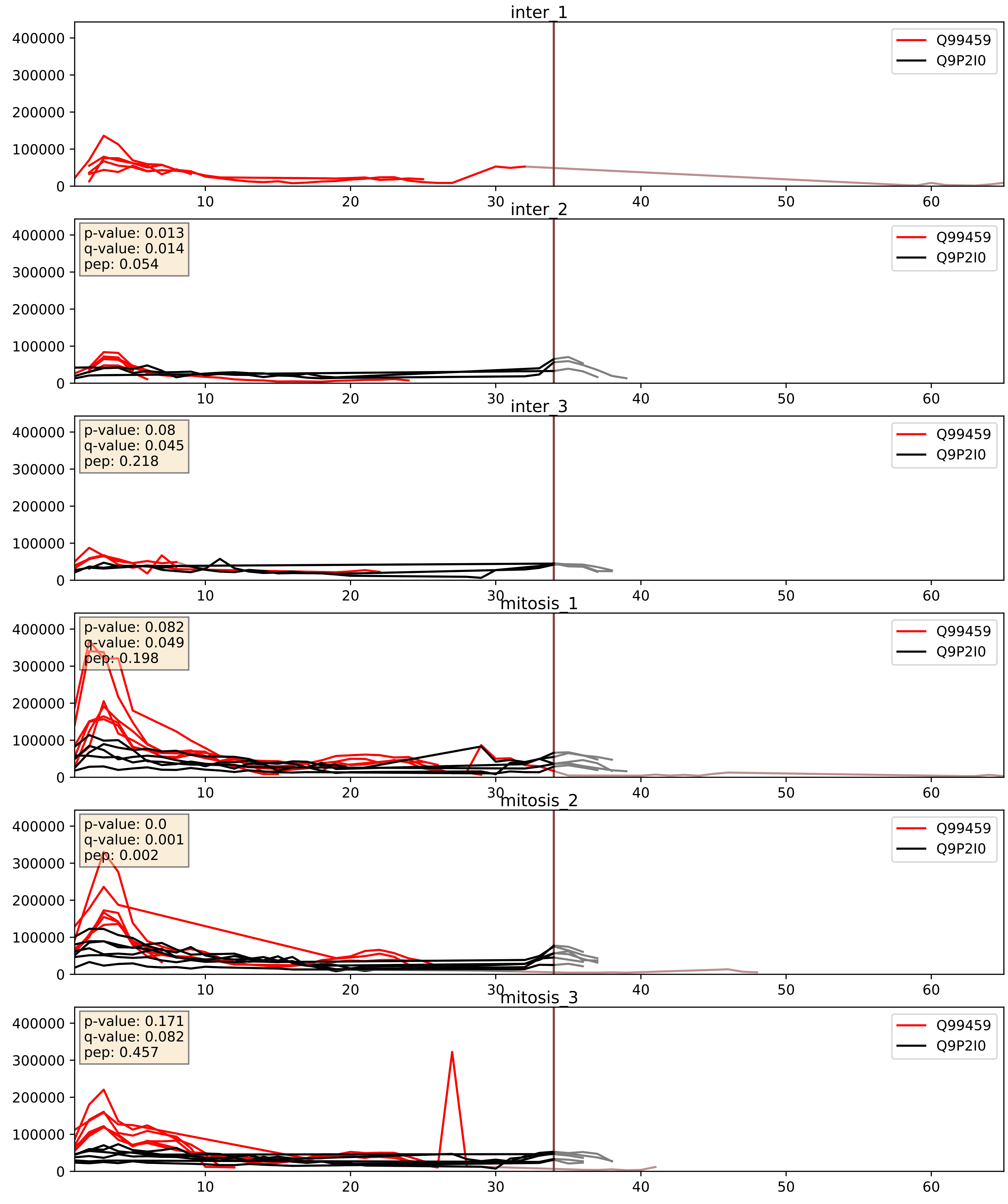


| level | condition_1 | condition_2 | pvalue | pvalue_adjusted |
|---|---|---|---|---|
| interactor_abundance | mitosis | inter | 0.0054405300799866795 | 0.06238571665731544 |
| complex_abundance | mitosis | inter | 0.009246753476483927 | 0.07106059027160733 |
| interactor_ratio | mitosis | inter | 0.011940403429490735 | 0.15773125517969241 |

Q99459_Q9UKF6
CDC5L_HUMAN vs CPSF3_HUMAN
p-value: 0.036 q-value: 0.025

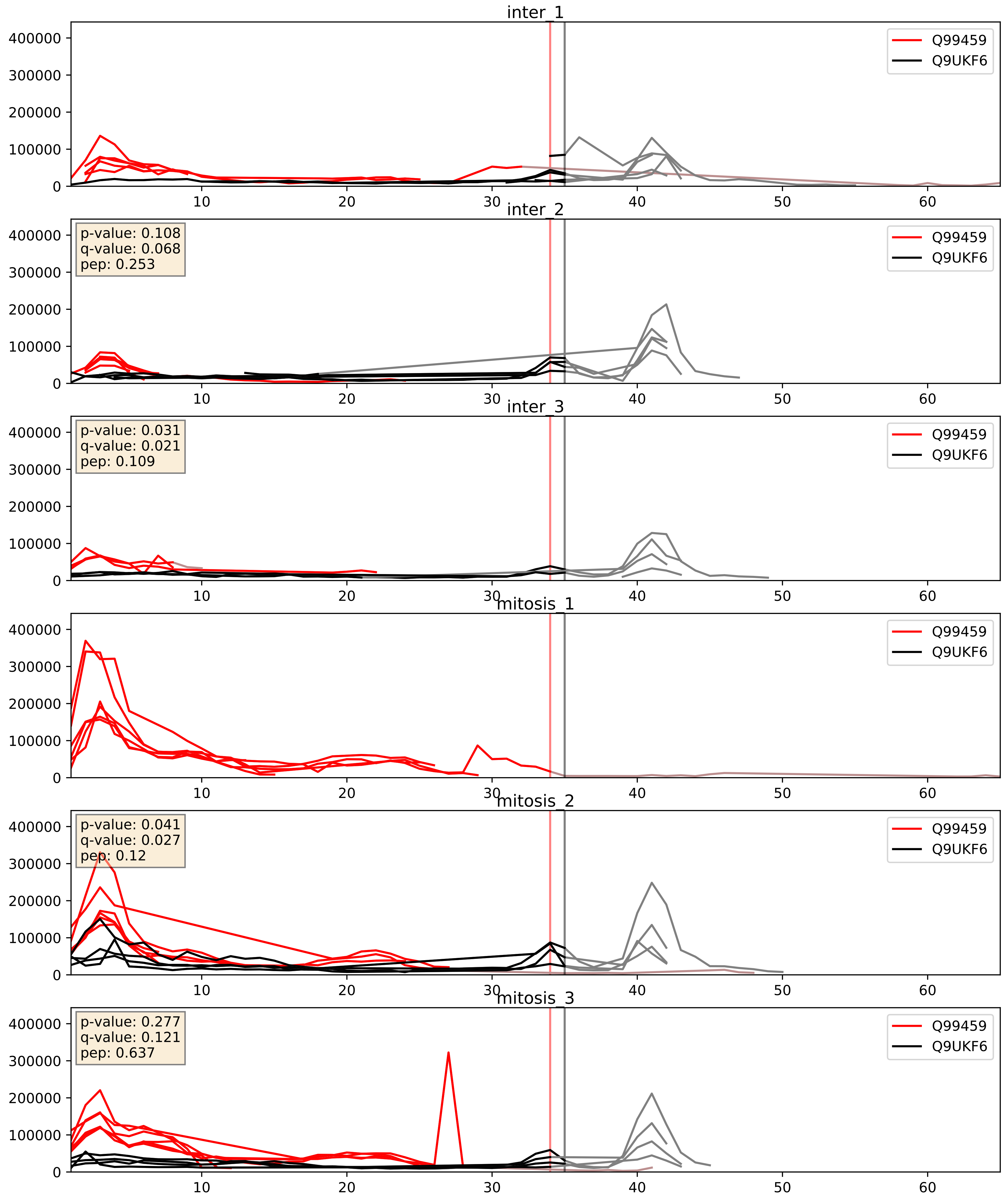


| level | condition_1 | condition_2 | pvalue | pvalue_adjusted |
|---|---|---|---|---|
| interactor_ratio | mitosis | inter | 0.10869958575448795 | 0.5473405393374283 |
| interactor_abundance | mitosis | inter | 0.5435347363676594 | 0.7002268557915816 |
| complex_abundance | mitosis | inter | 0.8315264262008489 | 0.890846834578131 |
