## Supplementary Data 2 for "SECAT: Quantifying differential protein-protein interaction states by network-centric analysis": hela_string_000030_O94906.pdf

O94906_O94906

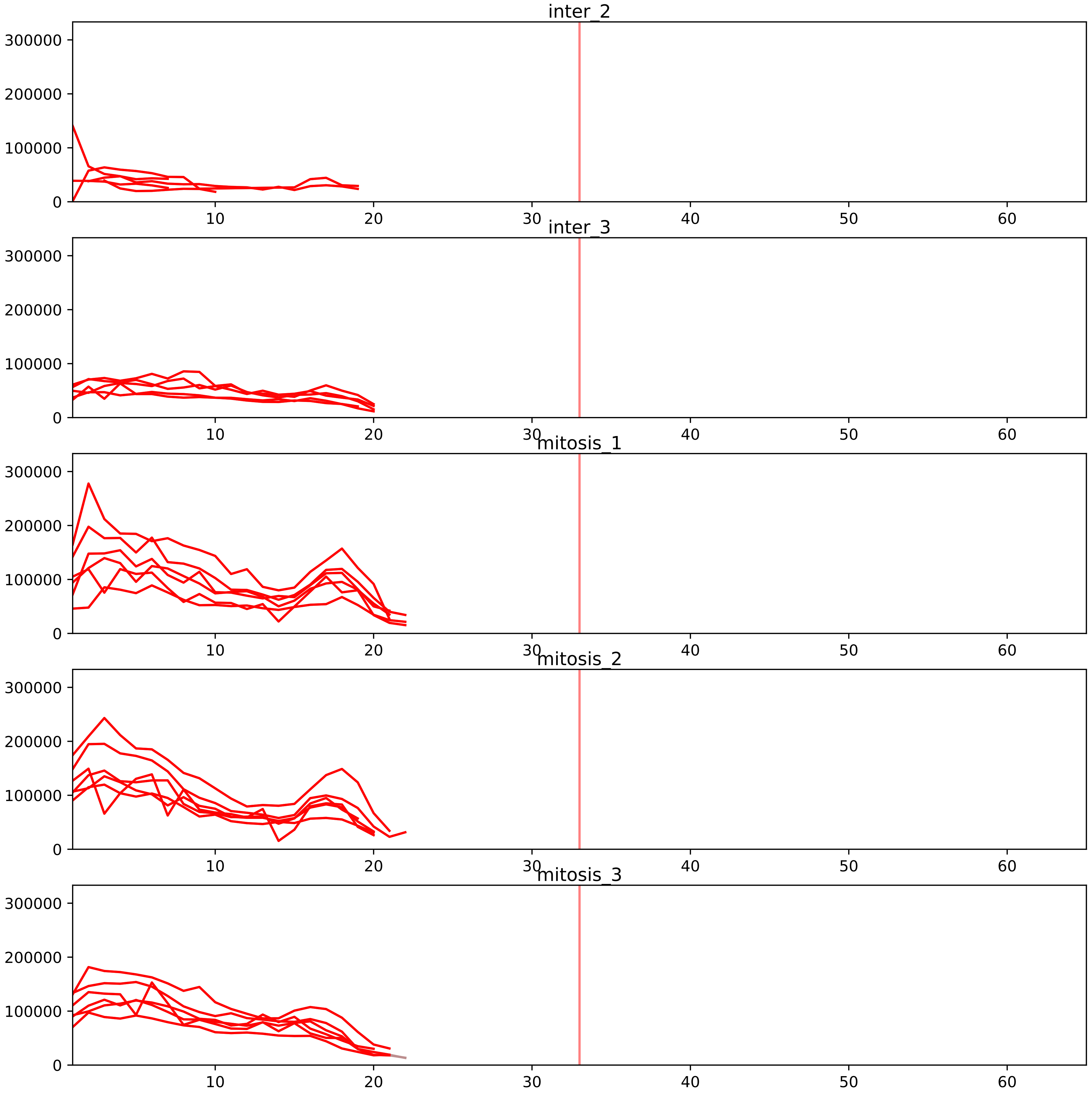


| level | condition_1 | condition_2 | pvalue | pvalue_adjusted |
|---|---|---|---|---|
| interactor_ratio | mitosis | inter | 8.087930149973278e-10 | 5.131652233086494e-08 |
| interactor_abundance | mitosis | inter | 8.293814753132651e-05 | 0.0020079762033900104 |
| complex_abundance | mitosis | inter | 0.00021243120657083453 | 0.003794887573692578 |
| total_abundance | mitosis | inter | 0.07383002876185442 | 0.33710195195210835 |
| assembled_abundance | mitosis | inter | 0.0862577503474618 | 0.34422210518028024 |

O43172_O94906
PRP4_HUMAN vs PRP6_HUMAN
p-value: 0.0 q-value: 0.001

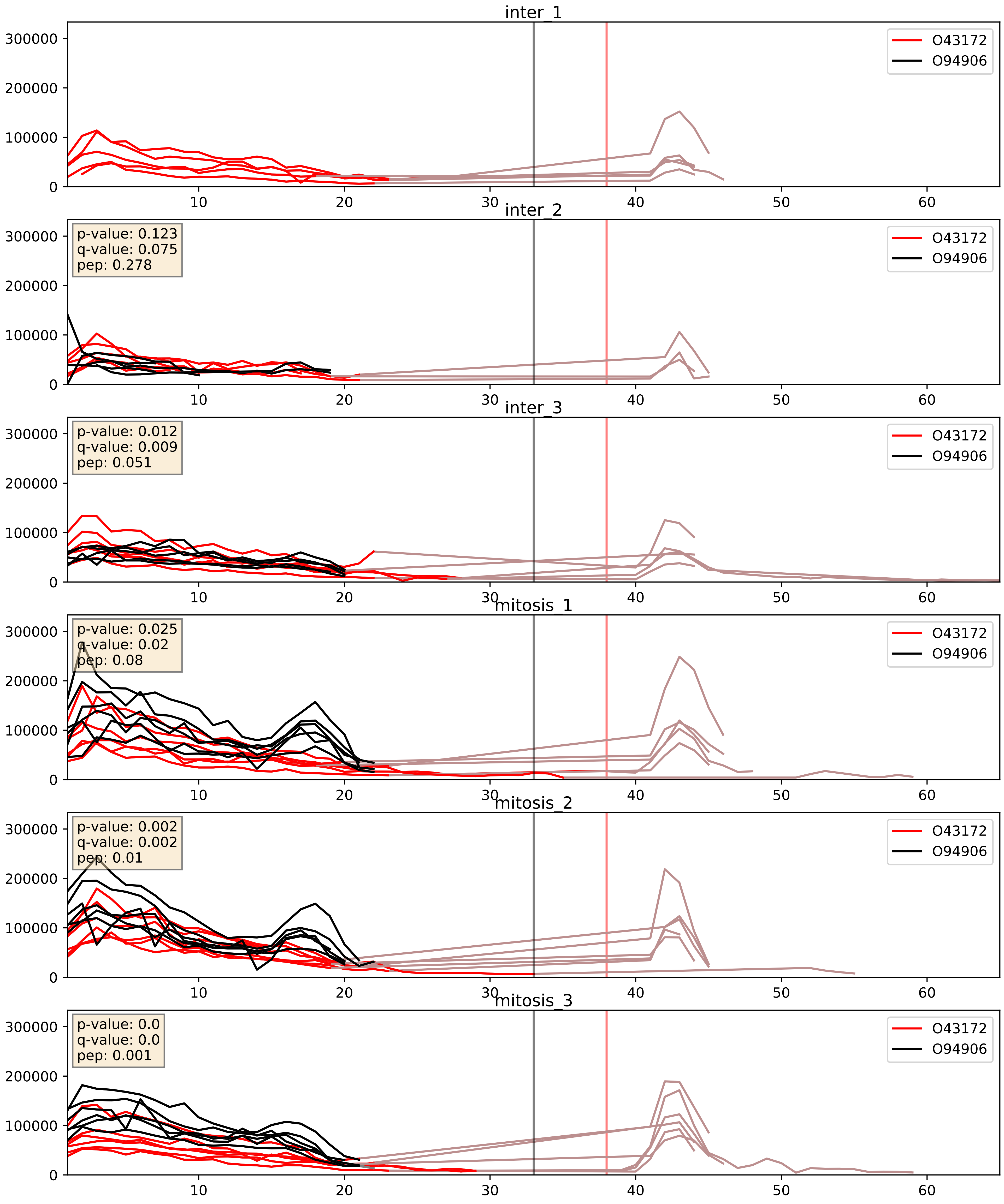


| level | condition_1 | condition_2 | pvalue | pvalue_adjusted |
|---|---|---|---|---|
| interactor_ratio | mitosis | inter | 0.013233768720388672 | 0.16610126135854406 |
| complex_abundance | mitosis | inter | 0.07775569399842704 | 0.20317116624741619 |
| interactor_abundance | mitosis | inter | 0.13641860484415502 | 0.32358874830969575 |

O43290_O94906
SNUT1_HUMAN vs PRP6_HUMAN
p-value: 0.024 q-value: 0.019

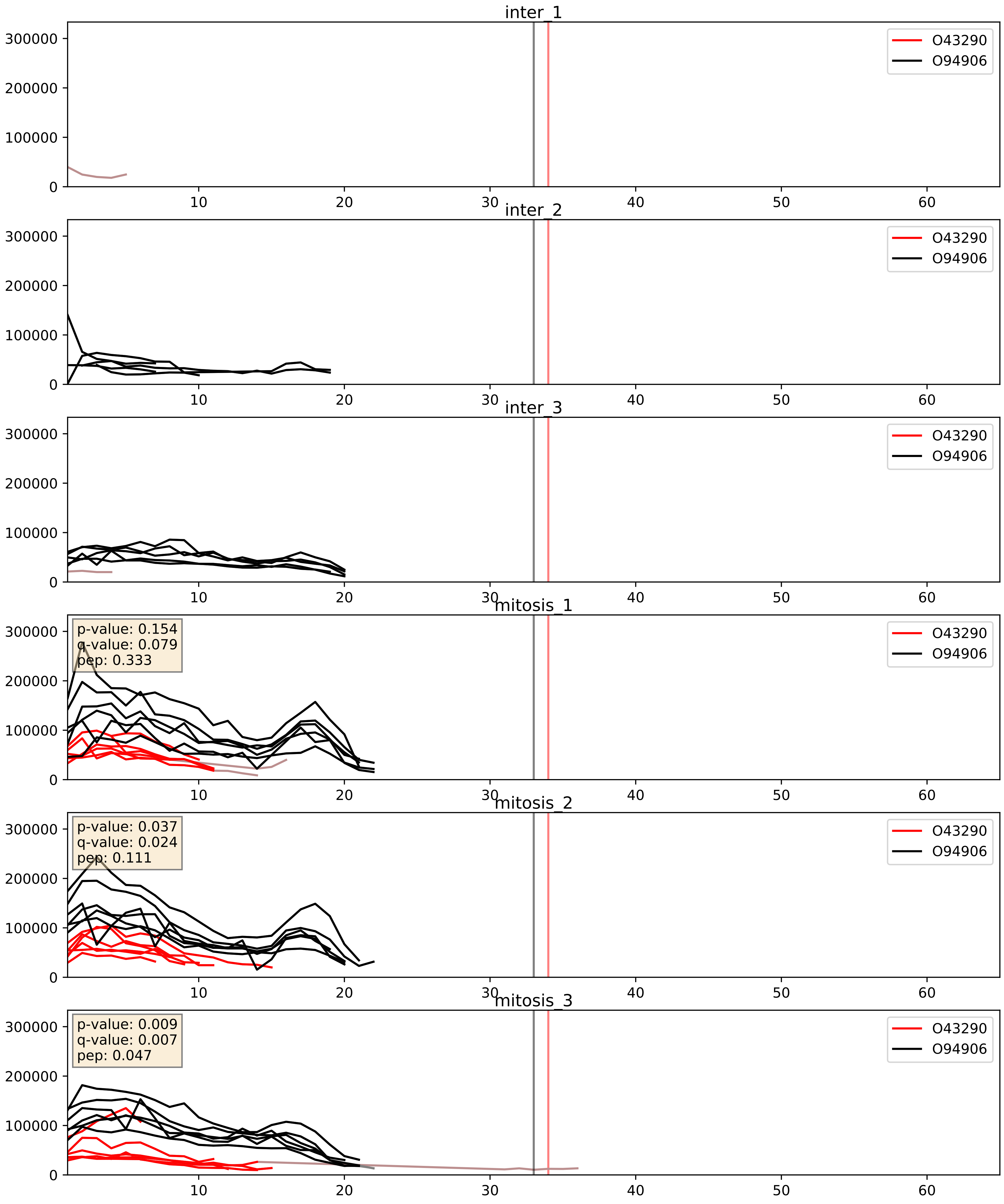


| level | condition_1 | condition_2 | pvalue | pvalue_adjusted |
|---|---|---|---|---|
| interactor_ratio | mitosis | inter | 0.00015386474196291182 | 0.012836278661937291 |
| complex_abundance | mitosis | inter | 0.0001611827964660317 | 0.008563834848633118 |
| interactor_abundance | mitosis | inter | 0.0010647499919498099 | 0.024049720606162197 |

O43395_O94906
PRPF3_HUMAN vs PRP6_HUMAN
p-value: 0.002 q-value: 0.003

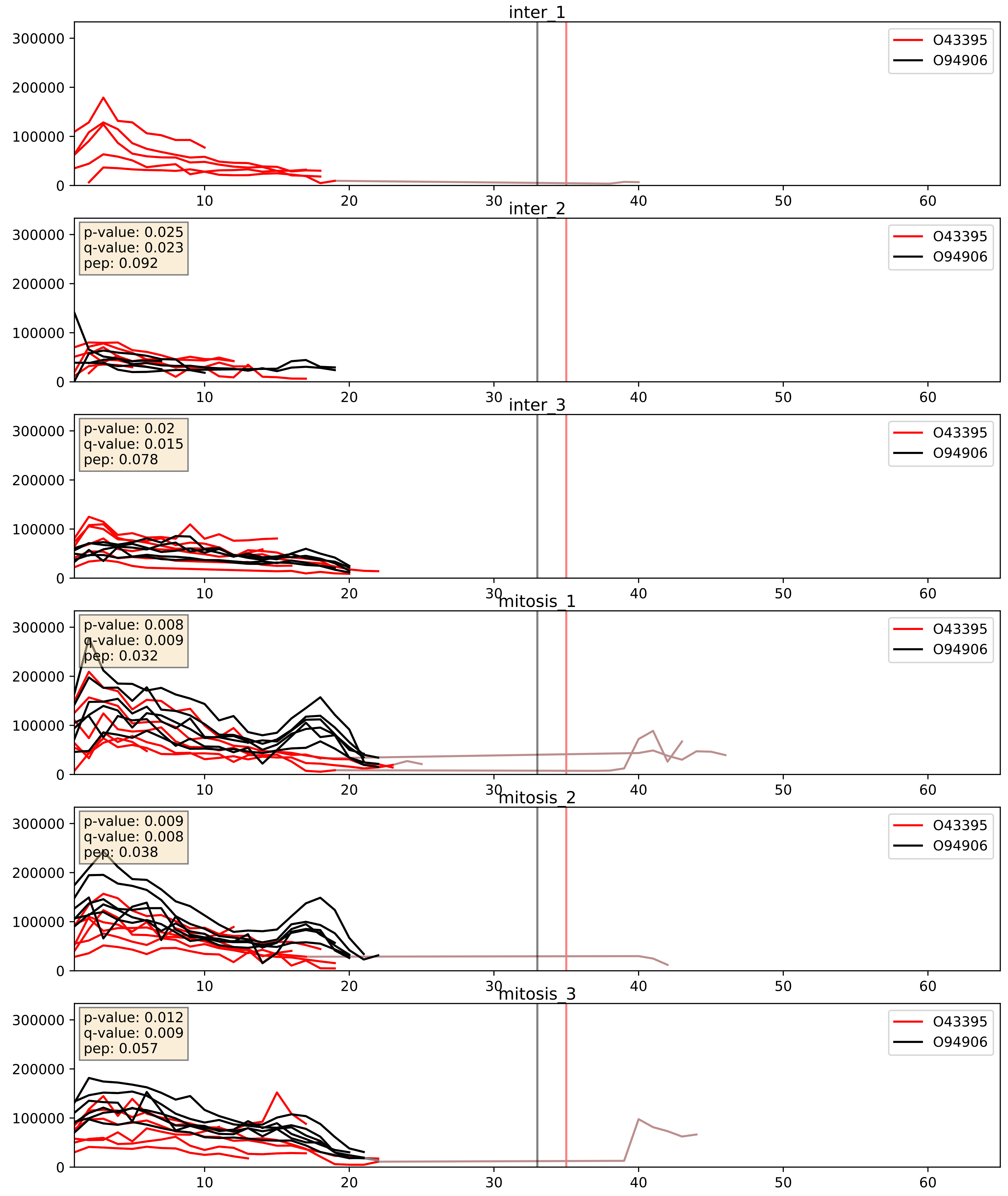


| level | condition_1 | condition_2 | pvalue | pvalue_adjusted |
|---|---|---|---|---|
| interactor_ratio | mitosis | inter | 0.006807476722737148 | 0.11448330205624752 |
| complex_abundance | mitosis | inter | 0.09721955987231426 | 0.23097403067083264 |
| interactor_abundance | mitosis | inter | 0.21396425439359726 | 0.409281344717138 |

O43447_O94906
PPIH_HUMAN vs PRP6_HUMAN
p-value: 0.042 q-value: 0.029

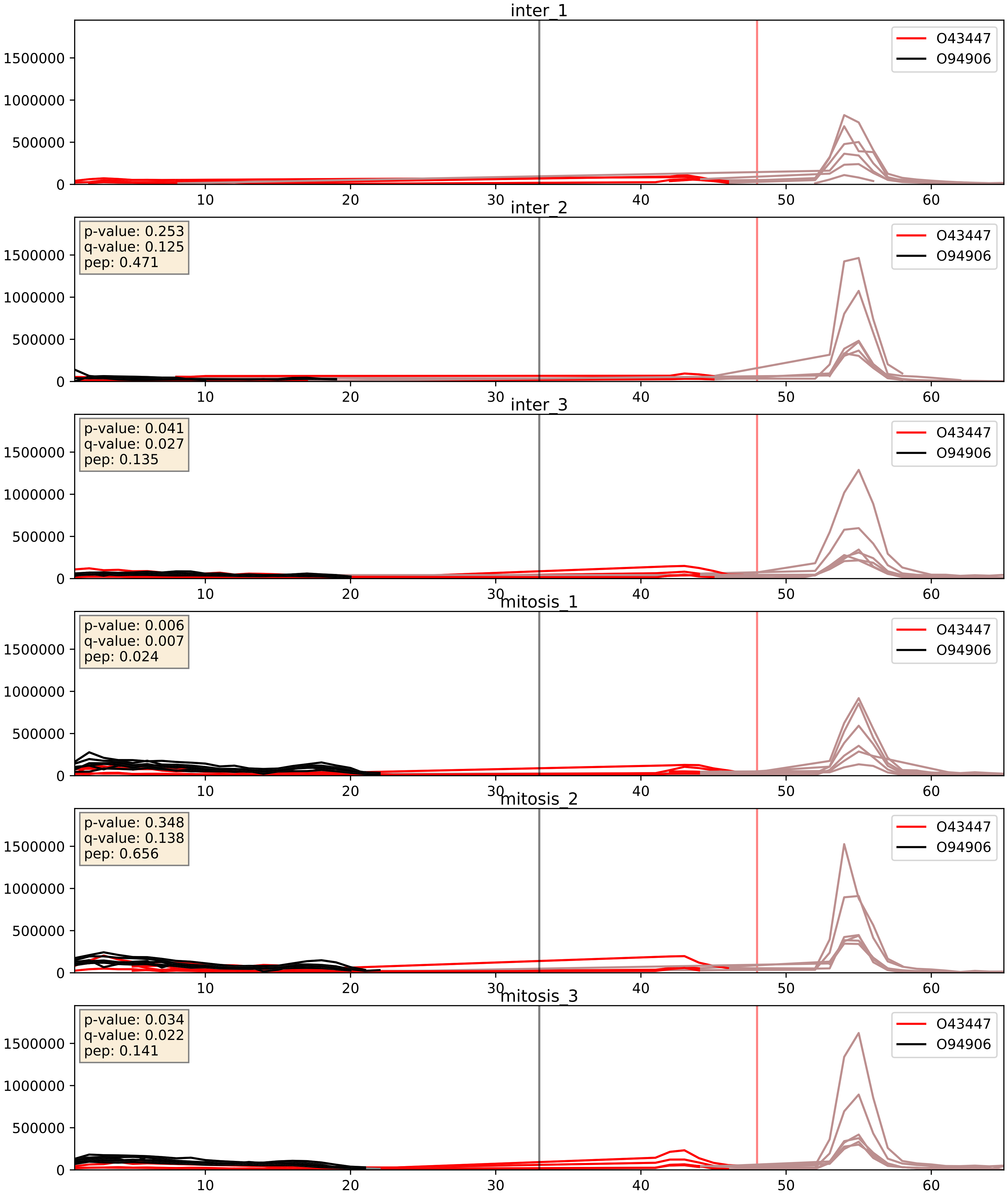


| level | condition_1 | condition_2 | pvalue | pvalue_adjusted |
|---|---|---|---|---|
| interactor_ratio | mitosis | inter | 0.040573208153838795 | 0.3329881704667882 |
| complex_abundance | mitosis | inter | 0.09551852194081667 | 0.22873067549909426 |
| interactor_abundance | mitosis | inter | 0.2972663313178642 | 0.49163725584376783 |

O94906_P55769
PRP6_HUMAN vs NH2L1_HUMAN
p-value: 0.005 q-value: 0.005

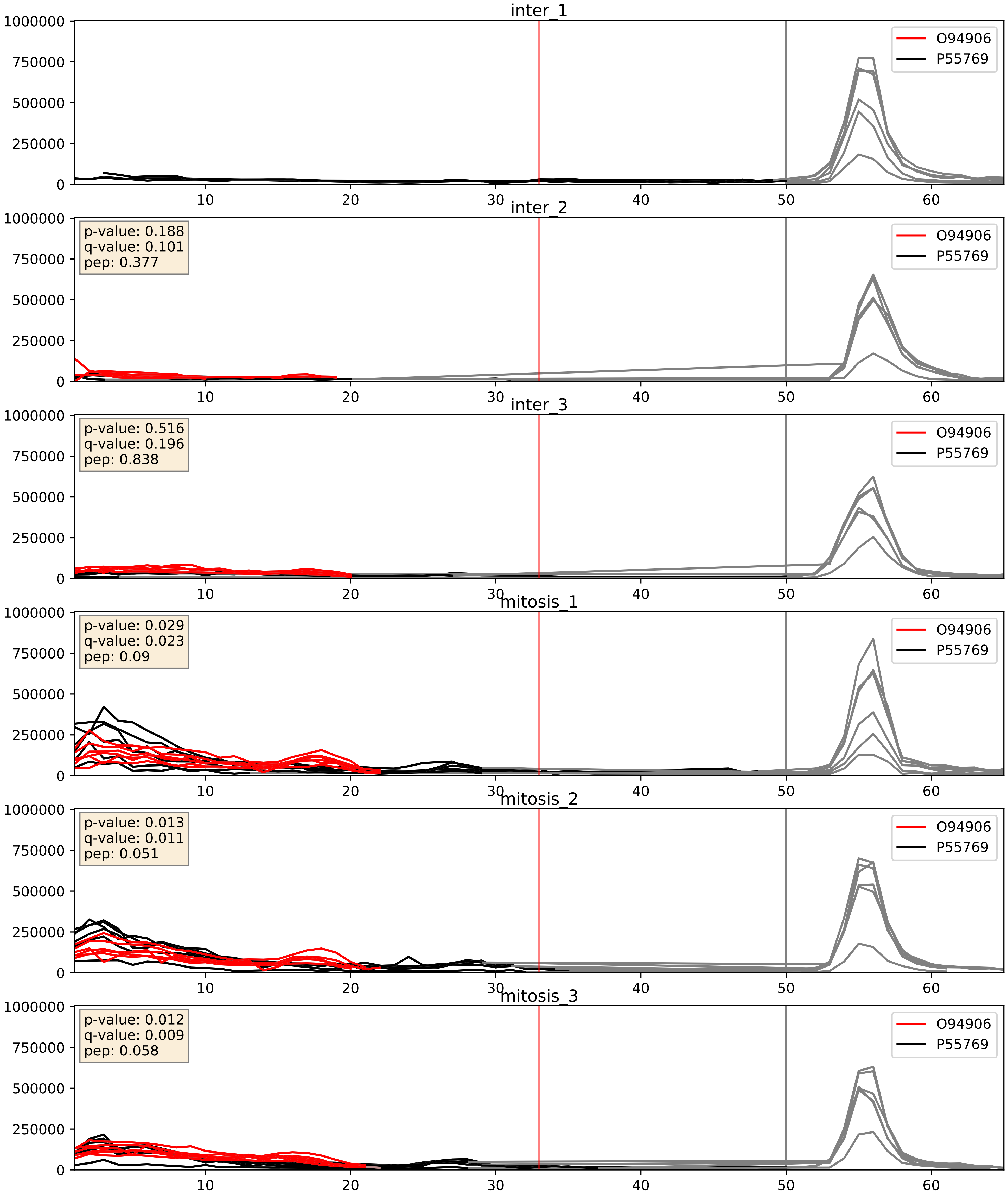


| level | condition_1 | condition_2 | pvalue | pvalue_adjusted |
|---|---|---|---|---|
| complex_abundance | mitosis | inter | 0.01249359904338843 | 0.08303199364239515 |
| interactor_abundance | mitosis | inter | 0.04665310860590202 | 0.1724311786124876 |
| interactor_ratio | mitosis | inter | 0.17781759270905903 | 0.6444111479450476 |

O94906_Q13123
PRP6_HUMAN vs RED_HUMAN
p-value: 0.042 q-value: 0.028

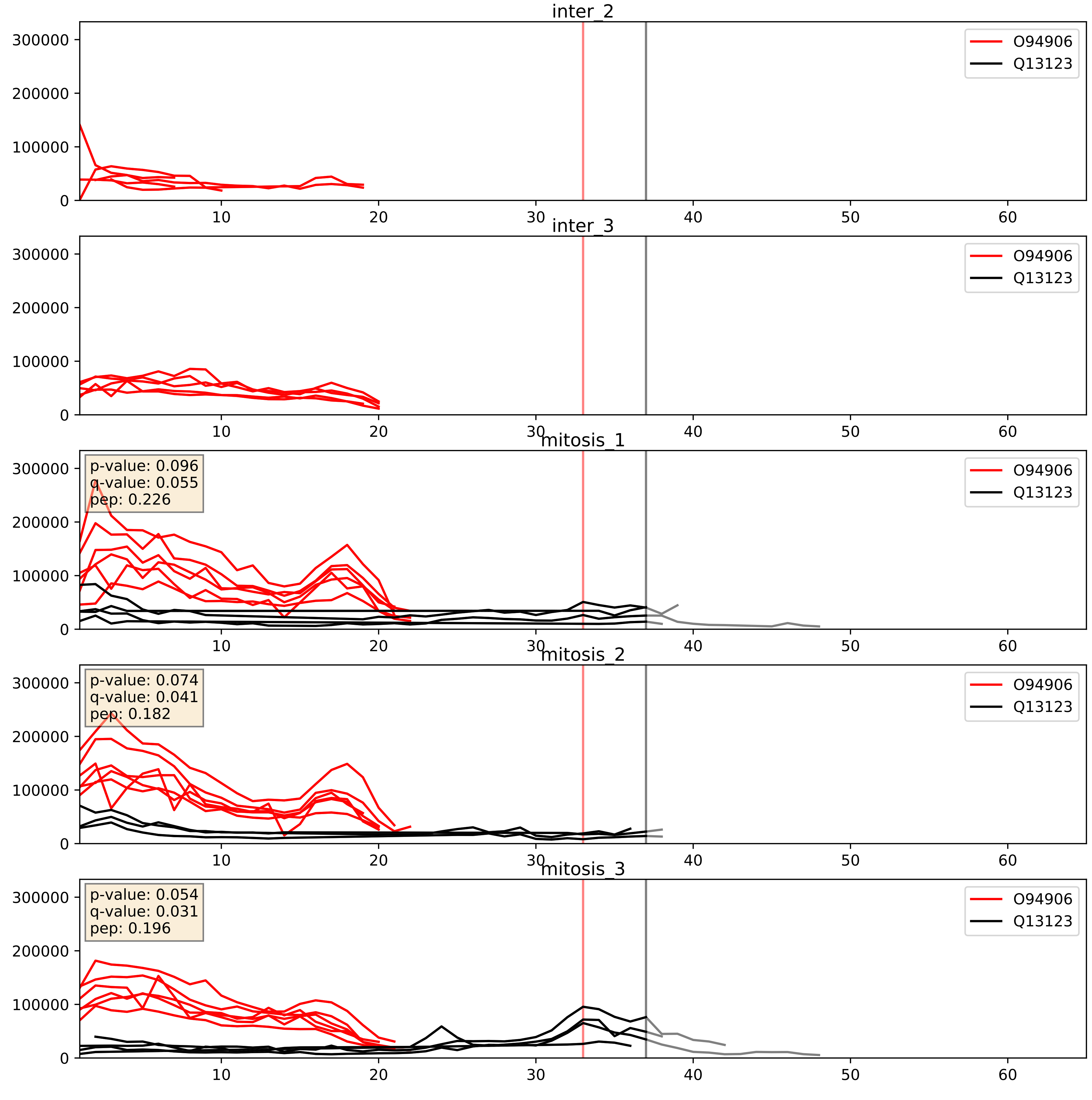


| level | condition_1 | condition_2 | pvalue | pvalue_adjusted |
|---|---|---|---|---|
| interactor_abundance | mitosis | inter | 0.0002447145305091371 | 0.011879056967425 26 |
| interactor_ratio | mitosis | inter | 0.0002458464698806514 | 0.016038997600655195 |
| complex_abundance | mitosis | inter | 0.00047785192447750004 | 0.01381896105921419 |

O94906_Q2TAY7
PRP6_HUMAN vs SMU1_HUMAN
p-value: 0.035 q-value: 0.025

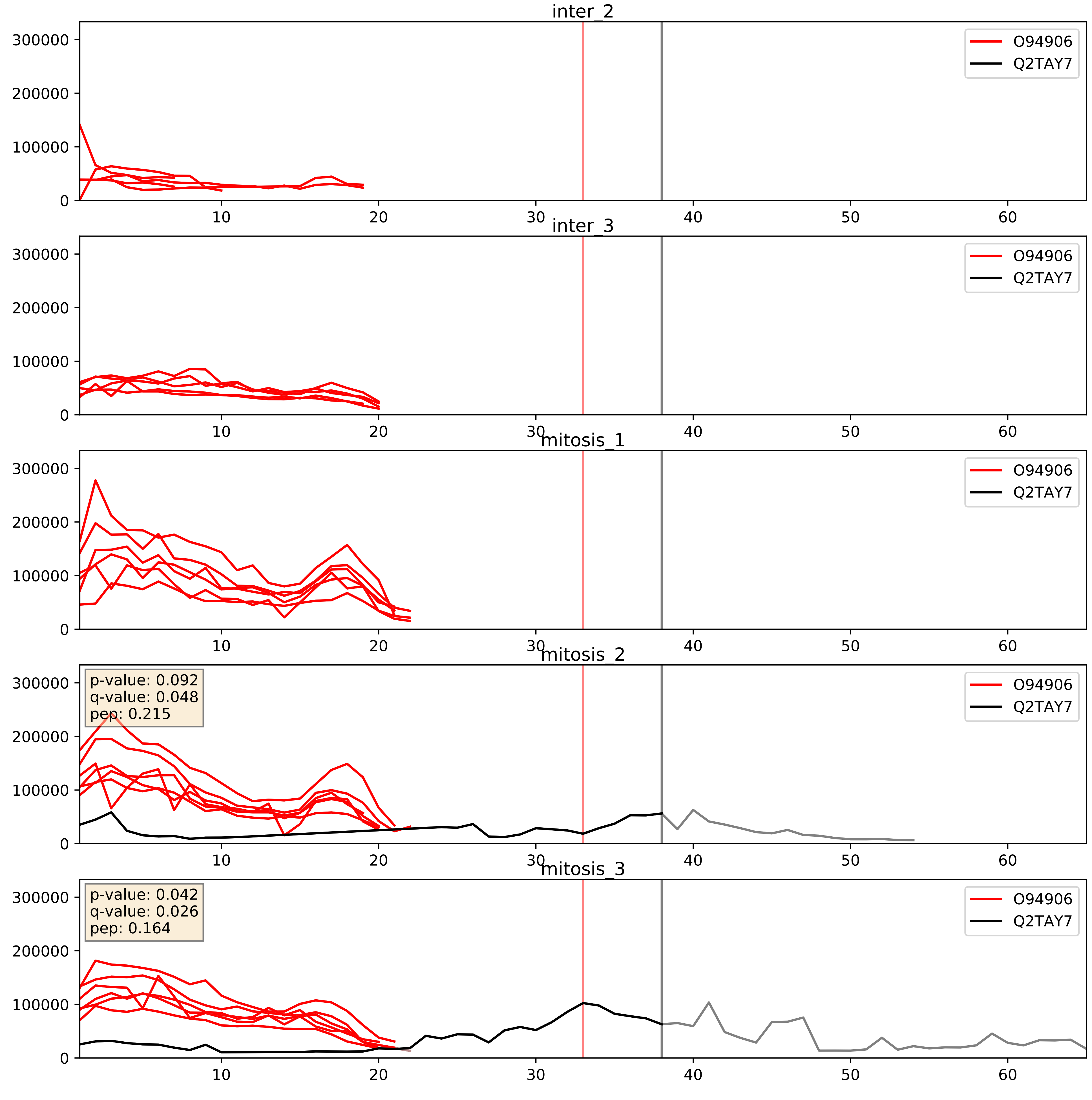


| level | condition_1 | condition_2 | pvalue | pvalue_adjusted |
|---|---|---|---|---|
| interactor_ratio | mitosis | inter | 0.14754299800444776 | 0.6019866839456972 |
| interactor_abundance | mitosis | inter | 0.16218201860998468 | 0.34963318876681726 |
| complex_abundance | mitosis | inter | 0.17584405082794036 | 0.32606993026301806 |

O94906_Q53GS9
PRP6_HUMAN vs SNUT2_HUMAN
p-value: 0.017 q-value: 0.014

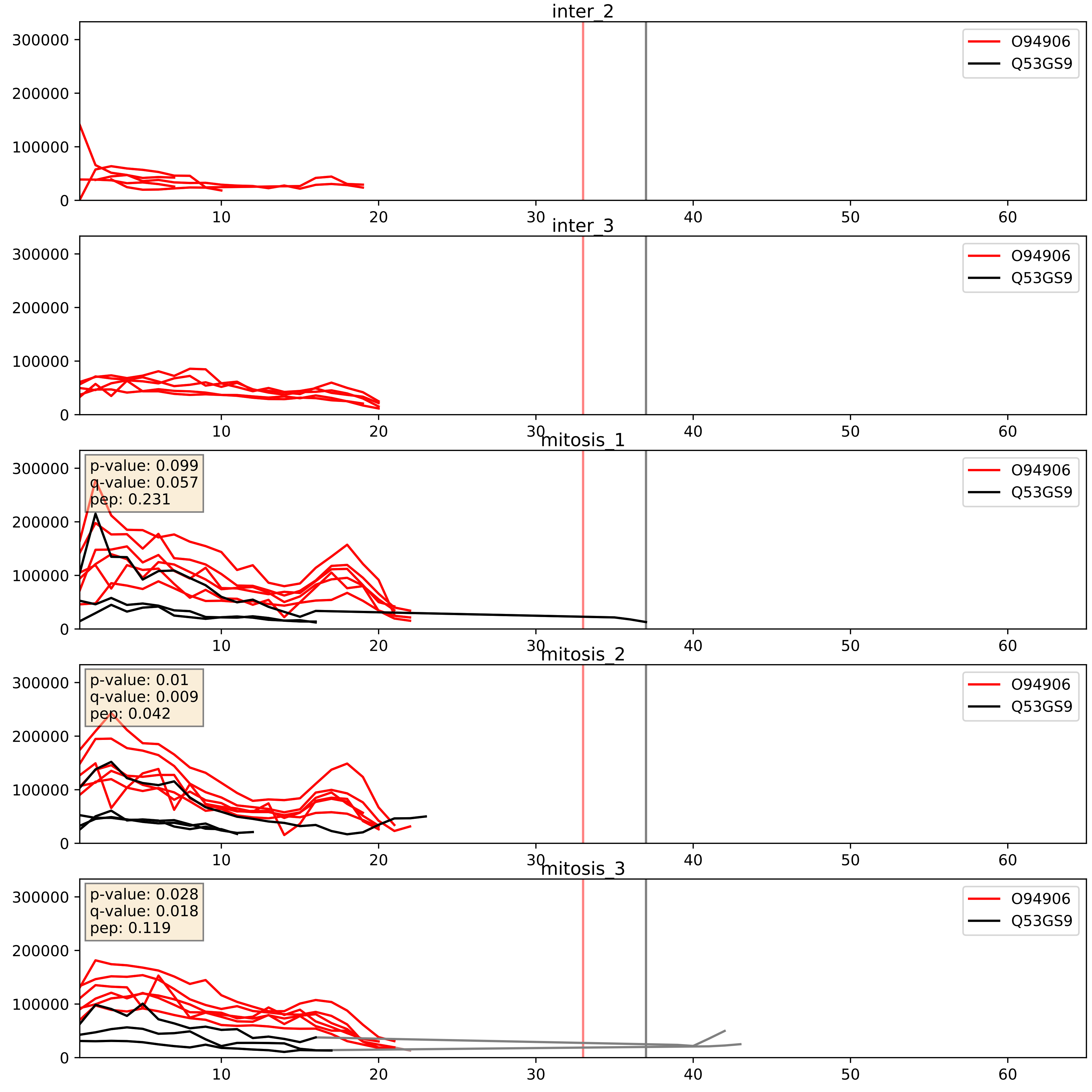


| level | condition_1 | condition_2 | pvalue | pvalue_adjusted |
|---|---|---|---|---|
| interactor_abundance | mitosis | inter | 3.439973865620678e-05 | 0.005164930486161031 |
| interactor_ratio | mitosis | inter | 9.480369998418357e-05 | 0.010510079284752772 |
| complex_abundance | mitosis | inter | 0.00015537452447804976 | 0.008563834848633118 |

O94906_Q99459
PRP6_HUMAN vs CDC5L_HUMAN
p-value: 0.019 q-value: 0.015

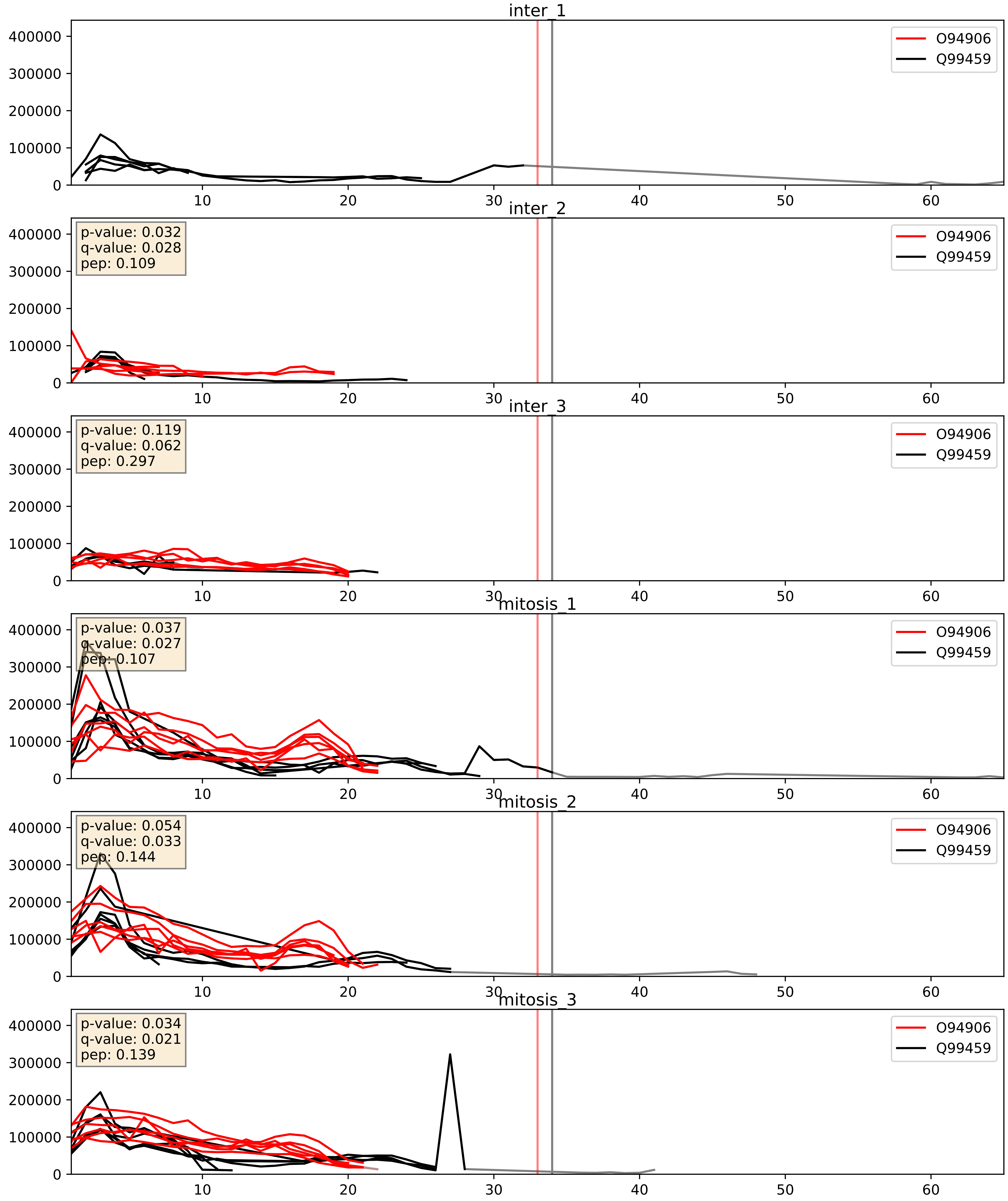


| level | condition_1 | condition_2 | pvalue | pvalue_adjusted |
|---|---|---|---|---|
| complex_abundance | mitosis | inter | 0.0063618354150012085 | 0.05948545934570088 |
| interactor_abundance | mitosis | inter | 0.014678960452509026 | 0.10605571190893752 |
| interactor_ratio | mitosis | inter | 0.9303107107087359 | 0.9878882449502933 |

O94906_Q9BUQ8
PRP6_HUMAN vs DDX23_HUMAN
p-value: 0.008 q-value: 0.007

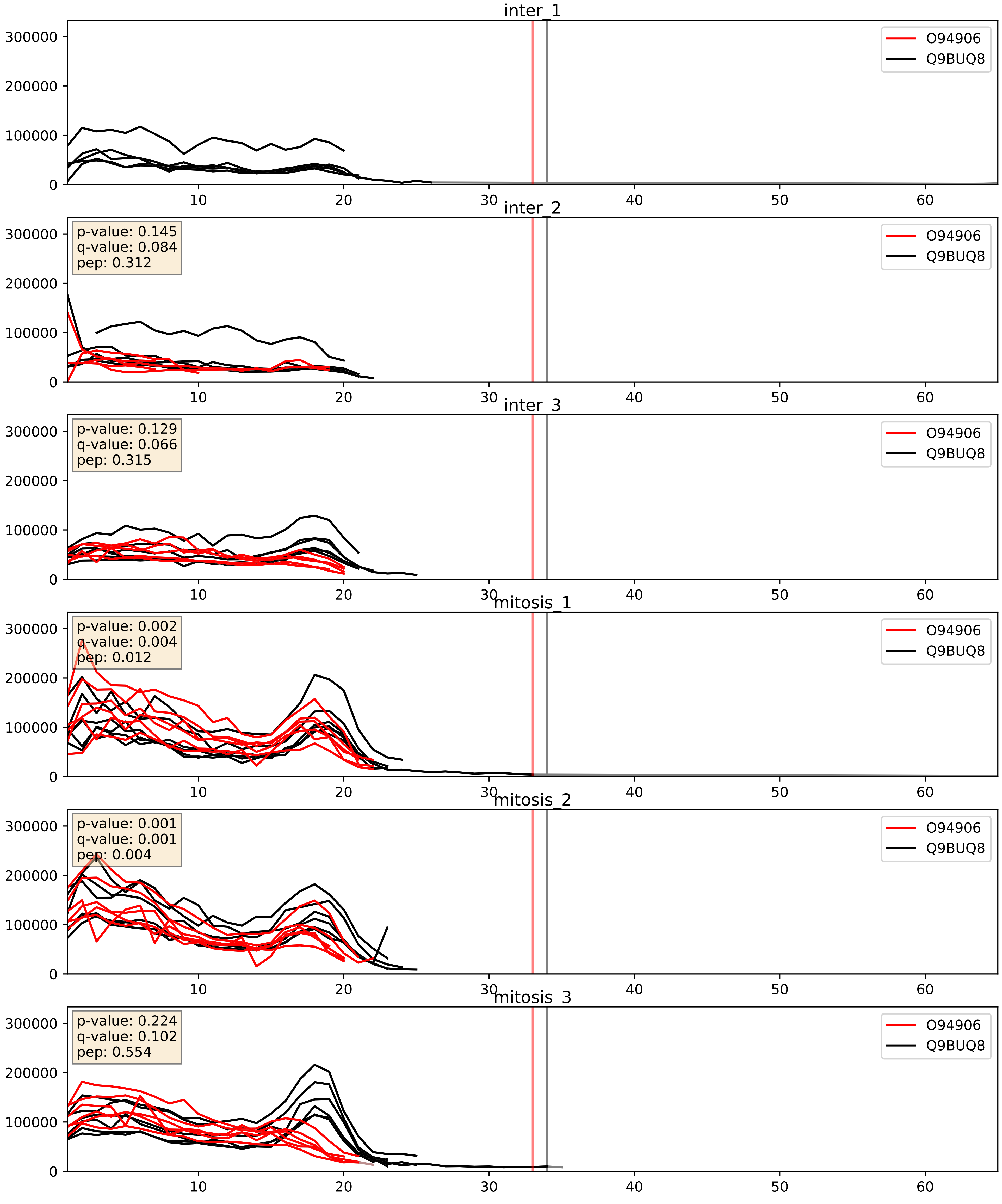


| level | condition_1 | condition_2 | pvalue | pvalue_adjusted |
|---|---|---|---|---|
| interactor_abundance | mitosis | inter | 0.04665310860590202 | 0.1724311786124876 |
| complex_abundance | mitosis | inter | 0.07150374961488644 | 0.19422056657737993 |
| interactor_ratio | mitosis | inter | 0.3936541743101139 | 0.8267123974716818 |

O94906_Q9UMS4
PRP6_HUMAN vs PRP19_HUMAN
p-value: 0.037 q-value: 0.026

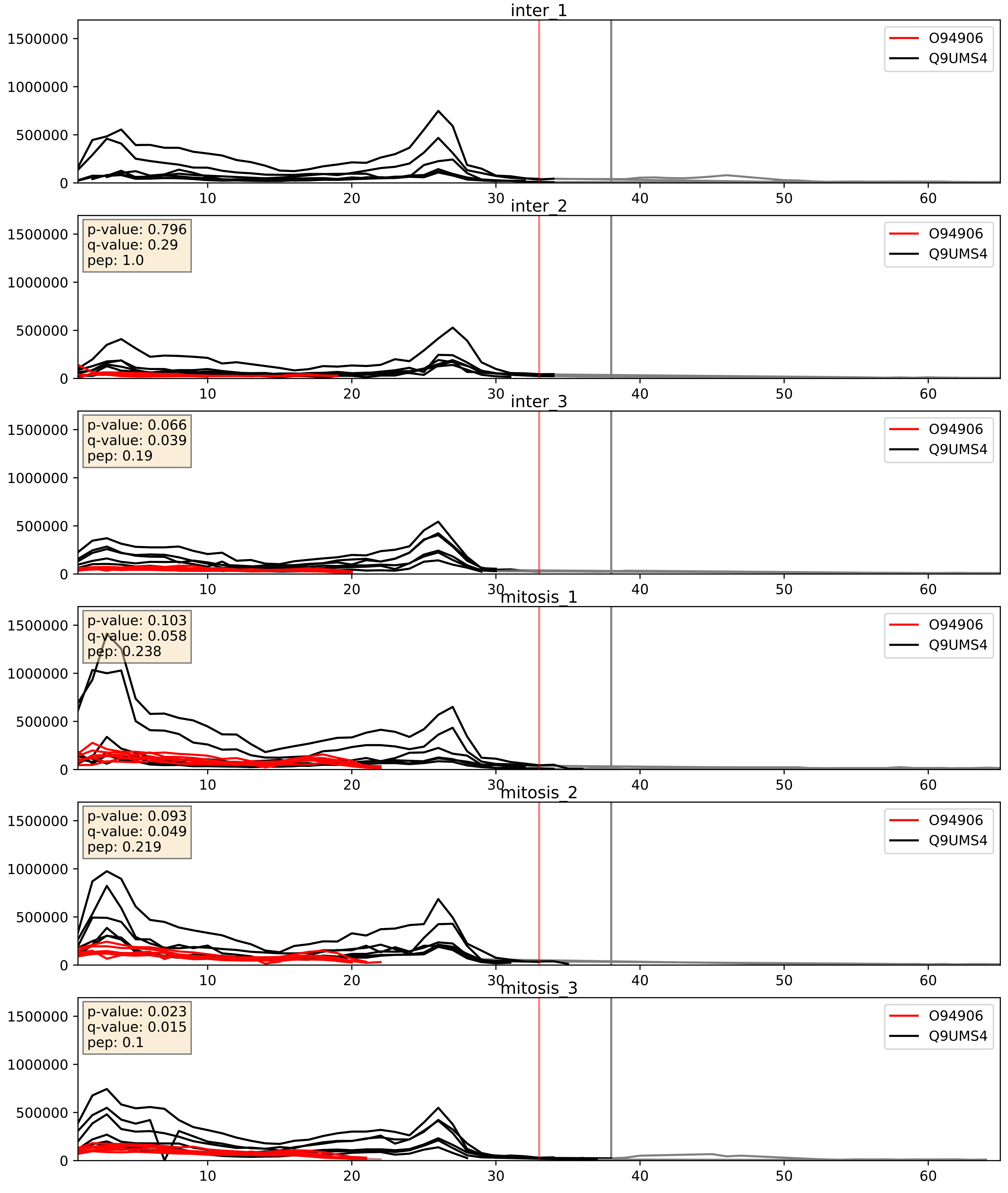


| level | condition_1 | condition_2 | pvalue | pvalue_adjusted |
|---|---|---|---|---|
| interactor_abundance | mitosis | inter | 0.04665310860590202 | 0.1724311786124876 |
| complex_abundance | mitosis | inter | 0.09555384294214499 | 0.22873067549909426 |
| interactor_ratio | mitosis | inter | 0.682256052881256 | 0.9349347226390419 |
