## Supplementary Data 2 for "SECAT: Quantifying differential protein-protein interaction states by network-centric analysis": hela_string_000031_P35232.pdf

P35232_P35232

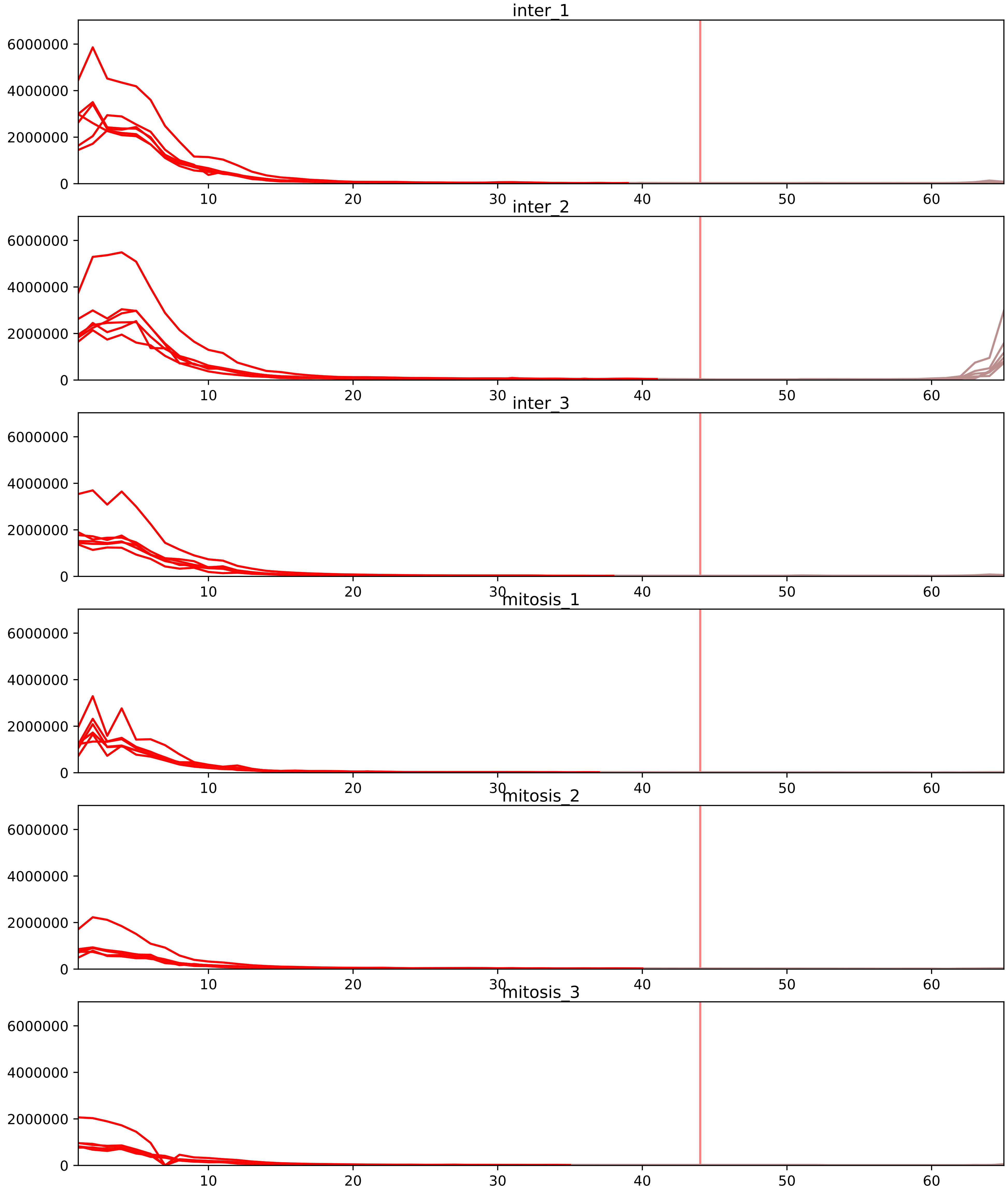


| level | condition_1 | condition_2 | pvalue | pvalue_adjusted |
|---|---|---|---|---|
| interactor_ratio | mitosis | inter | 1.945208564111527e-09 | 1.1930612526550698e-07 |
| interactor_abundance | mitosis | inter | 7.158437318226762e-05 | 0.0018479549585806118 |
| complex_abundance | mitosis | inter | 0.0003240085905083281 | 0.005229612338029155 |
| assembled_abundance | mitosis | inter | 0.0454848283401244 | 0.25437811405032534 |
| total_abundance | mitosis | inter | 0.04649794592168186 | 0.270064309647489 |
| monomer_abundance | mitosis | inter | 0.07350285819820047 | 0.37224850260048076 |

P00403_P35232
COX2_HUMAN vs PHB_HUMAN
p-value: 0.0 q-value: 0.022

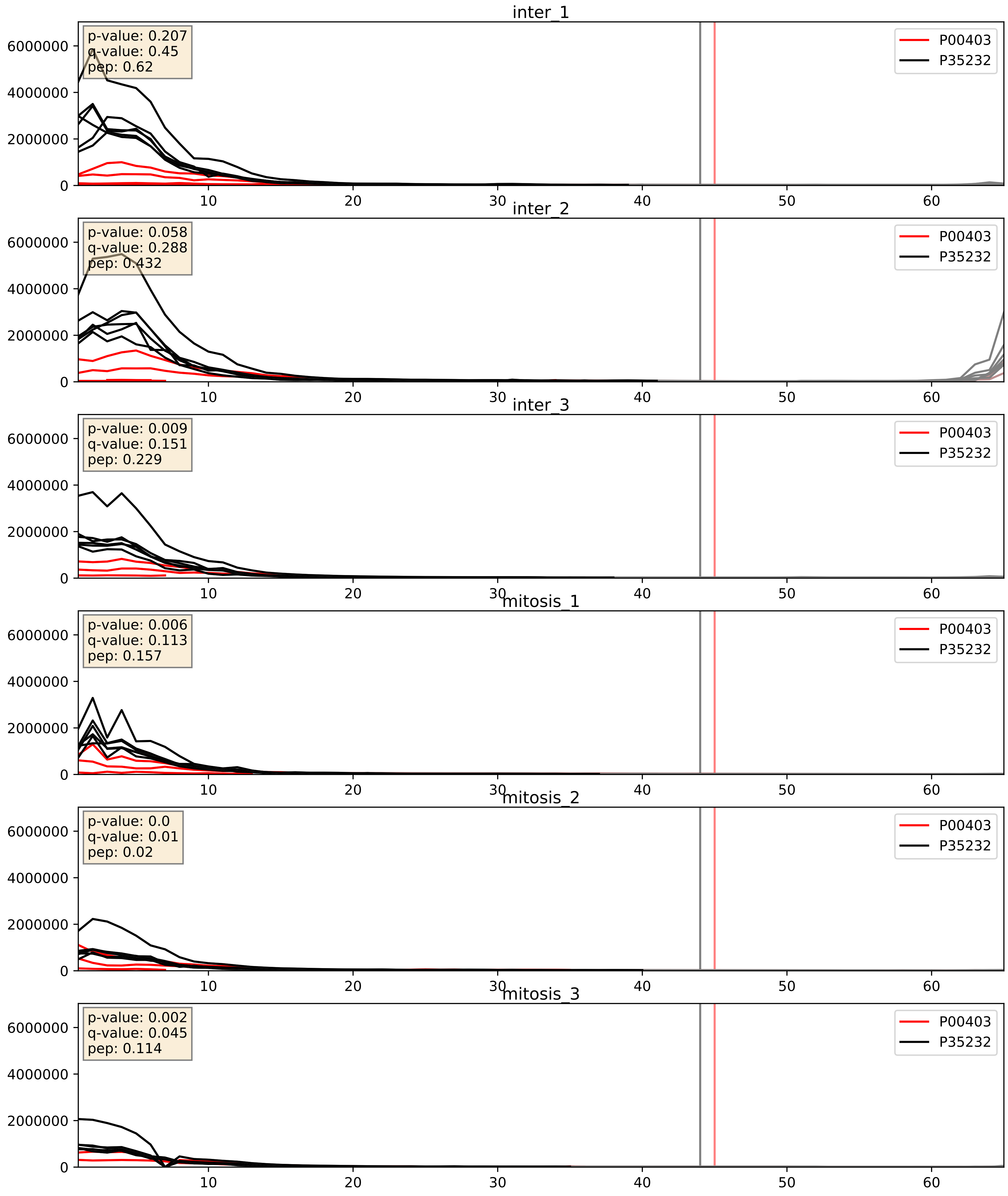


| level | condition_1 | condition_2 | pvalue | pvalue_adjusted |
|---|---|---|---|---|
| complex_abundance | mitosis | inter | 0.005505336892302833 | 0.055958215865660764 |
| interactor_ratio | mitosis | inter | 0.019625730779465705 | 0.21211648418210408 |
| interactor_abundance | mitosis | inter | 0.05021540360985647 | 0.17917626298473172 |

P18124_P35232
RL7_HUMAN vs PHB_HUMAN
p-value: 0.0 q-value: 0.016

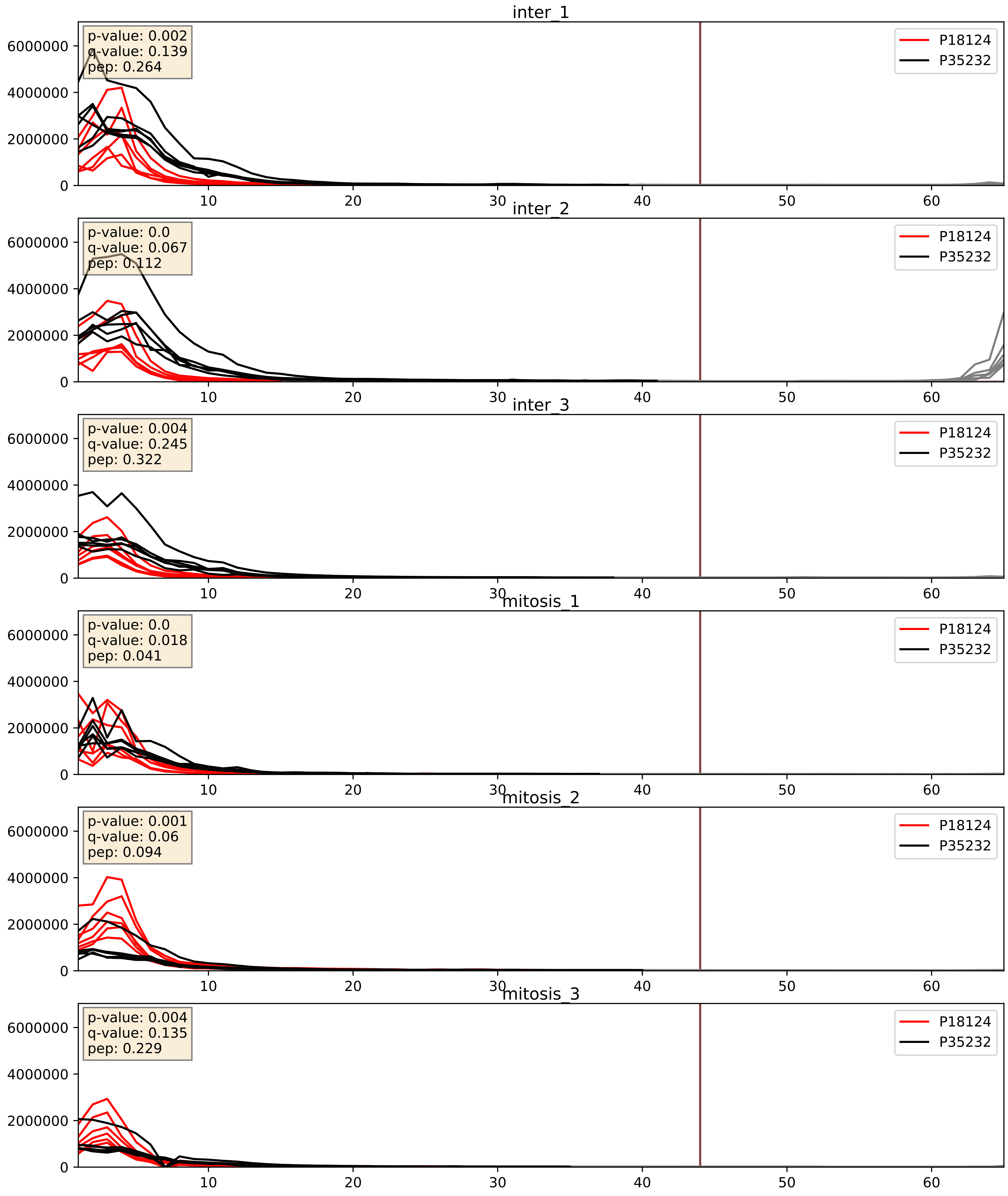


| level | condition_1 | condition_2 | pvalue | pvalue_adjusted |
|---|---|---|---|---|
| interactor_ratio | mitosis | inter | 0.002305266593070974 | 0.05997897275588917 |
| complex_abundance | mitosis | inter | 0.006501290201700229 | 0.05955783692156859 |
| interactor_abundance | mitosis | inter | 0.4927360017152748 | 0.6596528268193231 |

P19338_P35232
NUCL_HUMAN vs PHB_HUMAN
p-value: 0.0 q-value: 0.047

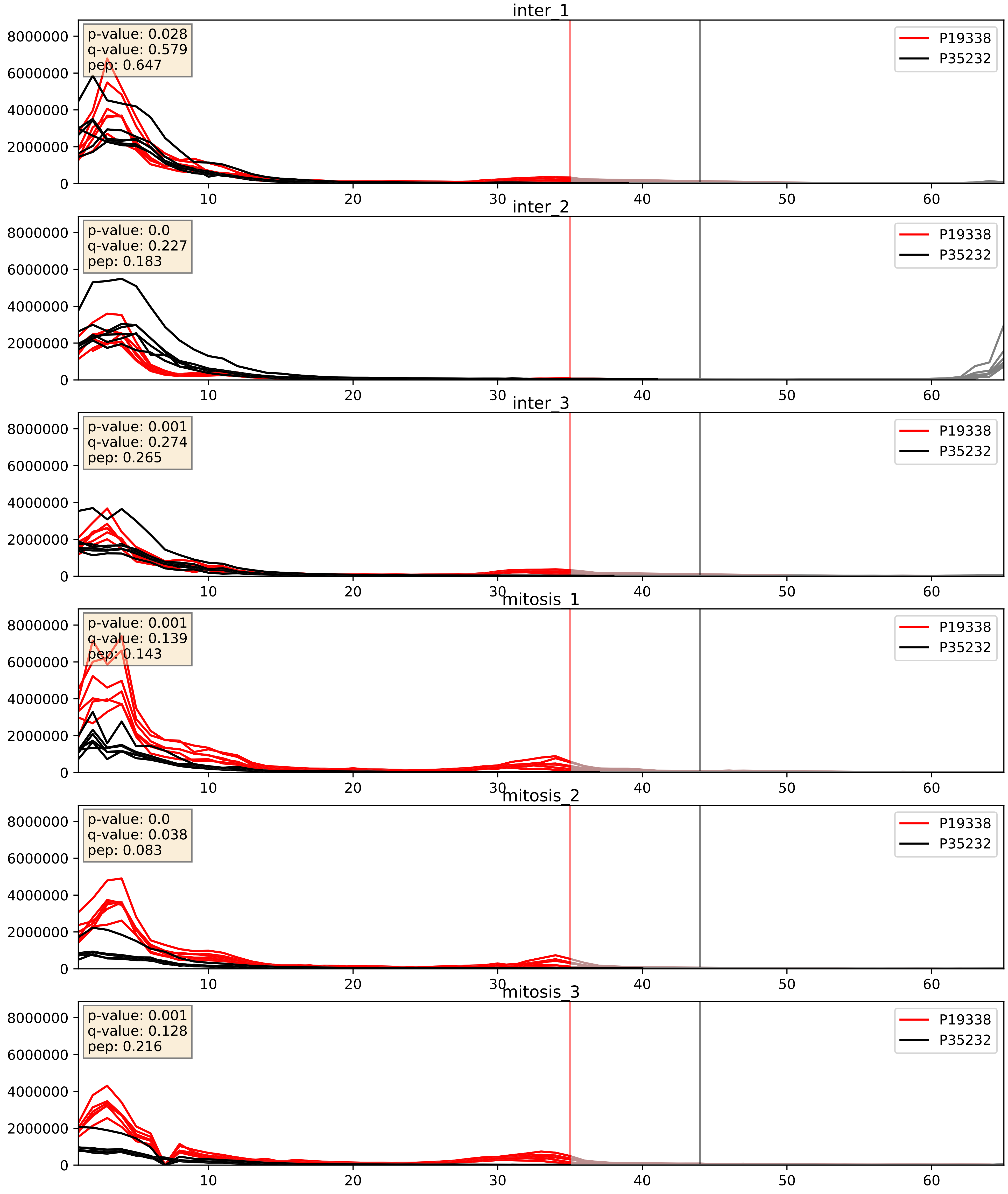


| level | condition_1 | condition_2 | pvalue | pvalue_adjusted |
|---|---|---|---|---|
| interactor_ratio | mitosis | inter | 0.00876770769960161 | 0.13236609860421478 |
| complex_abundance | mitosis | inter | 0.08977582856603068 | 0.2200690413875208 |
| interactor_abundance | mitosis | inter | 0.5623129846139611 | 0.7112621176332597 |

P35232_P61313
PHB_HUMAN vs RL15_HUMAN
p-value: 0.0 q-value: 0.026

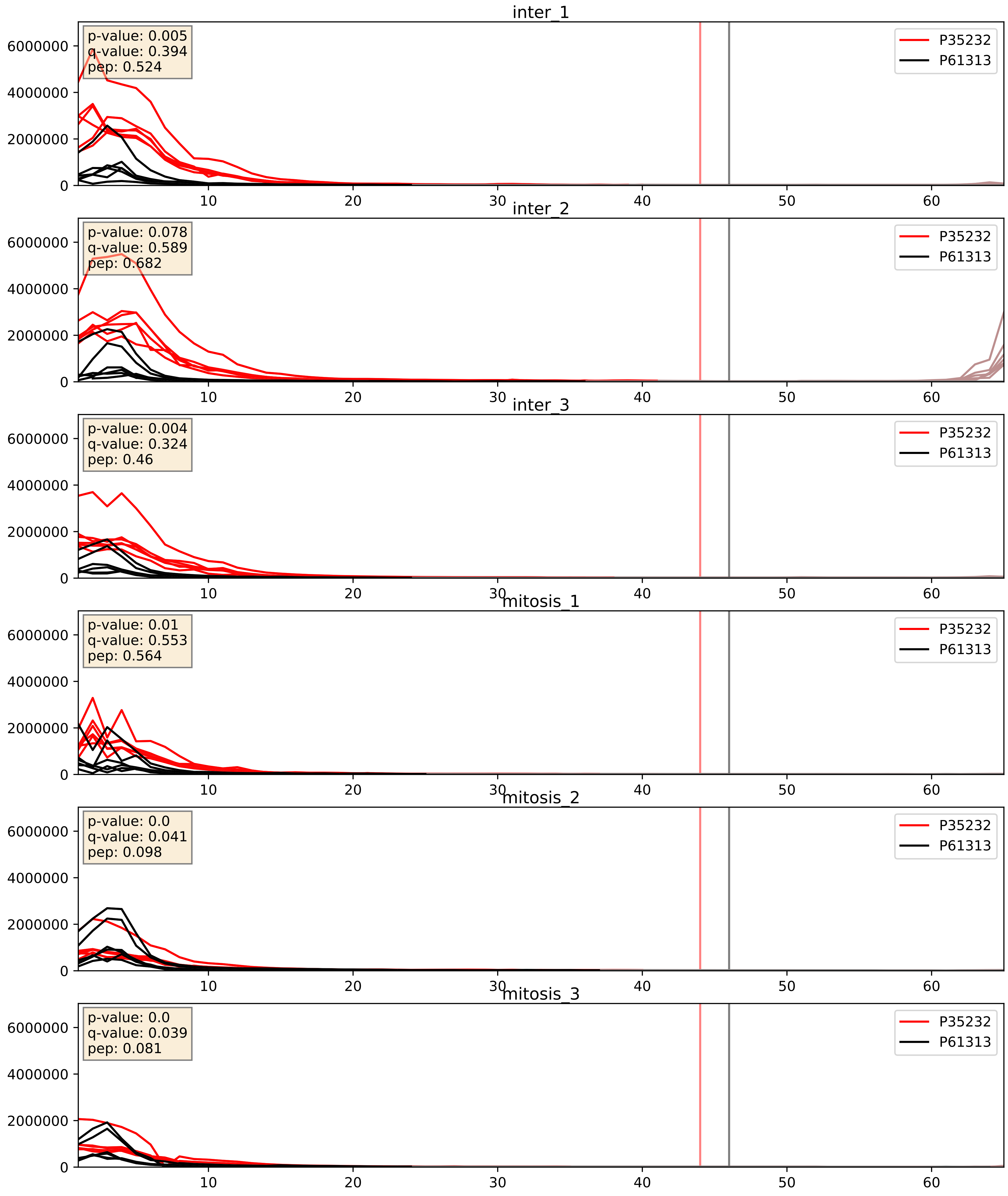


| level | condition_1 | condition_2 | pvalue | pvalue_adjusted |
|---|---|---|---|---|
| interactor_abundance | mitosis | inter | 0.0020095501929169087 | 0.03264187006238393 |
| complex_abundance | mitosis | inter | 0.002789280573146706 | 0.0383862406851058 |
| interactor_ratio | mitosis | inter | 0.0048106350951971025 | 0.09212312397066487 |

P35232_Q07020
PHB_HUMAN vs RL18_HUMAN
p-value: 0.0 q-value: 0.041

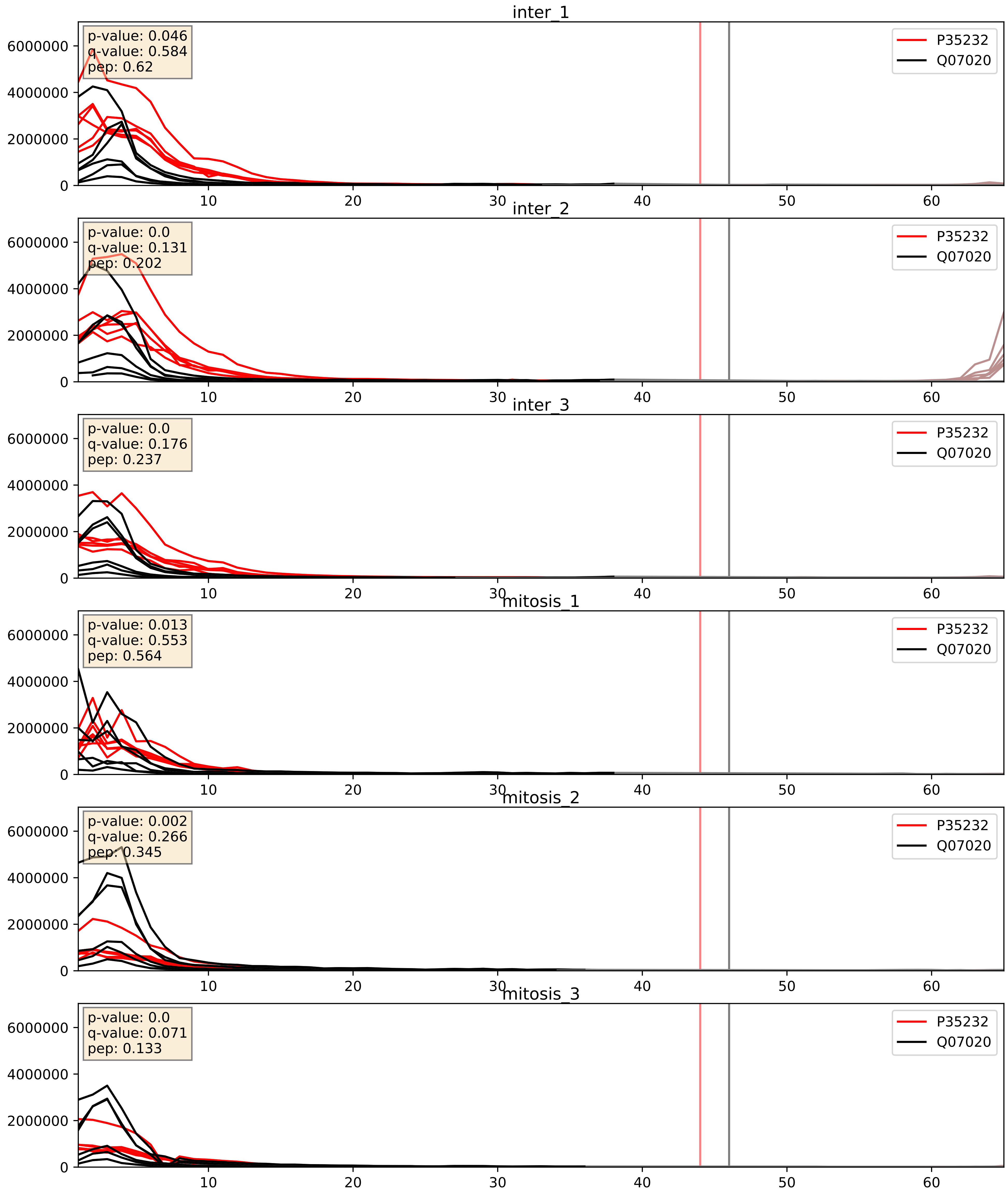


| level | condition_1 | condition_2 | pvalue | pvalue_adjusted |
|---|---|---|---|---|
| complex_abundance | mitosis | inter | 0.0010837200182553747 | 0.021833569470040306 |
| interactor_abundance | mitosis | inter | 0.002039835423541839 | 0.03279059385073829 |
| interactor_ratio | mitosis | inter | 0.016294221445469353 | 0.18848450753137524 |

P35232_Q07065
PHB_HUMAN vs CKAP4_HUMAN
p-value: 0.0 q-value: 0.04

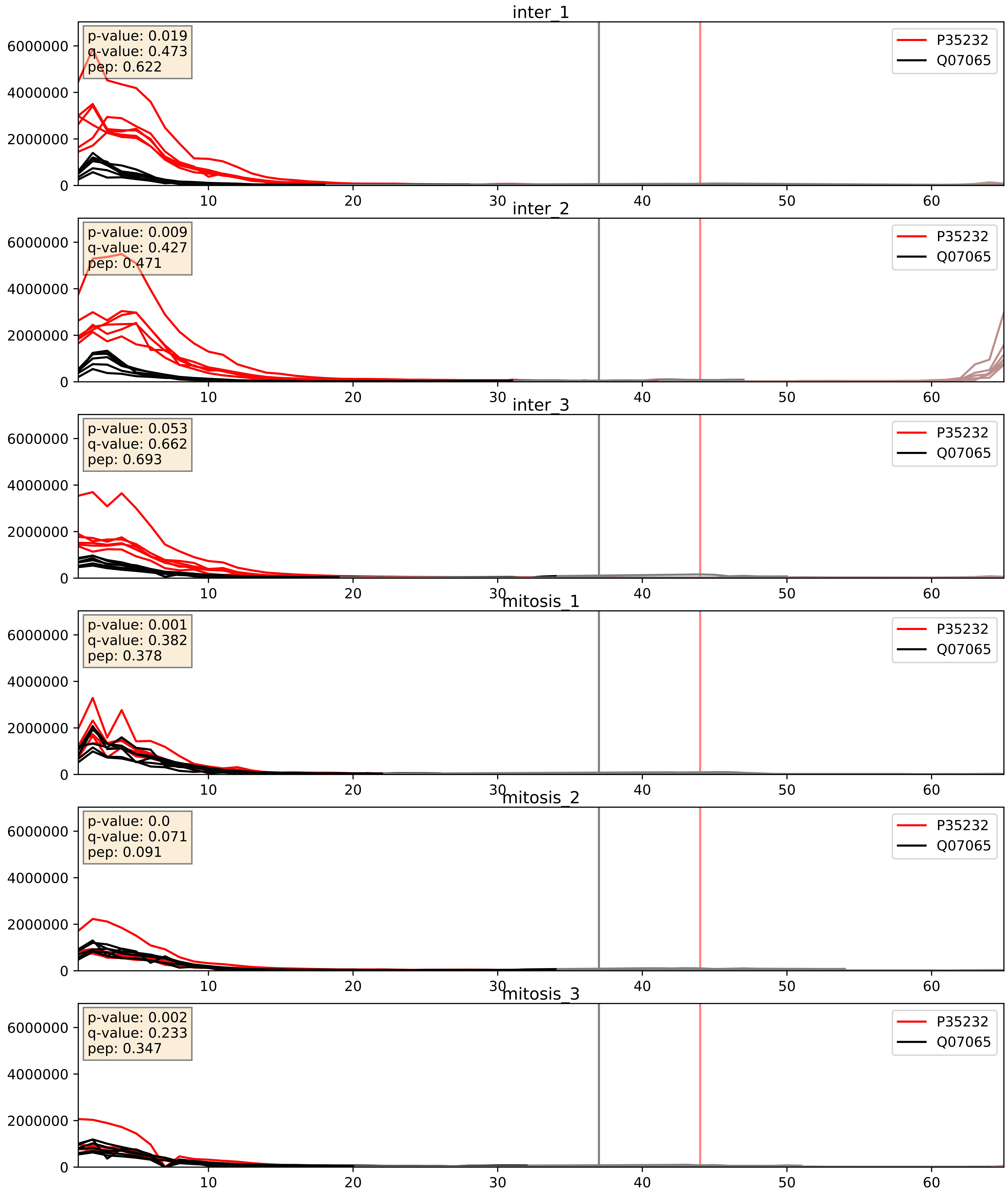


| level | condition_1 | condition_2 | pvalue | pvalue_adjusted |
|---|---|---|---|---|
| interactor_ratio | mitosis | inter | 5.502030197327859e-05 | 0.007596351369213947 |
| interactor_abundance | mitosis | inter | 0.002121211643892444 | 0.03376647511573598 |
| complex_abundance | mitosis | inter | 0.020268857034890116 | 0.10510110796643679 |

P35232_Q99623
PHB_HUMAN vs PHB2_HUMAN
p-value: 0.0 q-value: 0.0

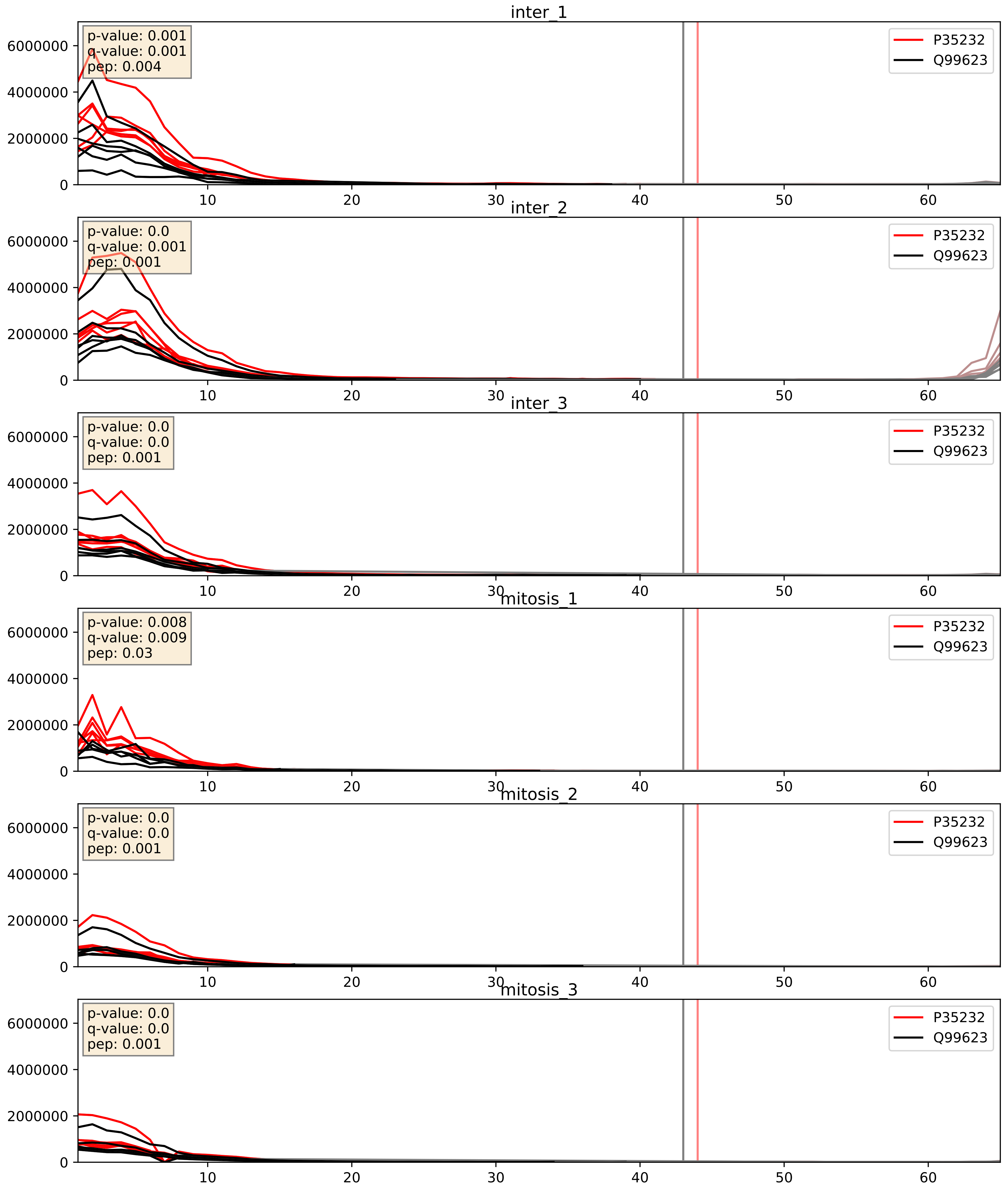


| level | condition_1 | condition_2 | pvalue | pvalue_adjusted |
|---|---|---|---|---|
| complex_abundance | mitosis | inter | 0.0005098332523952253 | 0.014318857255407222 |
| interactor_abundance | mitosis | inter | 0.0019578595900461063 | 0.032370554261591626 |
| interactor_ratio | mitosis | inter | 0.3720179690761096 | 0.8138388063636502 |
