## Supplementary Data 2 for "SECAT: Quantifying differential protein-protein interaction states by network-centric analysis": hela_string_000032_Q9NZE8.pdf

Q9NZE8_Q9NZE8

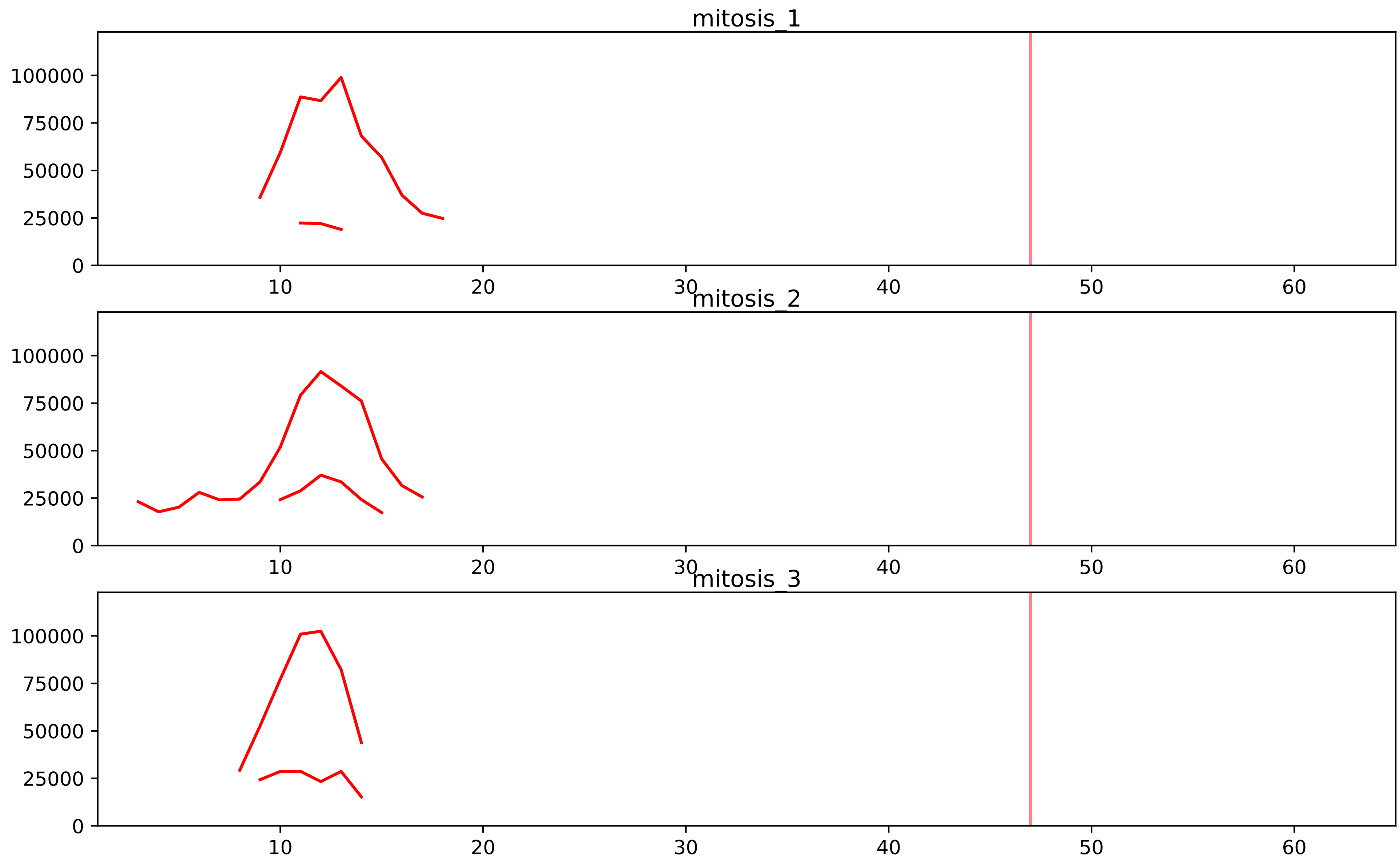


| level | condition_1 | condition_2 | pvalue | pvalue_adjusted |
|---|---|---|---|---|
| interactor_ratio | mitosis | inter | 2.68027092752261e-09 | 1.59087048601342e-07 |
| complex_abundance | mitosis | inter | 1.2761403113277732e-05 | 0.000572706871425147 |
| assembled_abundance | mitosis | inter | 0.0002644346611020986 | 0.016114995197788636 |
| total_abundance | mitosis | inter | 0.0011059578671142312 | 0.03653159157438113 |
| interactor_abundance | mitosis | inter | 0.0012131874289300728 | 0.013208667865274165 |

P09001_Q9NZE8
RM03_HUMAN vs RM35_HUMAN
p-value: 0.0 q-value: 0.0

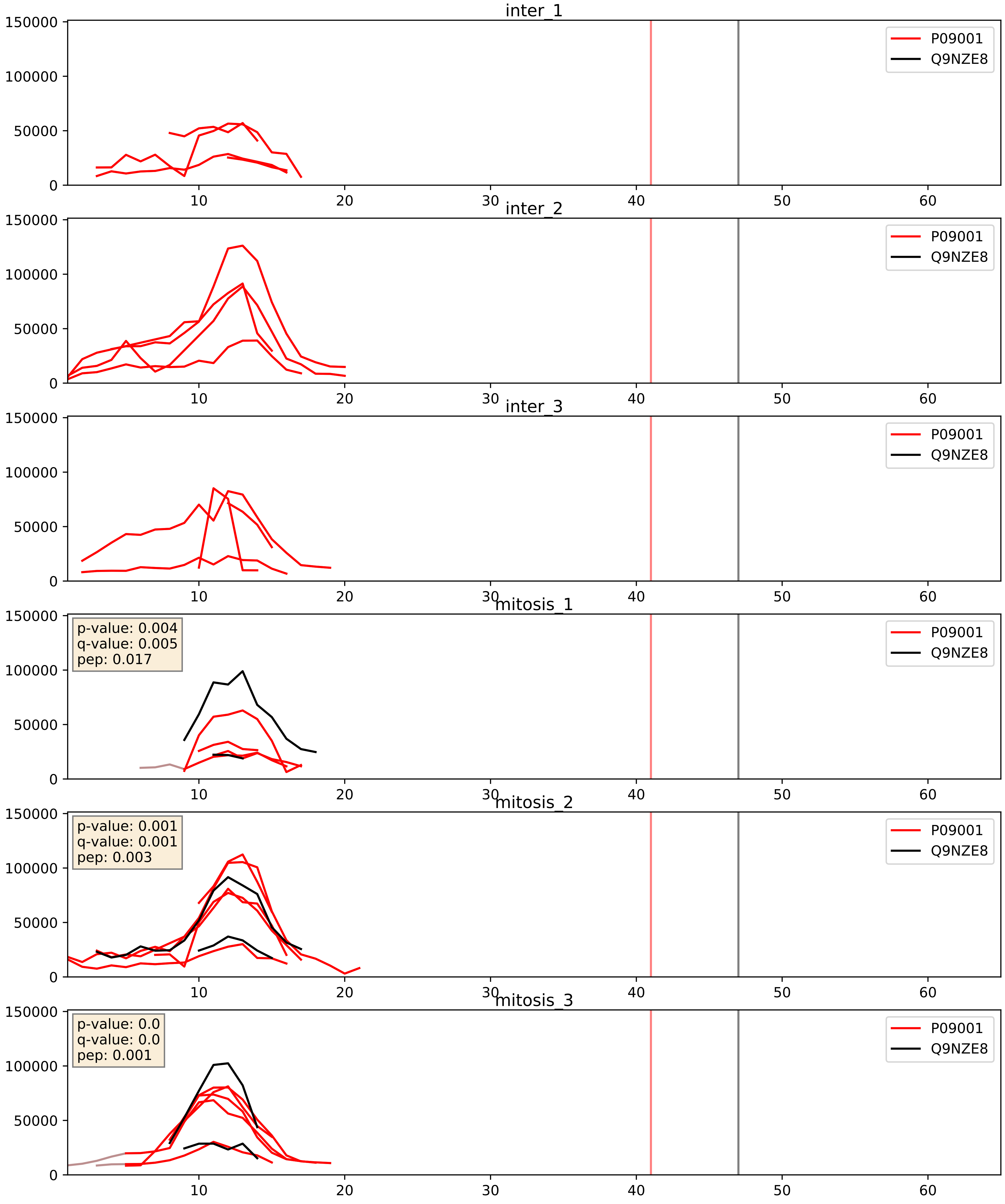


| level | condition_1 | condition_2 | pvalue | pvalue_adjusted |
|---|---|---|---|---|
| complex_abundance | mitosis | inter | 0.007516687712131053 | 0.06444768685385507 |
| interactor_abundance | mitosis | inter | 0.02167768374935654 | 0.12240169715995514 |
| interactor_ratio | mitosis | inter | 0.26072769036023485 | 0.7341204954098902 |

P49406_Q9NZE8
RM19_HUMAN vs RM35_HUMAN
p-value: 0.0 q-value: 0.001

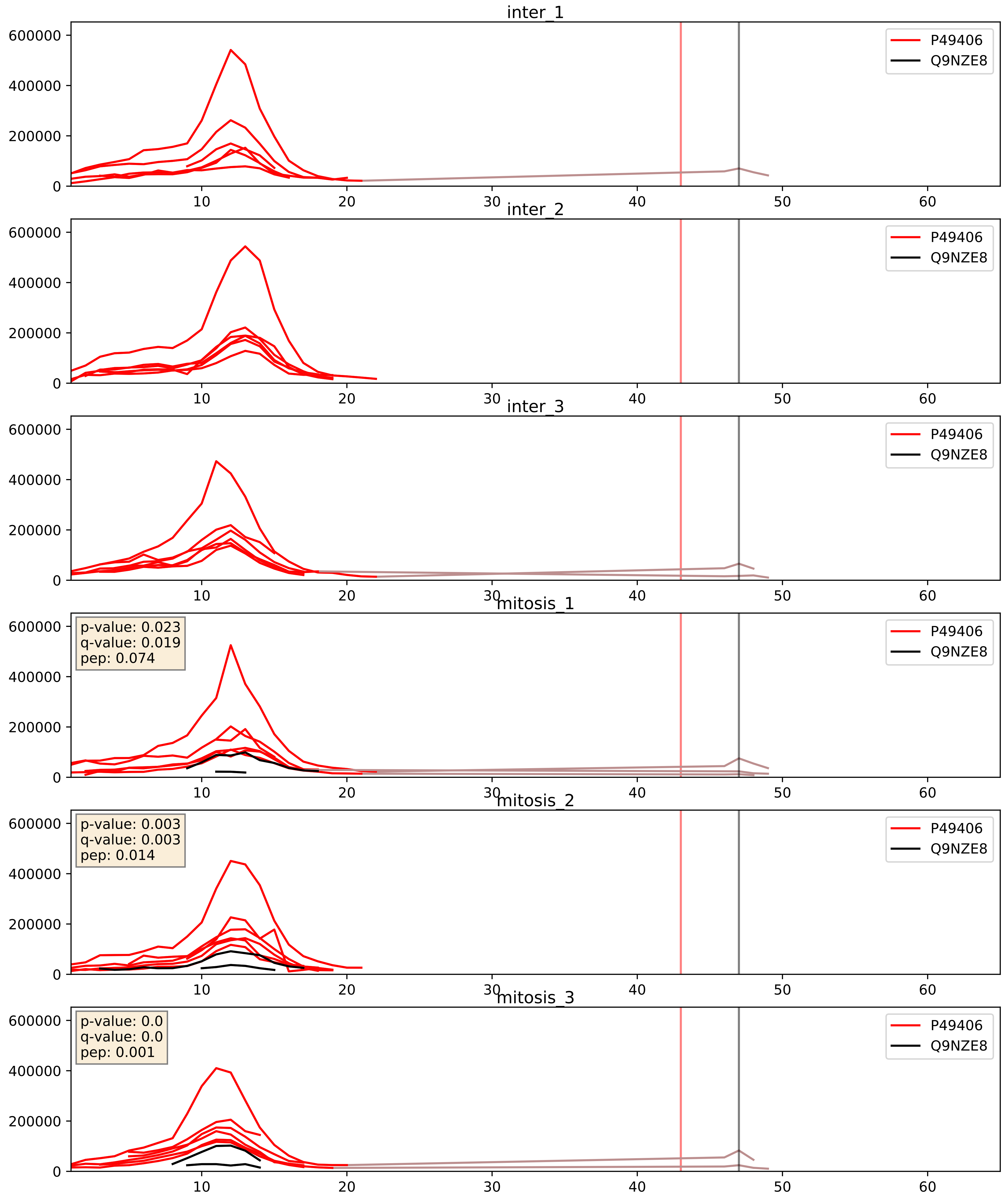


| level | condition_1 | condition_2 | pvalue | pvalue_adjusted |
|---|---|---|---|---|
| interactor_ratio | mitosis | inter | 4.703984768646224e-05 | 0.006815555856770076 |
| interactor_abundance | mitosis | inter | 0.0001364063589969551 | 0.00891327048102241 |
| complex_abundance | mitosis | inter | 0.00046429450983344415 | 0.013667862018171982 |

P82650_Q9NZE8
RT22_HUMAN vs RM35_HUMAN
p-value: 0.024 q-value: 0.018

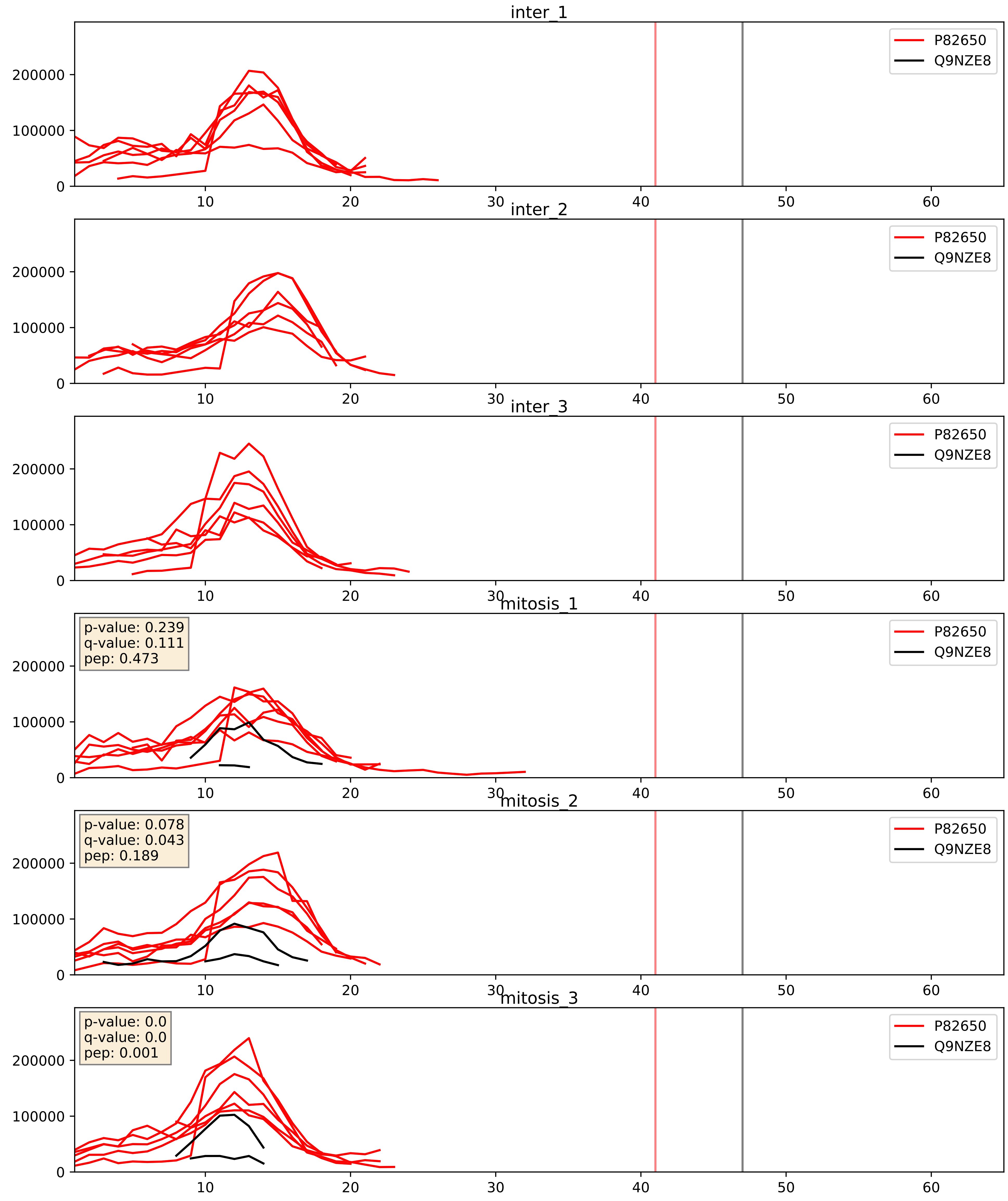


| level | condition_1 | condition_2 | pvalue | pvalue_adjusted |
|---|---|---|---|---|
| interactor_abundance | mitosis | inter | 0.00039383957194958876 | 0.013704336324749917 |
| interactor_ratio | mitosis | inter | 0.0006356762334973926 | 0.02925477719751441 |
| complex_abundance | mitosis | inter | 0.000663807866387126 | 0.016422529873623696 |

P82933_Q9NZE8
RT09_HUMAN vs RM35_HUMAN
p-value: 0.049 q-value: 0.032

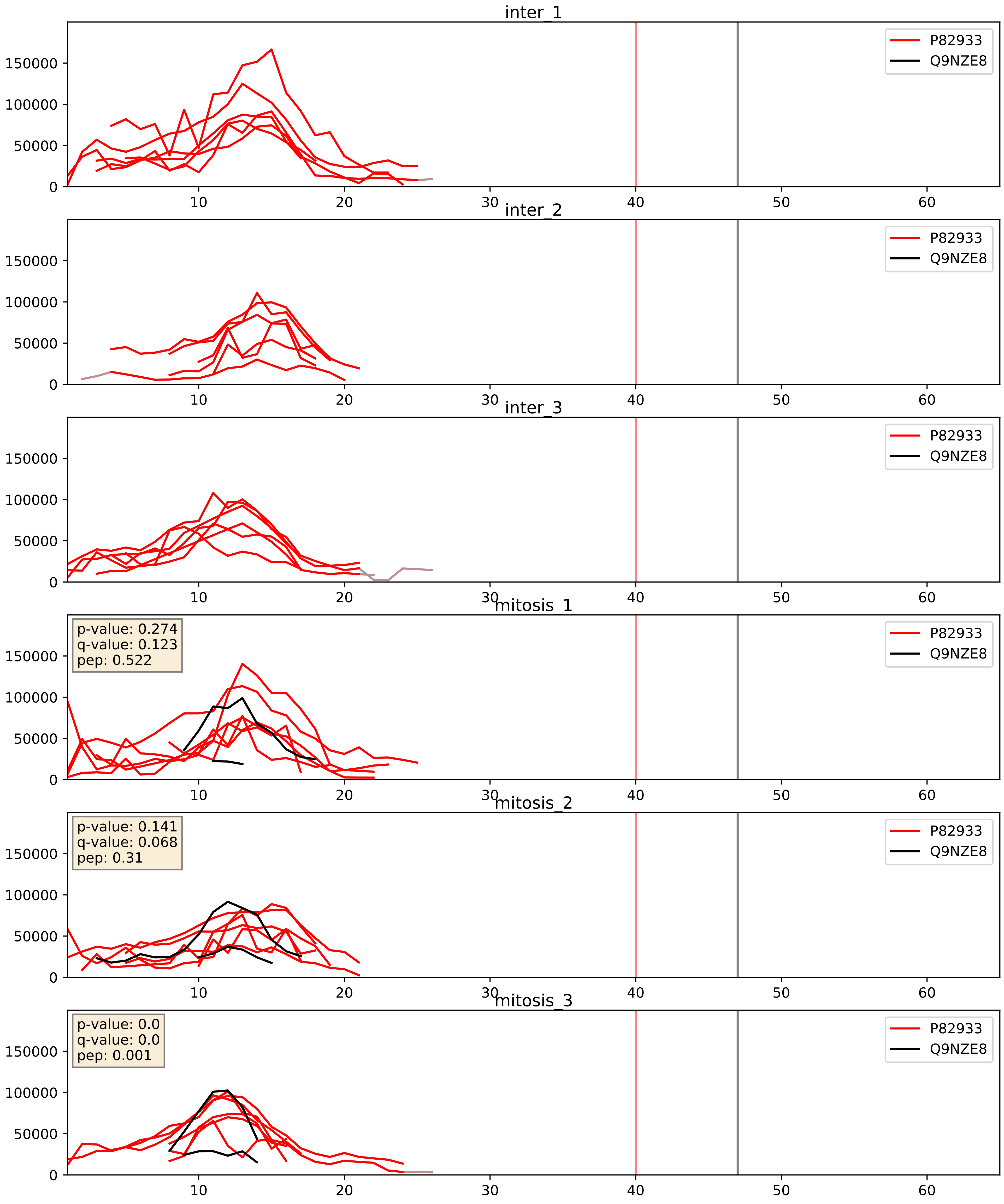


| level | condition_1 | condition_2 | pvalue | pvalue_adjusted |
|---|---|---|---|---|
| interactor_ratio | mitosis | inter | 0.004689685545013617 | 0.09159088101537749 |
| interactor_abundance | mitosis | inter | 0.006403949311252828 | 0.06847945796917453 |
| complex_abundance | mitosis | inter | 0.007787025536984597 | 0.06557020805367884 |

Q13084_Q9NZE8
RM28_HUMAN vs RM35_HUMAN
p-value: 0.0 q-value: 0.0

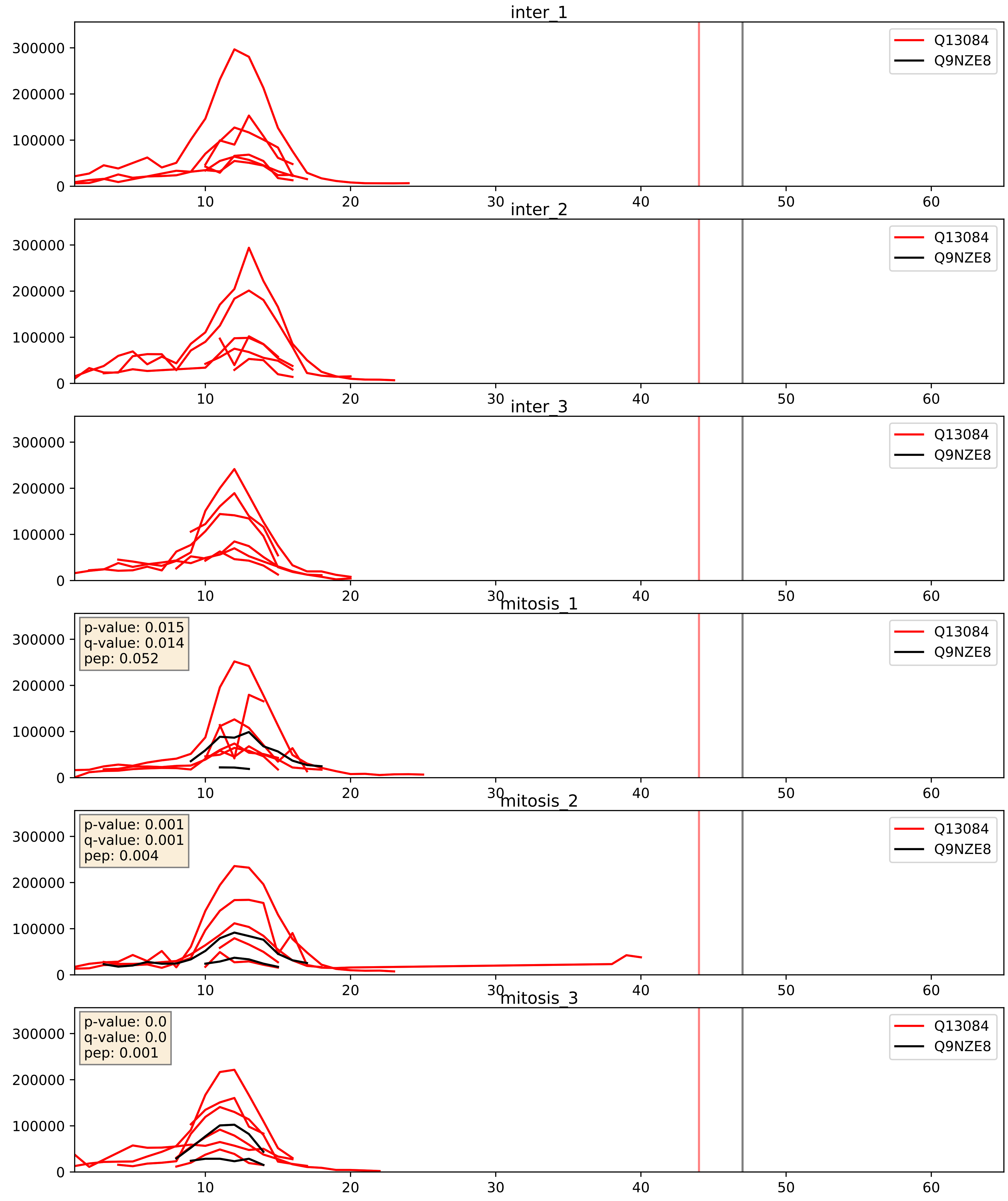


| level | condition_1 | condition_2 | pvalue | pvalue_adjusted |
|---|---|---|---|---|
| interactor_abundance | mitosis | inter | 0.0007778715672234191 | 0.02050037113506771 |
| interactor_ratio | mitosis | inter | 0.001073978392292625 | 0.038414698994273265 |
| complex_abundance | mitosis | inter | 0.0014060568822015964 | 0.025608184918395033 |

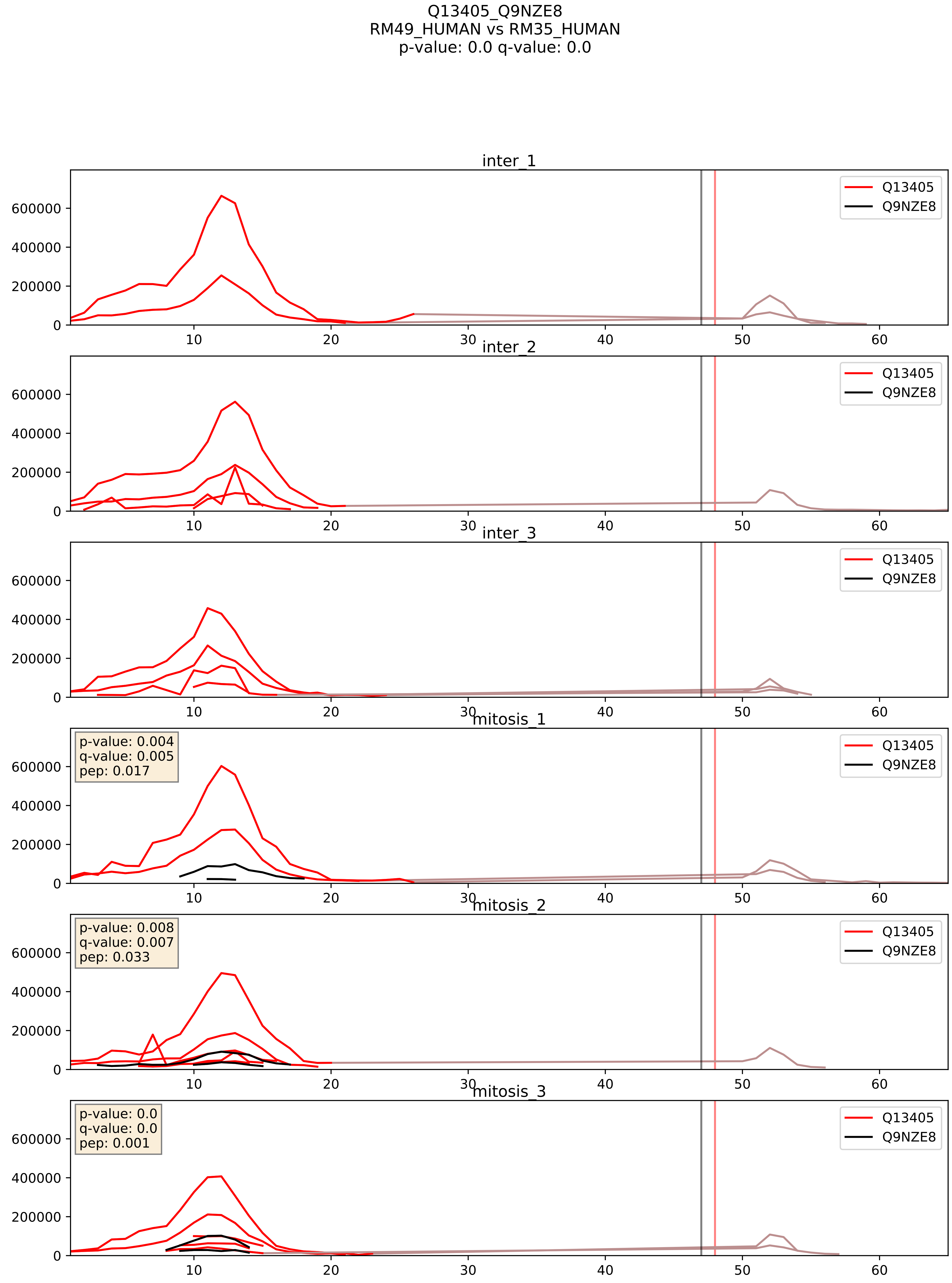


| level | condition_1 | condition_2 | pvalue | pvalue_adjusted |
|---|---|---|---|---|
| interactor_ratio | mitosis | inter | 3.435573560173287e-05 | 0.006185155691845816 |
| interactor_abundance | mitosis | inter | 0.0002456300330881158 | 0.01187905696742526 |
| complex_abundance | mitosis | inter | 0.0008334340525324971 | 0.019229637438485648 |

Q14197_Q9NZE8
ICT1_HUMAN vs RM35_HUMAN
p-value: 0.0 q-value: 0.003

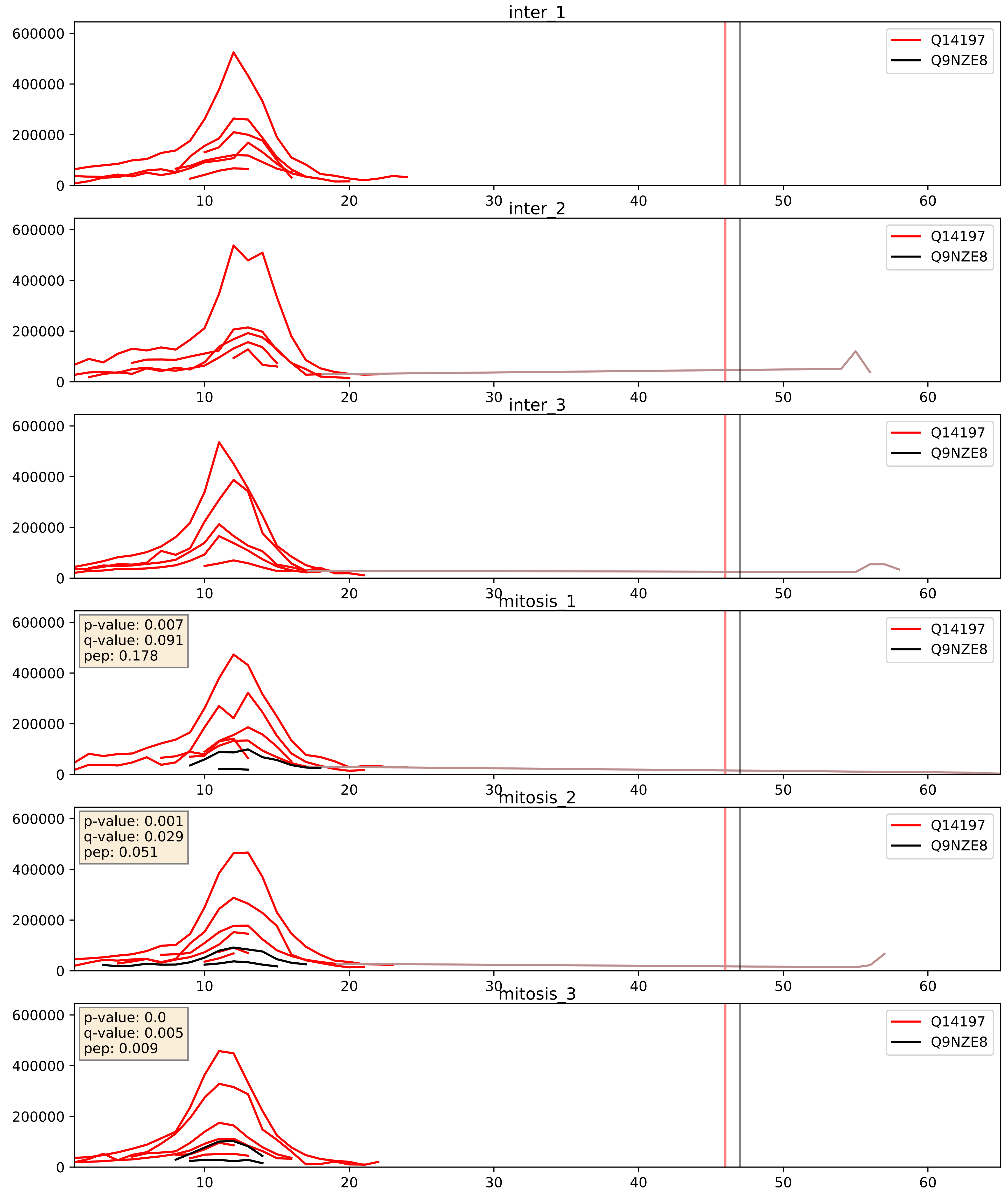


| level | condition_1 | condition_2 | pvalue | pvalue_adjusted |
|---|---|---|---|---|
| interactor_ratio | mitosis | inter | 1.202946967354636e-05 | 0.003960471554059879 |
| interactor_abundance | mitosis | inter | 6.029903163637628e-05 | 0.006459477818931243 |
| complex_abundance | mitosis | inter | 0.0002708323422165612 | 0.011039642139875067 |

Q16540_Q9NZE8
RM23_HUMAN vs RM35_HUMAN
p-value: 0.0 q-value: 0.008

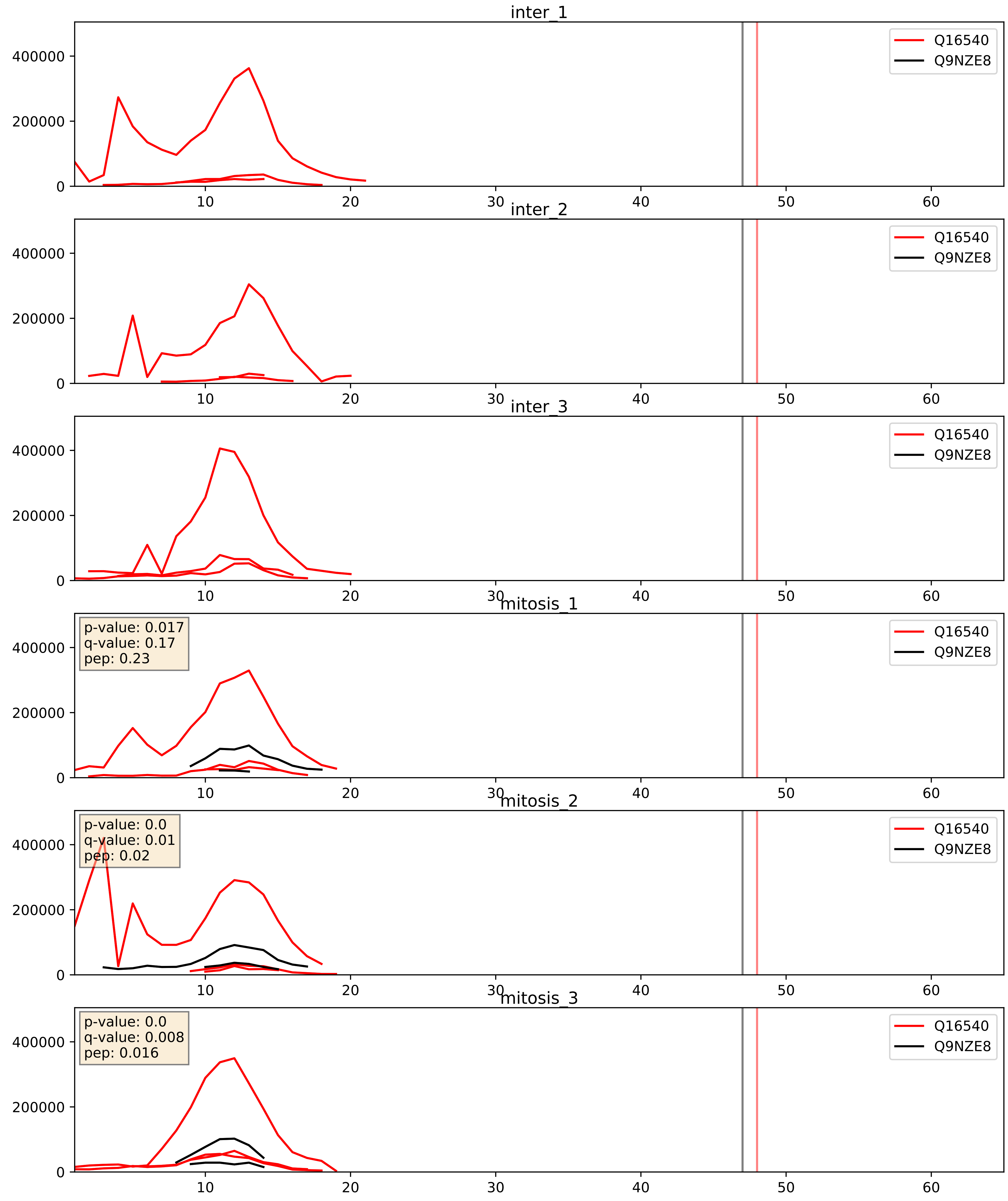


| level | condition_1 | condition_2 | pvalue | pvalue_adjusted |
|---|---|---|---|---|
| interactor_ratio | mitosis | inter | 0.001392665828104323 | 0.044212523606194314 |
| interactor_abundance | mitosis | inter | 0.0023128623936897216 | 0.035640147776748905 |
| complex_abundance | mitosis | inter | 0.003804959318430418 | 0.04558257253867179 |

Q5T653_Q9NZE8
RM02_HUMAN vs RM35_HUMAN
p-value: 0.002 q-value: 0.003

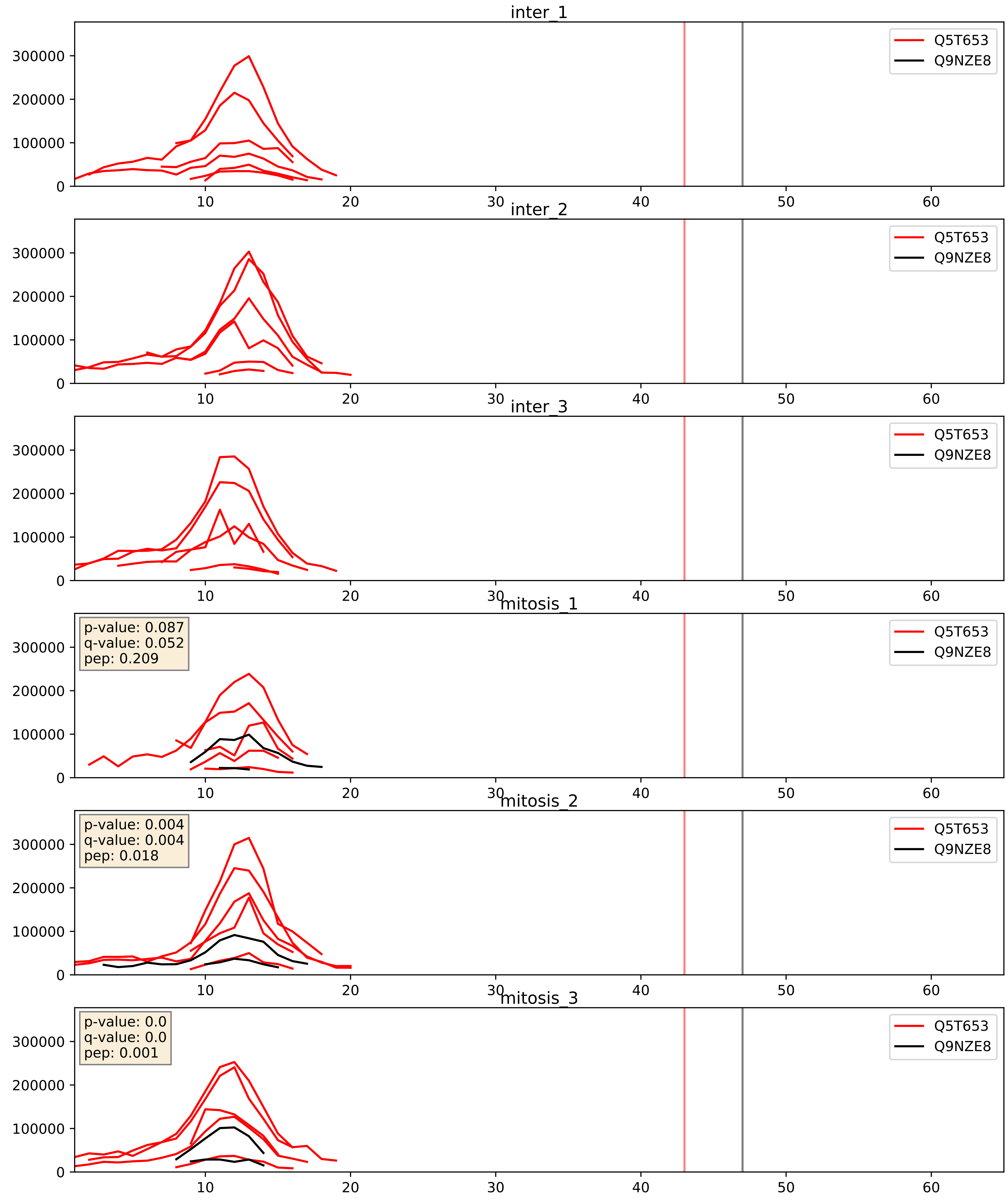


| level | condition_1 | condition_2 | pvalue | pvalue_adjusted |
|---|---|---|---|---|
| interactor_abundance | mitosis | inter | 0.000471319604376598 | 0.015253292300429789 |
| complex_abundance | mitosis | inter | 0.0006614826715483389 | 0.01642175463112936 |
| interactor_ratio | mitosis | inter | 0.0011684934404209454 | 0.04023858234363702 |

Q6P1L8_Q9NZE8
RM14_HUMAN vs RM35_HUMAN
p-value: 0.006 q-value: 0.006

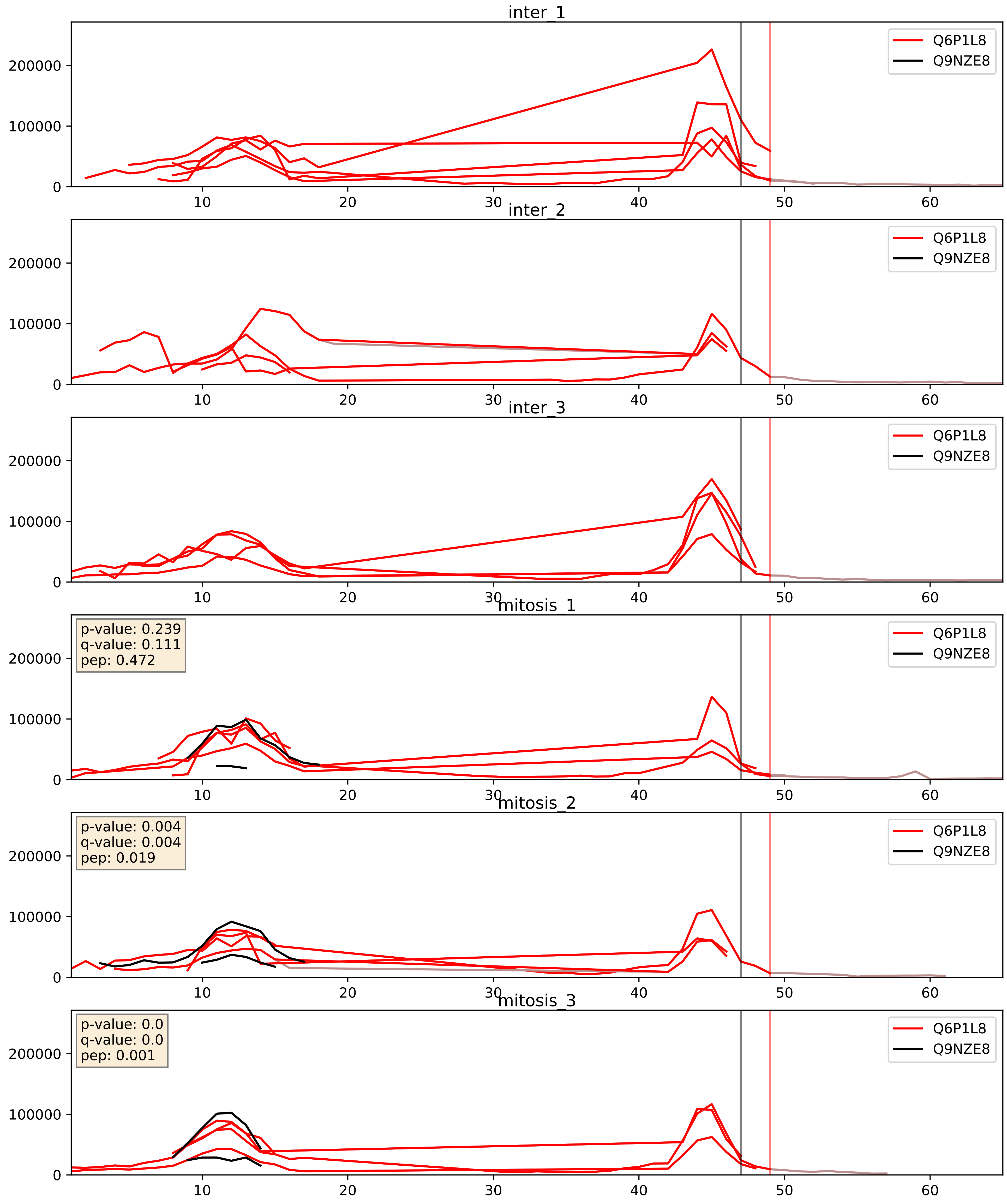


| level | condition_1 | condition_2 | pvalue | pvalue_adjusted |
|---|---|---|---|---|
| interactor_abundance | mitosis | inter | 0.013398151618360356 | 0.10304923015527873 |
| complex_abundance | mitosis | inter | 0.013554529540930305 | 0.0865877814270891 |
| interactor_ratio | mitosis | inter | 0.015739053098212975 | 0.1848097318528163 |

Q7Z2W9_Q9NZE8
RM21_HUMAN vs RM35_HUMAN
p-value: 0.0 q-value: 0.001

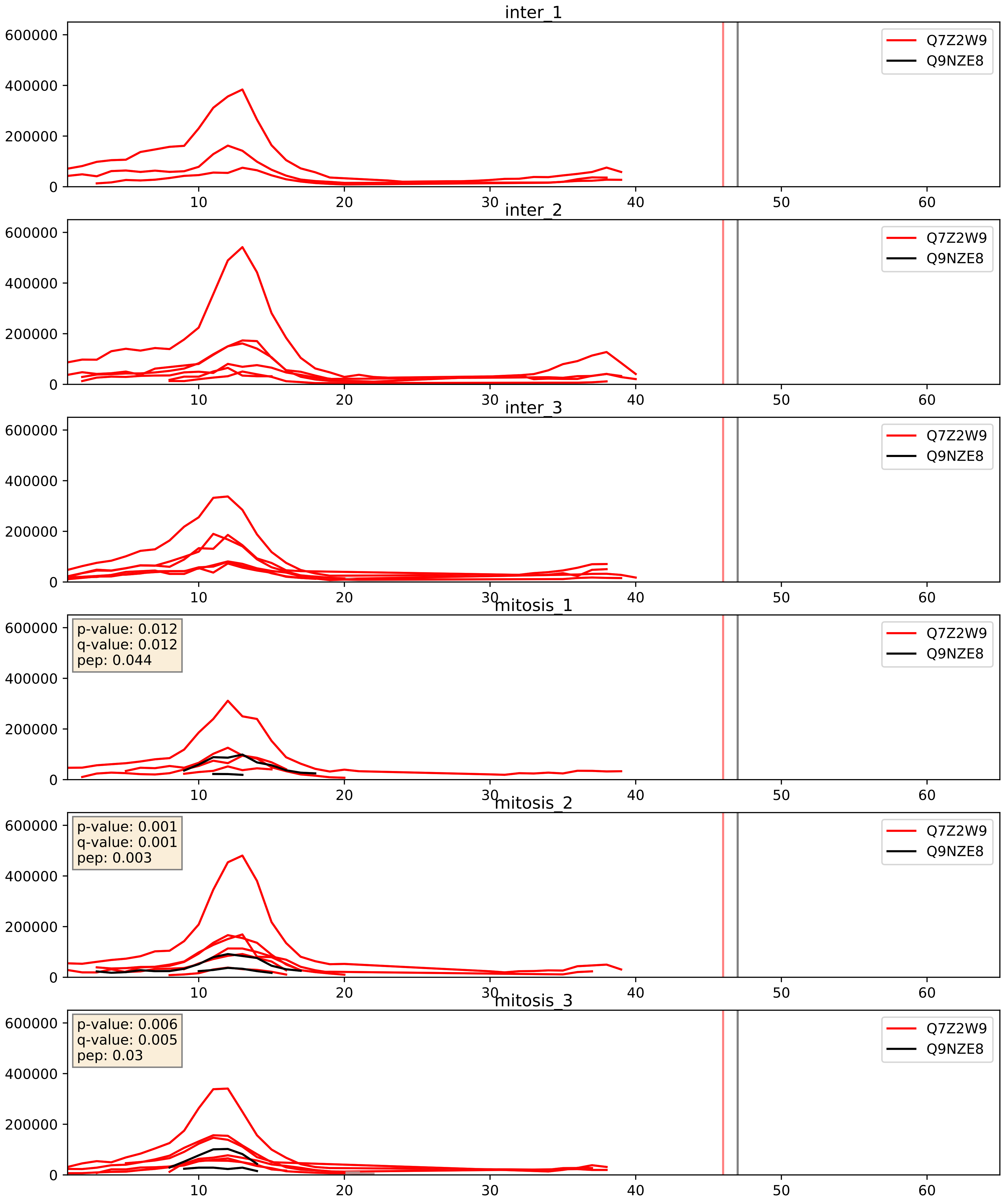


| level | condition_1 | condition_2 | pvalue | pvalue_adjusted |
|---|---|---|---|---|
| interactor_abundance | mitosis | inter | 0.00044538007768532095 | 0.014719897548209836 |
| complex_abundance | mitosis | inter | 0.0005987514747566535 | 0.01581504091436693 |
| interactor_ratio | mitosis | inter | 0.0012161139437632392 | 0.04130926729608463 |

Q7Z7F7_Q9NZE8
RM55_HUMAN vs RM35_HUMAN
p-value: 0.0 q-value: 0.0

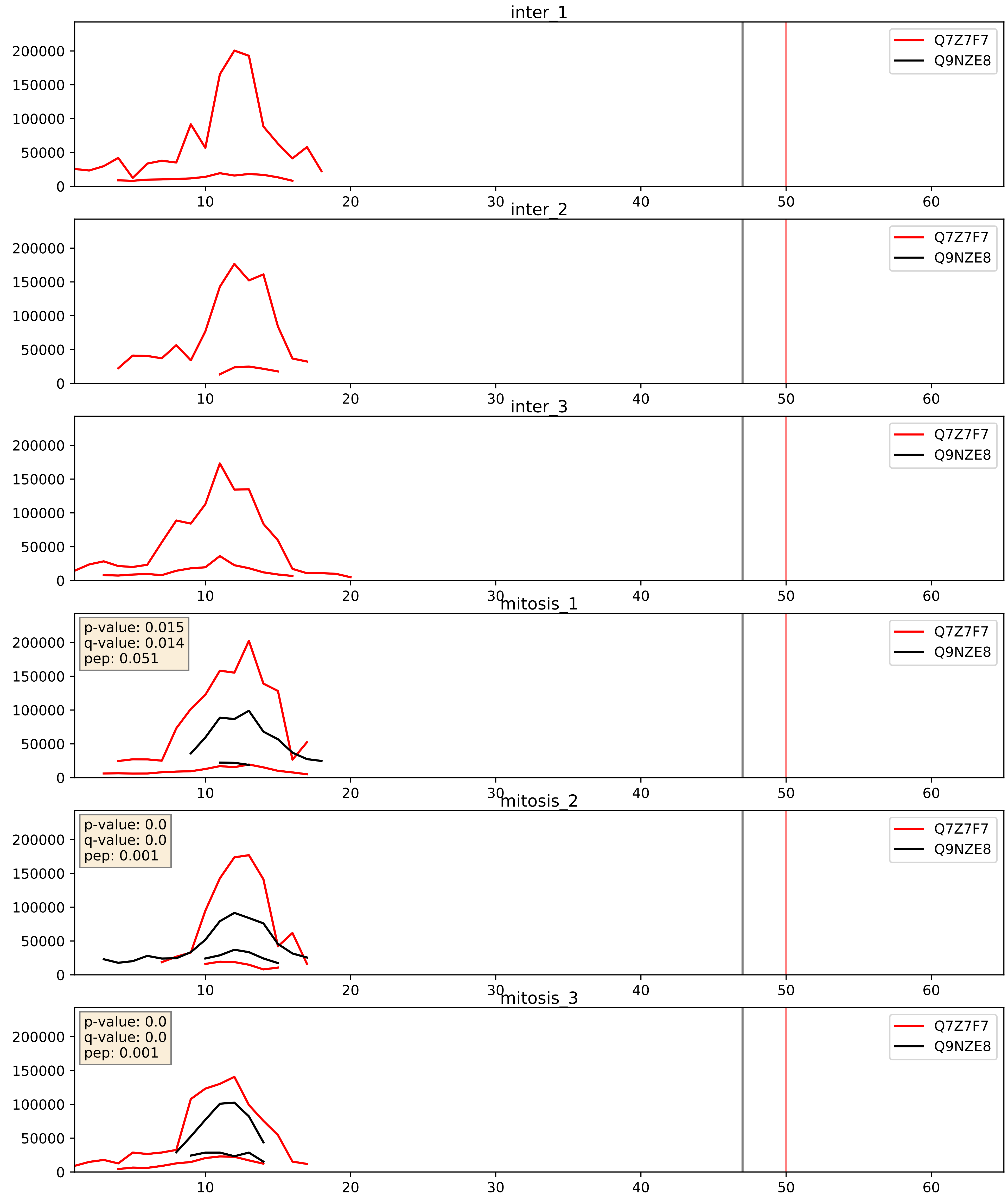


| level | condition_1 | condition_2 | pvalue | pvalue_adjusted |
|---|---|---|---|---|
| complex_abundance | mitosis | inter | 0.015683540735771637 | 0.09352166546417016 |
| interactor_abundance | mitosis | inter | 0.016193183296677703 | 0.11149217703460695 |
| interactor_ratio | mitosis | inter | 0.0423971923383099 | 0.3404502499211377 |

Q7Z7H8_Q9NZE8
RM10_HUMAN vs RM35_HUMAN
p-value: 0.003 q-value: 0.003

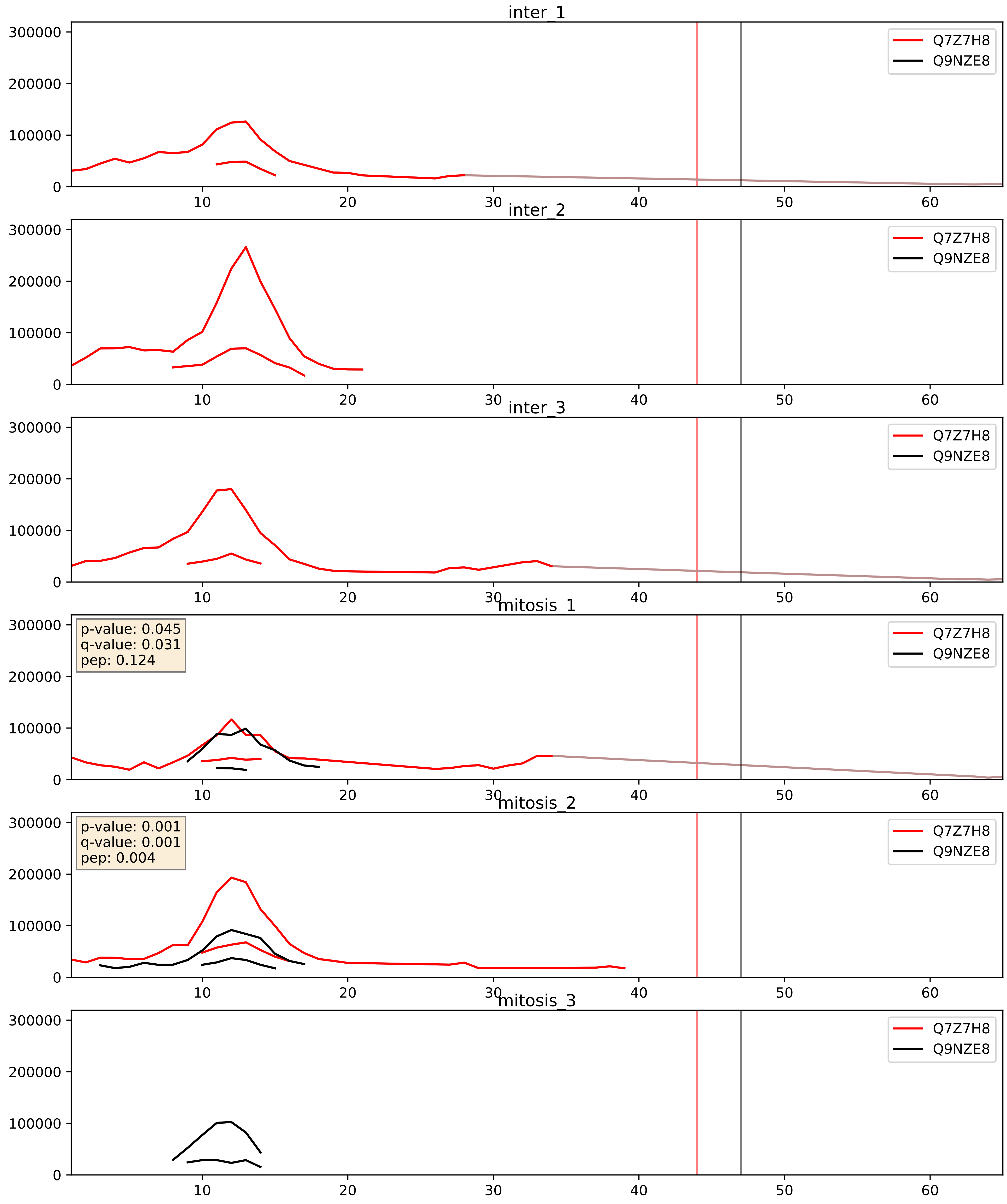


| level | condition_1 | condition_2 | pvalue | pvalue_adjusted |
|---|---|---|---|---|
| interactor_ratio | mitosis | inter | 0.22017207466890404 | 0.6977685891024875 |
| interactor_abundance | mitosis | inter | 0.4577203702037011 | 0.629609570279835 |
| complex_abundance | mitosis | inter | 0.5342414056390921 | 0.6696597499298035 |

Q8IXM3_Q9NZE8
RM41_HUMAN vs RM35_HUMAN
p-value: 0.0 q-value: 0.0

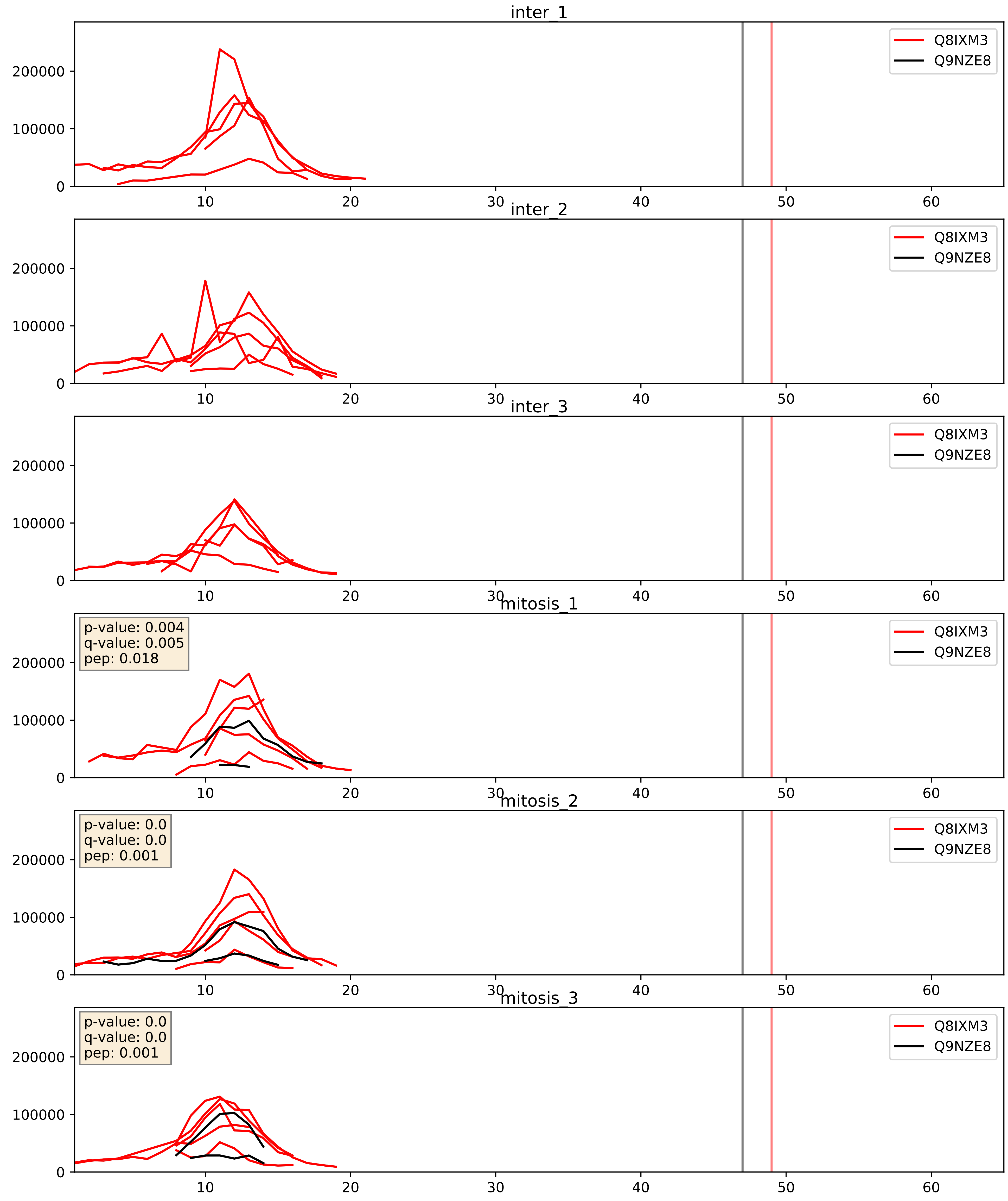


| level | condition_1 | condition_2 | pvalue | pvalue_adjusted |
|---|---|---|---|---|
| interactor_ratio | mitosis | inter | 0.0005304476854813565 | 0.02670960110423772 |
| interactor_abundance | mitosis | inter | 0.001815196307140194 | 0.03132677497806464 |
| complex_abundance | mitosis | inter | 0.0033037987621844524 | 0.04228056500010875 |

Q8N5N7_Q9NZE8
RM50_HUMAN vs RM35_HUMAN
p-value: 0.0 q-value: 0.001

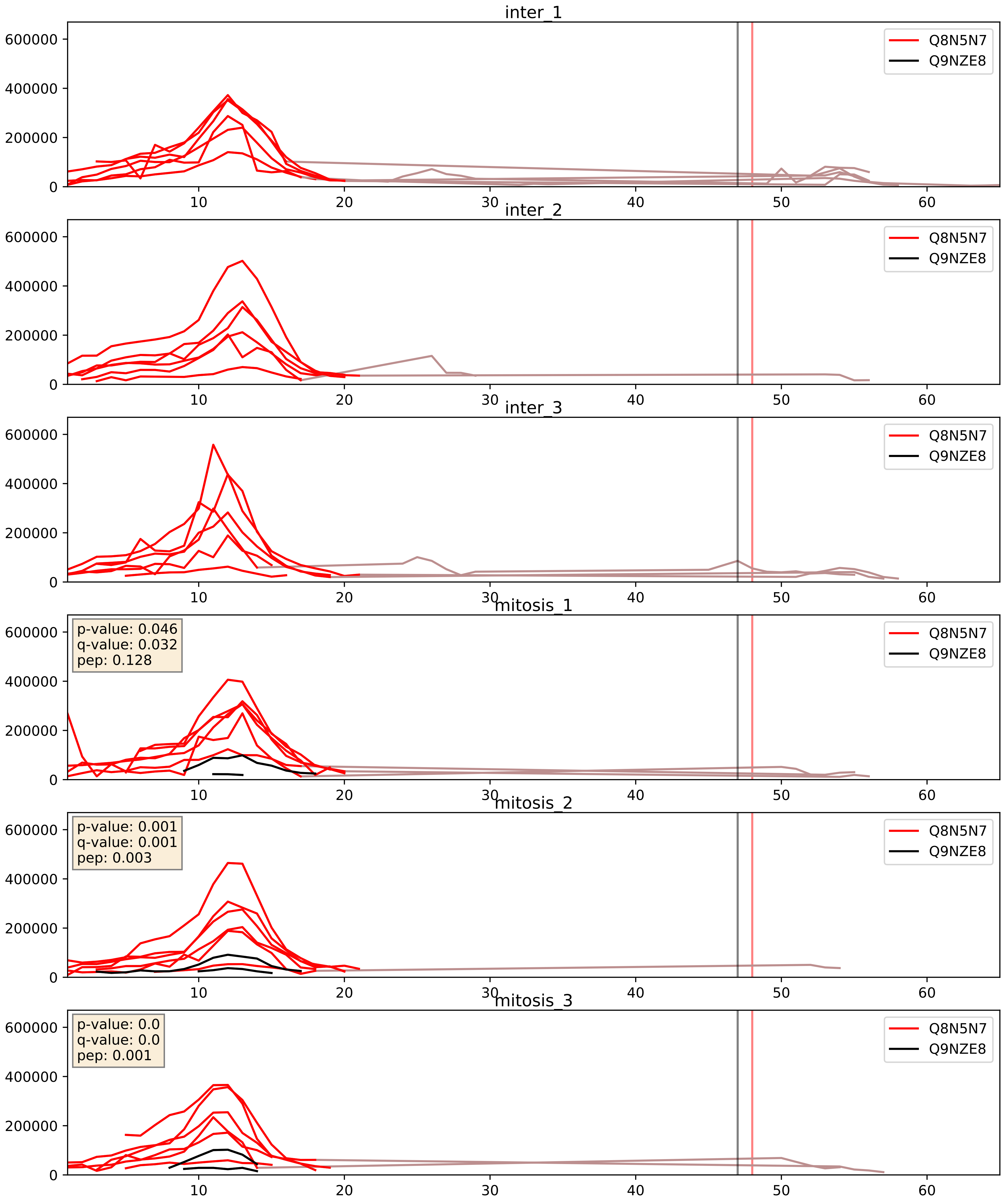


| level | condition_1 | condition_2 | pvalue | pvalue_adjusted |
|---|---|---|---|---|
| interactor_ratio | mitosis | inter | 1.5842048390654197e-05 | 0.004237747944499997 |
| interactor_abundance | mitosis | inter | 6.274450699580499e-05 | 0.006470999757639647 |
| complex_abundance | mitosis | inter | 0.00023742048328256082 | 0.010310695784295044 |

Q8N983_Q9NZE8
RM43_HUMAN vs RM35_HUMAN
p-value: 0.001 q-value: 0.002

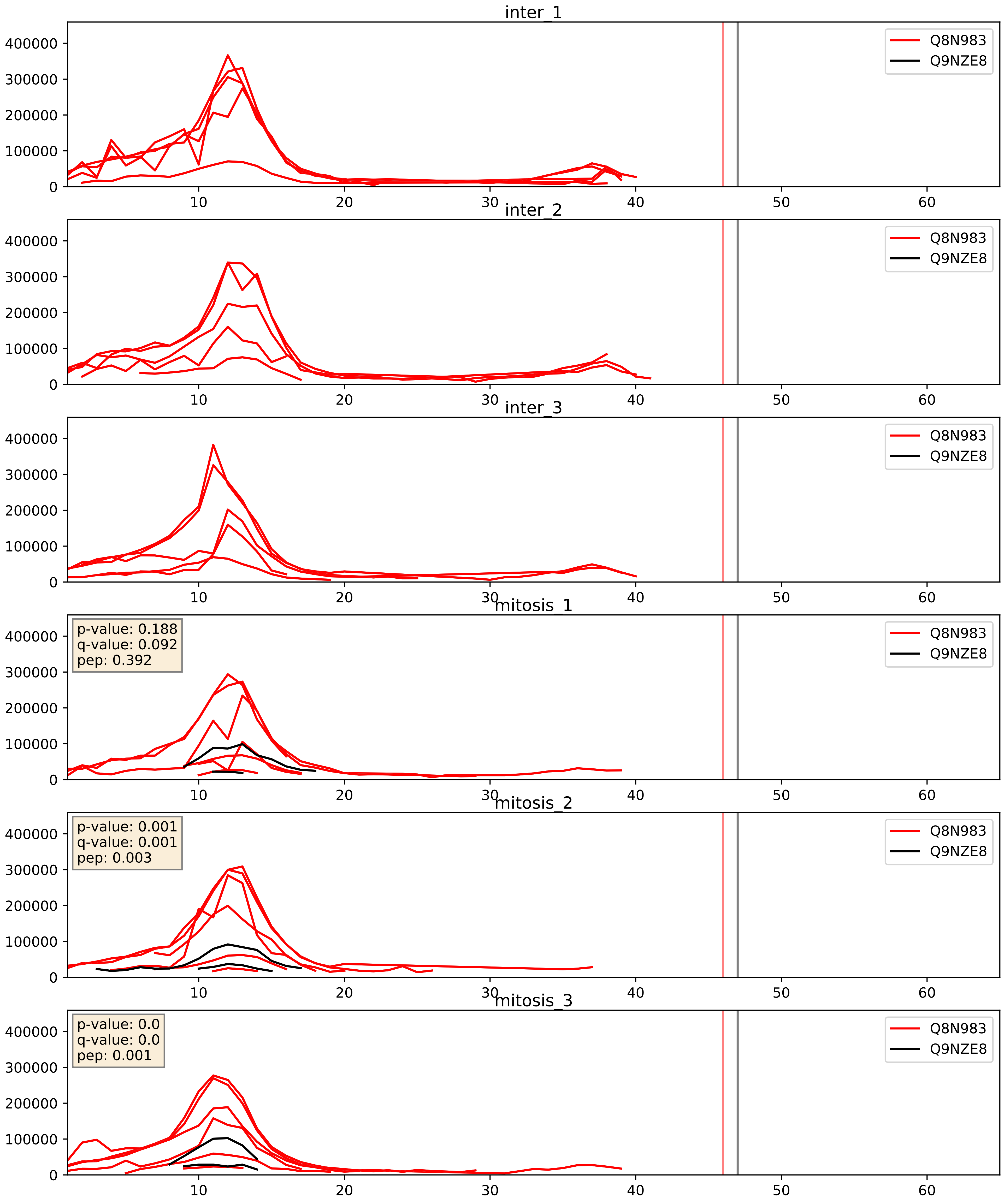


| level | condition_1 | condition_2 | pvalue | pvalue_adjusted |
|---|---|---|---|---|
| interactor_abundance | mitosis | inter | 0.000304688924010125 | 0.013073369371061003 |
| complex_abundance | mitosis | inter | 0.00043010621427248347 | 0.013091389064164169 |
| interactor_ratio | mitosis | inter | 0.0007653204461325218 | 0.03180166514026401 |

Q8TAE8_Q9NZE8
G45IP_HUMAN vs RM35_HUMAN
p-value: 0.001 q-value: 0.021

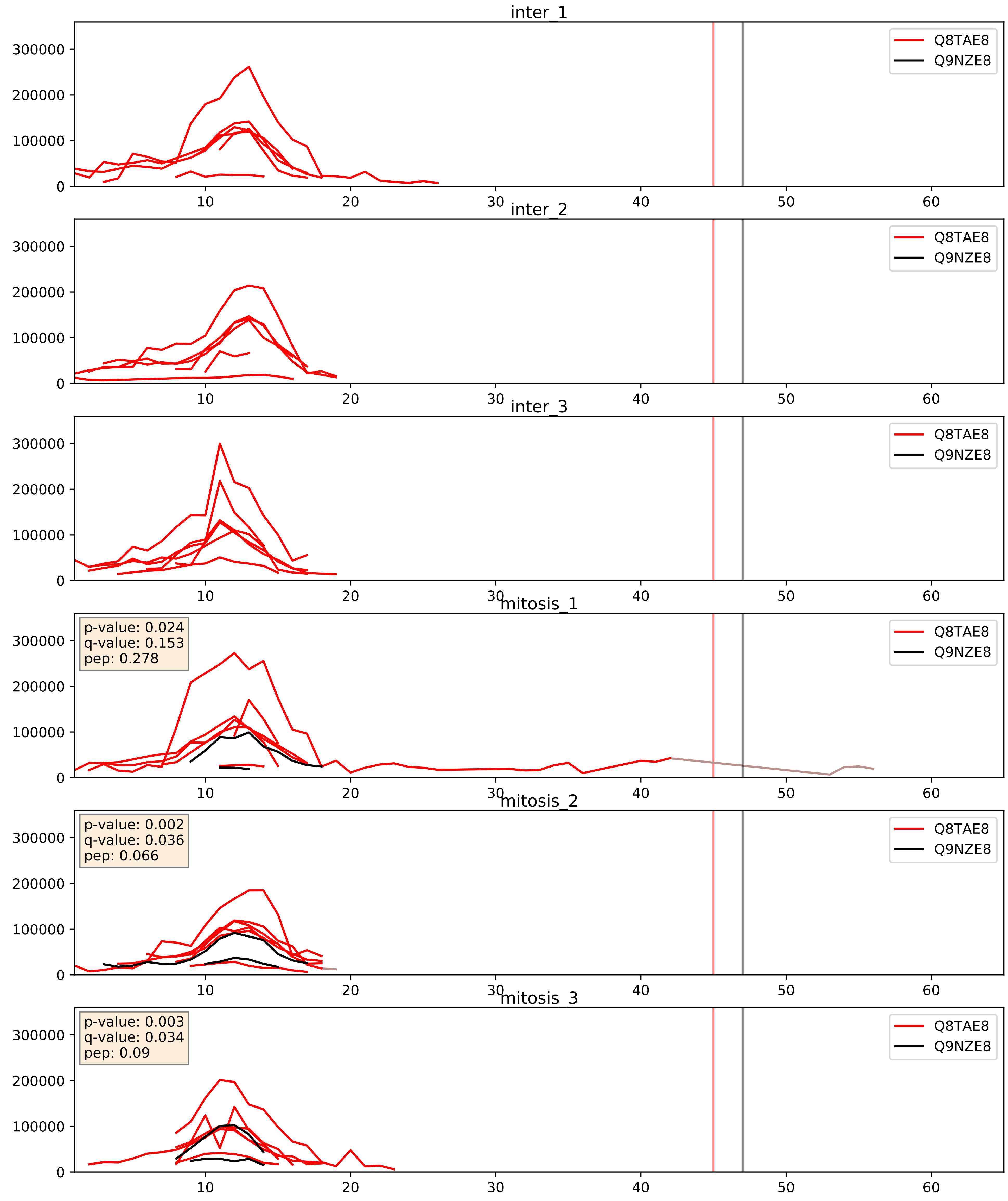


| level | condition_1 | condition_2 | pvalue | pvalue_adjusted |
|---|---|---|---|---|
| interactor_ratio | mitosis | inter | 0.0008167336421248371 | 0.03277774637652267 |
| interactor_abundance | mitosis | inter | 0.002081365263003045 | 0.03317781499312116 |
| complex_abundance | mitosis | inter | 0.003414130234741549 | 0.04302508960916016 |

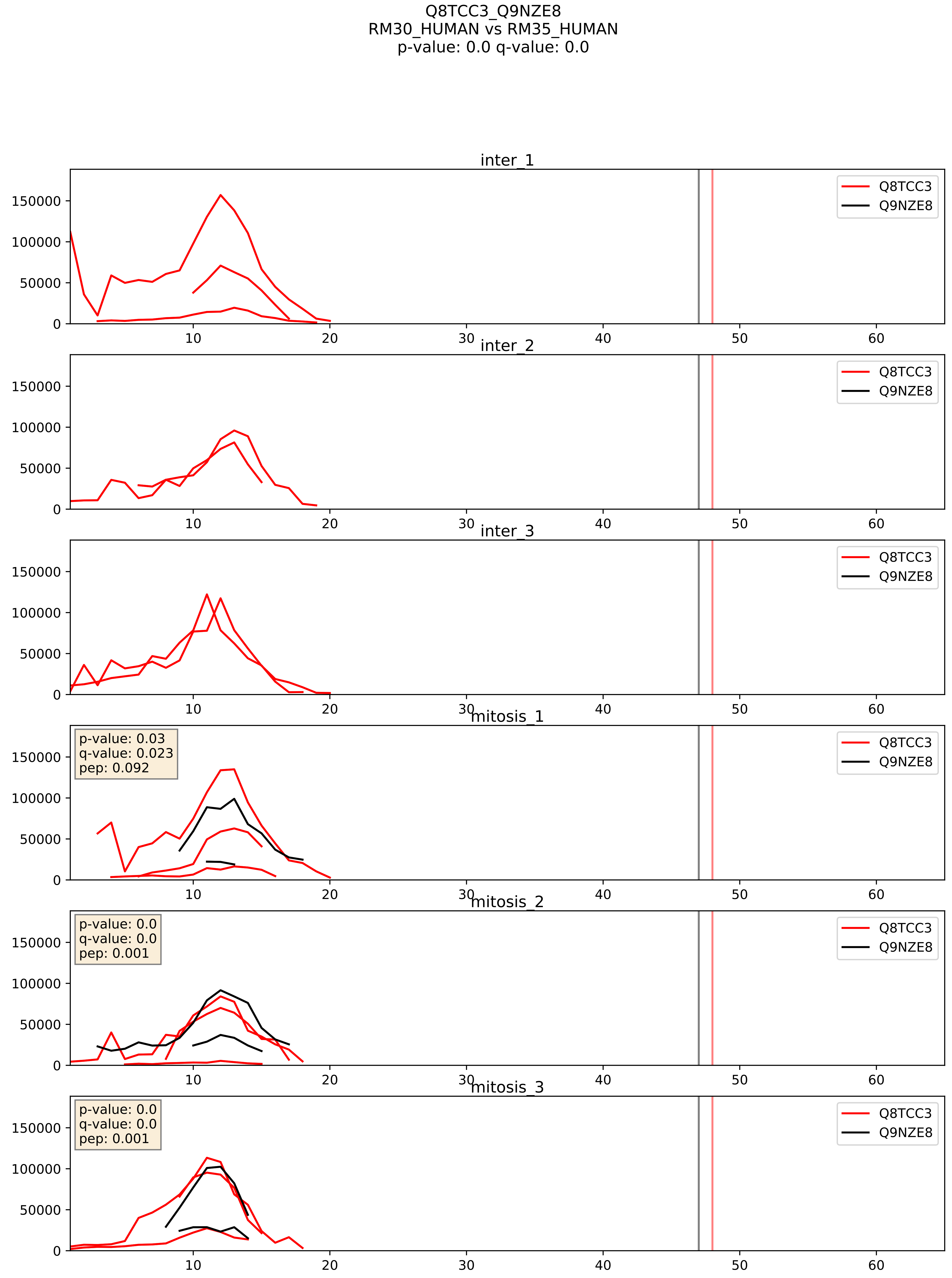


| level | condition_1 | condition_2 | pvalue | pvalue_adjusted |
|---|---|---|---|---|
| complex_abundance | mitosis | inter | 0.015724030307540635 | 0.09352166546417016 |
| interactor_abundance | mitosis | inter | 0.019249969920338075 | 0.11885079271397347 |
| interactor_ratio | mitosis | inter | 0.06015313210053977 | 0.40576108020537466 |

Q96A35_Q9NZE8
RM24_HUMAN vs RM35_HUMAN
p-value: 0.0 q-value: 0.0

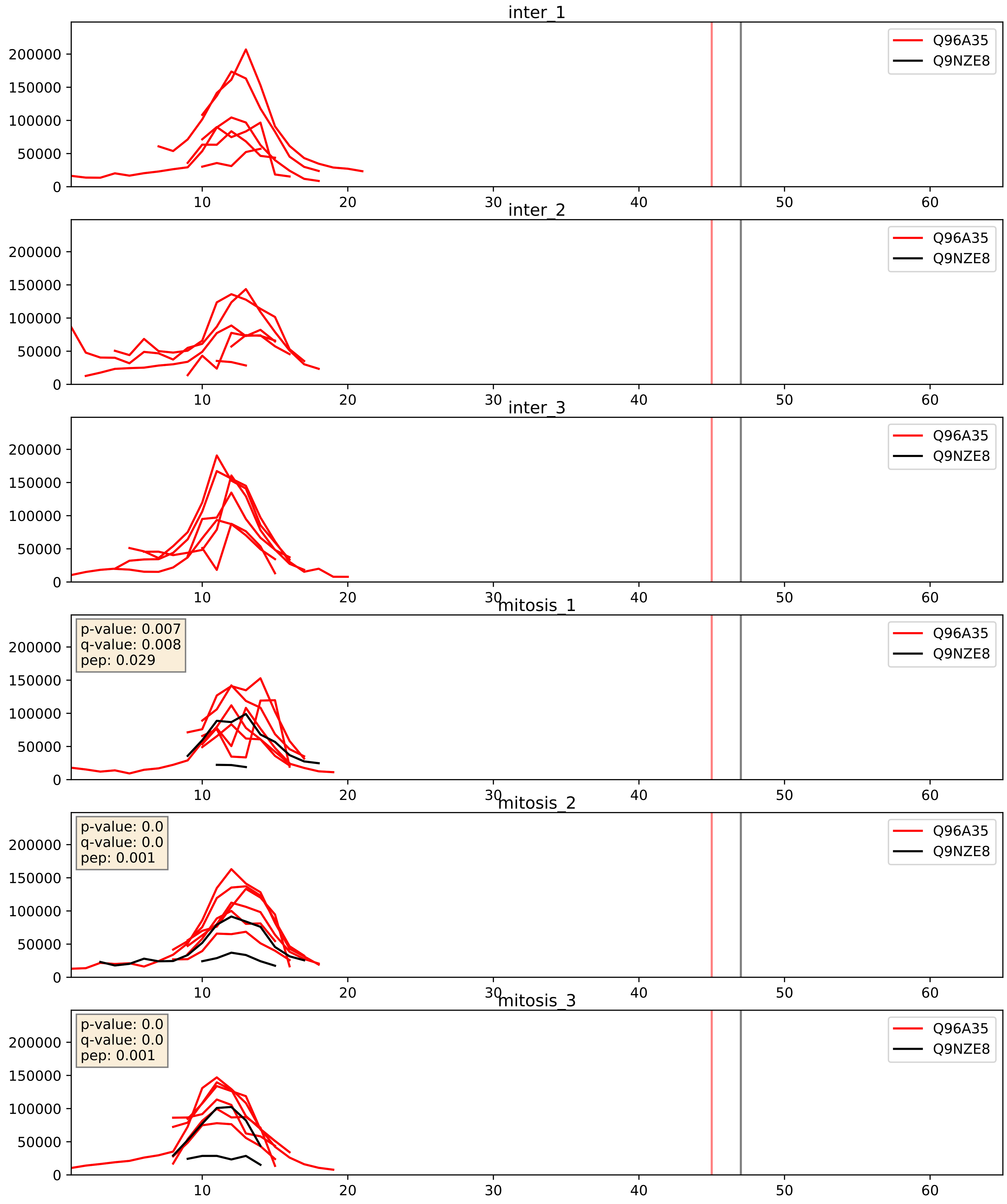


| level | condition_1 | condition_2 | pvalue | pvalue_adjusted |
|---|---|---|---|---|
| interactor_ratio | mitosis | inter | 3.468311602904196e-05 | 0.006185155691845816 |
| interactor_abundance | mitosis | inter | 0.0006188665072018961 | 0.01748938971196646 |
| complex_abundance | mitosis | inter | 0.0017767857790105326 | 0.029939539898287716 |

Q96DV4_Q9NZE8
RM38_HUMAN vs RM35_HUMAN
p-value: 0.0 q-value: 0.001

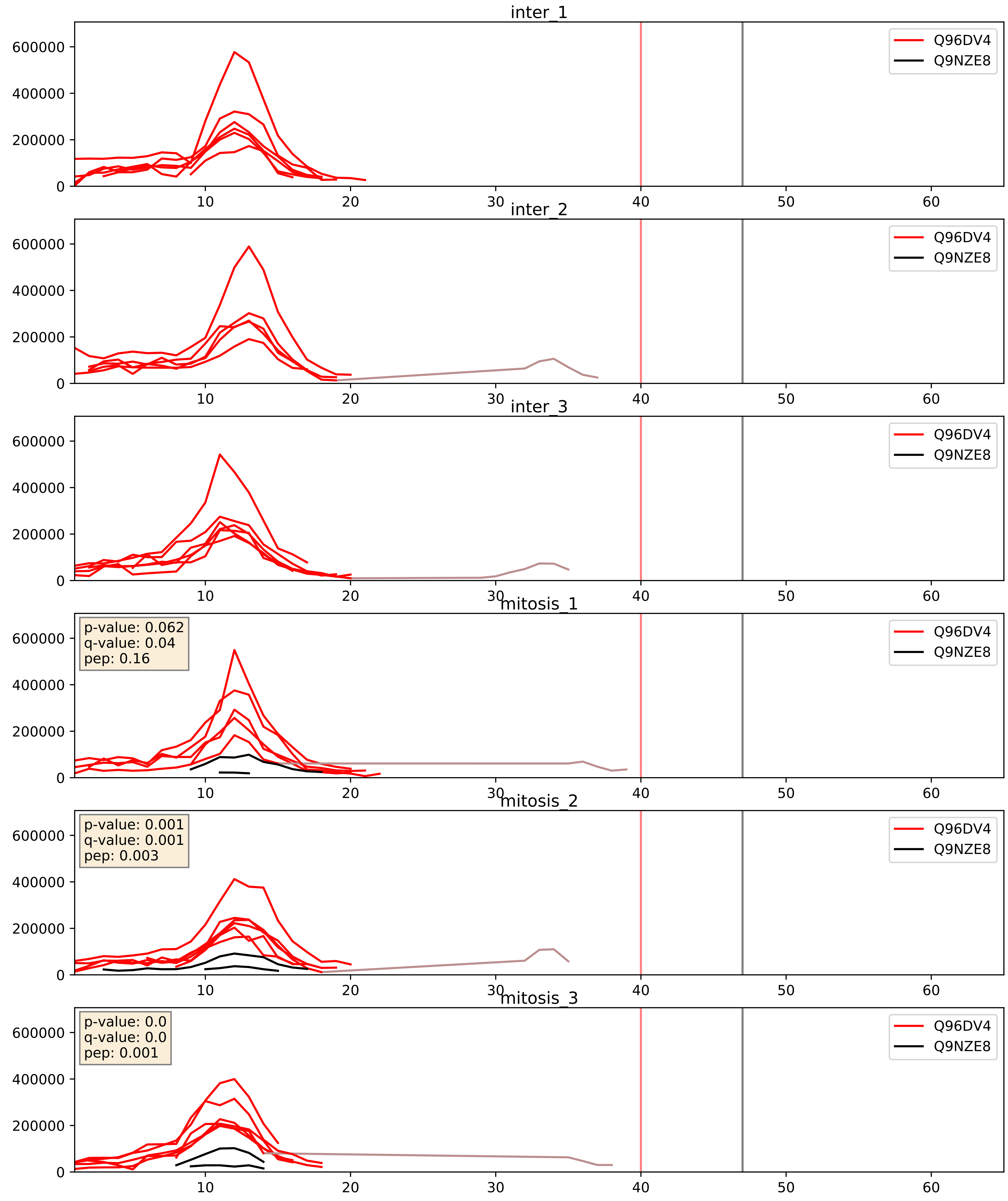


| level | condition_1 | condition_2 | pvalue | pvalue_adjusted |
|---|---|---|---|---|
| interactor_ratio | mitosis | inter | 3.97948876885048e-06 | 0.002520386835947235 |
| interactor_abundance | mitosis | inter | 8.740887541956166e-05 | 0.007396785432408699 |
| complex_abundance | mitosis | inter | 0.00035493161137333326 | 0.012039775462644836 |

Q96EL3_Q9NZE8
RM53_HUMAN vs RM35_HUMAN
p-value: 0.001 q-value: 0.002

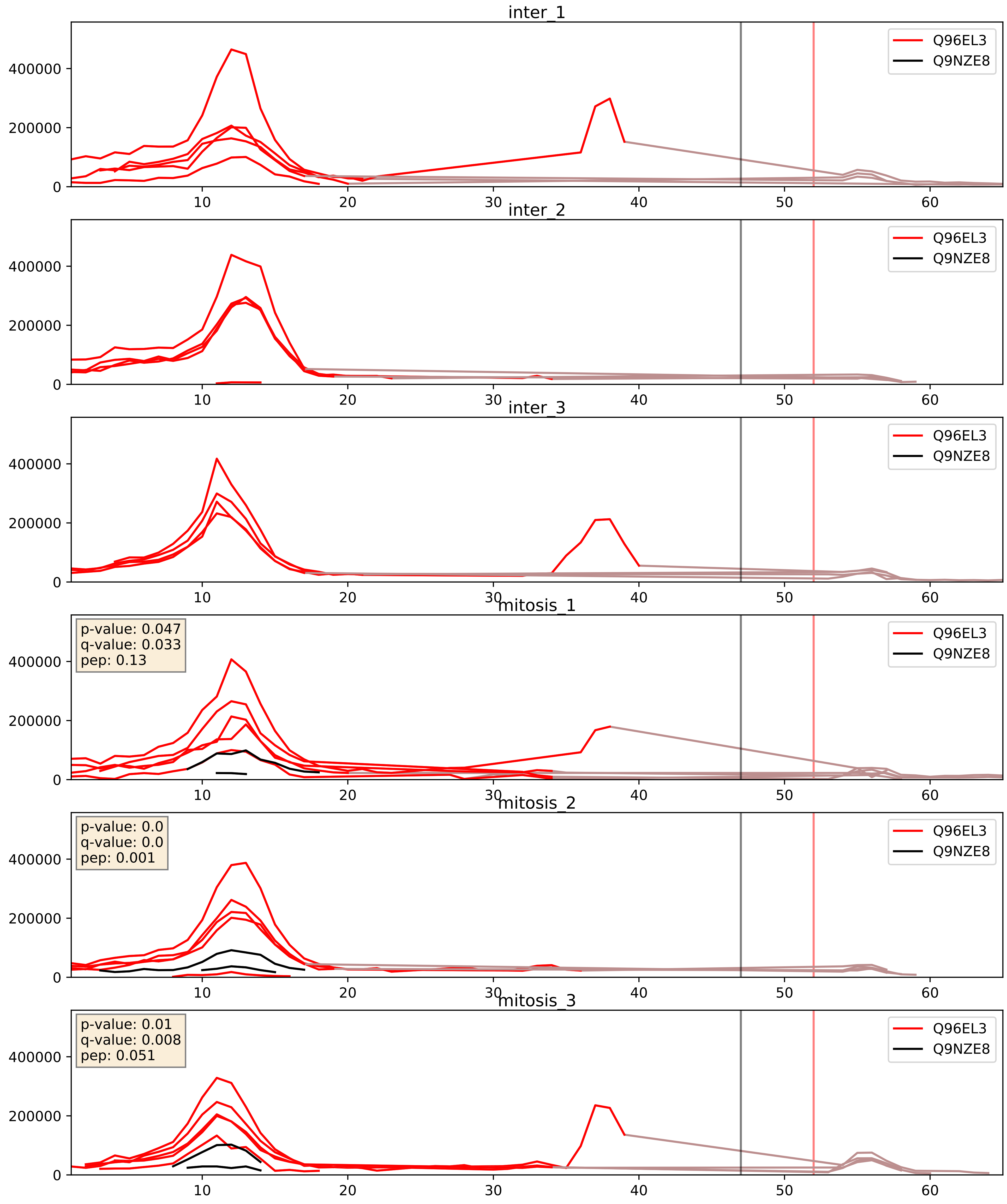


| level | condition_1 | condition_2 | pvalue | pvalue_adjusted |
|---|---|---|---|---|
| interactor_ratio | mitosis | inter | 0.00010644987613242638 | 0.010720128702277293 |
| interactor_abundance | mitosis | inter | 0.000203702071830833 | 0.010763516881925495 |
| complex_abundance | mitosis | inter | 0.0005163436775203946 | 0.014444123789459404 |

Q96GC5_Q9NZE8
RM48_HUMAN vs RM35_HUMAN
p-value: 0.0 q-value: 0.0

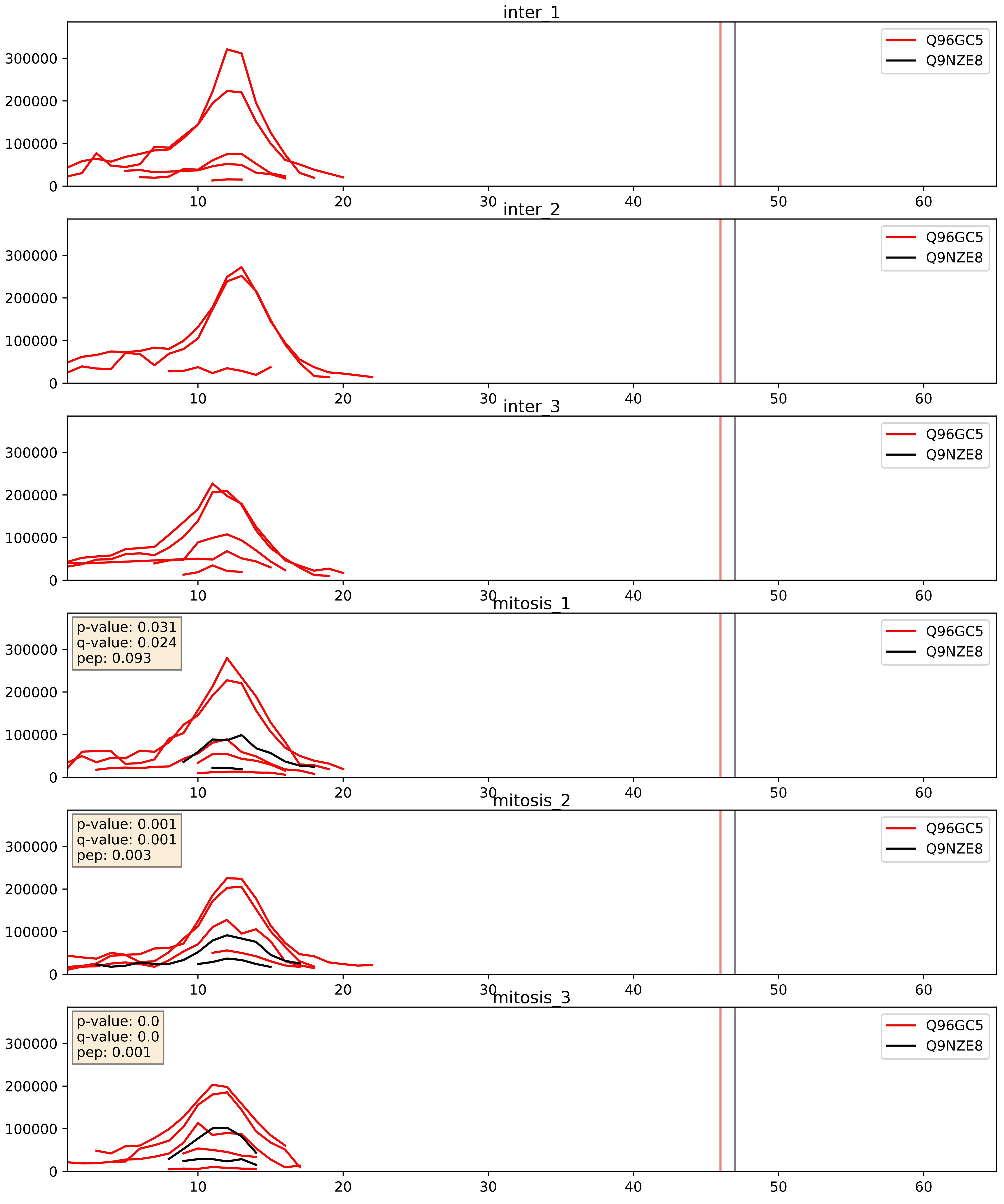


| level | condition_1 | condition_2 | pvalue | pvalue_adjusted |
|---|---|---|---|---|
| interactor_ratio | mitosis | inter | 0.00013707473310461304 | 0.012616771133069759 |
| interactor_abundance | mitosis | inter | 0.0004940523069066729 | 0.0157801781608997 |
| complex_abundance | mitosis | inter | 0.001217098331091602 | 0.023464778635459718 |

Q9BQ48_Q9NZE8
RM34_HUMAN vs RM35_HUMAN
p-value: 0.0 q-value: 0.0

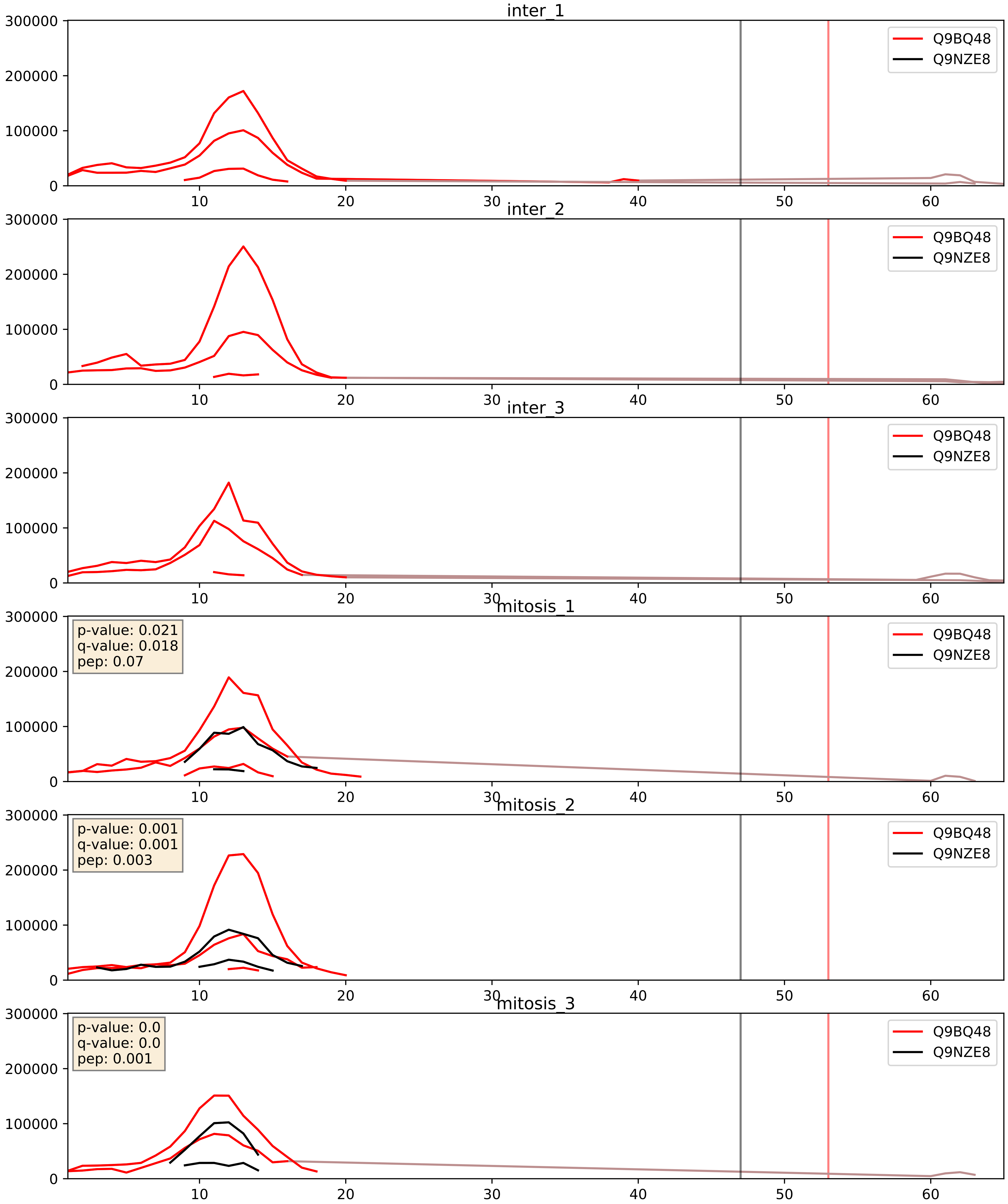


| level | condition_1 | condition_2 | pvalue | pvalue_adjusted |
|---|---|---|---|---|
| complex_abundance | mitosis | inter | 0.016194101194036054 | 0.09507647888953953 |
| interactor_abundance | mitosis | inter | 0.019774471828388575 | 0.1190514126751731 |
| interactor_ratio | mitosis | inter | 0.04217450998621126 | 0.3399226239910554 |

Q9BQC6_Q9NZE8
RT63_HUMAN vs RM35_HUMAN
p-value: 0.0 q-value: 0.003

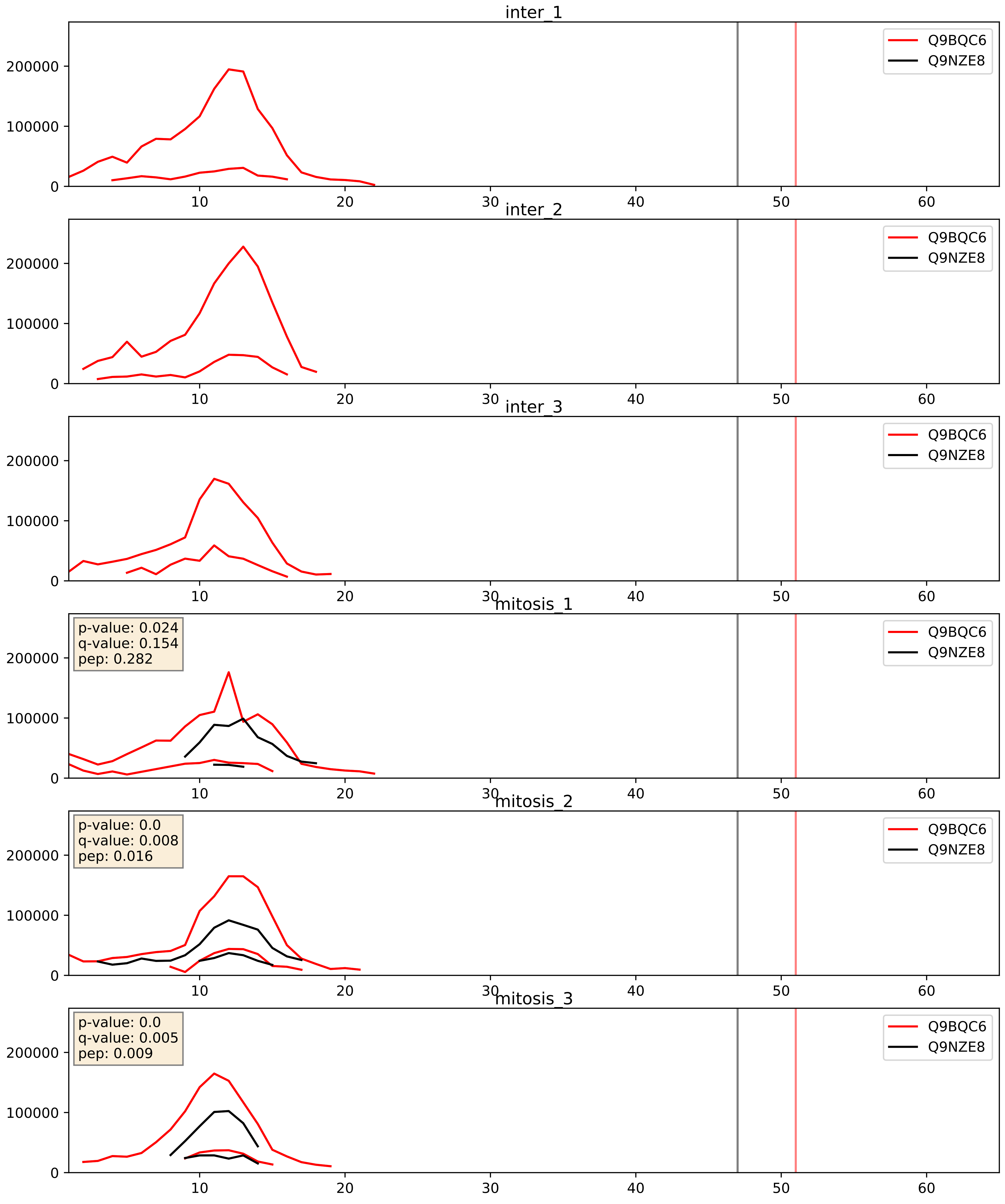


| level | condition_1 | condition_2 | pvalue | pvalue_adjusted |
|---|---|---|---|---|
| interactor_ratio | mitosis | inter | 0.0006732493598817932 | 0.029848225645041686 |
| interactor_abundance | mitosis | inter | 0.002403542935310808 | 0.03654409862568475 |
| complex_abundance | mitosis | inter | 0.005298968044648308 | 0.05478160200747526 |

Q9BYC8_Q9NZE8
RM32_HUMAN vs RM35_HUMAN
p-value: 0.0 q-value: 0.0

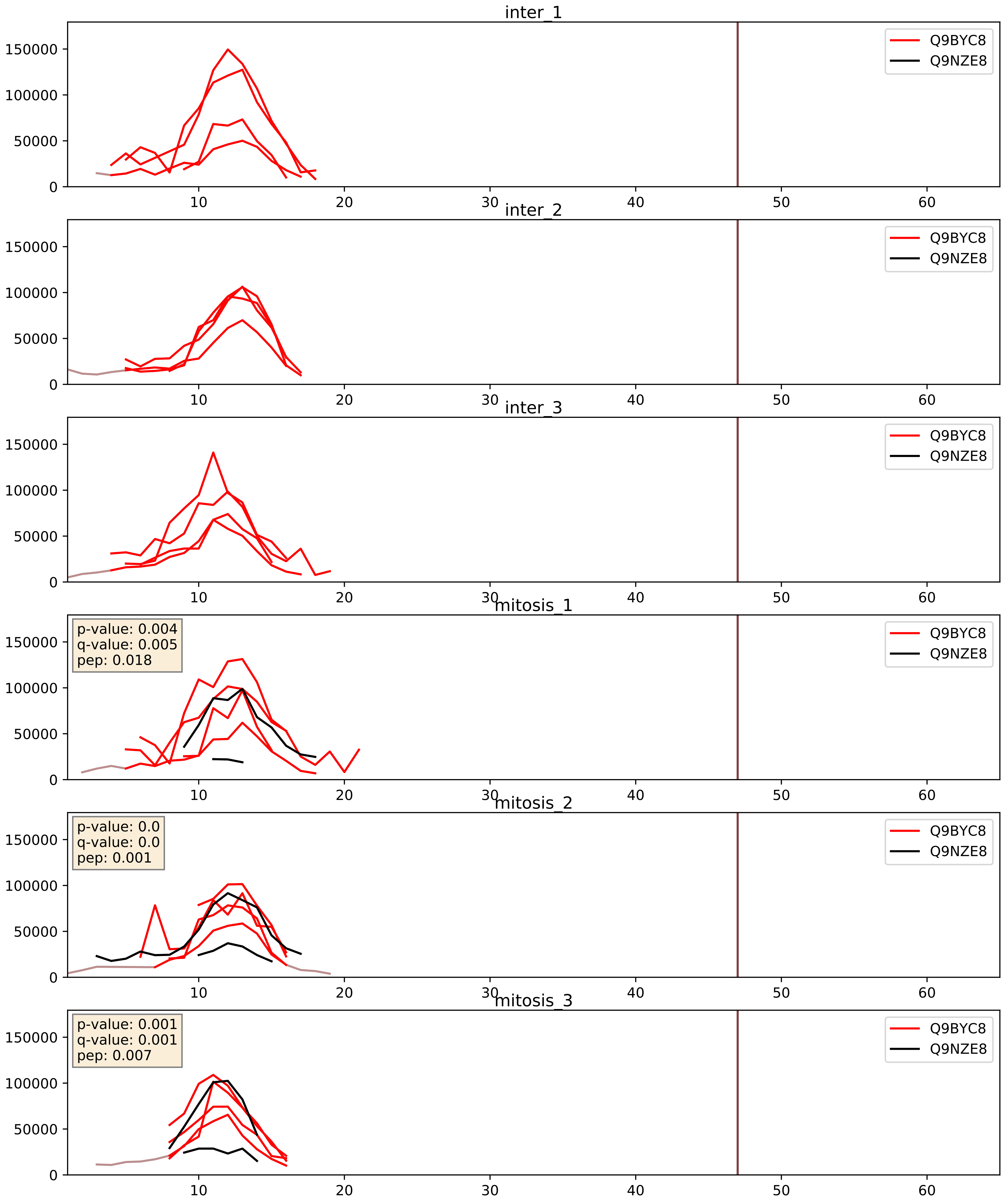


| level | condition_1 | condition_2 | pvalue | pvalue_adjusted |
|---|---|---|---|---|
| interactor_ratio | mitosis | inter | 0.002566068220898708 | 0.0651778770553566 |
| interactor_abundance | mitosis | inter | 0.006221004123186849 | 0.06721690780532076 |
| complex_abundance | mitosis | inter | 0.008606616325918843 | 0.06907915157215504 |

Q9BYC9_Q9NZE8
RM20_HUMAN vs RM35_HUMAN
p-value: 0.004 q-value: 0.004

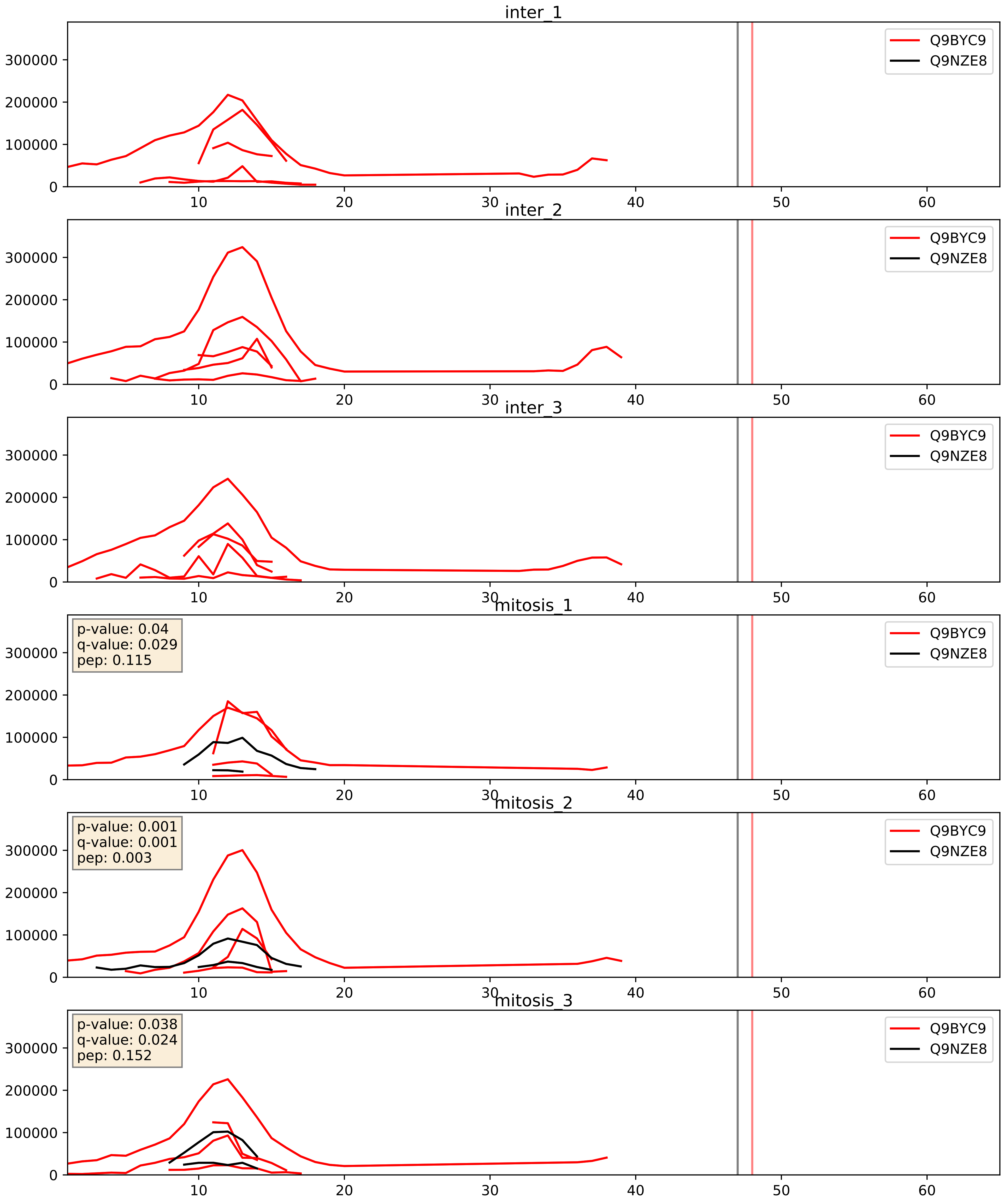


| level | condition_1 | condition_2 | pvalue | pvalue_adjusted |
|---|---|---|---|---|
| interactor_ratio | mitosis | inter | 0.0007447256434778975 | 0.03166103300210851 |
| interactor_abundance | mitosis | inter | 0.001307204373349643 | 0.026737561376040487 |
| complex_abundance | mitosis | inter | 0.0025052252042671035 | 0.03638642644133238 |

Q9BYD1_Q9NZE8
RM13_HUMAN vs RM35_HUMAN
p-value: 0.0 q-value: 0.0

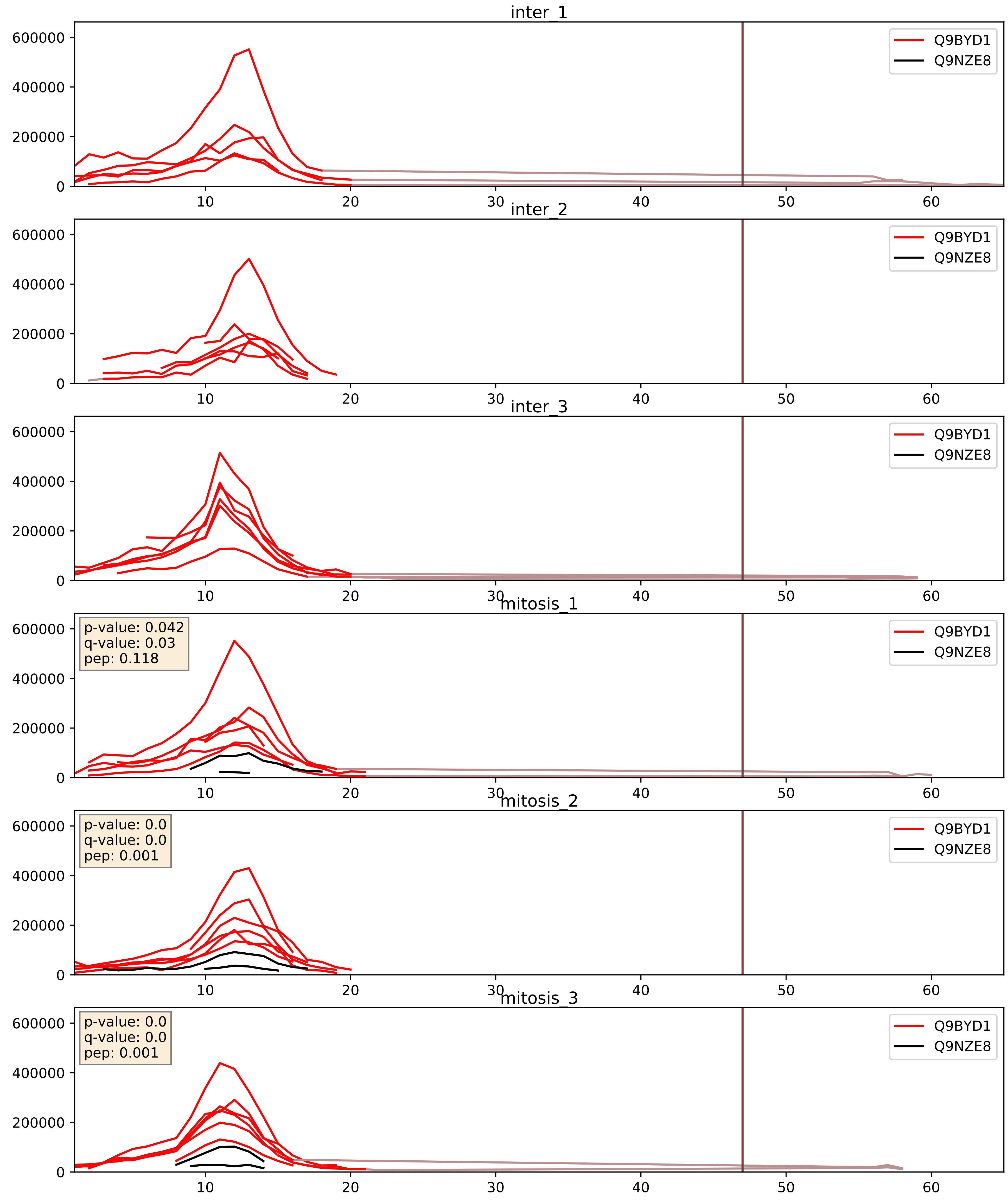


| level | condition_1 | condition_2 | pvalue | pvalue_adjusted |
|---|---|---|---|---|
| interactor_ratio | mitosis | inter | 1.1124146254019928e-05 | 0.003960471554059879 |
| interactor_abundance | mitosis | inter | 0.00017998465470889184 | 0.010140496760639767 |
| complex_abundance | mitosis | inter | 0.00057991351064485 | 0.01561025047521986 |

Q9BYD2_Q9NZE8
RM09_HUMAN vs RM35_HUMAN
p-value: 0.0 q-value: 0.0

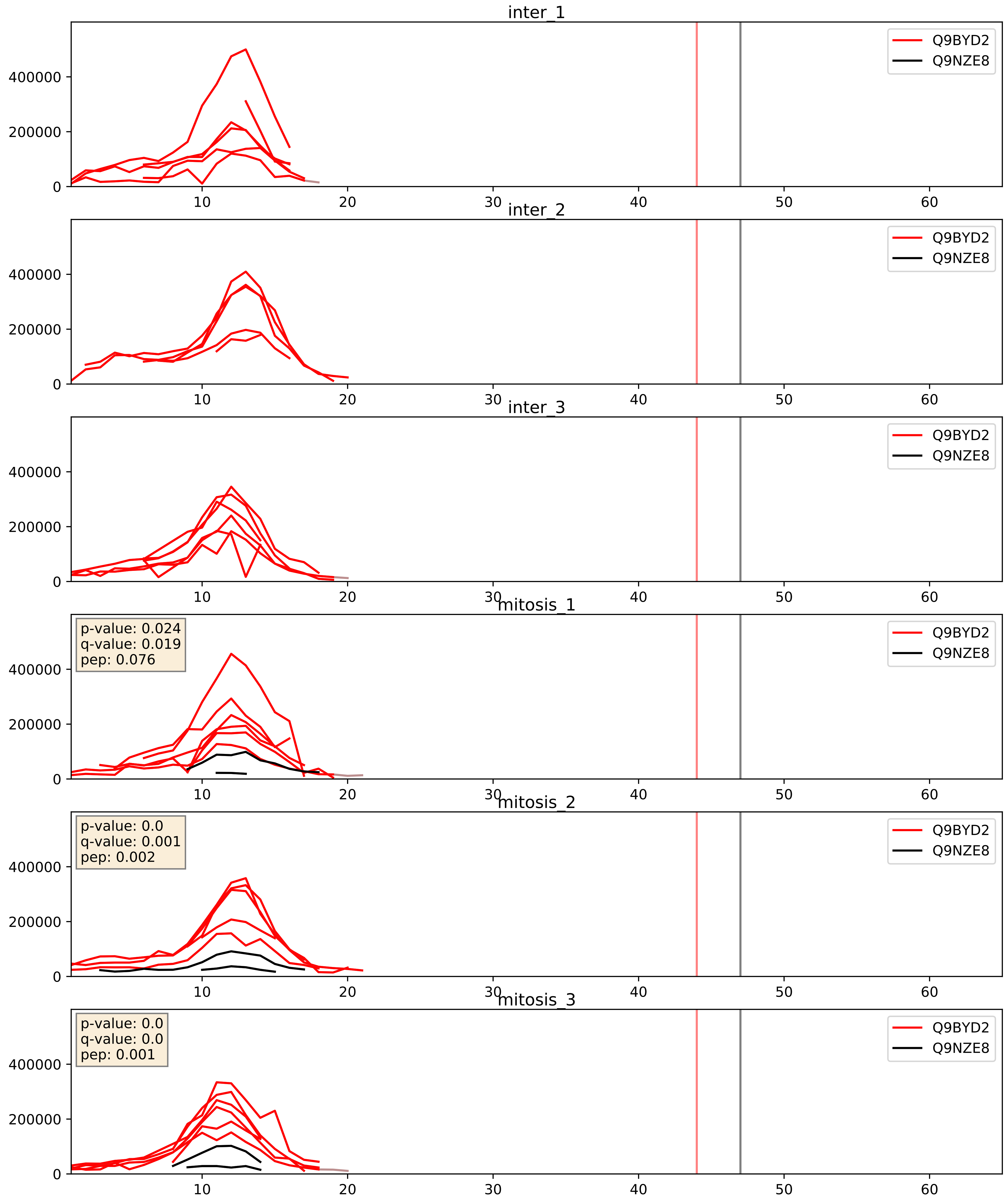


| level | condition_1 | condition_2 | pvalue | pvalue_adjusted |
|---|---|---|---|---|
| interactor_abundance | mitosis | inter | 0.00013025789724766 | 0.008677101948949181 |
| interactor_ratio | mitosis | inter | 0.00013578952627517755 | 0.012616771133069759 |
| complex_abundance | mitosis | inter | 0.0003167827076770335 | 0.01165207110963347 |

Q9BYD3_Q9NZE8
RM04_HUMAN vs RM35_HUMAN
p-value: 0.003 q-value: 0.003

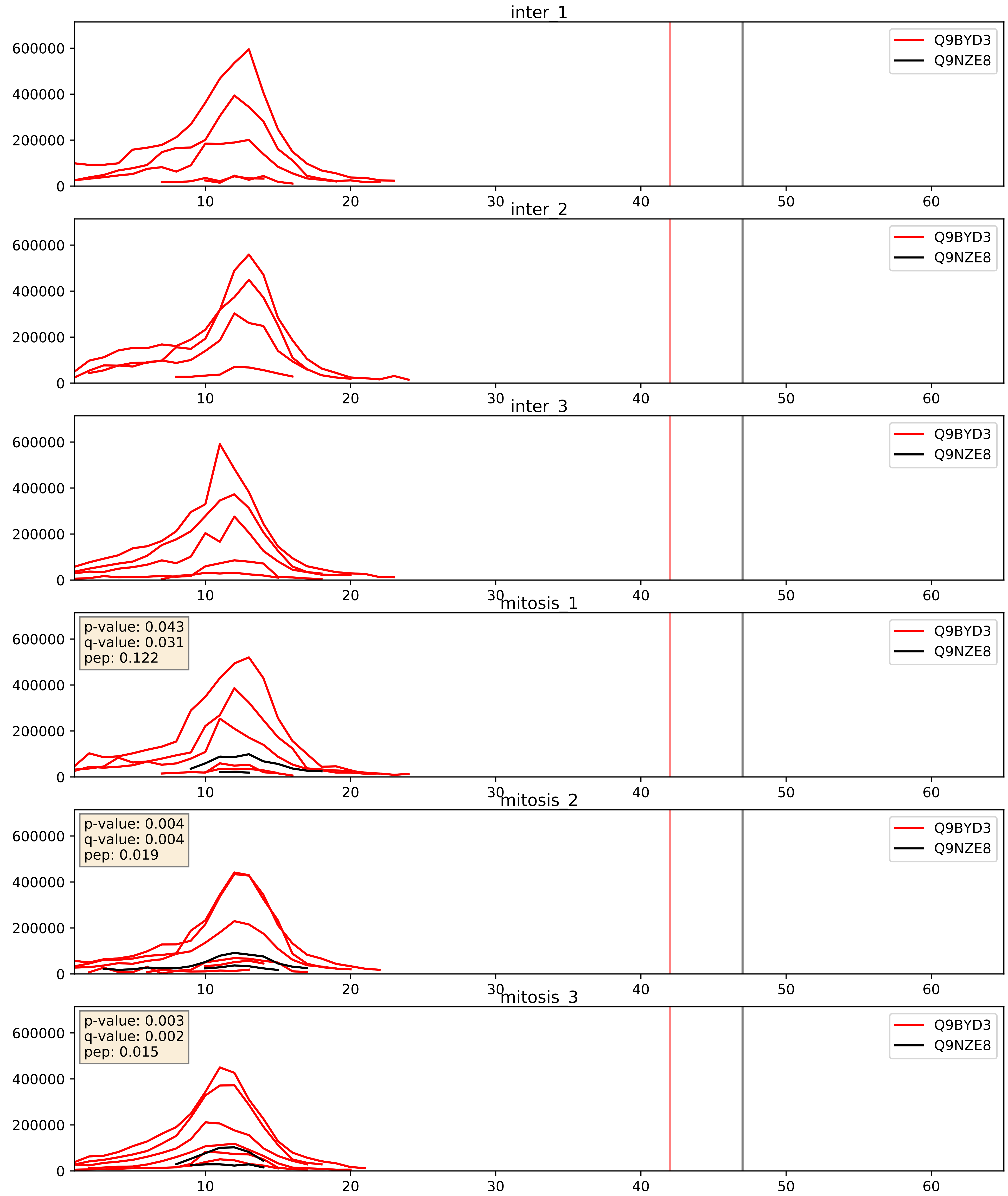


| level | condition_1 | condition_2 | pvalue | pvalue_adjusted |
|---|---|---|---|---|
| interactor_ratio | mitosis | inter | 1.3911120369424542e-05 | 0.003969306345409136 |
| interactor_abundance | mitosis | inter | 6.183318083340584e-05 | 0.006459477818931243 |
| complex_abundance | mitosis | inter | 0.0002425589183969702 | 0.010381521707390326 |

Q9BYD6_Q9NZE8
RM01_HUMAN vs RM35_HUMAN
p-value: 0.001 q-value: 0.015

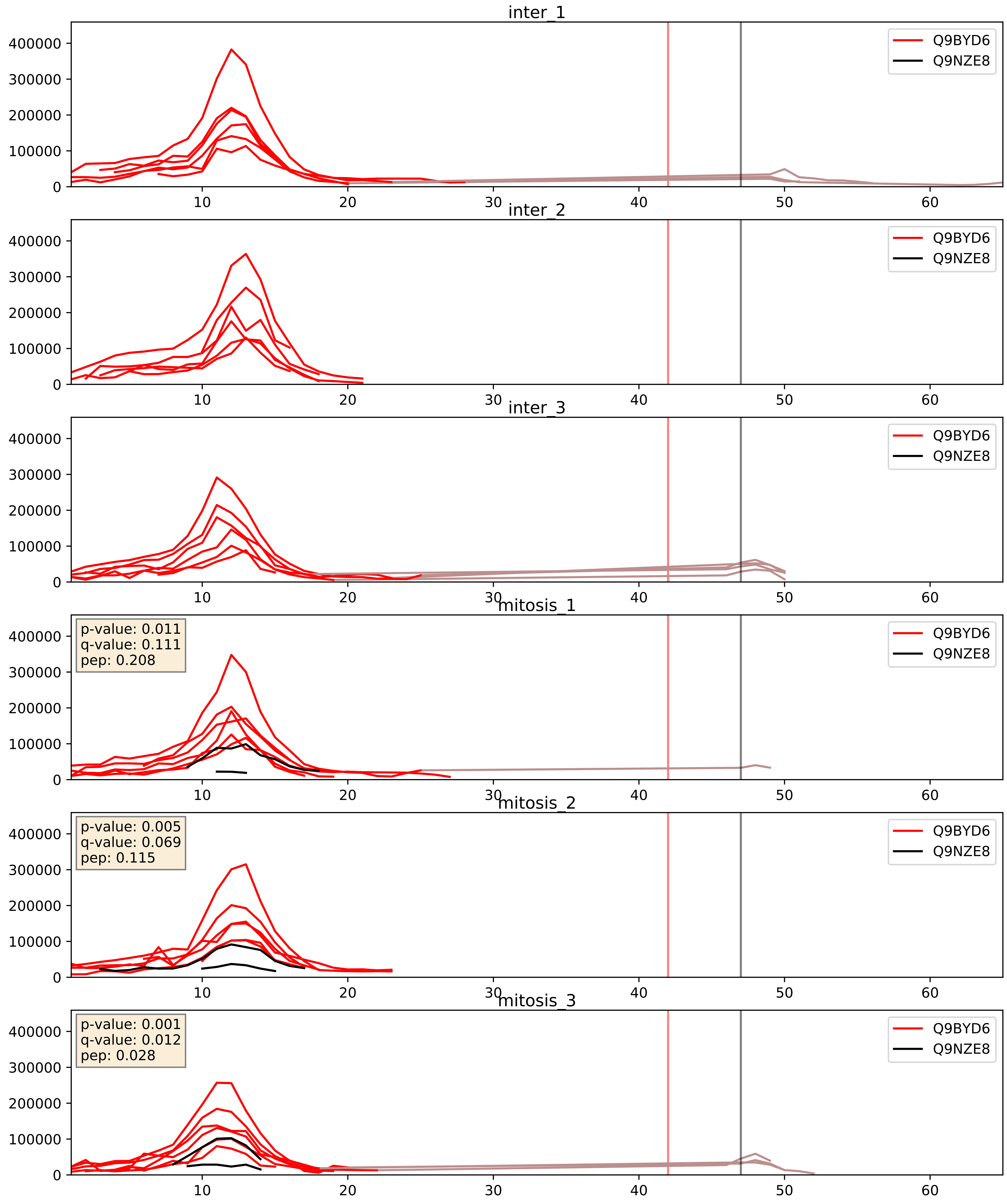


| level | condition_1 | condition_2 | pvalue | pvalue_adjusted |
|---|---|---|---|---|
| interactor_ratio | mitosis | inter | 0.00024286736594910028 | 0.016038997600655195 |
| interactor_abundance | mitosis | inter | 0.0004208994327649526 | 0.014156774634451844 |
| complex_abundance | mitosis | inter | 0.000941076245578817 | 0.020204817094612457 |

Q9BYN8_Q9NZE8
RT26_HUMAN vs RM35_HUMAN
p-value: 0.062 q-value: 0.039

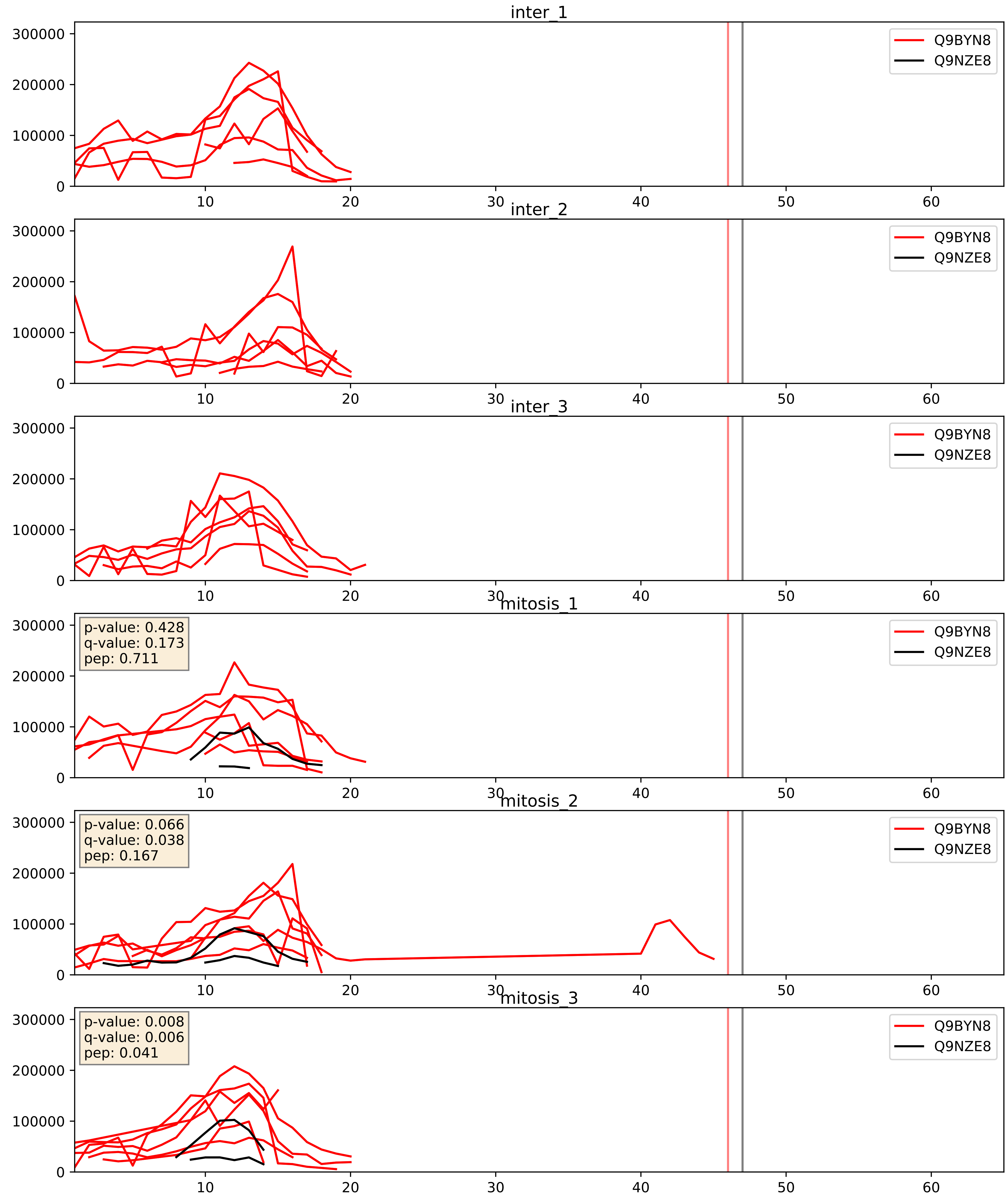


| level | condition_1 | condition_2 | pvalue | pvalue_adjusted |
|---|---|---|---|---|
| interactor_ratio | mitosis | inter | 4.2969370719897104e-05 | 0.006796360849516482 |
| interactor_abundance | mitosis | inter | 0.00046158732957023136 | 0.015023526772323879 |
| complex_abundance | mitosis | inter | 0.001216295247362876 | 0.023464778635459718 |

Q9BZE1_Q9NZE8
RM37_HUMAN vs RM35_HUMAN
p-value: 0.0 q-value: 0.003

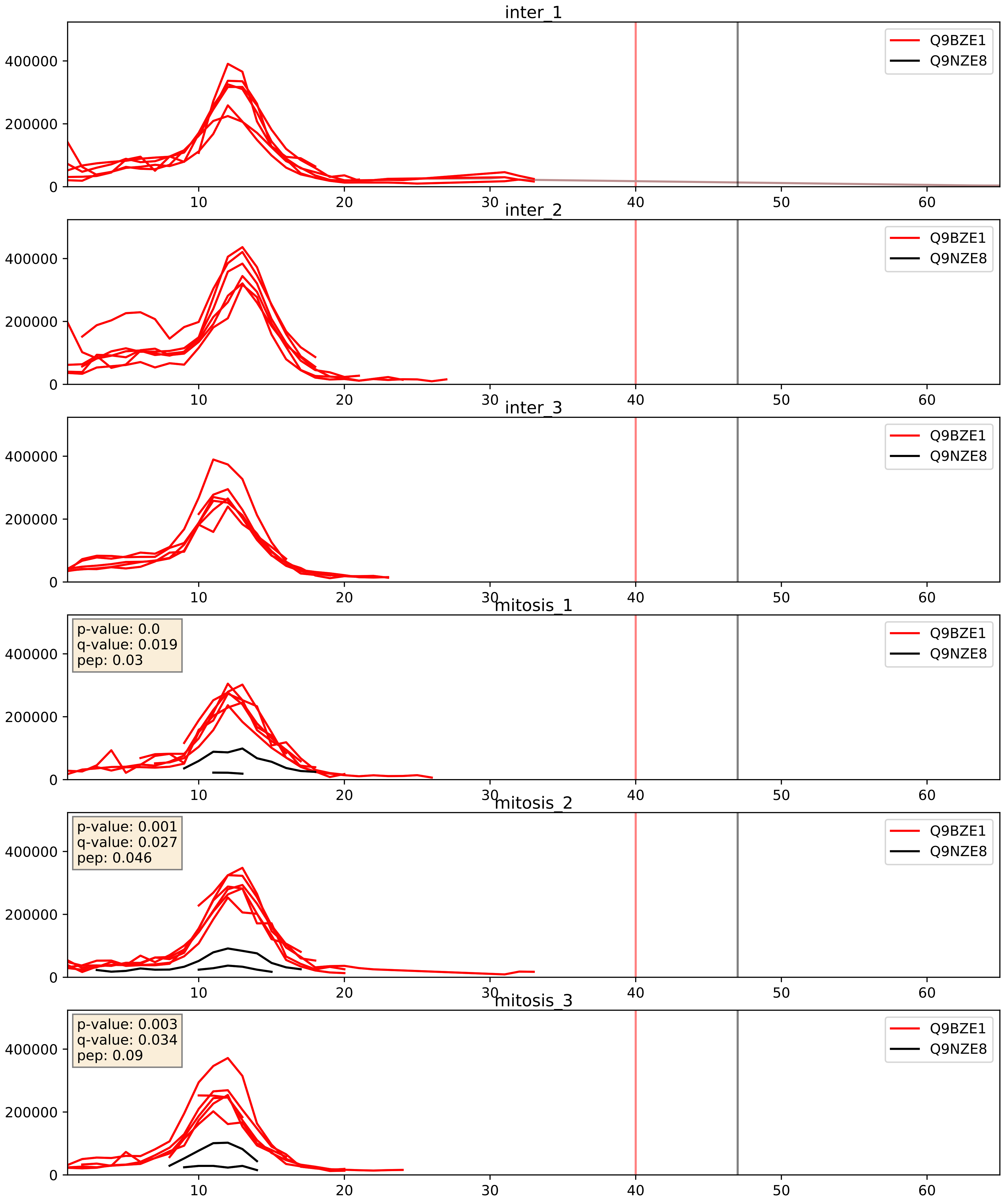


| level | condition_1 | condition_2 | pvalue | pvalue_adjusted |
|---|---|---|---|---|
| interactor_ratio | mitosis | inter | 2.3076733037133364e-05 | 0.004993481478971454 |
| interactor_abundance | mitosis | inter | 8.903101431711528e-05 | 0.007448207966538745 |
| complex_abundance | mitosis | inter | 0.000342001236020951 | 0.011998076148931723 |

Q9H0U6_Q9NZE8
RM18_HUMAN vs RM35_HUMAN
p-value: 0.001 q-value: 0.002

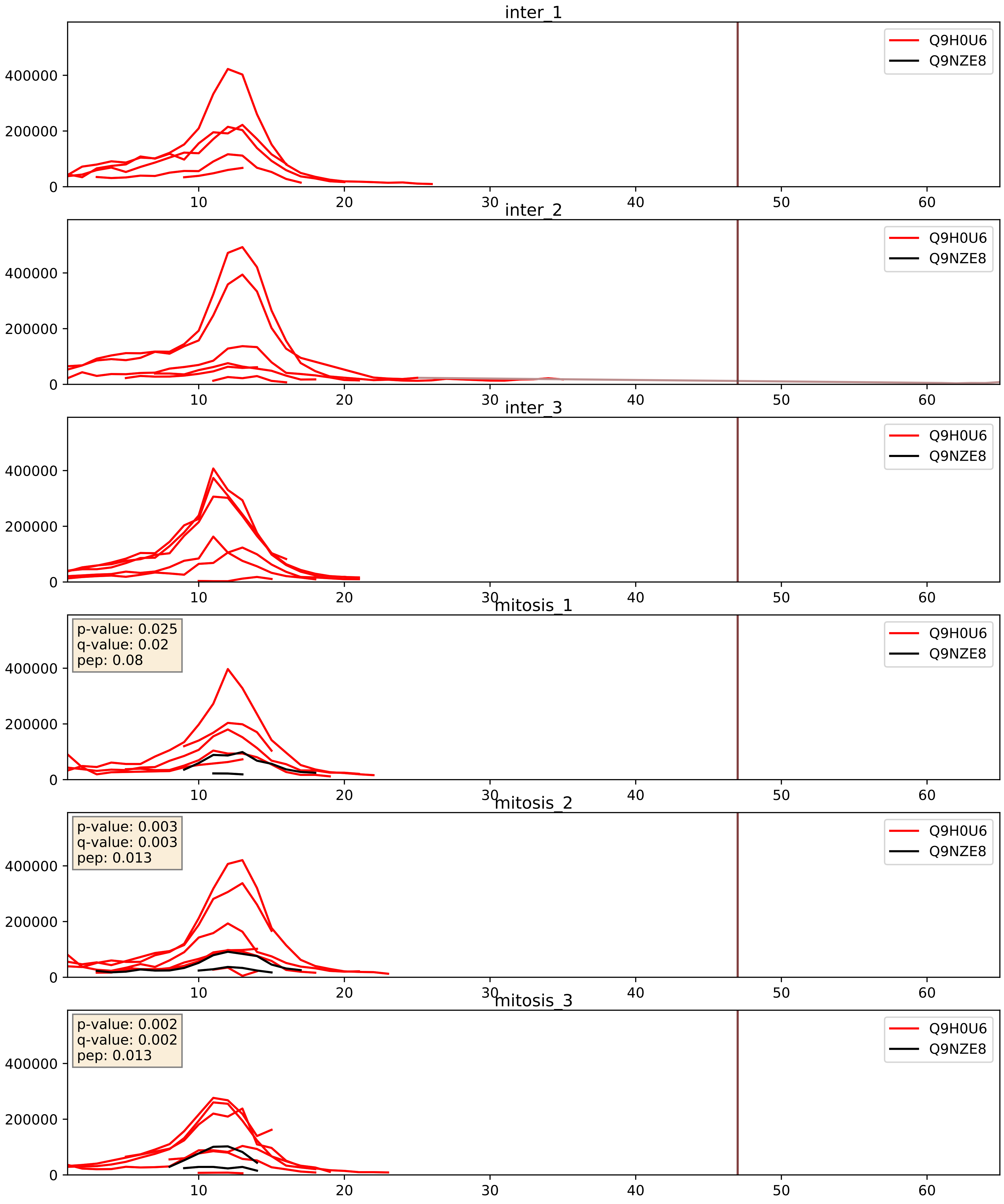


| level | condition_1 | condition_2 | pvalue | pvalue_adjusted |
|---|---|---|---|---|
| interactor_abundance | mitosis | inter | 0.0001467434459607821 | 0.009270287065861955 |
| interactor_ratio | mitosis | inter | 0.00016088856314043113 | 0.01311624857601991 |
| complex_abundance | mitosis | inter | 0.0003623610414143661 | 0.012039775462644836 |

Q9H2W6_Q9NZE8
RM46_HUMAN vs RM35_HUMAN
p-value: 0.0 q-value: 0.0

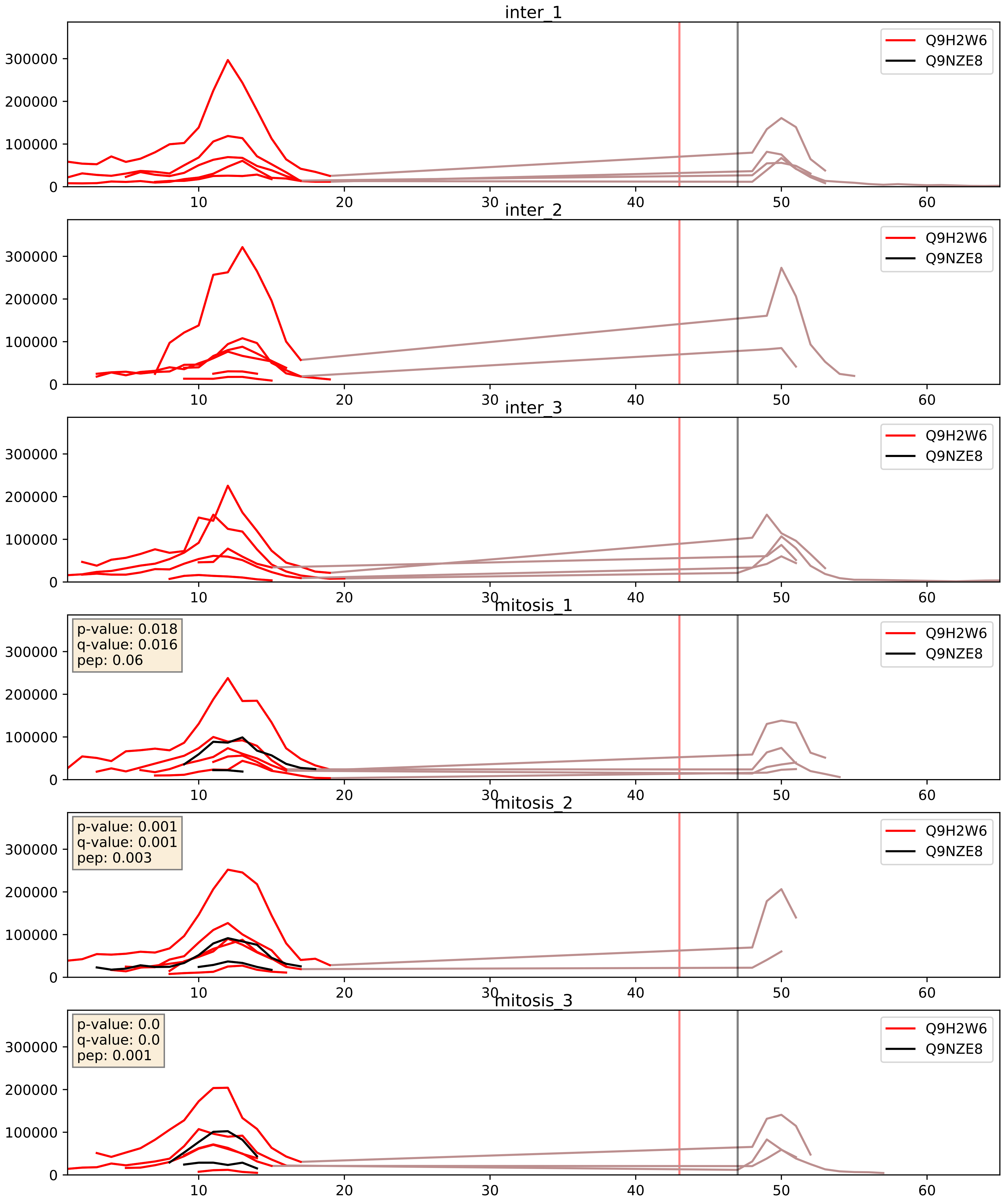


| level | condition_1 | condition_2 | pvalue | pvalue_adjusted |
|---|---|---|---|---|
| interactor_ratio | mitosis | inter | 0.0005259594433915967 | 0.026640312635692706 |
| interactor_abundance | mitosis | inter | 0.0010064124754765528 | 0.023667282390327723 |
| complex_abundance | mitosis | inter | 0.002063198559081843 | 0.03307299562872767 |

Q9H9J2_Q9NZE8
RM44_HUMAN vs RM35_HUMAN
p-value: 0.004 q-value: 0.004

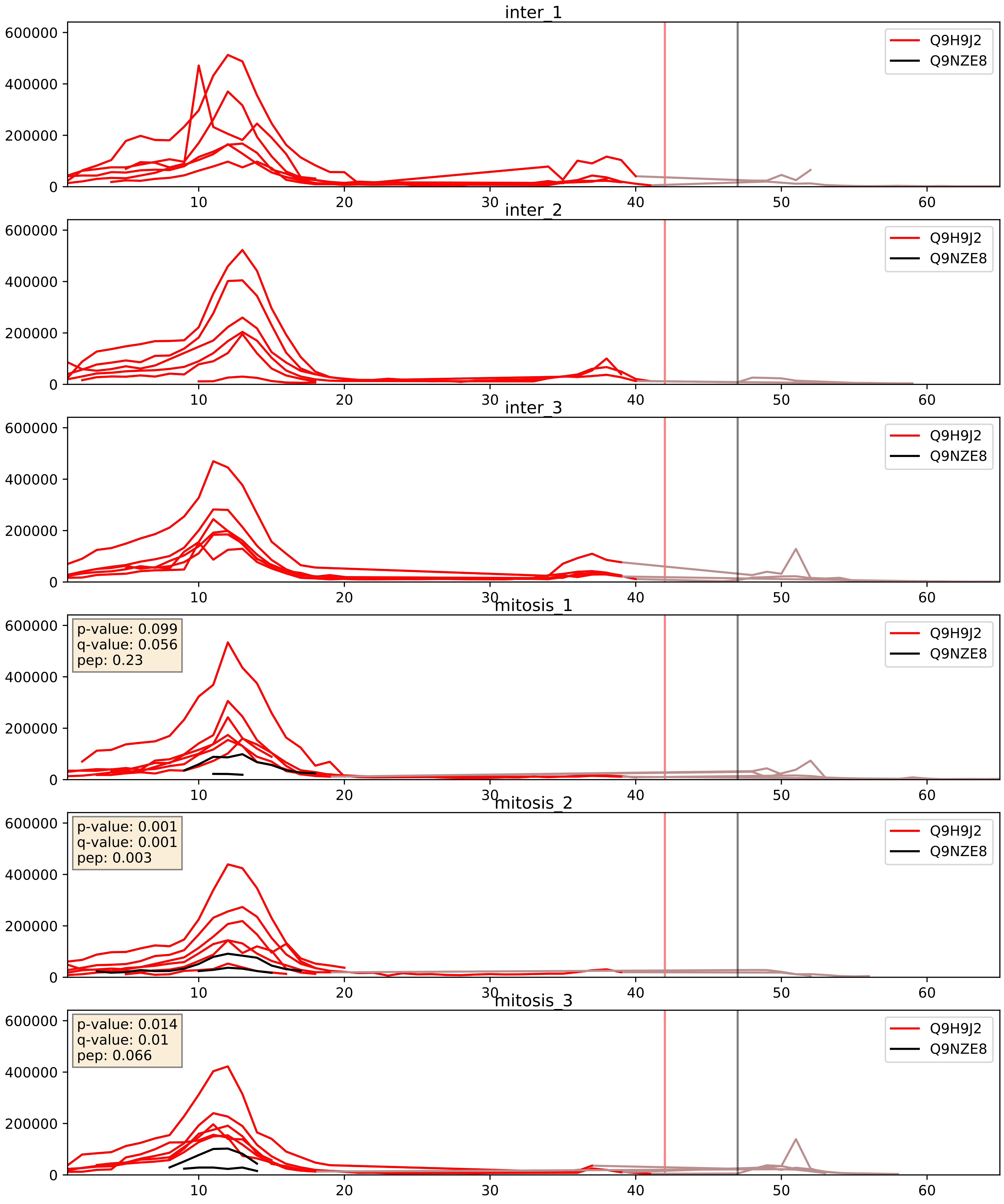


| level | condition_1 | condition_2 | pvalue | pvalue_adjusted |
|---|---|---|---|---|
| interactor_ratio | mitosis | inter | 9.74535639075859e-05 | 0.010559525405682725 |
| interactor_abundance | mitosis | inter | 0.00016603296423814968 | 0.009910743628815524 |
| complex_abundance | mitosis | inter | 0.0004300922958164677 | 0.013091389064164169 |

Q9HD33_Q9NZE8
RM47_HUMAN vs RM35_HUMAN
p-value: 0.0 q-value: 0.001

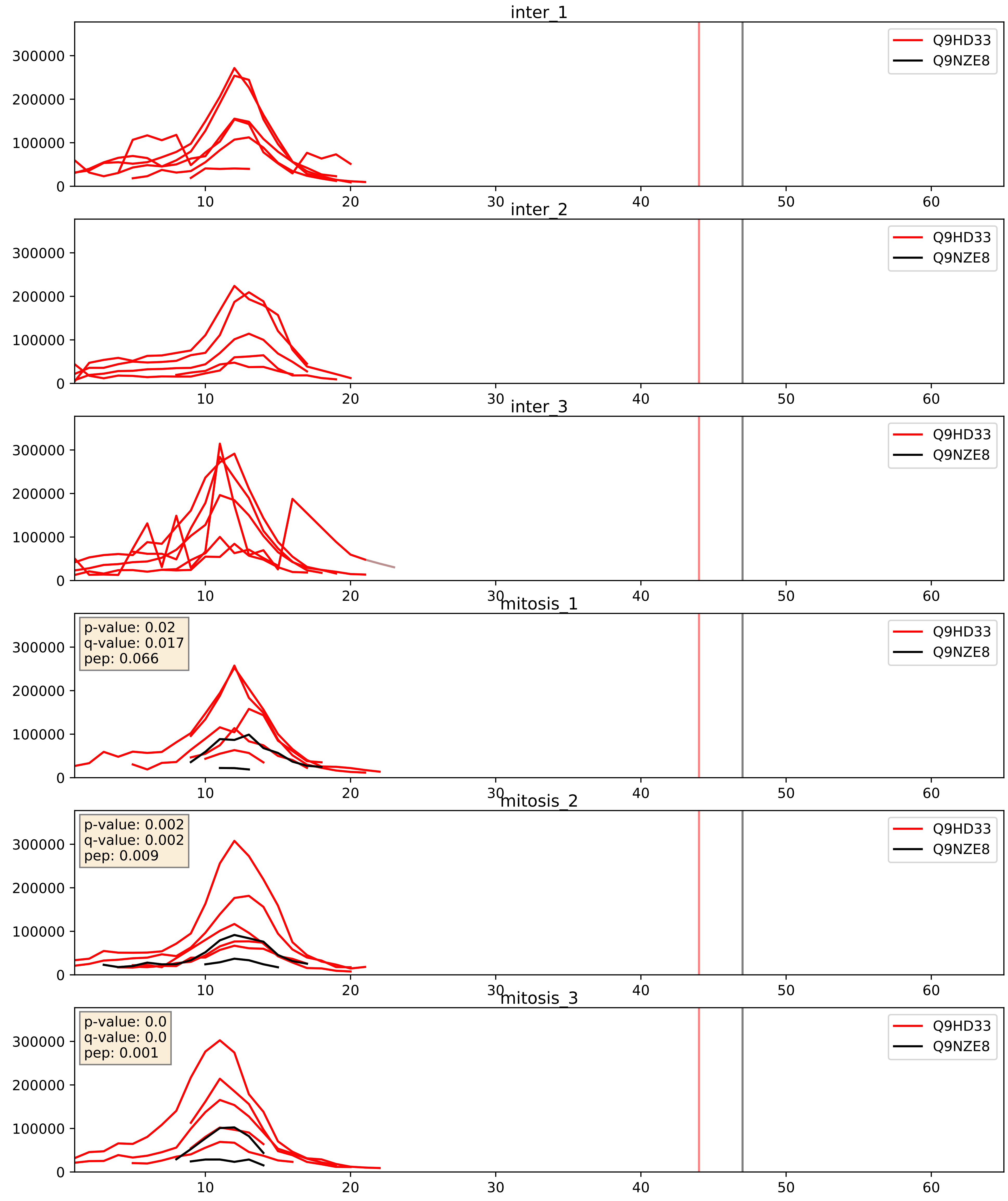


| level | condition_1 | condition_2 | pvalue | pvalue_adjusted |
|---|---|---|---|---|
| interactor_ratio | mitosis | inter | 1.525346211867344e-05 | 0.004211923733414343 |
| interactor_abundance | mitosis | inter | 0.00020639678666440104 | 0.01080585011527384 |
| complex_abundance | mitosis | inter | 0.0007389457172965143 | 0.017798602608495383 |

Q9NP92_Q9NZE8
RT30_HUMAN vs RM35_HUMAN
p-value: 0.004 q-value: 0.004

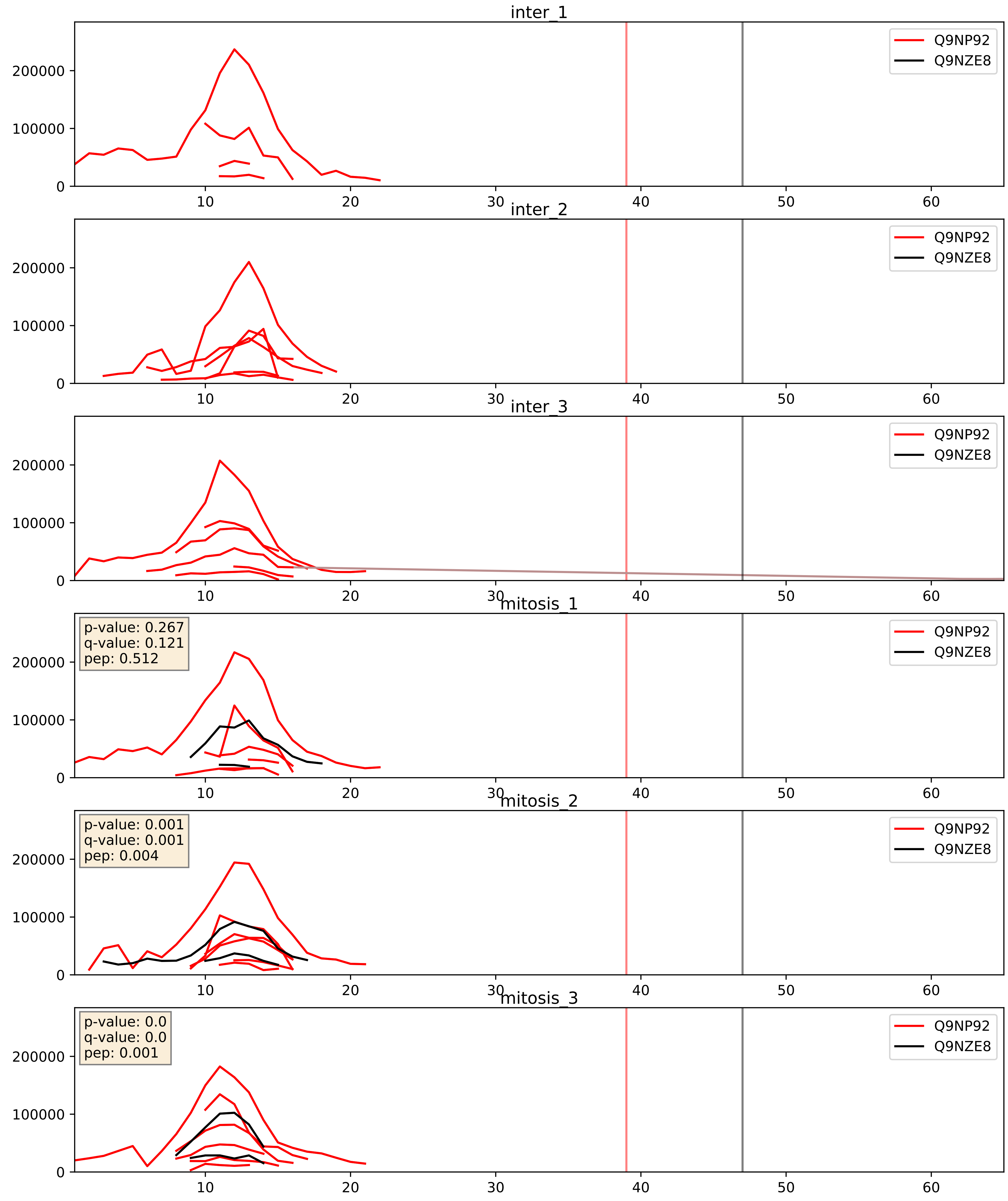


| level | condition_1 | condition_2 | pvalue | pvalue_adjusted |
|---|---|---|---|---|
| interactor_ratio | mitosis | inter | 9.479662648051581e-05 | 0.010510079284752772 |
| interactor_abundance | mitosis | inter | 0.001126614263499709 | 0.024137820492827596 |
| complex_abundance | mitosis | inter | 0.002708775939364819 | 0.03746658311311581 |

Q9NQ50_Q9NZE8
RM40_HUMAN vs RM35_HUMAN
p-value: 0.0 q-value: 0.0

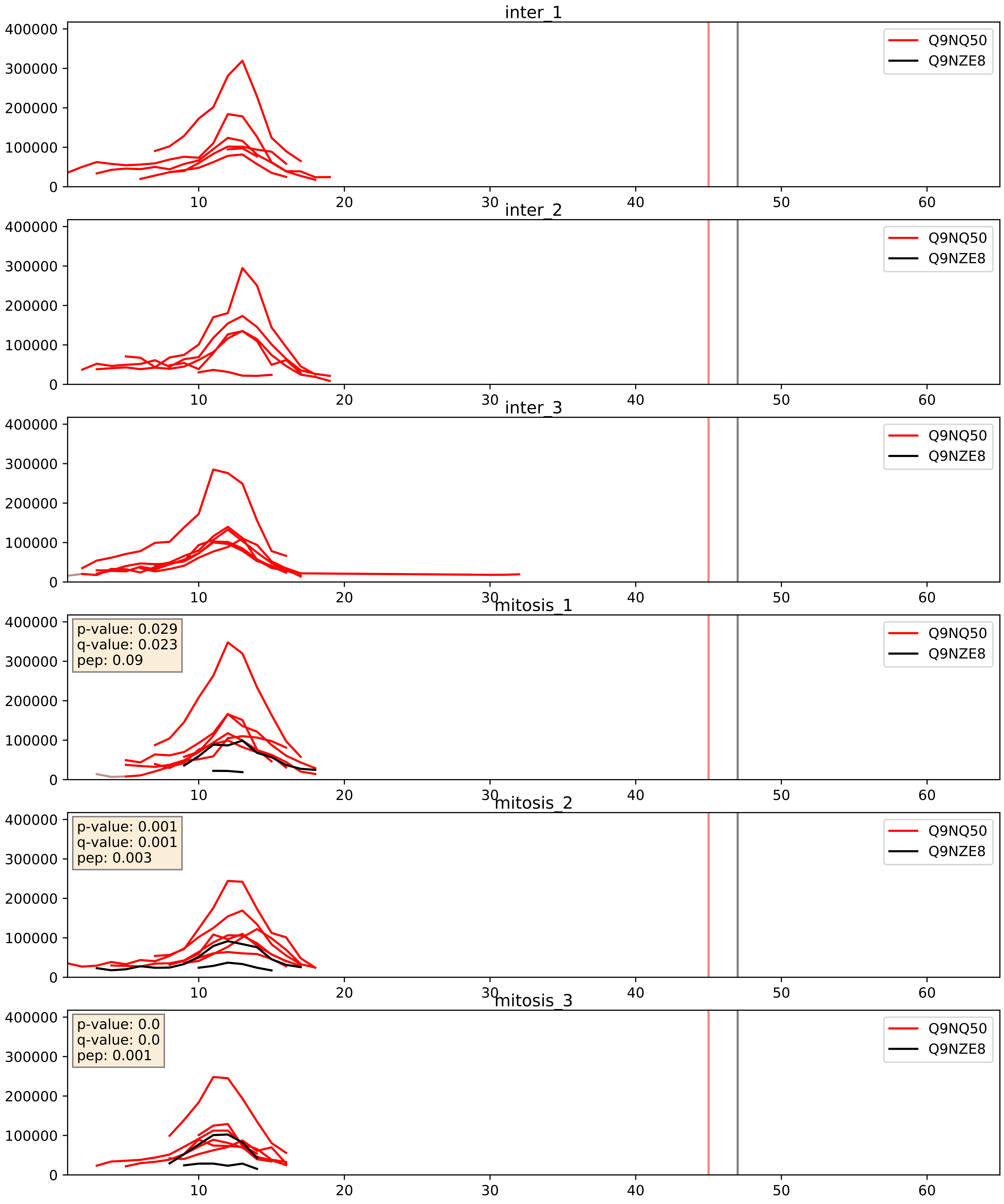


| level | condition_1 | condition_2 | pvalue | pvalue_adjusted |
|---|---|---|---|---|
| interactor_ratio | mitosis | inter | 0.0004512995297300989 | 0.02399455884776178 |
| interactor_abundance | mitosis | inter | 0.0012570895011381538 | 0.026054930096229046 |
| complex_abundance | mitosis | inter | 0.00232547312604243 | 0.035168510428347066 |

Q9NRX2_Q9NZE8
RM17_HUMAN vs RM35_HUMAN
p-value: 0.0 q-value: 0.0

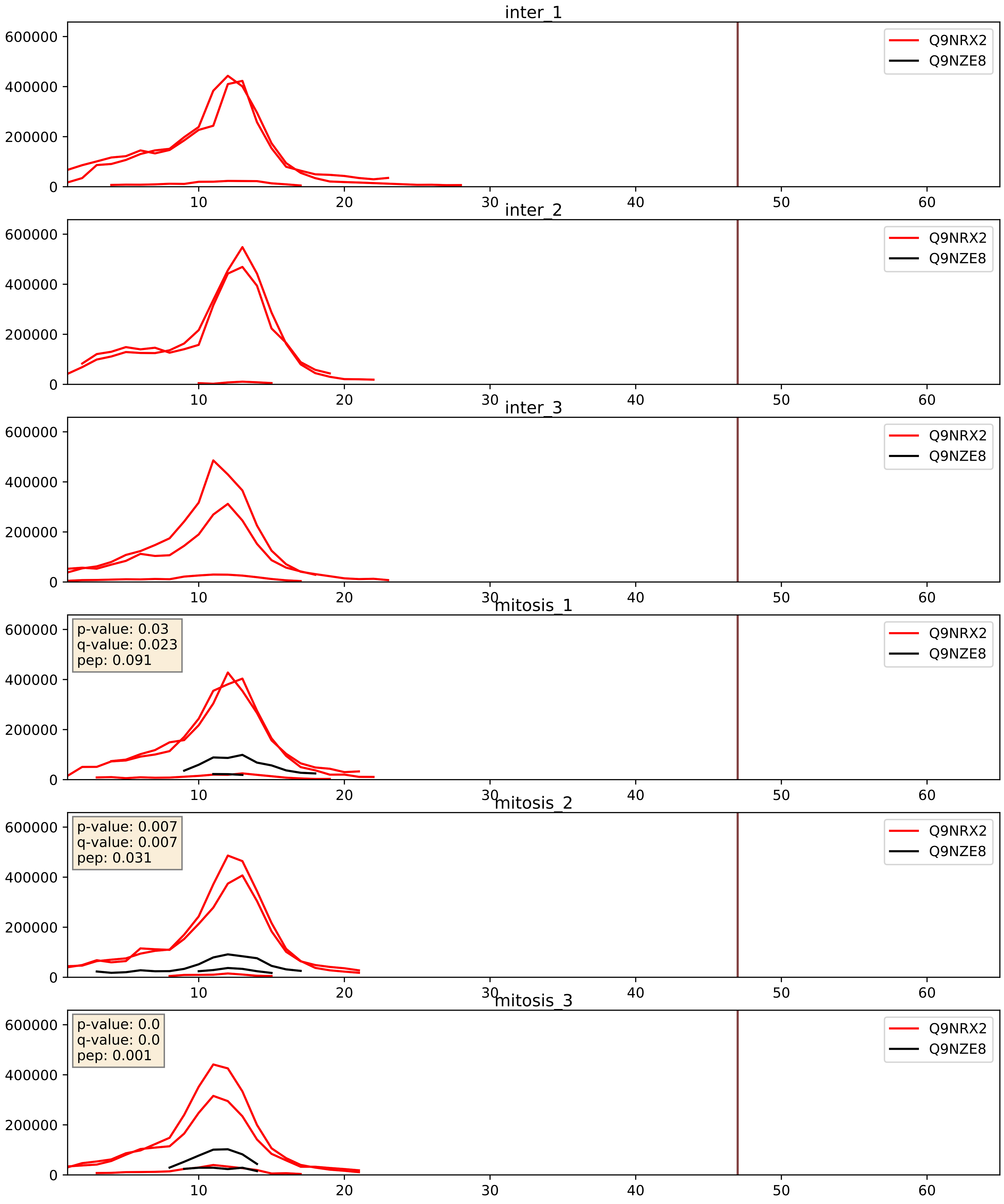


| level | condition_1 | condition_2 | pvalue | pvalue_adjusted |
|---|---|---|---|---|
| interactor_ratio | mitosis | inter | 6.434098341147061e-06 | 0.0028987306210641493 |
| interactor_abundance | mitosis | inter | 0.0002212146941997053 | 0.011215116209912415 |
| complex_abundance | mitosis | inter | 0.0007832167773801189 | 0.01846924411673228 |

Q9NVS2_Q9NZE8
RT18A_HUMAN vs RM35_HUMAN
p-value: 0.0 q-value: 0.0

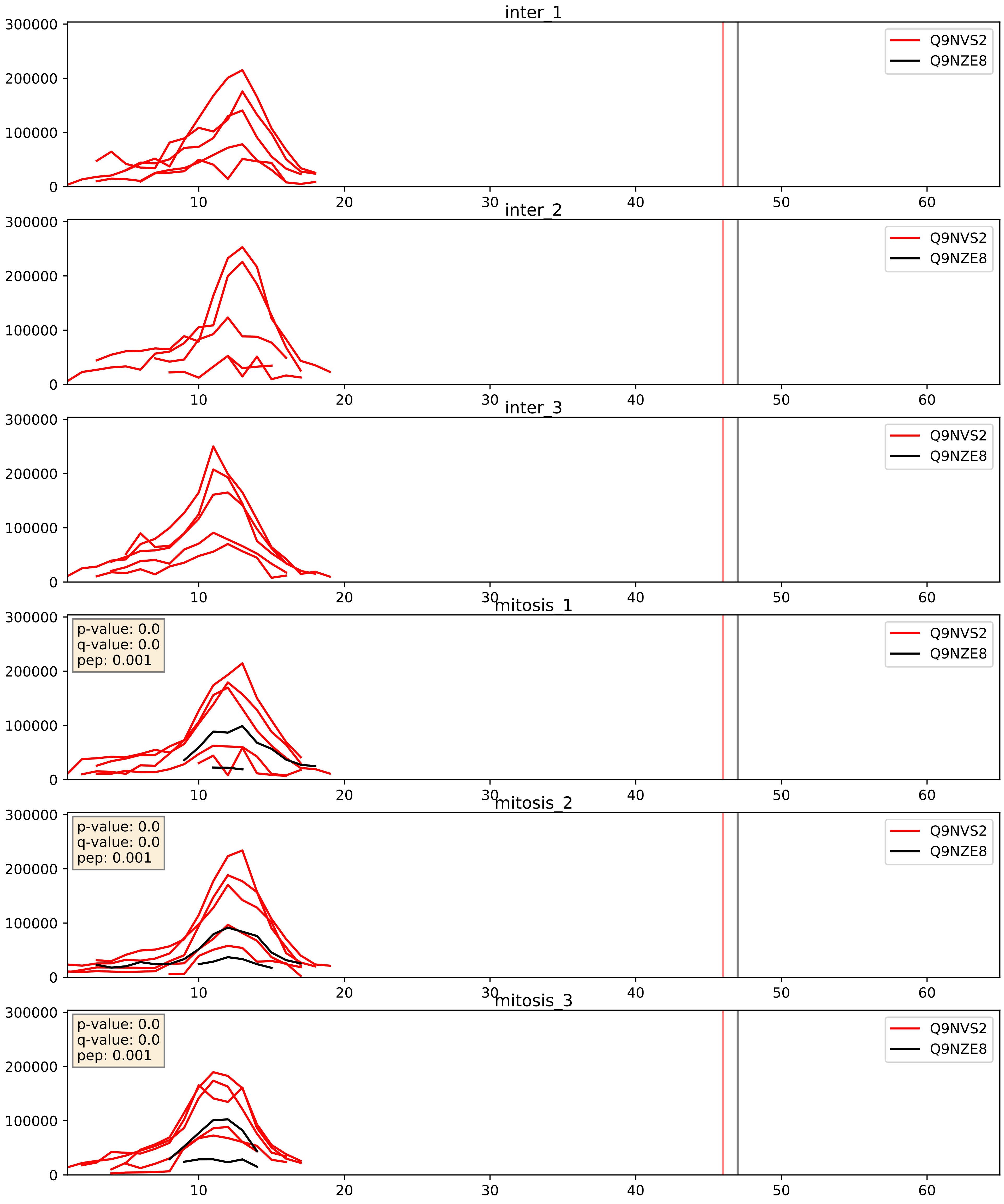


| level | condition_1 | condition_2 | pvalue | pvalue_adjusted |
|---|---|---|---|---|
| interactor_ratio | mitosis | inter | 4.7772587781098664e-05 | 0.006815555856770076 |
| interactor_abundance | mitosis | inter | 0.00030597083604629727 | 0.013095551782781524 |
| complex_abundance | mitosis | inter | 0.0009417899556951367 | 0.020204817094612457 |

Q9NWU5_Q9NZE8
RM22_HUMAN vs RM35_HUMAN
p-value: 0.001 q-value: 0.021

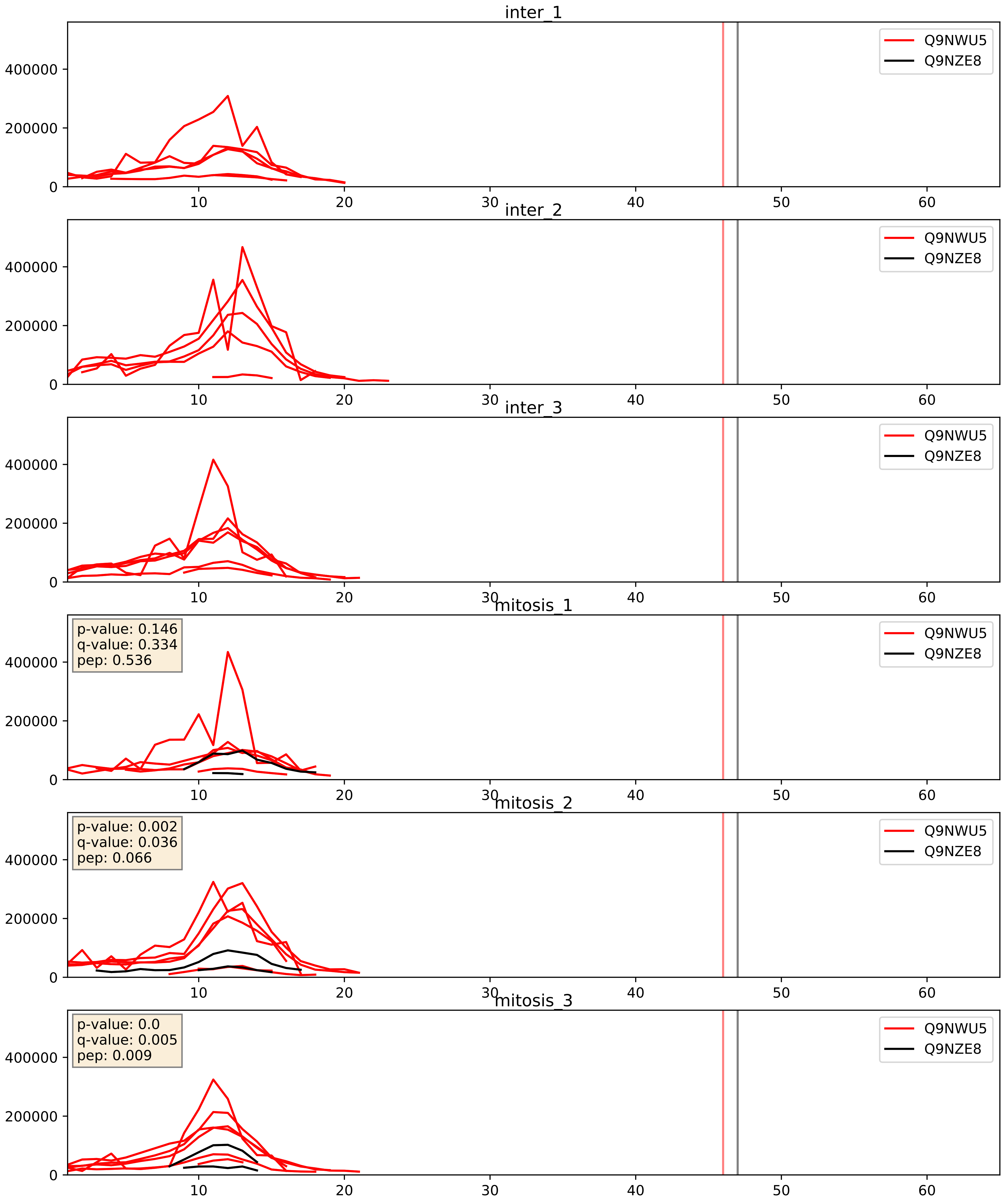


| level | condition_1 | condition_2 | pvalue | pvalue_adjusted |
|---|---|---|---|---|
| complex_abundance | mitosis | inter | 0.0010971091654939492 | 0.022013017310407657 |
| interactor_abundance | mitosis | inter | 0.0013581563127892076 | 0.02728785740910174 |
| interactor_ratio | mitosis | inter | 0.003858161780207222 | 0.08075090146433855 |

Q9NX20_Q9NZE8
RM16_HUMAN vs RM35_HUMAN
p-value: 0.005 q-value: 0.005

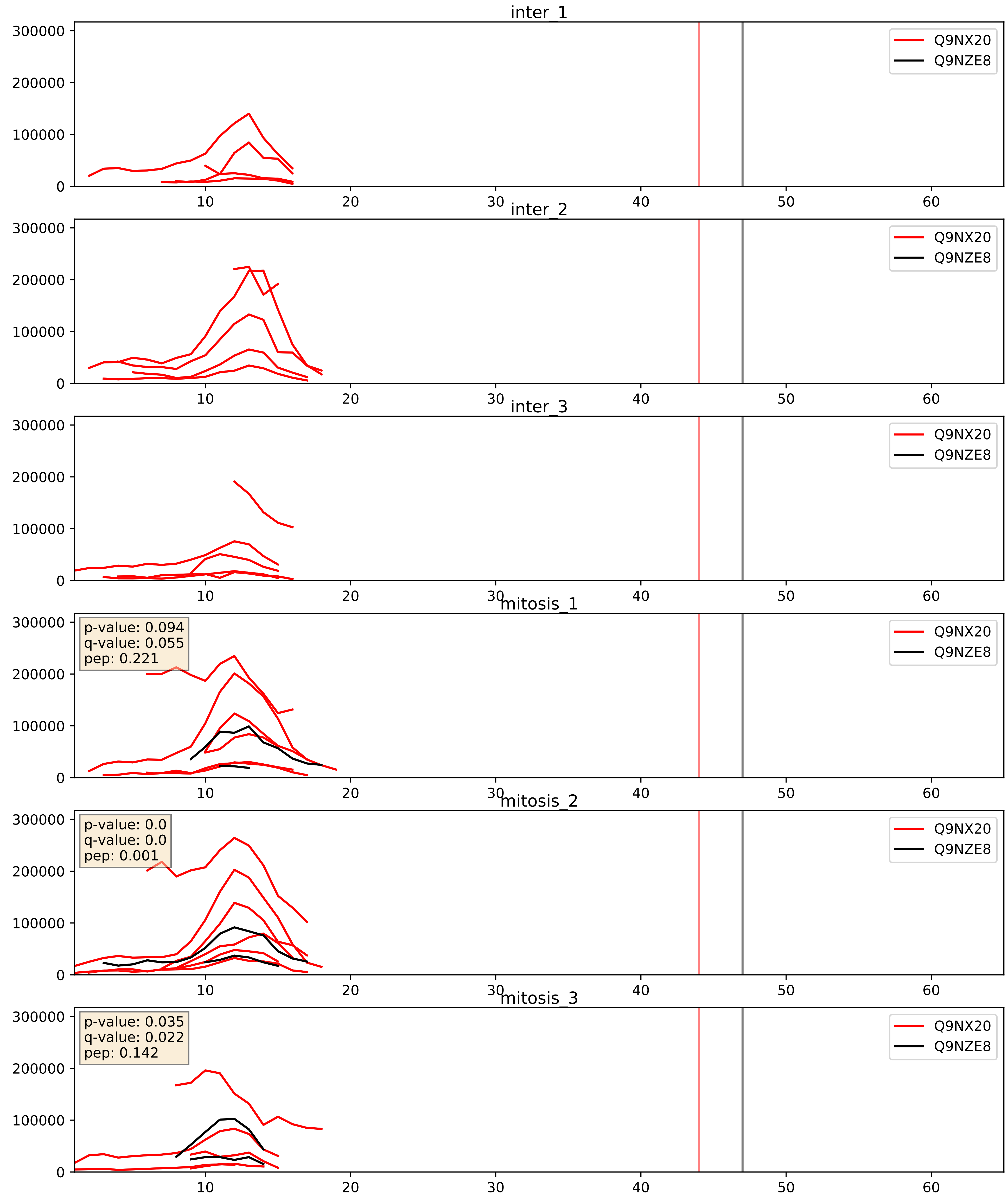


| level | condition_1 | condition_2 | pvalue | pvalue_adjusted |
|---|---|---|---|---|
| complex_abundance | mitosis | inter | 0.007325860091679865 | 0.06340877050320591 |
| interactor_abundance | mitosis | inter | 0.008635730724372138 | 0.0807312115073317 |
| interactor_ratio | mitosis | inter | 0.0149702995063588 | 0.17922484443976408 |

Q9NYK5_Q9NZE8
RM39_HUMAN vs RM35_HUMAN
p-value: 0.001 q-value: 0.015

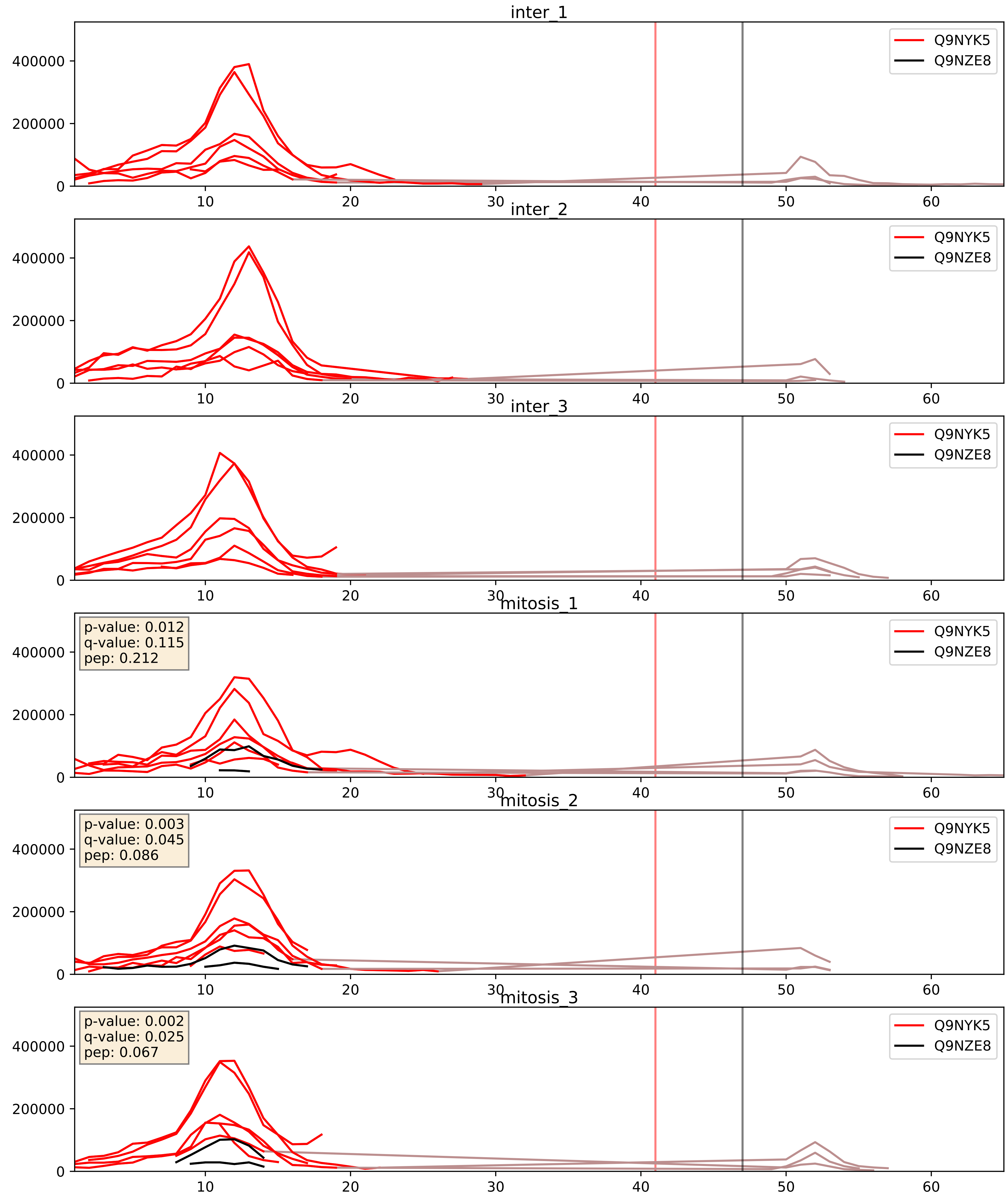


| level | condition_1 | condition_2 | pvalue | pvalue_adjusted |
|---|---|---|---|---|
| interactor_abundance | mitosis | inter | 0.00010544408643373516 | 0.007876481428251637 |
| interactor_ratio | mitosis | inter | 0.0001226960406114408 | 0.011800877613864419 |
| complex_abundance | mitosis | inter | 0.0002853446339259449 | 0.011360697983284132 |

Q9NZE8_Q9P015
RM35_HUMAN vs RM15_HUMAN
p-value: 0.0 q-value: 0.0

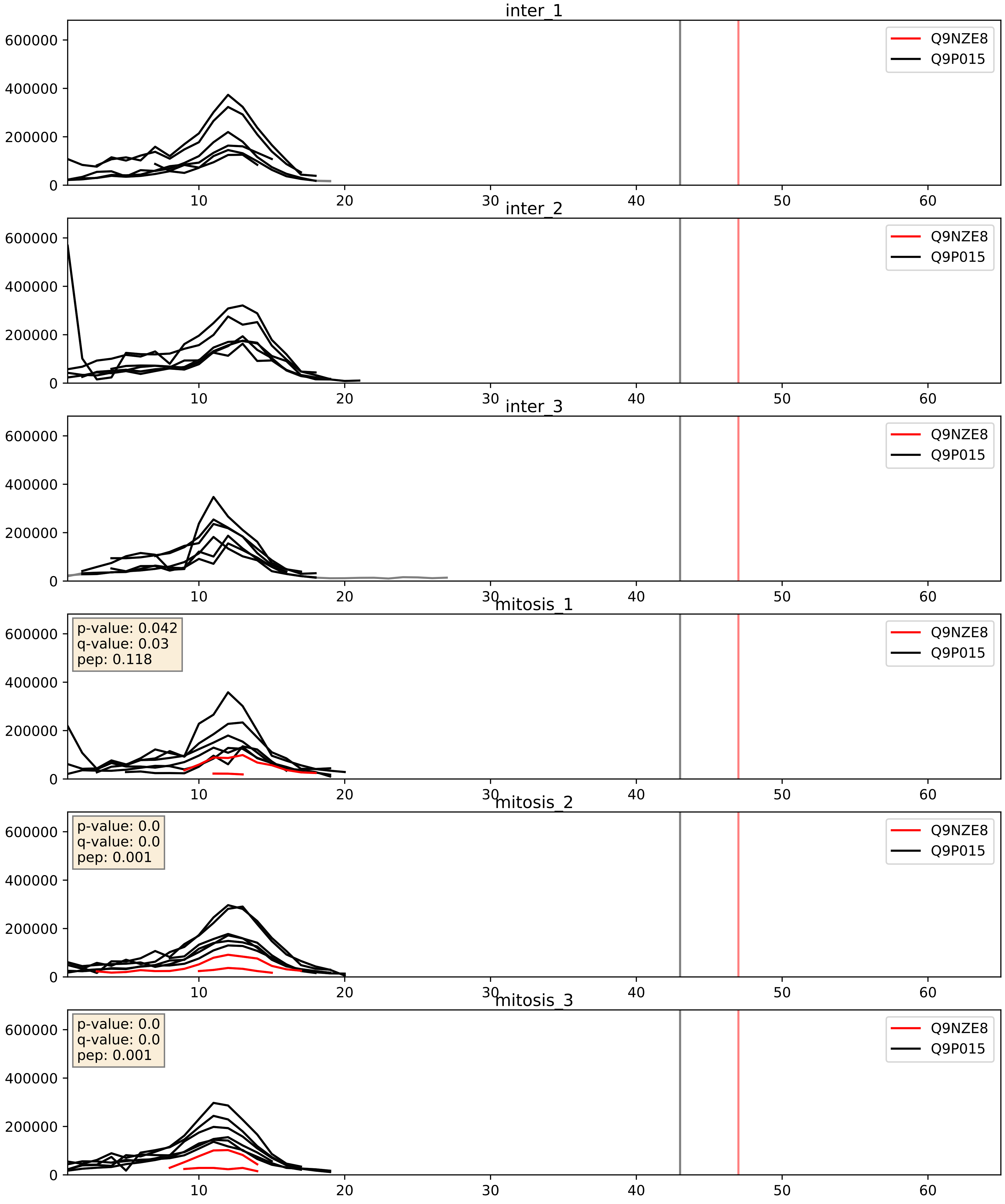


| level | condition_1 | condition_2 | pvalue | pvalue_adjusted |
|---|---|---|---|---|
| interactor_ratio | mitosis | inter | 0.00017291400110565987 | 0.013579301371233471 |
| complex_abundance | mitosis | inter | 0.00037669734334170777 | 0.012402035611557763 |
| interactor_abundance | mitosis | inter | 0.013555307611547178 | 0.10304923015527873 |

Q9NZE8_Q9P0M9
RM35_HUMAN vs RM27_HUMAN
p-value: 0.0 q-value: 0.0

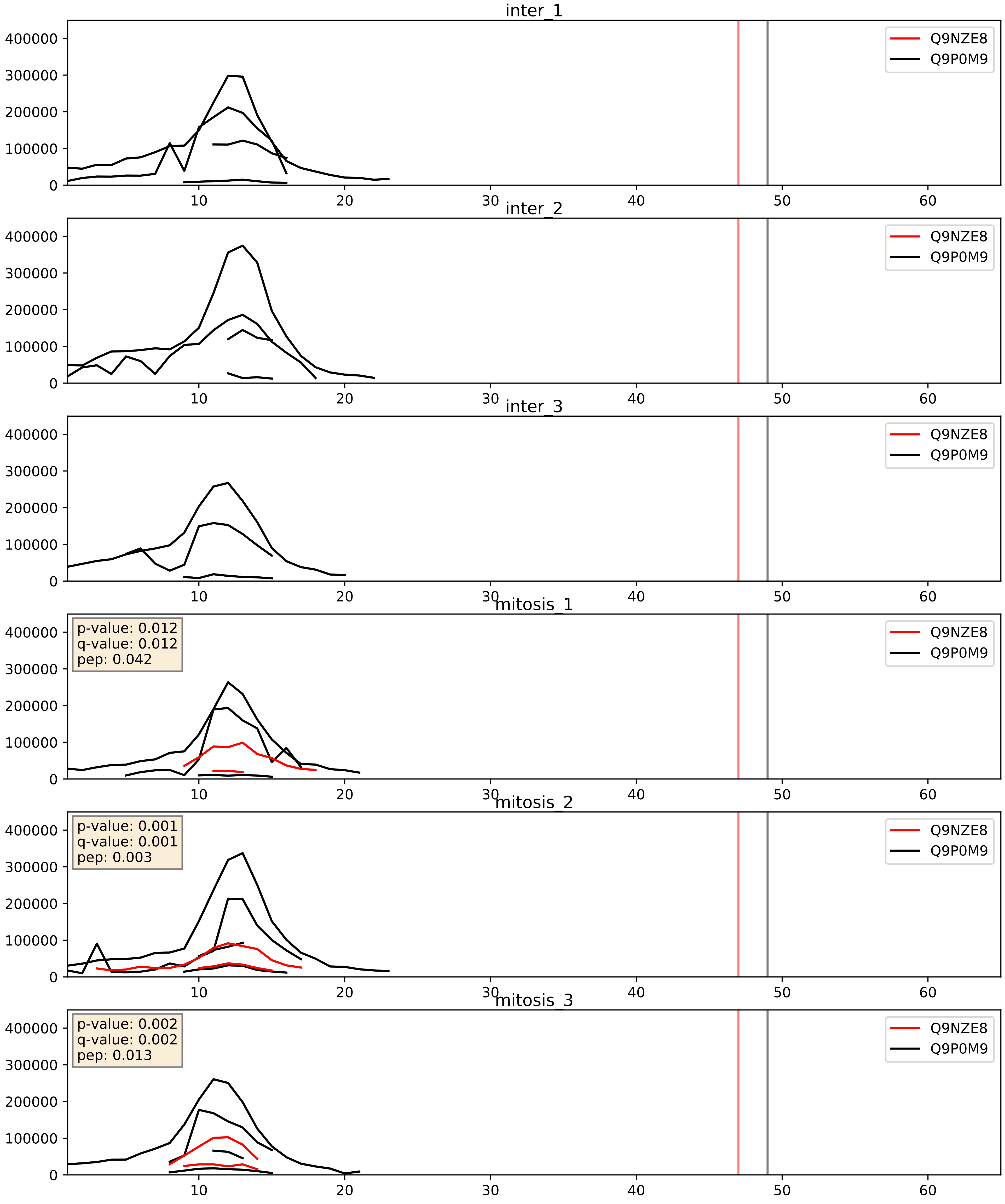


| level | condition_1 | condition_2 | pvalue | pvalue_adjusted |
|---|---|---|---|---|
| interactor_ratio | mitosis | inter | 0.000672938737816404 | 0.029848225645041686 |
| complex_abundance | mitosis | inter | 0.0009063075736871686 | 0.0197847007881119 |
| interactor_abundance | mitosis | inter | 0.013555307611547178 | 0.10304923015527873 |

Q9NZE8_Q9Y3B7
RM35_HUMAN vs RM11_HUMAN
p-value: 0.007 q-value: 0.006

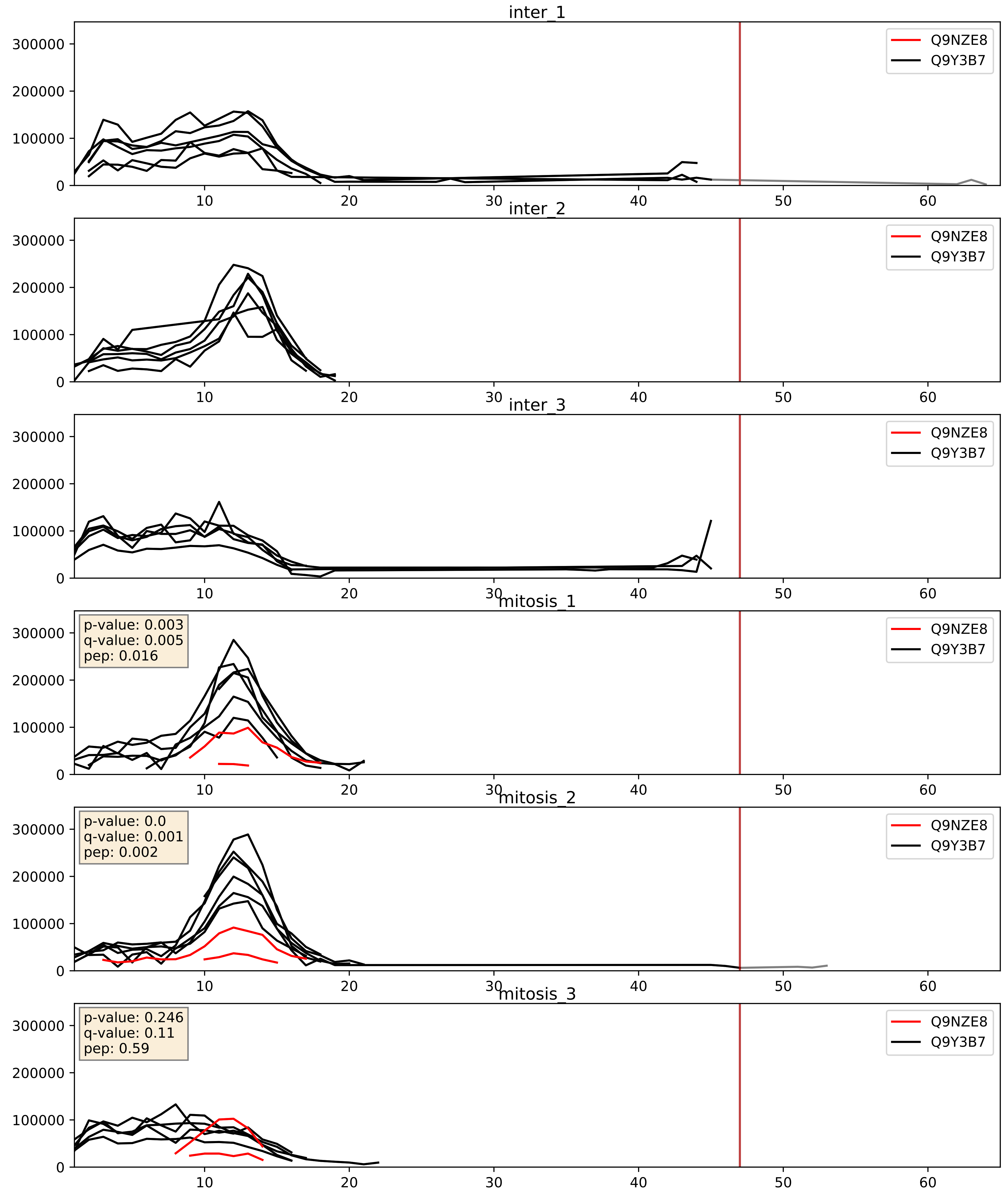


| level | condition_1 | condition_2 | pvalue | pvalue_adjusted |
|---|---|---|---|---|
| complex_abundance | mitosis | inter | 0.006448847346720888 | 0.05955783692156859 |
| interactor_ratio | mitosis | inter | 0.01026173625033943 | 0.14567240846252988 |
| interactor_abundance | mitosis | inter | 0.013555307611547178 | 0.10304923015527873 |

Q9NZE8_Q9Y3D3
RM35_HUMAN vs RT16_HUMAN
p-value: 0.0 q-value: 0.0

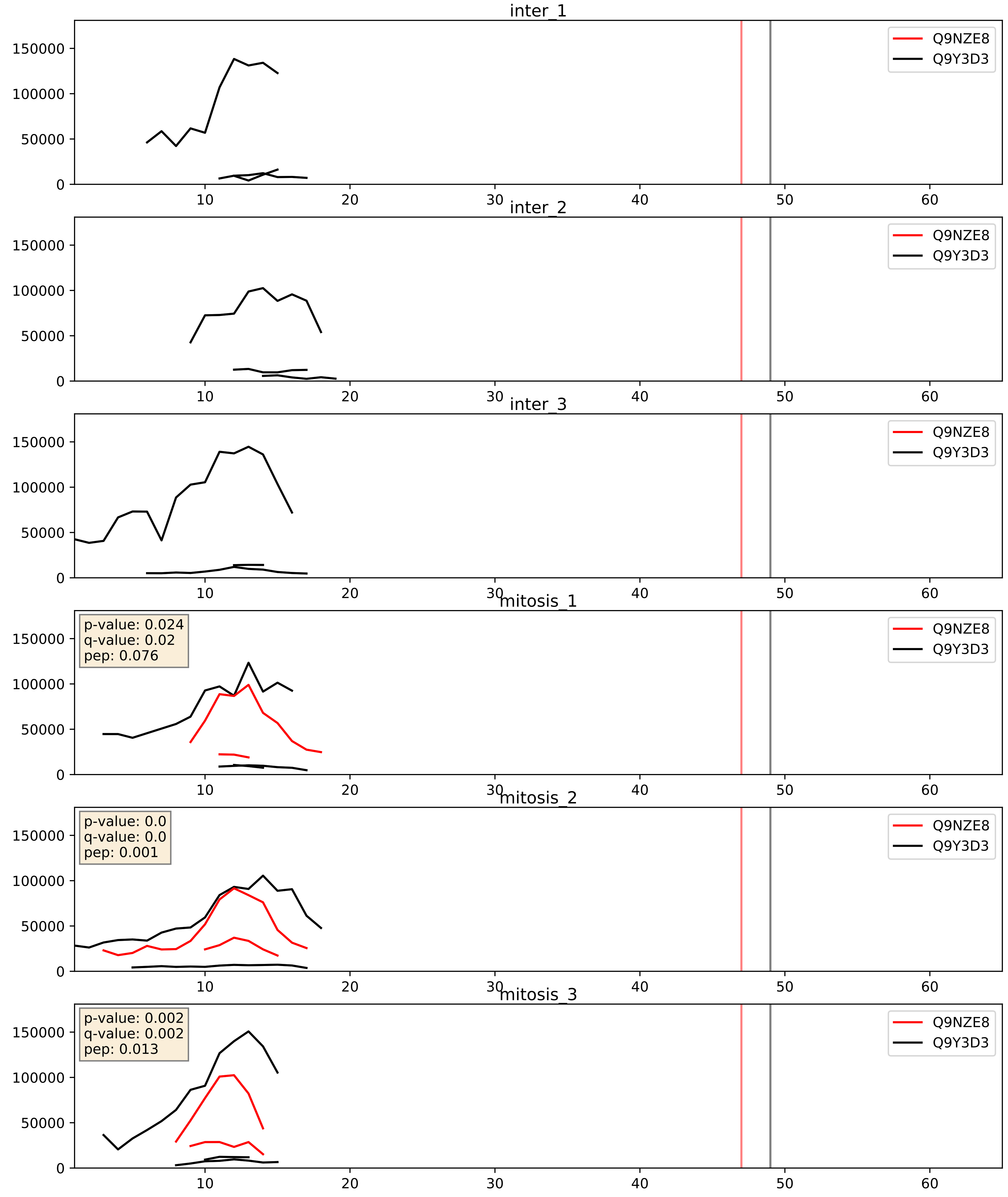


| level | condition_1 | condition_2 | pvalue | pvalue_adjusted |
|---|---|---|---|---|
| interactor_abundance | mitosis | inter | 0.012898459688029301 | 0.10167976967449956 |
| complex_abundance | mitosis | inter | 0.044967944465031065 | 0.15172471605071577 |
| interactor_ratio | mitosis | inter | 0.9139602152720305 | 0.9852905071335931 |
