## Supplementary Data 2 for "SECAT: Quantifying differential protein-protein interaction states by network-centric analysis": hela_string_000033_Q9BVP2.pdf

Q9BVP2_Q9BVP2

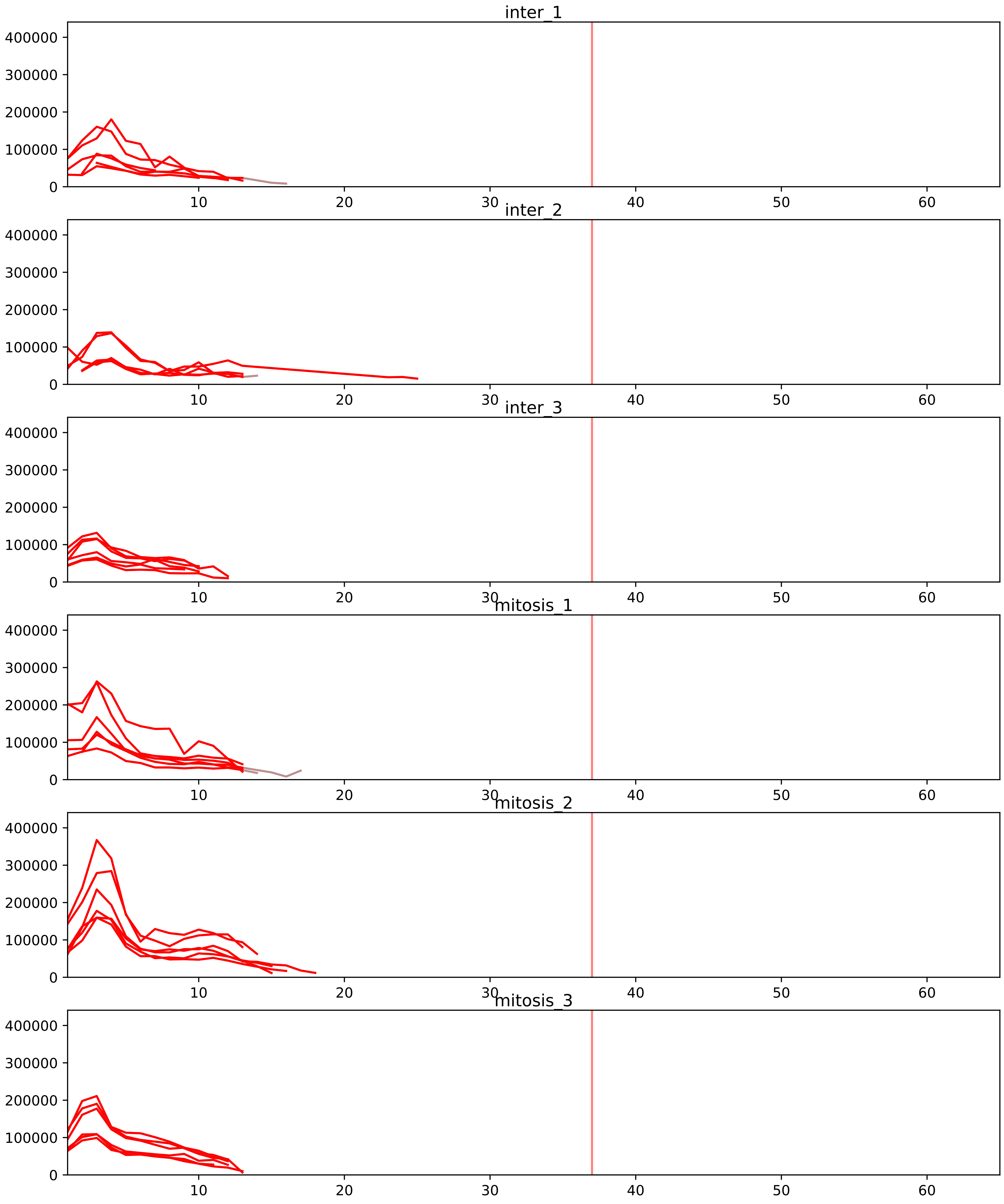


| level | condition_1 | condition_2 | pvalue | pvalue_adjusted |
|---|---|---|---|---|
| interactor_ratio | mitosis | inter | 7.594857695239889e-09 | 4.367043174762936e-07 |
| complex_abundance | mitosis | inter | 7.692067702793543e-05 | 0.0018871206097520157 |
| interactor_abundance | mitosis | inter | 0.00018316540539121243 | 0.0035106702699982386 |
| assembled_abundance | mitosis | inter | 0.019341807679005844 | 0.16491621447254345 |
| total_abundance | mitosis | inter | 0.021481863141287572 | 0.18835452556753943 |

O00567_Q9BVP2
NOP56_HUMAN vs GNL3_HUMAN
p-value: 0.0 q-value: 0.013

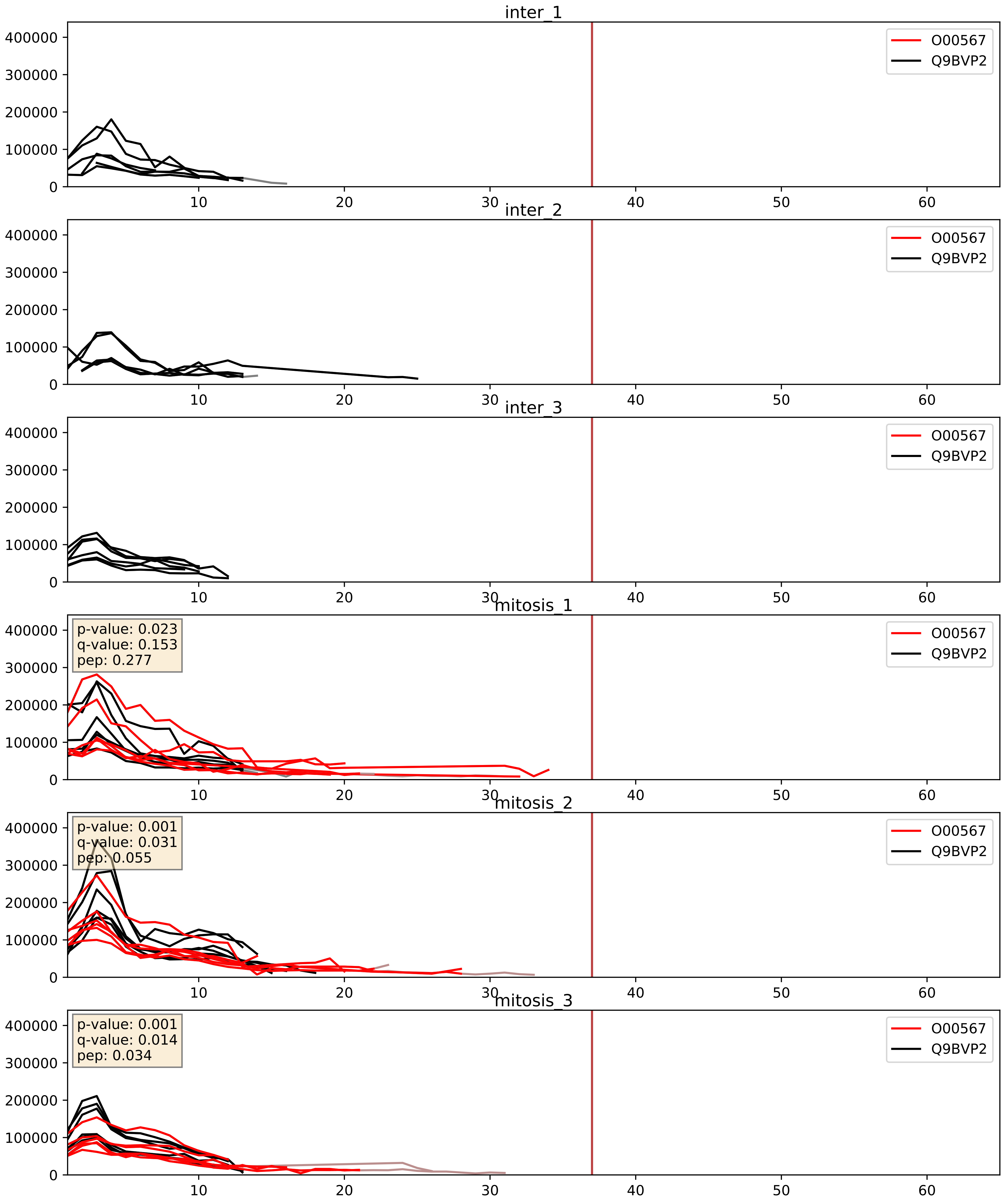


| level | condition_1 | condition_2 | pvalue | pvalue_adjusted |
|---|---|---|---|---|
| complex_abundance | mitosis | inter | 0.0003457116354080251 | 0.012015488415248196 |
| interactor_abundance | mitosis | inter | 0.0008877885552668727 | 0.02197564842852207 |
| interactor_ratio | mitosis | inter | 0.22699272553855118 | 0.7059145275954049 |

O76021_Q9BVP2
RL1D1_HUMAN vs GNL3_HUMAN
p-value: 0.0 q-value: 0.013

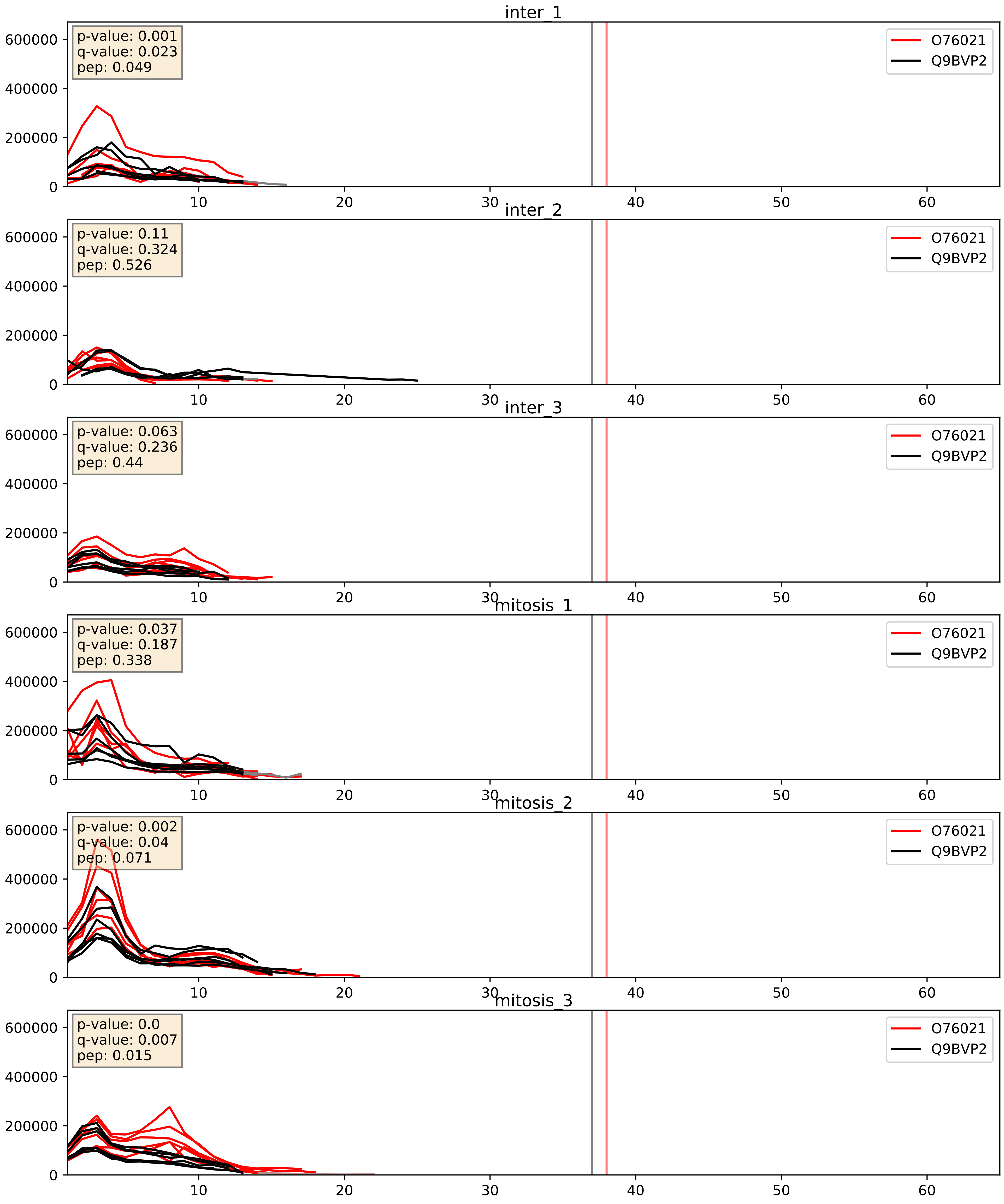


| level | condition_1 | condition_2 | pvalue | pvalue_adjusted |
|---|---|---|---|---|
| complex_abundance | mitosis | inter | 0.004581703881258023 | 0.05041052085291604 |
| interactor_abundance | mitosis | inter | 0.008085645358995957 | 0.077986618899161 |
| interactor_ratio | mitosis | inter | 0.3527656966303728 | 0.8022478353006512 |

P22087_Q9BVP2
FBRL_HUMAN vs GNL3_HUMAN
p-value: 0.001 q-value: 0.027

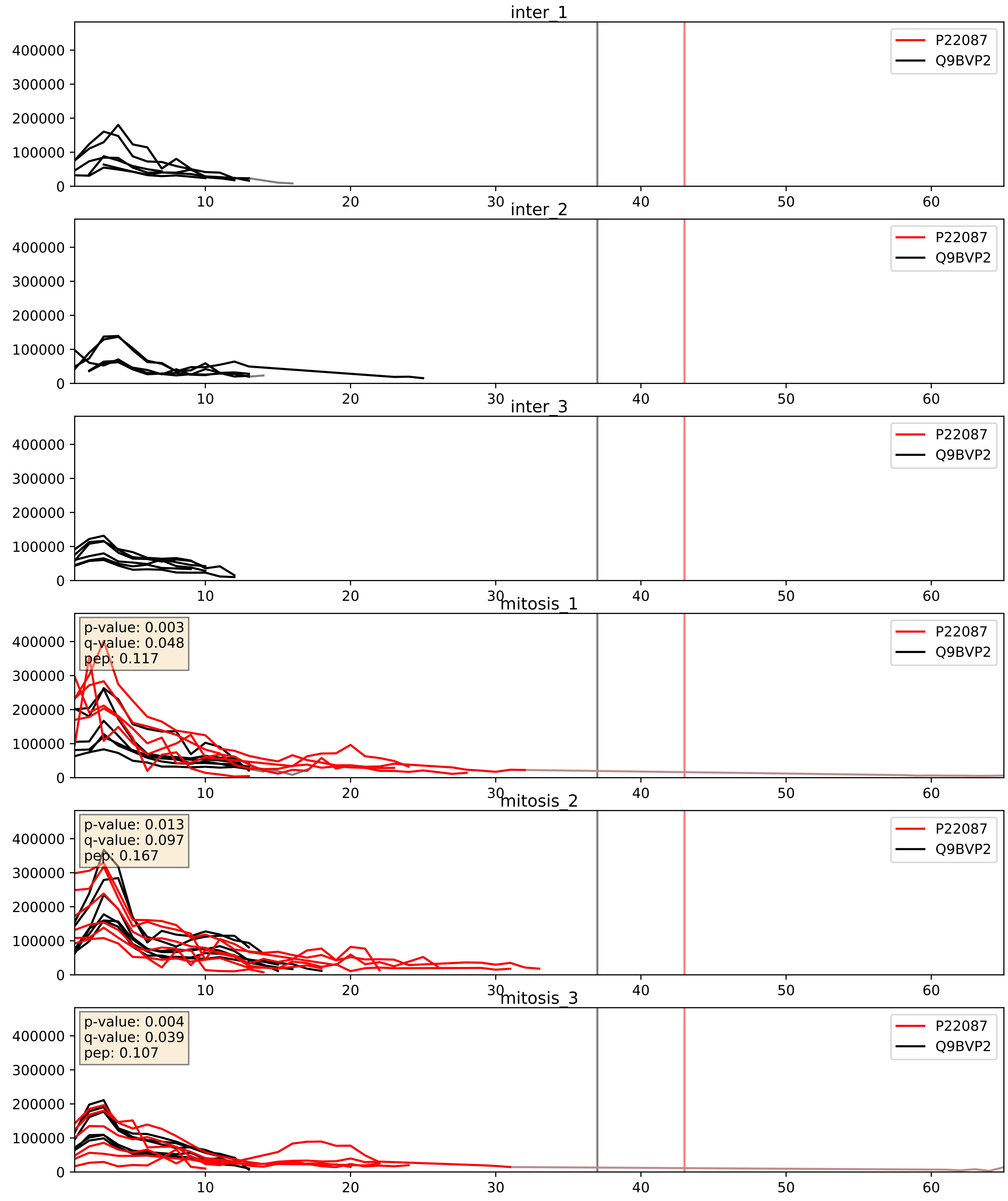


| level | condition_1 | condition_2 | pvalue | pvalue_adjusted |
|---|---|---|---|---|
| complex_abundance | mitosis | inter | 0.00030575611516796747 | 0.01165207110963347 |
| interactor_abundance | mitosis | inter | 0.0006136131553603794 | 0.017415659477552636 |
| interactor_ratio | mitosis | inter | 0.22854497040560845 | 0.7070495646135039 |

P46087_Q9BVP2
NOP2_HUMAN vs GNL3_HUMAN
p-value: 0.026 q-value: 0.019

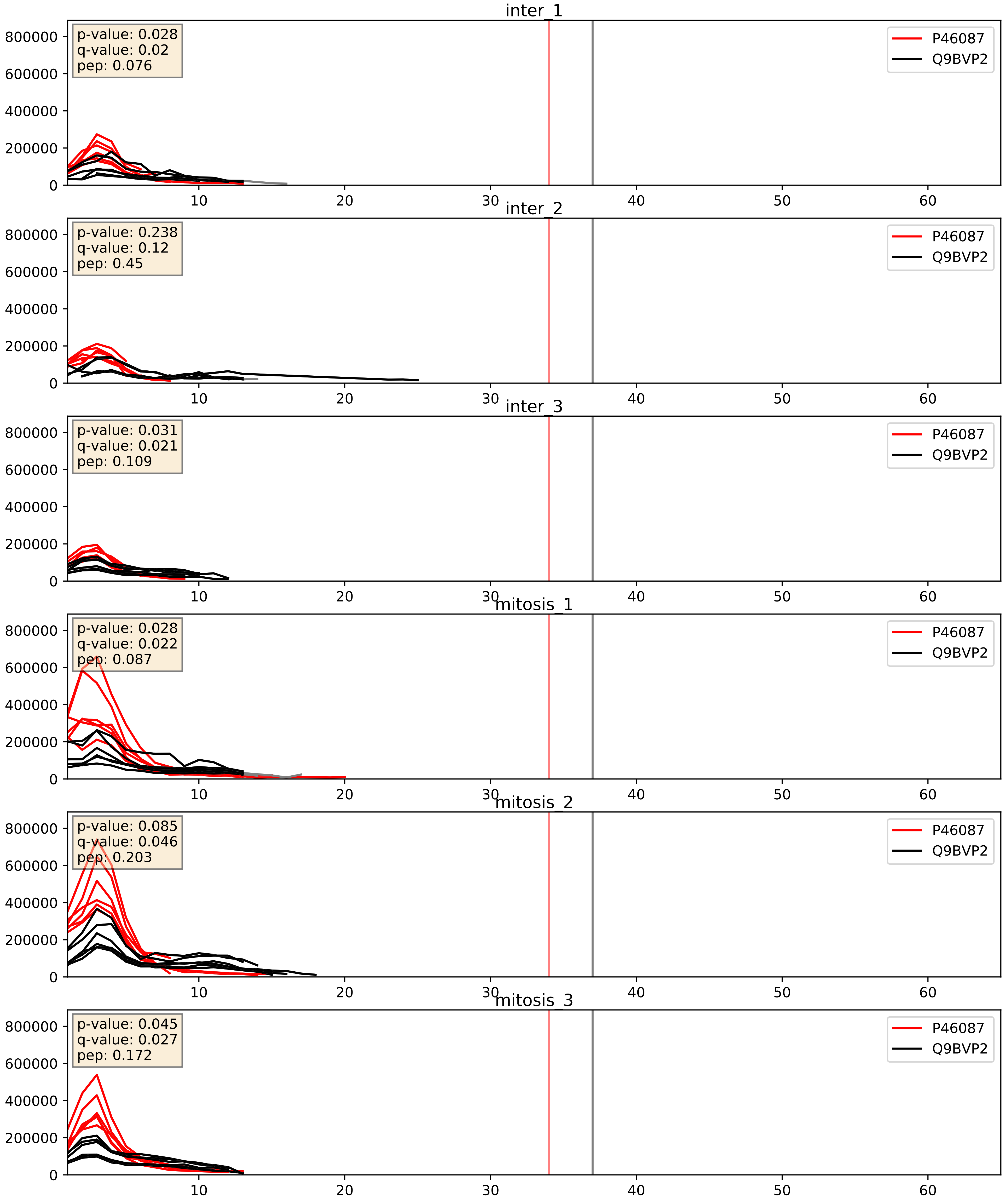


| level | condition_1 | condition_2 | pvalue | pvalue_adjusted |
|---|---|---|---|---|
| complex_abundance | mitosis | inter | 0.00781659329937401 | 0.0656624520536227 |
| interactor_abundance | mitosis | inter | 0.007951593035973885 | 0.07739128639901814 |
| interactor_ratio | mitosis | inter | 0.05096474016989619 | 0.373742437899162 |

P61513_Q9BVP2
RL37A_HUMAN vs GNL3_HUMAN
p-value: 0.001 q-value: 0.039

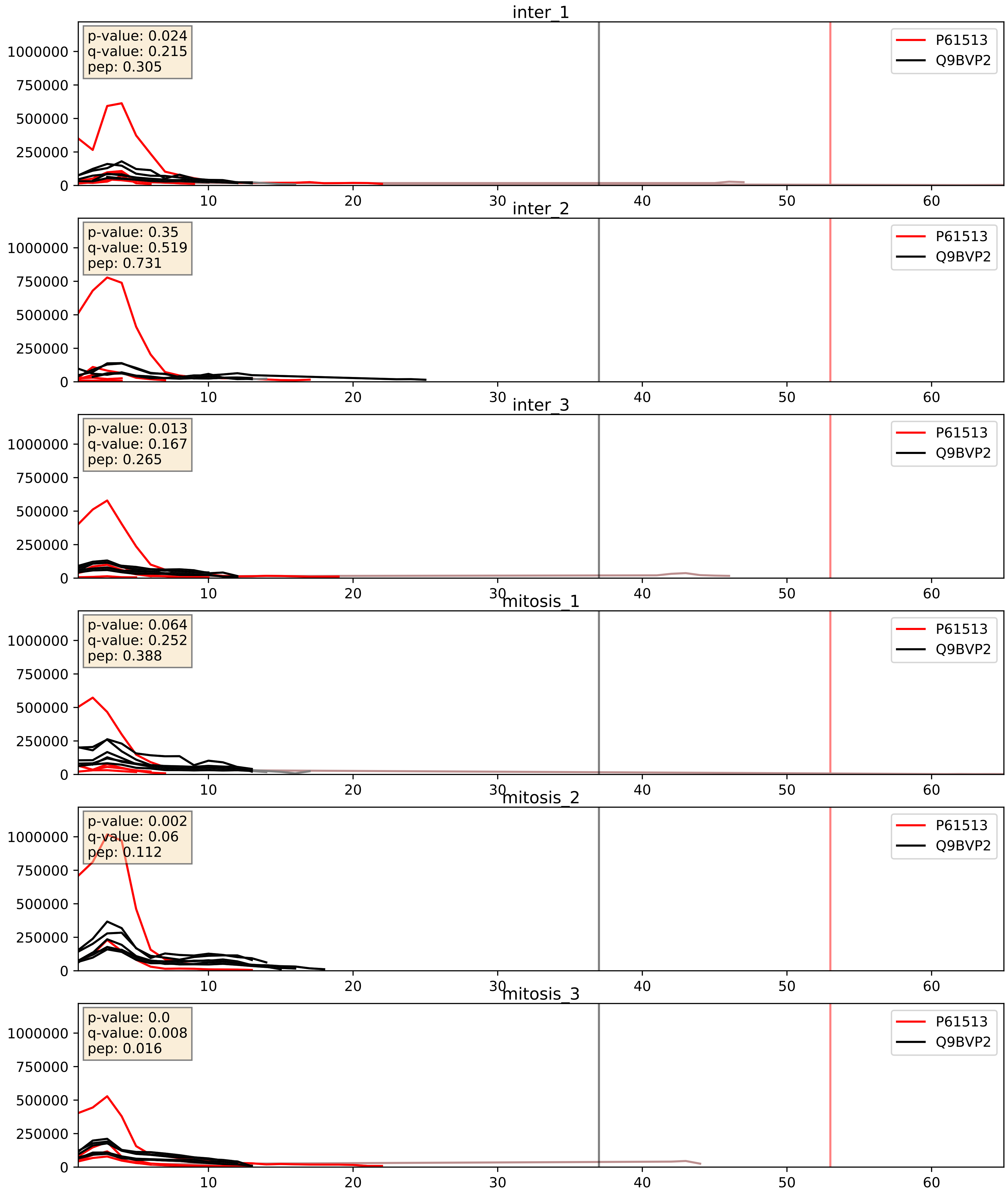


| level | condition_1 | condition_2 | pvalue | pvalue_adjusted |
|---|---|---|---|---|
| interactor_ratio | mitosis | inter | 0.02955716439929511 | 0.2722029531790272 |
| interactor_abundance | mitosis | inter | 0.041643683633610595 | 0.1640450676040988 |
| complex_abundance | mitosis | inter | 0.9739610278788303 | 0.9841938848592595 |

P62913_Q9BVP2
RL11_HUMAN vs GNL3_HUMAN
p-value: 0.0 q-value: 0.003

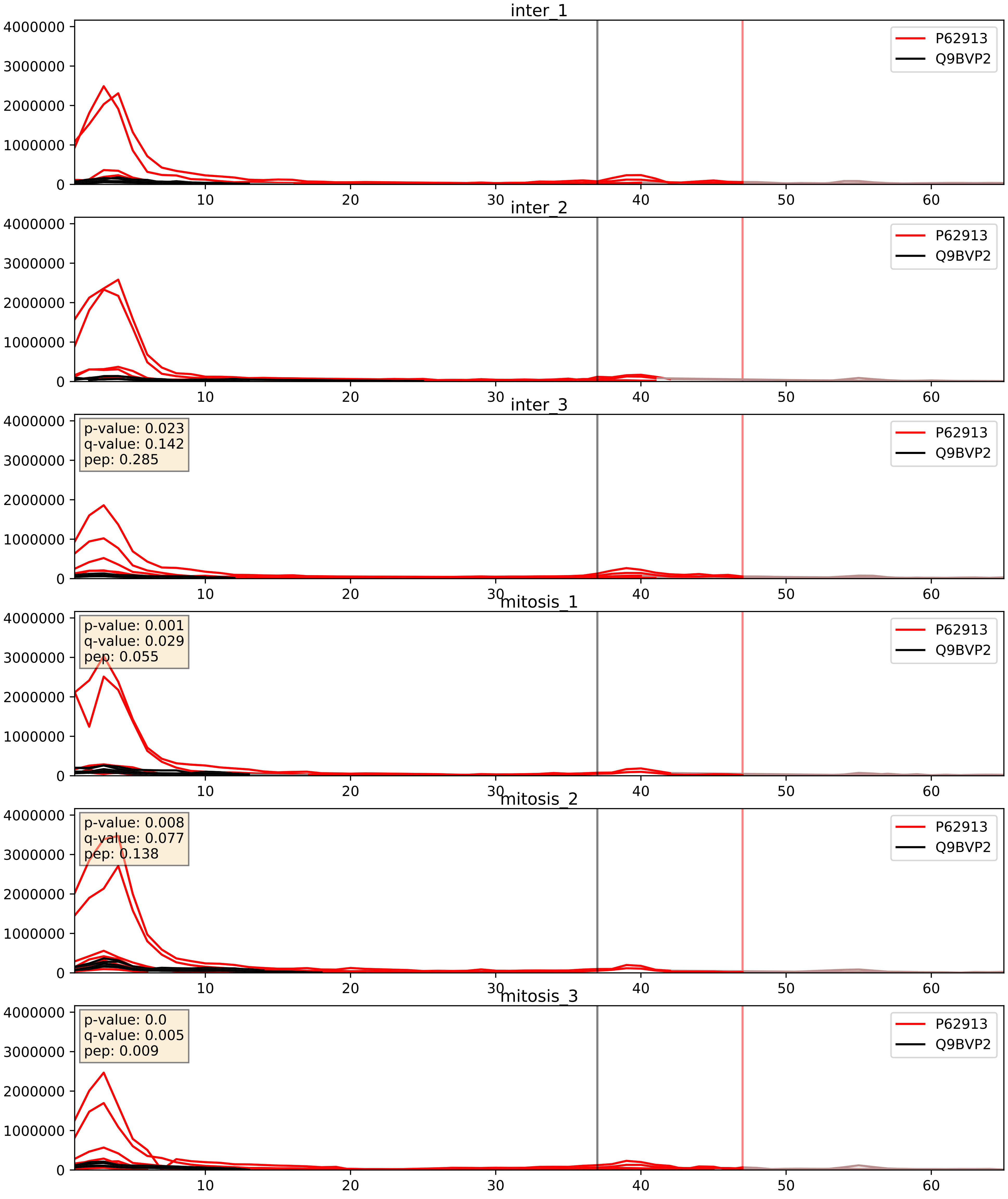


| level | condition_1 | condition_2 | pvalue | pvalue_adjusted |
|---|---|---|---|---|
| complex_abundance | mitosis | inter | 0.03208832444419898 | 0.13030173493469793 |
| interactor_ratio | mitosis | inter | 0.04593411677077528 | 0.35622692584467375 |
| interactor_abundance | mitosis | inter | 0.13052741730673784 | 0.31529718452669087 |

Q13823_Q9BVP2
NOG2_HUMAN vs GNL3_HUMAN
p-value: 0.034 q-value: 0.024

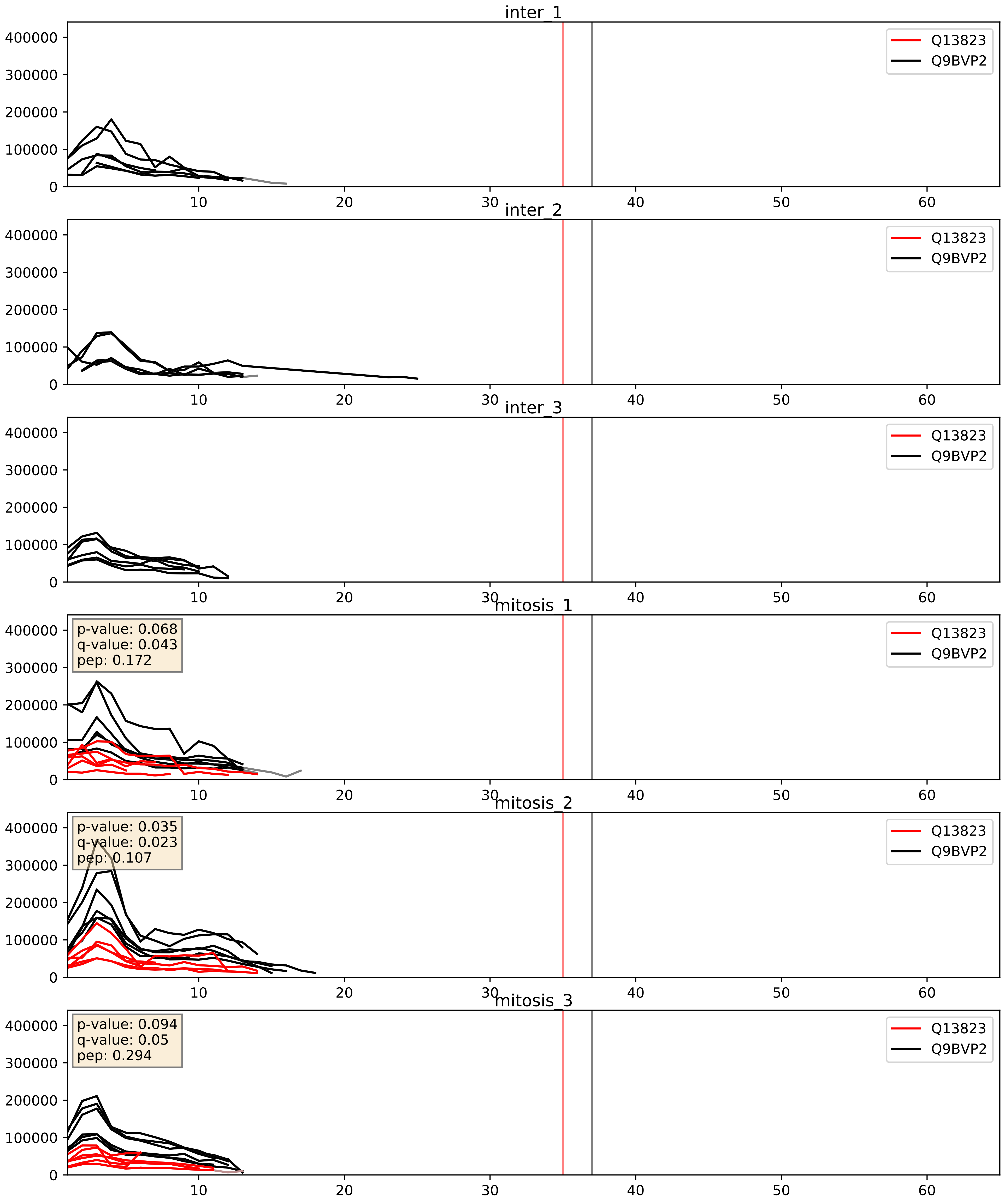


| level | condition_1 | condition_2 | pvalue | pvalue_adjusted |
|---|---|---|---|---|
| complex_abundance | mitosis | inter | 0.0009671264823373898 | 0.020644894485805625 |
| interactor_ratio | mitosis | inter | 0.0012027857202170427 | 0.041019305836884006 |
| interactor_abundance | mitosis | inter | 0.00933786289754201 | 0.08378627505551321 |

Q14137_Q9BVP2
BOP1_HUMAN vs GNL3_HUMAN
p-value: 0.042 q-value: 0.028

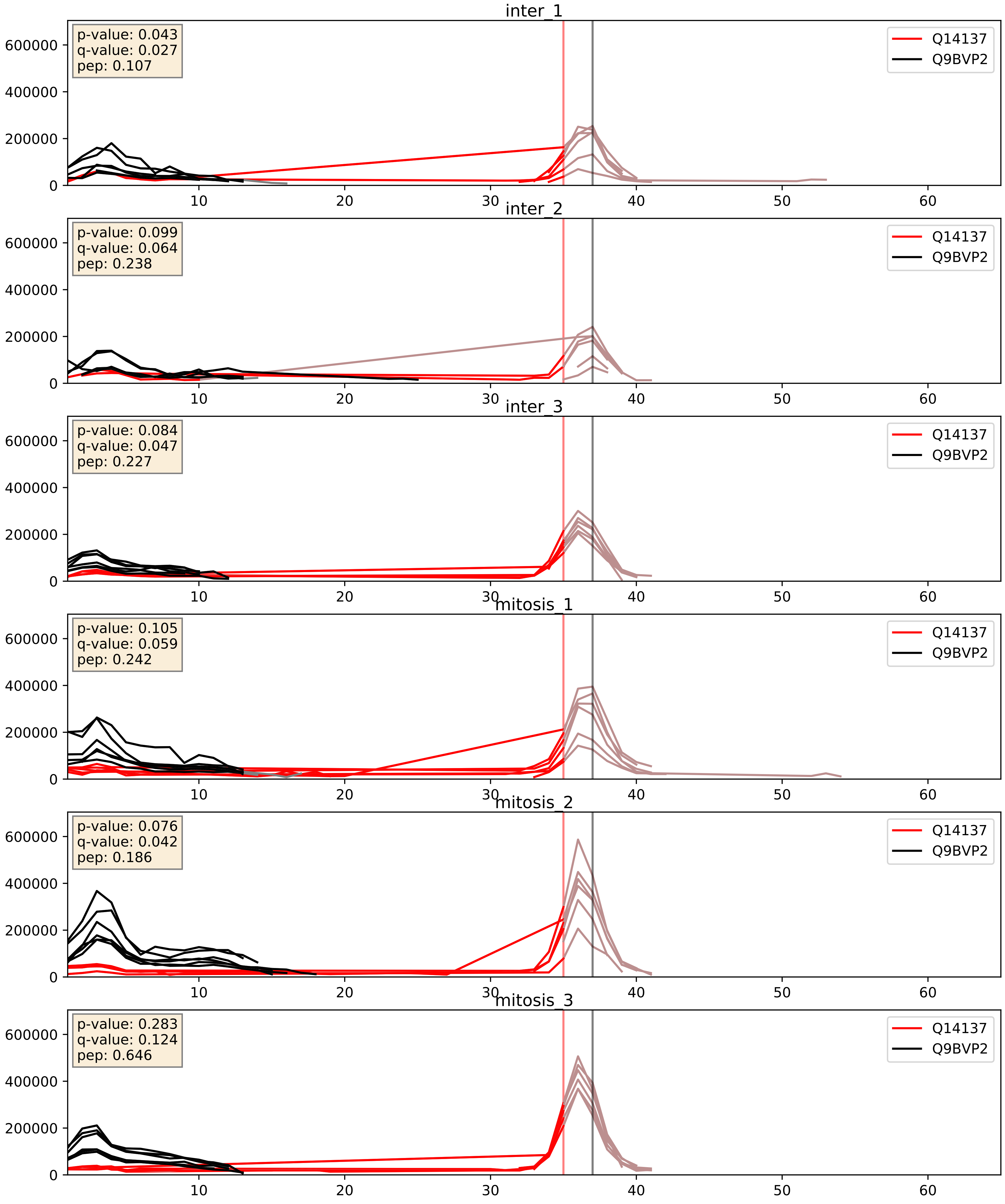


| level | condition_1 | condition_2 | pvalue | pvalue_adjusted |
|---|---|---|---|---|
| interactor_ratio | mitosis | inter | 0.011122406259462985 | 0.1518775136410363 |
| interactor_abundance | mitosis | inter | 0.09412782094612172 | 0.26266801867931605 |
| complex_abundance | mitosis | inter | 0.6045895512232708 | 0.7277860439419489 |

Q8IY81_Q9BVP2
SPB1_HUMAN vs GNL3_HUMAN
p-value: 0.027 q-value: 0.02

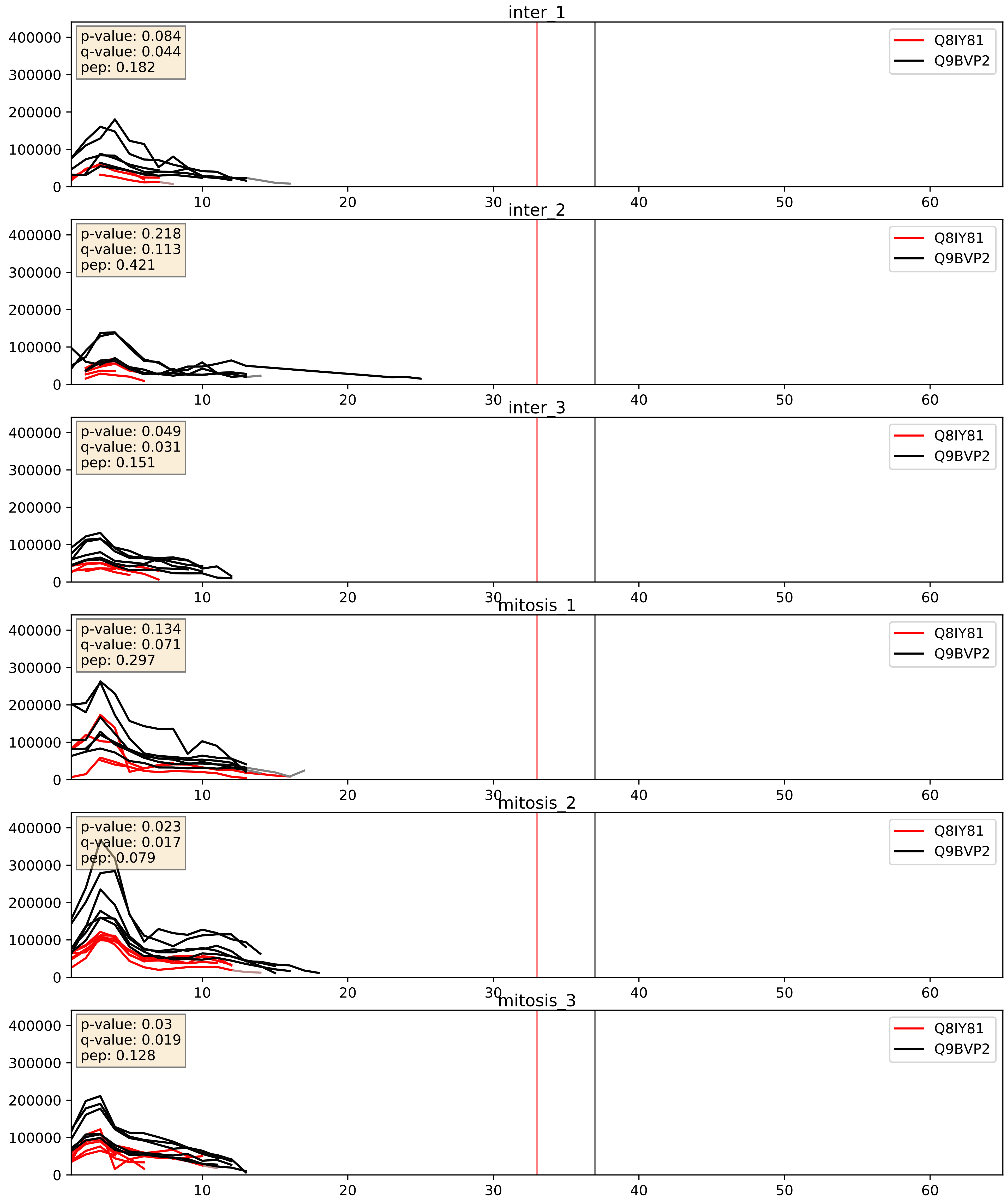


| level | condition_1 | condition_2 | pvalue | pvalue_adjusted |
|---|---|---|---|---|
| complex_abundance | mitosis | inter | 0.04433993120036613 | 0.15060165151763358 |
| interactor_ratio | mitosis | inter | 0.07249910086746936 | 0.44970456769966505 |
| interactor_abundance | mitosis | inter | 0.3789887745019212 | 0.5719072560134765 |

Q9BV38_Q9BVP2
WDR18_HUMAN vs GNL3_HUMAN
p-value: 0.003 q-value: 0.039

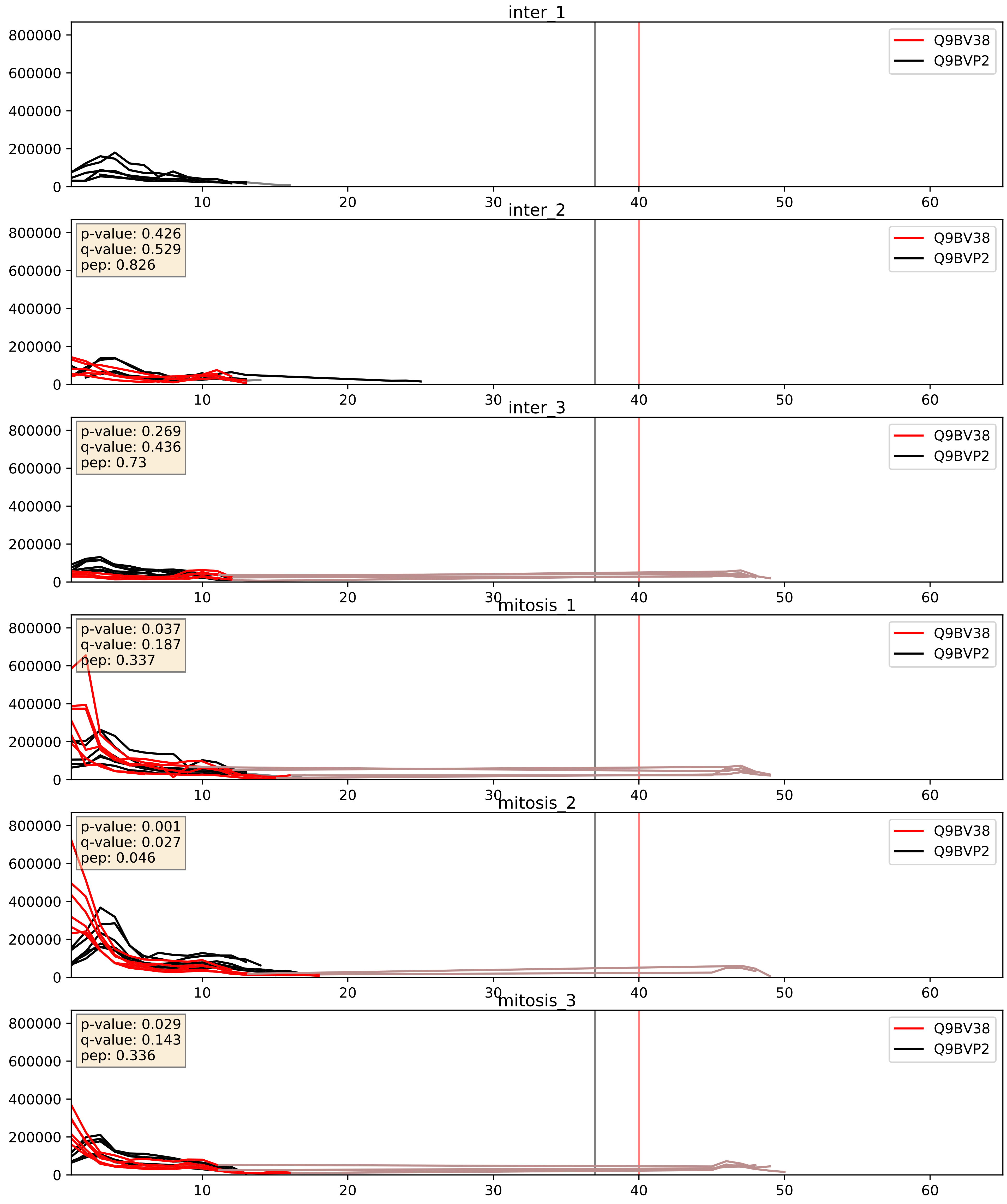


| level | condition_1 | condition_2 | pvalue | pvalue_adjusted |
|---|---|---|---|---|
| interactor_abundance | mitosis | inter | 0.03692462845525282 | 0.15802829520754763 |
| complex_abundance | mitosis | inter | 0.06145454048384596 | 0.17808086206557935 |
| interactor_ratio | mitosis | inter | 0.2008905695851433 | 0.6754215536719665 |

Q9BVP2_Q9BYG3
GNL3_HUMAN vs MK67I_HUMAN
p-value: 0.056 q-value: 0.036

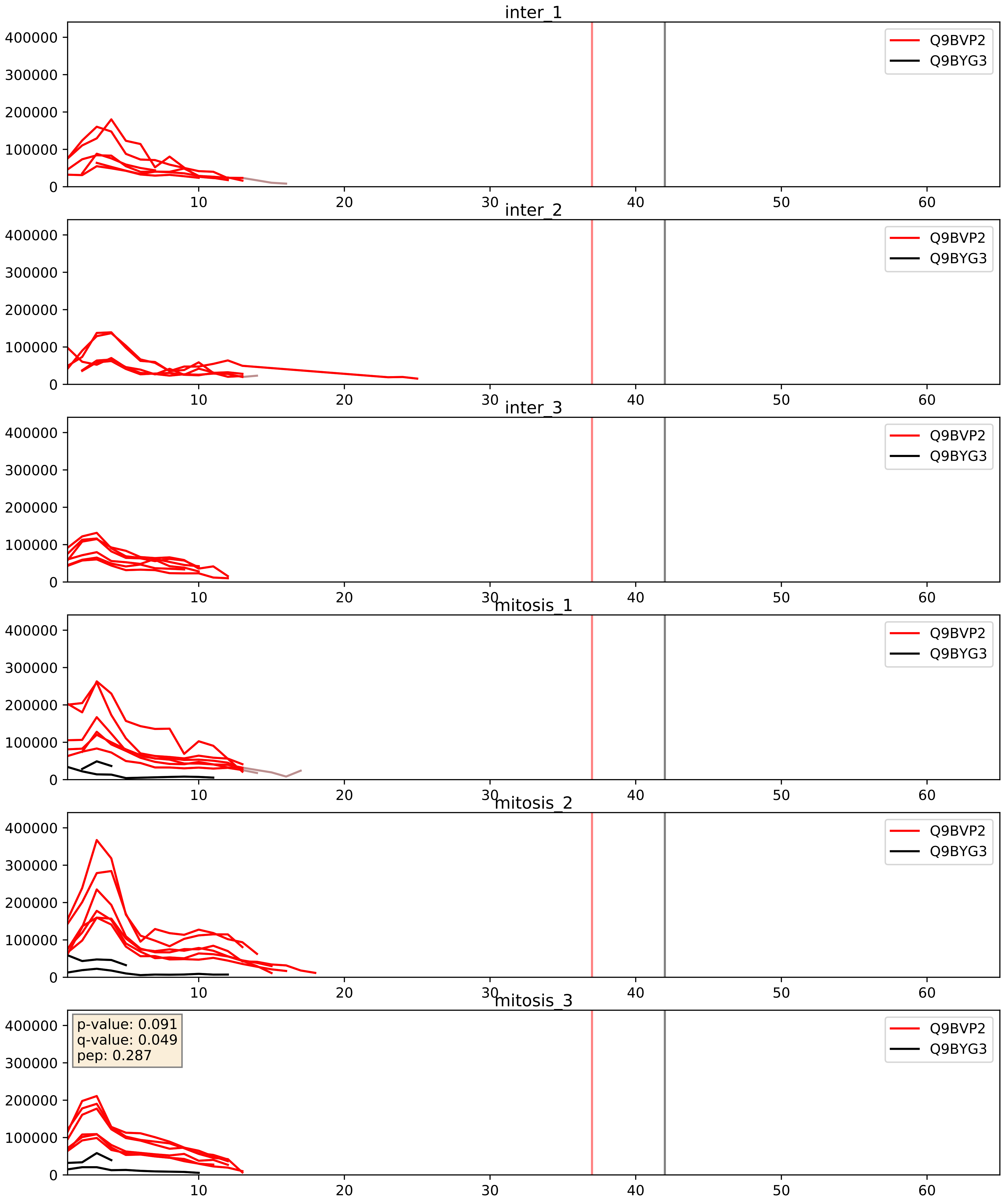


| level | condition_1 | condition_2 | pvalue | pvalue_adjusted |
|---|---|---|---|---|
| interactor_abundance | mitosis | inter | 0.00039498517428285436 | 0.01371632085947762 |
| interactor_ratio | mitosis | inter | 0.0007741550098460847 | 0.03185945617443502 |
| complex_abundance | mitosis | inter | 0.000841508846203478 | 0.019252661341274204 |

Q9BVP2_Q9BZE4
GNL3_HUMAN vs NOG1_HUMAN
p-value: 0.004 q-value: 0.004

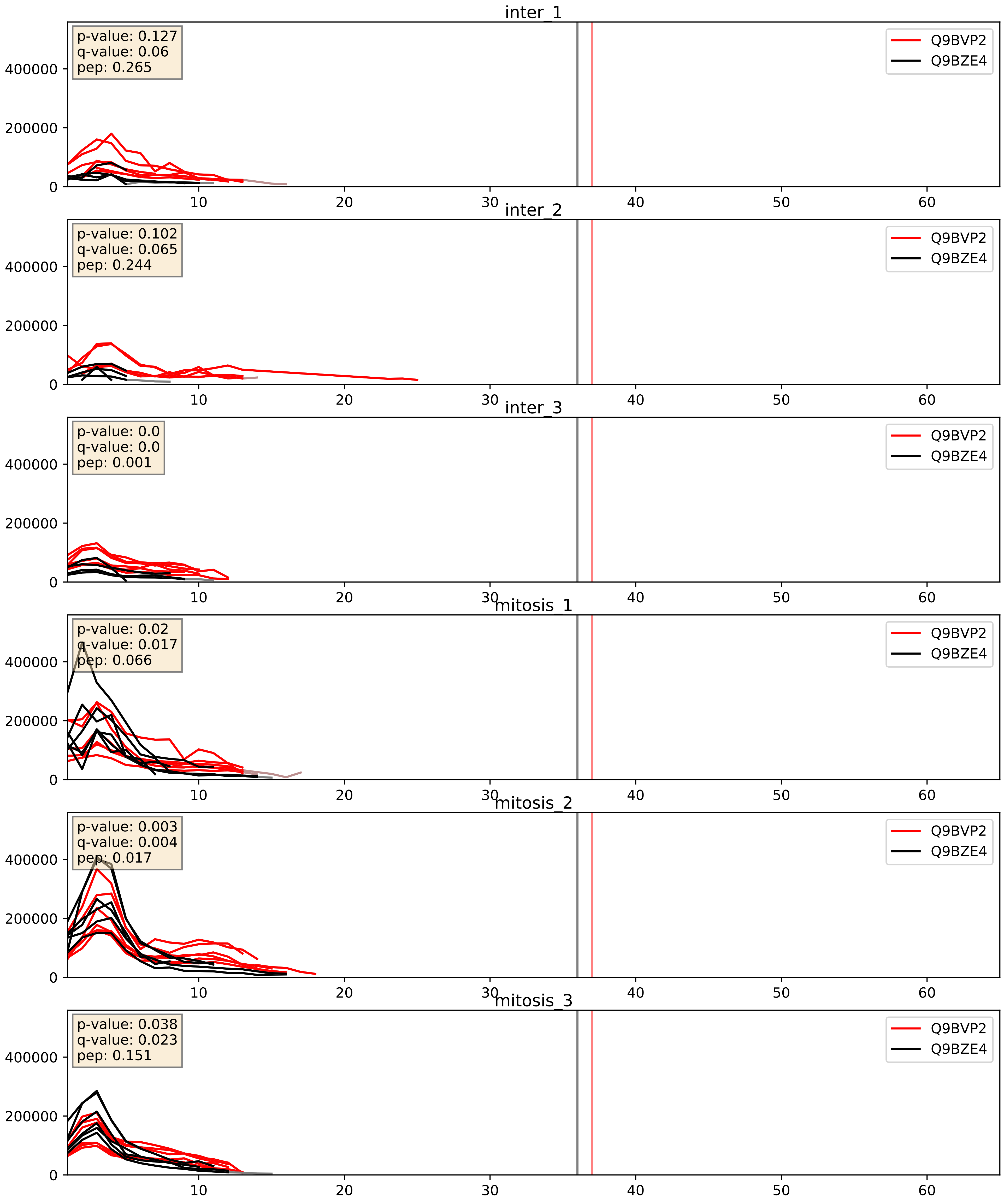


| level | condition_1 | condition_2 | pvalue | pvalue_adjusted |
|---|---|---|---|---|
| complex_abundance | mitosis | inter | 0.0013895549742411744 | 0.02552487248820698 |
| interactor_abundance | mitosis | inter | 0.006897488603137285 | 0.07148798621825644 |
| interactor_ratio | mitosis | inter | 0.011563542728058424 | 0.15494522140242886 |

Q9BVP2_Q9UKD2
GNL3_HUMAN vs MRT4_HUMAN
p-value: 0.019 q-value: 0.015

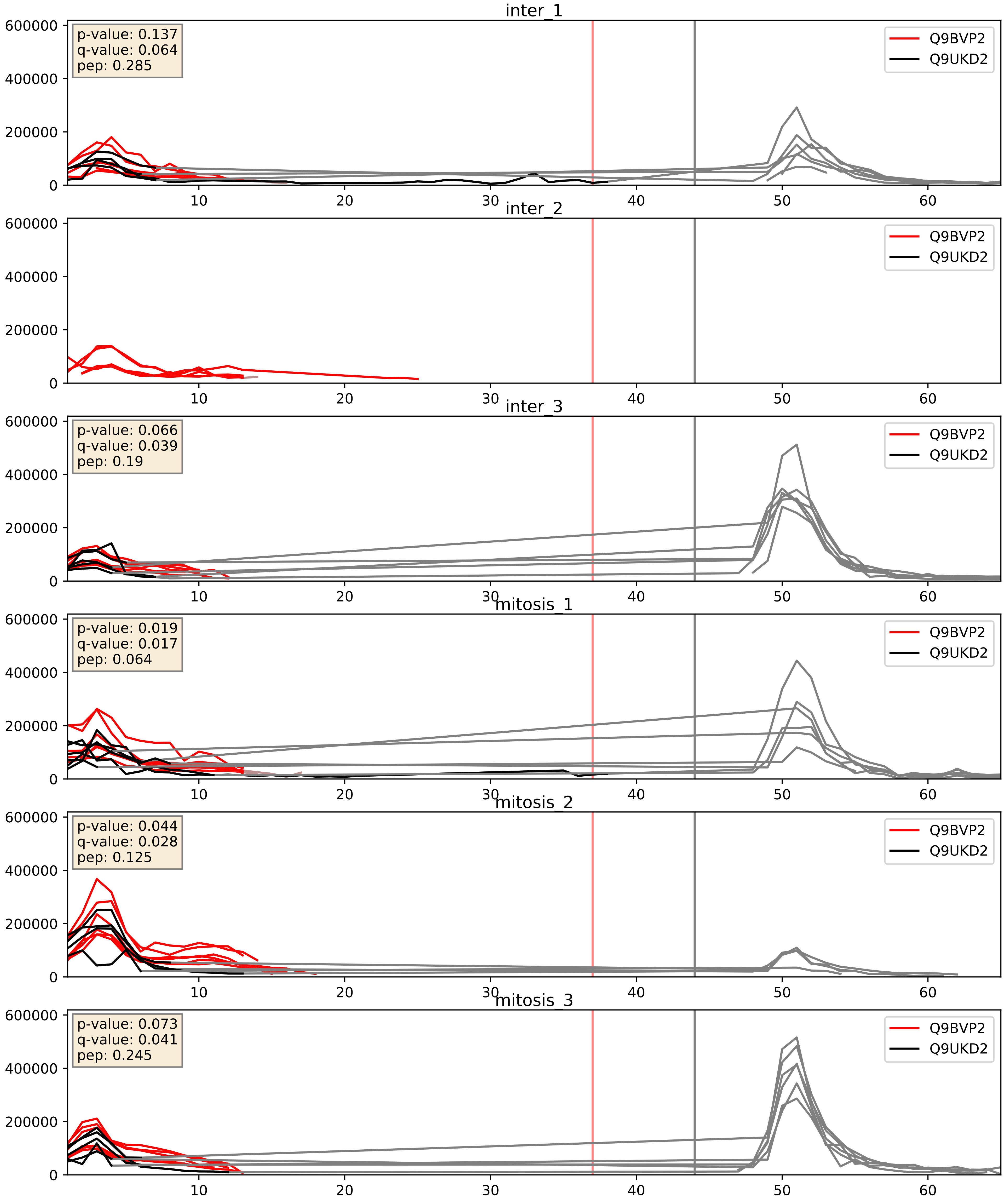


| level | condition_1 | condition_2 | pvalue | pvalue_adjusted |
|---|---|---|---|---|
| interactor_ratio | mitosis | inter | 0.03252851835389341 | 0.28824442764940744 |
| interactor_abundance | mitosis | inter | 0.0987233945687053 | 0.2701737135098137 |
| complex_abundance | mitosis | inter | 0.12841912353114857 | 0.27142412282139056 |

Q9BVP2_Q9Y2X3
GNL3_HUMAN vs NOP58_HUMAN
p-value: 0.002 q-value: 0.003

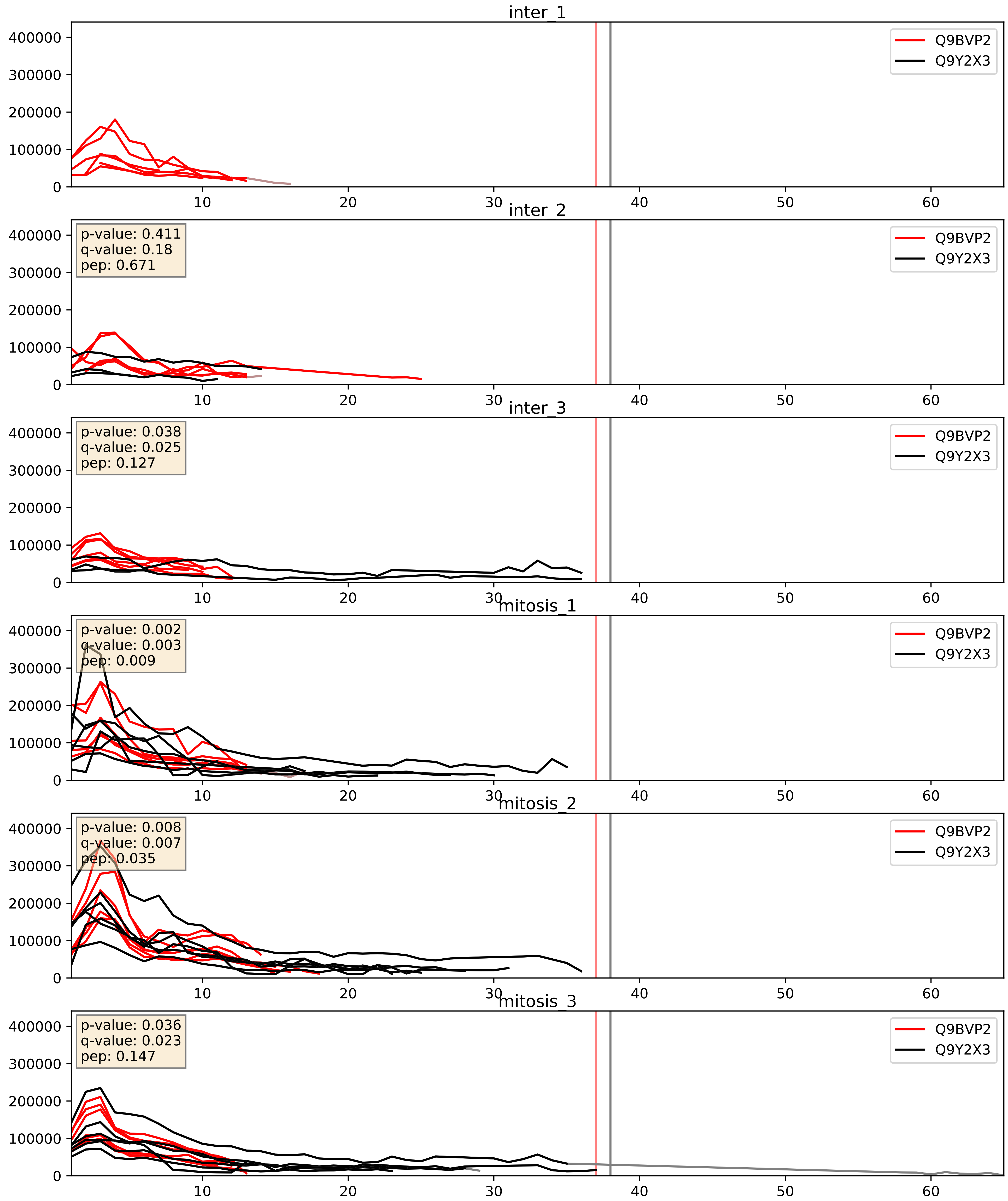


| level | condition_1 | condition_2 | pvalue | pvalue_adjusted |
|---|---|---|---|---|
| complex_abundance | mitosis | inter | 0.0274592583388612 | 0.12057983221353287 |
| interactor_ratio | mitosis | inter | 0.10574631048223244 | 0.5413806326123862 |
| interactor_abundance | mitosis | inter | 0.11740638492850289 | 0.29607210740171247 |
