## Supplementary Data 2 for "SECAT: Quantifying differential protein-protein interaction states by network-centric analysis": hela_string_000034_Q16629.pdf

Q16629_Q16629

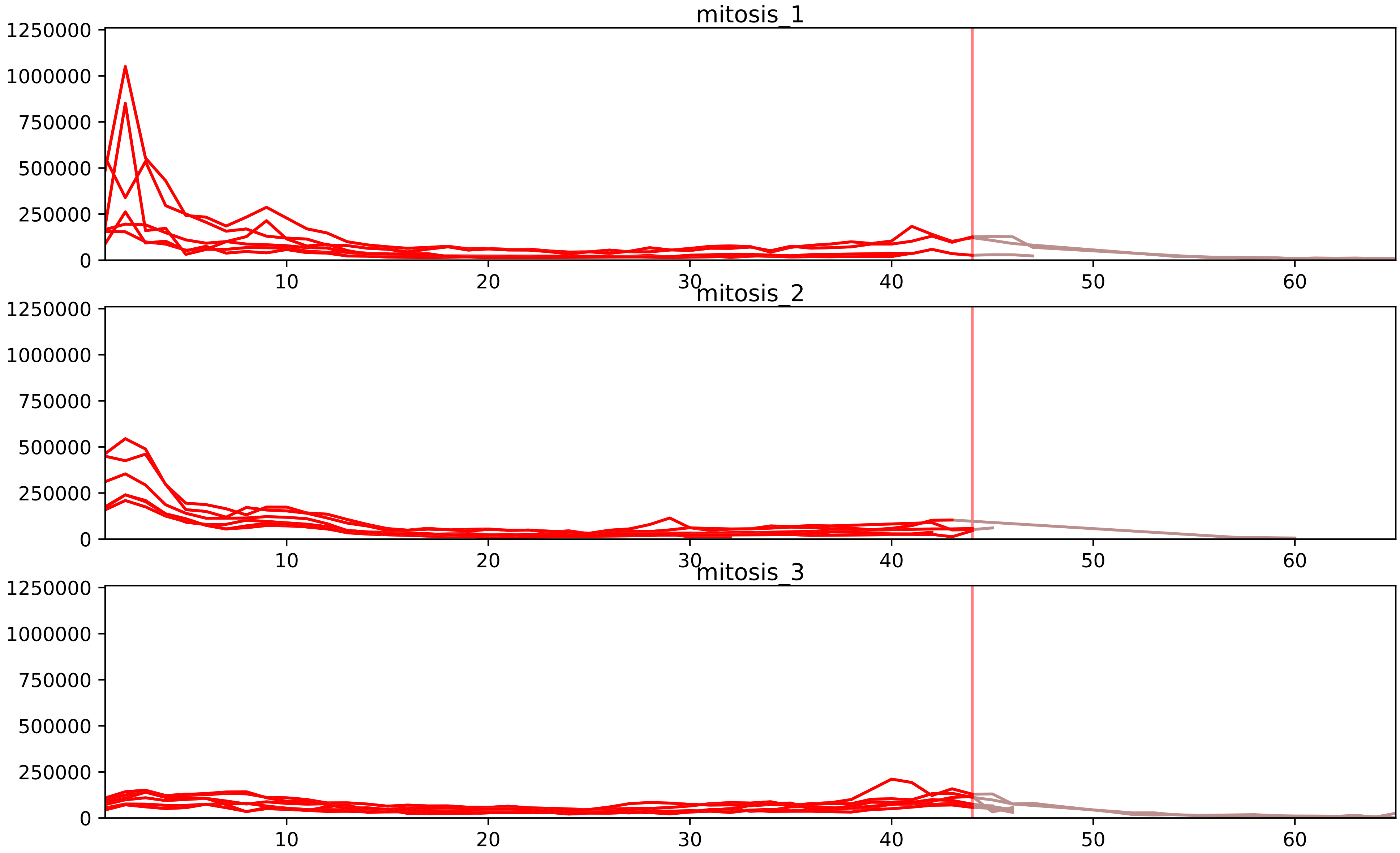


| level | condition_1 | condition_2 | pvalue | pvalue_adjusted |
|---|---|---|---|---|
| interactor_ratio | mitosis | inter | 8.563122365495420e-09 | 4.774589440155021e-07 |
| interactor_abundance | mitosis | inter | 3.972799678564088e-08 | 1.2183252347596537e-05 |
| complex_abundance | mitosis | inter | 4.5277608513934896e-08 | 1.4956025834220426e-05 |
| assembled_abundance | mitosis | inter | 0.0013466353772594625 | 0.03389032366102981 |
| total_abundance | mitosis | inter | 0.0015682485371086615 | 0.042983537408980685 |
| monomer_abundance | mitosis | inter | 0.012663085605216479 | 0.19546941175259344 |

P07910_Q16629
HNRPC_HUMAN vs SRSF7_HUMAN
p-value: 0.046 q-value: 0.031

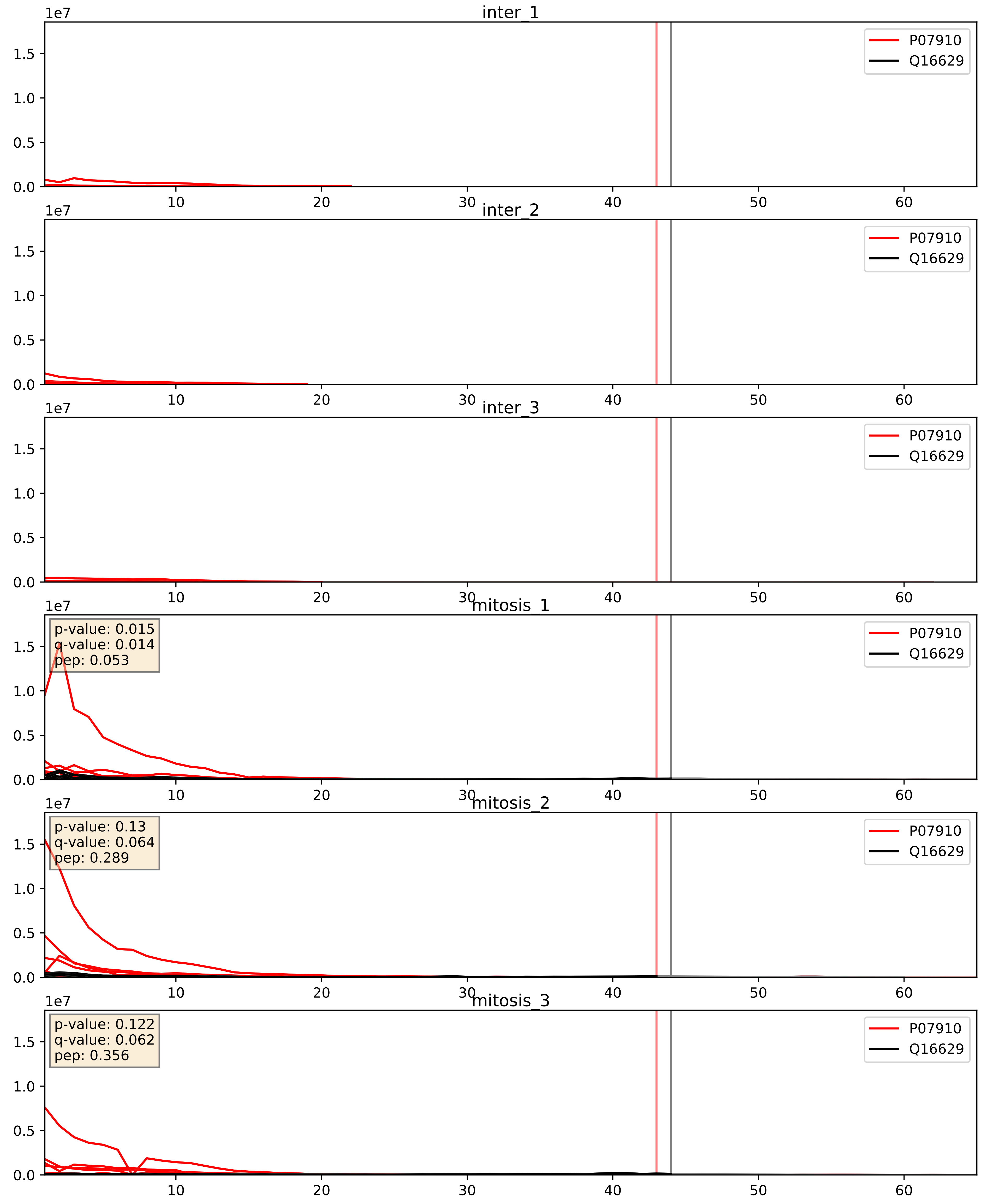


| level | condition_1 | condition_2 | pvalue | pvalue_adjusted |
|---|---|---|---|---|
| interactor_abundance | mitosis | inter | 2.525501836279769e-06 | 0.0033734327106189023 |
| complex_abundance | mitosis | inter | 5.288643111257365e-06 | 0.0056588481290453805 |
| interactor_ratio | mitosis | inter | 0.00024711905536094626 | 0.016038997600655195 |

P09651_Q16629
ROA1_HUMAN vs SRSF7_HUMAN
p-value: 0.008 q-value: 0.007

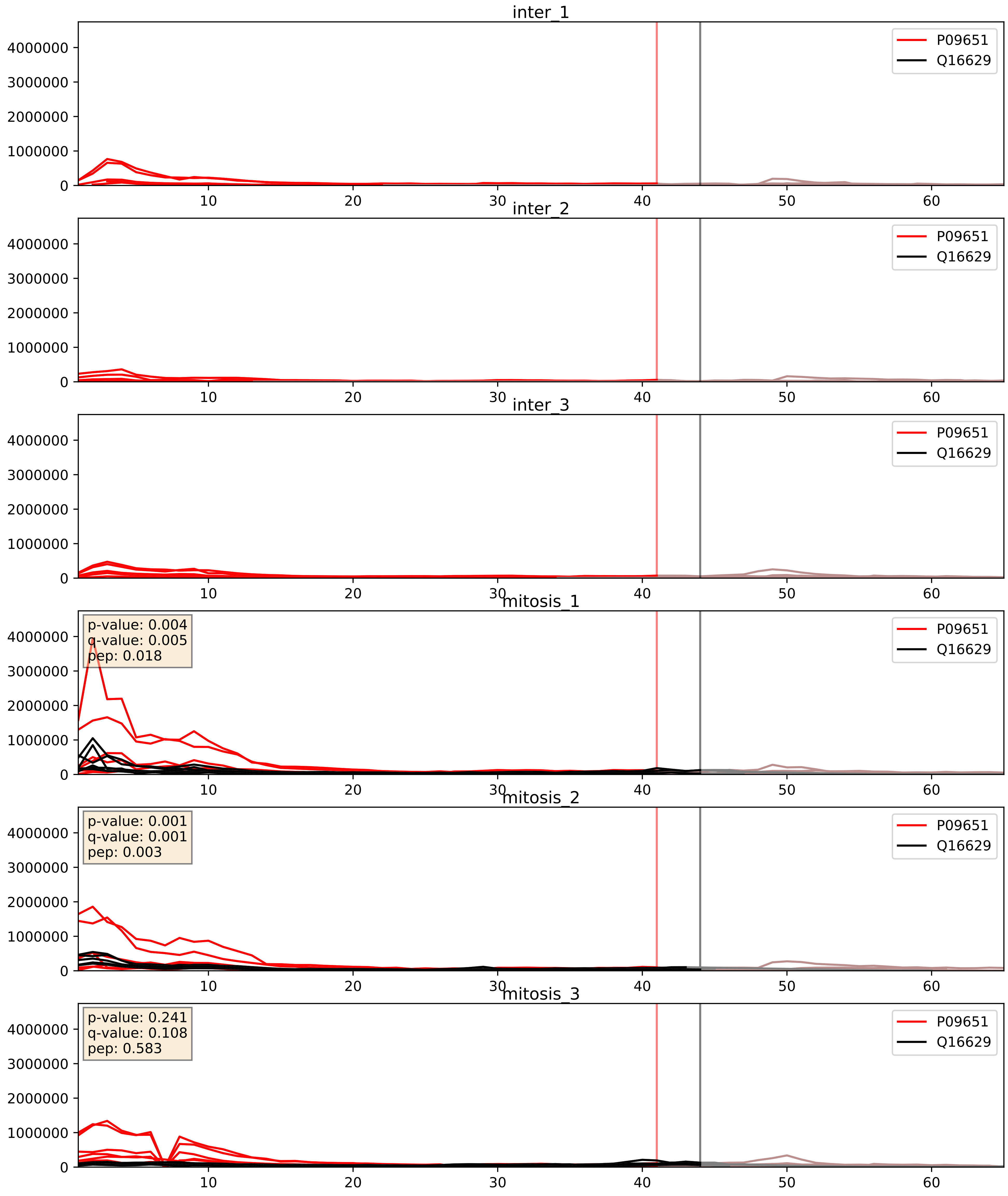


| level | condition_1 | condition_2 | pvalue | pvalue_adjusted |
|---|---|---|---|---|
| complex_abundance | mitosis | inter | 5.081957409361551e-05 | 0.005691857151652958 |
| interactor_abundance | mitosis | inter | 6.959157312257744e-05 | 0.006732838333317079 |
| interactor_ratio | mitosis | inter | 0.00173672861406567 | 0.05073855609693561 |

P14866_Q16629
HNRPL_HUMAN vs SRSF7_HUMAN
p-value: 0.003 q-value: 0.003

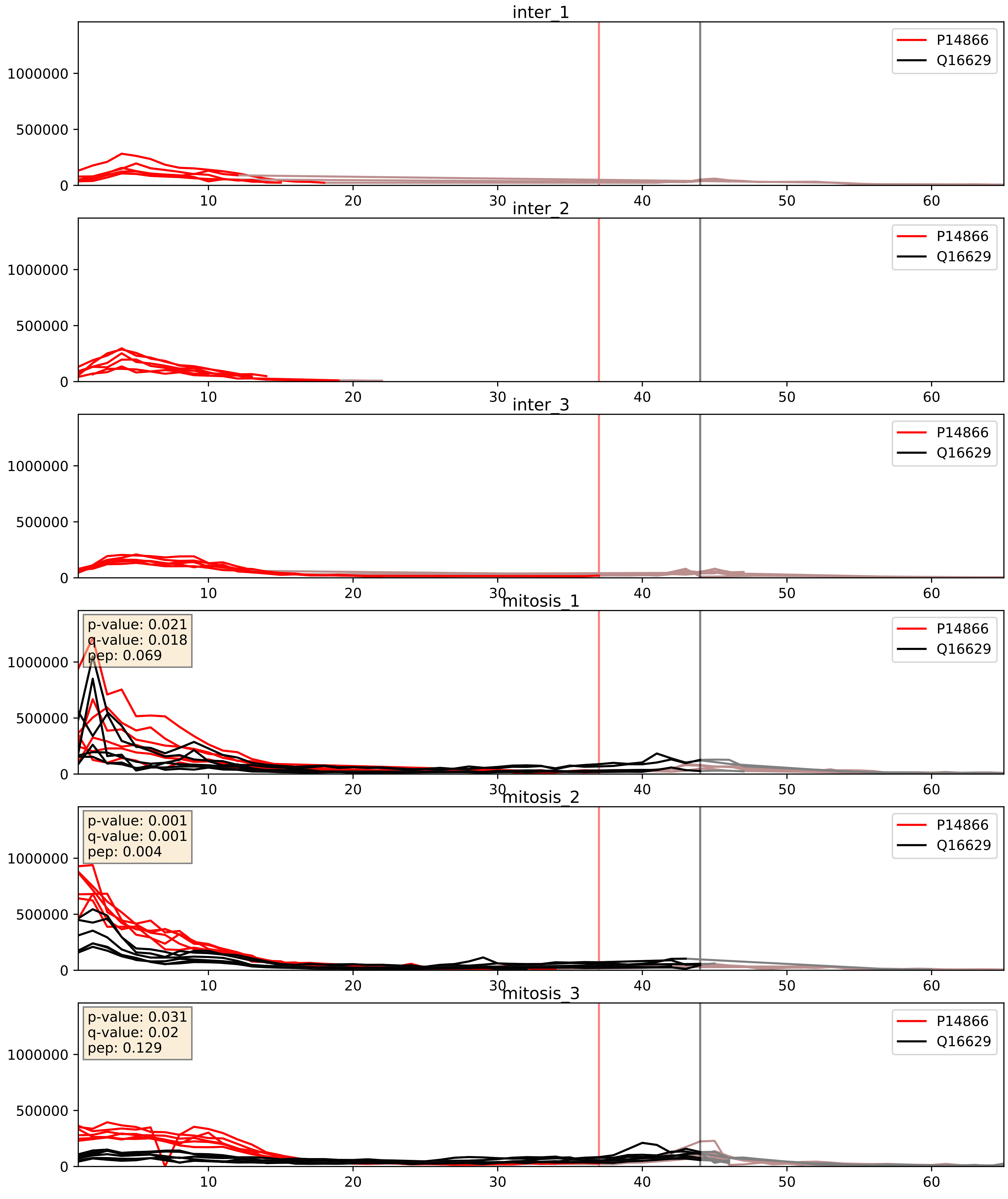


| level | condition_1 | condition_2 | pvalue | pvalue_adjusted |
|---|---|---|---|---|
| interactor_abundance | mitosis | inter | 1.2331621368333903e-05 | 0.003391165020149013 |
| complex_abundance | mitosis | inter | 3.0454949653656297e-05 | 0.005691857151652958 |
| interactor_ratio | mitosis | inter | 0.04207614494229382 | 0.3399226239910554 |

P17844_Q16629
DDX5_HUMAN vs SRSF7_HUMAN
p-value: 0.0 q-value: 0.01

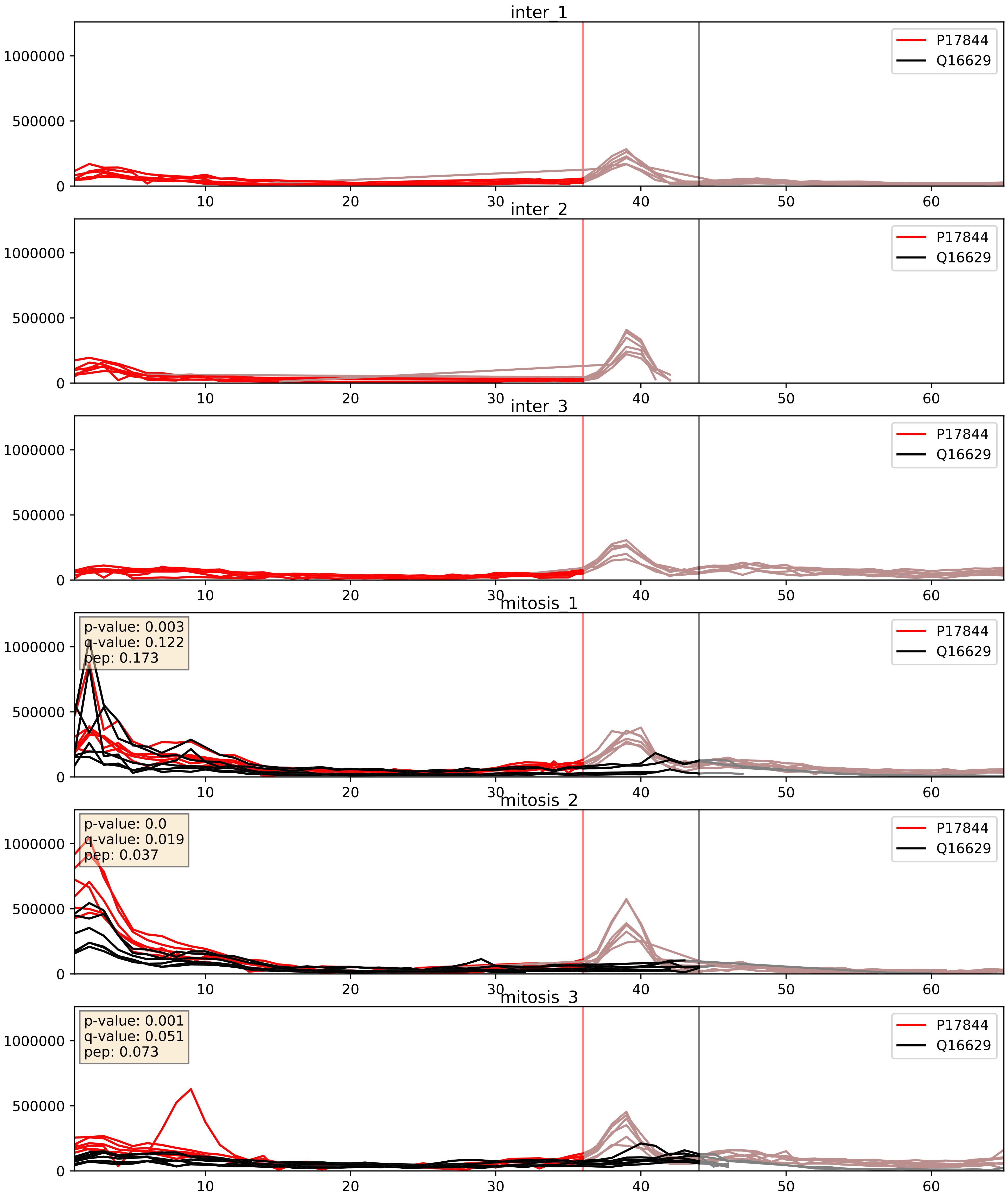


| level | condition_1 | condition_2 | pvalue | pvalue_adjusted |
|---|---|---|---|---|
| interactor_abundance | mitosis | inter | 2.9246981424557922e-05 | 0.004814503096042612 |
| complex_abundance | mitosis | inter | 4.447501338108865e-05 | 0.005691857151652958 |
| interactor_ratio | mitosis | inter | 0.15108645229943365 | 0.606604569117897 |

P22626_Q16629
ROA2_HUMAN vs SRSF7_HUMAN
p-value: 0.01 q-value: 0.009

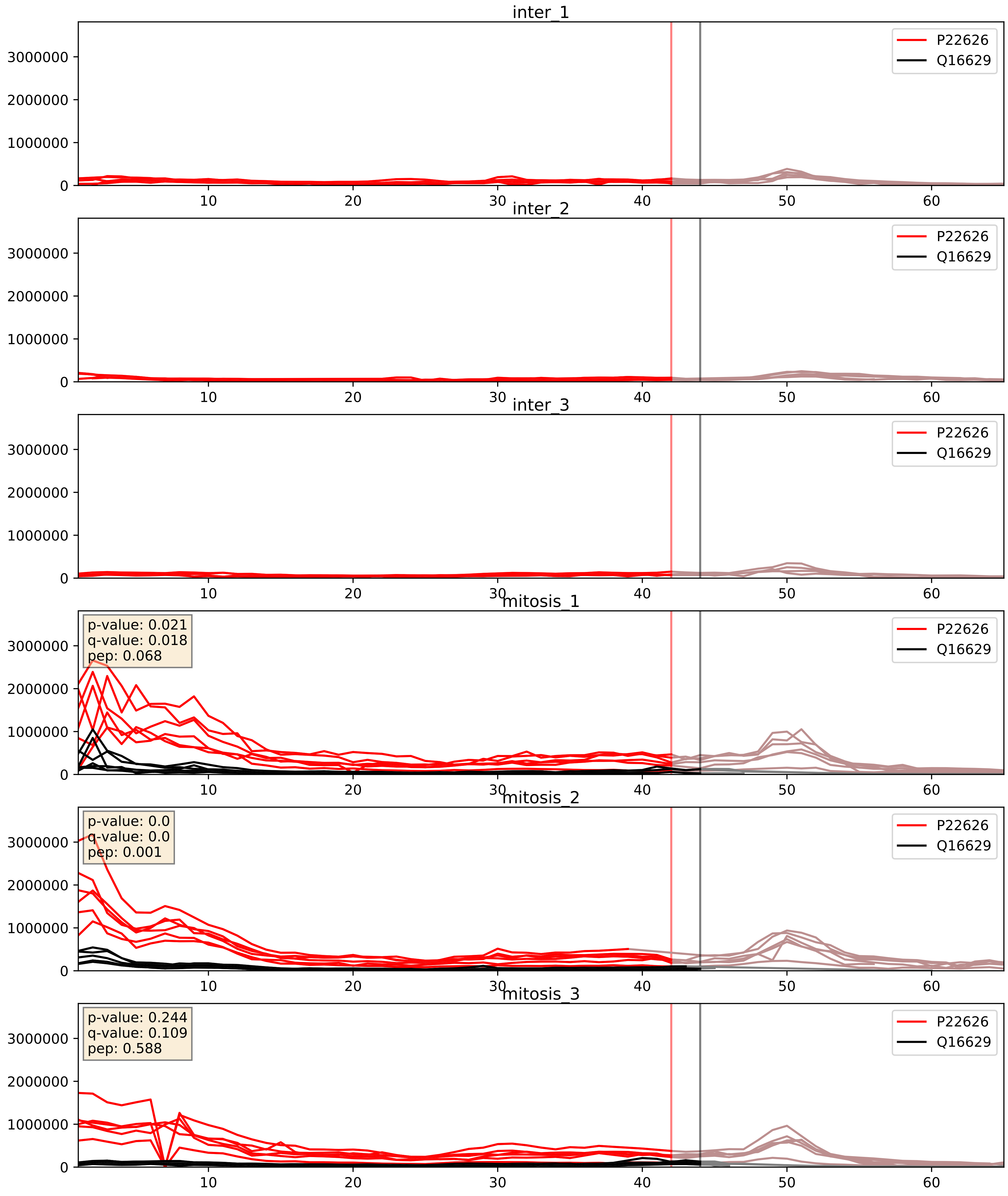


| level | condition_1 | condition_2 | pvalue | pvalue_adjusted |
|---|---|---|---|---|
| complex_abundance | mitosis | inter | 1.8097121496965717e-05 | 0.005691857151652958 |
| interactor_abundance | mitosis | inter | 2.0060878878898597e-05 | 0.004142545890653334 |
| interactor_ratio | mitosis | inter | 0.00021973308280332797 | 0.015168670877391028 |

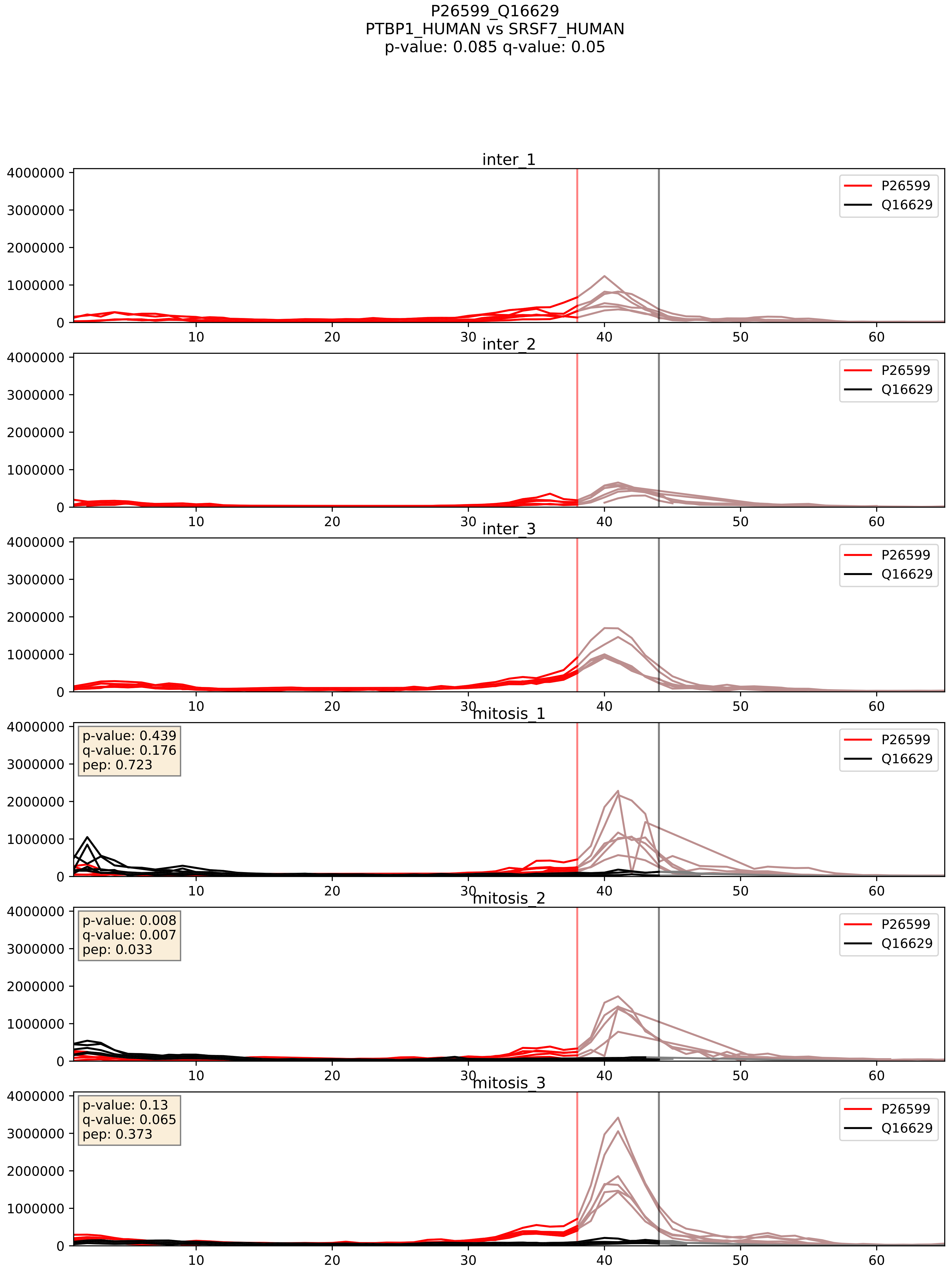


| level | condition_1 | condition_2 | pvalue | pvalue_adjusted |
|---|---|---|---|---|
| complex_abundance | mitosis | inter | 0.0006054510343287429 | 0.015834093001149168 |
| interactor_abundance | mitosis | inter | 0.00935589274653216 | 0.08390407743354142 |
| interactor_ratio | mitosis | inter | 0.4783463663699853 | 0.8660416446969278 |

P31943_Q16629
HNRH1_HUMAN vs SRSF7_HUMAN
p-value: 0.002 q-value: 0.003

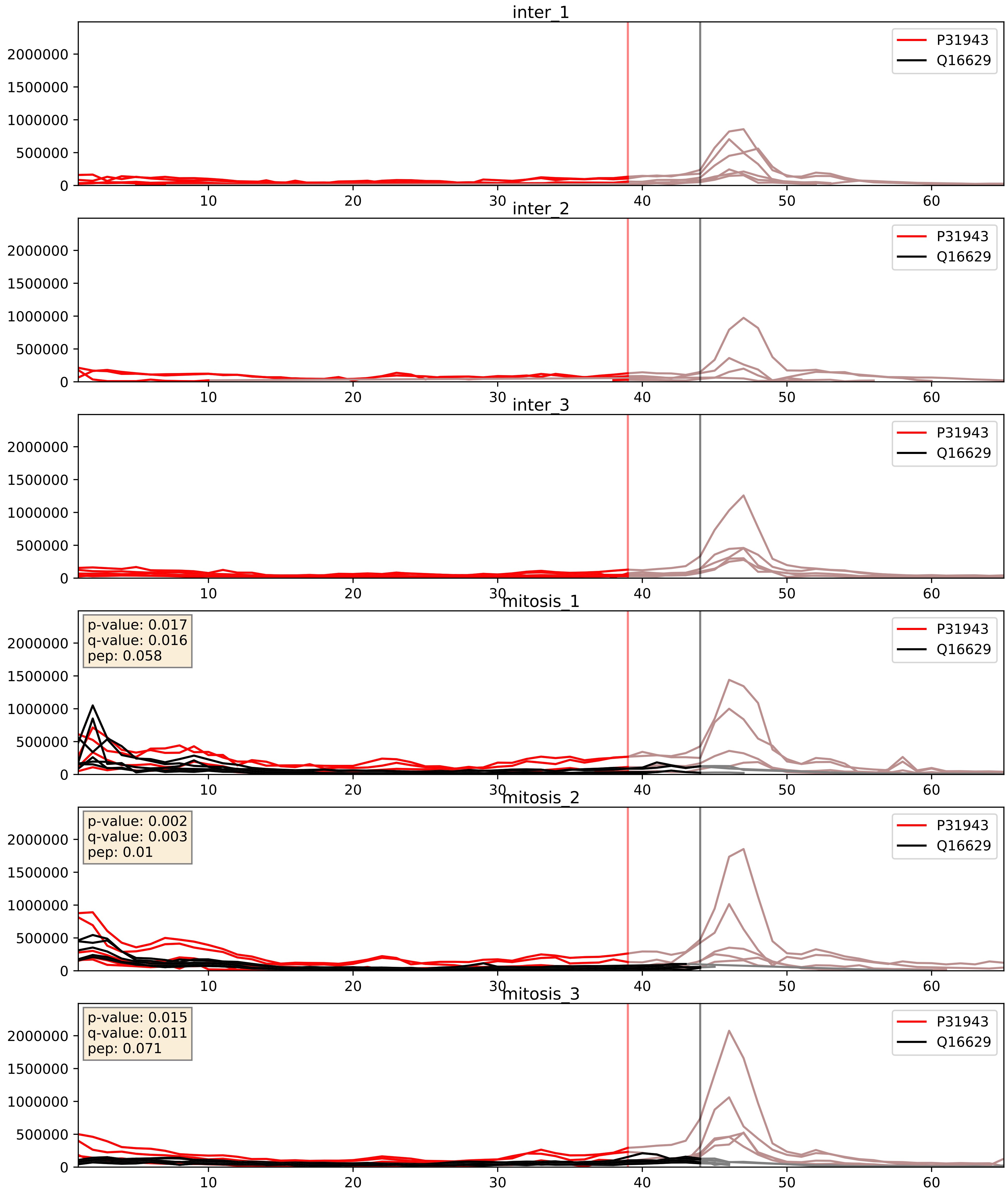


| level | condition_1 | condition_2 | pvalue | pvalue_adjusted |
|---|---|---|---|---|
| interactor_abundance | mitosis | inter | 2.510205397937702e-05 | 0.004476532959655569 |
| complex_abundance | mitosis | inter | 4.065873462401448e-05 | 0.005691857151652958 |
| interactor_ratio | mitosis | inter | 0.00017130672096930523 | 0.013577643810159747 |

P51991_Q16629
ROA3_HUMAN vs SRSF7_HUMAN
p-value: 0.0 q-value: 0.007

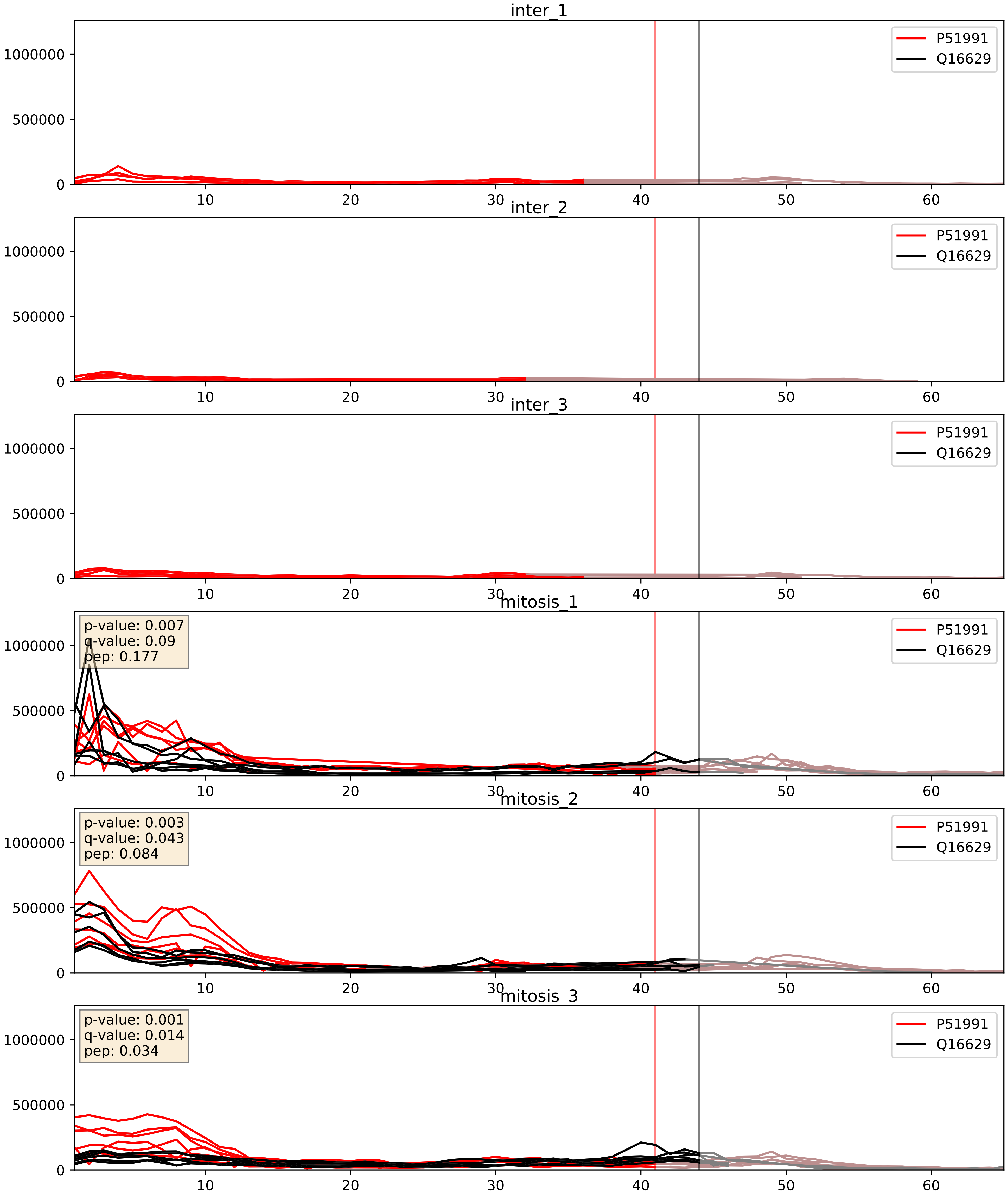


| level | condition_1 | condition_2 | pvalue | pvalue_adjusted |
|---|---|---|---|---|
| interactor_abundance | mitosis | inter | 1.150670630946923e-05 | 0.0033803141163598503 |
| complex_abundance | mitosis | inter | 2.3929388745179548e-05 | 0.005691857151652958 |
| interactor_ratio | mitosis | inter | 0.007352680833101884 | 0.11920255290028813 |

P52272_Q16629
HNRPM_HUMAN vs SRSF7_HUMAN
p-value: 0.025 q-value: 0.019

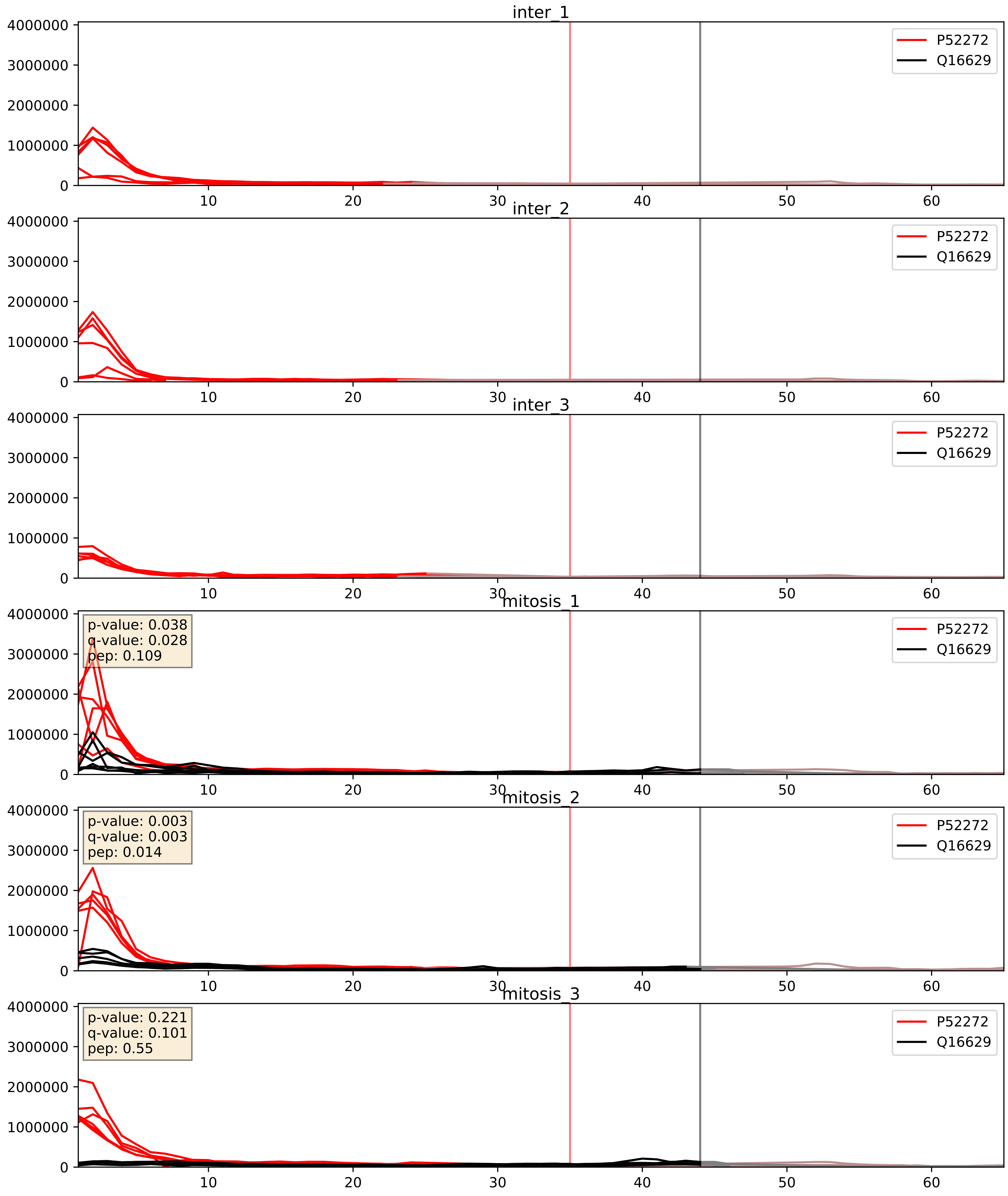


| level | condition_1 | condition_2 | pvalue | pvalue_adjusted |
|---|---|---|---|---|
| interactor_abundance | mitosis | inter | 2.2309750162464603e-05 | 0.0042438102531266 |
| complex_abundance | mitosis | inter | 3.735702009528526e-05 | 0.005691857151652958 |
| interactor_ratio | mitosis | inter | 0.0008463813255265196 | 0.033234055717922054 |

P52597_Q16629
HNRPF_HUMAN vs SRSF7_HUMAN
p-value: 0.0 q-value: 0.003

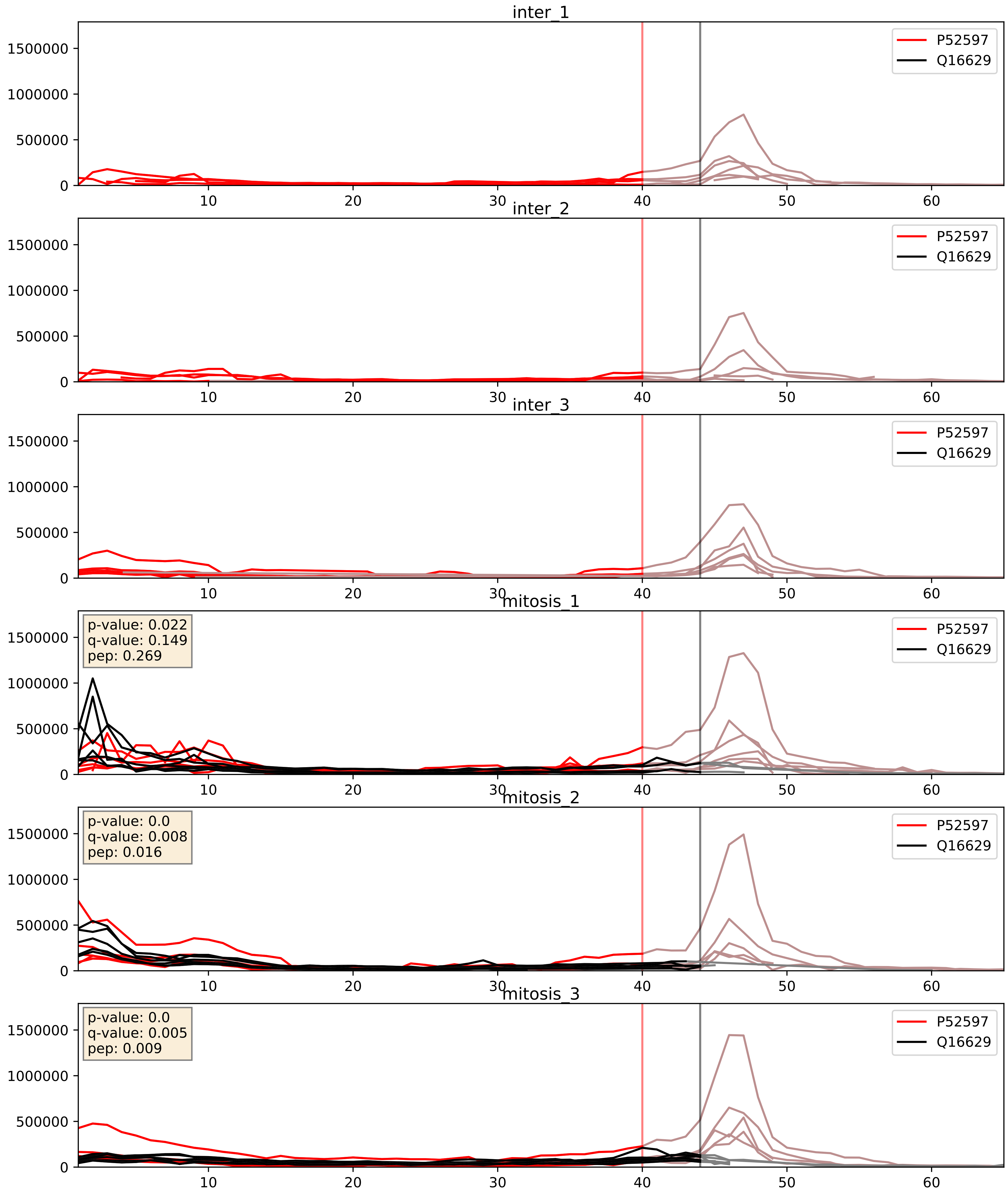


| level | condition_1 | condition_2 | pvalue | pvalue_adjusted |
|---|---|---|---|---|
| interactor_abundance | mitosis | inter | 4.447475947098241e-05 | 0.006058954526139308 |
| complex_abundance | mitosis | inter | 5.253696885978365e-05 | 0.005691857151652958 |
| interactor_ratio | mitosis | inter | 0.4299776830859134 | 0.8432094269126428 |

P55769_Q16629
NH2L1_HUMAN vs SRSF7_HUMAN
p-value: 0.001 q-value: 0.021

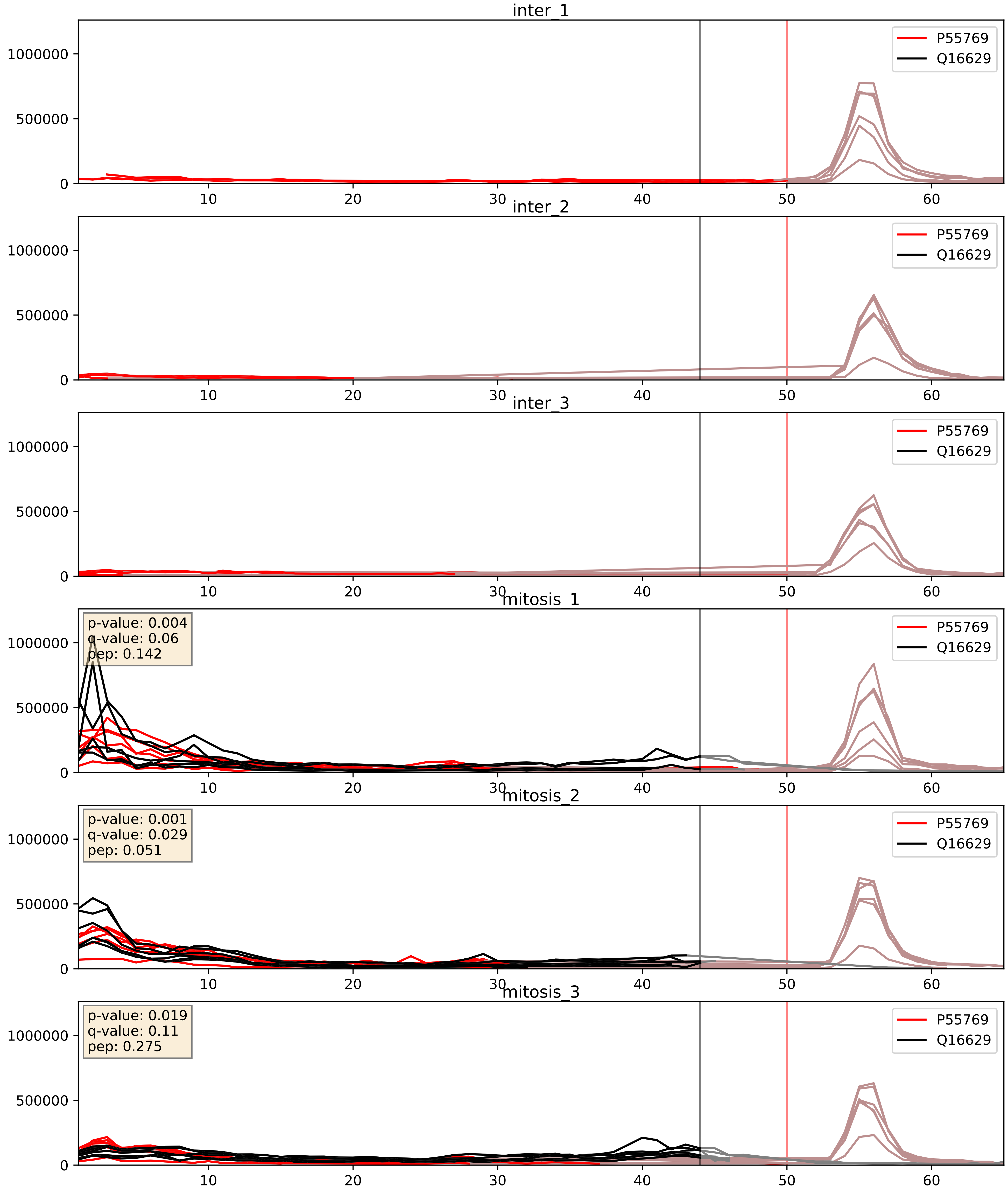


| level | condition_1 | condition_2 | pvalue | pvalue_adjusted |
|---|---|---|---|---|
| complex_abundance | mitosis | inter | 5.694619762543979e-05 | 0.005691857151652958 |
| interactor_abundance | mitosis | inter | 9.321731834197482e-05 | 0.007548664551888891 |
| interactor_ratio | mitosis | inter | 0.06548194300852817 | 0.42820888628953485 |

P61978_Q16629
HNRPK_HUMAN vs SRSF7_HUMAN
p-value: 0.049 q-value: 0.032

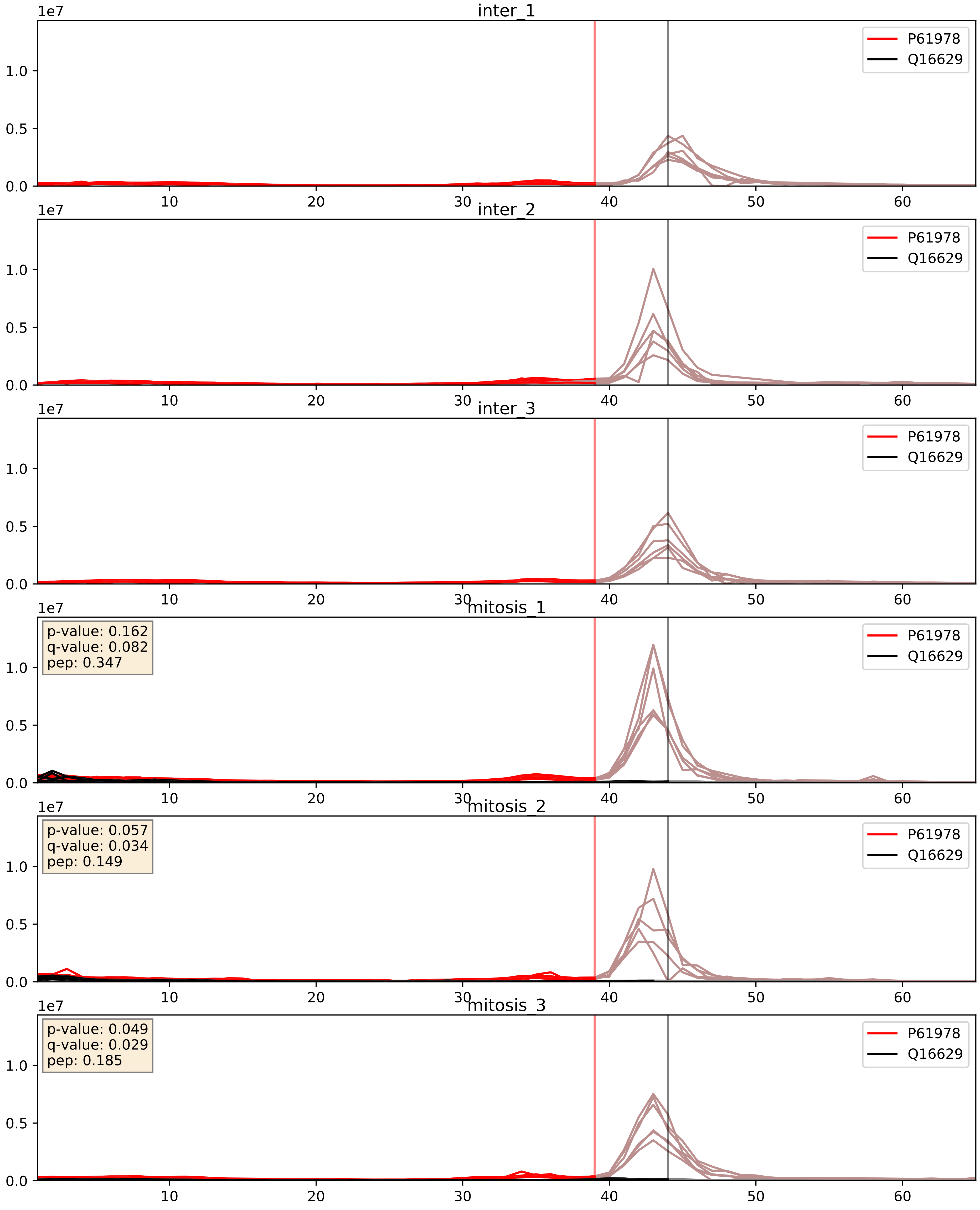


| level | condition_1 | condition_2 | pvalue | pvalue_adjusted |
|---|---|---|---|---|
| complex_abundance | mitosis | inter | 0.0007651916362873231 | 0.018128921229455204 |
| interactor_abundance | mitosis | inter | 0.004869975273961587 | 0.057420094139271606 |
| interactor_ratio | mitosis | inter | 0.4108932136976173 | 0.8340755186867097 |

P62995_Q16629
TRA2B_HUMAN vs SRSF7_HUMAN
p-value: 0.0 q-value: 0.001

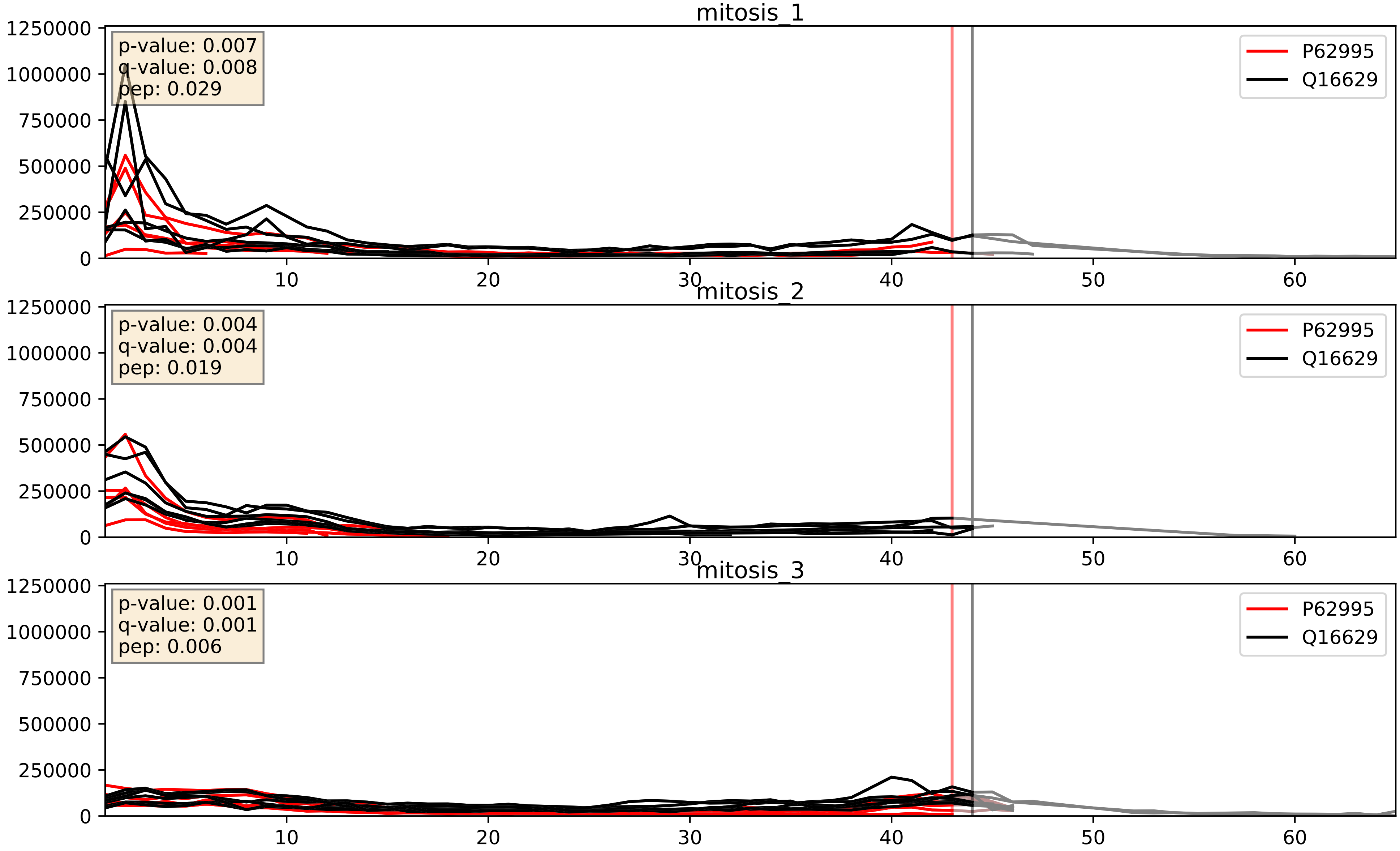


| level | condition_1 | condition_2 | pvalue | pvalue_adjusted |
|---|---|---|---|---|
| complex_abundance | mitosis | inter | 0.00017630276153620522 | 0.008807651911797798 |
| interactor_abundance | mitosis | inter | 0.0004584932841066445 | 0.014951247664582388 |
| interactor_ratio | mitosis | inter | 0.002039014647753622 | 0.05540941391990795 |

P84103_Q16629
SRSF3_HUMAN vs SRSF7_HUMAN
p-value: 0.001 q-value: 0.001

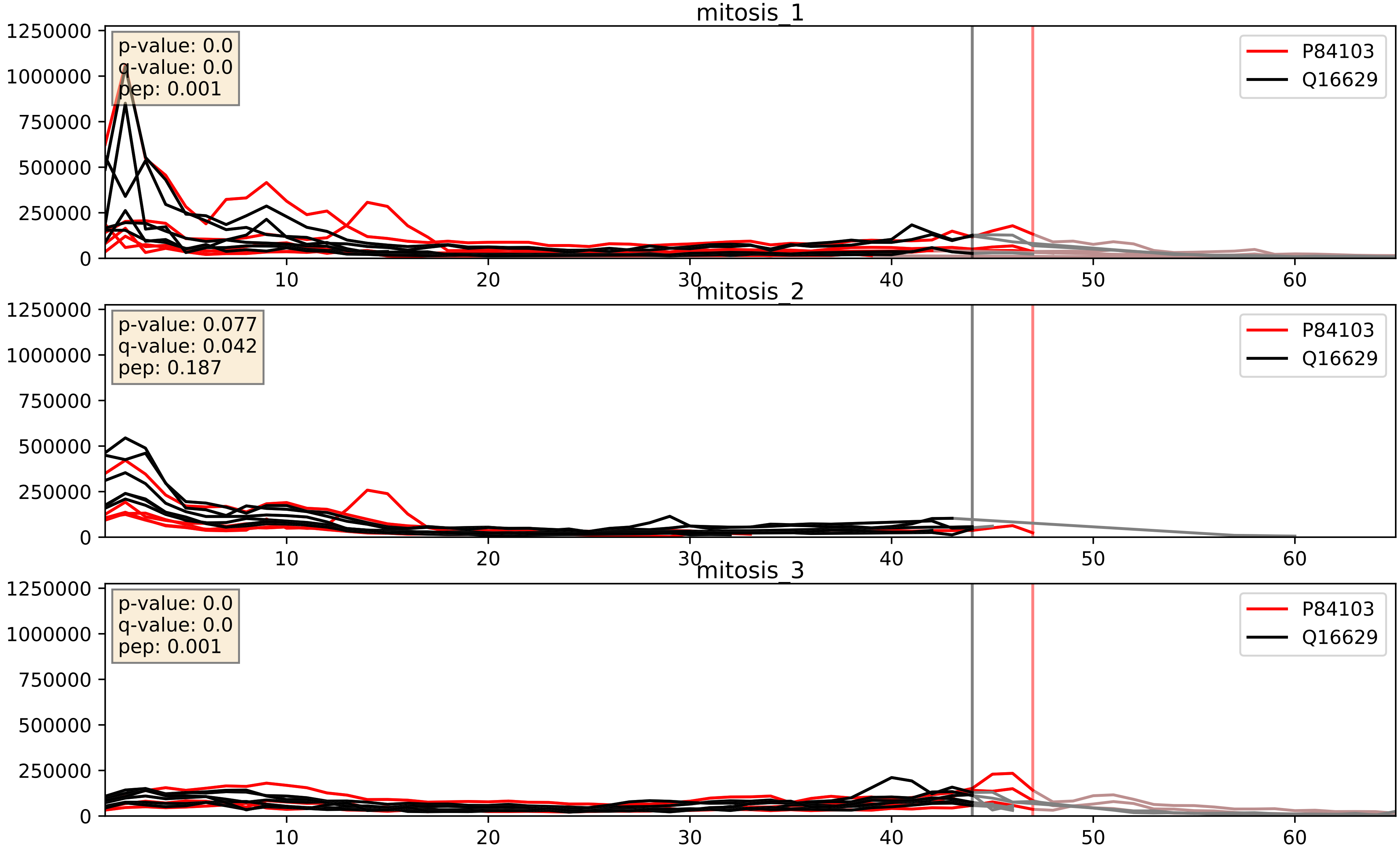


| level | condition_1 | condition_2 | pvalue | pvalue_adjusted |
|---|---|---|---|---|
| complex_abundance | mitosis | inter | 0.0003855844193721723 | 0.012502282688734073 |
| interactor_abundance | mitosis | inter | 0.002590890474122459 | 0.03806046117740333 |
| interactor_ratio | mitosis | inter | 0.026618515768801317 | 0.25614196664208316 |

Q05519_Q16629
SRS11_HUMAN vs SRSF7_HUMAN
p-value: 0.067 q-value: 0.041

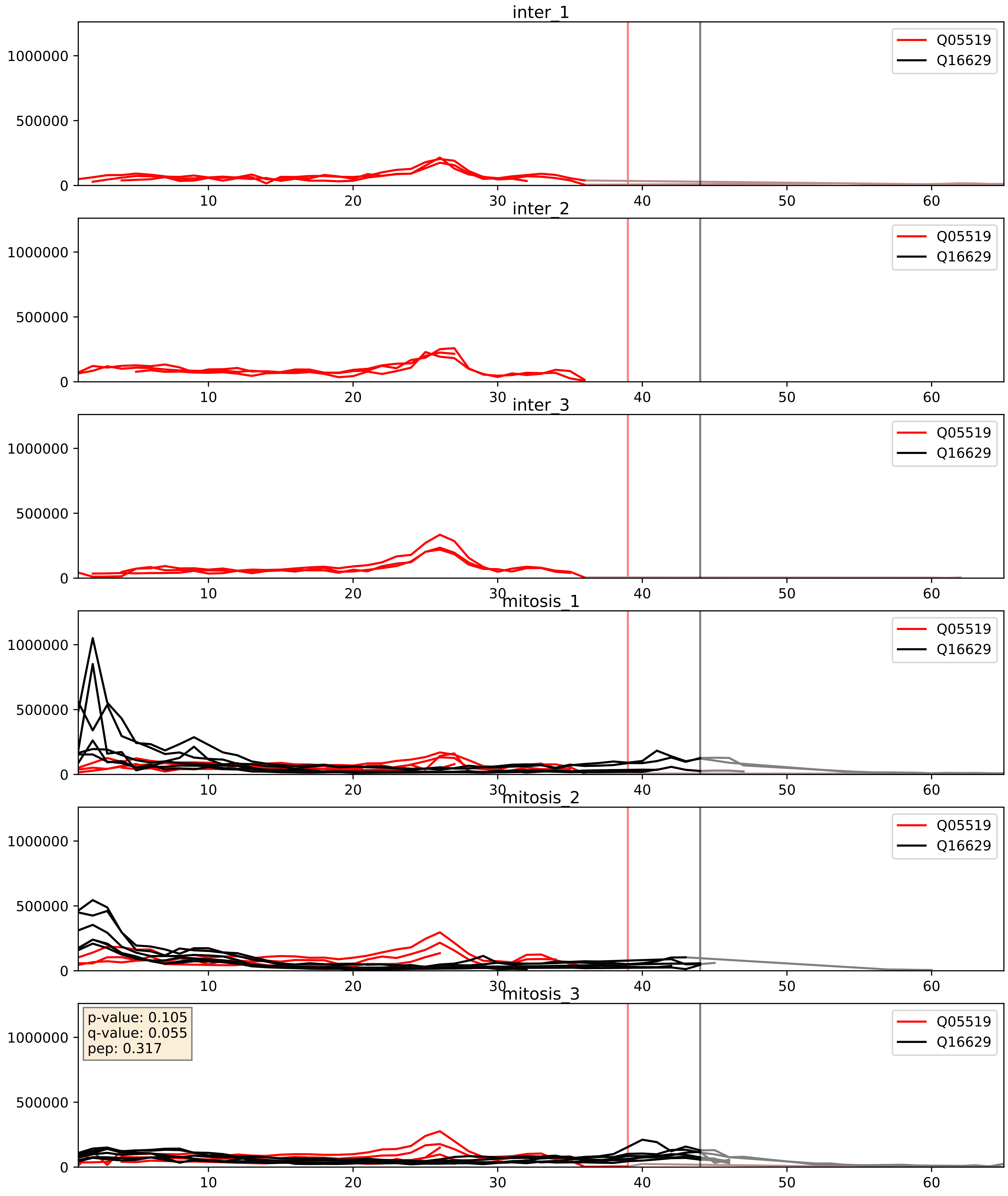


| level | condition_1 | condition_2 | pvalue | pvalue_adjusted |
|---|---|---|---|---|
| complex_abundance | mitosis | inter | 2.4413174450807402e-05 | 0.005691857151652958 |
| interactor_abundance | mitosis | inter | 6.634639281590001e-05 | 0.006642399093615253 |
| interactor_ratio | mitosis | inter | 0.05621811676799316 | 0.3926647223857623 |

Q07955_Q16629
SRSF1_HUMAN vs SRSF7_HUMAN
p-value: 0.0 q-value: 0.0

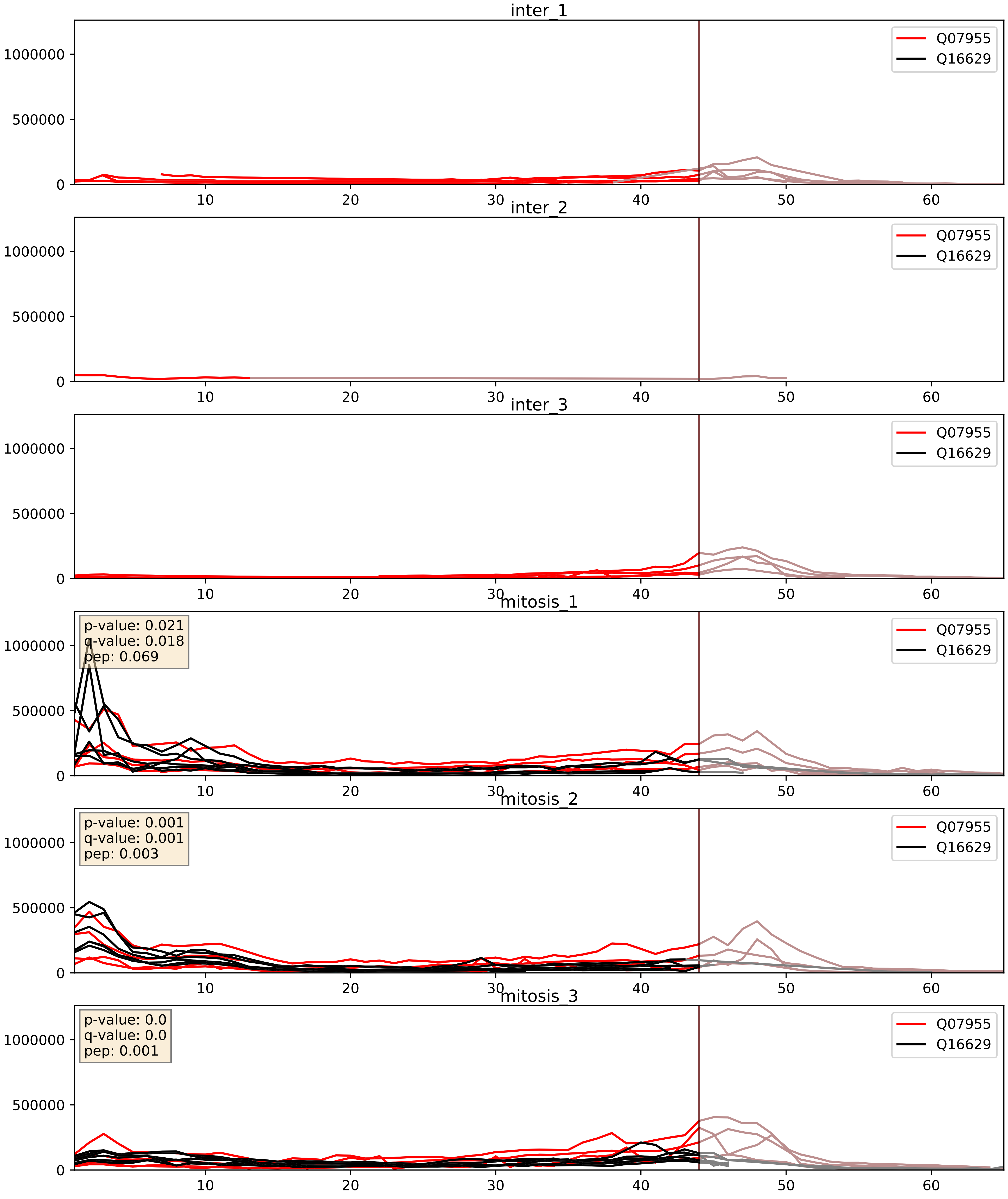


| level | condition_1 | condition_2 | pvalue | pvalue_adjusted |
|---|---|---|---|---|
| complex_abundance | mitosis | inter | 3.059501908721843e-05 | 0.005691857151652958 |
| interactor_abundance | mitosis | inter | 0.0006217246581139298 | 0.01750645747847118 |
| interactor_ratio | mitosis | inter | 0.4633387300269737 | 0.8588611490067614 |

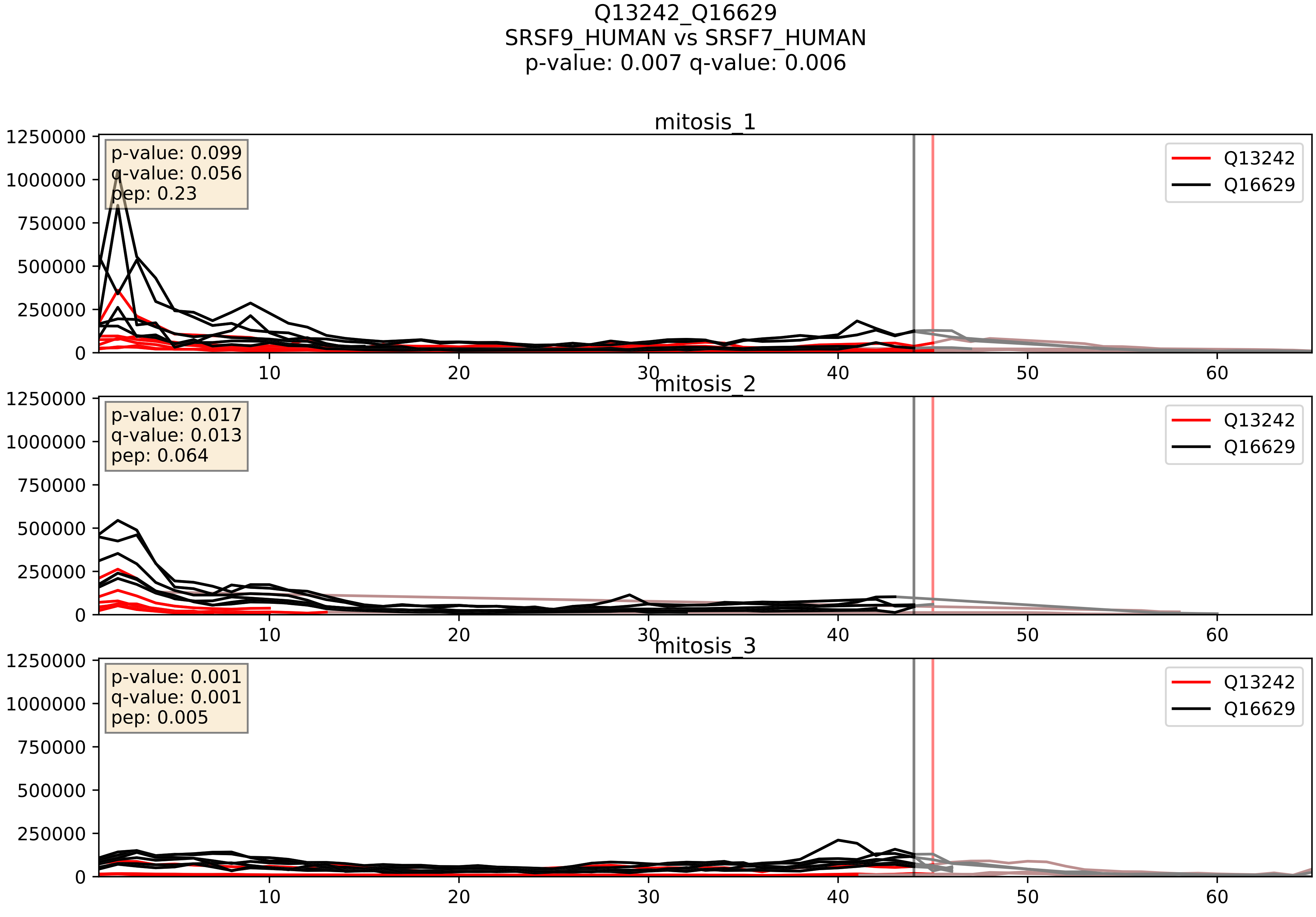


| level | condition_1 | condition_2 | pvalue | pvalue_adjusted |
|---|---|---|---|---|
| interactor_ratio | mitosis | inter | 0.0006742820733493766 | 0.029848225645041686 |
| complex_abundance | mitosis | inter | 0.0025654119015484406 | 0.03664546283589145 |
| interactor_abundance | mitosis | inter | 0.04867027237079332 | 0.175973614147409 |

Q13247_Q16629
SRSF6_HUMAN vs SRSF7_HUMAN
p-value: 0.001 q-value: 0.002

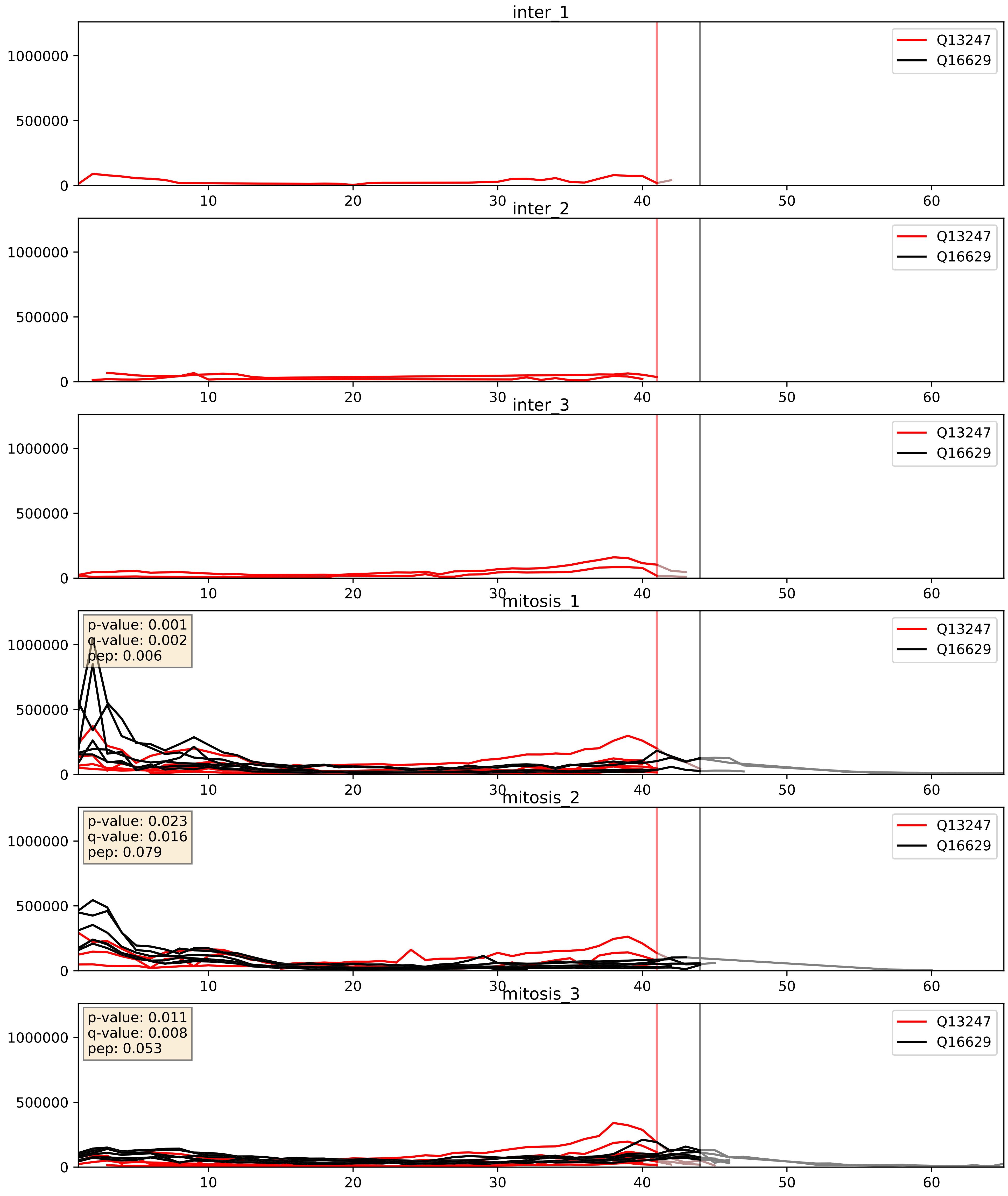


| level | condition_1 | condition_2 | pvalue | pvalue_adjusted |
|---|---|---|---|---|
| complex_abundance | mitosis | inter | 0.00033327015539205967 | 0.011886635542316795 |
| interactor_abundance | mitosis | inter | 0.00279726046188116 | 0.040175418714266324 |
| interactor_ratio | mitosis | inter | 0.11156417163406727 | 0.5532962393902757 |

Q15717_Q16629
ELAV1_HUMAN vs SRSF7_HUMAN
p-value: 0.005 q-value: 0.005

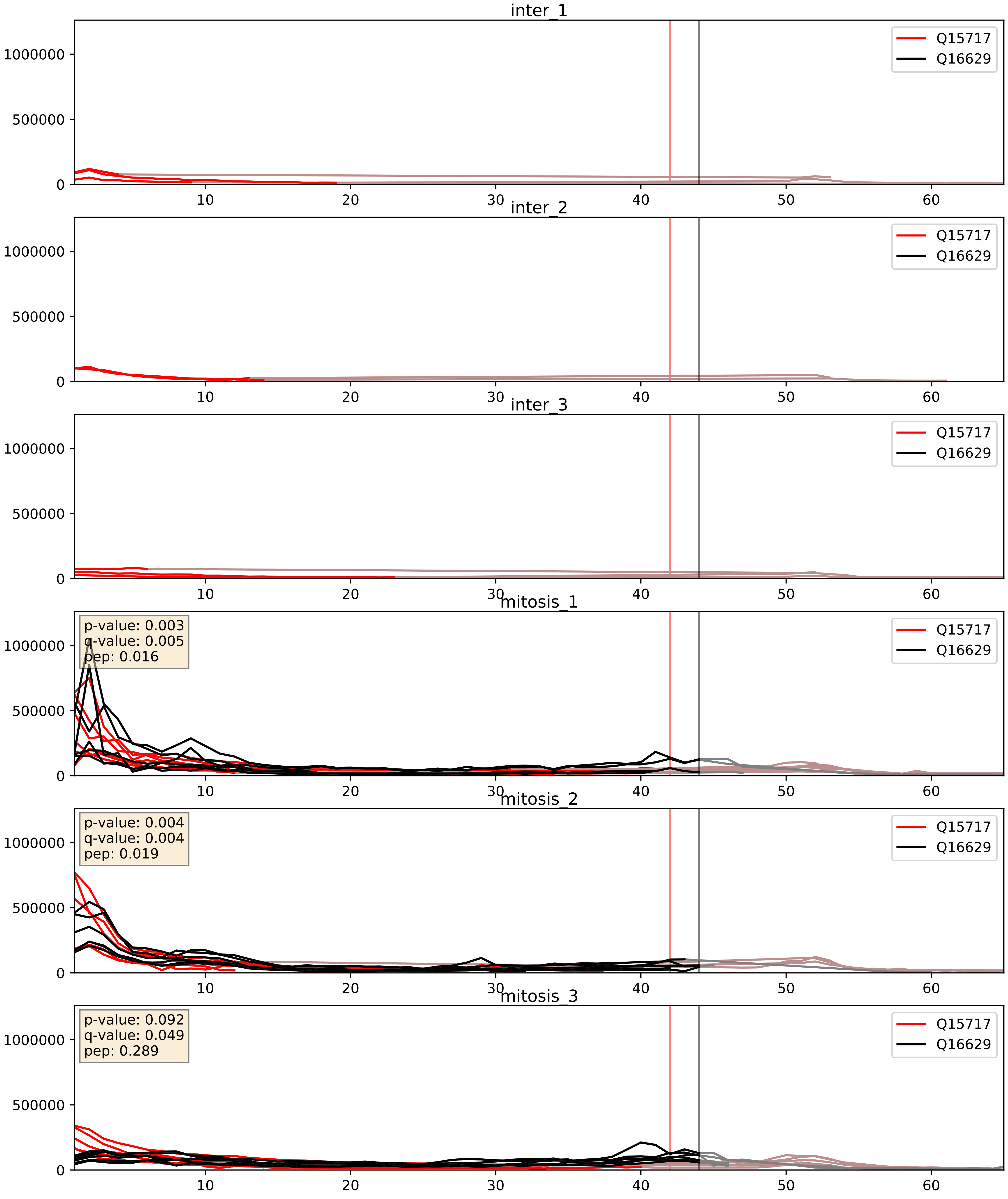


| level | condition_1 | condition_2 | pvalue | pvalue_adjusted |
|---|---|---|---|---|
| interactor_abundance | mitosis | inter | 6.059680363197429e-05 | 0.006459477818931243 |
| complex_abundance | mitosis | inter | 6.314734732233846e-05 | 0.005691857151652958 |
| interactor_ratio | mitosis | inter | 0.028137833796062196 | 0.2626606949774181 |

Q16629_Q99459
SRSF7_HUMAN vs CDC5L_HUMAN
p-value: 0.073 q-value: 0.044

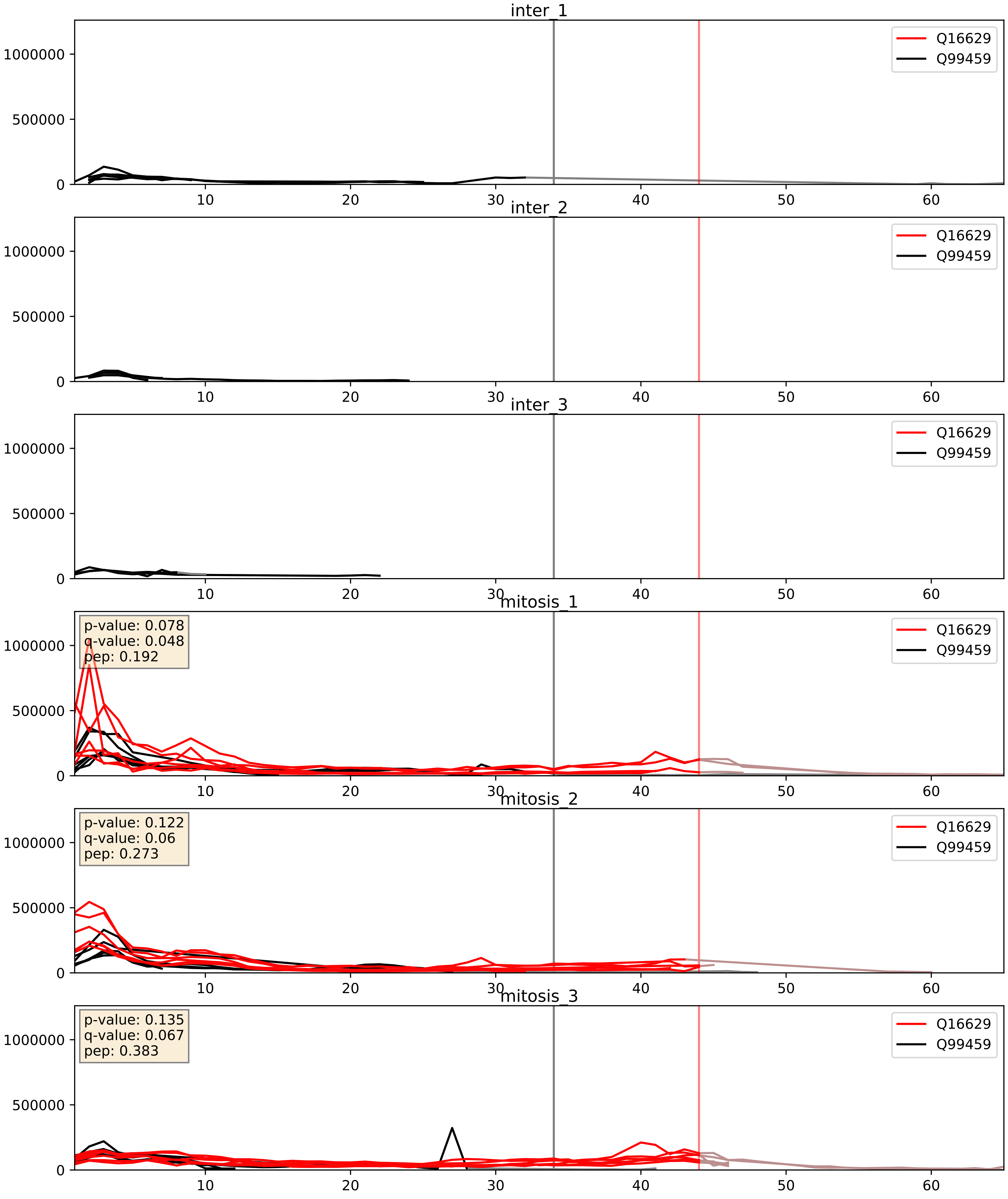


| level | condition_1 | condition_2 | pvalue | pvalue_adjusted |
|---|---|---|---|---|
| interactor_abundance | mitosis | inter | 0.00026344179809256095 | 0.012309909377725258 |
| complex_abundance | mitosis | inter | 0.0003628810828694355 | 0.012039775462644836 |
| interactor_ratio | mitosis | inter | 0.008617104652240045 | 0.13203195690147984 |
