## Supplementary Data 2 for "SECAT: Quantifying differential protein-protein interaction states by network-centric analysis": hela_string_000035_Q8NI36.pdf

# Q8NI36_Q8NI36

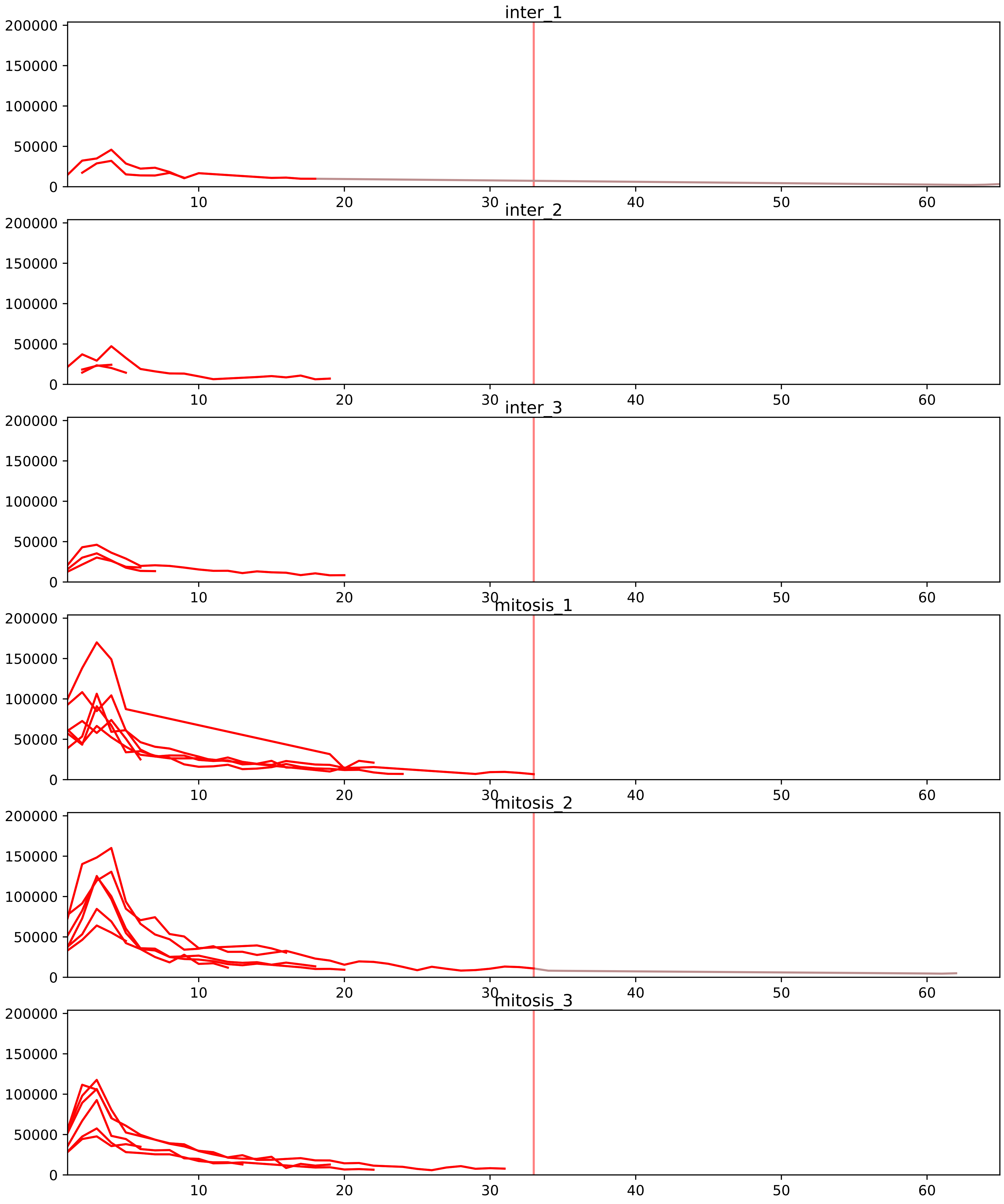


| level | condition_1 | condition_2 | pvalue | pvalue_adjusted |
|---|---|---|---|---|
| interactor_ratio | mitosis | inter | 1.1187930805991261e-08 | 6.05464490677174e-07 |
| complex_abundance | mitosis | inter | 2.7907264716360156e-05 | 0.0010068503348647586 |
| interactor_abundance | mitosis | inter | 3.109361200325882e-05 | 0.0011218087467842399 |
| total_abundance | mitosis | inter | 0.40254495777027205 | 0.7023154146704781 |
| assembled_abundance | mitosis | inter | 0.48614739509528304 | 0.7405323646082069 |
| monomer_abundance | mitosis | inter | 0.5278317490551343 | 0.7708772093497522 |

O00567_Q8NI36
NOP56_HUMAN vs WDR36_HUMAN
p-value: 0.005 q-value: 0.005

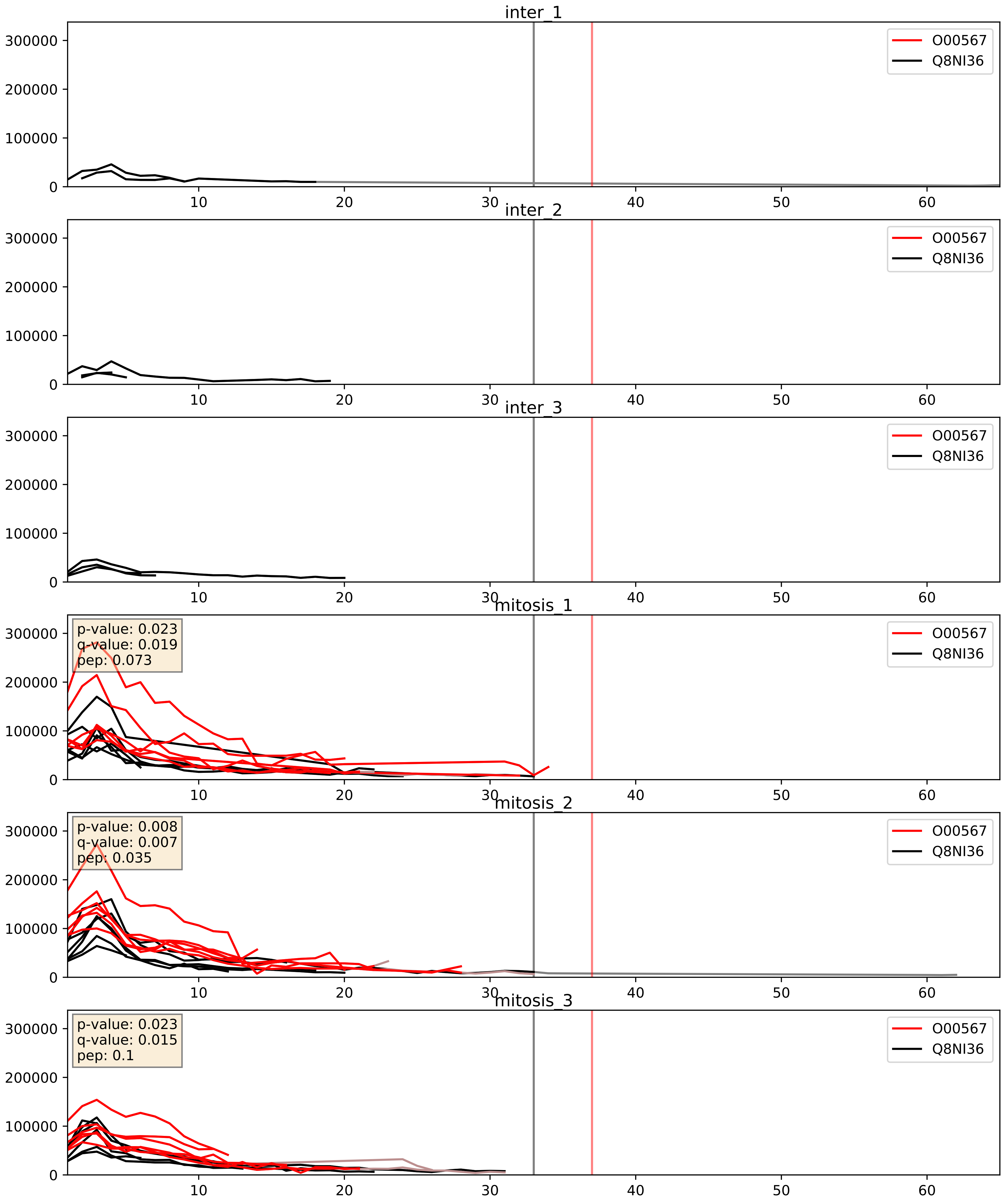


| level | condition_1 | condition_2 | pvalue | pvalue_adjusted |
|---|---|---|---|---|
| complex_abundance | mitosis | inter | 0.0007499876579213406 | 0.017895661700548352 |
| interactor_abundance | mitosis | inter | 0.0011052779688176132 | 0.024049720606162197 |
| interactor_ratio | mitosis | inter | 0.010958138448517993 | 0.15054322834266998 |

O60832_Q8NI36
DKC1_HUMAN vs WDR36_HUMAN
p-value: 0.004 q-value: 0.047

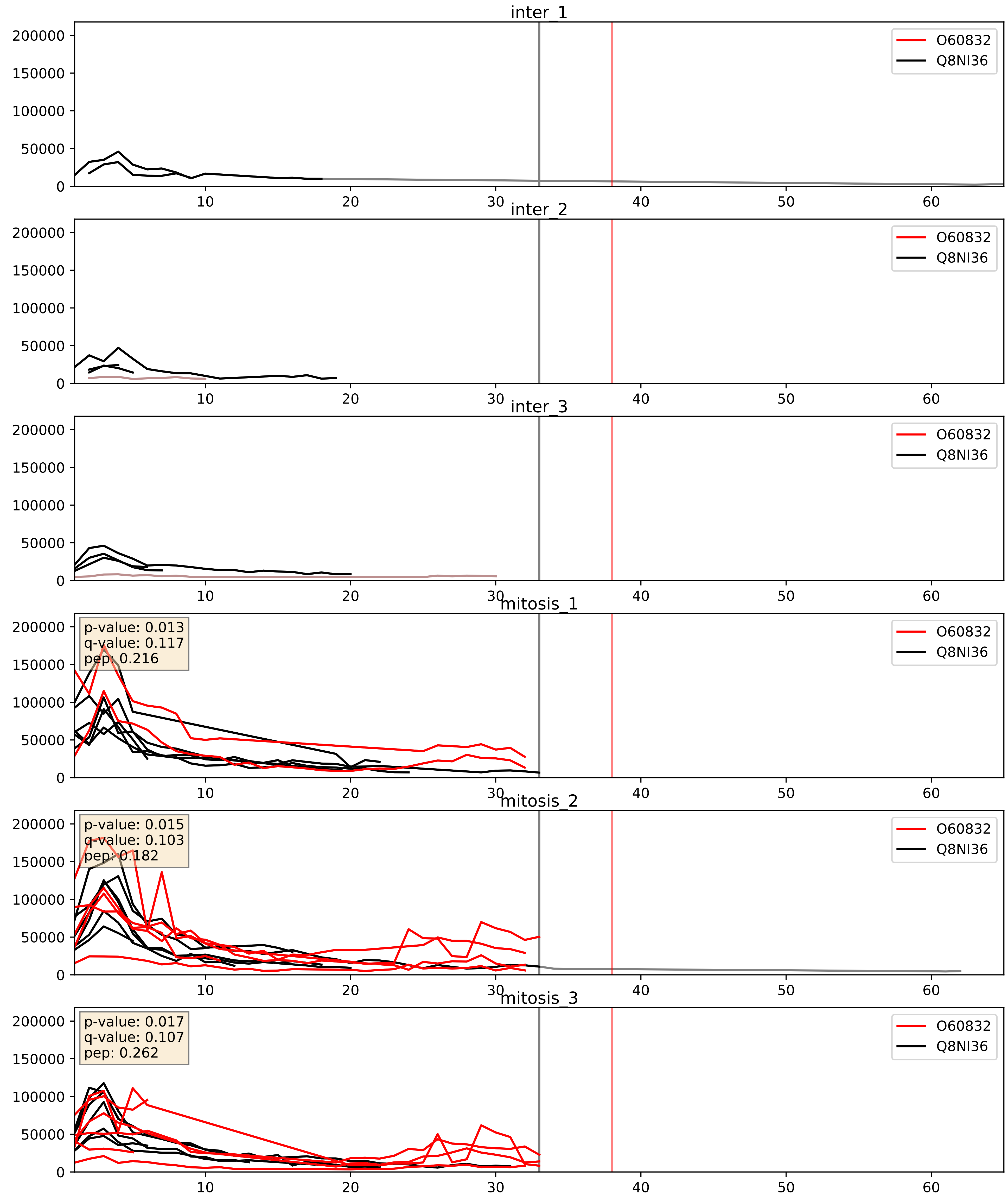


| level | condition_1 | condition_2 | pvalue | pvalue_adjusted |
|---|---|---|---|---|
| complex_abundance | mitosis | inter | 0.005657961196732316 | 0.05664578695208026 |
| interactor_abundance | mitosis | inter | 0.05267352363657362 | 0.18428385541163456 |
| interactor_ratio | mitosis | inter | 0.8878767344863951 | 0.9804177989881011 |

O76021_Q8NI36
RL1D1_HUMAN vs WDR36_HUMAN
p-value: 0.002 q-value: 0.003

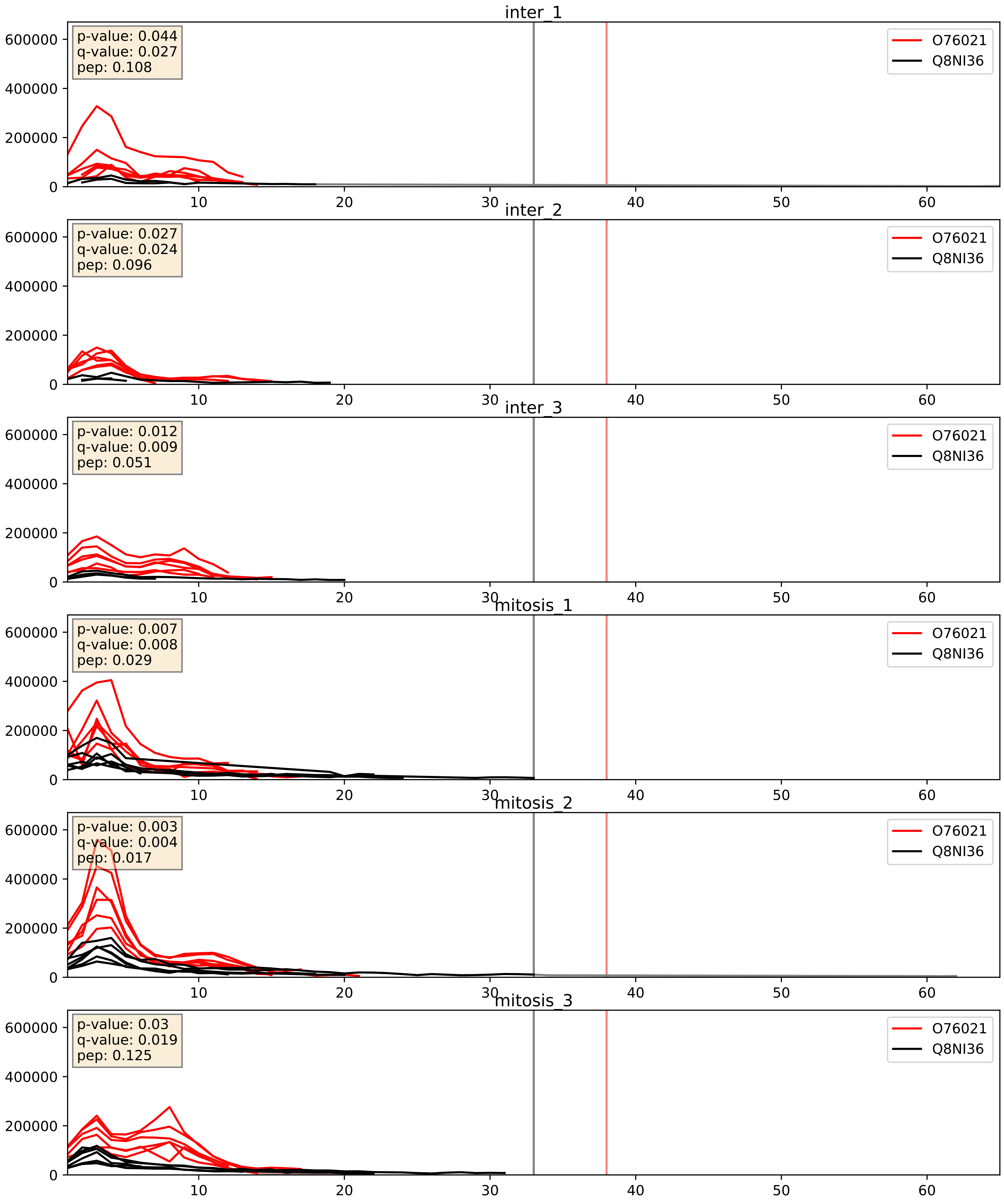


| level | condition_1 | condition_2 | pvalue | pvalue_adjusted |
|---|---|---|---|---|
| complex_abundance | mitosis | inter | 0.00046464344010374375 | 0.013667862018171982 |
| interactor_abundance | mitosis | inter | 0.005364620772517218 | 0.06171739355782327 |
| interactor_ratio | mitosis | inter | 0.9455967382350232 | 0.9912117205440141 |

P22087_Q8NI36
FBRL_HUMAN vs WDR36_HUMAN
p-value: 0.005 q-value: 0.005

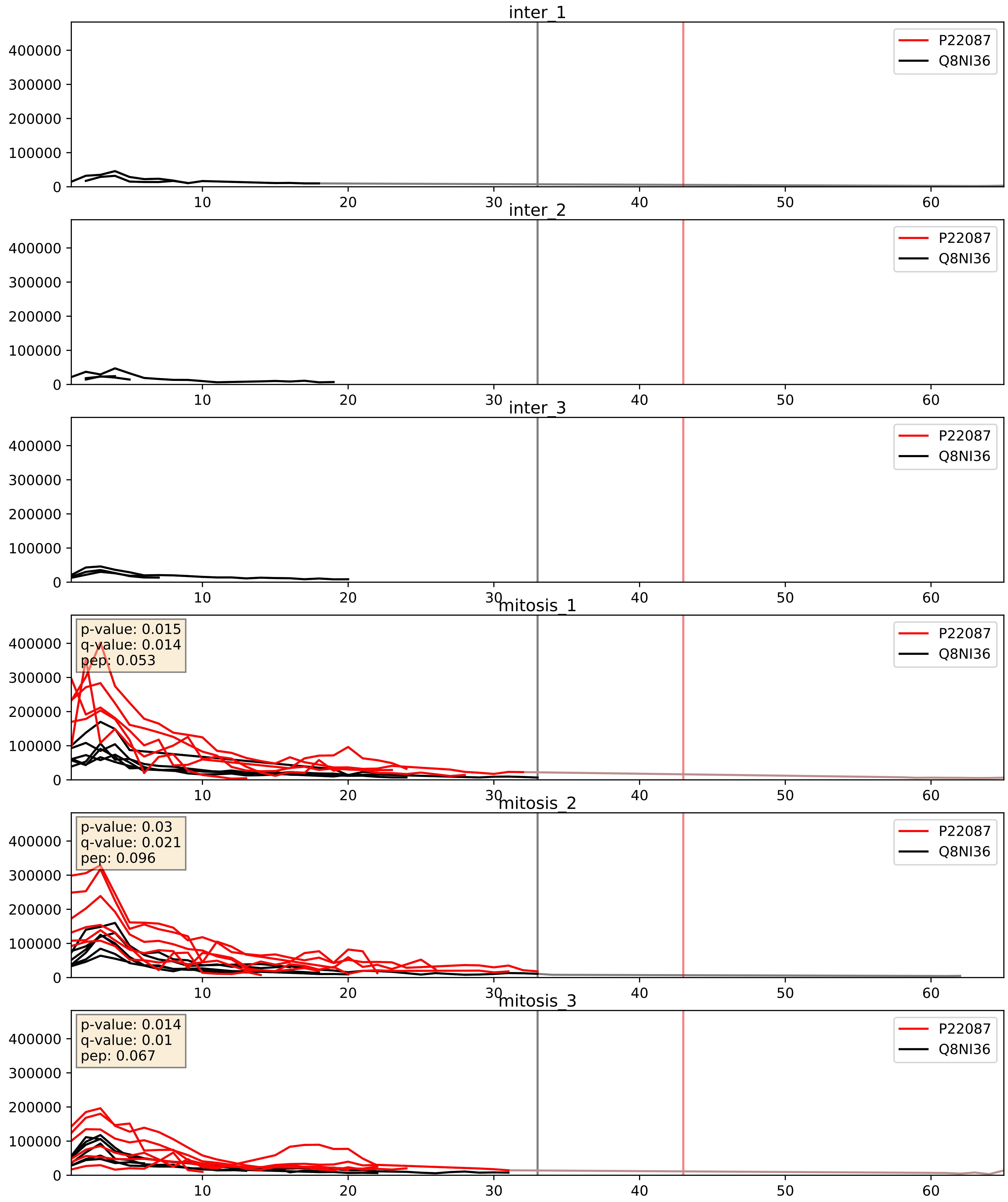


| level | condition_1 | condition_2 | pvalue | pvalue_adjusted |
|---|---|---|---|---|
| complex_abundance | mitosis | inter | 0.00034440768556571094 | 0.012015488415248196 |
| interactor_abundance | mitosis | inter | 0.0003496674327824524 | 0.01356427145975096 |
| interactor_ratio | mitosis | inter | 0.0010740795022735833 | 0.038414698994273265 |

P42285_Q8NI36
MTREX_HUMAN vs WDR36_HUMAN
p-value: 0.043 q-value: 0.029

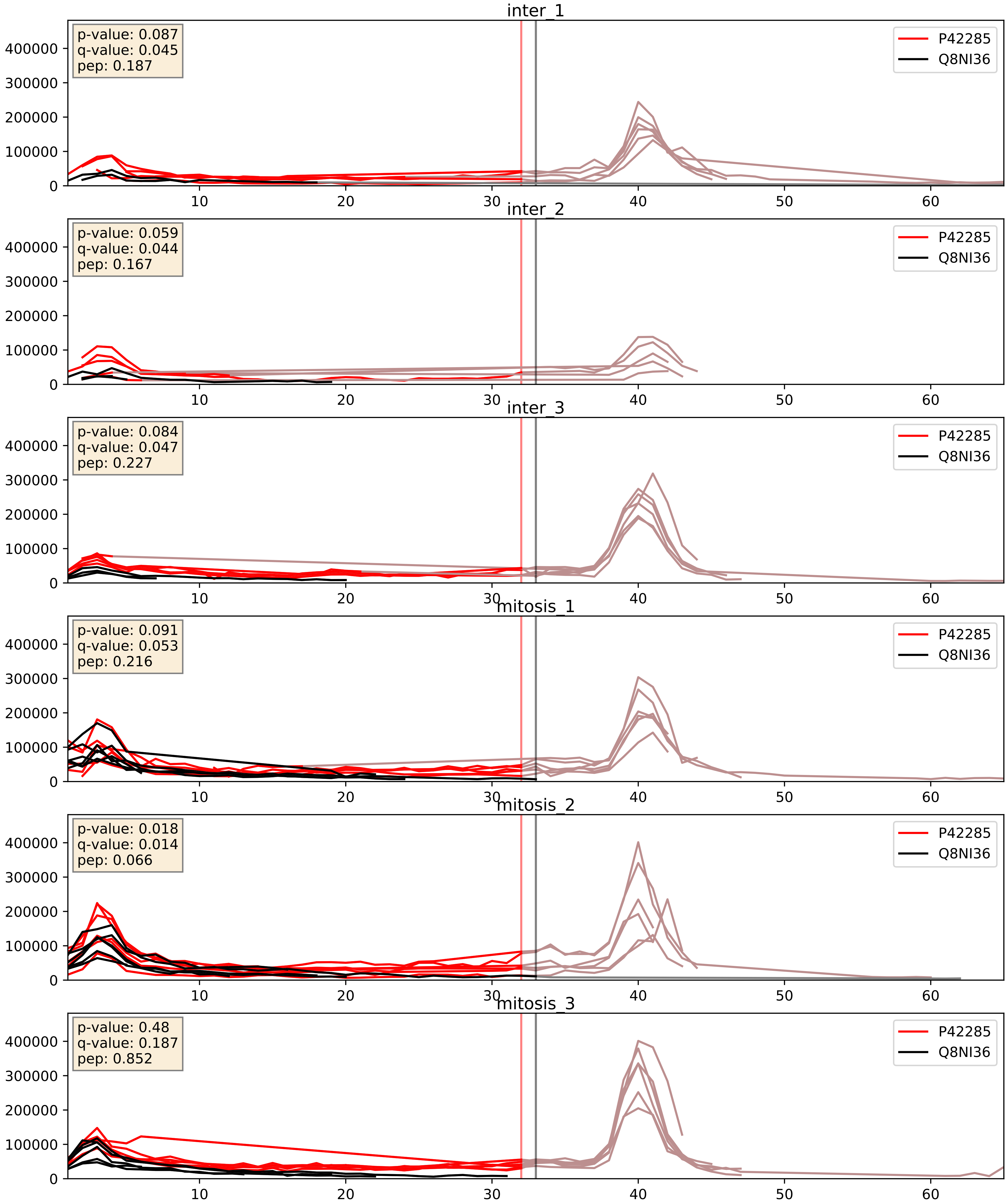


| level | condition_1 | condition_2 | pvalue | pvalue_adjusted |
|---|---|---|---|---|
| complex_abundance | mitosis | inter | 6.003680961344212e-05 | 0.005691857151652958 |
| interactor_abundance | mitosis | inter | 0.0011713522879652522 | 0.0247880731396355 |
| interactor_ratio | mitosis | inter | 0.5585467090945081 | 0.8918094730984788 |

P55769_Q8NI36
NH2L1_HUMAN vs WDR36_HUMAN
p-value: 0.017 q-value: 0.014

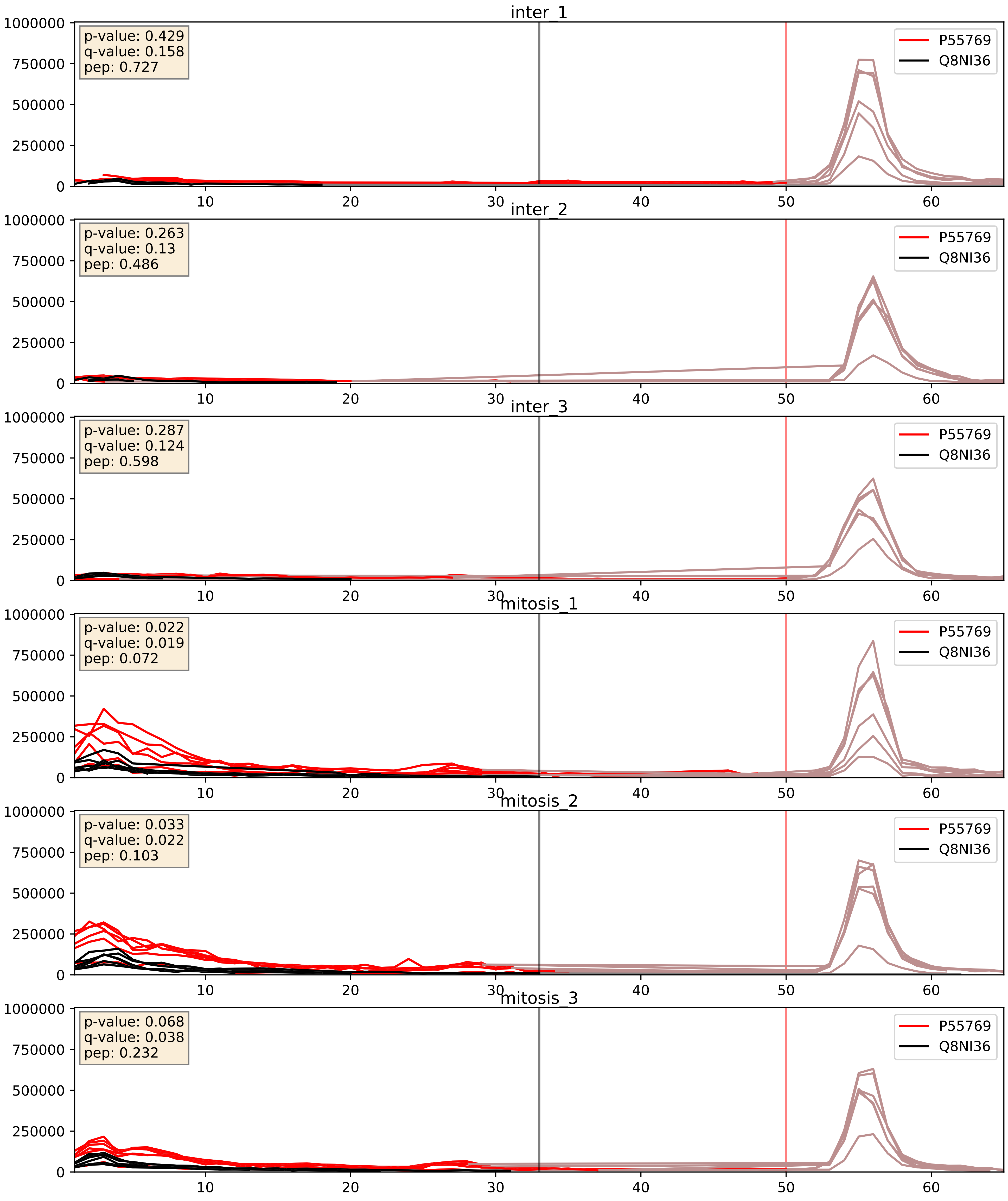


| level | condition_1 | condition_2 | pvalue | pvalue_adjusted |
|---|---|---|---|---|
| interactor_abundance | mitosis | inter | 0.0005601385574333299 | 0.017415659477552636 |
| complex_abundance | mitosis | inter | 0.0007027447438489493 | 0.01718712859242002 |
| interactor_ratio | mitosis | inter | 0.0015892366528110781 | 0.0480939461418057 |

P56182_Q8NI36
RRP1_HUMAN vs WDR36_HUMAN
p-value: 0.002 q-value: 0.033

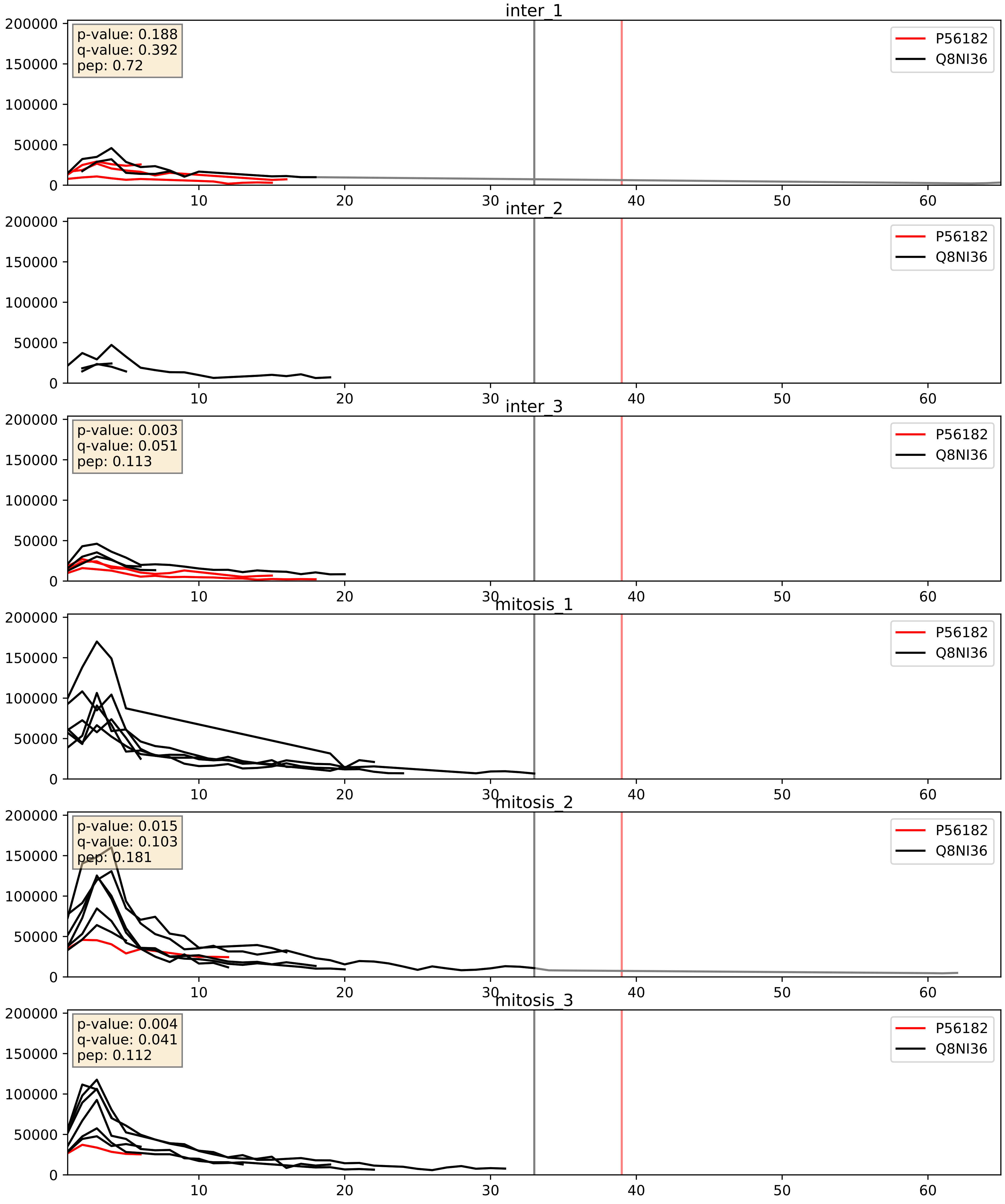


| level | condition_1 | condition_2 | pvalue | pvalue_adjusted |
|---|---|---|---|---|
| interactor_abundance | mitosis | inter | 0.029569223230101607 | 0.1428224596095078 |
| interactor_ratio | mitosis | inter | 0.09422286795448138 | 0.5094488449611217 |
| complex_abundance | mitosis | inter | 0.2277526150982002 | 0.38143173660824553 |

P62081_Q8NI36
RS7_HUMAN vs WDR36_HUMAN
p-value: 0.0 q-value: 0.013

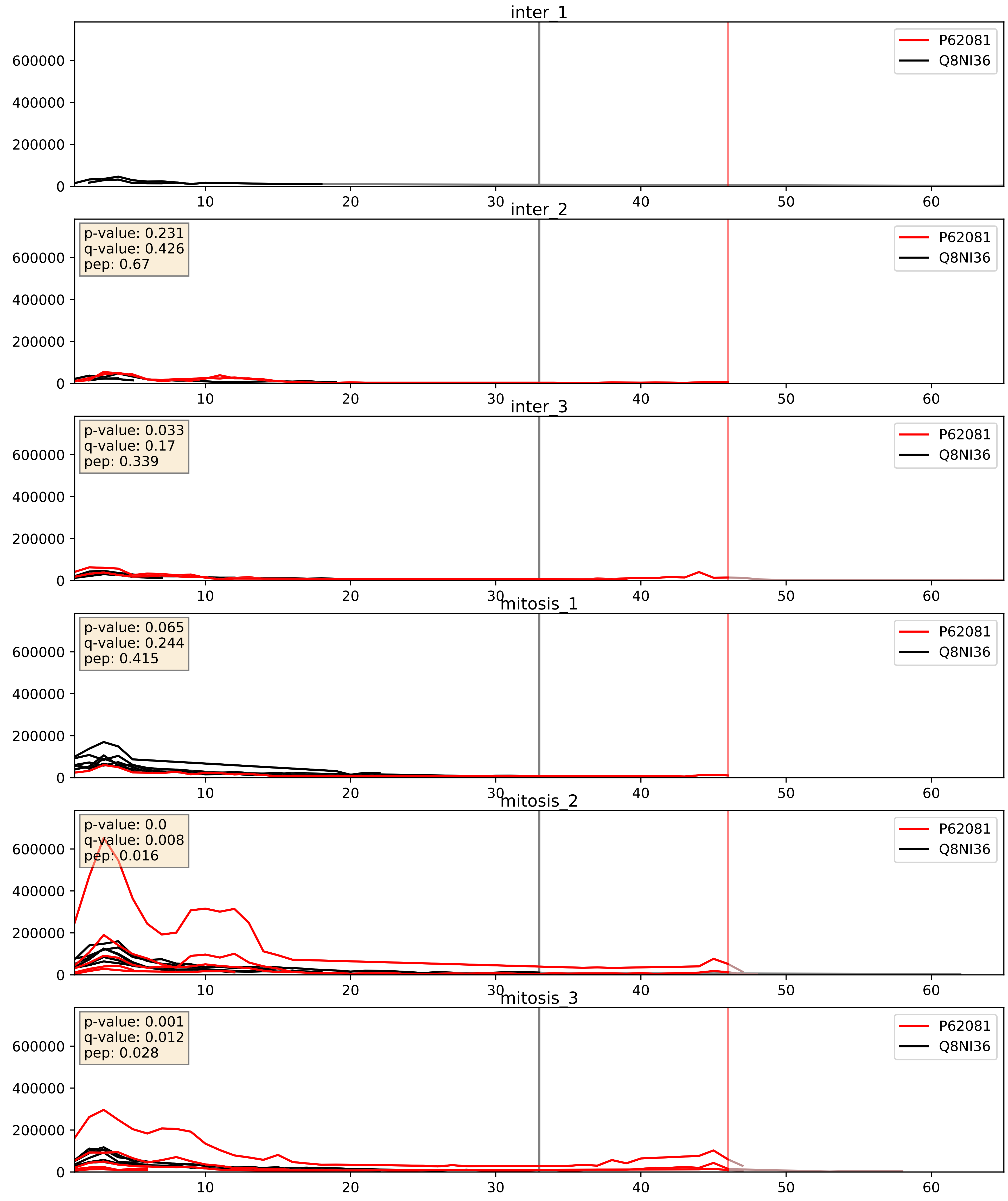


| level | condition_1 | condition_2 | pvalue | pvalue_adjusted |
|---|---|---|---|---|
| complex_abundance | mitosis | inter | 0.06040034320689314 | 0.17626794583586675 |
| interactor_abundance | mitosis | inter | 0.23996624546210205 | 0.4312235245182871 |
| interactor_ratio | mitosis | inter | 0.9280233883773915 | 0.987346851953133 |

P62280_Q8NI36
RS11_HUMAN vs WDR36_HUMAN
p-value: 0.004 q-value: 0.047

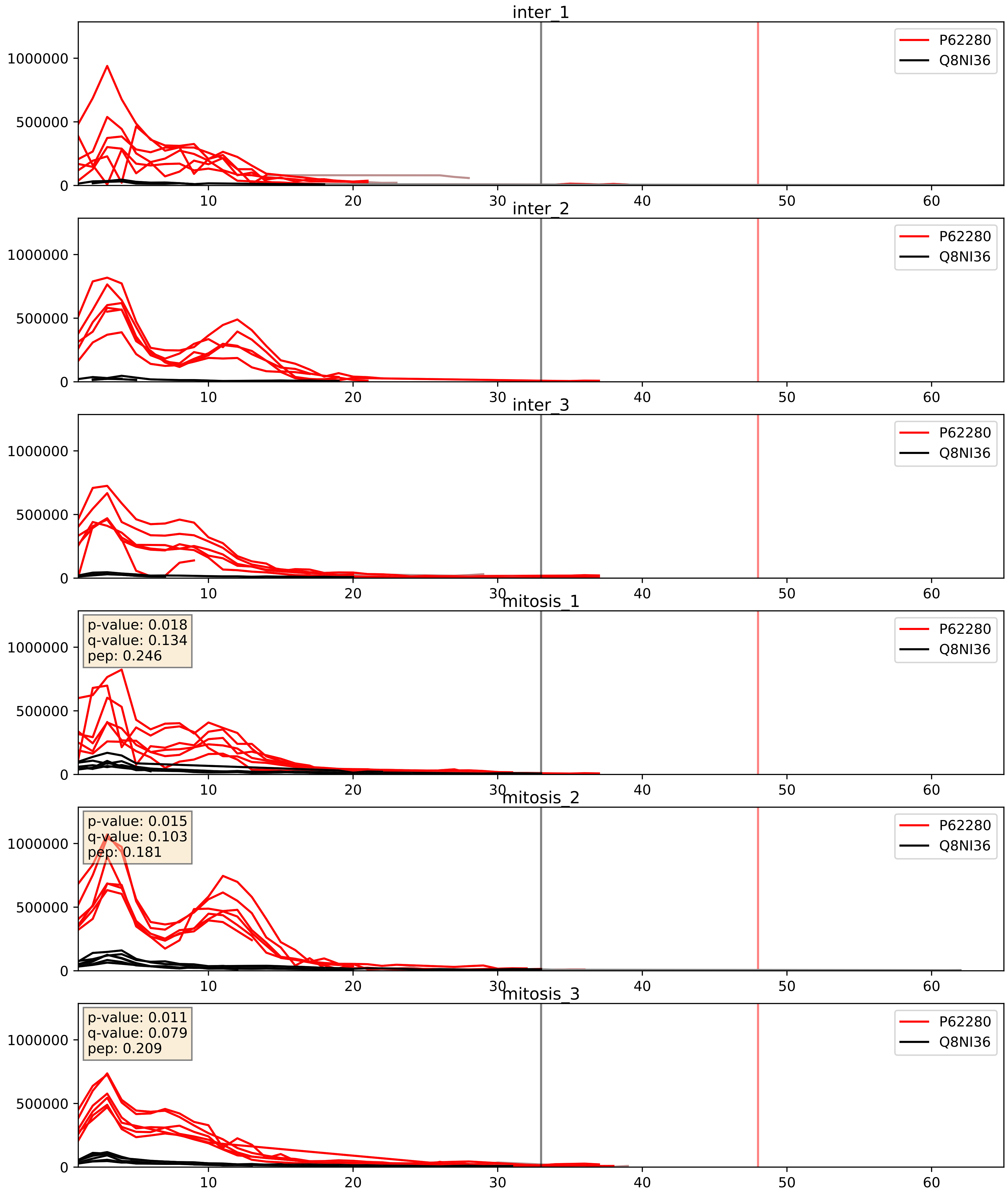


| level | condition_1 | condition_2 | pvalue | pvalue_adjusted |
|---|---|---|---|---|
| complex_abundance | mitosis | inter | 0.004925044309485646 | 0.052626777135169354 |
| interactor_ratio | mitosis | inter | 0.11241216105864874 | 0.5536525308757383 |
| interactor_abundance | mitosis | inter | 0.4465653089356643 | 0.6221171852045385 |

Q12788_Q8NI36
TBL3_HUMAN vs WDR36_HUMAN
p-value: 0.0 q-value: 0.001

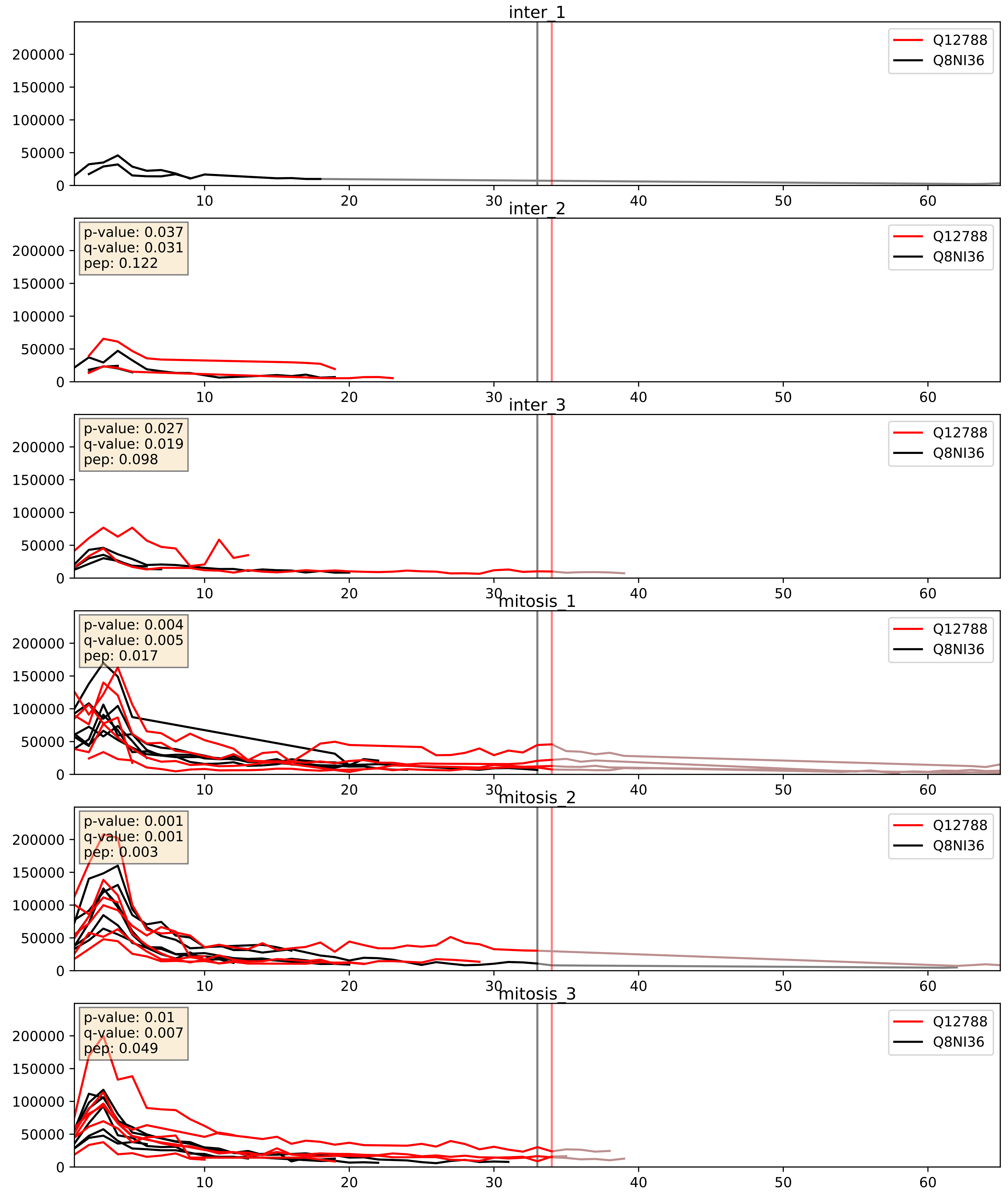


| level | condition_1 | condition_2 | pvalue | pvalue_adjusted |
|---|---|---|---|---|
| complex_abundance | mitosis | inter | 0.003360478542340517 | 0.04258563238370332 |
| interactor_abundance | mitosis | inter | 0.006662312924366897 | 0.06988896891247627 |
| interactor_ratio | mitosis | inter | 0.3561802533008551 | 0.8030936685700529 |

Q14137_Q8NI36
BOP1_HUMAN vs WDR36_HUMAN
p-value: 0.029 q-value: 0.021

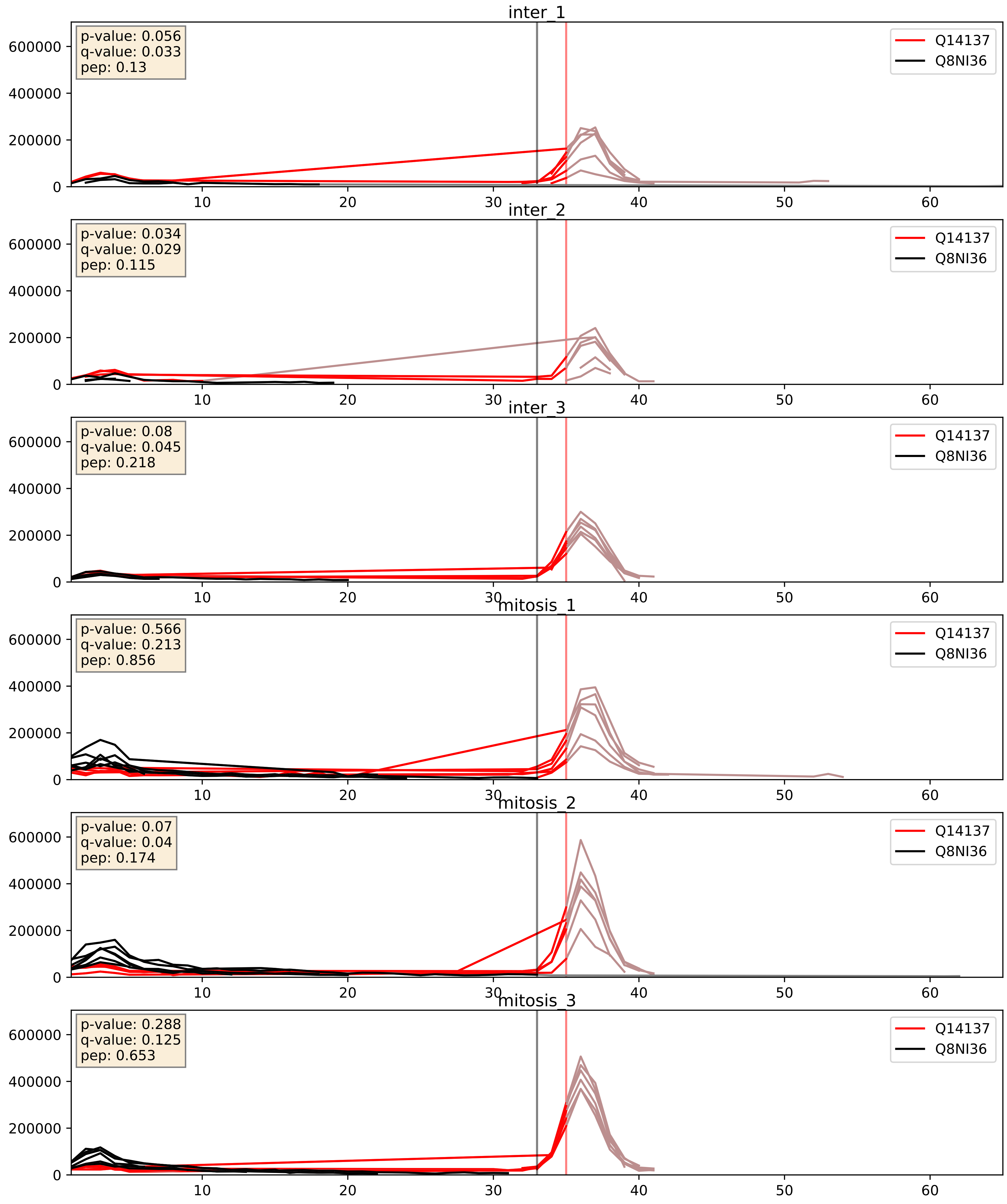


| level | condition_1 | condition_2 | pvalue | pvalue_adjusted |
|---|---|---|---|---|
| interactor_ratio | mitosis | inter | 0.0022534435788810043 | 0.05896690574521502 |
| interactor_abundance | mitosis | inter | 0.22206489816658428 | 0.4181424391346154 |
| complex_abundance | mitosis | inter | 0.5957970214043927 | 0.7204524977097283 |

Q15061_Q8NI36
WDR43_HUMAN vs WDR36_HUMAN
p-value: 0.08 q-value: 0.048

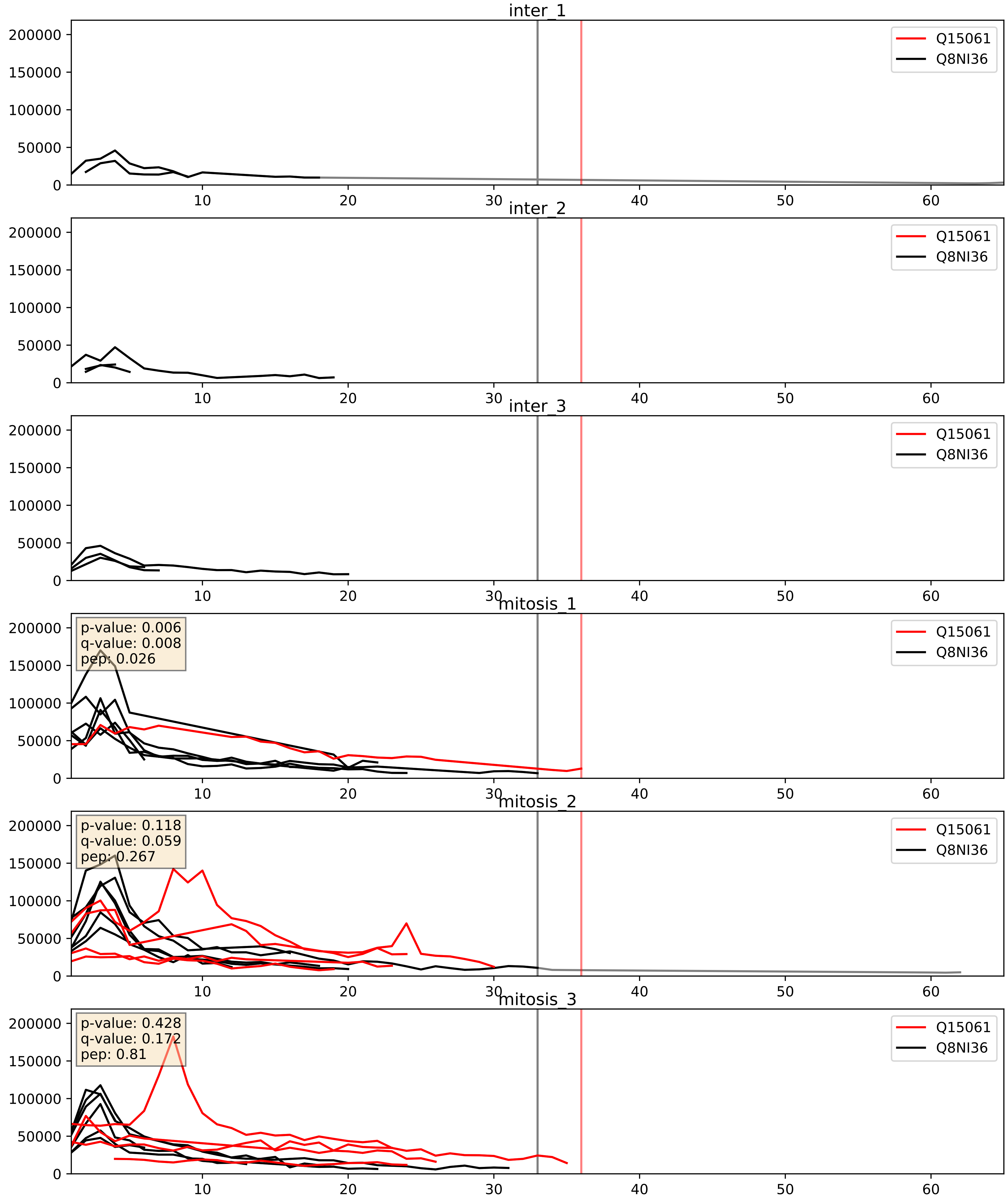


| level | condition_1 | condition_2 | pvalue | pvalue_adjusted |
|---|---|---|---|---|
| complex_abundance | mitosis | inter | 0.005860437094623565 | 0.05756820427608883 |
| interactor_abundance | mitosis | inter | 0.045200483218485674 | 0.16918064554011253 |
| interactor_ratio | mitosis | inter | 0.7727111896398332 | 0.9554532798105997 |

Q15269_Q8NI36
PWP2_HUMAN vs WDR36_HUMAN
p-value: 0.001 q-value: 0.001

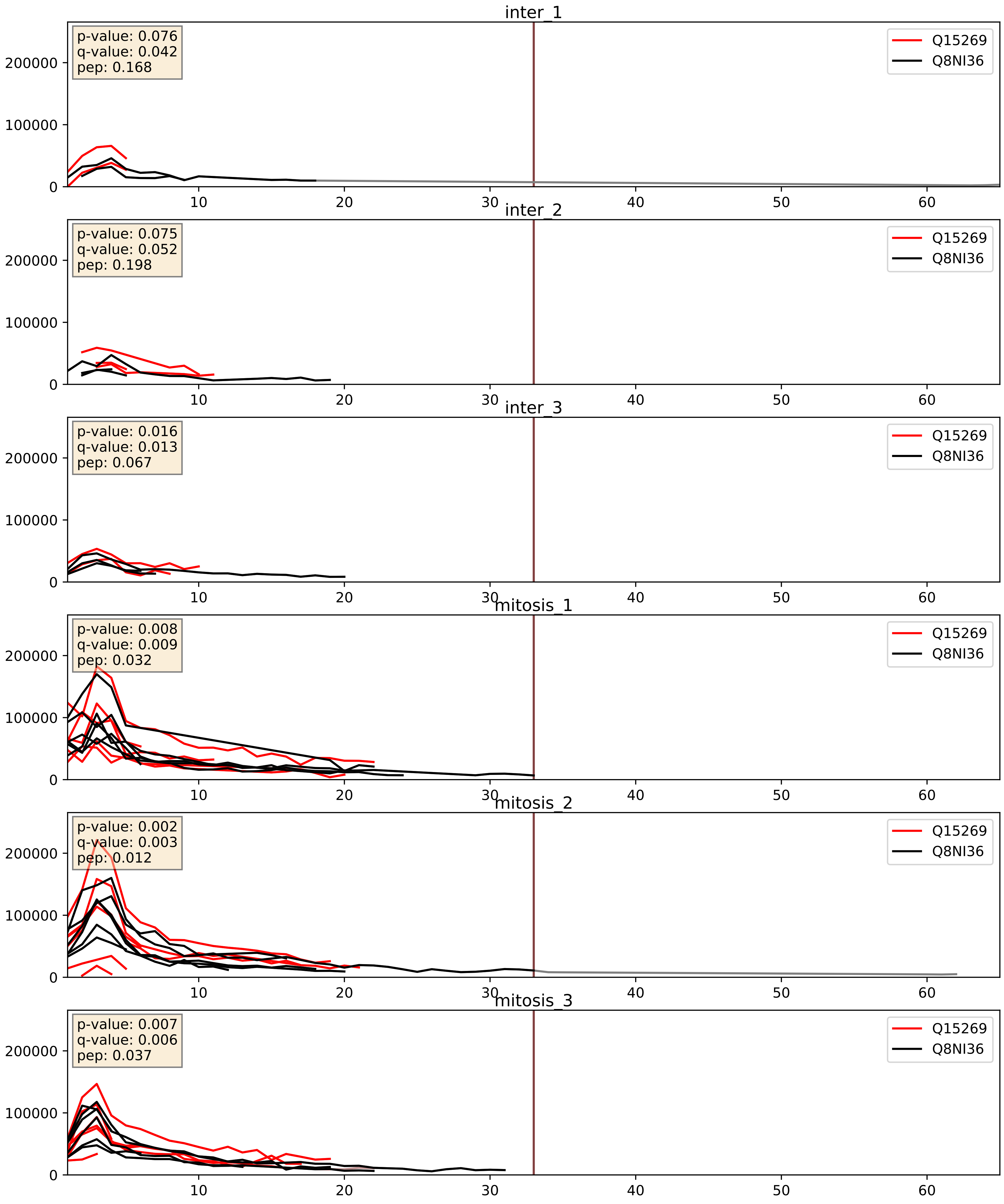


| level | condition_1 | condition_2 | pvalue | pvalue_adjusted |
|---|---|---|---|---|
| complex_abundance | mitosis | inter | 0.008082465810691268 | 0.06653038413698939 |
| interactor_abundance | mitosis | inter | 0.02134655497707949 | 0.1216554664472706 |
| interactor_ratio | mitosis | inter | 0.7936552315502841 | 0.9608508478447312 |

Q8IY81_Q8NI36
SPB1_HUMAN vs WDR36_HUMAN
p-value: 0.008 q-value: 0.008

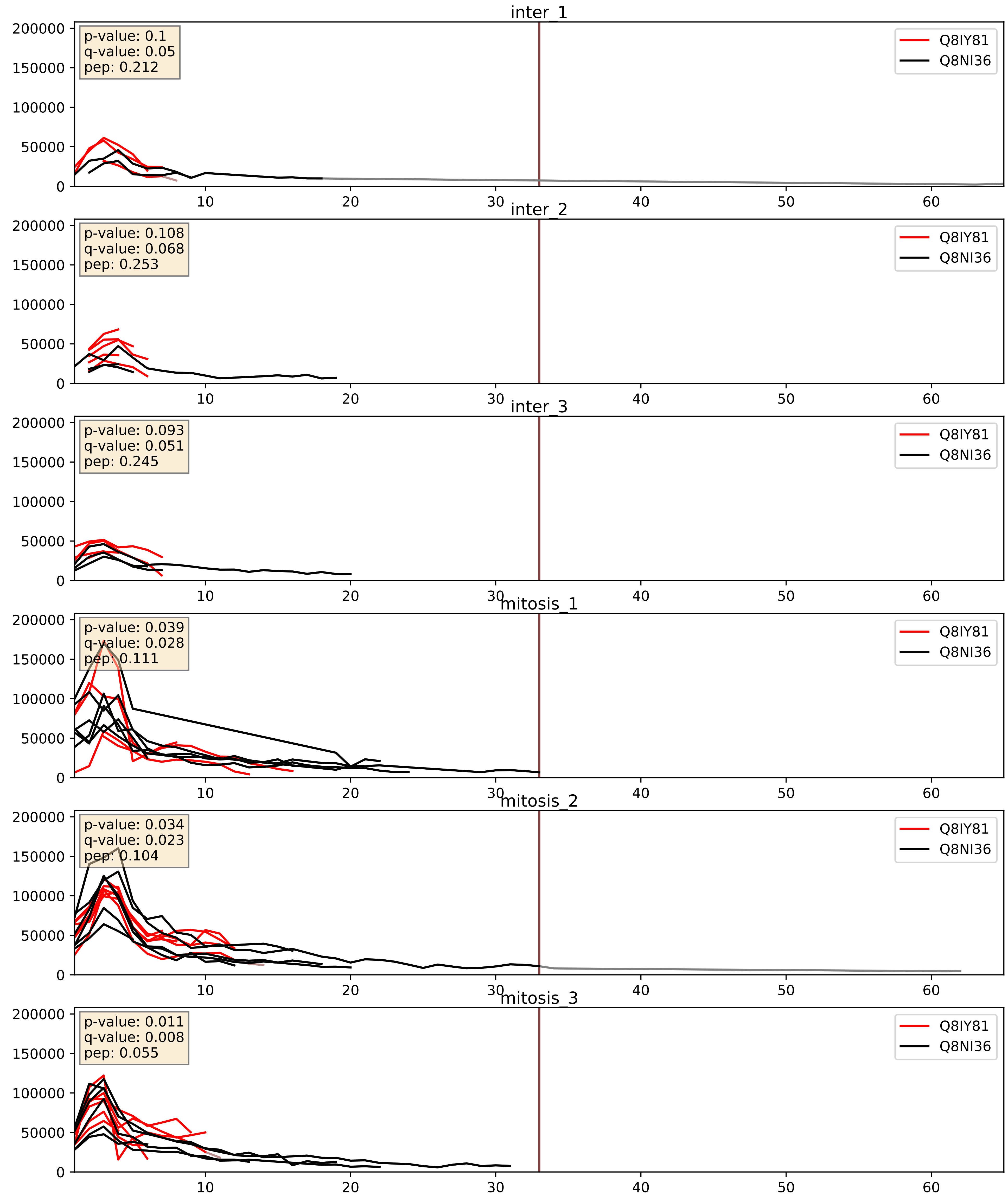


| level | condition_1 | condition_2 | pvalue | pvalue_adjusted |
|---|---|---|---|---|
| complex_abundance | mitosis | inter | 0.07129833141638876 | 0.1941068522539015 |
| interactor_ratio | mitosis | inter | 0.09852732059893228 | 0.5220660934699379 |
| interactor_abundance | mitosis | inter | 0.36653218238088725 | 0.5624025973314919 |

Q8NI36_Q8TED0
WDR36_HUMAN vs UTP15_HUMAN
p-value: 0.034 q-value: 0.024

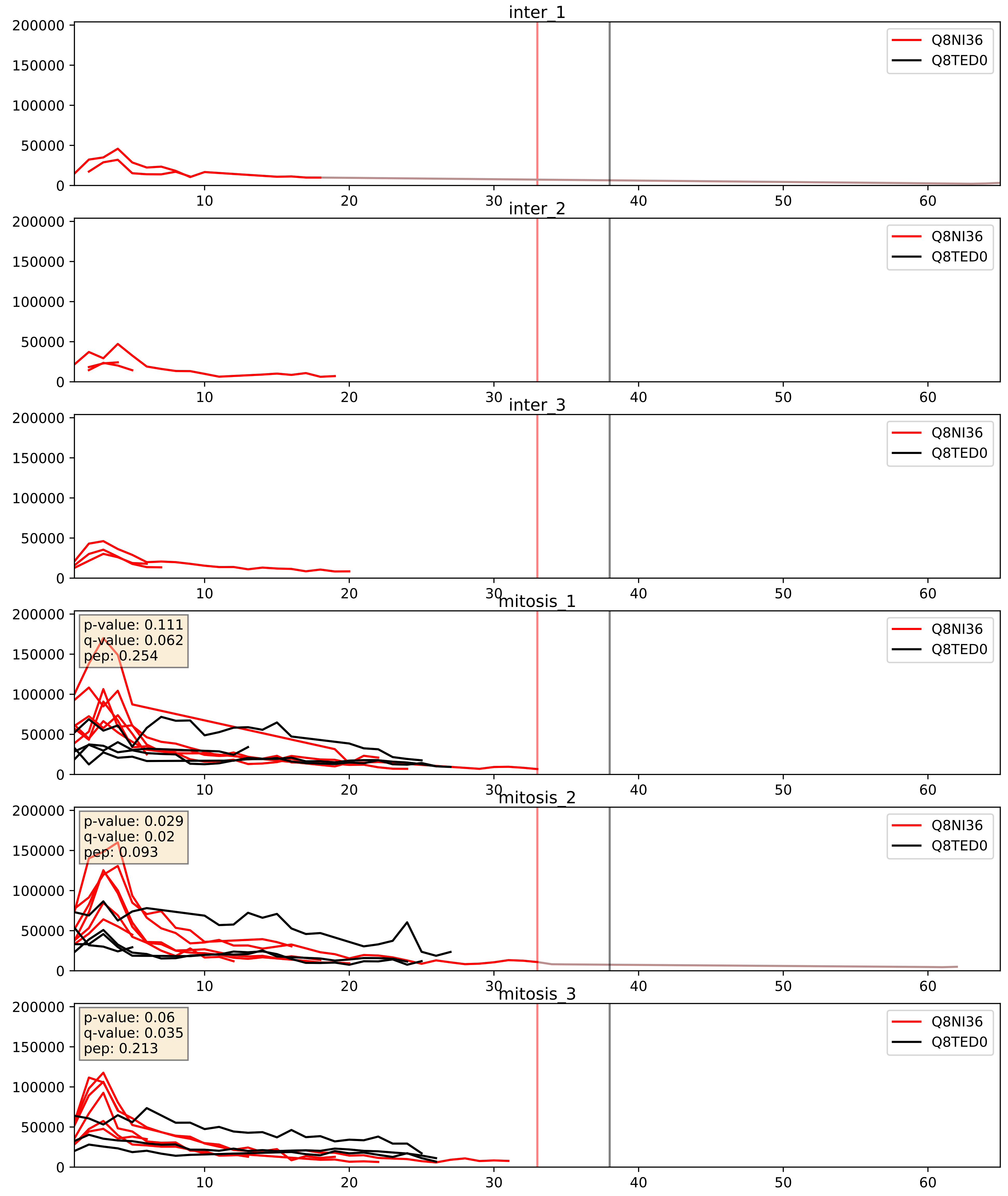


| level | condition_1 | condition_2 | pvalue | pvalue_adjusted |
|---|---|---|---|---|
| interactor_abundance | mitosis | inter | 0.0005649580704027154 | 0.017415659477552636 |
| complex_abundance | mitosis | inter | 0.0006846248474034801 | 0.016840197395901695 |
| interactor_ratio | mitosis | inter | 0.15977555659771123 | 0.616090239021059 |

Q8NI36_Q969X6
WDR36_HUMAN vs UTP4_HUMAN
p-value: 0.067 q-value: 0.041

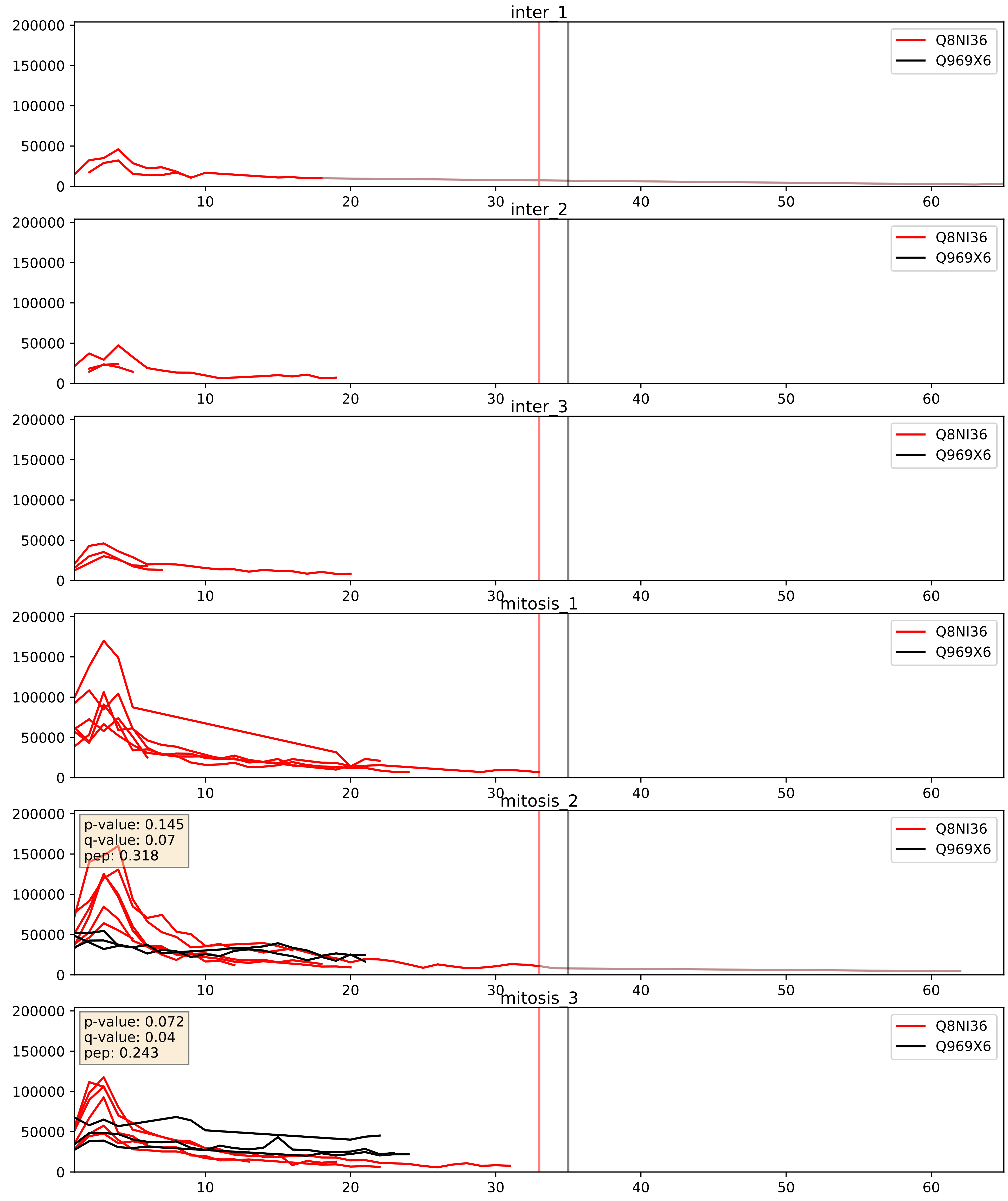


| level | condition_1 | condition_2 | pvalue | pvalue_adjusted |
|---|---|---|---|---|
| interactor_abundance | mitosis | inter | 0.20025917161989962 | 0.3948902347538219 |
| interactor_ratio | mitosis | inter | 0.21390918761331124 | 0.6906789033829601 |
| complex_abundance | mitosis | inter | 0.22057379217360717 | 0.3746808072497155 |

Q8NI36_Q9BQ67
WDR36_HUMAN vs GRWD1_HUMAN
p-value: 0.006 q-value: 0.006

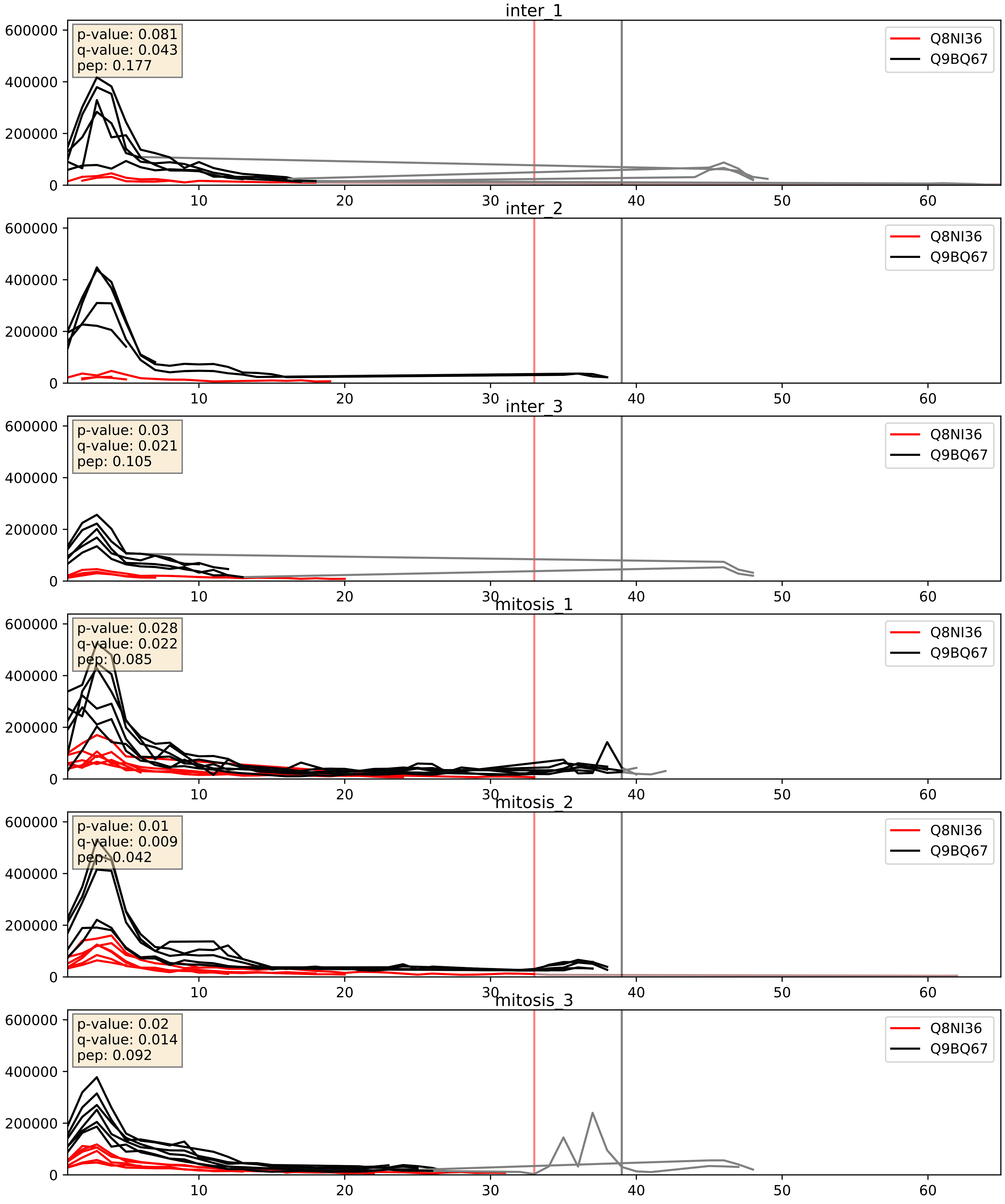


| level | condition_1 | condition_2 | pvalue | pvalue_adjusted |
|---|---|---|---|---|
| interactor_abundance | mitosis | inter | 0.004254336243081026 | 0.05203482881393319 |
| complex_abundance | mitosis | inter | 0.027185050581926013 | 0.12026048216087167 |
| interactor_ratio | mitosis | inter | 0.09811809124801148 | 0.5212210525862964 |

Q8NI36_Q9H0A0
WDR36_HUMAN vs NAT10_HUMAN
p-value: 0.008 q-value: 0.007

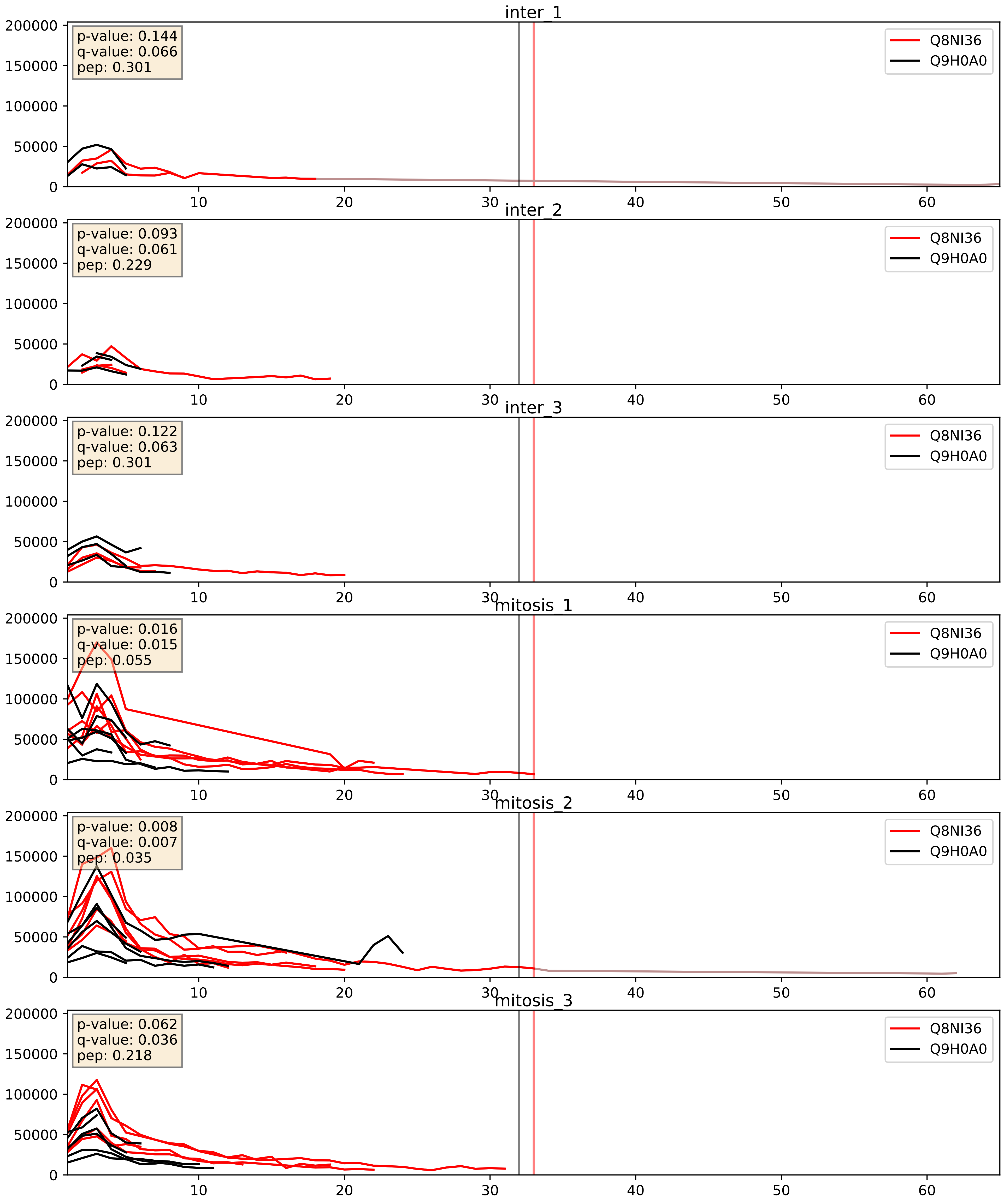


| level | condition_1 | condition_2 | pvalue | pvalue_adjusted |
|---|---|---|---|---|
| interactor_abundance | mitosis | inter | 0.00851704615394727 | 0.0803238047623781 |
| interactor_ratio | mitosis | inter | 0.018810868078371294 | 0.20643721891135677 |
| complex_abundance | mitosis | inter | 0.08899095423186006 | 0.2191545360681031 |

Q8NI36_Q9H0S4
WDR36_HUMAN vs DDX47_HUMAN
p-value: 0.004 q-value: 0.048

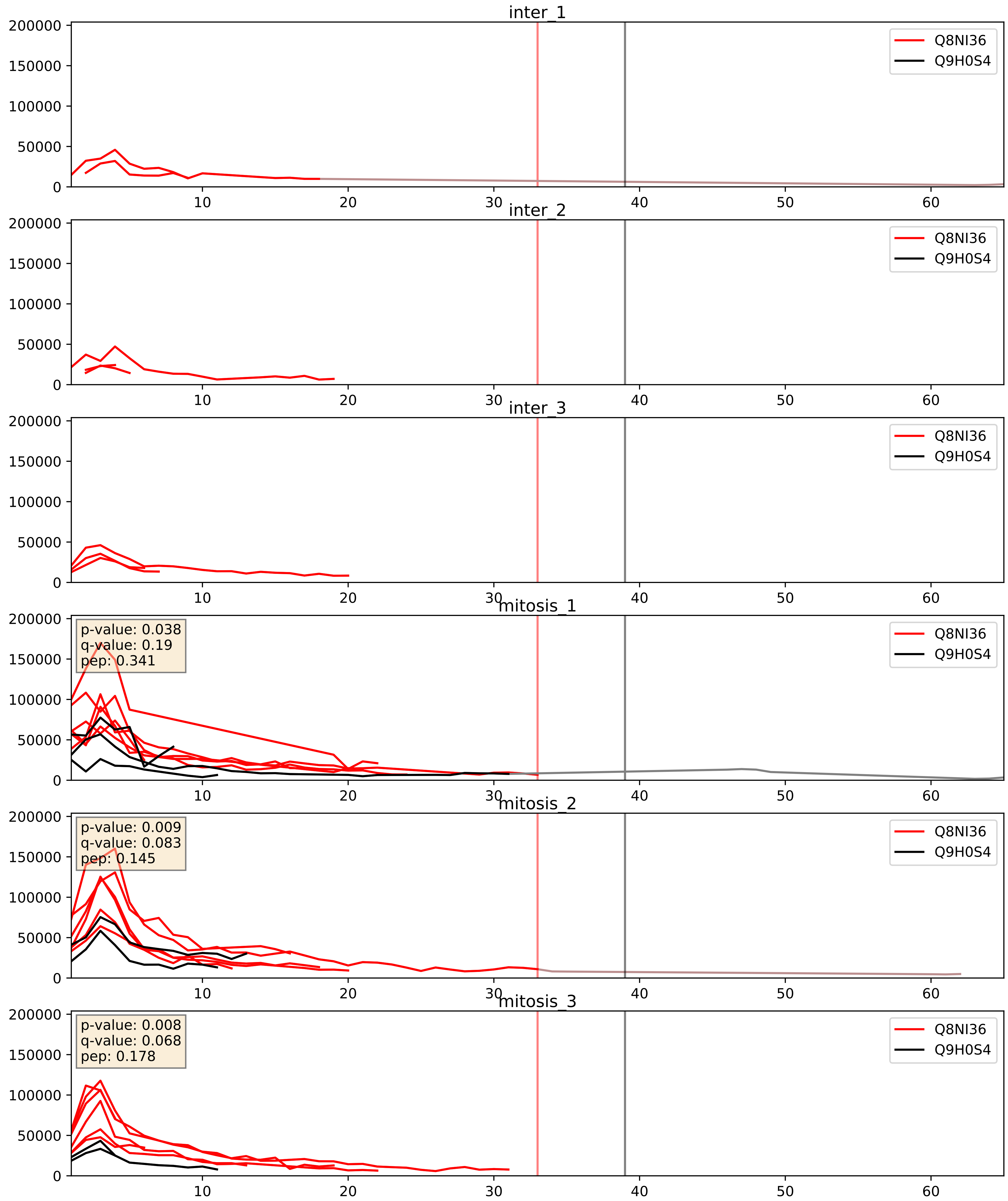


| level | condition_1 | condition_2 | pvalue | pvalue_adjusted |
|---|---|---|---|---|
| interactor_abundance | mitosis | inter | 0.001838446528961071 | 0.0315372791340817 |
| interactor_ratio | mitosis | inter | 0.009989406050391692 | 0.14263182888175546 |
| complex_abundance | mitosis | inter | 0.03375006447461892 | 0.13385180030241378 |

Q8NI36_Q9H583
WDR36_HUMAN vs HEAT1_HUMAN
p-value: 0.03 q-value: 0.022

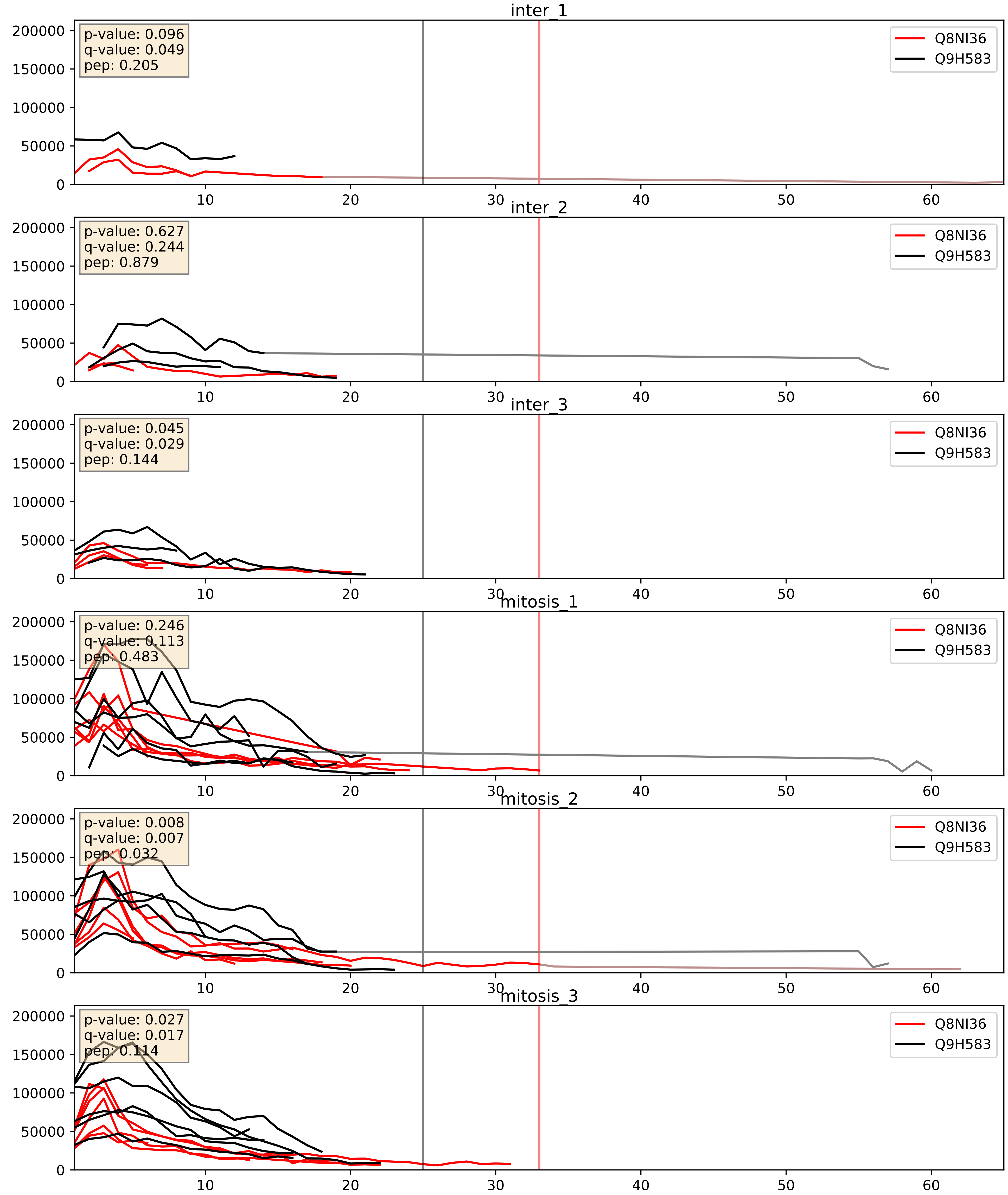


| level | condition_1 | condition_2 | pvalue | pvalue_adjusted |
|---|---|---|---|---|
| complex_abundance | mitosis | inter | 0.00031756796799436147 | 0.01165207110963347 |
| interactor_abundance | mitosis | inter | 0.0035552250741893126 | 0.04601489785035819 |
| interactor_ratio | mitosis | inter | 0.06616503681267413 | 0.4296400448702392 |

Q8NI36_Q9H6R4
WDR36_HUMAN vs NOL6_HUMAN
p-value: 0.032 q-value: 0.023

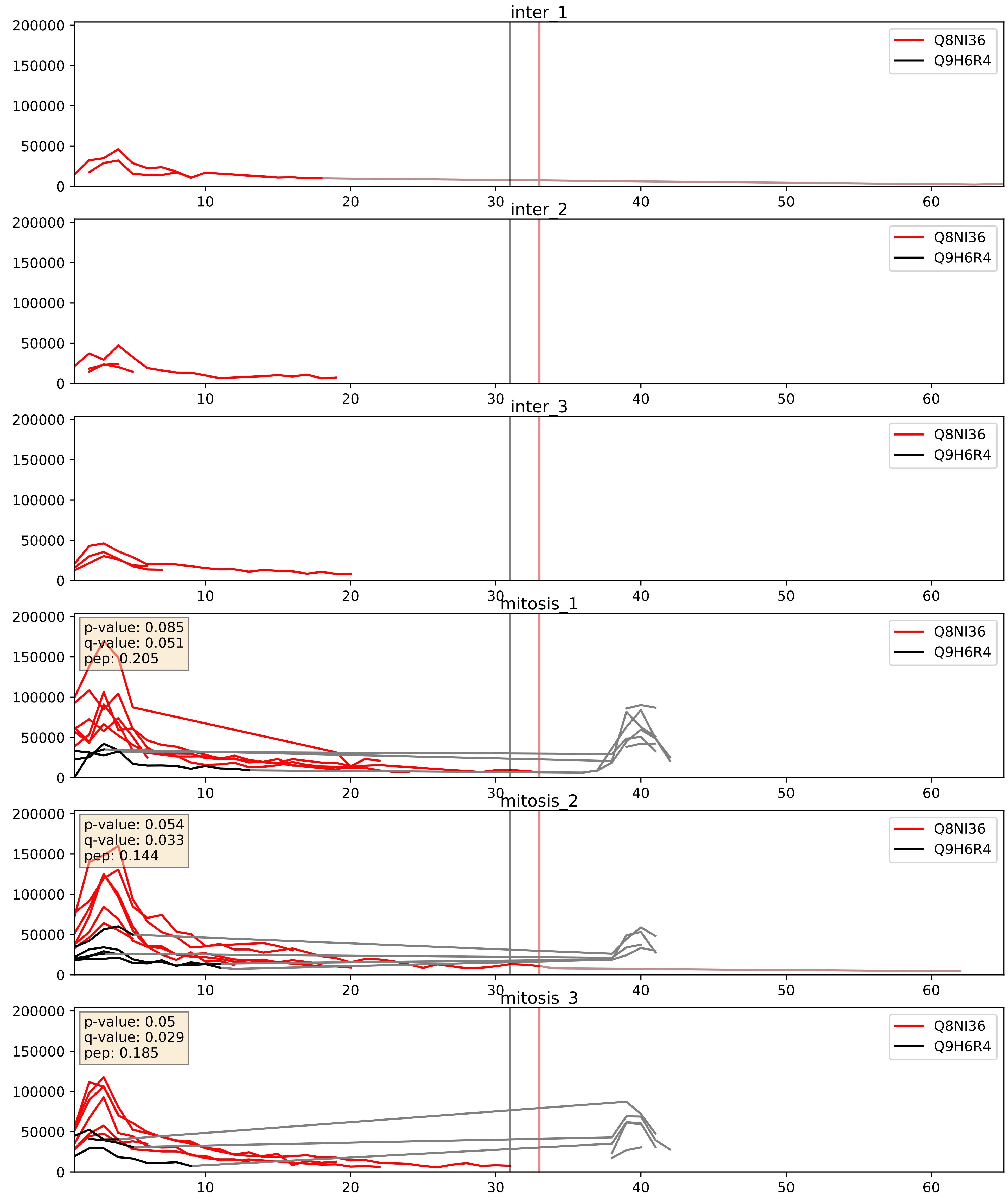


| level | condition_1 | condition_2 | pvalue | pvalue_adjusted |
|---|---|---|---|---|
| interactor_ratio | mitosis | inter | 0.0015565326714841756 | 0.04758542738537337 |
| interactor_abundance | mitosis | inter | 0.0022775002900272527 | 0.03523946087610491 |
| complex_abundance | mitosis | inter | 0.01595418682162696 | 0.09419630360707916 |

Q8NI36_Q9NVP1
WDR36_HUMAN vs DDX18_HUMAN
p-value: 0.034 q-value: 0.024

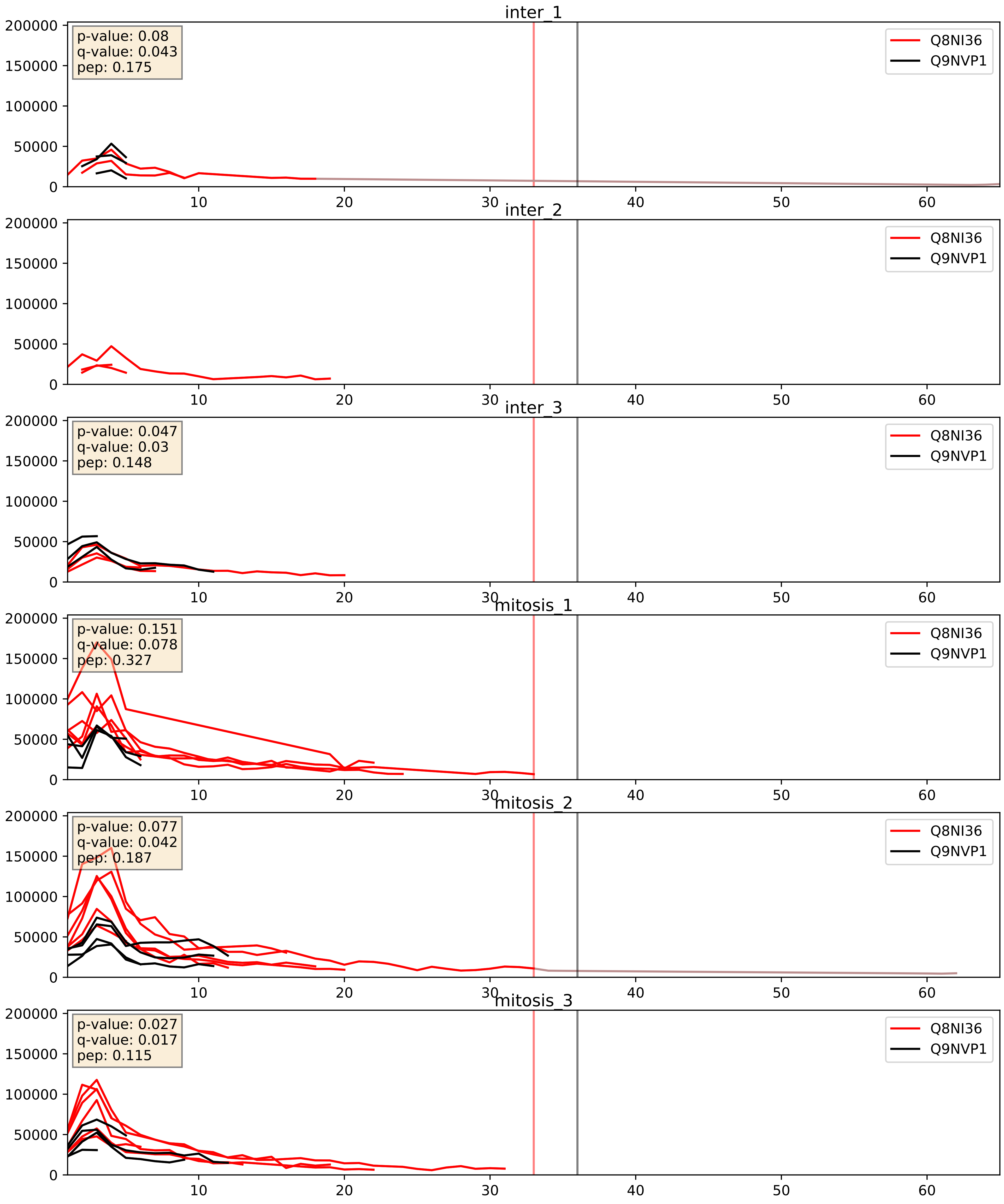


| level | condition_1 | condition_2 | pvalue | pvalue_adjusted |
|---|---|---|---|---|
| interactor_ratio | mitosis | inter | 0.009055793579369052 | 0.135757606023466 |
| interactor_abundance | mitosis | inter | 0.016895114881607966 | 0.11285383018850112 |
| complex_abundance | mitosis | inter | 0.13961312159519626 | 0.28468378070900036 |

Q8NI36_Q9NY93
WDR36_HUMAN vs DDX56_HUMAN
p-value: 0.062 q-value: 0.039

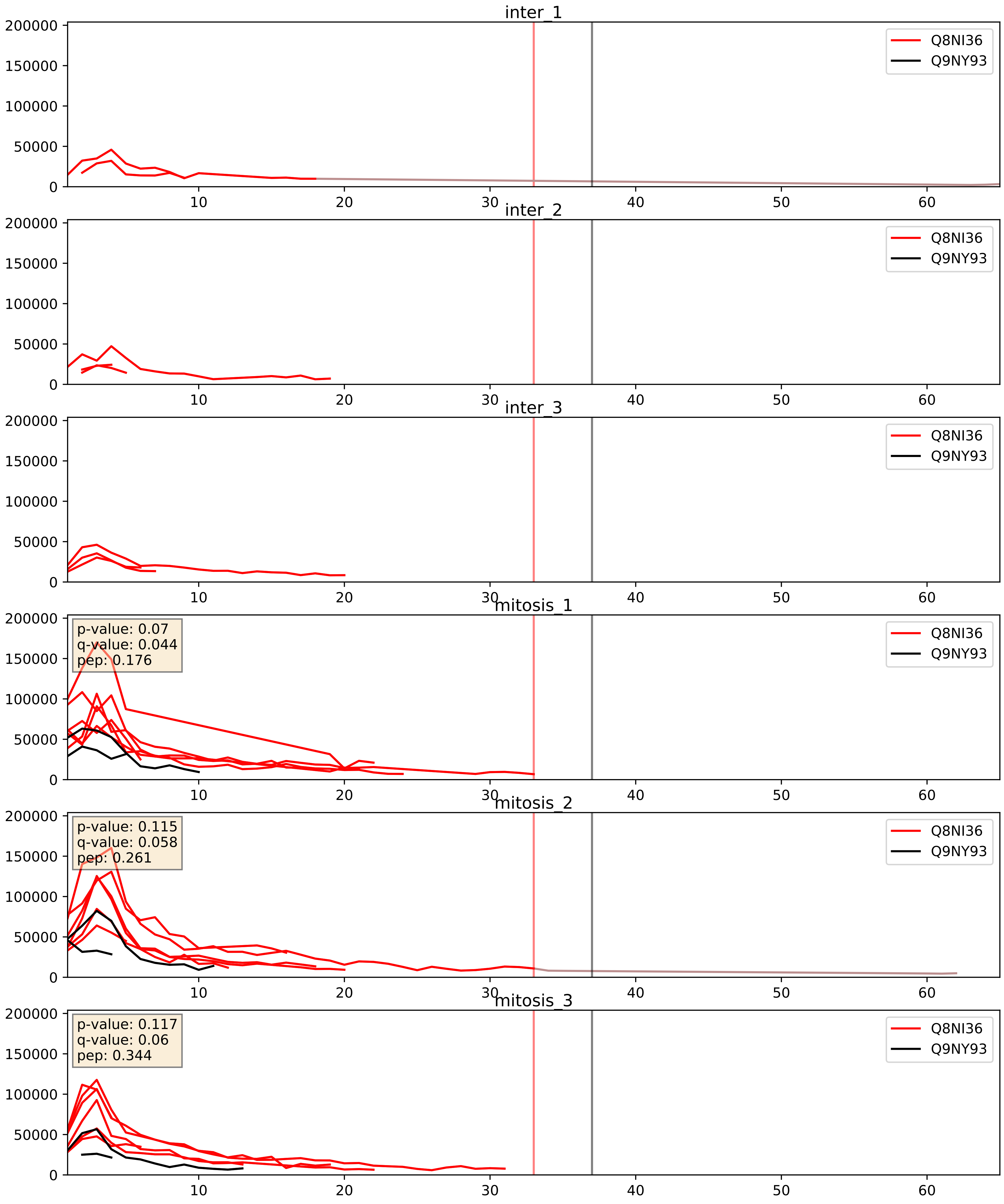


| level | condition_1 | condition_2 | pvalue | pvalue_adjusted |
|---|---|---|---|---|
| interactor_abundance | mitosis | inter | 0.000882265543690579 | 0.021954049575556266 |
| interactor_ratio | mitosis | inter | 0.0011153489338967212 | 0.03908562924515634 |
| complex_abundance | mitosis | inter | 0.004615842365066132 | 0.050629452773887605 |

Q8NI36_Q9UNX4
WDR36_HUMAN vs WDR3_HUMAN
p-value: 0.005 q-value: 0.005

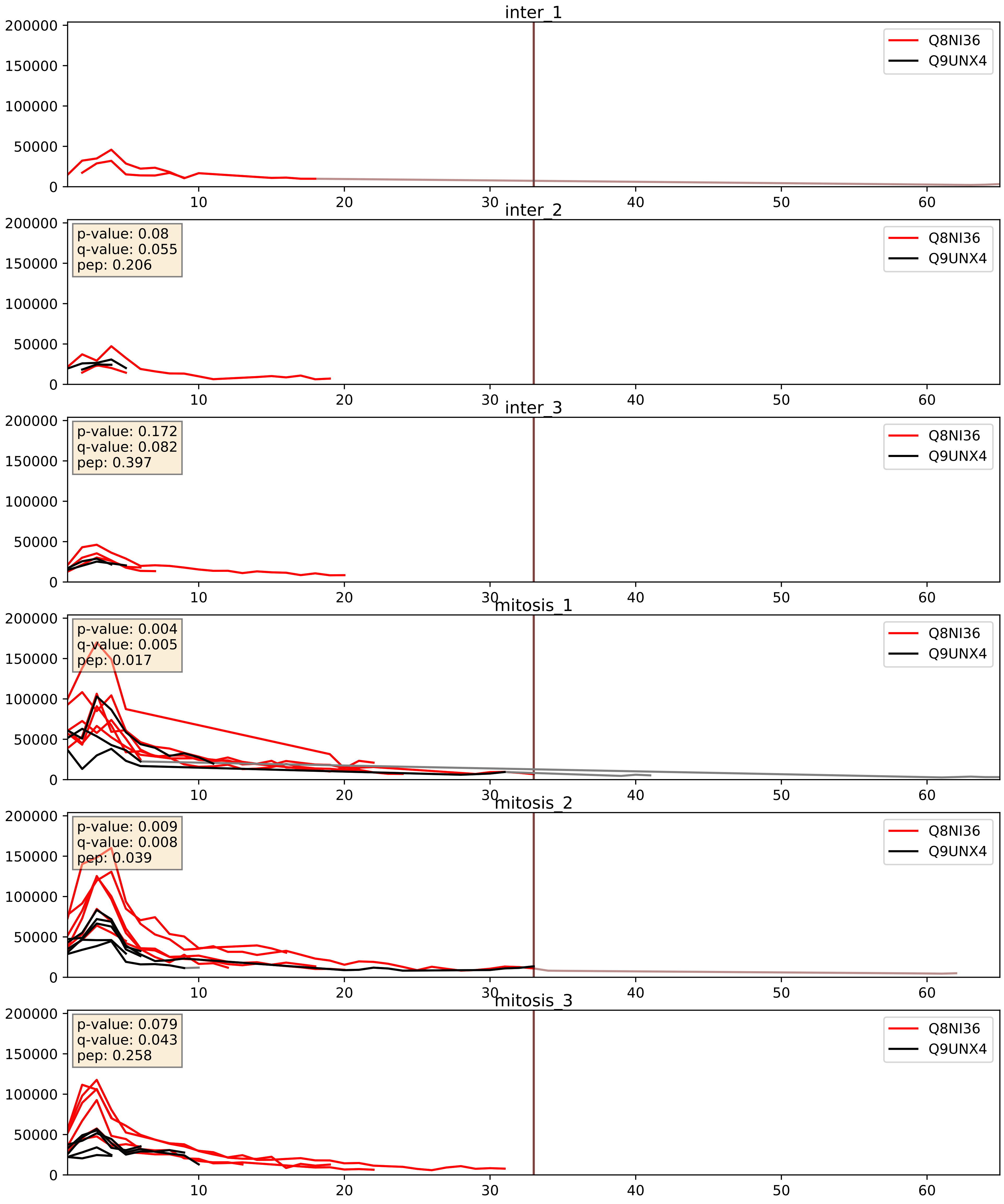


| level | condition_1 | condition_2 | pvalue | pvalue_adjusted |
|---|---|---|---|---|
| interactor_ratio | mitosis | inter | 0.0034412210964951434 | 0.07572939655107606 |
| interactor_abundance | mitosis | inter | 0.00922348089309837 | 0.08324021891678159 |
| complex_abundance | mitosis | inter | 0.03510041369517333 | 0.13586662078716946 |

Q8NI36_Q9Y221
WDR36_HUMAN vs NIP7_HUMAN
p-value: 0.038 q-value: 0.026

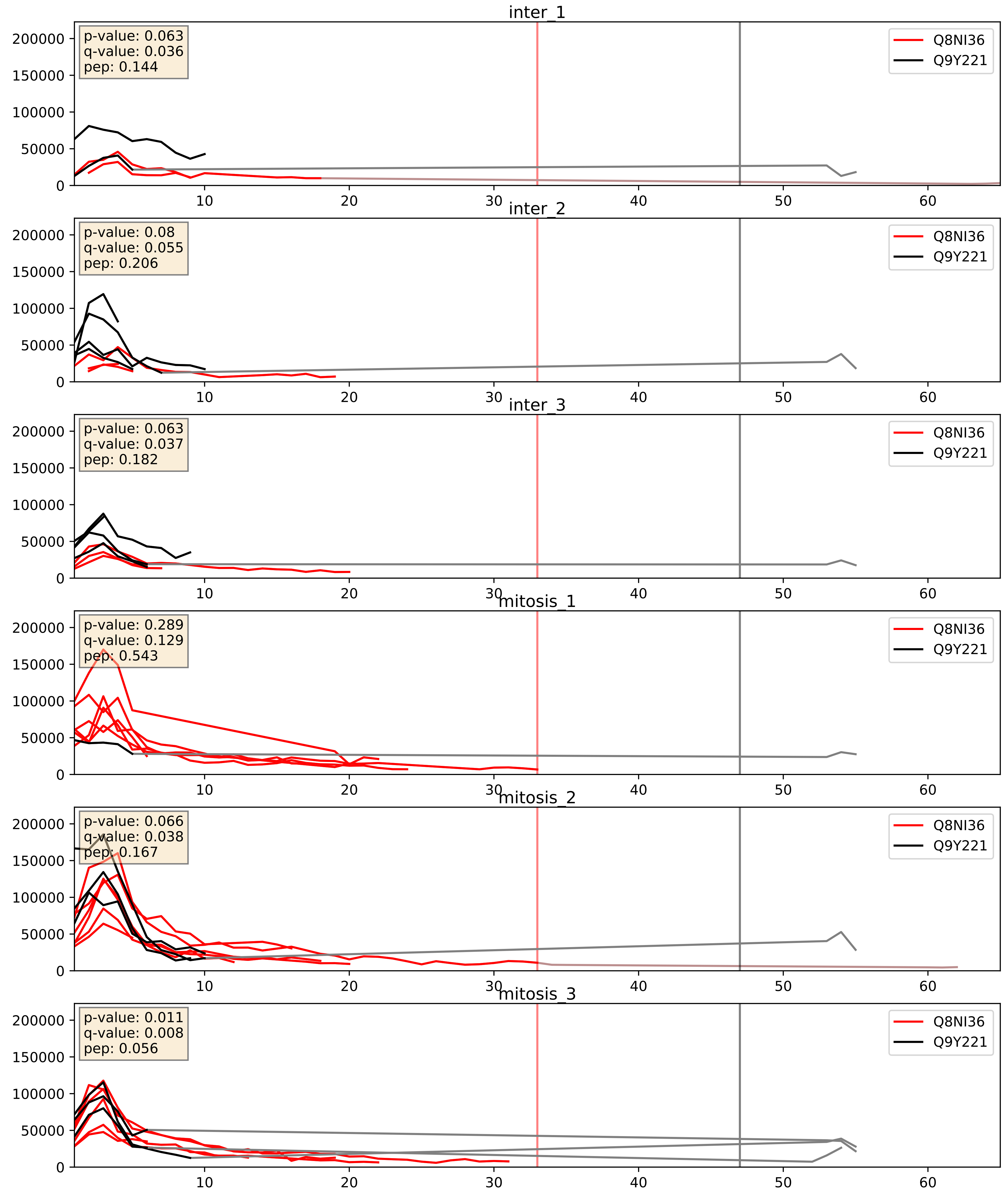


| level | condition_1 | condition_2 | pvalue | pvalue_adjusted |
|---|---|---|---|---|
| interactor_abundance | mitosis | inter | 0.039391110413609674 | 0.16083596262333216 |
| interactor_ratio | mitosis | inter | 0.08328109694013458 | 0.4841037832929192 |
| complex_abundance | mitosis | inter | 0.7940909055995328 | 0.8663545949441754 |

Q8NI36_Q9Y2X3
WDR36_HUMAN vs NOP58_HUMAN
p-value: 0.005 q-value: 0.005

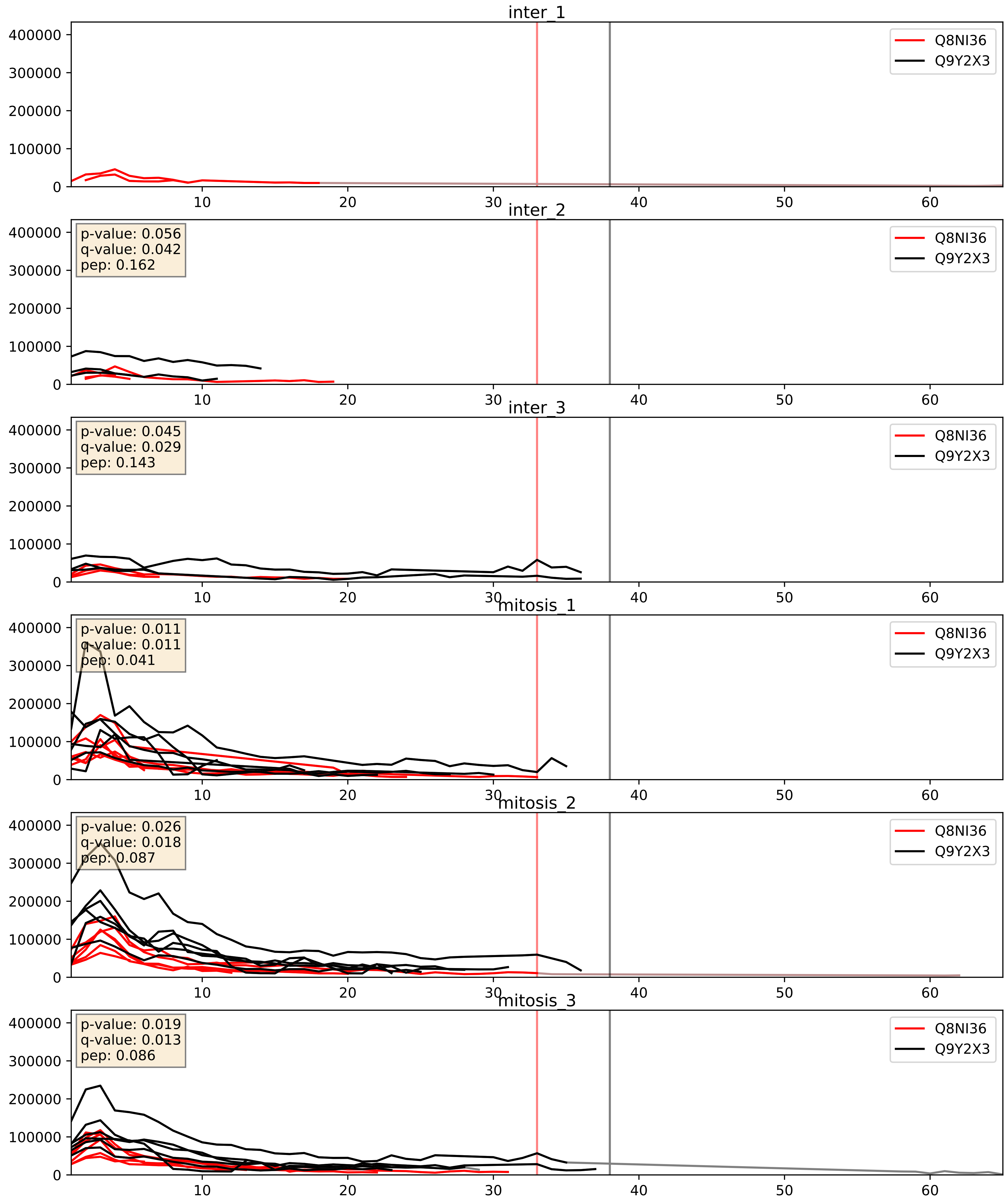


| level | condition_1 | condition_2 | pvalue | pvalue_adjusted |
|---|---|---|---|---|
| complex_abundance | mitosis | inter | 0.002522434141372613 | 0.036409709098919725 |
| interactor_abundance | mitosis | inter | 0.0036714075898180323 | 0.04704677989347658 |
| interactor_ratio | mitosis | inter | 0.003966749934868759 | 0.08201435780423773 |
