## Supplementary Data 2 for "SECAT: Quantifying differential protein-protein interaction states by network-centric analysis": hela_string_000036_Q9GZR7.pdf

Q9GZR7_Q9GZR7

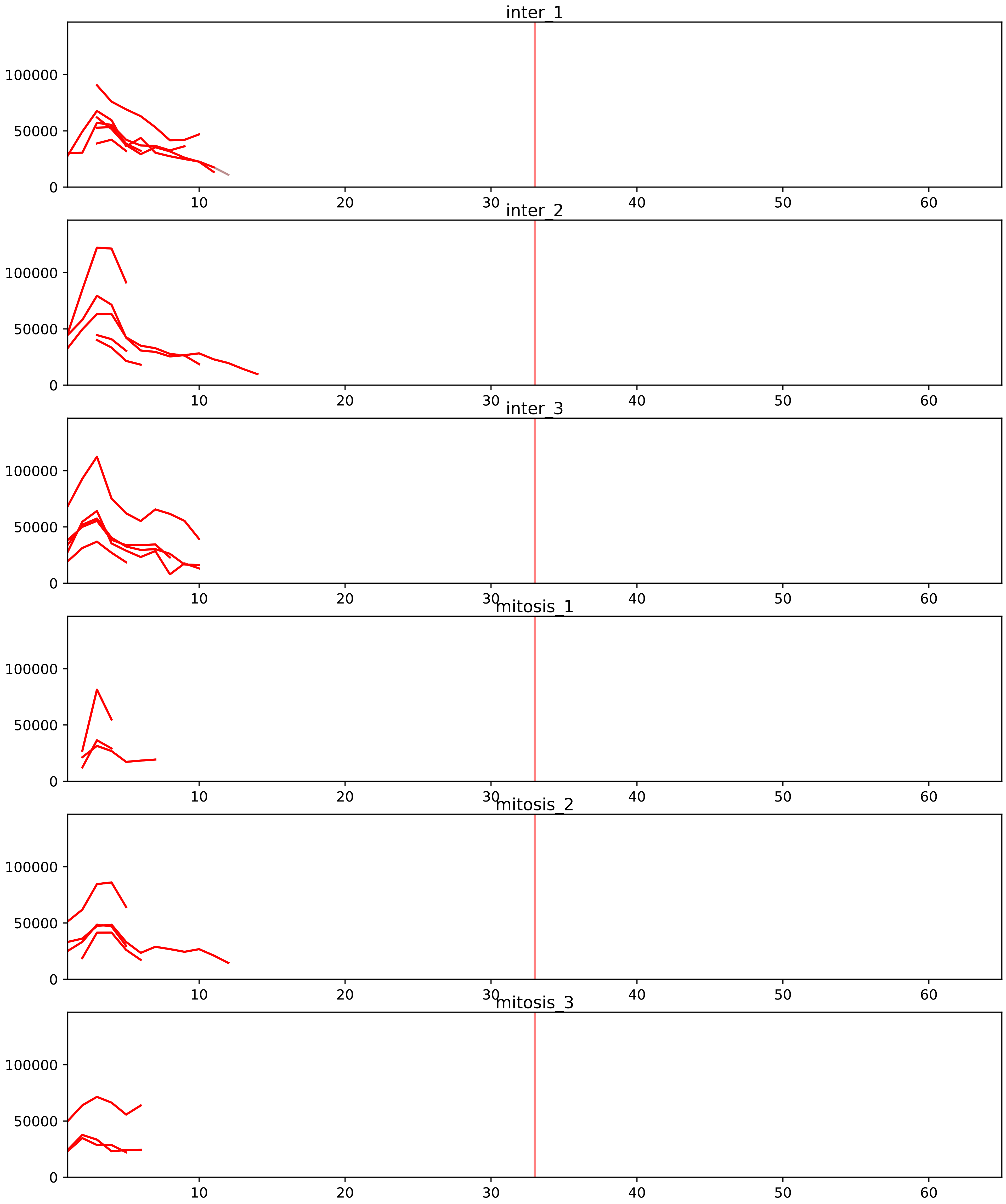


| level | condition_1 | condition_2 | pvalue | pvalue_adjusted |
|---|---|---|---|---|
| interactor_abundance | mitosis | inter | 1.2555544909371346e-08 | 6.3766625832553175e-06 |
| interactor_ratio | mitosis | inter | 1.7464865640666519e-06 | 4.943900427511753e-05 |
| assembled_abundance | mitosis | inter | 0.0232092602505601 | 0.18289991116645088 |
| total_abundance | mitosis | inter | 0.02381976610025858 | 0.19807504938328288 |
| complex_abundance | mitosis | inter | 0.02482496697611013 | 0.08041890710570887 |

P46087_Q9GZR7
NOP2_HUMAN vs DDX24_HUMAN
p-value: 0.039 q-value: 0.027

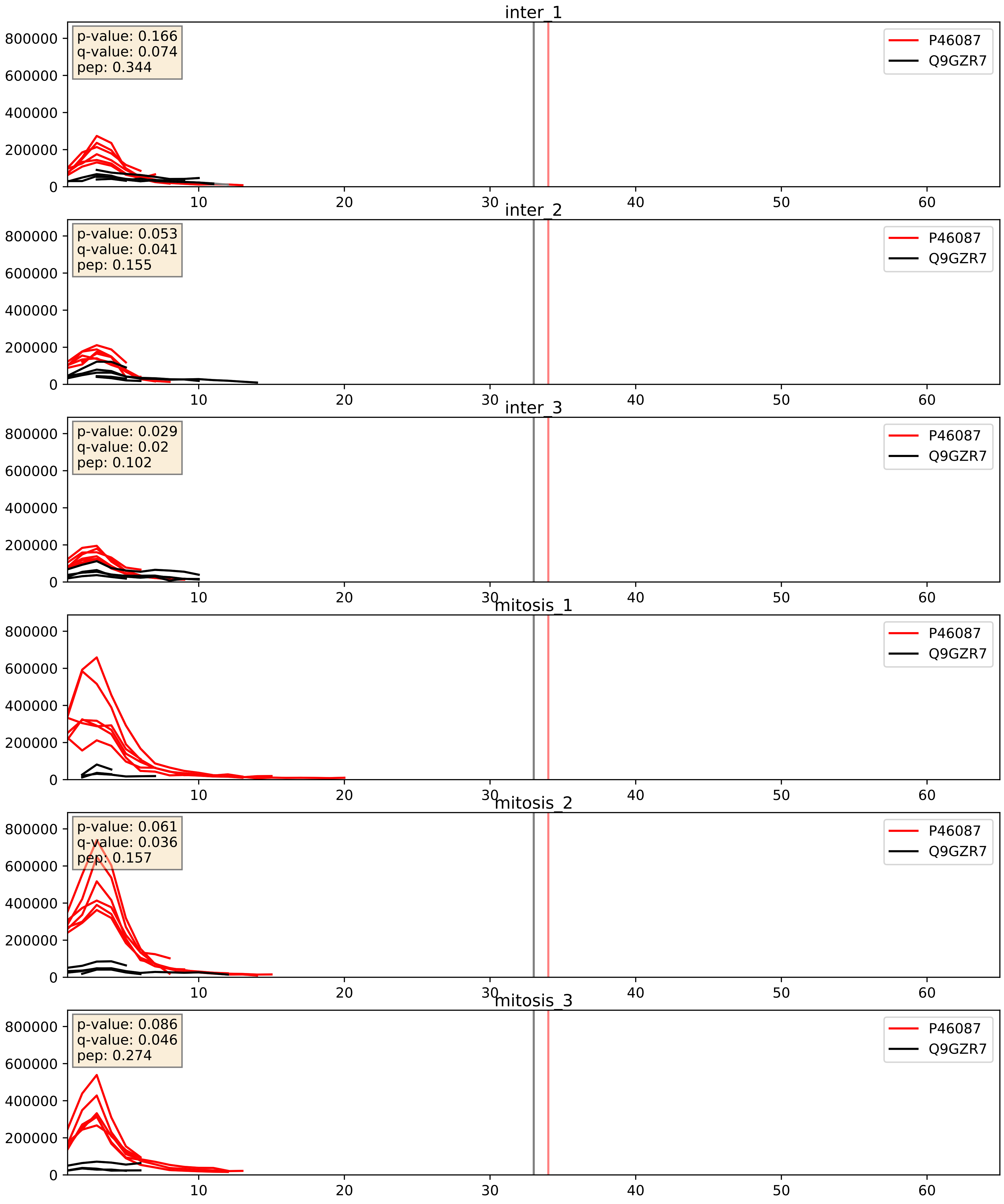


| level | condition_1 | condition_2 | pvalue | pvalue_adjusted |
|---|---|---|---|---|
| interactor_ratio | mitosis | inter | 0.0003942176979097222 | 0.021937350419330353 |
| interactor_abundance | mitosis | inter | 0.021324997778811212 | 0.1216554664472706 |
| complex_abundance | mitosis | inter | 0.8062023626534441 | 0.8745098220636244 |

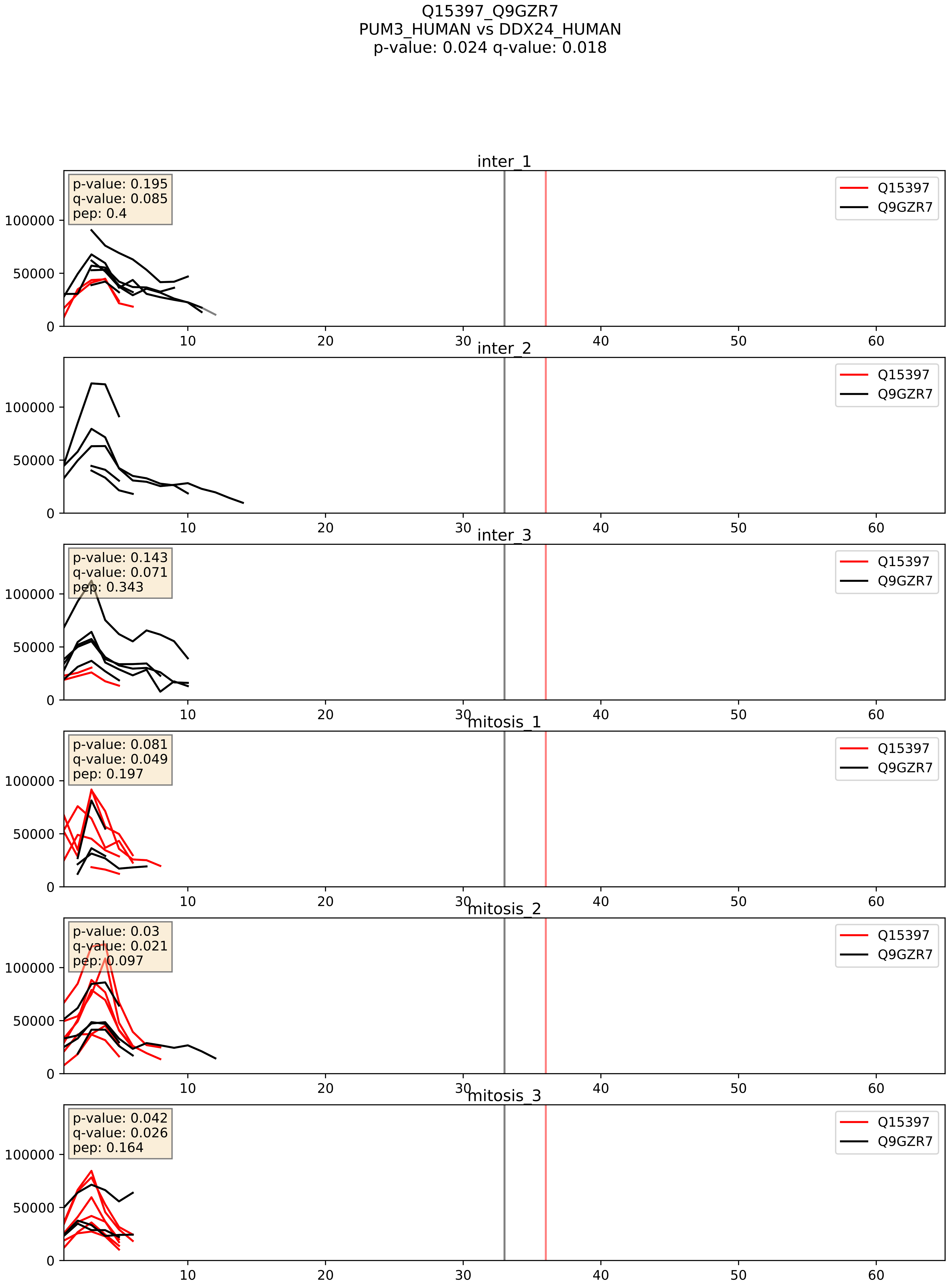


| level | condition_1 | condition_2 | pvalue | pvalue_adjusted |
|---|---|---|---|---|
| interactor_ratio | mitosis | inter | 0.04271903862311729 | 0.3405601671696144 |
| interactor_abundance | mitosis | inter | 0.30313654120020245 | 0.4984342667448584 |
| complex_abundance | mitosis | inter | 0.9077937729670189 | 0.9424158106875262 |

Q8TDN6_Q9GZR7
BRX1_HUMAN vs DDX24_HUMAN
p-value: 0.01 q-value: 0.009

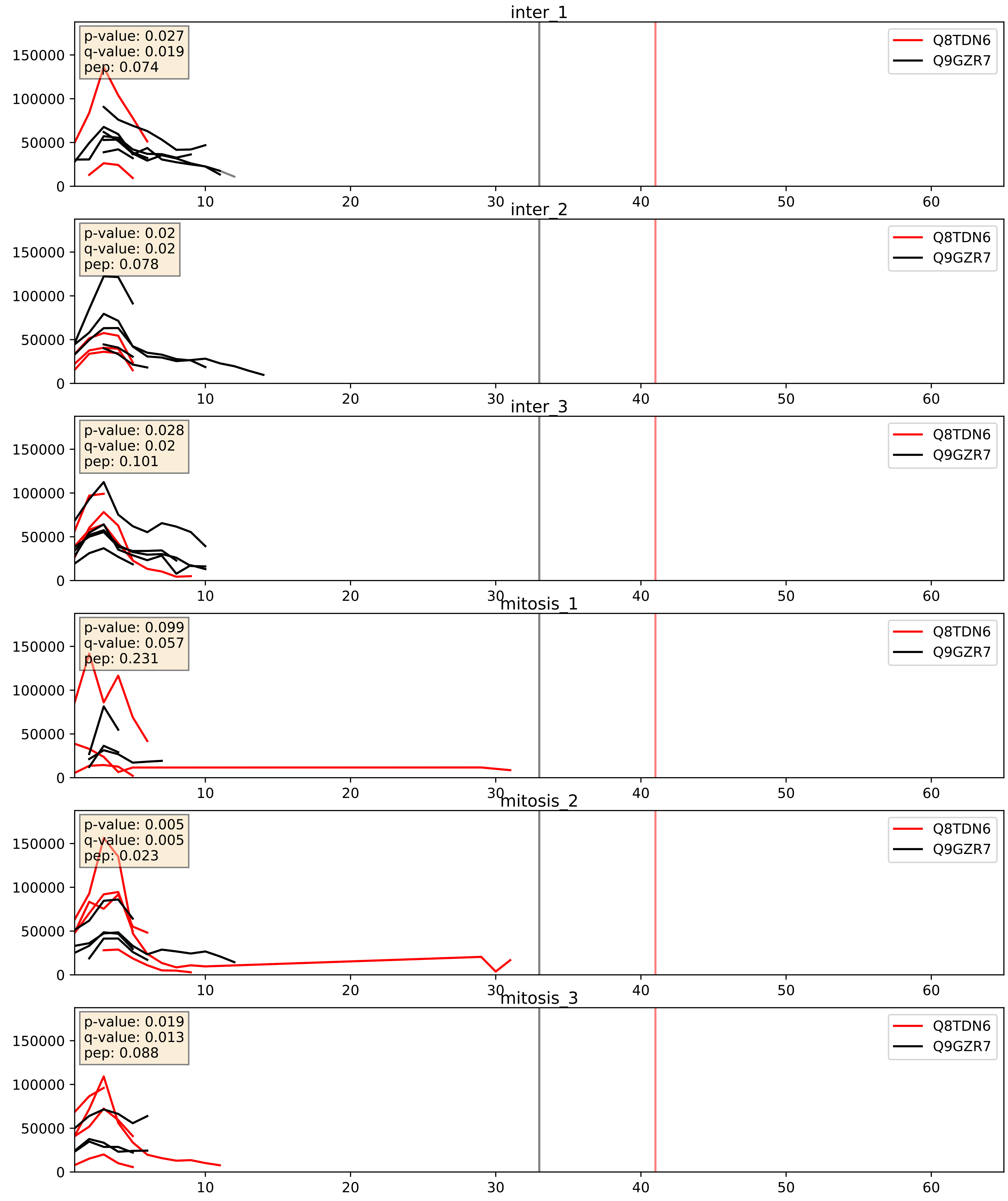


| level | condition_1 | condition_2 | pvalue | pvalue_adjusted |
|---|---|---|---|---|
| complex_abundance | mitosis | inter | 0.001617627551261813 | 0.02814408910325431 |
| interactor_ratio | mitosis | inter | 0.0017028547996455828 | 0.049919305085500645 |
| interactor_abundance | mitosis | inter | 0.04383239640025894 | 0.16685321787696922 |

Q9BYG3_Q9GZR7
MK67I_HUMAN vs DDX24_HUMAN
p-value: 0.024 q-value: 0.018

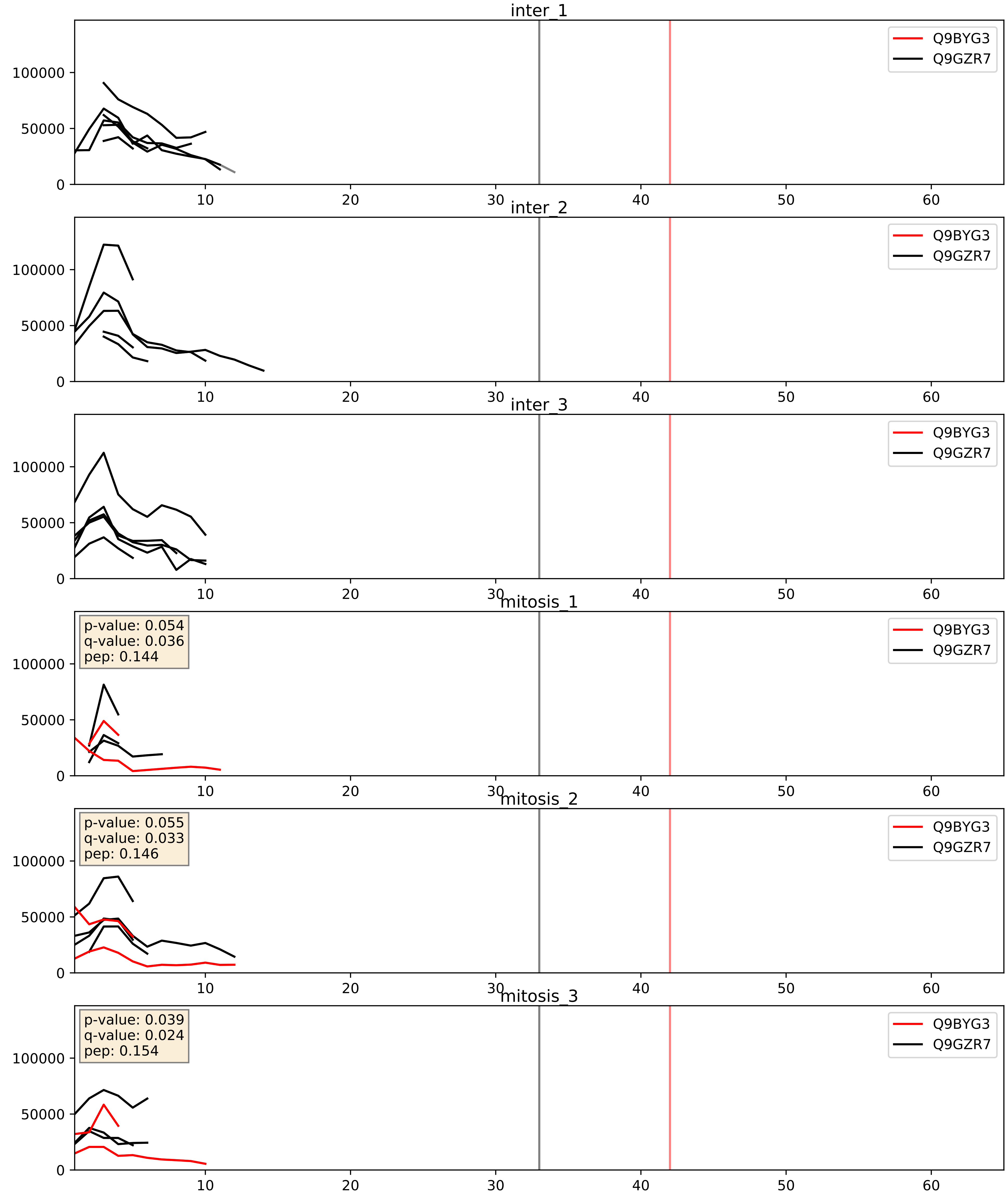


| level | condition_1 | condition_2 | pvalue | pvalue_adjusted |
|---|---|---|---|---|
| complex_abundance | mitosis | inter | 0.08265105601072668 | 0.21108552910002362 |
| interactor_ratio | mitosis | inter | 0.15249623026144915 | 0.6082794646029844 |
| interactor_abundance | mitosis | inter | 0.2535147171951053 | 0.4468626754113251 |

Q9BZE4_Q9GZR7
NOG1_HUMAN vs DDX24_HUMAN
p-value: 0.012 q-value: 0.01

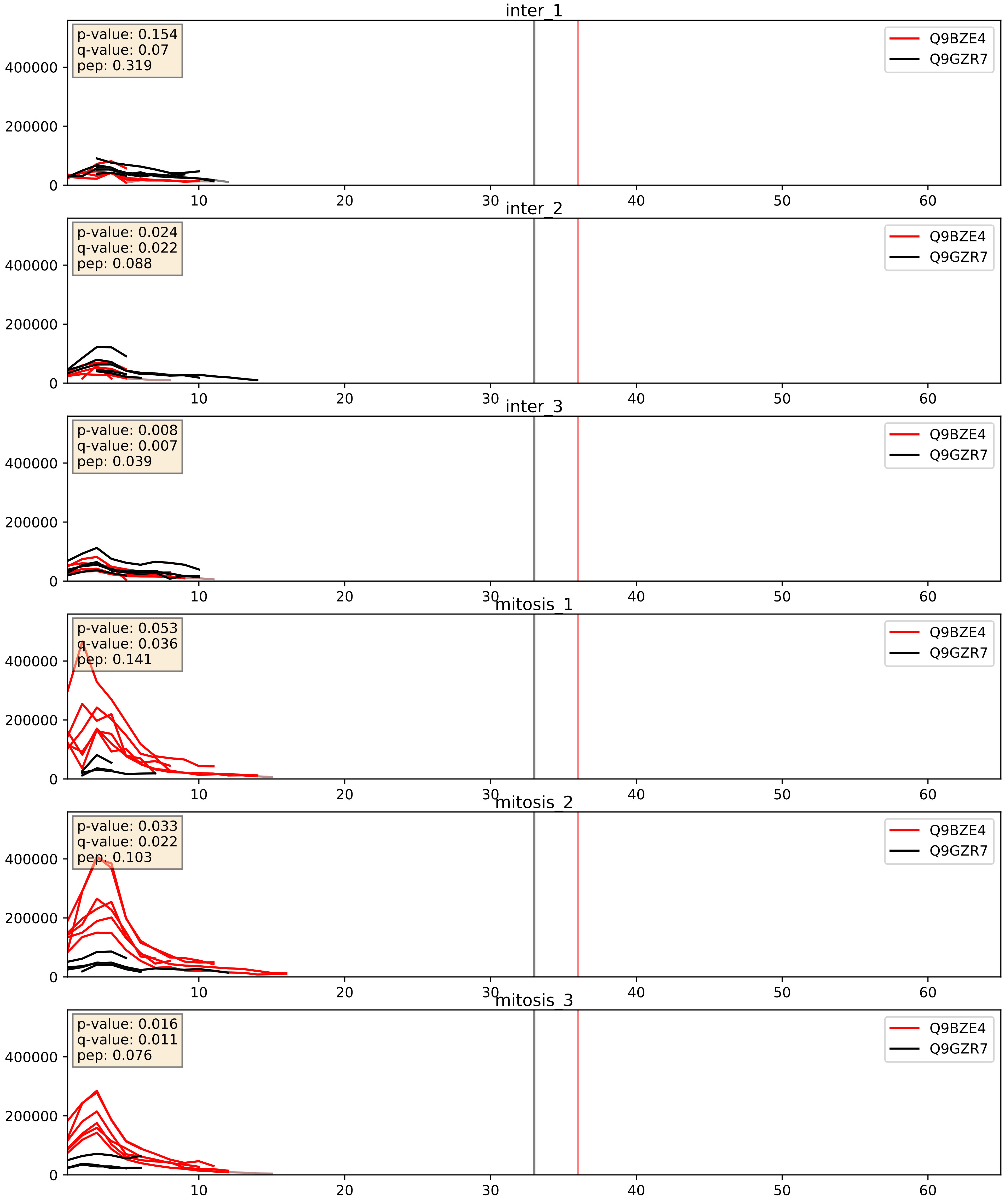


| level | condition_1 | condition_2 | pvalue | pvalue_adjusted |
|---|---|---|---|---|
| interactor_ratio | mitosis | inter | 0.00020599297265075348 | 0.014683183137172115 |
| interactor_abundance | mitosis | inter | 0.004638788434277854 | 0.05526517605764569 |
| complex_abundance | mitosis | inter | 0.2407443405492743 | 0.39200524160201405 |

Q9GZR7_Q9NY93
DDX24_HUMAN vs DDX56_HUMAN
p-value: 0.026 q-value: 0.02

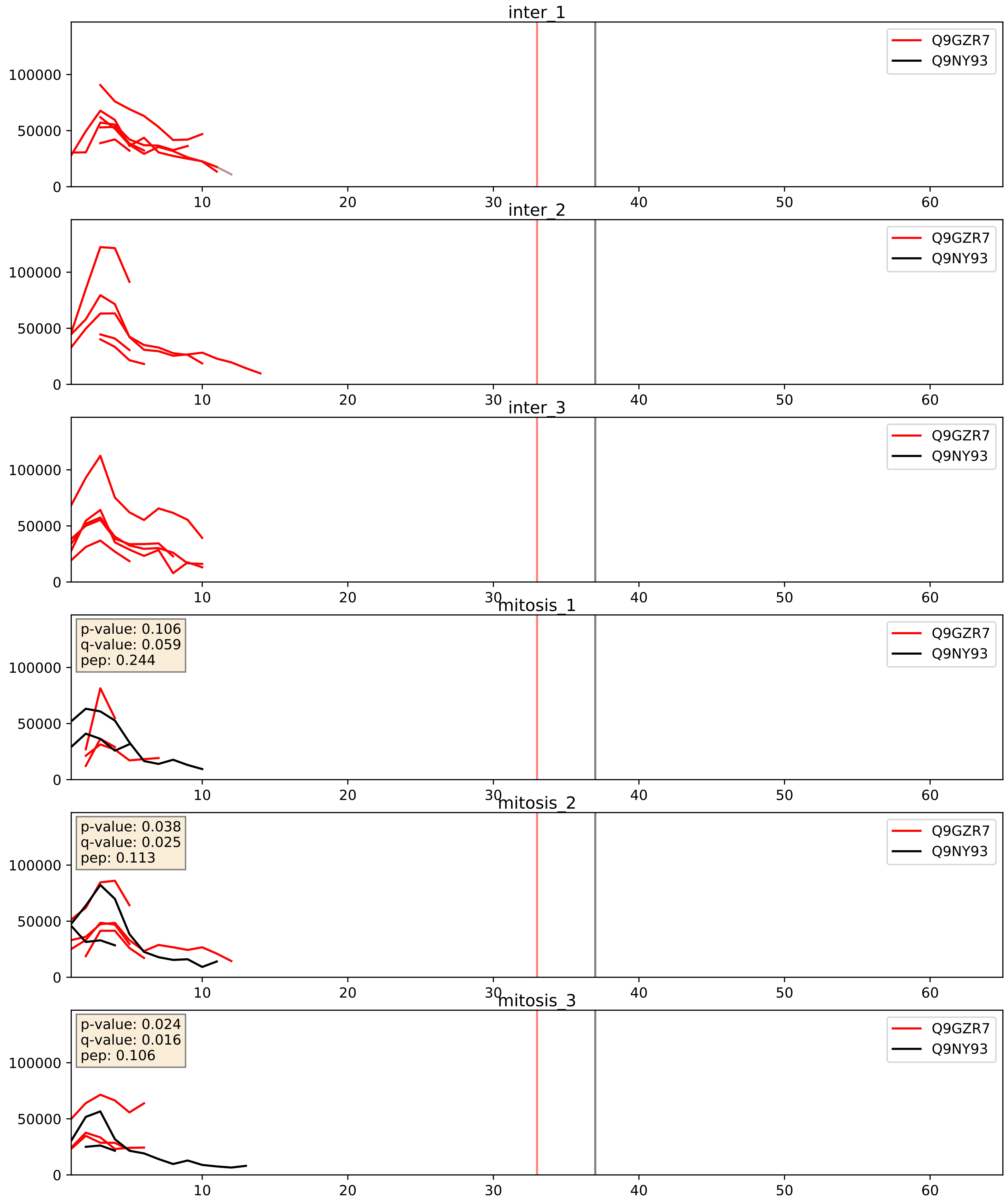


| level | condition_1 | condition_2 | pvalue | pvalue_adjusted |
|---|---|---|---|---|
| interactor_abundance | mitosis | inter | 0.04944767941813647 | 0.1776215425175192 |
| complex_abundance | mitosis | inter | 0.05943251262016525 | 0.1744639470382664 |
| interactor_ratio | mitosis | inter | 0.428561697848441 | 0.8432094269126428 |
