## Supplementary Data 2 for "SECAT: Quantifying differential protein-protein interaction states by network-centric analysis": hela_string_000037_Q9NY93.pdf

Q9NY93_Q9NY93

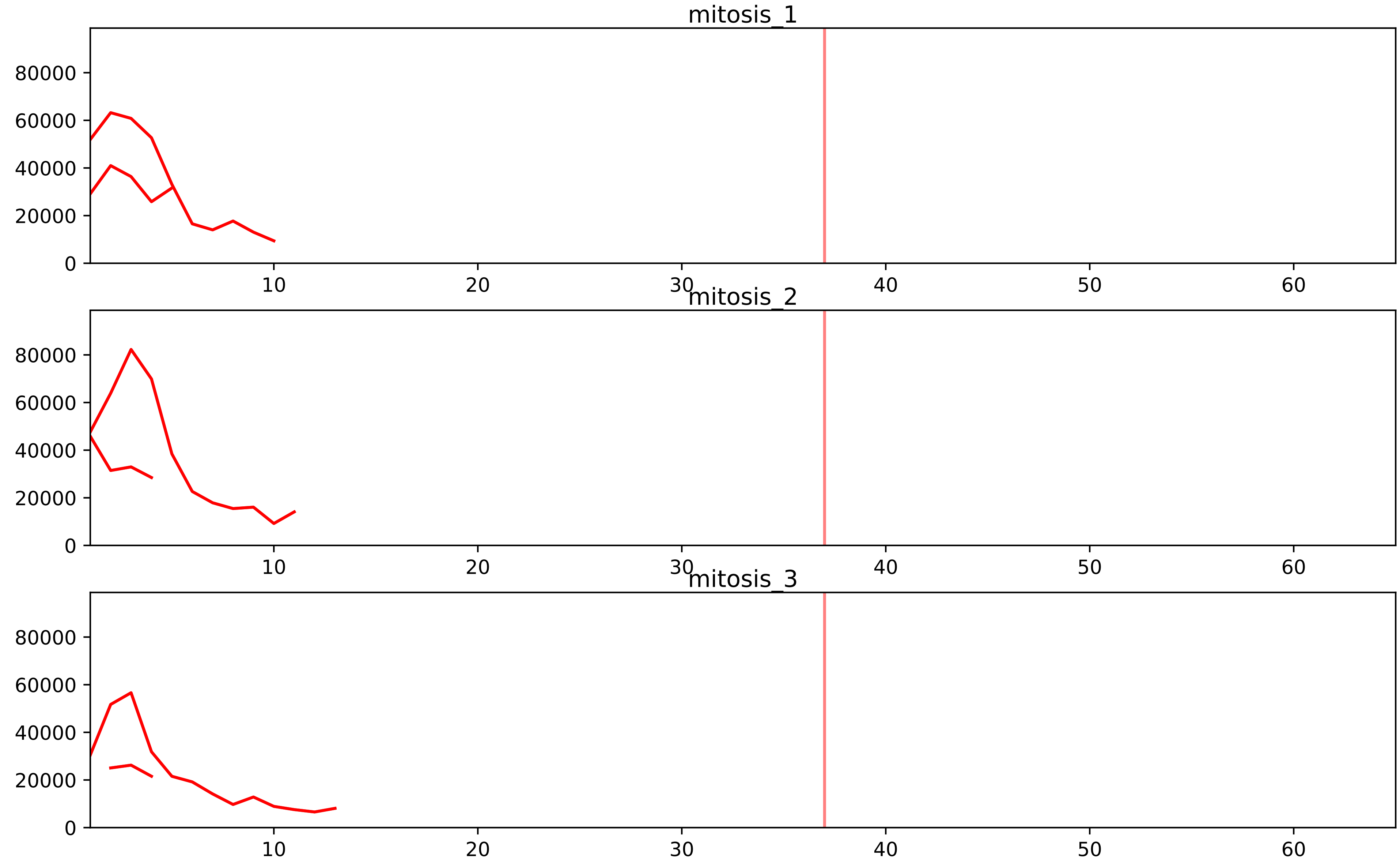


| level | condition_1 | condition_2 | pvalue | pvalue_adjusted |
|---|---|---|---|---|
| interactor_ratio | mitosis | inter | 1.329928828057854e-08 | 6.991625838932718e-07 |
| complex_abundance | mitosis | inter | 0.0007432445024254464 | 0.008938365257926937 |
| assembled_abundance | mitosis | inter | 0.0017716545054505001 | 0.039859774669462465 |
| total_abundance | mitosis | inter | 0.006877617881146651 | 0.09753190148445015 |
| interactor_abundance | mitosis | inter | 0.06748323189580736 | 0.17446085174586026 |

O00567_Q9NY93
NOP56_HUMAN vs DDX56_HUMAN
p-value: 0.032 q-value: 0.023

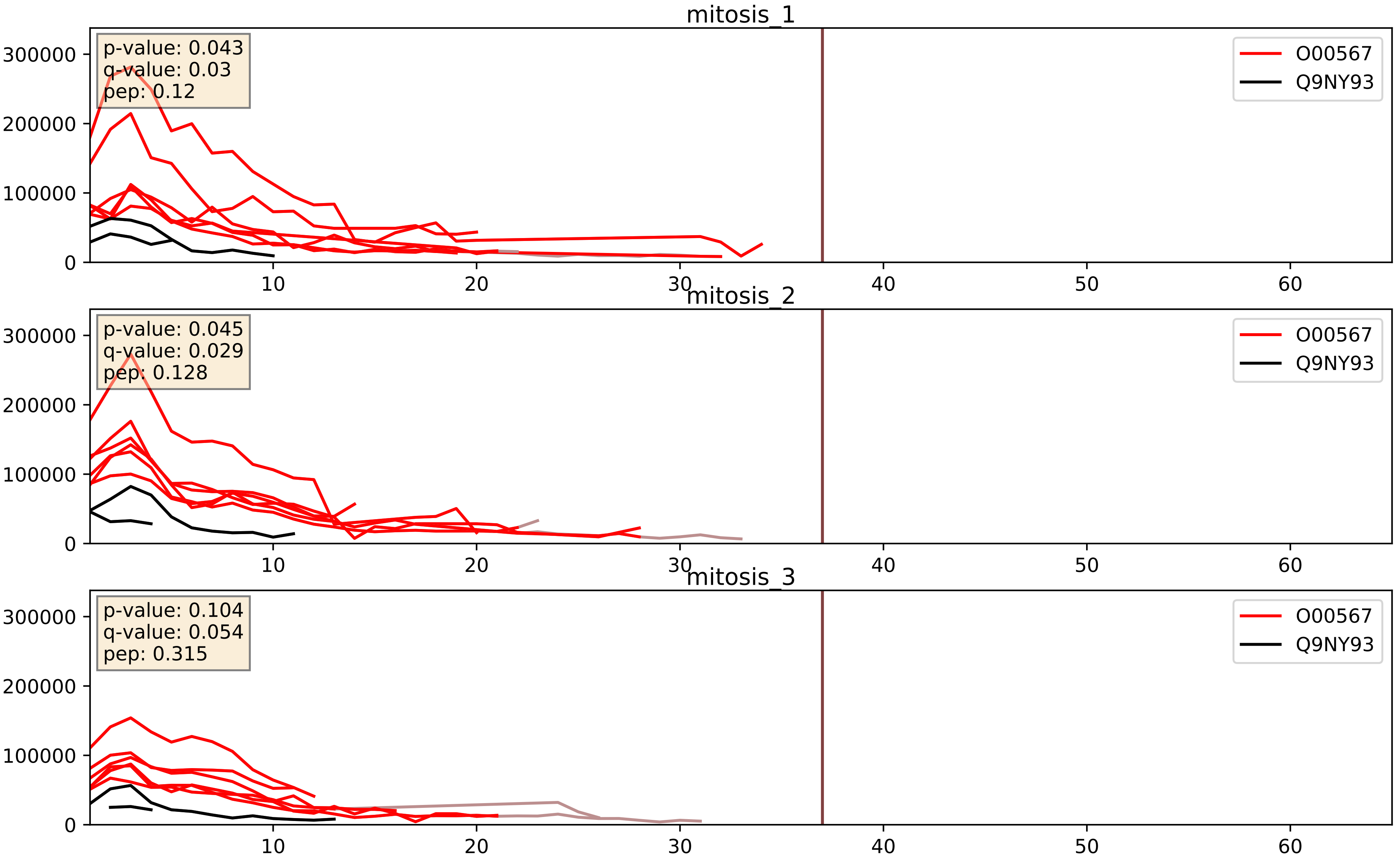


| level | condition_1 | condition_2 | pvalue | pvalue_adjusted |
|---|---|---|---|---|
| interactor_ratio | mitosis | inter | 2.3383393070343584e-05 | 0.004993481478971454 |
| interactor_abundance | mitosis | inter | 0.0005414629468052343 | 0.017071538948997444 |
| complex_abundance | mitosis | inter | 0.0026325520572434646 | 0.03679993741510981 |

P46087_Q9NY93
NOP2_HUMAN vs DDX56_HUMAN
p-value: 0.046 q-value: 0.031

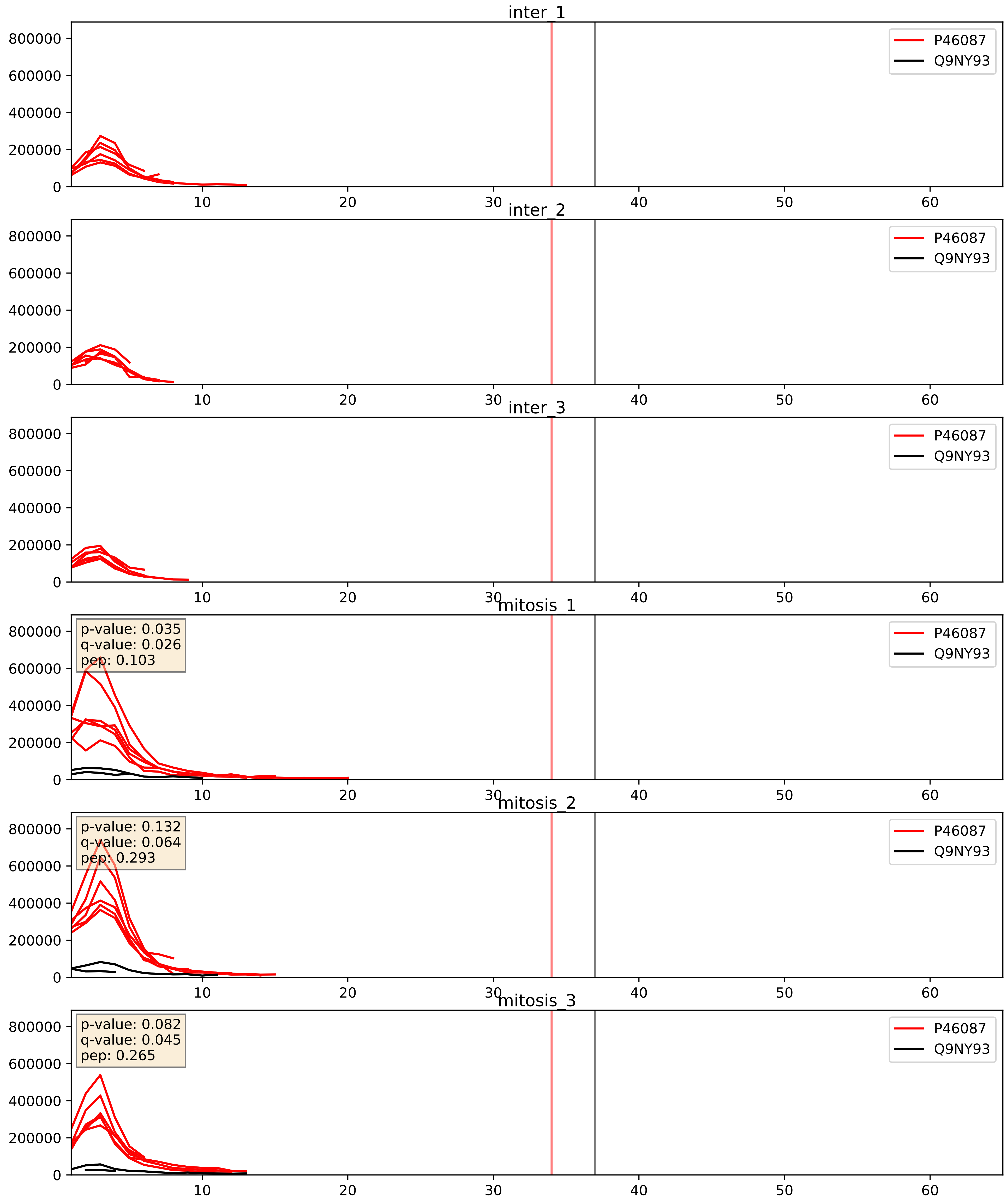


| level | condition_1 | condition_2 | pvalue | pvalue_adjusted |
|---|---|---|---|---|
| interactor_ratio | mitosis | inter | 2.6497983805864818e-05 | 0.005274947473911693 |
| interactor_abundance | mitosis | inter | 0.00012951041339078335 | 0.008661008895508637 |
| complex_abundance | mitosis | inter | 0.0006897093281200277 | 0.016916652861625893 |

Q03701_Q9NY93
CEBPZ_HUMAN vs DDX56_HUMAN
p-value: 0.011 q-value: 0.01

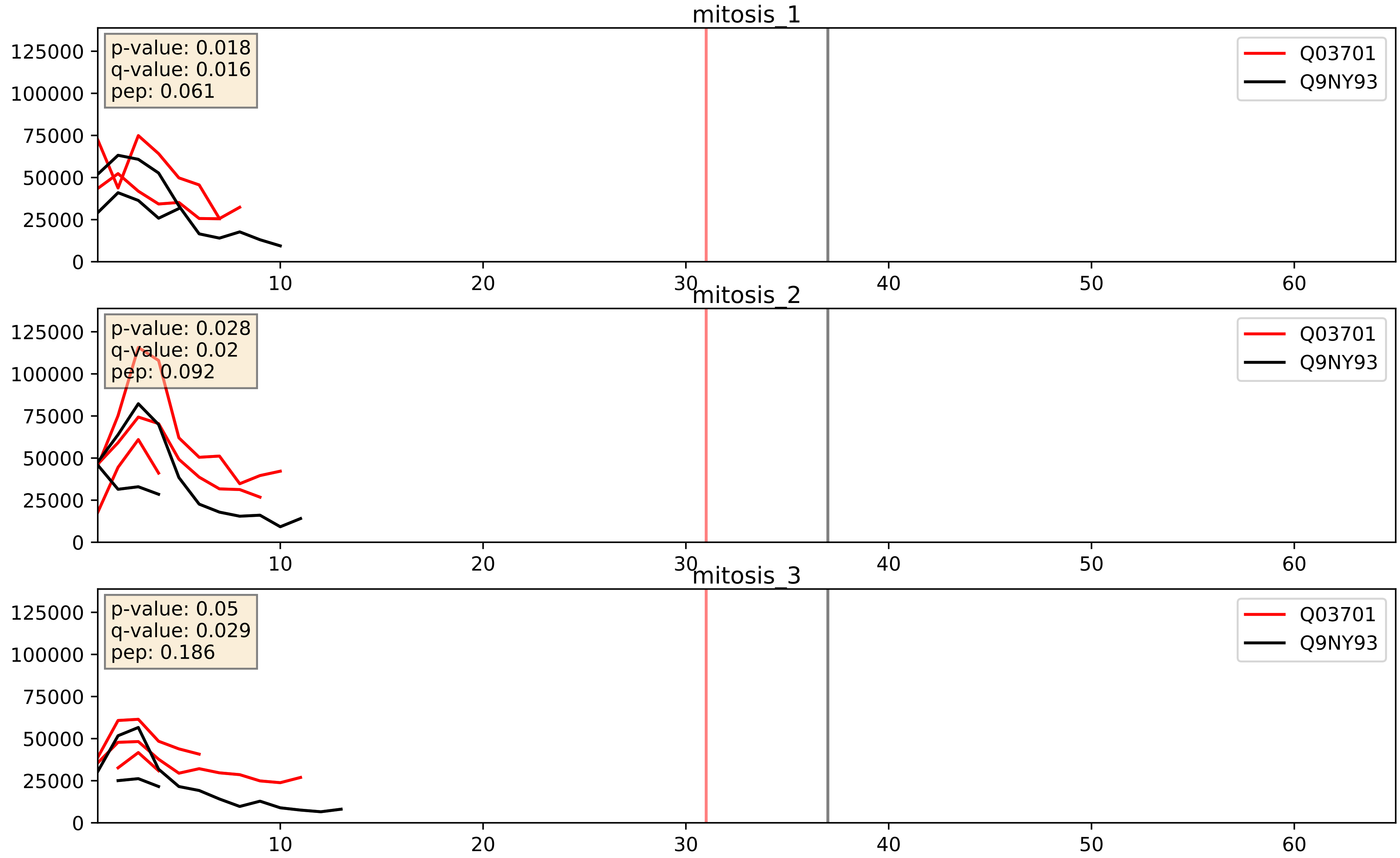


| level | condition_1 | condition_2 | pvalue | pvalue_adjusted |
|---|---|---|---|---|
| interactor_abundance | mitosis | inter | 0.019842839707373484 | 0.1190514126751731 |
| complex_abundance | mitosis | inter | 0.03402980185974759 | 0.1342992641398983 |
| interactor_ratio | mitosis | inter | 0.10349888889113636 | 0.5362896421962029 |

Q15050_Q9NY93
RRS1_HUMAN vs DDX56_HUMAN
p-value: 0.031 q-value: 0.023

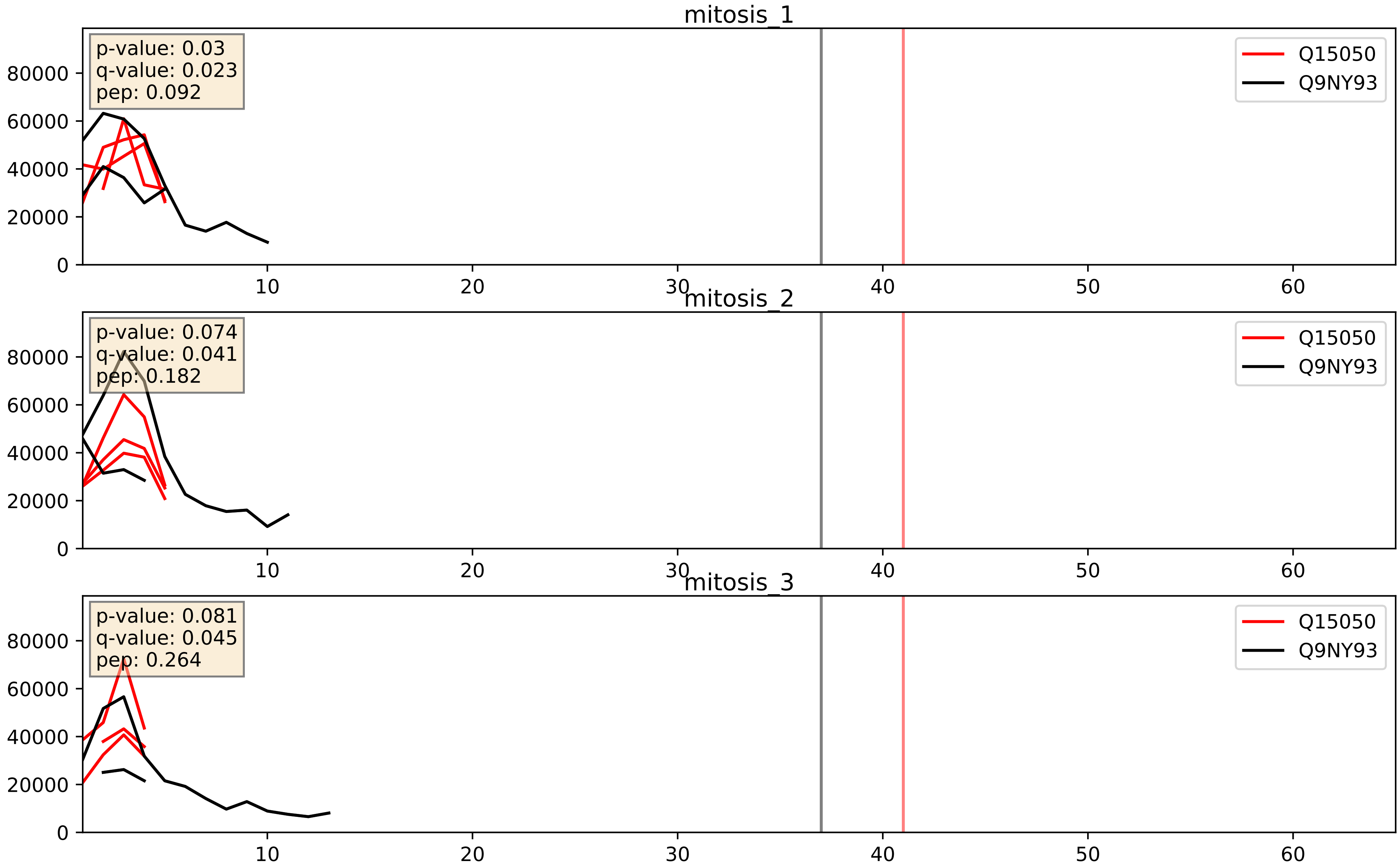


| level | condition_1 | condition_2 | pvalue | pvalue_adjusted |
|---|---|---|---|---|
| complex_abundance | mitosis | inter | 0.2143602303989282 | 0.36911920488133304 |
| interactor_abundance | mitosis | inter | 0.2162538244564617 | 0.4119120465837366 |
| interactor_ratio | mitosis | inter | 0.27768564921583294 | 0.7493660647186412 |

Q15397_Q9NY93
PUM3_HUMAN vs DDX56_HUMAN
p-value: 0.02 q-value: 0.016

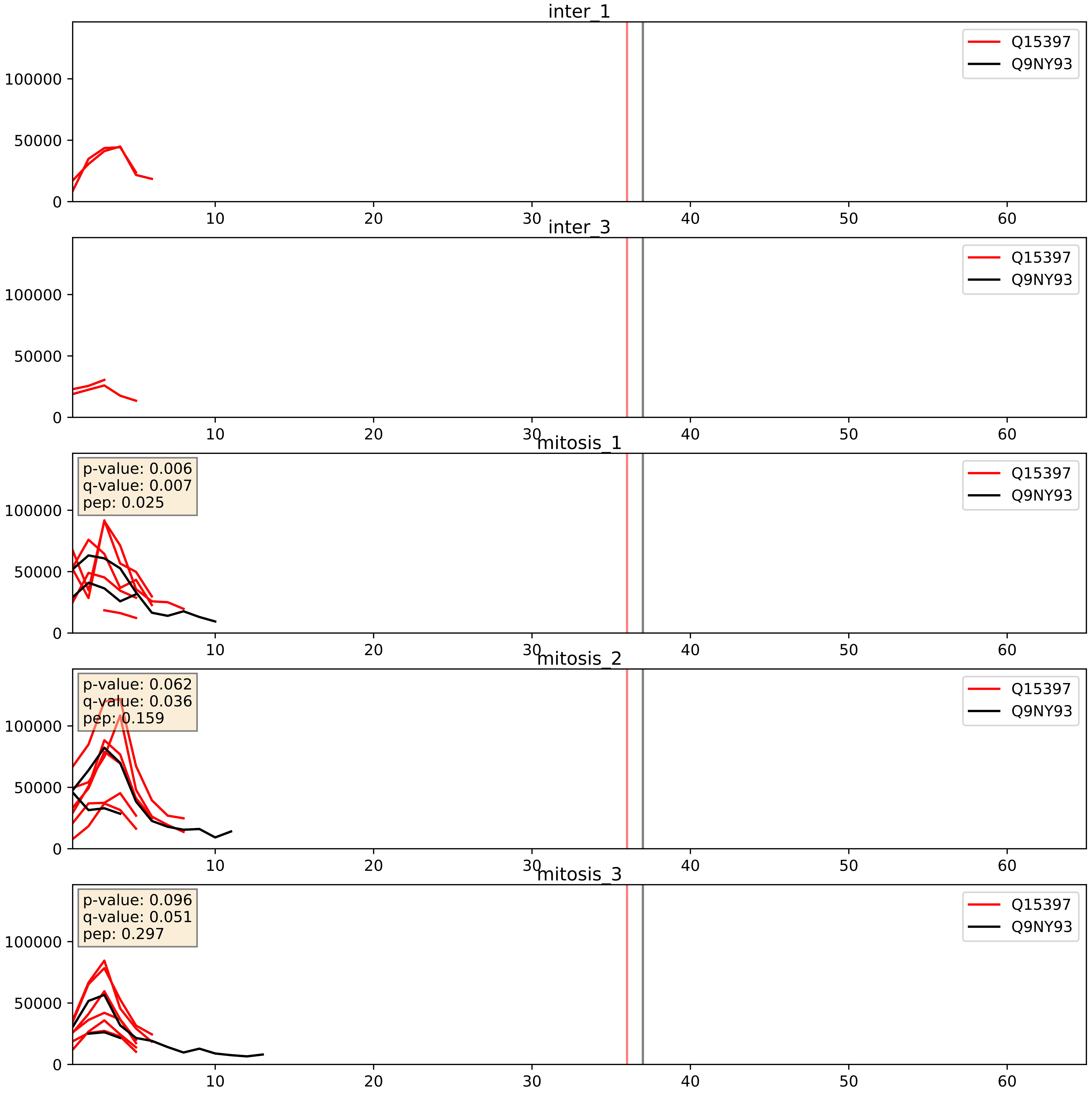


| level | condition_1 | condition_2 | pvalue | pvalue_adjusted |
|---|---|---|---|---|
| interactor_ratio | mitosis | inter | 0.0035194169325132555 | 0.07646419083591573 |
| interactor_abundance | mitosis | inter | 0.027567912416214223 | 0.13802761546789935 |
| complex_abundance | mitosis | inter | 0.05413600038038046 | 0.16599932138049858 |

Q8NI36_Q9NY93
WDR36_HUMAN vs DDX56_HUMAN
p-value: 0.062 q-value: 0.039

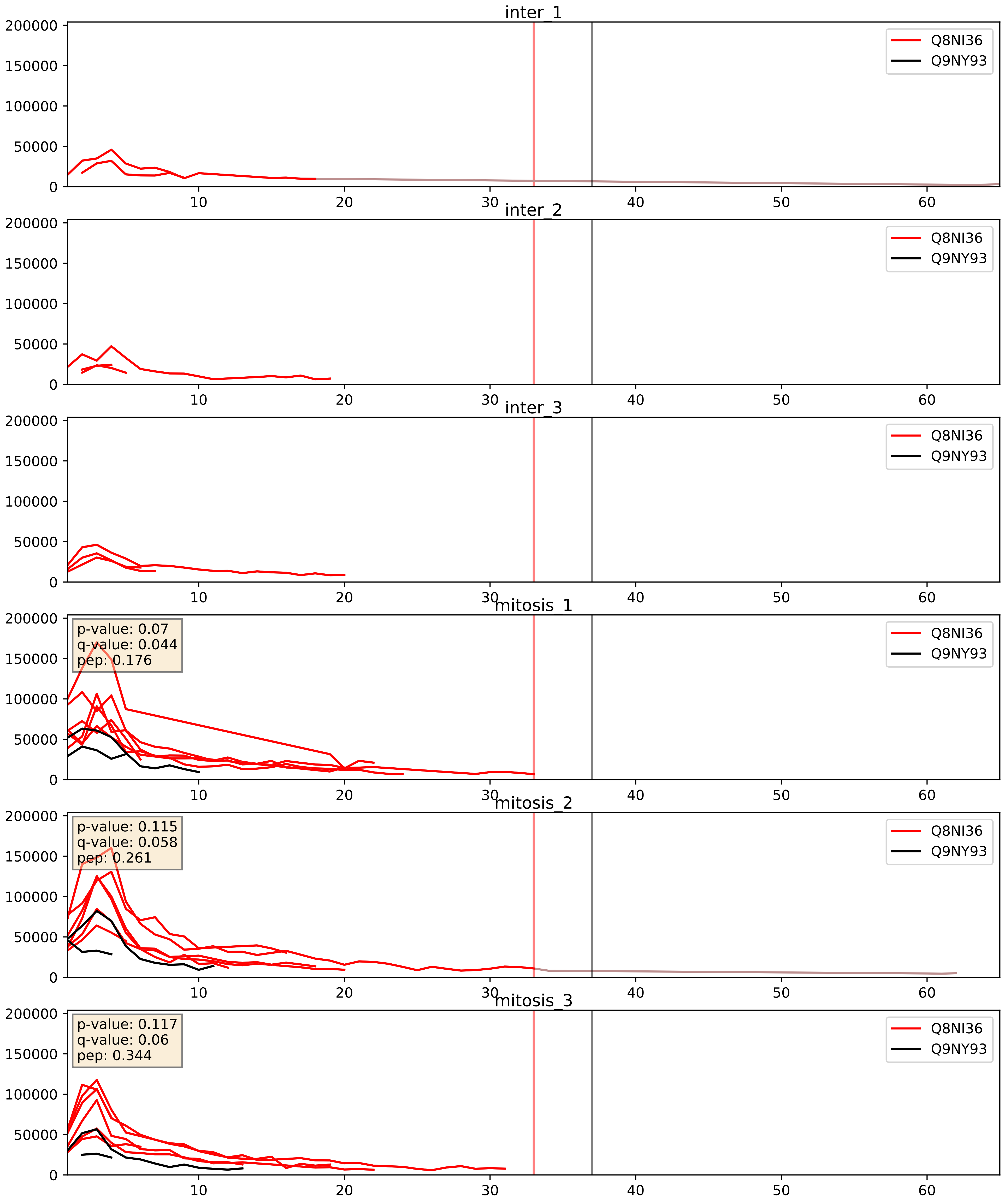


| level | condition_1 | condition_2 | pvalue | pvalue_adjusted |
|---|---|---|---|---|
| interactor_abundance | mitosis | inter | 0.000882265543690579 | 0.021954049575556266 |
| interactor_ratio | mitosis | inter | 0.0011153489338967212 | 0.03908562924515634 |
| complex_abundance | mitosis | inter | 0.004615842365066132 | 0.050629452773887605 |

Q8TDN6_Q9NY93
BRX1_HUMAN vs DDX56_HUMAN
p-value: 0.019 q-value: 0.015

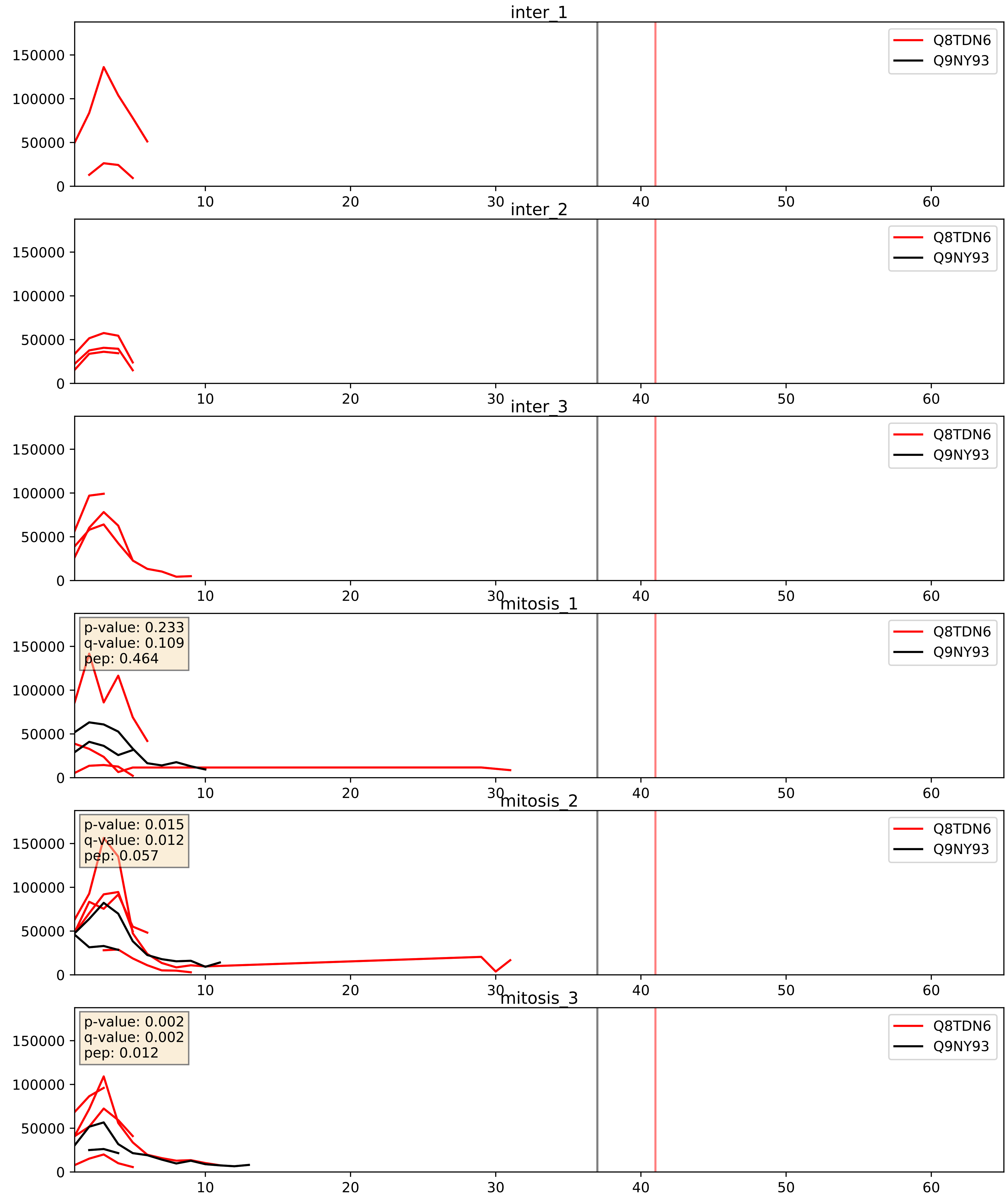


| level | condition_1 | condition_2 | pvalue | pvalue_adjusted |
|---|---|---|---|---|
| interactor_abundance | mitosis | inter | 0.013920743574672224 | 0.10466540623556807 |
| complex_abundance | mitosis | inter | 0.019947043532560785 | 0.10475257217099405 |
| interactor_ratio | mitosis | inter | 0.10974221992695789 | 0.5496158365704675 |

Q9BZE4_Q9NY93
NOG1_HUMAN vs DDX56_HUMAN
p-value: 0.051 q-value: 0.033

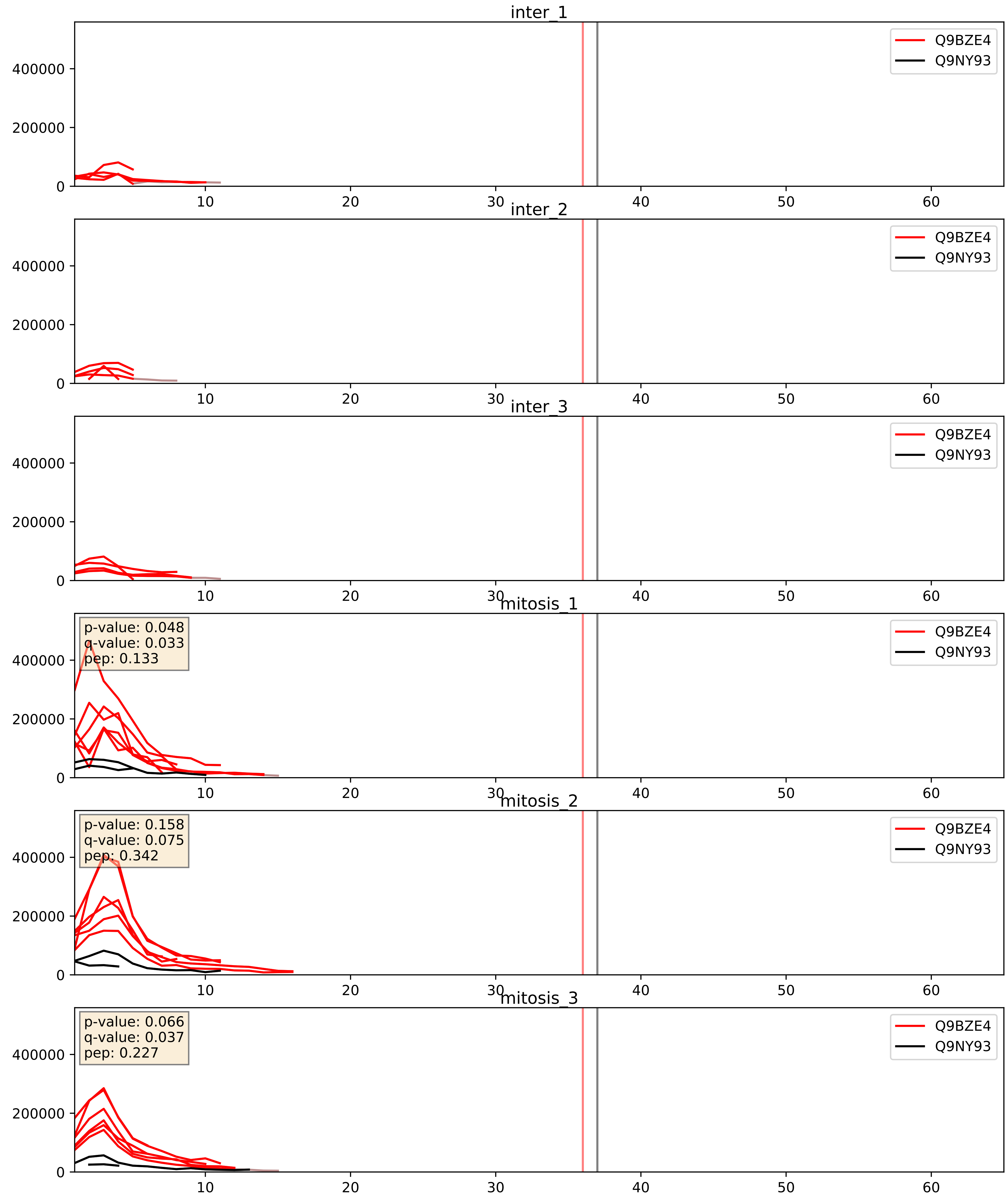


| level | condition_1 | condition_2 | pvalue | pvalue_adjusted |
|---|---|---|---|---|
| interactor_abundance | mitosis | inter | 0.0002829560238761811 | 0.012681170494136702 |
| interactor_ratio | mitosis | inter | 0.0002899670013636358 | 0.017729410940519448 |
| complex_abundance | mitosis | inter | 0.0011975320997738852 | 0.02329744266832831 |

Q9GZR7_Q9NY93
DDX24_HUMAN vs DDX56_HUMAN
p-value: 0.026 q-value: 0.02

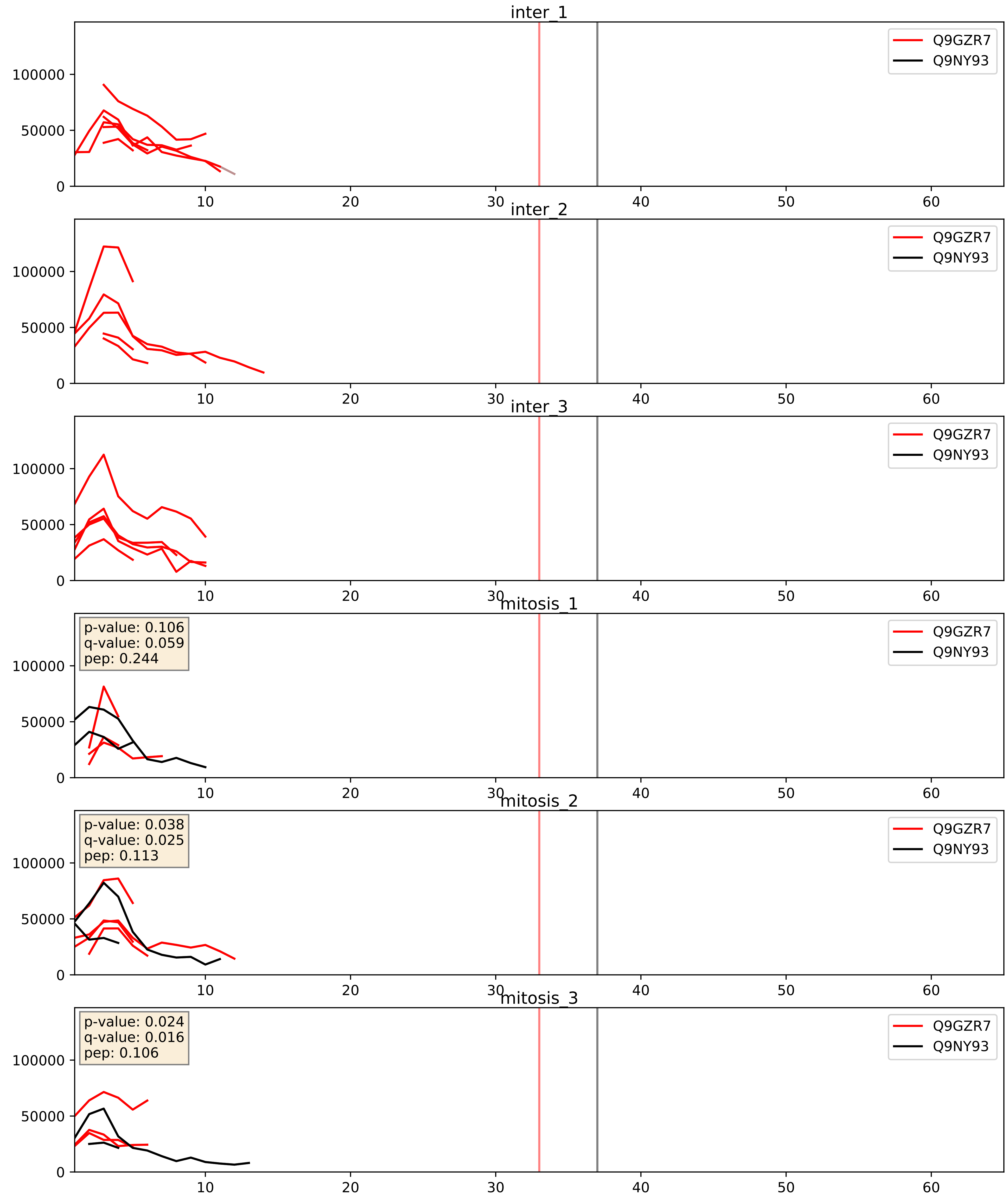


| level | condition_1 | condition_2 | pvalue | pvalue_adjusted |
|---|---|---|---|---|
| interactor_abundance | mitosis | inter | 0.04944767941813647 | 0.1776215425175192 |
| complex_abundance | mitosis | inter | 0.05943251262016525 | 0.1744639470382664 |
| interactor_ratio | mitosis | inter | 0.428561697848441 | 0.8432094269126428 |

Q9NVP1_Q9NY93
DDX18_HUMAN vs DDX56_HUMAN
p-value: 0.085 q-value: 0.05

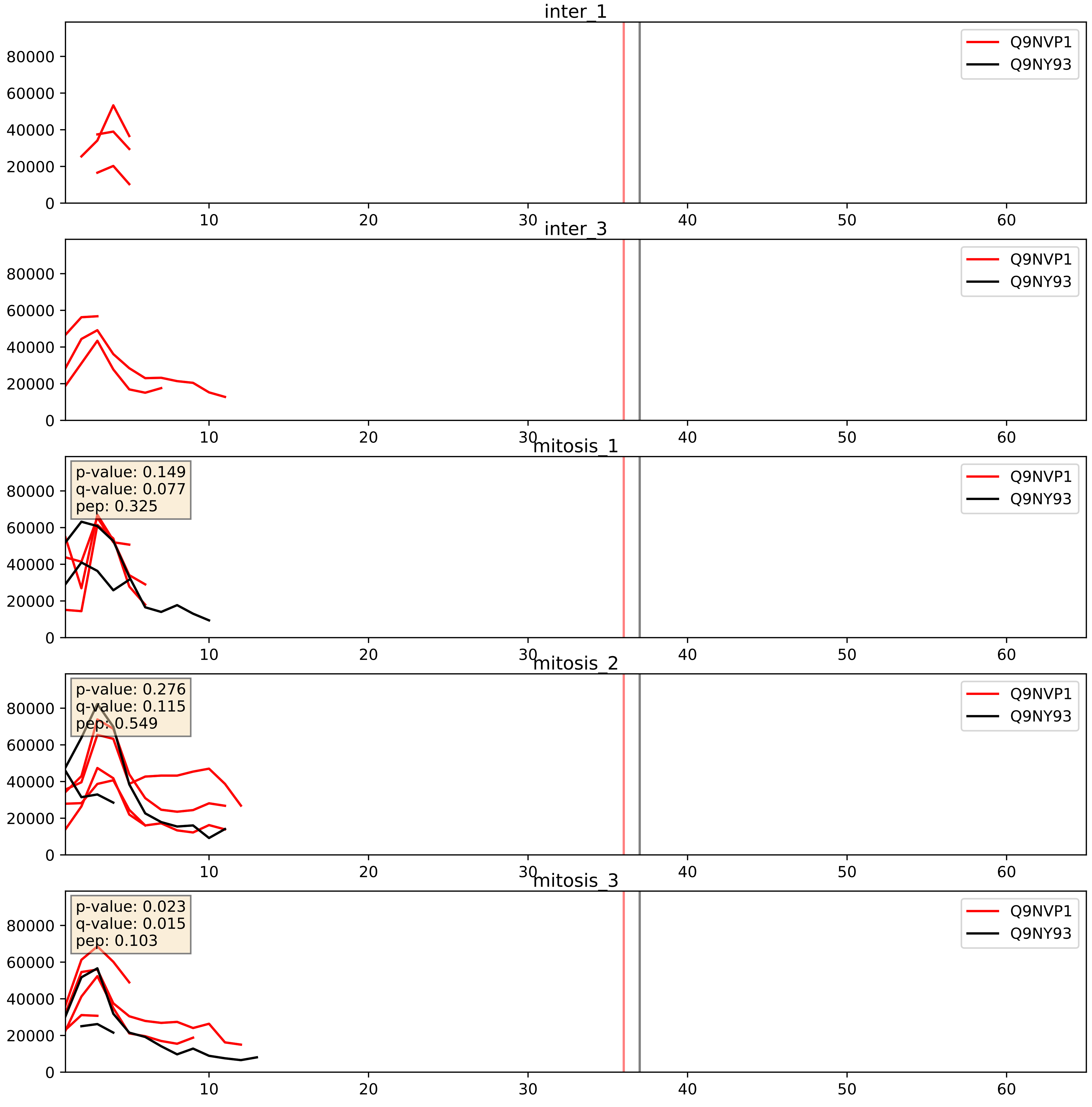


| level | condition_1 | condition_2 | pvalue | pvalue_adjusted |
|---|---|---|---|---|
| interactor_ratio | mitosis | inter | 0.003673014720945405 | 0.0787300451287945 |
| interactor_abundance | mitosis | inter | 0.010894066262585354 | 0.09160432928067842 |
| complex_abundance | mitosis | inter | 0.02682633767318529 | 0.11972879191111971 |

Q9NY93_Q9UKD2
DDX56_HUMAN vs MRT4_HUMAN
p-value: 0.018 q-value: 0.014

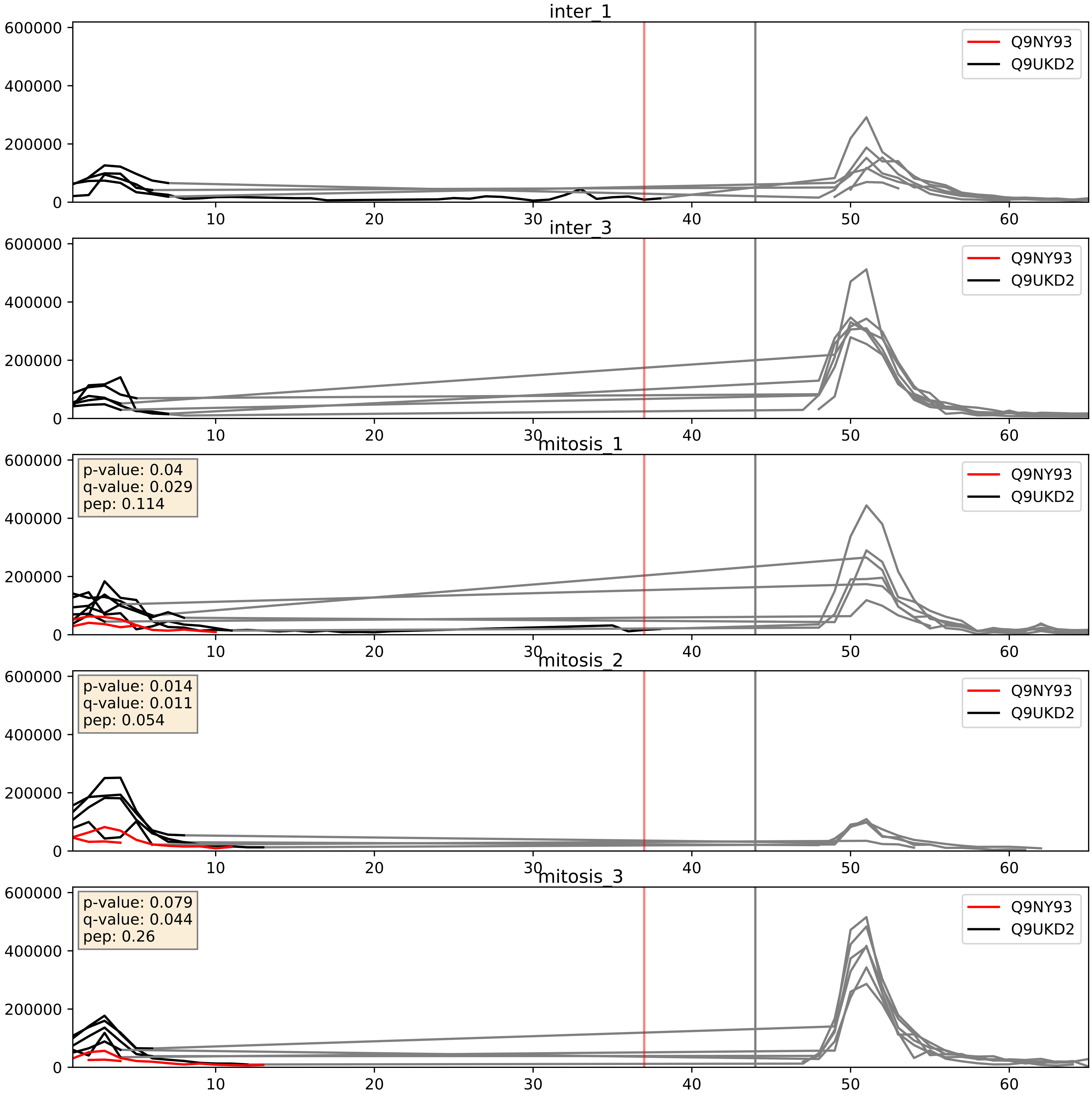


| level | condition_1 | condition_2 | pvalue | pvalue_adjusted |
|---|---|---|---|---|
| interactor_ratio | mitosis | inter | 0.0009603917975522726 | 0.03574327733498893 |
| complex_abundance | mitosis | inter | 0.003800492486110046 | 0.04558257253867179 |
| interactor_abundance | mitosis | inter | 0.10493275671361522 | 0.27679986972306914 |

Q9NY93_Q9Y2X3
DDX56_HUMAN vs NOP58_HUMAN
p-value: 0.021 q-value: 0.017

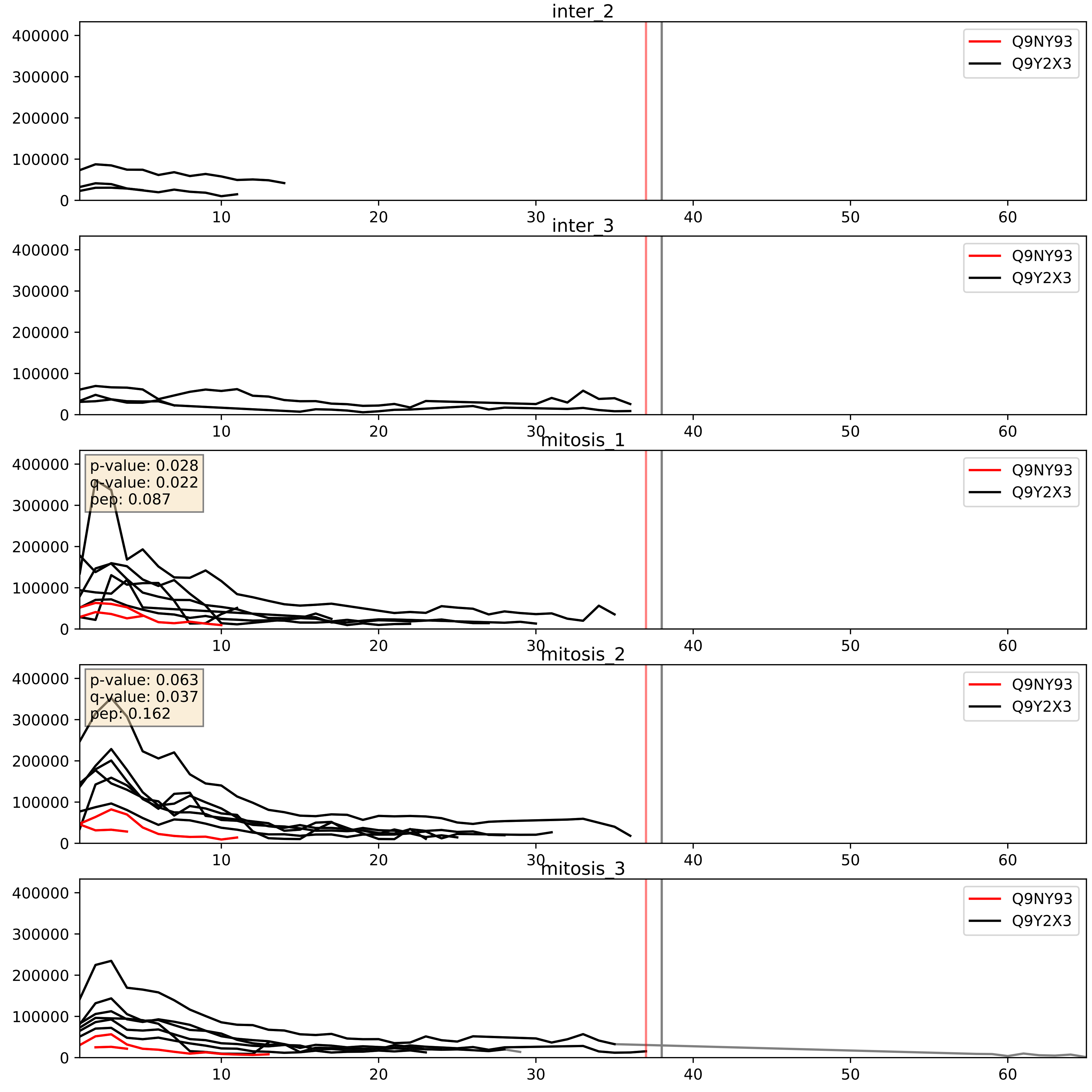


| level | condition_1 | condition_2 | pvalue | pvalue_adjusted |
|---|---|---|---|---|
| interactor_ratio | mitosis | inter | 0.00010063874199602321 | 0.010720128702277293 |
| complex_abundance | mitosis | inter | 0.0007327642214531893 | 0.017769013415408784 |
| interactor_abundance | mitosis | inter | 0.10159744920024559 | 0.27370850762629345 |
