## Supplementary Data 2 for "SECAT: Quantifying differential protein-protein interaction states by network-centric analysis": hela_string_000038_Q9H0S4.pdf

Q9H0S4_Q9H0S4

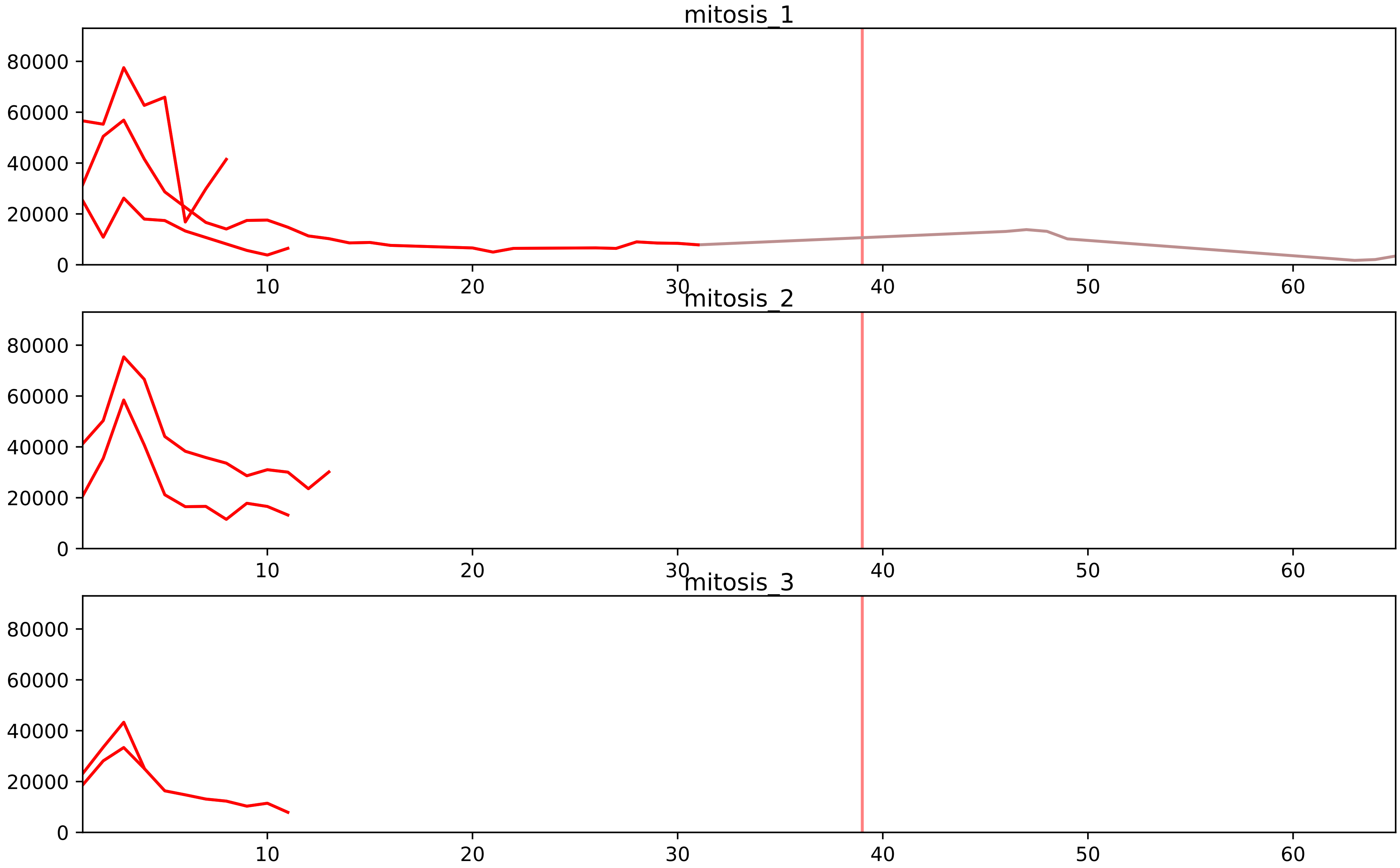


| level | condition_1 | condition_2 | pvalue | pvalue_adjusted |
|---|---|---|---|---|
| interactor_ratio | mitosis | inter | 1.5059582390526945e-08 | 7.69711988849155e-07 |
| complex_abundance | mitosis | inter | 0.00809167099222428 | 0.039077886156673694 |
| assembled_abundance | mitosis | inter | 0.03376955136076196 | 0.22091914204238025 |
| total_abundance | mitosis | inter | 0.08532322762517892 | 0.358776777060771 |
| monomer_abundance | mitosis | inter | 0.3942943740886353 | 0.6908227865123439 |
| interactor_abundance | mitosis | inter | 0.4746685095281915 | 0.6455211068232612 |

O00567_Q9H0S4
NOP56_HUMAN vs DDX47_HUMAN
p-value: 0.01 q-value: 0.009

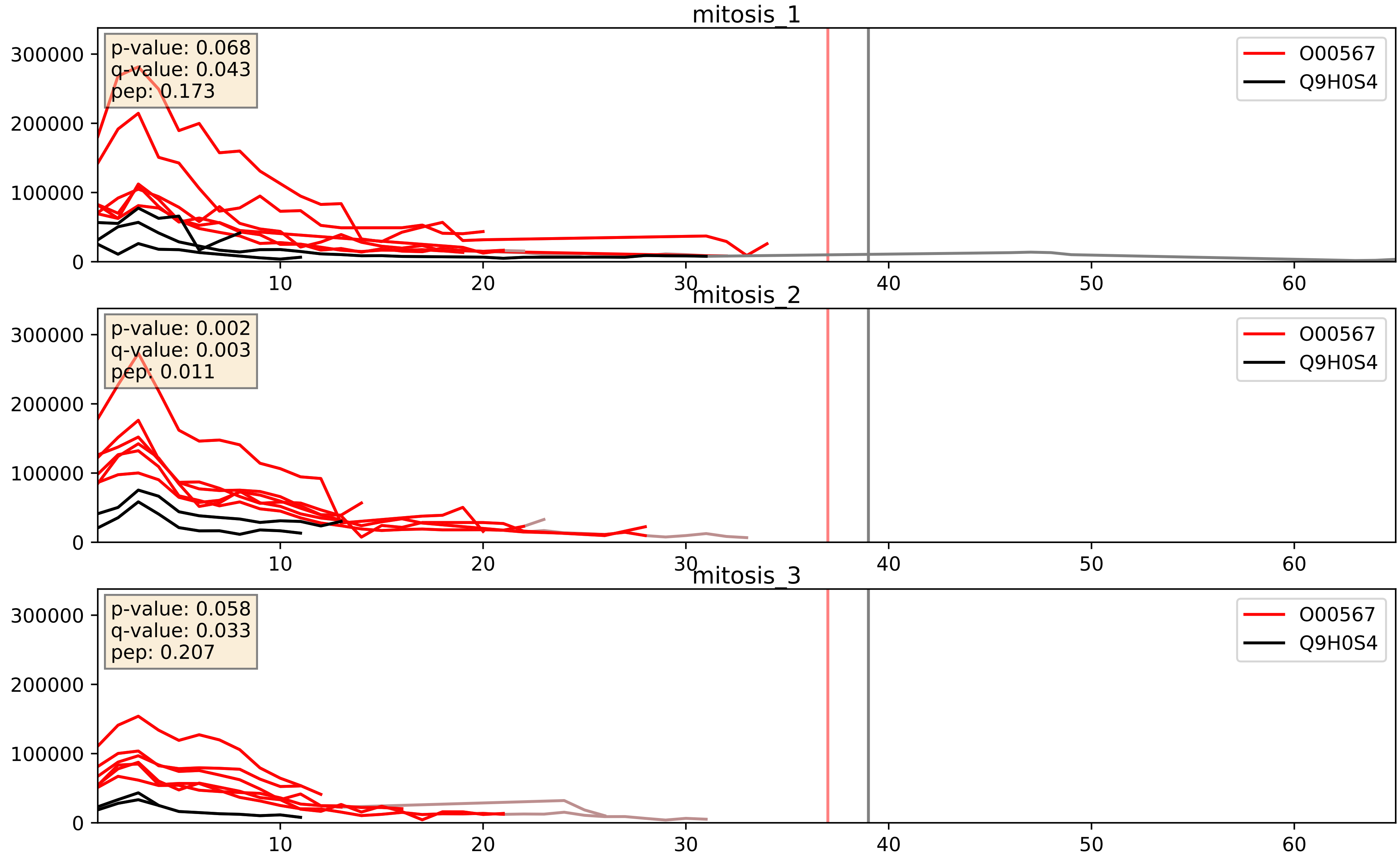


| level | condition_1 | condition_2 | pvalue | pvalue_adjusted |
|---|---|---|---|---|
| interactor_ratio | mitosis | inter | 0.0001712185653981663 | 0.013577643810159747 |
| interactor_abundance | mitosis | inter | 0.0014024757199137346 | 0.02753913185123736 |
| complex_abundance | mitosis | inter | 0.01641041278178478 | 0.09575537383236381 |

P22087_Q9H0S4
FBRL_HUMAN vs DDX47_HUMAN
p-value: 0.003 q-value: 0.042

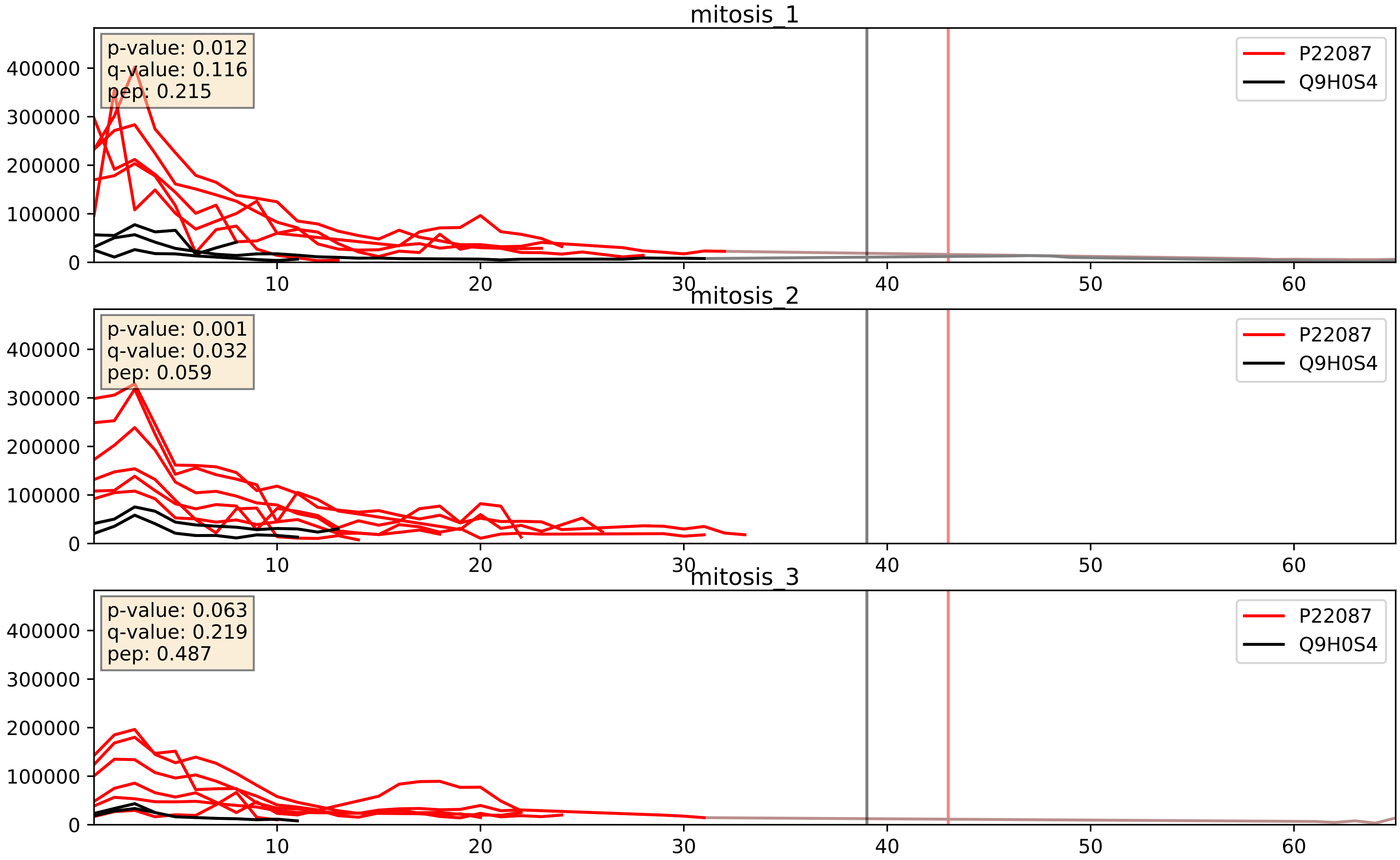


| level | condition_1 | condition_2 | pvalue | pvalue_adjusted |
|---|---|---|---|---|
| interactor_ratio | mitosis | inter | 6.650160997708512e-05 | 0.008371379138291892 |
| interactor_abundance | mitosis | inter | 0.0010898797318194015 | 0.024049720606162197 |
| complex_abundance | mitosis | inter | 0.011229383006423614 | 0.0785322863847926 |

Q12788_Q9H0S4
TBL3_HUMAN vs DDX47_HUMAN
p-value: 0.041 q-value: 0.028

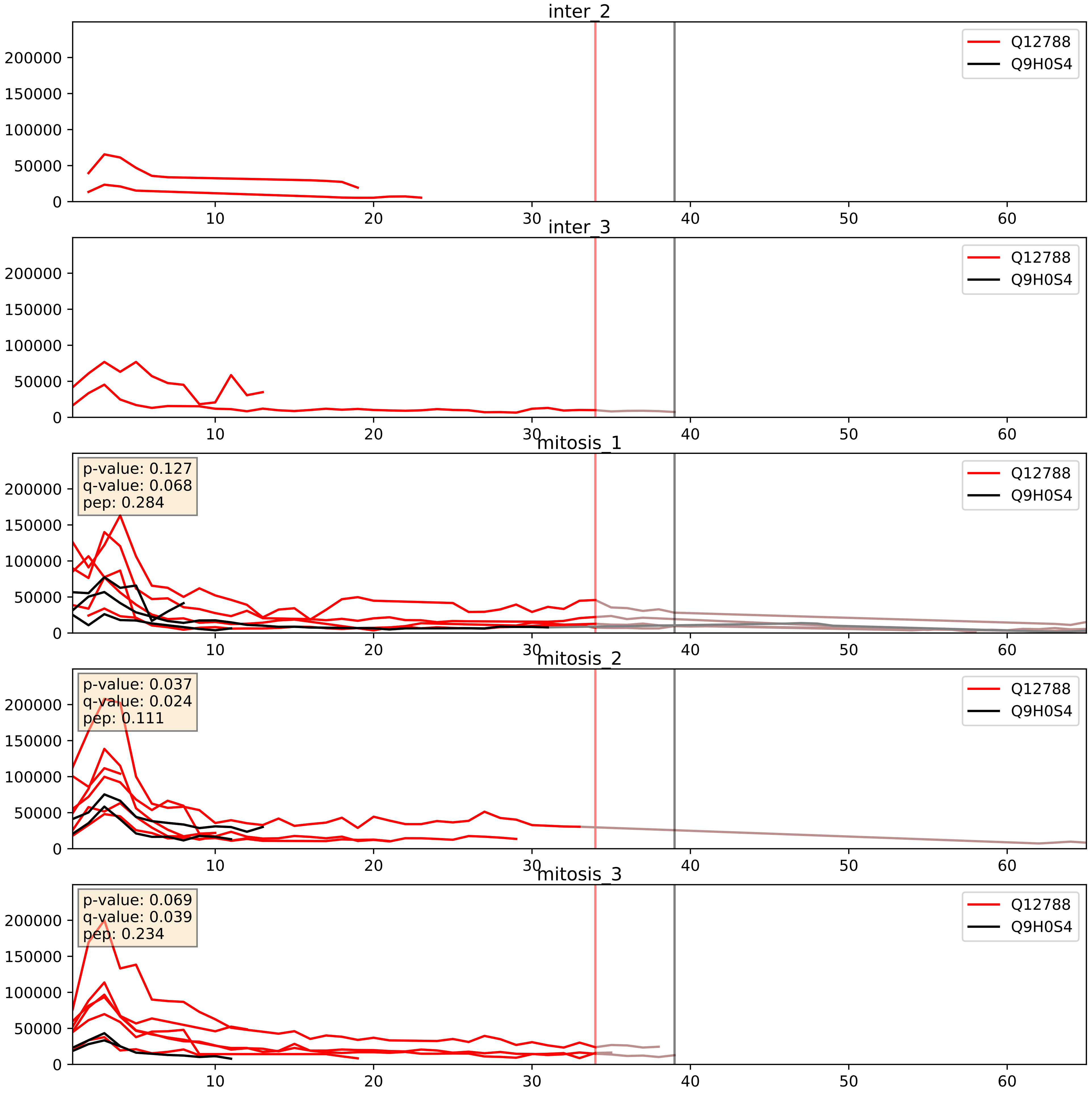


| level | condition_1 | condition_2 | pvalue | pvalue_adjusted |
|---|---|---|---|---|
| interactor_abundance | mitosis | inter | 0.0015075846392158566 | 0.02867761002597274 |
| interactor_ratio | mitosis | inter | 0.01308259178830462 | 0.16493001984594508 |
| complex_abundance | mitosis | inter | 0.02359161739620337 | 0.11181851877713225 |

Q8IY81_Q9H0S4
SPB1_HUMAN vs DDX47_HUMAN
p-value: 0.065 q-value: 0.04

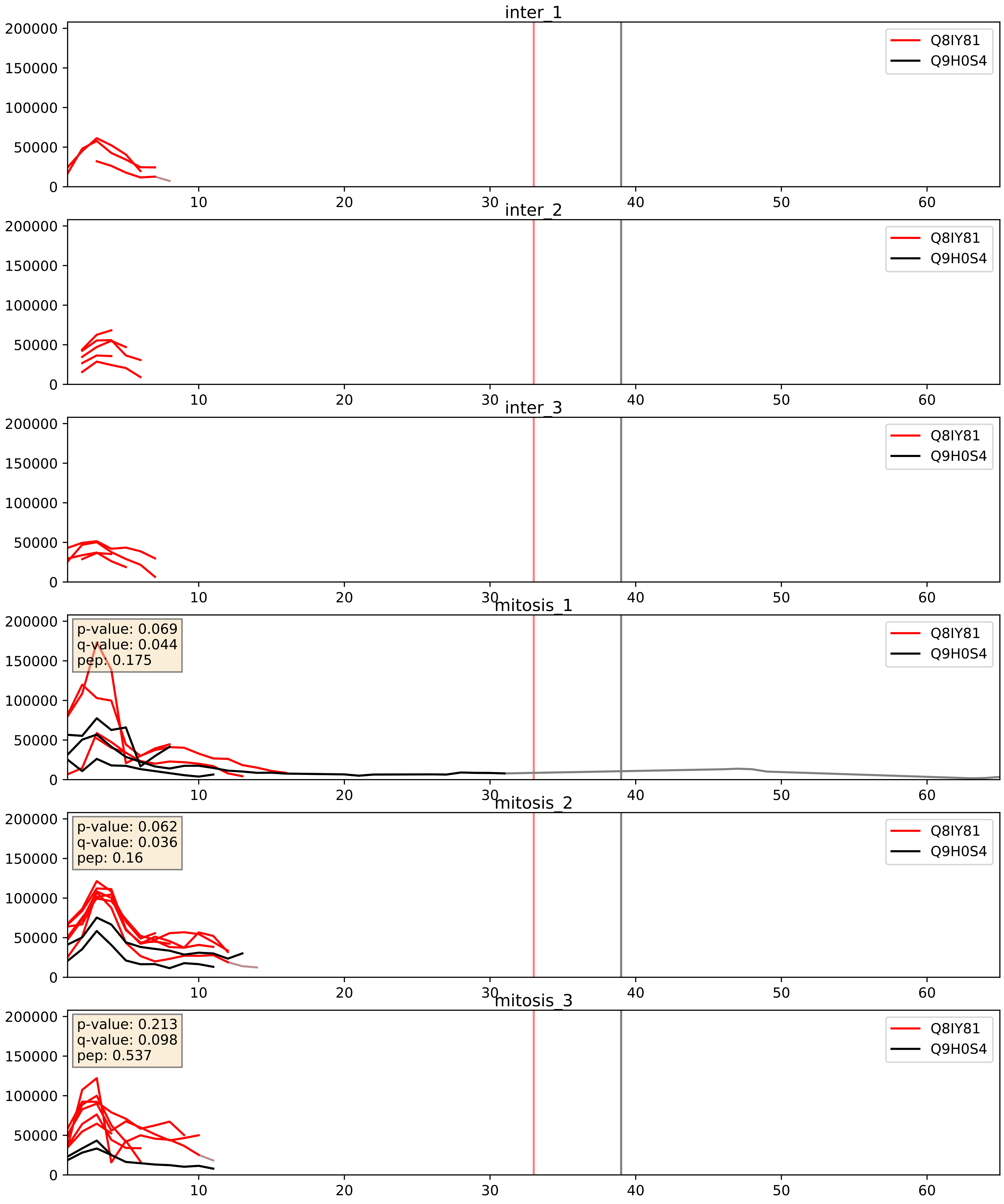


| level | condition_1 | condition_2 | pvalue | pvalue_adjusted |
|---|---|---|---|---|
| interactor_abundance | mitosis | inter | 0.0056588368861371875 | 0.06348577161904892 |
| complex_abundance | mitosis | inter | 0.03774864580132628 | 0.14178517247009784 |
| interactor_ratio | mitosis | inter | 0.10205027280289247 | 0.5320038582172714 |

Q8NI36_Q9H0S4
WDR36_HUMAN vs DDX47_HUMAN
p-value: 0.004 q-value: 0.048

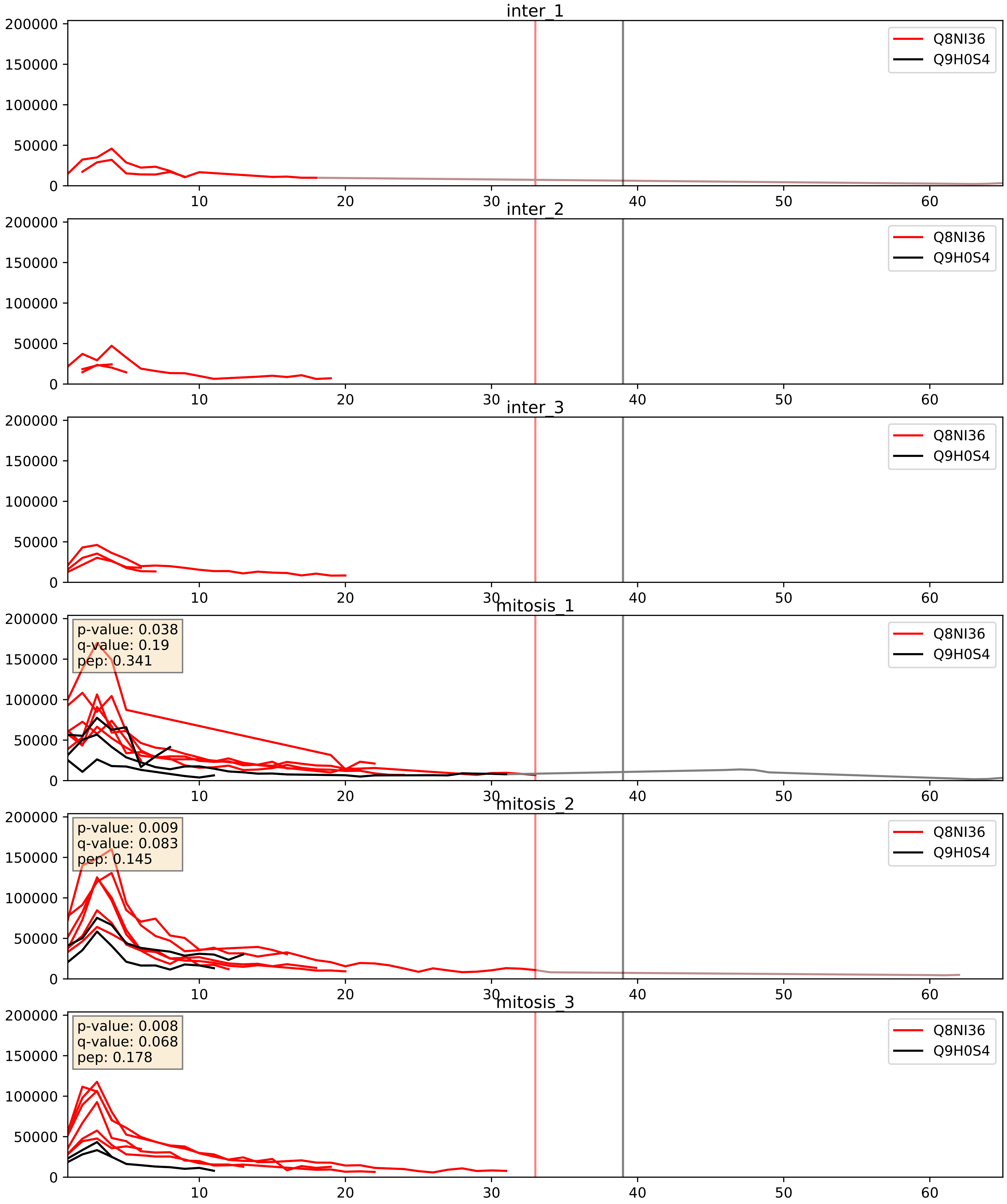


| level | condition_1 | condition_2 | pvalue | pvalue_adjusted |
|---|---|---|---|---|
| interactor_abundance | mitosis | inter | 0.001838446528961071 | 0.0315372791340817 |
| interactor_ratio | mitosis | inter | 0.009989406050391692 | 0.14263182888175546 |
| complex_abundance | mitosis | inter | 0.03375006447461892 | 0.13385180030241378 |

Q969X6_Q9H0S4
UTP4_HUMAN vs DDX47_HUMAN
p-value: 0.065 q-value: 0.04

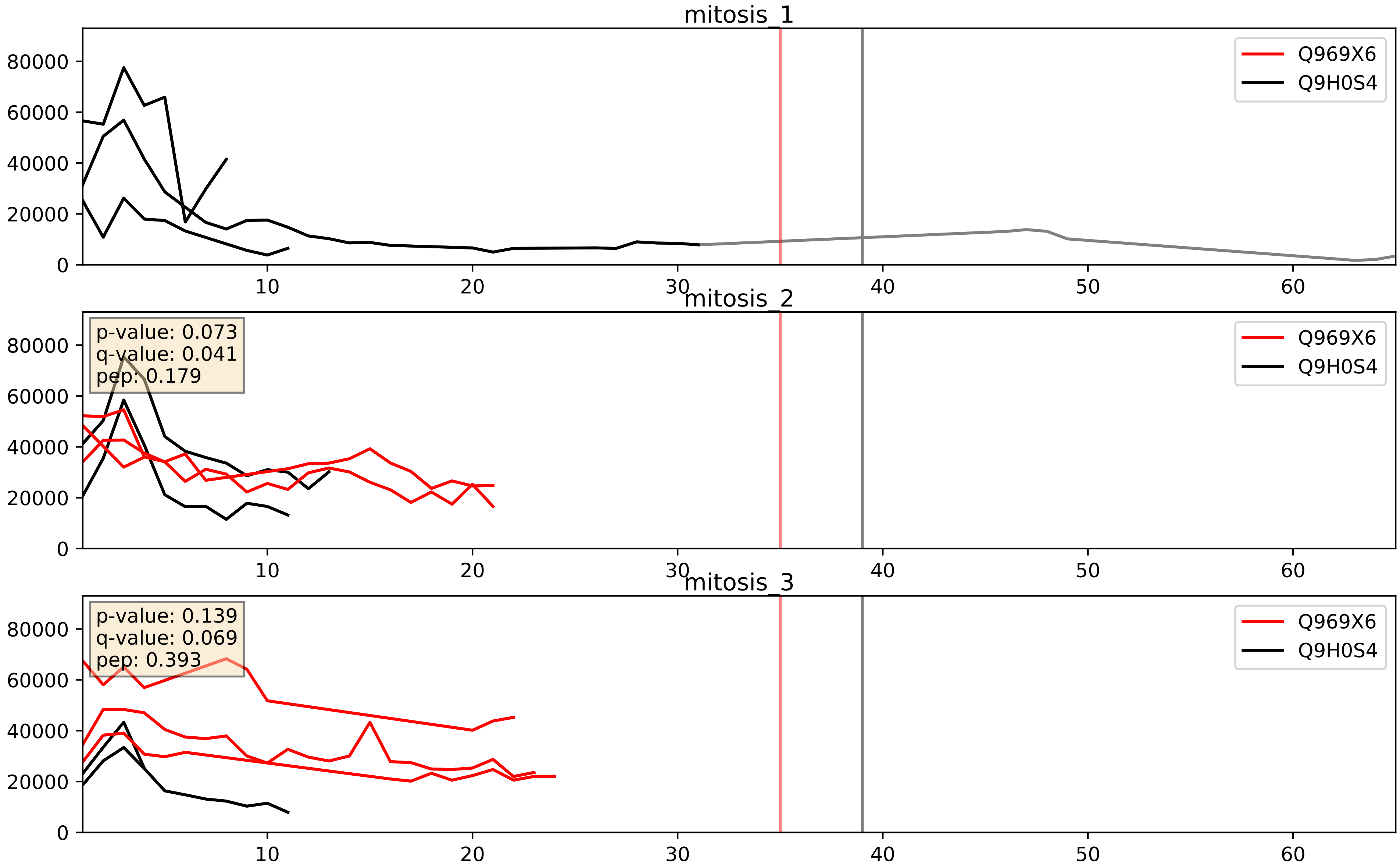


| level | condition_1 | condition_2 | pvalue | pvalue_adjusted |
|---|---|---|---|---|
| interactor_ratio | mitosis | inter | 0.18532727013006978 | 0.6555888709248134 |
| interactor_abundance | mitosis | inter | 0.3522158104571494 | 0.5503698117722354 |
| complex_abundance | mitosis | inter | 0.582834473306709 | 0.7111239938454823 |

Q9H0S4_Q9NW13
DDX47_HUMAN vs RBM28_HUMAN
p-value: 0.029 q-value: 0.021

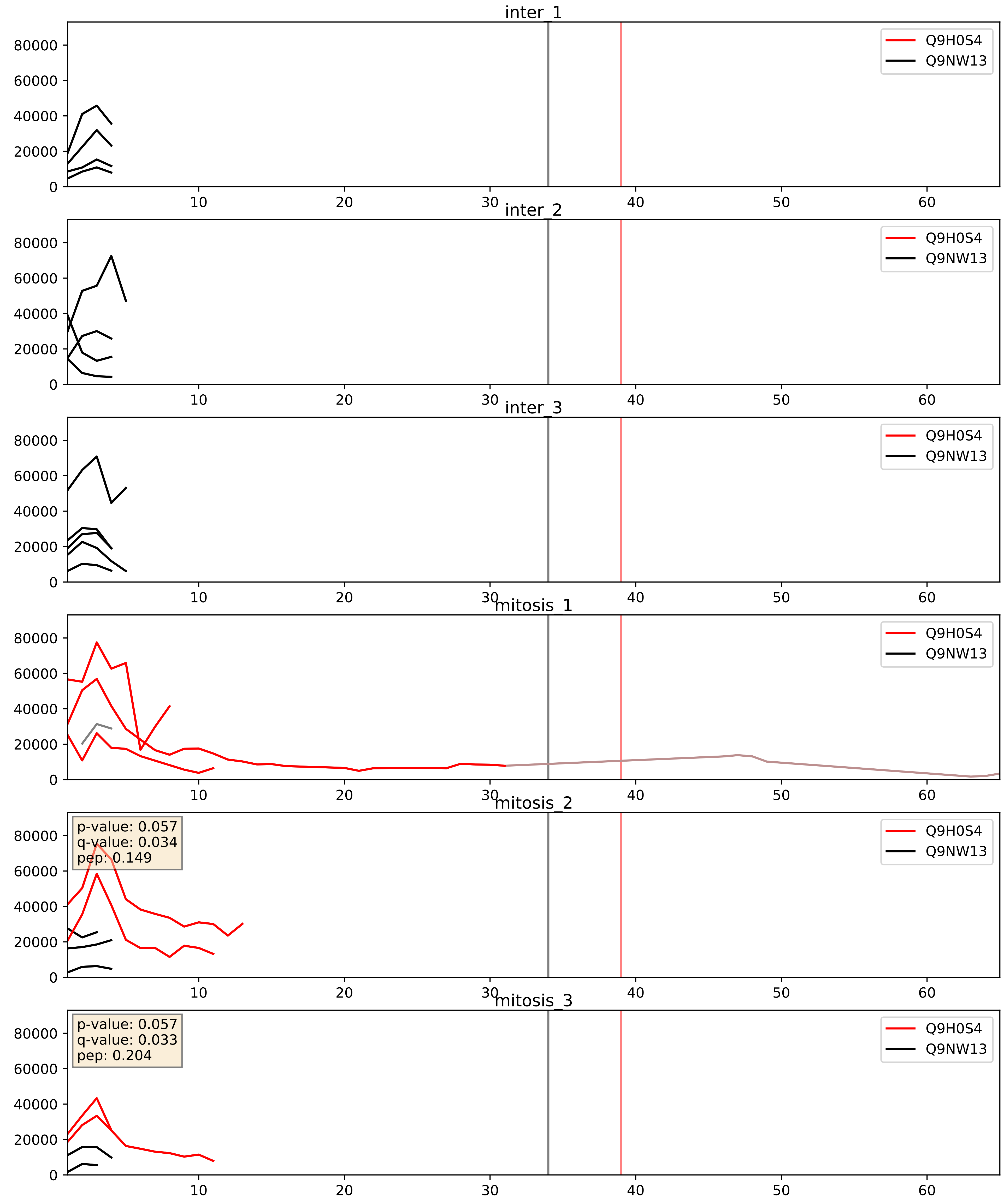


| level | condition_1 | condition_2 | pvalue | pvalue_adjusted |
|---|---|---|---|---|
| interactor_ratio | mitosis | inter | 0.18666469483853065 | 0.6572260773818791 |
| interactor_abundance | mitosis | inter | 0.2132781052453463 | 0.40861552381025035 |
| complex_abundance | mitosis | inter | 0.22958961967190683 | 0.38287300689490017 |

Q9H0S4_Q9Y221
DDX47_HUMAN vs NIP7_HUMAN
p-value: 0.044 q-value: 0.029

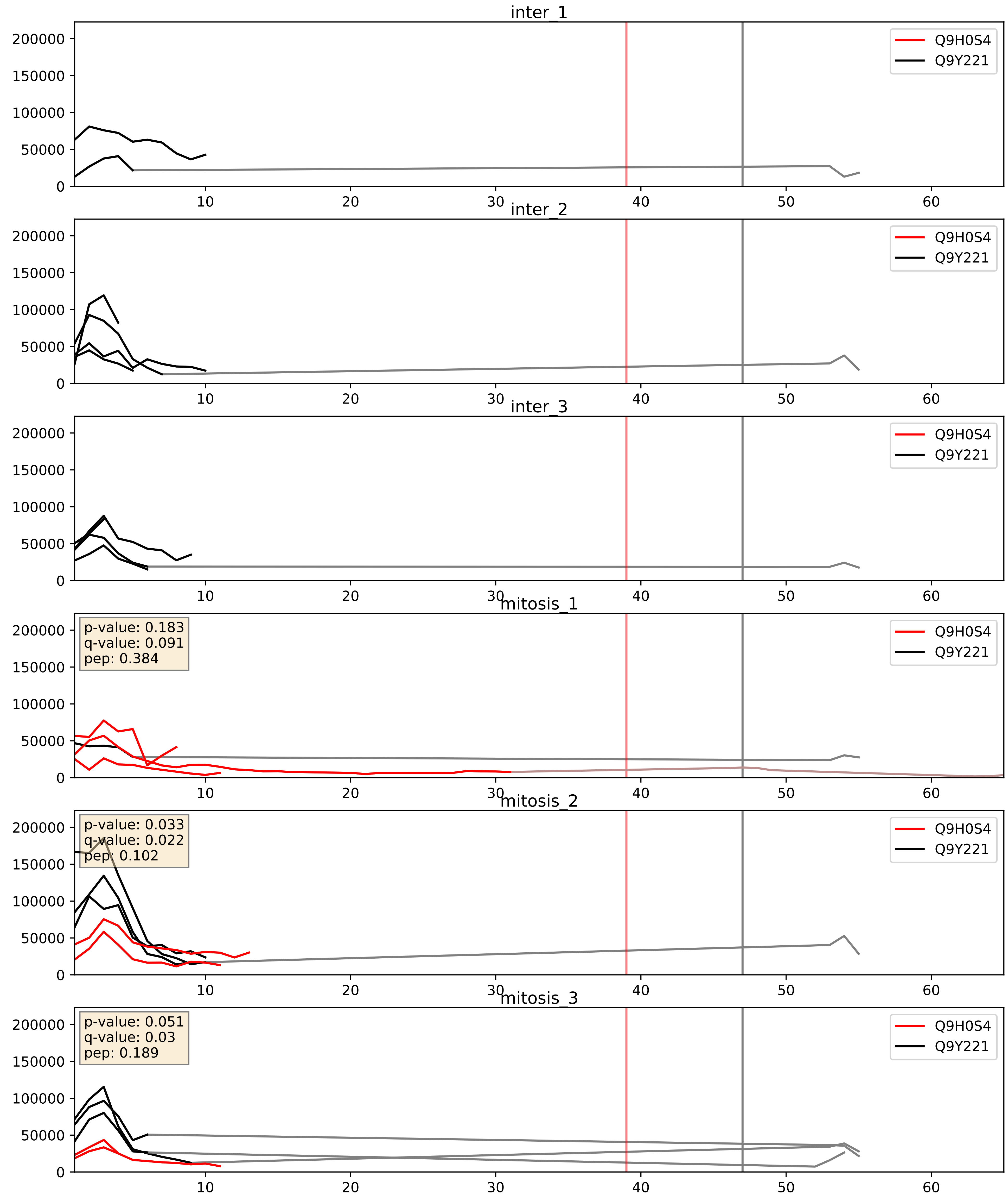


| level | condition_1 | condition_2 | pvalue | pvalue_adjusted |
|---|---|---|---|---|
| complex_abundance | mitosis | inter | 0.0477682613905785 | 0.15599228513863675 |
| interactor_ratio | mitosis | inter | 0.4503898579420976 | 0.8524836694963032 |
| interactor_abundance | mitosis | inter | 0.4816919099796174 | 0.6532967994019686 |

Q9H0S4_Q9Y2X3
DDX47_HUMAN vs NOP58_HUMAN
p-value: 0.001 q-value: 0.002

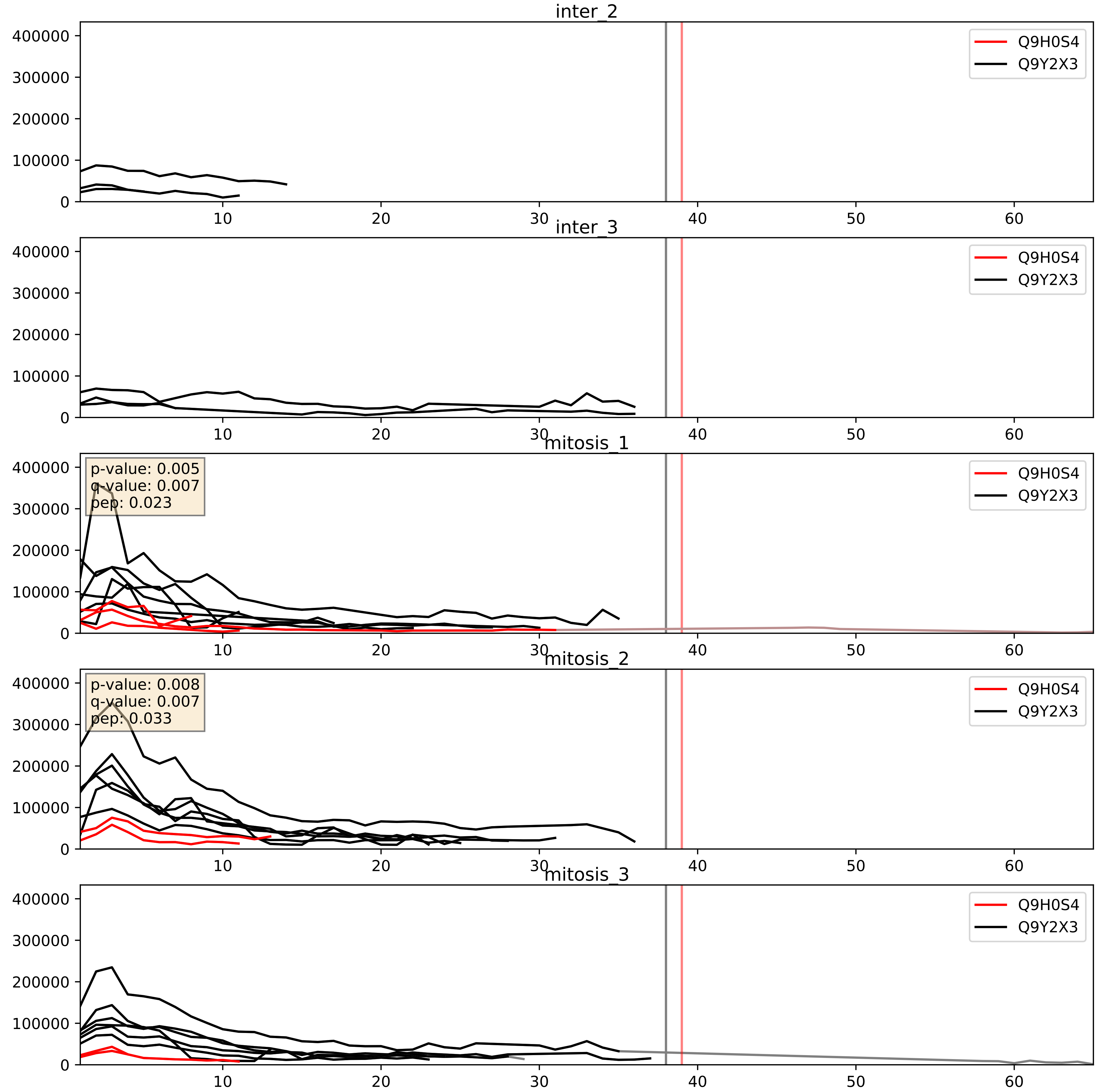


| level | condition_1 | condition_2 | pvalue | pvalue_adjusted |
| --- | --- | --- | --- | --- |
| interactor_ratio | mitosis | inter | 0.0006458171899288122 | 0.029562540886580918 |
| complex_abundance | mitosis | inter | 0.00802824854735332 | 0.0663337910862398 |
| interactor_abundance | mitosis | inter | 0.3974703705091973 | 0.588081647490922 |
