## Supplementary Data 2 for "SECAT: Quantifying differential protein-protein interaction states by network-centric analysis": hela_string_000039_O43390.pdf

O43390_O43390

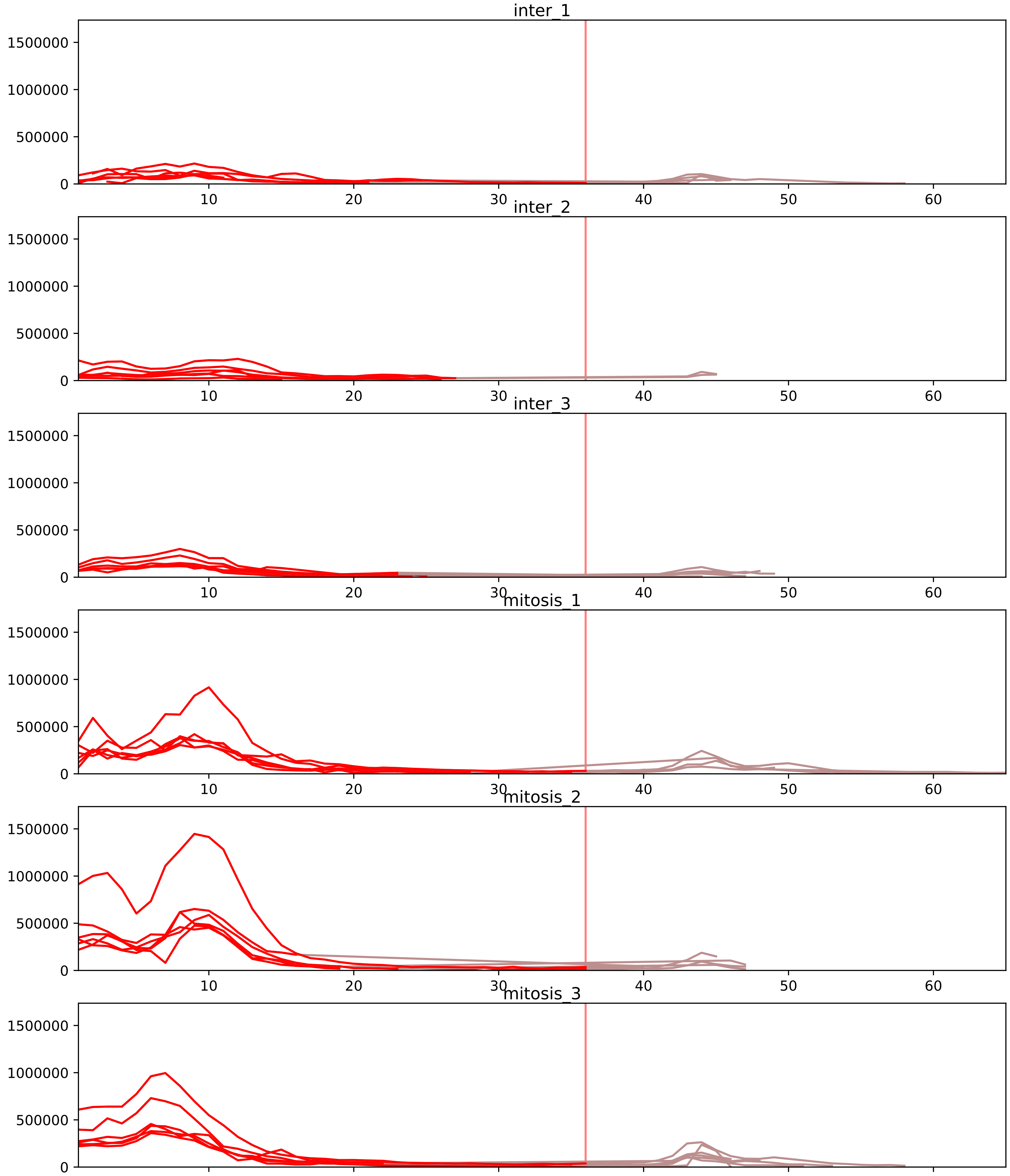


| level | condition_1 | condition_2 | pvalue | pvalue_adjusted |
|---|---|---|---|---|
| interactor_abundance | mitosis | inter | 1.514047685785222e-08 | 6.3766625832553175e-06 |
| complex_abundance | mitosis | inter | 2.6406100545791146e-08 | 1.2146806251063927e-05 |
| interactor_ratio | mitosis | inter | 0.0016698856546454126 | 0.016790107128675187 |
| assembled_abundance | mitosis | inter | 0.007407195109050039 | 0.09331793469809328 |
| total_abundance | mitosis | inter | 0.00888551053994586 | 0.11468713451692311 |
| monomer_abundance | mitosis | inter | 0.08668664683838957 | 0.38479695318022406 |

O43390_O60506
HNRPR_HUMAN vs HNRPQ_HUMAN
p-value: 0.002 q-value: 0.03

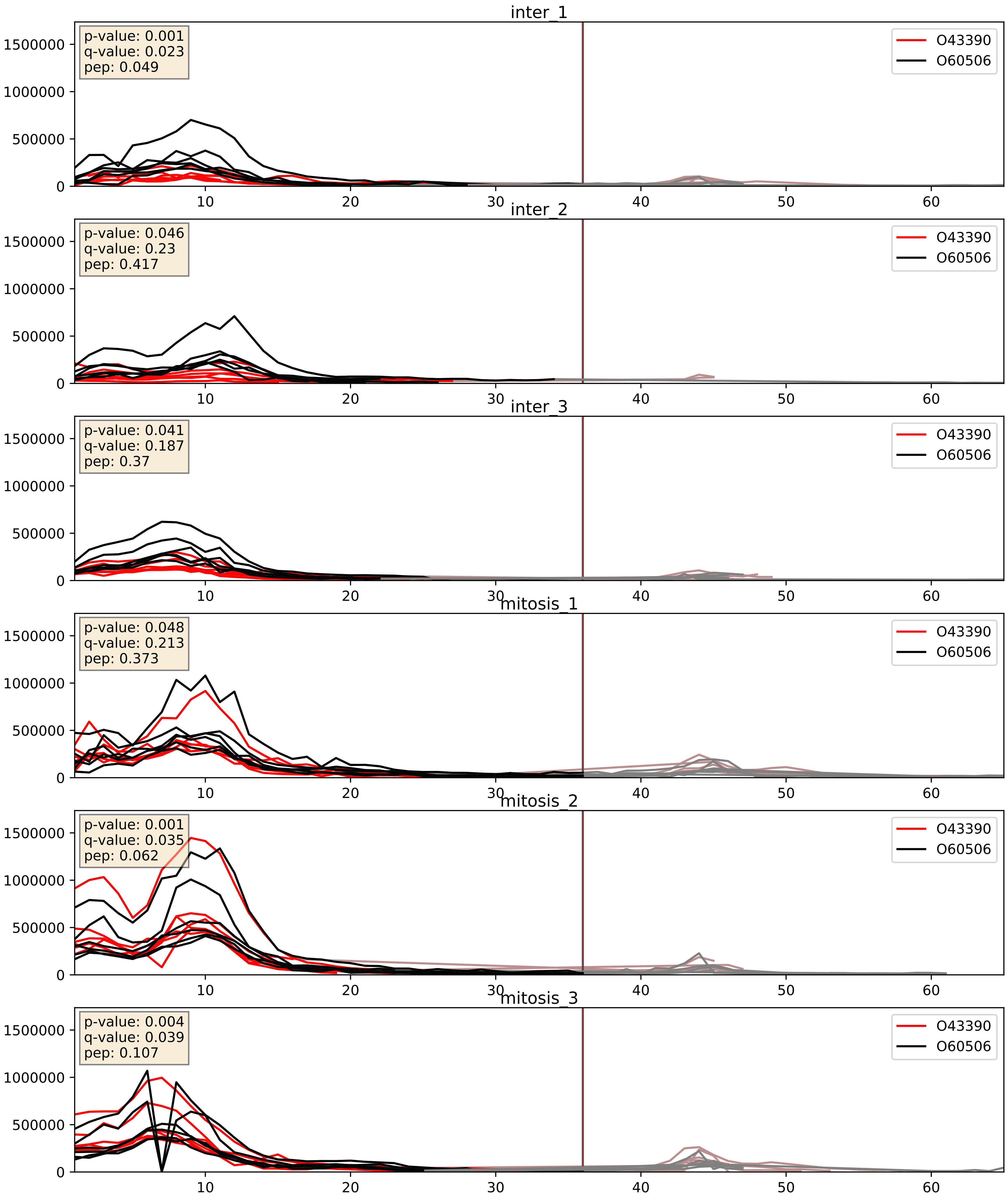


| level | condition_1 | condition_2 | pvalue | pvalue_adjusted |
|---|---|---|---|---|
| complex_abundance | mitosis | inter | 1.4707897835434114e-05 | 0.005691857151652958 |
| interactor_abundance | mitosis | inter | 0.00010796061997593195 | 0.007876481428251637 |
| interactor_ratio | mitosis | inter | 0.00949379078215129 | 0.13913572462569762 |

O43390_P09651
HNRPR_HUMAN vs ROA1_HUMAN
p-value: 0.065 q-value: 0.04

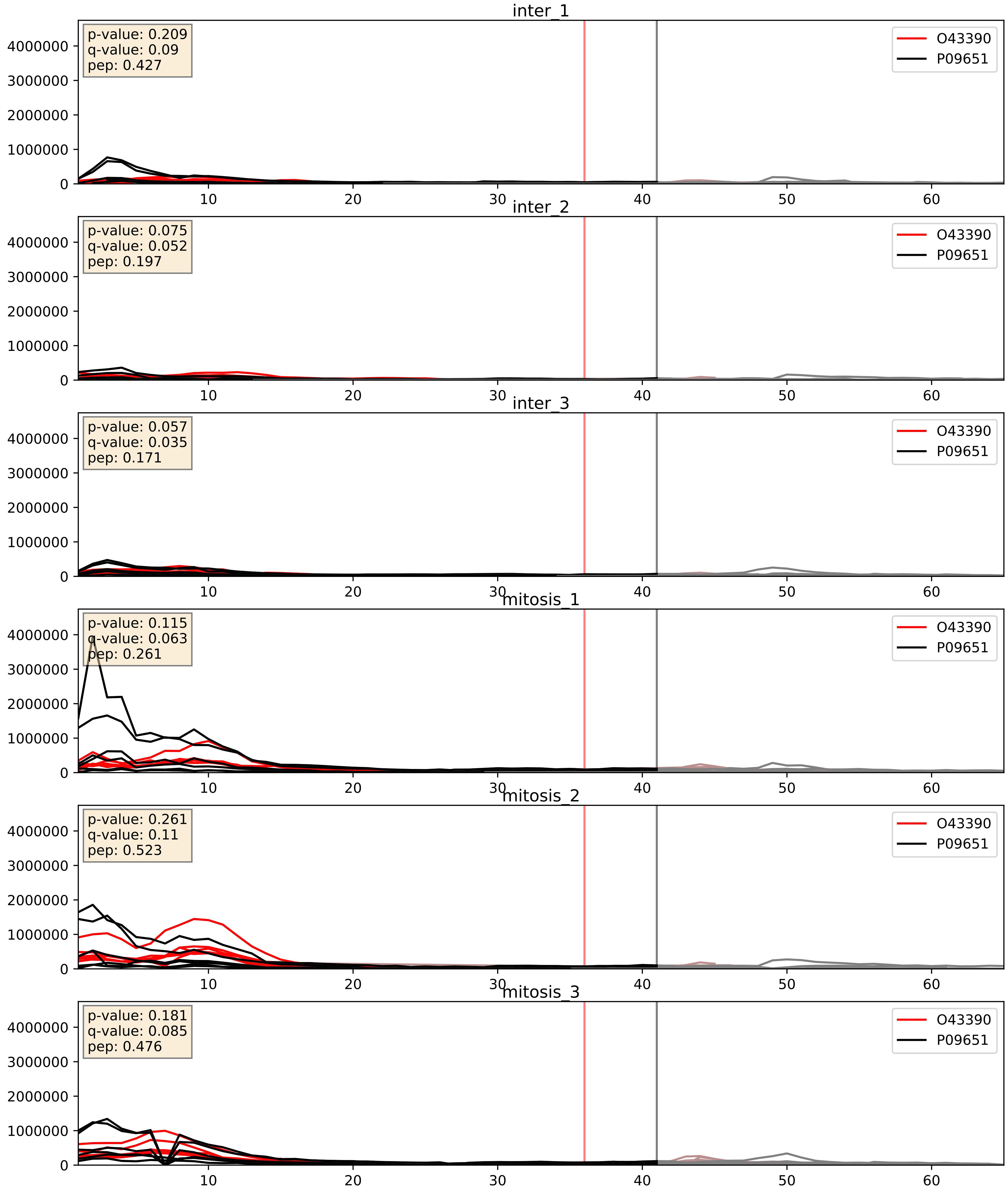


| level | condition_1 | condition_2 | pvalue | pvalue_adjusted |
|---|---|---|---|---|
| interactor_abundance | mitosis | inter | 0.00010796061997593195 | 0.007876481428251637 |
| complex_abundance | mitosis | inter | 0.0009535473068718876 | 0.020405912367058396 |
| interactor_ratio | mitosis | inter | 0.3502980208802063 | 0.8007410362093738 |

O43390_P14866
HNRPR_HUMAN vs HNRPL_HUMAN
p-value: 0.028 q-value: 0.021

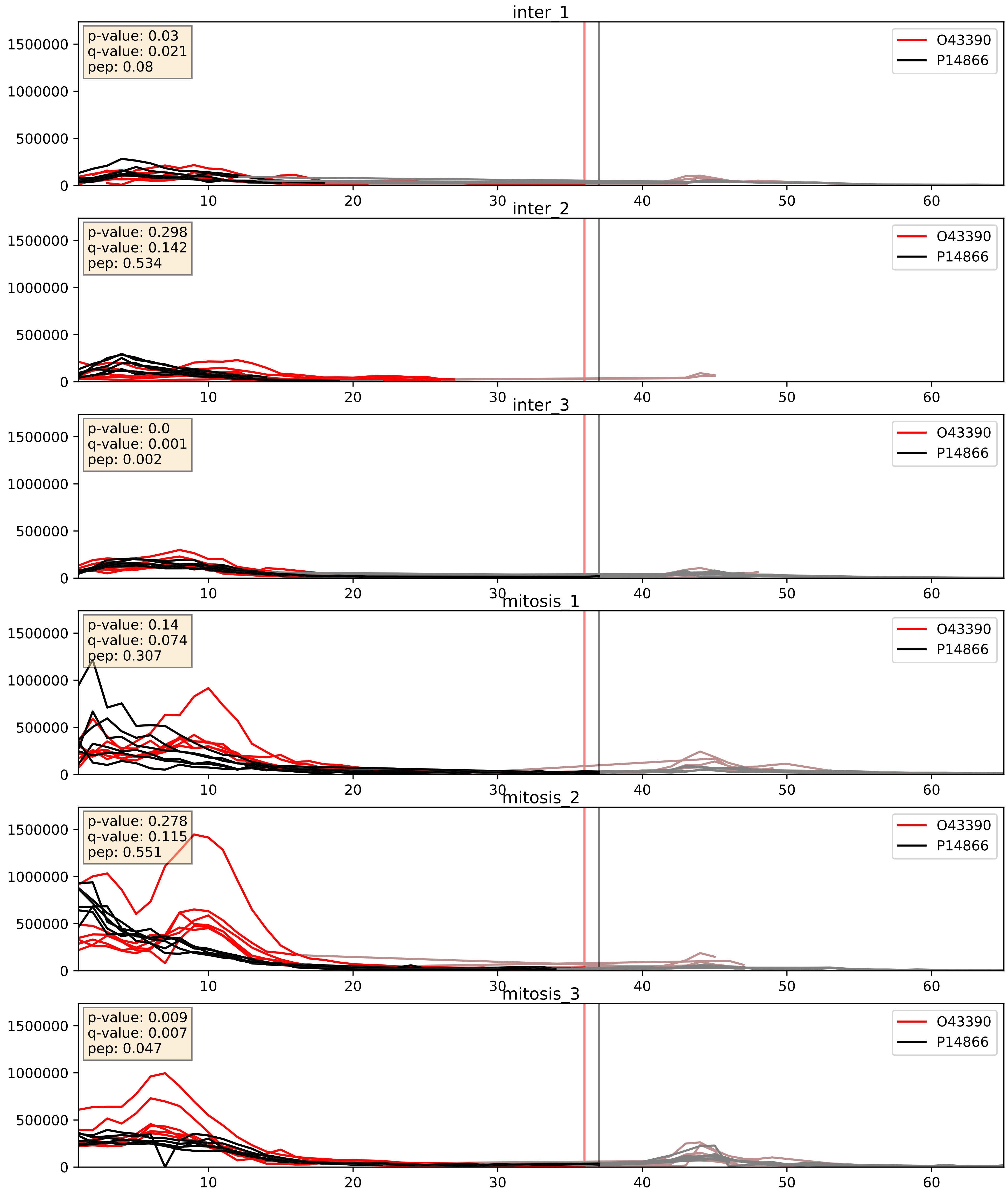


| level | condition_1 | condition_2 | pvalue | pvalue_adjusted |
|---|---|---|---|---|
| interactor_abundance | mitosis | inter | 5.893602312406929e-05 | 0.006459477818931243 |
| complex_abundance | mitosis | inter | 0.0011057804224422322 | 0.022115608448844644 |
| interactor_ratio | mitosis | inter | 0.7048469232789103 | 0.9378021986892063 |

O43390_P22626
HNRPR_HUMAN vs ROA2_HUMAN
p-value: 0.007 q-value: 0.007

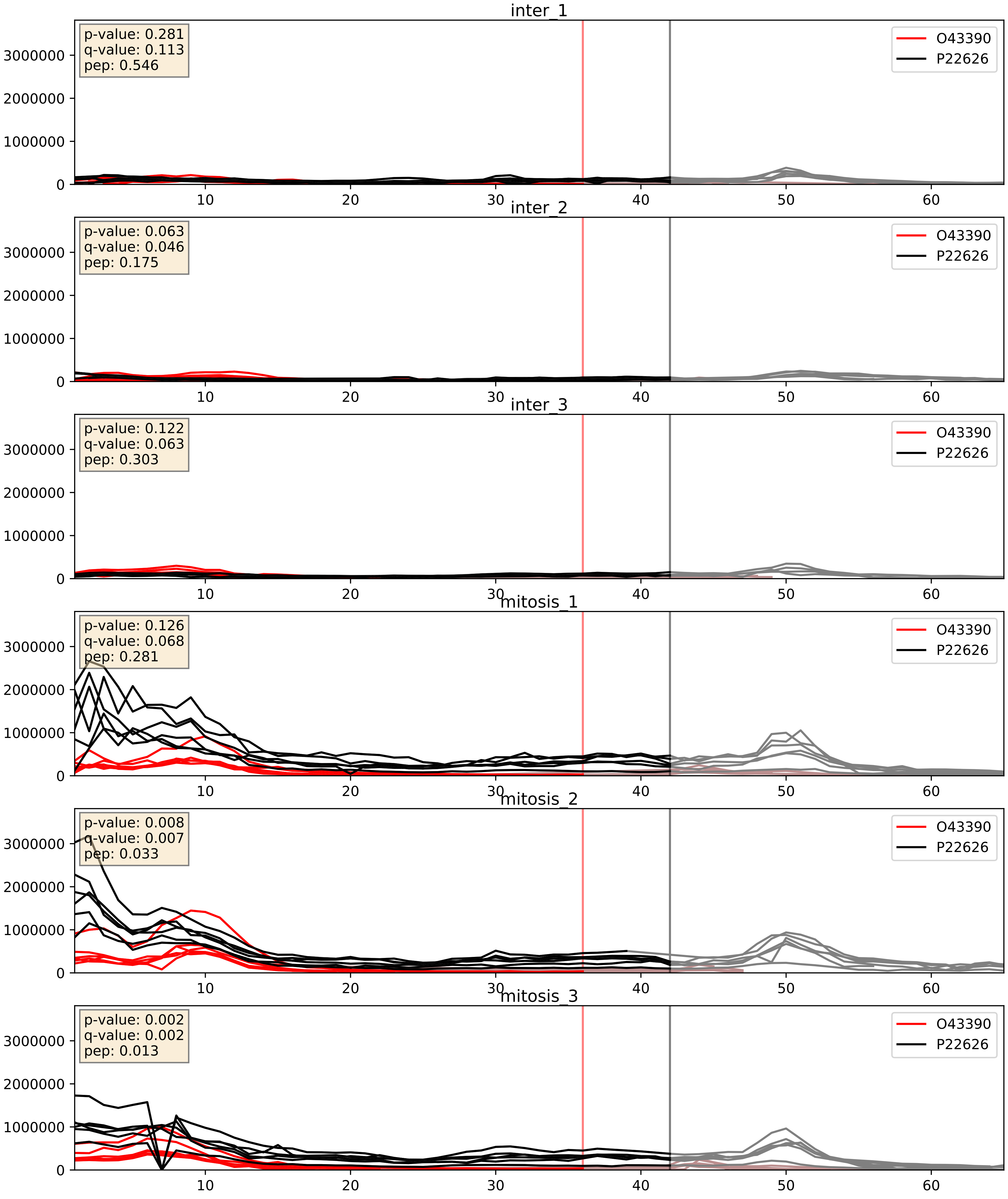


| level | condition_1 | condition_2 | pvalue | pvalue_adjusted |
|---|---|---|---|---|
| interactor_abundance | mitosis | inter | 0.00010796061997593195 | 0.007876481428251637 |
| complex_abundance | mitosis | inter | 0.0009205930867441553 | 0.019950067905139163 |
| interactor_ratio | mitosis | inter | 0.016834477256909862 | 0.19213750042553124 |

O43390_P31943
HNRPR_HUMAN vs HNRH1_HUMAN
p-value: 0.005 q-value: 0.005

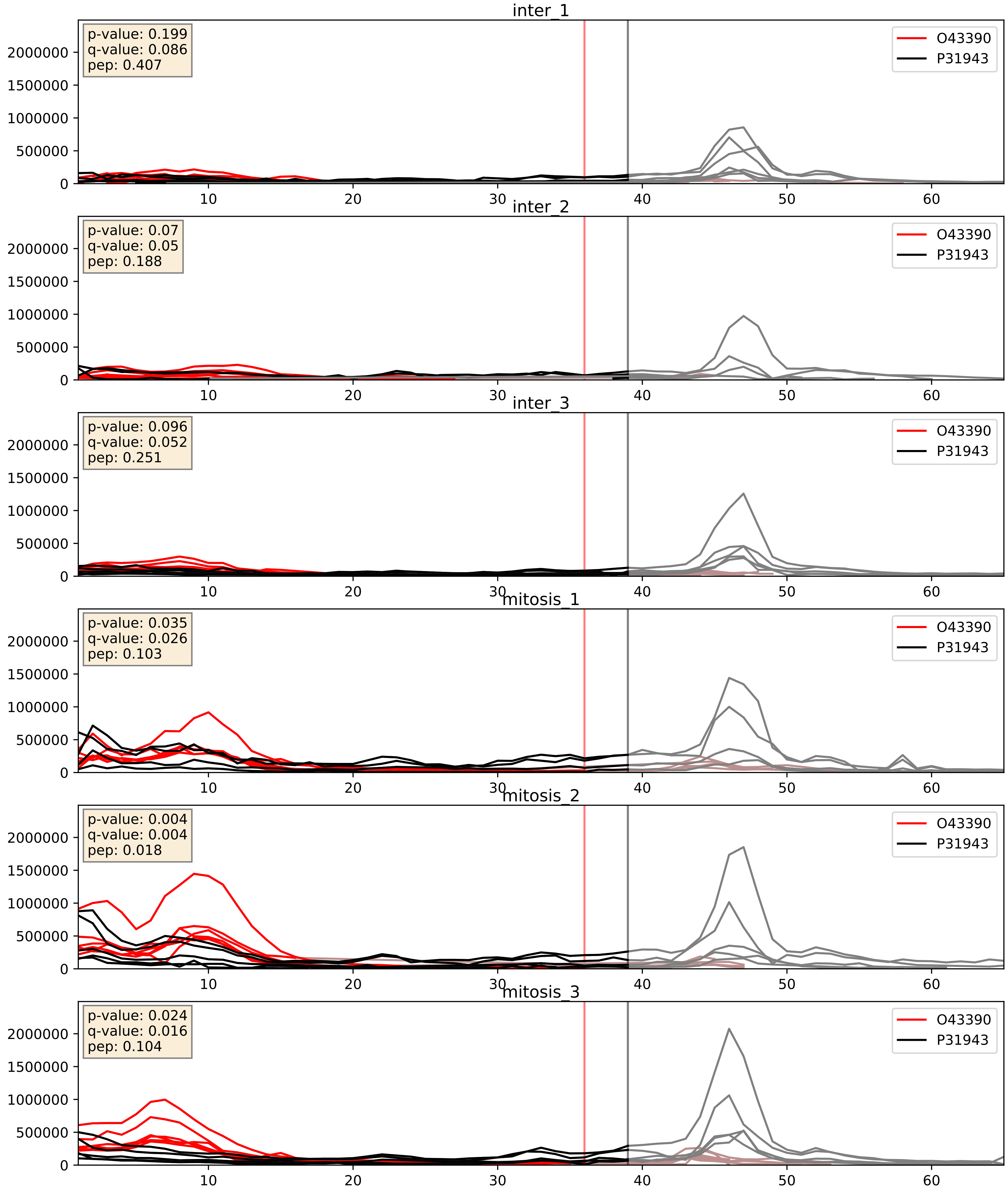


| level | condition_1 | condition_2 | pvalue | pvalue_adjusted |
|---|---|---|---|---|
| complex_abundance | mitosis | inter | 9.891557176946058e-05 | 0.006719978526560179 |
| interactor_abundance | mitosis | inter | 0.00010796061997593195 | 0.007876481428251637 |
| interactor_ratio | mitosis | inter | 0.4332742845030043 | 0.8435878041032946 |

O43390_P51991
HNRPR_HUMAN vs ROA3_HUMAN
p-value: 0.003 q-value: 0.041

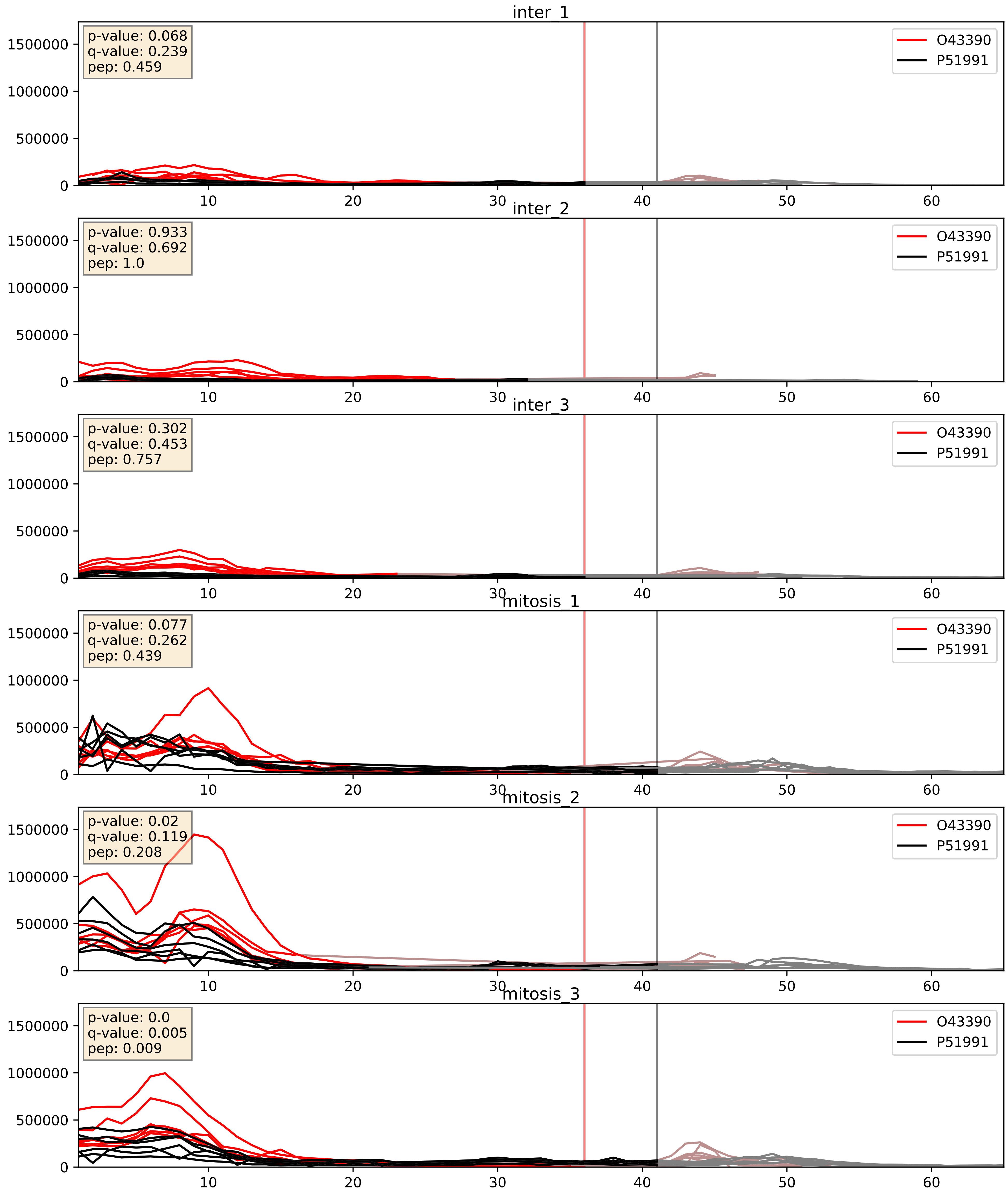


| level | condition_1 | condition_2 | pvalue | pvalue_adjusted |
|---|---|---|---|---|
| complex_abundance | mitosis | inter | 1.1164489343079466e-07 | 0.0009556802877676023 |
| interactor_abundance | mitosis | inter | 3.72135292356026e-05 | 0.005353744710197617 |
| interactor_ratio | mitosis | inter | 0.0004914306397162247 | 0.025464098294827937 |

O43390_P52272
HNRPR_HUMAN vs HNRPM_HUMAN
p-value: 0.058 q-value: 0.037

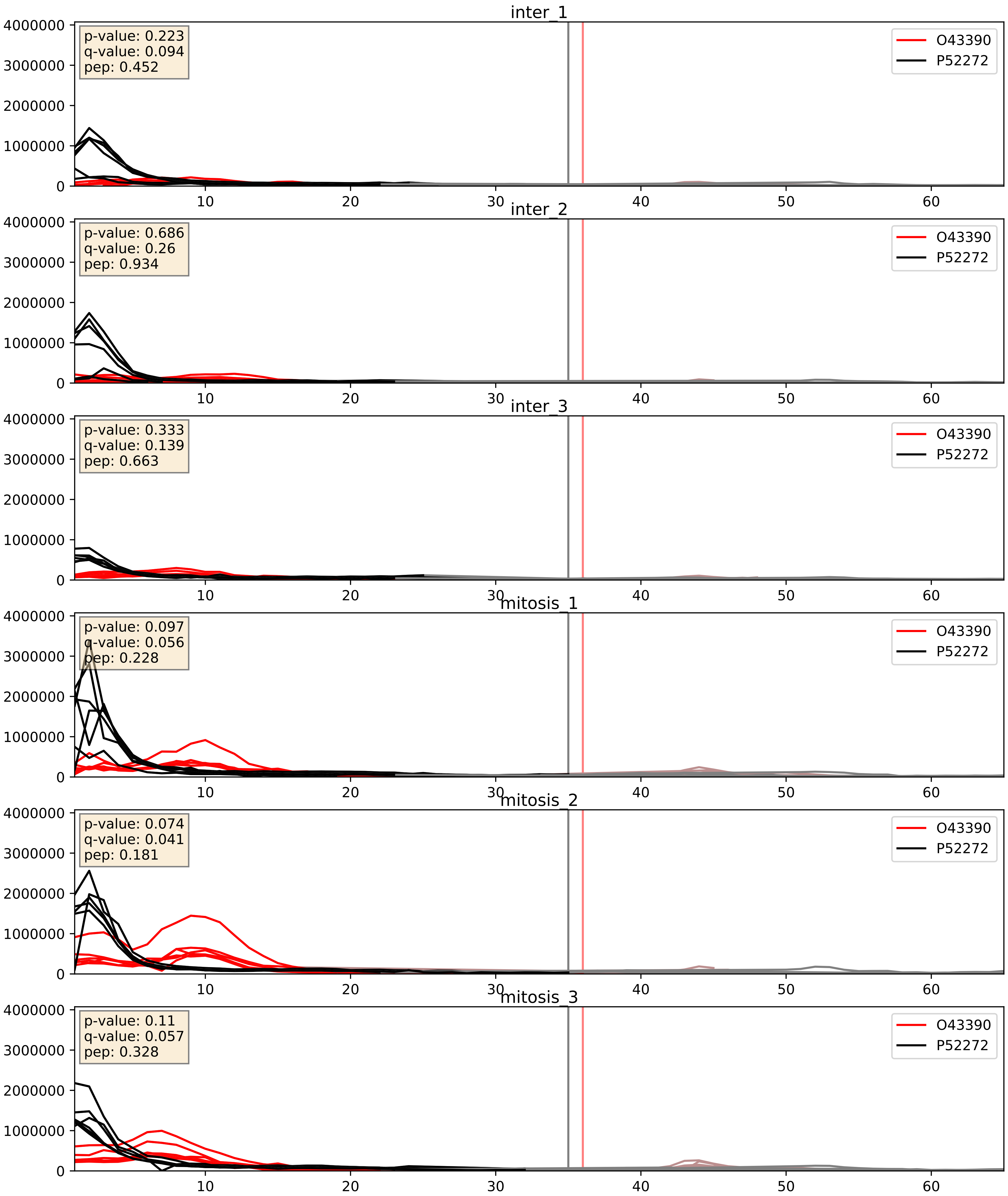


| level | condition_1 | condition_2 | pvalue | pvalue_adjusted |
|---|---|---|---|---|
| interactor_abundance | mitosis | inter | 6.140823405978327e-05 | 0.006459477818931243 |
| complex_abundance | mitosis | inter | 0.0025869915277018023 | 0.03664546283589145 |
| interactor_ratio | mitosis | inter | 0.04696217208361891 | 0.35838626305591875 |

O43390_P52597
HNRPR_HUMAN vs HNRPF_HUMAN
p-value: 0.003 q-value: 0.003

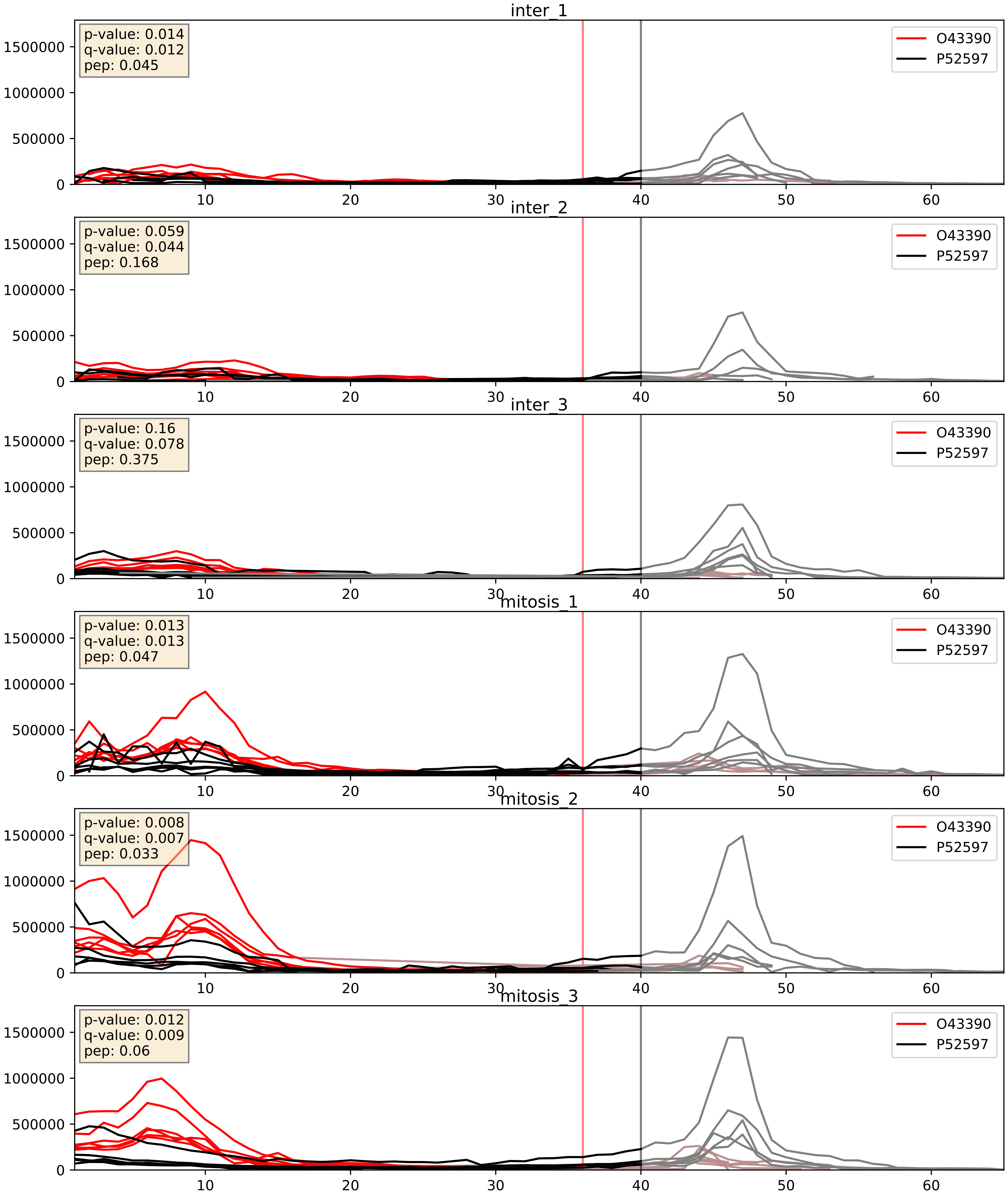


| level | condition_1 | condition_2 | pvalue | pvalue_adjusted |
|---|---|---|---|---|
| complex_abundance | mitosis | inter | 7.40732189271377e-05 | 0.005893640694944935 |
| interactor_abundance | mitosis | inter | 0.00010796061997593195 | 0.007876481428251637 |
| interactor_ratio | mitosis | inter | 0.29205419947420935 | 0.7616429249926009 |

O43390_P61978
HNRPR_HUMAN vs HNRPK_HUMAN
p-value: 0.058 q-value: 0.037

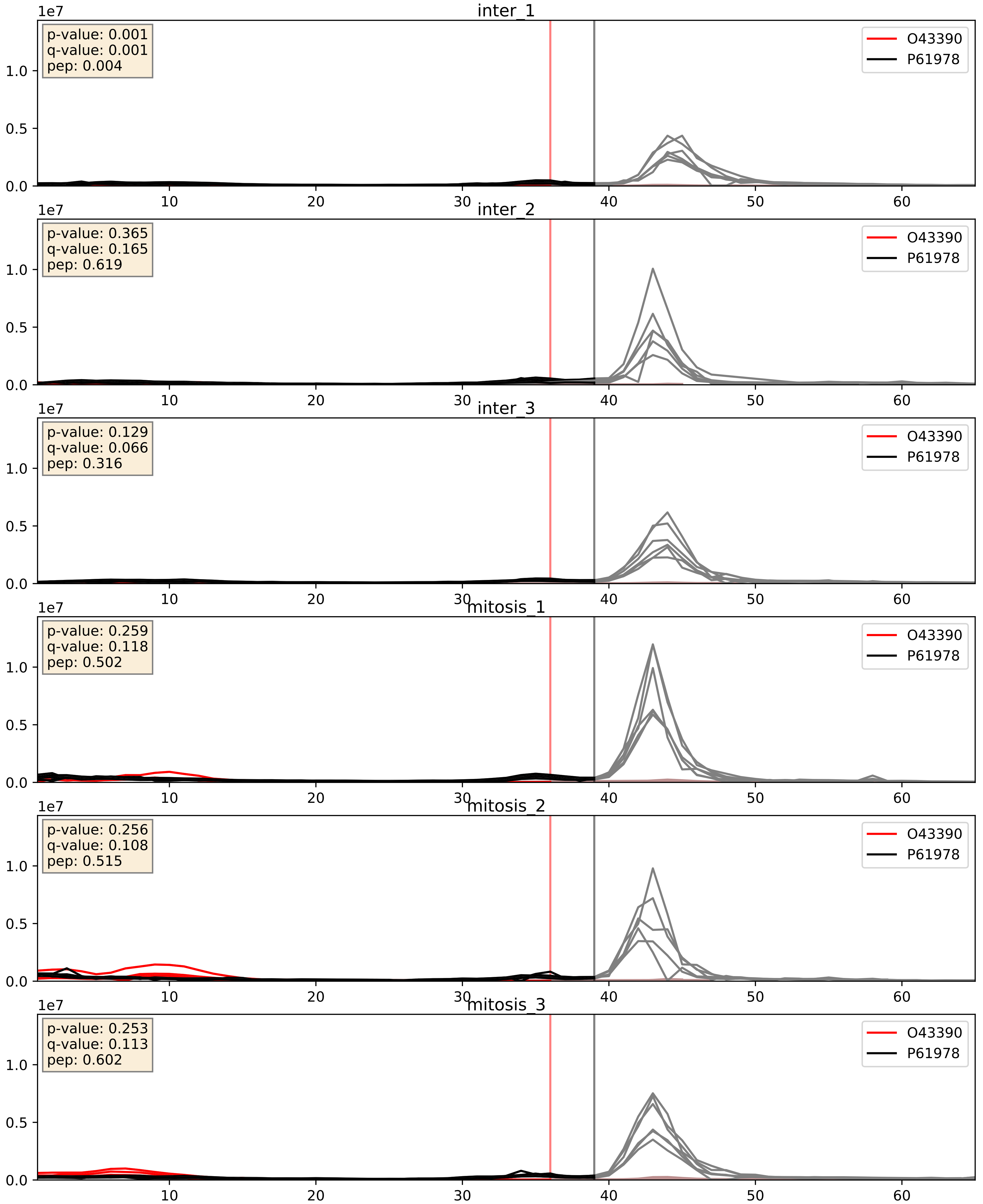


| level | condition_1 | condition_2 | pvalue | pvalue_adjusted |
|---|---|---|---|---|
| interactor_abundance | mitosis | inter | 0.00010796061997593195 | 0.007876481428251637 |
| complex_abundance | mitosis | inter | 0.10752277520932148 | 0.2443542206845082 |
| interactor_ratio | mitosis | inter | 0.272918022983022 | 0.7468898440591442 |

O43390_P67809
HNRPR_HUMAN vs YBOX1_HUMAN
p-value: 0.0 q-value: 0.005

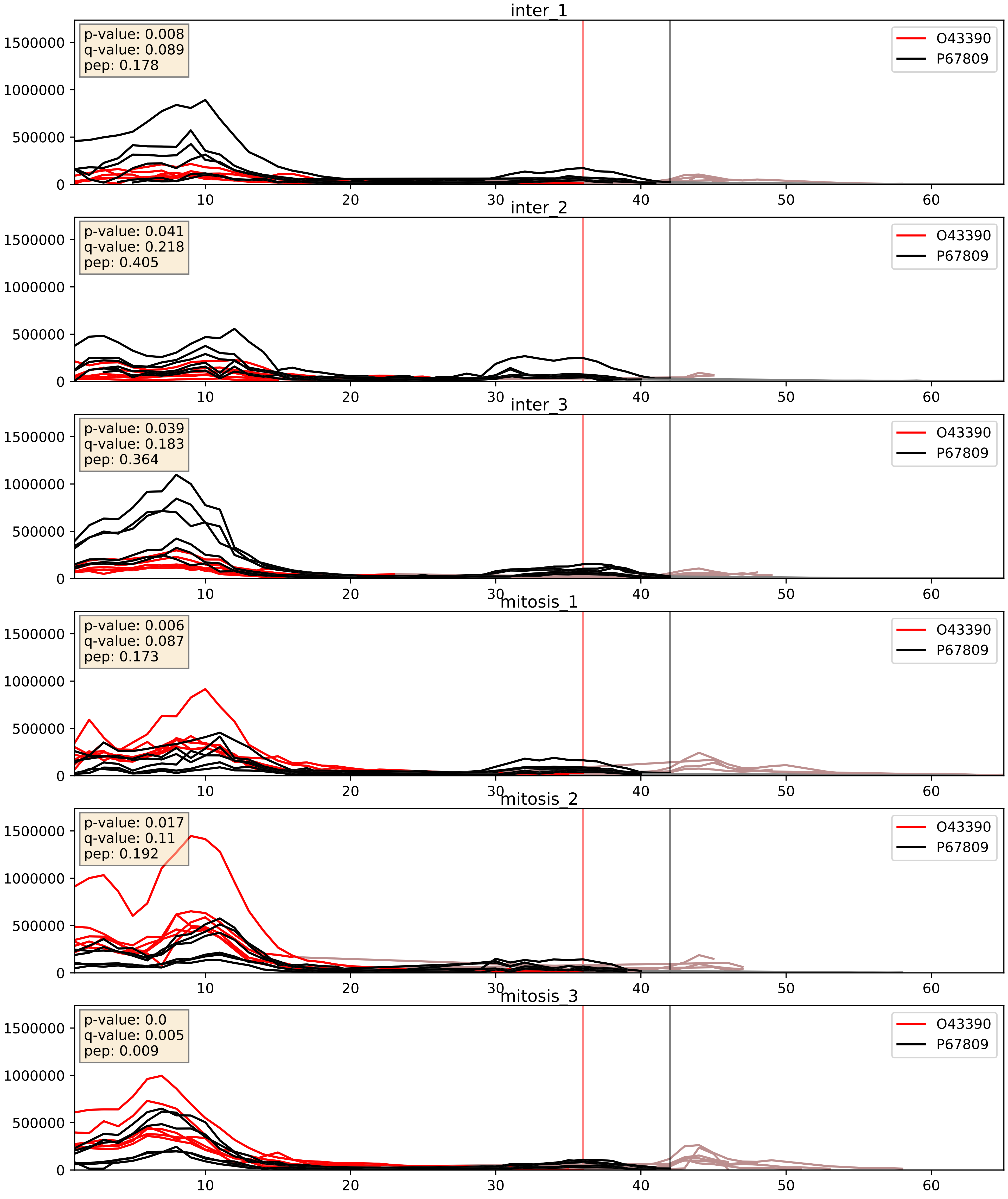


| level | condition_1 | condition_2 | pvalue | pvalue_adjusted |
|---|---|---|---|---|
| interactor_abundance | mitosis | inter | 0.00010796061997593195 | 0.007876481428251637 |
| interactor_ratio | mitosis | inter | 0.006983891252796053 | 0.11653432577764955 |
| complex_abundance | mitosis | inter | 0.03412451813730336 | 0.13430390263264738 |

O43390_Q07955
HNRPR_HUMAN vs SRSF1_HUMAN
p-value: 0.002 q-value: 0.002

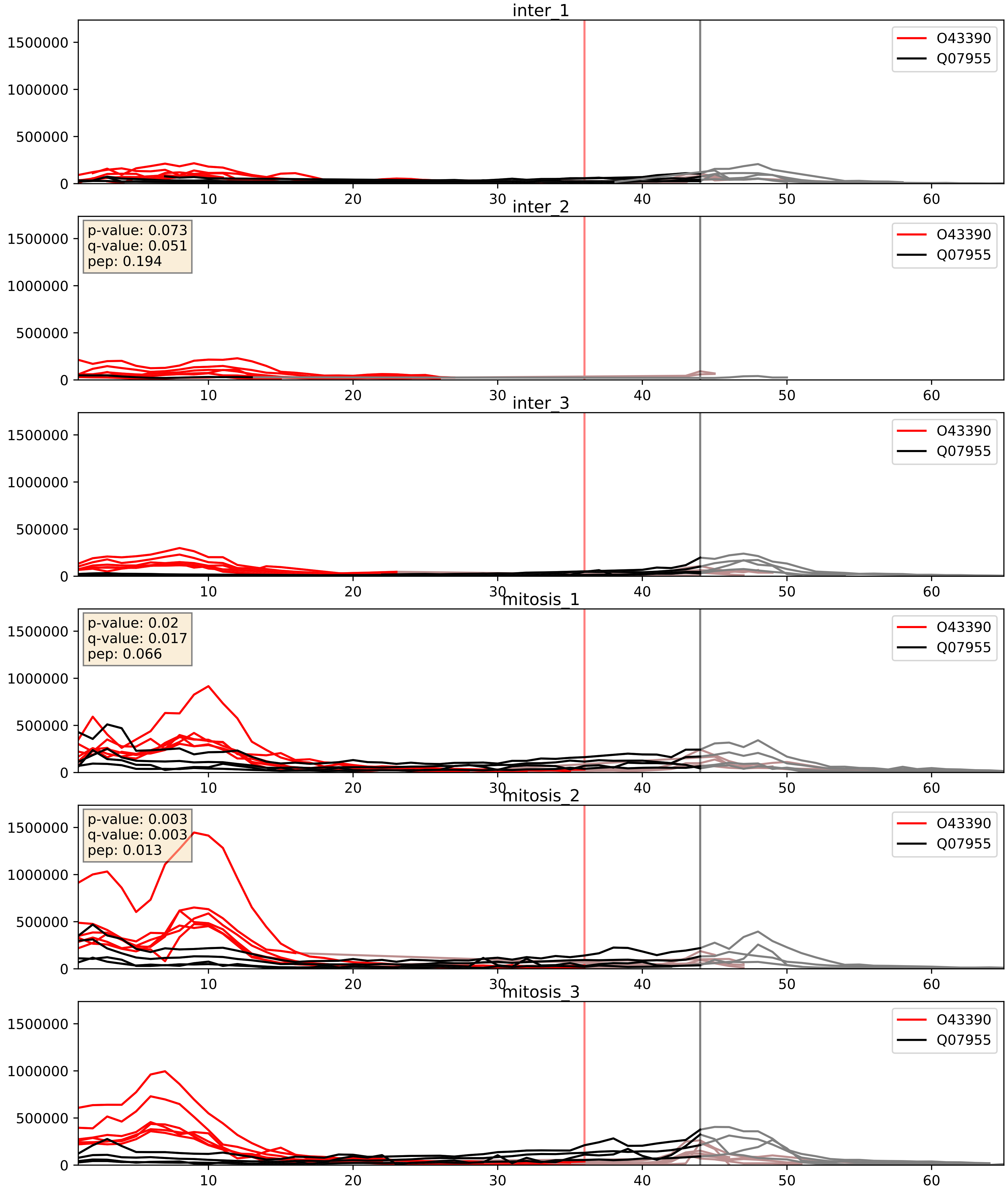


| level | condition_1 | condition_2 | pvalue | pvalue_adjusted |
|---|---|---|---|---|
| interactor_abundance | mitosis | inter | 0.0009696737069179261 | 0.02306454775731039 |
| complex_abundance | mitosis | inter | 0.004069476203020404 | 0.04732787841836958 |
| interactor_ratio | mitosis | inter | 0.16851302494823342 | 0.6310024031307429 |

O43390_Q08211
HNRPR_HUMAN vs DHX9_HUMAN
p-value: 0.004 q-value: 0.004

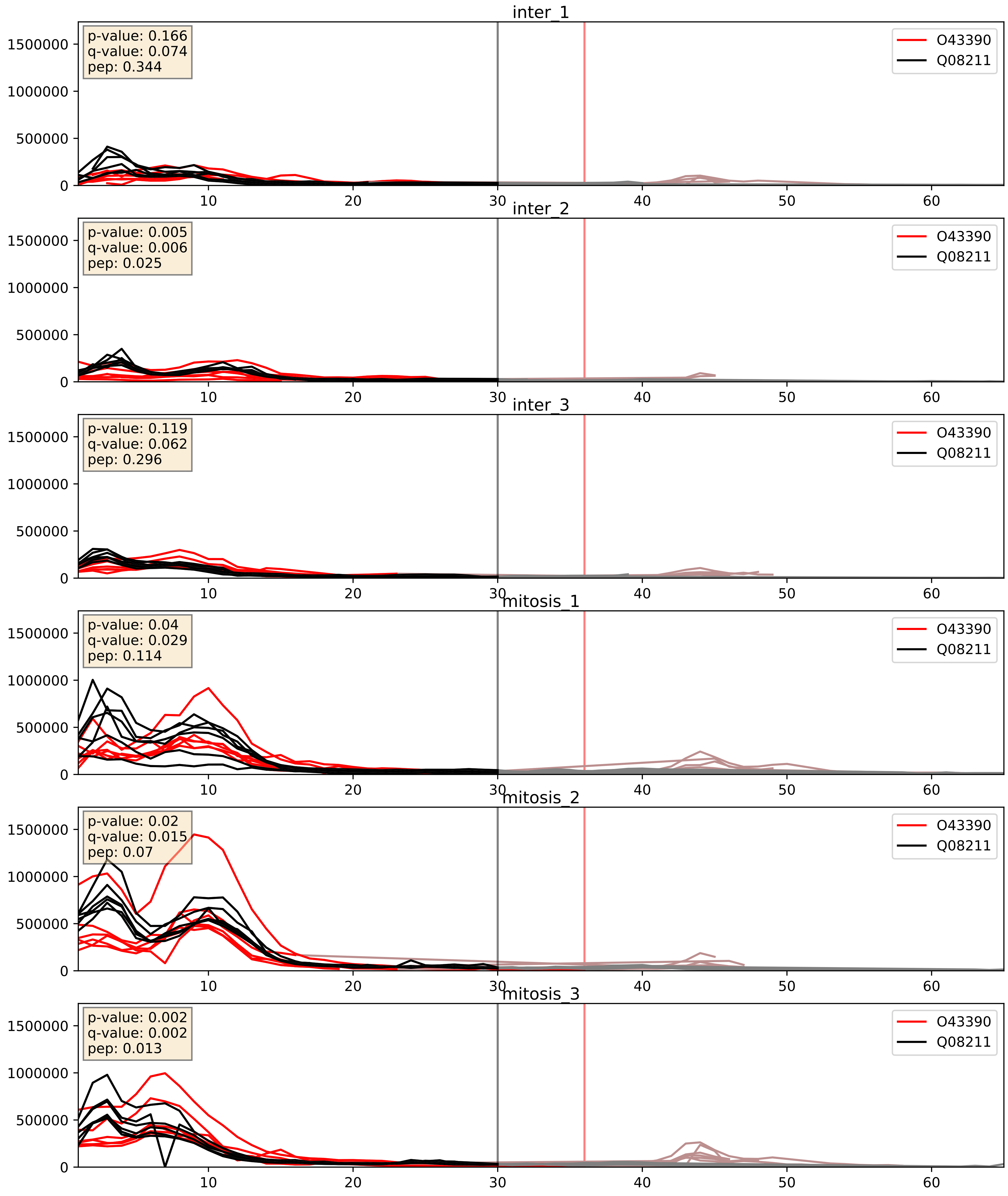


| level | condition_1 | condition_2 | pvalue | pvalue_adjusted |
|---|---|---|---|---|
| complex_abundance | mitosis | inter | 3.9289007550731724e-05 | 0.005691857151652958 |
| interactor_abundance | mitosis | inter | 0.00014528767019049324 | 0.009246561017328046 |
| interactor_ratio | mitosis | inter | 0.8637249543002753 | 0.9774585538872641 |

O43390_Q12905
HNRPR_HUMAN vs ILF2_HUMAN
p-value: 0.003 q-value: 0.041

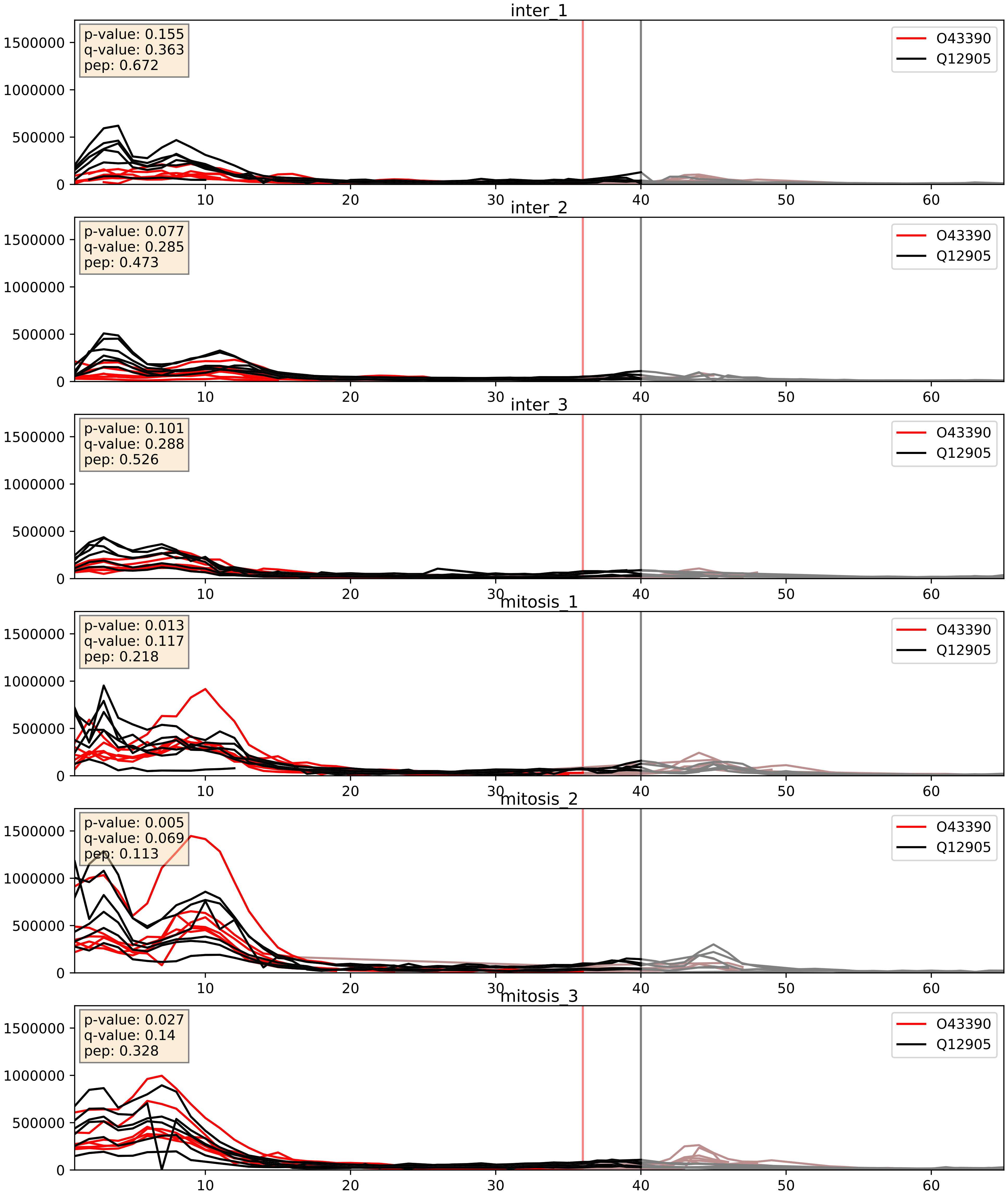


| level | condition_1 | condition_2 | pvalue | pvalue_adjusted |
|---|---|---|---|---|
| interactor_abundance | mitosis | inter | 0.00010796061997593195 | 0.007876481428251637 |
| complex_abundance | mitosis | inter | 0.0003909354311201745 | 0.012627952039202618 |
| interactor_ratio | mitosis | inter | 0.004324784485630598 | 0.08690177276290592 |

O43390_Q13151
HNRPR_HUMAN vs ROA0_HUMAN
p-value: 0.005 q-value: 0.005

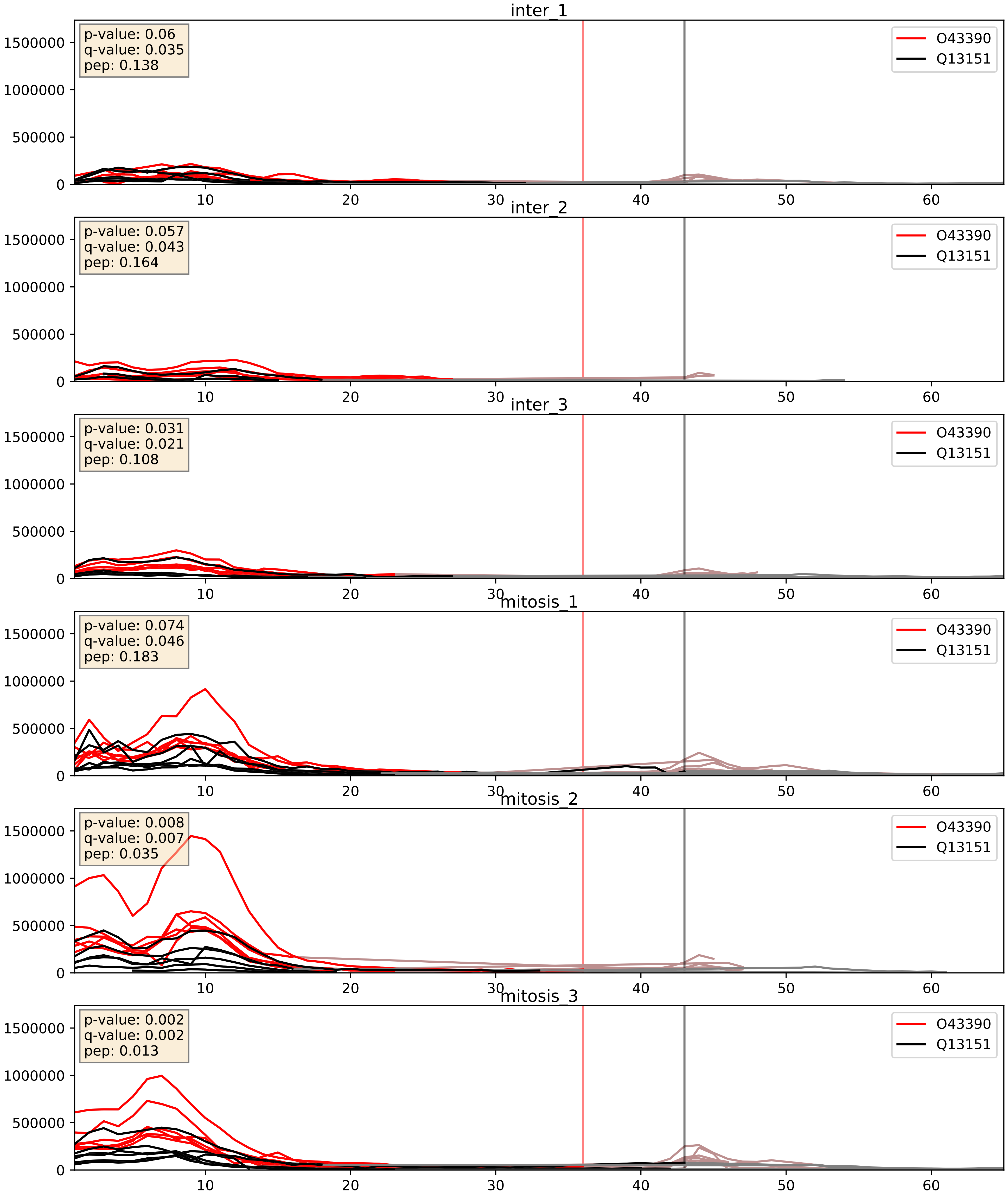


| level | condition_1 | condition_2 | pvalue | pvalue_adjusted |
|---|---|---|---|---|
| interactor_abundance | mitosis | inter | 3.1358423719273224e-05 | 0.004891461372878449 |
| complex_abundance | mitosis | inter | 0.0005214359642555288 | 0.01453706838573521 |
| interactor_ratio | mitosis | inter | 0.9691540655893558 | 0.9937294965826962 |

O43390_Q13247
HNRPR_HUMAN vs SRSF6_HUMAN
p-value: 0.0 q-value: 0.005

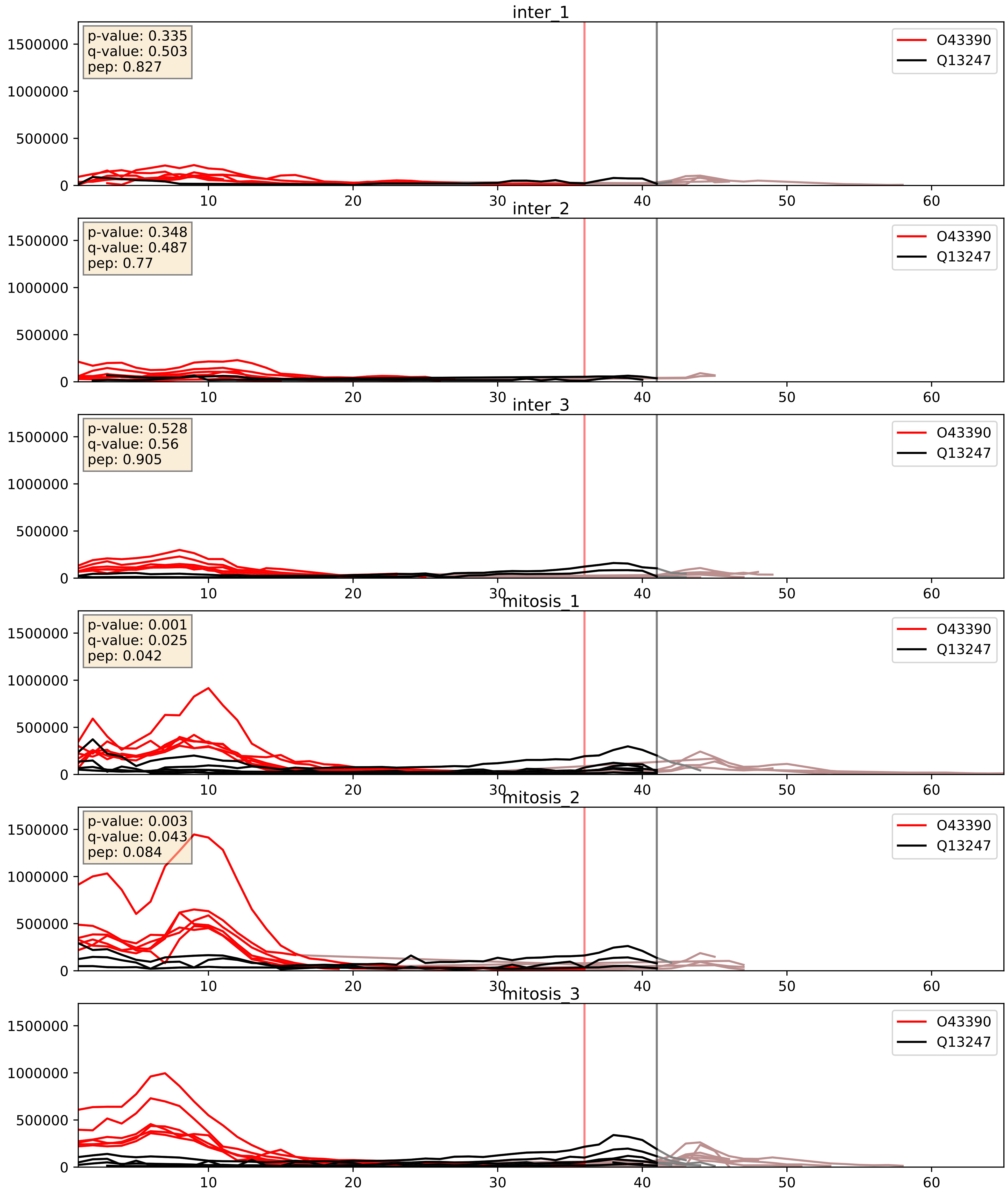


| level | condition_1 | condition_2 | pvalue | pvalue_adjusted |
|---|---|---|---|---|
| interactor_abundance | mitosis | inter | 0.0005707867388178207 | 0.017415659477552636 |
| complex_abundance | mitosis | inter | 0.0006004542403235107 | 0.01581504091436693 |
| interactor_ratio | mitosis | inter | 0.9652626032384787 | 0.9937294965826962 |
