## Supplementary Data 2 for "SECAT: Quantifying differential protein-protein interaction states by network-centric analysis": hela_string_000040_O43290.pdf

O43290_O43290

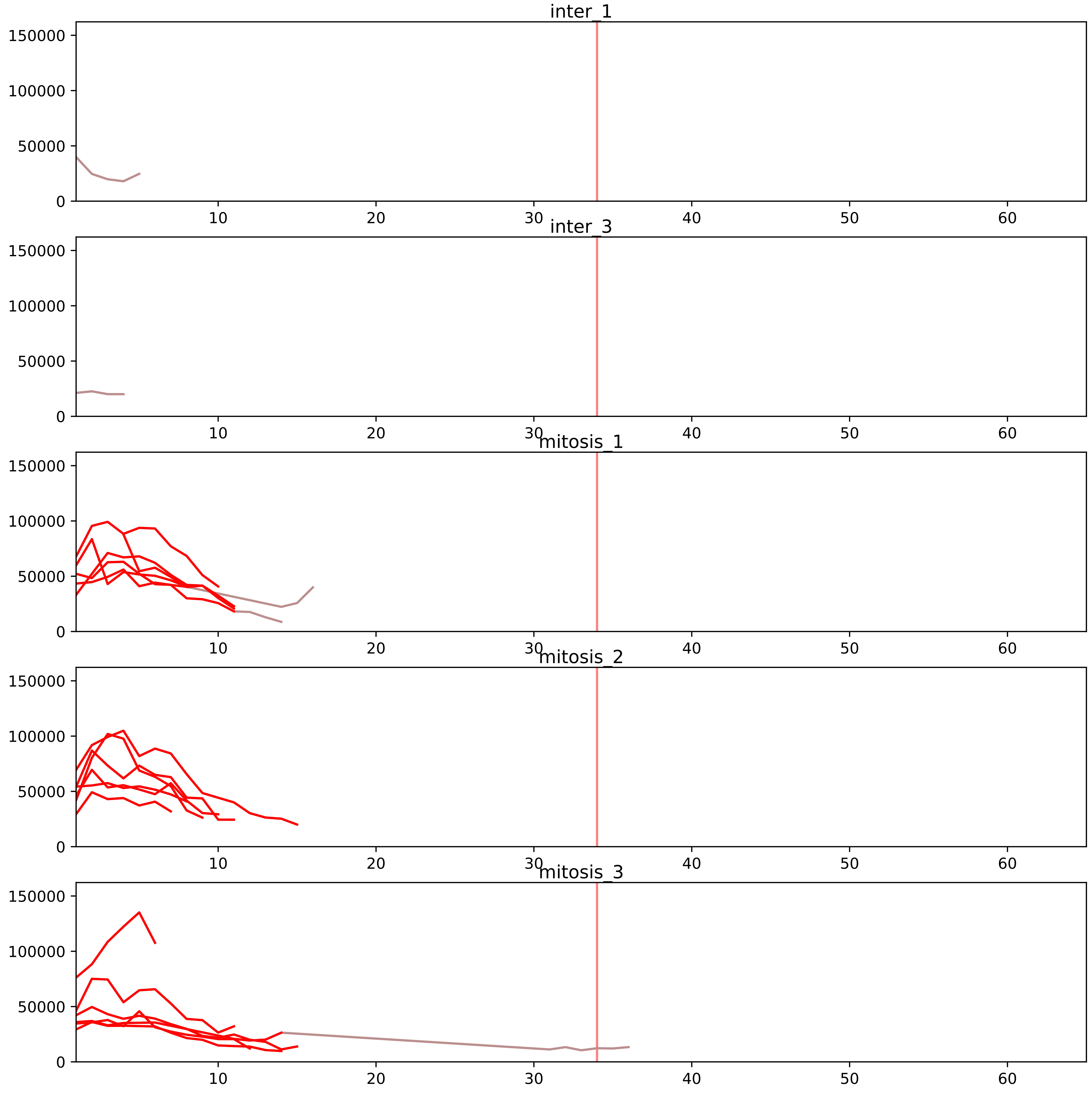


| level | condition_1 | condition_2 | pvalue | pvalue_adjusted |
|---|---|---|---|---|
| interactor_ratio | mitosis | inter | 1.7052039998574982e-08 | 8.479933404696749e-07 |
| complex_abundance | mitosis | inter | 1.7418937808938628e-05 | 0.0007284283083737971 |
| assembled_abundance | mitosis | inter | 7.932826583893875e-05 | 0.008635711916094704 |
| interactor_abundance | mitosis | inter | 0.00015653473084287266 | 0.0030970312338804915 |
| total_abundance | mitosis | inter | 0.00022447012640694944 | 0.01843189376673838 |

O43172_O43290
PRP4_HUMAN vs SNUT1_HUMAN
p-value: 0.032 q-value: 0.023

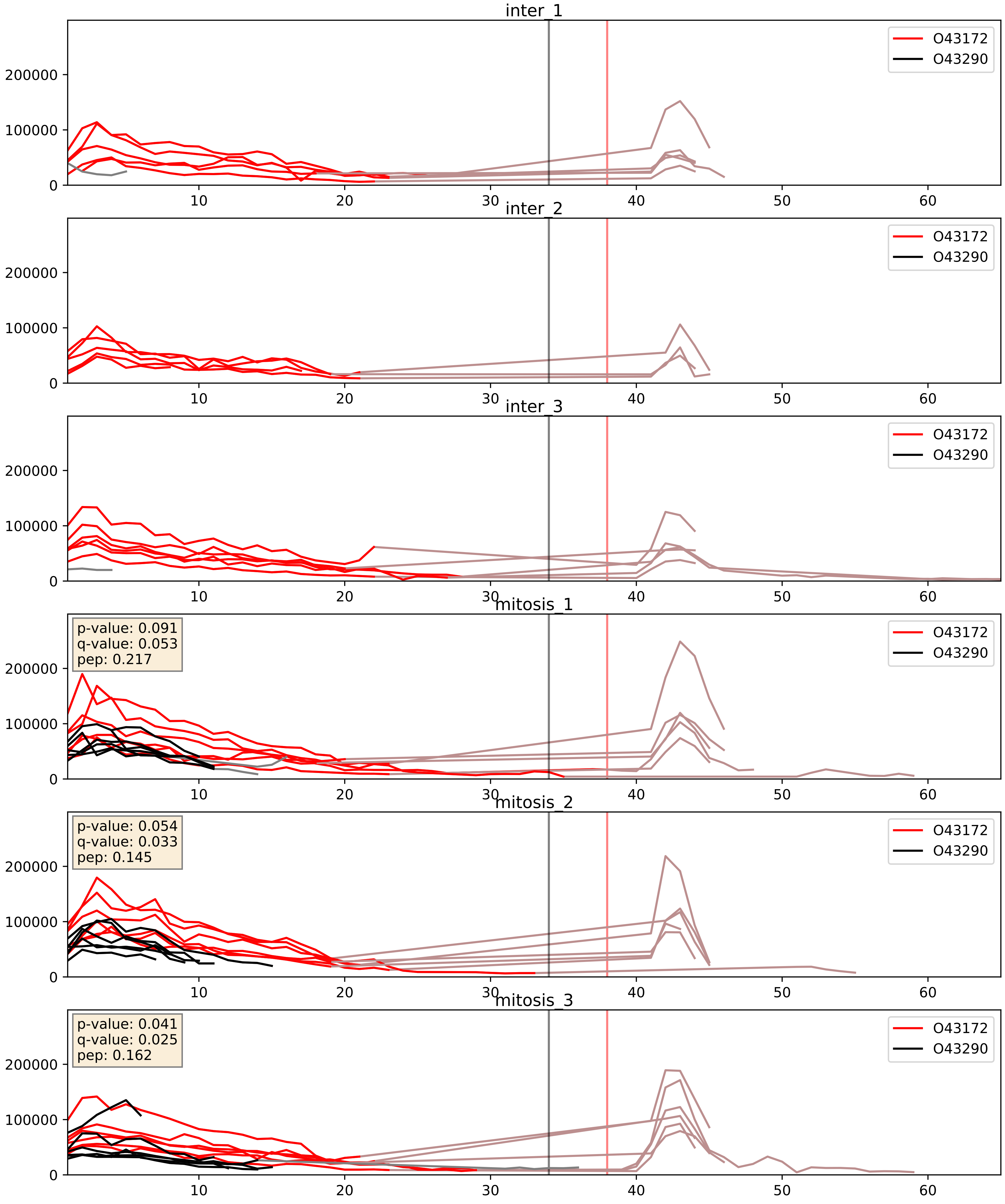


| level | condition_1 | condition_2 | pvalue | pvalue_adjusted |
|---|---|---|---|---|
| interactor_abundance | mitosis | inter | 0.00018106838416699807 | 0.010140496760639767 |
| complex_abundance | mitosis | inter | 0.0003207915922283504 | 0.011685004380743318 |
| interactor_ratio | mitosis | inter | 0.001184418604256362 | 0.04055449300973783 |

O43290_O43395
SNUT1_HUMAN vs PRPF3_HUMAN
p-value: 0.023 q-value: 0.018

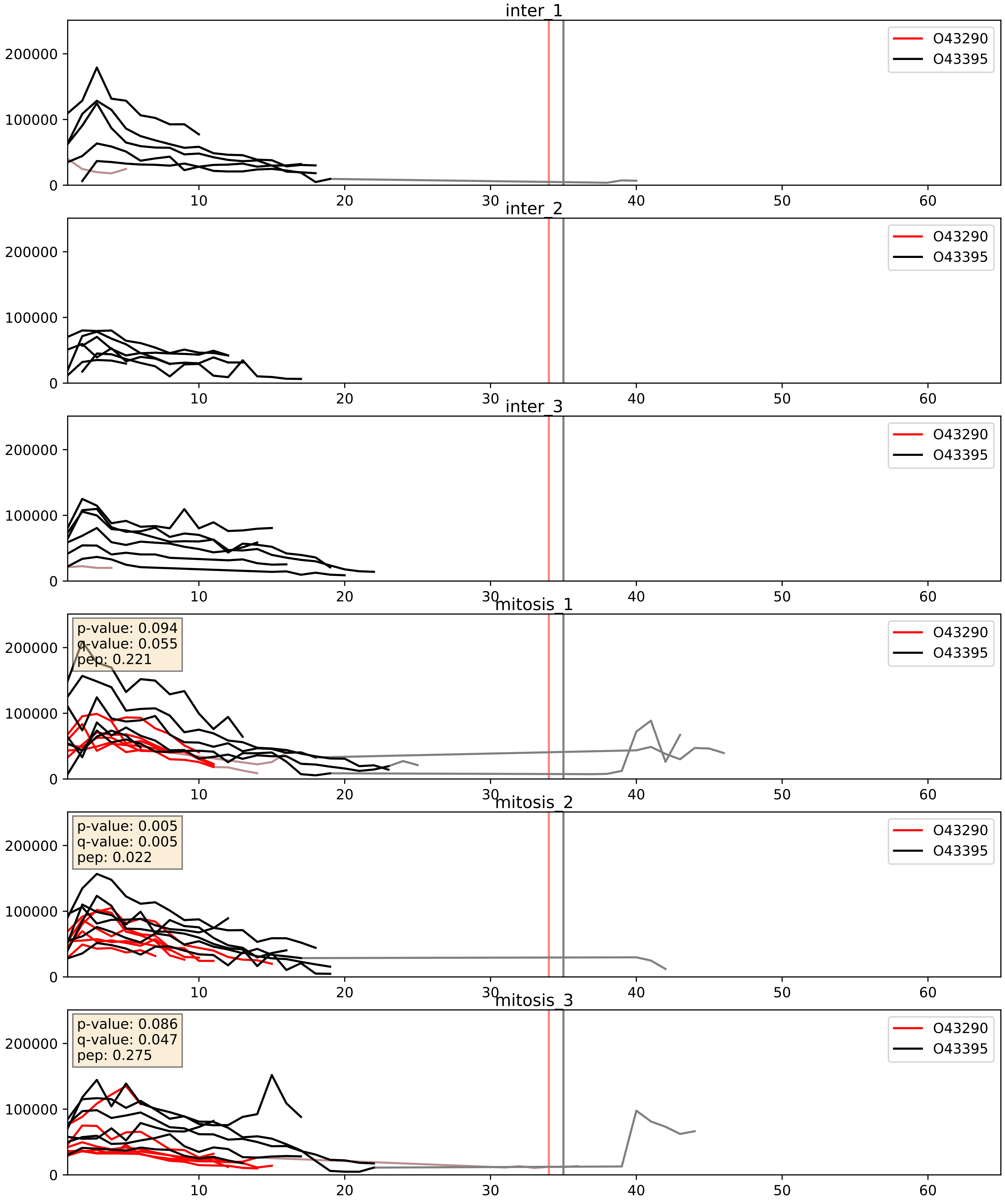


| level | condition_1 | condition_2 | pvalue | pvalue_adjusted |
|---|---|---|---|---|
| complex_abundance | mitosis | inter | 0.000327848443107248 | 0.011886635542316795 |
| interactor_abundance | mitosis | inter | 0.0010647499919498099 | 0.024049720606162197 |
| interactor_ratio | mitosis | inter | 0.007093762501146312 | 0.11727616584402438 |

O43290_O94906
SNUT1_HUMAN vs PRP6_HUMAN
p-value: 0.024 q-value: 0.019

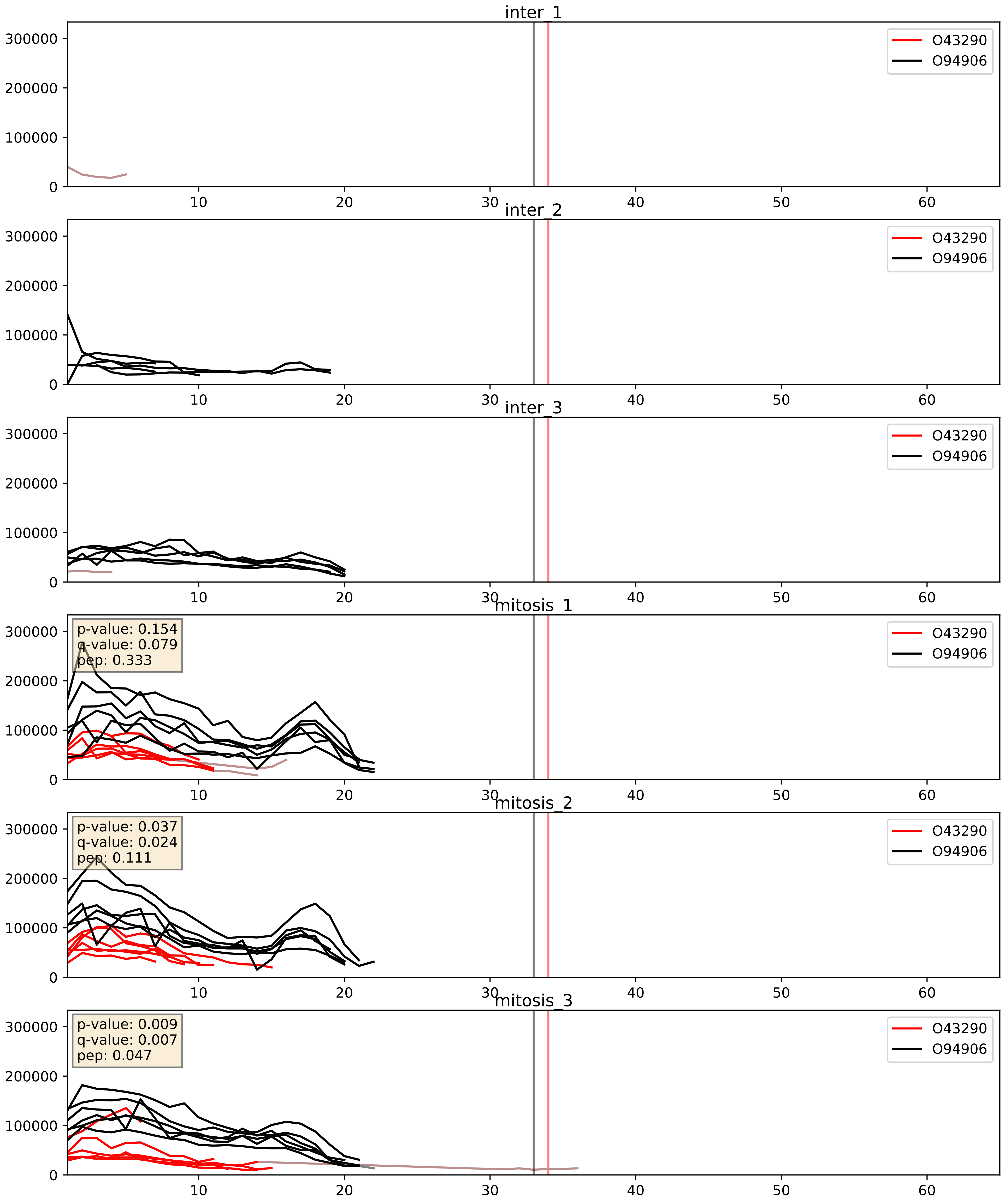


| level | condition_1 | condition_2 | pvalue | pvalue_adjusted |
|---|---|---|---|---|
| interactor_ratio | mitosis | inter | 0.00015386474196291182 | 0.012836278661937291 |
| complex_abundance | mitosis | inter | 0.0001611827964660317 | 0.008563834848633118 |
| interactor_abundance | mitosis | inter | 0.0010647499919498099 | 0.024049720606162197 |

O43290_Q2TAY7
SNUT1_HUMAN vs SMU1_HUMAN
p-value: 0.036 q-value: 0.025

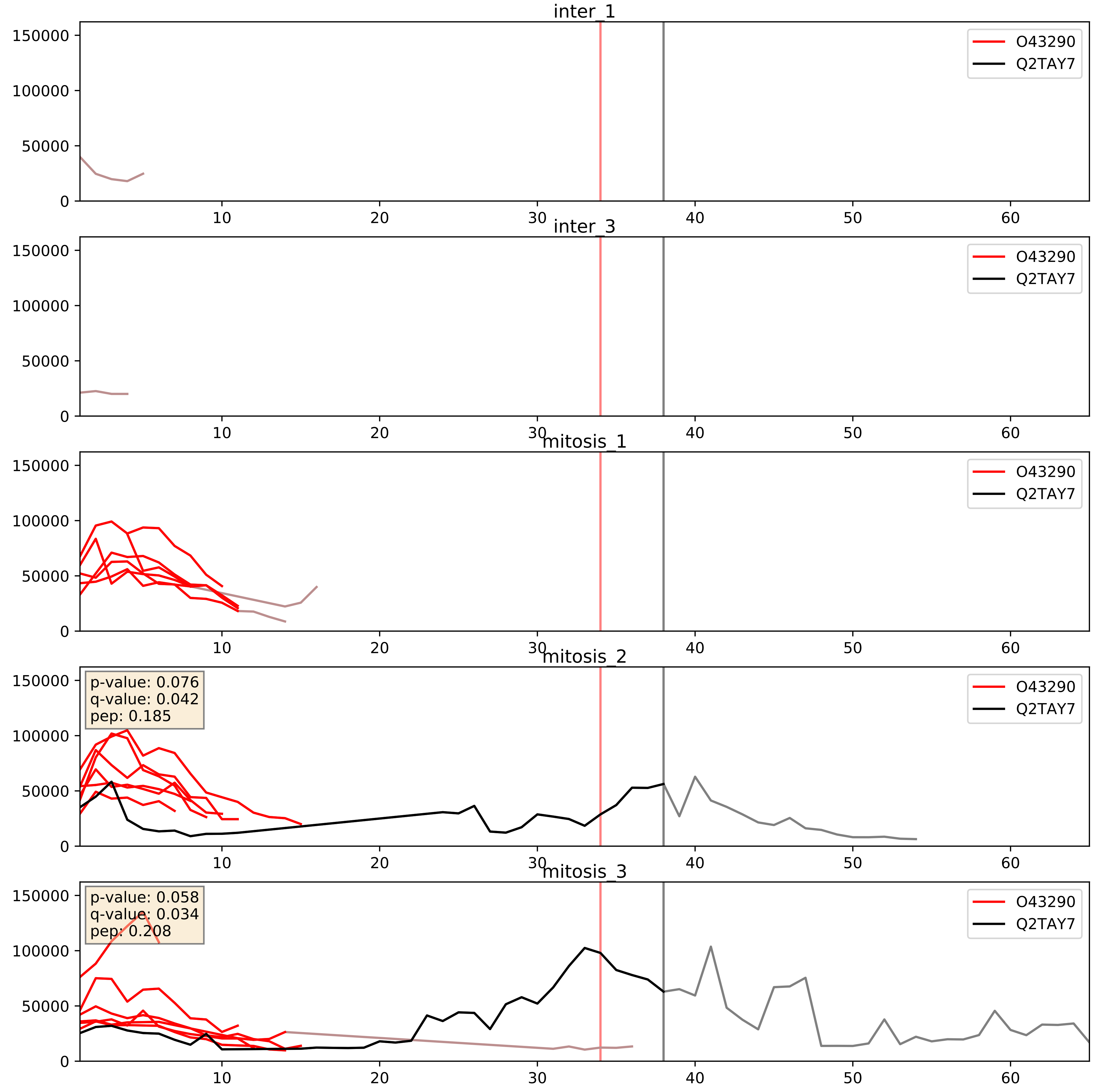


| level | condition_1 | condition_2 | pvalue | pvalue_adjusted |
|---|---|---|---|---|
| interactor_ratio | mitosis | inter | 0.194304615029616 | 0.667434793199644 |
| interactor_abundance | mitosis | inter | 0.2238361078991131 | 0.4190762663832913 |
| complex_abundance | mitosis | inter | 0.2490395466634567 | 0.3990599998950186 |

O43290_Q9BUQ8
SNUT1_HUMAN vs DDX23_HUMAN
p-value: 0.069 q-value: 0.042

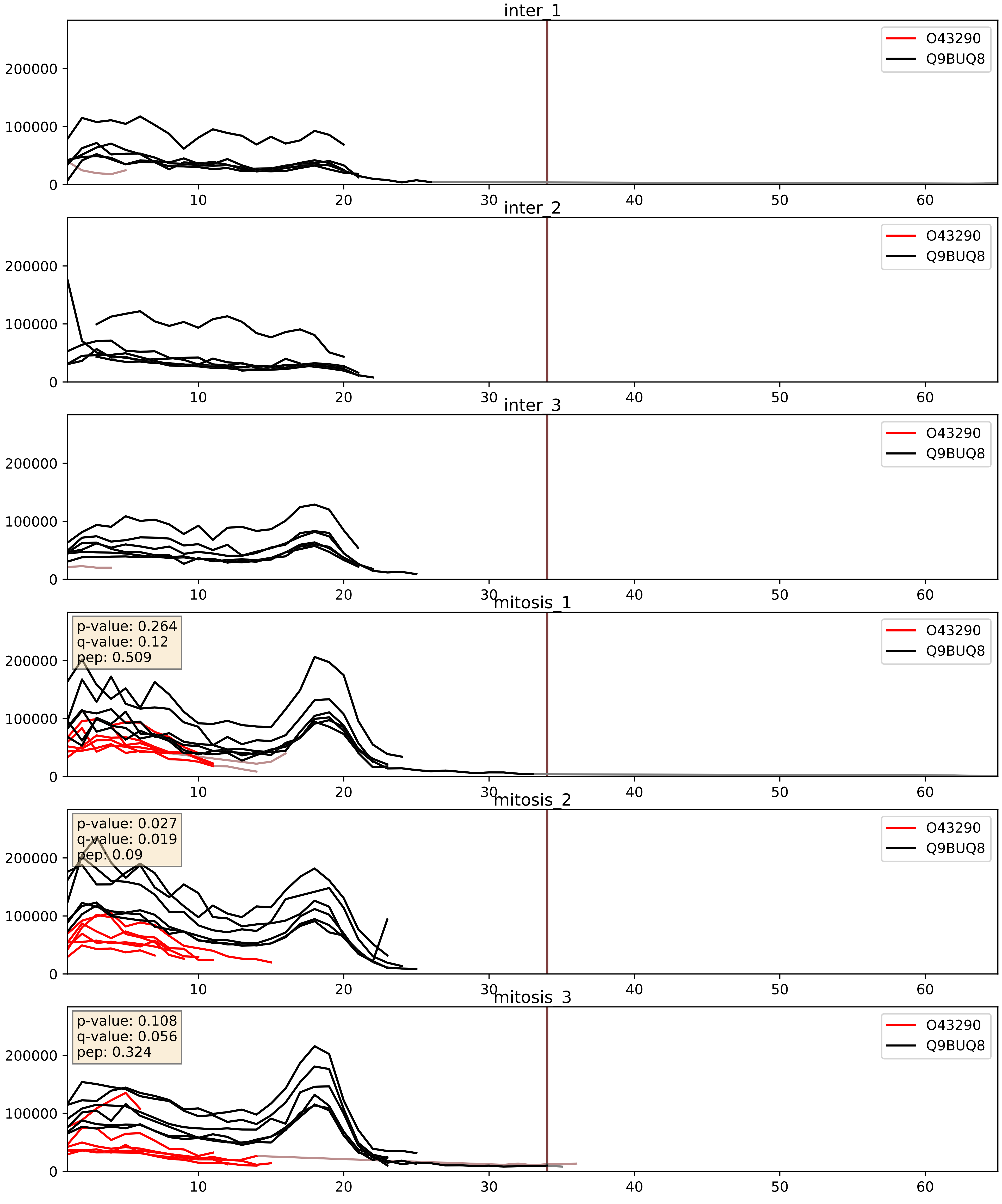


| level | condition_1 | condition_2 | pvalue | pvalue_adjusted |
|---|---|---|---|---|
| complex_abundance | mitosis | inter | 0.0002340547270430609 | 0.010221981956574496 |
| interactor_abundance | mitosis | inter | 0.0010647499919498099 | 0.024049720606162197 |
| interactor_ratio | mitosis | inter | 0.005120825250895749 | 0.09650938060012829 |

O43290_Q9BZJ0
SNUT1_HUMAN vs CRNL1_HUMAN
p-value: 0.047 q-value: 0.031

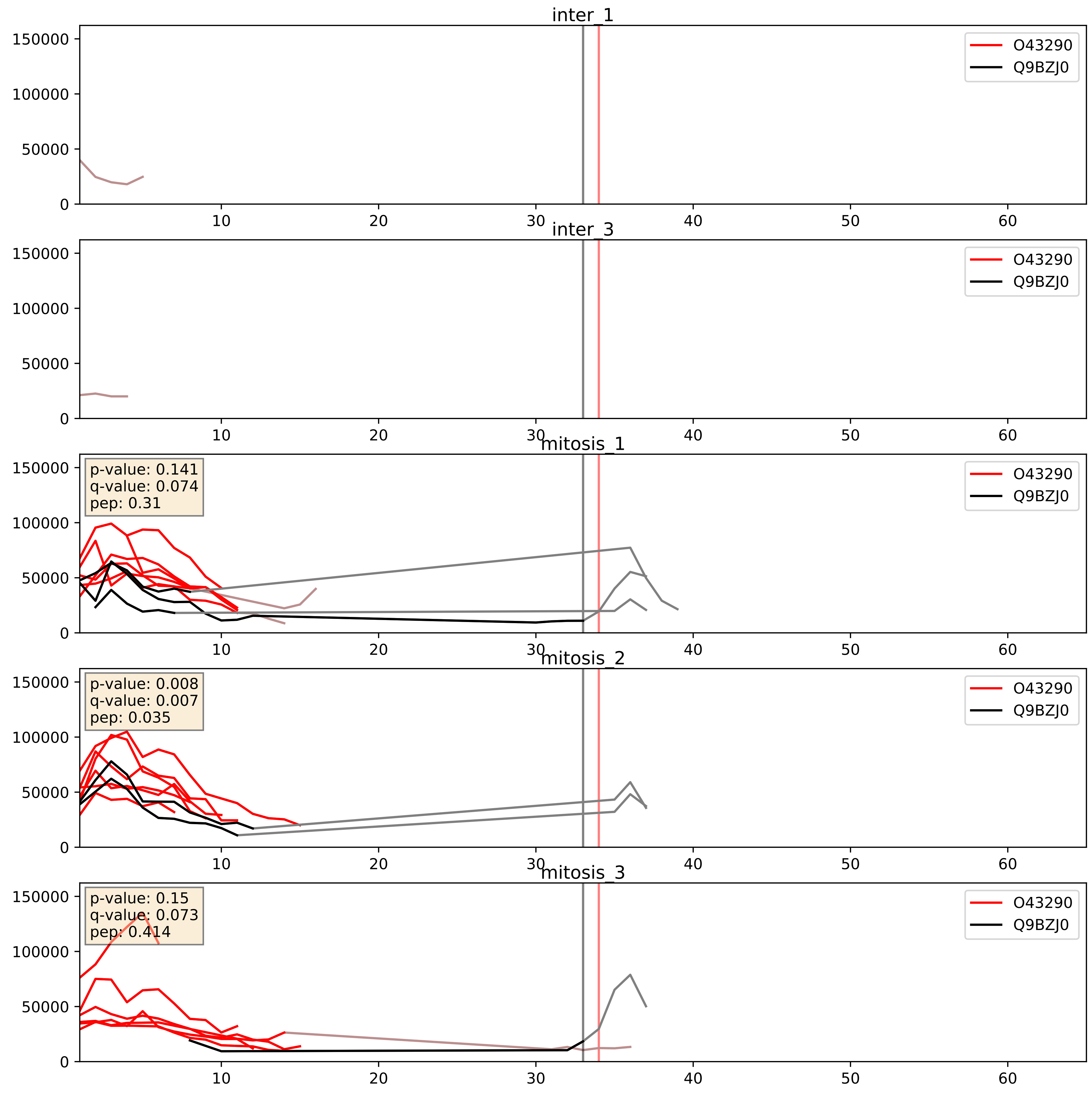


| level | condition_1 | condition_2 | pvalue | pvalue_adjusted |
|---|---|---|---|---|
| interactor_ratio | mitosis | inter | 0.0521009646673782 | 0.37764936302518703 |
| interactor_abundance | mitosis | inter | 0.2312435840999291 | 0.42318727300216413 |
| complex_abundance | mitosis | inter | 0.3821085485176027 | 0.5292636206004335 |
