## Supplementary Data 2 for "SECAT: Quantifying differential protein-protein interaction states by network-centric analysis": hela_string_000041_Q9BPX3.pdf

Q9BPX3_Q9BPX3

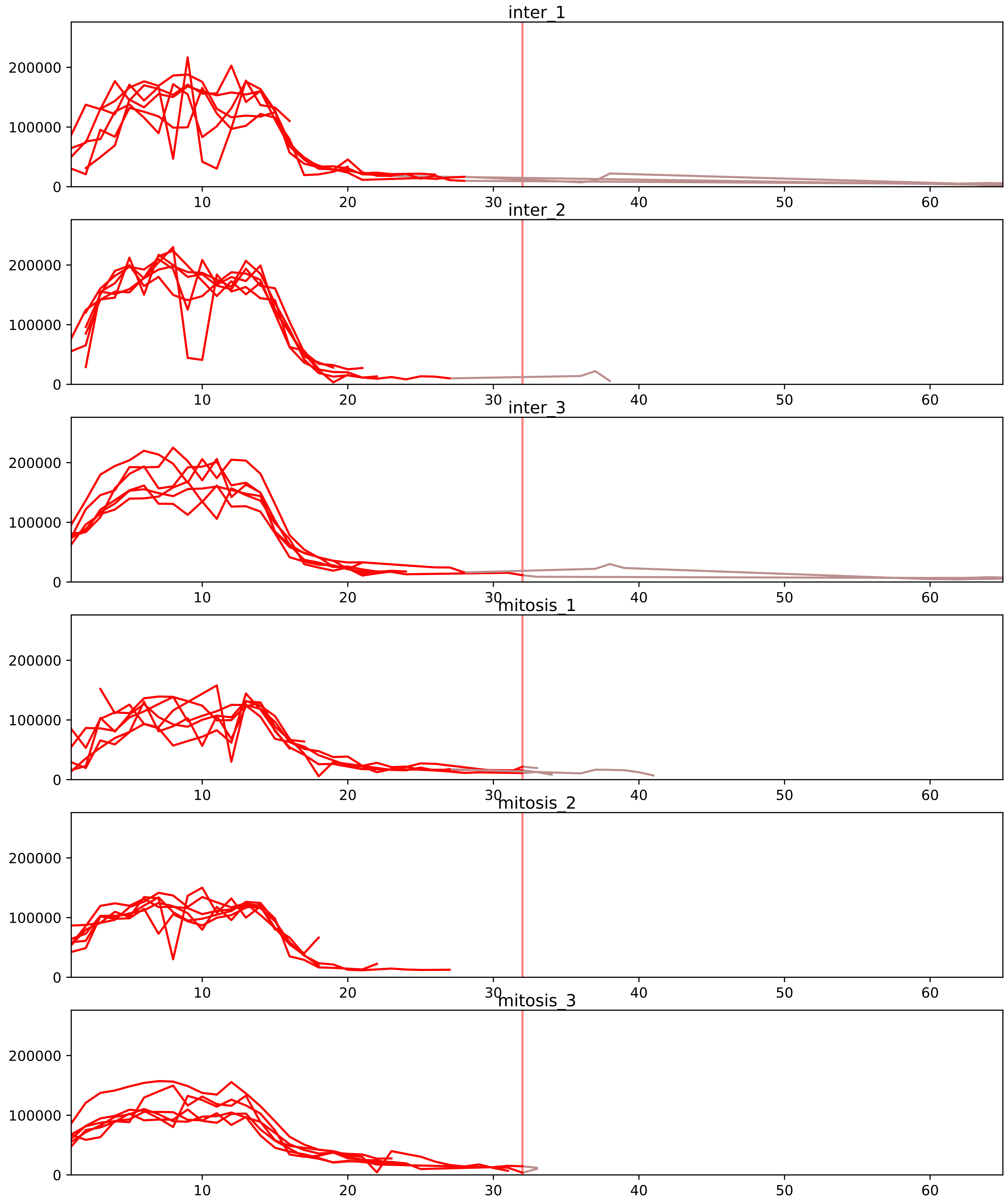


| level | condition_1 | condition_2 | pvalue | pvalue_adjusted |
|---|---|---|---|---|
| interactor_abundance | mitosis | inter | 1.7327887454498145e-08 | 6.3766625832553175e-06 |
| complex_abundance | mitosis | inter | 6.67266531213485e-08 | 1.5347130217910155e-05 |
| interactor_ratio | mitosis | inter | 0.003253396231383724 | 0.02797312647544884 |
| monomer_abundance | mitosis | inter | 0.4786370663152659 | 0.7397118297599564 |
| assembled_abundance | mitosis | inter | 0.6874720906675913 | 0.8693291740586497 |
| total_abundance | mitosis | inter | 0.711210875285267 | 0.8774393303188849 |

O43683_Q9BPX3
BUB1_HUMAN vs CND3_HUMAN
p-value: 0.075 q-value: 0.045

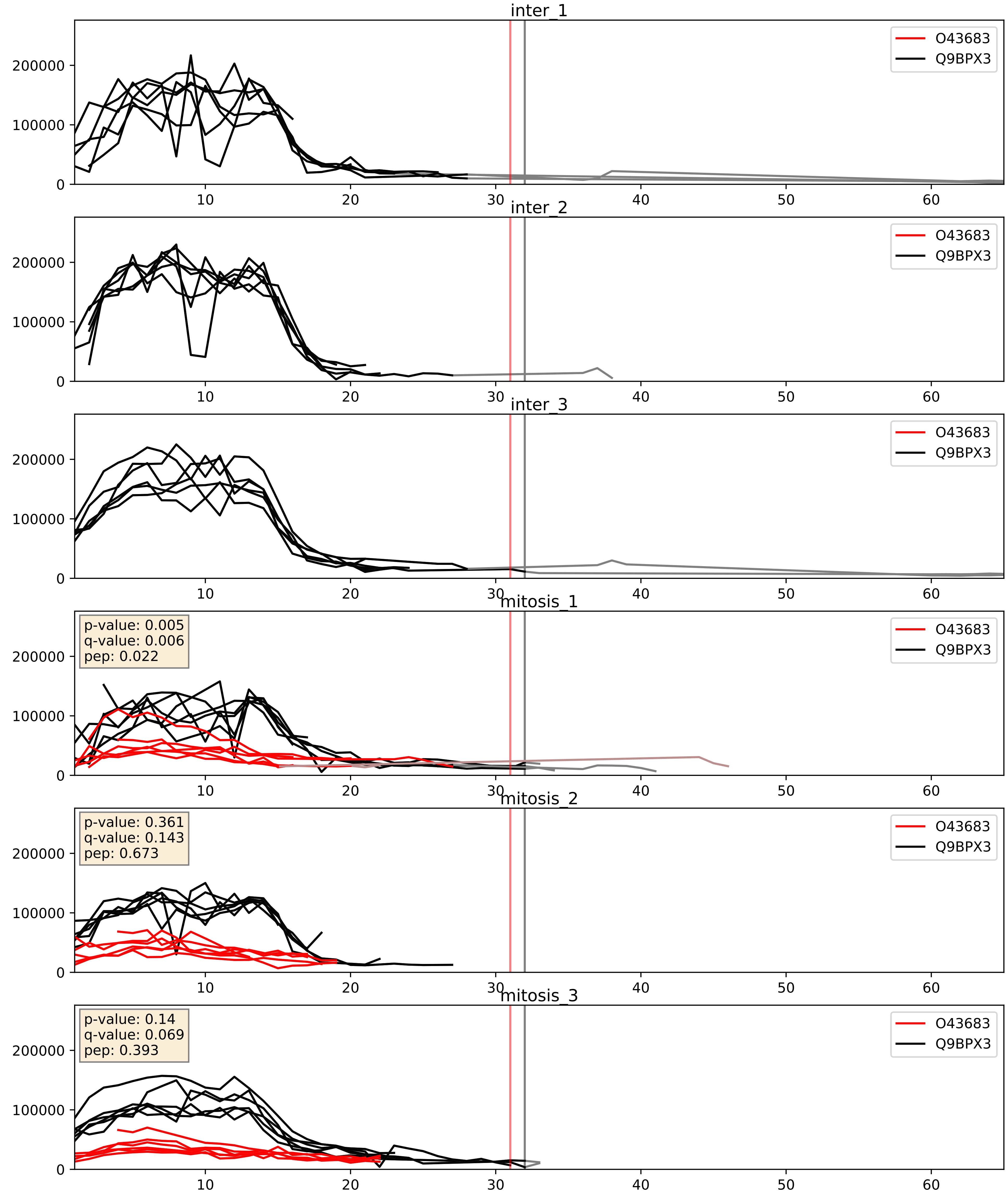


| level | condition_1 | condition_2 | pvalue | pvalue_adjusted |
|---|---|---|---|---|
| complex_abundance | mitosis | inter | 0.0004260739905454476 | 0.013091389064164169 |
| interactor_ratio | mitosis | inter | 0.0017705498104541042 | 0.05137595382199028 |
| interactor_abundance | mitosis | inter | 0.004465595680965726 | 0.05368749863632952 |

O95347_Q9BPX3
SMC2_HUMAN vs CND3_HUMAN
p-value: 0.013 q-value: 0.011

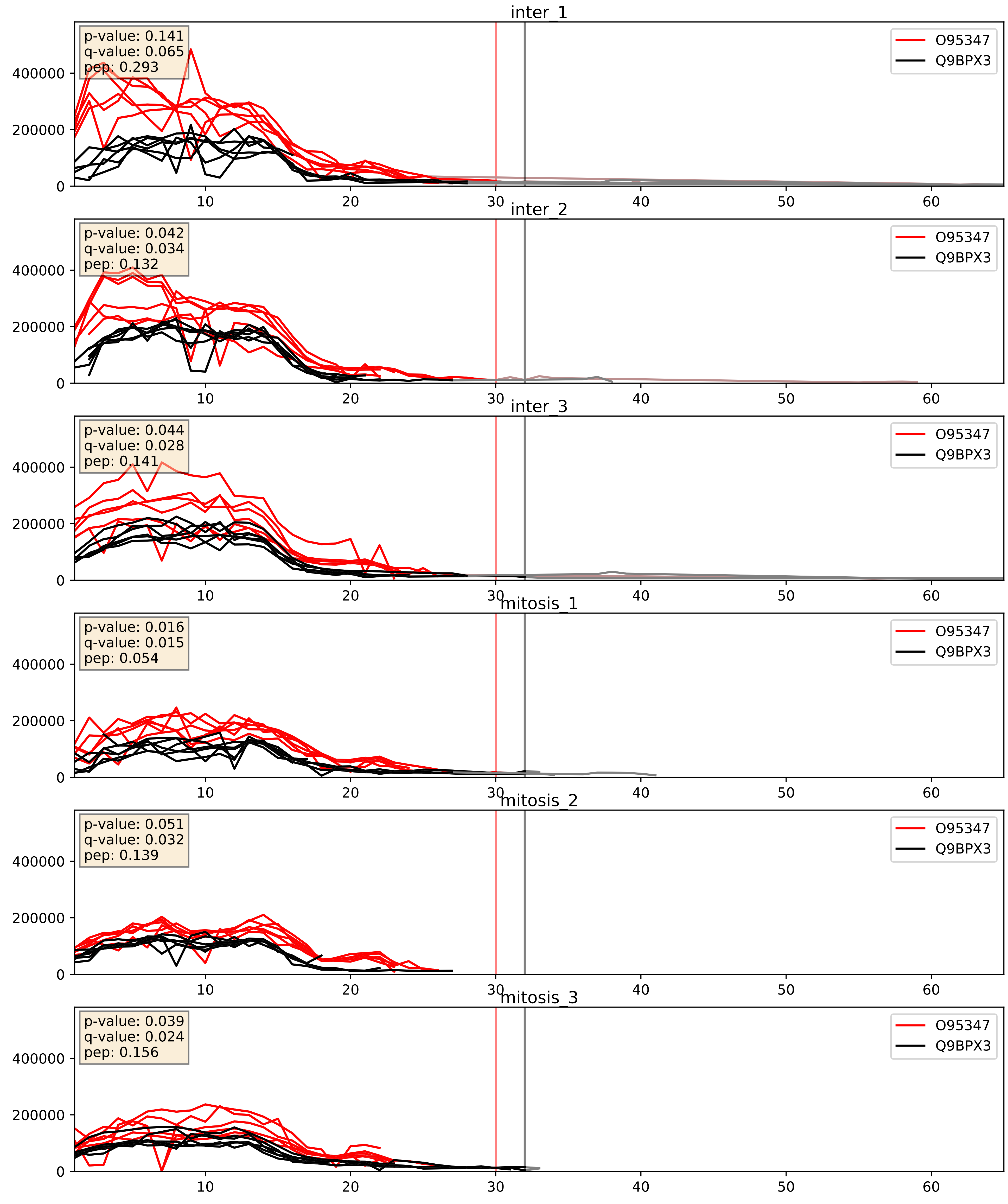


| level | condition_1 | condition_2 | pvalue | pvalue_adjusted |
|---|---|---|---|---|
| interactor_abundance | mitosis | inter | 0.0009422447006947195 | 0.02284551033392284 |
| complex_abundance | mitosis | inter | 0.001294321445799114 | 0.024459508269436846 |
| interactor_ratio | mitosis | inter | 0.012630388738437345 | 0.16210137600597313 |

P33981_Q9BPX3
TTK_HUMAN vs CND3_HUMAN
p-value: 0.051 q-value: 0.033

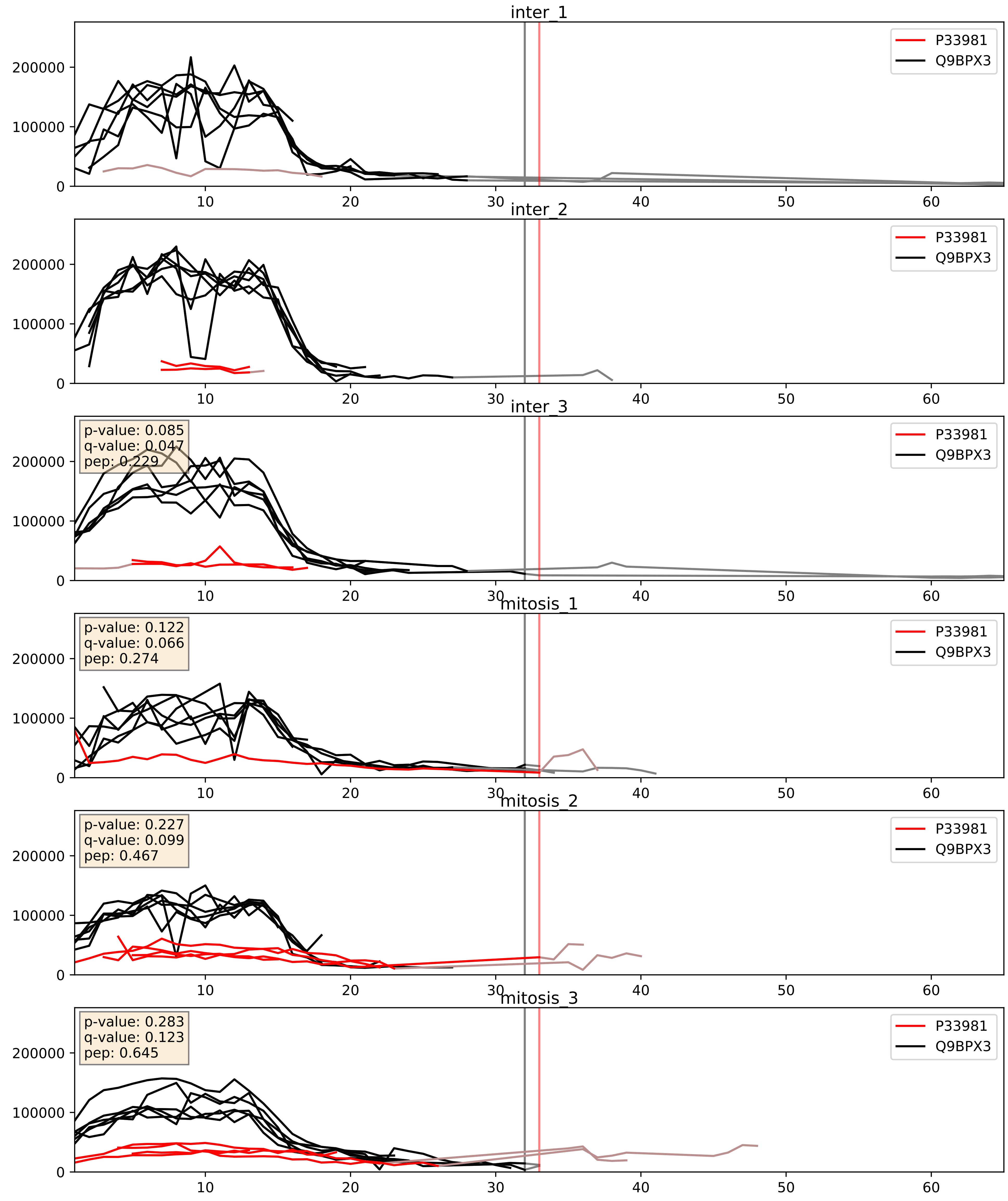


| level | condition_1 | condition_2 | pvalue | pvalue_adjusted |
|---|---|---|---|---|
| interactor_abundance | mitosis | inter | 0.08106202095833363 | 0.23762116958452006 |
| complex_abundance | mitosis | inter | 0.1393545669409426 | 0.28459313475568465 |
| interactor_ratio | mitosis | inter | 0.8458366137068474 | 0.9755634896635457 |

Q15003_Q9BPX3
CND2_HUMAN vs CND3_HUMAN
p-value: 0.003 q-value: 0.004

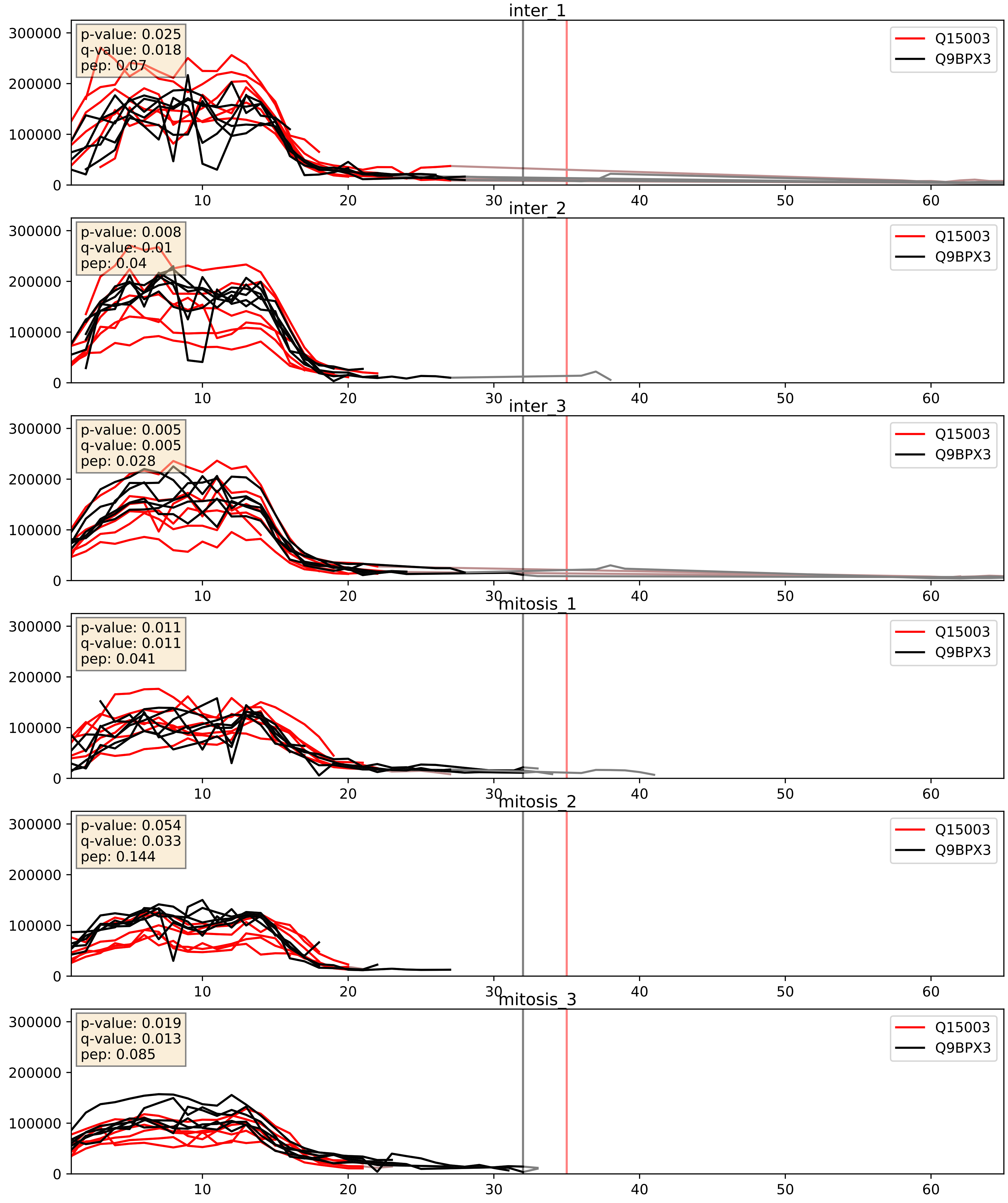


| level | condition_1 | condition_2 | pvalue | pvalue_adjusted |
|---|---|---|---|---|
| complex_abundance | mitosis | inter | 0.010946738379878694 | 0.07750544295431068 |
| interactor_abundance | mitosis | inter | 0.037758749960714884 | 0.1583513855205619 |
| interactor_ratio | mitosis | inter | 0.4893787185498002 | 0.8692763386835093 |

Q15021_Q9BPX3
CND1_HUMAN vs CND3_HUMAN
p-value: 0.004 q-value: 0.004

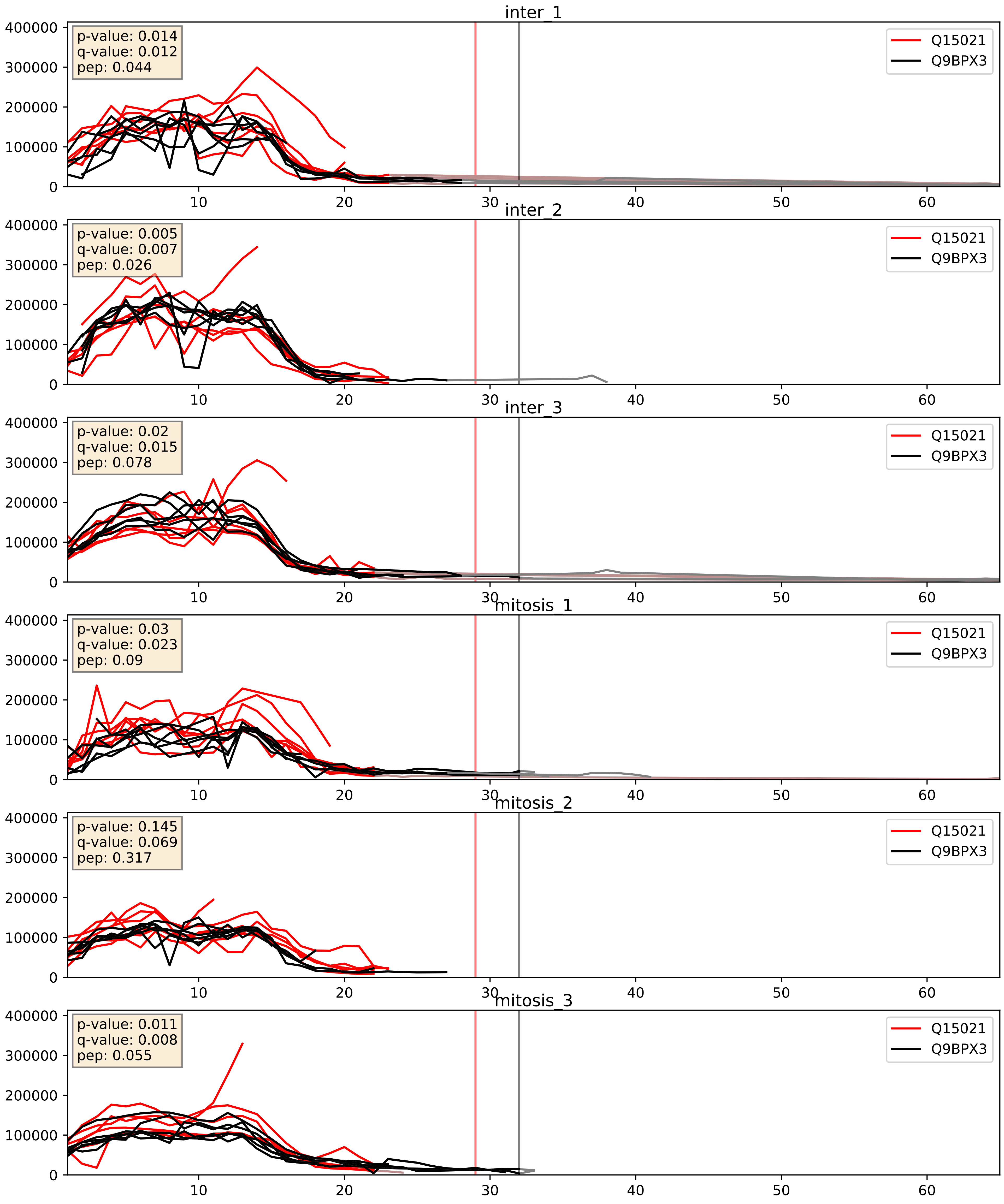


| level | condition_1 | condition_2 | pvalue | pvalue_adjusted |
|---|---|---|---|---|
| complex_abundance | mitosis | inter | 0.006059077368546302 | 0.05817678481497925 |
| interactor_abundance | mitosis | inter | 0.02229236740057479 | 0.12329937058940194 |
| interactor_ratio | mitosis | inter | 0.1652713568552461 | 0.6259835463189852 |

Q9BPX3_Q9NTJ3
CND3_HUMAN vs SMC4_HUMAN
p-value: 0.0 q-value: 0.001

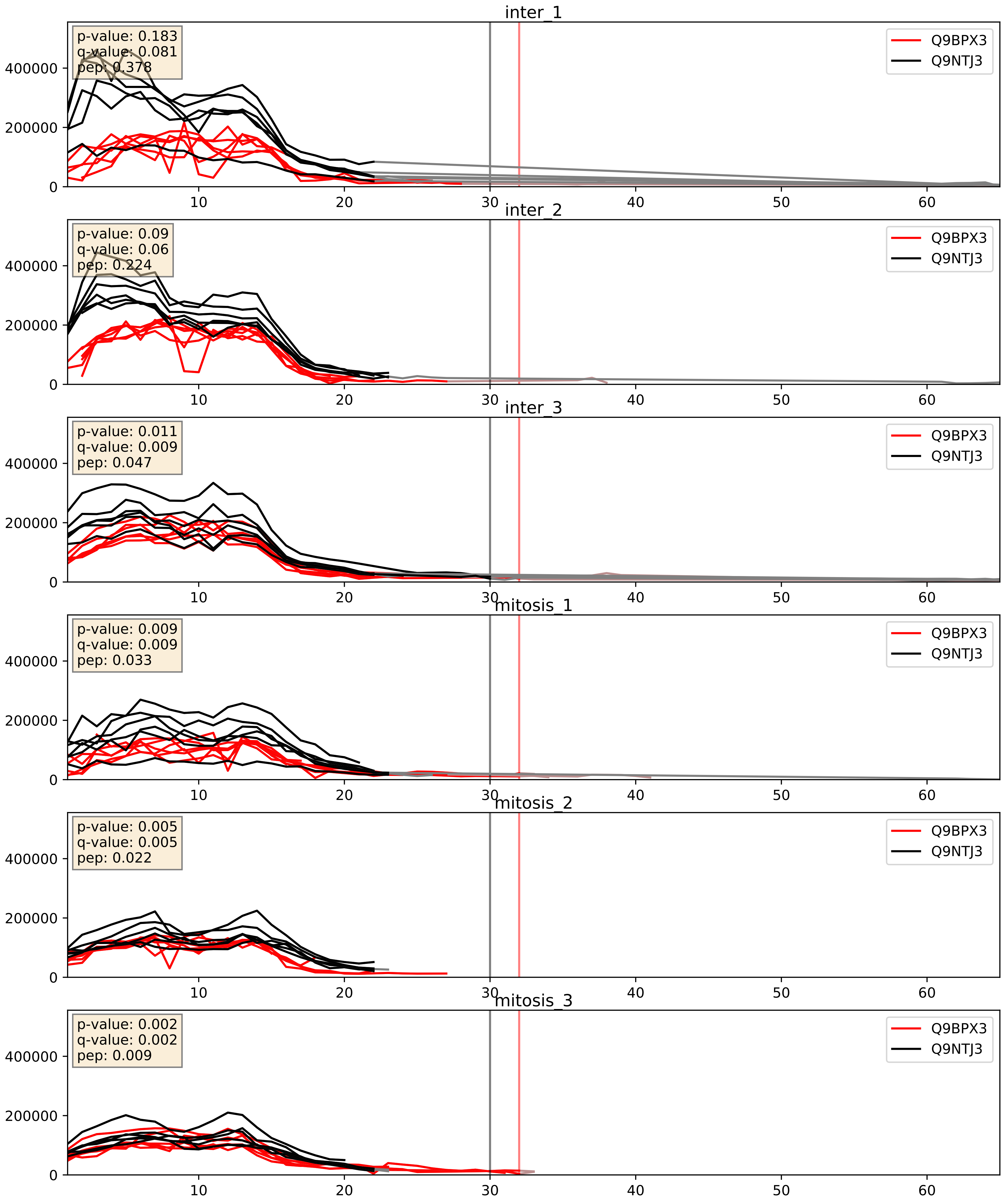


| level | condition_1 | condition_2 | pvalue | pvalue_adjusted |
|---|---|---|---|---|
| interactor_abundance | mitosis | inter | 0.0014070371627265526 | 0.027561185613133386 |
| complex_abundance | mitosis | inter | 0.004074531356902615 | 0.04732787841836958 |
| interactor_ratio | mitosis | inter | 0.24942022833694516 | 0.723755371092335 |
