## Supplementary Data 2 for "SECAT: Quantifying differential protein-protein interaction states by network-centric analysis": hela_string_000042_Q08211.pdf

Q08211_Q08211

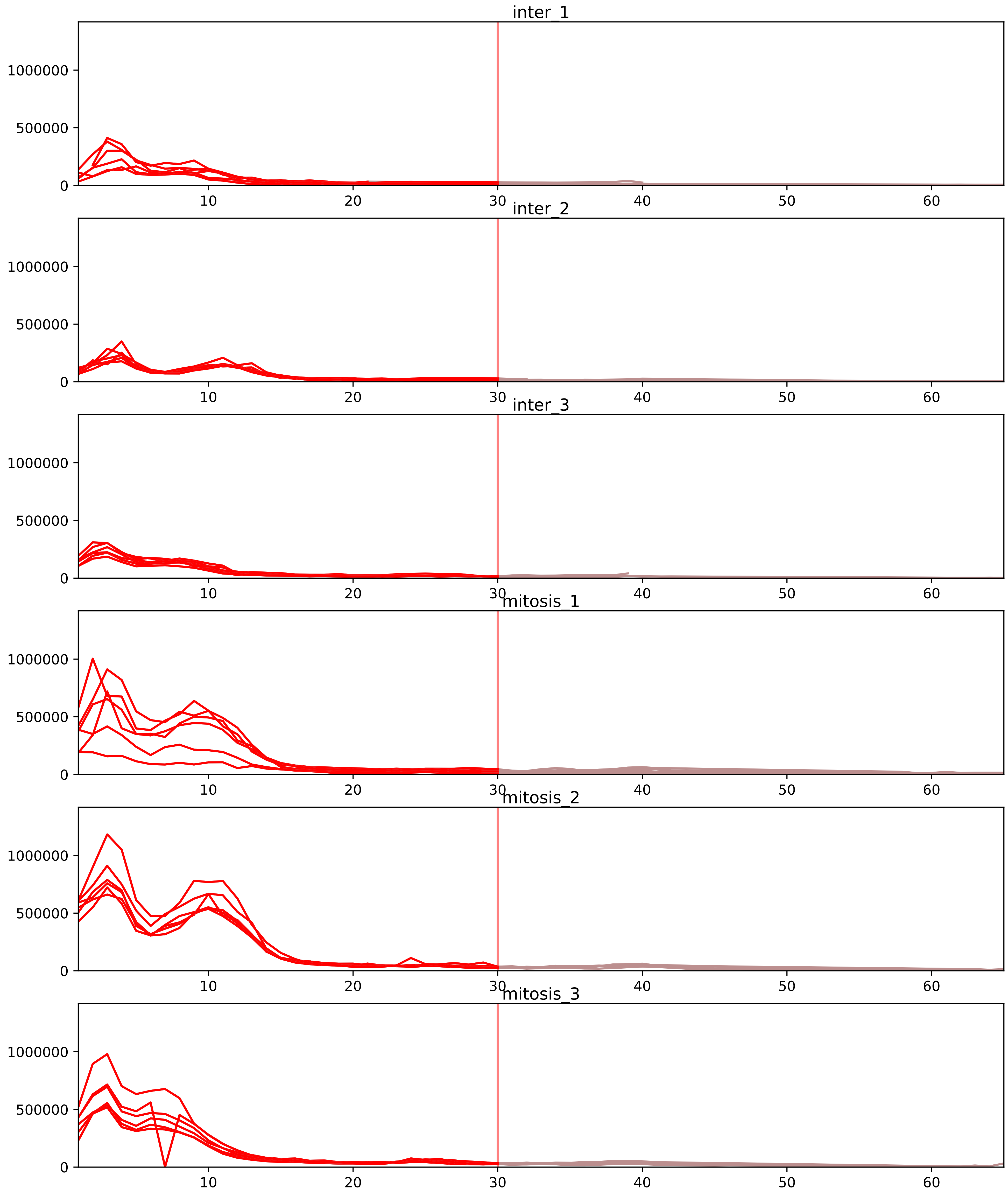


| level | condition_1 | condition_2 | pvalue | pvalue_adjusted |
|---|---|---|---|---|
| interactor_ratio | mitosis | inter | 1.79260803960625e-08 | 8.67999682335658e-07 |
| complex_abundance | mitosis | inter | 1.0645707909592191e-07 | 2.0418801137392286e-05 |
| interactor_abundance | mitosis | inter | 1.4275809298545349e-07 | 3.752498444189063e-05 |
| total_abundance | mitosis | inter | 0.0034808781952578058 | 0.0662034404709916 |
| assembled_abundance | mitosis | inter | 0.0036342999339008175 | 0.05995073678868365 |
| monomer_abundance | mitosis | inter | 0.005287723034330373 | 0.14516228843570478 |

O43390_Q08211
HNRPR_HUMAN vs DHX9_HUMAN
p-value: 0.004 q-value: 0.004

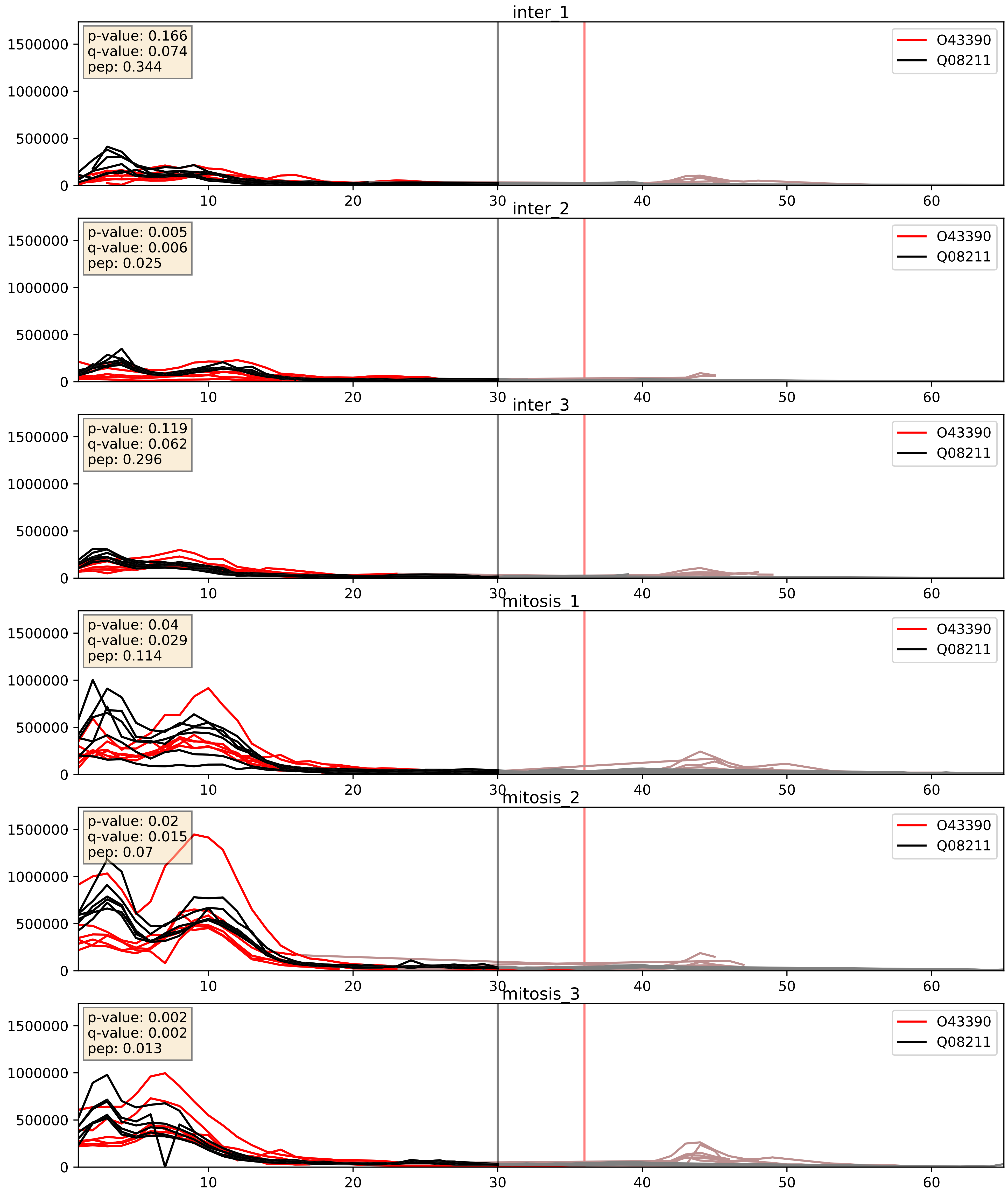


| level | condition_1 | condition_2 | pvalue | pvalue_adjusted |
|---|---|---|---|---|
| complex_abundance | mitosis | inter | 3.9289007550731724e-05 | 0.005691857151652958 |
| interactor_abundance | mitosis | inter | 0.00014528767019049324 | 0.009246561017328046 |
| interactor_ratio | mitosis | inter | 0.8637249543002753 | 0.9774585538872641 |

P07910_Q08211
HNRPC_HUMAN vs DHX9_HUMAN
p-value: 0.015 q-value: 0.012

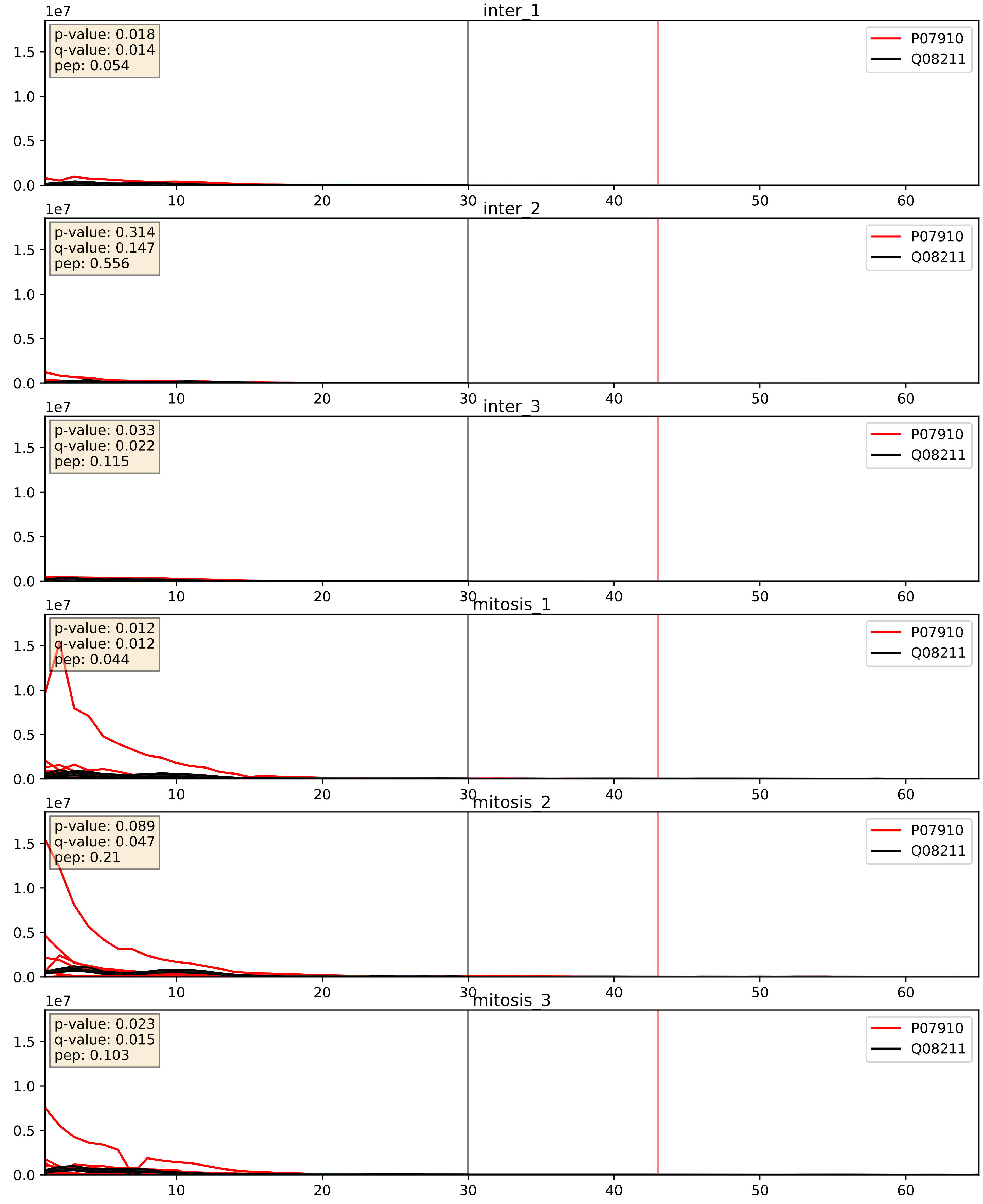


| level | condition_1 | condition_2 | pvalue | pvalue_adjusted |
|---|---|---|---|---|
| complex_abundance | mitosis | inter | 9.436874100373223e-06 | 0.005691857151652958 |
| interactor_abundance | mitosis | inter | 9.783708847801534e-06 | 0.0033734327106189023 |
| interactor_ratio | mitosis | inter | 8.825216175232504e-05 | 0.010208628440539222 |

P09651_Q08211
ROA1_HUMAN vs DHX9_HUMAN
p-value: 0.003 q-value: 0.004

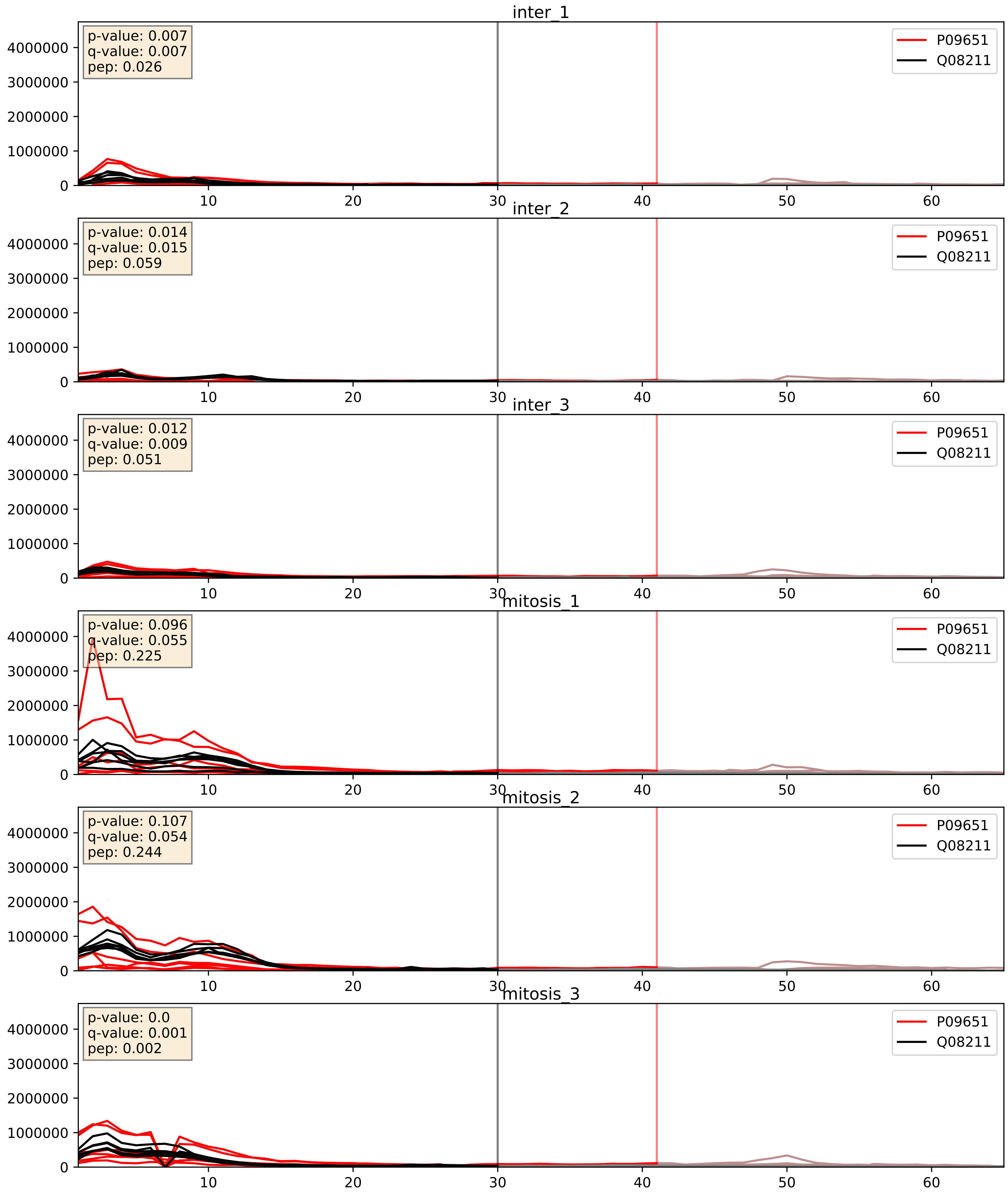


| level | condition_1 | condition_2 | pvalue | pvalue_adjusted |
|---|---|---|---|---|
| complex_abundance | mitosis | inter | 0.002467831850261896 | 0.03614494596242307 |
| interactor_abundance | mitosis | inter | 0.013517041923941197 | 0.10304923015527873 |
| interactor_ratio | mitosis | inter | 0.27952484058418636 | 0.7507789882022703 |

P14866_Q08211
HNRPL_HUMAN vs DHX9_HUMAN
p-value: 0.019 q-value: 0.015

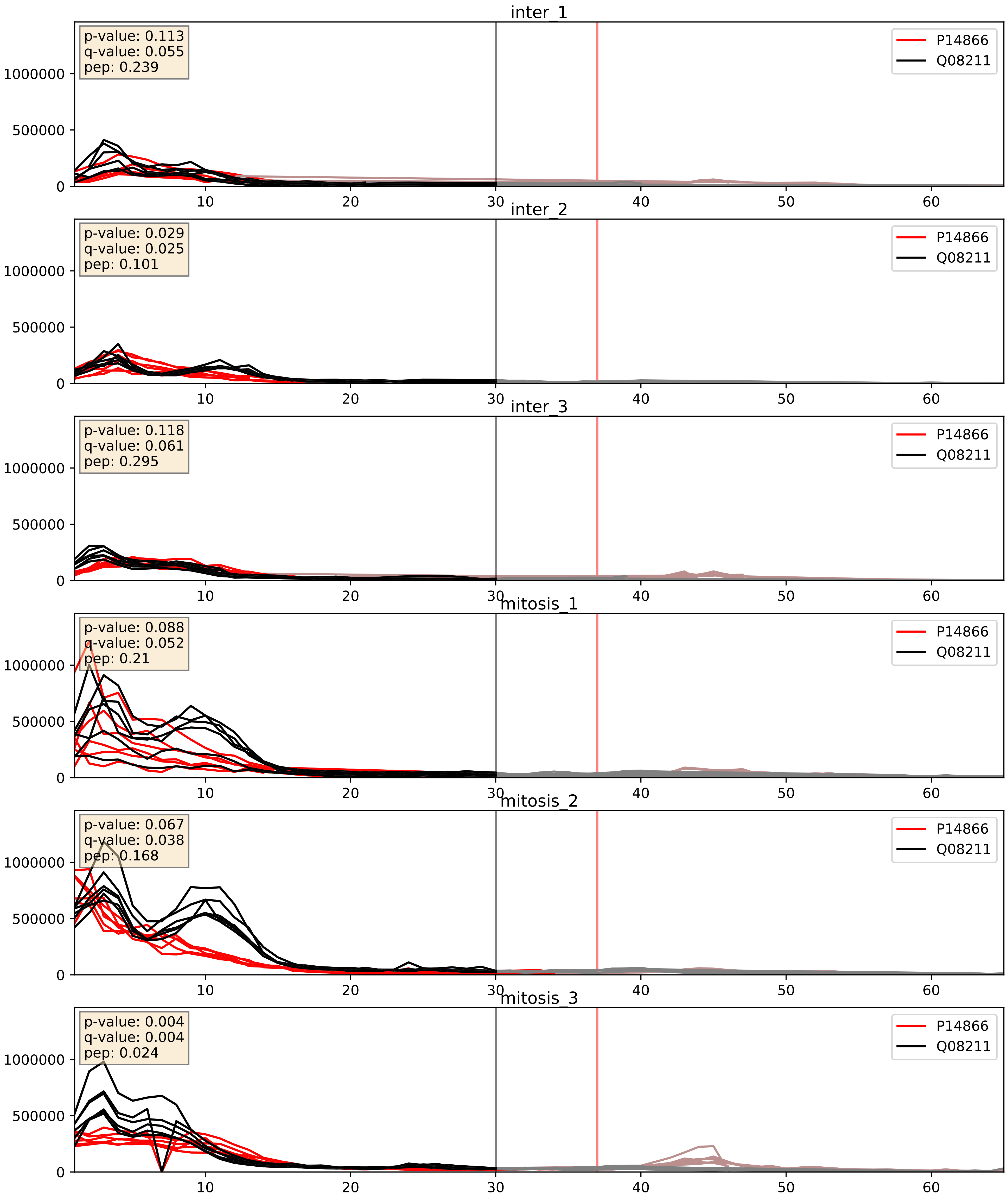


| level | condition_1 | condition_2 | pvalue | pvalue_adjusted |
|---|---|---|---|---|
| complex_abundance | mitosis | inter | 0.00043565000436869284 | 0.013137121901174406 |
| interactor_abundance | mitosis | inter | 0.010470812401431332 | 0.08918423299129571 |
| interactor_ratio | mitosis | inter | 0.9543473012403387 | 0.992614664476724 |

P17844_Q08211
DDX5_HUMAN vs DHX9_HUMAN
p-value: 0.004 q-value: 0.004

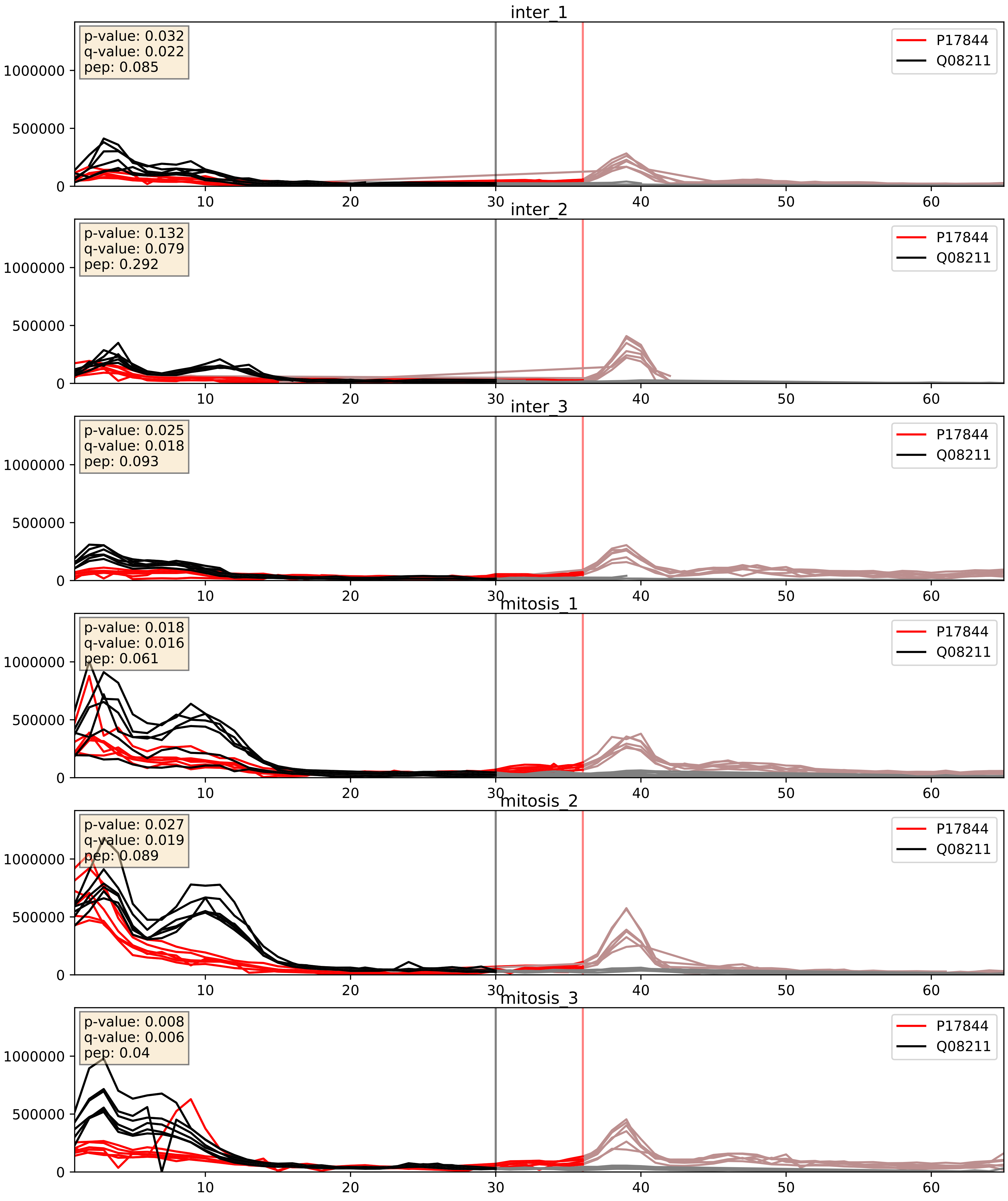


| level | condition_1 | condition_2 | pvalue | pvalue_adjusted |
|---|---|---|---|---|
| complex_abundance | mitosis | inter | 0.0004358577827025154 | 0.013137121901174406 |
| interactor_abundance | mitosis | inter | 0.0017533669129677133 | 0.030599022986755608 |
| interactor_ratio | mitosis | inter | 0.4374343152584505 | 0.8454166152138379 |

P22626_Q08211
ROA2_HUMAN vs DHX9_HUMAN
p-value: 0.007 q-value: 0.006

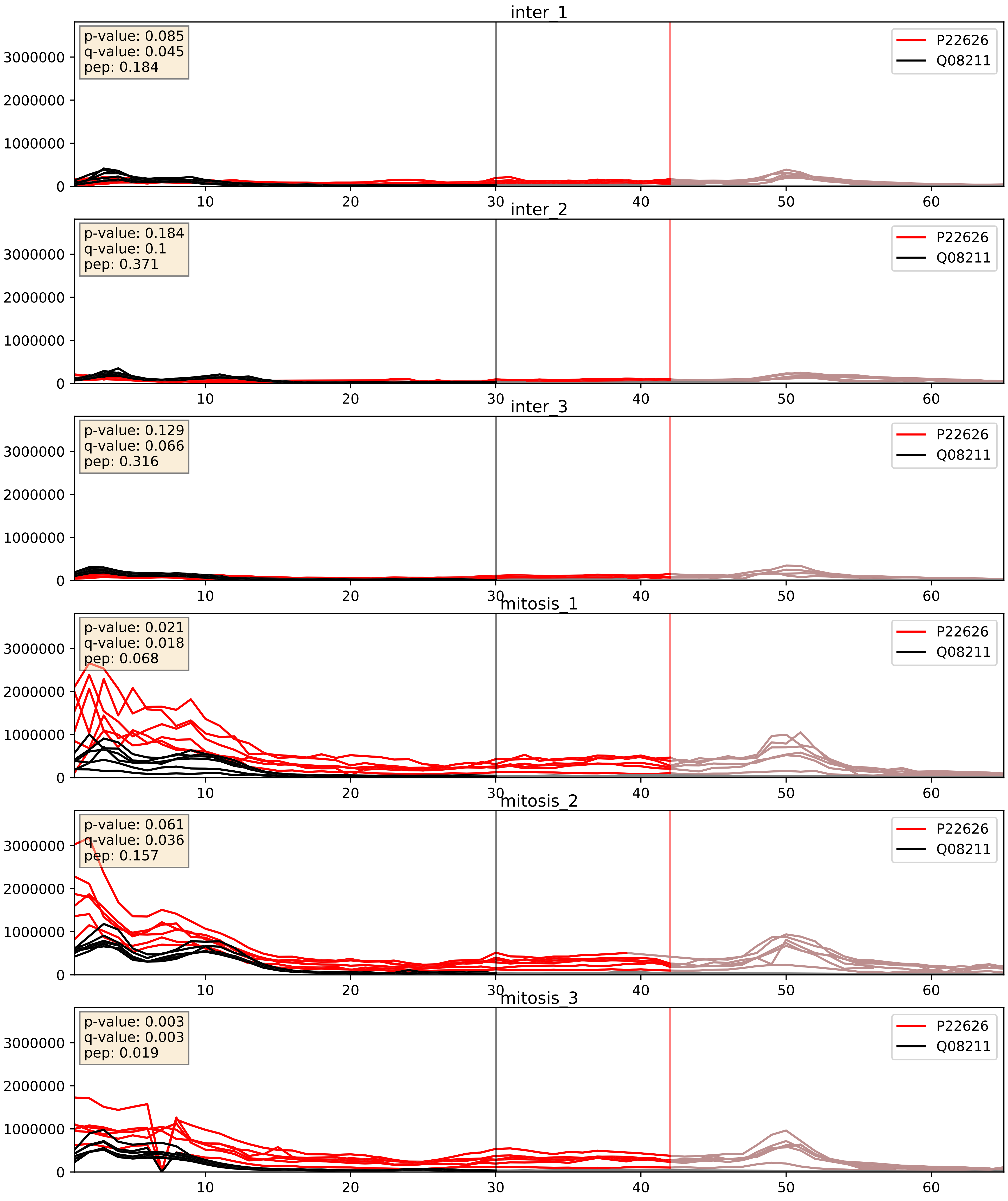


| level | condition_1 | condition_2 | pvalue | pvalue_adjusted |
|---|---|---|---|---|
| complex_abundance | mitosis | inter | 0.0009287786996386022 | 0.02007663047703645 |
| interactor_abundance | mitosis | inter | 0.0022087495902483206 | 0.034628015554076234 |
| interactor_ratio | mitosis | inter | 0.009213692076387548 | 0.13668839544866102 |

P51991_Q08211
ROA3_HUMAN vs DHX9_HUMAN
p-value: 0.003 q-value: 0.045

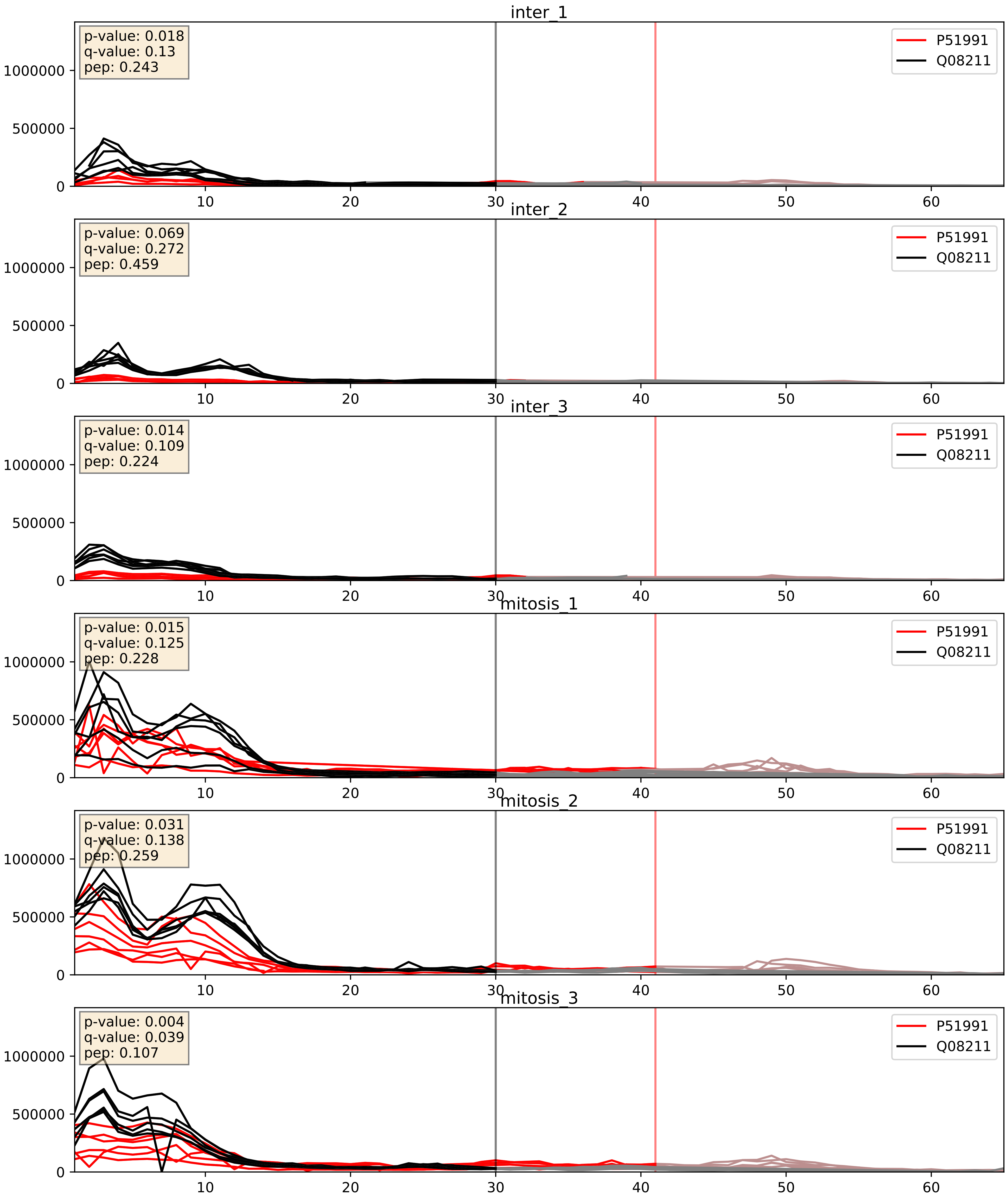


| level | condition_1 | condition_2 | pvalue | pvalue_adjusted |
|---|---|---|---|---|
| interactor_abundance | mitosis | inter | 4.591141207513425e-06 | 0.0033734327106189023 |
| complex_abundance | mitosis | inter | 7.4317189984560 32e-06 | 0.005691857151652958 |
| interactor_ratio | mitosis | inter | 0.0005736461993080531 | 0.0279000651481644 |

P52272_Q08211
HNRPM_HUMAN vs DHX9_HUMAN
p-value: 0.001 q-value: 0.002

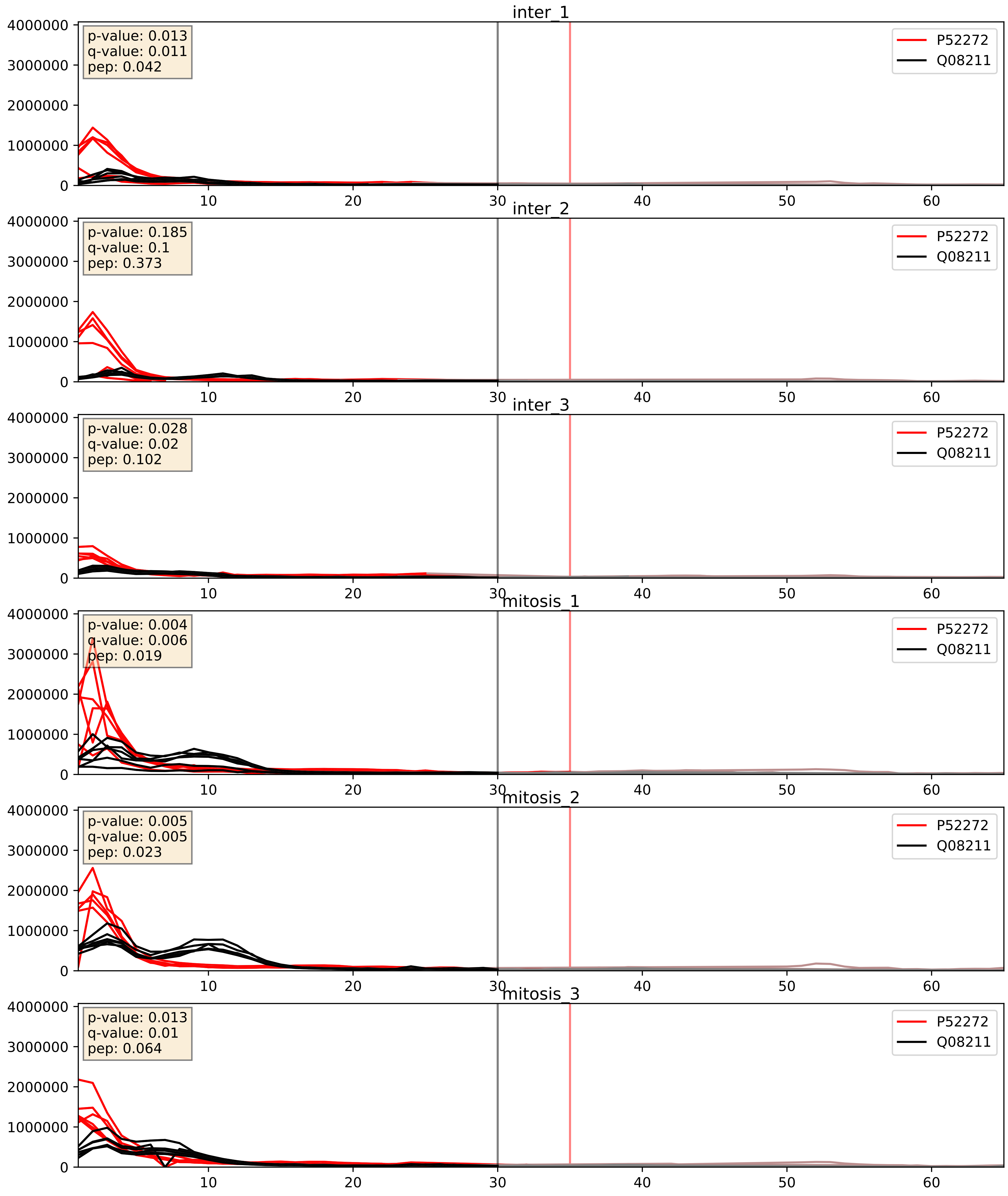


| level | condition_1 | condition_2 | pvalue | pvalue_adjusted |
|---|---|---|---|---|
| complex_abundance | mitosis | inter | 0.006357588474845068 | 0.05948545934570088 |
| interactor_ratio | mitosis | inter | 0.024442223536284675 | 0.2447639398105627 |
| interactor_abundance | mitosis | inter | 0.19956587550211832 | 0.39378164892295875 |

P52597_Q08211
HNRPF_HUMAN vs DHX9_HUMAN
p-value: 0.002 q-value: 0.03

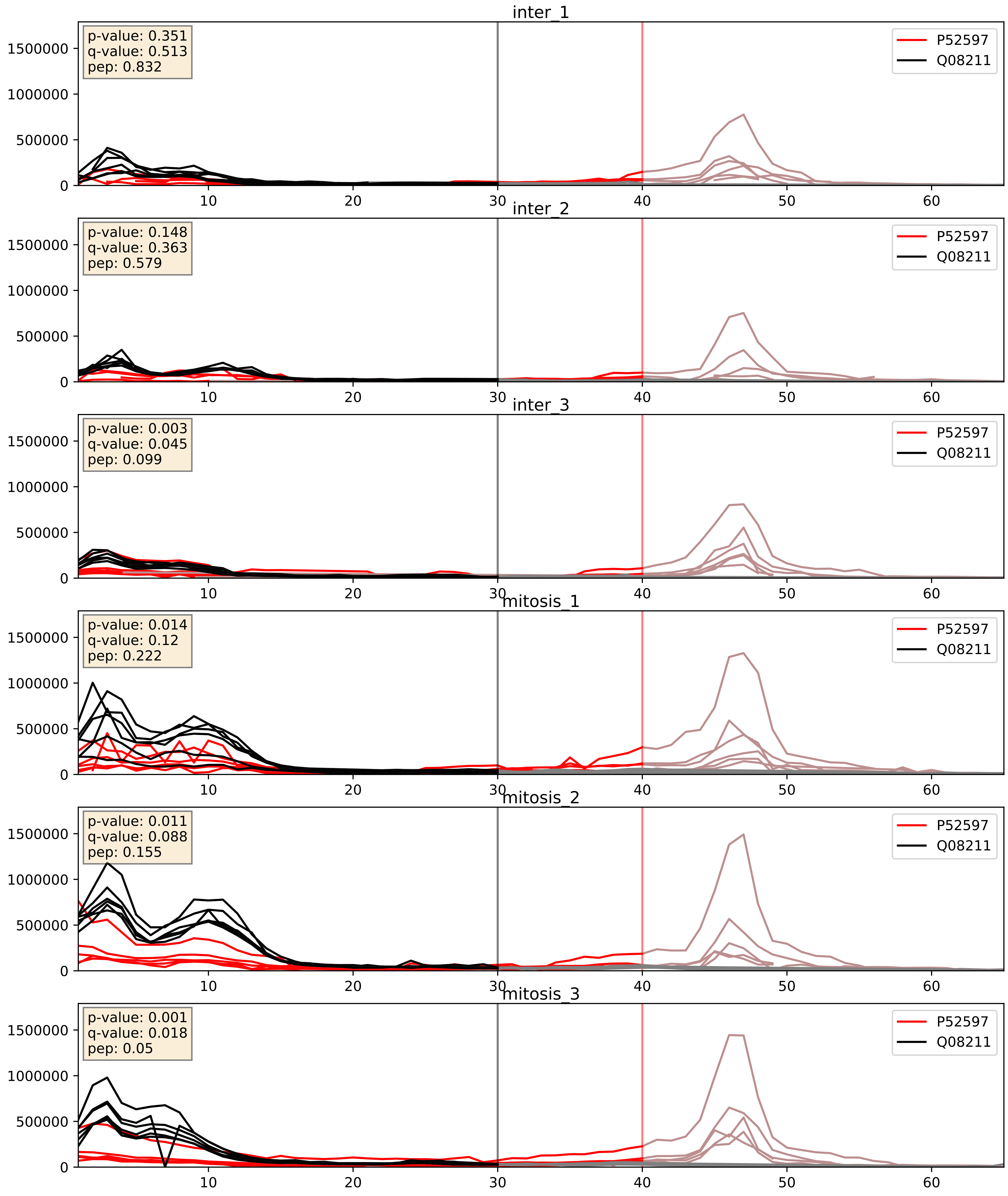


| level | condition_1 | condition_2 | pvalue | pvalue_adjusted |
|---|---|---|---|---|
| complex_abundance | mitosis | inter | 0.0005487925892557542 | 0.015047650716485379 |
| interactor_abundance | mitosis | inter | 0.007718766641450282 | 0.0762963538693007 |
| interactor_ratio | mitosis | inter | 0.1472602964791206 | 0.6016936218908221 |

P55769_Q08211
NH2L1_HUMAN vs DHX9_HUMAN
p-value: 0.001 q-value: 0.048

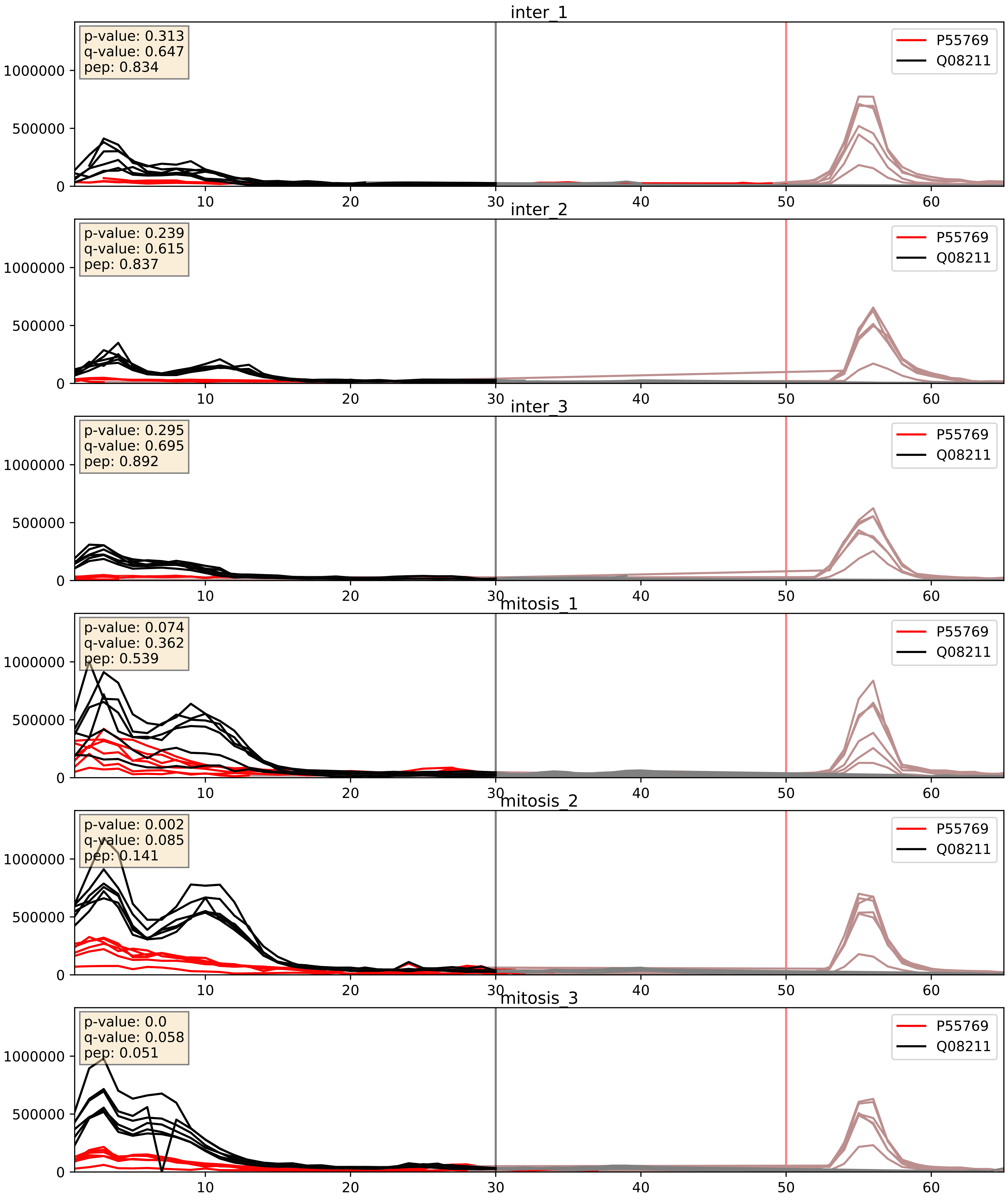


| level | condition_1 | condition_2 | pvalue | pvalue_adjusted |
|---|---|---|---|---|
| complex_abundance | mitosis | inter | 0.0007198582539638076 | 0.017505643903210 78 |
| interactor_abundance | mitosis | inter | 0.0013996461911011957 | 0.02753913185123736 |
| interactor_ratio | mitosis | inter | 0.012767744743810204 | 0.16336606129598708 |

P61978_Q08211
HNRPK_HUMAN vs DHX9_HUMAN
p-value: 0.023 q-value: 0.018

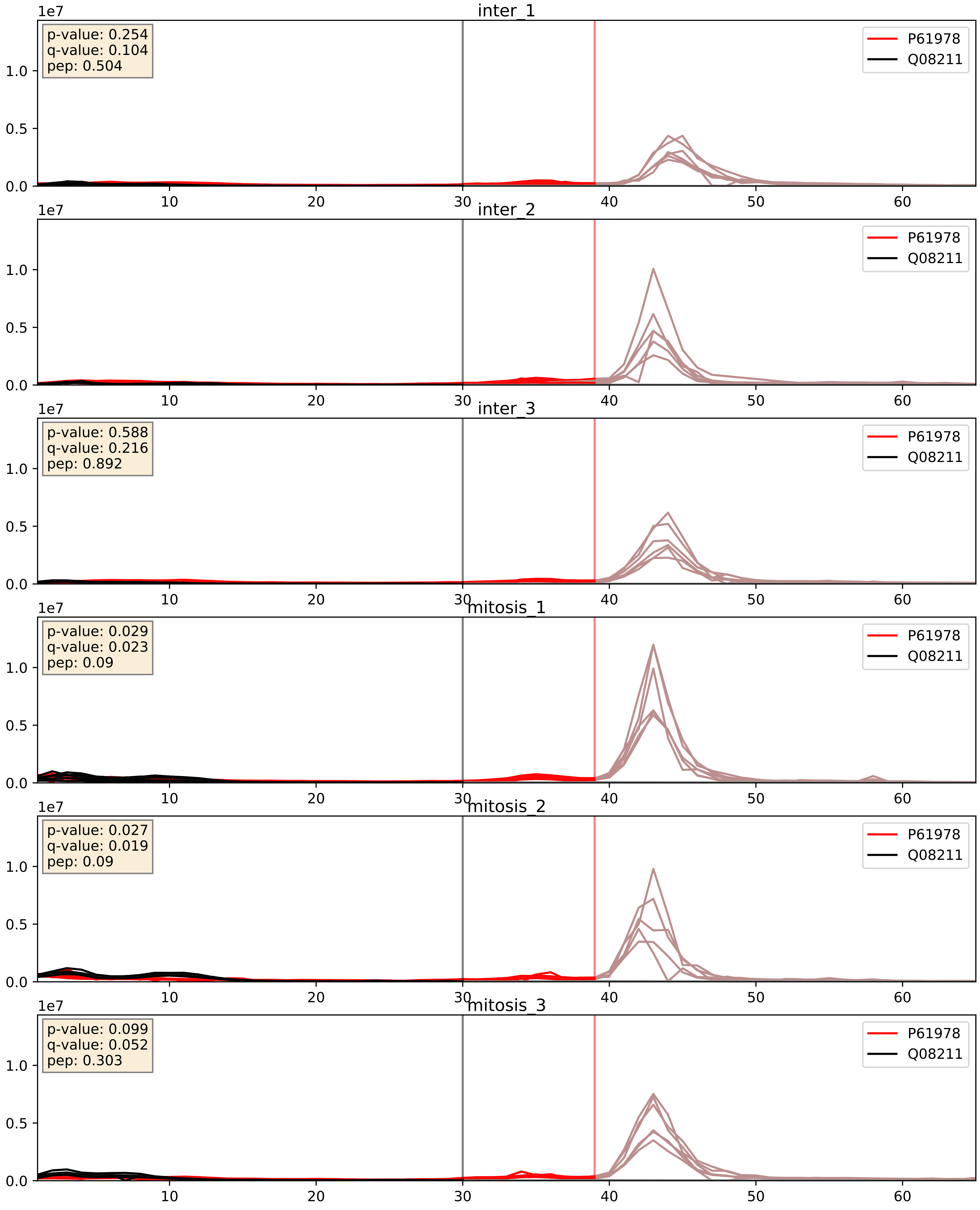


| level | condition_1 | condition_2 | pvalue | pvalue_adjusted |
|---|---|---|---|---|
| interactor_ratio | mitosis | inter | 0.11757582191833955 | 0.5644694535170984 |
| complex_abundance | mitosis | inter | 0.17223141055009858 | 0.32161886437802006 |
| interactor_abundance | mitosis | inter | 0.8464929270132362 | 0.915763596238016 |

Q07955_Q08211
SRSF1_HUMAN vs DHX9_HUMAN
p-value: 0.076 q-value: 0.046

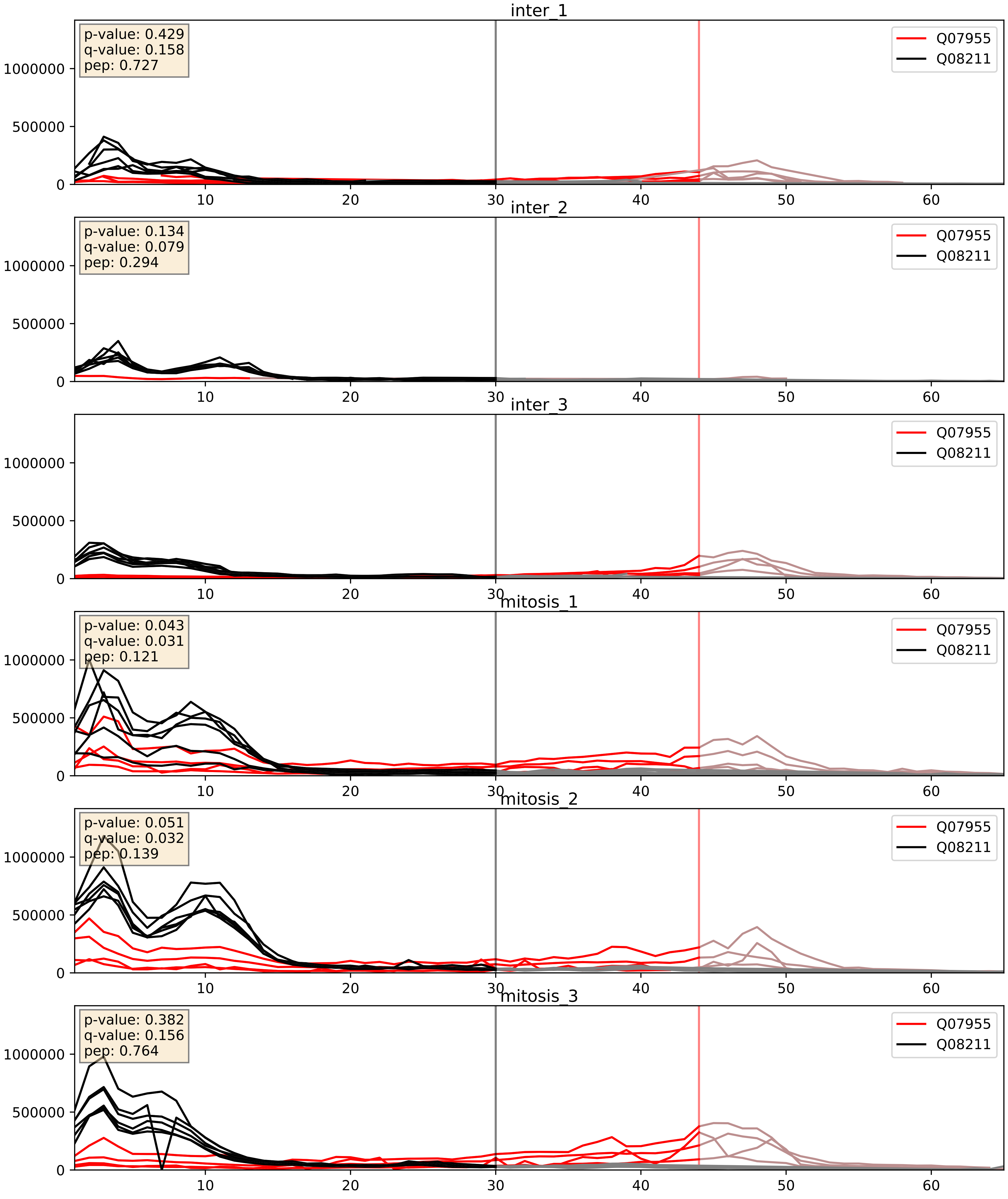


| level | condition_1 | condition_2 | pvalue | pvalue_adjusted |
|---|---|---|---|---|
| complex_abundance | mitosis | inter | 0.0014016180824241073 | 0.025581771397761962 |
| interactor_abundance | mitosis | inter | 0.0024364603316211677 | 0.03681201502126771 |
| interactor_ratio | mitosis | inter | 0.05437572514359929 | 0.38595042058806794 |

Q08211_Q12905
DHX9_HUMAN vs ILF2_HUMAN
p-value: 0.001 q-value: 0.039

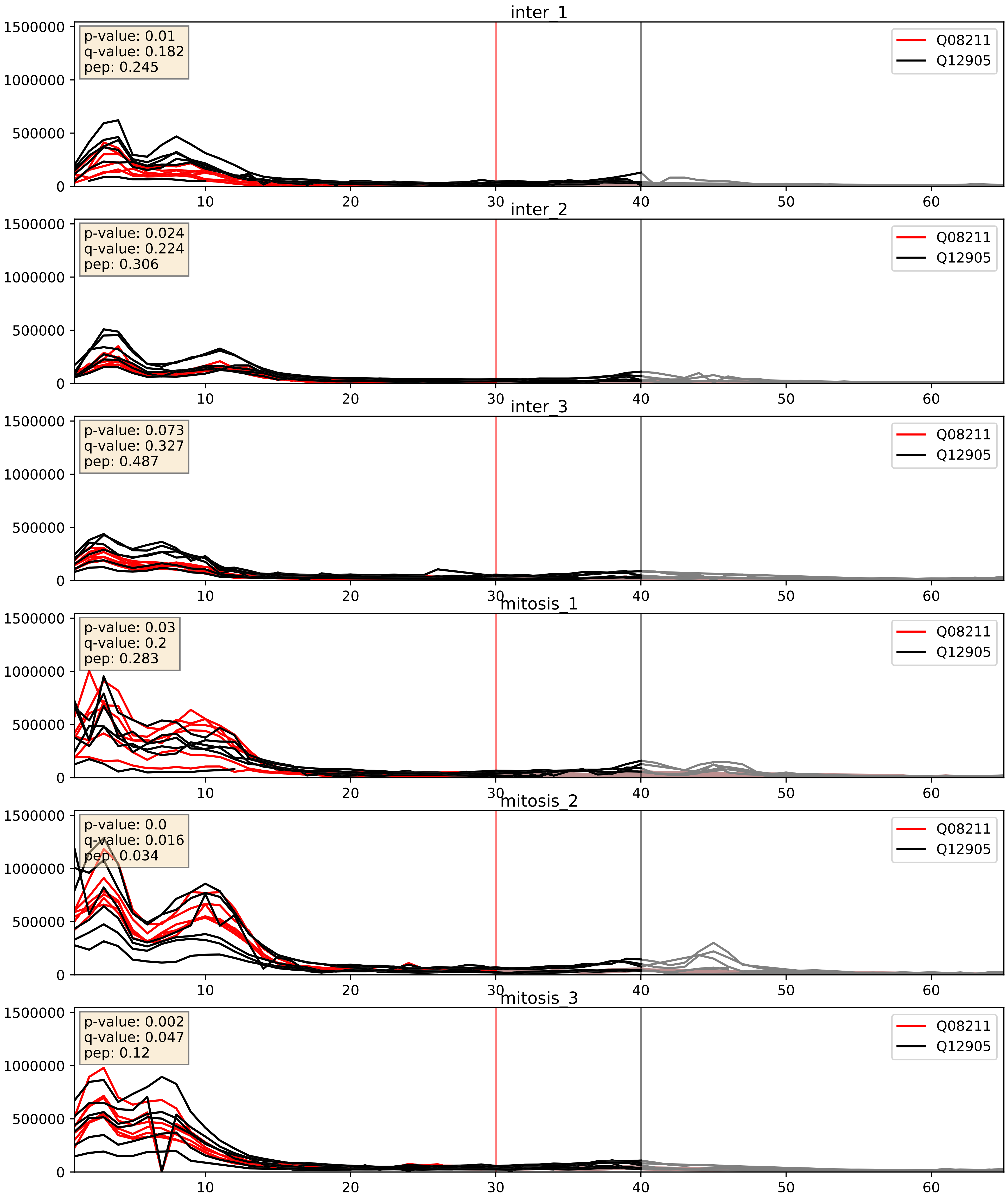


| level | condition_1 | condition_2 | pvalue | pvalue_adjusted |
|---|---|---|---|---|
| interactor_abundance | mitosis | inter | 0.0001655881985577161 | 0.009910743628815524 |
| complex_abundance | mitosis | inter | 0.00023397135977256306 | 0.010221981956574496 |
| interactor_ratio | mitosis | inter | 0.007958384649893444 | 0.1259990886723217 |
