## Supplementary Data 2 for "SECAT: Quantifying differential protein-protein interaction states by network-centric analysis": hela_string_000043_Q53GS9.pdf

Q53GS9_Q53GS9

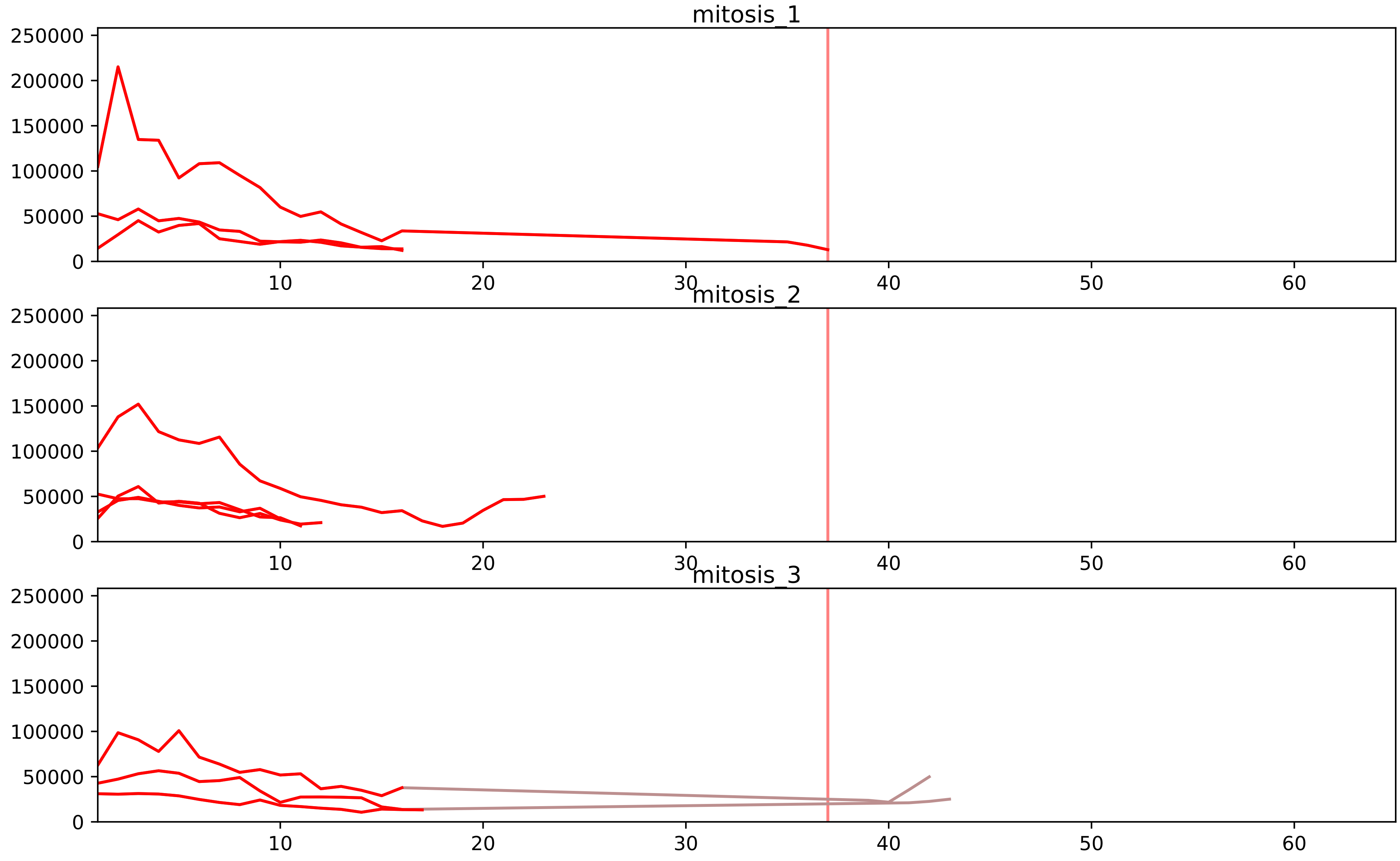


| level | condition_1 | condition_2 | pvalue | pvalue_adjusted |
|---|---|---|---|---|
| interactor_ratio | mitosis | inter | 2.9457902095187802e-08 | 1.3550634963786389e-06 |
| complex_abundance | mitosis | inter | 4.091948026604279e-06 | 0.00024287691512747981 |
| interactor_abundance | mitosis | inter | 3.710787342209604e-05 | 0.001219258698154584 |
| total_abundance | mitosis | inter | 0.020022510637017956 | 0.1794623268539761 |
| assembled_abundance | mitosis | inter | 0.020833909284398763 | 0.17079427208453696 |
| monomer_abundance | mitosis | inter | 0.19696945817626138 | 0.5688748940046396 |

O43172_Q53GS9
PRP4_HUMAN vs SNUT2_HUMAN
p-value: 0.003 q-value: 0.003

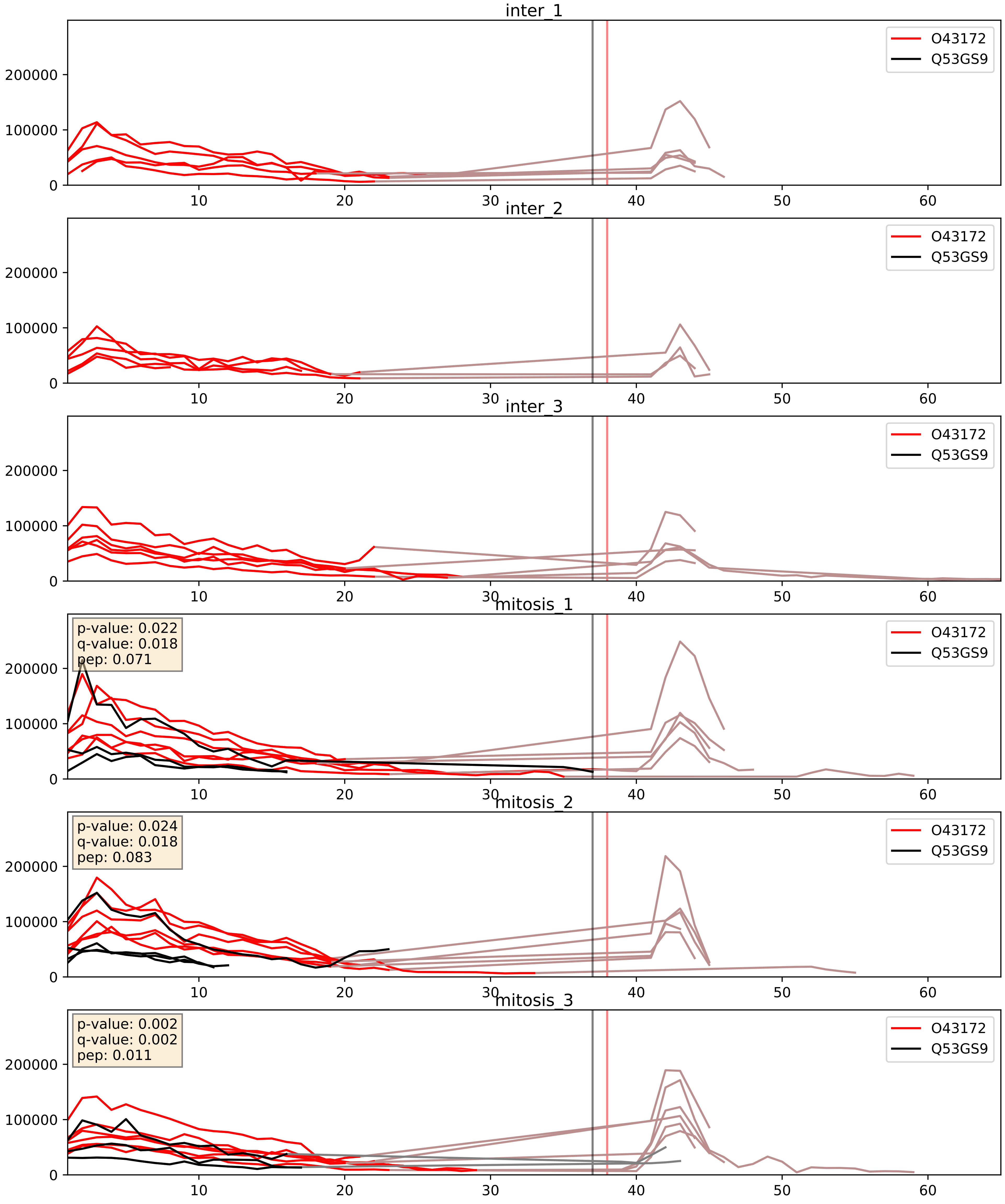


| level | condition_1 | condition_2 | pvalue | pvalue_adjusted |
|---|---|---|---|---|
| interactor_abundance | mitosis | inter | 0.00018211163019669088 | 0.010155541071554879 |
| complex_abundance | mitosis | inter | 0.0003182703109427418 | 0.01165207110963347 |
| interactor_ratio | mitosis | inter | 0.002963584722372241 | 0.07027225823685979 |

O43395_Q53GS9
PRPF3_HUMAN vs SNUT2_HUMAN
p-value: 0.004 q-value: 0.004

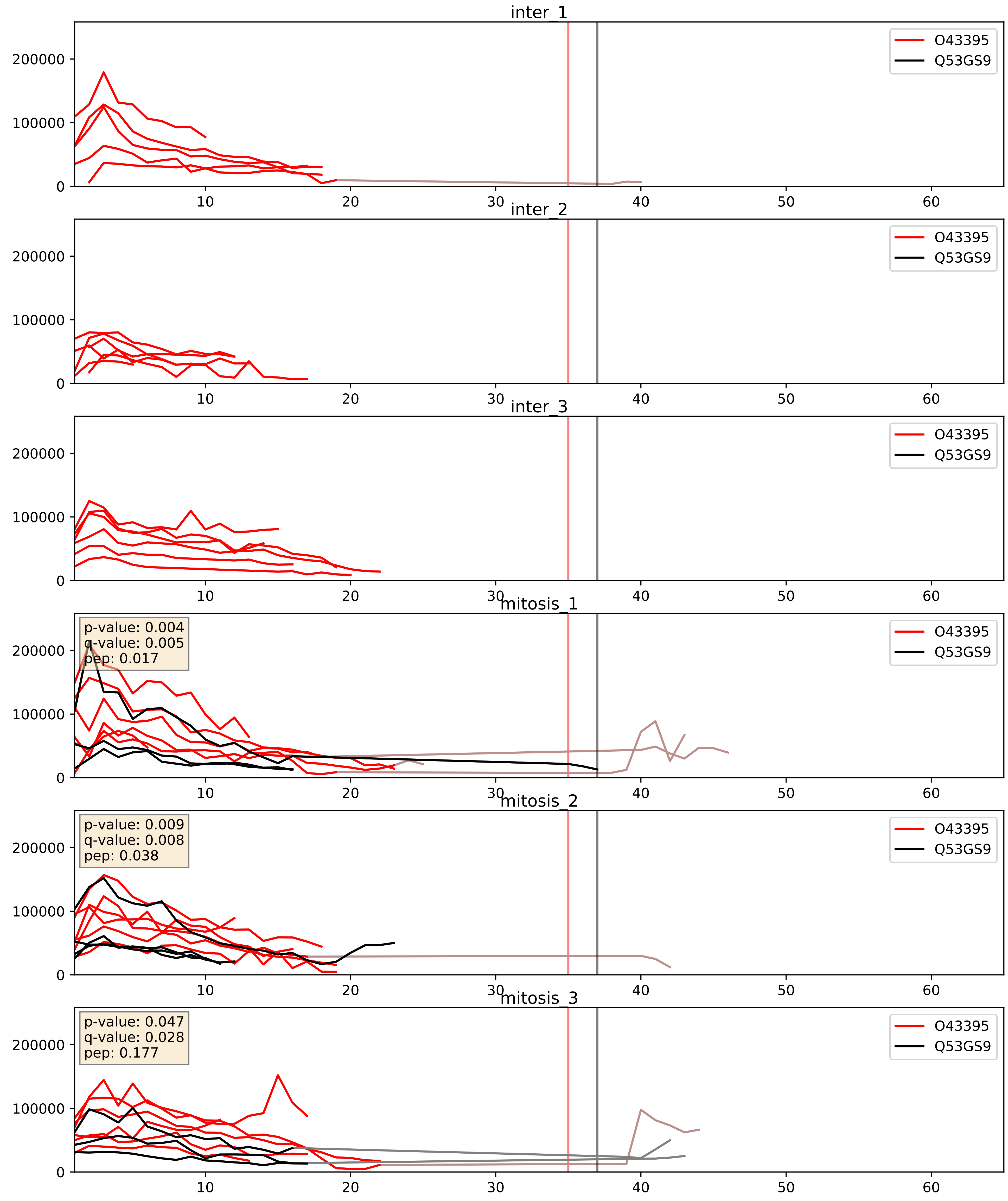


| level | condition_1 | condition_2 | pvalue | pvalue_adjusted |
|---|---|---|---|---|
| interactor_abundance | mitosis | inter | 0.00027487684997001897 | 0.01256848217931955 |
| complex_abundance | mitosis | inter | 0.00043128174253438033 | 0.013091389064164169 |
| interactor_ratio | mitosis | inter | 0.004718642351376341 | 0.09159088101537749 |

O94906_Q53GS9
PRP6_HUMAN vs SNUT2_HUMAN
p-value: 0.017 q-value: 0.014

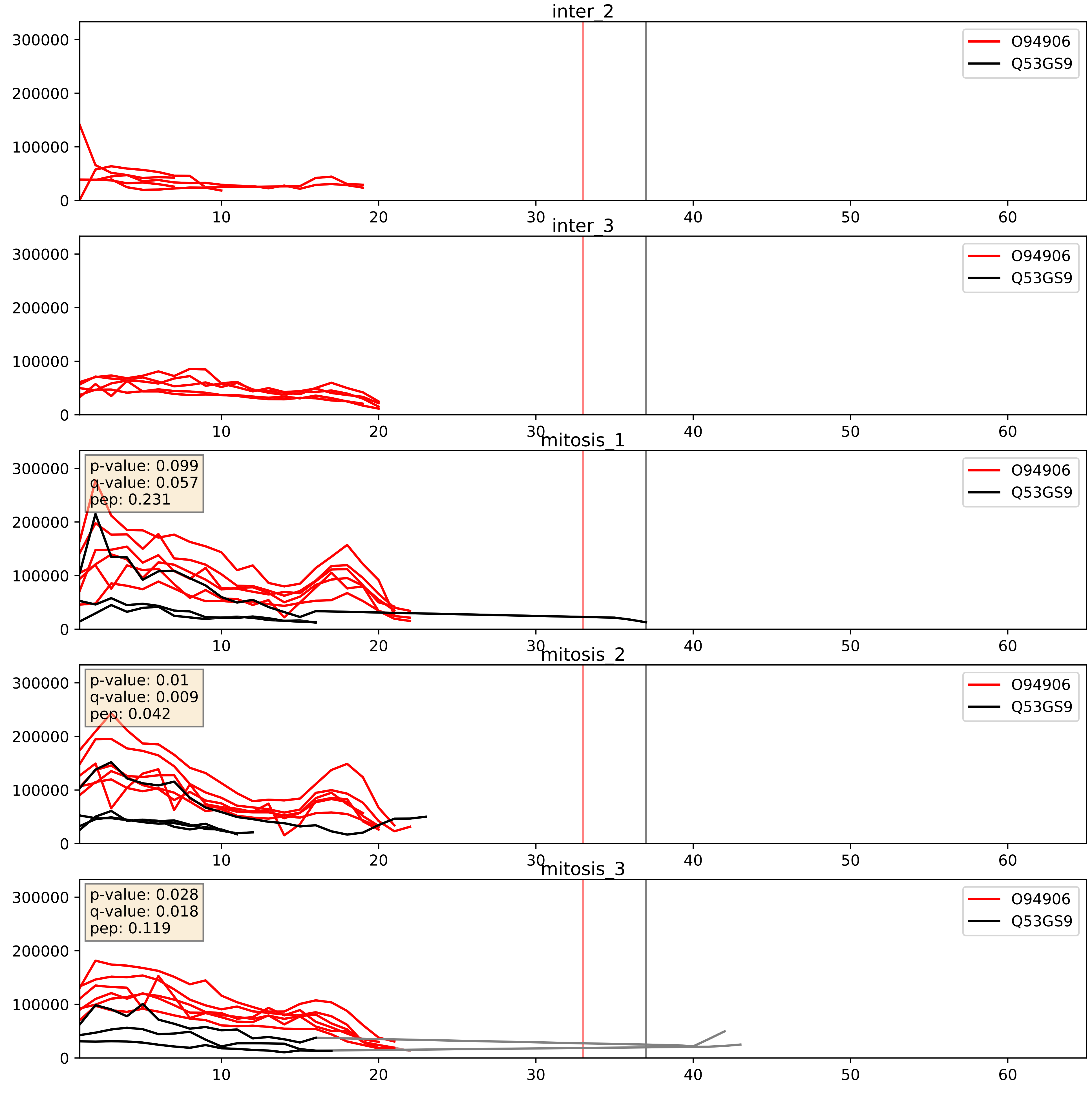


| level | condition_1 | condition_2 | pvalue | pvalue_adjusted |
|---|---|---|---|---|
| interactor_abundance | mitosis | inter | 3.439973865620678e-05 | 0.005164930486161031 |
| interactor_ratio | mitosis | inter | 9.480369998418357e-05 | 0.010510079284752772 |
| complex_abundance | mitosis | inter | 0.00015537452447804976 | 0.008563834848633118 |

P55769_Q53GS9
NH2L1_HUMAN vs SNUT2_HUMAN
p-value: 0.002 q-value: 0.002

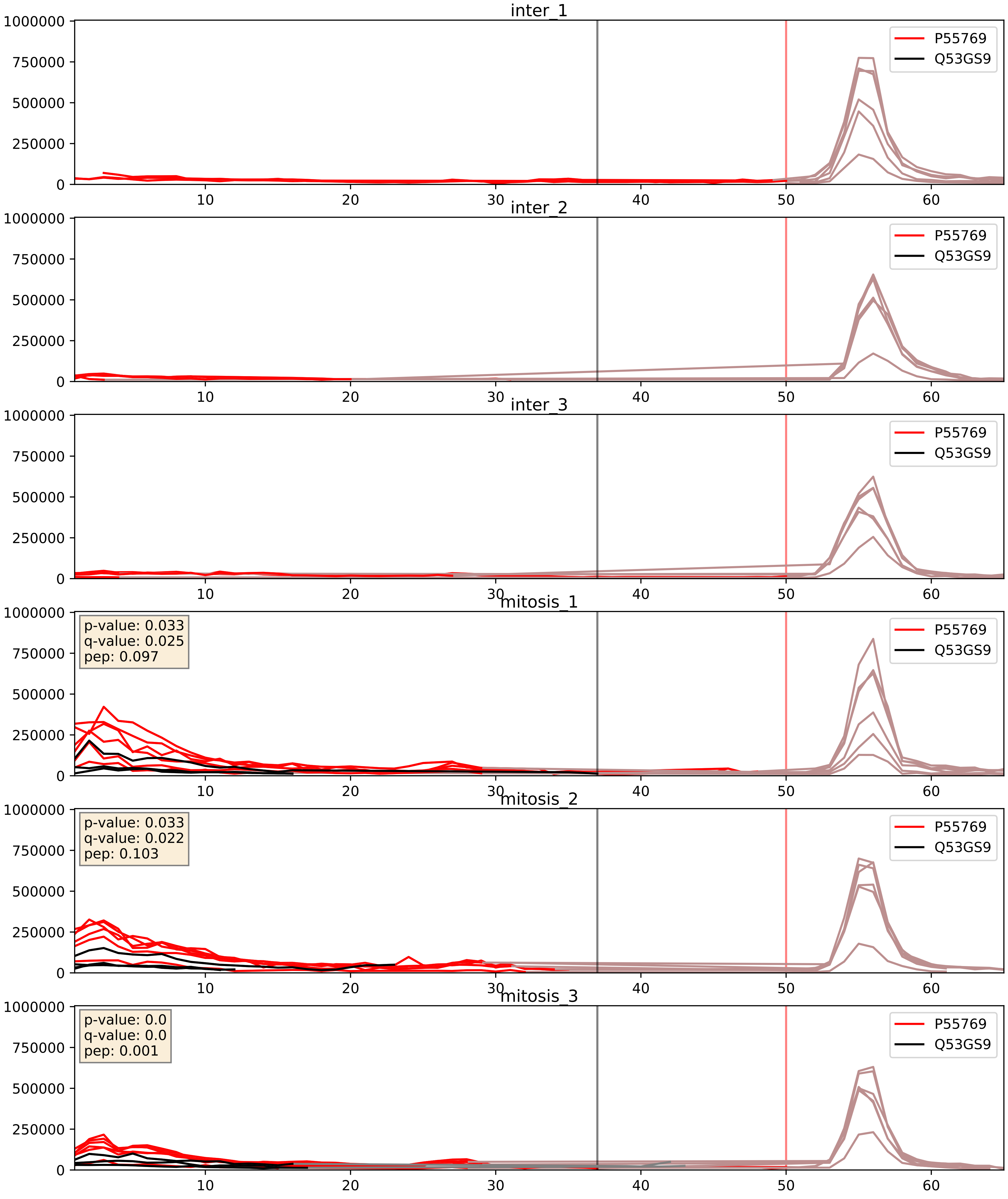


| level | condition_1 | condition_2 | pvalue | pvalue_adjusted |
|---|---|---|---|---|
| interactor_abundance | mitosis | inter | 0.00033269578556084927 | 0.013536301629984253 |
| complex_abundance | mitosis | inter | 0.0004200036215168646 | 0.013091389064164169 |
| interactor_ratio | mitosis | inter | 0.0011227623642383034 | 0.03908562924515634 |

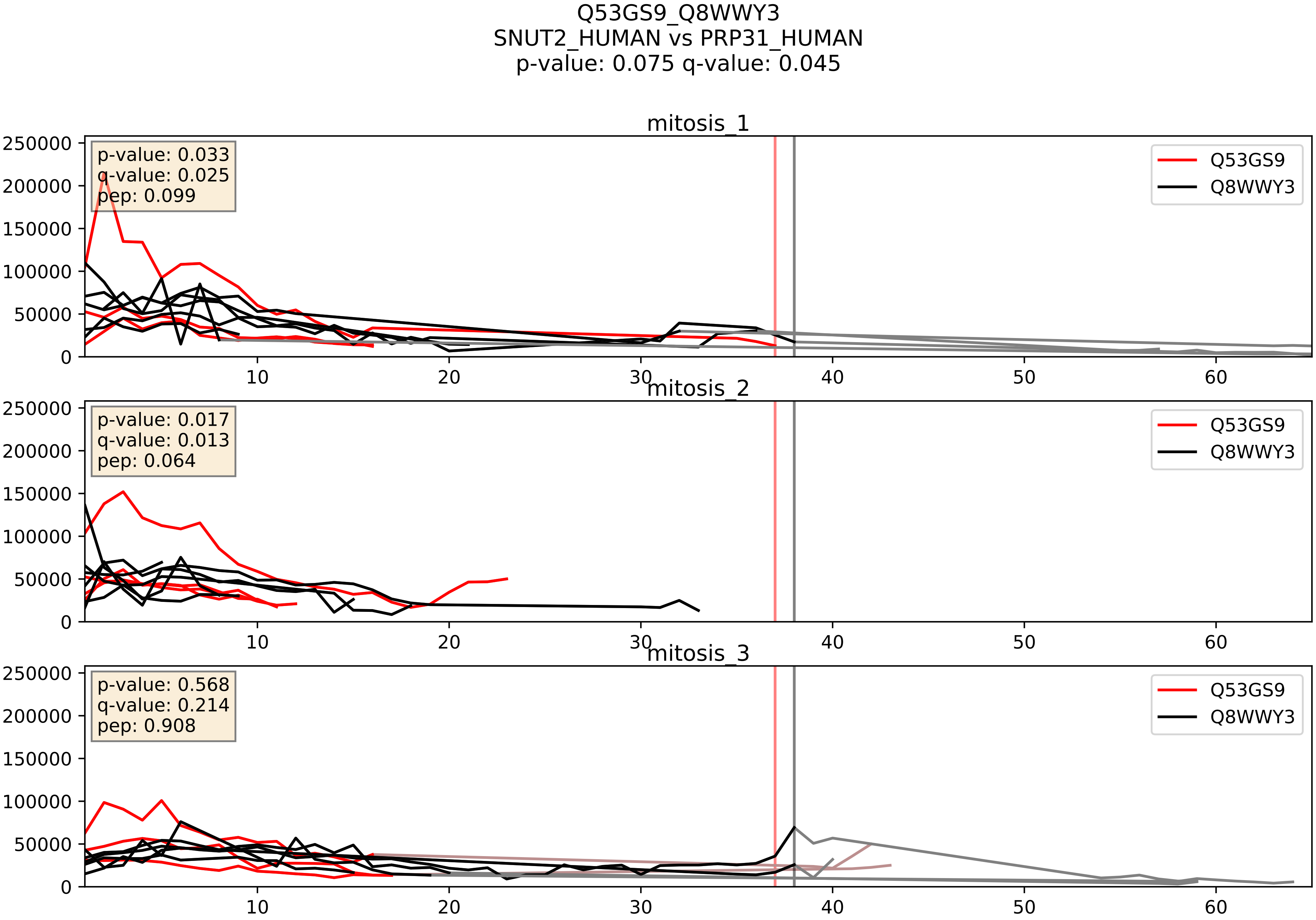


| level | condition_1 | condition_2 | pvalue | pvalue_adjusted |
|---|---|---|---|---|
| interactor_abundance | mitosis | inter | 0.0016940513754034087 | 0.030042528708242176 |
| complex_abundance | mitosis | inter | 0.0028637689480298383 | 0.03901194445107684 |
| interactor_ratio | mitosis | inter | 0.2851403101310909 | 0.755432081312949 |

Q53GS9_Q99459
SNUT2_HUMAN vs CDC5L_HUMAN
p-value: 0.037 q-value: 0.026

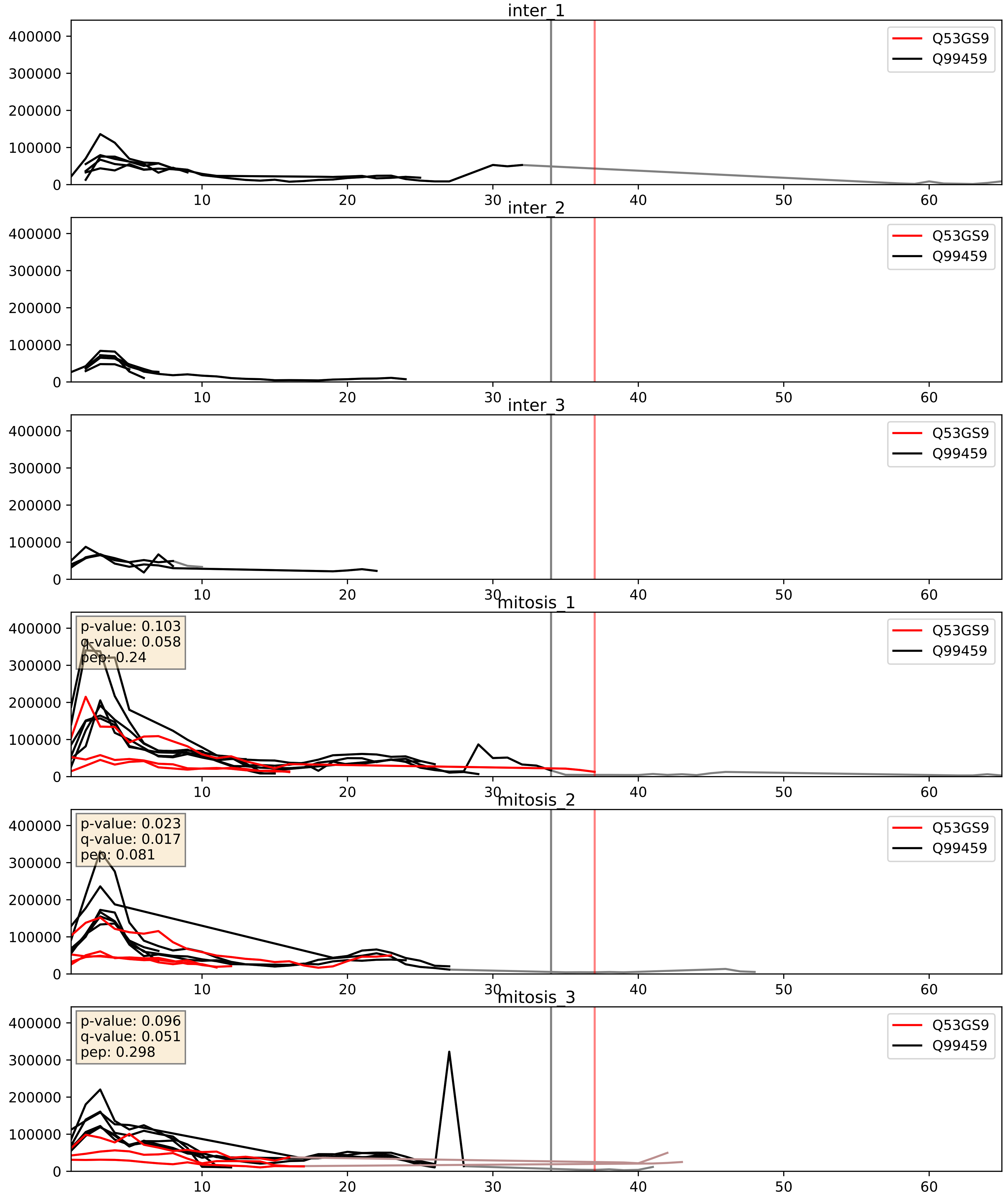


| level | condition_1 | condition_2 | pvalue | pvalue_adjusted |
|---|---|---|---|---|
| complex_abundance | mitosis | inter | 0.0010493796329760585 | 0.021612845224852777 |
| interactor_abundance | mitosis | inter | 0.0013451098335009928 | 0.027124005123129562 |
| interactor_ratio | mitosis | inter | 0.0020791470592695536 | 0.05614352942380876 |

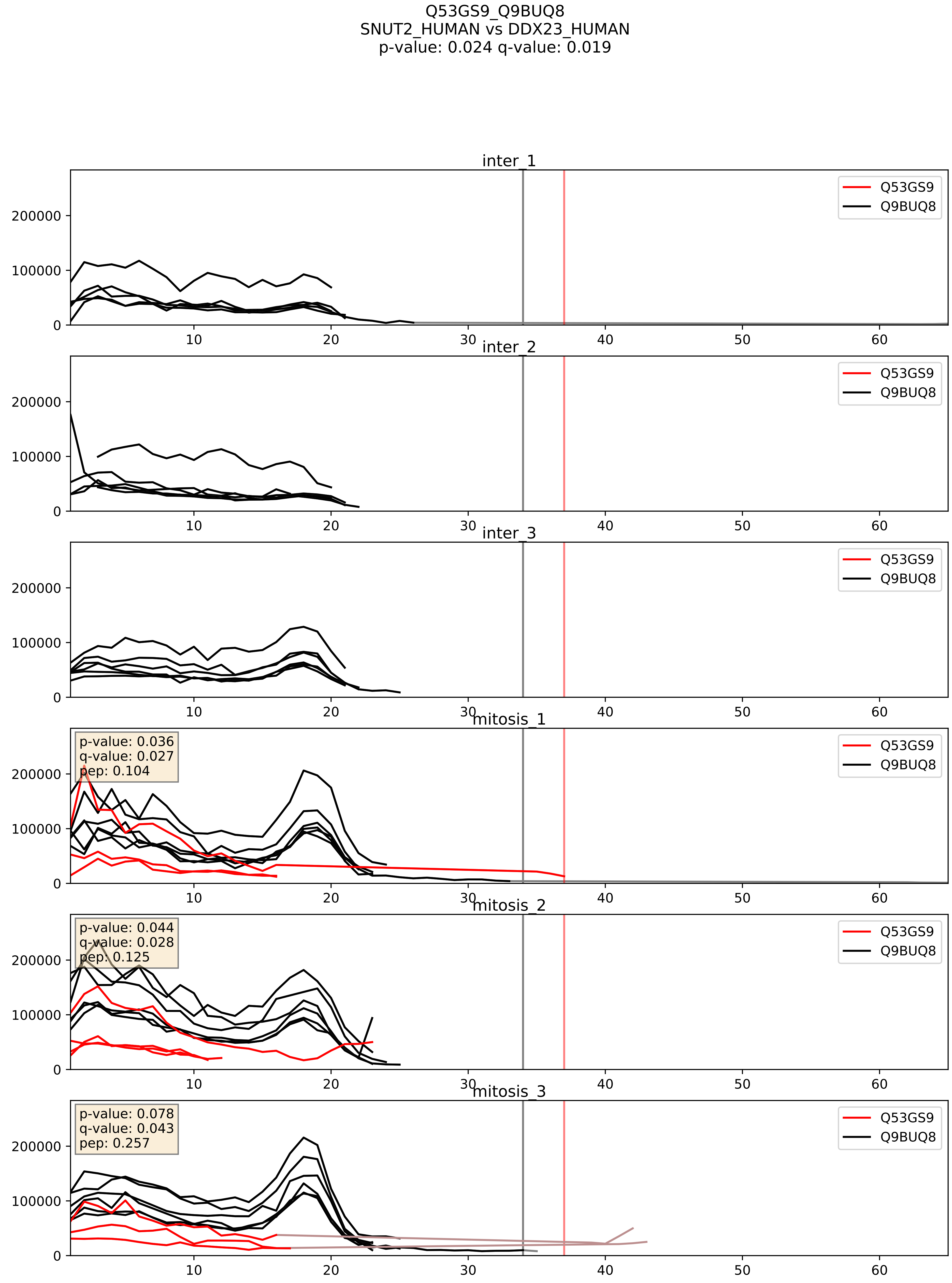


| level | condition_1 | condition_2 | pvalue | pvalue_adjusted |
|---|---|---|---|---|
| complex_abundance | mitosis | inter | 0.00014008468161613128 | 0.00812325774702682 |
| interactor_abundance | mitosis | inter | 0.0013451098335009928 | 0.027124005123129562 |
| interactor_ratio | mitosis | inter | 0.007320278961347636 | 0.11910596594407505 |
