## Supplementary Data 2 for "SECAT: Quantifying differential protein-protein interaction states by network-centric analysis": hela_string_000044_P06493.pdf

P06493_P06493

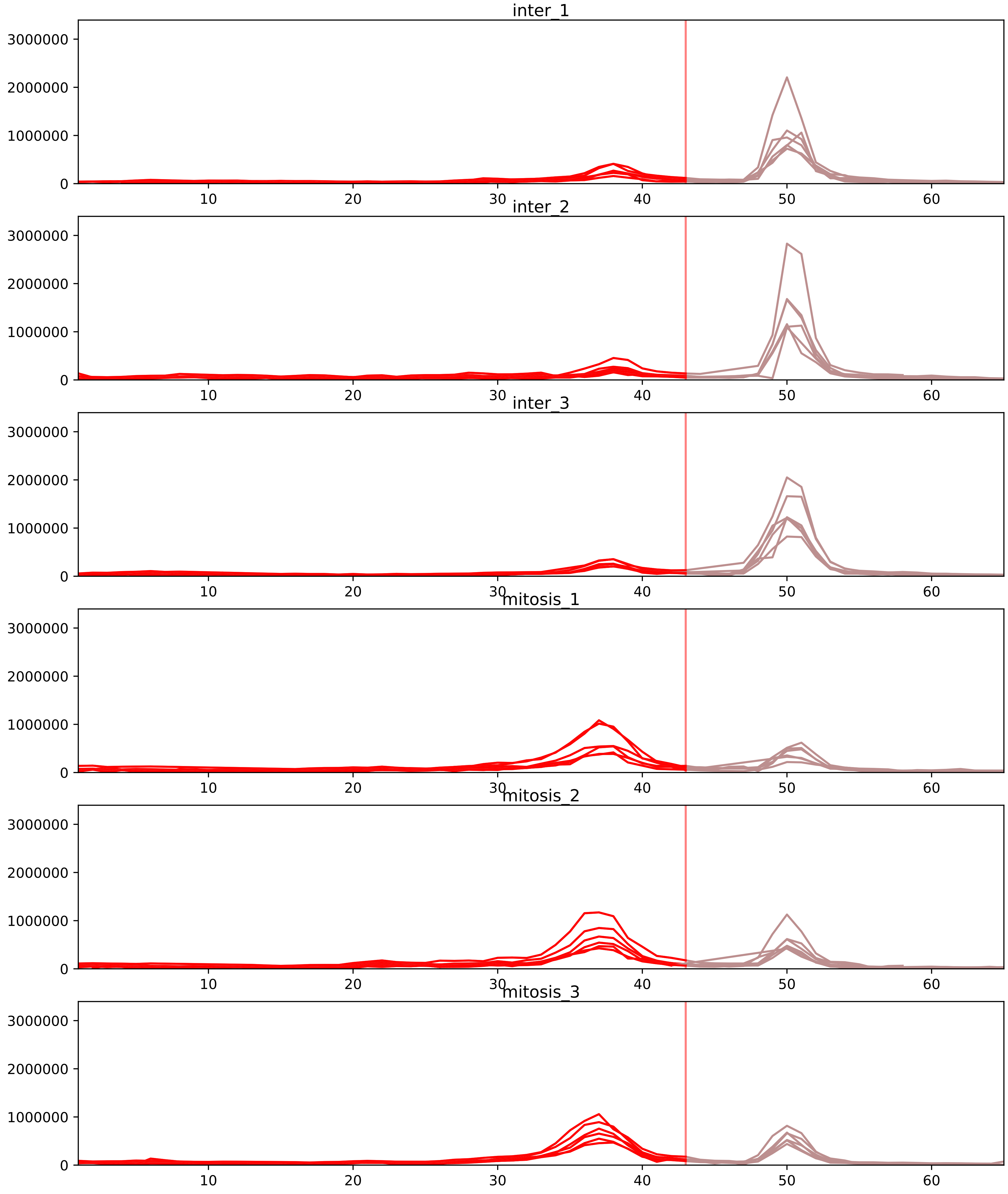


| level | condition_1 | condition_2 | pvalue | pvalue_adjusted |
|---|---|---|---|---|
| interactor_ratio | mitosis | inter | 4.885600014874697e-08 | 2.192561957894986e-06 |
| complex_abundance | mitosis | inter | 1.109717453119146e-07 | 2.0418801137392286e-05 |
| interactor_abundance | mitosis | inter | 2.7822022144841254e-07 | 4.6538655224098096e-05 |
| assembled_abundance | mitosis | inter | 0.0036712197393097236 | 0.0601054752900627 |
| monomer_abundance | mitosis | inter | 0.007789245288793864 | 0.16833884898068865 |
| total_abundance | mitosis | inter | 0.4790822483745524 | 0.7481618792867626 |

O43684_P06493
BUB3_HUMAN vs CDK1_HUMAN
p-value: 0.04 q-value: 0.027

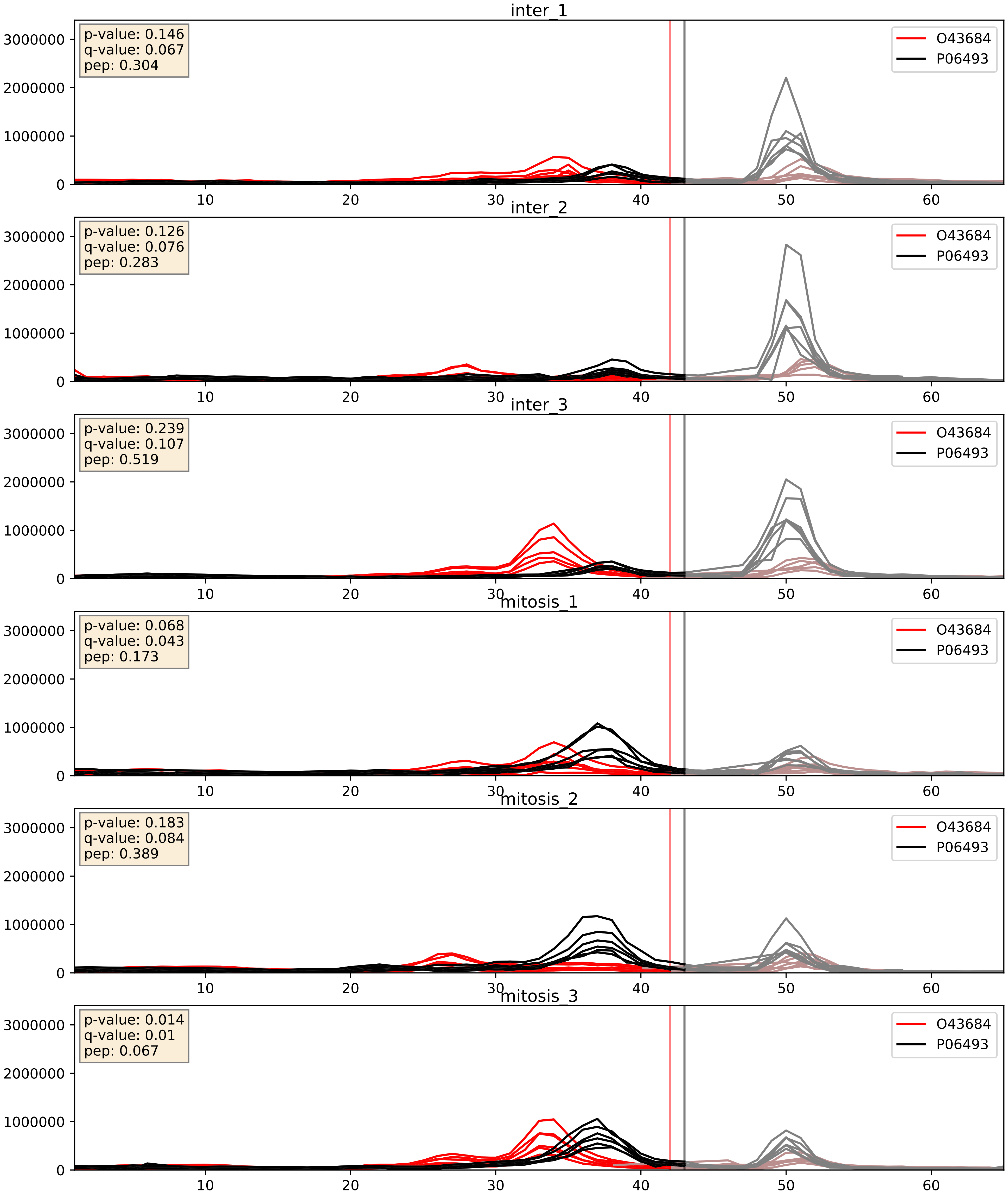


| level | condition_1 | condition_2 | pvalue | pvalue_adjusted |
|---|---|---|---|---|
| complex_abundance | mitosis | inter | 0.07570168337064632 | 0.1998767548072416 |
| interactor_ratio | mitosis | inter | 0.17543155593512502 | 0.6412015878756064 |
| interactor_abundance | mitosis | inter | 0.8856843168093265 | 0.9361222224987428 |

O95067_P06493
CCNB2_HUMAN vs CDK1_HUMAN
p-value: 0.063 q-value: 0.039

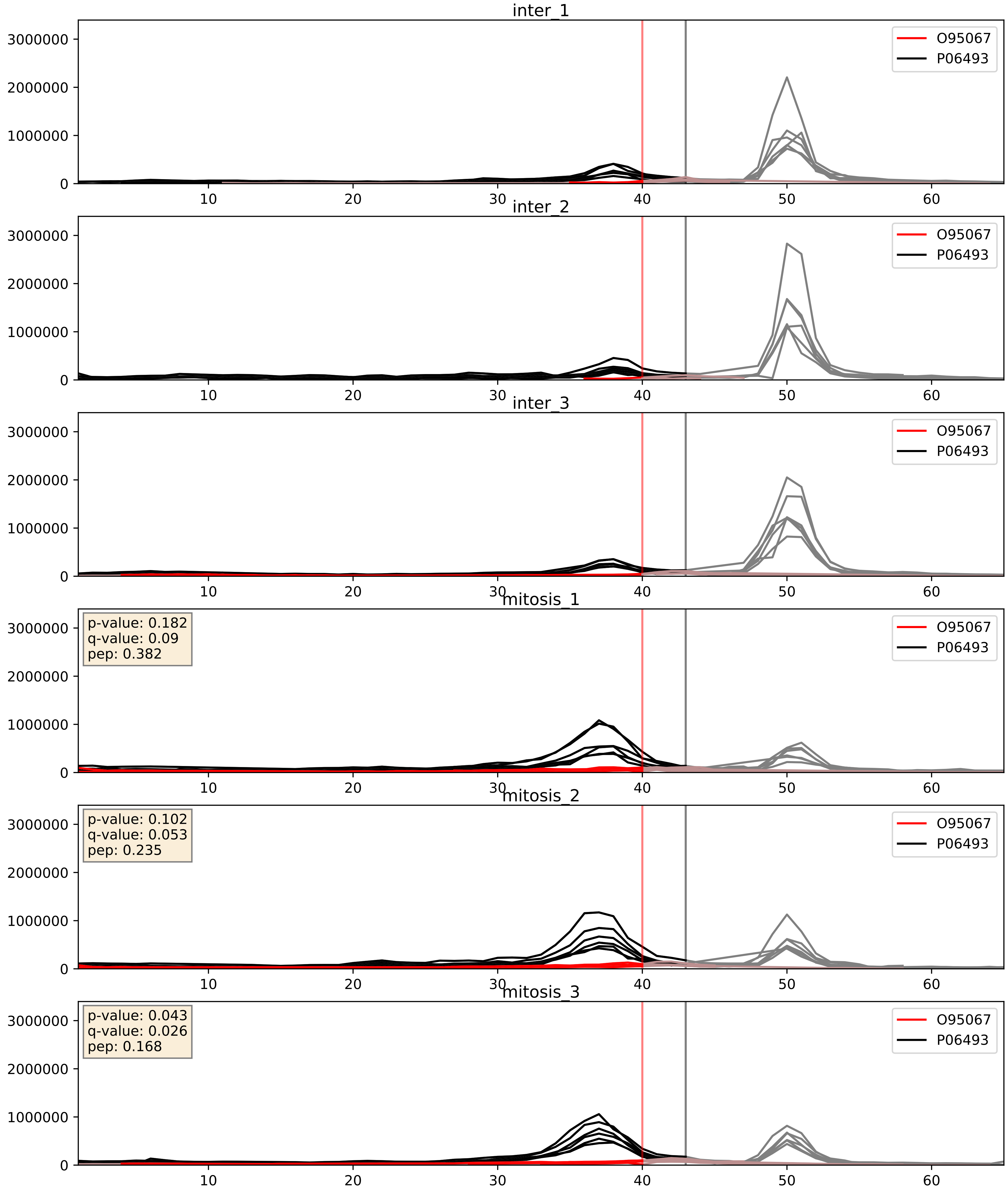


| level | condition_1 | condition_2 | pvalue | pvalue_adjusted |
|---|---|---|---|---|
| complex_abundance | mitosis | inter | 0.00017800511457254897 | 0.008807651911797798 |
| interactor_abundance | mitosis | inter | 0.0031005769388121997 | 0.04271207144514066 |
| interactor_ratio | mitosis | inter | 0.005268855769537838 | 0.0974780445871482 |

P05455_P06493
LA_HUMAN vs CDK1_HUMAN
p-value: 0.0 q-value: 0.042

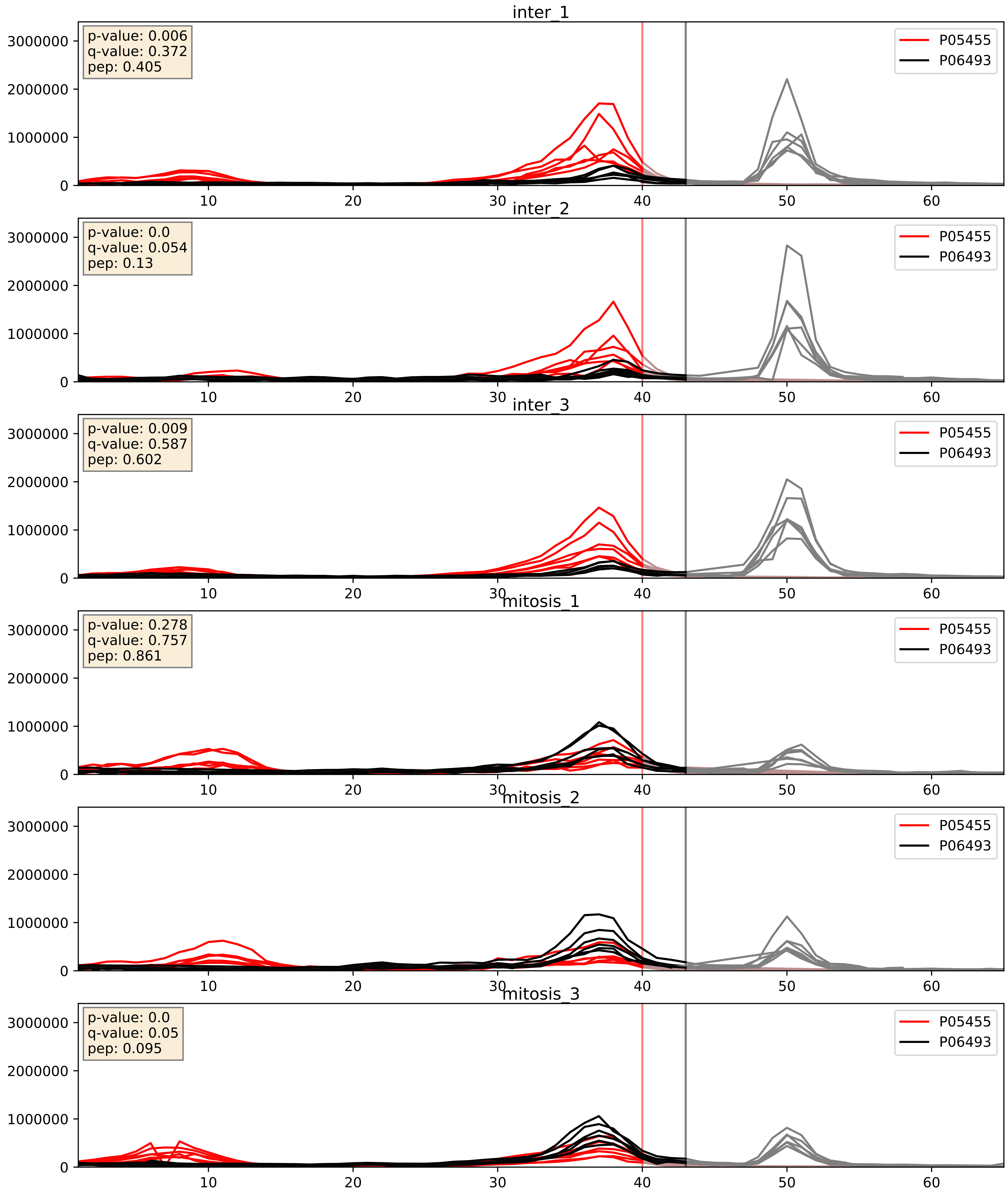


| level | condition_1 | condition_2 | pvalue | pvalue_adjusted |
|---|---|---|---|---|
| interactor_ratio | mitosis | inter | 0.013082649938714568 | 0.16493001984594508 |
| interactor_abundance | mitosis | inter | 0.049423900646237605 | 0.17757338490316638 |
| complex_abundance | mitosis | inter | 0.20200239193080372 | 0.3549867532185752 |

P06493_P14635
CDK1_HUMAN vs CCNB1_HUMAN
p-value: 0.0 q-value: 0.0

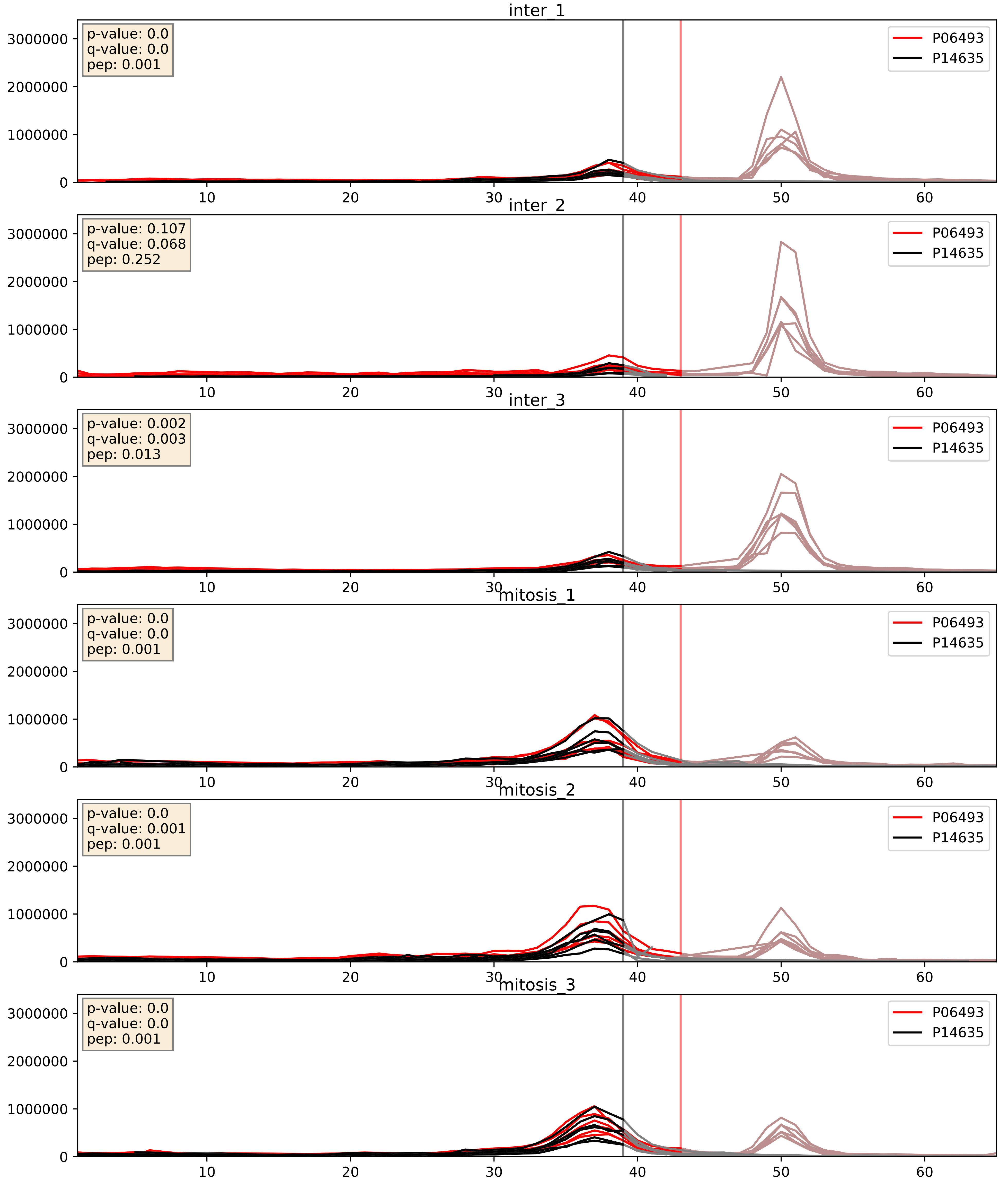


| level | condition_1 | condition_2 | pvalue | pvalue_adjusted |
|---|---|---|---|---|
| interactor_abundance | mitosis | inter | 0.0002579902051684421 | 0.012234881752032491 |
| complex_abundance | mitosis | inter | 0.0009312502745939036 | 0.020079351008876107 |
| interactor_ratio | mitosis | inter | 0.0302146448575311 | 0.2765852972945998 |

P06493_P24941
CDK1_HUMAN vs CDK2_HUMAN
p-value: 0.001 q-value: 0.001

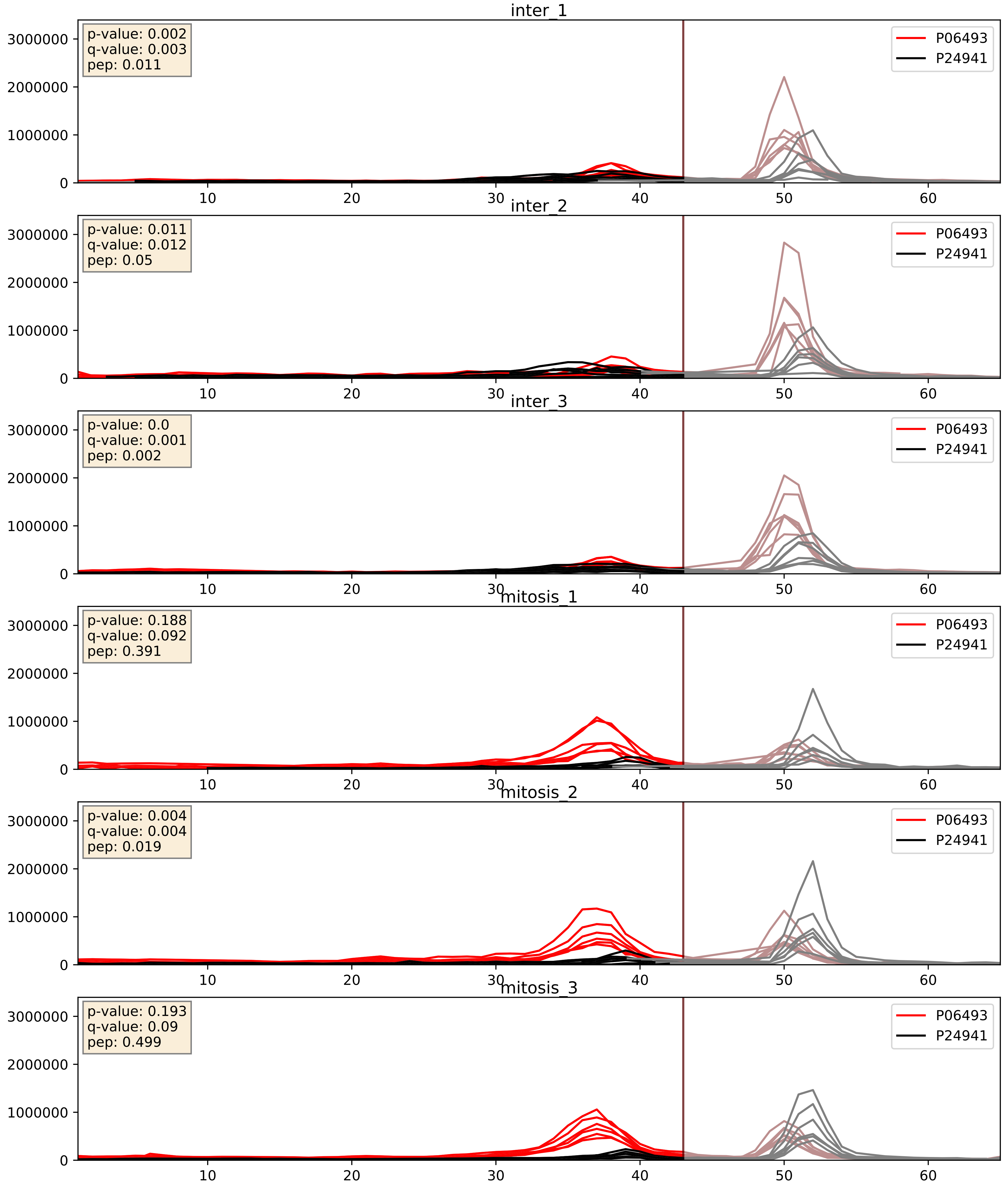


| level | condition_1 | condition_2 | pvalue | pvalue_adjusted |
|---|---|---|---|---|
| interactor_ratio | mitosis | inter | 0.00019753625235696597 | 0.014576813104962318 |
| interactor_abundance | mitosis | inter | 0.017640941221411546 | 0.11492120004207217 |
| complex_abundance | mitosis | inter | 0.34351583213978315 | 0.4915572589629796 |

P06493_P31350
CDK1_HUMAN vs RIR2_HUMAN
p-value: 0.039 q-value: 0.027

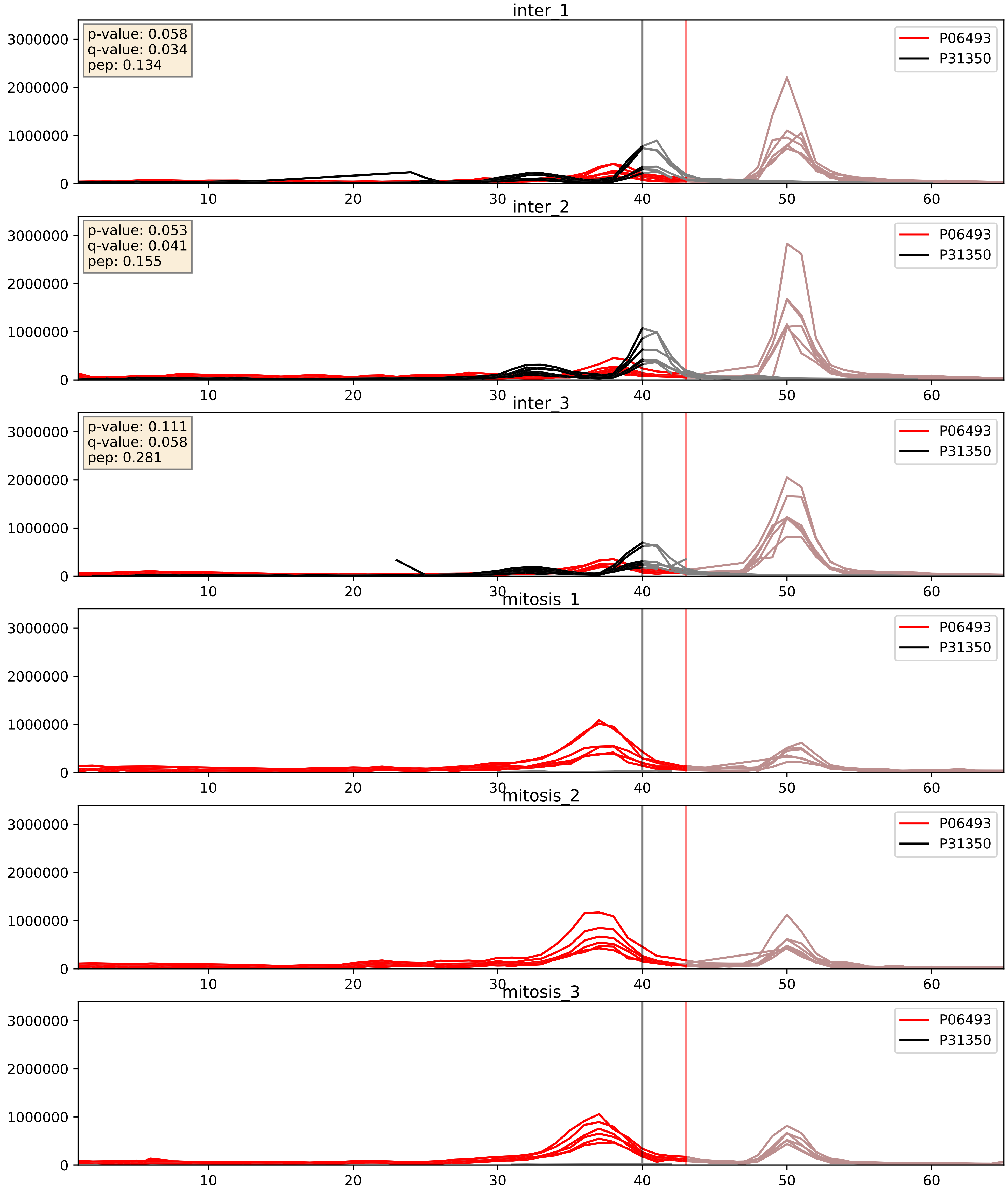


| level | condition_1 | condition_2 | pvalue | pvalue_adjusted |
|---|---|---|---|---|
| complex_abundance | mitosis | inter | 5.655065103525419e-05 | 0.005691857151652958 |
| interactor_abundance | mitosis | inter | 8.436905755951776e-05 | 0.007227629947416683 |
| interactor_ratio | mitosis | inter | 0.11337985380266258 | 0.553985317646331 |

P06493_P33552
CDK1_HUMAN vs CKS2_HUMAN
p-value: 0.031 q-value: 0.022

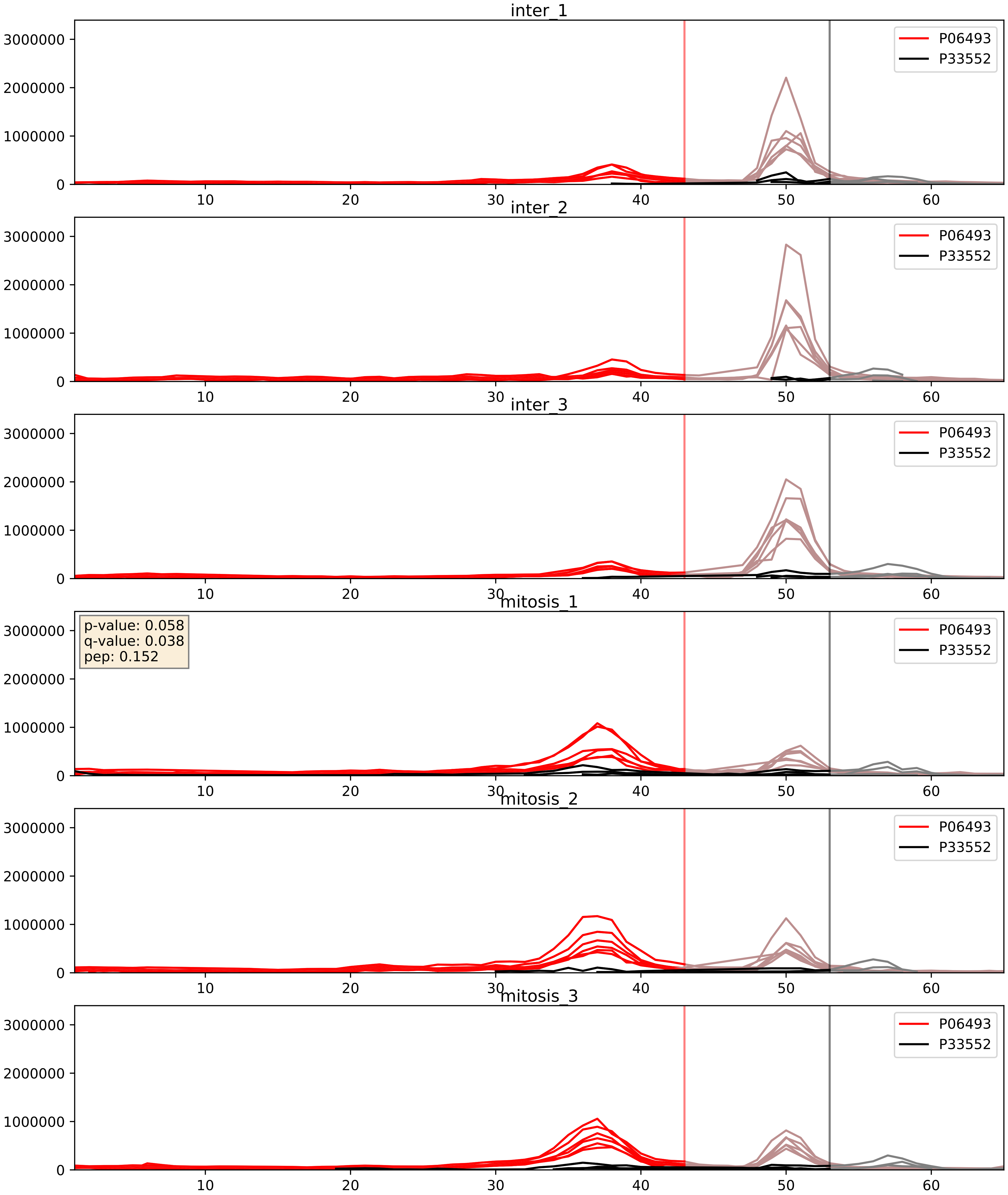


| level | condition_1 | condition_2 | pvalue | pvalue_adjusted |
|---|---|---|---|---|
| interactor_abundance | mitosis | inter | 0.014409321007444237 | 0.10605571190893752 |
| complex_abundance | mitosis | inter | 0.019583703214056107 | 0.10412204938653433 |
| interactor_ratio | mitosis | inter | 0.047321467362668845 | 0.3600637872217292 |

P06493_P63151
CDK1_HUMAN vs 2ABA_HUMAN
p-value: 0.0 q-value: 0.009

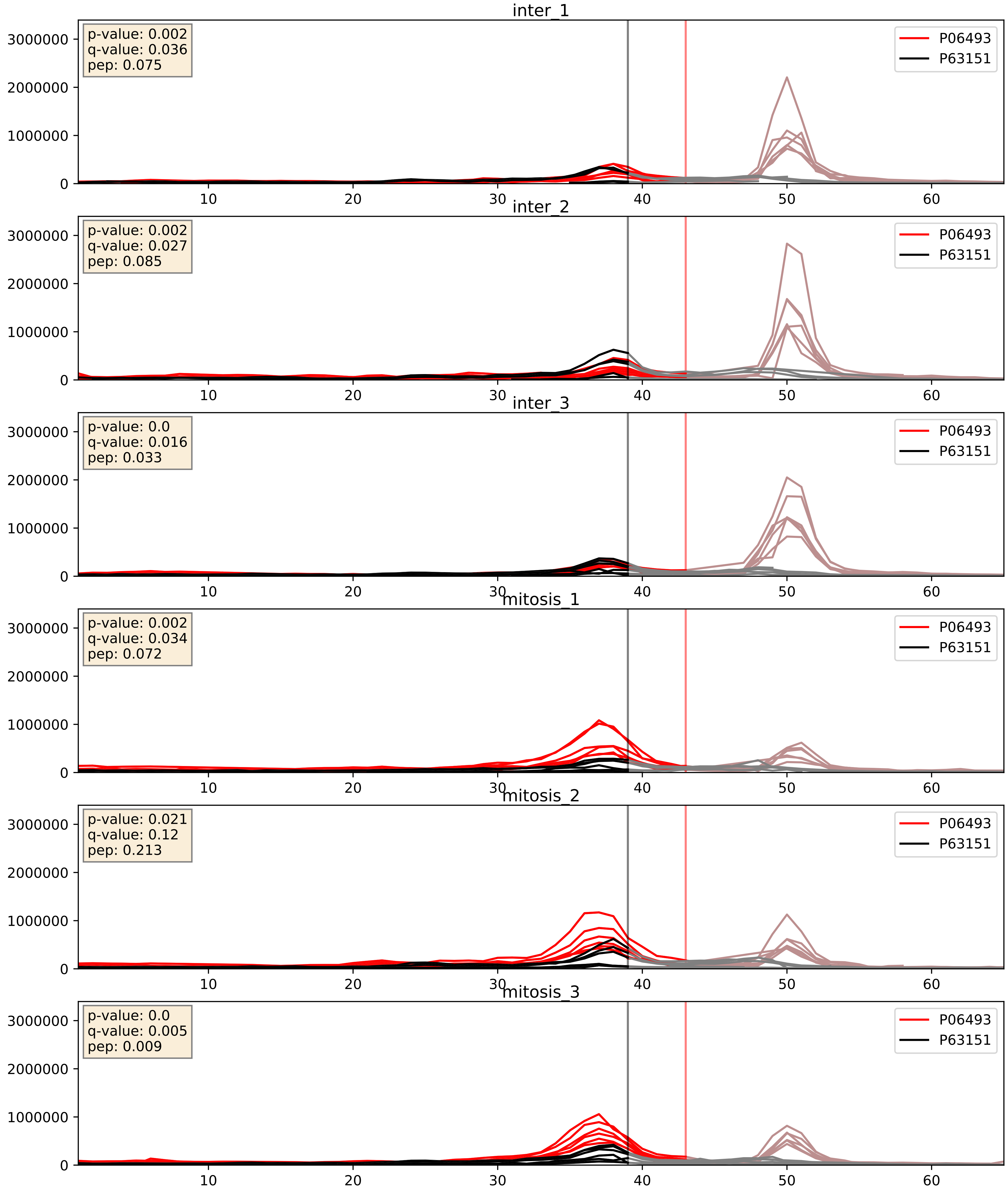


| level | condition_1 | condition_2 | pvalue | pvalue_adjusted |
|---|---|---|---|---|
| interactor_ratio | mitosis | inter | 0.011824811140129647 | 0.15690780467250182 |
| interactor_abundance | mitosis | inter | 0.014367474382127028 | 0.10605571190893752 |
| complex_abundance | mitosis | inter | 0.027827045223281145 | 0.12135715598727036 |
