## Supplementary Data 2 for "SECAT: Quantifying differential protein-protein interaction states by network-centric analysis": hela_string_000045_Q15717.pdf

Q15717_Q15717

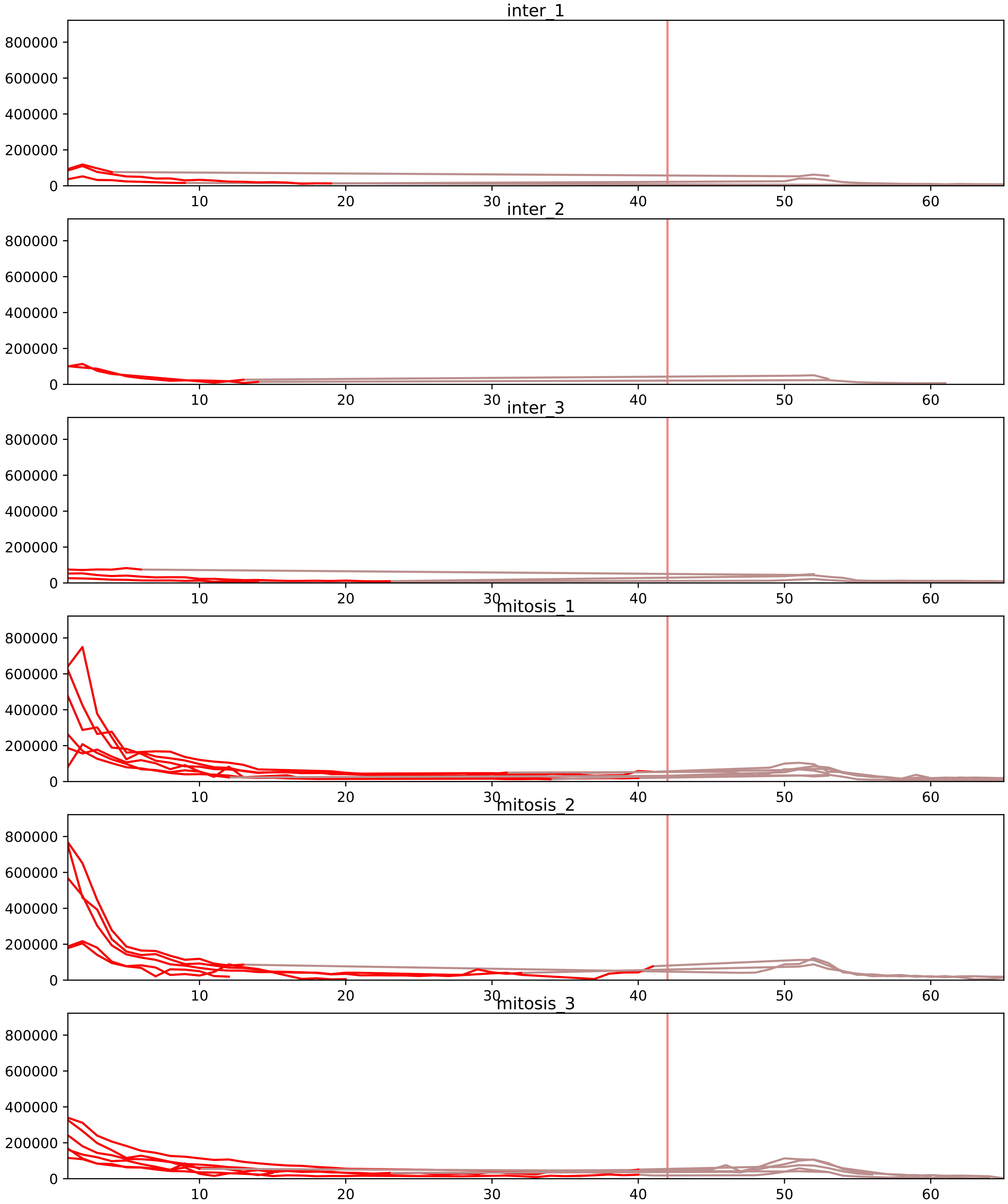


| level | condition_1 | condition_2 | pvalue | pvalue_adjusted |
|---|---|---|---|---|
| interactor_ratio | mitosis | inter | 5.118782640151508e-08 | 2.2425142994949464e-06 |
| complex_abundance | mitosis | inter | 5.6110678620008565e-08 | 1.4956025834220426e-05 |
| interactor_abundance | mitosis | inter | 2.1578100855933622e-07 | 4.41152284165754e-05 |
| monomer_abundance | mitosis | inter | 1.0251083213496417e-05 | 0.020825075548217973 |
| total_abundance | mitosis | inter | 2.26652545453519e-05 | 0.005915549590004797 |
| assembled_abundance | mitosis | inter | 5.496351324193898e-05 | 0.007270487507555603 |

P07910_Q15717
HNRPC_HUMAN vs ELAV1_HUMAN
p-value: 0.026 q-value: 0.02

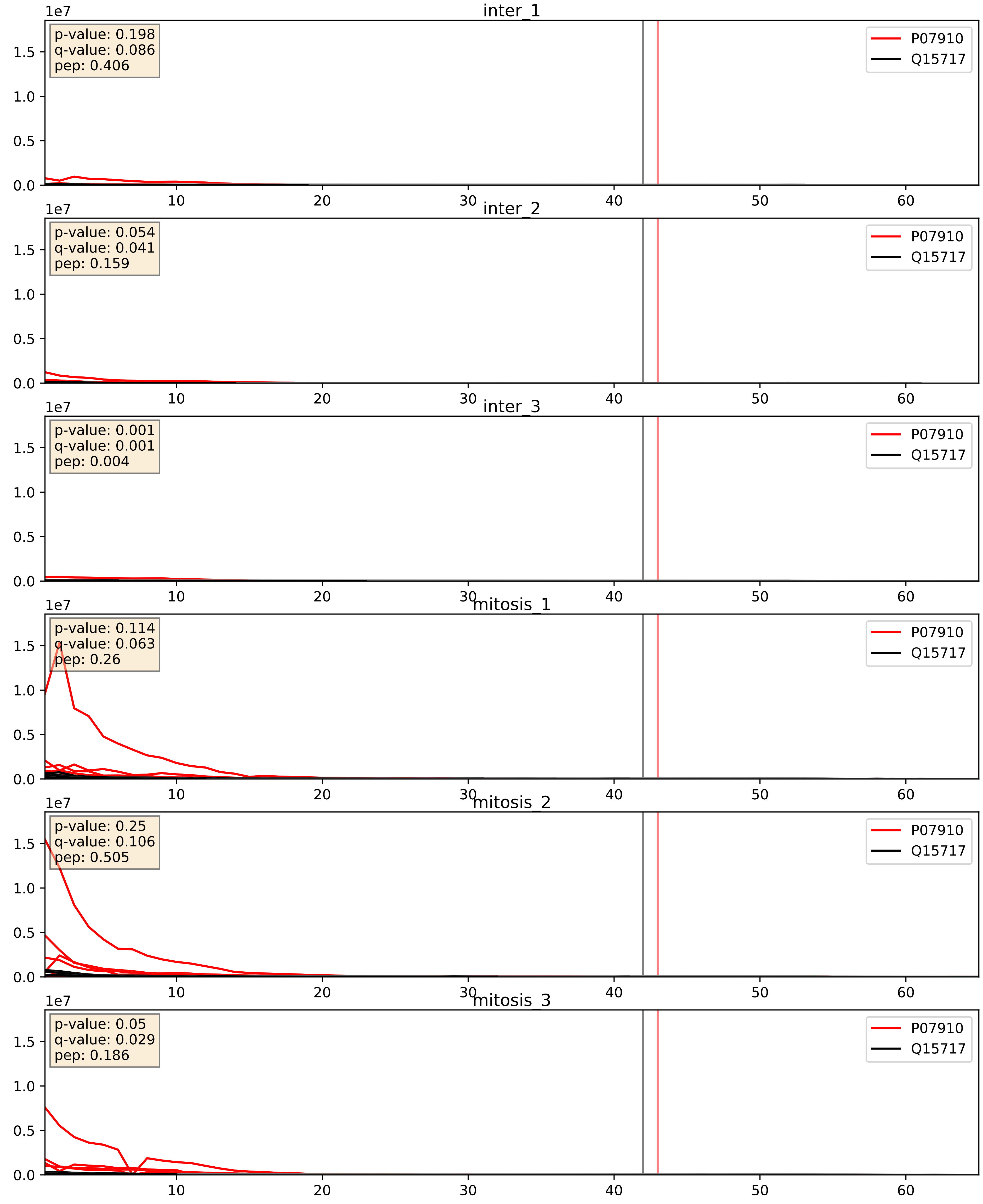


| level | condition_1 | condition_2 | pvalue | pvalue_adjusted |
|---|---|---|---|---|
| interactor_abundance | mitosis | inter | 5.855680651902104e-06 | 0.0033734327106189023 |
| complex_abundance | mitosis | inter | 1.6805290568155707e-05 | 0.005691857151652958 |
| interactor_ratio | mitosis | inter | 0.0014640241868224817 | 0.04557108014254707 |

P09651_Q15717
ROA1_HUMAN vs ELAV1_HUMAN
p-value: 0.006 q-value: 0.005

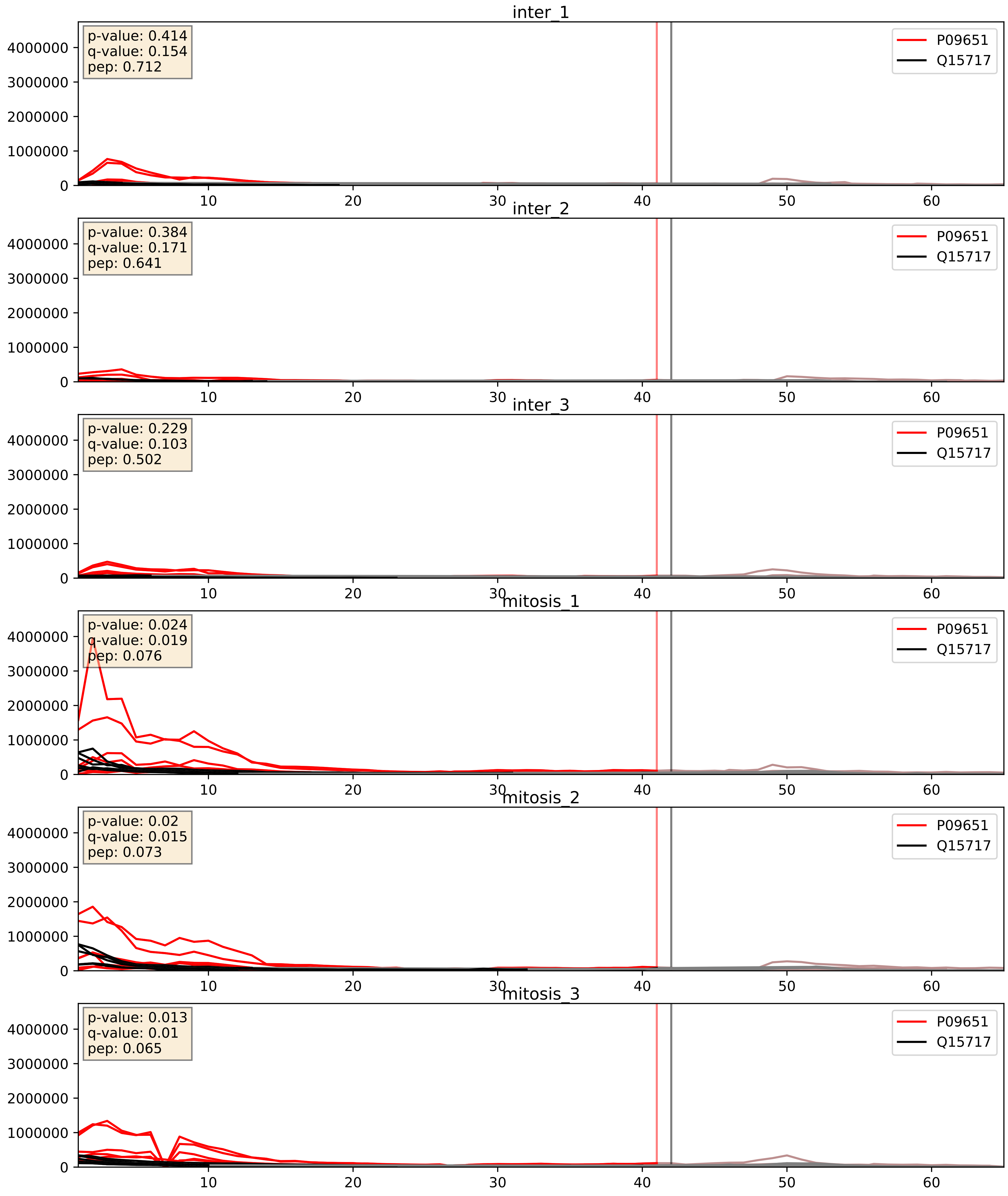


| level | condition_1 | condition_2 | pvalue | pvalue_adjusted |
|---|---|---|---|---|
| complex_abundance | mitosis | inter | 0.0009095527070255489 | 0.0197847007881119 |
| interactor_abundance | mitosis | inter | 0.012699265612423818 | 0.10046738783950822 |
| interactor_ratio | mitosis | inter | 0.4221604689601121 | 0.8388785723072948 |

P14866_Q15717
HNRPL_HUMAN vs ELAV1_HUMAN
p-value: 0.007 q-value: 0.007

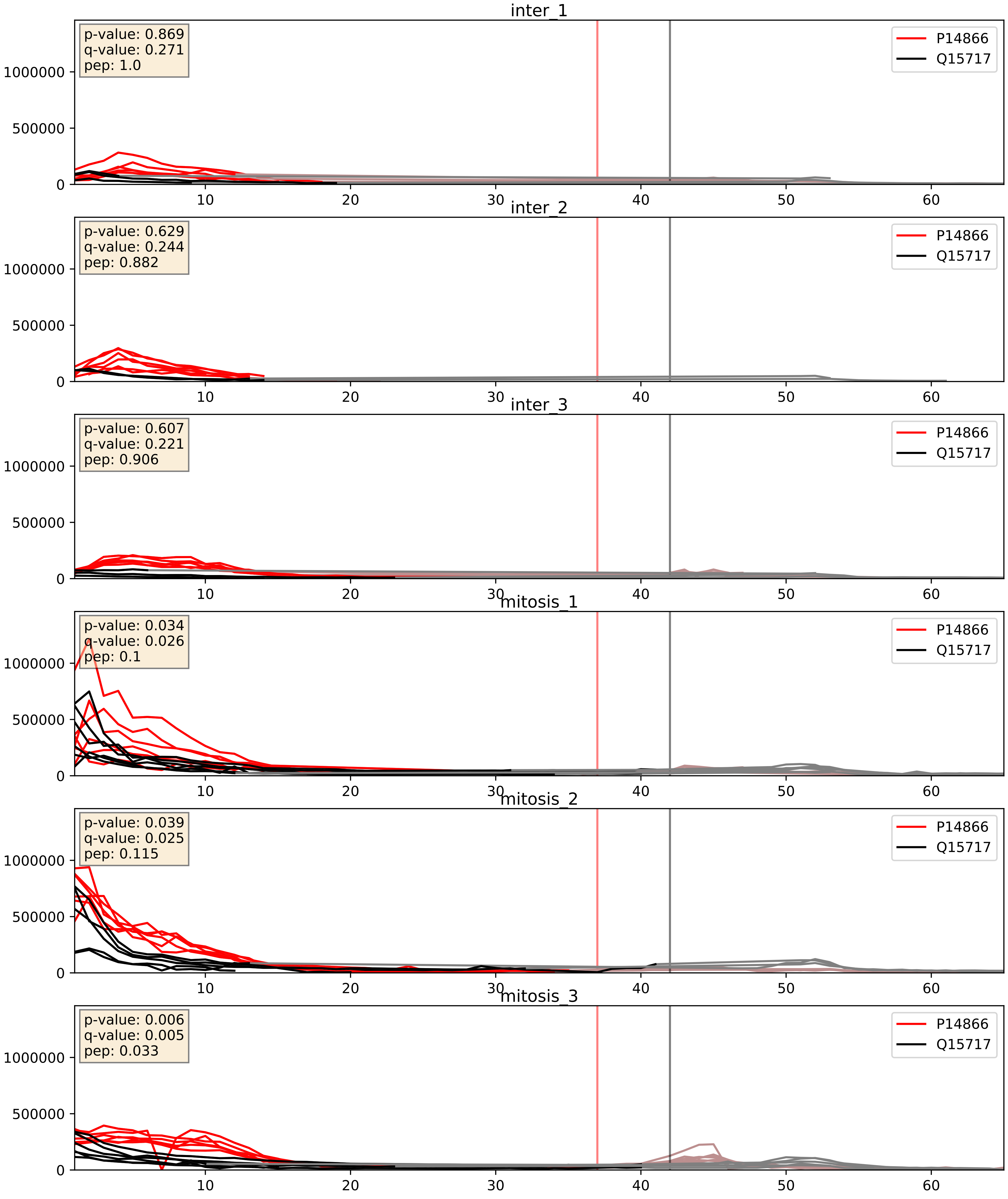


| level | condition_1 | condition_2 | pvalue | pvalue_adjusted |
|---|---|---|---|---|
| complex_abundance | mitosis | inter | 0.00015412022185430893 | 0.008563834848633118 |
| interactor_abundance | mitosis | inter | 0.009449586988451214 | 0.08425881731368999 |
| interactor_ratio | mitosis | inter | 0.04986156517554219 | 0.36889801028750313 |

P22626_Q15717
ROA2_HUMAN vs ELAV1_HUMAN
p-value: 0.019 q-value: 0.015

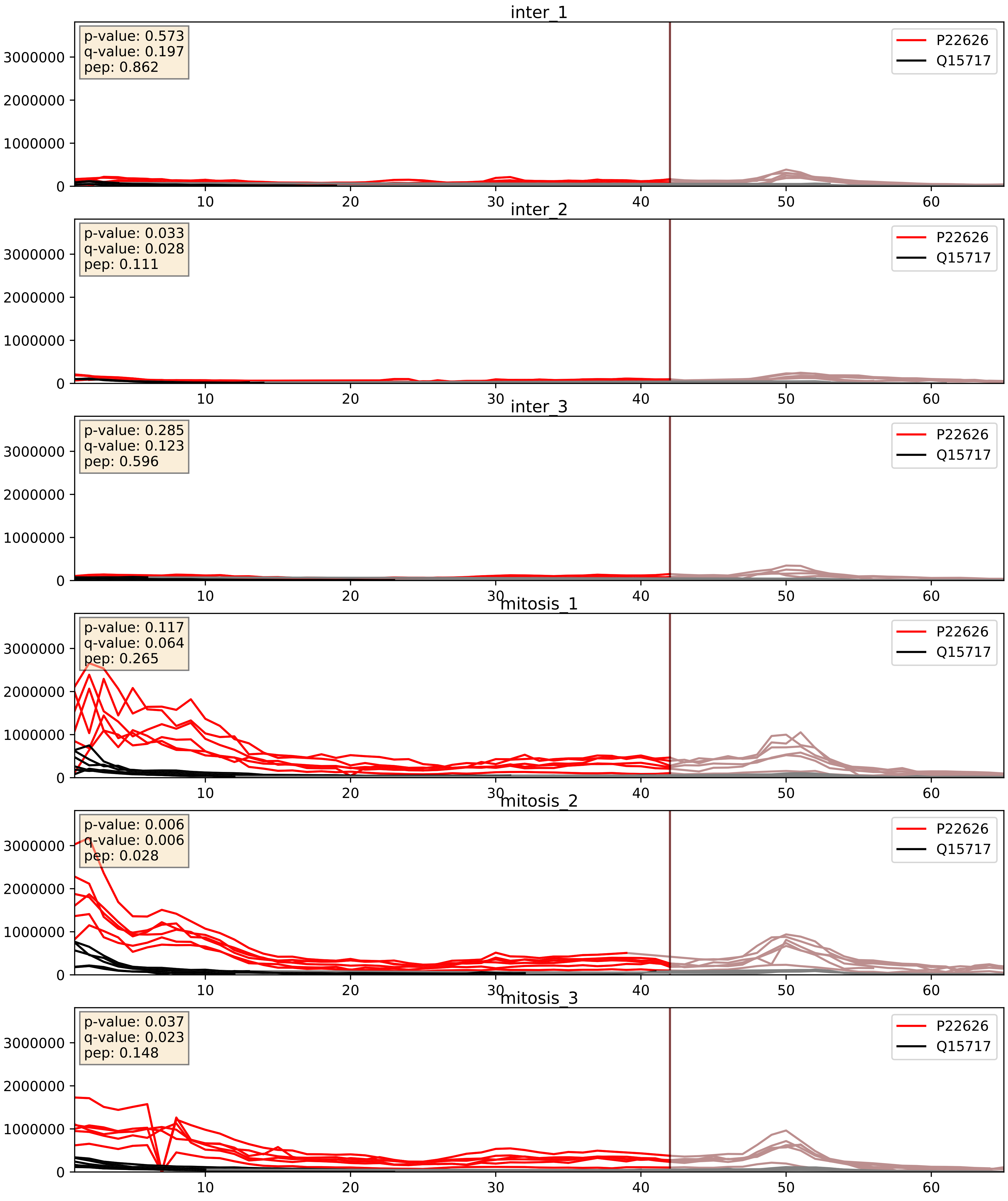


| level | condition_1 | condition_2 | pvalue | pvalue_adjusted |
|---|---|---|---|---|
| complex_abundance | mitosis | inter | 0.00036263180025352423 | 0.012039775462644836 |
| interactor_abundance | mitosis | inter | 0.0009434448484861828 | 0.02284551033392284 |
| interactor_ratio | mitosis | inter | 0.013987766740221716 | 0.17403384200043295 |

P31943_Q15717
HNRH1_HUMAN vs ELAV1_HUMAN
p-value: 0.006 q-value: 0.006

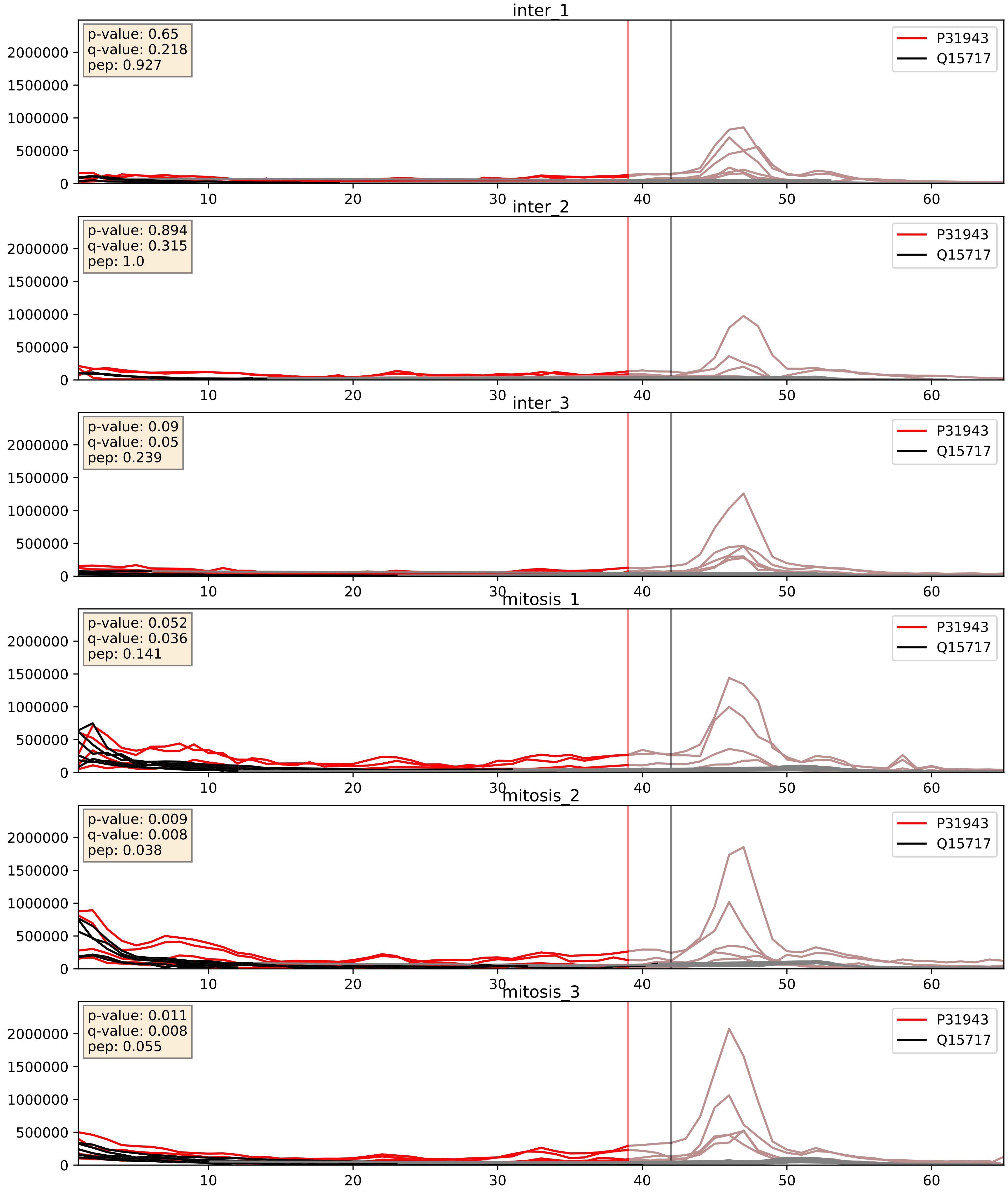


| level | condition_1 | condition_2 | pvalue | pvalue_adjusted |
|---|---|---|---|---|
| complex_abundance | mitosis | inter | 0.00017736744901018607 | 0.008807651911797798 |
| interactor_abundance | mitosis | inter | 0.0014904490749389305 | 0.0285056509549173 |
| interactor_ratio | mitosis | inter | 0.12665350951524584 | 0.5750062399685051 |

P38159_Q15717
RBMX_HUMAN vs ELAV1_HUMAN
p-value: 0.0 q-value: 0.0

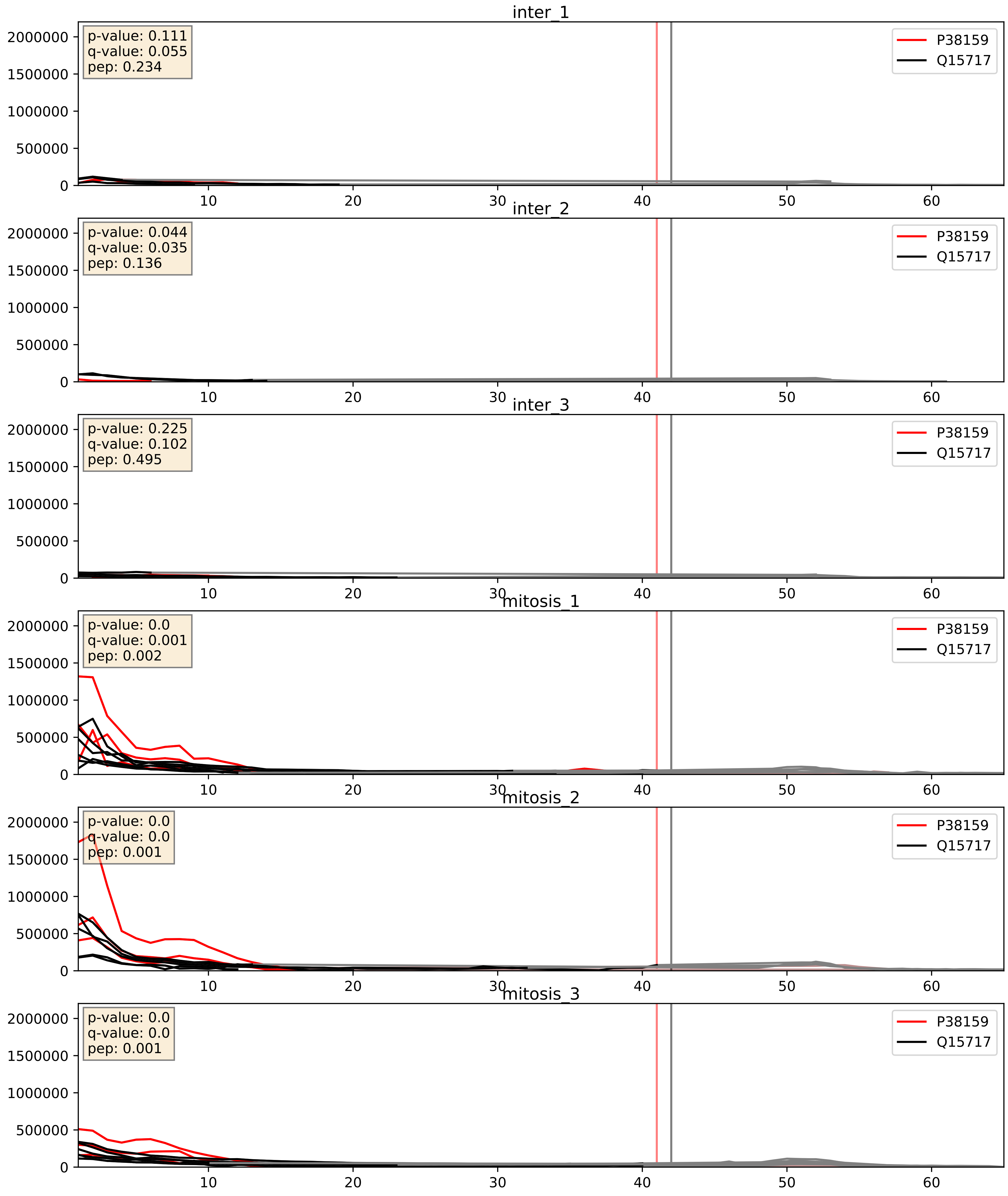


| level | condition_1 | condition_2 | pvalue | pvalue_adjusted |
|---|---|---|---|---|
| complex_abundance | mitosis | inter | 0.0004288806060394006 | 0.013091389064164169 |
| interactor_abundance | mitosis | inter | 0.0010979128258399222 | 0.024049720606162197 |
| interactor_ratio | mitosis | inter | 0.3588347875269688 | 0.8049703690863734 |

P51991_Q15717
ROA3_HUMAN vs ELAV1_HUMAN
p-value: 0.001 q-value: 0.024

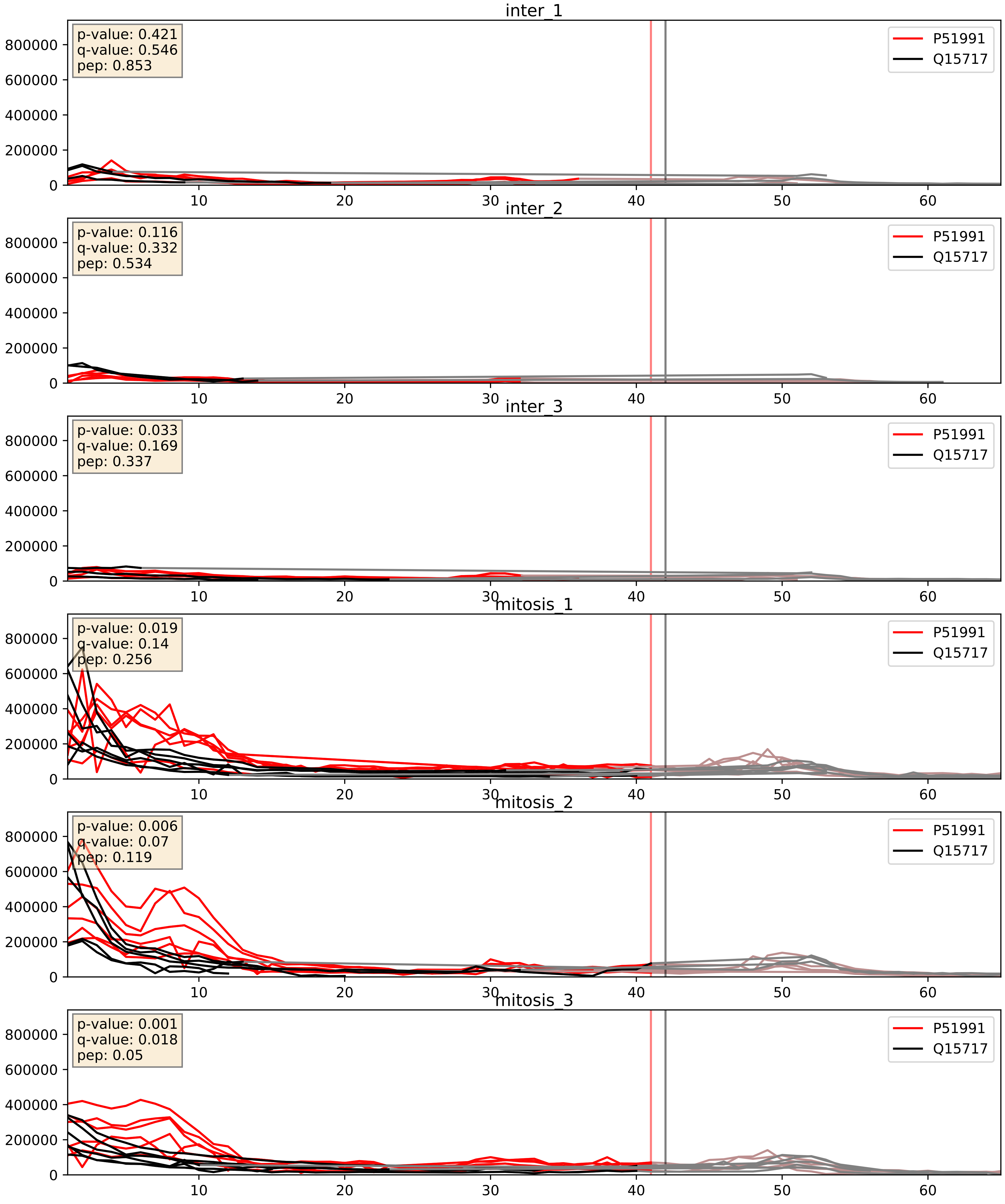


| level | condition_1 | condition_2 | pvalue | pvalue_adjusted |
|---|---|---|---|---|
| interactor_abundance | mitosis | inter | 3.736100323192025e-06 | 0.0033734327106189023 |
| complex_abundance | mitosis | inter | 1.684569173590749e-05 | 0.005691857151652958 |
| interactor_ratio | mitosis | inter | 0.24176845376887507 | 0.7200661897289671 |

P52272_Q15717
HNRPM_HUMAN vs ELAV1_HUMAN
p-value: 0.001 q-value: 0.002

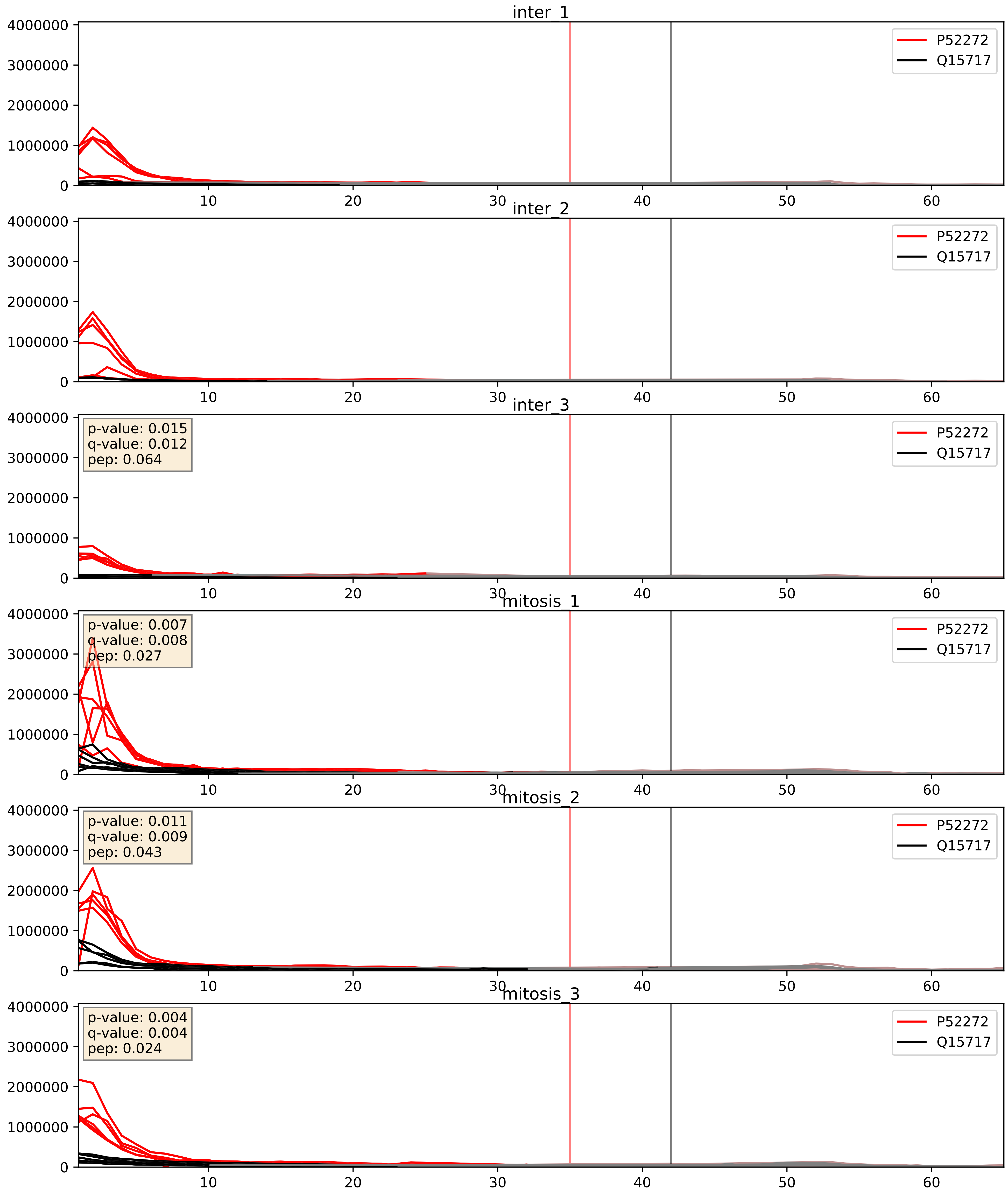


| level | condition_1 | condition_2 | pvalue | pvalue_adjusted |
|---|---|---|---|---|
| interactor_ratio | mitosis | inter | 0.0003243129433665509 | 0.019014512296011477 |
| complex_abundance | mitosis | inter | 0.002303087840743302 | 0.03502731659907978 |
| interactor_abundance | mitosis | inter | 0.1314788701529947 | 0.31658484627556527 |

P52597_Q15717
HNRPF_HUMAN vs ELAV1_HUMAN
p-value: 0.005 q-value: 0.005

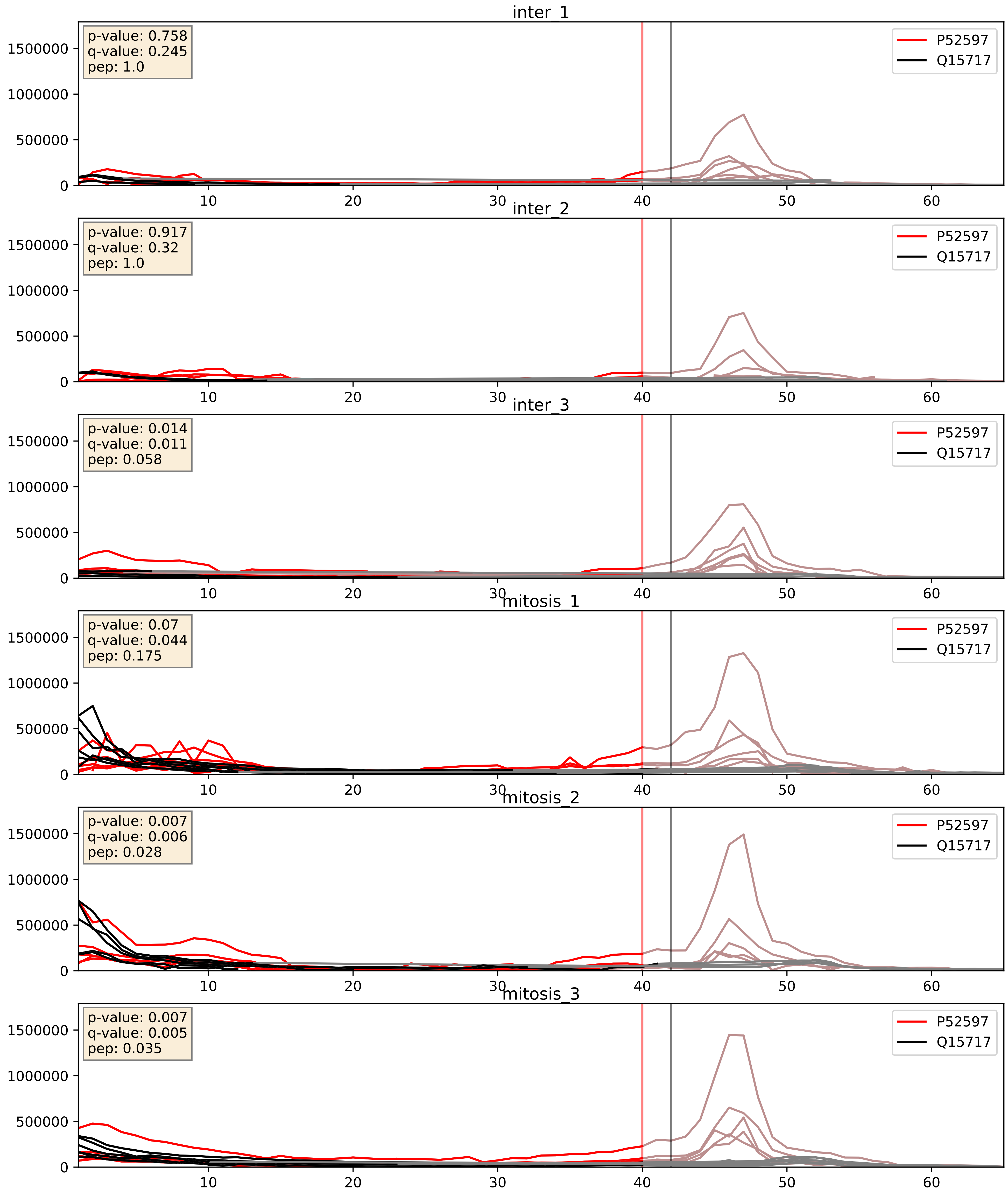


| level | condition_1 | condition_2 | pvalue | pvalue_adjusted |
|---|---|---|---|---|
| complex_abundance | mitosis | inter | 0.00016915016388839636 | 0.008722442186052247 |
| interactor_abundance | mitosis | inter | 0.002413019482392022 | 0.036558312865974704 |
| interactor_ratio | mitosis | inter | 0.014023604107891958 | 0.1741671265118786 |

P62304_Q15717
RUXE_HUMAN vs ELAV1_HUMAN
p-value: 0.082 q-value: 0.048

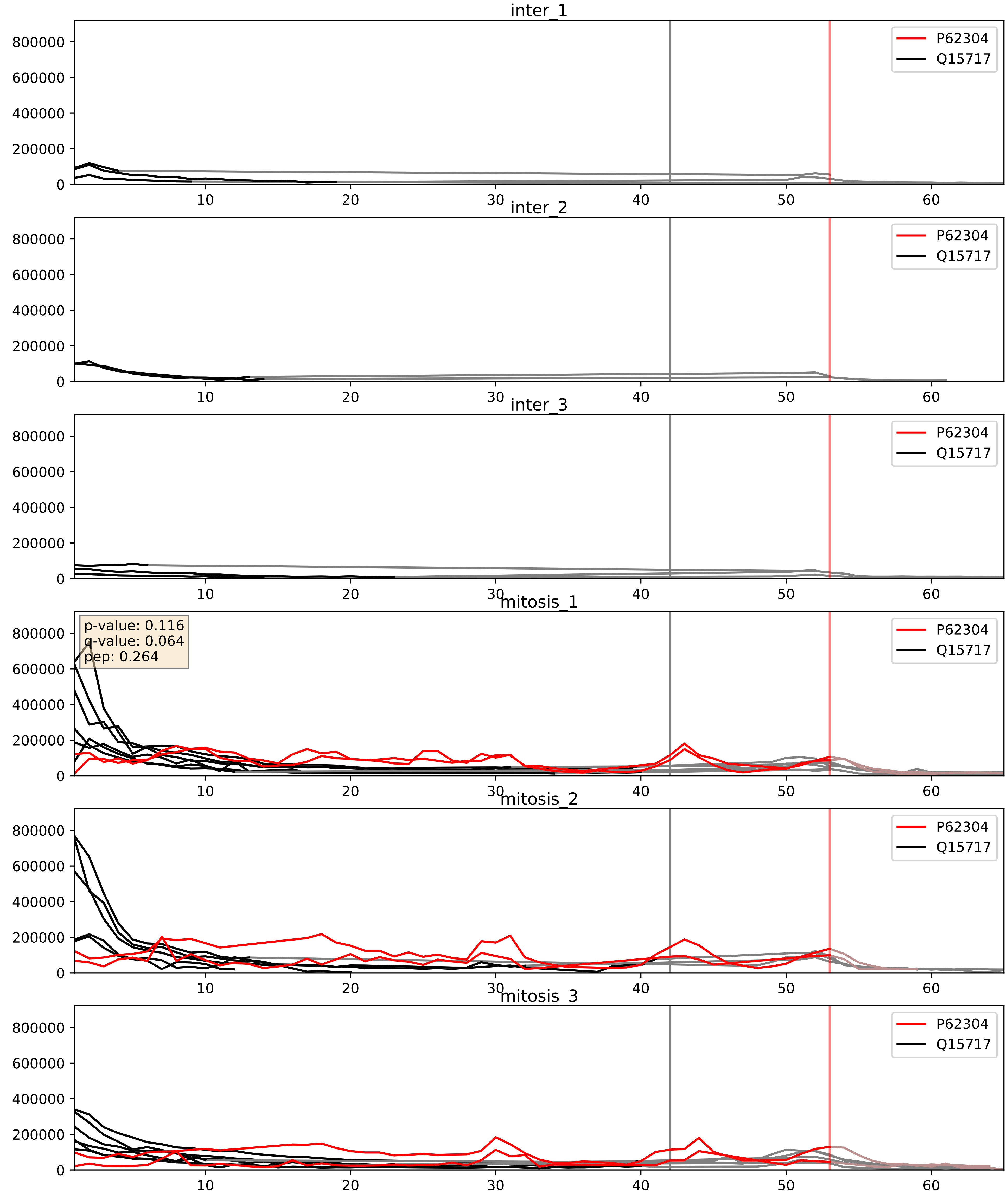


| level | condition_1 | condition_2 | pvalue | pvalue_adjusted |
|---|---|---|---|---|
| interactor_ratio | mitosis | inter | 1.3829541673086077e-05 | 0.003969306345409136 |
| complex_abundance | mitosis | inter | 9.823710518782713e-05 | 0.006719978526560179 |
| interactor_abundance | mitosis | inter | 0.00019797887174147875 | 0.01052608162799415 |

P62995_Q15717
TRA2B_HUMAN vs ELAV1_HUMAN
p-value: 0.0 q-value: 0.0

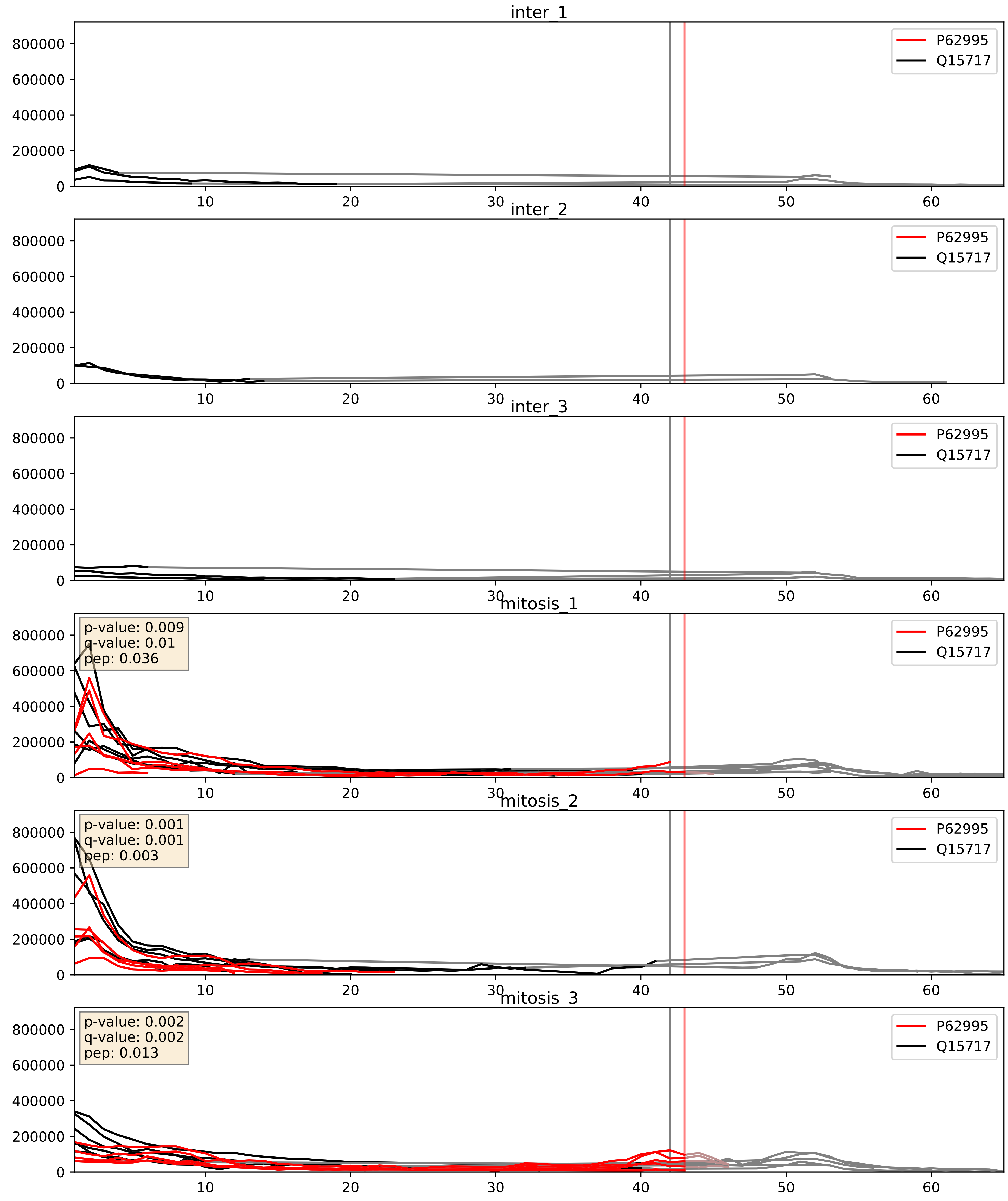


| level | condition_1 | condition_2 | pvalue | pvalue_adjusted |
|---|---|---|---|---|
| complex_abundance | mitosis | inter | 0.0001242130151056852 | 0.007704807313801923 |
| interactor_abundance | mitosis | inter | 0.0004121602900123102 | 0.014019253191624928 |
| interactor_ratio | mitosis | inter | 0.01618022122043033 | 0.18789564671895648 |

P84103_Q15717
SRSF3_HUMAN vs ELAV1_HUMAN
p-value: 0.04 q-value: 0.028

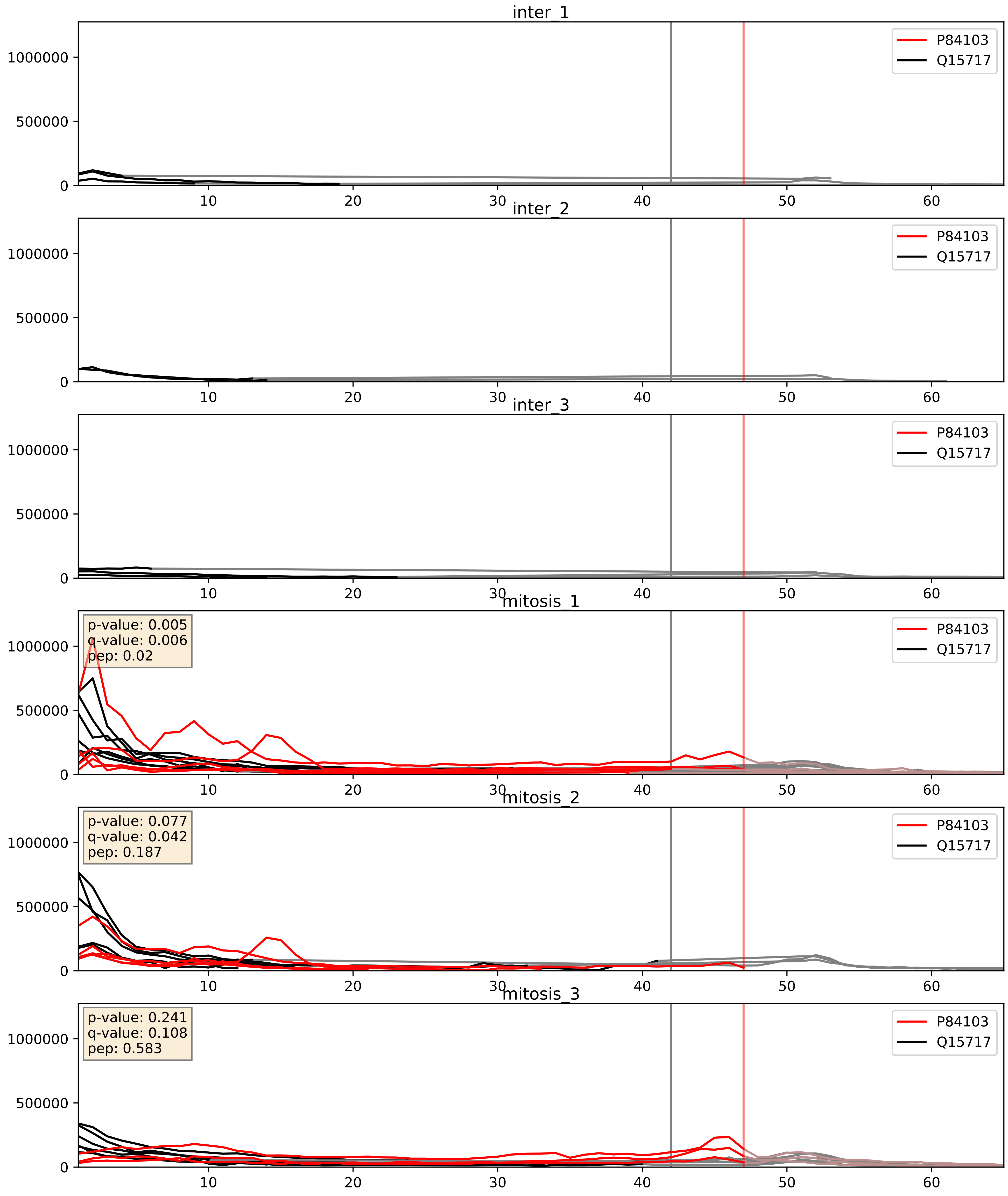


| level | condition_1 | condition_2 | pvalue | pvalue_adjusted |
|---|---|---|---|---|
| complex_abundance | mitosis | inter | 0.0005540143380544626 | 0.015065929247636585 |
| interactor_abundance | mitosis | inter | 0.0033330748758513776 | 0.04454507562418079 |
| interactor_ratio | mitosis | inter | 0.05931613769021241 | 0.40293896347252833 |

Q07955_Q15717
SRSF1_HUMAN vs ELAV1_HUMAN
p-value: 0.011 q-value: 0.01

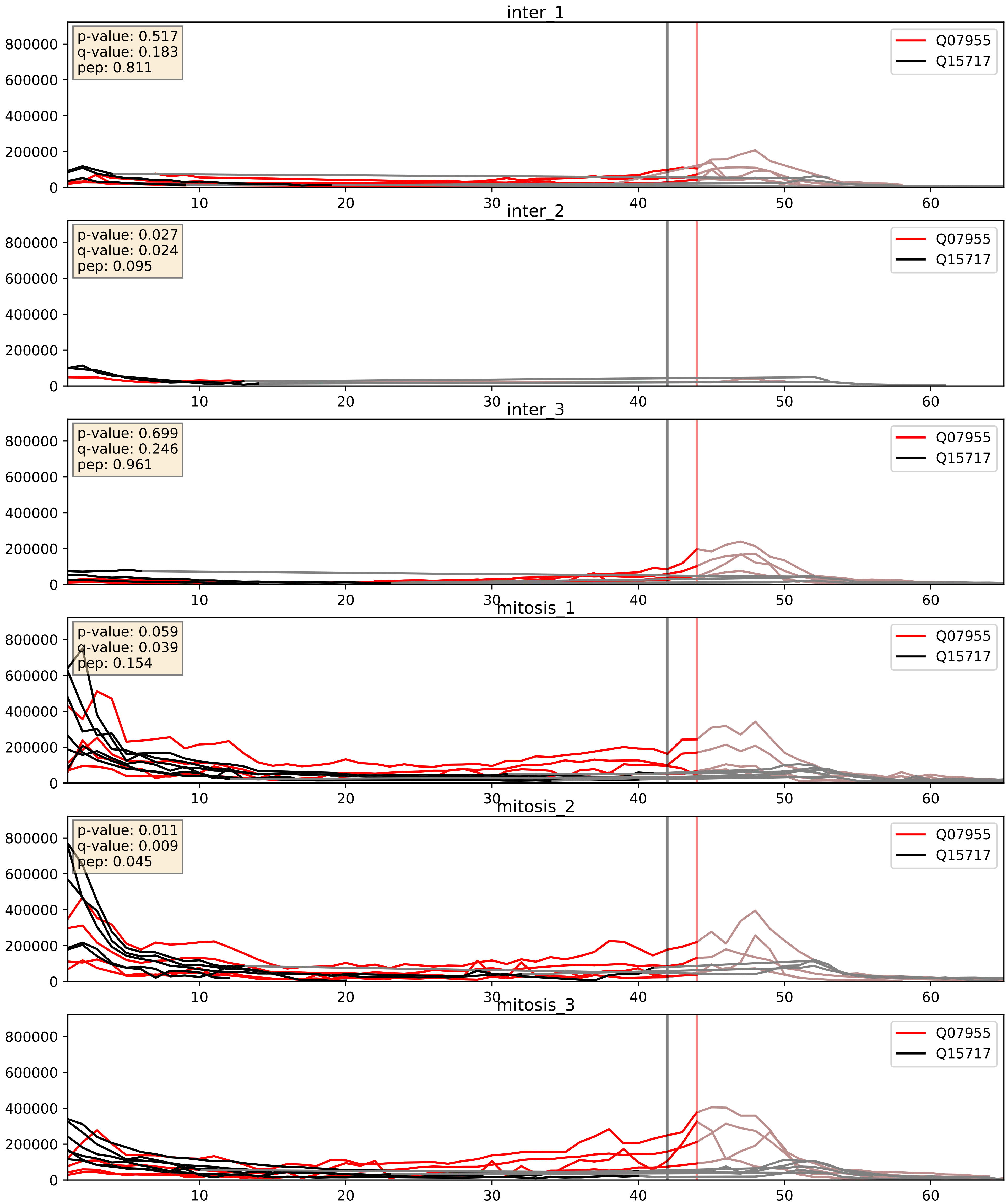


| level | condition_1 | condition_2 | pvalue | pvalue_adjusted |
|---|---|---|---|---|
| complex_abundance | mitosis | inter | 0.0001877702555432865 | 0.009111526782692203 |
| interactor_abundance | mitosis | inter | 0.001583237558275299 | 0.029025269221018554 |
| interactor_ratio | mitosis | inter | 0.8402338042736714 | 0.9741841208970103 |

Q12905_Q15717
ILF2_HUMAN vs ELAV1_HUMAN
p-value: 0.004 q-value: 0.047

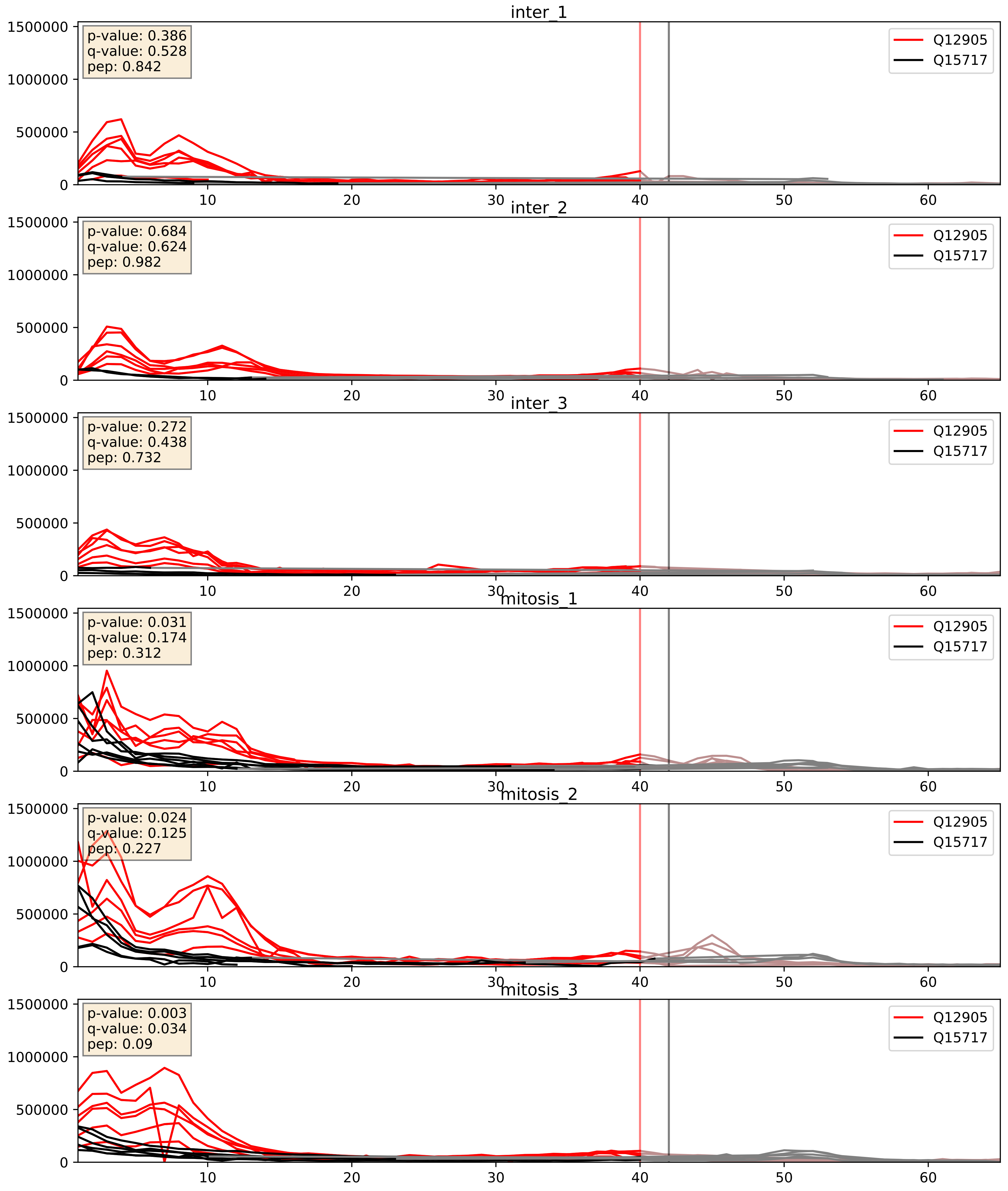


| level | condition_1 | condition_2 | pvalue | pvalue_adjusted |
|---|---|---|---|---|
| complex_abundance | mitosis | inter | 4.5501477611336626e-05 | 0.005691857151652958 |
| interactor_abundance | mitosis | inter | 0.0015109761494022987 | 0.028710223837699618 |
| interactor_ratio | mitosis | inter | 0.003190844185944654 | 0.07382061143698984 |

Q13151_Q15717
ROA0_HUMAN vs ELAV1_HUMAN
p-value: 0.026 q-value: 0.019

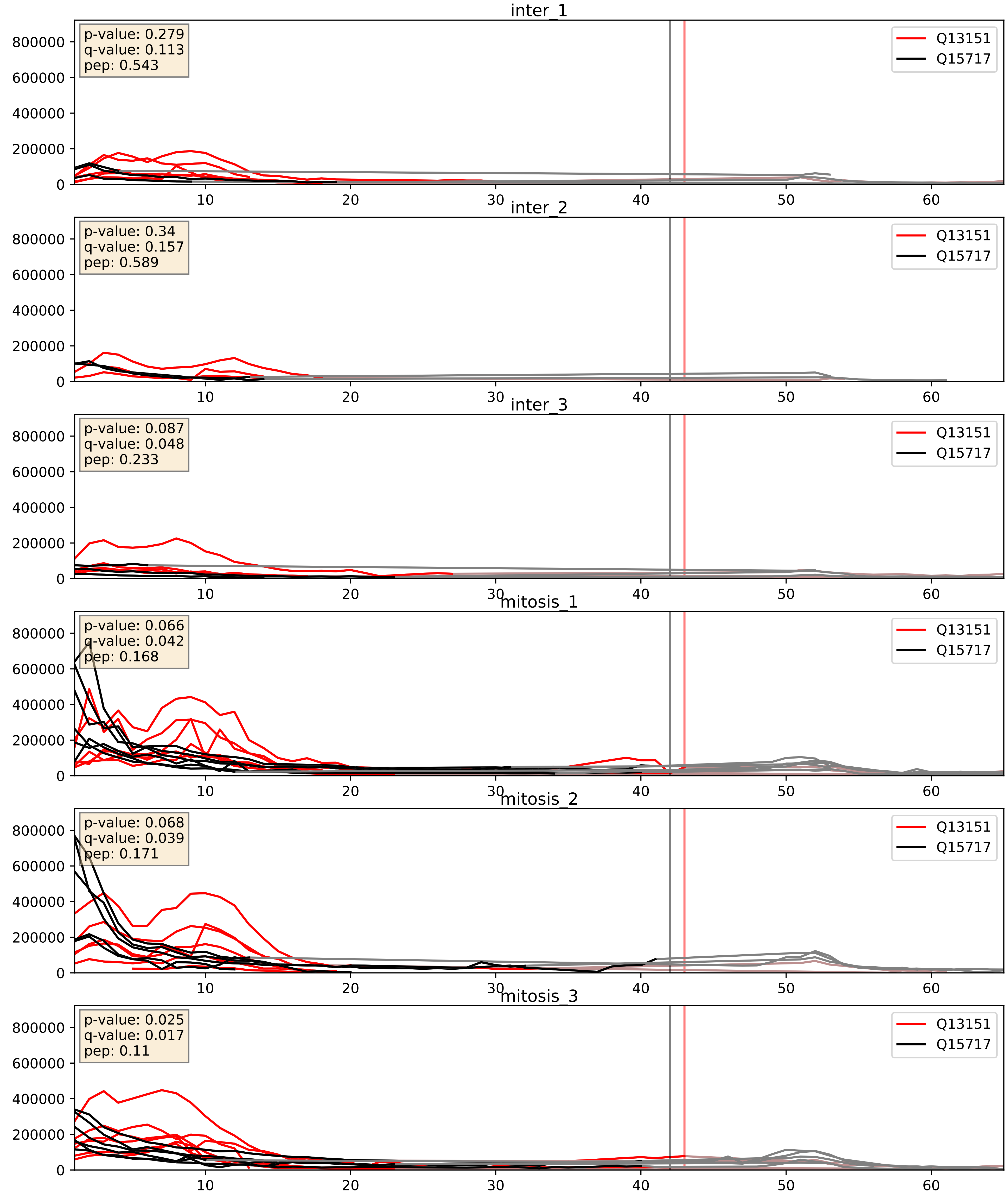


| level | condition_1 | condition_2 | pvalue | pvalue_adjusted |
|---|---|---|---|---|
| complex_abundance | mitosis | inter | 0.0008222134400985702 | 0.01907356923372293 |
| interactor_abundance | mitosis | inter | 0.006871344350993082 | 0.0713363448150053 |
| interactor_ratio | mitosis | inter | 0.009973473593403892 | 0.14263182888175546 |

Q15366_Q15717
PCBP2_HUMAN vs ELAV1_HUMAN
p-value: 0.068 q-value: 0.042

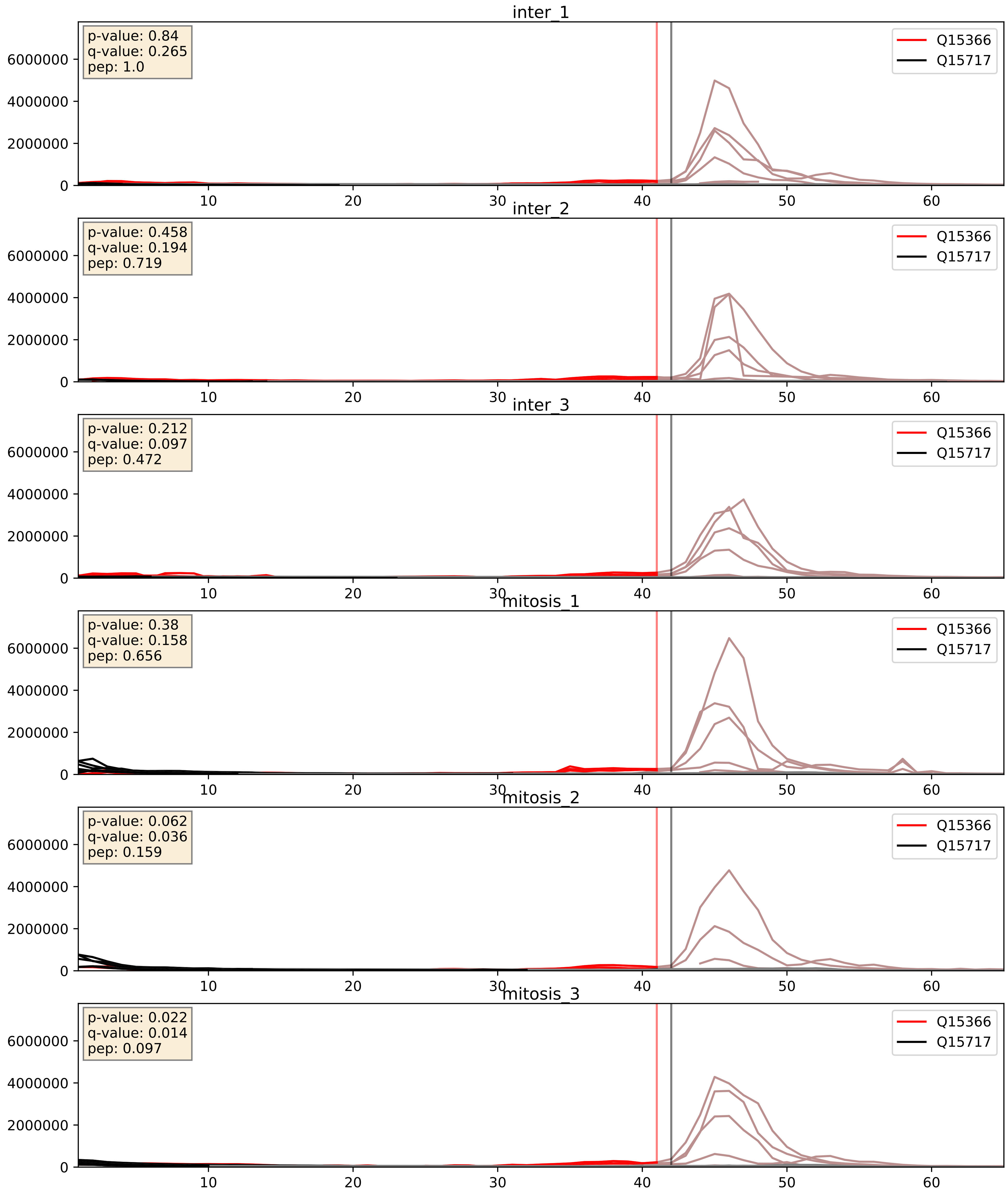


| level | condition_1 | condition_2 | pvalue | pvalue_adjusted |
|---|---|---|---|---|
| complex_abundance | mitosis | inter | 0.012844837756186694 | 0.08405635484106207 |
| interactor_ratio | mitosis | inter | 0.05766431446787582 | 0.3961608945972482 |
| interactor_abundance | mitosis | inter | 0.5222800748346491 | 0.683447551765955 |

Q15717_Q16629
ELAV1_HUMAN vs SRSF7_HUMAN
p-value: 0.005 q-value: 0.005

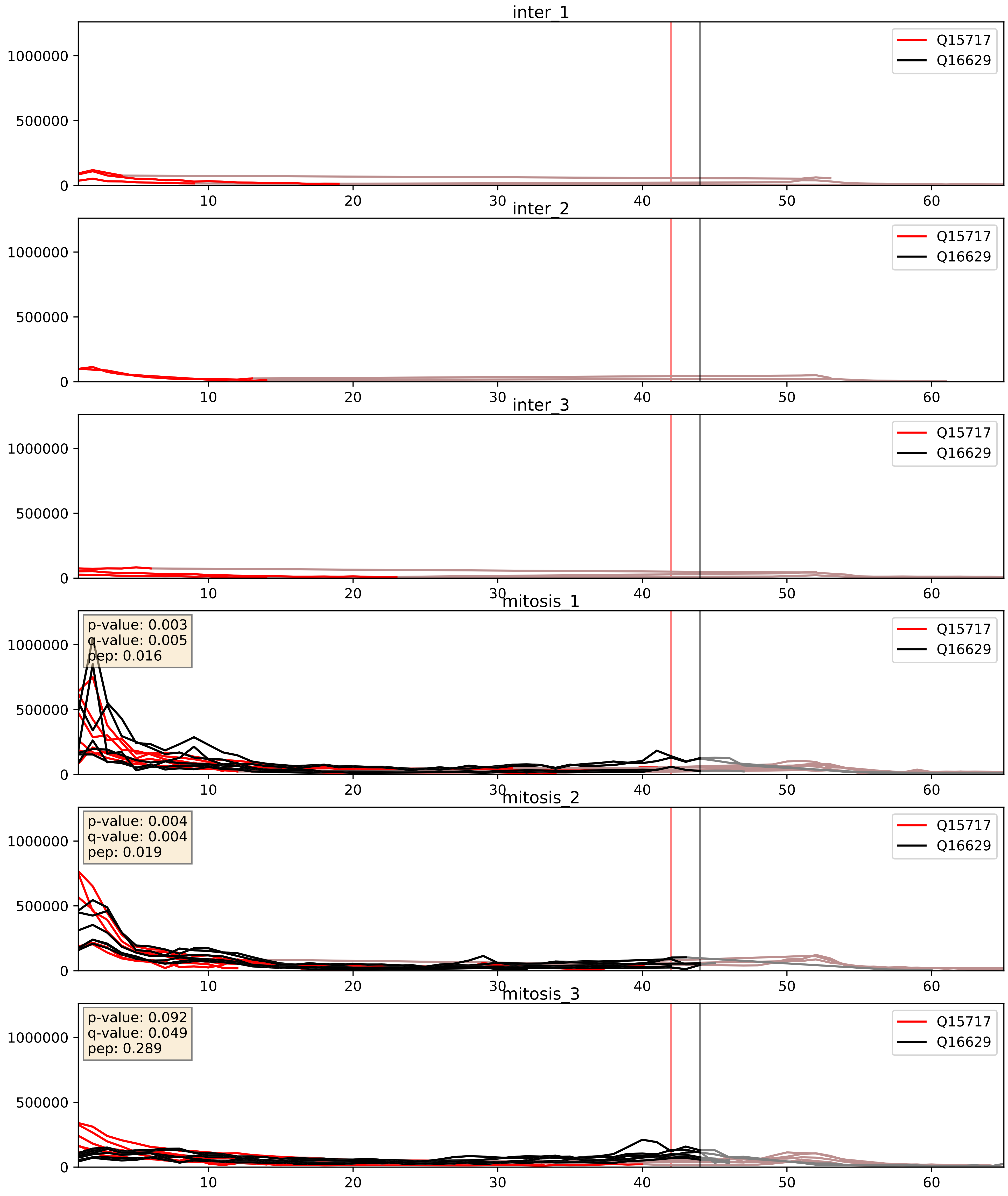


| level | condition_1 | condition_2 | pvalue | pvalue_adjusted |
|---|---|---|---|---|
| interactor_abundance | mitosis | inter | 6.059680363197429e-05 | 0.006459477818931243 |
| complex_abundance | mitosis | inter | 6.314734732233846e-05 | 0.005691857151652958 |
| interactor_ratio | mitosis | inter | 0.028137833796062196 | 0.2626606949774181 |

Q15717_Q99459
ELAV1_HUMAN vs CDC5L_HUMAN
p-value: 0.029 q-value: 0.021

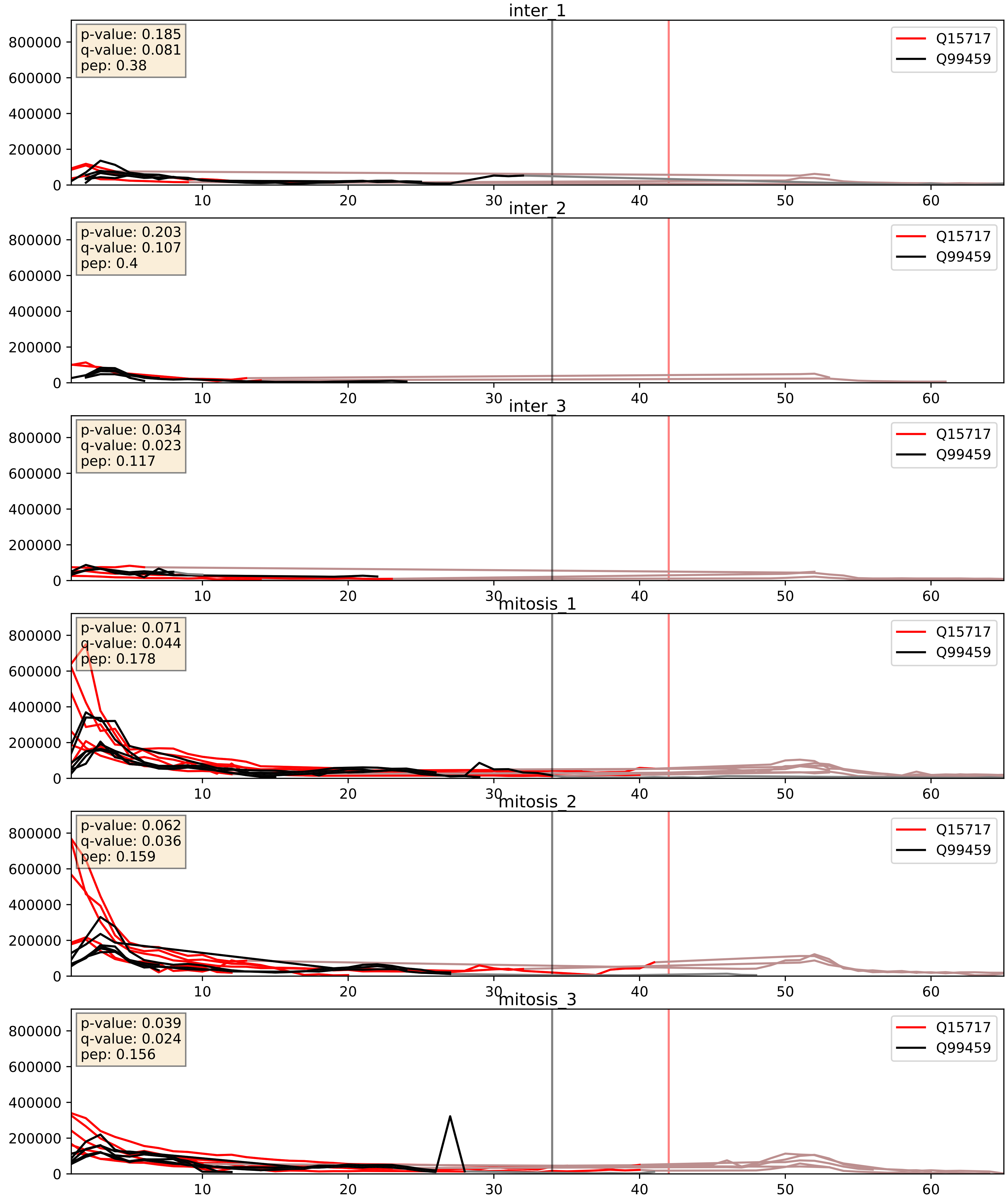


| level | condition_1 | condition_2 | pvalue | pvalue_adjusted |
|---|---|---|---|---|
| interactor_abundance | mitosis | inter | 0.00036534728597032297 | 0.01356427145975096 |
| complex_abundance | mitosis | inter | 0.002092462180363407 | 0.03329270681024306 |
| interactor_ratio | mitosis | inter | 0.046977446550768735 | 0.35838626305591875 |
