## Supplementary Data 2 for "SECAT: Quantifying differential protein-protein interaction states by network-centric analysis": hela_string_000046_P14866.pdf

P14866_P14866

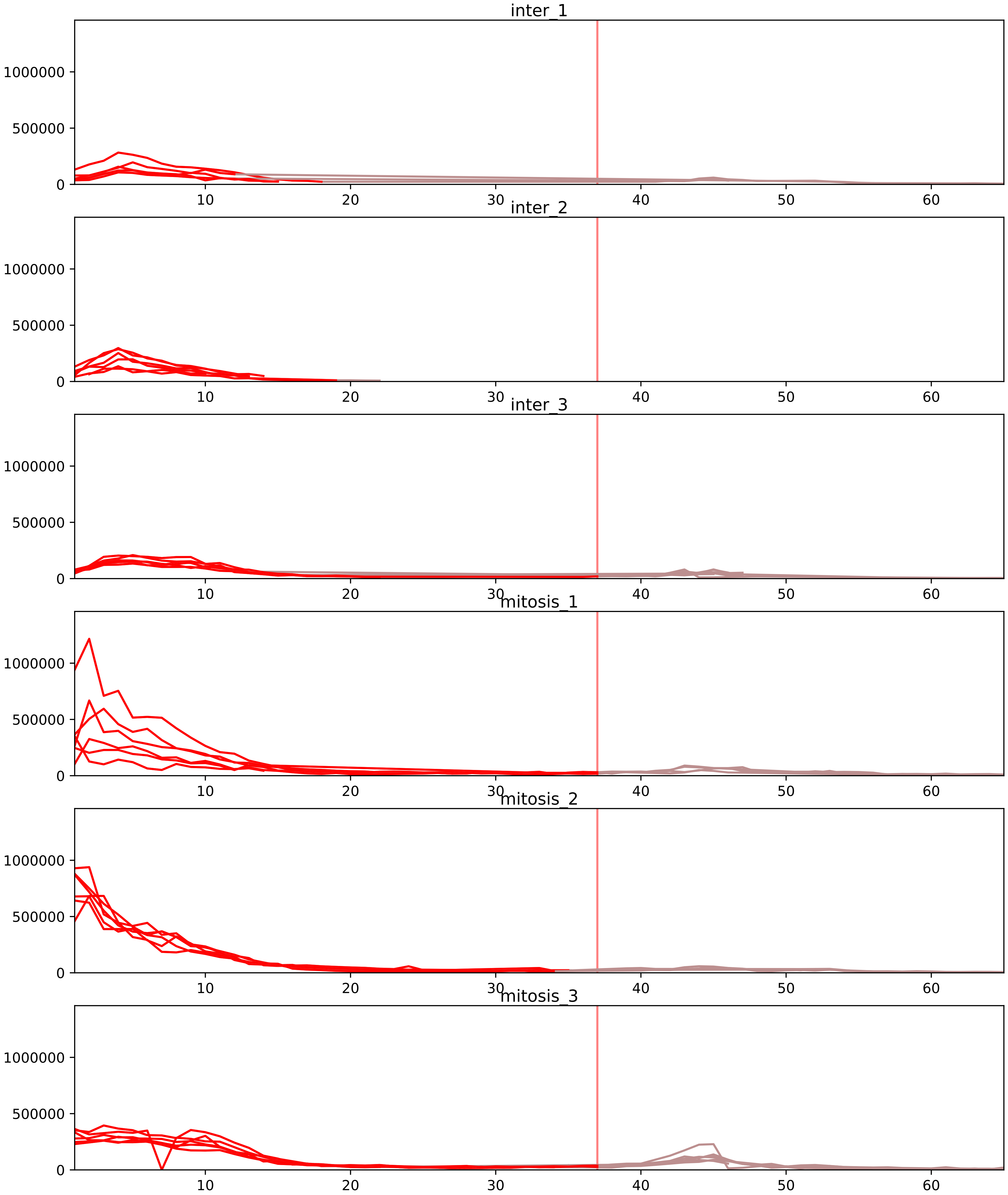


| level | condition_1 | condition_2 | pvalue | pvalue_adjusted |
|---|---|---|---|---|
| interactor_ratio | mitosis | inter | 6.118649057236363e-08 | 2.6182126198406763e-06 |
| complex_abundance | mitosis | inter | 1.4938350993050998e-07 | 2.3521134003385493e-05 |
| interactor_abundance | mitosis | inter | 2.3740192659613475e-05 | 0.0009294032870997615 |
| total_abundance | mitosis | inter | 0.015137367952719655 | 0.15537165372438663 |
| assembled_abundance | mitosis | inter | 0.02002798757353147 | 0.16728861540201573 |
| monomer_abundance | mitosis | inter | 0.06060822866273875 | 0.3544344065380258 |

O14979_P14866
HNRDL_HUMAN vs HNRPL_HUMAN
p-value: 0.003 q-value: 0.044

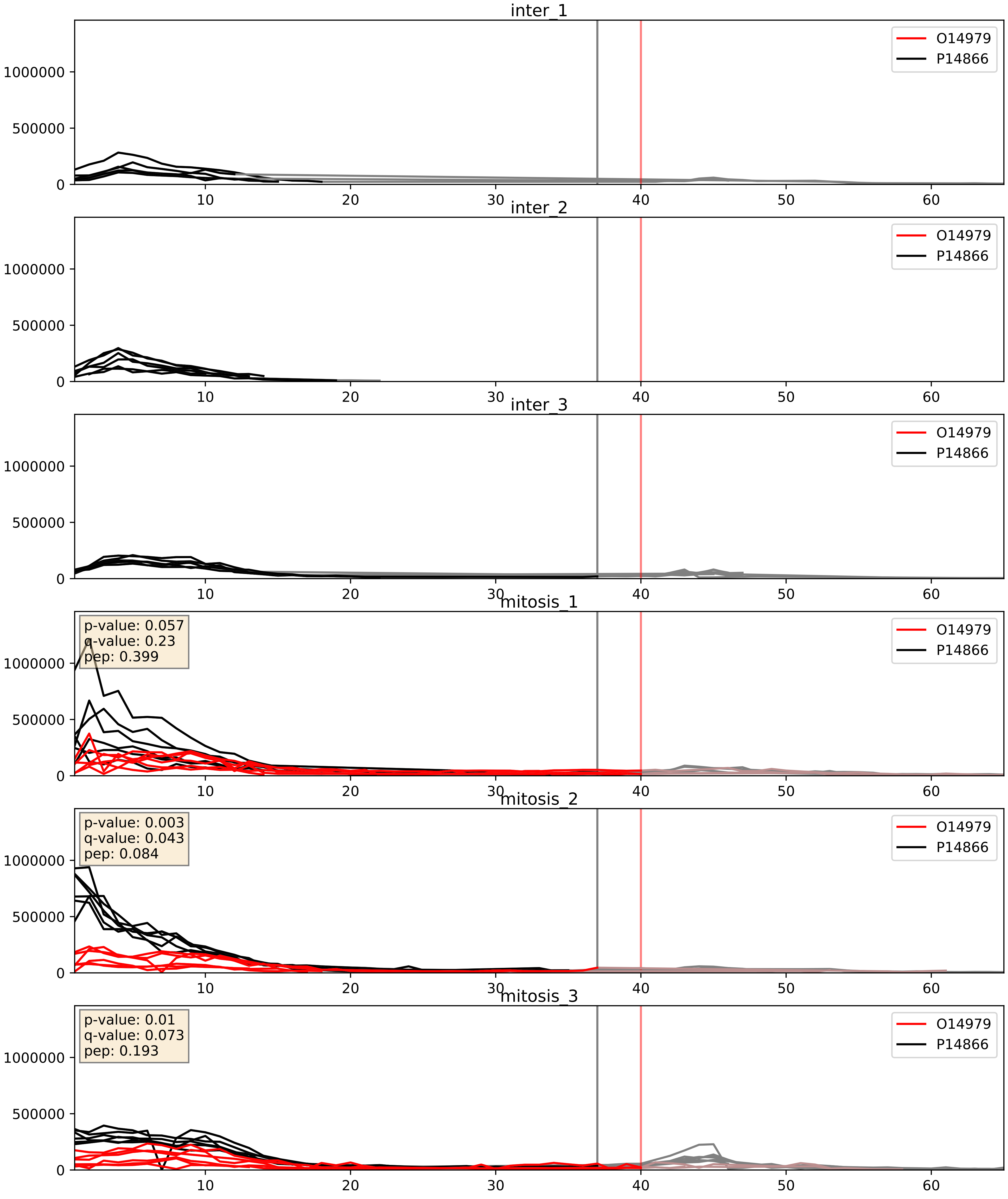


| level | condition_1 | condition_2 | pvalue | pvalue_adjusted |
|---|---|---|---|---|
| complex_abundance | mitosis | inter | 9.401708878998684e-05 | 0.006596608852805635 |
| interactor_abundance | mitosis | inter | 0.0007528746798464765 | 0.020141995013343572 |
| interactor_ratio | mitosis | inter | 0.012414827969505712 | 0.16080750275117184 |

O43390_P14866
HNRPR_HUMAN vs HNRPL_HUMAN
p-value: 0.028 q-value: 0.021

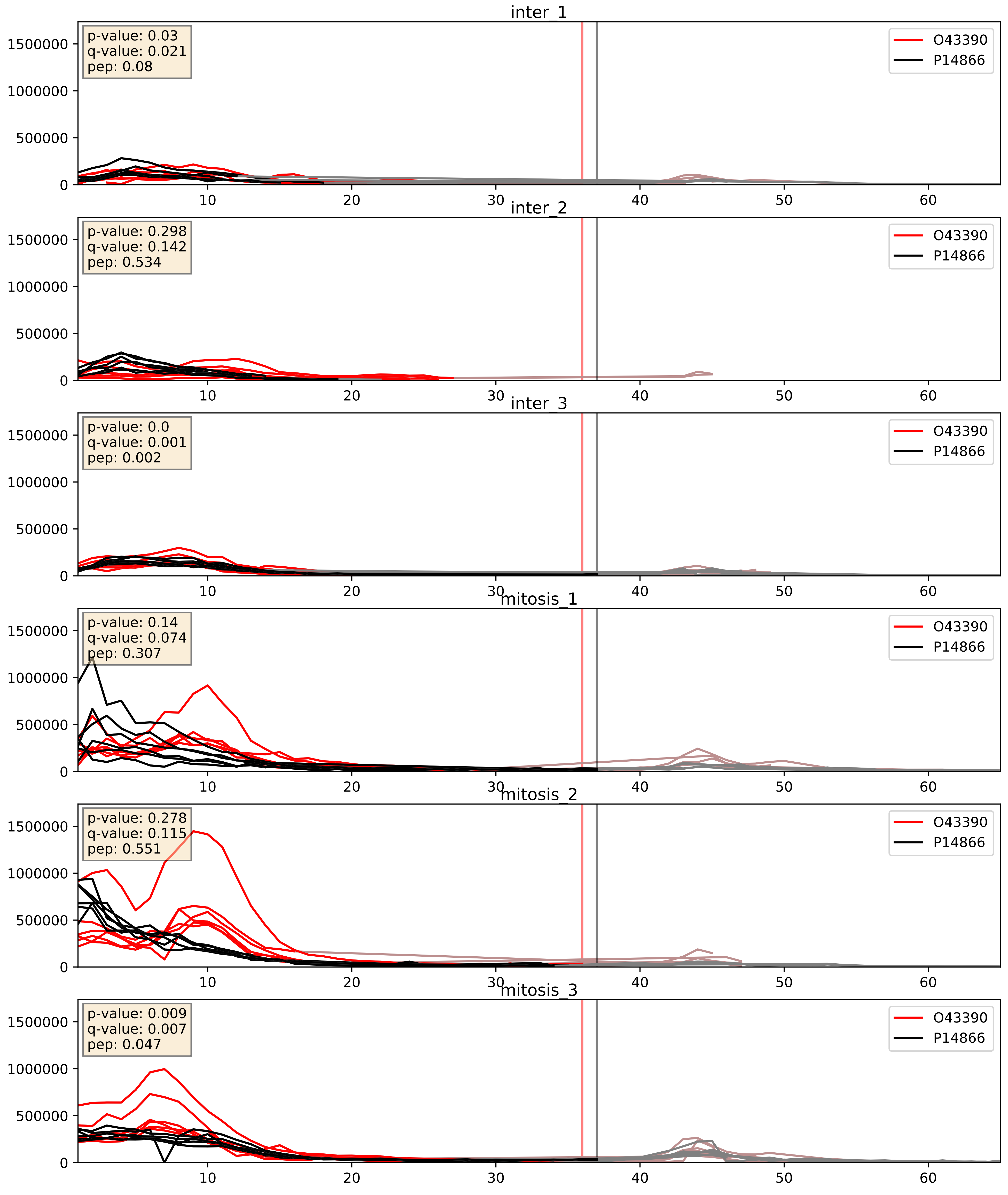


| level | condition_1 | condition_2 | pvalue | pvalue_adjusted |
|---|---|---|---|---|
| interactor_abundance | mitosis | inter | 5.893602312406929e-05 | 0.006459477818931243 |
| complex_abundance | mitosis | inter | 0.0011057804224422322 | 0.022115608448844644 |
| interactor_ratio | mitosis | inter | 0.7048469232789103 | 0.9378021986892063 |

P07910_P14866
HNRPC_HUMAN vs HNRPL_HUMAN
p-value: 0.011 q-value: 0.01

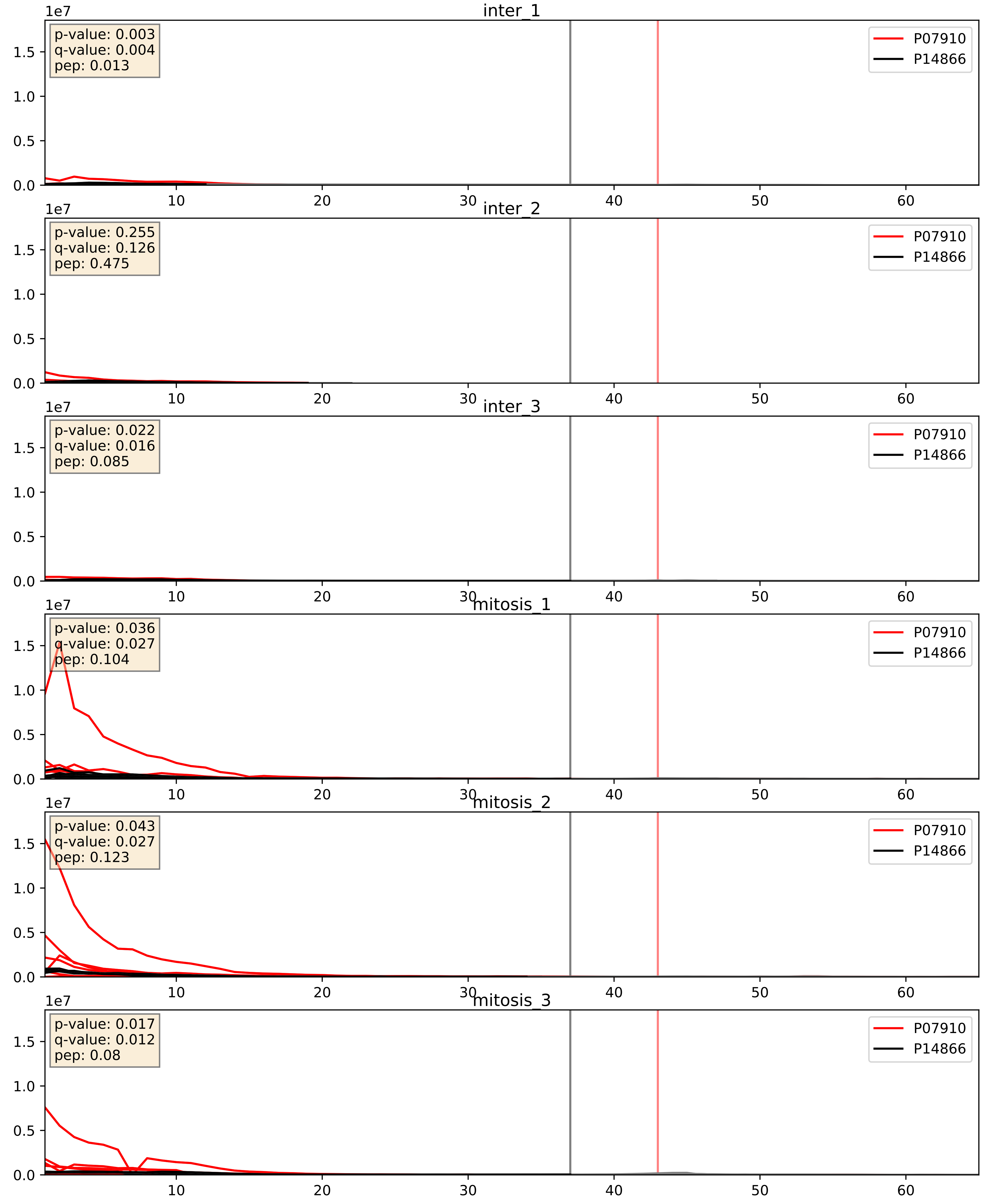


| level | condition_1 | condition_2 | pvalue | pvalue_adjusted |
|---|---|---|---|---|
| interactor_abundance | mitosis | inter | 8.645329666757008e-06 | 0.0033734327106189023 |
| complex_abundance | mitosis | inter | 3.986857495835857e-05 | 0.005691857151652958 |
| interactor_ratio | mitosis | inter | 0.0022228260684612118 | 0.05890833172144883 |

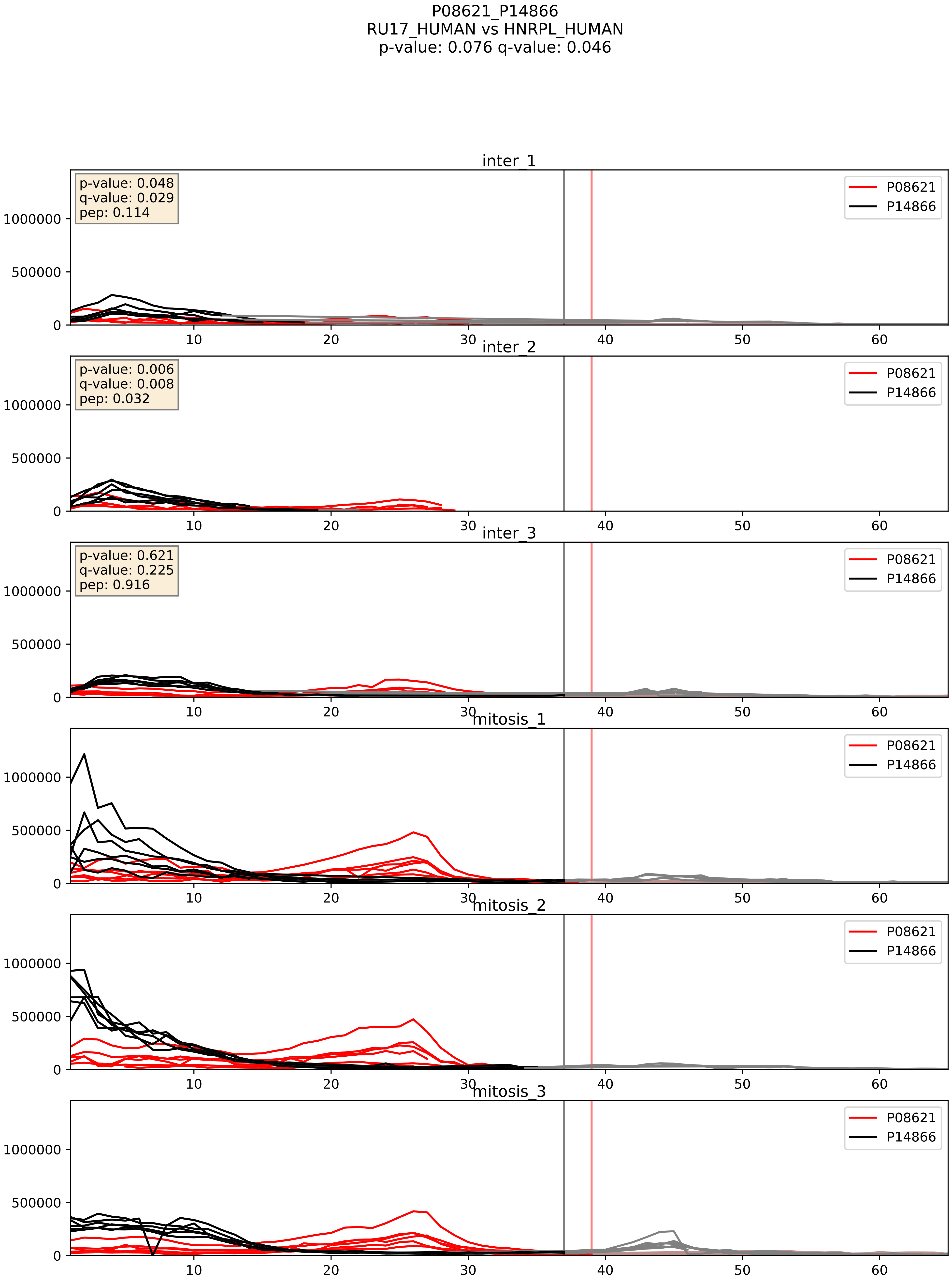


| level | condition_1 | condition_2 | pvalue | pvalue_adjusted |
|---|---|---|---|---|
| interactor_abundance | mitosis | inter | 1.057401869558351e-05 | 0.0033734327106189023 |
| complex_abundance | mitosis | inter | 0.0003837667920089791 | 0.01249066060683217 |
| interactor_ratio | mitosis | inter | 0.14042639881275756 | 0.5907826388074301 |

P09651_P14866
ROA1_HUMAN vs HNRPL_HUMAN
p-value: 0.005 q-value: 0.005

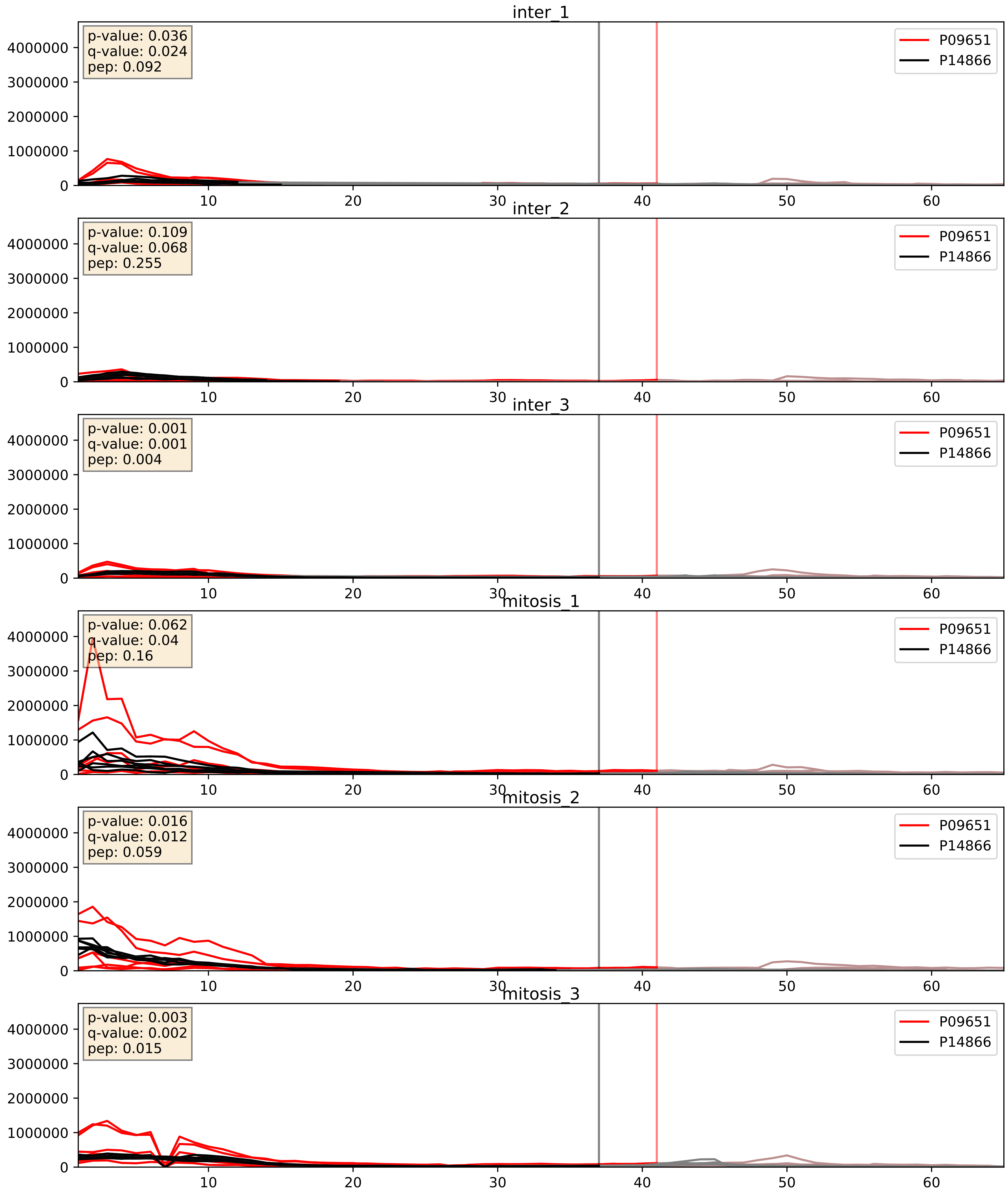


| level | condition_1 | condition_2 | pvalue | pvalue_adjusted |
|---|---|---|---|---|
| complex_abundance | mitosis | inter | 0.0022461160486105898 | 0.03476808928771546 |
| interactor_abundance | mitosis | inter | 0.010916100555075968 | 0.0916545568920552 |
| interactor_ratio | mitosis | inter | 0.343200993452667 | 0.7955051459395694 |

P14866_P17844
HNRPL_HUMAN vs DDX5_HUMAN
p-value: 0.0 q-value: 0.01

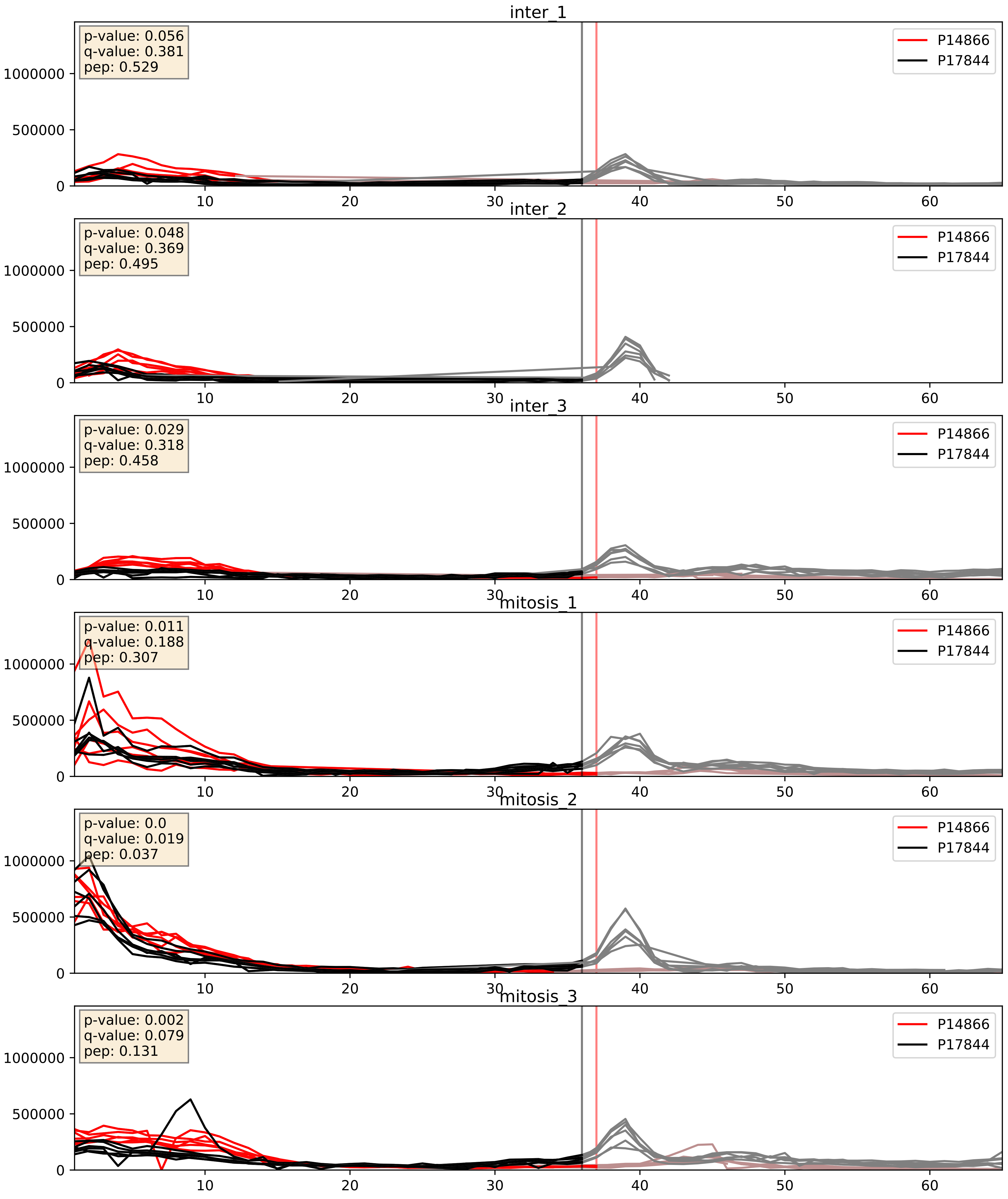


| level | condition_1 | condition_2 | pvalue | pvalue_adjusted |
| --- | --- | --- | --- | --- |
| complex_abundance | mitosis | inter | 0.0013977644692216144 | 0.025565948411403885 |
| interactor_abundance | mitosis | inter | 0.009741046732019873 | 0.08547756025227074 |
| interactor_ratio | mitosis | inter | 0.7081208830132038 | 0.9391051699295014 |

P14866_P22626
HNRPL_HUMAN vs ROA2_HUMAN
p-value: 0.012 q-value: 0.01

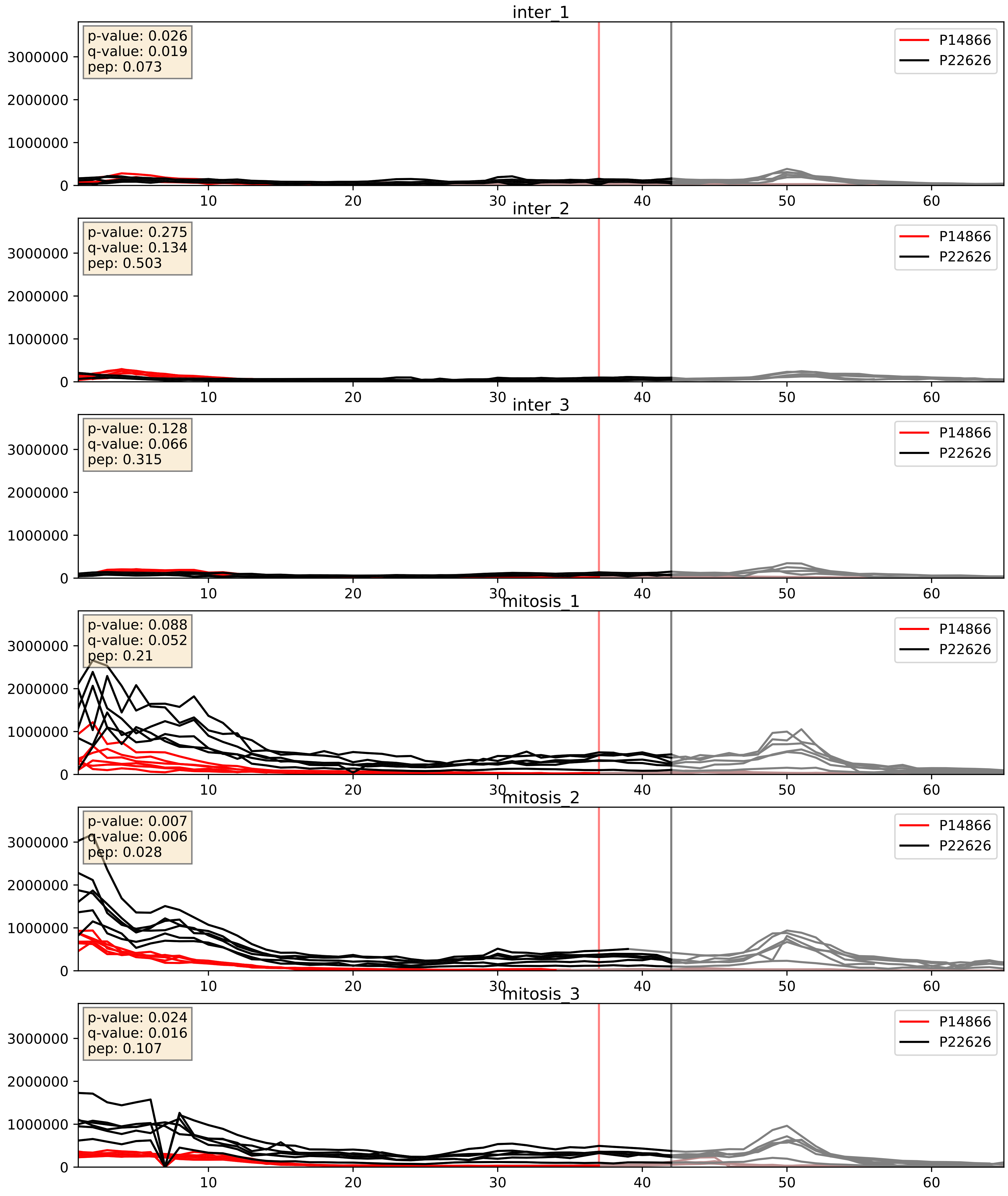


| level | condition_1 | condition_2 | pvalue | pvalue_adjusted |
|---|---|---|---|---|
| complex_abundance | mitosis | inter | 9.663556889611487e-05 | 0.00670982287146329 |
| interactor_abundance | mitosis | inter | 0.009697640385696541 | 0.08525122834595471 |
| interactor_ratio | mitosis | inter | 0.0163886342509947 | 0.1887653035437297 |

P14866_P26599
HNRPL_HUMAN vs PTBP1_HUMAN
p-value: 0.076 q-value: 0.046

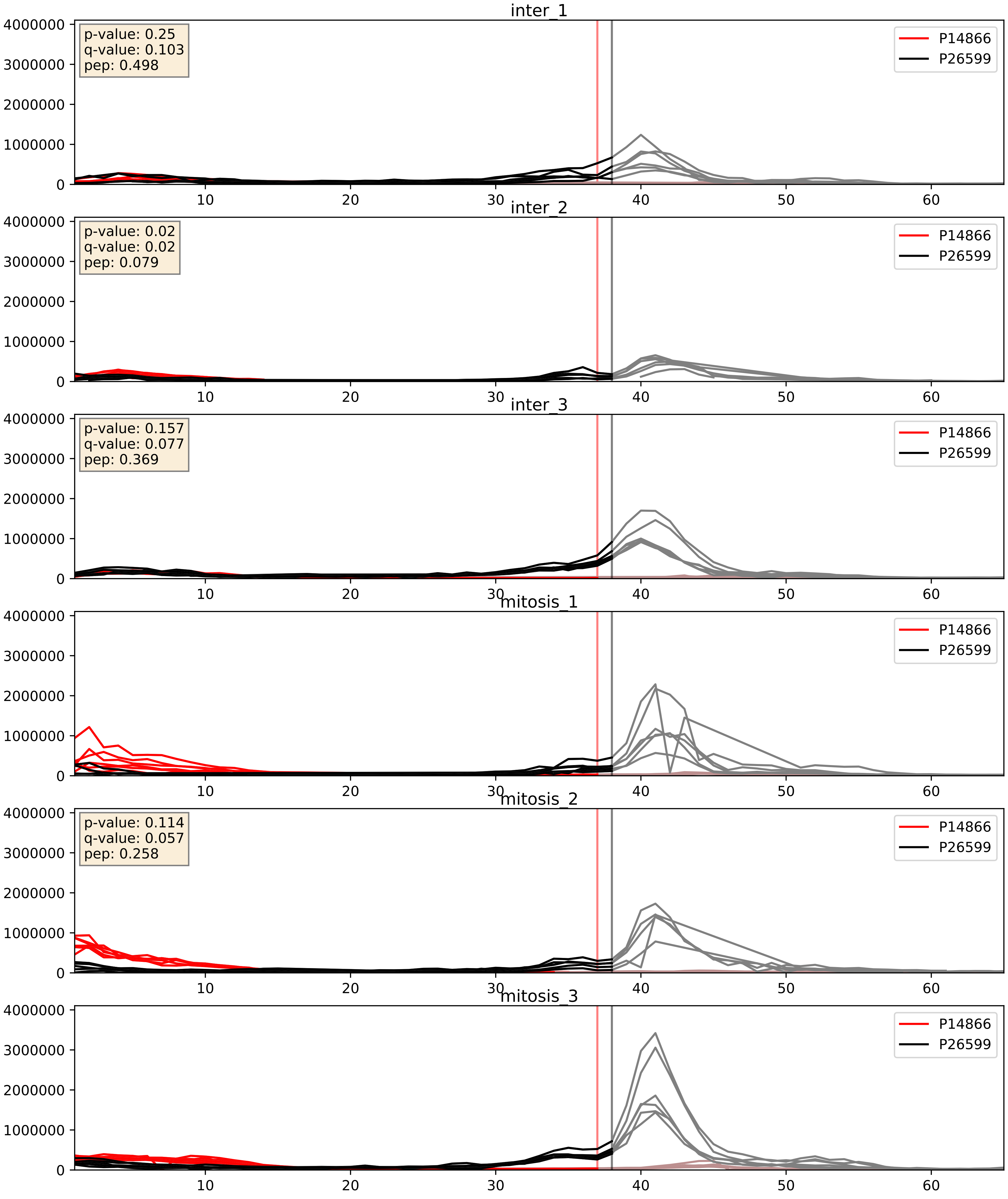


| level | condition_1 | condition_2 | pvalue | pvalue_adjusted |
|---|---|---|---|---|
| interactor_abundance | mitosis | inter | 0.009524831403908897 | 0.08470914994021835 |
| complex_abundance | mitosis | inter | 0.14431260111389285 | 0.29021207390906123 |
| interactor_ratio | mitosis | inter | 0.1898514385945017 | 0.6619009156224659 |

P14866_P31943
HNRPL_HUMAN vs HNRH1_HUMAN
p-value: 0.001 q-value: 0.002

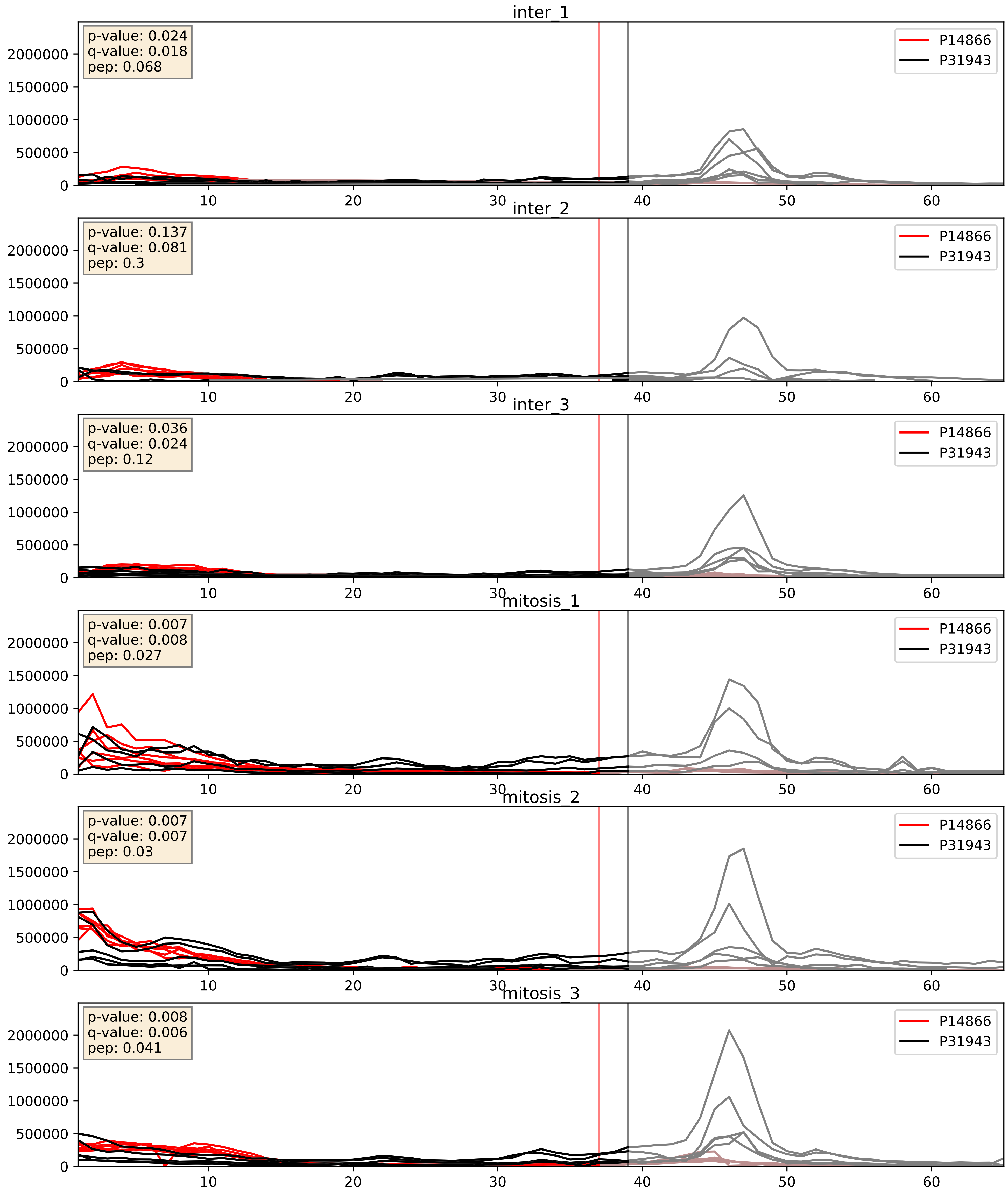


| level | condition_1 | condition_2 | pvalue | pvalue_adjusted |
|---|---|---|---|---|
| complex_abundance | mitosis | inter | 0.0010805657853290274 | 0.021833569470040306 |
| interactor_abundance | mitosis | inter | 0.009697640385696541 | 0.08525122834595471 |
| interactor_ratio | mitosis | inter | 0.20279183069839515 | 0.678106014366403 |

P14866_P38159
HNRPL_HUMAN vs RBMX_HUMAN
p-value: 0.008 q-value: 0.008

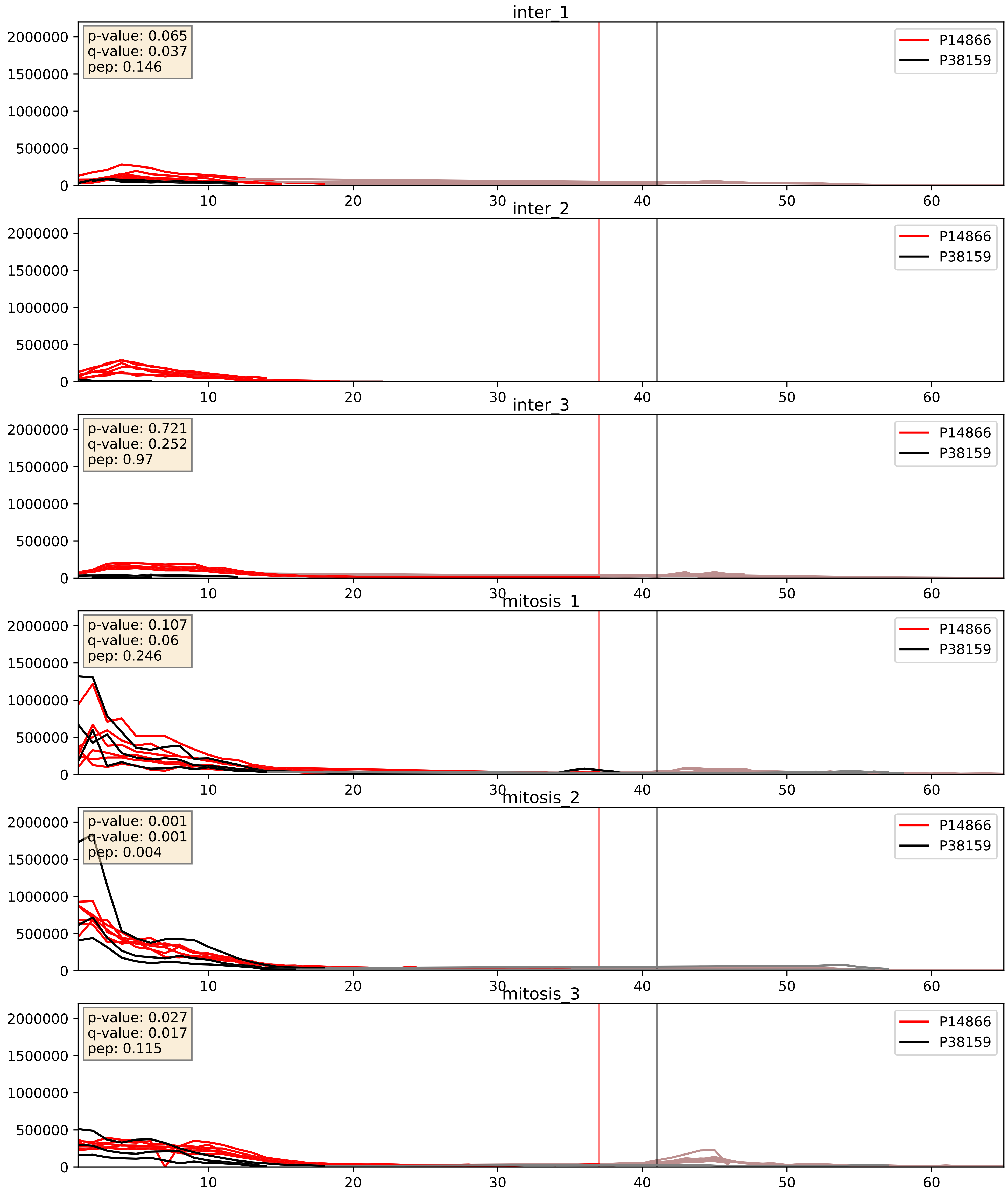


| level | condition_1 | condition_2 | pvalue | pvalue_adjusted |
|---|---|---|---|---|
| complex_abundance | mitosis | inter | 0.00042915958677650936 | 0.013091389064164169 |
| interactor_abundance | mitosis | inter | 0.002222136362133671 | 0.03466841143289118 |
| interactor_ratio | mitosis | inter | 0.04602651635329546 | 0.35622692584467375 |

P14866_P51991
HNRPL_HUMAN vs ROA3_HUMAN
p-value: 0.001 q-value: 0.002

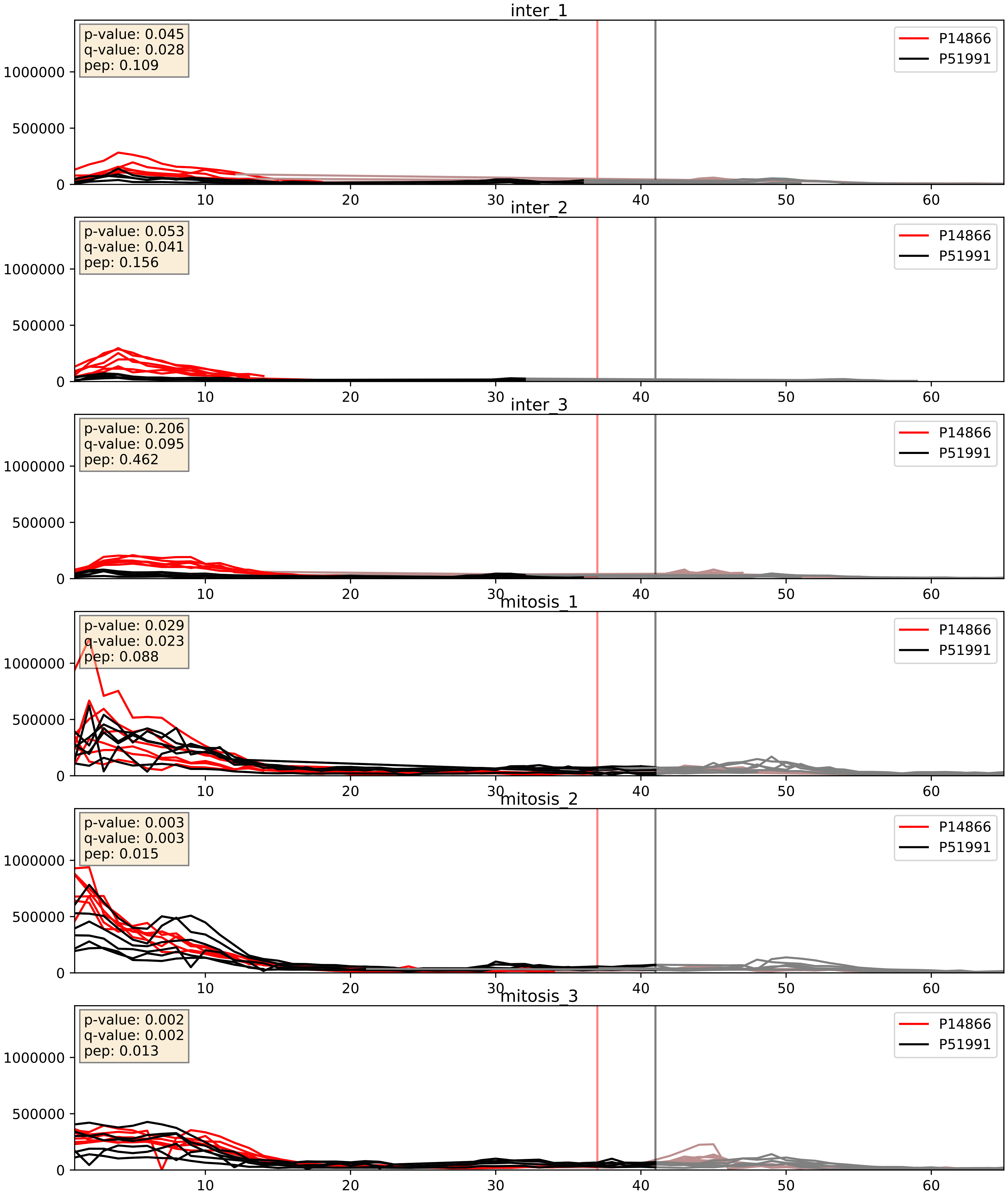


| level | condition_1 | condition_2 | pvalue | pvalue_adjusted |
|---|---|---|---|---|
| complex_abundance | mitosis | inter | 0.000306389187647456 | 0.01165207110963347 |
| interactor_ratio | mitosis | inter | 0.005592202135892223 | 0.10141790314245218 |
| interactor_abundance | mitosis | inter | 0.009697640385696541 | 0.08525122834595471 |

P14866_P52272
HNRPL_HUMAN vs HNRPM_HUMAN
p-value: 0.01 q-value: 0.009

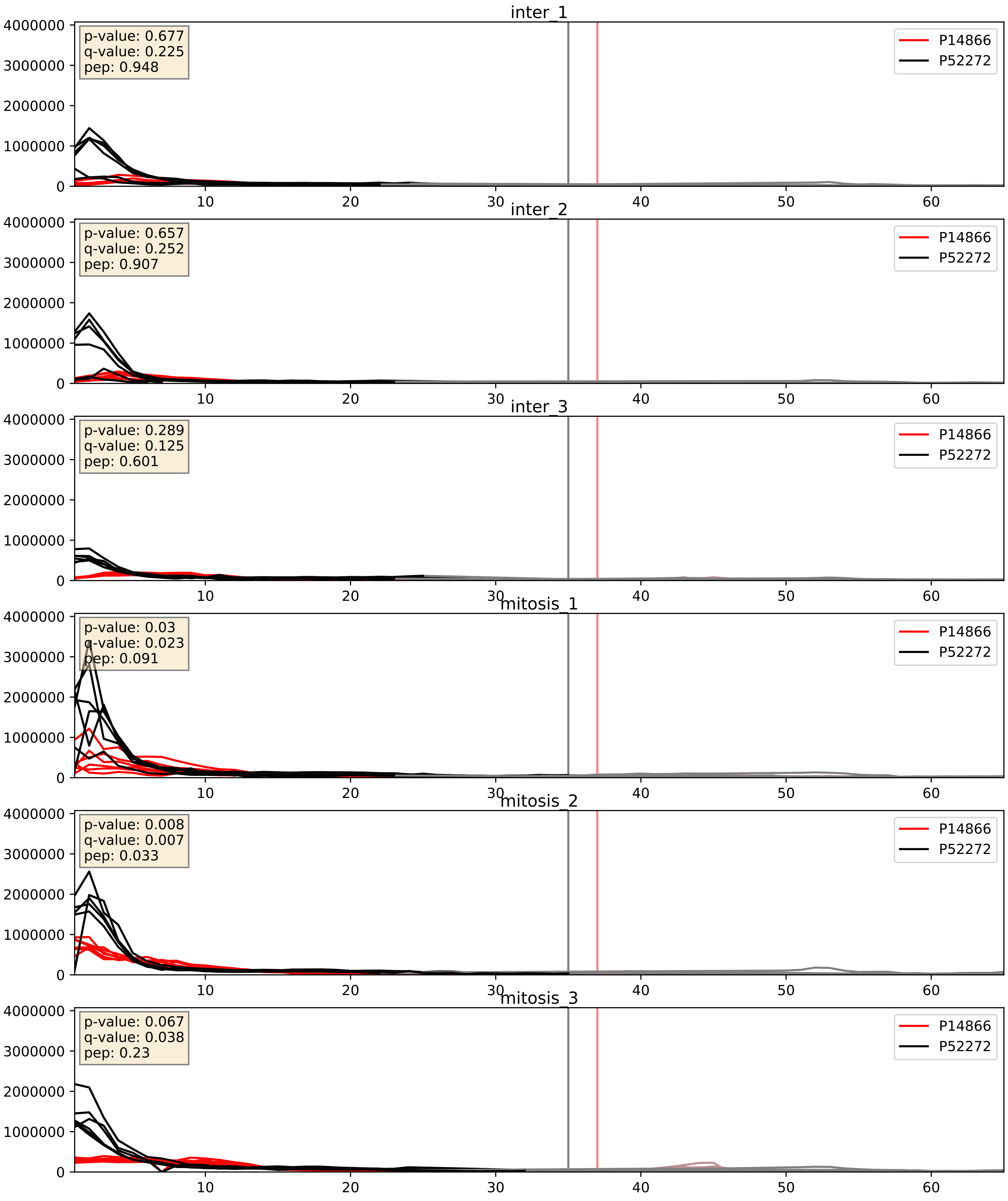


| level | condition_1 | condition_2 | pvalue | pvalue_adjusted |
|---|---|---|---|---|
| complex_abundance | mitosis | inter | 0.00290128716918596 | 0.03923383596877065 |
| interactor_abundance | mitosis | inter | 0.00986072689417613 | 0.08626246521629807 |
| interactor_ratio | mitosis | inter | 0.16723764333693136 | 0.6293444626080091 |

P14866_P52597
HNRPL_HUMAN vs HNRPF_HUMAN
p-value: 0.001 q-value: 0.002

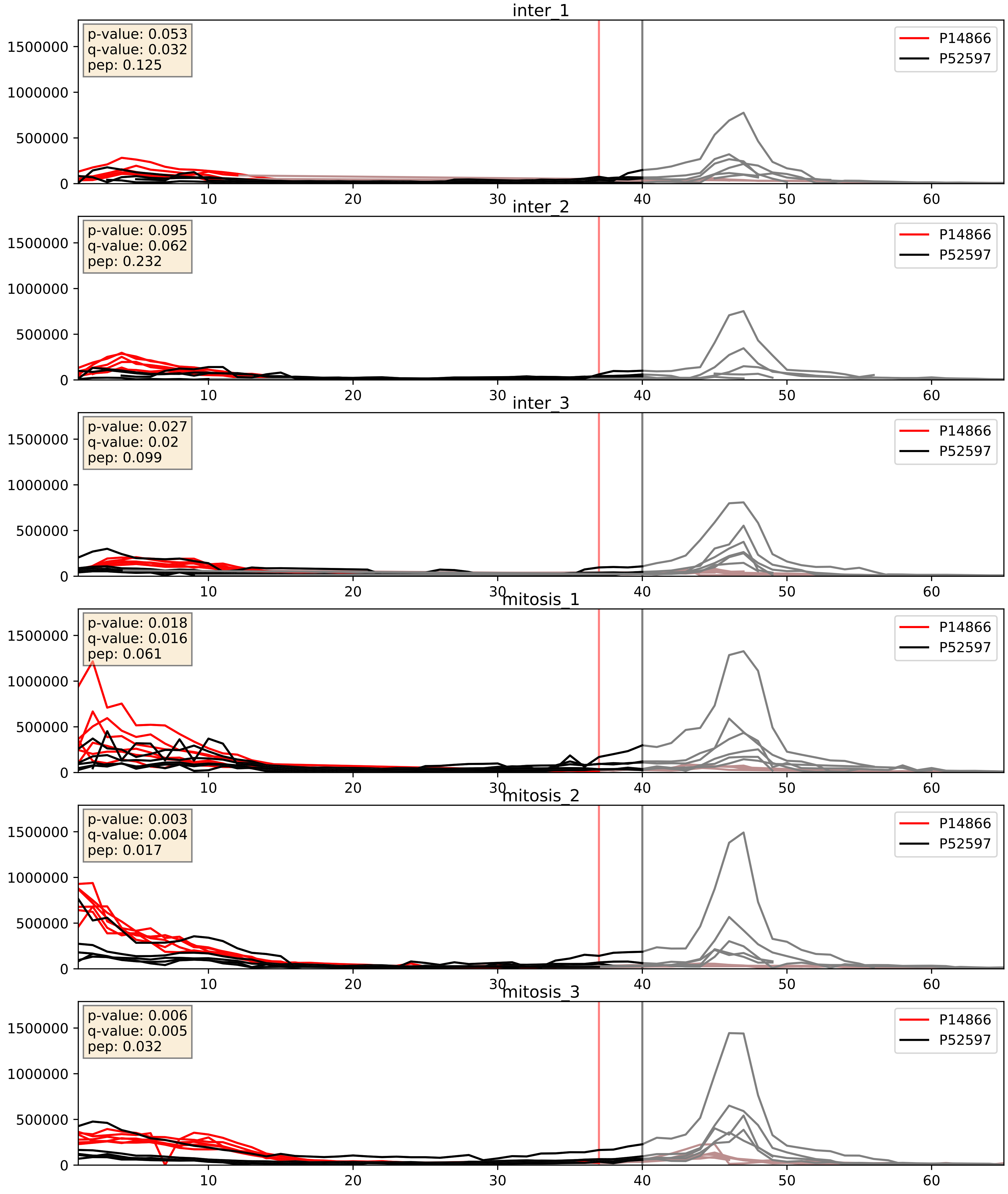


| level | condition_1 | condition_2 | pvalue | pvalue_adjusted |
|---|---|---|---|---|
| complex_abundance | mitosis | inter | 0.005276617137903093 | 0.054740033961712564 |
| interactor_abundance | mitosis | inter | 0.009697640385696541 | 0.08525122834595471 |
| interactor_ratio | mitosis | inter | 0.48618414815202177 | 0.8688384776996464 |

P14866_P62306
HNRPL_HUMAN vs RUXF_HUMAN
p-value: 0.069 q-value: 0.042

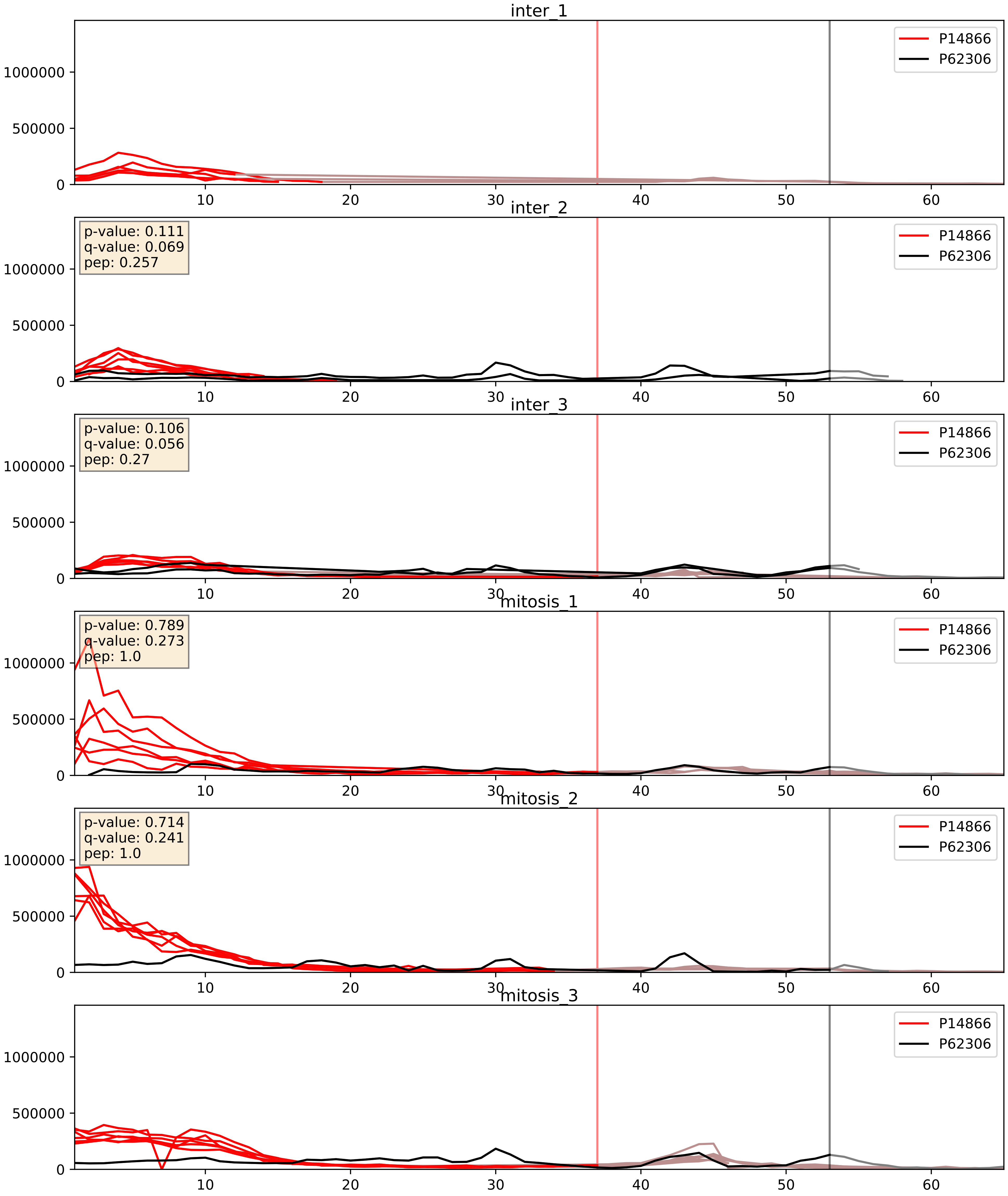


| level | condition_1 | condition_2 | pvalue | pvalue_adjusted |
| --- | --- | --- | --- | --- |
| interactor_ratio | mitosis | inter | 0.01297337334516381 | 0.16452159382904033 |
| interactor_abundance | mitosis | inter | 0.10508626811479461 | 0.2769210432287962 |
| complex_abundance | mitosis | inter | 0.2692733160070283 | 0.41878262809232597 |

P14866_P62316
HNRPL_HUMAN vs SMD2_HUMAN
p-value: 0.073 q-value: 0.044

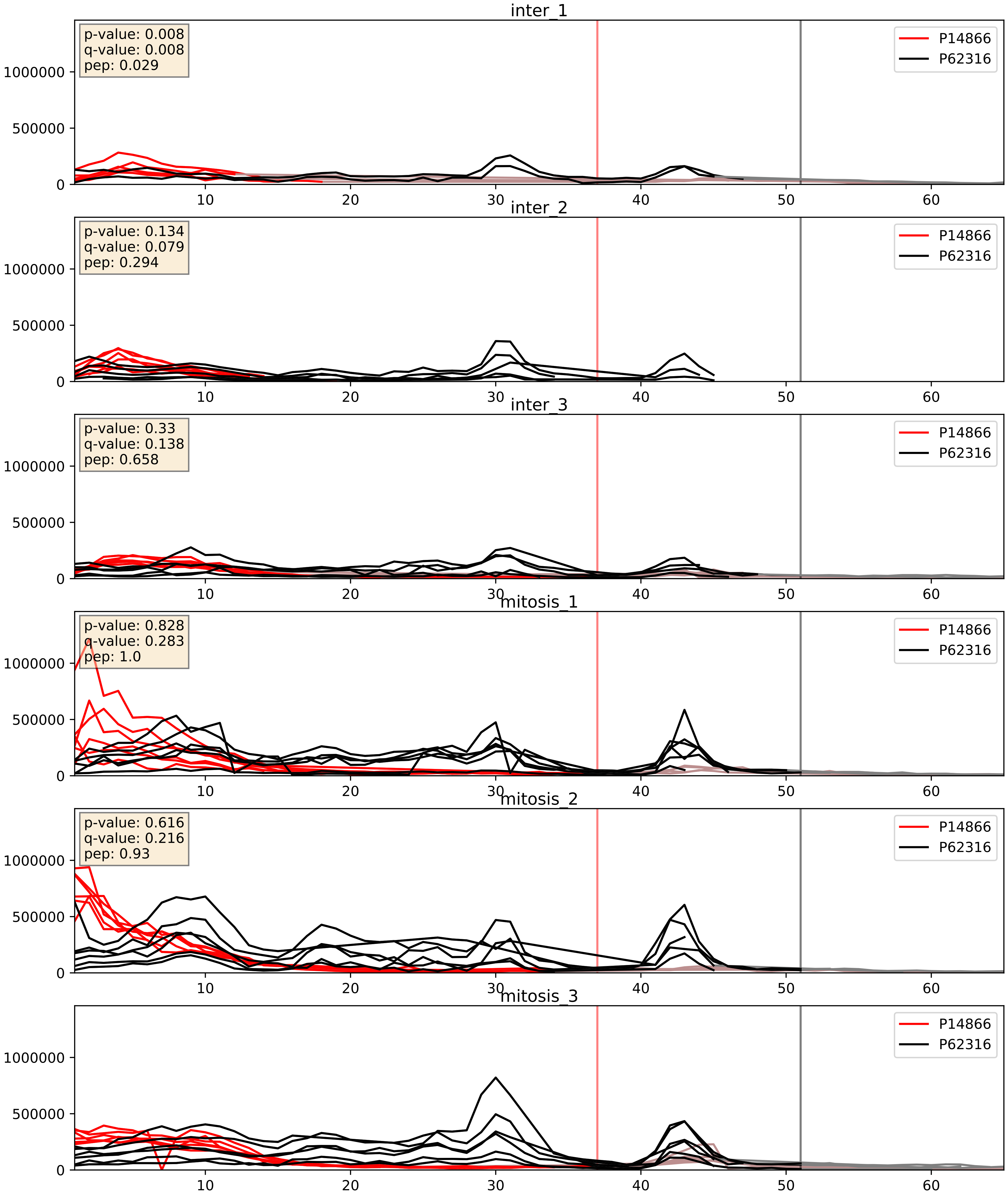


| level | condition_1 | condition_2 | pvalue | pvalue_adjusted |
|---|---|---|---|---|
| complex_abundance | mitosis | inter | 0.000861046445581631 | 0.019513211095311817 |
| interactor_abundance | mitosis | inter | 0.009697640385696541 | 0.08525122834595471 |
| interactor_ratio | mitosis | inter | 0.009765737212506018 | 0.14216787506641412 |

P14866_P62995
HNRPL_HUMAN vs TRA2B_HUMAN
p-value: 0.007 q-value: 0.007

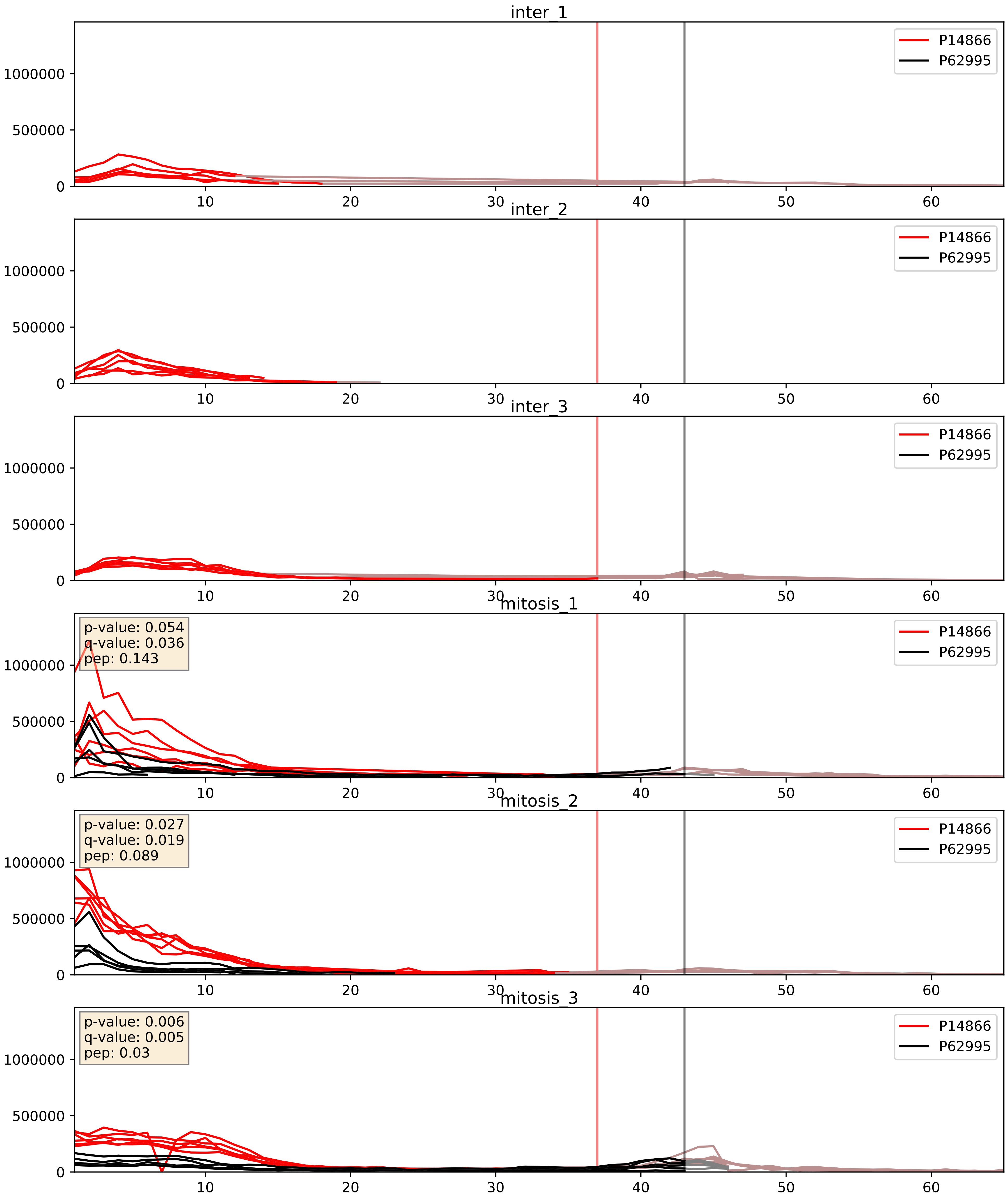


| level | condition_1 | condition_2 | pvalue | pvalue_adjusted |
|---|---|---|---|---|
| interactor_abundance | mitosis | inter | 1.2674740761041976e-05 | 0.003391165020149013 |
| complex_abundance | mitosis | inter | 5.6324767565087076e-05 | 0.005691857151652958 |
| interactor_ratio | mitosis | inter | 0.000967869032068439 | 0.03586562300651878 |

P14866_P84103
HNRPL_HUMAN vs SRSF3_HUMAN
p-value: 0.039 q-value: 0.027

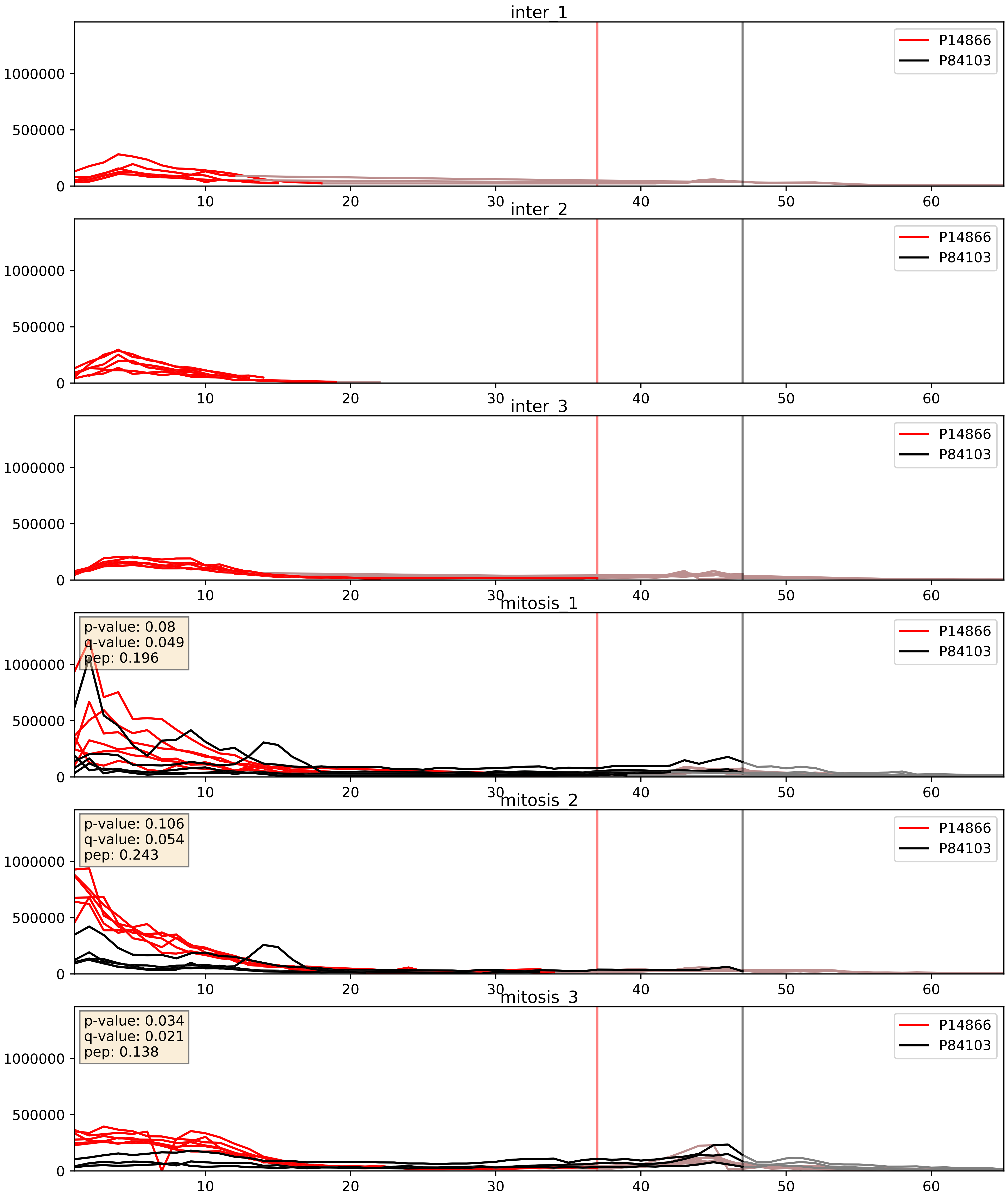


| level | condition_1 | condition_2 | pvalue | pvalue_adjusted |
|---|---|---|---|---|
| interactor_abundance | mitosis | inter | 1.2564597628871335e-05 | 0.003391165020149013 |
| complex_abundance | mitosis | inter | 0.00025225809034102616 | 0.010637089917828493 |
| interactor_ratio | mitosis | inter | 0.02957345168884291 | 0.2722029531790272 |

P14866_Q07955
HNRPL_HUMAN vs SRSF1_HUMAN
p-value: 0.005 q-value: 0.005

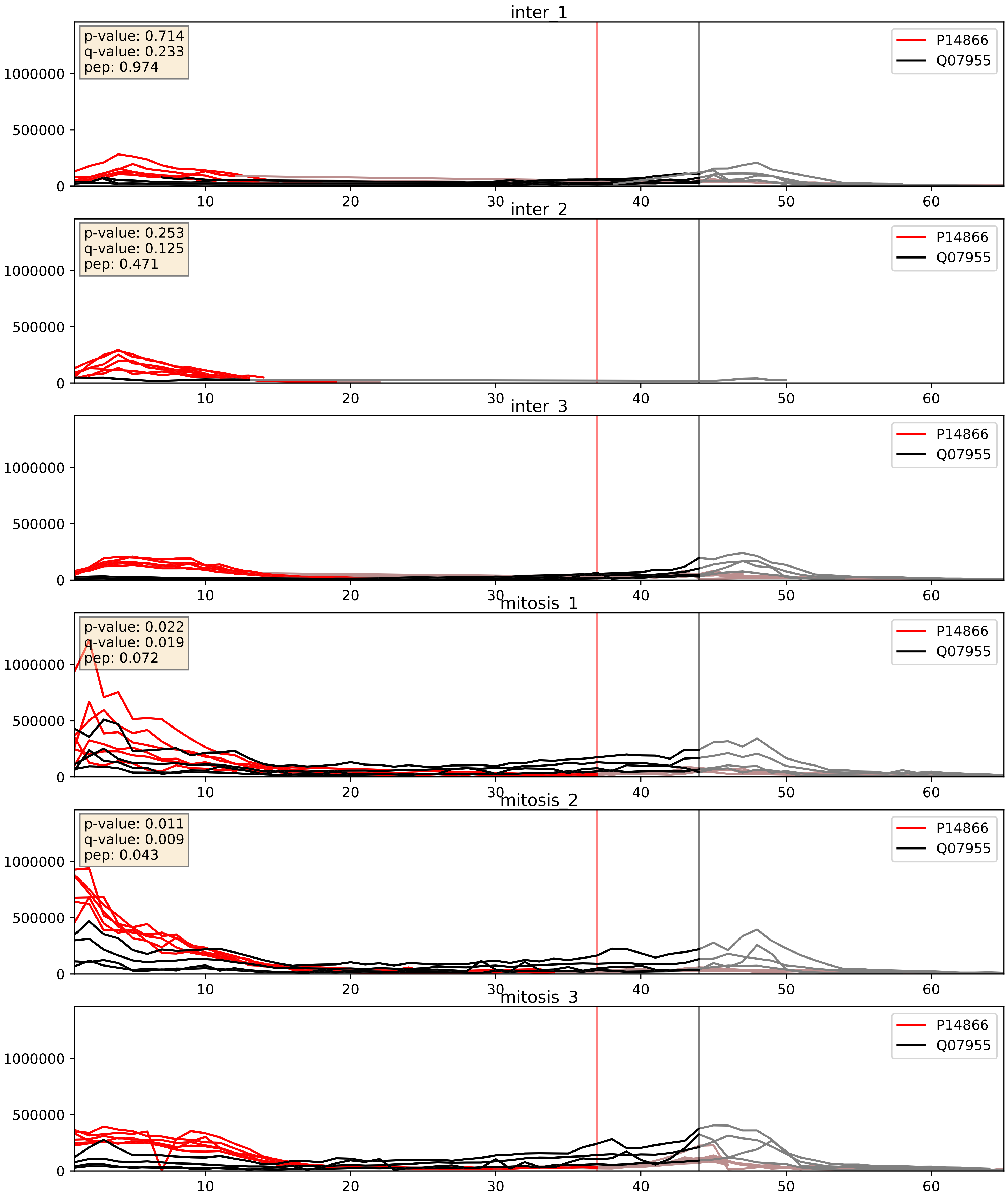


| level | condition_1 | condition_2 | pvalue | pvalue_adjusted |
|---|---|---|---|---|
| complex_abundance | mitosis | inter | 4.997121611729198e-05 | 0.005691857151652958 |
| interactor_abundance | mitosis | inter | 0.0019001893143151473 | 0.03196215383917848 |
| interactor_ratio | mitosis | inter | 0.15454422094563686 | 0.61160357433872 |

P14866_Q08211
HNRPL_HUMAN vs DHX9_HUMAN
p-value: 0.019 q-value: 0.015

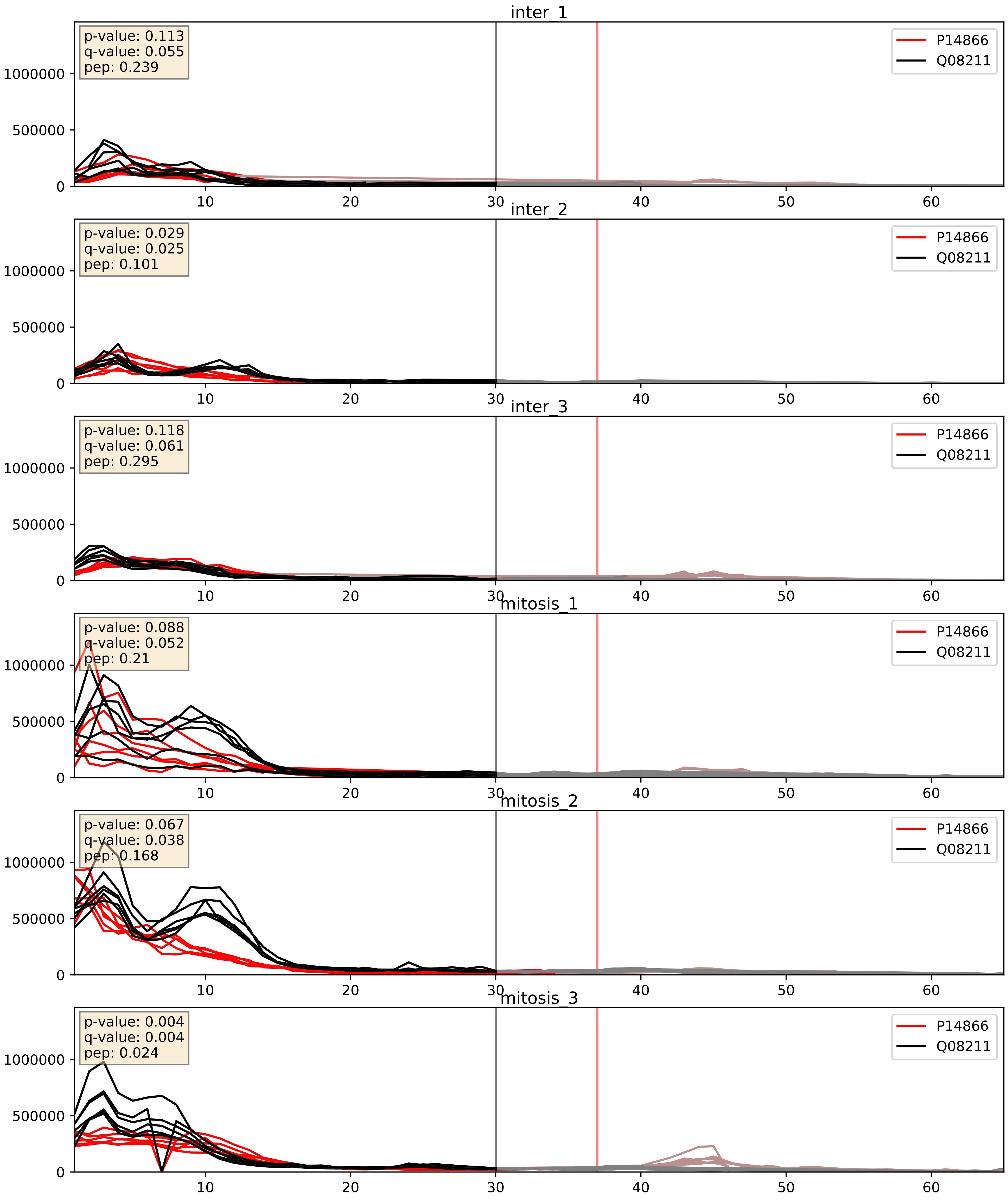


| level | condition_1 | condition_2 | pvalue | pvalue_adjusted |
|---|---|---|---|---|
| complex_abundance | mitosis | inter | 0.00043565000436869284 | 0.013137121901174406 |
| interactor_abundance | mitosis | inter | 0.010470812401431332 | 0.08918423299129571 |
| interactor_ratio | mitosis | inter | 0.9543473012403387 | 0.992614664476724 |

P14866_Q12906
HNRPL_HUMAN vs ILF3_HUMAN
p-value: 0.0 q-value: 0.001

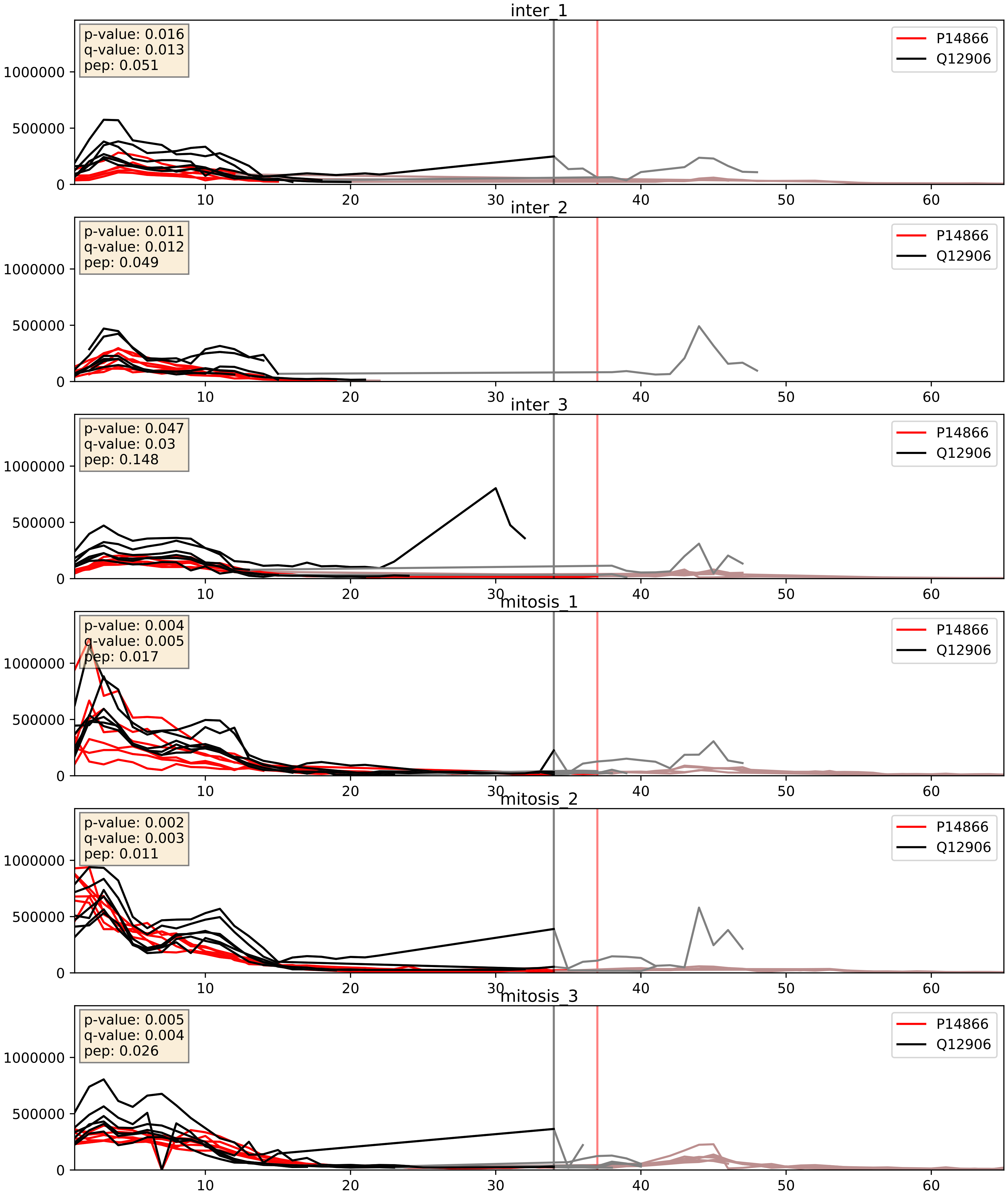


| level | condition_1 | condition_2 | pvalue | pvalue_adjusted |
|---|---|---|---|---|
| complex_abundance | mitosis | inter | 0.0028666487219307637 | 0.03901194445107684 |
| interactor_abundance | mitosis | inter | 0.010616908323401677 | 0.0901594595717444 |
| interactor_ratio | mitosis | inter | 0.08297559701664442 | 0.4838359063095887 |

P14866_Q13151
HNRPL_HUMAN vs ROA0_HUMAN
p-value: 0.014 q-value: 0.012

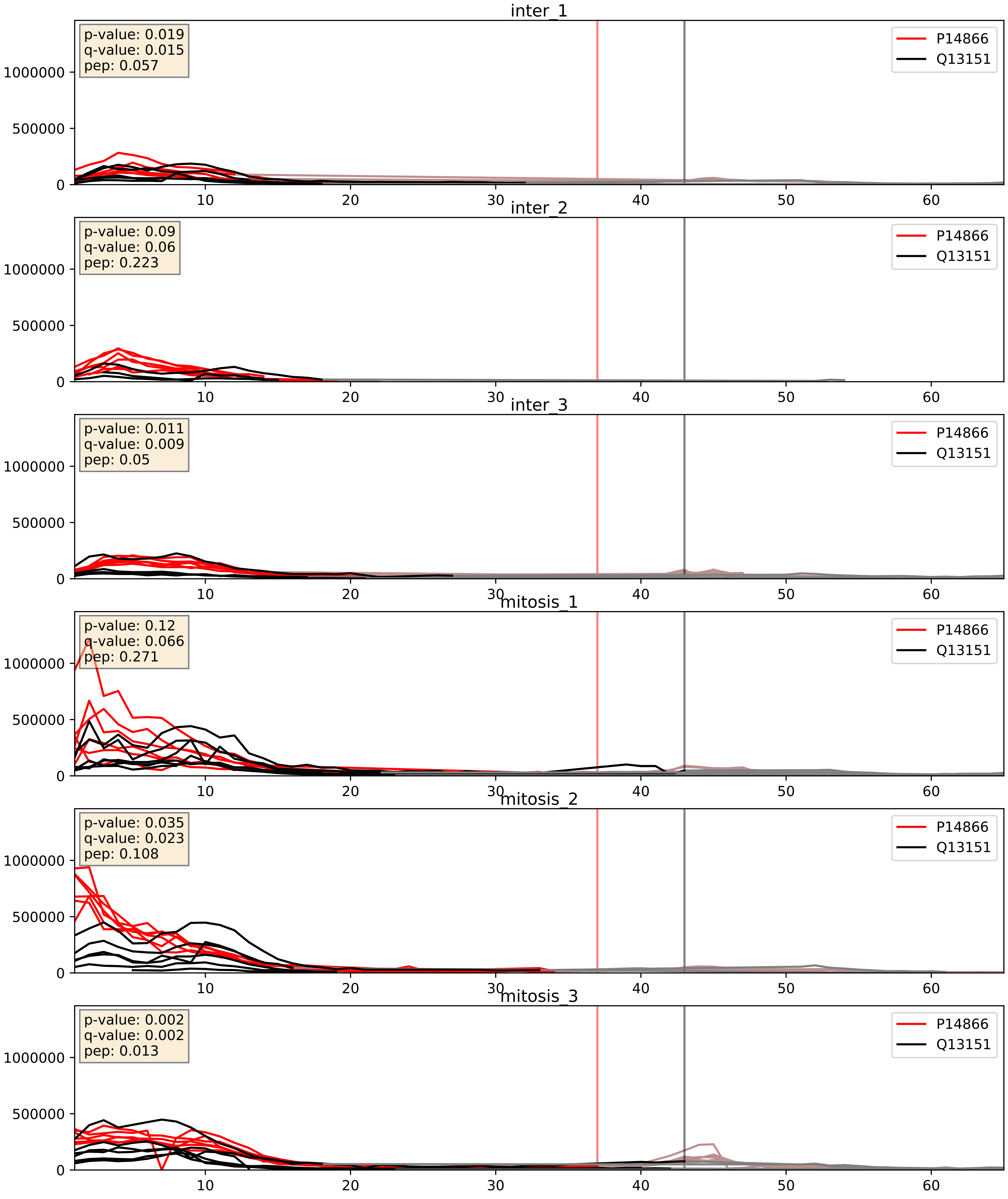


| level | condition_1 | condition_2 | pvalue | pvalue_adjusted |
|---|---|---|---|---|
| complex_abundance | mitosis | inter | 2.5979691148047346e-05 | 0.005691857151652958 |
| interactor_abundance | mitosis | inter | 0.00984142502181702 | 0.08622579138869364 |
| interactor_ratio | mitosis | inter | 0.8500340674765395 | 0.9755634896635457 |

P14866_Q15365
HNRPL_HUMAN vs PCBP1_HUMAN
p-value: 0.037 q-value: 0.026

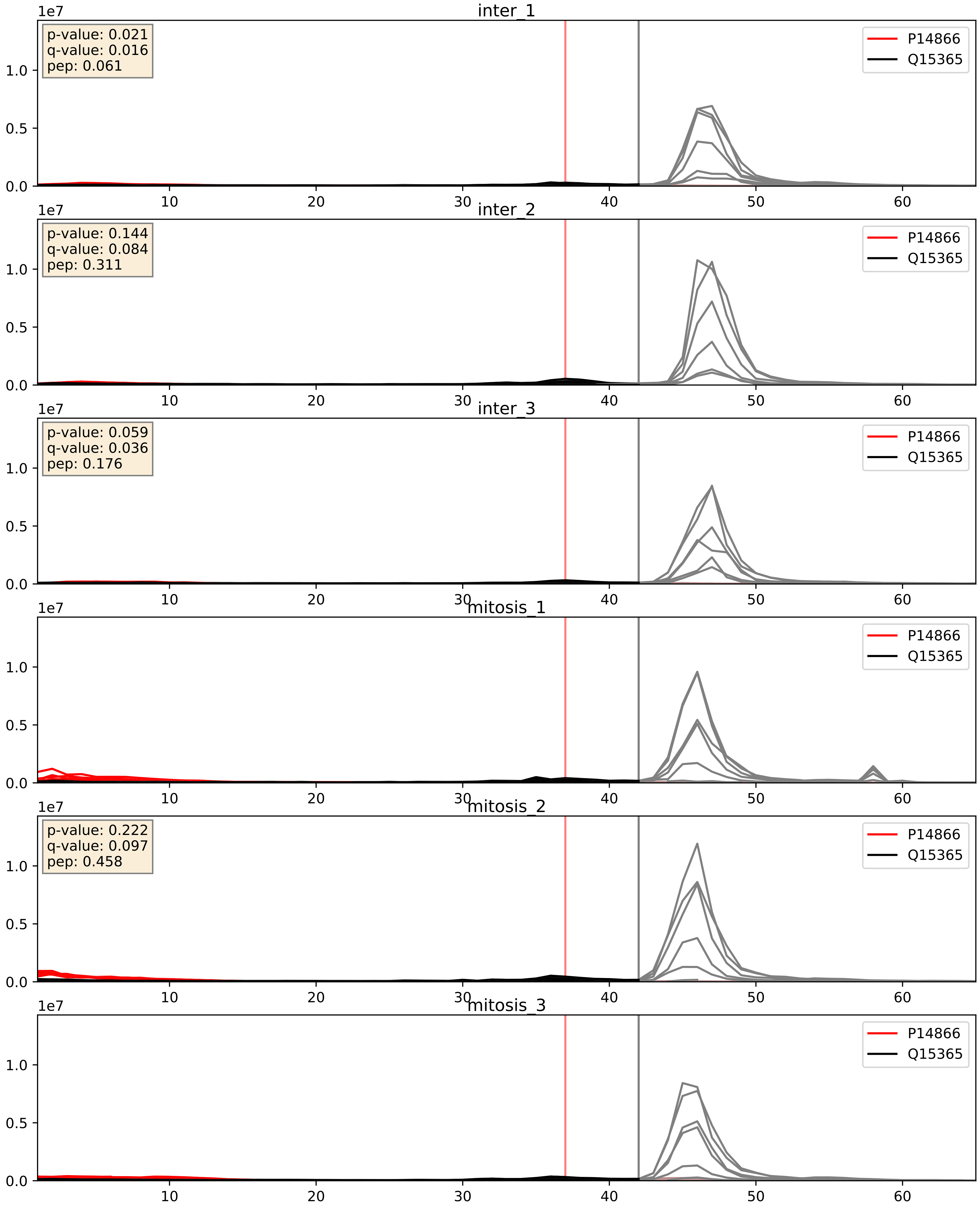


| level | condition_1 | condition_2 | pvalue | pvalue_adjusted |
|---|---|---|---|---|
| complex_abundance | mitosis | inter | 0.006117488702226094 | 0.05820202463873419 |
| interactor_abundance | mitosis | inter | 0.009697640385696541 | 0.08525122834595471 |
| interactor_ratio | mitosis | inter | 0.04505144601463991 | 0.35282742715948545 |

P14866_Q15717
HNRPL_HUMAN vs ELAV1_HUMAN
p-value: 0.007 q-value: 0.007

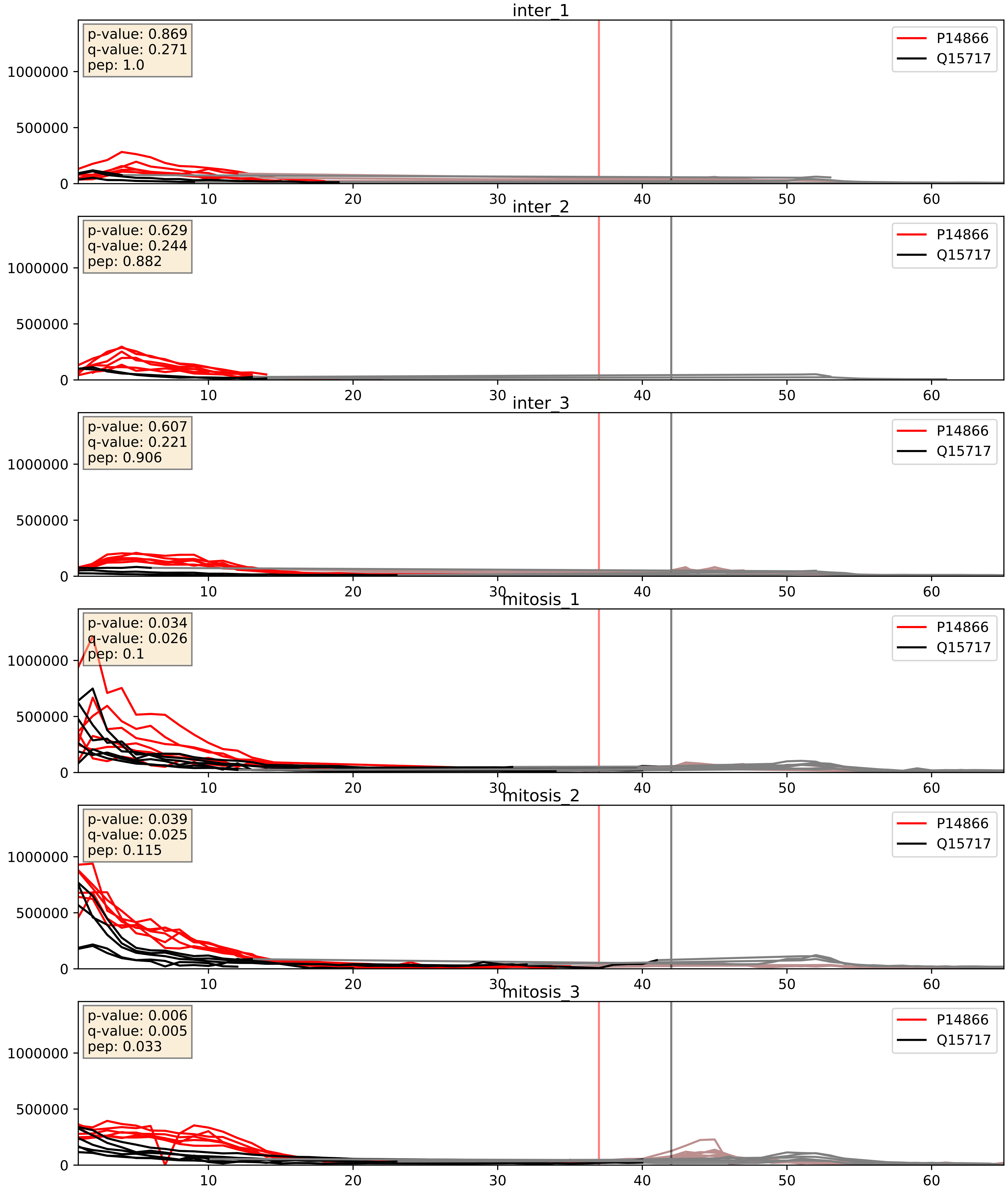


| level | condition_1 | condition_2 | pvalue | pvalue_adjusted |
|---|---|---|---|---|
| complex_abundance | mitosis | inter | 0.00015412022185430893 | 0.008563834848633118 |
| interactor_abundance | mitosis | inter | 0.009449586988451214 | 0.08425881731368999 |
| interactor_ratio | mitosis | inter | 0.04986156517554219 | 0.36889801028750313 |

P14866_Q16629
HNRPL_HUMAN vs SRSF7_HUMAN
p-value: 0.003 q-value: 0.003

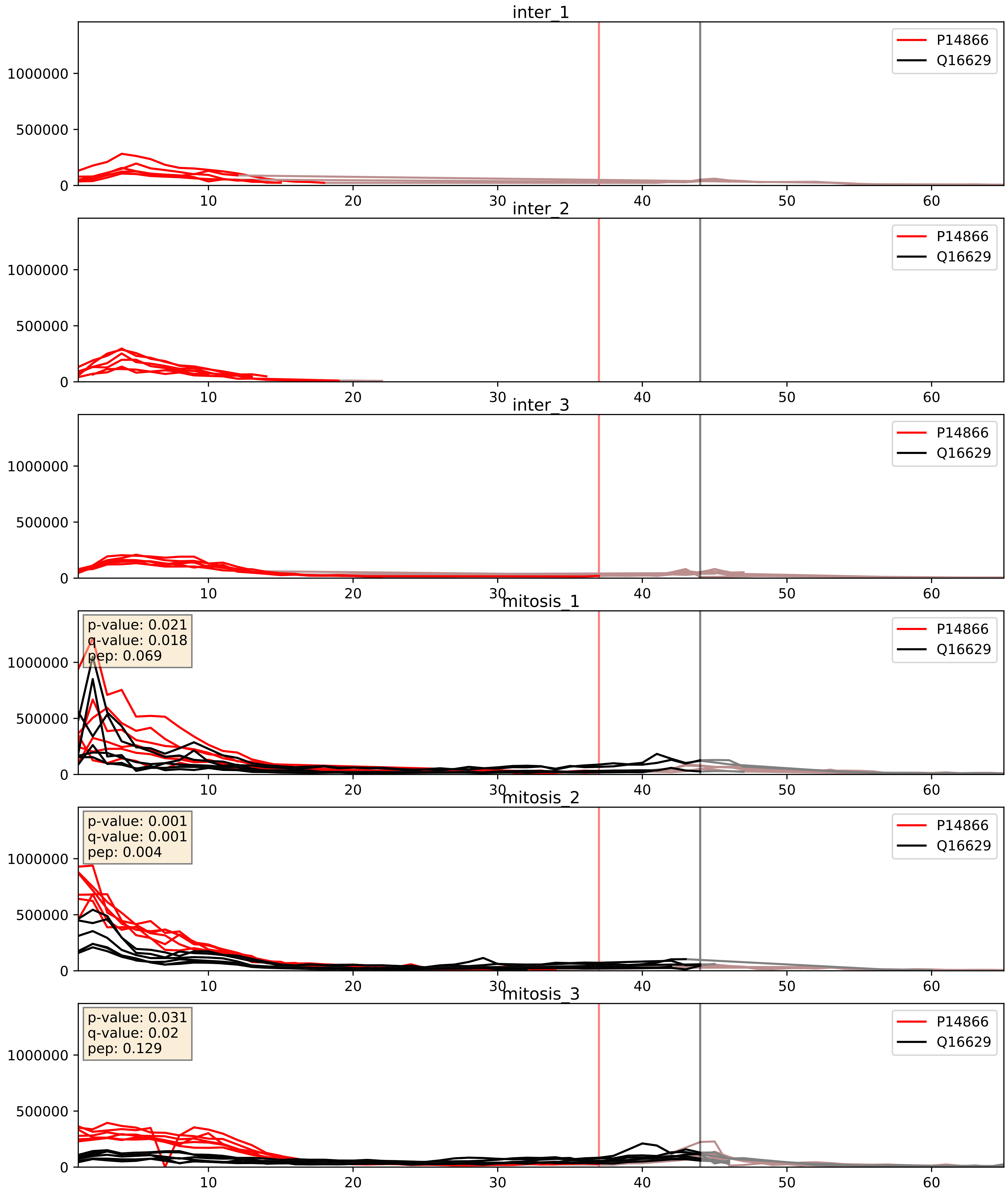


| level | condition_1 | condition_2 | pvalue | pvalue_adjusted |
|---|---|---|---|---|
| interactor_abundance | mitosis | inter | 1.2331621368333903e-05 | 0.003391165020149013 |
| complex_abundance | mitosis | inter | 3.0454949653656297e-05 | 0.005691857151652958 |
| interactor_ratio | mitosis | inter | 0.04207614494229382 | 0.3399226239910554 |

P14866_Q9UMS4
HNRPL_HUMAN vs PRP19_HUMAN
p-value: 0.006 q-value: 0.006

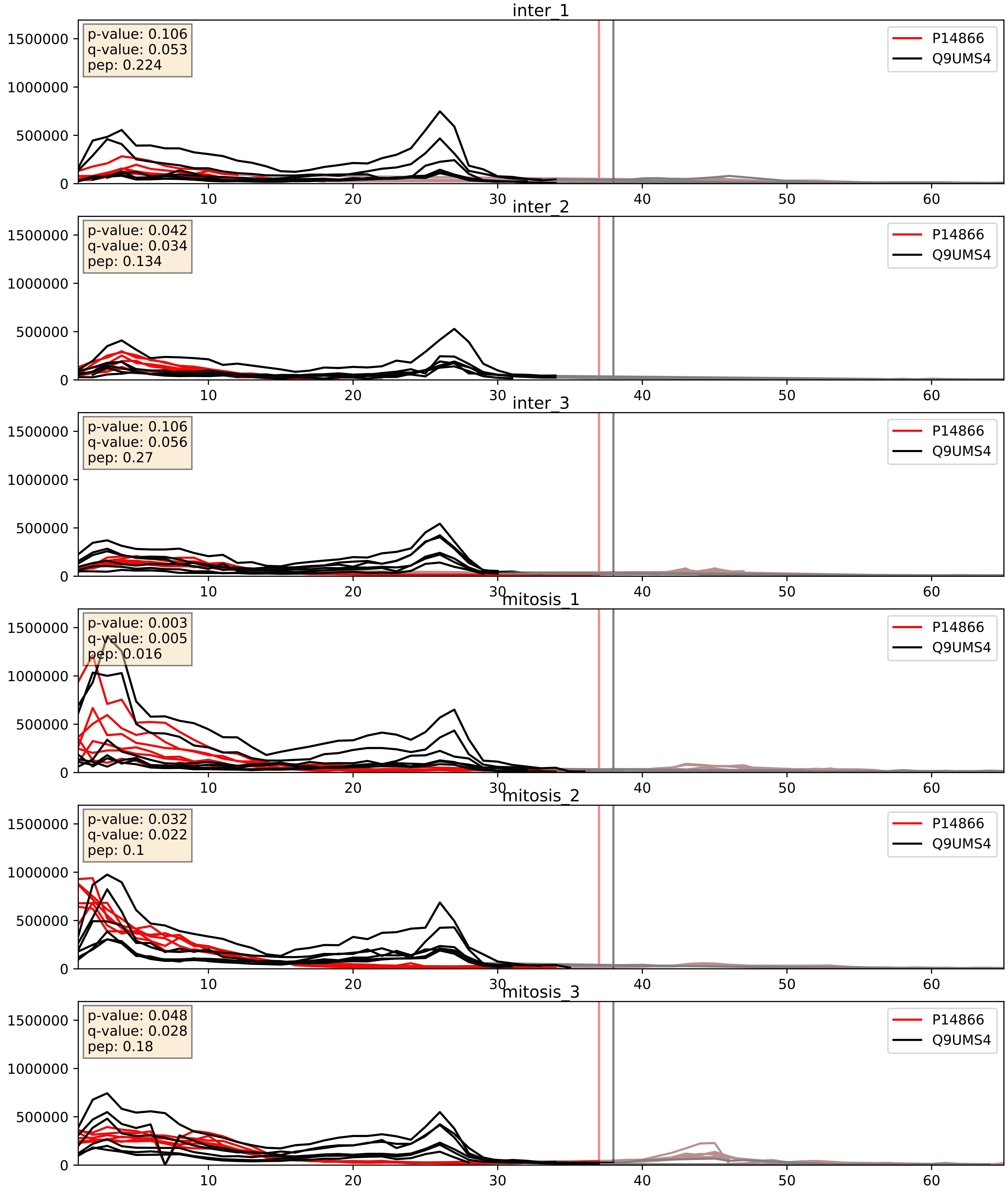


| level | condition_1 | condition_2 | pvalue | pvalue_adjusted |
|---|---|---|---|---|
| complex_abundance | mitosis | inter | 0.0034585900486918215 | 0.04302508960916016 |
| interactor_abundance | mitosis | inter | 0.009606390243086767 | 0.08507272274138435 |
| interactor_ratio | mitosis | inter | 0.7612085472999965 | 0.9536594850393604 |
