## Supplementary Data 2 for "SECAT: Quantifying differential protein-protein interaction states by network-centric analysis": hela_string_000047_P22626.pdf

# P22626_P22626

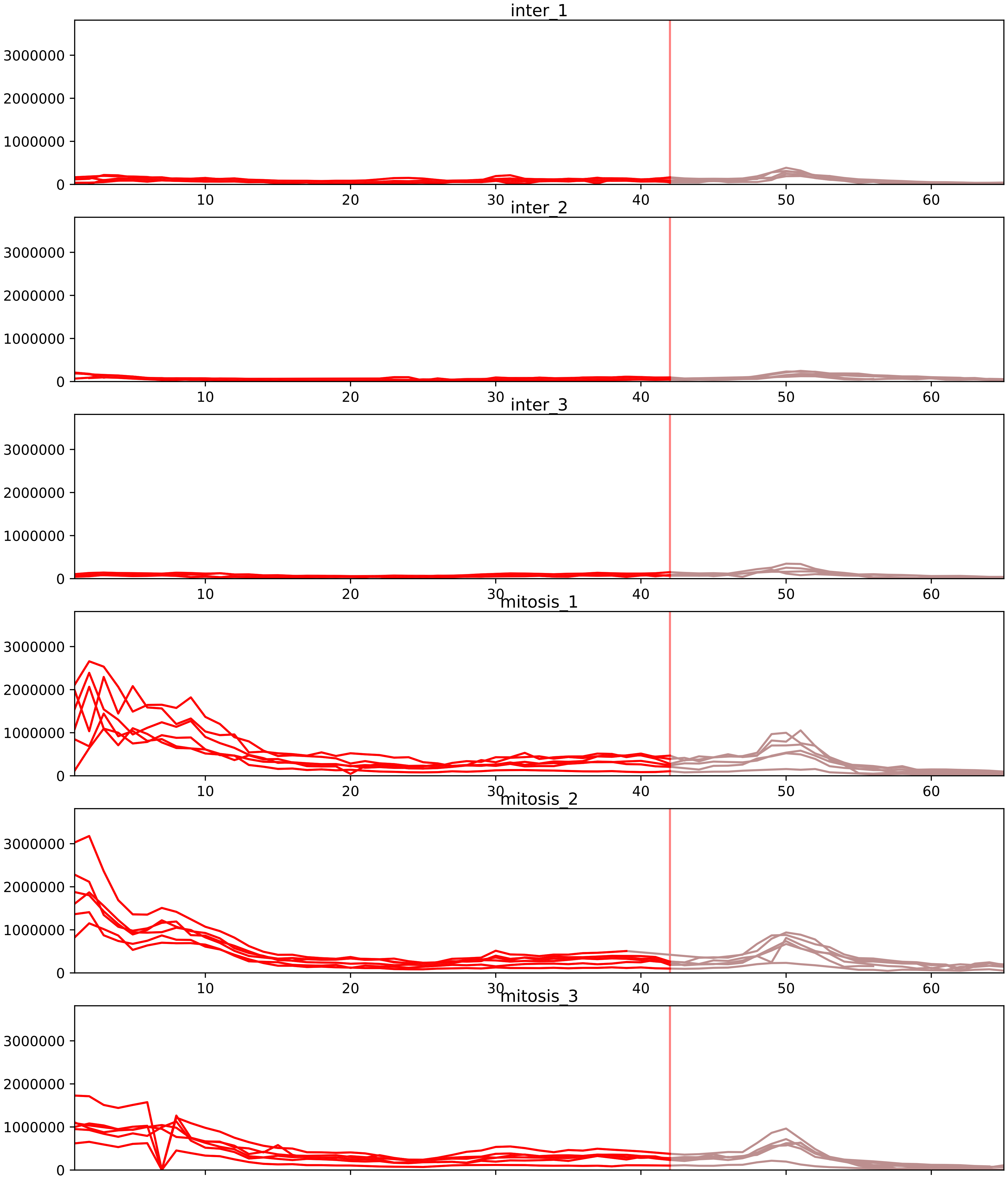


| level | condition_1 | condition_2 | pvalue | pvalue_adjusted |
|---|---|---|---|---|
| interactor_ratio | mitosis | inter | 1.0010748619776324e-07 | 4.18631305917919e-06 |
| complex_abundance | mitosis | inter | 8.091877387277929e-06 | 0.00041358484442386497 |
| interactor_abundance | mitosis | inter | 1.4399495152893553e-05 | 0.0006689893096955349 |
| assembled_abundance | mitosis | inter | 0.007443493909955384 | 0.09331793469809328 |
| total_abundance | mitosis | inter | 0.012632435096449722 | 0.14134445511214405 |
| monomer_abundance | mitosis | inter | 0.03804224057878451 | 0.3150273663293861 |

O43390_P22626
HNRPR_HUMAN vs ROA2_HUMAN
p-value: 0.007 q-value: 0.007

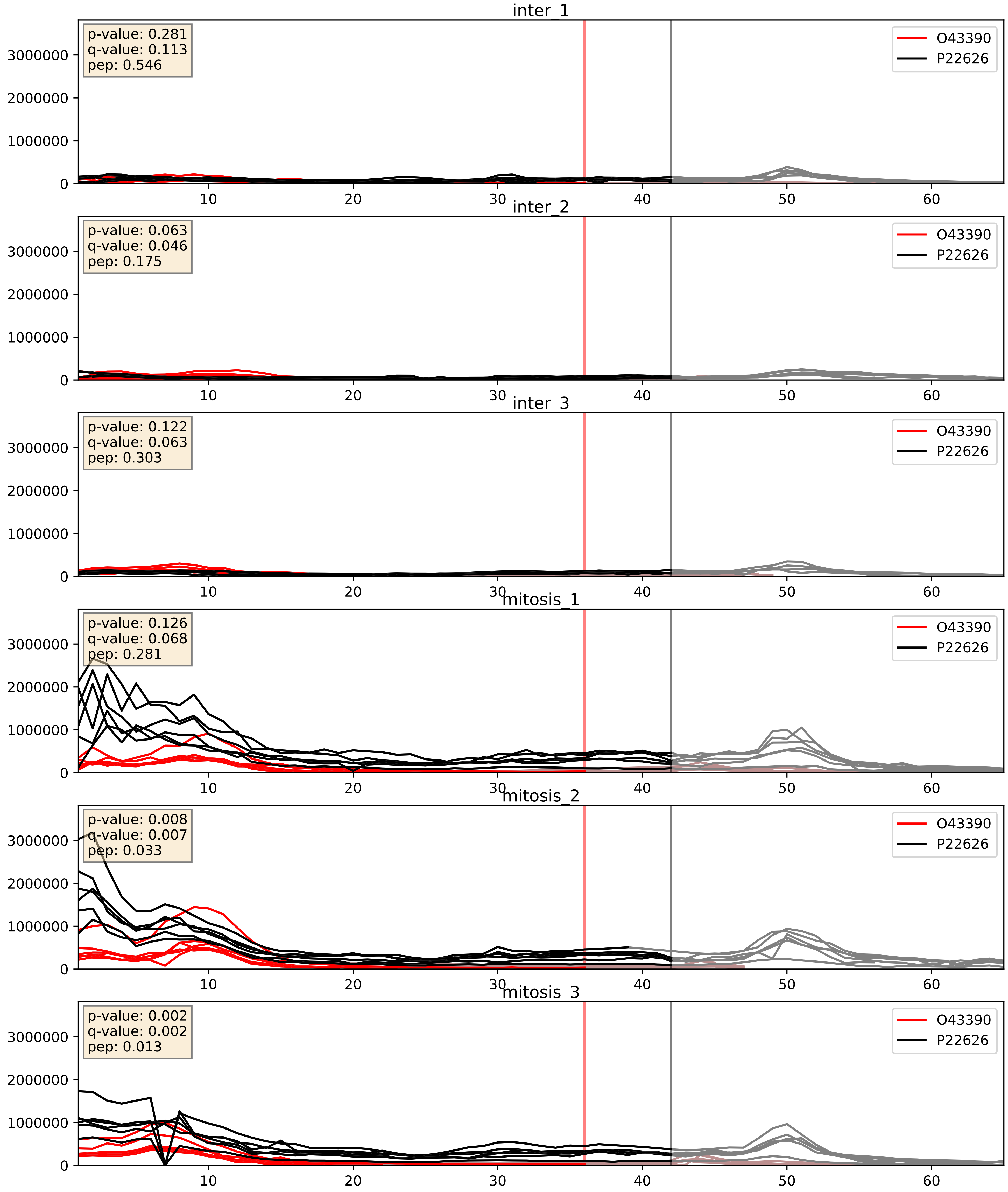


| level | condition_1 | condition_2 | pvalue | pvalue_adjusted |
|---|---|---|---|---|
| interactor_abundance | mitosis | inter | 0.00010796061997593195 | 0.007876481428251637 |
| complex_abundance | mitosis | inter | 0.0009205930867441553 | 0.019950067905139163 |
| interactor_ratio | mitosis | inter | 0.016834477256909862 | 0.19213750042553124 |

P07910_P22626
HNRPC_HUMAN vs ROA2_HUMAN
p-value: 0.0 q-value: 0.0

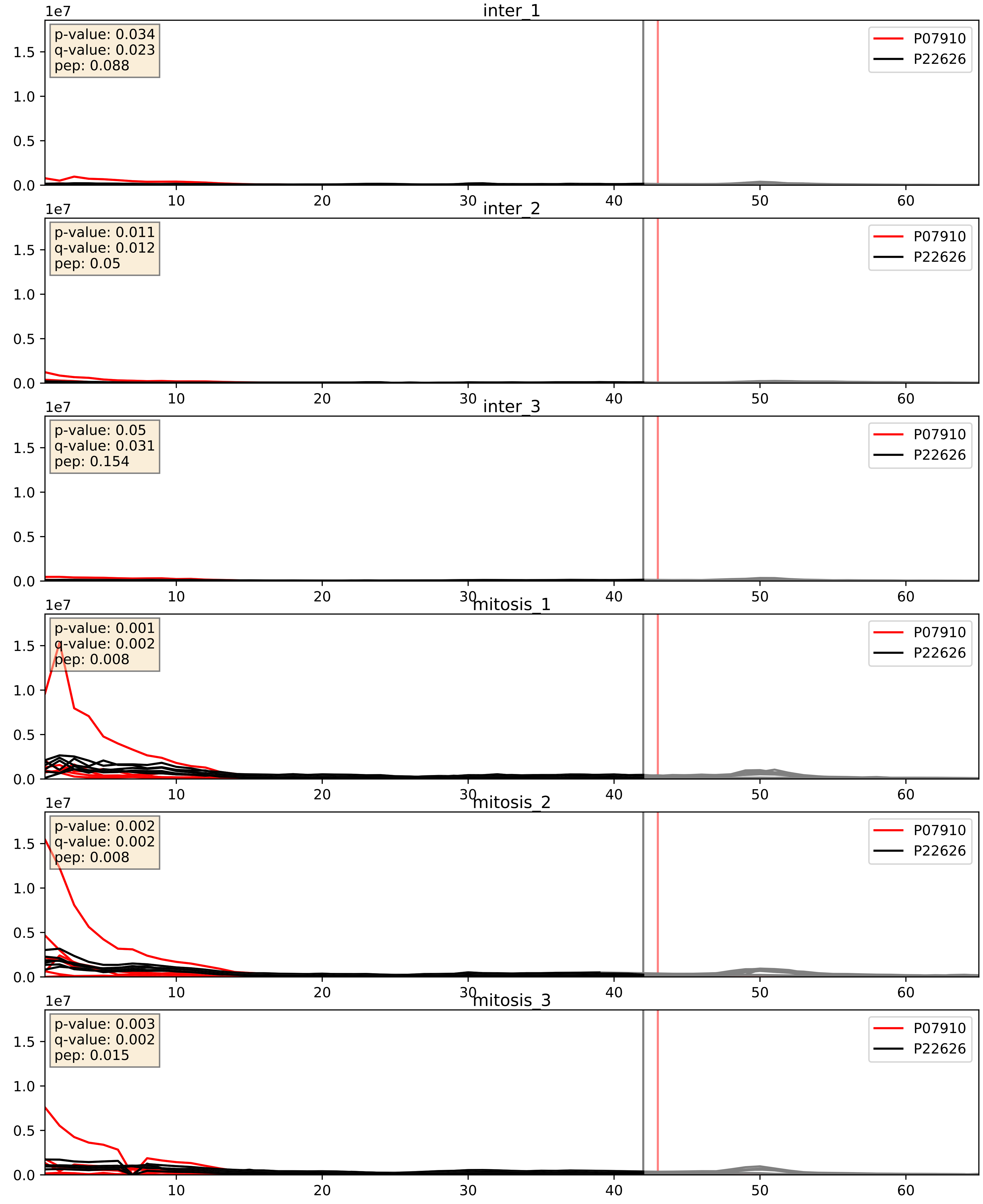


| level | condition_1 | condition_2 | pvalue | pvalue_adjusted |
|---|---|---|---|---|
| interactor_abundance | mitosis | inter | 9.148564965046394e-06 | 0.0033734327106189023 |
| complex_abundance | mitosis | inter | 0.0001404488488971927 | 0.00812325774702682 |
| interactor_ratio | mitosis | inter | 0.417882469042839 | 0.8387043223931305 |

P09651_P22626
ROA1_HUMAN vs ROA2_HUMAN
p-value: 0.0 q-value: 0.0

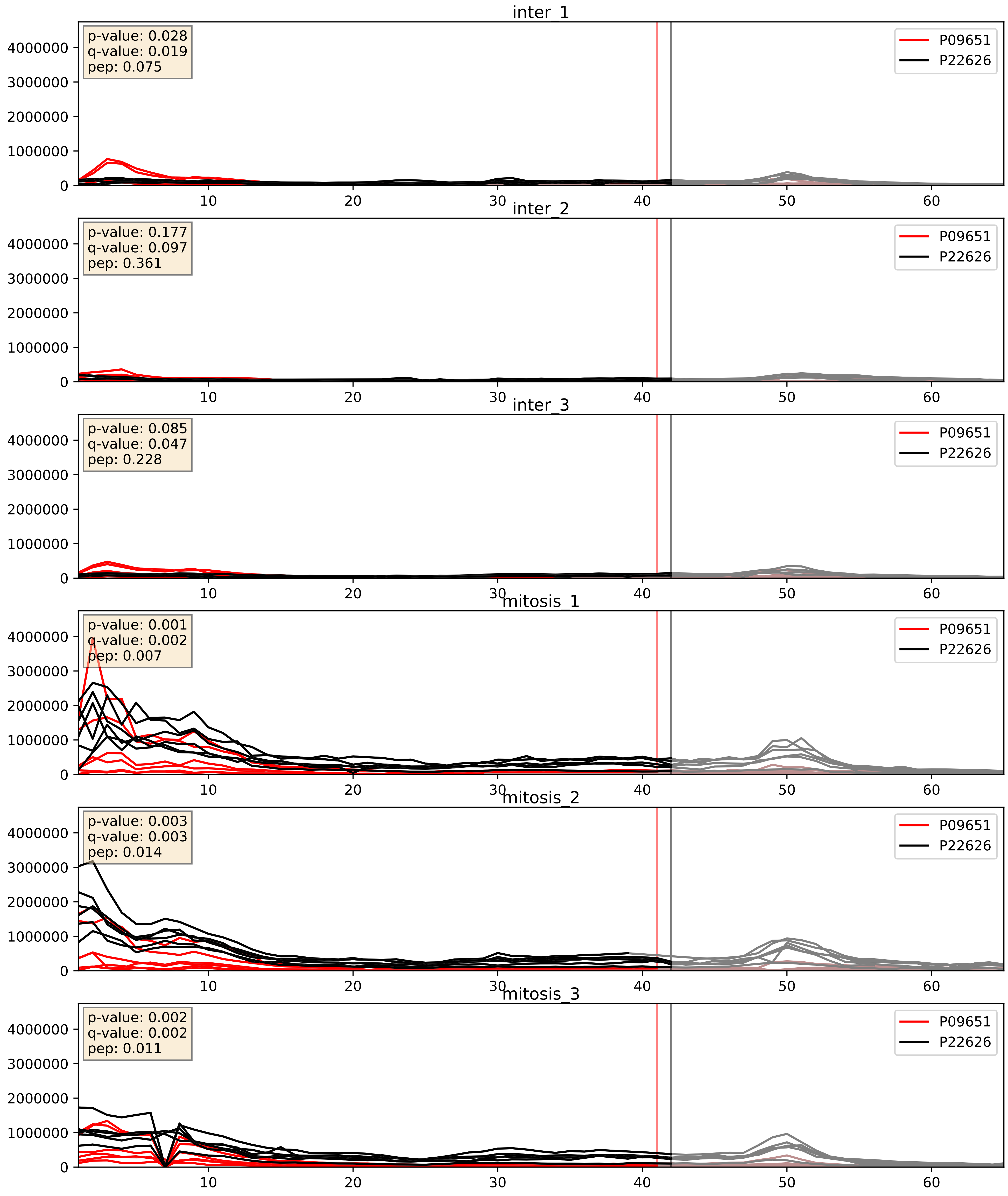


| level | condition_1 | condition_2 | pvalue | pvalue_adjusted |
|---|---|---|---|---|
| complex_abundance | mitosis | inter | 0.0022885483088337477 | 0.03502731659907978 |
| interactor_abundance | mitosis | inter | 0.016856414137678265 | 0.11277132084292767 |
| interactor_ratio | mitosis | inter | 0.066322140233217 | 0.4296400448702392 |

P14866_P22626
HNRPL_HUMAN vs ROA2_HUMAN
p-value: 0.012 q-value: 0.01

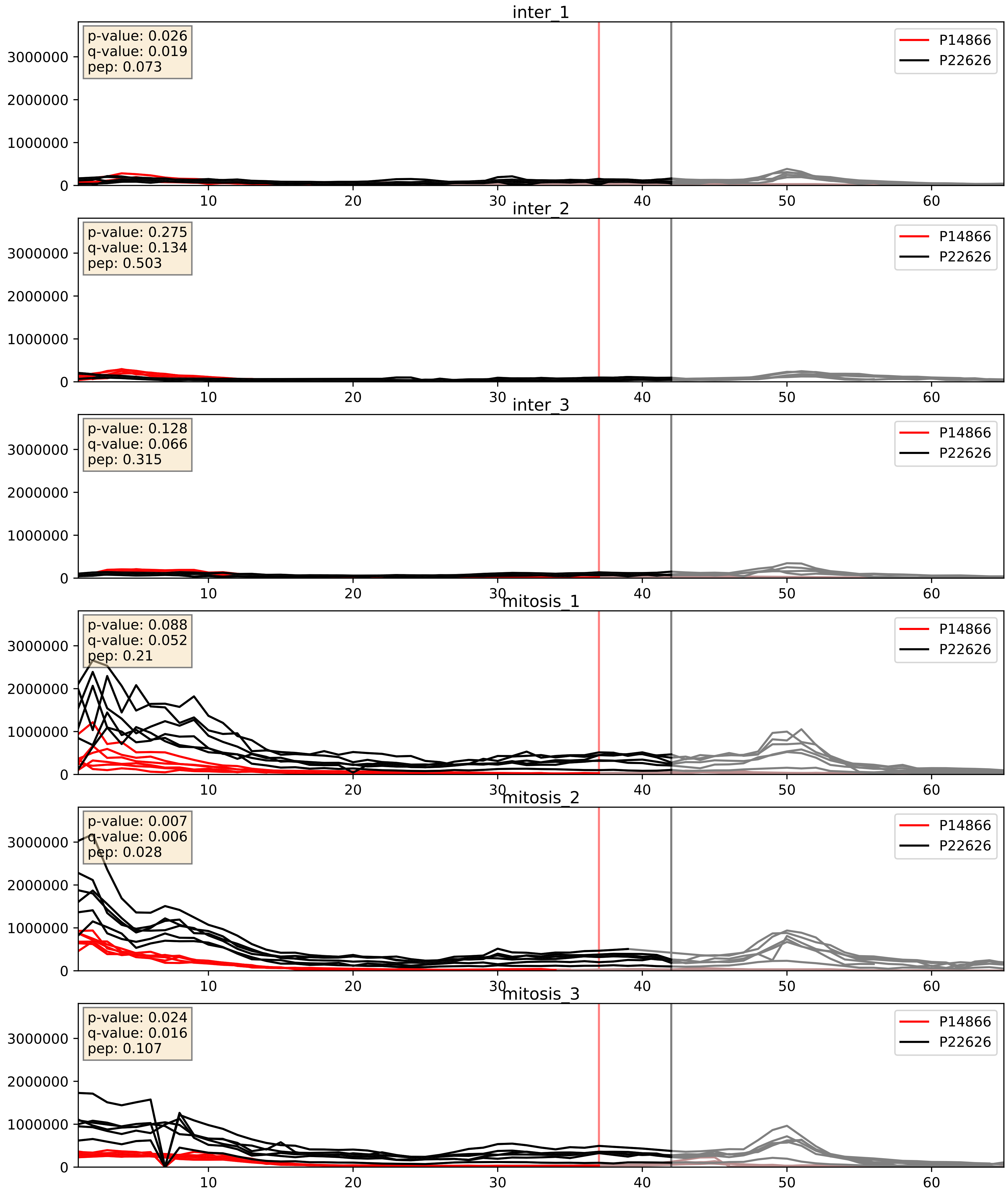


| level | condition_1 | condition_2 | pvalue | pvalue_adjusted |
|---|---|---|---|---|
| complex_abundance | mitosis | inter | 9.663556889611487e-05 | 0.00670982287146329 |
| interactor_abundance | mitosis | inter | 0.009697640385696541 | 0.08525122834595471 |
| interactor_ratio | mitosis | inter | 0.0163886342509947 | 0.1887653035437297 |

P15880_P22626
RS2_HUMAN vs ROA2_HUMAN
p-value: 0.0 q-value: 0.041

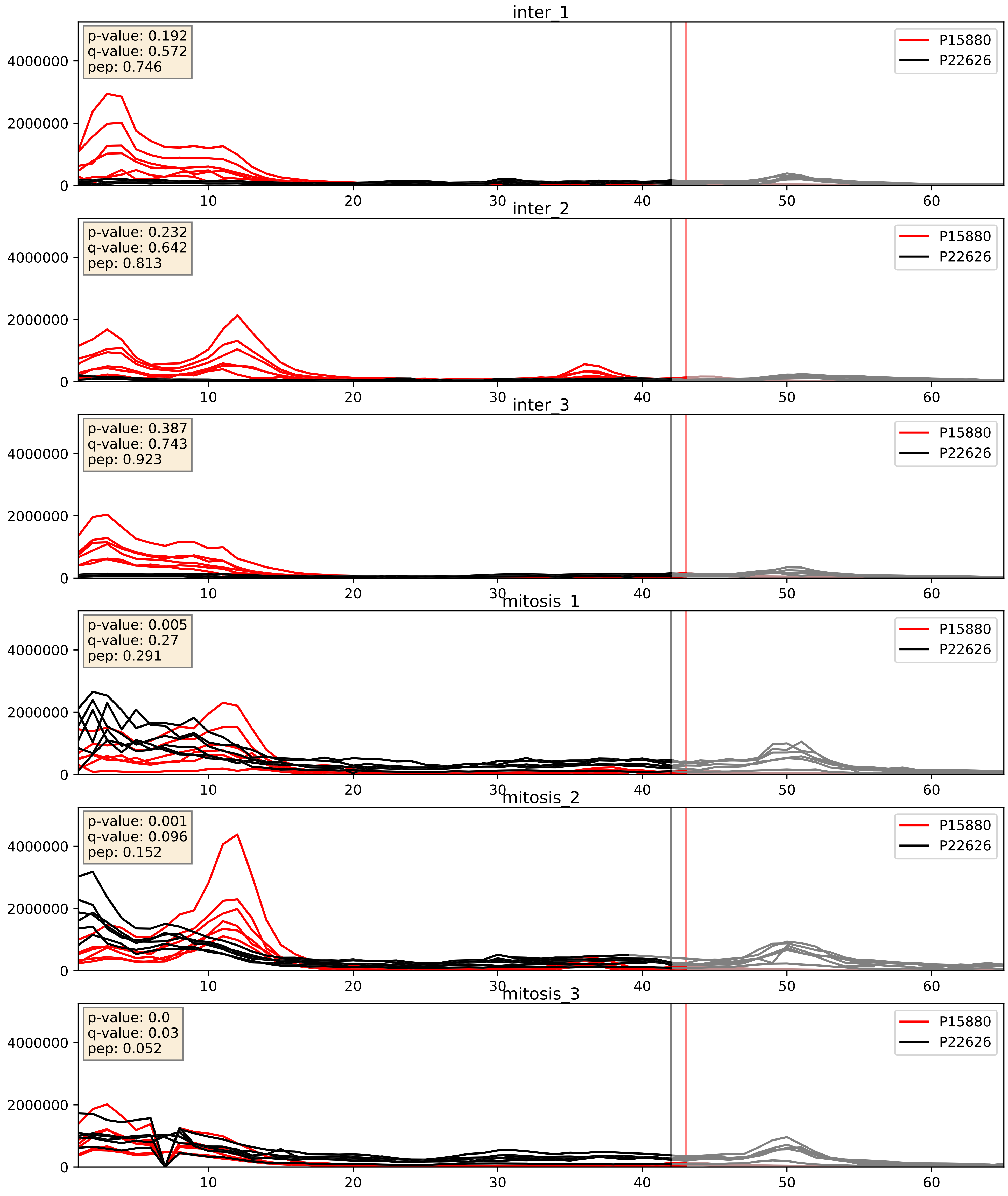


| level | condition_1 | condition_2 | pvalue | pvalue_adjusted |
|---|---|---|---|---|
| complex_abundance | mitosis | inter | 0.002574269466773442 | 0.03664546283589145 |
| interactor_ratio | mitosis | inter | 0.0049215113311436845 | 0.09361808221019986 |
| interactor_abundance | mitosis | inter | 0.294518135901904 | 0.48914925171135015 |

P17844_P22626
DDX5_HUMAN vs ROA2_HUMAN
p-value: 0.002 q-value: 0.033

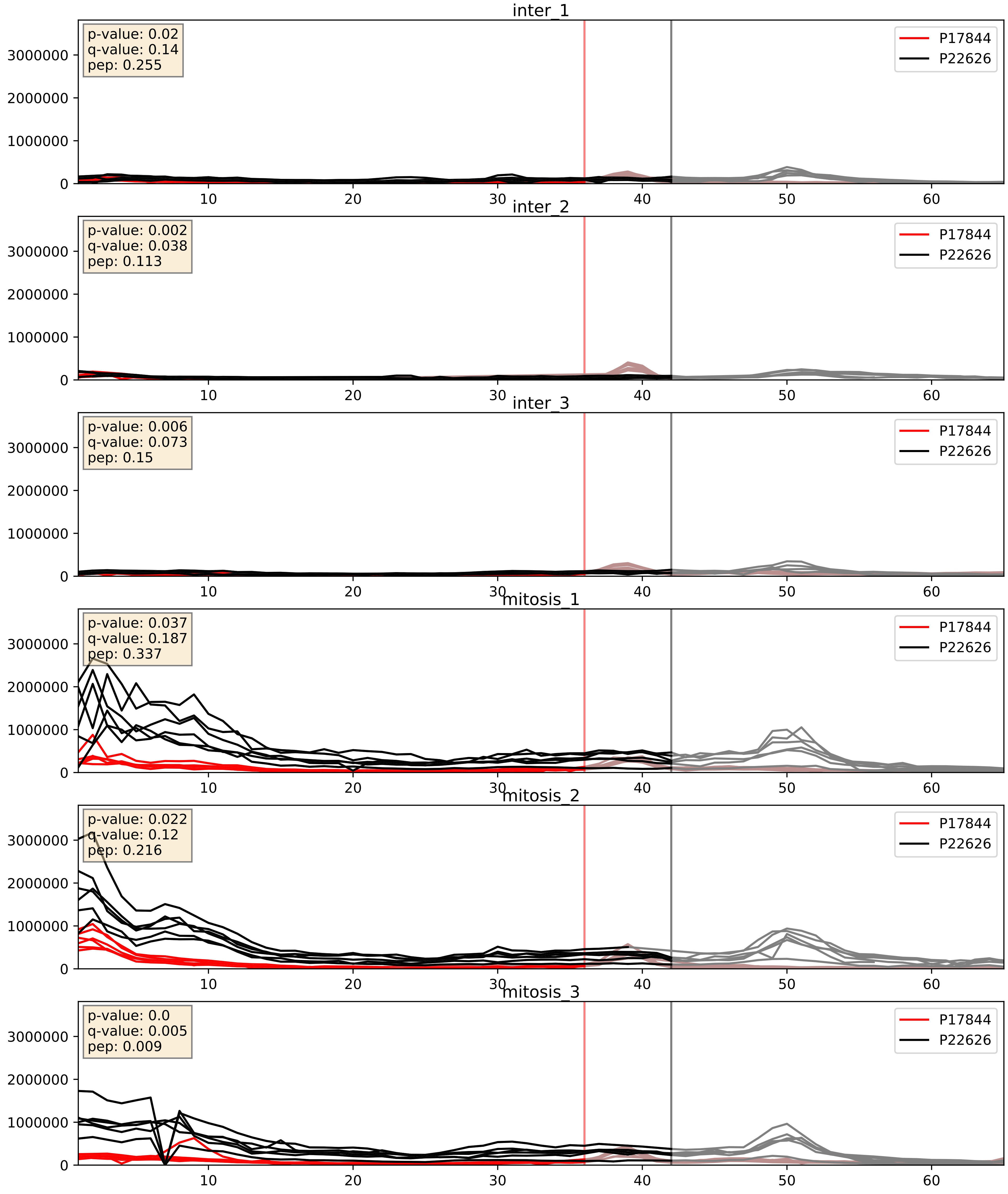


| level | condition_1 | condition_2 | pvalue | pvalue_adjusted |
|---|---|---|---|---|
| complex_abundance | mitosis | inter | 0.0004483662606115384 | 0.013372875229389437 |
| interactor_abundance | mitosis | inter | 0.0021426459467421676 | 0.033870820506210444 |
| interactor_ratio | mitosis | inter | 0.017003414810485822 | 0.1935495090129769 |

P22626_P26599
ROA2_HUMAN vs PTBP1_HUMAN
p-value: 0.007 q-value: 0.006

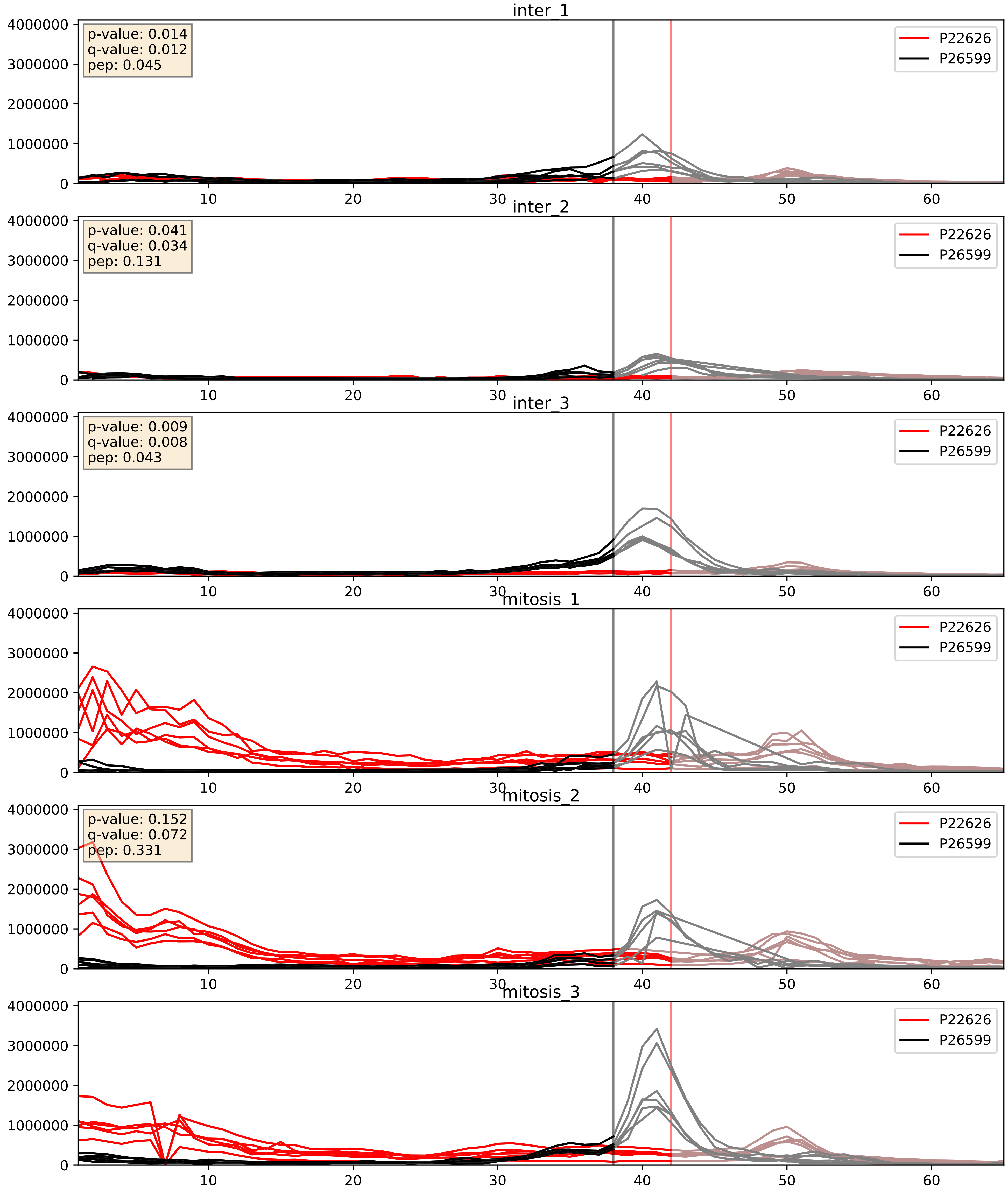


| level | condition_1 | condition_2 | pvalue | pvalue_adjusted |
|---|---|---|---|---|
| interactor_abundance | mitosis | inter | 0.00287274975376958 | 0.04063687618579503 |
| interactor_ratio | mitosis | inter | 0.00514591084778699 | 0.09659867731810666 |
| complex_abundance | mitosis | inter | 0.05656779343510957 | 0.16946458813160903 |

P22626_P31943
ROA2_HUMAN vs HNRH1_HUMAN
p-value: 0.003 q-value: 0.003

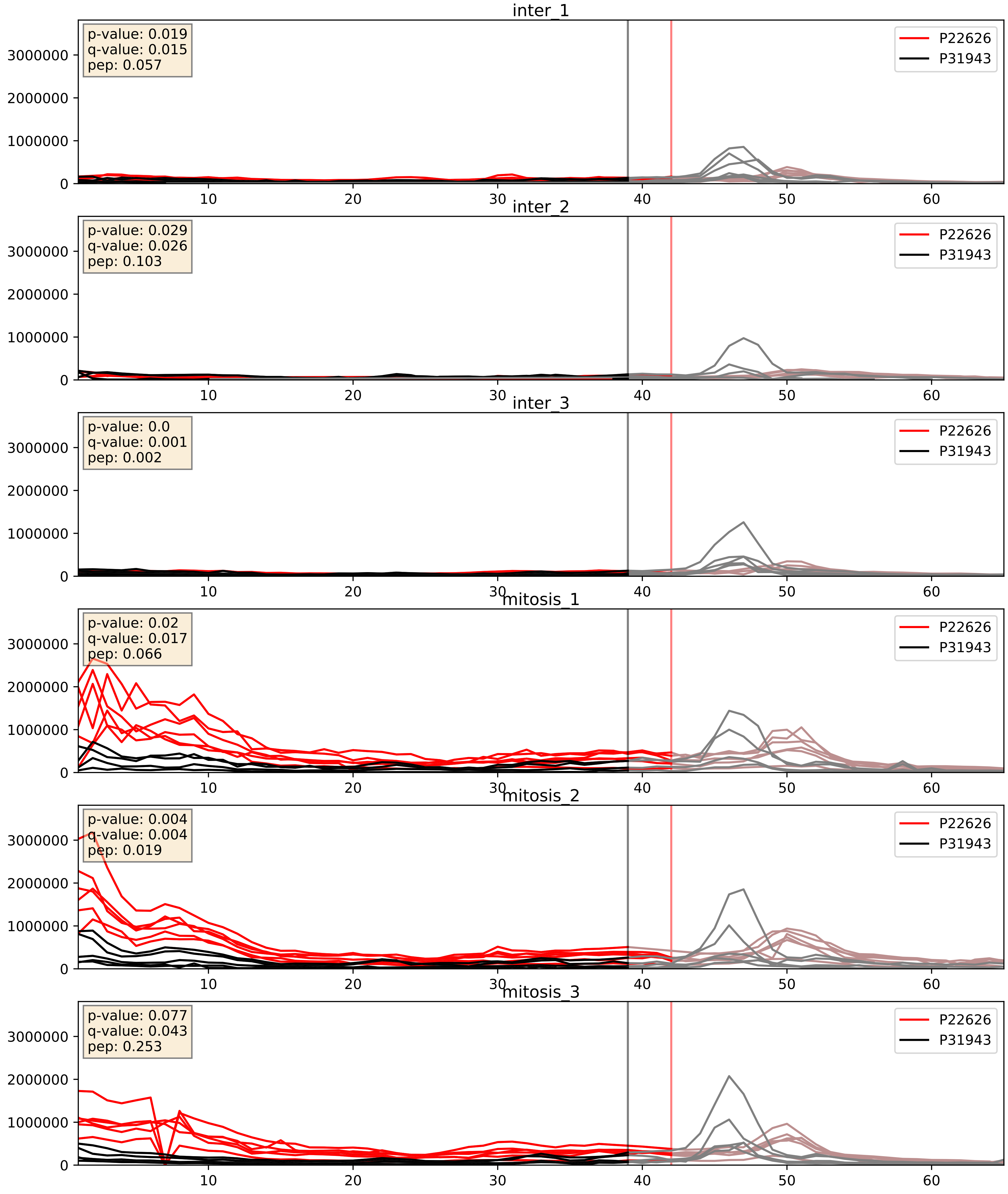


| level | condition_1 | condition_2 | pvalue | pvalue_adjusted |
|---|---|---|---|---|
| complex_abundance | mitosis | inter | 0.0008688599728642934 | 0.019513211095311817 |
| interactor_abundance | mitosis | inter | 0.003279687783791118 | 0.0442112242980346 |
| interactor_ratio | mitosis | inter | 0.02932575485422149 | 0.27108905135219863 |

P22626_P38159
ROA2_HUMAN vs RBMX_HUMAN
p-value: 0.011 q-value: 0.009

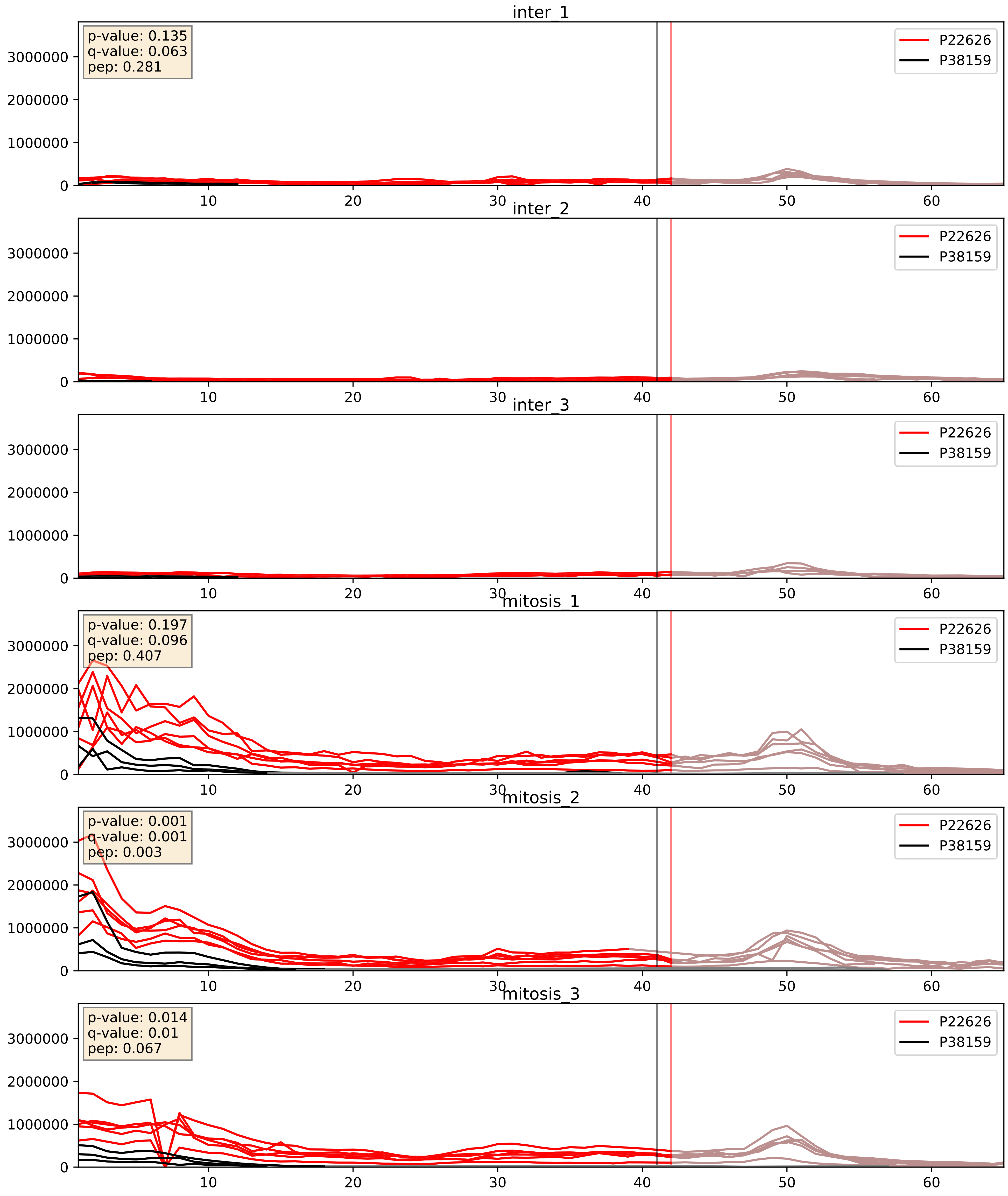


| level | condition_1 | condition_2 | pvalue | pvalue_adjusted |
|---|---|---|---|---|
| complex_abundance | mitosis | inter | 0.0005850625769621449 | 0.015650423933737376 |
| interactor_abundance | mitosis | inter | 0.0012988409915939223 | 0.02663012907315922 |
| interactor_ratio | mitosis | inter | 0.05283895661774645 | 0.3800852677713526 |

P22626_P51991
ROA2_HUMAN vs ROA3_HUMAN
p-value: 0.0 q-value: 0.001

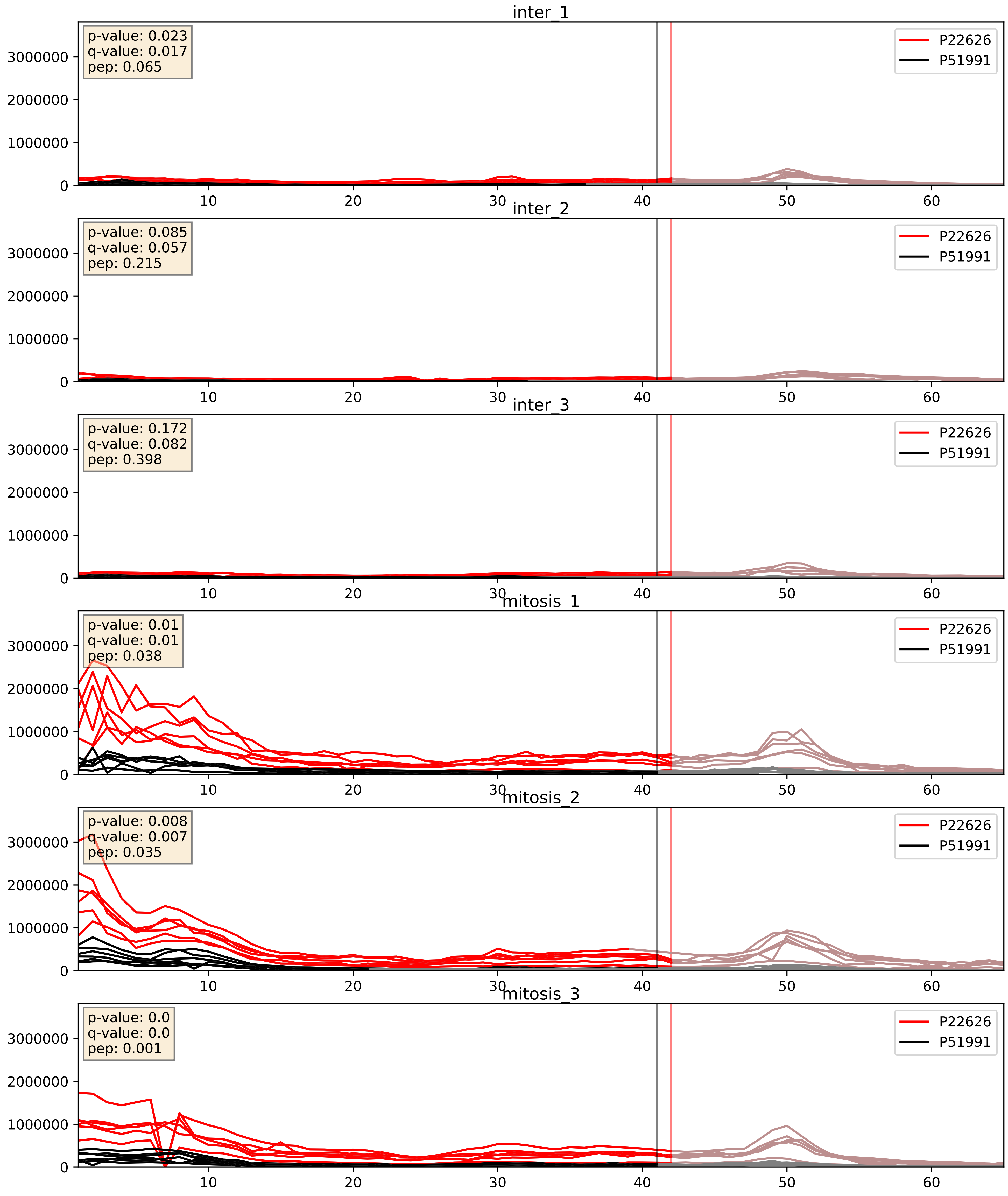


| level | condition_1 | condition_2 | pvalue | pvalue_adjusted |
|---|---|---|---|---|
| complex_abundance | mitosis | inter | 0.00023249210353232296 | 0.010221981956574496 |
| interactor_abundance | mitosis | inter | 0.0019123553121689623 | 0.031986606092438766 |
| interactor_ratio | mitosis | inter | 0.08015668685278068 | 0.47352742543809706 |

P22626_P52272
ROA2_HUMAN vs HNRPM_HUMAN
p-value: 0.003 q-value: 0.003

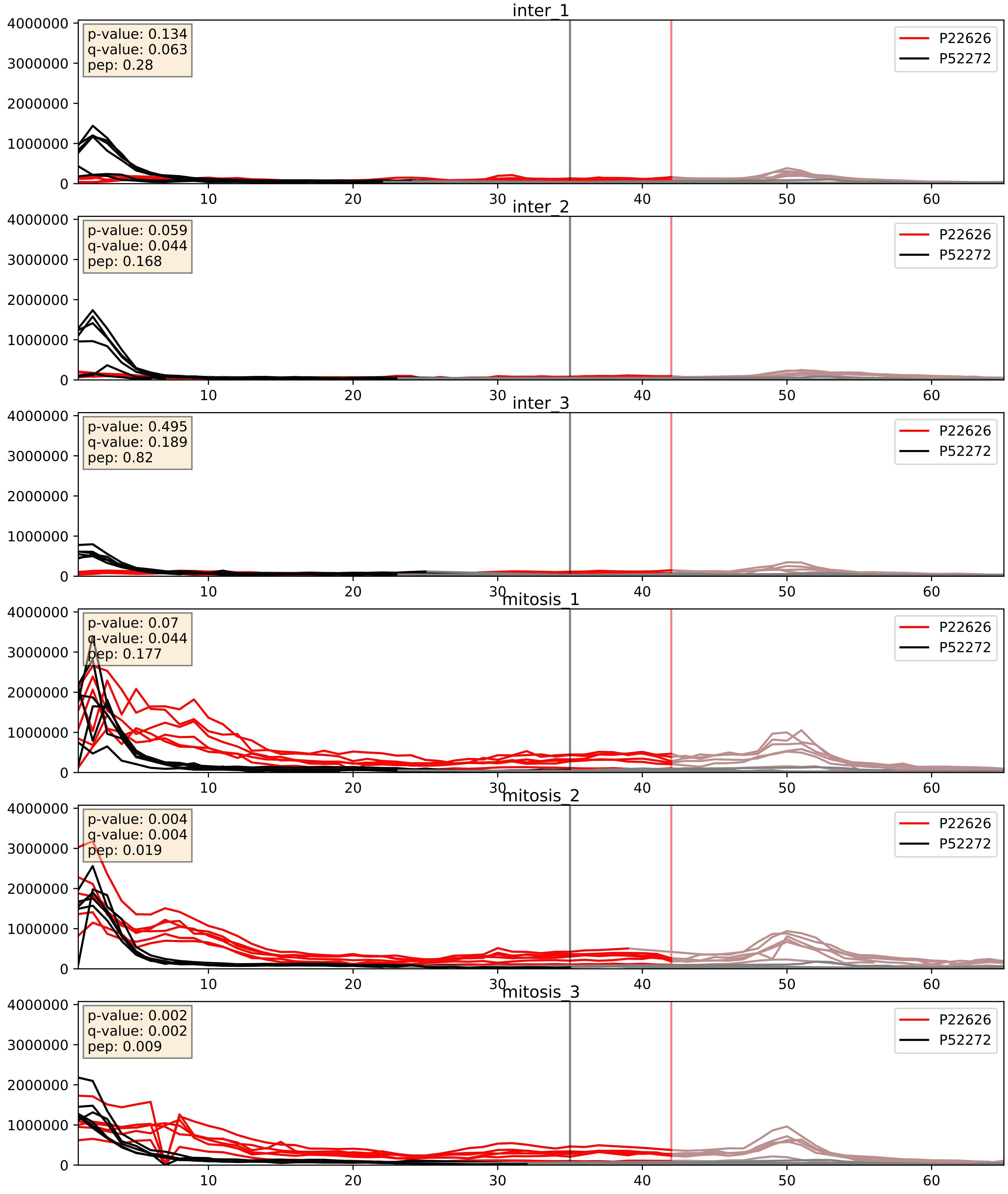


| level | condition_1 | condition_2 | pvalue | pvalue_adjusted |
|---|---|---|---|---|
| interactor_ratio | mitosis | inter | 0.0006314989468148049 | 0.029219626944512057 |
| interactor_abundance | mitosis | inter | 0.0029874185243059705 | 0.04171664366730687 |
| complex_abundance | mitosis | inter | 0.007981194454066766 | 0.06600871934957633 |

P22626_P52597
ROA2_HUMAN vs HNRPF_HUMAN
p-value: 0.006 q-value: 0.006

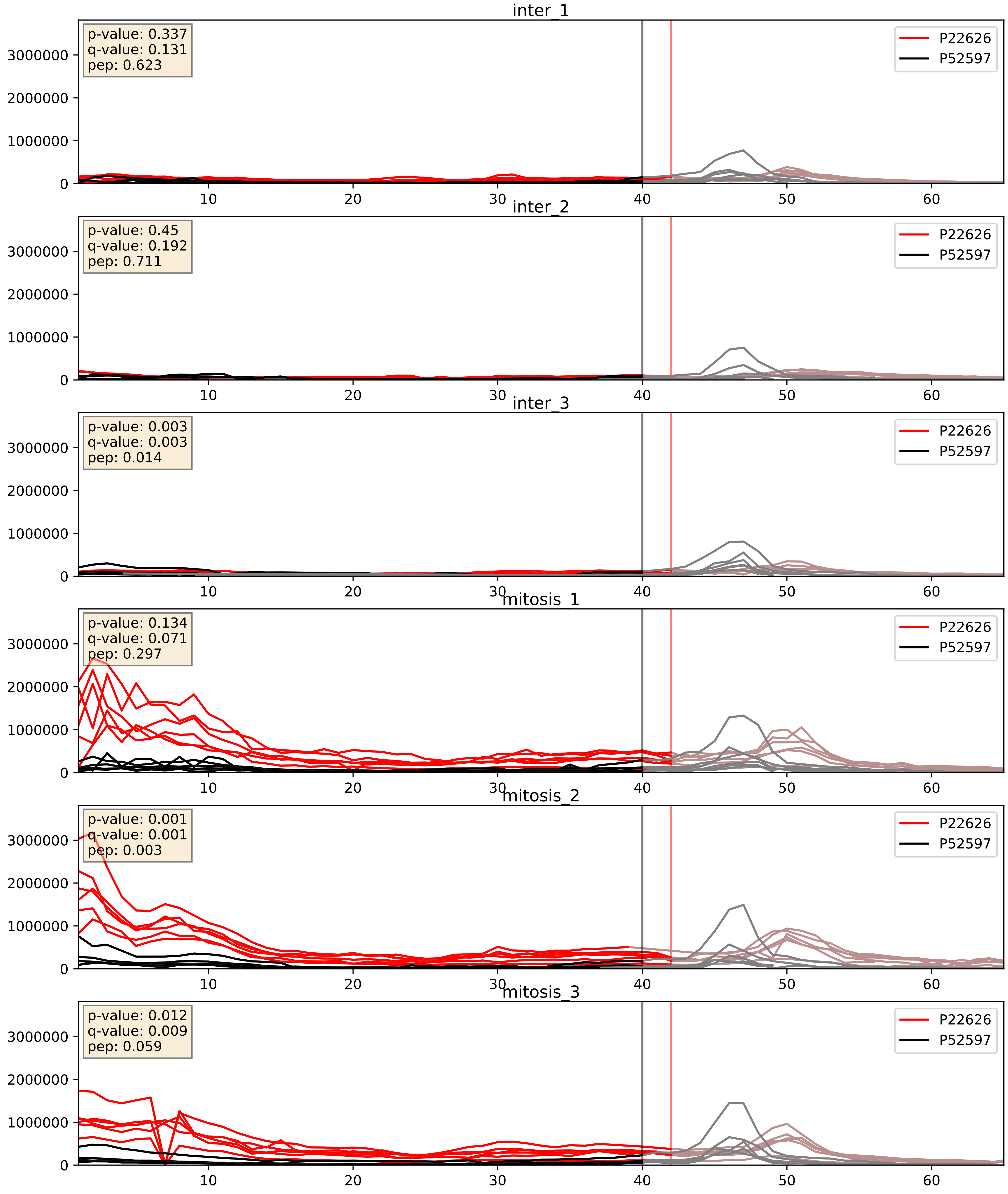


| level | condition_1 | condition_2 | pvalue | pvalue_adjusted |
|---|---|---|---|---|
| complex_abundance | mitosis | inter | 0.0010673061249328343 | 0.021804631096479864 |
| interactor_abundance | mitosis | inter | 0.003361167395046095 | 0.04478068934100323 |
| interactor_ratio | mitosis | inter | 0.01572743448355028 | 0.1848097318528163 |

P22626_P61978
ROA2_HUMAN vs HNRPK_HUMAN
p-value: 0.004 q-value: 0.004

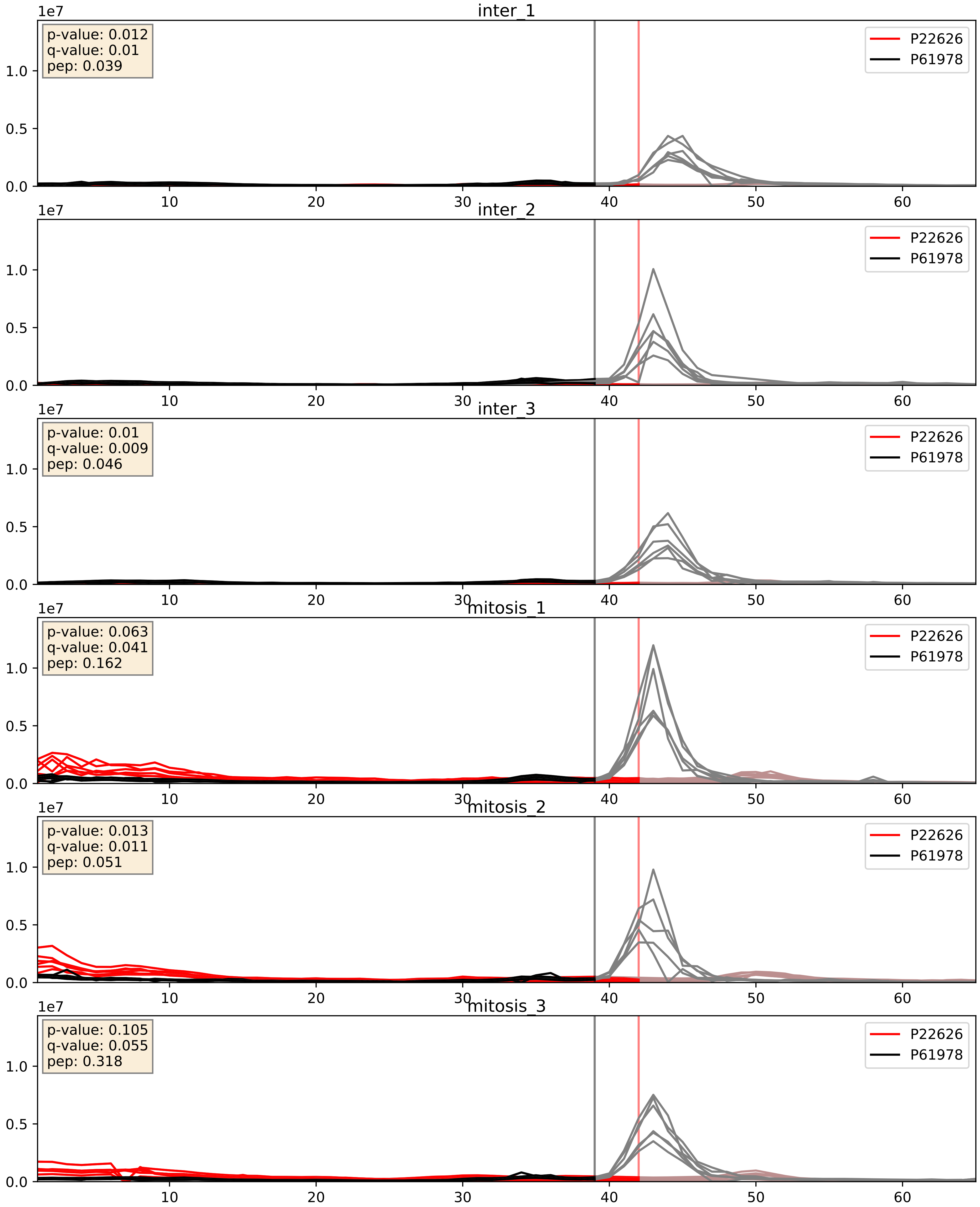


| level | condition_1 | condition_2 | pvalue | pvalue_adjusted |
|---|---|---|---|---|
| interactor_abundance | mitosis | inter | 0.003279687783791118 | 0.0442112242980346 |
| interactor_ratio | mitosis | inter | 0.009137891625288207 | 0.13627239078827014 |
| complex_abundance | mitosis | inter | 0.043583982673943734 | 0.14954839107468462 |

P22626_P62314
ROA2_HUMAN vs SMD1_HUMAN
p-value: 0.0 q-value: 0.001

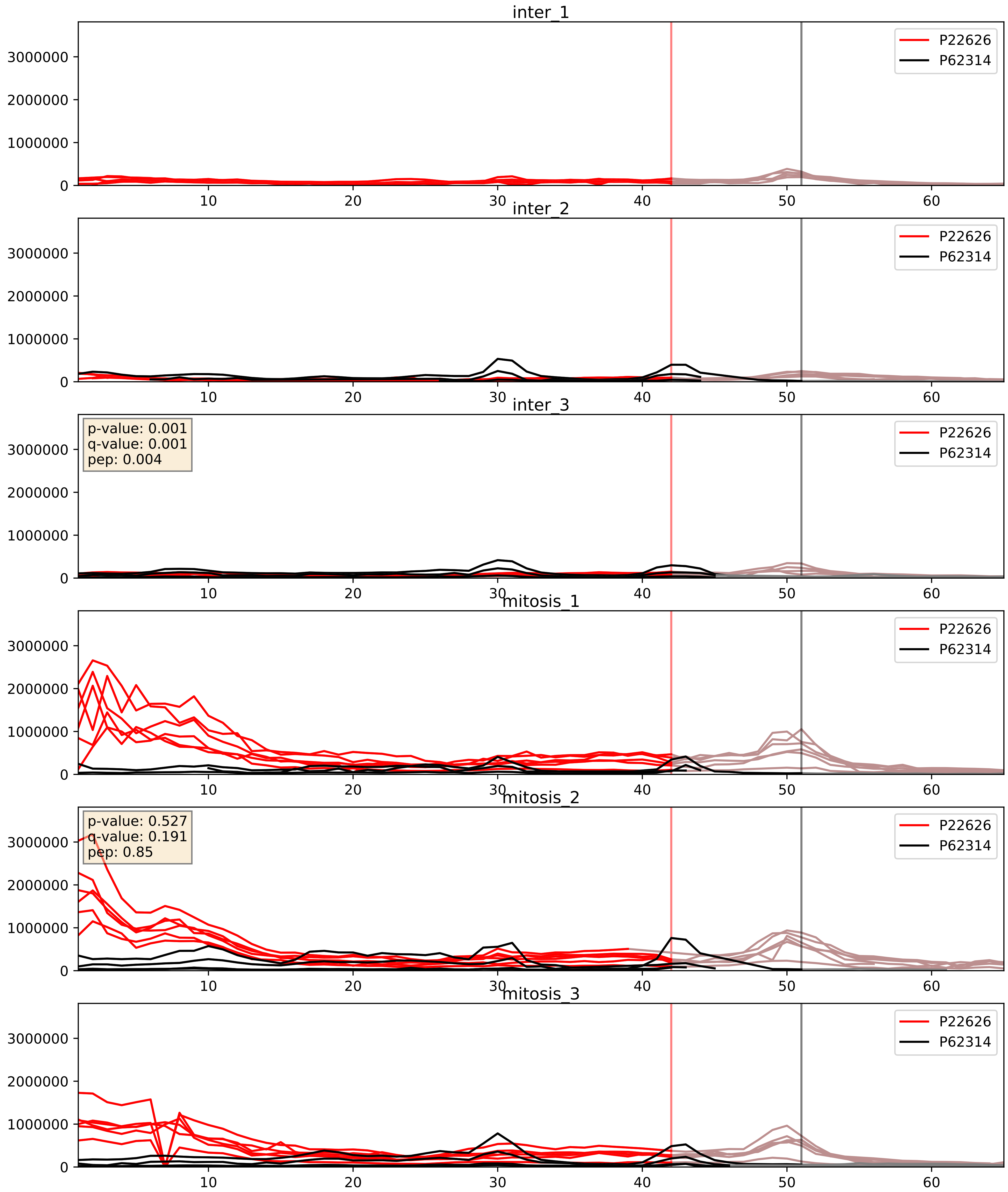


| level | condition_1 | condition_2 | pvalue | pvalue_adjusted |
|---|---|---|---|---|
| interactor_abundance | mitosis | inter | 0.010521951914376396 | 0.08948624777651461 |
| interactor_ratio | mitosis | inter | 0.010795815399431462 | 0.14905190293408602 |
| complex_abundance | mitosis | inter | 0.04151925020286664 | 0.14668491551379267 |

P22626_P62316
ROA2_HUMAN vs SMD2_HUMAN
p-value: 0.0 q-value: 0.001

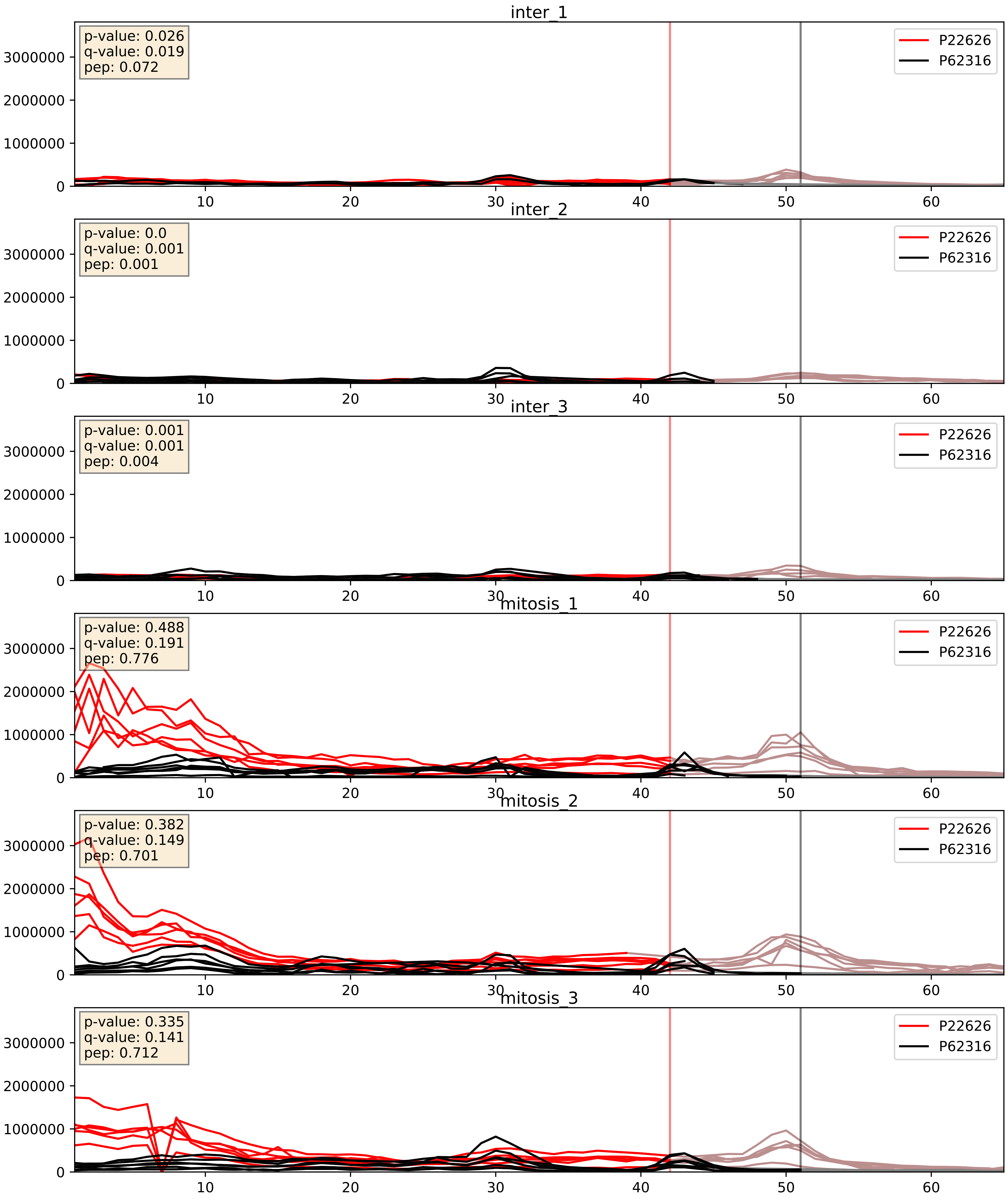


| level | condition_1 | condition_2 | pvalue | pvalue_adjusted |
|---|---|---|---|---|
| complex_abundance | mitosis | inter | 0.0008978115482066242 | 0.01975647005822289 |
| interactor_abundance | mitosis | inter | 0.0034466655543236775 | 0.04511548235131213 |
| interactor_ratio | mitosis | inter | 0.0603049372865427 | 0.40594115513296747 |

P22626_P62318
ROA2_HUMAN vs SMD3_HUMAN
p-value: 0.036 q-value: 0.025

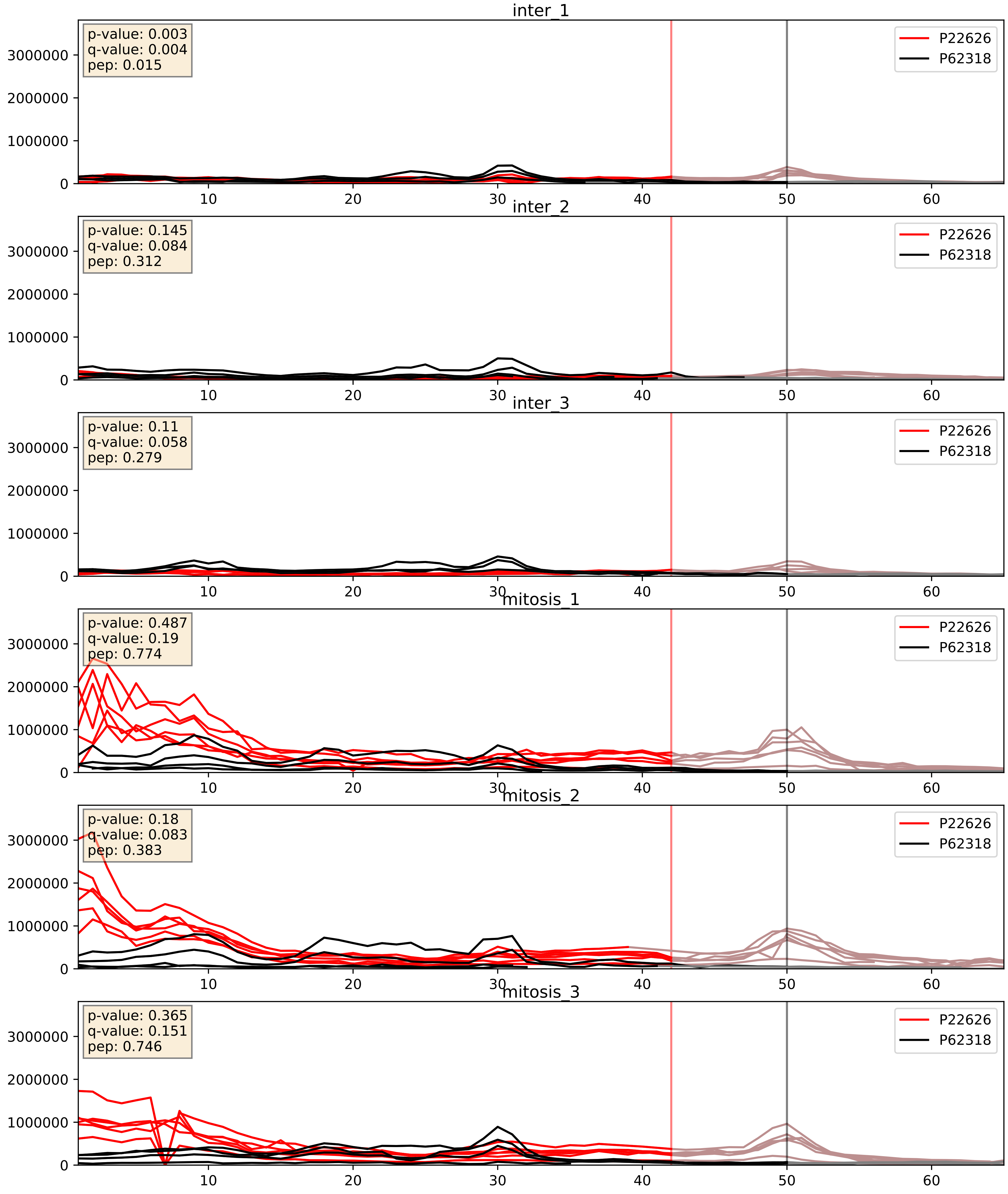


| level | condition_1 | condition_2 | pvalue | pvalue_adjusted |
|---|---|---|---|---|
| interactor_abundance | mitosis | inter | 0.0034466655543236775 | 0.04511548235131213 |
| complex_abundance | mitosis | inter | 0.015254221642728273 | 0.09174685245214603 |
| interactor_ratio | mitosis | inter | 0.052785200781211176 | 0.38001792993033445 |

P22626_P62753
ROA2_HUMAN vs RS6_HUMAN
p-value: 0.0 q-value: 0.018

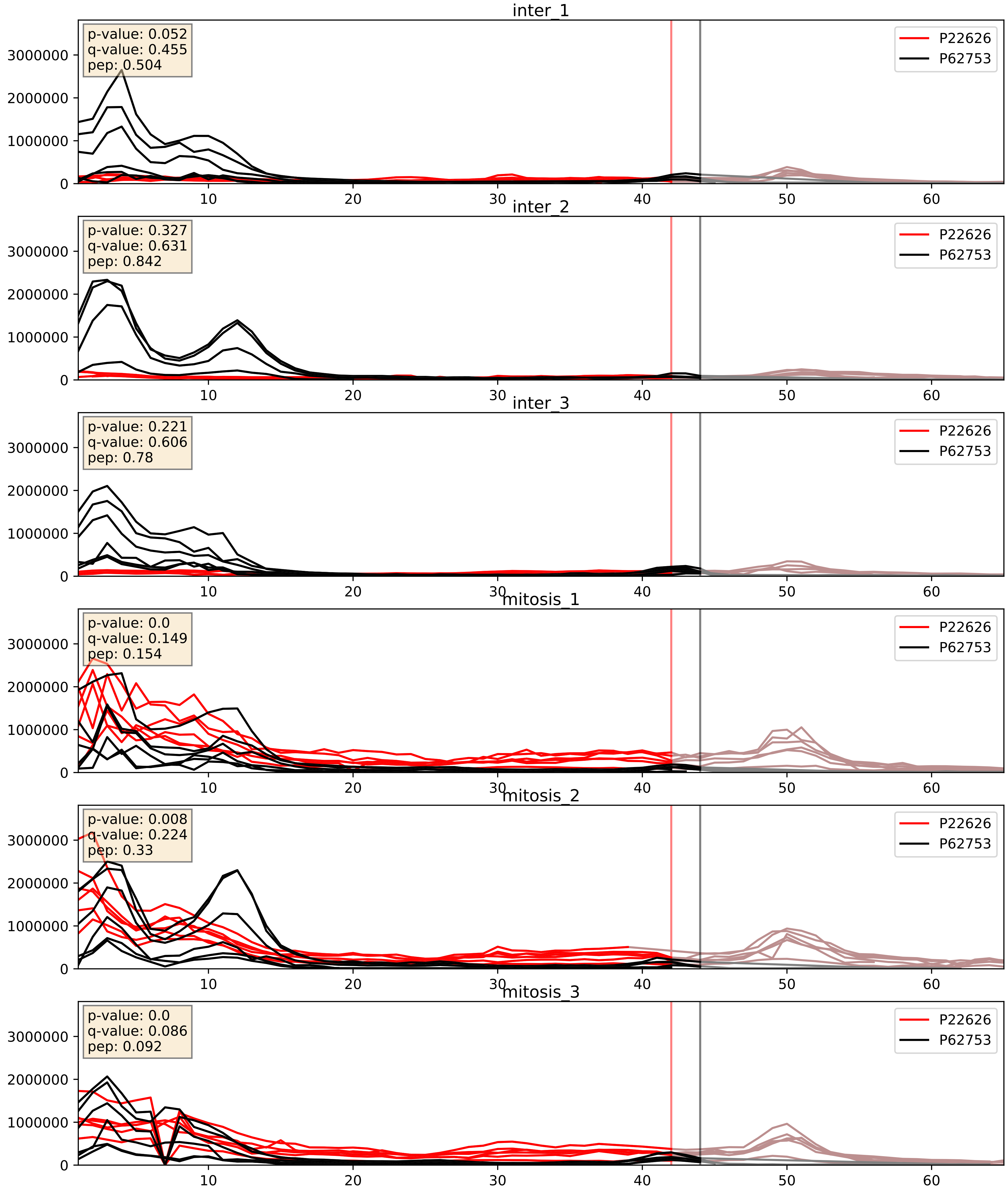


| level | condition_1 | condition_2 | pvalue | pvalue_adjusted |
|---|---|---|---|---|
| complex_abundance | mitosis | inter | 0.0020990721188606045 | 0.033335913427545036 |
| interactor_abundance | mitosis | inter | 0.0034466655543236775 | 0.04511548235131213 |
| interactor_ratio | mitosis | inter | 0.006351778787069153 | 0.11006321137107682 |

P22626_P84103
ROA2_HUMAN vs SRSF3_HUMAN
p-value: 0.015 q-value: 0.012

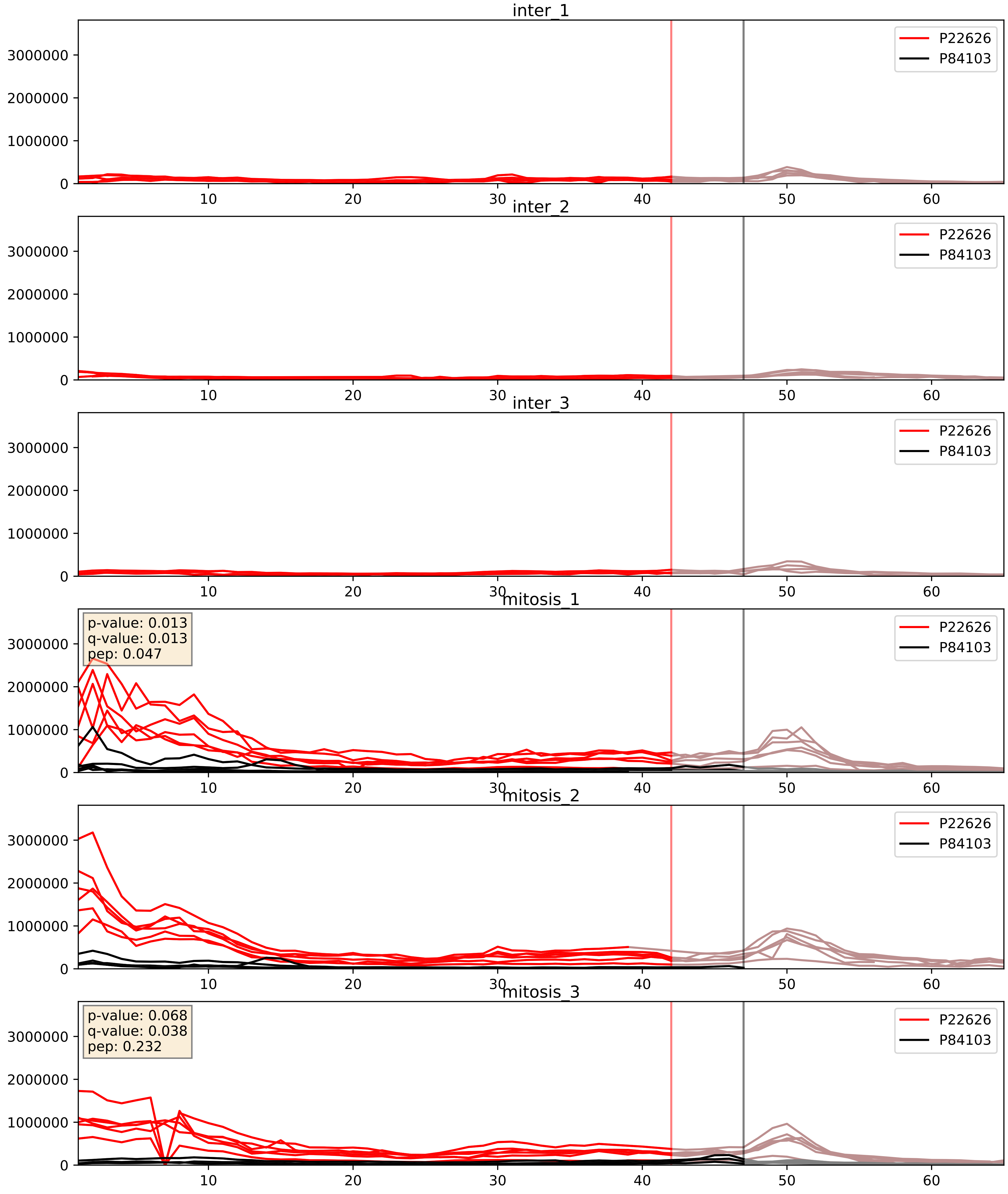


| level | condition_1 | condition_2 | pvalue | pvalue_adjusted |
|---|---|---|---|---|
| interactor_abundance | mitosis | inter | 3.003313050728667e-05 | 0.004850633908346677 |
| complex_abundance | mitosis | inter | 6.433903659397418e-05 | 0.005691857151652958 |
| interactor_ratio | mitosis | inter | 0.0014422267222096987 | 0.045221467919835244 |

P22626_Q07955
ROA2_HUMAN vs SRSF1_HUMAN
p-value: 0.028 q-value: 0.021

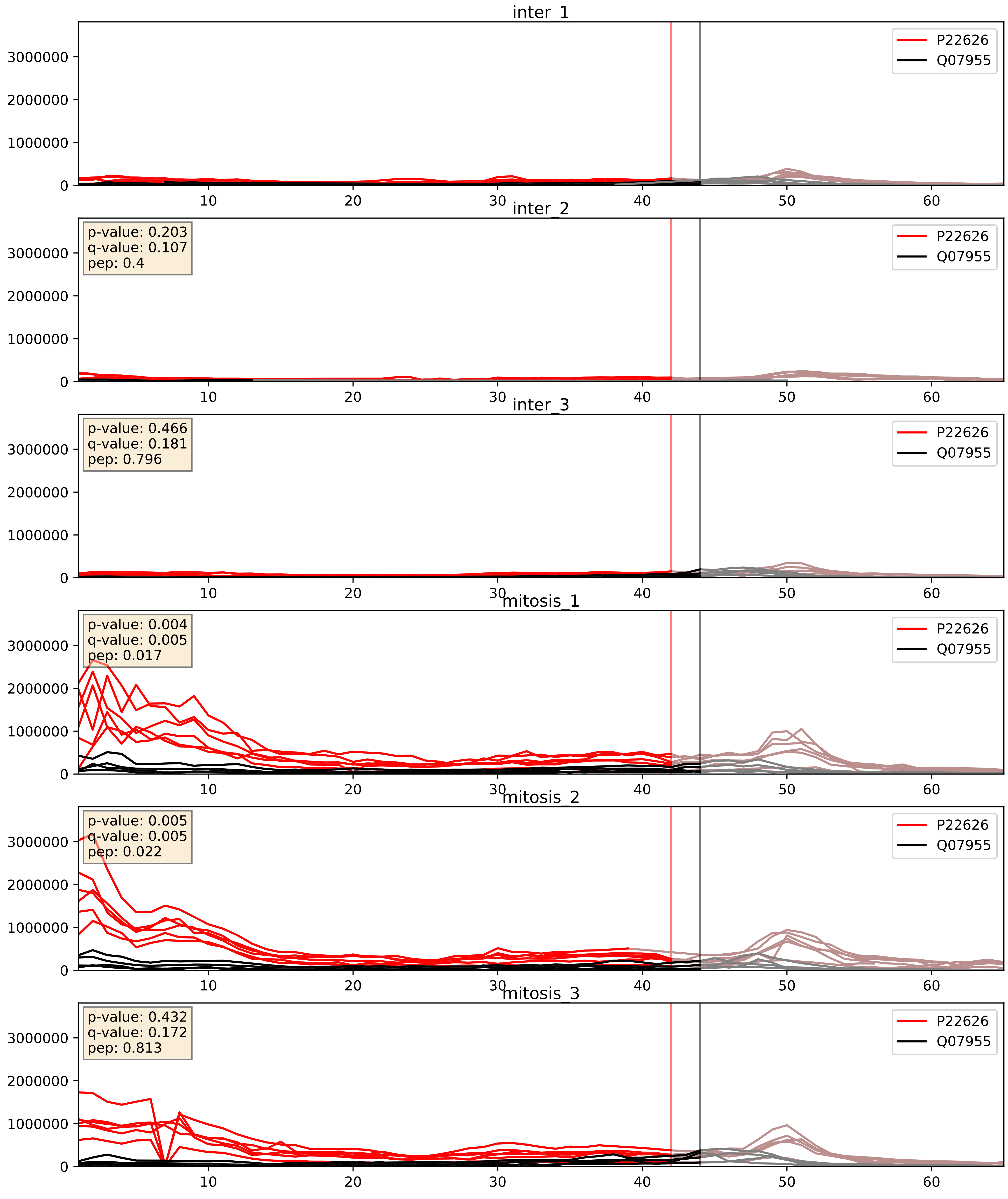


| level | condition_1 | condition_2 | pvalue | pvalue_adjusted |
|---|---|---|---|---|
| interactor_abundance | mitosis | inter | 0.0036653640621034087 | 0.0470045189087718 |
| complex_abundance | mitosis | inter | 0.004020555198830934 | 0.04695218622372823 |
| interactor_ratio | mitosis | inter | 0.03180859881707865 | 0.28451578461253213 |

P22626_Q08211
ROA2_HUMAN vs DHX9_HUMAN
p-value: 0.007 q-value: 0.006

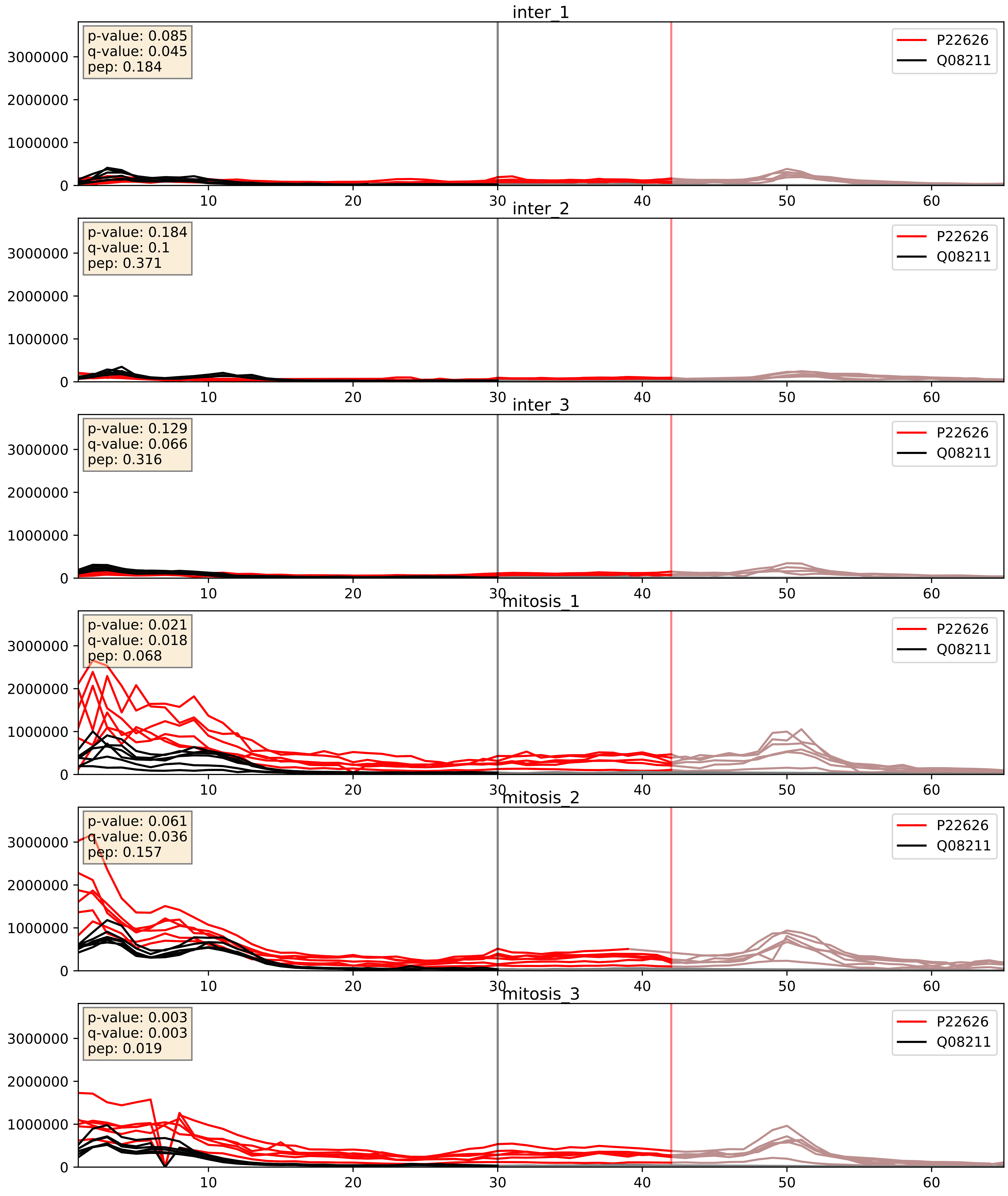


| level | condition_1 | condition_2 | pvalue | pvalue_adjusted |
|---|---|---|---|---|
| complex_abundance | mitosis | inter | 0.0009287786996386022 | 0.02007663047703645 |
| interactor_abundance | mitosis | inter | 0.0022087495902483206 | 0.034628015554076234 |
| interactor_ratio | mitosis | inter | 0.009213692076387548 | 0.13668839544866102 |

P22626_Q12905
ROA2_HUMAN vs ILF2_HUMAN
p-value: 0.002 q-value: 0.035

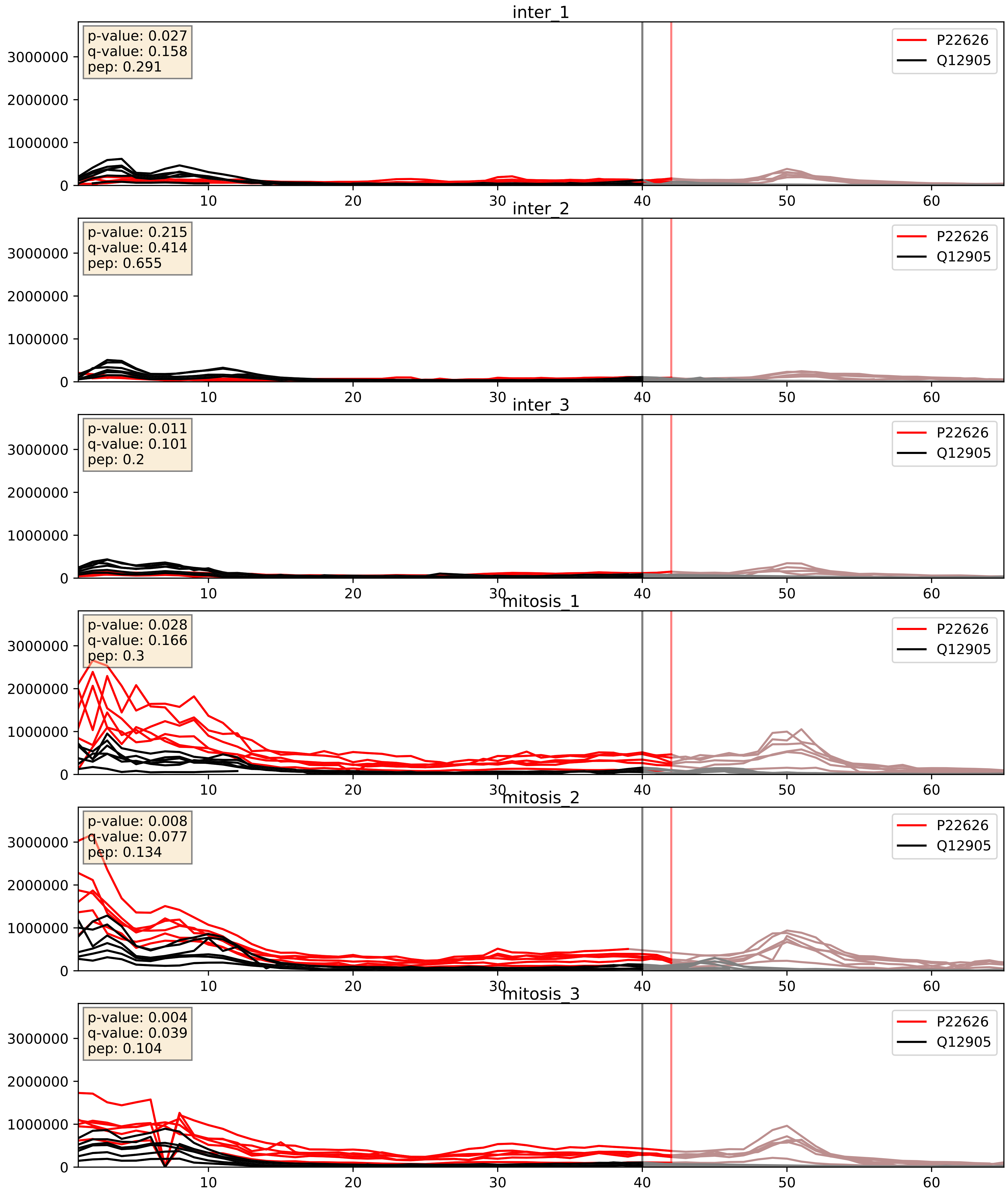


| level | condition_1 | condition_2 | pvalue | pvalue_adjusted |
|---|---|---|---|---|
| complex_abundance | mitosis | inter | 0.0022575804114793045 | 0.034819618598671796 |
| interactor_abundance | mitosis | inter | 0.003361167395046095 | 0.04478068934100323 |
| interactor_ratio | mitosis | inter | 0.006434293228359545 | 0.11104344765072117 |

P22626_Q12906
ROA2_HUMAN vs ILF3_HUMAN
p-value: 0.003 q-value: 0.042

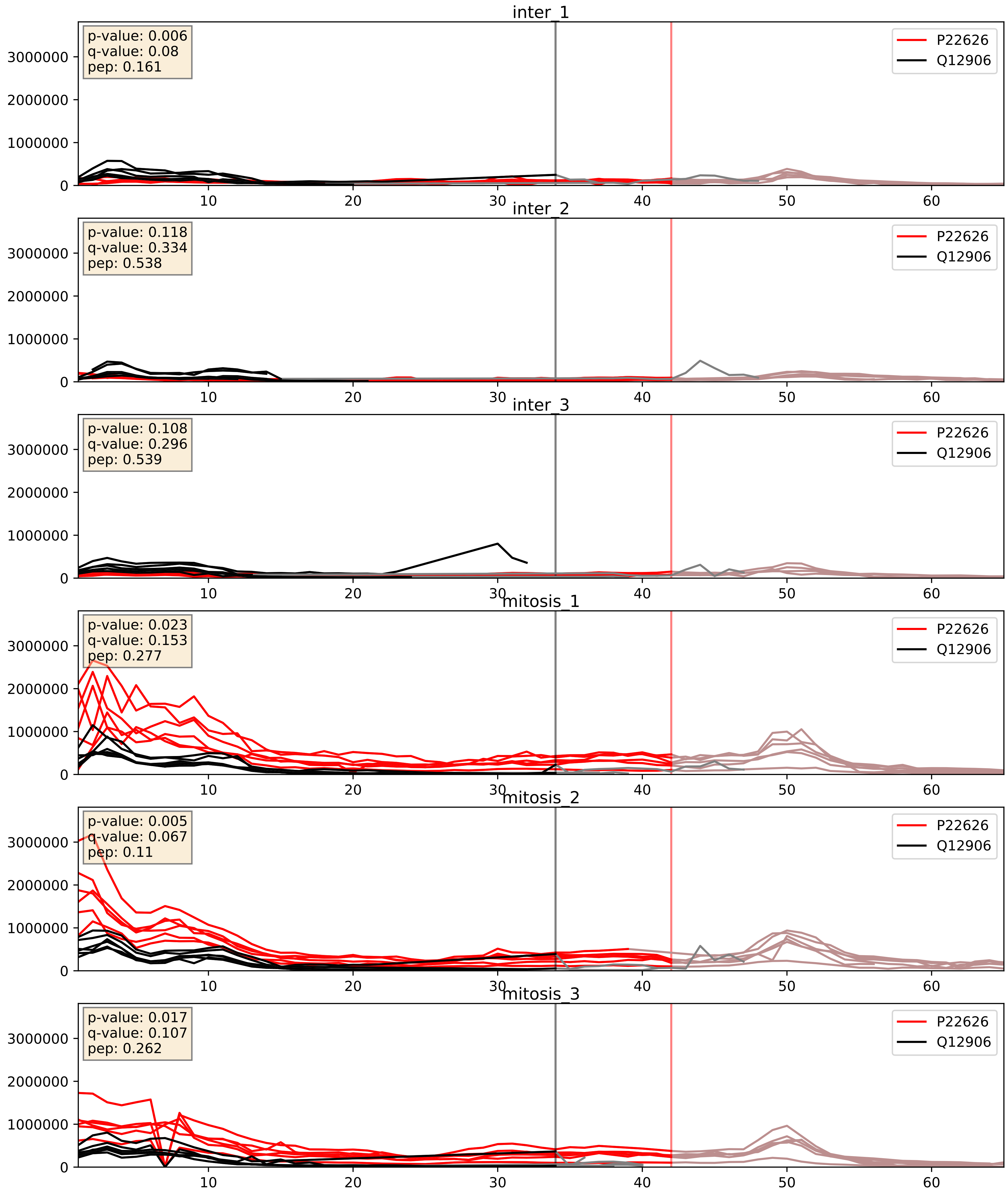


| level | condition_1 | condition_2 | pvalue | pvalue_adjusted |
|---|---|---|---|---|
| complex_abundance | mitosis | inter | 0.0006580872733274757 | 0.01642175463112936 |
| interactor_abundance | mitosis | inter | 0.0011041441288093116 | 0.024049720606162197 |
| interactor_ratio | mitosis | inter | 0.002740523287269679 | 0.06759600533681397 |

P22626_Q13151
ROA2_HUMAN vs ROA0_HUMAN
p-value: 0.045 q-value: 0.03

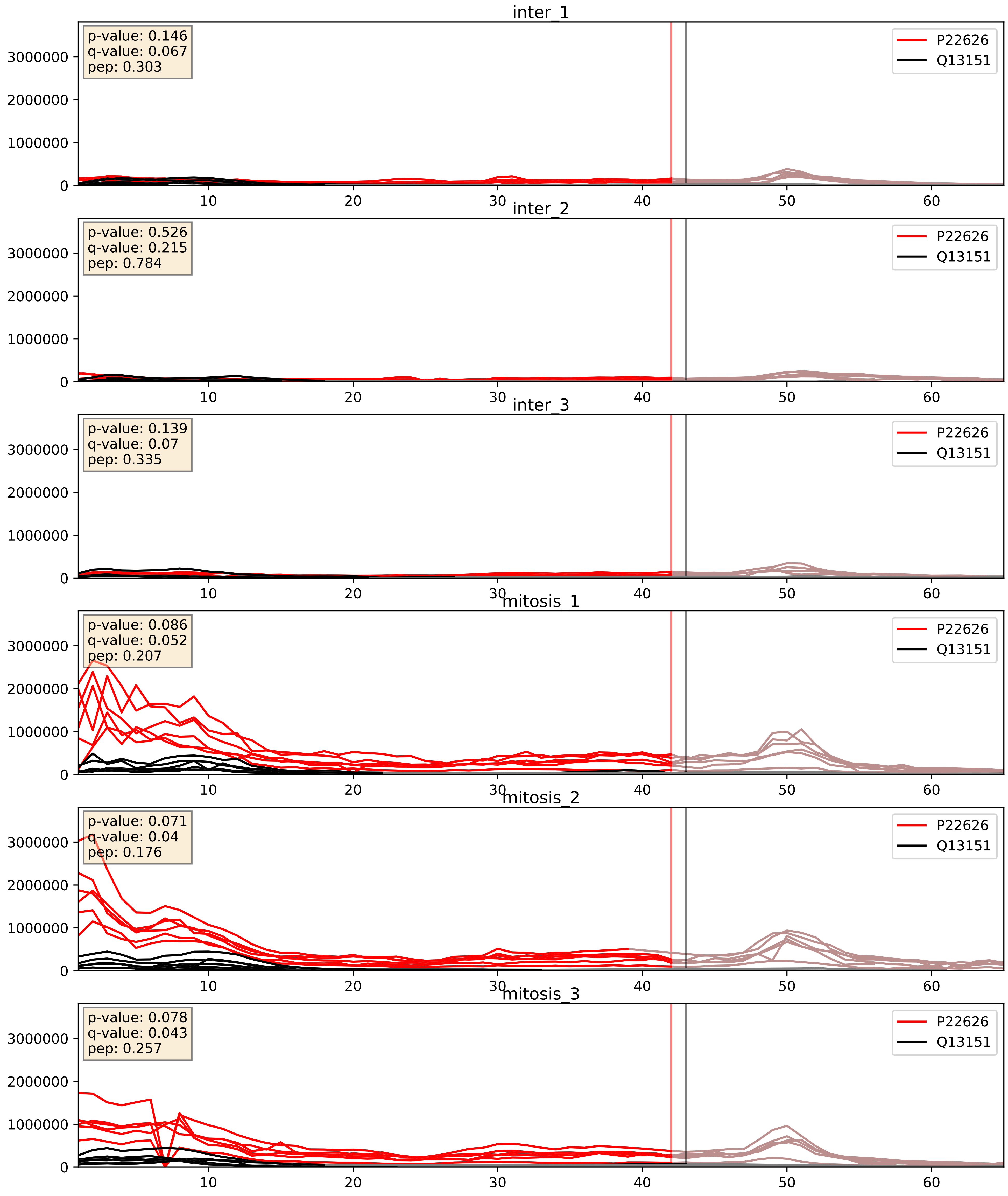


| level | condition_1 | condition_2 | pvalue | pvalue_adjusted |
|---|---|---|---|---|
| interactor_ratio | mitosis | inter | 0.0031237624803896113 | 0.0729098066746082 |
| interactor_abundance | mitosis | inter | 0.0034862800934623243 | 0.04542246210051369 |
| complex_abundance | mitosis | inter | 0.0041541914710077335 | 0.04792436521809461 |

P22626_Q14103
ROA2_HUMAN vs HNRPD_HUMAN
p-value: 0.036 q-value: 0.025

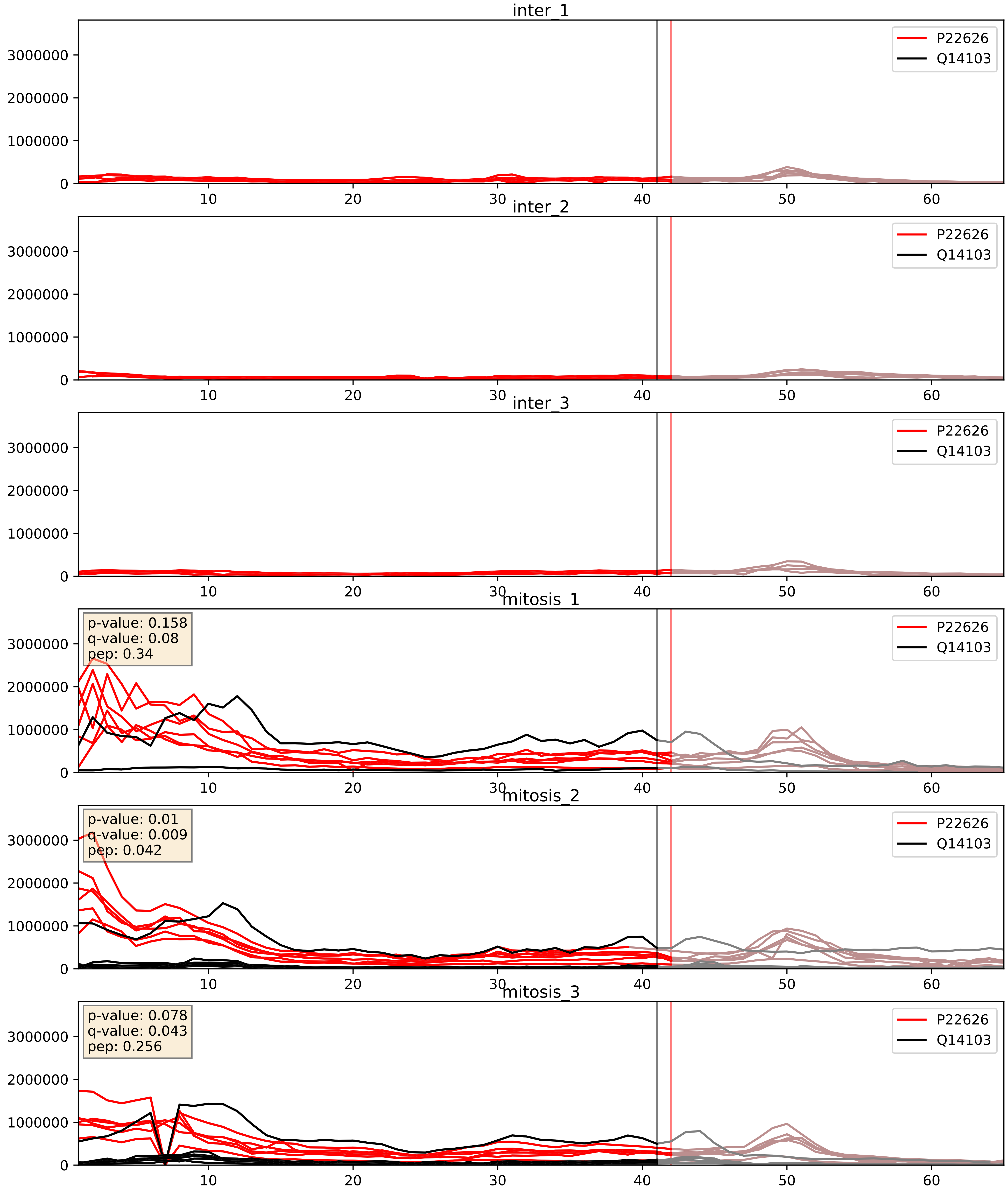


| level | condition_1 | condition_2 | pvalue | pvalue_adjusted |
|---|---|---|---|---|
| interactor_abundance | mitosis | inter | 2.1731713857435016e-05 | 0.004180302710553792 |
| complex_abundance | mitosis | inter | 0.0002607874150841398 | 0.010784252527150902 |
| interactor_ratio | mitosis | inter | 0.026748360525100004 | 0.2569763929235197 |

P22626_Q15365
ROA2_HUMAN vs PCBP1_HUMAN
p-value: 0.0 q-value: 0.0

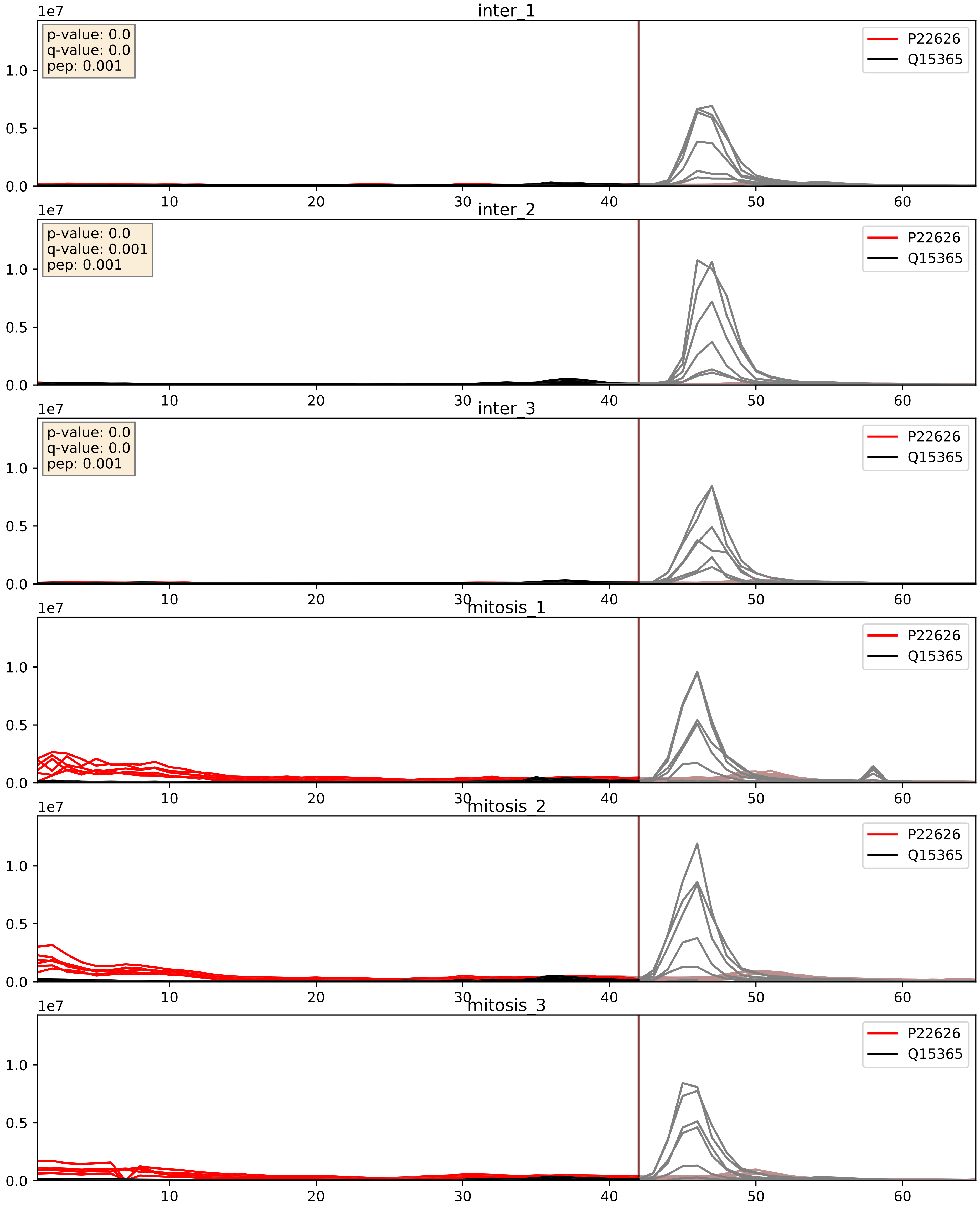


| level | condition_1 | condition_2 | pvalue | pvalue_adjusted |
|---|---|---|---|---|
| interactor_ratio | mitosis | inter | 0.0018969253504078144 | 0.052986711193083874 |
| interactor_abundance | mitosis | inter | 0.0034466655543236775 | 0.04511548235131213 |
| complex_abundance | mitosis | inter | 0.006247593176456641 | 0.05896295213943644 |

P22626_Q15366
ROA2_HUMAN vs PCBP2_HUMAN
p-value: 0.001 q-value: 0.029

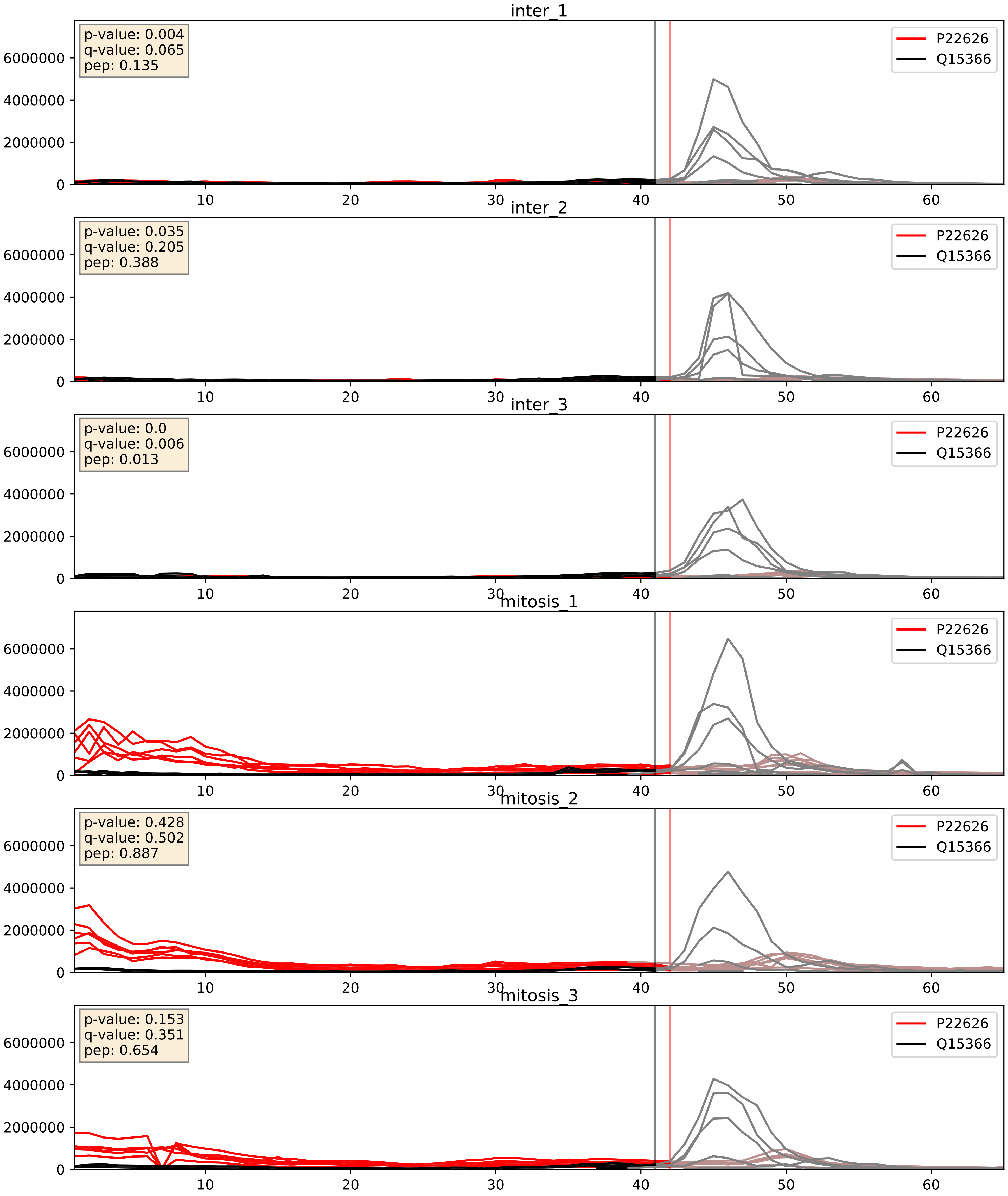


| level | condition_1 | condition_2 | pvalue | pvalue_adjusted |
|---|---|---|---|---|
| interactor_abundance | mitosis | inter | 0.0034576483826077656 | 0.045152509771353884 |
| interactor_ratio | mitosis | inter | 0.0042911301440727335 | 0.08642840949002965 |
| complex_abundance | mitosis | inter | 0.038506417698962146 | 0.14343034002958538 |

P22626_Q15717
ROA2_HUMAN vs ELAV1_HUMAN
p-value: 0.019 q-value: 0.015

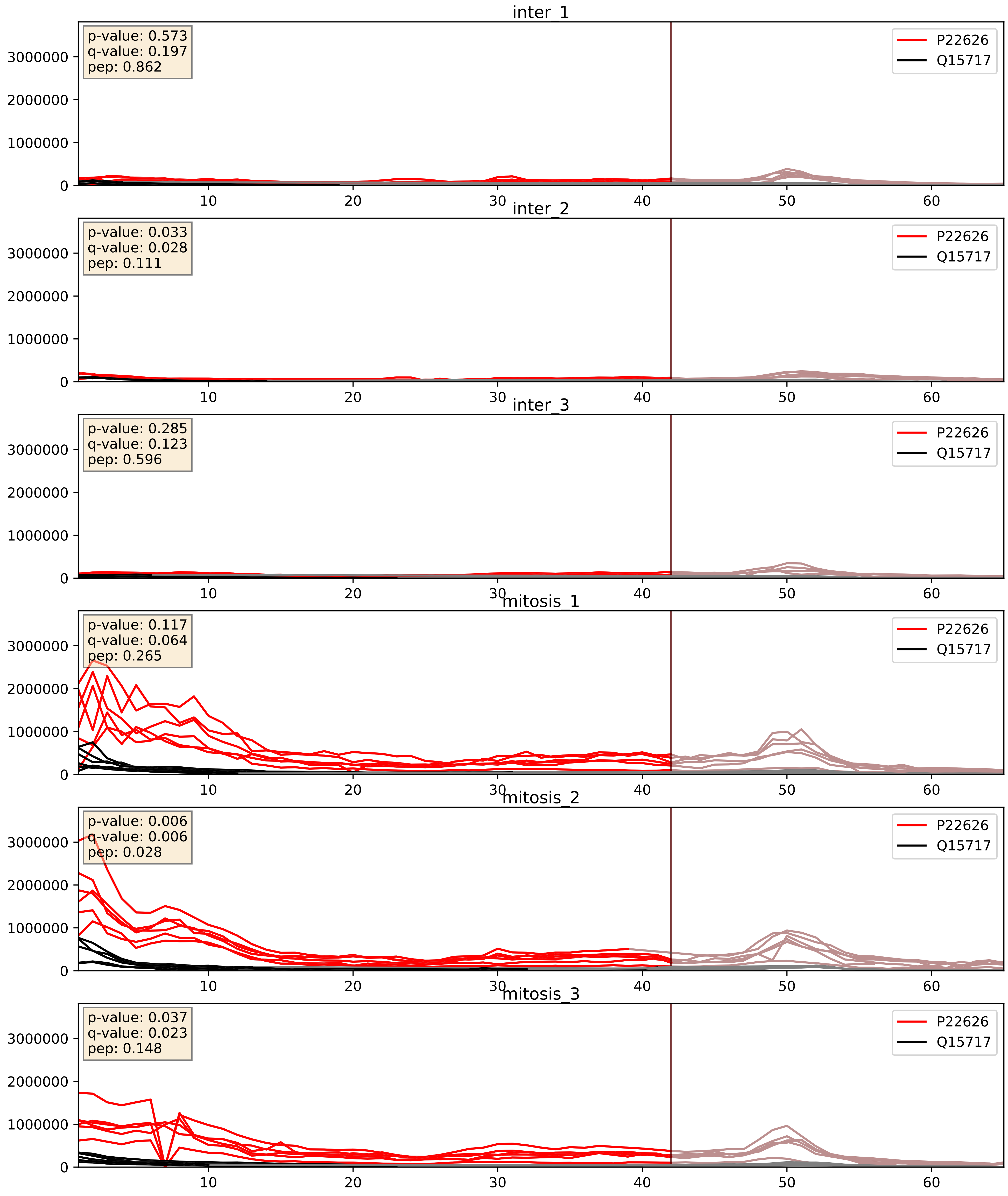


| level | condition_1 | condition_2 | pvalue | pvalue_adjusted |
|---|---|---|---|---|
| complex_abundance | mitosis | inter | 0.00036263180025352423 | 0.012039775462644836 |
| interactor_abundance | mitosis | inter | 0.0009434448484861828 | 0.02284551033392284 |
| interactor_ratio | mitosis | inter | 0.013987766740221716 | 0.17403384200043295 |

P22626_Q16629
ROA2_HUMAN vs SRSF7_HUMAN
p-value: 0.01 q-value: 0.009

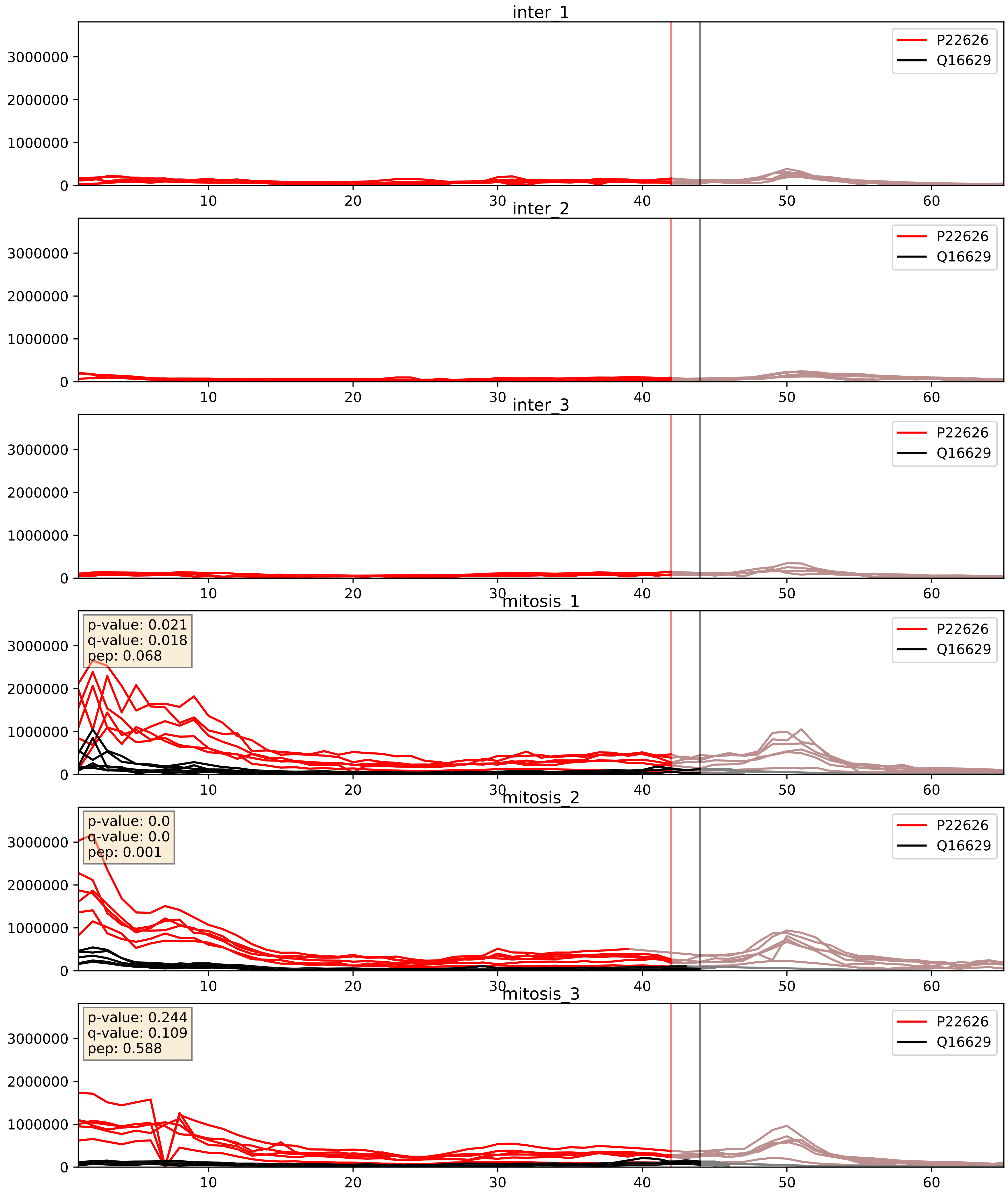


| level | condition_1 | condition_2 | pvalue | pvalue_adjusted |
|---|---|---|---|---|
| complex_abundance | mitosis | inter | 1.8097121496965717e-05 | 0.005691857151652958 |
| interactor_abundance | mitosis | inter | 2.0060878878898597e-05 | 0.004142545890653334 |
| interactor_ratio | mitosis | inter | 0.00021973308280332797 | 0.015168670877391028 |

P22626_Q9UMS4
ROA2_HUMAN vs PRP19_HUMAN
p-value: 0.051 q-value: 0.033

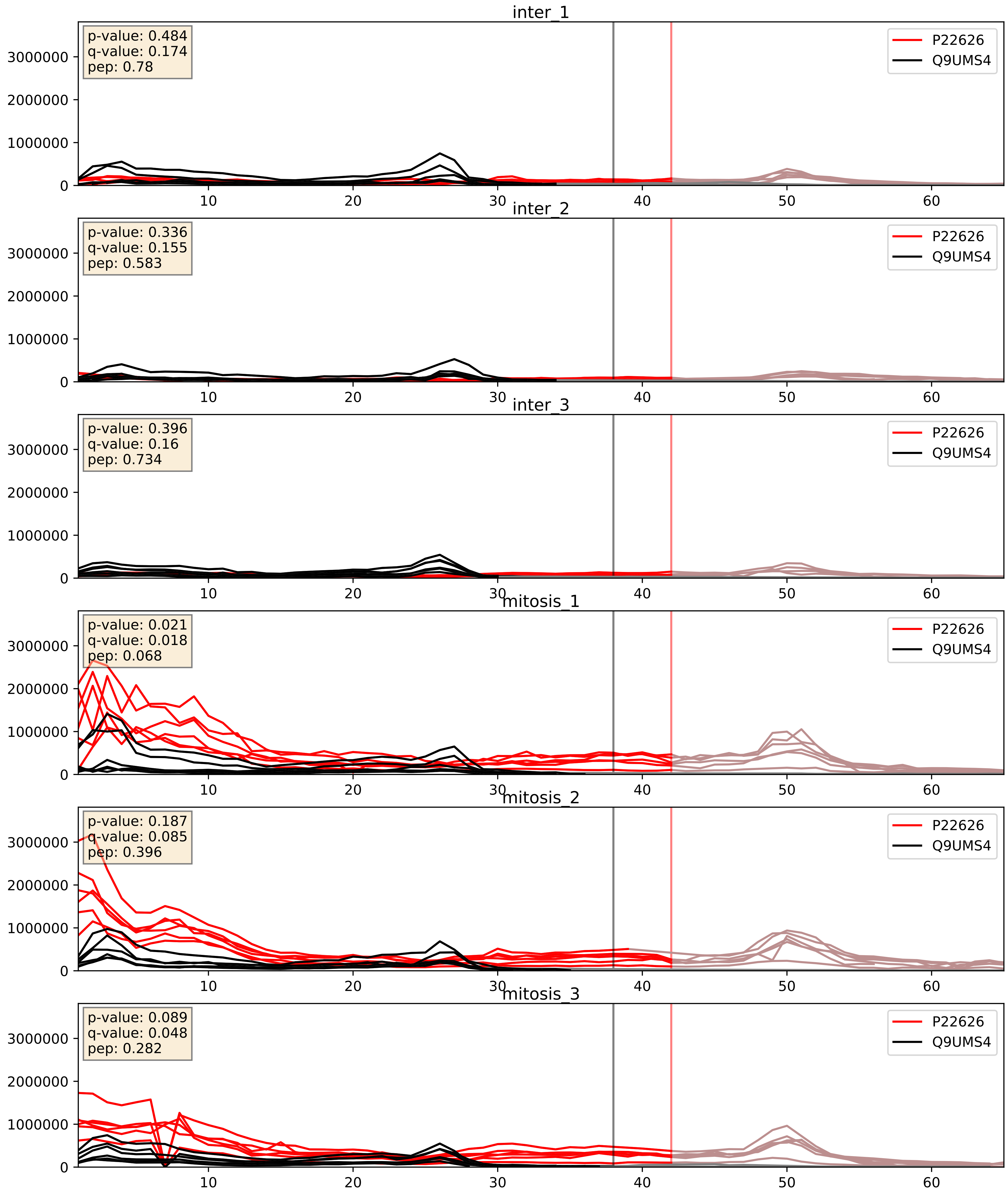


| level | condition_1 | condition_2 | pvalue | pvalue_adjusted |
|---|---|---|---|---|
| interactor_abundance | mitosis | inter | 0.0029005306755569248 | 0.040802863735032495 |
| complex_abundance | mitosis | inter | 0.0031675030266533457 | 0.041269141412713296 |
| interactor_ratio | mitosis | inter | 0.004793317480929459 | 0.09212312397066487 |
