## Supplementary Data 2 for "SECAT: Quantifying differential protein-protein interaction states by network-centric analysis": hela_string_000048_P62306.pdf

P62306_P62306

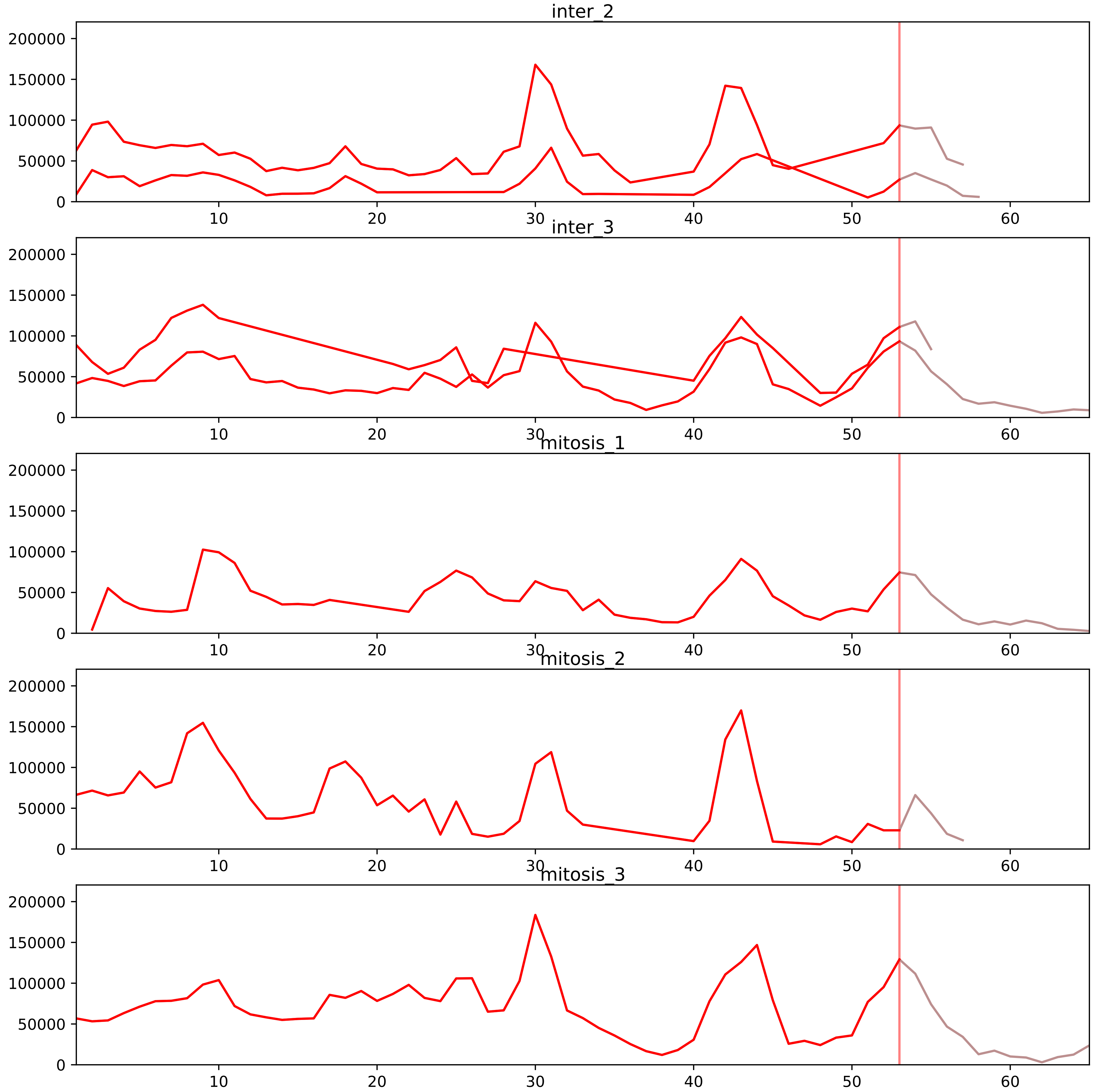


| level | condition_1 | condition_2 | pvalue | pvalue_adjusted |
|---|---|---|---|---|
| interactor_ratio | mitosis | inter | 1.0750077767238586e-07 | 4.395587353715333e-06 |
| complex_abundance | mitosis | inter | 0.03511770299781572 | 0.09941011310150911 |
| interactor_abundance | mitosis | inter | 0.10600403103999607 | 0.2355645134222135 |
| total_abundance | mitosis | inter | 0.6310445487946666 | 0.8348876813704906 |
| assembled_abundance | mitosis | inter | 0.6427882695008473 | 0.8380560544698799 |
| monomer_abundance | mitosis | inter | 0.7287897941939578 | 0.8860182327378967 |

P08579_P62306
RU2B_HUMAN vs RUXF_HUMAN
p-value: 0.017 q-value: 0.014

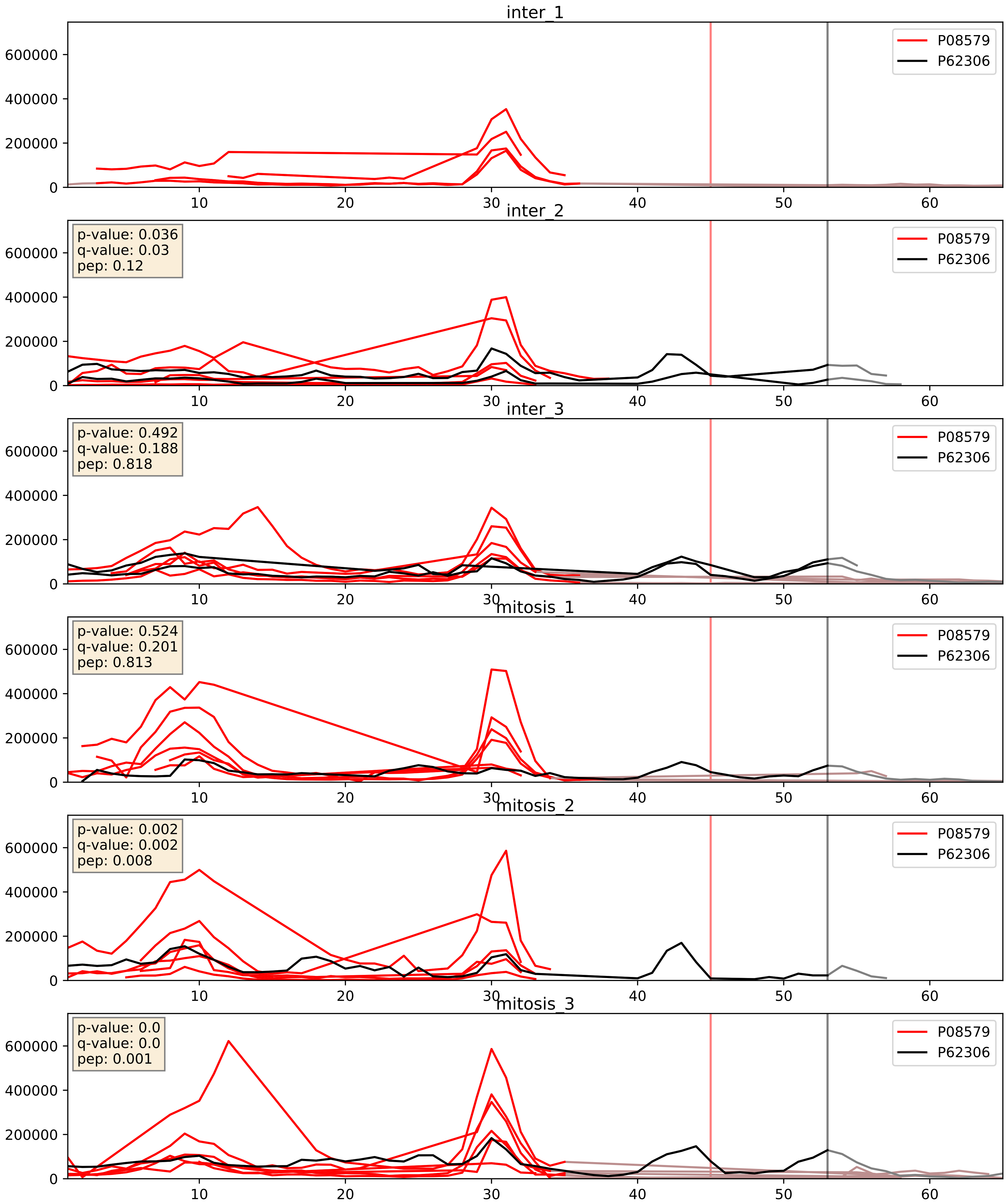


| level | condition_1 | condition_2 | pvalue | pvalue_adjusted |
|---|---|---|---|---|
| interactor_ratio | mitosis | inter | 0.02755407519938913 | 0.2597608851396156 |
| interactor_abundance | mitosis | inter | 0.28440697634059425 | 0.4784643835424518 |
| complex_abundance | mitosis | inter | 0.6120203817518135 | 0.733533249481311 |

P14866_P62306
HNRPL_HUMAN vs RUXF_HUMAN
p-value: 0.069 q-value: 0.042

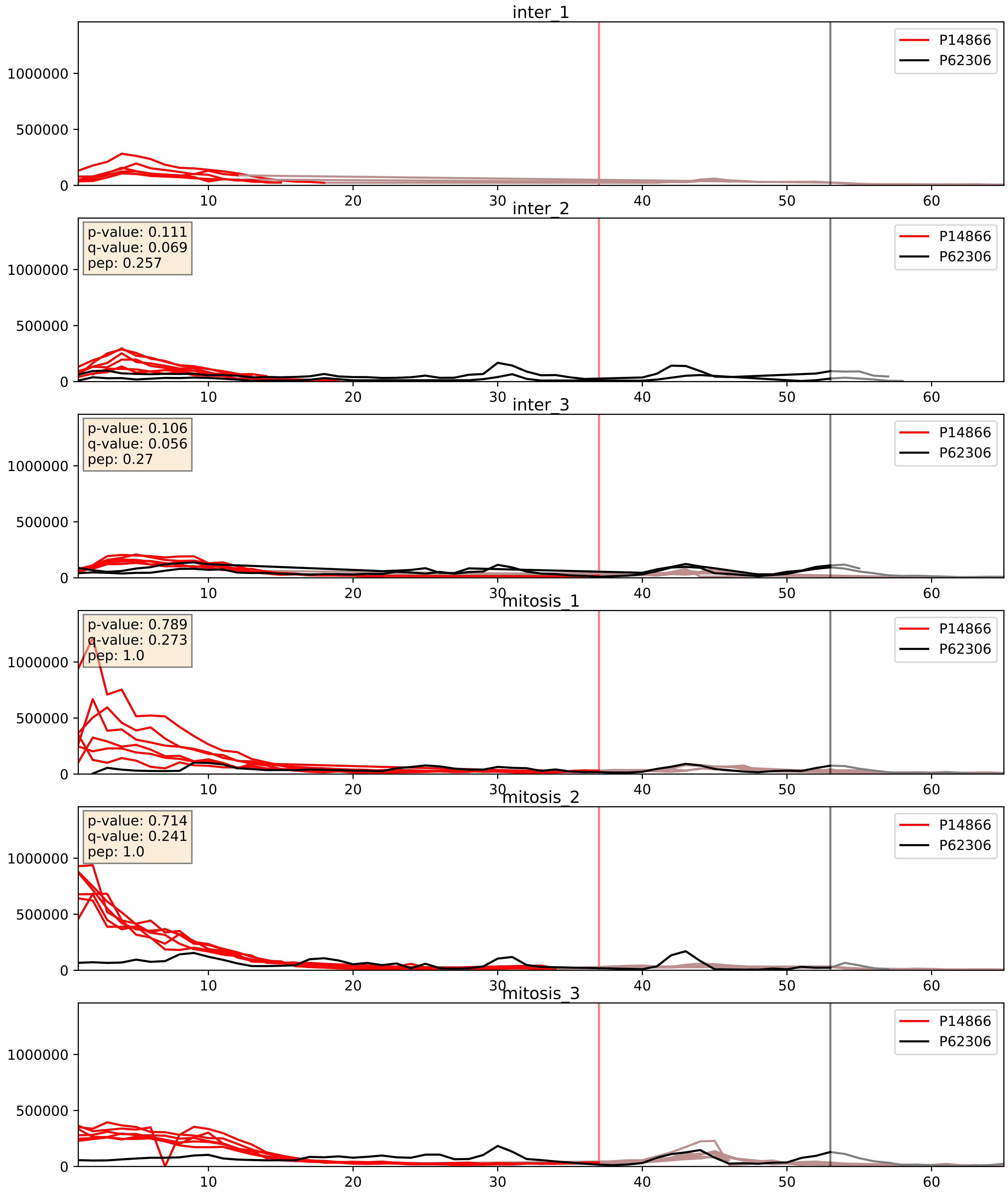


| level | condition_1 | condition_2 | pvalue | pvalue_adjusted |
|---|---|---|---|---|
| interactor_ratio | mitosis | inter | 0.01297337334516381 | 0.16452159382904033 |
| interactor_abundance | mitosis | inter | 0.10508626811479461 | 0.2769210432287962 |
| complex_abundance | mitosis | inter | 0.2692733160070283 | 0.41878262809232597 |

P62304_P62306
RUXE_HUMAN vs RUXF_HUMAN
p-value: 0.0 q-value: 0.0

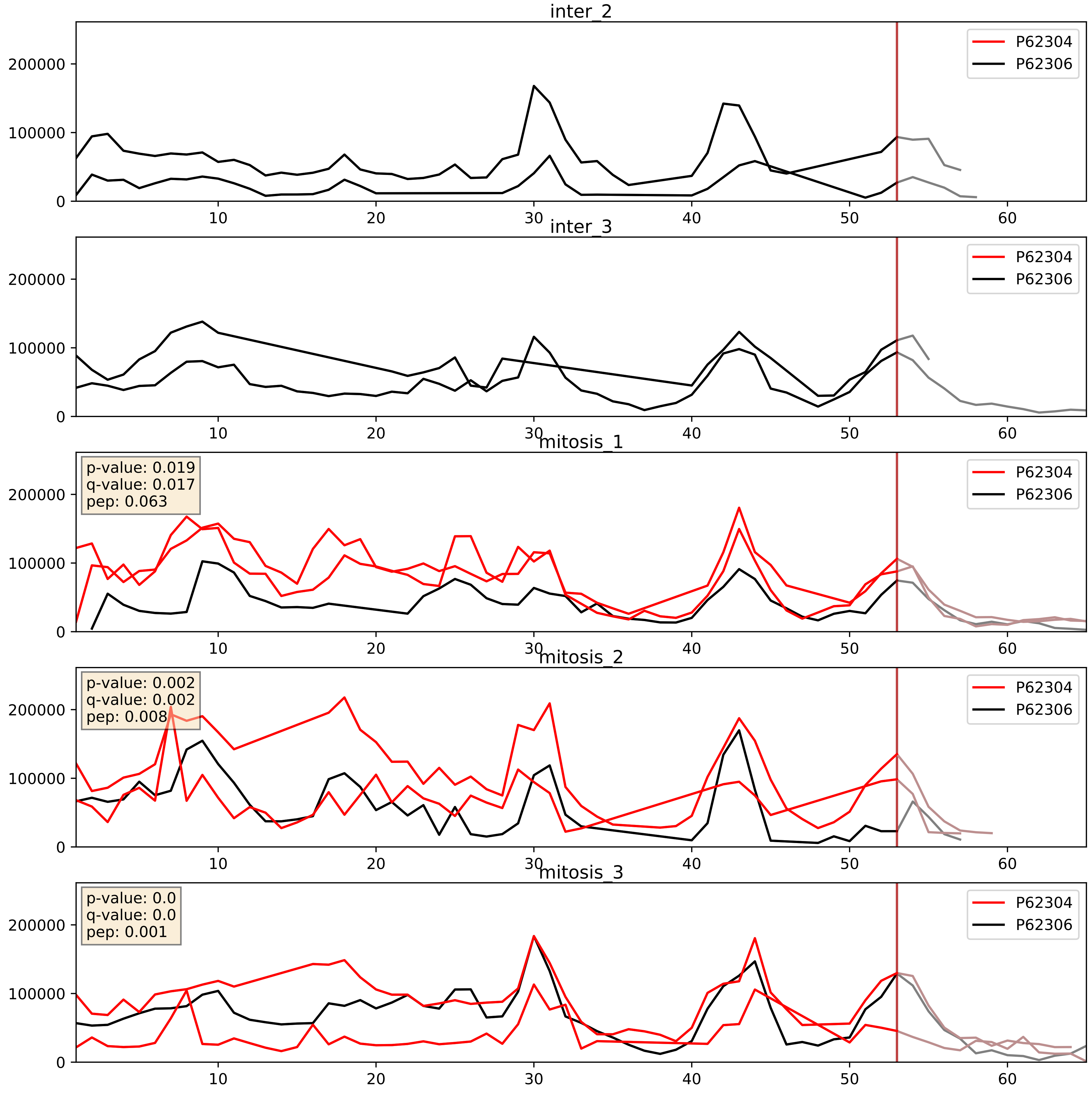


| level | condition_1 | condition_2 | pvalue | pvalue_adjusted |
|---|---|---|---|---|
| complex_abundance | mitosis | inter | 1.9058781942551378e-05 | 0.005691857151652958 |
| interactor_abundance | mitosis | inter | 5.613578522364216e-05 | 0.006414334379145961 |
| interactor_ratio | mitosis | inter | 0.0016271261094656015 | 0.048854197059354385 |

P62306_P62314
RUXF_HUMAN vs SMD1_HUMAN
p-value: 0.0 q-value: 0.001

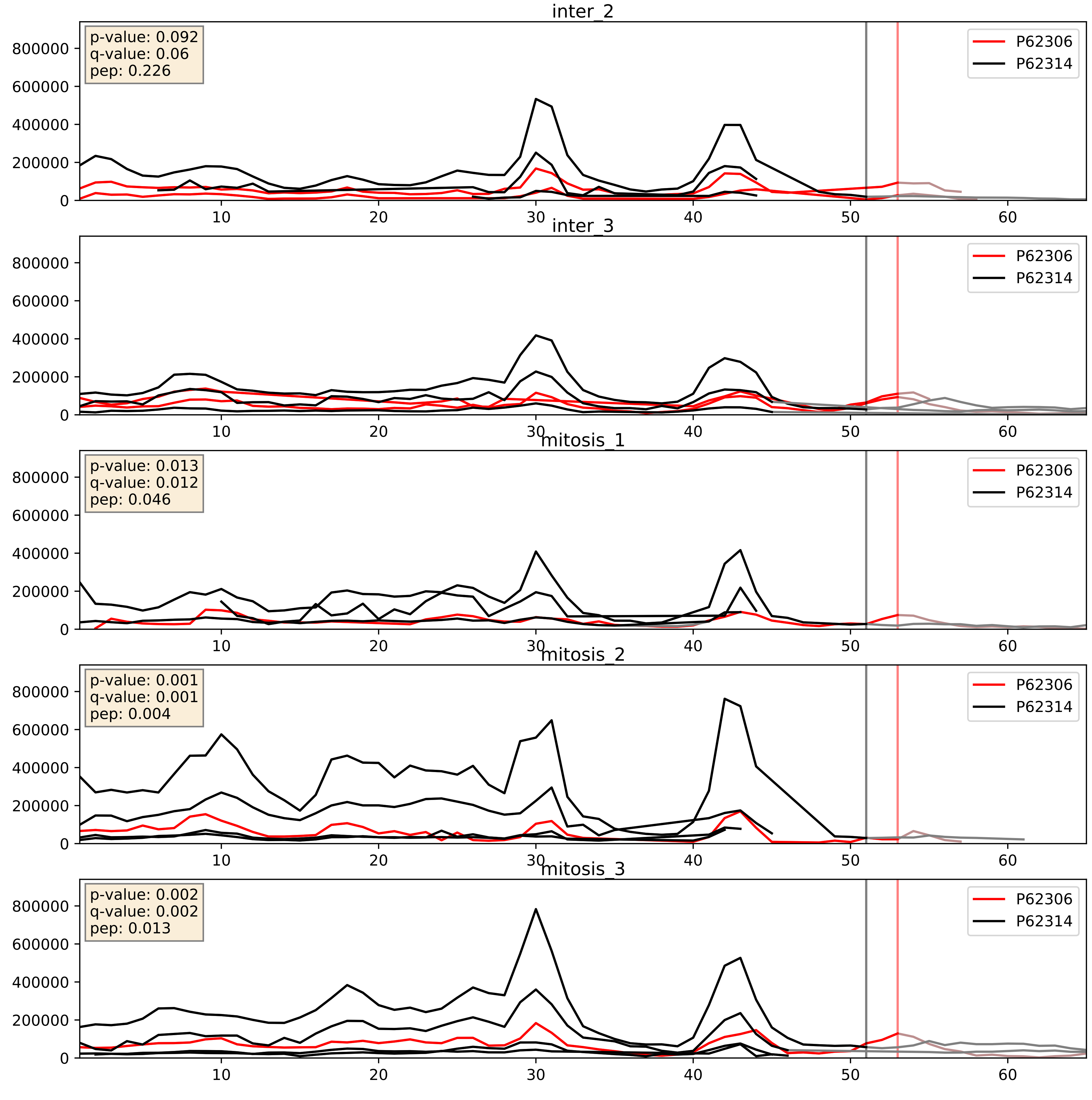


| level | condition_1 | condition_2 | pvalue | pvalue_adjusted |
|---|---|---|---|---|
| interactor_ratio | mitosis | inter | 0.06715288545104708 | 0.43118262177163247 |
| interactor_abundance | mitosis | inter | 0.4772754767470721 | 0.6486945190465128 |
| complex_abundance | mitosis | inter | 0.6946400609259482 | 0.7941924564613486 |

P62306_P62316
RUXF_HUMAN vs SMD2_HUMAN
p-value: 0.0 q-value: 0.0

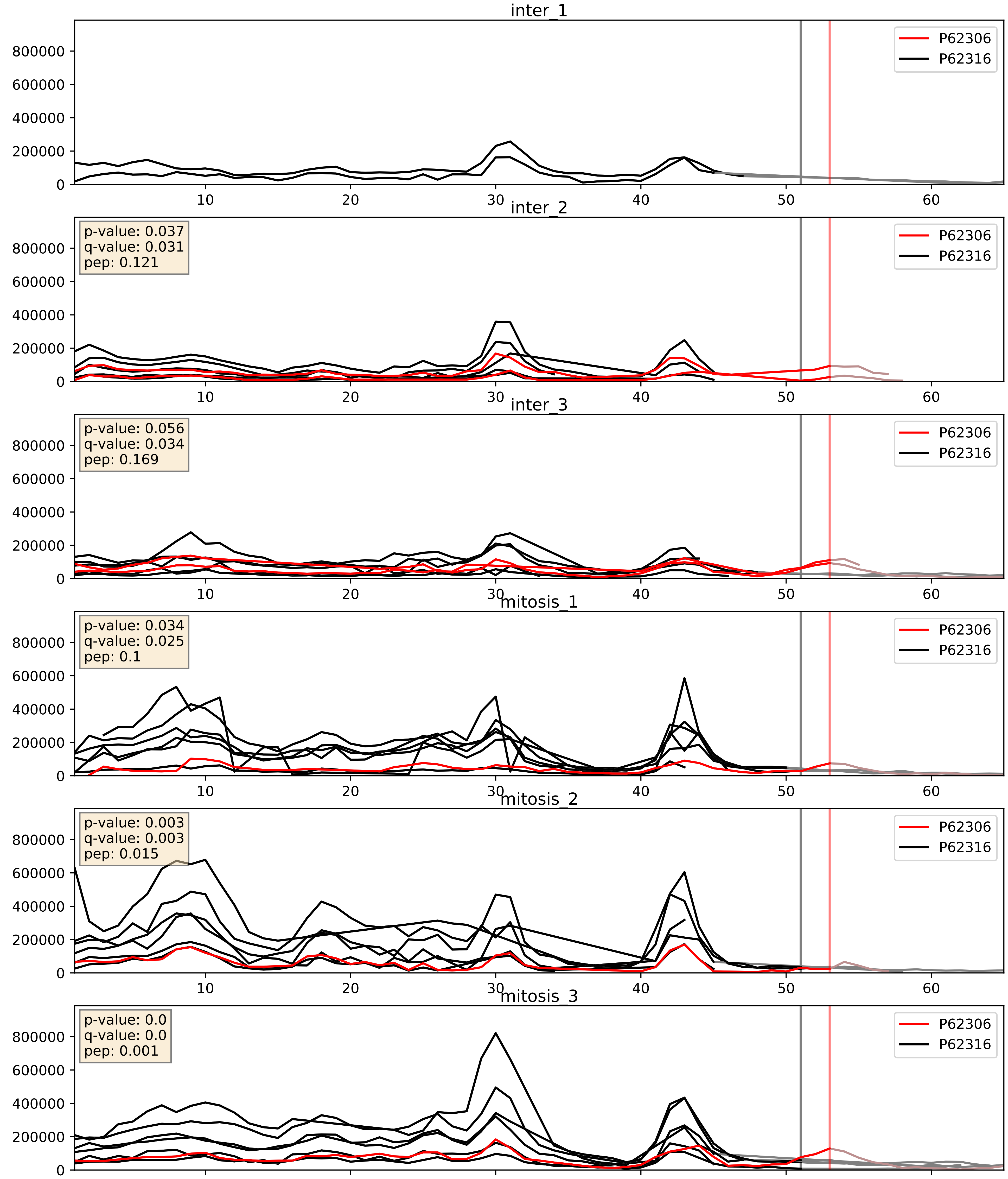


| level | condition_1 | condition_2 | pvalue | pvalue_adjusted |
|---|---|---|---|---|
| interactor_ratio | mitosis | inter | 0.00031290593502055766 | 0.018787491549285 96 |
| complex_abundance | mitosis | inter | 0.18206089764979508 | 0.33292913562961884 |
| interactor_abundance | mitosis | inter | 0.4800162238922761 | 0.6514886438113023 |

P62306_P62318
RUXF_HUMAN vs SMD3_HUMAN
p-value: 0.03 q-value: 0.022

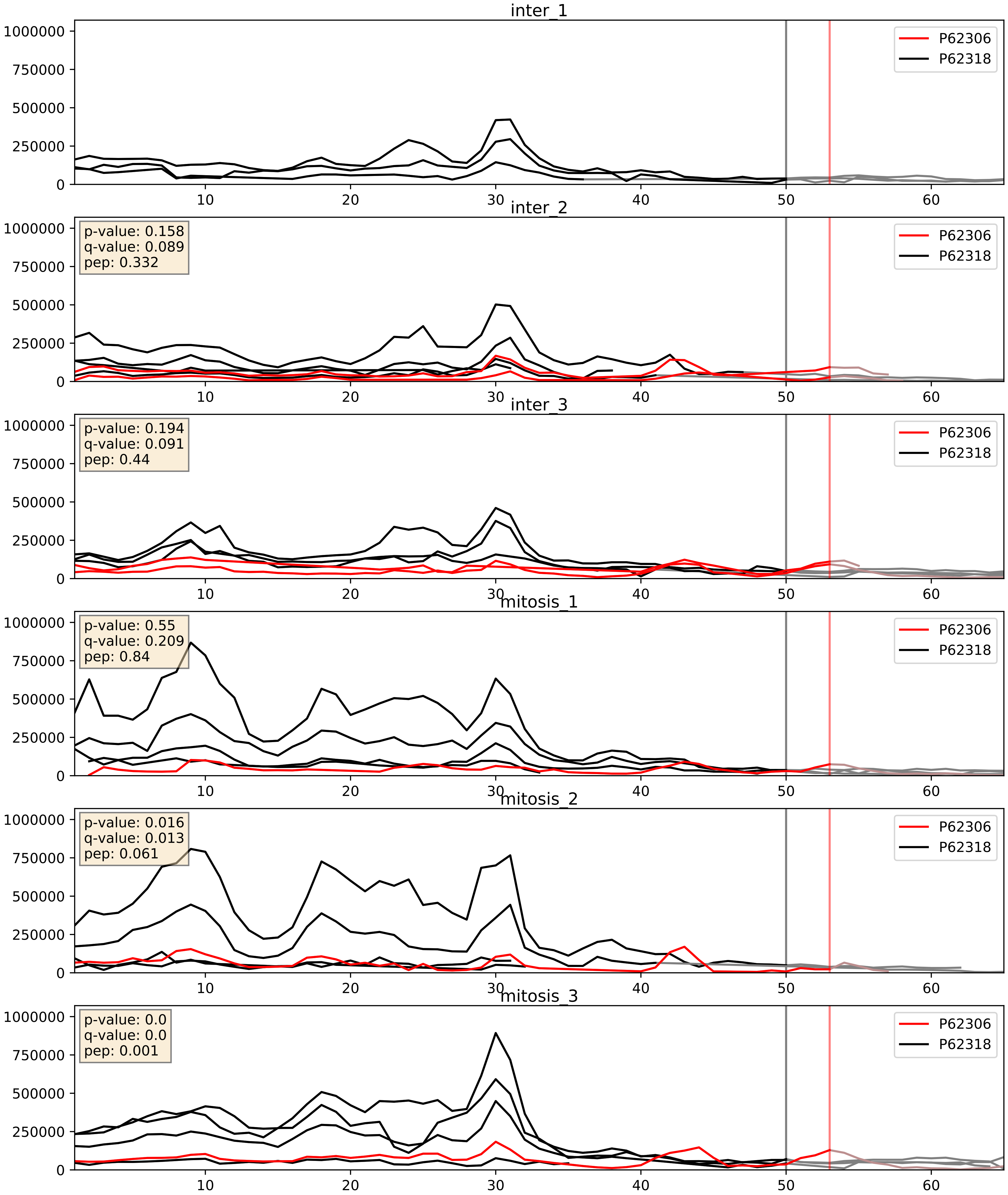


| level | condition_1 | condition_2 | pvalue | pvalue_adjusted |
|---|---|---|---|---|
| interactor_ratio | mitosis | inter | 0.12209918771763113 | 0.5708187039120276 |
| interactor_abundance | mitosis | inter | 0.4793717894680107 | 0.6509752546753663 |
| complex_abundance | mitosis | inter | 0.6668080091316944 | 0.7782580688435843 |

P62306_Q15428
RUXF_HUMAN vs SF3A2_HUMAN
p-value: 0.063 q-value: 0.039

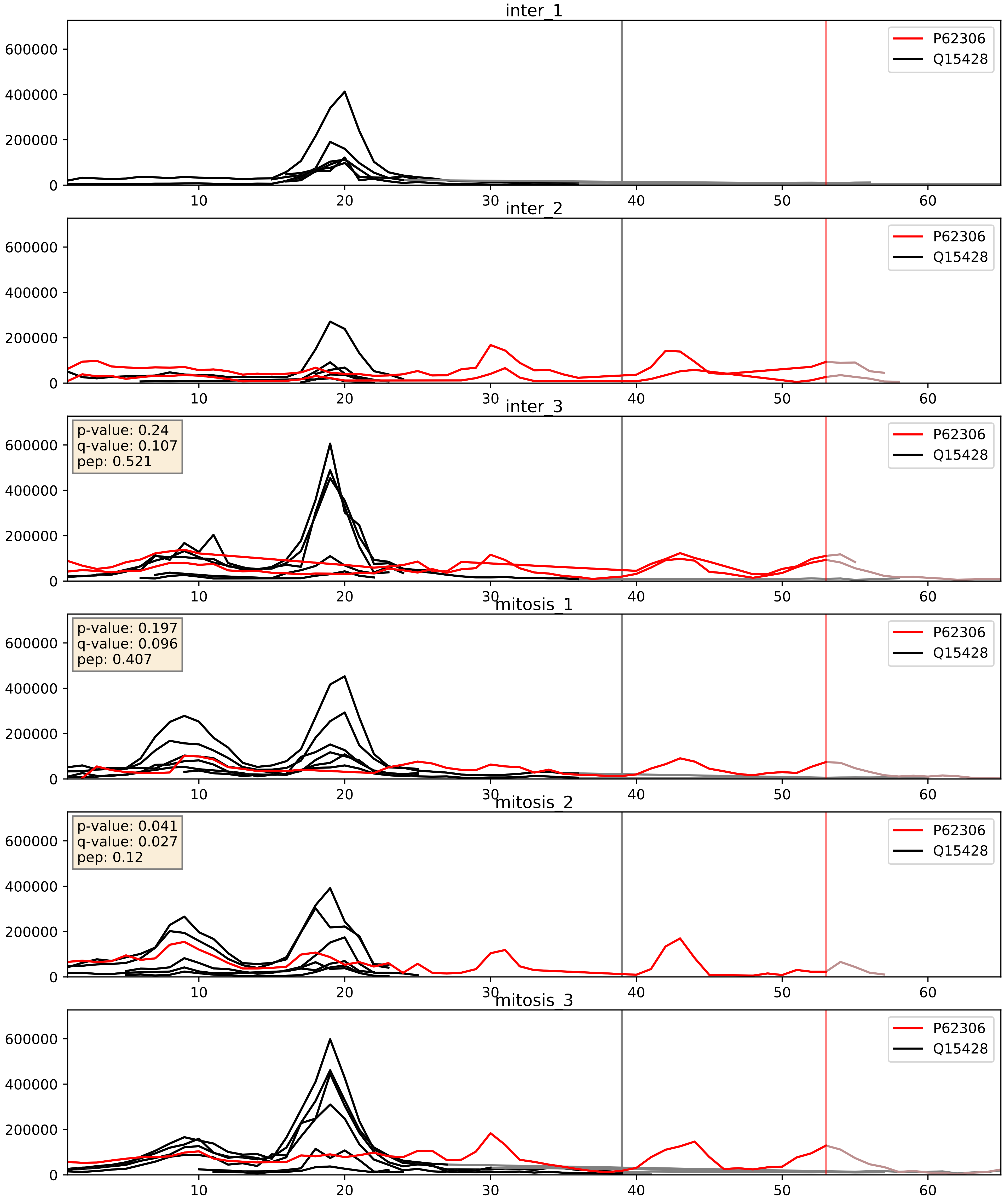


| level | condition_1 | condition_2 | pvalue | pvalue_adjusted |
|---|---|---|---|---|
| interactor_ratio | mitosis | inter | 0.05627708430605203 | 0.3926647223857623 |
| interactor_abundance | mitosis | inter | 0.5441496332187279 | 0.7009135295090378 |
| complex_abundance | mitosis | inter | 0.5509767597282238 | 0.6846220152814045 |
