## Supplementary Data 2 for "SECAT: Quantifying differential protein-protein interaction states by network-centric analysis": hela_string_000049_P17844.pdf

P17844_P17844

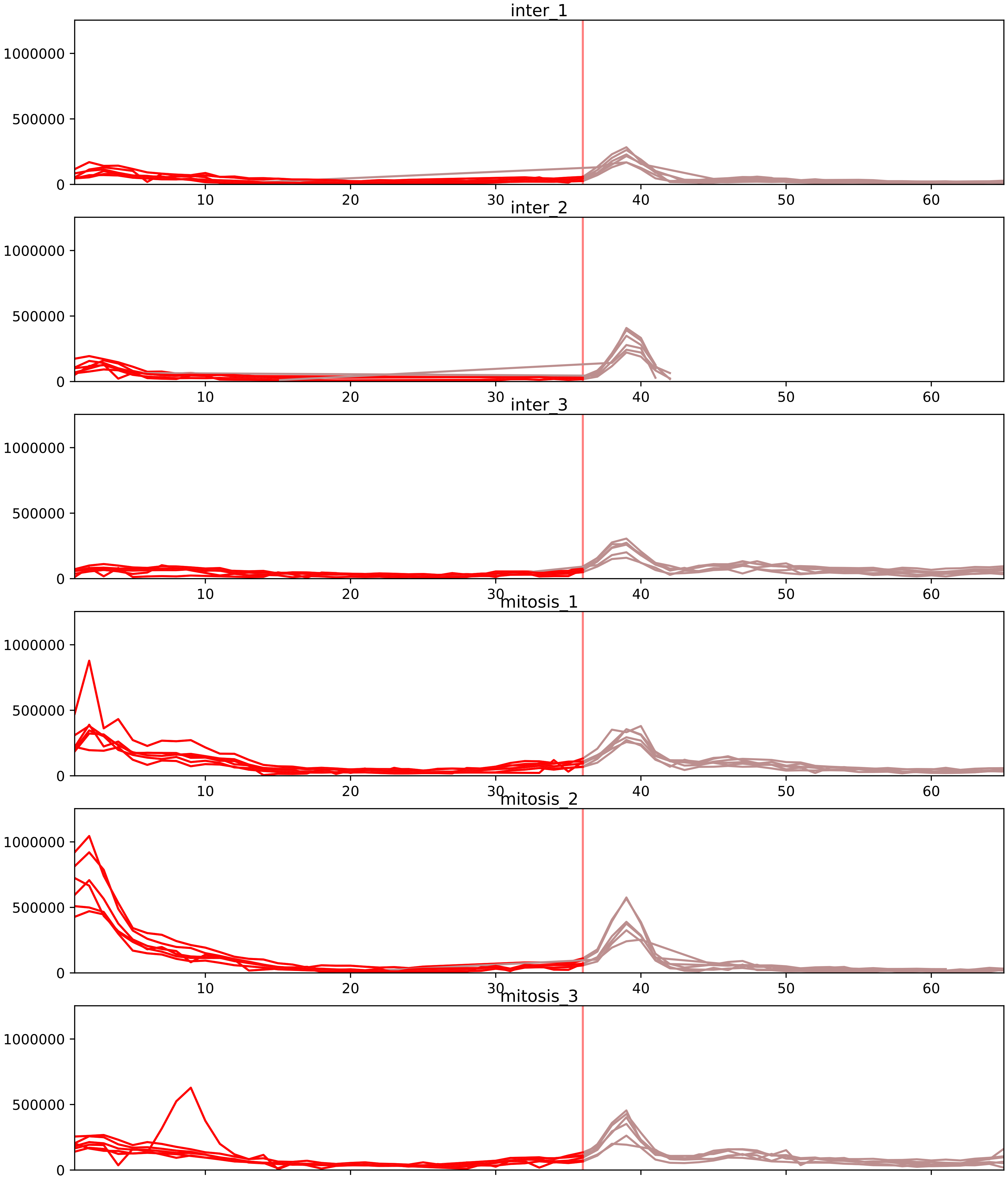


| level | condition_1 | condition_2 | pvalue | pvalue_adjusted |
|---|---|---|---|---|
| interactor_ratio | mitosis | inter | 1.3282135748001743e-07 | 5.312854299200697e-06 |
| interactor_abundance | mitosis | inter | 1.1761939061874725e-06 | 0.00013353840650924488 |
| complex_abundance | mitosis | inter | 1.7550642199007419e-06 | 0.00014678718930078932 |
| assembled_abundance | mitosis | inter | 0.003916771717355212 | 0.06212029154760582 |
| total_abundance | mitosis | inter | 0.017124185641070974 | 0.1657399792750805 |
| monomer_abundance | mitosis | inter | 0.14643285824768854 | 0.504947747149949 |

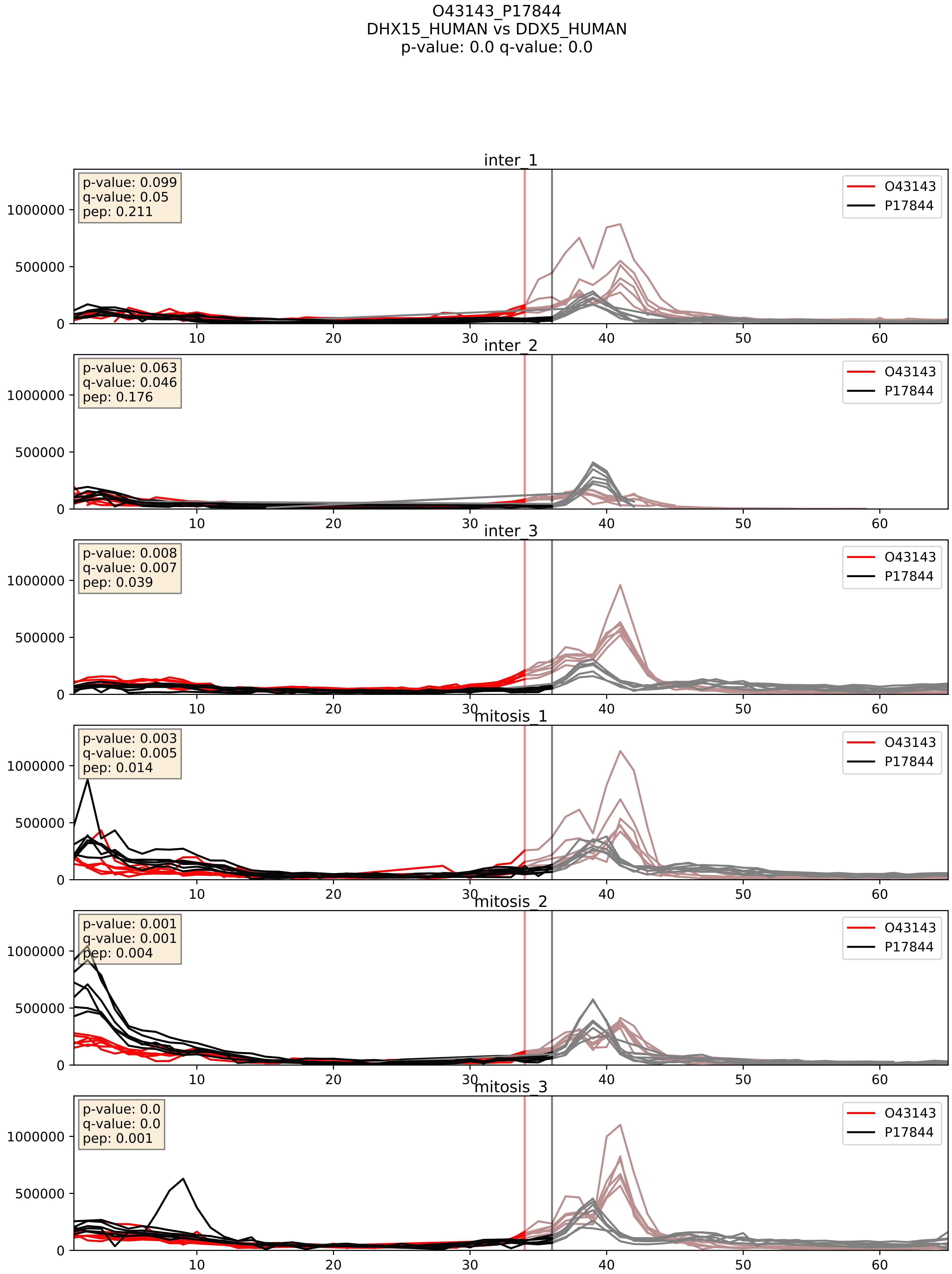


| level | condition_1 | condition_2 | pvalue | pvalue_adjusted |
|---|---|---|---|---|
| complex_abundance | mitosis | inter | 0.021707386299522128 | 0.10853693149761064 |
| interactor_ratio | mitosis | inter | 0.06299601081169943 | 0.4167278613200519 |
| interactor_abundance | mitosis | inter | 0.29892011664777984 | 0.49344303591965044 |

P07910_P17844
HNRPC_HUMAN vs DDX5_HUMAN
p-value: 0.002 q-value: 0.034

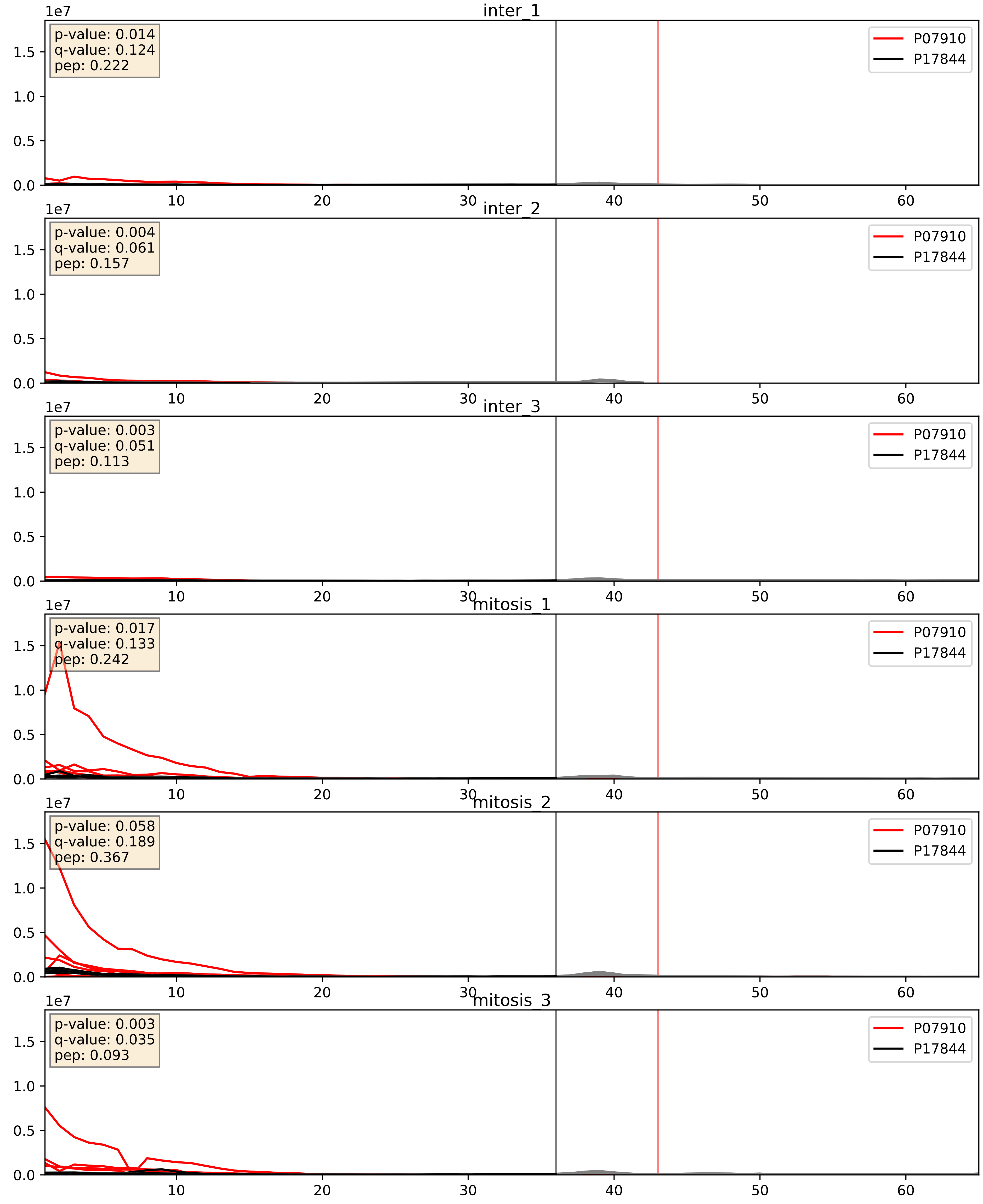


| level | condition_1 | condition_2 | pvalue | pvalue_adjusted |
|---|---|---|---|---|
| complex_abundance | mitosis | inter | 8.901605986839528e-06 | 0.005691857151652958 |
| interactor_abundance | mitosis | inter | 1.005842017873748e-05 | 0.0033734327106189023 |
| interactor_ratio | mitosis | inter | 0.00013562784083812567 | 0.012616771133069759 |

P09651_P17844
ROA1_HUMAN vs DDX5_HUMAN
p-value: 0.001 q-value: 0.015

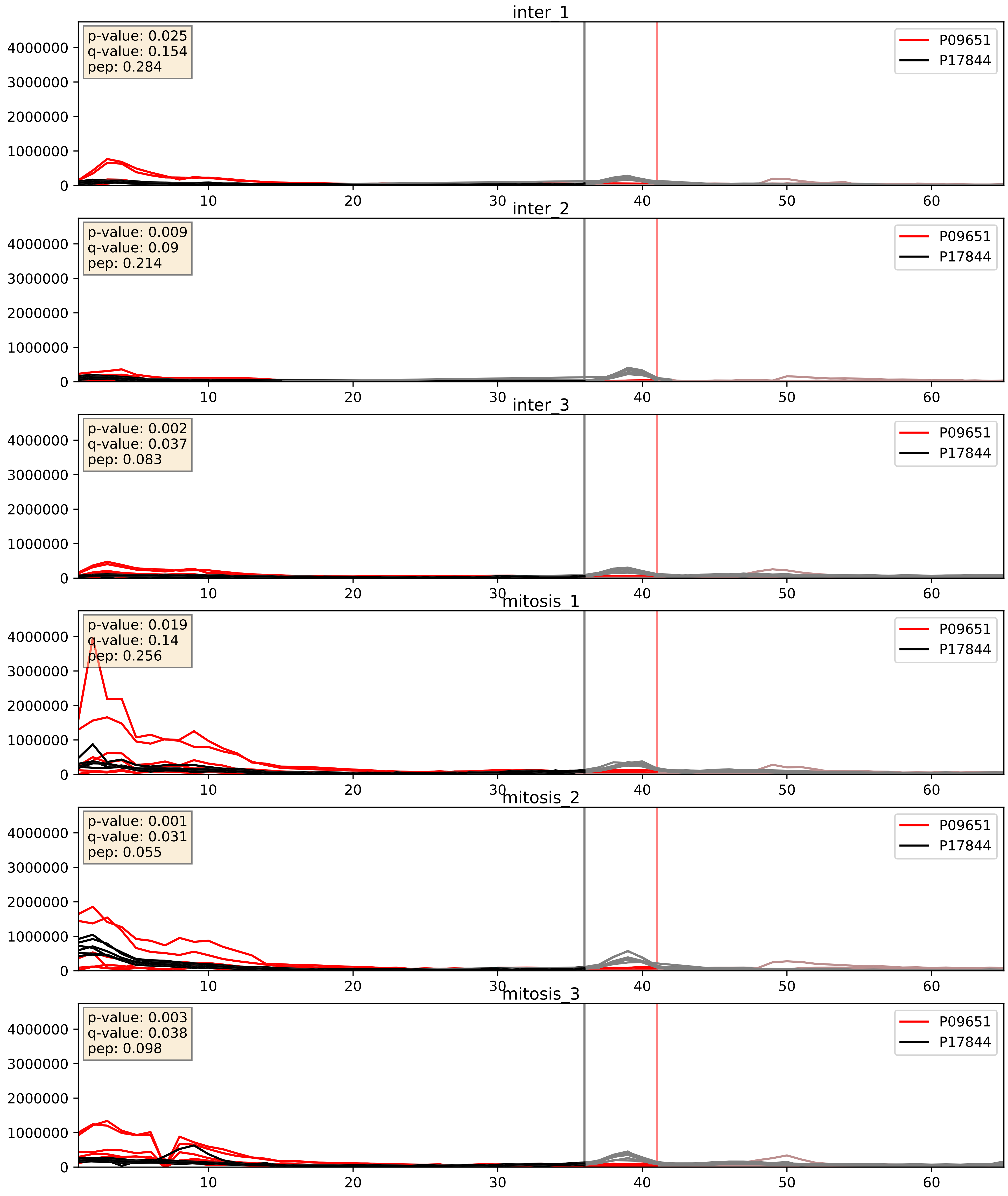


| level | condition_1 | condition_2 | pvalue | pvalue_adjusted |
|---|---|---|---|---|
| complex_abundance | mitosis | inter | 0.005592113315961333 | 0.056382202573179045 |
| interactor_abundance | mitosis | inter | 0.017017205188648715 | 0.11334462047152971 |
| interactor_ratio | mitosis | inter | 0.1872358107783095 | 0.657442464198634 |

P11940_P17844
PABP1_HUMAN vs DDX5_HUMAN
p-value: 0.0 q-value: 0.016

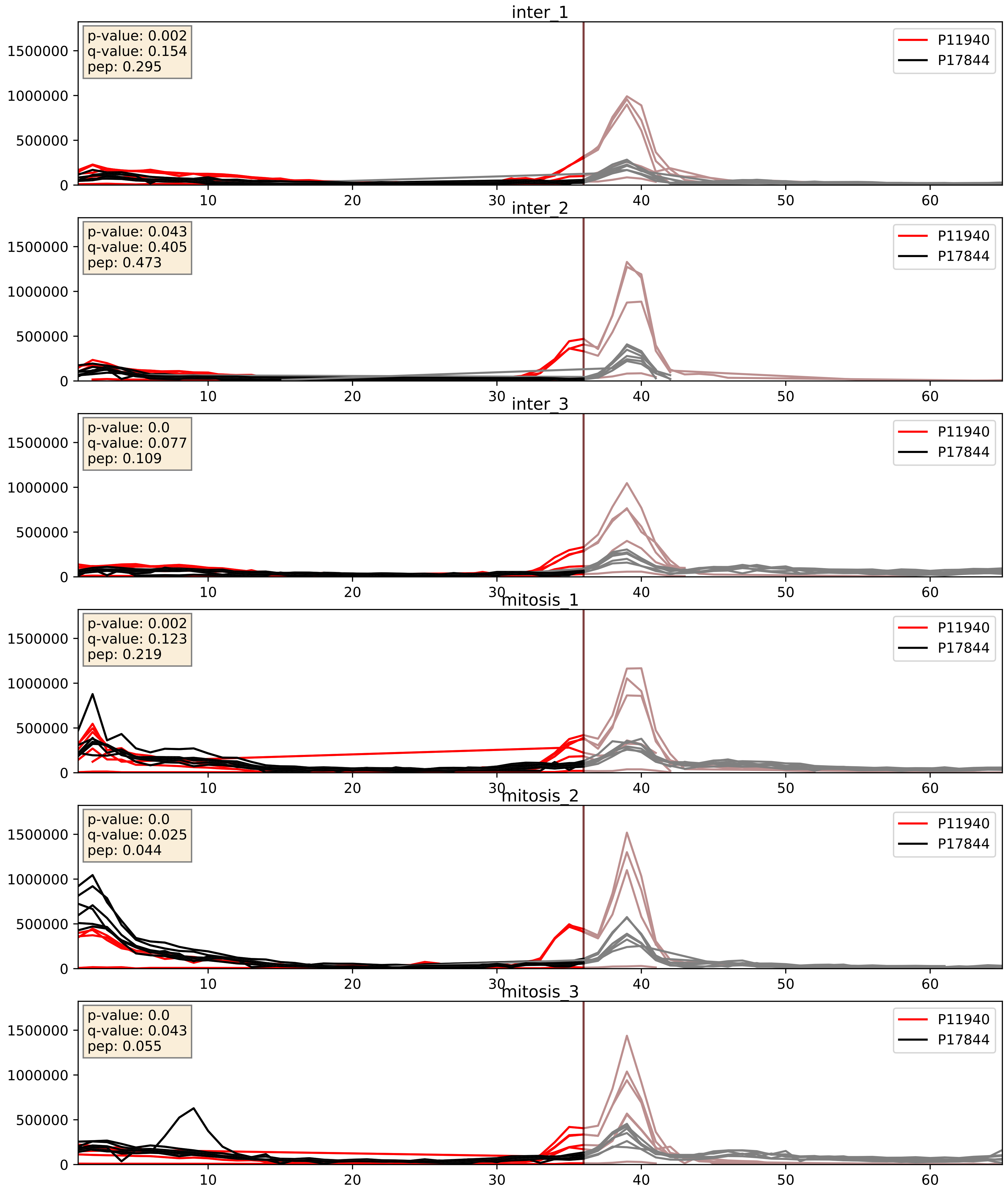


| level | condition_1 | condition_2 | pvalue | pvalue_adjusted |
|---|---|---|---|---|
| complex_abundance | mitosis | inter | 0.003636591319936893 | 0.04428054295684183 |
| interactor_ratio | mitosis | inter | 0.006113461516559377 | 0.10745632562987323 |
| interactor_abundance | mitosis | inter | 0.13490704099728298 | 0.3215381514511325 |

P14866_P17844
HNRPL_HUMAN vs DDX5_HUMAN
p-value: 0.0 q-value: 0.01

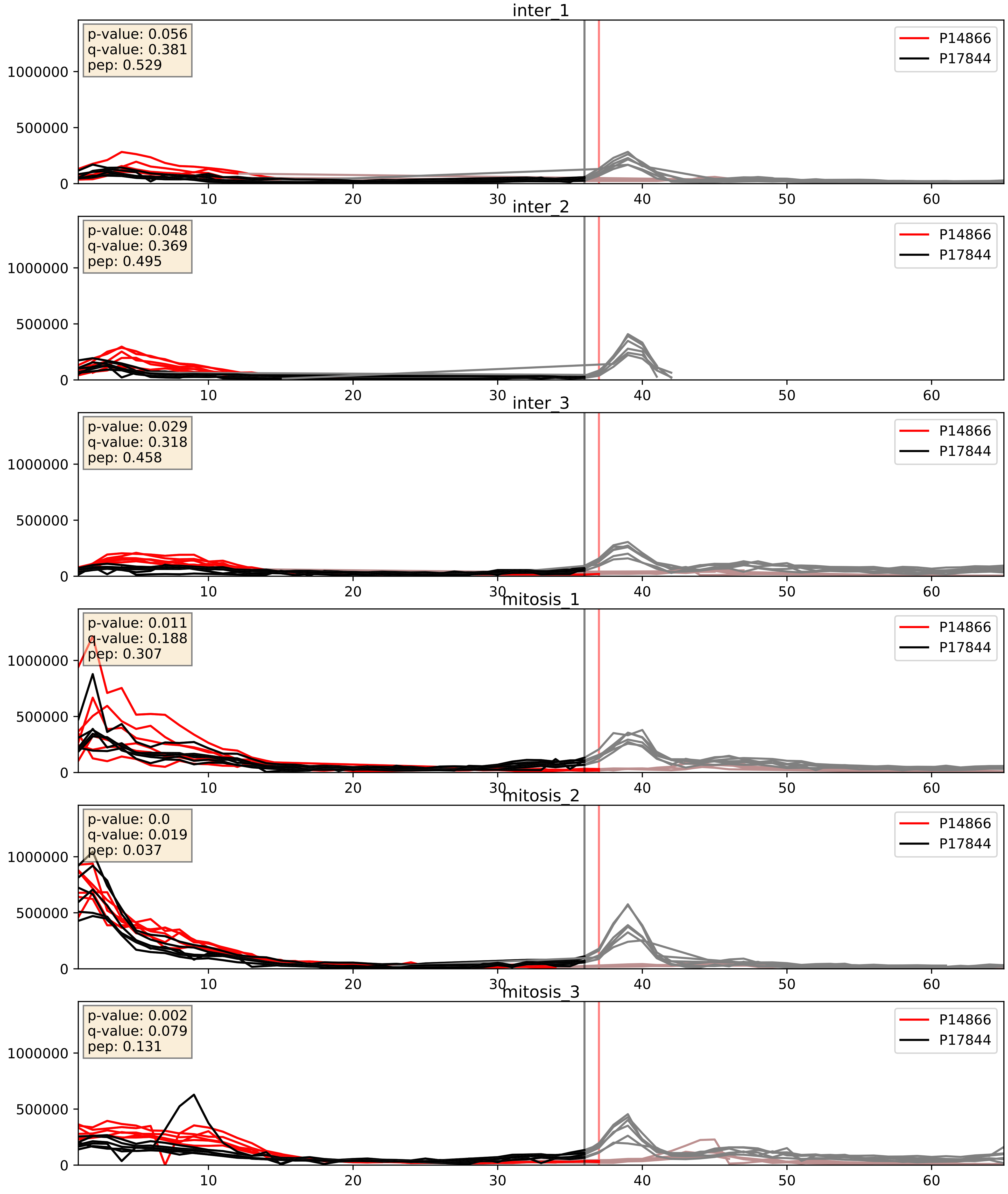


| level | condition_1 | condition_2 | pvalue | pvalue_adjusted |
|---|---|---|---|---|
| complex_abundance | mitosis | inter | 0.0013977644692216144 | 0.025565948411403885 |
| interactor_abundance | mitosis | inter | 0.009741046732019873 | 0.08547756025227074 |
| interactor_ratio | mitosis | inter | 0.7081208830132038 | 0.9391051699295014 |

P17844_P22626
DDX5_HUMAN vs ROA2_HUMAN
p-value: 0.002 q-value: 0.033

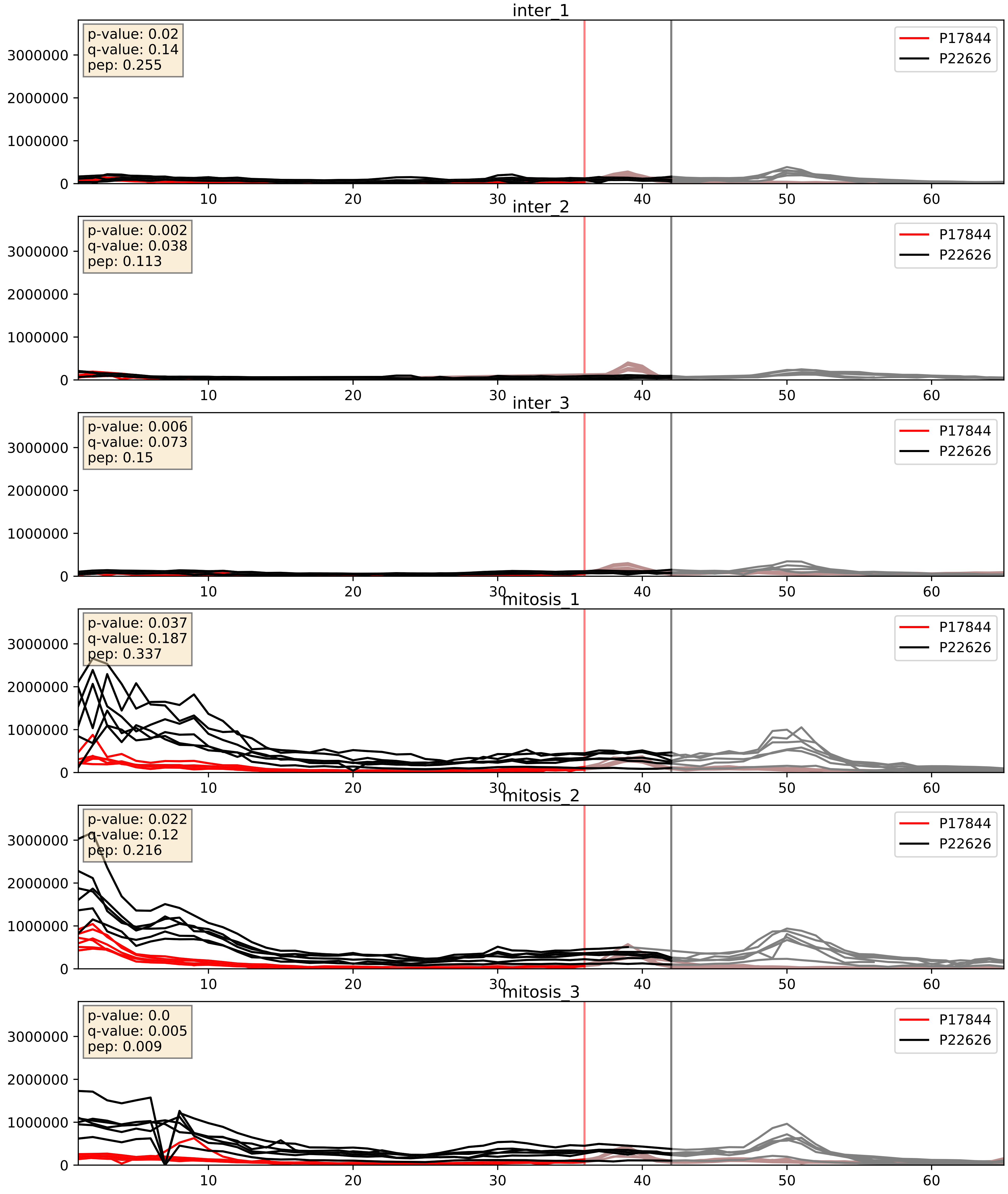


| level | condition_1 | condition_2 | pvalue | pvalue_adjusted |
|---|---|---|---|---|
| complex_abundance | mitosis | inter | 0.0004483662606115384 | 0.013372875229389437 |
| interactor_abundance | mitosis | inter | 0.0021426459467421676 | 0.033870820506210444 |
| interactor_ratio | mitosis | inter | 0.017003414810485822 | 0.1935495090129769 |

P17844_P31942
DDX5_HUMAN vs HNRH3_HUMAN
p-value: 0.0 q-value: 0.043

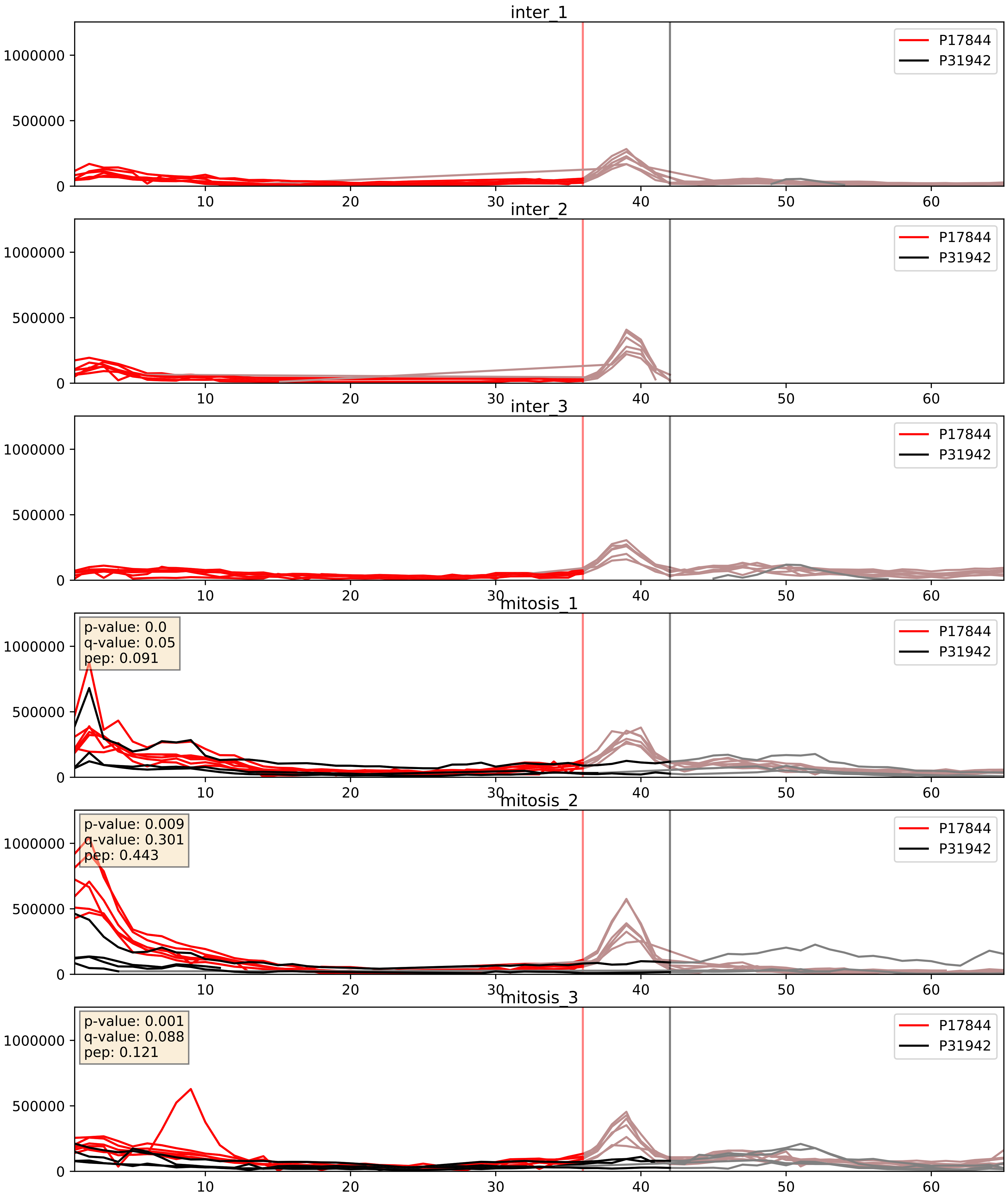


| level | condition_1 | condition_2 | pvalue | pvalue_adjusted |
|---|---|---|---|---|
| interactor_abundance | mitosis | inter | 2.8004503716417208e-05 | 0.004746902016089729 |
| complex_abundance | mitosis | inter | 5.883087896907052e-05 | 0.005691857151652958 |
| interactor_ratio | mitosis | inter | 0.016560873095316366 | 0.18977386036935487 |

P17844_P31943
DDX5_HUMAN vs HNRH1_HUMAN
p-value: 0.0 q-value: 0.0

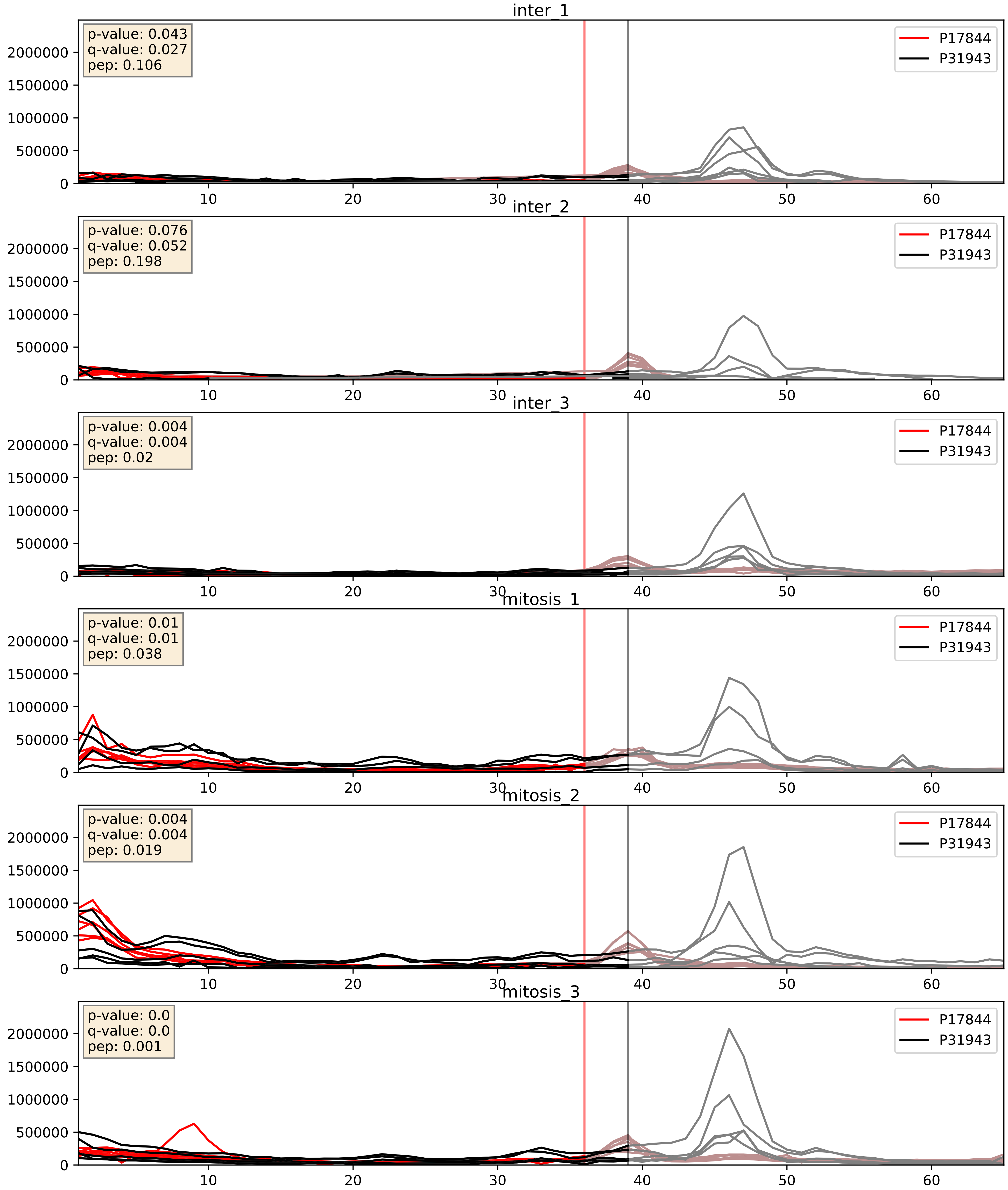


| level | condition_1 | condition_2 | pvalue | pvalue_adjusted |
|---|---|---|---|---|
| interactor_abundance | mitosis | inter | 0.0021426459467421676 | 0.033870820506210444 |
| complex_abundance | mitosis | inter | 0.0032360173229818456 | 0.04165459892439789 |
| interactor_ratio | mitosis | inter | 0.12617781079044613 | 0.5750062399685051 |

P17844_P38159
DDX5_HUMAN vs RBMX_HUMAN
p-value: 0.0 q-value: 0.003

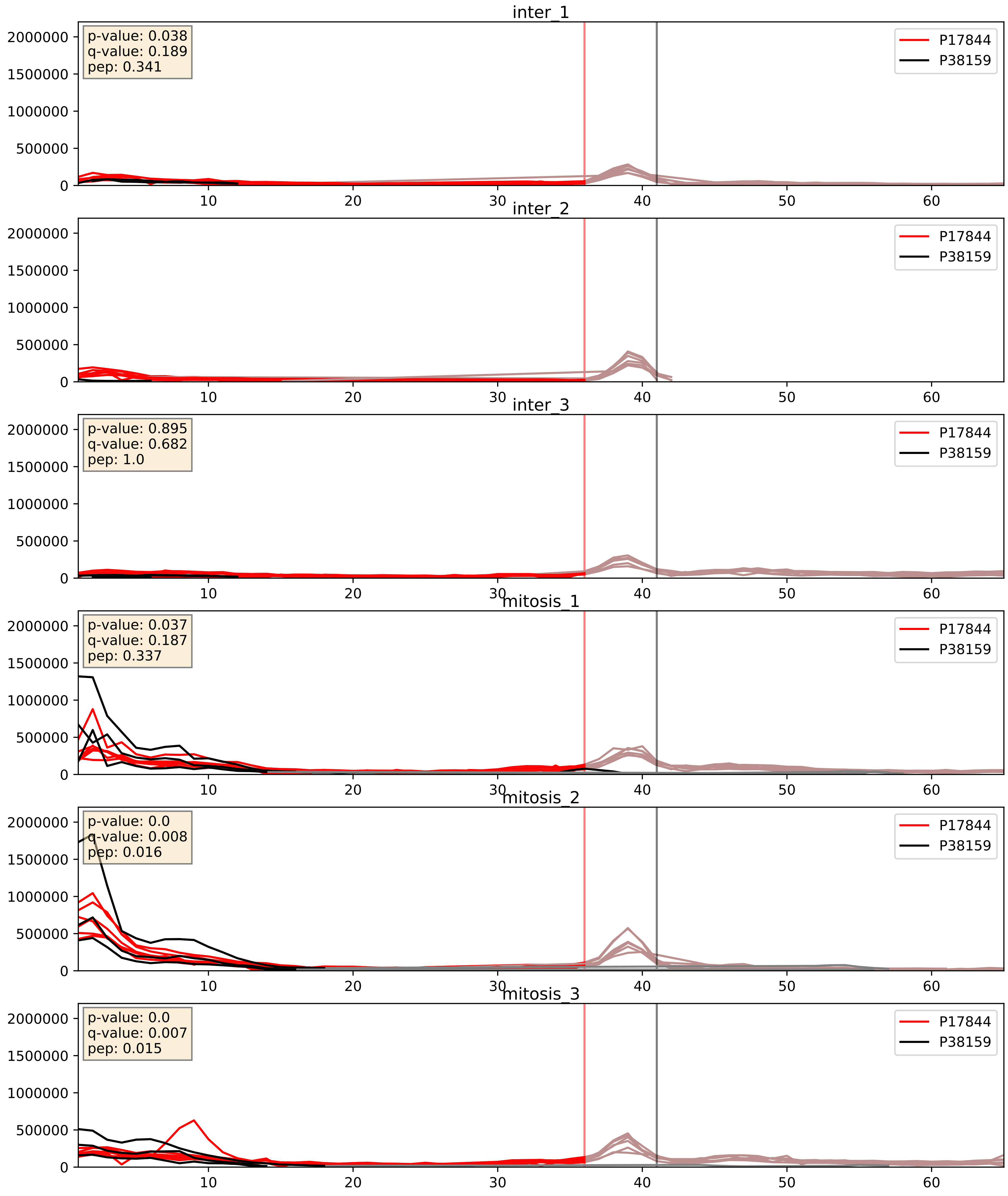


| level | condition_1 | condition_2 | pvalue | pvalue_adjusted |
|---|---|---|---|---|
| complex_abundance | mitosis | inter | 0.0010528687451826645 | 0.021612845224852777 |
| interactor_abundance | mitosis | inter | 0.0016025115124044603 | 0.029124200734994013 |
| interactor_ratio | mitosis | inter | 0.005020220566009392 | 0.09528400896904744 |

P17844_P42285
DDX5_HUMAN vs MTREX_HUMAN
p-value: 0.047 q-value: 0.031

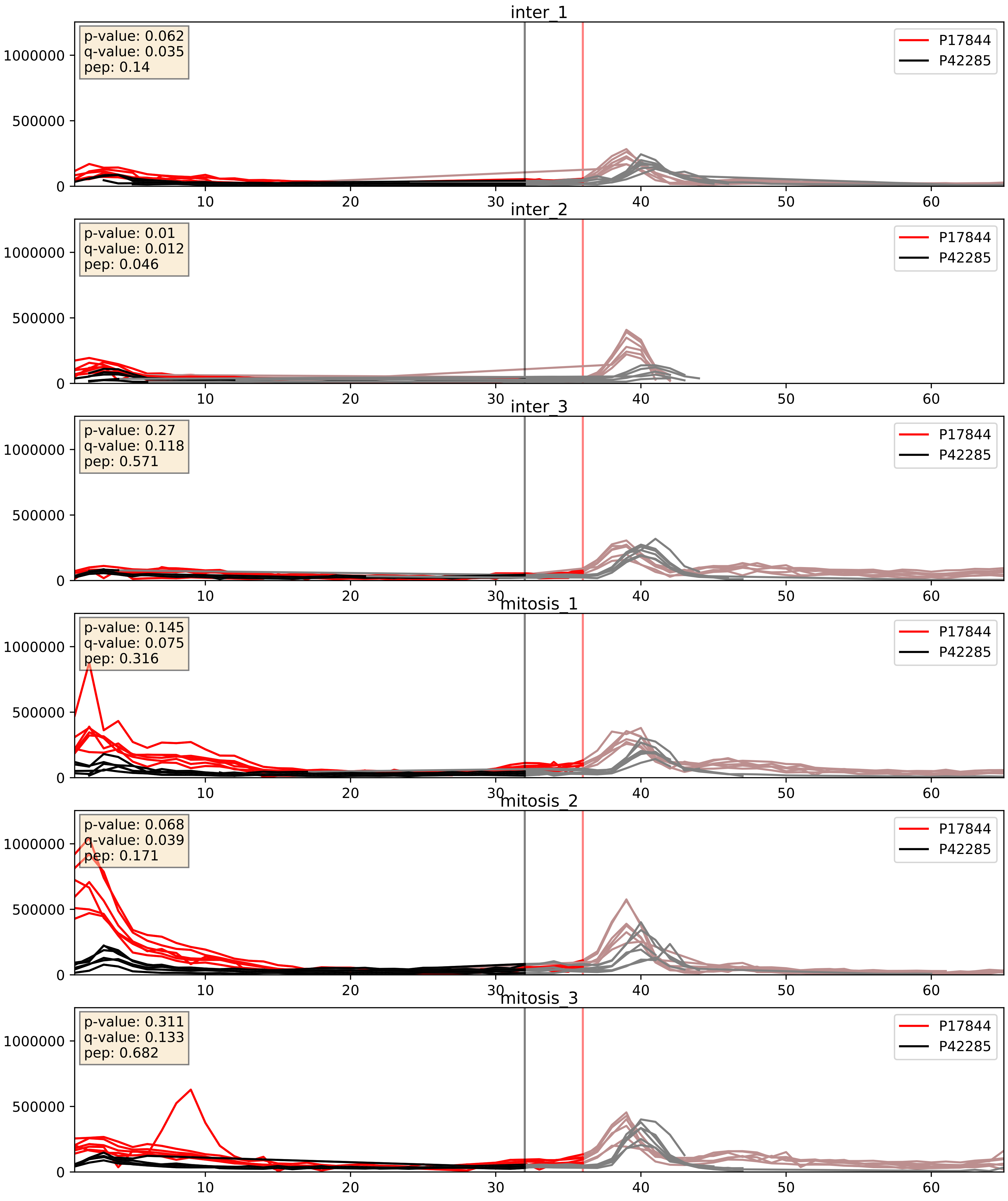


| level | condition_1 | condition_2 | pvalue | pvalue_adjusted |
|---|---|---|---|---|
| interactor_abundance | mitosis | inter | 0.0019402894813686398 | 0.032281589816356766 |
| complex_abundance | mitosis | inter | 0.0026967538514306786 | 0.03746658311311581 |
| interactor_ratio | mitosis | inter | 0.012520475632328332 | 0.1615861374955141 |

P17844_P52272
DDX5_HUMAN vs HNRPM_HUMAN
p-value: 0.0 q-value: 0.005

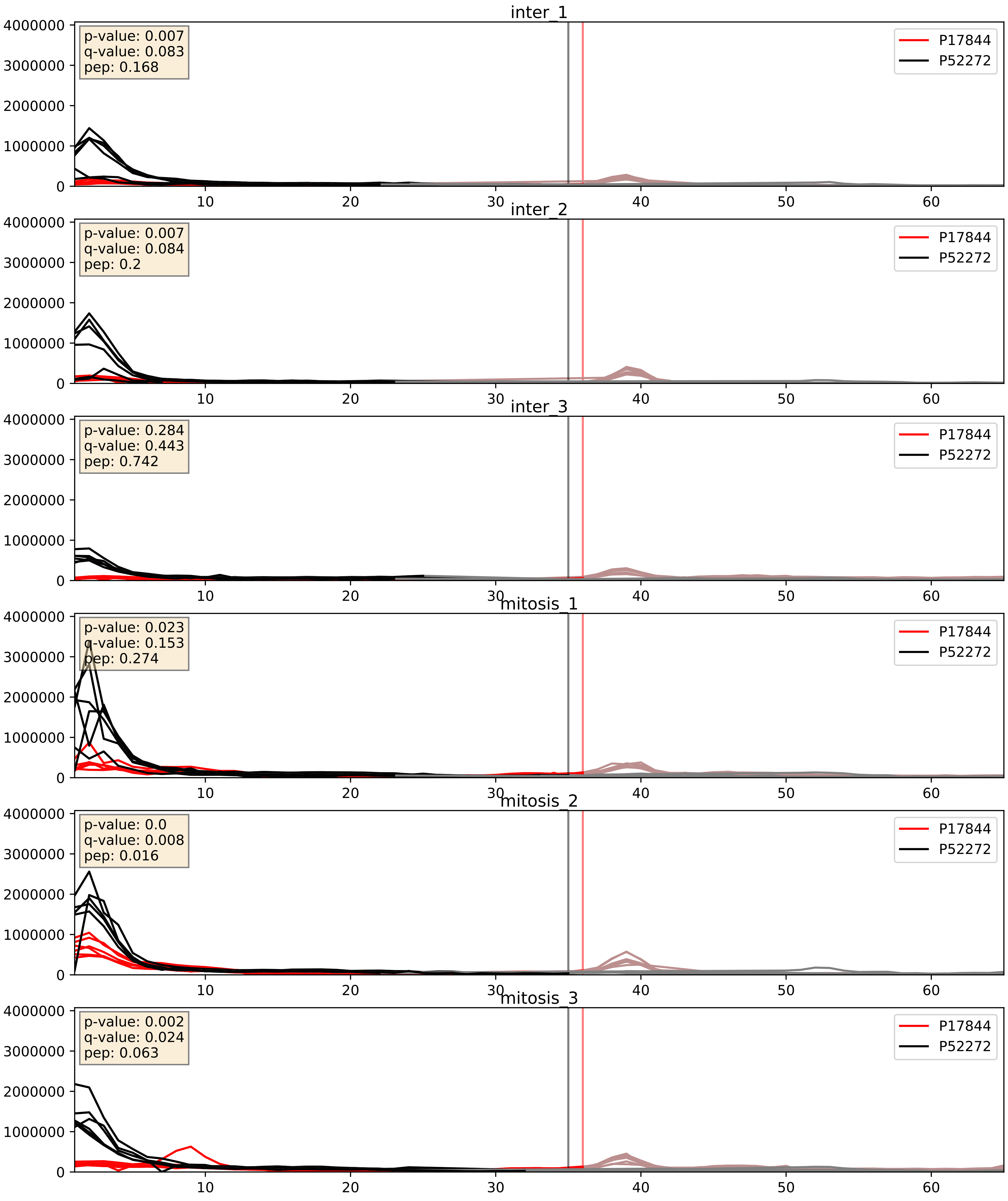


| level | condition_1 | condition_2 | pvalue | pvalue_adjusted |
|---|---|---|---|---|
| interactor_abundance | mitosis | inter | 0.0008880993781604356 | 0.02197564842852207 |
| complex_abundance | mitosis | inter | 0.003908349803170464 | 0.0462092186673193 |
| interactor_ratio | mitosis | inter | 0.08332897328355368 | 0.4841037832929192 |

P17844_P52597
DDX5_HUMAN vs HNRPF_HUMAN
p-value: 0.0 q-value: 0.005

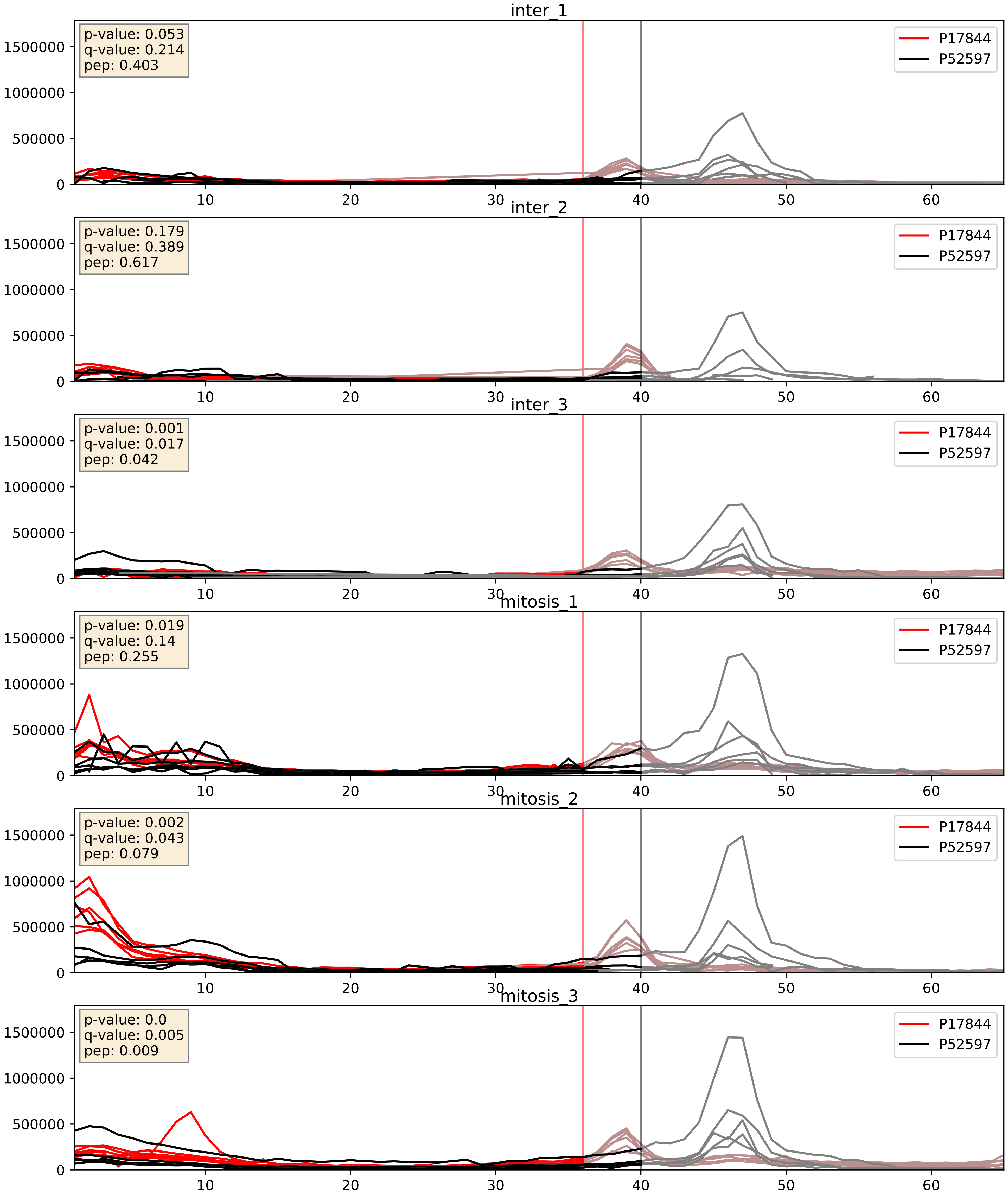


| level | condition_1 | condition_2 | pvalue | pvalue_adjusted |
|---|---|---|---|---|
| interactor_abundance | mitosis | inter | 0.0021426459467421676 | 0.033870820506210444 |
| complex_abundance | mitosis | inter | 0.002394620564026592 | 0.035463584823646414 |
| interactor_ratio | mitosis | inter | 0.413906743817922 | 0.8357726189348997 |

P17844_P61978
DDX5_HUMAN vs HNRPK_HUMAN
p-value: 0.004 q-value: 0.047

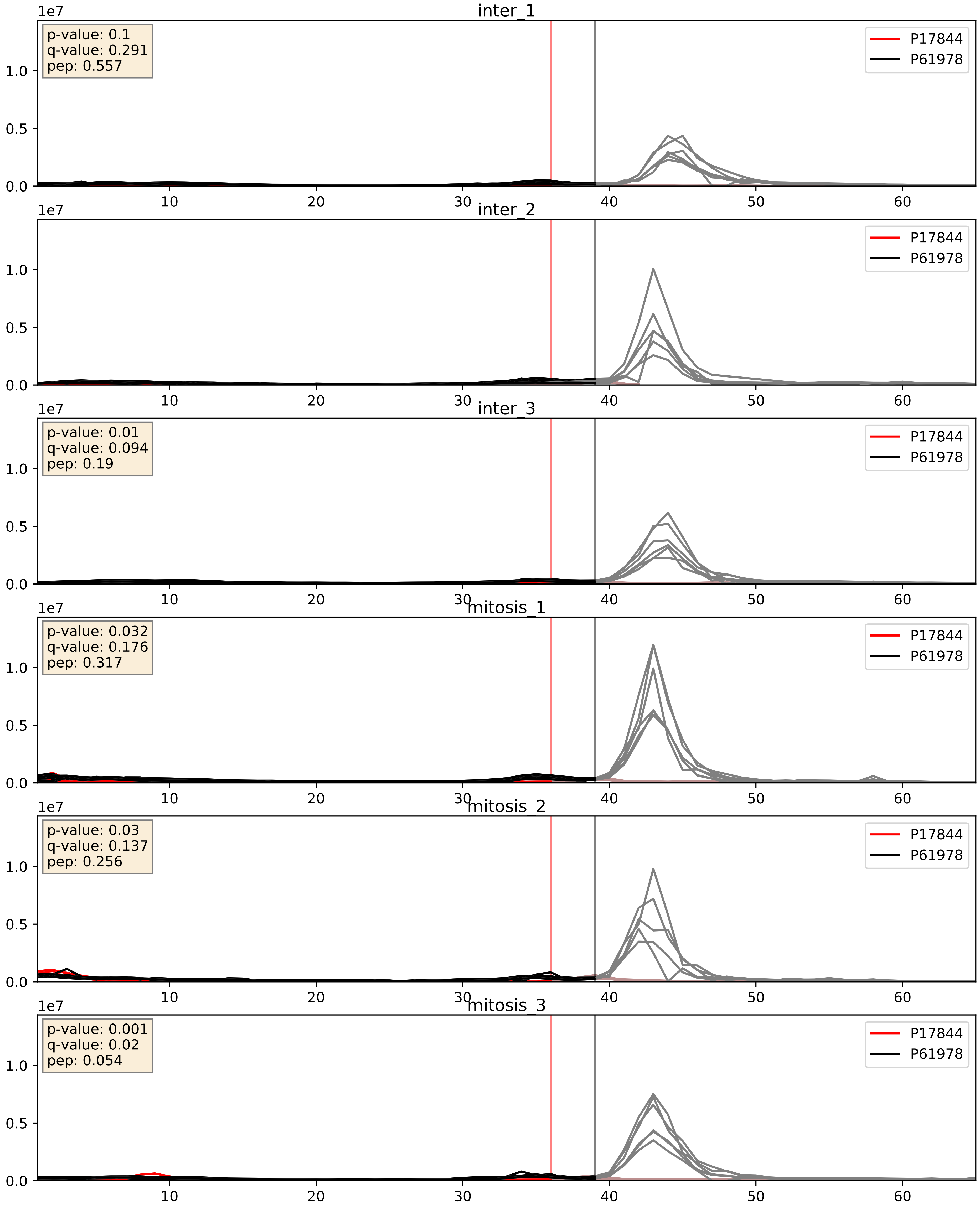


| level | condition_1 | condition_2 | pvalue | pvalue_adjusted |
|---|---|---|---|---|
| interactor_abundance | mitosis | inter | 0.0021426459467421676 | 0.033870820506210444 |
| interactor_ratio | mitosis | inter | 0.20336790116763848 | 0.6786712771498306 |
| complex_abundance | mitosis | inter | 0.2639866365284098 | 0.4140966847504468 |

P17844_Q07955
DDX5_HUMAN vs SRSF1_HUMAN
p-value: 0.001 q-value: 0.015

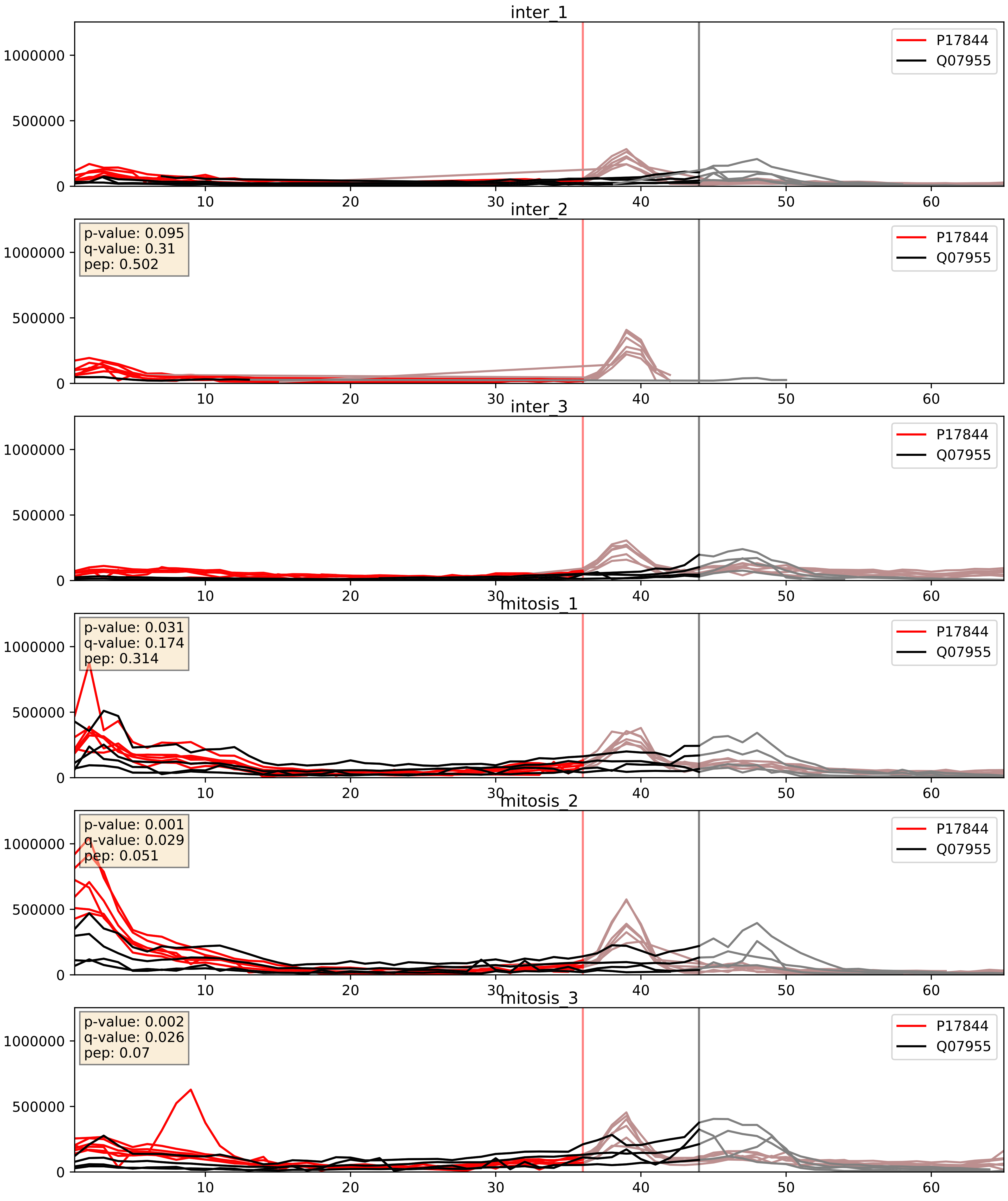


| level | condition_1 | condition_2 | pvalue | pvalue_adjusted |
|---|---|---|---|---|
| interactor_abundance | mitosis | inter | 0.0007813490254633644 | 0.02050037113506771 |
| complex_abundance | mitosis | inter | 0.0024237962141353607 | 0.035648961499997744 |
| interactor_ratio | mitosis | inter | 0.14930404490866453 | 0.6039771666075939 |

P17844_Q08211
DDX5_HUMAN vs DHX9_HUMAN
p-value: 0.004 q-value: 0.004

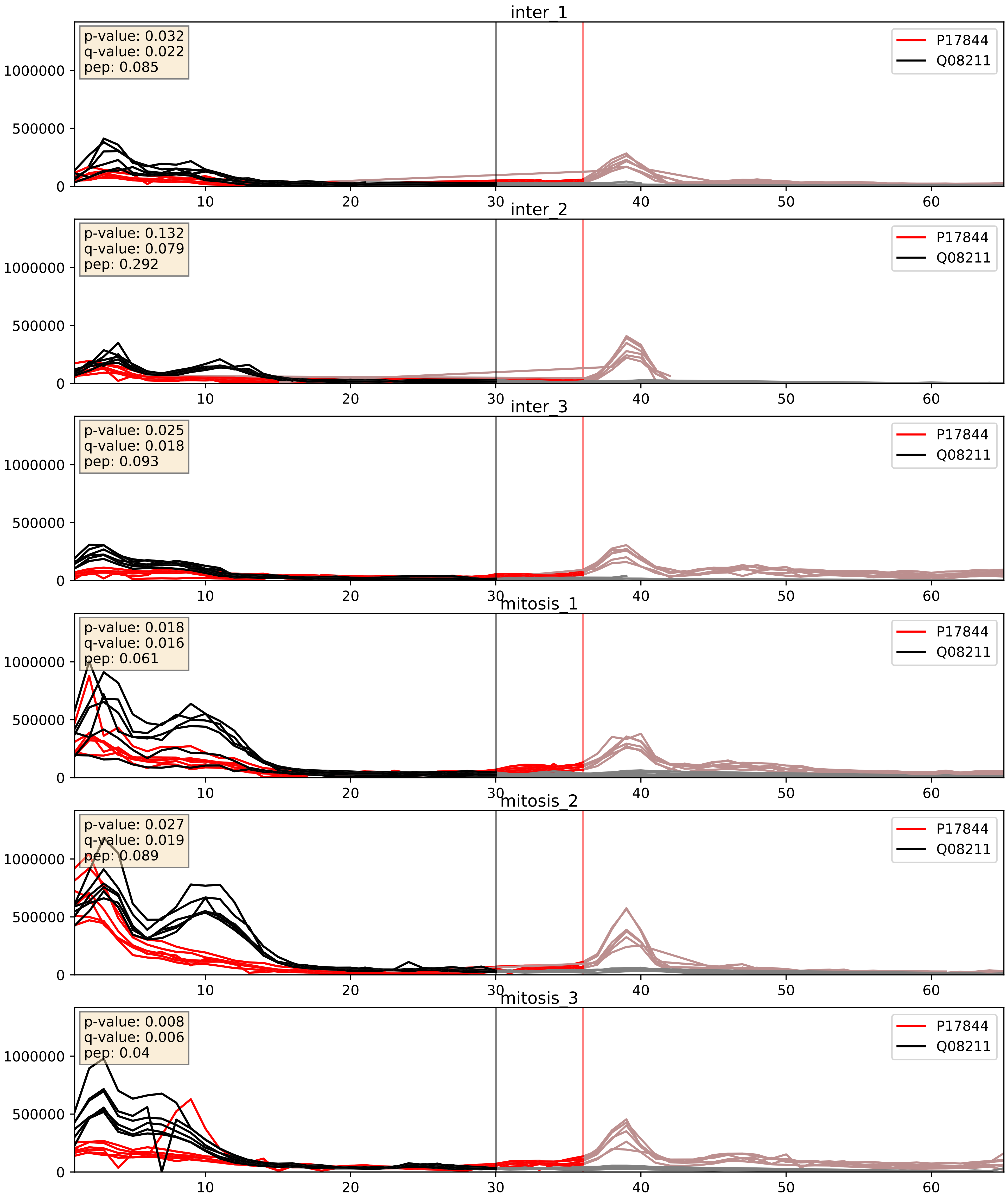


| level | condition_1 | condition_2 | pvalue | pvalue_adjusted |
|---|---|---|---|---|
| complex_abundance | mitosis | inter | 0.0004358577827025154 | 0.013137121901174406 |
| interactor_abundance | mitosis | inter | 0.0017533669129677133 | 0.030599022986755608 |
| interactor_ratio | mitosis | inter | 0.4374343152584505 | 0.8454166152138379 |

P17844_Q13247
DDX5_HUMAN vs SRSF6_HUMAN
p-value: 0.0 q-value: 0.005

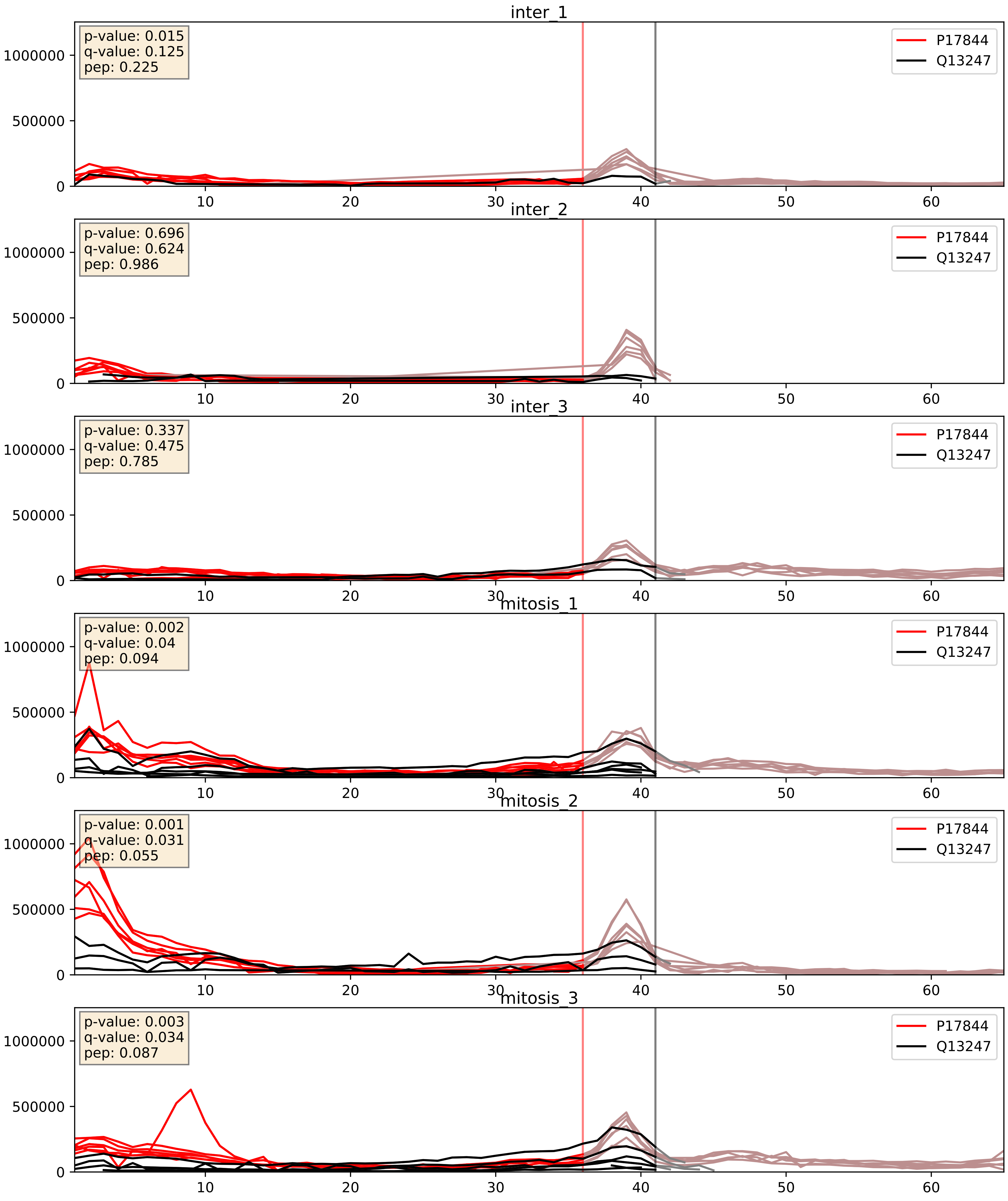


| level | condition_1 | condition_2 | pvalue | pvalue_adjusted |
|---|---|---|---|---|
| complex_abundance | mitosis | inter | 0.002718775066662734 | 0.03753663640424678 |
| interactor_abundance | mitosis | inter | 0.003078864127685547 | 0.042536422819834926 |
| interactor_ratio | mitosis | inter | 0.6545547922536914 | 0.9236711212811735 |

P17844_Q16629
DDX5_HUMAN vs SRSF7_HUMAN
p-value: 0.0 q-value: 0.01

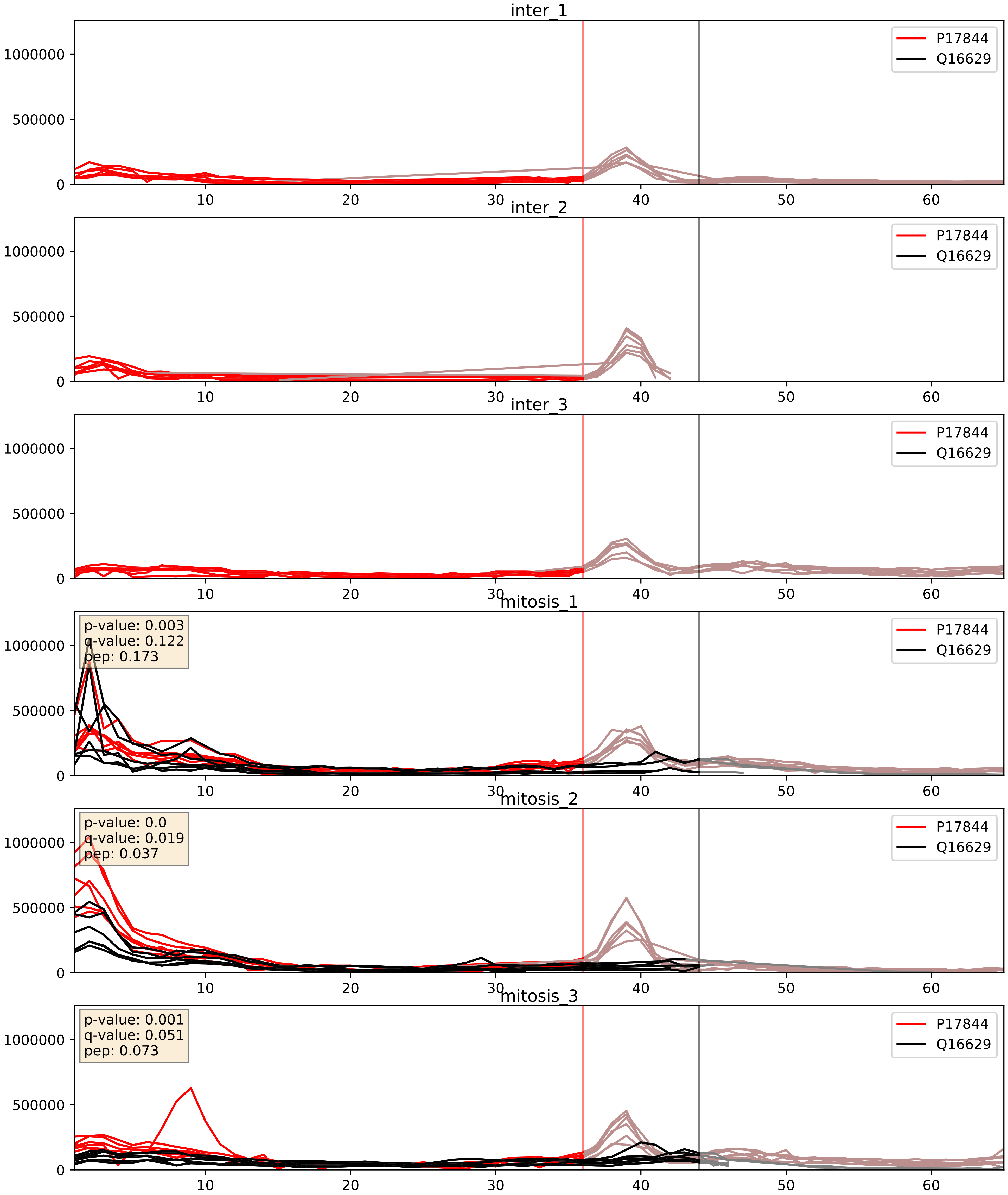


| level | condition_1 | condition_2 | pvalue | pvalue_adjusted |
|---|---|---|---|---|
| interactor_abundance | mitosis | inter | 2.9246981424557922e-05 | 0.004814503096042612 |
| complex_abundance | mitosis | inter | 4.447501338108865e-05 | 0.005691857151652958 |
| interactor_ratio | mitosis | inter | 0.15108645229943365 | 0.606604569117897 |

P17844_Q99459
DDX5_HUMAN vs CDC5L_HUMAN
p-value: 0.007 q-value: 0.006

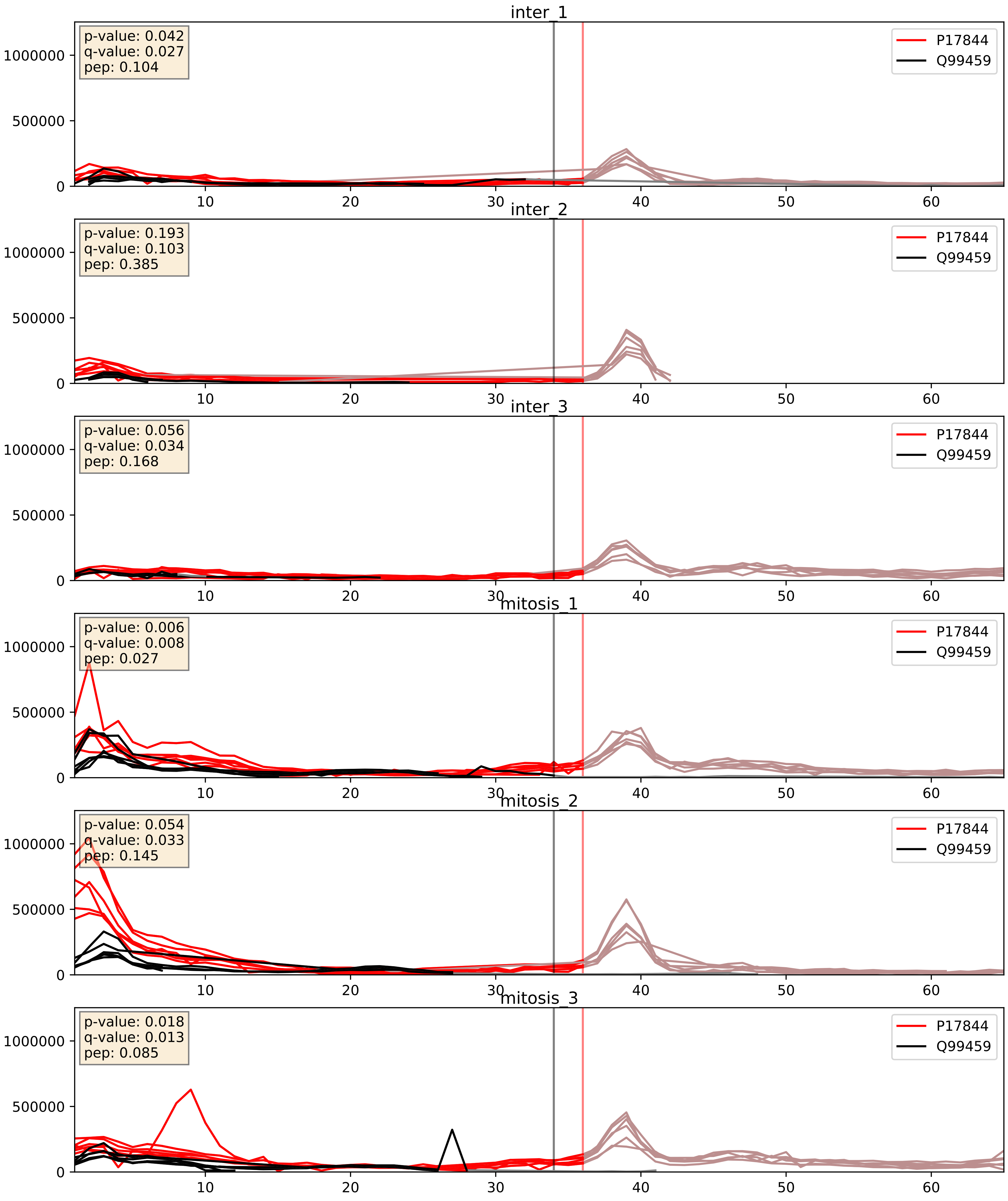


| level | condition_1 | condition_2 | pvalue | pvalue_adjusted |
|---|---|---|---|---|
| interactor_abundance | mitosis | inter | 0.006359400241119042 | 0.06830171400750187 |
| complex_abundance | mitosis | inter | 0.008211982545060673 | 0.06720322235728429 |
| interactor_ratio | mitosis | inter | 0.4083540240816134 | 0.8335535006745441 |

P17844_Q9UMS4
DDX5_HUMAN vs PRP19_HUMAN
p-value: 0.003 q-value: 0.041

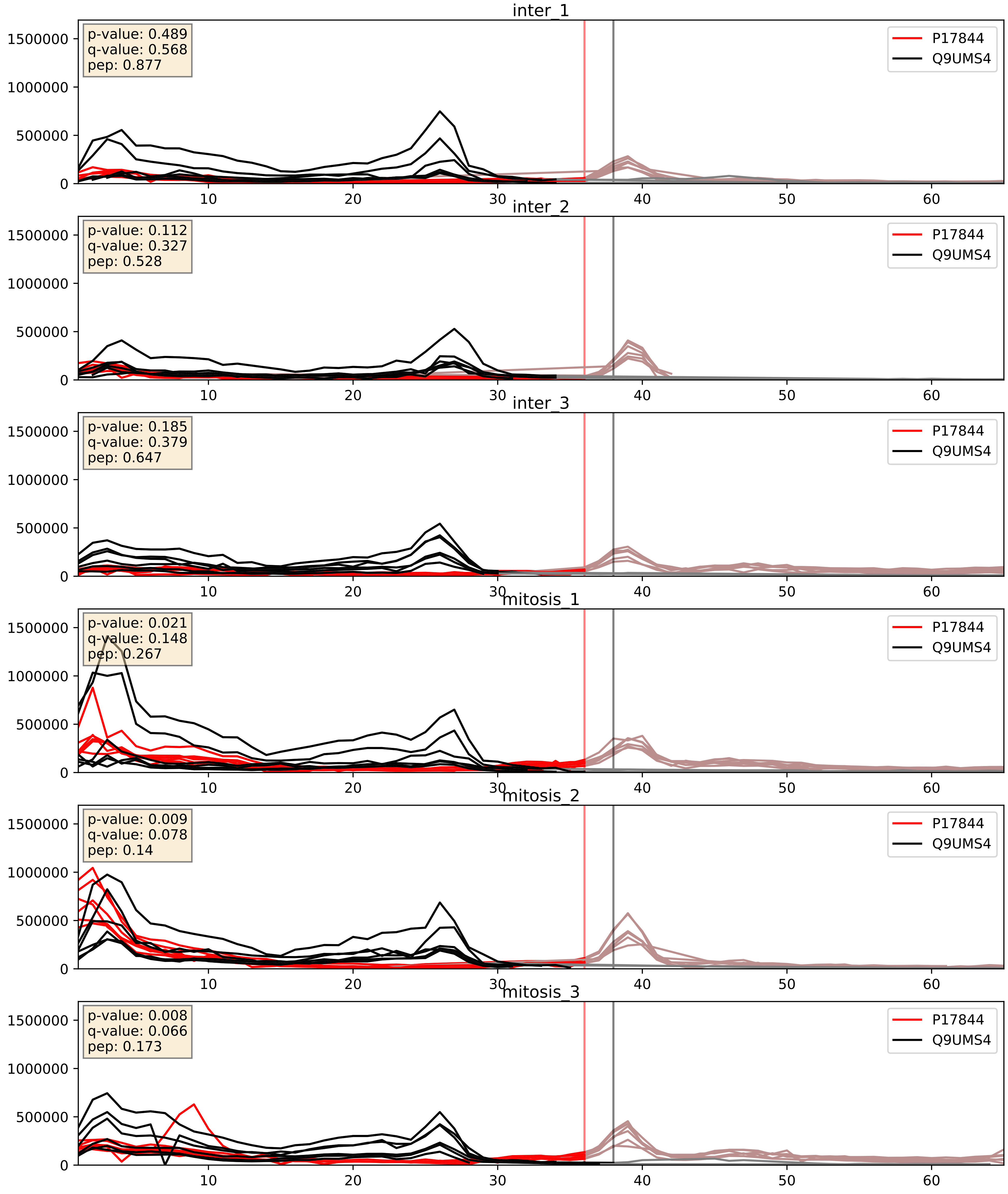


| level | condition_1 | condition_2 | pvalue | pvalue_adjusted |
|---|---|---|---|---|
| interactor_abundance | mitosis | inter | 0.0013659136984398564 | 0.027354532106093617 |
| complex_abundance | mitosis | inter | 0.01277054821565052 | 0.08405635484106207 |
| interactor_ratio | mitosis | inter | 0.33717069446515985 | 0.7922357565214662 |
