## Supplementary Data 2 for "SECAT: Quantifying differential protein-protein interaction states by network-centric analysis": hela_string_000050_Q9BYG3.pdf

Q9BYG3_Q9BYG3

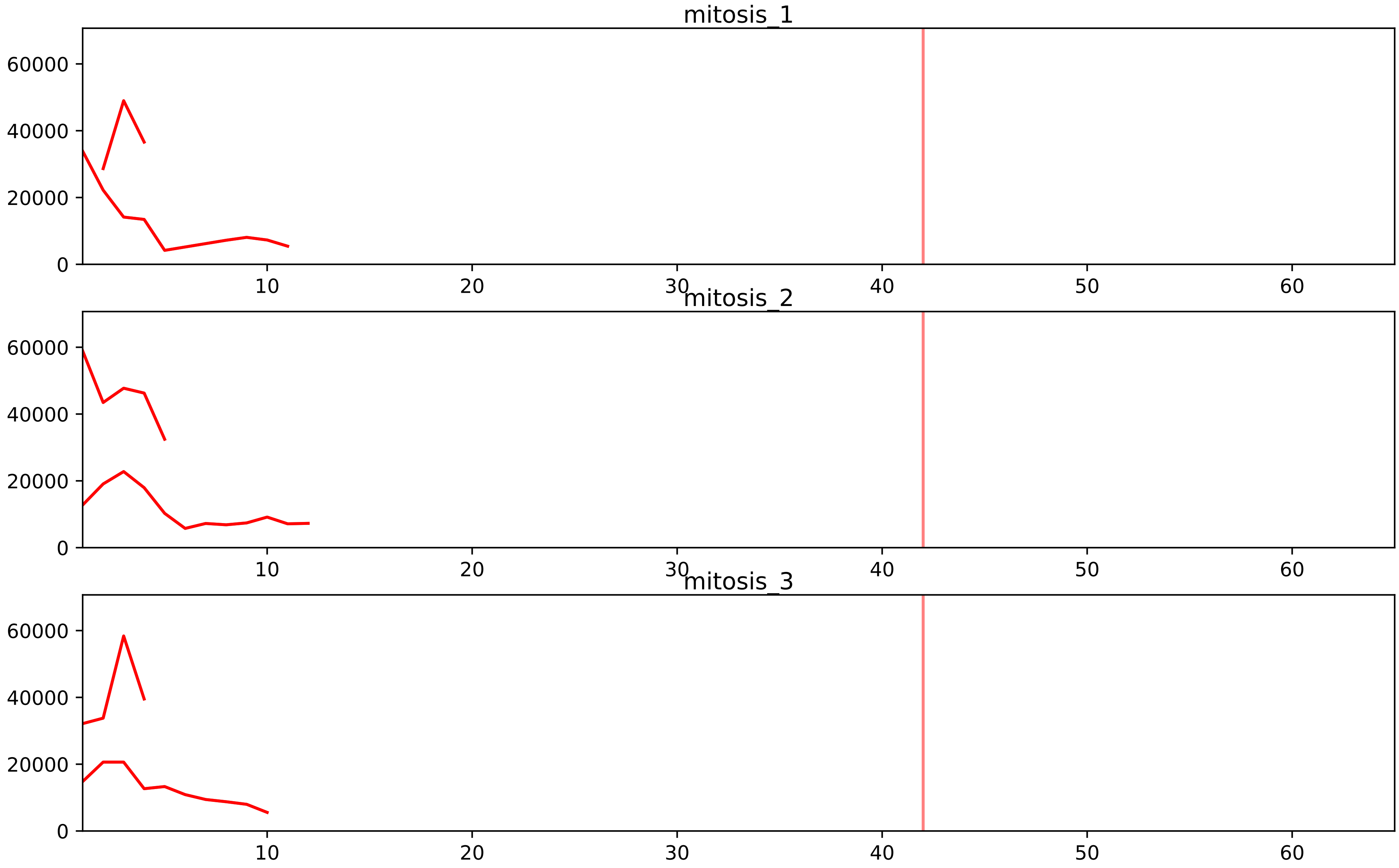


| level | condition_1 | condition_2 | pvalue | pvalue_adjusted |
|---|---|---|---|---|
| interactor_ratio | mitosis | inter | 1.4013184535026228e-07 | 5.371720738426721e-06 |
| assembled_abundance | mitosis | inter | 0.0010577054243156021 | 0.029826018621815265 |
| complex_abundance | mitosis | inter | 0.003380456956523474 | 0.02338361203008719 |
| total_abundance | mitosis | inter | 0.009216139518524269 | 0.11549069945635781 |
| interactor_abundance | mitosis | inter | 0.2464823153220926 | 0.4056596245014762 |

O00567_Q9BYG3
NOP56_HUMAN vs MK67I_HUMAN
p-value: 0.03 q-value: 0.022

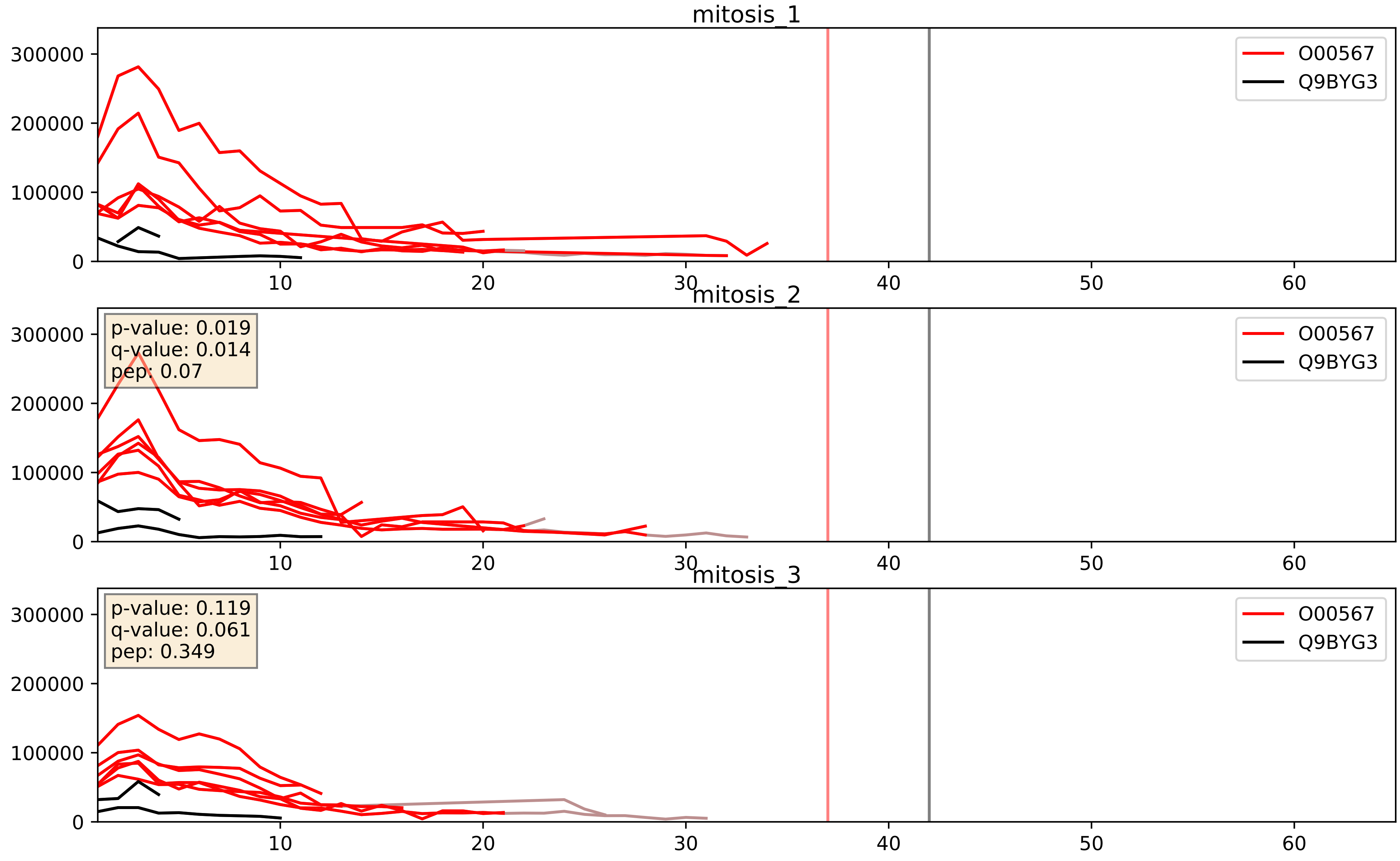


| level | condition_1 | condition_2 | pvalue | pvalue_adjusted |
|---|---|---|---|---|
| interactor_ratio | mitosis | inter | 0.00025624310259385303 | 0.016247710801506533 |
| interactor_abundance | mitosis | inter | 0.000695734930284271 | 0.019073880342257484 |
| complex_abundance | mitosis | inter | 0.002507942943970339 | 0.03638642644133238 |

O75683_Q9BYG3
SURF6_HUMAN vs MK67I_HUMAN
p-value: 0.019 q-value: 0.015

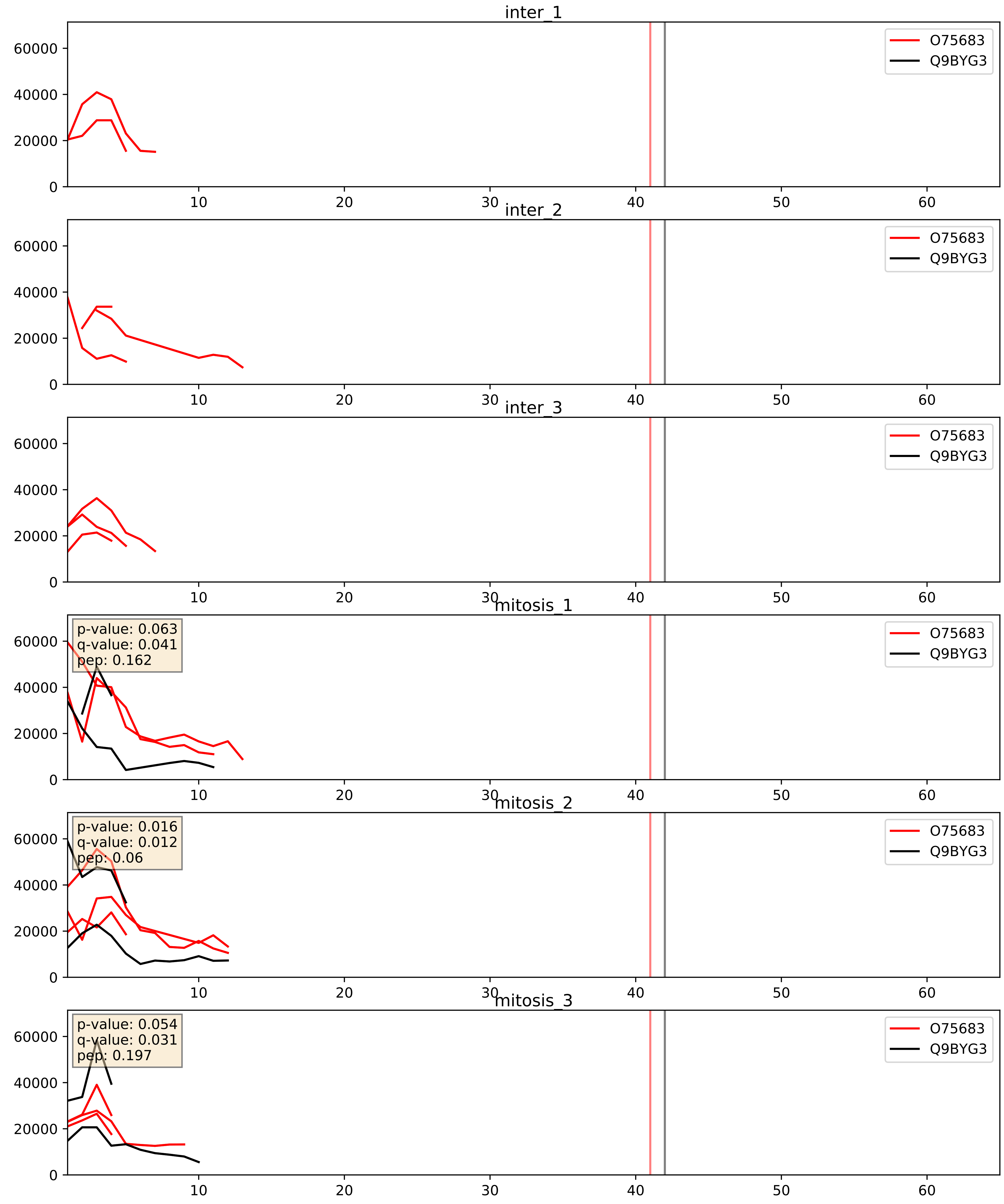


| level | condition_1 | condition_2 | pvalue | pvalue_adjusted |
|---|---|---|---|---|
| complex_abundance | mitosis | inter | 0.21693975168529286 | 0.3715494746750915 |
| interactor_abundance | mitosis | inter | 0.22189639792716673 | 0.4180090594754725 |
| interactor_ratio | mitosis | inter | 0.33369938135990024 | 0.7906537175374388 |

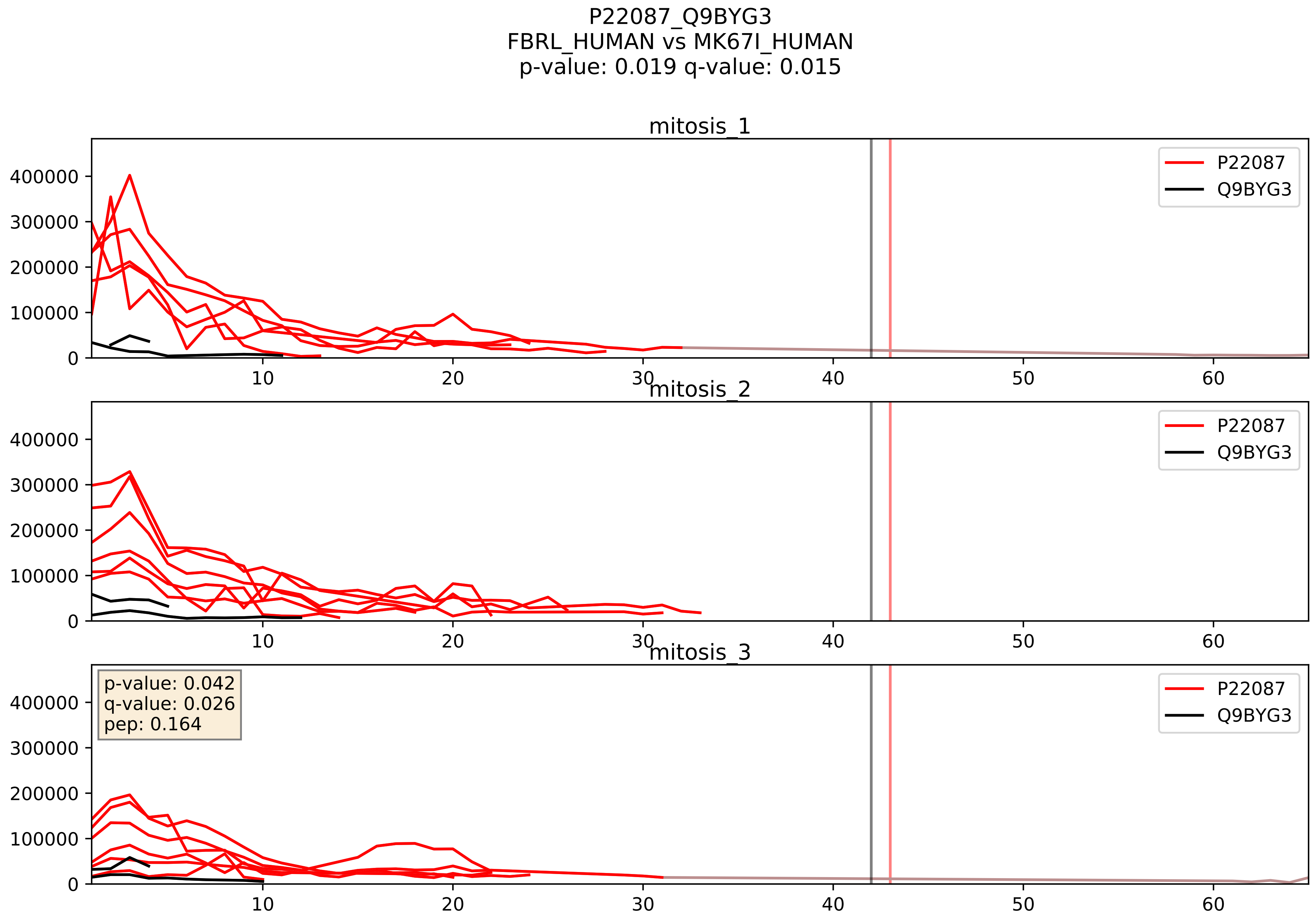


| level | condition_1 | condition_2 | pvalue | pvalue_adjusted |
|---|---|---|---|---|
| interactor_ratio | mitosis | inter | 0.00022528201190776103 | 0.01529848062979237 |
| interactor_abundance | mitosis | inter | 0.0005233714416599235 | 0.016562142479145826 |
| complex_abundance | mitosis | inter | 0.0016144918603450086 | 0.02814408910325431 |

Q15050_Q9BYG3
RRS1_HUMAN vs MK67I_HUMAN
p-value: 0.033 q-value: 0.023

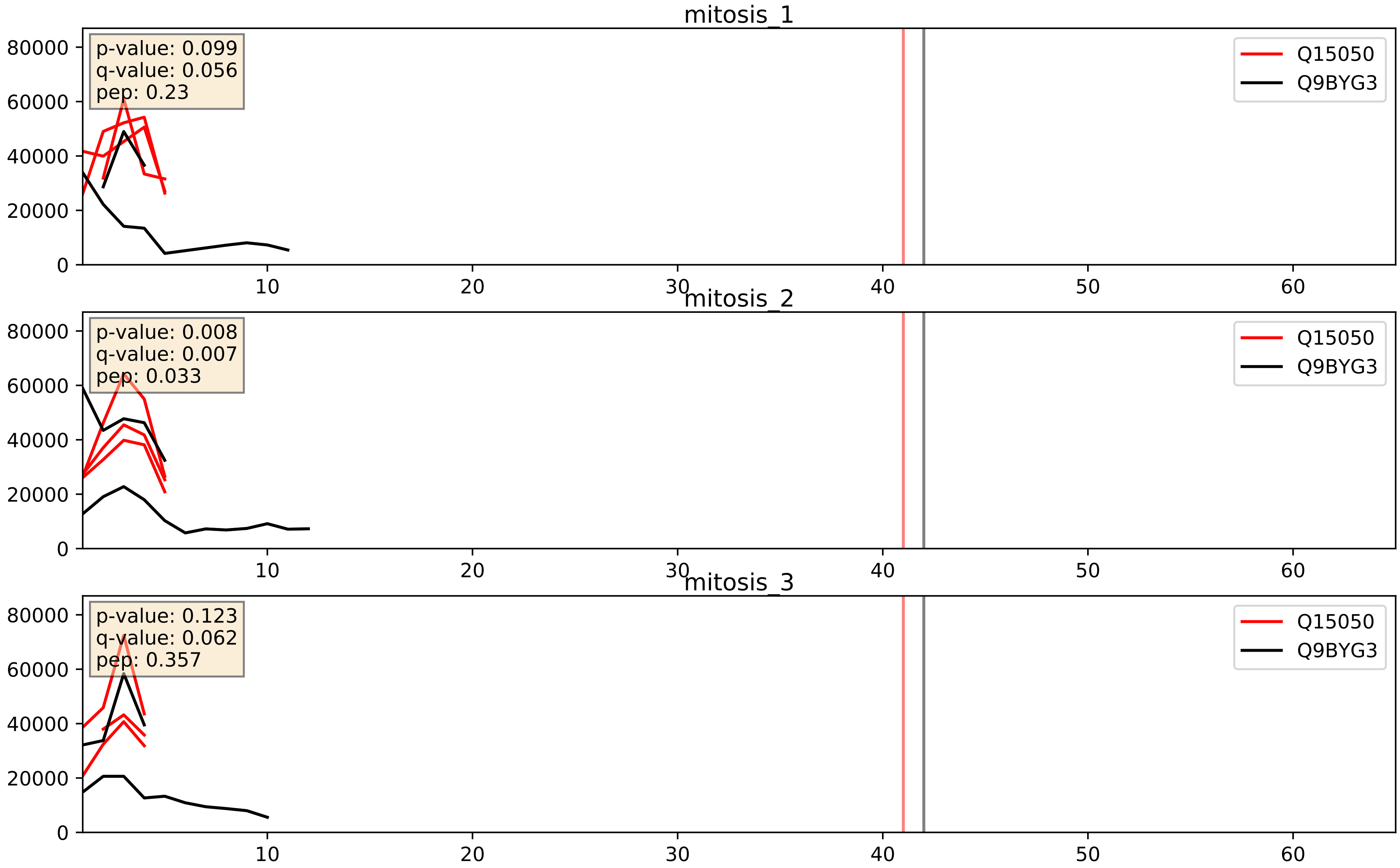


| level | condition_1 | condition_2 | pvalue | pvalue_adjusted |
|---|---|---|---|---|
| interactor_ratio | mitosis | inter | 0.12735748141619438 | 0.5750062399685051 |
| interactor_abundance | mitosis | inter | 0.2162538244564617 | 0.4119120465837366 |
| complex_abundance | mitosis | inter | 0.251694037814913 | 0.40201339026612704 |

Q15397_Q9BYG3
PUM3_HUMAN vs MK67I_HUMAN
p-value: 0.052 q-value: 0.033

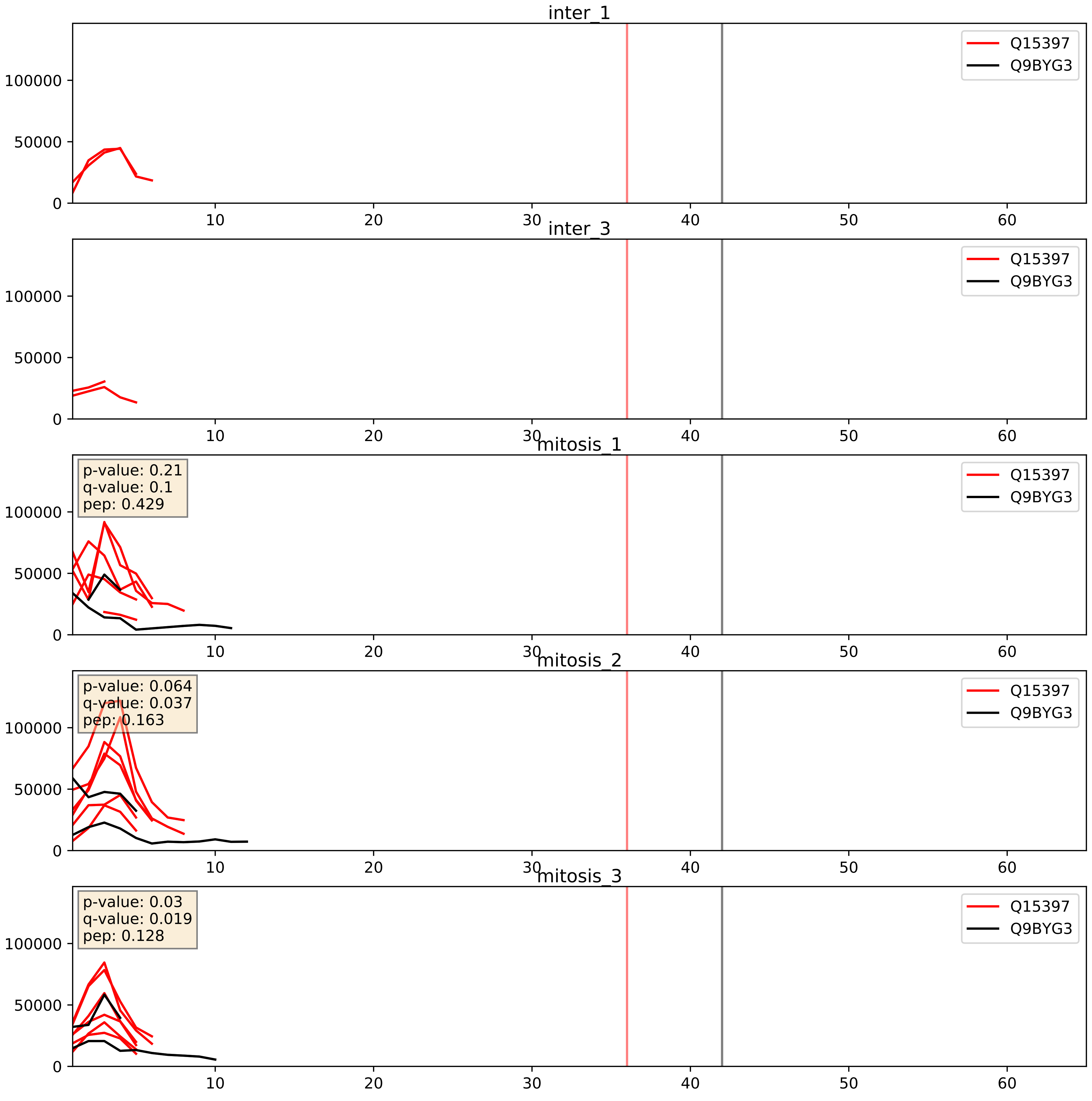


| level | condition_1 | condition_2 | pvalue | pvalue_adjusted |
|---|---|---|---|---|
| interactor_ratio | mitosis | inter | 0.004748651113544903 | 0.09196482699534925 |
| interactor_abundance | mitosis | inter | 0.024994758678114595 | 0.1308194034146505 |
| complex_abundance | mitosis | inter | 0.05783440640842851 | 0.1717178351911717 |

Q8TDN6_Q9BYG3
BRX1_HUMAN vs MK67I_HUMAN
p-value: 0.044 q-value: 0.029

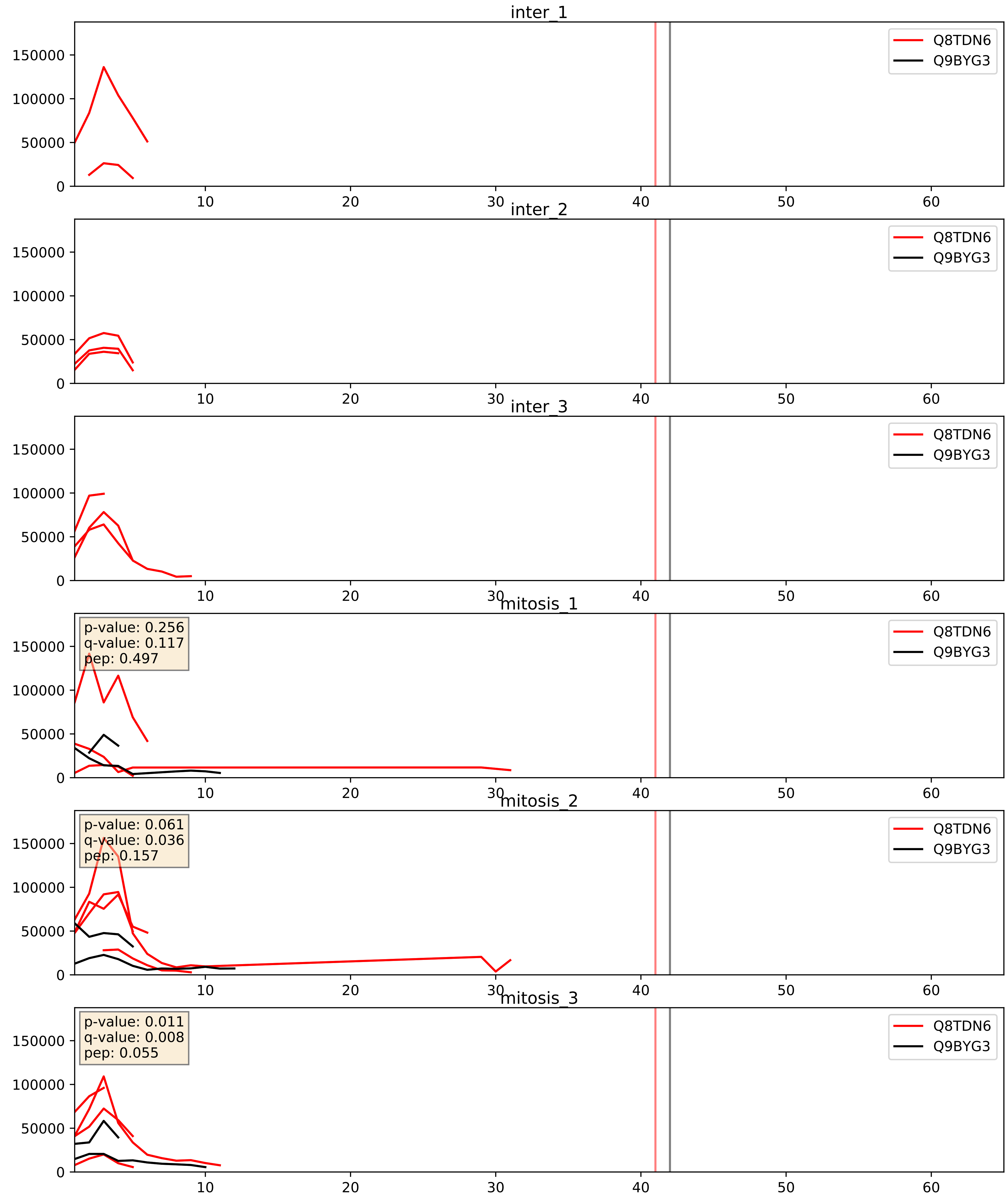


| level | condition_1 | condition_2 | pvalue | pvalue_adjusted |
|---|---|---|---|---|
| interactor_abundance | mitosis | inter | 0.015356739953974529 | 0.10851551243015083 |
| complex_abundance | mitosis | inter | 0.02656820842384979 | 0.11938260583105208 |
| interactor_ratio | mitosis | inter | 0.045559994696426284 | 0.3548621970895441 |

Q96GQ7_Q9BYG3
DDX27_HUMAN vs MK67I_HUMAN
p-value: 0.024 q-value: 0.018

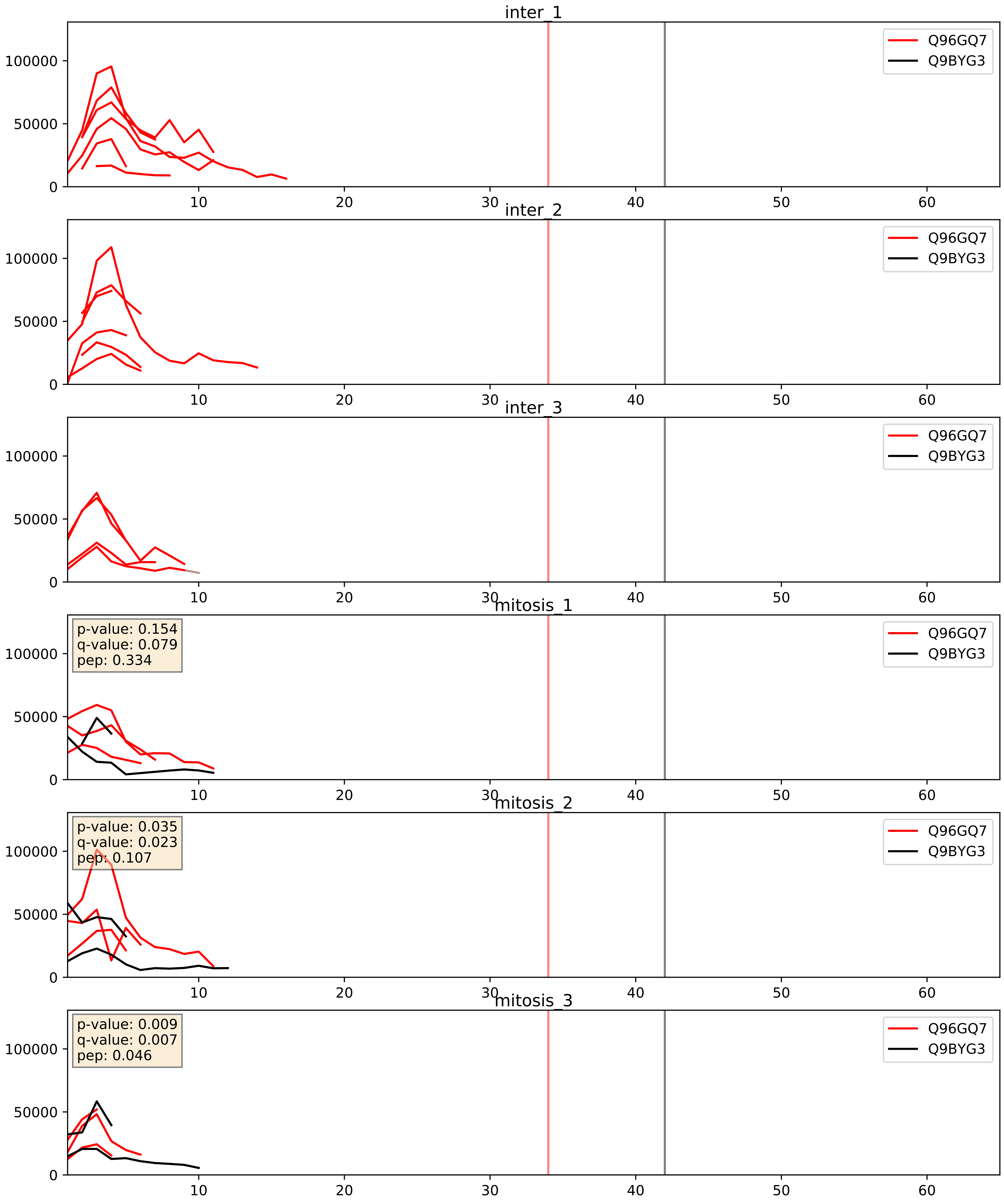


| level | condition_1 | condition_2 | pvalue | pvalue_adjusted |
|---|---|---|---|---|
| interactor_ratio | mitosis | inter | 0.13162543328171866 | 0.5797318033557721 |
| interactor_abundance | mitosis | inter | 0.17040407116164052 | 0.3598133054863397 |
| complex_abundance | mitosis | inter | 0.1935502198005397 | 0.34516455864429585 |

Q9BVP2_Q9BYG3
GNL3_HUMAN vs MK67I_HUMAN
p-value: 0.056 q-value: 0.036

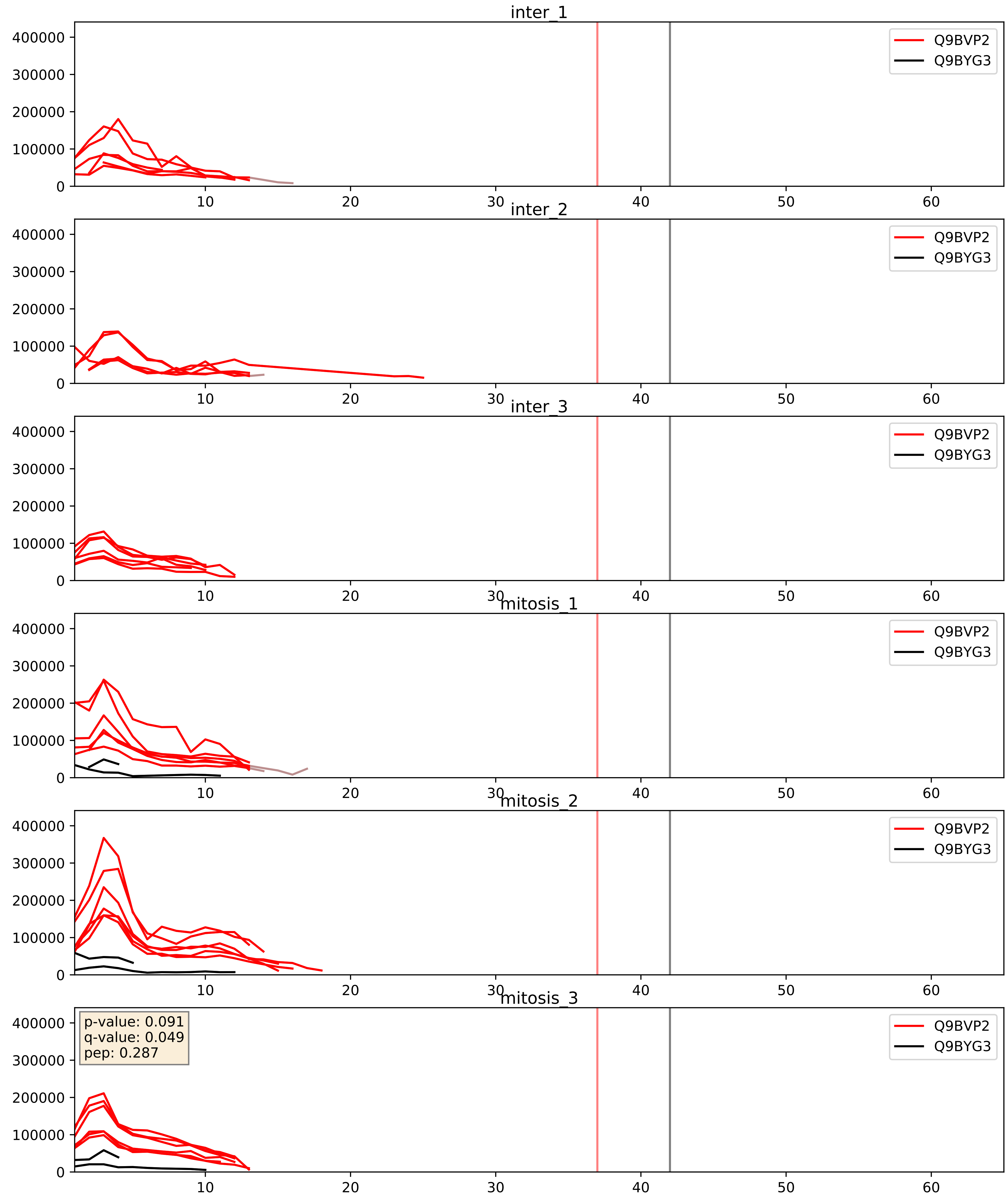


| level | condition_1 | condition_2 | pvalue | pvalue_adjusted |
|---|---|---|---|---|
| interactor_abundance | mitosis | inter | 0.00039498517428285436 | 0.01371632085947762 |
| interactor_ratio | mitosis | inter | 0.0007741550098460847 | 0.03185945617443502 |
| complex_abundance | mitosis | inter | 0.000841508846203478 | 0.019252661341274204 |

Q9BYG3_Q9BZE4
MK67I_HUMAN vs NOG1_HUMAN
p-value: 0.068 q-value: 0.041

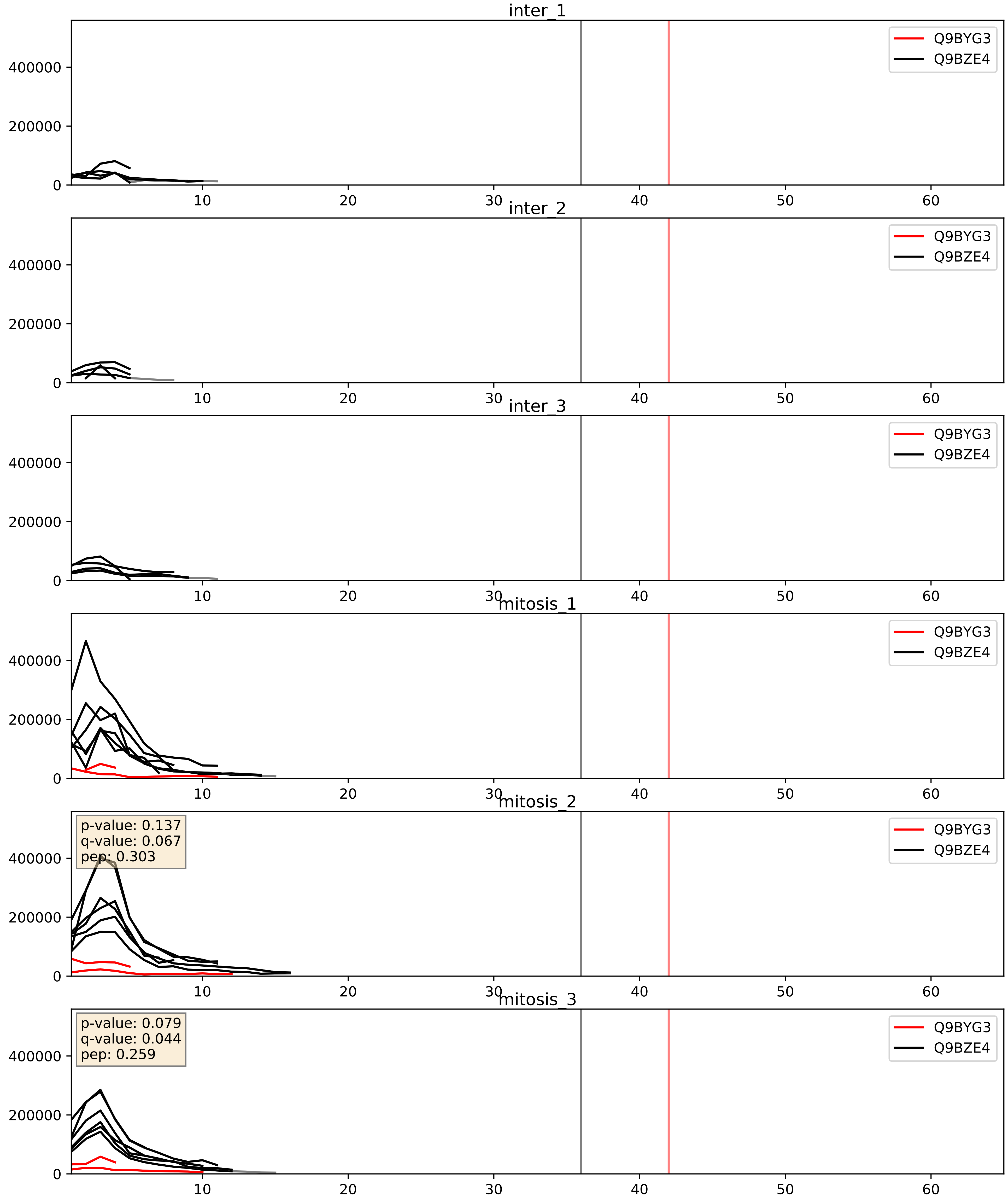


| level | condition_1 | condition_2 | pvalue | pvalue_adjusted |
|---|---|---|---|---|
| interactor_ratio | mitosis | inter | 0.0006141803584999456 | 0.029219626944512057 |
| complex_abundance | mitosis | inter | 0.0011171790006910644 | 0.022133971147621622 |
| interactor_abundance | mitosis | inter | 0.21337749899904426 | 0.40861552381025035 |

Q9BYG3_Q9GZR7
MK67I_HUMAN vs DDX24_HUMAN
p-value: 0.024 q-value: 0.018

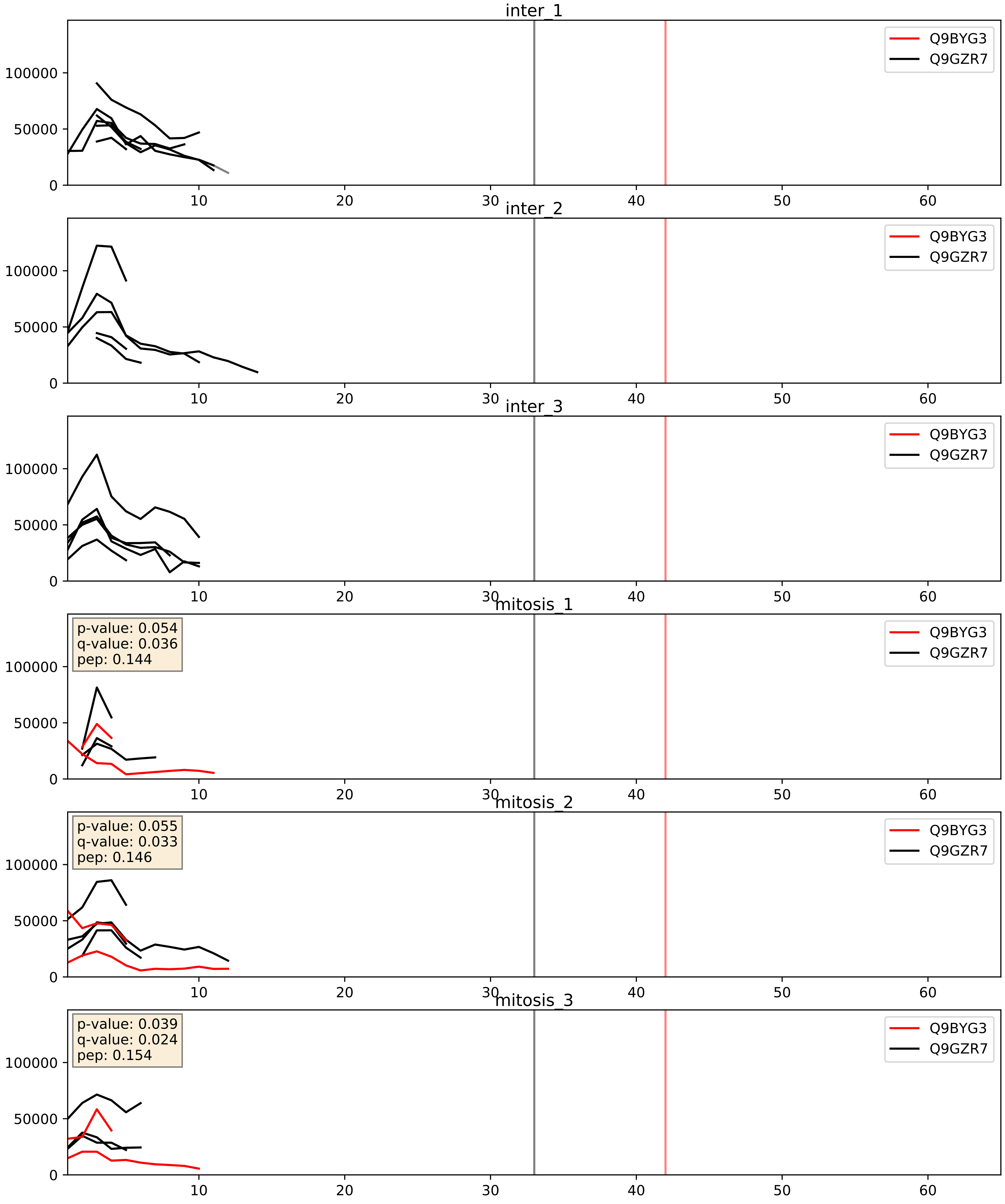


| level | condition_1 | condition_2 | pvalue | pvalue_adjusted |
|---|---|---|---|---|
| complex_abundance | mitosis | inter | 0.08265105601072668 | 0.21108552910002362 |
| interactor_ratio | mitosis | inter | 0.15249623026144915 | 0.6082794646029844 |
| interactor_abundance | mitosis | inter | 0.2535147171951053 | 0.4468626754113251 |

Q9BYG3_Q9H7B2
MK67I_HUMAN vs RPF2_HUMAN
p-value: 0.036 q-value: 0.025

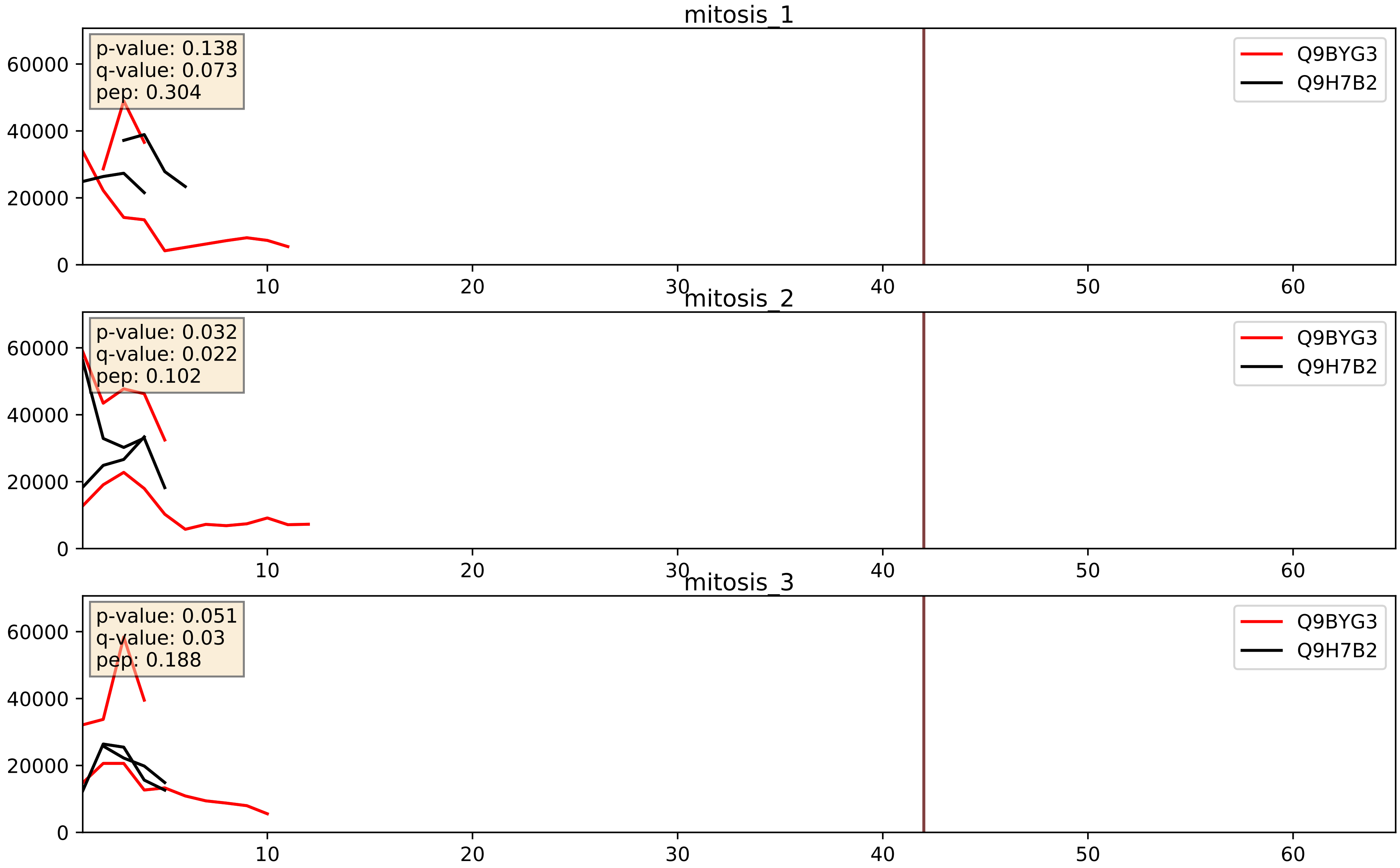


| level | condition_1 | condition_2 | pvalue | pvalue_adjusted |
|---|---|---|---|---|
| interactor_ratio | mitosis | inter | 0.15944016446652348 | 0.616090239021059 |
| interactor_abundance | mitosis | inter | 0.3187687602568554 | 0.5167917779921747 |
| complex_abundance | mitosis | inter | 0.3970451119903028 | 0.5437591915331307 |

Q9BYG3_Q9NW13
MK67I_HUMAN vs RBM28_HUMAN
p-value: 0.054 q-value: 0.035

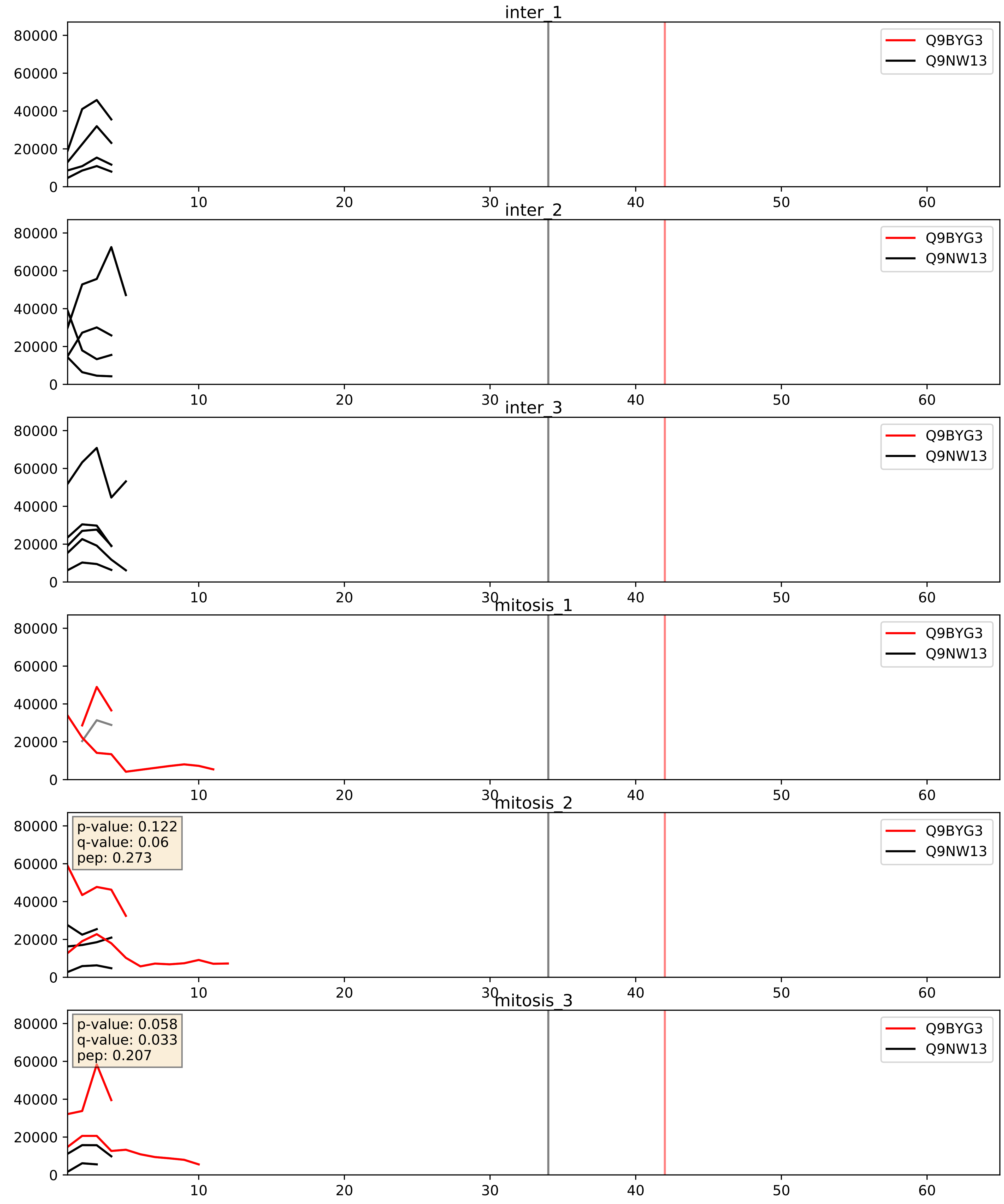


| level | condition_1 | condition_2 | pvalue | pvalue_adjusted |
|---|---|---|---|---|
| complex_abundance | mitosis | inter | 0.5935004295074089 | 0.7184788115660331 |
| interactor_ratio | mitosis | inter | 0.7413797794565655 | 0.9491588884573259 |
| interactor_abundance | mitosis | inter | 0.7811482656816368 | 0.8726814827254089 |

Q9BYG3_Q9UKD2
MK67I_HUMAN vs MRT4_HUMAN
p-value: 0.081 q-value: 0.048

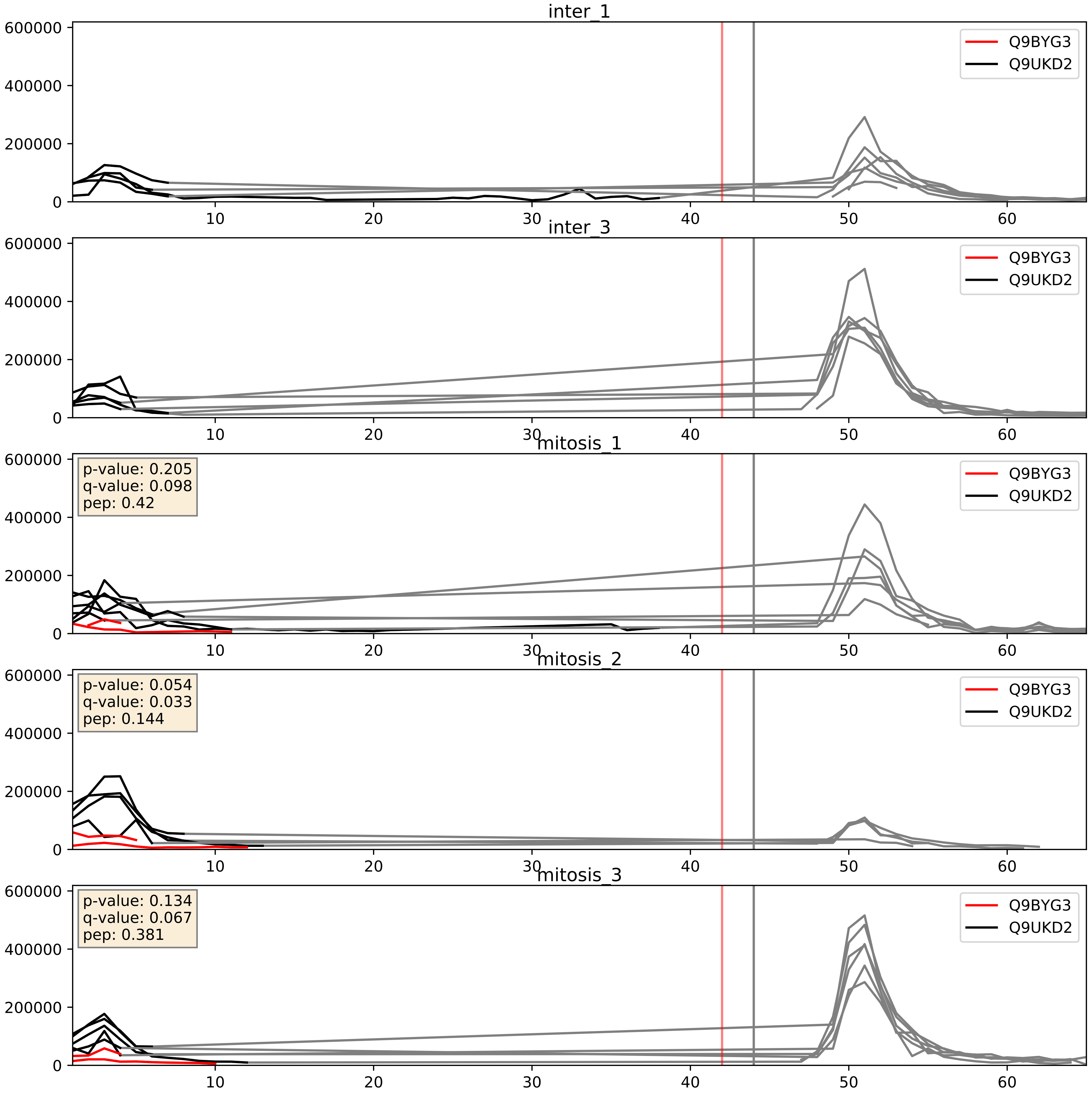


| level | condition_1 | condition_2 | pvalue | pvalue_adjusted |
|---|---|---|---|---|
| interactor_ratio | mitosis | inter | 0.0017848561268288656 | 0.051461731073100085 |
| complex_abundance | mitosis | inter | 0.0034654008381990245 | 0.04302508960916016 |
| interactor_abundance | mitosis | inter | 0.21337749899904426 | 0.40861552381025035 |

Q9BYG3_Q9Y221
MK67I_HUMAN vs NIP7_HUMAN
p-value: 0.042 q-value: 0.028

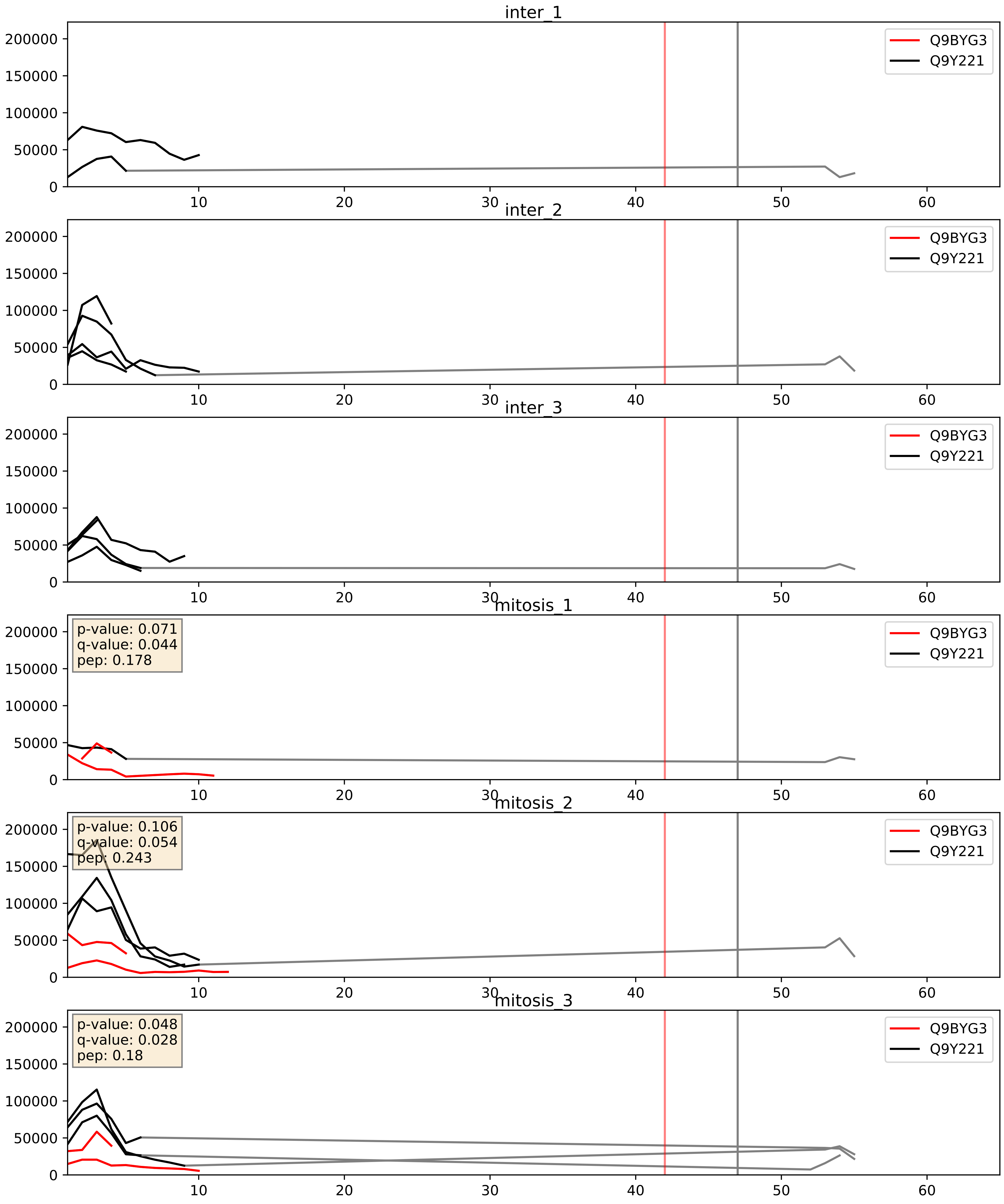


| level | condition_1 | condition_2 | pvalue | pvalue_adjusted |
|---|---|---|---|---|
| complex_abundance | mitosis | inter | 0.05436776342416502 | 0.16632882591524398 |
| interactor_abundance | mitosis | inter | 0.22797553195418468 | 0.4210333251045526 |
| interactor_ratio | mitosis | inter | 0.26800099815205625 | 0.7435173930604346 |
