## Supplementary Data 2 for "SECAT: Quantifying differential protein-protein interaction states by network-centric analysis": hela_string_000051_Q9BV38.pdf

Q9BV38_Q9BV38

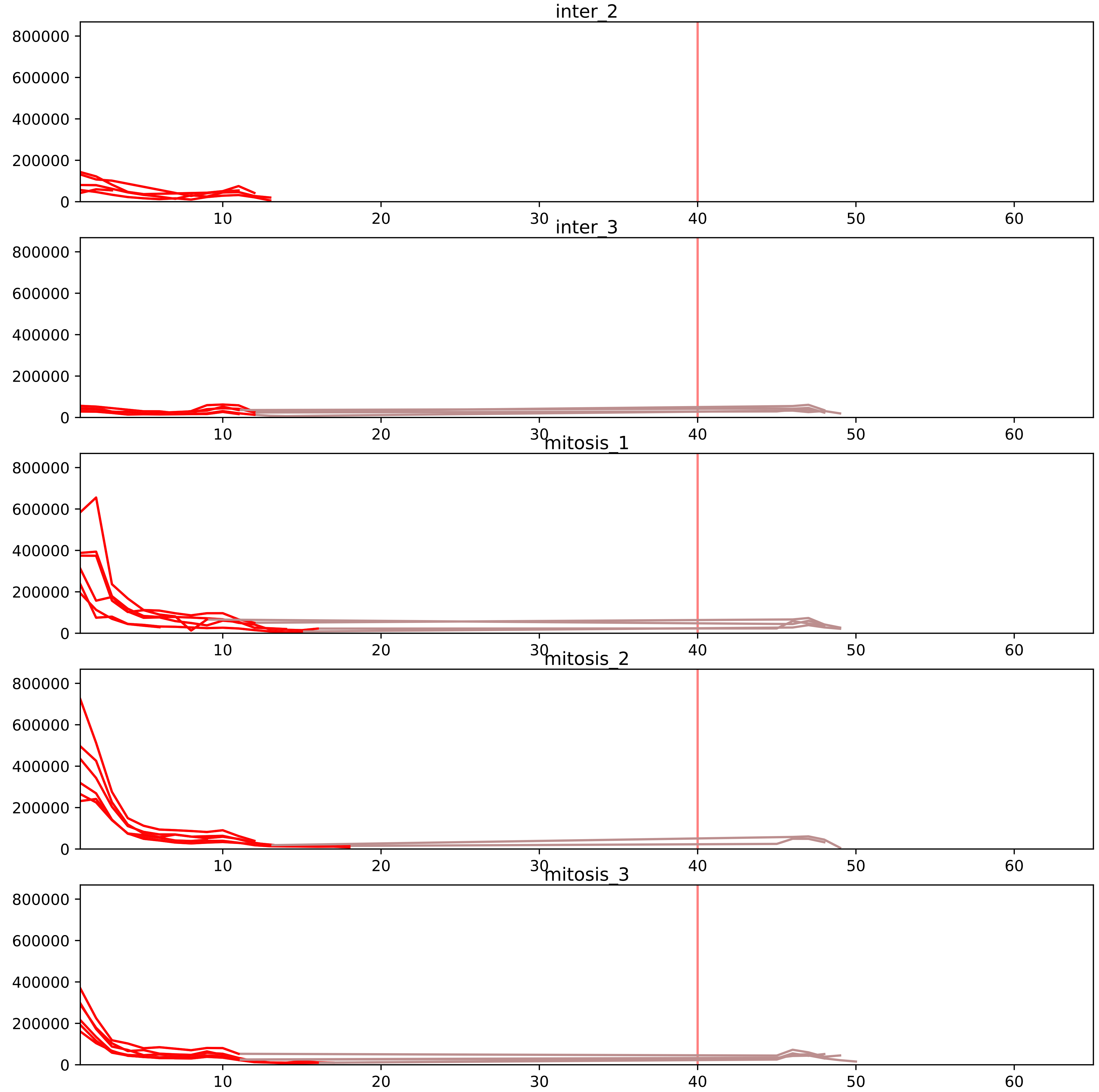


| level | condition_1 | condition_2 | pvalue | pvalue_adjusted |
|---|---|---|---|---|
| interactor_ratio | mitosis | inter | 1.7019040626548957e-07 | 6.390823418948996e-06 |
| interactor_abundance | mitosis | inter | 0.00297728379962917 | 0.024892462452794143 |
| complex_abundance | mitosis | inter | 0.011440001815532655 | 0.048838986869095324 |
| total_abundance | mitosis | inter | 0.12544562938414316 | 0.43953454865428276 |
| monomer_abundance | mitosis | inter | 0.13142496679295312 | 0.4797660737464228 |
| assembled_abundance | mitosis | inter | 0.14515258116211718 | 0.45056978277179743 |

O43818_Q9BV38
U3IP2_HUMAN vs WDR18_HUMAN
p-value: 0.08 q-value: 0.047

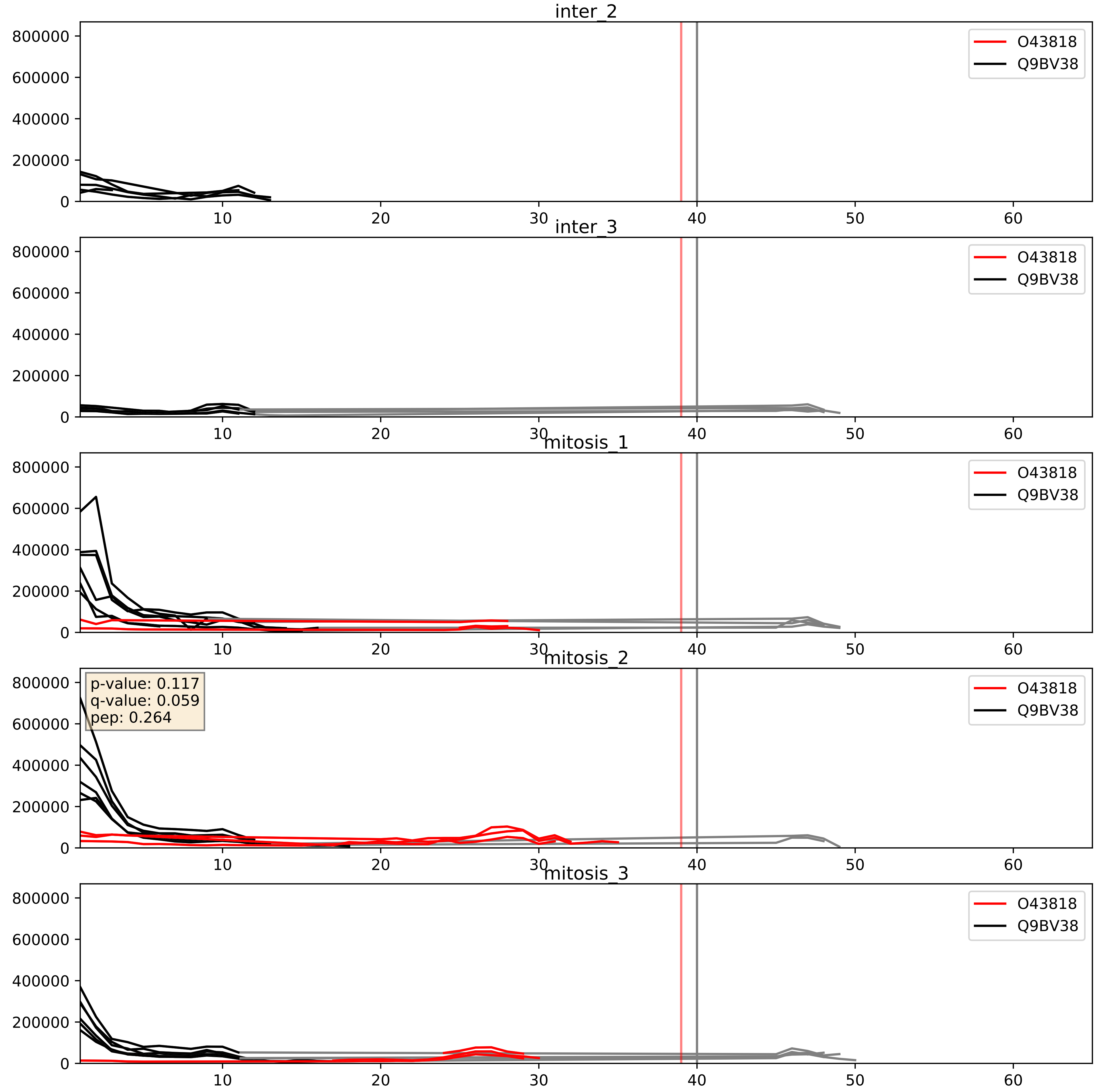


| level | condition_1 | condition_2 | pvalue | pvalue_adjusted |
|---|---|---|---|---|
| interactor_ratio | mitosis | inter | 2.0420331298463077e-05 | 0.004993481478971454 |
| complex_abundance | mitosis | inter | 0.021073640860237986 | 0.1071838180413768 |
| interactor_abundance | mitosis | inter | 0.6958430252720476 | 0.7990899243800278 |

Q14137_Q9BV38
BOP1_HUMAN vs WDR18_HUMAN
p-value: 0.073 q-value: 0.044

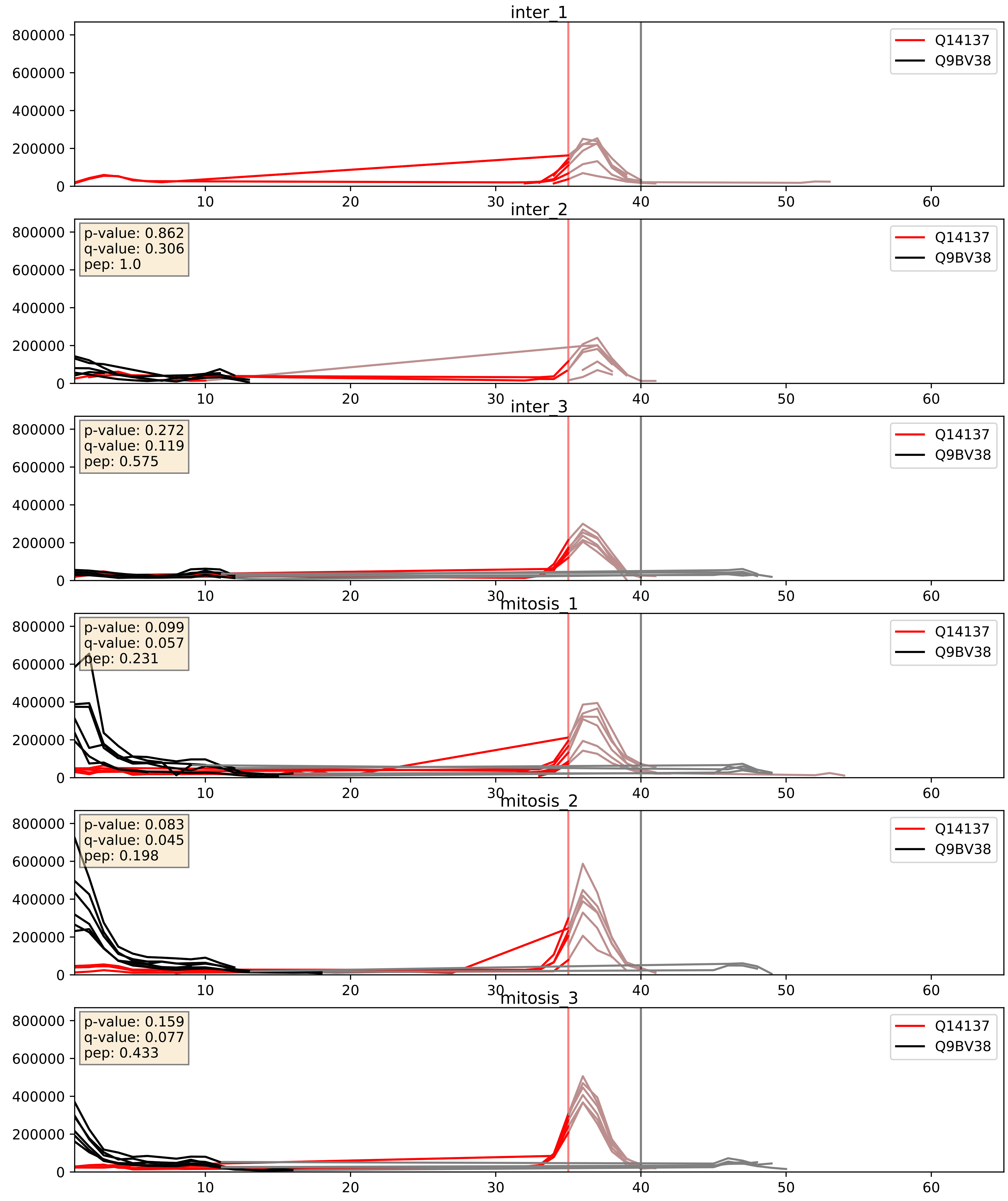


| level | condition_1 | condition_2 | pvalue | pvalue_adjusted |
|---|---|---|---|---|
| interactor_ratio | mitosis | inter | 0.001869302341360184 | 0.0524630427607973 |
| complex_abundance | mitosis | inter | 0.21507677809306397 | 0.3695903124243094 |
| interactor_abundance | mitosis | inter | 0.5302080199241265 | 0.689353305367806 |

Q5SY16_Q9BV38
NOL9_HUMAN vs WDR18_HUMAN
p-value: 0.01 q-value: 0.009

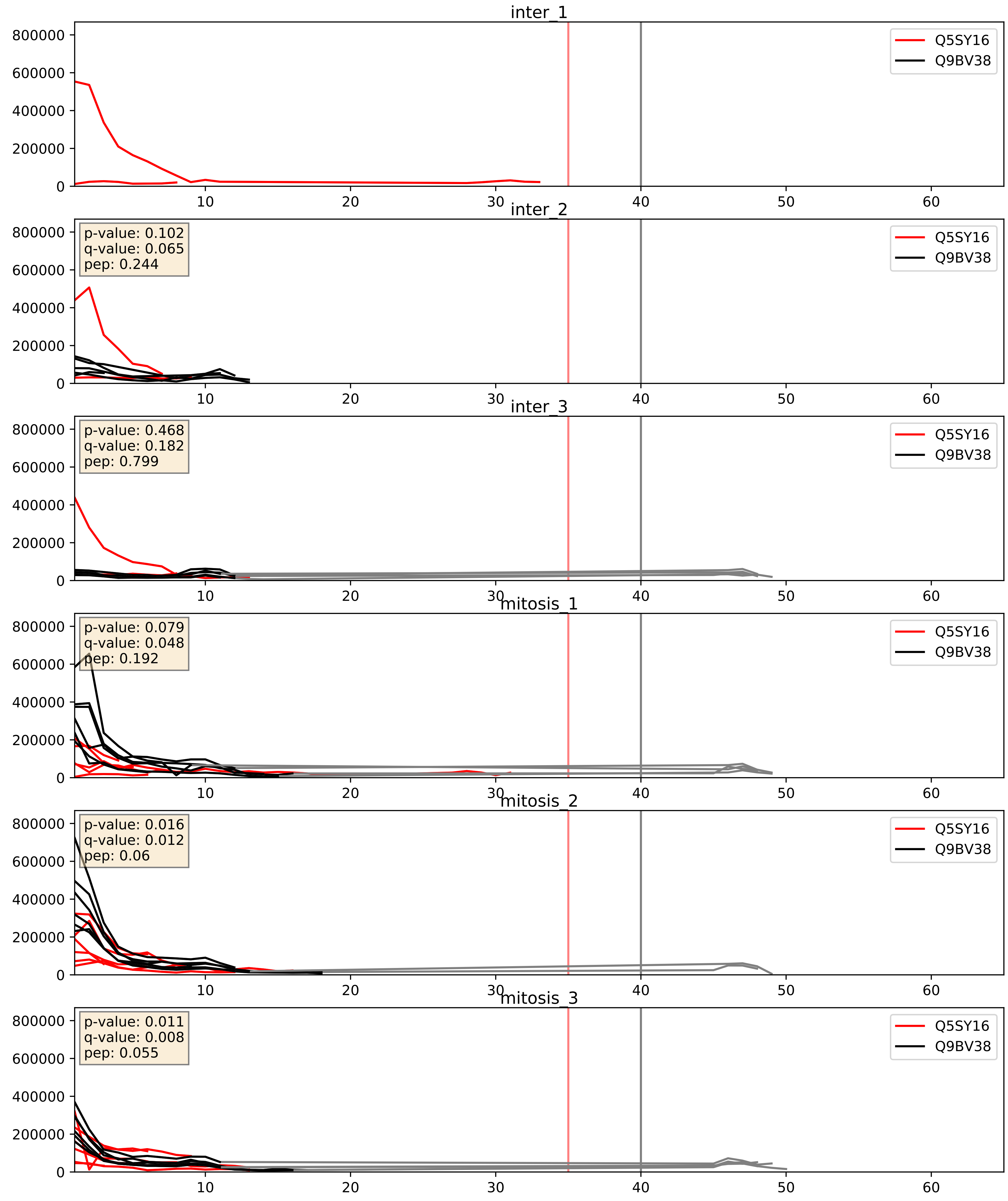


| level | condition_1 | condition_2 | pvalue | pvalue_adjusted |
|---|---|---|---|---|
| complex_abundance | mitosis | inter | 0.046844300591380464 | 0.15476511311934504 |
| interactor_ratio | mitosis | inter | 0.1031107732984949 | 0.5348944473901304 |
| interactor_abundance | mitosis | inter | 0.23255993354729296 | 0.423826491625469 |

Q9BV38_Q9BVP2
WDR18_HUMAN vs GNL3_HUMAN
p-value: 0.003 q-value: 0.039

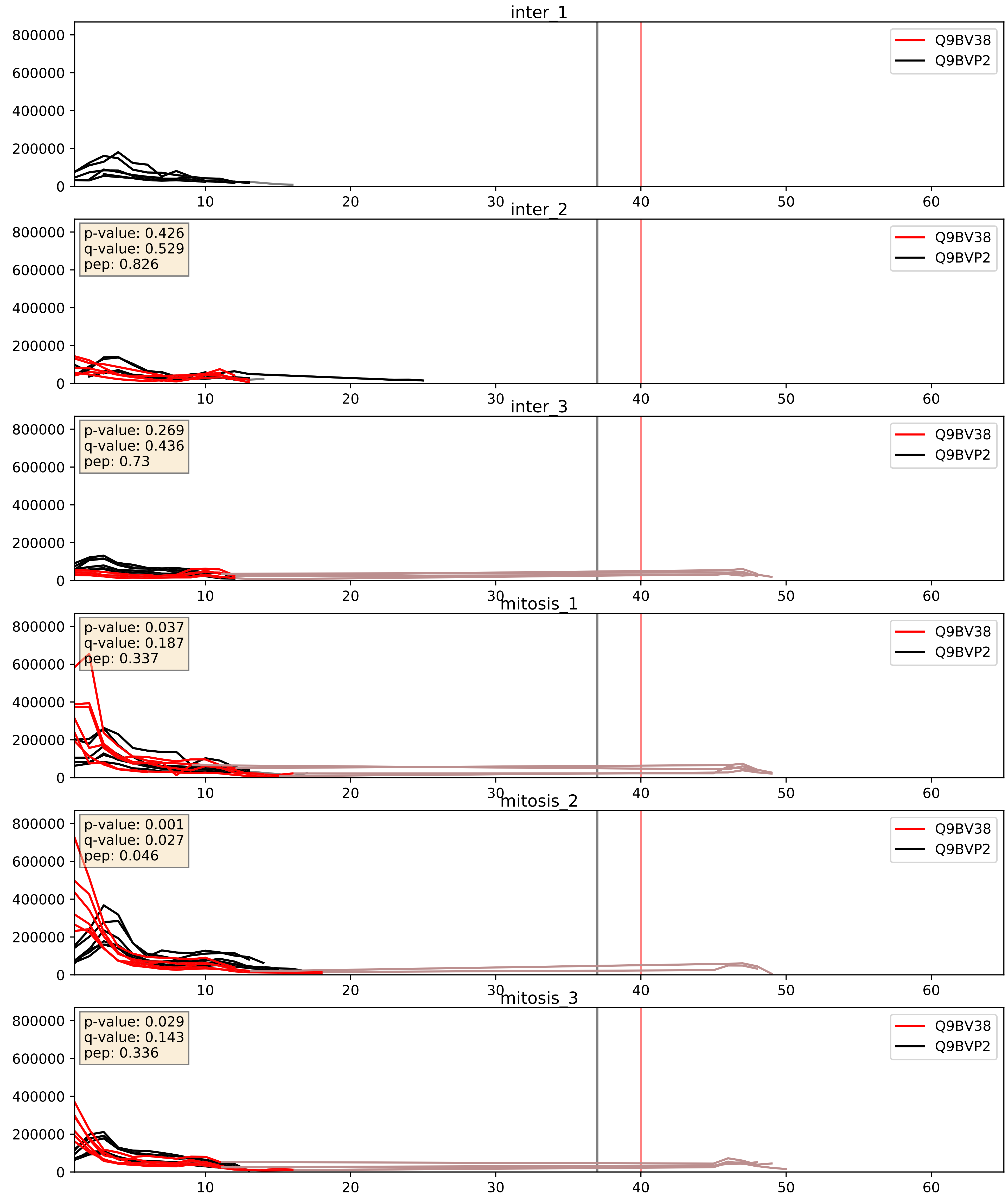


| level | condition_1 | condition_2 | pvalue | pvalue_adjusted |
|---|---|---|---|---|
| interactor_abundance | mitosis | inter | 0.03692462845525282 | 0.15802829520754763 |
| complex_abundance | mitosis | inter | 0.06145454048384596 | 0.17808086206557935 |
| interactor_ratio | mitosis | inter | 0.2008905695851433 | 0.6754215536719665 |

Q9BV38_Q9H4L4
WDR18_HUMAN vs SENP3_HUMAN
p-value: 0.028 q-value: 0.021

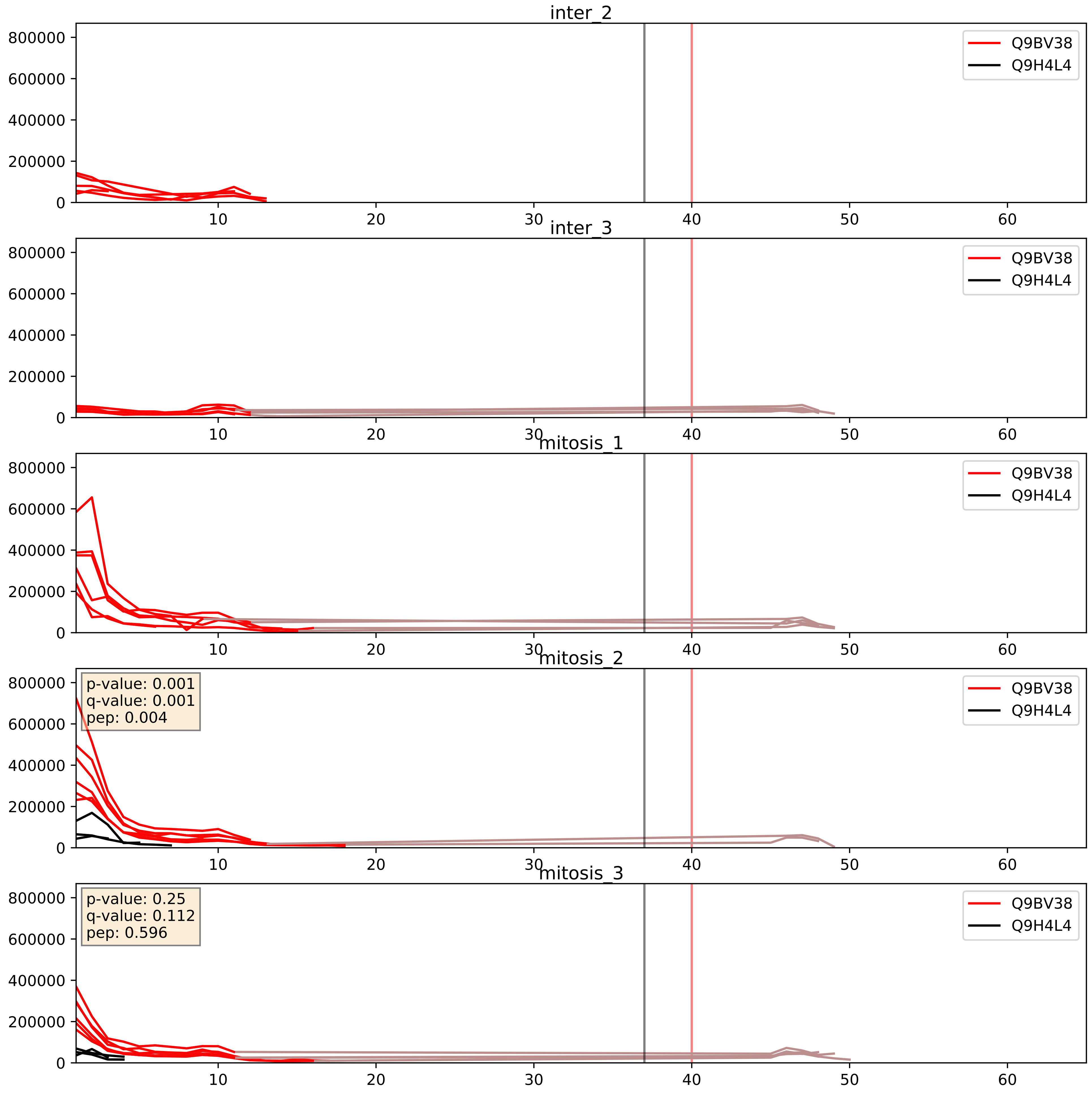


| level | condition_1 | condition_2 | pvalue | pvalue_adjusted |
|---|---|---|---|---|
| interactor_ratio | mitosis | inter | 0.1307547102105852 | 0.5784291056344235 |
| interactor_abundance | mitosis | inter | 0.19698727133955546 | 0.39091481225607855 |
| complex_abundance | mitosis | inter | 0.25744030055041556 | 0.40786681829759663 |

Q9BV38_Q9NVX2
WDR18_HUMAN vs NLE1_HUMAN
p-value: 0.048 q-value: 0.032

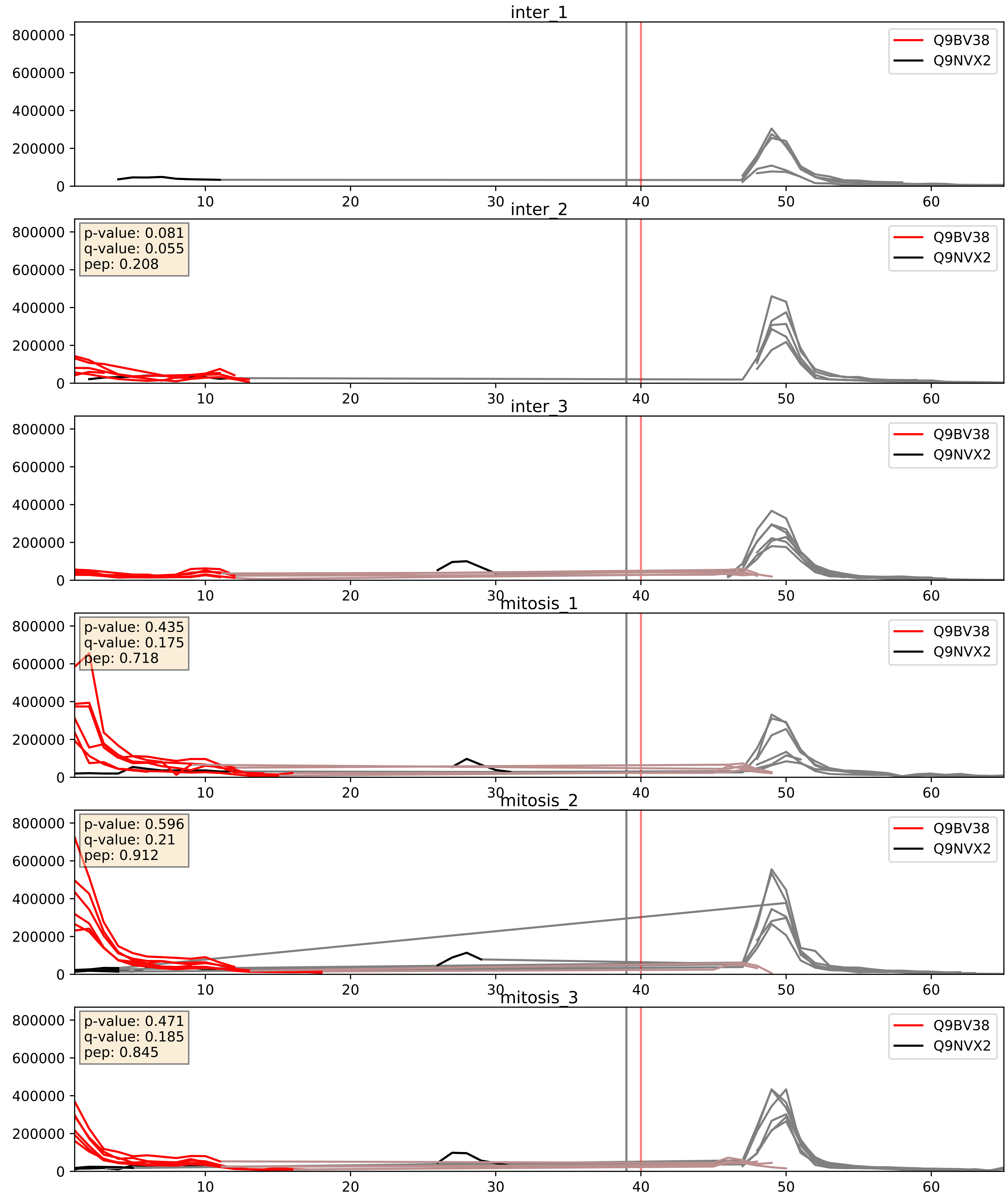


| level | condition_1 | condition_2 | pvalue | pvalue_adjusted |
|---|---|---|---|---|
| interactor_ratio | mitosis | inter | 0.008126153594009871 | 0.12739903803063093 |
| interactor_abundance | mitosis | inter | 0.027671848141419618 | 0.13815894675452584 |
| complex_abundance | mitosis | inter | 0.061232504518269844 | 0.1778907646484952 |

Q9BV38_Q9NXF1
WDR18_HUMAN vs TEX10_HUMAN
p-value: 0.005 q-value: 0.005

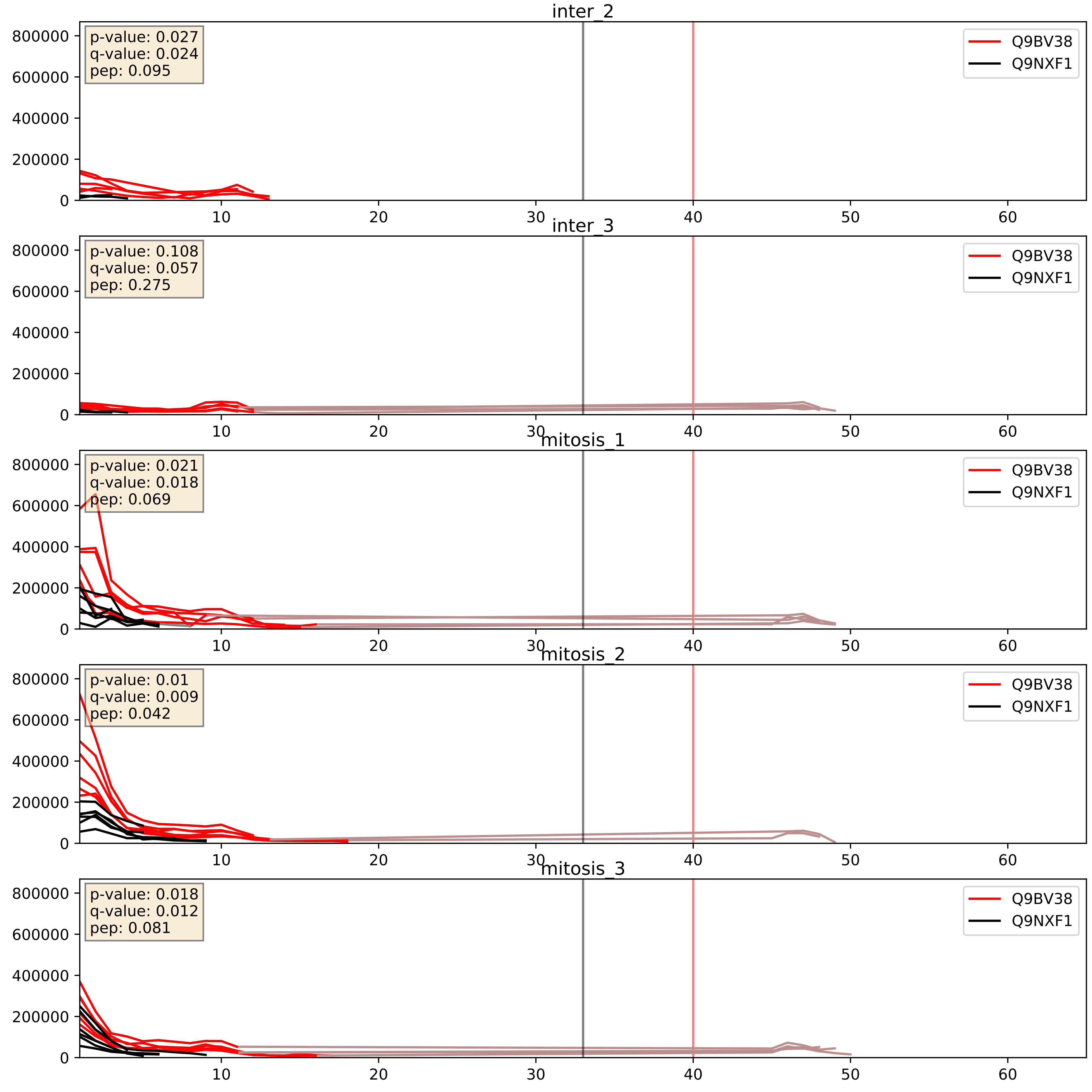


| level | condition_1 | condition_2 | pvalue | pvalue_adjusted |
|---|---|---|---|---|
| complex_abundance | mitosis | inter | 0.0064532229320805835 | 0.05955783692156859 |
| interactor_abundance | mitosis | inter | 0.015148920743567303 | 0.10774803619853436 |
| interactor_ratio | mitosis | inter | 0.10491269377910745 | 0.5400196384540948 |

Q9BV38_Q9Y4W2
WDR18_HUMAN vs LAS1L_HUMAN
p-value: 0.003 q-value: 0.003

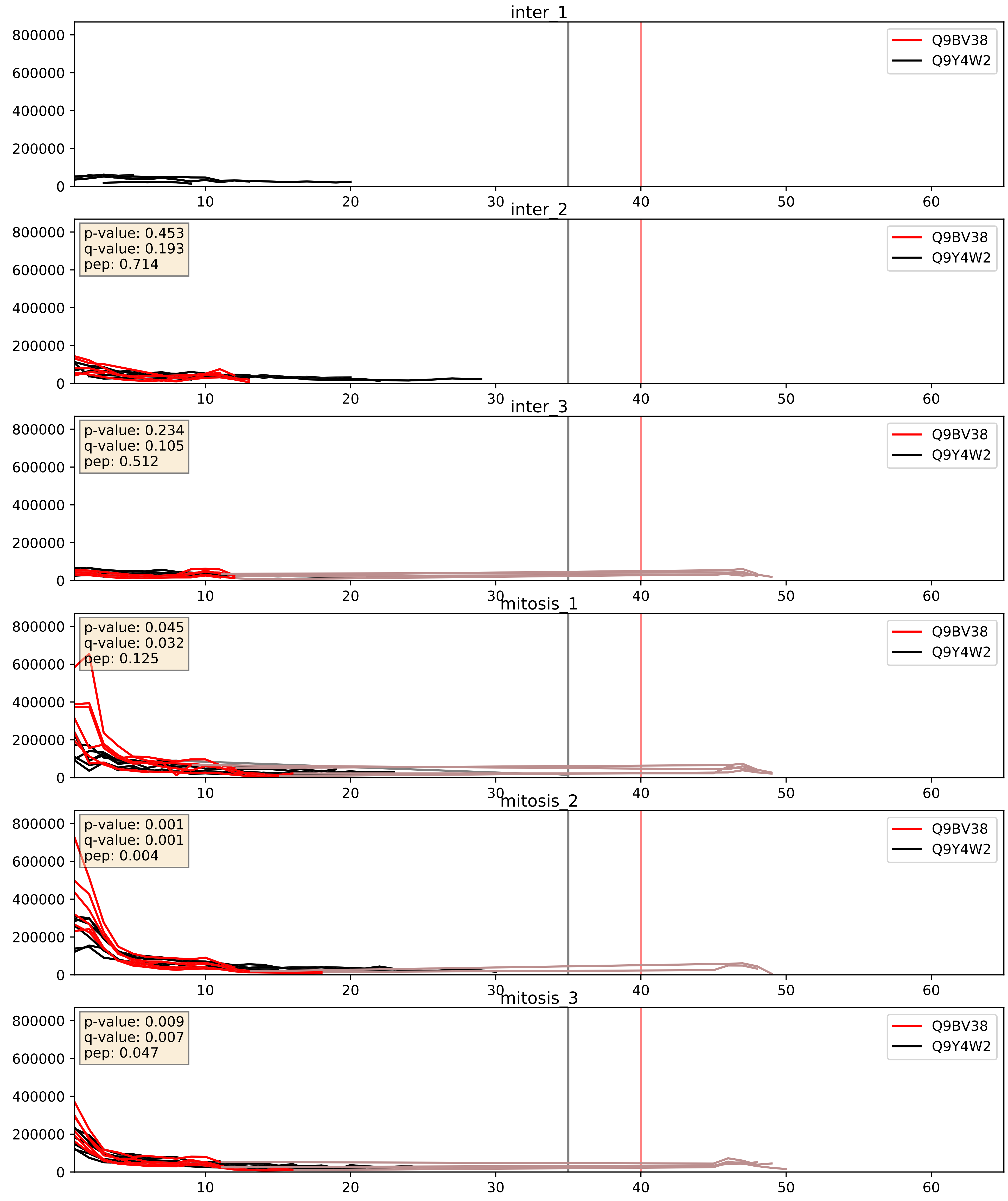


| level | condition_1 | condition_2 | pvalue | pvalue_adjusted |
|---|---|---|---|---|
| interactor_abundance | mitosis | inter | 0.03368708868945364 | 0.1532530798125457 |
| complex_abundance | mitosis | inter | 0.05042502252144245 | 0.15980680961997312 |
| interactor_ratio | mitosis | inter | 0.251857865158055 | 0.7251677296572153 |
