## Supplementary Data 2 for "SECAT: Quantifying differential protein-protein interaction states by network-centric analysis": hela_string_000052_Q13242.pdf

Q13242_Q13242

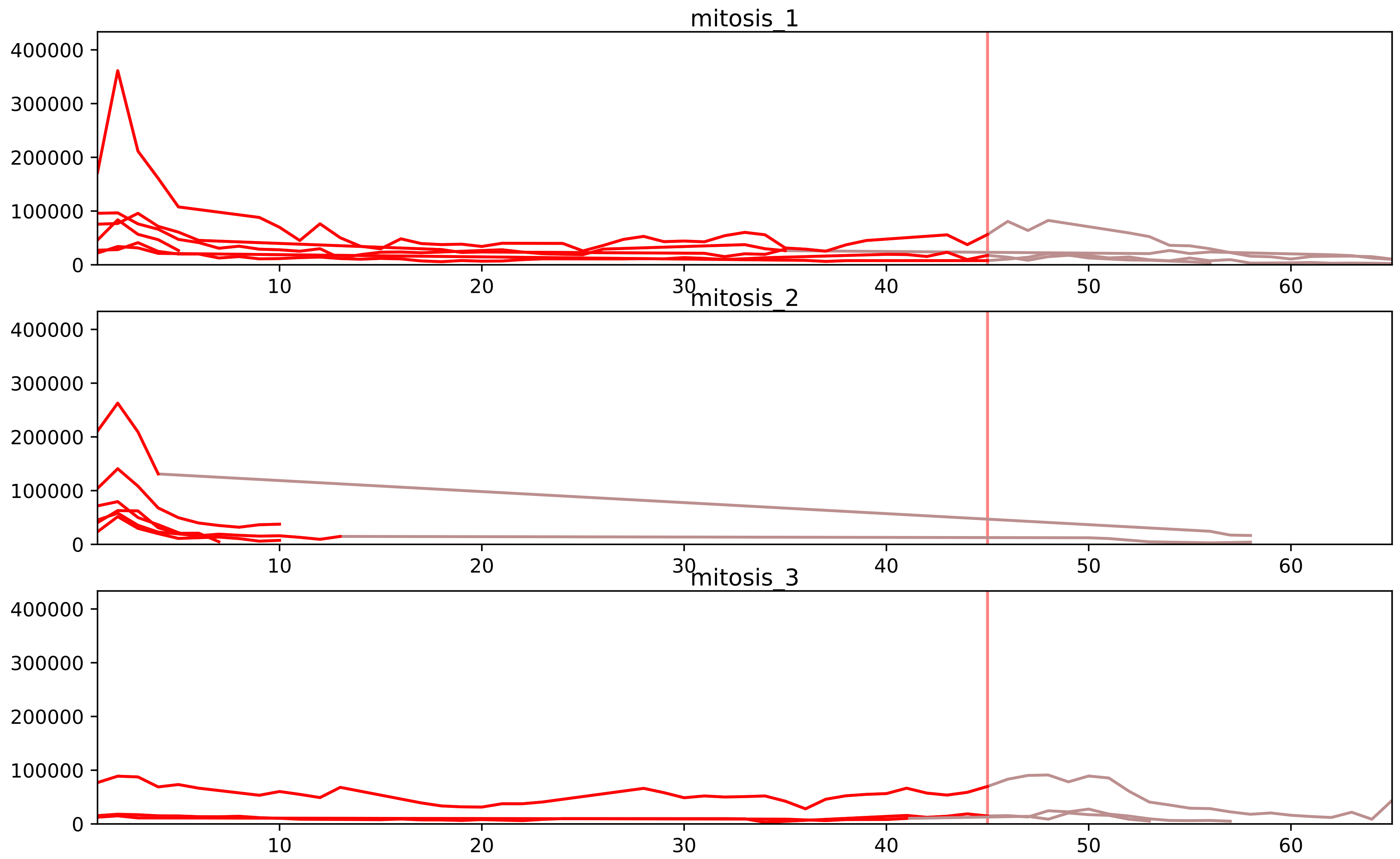


| level | condition_1 | condition_2 | pvalue | pvalue_adjusted |
|---|---|---|---|---|
| interactor_ratio | mitosis | inter | 1.829397131985465e-07 | 6.732181445706511e-06 |
| complex_abundance | mitosis | inter | 0.0005006804788283428 | 0.00687501553018023 |
| monomer_abundance | mitosis | inter | 0.011554247313514999 | 0.1902553554136748 |
| assembled_abundance | mitosis | inter | 0.02076233762522784 | 0.1705061446029676 |
| total_abundance | mitosis | inter | 0.021856082489921543 | 0.18835452556753943 |
| interactor_abundance | mitosis | inter | 0.028160152613427185 | 0.09964361693981928 |

P38159_Q13242
RBMX_HUMAN vs SRSF9_HUMAN
p-value: 0.042 q-value: 0.028

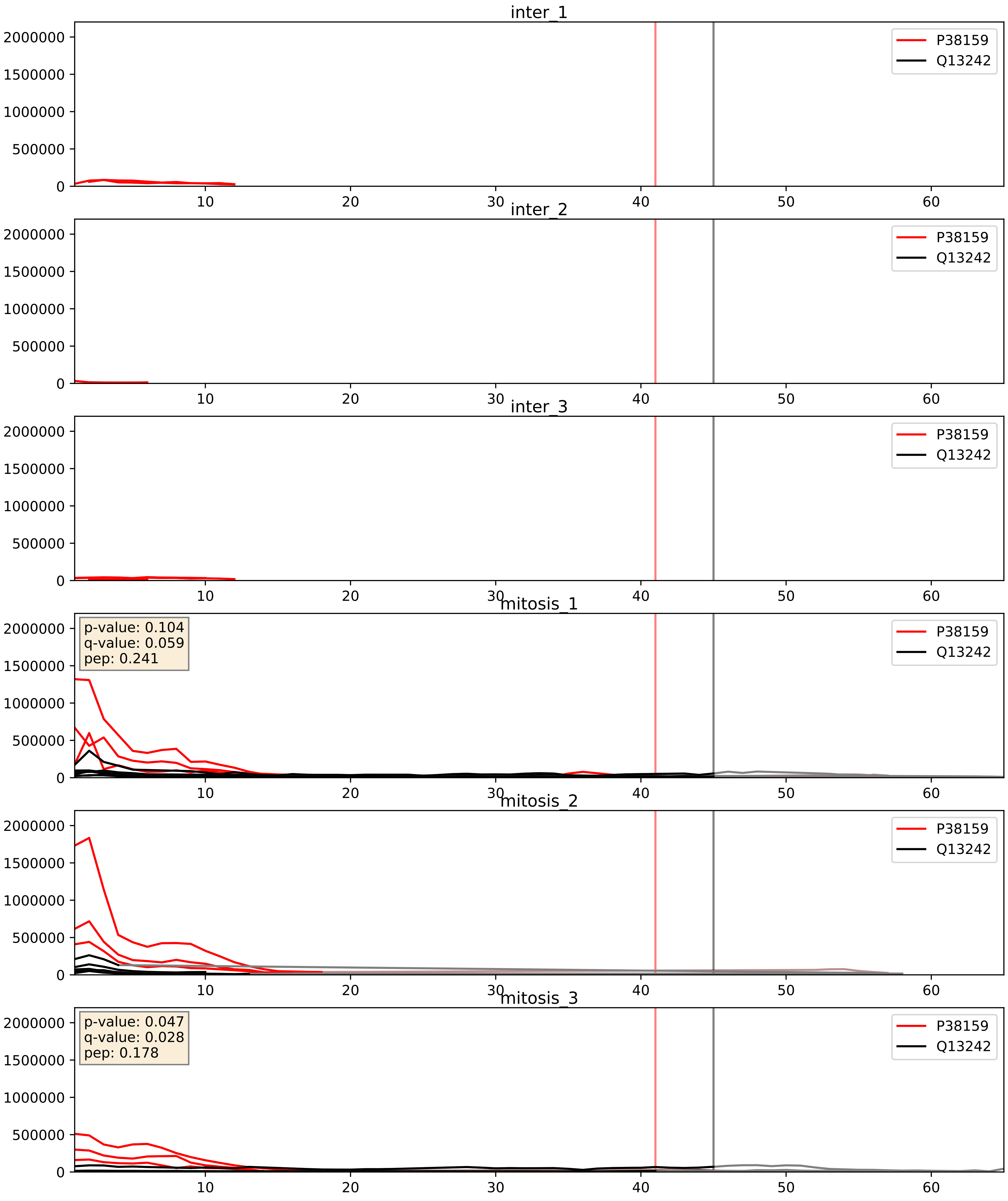


| level | condition_1 | condition_2 | pvalue | pvalue_adjusted |
|---|---|---|---|---|
| interactor_abundance | mitosis | inter | 0.00016138549462100720 | 0.009910743628815524 |
| interactor_ratio | mitosis | inter | 0.0022484423567735888 | 0.05896690574521502 |
| complex_abundance | mitosis | inter | 0.0029712233791559066 | 0.03988944941488032 |

P62995_Q13242
TRA2B_HUMAN vs SRSF9_HUMAN
p-value: 0.001 q-value: 0.002

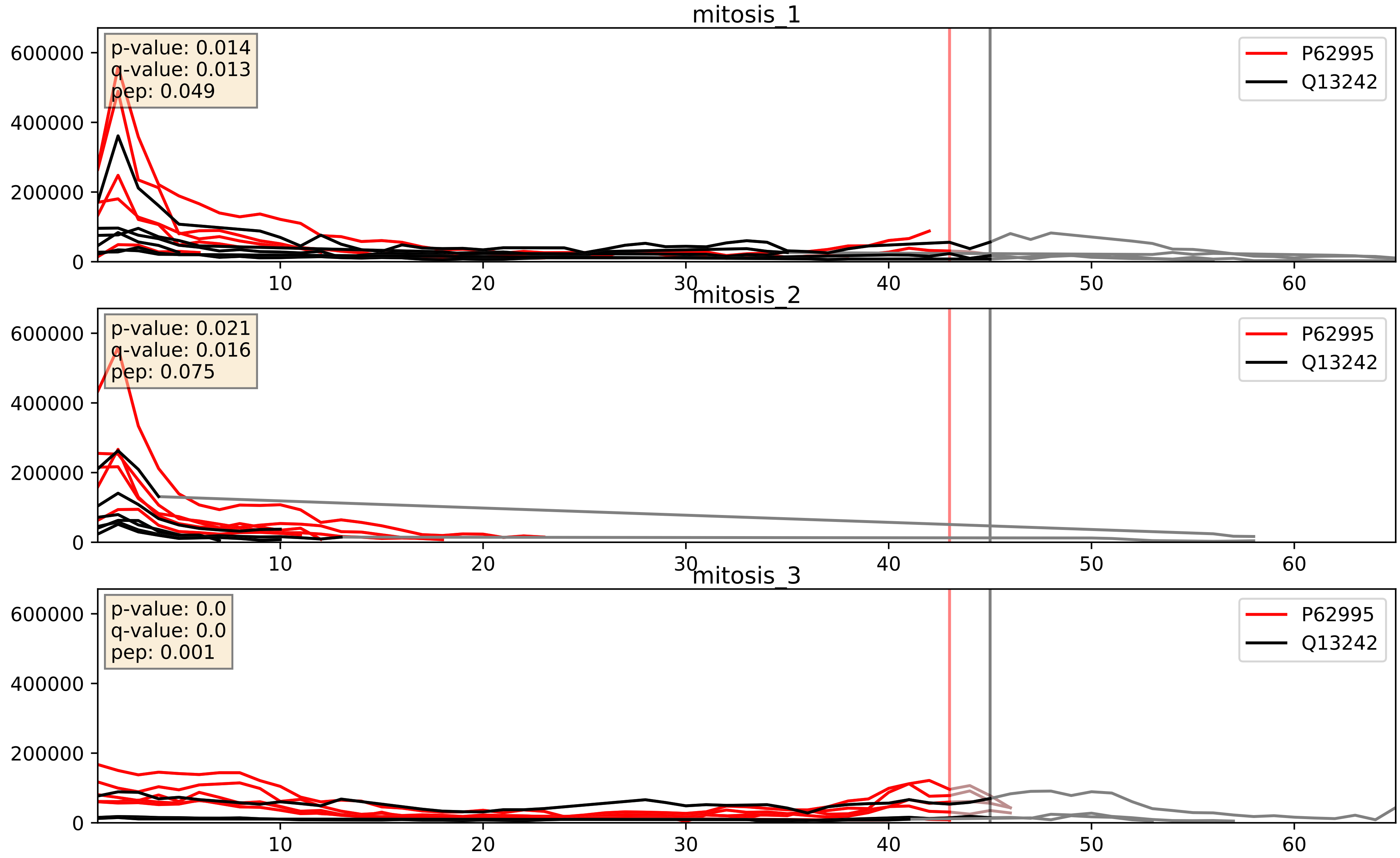


| level | condition_1 | condition_2 | pvalue | pvalue_adjusted |
|---|---|---|---|---|
| interactor_abundance | mitosis | inter | 0.0006196284287297792 | 0.01748938971196646 |
| complex_abundance | mitosis | inter | 0.004301196506507519 | 0.04889540782962067 |
| interactor_ratio | mitosis | inter | 0.008613258867848741 | 0.13203195690147984 |

P84103_Q13242
SRSF3_HUMAN vs SRSF9_HUMAN
p-value: 0.059 q-value: 0.037

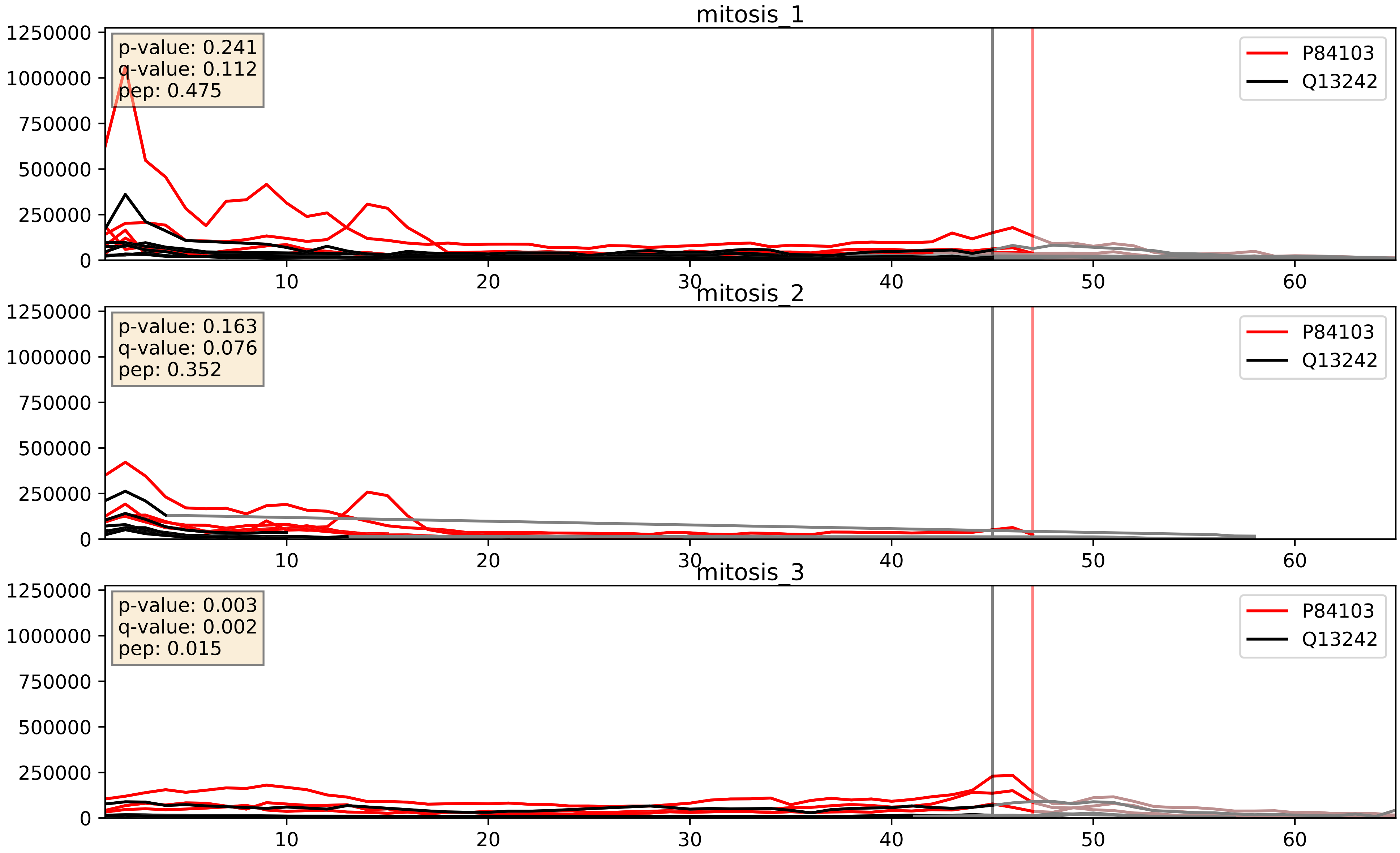


| level | condition_1 | condition_2 | pvalue | pvalue_adjusted |
|---|---|---|---|---|
| interactor_ratio | mitosis | inter | 0.00018729042781272996 | 0.014134487523529912 |
| interactor_abundance | mitosis | inter | 0.004192456982592769 | 0.05145151508386251 |
| complex_abundance | mitosis | inter | 0.012111824621720255 | 0.0818150085787963 |

Q13242_Q16629
SRSF9_HUMAN vs SRSF7_HUMAN
p-value: 0.007 q-value: 0.006

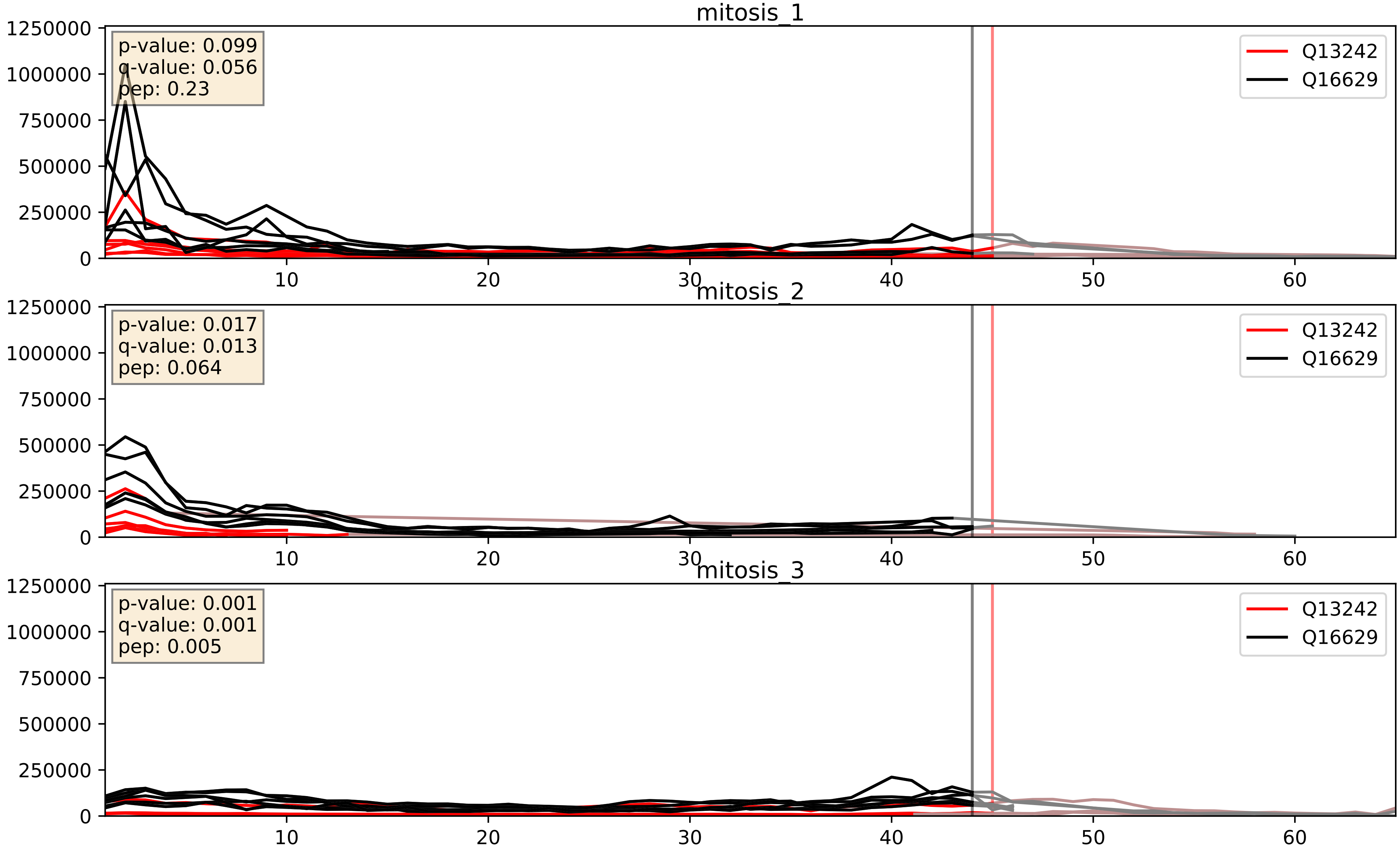


| level | condition_1 | condition_2 | pvalue | pvalue_adjusted |
|---|---|---|---|---|
| interactor_ratio | mitosis | inter | 0.0006742820733493766 | 0.029848225645041686 |
| complex_abundance | mitosis | inter | 0.0025654119015484406 | 0.03664546283589145 |
| interactor_abundance | mitosis | inter | 0.04867027237079332 | 0.175973614147409 |
