## Supplementary Data 2 for "SECAT: Quantifying differential protein-protein interaction states by network-centric analysis": hela_string_000053_P12236.pdf

P12236_P12236

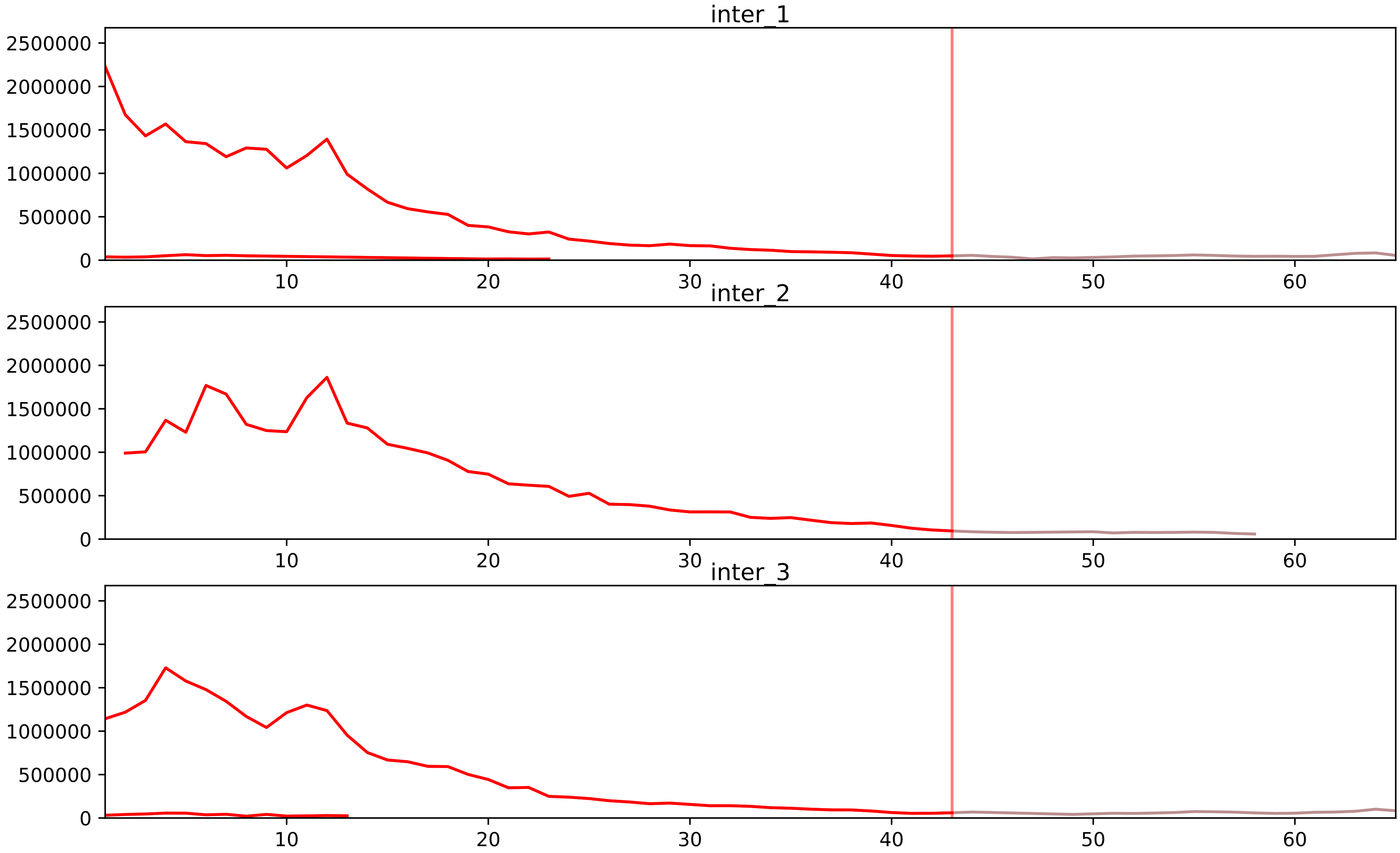


| level | condition_1 | condition_2 | pvalue | pvalue_adjusted |
|---|---|---|---|---|
| assembled_abundance | mitosis | inter | 1.8679667460981593e-07 | 0.0008743952338485483 |
| total_abundance | mitosis | inter | 2.943906534735538e-07 | 0.0014987428168338624 |
| complex_abundance | mitosis | inter | 7.987306921555082e-06 | 0.00041358484442386497 |
| interactor_abundance | mitosis | inter | 9.11661826607866e-06 | 0.000492637777039522 |
| monomer_abundance | mitosis | inter | 3.148634826898521e-05 | 0.0267281636508792 |
| interactor_ratio | mitosis | inter | 0.00019054730348443444 | 0.002873828183699666 |

P12236_P15880
ADT3_HUMAN vs RS2_HUMAN
p-value: 0.0 q-value: 0.018

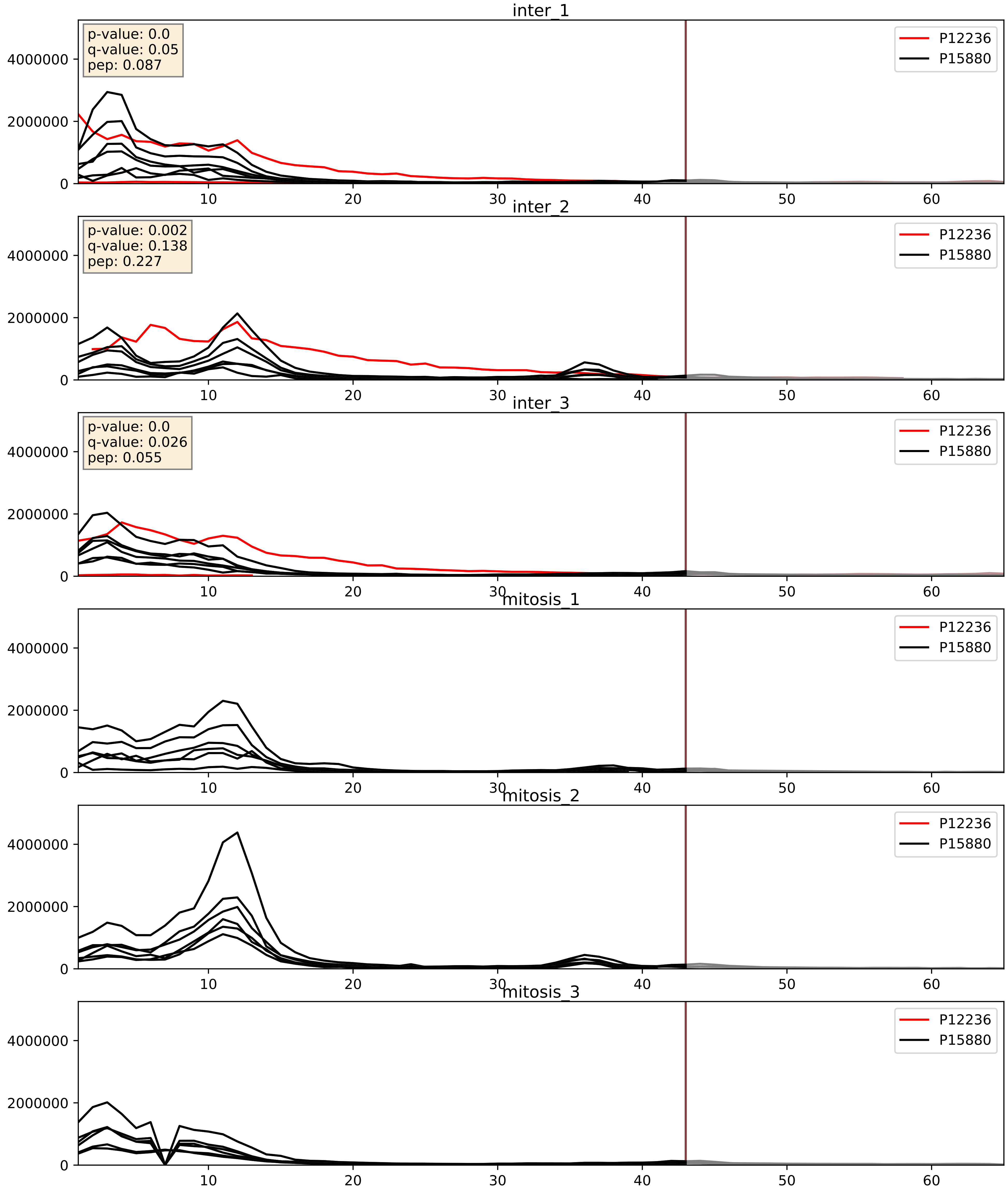


| level | condition_1 | condition_2 | pvalue | pvalue_adjusted |
|---|---|---|---|---|
| interactor_ratio | mitosis | inter | 4.0034237732346944e-06 | 0.002520386835947235 |
| complex_abundance | mitosis | inter | 1.3908973094562453e-05 | 0.005691857151652958 |
| interactor_abundance | mitosis | inter | 6.587220104638708e-05 | 0.006633718128906747 |

P12236_P63220
ADT3_HUMAN vs RS21_HUMAN
p-value: 0.0 q-value: 0.042

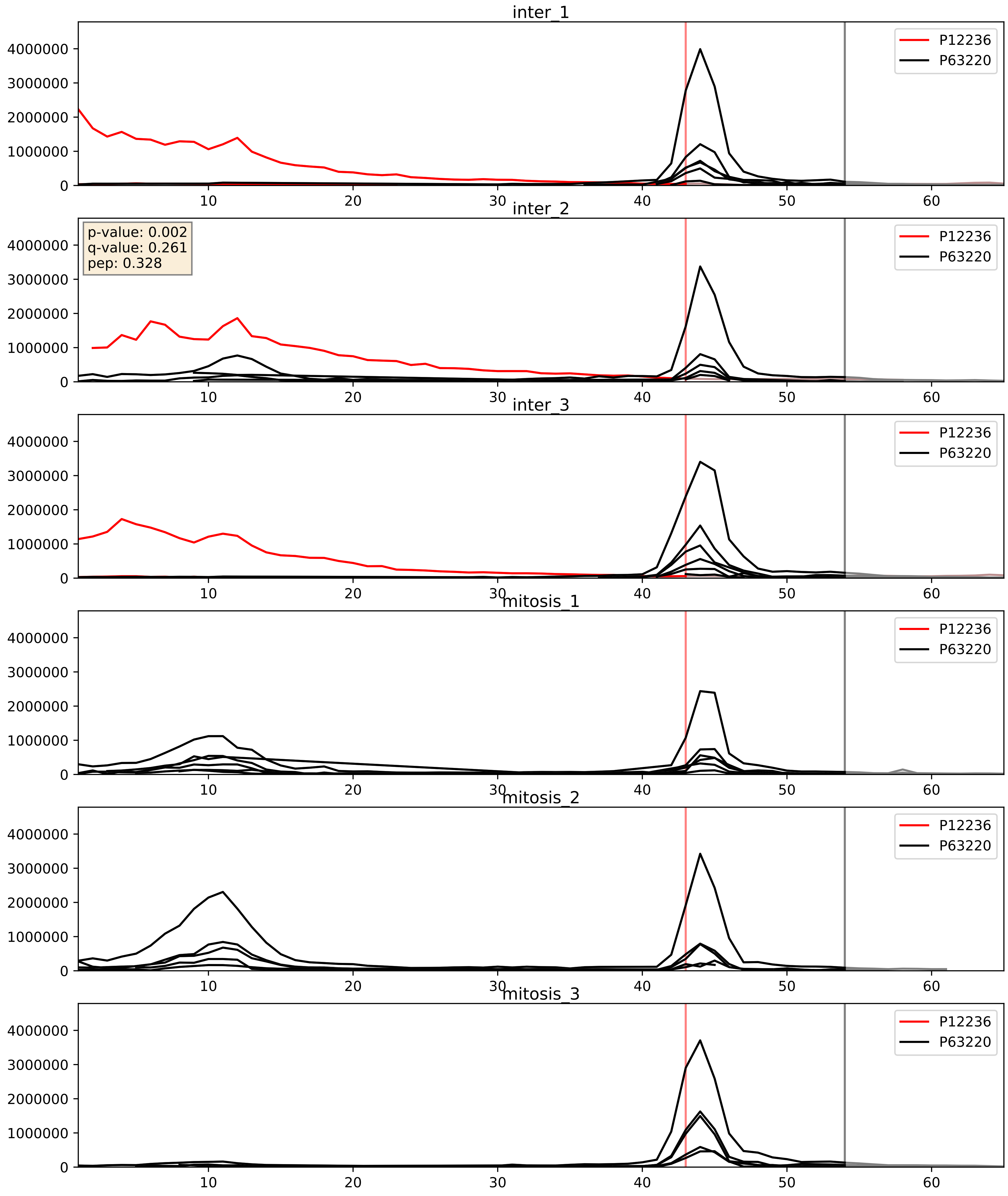


| level | condition_1 | condition_2 | pvalue | pvalue_adjusted |
|---|---|---|---|---|
| interactor_abundance | mitosis | inter | 0.0001795330340236242 | 0.010140496760639767 |
| complex_abundance | mitosis | inter | 0.0007998830179067109 | 0.018707646539020344 |
| interactor_ratio | mitosis | inter | 0.5219840227956803 | 0.8786895401167326 |
