## Supplementary Data 2 for "SECAT: Quantifying differential protein-protein interaction states by network-centric analysis": hela_string_000054_P14635.pdf

# P14635_P14635

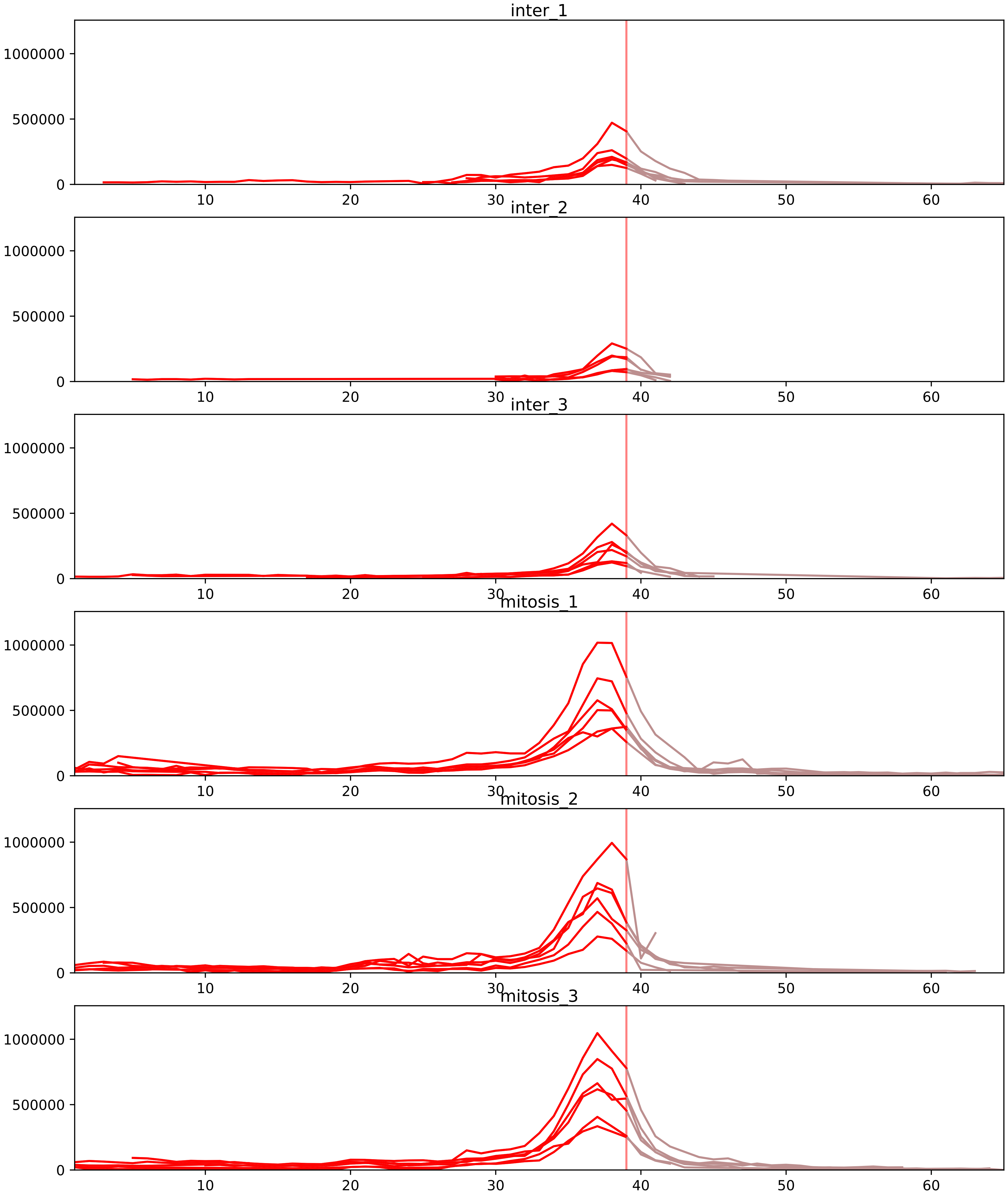


| level | condition_1 | condition_2 | pvalue | pvalue_adjusted |
|---|---|---|---|---|
| interactor_ratio | mitosis | inter | 2.1777427908411636e-07 | 7.84365637877767e-06 |
| interactor_abundance | mitosis | inter | 0.0001320465818999694 | 0.0028245321848403312 |
| complex_abundance | mitosis | inter | 0.0006016033277756815 | 0.007795423402163761 |
| assembled_abundance | mitosis | inter | 0.0018742887641910978 | 0.04067828623906237 |
| total_abundance | mitosis | inter | 0.0019659457427467046 | 0.04766014179201654 |
| monomer_abundance | mitosis | inter | 0.005753397967977147 | 0.14889207607573982 |

O95067_P14635
CCNB2_HUMAN vs CCNB1_HUMAN
p-value: 0.01 q-value: 0.008

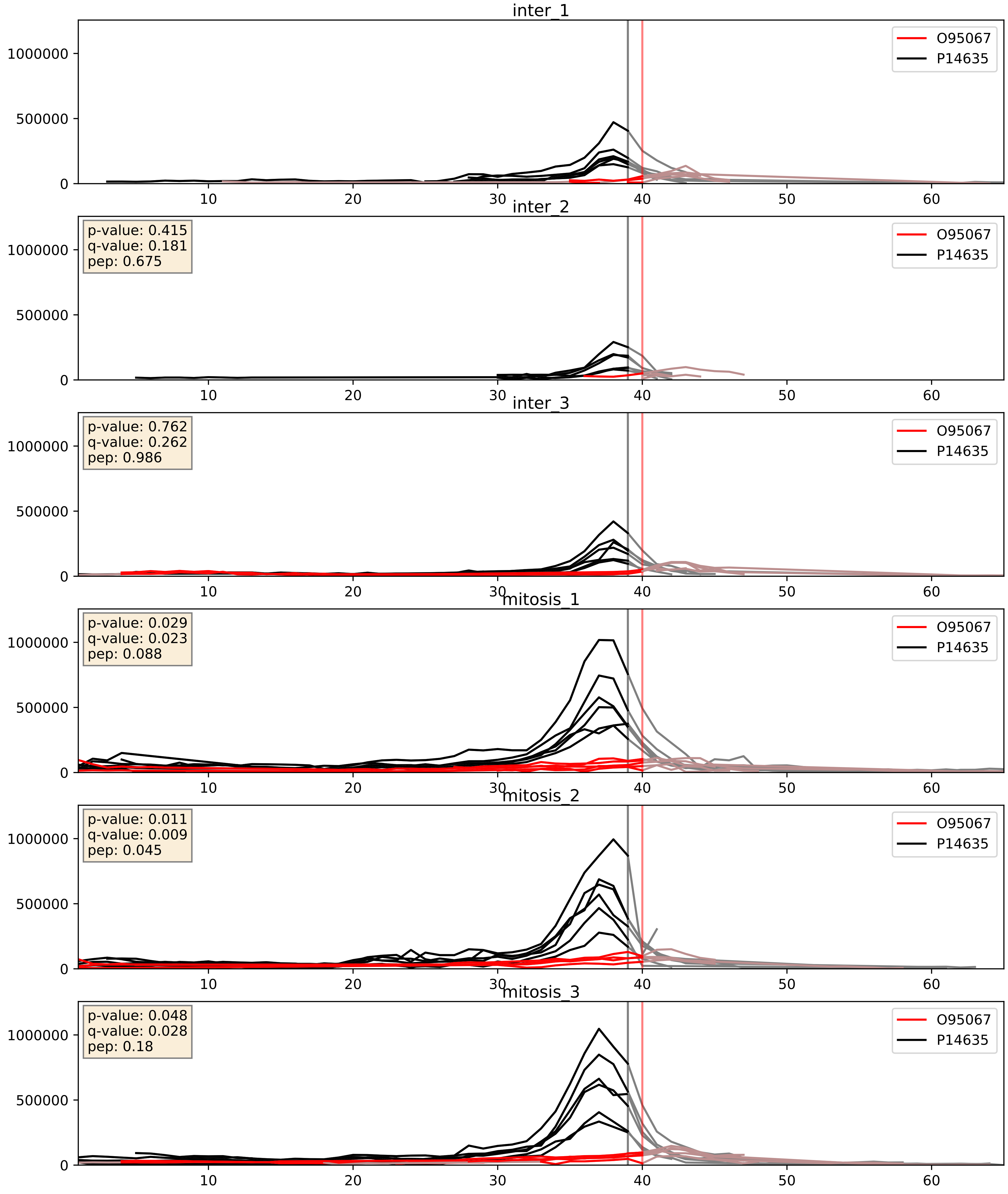


| level | condition_1 | condition_2 | pvalue | pvalue_adjusted |
|---|---|---|---|---|
| complex_abundance | mitosis | inter | 0.000540651241243441 | 0.014880947347407894 |
| interactor_abundance | mitosis | inter | 0.0025820700836980265 | 0.03806046117740333 |
| interactor_ratio | mitosis | inter | 0.003698695469143503 | 0.07875829158176215 |

P06493_P14635
CDK1_HUMAN vs CCNB1_HUMAN
p-value: 0.0 q-value: 0.0

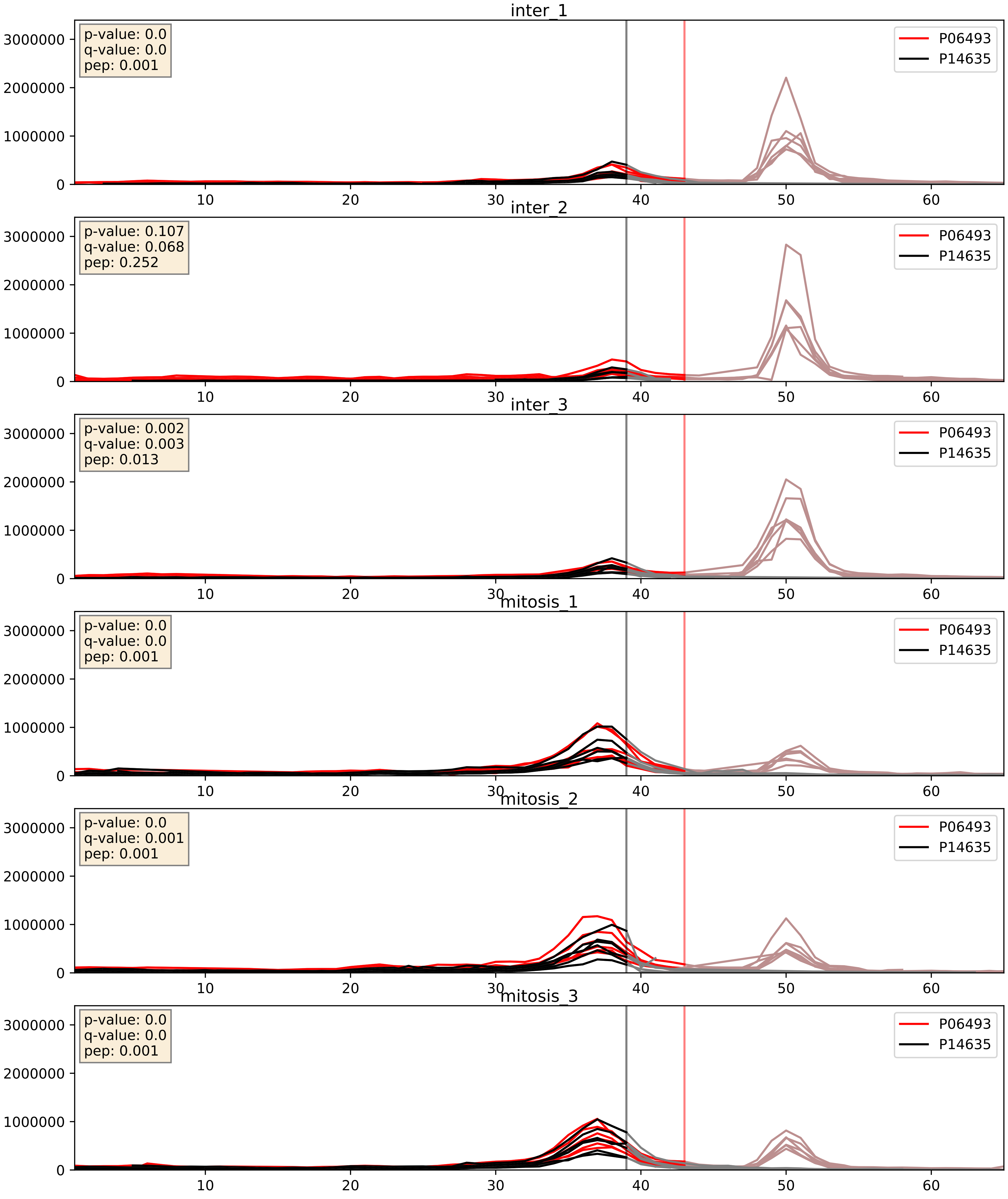


| level | condition_1 | condition_2 | pvalue | pvalue_adjusted |
|---|---|---|---|---|
| interactor_abundance | mitosis | inter | 0.0002579902051684421 | 0.012234881752032491 |
| complex_abundance | mitosis | inter | 0.0009312502745939036 | 0.020079351008876107 |
| interactor_ratio | mitosis | inter | 0.0302146448575311 | 0.2765852972945998 |

P14635_P24941
CCNB1_HUMAN vs CDK2_HUMAN
p-value: 0.023 q-value: 0.018

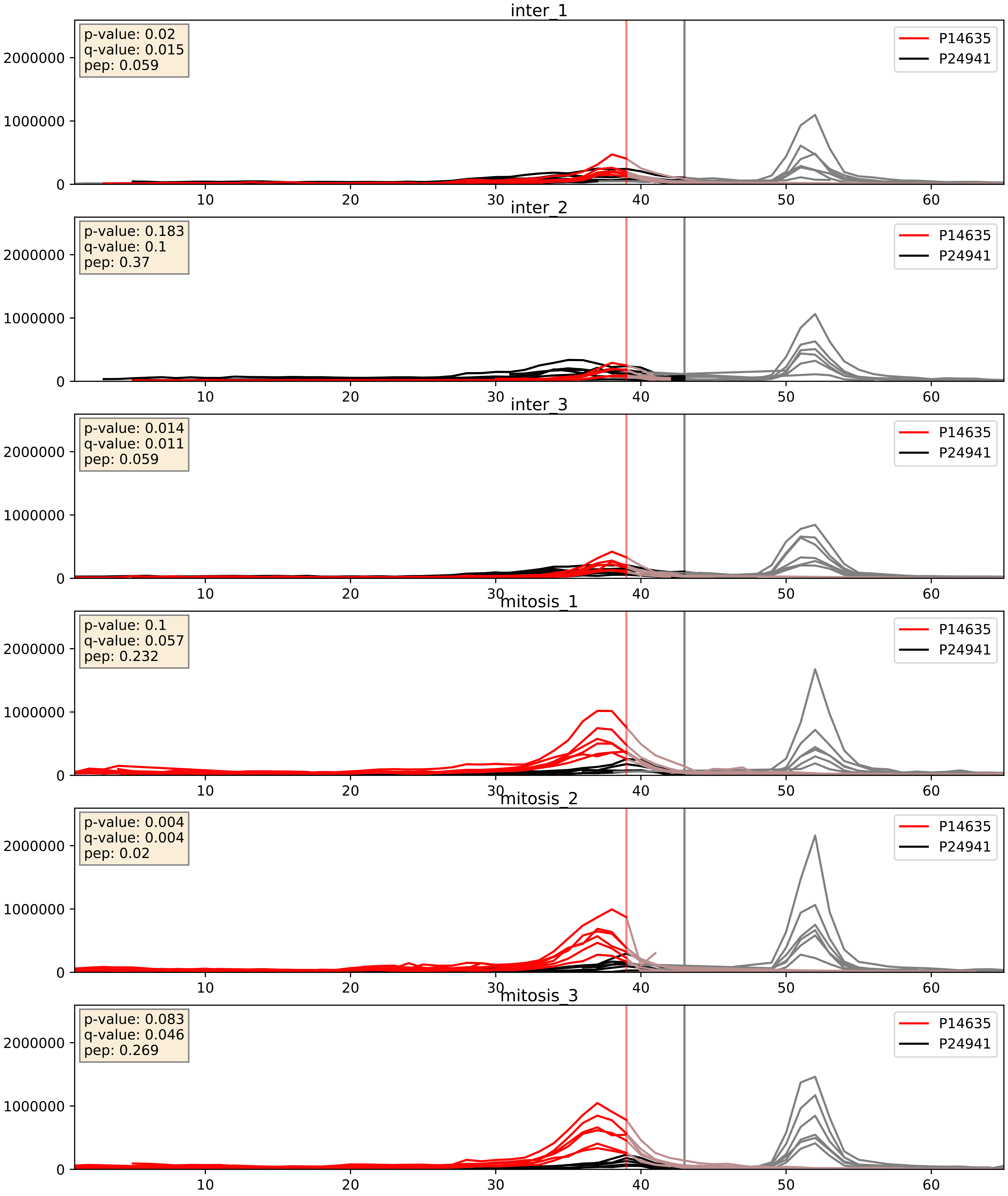


| level | condition_1 | condition_2 | pvalue | pvalue_adjusted |
|---|---|---|---|---|
| interactor_ratio | mitosis | inter | 0.0008074983906906686 | 0.032604652001472285 |
| interactor_abundance | mitosis | inter | 0.002332060110087462 | 0.03586830170101165 |
| complex_abundance | mitosis | inter | 0.0425001476113568 | 0.1476929246638287 |

P14635_P30307
CCNB1_HUMAN vs MPIP3_HUMAN
p-value: 0.013 q-value: 0.011

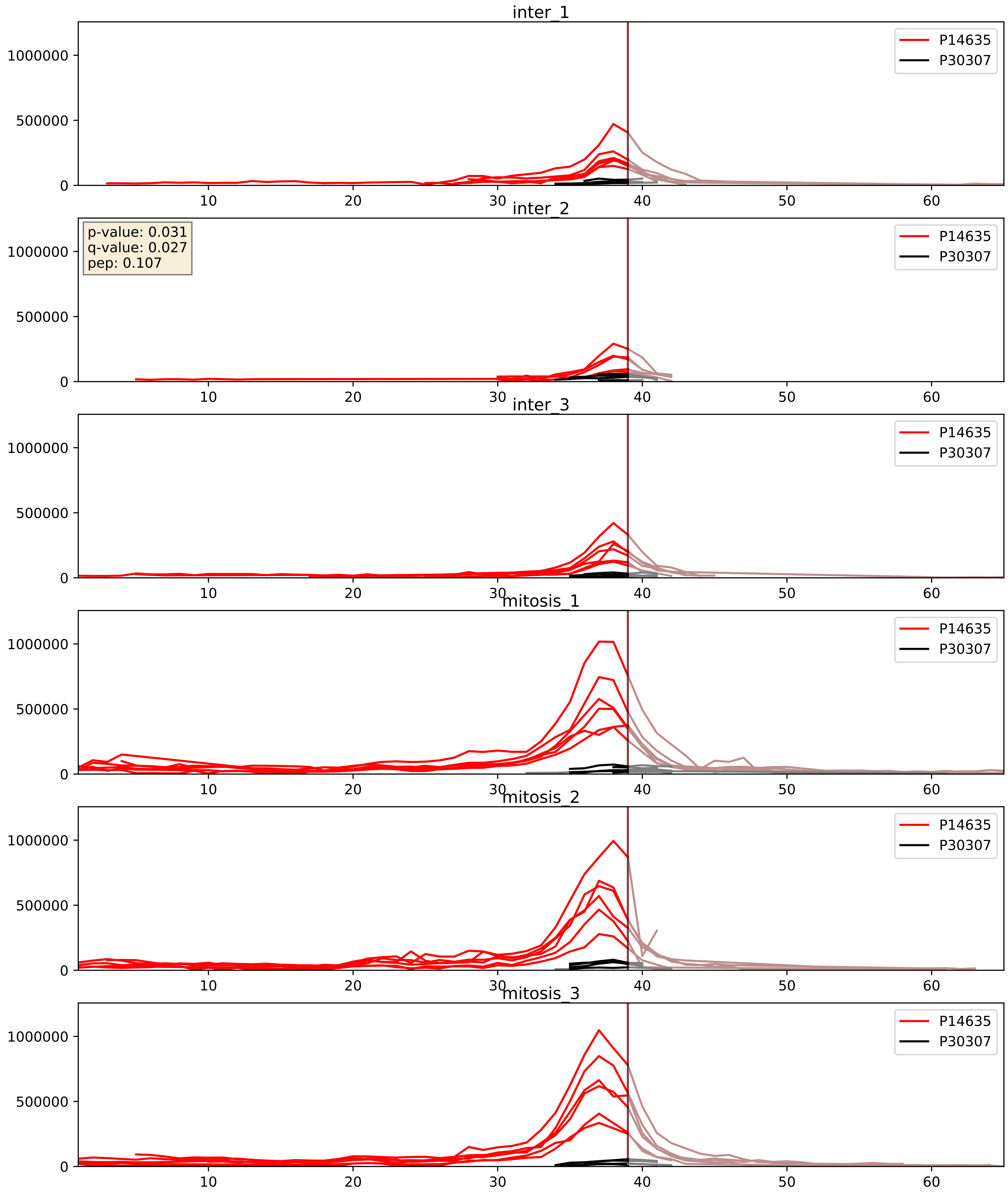


| level | condition_1 | condition_2 | pvalue | pvalue_adjusted |
|---|---|---|---|---|
| interactor_abundance | mitosis | inter | 0.0020507322378847045 | 0.03290396992744718 |
| interactor_ratio | mitosis | inter | 0.005842696997531102 | 0.10441228872414662 |
| complex_abundance | mitosis | inter | 0.012644321240594406 | 0.08351496128046922 |

P14635_P33552
CCNB1_HUMAN vs CKS2_HUMAN
p-value: 0.037 q-value: 0.026

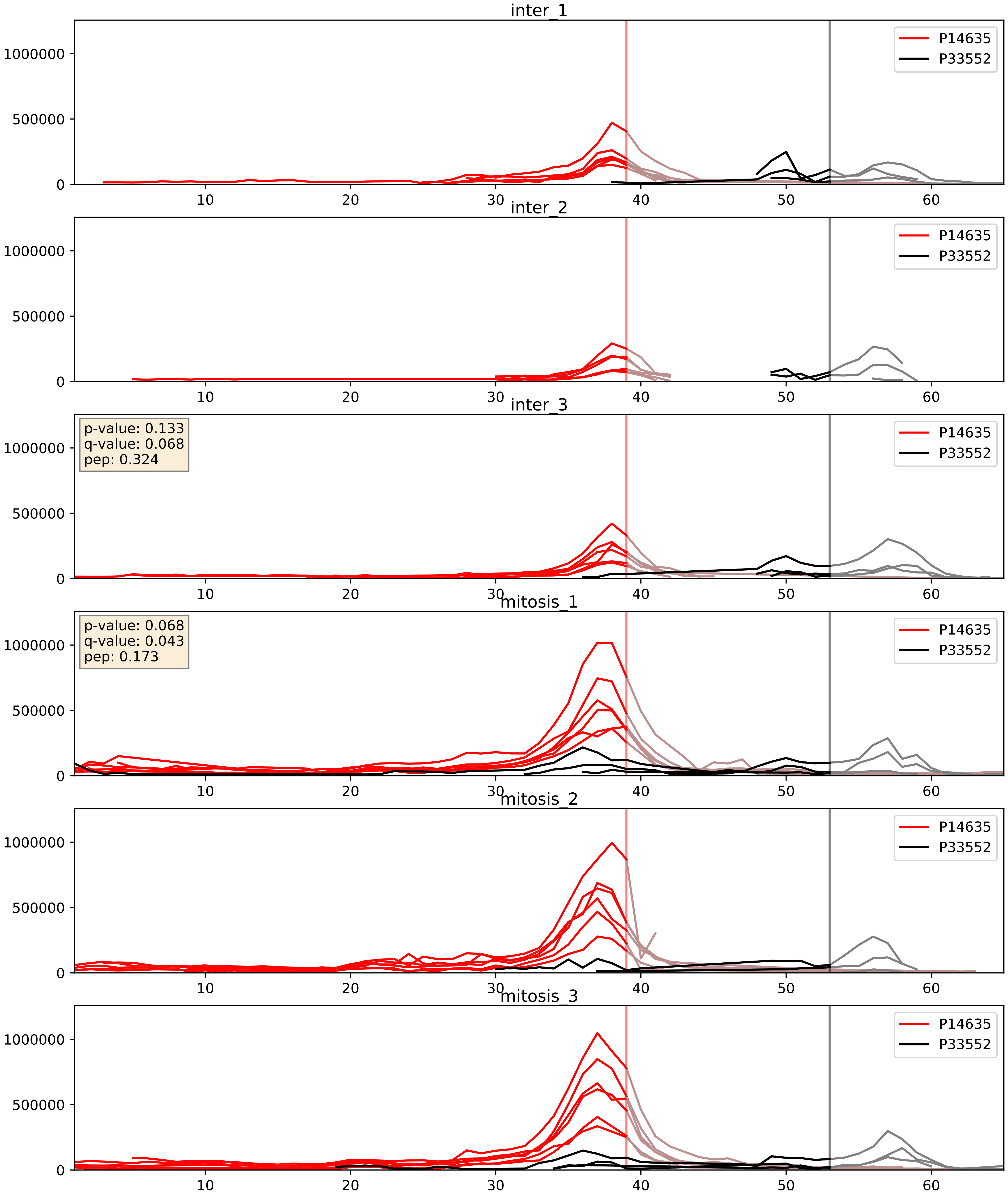


| level | condition_1 | condition_2 | pvalue | pvalue_adjusted |
|---|---|---|---|---|
| interactor_abundance | mitosis | inter | 0.006463740447287108 | 0.0687067394609051 |
| complex_abundance | mitosis | inter | 0.01479460537239751 | 0.09014873100884367 |
| interactor_ratio | mitosis | inter | 0.01952895271025558 | 0.21160485468327564 |

P14635_P61024
CCNB1_HUMAN vs CKS1_HUMAN
p-value: 0.041 q-value: 0.028

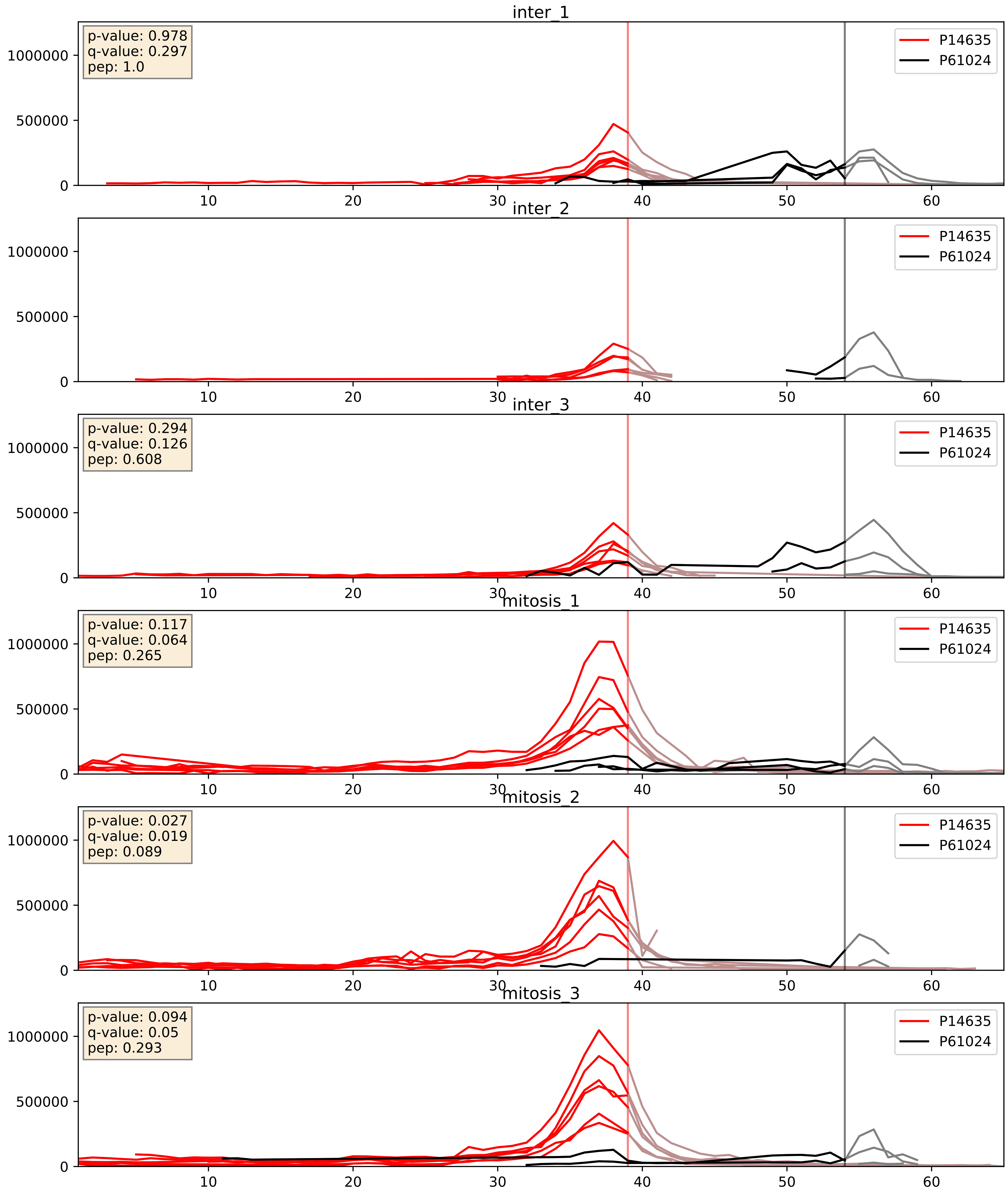


| level | condition_1 | condition_2 | pvalue | pvalue_adjusted |
|---|---|---|---|---|
| interactor_ratio | mitosis | inter | 0.025387313557422072 | 0.25152245839297793 |
| interactor_abundance | mitosis | inter | 0.0437690238659883 | 0.16685321787696922 |
| complex_abundance | mitosis | inter | 0.134842963119675 | 0.2798195792253135 |
