## Supplementary Data 2 for "SECAT: Quantifying differential protein-protein interaction states by network-centric analysis": hela_string_000055_Q99547.pdf

Q99547_Q99547

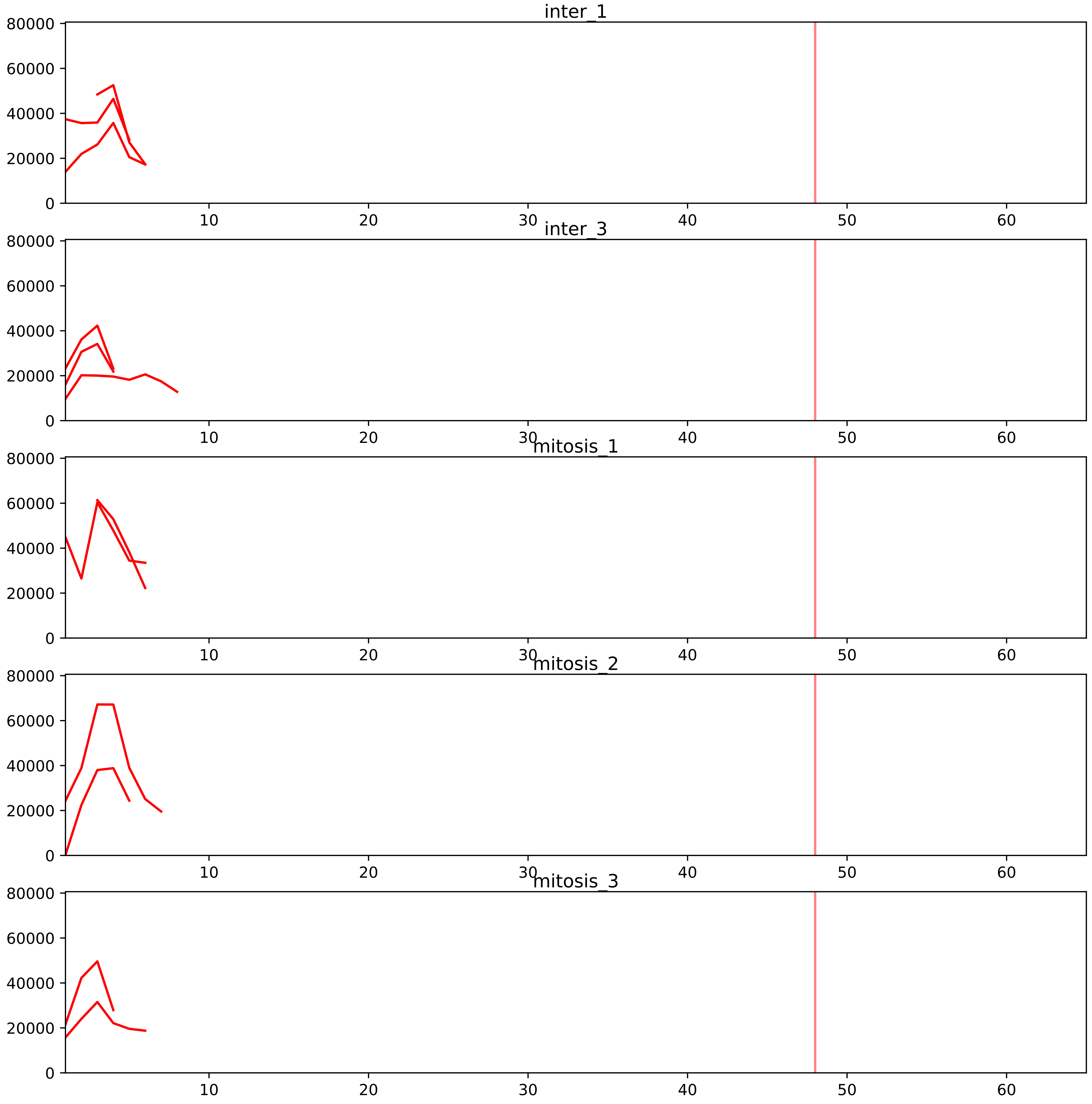


| level | condition_1 | condition_2 | pvalue | pvalue_adjusted |
|---|---|---|---|---|
| interactor_ratio | mitosis | inter | 2.2166854983502107e-07 | 7.84365637877767e-06 |
| interactor_abundance | mitosis | inter | 0.13544338987855314 | 0.27629250263474253 |
| total_abundance | mitosis | inter | 0.4998178241775279 | 0.762128537619219 |
| complex_abundance | mitosis | inter | 0.5821049112792518 | 0.681826148855068 |
| assembled_abundance | mitosis | inter | 0.6049895414385338 | 0.8154206862867195 |

Q01780_Q99547
EXOSX_HUMAN vs MPH6_HUMAN
p-value: 0.024 q-value: 0.018

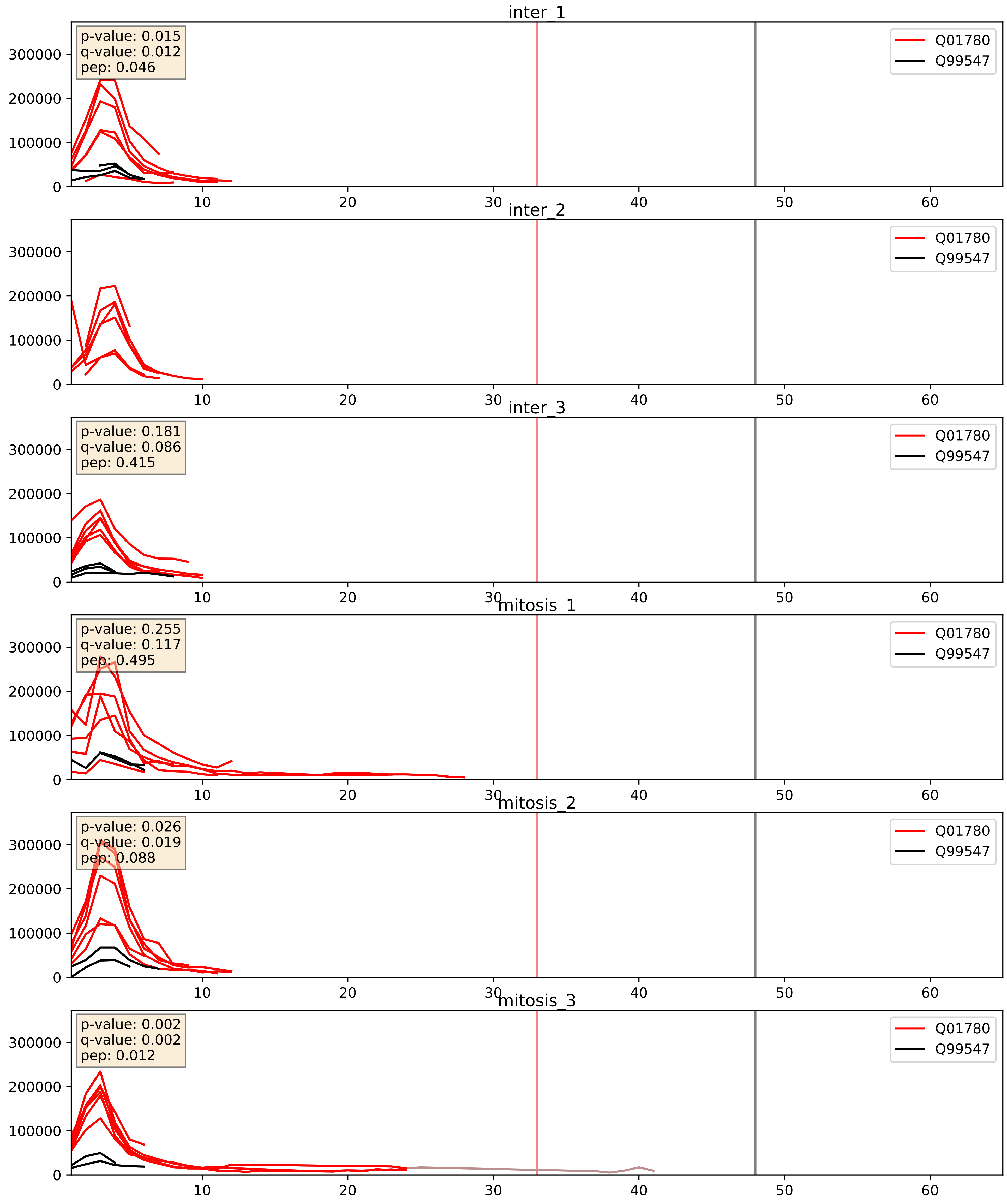


| level | condition_1 | condition_2 | pvalue | pvalue_adjusted |
|---|---|---|---|---|
| interactor_ratio | mitosis | inter | 0.027135379938690897 | 0.2577730310940899 |
| interactor_abundance | mitosis | inter | 0.3953301978901166 | 0.5864355764560087 |
| complex_abundance | mitosis | inter | 0.9932760166578614 | 0.9963021681030341 |

Q15024_Q99547
EXOS7_HUMAN vs MPH6_HUMAN
p-value: 0.0 q-value: 0.001

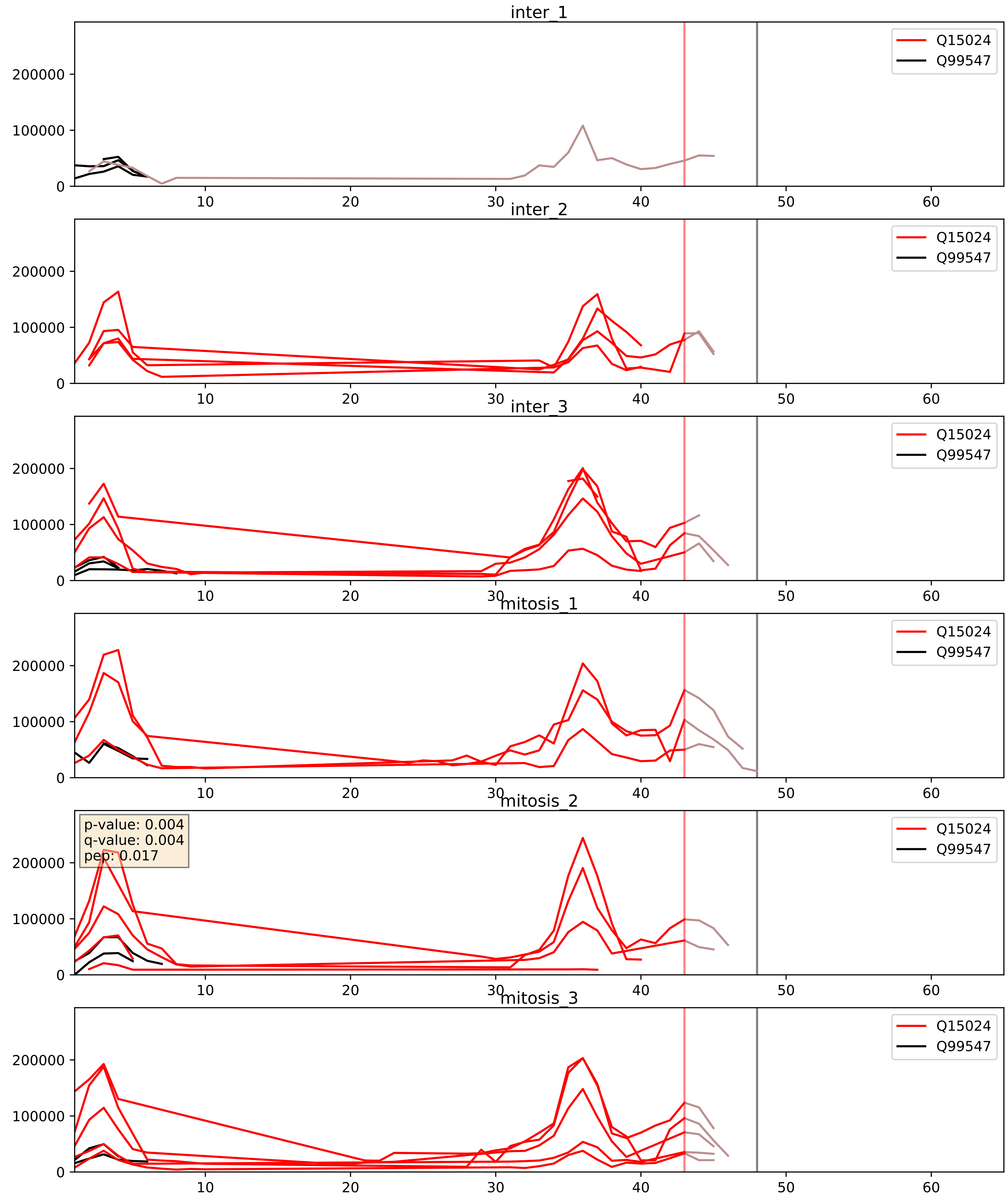


| level | condition_1 | condition_2 | pvalue | pvalue_adjusted |
|---|---|---|---|---|
| interactor_ratio | mitosis | inter | 0.04555942657221424 | 0.3548621970895441 |
| interactor_abundance | mitosis | inter | 0.13020836346228779 | 0.3149430887926486 |
| complex_abundance | mitosis | inter | 0.4625290419485175 | 0.6037280571941613 |

Q5RKV6_Q99547
EXOS6_HUMAN vs MPH6_HUMAN
p-value: 0.023 q-value: 0.018

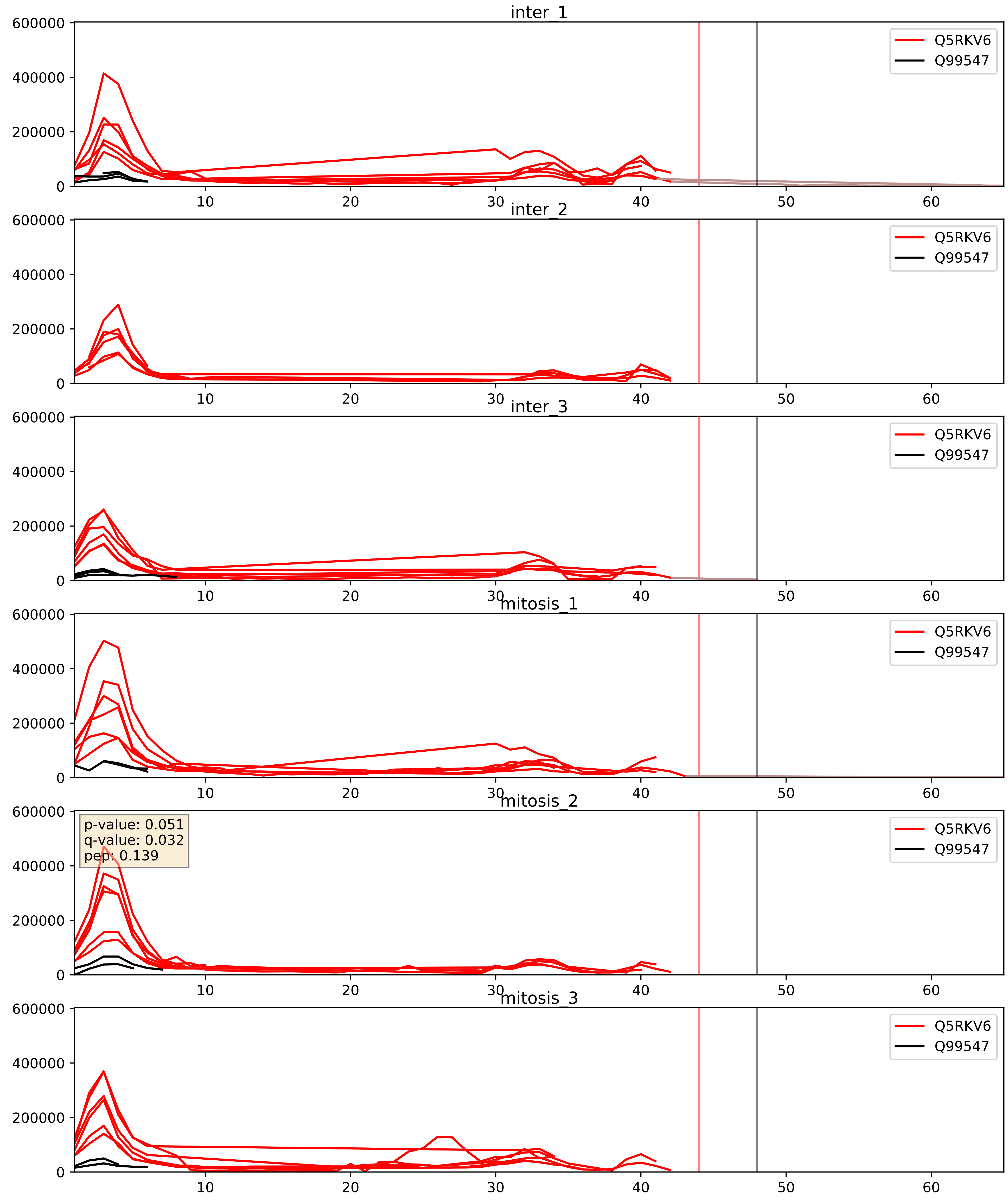


| level | condition_1 | condition_2 | pvalue | pvalue_adjusted |
|---|---|---|---|---|
| interactor_ratio | mitosis | inter | 0.02063381208485944 | 0.21698455951645798 |
| interactor_abundance | mitosis | inter | 0.24623688817456618 | 0.4397469390695545 |
| complex_abundance | mitosis | inter | 0.7726168130820598 | 0.8500771105375876 |

Q96B26_Q99547
EXOS8_HUMAN vs MPH6_HUMAN
p-value: 0.009 q-value: 0.008

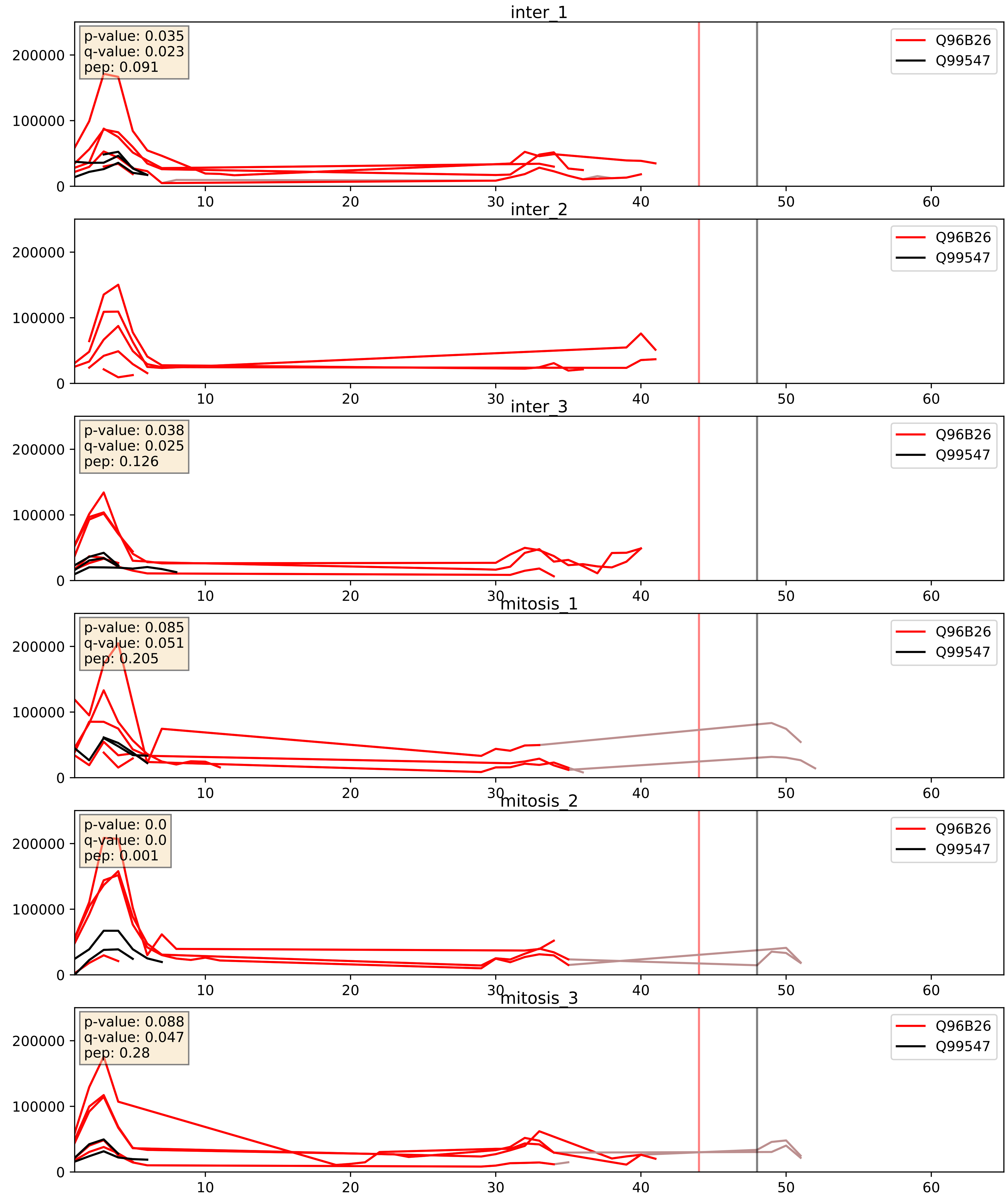


| level | condition_1 | condition_2 | pvalue | pvalue_adjusted |
|---|---|---|---|---|
| interactor_ratio | mitosis | inter | 0.020164844675413888 | 0.21442368996464944 |
| interactor_abundance | mitosis | inter | 0.6106227564157307 | 0.7445173329393203 |
| complex_abundance | mitosis | inter | 0.7029795413662915 | 0.8007325181763746 |

Q99547_Q9NQT4
MPH6_HUMAN vs EXOS5_HUMAN
p-value: 0.019 q-value: 0.015

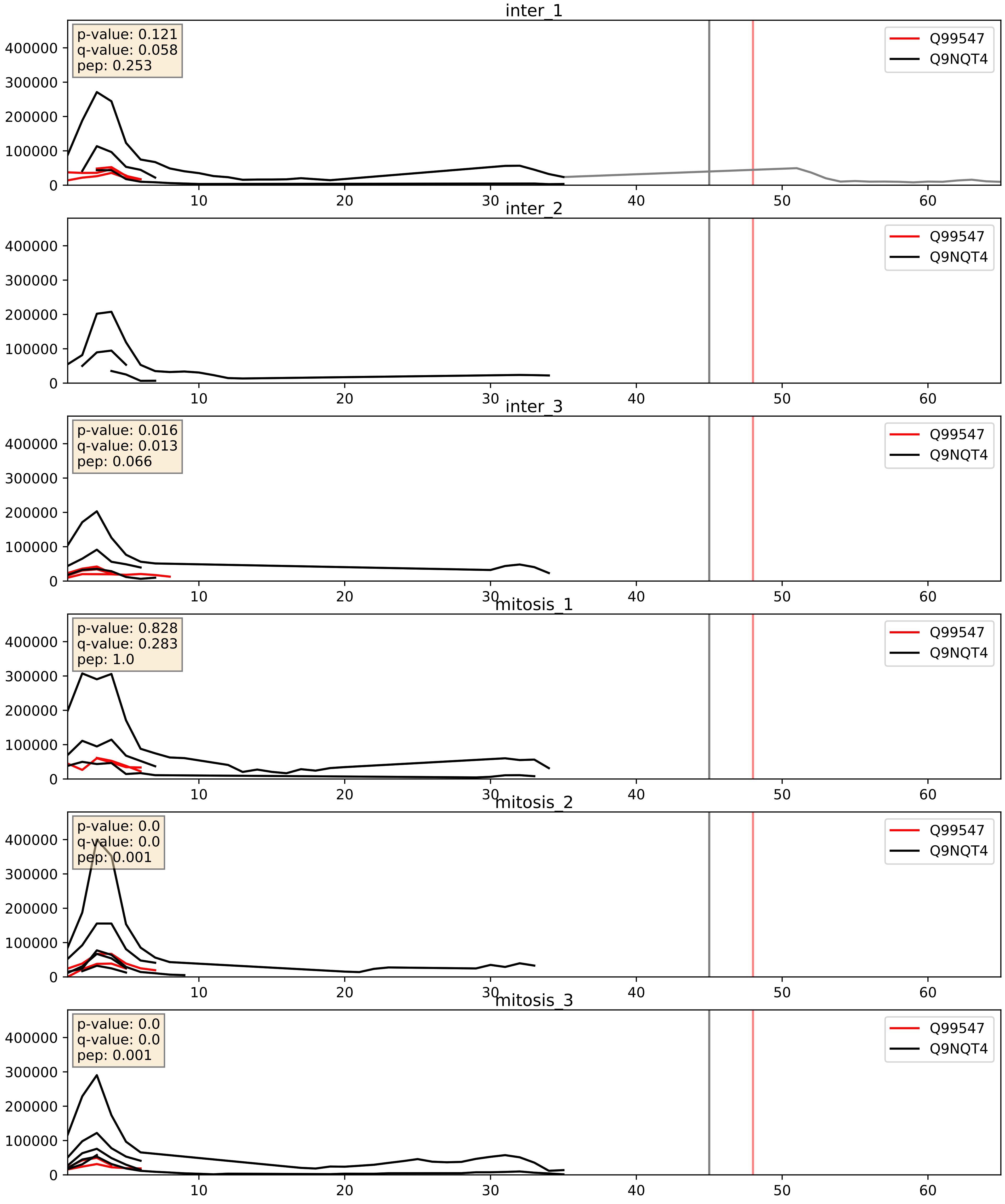


| level | condition_1 | condition_2 | pvalue | pvalue_adjusted |
|---|---|---|---|---|
| interactor_ratio | mitosis | inter | 0.009648639079298532 | 0.14070247107120176 |
| interactor_abundance | mitosis | inter | 0.19800317018799962 | 0.39175017607980506 |
| complex_abundance | mitosis | inter | 0.7793227980672002 | 0.8553664766579349 |

Q99547_Q9NQT5
MPH6_HUMAN vs EXOS3_HUMAN
p-value: 0.004 q-value: 0.004

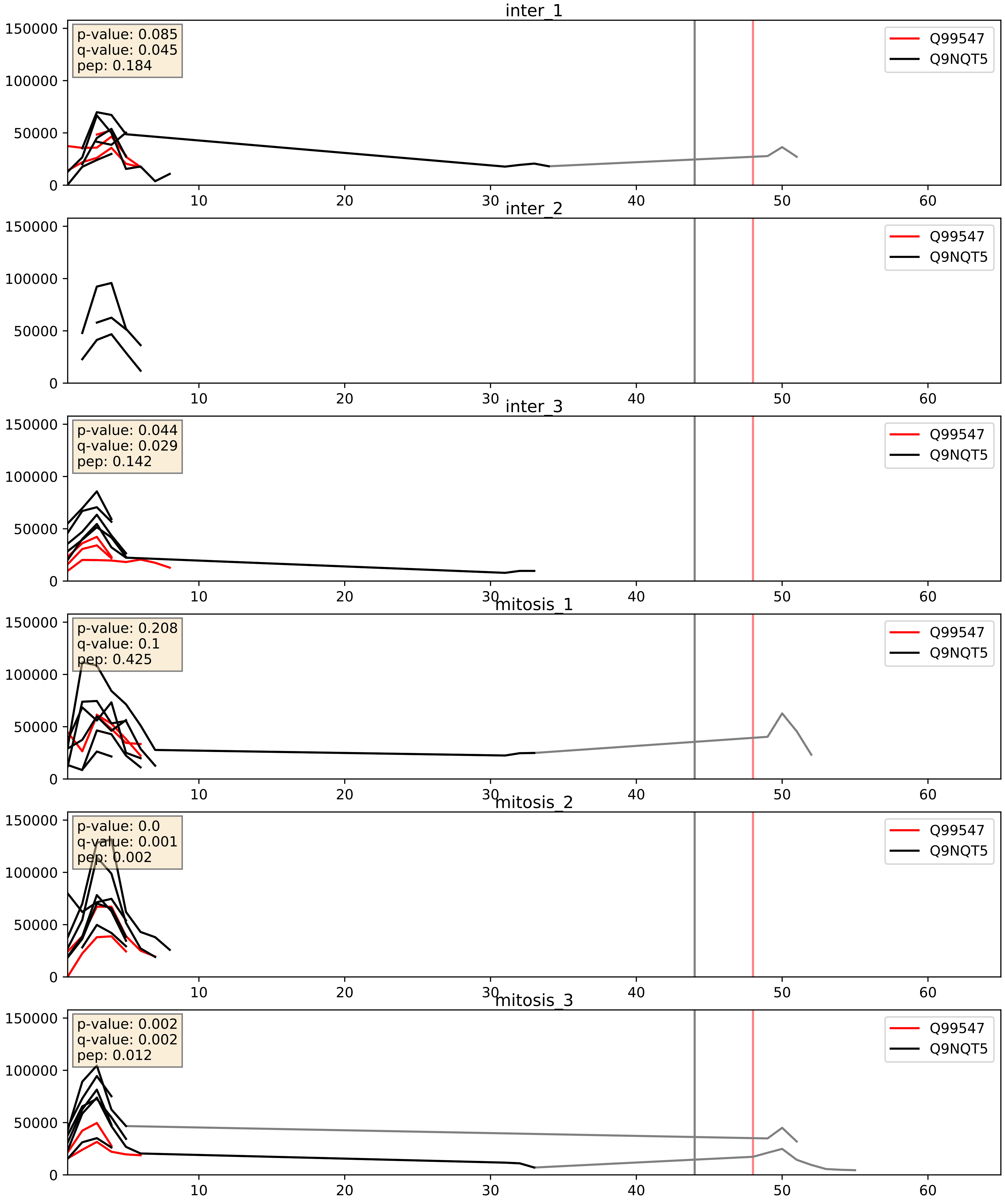


| level | condition_1 | condition_2 | pvalue | pvalue_adjusted |
|---|---|---|---|---|
| interactor_ratio | mitosis | inter | 0.005876576927210164 | 0.10458107795617257 |
| interactor_abundance | mitosis | inter | 0.20467957343369572 | 0.40102017591953204 |
| complex_abundance | mitosis | inter | 0.5525719831294813 | 0.6855976776987645 |

Q99547_Q9Y2L1
MPH6_HUMAN vs RRP44_HUMAN
p-value: 0.047 q-value: 0.031

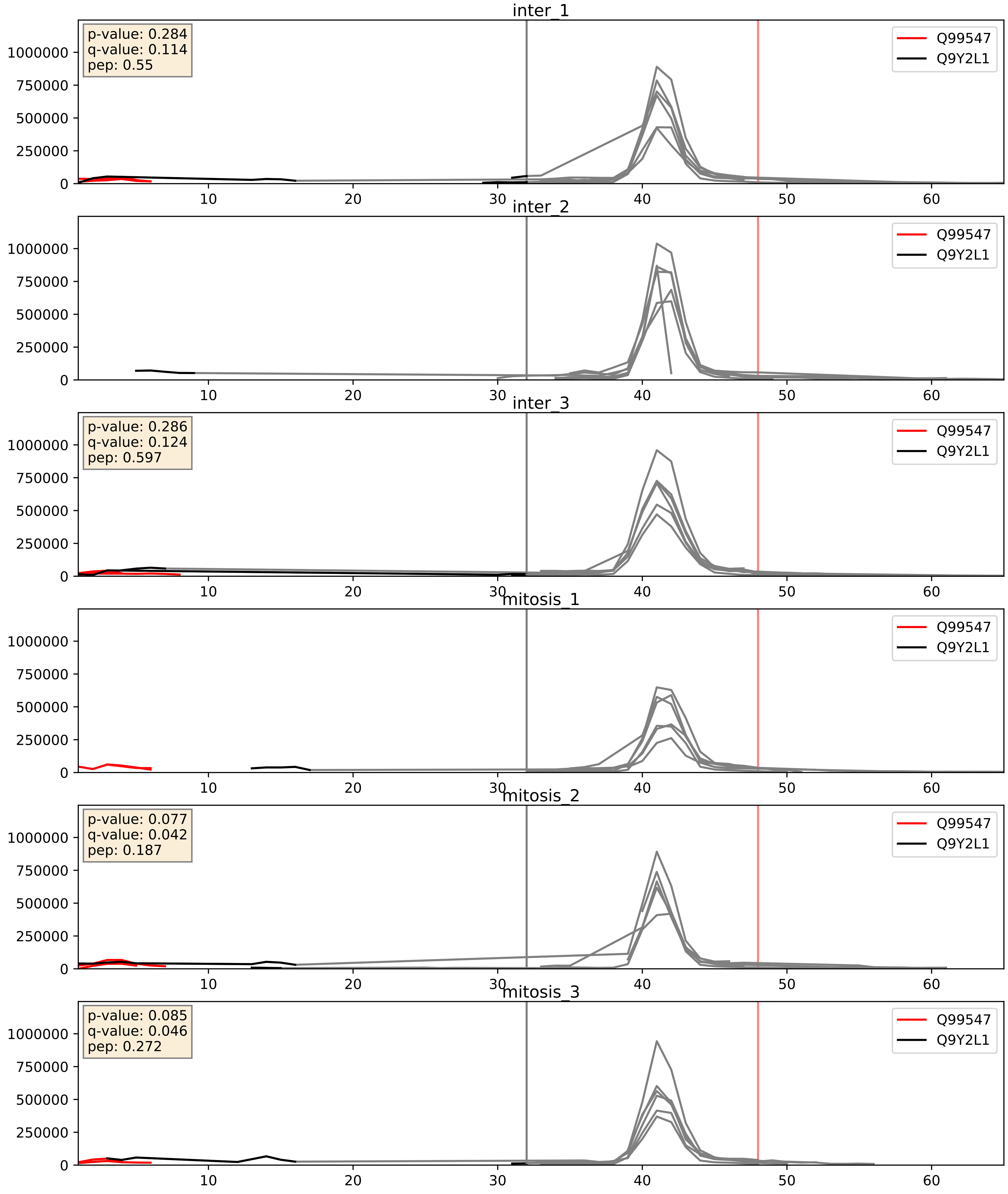


| level | condition_1 | condition_2 | pvalue | pvalue_adjusted |
|---|---|---|---|---|
| complex_abundance | mitosis | inter | 0.02310054738443675 | 0.11091884921279084 |
| interactor_abundance | mitosis | inter | 0.03625634554697996 | 0.15802829520754763 |
| interactor_ratio | mitosis | inter | 0.14941800190579227 | 0.6039771666075939 |

Q99547_Q9Y3B2
MPH6_HUMAN vs EXOS1_HUMAN
p-value: 0.012 q-value: 0.01

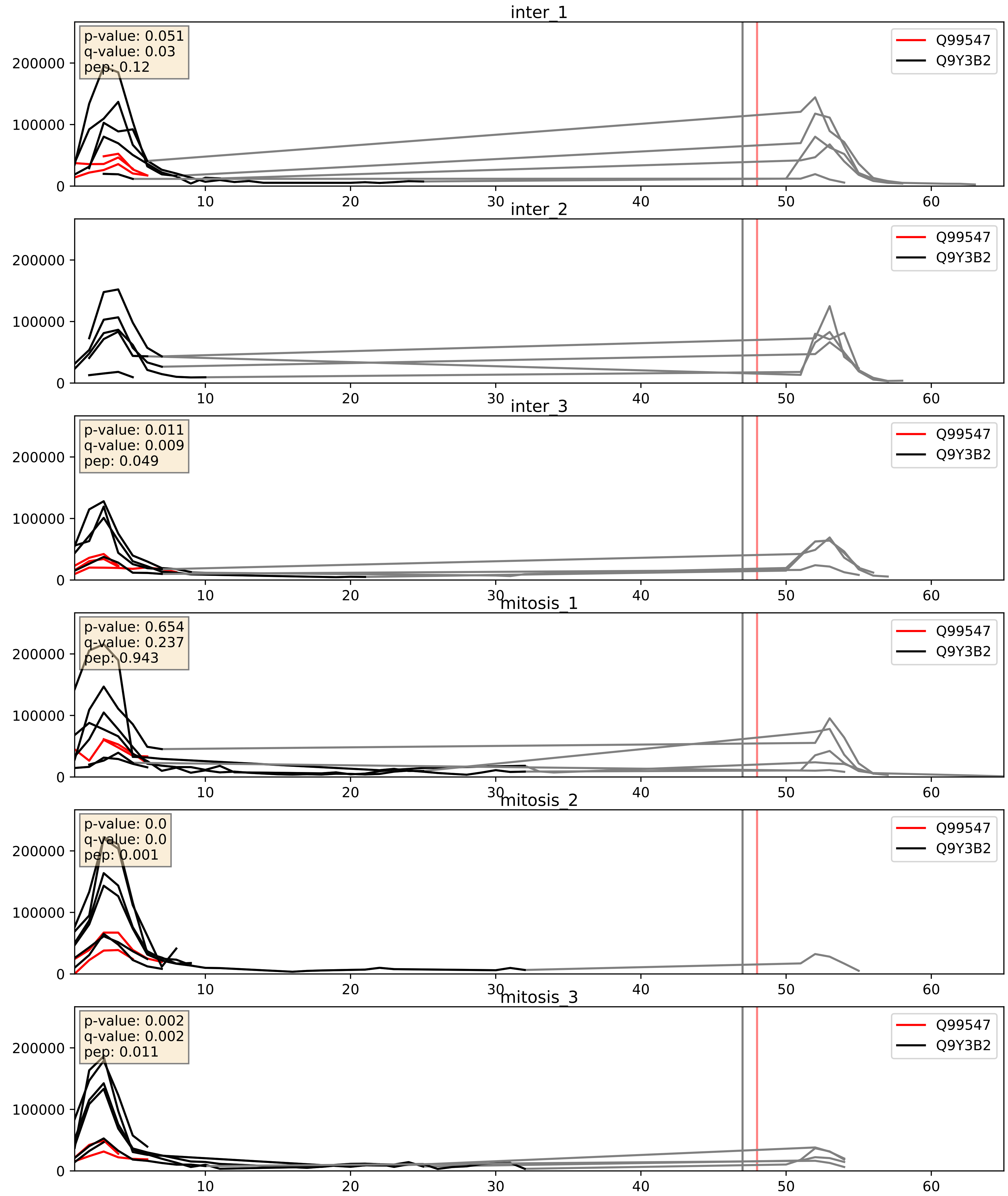


| level | condition_1 | condition_2 | pvalue | pvalue_adjusted |
|---|---|---|---|---|
| interactor_ratio | mitosis | inter | 0.007134109691818039 | 0.11743842108069695 |
| interactor_abundance | mitosis | inter | 0.19633954672645476 | 0.38990059621353734 |
| complex_abundance | mitosis | inter | 0.9599822384690955 | 0.9765238219008268 |
