## Supplementary Data 2 for "SECAT: Quantifying differential protein-protein interaction states by network-centric analysis": hela_string_000056_O15160.pdf

# O15160_O15160

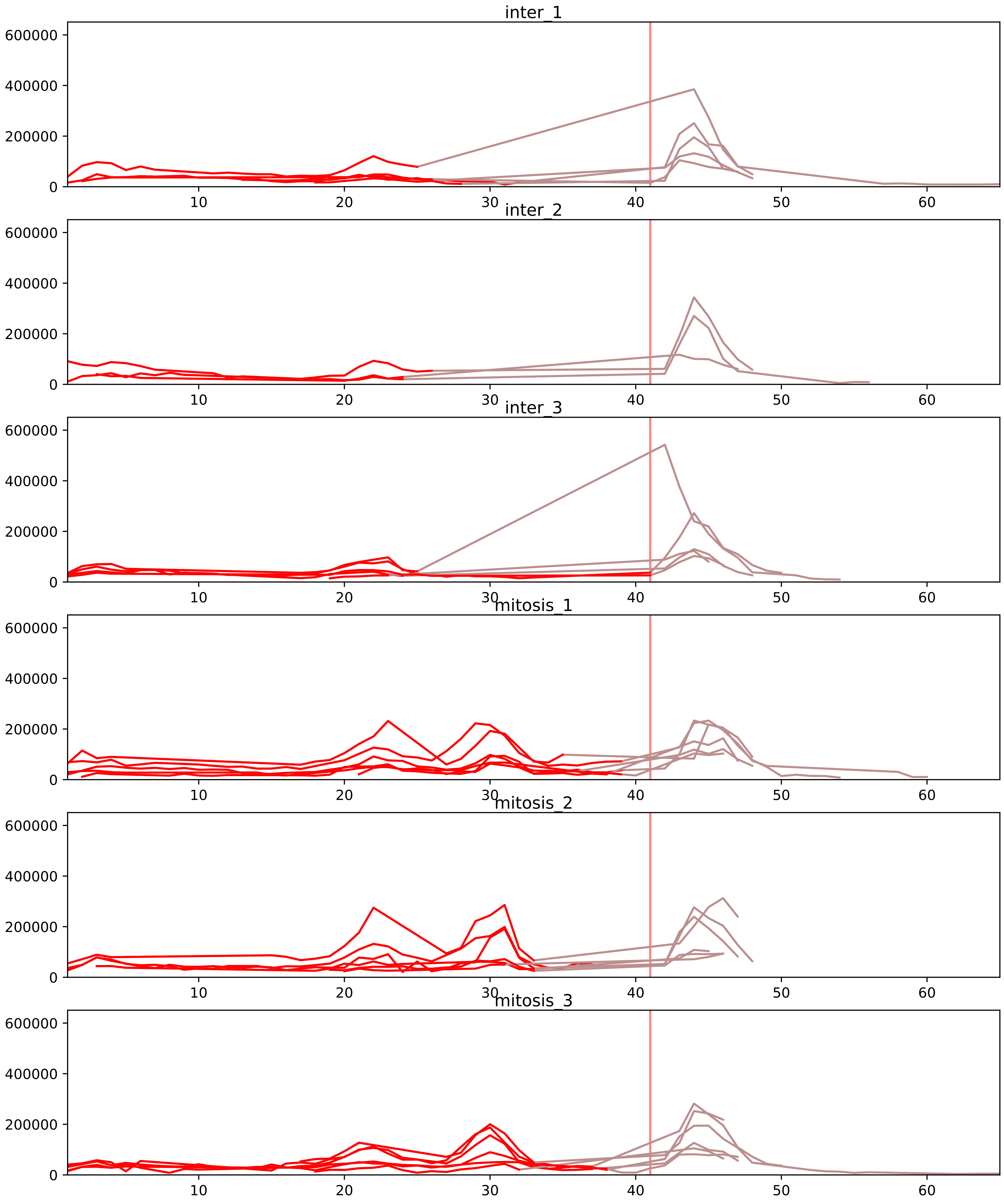


| level | condition_1 | condition_2 | pvalue | pvalue_adjusted |
|---|---|---|---|---|
| interactor_ratio | mitosis | inter | 2.598890731785029e-07 | 9.022564049970665e-06 |
| total_abundance | mitosis | inter | 6.495359447648242e-05 | 0.011022624982659067 |
| interactor_abundance | mitosis | inter | 9.185095985587951e-05 | 0.002139313495377447 |
| assembled_abundance | mitosis | inter | 0.00028310530753734625 | 0.016497717306915387 |
| monomer_abundance | mitosis | inter | 0.00037427250433783737 | 0.05243686845257356 |
| complex_abundance | mitosis | inter | 0.00840073756588944 | 0.03953288266300913 |

O14802_O15160
RPC1_HUMAN vs RPAC1_HUMAN
p-value: 0.073 q-value: 0.044

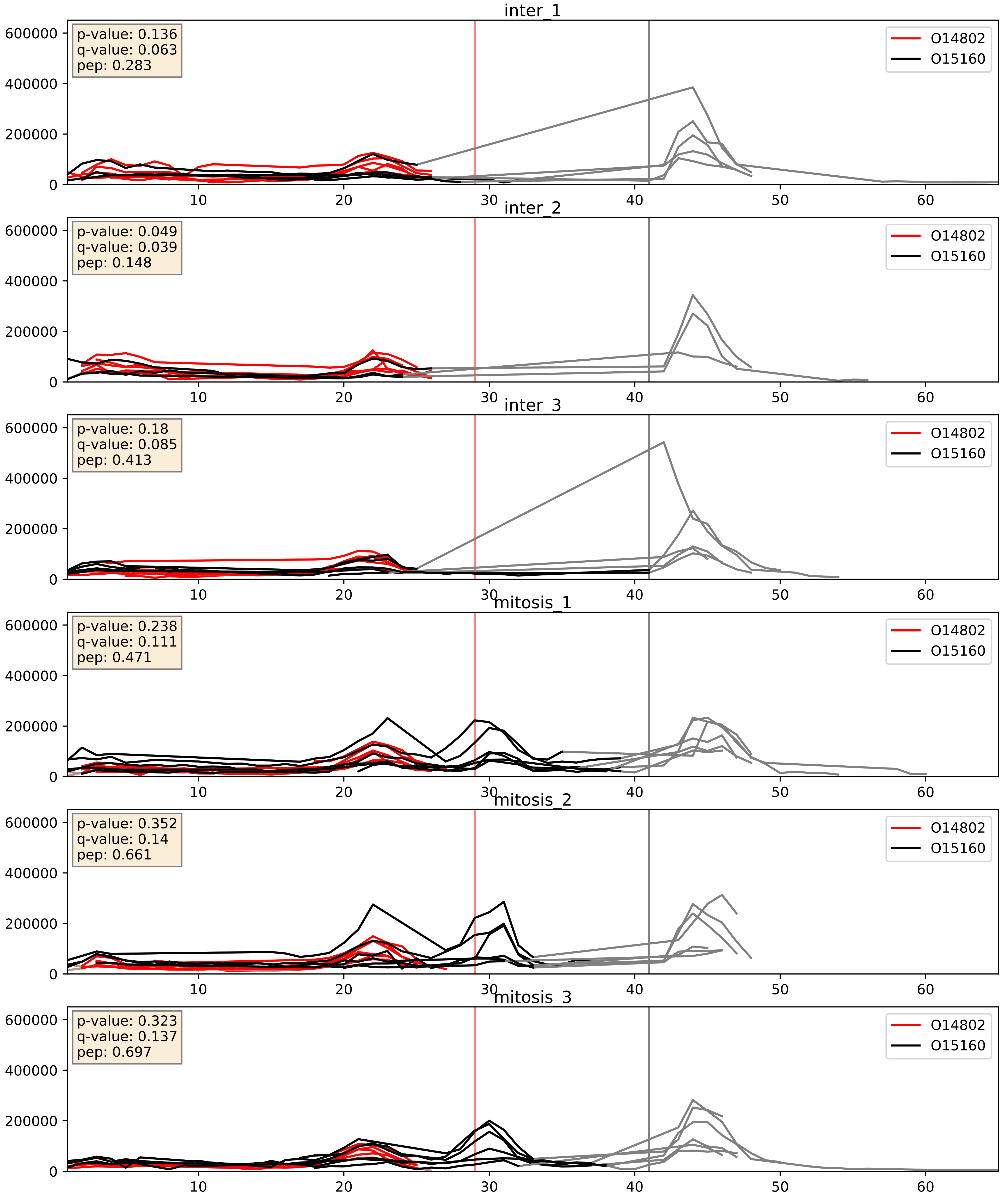


| level | condition_1 | condition_2 | pvalue | pvalue_adjusted |
|---|---|---|---|---|
| interactor_ratio | mitosis | inter | 0.0016630222471381159 | 0.04925768316782793 |
| interactor_abundance | mitosis | inter | 0.06556300075007898 | 0.21218120469590776 |
| complex_abundance | mitosis | inter | 0.8559028069573468 | 0.9086603035538743 |

O15160_P05423
RPAC1_HUMAN vs RPC4_HUMAN
p-value: 0.013 q-value: 0.011

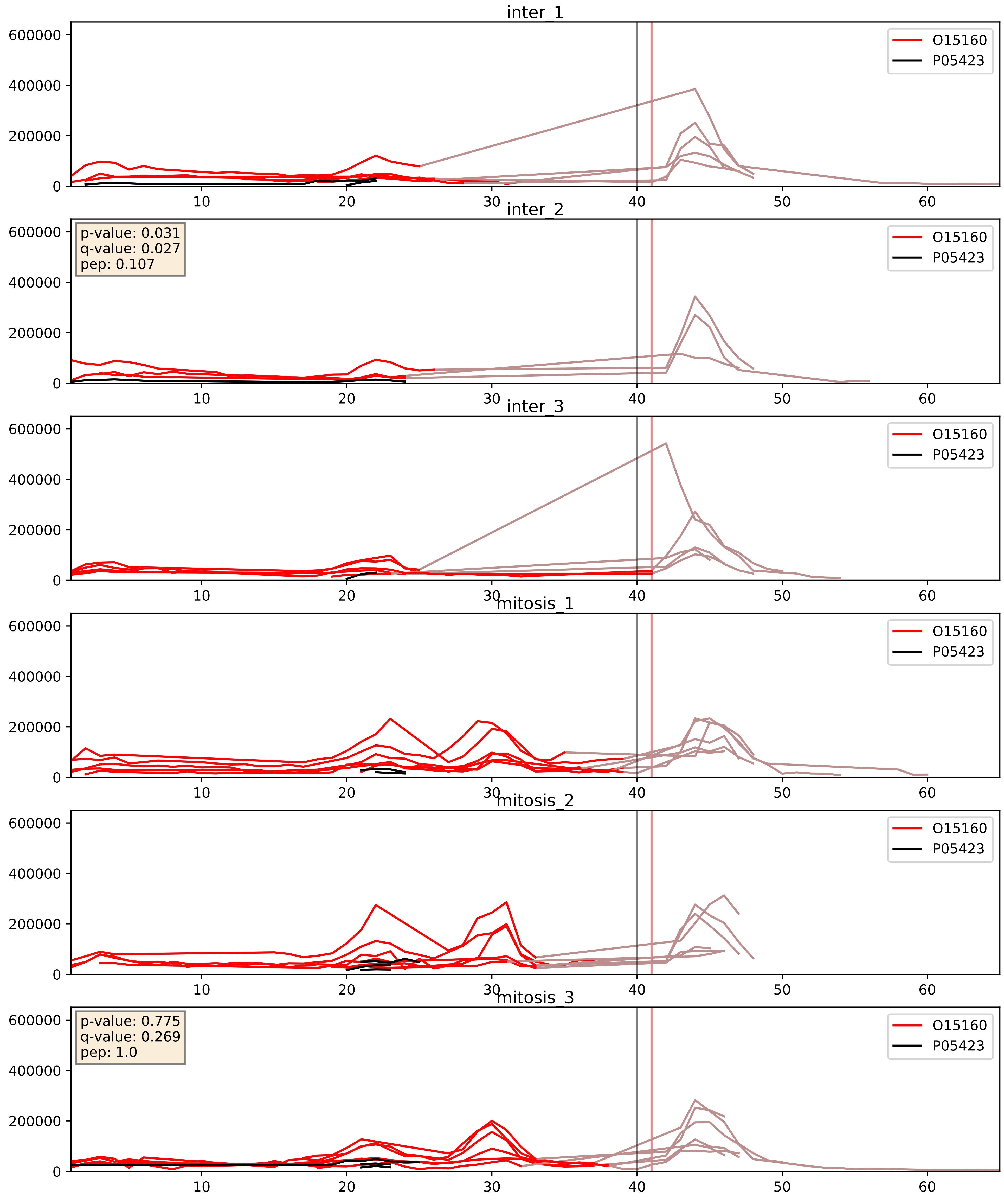


| level | condition_1 | condition_2 | pvalue | pvalue_adjusted |
|---|---|---|---|---|
| interactor_ratio | mitosis | inter | 0.00993696687049424 | 0.14263182888175546 |
| complex_abundance | mitosis | inter | 0.5549211692552949 | 0.6869692244436636 |
| interactor_abundance | mitosis | inter | 0.6713569734985754 | 0.7847089087386913 |

O15160_P19388
RPAC1_HUMAN vs RPAB1_HUMAN
p-value: 0.036 q-value: 0.025

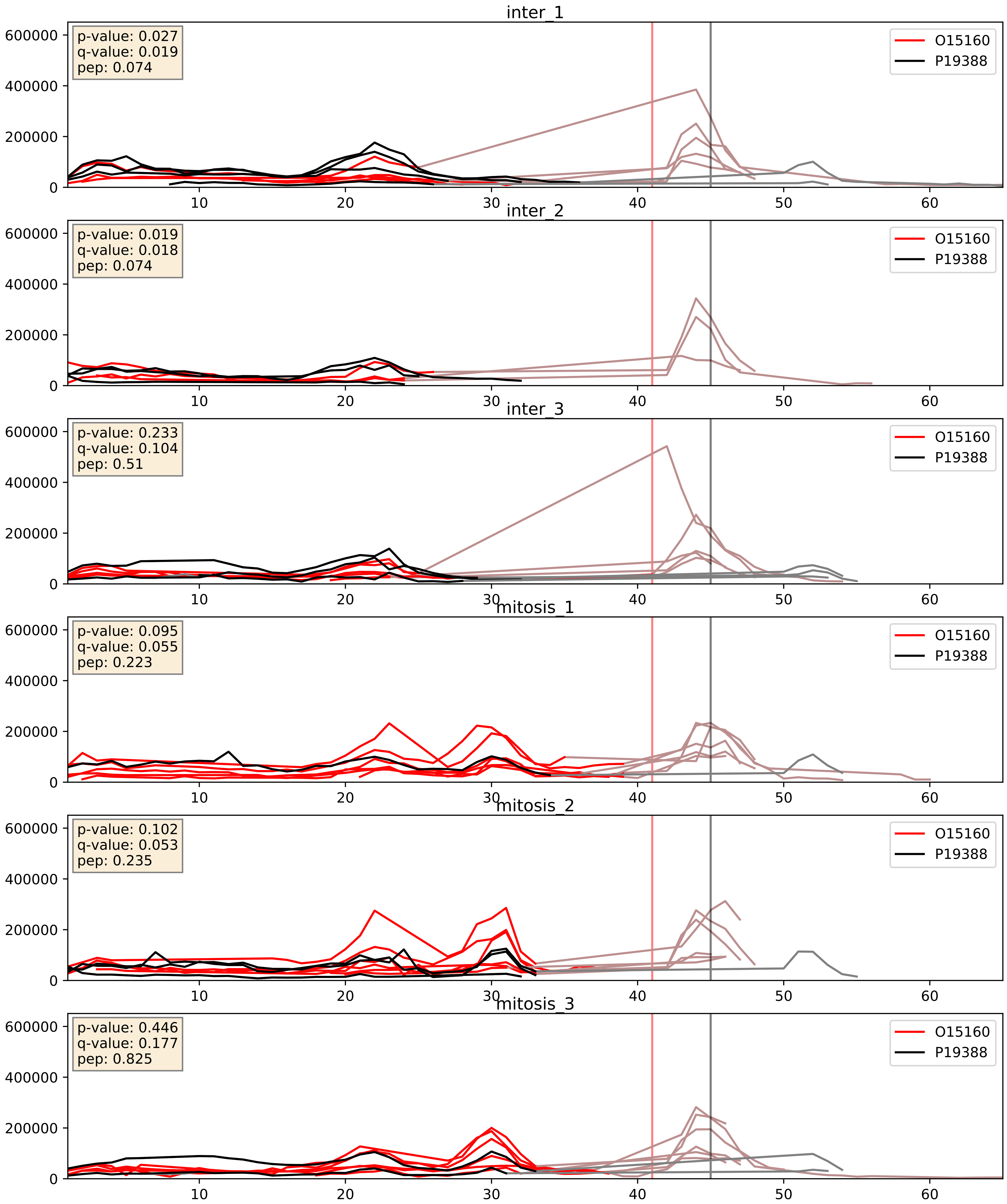


| level | condition_1 | condition_2 | pvalue | pvalue_adjusted |
|---|---|---|---|---|
| interactor_abundance | mitosis | inter | 0.0013185480112738036 | 0.026872298192663256 |
| interactor_ratio | mitosis | inter | 0.004474652467414517 | 0.0890768026071355 |
| complex_abundance | mitosis | inter | 0.6726891184550716 | 0.7819417237880789 |

O15160_P52434
RPAC1_HUMAN vs RPAB3_HUMAN
p-value: 0.032 q-value: 0.023

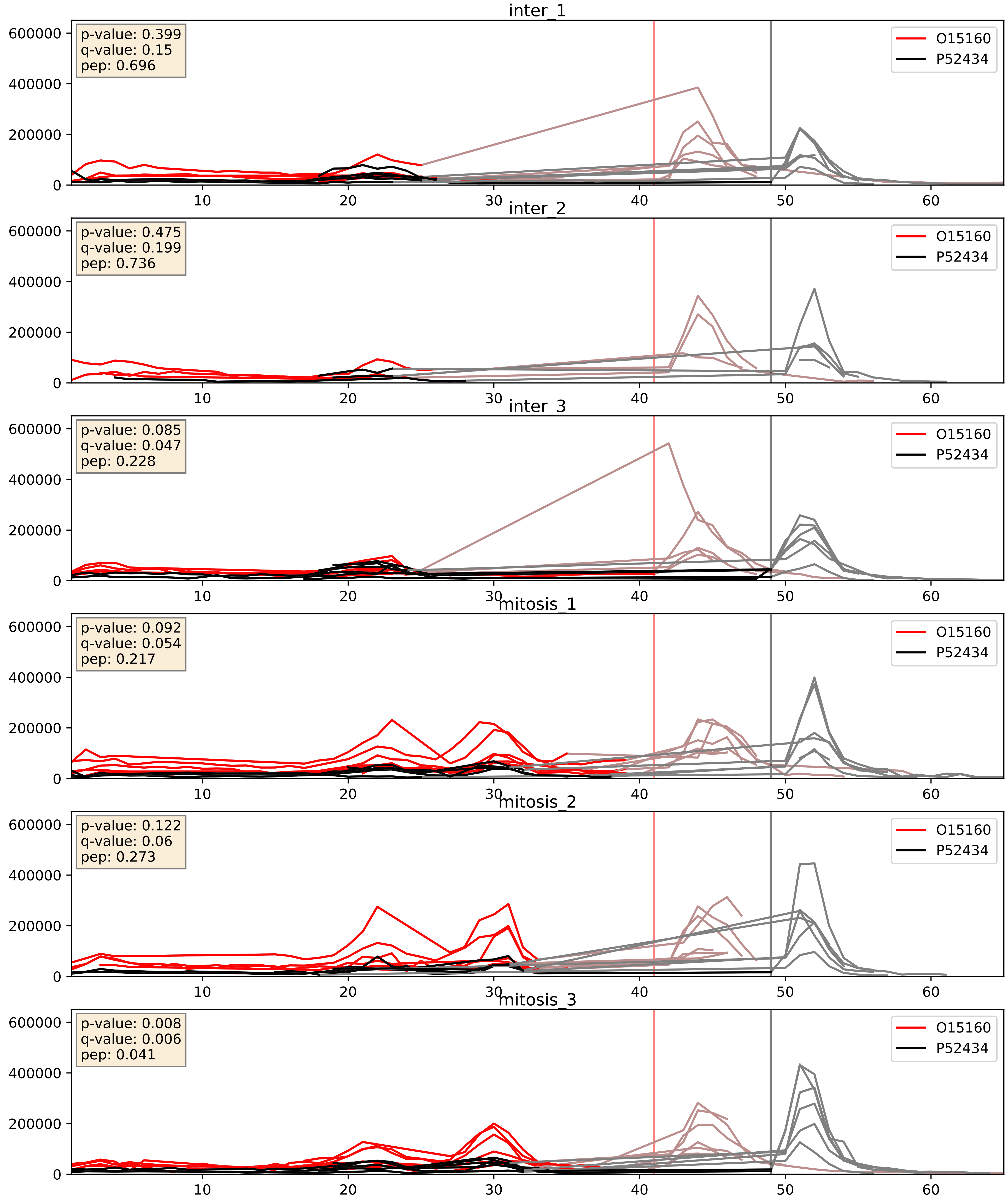


| level | condition_1 | condition_2 | pvalue | pvalue_adjusted |
|---|---|---|---|---|
| interactor_ratio | mitosis | inter | 0.0007963931047073435 | 0.03250474790144618 |
| interactor_abundance | mitosis | inter | 0.0015023531794424757 | 0.028609884796501873 |
| complex_abundance | mitosis | inter | 0.019617192368517836 | 0.1042142626599879 |

O15160_Q9BUI4
RPAC1_HUMAN vs RPC3_HUMAN
p-value: 0.027 q-value: 0.02

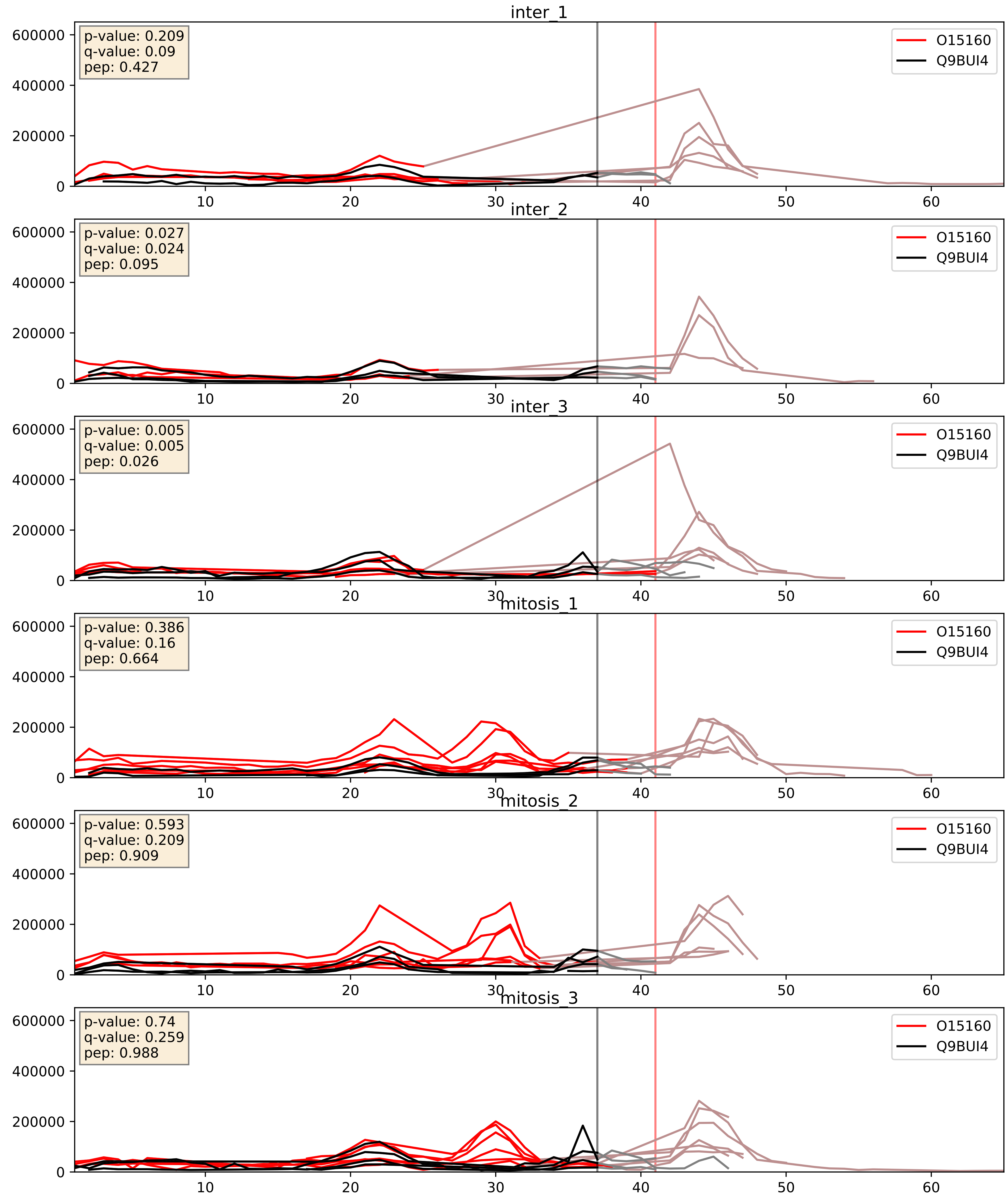


| level | condition_1 | condition_2 | pvalue | pvalue_adjusted |
|---|---|---|---|---|
| interactor_abundance | mitosis | inter | 0.00038216274814687997 | 0.013573913378163039 |
| complex_abundance | mitosis | inter | 0.0010575394663l5369 | 0.021656789070955883 |
| interactor_ratio | mitosis | inter | 0.00709215084655209 | 0.11727616584402438 |

O15160_Q9H1D9
RPAC1_HUMAN vs RPC6_HUMAN
p-value: 0.026 q-value: 0.02

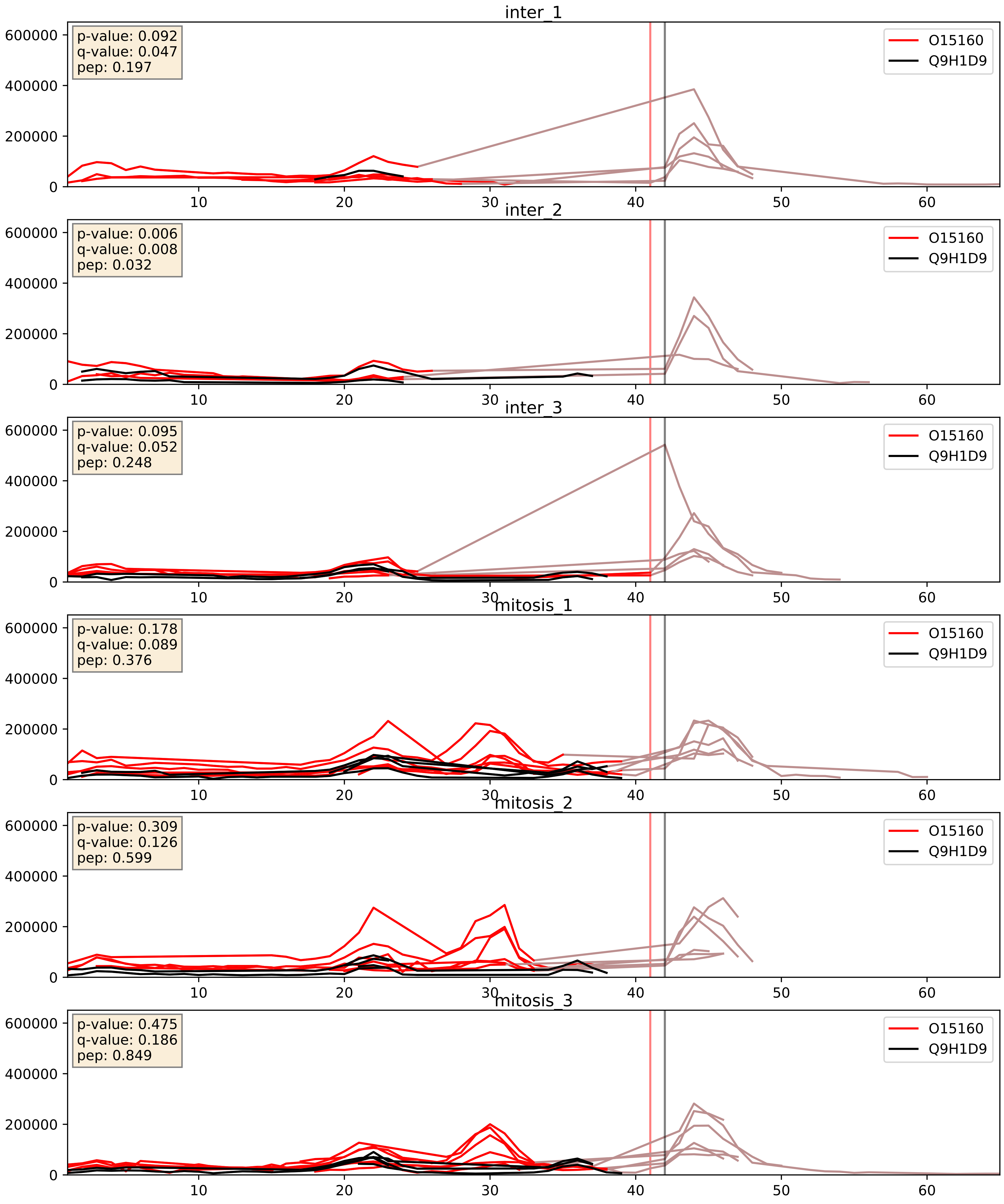


| level | condition_1 | condition_2 | pvalue | pvalue_adjusted |
|---|---|---|---|---|
| interactor_ratio | mitosis | inter | 0.0020583575769718685 | 0.055758040692655685 |
| interactor_abundance | mitosis | inter | 0.0025922019703768858 | 0.03806046117740333 |
| complex_abundance | mitosis | inter | 0.022906312351345822 | 0.11090386523049788 |
