## Supplementary Data 2 for "SECAT: Quantifying differential protein-protein interaction states by network-centric analysis": hela_string_000057_P08621.pdf

P08621_P08621

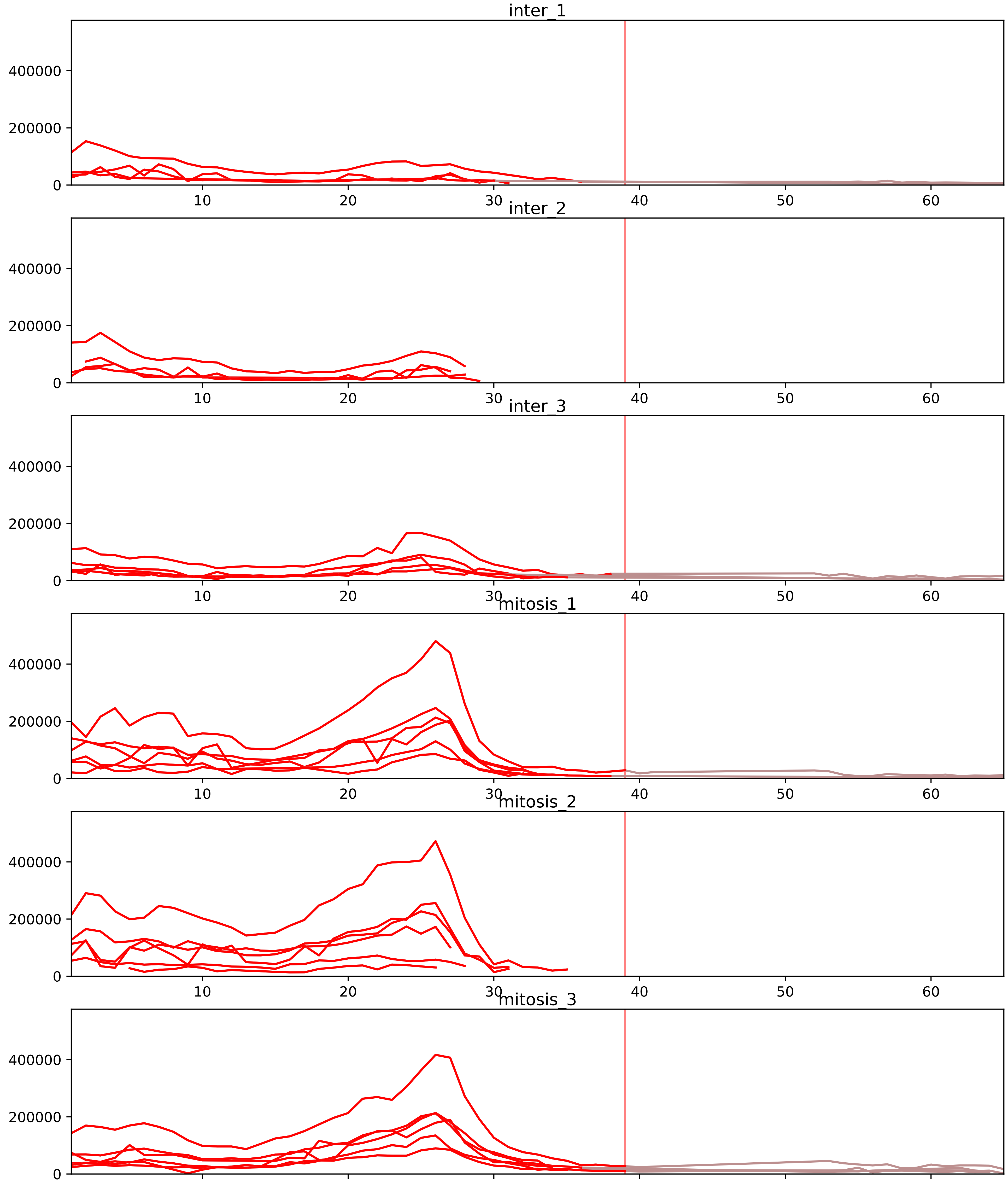


| level | condition_1 | condition_2 | pvalue | pvalue_adjusted |
|---|---|---|---|---|
| interactor_abundance | mitosis | inter | 2.726107437283065e-07 | 4.6538655224098096e-05 |
| complex_abundance | mitosis | inter | 3.7299027817934944e-07 | 4.575347412333353e-05 |
| interactor_ratio | mitosis | inter | 0.00024234752088816522 | 0.003483745612767375 |
| total_abundance | mitosis | inter | 0.30218610925117295 | 0.6294719648926848 |
| assembled_abundance | mitosis | inter | 0.31925768911536245 | 0.6097287812113471 |
| monomer_abundance | mitosis | inter | 0.5165554110025651 | 0.7637425891206047 |

P07910_P08621
HNRPC_HUMAN vs RU17_HUMAN
p-value: 0.031 q-value: 0.022

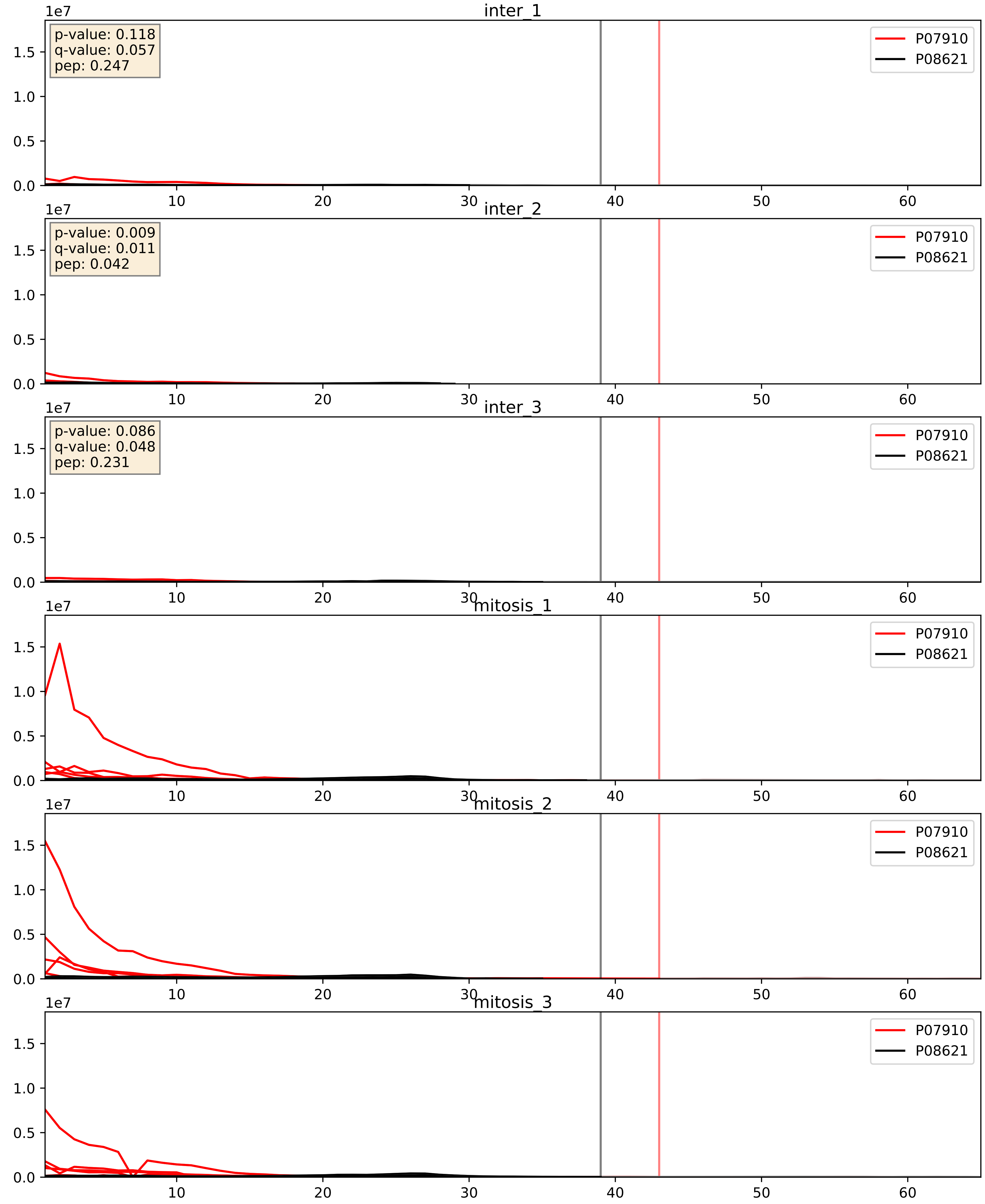


| level | condition_1 | condition_2 | pvalue | pvalue_adjusted |
|---|---|---|---|---|
| interactor_abundance | mitosis | inter | 9.496688410058796e-06 | 0.0033734327106189023 |
| complex_abundance | mitosis | inter | 1.5065308283938132e-05 | 0.005691857151652958 |
| interactor_ratio | mitosis | inter | 0.0001285712217163946 | 0.01222855175435931 |

P08621_P09012
RU17_HUMAN vs SNRPA_HUMAN
p-value: 0.012 q-value: 0.01

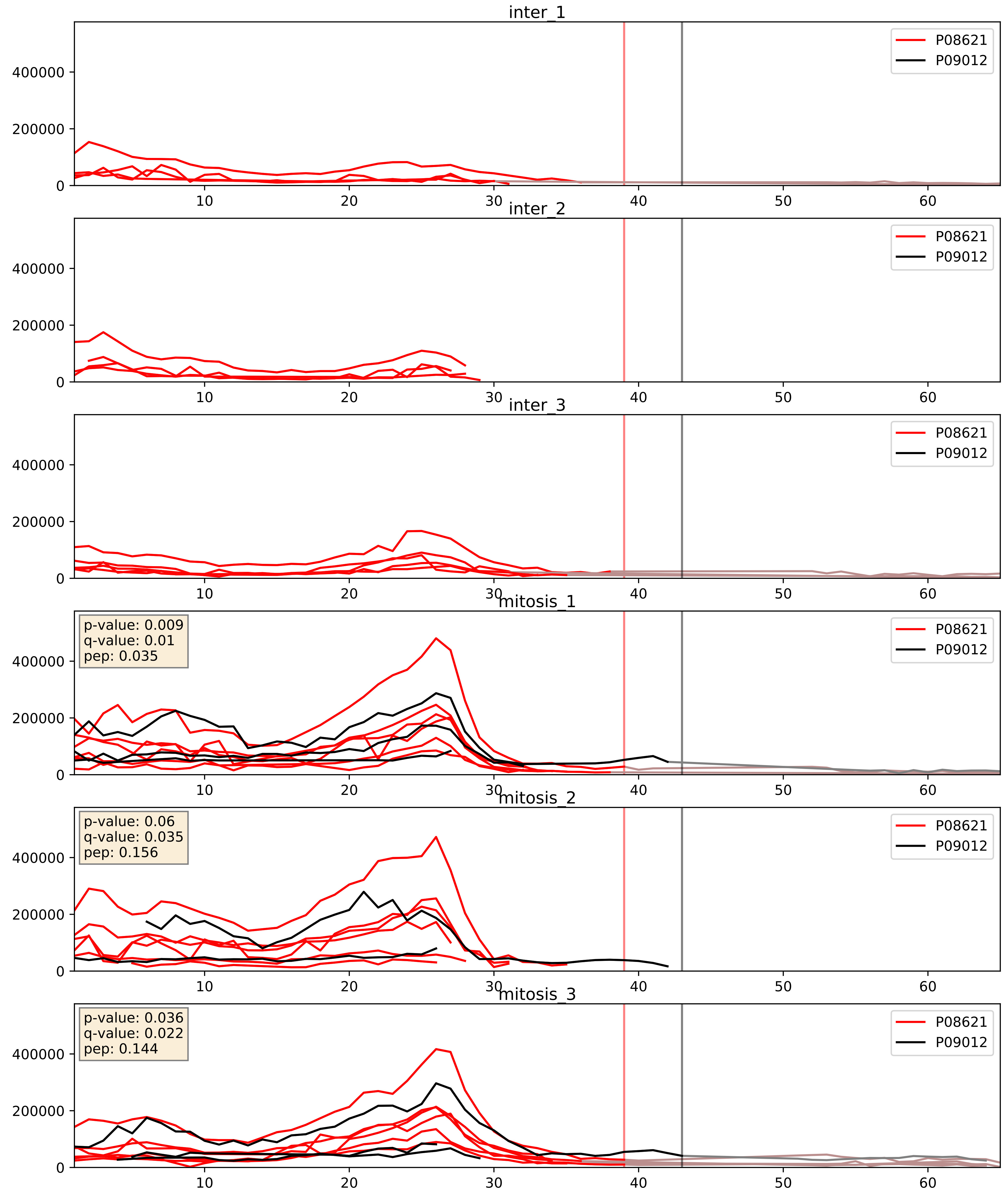


| level | condition_1 | condition_2 | pvalue | pvalue_adjusted |
|---|---|---|---|---|
| interactor_abundance | mitosis | inter | 1.4732188876160835e-05 | 0.0037089335794386596 |
| complex_abundance | mitosis | inter | 0.00033999865555559715 | 0.011976907372671259 |
| interactor_ratio | mitosis | inter | 0.026390044015689475 | 0.25543205403291264 |

P08621_P14866
RU17_HUMAN vs HNRPL_HUMAN
p-value: 0.076 q-value: 0.046

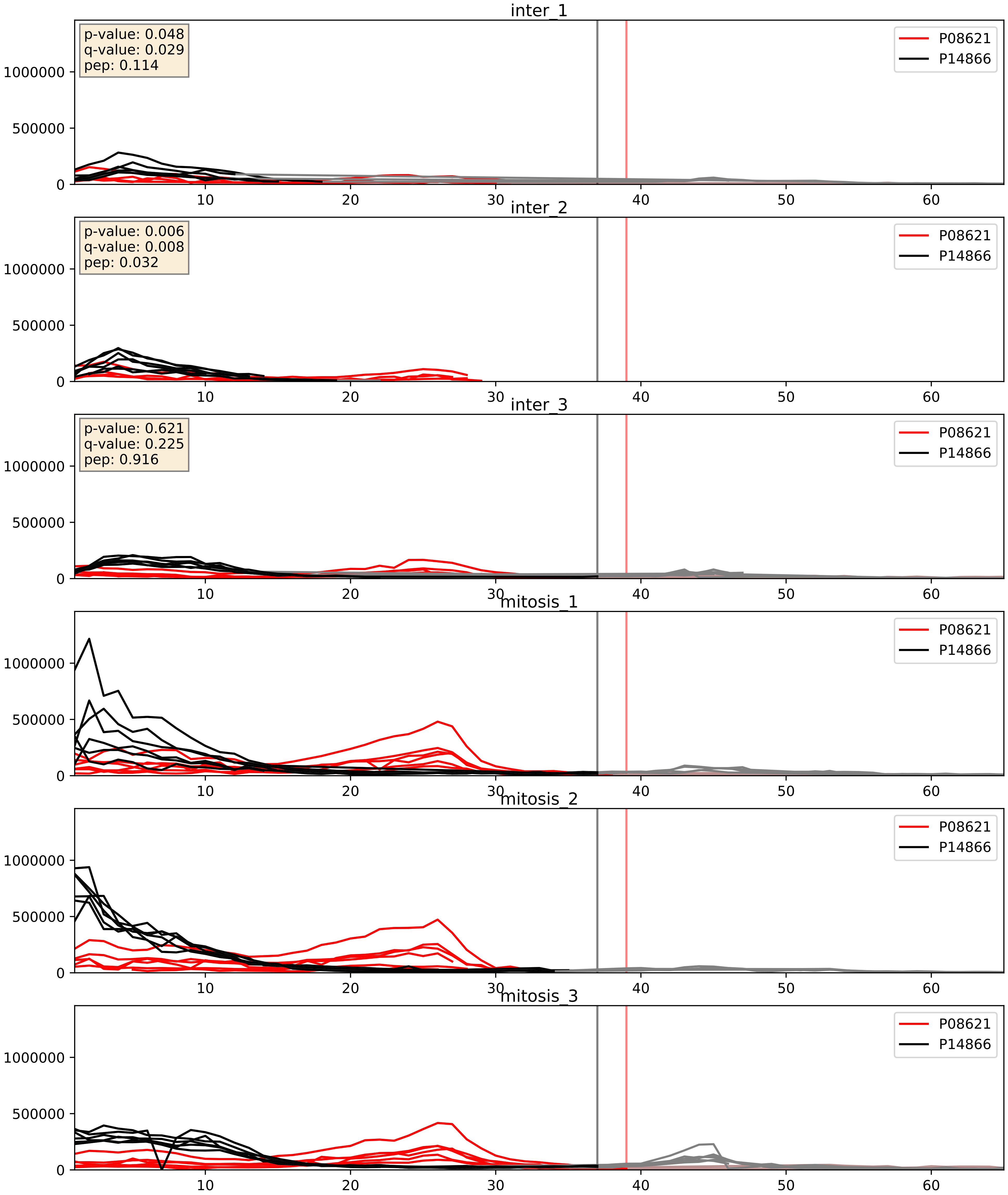


| level | condition_1 | condition_2 | pvalue | pvalue_adjusted |
|---|---|---|---|---|
| interactor_abundance | mitosis | inter | 1.057401869558351e-05 | 0.0033734327106189023 |
| complex_abundance | mitosis | inter | 0.0003837667920089791 | 0.01249066060683217 |
| interactor_ratio | mitosis | inter | 0.14042639881275756 | 0.5907826388074301 |
