## Supplementary Data 2 for "SECAT: Quantifying differential protein-protein interaction states by network-centric analysis": hela_string_000058_P62316.pdf

P62316_P62316

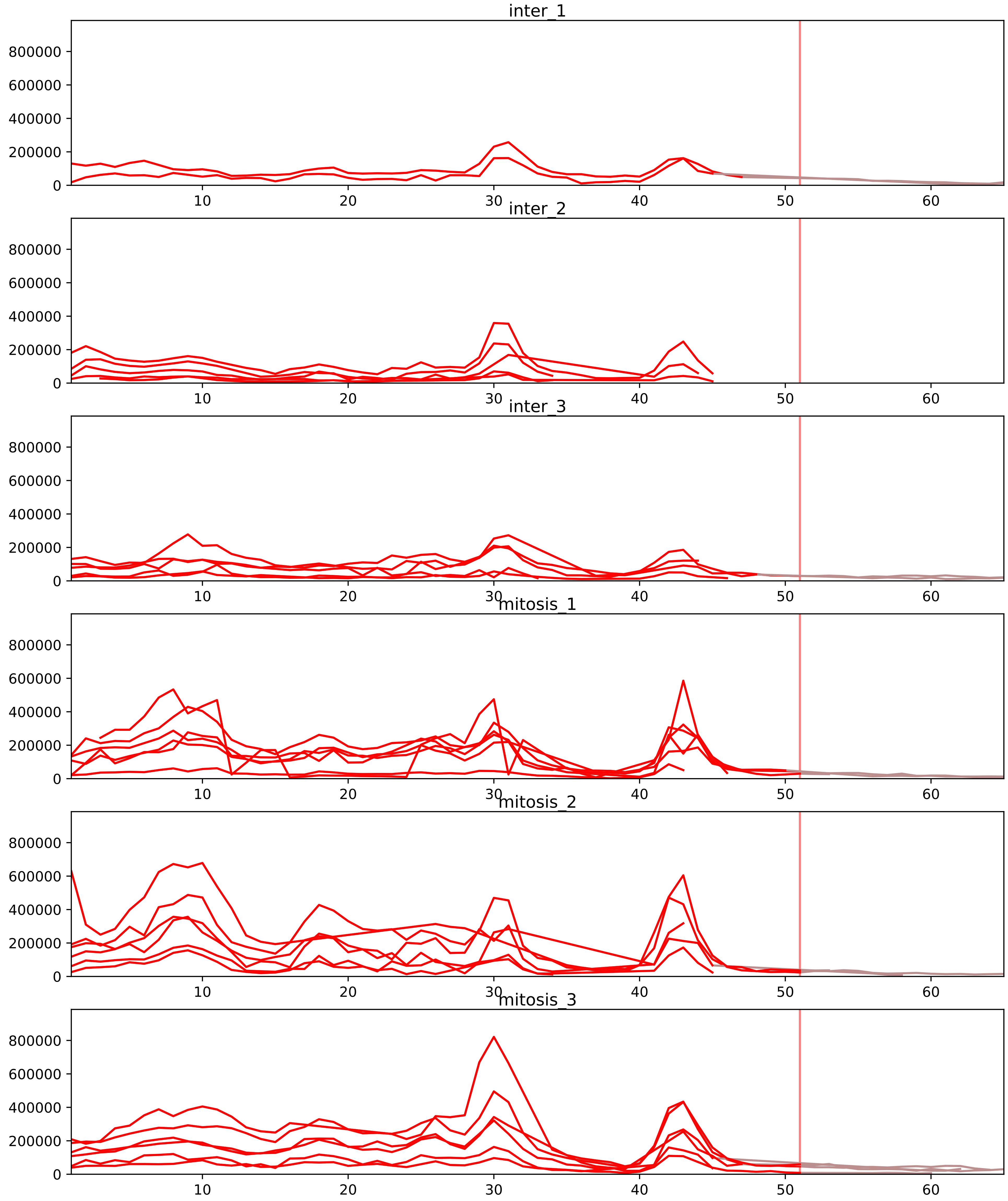


| level | condition_1 | condition_2 | pvalue | pvalue_adjusted |
|---|---|---|---|---|
| interactor_ratio | mitosis | inter | 2.8591201935579615e-07 | 9.742187326197498e-06 |
| interactor_abundance | mitosis | inter | 3.472881460334201e-06 | 0.00026625424529228876 |
| complex_abundance | mitosis | inter | 2.3002549277501173e-05 | 0.0008637691973592277 |
| monomer_abundance | mitosis | inter | 0.040789925219609294 | 0.3157957803087734 |
| total_abundance | mitosis | inter | 0.10041792064032158 | 0.3897285356427332 |
| assembled_abundance | mitosis | inter | 0.10569750250809032 | 0.38175503685879025 |

P08579_P62316
RU2B_HUMAN vs SMD2_HUMAN
p-value: 0.01 q-value: 0.008

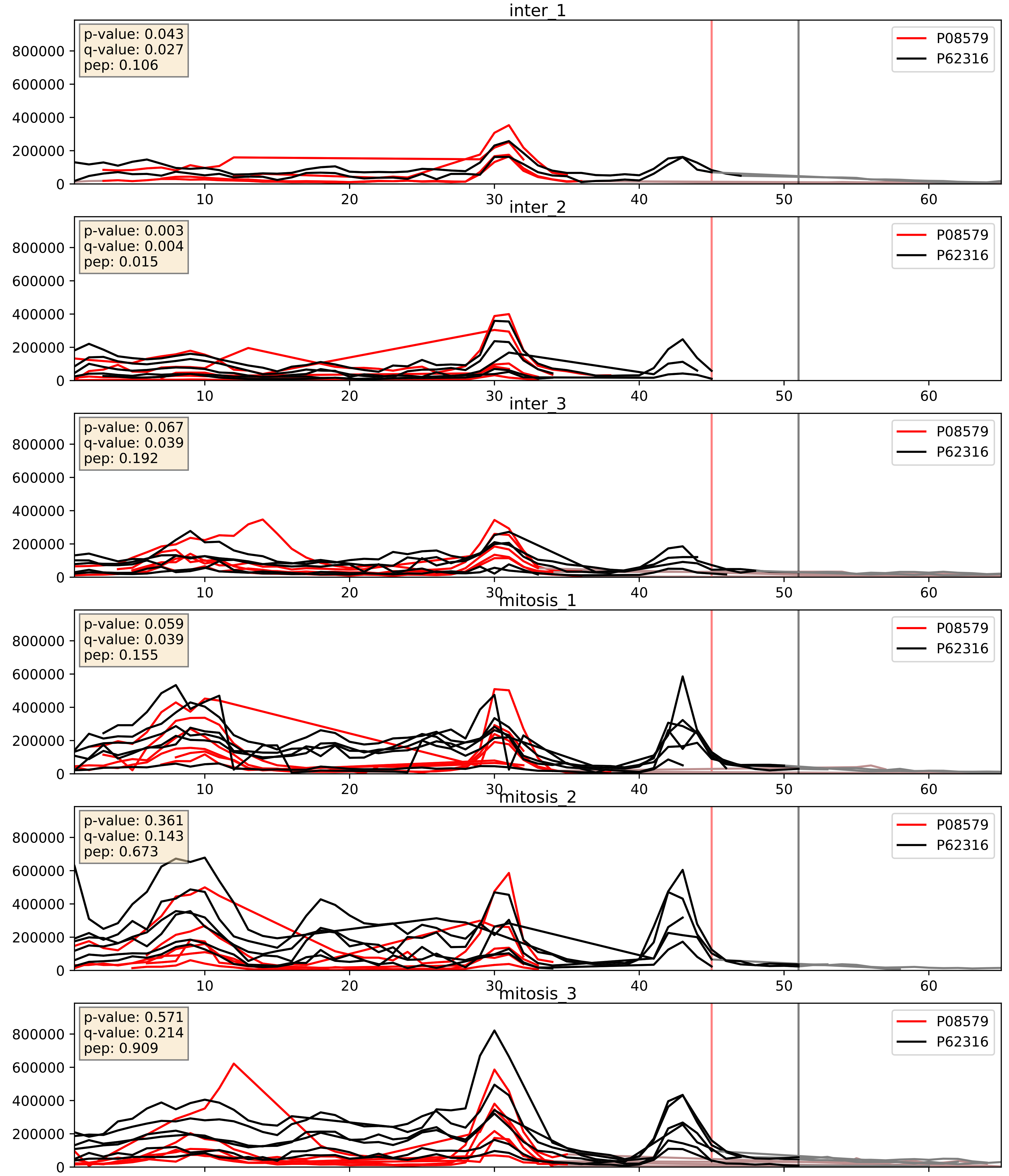


| level | condition_1 | condition_2 | pvalue | pvalue_adjusted |
|---|---|---|---|---|
| complex_abundance | mitosis | inter | 0.0025128797738631346 | 0.03639636356052189 |
| interactor_ratio | mitosis | inter | 0.004773103757202409 | 0.09212312397066487 |
| interactor_abundance | mitosis | inter | 0.18070678199639797 | 0.37336472456895164 |

P09661_P62316
RU2A_HUMAN vs SMD2_HUMAN
p-value: 0.003 q-value: 0.003

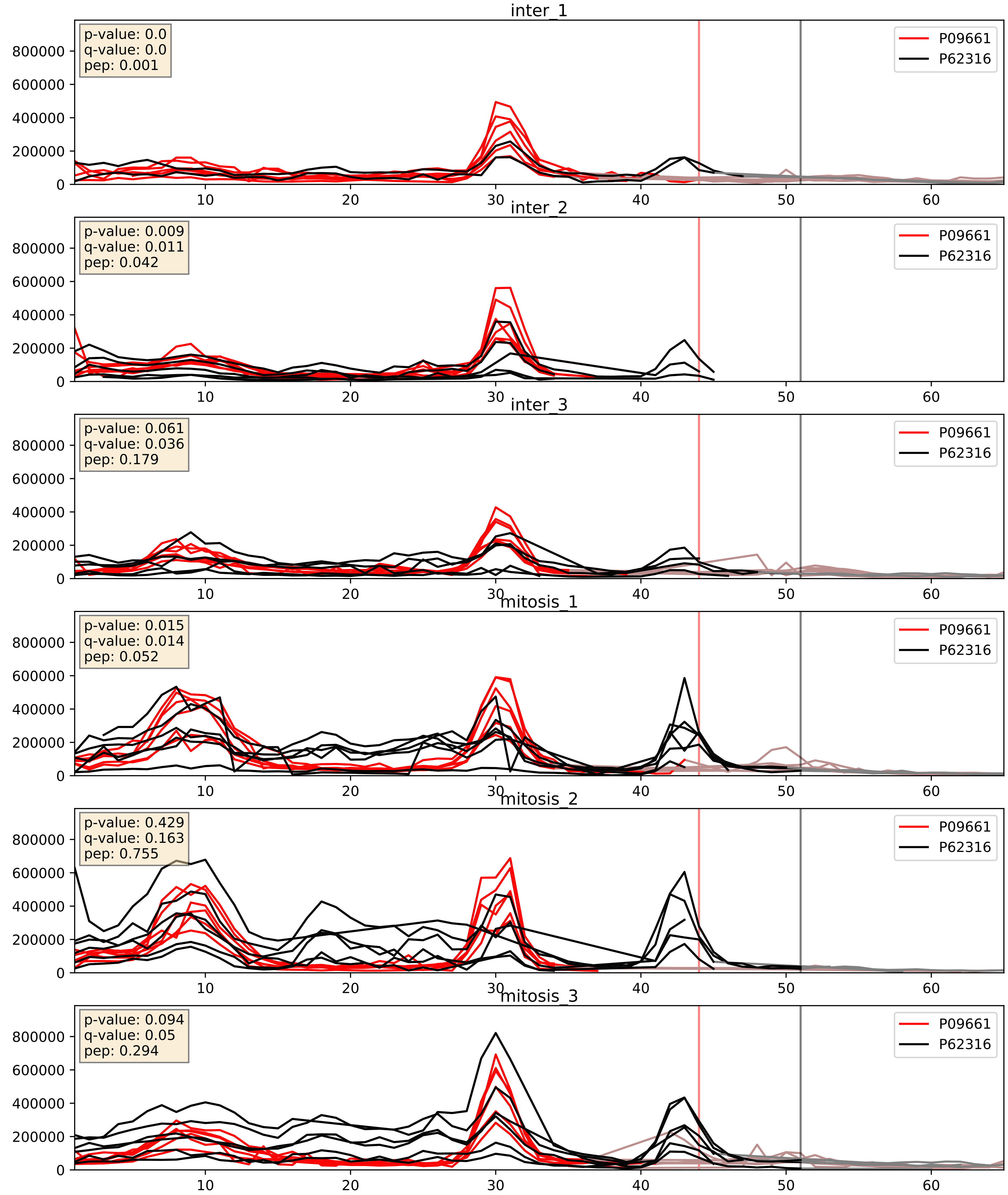


| level | condition_1 | condition_2 | pvalue | pvalue_adjusted |
|---|---|---|---|---|
| complex_abundance | mitosis | inter | 0.0005502236768995238 | 0.015047650716485379 |
| interactor_ratio | mitosis | inter | 0.003244967118469916 | 0.07405554735675686 |
| interactor_abundance | mitosis | inter | 0.21742217104749603 | 0.41312625619679605 |

P14866_P62316
HNRPL_HUMAN vs SMD2_HUMAN
p-value: 0.073 q-value: 0.044

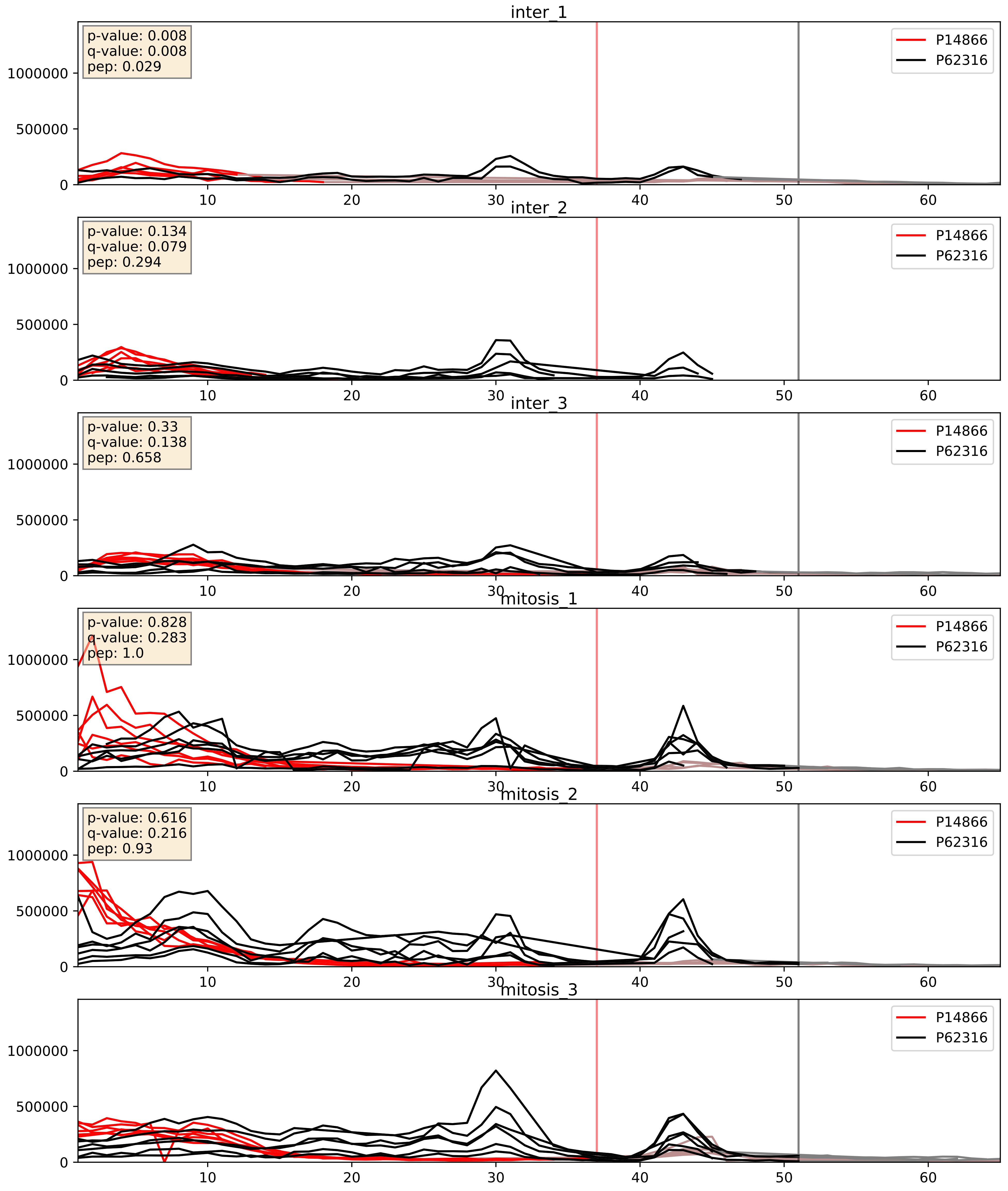


| level | condition_1 | condition_2 | pvalue | pvalue_adjusted |
|---|---|---|---|---|
| complex_abundance | mitosis | inter | 0.000861046445581631 | 0.019513211095311817 |
| interactor_abundance | mitosis | inter | 0.009697640385696541 | 0.08525122834595471 |
| interactor_ratio | mitosis | inter | 0.009765737212506018 | 0.14216787506641412 |

P22626_P62316
ROA2_HUMAN vs SMD2_HUMAN
p-value: 0.0 q-value: 0.001

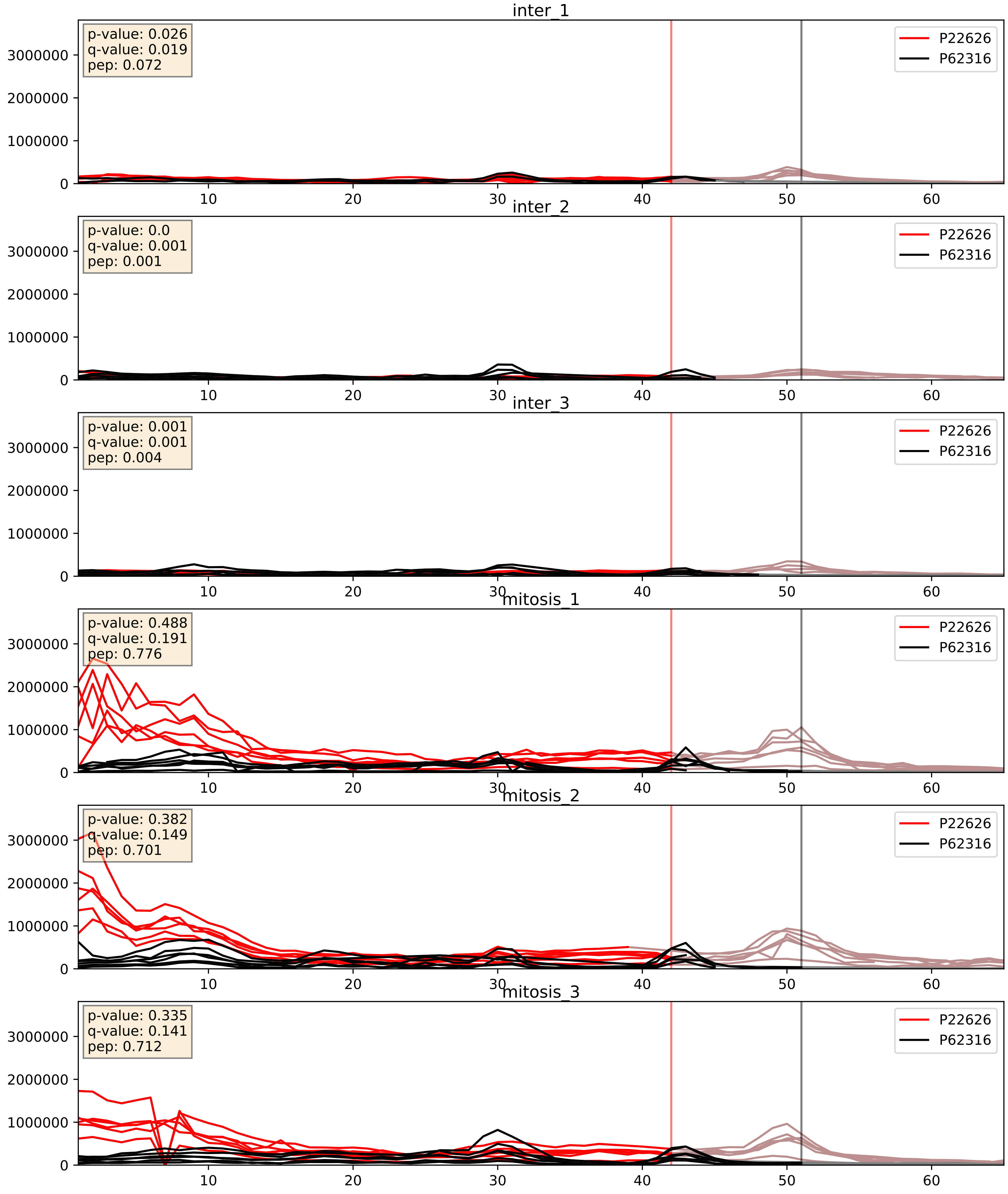


| level | condition_1 | condition_2 | pvalue | pvalue_adjusted |
|---|---|---|---|---|
| complex_abundance | mitosis | inter | 0.0008978115482066242 | 0.01975647005822289 |
| interactor_abundance | mitosis | inter | 0.0034466655543236775 | 0.04511548235131213 |
| interactor_ratio | mitosis | inter | 0.0603049372865427 | 0.40594115513296747 |

P62304_P62316
RUXE_HUMAN vs SMD2_HUMAN
p-value: 0.001 q-value: 0.002

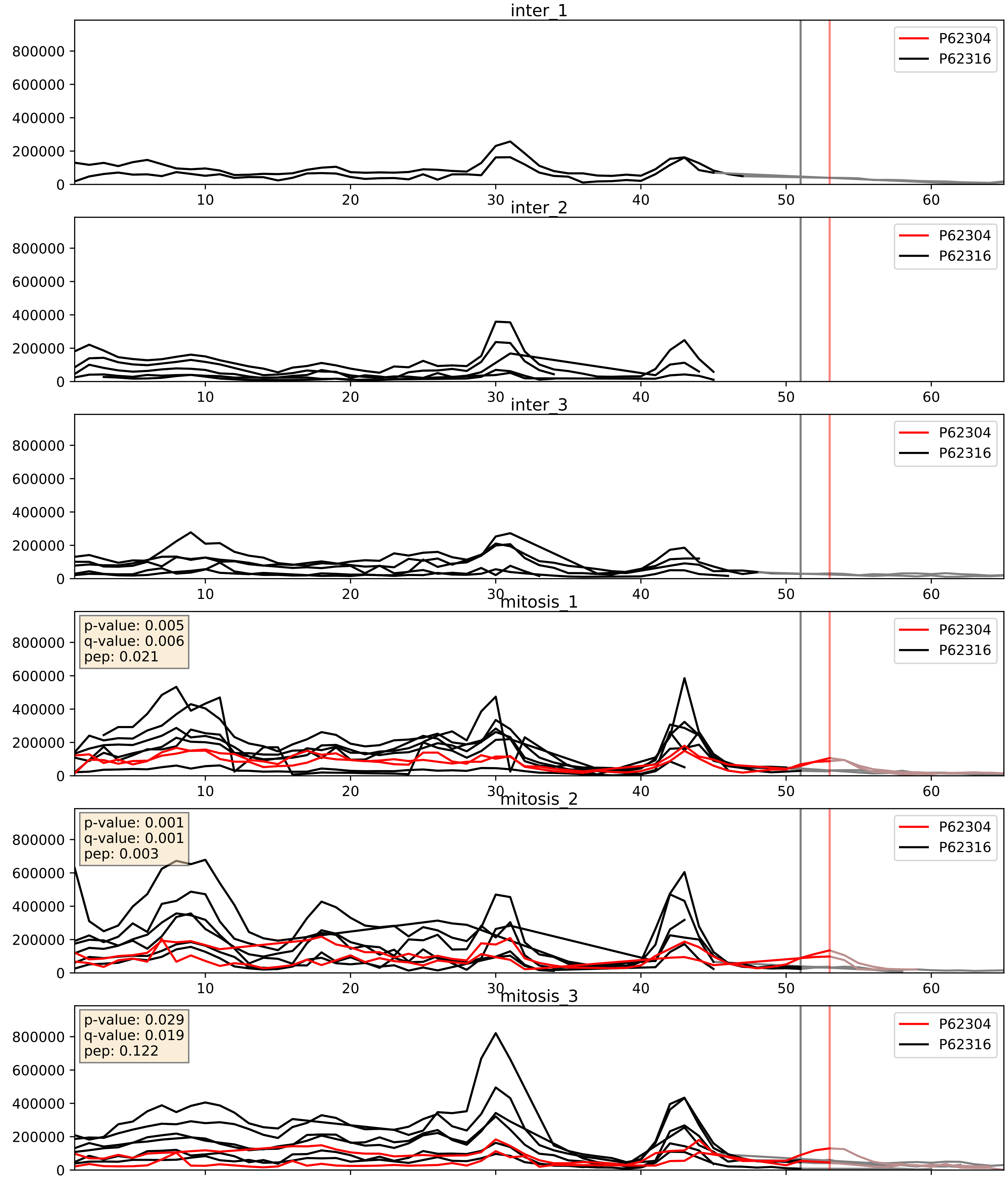


| level | condition_1 | condition_2 | pvalue | pvalue_adjusted |
|---|---|---|---|---|
| complex_abundance | mitosis | inter | 1.149781427818535e-05 | 0.005691857151652958 |
| interactor_abundance | mitosis | inter | 0.00011451049632638166 | 0.008168415404615225 |
| interactor_ratio | mitosis | inter | 0.001270295386270424 | 0.0423373831474418 |

P62306_P62316
RUXF_HUMAN vs SMD2_HUMAN
p-value: 0.0 q-value: 0.0

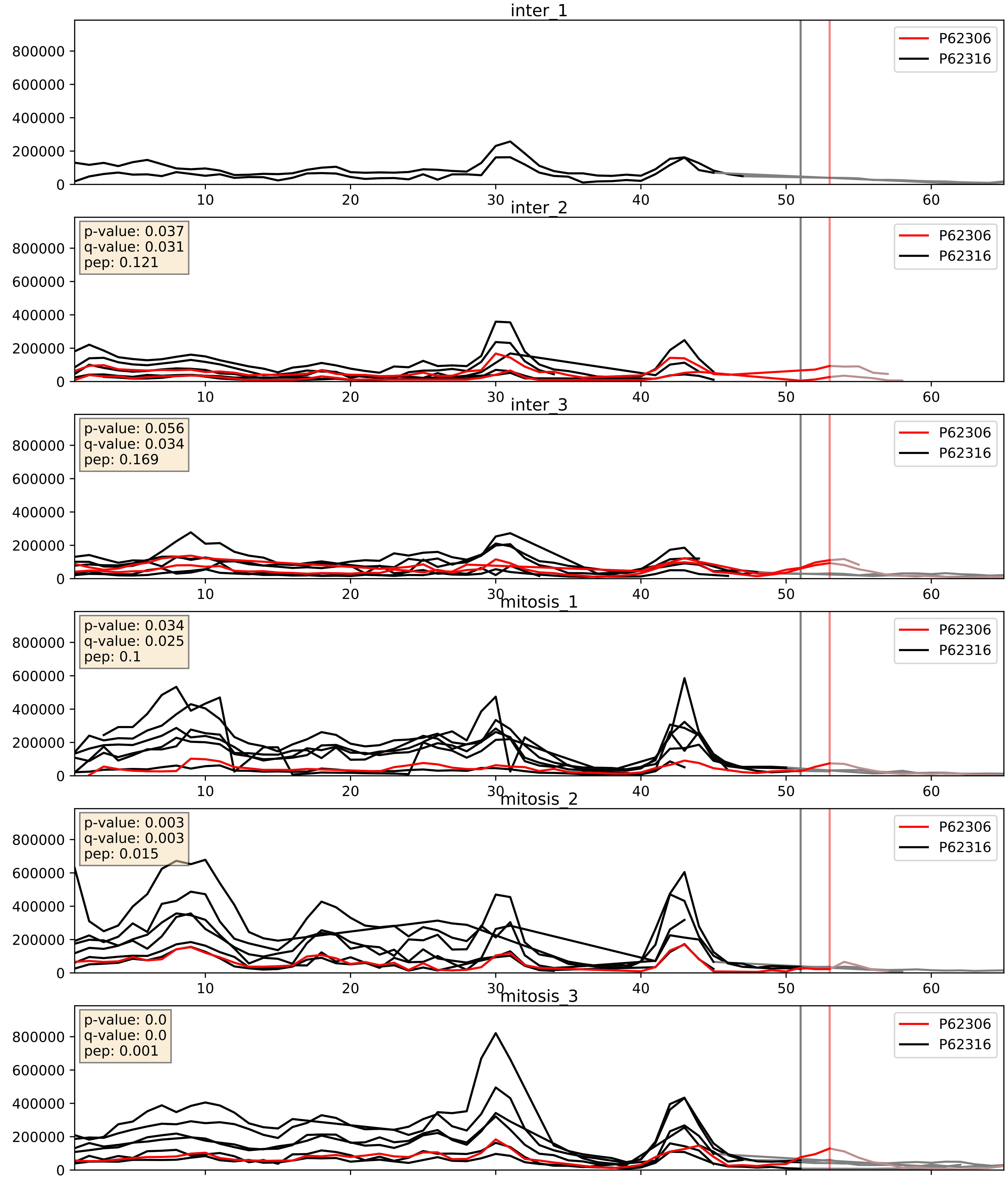


| level | condition_1 | condition_2 | pvalue | pvalue_adjusted |
|---|---|---|---|---|
| interactor_ratio | mitosis | inter | 0.00031290593502055766 | 0.01878749154928596 |
| complex_abundance | mitosis | inter | 0.18206089764979508 | 0.33292913562961884 |
| interactor_abundance | mitosis | inter | 0.4800162238922761 | 0.6514886438113023 |

P62314_P62316
SMD1_HUMAN vs SMD2_HUMAN
p-value: 0.007 q-value: 0.006

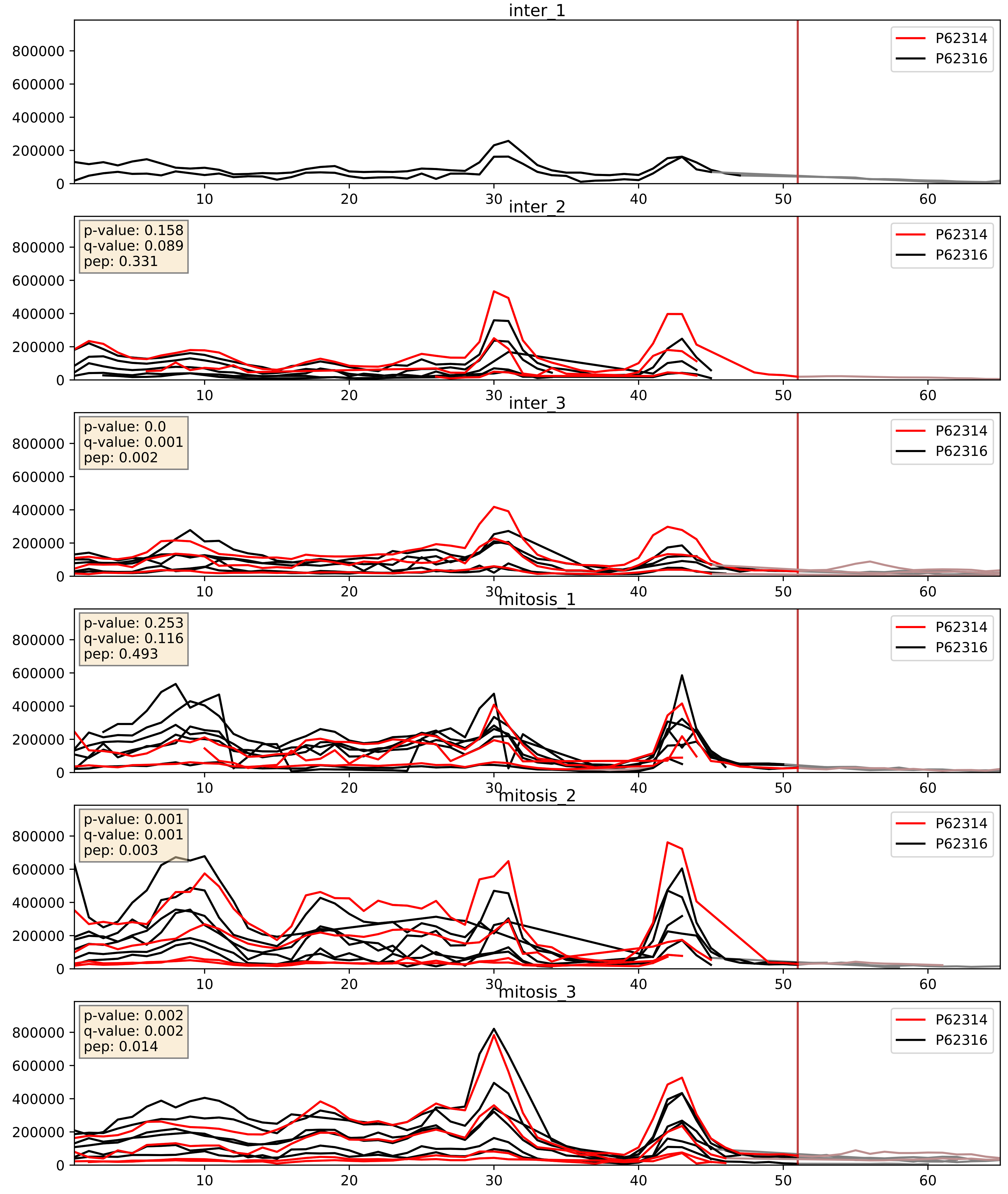


| level | condition_1 | condition_2 | pvalue | pvalue_adjusted |
|---|---|---|---|---|
| interactor_ratio | mitosis | inter | 0.030400220361707538 | 0.27742631801302403 |
| complex_abundance | mitosis | inter | 0.1044510172596175 | 0.2402721956553067 |
| interactor_abundance | mitosis | inter | 0.3411573116383529 | 0.5396541039907444 |

P62316_P62318
SMD2_HUMAN vs SMD3_HUMAN
p-value: 0.001 q-value: 0.001

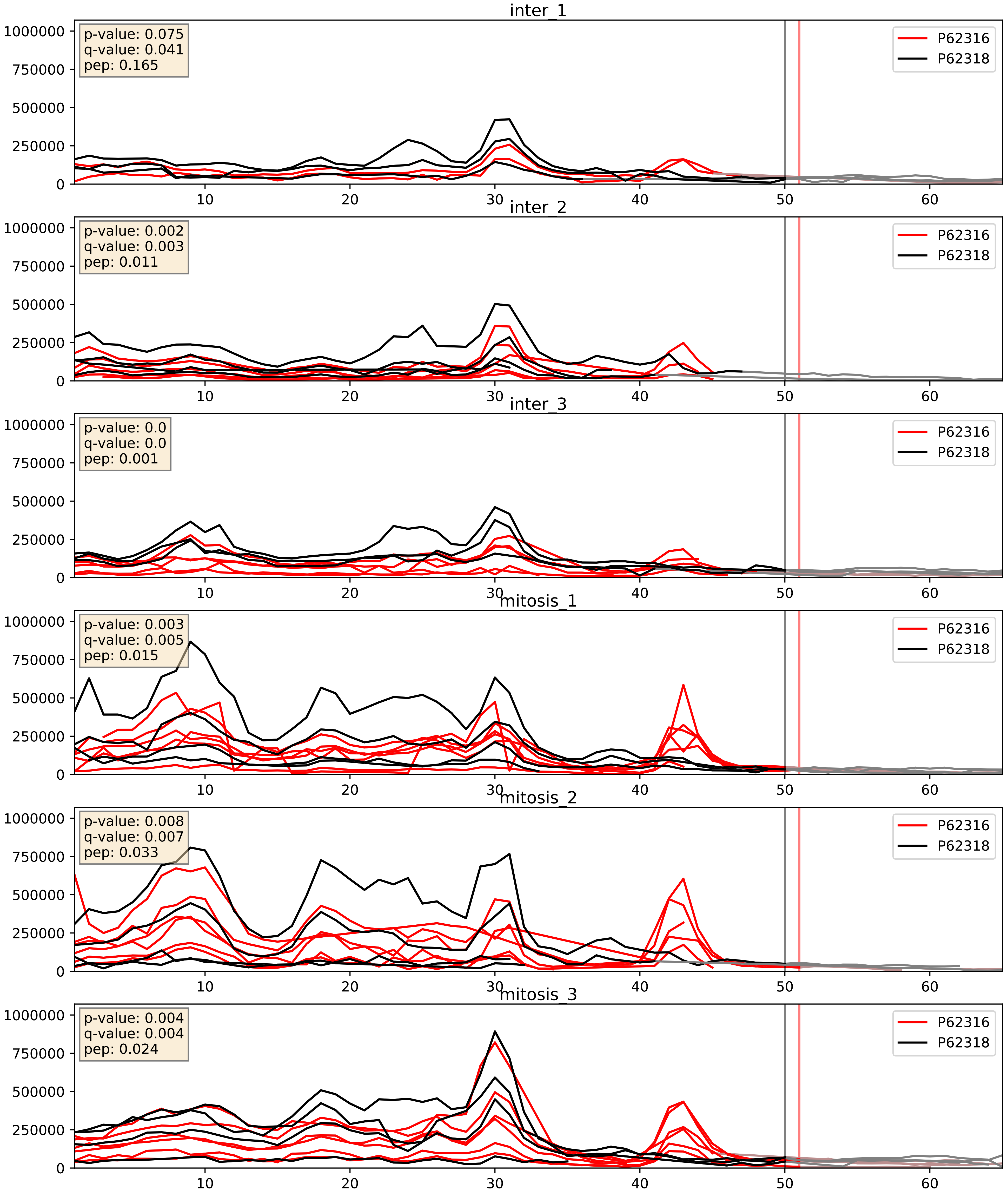


| level | condition_1 | condition_2 | pvalue | pvalue_adjusted |
|---|---|---|---|---|
| interactor_abundance | mitosis | inter | 0.0003865021250090336 | 0.013615054280153612 |
| complex_abundance | mitosis | inter | 0.05118009301339294 | 0.161227792796871 |
| interactor_ratio | mitosis | inter | 0.1220440593729674 | 0.5708187039120276 |
