## Supplementary Data 2 for "SECAT: Quantifying differential protein-protein interaction states by network-centric analysis": hela_string_000059_Q9UKD2.pdf

Q9UKD2_Q9UKD2

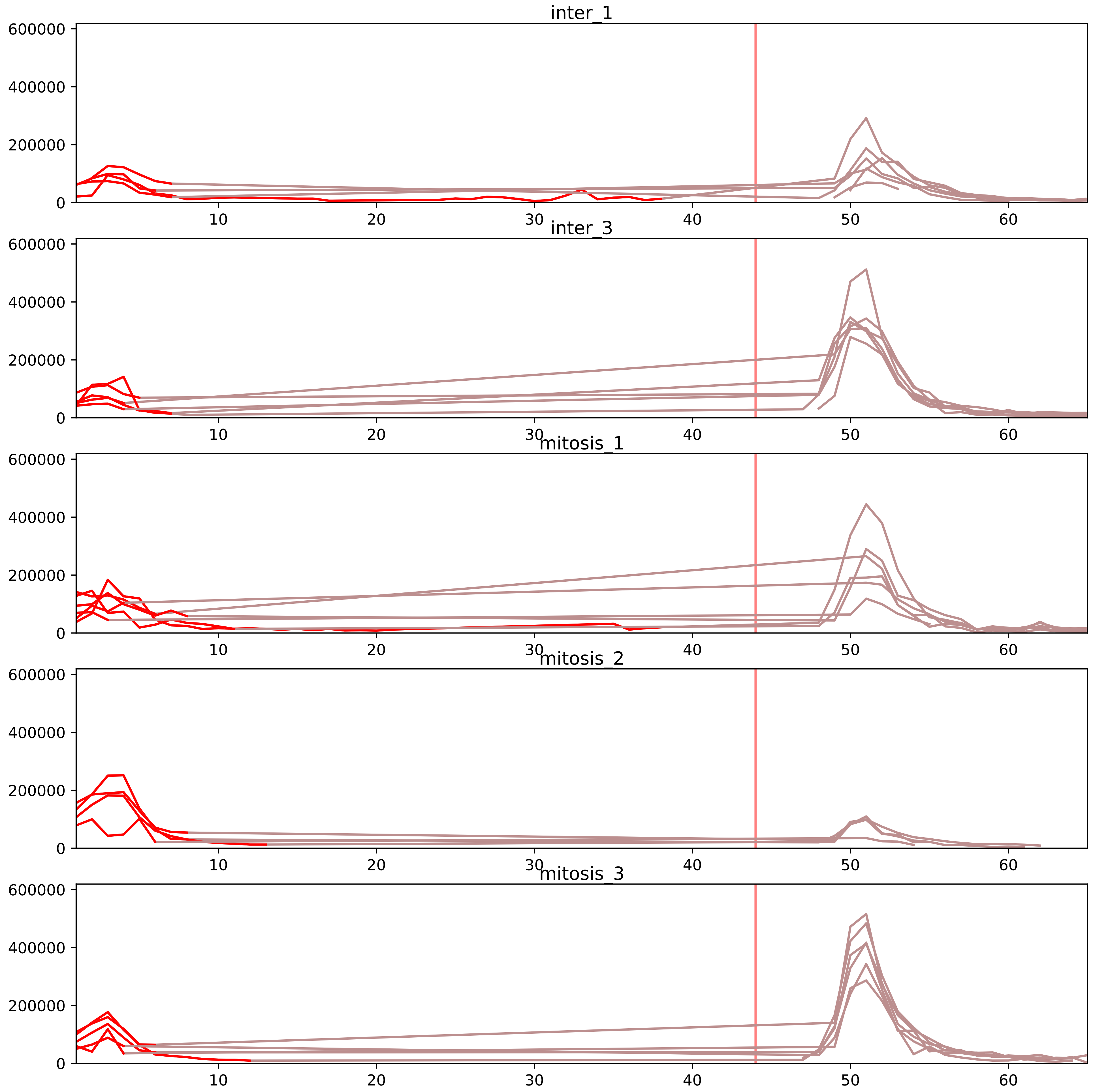


| level | condition_1 | condition_2 | pvalue | pvalue_adjusted |
|---|---|---|---|---|
| interactor_ratio | mitosis | inter | 3.133644431089284e-07 | 1.0483465005825969e-05 |
| interactor_abundance | mitosis | inter | 0.035827861245746995 | 0.11425175856529371 |
| complex_abundance | mitosis | inter | 0.03777799560120493 | 0.10320748069892423 |
| assembled_abundance | mitosis | inter | 0.1652042174681168 | 0.4775952195267642 |
| total_abundance | mitosis | inter | 0.3389117620190077 | 0.6566203004936698 |
| monomer_abundance | mitosis | inter | 0.49092934063814164 | 0.750406864904721 |

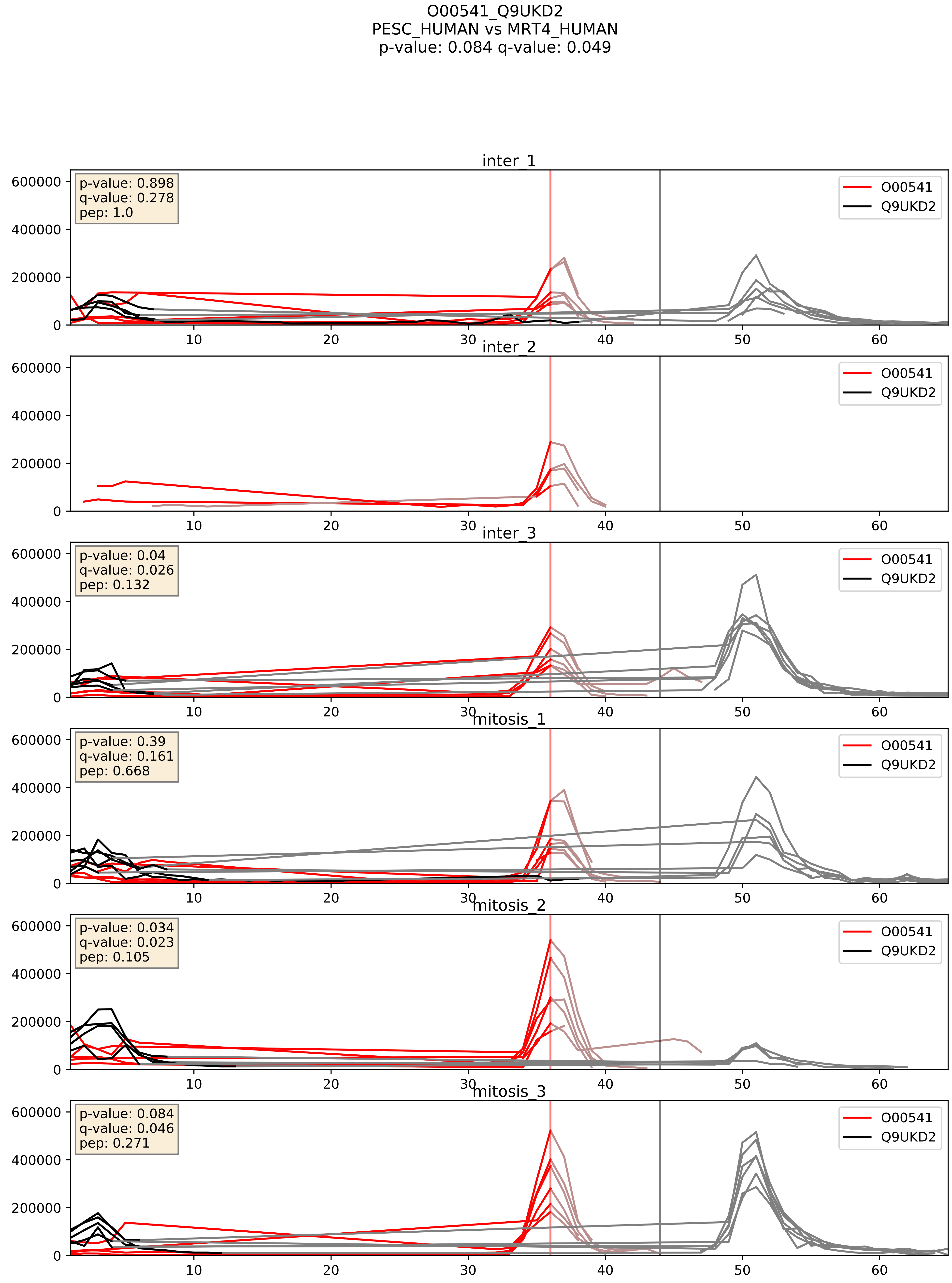


| level | condition_1 | condition_2 | pvalue | pvalue_adjusted |
|---|---|---|---|---|
| interactor_ratio | mitosis | inter | 0.039784780596399914 | 0.3296783367910777 |
| complex_abundance | mitosis | inter | 0.6585396611988352 | 0.7711057603735534 |
| interactor_abundance | mitosis | inter | 0.7805251561649456 | 0.87240260322151 |

O43159_Q9UKD2
RRP8_HUMAN vs MRT4_HUMAN
p-value: 0.001 q-value: 0.032

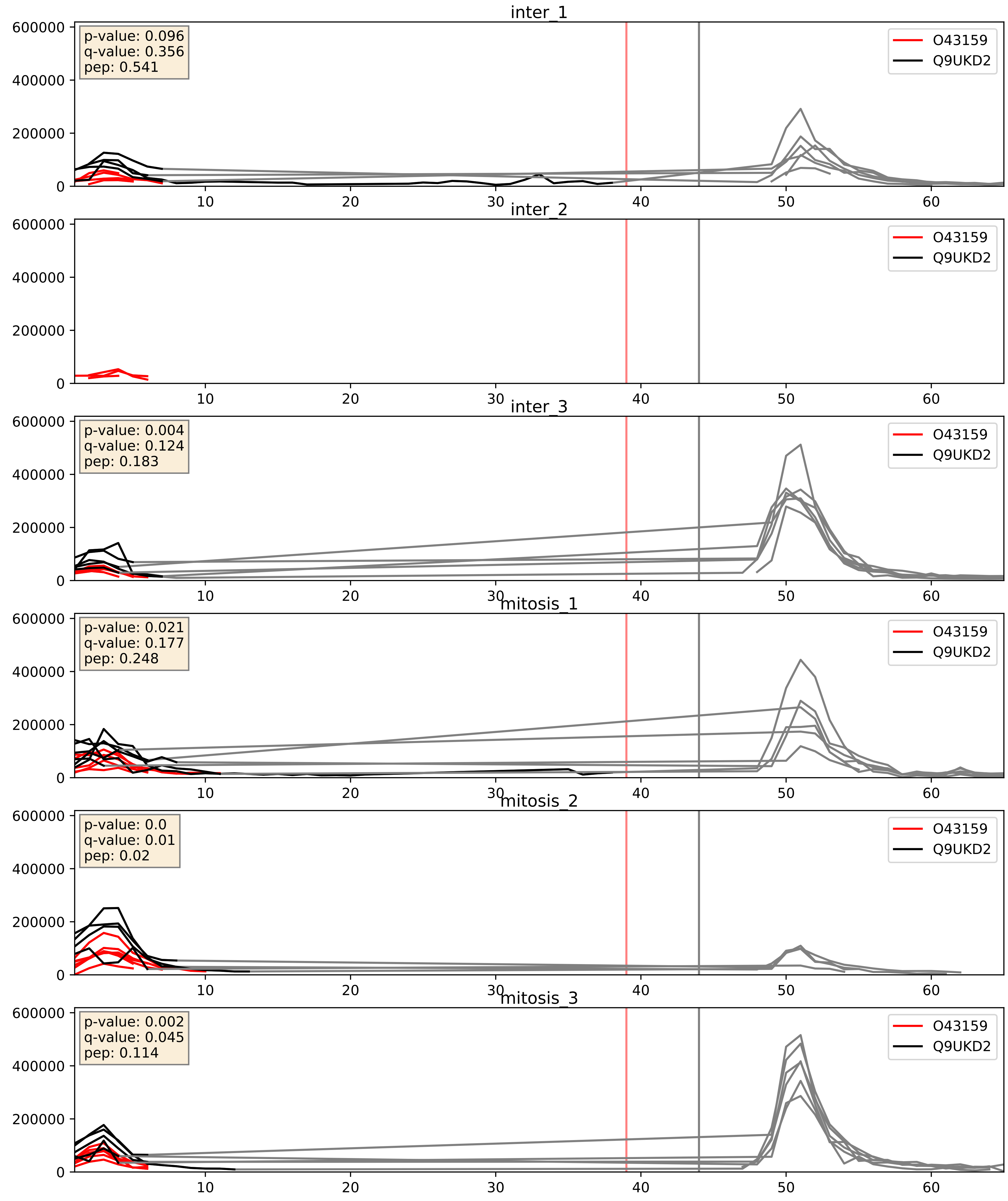


| level | condition_1 | condition_2 | pvalue | pvalue_adjusted |
|---|---|---|---|---|
| interactor_ratio | mitosis | inter | 0.07462663170364646 | 0.4589108961086305 |
| complex_abundance | mitosis | inter | 0.26740809740739513 | 0.4169423158118948 |
| interactor_abundance | mitosis | inter | 0.4228129447317011 | 0.6037162313433464 |

P18077_Q9UKD2
RL35A_HUMAN vs MRT4_HUMAN
p-value: 0.0 q-value: 0.023

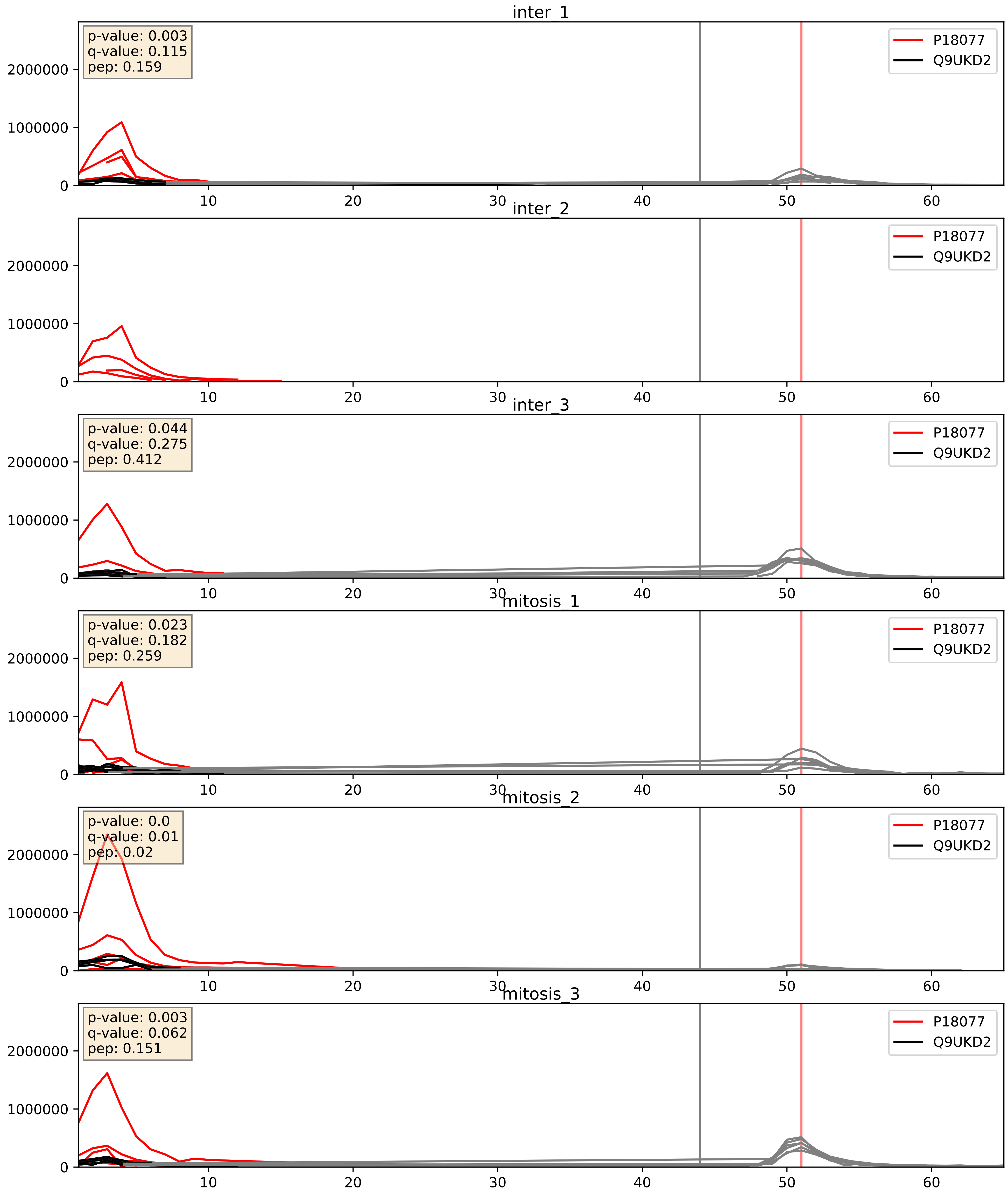


| level | condition_1 | condition_2 | pvalue | pvalue_adjusted |
|---|---|---|---|---|
| complex_abundance | mitosis | inter | 0.2786450579946821 | 0.4272258098575101 |
| interactor_abundance | mitosis | inter | 0.3379896833011881 | 0.5374183503405164 |
| interactor_ratio | mitosis | inter | 0.5741043006239598 | 0.894979568993097 |

P22087_Q9UKD2
FBRL_HUMAN vs MRT4_HUMAN
p-value: 0.002 q-value: 0.034

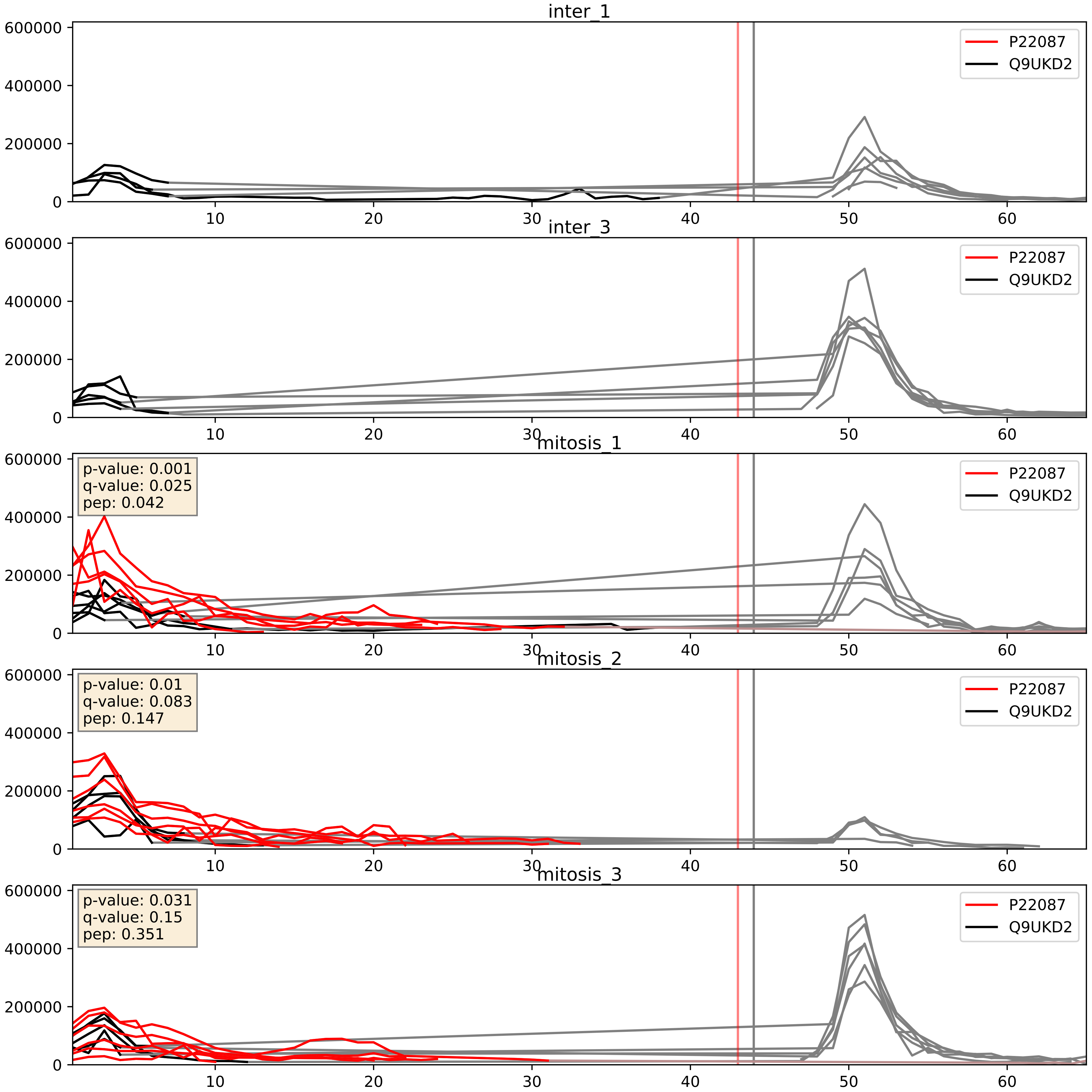


| level | condition_1 | condition_2 | pvalue | pvalue_adjusted |
|---|---|---|---|---|
| complex_abundance | mitosis | inter | 0.0008743237108210411 | 0.019513211095311817 |
| interactor_abundance | mitosis | inter | 0.0009724779092061511 | 0.023091292379485866 |
| interactor_ratio | mitosis | inter | 0.004228737105466719 | 0.08567853202577967 |

P32969_Q9UKD2
RL9_HUMAN vs MRT4_HUMAN
p-value: 0.0 q-value: 0.015

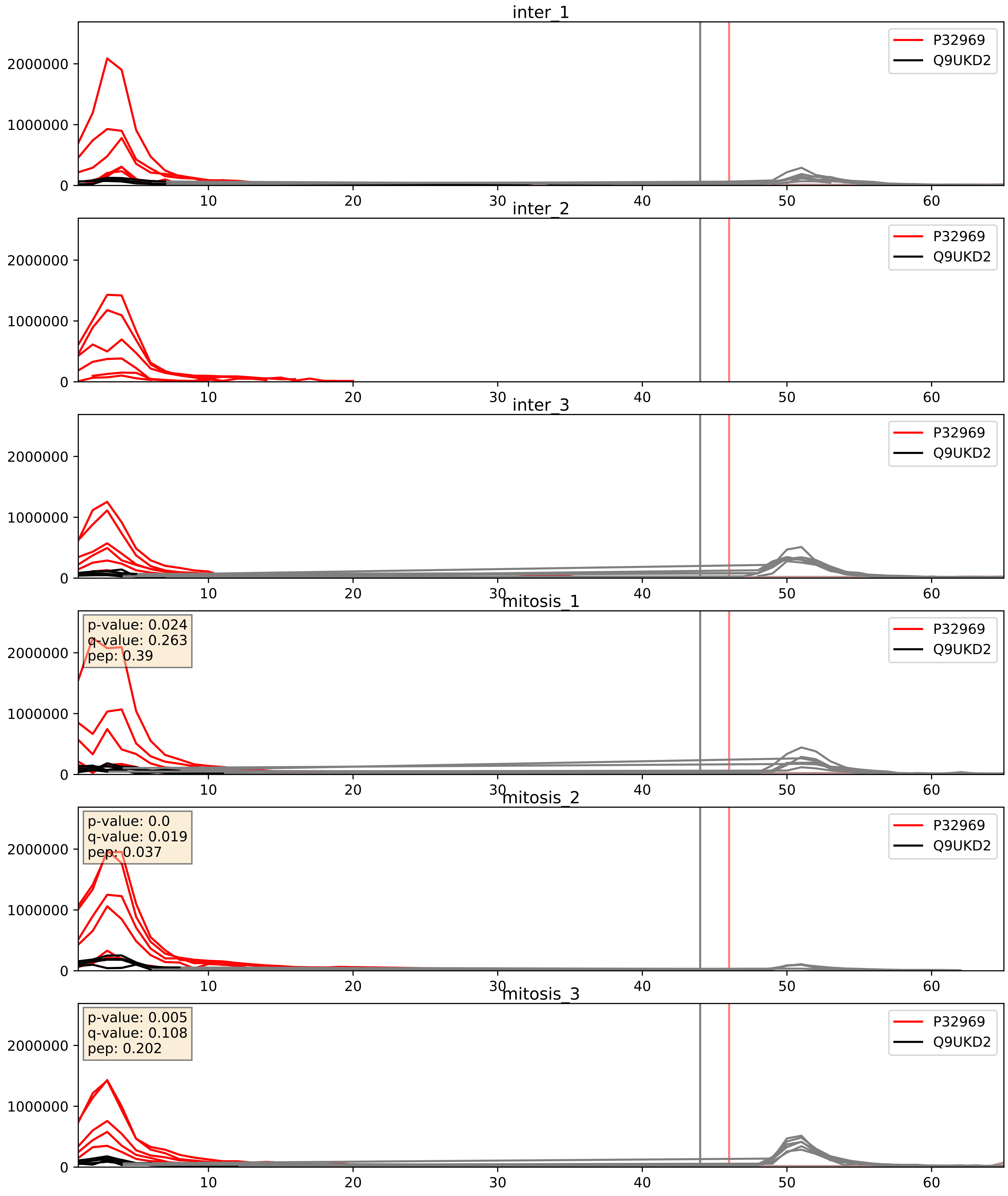


| level | condition_1 | condition_2 | pvalue | pvalue_adjusted |
|---|---|---|---|---|
| complex_abundance | mitosis | inter | 0.26136949740300996 | 0.4114238502702768 |
| interactor_abundance | mitosis | inter | 0.30057899301007135 | 0.49537084716330587 |
| interactor_ratio | mitosis | inter | 0.4042759720104136 | 0.8313271231500527 |

P40429_Q9UKD2
RL13A_HUMAN vs MRT4_HUMAN
p-value: 0.0 q-value: 0.017

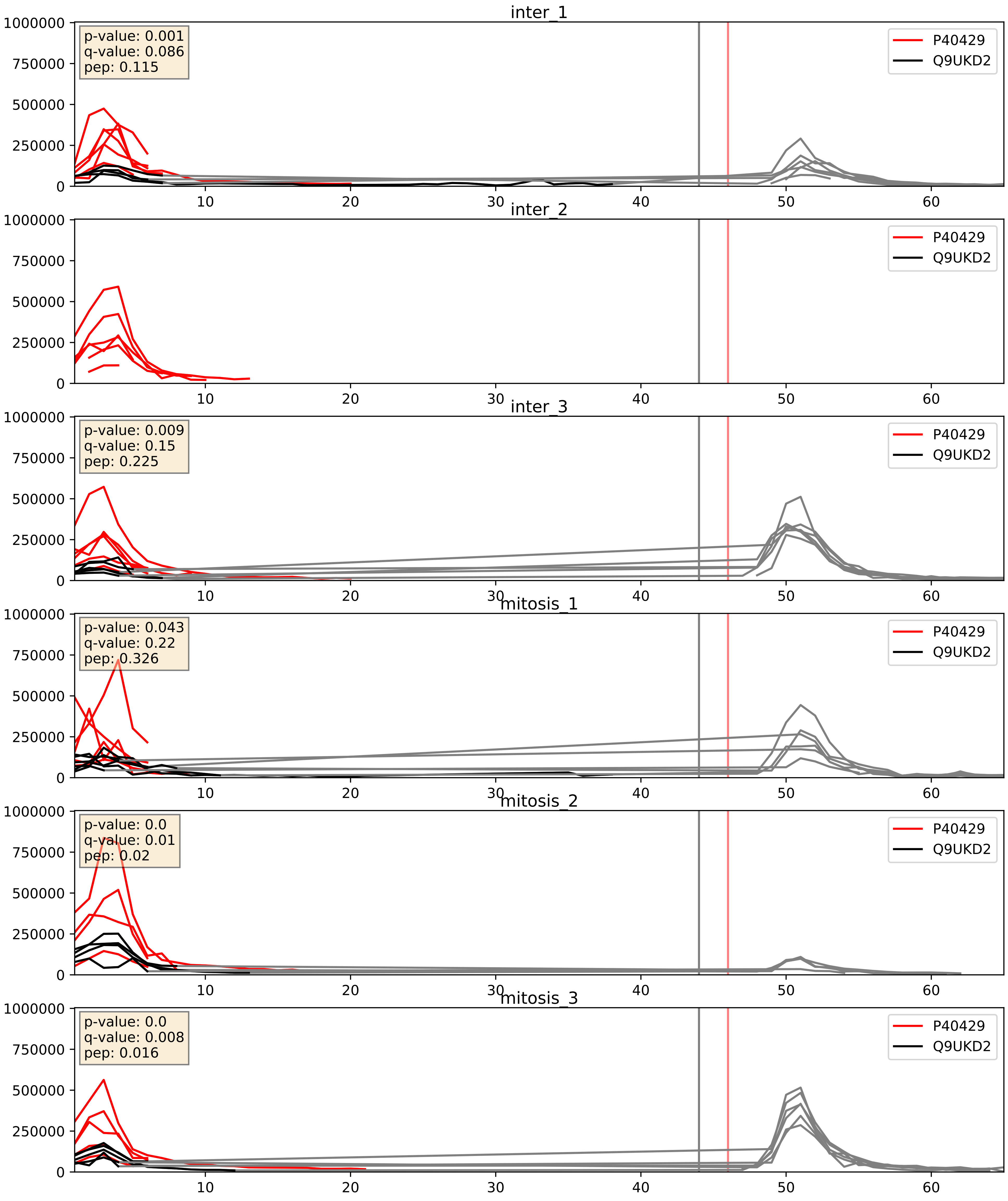


| level | condition_1 | condition_2 | pvalue | pvalue_adjusted |
|---|---|---|---|---|
| complex_abundance | mitosis | inter | 0.27422769777551415 | 0.4234662136453529 |
| interactor_abundance | mitosis | inter | 0.3433035485448669 | 0.5420201629725344 |
| interactor_ratio | mitosis | inter | 0.6842214633715376 | 0.9349347226390419 |

P46087_Q9UKD2
NOP2_HUMAN vs MRT4_HUMAN
p-value: 0.002 q-value: 0.003

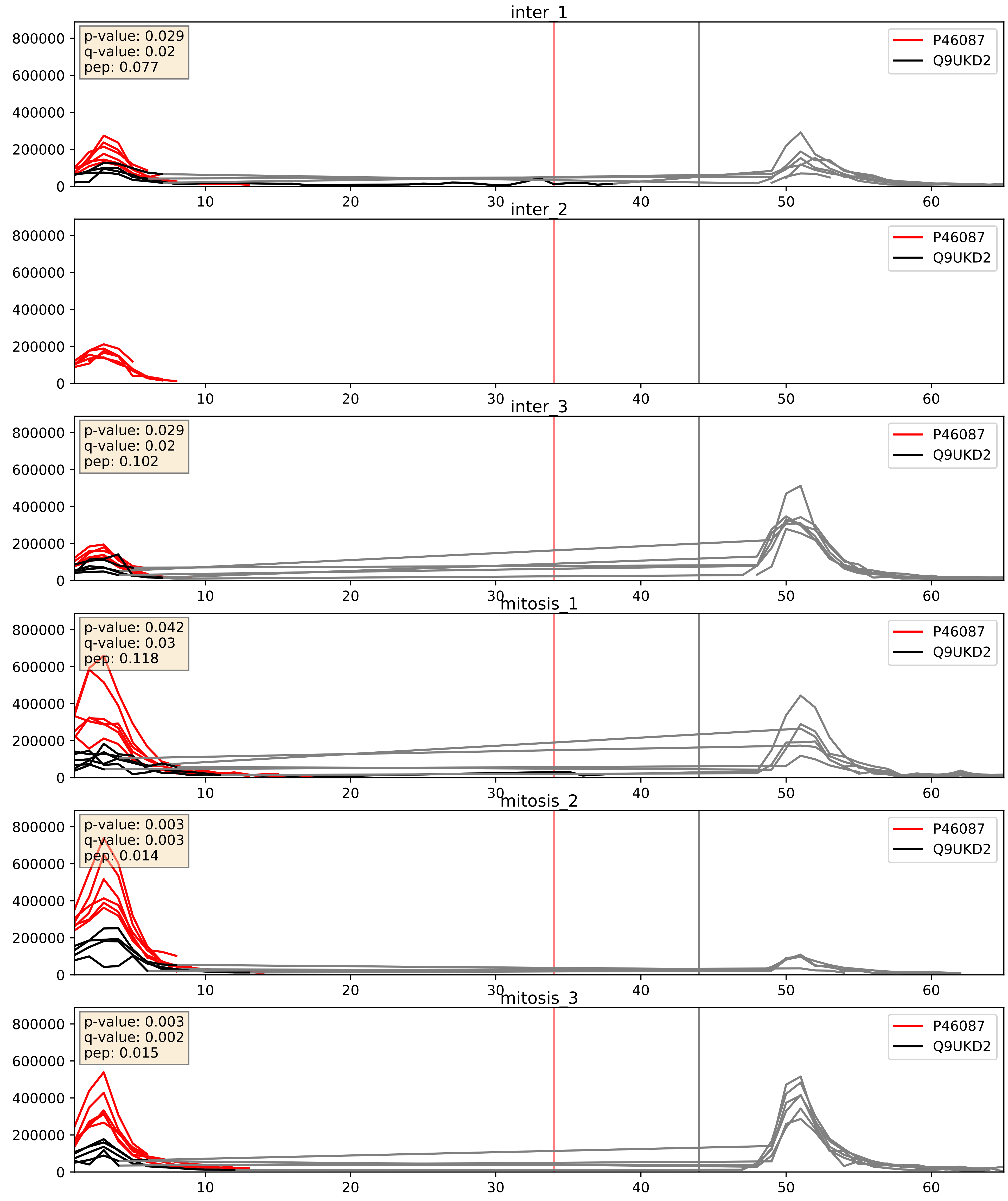


| level | condition_1 | condition_2 | pvalue | pvalue_adjusted |
|---|---|---|---|---|
| interactor_ratio | mitosis | inter | 0.008169453022261623 | 0.12756688776860511 |
| interactor_abundance | mitosis | inter | 0.05561217096474677 | 0.1913344788819262 |
| complex_abundance | mitosis | inter | 0.08848513121730248 | 0.21834324682044082 |

P46779_Q9UKD2
RL28_HUMAN vs MRT4_HUMAN
p-value: 0.0 q-value: 0.008

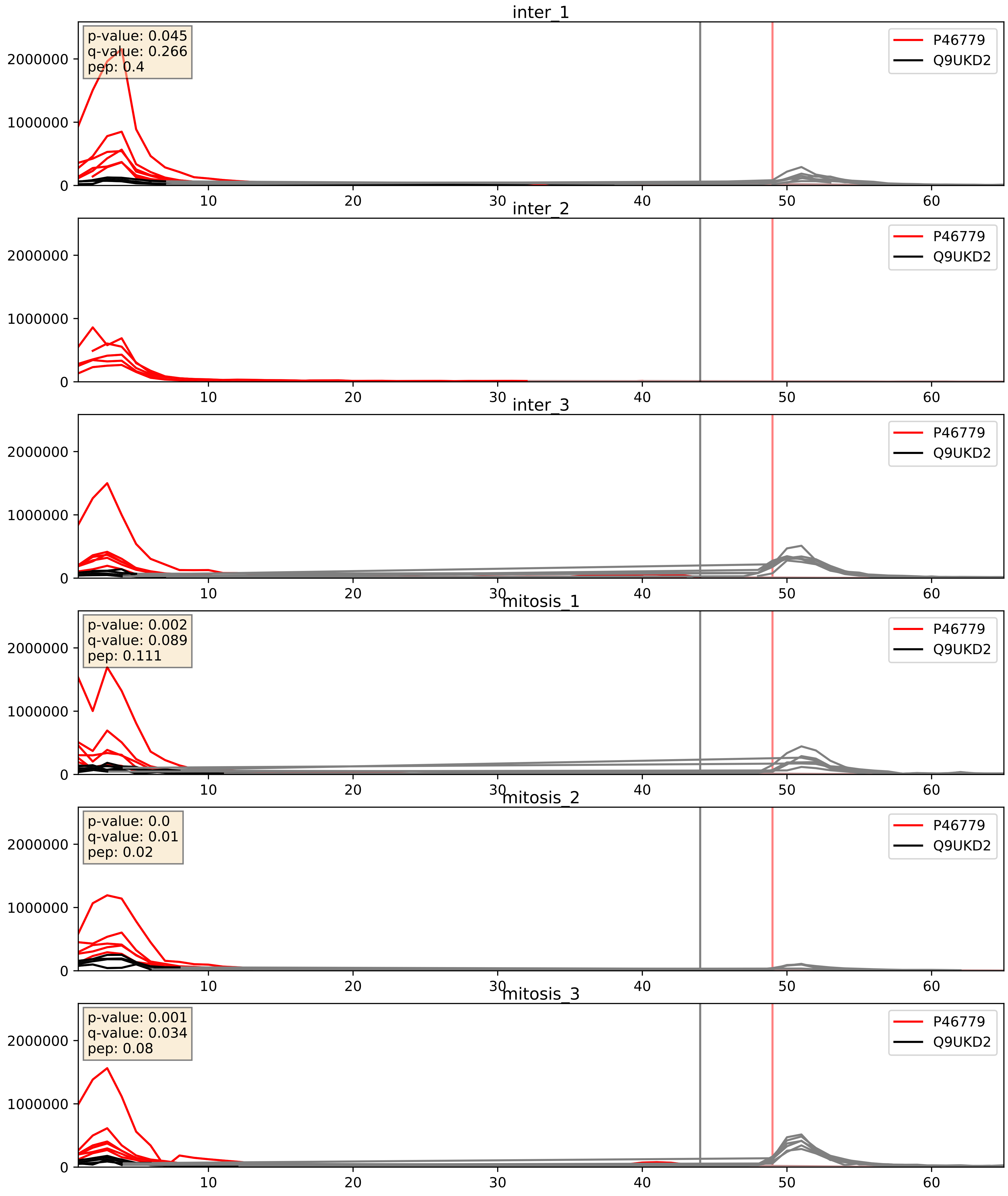


| level | condition_1 | condition_2 | pvalue | pvalue_adjusted |
|---|---|---|---|---|
| complex_abundance | mitosis | inter | 0.3564277459855168 | 0.5039678734119629 |
| interactor_abundance | mitosis | inter | 0.44751503442150253 | 0.6226032671235112 |
| interactor_ratio | mitosis | inter | 0.7406373205653809 | 0.9491588884573259 |

P49207_Q9UKD2
RL34_HUMAN vs MRT4_HUMAN
p-value: 0.0 q-value: 0.017

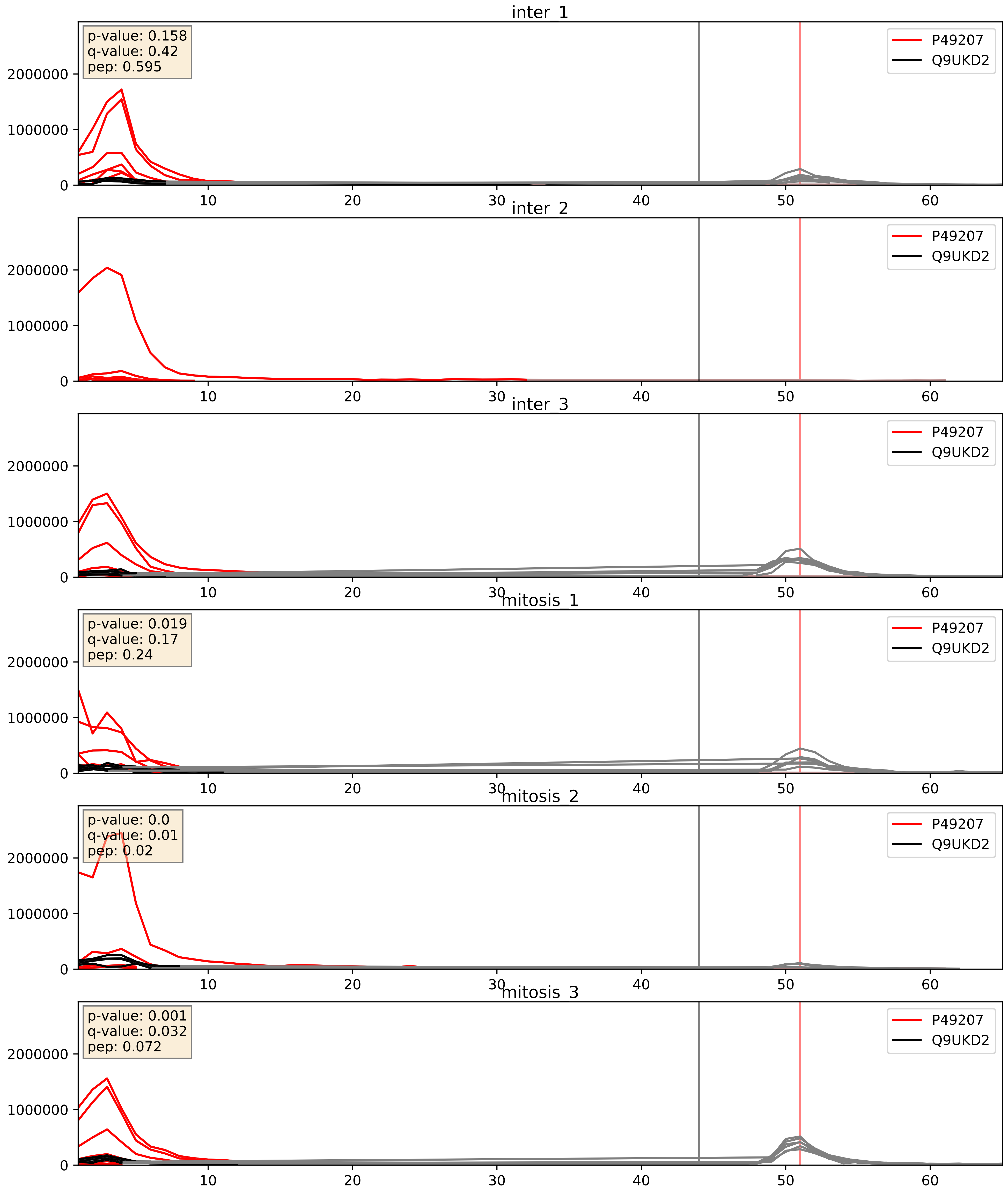


| level | condition_1 | condition_2 | pvalue | pvalue_adjusted |
|---|---|---|---|---|
| complex_abundance | mitosis | inter | 0.5248532931740976 | 0.6603548474405384 |
| interactor_abundance | mitosis | inter | 0.7240257921696673 | 0.8244310982337681 |
| interactor_ratio | mitosis | inter | 0.897637502885408 | 0.9830084986132549 |

P55769_Q9UKD2
NH2L1_HUMAN vs MRT4_HUMAN
p-value: 0.001 q-value: 0.029

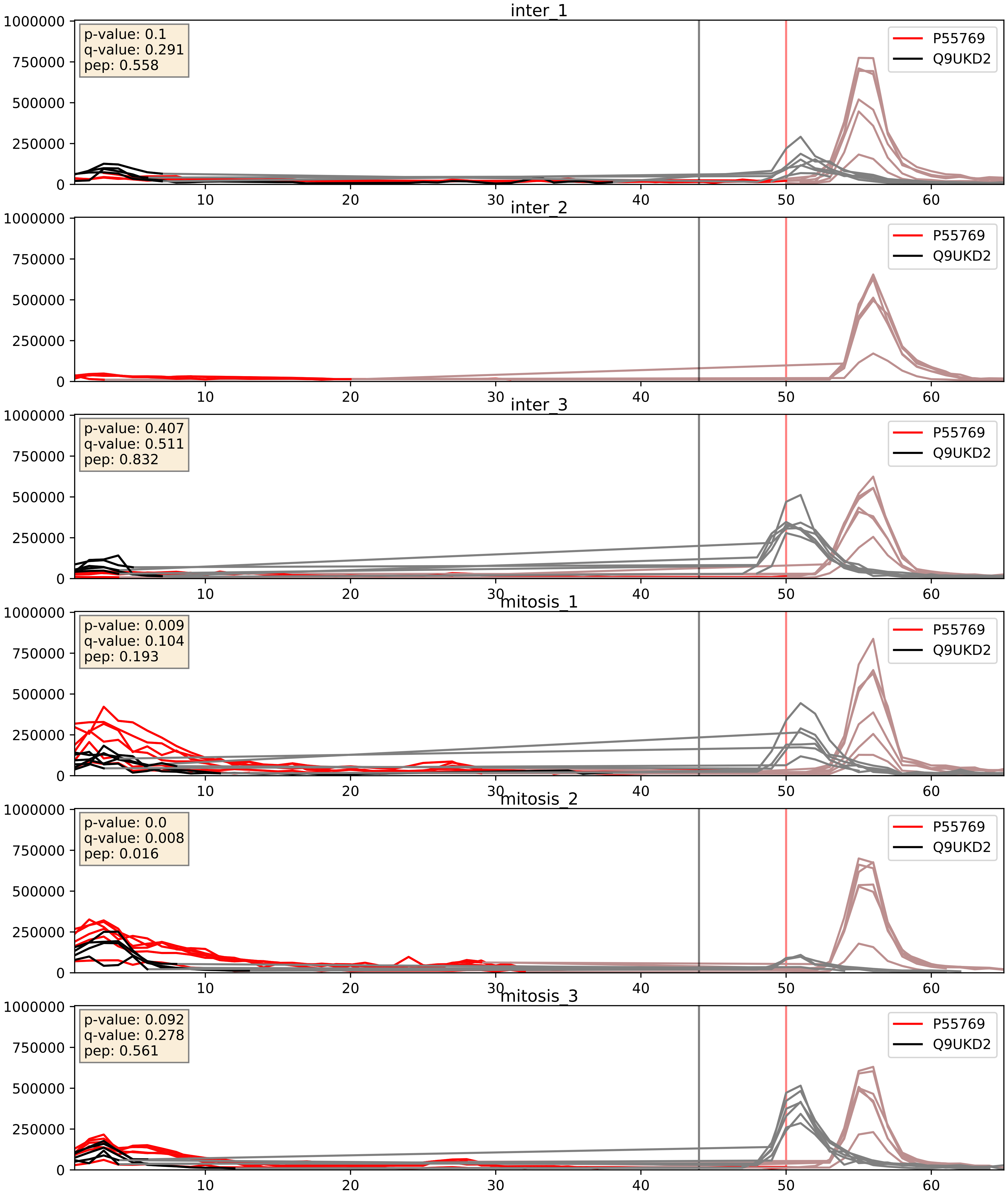


| level | condition_1 | condition_2 | pvalue | pvalue_adjusted |
|---|---|---|---|---|
| interactor_ratio | mitosis | inter | 0.0025457301912372872 | 0.06485550725294995 |
| interactor_abundance | mitosis | inter | 0.019320041195118556 | 0.1190514126751731 |
| complex_abundance | mitosis | inter | 0.051399232483414346 | 0.16145960736074377 |

P56537_Q9UKD2
IF6_HUMAN vs MRT4_HUMAN
p-value: 0.002 q-value: 0.035

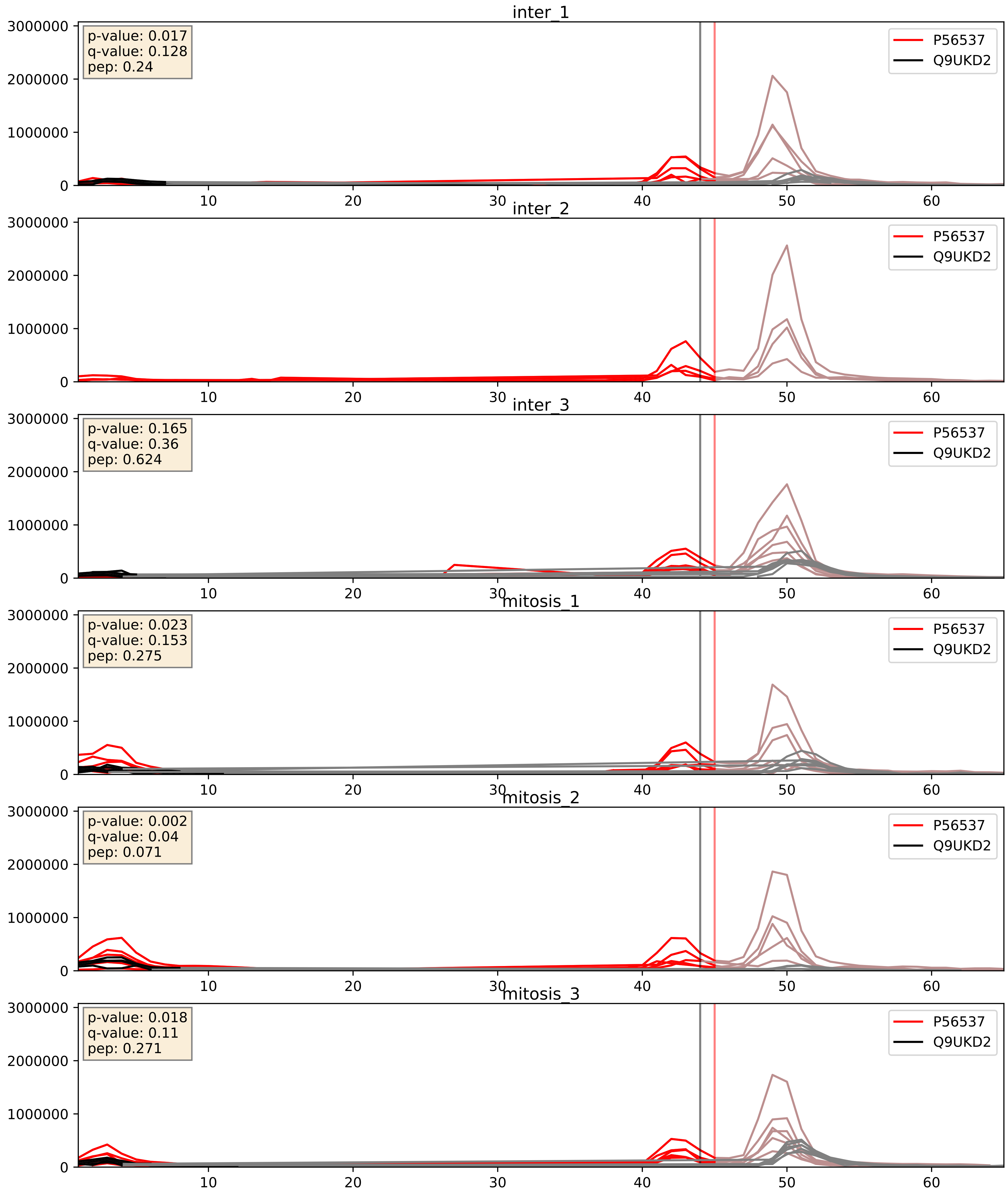


| level | condition_1 | condition_2 | pvalue | pvalue_adjusted |
|---|---|---|---|---|
| interactor_ratio | mitosis | inter | 0.0007143459035063286 | 0.03088283300007158 |
| interactor_abundance | mitosis | inter | 0.025852706349435345 | 0.13331275081395577 |
| complex_abundance | mitosis | inter | 0.060938893075678645 | 0.17730690847308267 |

P61353_Q9UKD2
RL27_HUMAN vs MRT4_HUMAN
p-value: 0.0 q-value: 0.014

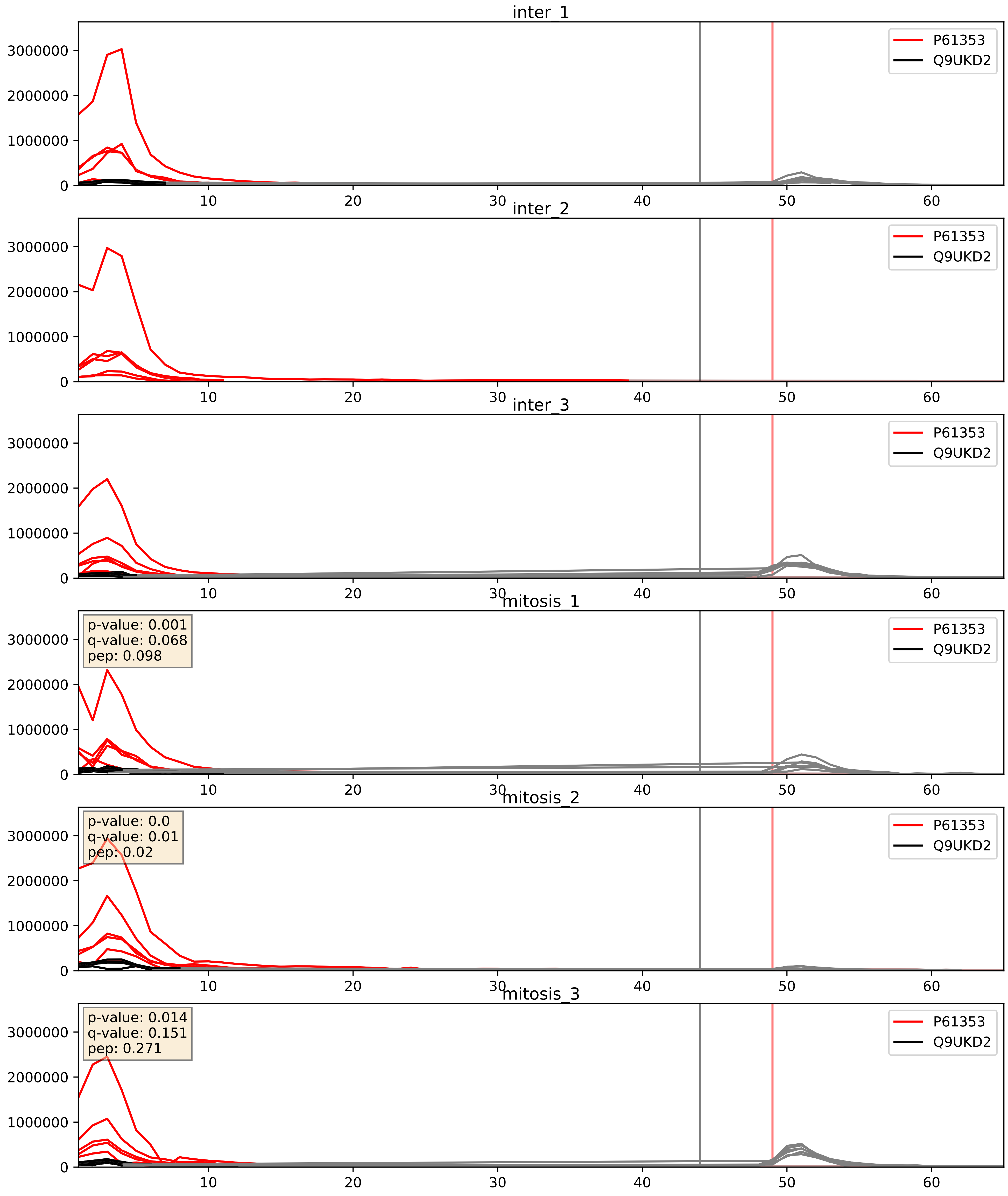


| level | condition_1 | condition_2 | pvalue | pvalue_adjusted |
|---|---|---|---|---|
| complex_abundance | mitosis | inter | 0.3247354238587871 | 0.4731464218265902 |
| interactor_abundance | mitosis | inter | 0.38578530080827556 | 0.5780812462969932 |
| interactor_ratio | mitosis | inter | 0.5443881917068102 | 0.8859730852936224 |

P62910_Q9UKD2
RL32_HUMAN vs MRT4_HUMAN
p-value: 0.0 q-value: 0.008

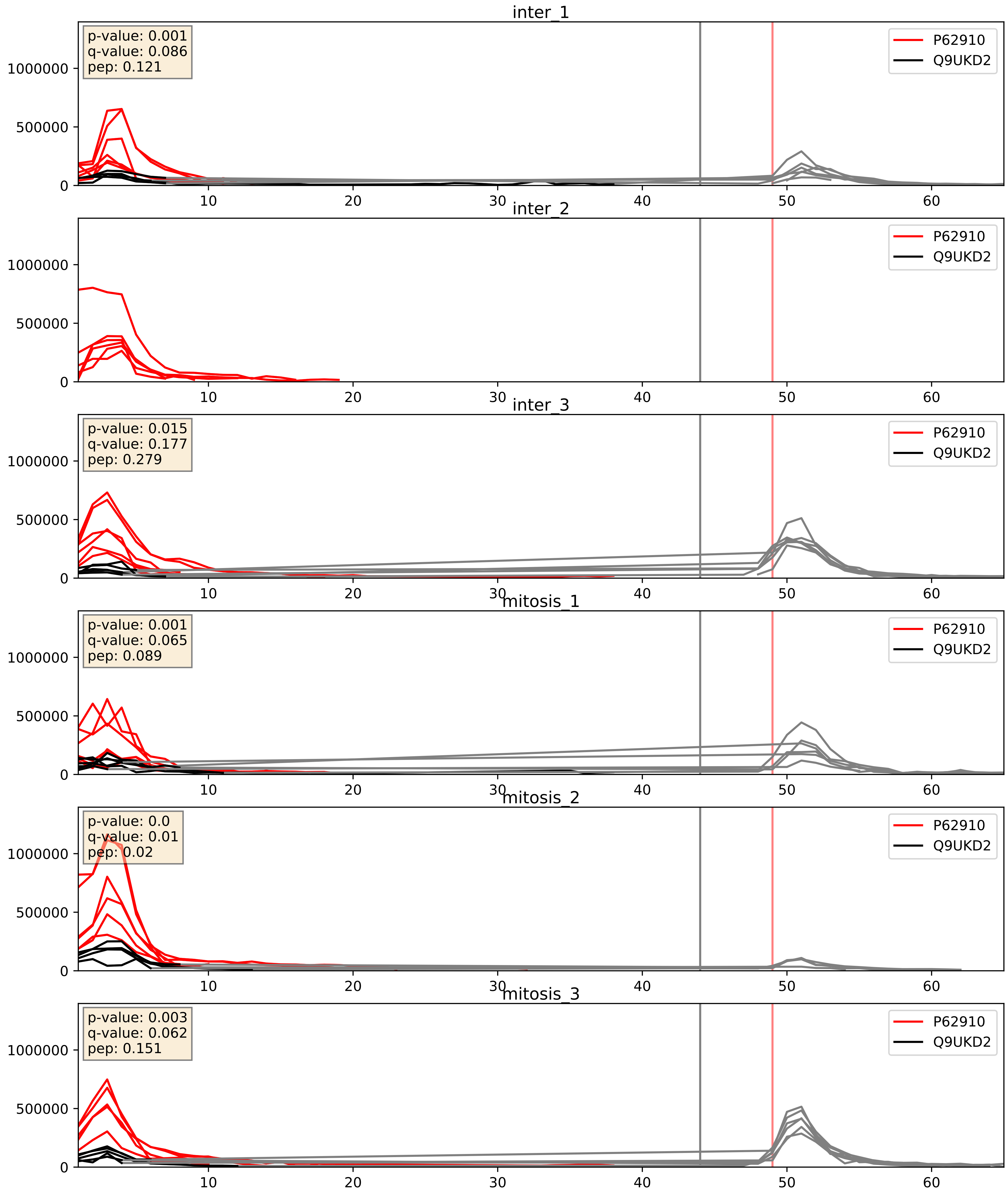


| level | condition_1 | condition_2 | pvalue | pvalue_adjusted |
|---|---|---|---|---|
| complex_abundance | mitosis | inter | 0.24869334211723995 | 0.39890741648141304 |
| interactor_abundance | mitosis | inter | 0.3056049316226747 | 0.5012893005059108 |
| interactor_ratio | mitosis | inter | 0.5178375704684913 | 0.8775869903454201 |

Q02543_Q9UKD2
RL18A_HUMAN vs MRT4_HUMAN
p-value: 0.0 q-value: 0.049

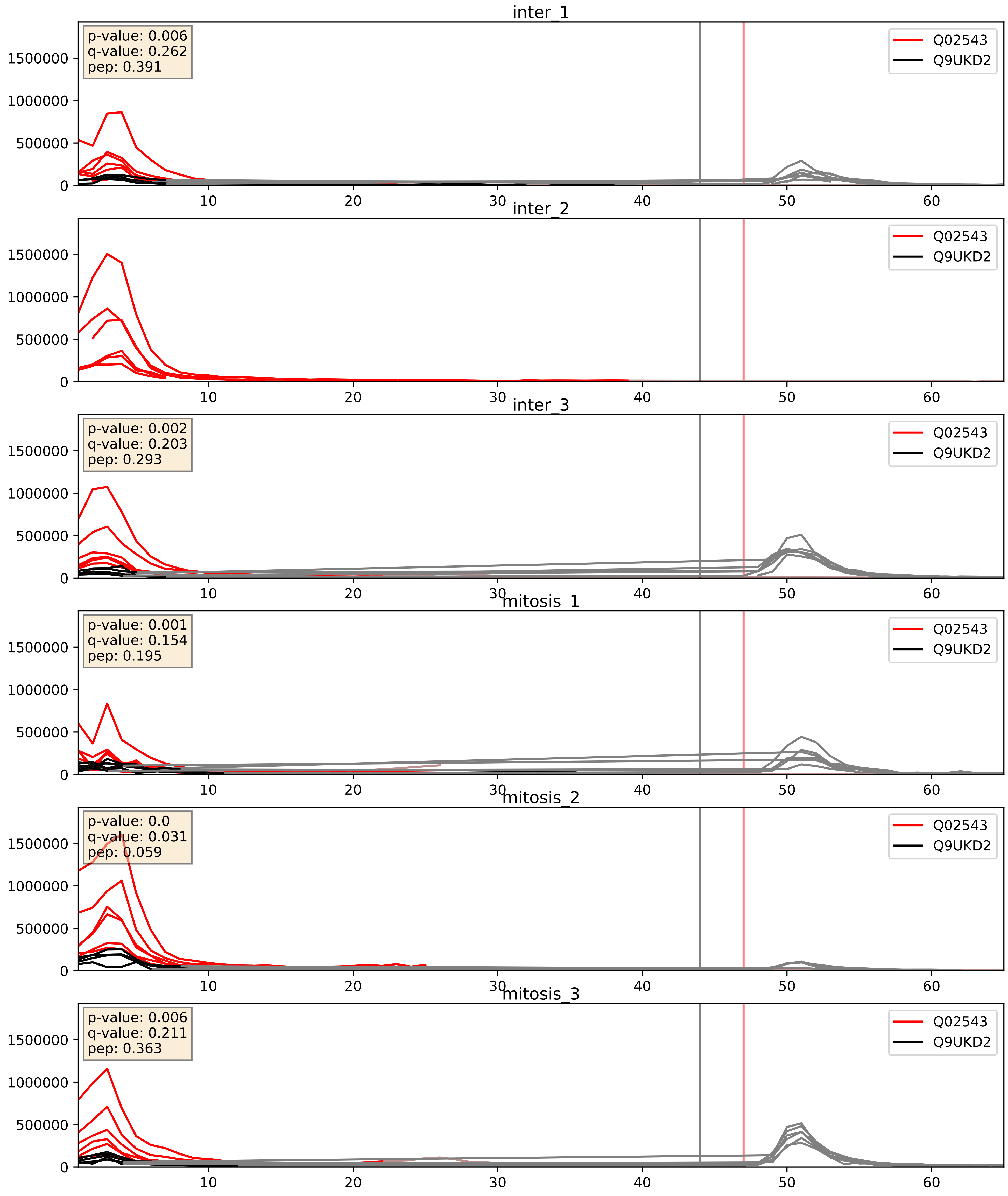


| level | condition_1 | condition_2 | pvalue | pvalue_adjusted |
|---|---|---|---|---|
| complex_abundance | mitosis | inter | 0.2749810168402923 | 0.4241912964773657 |
| interactor_abundance | mitosis | inter | 0.3497635812834094 | 0.548849909401647 |
| interactor_ratio | mitosis | inter | 0.6073602492652684 | 0.9055803577343394 |

Q13823_Q9UKD2
NOG2_HUMAN vs MRT4_HUMAN
p-value: 0.01 q-value: 0.008

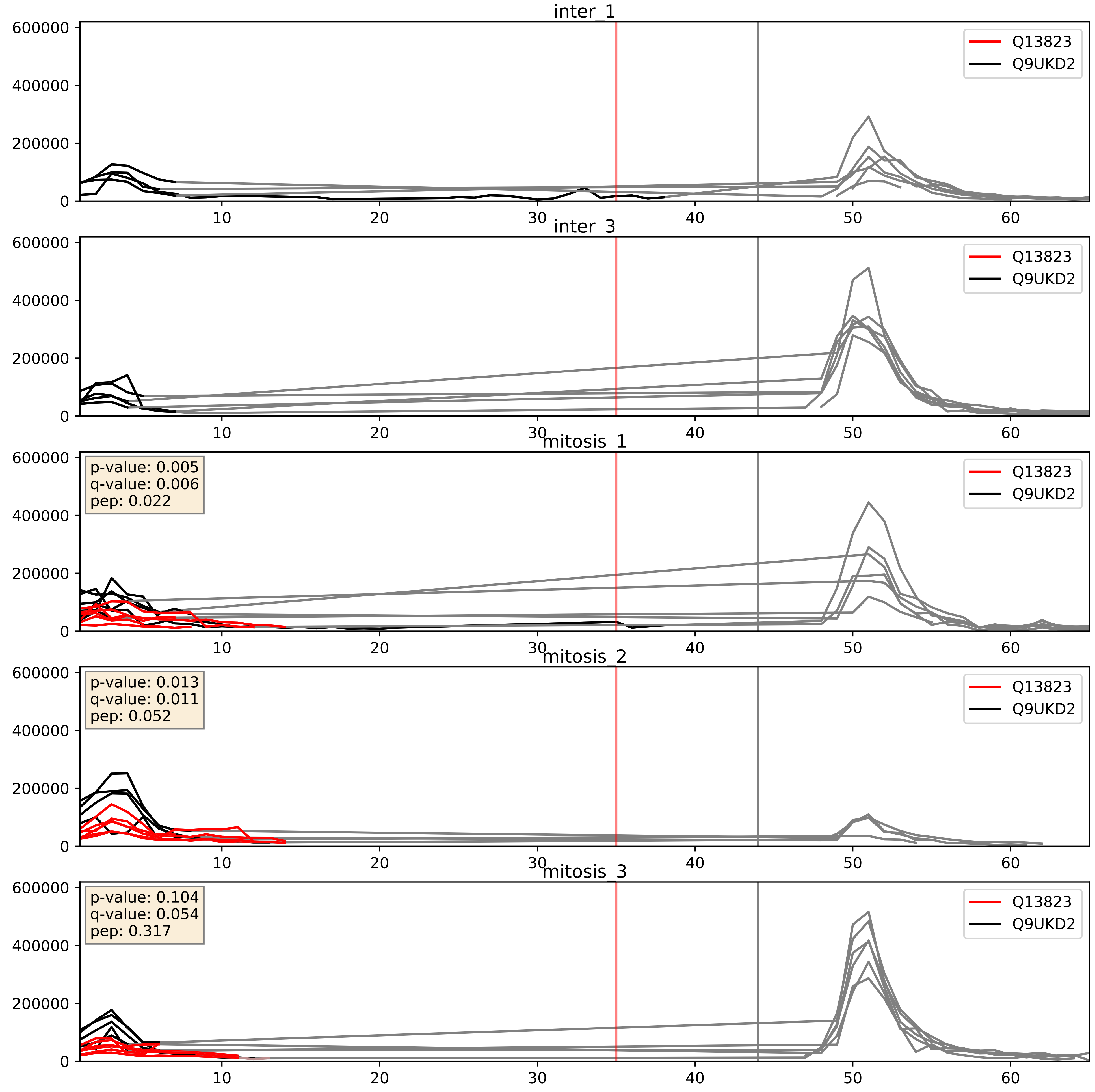


| level | condition_1 | condition_2 | pvalue | pvalue_adjusted |
|---|---|---|---|---|
| complex_abundance | mitosis | inter | 0.0028361034104119874 | 0.038719370323965885 |
| interactor_abundance | mitosis | inter | 0.009614702921018391 | 0.08507272274138435 |
| interactor_ratio | mitosis | inter | 0.012338641662349704 | 0.16051485202084112 |

Q15397_Q9UKD2
PUM3_HUMAN vs MRT4_HUMAN
p-value: 0.003 q-value: 0.041

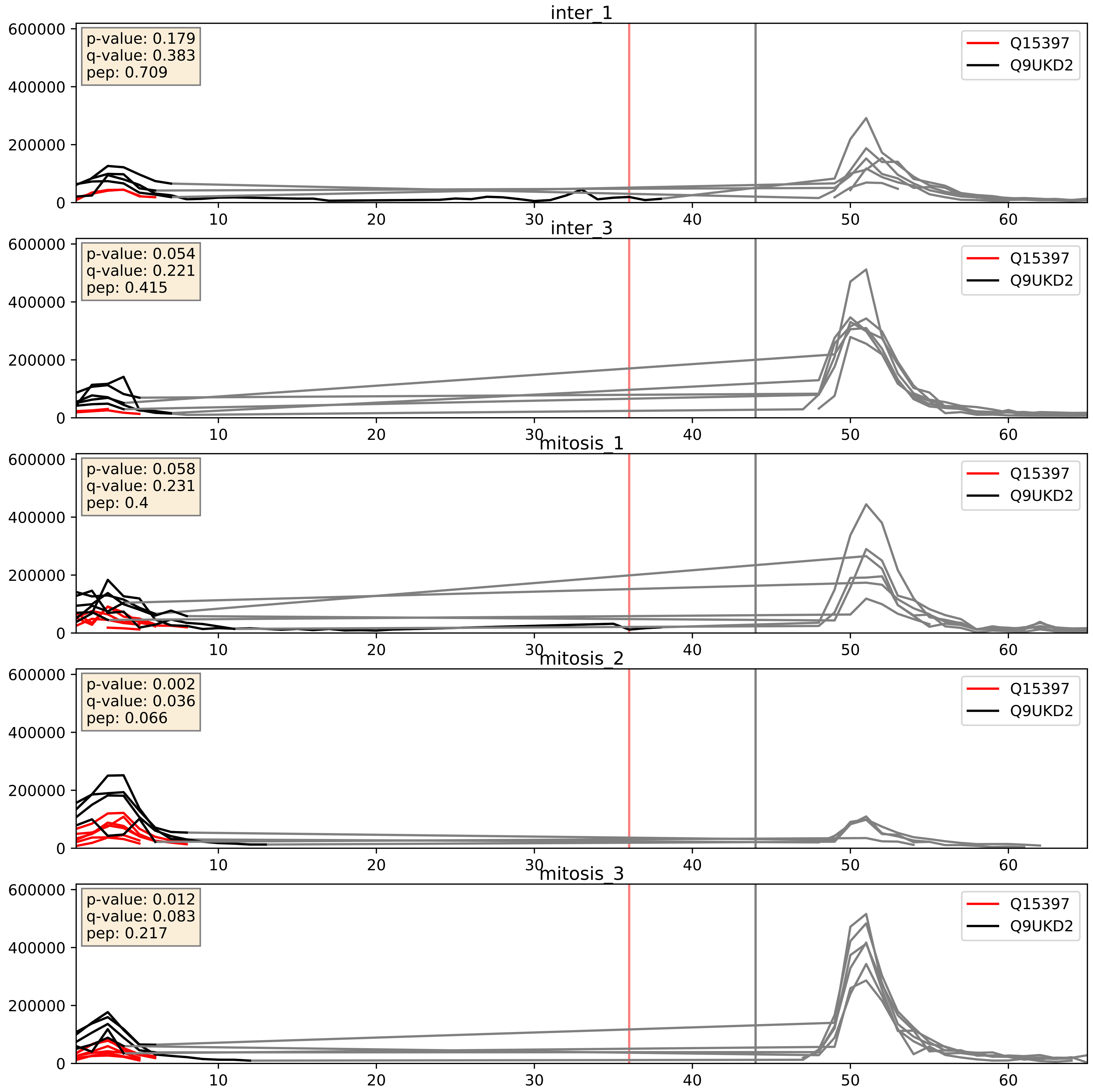


| level | condition_1 | condition_2 | pvalue | pvalue_adjusted |
|---|---|---|---|---|
| complex_abundance | mitosis | inter | 0.19909423997113293 | 0.3518989663747466 |
| interactor_ratio | mitosis | inter | 0.27205097674397627 | 0.7468898440591442 |
| interactor_abundance | mitosis | inter | 0.28137845341557466 | 0.4746944346151595 |

Q8IY81_Q9UKD2
SPB1_HUMAN vs MRT4_HUMAN
p-value: 0.024 q-value: 0.018

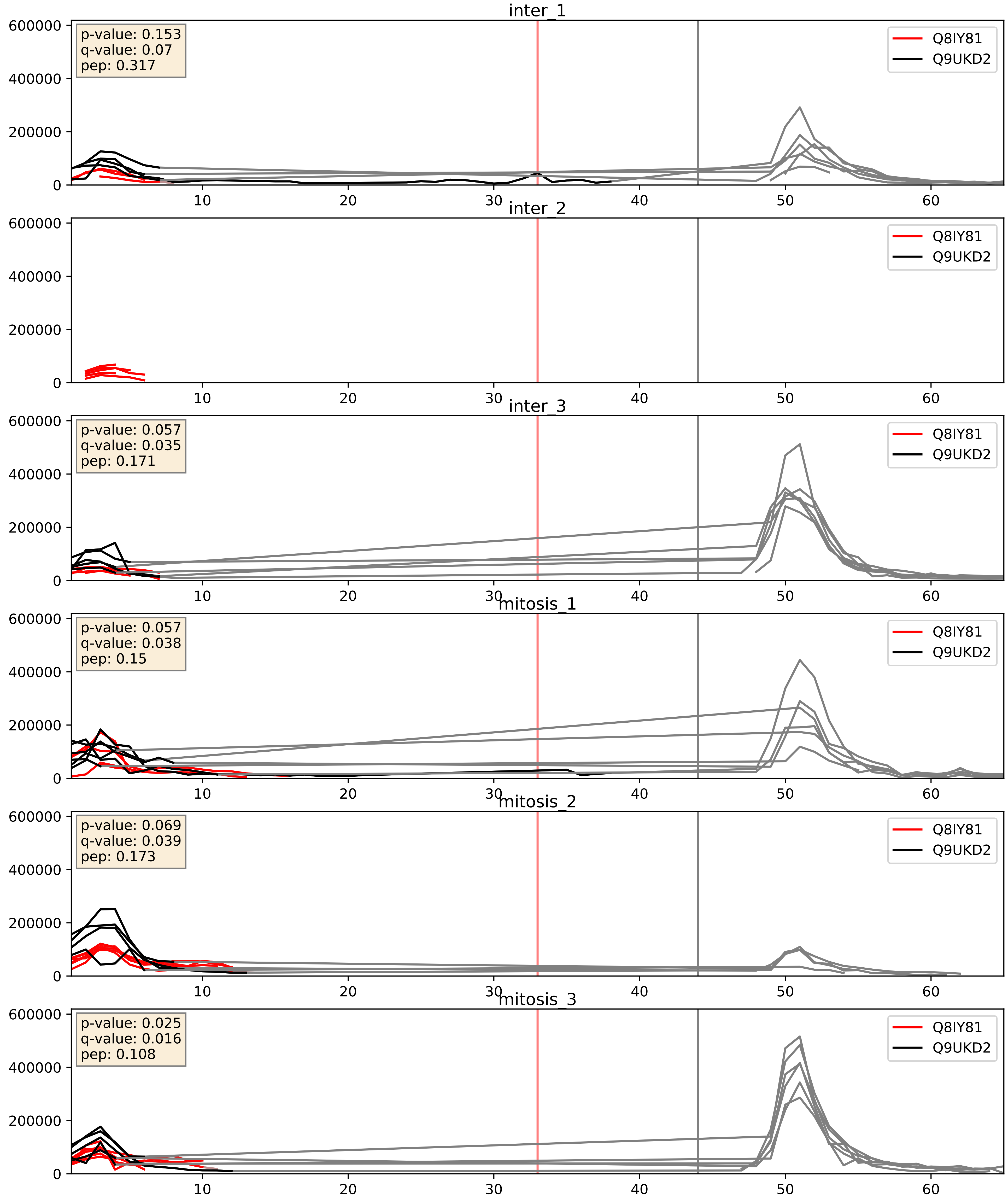


| level | condition_1 | condition_2 | pvalue | pvalue_adjusted |
|---|---|---|---|---|
| interactor_abundance | mitosis | inter | 0.10319084594938516 | 0.2756657235420093 |
| complex_abundance | mitosis | inter | 0.1365404073747575 | 0.28185013765792694 |
| interactor_ratio | mitosis | inter | 0.5766472200424113 | 0.8958439570894811 |

Q8TDN6_Q9UKD2
BRX1_HUMAN vs MRT4_HUMAN
p-value: 0.026 q-value: 0.019

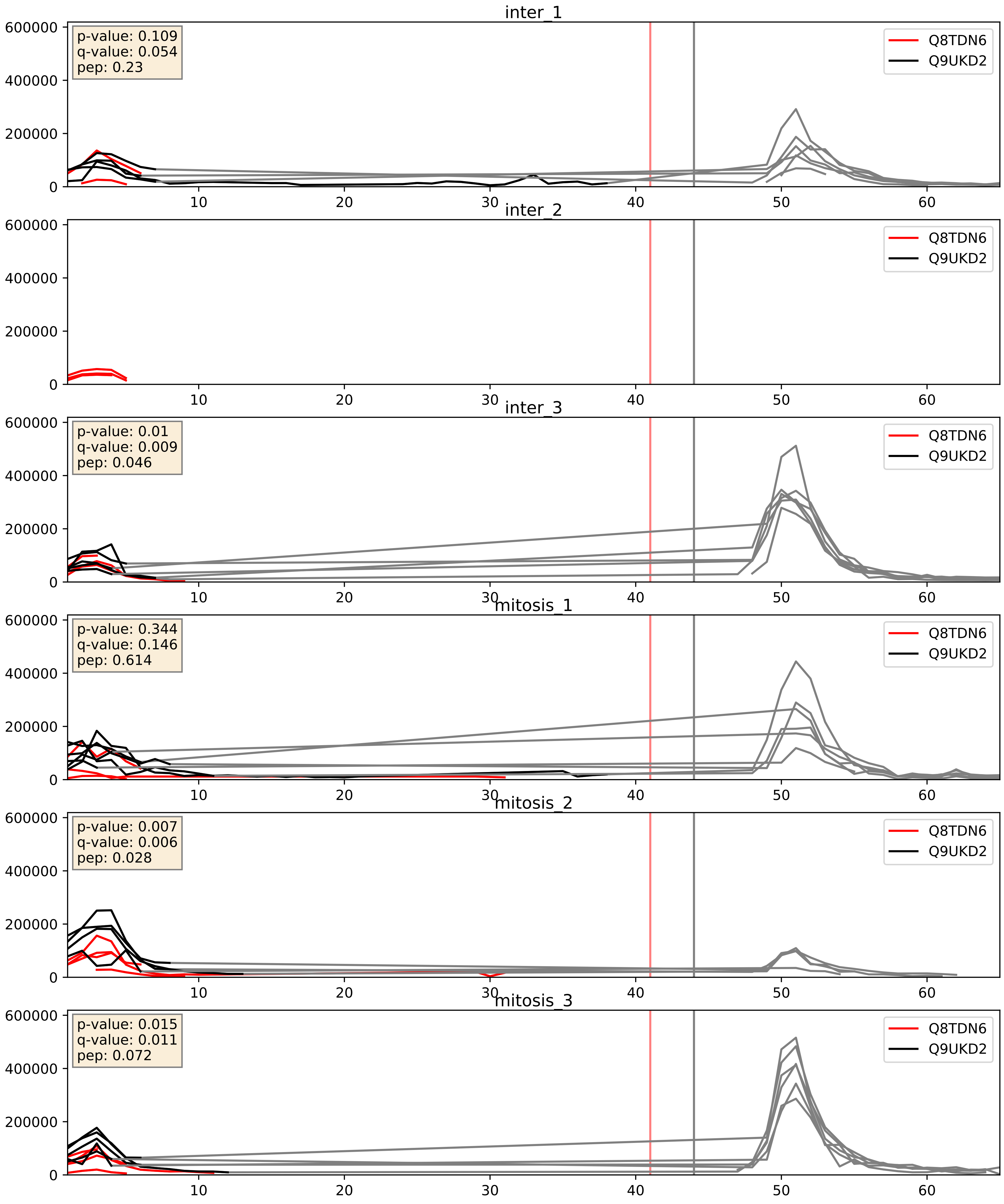


| level | condition_1 | condition_2 | pvalue | pvalue_adjusted |
|---|---|---|---|---|
| interactor_ratio | mitosis | inter | 0.026427216938707084 | 0.25543205403291264 |
| complex_abundance | mitosis | inter | 0.3724276336005471 | 0.5196382304190192 |
| interactor_abundance | mitosis | inter | 0.9044560948065035 | 0.9461832168094921 |

Q9BRT6_Q9UKD2
LLPH_HUMAN vs MRT4_HUMAN
p-value: 0.0 q-value: 0.041

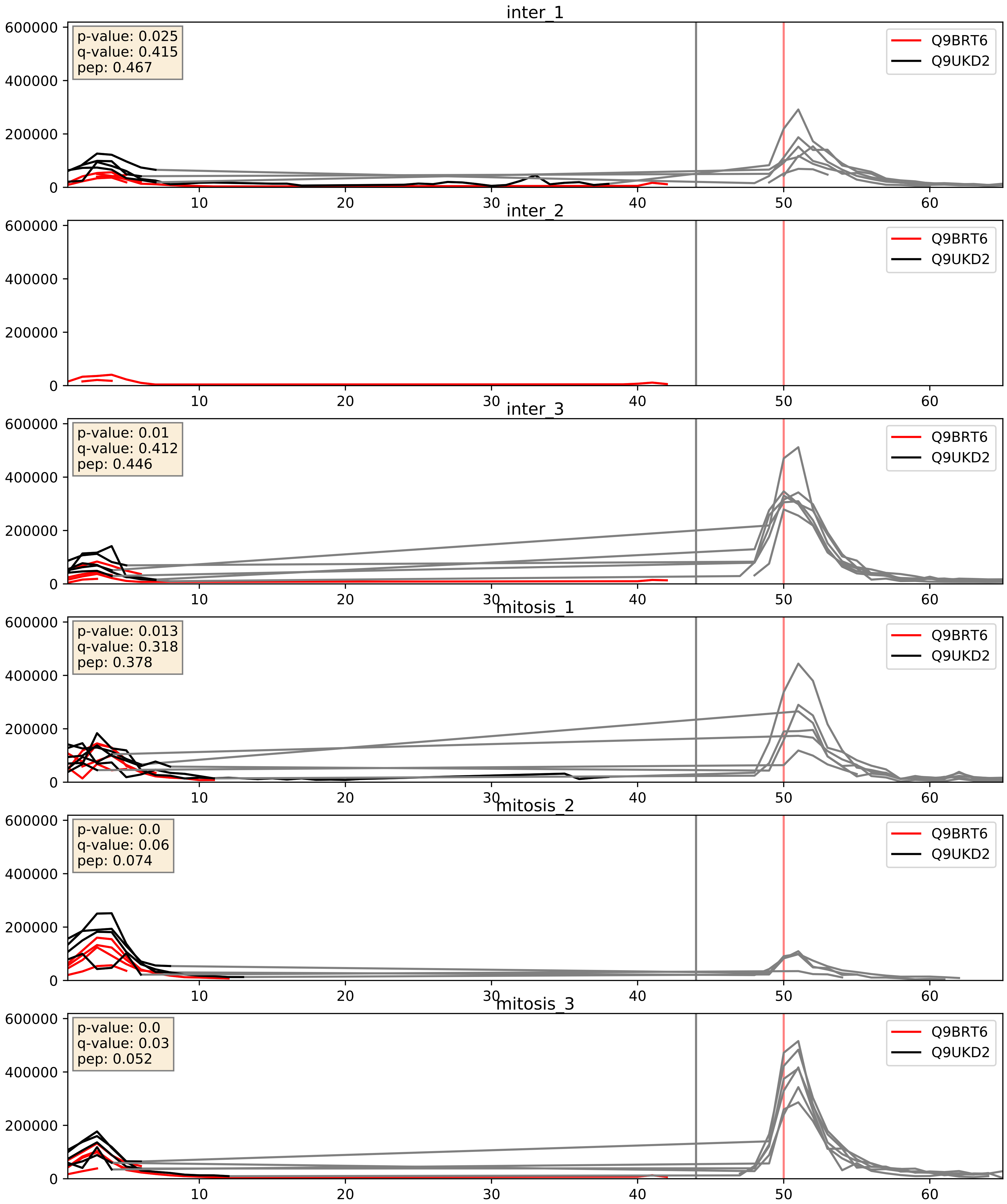


| level | condition_1 | condition_2 | pvalue | pvalue_adjusted |
|---|---|---|---|---|
| interactor_ratio | mitosis | inter | 0.17657394314745203 | 0.6420034343641515 |
| complex_abundance | mitosis | inter | 0.2156689053570195 | 0.3703361744947015 |
| interactor_abundance | mitosis | inter | 0.26832126808793094 | 0.4602233664765775 |

Q9BVP2_Q9UKD2
GNL3_HUMAN vs MRT4_HUMAN
p-value: 0.019 q-value: 0.015

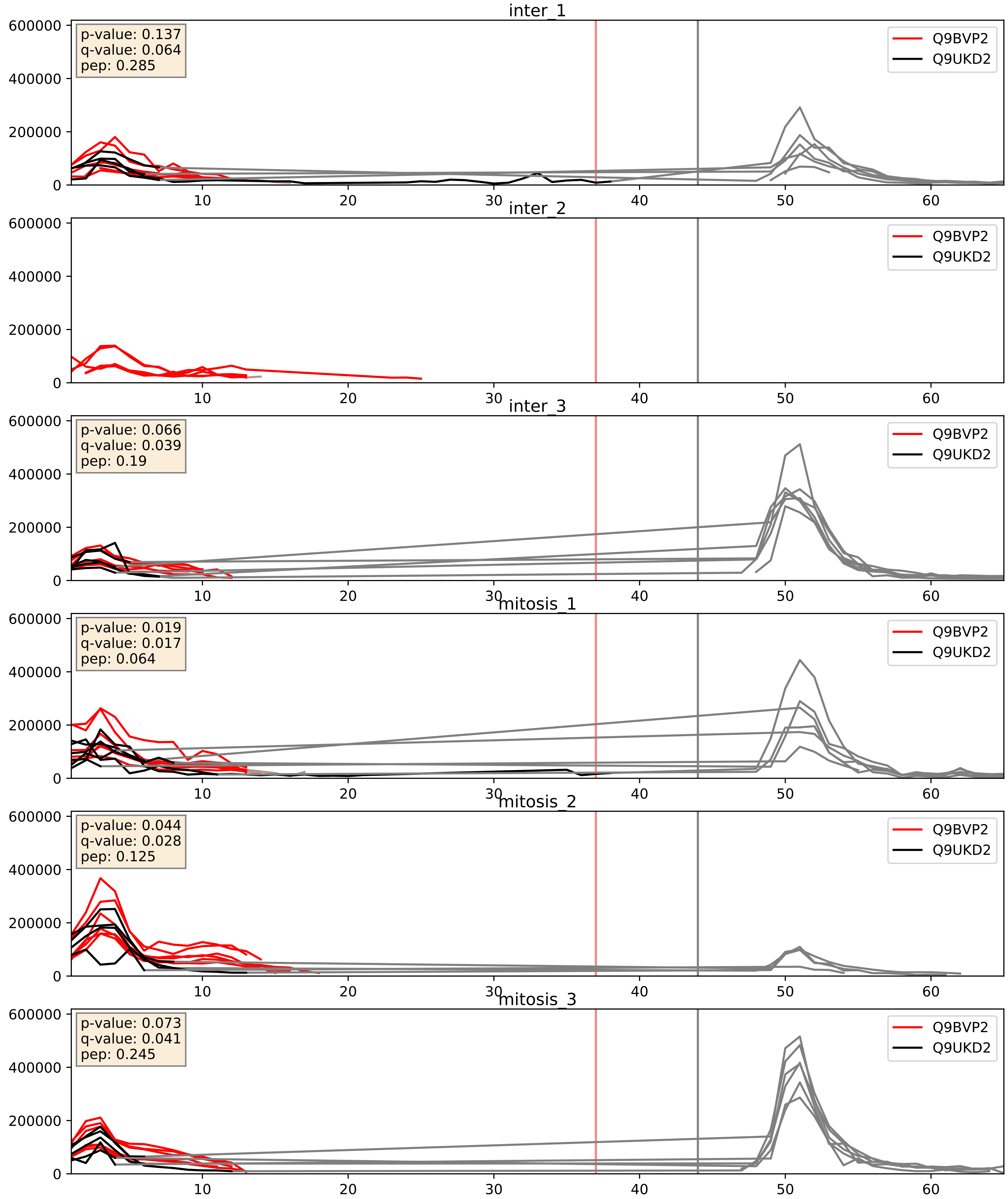


| level | condition_1 | condition_2 | pvalue | pvalue_adjusted |
|---|---|---|---|---|
| interactor_ratio | mitosis | inter | 0.03252851835389341 | 0.28824442764940744 |
| interactor_abundance | mitosis | inter | 0.0987233945687053 | 0.2701737135098137 |
| complex_abundance | mitosis | inter | 0.12841912353114857 | 0.27142412282139056 |

Q9BYG3_Q9UKD2
MK67I_HUMAN vs MRT4_HUMAN
p-value: 0.081 q-value: 0.048

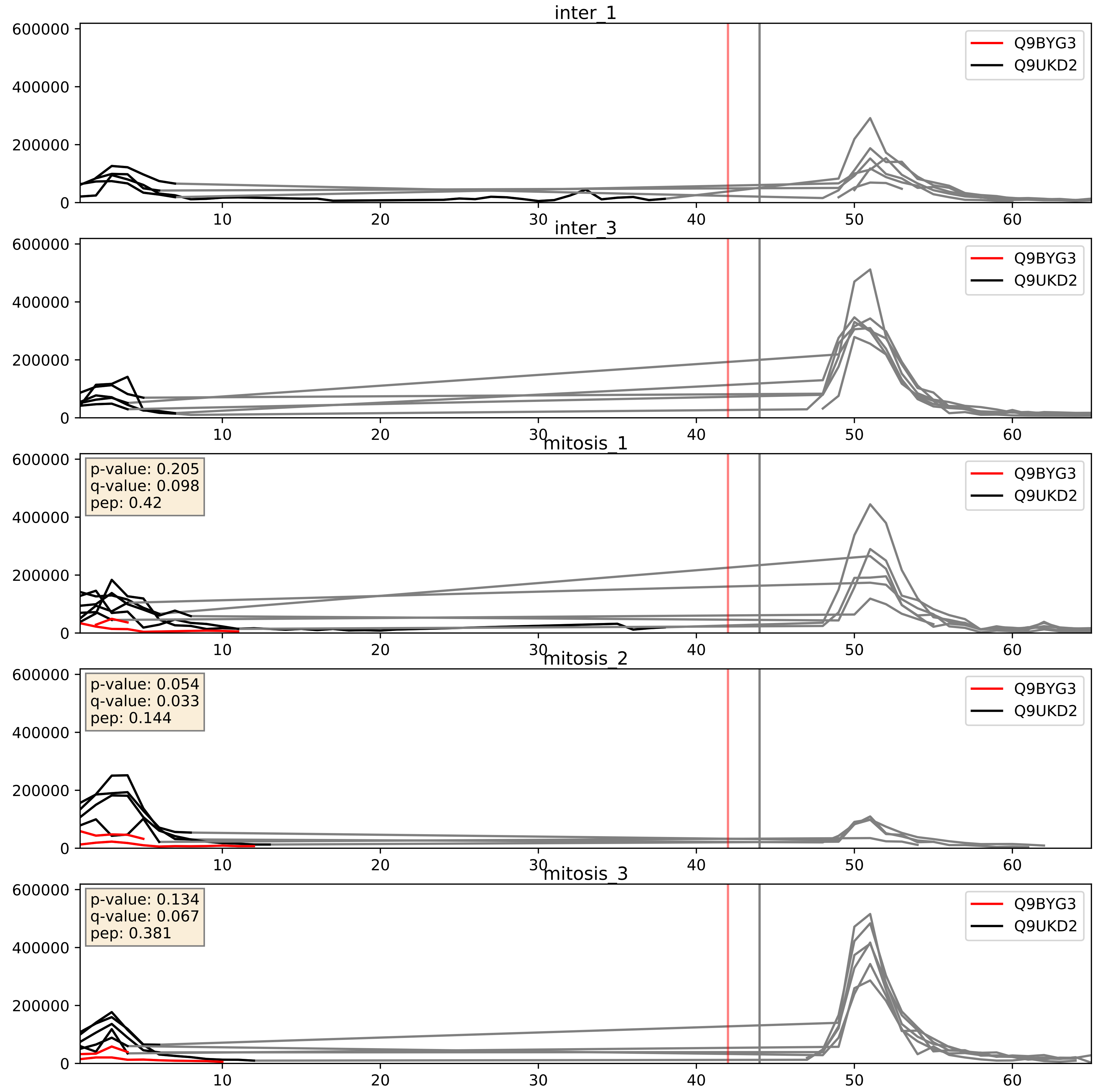


| level | condition_1 | condition_2 | pvalue | pvalue_adjusted |
|---|---|---|---|---|
| interactor_ratio | mitosis | inter | 0.0017848561268288656 | 0.051461731073100085 |
| complex_abundance | mitosis | inter | 0.0034654008381990245 | 0.04302508960916016 |
| interactor_abundance | mitosis | inter | 0.21337749899904426 | 0.40861552381025035 |

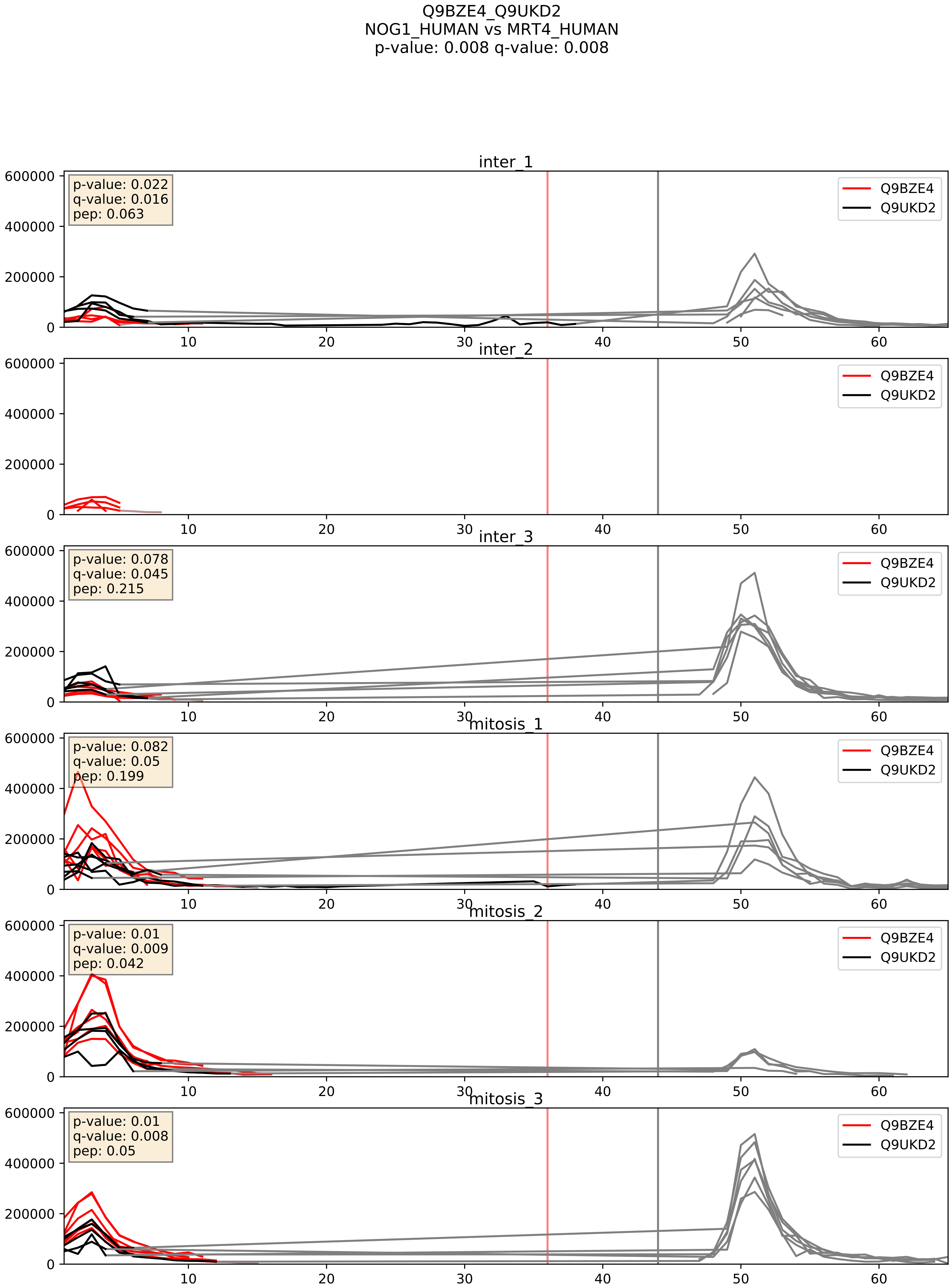


| level | condition_1 | condition_2 | pvalue | pvalue_adjusted |
|---|---|---|---|---|
| interactor_abundance | mitosis | inter | 0.0030352039949890198 | 0.0421728093362545 |
| interactor_ratio | mitosis | inter | 0.0047183833657339245 | 0.09159088101537749 |
| complex_abundance | mitosis | inter | 0.02221883361752925 | 0.10952717692621856 |

Q9NY93_Q9UKD2
DDX56_HUMAN vs MRT4_HUMAN
p-value: 0.018 q-value: 0.014

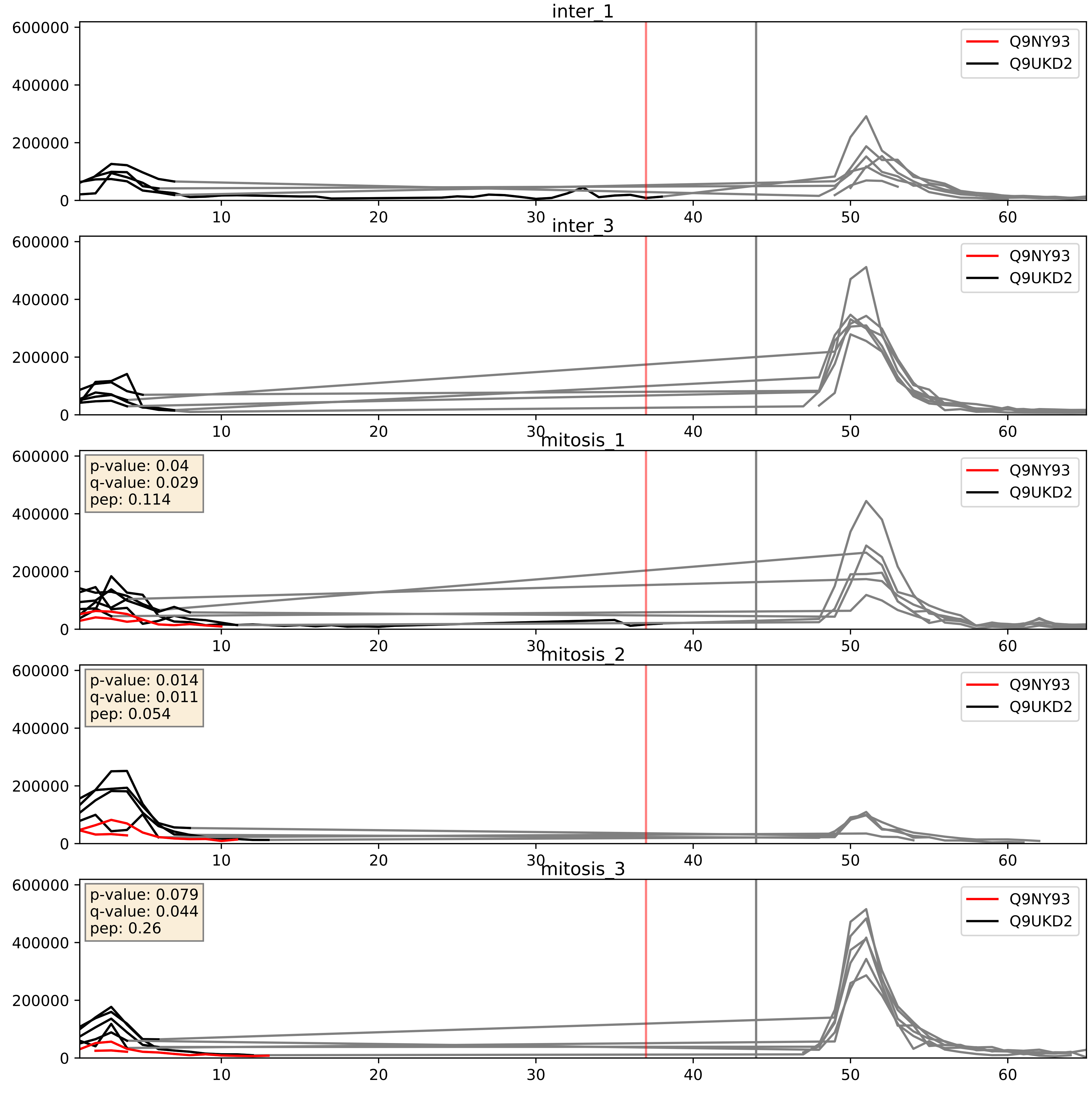


| level | condition_1 | condition_2 | pvalue | pvalue_adjusted |
|---|---|---|---|---|
| interactor_ratio | mitosis | inter | 0.0009603917975522726 | 0.03574327733498893 |
| complex_abundance | mitosis | inter | 0.003800492486110046 | 0.04558257253867179 |
| interactor_abundance | mitosis | inter | 0.10493275671361522 | 0.27679986972306914 |

Q9UKD2_Q9Y221
MRT4_HUMAN vs NIP7_HUMAN
p-value: 0.007 q-value: 0.006

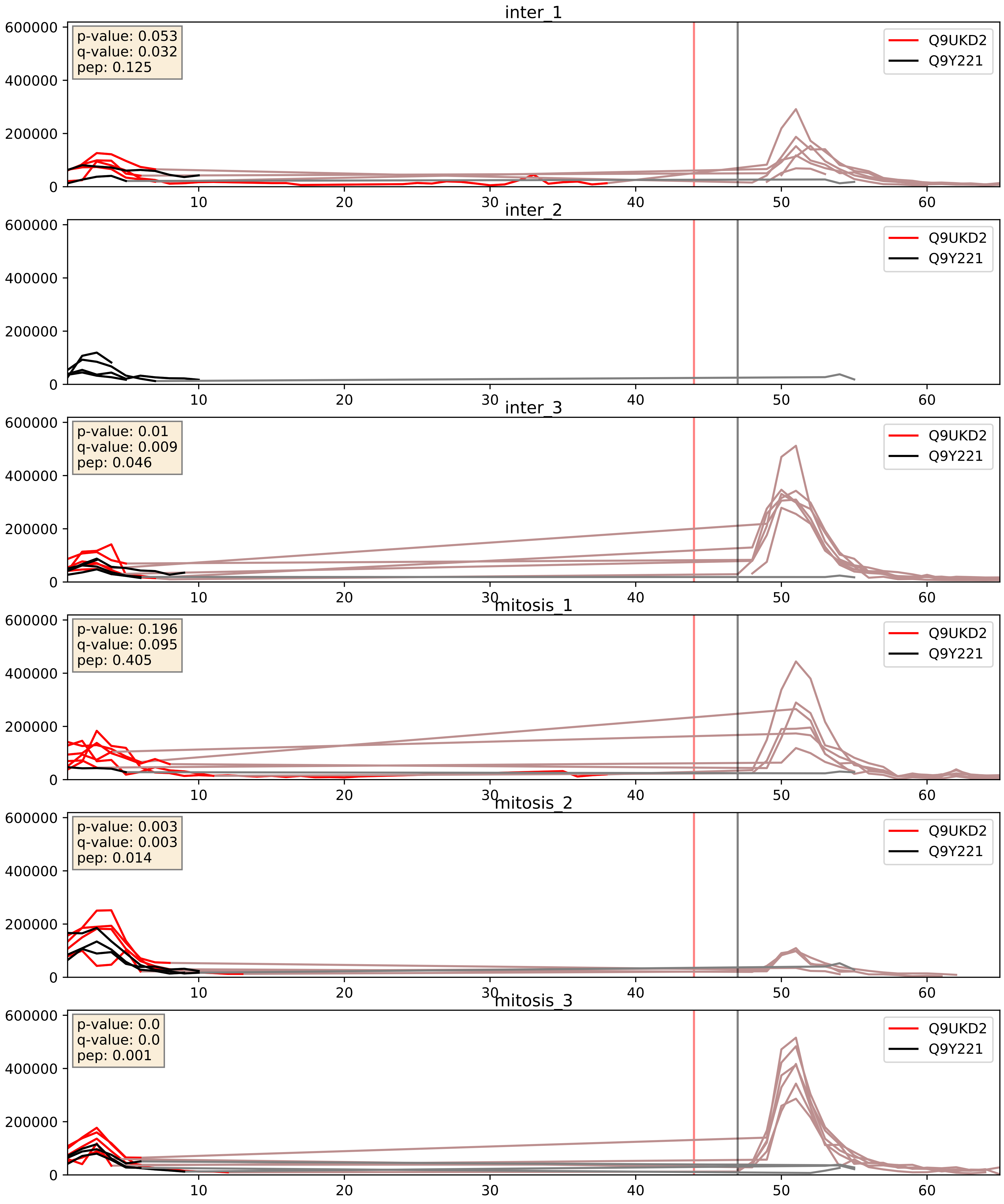


| level | condition_1 | condition_2 | pvalue | pvalue_adjusted |
|---|---|---|---|---|
| interactor_ratio | mitosis | inter | 0.1631762817819746 | 0.6224549786335574 |
| interactor_abundance | mitosis | inter | 0.23967720596818032 | 0.43092562131645107 |
| complex_abundance | mitosis | inter | 0.45234766825486494 | 0.5944163667216406 |

Q9UKD2_Q9Y3C1
MRT4_HUMAN vs NOP16_HUMAN
p-value: 0.002 q-value: 0.036

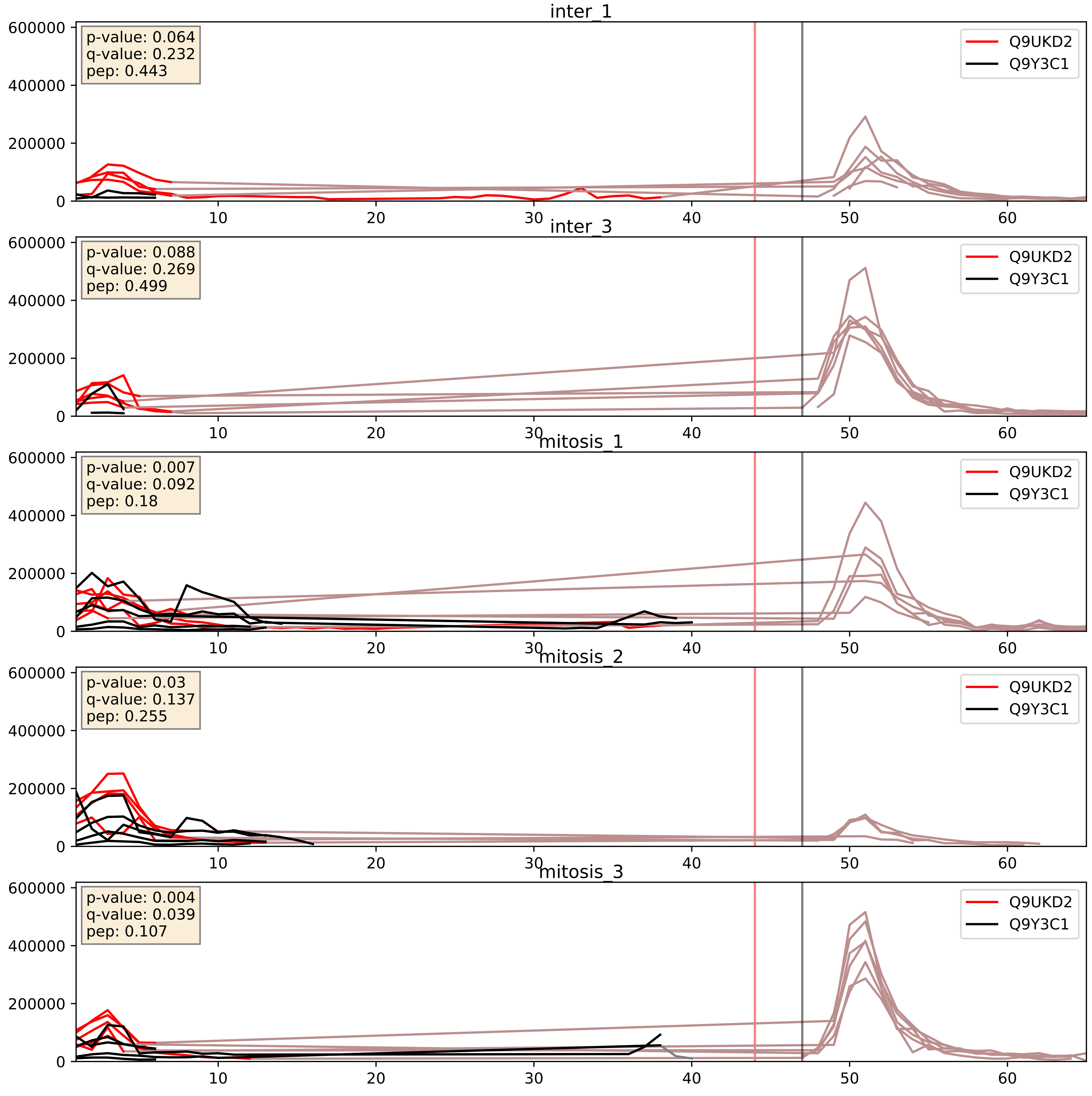


| level | condition_1 | condition_2 | pvalue | pvalue_adjusted |
|---|---|---|---|---|
| complex_abundance | mitosis | inter | 0.042034973880429934 | 0.14726323635770058 |
| interactor_abundance | mitosis | inter | 0.14892390476455458 | 0.3352060543740697 |
| interactor_ratio | mitosis | inter | 0.3243789503904871 | 0.7829618581024819 |

Q9UKD2_Q9Y3U8
MRT4_HUMAN vs RL36_HUMAN
p-value: 0.0 q-value: 0.023

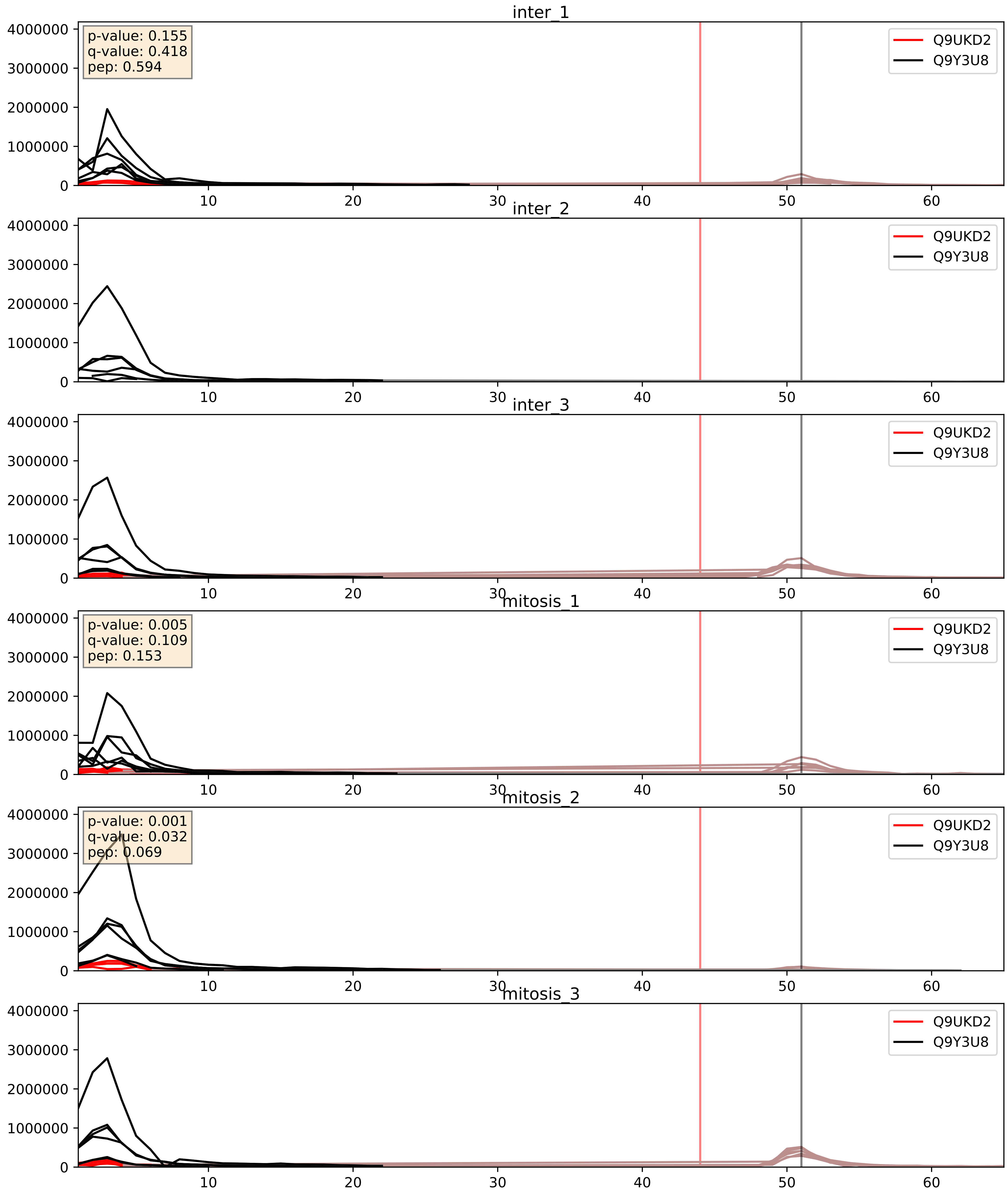


| level | condition_1 | condition_2 | pvalue | pvalue_adjusted |
|---|---|---|---|---|
| interactor_abundance | mitosis | inter | 0.1951908537681602 | 0.3882076736204408 |
| complex_abundance | mitosis | inter | 0.2862553181621983 | 0.434779465129125 |
| interactor_ratio | mitosis | inter | 0.4841284538060675 | 0.8678583660809296 |
