## Supplementary Data 2 for "SECAT: Quantifying differential protein-protein interaction states by network-centric analysis": hela_string_000060_P38159.pdf

P38159_P38159

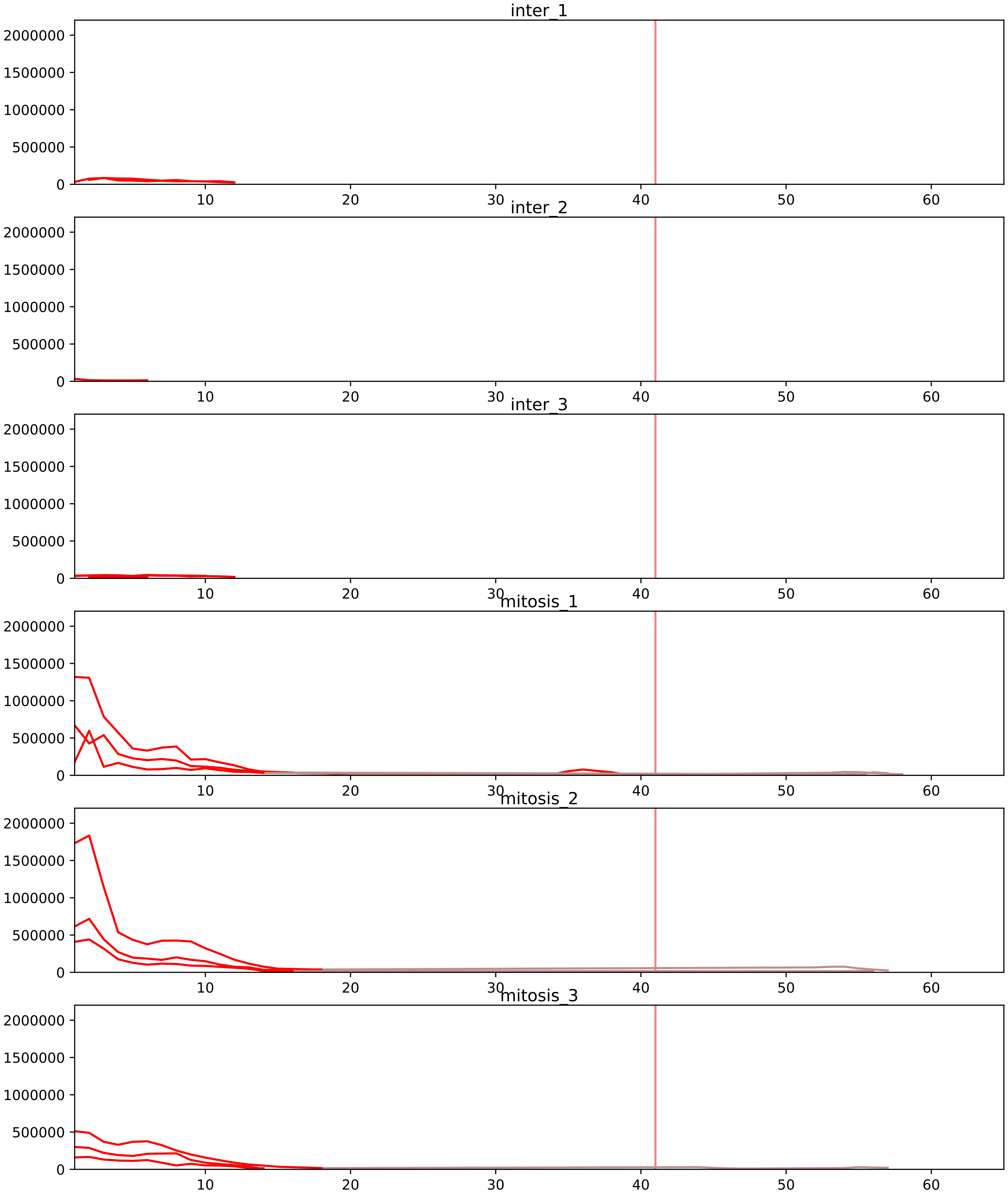


| level | condition_1 | condition_2 | pvalue | pvalue_adjusted |
|---|---|---|---|---|
| interactor_ratio | mitosis | inter | 5.0275180233444e-07 | 1.622918098763806e-05 |
| complex_abundance | mitosis | inter | 2.0145910149299308e-06 | 0.00014827389869884292 |
| interactor_abundance | mitosis | inter | 4.439460271806345e-06 | 0.000300834717183481 |
| monomer_abundance | mitosis | inter | 0.053487481824493845 | 0.34111396383483944 |
| total_abundance | mitosis | inter | 0.12747123038640995 | 0.4419285945231154 |
| assembled_abundance | mitosis | inter | 0.1440999257091801 | 0.4490802322359216 |

P07910_P38159
HNRPC_HUMAN vs RBMX_HUMAN
p-value: 0.022 q-value: 0.017

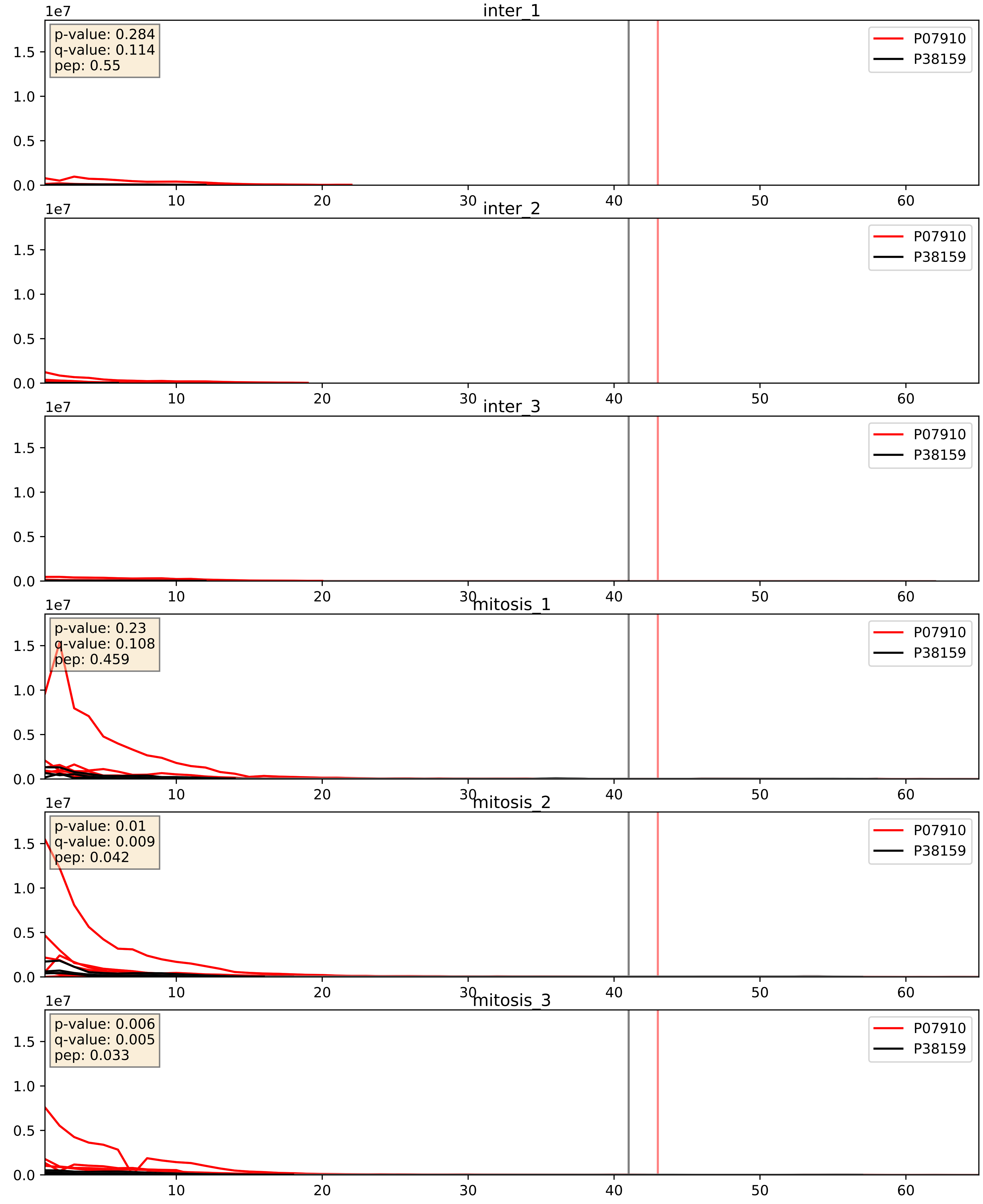


| level | condition_1 | condition_2 | pvalue | pvalue_adjusted |
|---|---|---|---|---|
| interactor_abundance | mitosis | inter | 2.354035126084445e-06 | 0.0033734327106189023 |
| complex_abundance | mitosis | inter | 4.105555000846496e-05 | 0.005691857151652958 |
| interactor_ratio | mitosis | inter | 0.018442867517554088 | 0.20476127879411543 |

P09651_P38159
ROA1_HUMAN vs RBMX_HUMAN
p-value: 0.004 q-value: 0.004

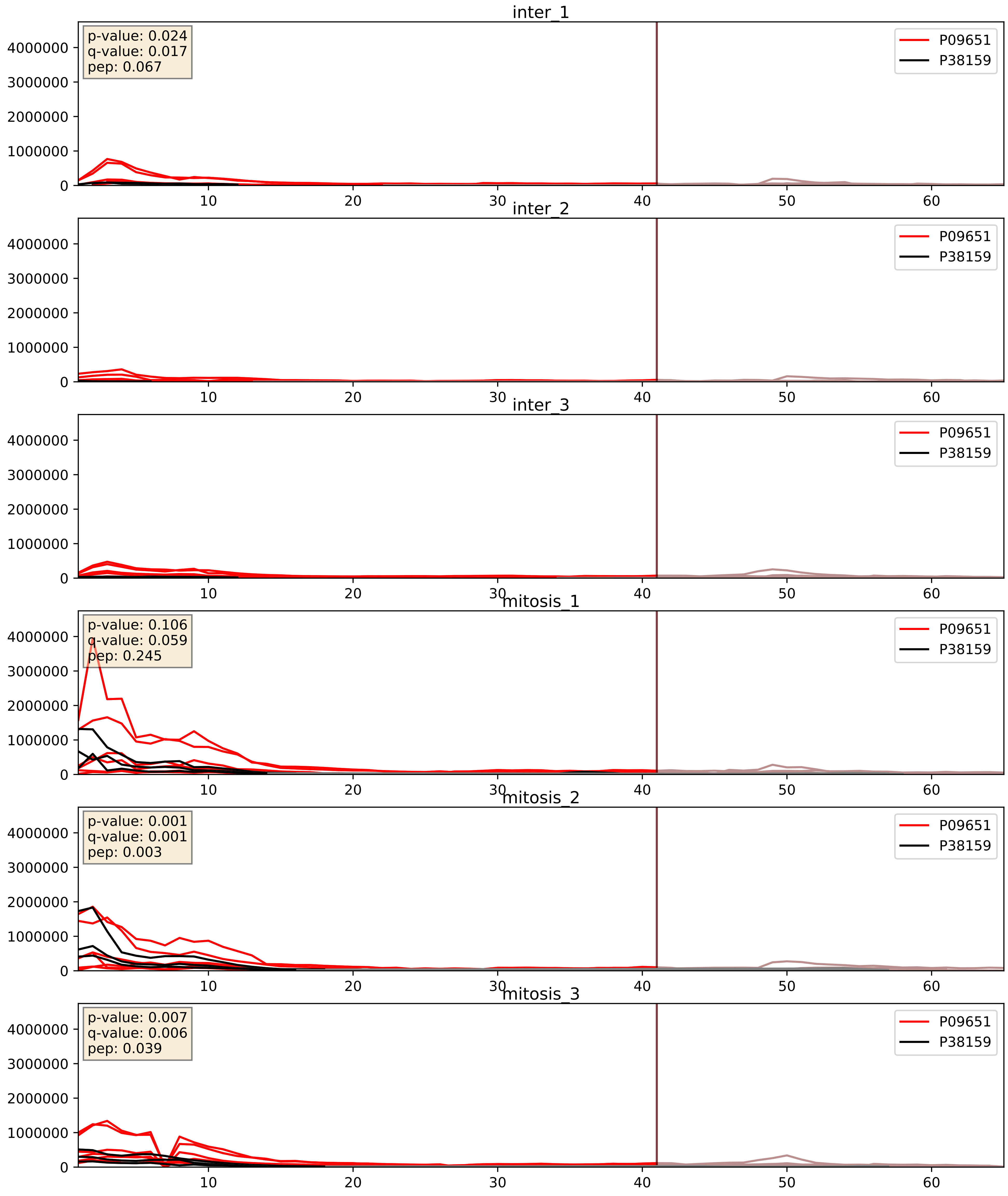


| level | condition_1 | condition_2 | pvalue | pvalue_adjusted |
|---|---|---|---|---|
| complex_abundance | mitosis | inter | 0.003021295973201694 | 0.04033974863104599 |
| interactor_abundance | mitosis | inter | 0.013856881209314718 | 0.10436859054266079 |
| interactor_ratio | mitosis | inter | 0.2375025223286475 | 0.7164022336563056 |

P14866_P38159
HNRPL_HUMAN vs RBMX_HUMAN
p-value: 0.008 q-value: 0.008

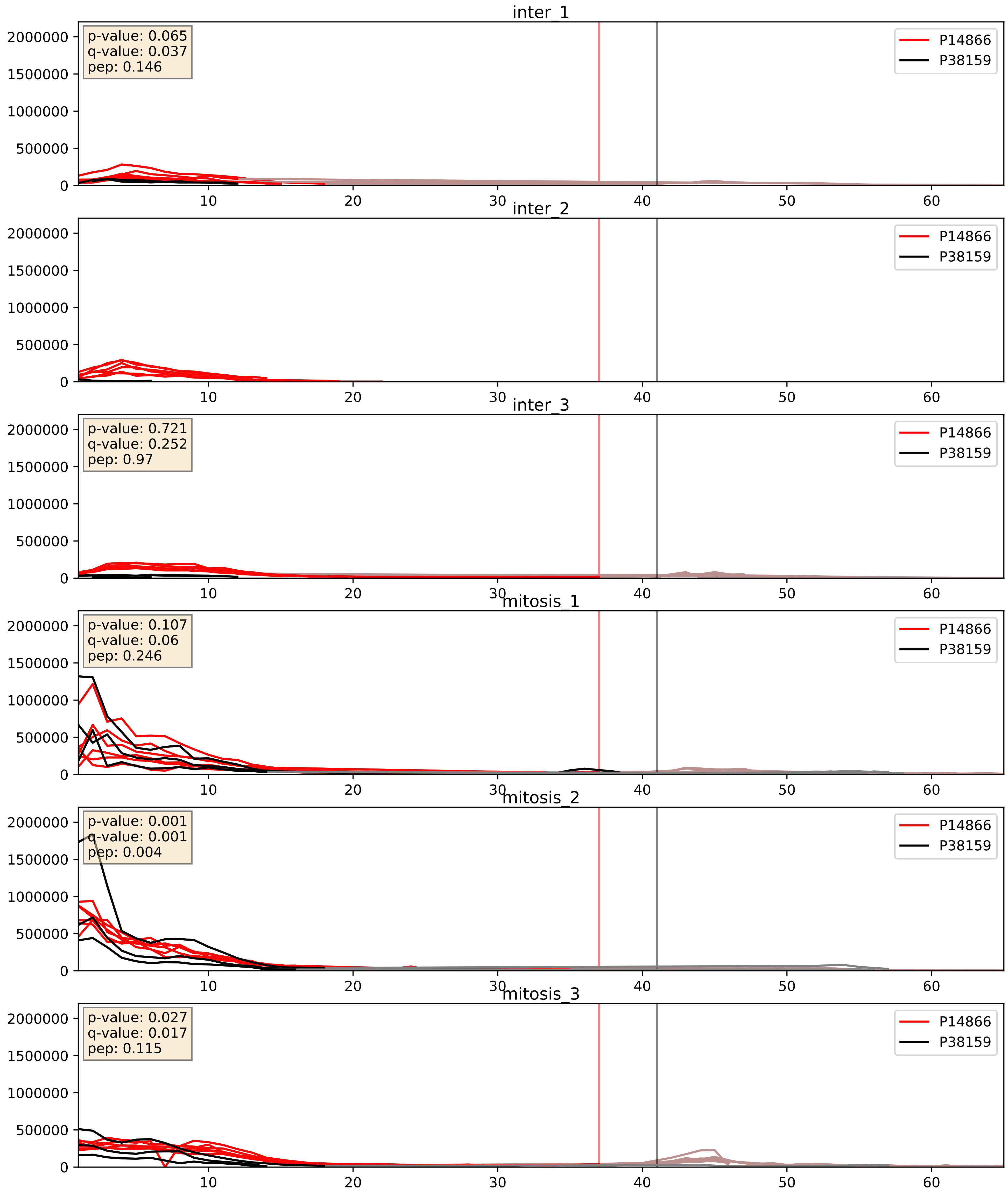


| level | condition_1 | condition_2 | pvalue | pvalue_adjusted |
|---|---|---|---|---|
| complex_abundance | mitosis | inter | 0.00042915958677650936 | 0.013091389064164169 |
| interactor_abundance | mitosis | inter | 0.002222136362133671 | 0.03466841143289118 |
| interactor_ratio | mitosis | inter | 0.04602651635329546 | 0.35622692584467375 |

P17844_P38159
DDX5_HUMAN vs RBMX_HUMAN
p-value: 0.0 q-value: 0.003

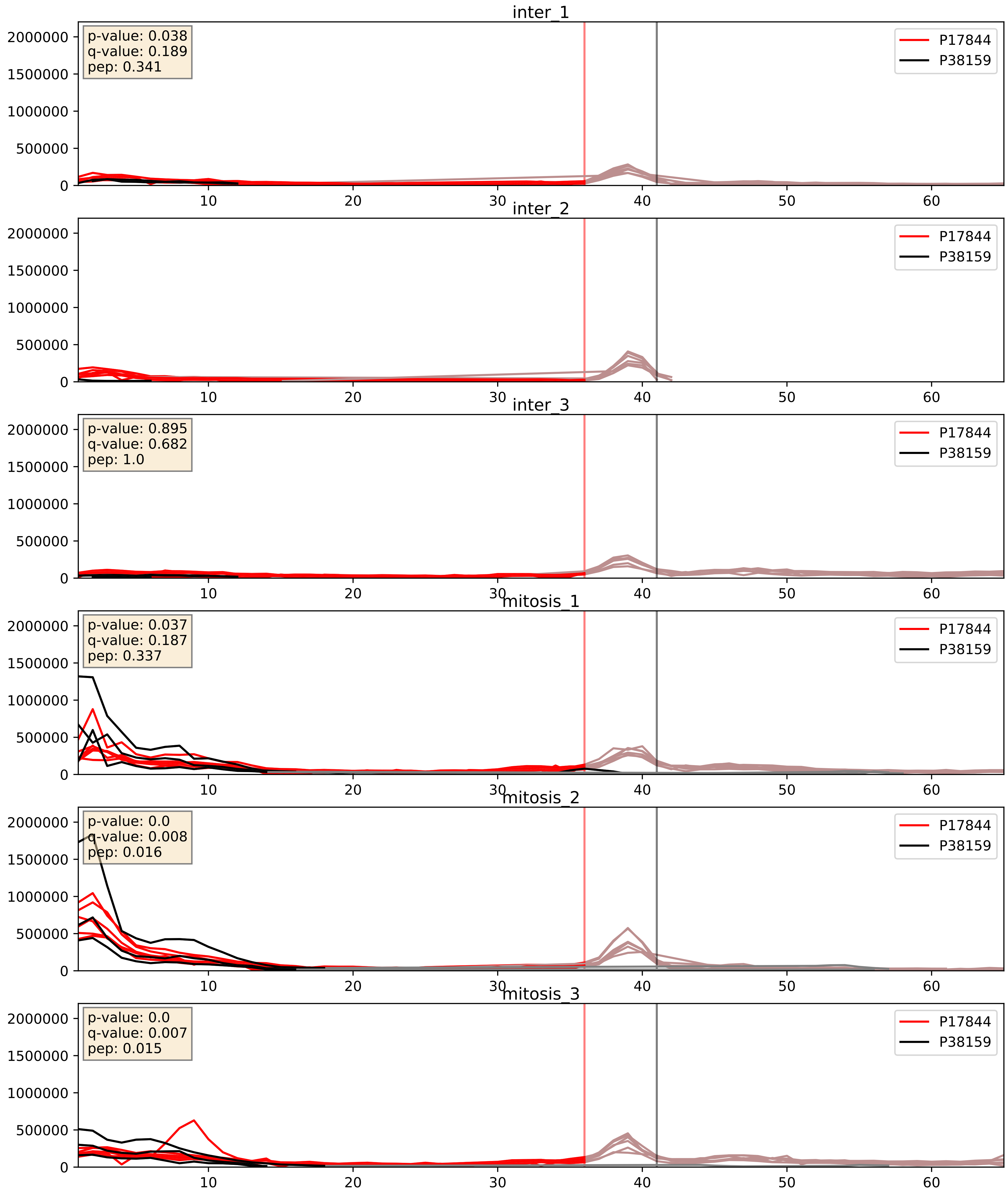


| level | condition_1 | condition_2 | pvalue | pvalue_adjusted |
|---|---|---|---|---|
| complex_abundance | mitosis | inter | 0.0010528687451826645 | 0.021612845224852777 |
| interactor_abundance | mitosis | inter | 0.0016025115124044603 | 0.029124200734994013 |
| interactor_ratio | mitosis | inter | 0.005020220566009392 | 0.09528400896904744 |

P22626_P38159
ROA2_HUMAN vs RBMX_HUMAN
p-value: 0.011 q-value: 0.009

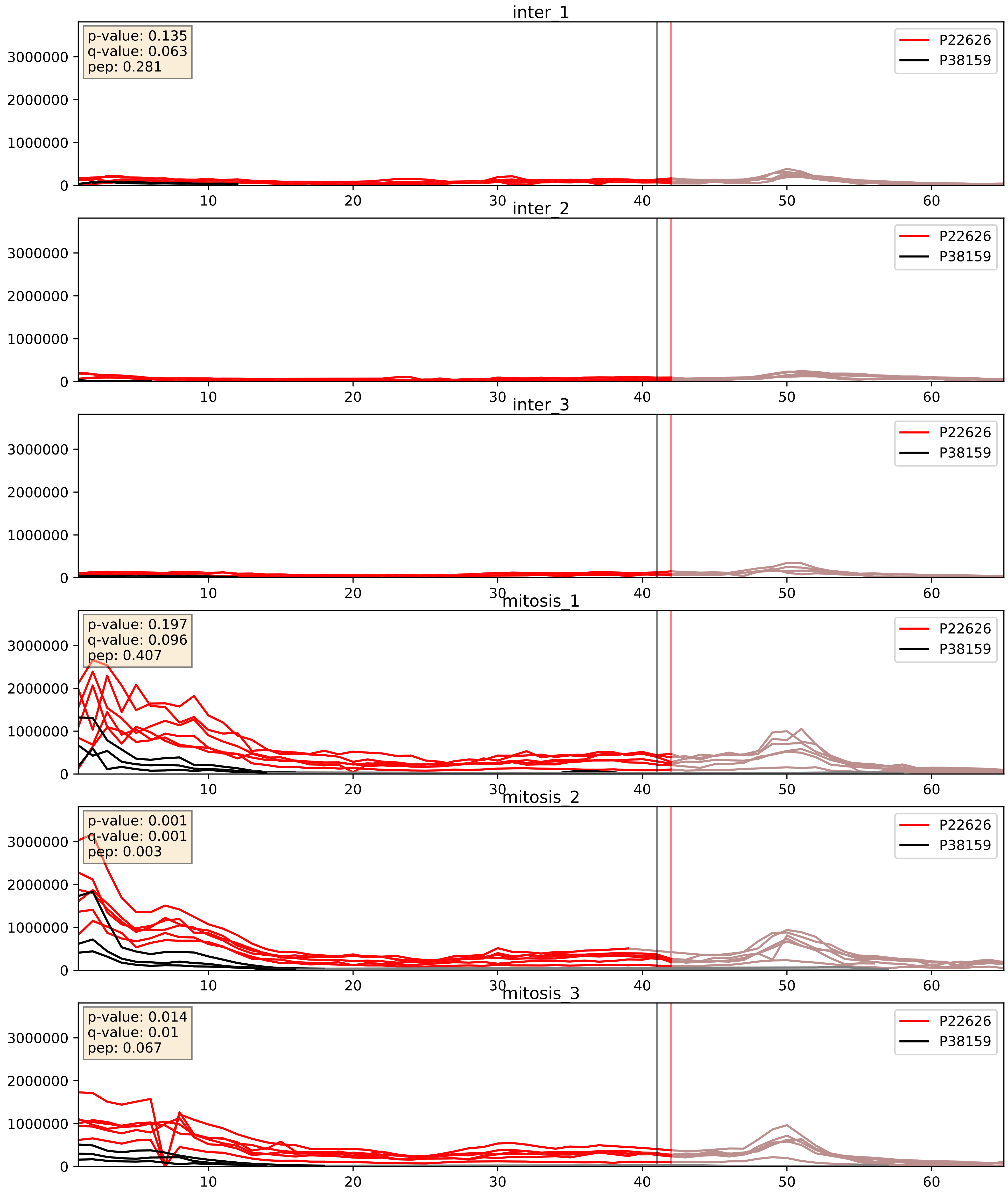


| level | condition_1 | condition_2 | pvalue | pvalue_adjusted |
|---|---|---|---|---|
| complex_abundance | mitosis | inter | 0.0005850625769621449 | 0.015650423933737376 |
| interactor_abundance | mitosis | inter | 0.0012988409915939223 | 0.02663012907315922 |
| interactor_ratio | mitosis | inter | 0.05283895661774645 | 0.3800852677713526 |

P31943_P38159
HNRH1_HUMAN vs RBMX_HUMAN
p-value: 0.004 q-value: 0.004

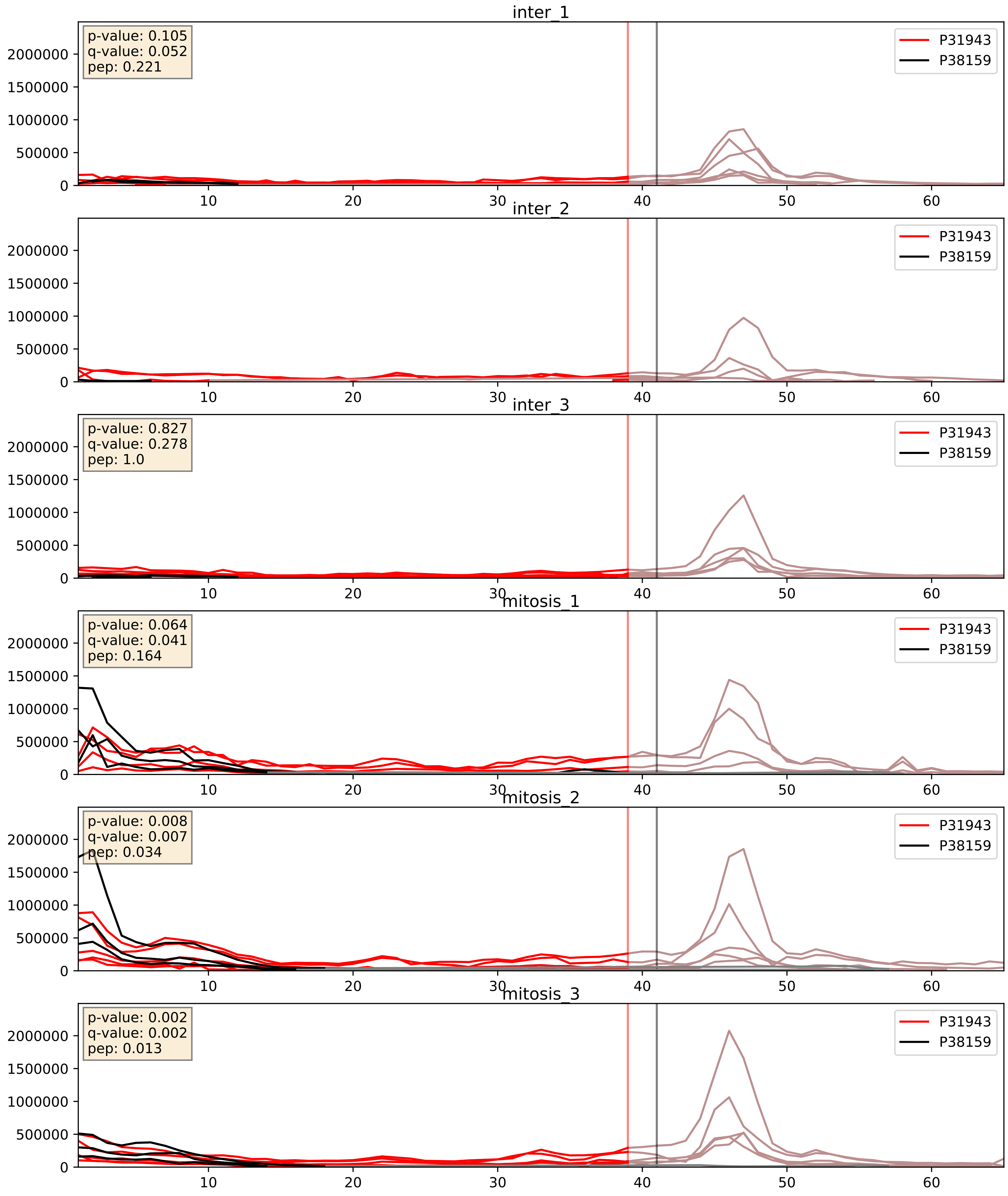


| level | condition_1 | condition_2 | pvalue | pvalue_adjusted |
|---|---|---|---|---|
| complex_abundance | mitosis | inter | 0.002318702437818041 | 0.03512936790747333 |
| interactor_ratio | mitosis | inter | 0.006225715441810617 | 0.10853793112403032 |
| interactor_abundance | mitosis | inter | 0.0062904982907834375 | 0.06768908280214485 |

P38159_P51991
RBMX_HUMAN vs ROA3_HUMAN
p-value: 0.003 q-value: 0.039

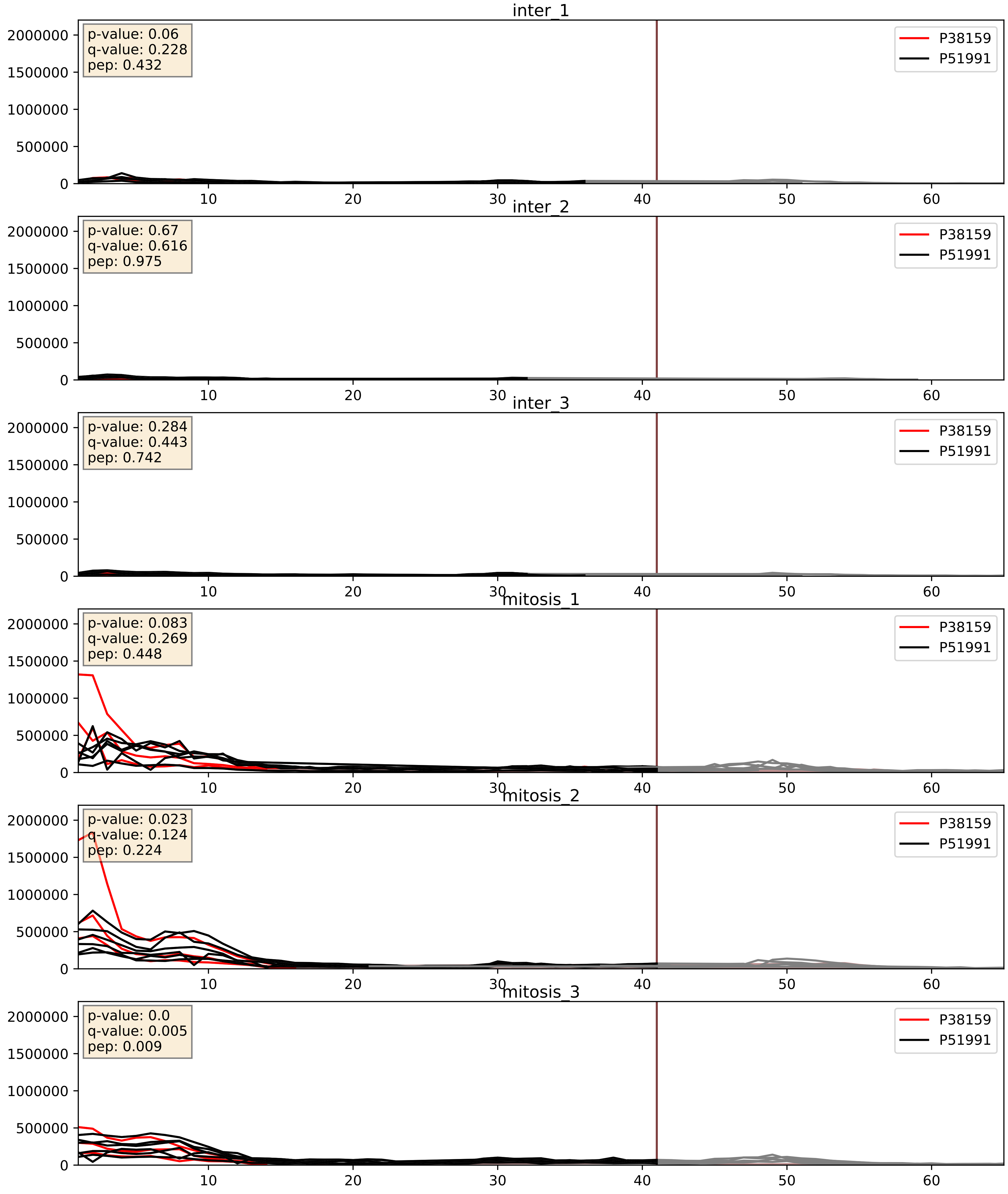


| level | condition_1 | condition_2 | pvalue | pvalue_adjusted |
|---|---|---|---|---|
| complex_abundance | mitosis | inter | 0.000160188868584408 | 0.008563834848633118 |
| interactor_abundance | mitosis | inter | 0.0010979128258399222 | 0.024049720606162197 |
| interactor_ratio | mitosis | inter | 0.9349452147444461 | 0.9881103513214468 |

P38159_P52272
RBMX_HUMAN vs HNRPM_HUMAN
p-value: 0.001 q-value: 0.002

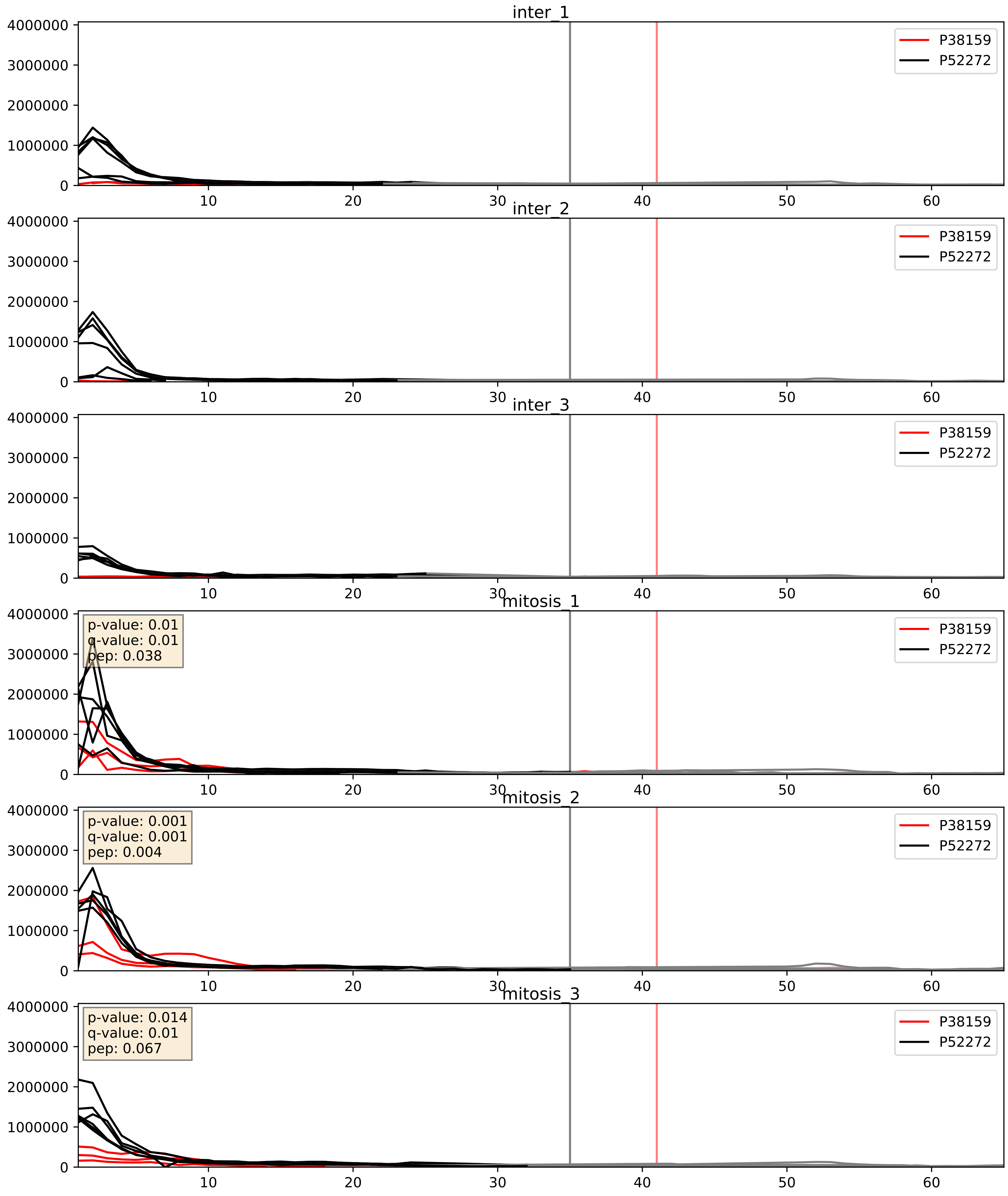


| level | condition_1 | condition_2 | pvalue | pvalue_adjusted |
|---|---|---|---|---|
| interactor_abundance | mitosis | inter | 0.0010812776417605033 | 0.024049720606162197 |
| complex_abundance | mitosis | inter | 0.002146905767739123 | 0.033596916584729235 |
| interactor_ratio | mitosis | inter | 0.00772024973379348 | 0.12375531408477938 |

P38159_P52597
RBMX_HUMAN vs HNRPF_HUMAN
p-value: 0.005 q-value: 0.005

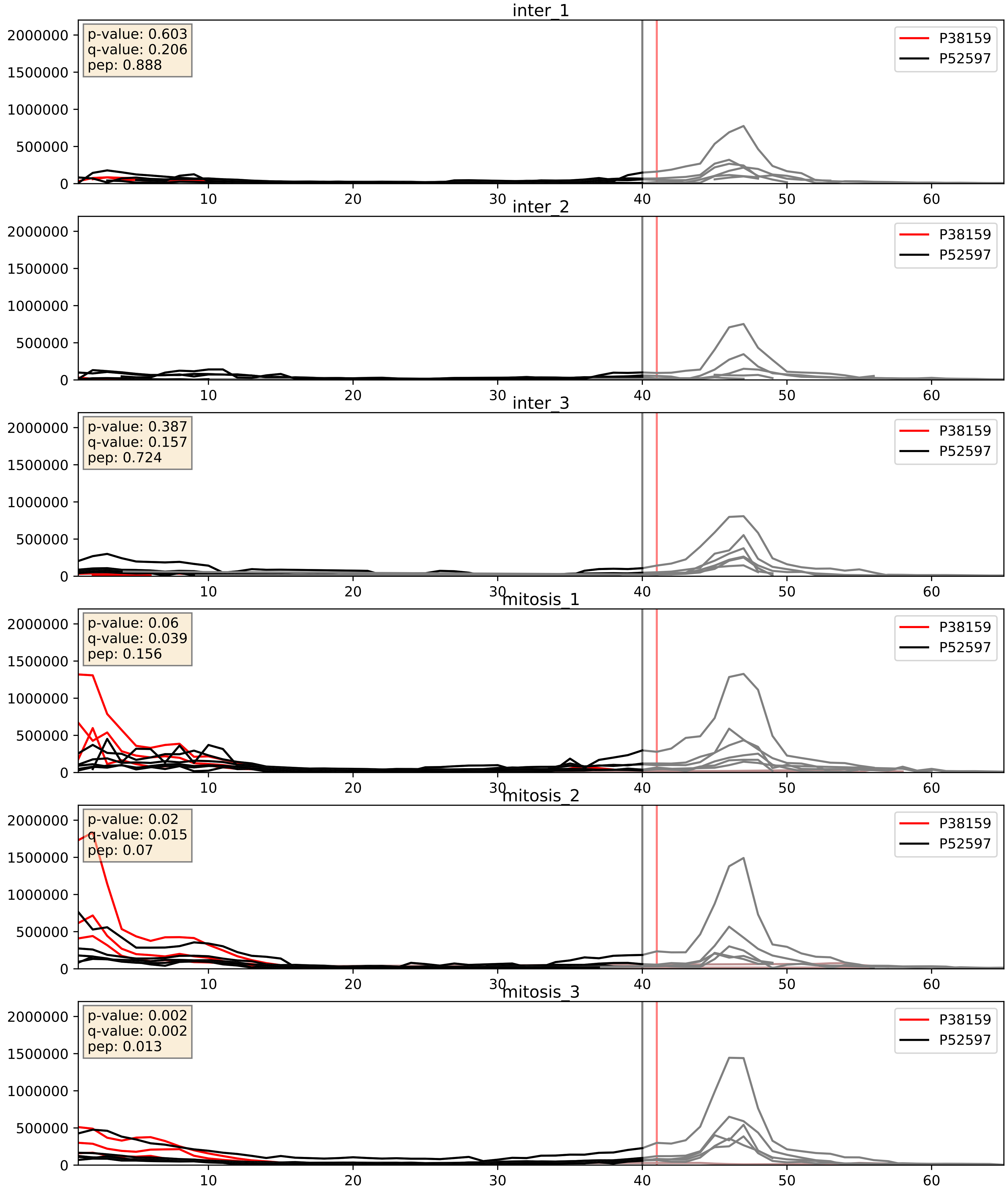


| level | condition_1 | condition_2 | pvalue | pvalue_adjusted |
|---|---|---|---|---|
| interactor_abundance | mitosis | inter | 0.0010973699494020942 | 0.024049720606162197 |
| complex_abundance | mitosis | inter | 0.0032164973285077132 | 0.0416428285245195 |
| interactor_ratio | mitosis | inter | 0.00676965061563321 | 0.11425781573990595 |

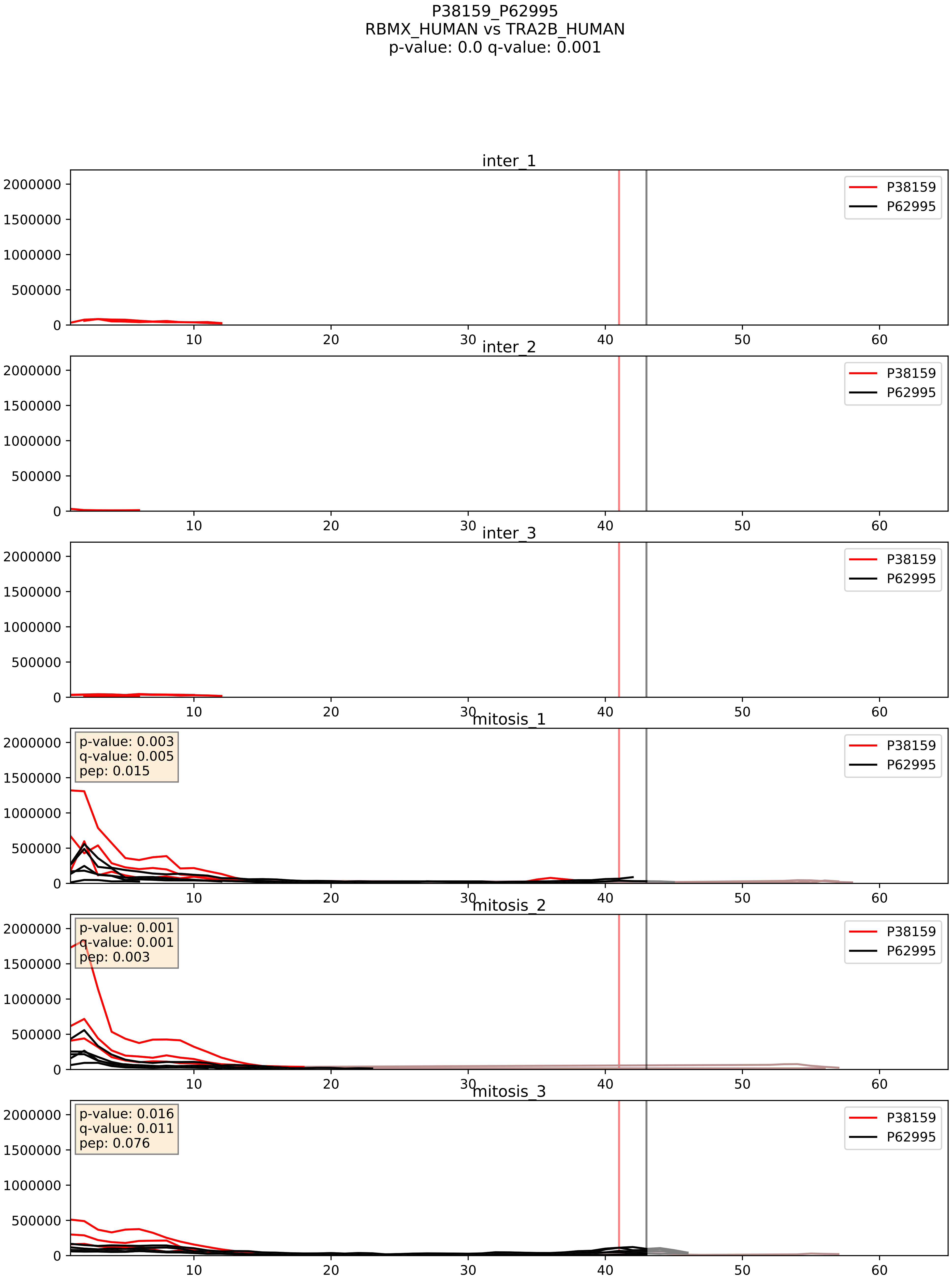


| level | condition_1 | condition_2 | pvalue | pvalue_adjusted |
|---|---|---|---|---|
| interactor_abundance | mitosis | inter | 0.0001647383149740902 | 0.009910743628815524 |
| complex_abundance | mitosis | inter | 0.00022597518718696217 | 0.010219389159334393 |
| interactor_ratio | mitosis | inter | 0.011485414155784251 | 0.15494522140242886 |

P38159_Q07955
RBMX_HUMAN vs SRSF1_HUMAN
p-value: 0.074 q-value: 0.045

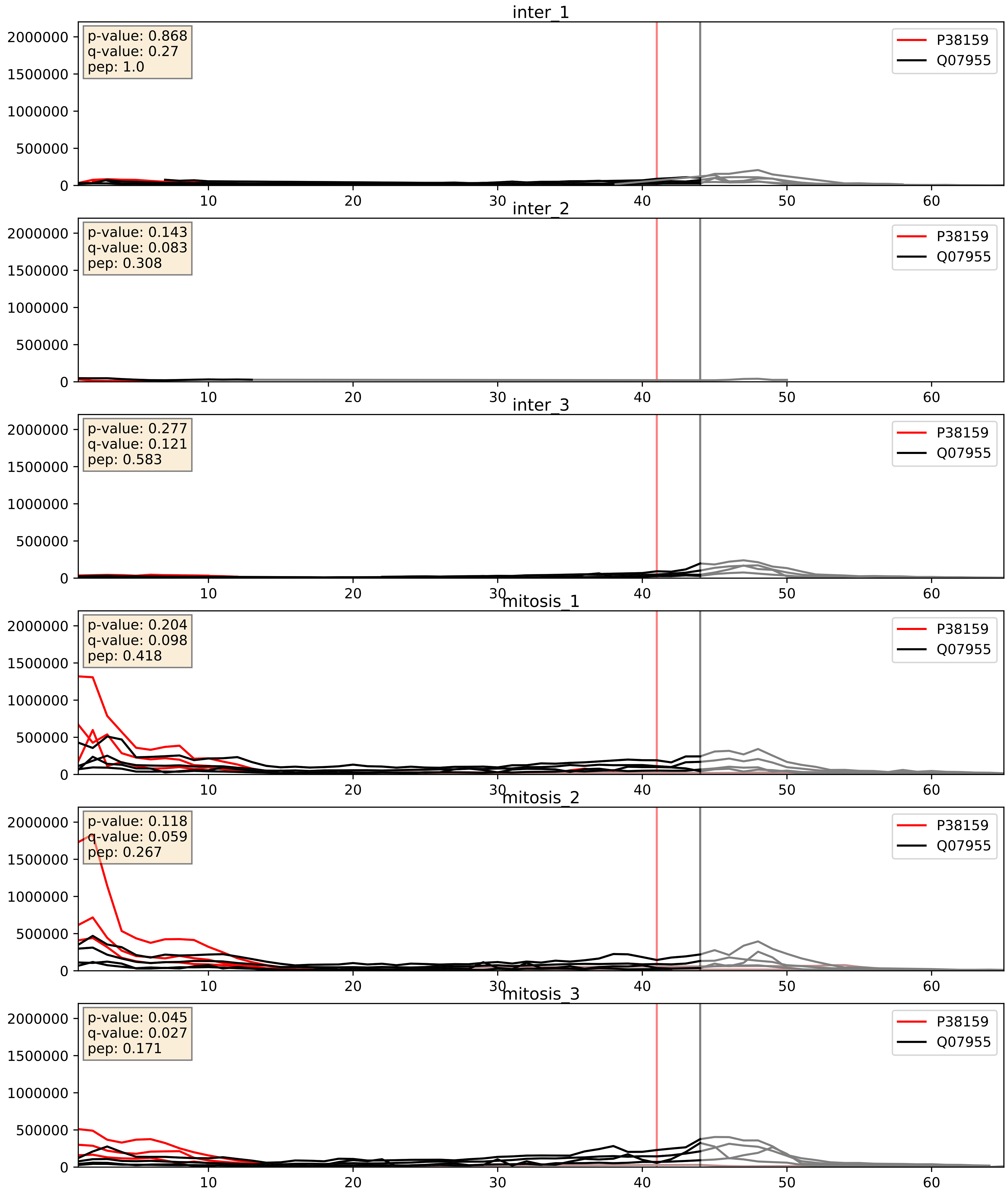


| level | condition_1 | condition_2 | pvalue | pvalue_adjusted |
|---|---|---|---|---|
| interactor_abundance | mitosis | inter | 0.0010393306972951088 | 0.024049720606162197 |
| complex_abundance | mitosis | inter | 0.0014295676213636915 | 0.02592605686201949 |
| interactor_ratio | mitosis | inter | 0.05120606381407092 | 0.37414961988671497 |

P38159_Q13151
RBMX_HUMAN vs ROA0_HUMAN
p-value: 0.017 q-value: 0.014

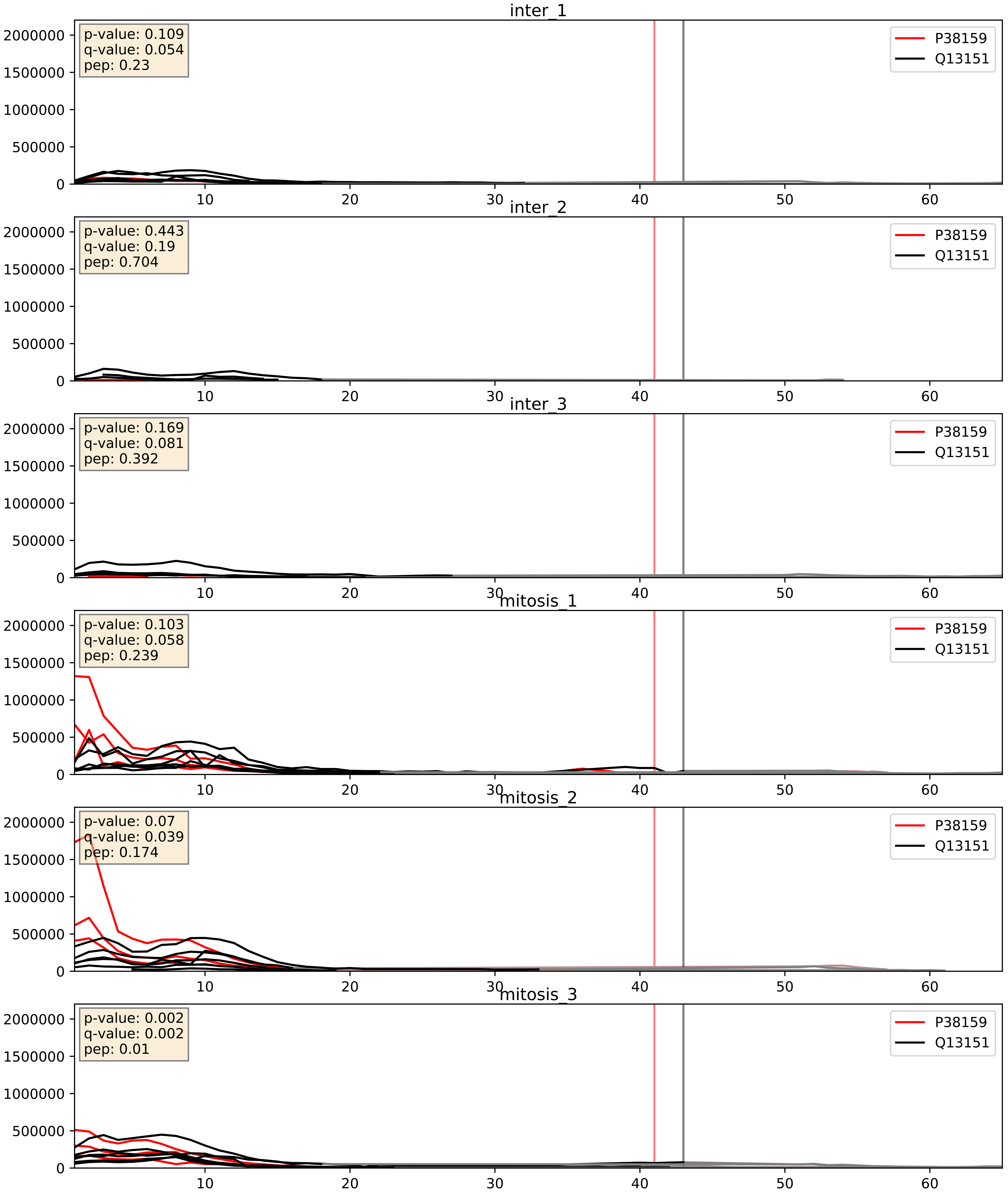


| level | condition_1 | condition_2 | pvalue | pvalue_adjusted |
|---|---|---|---|---|
| interactor_abundance | mitosis | inter | 0.0010950989897773285 | 0.024049720606162197 |
| complex_abundance | mitosis | inter | 0.001785591685100528 | 0.030007039692728107 |
| interactor_ratio | mitosis | inter | 0.02637445981207413 | 0.25543205403291264 |

P38159_Q13242
RBMX_HUMAN vs SRSF9_HUMAN
p-value: 0.042 q-value: 0.028

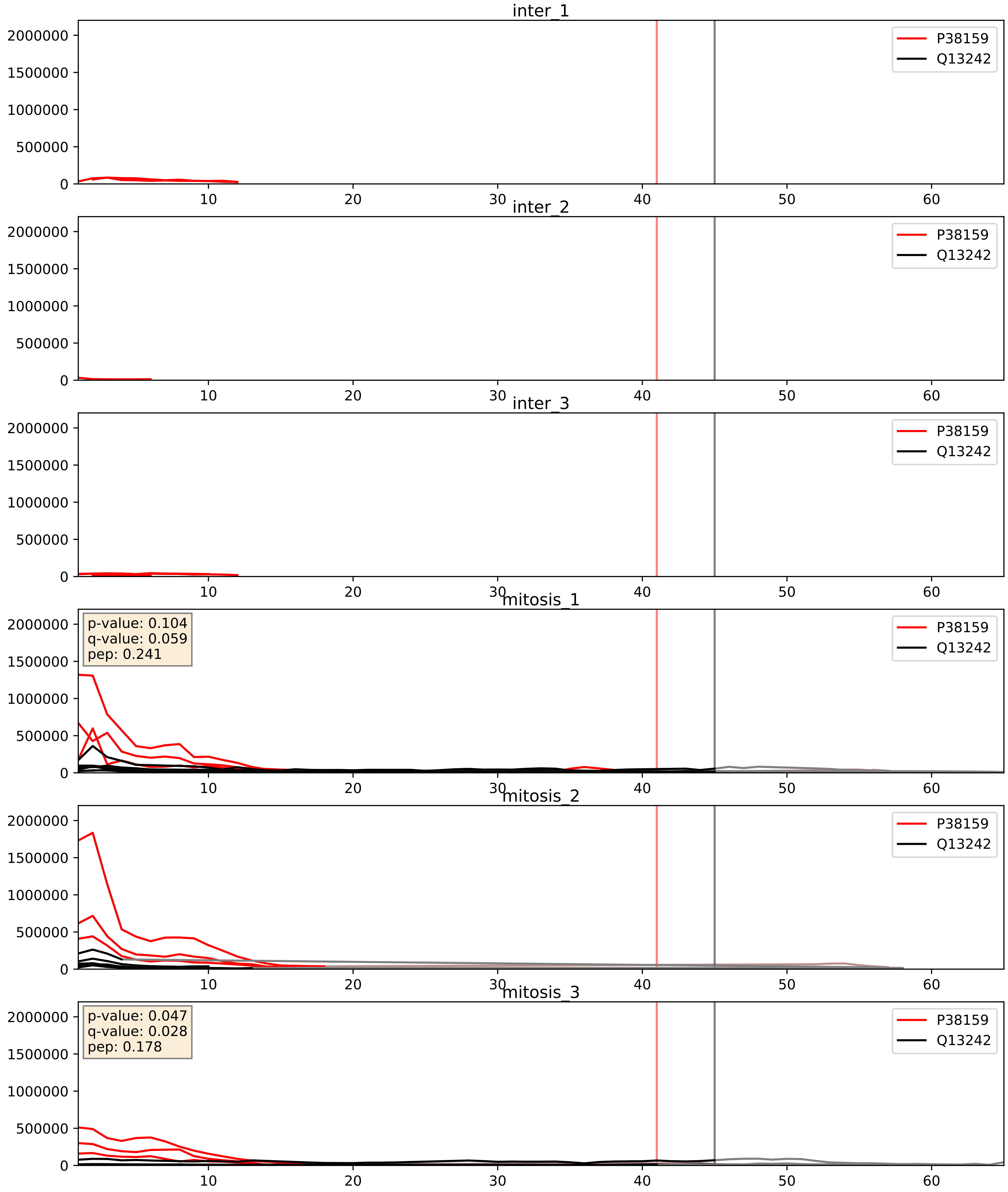


| level | condition_1 | condition_2 | pvalue | pvalue_adjusted |
|---|---|---|---|---|
| interactor_abundance | mitosis | inter | 0.0001613854946210072 | 0.009910743628815524 |
| interactor_ratio | mitosis | inter | 0.0022484423567735888 | 0.05896690574521502 |
| complex_abundance | mitosis | inter | 0.0029712233791559066 | 0.03988944941488032 |

P38159_Q15717
RBMX_HUMAN vs ELAV1_HUMAN
p-value: 0.0 q-value: 0.0

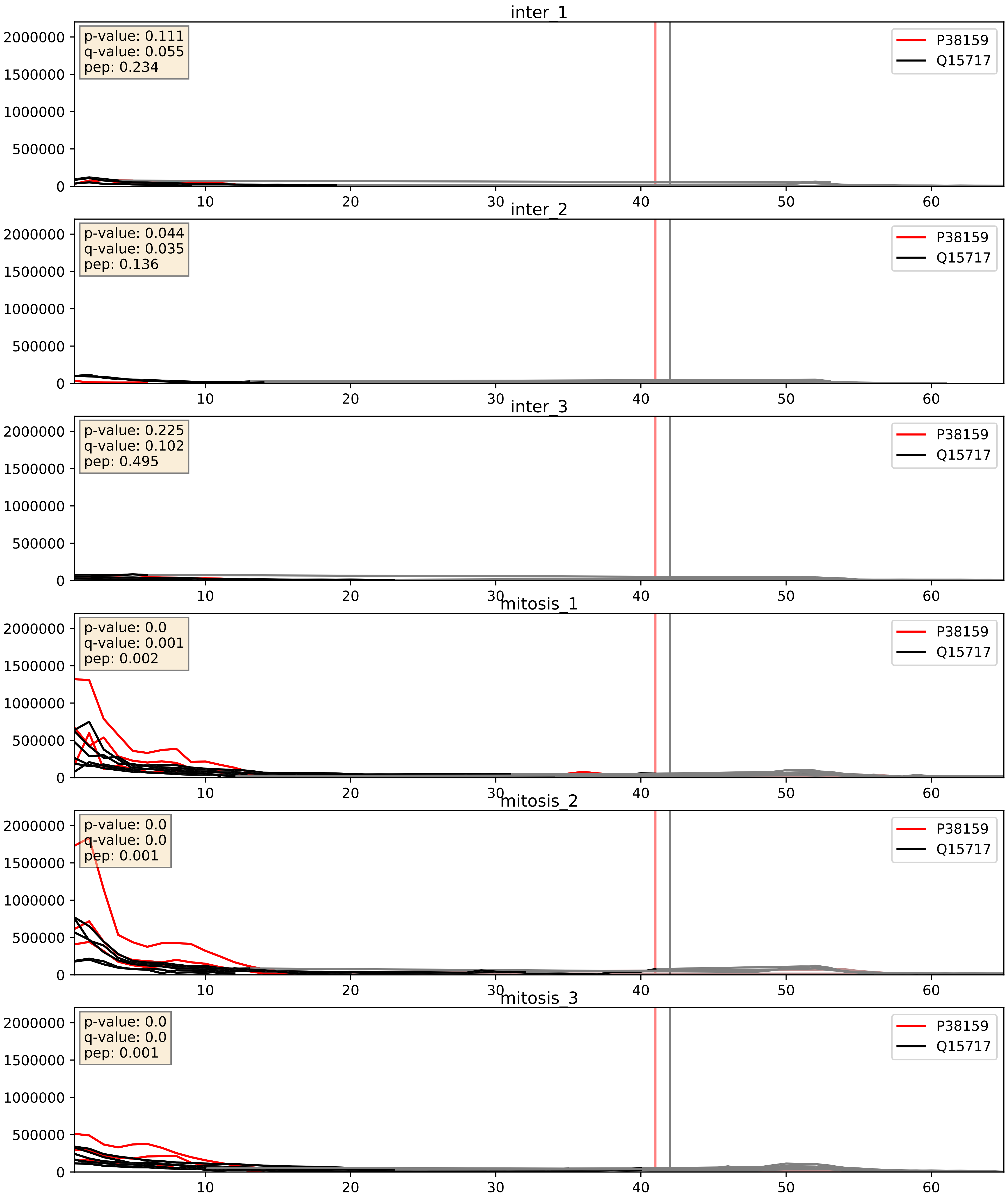


| level | condition_1 | condition_2 | pvalue | pvalue_adjusted |
|---|---|---|---|---|
| complex_abundance | mitosis | inter | 0.0004288806060394006 | 0.013091389064164169 |
| interactor_abundance | mitosis | inter | 0.0010979128258399222 | 0.024049720606162197 |
| interactor_ratio | mitosis | inter | 0.3588347875269688 | 0.8049703690863734 |

P38159_Q99459
RBMX_HUMAN vs CDC5L_HUMAN
p-value: 0.036 q-value: 0.025

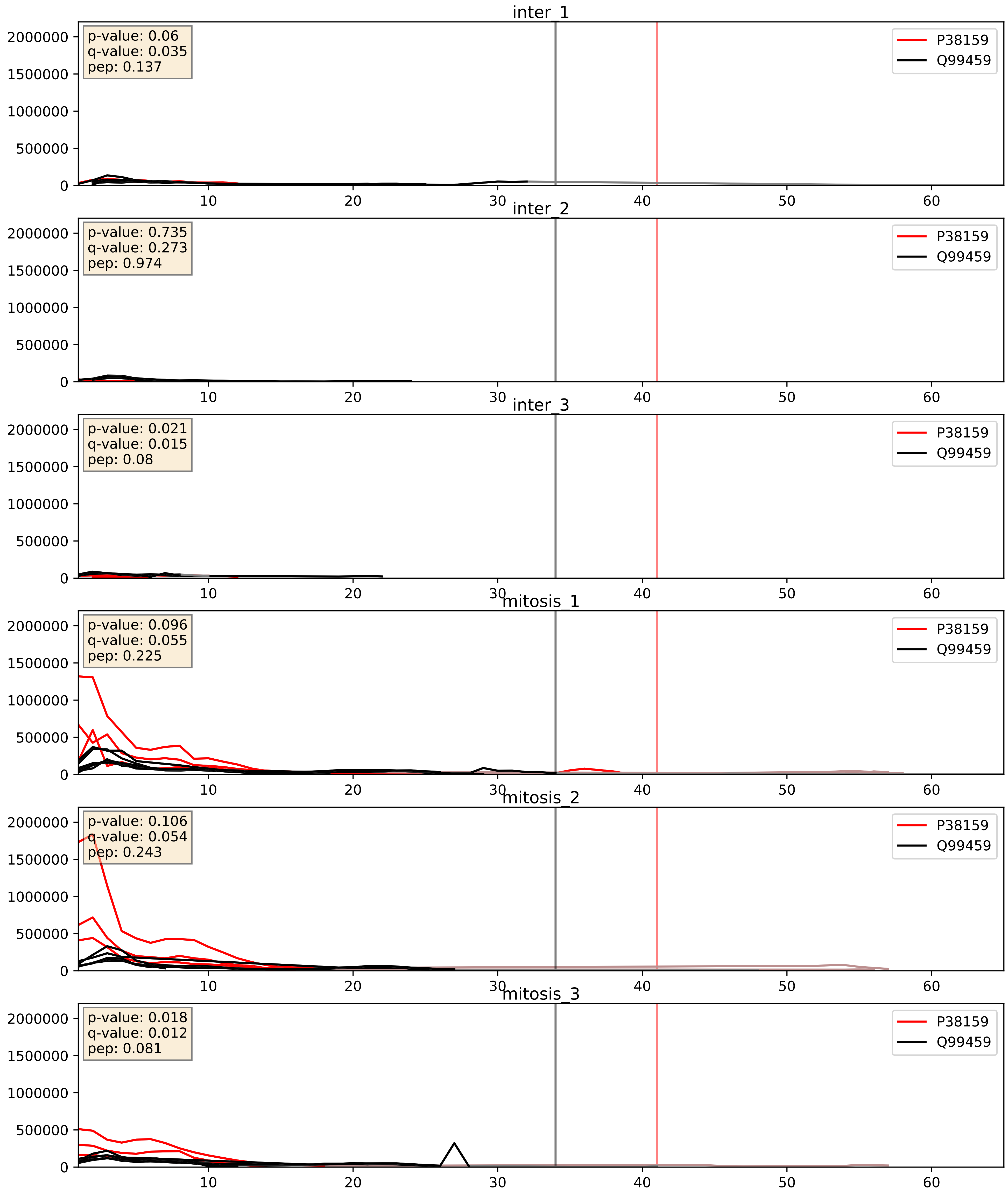


| level | condition_1 | condition_2 | pvalue | pvalue_adjusted |
|---|---|---|---|---|
| interactor_abundance | mitosis | inter | 0.0010224373165272904 | 0.0238801184978816 |
| complex_abundance | mitosis | inter | 0.0023534385428367085 | 0.035168510428347066 |
| interactor_ratio | mitosis | inter | 0.05399913247773674 | 0.38487308410443505 |
