## Supplementary Data 2 for "SECAT: Quantifying differential protein-protein interaction states by network-centric analysis": hela_string_000061_P84103.pdf

P84103_P84103

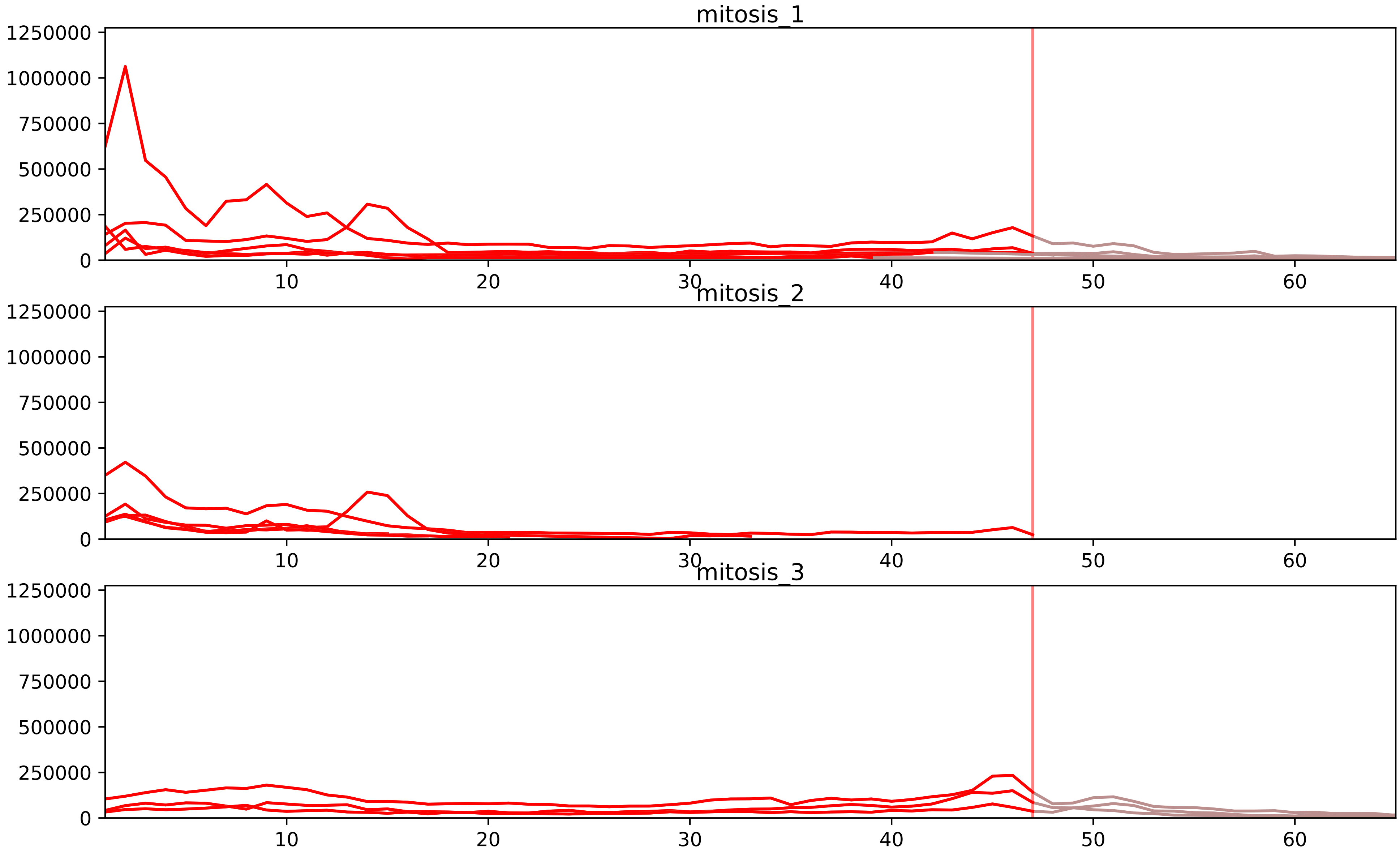


| level | condition_1 | condition_2 | pvalue | pvalue_adjusted |
|---|---|---|---|---|
| interactor_ratio | mitosis | inter | 5.478163290116805e-07 | 1.737900078243952e-05 |
| complex_abundance | mitosis | inter | 1.1406230345760348e-06 | 0.00010493731918099521 |
| interactor_abundance | mitosis | inter | 8.640934153678481e-05 | 0.0020648466029569356 |
| assembled_abundance | mitosis | inter | 0.03000502481589873 | 0.20994547259076526 |
| total_abundance | mitosis | inter | 0.03116287888907052 | 0.22458279983311685 |
| monomer_abundance | mitosis | inter | 0.04076279511363155 | 0.3157957803087734 |

P07910_P84103
HNRPC_HUMAN vs SRSF3_HUMAN
p-value: 0.043 q-value: 0.029

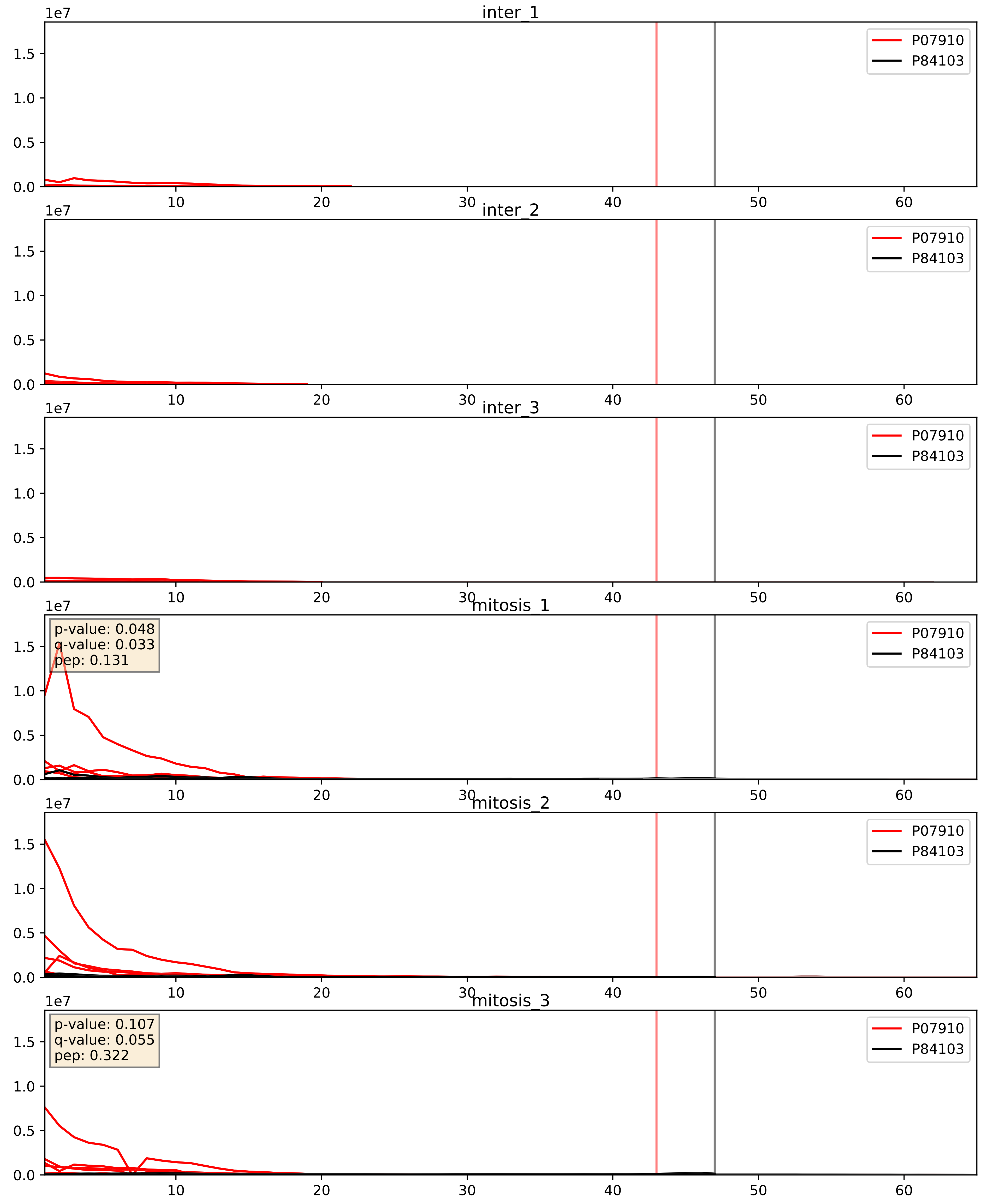


| level | condition_1 | condition_2 | pvalue | pvalue_adjusted |
|---|---|---|---|---|
| interactor_abundance | mitosis | inter | 2.2213691150803744e-06 | 0.0033734327106189023 |
| complex_abundance | mitosis | inter | 4.449045517390577e-05 | 0.005691857151652958 |
| interactor_ratio | mitosis | inter | 0.0013366807319858036 | 0.04334578357589157 |

P09651_P84103
ROA1_HUMAN vs SRSF3_HUMAN
p-value: 0.051 q-value: 0.033

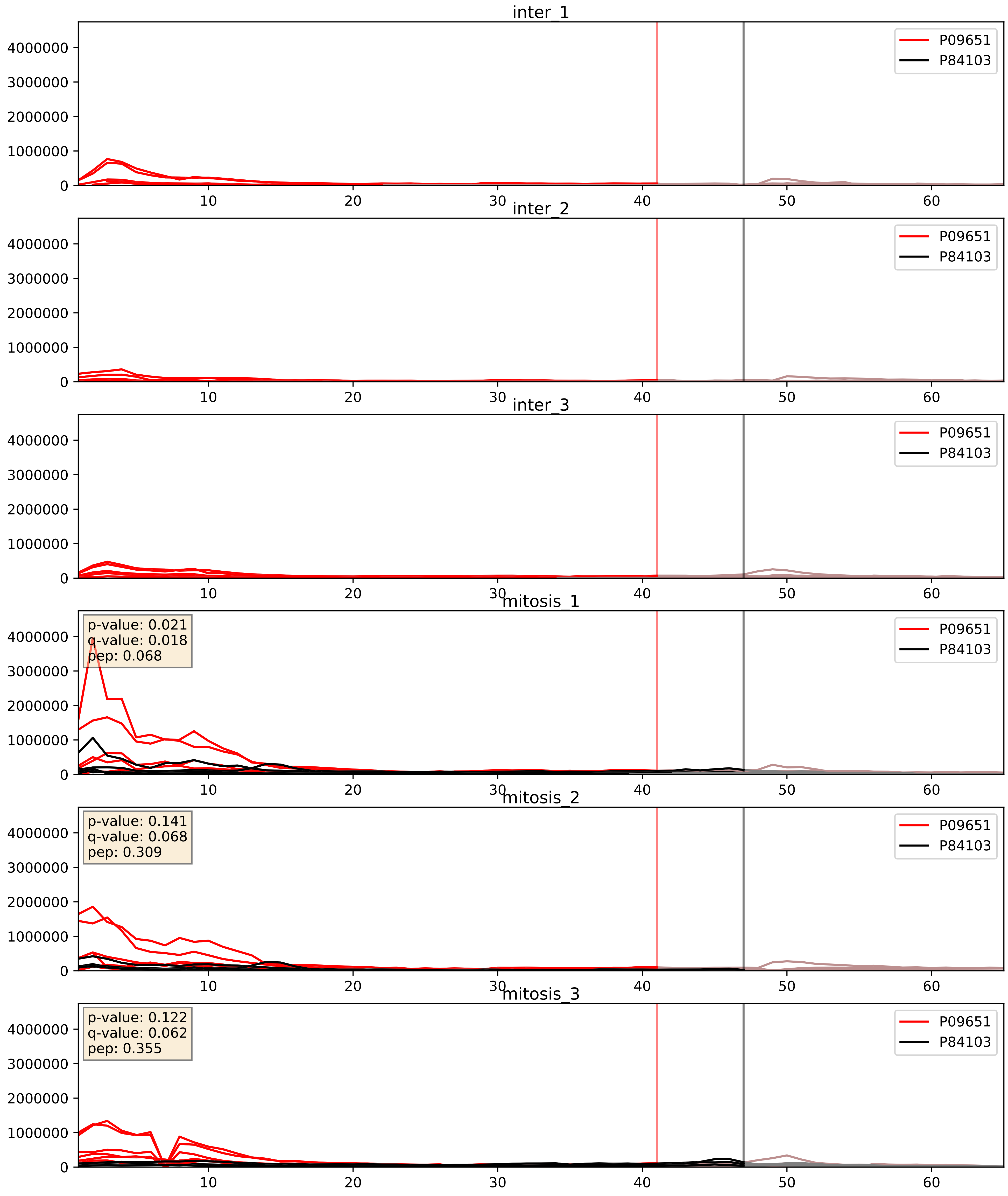


| level | condition_1 | condition_2 | pvalue | pvalue_adjusted |
|---|---|---|---|---|
| interactor_abundance | mitosis | inter | 7.226077128975986e-05 | 0.006834830963981706 |
| complex_abundance | mitosis | inter | 0.0002291506843386366 | 0.010219389159334393 |
| interactor_ratio | mitosis | inter | 0.004188942000443196 | 0.08517183734867877 |

P14866_P84103
HNRPL_HUMAN vs SRSF3_HUMAN
p-value: 0.039 q-value: 0.027

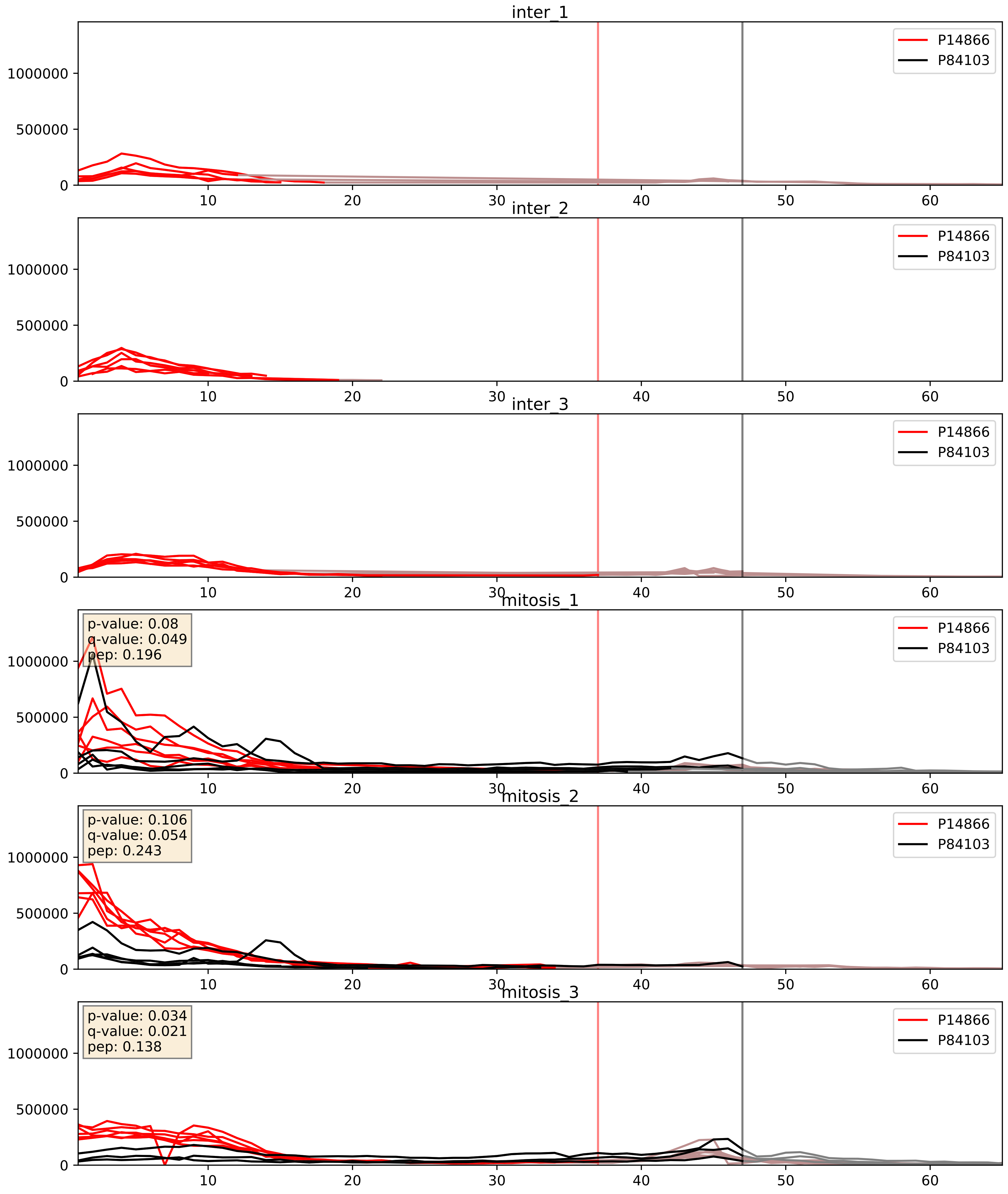


| level | condition_1 | condition_2 | pvalue | pvalue_adjusted |
|---|---|---|---|---|
| interactor_abundance | mitosis | inter | 1.2564597628871335e-05 | 0.003391165020149013 |
| complex_abundance | mitosis | inter | 0.00025225809034102616 | 0.010637089917828493 |
| interactor_ratio | mitosis | inter | 0.02957345168884291 | 0.2722029531790272 |

P22626_P84103
ROA2_HUMAN vs SRSF3_HUMAN
p-value: 0.015 q-value: 0.012

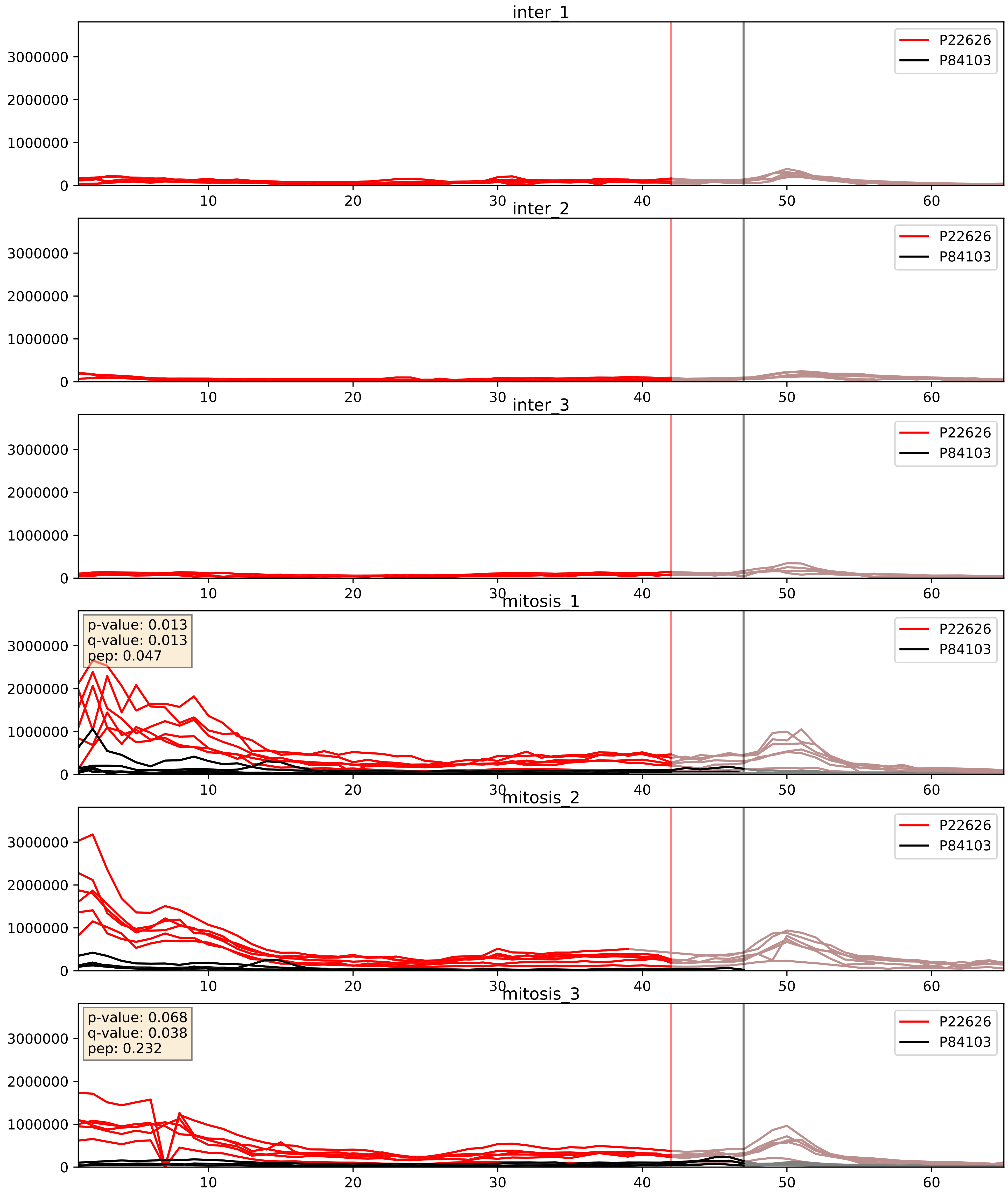


| level | condition_1 | condition_2 | pvalue | pvalue_adjusted |
|---|---|---|---|---|
| interactor_abundance | mitosis | inter | 3.003313050728667e-05 | 0.004850633908346677 |
| complex_abundance | mitosis | inter | 6.433903659397418e-05 | 0.005691857151652958 |
| interactor_ratio | mitosis | inter | 0.0014422267222096987 | 0.045221467919835244 |

P26599_P84103
PTBP1_HUMAN vs SRSF3_HUMAN
p-value: 0.048 q-value: 0.032

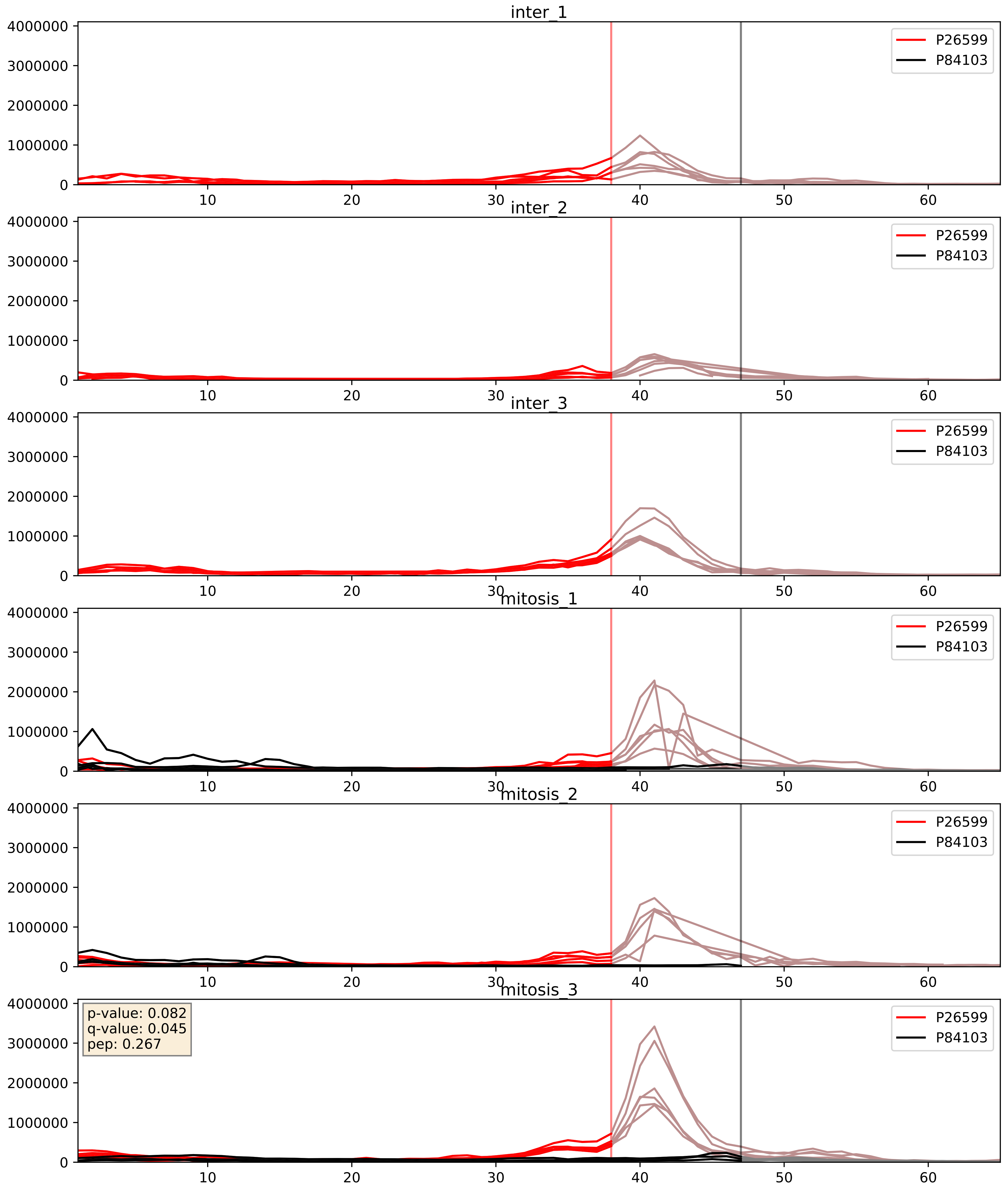


| level | condition_1 | condition_2 | pvalue | pvalue_adjusted |
|---|---|---|---|---|
| complex_abundance | mitosis | inter | 0.0008711643078724355 | 0.019513211095311817 |
| interactor_abundance | mitosis | inter | 0.010211870122103946 | 0.0879625269700301 |
| interactor_ratio | mitosis | inter | 0.7275771855558608 | 0.947216680824468 |

P31943_P84103
HNRH1_HUMAN vs SRSF3_HUMAN
p-value: 0.005 q-value: 0.005

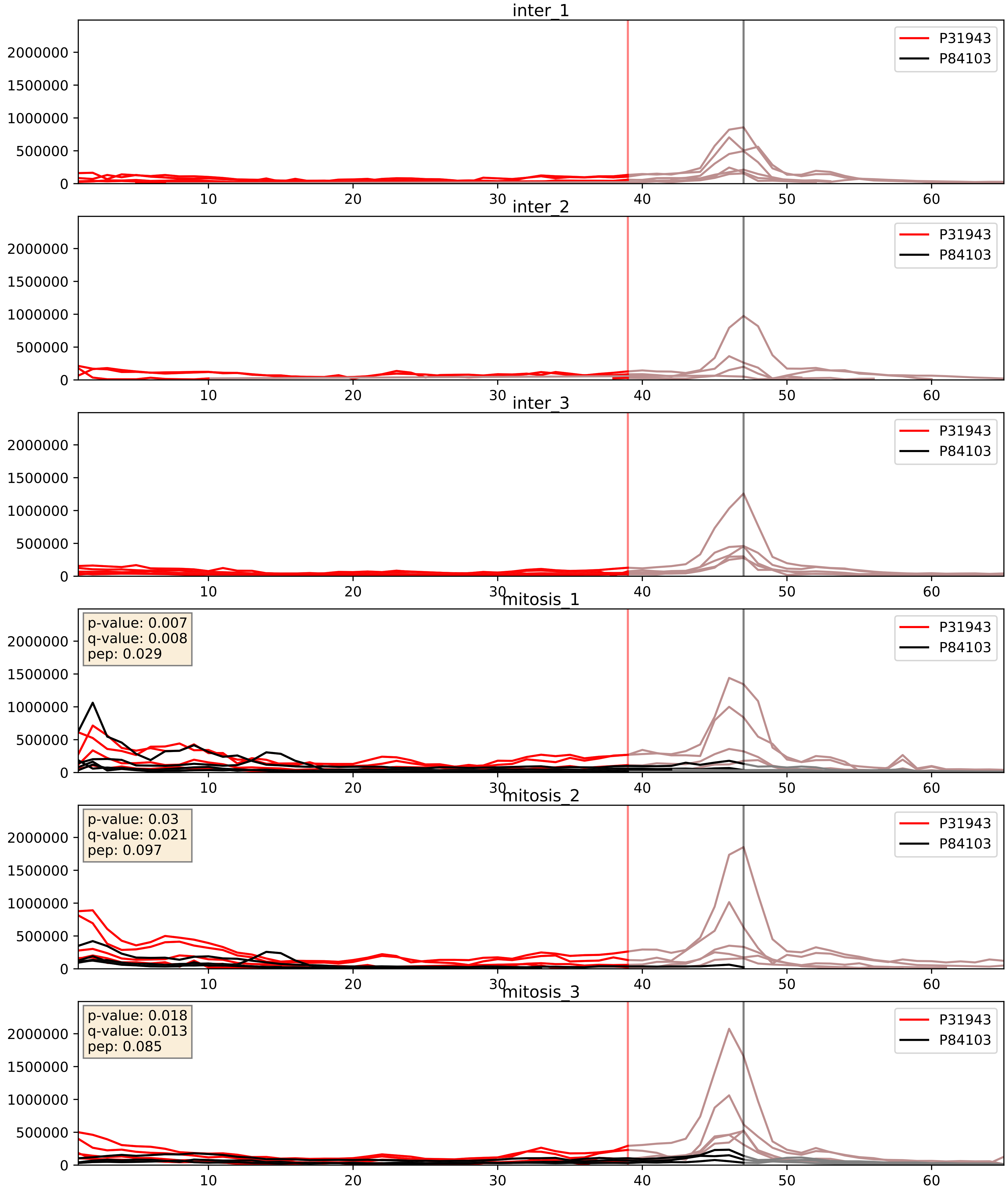


| level | condition_1 | condition_2 | pvalue | pvalue_adjusted |
|---|---|---|---|---|
| interactor_abundance | mitosis | inter | 2.5609271594898103e-05 | 0.004519904429944902 |
| complex_abundance | mitosis | inter | 0.0003358253138333869 | 0.011893558089955385 |
| interactor_ratio | mitosis | inter | 0.005707332319897169 | 0.10328702887594031 |

P52597_P84103
HNRPF_HUMAN vs SRSF3_HUMAN
p-value: 0.001 q-value: 0.002

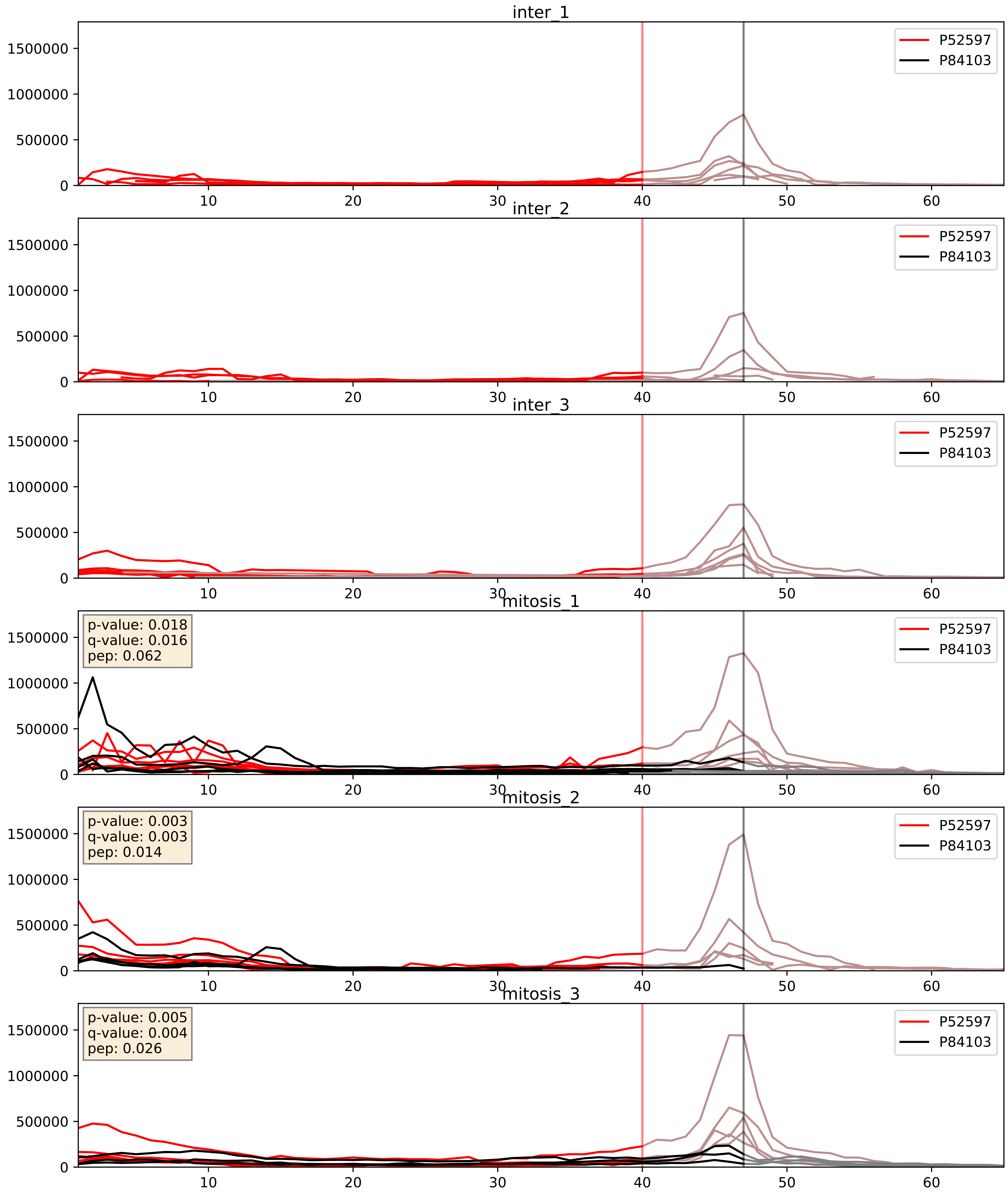


| level | condition_1 | condition_2 | pvalue | pvalue_adjusted |
|---|---|---|---|---|
| interactor_abundance | mitosis | inter | 4.920255045596975e-05 | 0.006283598333582311 |
| complex_abundance | mitosis | inter | 0.00044192382616051125 | 0.013273220883978866 |
| interactor_ratio | mitosis | inter | 0.0357560041616608 | 0.30712516692473574 |

P62995_P84103
TRA2B_HUMAN vs SRSF3_HUMAN
p-value: 0.008 q-value: 0.008

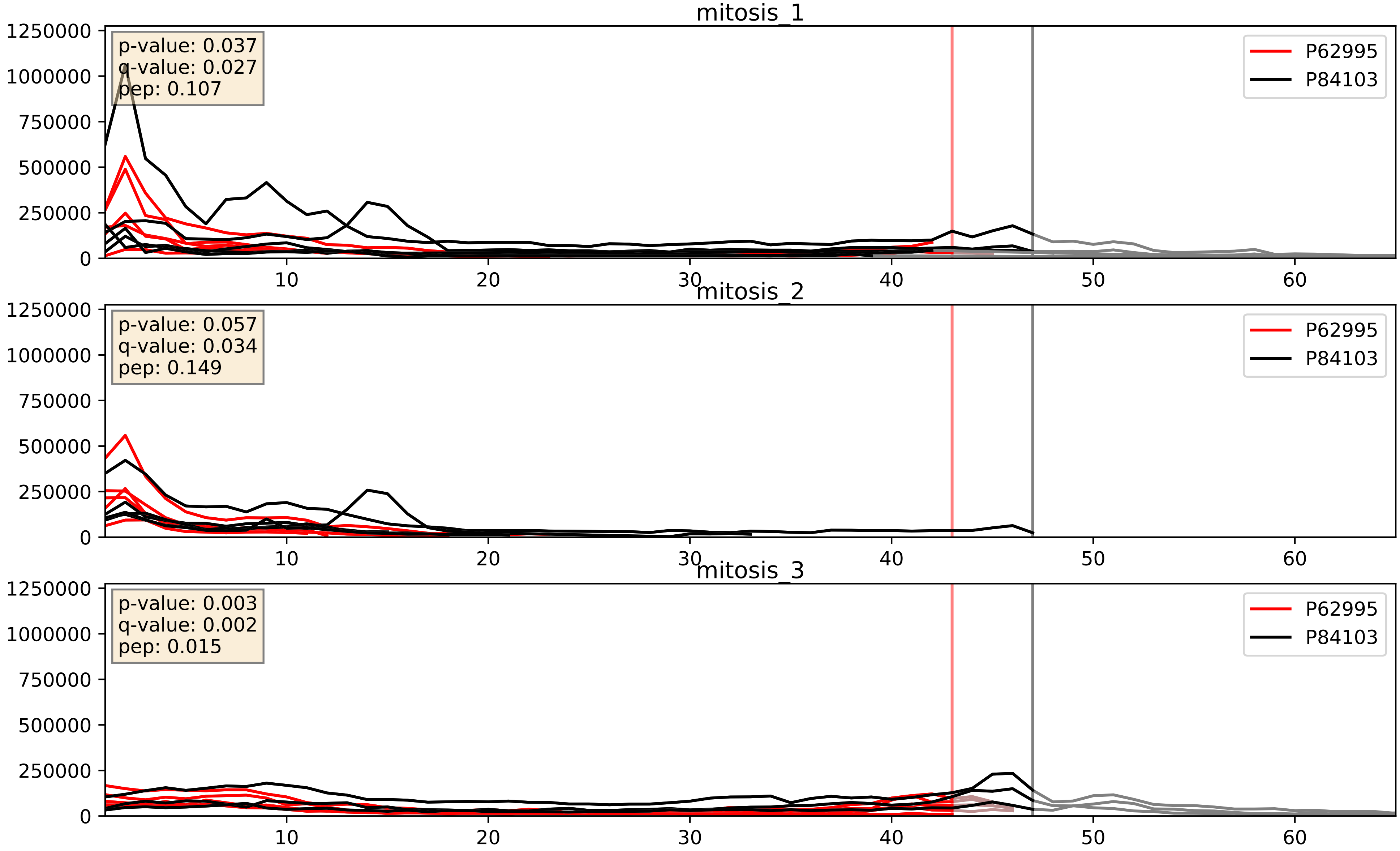


| level | condition_1 | condition_2 | pvalue | pvalue_adjusted |
|---|---|---|---|---|
| interactor_abundance | mitosis | inter | 0.0004584932841066445 | 0.014951247664582388 |
| complex_abundance | mitosis | inter | 0.000642767496886288 | 0.016393529992559892 |
| interactor_ratio | mitosis | inter | 0.8736089821848678 | 0.980069417404396 |

P84103_Q07955
SRSF3_HUMAN vs SRSF1_HUMAN
p-value: 0.054 q-value: 0.035

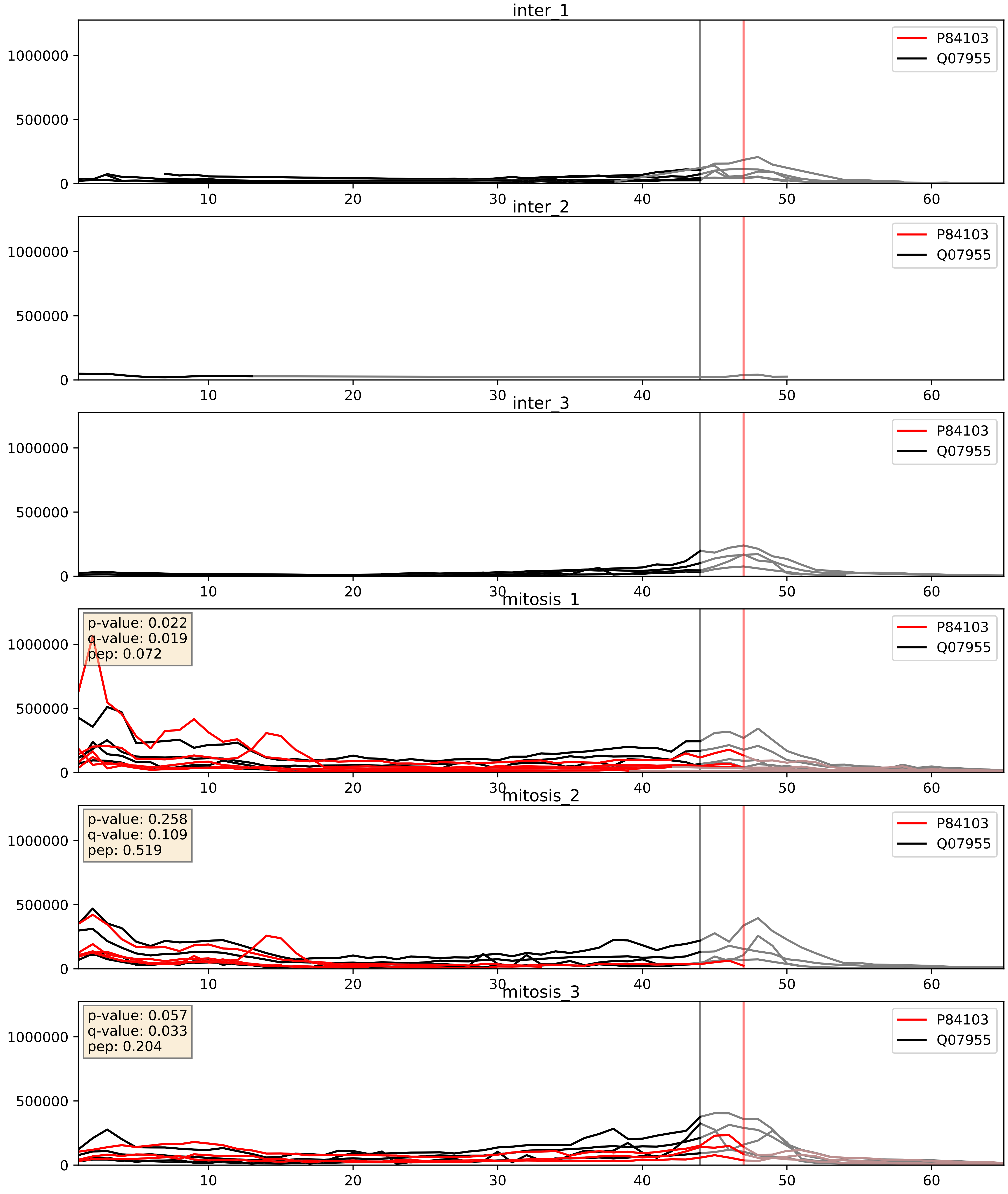


| level | condition_1 | condition_2 | pvalue | pvalue_adjusted |
|---|---|---|---|---|
| complex_abundance | mitosis | inter | 3.7310233198972706e-05 | 0.005691857151652958 |
| interactor_abundance | mitosis | inter | 0.002590890474122459 | 0.03806046117740333 |
| interactor_ratio | mitosis | inter | 0.4549089543182232 | 0.8545518508495171 |

P84103_Q13151
SRSF3_HUMAN vs ROA0_HUMAN
p-value: 0.009 q-value: 0.008

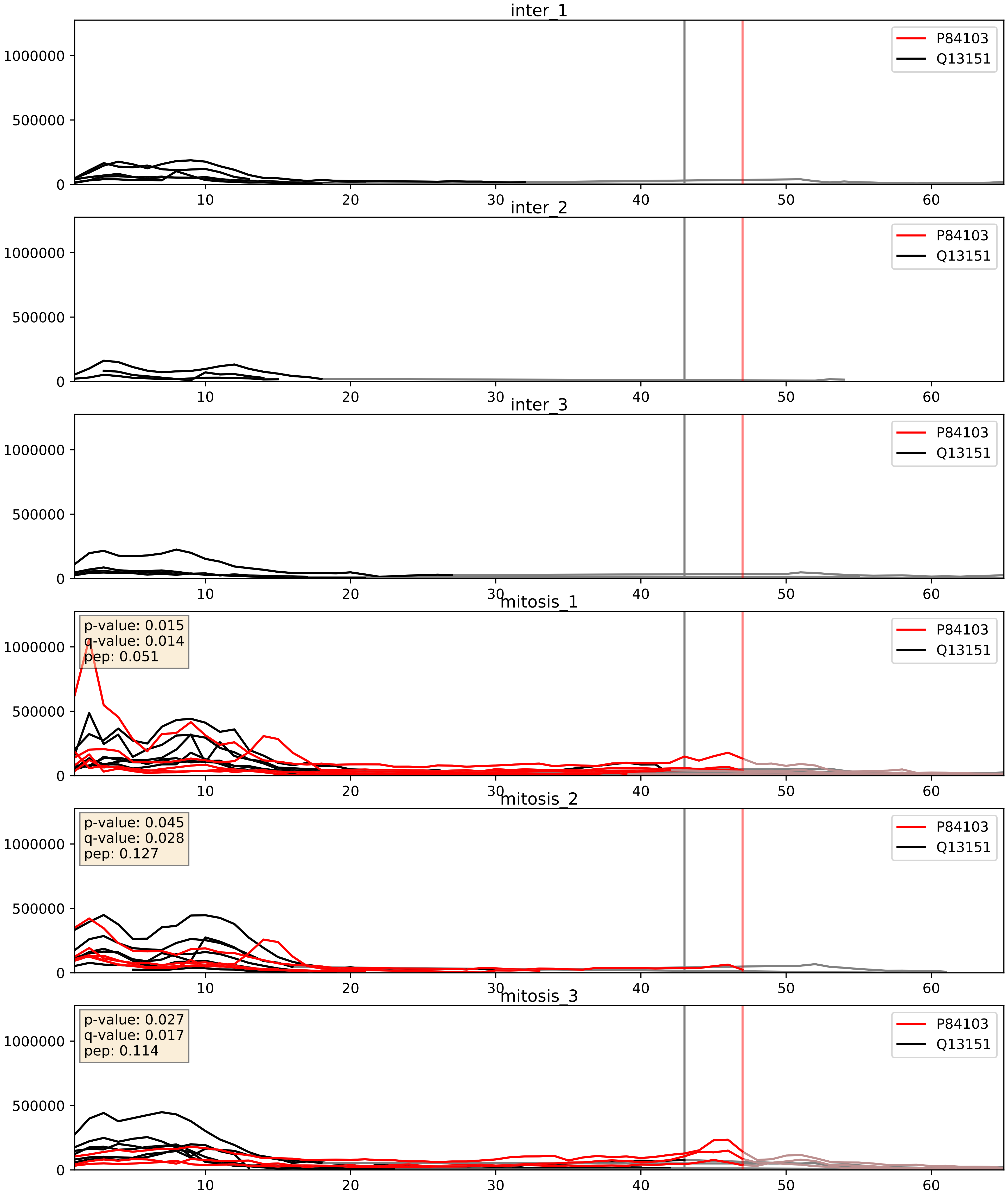


| level | condition_1 | condition_2 | pvalue | pvalue_adjusted |
|---|---|---|---|---|
| complex_abundance | mitosis | inter | 0.0002598167279666148 | 0.010784252527150902 |
| interactor_abundance | mitosis | inter | 0.0026696686527179834 | 0.038732819775027016 |
| interactor_ratio | mitosis | inter | 0.11994976755506524 | 0.5692297679773661 |

P84103_Q13242
SRSF3_HUMAN vs SRSF9_HUMAN
p-value: 0.059 q-value: 0.037

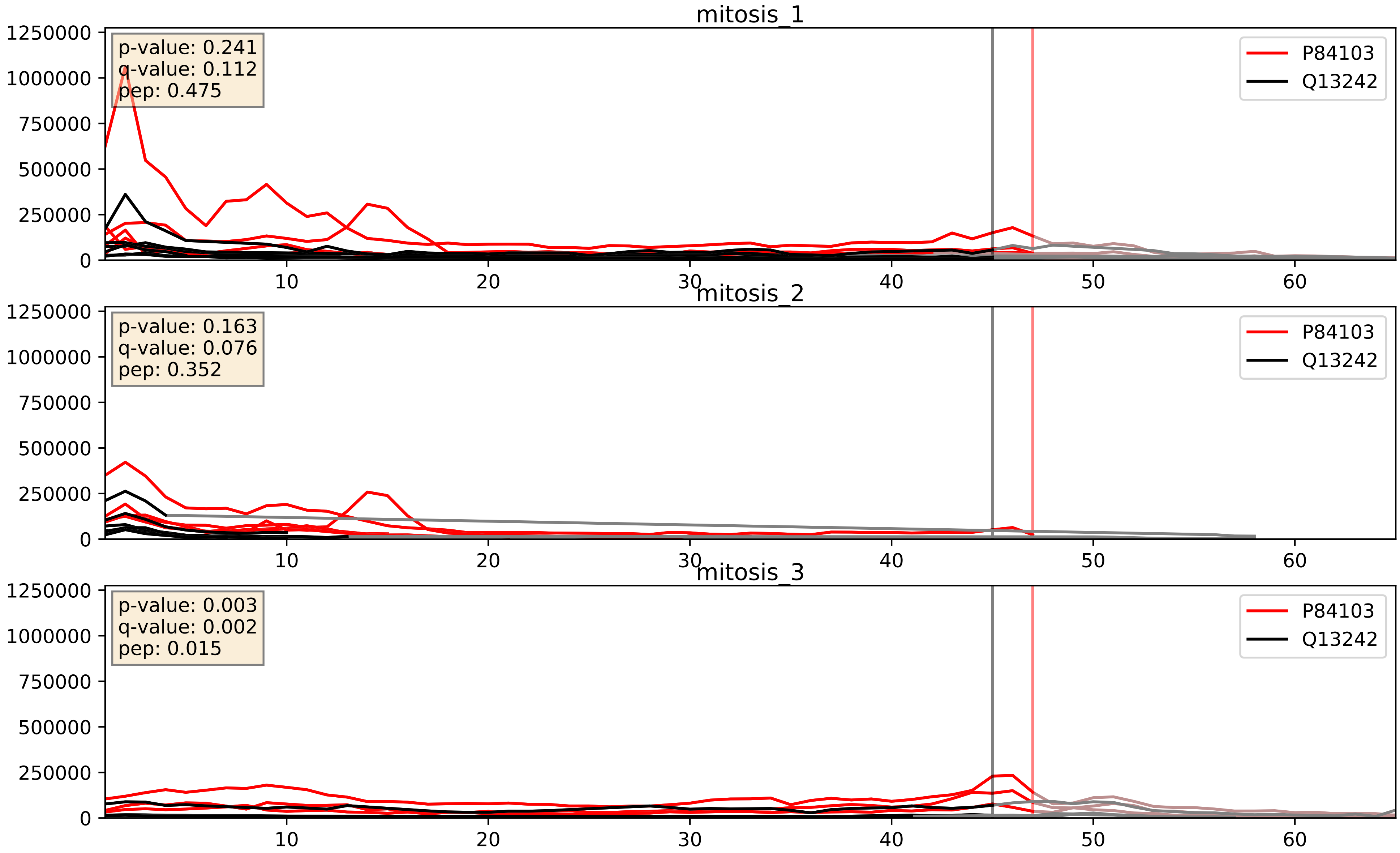


| level | condition_1 | condition_2 | pvalue | pvalue_adjusted |
|---|---|---|---|---|
| interactor_ratio | mitosis | inter | 0.00018729042781272996 | 0.014134487523529912 |
| interactor_abundance | mitosis | inter | 0.004192456982592769 | 0.05145151508386251 |
| complex_abundance | mitosis | inter | 0.012111824621720255 | 0.0818150085787963 |

P84103_Q15365
SRSF3_HUMAN vs PCBP1_HUMAN
p-value: 0.002 q-value: 0.03

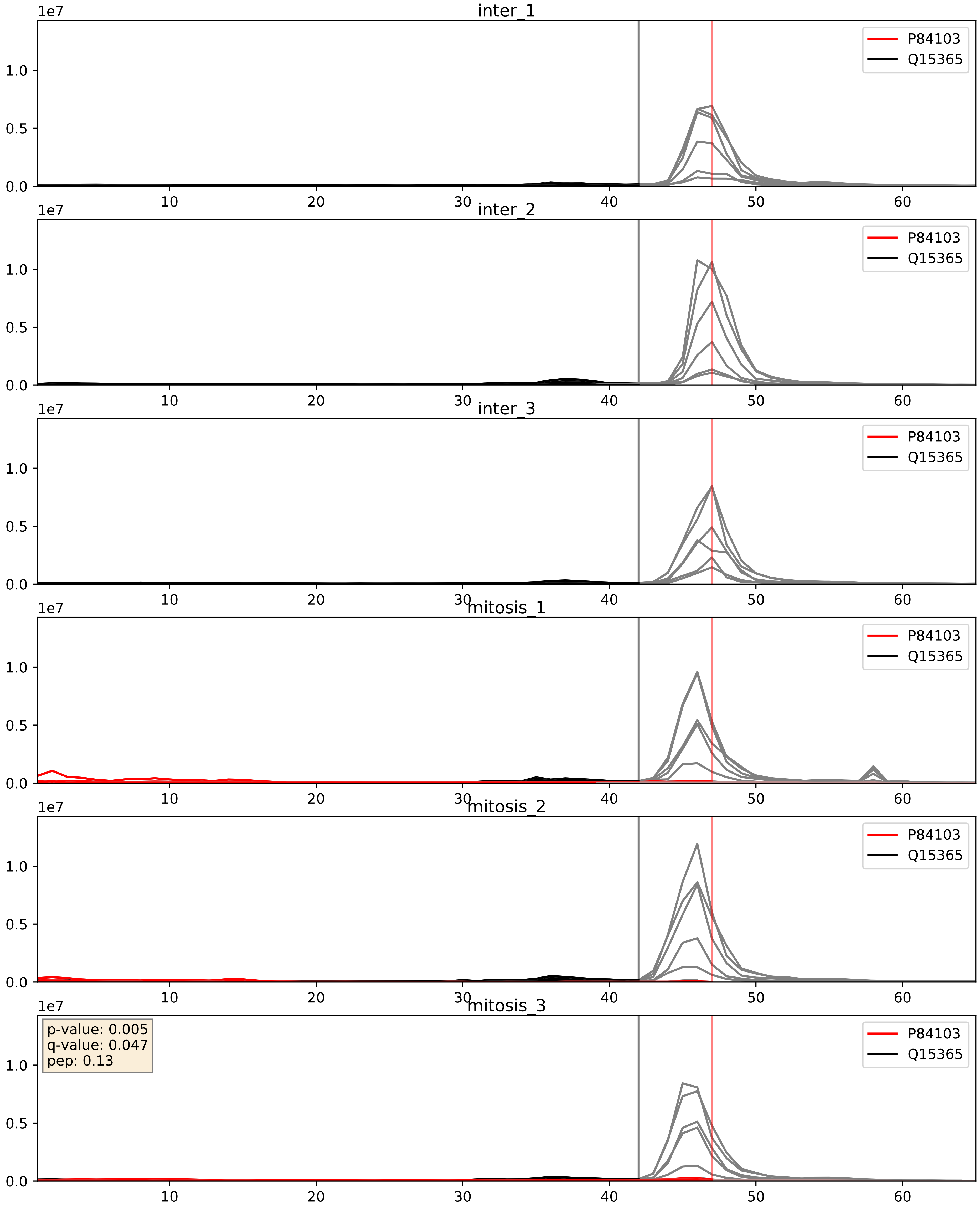


| level | condition_1 | condition_2 | pvalue | pvalue_adjusted |
|---|---|---|---|---|
| complex_abundance | mitosis | inter | 0.00026638034582209696 | 0.010910123254723205 |
| interactor_abundance | mitosis | inter | 0.00295284789649354 | 0.041301271232001155 |
| interactor_ratio | mitosis | inter | 0.041404094351652806 | 0.33818611416998856 |

P84103_Q15366
SRSF3_HUMAN vs PCBP2_HUMAN
p-value: 0.0 q-value: 0.0

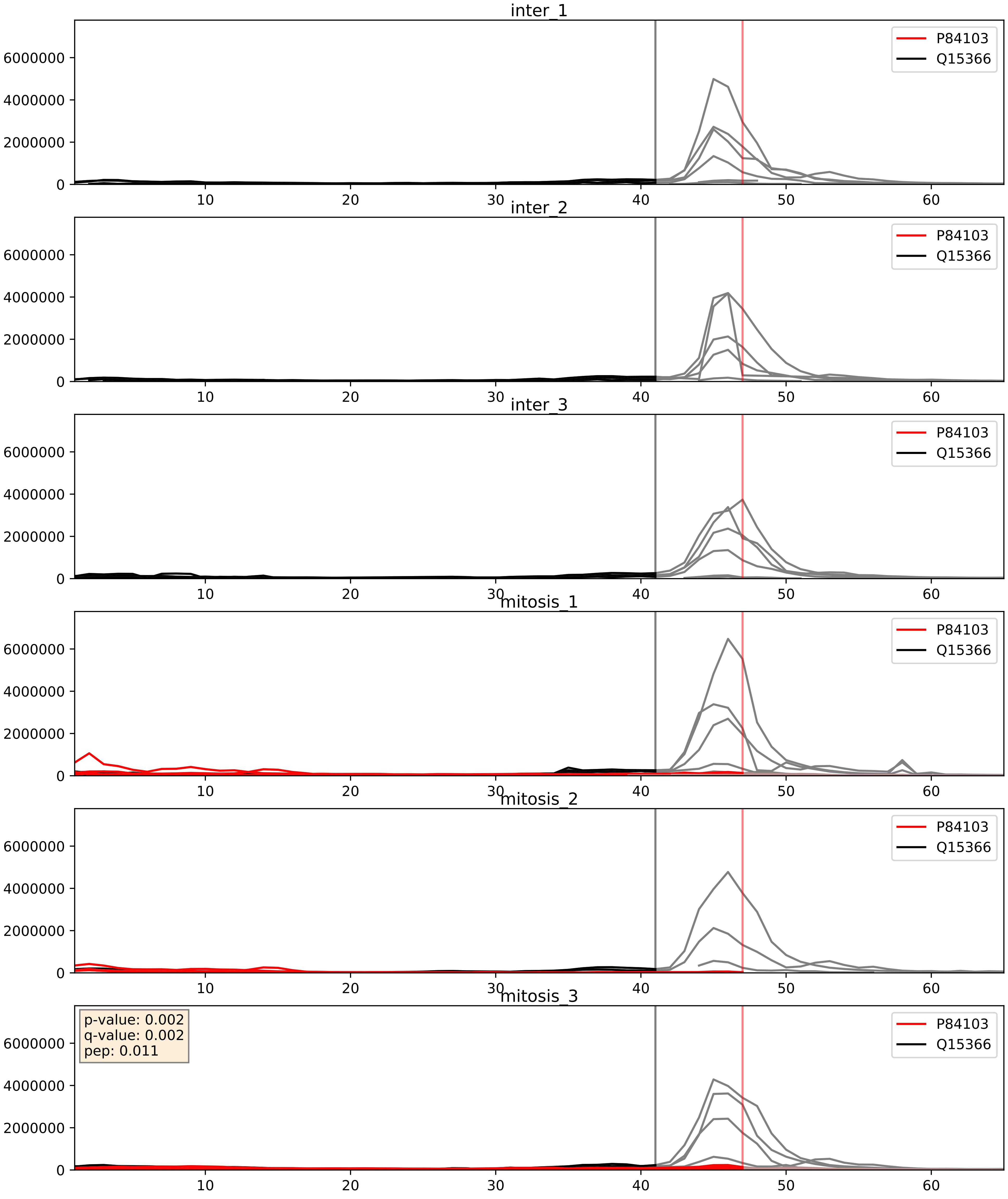


| level | condition_1 | condition_2 | pvalue | pvalue_adjusted |
|---|---|---|---|---|
| complex_abundance | mitosis | inter | 0.0015689875803000727 | 0.027691822035811595 |
| interactor_abundance | mitosis | inter | 0.003039720594488742 | 0.0421728093362545 |
| interactor_ratio | mitosis | inter | 0.6785452969895303 | 0.9339333657958224 |

P84103_Q15717
SRSF3_HUMAN vs ELAV1_HUMAN
p-value: 0.04 q-value: 0.028

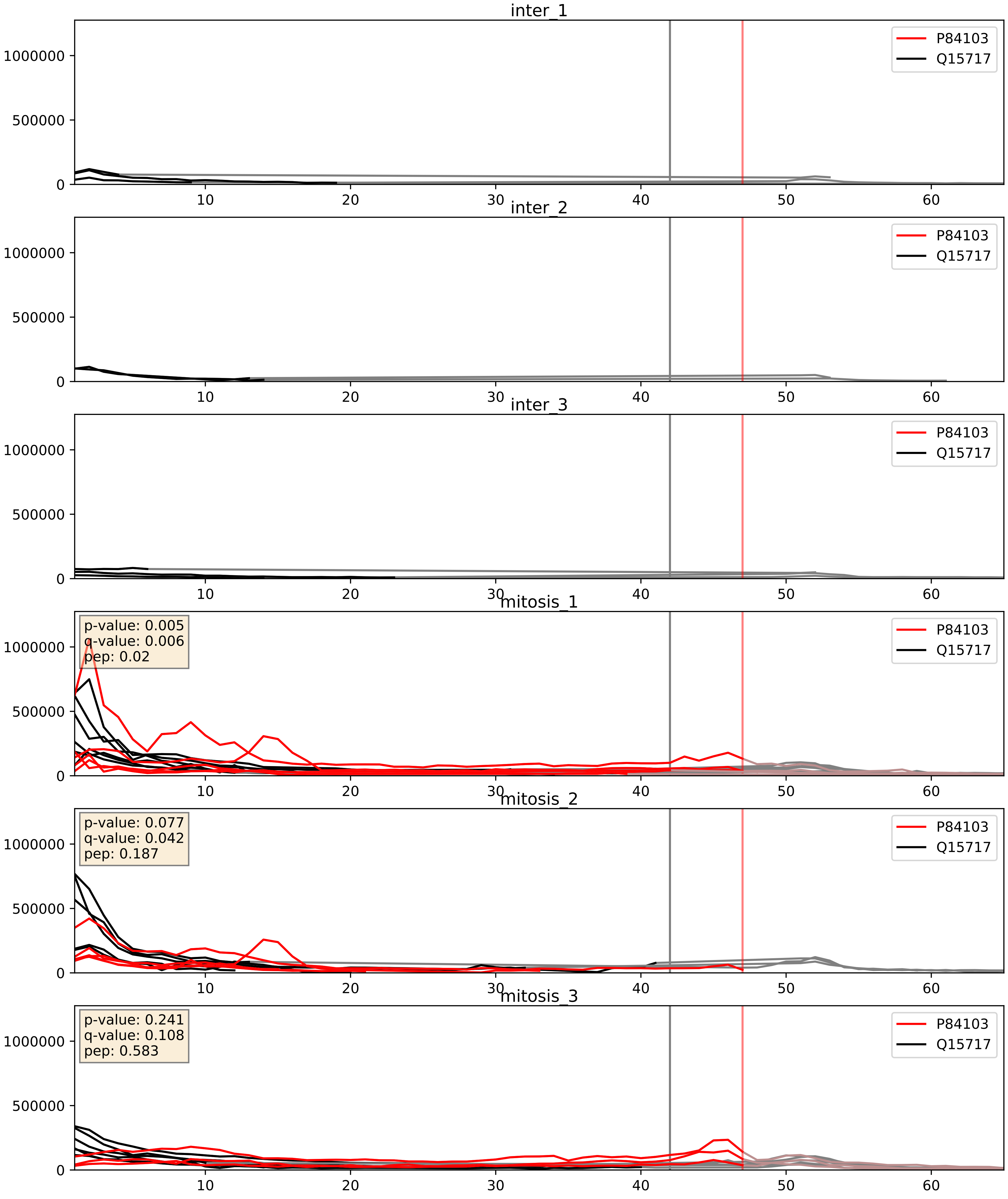


| level | condition_1 | condition_2 | pvalue | pvalue_adjusted |
|---|---|---|---|---|
| complex_abundance | mitosis | inter | 0.0005540143380544626 | 0.015065929247636585 |
| interactor_abundance | mitosis | inter | 0.0033330748758513776 | 0.04454507562418079 |
| interactor_ratio | mitosis | inter | 0.05931613769021241 | 0.40293896347252833 |

P84103_Q16629
SRSF3_HUMAN vs SRSF7_HUMAN
p-value: 0.001 q-value: 0.001

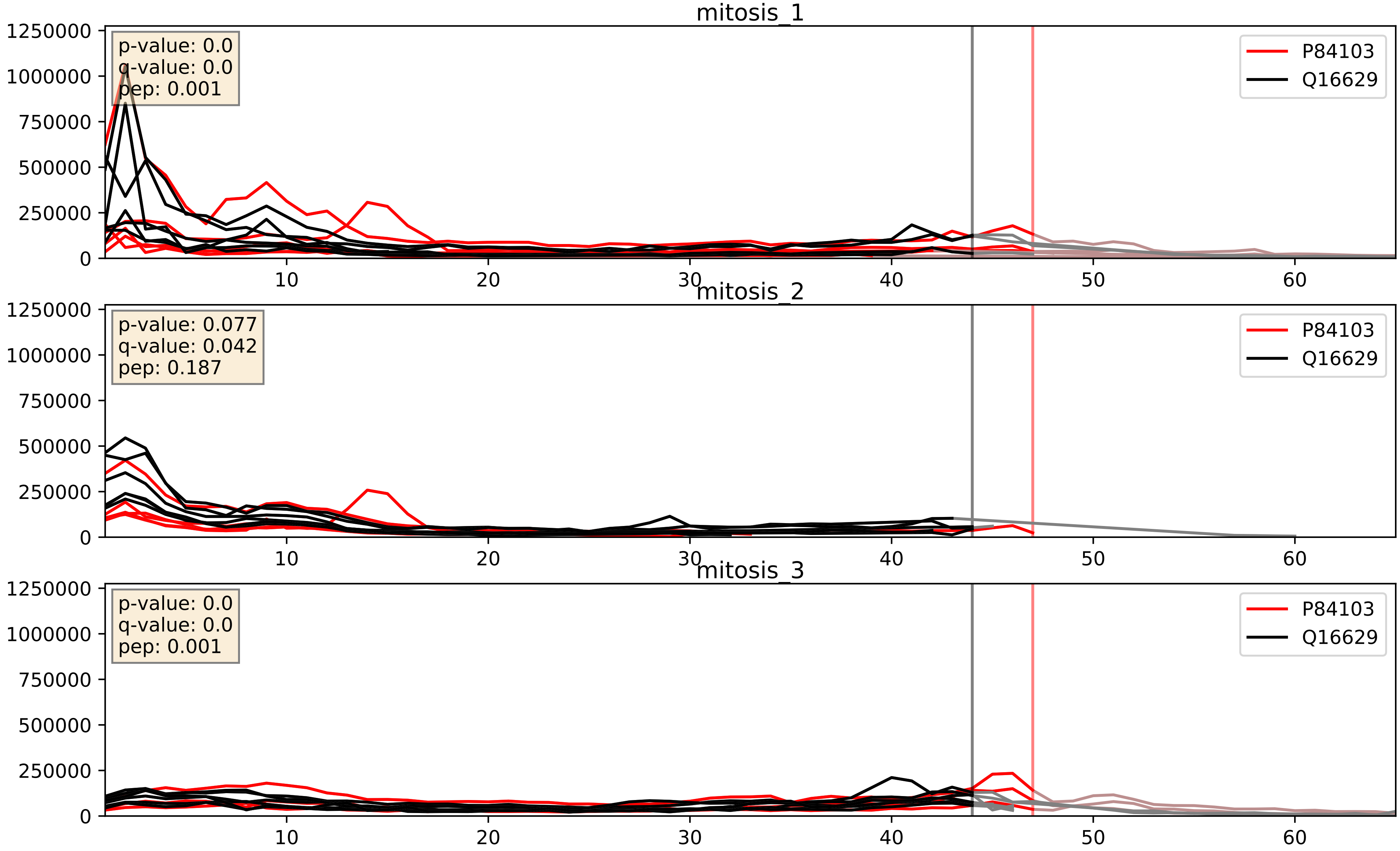


| level | condition_1 | condition_2 | pvalue | pvalue_adjusted |
|---|---|---|---|---|
| complex_abundance | mitosis | inter | 0.0003855844193721723 | 0.012502282688734073 |
| interactor_abundance | mitosis | inter | 0.002590890474122459 | 0.03806046117740333 |
| interactor_ratio | mitosis | inter | 0.026618515768801317 | 0.25614196664208316 |
