## Supplementary Data 2 for "SECAT: Quantifying differential protein-protein interaction states by network-centric analysis": hela_string_000062_P16949.pdf

P16949_P16949

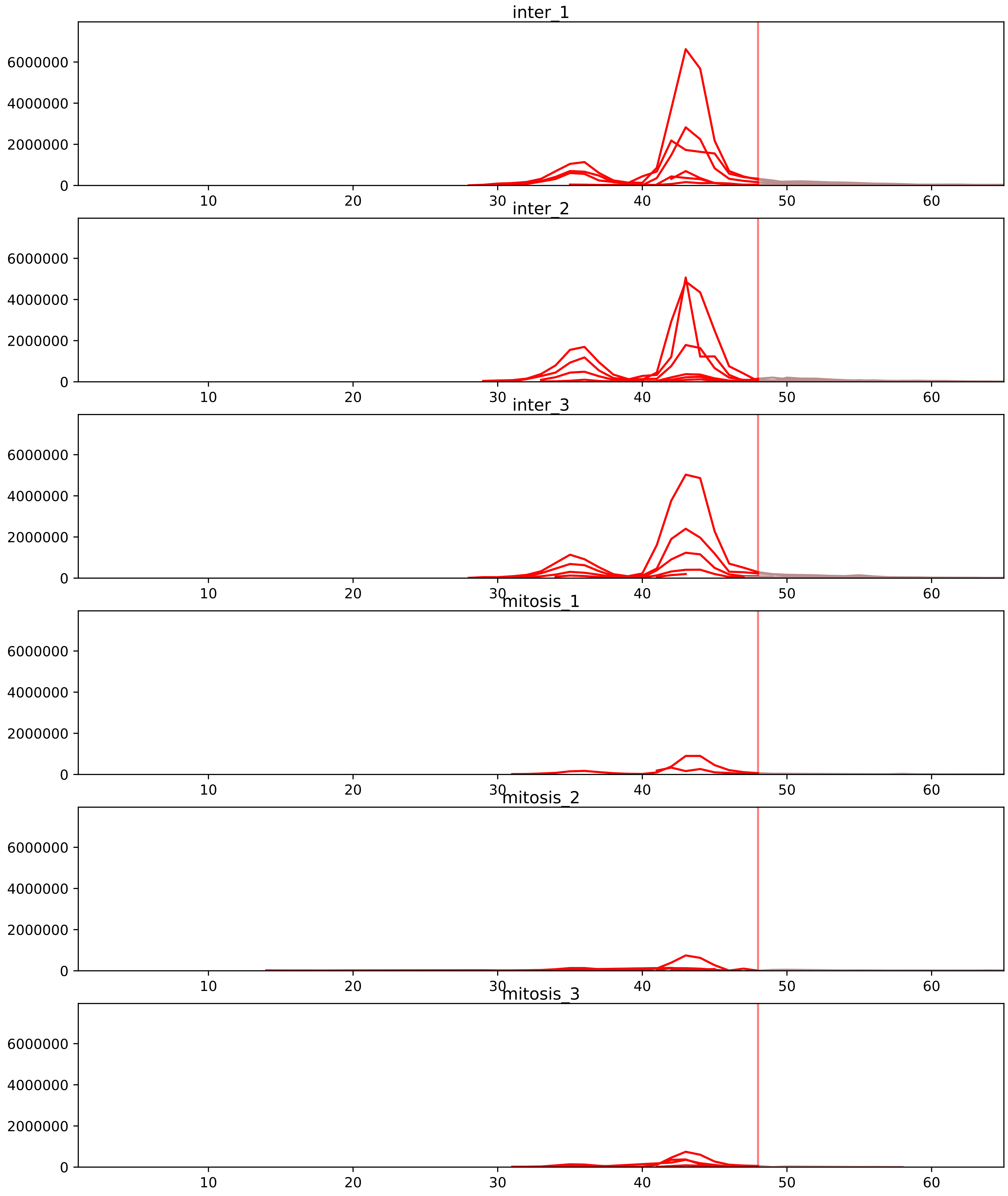


| level | condition_1 | condition_2 | pvalue | pvalue_adjusted |
|---|---|---|---|---|
| interactor_abundance | mitosis | inter | 5.684436968655596e-07 | 8.04566463255869e-05 |
| assembled_abundance | mitosis | inter | 1.952523690631252e-06 | 0.0023535084180592012 |
| interactor_ratio | mitosis | inter | 2.7221622757980846e-06 | 7.365850863924229e-05 |
| total_abundance | mitosis | inter | 4.605053659334374e-06 | 0.003540864511957668 |
| complex_abundance | mitosis | inter | 1.9182955666344685e-05 | 0.0007509923069377493 |
| monomer_abundance | mitosis | inter | 0.0006120182014698664 | 0.06224773594363797 |

P16949_Q99497
STMN1_HUMAN vs PARK7_HUMAN
p-value: 0.0 q-value: 0.038

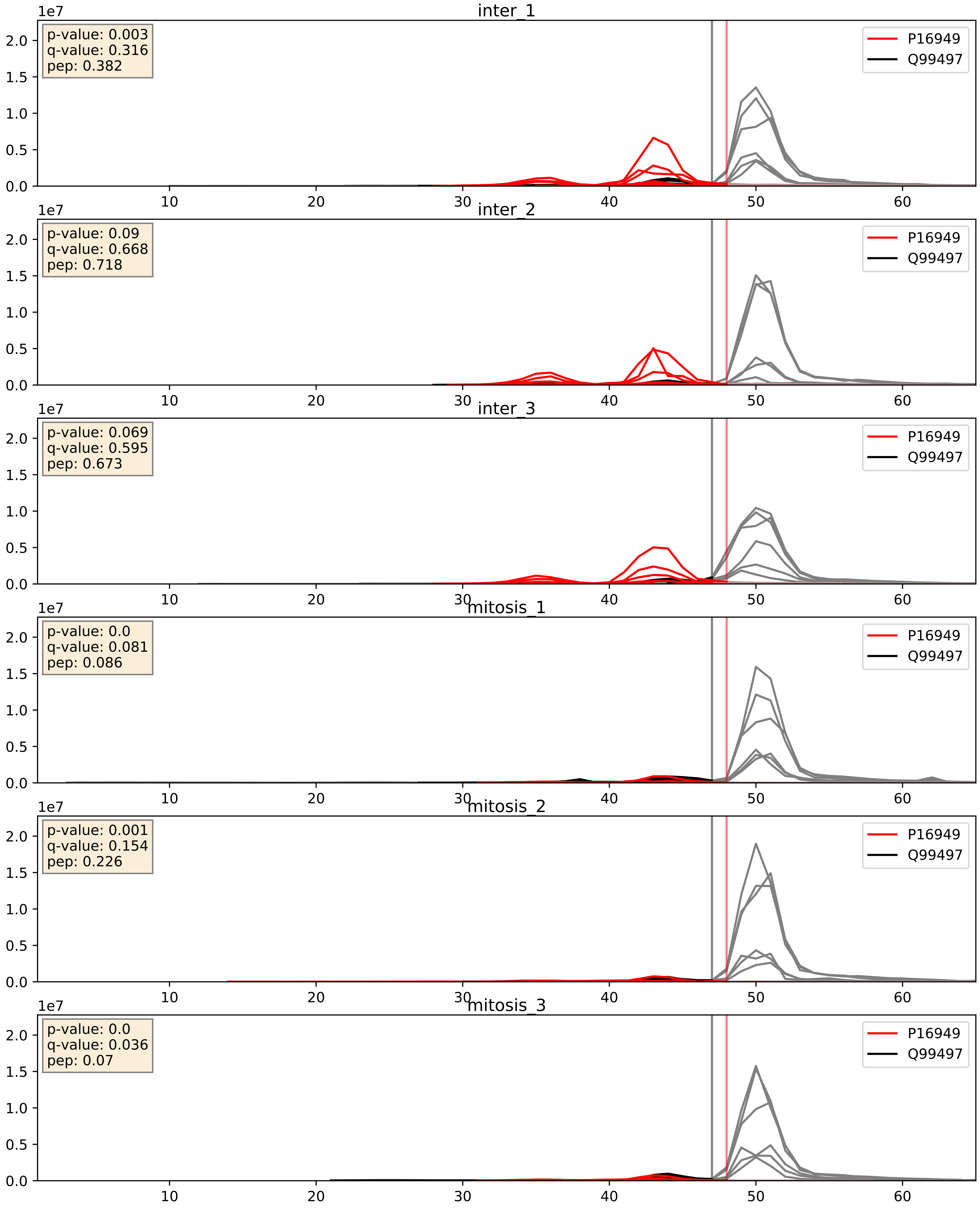


| level | condition_1 | condition_2 | pvalue | pvalue_adjusted |
|---|---|---|---|---|
| interactor_abundance | mitosis | inter | 1.1380048322495522e-05 | 0.0033803141163598503 |
| interactor_ratio | mitosis | inter | 7.190770405545085e-05 | 0.008863126489088583 |
| complex_abundance | mitosis | inter | 0.0003066083054506631 | 0.01165207110963347 |

P16949_Q9H910
STMN1_HUMAN vs JUPI2_HUMAN
p-value: 0.001 q-value: 0.045

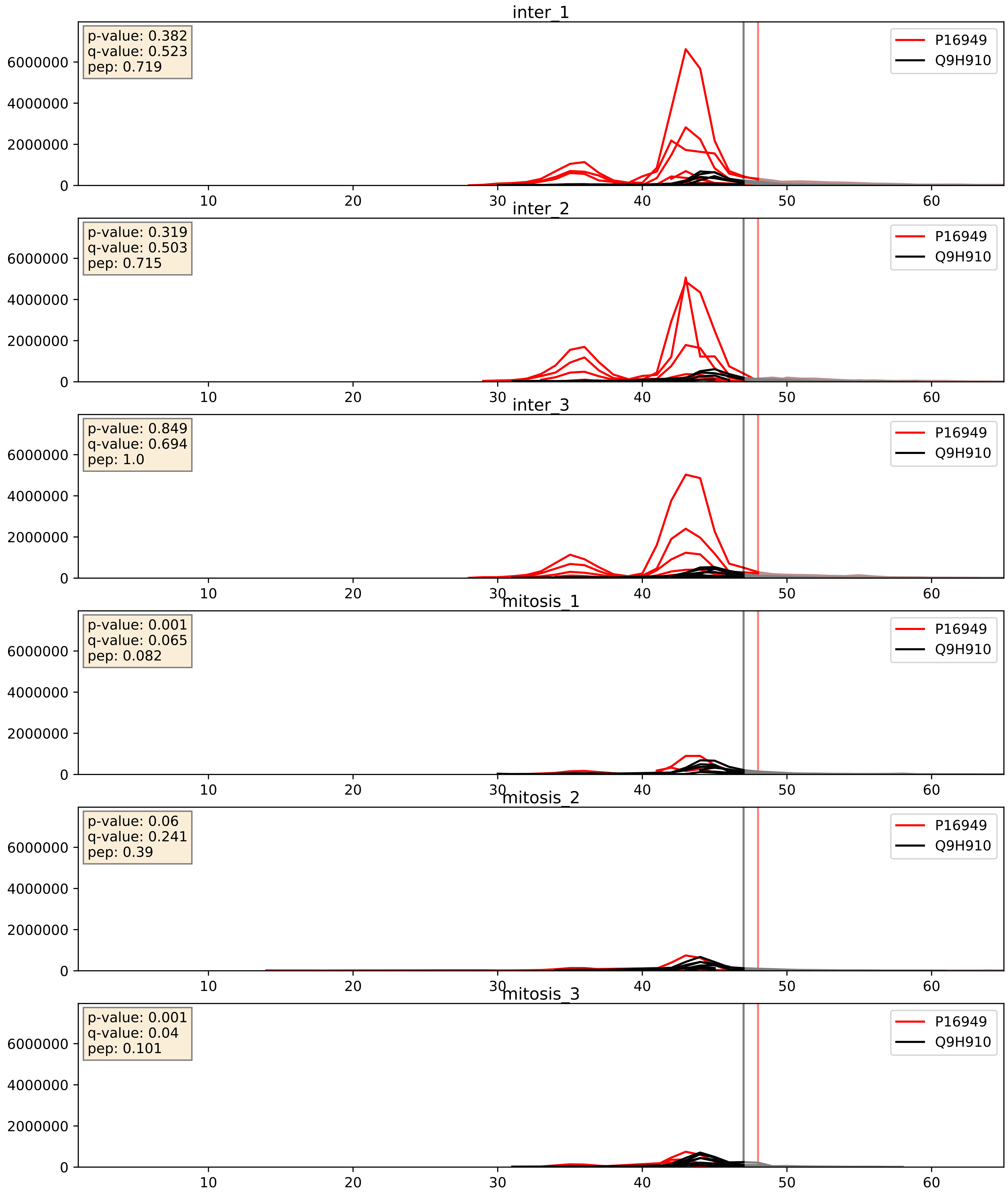


| level | condition_1 | condition_2 | pvalue | pvalue_adjusted |
|---|---|---|---|---|
| interactor_abundance | mitosis | inter | 1.5772574337866112e-05 | 0.0037089335794386596 |
| interactor_ratio | mitosis | inter | 3.7446357313653444e-05 | 0.00641081637209747 |
| complex_abundance | mitosis | inter | 0.00010635052484114434 | 0.0070027730203091965 |
