## Supplementary Data 2 for "SECAT: Quantifying differential protein-protein interaction states by network-centric analysis": hela_string_000063_O76021.pdf

O76021_O76021

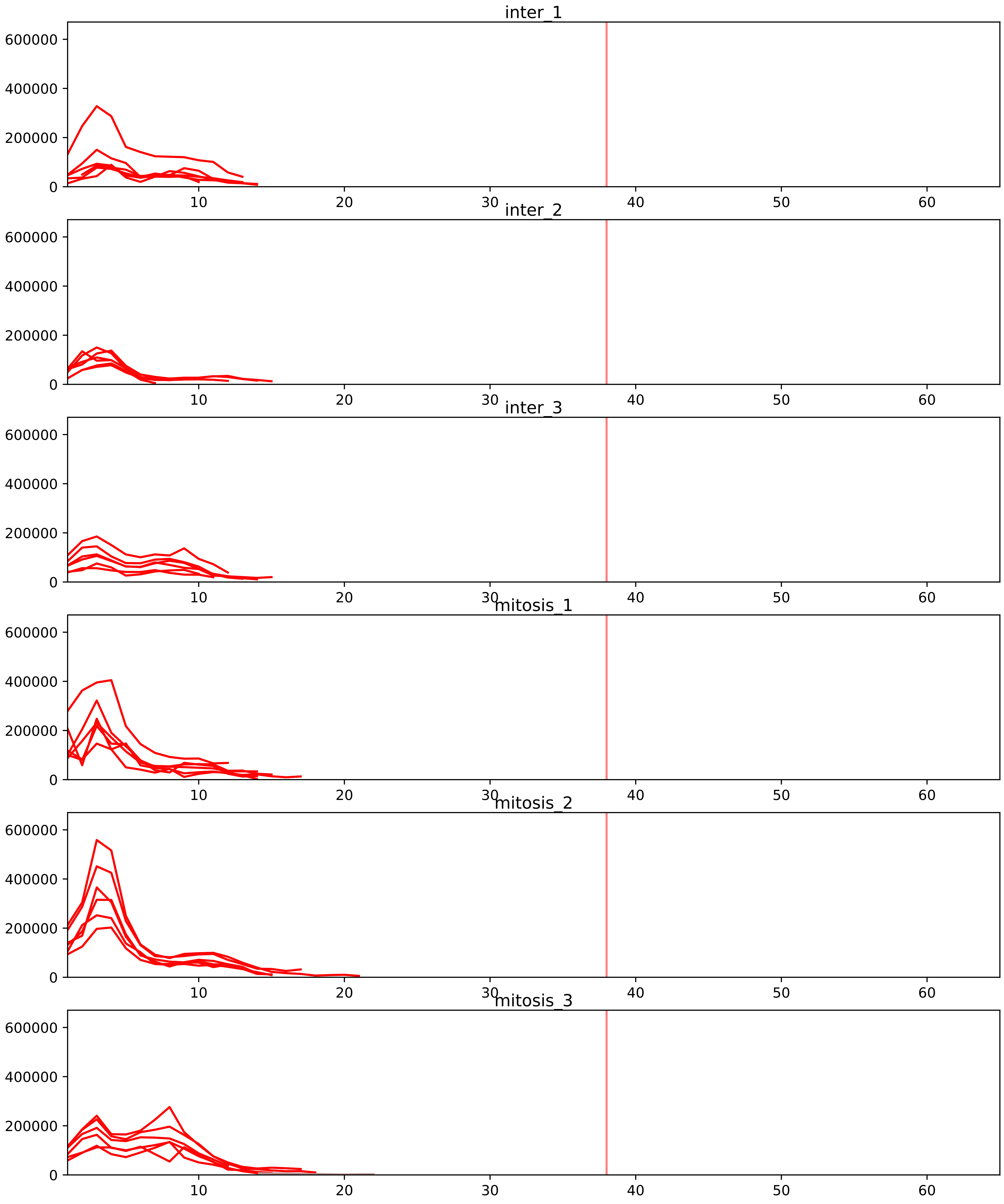


| level | condition_1 | condition_2 | pvalue | pvalue_adjusted |
|---|---|---|---|---|
| interactor_ratio | mitosis | inter | 6.070063786131054e-07 | 1.8930368417764642e-05 |
| complex_abundance | mitosis | inter | 2.816744772302714e-06 | 0.00018666475235760692 |
| interactor_abundance | mitosis | inter | 2.0151461781199695e-05 | 0.0008828259447001771 |
| total_abundance | mitosis | inter | 0.006773744646397593 | 0.09686835391800604 |
| assembled_abundance | mitosis | inter | 0.00794319085537495 | 0.09552127543439592 |

O00567_O76021
NOP56_HUMAN vs RL1D1_HUMAN
p-value: 0.001 q-value: 0.001

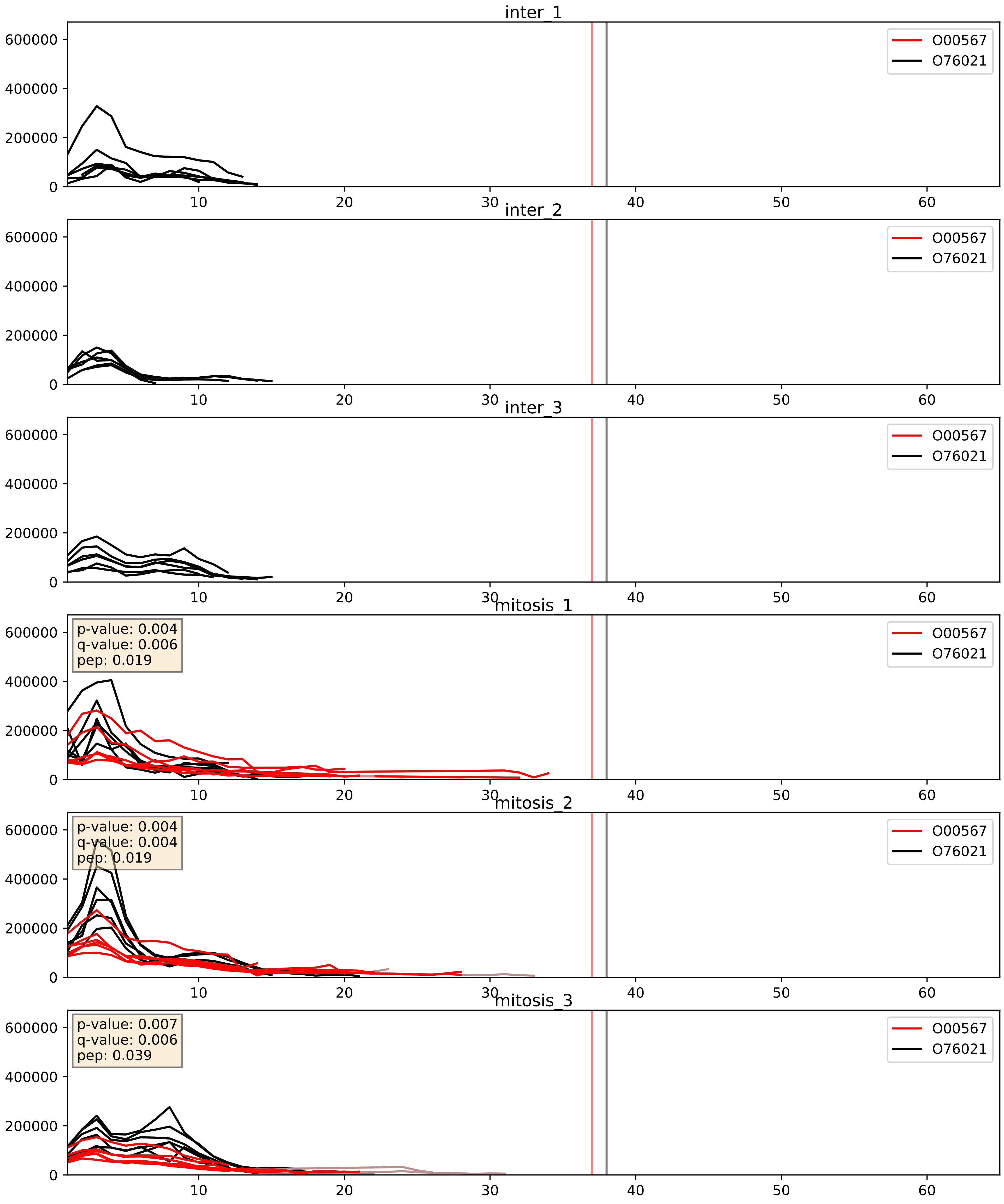


| level | condition_1 | condition_2 | pvalue | pvalue_adjusted |
|---|---|---|---|---|
| complex_abundance | mitosis | inter | 0.00010088212909086971 | 0.006746492382951911 |
| interactor_abundance | mitosis | inter | 0.0009352754344708583 | 0.022711936791689494 |
| interactor_ratio | mitosis | inter | 0.13519290058957723 | 0.5858060166978465 |

O75683_O76021
SURF6_HUMAN vs RL1D1_HUMAN
p-value: 0.0 q-value: 0.023

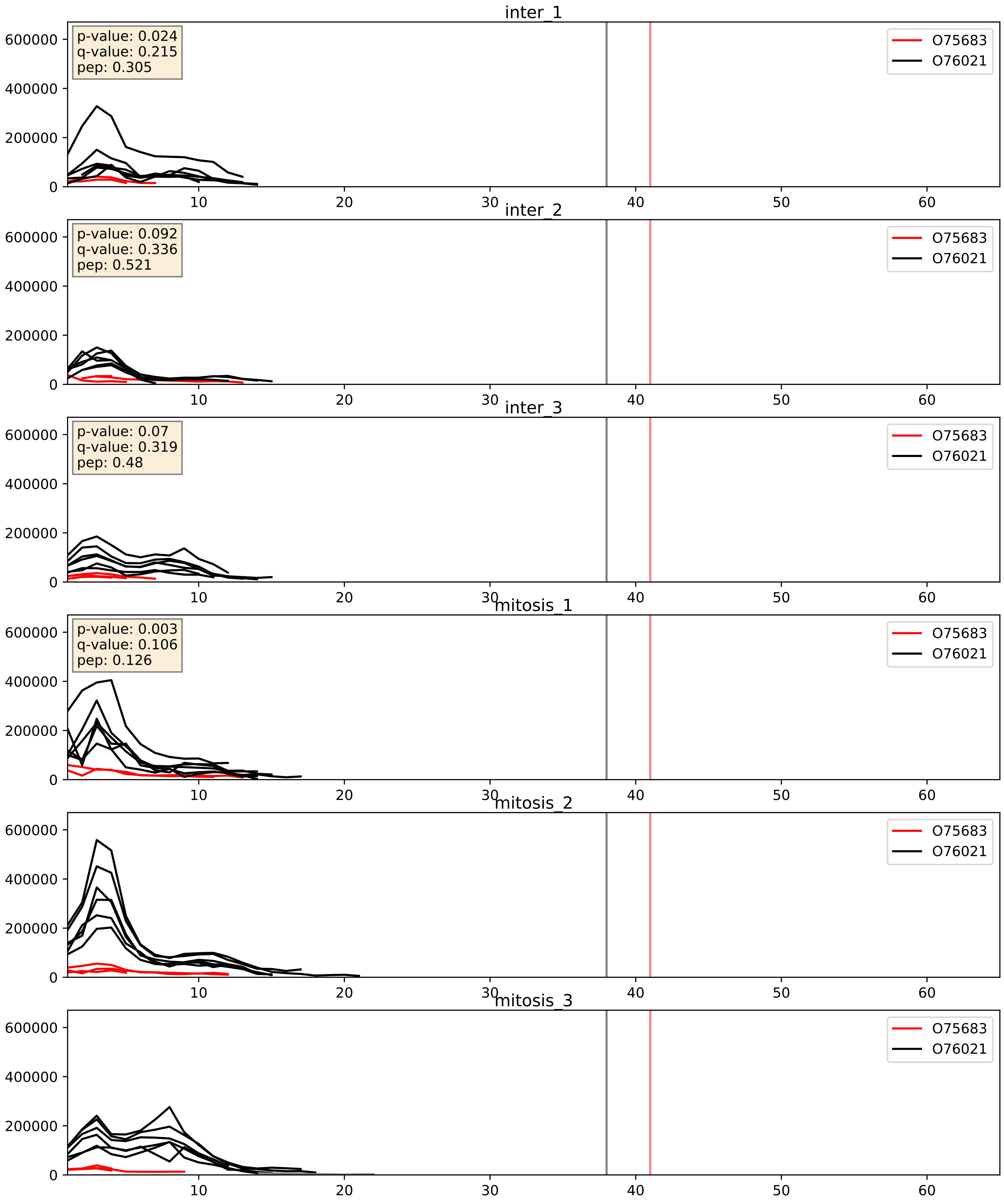


| level | condition_1 | condition_2 | pvalue | pvalue_adjusted |
|---|---|---|---|---|
| interactor_ratio | mitosis | inter | 0.0031769021054309015 | 0.07369724125335642 |
| interactor_abundance | mitosis | inter | 0.06023012635531873 | 0.20080618562863808 |
| complex_abundance | mitosis | inter | 0.11262448179668066 | 0.25199910682811305 |

O76021_P22087
RL1D1_HUMAN vs FBRL_HUMAN
p-value: 0.003 q-value: 0.004

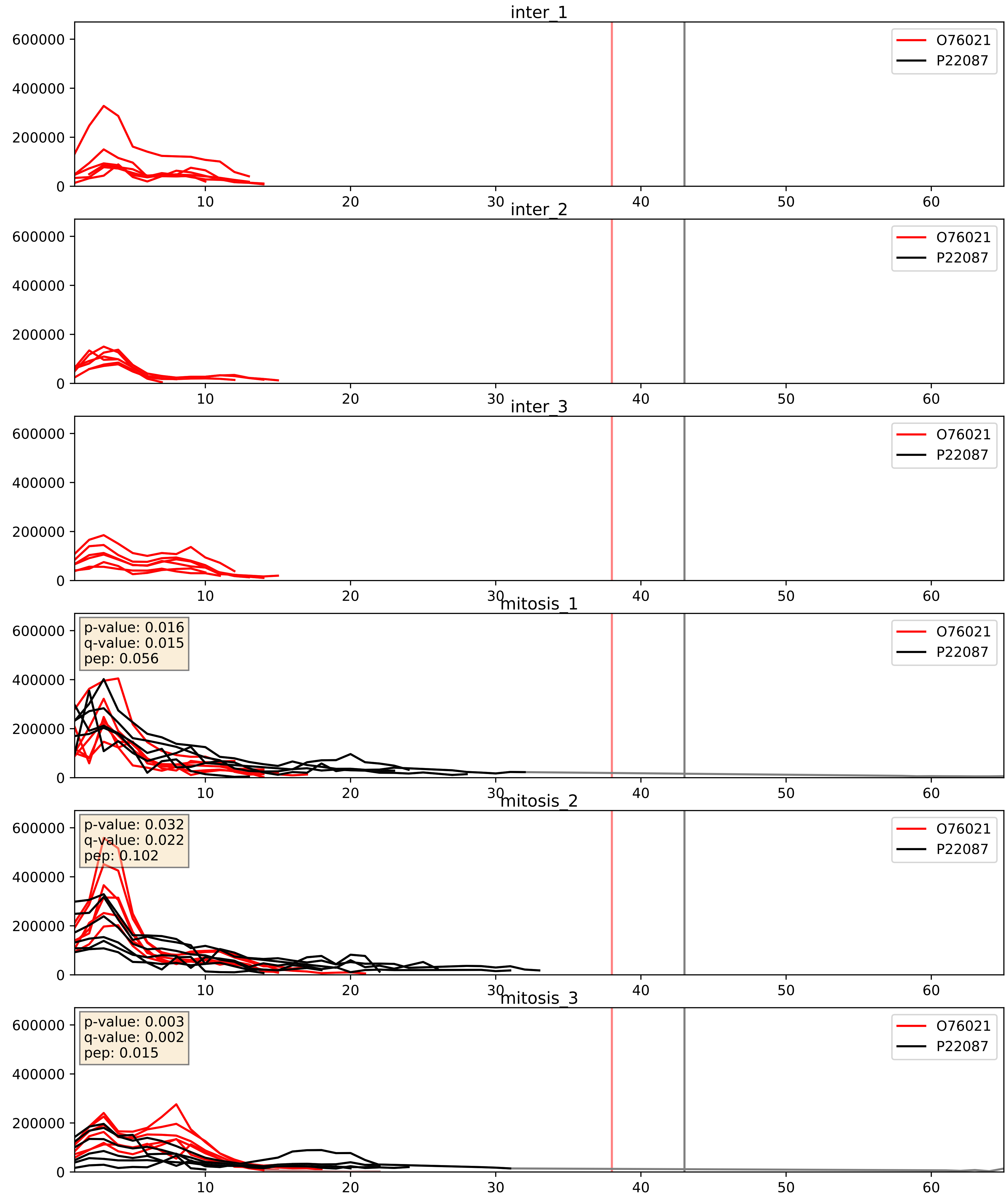


| level | condition_1 | condition_2 | pvalue | pvalue_adjusted |
|---|---|---|---|---|
| complex_abundance | mitosis | inter | 7.351496278048426e-05 | 0.005893640694944935 |
| interactor_abundance | mitosis | inter | 8.109392026765088e-05 | 0.007119630332419645 |
| interactor_ratio | mitosis | inter | 0.9974833049507463 | 0.9996442035996809 |

O76021_P42766
RL1D1_HUMAN vs RL35_HUMAN
p-value: 0.0 q-value: 0.041

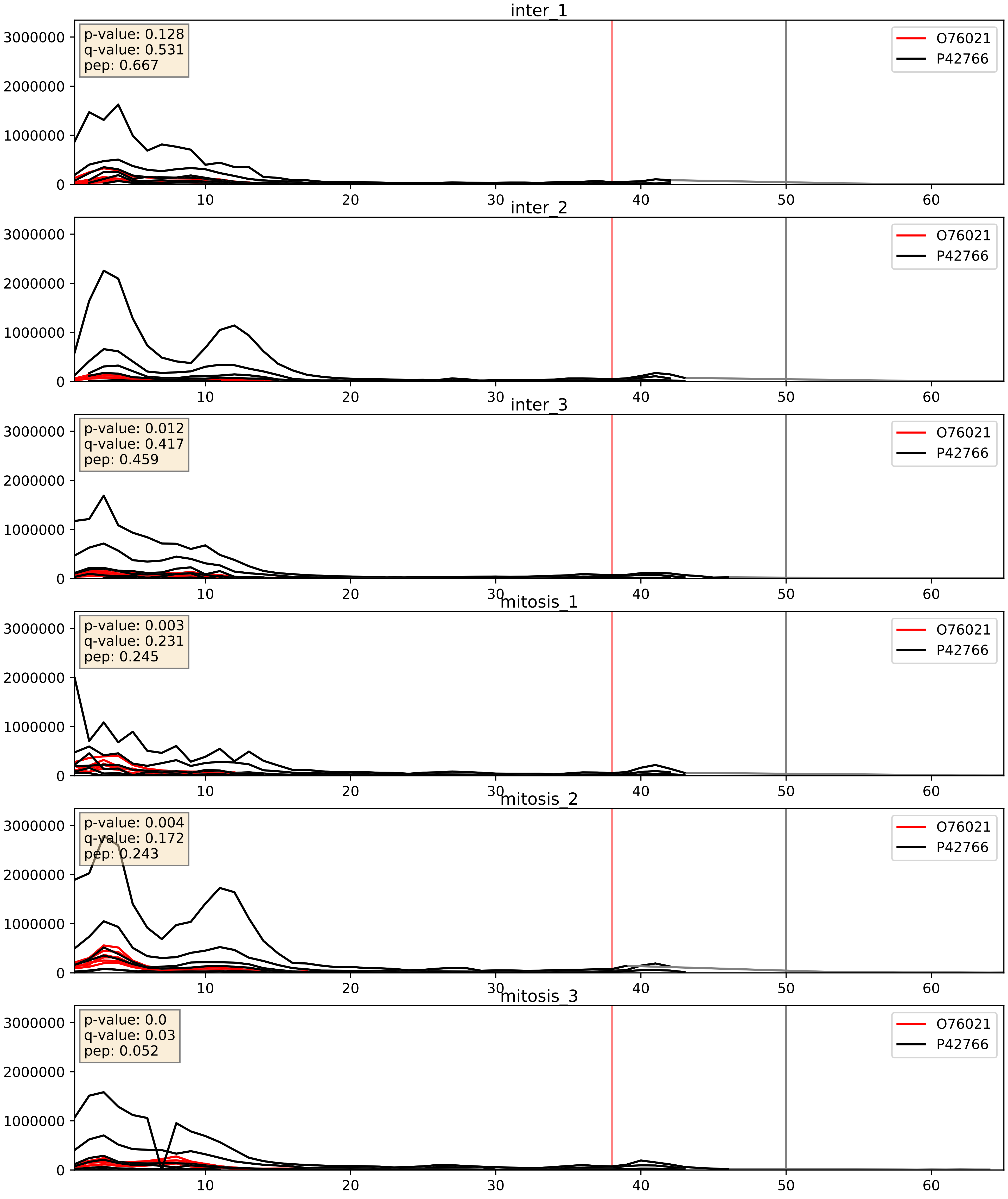


| level | condition_1 | condition_2 | pvalue | pvalue_adjusted |
|---|---|---|---|---|
| interactor_abundance | mitosis | inter | 0.007302544007558834 | 0.07362753439894419 |
| interactor_ratio | mitosis | inter | 0.028800205157091852 | 0.2673858526515252 |
| complex_abundance | mitosis | inter | 0.04911757511194777 | 0.15812201690796274 |

O76021_P46087
RL1D1_HUMAN vs NOP2_HUMAN
p-value: 0.018 q-value: 0.014

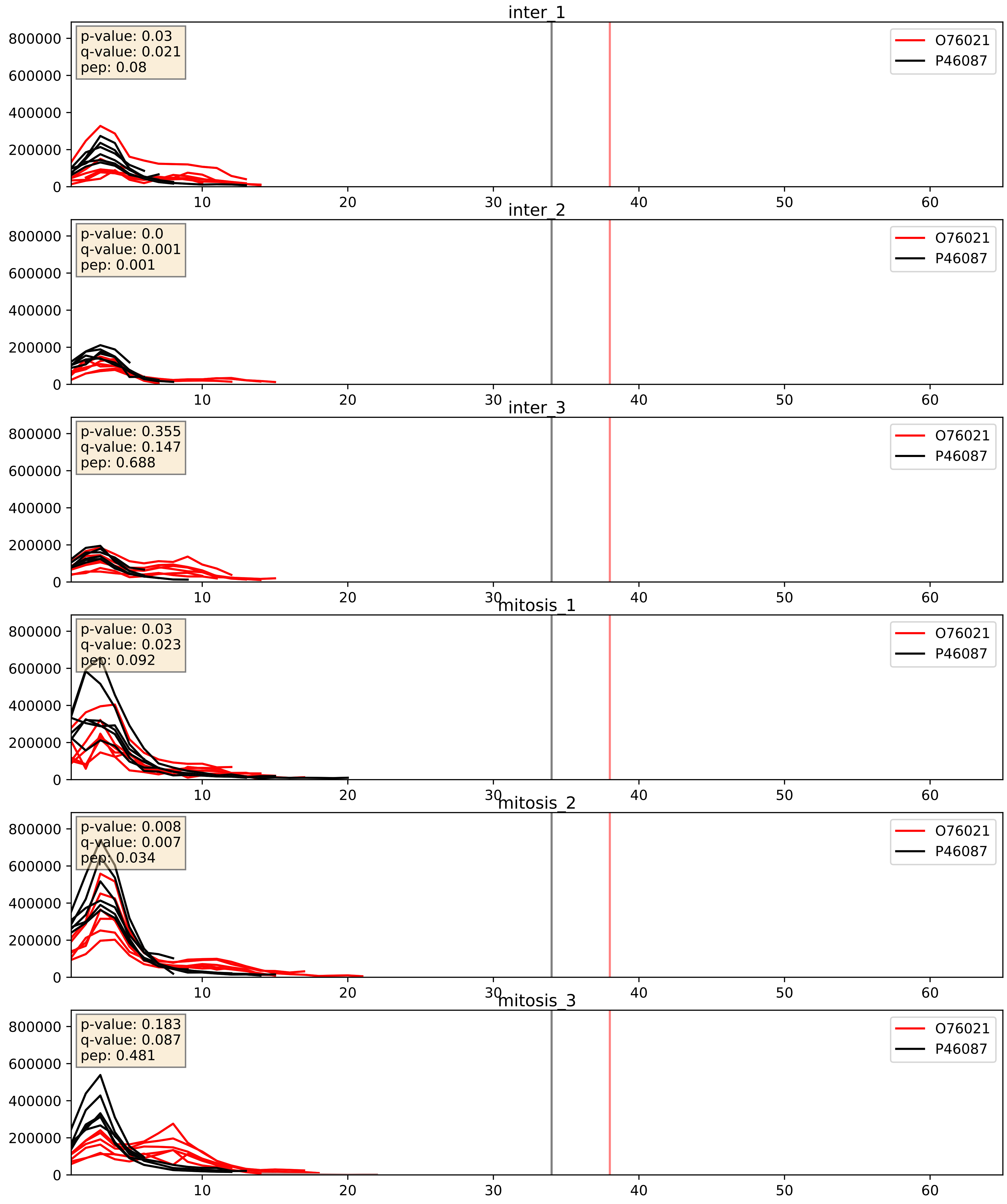


| level | condition_1 | condition_2 | pvalue | pvalue_adjusted |
|---|---|---|---|---|
| complex_abundance | mitosis | inter | 0.0026377819363957414 | 0.03679993741510981 |
| interactor_abundance | mitosis | inter | 0.007992044606047556 | 0.07749253525281795 |
| interactor_ratio | mitosis | inter | 0.4410474716757986 | 0.8466845385837265 |

O76021_P46776
RL1D1_HUMAN vs RL27A_HUMAN
p-value: 0.0 q-value: 0.043

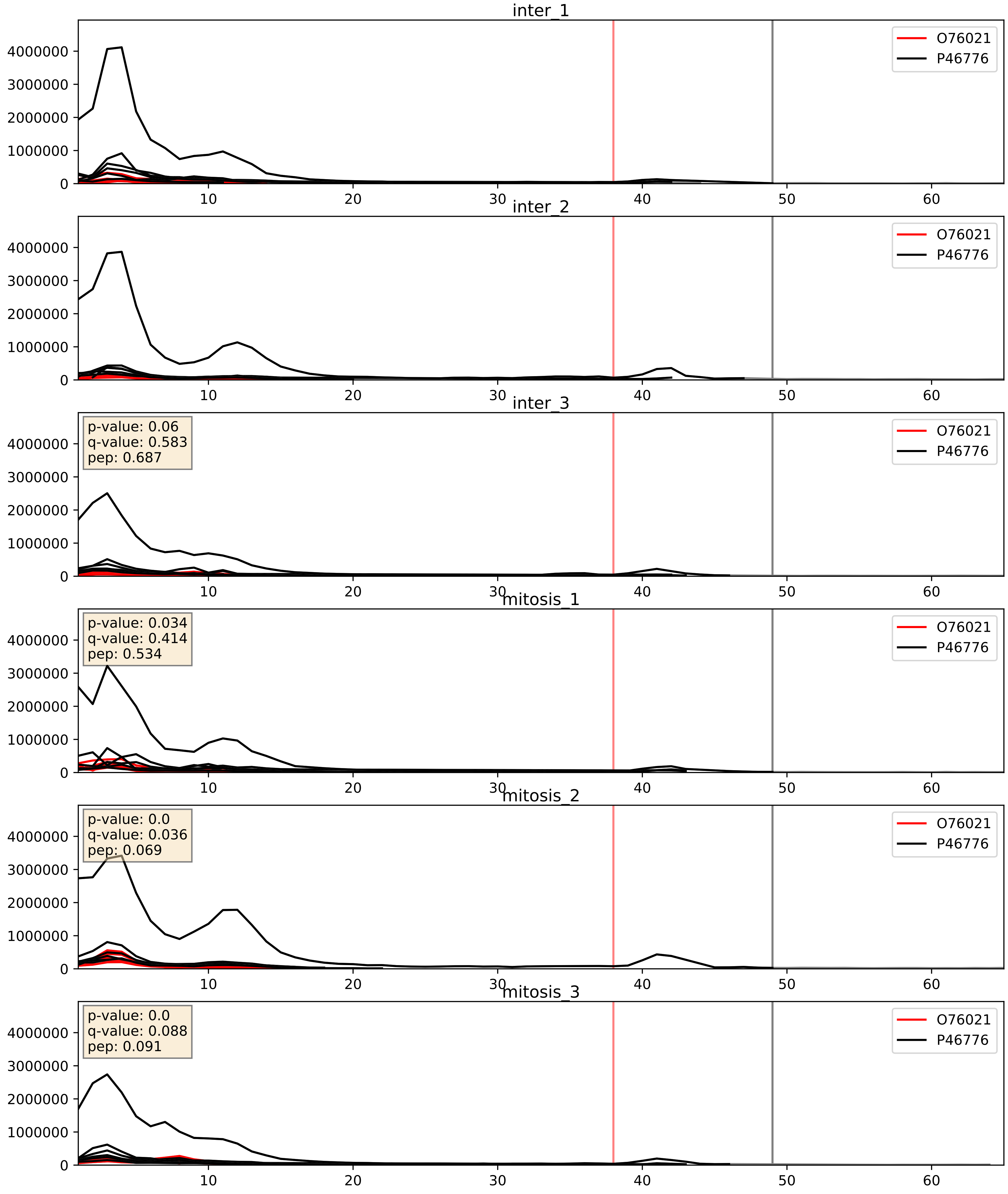


| level | condition_1 | condition_2 | pvalue | pvalue_adjusted |
|---|---|---|---|---|
| interactor_abundance | mitosis | inter | 0.007302544007558834 | 0.07362753439894419 |
| interactor_ratio | mitosis | inter | 0.010401885614920456 | 0.14681567707444748 |
| complex_abundance | mitosis | inter | 0.038069591788847876 | 0.1424797457329857 |

O76021_P55769
RL1D1_HUMAN vs NH2L1_HUMAN
p-value: 0.0 q-value: 0.013

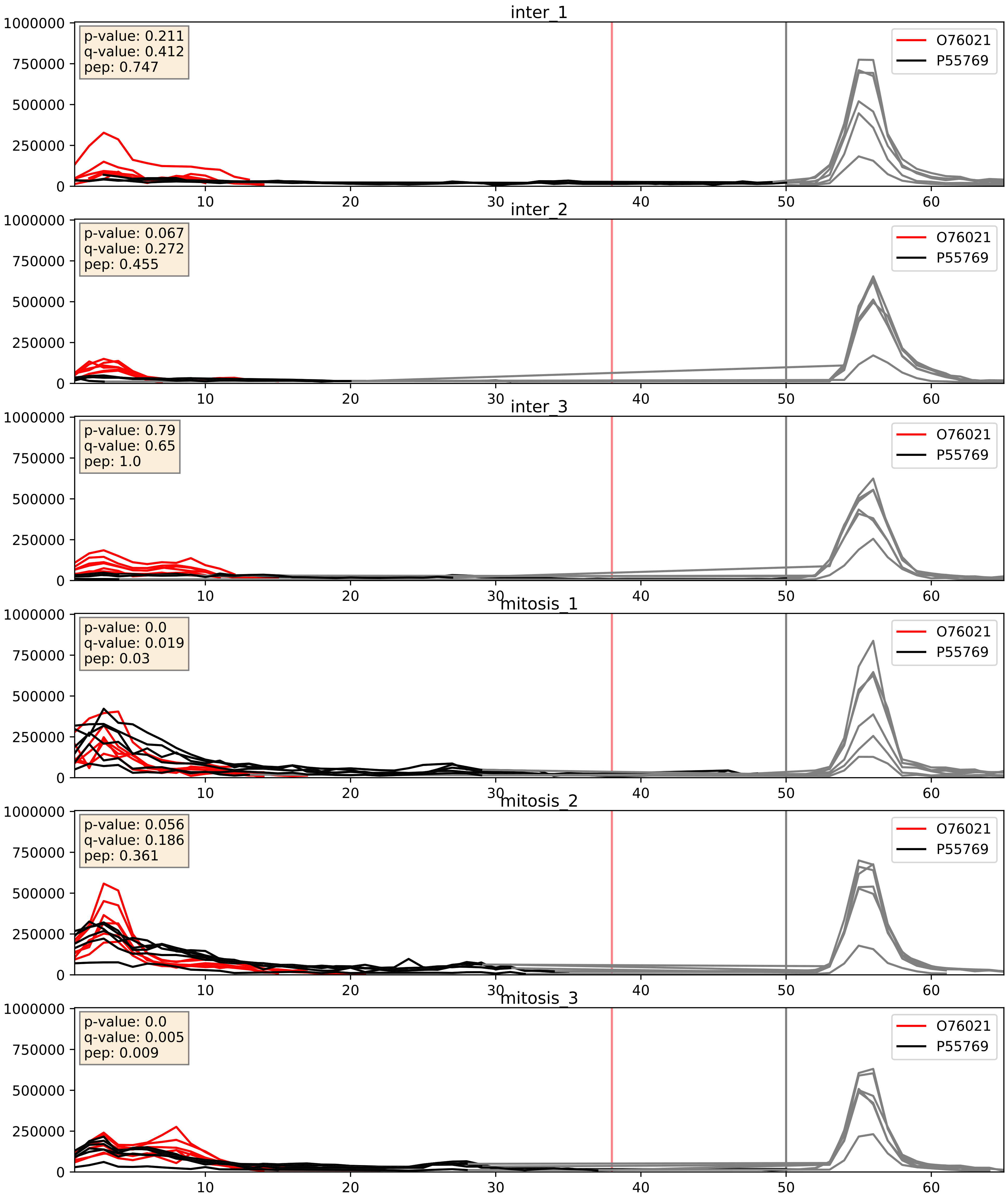


| level | condition_1 | condition_2 | pvalue | pvalue_adjusted |
|---|---|---|---|---|
| complex_abundance | mitosis | inter | 0.0010308703241052963 | 0.021560581032375372 |
| interactor_abundance | mitosis | inter | 0.007302544007558834 | 0.07362753439894419 |
| interactor_ratio | mitosis | inter | 0.018751790484042963 | 0.20631790044139817 |

O76021_P56537
RL1D1_HUMAN vs IF6_HUMAN
p-value: 0.001 q-value: 0.039

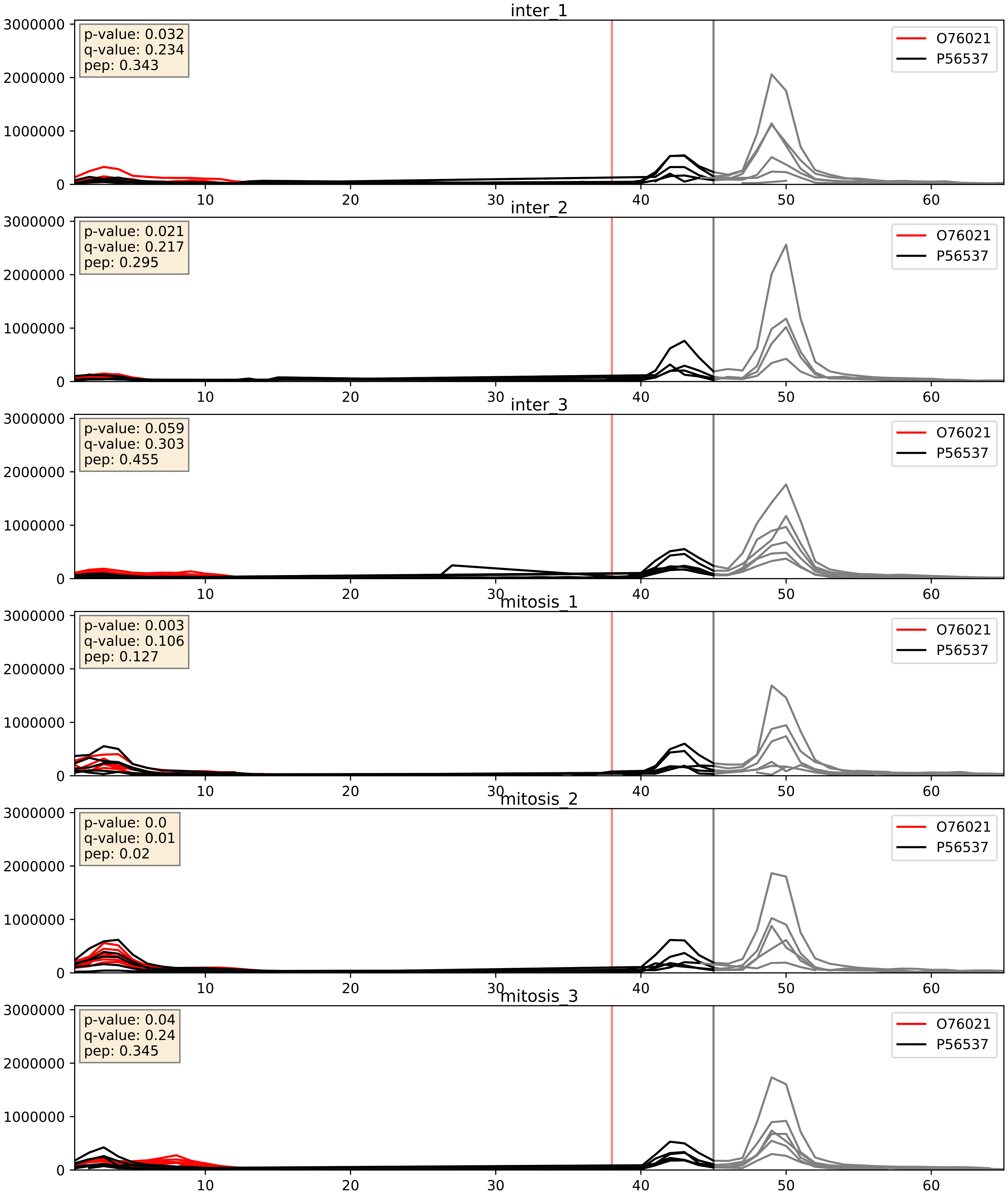


| level | condition_1 | condition_2 | pvalue | pvalue_adjusted |
|---|---|---|---|---|
| complex_abundance | mitosis | inter | 0.001587159867408426 | 0.027897512248493073 |
| interactor_abundance | mitosis | inter | 0.007302544007558834 | 0.07362753439894419 |
| interactor_ratio | mitosis | inter | 0.17049014932935702 | 0.6336933036297422 |

O76021_P62081
RL1D1_HUMAN vs RS7_HUMAN
p-value: 0.002 q-value: 0.03

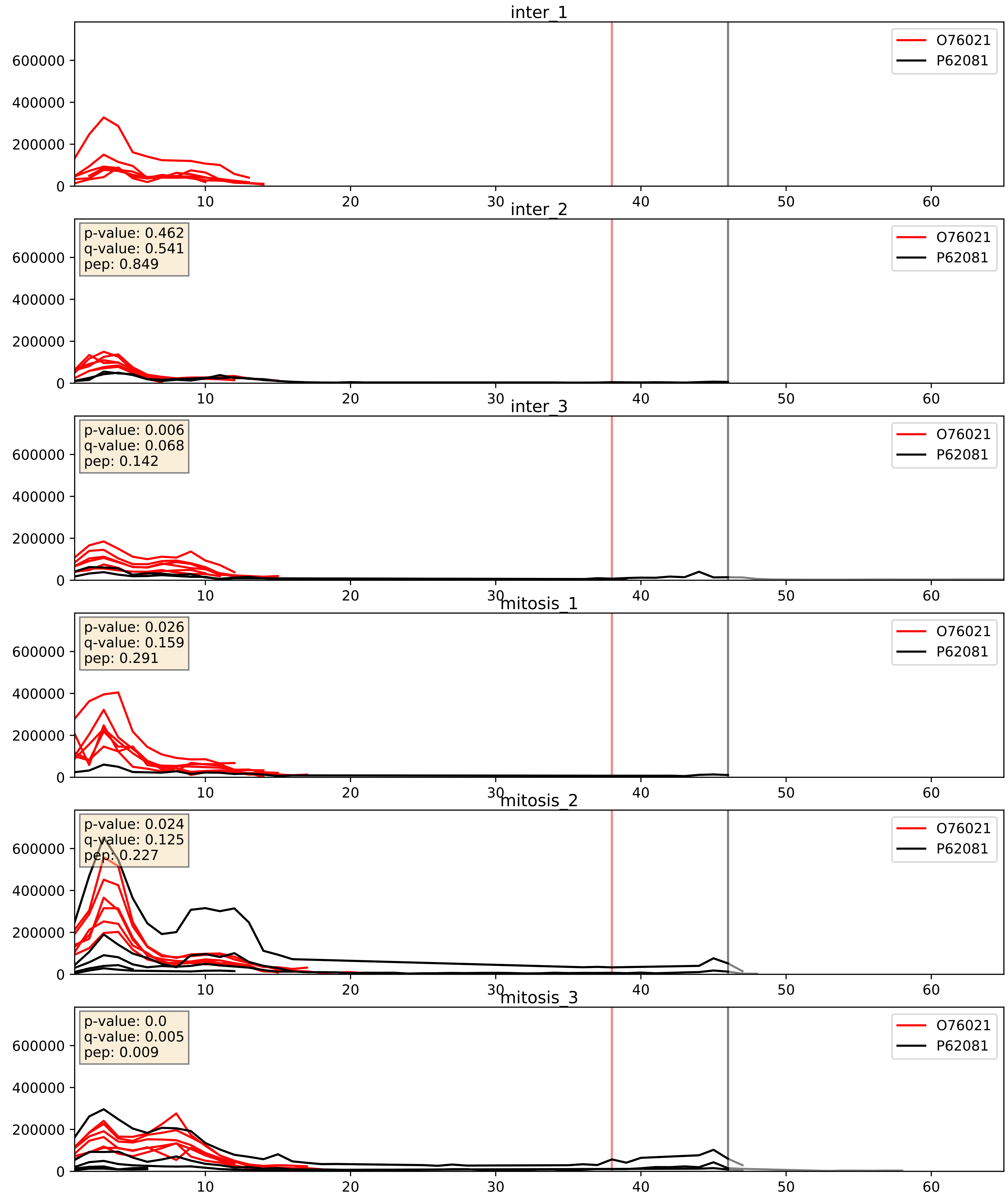


| level | condition_1 | condition_2 | pvalue | pvalue_adjusted |
|---|---|---|---|---|
| interactor_abundance | mitosis | inter | 0.08189804567341344 | 0.23942871276107208 |
| complex_abundance | mitosis | inter | 0.0997567739797323 | 0.2344092031621868 |
| interactor_ratio | mitosis | inter | 0.5421276788780396 | 0.8859730852936224 |

O76021_P62266
RL1D1_HUMAN vs RS23_HUMAN
p-value: 0.0 q-value: 0.017

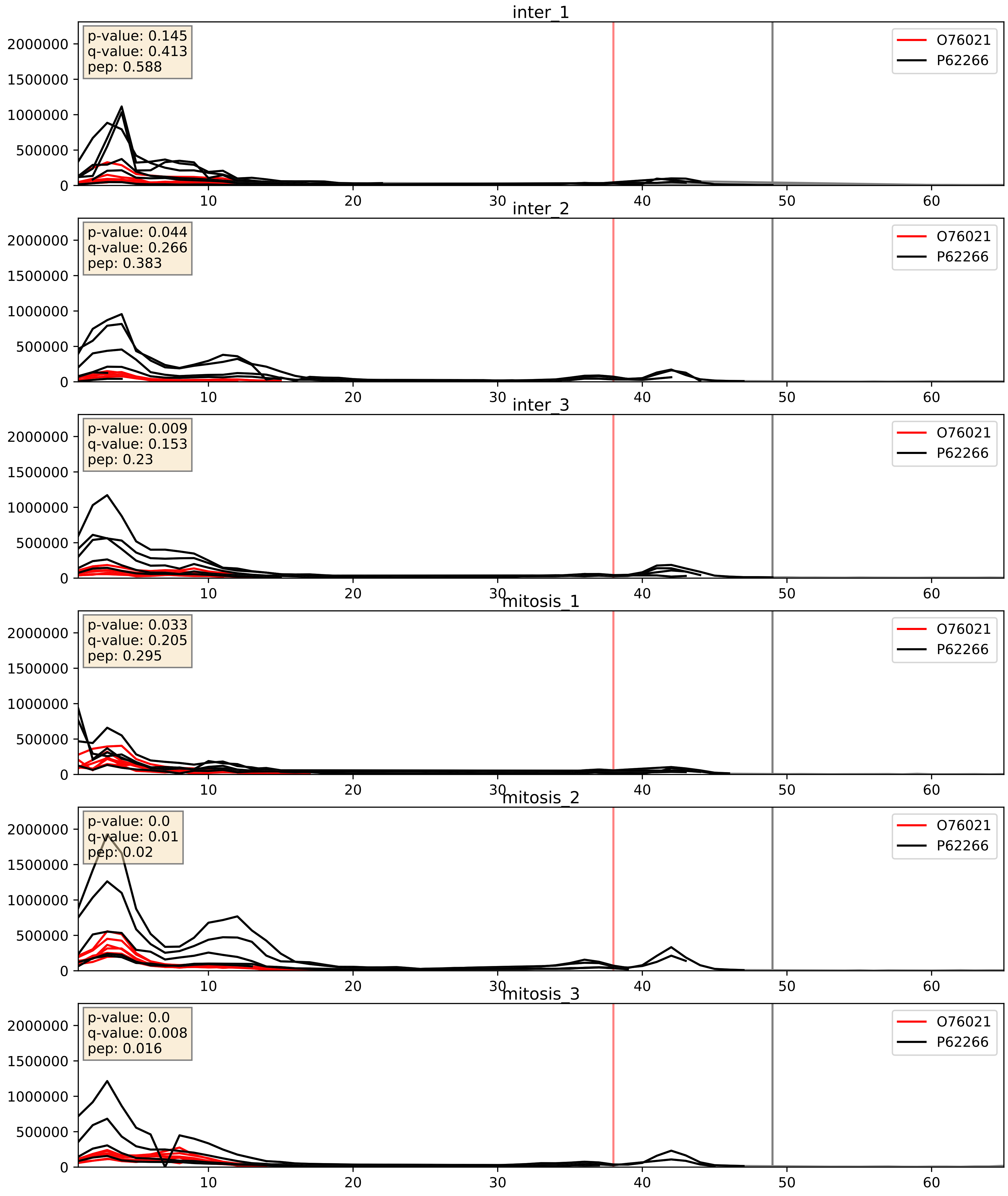


| level | condition_1 | condition_2 | pvalue | pvalue_adjusted |
|---|---|---|---|---|
| interactor_abundance | mitosis | inter | 0.007302544007558834 | 0.07362753439894419 |
| interactor_ratio | mitosis | inter | 0.07994343343495322 | 0.4732286766833245 |
| complex_abundance | mitosis | inter | 0.8426589559678248 | 0.8997331499419459 |

O76021_P62750
RL1D1_HUMAN vs RL23A_HUMAN
p-value: 0.0 q-value: 0.022

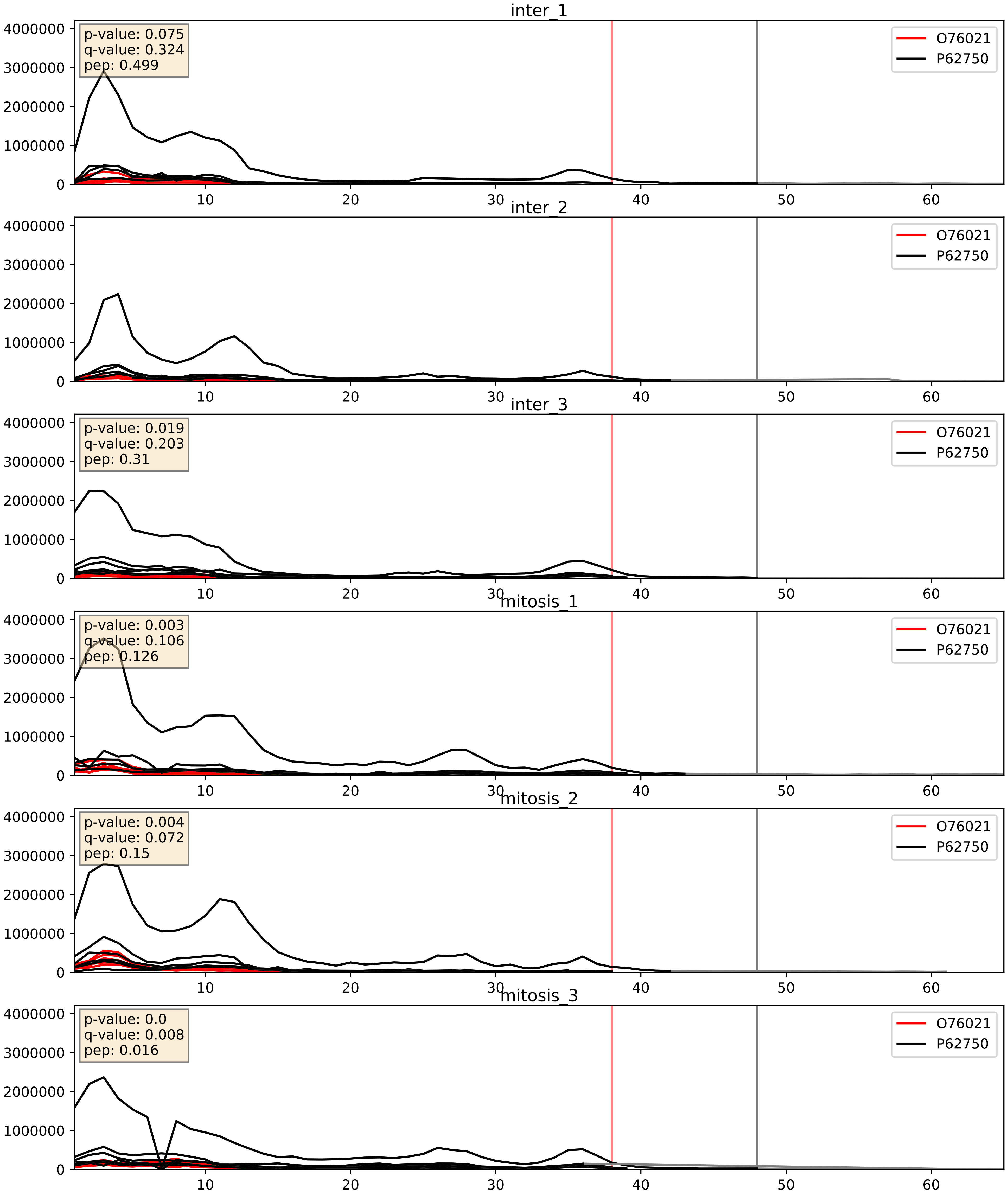


| level | condition_1 | condition_2 | pvalue | pvalue_adjusted |
|---|---|---|---|---|
| interactor_abundance | mitosis | inter | 0.007302544007558834 | 0.07362753439894419 |
| complex_abundance | mitosis | inter | 0.019907099219111473 | 0.10473556811038366 |
| interactor_ratio | mitosis | inter | 0.04588133509498198 | 0.35622692584467375 |

O76021_P62829
RL1D1_HUMAN vs RL23_HUMAN
p-value: 0.0 q-value: 0.041

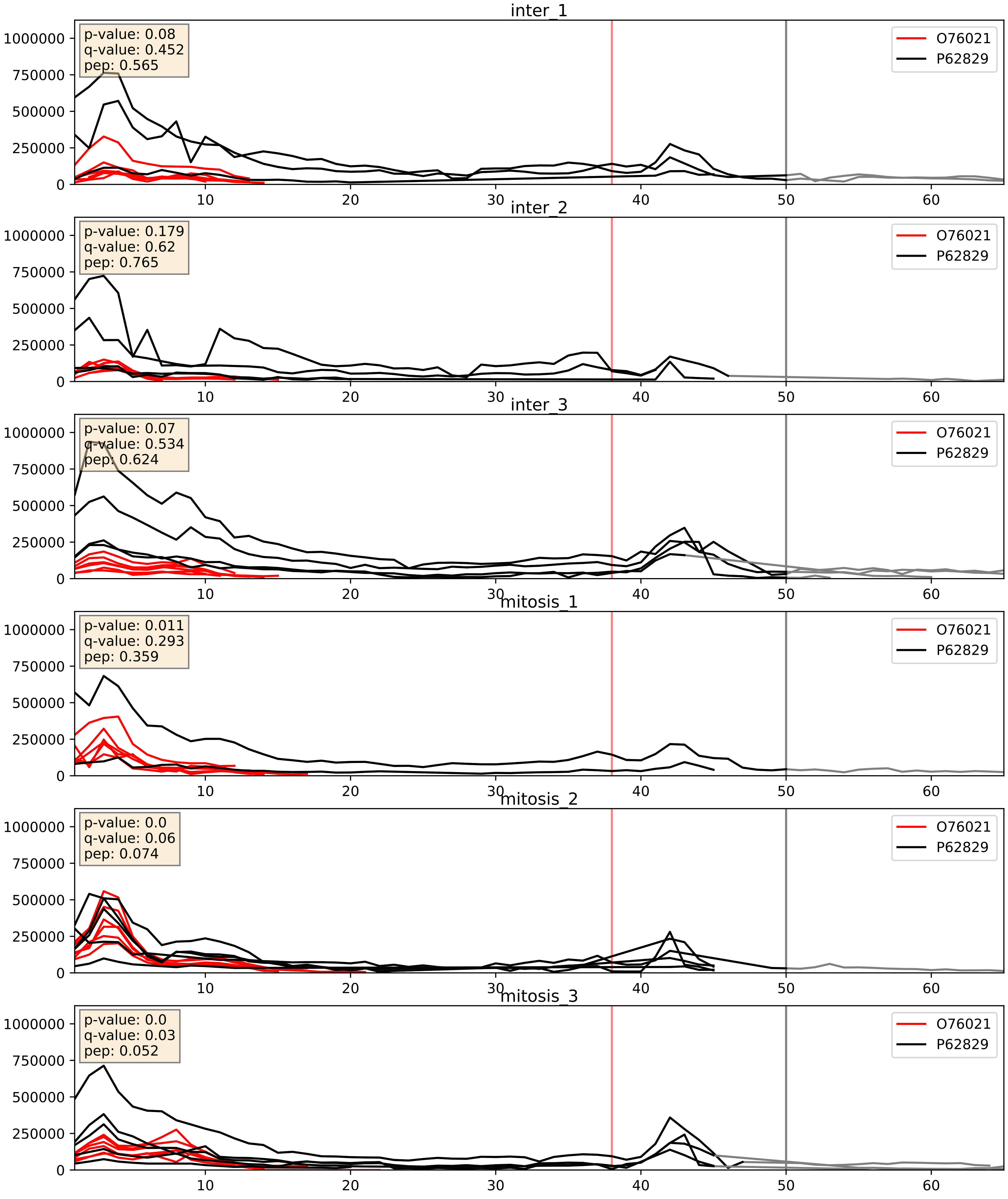


| level | condition_1 | condition_2 | pvalue | pvalue_adjusted |
|---|---|---|---|---|
| interactor_ratio | mitosis | inter | 0.0003092702819777167 | 0.018775557544179113 |
| interactor_abundance | mitosis | inter | 0.007302544007558834 | 0.07362753439894419 |
| complex_abundance | mitosis | inter | 0.086333506104879 | 0.2153229384262426 |

O76021_Q12788
RL1D1_HUMAN vs TBL3_HUMAN
p-value: 0.008 q-value: 0.007

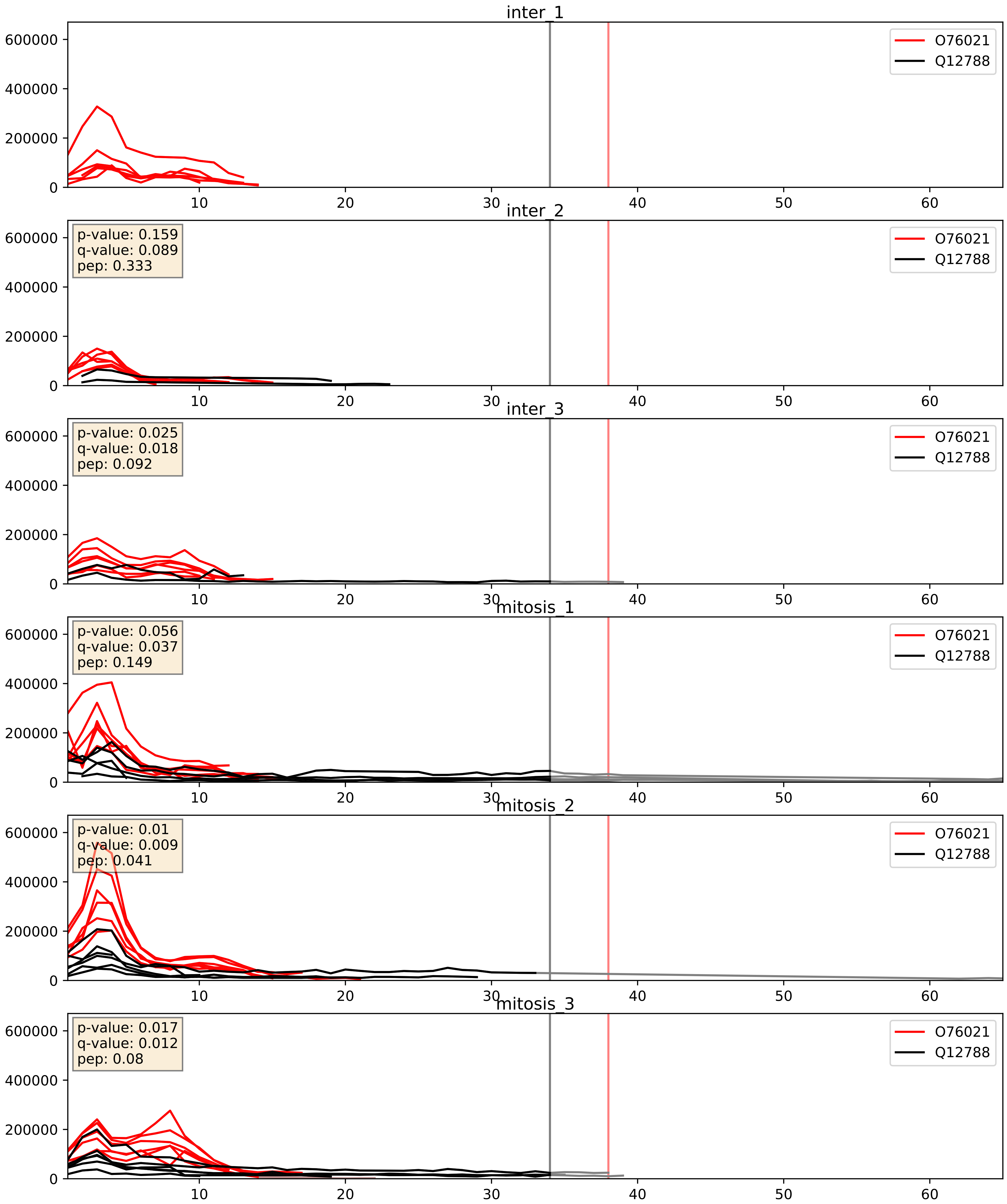


| level | condition_1 | condition_2 | pvalue | pvalue_adjusted |
|---|---|---|---|---|
| complex_abundance | mitosis | inter | 0.028907339226821576 | 0.12366158110024622 |
| interactor_abundance | mitosis | inter | 0.06155864533413795 | 0.2029091986005969 |
| interactor_ratio | mitosis | inter | 0.37222432746257683 | 0.8138388063636502 |

O76021_Q14137
RL1D1_HUMAN vs BOP1_HUMAN
p-value: 0.003 q-value: 0.003

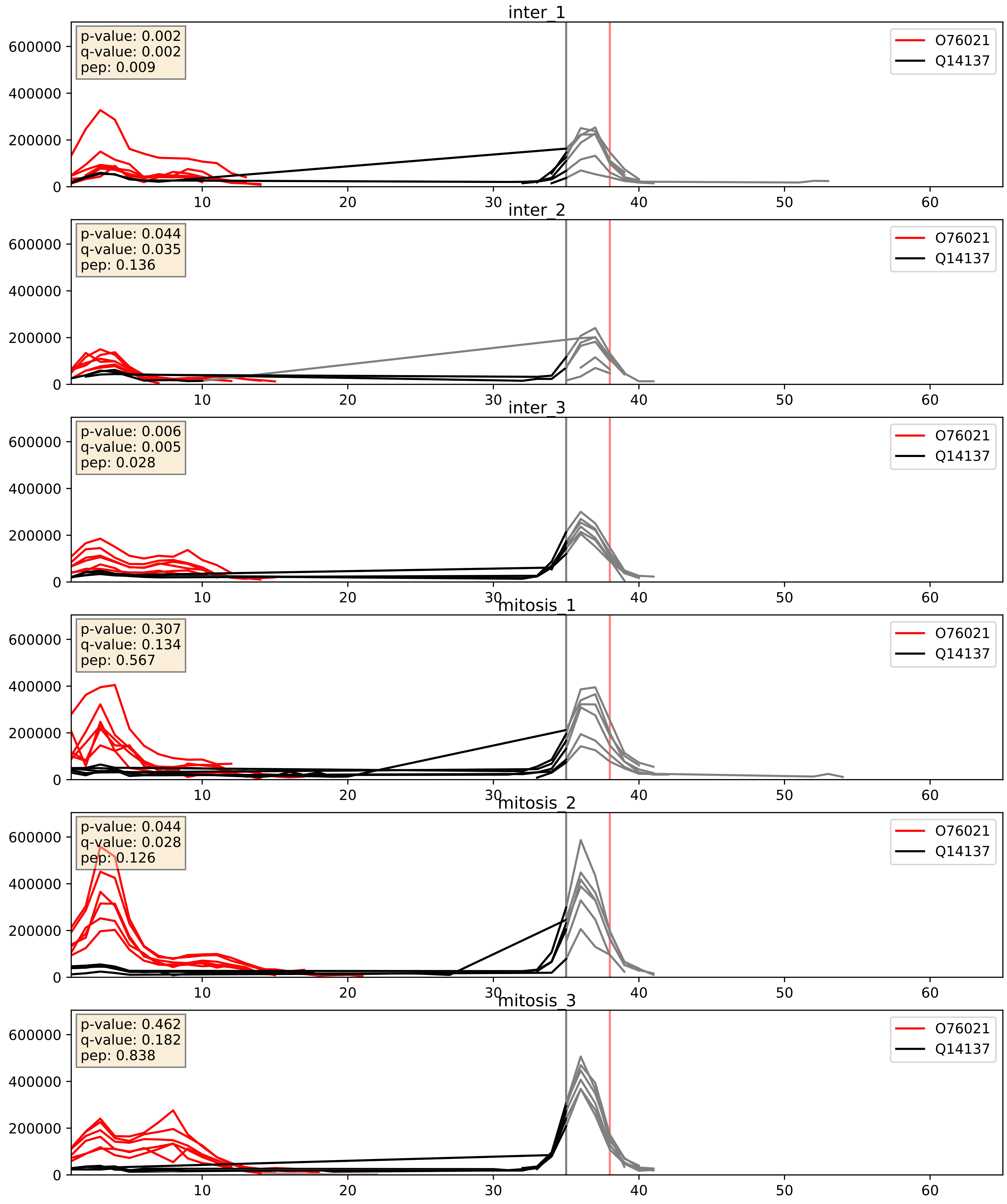


| level | condition_1 | condition_2 | pvalue | pvalue_adjusted |
|---|---|---|---|---|
| interactor_ratio | mitosis | inter | 0.014429314087919975 | 0.17619818629471468 |
| interactor_abundance | mitosis | inter | 0.04106316041394885 | 0.16296731794217187 |
| complex_abundance | mitosis | inter | 0.7782942865568719 | 0.8546759580406446 |

O76021_Q15050
RL1D1_HUMAN vs RRS1_HUMAN
p-value: 0.056 q-value: 0.036

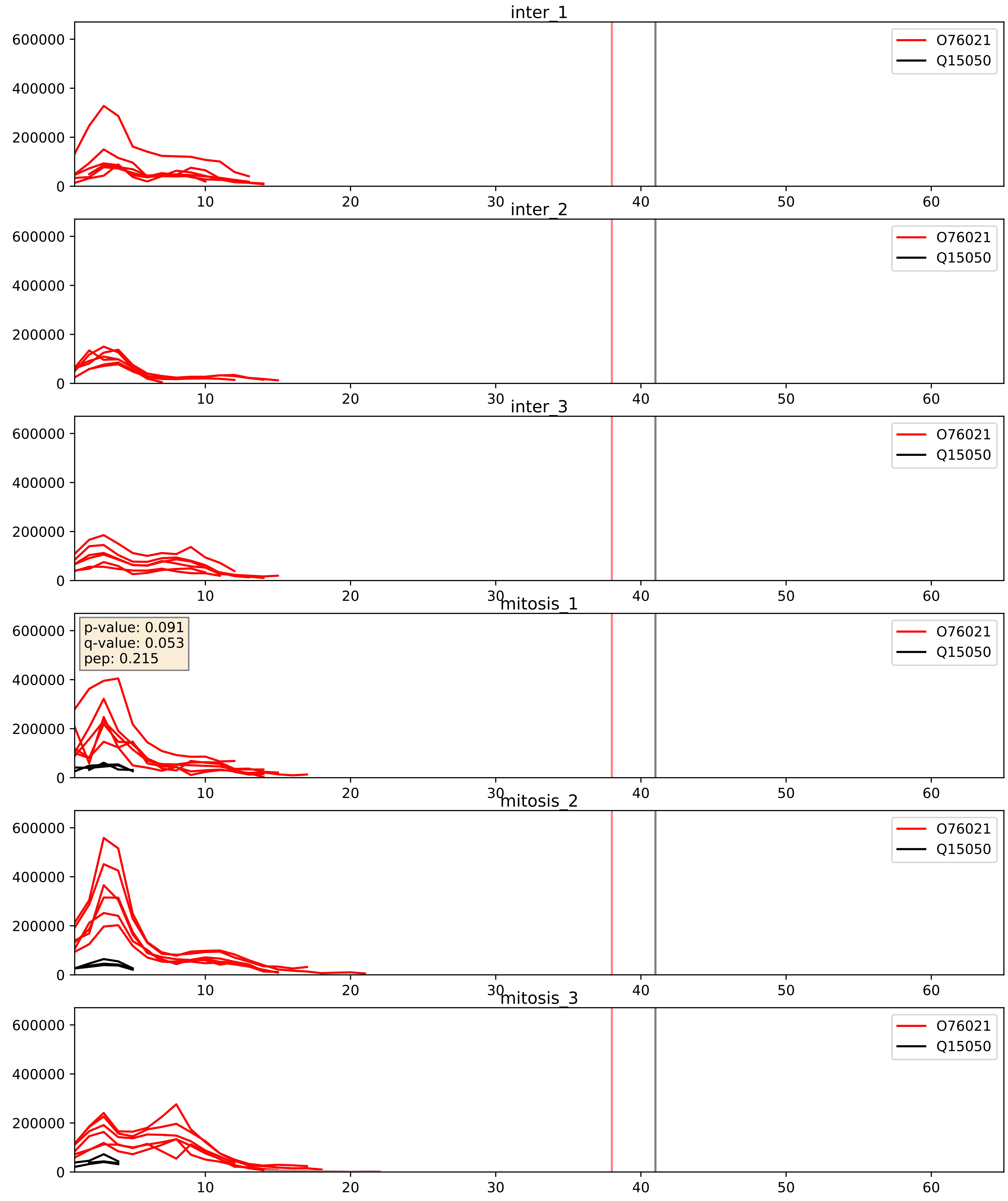


| level | condition_1 | condition_2 | pvalue | pvalue_adjusted |
|---|---|---|---|---|
| interactor_abundance | mitosis | inter | 0.002514210492437318 | 0.03746151752004081 |
| interactor_ratio | mitosis | inter | 0.003845030598611075 | 0.08075090146433855 |
| complex_abundance | mitosis | inter | 0.0081844105880903 | 0.06710589524334576 |

O76021_Q15397
RL1D1_HUMAN vs PUM3_HUMAN
p-value: 0.003 q-value: 0.039

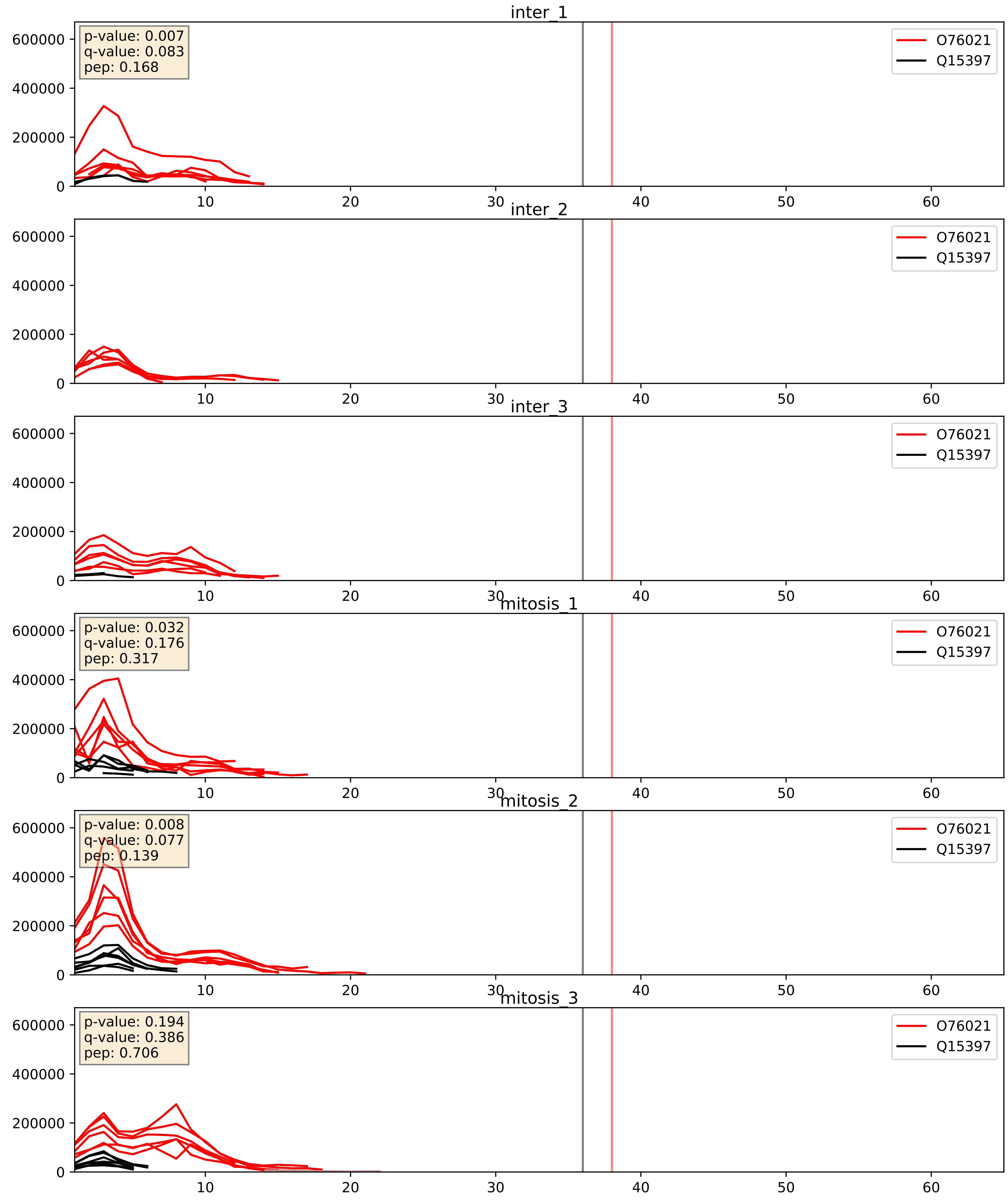


| level | condition_1 | condition_2 | pvalue | pvalue_adjusted |
|---|---|---|---|---|
| interactor_ratio | mitosis | inter | 0.09439087244256297 | 0.5094488449611217 |
| interactor_abundance | mitosis | inter | 0.09527655722192445 | 0.264708643239102 |
| complex_abundance | mitosis | inter | 0.1206192827943469 | 0.2626560826048358 |

O76021_Q8NI36
RL1D1_HUMAN vs WDR36_HUMAN
p-value: 0.002 q-value: 0.003

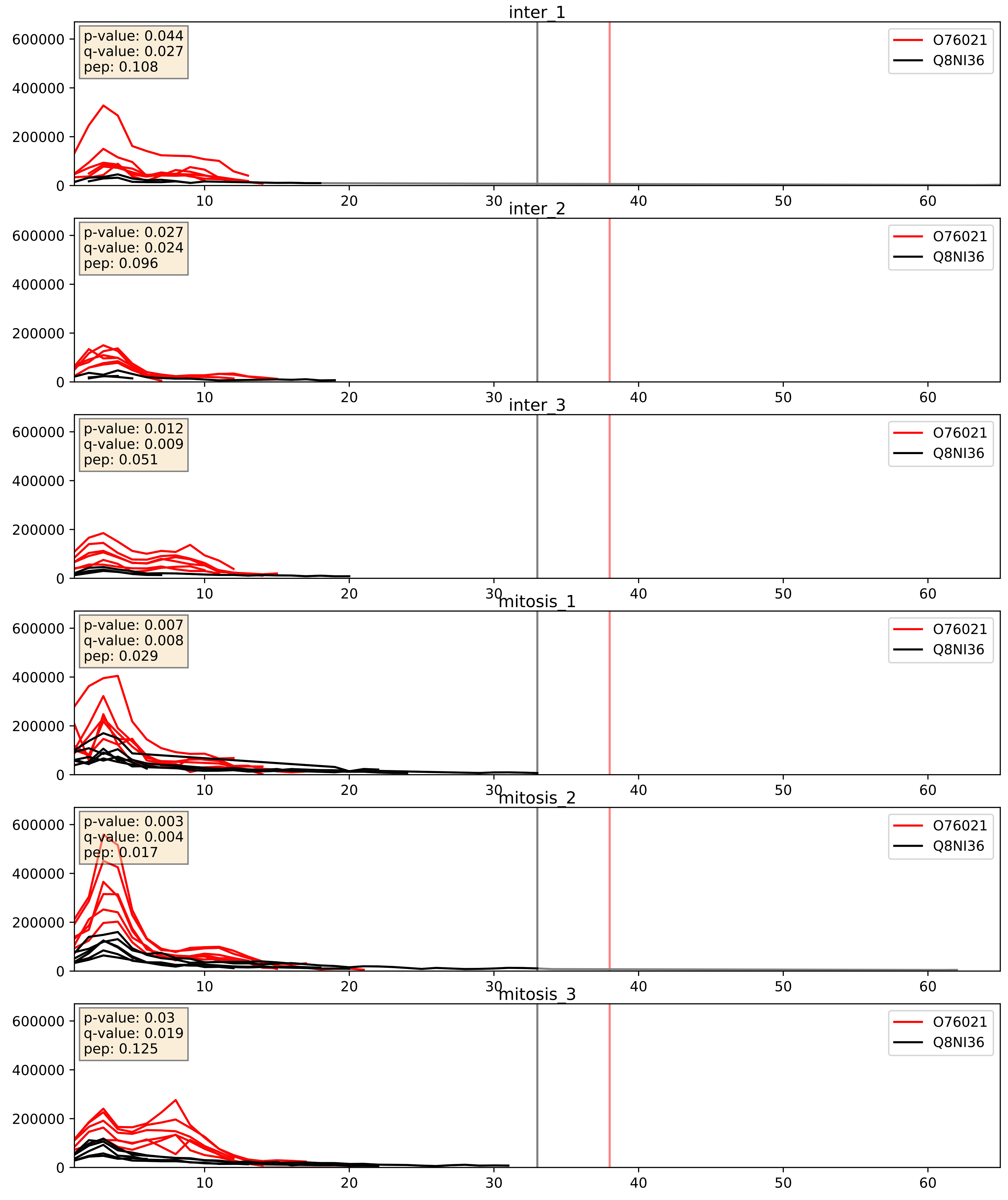


| level | condition_1 | condition_2 | pvalue | pvalue_adjusted |
|---|---|---|---|---|
| complex_abundance | mitosis | inter | 0.00046464344010374375 | 0.013667862018171982 |
| interactor_abundance | mitosis | inter | 0.005364620772517218 | 0.06171739355782327 |
| interactor_ratio | mitosis | inter | 0.9455967382350232 | 0.9912117205440141 |

O76021_Q8TED0
RL1D1_HUMAN vs UTP15_HUMAN
p-value: 0.066 q-value: 0.041

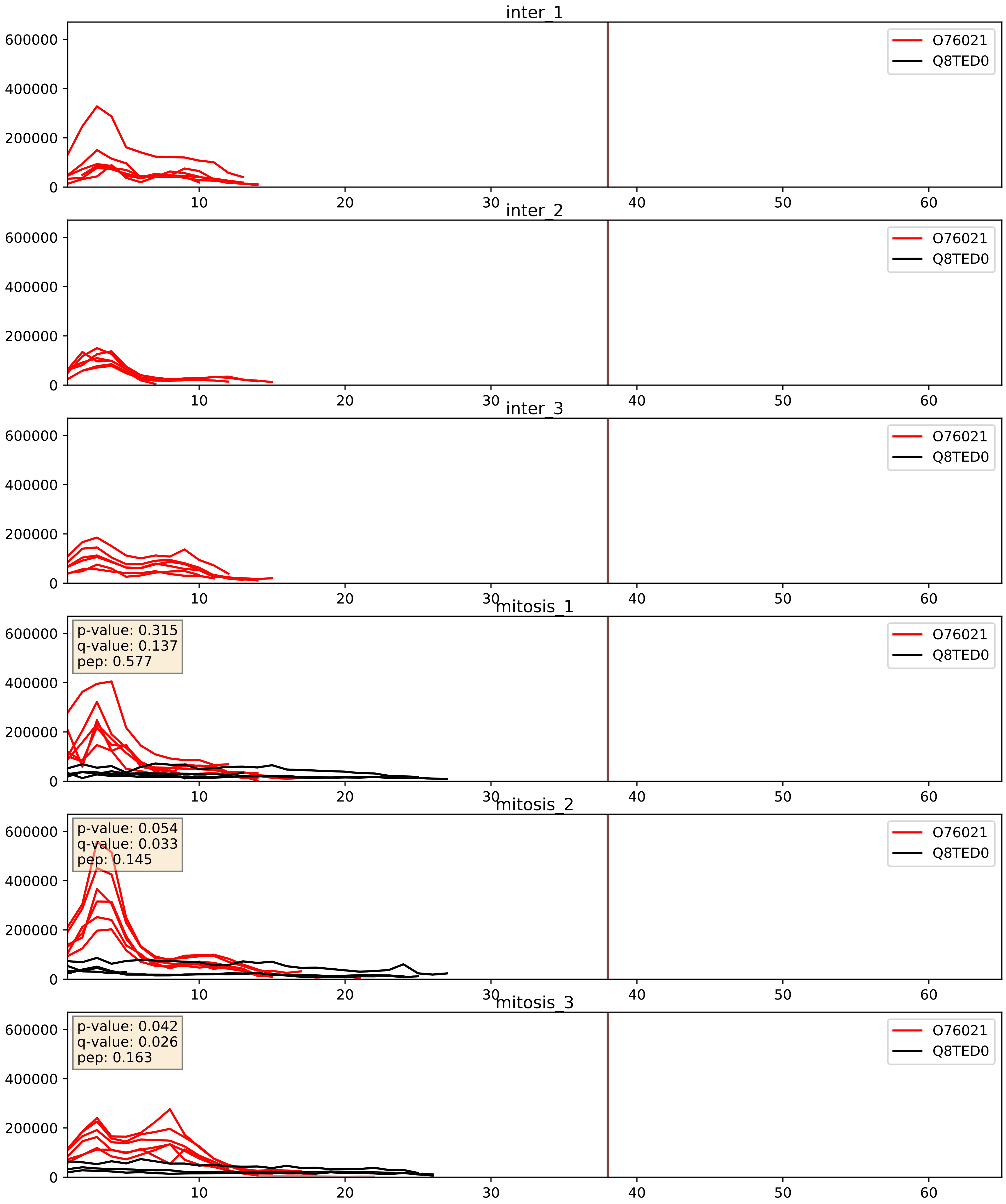


| level | condition_1 | condition_2 | pvalue | pvalue_adjusted |
|---|---|---|---|---|
| interactor_abundance | mitosis | inter | 8.109392026765088e-05 | 0.007119630332419645 |
| complex_abundance | mitosis | inter | 0.00016019902877882495 | 0.008563834848633118 |
| interactor_ratio | mitosis | inter | 0.0006707240247119572 | 0.029848225645041686 |

O76021_Q9BVP2
RL1D1_HUMAN vs GNL3_HUMAN
p-value: 0.0 q-value: 0.013

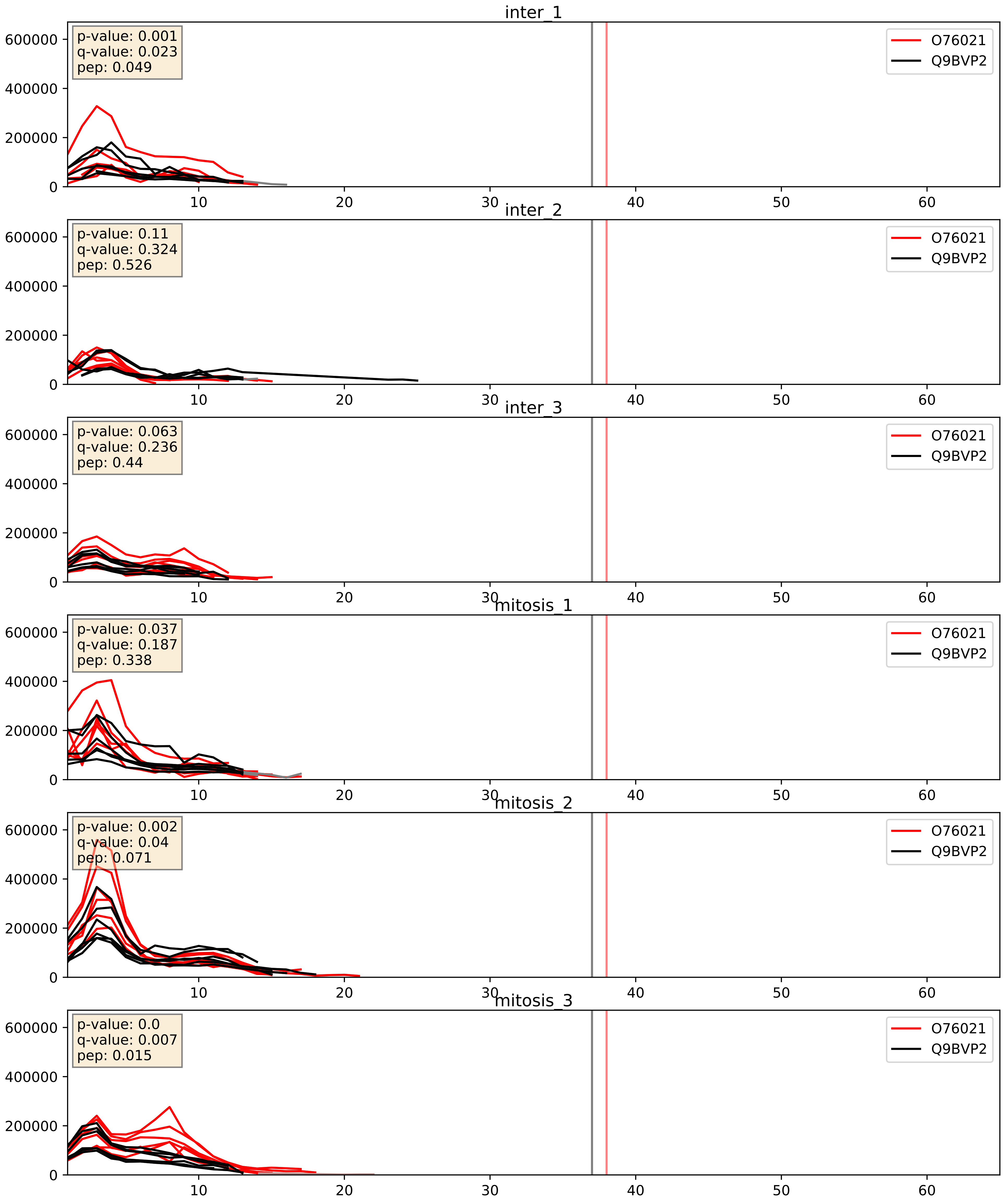


| level | condition_1 | condition_2 | pvalue | pvalue_adjusted |
|---|---|---|---|---|
| complex_abundance | mitosis | inter | 0.004581703881258023 | 0.05041052085291604 |
| interactor_abundance | mitosis | inter | 0.008085645358995957 | 0.077986618899161 |
| interactor_ratio | mitosis | inter | 0.3527656966303728 | 0.8022478353006512 |

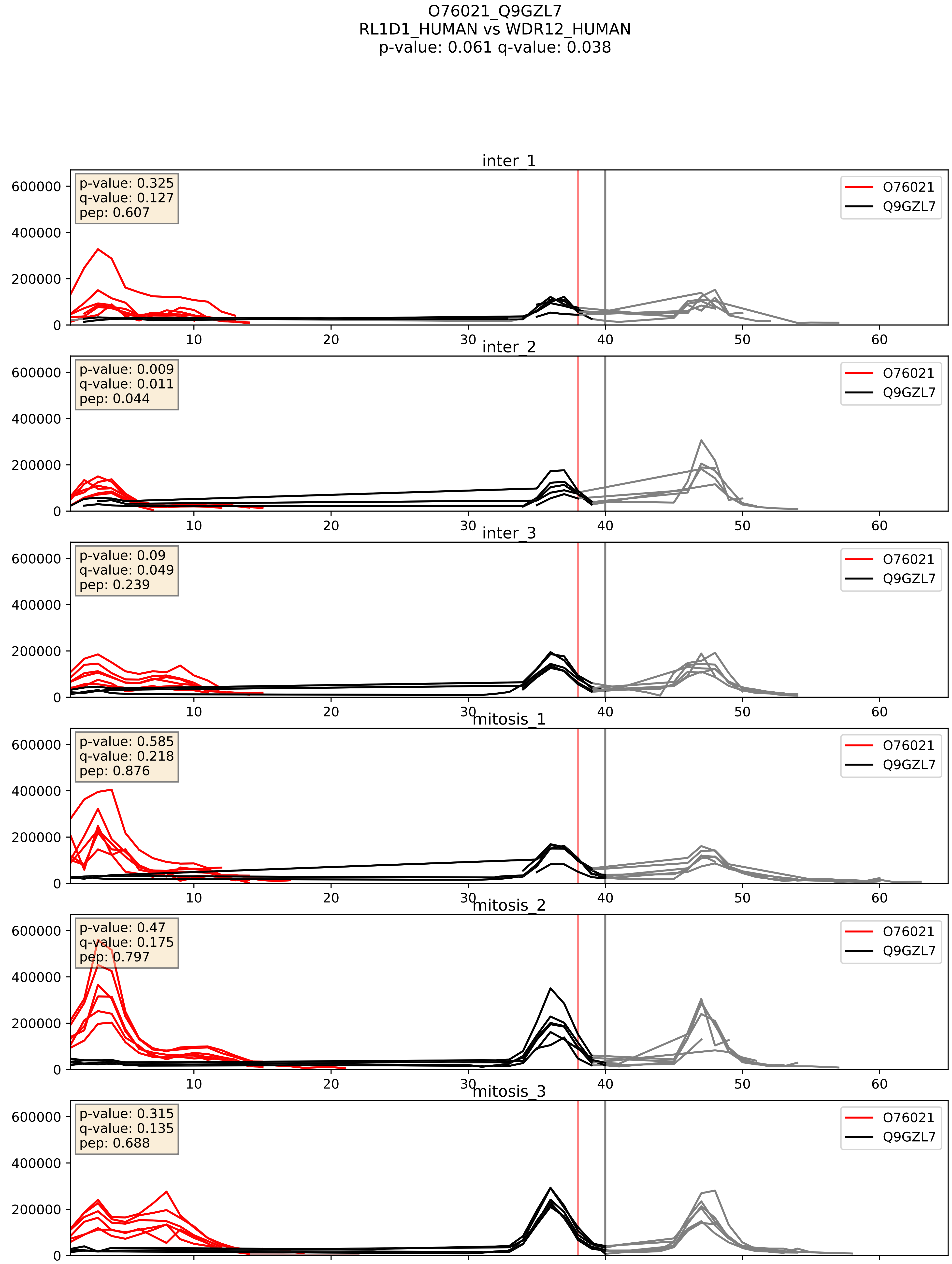


| level | condition_1 | condition_2 | pvalue | pvalue_adjusted |
|---|---|---|---|---|
| interactor_ratio | mitosis | inter | 0.033075047598448176 | 0.29157817450331247 |
| complex_abundance | mitosis | inter | 0.4134197061604915 | 0.5587105596359025 |
| interactor_abundance | mitosis | inter | 0.42772784767610067 | 0.6080462303591168 |

O76021_Q9H0A0
RL1D1_HUMAN vs NAT10_HUMAN
p-value: 0.078 q-value: 0.047

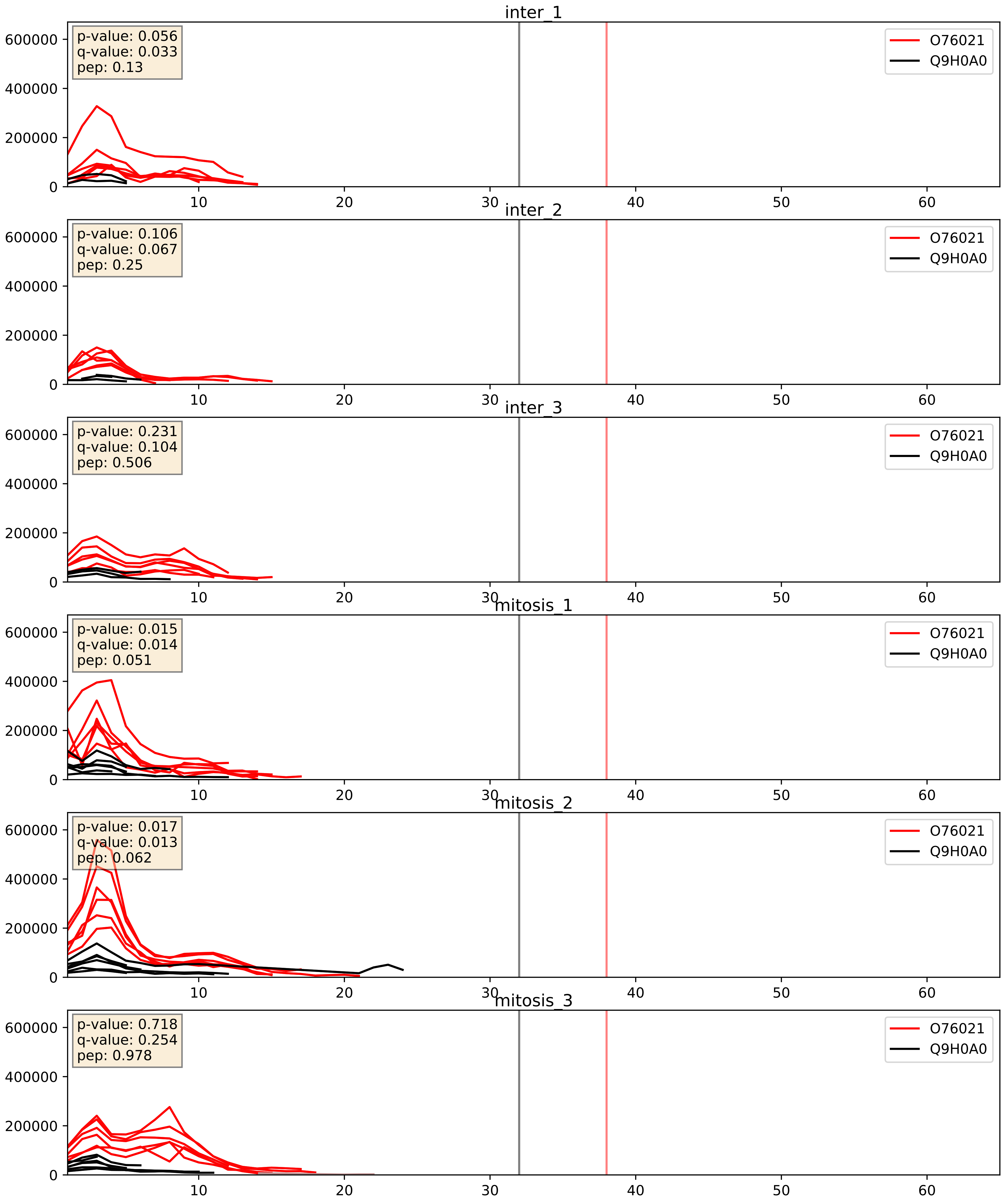


| level | condition_1 | condition_2 | pvalue | pvalue_adjusted |
|---|---|---|---|---|
| interactor_abundance | mitosis | inter | 0.0004232024324935983 | 0.014167094422781207 |
| complex_abundance | mitosis | inter | 0.002537322887836132 | 0.036409709098919725 |
| interactor_ratio | mitosis | inter | 0.02791082622771571 | 0.26184787474031573 |

O76021_Q9H583
RL1D1_HUMAN vs HEAT1_HUMAN
p-value: 0.022 q-value: 0.017

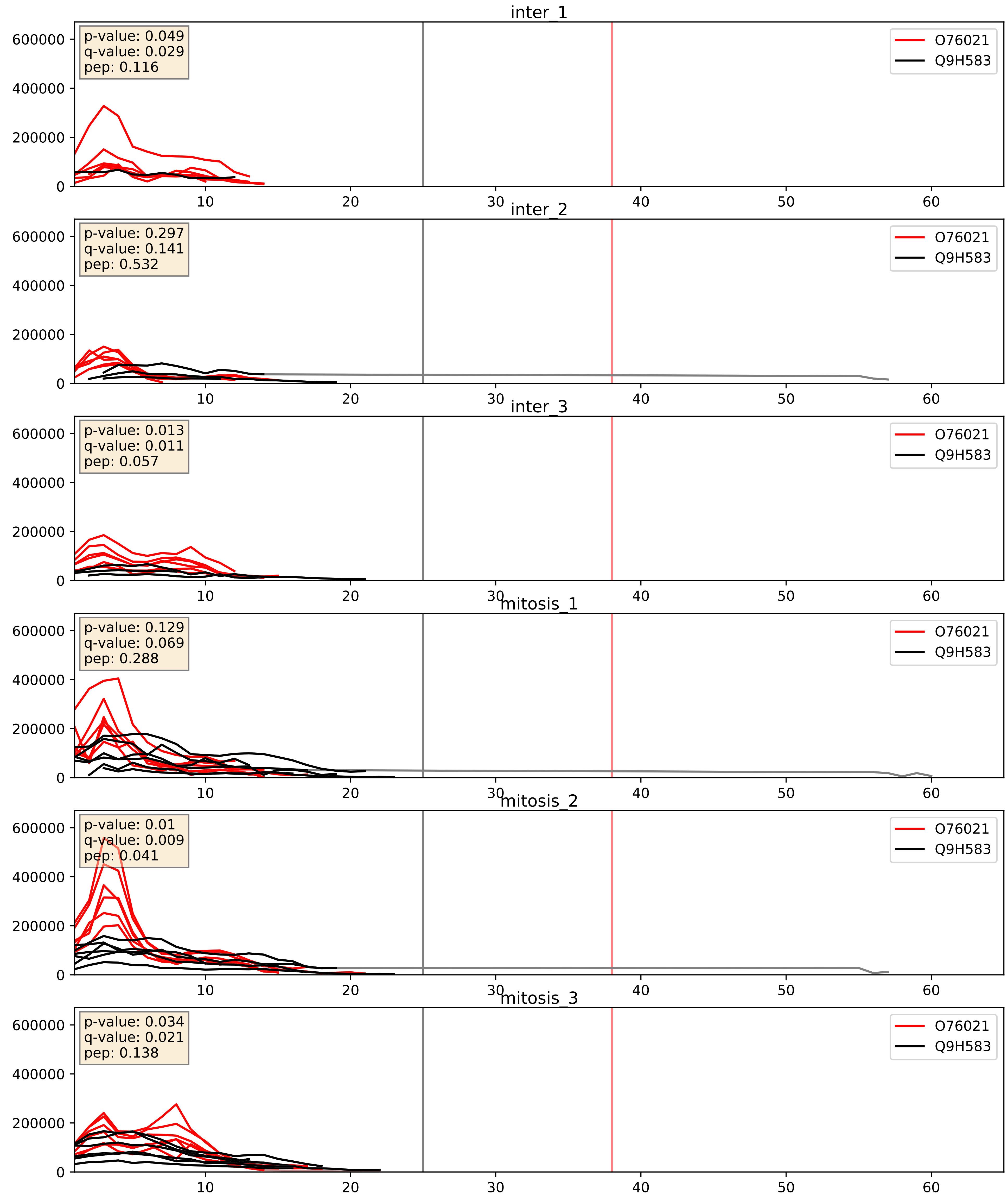


| level | condition_1 | condition_2 | pvalue | pvalue_adjusted |
|---|---|---|---|---|
| complex_abundance | mitosis | inter | 0.00022168473140128019 | 0.010113501428602107 |
| interactor_abundance | mitosis | inter | 0.009409188518133108 | 0.08407375126849626 |
| interactor_ratio | mitosis | inter | 0.1385553890354226 | 0.5876014454413842 |

O76021_Q9H6R4
RL1D1_HUMAN vs NOL6_HUMAN
p-value: 0.02 q-value: 0.016

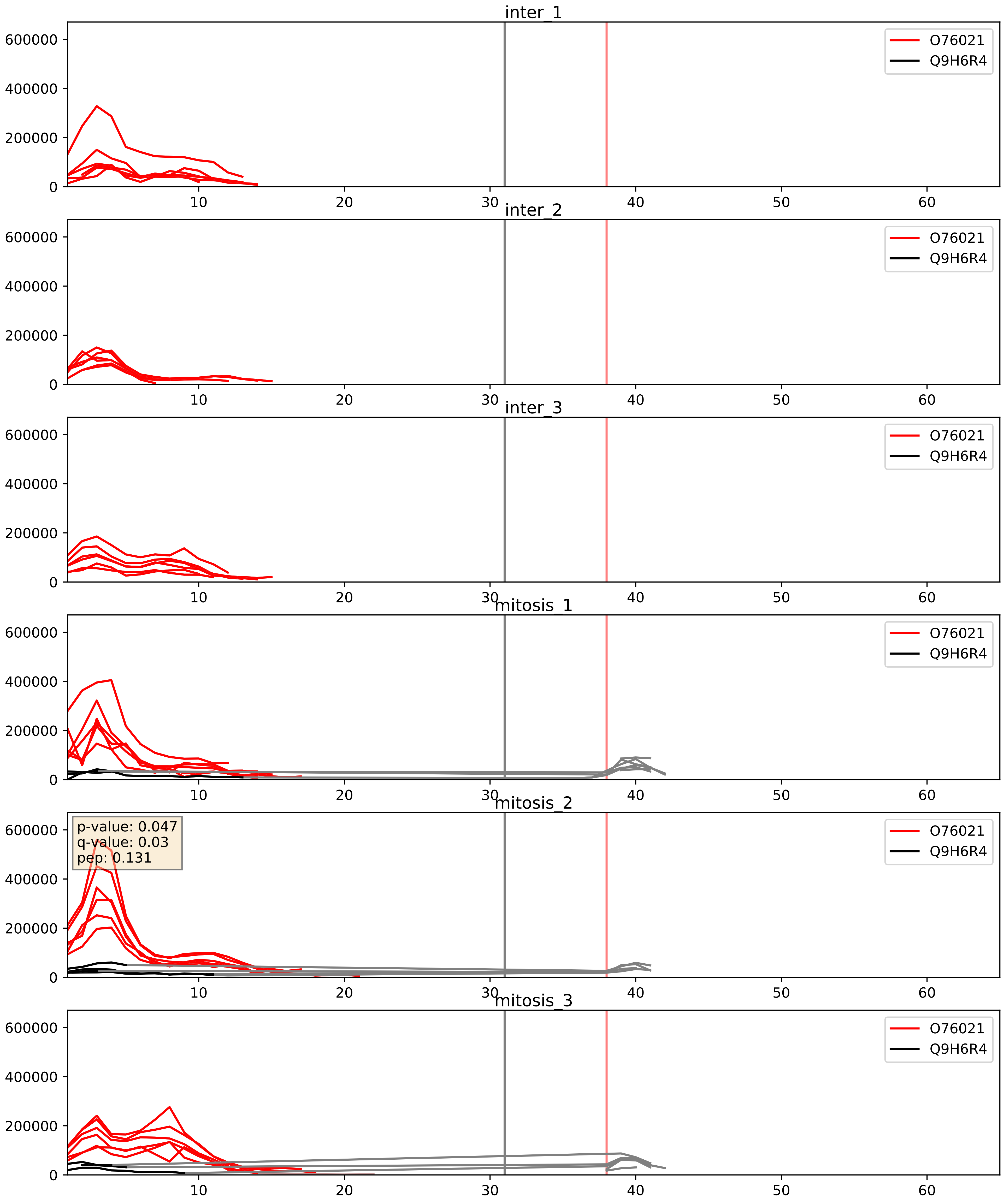


| level | condition_1 | condition_2 | pvalue | pvalue_adjusted |
|---|---|---|---|---|
| interactor_abundance | mitosis | inter | 6.462019261636246e-05 | 0.0065503332782105754 |
| interactor_ratio | mitosis | inter | 0.0002037188630223042 | 0.014683183137172115 |
| complex_abundance | mitosis | inter | 0.0007402222580168641 | 0.017798602608495383 |

O76021_Q9UNX4
RL1D1_HUMAN vs WDR3_HUMAN
p-value: 0.014 q-value: 0.011

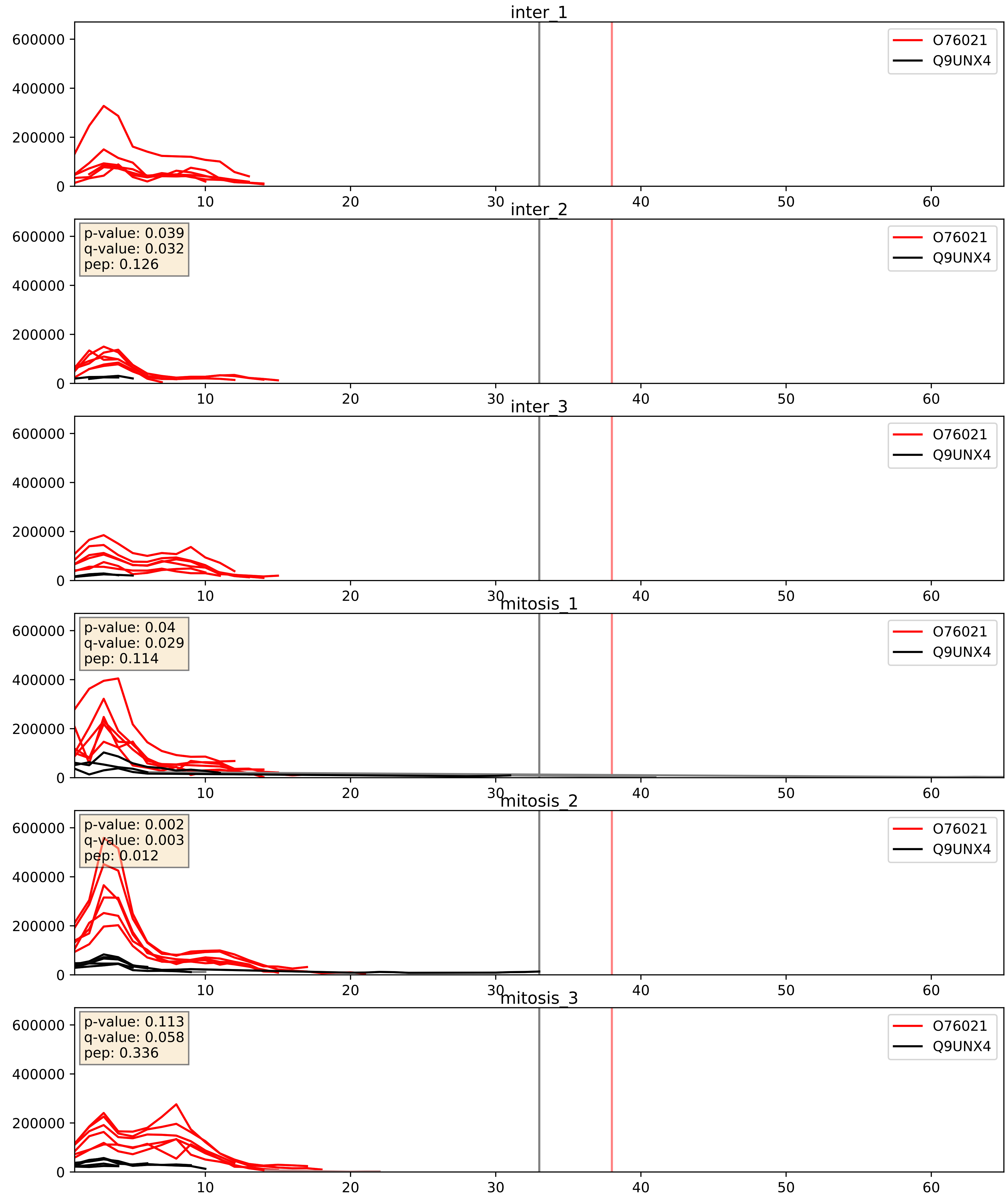


| level | condition_1 | condition_2 | pvalue | pvalue_adjusted |
|---|---|---|---|---|
| interactor_abundance | mitosis | inter | 0.022557396454085495 | 0.12329937058940194 |
| complex_abundance | mitosis | inter | 0.02766391673322131 | 0.12094133158139654 |
| interactor_ratio | mitosis | inter | 0.033938259800389954 | 0.296589834089654 |

O76021_Q9Y2X3
RL1D1_HUMAN vs NOP58_HUMAN
p-value: 0.001 q-value: 0.002

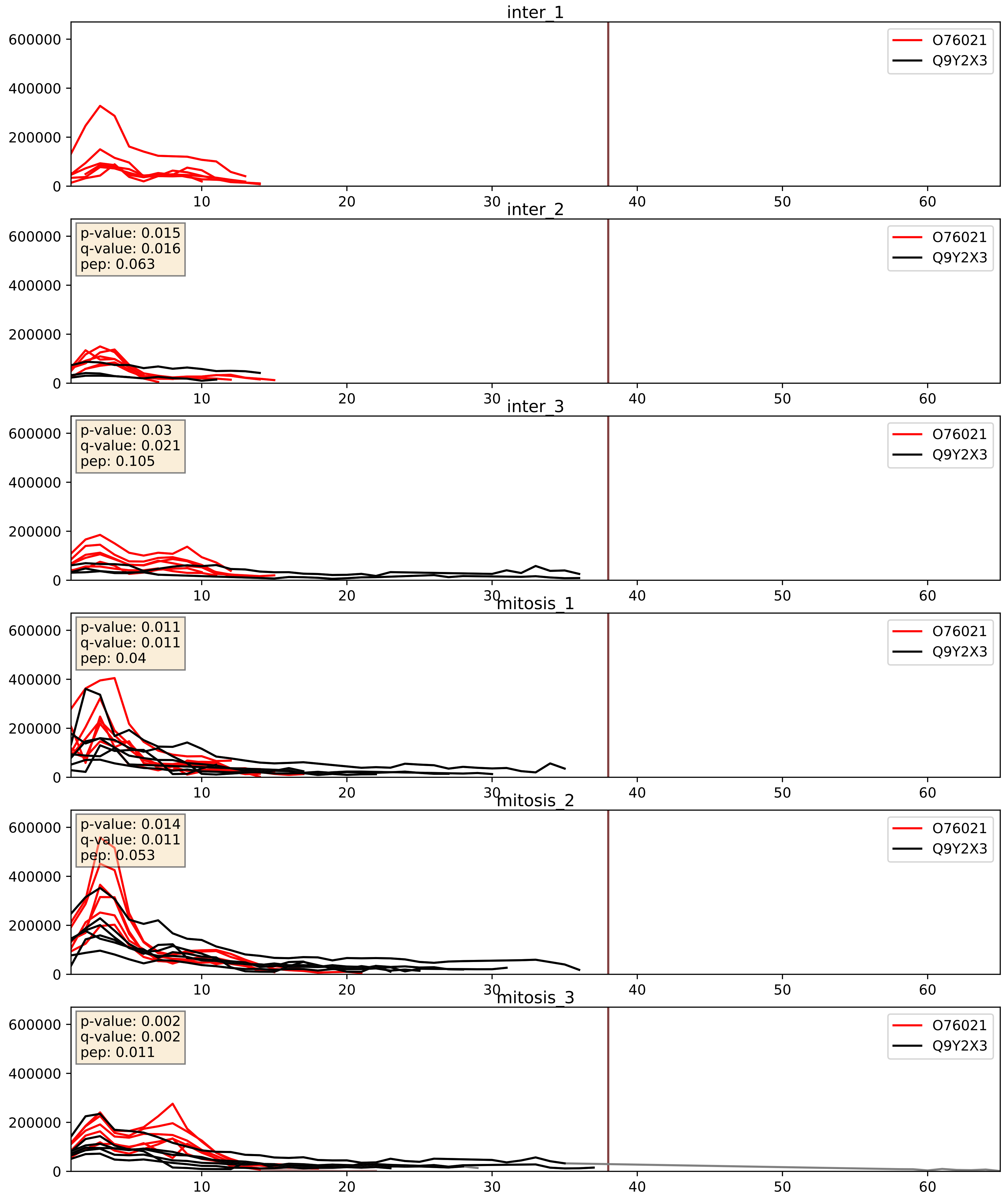


| level | condition_1 | condition_2 | pvalue | pvalue_adjusted |
|---|---|---|---|---|
| complex_abundance | mitosis | inter | 0.01952163921039481 | 0.10409657315782883 |
| interactor_abundance | mitosis | inter | 0.0810993279136887 | 0.23762116958452006 |
| interactor_ratio | mitosis | inter | 0.3037548150127056 | 0.7719380618221225 |

O76021_Q9Y5J1
RL1D1_HUMAN vs UTP18_HUMAN
p-value: 0.068 q-value: 0.042

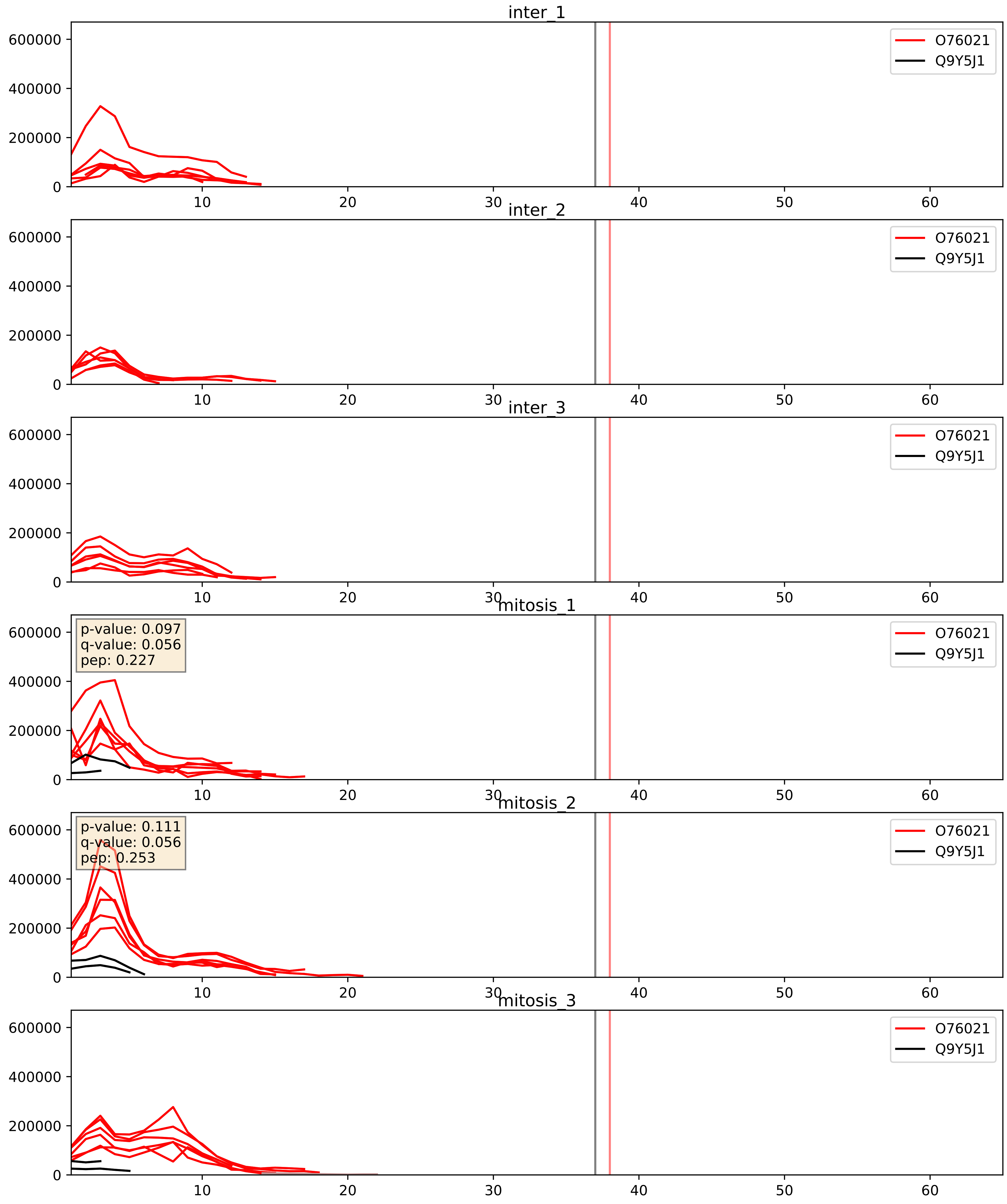


| level | condition_1 | condition_2 | pvalue | pvalue_adjusted |
|---|---|---|---|---|
| interactor_ratio | mitosis | inter | 0.0008389471802542103 | 0.03318433890476037 |
| interactor_abundance | mitosis | inter | 0.0012303243090360165 | 0.02571813451855507 |
| complex_abundance | mitosis | inter | 0.0035472690192235807 | 0.043600374082146835 |
