## Supplementary Data 2 for "SECAT: Quantifying differential protein-protein interaction states by network-centric analysis": hela_string_000064_Q13823.pdf

Q13823_Q13823

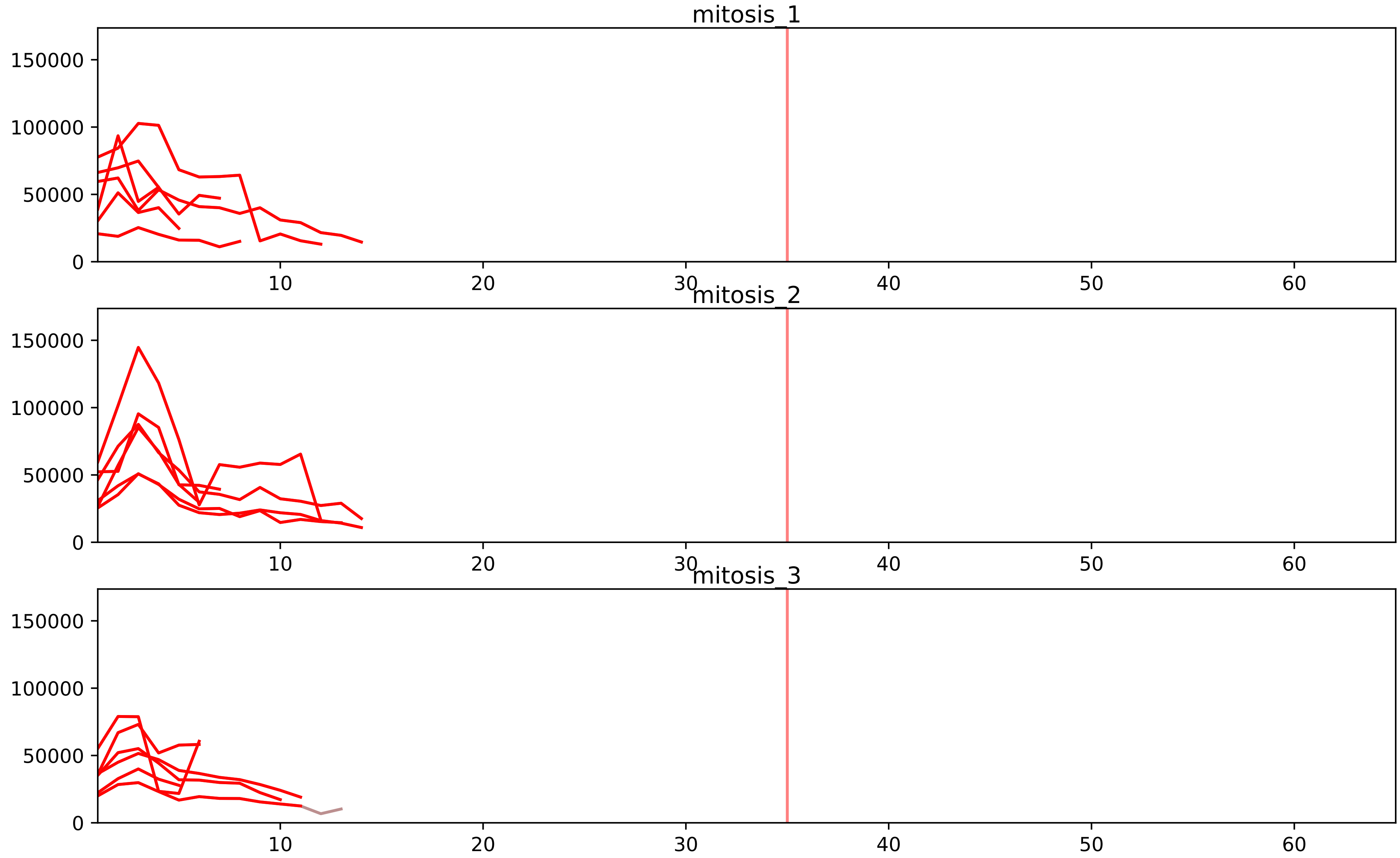


| level | condition_1 | condition_2 | pvalue | pvalue_adjusted |
|---|---|---|---|---|
| interactor_ratio | mitosis | inter | 6.671228152205581e-07 | 2.0458433000097115e-05 |
| complex_abundance | mitosis | inter | 0.0004578837743306342 | 0.006531055385801294 |
| assembled_abundance | mitosis | inter | 0.0007253582786950801 | 0.024427353255911294 |
| total_abundance | mitosis | inter | 0.0020798672876501767 | 0.04923864988533268 |
| interactor_abundance | mitosis | inter | 0.0027583244969317665 | 0.023716434926889955 |

Q13823_Q9BVP2
NOG2_HUMAN vs GNL3_HUMAN
p-value: 0.034 q-value: 0.024

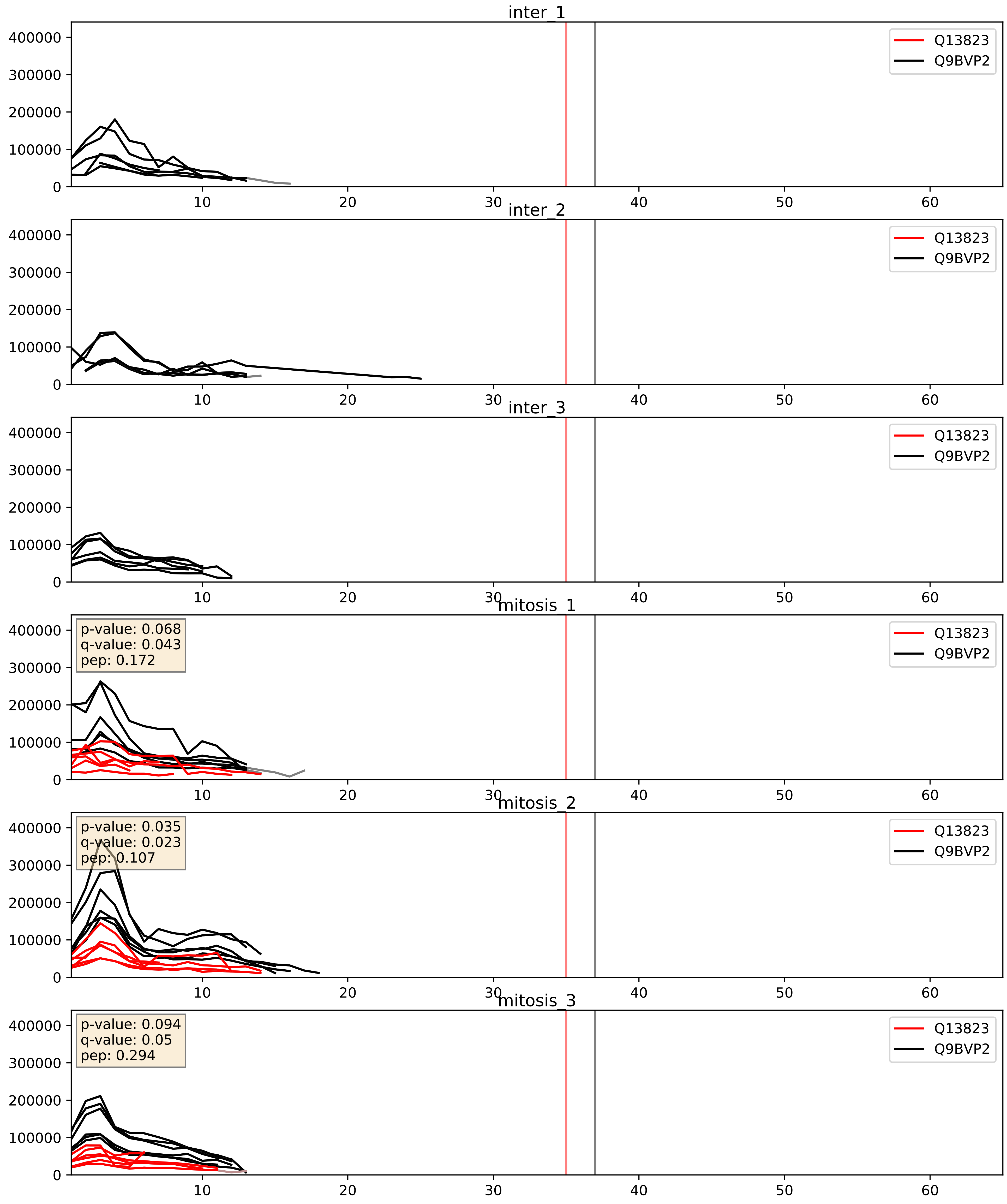


| level | condition_1 | condition_2 | pvalue | pvalue_adjusted |
|---|---|---|---|---|
| complex_abundance | mitosis | inter | 0.0009671264823373898 | 0.020644894485805625 |
| interactor_ratio | mitosis | inter | 0.0012027857202170427 | 0.041019305836884006 |
| interactor_abundance | mitosis | inter | 0.00933786289754201 | 0.08378627505551321 |

Q13823_Q9BZE4
NOG2_HUMAN vs NOG1_HUMAN
p-value: 0.007 q-value: 0.006

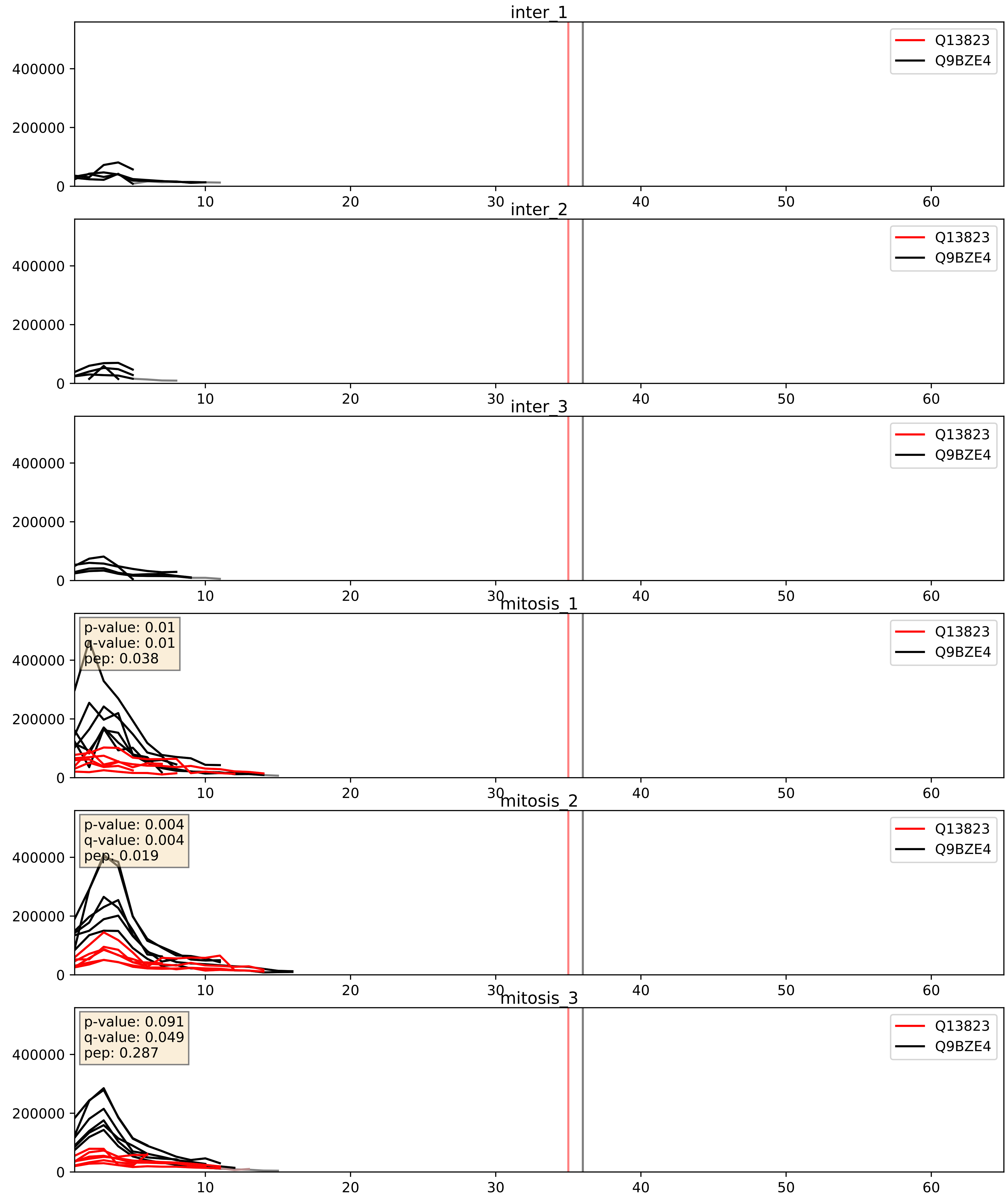


| level | condition_1 | condition_2 | pvalue | pvalue_adjusted |
|---|---|---|---|---|
| interactor_ratio | mitosis | inter | 0.0009758498273769684 | 0.03599129628116289 |
| complex_abundance | mitosis | inter | 0.0012618827871512452 | 0.024110974683068435 |
| interactor_abundance | mitosis | inter | 0.009576535749111607 | 0.08503645851908231 |

Q13823_Q9H7B2
NOG2_HUMAN vs RPF2_HUMAN
p-value: 0.049 q-value: 0.032

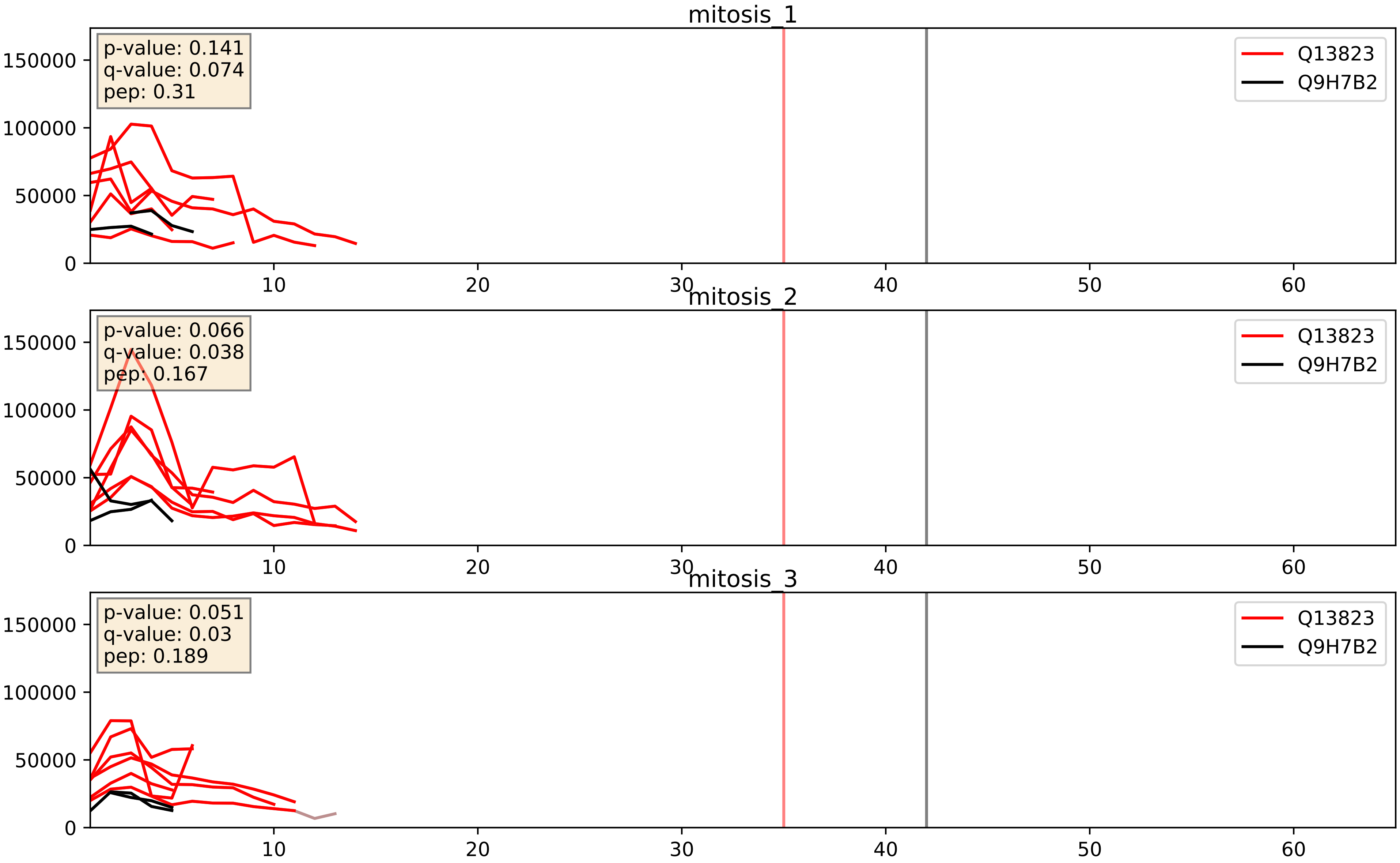


| level | condition_1 | condition_2 | pvalue | pvalue_adjusted |
|---|---|---|---|---|
| interactor_ratio | mitosis | inter | 0.0008412384979360982 | 0.03318433890476037 |
| interactor_abundance | mitosis | inter | 0.03007008855576401 | 0.14418523491160162 |
| complex_abundance | mitosis | inter | 0.08703996497906602 | 0.21596002904950873 |

Q13823_Q9UKD2
NOG2_HUMAN vs MRT4_HUMAN
p-value: 0.01 q-value: 0.008

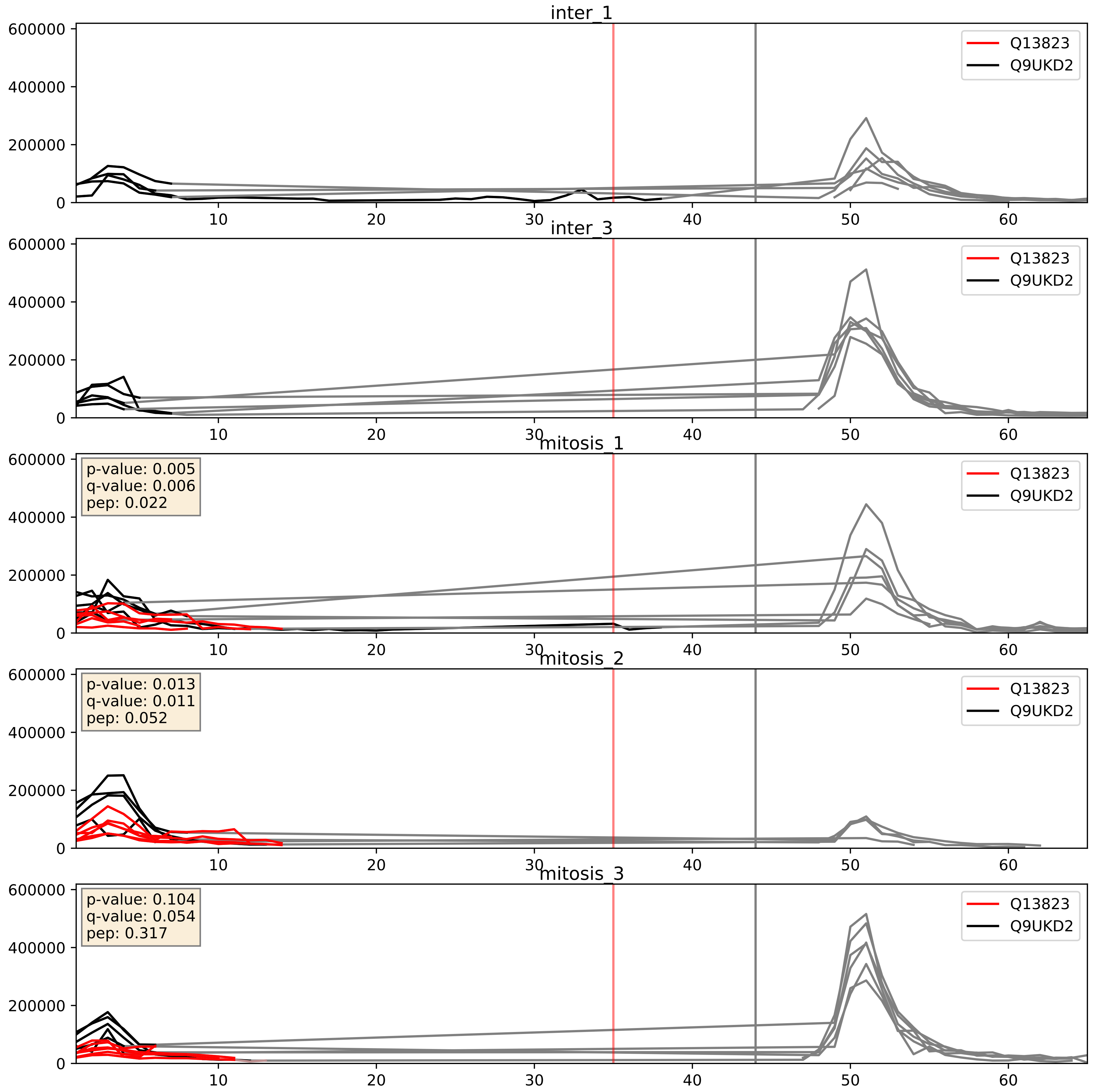


| level | condition_1 | condition_2 | pvalue | pvalue_adjusted |
|---|---|---|---|---|
| complex_abundance | mitosis | inter | 0.0028361034104119874 | 0.038719370323965885 |
| interactor_abundance | mitosis | inter | 0.009614702921018391 | 0.08507272274138435 |
| interactor_ratio | mitosis | inter | 0.012338641662349704 | 0.16051485202084112 |
