## Supplementary Data 2 for "SECAT: Quantifying differential protein-protein interaction states by network-centric analysis": hela_string_000065_Q9H583.pdf

Q9H583_Q9H583

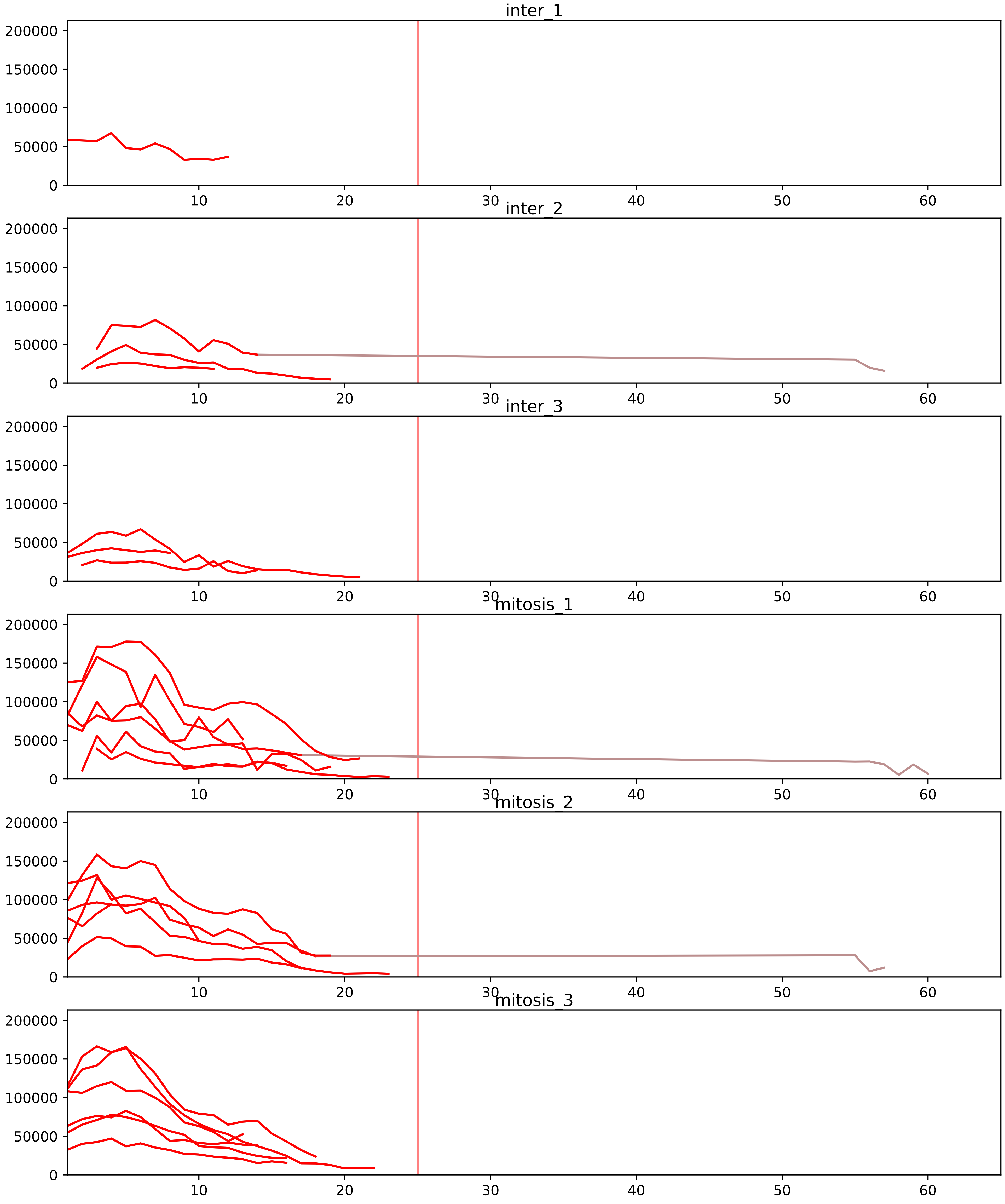


| level | condition_1 | condition_2 | pvalue | pvalue_adjusted |
|---|---|---|---|---|
| complex_abundance | mitosis | inter | 6.735402775045637e-07 | 7.290083003578808e-05 |
| interactor_abundance | mitosis | inter | 1.2719514669764521e-06 | 0.00013353840650924488 |
| interactor_ratio | mitosis | inter | 0.0014822219198476802 | 0.015236247667707998 |
| total_abundance | mitosis | inter | 0.4813484227093995 | 0.7510097517663356 |
| assembled_abundance | mitosis | inter | 0.5150797151285518 | 0.7572290843490062 |
| monomer_abundance | mitosis | inter | 0.6073888843499315 | 0.8144623884863933 |

O00567_Q9H583
NOP56_HUMAN vs HEAT1_HUMAN
p-value: 0.012 q-value: 0.01

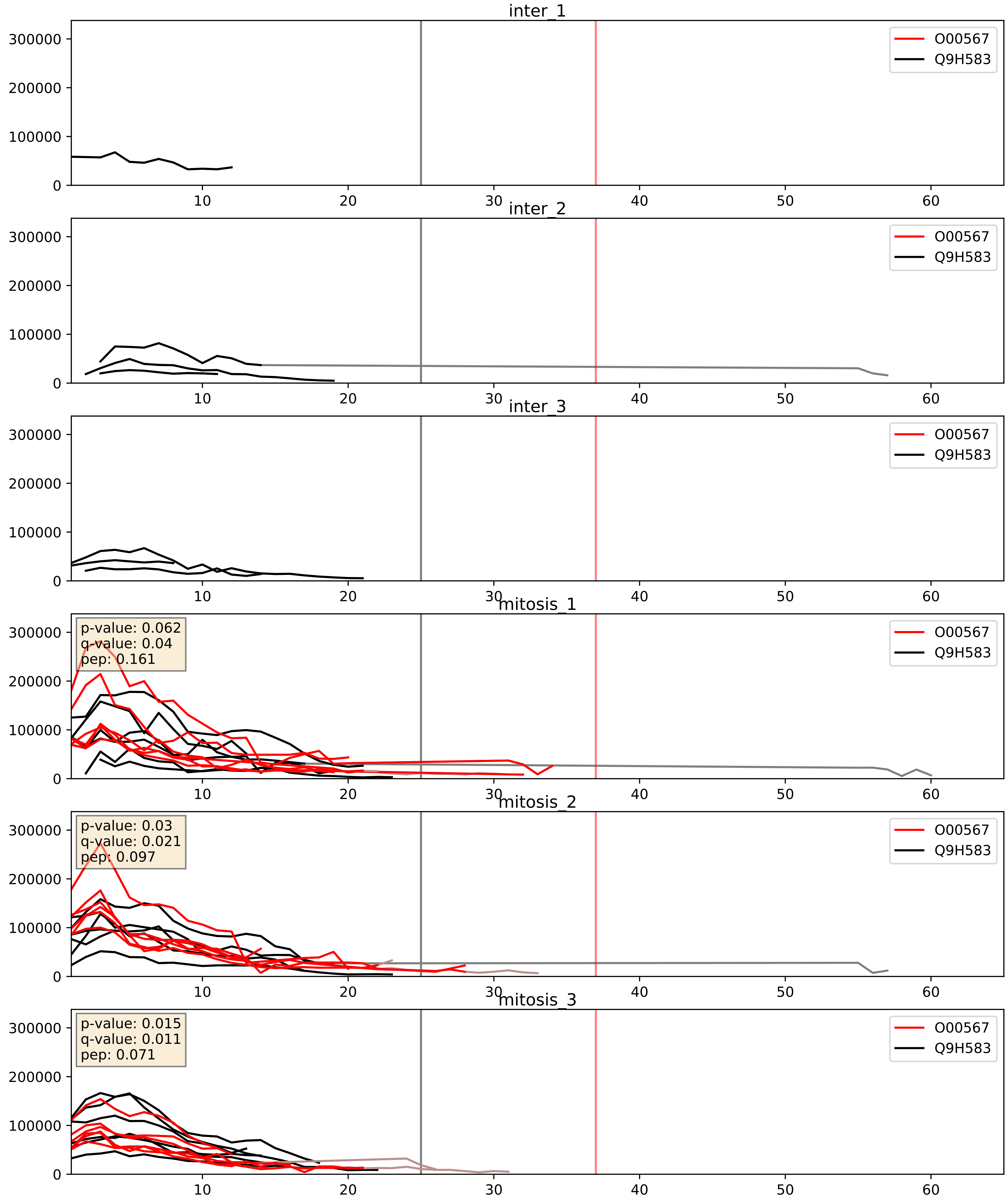


| level | condition_1 | condition_2 | pvalue | pvalue_adjusted |
|---|---|---|---|---|
| complex_abundance | mitosis | inter | 0.00016025150516764934 | 0.008563834848633118 |
| interactor_abundance | mitosis | inter | 0.001056251194238277 | 0.024049720606162197 |
| interactor_ratio | mitosis | inter | 0.8485851427407834 | 0.9755634896635457 |

O76021_Q9H583
RL1D1_HUMAN vs HEAT1_HUMAN
p-value: 0.022 q-value: 0.017

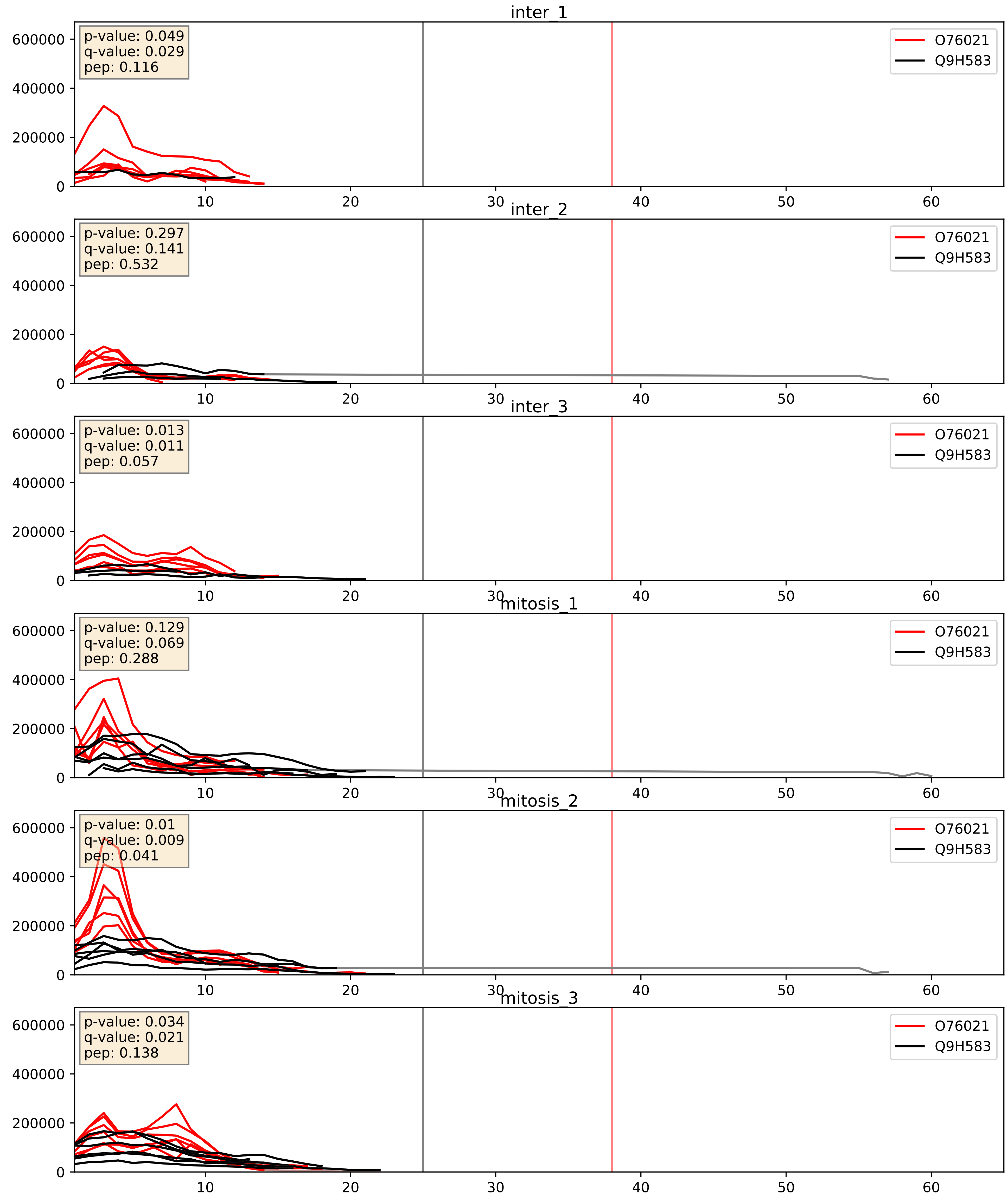


| level | condition_1 | condition_2 | pvalue | pvalue_adjusted |
|---|---|---|---|---|
| complex_abundance | mitosis | inter | 0.00022168473140128019 | 0.010113501428602107 |
| interactor_abundance | mitosis | inter | 0.009409188518133108 | 0.08407375126849626 |
| interactor_ratio | mitosis | inter | 0.1385553890354226 | 0.5876014454413842 |

P22087_Q9H583
FBRL_HUMAN vs HEAT1_HUMAN
p-value: 0.069 q-value: 0.042

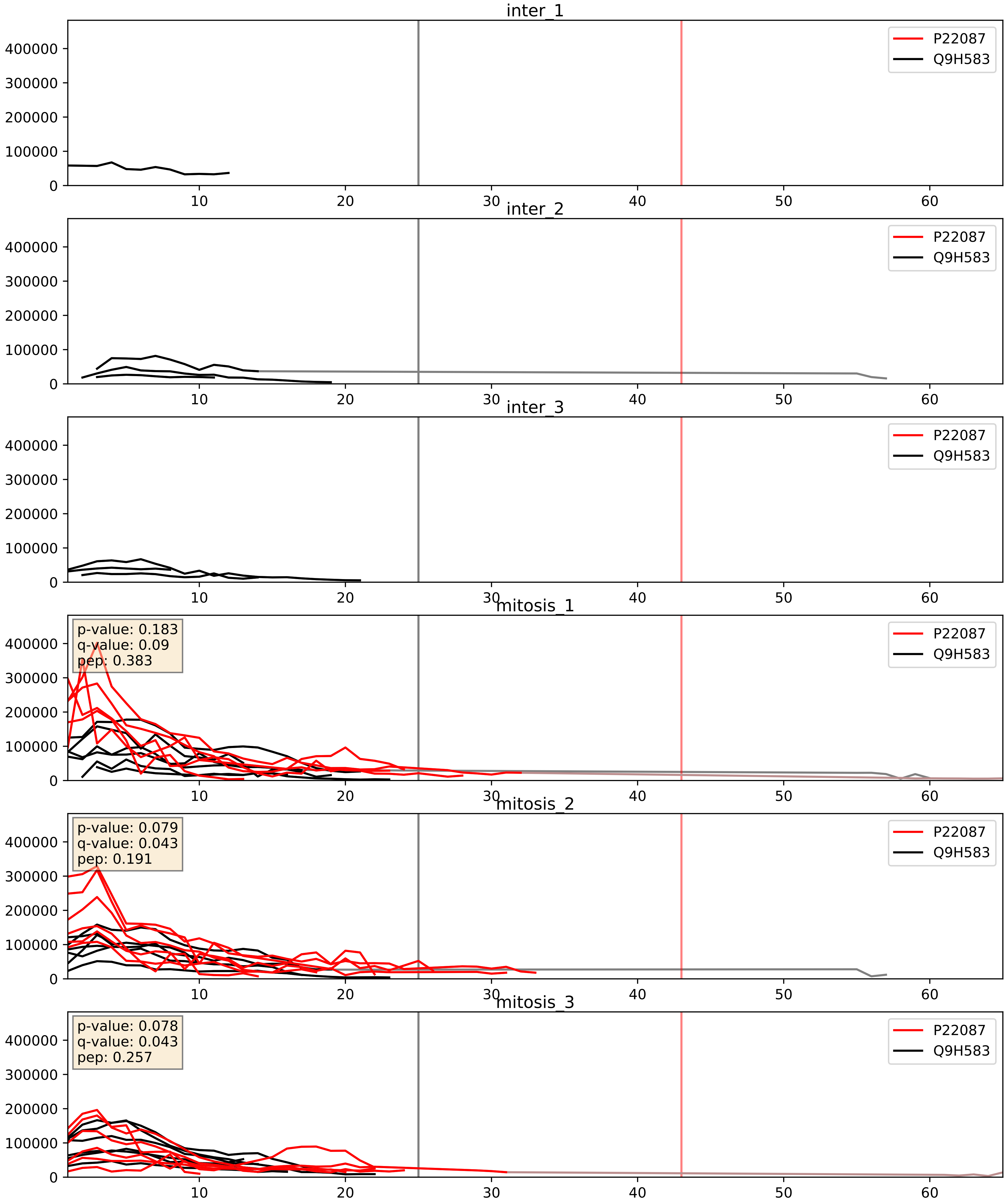


| level | condition_1 | condition_2 | pvalue | pvalue_adjusted |
|---|---|---|---|---|
| complex_abundance | mitosis | inter | 9.233901065022553e-05 | 0.006532412654263889 |
| interactor_abundance | mitosis | inter | 0.00038515934981581823 | 0.013595727977003728 |
| interactor_ratio | mitosis | inter | 0.08842363453252496 | 0.4953575337685953 |

P42285_Q9H583
MTREX_HUMAN vs HEAT1_HUMAN
p-value: 0.032 q-value: 0.023

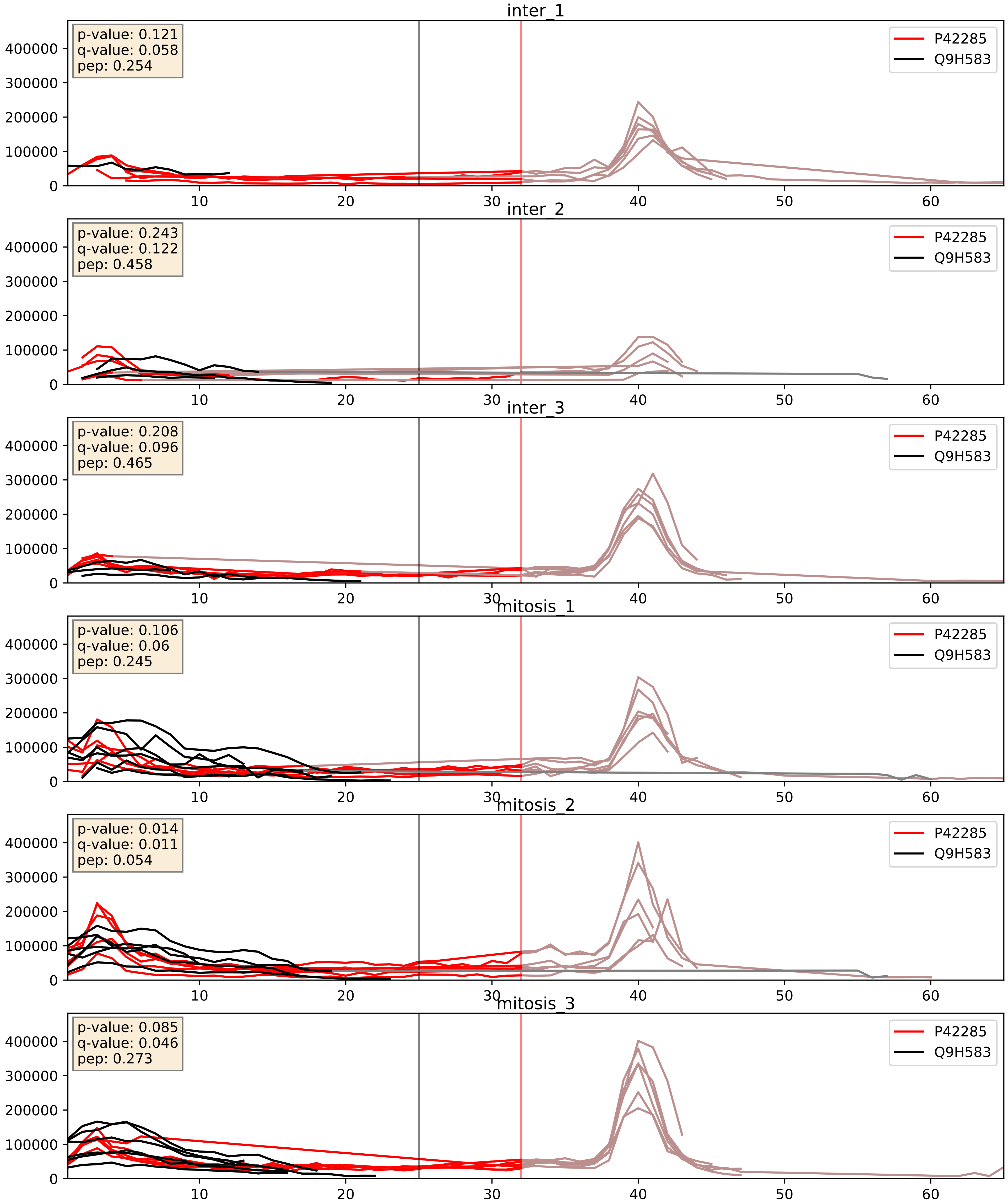


| level | condition_1 | condition_2 | pvalue | pvalue_adjusted |
|---|---|---|---|---|
| complex_abundance | mitosis | inter | 0.00024714820281548363 | 0.010525316498012636 |
| interactor_abundance | mitosis | inter | 0.00721018643363647 | 0.07343152394042615 |
| interactor_ratio | mitosis | inter | 0.11491151406247362 | 0.559317550599762 |

P61247_Q9H583
RS3A_HUMAN vs HEAT1_HUMAN
p-value: 0.071 q-value: 0.043

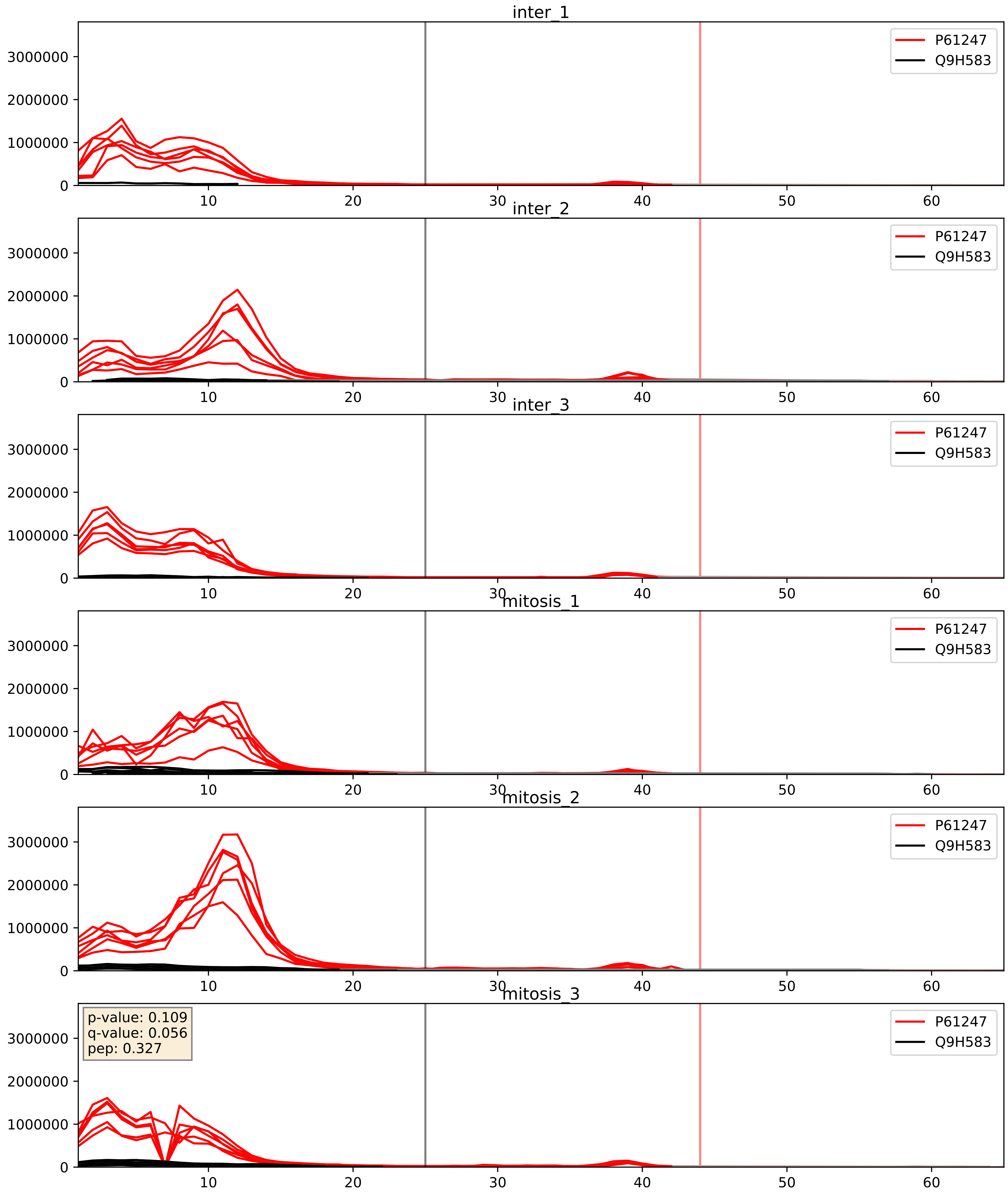


| level | condition_1 | condition_2 | pvalue | pvalue_adjusted |
|---|---|---|---|---|
| complex_abundance | mitosis | inter | 0.00423876784503886 | 0.0484430610861584 |
| interactor_ratio | mitosis | inter | 0.016270844431761702 | 0.18846877988617075 |
| interactor_abundance | mitosis | inter | 0.4025920410924137 | 0.5901502794223283 |

Q12788_Q9H583
TBL3_HUMAN vs HEAT1_HUMAN
p-value: 0.023 q-value: 0.017

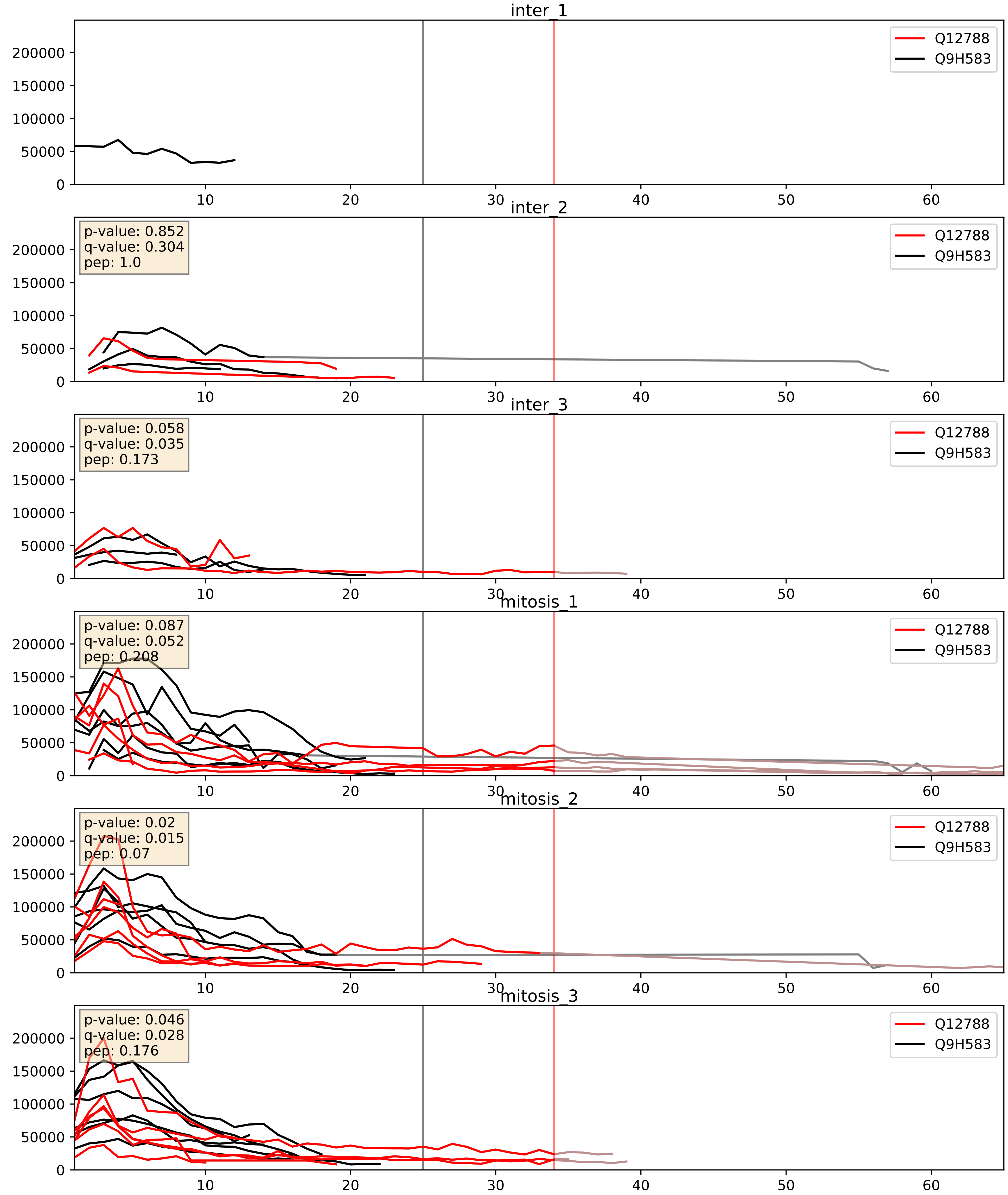


| level | condition_1 | condition_2 | pvalue | pvalue_adjusted |
|---|---|---|---|---|
| interactor_ratio | mitosis | inter | 0.0032358533534543603 | 0.07405554735675686 |
| complex_abundance | mitosis | inter | 0.004224521690450446 | 0.048345498563686945 |
| interactor_abundance | mitosis | inter | 0.009154209095090269 | 0.08324021891678159 |

Q15061_Q9H583
WDR43_HUMAN vs HEAT1_HUMAN
p-value: 0.034 q-value: 0.024

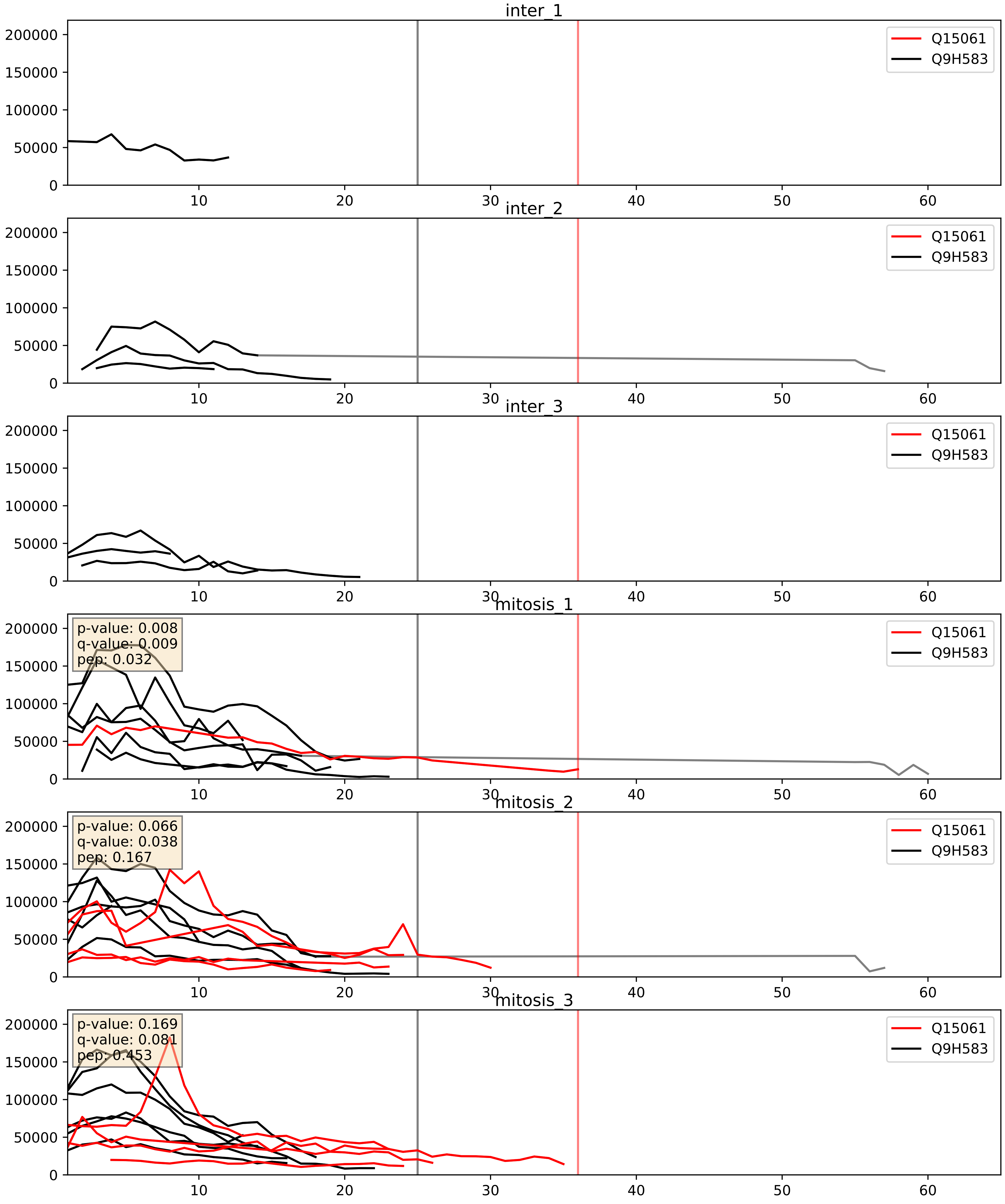


| level | condition_1 | condition_2 | pvalue | pvalue_adjusted |
|---|---|---|---|---|
| complex_abundance | mitosis | inter | 0.005287778237647574 | 0.054740033961712564 |
| interactor_ratio | mitosis | inter | 0.03948312491726528 | 0.32783954136208876 |
| interactor_abundance | mitosis | inter | 0.04327794470388063 | 0.16685321787696922 |

Q15269_Q9H583
PWP2_HUMAN vs HEAT1_HUMAN
p-value: 0.029 q-value: 0.021

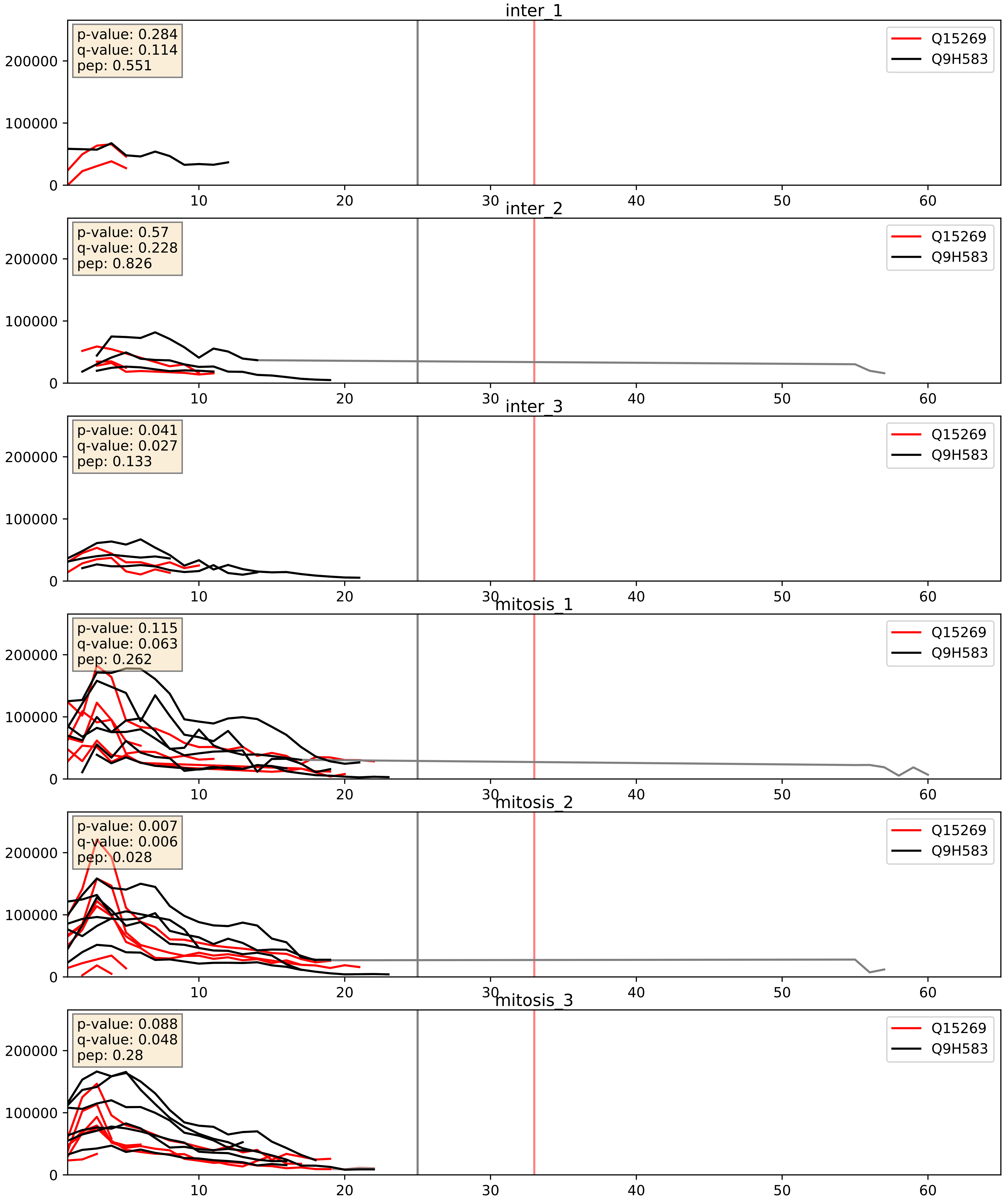


| level | condition_1 | condition_2 | pvalue | pvalue_adjusted |
|---|---|---|---|---|
| complex_abundance | mitosis | inter | 0.0013443121273097713 | 0.025015895238634005 |
| interactor_abundance | mitosis | inter | 0.02134655497707949 | 0.1216554664472706 |
| interactor_ratio | mitosis | inter | 0.06289218988803522 | 0.4166850970910073 |

Q8NI36_Q9H583
WDR36_HUMAN vs HEAT1_HUMAN
p-value: 0.03 q-value: 0.022

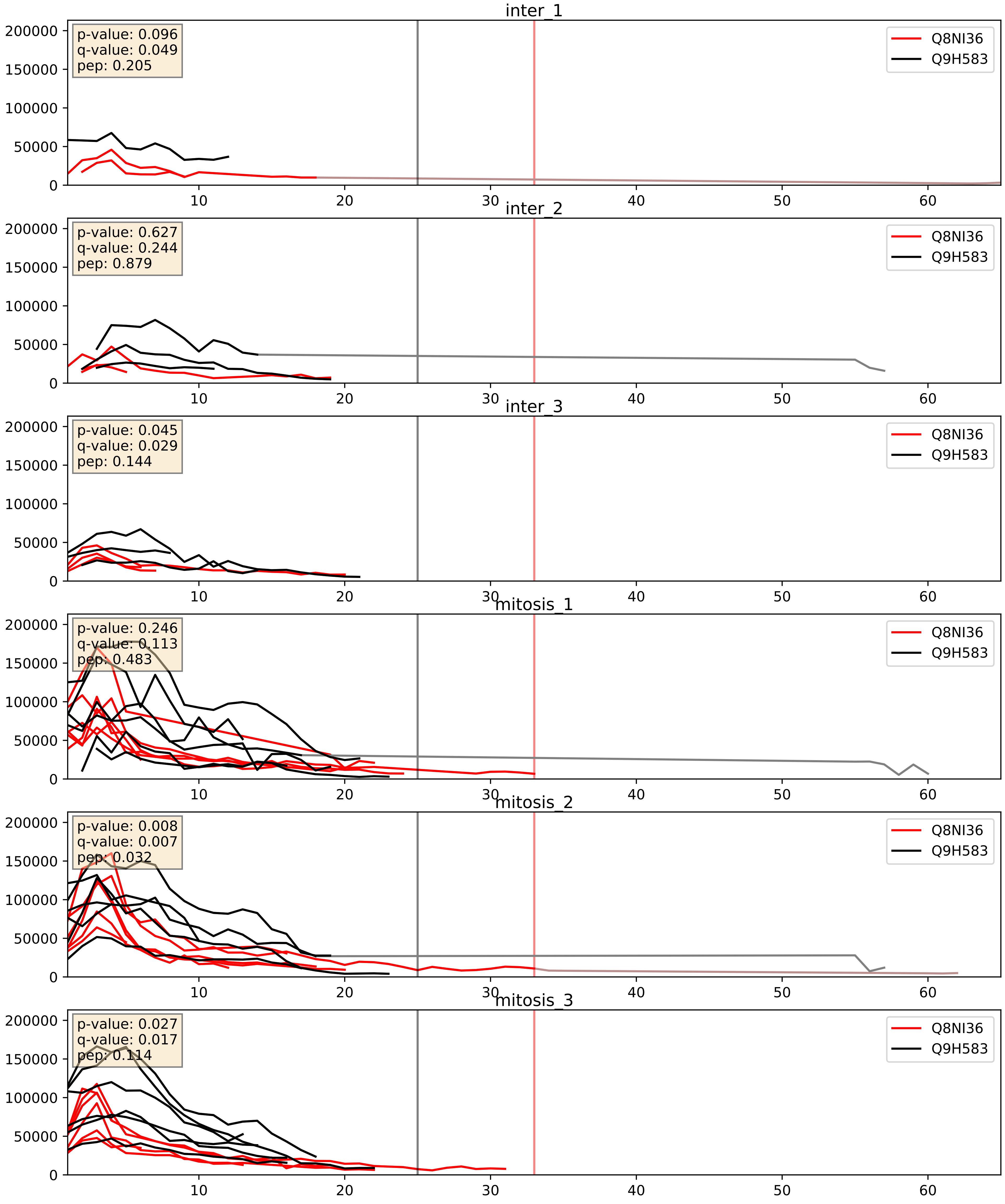


| level | condition_1 | condition_2 | pvalue | pvalue_adjusted |
|---|---|---|---|---|
| complex_abundance | mitosis | inter | 0.00031756796799436147 | 0.011652071109663347 |
| interactor_abundance | mitosis | inter | 0.0035552250741893126 | 0.04601489785035819 |
| interactor_ratio | mitosis | inter | 0.06616503681267413 | 0.4296400448702392 |

Q8TED0_Q9H583
UTP15_HUMAN vs HEAT1_HUMAN
p-value: 0.048 q-value: 0.032

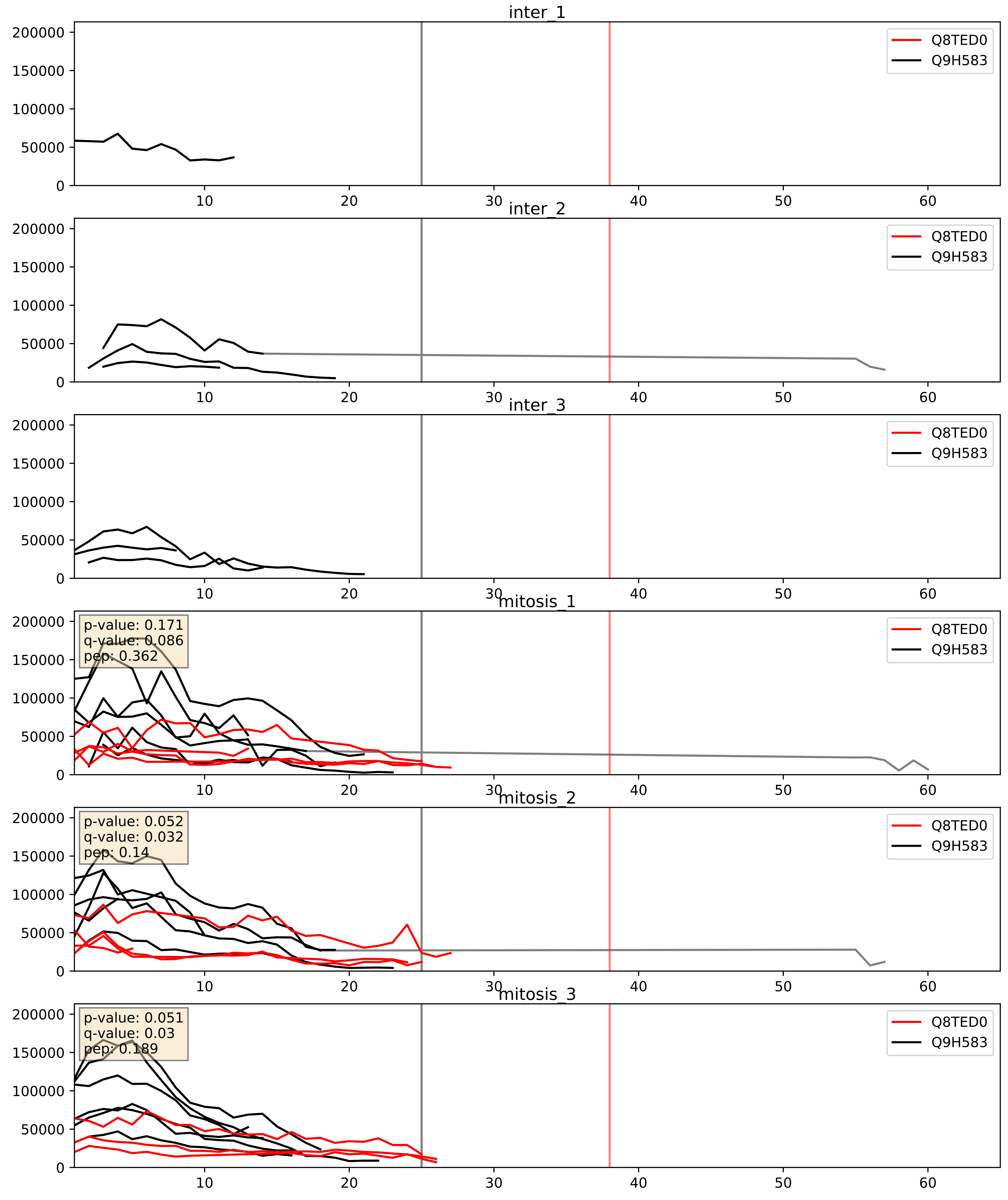


| level | condition_1 | condition_2 | pvalue | pvalue_adjusted |
|---|---|---|---|---|
| complex_abundance | mitosis | inter | 0.00031488057374661957 | 0.01165207110963347 |
| interactor_abundance | mitosis | inter | 0.0015696169880338438 | 0.029025269221018554 |
| interactor_ratio | mitosis | inter | 0.003125922786166029 | 0.0729098066746082 |

Q9BQG0_Q9H583
MBB1A_HUMAN vs HEAT1_HUMAN
p-value: 0.036 q-value: 0.025

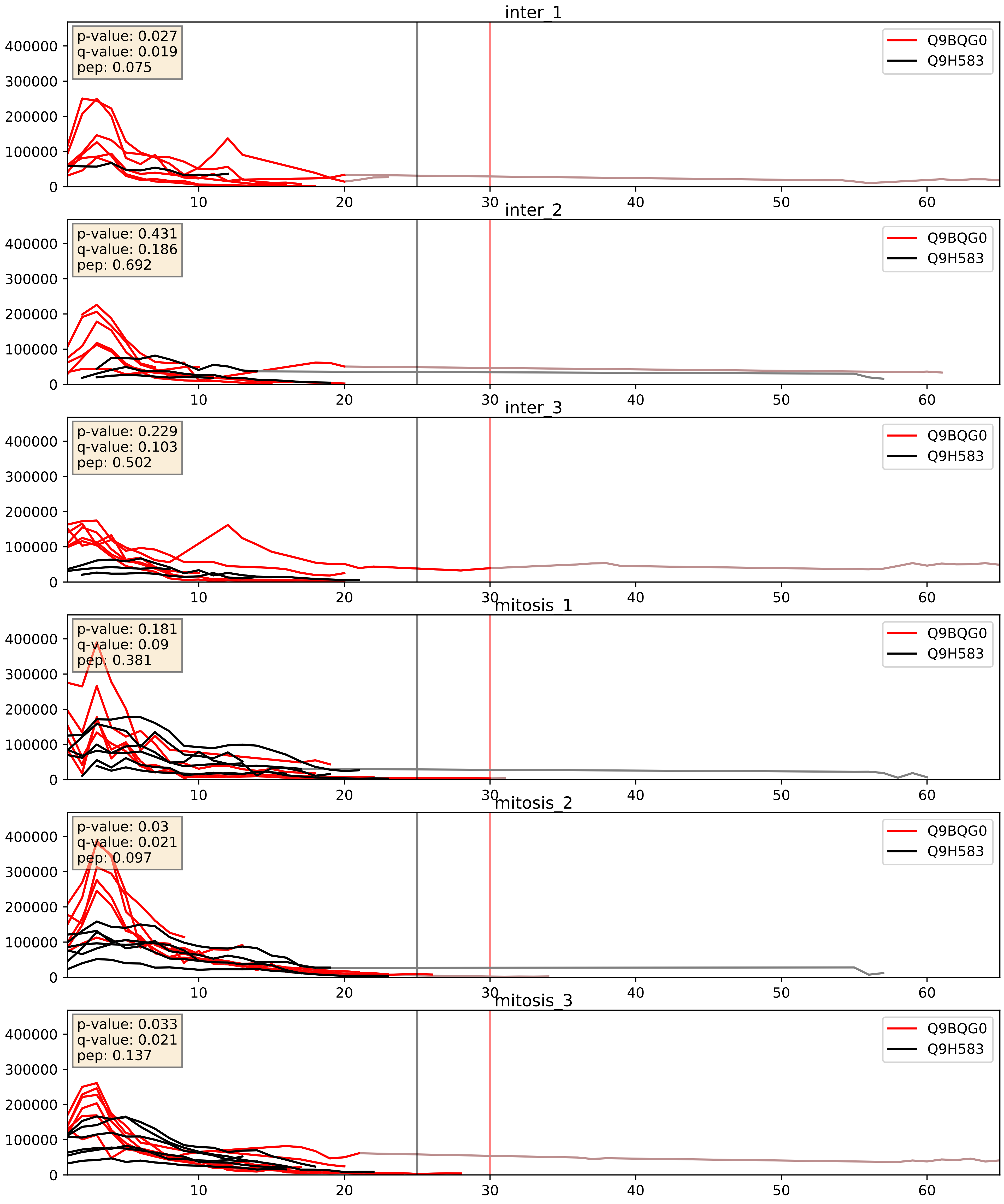


| level | condition_1 | condition_2 | pvalue | pvalue_adjusted |
|---|---|---|---|---|
| complex_abundance | mitosis | inter | 0.0012715449926405742 | 0.024187611415562922 |
| interactor_ratio | mitosis | inter | 0.012032971951948402 | 0.15846498447488974 |
| interactor_abundance | mitosis | inter | 0.03368869821852597 | 0.1532530798125457 |

Q9H583_Q9Y2X3
HEAT1_HUMAN vs NOP58_HUMAN
p-value: 0.08 q-value: 0.047

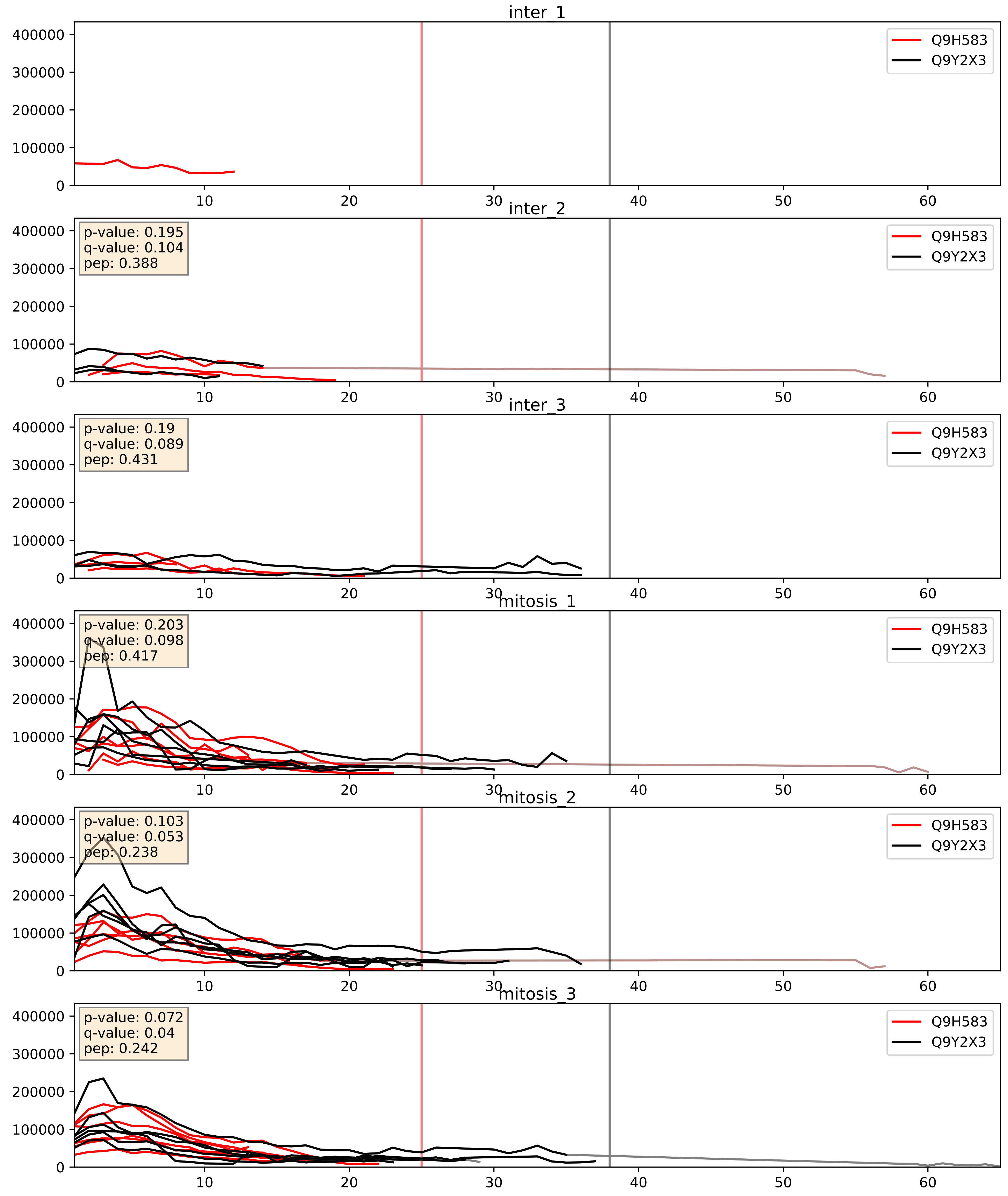


| level | condition_1 | condition_2 | pvalue | pvalue_adjusted |
|---|---|---|---|---|
| complex_abundance | mitosis | inter | 0.003745309179790294 | 0.04515471349155622 |
| interactor_abundance | mitosis | inter | 0.006814646959063384 | 0.07092204008459886 |
| interactor_ratio | mitosis | inter | 0.05599641218001745 | 0.3926647223857623 |
