## Supplementary Data 2 for "SECAT: Quantifying differential protein-protein interaction states by network-centric analysis": hela_string_000066_P53350.pdf

P53350_P53350

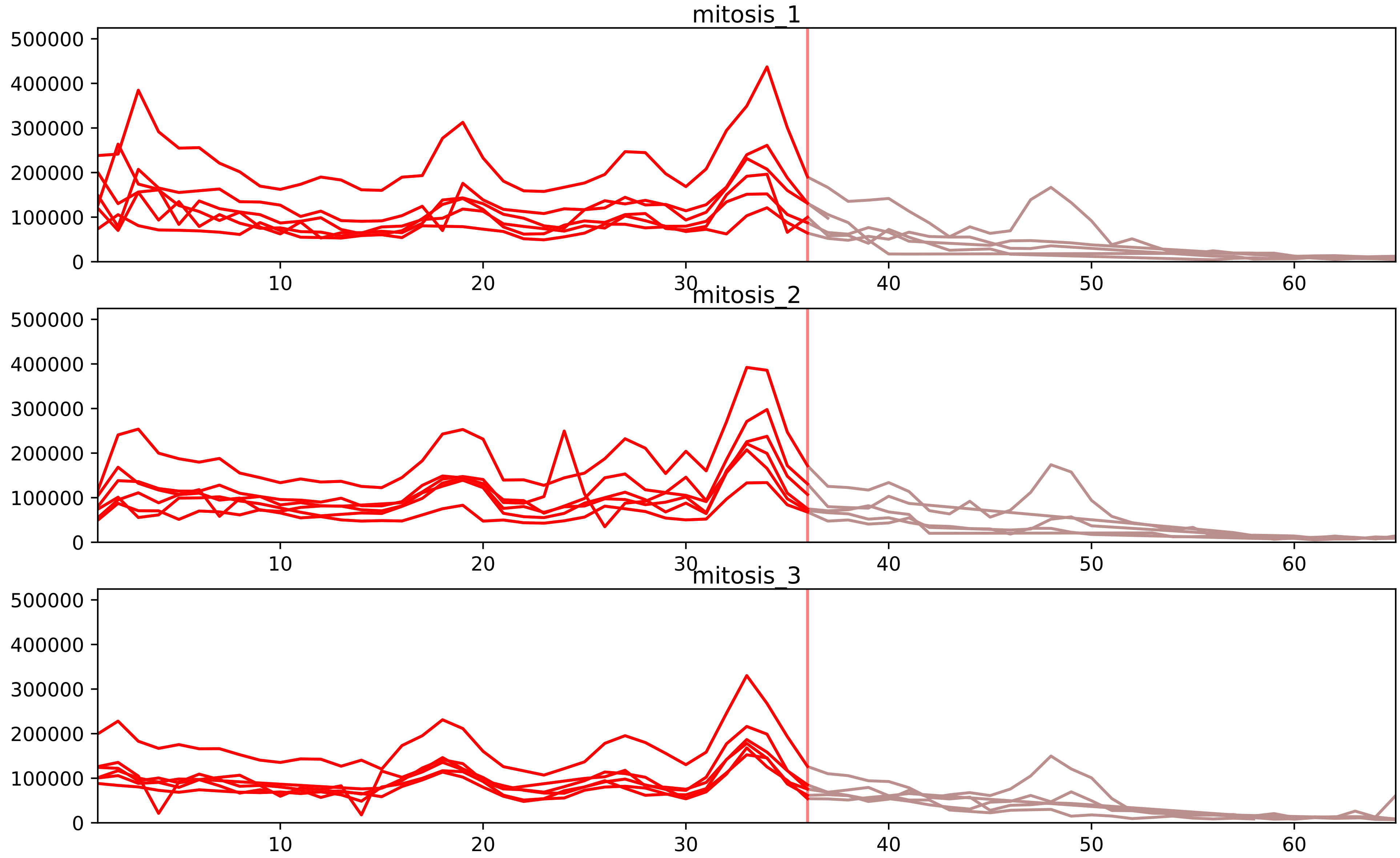


| level | condition_1 | condition_2 | pvalue | pvalue_adjusted |
|---|---|---|---|---|
| complex_abundance | mitosis | inter | 7.969921489367675e-07 | 8.147030855798067e-05 |
| interactor_abundance | mitosis | inter | 8.703069782027219e-07 | 0.00011438320284950059 |
| total_abundance | mitosis | inter | 2.4403327490792538e-06 | 0.003540864511957668 |
| assembled_abundance | mitosis | inter | 2.5138949135432614e-06 | 0.0023535084180592012 |
| monomer_abundance | mitosis | inter | 9.632706566599342e-05 | 0.03261473898341094 |
| interactor_ratio | mitosis | inter | 0.0015349348721193973 | 0.015690445359442726 |

O43684_P53350
BUB3_HUMAN vs PLK1_HUMAN
p-value: 0.051 q-value: 0.033

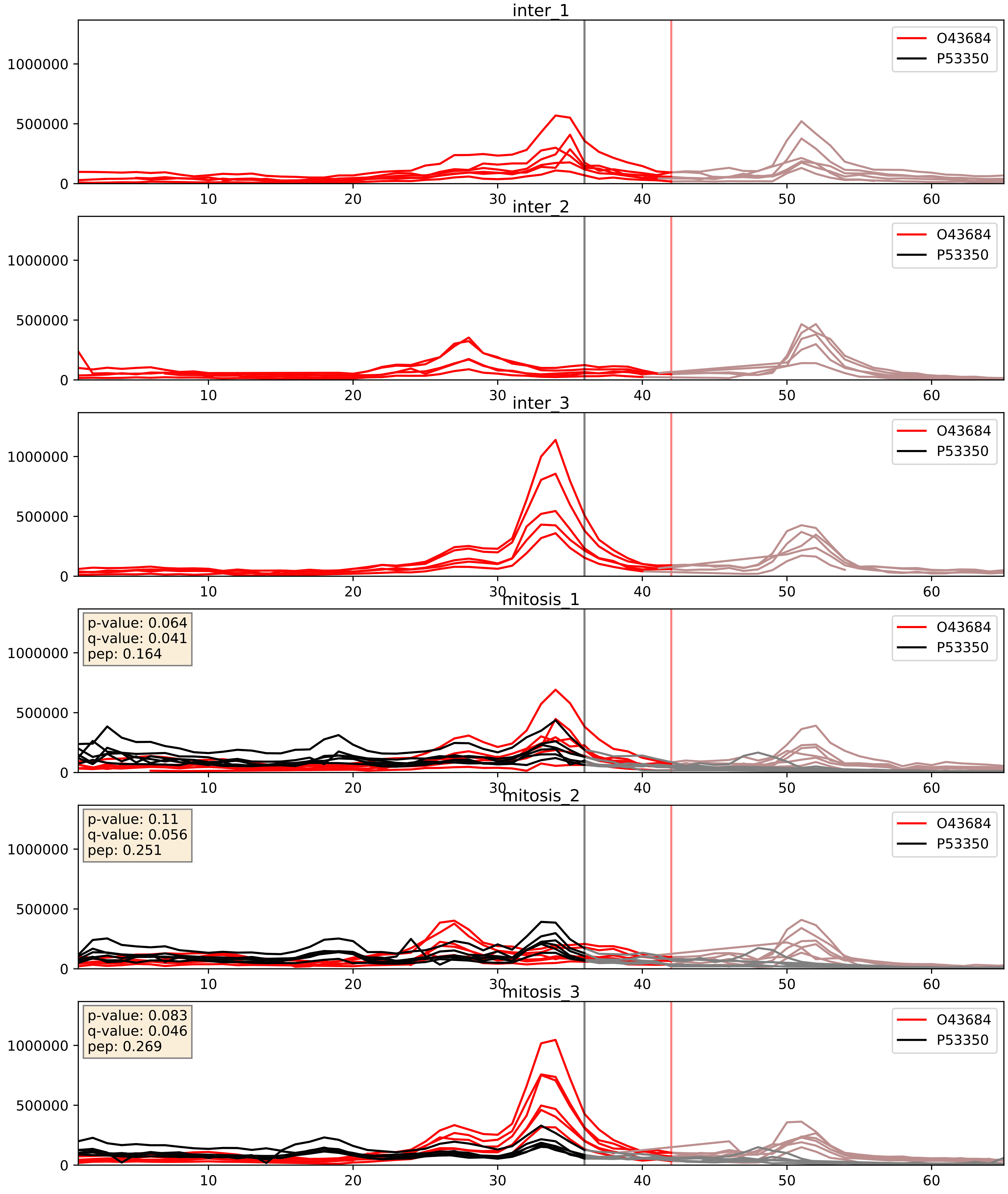


| level | condition_1 | condition_2 | pvalue | pvalue_adjusted |
|---|---|---|---|---|
| complex_abundance | mitosis | inter | 2.6656409536998423e-05 | 0.005691857151652958 |
| interactor_abundance | mitosis | inter | 7.51921713288653e-05 | 0.00688390359973355 |
| interactor_ratio | mitosis | inter | 0.9157420255633383 | 0.9855669004419936 |

P53350_Q9Y266
PLK1_HUMAN vs NUDC_HUMAN
p-value: 0.073 q-value: 0.044

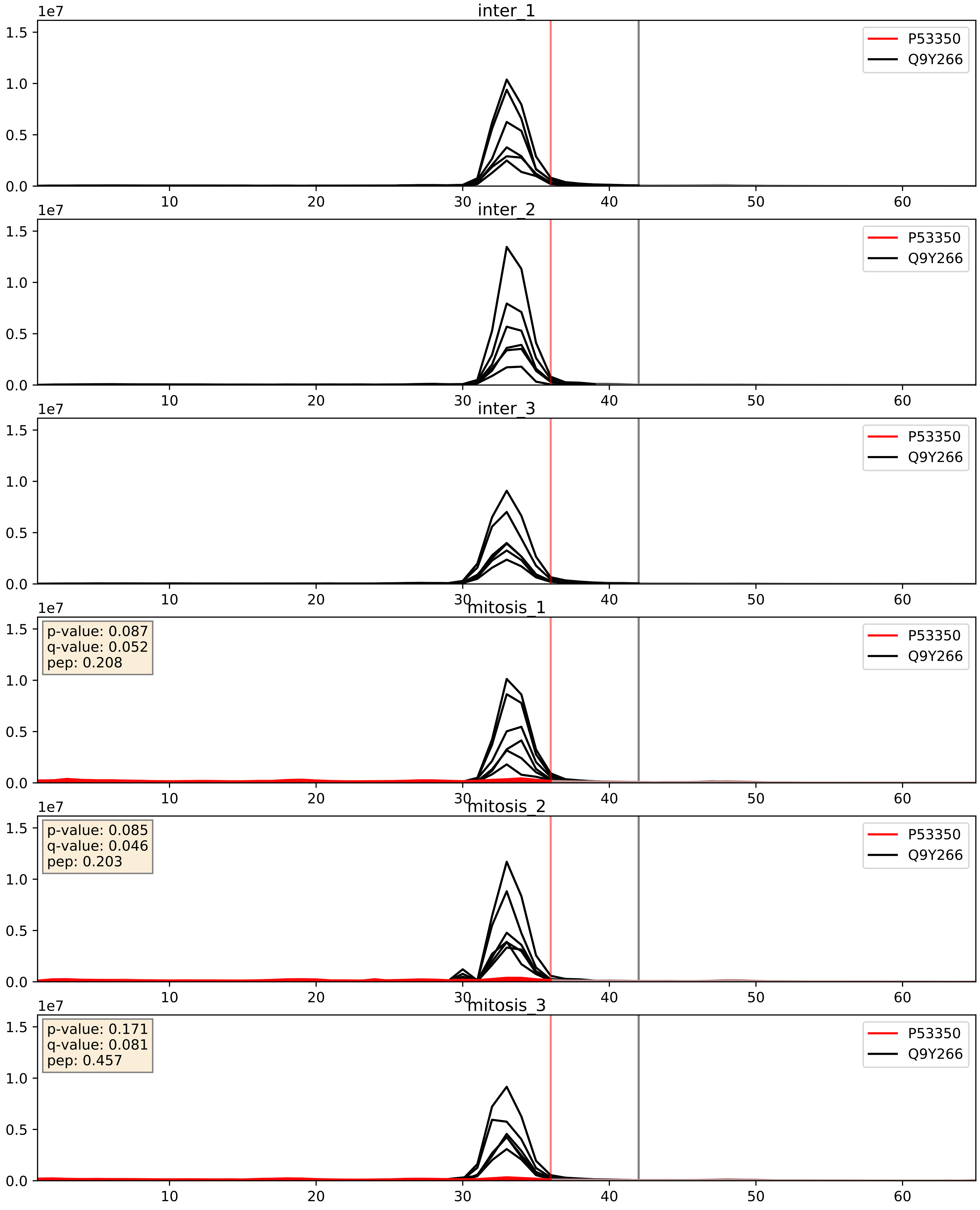


| level | condition_1 | condition_2 | pvalue | pvalue_adjusted |
|---|---|---|---|---|
| complex_abundance | mitosis | inter | 1.1936559716790173e-05 | 0.005691857151652958 |
| interactor_abundance | mitosis | inter | 1.8795469290468416e-05 | 0.004081879629435812 |
| interactor_ratio | mitosis | inter | 0.0001559547874814811 | 0.012836278661937291 |
