## Supplementary Data 2 for "SECAT: Quantifying differential protein-protein interaction states by network-centric analysis": hela_string_000067_Q1KMD3.pdf

Q1KMD3_Q1KMD3

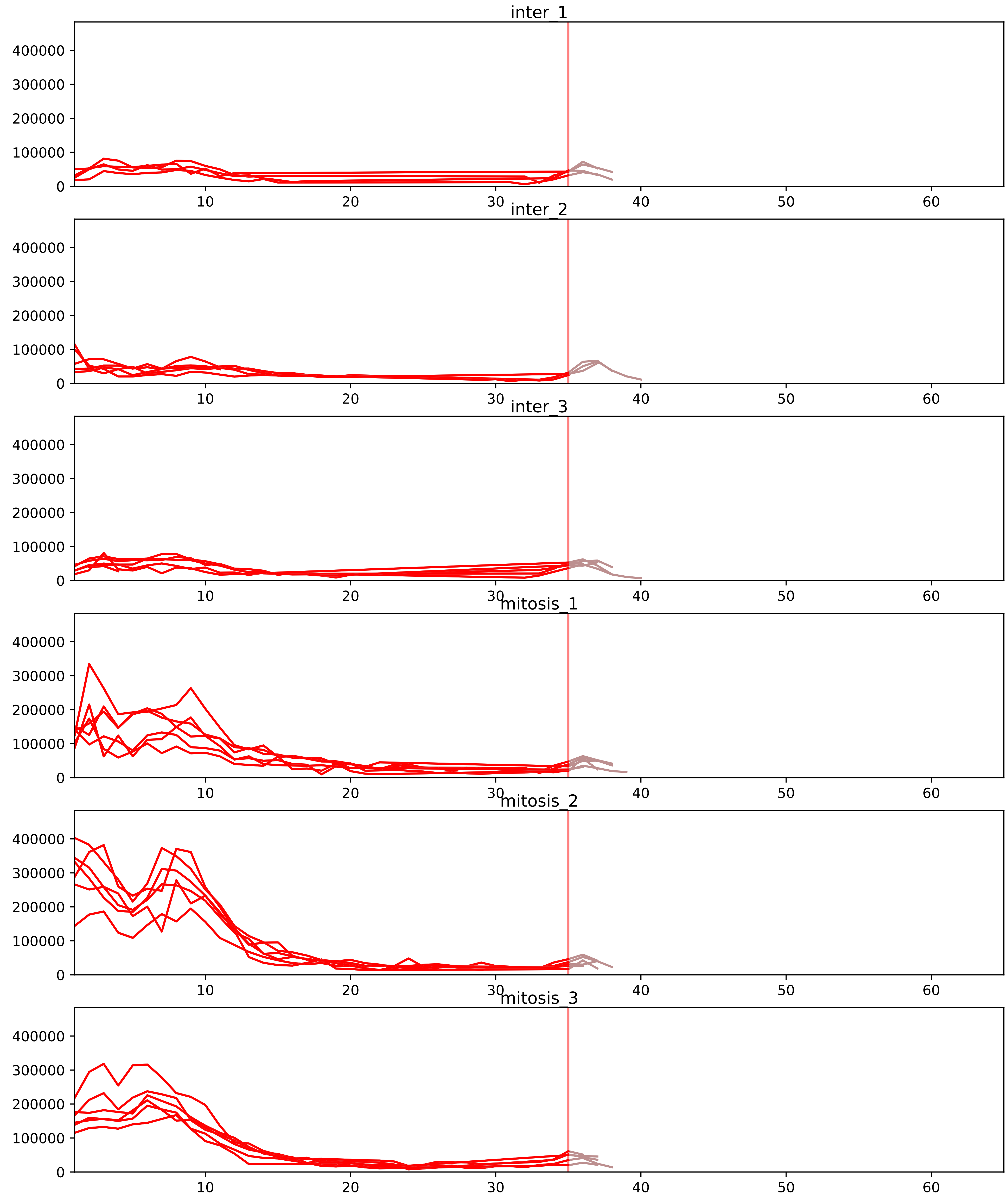


| level | condition_1 | condition_2 | pvalue | pvalue_adjusted |
|---|---|---|---|---|
| complex_abundance | mitosis | inter | 8.668080290568026e-07 | 8.39435143928693e-05 |
| interactor_abundance | mitosis | inter | 0.00012634120272096616 | 0.002734915447136209 |
| total_abundance | mitosis | inter | 0.0028083301637058557 | 0.05744188418058572 |
| assembled_abundance | mitosis | inter | 0.003004915549823413 | 0.053918527981348933 |
| interactor_ratio | mitosis | inter | 0.005567869221764732 | 0.04268699736686295 |
| monomer_abundance | mitosis | inter | 0.8767857745578896 | 0.9675123851245806 |

P51991_Q1KMD3
ROA3_HUMAN vs HNRL2_HUMAN
p-value: 0.0 q-value: 0.047

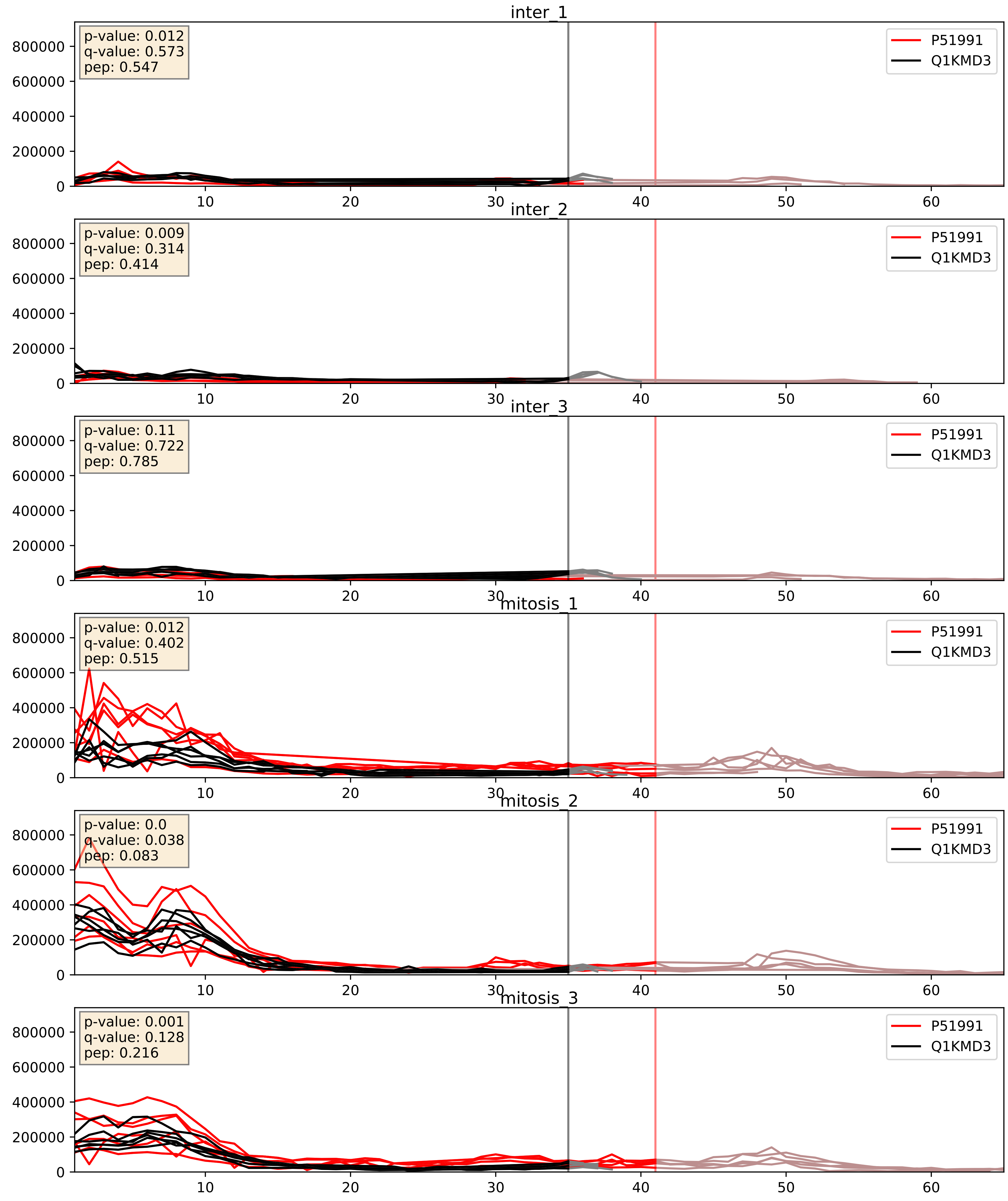


| level | condition_1 | condition_2 | pvalue | pvalue_adjusted |
|---|---|---|---|---|
| complex_abundance | mitosis | inter | 8.668080290568026e-07 | 0.0024732922429087437 |
| interactor_abundance | mitosis | inter | 3.1946371197007753e-06 | 0.0033734327106189023 |
| interactor_ratio | mitosis | inter | 0.005567869221764732 | 0.10119099901975818 |
