## Supplementary Data 2 for "SECAT: Quantifying differential protein-protein interaction states by network-centric analysis": hela_string_000068_O43395.pdf

O43395_O43395

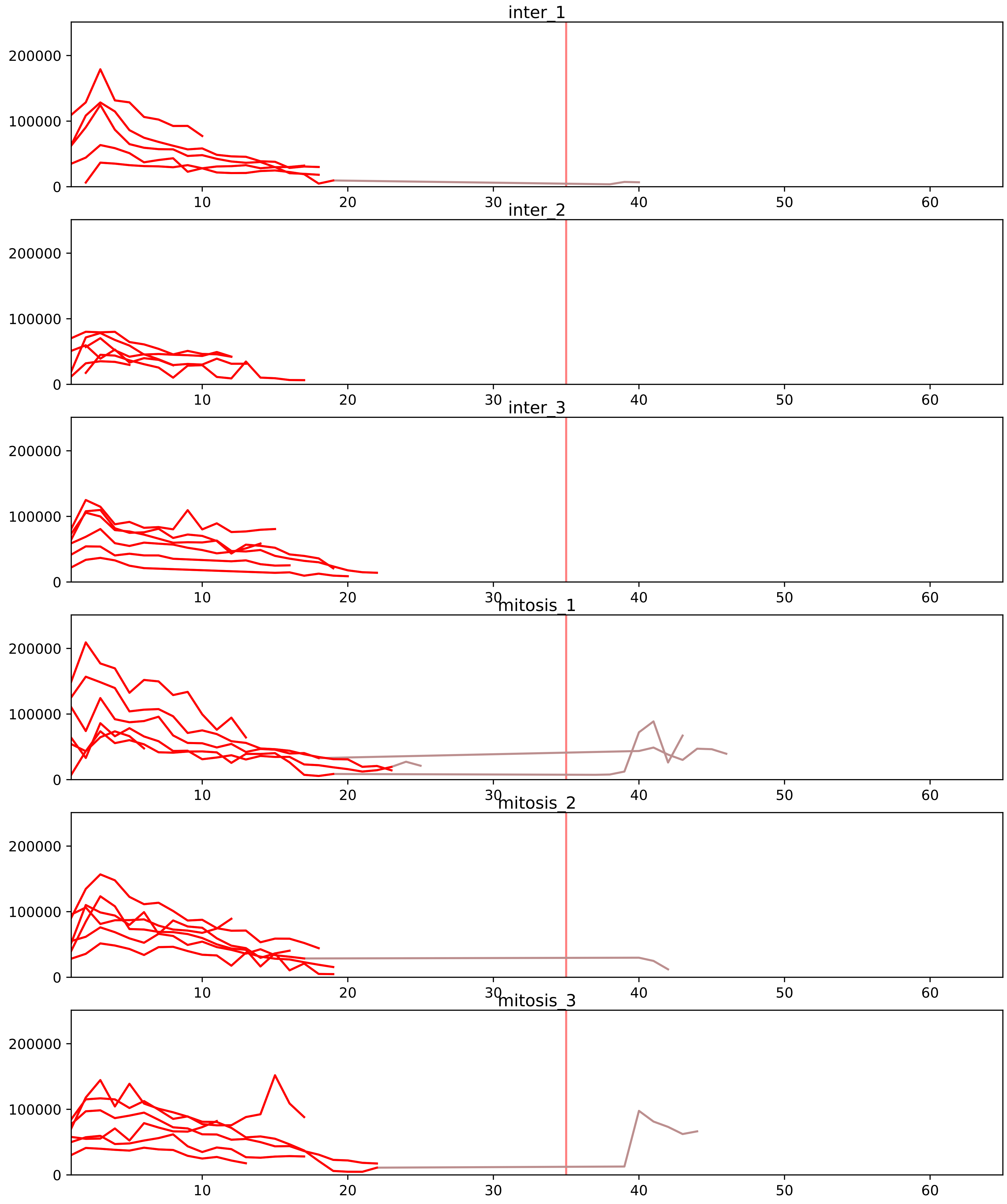


| level | condition_1 | condition_2 | pvalue | pvalue_adjusted |
|---|---|---|---|---|
| interactor_ratio | mitosis | inter | 1.0589666635536282e-06 | 3.1942600998994685e-05 |
| complex_abundance | mitosis | inter | 0.0006751661667077167 | 0.0083939577482581 |
| interactor_abundance | mitosis | inter | 0.0017968759079445135 | 0.017775546616225296 |
| monomer_abundance | mitosis | inter | 0.13779225242897966 | 0.4888918124193968 |
| assembled_abundance | mitosis | inter | 0.641886138035716 | 0.8374216867740208 |
| total_abundance | mitosis | inter | 0.6988766237598585 | 0.8697093355075628 |

O43172_O43395
PRP4_HUMAN vs PRPF3_HUMAN
p-value: 0.001 q-value: 0.001

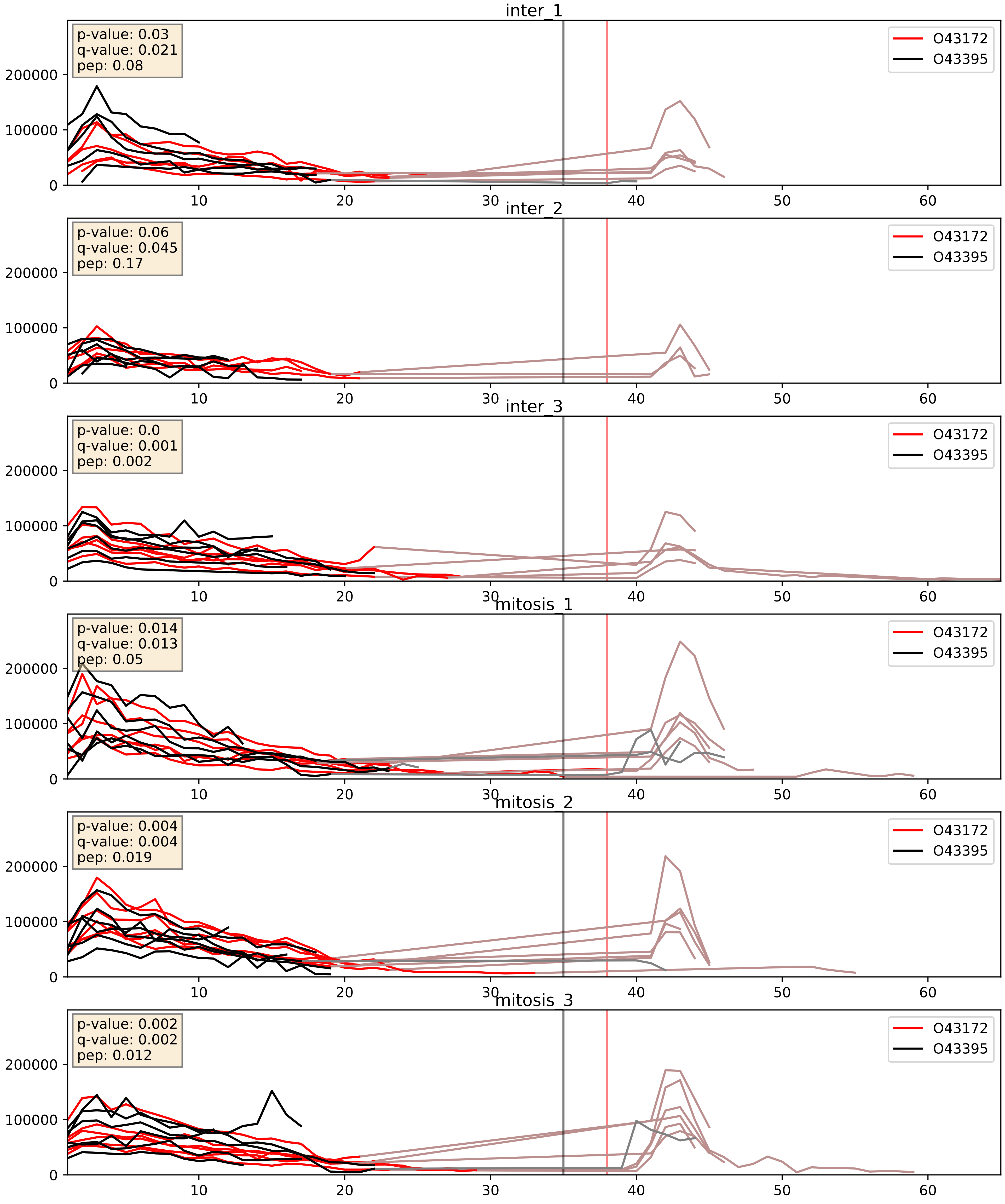


| level | condition_1 | condition_2 | pvalue | pvalue_adjusted |
|---|---|---|---|---|
| interactor_abundance | mitosis | inter | 0.2011910805631144 | 0.39604361265269844 |
| complex_abundance | mitosis | inter | 0.21998527108379787 | 0.3743685726595049 |
| interactor_ratio | mitosis | inter | 0.5720390309274075 | 0.8945917155252349 |

O43290_O43395
SNUT1_HUMAN vs PRPF3_HUMAN
p-value: 0.023 q-value: 0.018

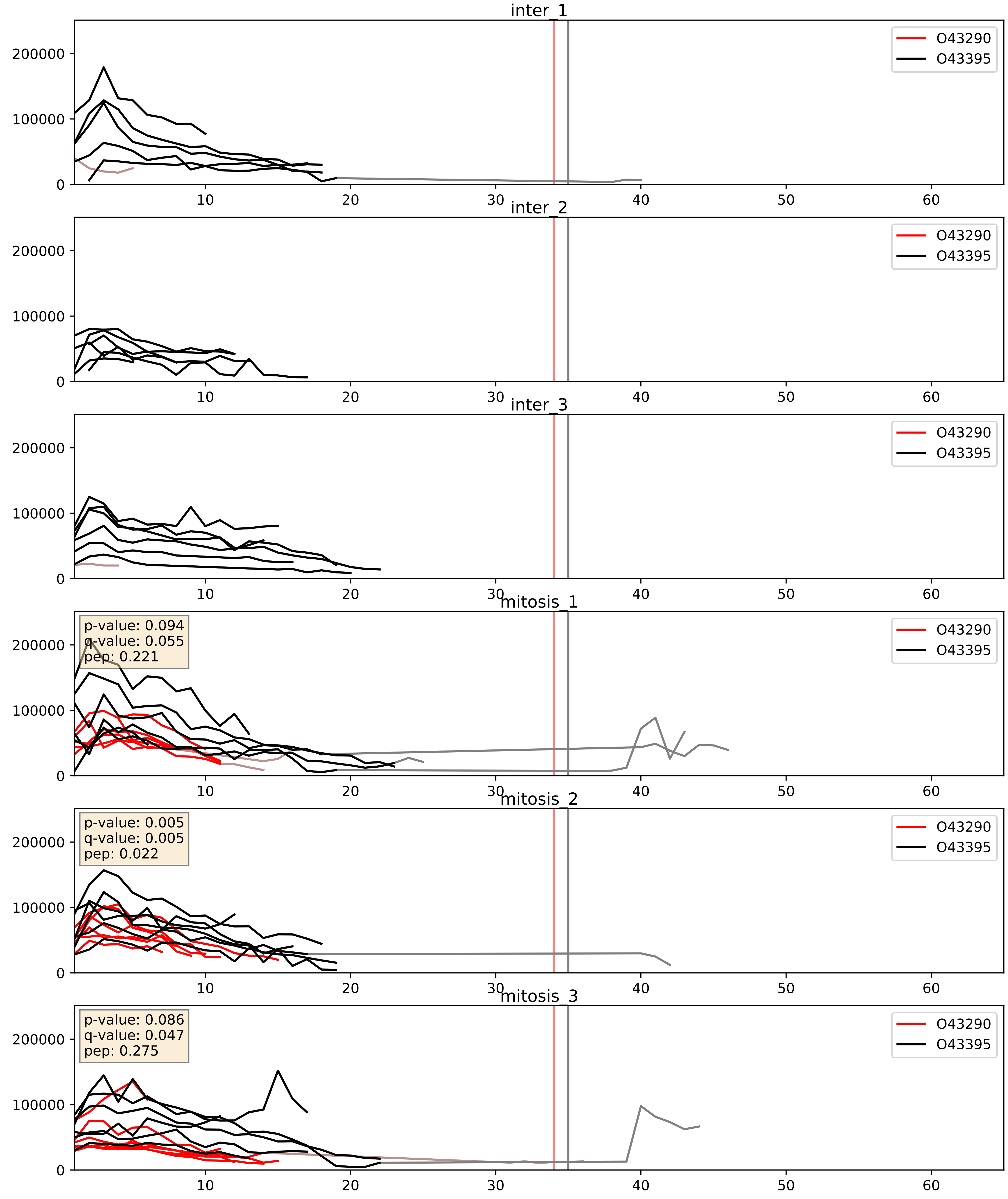


| level | condition_1 | condition_2 | pvalue | pvalue_adjusted |
|---|---|---|---|---|
| complex_abundance | mitosis | inter | 0.000327848443107248 | 0.011886635542316795 |
| interactor_abundance | mitosis | inter | 0.0010647499919498099 | 0.024049720606162197 |
| interactor_ratio | mitosis | inter | 0.007093762501146312 | 0.11727616584402438 |

O43395_O43447
PRPF3_HUMAN vs PPIH_HUMAN
p-value: 0.028 q-value: 0.021

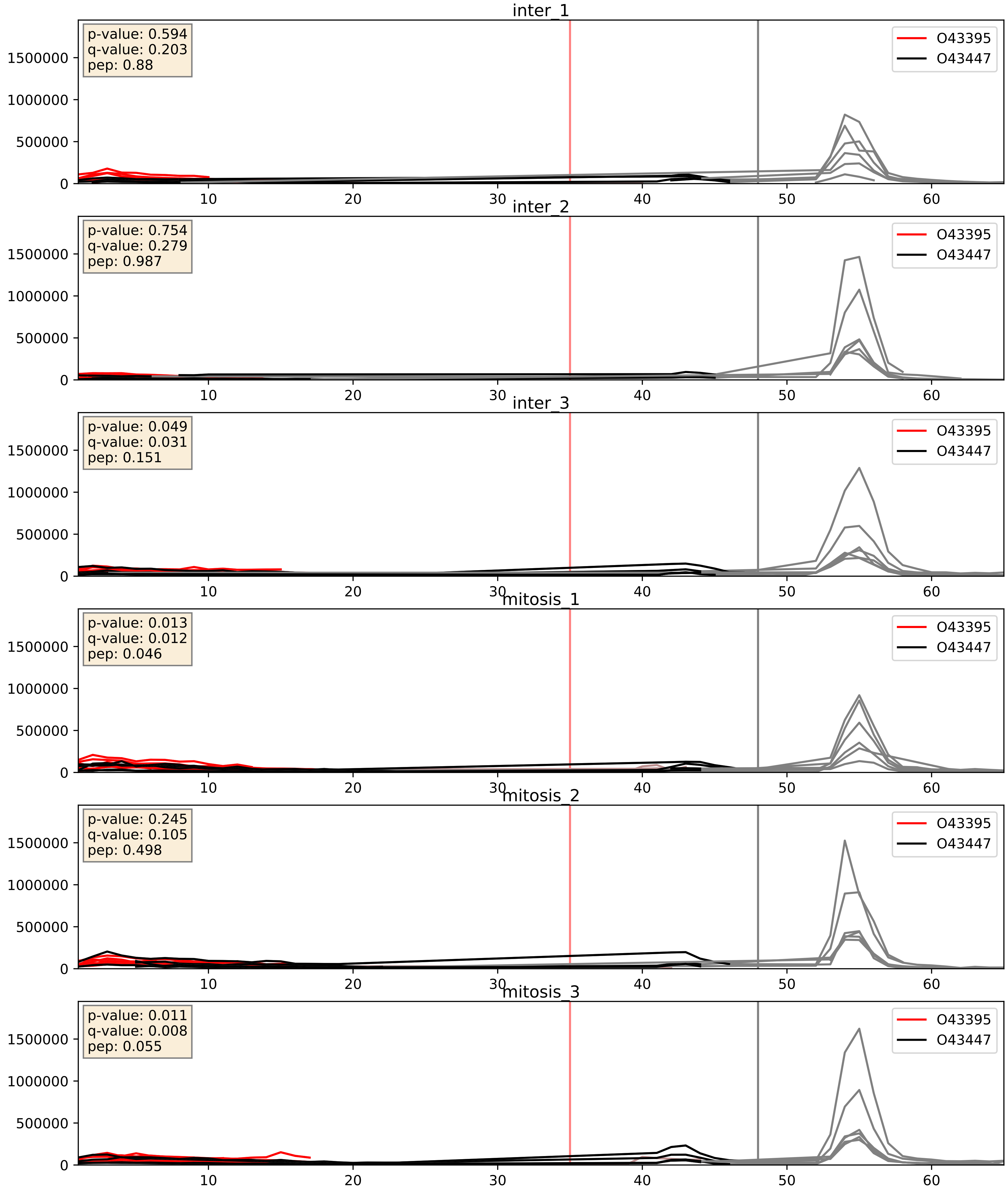


| level | condition_1 | condition_2 | pvalue | pvalue_adjusted |
|---|---|---|---|---|
| complex_abundance | mitosis | inter | 0.3081016439164507 | 0.45796466918390843 |
| interactor_abundance | mitosis | inter | 0.3100367719395898 | 0.5063273428985764 |
| interactor_ratio | mitosis | inter | 0.8500431886764133 | 0.9755634896635457 |

O43395_O75934
PRPF3_HUMAN vs SPF27_HUMAN
p-value: 0.017 q-value: 0.014

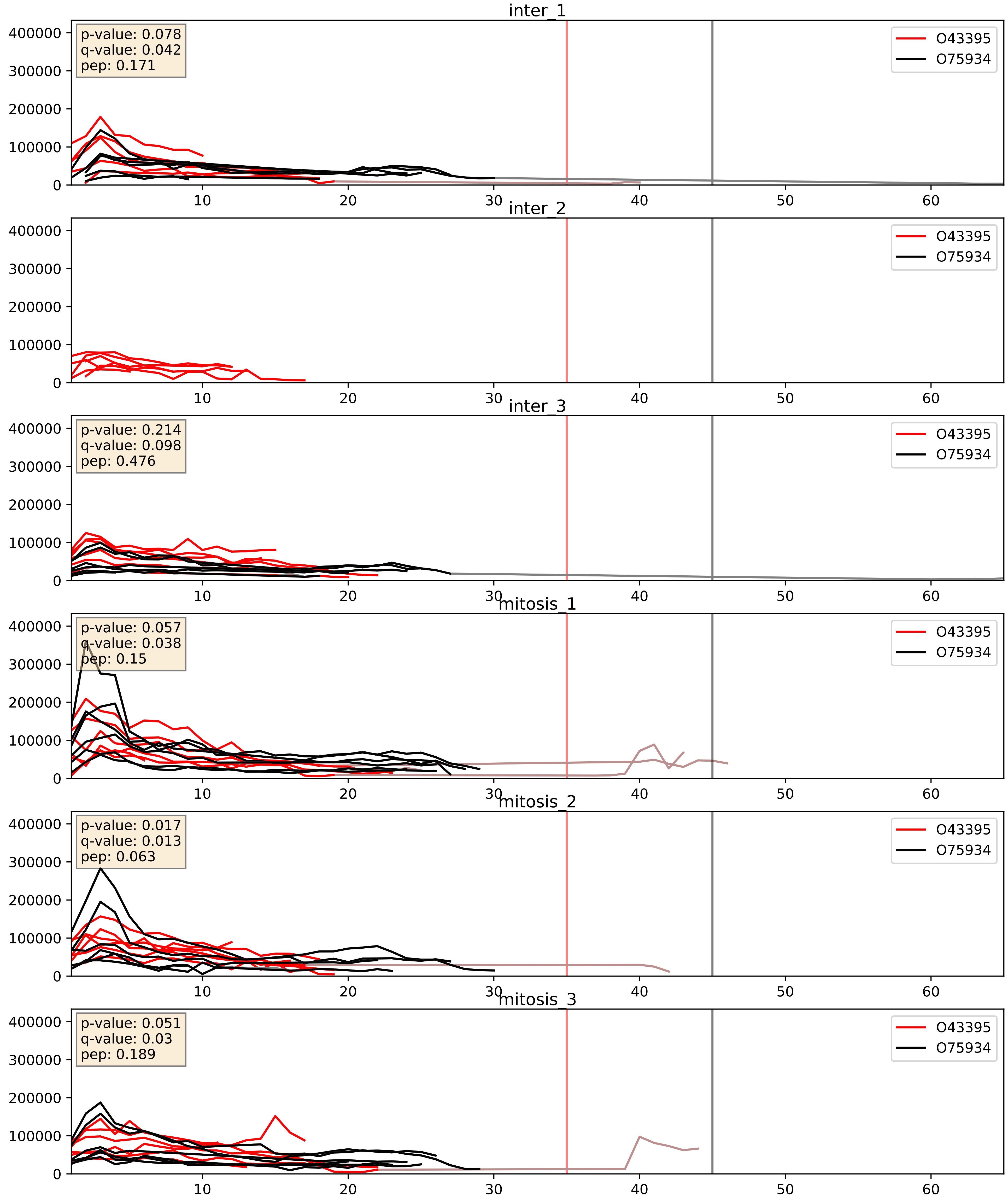


| level | condition_1 | condition_2 | pvalue | pvalue_adjusted |
|---|---|---|---|---|
| complex_abundance | mitosis | inter | 0.1851475550568205 | 0.336615734724054 |
| interactor_abundance | mitosis | inter | 0.29173386071592 | 0.4862448032696506 |
| interactor_ratio | mitosis | inter | 0.3146050758170687 | 0.7783421708721073 |

# O43395_O94906
PRPF3_HUMAN vs PRP6_HUMAN
p-value: 0.002 q-value: 0.003

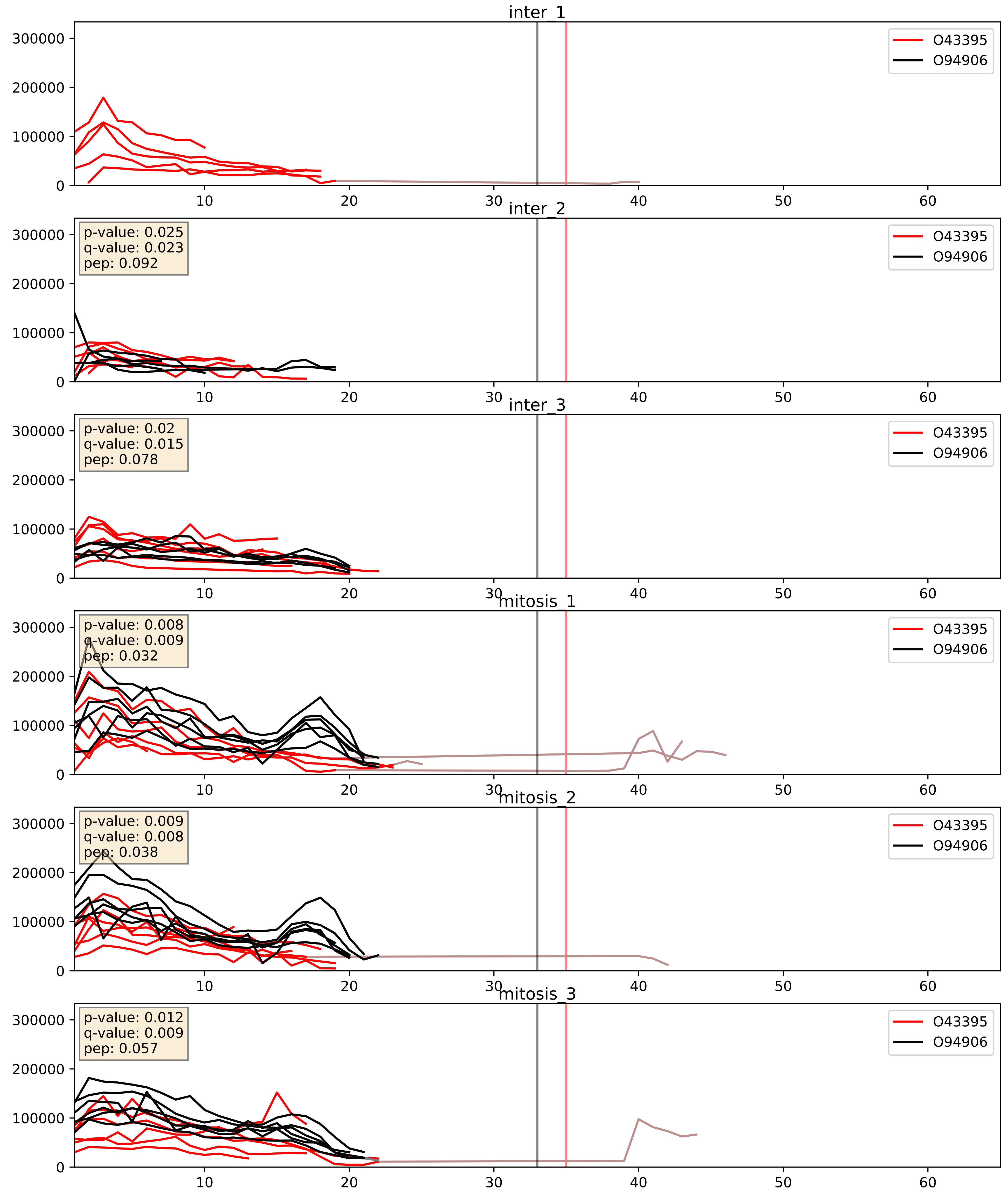


| level | condition_1 | condition_2 | pvalue | pvalue_adjusted |
|---|---|---|---|---|
| interactor_ratio | mitosis | inter | 0.006807476722737148 | 0.11448330205624752 |
| complex_abundance | mitosis | inter | 0.09721955987231426 | 0.23097403067083264 |
| interactor_abundance | mitosis | inter | 0.21396425439359726 | 0.409281344717138 |

O43395_P55769
PRPF3_HUMAN vs NH2L1_HUMAN
p-value: 0.066 q-value: 0.041

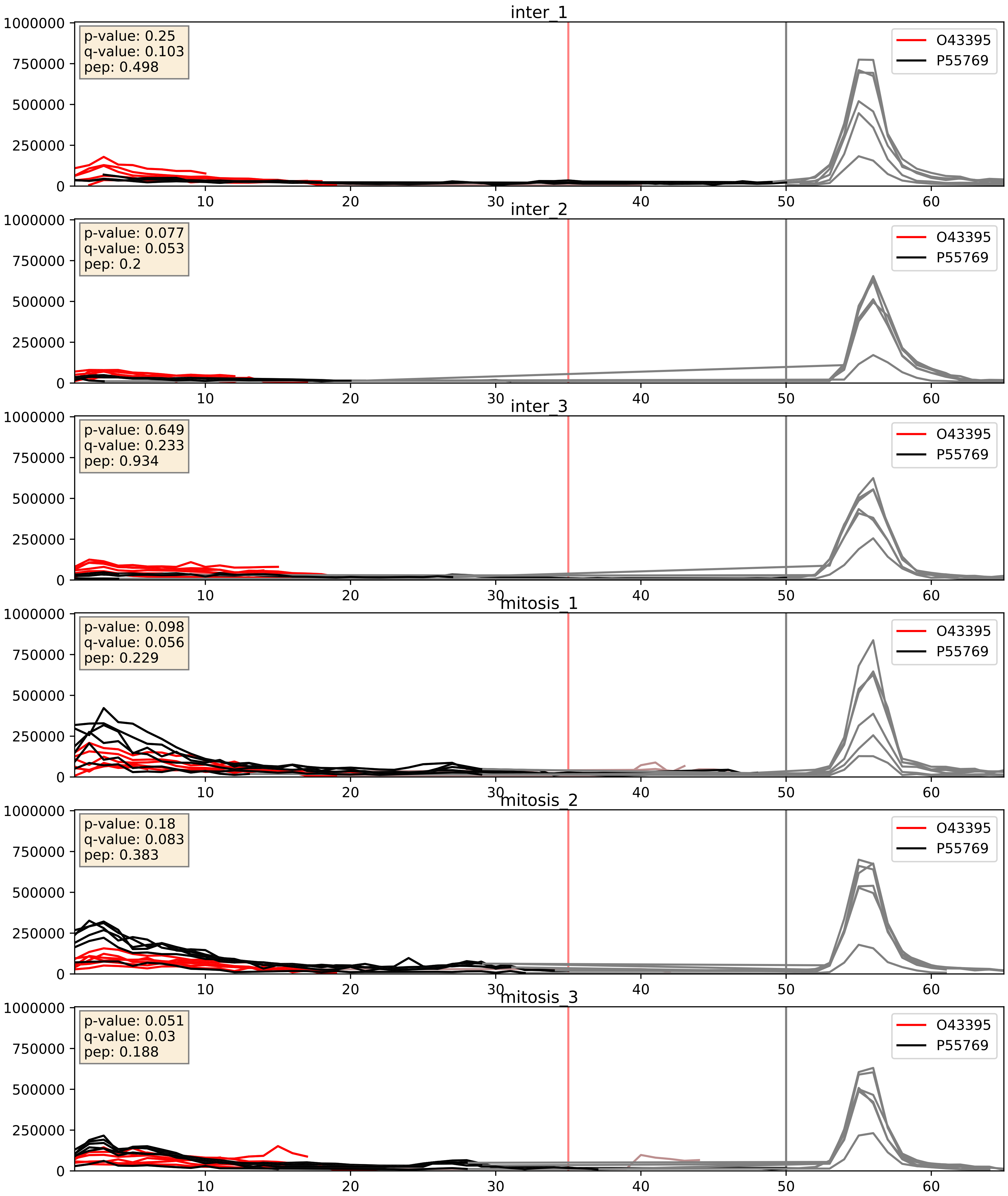


| level | condition_1 | condition_2 | pvalue | pvalue_adjusted |
|---|---|---|---|---|
| complex_abundance | mitosis | inter | 0.0049893793514525385 | 0.05279244406481302 |
| interactor_ratio | mitosis | inter | 0.016058636694013865 | 0.18702303415069207 |
| interactor_abundance | mitosis | inter | 0.3100367719395898 | 0.5063273428985764 |

O43395_Q53GS9
PRPF3_HUMAN vs SNUT2_HUMAN
p-value: 0.004 q-value: 0.004

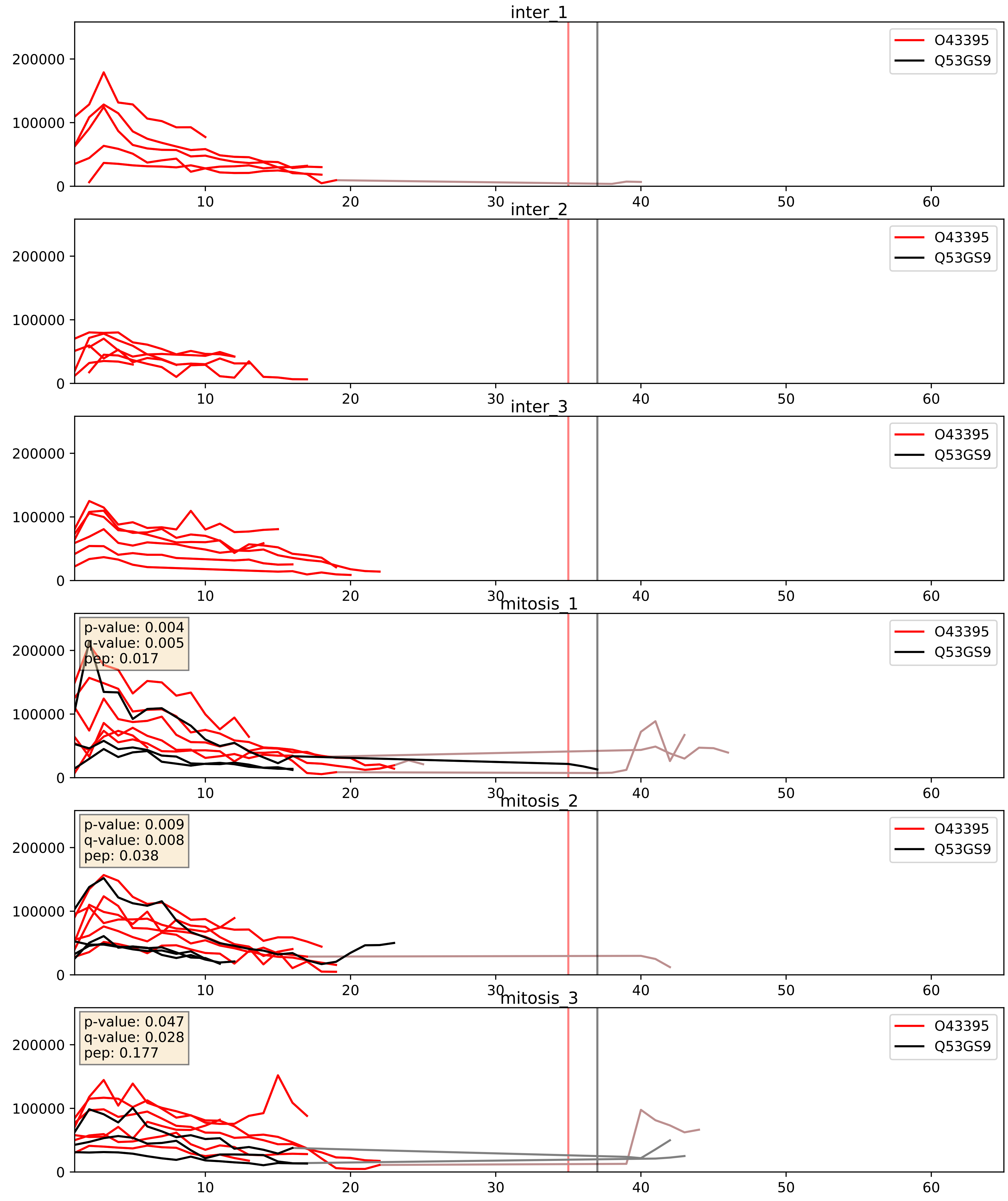


| level | condition_1 | condition_2 | pvalue | pvalue_adjusted |
|---|---|---|---|---|
| interactor_abundance | mitosis | inter | 0.00027487684997001897 | 0.012568482179319555 |
| complex_abundance | mitosis | inter | 0.00043128174253438033 | 0.013091389064164169 |
| interactor_ratio | mitosis | inter | 0.004718642351376341 | 0.09159088101537749 |

O43395_Q8WWY3
PRPF3_HUMAN vs PRP31_HUMAN
p-value: 0.036 q-value: 0.025

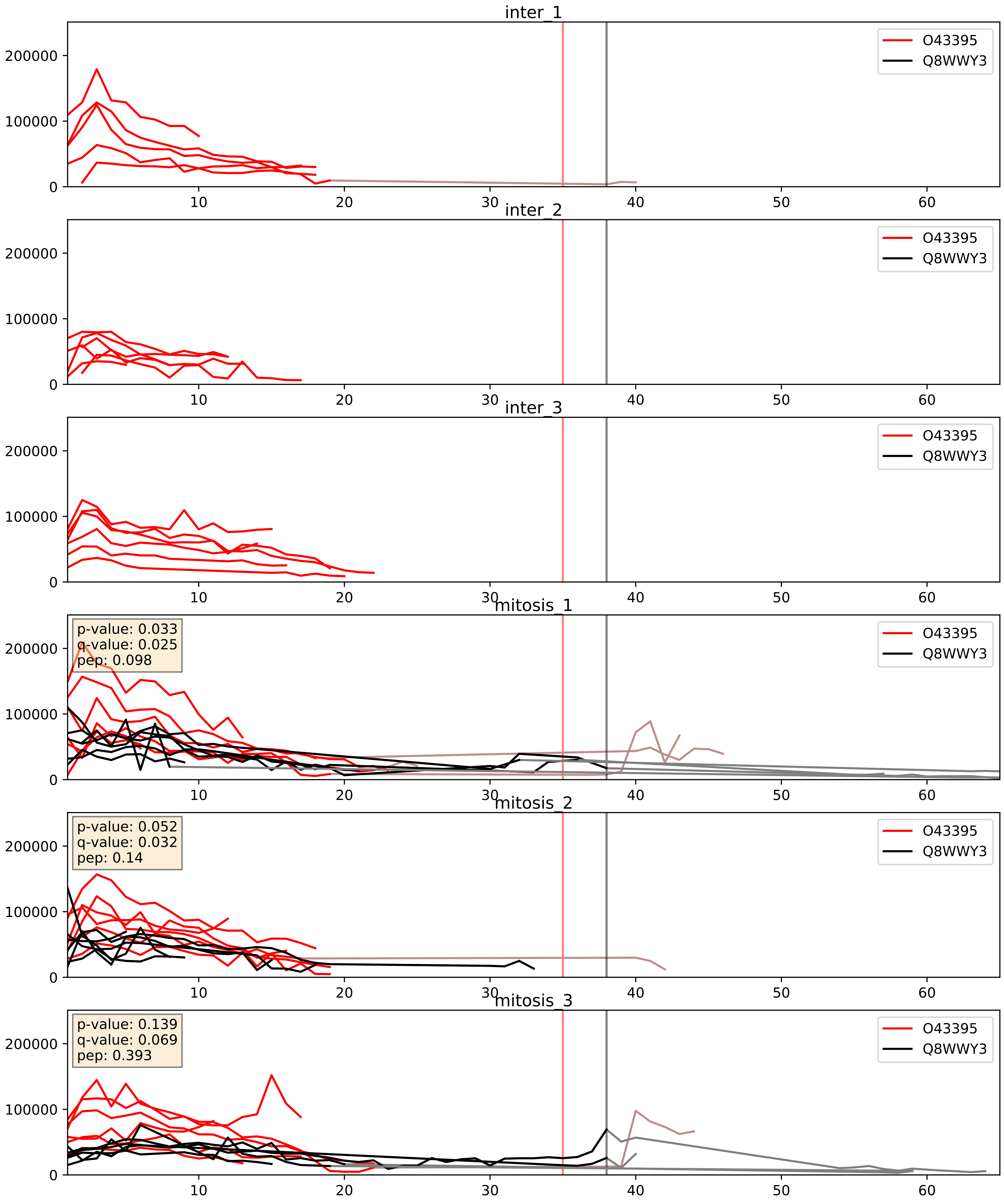


| level | condition_1 | condition_2 | pvalue | pvalue_adjusted |
|---|---|---|---|---|
| interactor_abundance | mitosis | inter | 0.0003488542111186024 | 0.01356427145975096 |
| complex_abundance | mitosis | inter | 0.0006558599407001601 | 0.01642175463112936 |
| interactor_ratio | mitosis | inter | 0.011383459471981348 | 0.15421453104903593 |

O43395_Q99459
PRPF3_HUMAN vs CDC5L_HUMAN
p-value: 0.003 q-value: 0.004

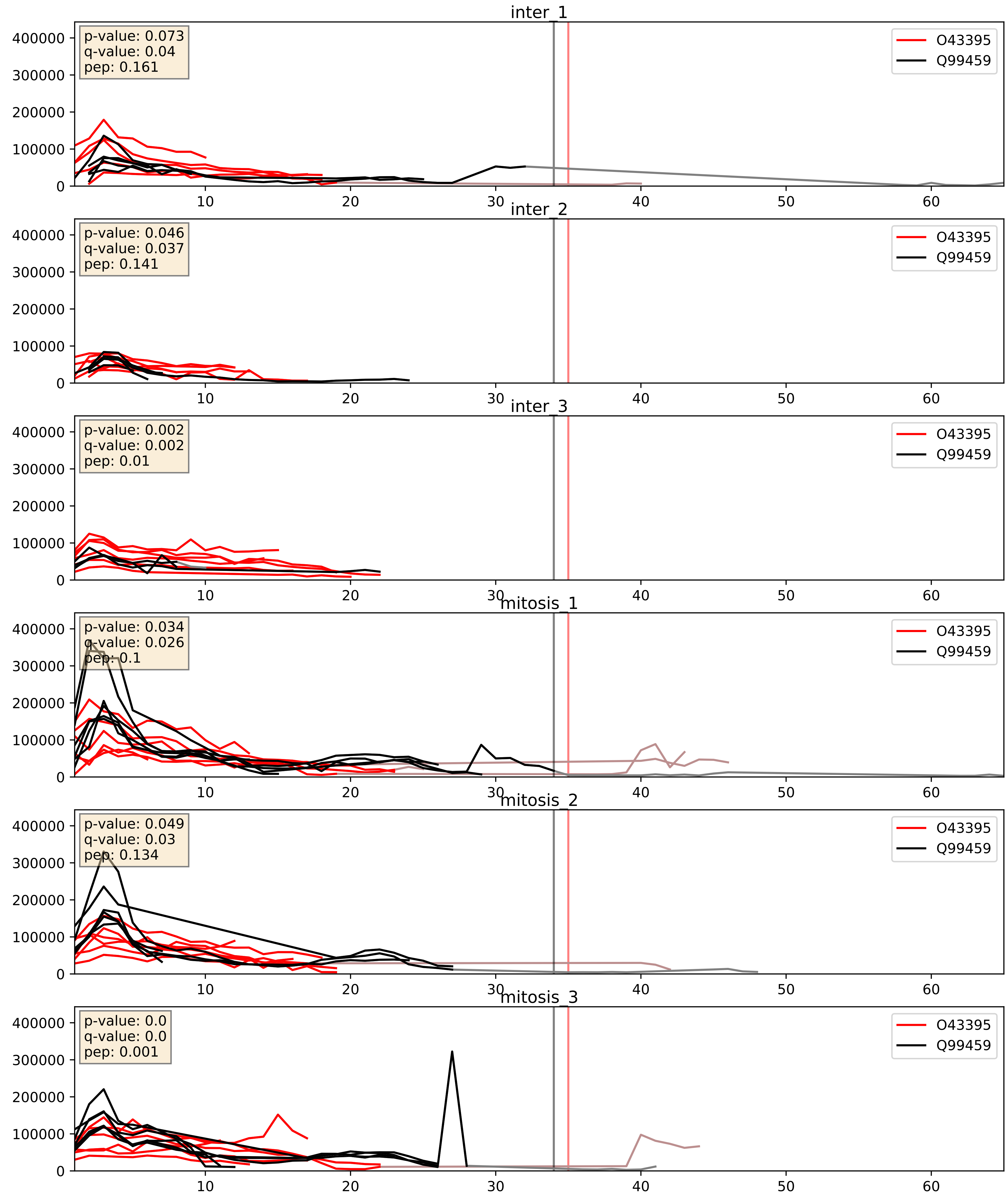


| level | condition_1 | condition_2 | pvalue | pvalue_adjusted |
|---|---|---|---|---|
| complex_abundance | mitosis | inter | 0.020735988987782956 | 0.1061724104661514 |
| interactor_ratio | mitosis | inter | 0.05638110929547731 | 0.3926647223857623 |
| interactor_abundance | mitosis | inter | 0.06577503851832661 | 0.21269195540886726 |

O43395_Q9BZJ0
PRPF3_HUMAN vs CRNL1_HUMAN
p-value: 0.022 q-value: 0.017

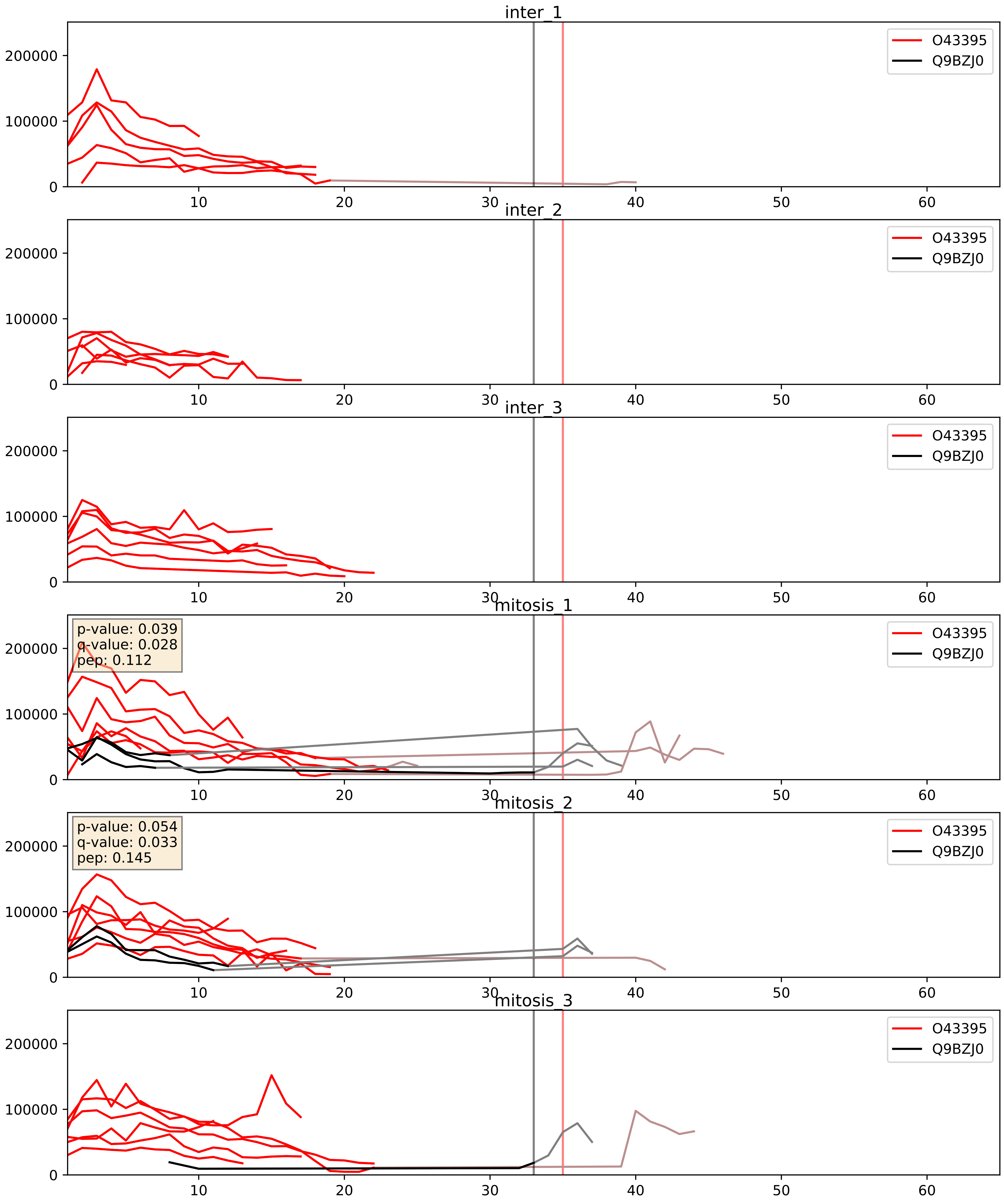


| level | condition_1 | condition_2 | pvalue | pvalue_adjusted |
|---|---|---|---|---|
| interactor_ratio | mitosis | inter | 0.0010612977904857902 | 0.038414698994273265 |
| interactor_abundance | mitosis | inter | 0.050901463188763756 | 0.18087028845820577 |
| complex_abundance | mitosis | inter | 0.16968315397865294 | 0.31873042673431795 |

O43395_Q9UMS4
PRPF3_HUMAN vs PRP19_HUMAN
p-value: 0.007 q-value: 0.007

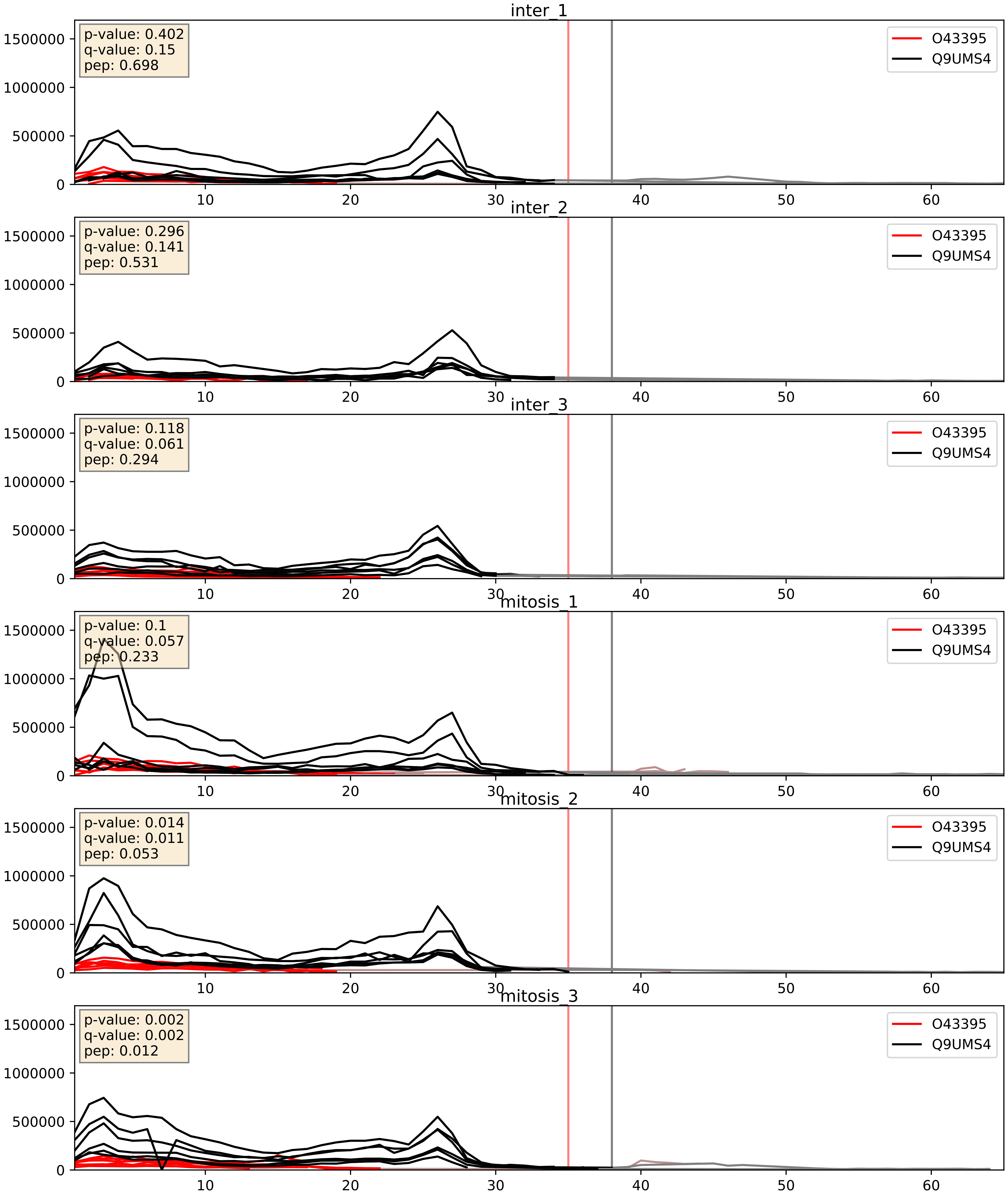


| level | condition_1 | condition_2 | pvalue | pvalue_adjusted |
|---|---|---|---|---|
| interactor_ratio | mitosis | inter | 0.007326432304467806 | 0.11910596594407505 |
| complex_abundance | mitosis | inter | 0.11767997417224624 | 0.25891907449070617 |
| interactor_abundance | mitosis | inter | 0.3100367719395898 | 0.5063273428985764 |

O43395_Q9Y4Z0
PRPF3_HUMAN vs LSM4_HUMAN
p-value: 0.043 q-value: 0.029

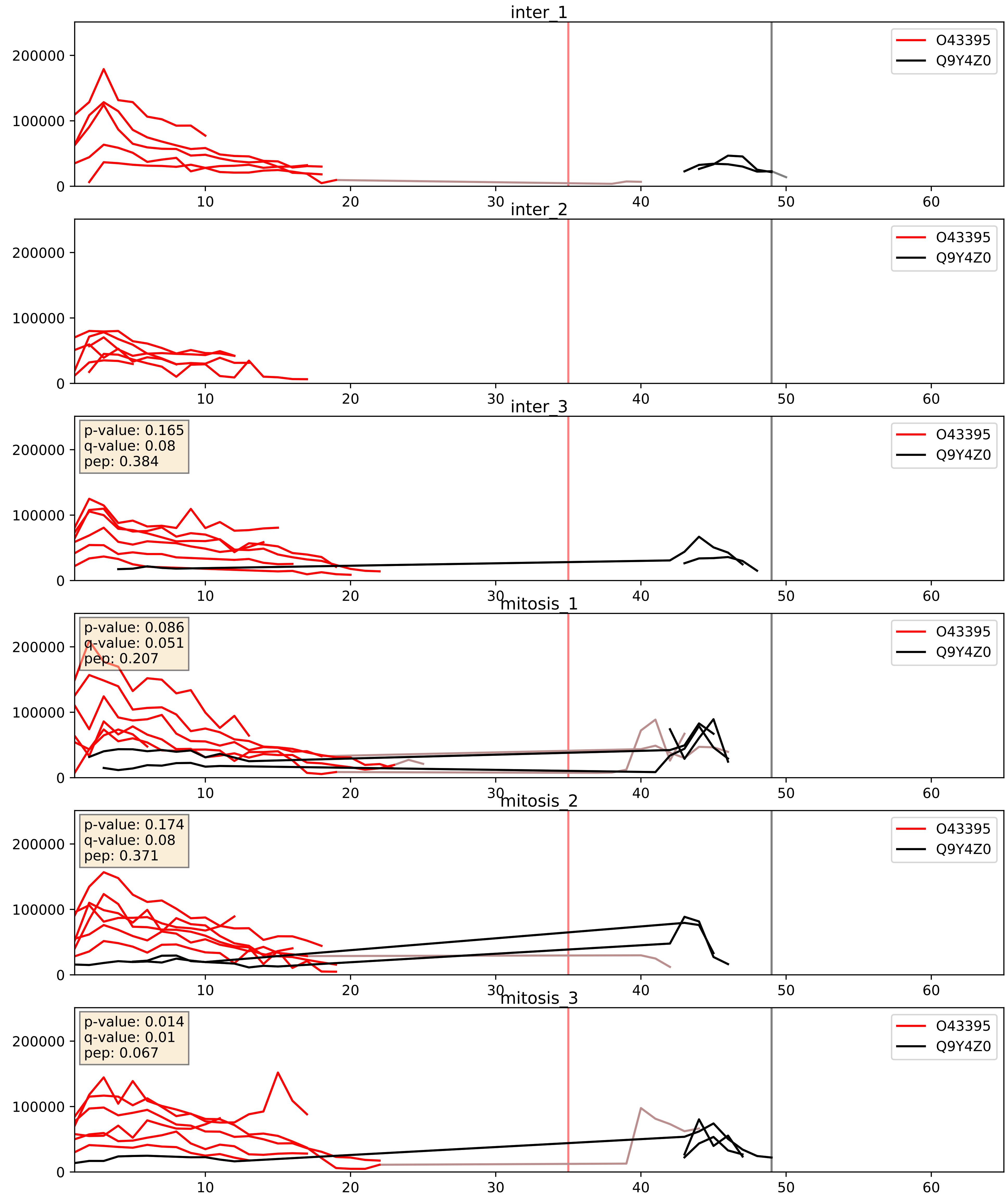


| level | condition_1 | condition_2 | pvalue | pvalue_adjusted |
|---|---|---|---|---|
| interactor_abundance | mitosis | inter | 0.011098725742666608 | 0.09286910298848645 |
| interactor_ratio | mitosis | inter | 0.019191571727531852 | 0.20874187291953322 |
| complex_abundance | mitosis | inter | 0.023033384633034417 | 0.11091884921279084 |
