## Supplementary Data 2 for "SECAT: Quantifying differential protein-protein interaction states by network-centric analysis": hela_string_000069_Q15365.pdf

Q15365_Q15365

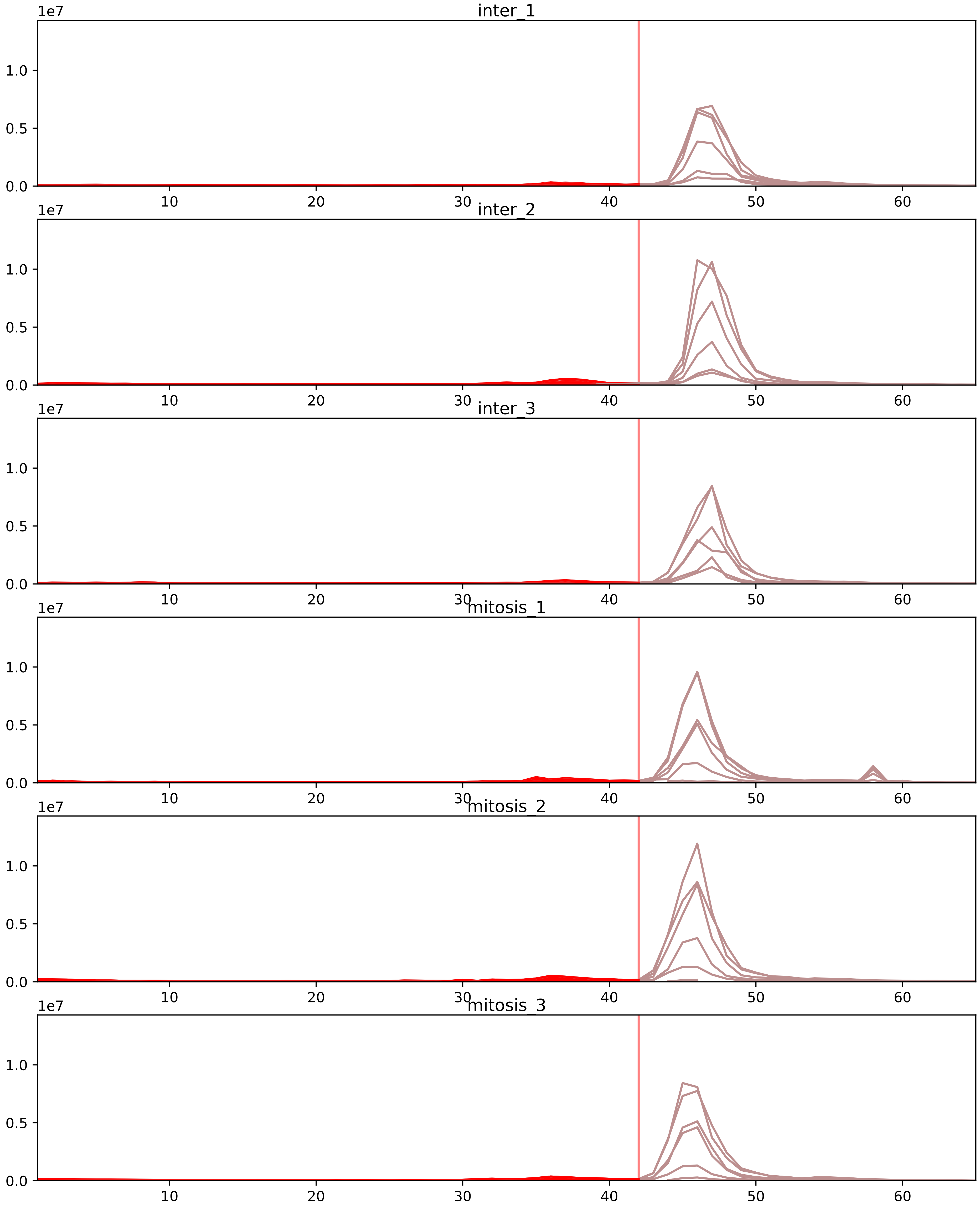


| level | condition_1 | condition_2 | pvalue | pvalue_adjusted |
|---|---|---|---|---|
| interactor_ratio | mitosis | inter | 1.23175751843519e-06 | 3.6555384418076606e-05 |
| complex_abundance | mitosis | inter | 1.6972935074249184e-05 | 0.0007262837334097326 |
| interactor_abundance | mitosis | inter | 0.007579964161305776 | 0.04385891212831015 |
| assembled_abundance | mitosis | inter | 0.2187000587398937 | 0.5256806699531161 |
| monomer_abundance | mitosis | inter | 0.4370577229868118 | 0.718778588275368 |
| total_abundance | mitosis | inter | 0.4786045520874924 | 0.7479754392421735 |

P07910_Q15365
HNRPC_HUMAN vs PCBP1_HUMAN
p-value: 0.026 q-value: 0.019

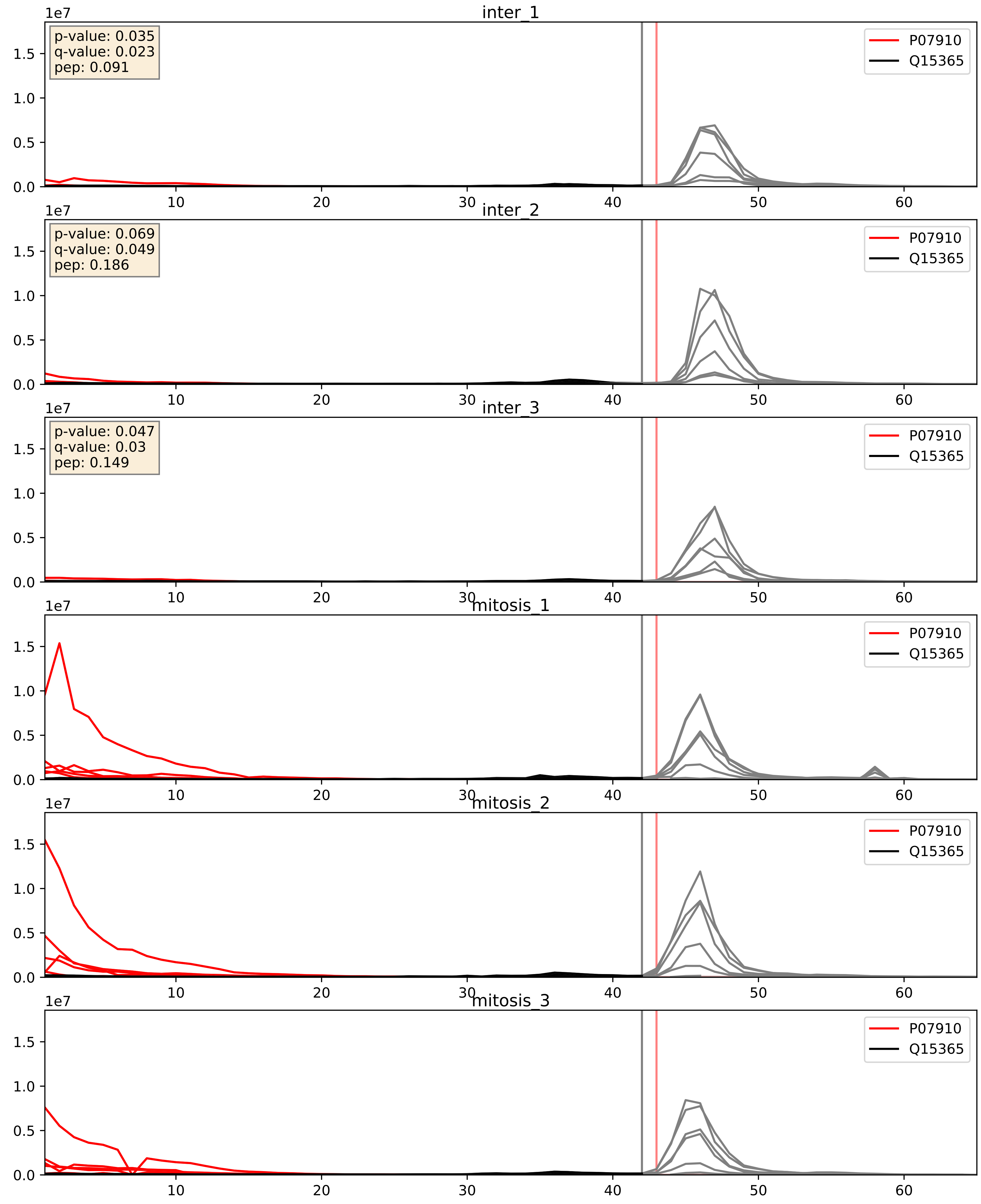


| level | condition_1 | condition_2 | pvalue | pvalue_adjusted |
|---|---|---|---|---|
| interactor_abundance | mitosis | inter | 9.148564965046394e-06 | 0.0033734327106189023 |
| complex_abundance | mitosis | inter | 1.030468151281165e-05 | 0.005691857151652958 |
| interactor_ratio | mitosis | inter | 4.476005324973264e-05 | 0.006796360849516482 |

P11940_Q15365
PABP1_HUMAN vs PCBP1_HUMAN
p-value: 0.0 q-value: 0.023

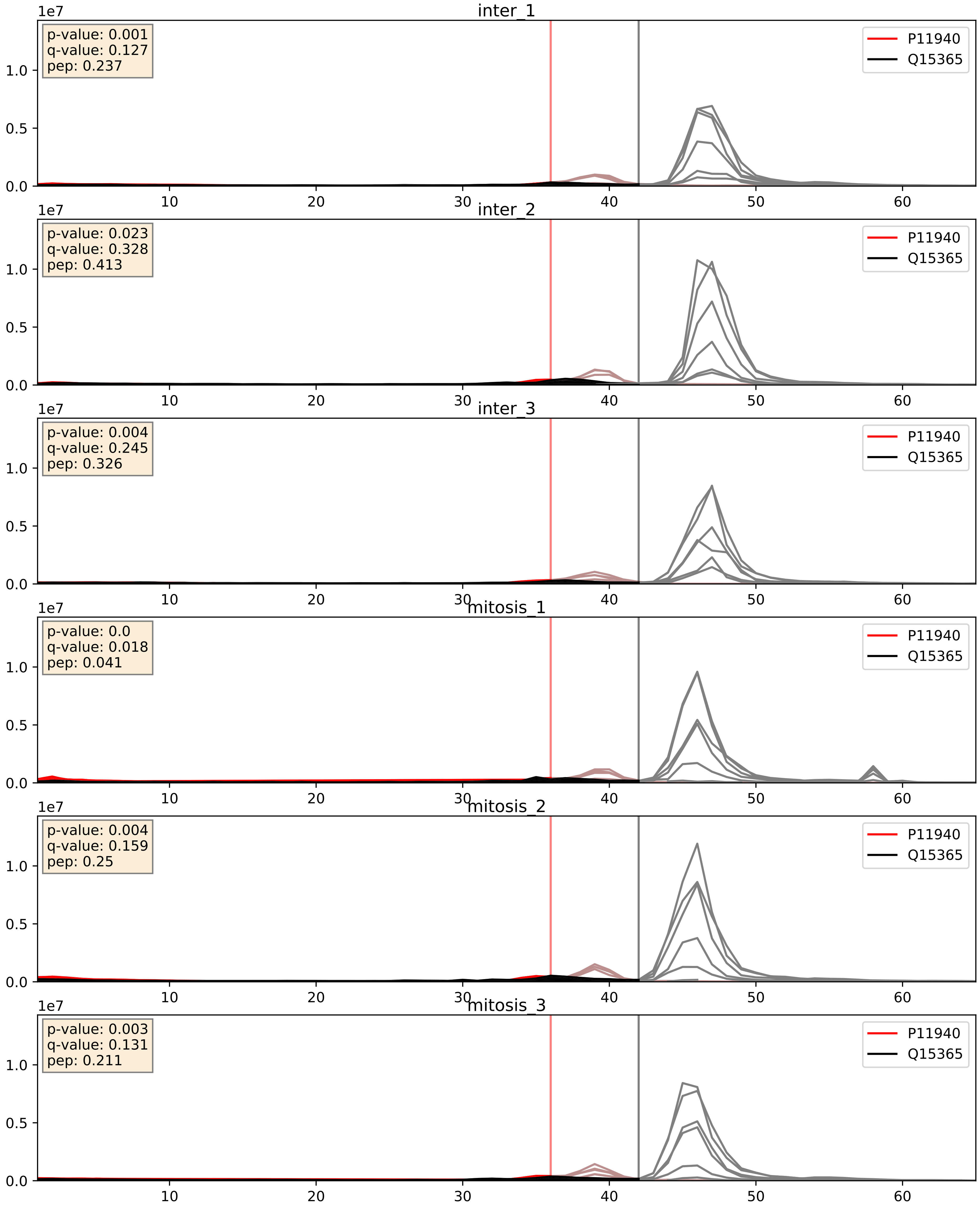


| level | condition_1 | condition_2 | pvalue | pvalue_adjusted |
|---|---|---|---|---|
| interactor_ratio | mitosis | inter | 0.06976990871901345 | 0.4375075131827156 |
| interactor_abundance | mitosis | inter | 0.15090604372175434 | 0.33732834796845007 |
| complex_abundance | mitosis | inter | 0.29680960341908563 | 0.4463615961467627 |

P14866_Q15365
HNRPL_HUMAN vs PCBP1_HUMAN
p-value: 0.037 q-value: 0.026

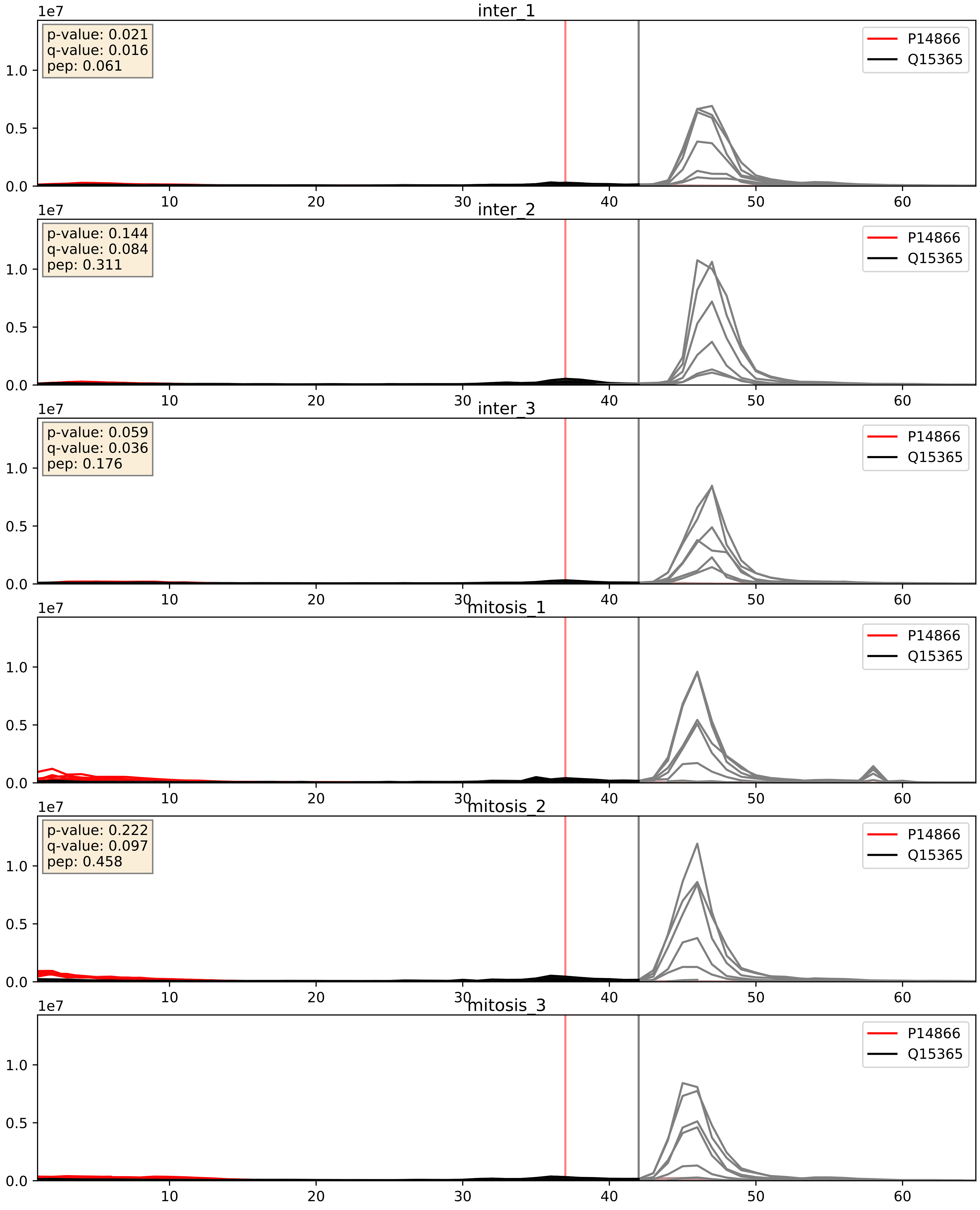


| level | condition_1 | condition_2 | pvalue | pvalue_adjusted |
|---|---|---|---|---|
| complex_abundance | mitosis | inter | 0.006117488702226094 | 0.05820202463873419 |
| interactor_abundance | mitosis | inter | 0.009697640385696541 | 0.08525122834595471 |
| interactor_ratio | mitosis | inter | 0.04505144601463991 | 0.35282742715948545 |

P22626_Q15365
ROA2_HUMAN vs PCBP1_HUMAN
p-value: 0.0 q-value: 0.0

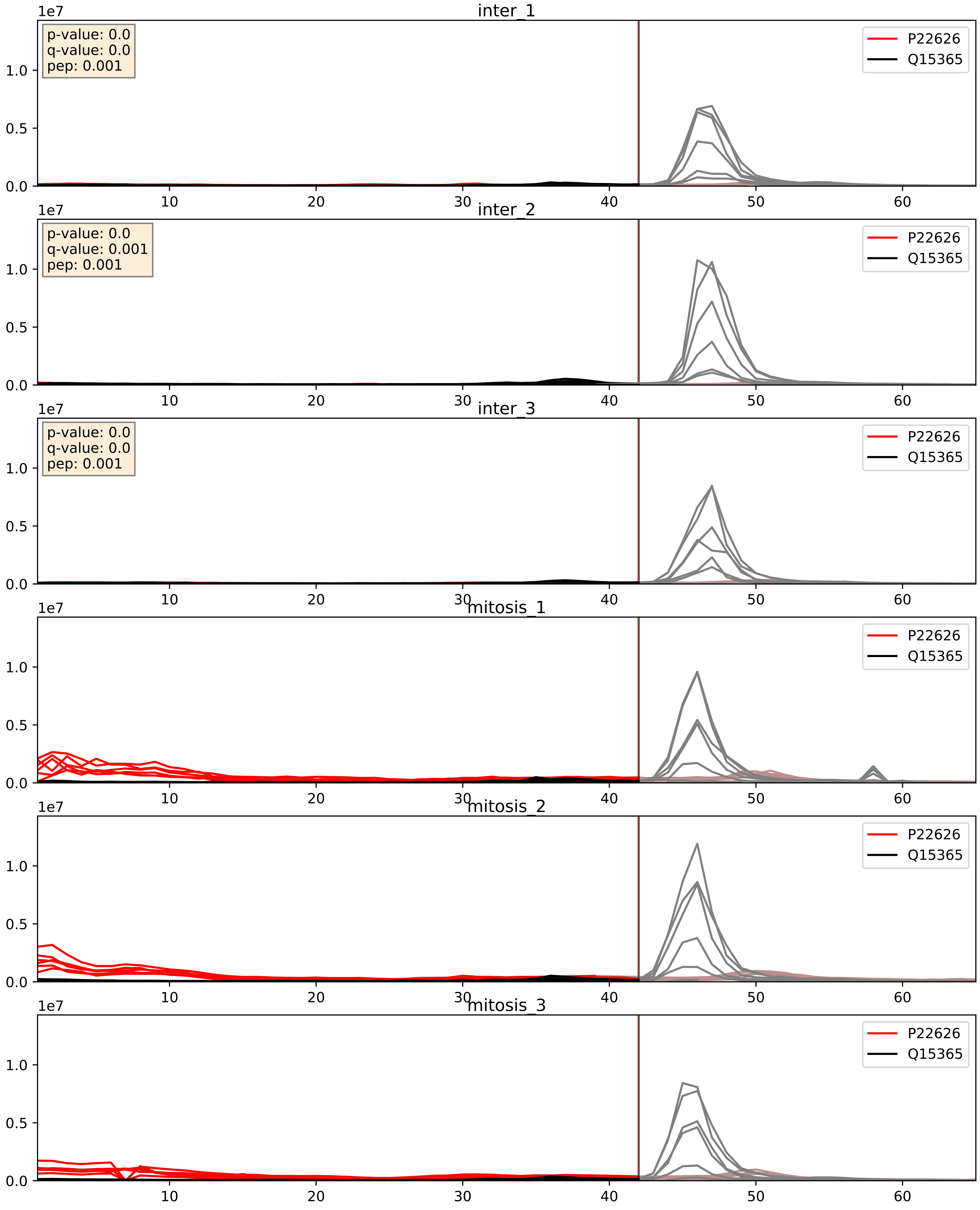


| level | condition_1 | condition_2 | pvalue | pvalue_adjusted |
|---|---|---|---|---|
| interactor_ratio | mitosis | inter | 0.0018969253504078144 | 0.052986711193083874 |
| interactor_abundance | mitosis | inter | 0.0034466655543236775 | 0.04511548235131213 |
| complex_abundance | mitosis | inter | 0.006247593176456641 | 0.05896295213943644 |

P26599_Q15365
PTBP1_HUMAN vs PCBP1_HUMAN
p-value: 0.0 q-value: 0.0

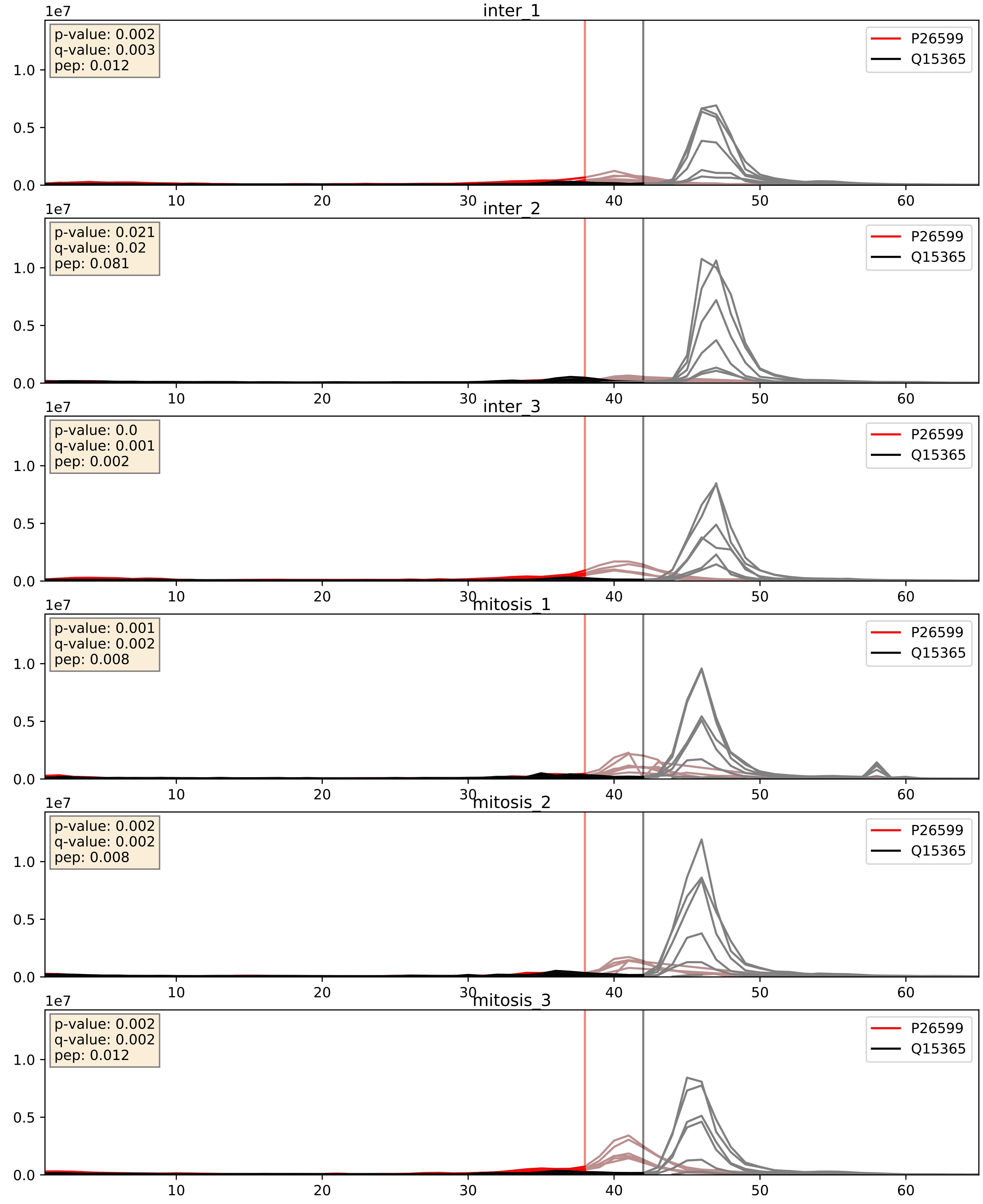


| level | condition_1 | condition_2 | pvalue | pvalue_adjusted |
|---|---|---|---|---|
| interactor_abundance | mitosis | inter | 0.3985986073620091 | 0.5886697292275717 |
| complex_abundance | mitosis | inter | 0.3987492173530314 | 0.5452545208533465 |
| interactor_ratio | mitosis | inter | 0.4019463103853237 | 0.8311893393797354 |

P31943_Q15365
HNRH1_HUMAN vs PCBP1_HUMAN
p-value: 0.028 q-value: 0.021

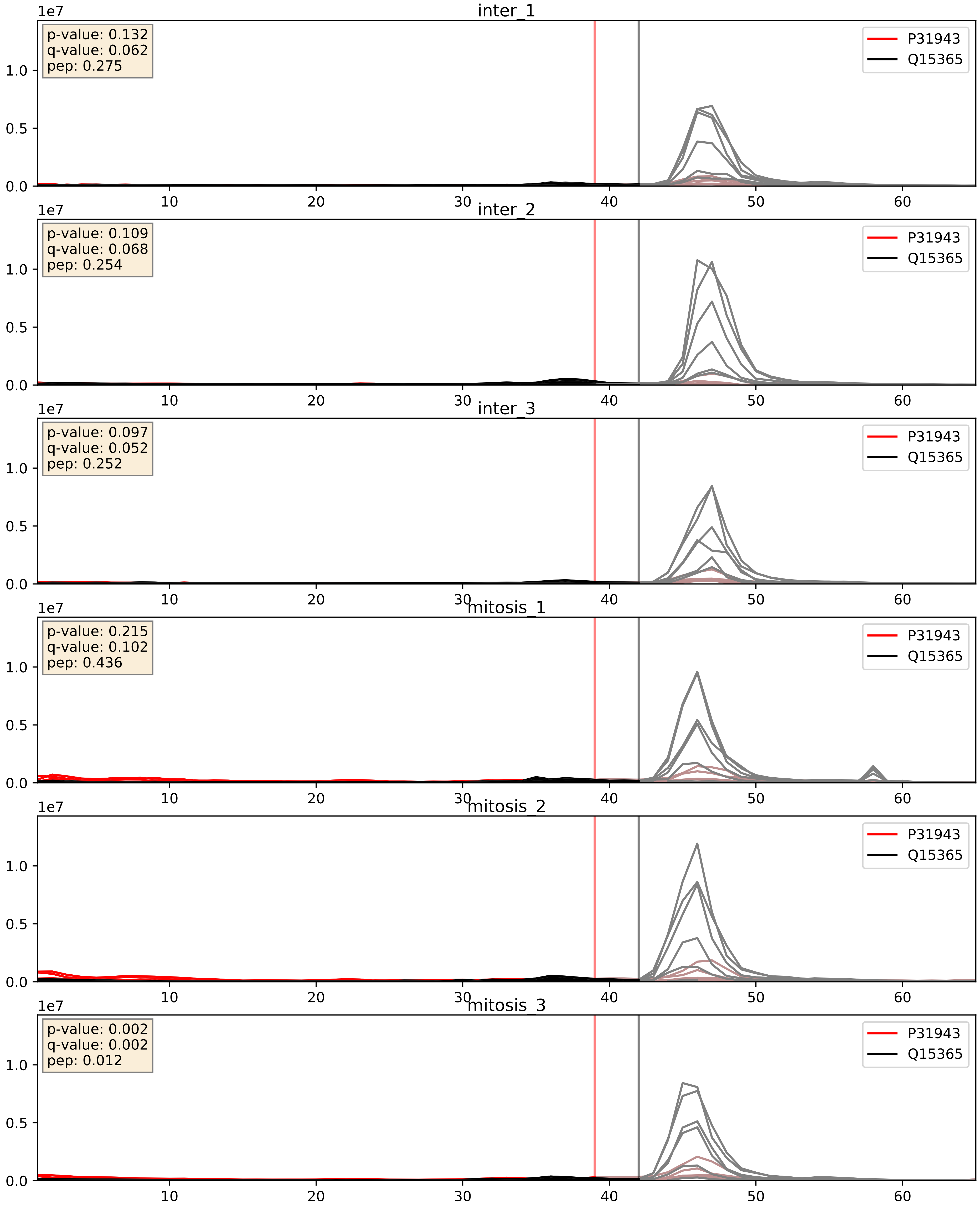


| level | condition_1 | condition_2 | pvalue | pvalue_adjusted |
|---|---|---|---|---|
| interactor_abundance | mitosis | inter | 0.0035613165684418575 | 0.04601489785035819 |
| complex_abundance | mitosis | inter | 0.004390077064485453 | 0.04944613114736247 |
| interactor_ratio | mitosis | inter | 0.005505267863595088 | 0.1004799422438677 |

P52597_Q15365
HNRPF_HUMAN vs PCBP1_HUMAN
p-value: 0.002 q-value: 0.035

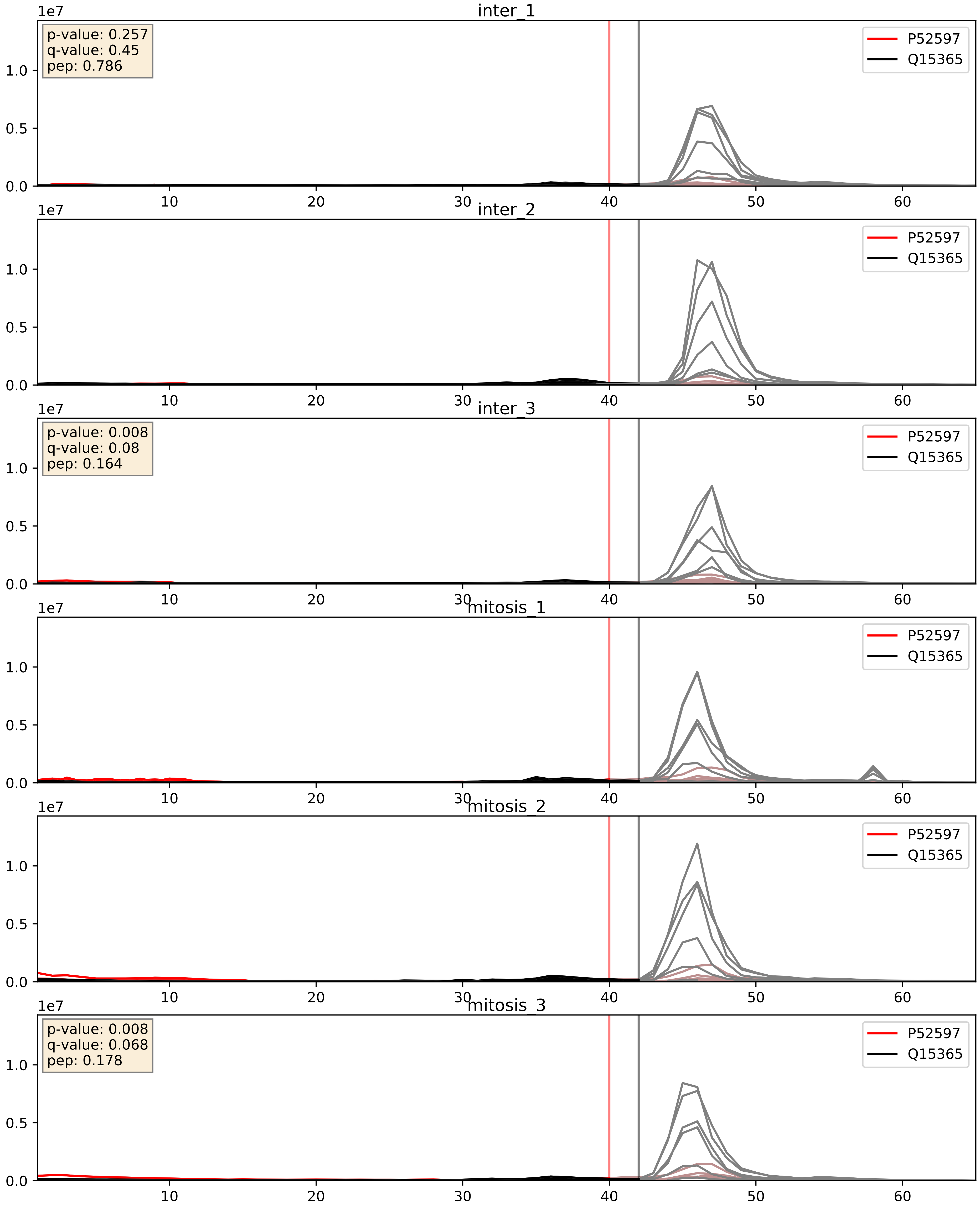


| level | condition_1 | condition_2 | pvalue | pvalue_adjusted |
|---|---|---|---|---|
| interactor_ratio | mitosis | inter | 0.0031499845140135186 | 0.07327137891292315 |
| interactor_abundance | mitosis | inter | 0.003829085421829145 | 0.04830798999389459 |
| complex_abundance | mitosis | inter | 0.01784052834109063 | 0.1000532792033756 |

P61978_Q15365
HNRPK_HUMAN vs PCBP1_HUMAN
p-value: 0.026 q-value: 0.019

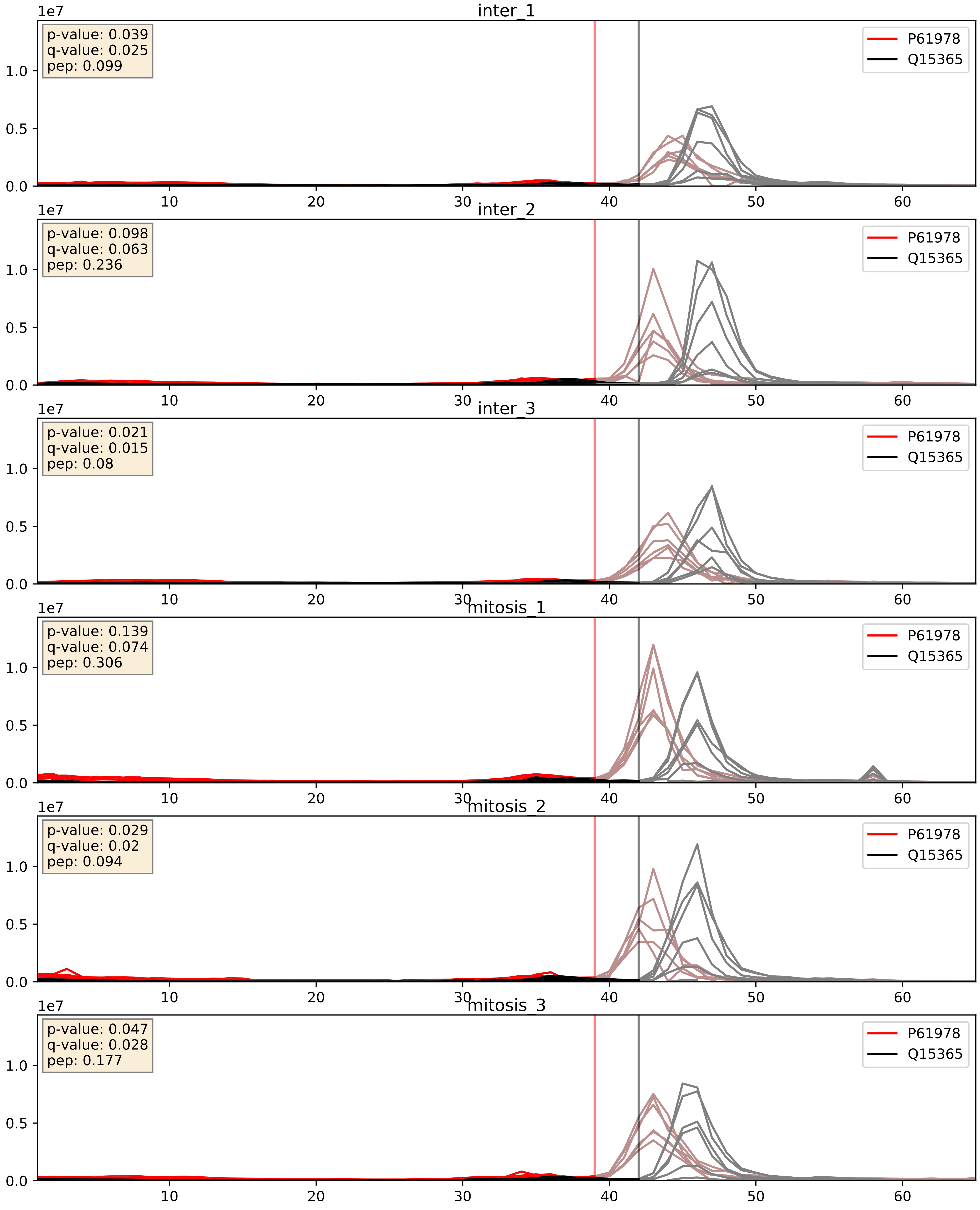


| level | condition_1 | condition_2 | pvalue | pvalue_adjusted |
|---|---|---|---|---|
| complex_abundance | mitosis | inter | 0.7714308414862175 | 0.8494551363838663 |
| interactor_abundance | mitosis | inter | 0.8366334366207034 | 0.9107372311913552 |
| interactor_ratio | mitosis | inter | 0.9089820075695871 | 0.9852158746906624 |

P84103_Q15365
SRSF3_HUMAN vs PCBP1_HUMAN
p-value: 0.002 q-value: 0.03

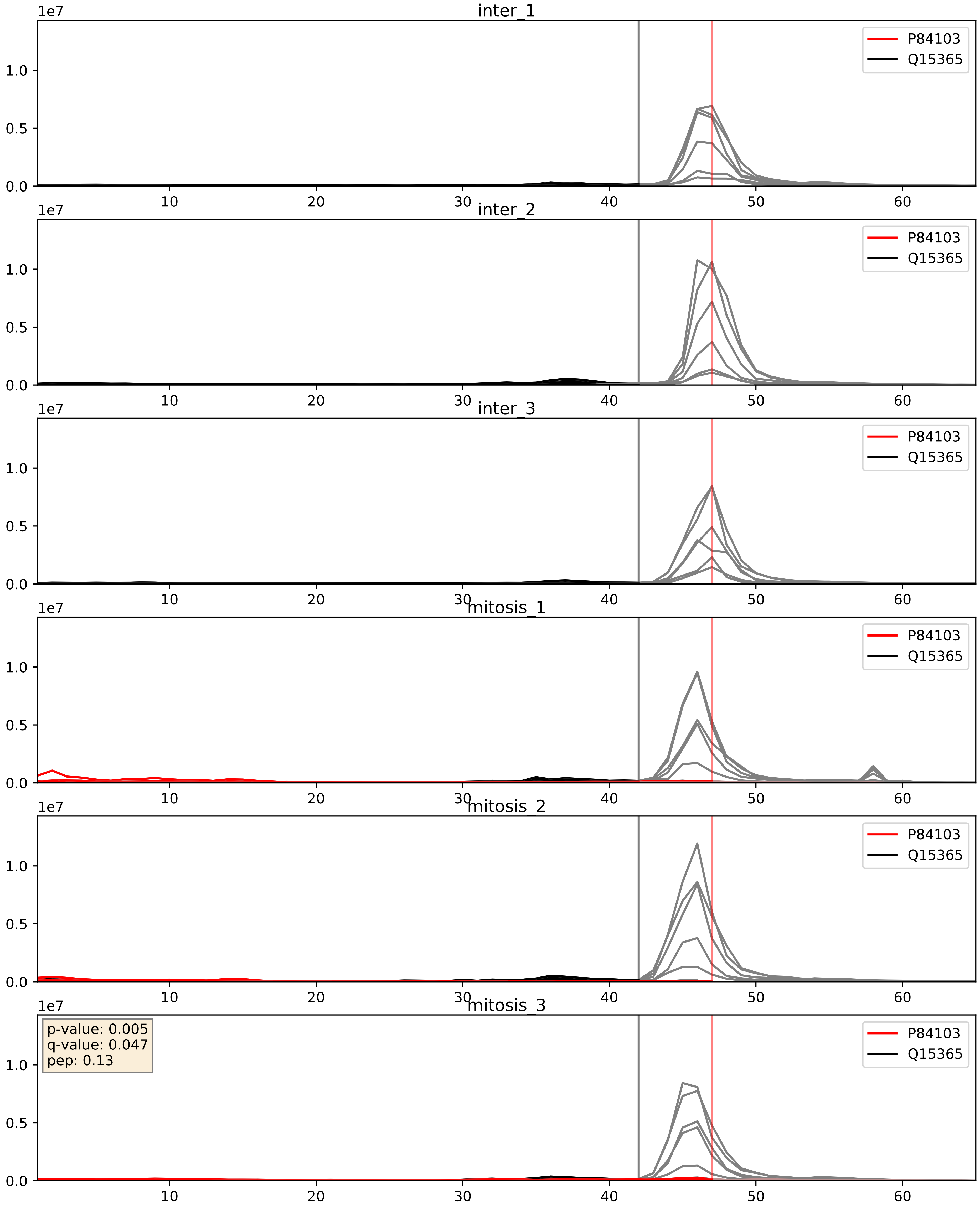


| level | condition_1 | condition_2 | pvalue | pvalue_adjusted |
|---|---|---|---|---|
| complex_abundance | mitosis | inter | 0.00026638034582209696 | 0.010910123254723205 |
| interactor_abundance | mitosis | inter | 0.00295284789649354 | 0.041301271232001155 |
| interactor_ratio | mitosis | inter | 0.041404094351652806 | 0.33818611416998856 |

Q07955_Q15365
SRSF1_HUMAN vs PCBP1_HUMAN
p-value: 0.0 q-value: 0.009

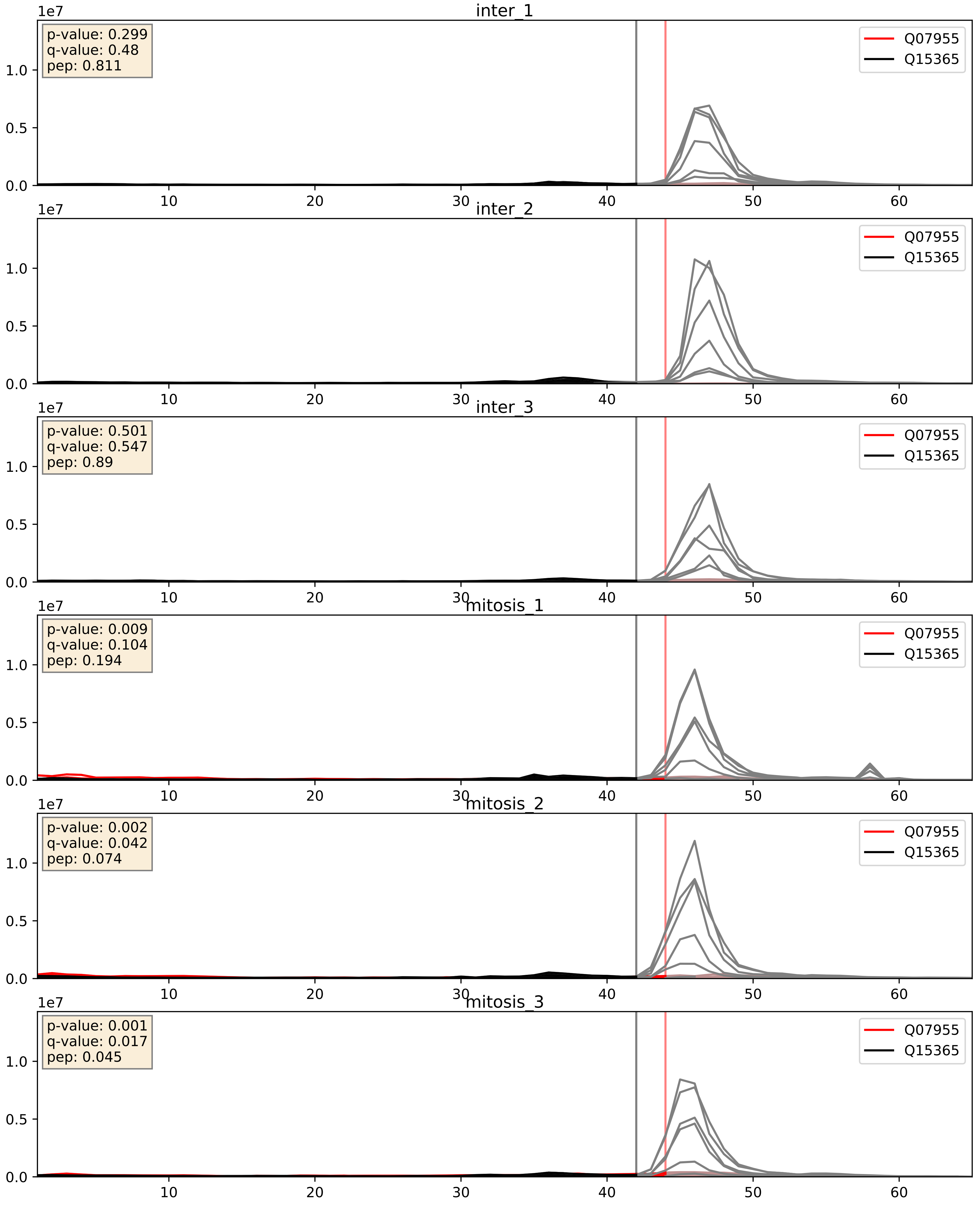


| level | condition_1 | condition_2 | pvalue | pvalue_adjusted |
|---|---|---|---|---|
| interactor_ratio | mitosis | inter | 0.014640895252484376 | 0.17726458749825497 |
| interactor_abundance | mitosis | inter | 0.015951526618393257 | 0.11088021938056777 |
| complex_abundance | mitosis | inter | 0.042812804870910104 | 0.1481949160937304 |

Q15365_Q15366
PCBP1_HUMAN vs PCBP2_HUMAN
p-value: 0.0 q-value: 0.003

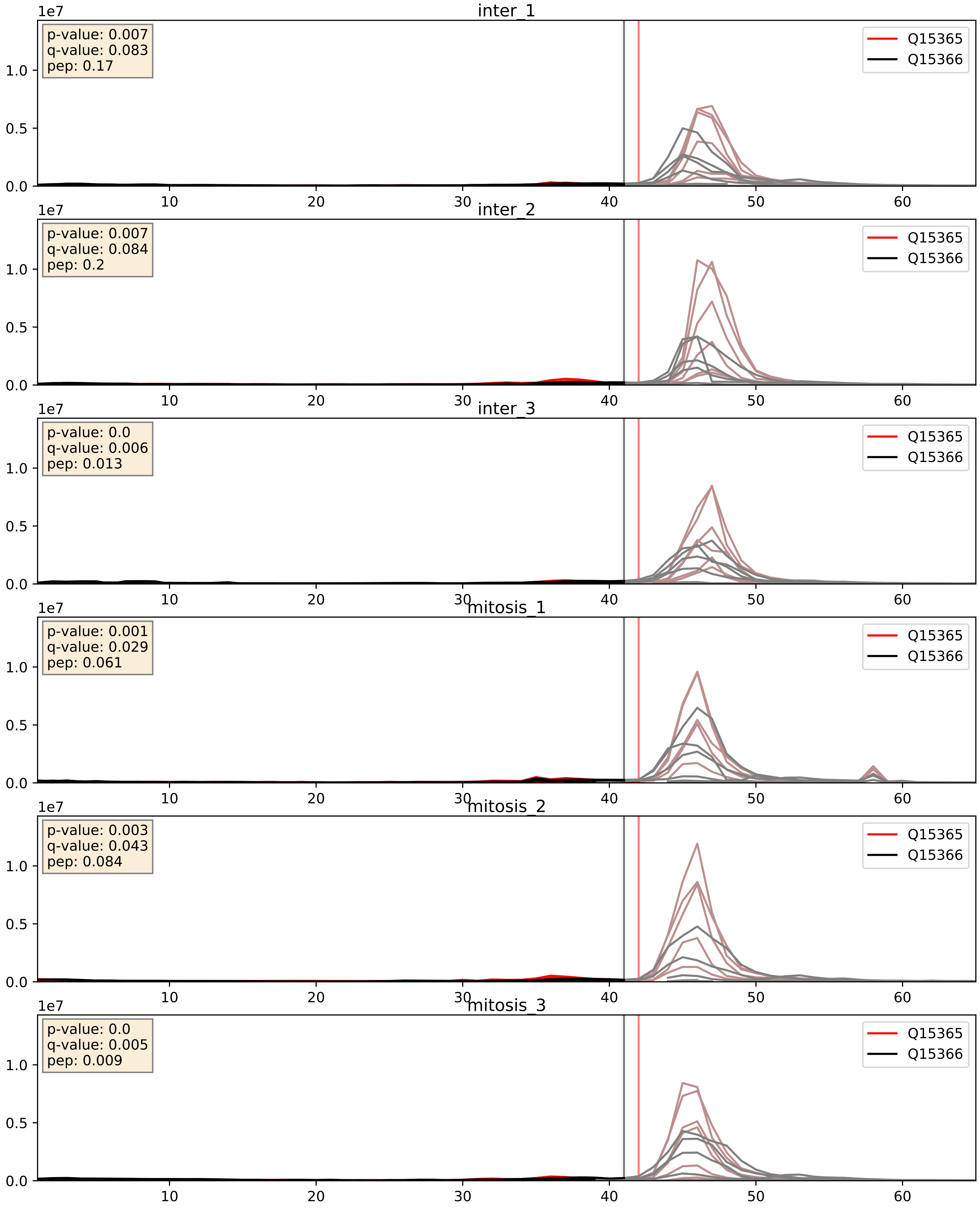


| level | condition_1 | condition_2 | pvalue | pvalue_adjusted |
|---|---|---|---|---|
| complex_abundance | mitosis | inter | 0.4033790850730204 | 0.5486684730185443 |
| interactor_abundance | mitosis | inter | 0.46548403340294203 | 0.6375269321486694 |
| interactor_ratio | mitosis | inter | 0.511381421849717 | 0.8744356714010344 |
