## Supplementary Data 2 for "SECAT: Quantifying differential protein-protein interaction states by network-centric analysis": hela_string_000070_P15954.pdf

# P15954_P15954

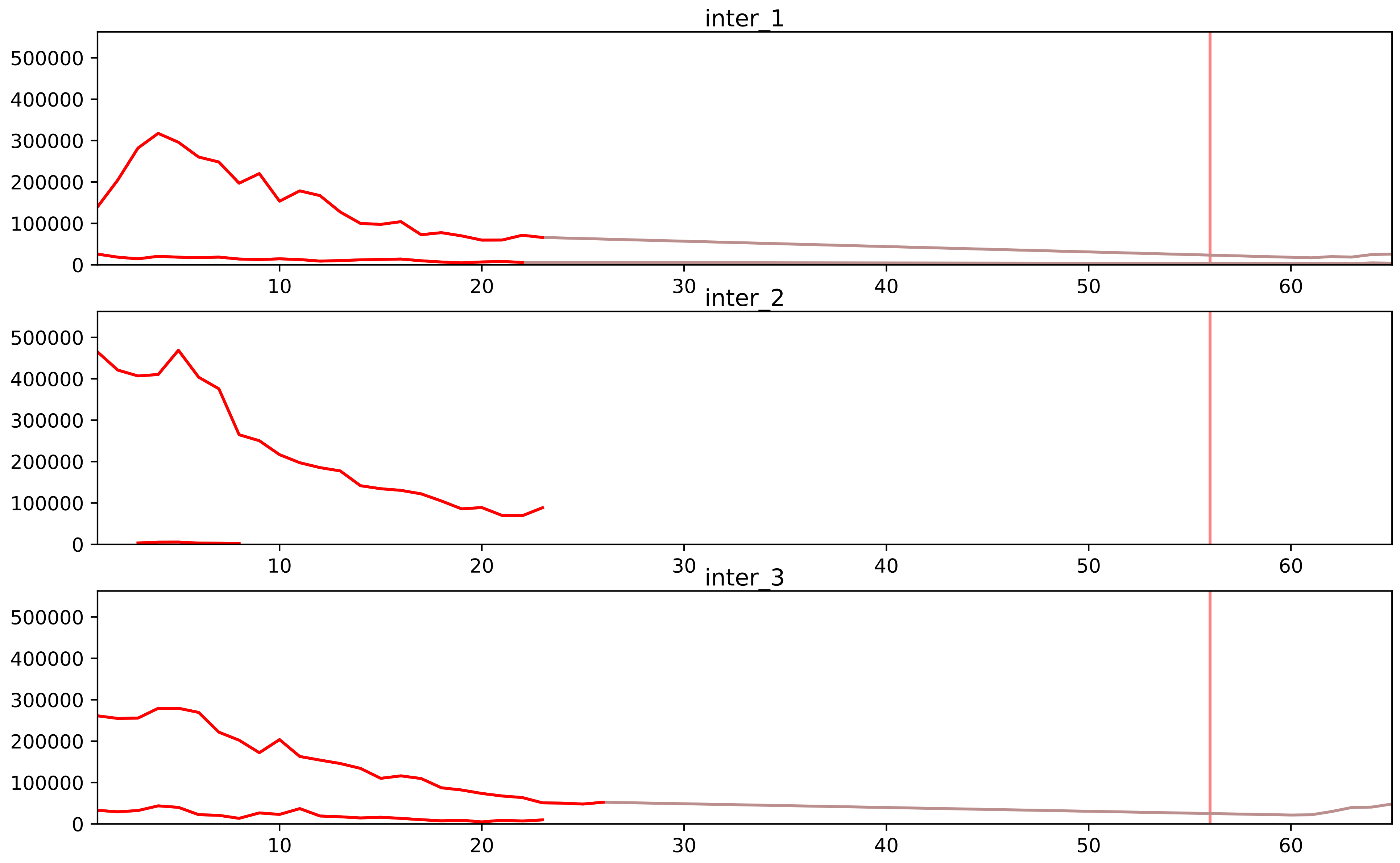


| level | condition_1 | condition_2 | pvalue | pvalue_adjusted |
|---|---|---|---|---|
| interactor_abundance | mitosis | inter | 1.2856791442740717e-06 | 0.00013353840650924488 |
| complex_abundance | mitosis | inter | 1.8370053612633993e-06 | 0.00014696042890107193 |
| interactor_ratio | mitosis | inter | 0.00011008615106774324 | 0.0018085581961129246 |
| total_abundance | mitosis | inter | 0.0005020398266295374 | 0.027376525239428108 |
| assembled_abundance | mitosis | inter | 0.0006947865327672442 | 0.024360801173422066 |
| monomer_abundance | mitosis | inter | 0.03277868081513594 | 0.30153627455136073 |

O14548_P15954
COX7R_HUMAN vs COX7C_HUMAN
p-value: 0.001 q-value: 0.025

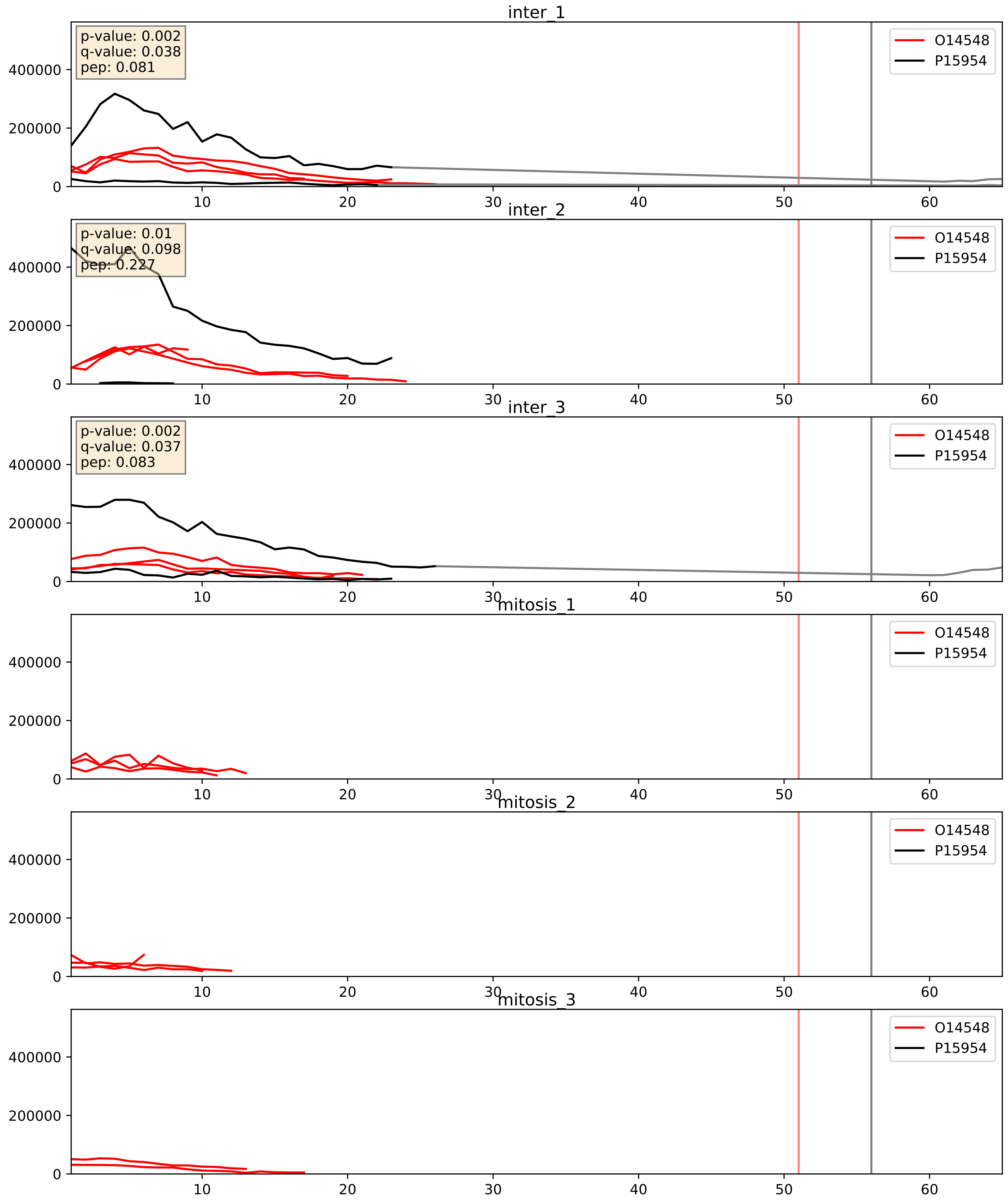


| level | condition_1 | condition_2 | pvalue | pvalue_adjusted |
|---|---|---|---|---|
| complex_abundance | mitosis | inter | 0.00021884711152757414 | 0.010087086090925927 |
| interactor_abundance | mitosis | inter | 0.0004706587247677187 | 0.015253292300429789 |
| interactor_ratio | mitosis | inter | 0.014546406987366608 | 0.17686306829592754 |

O14949_P15954
QCR8_HUMAN vs COX7C_HUMAN
p-value: 0.0 q-value: 0.0

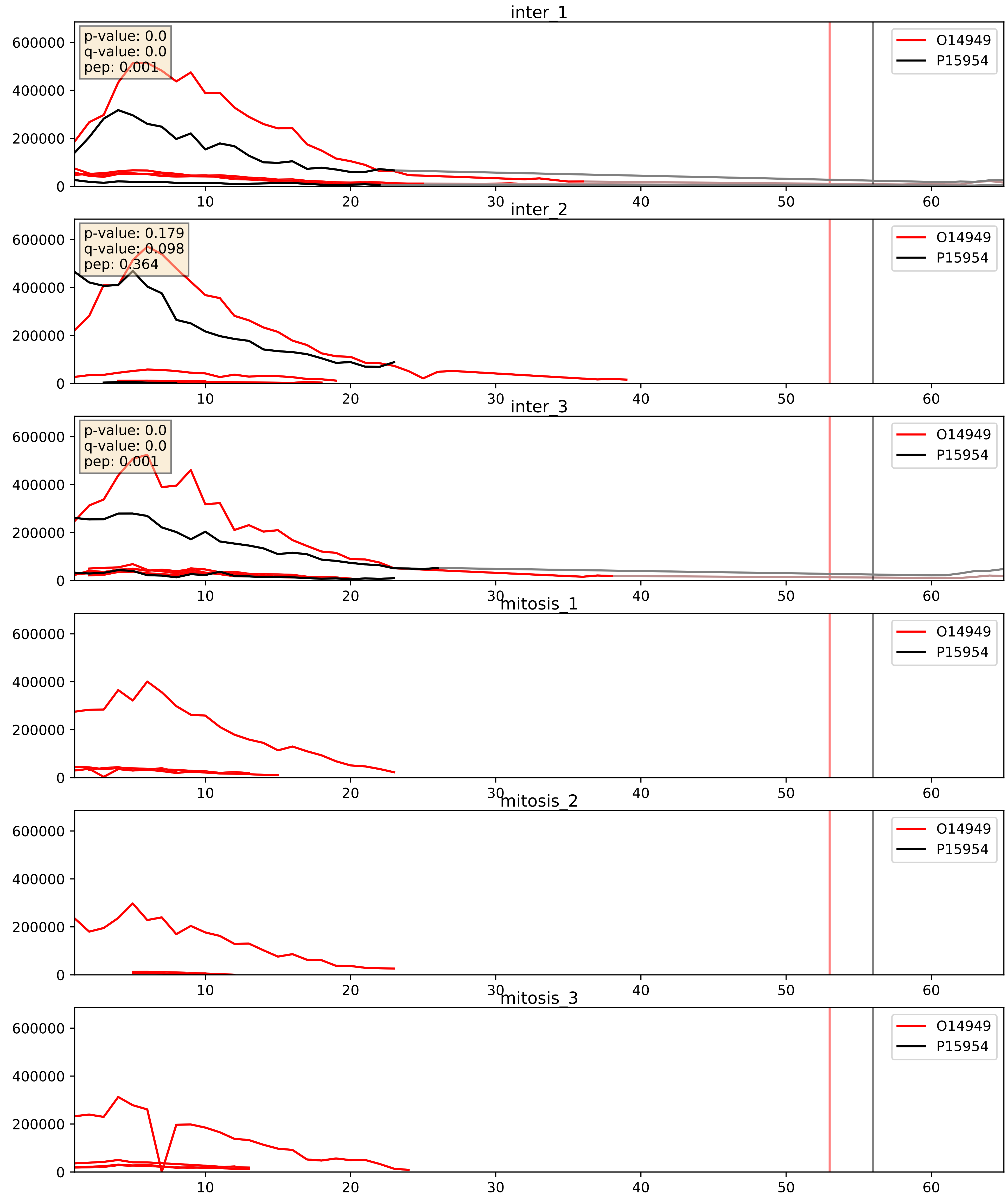


| level | condition_1 | condition_2 | pvalue | pvalue_adjusted |
|---|---|---|---|---|
| complex_abundance | mitosis | inter | 0.00019629554854570515 | 0.00938709438855439 |
| interactor_abundance | mitosis | inter | 0.00044896370615175136 | 0.014781266633303813 |
| interactor_ratio | mitosis | inter | 0.011007868718762004 | 0.1507637699721644 |

P00403_P15954
COX2_HUMAN vs COX7C_HUMAN
p-value: 0.019 q-value: 0.015

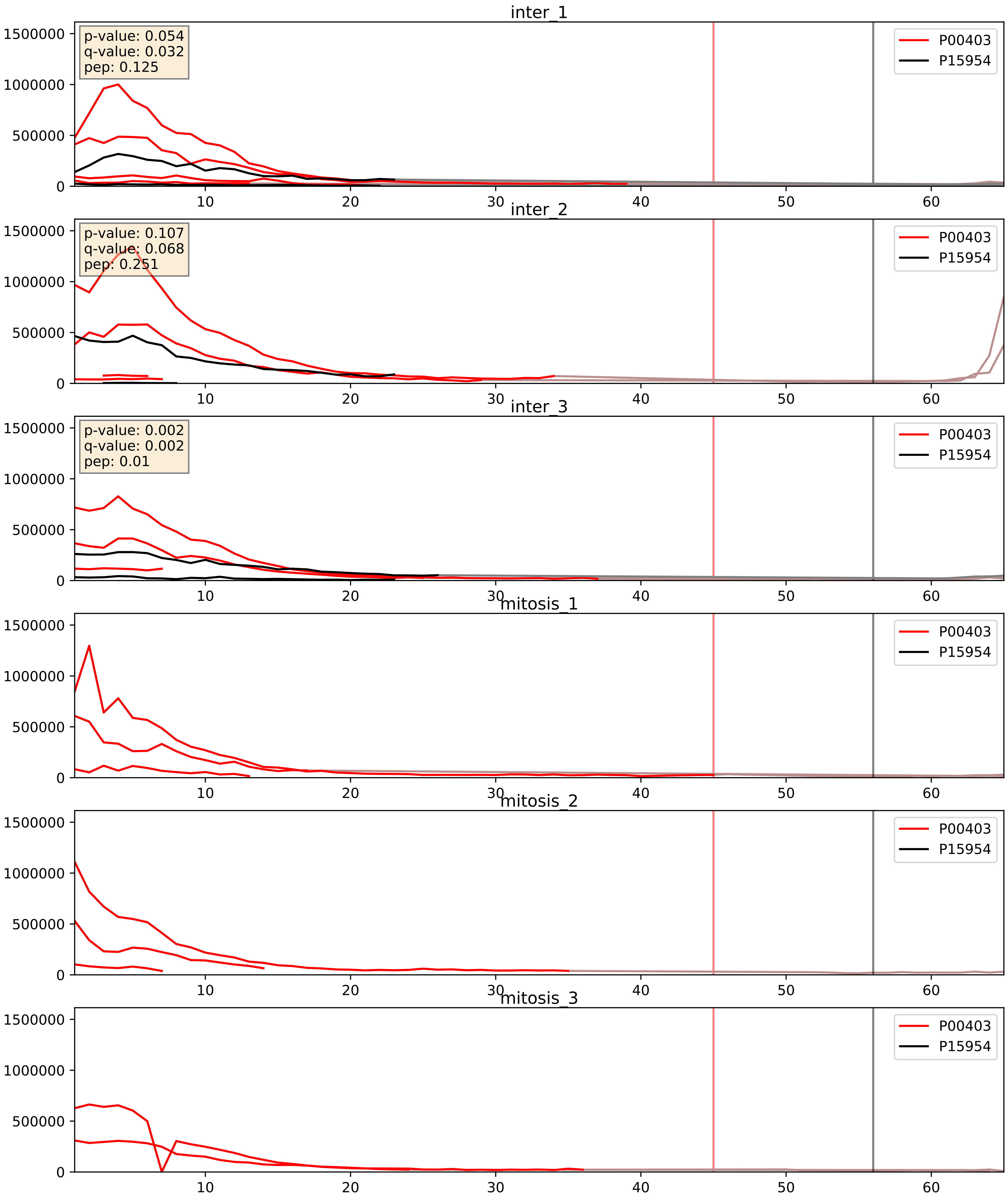


| level | condition_1 | condition_2 | pvalue | pvalue_adjusted |
|---|---|---|---|---|
| interactor_abundance | mitosis | inter | 4.4946683692739026e-05 | 0.006058954526139308 |
| complex_abundance | mitosis | inter | 5.2983607009485405e-05 | 0.005691857151652958 |
| interactor_ratio | mitosis | inter | 5.4319379841399415e-05 | 0.007596351369213947 |

P09669_P15954
COX6C_HUMAN vs COX7C_HUMAN
p-value: 0.003 q-value: 0.003

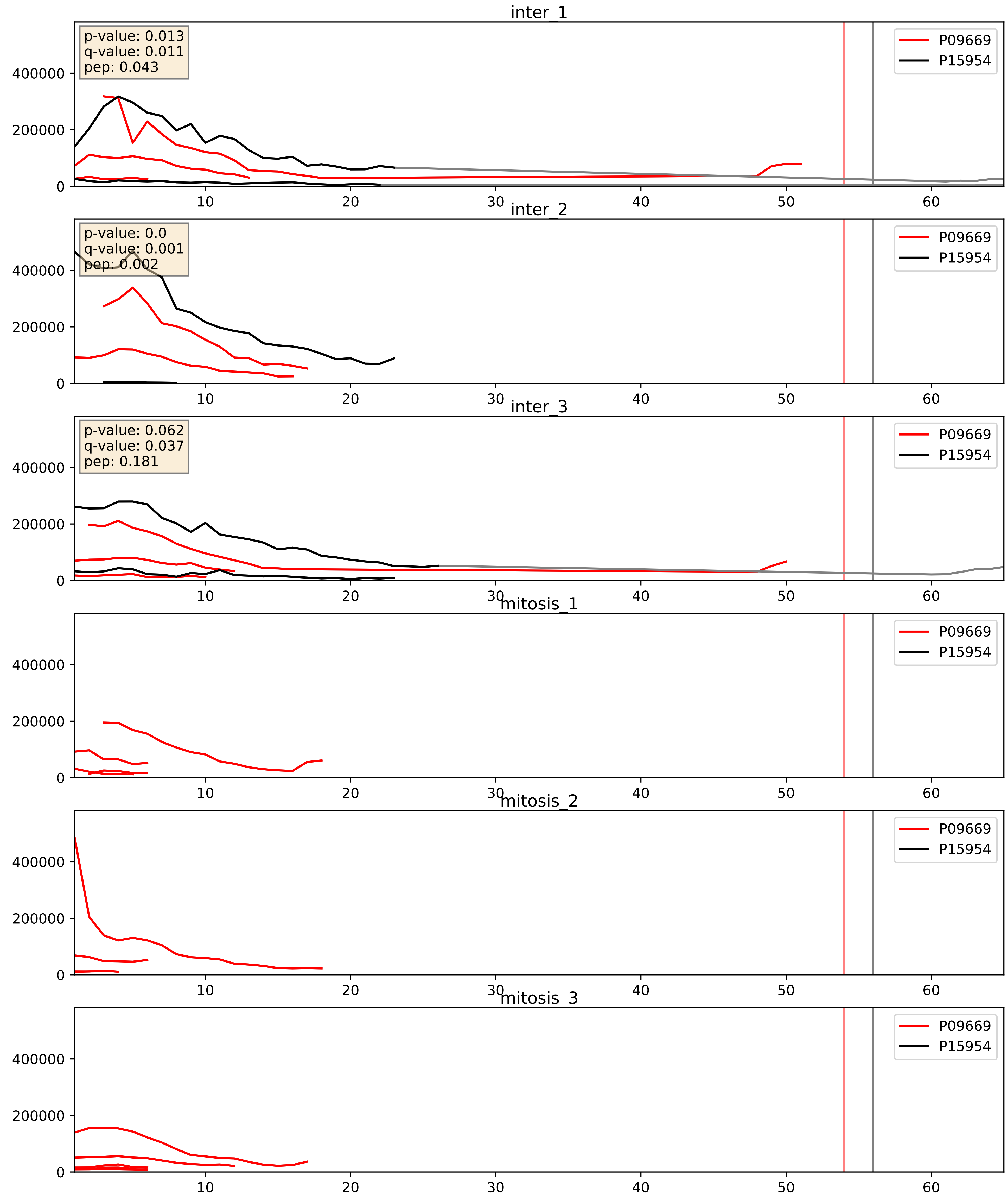


| level | condition_1 | condition_2 | pvalue | pvalue_adjusted |
|---|---|---|---|---|
| complex_abundance | mitosis | inter | 0.0002490403582267437 | 0.010553393398123397 |
| interactor_abundance | mitosis | inter | 0.00047503684757209603 | 0.015315688946203925 |
| interactor_ratio | mitosis | inter | 0.019164620158193375 | 0.20874187291953322 |

P10606_P15954
COX5B_HUMAN vs COX7C_HUMAN
p-value: 0.0 q-value: 0.0

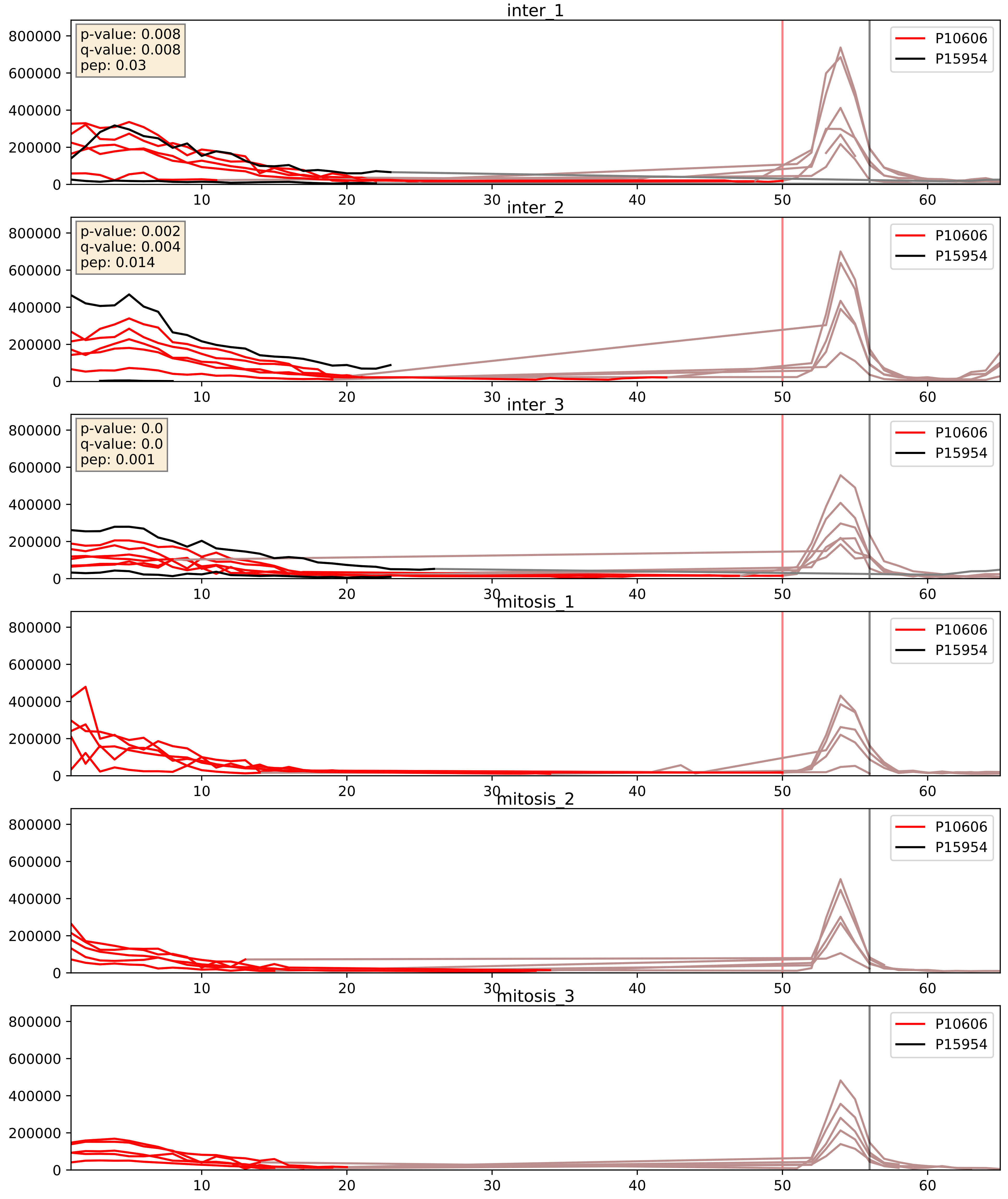


| level | condition_1 | condition_2 | pvalue | pvalue_adjusted |
|---|---|---|---|---|
| complex_abundance | mitosis | inter | 0.0001127294461587089 | 0.007310333781201123 |
| interactor_abundance | mitosis | inter | 0.00018045141508985833 | 0.010140496760639767 |
| interactor_ratio | mitosis | inter | 0.0011303731424321664 | 0.039174065179025684 |

P14927_P15954
QCR7_HUMAN vs COX7C_HUMAN
p-value: 0.003 q-value: 0.004

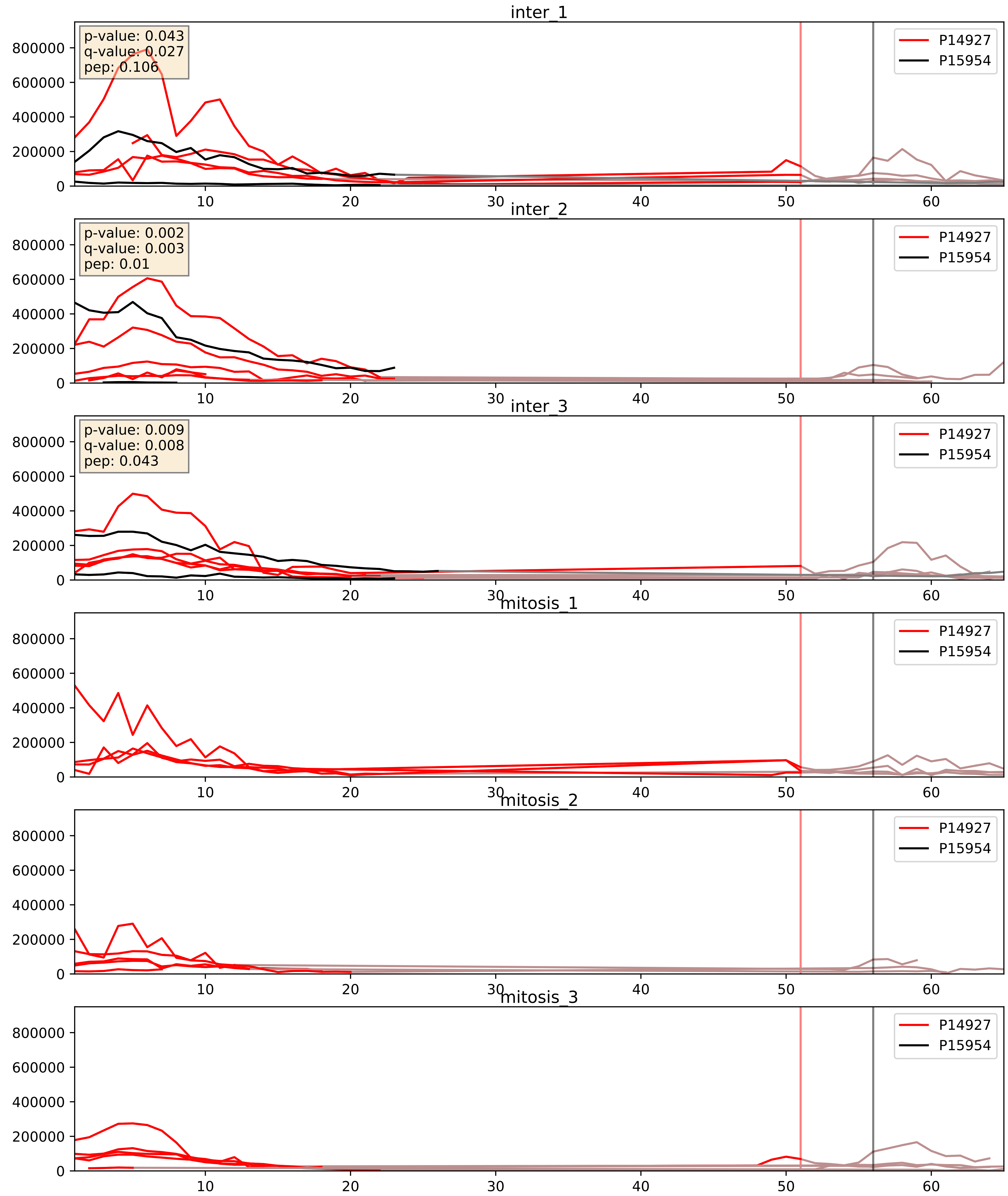


| level | condition_1 | condition_2 | pvalue | pvalue_adjusted |
|---|---|---|---|---|
| complex_abundance | mitosis | inter | 6.715859489683981e-05 | 0.005691857151652958 |
| interactor_abundance | mitosis | inter | 6.877853573801889e-05 | 0.006732838333317079 |
| interactor_ratio | mitosis | inter | 0.00015319123143554144 | 0.012836278661937291 |

P15954_P47985
COX7C_HUMAN vs UCRI_HUMAN
p-value: 0.034 q-value: 0.024

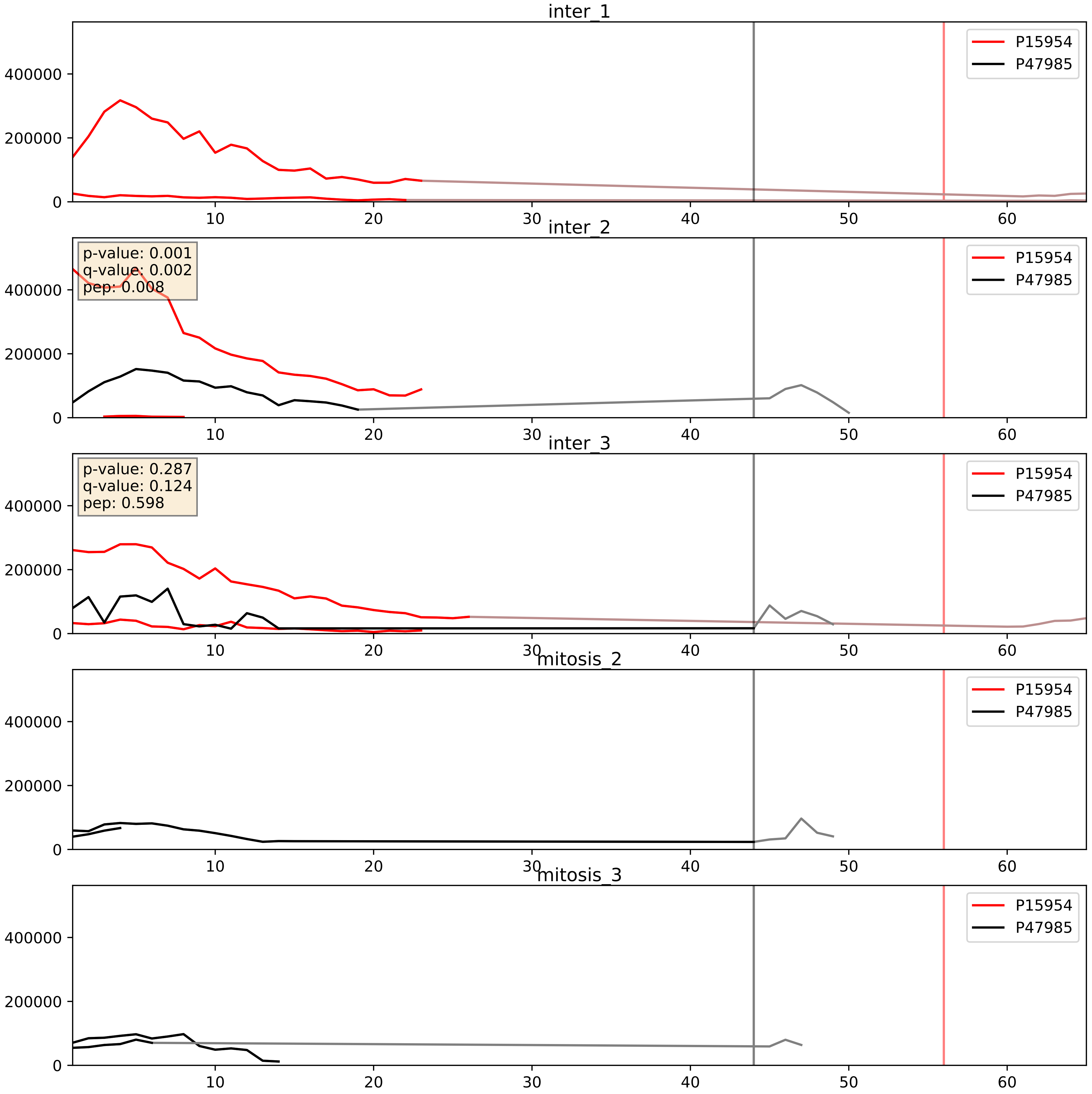


| level | condition_1 | condition_2 | pvalue | pvalue_adjusted |
|---|---|---|---|---|
| interactor_abundance | mitosis | inter | 0.04105574154351291 | 0.16296731794217187 |
| complex_abundance | mitosis | inter | 0.04203040597573723 | 0.14726323635770058 |
| interactor_ratio | mitosis | inter | 0.04220804271295535 | 0.3399226239910554 |
