## Supplementary Data 2 for "SECAT: Quantifying differential protein-protein interaction states by network-centric analysis": hela_string_000071_Q16795.pdf

# Q16795_Q16795

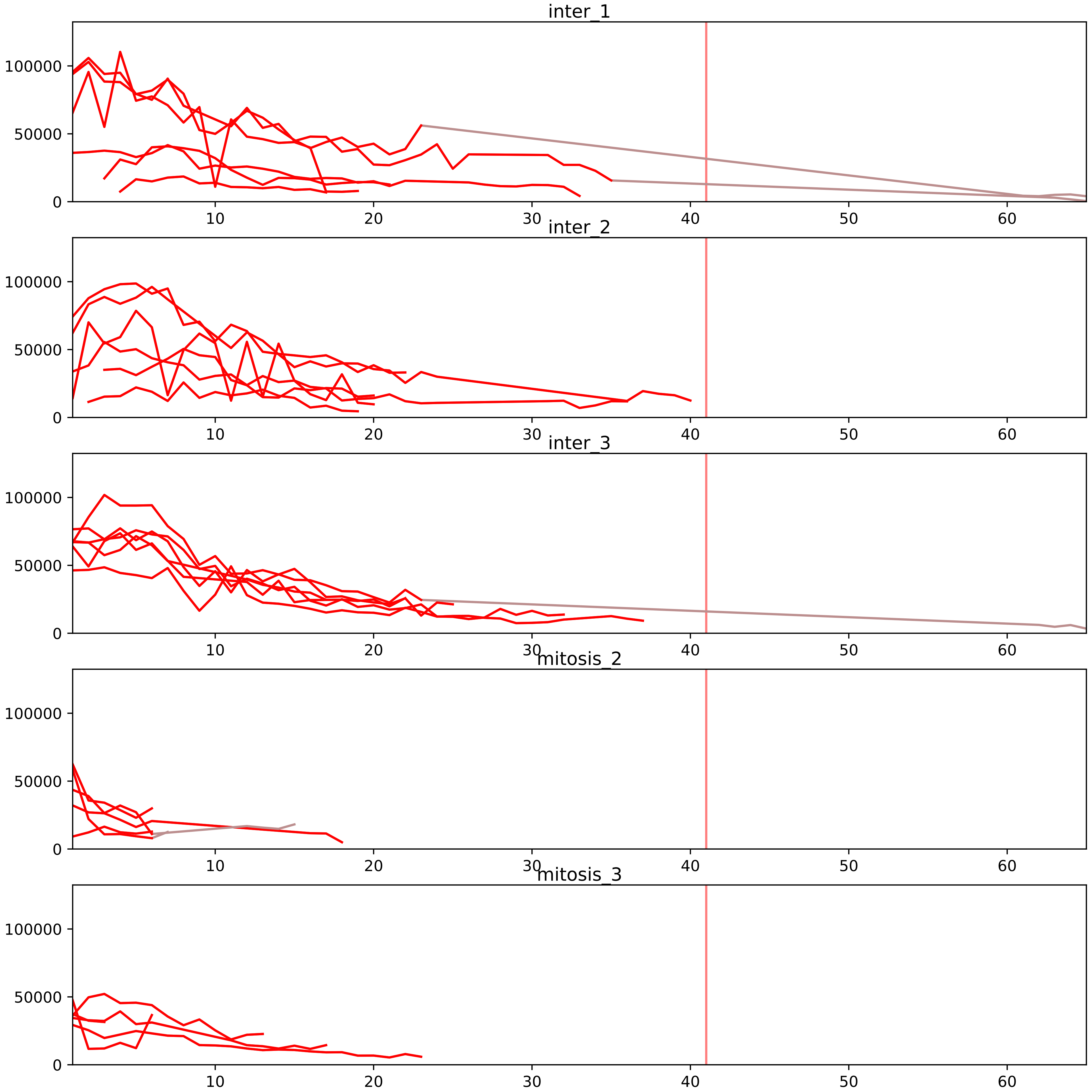


| level | condition_1 | condition_2 | pvalue | pvalue_adjusted |
|---|---|---|---|---|
| interactor_abundance | mitosis | inter | 1.378929197649811e-06 | 0.00013353840650924488 |
| complex_abundance | mitosis | inter | 0.0002502546759737164 | 0.004330299595990477 |
| interactor_ratio | mitosis | inter | 0.007278869607861483 | 0.04978855047756554 |
| total_abundance | mitosis | inter | 0.06399309673362598 | 0.3141941206356268 |
| monomer_abundance | mitosis | inter | 0.07303024035281172 | 0.37224850260048076 |
| assembled_abundance | mitosis | inter | 0.08099785778057741 | 0.33692014801381864 |

O00217_Q16795
NDUS8_HUMAN vs NDUA9_HUMAN
p-value: 0.015 q-value: 0.012

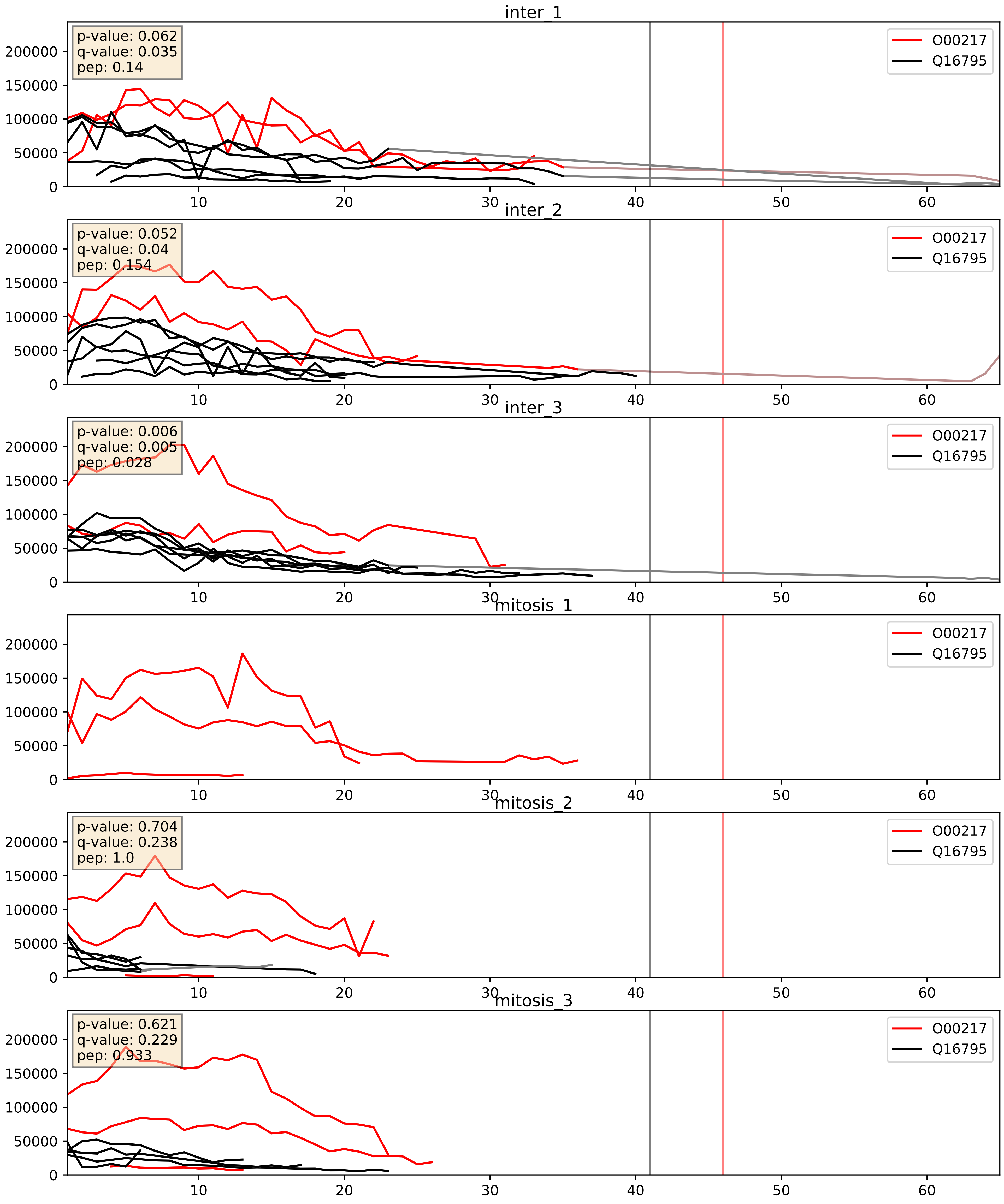


| level | condition_1 | condition_2 | pvalue | pvalue_adjusted |
|---|---|---|---|---|
| complex_abundance | mitosis | inter | 0.008621820707429922 | 0.06907915157215504 |
| interactor_abundance | mitosis | inter | 0.12144296724439234 | 0.30272329633430356 |
| interactor_ratio | mitosis | inter | 0.1440128515978898 | 0.5958192410236522 |

O43674_Q16795
NDUB5_HUMAN vs NDUA9_HUMAN
p-value: 0.009 q-value: 0.008

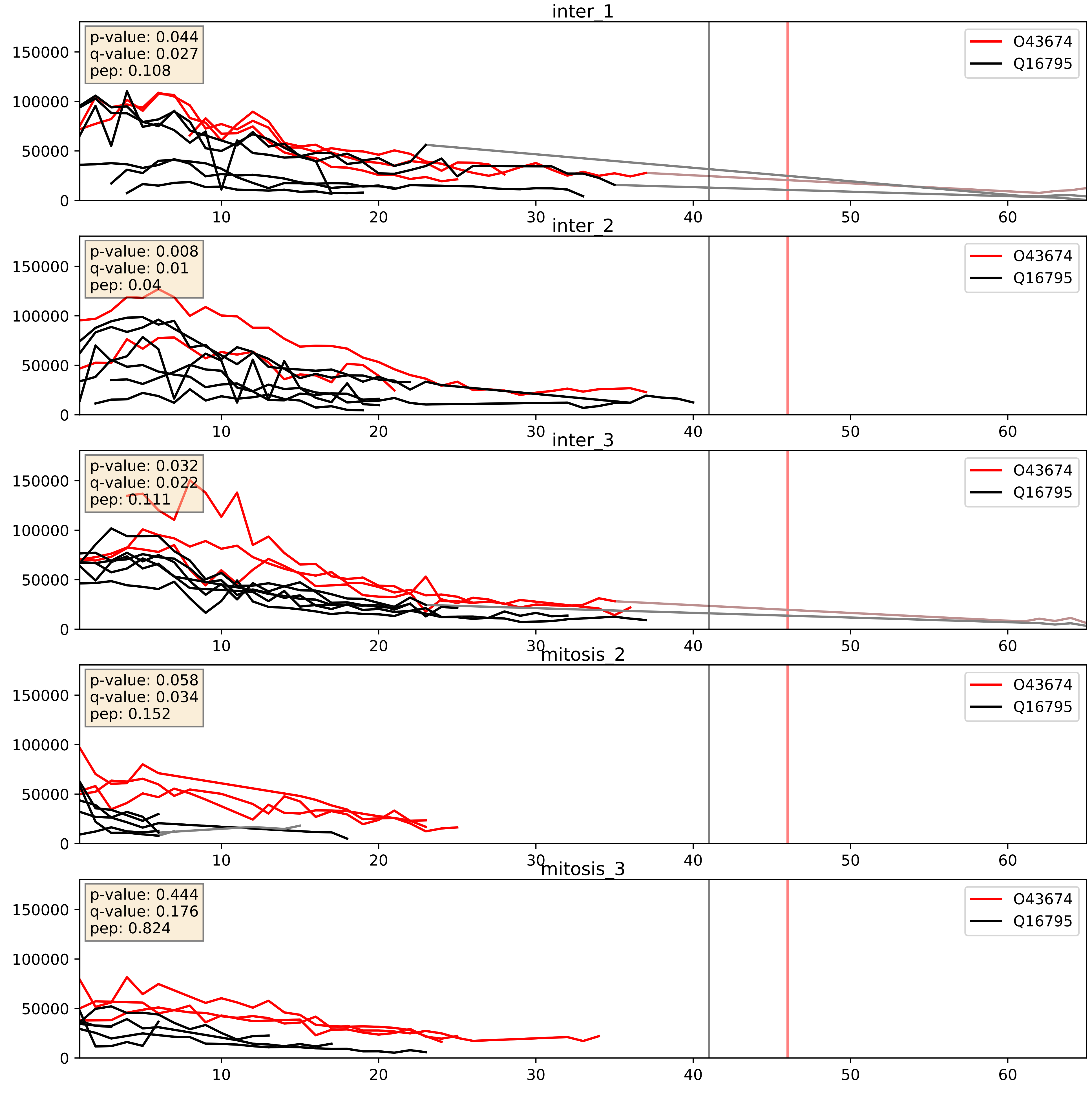


| level | condition_1 | condition_2 | pvalue | pvalue_adjusted |
|---|---|---|---|---|
| complex_abundance | mitosis | inter | 0.004012905502463791 | 0.04692687308892084 |
| interactor_abundance | mitosis | inter | 0.060074346990281614 | 0.20049308530665713 |
| interactor_ratio | mitosis | inter | 0.3349993602569867 | 0.7914334552464135 |

O75251_Q16795
NDUS7_HUMAN vs NDUA9_HUMAN
p-value: 0.005 q-value: 0.005

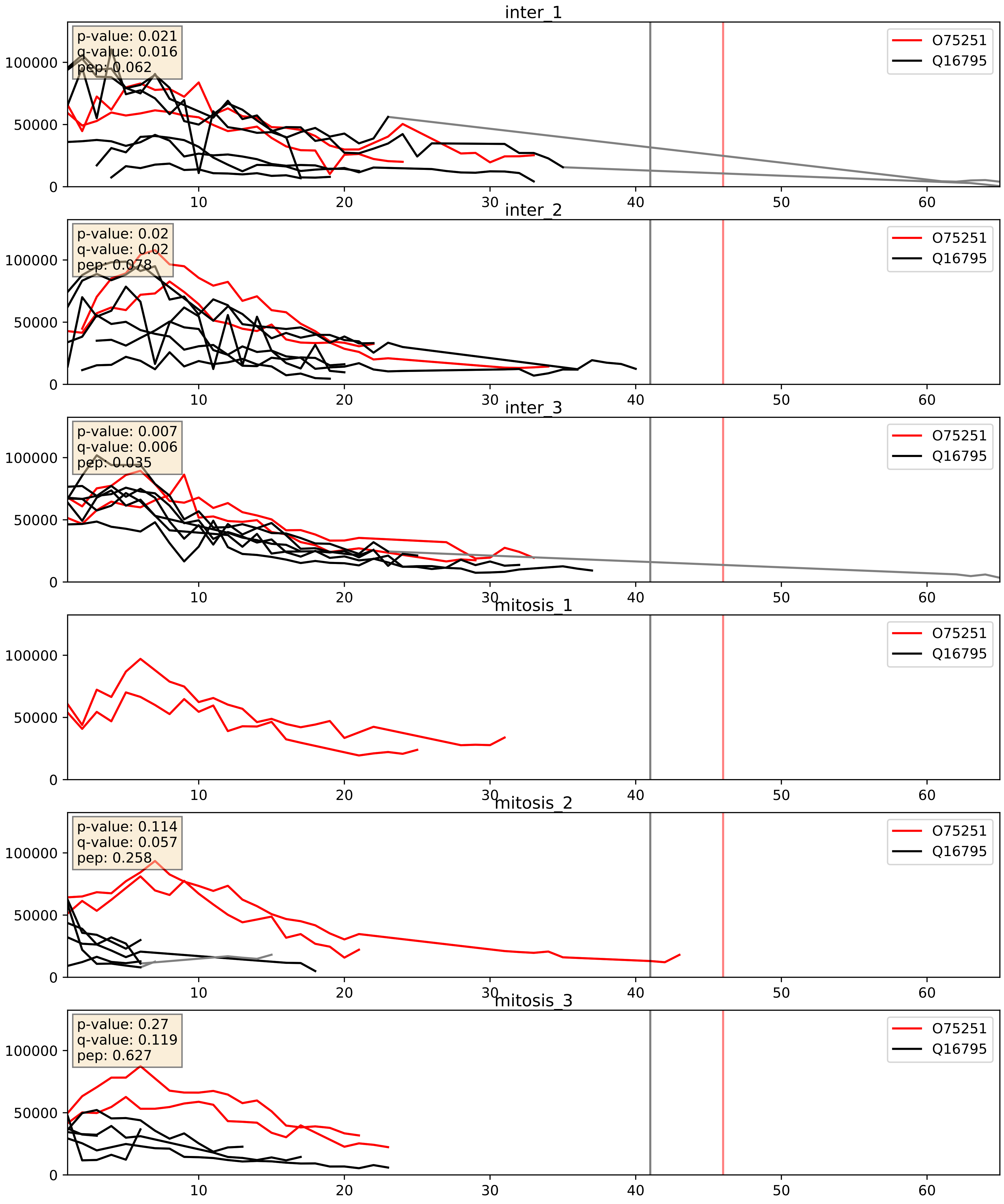


| level | condition_1 | condition_2 | pvalue | pvalue_adjusted |
|---|---|---|---|---|
| complex_abundance | mitosis | inter | 0.0034451756310002806 | 0.04302508960916016 |
| interactor_ratio | mitosis | inter | 0.03225972896418006 | 0.2870512265419764 |
| interactor_abundance | mitosis | inter | 0.08152270129016821 | 0.23860533505399845 |

O75306_Q16795
NDUS2_HUMAN vs NDUA9_HUMAN
p-value: 0.003 q-value: 0.003

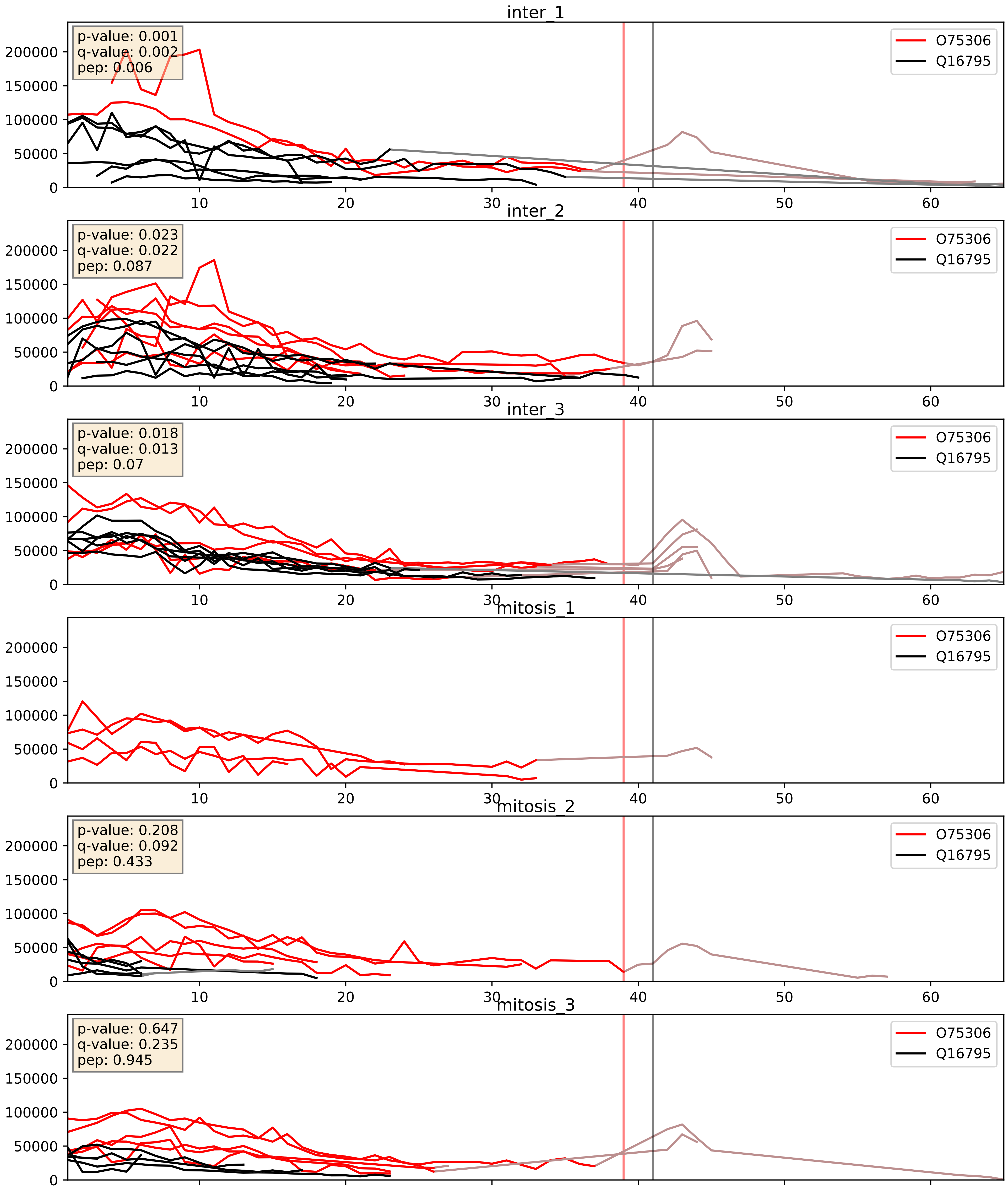


| level | condition_1 | condition_2 | pvalue | pvalue_adjusted |
|---|---|---|---|---|
| complex_abundance | mitosis | inter | 0.01920368528974783 | 0.10370391544427712 |
| interactor_abundance | mitosis | inter | 0.2467425181794483 | 0.4402075772438678 |
| interactor_ratio | mitosis | inter | 0.3089239106644241 | 0.7754077999721584 |

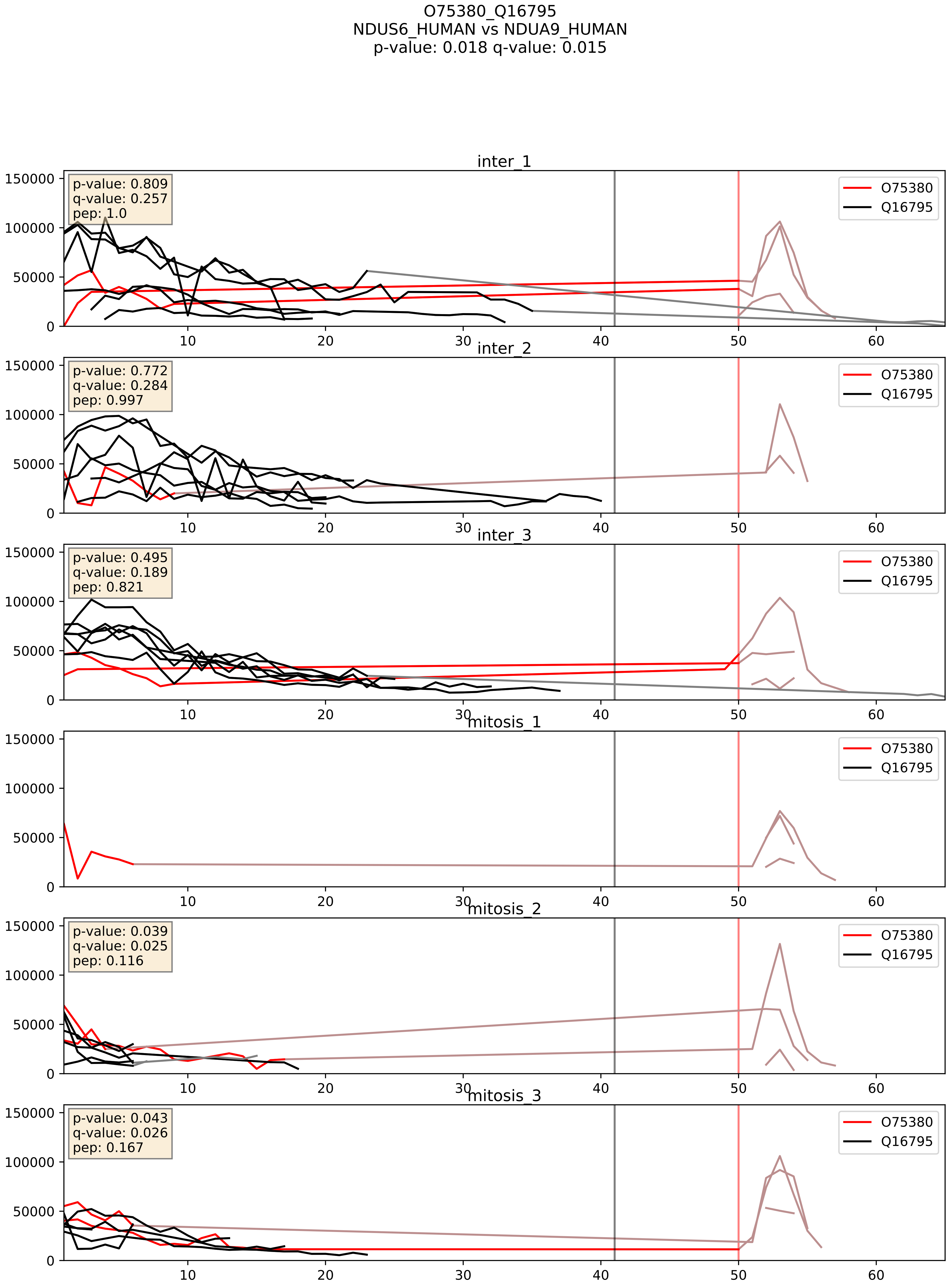


| level | condition_1 | condition_2 | pvalue | pvalue_adjusted |
|---|---|---|---|---|
| complex_abundance | mitosis | inter | 0.001864634710170751 | 0.030813268569617046 |
| interactor_ratio | mitosis | inter | 0.002026444247215246 | 0.0552431934909634 |
| interactor_abundance | mitosis | inter | 0.03282739912654744 | 0.15091435903504086 |

O75489_Q16795
NDUS3_HUMAN vs NDUA9_HUMAN
p-value: 0.016 q-value: 0.013

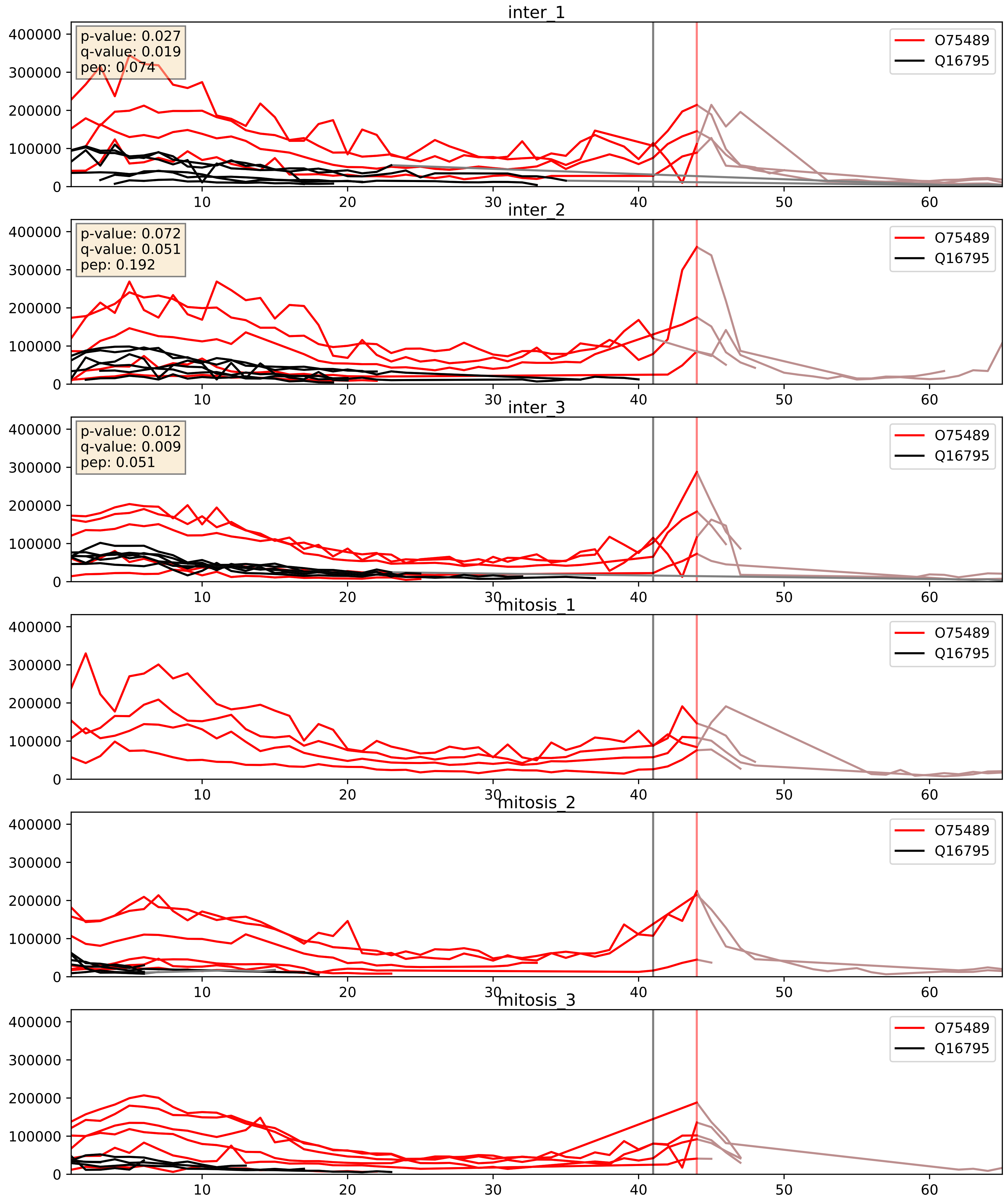


| level | condition_1 | condition_2 | pvalue | pvalue_adjusted |
|---|---|---|---|---|
| complex_abundance | mitosis | inter | 0.014394929509635883 | 0.08935503742021984 |
| interactor_abundance | mitosis | inter | 0.09482980261171356 | 0.26385279062449796 |
| interactor_ratio | mitosis | inter | 0.9224742852556895 | 0.9869241197086243 |

O95168_Q16795
NDUB4_HUMAN vs NDUA9_HUMAN
p-value: 0.005 q-value: 0.005

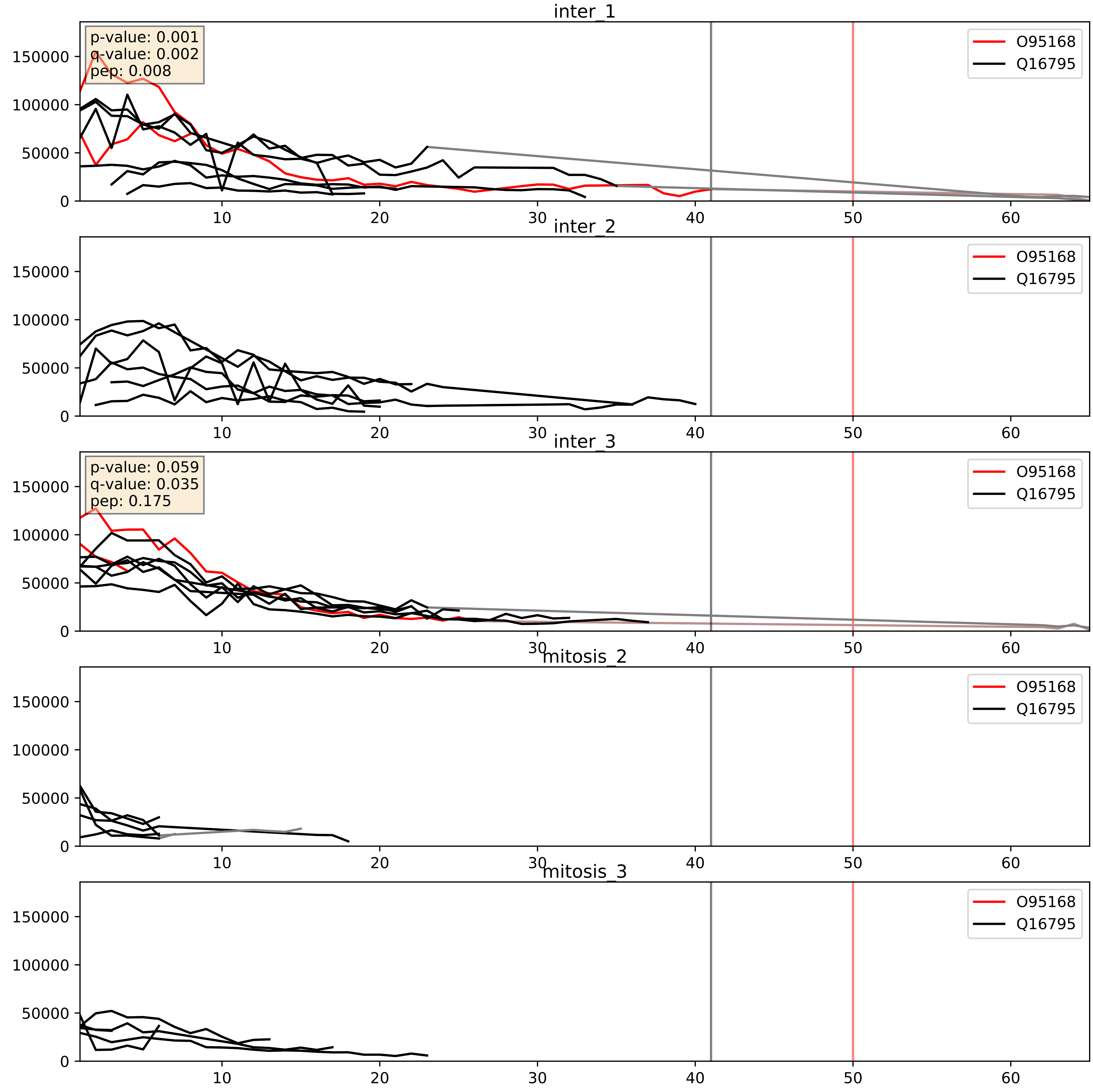


| level | condition_1 | condition_2 | pvalue | pvalue_adjusted |
|---|---|---|---|---|
| interactor_abundance | mitosis | inter | 0.04013928536710259 | 0.16166968180878097 |
| complex_abundance | mitosis | inter | 0.04204256652689513 | 0.14726323635770058 |
| interactor_ratio | mitosis | inter | 0.04903597586755165 | 0.3653158863587834 |

O95299_Q16795
NDUAA_HUMAN vs NDUA9_HUMAN
p-value: 0.007 q-value: 0.006

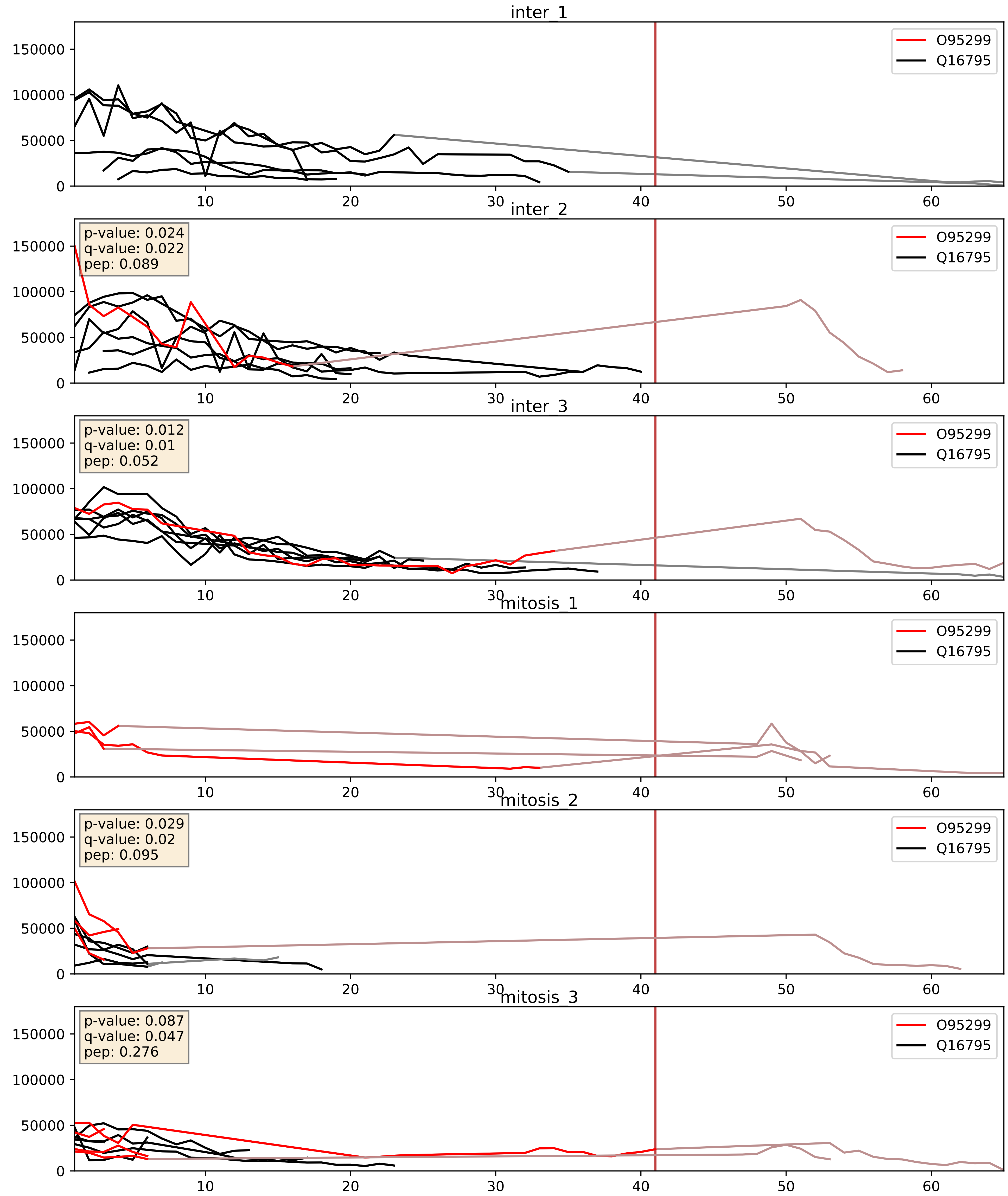


| level | condition_1 | condition_2 | pvalue | pvalue_adjusted |
|---|---|---|---|---|
| complex_abundance | mitosis | inter | 0.05404699082966155 | 0.16588104750874968 |
| interactor_ratio | mitosis | inter | 0.07816682136310352 | 0.46754569965771314 |
| interactor_abundance | mitosis | inter | 0.08765046374938844 | 0.2508734781283528 |

O96000_Q16795
NDUBA_HUMAN vs NDUA9_HUMAN
p-value: 0.016 q-value: 0.014

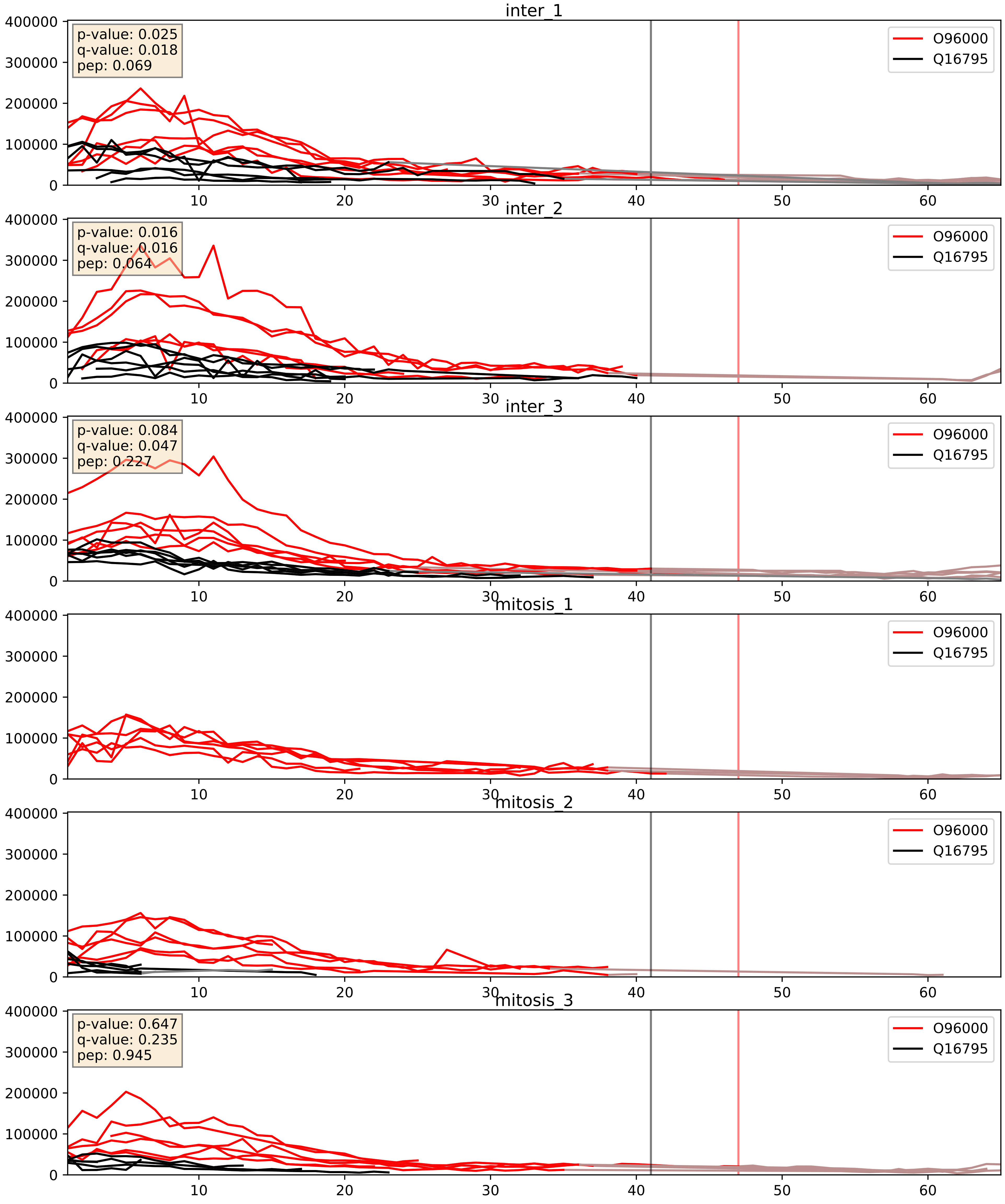


| level | condition_1 | condition_2 | pvalue | pvalue_adjusted |
|---|---|---|---|---|
| complex_abundance | mitosis | inter | 0.005866859048392097 | 0.05756820427608883 |
| interactor_abundance | mitosis | inter | 0.038734556862202 | 0.16006169767822792 |
| interactor_ratio | mitosis | inter | 0.8425742511809455 | 0.9753124530235151 |

P10606_Q16795
COX5B_HUMAN vs NDUA9_HUMAN
p-value: 0.001 q-value: 0.041

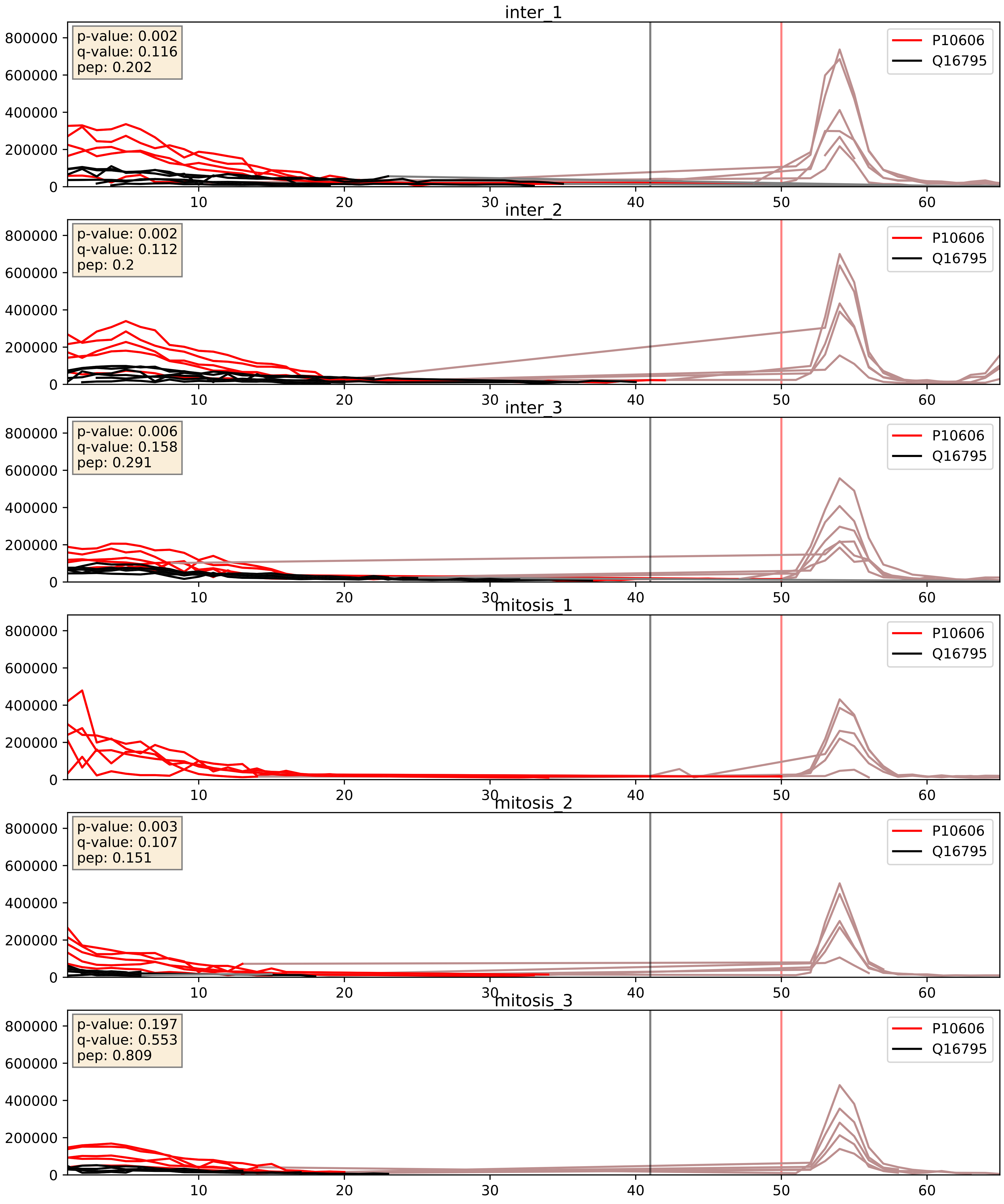


| level | condition_1 | condition_2 | pvalue | pvalue_adjusted |
|---|---|---|---|---|
| complex_abundance | mitosis | inter | 0.007677520966252494 | 0.06519315066658214 |
| interactor_abundance | mitosis | inter | 0.056456994347416016 | 0.19306244235182227 |
| interactor_ratio | mitosis | inter | 0.8094301625805784 | 0.9652518042298145 |

P17568_Q16795
NDUB7_HUMAN vs NDUA9_HUMAN
p-value: 0.034 q-value: 0.024

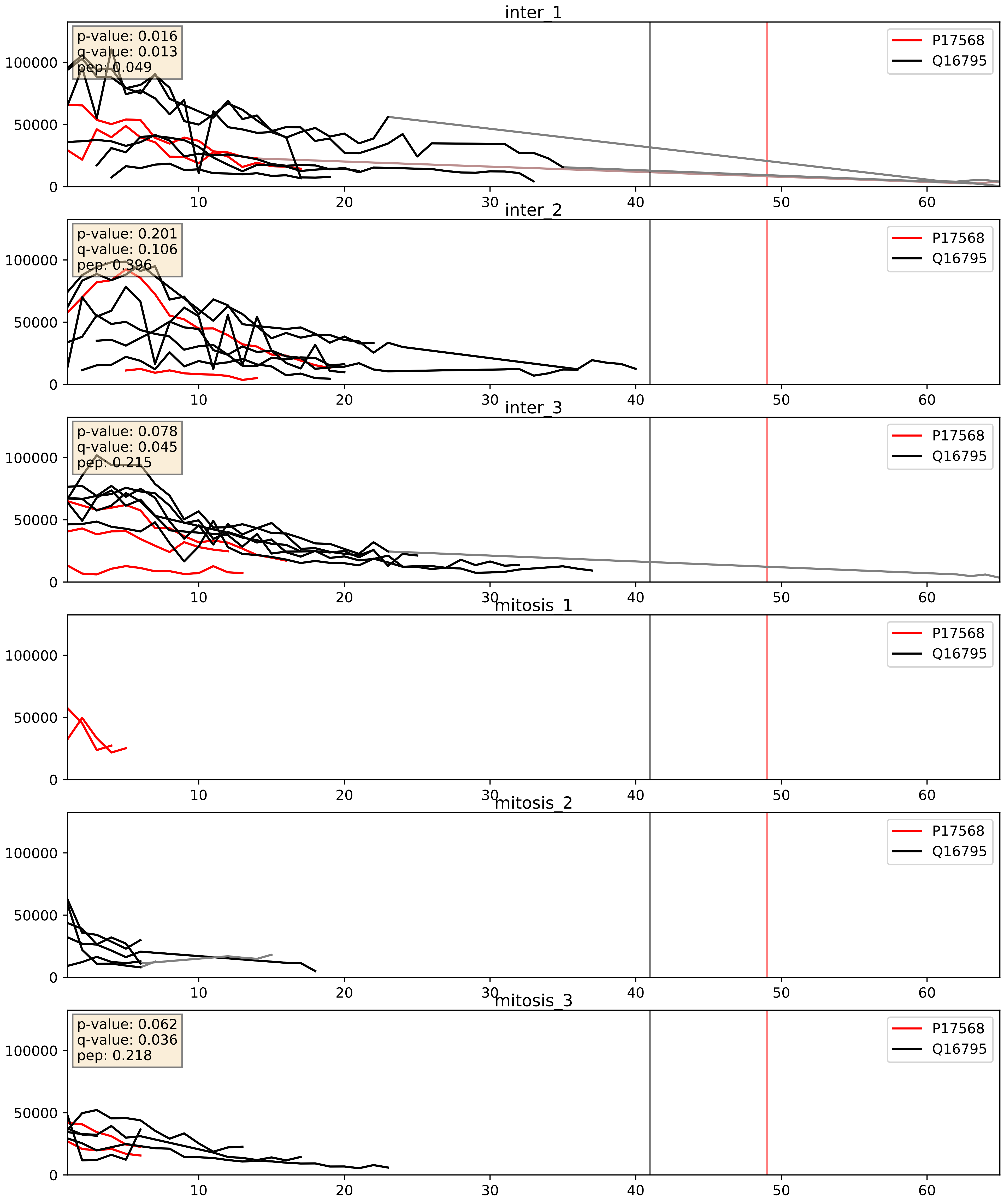


| level | condition_1 | condition_2 | pvalue | pvalue_adjusted |
|---|---|---|---|---|
| complex_abundance | mitosis | inter | 0.0011818146794331647 | 0.023096652182529428 |
| interactor_abundance | mitosis | inter | 0.002814834052395079 | 0.04032632550376883 |
| interactor_ratio | mitosis | inter | 0.007845783748905964 | 0.12516692290607134 |

P19404_Q16795
NDUV2_HUMAN vs NDUA9_HUMAN
p-value: 0.02 q-value: 0.016

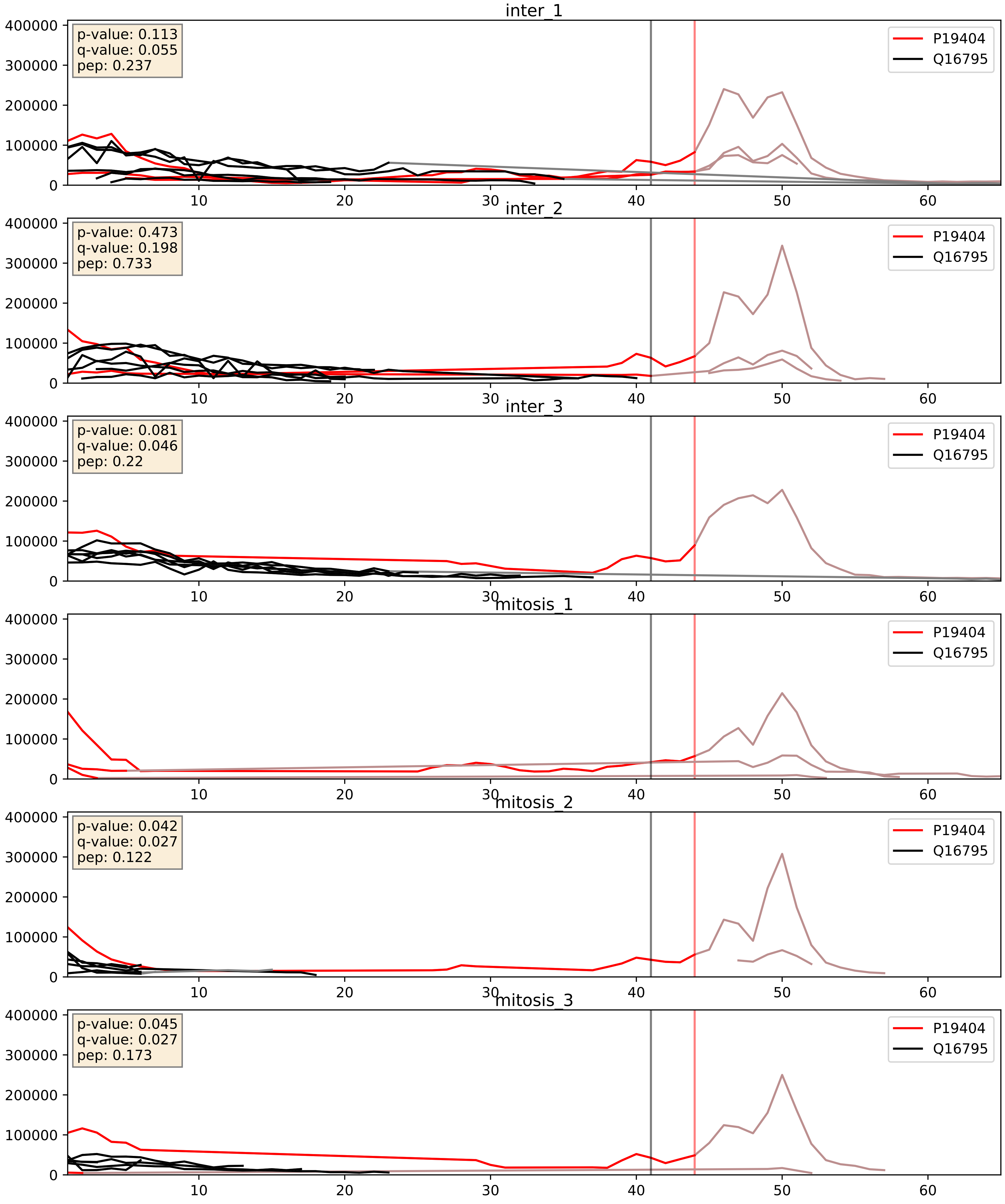


| level | condition_1 | condition_2 | pvalue | pvalue_adjusted |
|---|---|---|---|---|
| interactor_ratio | mitosis | inter | 0.002393274965868345 | 0.06170612562600311 |
| complex_abundance | mitosis | inter | 0.0044378730591735594 | 0.049683977369634376 |
| interactor_abundance | mitosis | inter | 0.014079905488348197 | 0.10514879448915647 |

P22695_Q16795
QCR2_HUMAN vs NDUA9_HUMAN
p-value: 0.059 q-value: 0.037

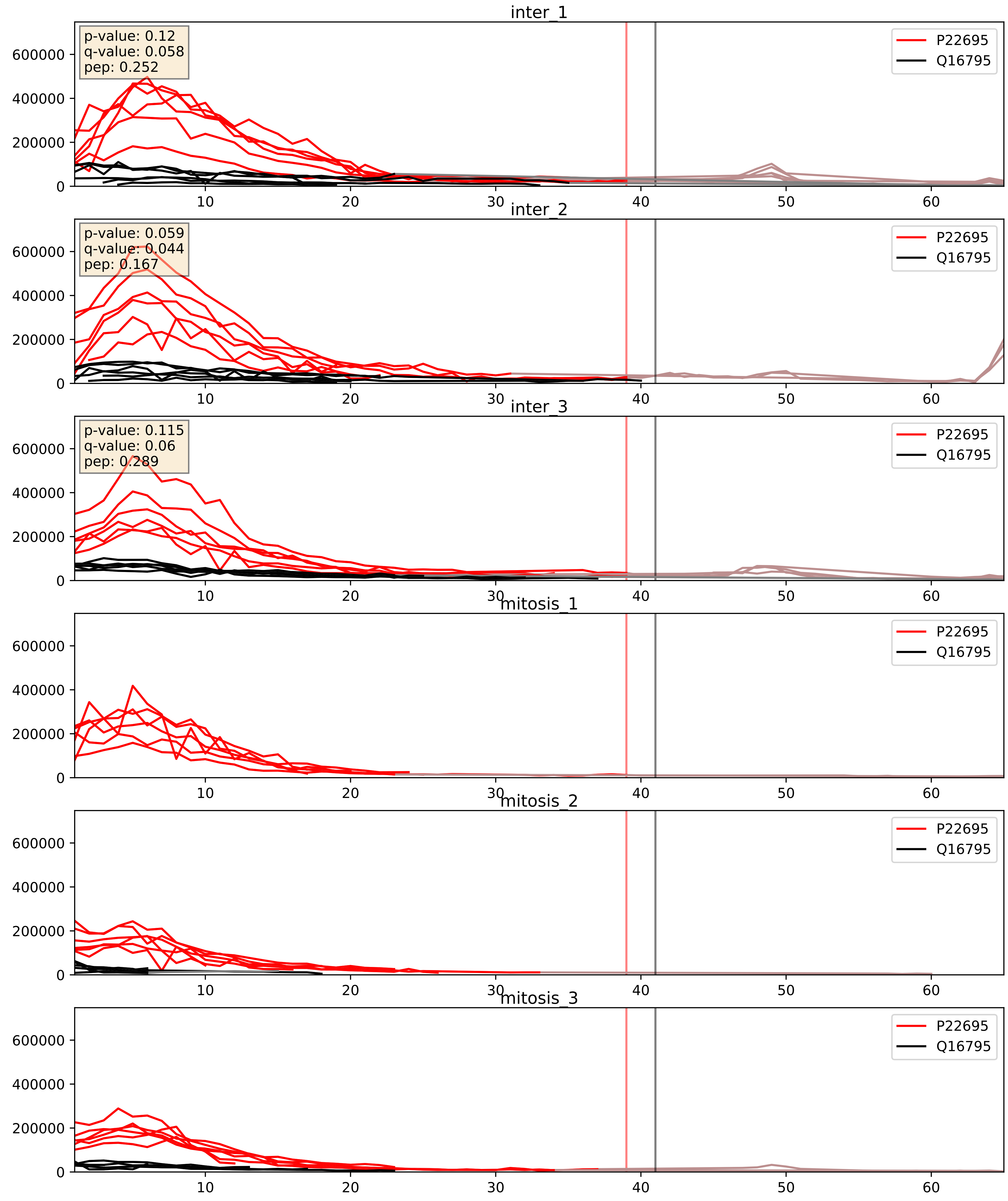


| level | condition_1 | condition_2 | pvalue | pvalue_adjusted |
|---|---|---|---|---|
| complex_abundance | mitosis | inter | 0.010152476802326144 | 0.07446889582511722 |
| interactor_abundance | mitosis | inter | 0.05984572023708979 | 0.19991389862614187 |
| interactor_ratio | mitosis | inter | 0.7776608823156874 | 0.9571917866015084 |

P28331_Q16795
NDUS1_HUMAN vs NDUA9_HUMAN
p-value: 0.032 q-value: 0.023

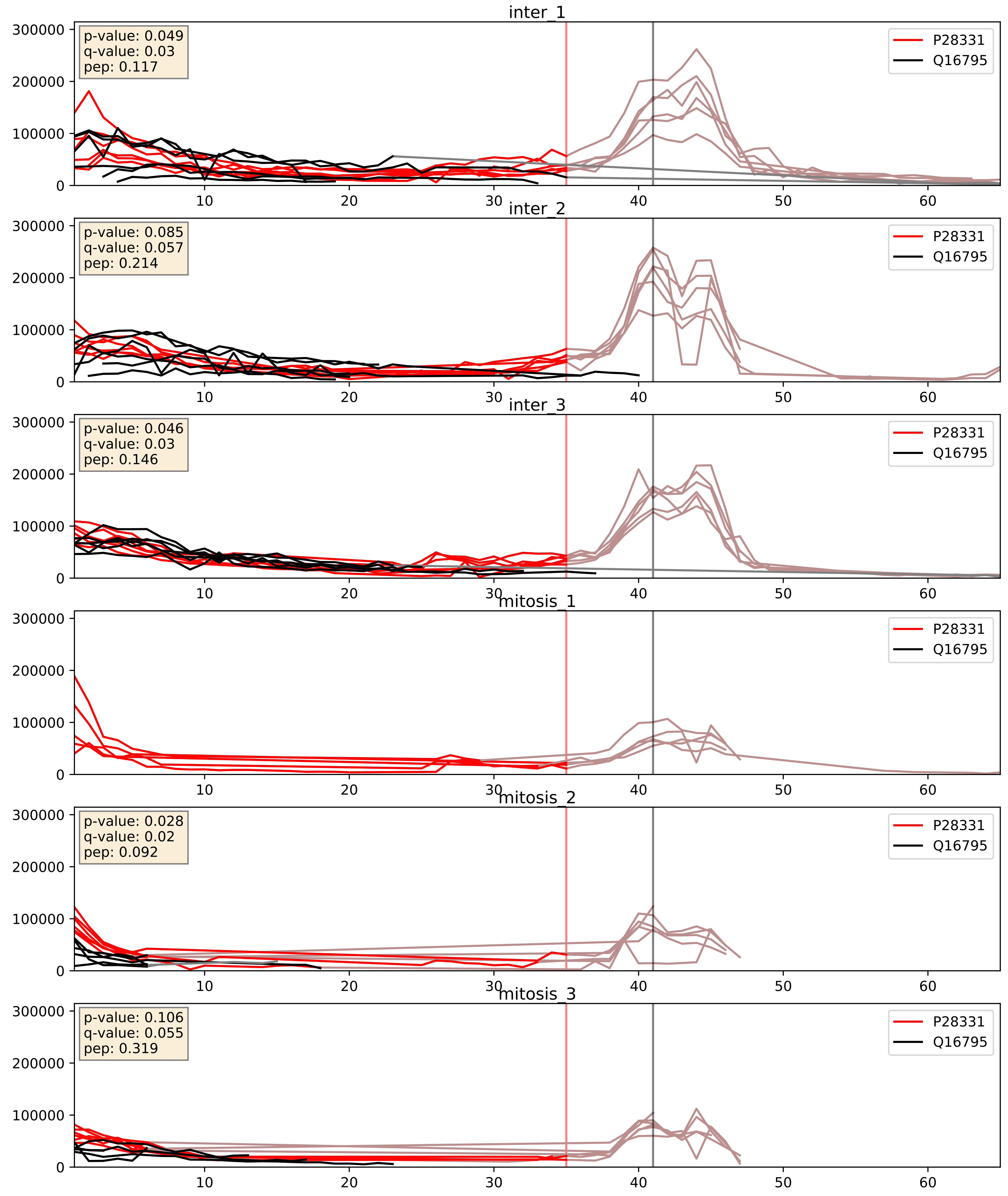


| level | condition_1 | condition_2 | pvalue | pvalue_adjusted |
|---|---|---|---|---|
| complex_abundance | mitosis | inter | 0.0004818900669497129 | 0.013833744180327739 |
| interactor_abundance | mitosis | inter | 0.005524204947292736 | 0.06317594435380872 |
| interactor_ratio | mitosis | inter | 0.15636129142680918 | 0.6134063495020562 |

P47985_Q16795
UCRI_HUMAN vs NDUA9_HUMAN
p-value: 0.011 q-value: 0.01

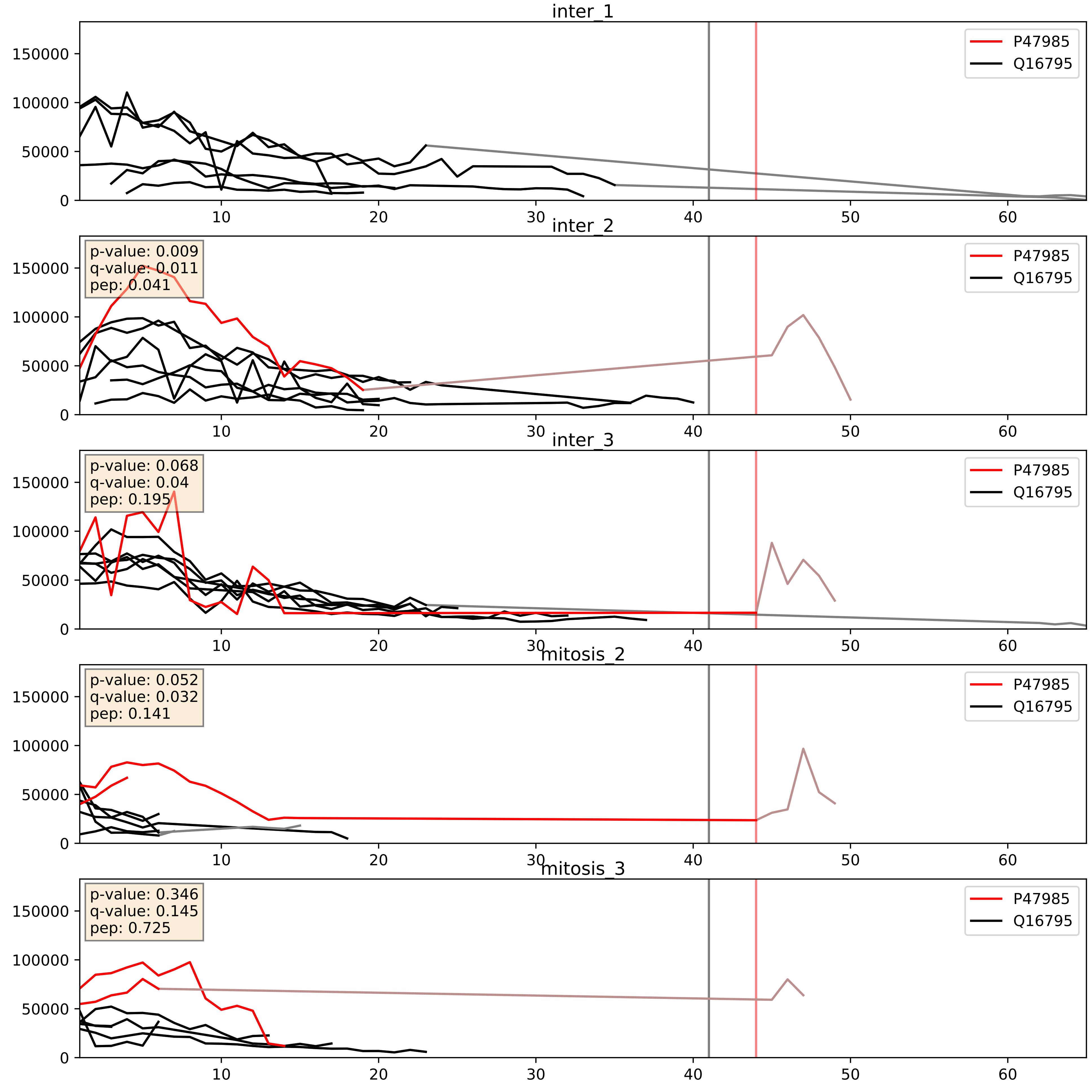


| level | condition_1 | condition_2 | pvalue | pvalue_adjusted |
| --- | --- | --- | --- | --- |
| interactor_ratio | mitosis | inter | 0.04227297338141573 | 0.3399226239910554 |
| complex_abundance | mitosis | inter | 0.13944074790777403 | 0.28459313475568465 |
| interactor_abundance | mitosis | inter | 0.654921269108097 | 0.7704956107154083 |

P49821_Q16795
NDUV1_HUMAN vs NDUA9_HUMAN
p-value: 0.031 q-value: 0.022

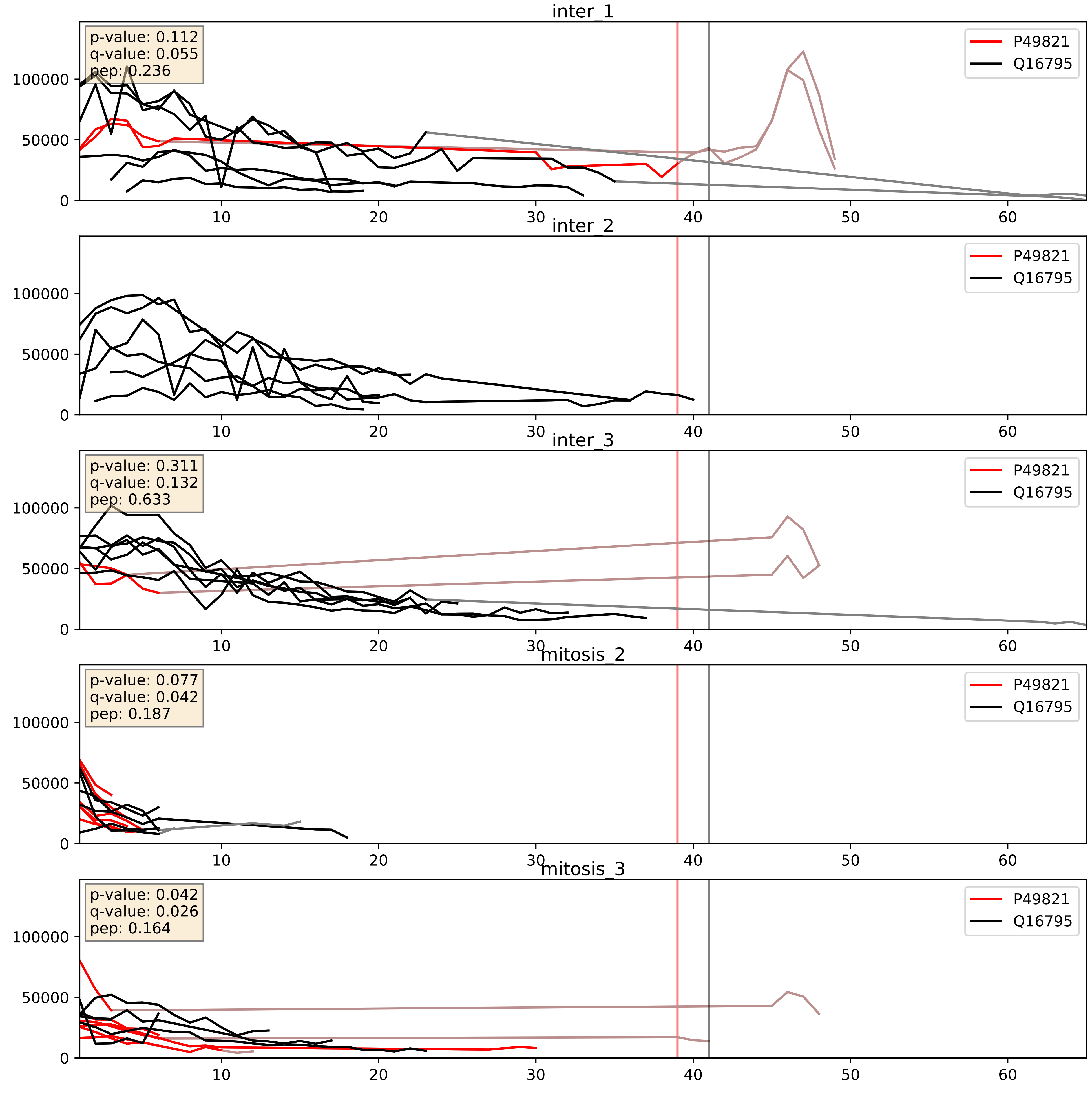


| level | condition_1 | condition_2 | pvalue | pvalue_adjusted |
|---|---|---|---|---|
| interactor_ratio | mitosis | inter | 0.05423293268660425 | 0.38557633205758507 |
| complex_abundance | mitosis | inter | 0.08829398621743838 | 0.21793440658052843 |
| interactor_abundance | mitosis | inter | 0.17744486878959026 | 0.36863408771116224 |

Q16718_Q16795
NDUA5_HUMAN vs NDUA9_HUMAN
p-value: 0.026 q-value: 0.02

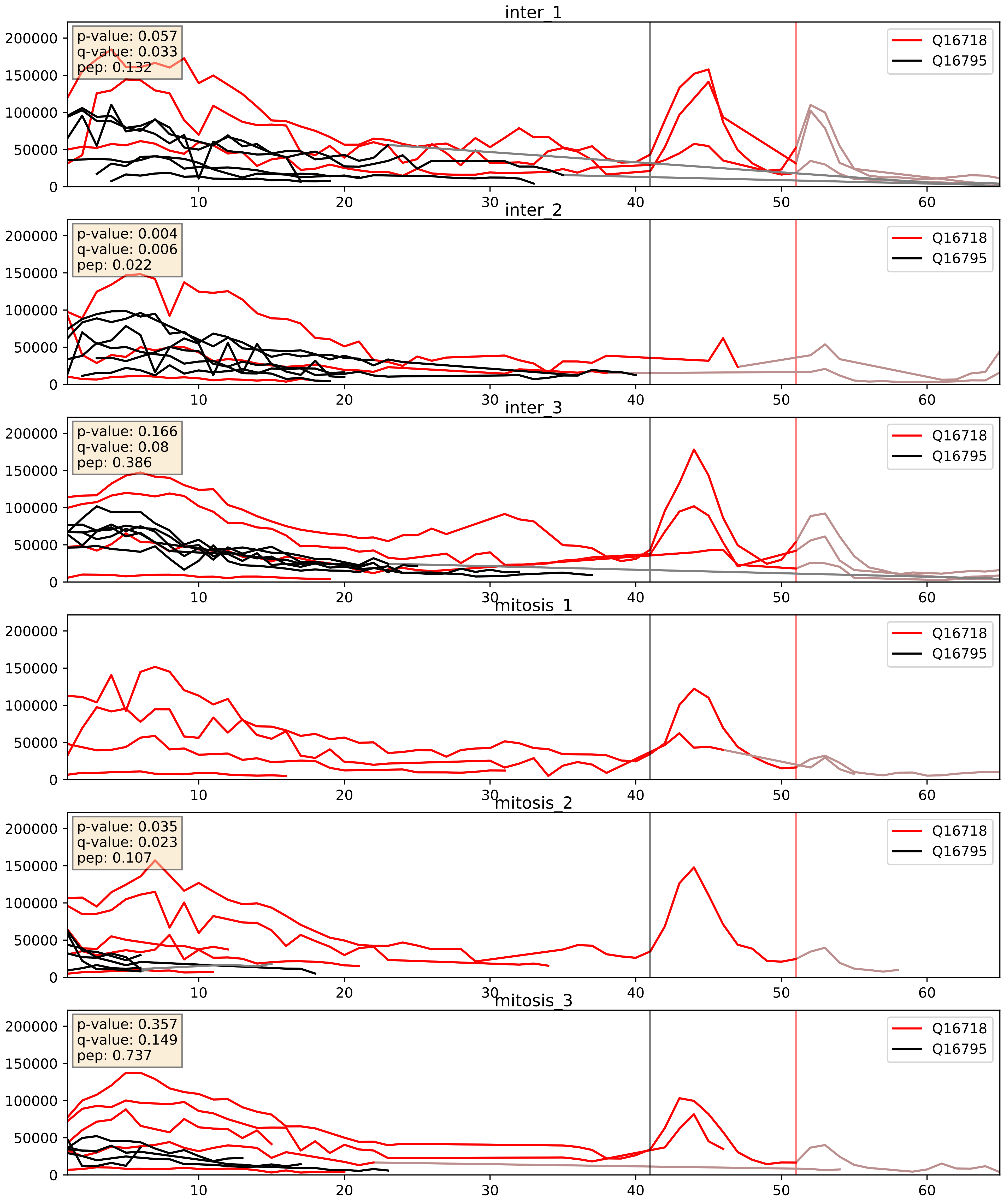


| level | condition_1 | condition_2 | pvalue | pvalue_adjusted |
|---|---|---|---|---|
| complex_abundance | mitosis | inter | 0.011637535380455984 | 0.08007821773046883 |
| interactor_abundance | mitosis | inter | 0.1350462351227089 | 0.3218140164115769 |
| interactor_ratio | mitosis | inter | 0.3856970585520332 | 0.8223080501134257 |

Q16795_Q9P0J0
NDUA9_HUMAN vs NDUAD_HUMAN
p-value: 0.006 q-value: 0.006

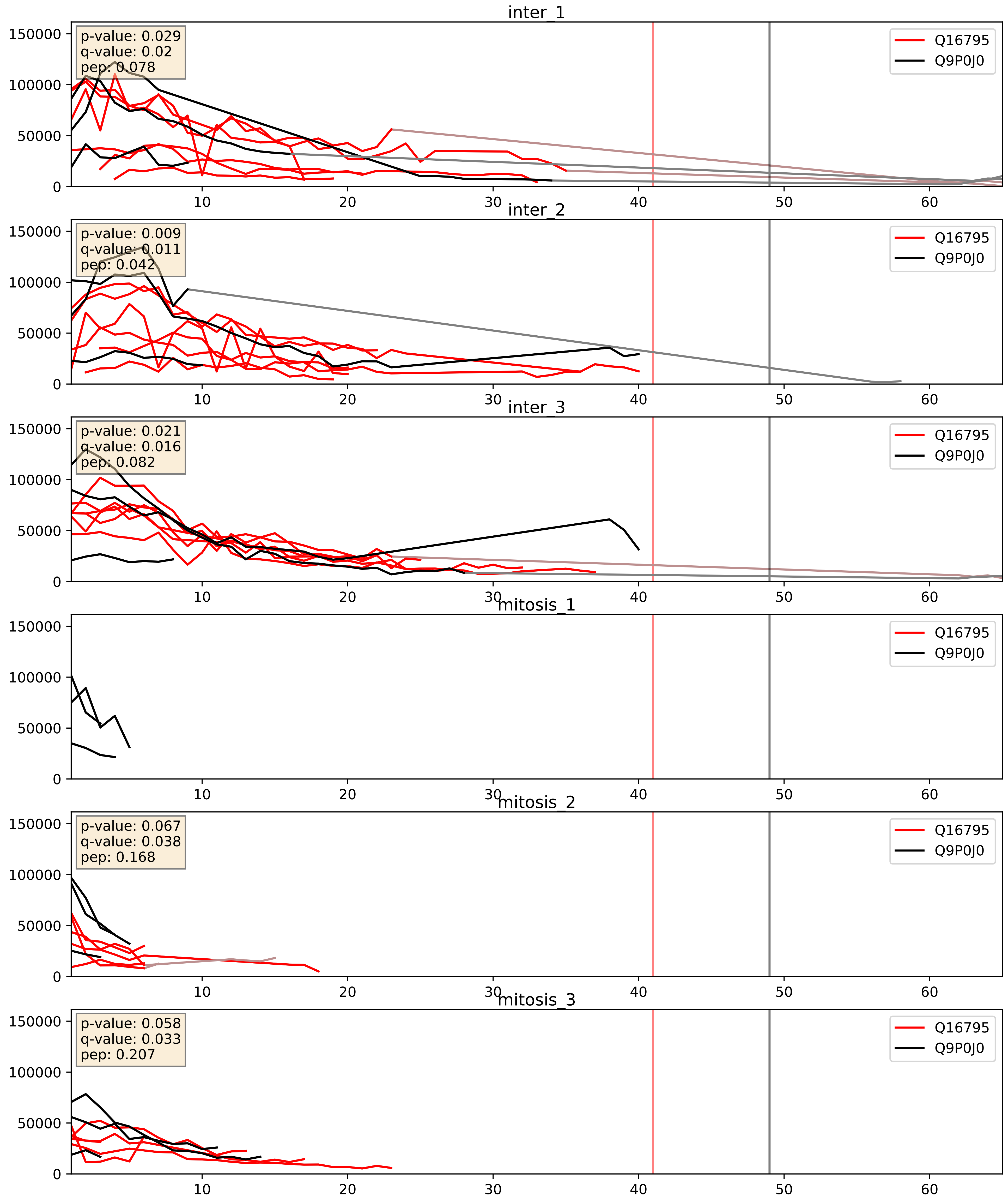


| level | condition_1 | condition_2 | pvalue | pvalue_adjusted |
|---|---|---|---|---|
| interactor_abundance | mitosis | inter | 5.119480063602888e-05 | 0.006283598333582311 |
| complex_abundance | mitosis | inter | 0.0004070925892423535 | 0.012858717948024156 |
| interactor_ratio | mitosis | inter | 0.040201029565406006 | 0.33216294698829674 |
