## Supplementary Data 2 for "SECAT: Quantifying differential protein-protein interaction states by network-centric analysis": hela_string_000072_P62310.pdf

P62310_P62310

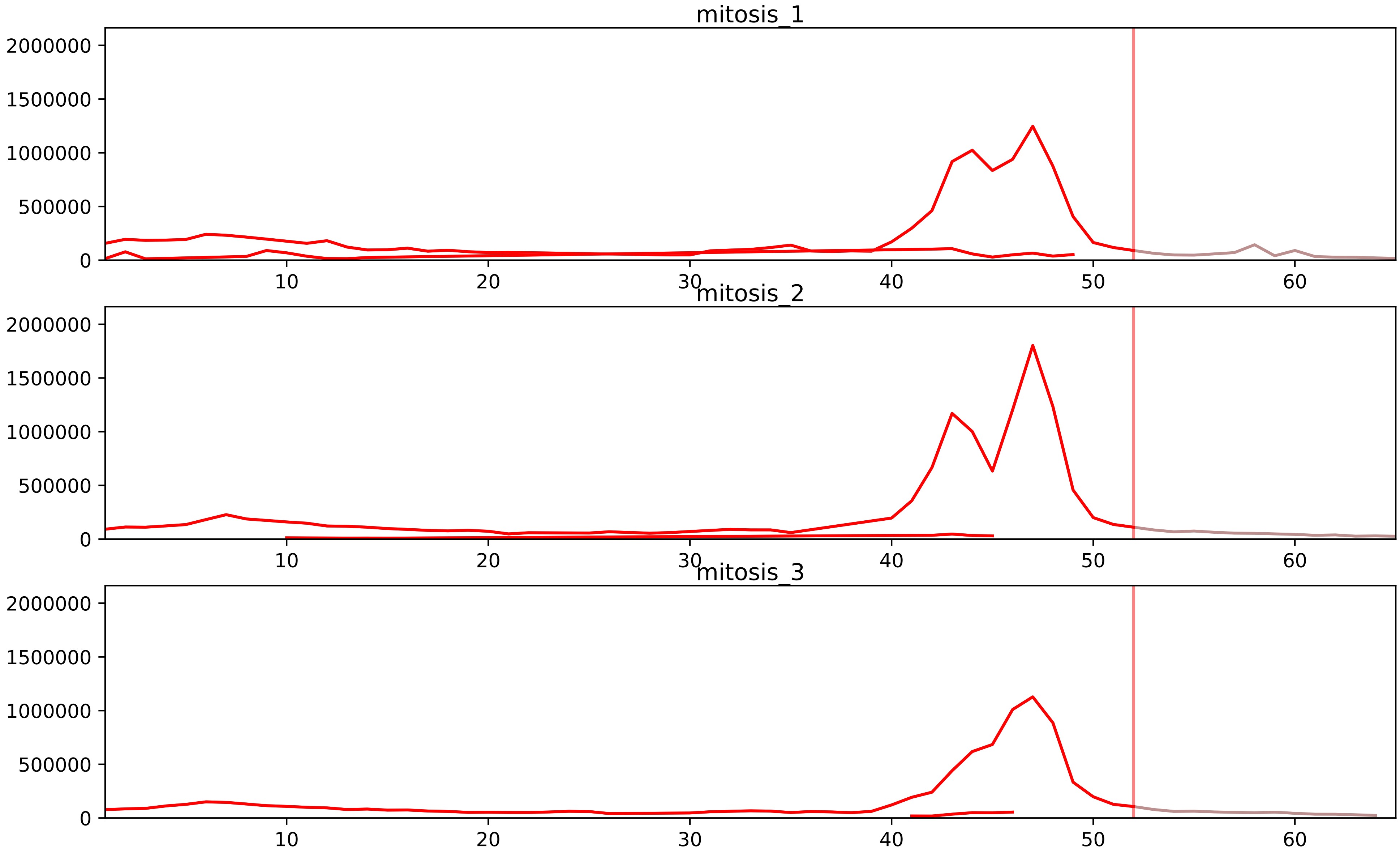


| level | condition_1 | condition_2 | pvalue | pvalue_adjusted |
|---|---|---|---|---|
| interactor_ratio | mitosis | inter | 1.5345480972589888e-06 | 4.481854760248476e-05 |
| assembled_abundance | mitosis | inter | 2.2094447180164275e-06 | 0.0023535084180592012 |
| total_abundance | mitosis | inter | 3.188054062278942e-06 | 0.003540864511957668 |
| complex_abundance | mitosis | inter | 2.0950924648982223e-05 | 0.0008031187782109853 |
| monomer_abundance | mitosis | inter | 0.00015110494156855934 | 0.038277056377303 |
| interactor_abundance | mitosis | inter | 0.00025196942887565314 | 0.004457920664723094 |

O95777_P62310
LSM8_HUMAN vs LSM3_HUMAN
p-value: 0.033 q-value: 0.024

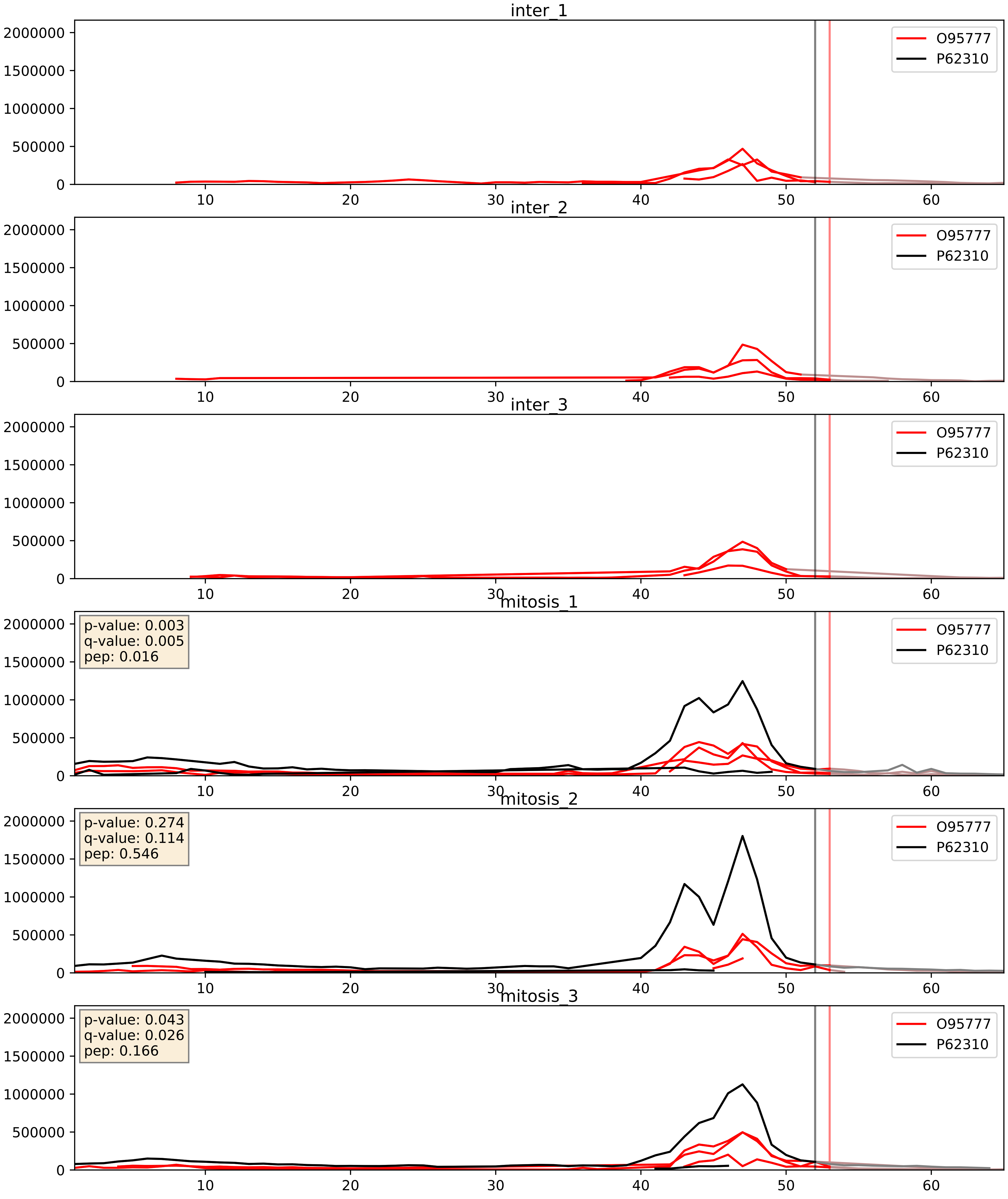


| level | condition_1 | condition_2 | pvalue | pvalue_adjusted |
|---|---|---|---|---|
| complex_abundance | mitosis | inter | 0.0007661190734198726 | 0.018128921229455204 |
| interactor_abundance | mitosis | inter | 0.0007667676877166997 | 0.02034469828315081 |
| interactor_ratio | mitosis | inter | 0.0036881715066175926 | 0.0787300451287945 |

P62310_Q13435
LSM3_HUMAN vs SF3B2_HUMAN
p-value: 0.018 q-value: 0.014

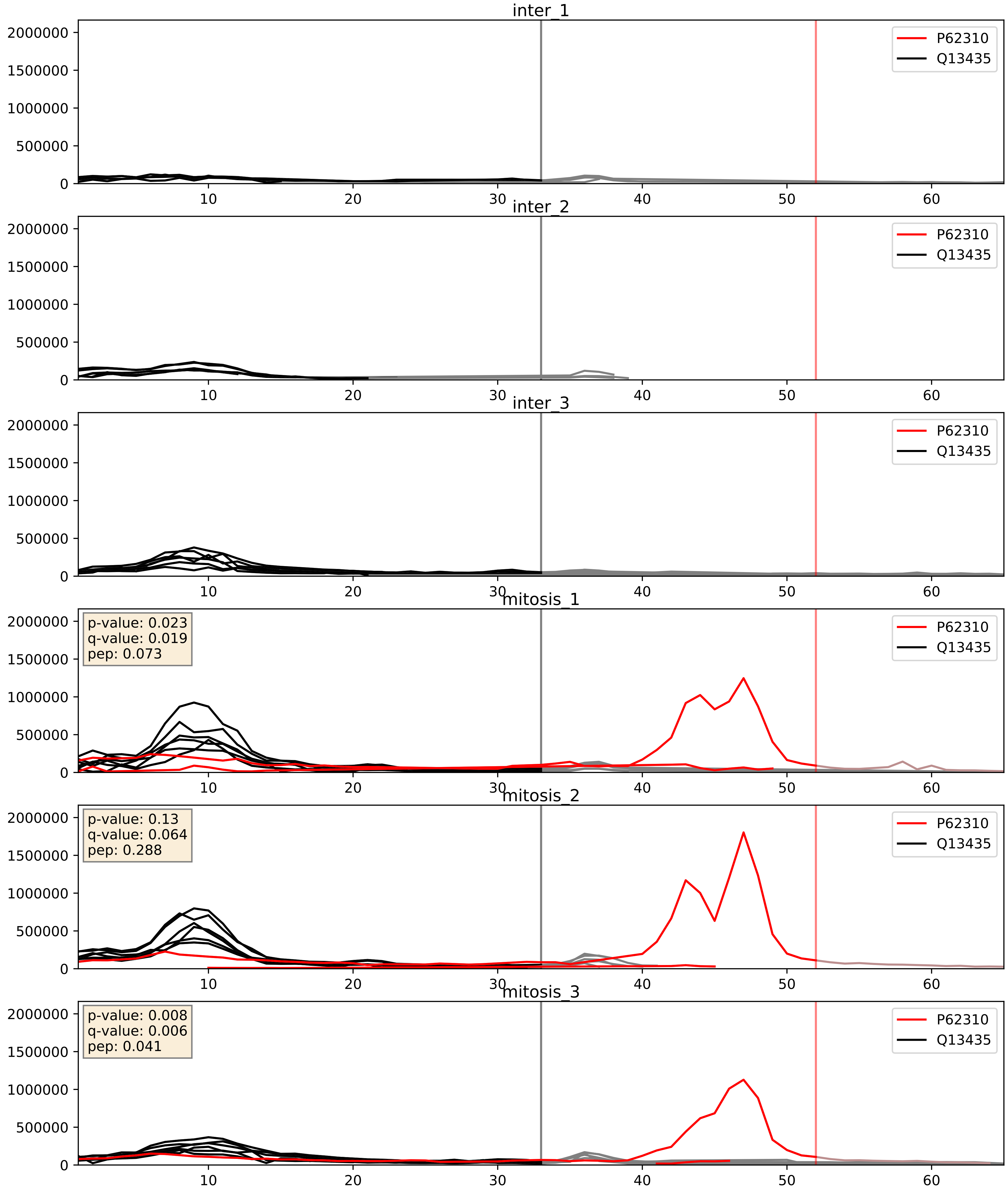


| level | condition_1 | condition_2 | pvalue | pvalue_adjusted |
|---|---|---|---|---|
| interactor_ratio | mitosis | inter | 4.374856816950016e-06 | 0.002520386835947235 |
| complex_abundance | mitosis | inter | 0.00047615701417451177 | 0.013816623868928207 |
| interactor_abundance | mitosis | inter | 0.011614432178085941 | 0.0950928163026453 |

P62310_Q9Y333
LSM3_HUMAN vs LSM2_HUMAN
p-value: 0.063 q-value: 0.039

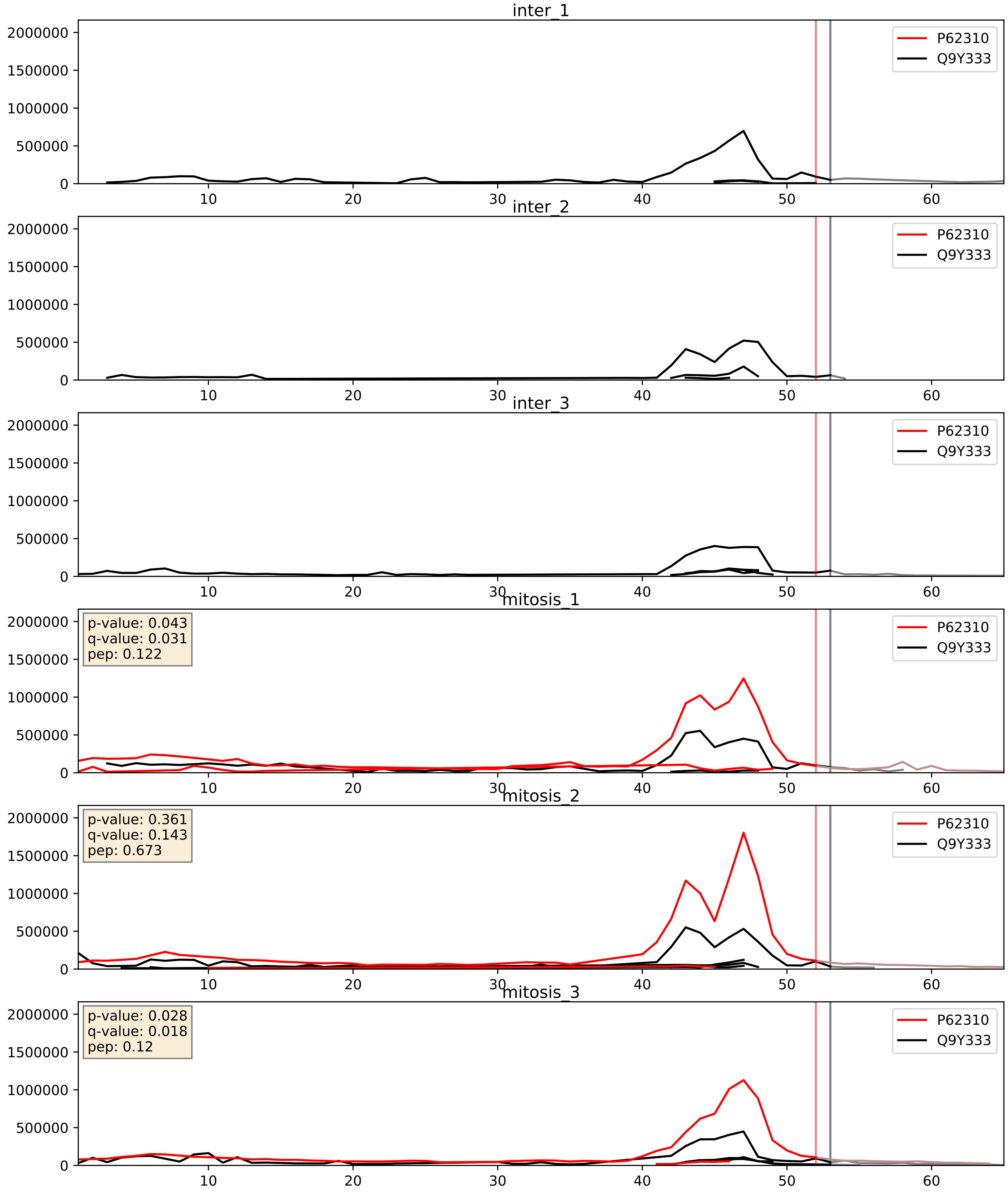


| level | condition_1 | condition_2 | pvalue | pvalue_adjusted |
|---|---|---|---|---|
| complex_abundance | mitosis | inter | 0.0002191820108542316 | 0.010087086090925927 |
| interactor_abundance | mitosis | inter | 0.0008888344561131203 | 0.02197564842852207 |
| interactor_ratio | mitosis | inter | 0.49118645642073633 | 0.8696082868586356 |
