## Supplementary Data 2 for "SECAT: Quantifying differential protein-protein interaction states by network-centric analysis": hela_string_000073_Q07955.pdf

Q07955_Q07955

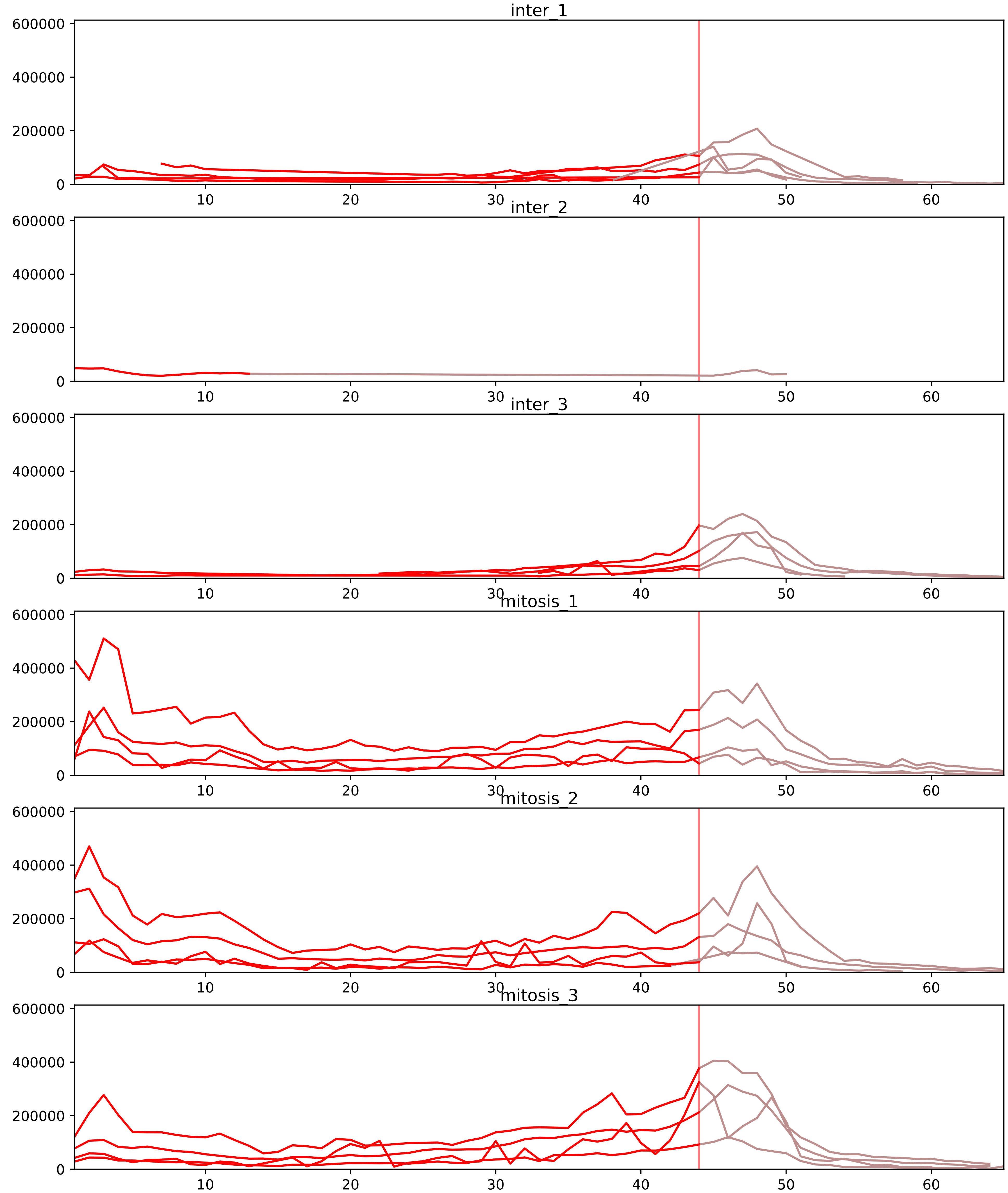


| level | condition_1 | condition_2 | pvalue | pvalue_adjusted |
|---|---|---|---|---|
| interactor_ratio | mitosis | inter | 1.6299757380016584e-06 | 4.686180246754768e-05 |
| complex_abundance | mitosis | inter | 1.54436244689788e-05 | 0.0006765778338790712 |
| interactor_abundance | mitosis | inter | 0.00010059875386324417 | 0.002285206260597151 |
| assembled_abundance | mitosis | inter | 0.04235561470036489 | 0.2467295239387945 |
| total_abundance | mitosis | inter | 0.12768901995003928 | 0.4419285945231154 |
| monomer_abundance | mitosis | inter | 0.3299864236890308 | 0.6718717495121003 |

O43390_Q07955
HNRPR_HUMAN vs SRSF1_HUMAN
p-value: 0.002 q-value: 0.002

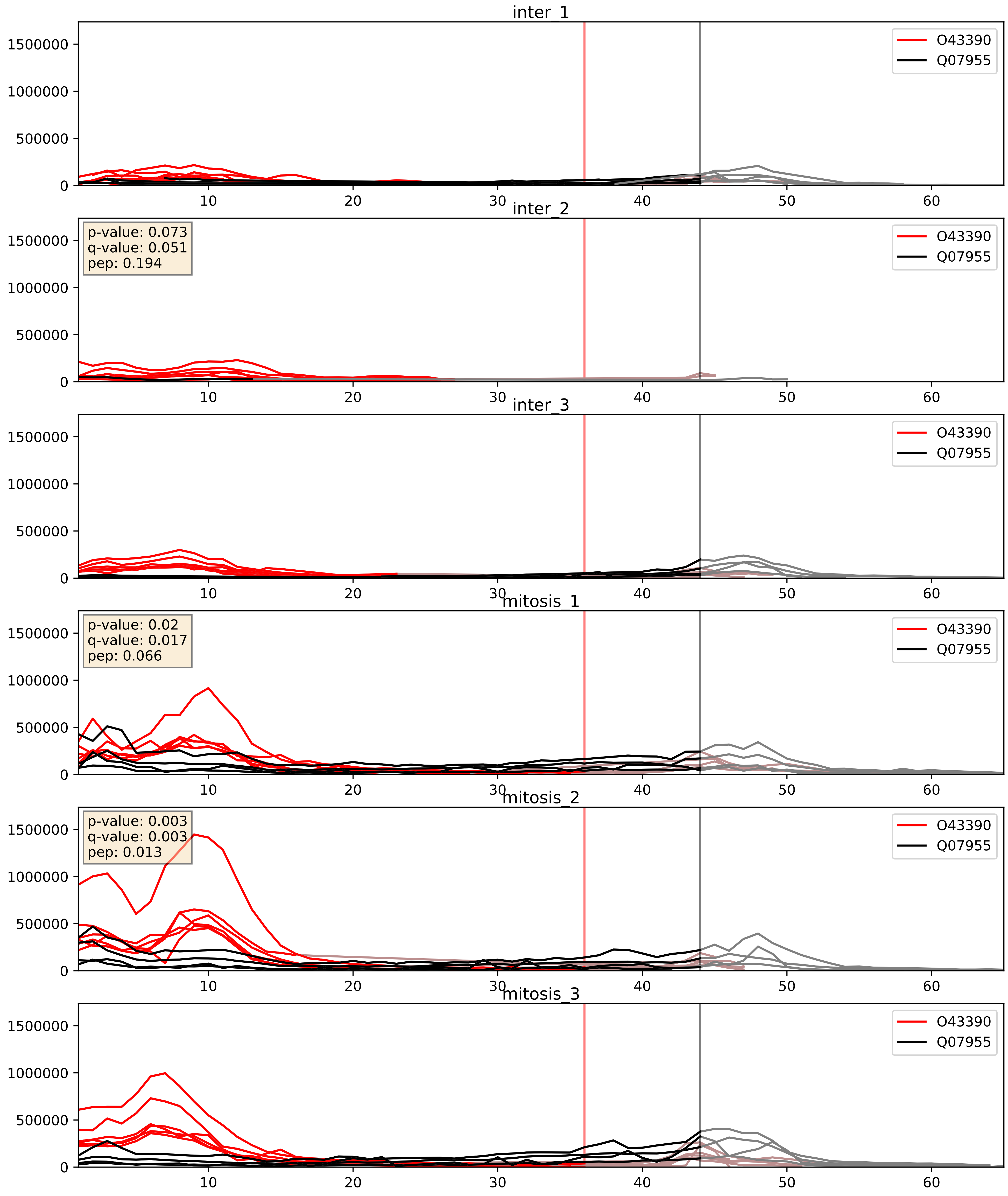


| level | condition_1 | condition_2 | pvalue | pvalue_adjusted |
|---|---|---|---|---|
| interactor_abundance | mitosis | inter | 0.0009696737069179261 | 0.02306454775731039 |
| complex_abundance | mitosis | inter | 0.004069476203020404 | 0.04732787841836958 |
| interactor_ratio | mitosis | inter | 0.16851302494823342 | 0.6310024031307429 |

P07910_Q07955
HNRPC_HUMAN vs SRSF1_HUMAN
p-value: 0.038 q-value: 0.026

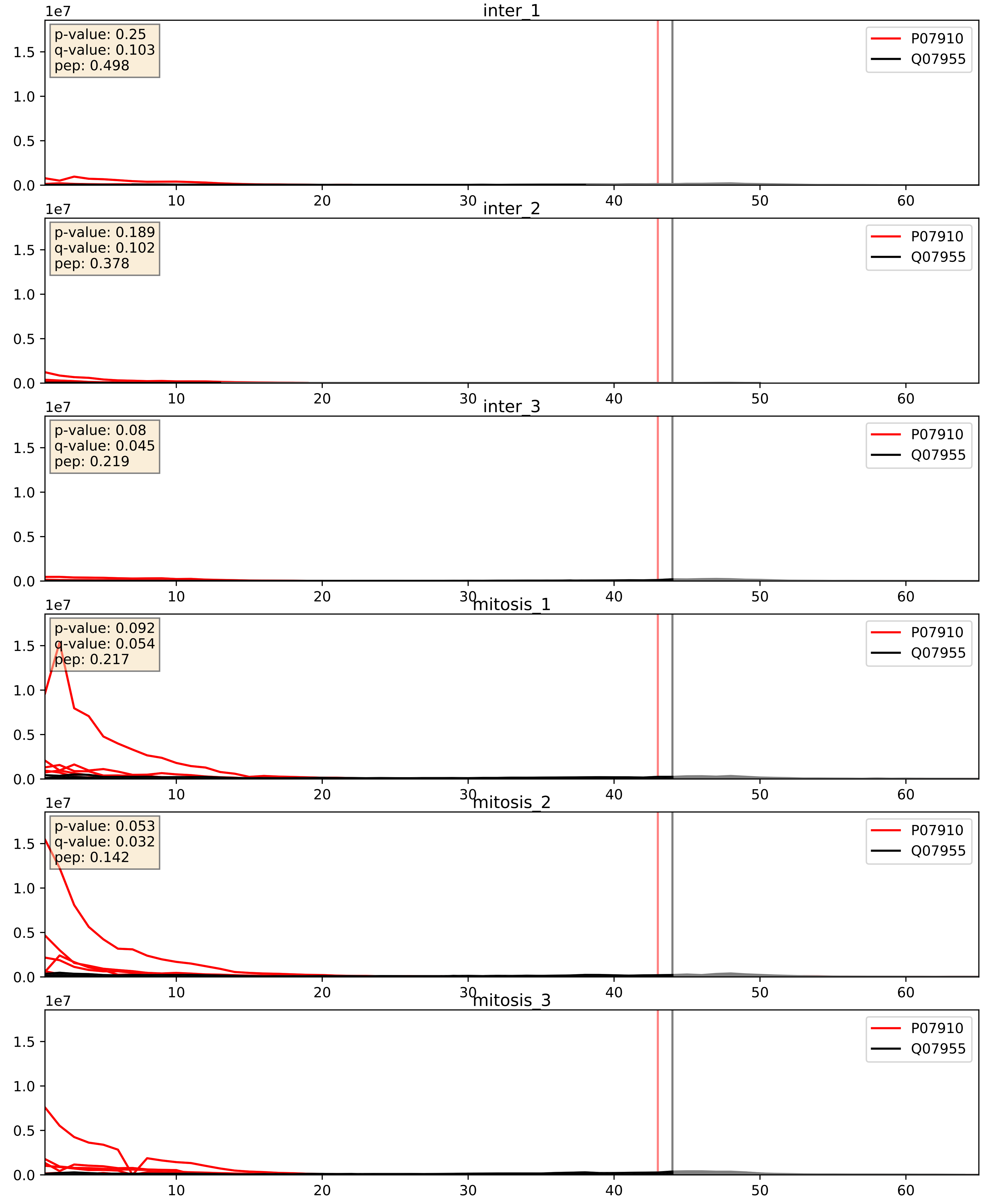


| level | condition_1 | condition_2 | pvalue | pvalue_adjusted |
|---|---|---|---|---|
| interactor_abundance | mitosis | inter | 4.923890450505053e-05 | 0.006283598333582311 |
| complex_abundance | mitosis | inter | 0.00034811228118943375 | 0.012015488415248196 |
| interactor_ratio | mitosis | inter | 0.01880601904879865 | 0.20643721891135677 |

P09651_Q07955
ROA1_HUMAN vs SRSF1_HUMAN
p-value: 0.0 q-value: 0.0

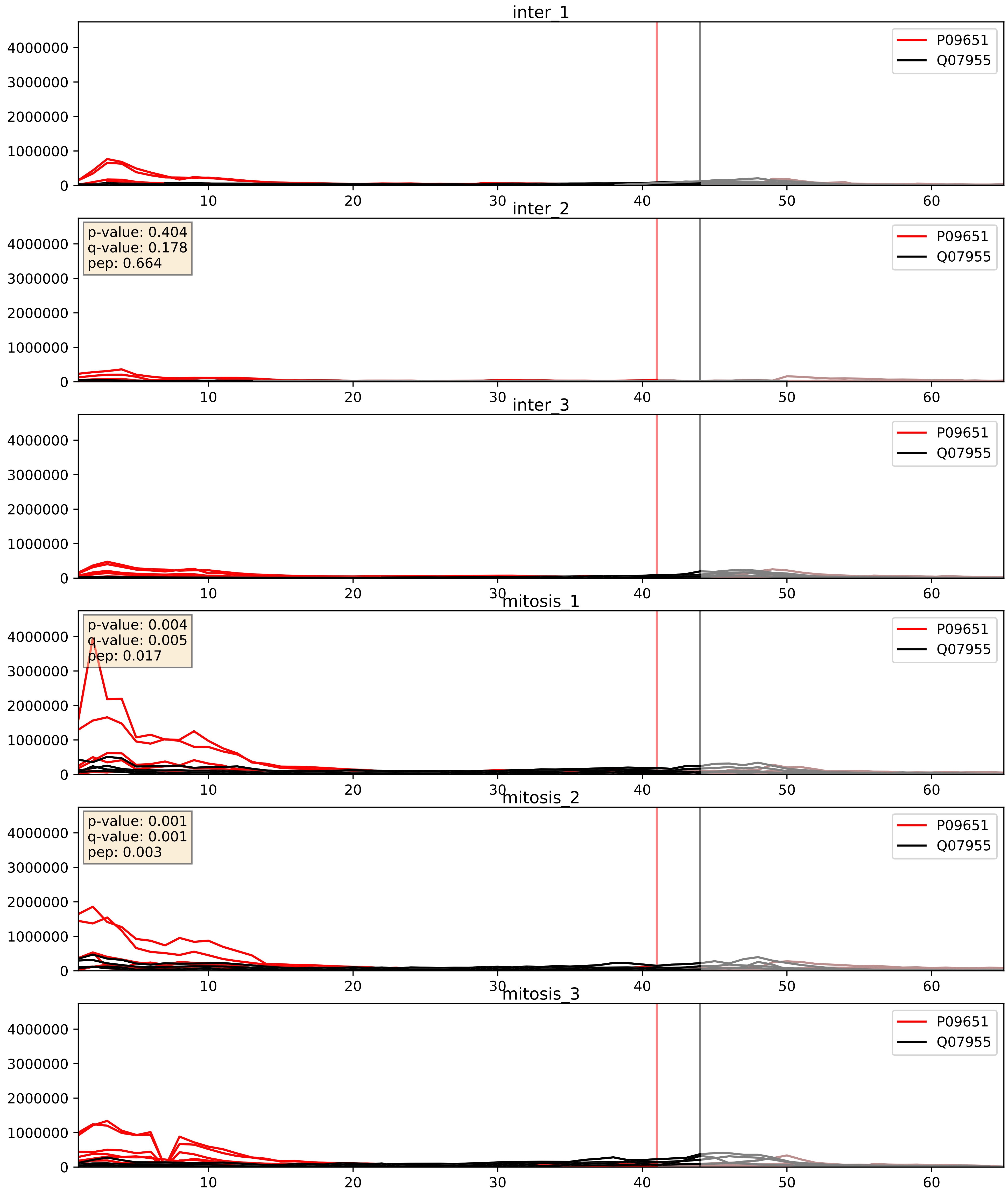


| level | condition_1 | condition_2 | pvalue | pvalue_adjusted |
|---|---|---|---|---|
| complex_abundance | mitosis | inter | 0.008509151419512374 | 0.0686506467021922 |
| interactor_abundance | mitosis | inter | 0.016490670357221612 | 0.11276964141946393 |
| interactor_ratio | mitosis | inter | 0.9147749642922947 | 0.9855669004419936 |

P14866_Q07955
HNRPL_HUMAN vs SRSF1_HUMAN
p-value: 0.005 q-value: 0.005

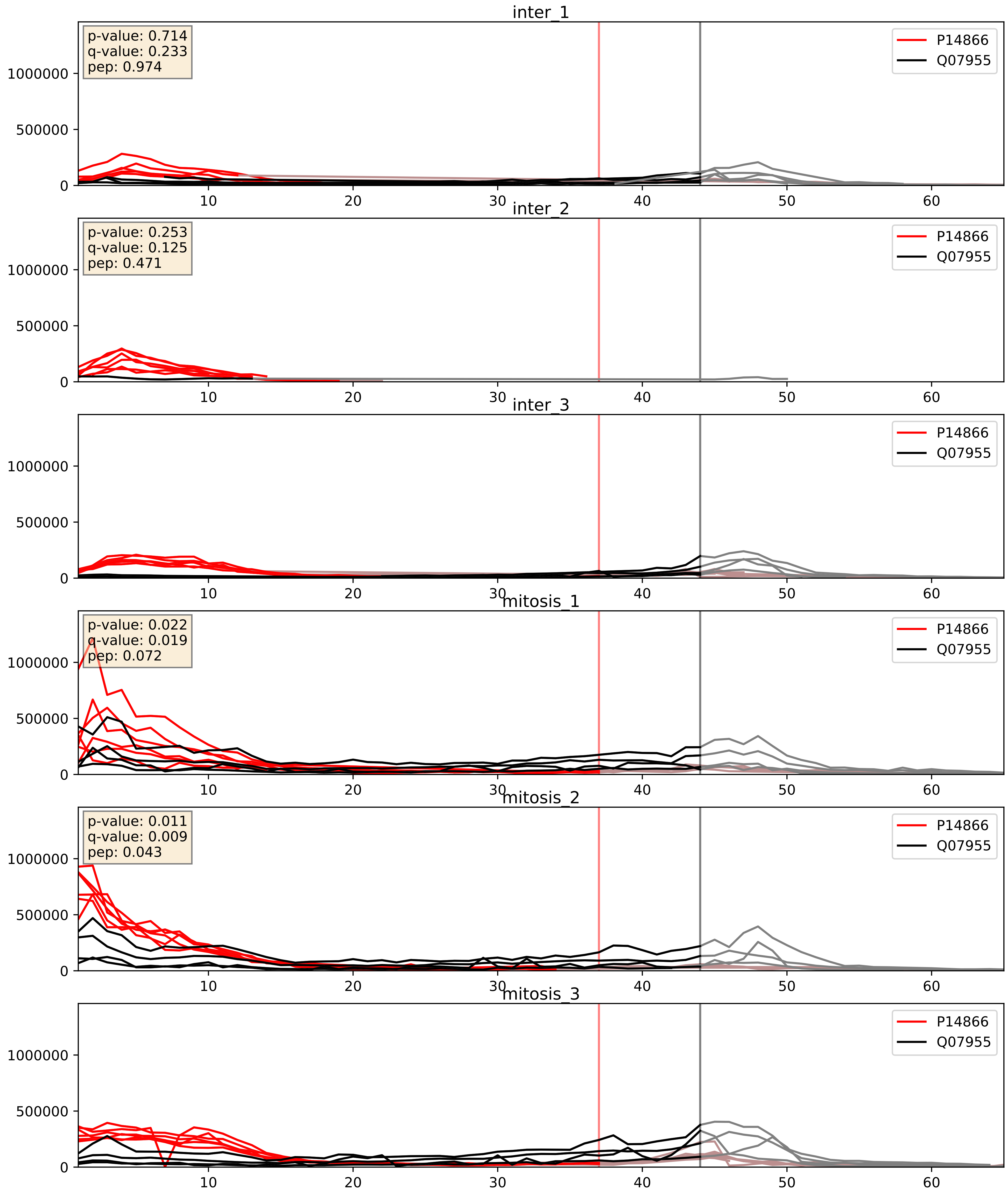


| level | condition_1 | condition_2 | pvalue | pvalue_adjusted |
|---|---|---|---|---|
| complex_abundance | mitosis | inter | 4.997121611729198e-05 | 0.005691857151652958 |
| interactor_abundance | mitosis | inter | 0.0019001893143151473 | 0.03196215383917848 |
| interactor_ratio | mitosis | inter | 0.15454422094563686 | 0.61160357433872 |

P17844_Q07955
DDX5_HUMAN vs SRSF1_HUMAN
p-value: 0.001 q-value: 0.015

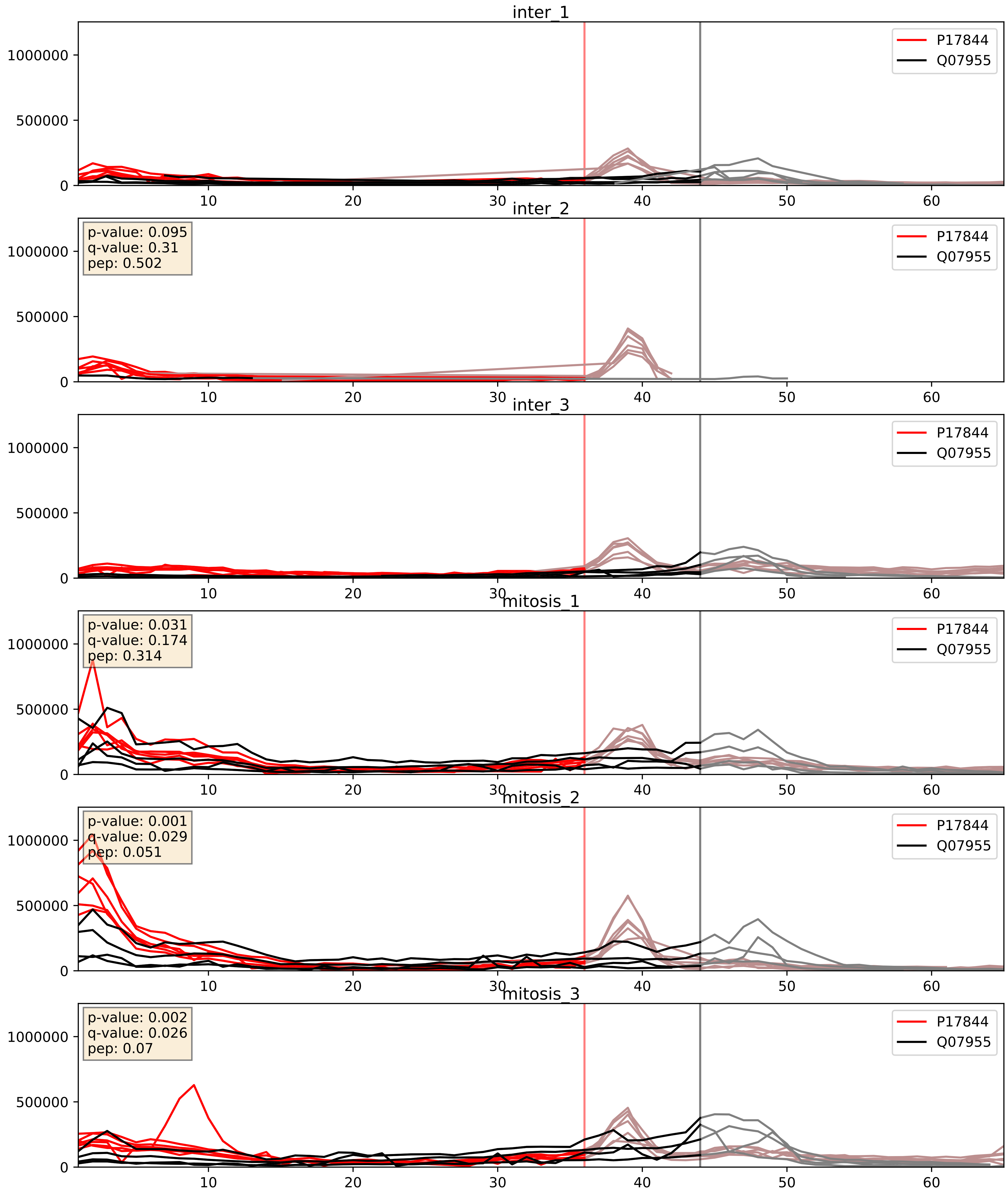


| level | condition_1 | condition_2 | pvalue | pvalue_adjusted |
|---|---|---|---|---|
| interactor_abundance | mitosis | inter | 0.0007813490254633644 | 0.02050037113506771 |
| complex_abundance | mitosis | inter | 0.0024237962141353607 | 0.035648961499997744 |
| interactor_ratio | mitosis | inter | 0.14930404490866453 | 0.6039771666075939 |

P22626_Q07955
ROA2_HUMAN vs SRSF1_HUMAN
p-value: 0.028 q-value: 0.021

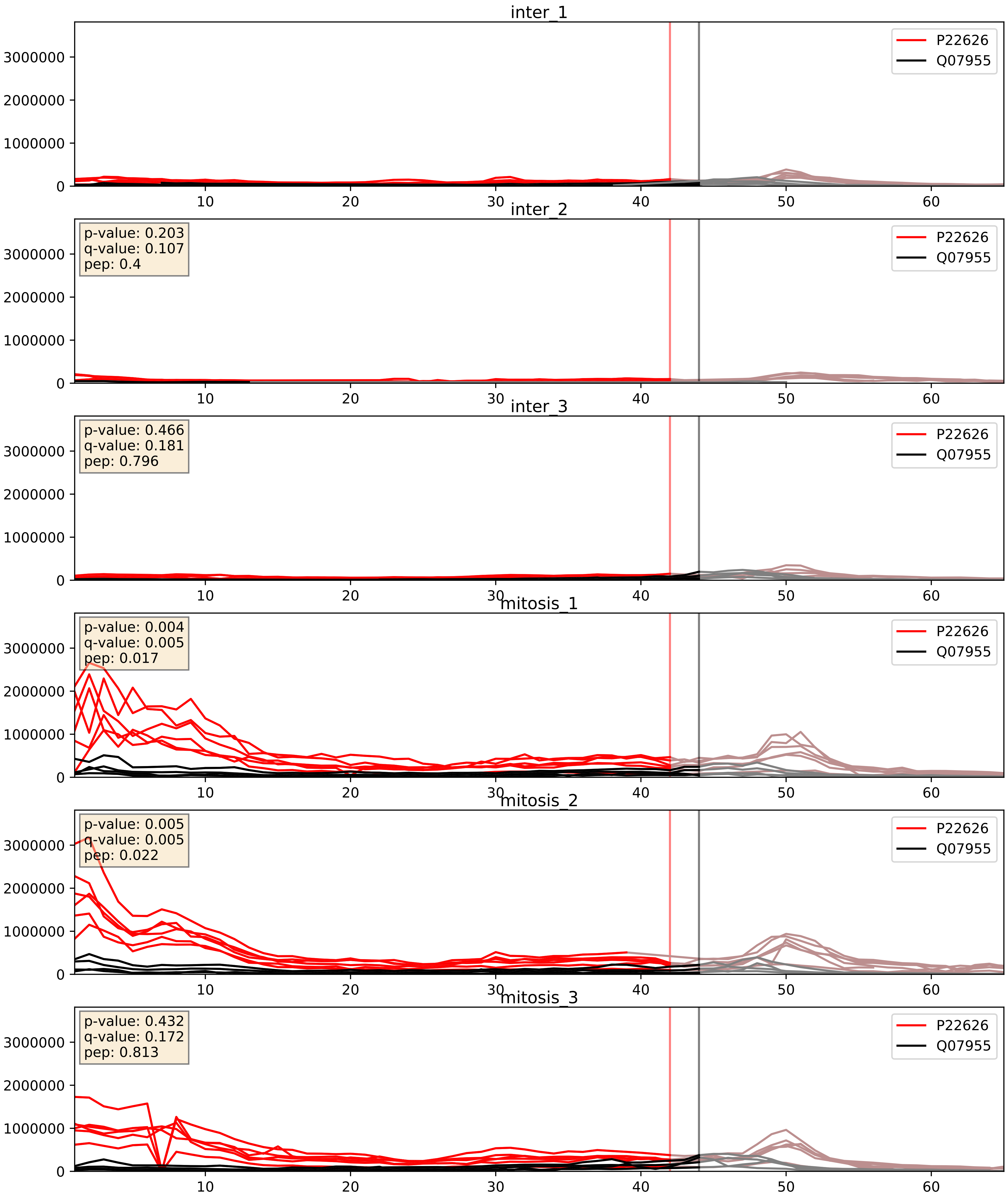


| level | condition_1 | condition_2 | pvalue | pvalue_adjusted |
|---|---|---|---|---|
| interactor_abundance | mitosis | inter | 0.0036653640621034087 | 0.0470045189087718 |
| complex_abundance | mitosis | inter | 0.004020555198830934 | 0.04695218622372823 |
| interactor_ratio | mitosis | inter | 0.03180859881707865 | 0.28451578461253213 |

P26599_Q07955
PTBP1_HUMAN vs SRSF1_HUMAN
p-value: 0.061 q-value: 0.038

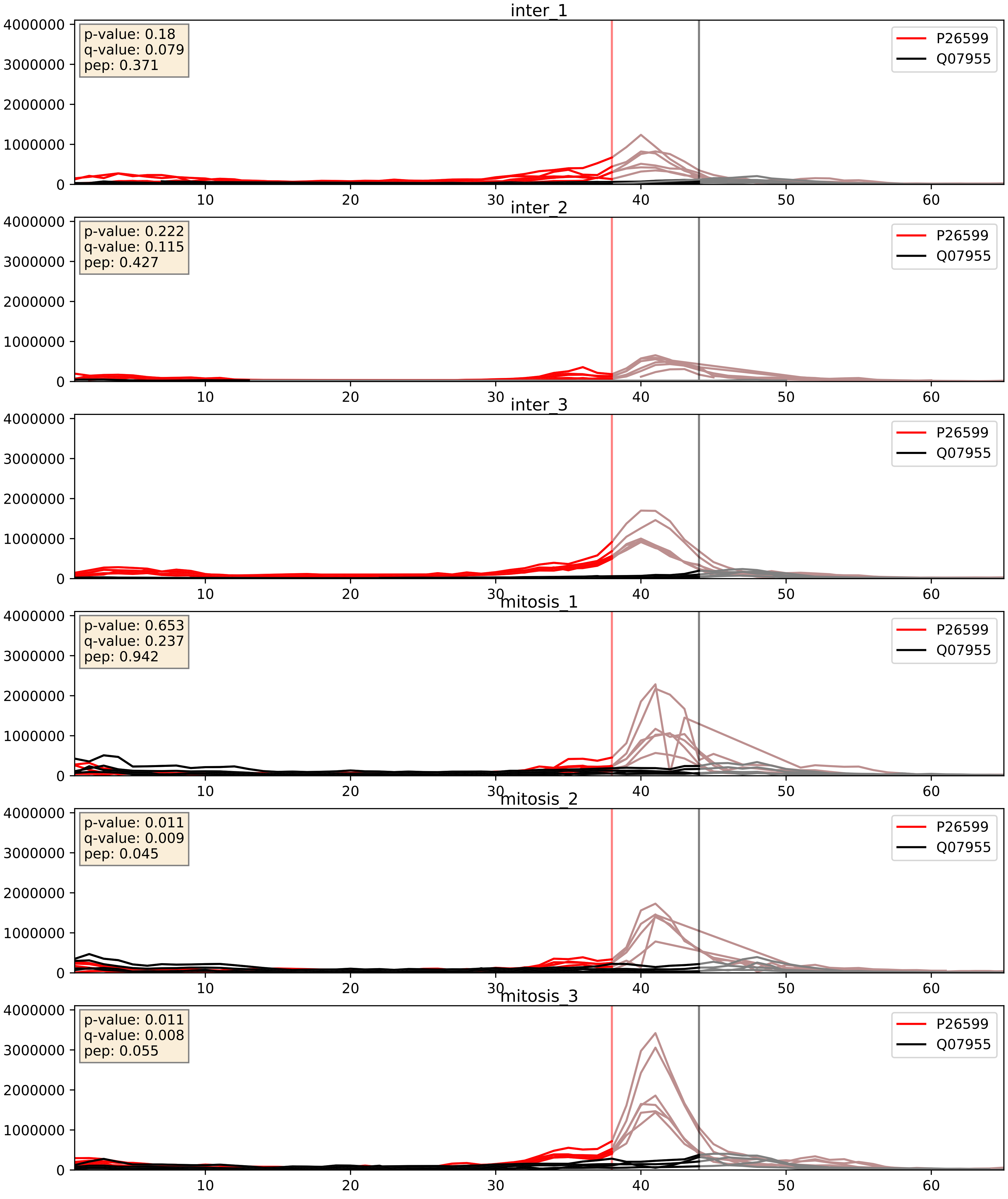


| level | condition_1 | condition_2 | pvalue | pvalue_adjusted |
|---|---|---|---|---|
| interactor_ratio | mitosis | inter | 0.0406622192886054 | 0.3333990393778374 |
| complex_abundance | mitosis | inter | 0.2472052067521012 | 0.3977727299820493 |
| interactor_abundance | mitosis | inter | 0.6816772227862057 | 0.7933102642712379 |

P31943_Q07955
HNRH1_HUMAN vs SRSF1_HUMAN
p-value: 0.001 q-value: 0.002

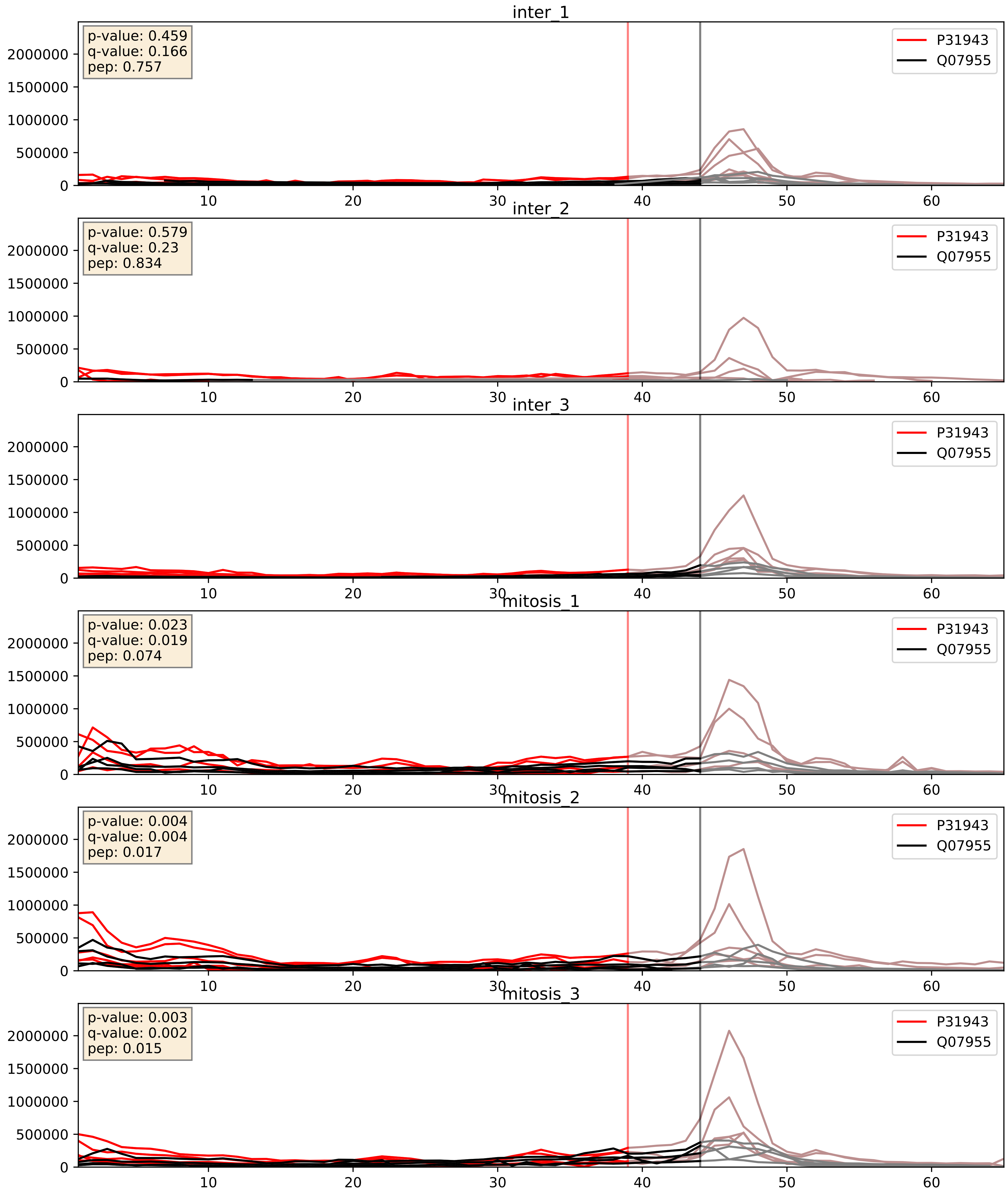


| level | condition_1 | condition_2 | pvalue | pvalue_adjusted |
|---|---|---|---|---|
| complex_abundance | mitosis | inter | 0.007690789144795531 | 0.06519315066658214 |
| interactor_abundance | mitosis | inter | 0.013391224463716843 | 0.10304923015527873 |
| interactor_ratio | mitosis | inter | 0.43412575717983676 | 0.8435878041032946 |

P38159_Q07955
RBMX_HUMAN vs SRSF1_HUMAN
p-value: 0.074 q-value: 0.045

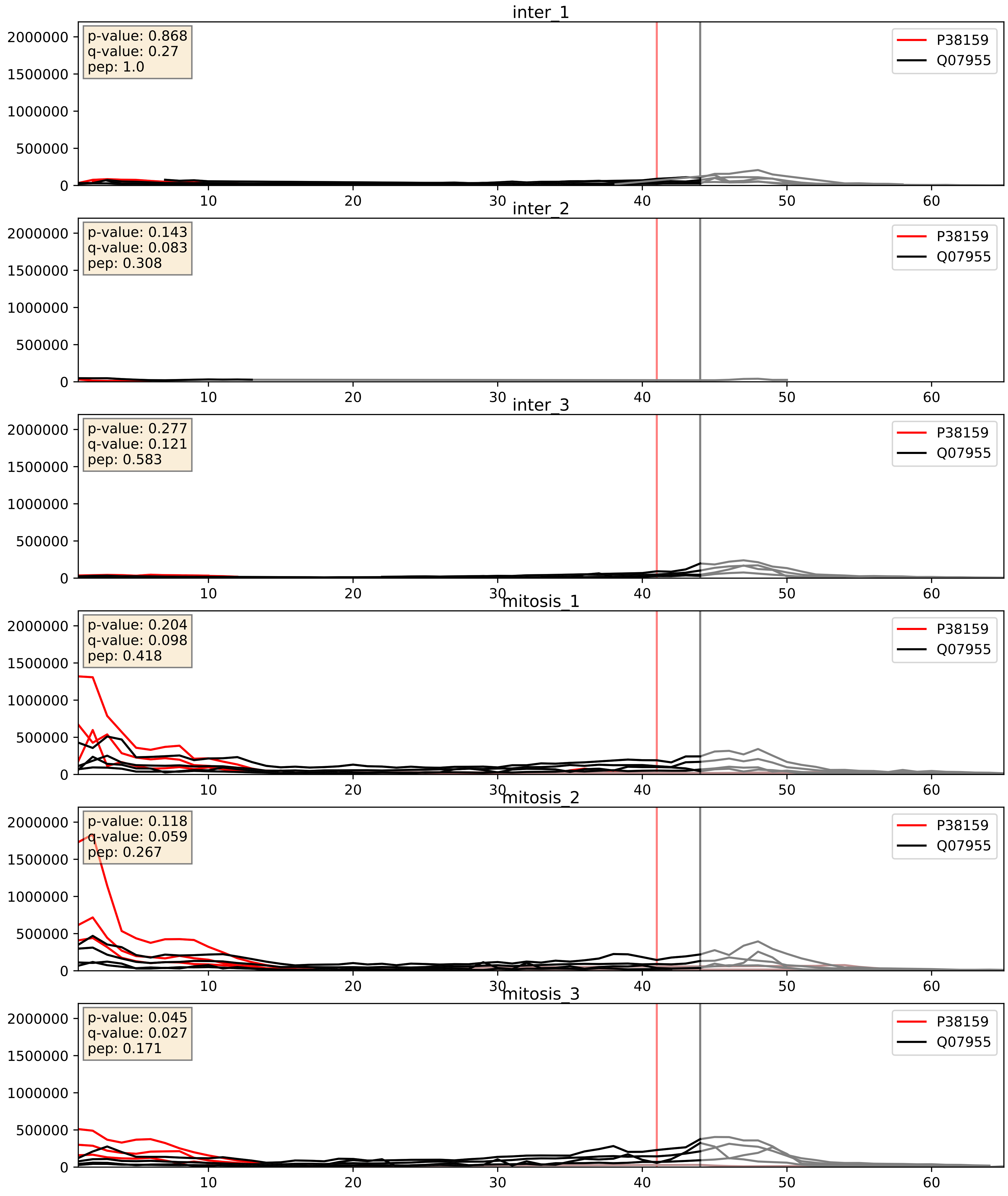


| level | condition_1 | condition_2 | pvalue | pvalue_adjusted |
|---|---|---|---|---|
| interactor_abundance | mitosis | inter | 0.0010393306972951088 | 0.024049720606162197 |
| complex_abundance | mitosis | inter | 0.0014295676213636915 | 0.02592605686201949 |
| interactor_ratio | mitosis | inter | 0.05120606381407092 | 0.37414961988671497 |

P51991_Q07955
ROA3_HUMAN vs SRSF1_HUMAN
p-value: 0.0 q-value: 0.0

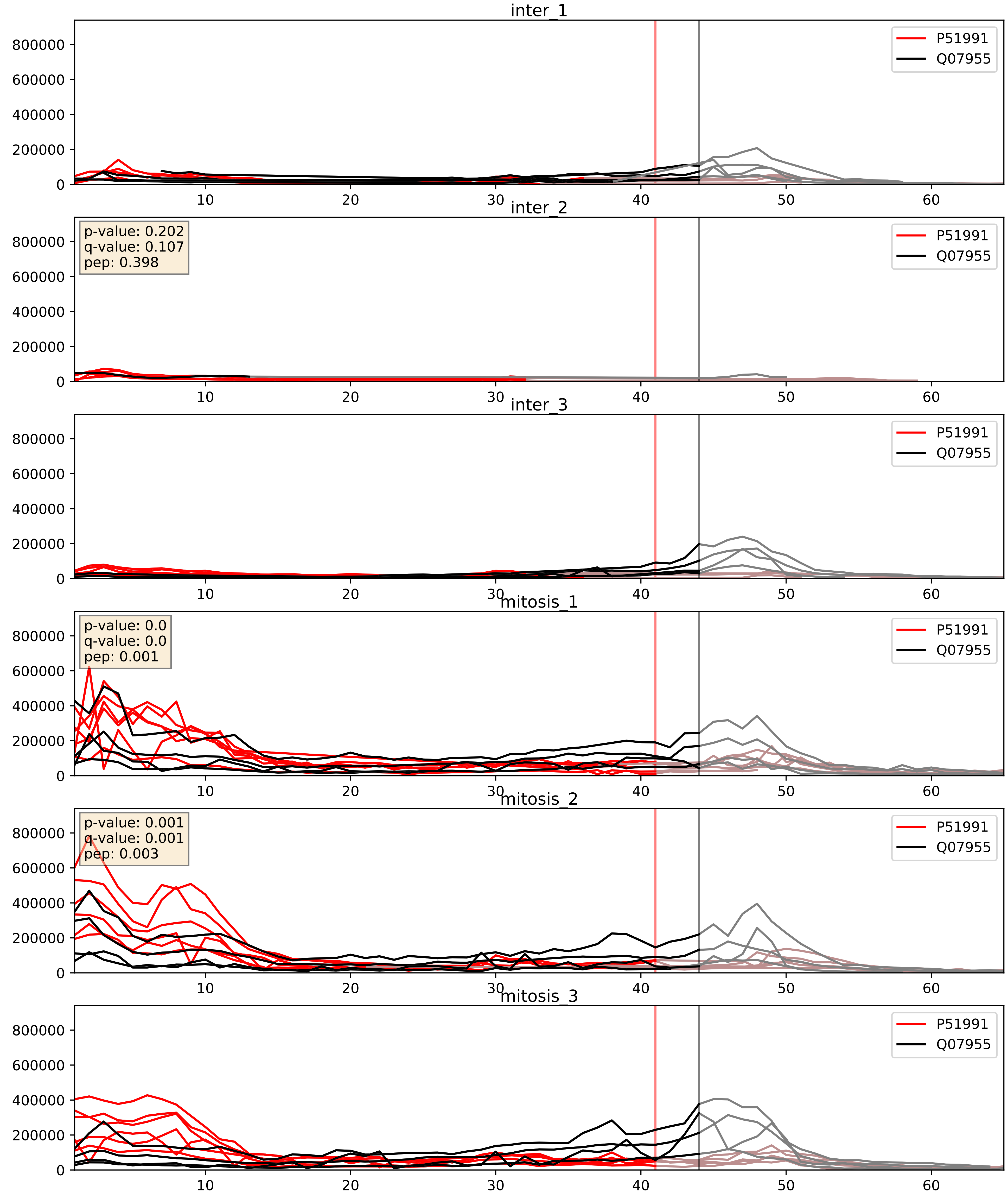


| level | condition_1 | condition_2 | pvalue | pvalue_adjusted |
|---|---|---|---|---|
| interactor_abundance | mitosis | inter | 4.395510848628261e-06 | 0.0033734327106189023 |
| complex_abundance | mitosis | inter | 0.0004743031873480036 | 0.013816623868928207 |
| interactor_ratio | mitosis | inter | 0.39080113396936994 | 0.825249705827432 |

P52272_Q07955
HNRPM_HUMAN vs SRSF1_HUMAN
p-value: 0.003 q-value: 0.004

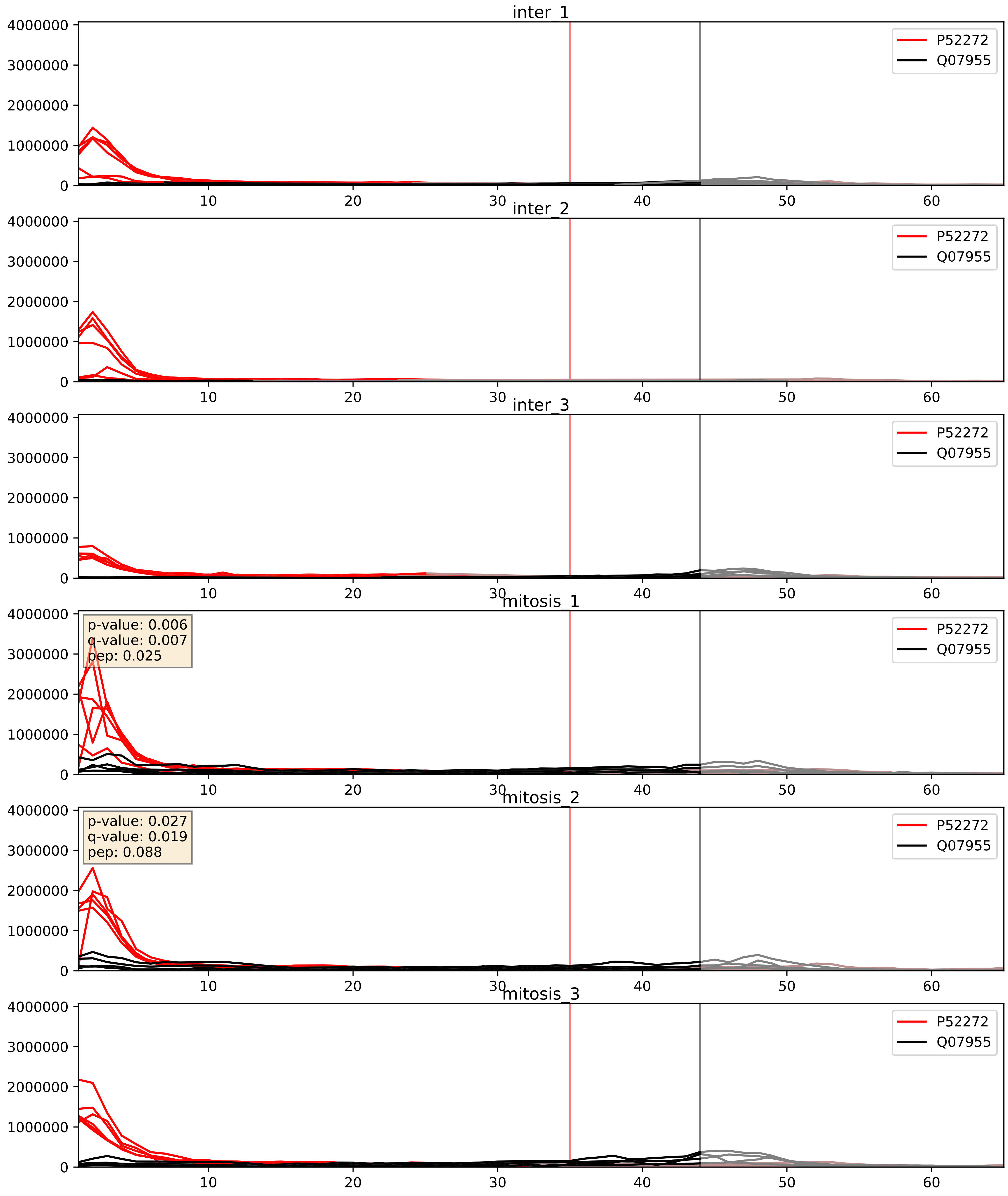


| level | condition_1 | condition_2 | pvalue | pvalue_adjusted |
|---|---|---|---|---|
| complex_abundance | mitosis | inter | 0.015294076257498249 | 0.09174685245214603 |
| interactor_ratio | mitosis | inter | 0.026601540420708367 | 0.25614196664208316 |
| interactor_abundance | mitosis | inter | 0.18545035871799714 | 0.37784844239126014 |

P52597_Q07955
HNRPF_HUMAN vs SRSF1_HUMAN
p-value: 0.0 q-value: 0.0

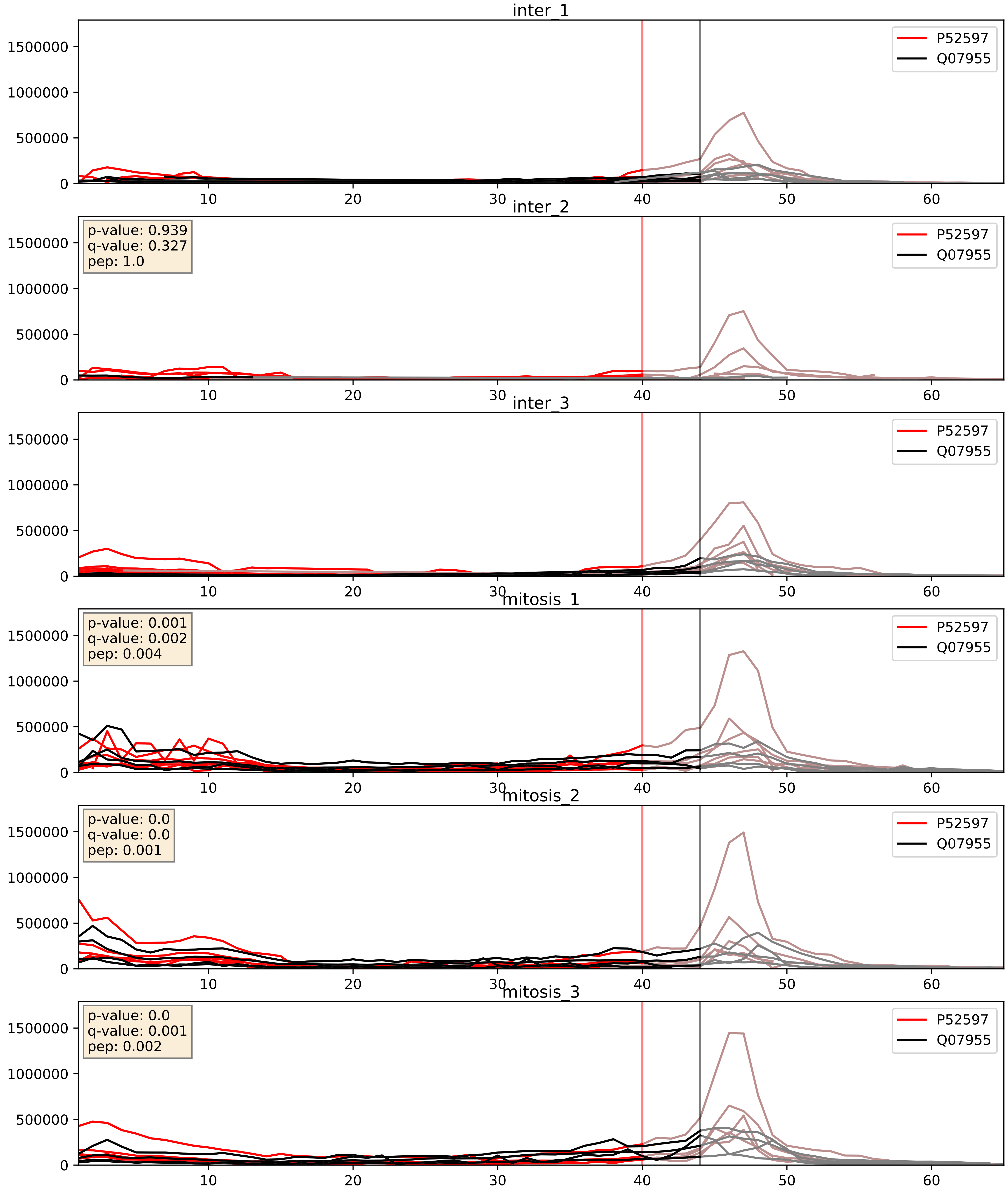


| level | condition_1 | condition_2 | pvalue | pvalue_adjusted |
|---|---|---|---|---|
| interactor_abundance | mitosis | inter | 0.0030437040272975205 | 0.042192884977598016 |
| complex_abundance | mitosis | inter | 0.003478199315974476 | 0.043087389500349516 |
| interactor_ratio | mitosis | inter | 0.17421734291907873 | 0.6394941918470471 |

P61978_Q07955
HNRPK_HUMAN vs SRSF1_HUMAN
p-value: 0.017 q-value: 0.014

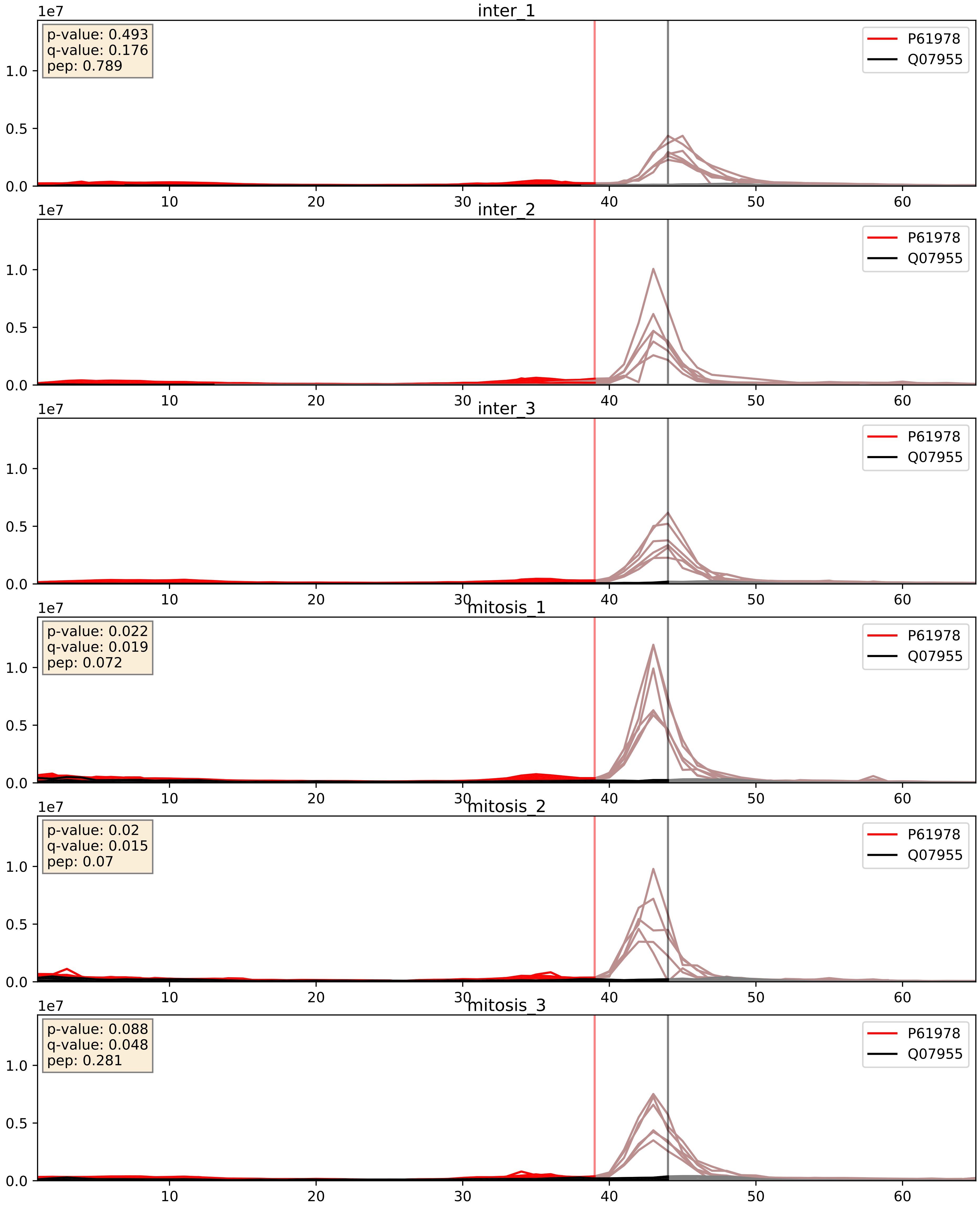


| level | condition_1 | condition_2 | pvalue | pvalue_adjusted |
|---|---|---|---|---|
| complex_abundance | mitosis | inter | 0.11800608492463315 | 0.259340715521145 |
| interactor_ratio | mitosis | inter | 0.20258983323506588 | 0.678106014366403 |
| interactor_abundance | mitosis | inter | 0.6678421614254995 | 0.7821493116196215 |

P62995_Q07955
TRA2B_HUMAN vs SRSF1_HUMAN
p-value: 0.019 q-value: 0.015

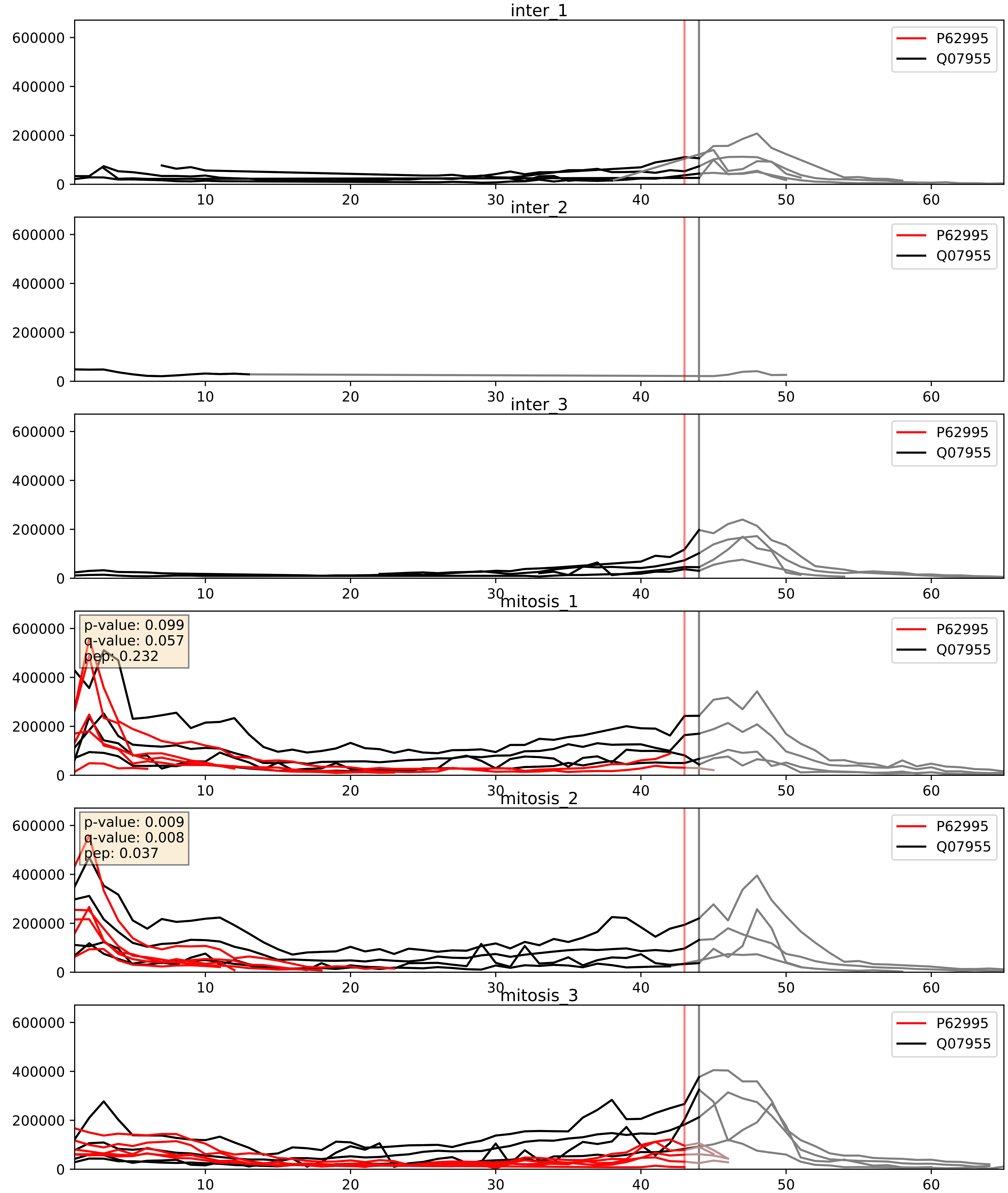


| level | condition_1 | condition_2 | pvalue | pvalue_adjusted |
|---|---|---|---|---|
| complex_abundance | mitosis | inter | 0.0002950478593069016 | 0.011617922212881656 |
| interactor_abundance | mitosis | inter | 0.0004584932841066445 | 0.014951247664582388 |
| interactor_ratio | mitosis | inter | 0.4511140392976068 | 0.8525838327540917 |

P67809_Q07955
YBOX1_HUMAN vs SRSF1_HUMAN
p-value: 0.0 q-value: 0.013

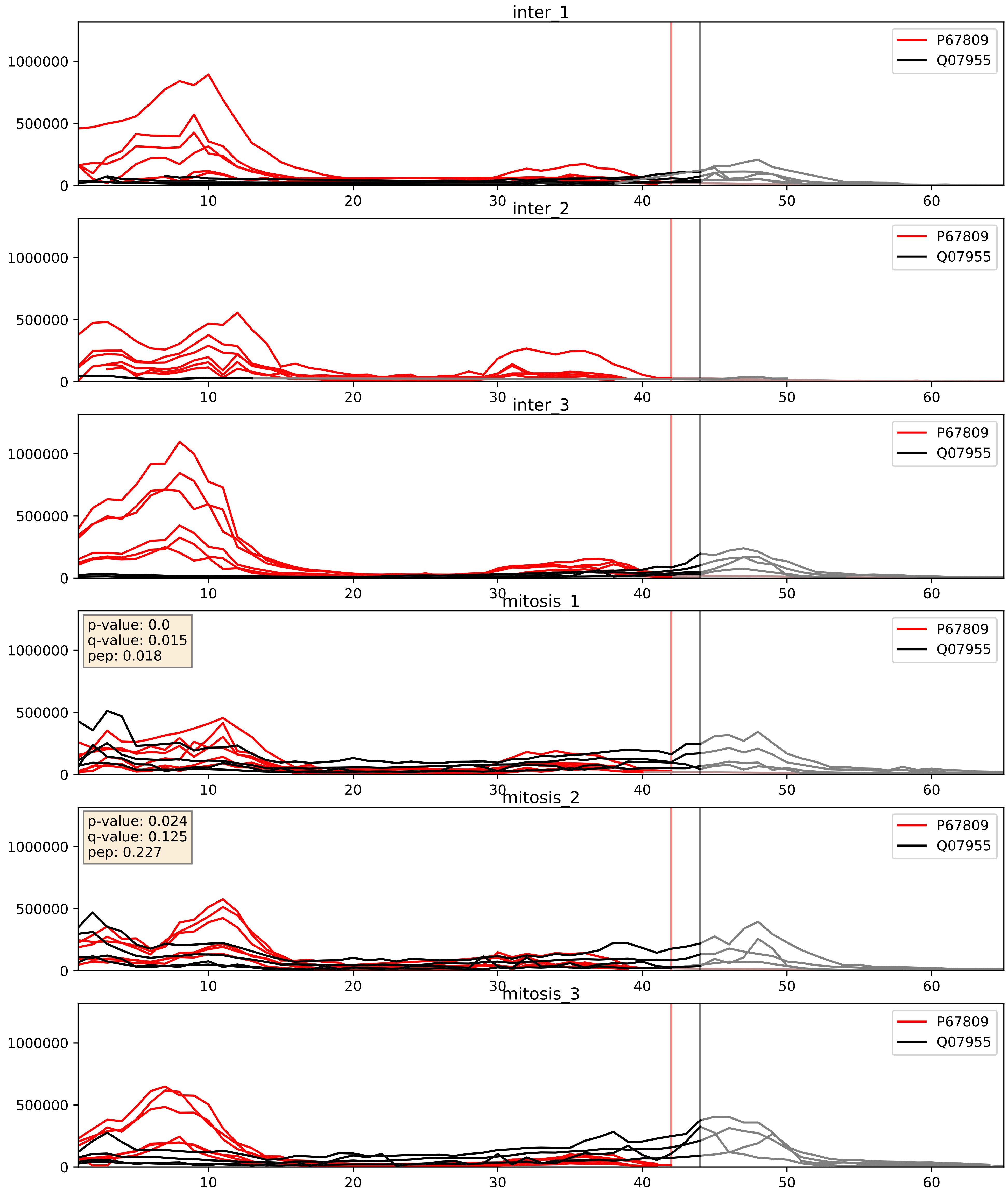


| level | condition_1 | condition_2 | pvalue | pvalue_adjusted |
|---|---|---|---|---|
| interactor_ratio | mitosis | inter | 0.0033517438699698825 | 0.07550244086037419 |
| complex_abundance | mitosis | inter | 0.06885530639151455 | 0.1900061443022207 |
| interactor_abundance | mitosis | inter | 0.44114345823369483 | 0.6185402133465074 |

P84103_Q07955
SRSF3_HUMAN vs SRSF1_HUMAN
p-value: 0.054 q-value: 0.035

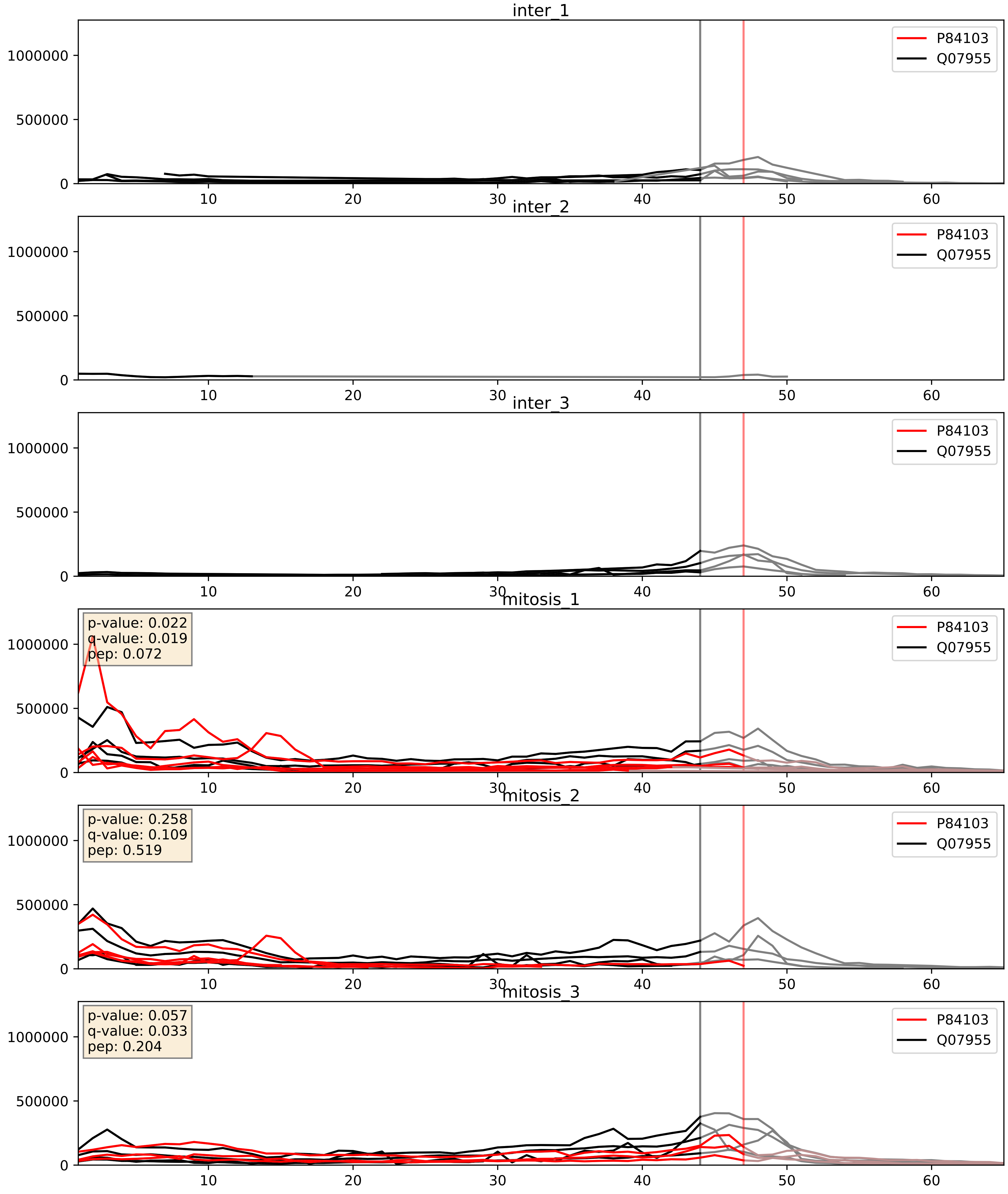


| level | condition_1 | condition_2 | pvalue | pvalue_adjusted |
|---|---|---|---|---|
| complex_abundance | mitosis | inter | 3.7310233198972706e-05 | 0.005691857151652958 |
| interactor_abundance | mitosis | inter | 0.002590890474122459 | 0.03806046117740333 |
| interactor_ratio | mitosis | inter | 0.4549089543182232 | 0.8545518508495171 |

Q07955_Q08211
SRSF1_HUMAN vs DHX9_HUMAN
p-value: 0.076 q-value: 0.046

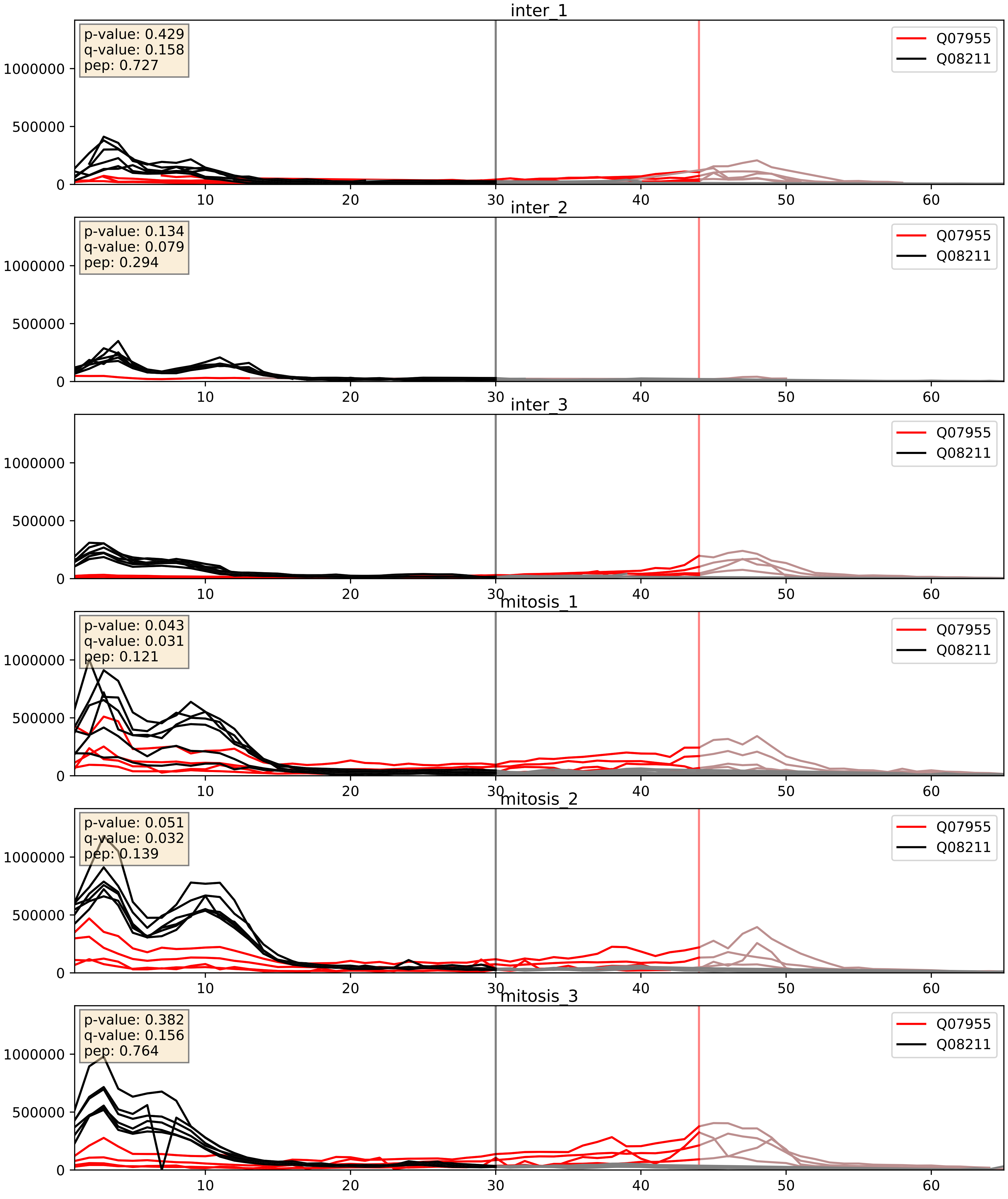


| level | condition_1 | condition_2 | pvalue | pvalue_adjusted |
|---|---|---|---|---|
| complex_abundance | mitosis | inter | 0.0014016180824241073 | 0.025581771397761962 |
| interactor_abundance | mitosis | inter | 0.0024364603316211677 | 0.03681201502126771 |
| interactor_ratio | mitosis | inter | 0.05437572514359929 | 0.38595042058806794 |

Q07955_Q13151
SRSF1_HUMAN vs ROA0_HUMAN
p-value: 0.007 q-value: 0.006

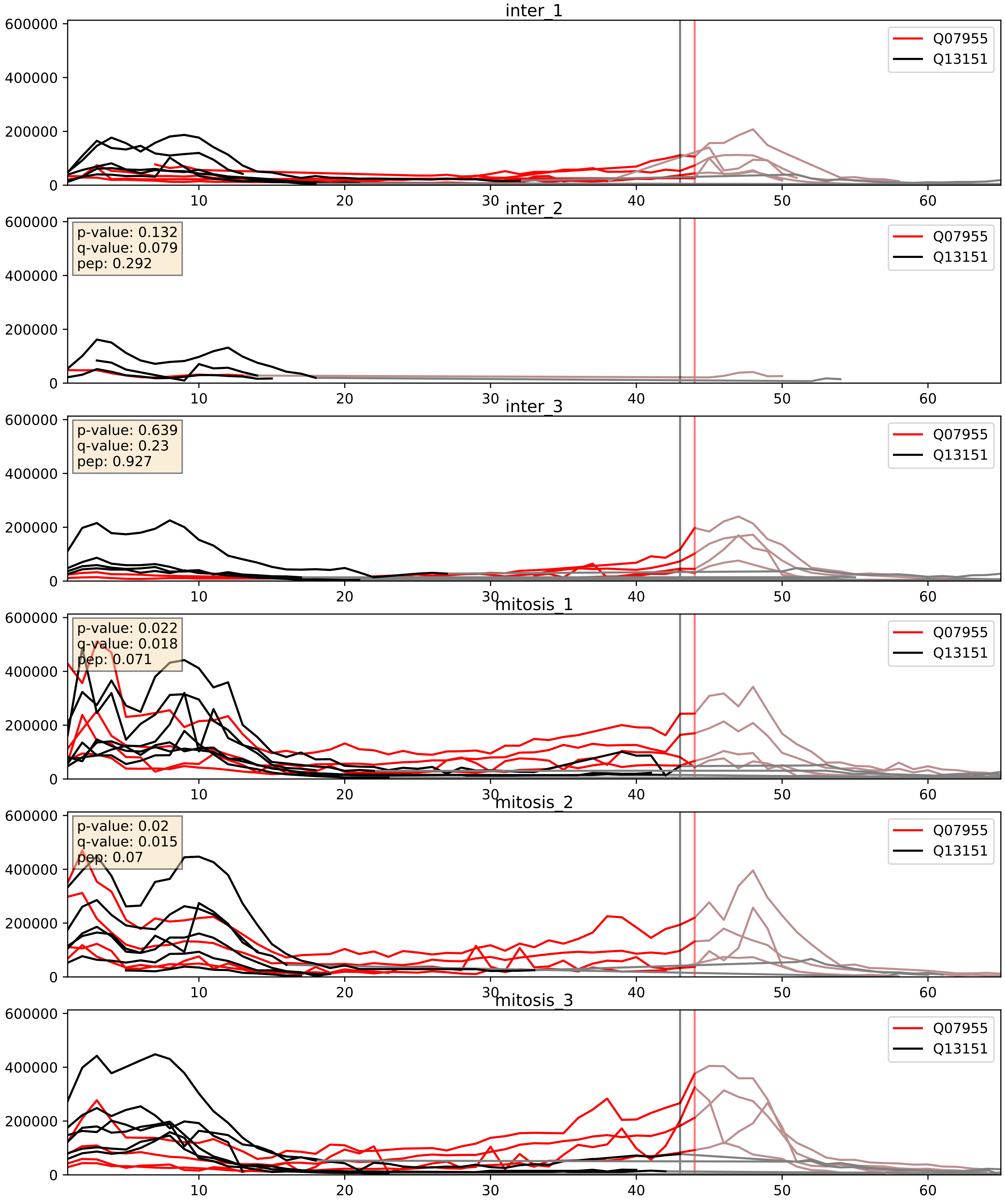


| level | condition_1 | condition_2 | pvalue | pvalue_adjusted |
|---|---|---|---|---|
| complex_abundance | mitosis | inter | 0.007513005302259508 | 0.06444768685385507 |
| interactor_abundance | mitosis | inter | 0.007752548845619896 | 0.07645370750979989 |
| interactor_ratio | mitosis | inter | 0.08841558352285728 | 0.4953575337685953 |

Q07955_Q13247
SRSF1_HUMAN vs SRSF6_HUMAN
p-value: 0.0 q-value: 0.0

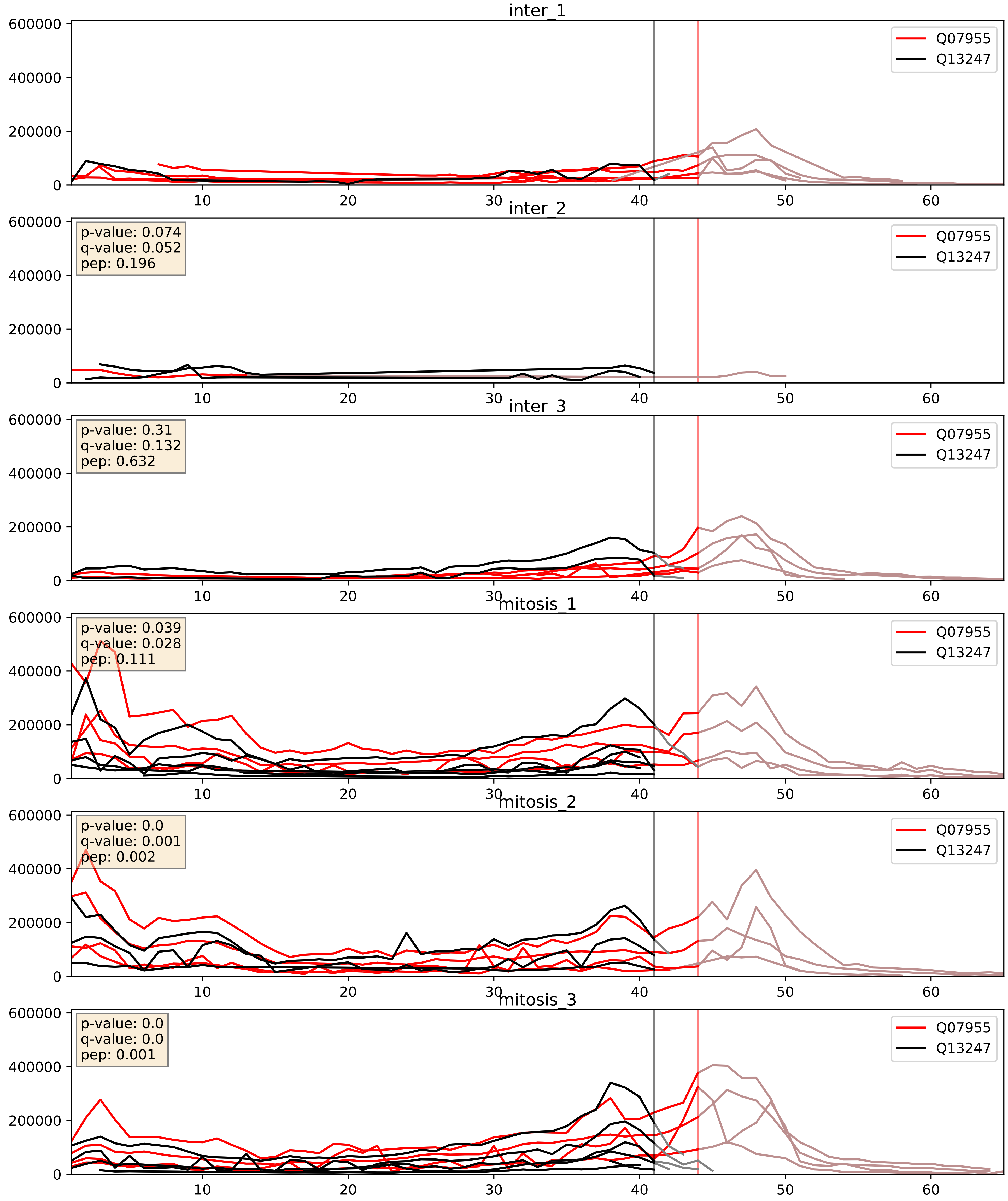


| level | condition_1 | condition_2 | pvalue | pvalue_adjusted |
|---|---|---|---|---|
| complex_abundance | mitosis | inter | 0.0021933030847482732 | 0.034197949736694384 |
| interactor_abundance | mitosis | inter | 0.009132210010912856 | 0.08324021891678159 |
| interactor_ratio | mitosis | inter | 0.6314956834780092 | 0.9183953086181231 |

Q07955_Q15365
SRSF1_HUMAN vs PCBP1_HUMAN
p-value: 0.0 q-value: 0.009

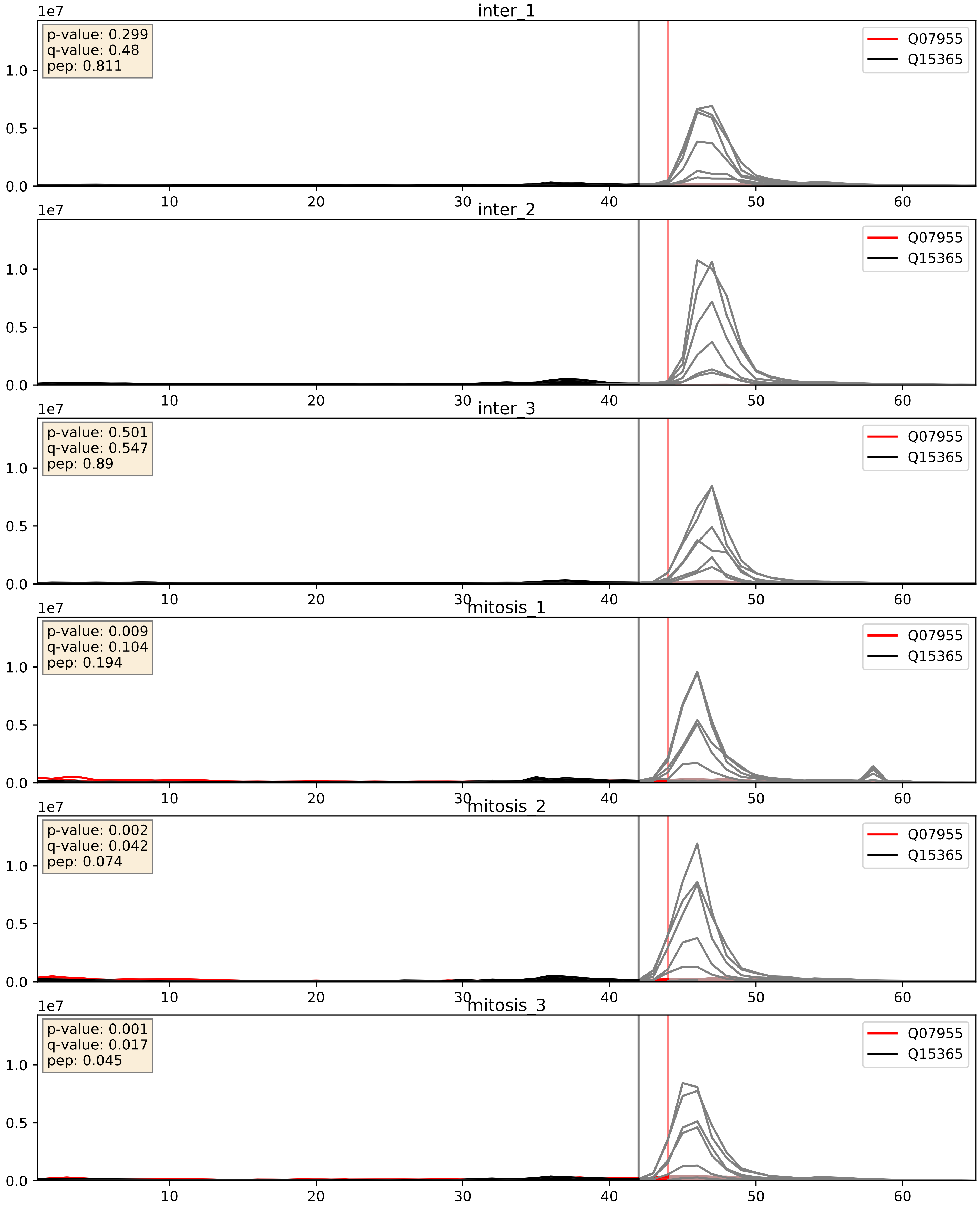


| level | condition_1 | condition_2 | pvalue | pvalue_adjusted |
|---|---|---|---|---|
| interactor_ratio | mitosis | inter | 0.014640895252484376 | 0.17726458749825497 |
| interactor_abundance | mitosis | inter | 0.015951526618393257 | 0.11088021938056777 |
| complex_abundance | mitosis | inter | 0.042812804870910104 | 0.1481949160937304 |

Q07955_Q15366
SRSF1_HUMAN vs PCBP2_HUMAN
p-value: 0.004 q-value: 0.047

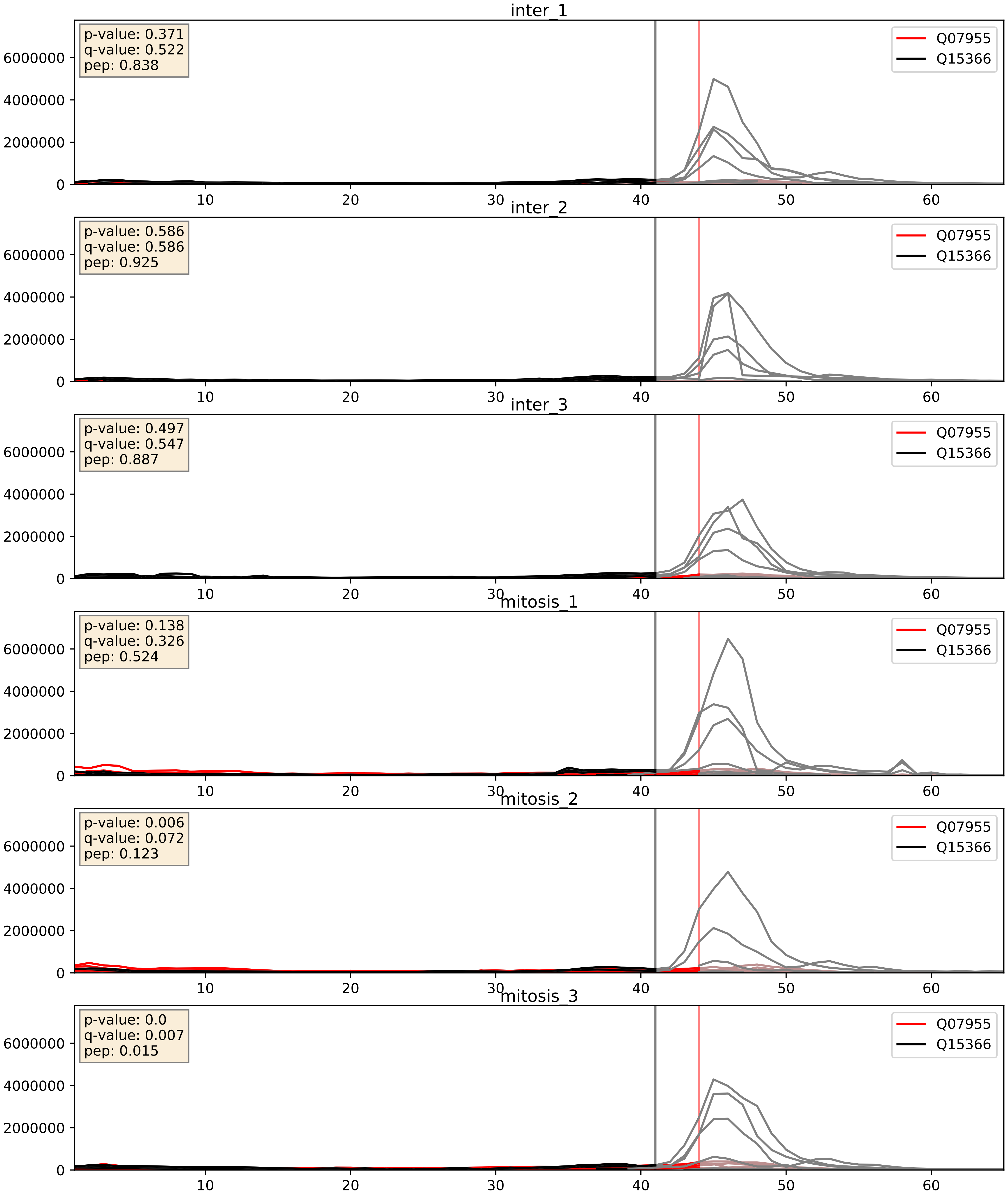


| level | condition_1 | condition_2 | pvalue | pvalue_adjusted |
|---|---|---|---|---|
| interactor_abundance | mitosis | inter | 0.013960395614656259 | 0.10473355518094442 |
| interactor_ratio | mitosis | inter | 0.06345390122101714 | 0.4184633239228866 |
| complex_abundance | mitosis | inter | 0.1106354971404223 | 0.24912488454653373 |

Q07955_Q15717
SRSF1_HUMAN vs ELAV1_HUMAN
p-value: 0.011 q-value: 0.01

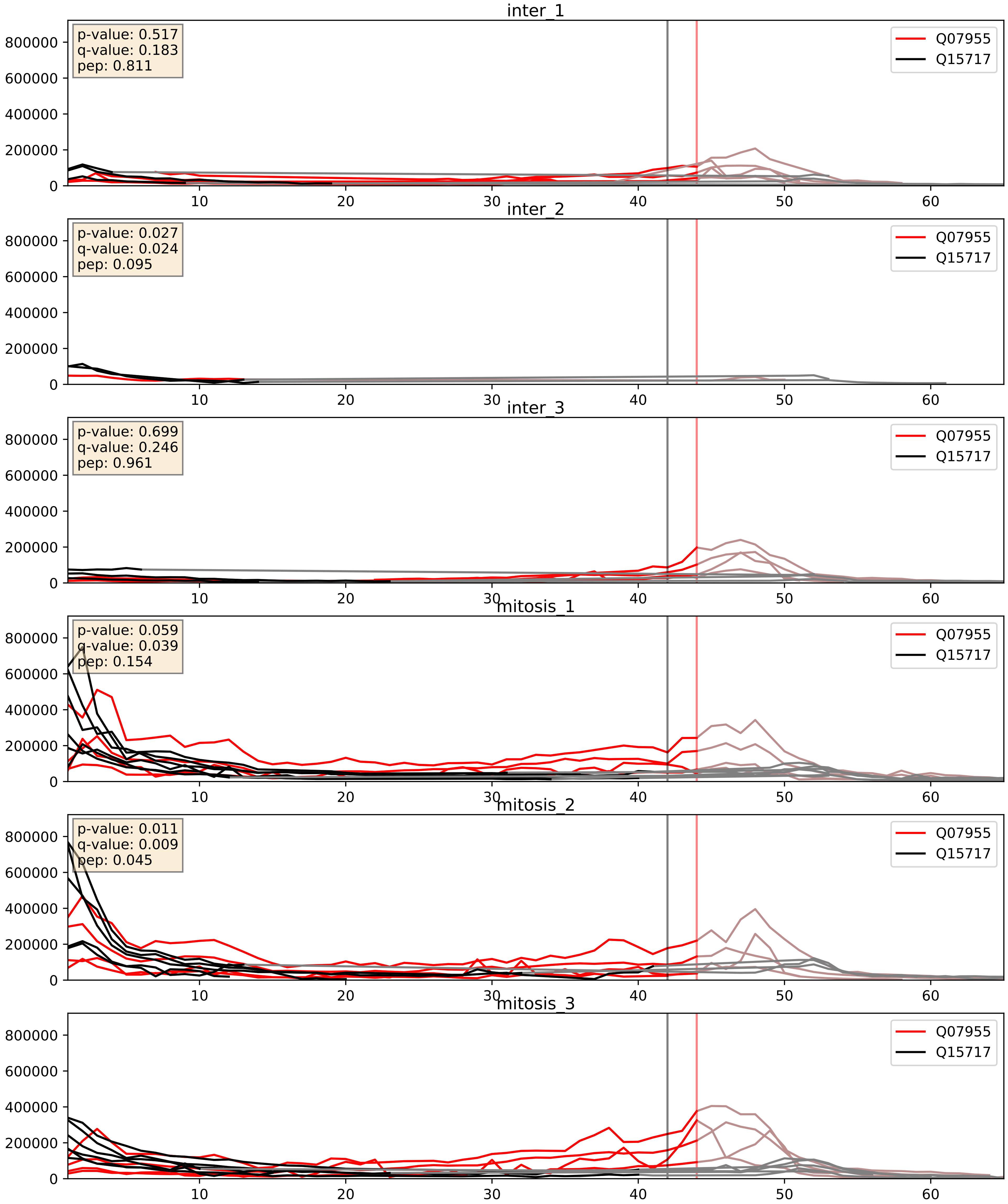


| level | condition_1 | condition_2 | pvalue | pvalue_adjusted |
|---|---|---|---|---|
| complex_abundance | mitosis | inter | 0.0001877702555432865 | 0.009111526782692203 |
| interactor_abundance | mitosis | inter | 0.0015832375582752990 | 0.029025269221018554 |
| interactor_ratio | mitosis | inter | 0.8402338042736714 | 0.9741841208970103 |

Q07955_Q16629
SRSF1_HUMAN vs SRSF7_HUMAN
p-value: 0.0 q-value: 0.0

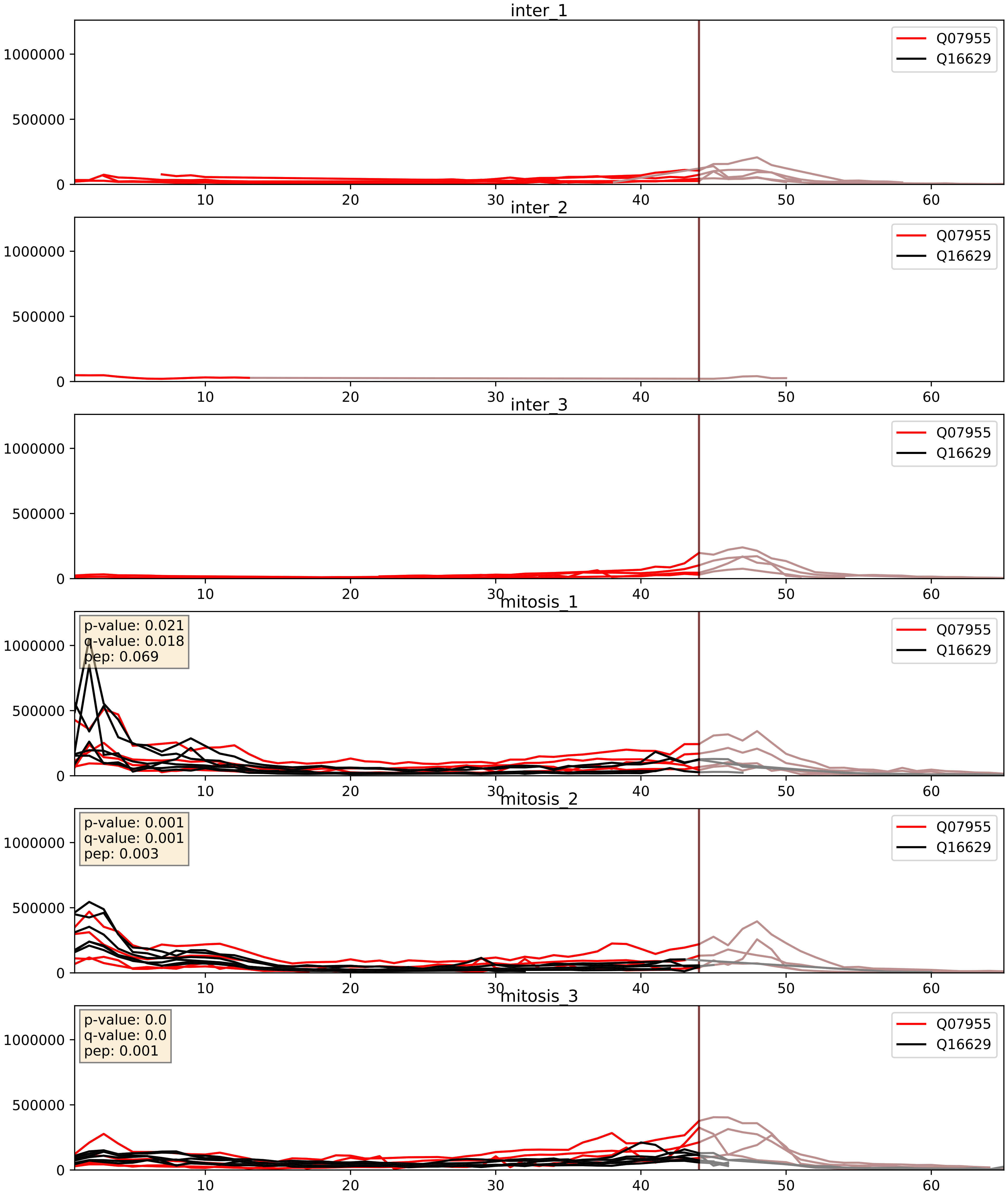


| level | condition_1 | condition_2 | pvalue | pvalue_adjusted |
|---|---|---|---|---|
| complex_abundance | mitosis | inter | 3.059501908721843e-05 | 0.005691857151652958 |
| interactor_abundance | mitosis | inter | 0.0006217246581139298 | 0.01750645747847118 |
| interactor_ratio | mitosis | inter | 0.4633387300269737 | 0.8588611490067614 |

Q07955_Q96SB4
SRSF1_HUMAN vs SRPK1_HUMAN
p-value: 0.074 q-value: 0.045

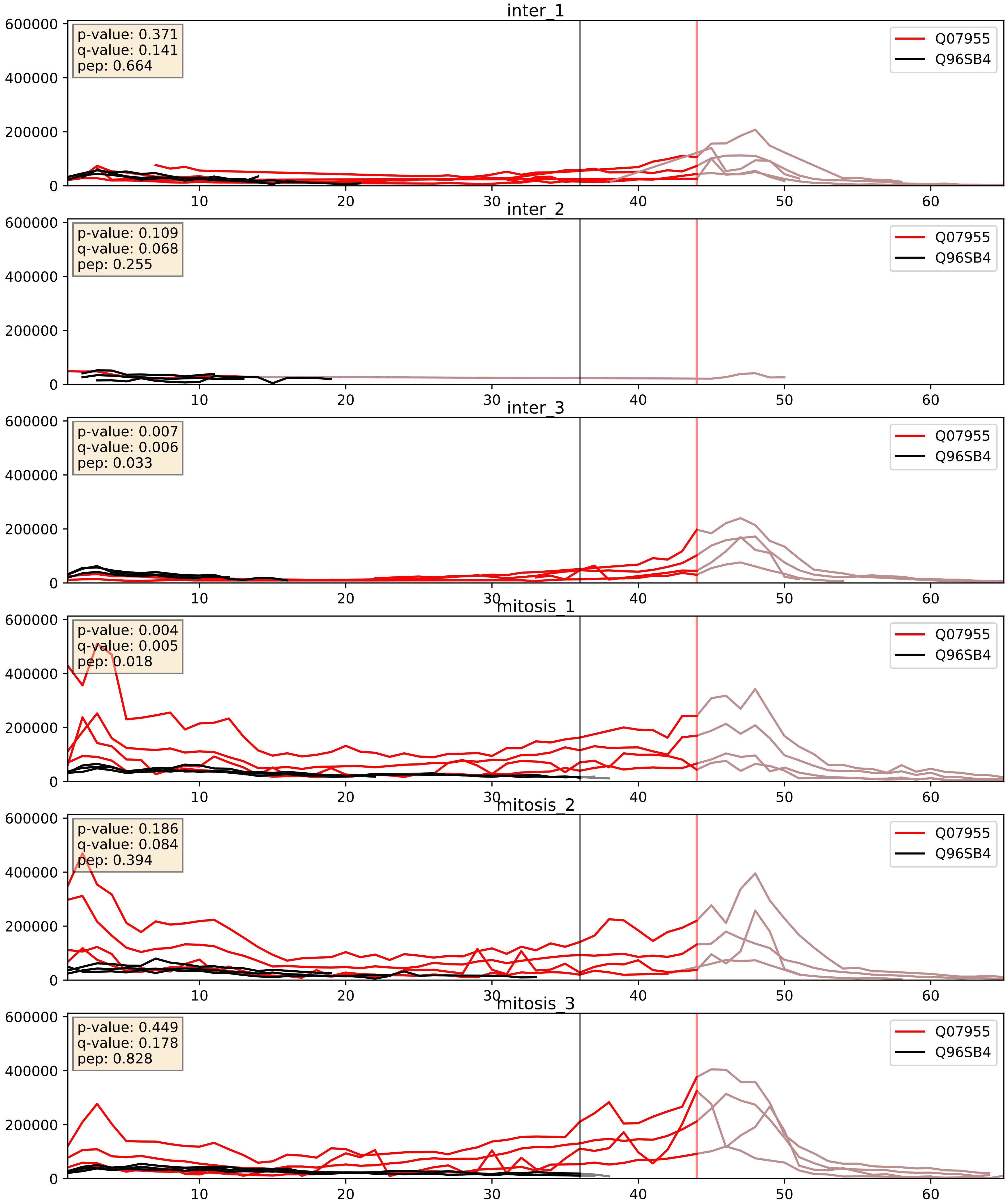


| level | condition_1 | condition_2 | pvalue | pvalue_adjusted |
|---|---|---|---|---|
| interactor_abundance | mitosis | inter | 0.0025420288426528524 | 0.03777170158161676 |
| complex_abundance | mitosis | inter | 0.00482263108415302 | 0.052189282023198294 |
| interactor_ratio | mitosis | inter | 0.00581869698400571 | 0.10426786862612165 |
