## Supplementary Data 2 for "SECAT: Quantifying differential protein-protein interaction states by network-centric analysis": hela_string_000074_Q5J8M3.pdf

Q5J8M3_Q5J8M3

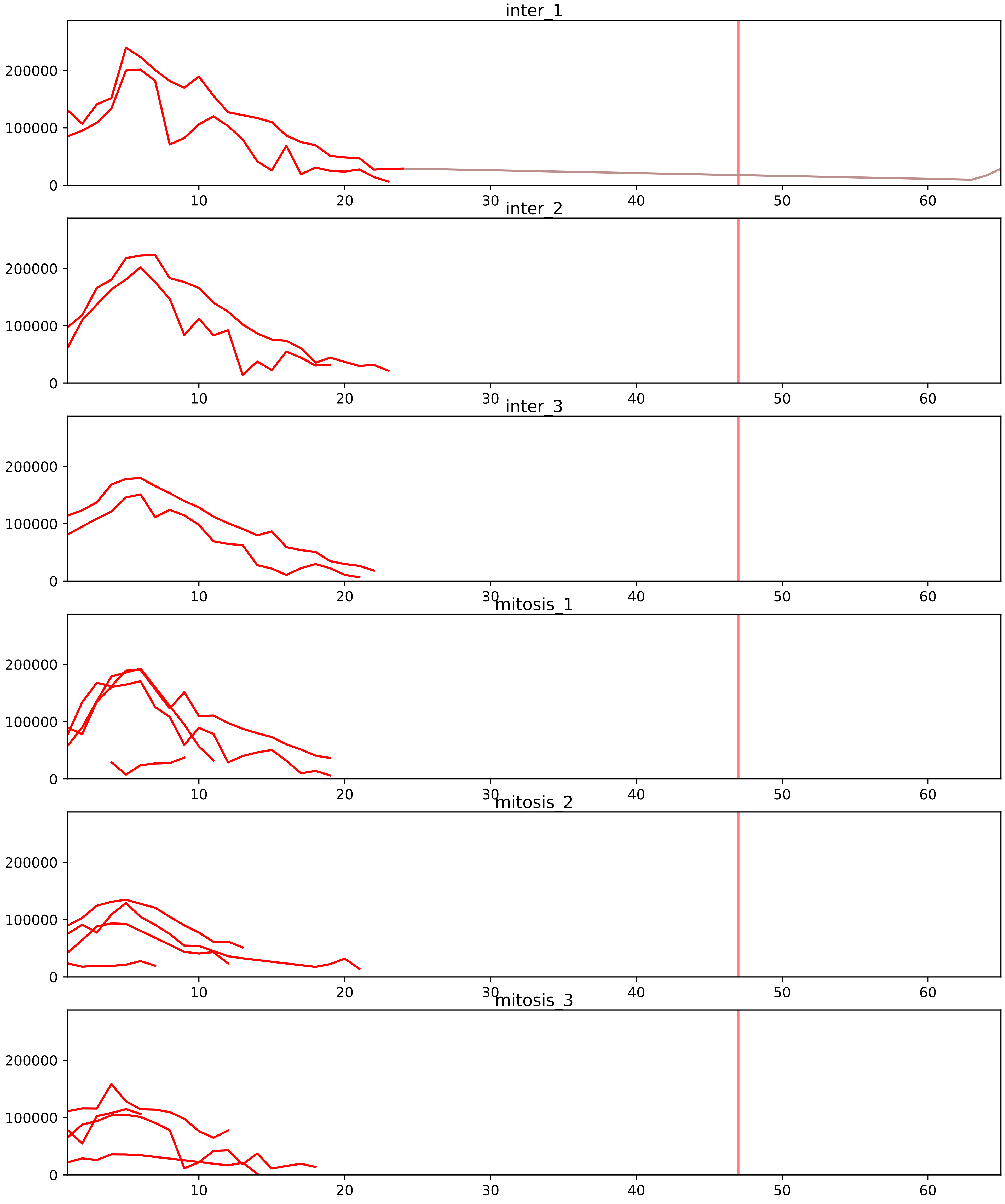


| level | condition_1 | condition_2 | pvalue | pvalue_adjusted |
|---|---|---|---|---|
| interactor_ratio | mitosis | inter | 1.874963828530872e-06 | 5.227171885601219e-05 |
| complex_abundance | mitosis | inter | 0.08631593278002163 | 0.1776524790998208 |
| assembled_abundance | mitosis | inter | 0.11952457844536159 | 0.40660941257466393 |
| total_abundance | mitosis | inter | 0.14902579470173236 | 0.47756083052049103 |
| monomer_abundance | mitosis | inter | 0.30774339273970164 | 0.6718670942432774 |
| interactor_abundance | mitosis | inter | 0.8348967656500583 | 0.9095382171676183 |

O43402_Q5J8M3
EMC8_HUMAN vs EMC4_HUMAN
p-value: 0.0 q-value: 0.0

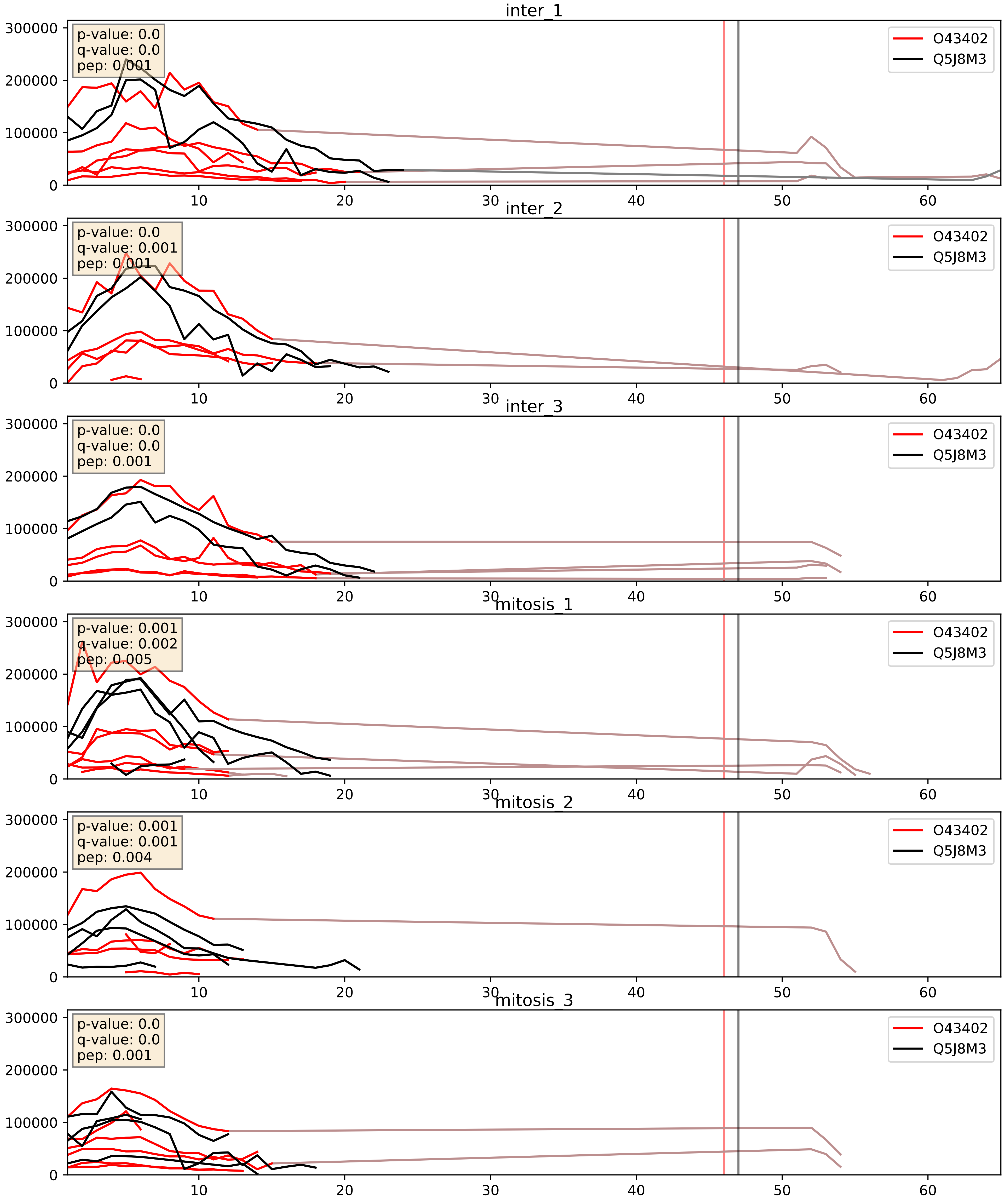


| level | condition_1 | condition_2 | pvalue | pvalue_adjusted |
|---|---|---|---|---|
| interactor_abundance | mitosis | inter | 0.014260968788348653 | 0.10550702235457024 |
| interactor_ratio | mitosis | inter | 0.015594769379646265 | 0.1843379164928052 |
| complex_abundance | mitosis | inter | 0.12383569520881246 | 0.26577856228555974 |

O75947_Q5J8M3
ATP5H_HUMAN vs EMC4_HUMAN
p-value: 0.0 q-value: 0.047

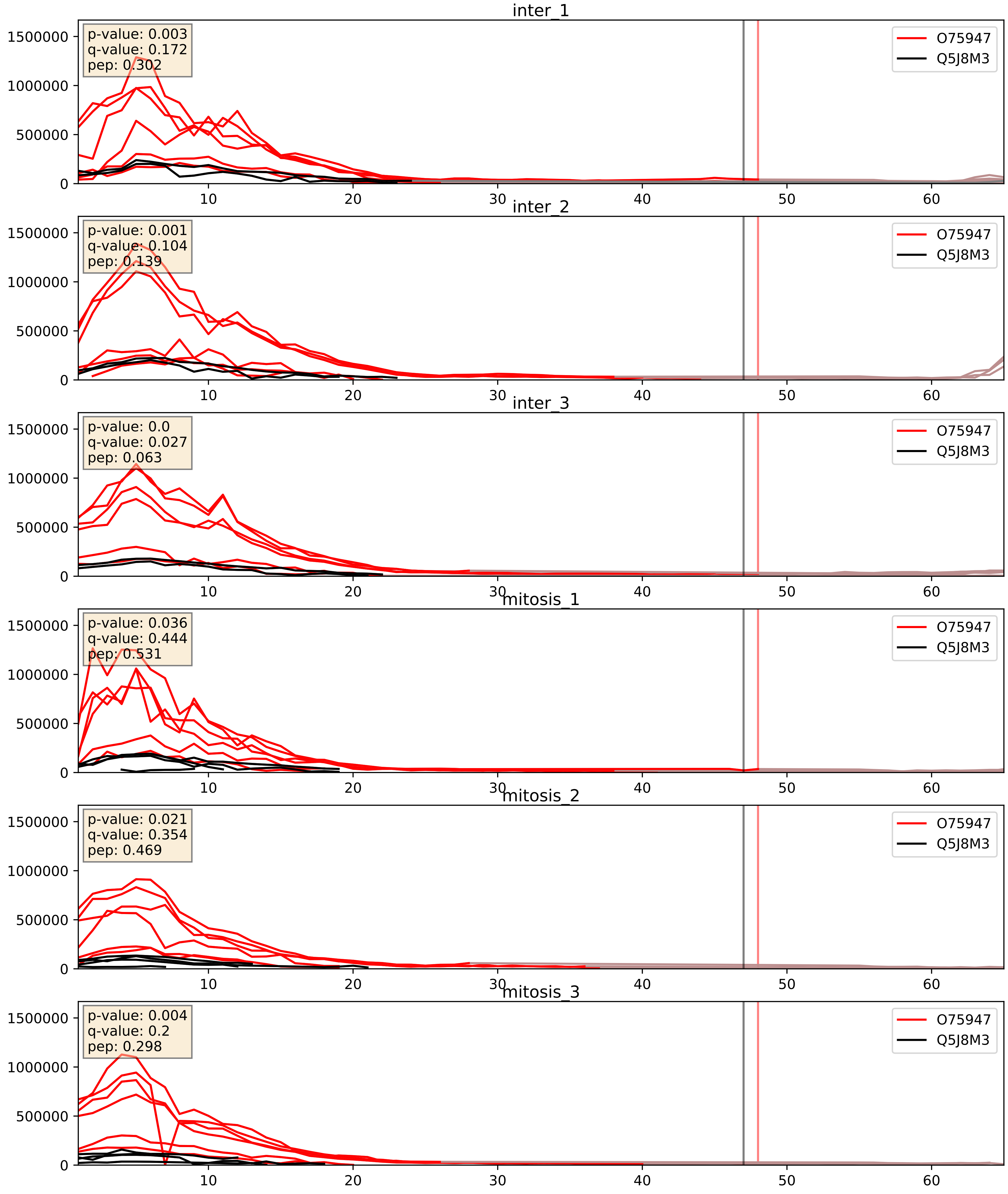


| level | condition_1 | condition_2 | pvalue | pvalue_adjusted |
|---|---|---|---|---|
| interactor_abundance | mitosis | inter | 0.011134689743319312 | 0.0930899374593446 |
| interactor_ratio | mitosis | inter | 0.03502169474477365 | 0.30342682896281625 |
| complex_abundance | mitosis | inter | 0.12225804843489237 | 0.2644085130375641 |

Q15006_Q5J8M3
EMC2_HUMAN vs EMC4_HUMAN
p-value: 0.0 q-value: 0.0

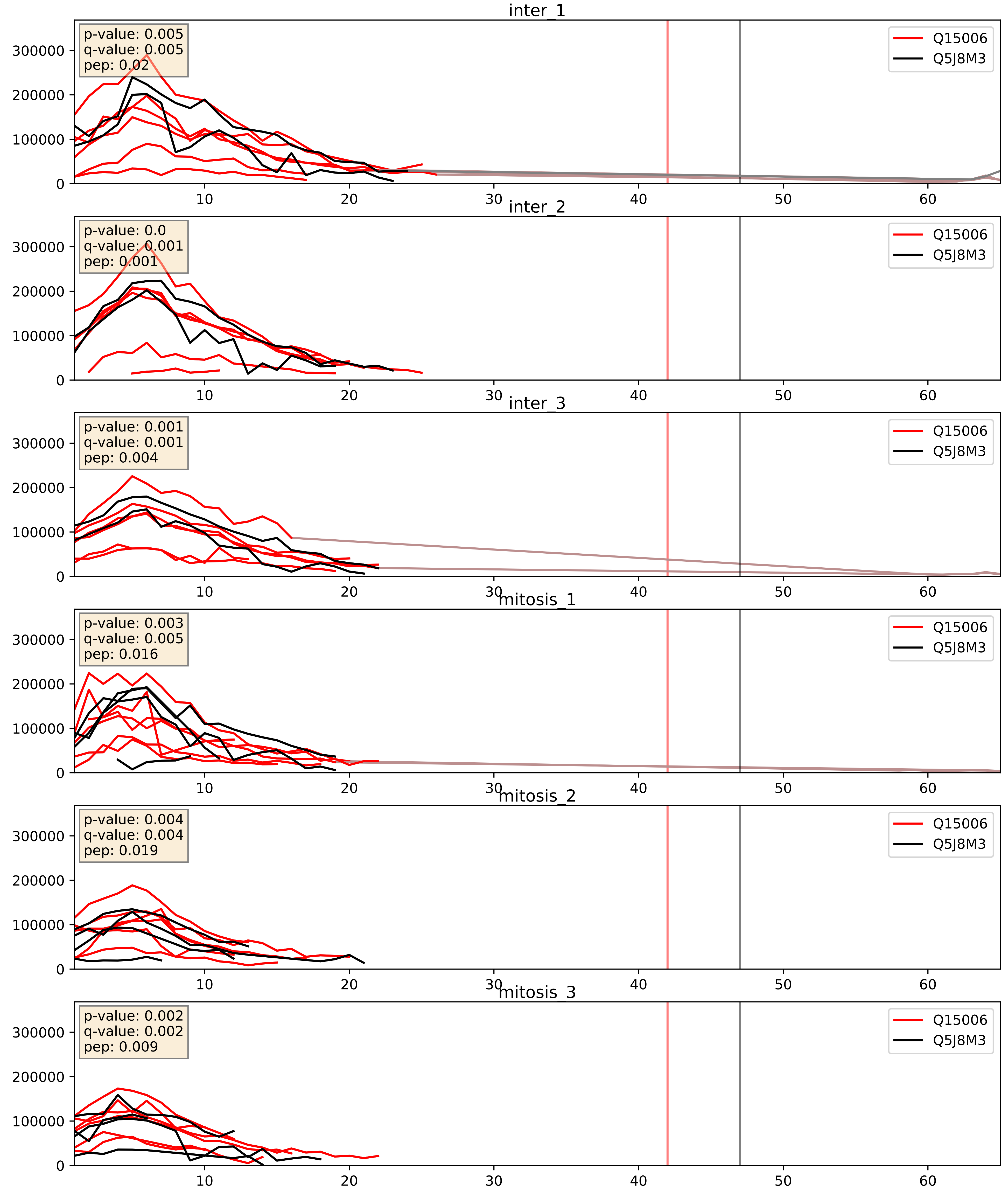


| level | condition_1 | condition_2 | pvalue | pvalue_adjusted |
|---|---|---|---|---|
| interactor_ratio | mitosis | inter | 0.004420734671358082 | 0.08831792861907116 |
| interactor_abundance | mitosis | inter | 0.02338918674609693 | 0.12556564873974368 |
| complex_abundance | mitosis | inter | 0.16992970498594384 | 0.31873042673431795 |

Q5J8M3_Q5UCC4
EMC4_HUMAN vs EMC10_HUMAN
p-value: 0.001 q-value: 0.015

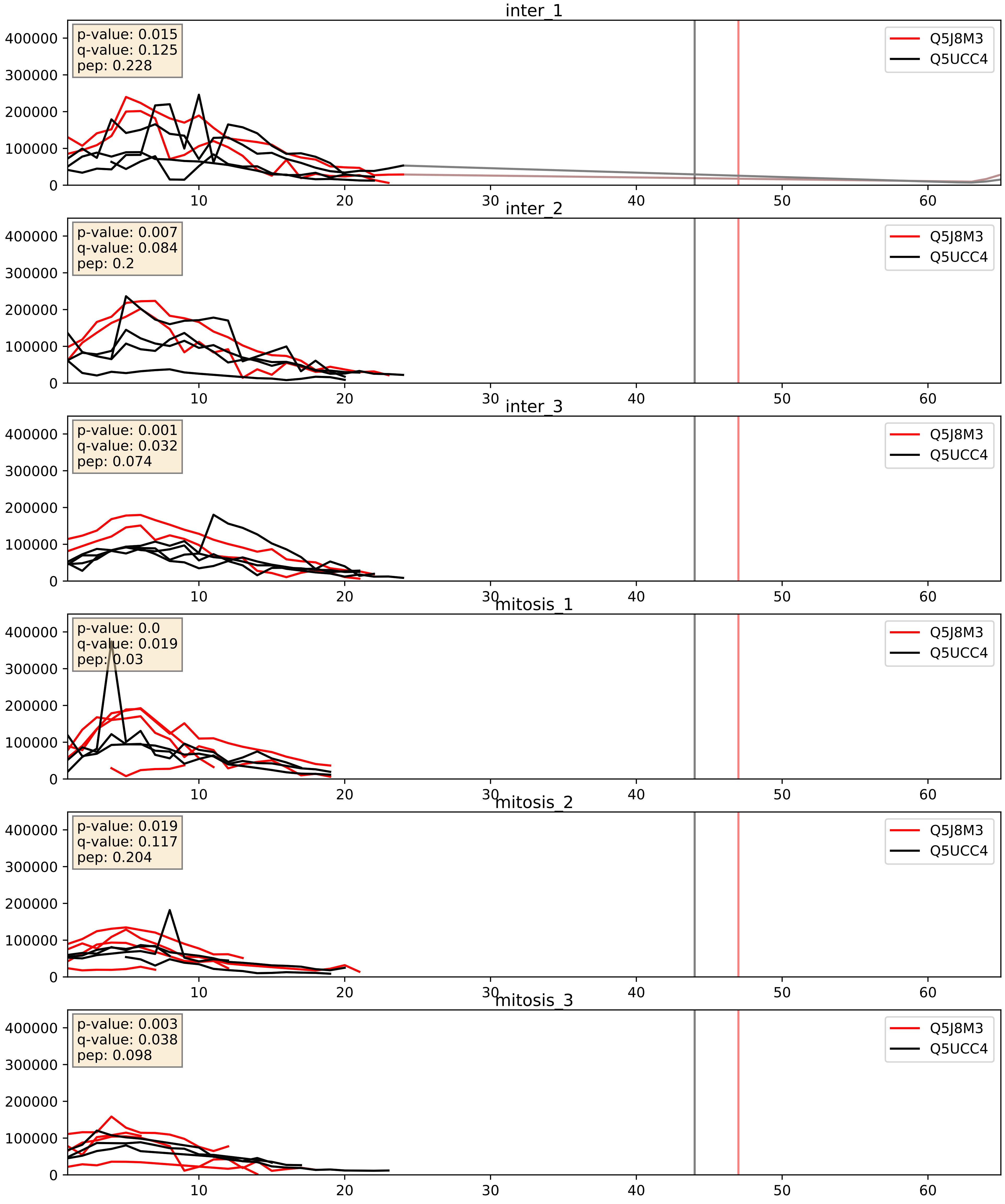


| level | condition_1 | condition_2 | pvalue | pvalue_adjusted |
|---|---|---|---|---|
| interactor_ratio | mitosis | inter | 0.009885240731487515 | 0.14263182888175546 |
| complex_abundance | mitosis | inter | 0.18882019288712634 | 0.3406324238385251 |
| interactor_abundance | mitosis | inter | 0.8457769001313749 | 0.9152981529527123 |

Q5J8M3_Q8N4V1
EMC4_HUMAN vs MMGT1_HUMAN
p-value: 0.0 q-value: 0.0

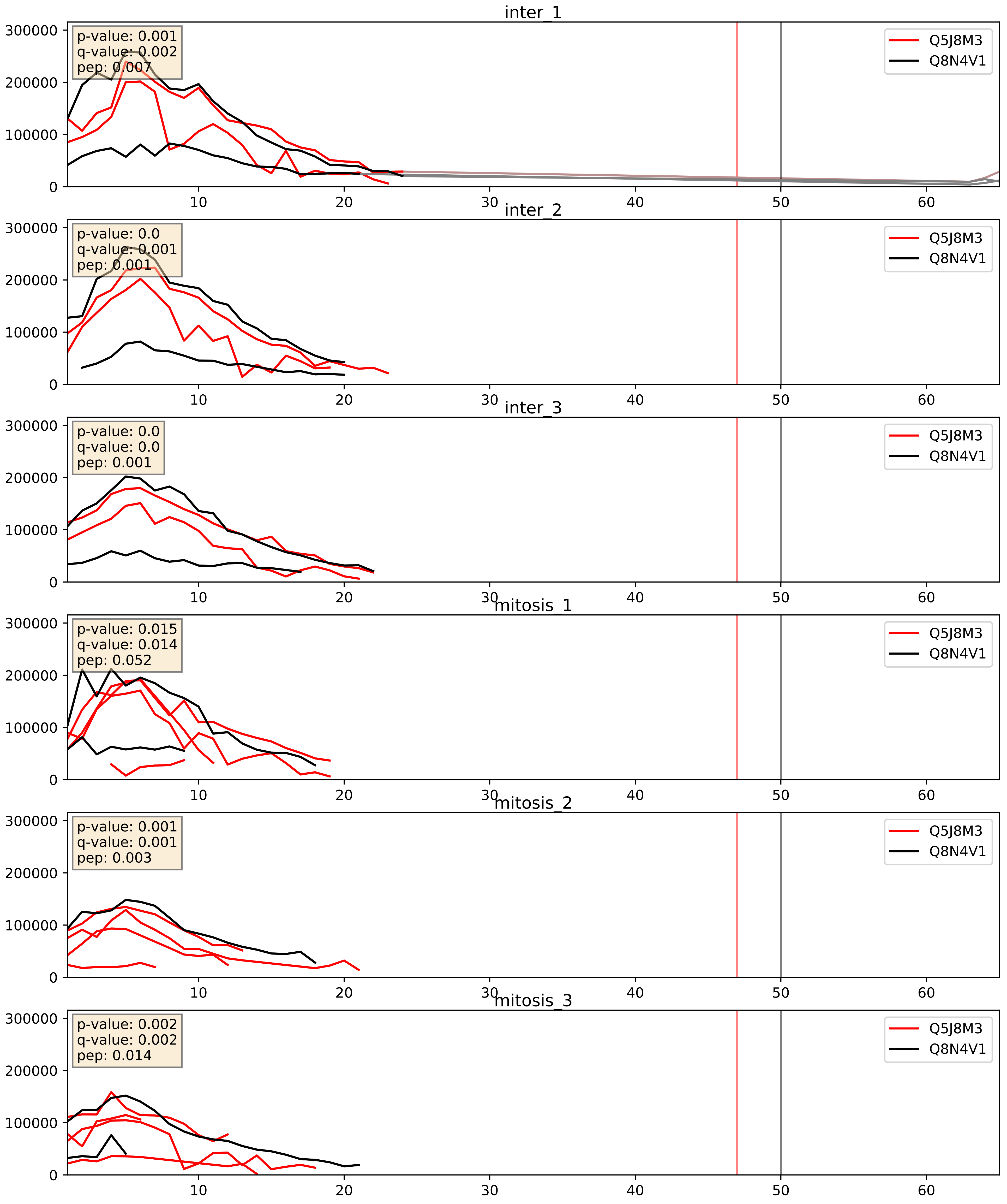


| level | condition_1 | condition_2 | pvalue | pvalue_adjusted |
|---|---|---|---|---|
| interactor_ratio | mitosis | inter | 0.0029469094962753088 | 0.07007095913365735 |
| complex_abundance | mitosis | inter | 0.21591933035717598 | 0.3706234327307472 |
| interactor_abundance | mitosis | inter | 0.8261440897441086 | 0.9033395169201723 |

Q5J8M3_Q8N766
EMC4_HUMAN vs EMC1_HUMAN
p-value: 0.011 q-value: 0.01

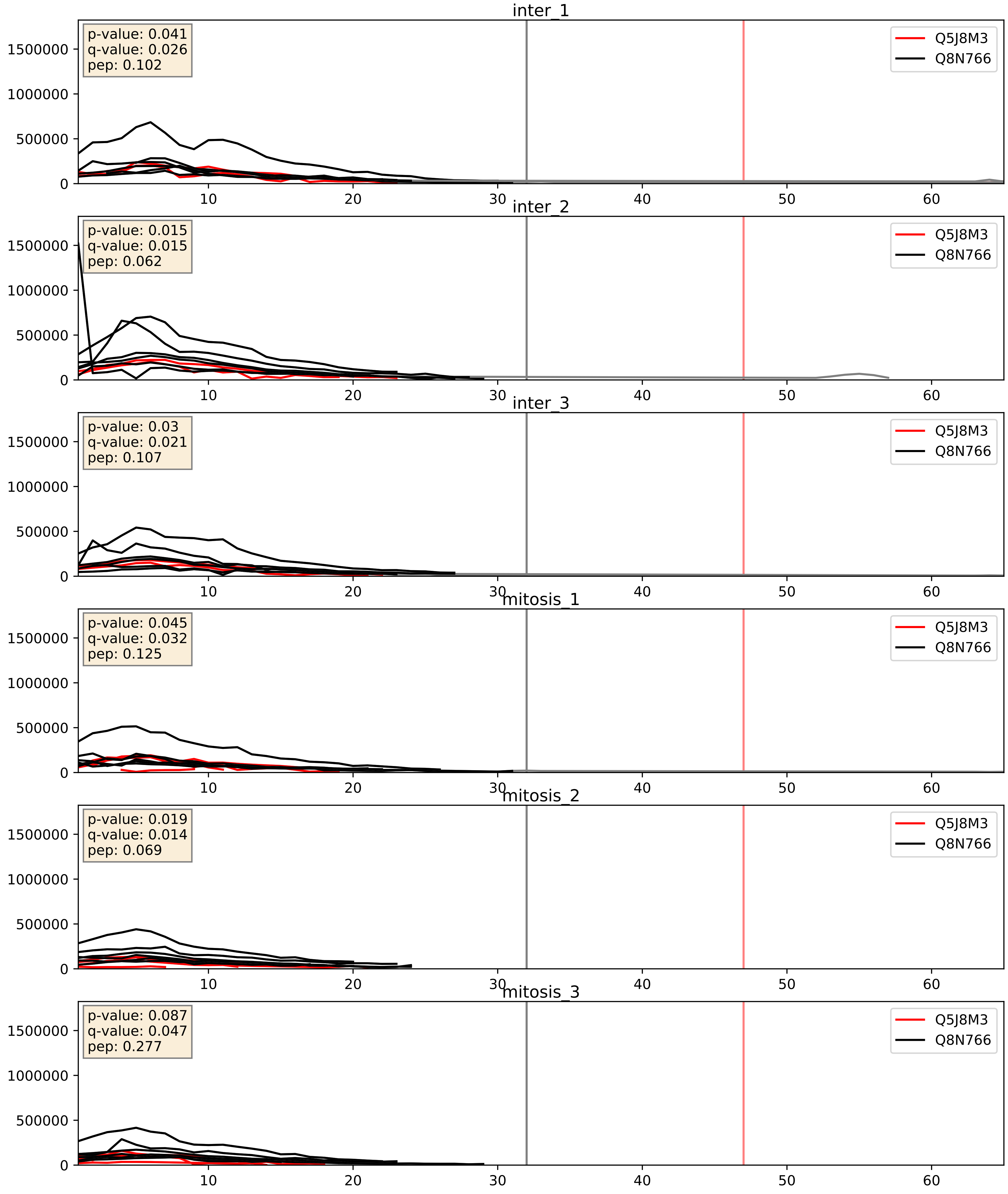


| level | condition_1 | condition_2 | pvalue | pvalue_adjusted |
|---|---|---|---|---|
| interactor_ratio | mitosis | inter | 0.07380800269676804 | 0.4548570936532285 |
| complex_abundance | mitosis | inter | 0.07961652324776375 | 0.20620799969768766 |
| interactor_abundance | mitosis | inter | 0.8539580253675086 | 0.9216480512950259 |

Q5J8M3_Q9BV81
EMC4_HUMAN vs EMC6_HUMAN
p-value: 0.0 q-value: 0.0

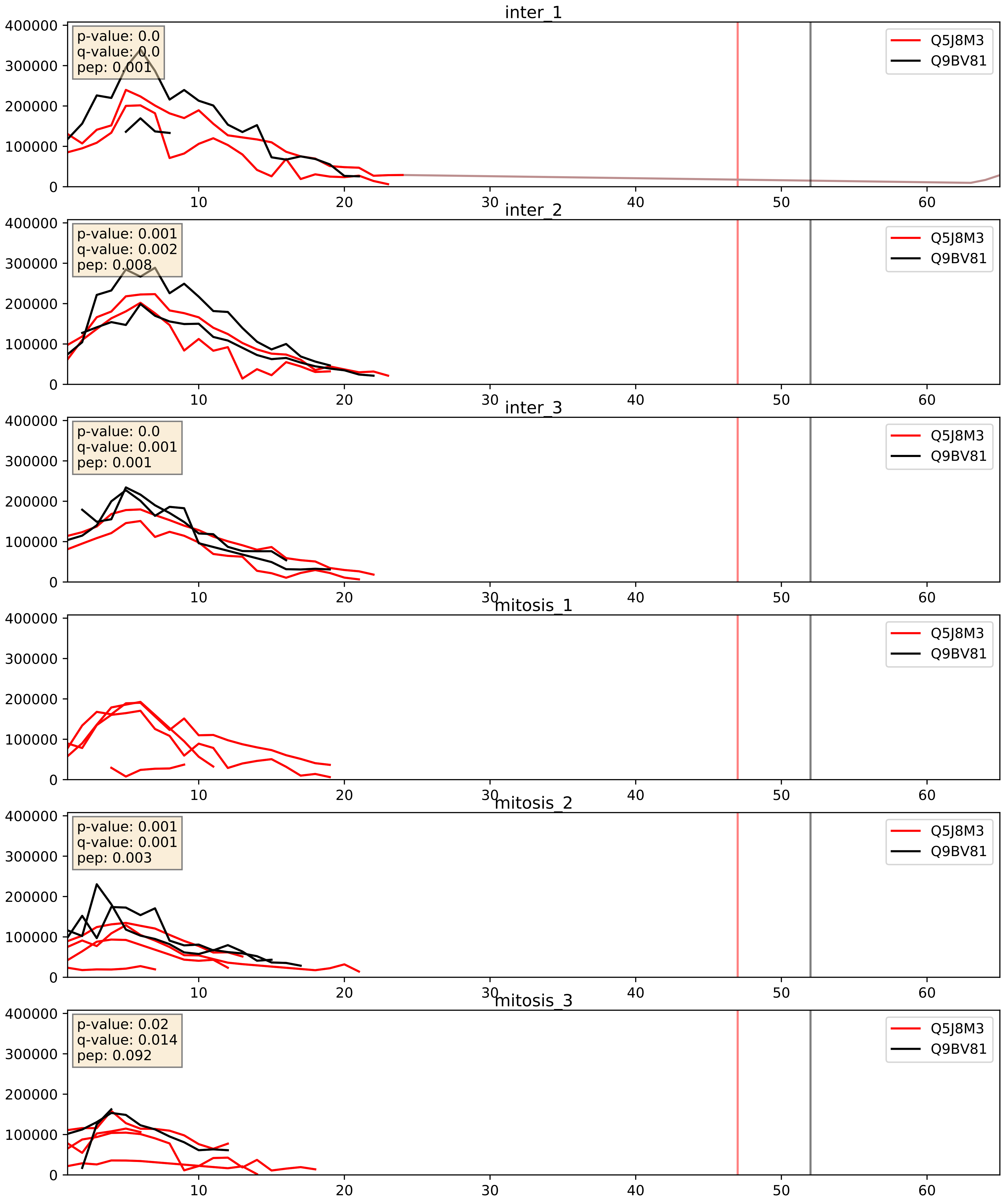


| level | condition_1 | condition_2 | pvalue | pvalue_adjusted |
|---|---|---|---|---|
| complex_abundance | mitosis | inter | 0.11404567393205839 | 0.25396227077482303 |
| interactor_abundance | mitosis | inter | 0.19796297502201385 | 0.39175017607980506 |
| interactor_ratio | mitosis | inter | 0.4801292005854926 | 0.8668753814420835 |

Q5J8M3_Q9NPA0
EMC4_HUMAN vs EMC7_HUMAN
p-value: 0.0 q-value: 0.0

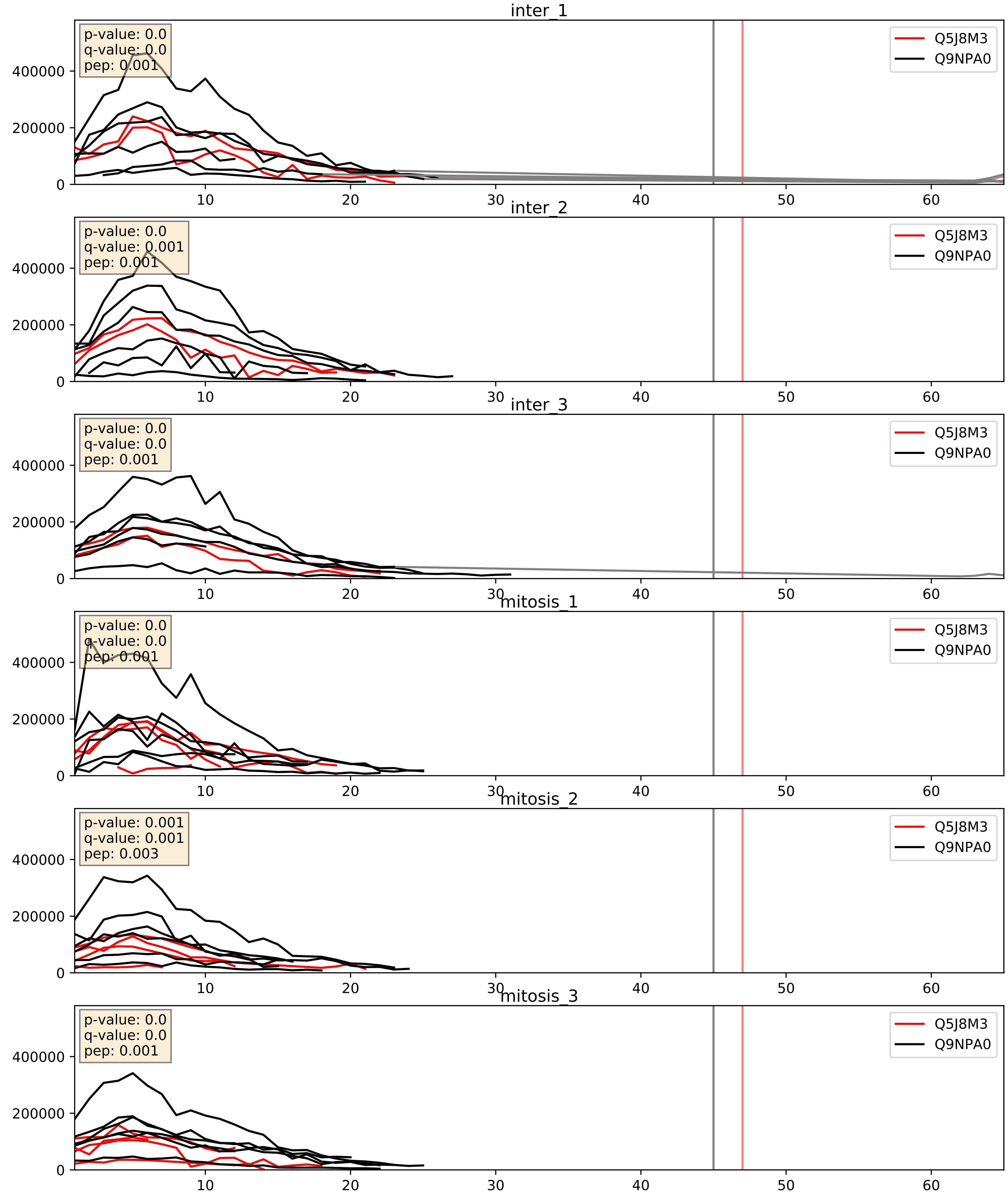


| level | condition_1 | condition_2 | pvalue | pvalue_adjusted |
|---|---|---|---|---|
| interactor_ratio | mitosis | inter | 0.008177226250630613 | 0.12756688776860511 |
| complex_abundance | mitosis | inter | 0.2177659373008419 | 0.37236844252800777 |
| interactor_abundance | mitosis | inter | 0.8539580253675086 | 0.9216480512950259 |

Q5J8M3_Q9P0I2
EMC4_HUMAN vs EMC3_HUMAN
p-value: 0.0 q-value: 0.0

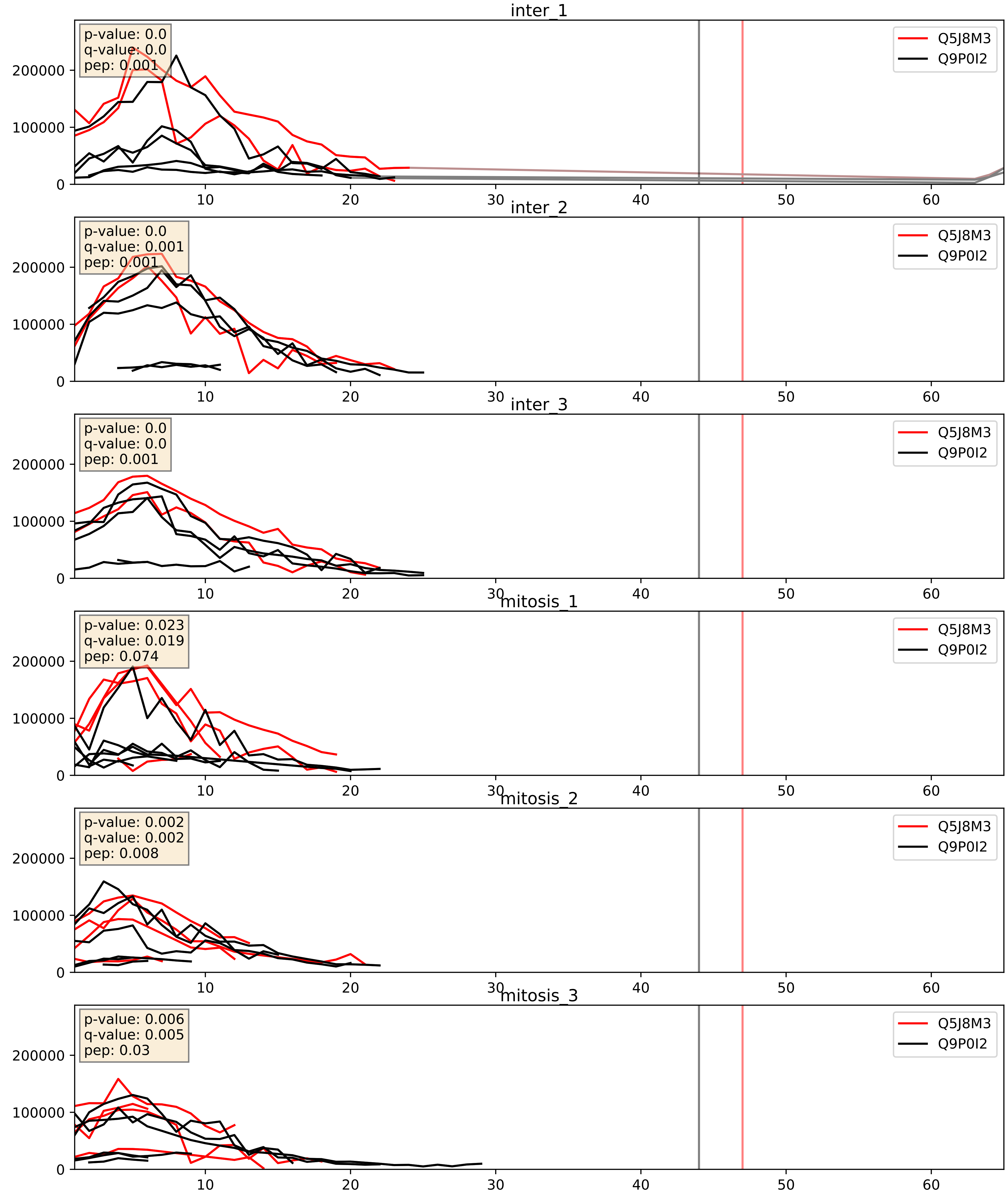


| level | condition_1 | condition_2 | pvalue | pvalue_adjusted |
|---|---|---|---|---|
| complex_abundance | mitosis | inter | 0.03814173657411883 | 0.14263576455852214 |
| interactor_ratio | mitosis | inter | 0.11942405731474313 | 0.5684776613996082 |
| interactor_abundance | mitosis | inter | 0.8556910364096157 | 0.9226244201620243 |

Q5J8M3_Q9UBU6
EMC4_HUMAN vs FA8A1_HUMAN
p-value: 0.0 q-value: 0.018

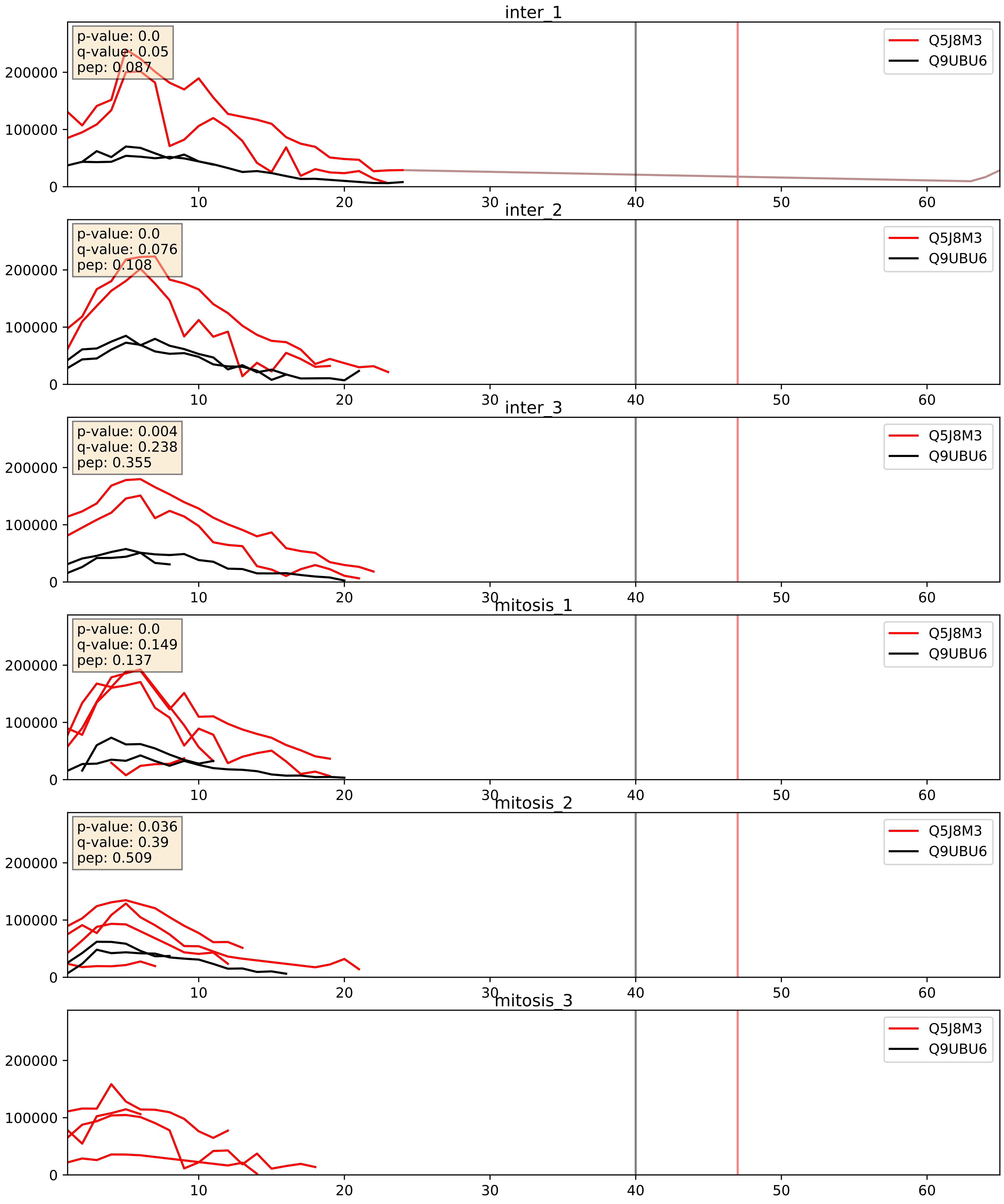


| level | condition_1 | condition_2 | pvalue | pvalue_adjusted |
|---|---|---|---|---|
| complex_abundance | mitosis | inter | 0.28368070908304627 | 0.4321599697011703 |
| interactor_abundance | mitosis | inter | 0.41235768611693535 | 0.5955929795260215 |
| interactor_ratio | mitosis | inter | 0.9046300761879054 | 0.9852158746906624 |

Q5J8M3_Q9Y3B3
EMC4_HUMAN vs TMED7_HUMAN
p-value: 0.0 q-value: 0.038

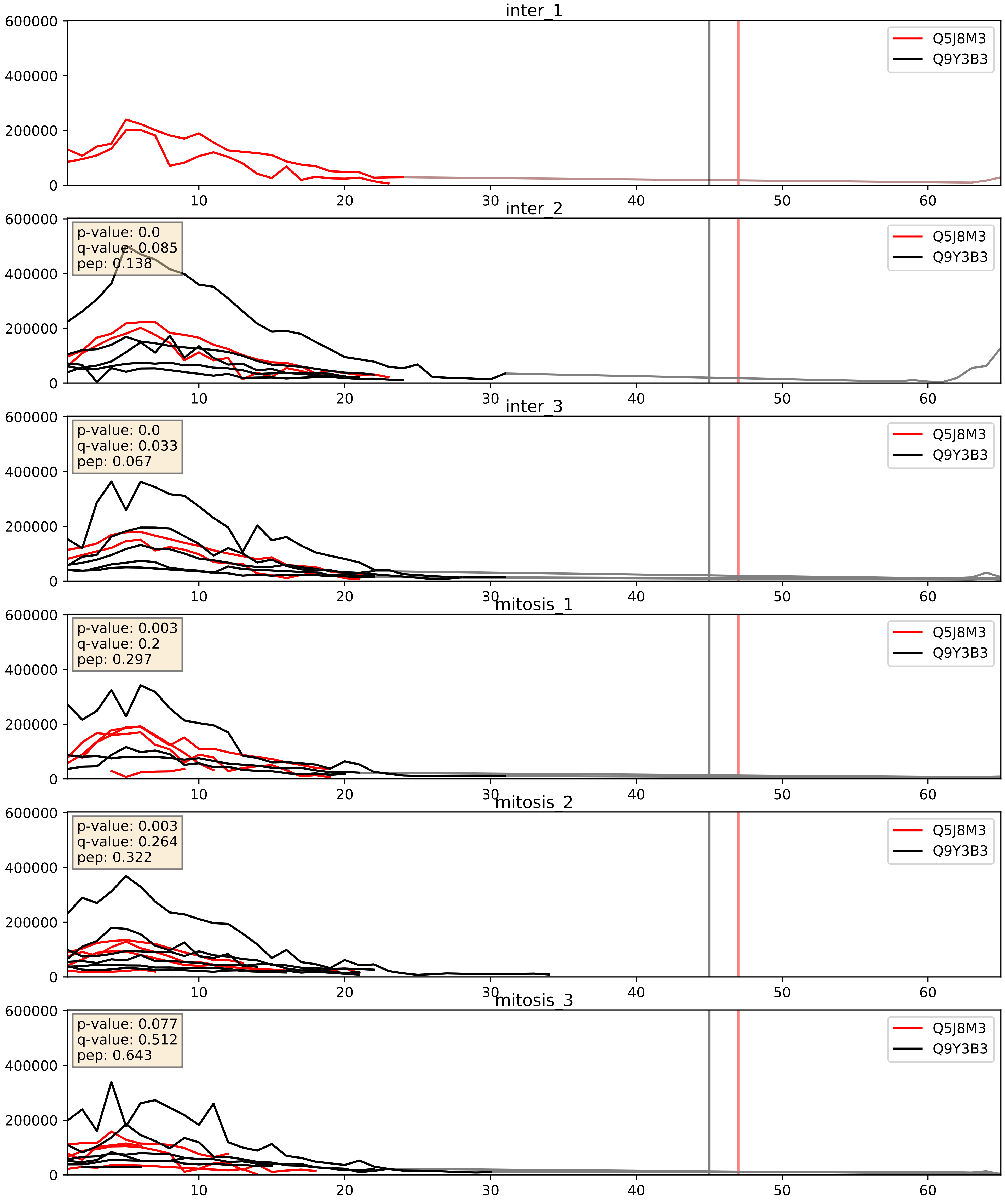


| level | condition_1 | condition_2 | pvalue | pvalue_adjusted |
|---|---|---|---|---|
| interactor_abundance | mitosis | inter | 0.4055903846935091 | 0.5919614139772273 |
| complex_abundance | mitosis | inter | 0.5901790048787492 | 0.7159767973018839 |
| interactor_ratio | mitosis | inter | 0.652053207023245 | 0.9220833993116596 |
