## Supplementary Data 2 for "SECAT: Quantifying differential protein-protein interaction states by network-centric analysis": hela_string_000075_O14979.pdf

O14979_O14979

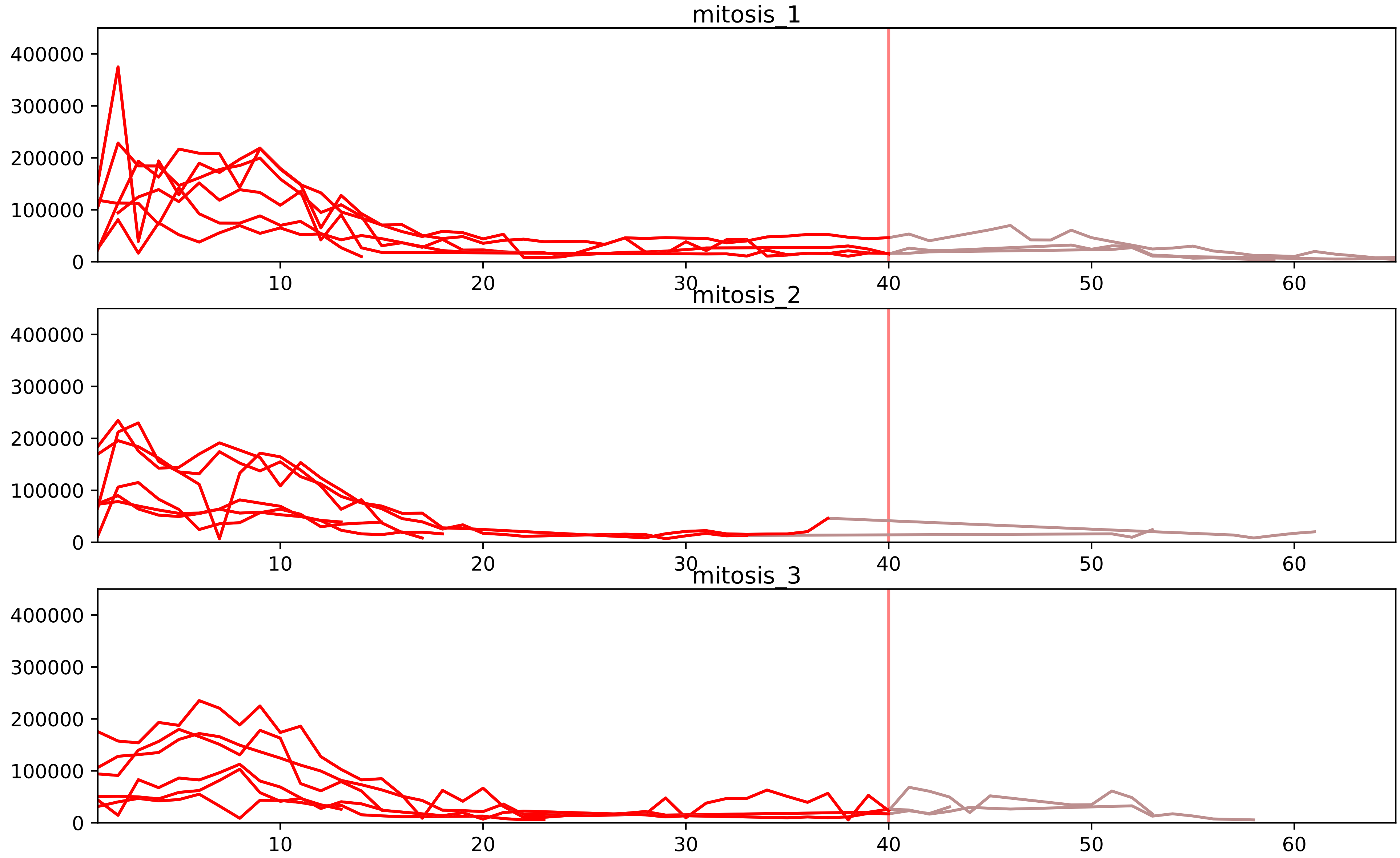


| level | condition_1 | condition_2 | pvalue | pvalue_adjusted |
|---|---|---|---|---|
| complex_abundance | mitosis | inter | 1.925630294141309e-06 | 0.00014763165588416703 |
| interactor_abundance | mitosis | inter | 2.1427679895569332e-05 | 0.0008896139540965403 |
| assembled_abundance | mitosis | inter | 0.004947683974771761 | 0.07135056685689445 |
| total_abundance | mitosis | inter | 0.005753070847610782 | 0.08742950353787012 |
| monomer_abundance | mitosis | inter | 0.014093359256851295 | 0.20378095477374408 |
| interactor_ratio | mitosis | inter | 0.022147563448601425 | 0.1138310523615269 |

O14979_P14866
HNRDL_HUMAN vs HNRPL_HUMAN
p-value: 0.003 q-value: 0.044

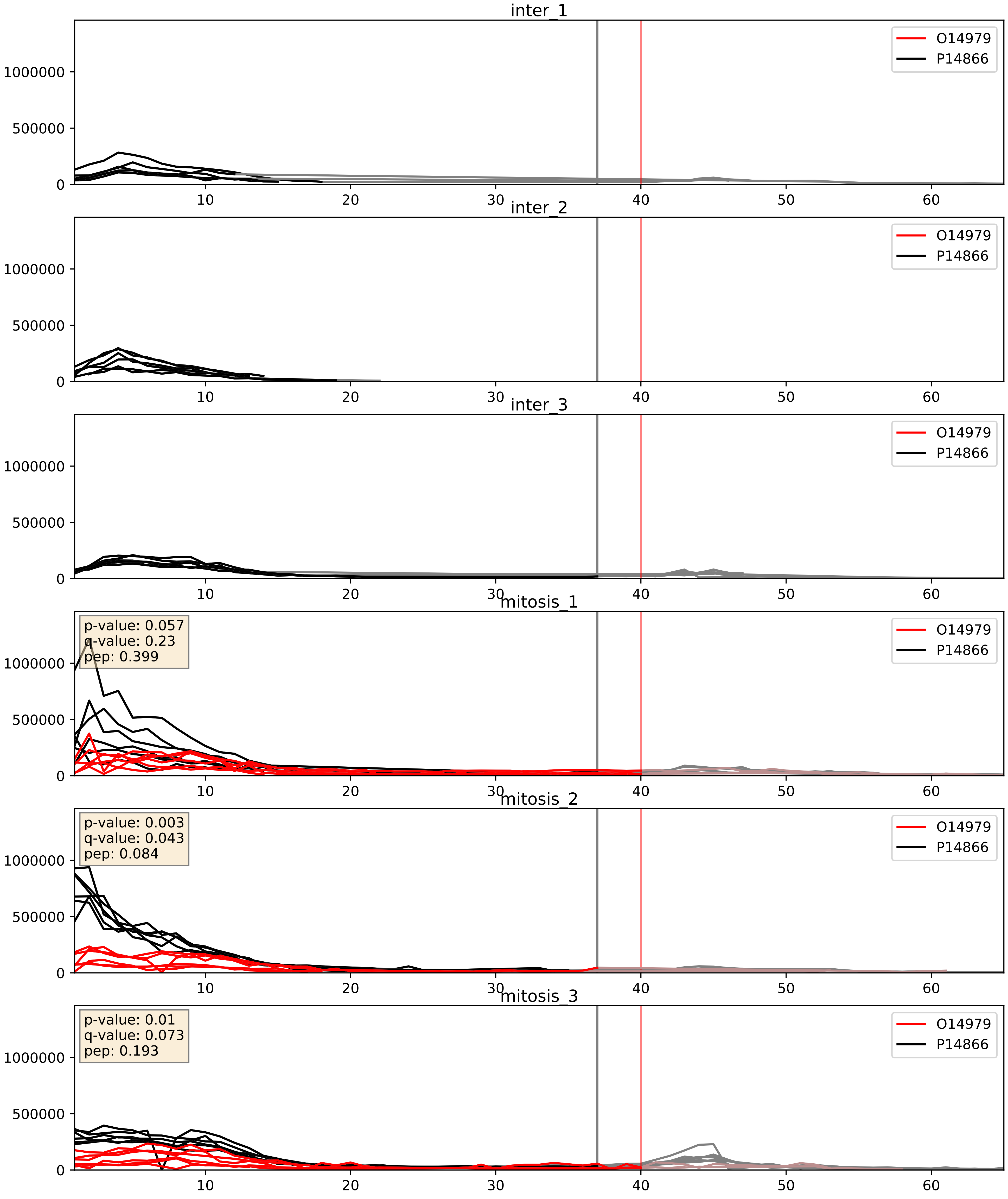


| level | condition_1 | condition_2 | pvalue | pvalue_adjusted |
|---|---|---|---|---|
| complex_abundance | mitosis | inter | 9.401708878998684e-05 | 0.006596608852805635 |
| interactor_abundance | mitosis | inter | 0.0007528746798464765 | 0.020141995013343572 |
| interactor_ratio | mitosis | inter | 0.012414827969505712 | 0.16080750275117184 |

O14979_P31942
HNRDL_HUMAN vs HNRH3_HUMAN
p-value: 0.004 q-value: 0.047

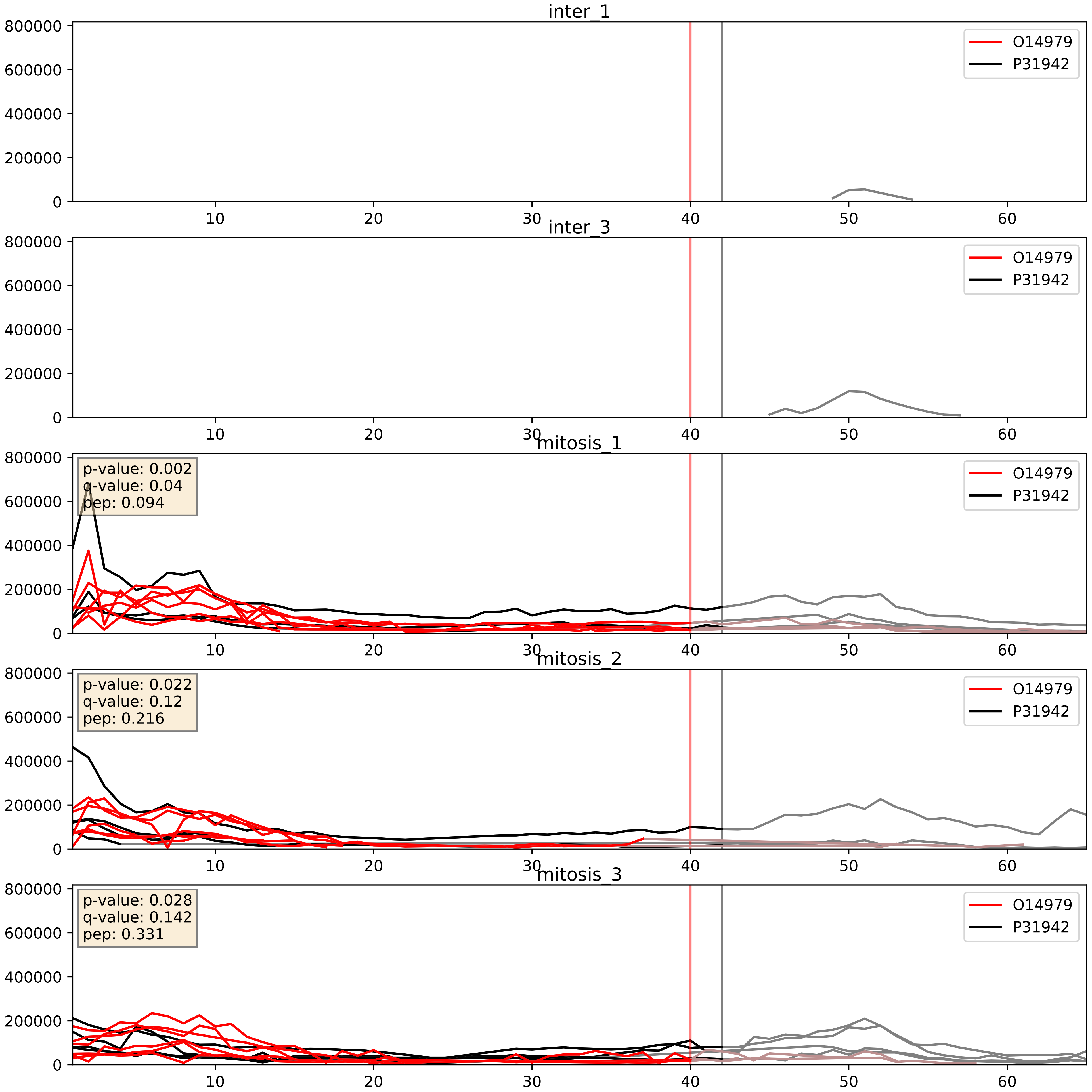


| level | condition_1 | condition_2 | pvalue | pvalue_adjusted |
|---|---|---|---|---|
| complex_abundance | mitosis | inter | 0.0004757589173787532 | 0.013816623868928207 |
| interactor_abundance | mitosis | inter | 0.0008235085849705327 | 0.02107394166621154 |
| interactor_ratio | mitosis | inter | 0.2439500030606306 | 0.7200661897289671 |

O14979_P52597
HNRDL_HUMAN vs HNRPF_HUMAN
p-value: 0.001 q-value: 0.024

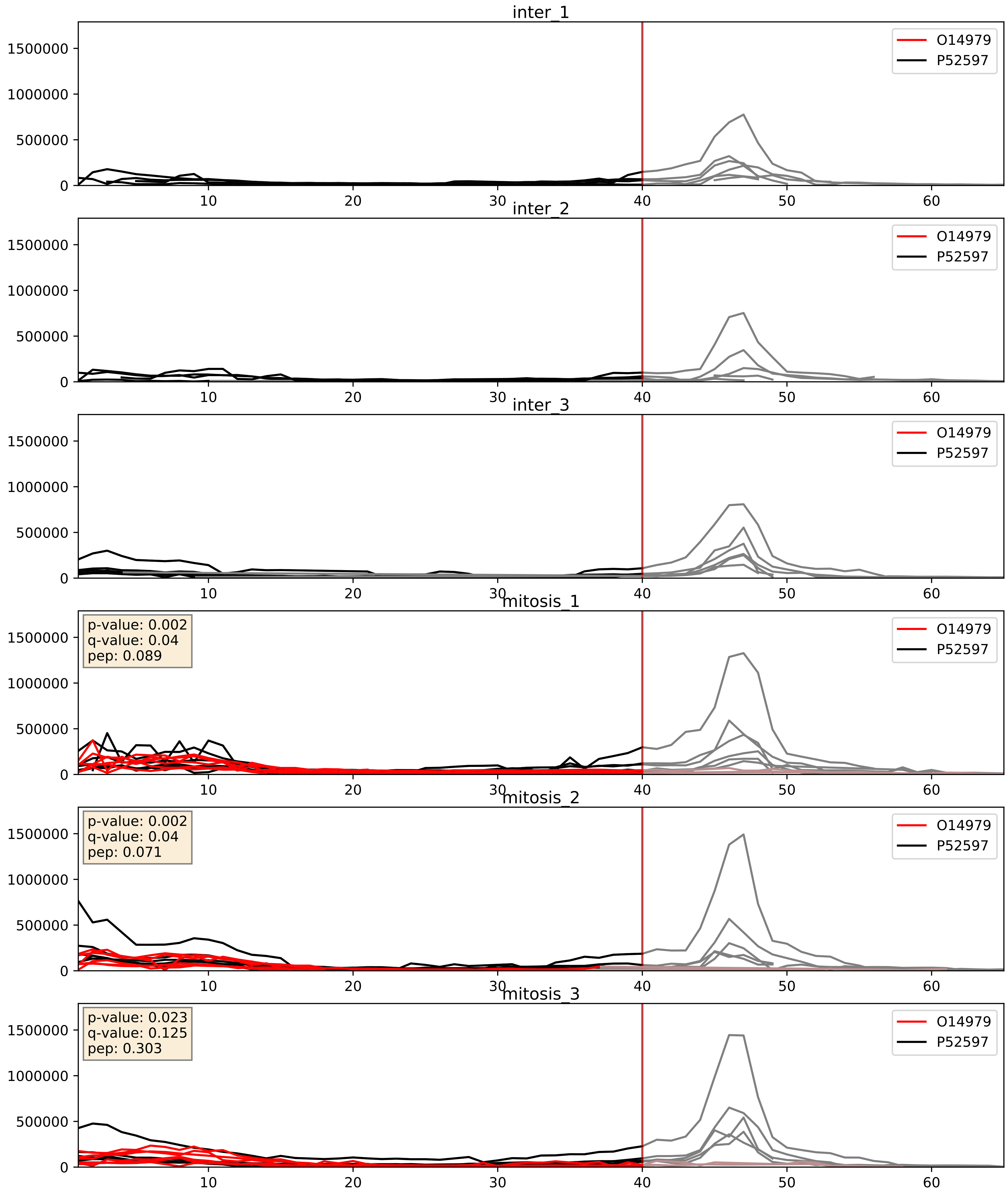


| level | condition_1 | condition_2 | pvalue | pvalue_adjusted |
|---|---|---|---|---|
| complex_abundance | mitosis | inter | 0.00023969958657414883 | 0.010310695784295044 |
| interactor_abundance | mitosis | inter | 0.0008164873709066391 | 0.021004694279574423 |
| interactor_ratio | mitosis | inter | 0.009098125486910772 | 0.13591615038037733 |

O14979_Q13151
HNRDL_HUMAN vs ROA0_HUMAN
p-value: 0.0 q-value: 0.026

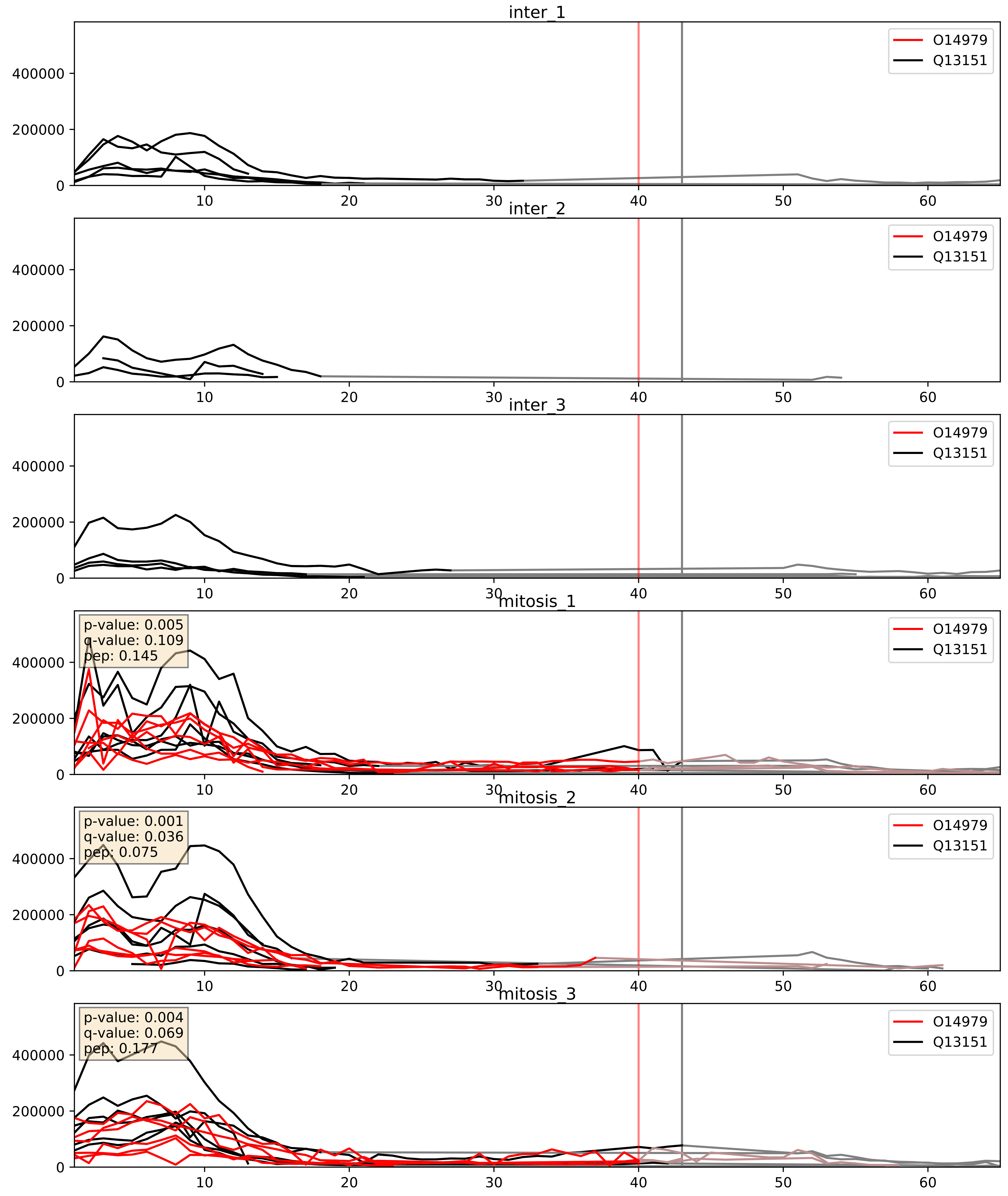


| level | condition_1 | condition_2 | pvalue | pvalue_adjusted |
|---|---|---|---|---|
| complex_abundance | mitosis | inter | 0.00015633998550700518 | 0.008563834848633118 |
| interactor_abundance | mitosis | inter | 0.0008316218584732691 | 0.021249800323973684 |
| interactor_ratio | mitosis | inter | 0.08960471804200747 | 0.4993680014453239 |

O14979_Q9BUJ2
HNRDL_HUMAN vs HNRL1_HUMAN
p-value: 0.0 q-value: 0.041

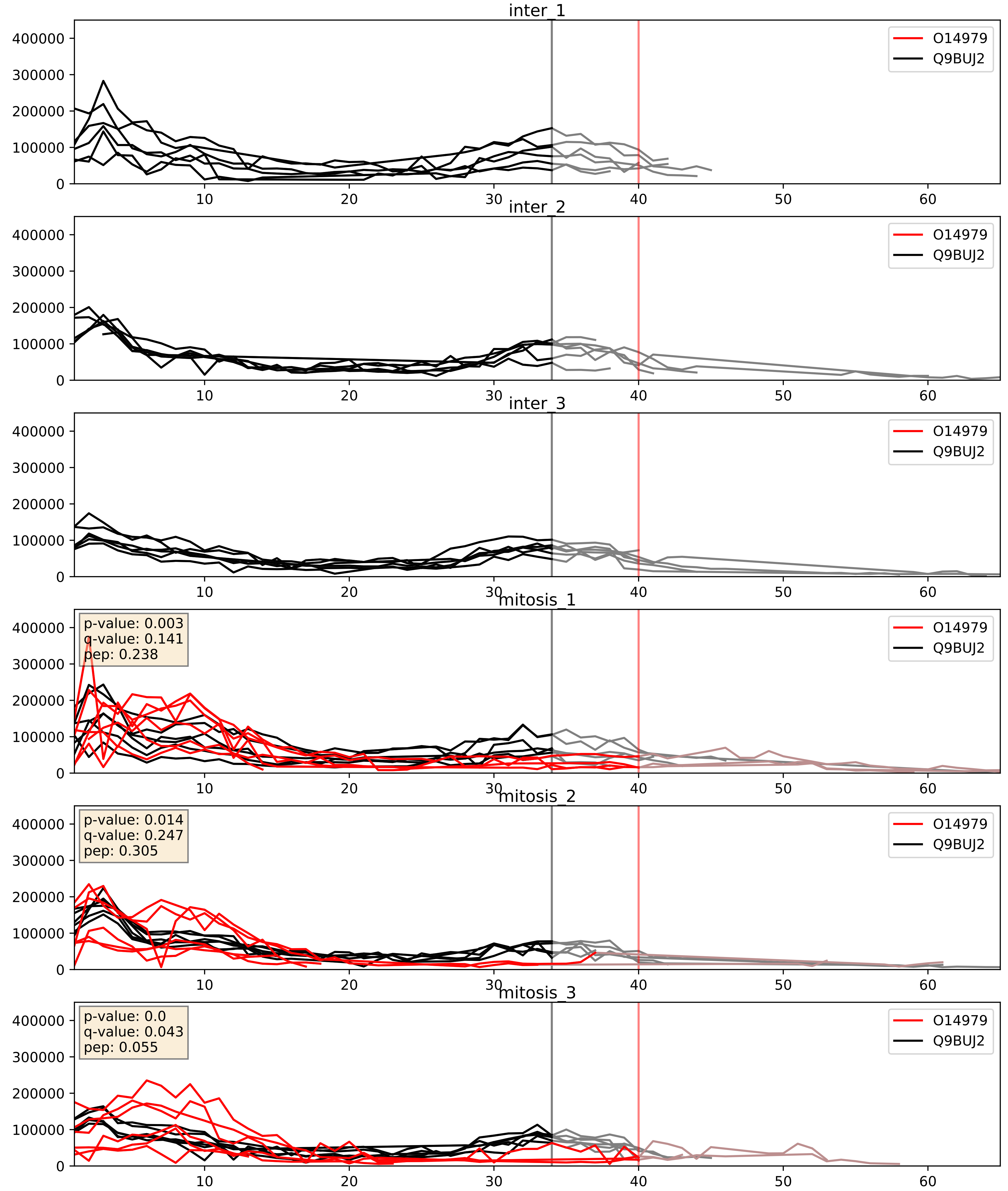


| level | condition_1 | condition_2 | pvalue | pvalue_adjusted |
|---|---|---|---|---|
| complex_abundance | mitosis | inter | 0.00016207257540637443 | 0.008563834848633118 |
| interactor_abundance | mitosis | inter | 0.0007200751823208013 | 0.019561481000604793 |
| interactor_ratio | mitosis | inter | 0.5211793406809895 | 0.8786895401167326 |
