## Supplementary Data 2 for "SECAT: Quantifying differential protein-protein interaction states by network-centric analysis": hela_string_000076_Q13151.pdf

Q13151_Q13151

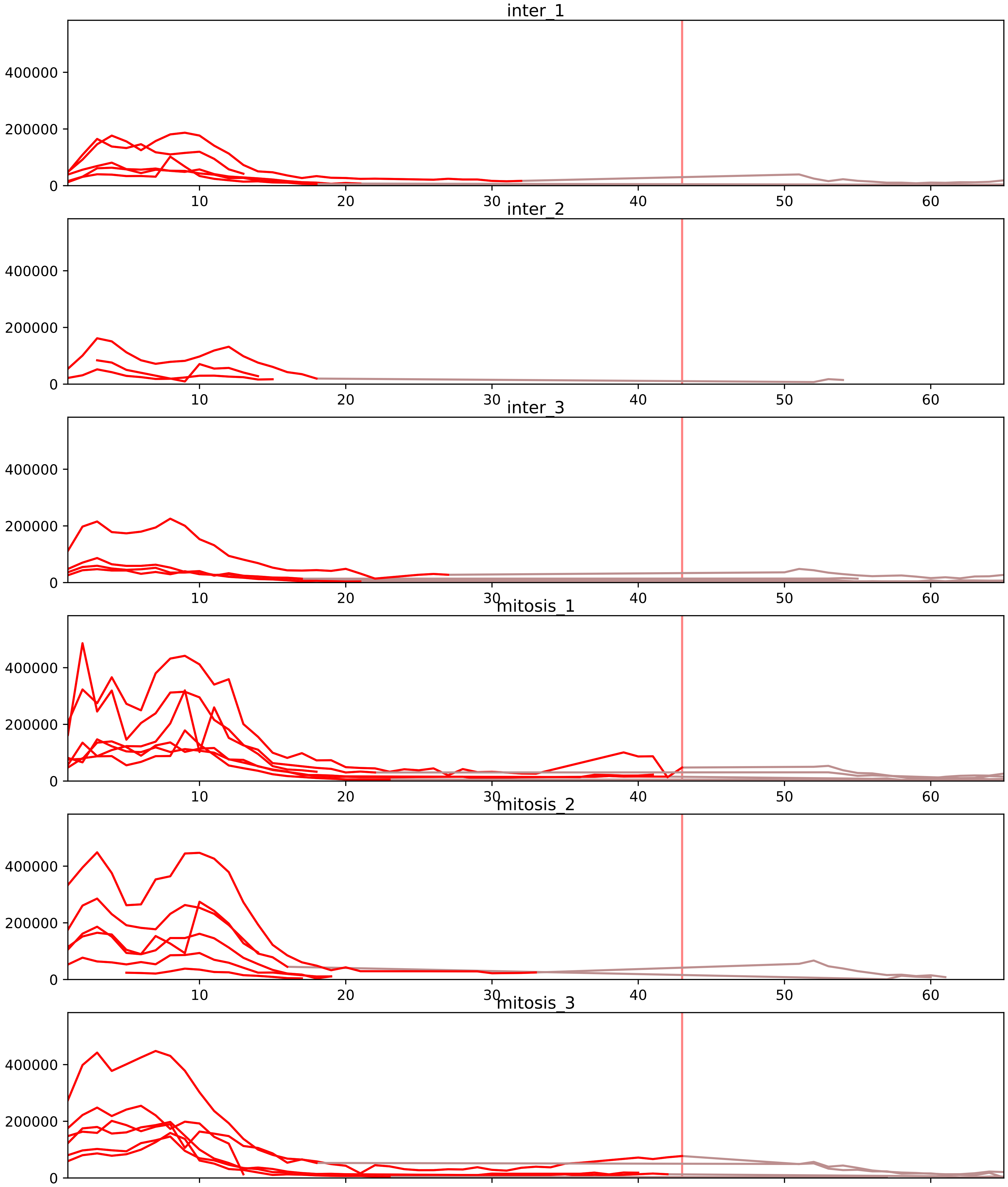


| level | condition_1 | condition_2 | pvalue | pvalue_adjusted |
|---|---|---|---|---|
| interactor_ratio | mitosis | inter | 2.0162738161111464e-06 | 5.537229584544044e-05 |
| complex_abundance | mitosis | inter | 2.840550579354888e-06 | 0.00018666475235760692 |
| interactor_abundance | mitosis | inter | 3.549540757007389e-05 | 0.0012094731468321473 |
| assembled_abundance | mitosis | inter | 0.611092850073818 | 0.8184750844848916 |
| total_abundance | mitosis | inter | 0.6122584010111752 | 0.8253485937924925 |
| monomer_abundance | mitosis | inter | 0.6435877567248027 | 0.8389146793624875 |

O14979_Q13151
HNRDL_HUMAN vs ROA0_HUMAN
p-value: 0.0 q-value: 0.026

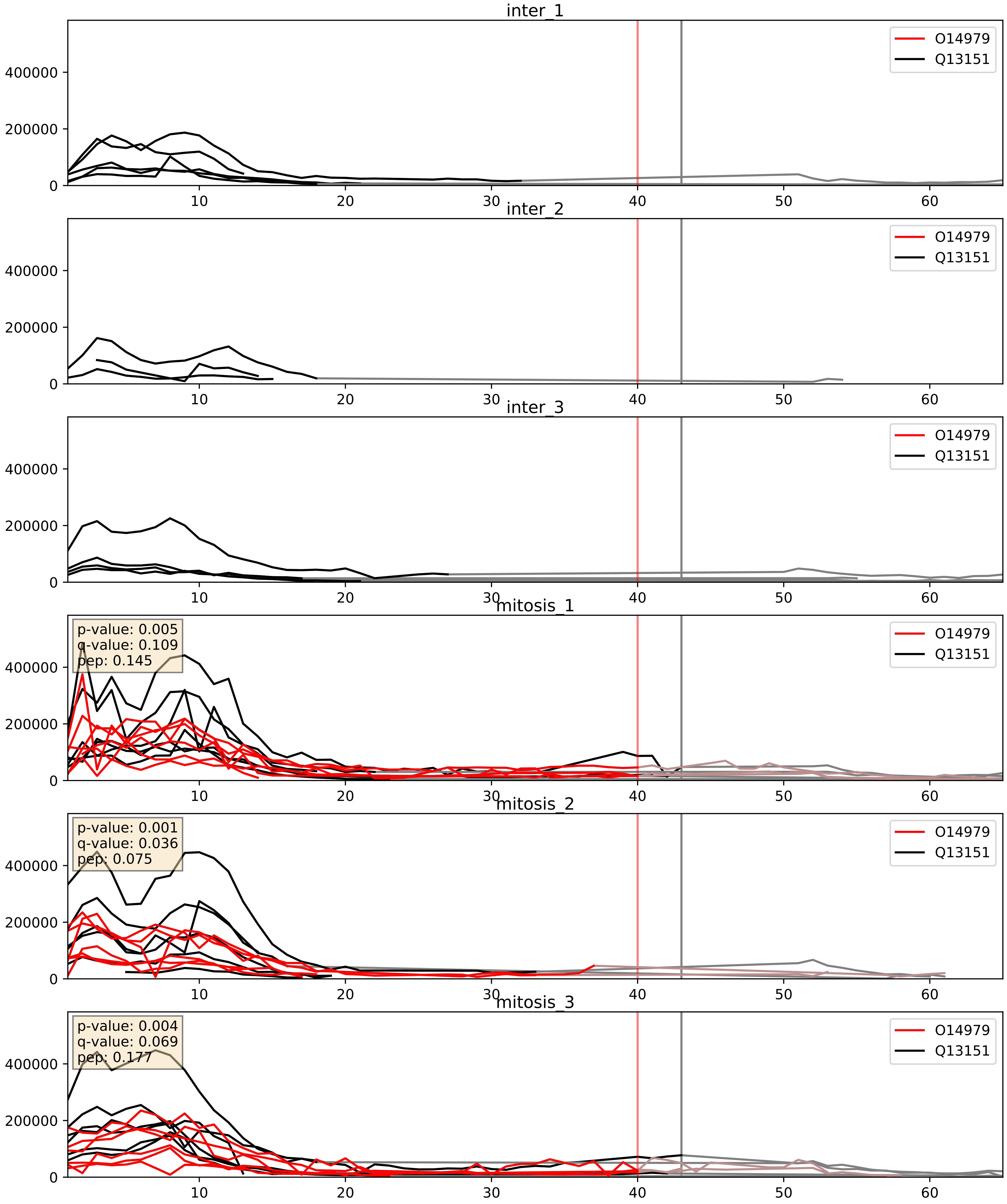


| level | condition_1 | condition_2 | pvalue | pvalue_adjusted |
|---|---|---|---|---|
| complex_abundance | mitosis | inter | 0.00015633998550700518 | 0.008563834848633118 |
| interactor_abundance | mitosis | inter | 0.0008316218584732691 | 0.021249800323973684 |
| interactor_ratio | mitosis | inter | 0.08960471804200747 | 0.4993680014453239 |

O43390_Q13151
HNRPR_HUMAN vs ROA0_HUMAN
p-value: 0.005 q-value: 0.005

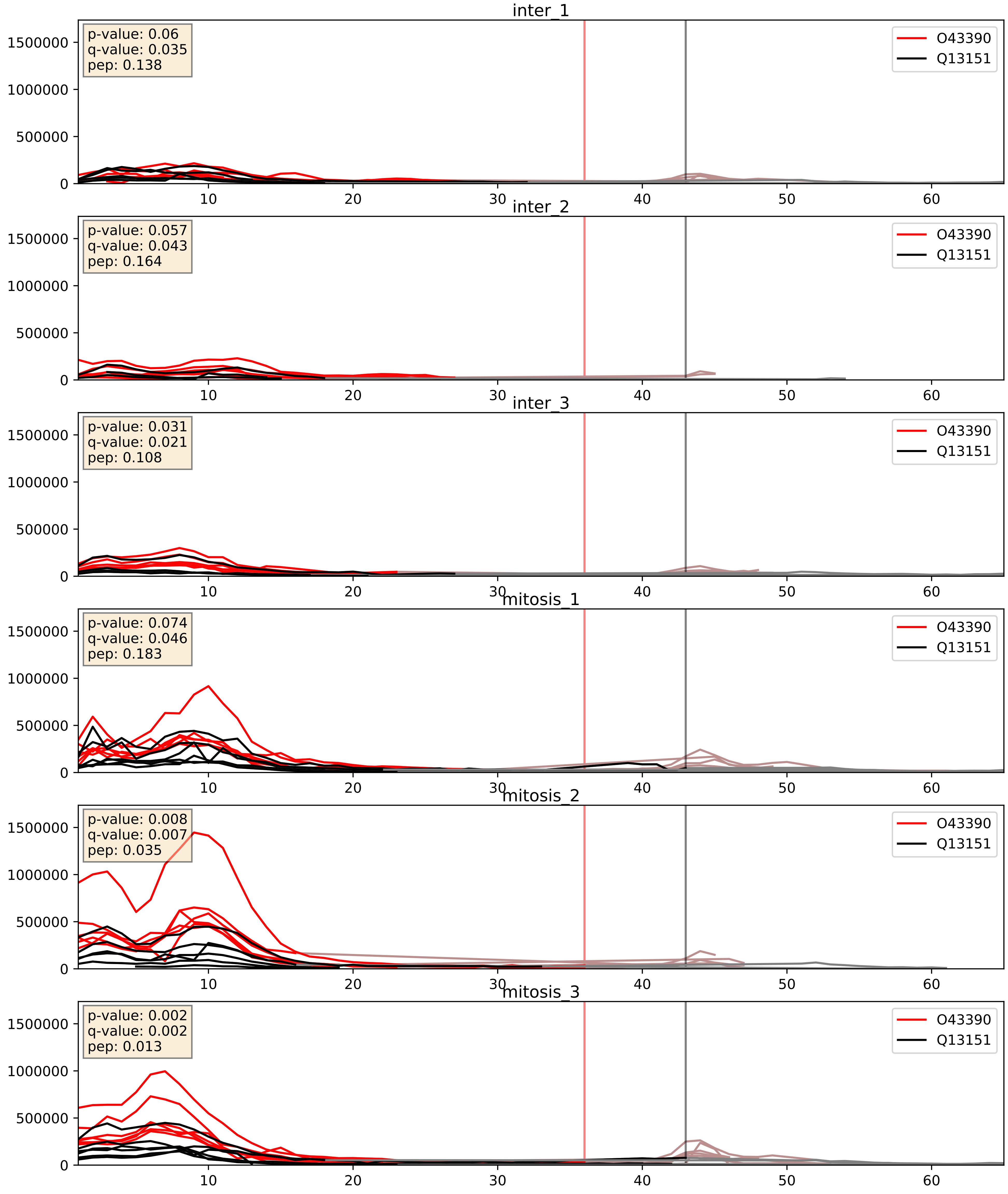


| level | condition_1 | condition_2 | pvalue | pvalue_adjusted |
|---|---|---|---|---|
| interactor_abundance | mitosis | inter | 3.1358423719273224e-05 | 0.004891461372878449 |
| complex_abundance | mitosis | inter | 0.0005214359642555288 | 0.01453706838573521 |
| interactor_ratio | mitosis | inter | 0.9691540655893558 | 0.9937294965826962 |

P07910_Q13151
HNRPC_HUMAN vs ROA0_HUMAN
p-value: 0.048 q-value: 0.032

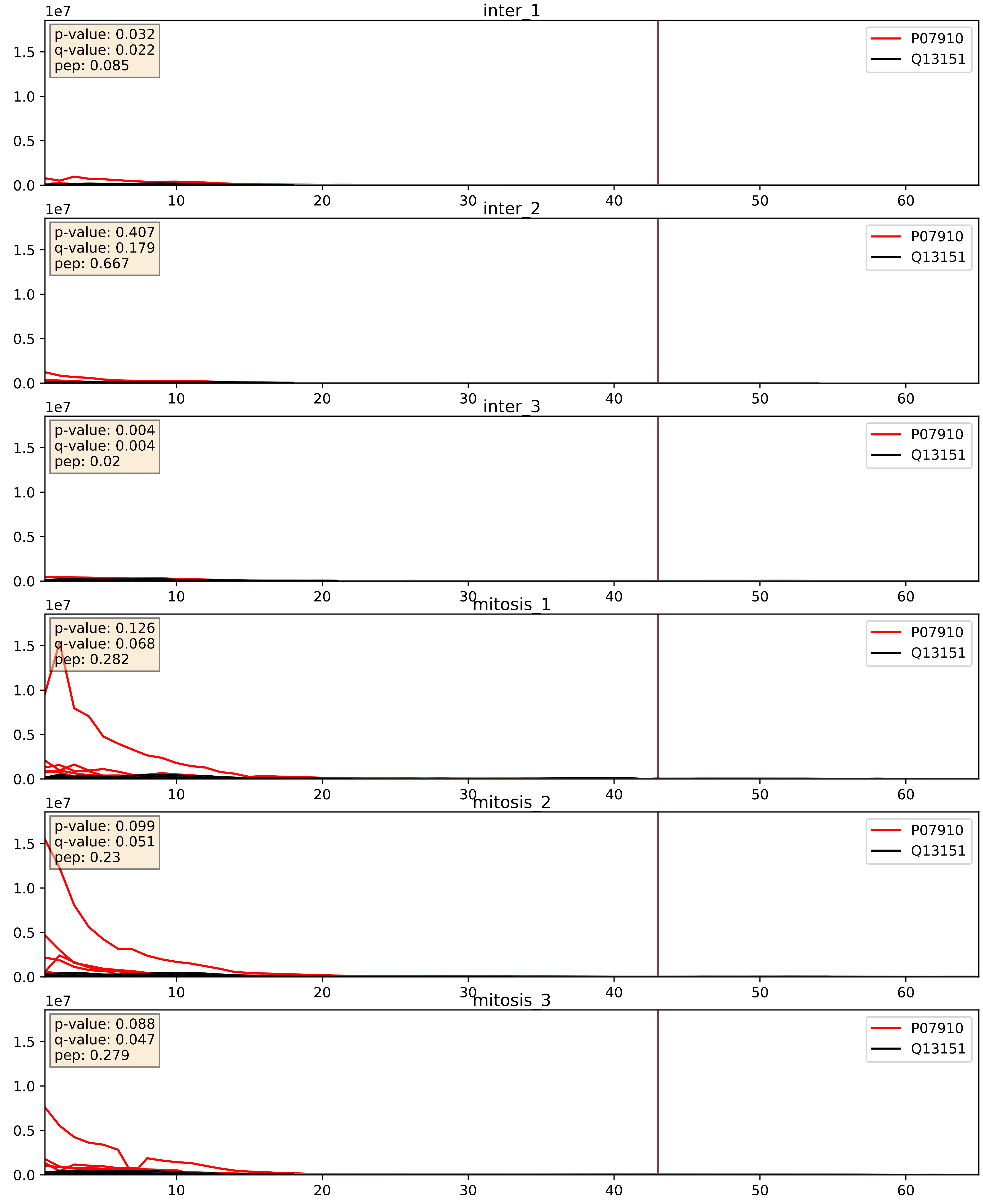


| level | condition_1 | condition_2 | pvalue | pvalue_adjusted |
|---|---|---|---|---|
| interactor_abundance | mitosis | inter | 6.849684494635132e-06 | 0.0033734327106189023 |
| complex_abundance | mitosis | inter | 0.00014564121967907843 | 0.008317961743055323 |
| interactor_ratio | mitosis | inter | 0.0007352419354070836 | 0.03146835483542318 |

P09651_Q13151
ROA1_HUMAN vs ROA0_HUMAN
p-value: 0.019 q-value: 0.015

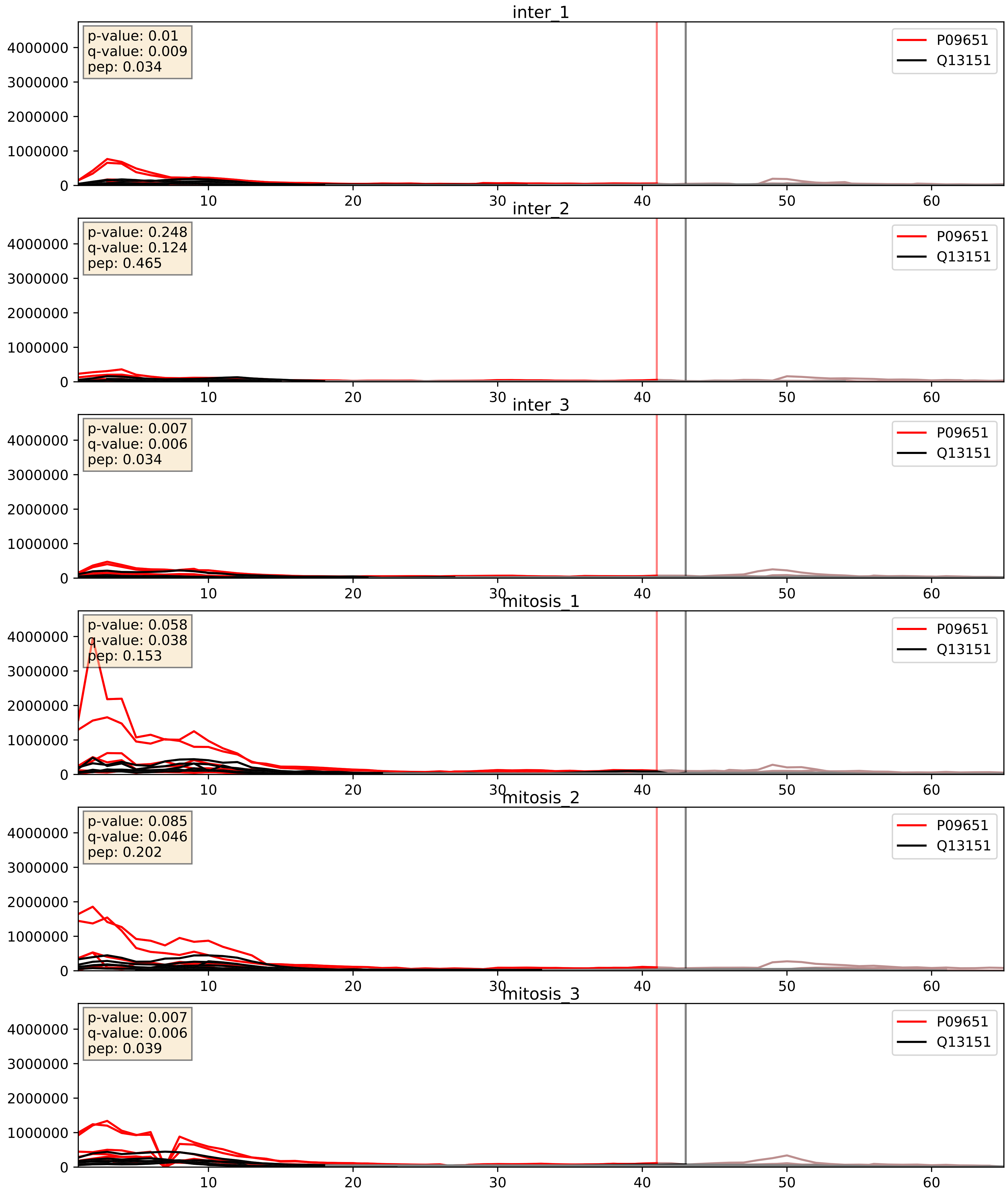


| level | condition_1 | condition_2 | pvalue | pvalue_adjusted |
|---|---|---|---|---|
| complex_abundance | mitosis | inter | 0.0063288454482083225 | 0.05946752693376865 |
| interactor_abundance | mitosis | inter | 0.014789027585019647 | 0.10635646274438217 |
| interactor_ratio | mitosis | inter | 0.3390820858283831 | 0.7924528012231136 |

P14866_Q13151
HNRPL_HUMAN vs ROA0_HUMAN
p-value: 0.014 q-value: 0.012

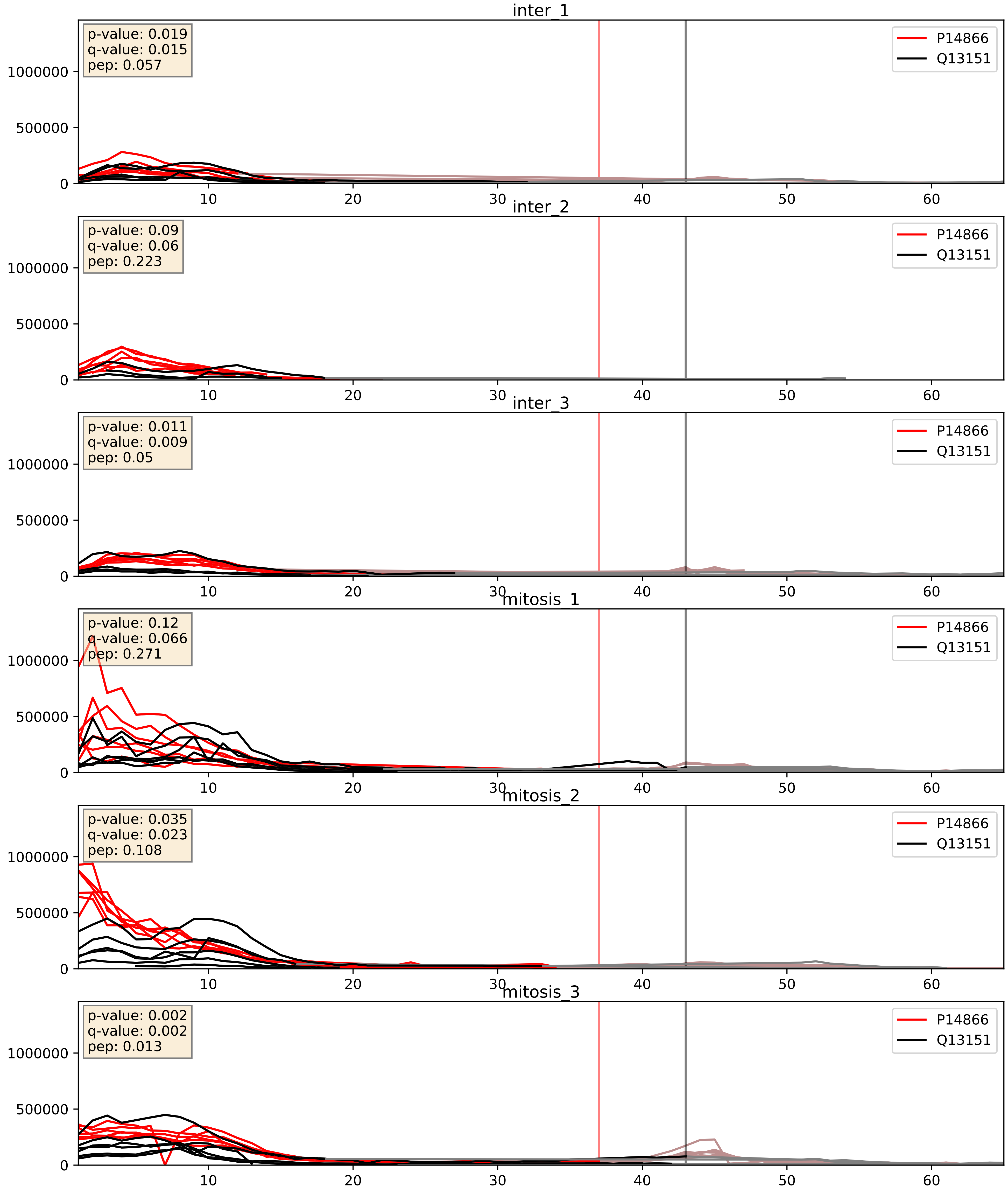


| level | condition_1 | condition_2 | pvalue | pvalue_adjusted |
|---|---|---|---|---|
| complex_abundance | mitosis | inter | 2.5979691148047346e-05 | 0.005691857151652958 |
| interactor_abundance | mitosis | inter | 0.00984142502181702 | 0.08622579138869364 |
| interactor_ratio | mitosis | inter | 0.8500340674765395 | 0.9755634896635457 |

P22626_Q13151
ROA2_HUMAN vs ROA0_HUMAN
p-value: 0.045 q-value: 0.03

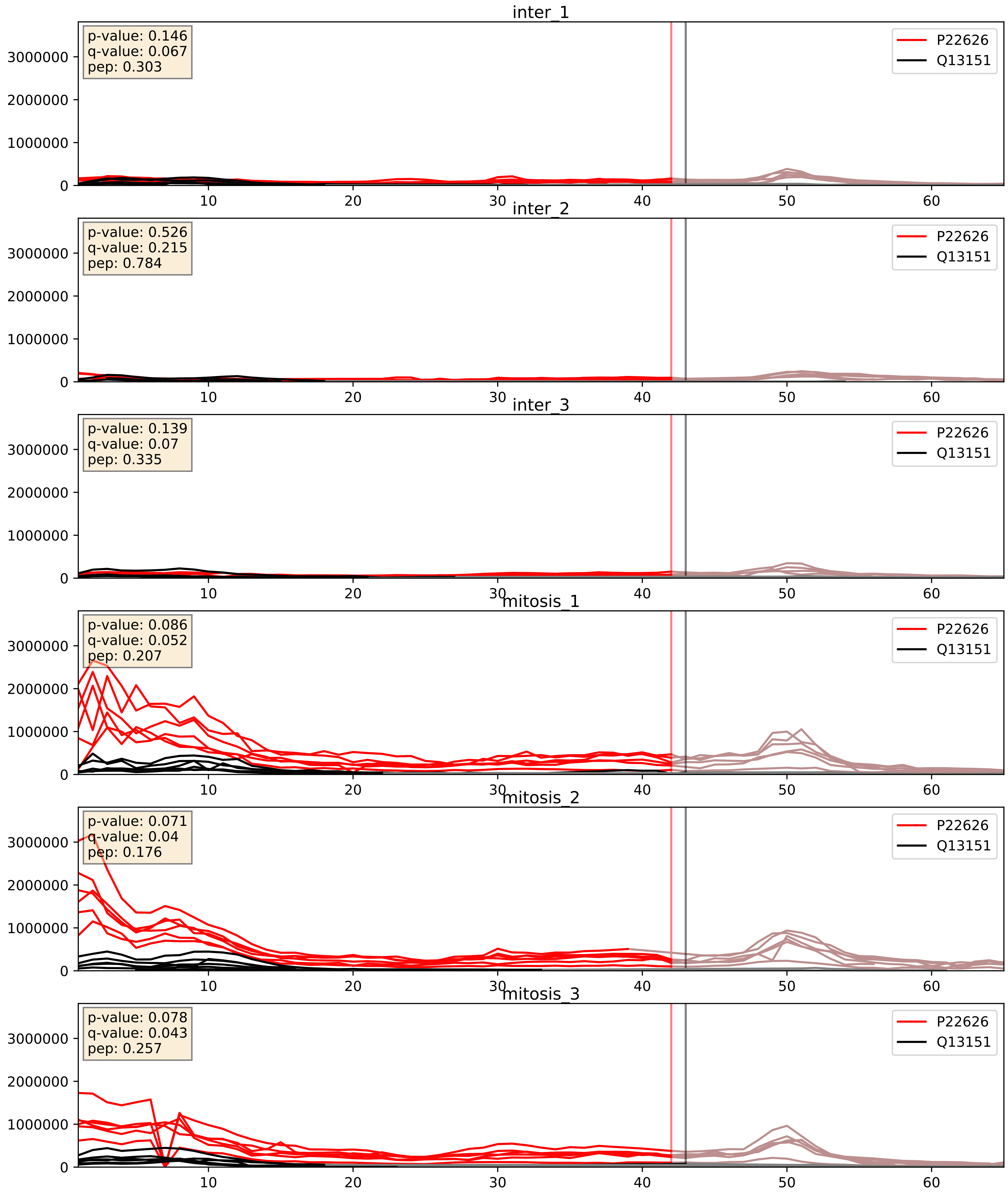


| level | condition_1 | condition_2 | pvalue | pvalue_adjusted |
|---|---|---|---|---|
| interactor_ratio | mitosis | inter | 0.0031237624803896113 | 0.0729098066746082 |
| interactor_abundance | mitosis | inter | 0.0034862800934623243 | 0.04542246210051369 |
| complex_abundance | mitosis | inter | 0.0041541914710077335 | 0.04792436521809461 |

P31943_Q13151
HNRH1_HUMAN vs ROA0_HUMAN
p-value: 0.021 q-value: 0.016

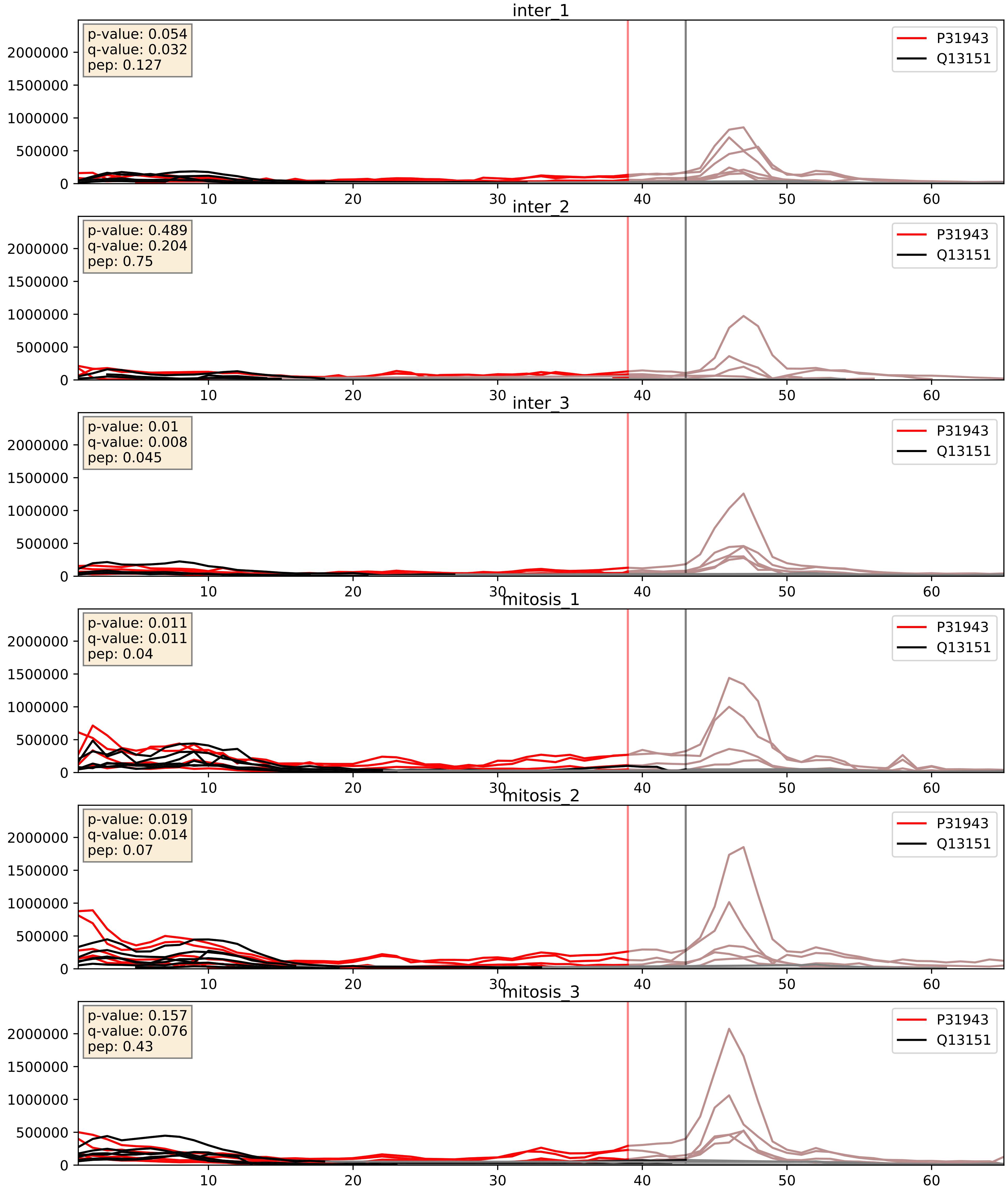


| level | condition_1 | condition_2 | pvalue | pvalue_adjusted |
|---|---|---|---|---|
| complex_abundance | mitosis | inter | 0.002219411769080441 | 0.03446495495307955 |
| interactor_abundance | mitosis | inter | 0.0024193513526238088 | 0.0366218347983374 |
| interactor_ratio | mitosis | inter | 0.413220111473216 | 0.8352217601442099 |

P38159_Q13151
RBMX_HUMAN vs ROA0_HUMAN
p-value: 0.017 q-value: 0.014

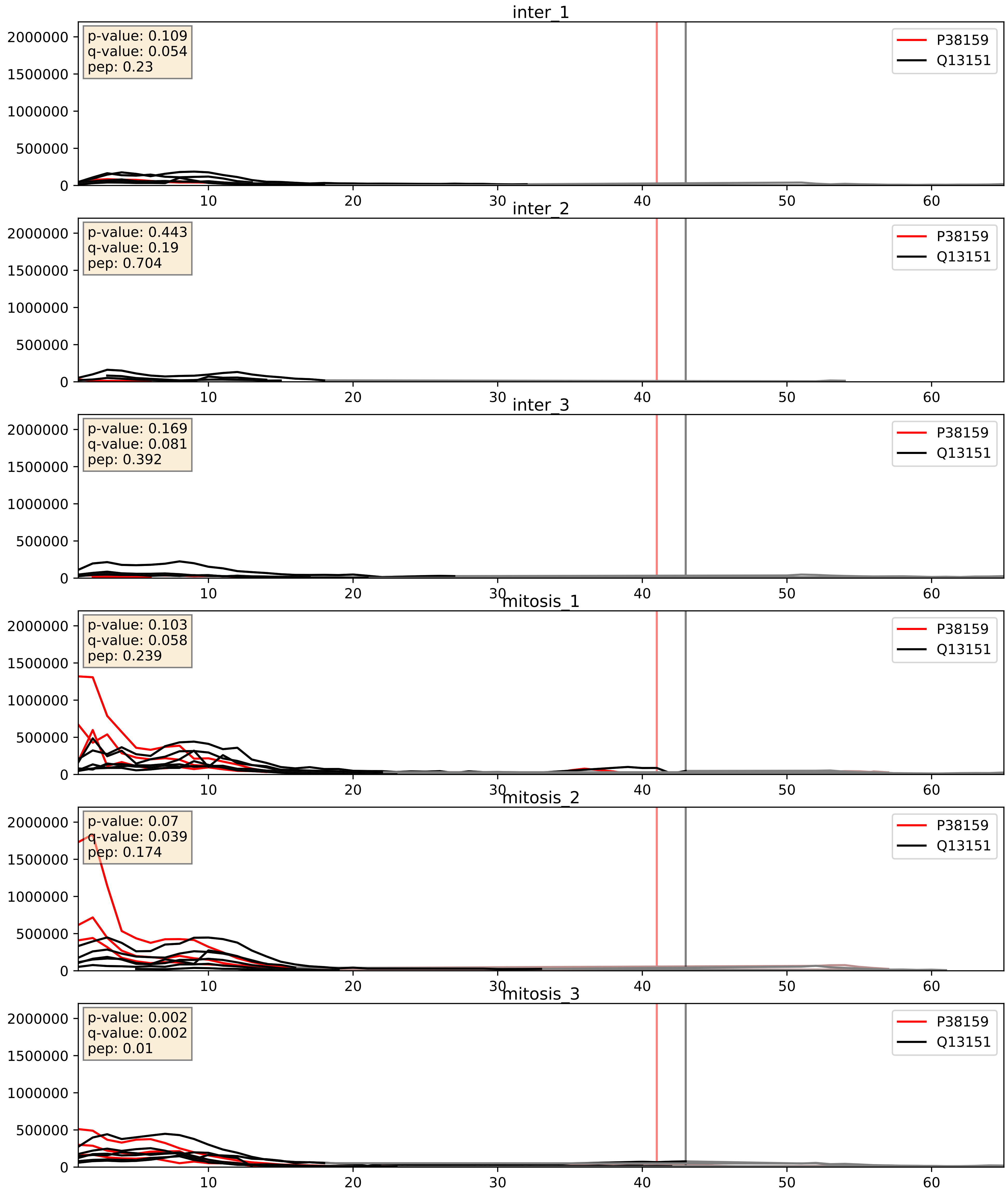


| level | condition_1 | condition_2 | pvalue | pvalue_adjusted |
|---|---|---|---|---|
| interactor_abundance | mitosis | inter | 0.0010950989897773285 | 0.024049720606162197 |
| complex_abundance | mitosis | inter | 0.001785591685100528 | 0.030007039692728107 |
| interactor_ratio | mitosis | inter | 0.02637445981207413 | 0.25543205403291264 |

P51991_Q13151
ROA3_HUMAN vs ROA0_HUMAN
p-value: 0.0 q-value: 0.005

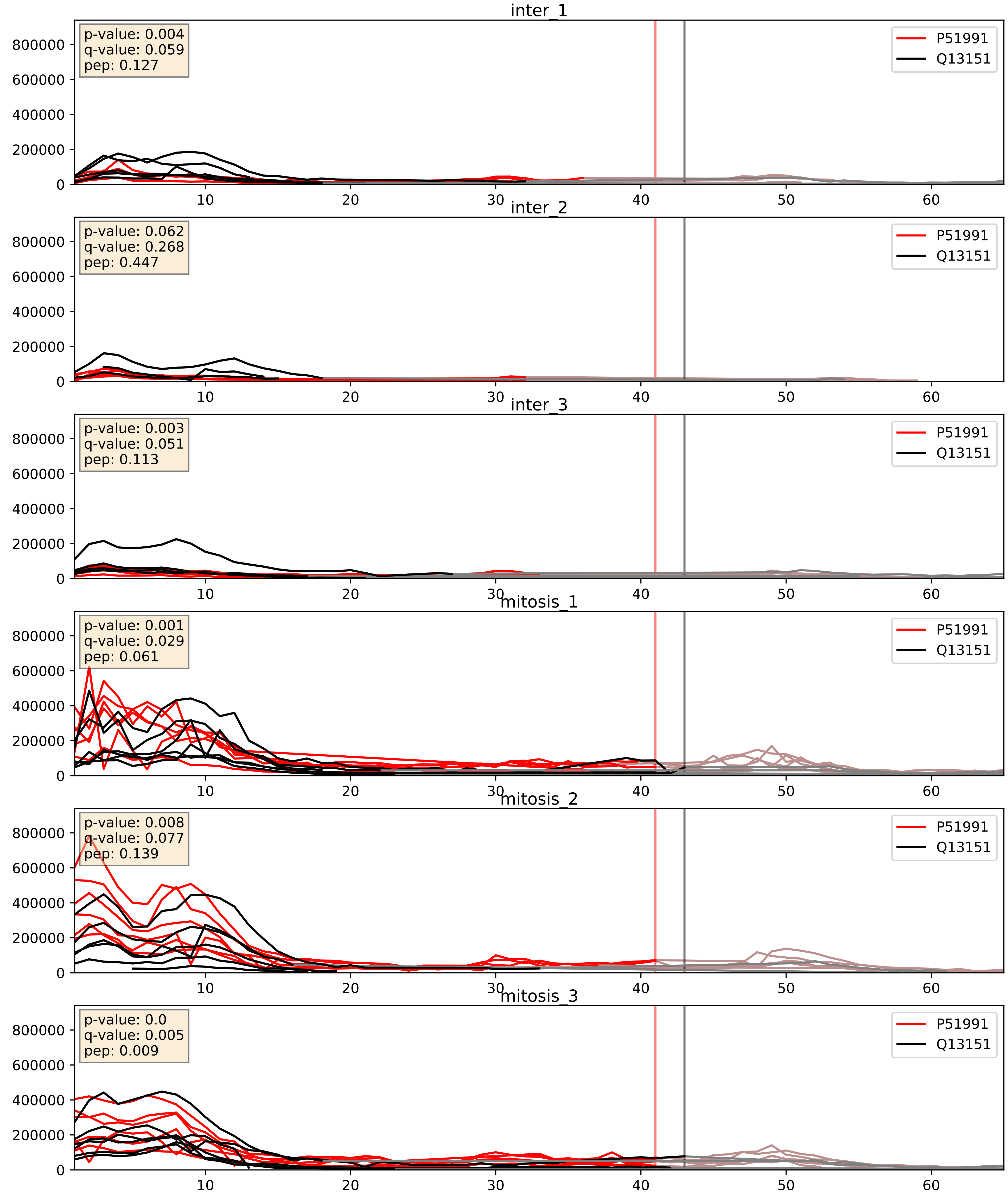


| level | condition_1 | condition_2 | pvalue | pvalue_adjusted |
|---|---|---|---|---|
| interactor_abundance | mitosis | inter | 4.098374668489626e-06 | 0.0033734327106189023 |
| complex_abundance | mitosis | inter | 0.00021831661319998973 | 0.010087086090925927 |
| interactor_ratio | mitosis | inter | 0.014115225863805373 | 0.17445264757370998 |

P52597_Q13151
HNRPF_HUMAN vs ROA0_HUMAN
p-value: 0.005 q-value: 0.005

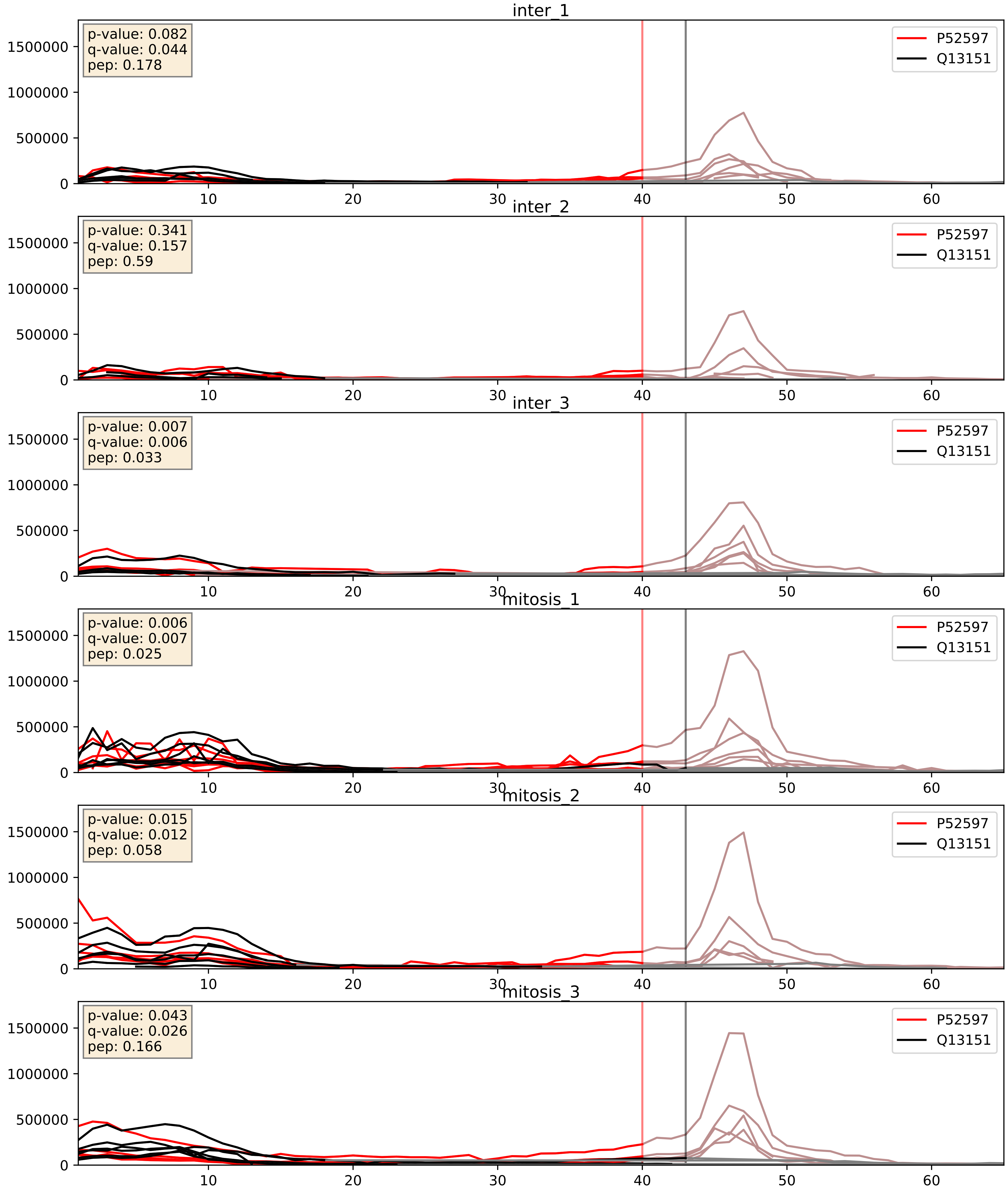


| level | condition_1 | condition_2 | pvalue | pvalue_adjusted |
|---|---|---|---|---|
| complex_abundance | mitosis | inter | 0.001076328295497409 | 0.021833569470040306 |
| interactor_abundance | mitosis | inter | 0.003070826023648689 | 0.042465703978082026 |
| interactor_ratio | mitosis | inter | 0.5732357411651294 | 0.8946257776299471 |

P55769_Q13151
NH2L1_HUMAN vs ROA0_HUMAN
p-value: 0.001 q-value: 0.041

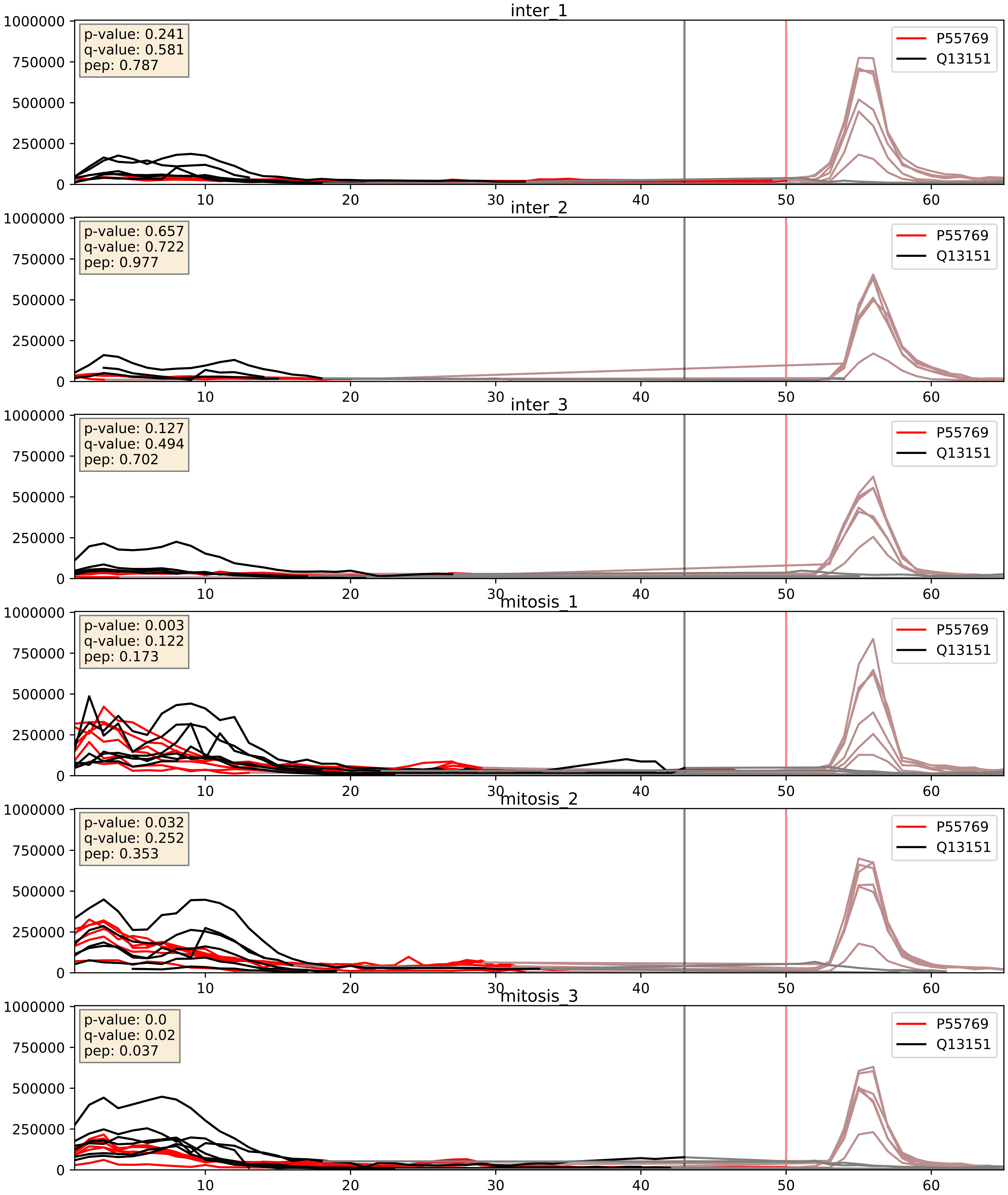


| level | condition_1 | condition_2 | pvalue | pvalue_adjusted |
|---|---|---|---|---|
| interactor_abundance | mitosis | inter | 0.0012966100224587672 | 0.026616263290760302 |
| complex_abundance | mitosis | inter | 0.002280148765337147 | 0.03502731659907978 |
| interactor_ratio | mitosis | inter | 0.06995079943241257 | 0.4380240257069873 |

P62995_Q13151
TRA2B_HUMAN vs ROA0_HUMAN
p-value: 0.005 q-value: 0.005

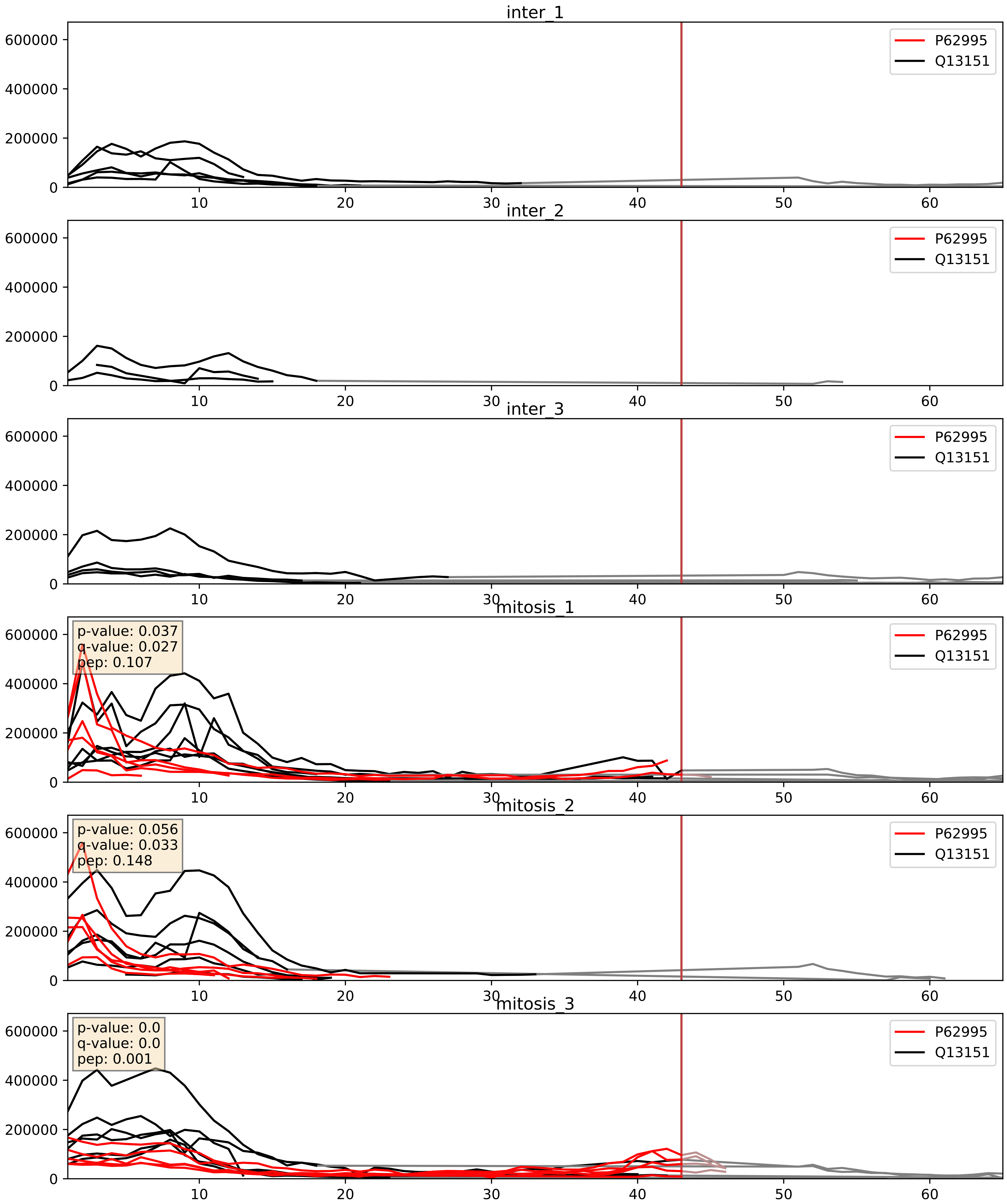


| level | condition_1 | condition_2 | pvalue | pvalue_adjusted |
|---|---|---|---|---|
| complex_abundance | mitosis | inter | 0.00011818315124916703 | 0.007493687219947184 |
| interactor_abundance | mitosis | inter | 0.0004573130702955831 | 0.014951247664582388 |
| interactor_ratio | mitosis | inter | 0.008403991269352195 | 0.13008709812957467 |

P84103_Q13151
SRSF3_HUMAN vs ROA0_HUMAN
p-value: 0.009 q-value: 0.008

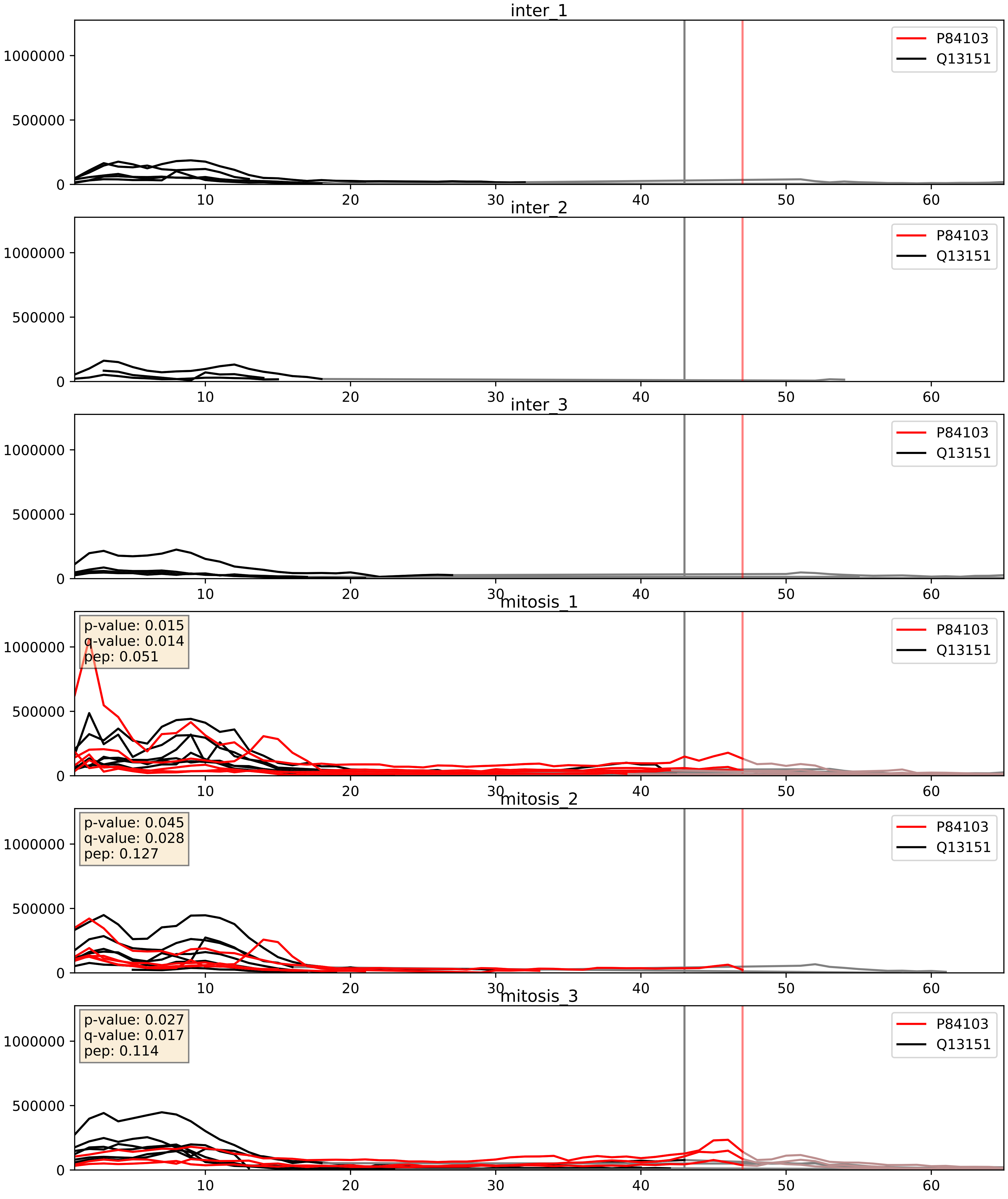


| level | condition_1 | condition_2 | pvalue | pvalue_adjusted |
|---|---|---|---|---|
| complex_abundance | mitosis | inter | 0.0002598167279666148 | 0.010784252527150902 |
| interactor_abundance | mitosis | inter | 0.0026696686527179834 | 0.038732819775027016 |
| interactor_ratio | mitosis | inter | 0.11994976755506524 | 0.5692297679773661 |

Q07955_Q13151
SRSF1_HUMAN vs ROA0_HUMAN
p-value: 0.007 q-value: 0.006

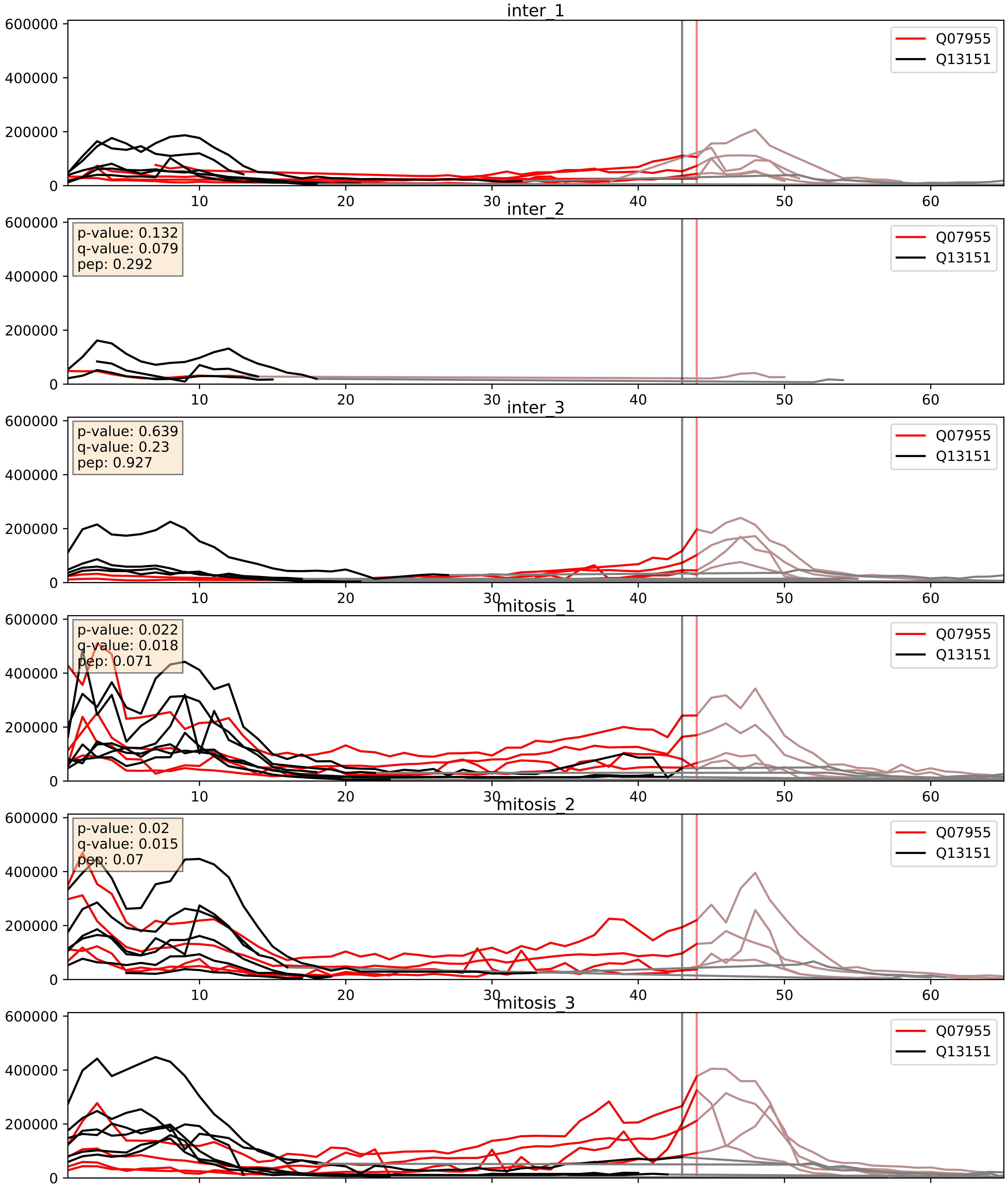


| level | condition_1 | condition_2 | pvalue | pvalue_adjusted |
|---|---|---|---|---|
| complex_abundance | mitosis | inter | 0.007513005302259508 | 0.06444768685385507 |
| interactor_abundance | mitosis | inter | 0.007752548845619896 | 0.07645370750979989 |
| interactor_ratio | mitosis | inter | 0.08841558352285728 | 0.4953575337685953 |

Q13151_Q14103
ROA0_HUMAN vs HNRPD_HUMAN
p-value: 0.076 q-value: 0.046

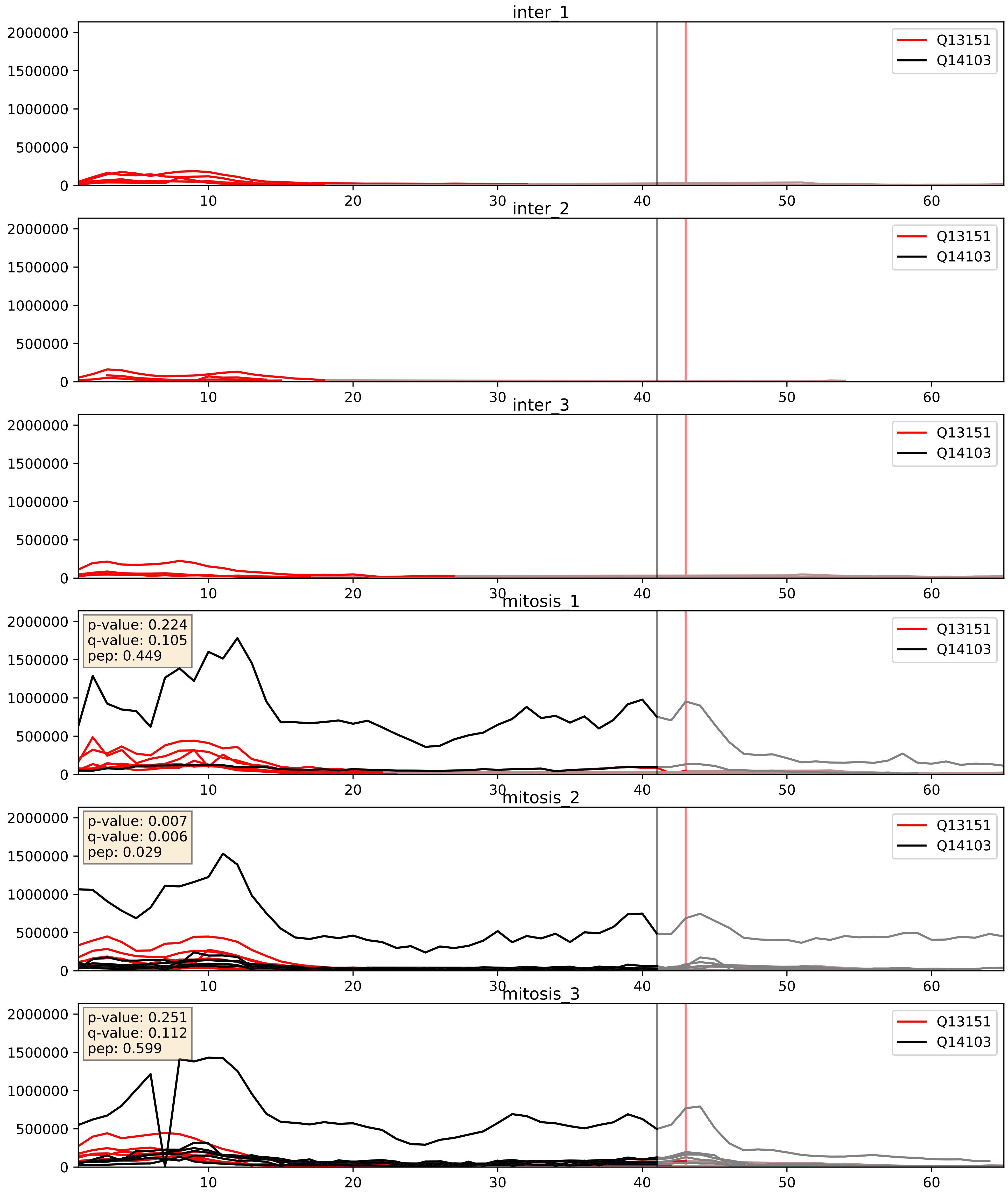


| level | condition_1 | condition_2 | pvalue | pvalue_adjusted |
|---|---|---|---|---|
| interactor_abundance | mitosis | inter | 2.9704194402271474e-05 | 0.0048431981730179775 |
| complex_abundance | mitosis | inter | 0.0009732578966867303 | 0.020686613267949534 |
| interactor_ratio | mitosis | inter | 0.7783913756132336 | 0.9574694891865612 |

Q13151_Q15717
ROA0_HUMAN vs ELAV1_HUMAN
p-value: 0.026 q-value: 0.019

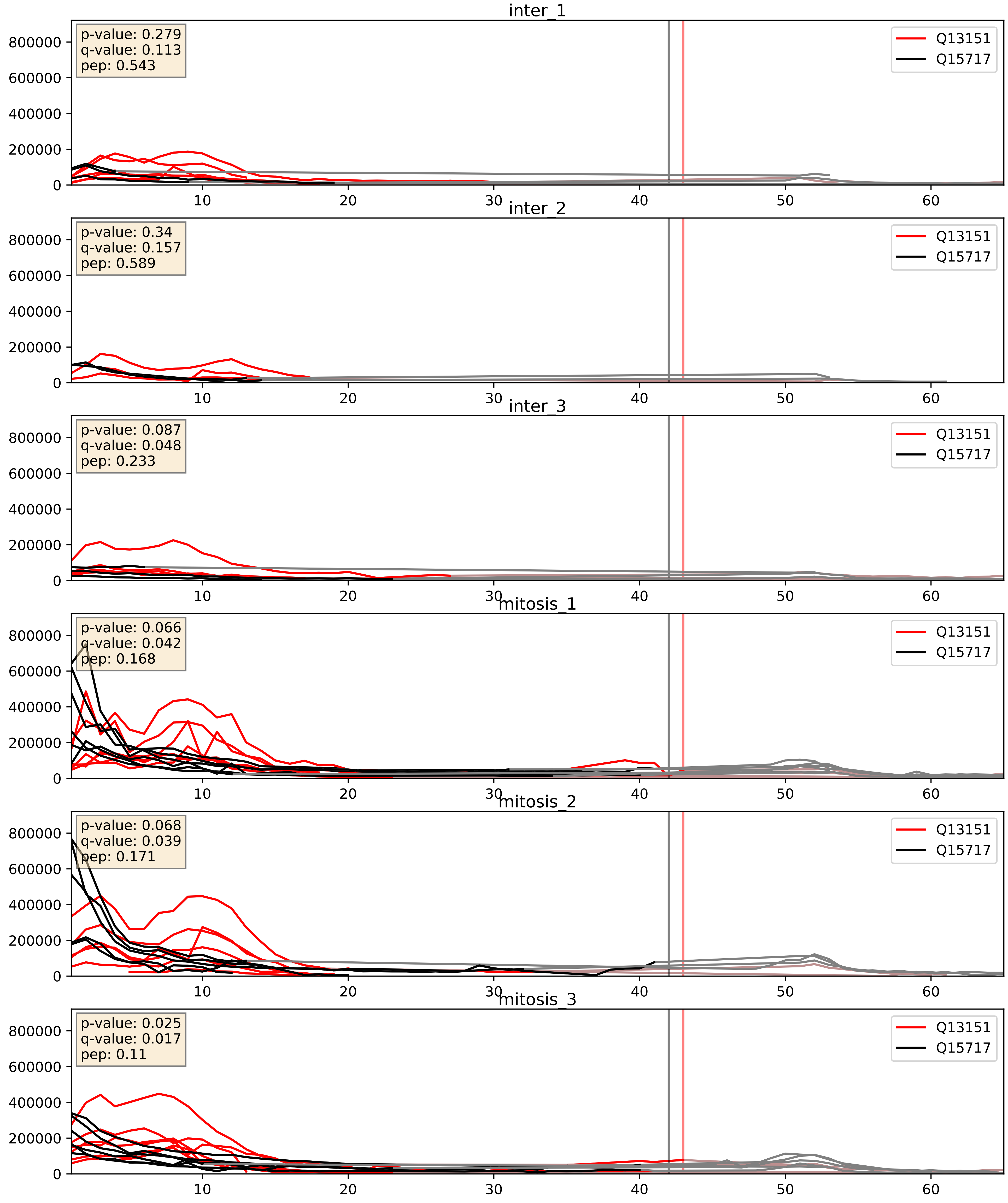


| level | condition_1 | condition_2 | pvalue | pvalue_adjusted |
|---|---|---|---|---|
| complex_abundance | mitosis | inter | 0.0008222134400985702 | 0.01907356923372293 |
| interactor_abundance | mitosis | inter | 0.006871344350993082 | 0.0713363448150053 |
| interactor_ratio | mitosis | inter | 0.009973473593403892 | 0.14263182888175546 |

Q13151_Q9BUJ2
ROA0_HUMAN vs HNRL1_HUMAN
p-value: 0.002 q-value: 0.035

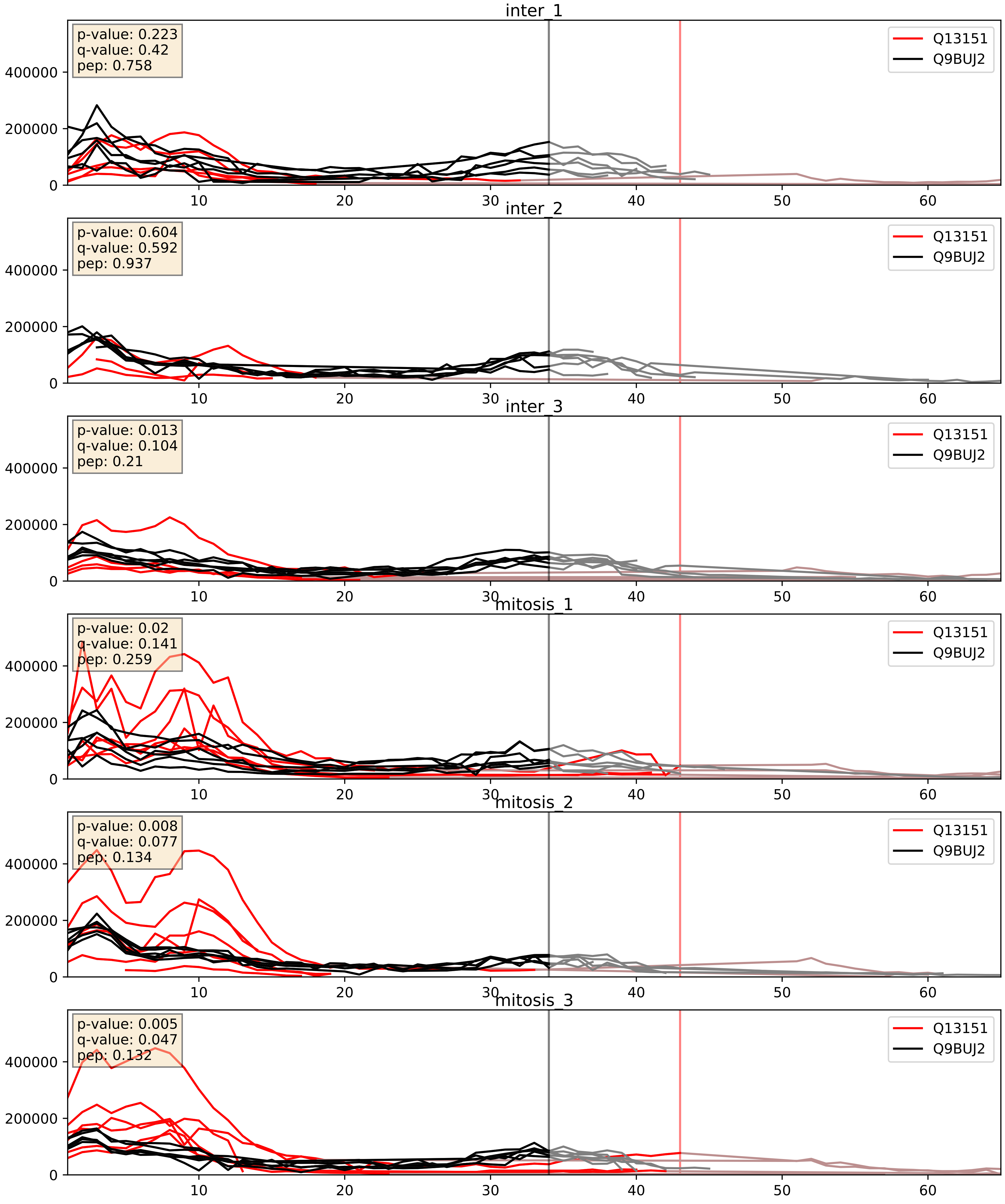


| level | condition_1 | condition_2 | pvalue | pvalue_adjusted |
|---|---|---|---|---|
| interactor_ratio | mitosis | inter | 0.002095252898243131 | 0.05622371413467461 |
| interactor_abundance | mitosis | inter | 0.008471091441998648 | 0.08017189393261118 |
| complex_abundance | mitosis | inter | 0.02812116174369059 | 0.1220004556068812 |
