## Supplementary Data 2 for "SECAT: Quantifying differential protein-protein interaction states by network-centric analysis": hela_string_000077_Q9UNL2.pdf

### Q9UNL2_Q9UNL2

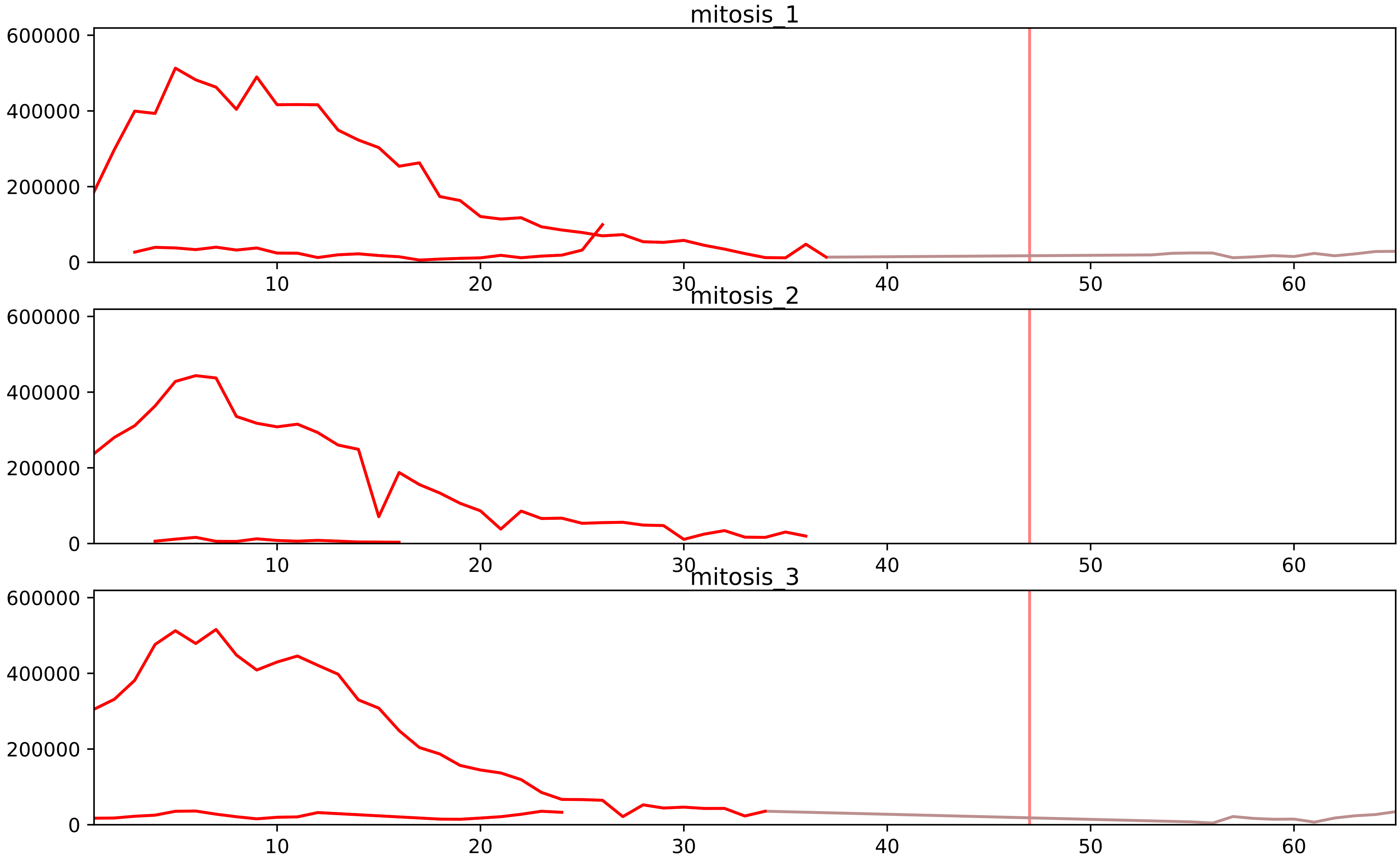


| level | condition_1 | condition_2 | pvalue | pvalue_adjusted |
|---|---|---|---|---|
| complex_abundance | mitosis | inter | 2.107790901493575e-06 | 0.00014916674072108375 |
| interactor_ratio | mitosis | inter | 8.909942089051232e-06 | 0.00020734899693854836 |
| total_abundance | mitosis | inter | 0.00011262652682320049 | 0.012743457225227978 |
| interactor_abundance | mitosis | inter | 0.00014260316316279663 | 0.002948200227185908 |
| assembled_abundance | mitosis | inter | 0.00016610493895286832 | 0.013884593200685297 |
| monomer_abundance | mitosis | inter | 0.0787687166946028 | 0.37600939822766133 |

P39656_Q9UNL2
OST48_HUMAN vs SSRG_HUMAN
p-value: 0.052 q-value: 0.034

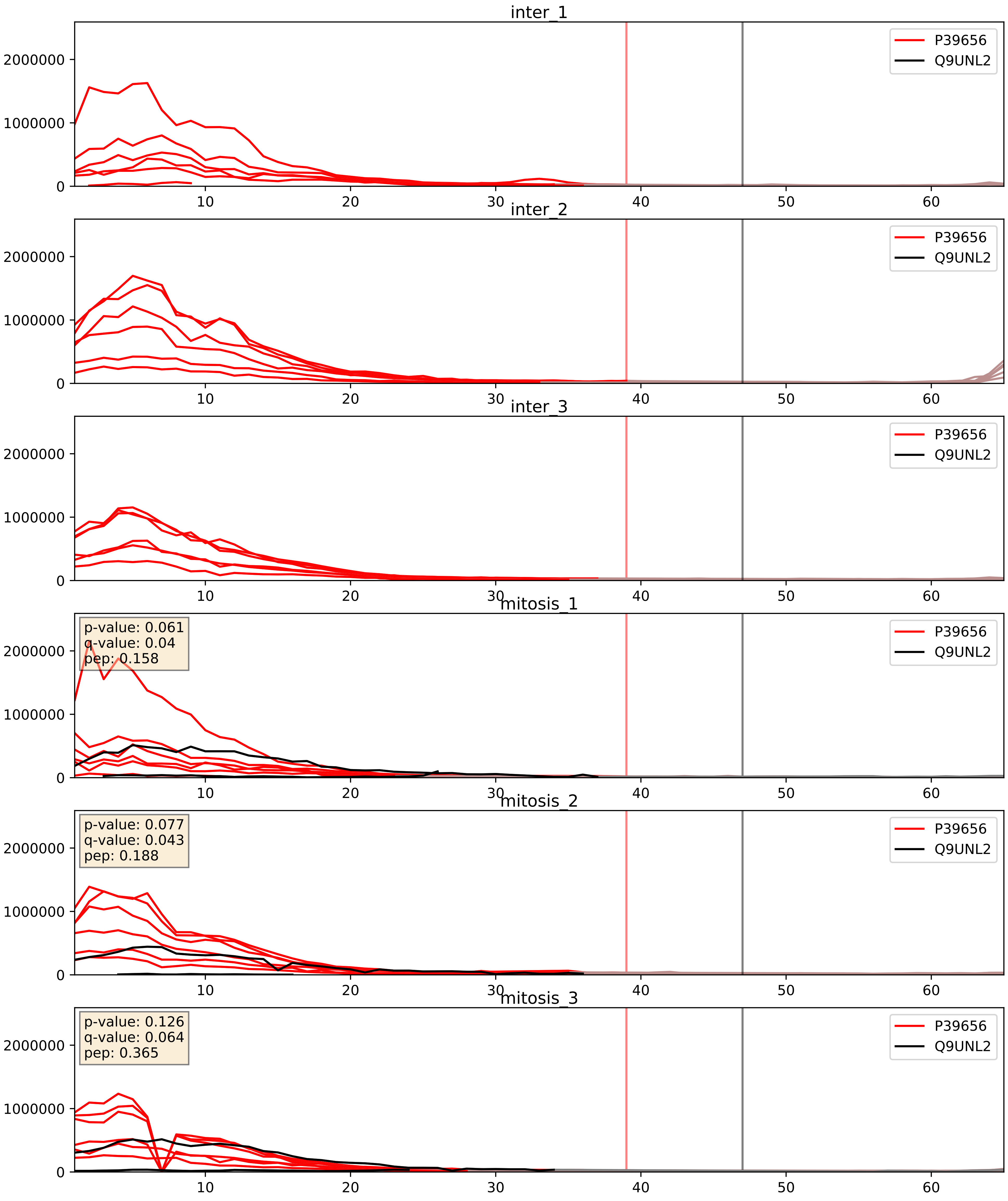


| level | condition_1 | condition_2 | pvalue | pvalue_adjusted |
|---|---|---|---|---|
| interactor_abundance | mitosis | inter | 1.0306611770522181e-05 | 0.0033734327106189023 |
| complex_abundance | mitosis | inter | 4.283591206693643e-05 | 0.005691857151652958 |
| interactor_ratio | mitosis | inter | 0.00020211075575767758 | 0.014683183137172115 |

P43307_Q9UNL2
SSRA_HUMAN vs SSRG_HUMAN
p-value: 0.01 q-value: 0.008

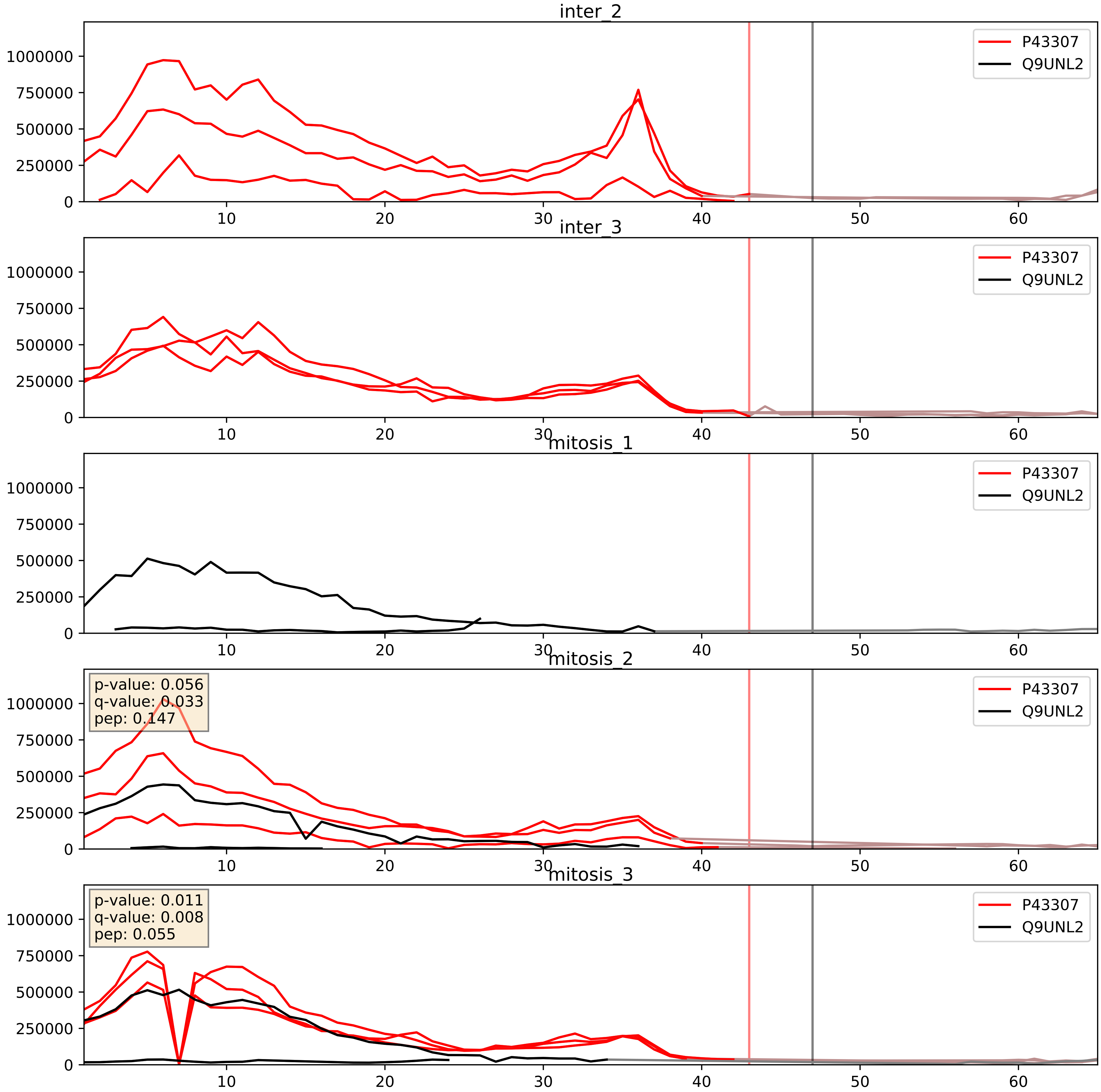


| level | condition_1 | condition_2 | pvalue | pvalue_adjusted |
|---|---|---|---|---|
| interactor_ratio | mitosis | inter | 0.1270531195040438 | 0.5750062399685051 |
| interactor_abundance | mitosis | inter | 0.14087666296675366 | 0.32795872586222774 |
| complex_abundance | mitosis | inter | 0.14883577768785208 | 0.29477886557334887 |

P51571_Q9UNL2
SSRD_HUMAN vs SSRG_HUMAN
p-value: 0.002 q-value: 0.003

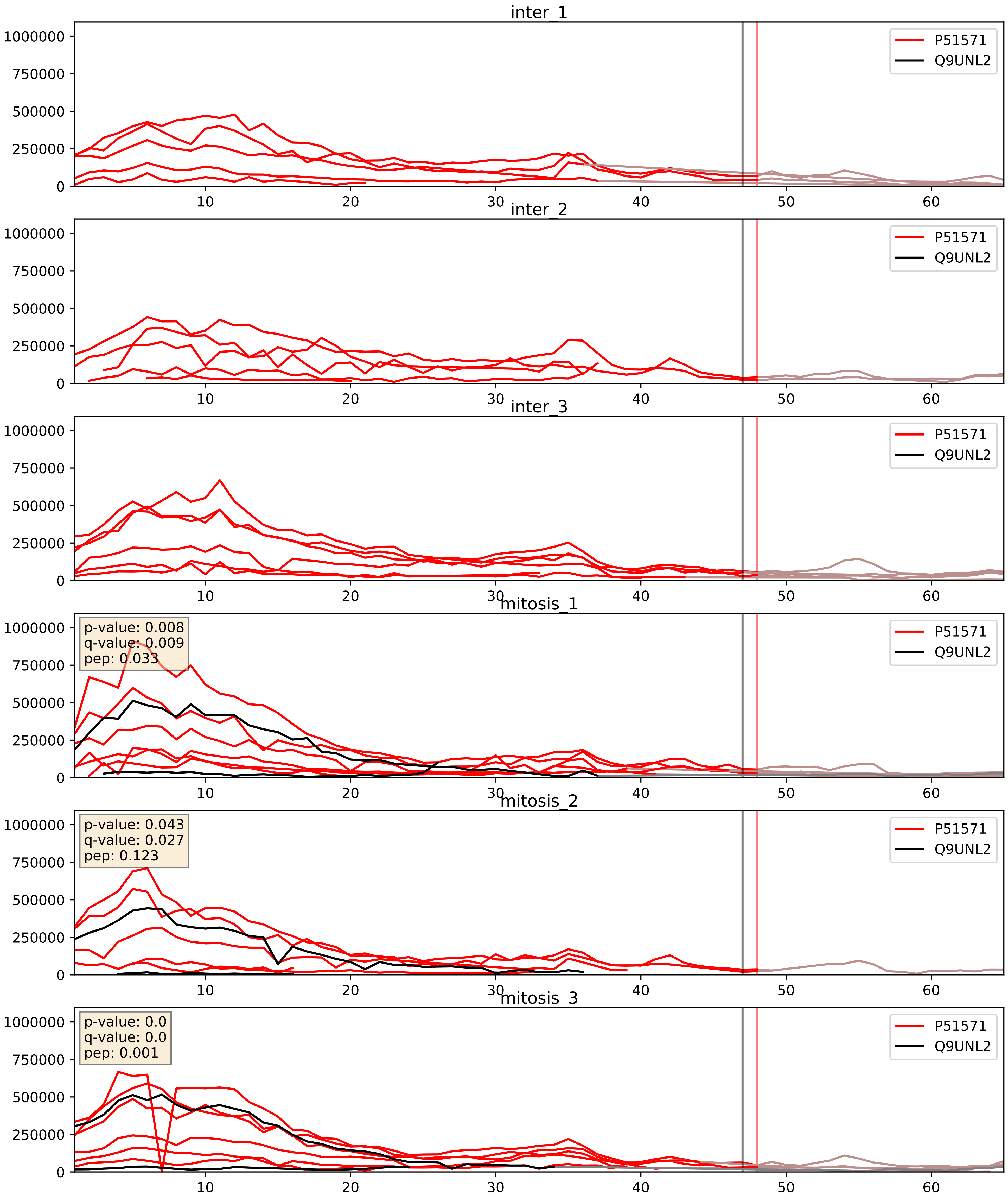


| level | condition_1 | condition_2 | pvalue | pvalue_adjusted |
|---|---|---|---|---|
| interactor_ratio | mitosis | inter | 3.068891572697528e-06 | 0.002520386835947235 |
| interactor_abundance | mitosis | inter | 3.390482672649808e-05 | 0.005164930486161031 |
| complex_abundance | mitosis | inter | 0.00016706521830134896 | 0.00871998944304602 |

P60468_Q9UNL2
SC61B_HUMAN vs SSRG_HUMAN
p-value: 0.003 q-value: 0.003

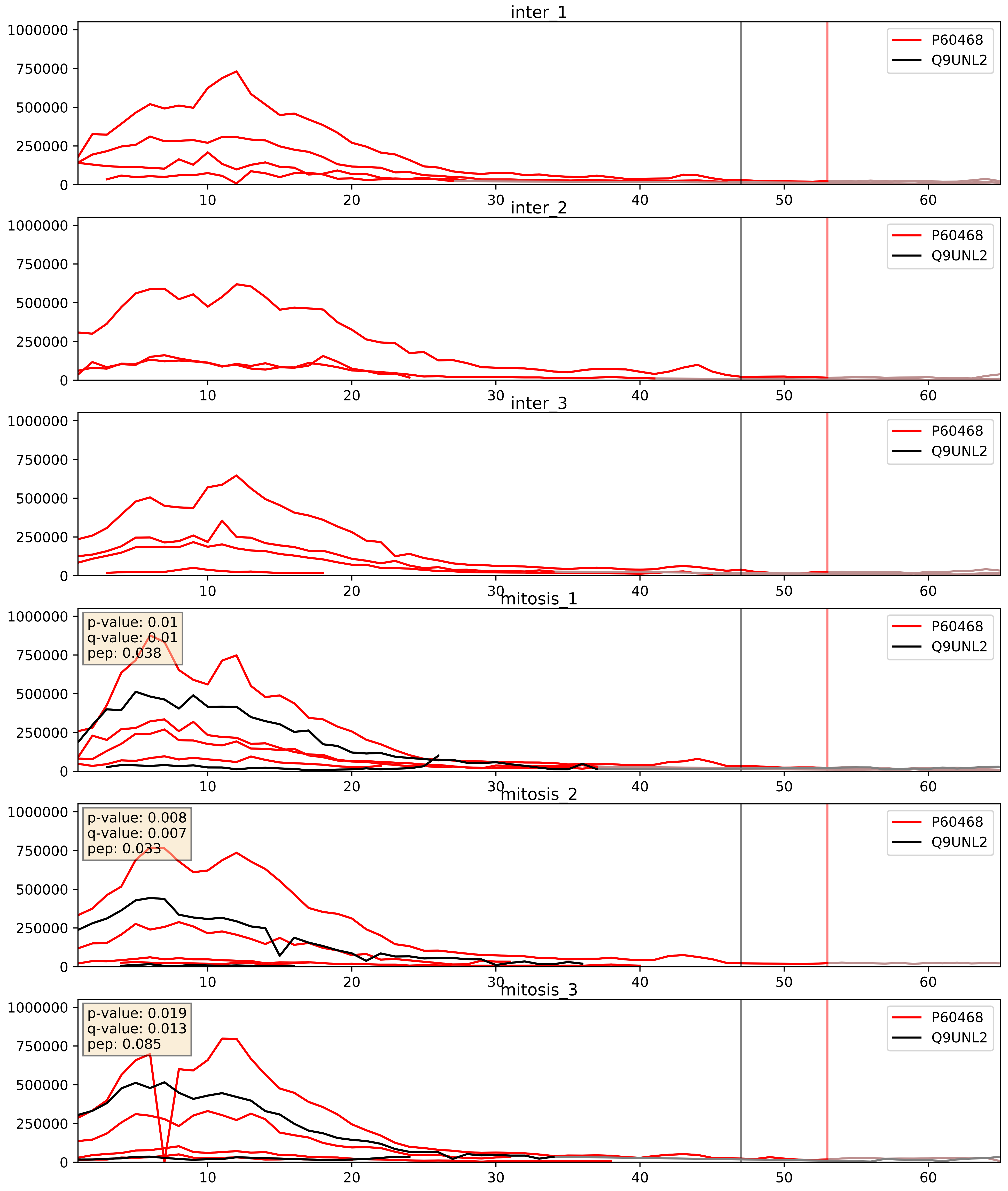


| level | condition_1 | condition_2 | pvalue | pvalue_adjusted |
|---|---|---|---|---|
| interactor_ratio | mitosis | inter | 4.323183886814836e-05 | 0.006796360849516482 |
| interactor_abundance | mitosis | inter | 0.0002380601724361412 | 0.011847645791007956 |
| complex_abundance | mitosis | inter | 0.0005101929279087853 | 0.014318857255407222 |

P61619_Q9UNL2
S61A1_HUMAN vs SSRG_HUMAN
p-value: 0.027 q-value: 0.02

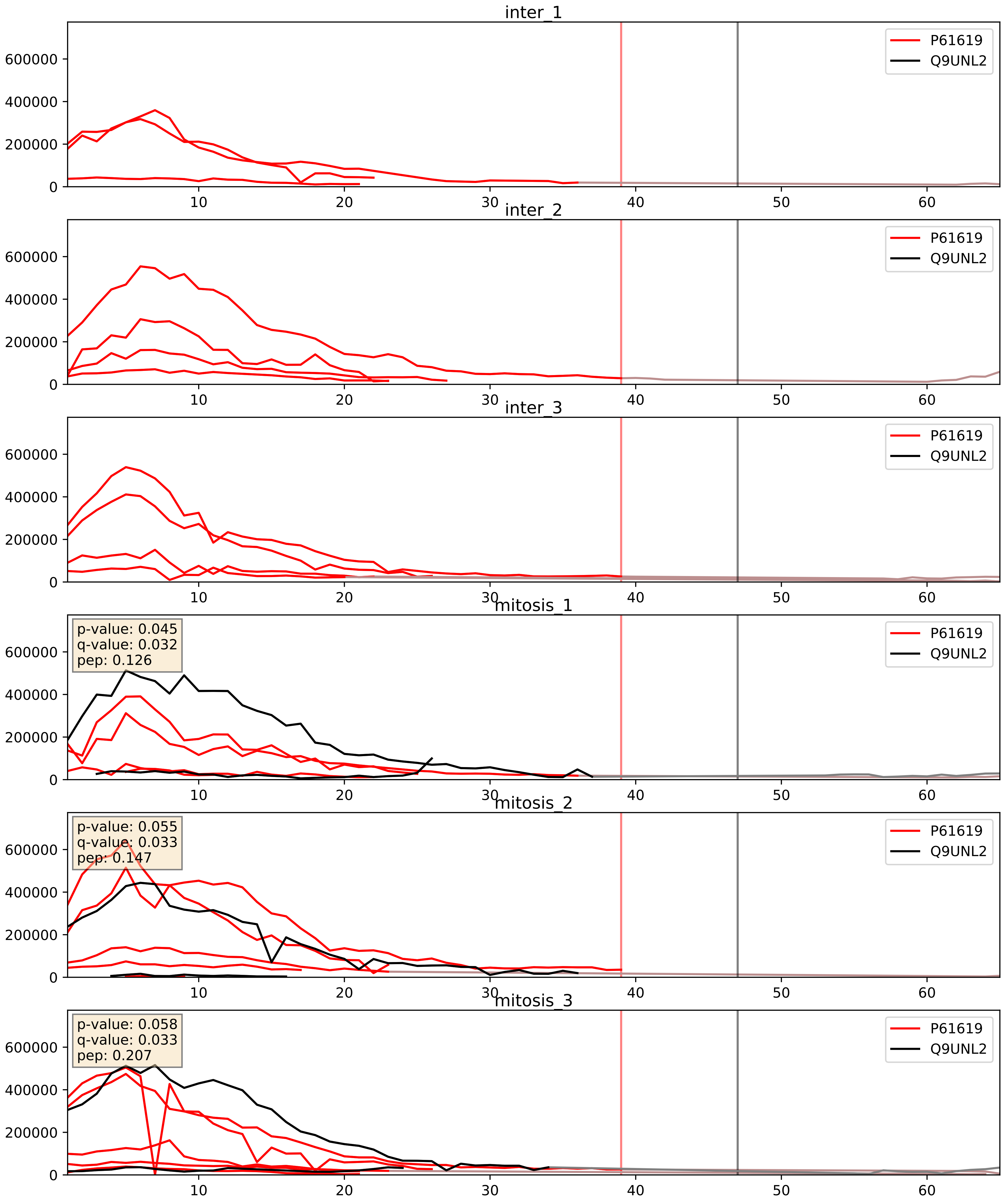


| level | condition_1 | condition_2 | pvalue | pvalue_adjusted |
|---|---|---|---|---|
| complex_abundance | mitosis | inter | 6.590674530976401e-05 | 0.005691857151652958 |
| interactor_abundance | mitosis | inter | 0.00017352230478394034 | 0.01006849049698354 |
| interactor_ratio | mitosis | inter | 0.012973195949404931 | 0.16452159382904033 |

P67812_Q9UNL2
SC11A_HUMAN vs SSRG_HUMAN
p-value: 0.002 q-value: 0.038

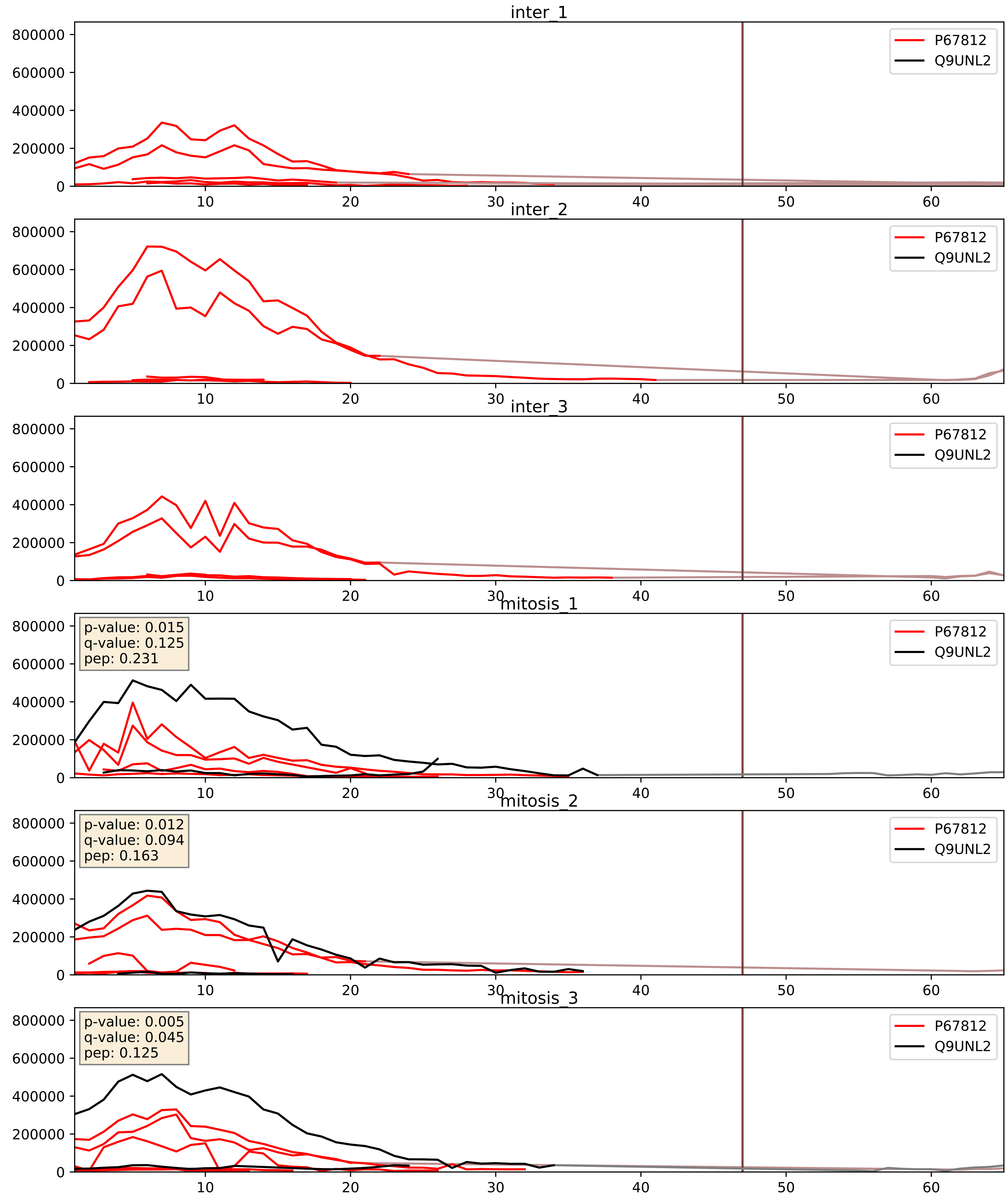


| level | condition_1 | condition_2 | pvalue | pvalue_adjusted |
|---|---|---|---|---|
| interactor_abundance | mitosis | inter | 0.00020875019033178539 | 0.010866247963489511 |
| complex_abundance | mitosis | inter | 0.00027778486441133314 | 0.011216219053589679 |
| interactor_ratio | mitosis | inter | 0.010642534290059028 | 0.14788976221250855 |

Q15005_Q9UNL2
SPCS2_HUMAN vs SSRG_HUMAN
p-value: 0.002 q-value: 0.003

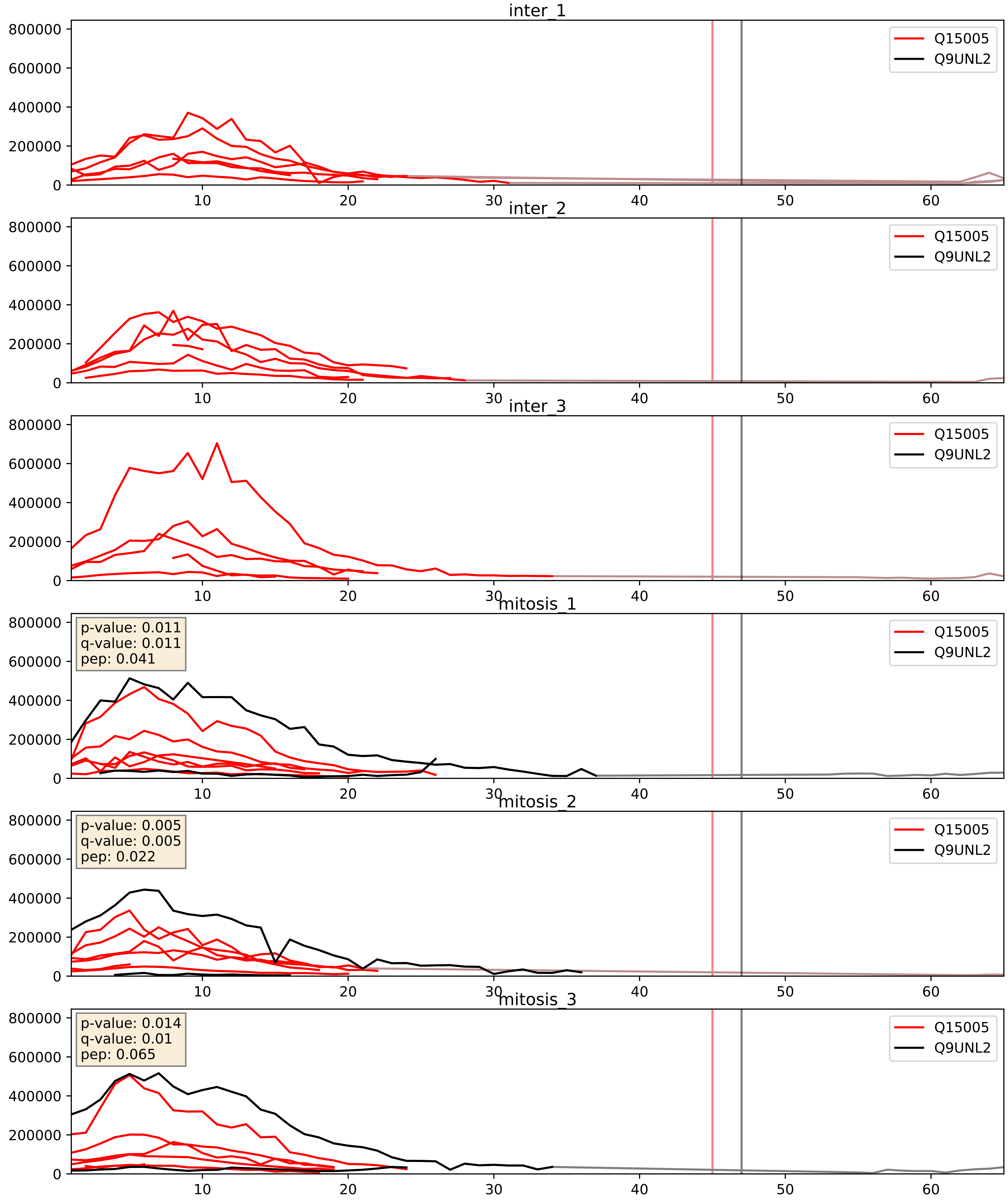


| level | condition_1 | condition_2 | pvalue | pvalue_adjusted |
|---|---|---|---|---|
| interactor_abundance | mitosis | inter | 4.11755258236053e-05 | 0.005825826463637378 |
| complex_abundance | mitosis | inter | 0.00023041379763452546 | 0.010219389159334393 |
| interactor_ratio | mitosis | inter | 0.0006764667961609915 | 0.029848225645041686 |
