## Supplementary Data 2 for "SECAT: Quantifying differential protein-protein interaction states by network-centric analysis": hela_string_000078_P31350.pdf

P31350_P31350

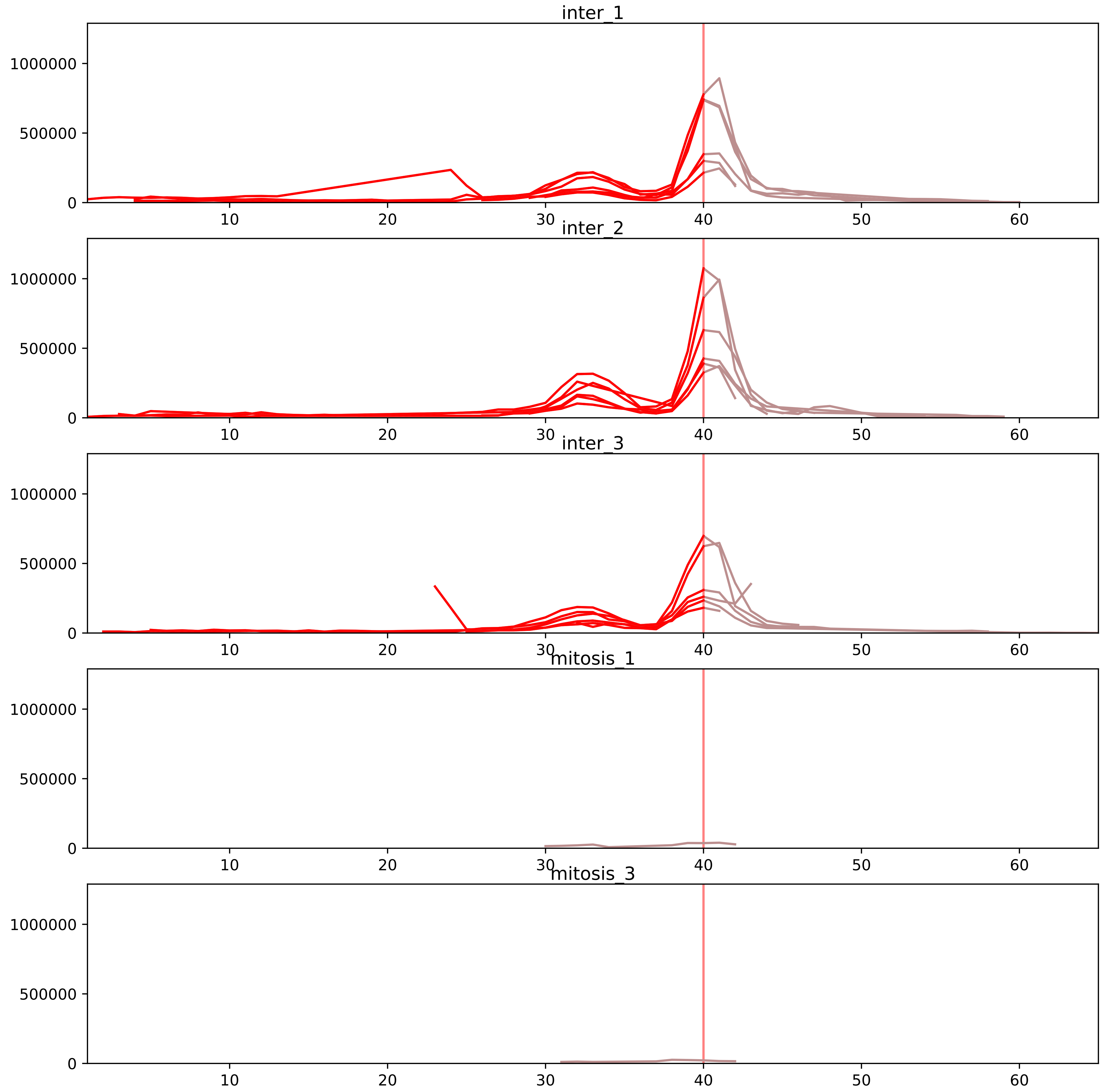


| level | condition_1 | condition_2 | pvalue | pvalue_adjusted |
|---|---|---|---|---|
| assembled_abundance | mitosis | inter | 2.2502033865927023e-06 | 0.0023535084180592012 |
| total_abundance | mitosis | inter | 5.3556651471354305e-06 | 0.003540864511957668 |
| monomer_abundance | mitosis | inter | 5.262744504234152e-05 | 0.0267281636508792 |
| complex_abundance | mitosis | inter | 5.655065103525419e-05 | 0.0016008184293056572 |
| interactor_abundance | mitosis | inter | 7.444159821538425e-05 | 0.0018763361741959866 |
| interactor_ratio | mitosis | inter | 0.11337985380266258 | 0.3535914084693206 |

P06493_P31350
CDK1_HUMAN vs RIR2_HUMAN
p-value: 0.039 q-value: 0.027

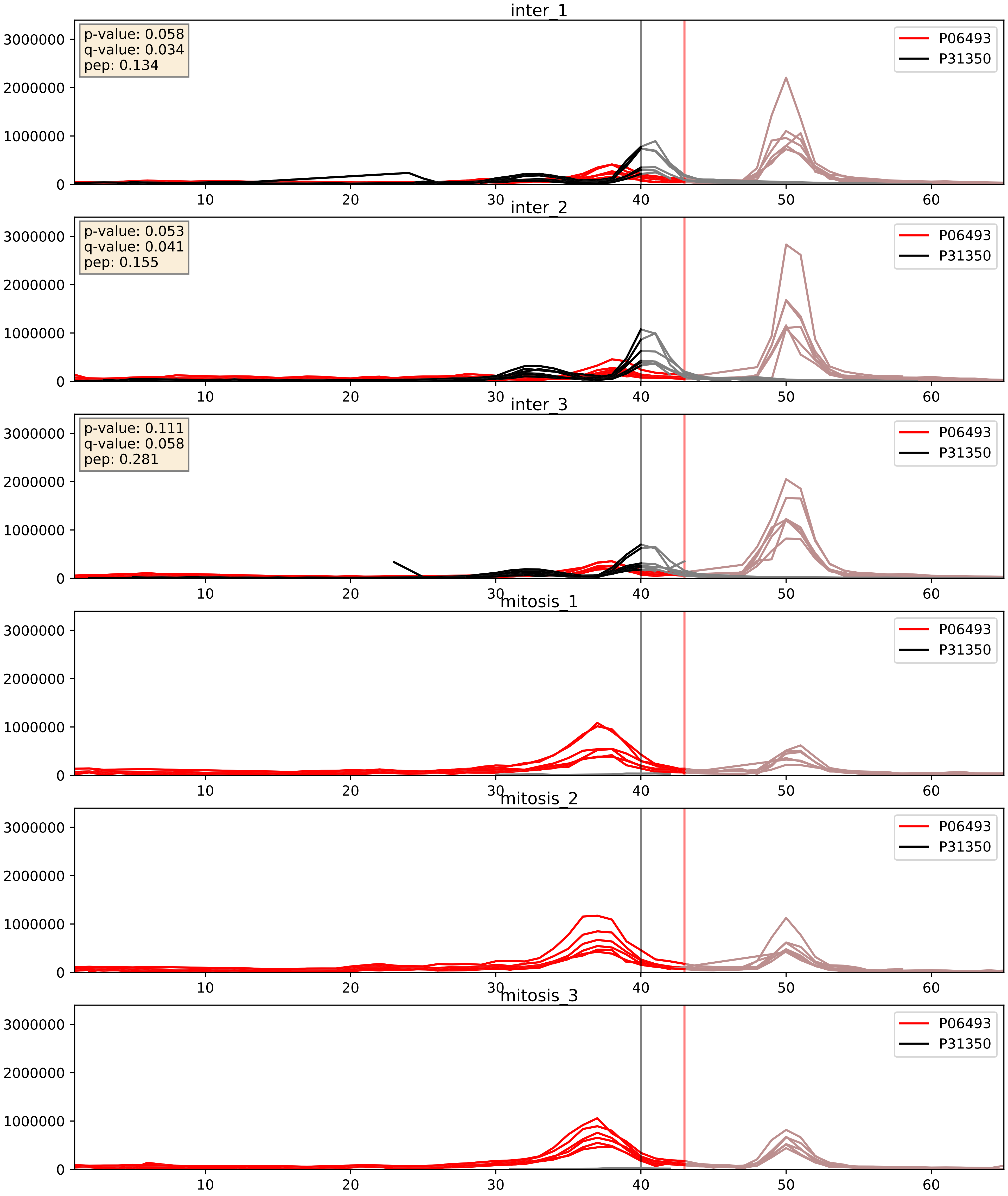


| level | condition_1 | condition_2 | pvalue | pvalue_adjusted |
|---|---|---|---|---|
| complex_abundance | mitosis | inter | 5.655065103525419e-05 | 0.005691857151652958 |
| interactor_abundance | mitosis | inter | 8.436905755951776e-05 | 0.007227629947416683 |
| interactor_ratio | mitosis | inter | 0.11337985380266258 | 0.553985317646331 |
