## Supplementary Data 2 for "SECAT: Quantifying differential protein-protein interaction states by network-centric analysis": hela_string_000079_Q92979.pdf

Q92979_Q92979

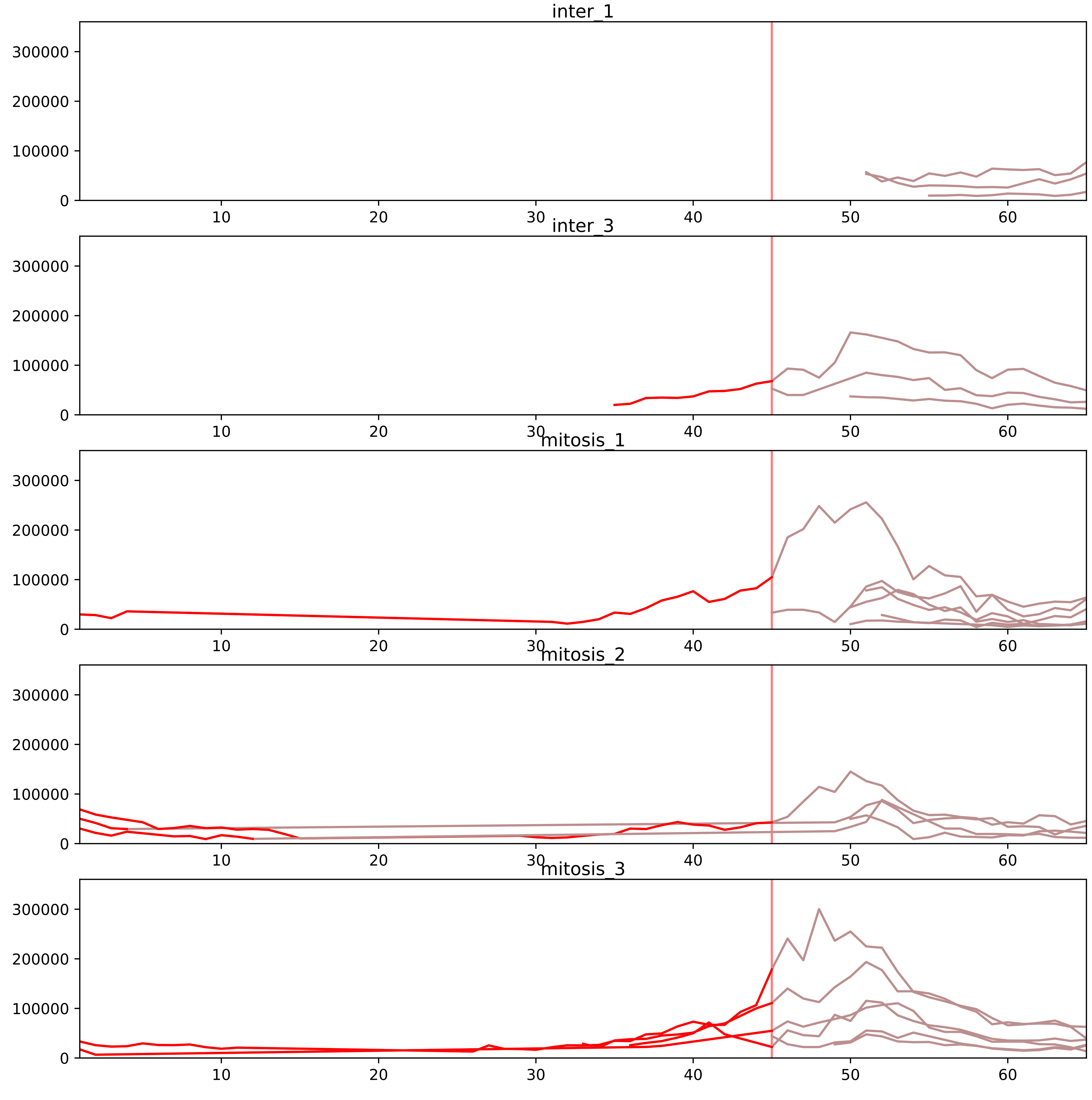


| level | condition_1 | condition_2 | pvalue | pvalue_adjusted |
|---|---|---|---|---|
| interactor_ratio | mitosis | inter | 3.410060089354032e-06 | 8.96358652058774e-05 |
| complex_abundance | mitosis | inter | 0.002367094680566051 | 0.01868585139522817 |
| monomer_abundance | mitosis | inter | 0.033937249516573814 | 0.30564884234321027 |
| total_abundance | mitosis | inter | 0.04291260230984743 | 0.25874247110331355 |
| assembled_abundance | mitosis | inter | 0.06903614956737186 | 0.31192878004330854 |
| interactor_abundance | mitosis | inter | 0.5878029340877351 | 0.7301006101642894 |

O00567_Q92979
NOP56_HUMAN vs NEP1_HUMAN
p-value: 0.068 q-value: 0.042

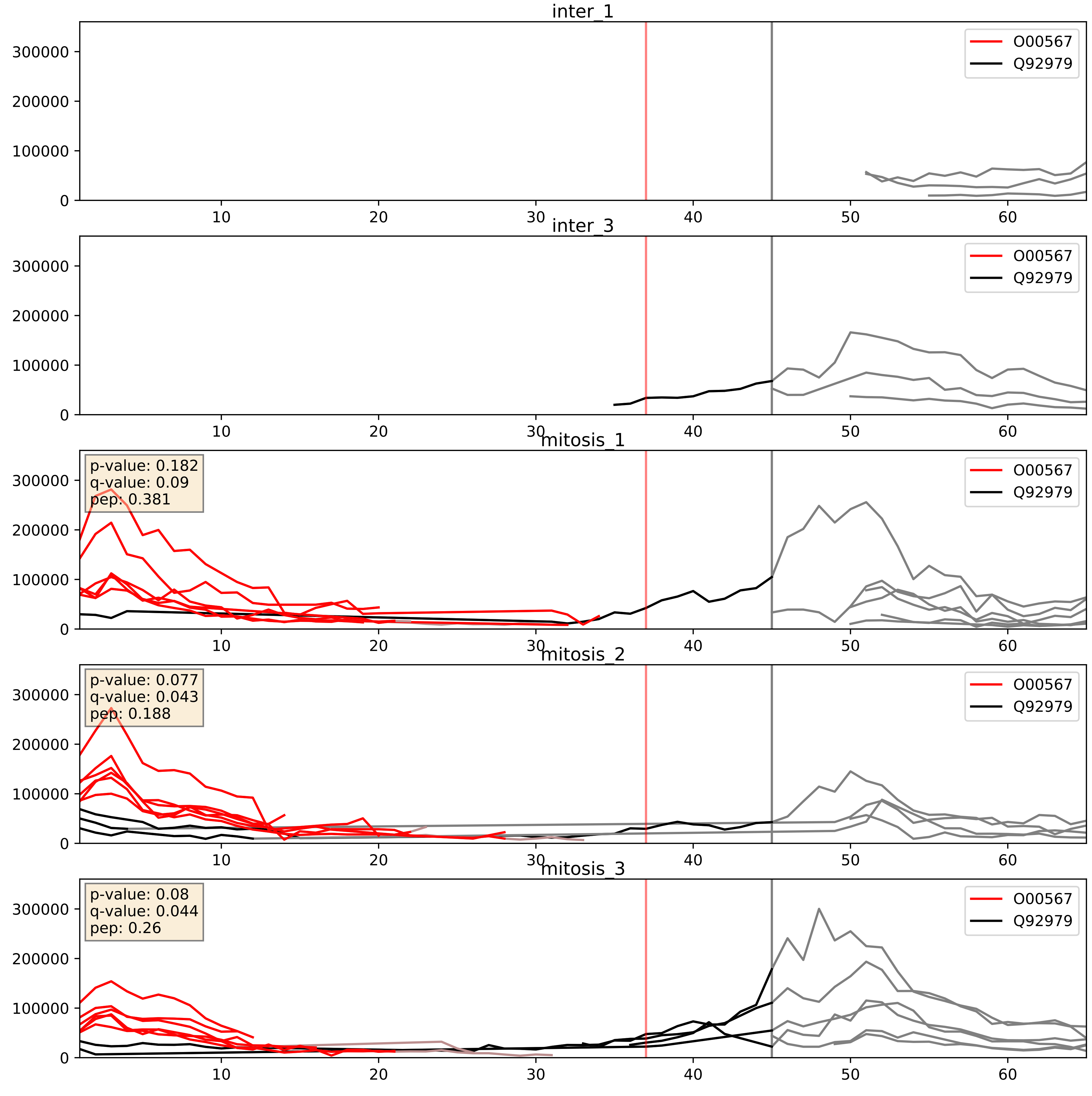


| level | condition_1 | condition_2 | pvalue | pvalue_adjusted |
|---|---|---|---|---|
| interactor_ratio | mitosis | inter | 0.00021846409665752416 | 0.015168670877391028 |
| interactor_abundance | mitosis | inter | 0.0013322765484747097 | 0.026992395869688792 |
| complex_abundance | mitosis | inter | 0.0060296068177722595 | 0.05806974312565789 |

O75937_Q92979
DNJC8_HUMAN vs NEP1_HUMAN
p-value: 0.0 q-value: 0.044

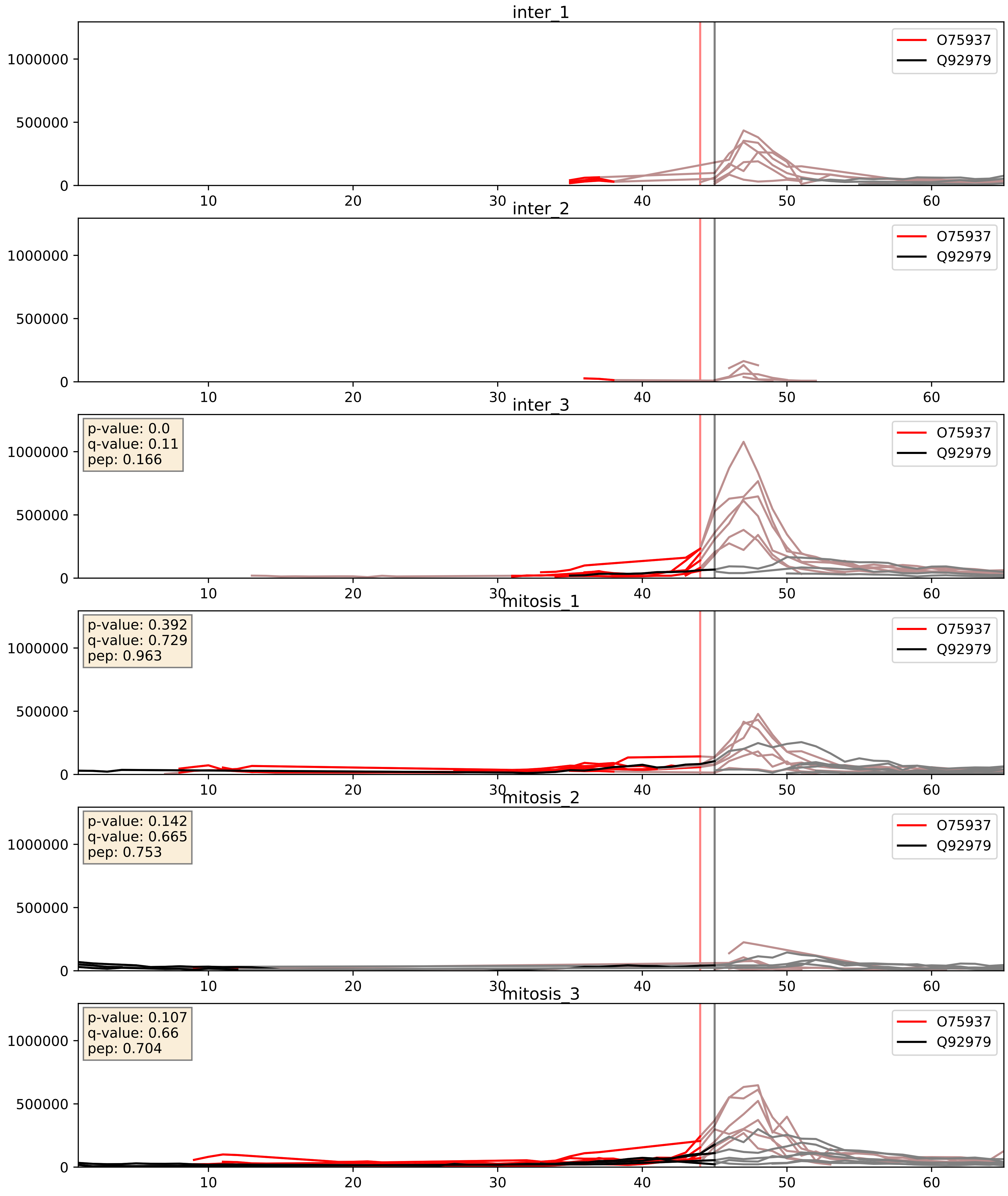


| level | condition_1 | condition_2 | pvalue | pvalue_adjusted |
|---|---|---|---|---|
| interactor_ratio | mitosis | inter | 0.6465451377513514 | 0.9217731067492805 |
| interactor_abundance | mitosis | inter | 0.6548108575870751 | 0.7704956107154083 |
| complex_abundance | mitosis | inter | 0.6918925155865954 | 0.7929571389337764 |

P22087_Q92979
FBRL_HUMAN vs NEP1_HUMAN
p-value: 0.035 q-value: 0.024

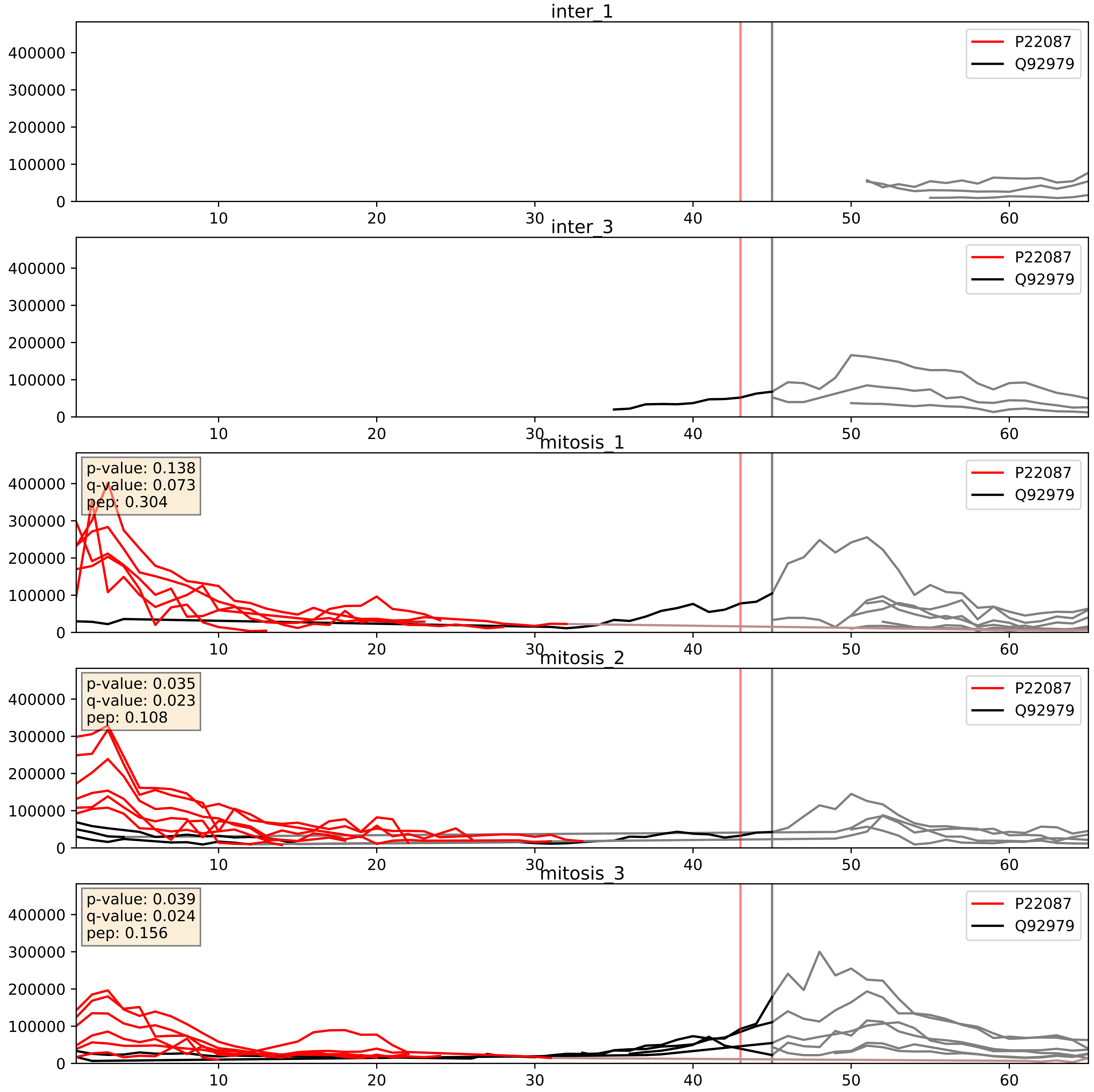


| level | condition_1 | condition_2 | pvalue | pvalue_adjusted |
|---|---|---|---|---|
| interactor_ratio | mitosis | inter | 9.565040877399547e-05 | 0.010510079284752772 |
| interactor_abundance | mitosis | inter | 0.0010078483561365378 | 0.023668537526827883 |
| complex_abundance | mitosis | inter | 0.0040748418684974745 | 0.04732787841836958 |
