## Supplementary Data 2 for "SECAT: Quantifying differential protein-protein interaction states by network-centric analysis": hela_string_000080_Q15773.pdf

Q15773_Q15773

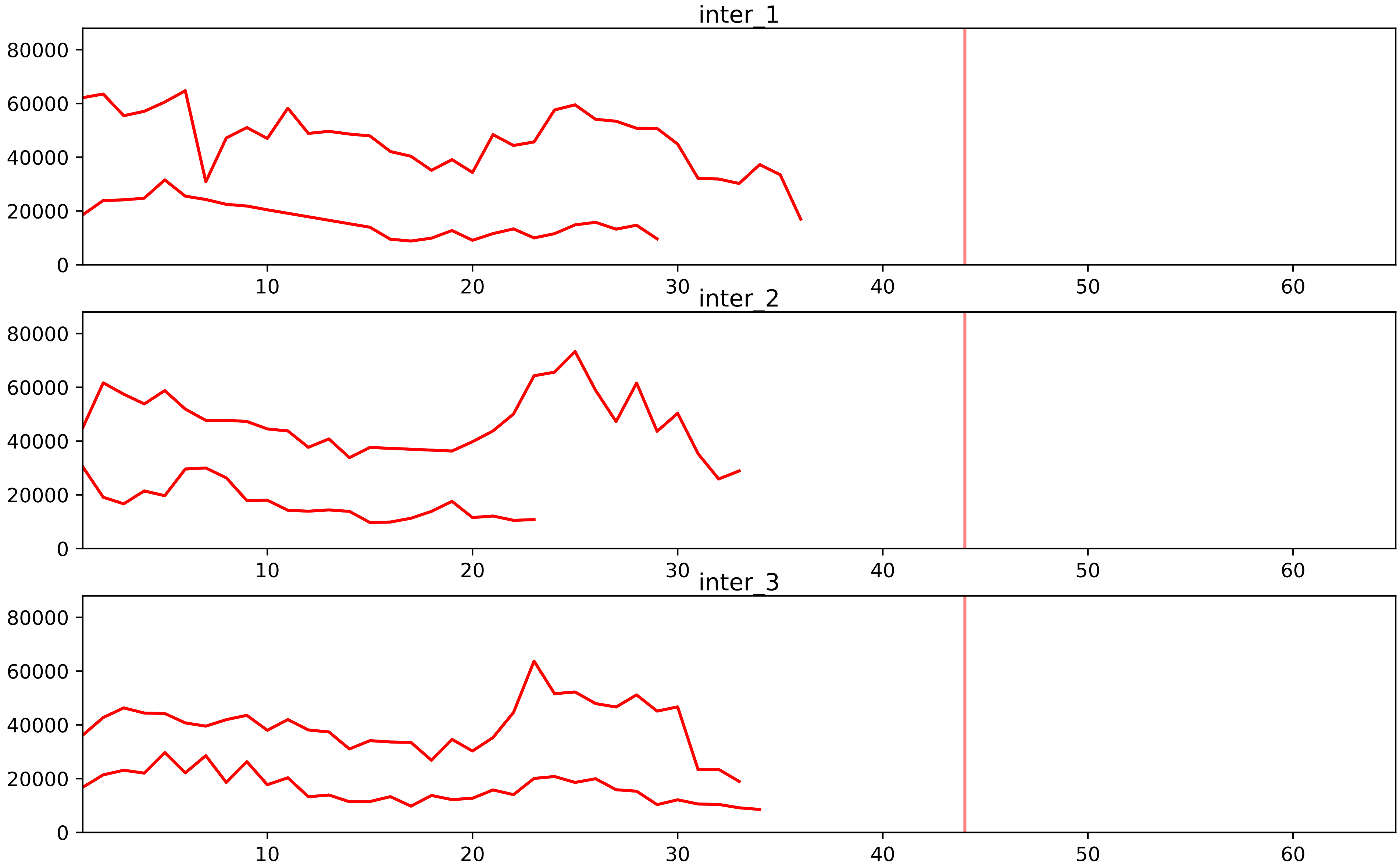


| level | condition_1 | condition_2 | pvalue | pvalue_adjusted |
|---|---|---|---|---|
| assembled_abundance | mitosis | inter | 3.6184745119491586e-06 | 0.0028230131984056687 |
| total_abundance | mitosis | inter | 8.219675450700892e-06 | 0.004123130635959455 |
