## Supplementary Data 2 for "SECAT: Quantifying differential protein-protein interaction states by network-centric analysis": hela_string_000081_P31942.pdf

P31942_P31942

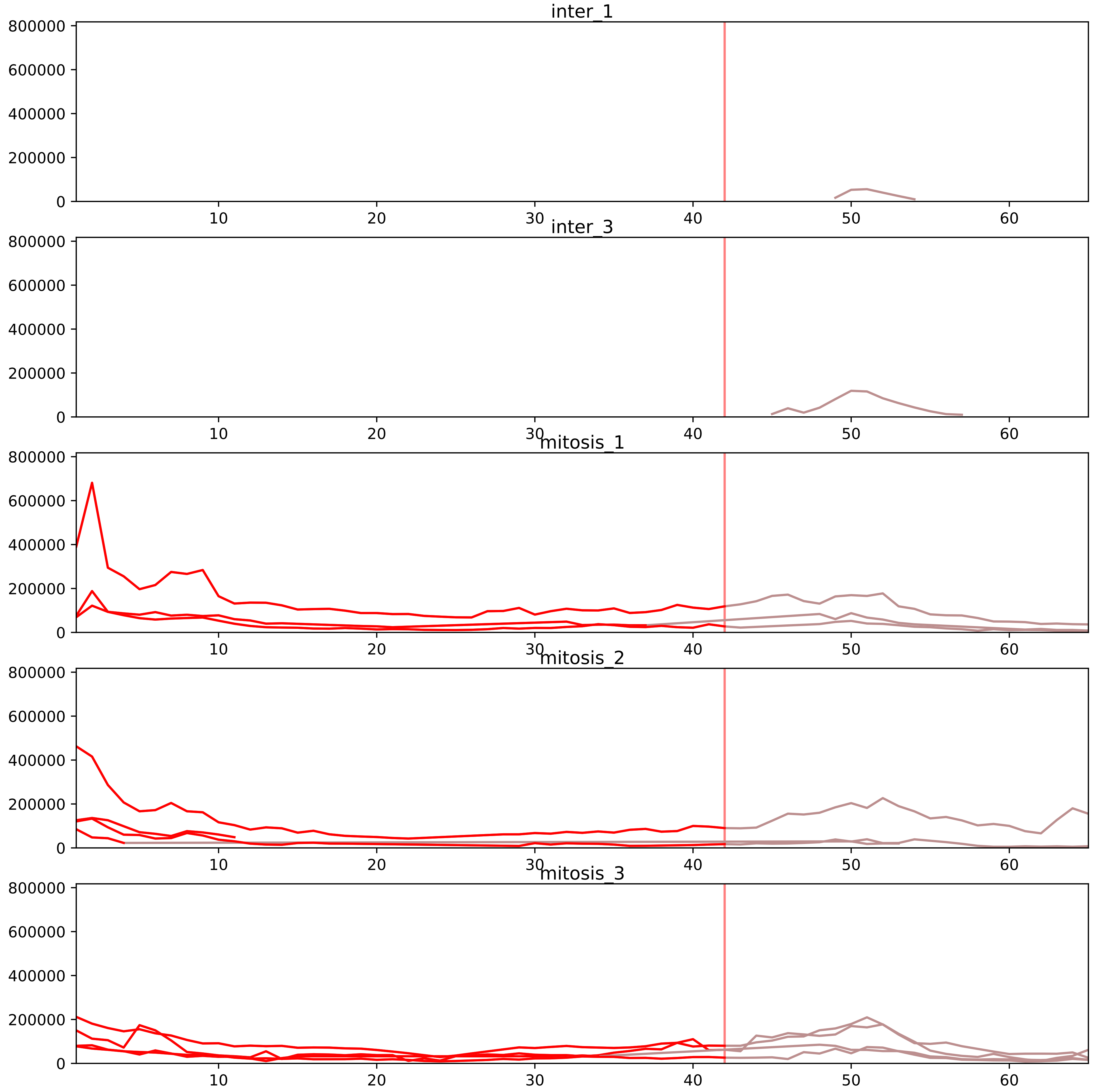


| level | condition_1 | condition_2 | pvalue | pvalue_adjusted |
|---|---|---|---|---|
| complex_abundance | mitosis | inter | 3.6268297864037747e-06 | 0.00022244556023276484 |
| interactor_abundance | mitosis | inter | 0.00023998795649377193 | 0.004329194509299415 |
| assembled_abundance | mitosis | inter | 0.0007887940115821994 | 0.02511799162051888 |
| total_abundance | mitosis | inter | 0.0021693710656859877 | 0.05005132055560529 |
| interactor_ratio | mitosis | inter | 0.0026000076695202583 | 0.02368323817780829 |
| monomer_abundance | mitosis | inter | 0.0055310707513959384 | 0.147846976729748 |

O14979_P31942
HNRDL_HUMAN vs HNRH3_HUMAN
p-value: 0.004 q-value: 0.047

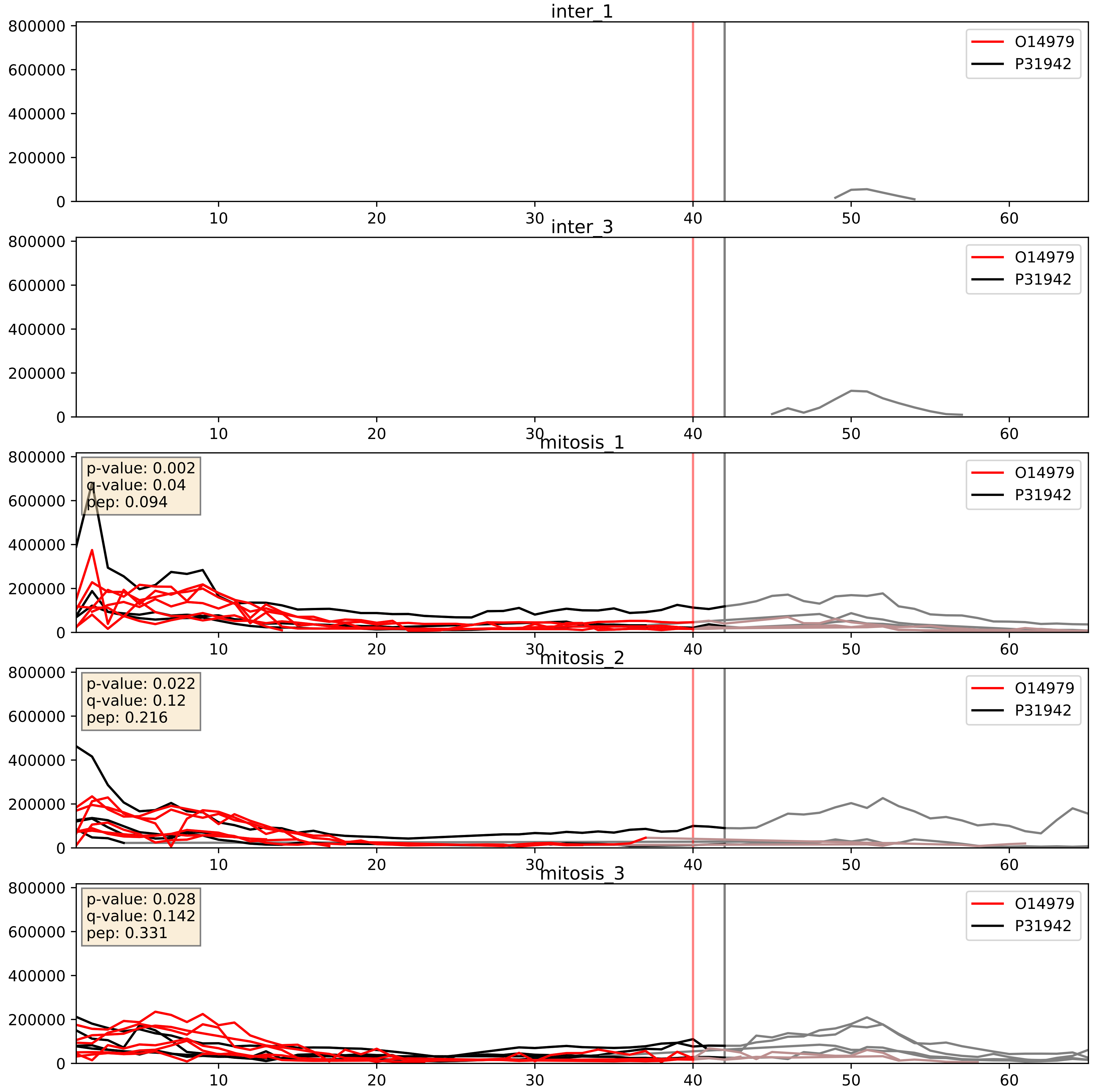


| level | condition_1 | condition_2 | pvalue | pvalue_adjusted |
|---|---|---|---|---|
| complex_abundance | mitosis | inter | 0.0004757589173787532 | 0.013816623868928207 |
| interactor_abundance | mitosis | inter | 0.0008235085849705327 | 0.02107394166621154 |
| interactor_ratio | mitosis | inter | 0.2439500030606306 | 0.7200661897289671 |

P17844_P31942
DDX5_HUMAN vs HNRH3_HUMAN
p-value: 0.0 q-value: 0.043

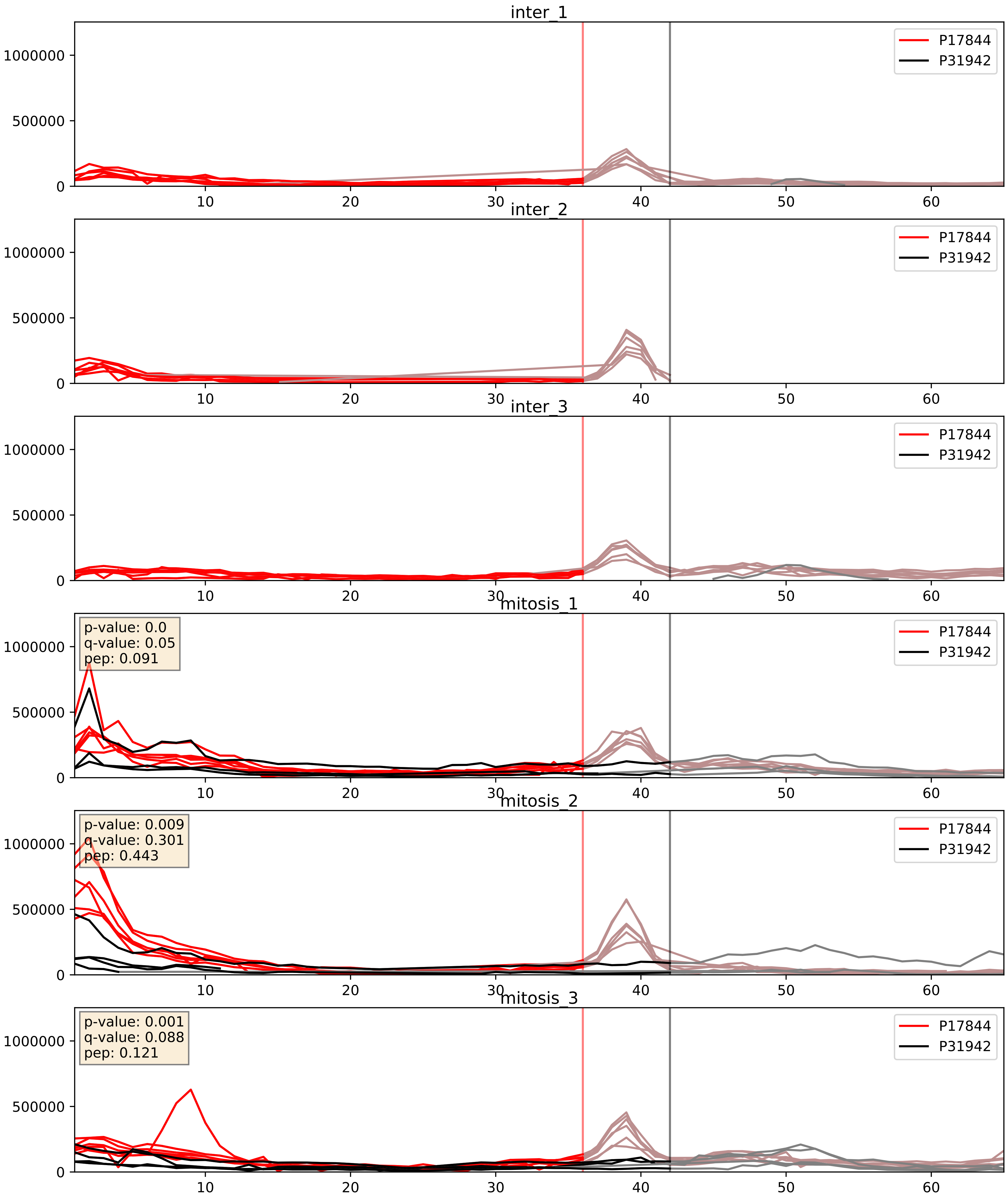


| level | condition_1 | condition_2 | pvalue | pvalue_adjusted |
|---|---|---|---|---|
| interactor_abundance | mitosis | inter | 2.8004503716417208e-05 | 0.004746902016089729 |
| complex_abundance | mitosis | inter | 5.883087896907052e-05 | 0.005691857151652958 |
| interactor_ratio | mitosis | inter | 0.016560873095316366 | 0.18977386036935487 |

P31942_P31943
HNRH3_HUMAN vs HNRH1_HUMAN
p-value: 0.002 q-value: 0.03

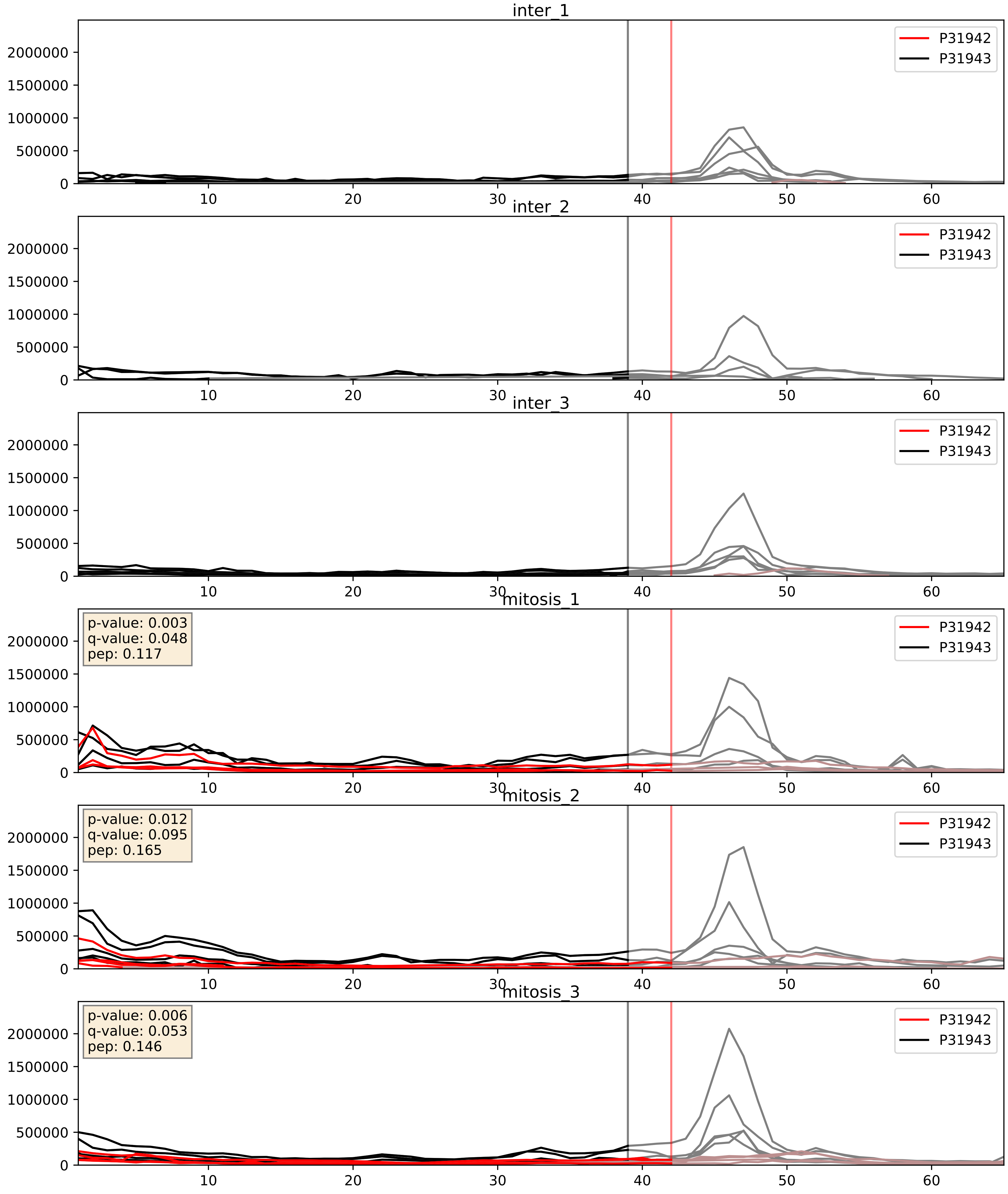


| level | condition_1 | condition_2 | pvalue | pvalue_adjusted |
|---|---|---|---|---|
| complex_abundance | mitosis | inter | 7.183772469446166e-05 | 0.005893640694944935 |
| interactor_abundance | mitosis | inter | 0.002362609793349813 | 0.03614645188753243 |
| interactor_ratio | mitosis | inter | 0.008559803010908383 | 0.13203195690147984 |

P31942_P62995
HNRH3_HUMAN vs TRA2B_HUMAN
p-value: 0.0 q-value: 0.023

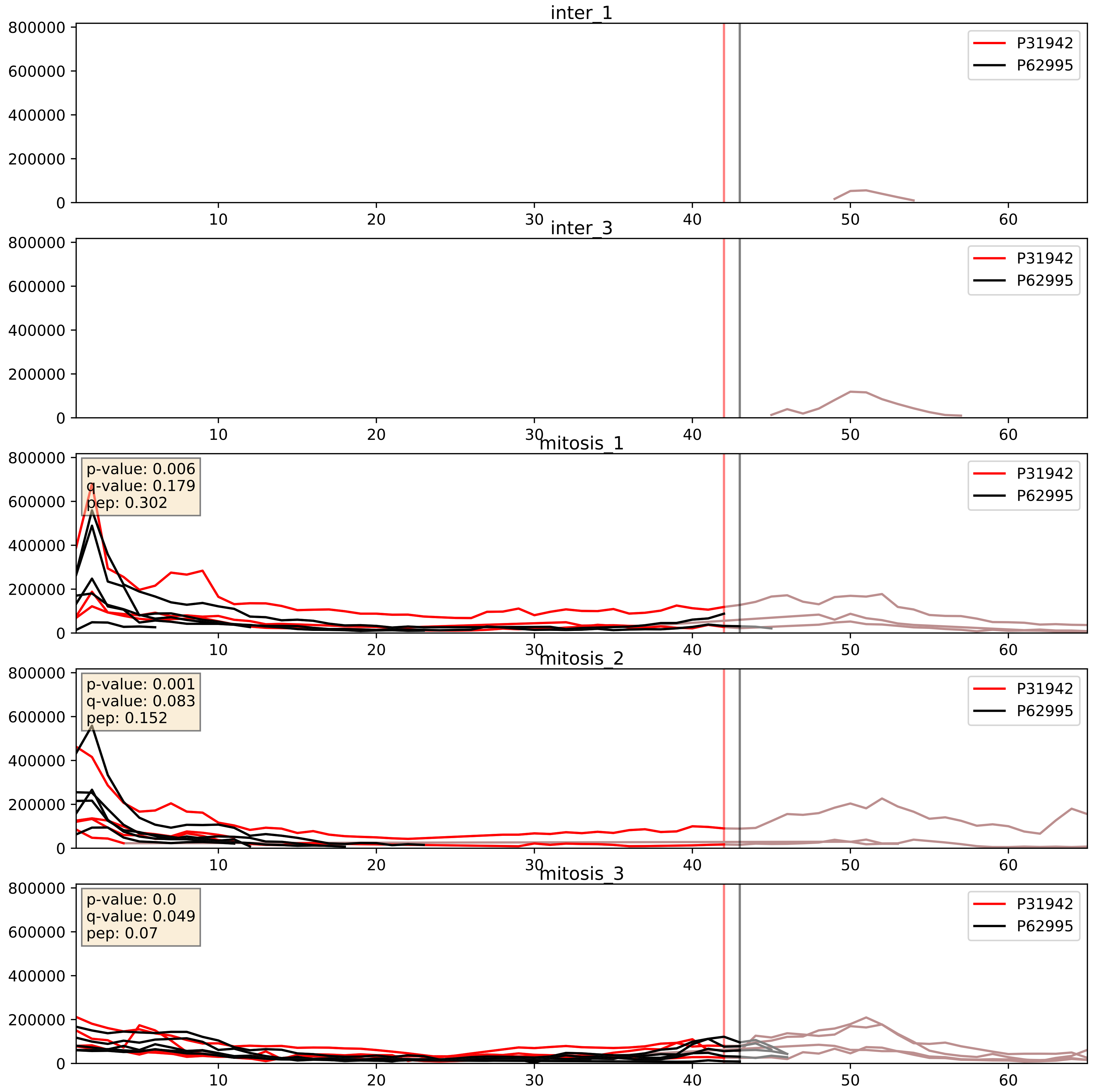


| level | condition_1 | condition_2 | pvalue | pvalue_adjusted |
|---|---|---|---|---|
| complex_abundance | mitosis | inter | 0.0010827209575059045 | 0.021833569470040306 |
| interactor_abundance | mitosis | inter | 0.004530454063144402 | 0.05416297036384927 |
| interactor_ratio | mitosis | inter | 0.20927687488332142 | 0.686697008467737 |
