## Supplementary Data 2 for "SECAT: Quantifying differential protein-protein interaction states by network-centric analysis": hela_string_000082_Q86U86.pdf

Q86U86_Q86U86

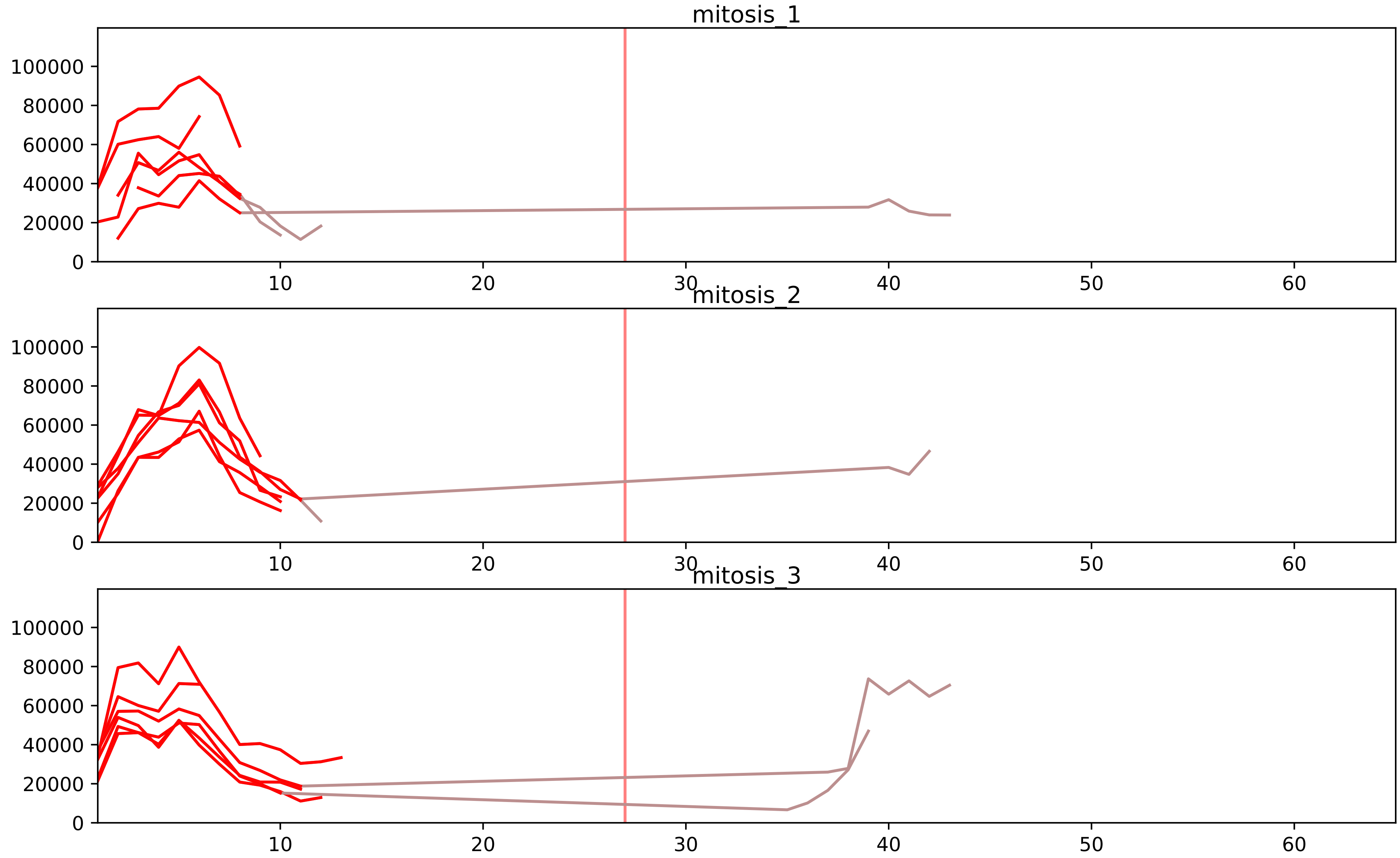


| level | condition_1 | condition_2 | pvalue | pvalue_adjusted |
|---|---|---|---|---|
| interactor_ratio | mitosis | inter | 3.6869938937361786e-06 | 9.55502642883742e-05 |
| interactor_abundance | mitosis | inter | 0.0007910613730607635 | 0.009703686176212032 |
| complex_abundance | mitosis | inter | 0.003365051311760884 | 0.023364884579773686 |
| total_abundance | mitosis | inter | 0.08678018567908966 | 0.36153676374160837 |
| assembled_abundance | mitosis | inter | 0.13990672372168597 | 0.44310106477754535 |
| monomer_abundance | mitosis | inter | 0.17133968195827404 | 0.5363275252669241 |

Q68CP9_Q86U86
ARID2_HUMAN vs PB1_HUMAN
p-value: 0.002 q-value: 0.002

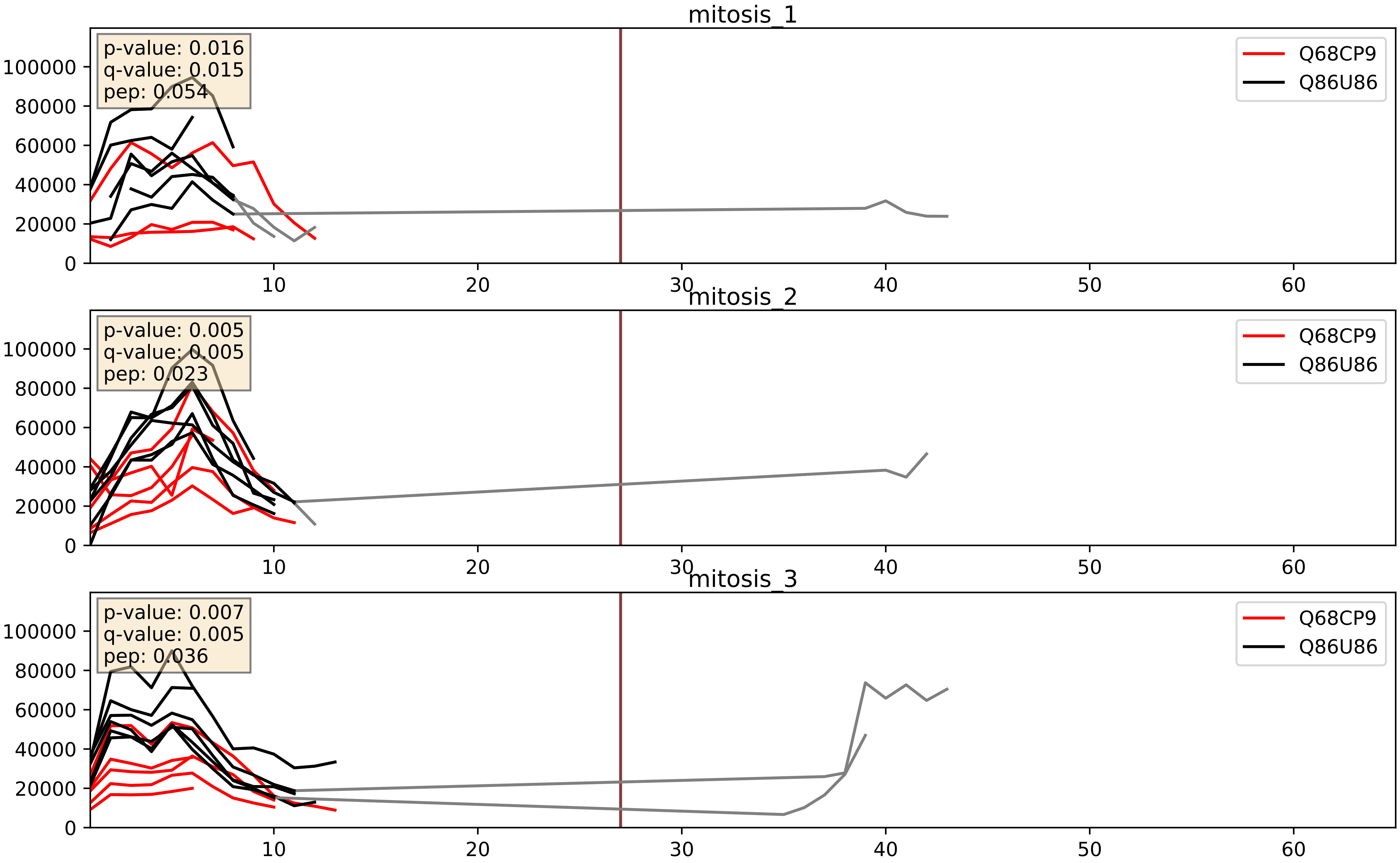


| level | condition_1 | condition_2 | pvalue | pvalue_adjusted |
|---|---|---|---|---|
| interactor_ratio | mitosis | inter | 0.00031824606012225044 | 0.01878749154928596 |
| complex_abundance | mitosis | inter | 0.004464316990302555 | 0.049788645279664145 |
| interactor_abundance | mitosis | inter | 0.018643676110411646 | 0.11769164270289358 |

Q86U86_Q9NPI1
PB1_HUMAN vs BRD7_HUMAN
p-value: 0.039 q-value: 0.027

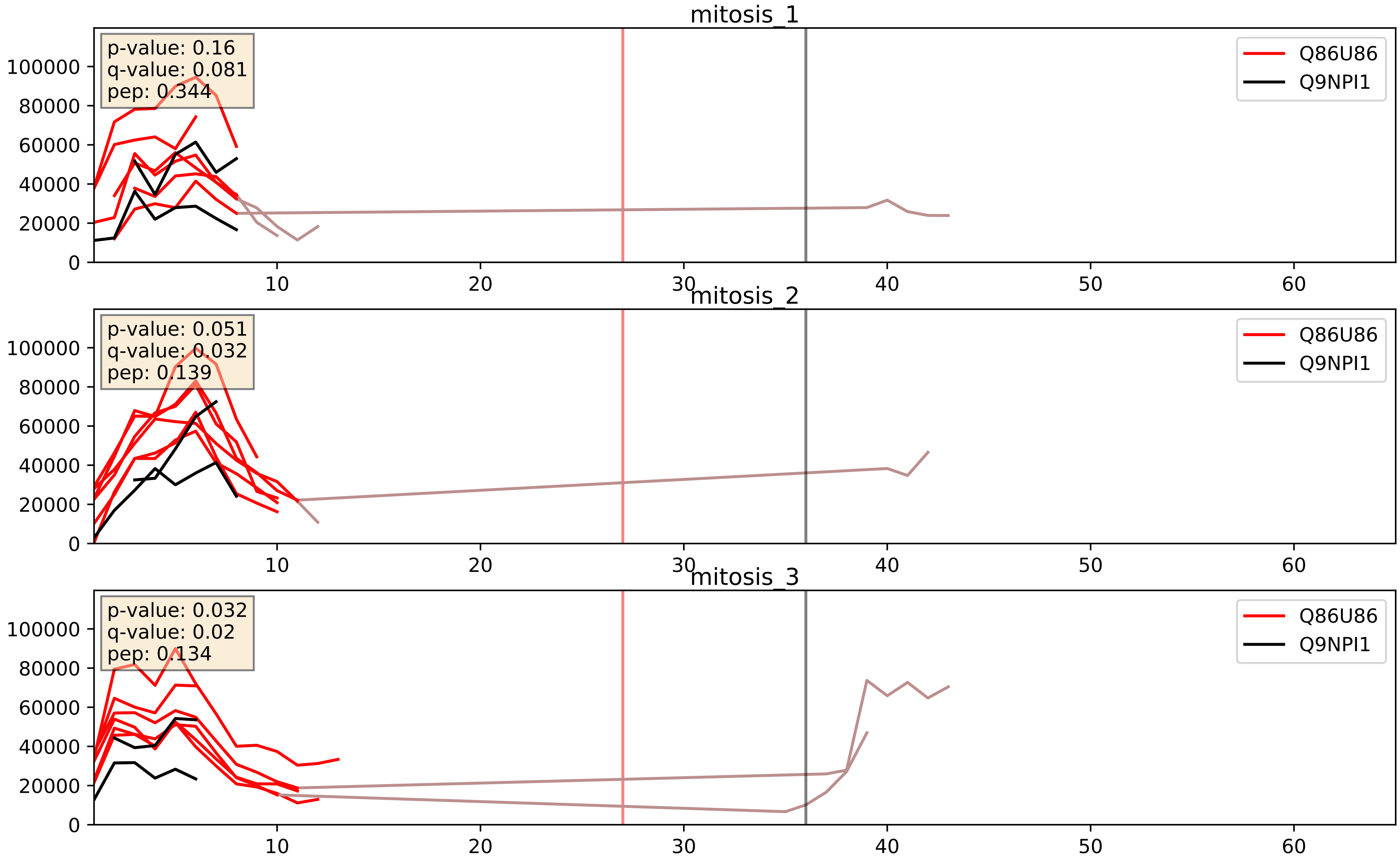


| level | condition_1 | condition_2 | pvalue | pvalue_adjusted |
|---|---|---|---|---|
| interactor_ratio | mitosis | inter | 0.0007106714449763887 | 0.030879936898466437 |
| interactor_abundance | mitosis | inter | 0.005336126744332609 | 0.06160113948953086 |
| complex_abundance | mitosis | inter | 0.016751616487354203 | 0.09662657488662532 |
