## Supplementary Data 2 for "SECAT: Quantifying differential protein-protein interaction states by network-centric analysis": hela_string_000083_O95067.pdf

O95067_O95067

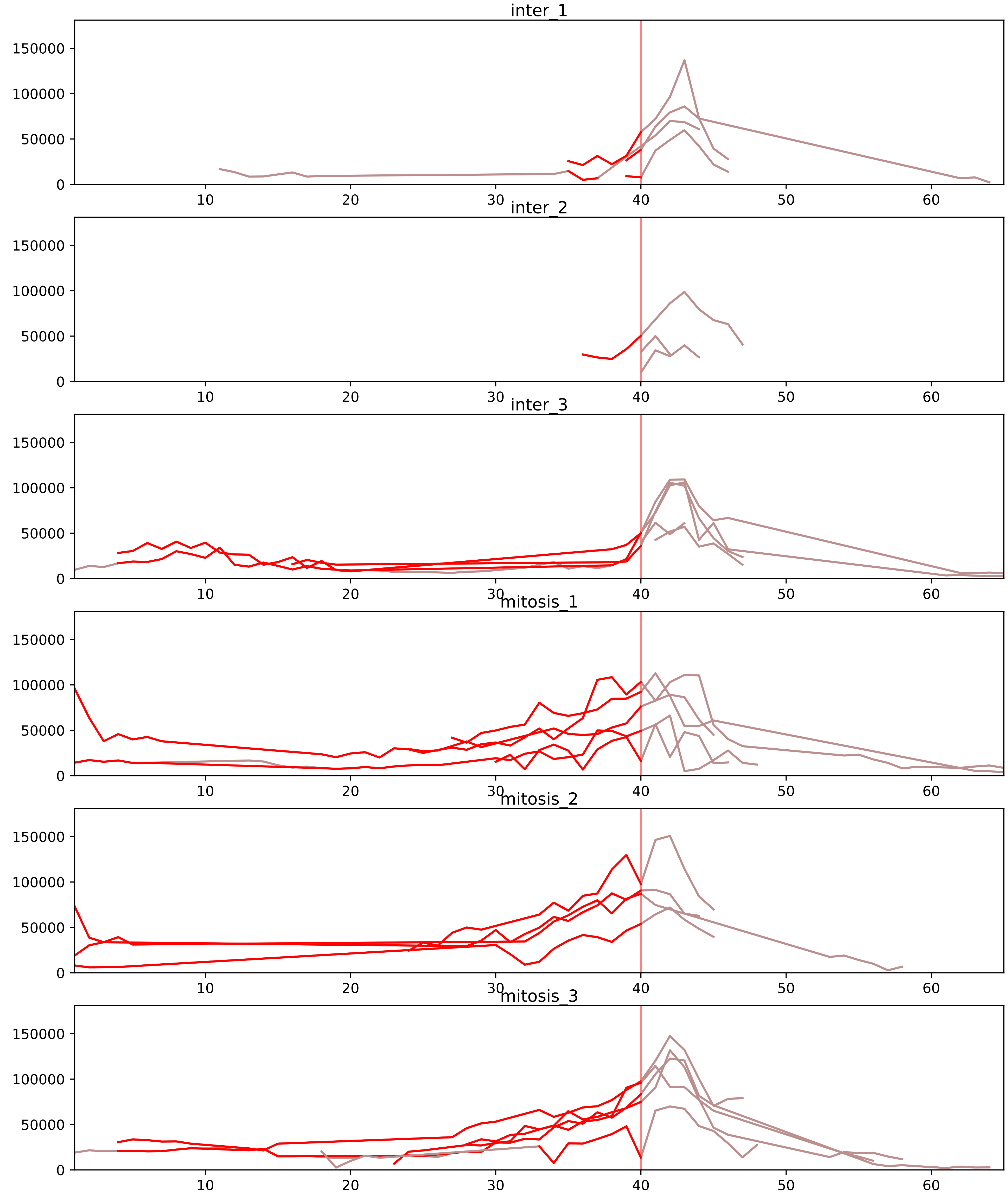


| level | condition_1 | condition_2 | pvalue | pvalue_adjusted |
|---|---|---|---|---|
| interactor_ratio | mitosis | inter | 3.993680251547266e-06 | 0.00010206071753954125 |
| complex_abundance | mitosis | inter | 0.0002616716573180596 | 0.004377053176956634 |
| assembled_abundance | mitosis | inter | 0.000265083236537006 | 0.016114995197788636 |
| interactor_abundance | mitosis | inter | 0.0002915976523666997 | 0.005027899116641351 |
| total_abundance | mitosis | inter | 0.003141515632191738 | 0.06263245817916394 |
| monomer_abundance | mitosis | inter | 0.037714679231998054 | 0.3150273663293861 |

O95067_P06493
CCNB2_HUMAN vs CDK1_HUMAN
p-value: 0.063 q-value: 0.039

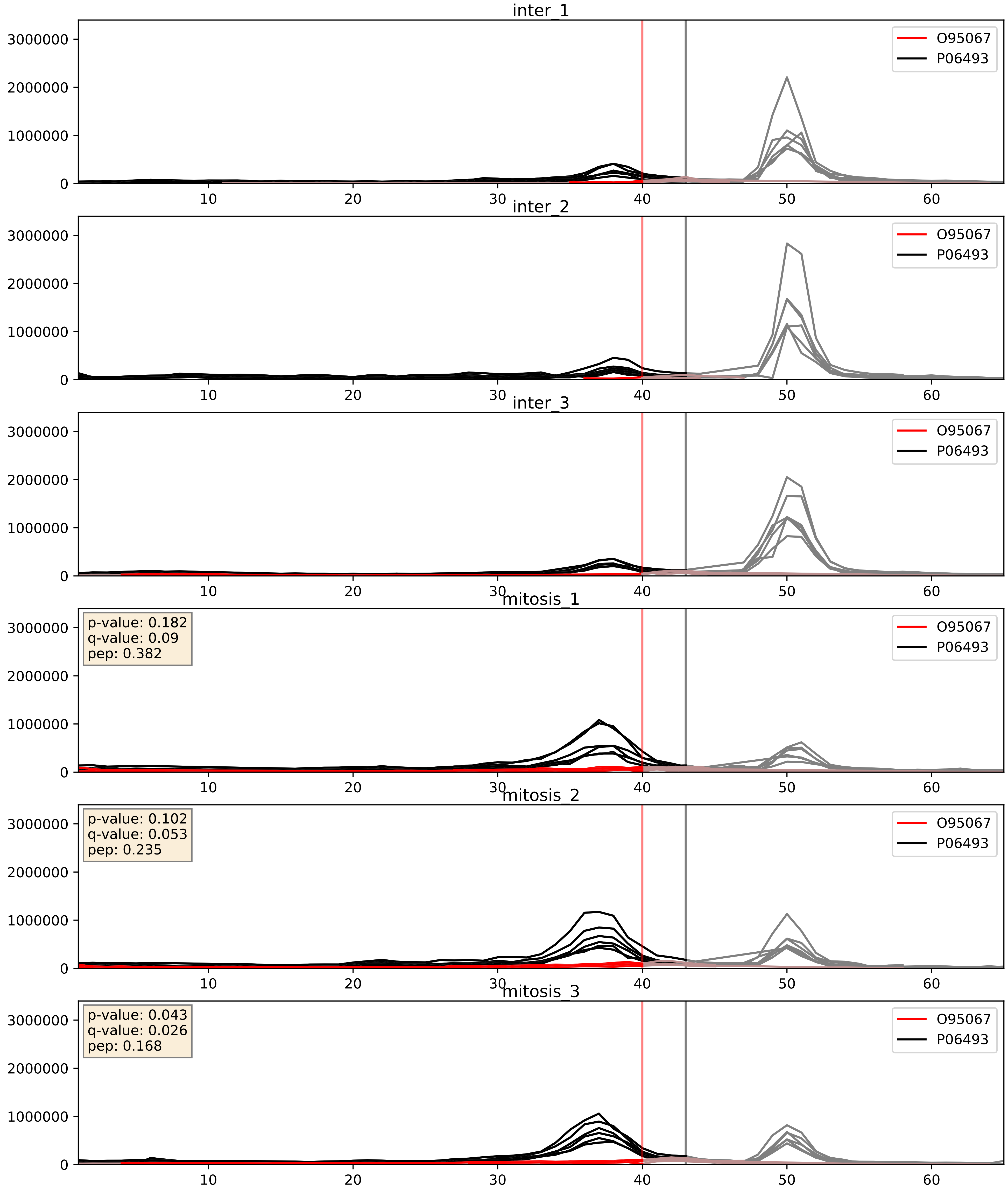


| level | condition_1 | condition_2 | pvalue | pvalue_adjusted |
|---|---|---|---|---|
| complex_abundance | mitosis | inter | 0.00017800511457254897 | 0.008807651911797798 |
| interactor_abundance | mitosis | inter | 0.0031005769388121997 | 0.04271207144514066 |
| interactor_ratio | mitosis | inter | 0.005268855769537838 | 0.0974780445871482 |

O95067_P14635
CCNB2_HUMAN vs CCNB1_HUMAN
p-value: 0.01 q-value: 0.008

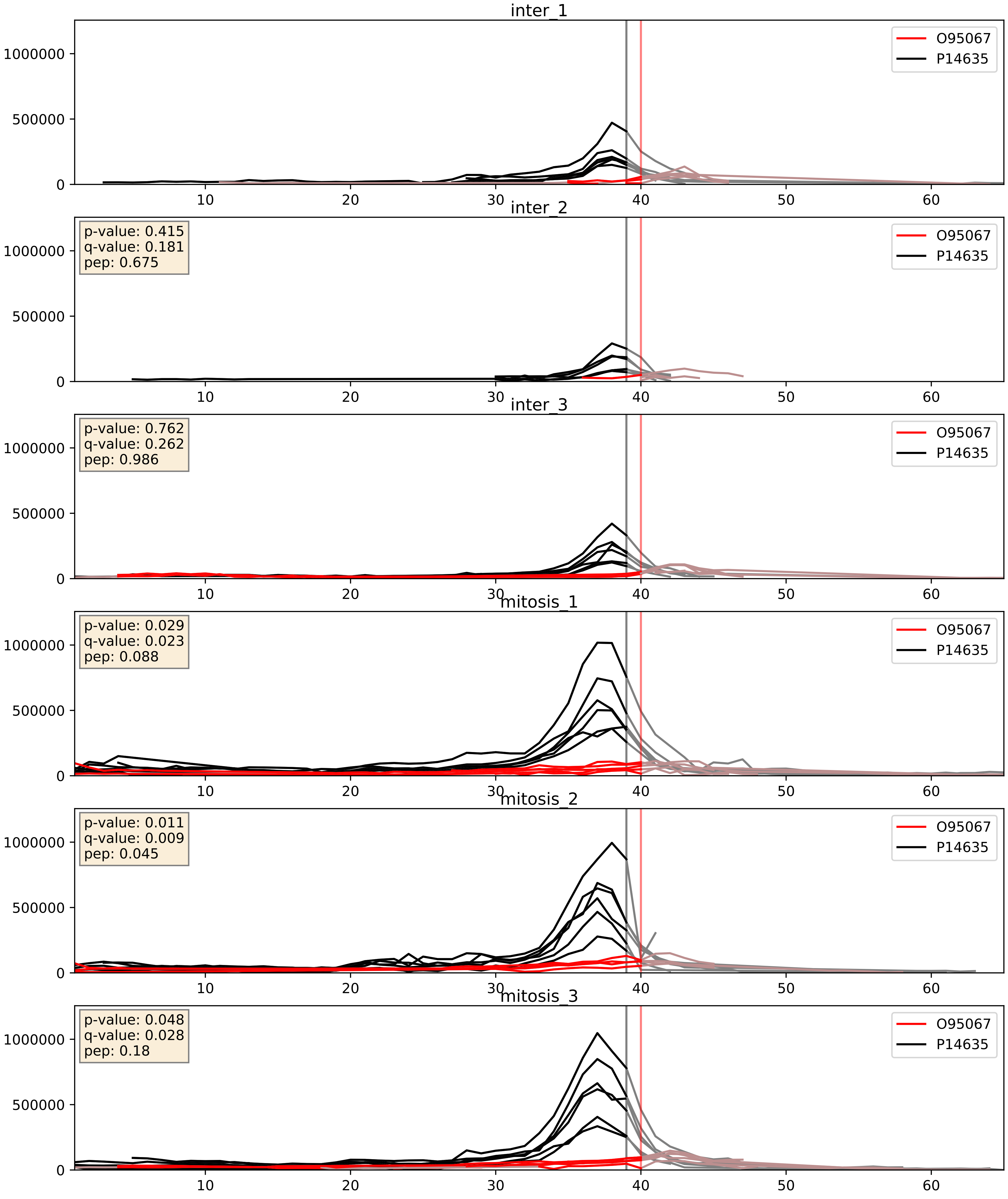


| level | condition_1 | condition_2 | pvalue | pvalue_adjusted |
|---|---|---|---|---|
| complex_abundance | mitosis | inter | 0.000540651241243441 | 0.014880947347407894 |
| interactor_abundance | mitosis | inter | 0.0025820700836980265 | 0.03806046117740333 |
| interactor_ratio | mitosis | inter | 0.003698695469143503 | 0.07875829158176215 |

O95067_P24941
CCNB2_HUMAN vs CDK2_HUMAN
p-value: 0.004 q-value: 0.004

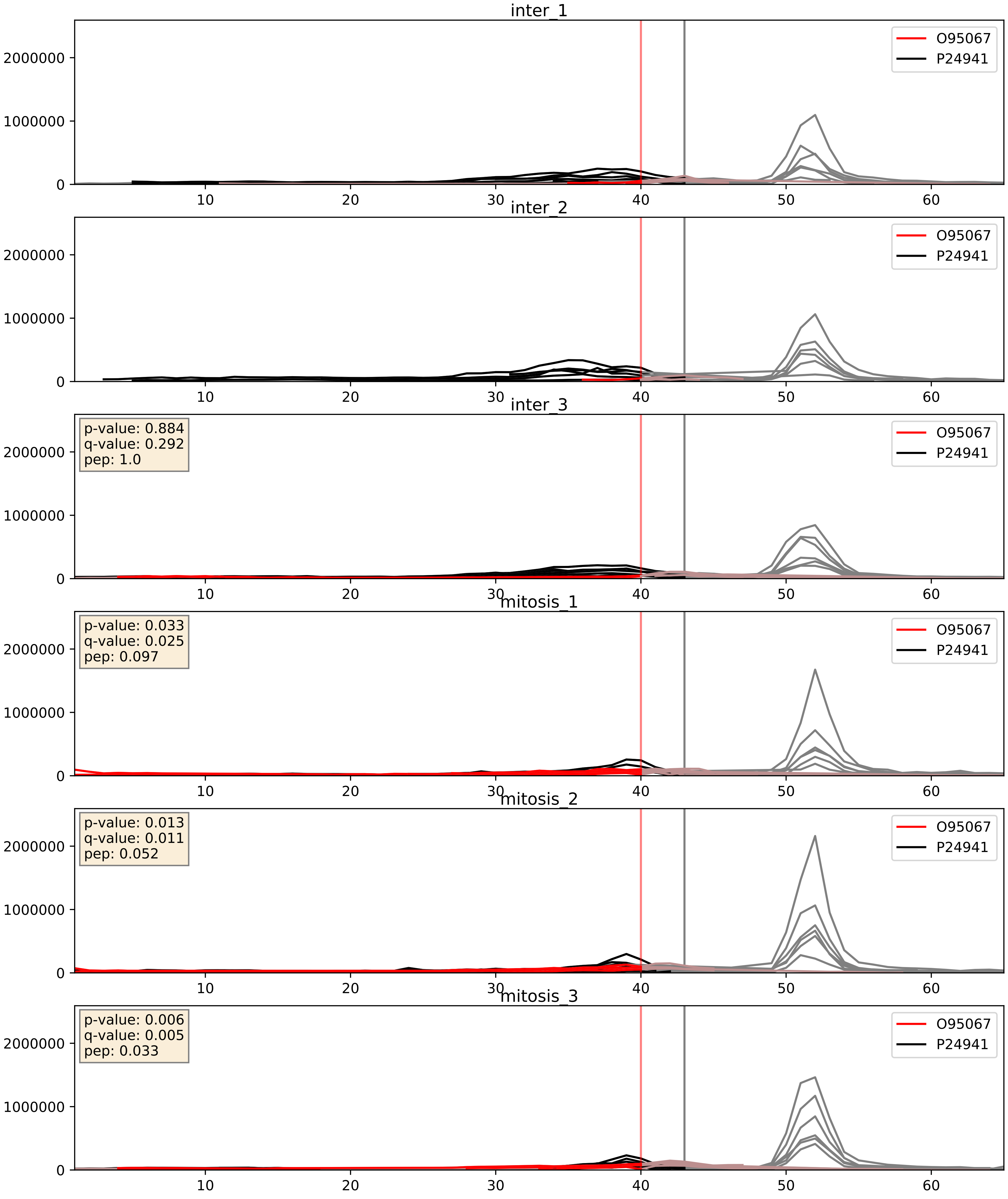


| level | condition_1 | condition_2 | pvalue | pvalue_adjusted |
|---|---|---|---|---|
| interactor_abundance | mitosis | inter | 0.001443970468219757 | 0.027996347016899476 |
| interactor_ratio | mitosis | inter | 0.011385932666237232 | 0.15421453104903593 |
| complex_abundance | mitosis | inter | 0.04027580454763698 | 0.14508340529050803 |

O95067_P30307
CCNB2_HUMAN vs MPIP3_HUMAN
p-value: 0.074 q-value: 0.045

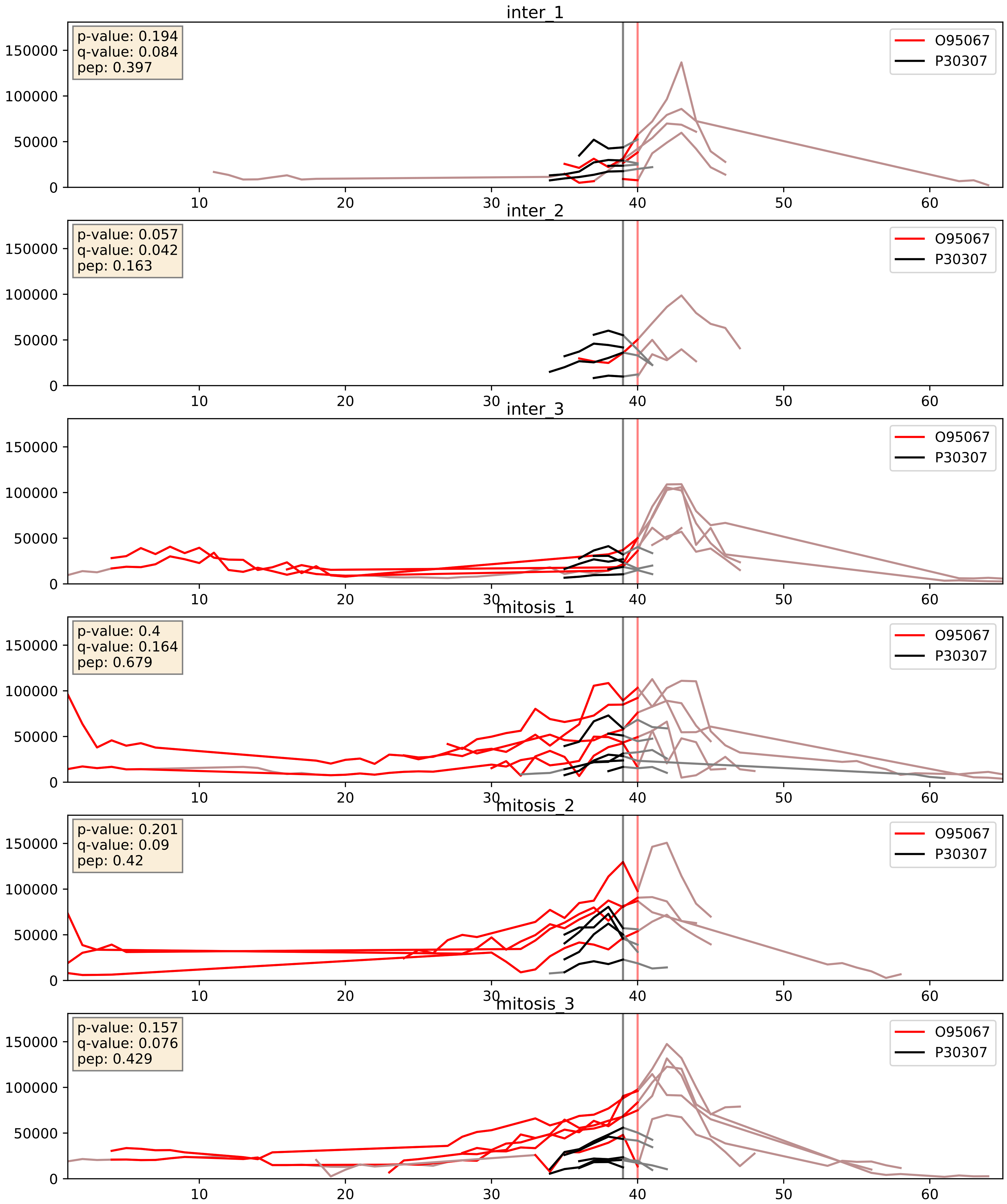


| level | condition_1 | condition_2 | pvalue | pvalue_adjusted |
|---|---|---|---|---|
| interactor_ratio | mitosis | inter | 0.00775417243450131 | 0.12406675895202096 |
| interactor_abundance | mitosis | inter | 0.08453425855201002 | 0.24442264928397425 |
| complex_abundance | mitosis | inter | 0.8738368412836998 | 0.9213010668048368 |

O95067_Q96FF9
CCNB2_HUMAN vs CDCA5_HUMAN
p-value: 0.034 q-value: 0.024

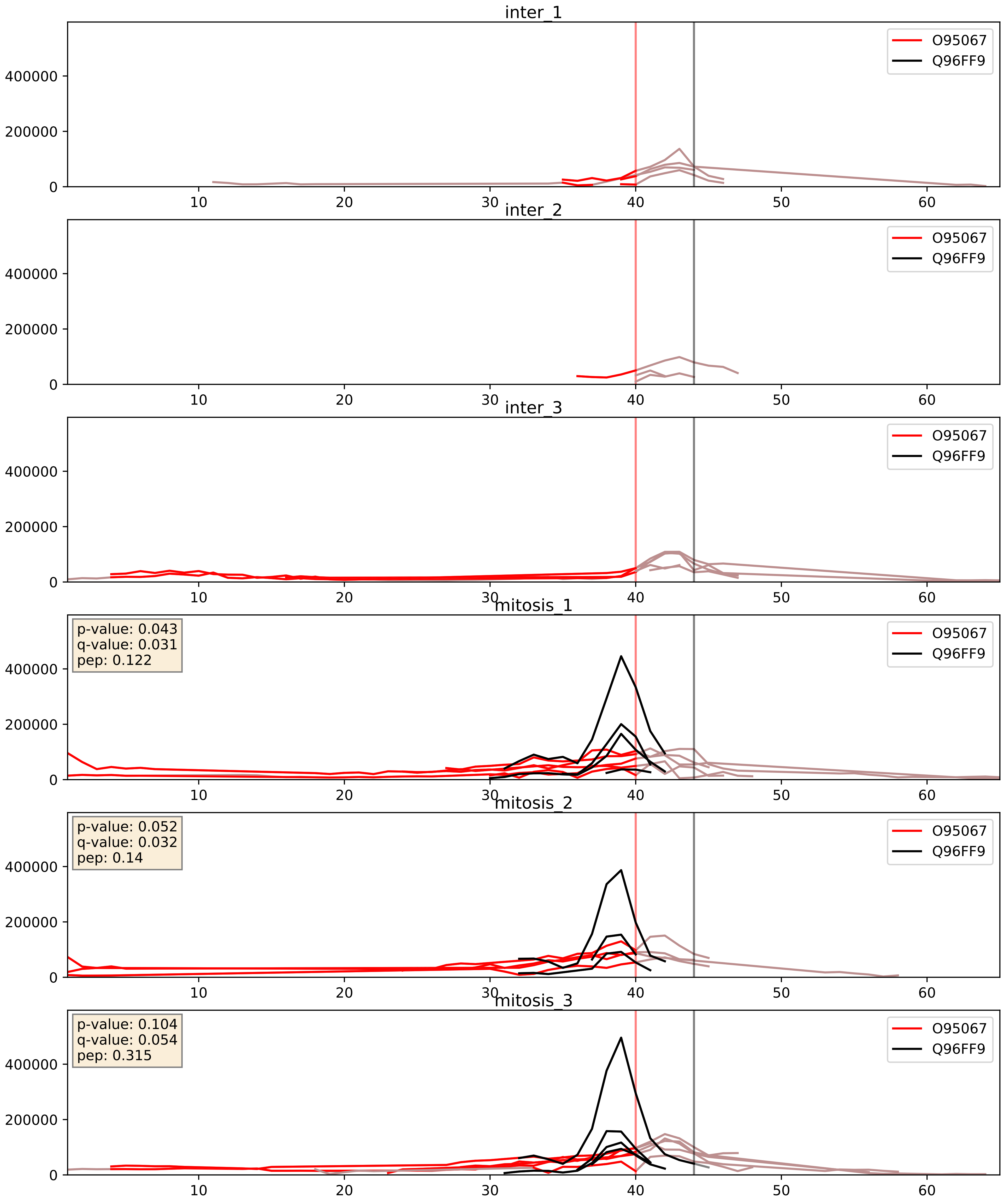


| level | condition_1 | condition_2 | pvalue | pvalue_adjusted |
|---|---|---|---|---|
| interactor_abundance | mitosis | inter | 0.002773496431292935 | 0.03993461640347775 |
| complex_abundance | mitosis | inter | 0.003336551402585707 | 0.04242071705738532 |
| interactor_ratio | mitosis | inter | 0.20486220931689902 | 0.6802724376270011 |
