## Supplementary Data 2 for "SECAT: Quantifying differential protein-protein interaction states by network-centric analysis": hela_string_000084_P11172.pdf

# P11172_P11172

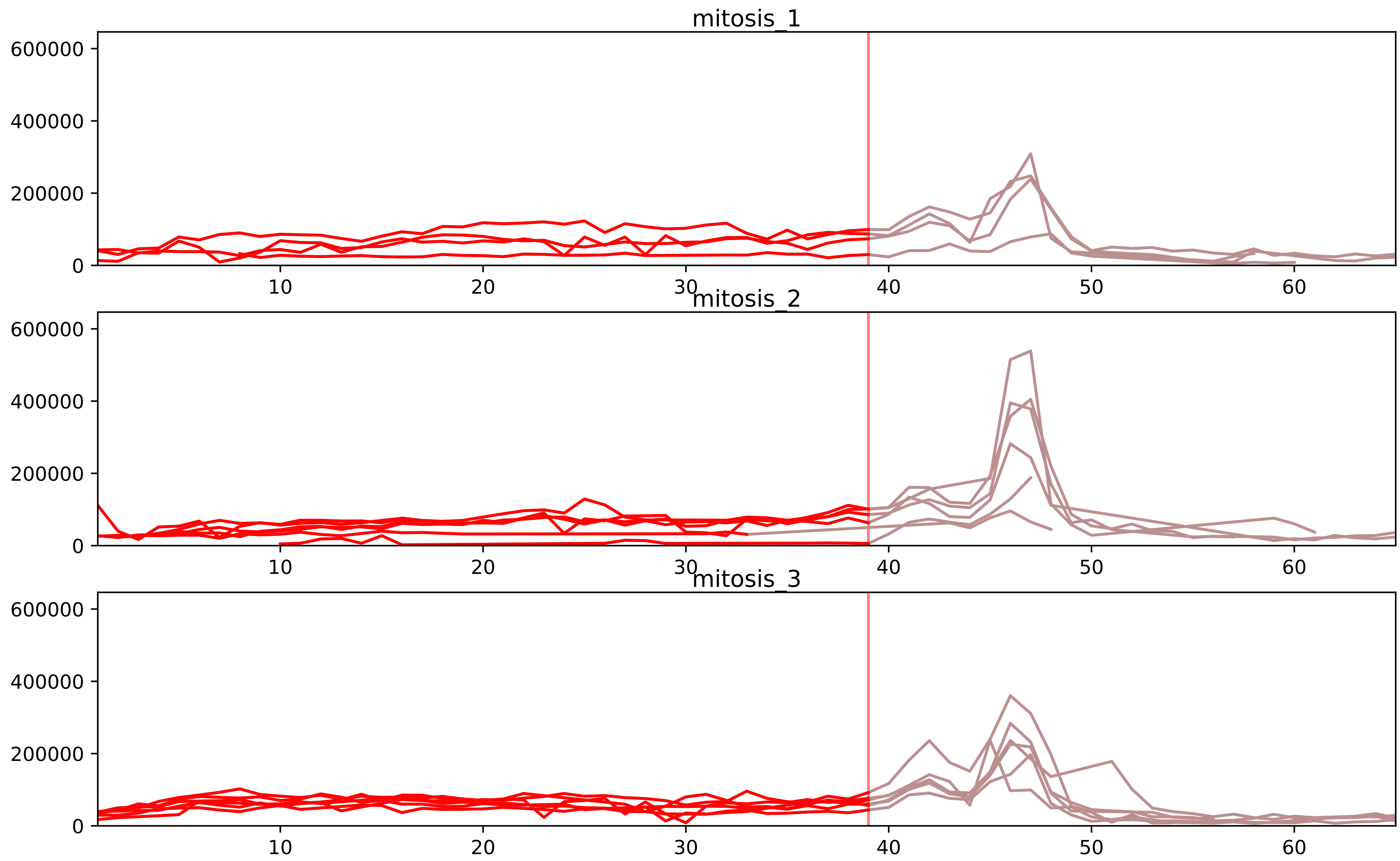


| level | condition_1 | condition_2 | pvalue | pvalue_adjusted |
|---|---|---|---|---|
| assembled_abundance | mitosis | inter | 4.5700501484849455e-06 | 0.002899782131414633 |
| total_abundance | mitosis | inter | 5.079406039091716e-06 | 0.003540864511957668 |
| monomer_abundance | mitosis | inter | 8.100837042744166e-05 | 0.03261473898341094 |
