## Supplementary Data 2 for "SECAT: Quantifying differential protein-protein interaction states by network-centric analysis": hela_string_000085_O95235.pdf

O95235_O95235

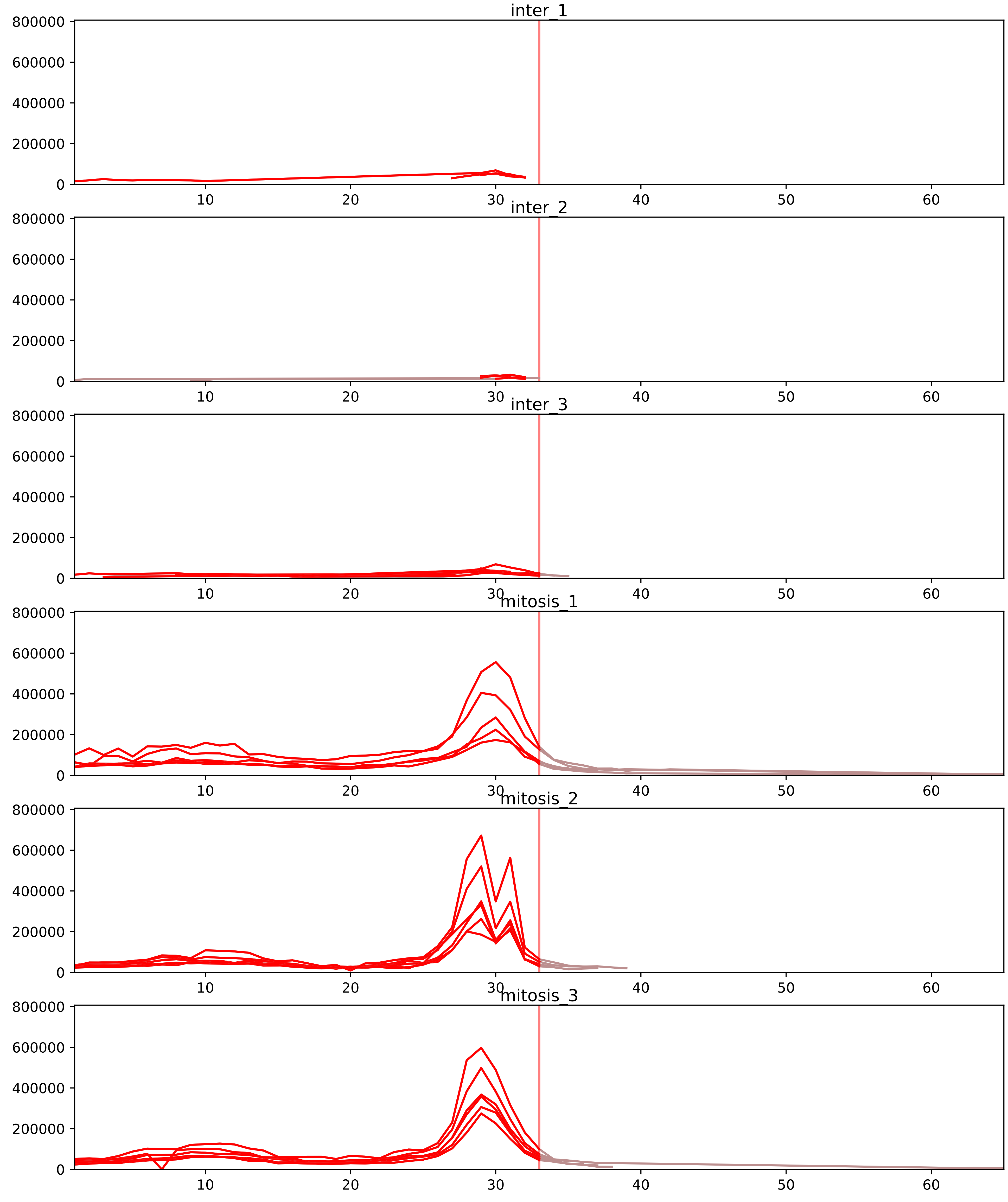


| level | condition_1 | condition_2 | pvalue | pvalue_adjusted |
|---|---|---|---|---|
| total_abundance | mitosis | inter | 4.740361078185997e-06 | 0.003540864511957668 |
| assembled_abundance | mitosis | inter | 7.7508278112276e-06 | 0.0032983295440323994 |
| interactor_abundance | mitosis | inter | 0.002491334705596794 | 0.022126626540542434 |
| monomer_abundance | mitosis | inter | 0.0033373251333708043 | 0.11999603554765999 |
| interactor_ratio | mitosis | inter | 0.009413807097681002 | 0.05993565764613509 |
| complex_abundance | mitosis | inter | 0.040808984966054784 | 0.10835286051593189 |

O95235_Q99661
KI20A_HUMAN vs KIF2C_HUMAN
p-value: 0.062 q-value: 0.039

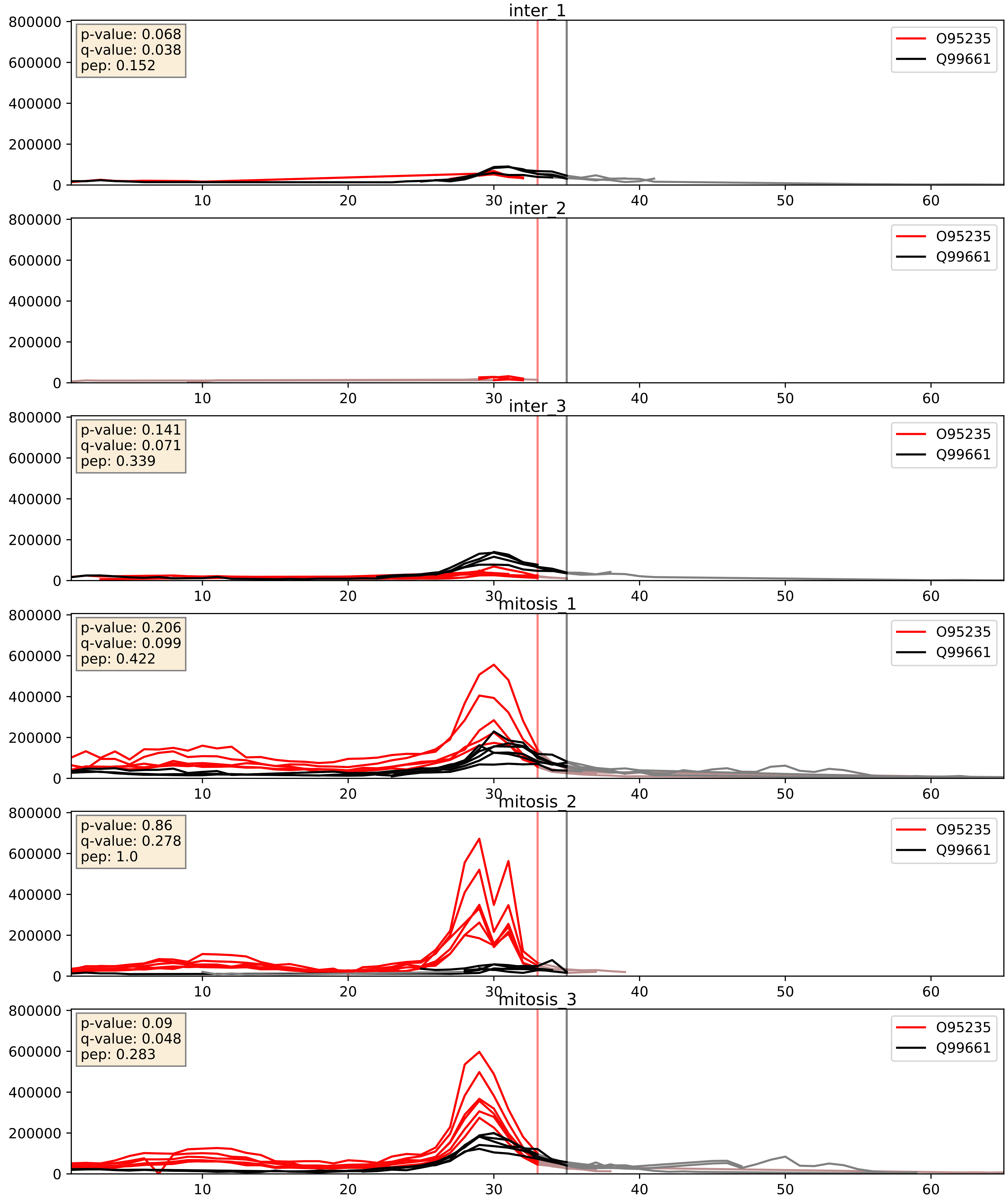


| level | condition_1 | condition_2 | pvalue | pvalue_adjusted |
|---|---|---|---|---|
| interactor_abundance | mitosis | inter | 0.002491334705596794 | 0.0373482050436227 |
| interactor_ratio | mitosis | inter | 0.009413807097681002 | 0.13869567772142752 |
| complex_abundance | mitosis | inter | 0.040808984966054784 | 0.14549142495186546 |
