## Supplementary Data 2 for "SECAT: Quantifying differential protein-protein interaction states by network-centric analysis": hela_string_000086_P19022.pdf

P19022_P19022

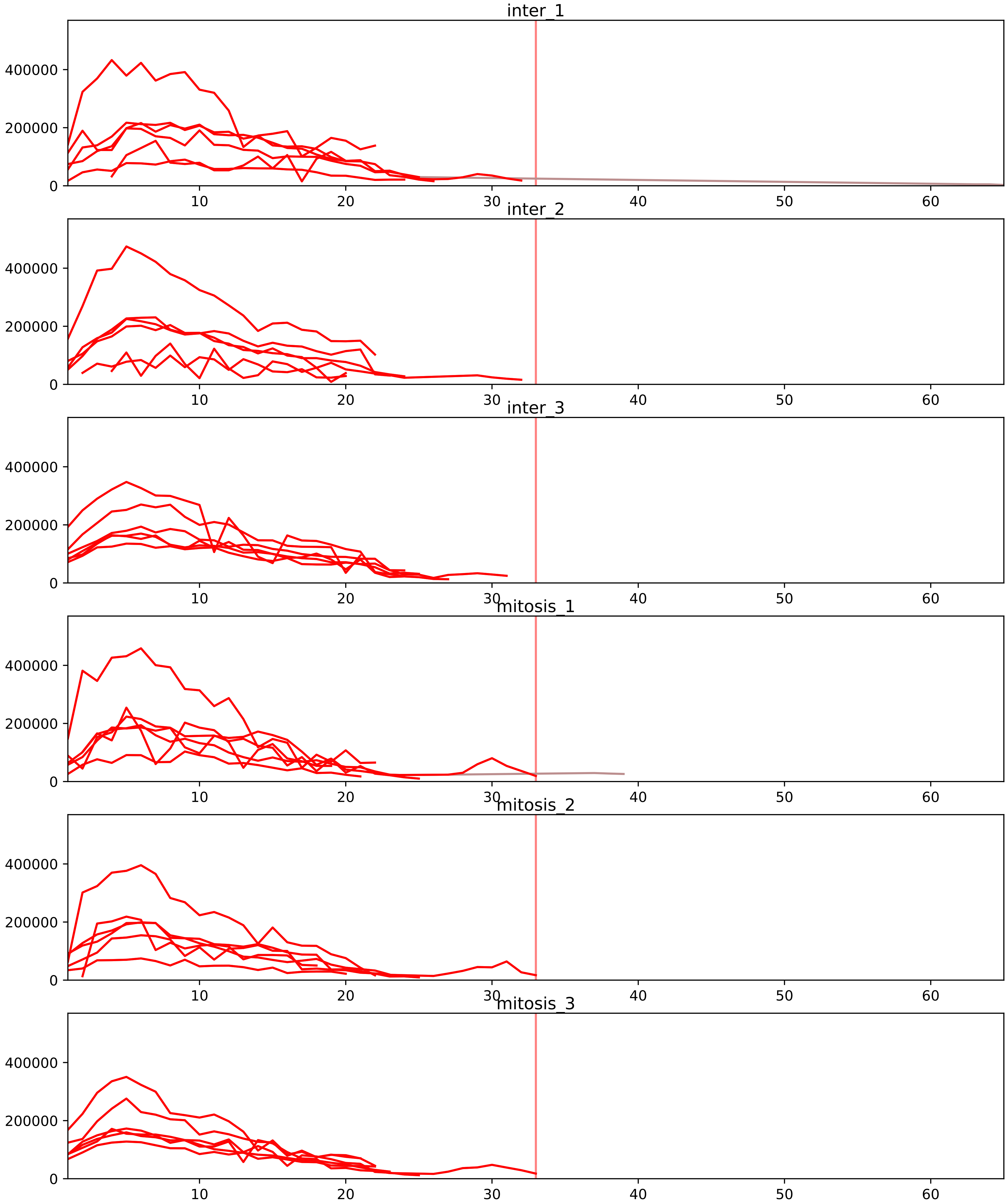


| level | condition_1 | condition_2 | pvalue | pvalue_adjusted |
|---|---|---|---|---|
| assembled_abundance | mitosis | inter | 4.95583359353067e-06 | 0.002899782131414633 |
| total_abundance | mitosis | inter | 9.718634380576206e-06 | 0.004123130635959455 |
| complex_abundance | mitosis | inter | 0.024128167685714394 | 0.07942008683669854 |
| monomer_abundance | mitosis | inter | 0.03320675671732449 | 0.30183233230981965 |
| interactor_abundance | mitosis | inter | 0.03531537796760526 | 0.11380086770646879 |
| interactor_ratio | mitosis | inter | 0.17348866454434866 | 0.4534362823318204 |

P14923_P19022
PLAK_HUMAN vs CADH2_HUMAN
p-value: 0.002 q-value: 0.003

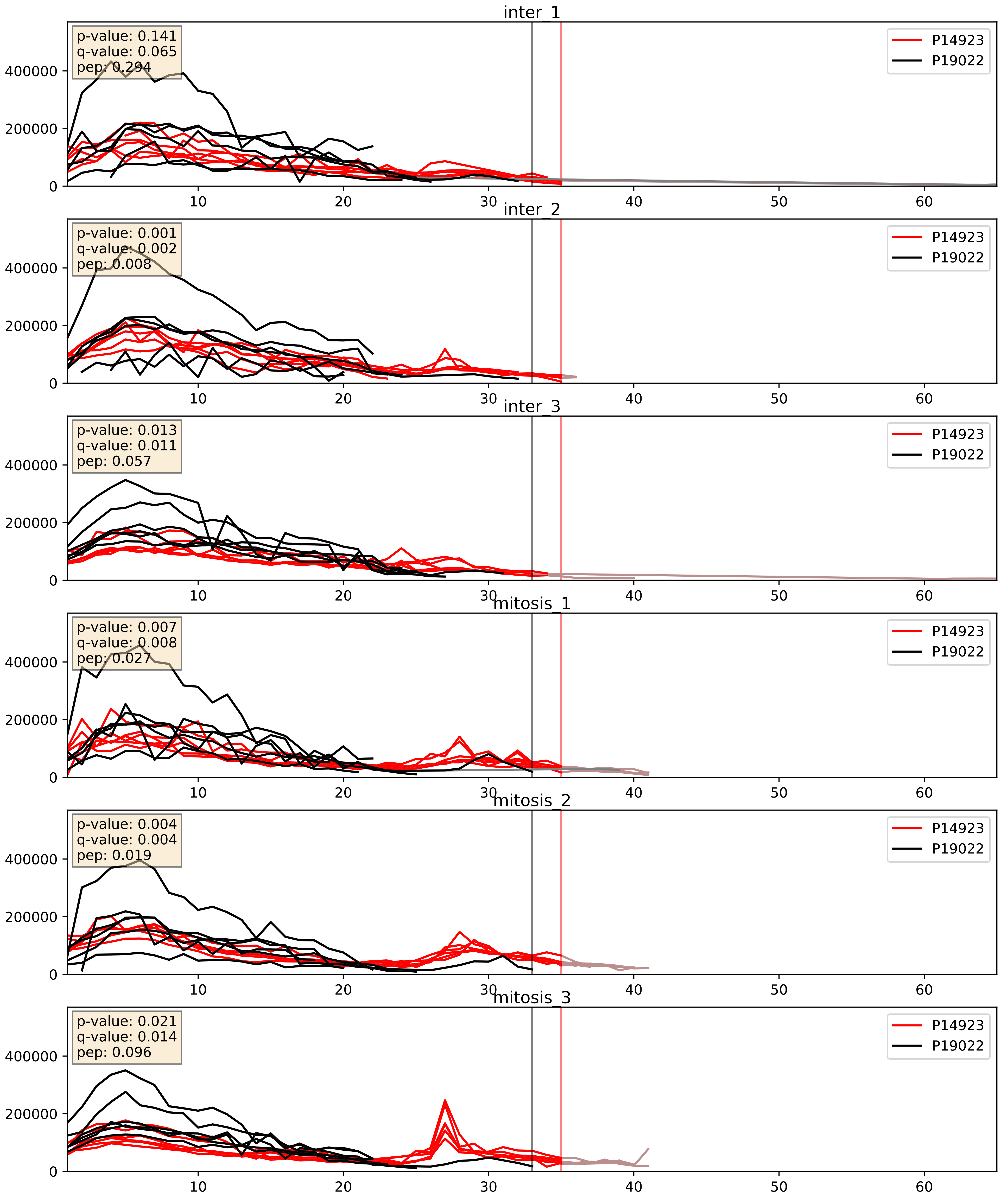


| level | condition_1 | condition_2 | pvalue | pvalue_adjusted |
|---|---|---|---|---|
| complex_abundance | mitosis | inter | 0.07084888972786241 | 0.19345012392626001 |
| interactor_ratio | mitosis | inter | 0.0744644247556564 | 0.45824261388096243 |
| interactor_abundance | mitosis | inter | 0.12013392513199599 | 0.30029279311852347 |

P19022_P35221
CADH2_HUMAN vs CTNA1_HUMAN
p-value: 0.004 q-value: 0.004

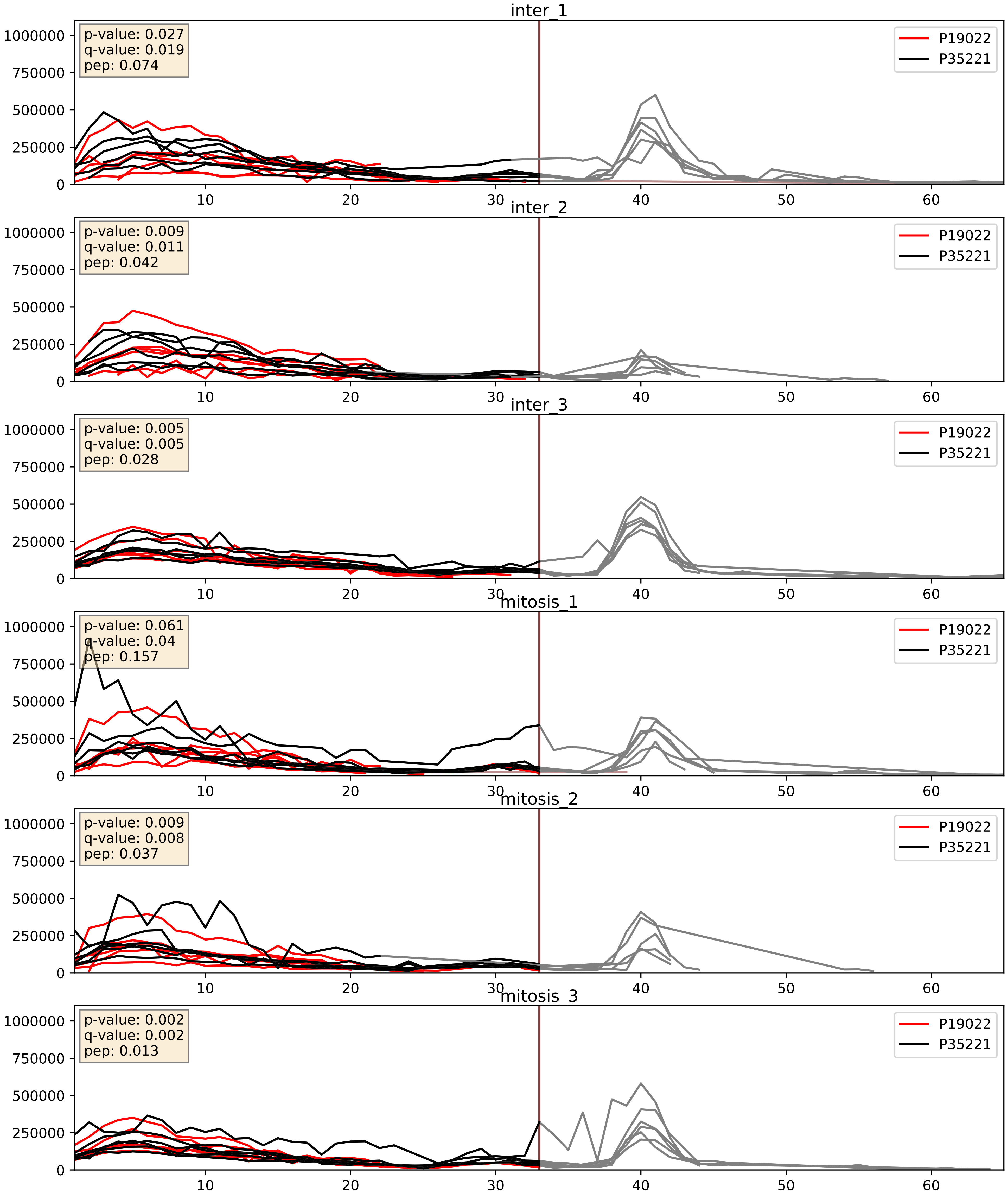


| level | condition_1 | condition_2 | pvalue | pvalue_adjusted |
|---|---|---|---|---|
| interactor_abundance | mitosis | inter | 0.061915759789182935 | 0.20357169341094908 |
| complex_abundance | mitosis | inter | 0.09322618704752257 | 0.22532498068132825 |
| interactor_ratio | mitosis | inter | 0.21091091773910894 | 0.6885573820925905 |

P19022_P35222
CADH2_HUMAN vs CTNB1_HUMAN
p-value: 0.003 q-value: 0.003

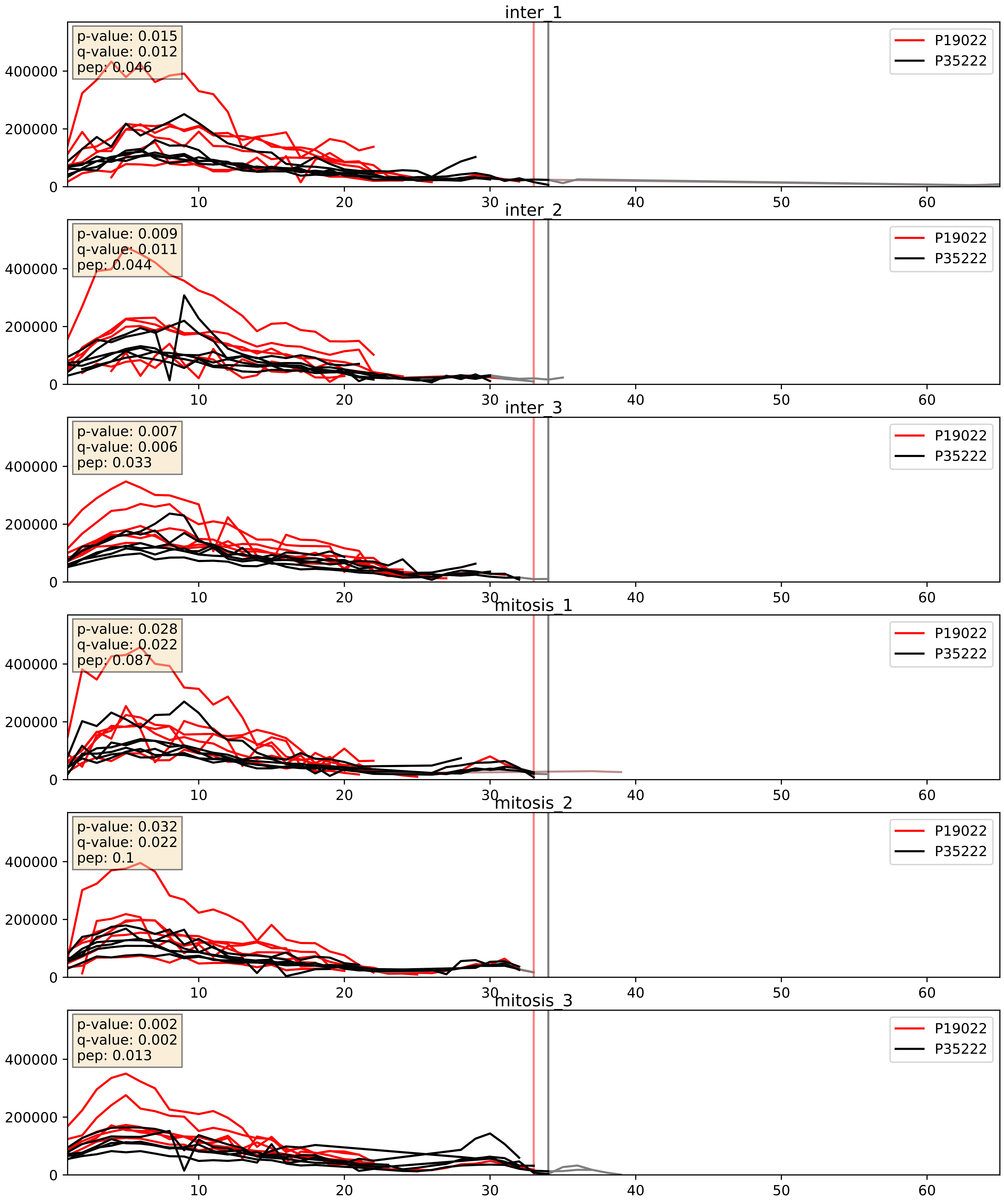


| level | condition_1 | condition_2 | pvalue | pvalue_adjusted |
|---|---|---|---|---|
| complex_abundance | mitosis | inter | 0.016394450140038135 | 0.09572748512873563 |
| interactor_abundance | mitosis | inter | 0.059200885939813894 | 0.1984691235397679 |
| interactor_ratio | mitosis | inter | 0.7067757583156111 | 0.9386176777228961 |
