## Supplementary Data 2 for "SECAT: Quantifying differential protein-protein interaction states by network-centric analysis": hela_string_000087_Q9Y2X3.pdf

Q9Y2X3_Q9Y2X3

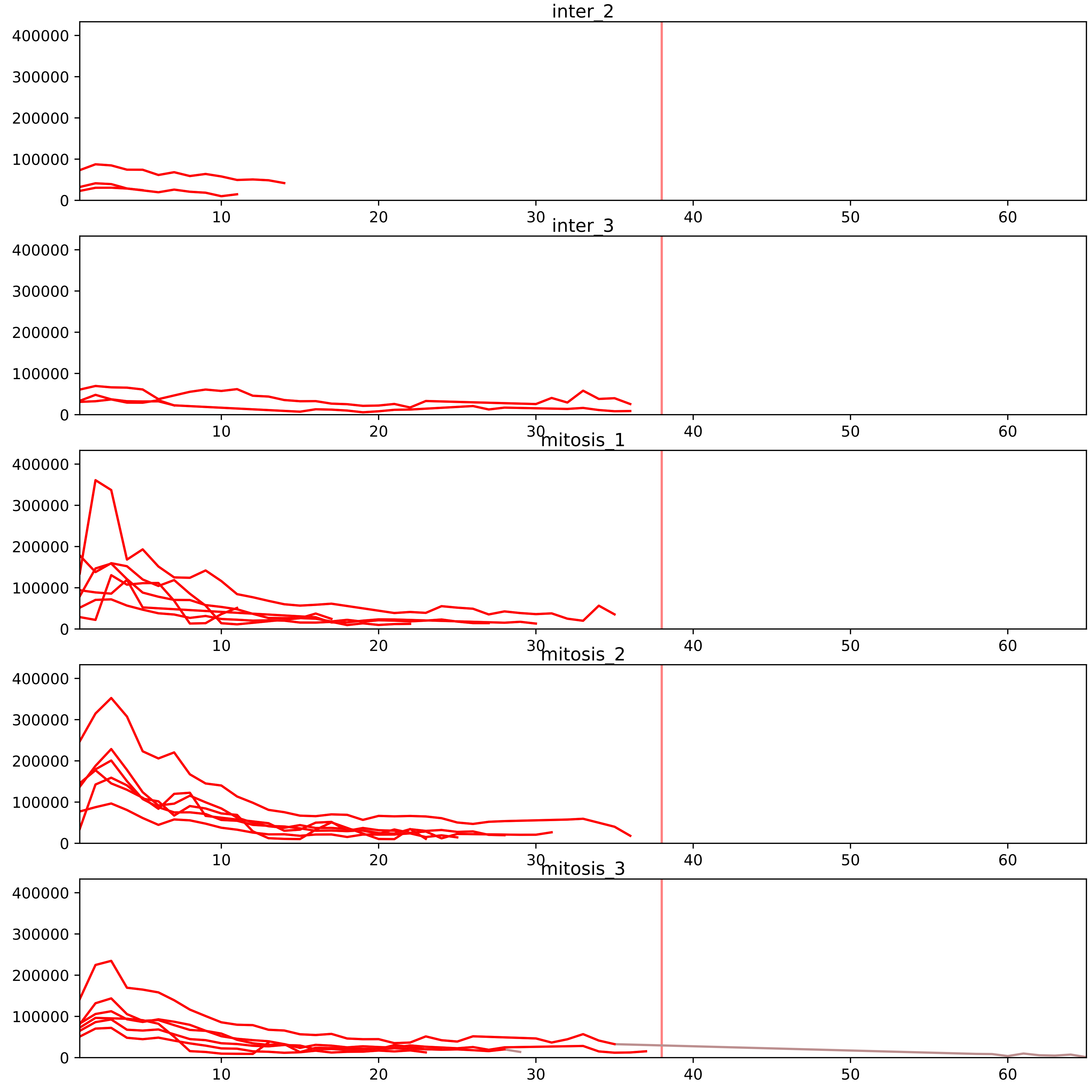


| level | condition_1 | condition_2 | pvalue | pvalue_adjusted |
|---|---|---|---|---|
| interactor_abundance | mitosis | inter | 5.0170917506979904e-06 | 0.00030771496070947676 |
| interactor_ratio | mitosis | inter | 5.958503231955422e-05 | 0.0011074389845250482 |
| complex_abundance | mitosis | inter | 0.0006151714388822871 | 0.00780631343133385 |
| total_abundance | mitosis | inter | 0.17580994545726875 | 0.5107045870665315 |
| assembled_abundance | mitosis | inter | 0.1820924595222927 | 0.49071934386920535 |
| monomer_abundance | mitosis | inter | 0.40512756950638223 | 0.6978579466069973 |

O00567_Q9Y2X3
NOP56_HUMAN vs NOP58_HUMAN
p-value: 0.001 q-value: 0.001

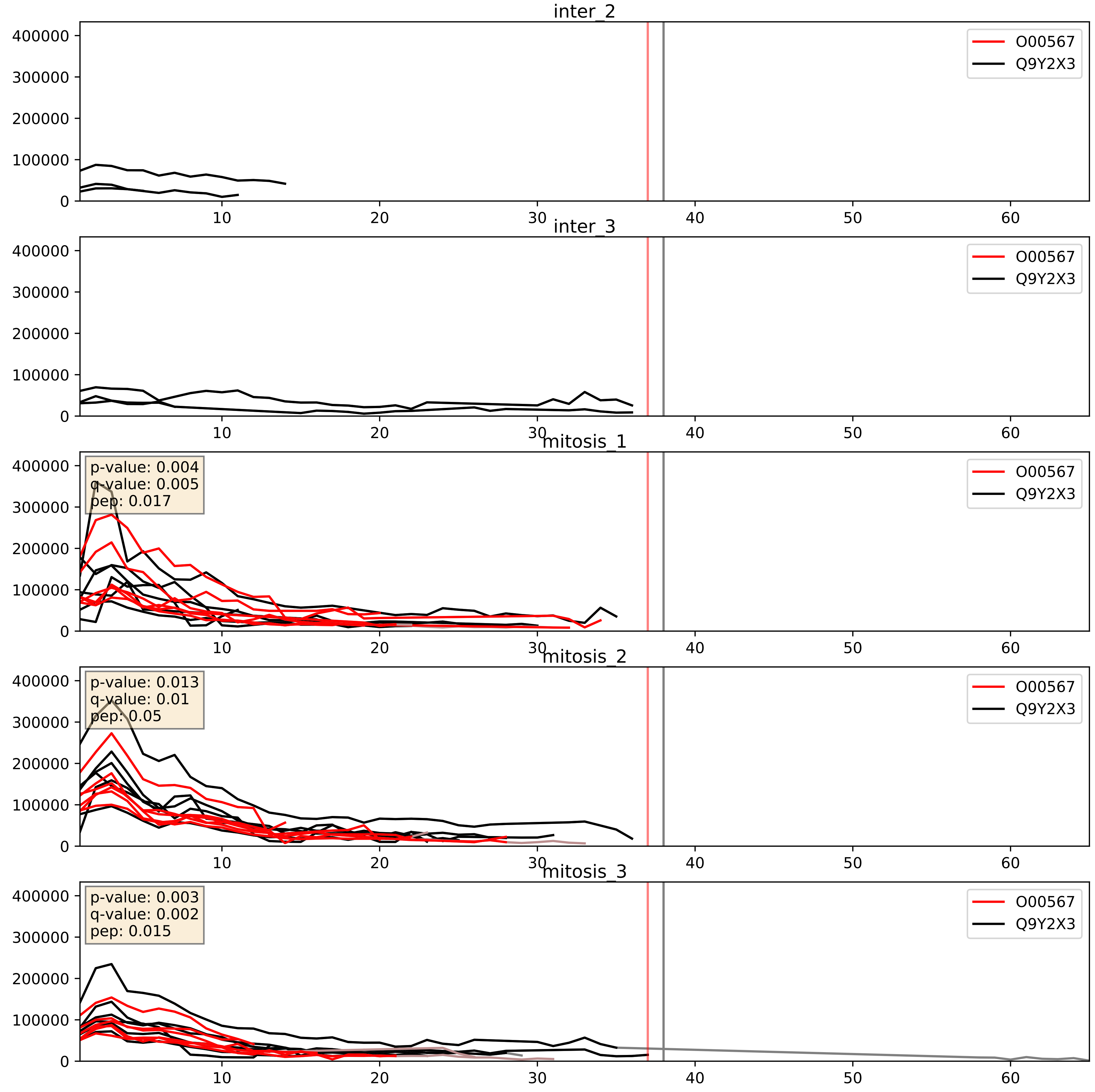


| level | condition_1 | condition_2 | pvalue | pvalue_adjusted |
|---|---|---|---|---|
| complex_abundance | mitosis | inter | 0.00028226689255234446 | 0.011311068978518664 |
| interactor_abundance | mitosis | inter | 0.0011181996263145508 | 0.024049720606162197 |
| interactor_ratio | mitosis | inter | 0.12583682042012595 | 0.5750062399685051 |

O60832_Q9Y2X3
DKC1_HUMAN vs NOP58_HUMAN
p-value: 0.01 q-value: 0.009

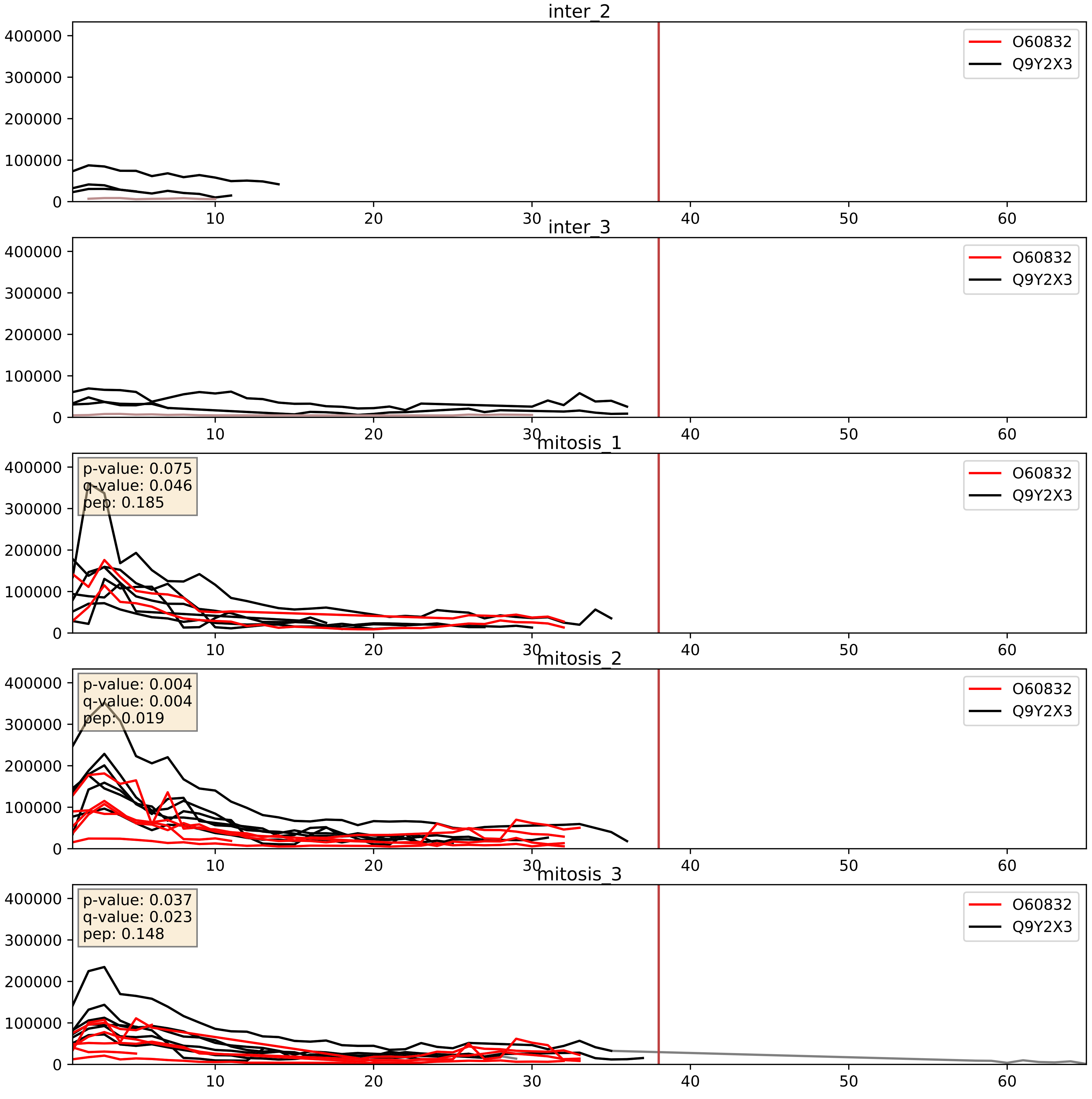


| level | condition_1 | condition_2 | pvalue | pvalue_adjusted |
|---|---|---|---|---|
| complex_abundance | mitosis | inter | 0.002344505735626372 | 0.035168510428347066 |
| interactor_abundance | mitosis | inter | 0.04900154453194635 | 0.17683525345424148 |
| interactor_ratio | mitosis | inter | 0.05935818141809091 | 0.40293896347252833 |

O76021_Q9Y2X3
RL1D1_HUMAN vs NOP58_HUMAN
p-value: 0.001 q-value: 0.002

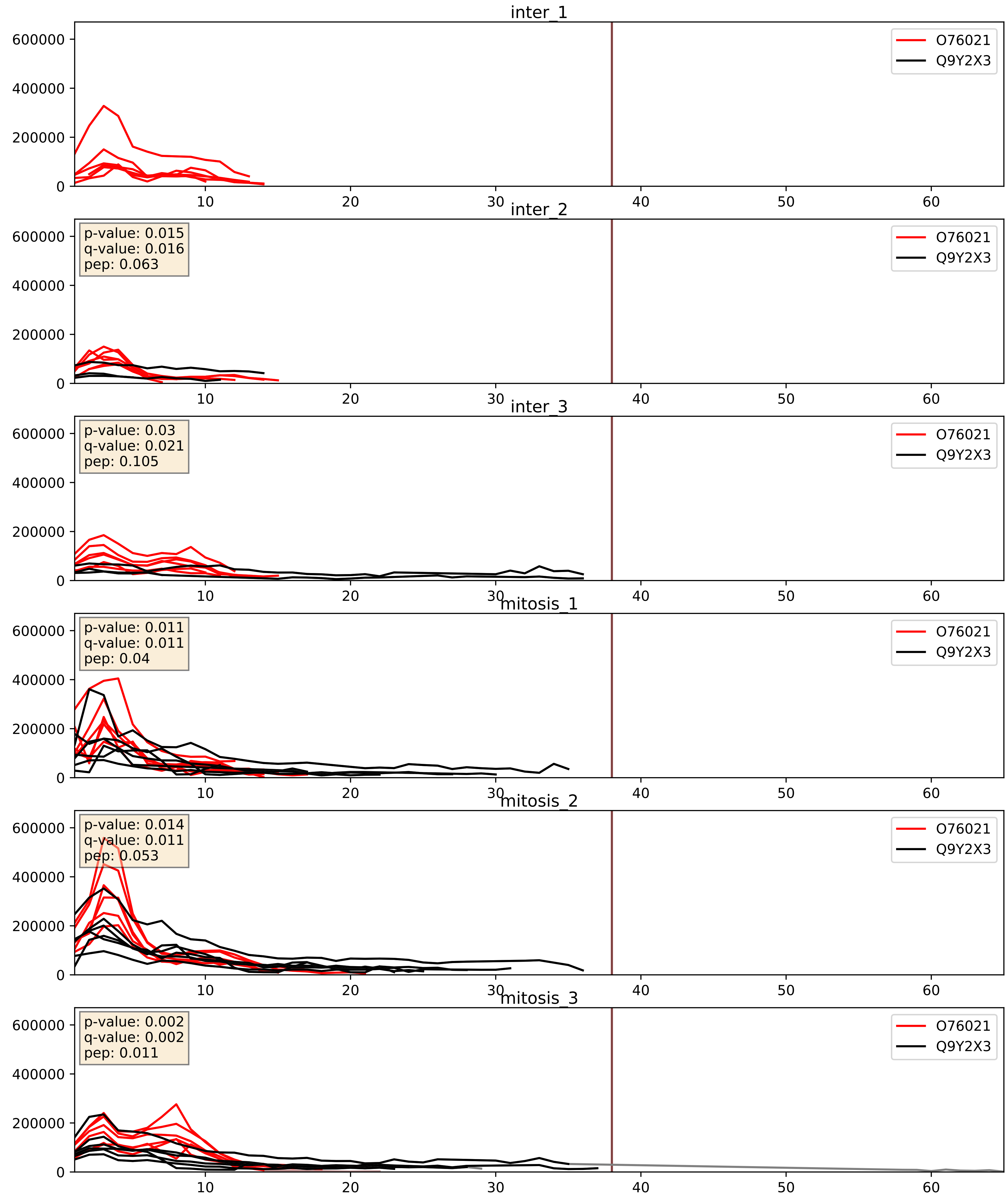


| level | condition_1 | condition_2 | pvalue | pvalue_adjusted |
|---|---|---|---|---|
| complex_abundance | mitosis | inter | 0.01952163921039481 | 0.10409657315782883 |
| interactor_abundance | mitosis | inter | 0.0810993279136887 | 0.23762116958452006 |
| interactor_ratio | mitosis | inter | 0.3037548150127056 | 0.7719380618221225 |

P18124_Q9Y2X3
RL7_HUMAN vs NOP58_HUMAN
p-value: 0.001 q-value: 0.039

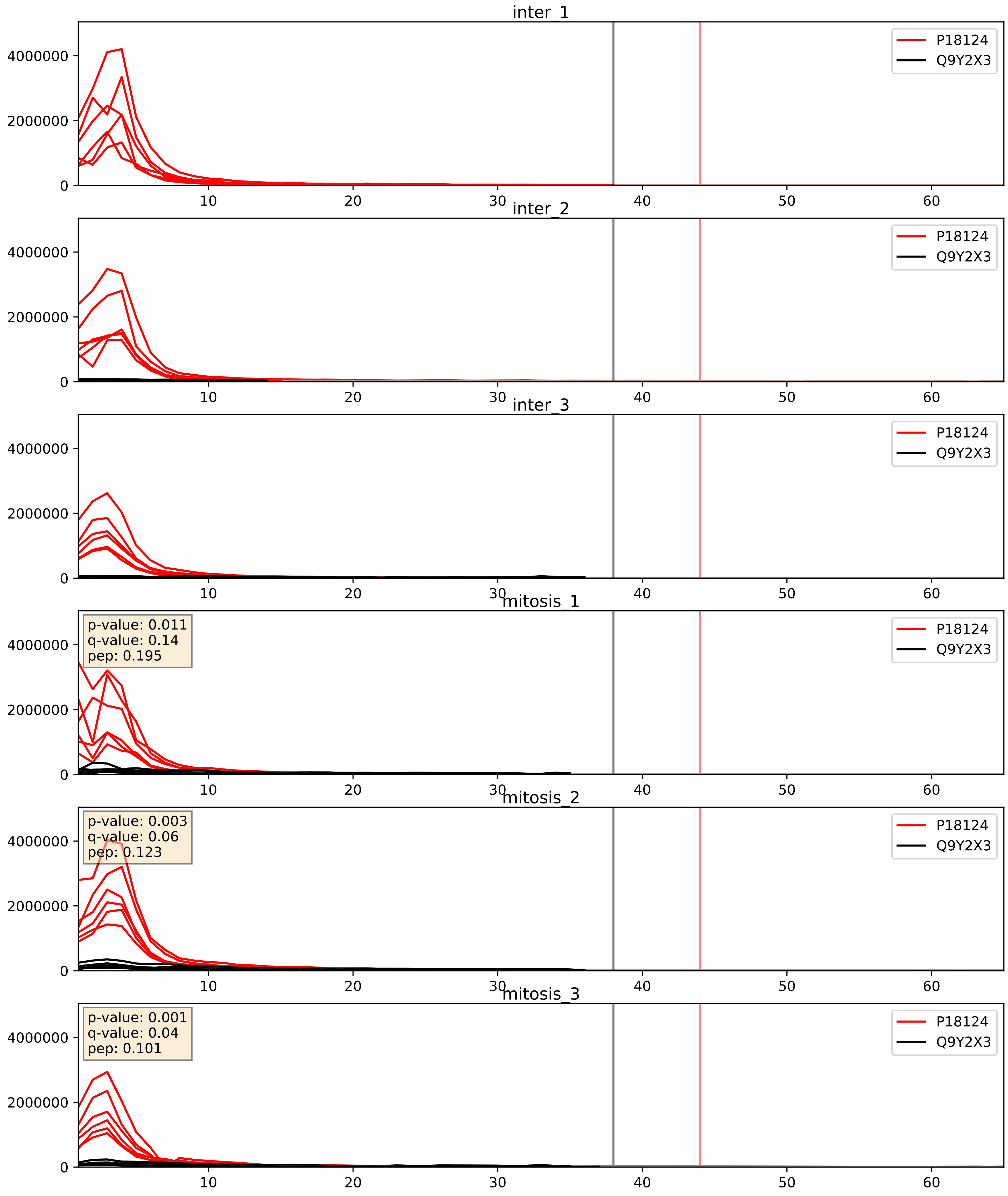


| level | condition_1 | condition_2 | pvalue | pvalue_adjusted |
|---|---|---|---|---|
| complex_abundance | mitosis | inter | 0.13604284374140788 | 0.2812866527600124 |
| interactor_abundance | mitosis | inter | 0.35877726225927037 | 0.55606253212735 |
| interactor_ratio | mitosis | inter | 0.8749925269452397 | 0.9802747470617126 |

P22087_Q9Y2X3
FBRL_HUMAN vs NOP58_HUMAN
p-value: 0.0 q-value: 0.0

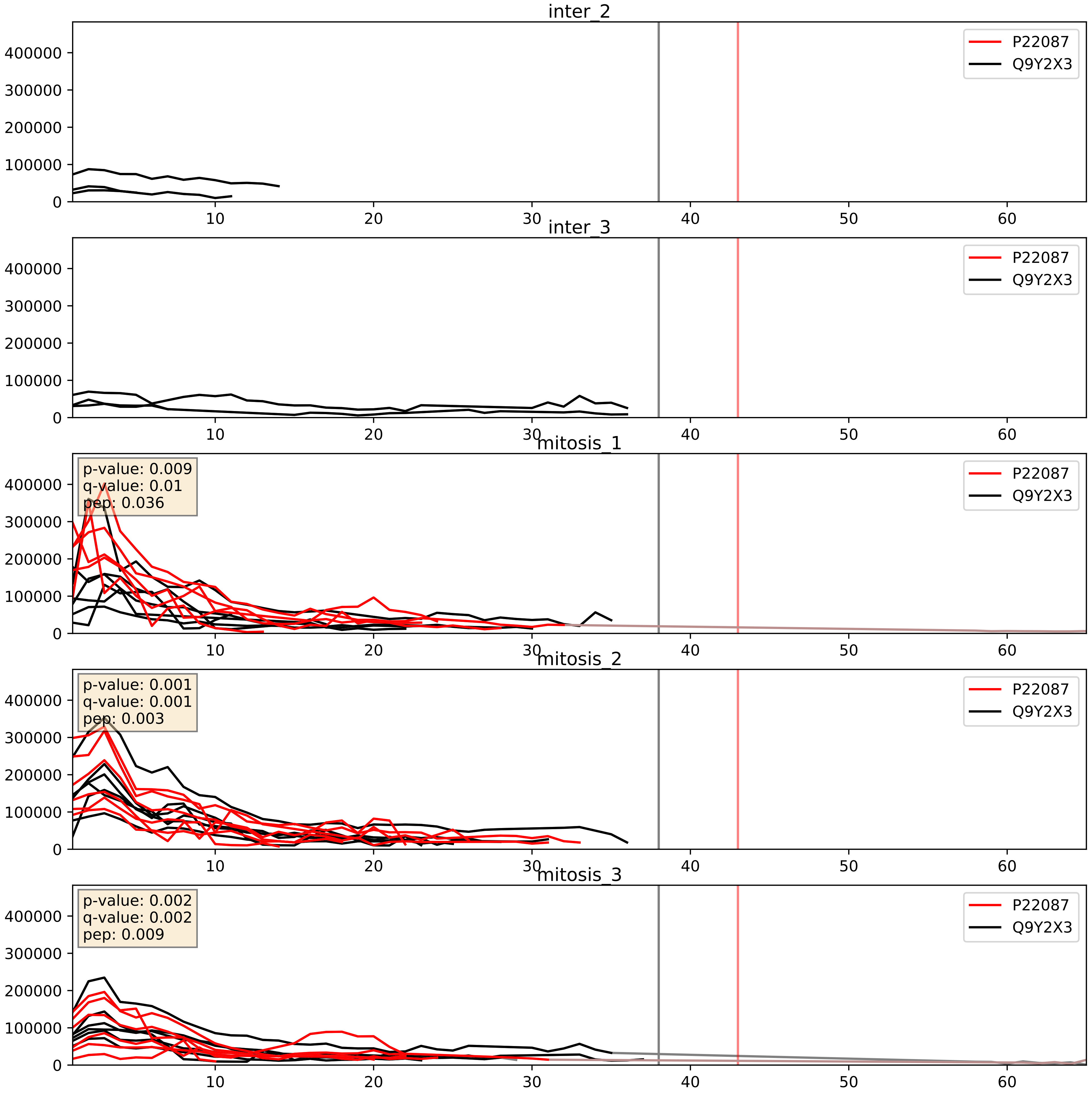


| level | condition_1 | condition_2 | pvalue | pvalue_adjusted |
|---|---|---|---|---|
| complex_abundance | mitosis | inter | 0.00015183324738868412 | 0.008550609195046946 |
| interactor_abundance | mitosis | inter | 0.00038188998169085025 | 0.013573913378163039 |
| interactor_ratio | mitosis | inter | 0.42563529182084064 | 0.841246386050888 |

P42285_Q9Y2X3
MTREX_HUMAN vs NOP58_HUMAN
p-value: 0.061 q-value: 0.038

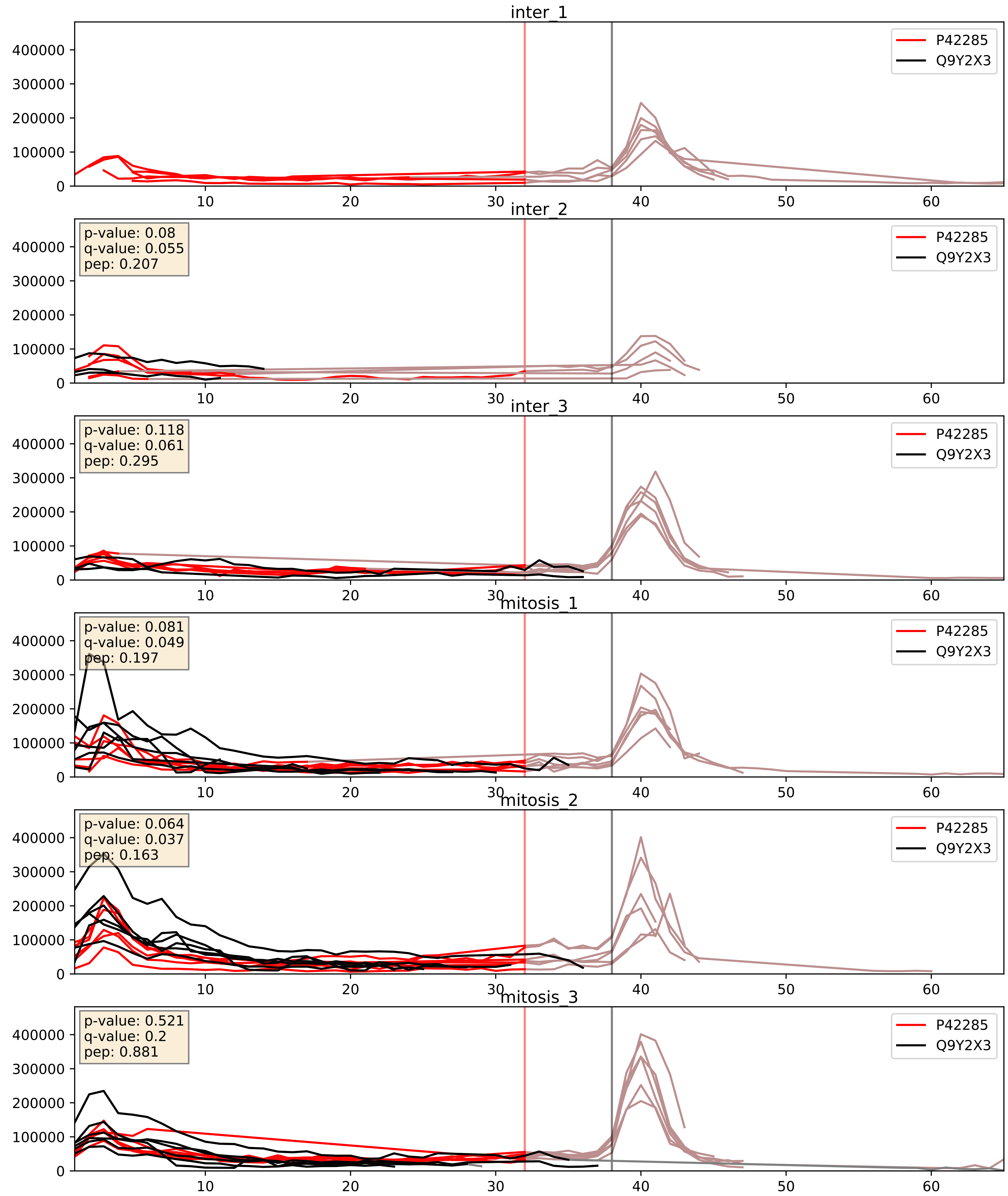


| level | condition_1 | condition_2 | pvalue | pvalue_adjusted |
|---|---|---|---|---|
| complex_abundance | mitosis | inter | 0.010746925639457542 | 0.07672534067869605 |
| interactor_ratio | mitosis | inter | 0.03074091262697153 | 0.2793441741898899 |
| interactor_abundance | mitosis | inter | 0.04261221300826687 | 0.16636566759504254 |

P46087_Q9Y2X3
NOP2_HUMAN vs NOP58_HUMAN
p-value: 0.014 q-value: 0.011

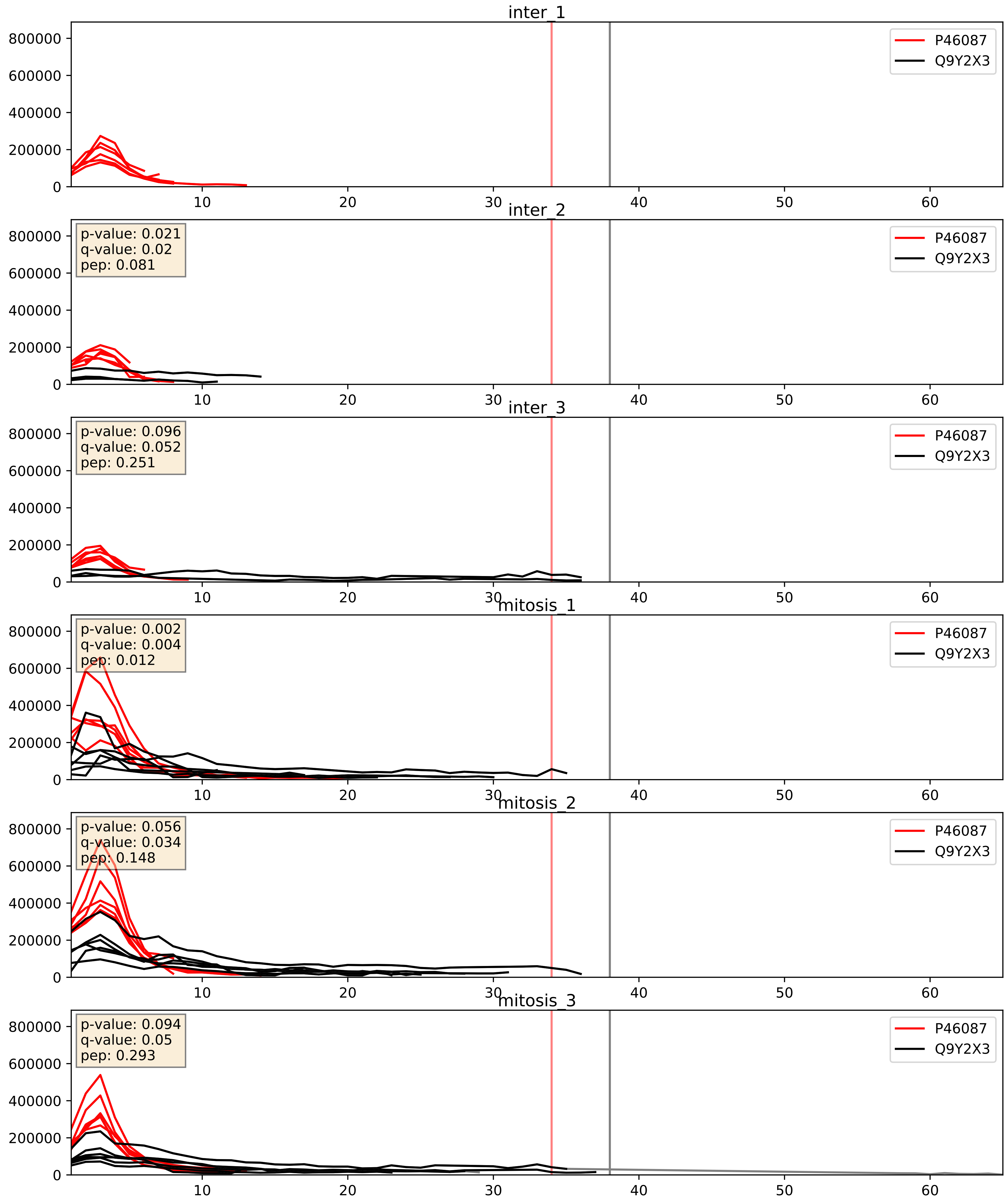


| level | condition_1 | condition_2 | pvalue | pvalue_adjusted |
|---|---|---|---|---|
| complex_abundance | mitosis | inter | 0.012113264791281528 | 0.0818150085787963 |
| interactor_abundance | mitosis | inter | 0.03839338005658009 | 0.15936046986653518 |
| interactor_ratio | mitosis | inter | 0.6680032234742932 | 0.9302273617927362 |

P55769_Q9Y2X3
NH2L1_HUMAN vs NOP58_HUMAN
p-value: 0.0 q-value: 0.0

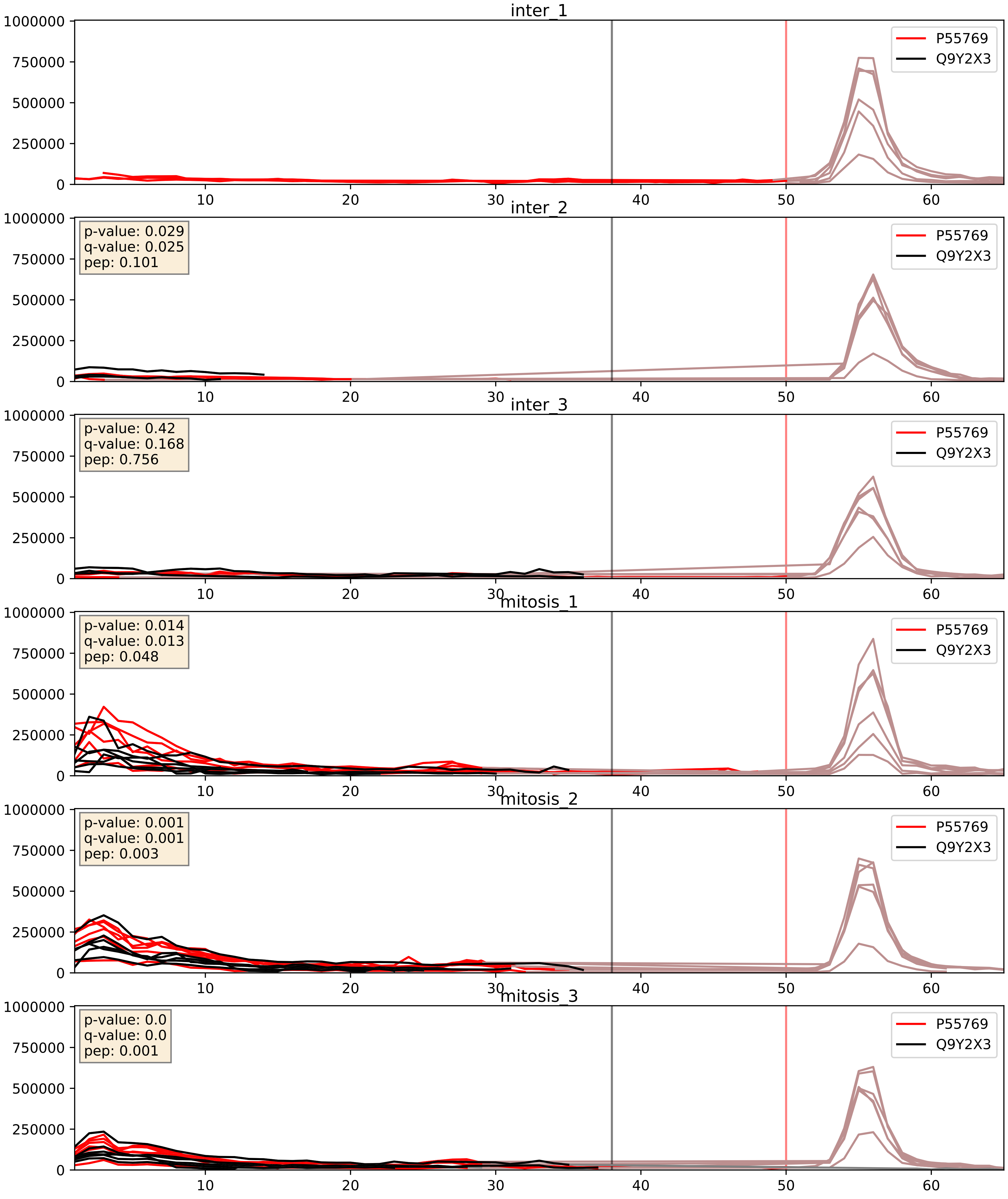


| level | condition_1 | condition_2 | pvalue | pvalue_adjusted |
|---|---|---|---|---|
| interactor_abundance | mitosis | inter | 0.0024610293995718582 | 0.037056133087660696 |
| complex_abundance | mitosis | inter | 0.0024997379458740598 | 0.03638642644133238 |
| interactor_ratio | mitosis | inter | 0.04568692907471442 | 0.35552737534505047 |

P56537_Q9Y2X3
IF6_HUMAN vs NOP58_HUMAN
p-value: 0.002 q-value: 0.03

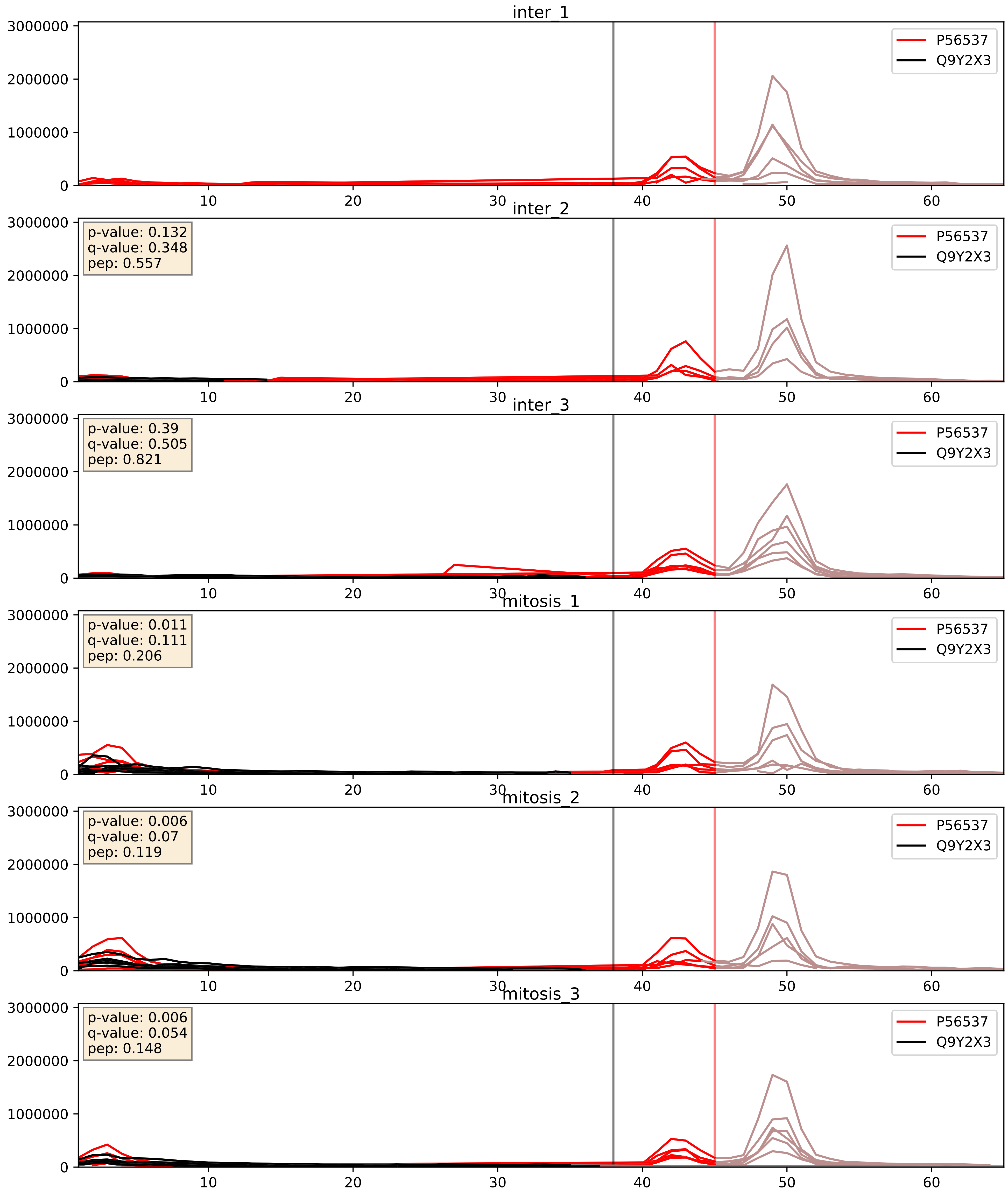


| level | condition_1 | condition_2 | pvalue | pvalue_adjusted |
|---|---|---|---|---|
| complex_abundance | mitosis | inter | 0.008291925197678492 | 0.06742444735376033 |
| interactor_abundance | mitosis | inter | 0.018773691505101508 | 0.11803363884220999 |
| interactor_ratio | mitosis | inter | 0.2376377875942226 | 0.7165126670681738 |

P61247_Q9Y2X3
RS3A_HUMAN vs NOP58_HUMAN
p-value: 0.084 q-value: 0.049

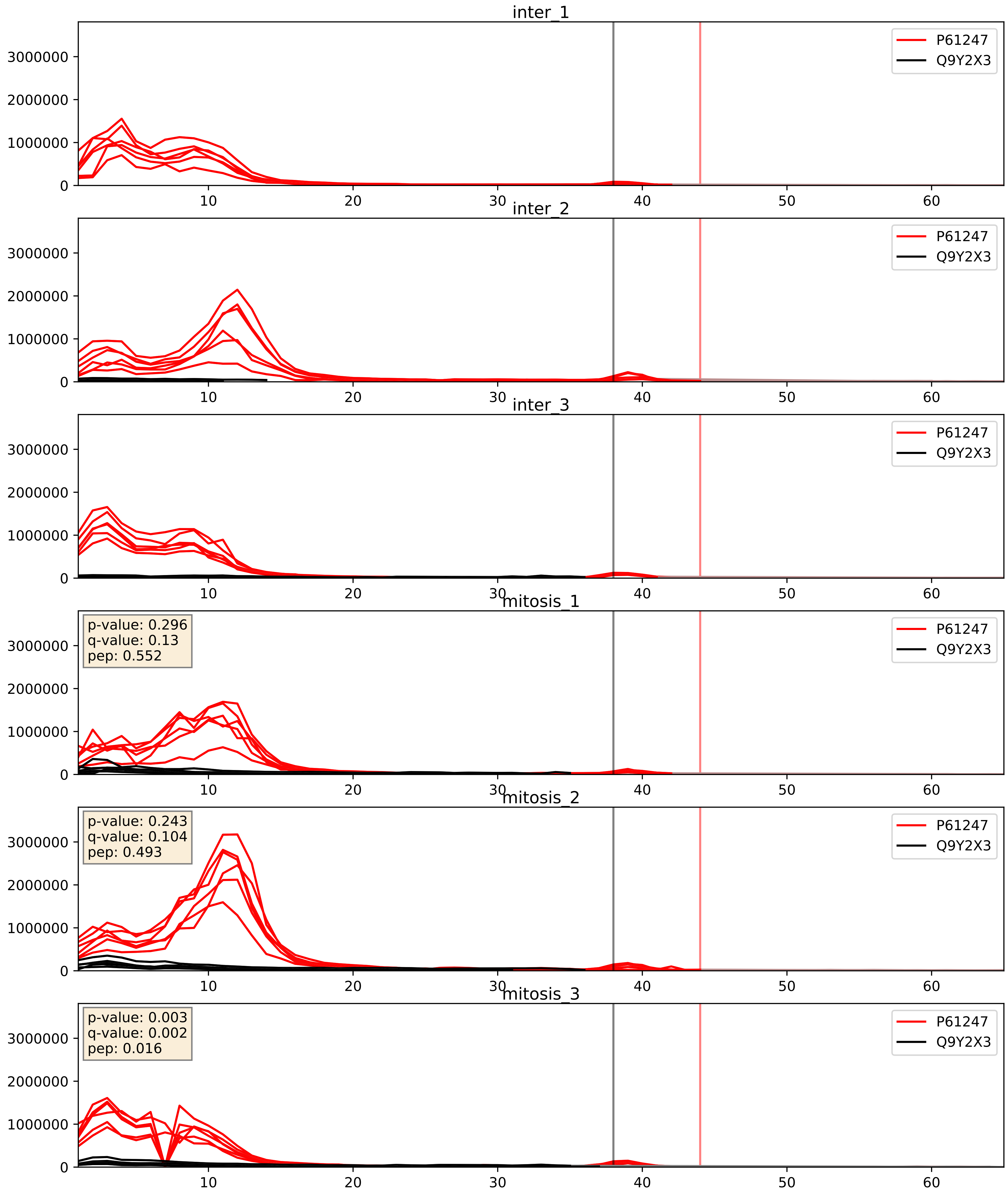


| level | condition_1 | condition_2 | pvalue | pvalue_adjusted |
|---|---|---|---|---|
| complex_abundance | mitosis | inter | 0.13929258382575133 | 0.28456909726692875 |
| interactor_abundance | mitosis | inter | 0.3433868839104531 | 0.5420201629725344 |
| interactor_ratio | mitosis | inter | 0.9892629882678046 | 0.9981863229331772 |

P62081_Q9Y2X3
RS7_HUMAN vs NOP58_HUMAN
p-value: 0.005 q-value: 0.005

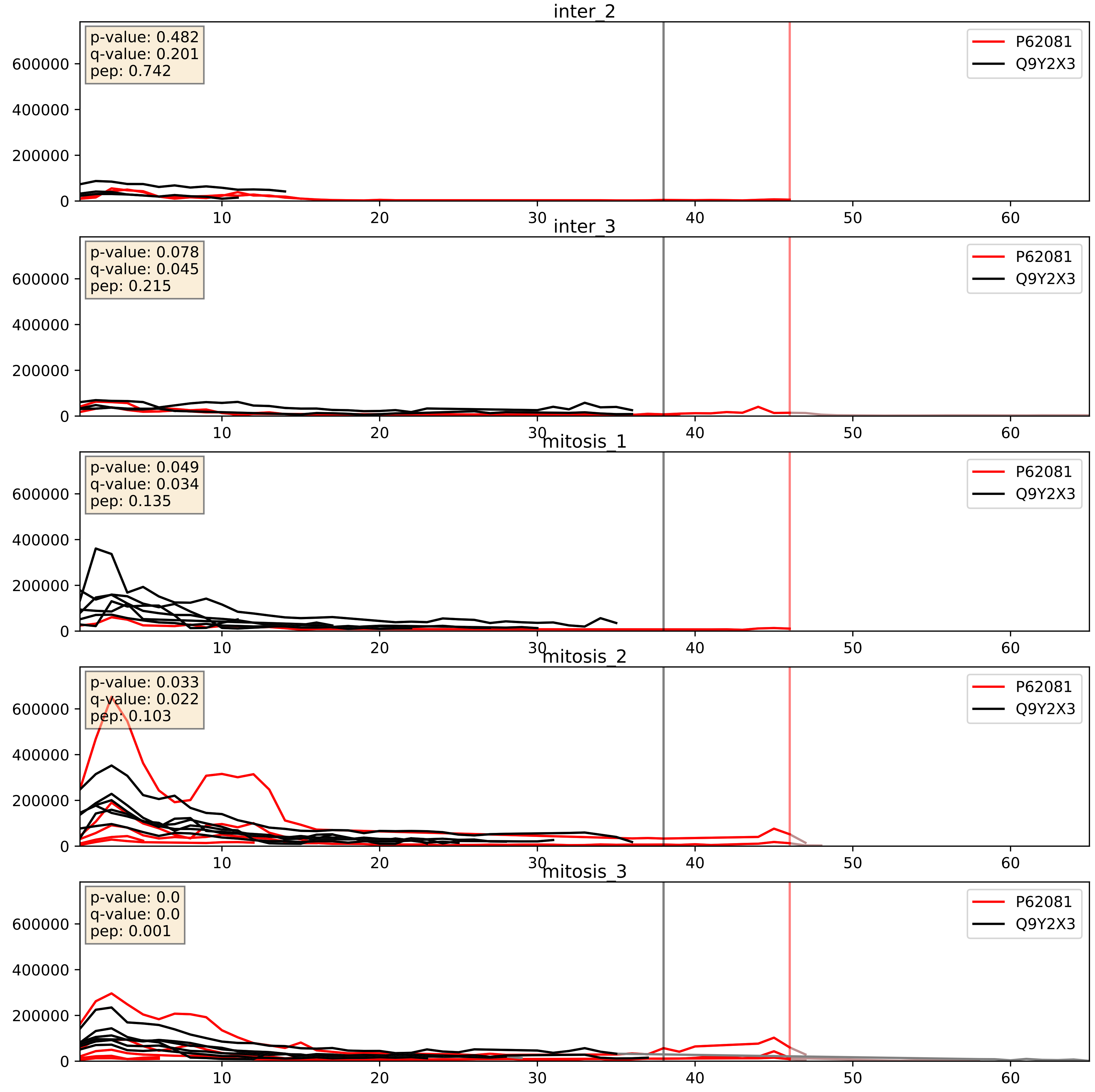


| level | condition_1 | condition_2 | pvalue | pvalue_adjusted |
|---|---|---|---|---|
| complex_abundance | mitosis | inter | 0.03412511544696356 | 0.13430390263264738 |
| interactor_ratio | mitosis | inter | 0.1688476843199037 | 0.6317028749031363 |
| interactor_abundance | mitosis | inter | 0.24658376608834062 | 0.4400220317547362 |

P62241_Q9Y2X3
RS8_HUMAN vs NOP58_HUMAN
p-value: 0.001 q-value: 0.001

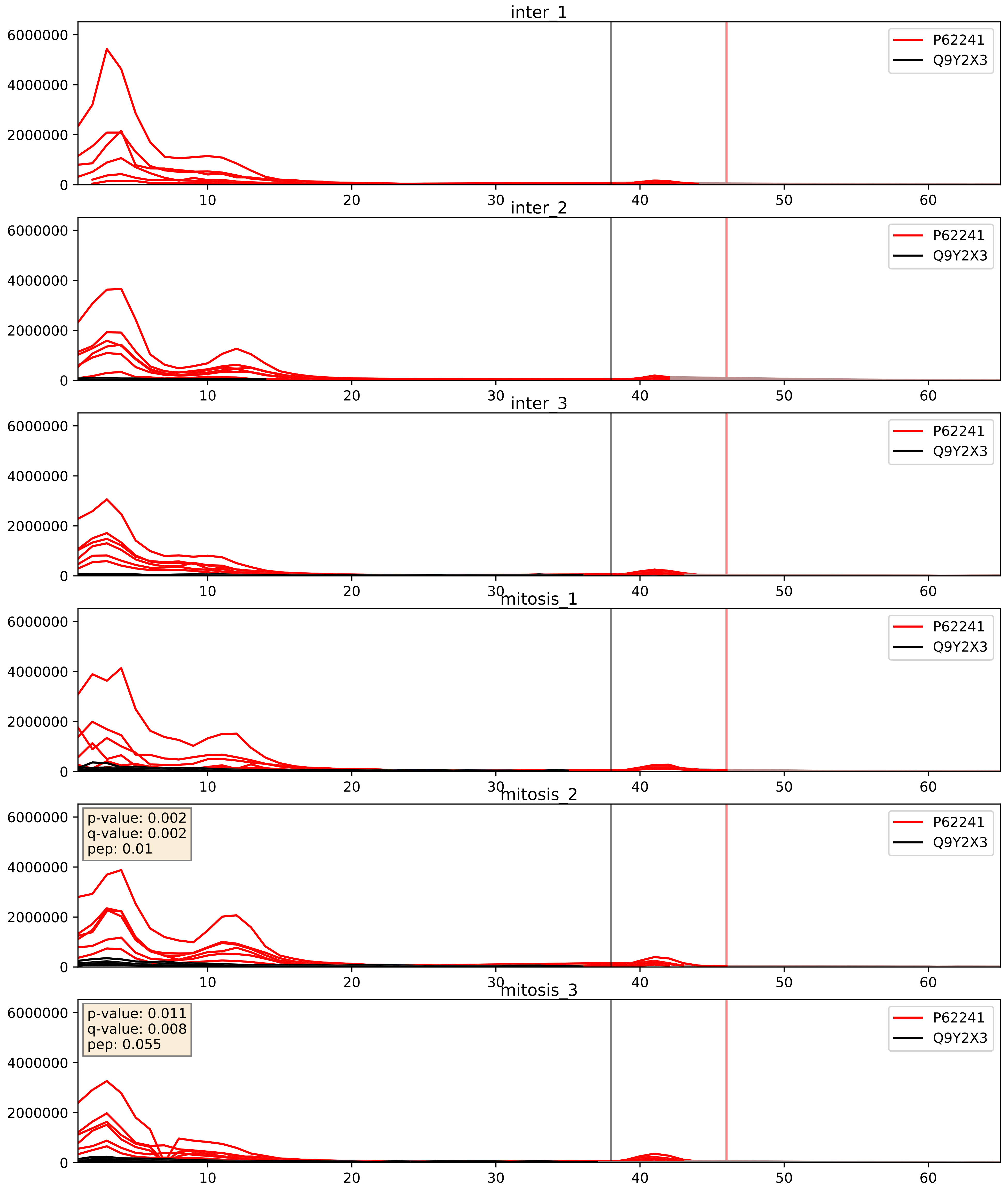


| level | condition_1 | condition_2 | pvalue | pvalue_adjusted |
|---|---|---|---|---|
| complex_abundance | mitosis | inter | 0.15225454898690197 | 0.29878471786517213 |
| interactor_abundance | mitosis | inter | 0.37055777769805676 | 0.5647325887398945 |
| interactor_ratio | mitosis | inter | 0.9670504853360298 | 0.9937294965826962 |

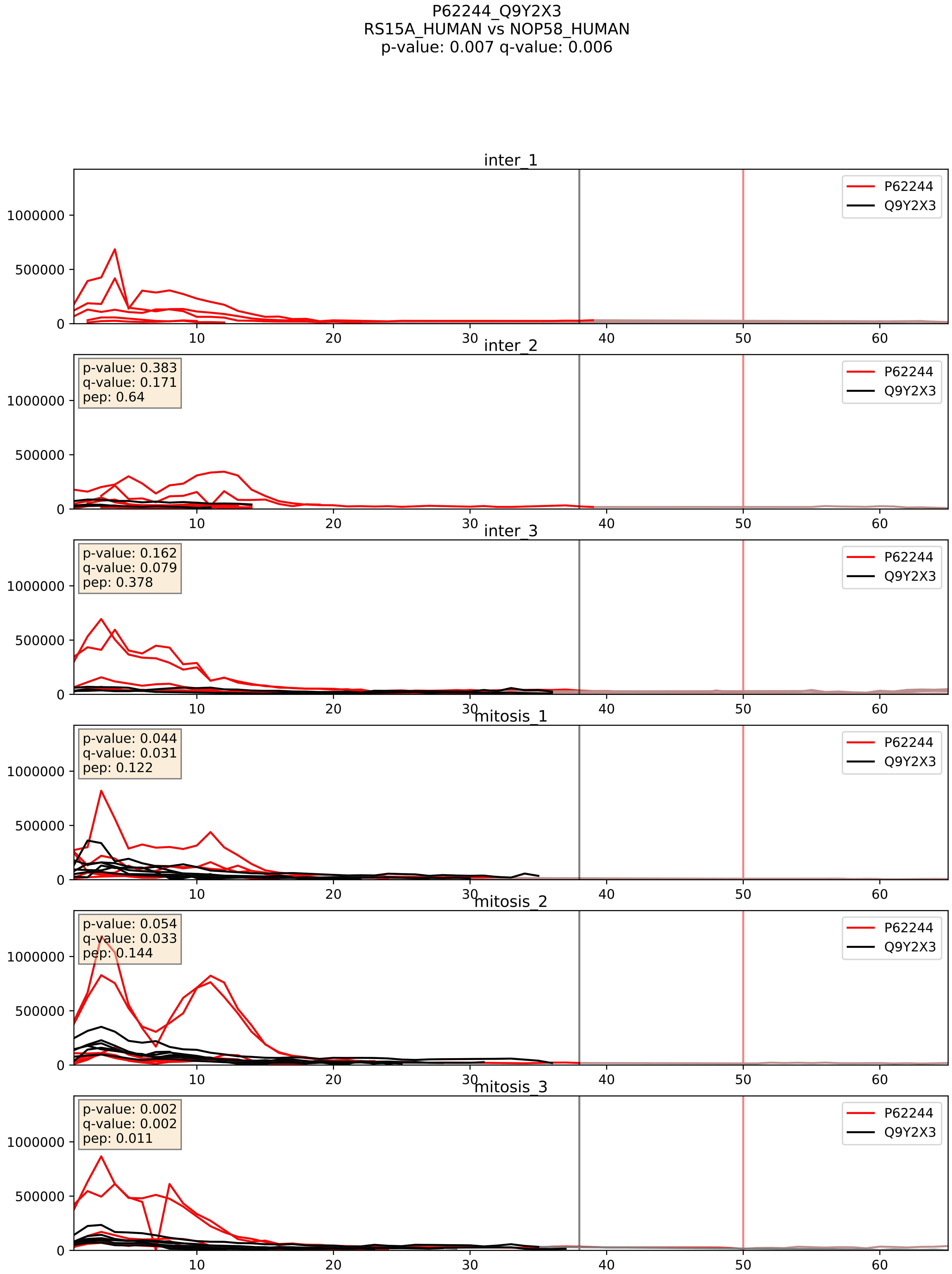


| level | condition_1 | condition_2 | pvalue | pvalue_adjusted |
|---|---|---|---|---|
| complex_abundance | mitosis | inter | 0.05590404049206118 | 0.16855885403735246 |
| interactor_abundance | mitosis | inter | 0.2149769649605422 | 0.4103931356071011 |
| interactor_ratio | mitosis | inter | 0.45891290366799514 | 0.8567708735873585 |

P62249_Q9Y2X3
RS16_HUMAN vs NOP58_HUMAN
p-value: 0.004 q-value: 0.047

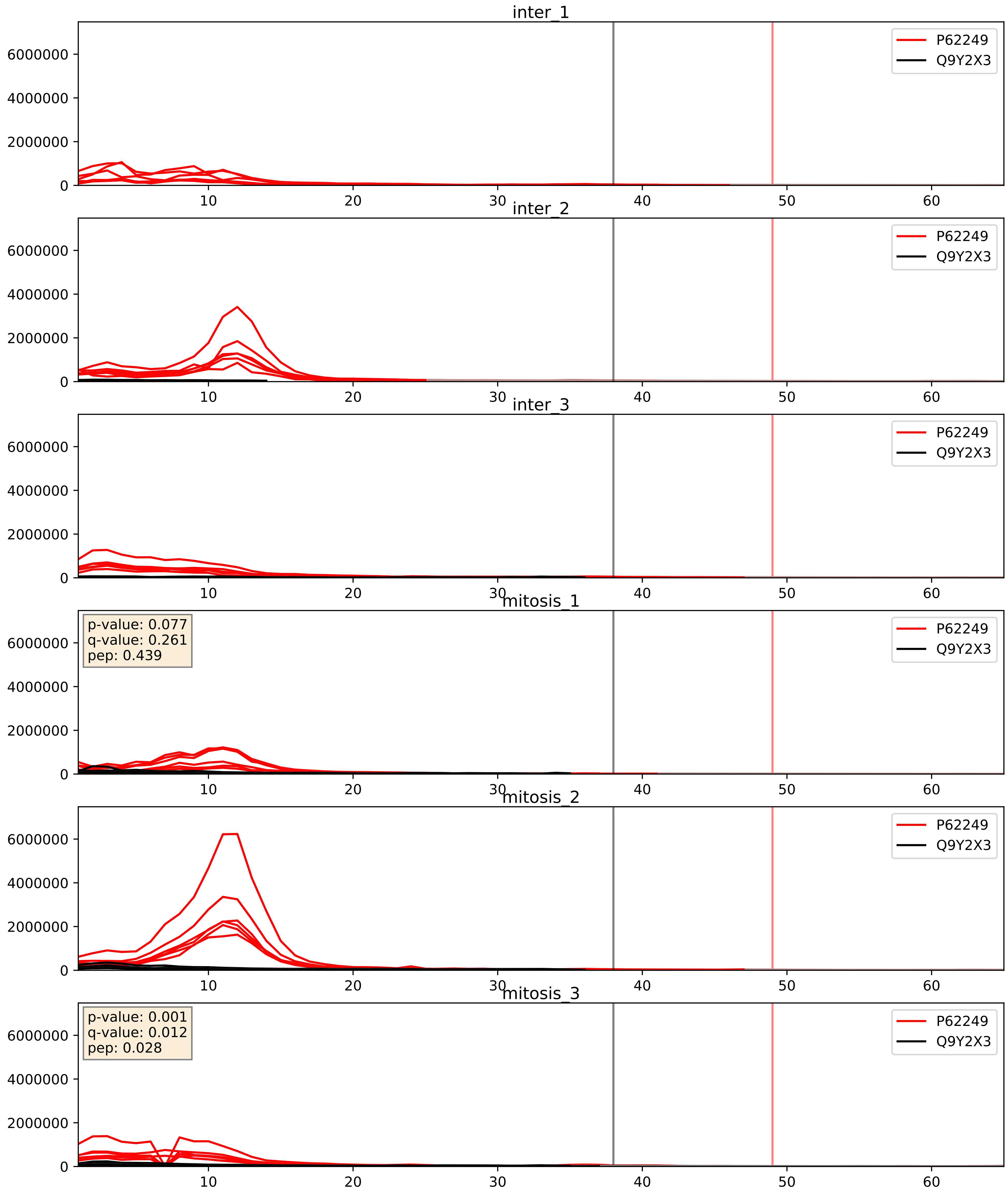


| level | condition_1 | condition_2 | pvalue | pvalue_adjusted |
|---|---|---|---|---|
| complex_abundance | mitosis | inter | 0.12973014762503704 | 0.273183287495773 |
| interactor_abundance | mitosis | inter | 0.3253739965644165 | 0.5246652847983095 |
| interactor_ratio | mitosis | inter | 0.9919370470547214 | 0.9981863229331772 |

P62263_Q9Y2X3
RS14_HUMAN vs NOP58_HUMAN
p-value: 0.006 q-value: 0.006

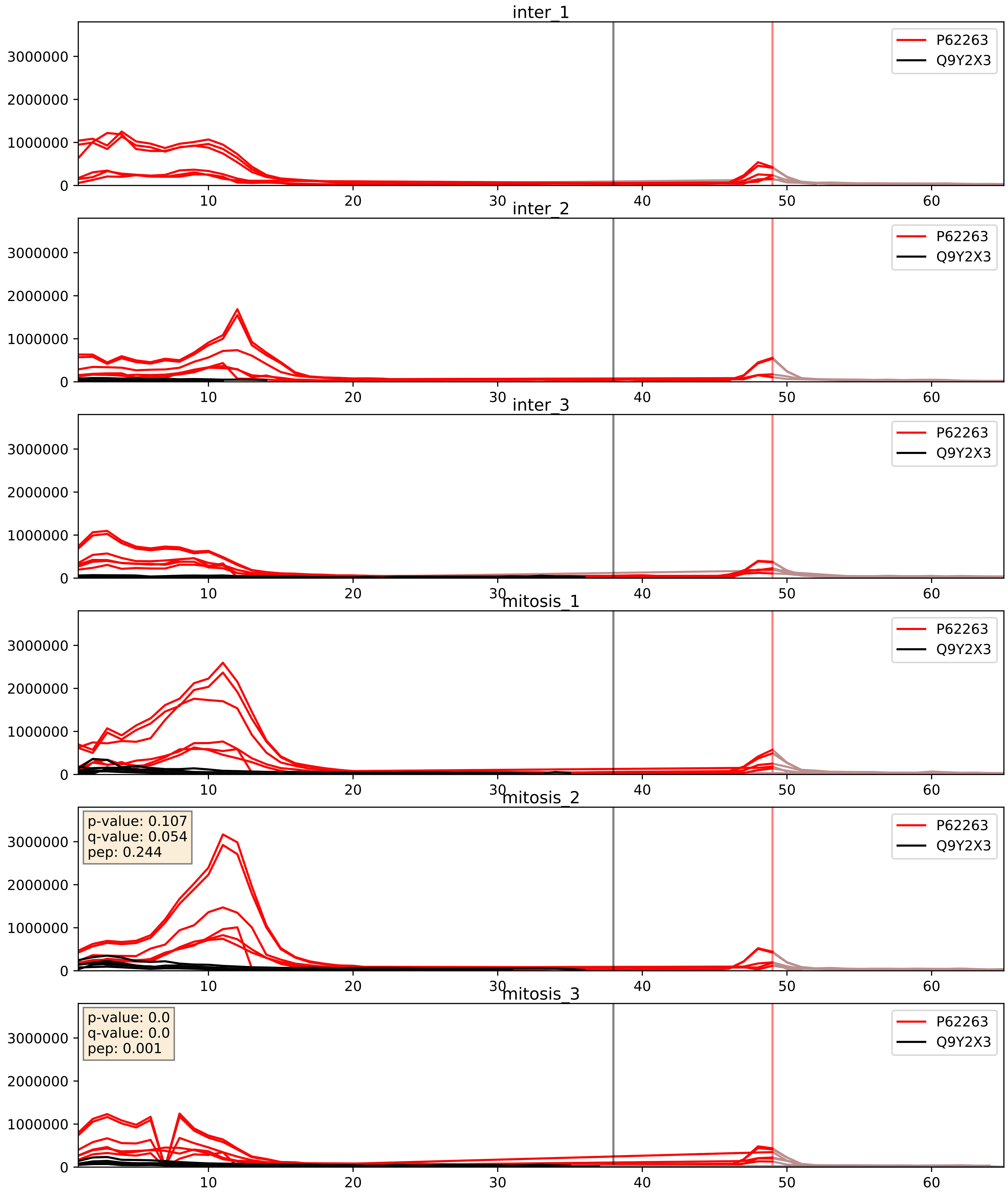


| level | condition_1 | condition_2 | pvalue | pvalue_adjusted |
|---|---|---|---|---|
| complex_abundance | mitosis | inter | 0.0891323399197668 | 0.21924506600954133 |
| interactor_abundance | mitosis | inter | 0.22504117249951913 | 0.4190762663832913 |
| interactor_ratio | mitosis | inter | 0.8164187265788692 | 0.9666192201555632 |

P62266_Q9Y2X3
RS23_HUMAN vs NOP58_HUMAN
p-value: 0.0 q-value: 0.003

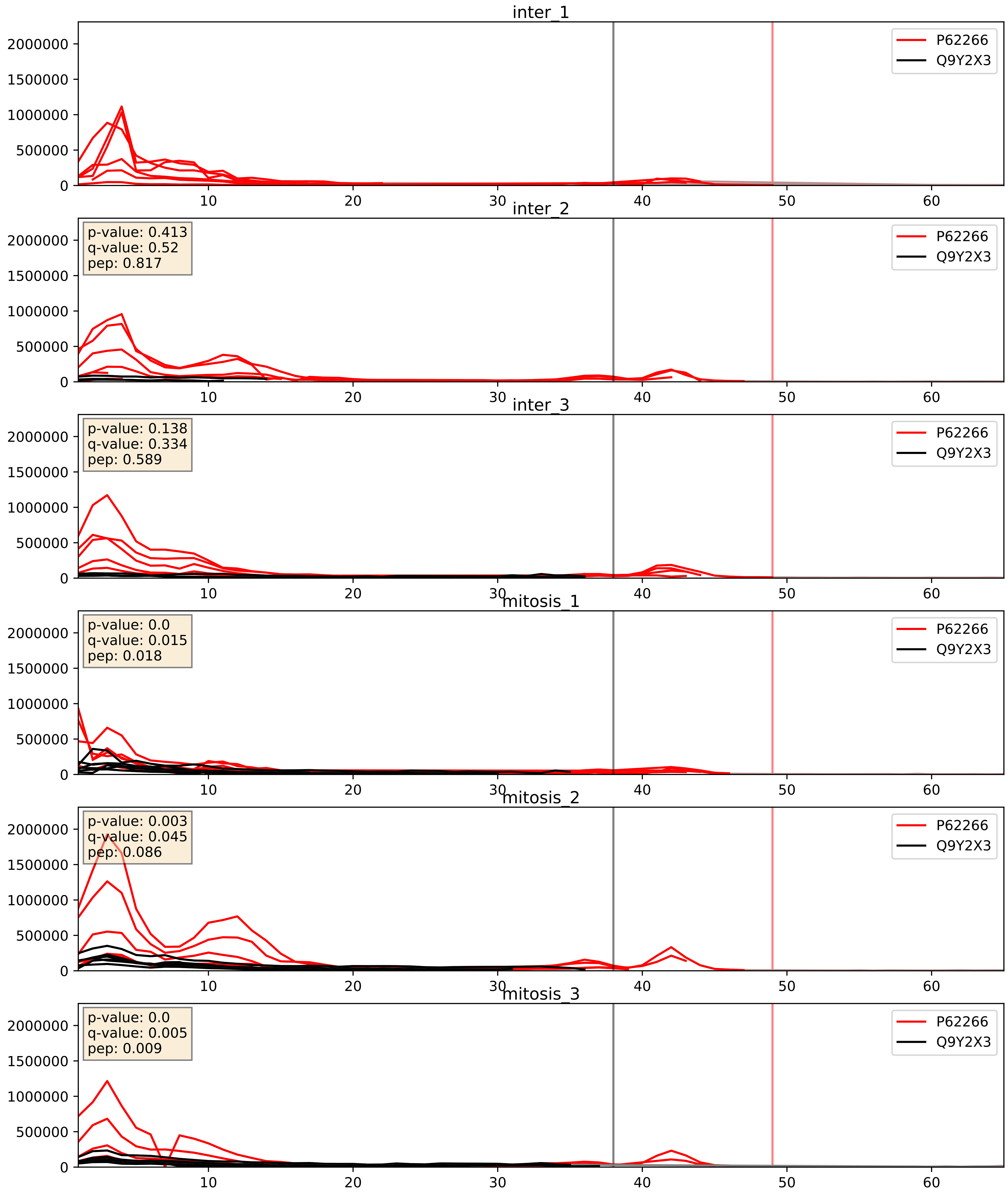


| level | condition_1 | condition_2 | pvalue | pvalue_adjusted |
|---|---|---|---|---|
| complex_abundance | mitosis | inter | 0.19107337617599943 | 0.34251727980964264 |
| interactor_ratio | mitosis | inter | 0.33247181184909896 | 0.7893553151713977 |
| interactor_abundance | mitosis | inter | 0.6259298185744482 | 0.7540727521353039 |

P62269_Q9Y2X3
RS18_HUMAN vs NOP58_HUMAN
p-value: 0.0 q-value: 0.024

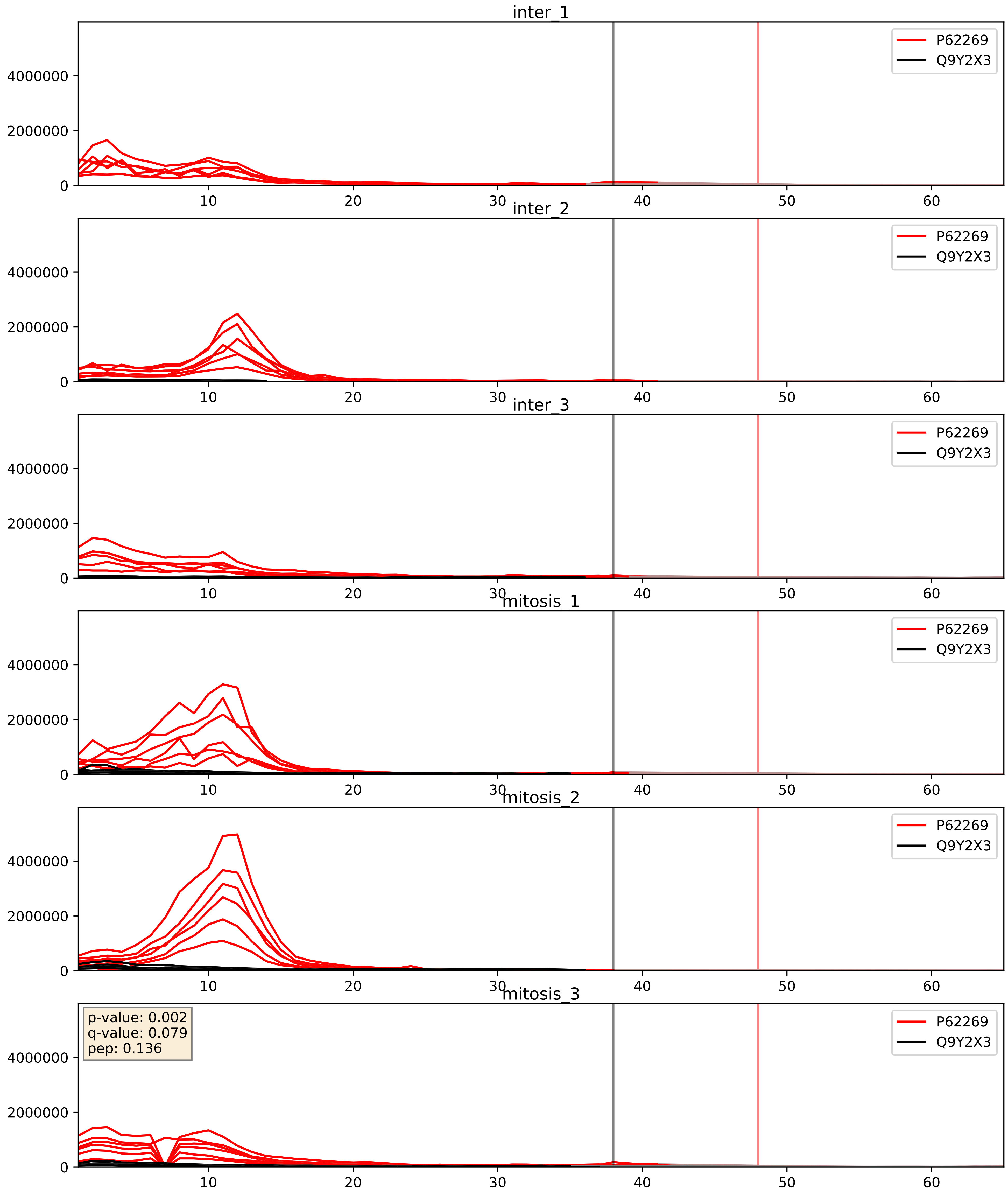


| level | condition_1 | condition_2 | pvalue | pvalue_adjusted |
|---|---|---|---|---|
| complex_abundance | mitosis | inter | 0.09773888757235956 | 0.23169340283007417 |
| interactor_abundance | mitosis | inter | 0.22976059832030665 | 0.42246864822439595 |
| interactor_ratio | mitosis | inter | 0.7394020622773316 | 0.9491588884573259 |

P62277_Q9Y2X3
RS13_HUMAN vs NOP58_HUMAN
p-value: 0.007 q-value: 0.007

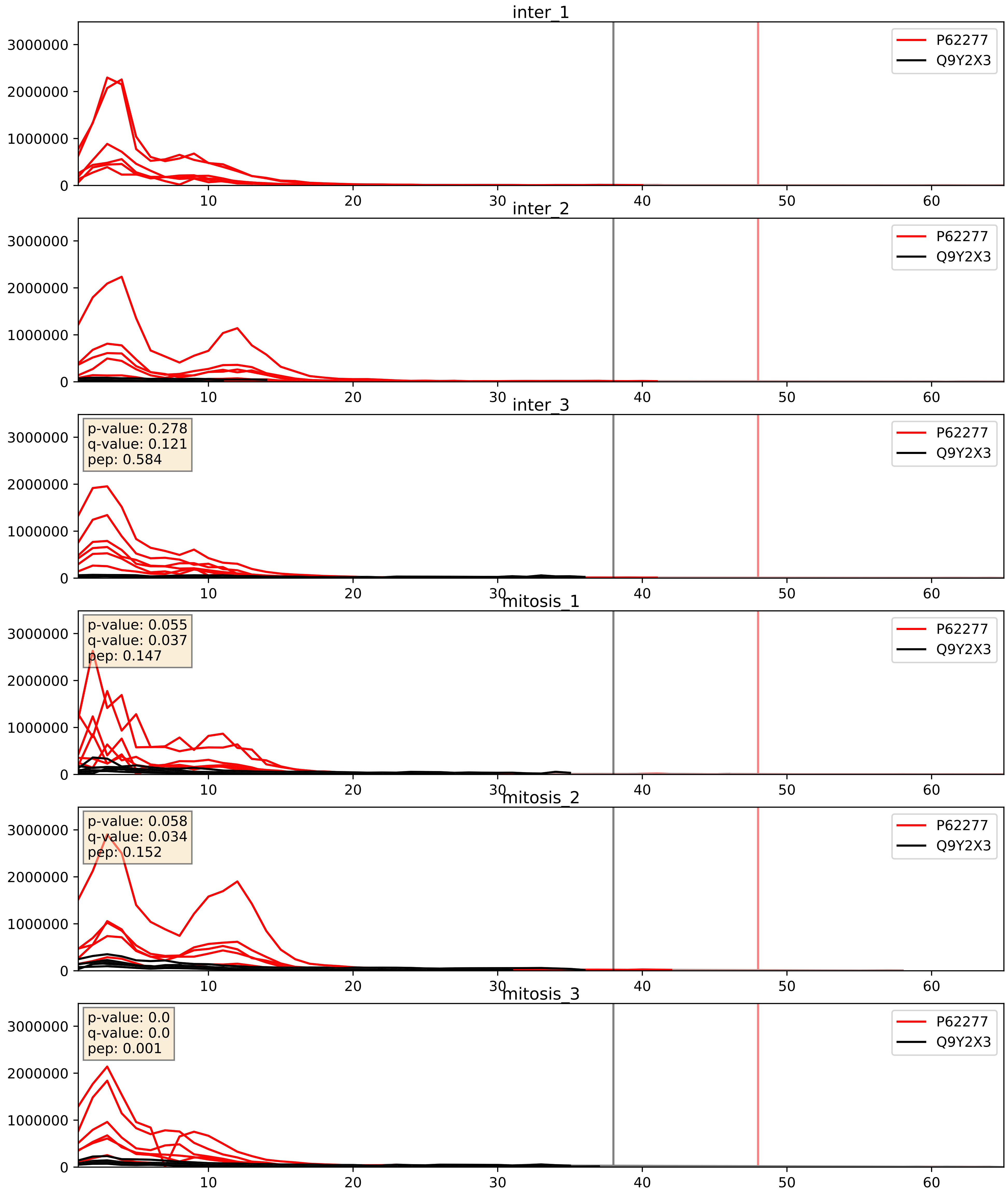


| level | condition_1 | condition_2 | pvalue | pvalue_adjusted |
|---|---|---|---|---|
| complex_abundance | mitosis | inter | 0.11815695899424583 | 0.2595894753207492 |
| interactor_abundance | mitosis | inter | 0.3333431452470456 | 0.5323539782303565 |
| interactor_ratio | mitosis | inter | 0.824285040248311 | 0.9690811625498615 |

P62280_Q9Y2X3
RS11_HUMAN vs NOP58_HUMAN
p-value: 0.001 q-value: 0.001

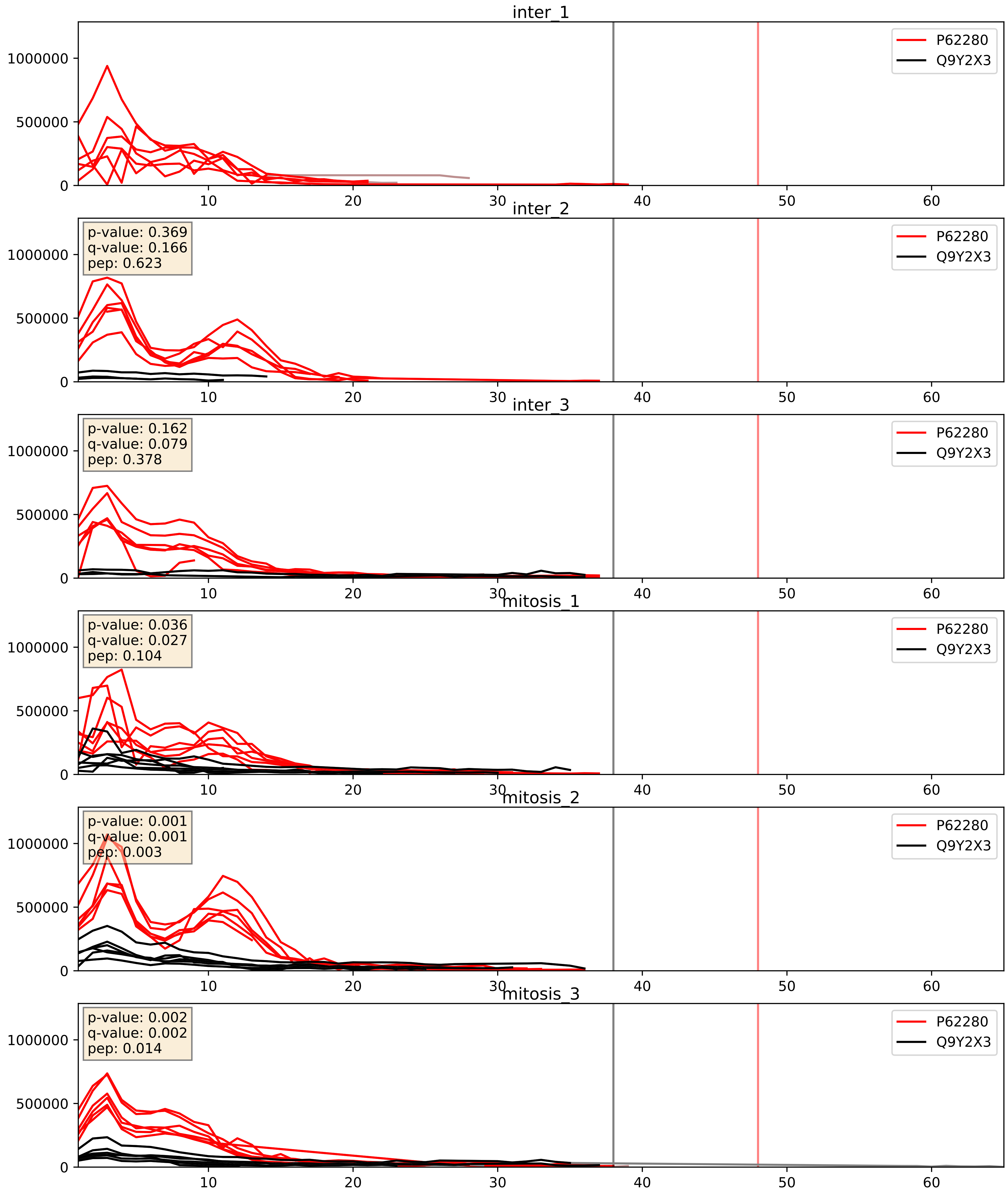


| level | condition_1 | condition_2 | pvalue | pvalue_adjusted |
|---|---|---|---|---|
| complex_abundance | mitosis | inter | 0.1201984451409299 | 0.26253351312960976 |
| interactor_abundance | mitosis | inter | 0.3892272604350498 | 0.5812098297992196 |
| interactor_ratio | mitosis | inter | 0.5589006890709313 | 0.8918094730984788 |

P62701_Q9Y2X3
RS4X_HUMAN vs NOP58_HUMAN
p-value: 0.015 q-value: 0.012

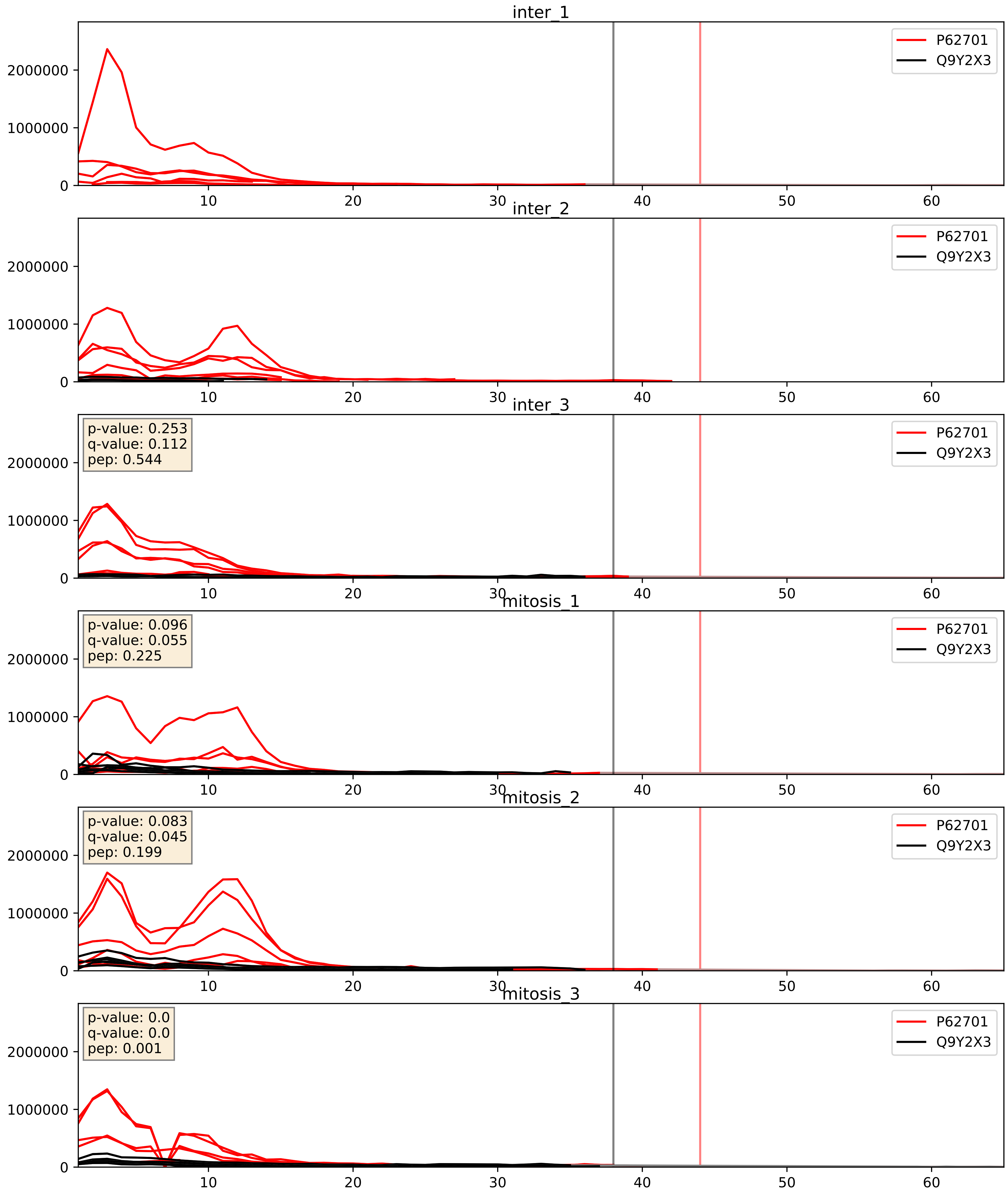


| level | condition_1 | condition_2 | pvalue | pvalue_adjusted |
|---|---|---|---|---|
| complex_abundance | mitosis | inter | 0.12463106614190488 | 0.26657719294720283 |
| interactor_abundance | mitosis | inter | 0.3601356061740694 | 0.5572597232194567 |
| interactor_ratio | mitosis | inter | 0.741357642697343 | 0.9491588884573259 |

P62753_Q9Y2X3
RS6_HUMAN vs NOP58_HUMAN
p-value: 0.043 q-value: 0.029

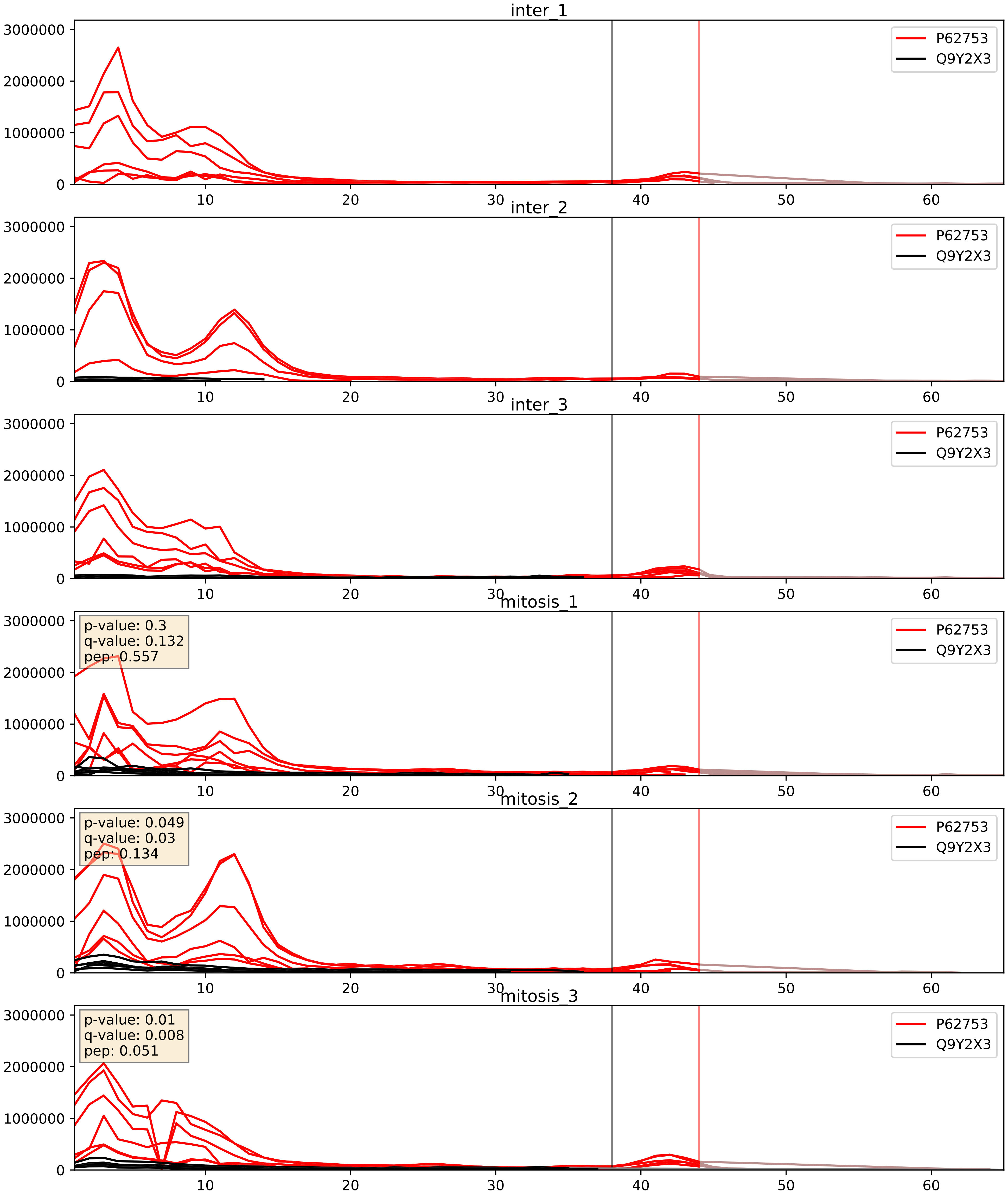


| level | condition_1 | condition_2 | pvalue | pvalue_adjusted |
|---|---|---|---|---|
| complex_abundance | mitosis | inter | 0.15629465876211507 | 0.3031684294139372 |
| interactor_abundance | mitosis | inter | 0.3758912235687873 | 0.5690889412360841 |
| interactor_ratio | mitosis | inter | 0.9662225531731017 | 0.9937294965826962 |

P62829_Q9Y2X3
RL23_HUMAN vs NOP58_HUMAN
p-value: 0.0 q-value: 0.047

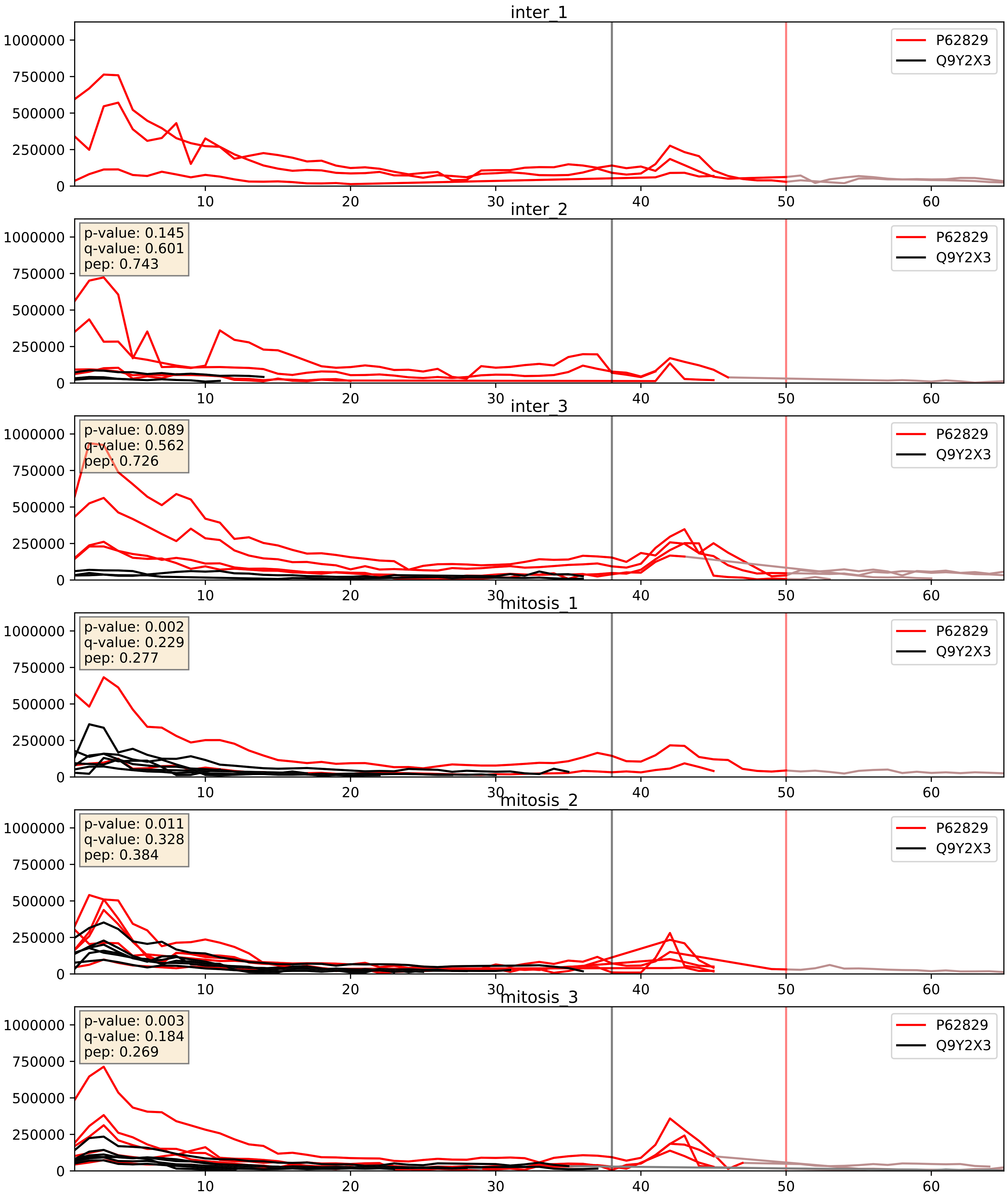


| level | condition_1 | condition_2 | pvalue | pvalue_adjusted |
|---|---|---|---|---|
| interactor_ratio | mitosis | inter | 0.08336086174927138 | 0.4841037832929192 |
| complex_abundance | mitosis | inter | 0.36594266613487514 | 0.5137722194709745 |
| interactor_abundance | mitosis | inter | 0.7978512935014155 | 0.8820363001901222 |

P62847_Q9Y2X3
RS24_HUMAN vs NOP58_HUMAN
p-value: 0.016 q-value: 0.013

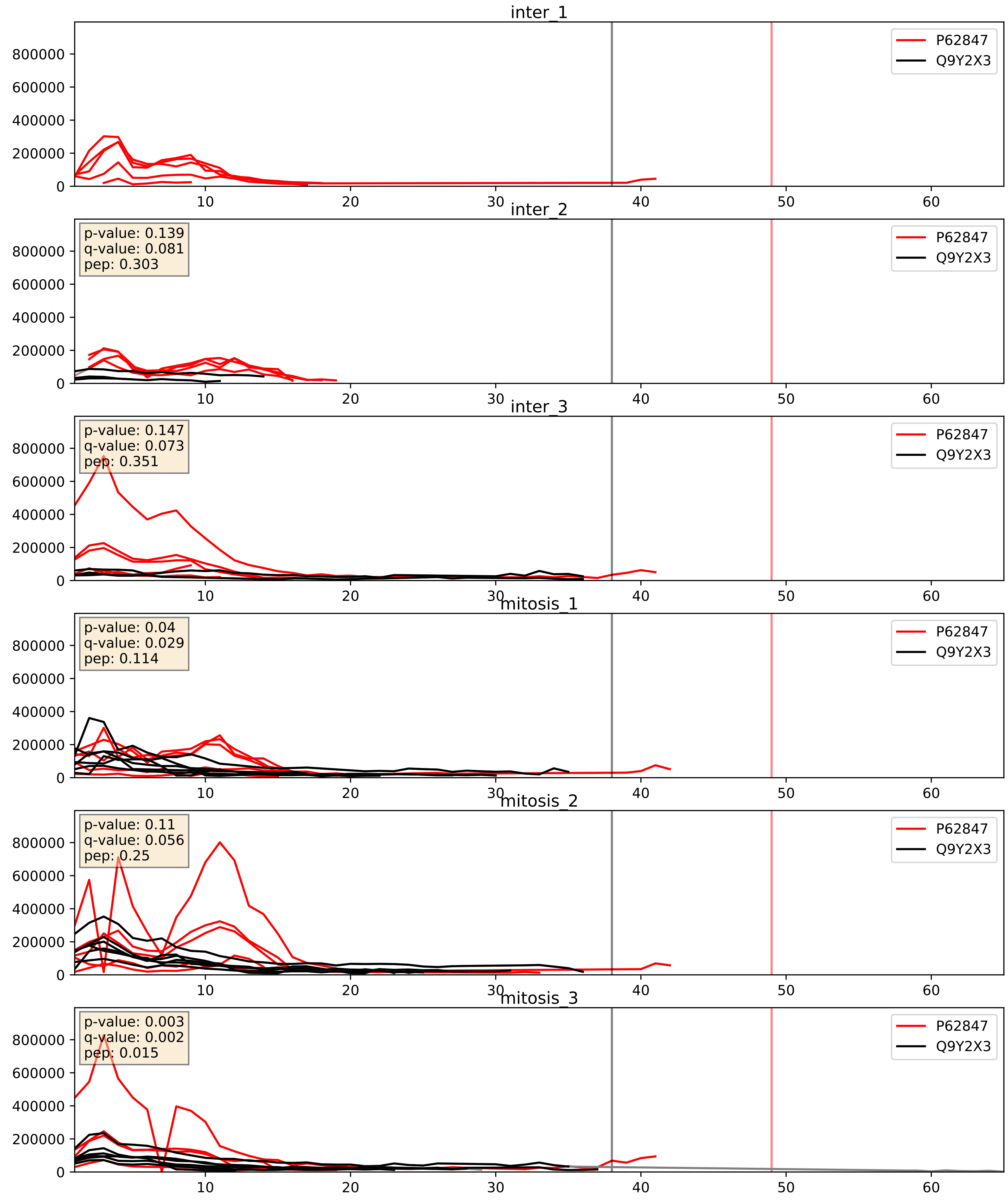


| level | condition_1 | condition_2 | pvalue | pvalue_adjusted |
|---|---|---|---|---|
| complex_abundance | mitosis | inter | 0.060269745015100636 | 0.17607816291101072 |
| interactor_abundance | mitosis | inter | 0.24539121493621377 | 0.4386653022562368 |
| interactor_ratio | mitosis | inter | 0.38884016459443194 | 0.8251906958403389 |

P62888_Q9Y2X3
RL30_HUMAN vs NOP58_HUMAN
p-value: 0.0 q-value: 0.023

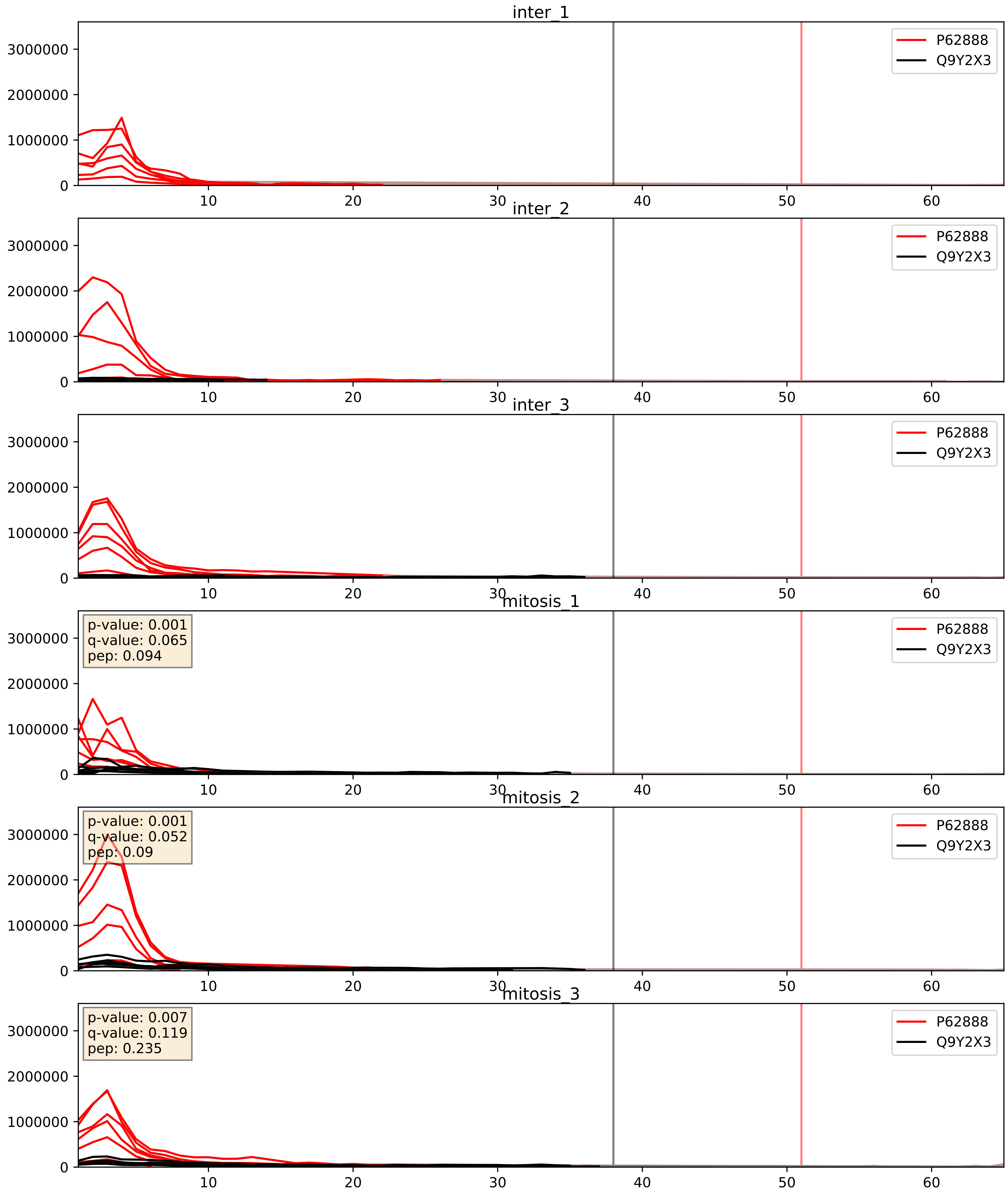


| level | condition_1 | condition_2 | pvalue | pvalue_adjusted |
|---|---|---|---|---|
| complex_abundance | mitosis | inter | 0.16679031270440656 | 0.31569245893417724 |
| interactor_abundance | mitosis | inter | 0.5030374445257547 | 0.6676685514632366 |
| interactor_ratio | mitosis | inter | 0.5386671452901584 | 0.8851272942373031 |

P62913_Q9Y2X3
RL11_HUMAN vs NOP58_HUMAN
p-value: 0.0 q-value: 0.034

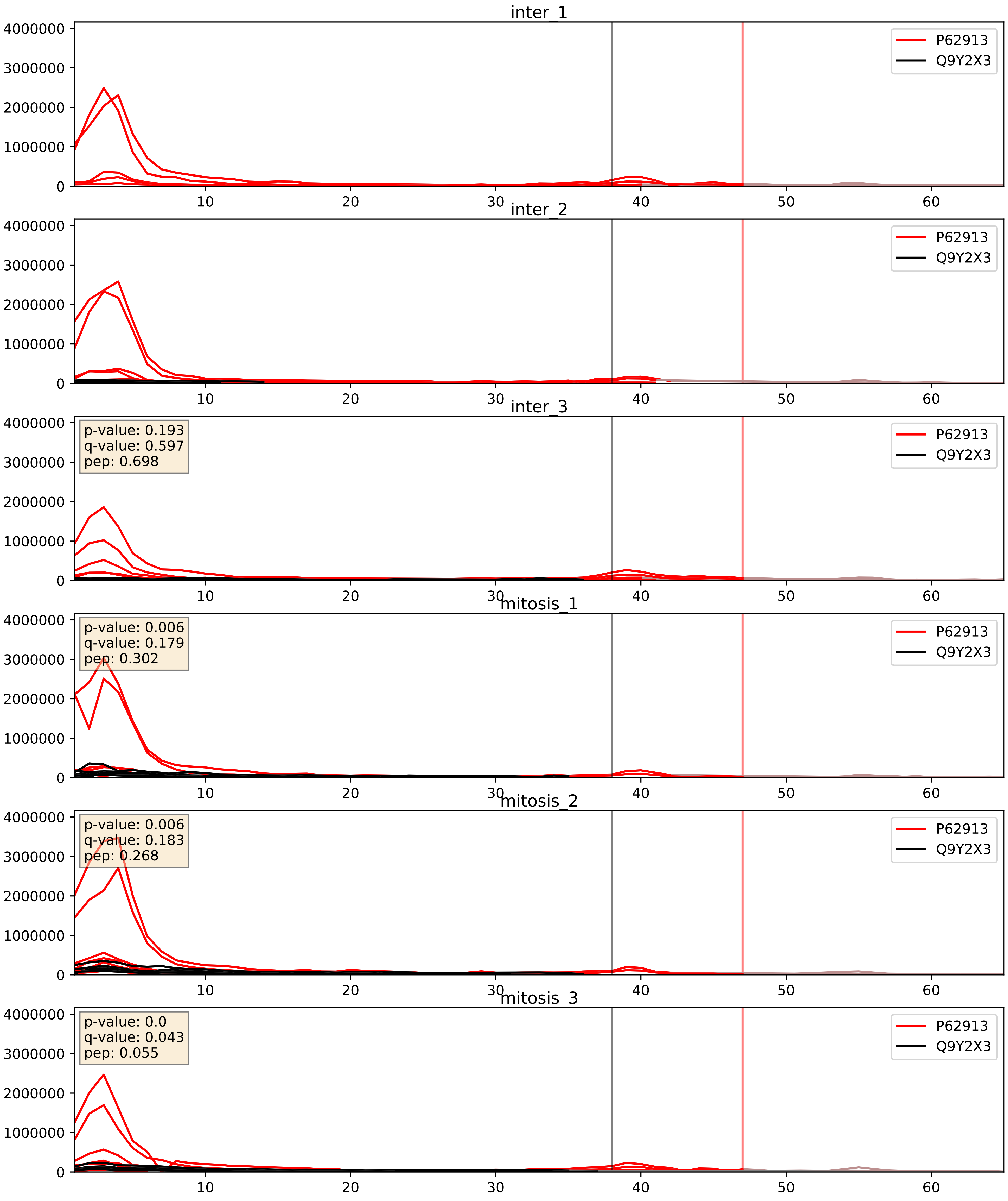


| level | condition_1 | condition_2 | pvalue | pvalue_adjusted |
|---|---|---|---|---|
| complex_abundance | mitosis | inter | 0.10070105674231268 | 0.2354669386569353 |
| interactor_abundance | mitosis | inter | 0.30531849198166233 | 0.5011555688136202 |
| interactor_ratio | mitosis | inter | 0.7510161906806581 | 0.9497901653572102 |

Q02543_Q9Y2X3
RL18A_HUMAN vs NOP58_HUMAN
p-value: 0.0 q-value: 0.023

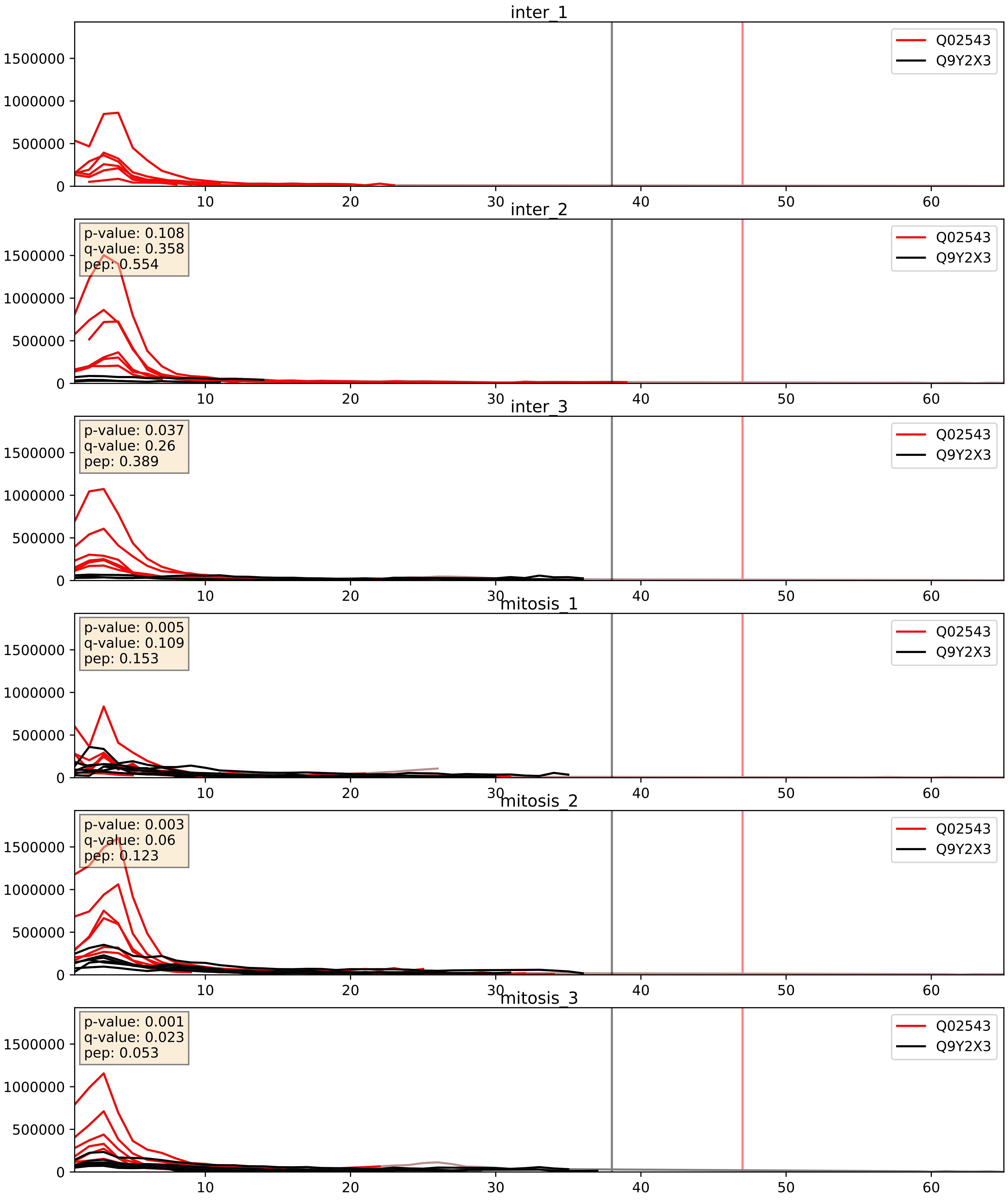


| level | condition_1 | condition_2 | pvalue | pvalue_adjusted |
|---|---|---|---|---|
| complex_abundance | mitosis | inter | 0.1564649350975027 | 0.30336123316752506 |
| interactor_ratio | mitosis | inter | 0.2844825492116298 | 0.7553021039341069 |
| interactor_abundance | mitosis | inter | 0.5756627867670725 | 0.7211739103273256 |

Q03701_Q9Y2X3
CEBPZ_HUMAN vs NOP58_HUMAN
p-value: 0.076 q-value: 0.046

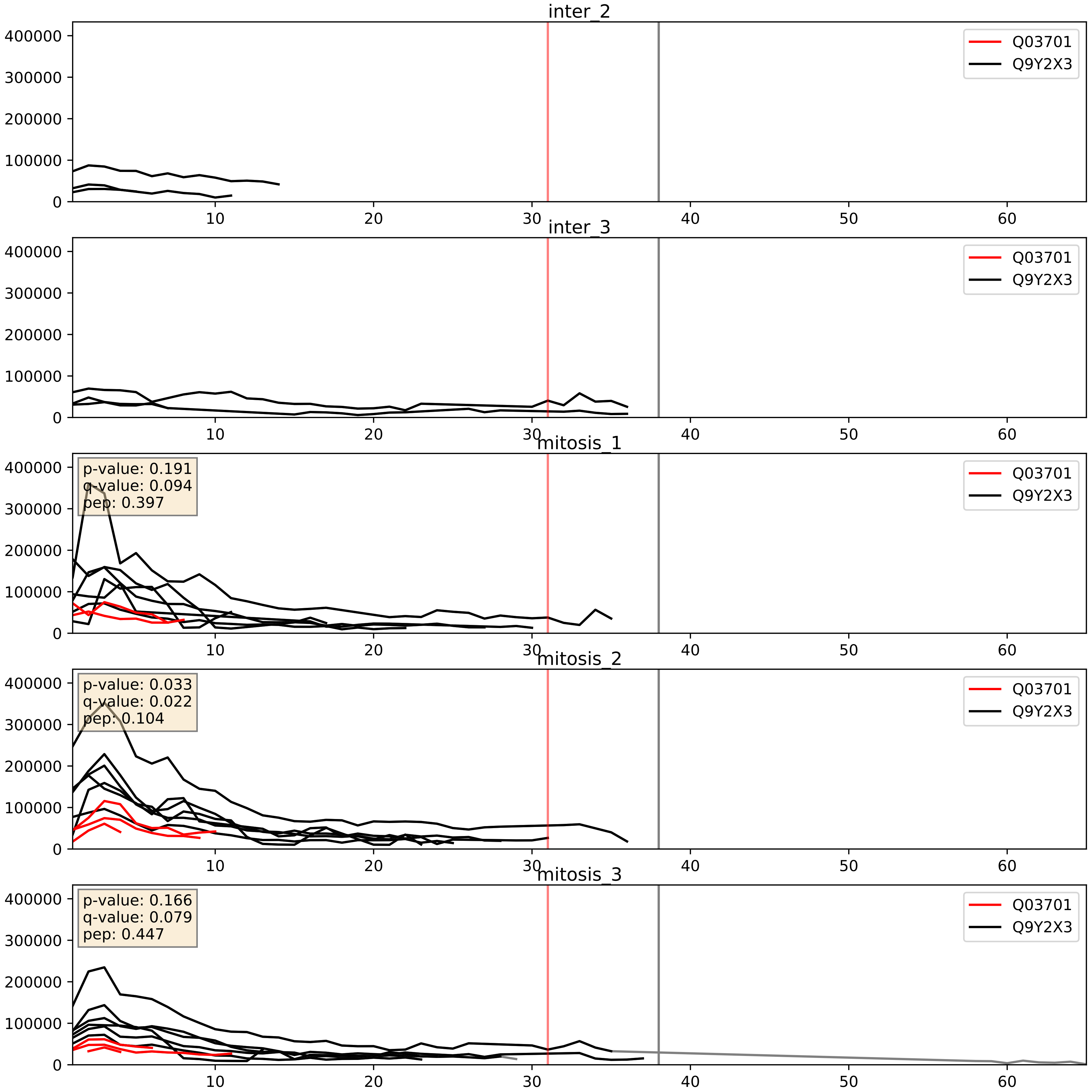


| level | condition_1 | condition_2 | pvalue | pvalue_adjusted |
|---|---|---|---|---|
| interactor_ratio | mitosis | inter | 1.052988896845605e-07 | 0.0004506792478499189 |
| complex_abundance | mitosis | inter | 0.000883253249181854 | 0.019570164268135633 |
| interactor_abundance | mitosis | inter | 0.019842839707373484 | 0.1190514126751731 |

Q12788_Q9Y2X3
TBL3_HUMAN vs NOP58_HUMAN
p-value: 0.001 q-value: 0.002

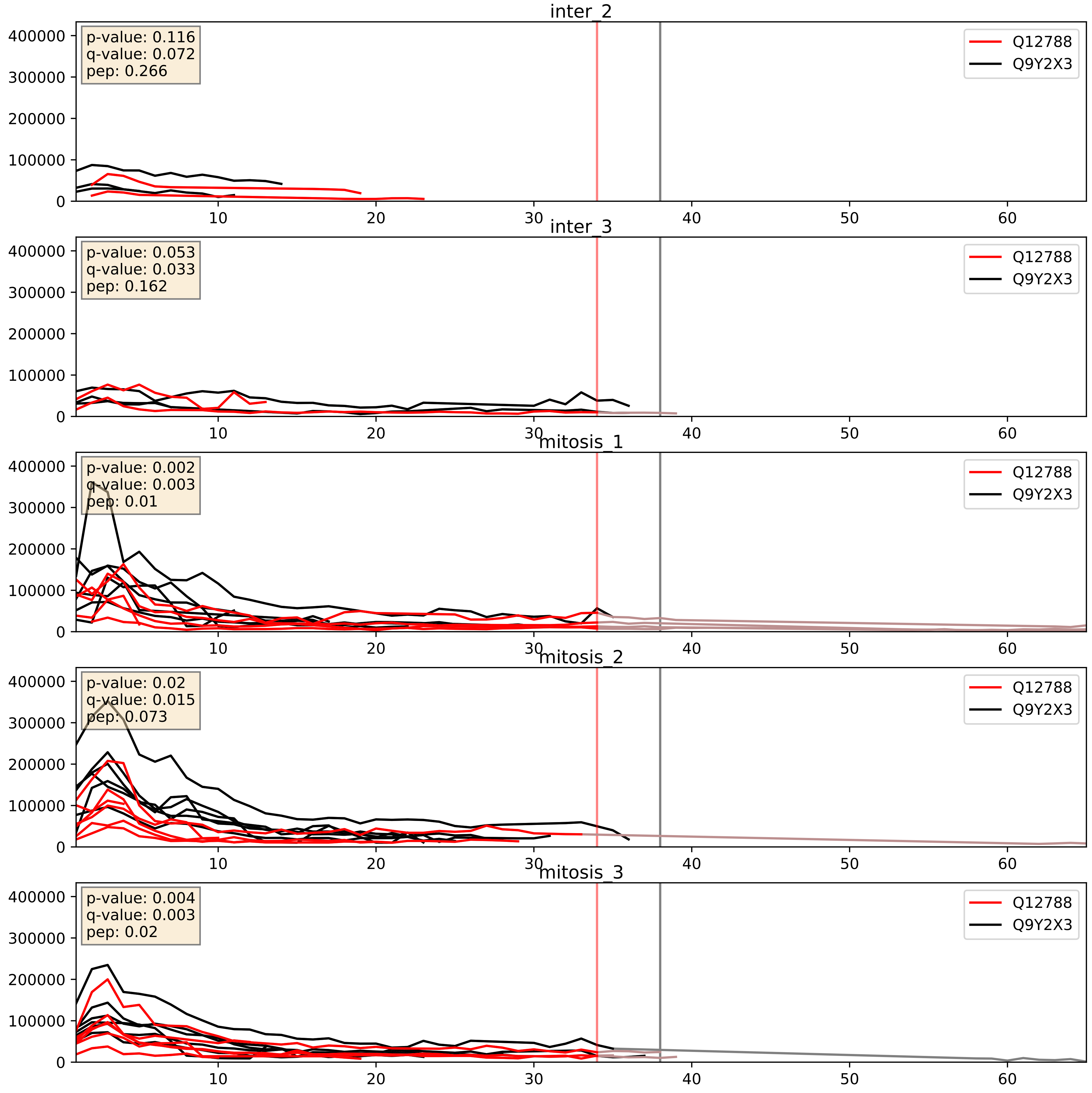


| level | condition_1 | condition_2 | pvalue | pvalue_adjusted |
|---|---|---|---|---|
| complex_abundance | mitosis | inter | 0.002118335372975017 | 0.033489340249041655 |
| interactor_abundance | mitosis | inter | 0.005521190842296777 | 0.06317594435380872 |
| interactor_ratio | mitosis | inter | 0.029449555755858125 | 0.27164676429972584 |

Q15269_Q9Y2X3
PWP2_HUMAN vs NOP58_HUMAN
p-value: 0.017 q-value: 0.014

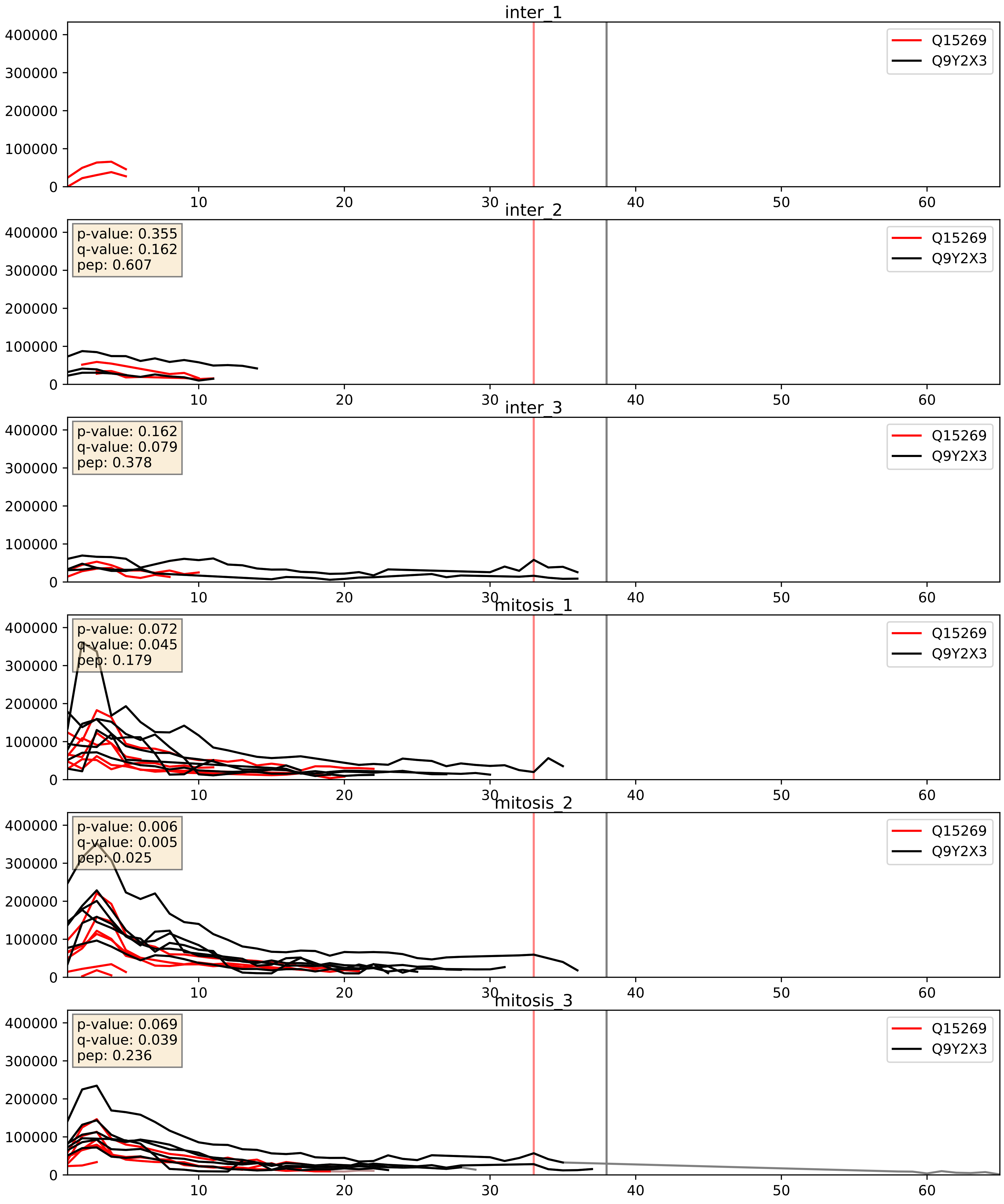


| level | condition_1 | condition_2 | pvalue | pvalue_adjusted |
|---|---|---|---|---|
| interactor_ratio | mitosis | inter | 0.0006251497771619007 | 0.029219626944512057 |
| complex_abundance | mitosis | inter | 0.0053638205370739305 | 0.055047043516043566 |
| interactor_abundance | mitosis | inter | 0.024889437139602743 | 0.13047955794942243 |

Q15397_Q9Y2X3
PUM3_HUMAN vs NOP58_HUMAN
p-value: 0.067 q-value: 0.041

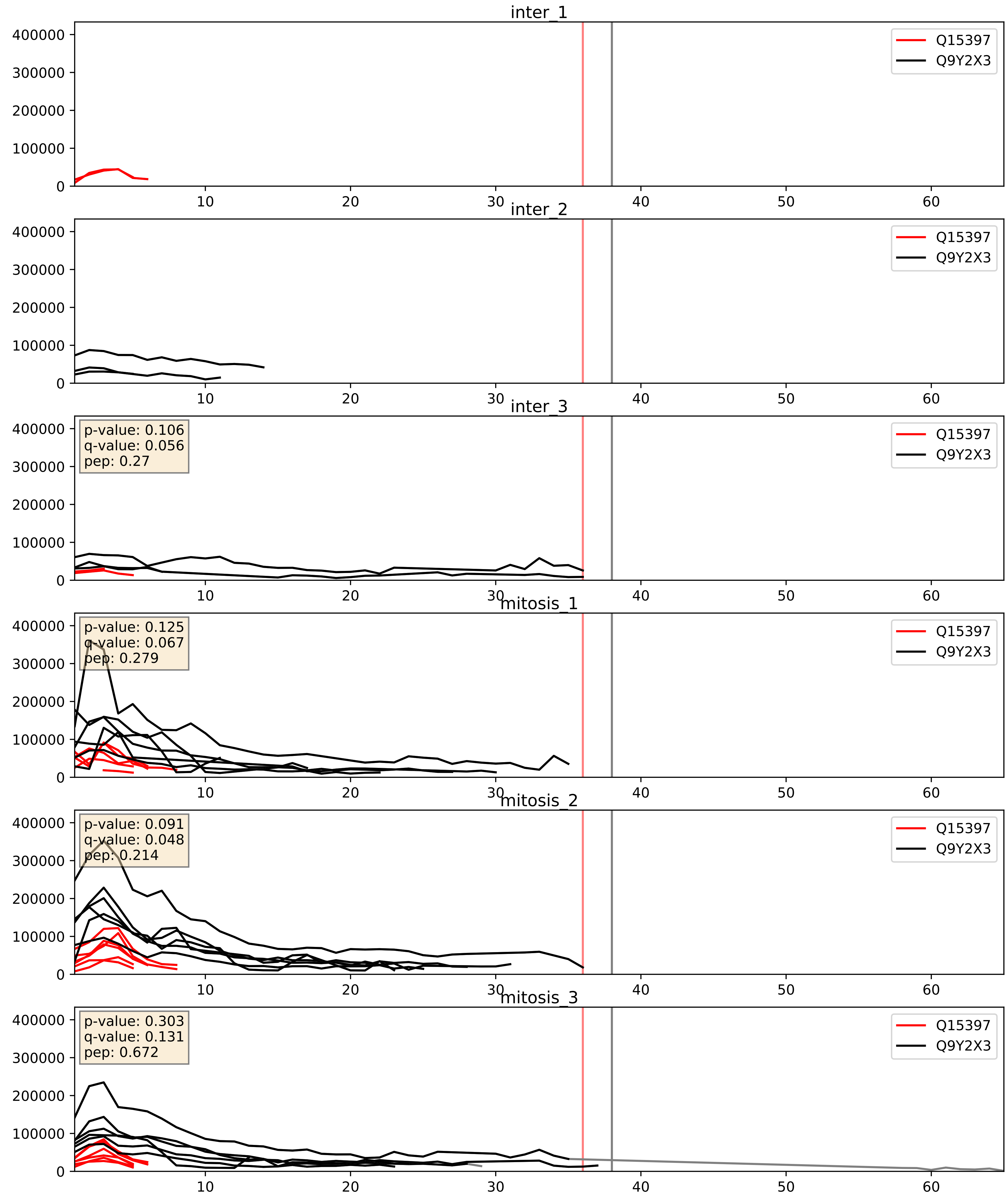


| level | condition_1 | condition_2 | pvalue | pvalue_adjusted |
|---|---|---|---|---|
| interactor_ratio | mitosis | inter | 0.0005521647453949105 | 0.02716396678494502 |
| complex_abundance | mitosis | inter | 0.004349412054734241 | 0.049247311095932673 |
| interactor_abundance | mitosis | inter | 0.02975596763925335 | 0.1434587907586644 |

Q5JTH9_Q9Y2X3
RRP12_HUMAN vs NOP58_HUMAN
p-value: 0.019 q-value: 0.015

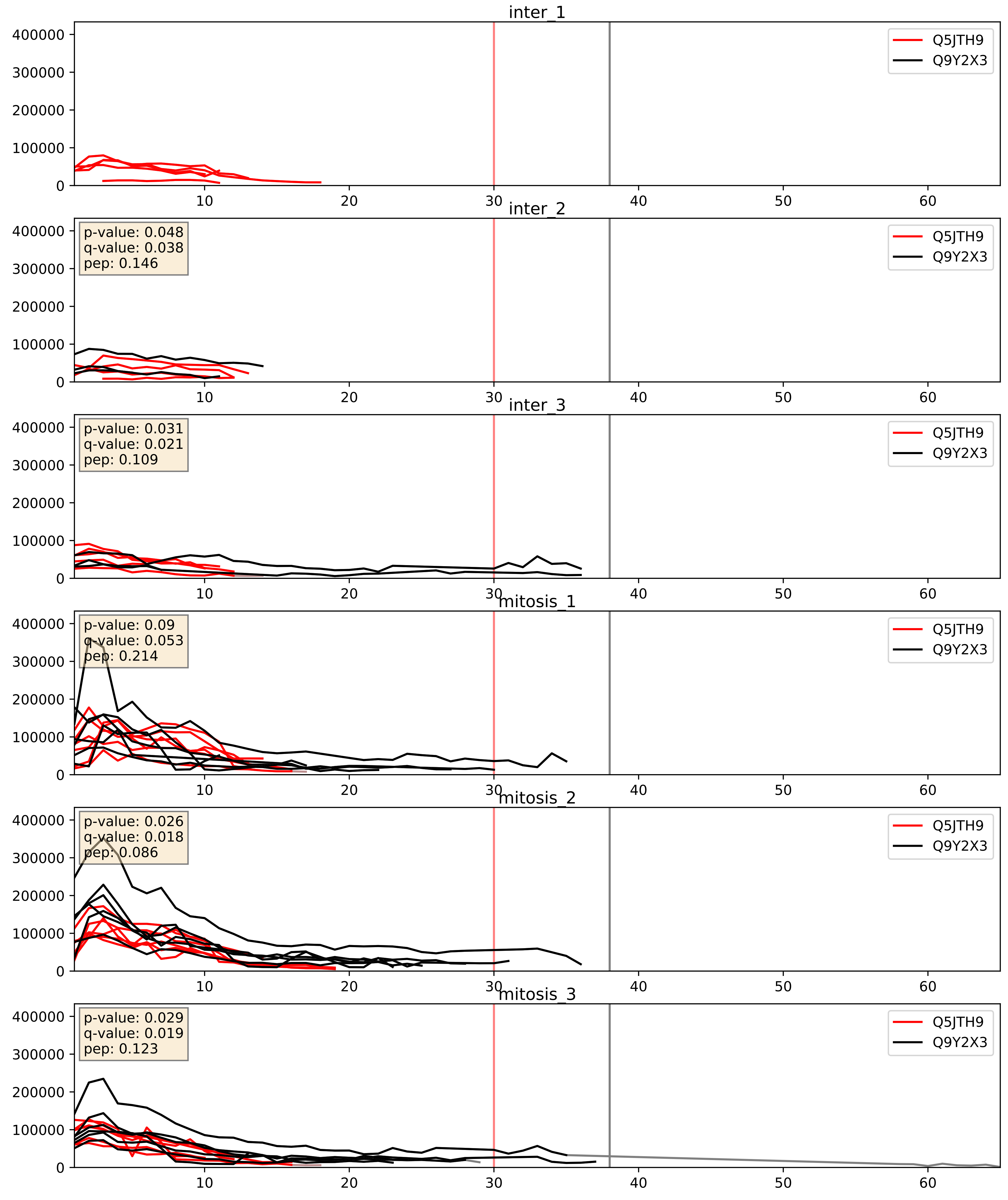


| level | condition_1 | condition_2 | pvalue | pvalue_adjusted |
|---|---|---|---|---|
| complex_abundance | mitosis | inter | 0.018047474157536936 | 0.10031583038215336 |
| interactor_ratio | mitosis | inter | 0.024352033719538677 | 0.2447639398105627 |
| interactor_abundance | mitosis | inter | 0.11040558256739492 | 0.286081969661541 |

Q8IY81_Q9Y2X3
SPB1_HUMAN vs NOP58_HUMAN
p-value: 0.063 q-value: 0.039

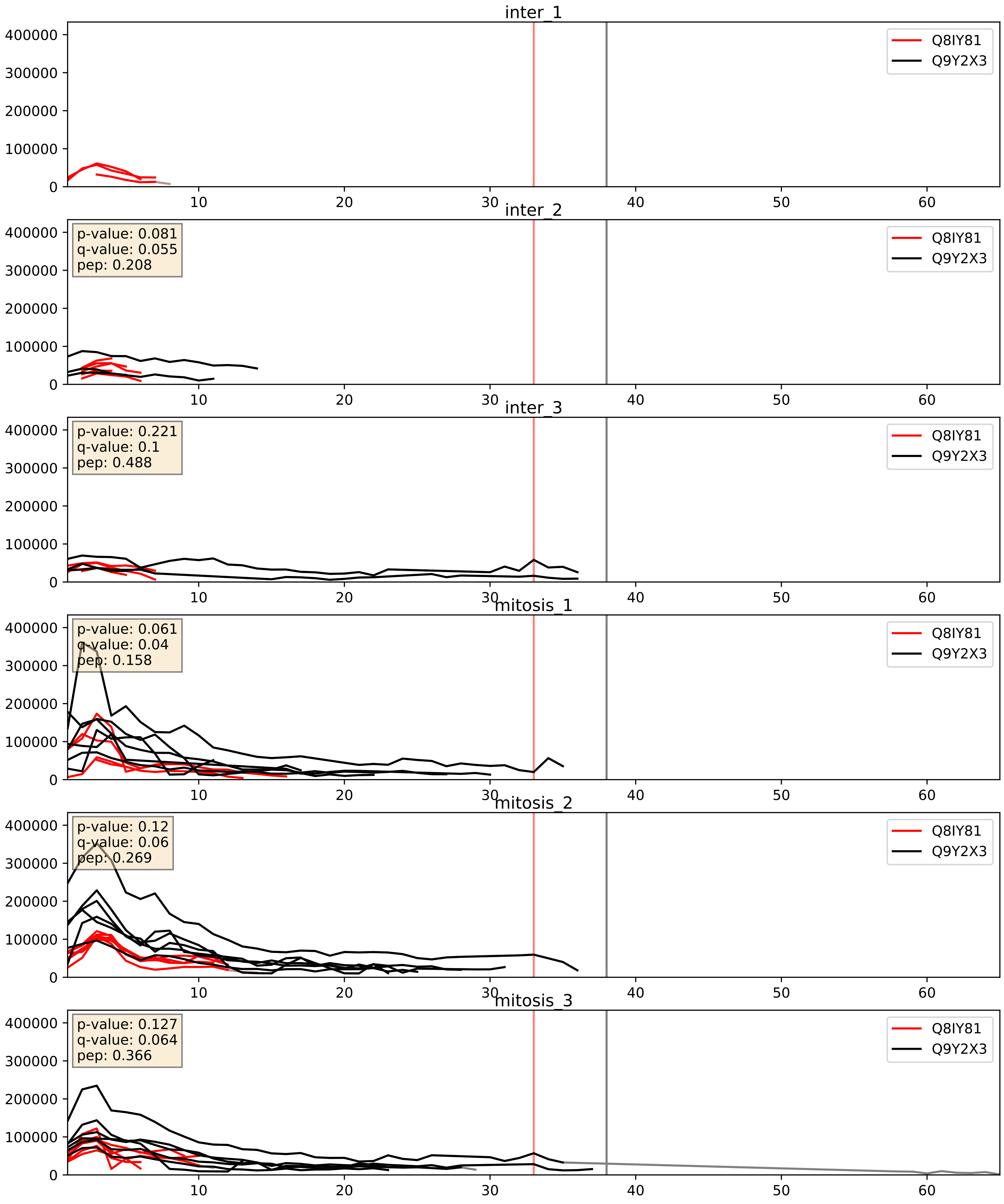


| level | condition_1 | condition_2 | pvalue | pvalue_adjusted |
|---|---|---|---|---|
| interactor_ratio | mitosis | inter | 0.0019003411607799942 | 0.052986711193083874 |
| complex_abundance | mitosis | inter | 0.01284951872274085 | 0.08405635484106207 |
| interactor_abundance | mitosis | inter | 0.18344633002469837 | 0.3757436970203683 |

Q8NI36_Q9Y2X3
WDR36_HUMAN vs NOP58_HUMAN
p-value: 0.005 q-value: 0.005

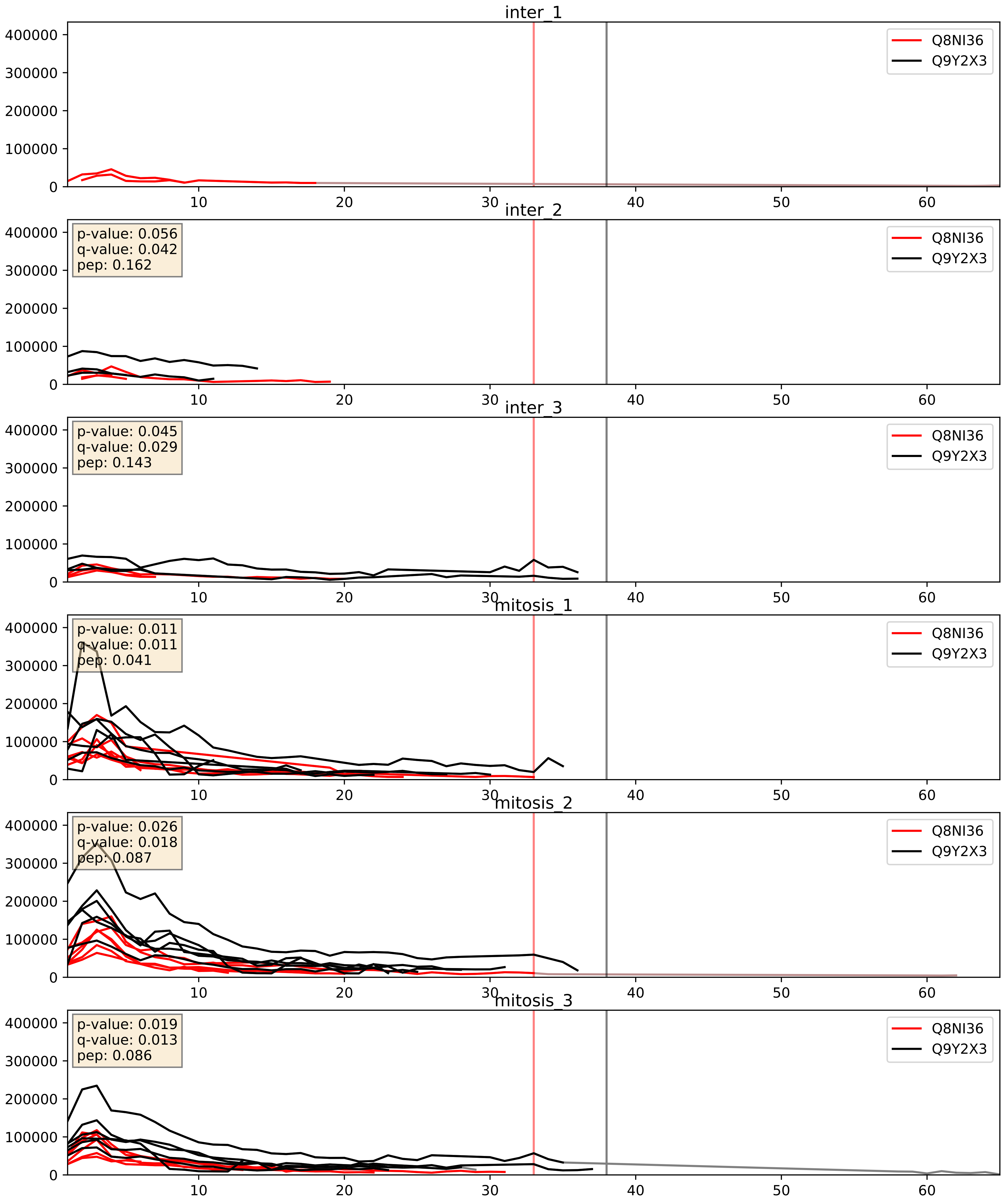


| level | condition_1 | condition_2 | pvalue | pvalue_adjusted |
|---|---|---|---|---|
| complex_abundance | mitosis | inter | 0.002522434141372613 | 0.036409709098919725 |
| interactor_abundance | mitosis | inter | 0.0036714075898180323 | 0.04704677989347658 |
| interactor_ratio | mitosis | inter | 0.003966749934868759 | 0.08201435780423773 |

Q8TDN6_Q9Y2X3
BRX1_HUMAN vs NOP58_HUMAN
p-value: 0.043 q-value: 0.029

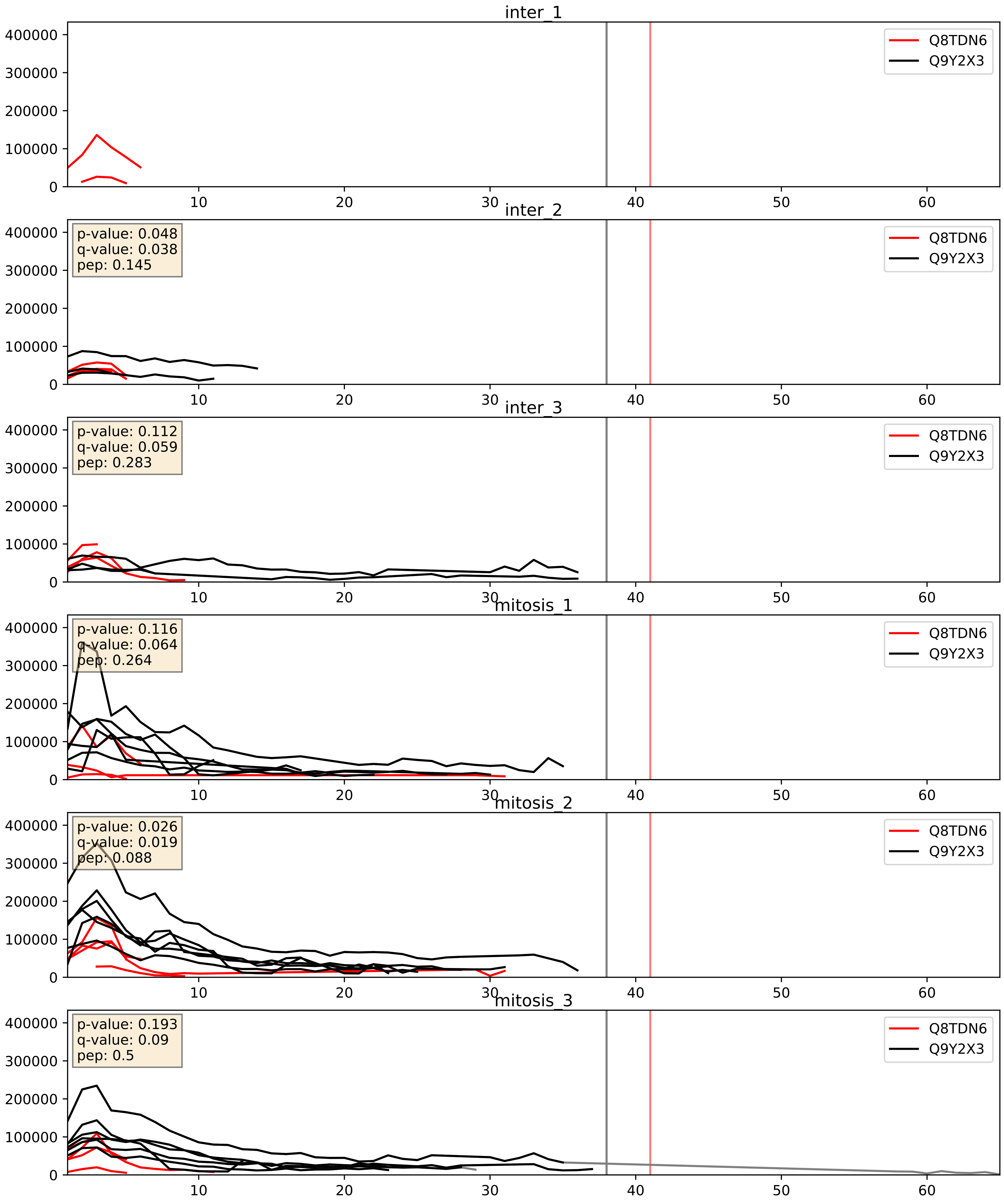


| level | condition_1 | condition_2 | pvalue | pvalue_adjusted |
|---|---|---|---|---|
| interactor_ratio | mitosis | inter | 4.525614117084573e-05 | 0.006796360849516482 |
| complex_abundance | mitosis | inter | 0.03832260841992085 | 0.14299979427834458 |
| interactor_abundance | mitosis | inter | 0.9343585822556804 | 0.962096690999235 |

Q9BVI4_Q9Y2X3
NOC4L_HUMAN vs NOP58_HUMAN
p-value: 0.081 q-value: 0.048

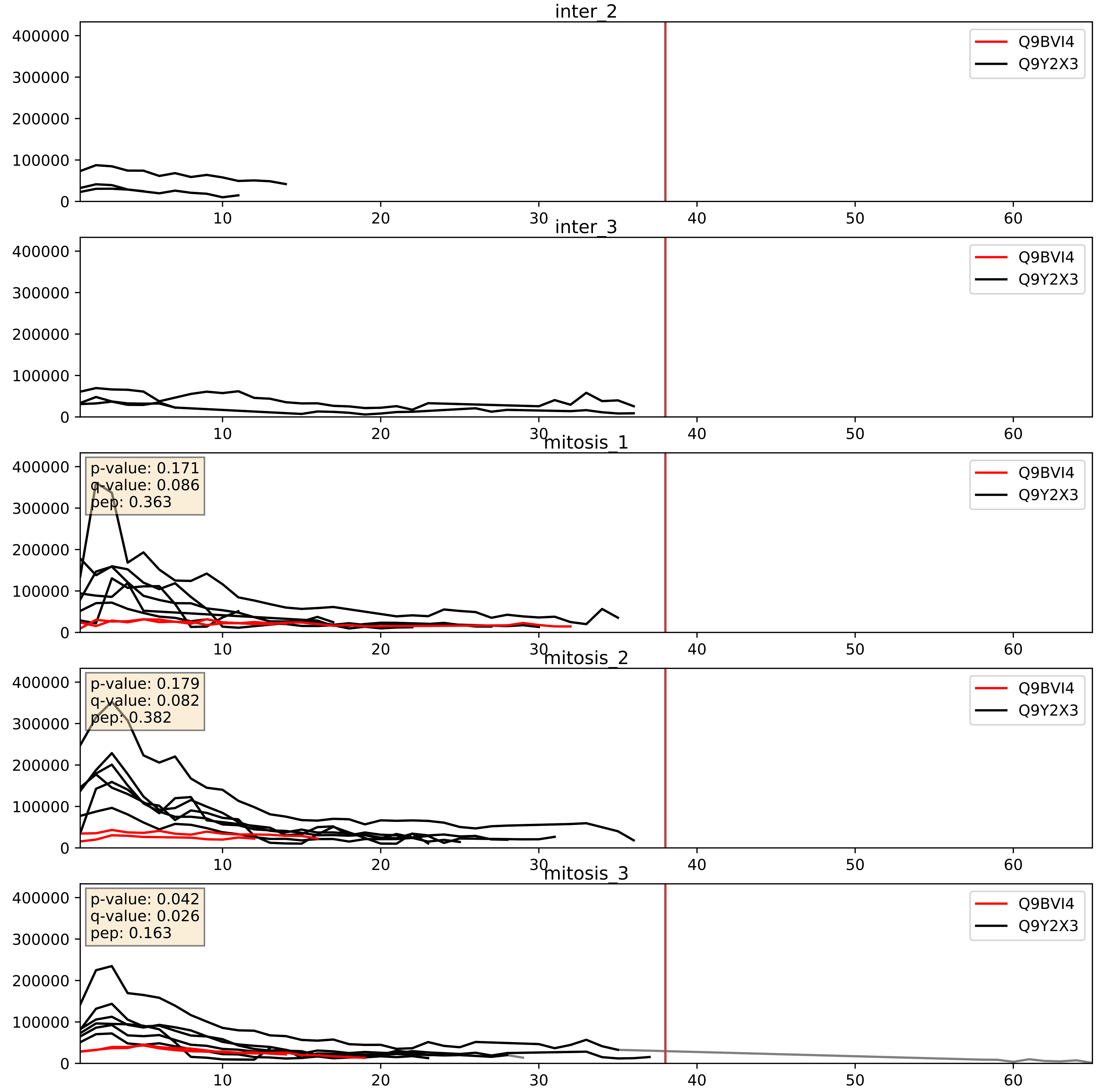


| level | condition_1 | condition_2 | pvalue | pvalue_adjusted |
|---|---|---|---|---|
| interactor_ratio | mitosis | inter | 0.00017598563399961114 | 0.01369488206396974 |
| complex_abundance | mitosis | inter | 0.0002596632330815969 | 0.010784252527150902 |
| interactor_abundance | mitosis | inter | 0.009419913204717079 | 0.08408839833172471 |

Q9BVP2_Q9Y2X3
GNL3_HUMAN vs NOP58_HUMAN
p-value: 0.002 q-value: 0.003

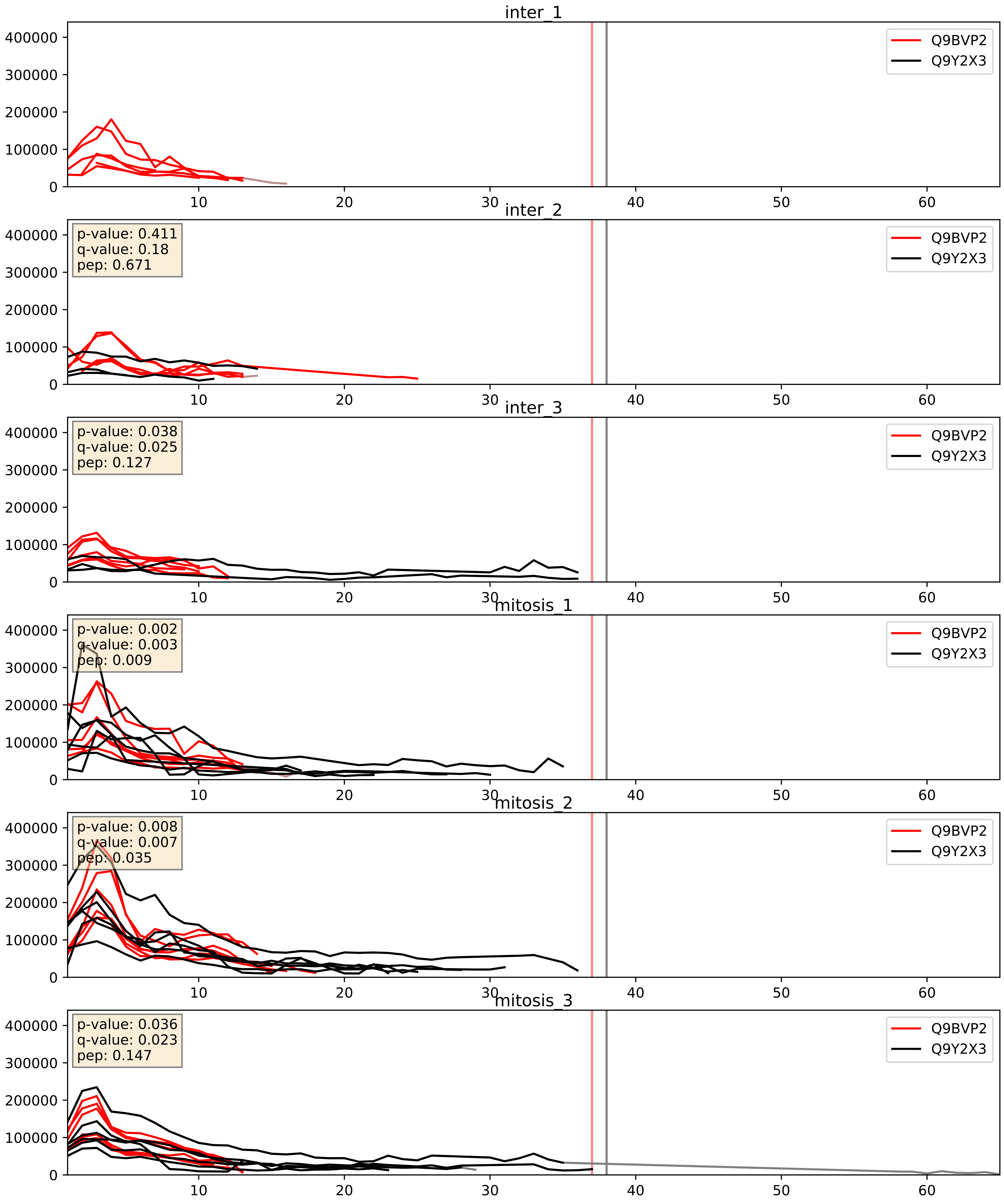


| level | condition_1 | condition_2 | pvalue | pvalue_adjusted |
|---|---|---|---|---|
| complex_abundance | mitosis | inter | 0.0274592583388612 | 0.12057983221353287 |
| interactor_ratio | mitosis | inter | 0.10574631048223244 | 0.5413806326123862 |
| interactor_abundance | mitosis | inter | 0.11740638492850289 | 0.29607210740171247 |

Q9BZE4_Q9Y2X3
NOG1_HUMAN vs NOP58_HUMAN
p-value: 0.034 q-value: 0.024

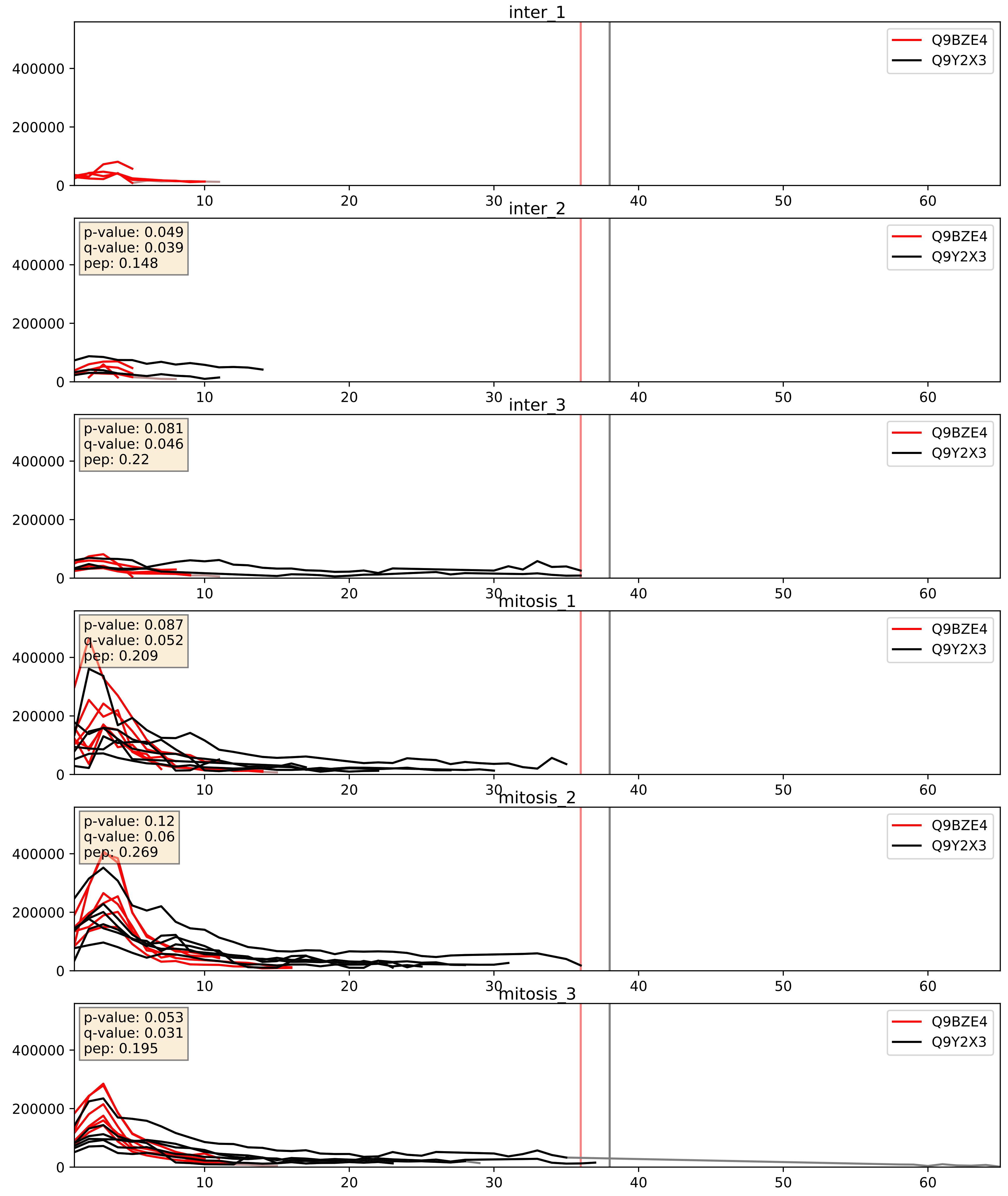


| level | condition_1 | condition_2 | pvalue | pvalue_adjusted |
|---|---|---|---|---|
| complex_abundance | mitosis | inter | 0.0015925256220138068 | 0.027934465828766776 |
| interactor_abundance | mitosis | inter | 0.001698666342849207 | 0.030042528708242176 |
| interactor_ratio | mitosis | inter | 0.04024854859360579 | 0.33223488520854927 |

Q9GZL7_Q9Y2X3
WDR12_HUMAN vs NOP58_HUMAN
p-value: 0.032 q-value: 0.023

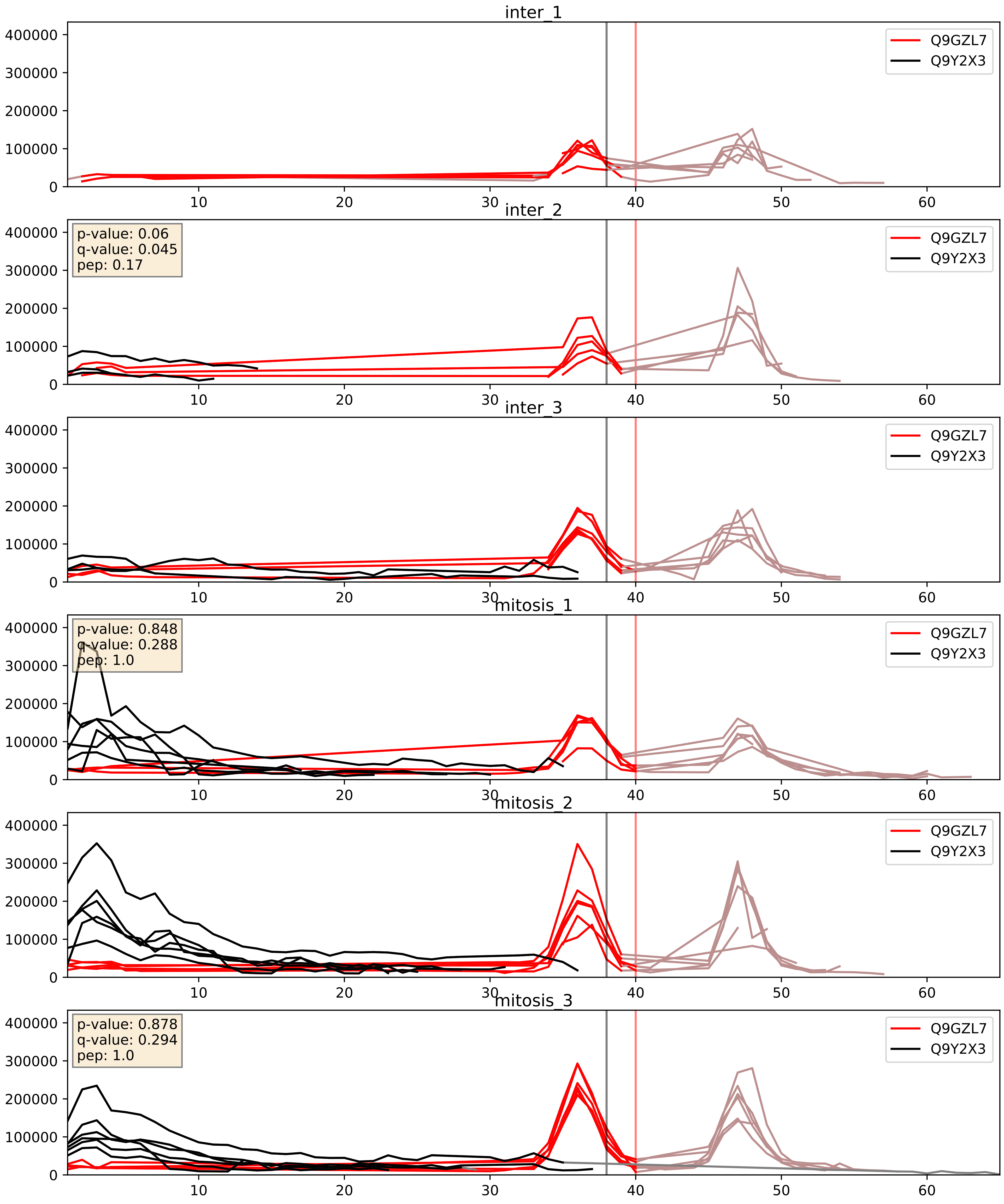


| level | condition_1 | condition_2 | pvalue | pvalue_adjusted |
|---|---|---|---|---|
| complex_abundance | mitosis | inter | 0.04103193725670029 | 0.1456795449677953 |
| interactor_ratio | mitosis | inter | 0.2261507288045601 | 0.7051563050171857 |
| interactor_abundance | mitosis | inter | 0.2603885476978522 | 0.4522065263326465 |

Q9H0A0_Q9Y2X3
NAT10_HUMAN vs NOP58_HUMAN
p-value: 0.055 q-value: 0.035

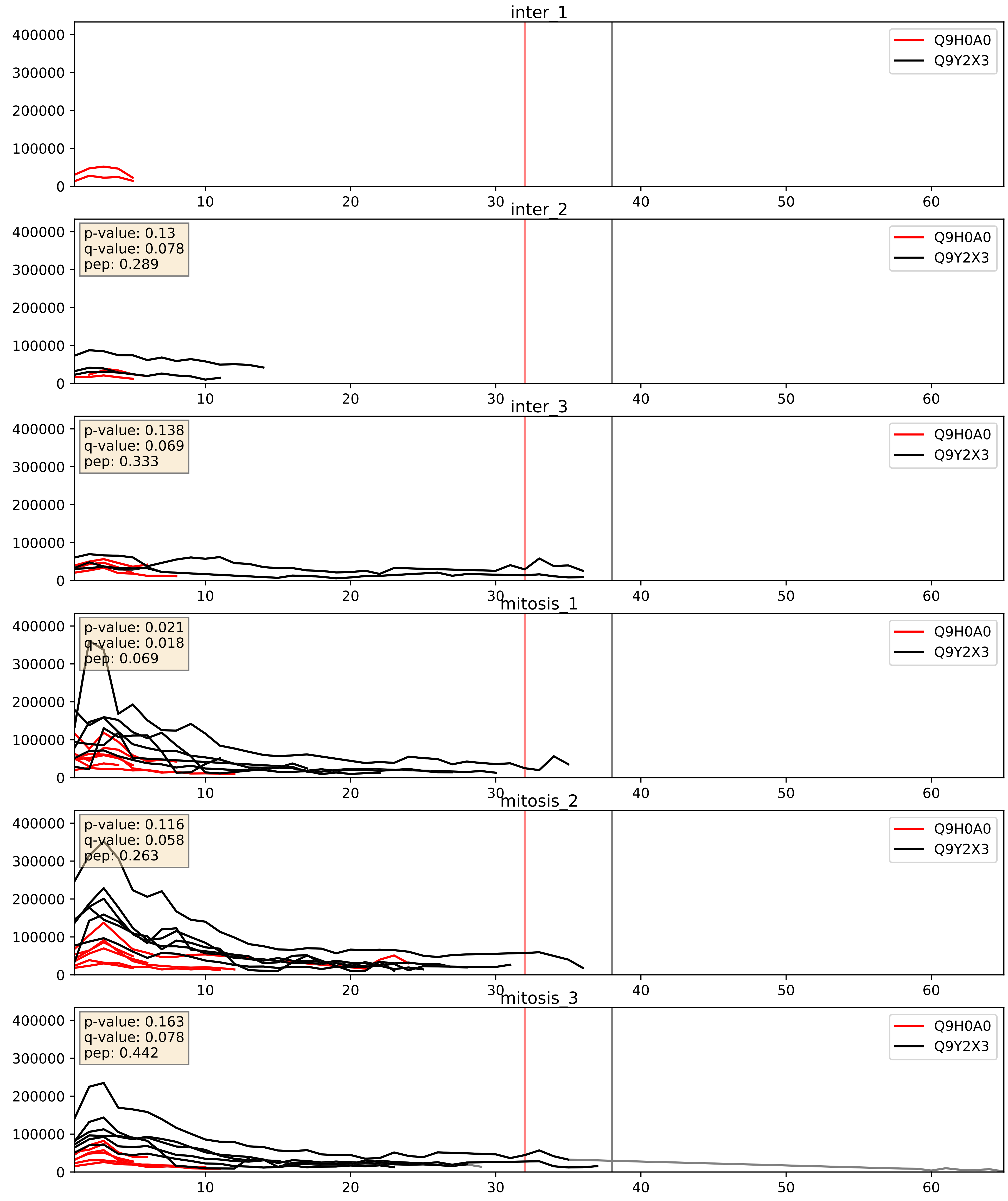


| level | condition_1 | condition_2 | pvalue | pvalue_adjusted |
|---|---|---|---|---|
| interactor_ratio | mitosis | inter | 0.0002532636495883736 | 0.016178633137884164 |
| complex_abundance | mitosis | inter | 0.02044007375838056 | 0.10540182612755276 |
| interactor_abundance | mitosis | inter | 0.45474744236861303 | 0.6283515910694637 |

Q9H0S4_Q9Y2X3
DDX47_HUMAN vs NOP58_HUMAN
p-value: 0.001 q-value: 0.002

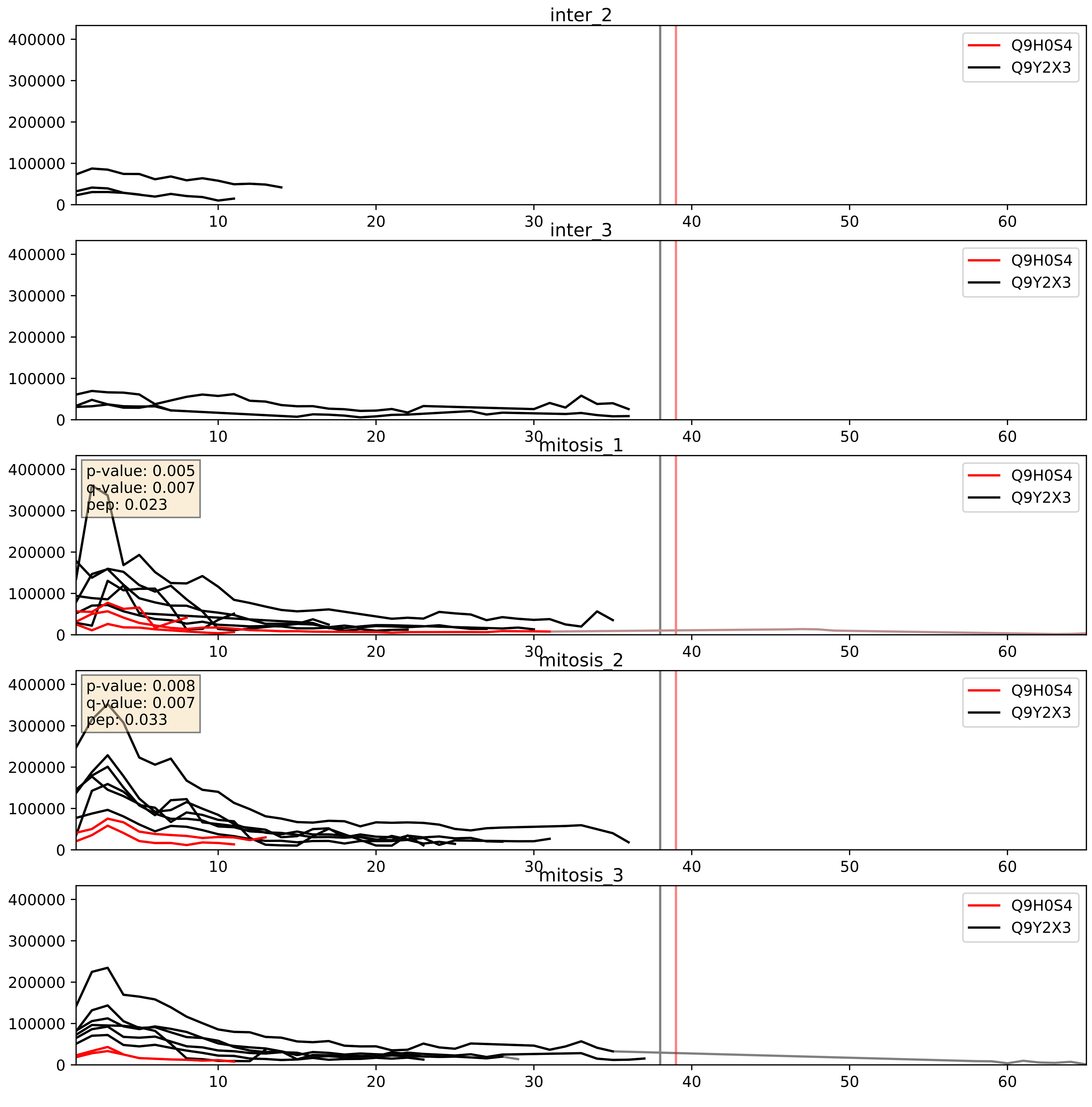


| level | condition_1 | condition_2 | pvalue | pvalue_adjusted |
|---|---|---|---|---|
| interactor_ratio | mitosis | inter | 0.0006458171899288122 | 0.029562540886580918 |
| complex_abundance | mitosis | inter | 0.00802824854735332 | 0.0663337910862398 |
| interactor_abundance | mitosis | inter | 0.3974703705091973 | 0.588081647490922 |

Q9H583_Q9Y2X3
HEAT1_HUMAN vs NOP58_HUMAN
p-value: 0.08 q-value: 0.047

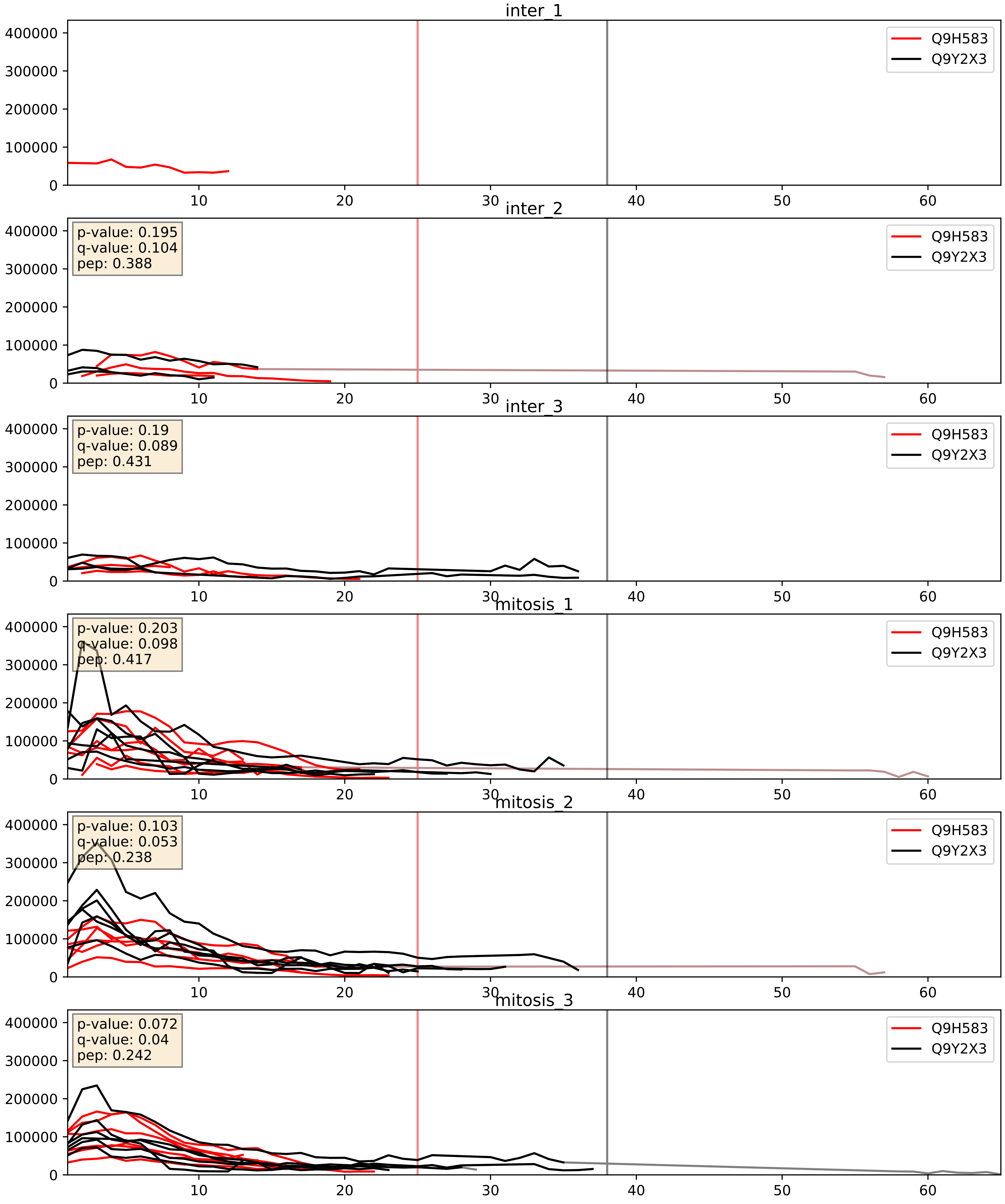


| level | condition_1 | condition_2 | pvalue | pvalue_adjusted |
|---|---|---|---|---|
| complex_abundance | mitosis | inter | 0.003745309179790294 | 0.04515471349155622 |
| interactor_abundance | mitosis | inter | 0.006814646959063384 | 0.07092204008459886 |
| interactor_ratio | mitosis | inter | 0.05599641218001745 | 0.3926647223857623 |

Q9NVP1_Q9Y2X3
DDX18_HUMAN vs NOP58_HUMAN
p-value: 0.049 q-value: 0.032

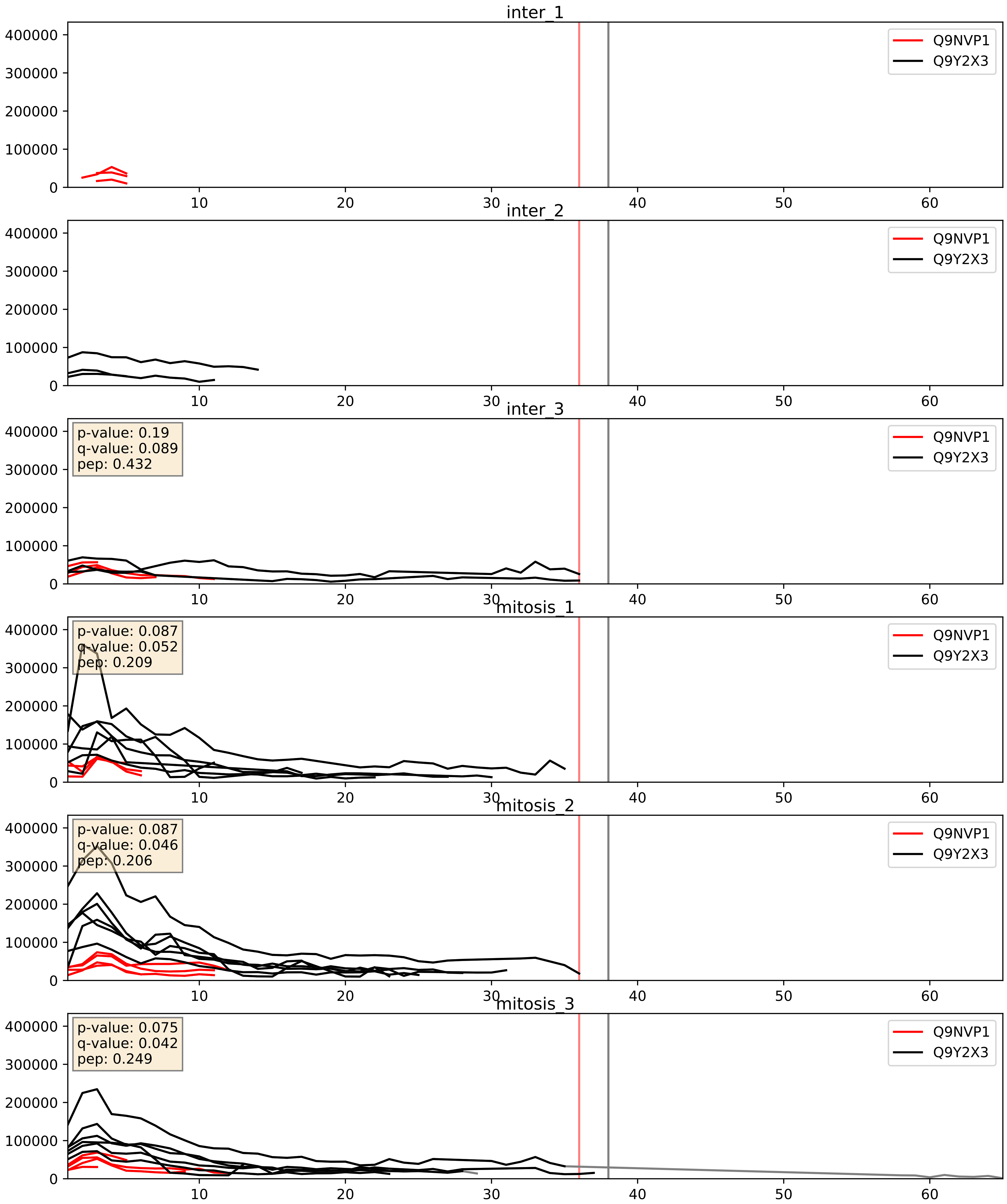


| level | condition_1 | condition_2 | pvalue | pvalue_adjusted |
|---|---|---|---|---|
| interactor_ratio | mitosis | inter | 0.0012725532055782418 | 0.0423373831474418 |
| complex_abundance | mitosis | inter | 0.006119371749399622 | 0.05820202463873419 |
| interactor_abundance | mitosis | inter | 0.16570946498784456 | 0.3537339202733041 |

Q9NX24_Q9Y2X3
NHP2_HUMAN vs NOP58_HUMAN
p-value: 0.001 q-value: 0.001

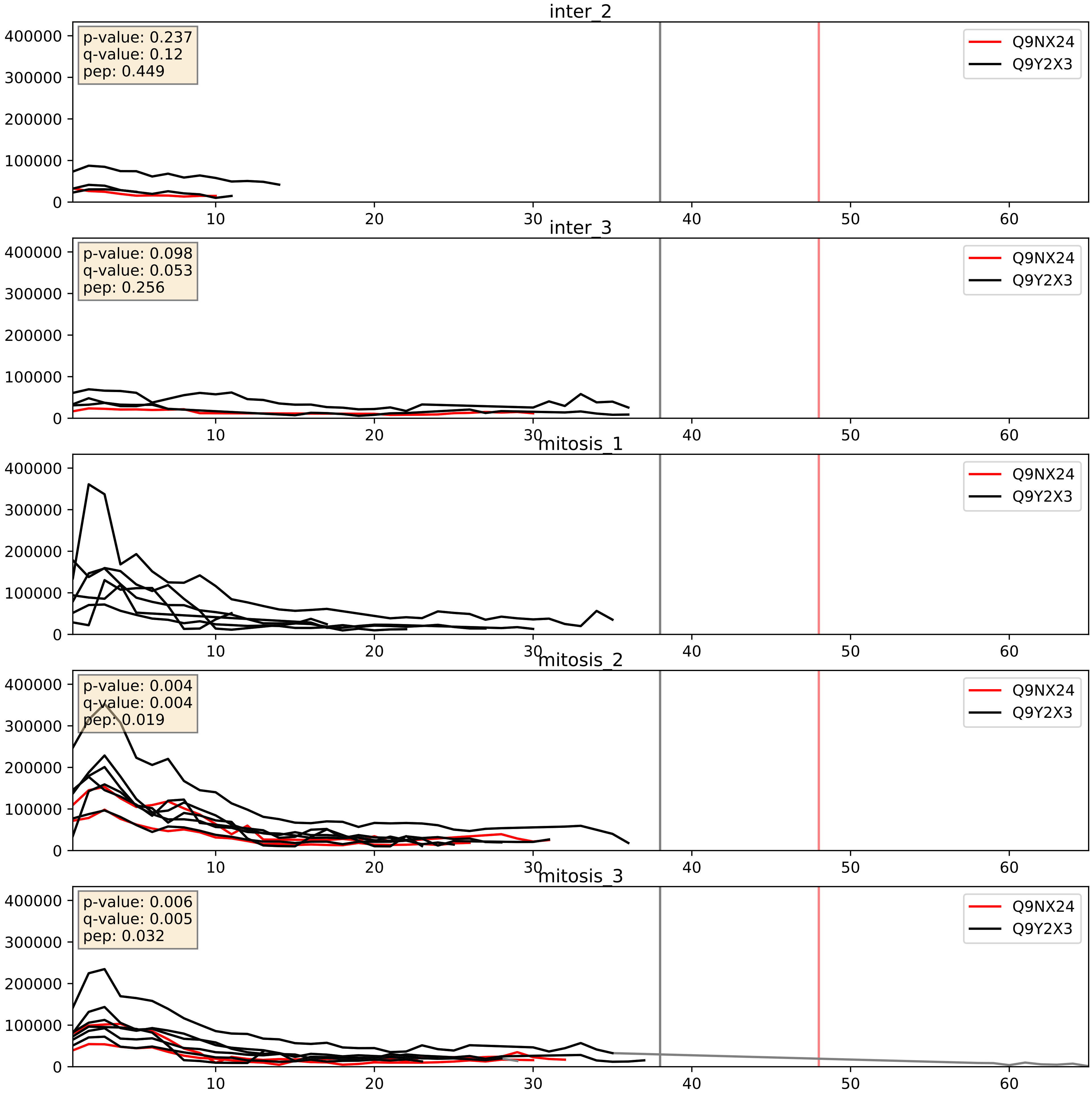


| level | condition_1 | condition_2 | pvalue | pvalue_adjusted |
|---|---|---|---|---|
| interactor_abundance | mitosis | inter | 0.3322629281627048 | 0.5313723802097624 |
| complex_abundance | mitosis | inter | 0.34406044365467237 | 0.4920075839766113 |
| interactor_ratio | mitosis | inter | 0.370927683823078 | 0.8138388063636502 |

Q9NY93_Q9Y2X3
DDX56_HUMAN vs NOP58_HUMAN
p-value: 0.021 q-value: 0.017

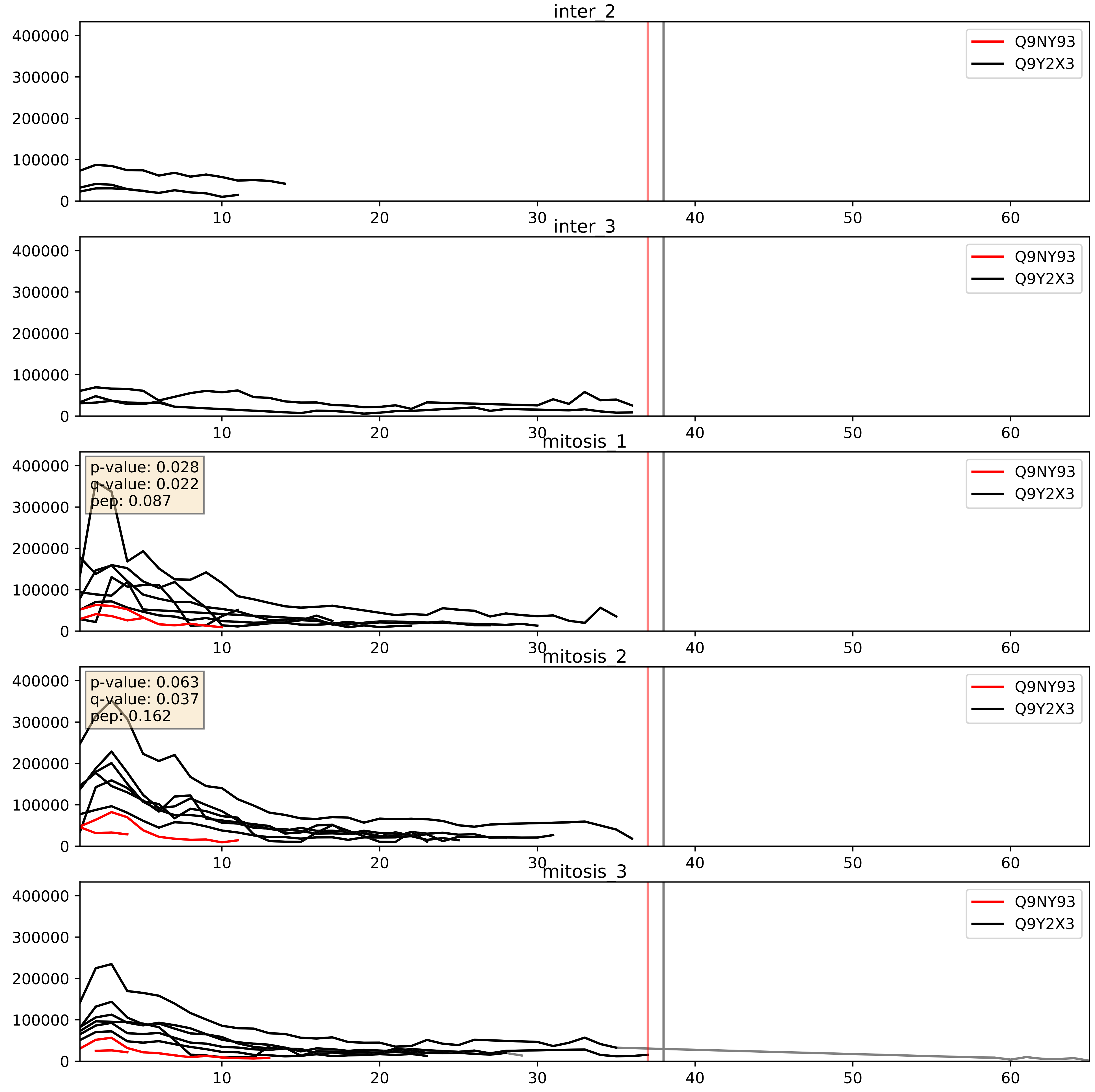


| level | condition_1 | condition_2 | pvalue | pvalue_adjusted |
|---|---|---|---|---|
| interactor_ratio | mitosis | inter | 0.00010063874199602321 | 0.010720128702277293 |
| complex_abundance | mitosis | inter | 0.0007327642214531893 | 0.017769013415408784 |
| interactor_abundance | mitosis | inter | 0.10159744920024559 | 0.27370850762629345 |

Q9UNX4_Q9Y2X3
WDR3_HUMAN vs NOP58_HUMAN
p-value: 0.052 q-value: 0.034

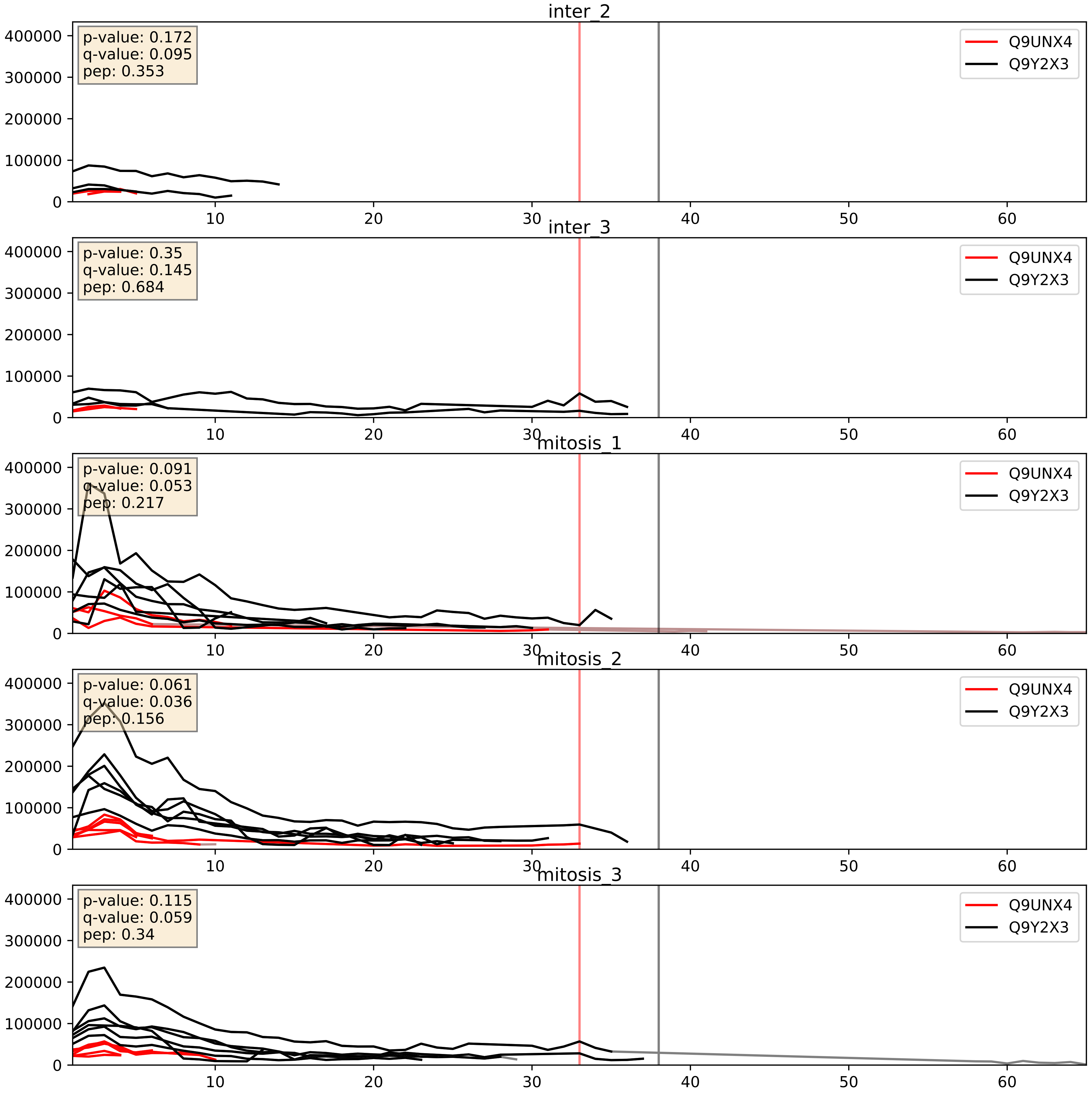


| level | condition_1 | condition_2 | pvalue | pvalue_adjusted |
|---|---|---|---|---|
| interactor_ratio | mitosis | inter | 0.0008632207210684006 | 0.033740499417102776 |
| complex_abundance | mitosis | inter | 0.00703579685561239 | 0.06202515044700521 |
| interactor_abundance | mitosis | inter | 0.15972059028570332 | 0.34604106627325243 |

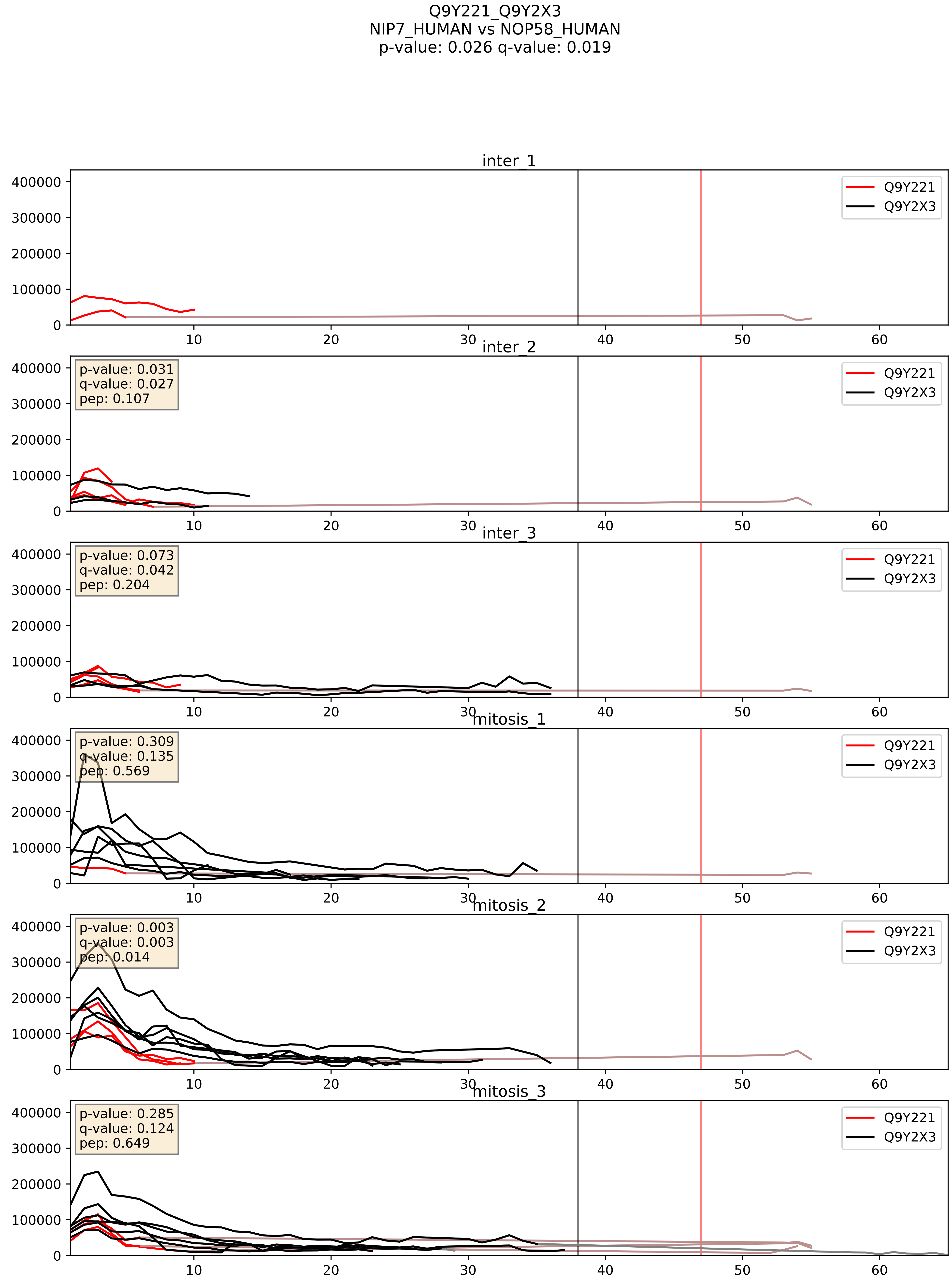


| level | condition_1 | condition_2 | pvalue | pvalue_adjusted |
|---|---|---|---|---|
| interactor_ratio | mitosis | inter | 0.0028583564607708 | 0.069510032114199 |
| complex_abundance | mitosis | inter | 0.13950383100874447 | 0.28459313475568465 |
| interactor_abundance | mitosis | inter | 0.9311568103729986 | 0.9610620620778482 |

Q9Y2X3_Q9Y3T9
NOP58_HUMAN vs NOC2L_HUMAN
p-value: 0.04 q-value: 0.027

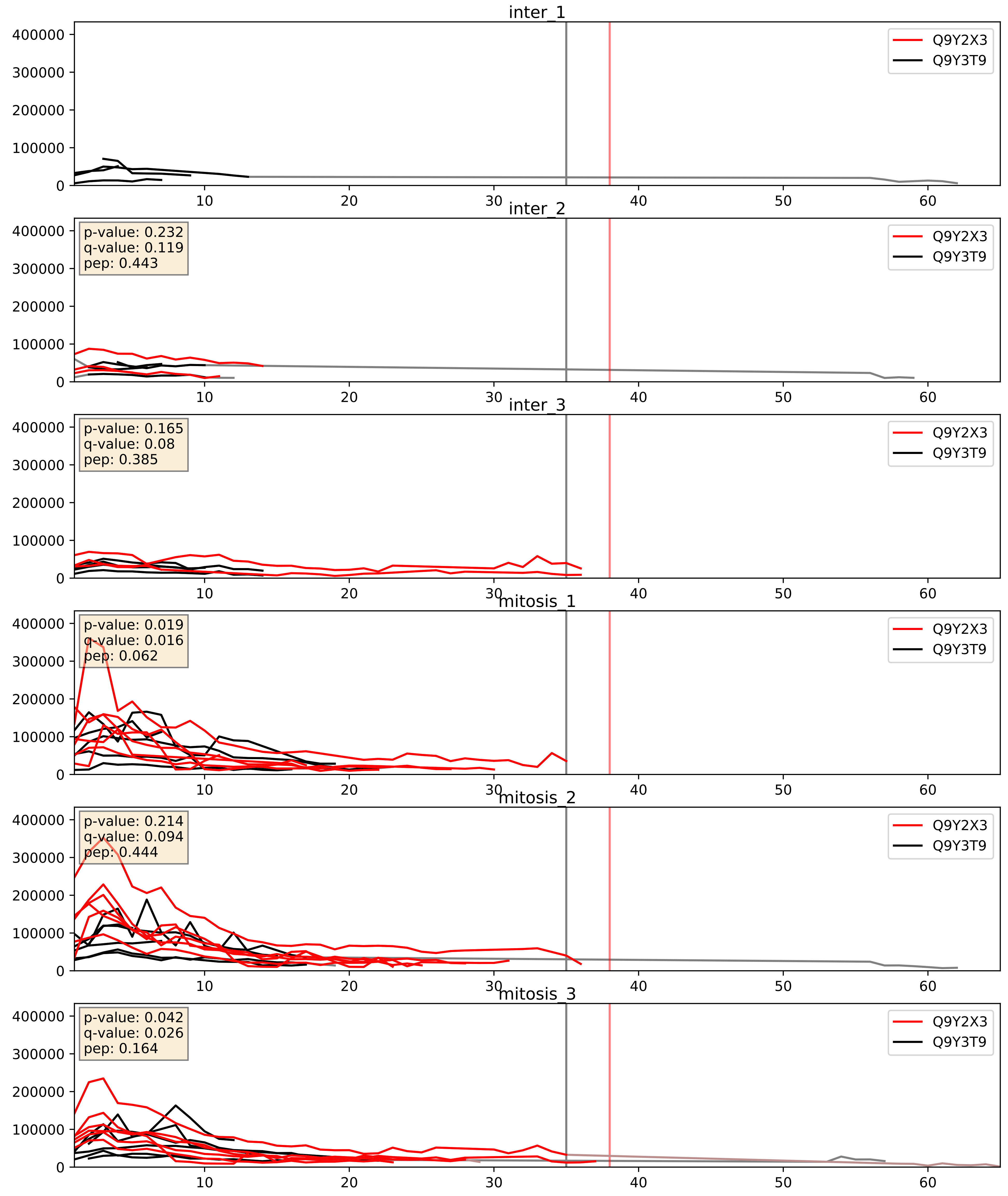


| level | condition_1 | condition_2 | pvalue | pvalue_adjusted |
|---|---|---|---|---|
| interactor_abundance | mitosis | inter | 0.0015817191617625384 | 0.029025269221018554 |
| complex_abundance | mitosis | inter | 0.0023478262388268484 | 0.035168510428347066 |
| interactor_ratio | mitosis | inter | 0.03146494532124448 | 0.28316826136323564 |
