## Supplementary Data 2 for "SECAT: Quantifying differential protein-protein interaction states by network-centric analysis": hela_string_000088_P08670.pdf

P08670_P08670

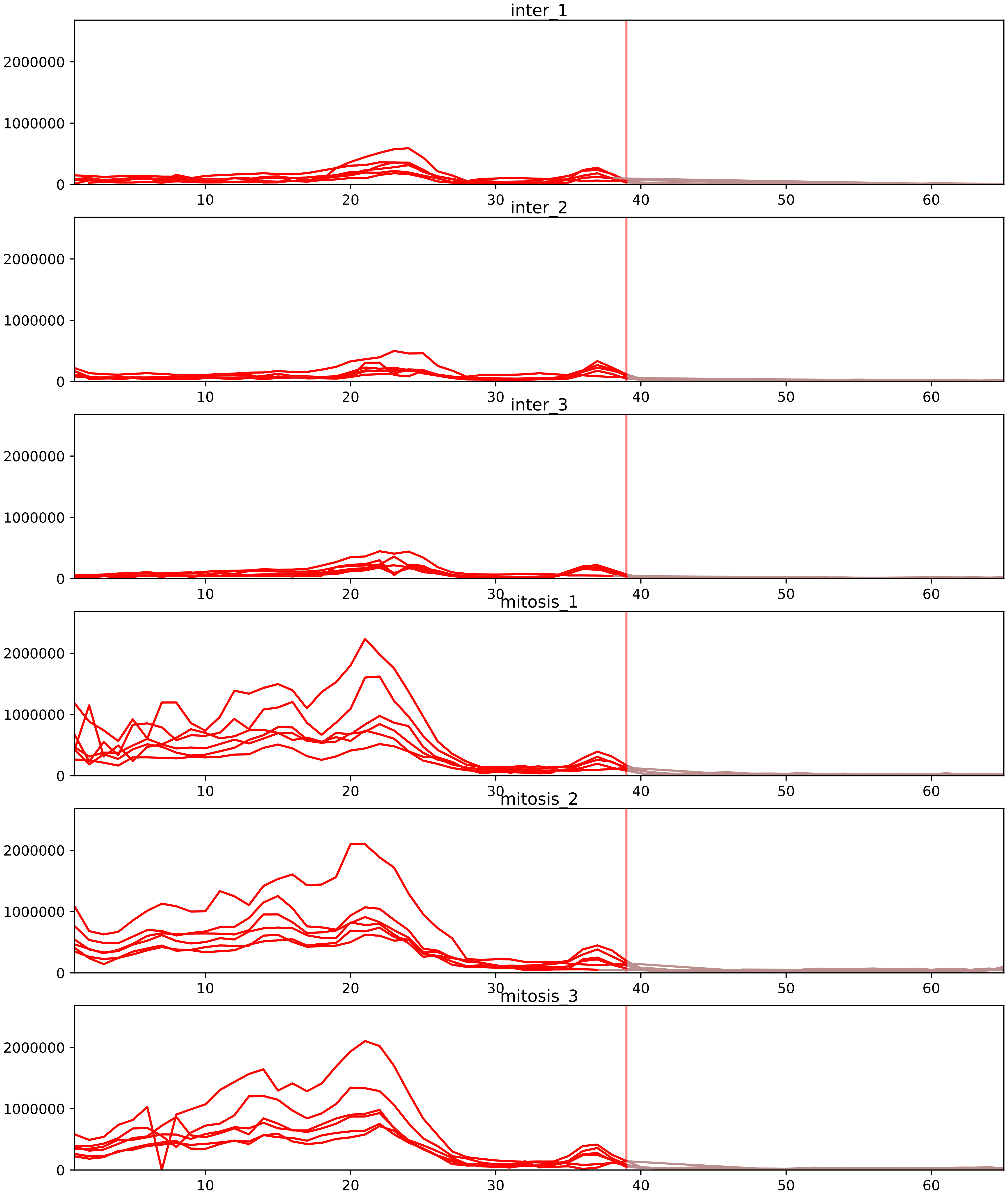


| level | condition_1 | condition_2 | pvalue | pvalue_adjusted |
|---|---|---|---|---|
| total_abundance | mitosis | inter | 5.564116302428078e-06 | 0.003540864511957668 |
| assembled_abundance | mitosis | inter | 5.777758293437989e-06 | 0.0030050762857314695 |
| monomer_abundance | mitosis | inter | 0.058025257644455895 | 0.3497872727142794 |
