## Supplementary Data 2 for "SECAT: Quantifying differential protein-protein interaction states by network-centric analysis": hela_string_000089_Q9BQ67.pdf

Q9BQ67_Q9BQ67

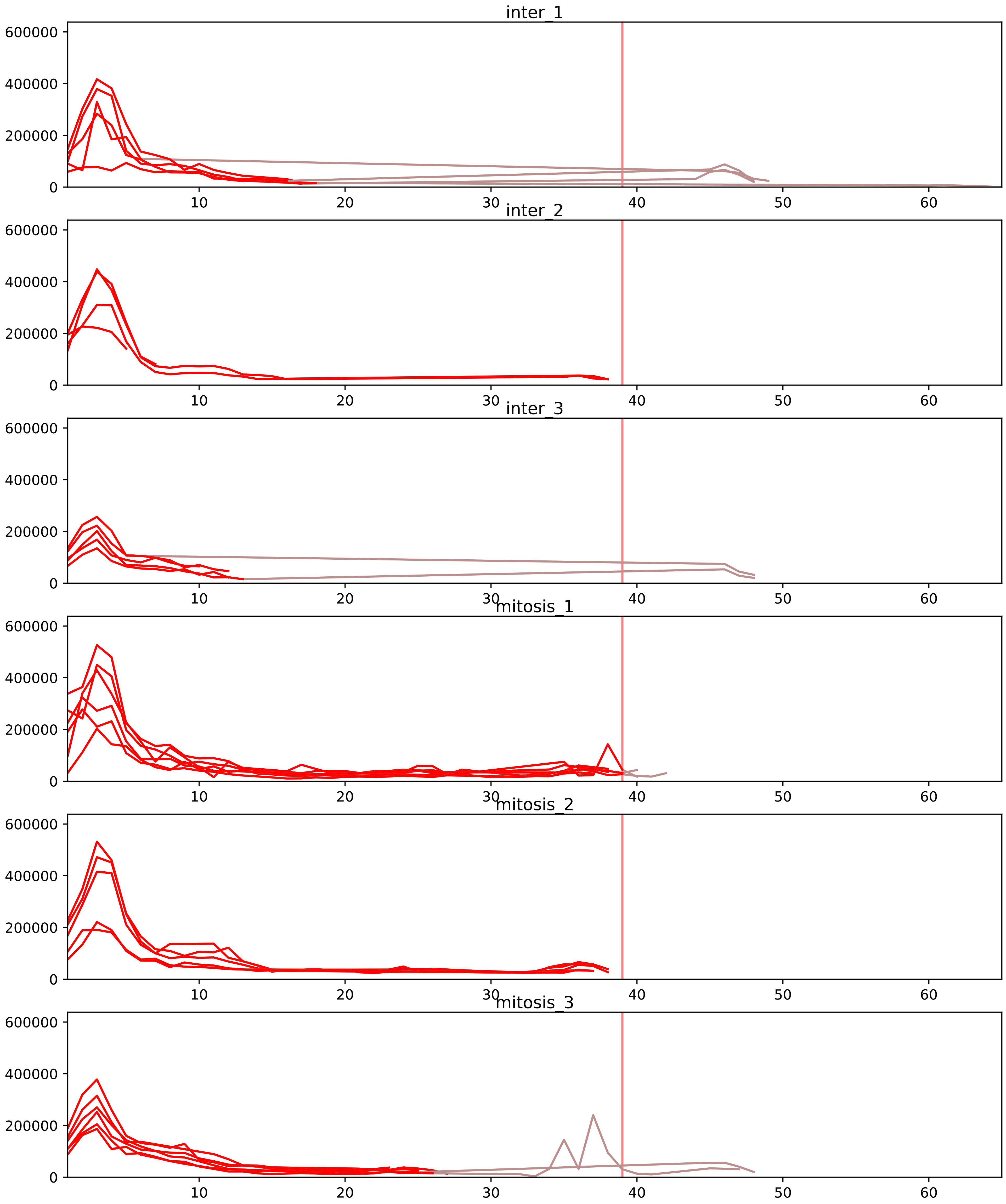


| level | condition_1 | condition_2 | pvalue | pvalue_adjusted |
|---|---|---|---|---|
| interactor_ratio | mitosis | inter | 5.976568051157773e-06 | 0.0001486065569477068 |
| complex_abundance | mitosis | inter | 0.0038480681352297524 | 0.02425761451640483 |
| interactor_abundance | mitosis | inter | 0.018330301400308575 | 0.07478437821855383 |
| monomer_abundance | mitosis | inter | 0.26949302476518566 | 0.6488006005486777 |
| total_abundance | mitosis | inter | 0.709124795526952 | 0.876251051948474 |
| assembled_abundance | mitosis | inter | 0.7279873687768754 | 0.8892768458362614 |

P22087_Q9BQ67
FBRL_HUMAN vs GRWD1_HUMAN
p-value: 0.001 q-value: 0.029

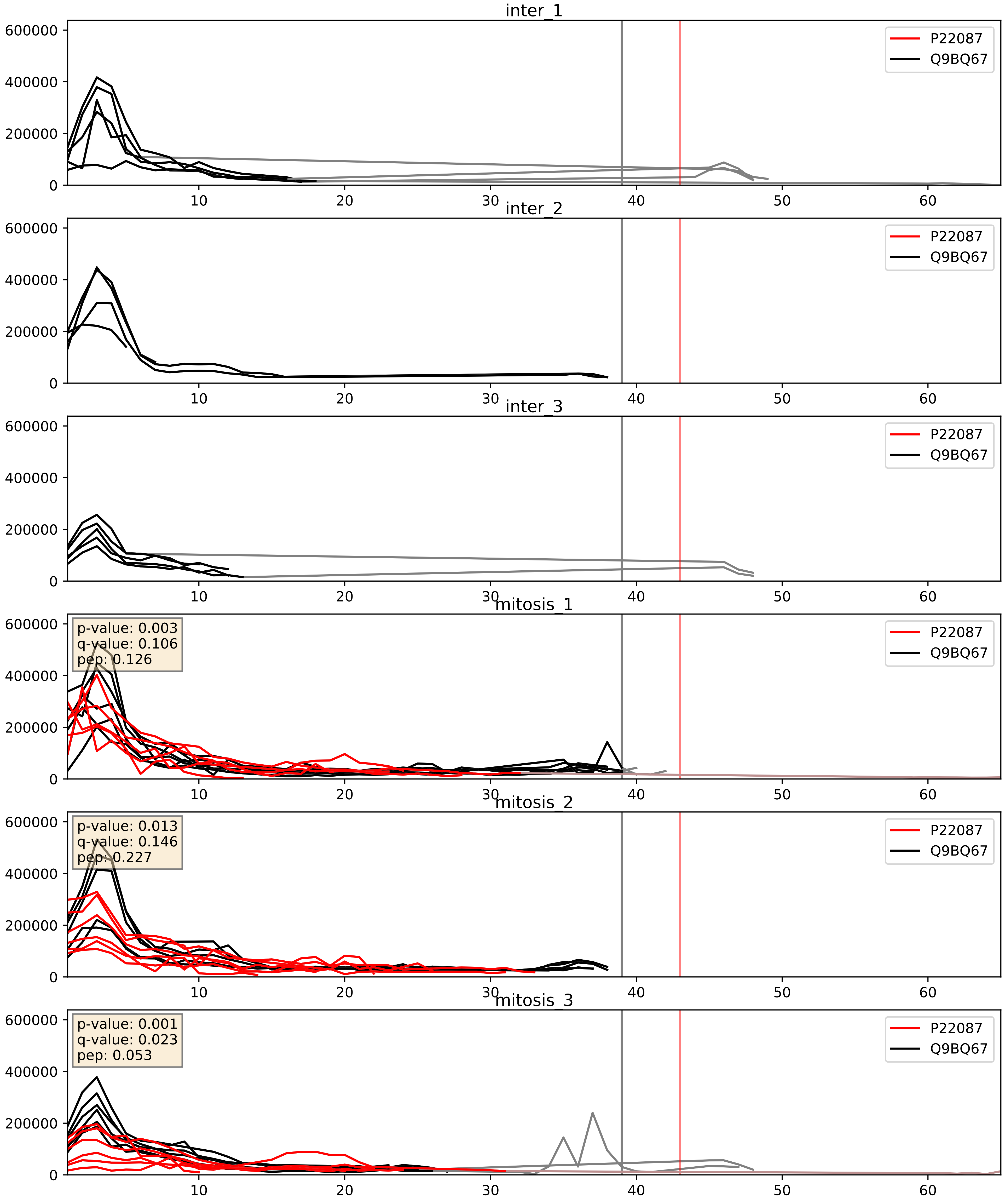


| level | condition_1 | condition_2 | pvalue | pvalue_adjusted |
|---|---|---|---|---|
| complex_abundance | mitosis | inter | 0.00022948761081750734 | 0.010219389159334393 |
| interactor_abundance | mitosis | inter | 0.0003900245617774174 | 0.01362698060740691 |
| interactor_ratio | mitosis | inter | 0.011537958352666234 | 0.15494522140242886 |

P30050_Q9BQ67
RL12_HUMAN vs GRWD1_HUMAN
p-value: 0.0 q-value: 0.042

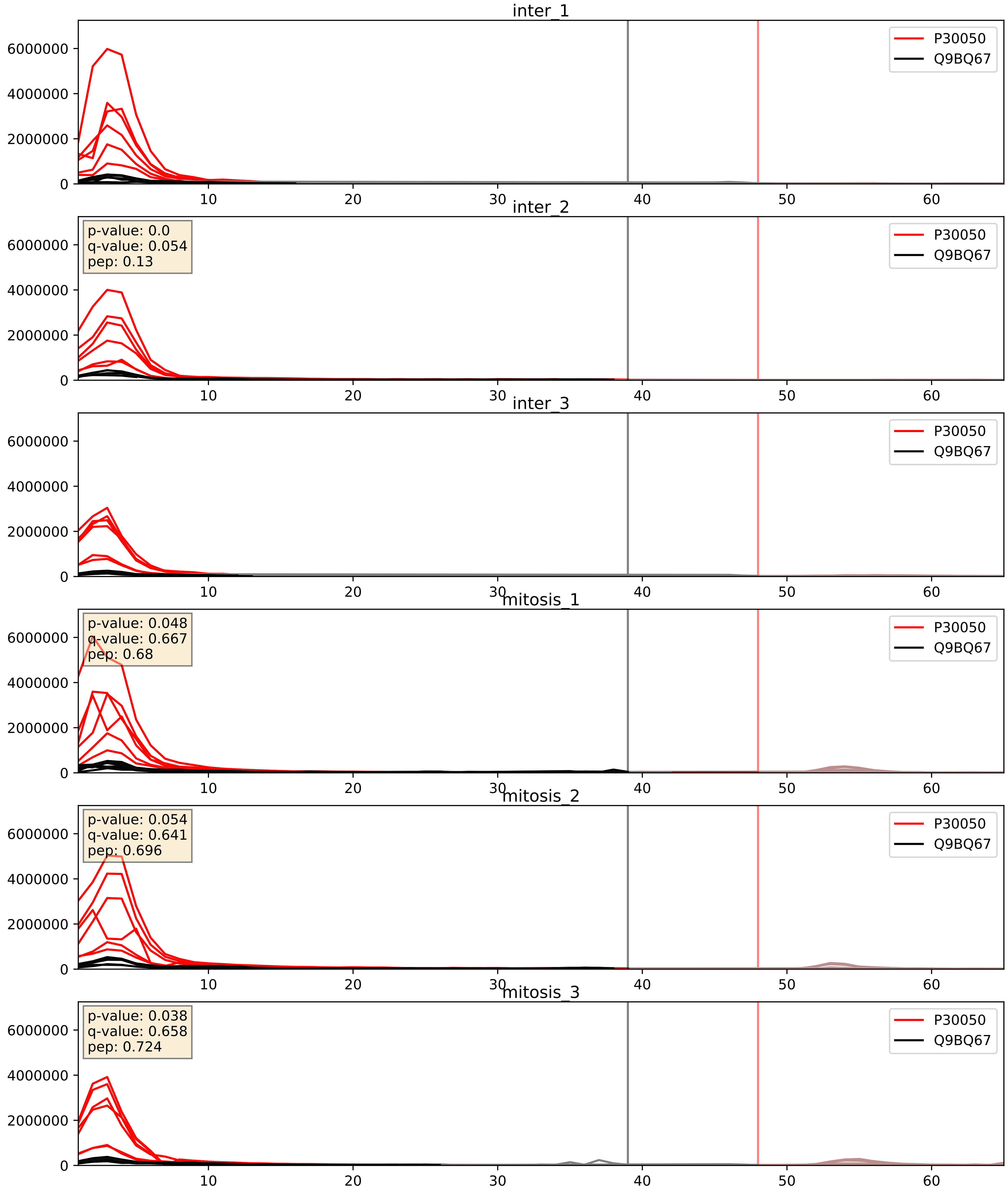


| level | condition_1 | condition_2 | pvalue | pvalue_adjusted |
|---|---|---|---|---|
| complex_abundance | mitosis | inter | 0.23344374732029655 | 0.3853960418634018 |
| interactor_ratio | mitosis | inter | 0.24945930068401068 | 0.723755371092335 |
| interactor_abundance | mitosis | inter | 0.8353553883695714 | 0.9101925168311928 |

P39023_Q9BQ67
RL3_HUMAN vs GRWD1_HUMAN
p-value: 0.001 q-value: 0.045

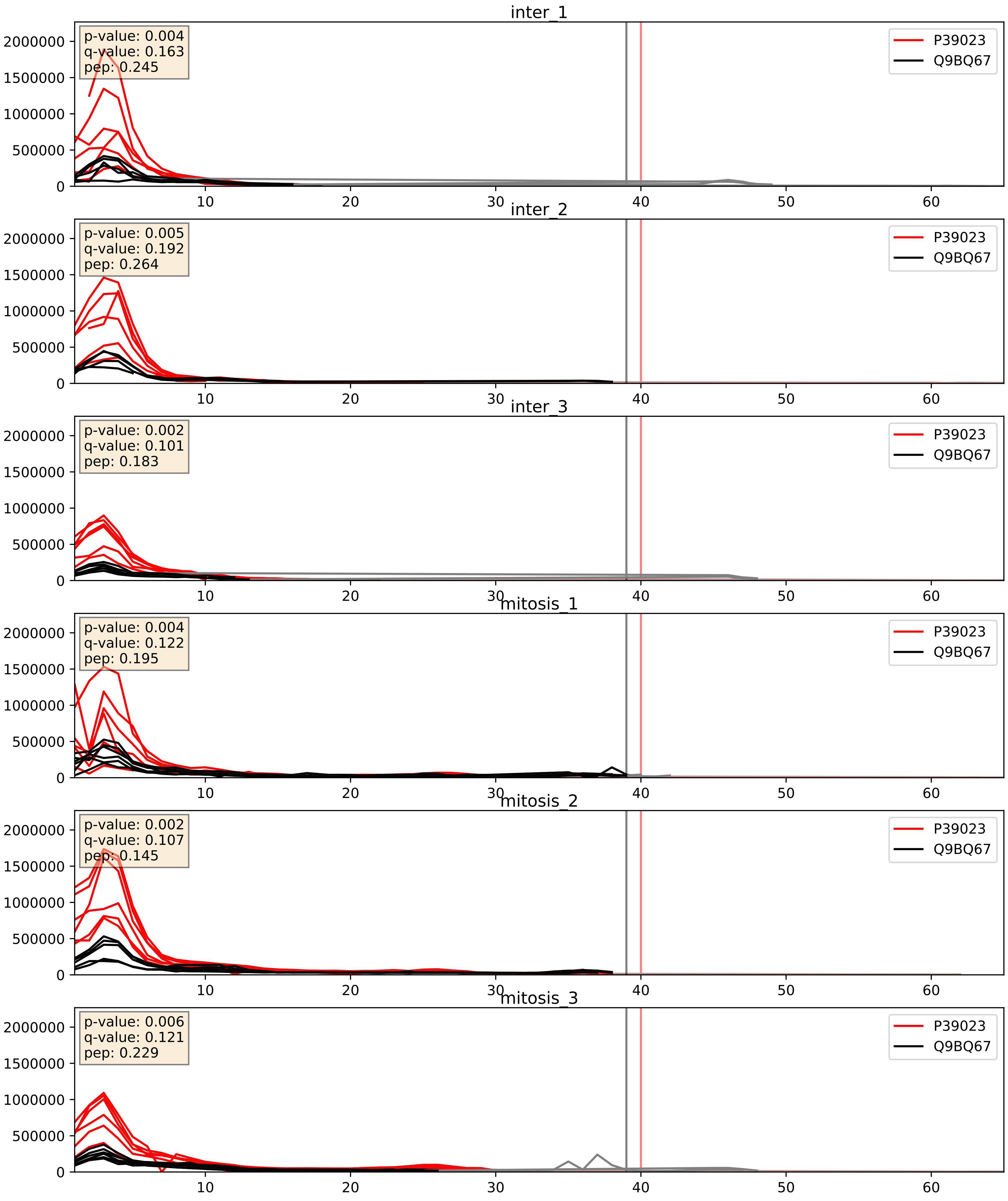


| level | condition_1 | condition_2 | pvalue | pvalue_adjusted |
|---|---|---|---|---|
| interactor_ratio | mitosis | inter | 0.1590076783003557 | 0.616090239021059 |
| complex_abundance | mitosis | inter | 0.18192039652865455 | 0.33281440356599334 |
| interactor_abundance | mitosis | inter | 0.45576812354633084 | 0.6293625806063166 |

P56537_Q9BQ67
IF6_HUMAN vs GRWD1_HUMAN
p-value: 0.0 q-value: 0.046

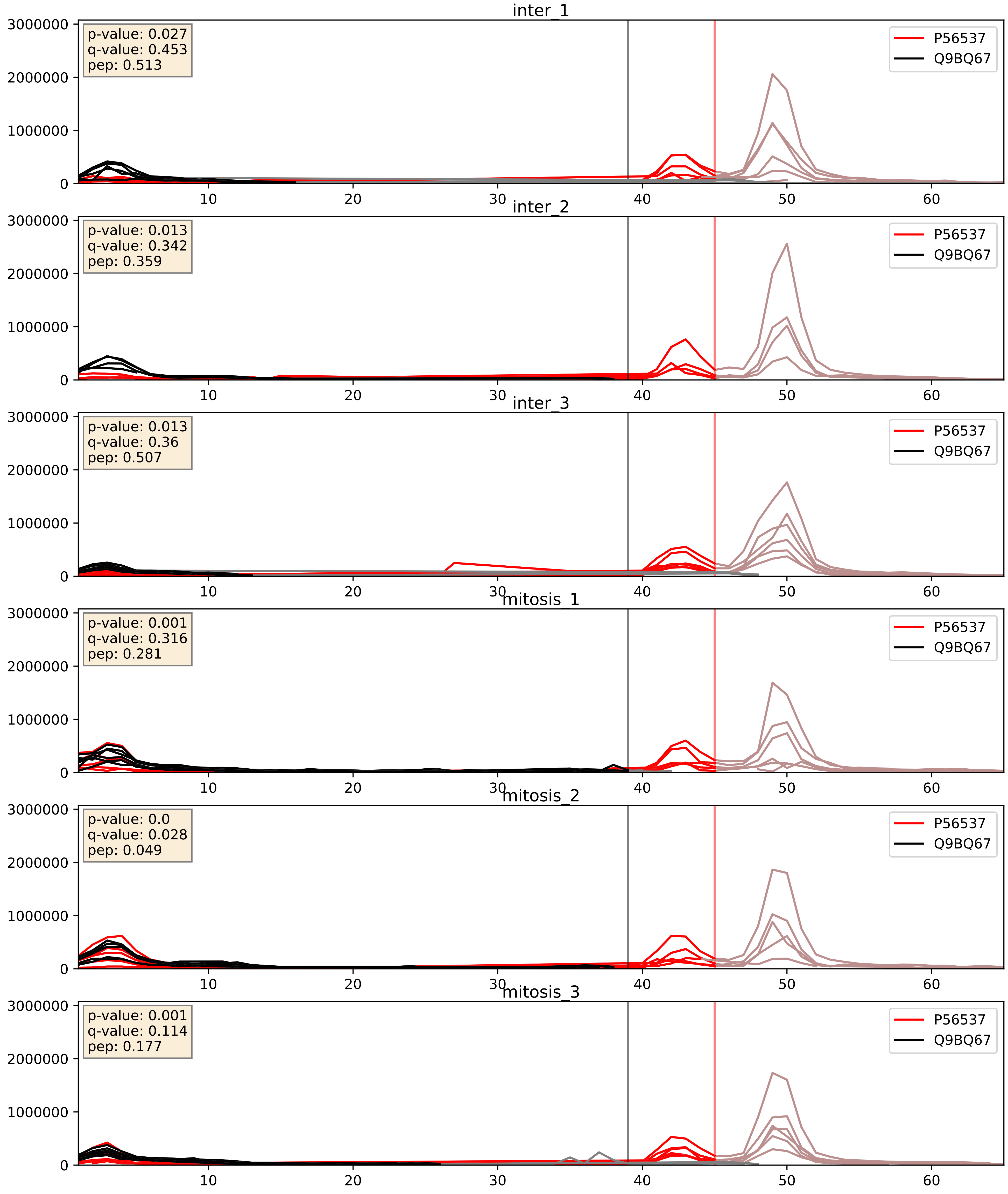


| level | condition_1 | condition_2 | pvalue | pvalue_adjusted |
|---|---|---|---|---|
| interactor_ratio | mitosis | inter | 0.0052956920235738775 | 0.0974780445871482 |
| interactor_abundance | mitosis | inter | 0.007702330616543858 | 0.07617787415091325 |
| complex_abundance | mitosis | inter | 0.03073328011425226 | 0.12751204260889337 |

Q8NI36_Q9BQ67
WDR36_HUMAN vs GRWD1_HUMAN
p-value: 0.006 q-value: 0.006

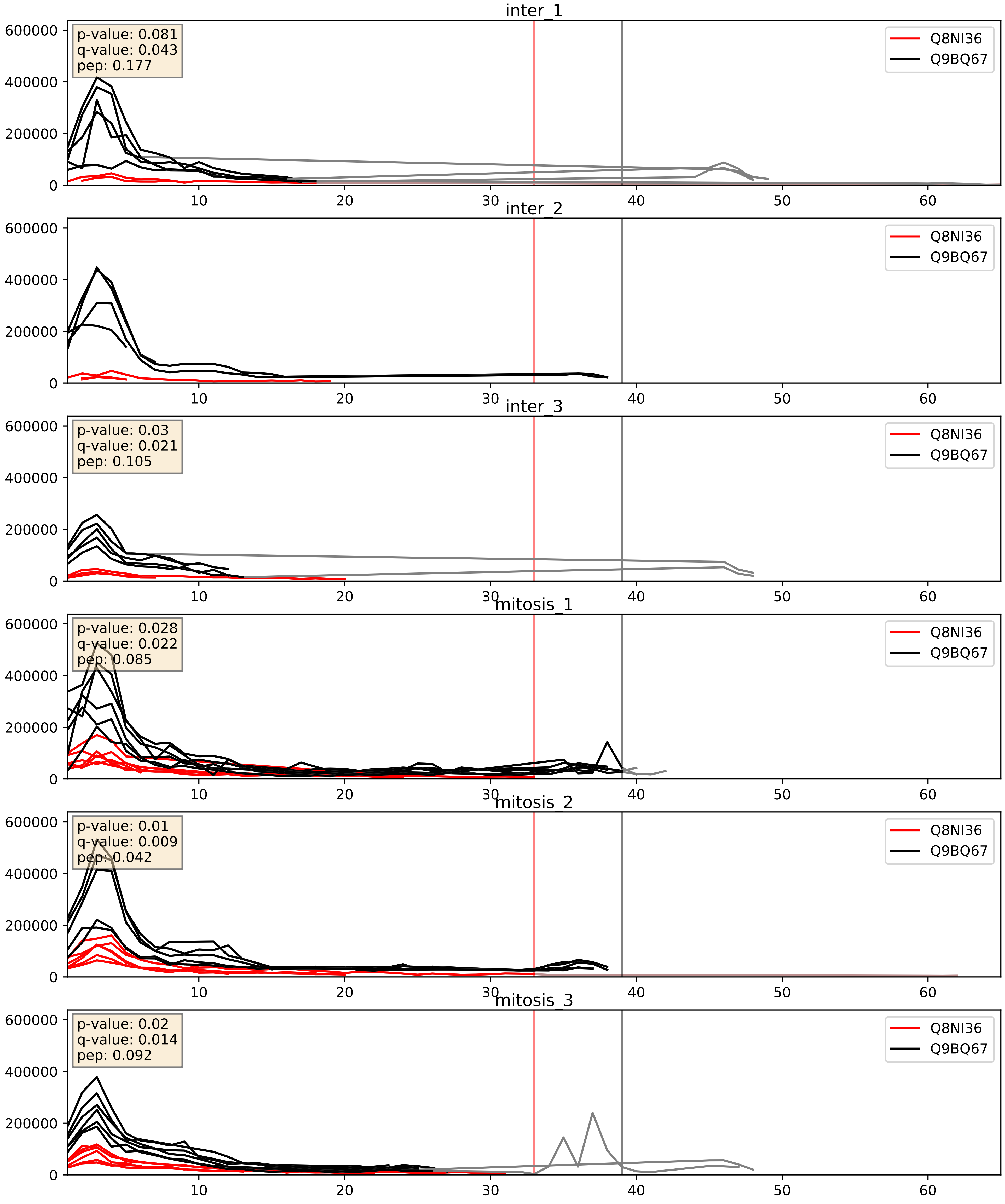


| level | condition_1 | condition_2 | pvalue | pvalue_adjusted |
|---|---|---|---|---|
| interactor_abundance | mitosis | inter | 0.004254336243081026 | 0.05203482881393319 |
| complex_abundance | mitosis | inter | 0.027185050581926013 | 0.12026048216087167 |
| interactor_ratio | mitosis | inter | 0.09811809124801148 | 0.5212210525862964 |

Q9BQ67_Q9BQG0
GRWD1_HUMAN vs MBB1A_HUMAN
p-value: 0.001 q-value: 0.047

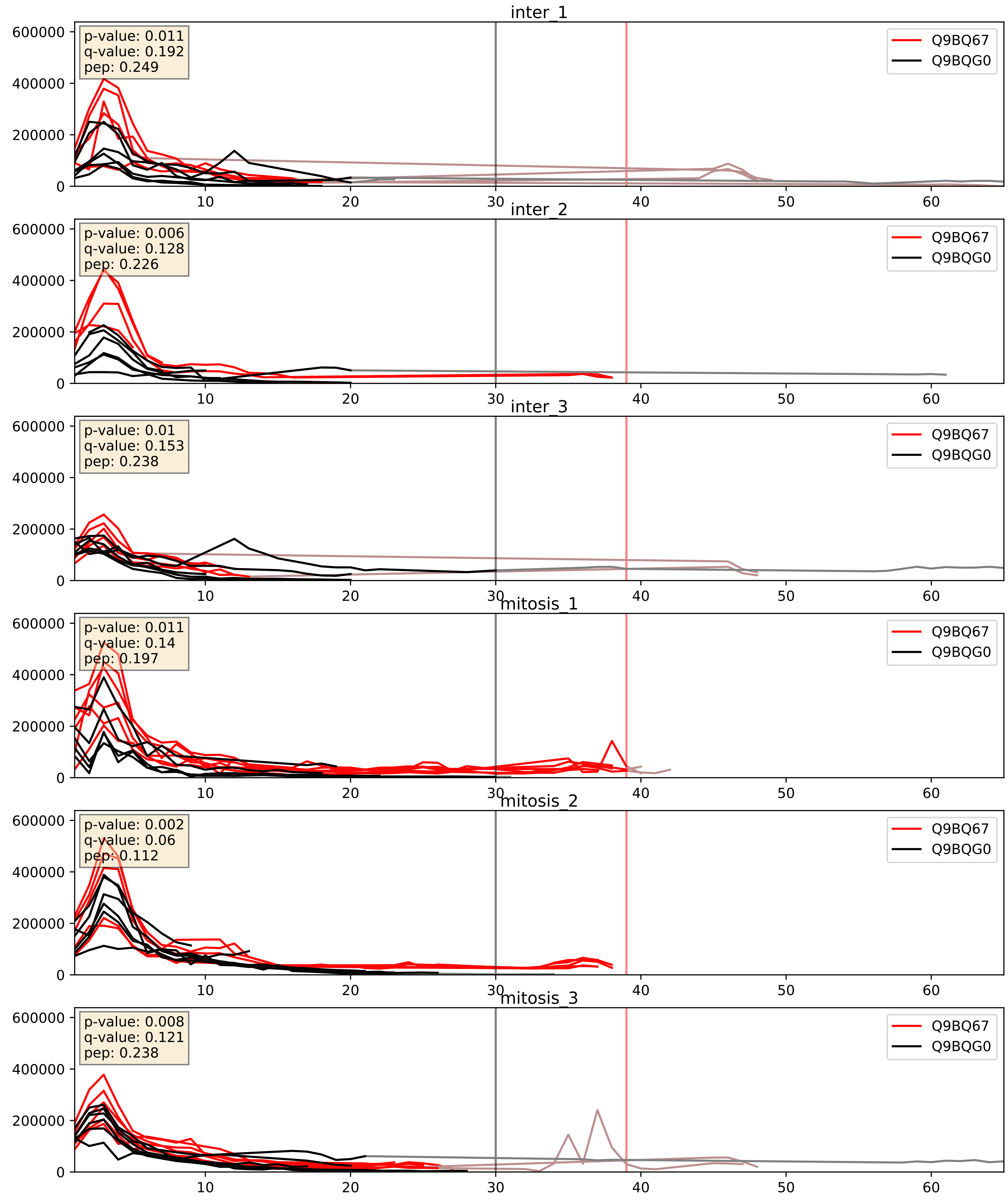


| level | condition_1 | condition_2 | pvalue | pvalue_adjusted |
|---|---|---|---|---|
| complex_abundance | mitosis | inter | 0.054598838712082653 | 0.16644090433597847 |
| interactor_abundance | mitosis | inter | 0.16213543266863736 | 0.34963318876681726 |
| interactor_ratio | mitosis | inter | 0.9244852212565758 | 0.9871879493798719 |

Q9BQ67_Q9BZE4
GRWD1_HUMAN vs NOG1_HUMAN
p-value: 0.003 q-value: 0.004

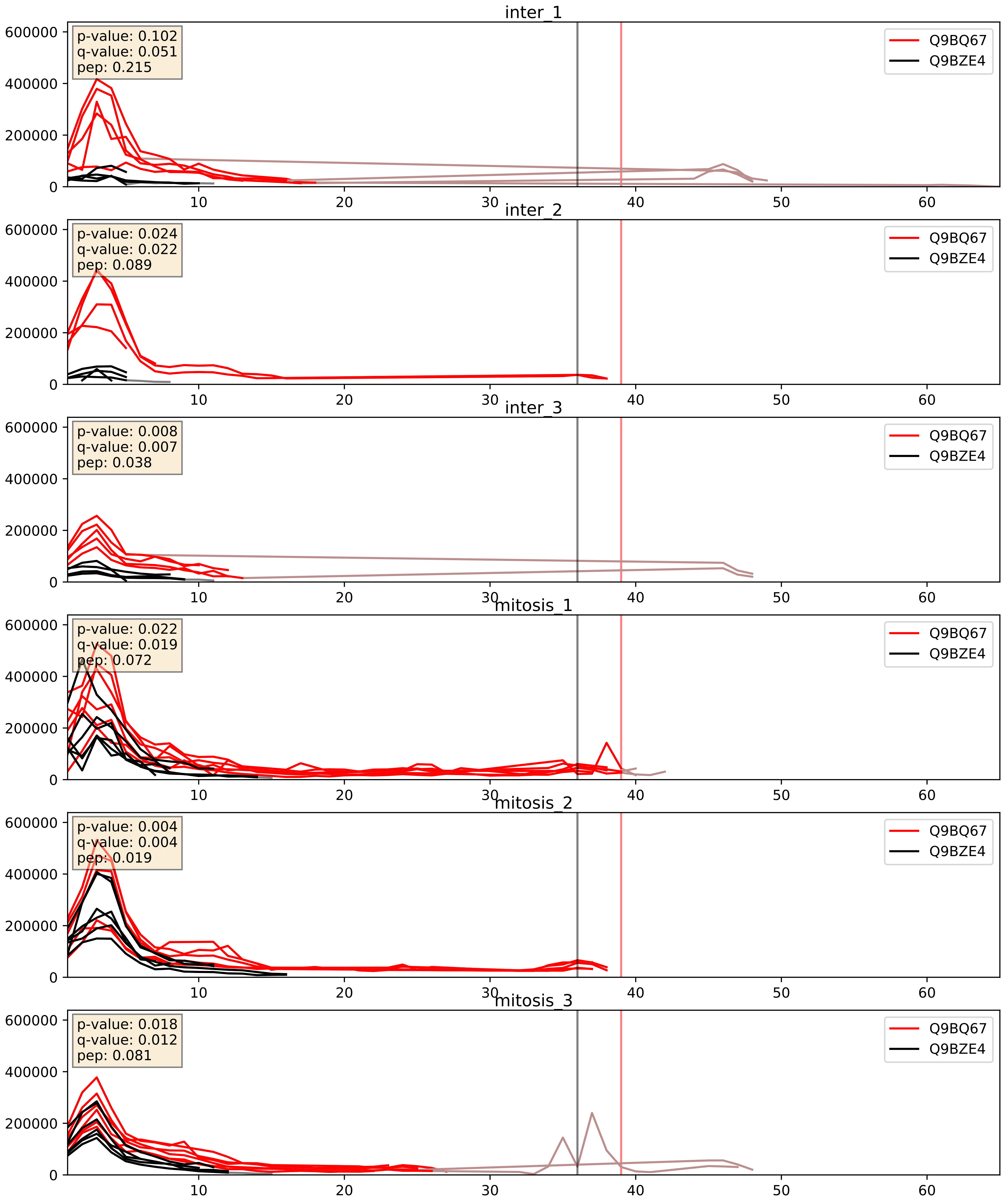


| level | condition_1 | condition_2 | pvalue | pvalue_adjusted |
|---|---|---|---|---|
| interactor_ratio | mitosis | inter | 0.0016120197660940117 | 0.048587638020298377 |
| complex_abundance | mitosis | inter | 0.007234422190094093 | 0.06298977015239762 |
| interactor_abundance | mitosis | inter | 0.08268426471817841 | 0.24090473795481276 |

Q9BQ67_Q9NVP1
GRWD1_HUMAN vs DDX18_HUMAN
p-value: 0.019 q-value: 0.015

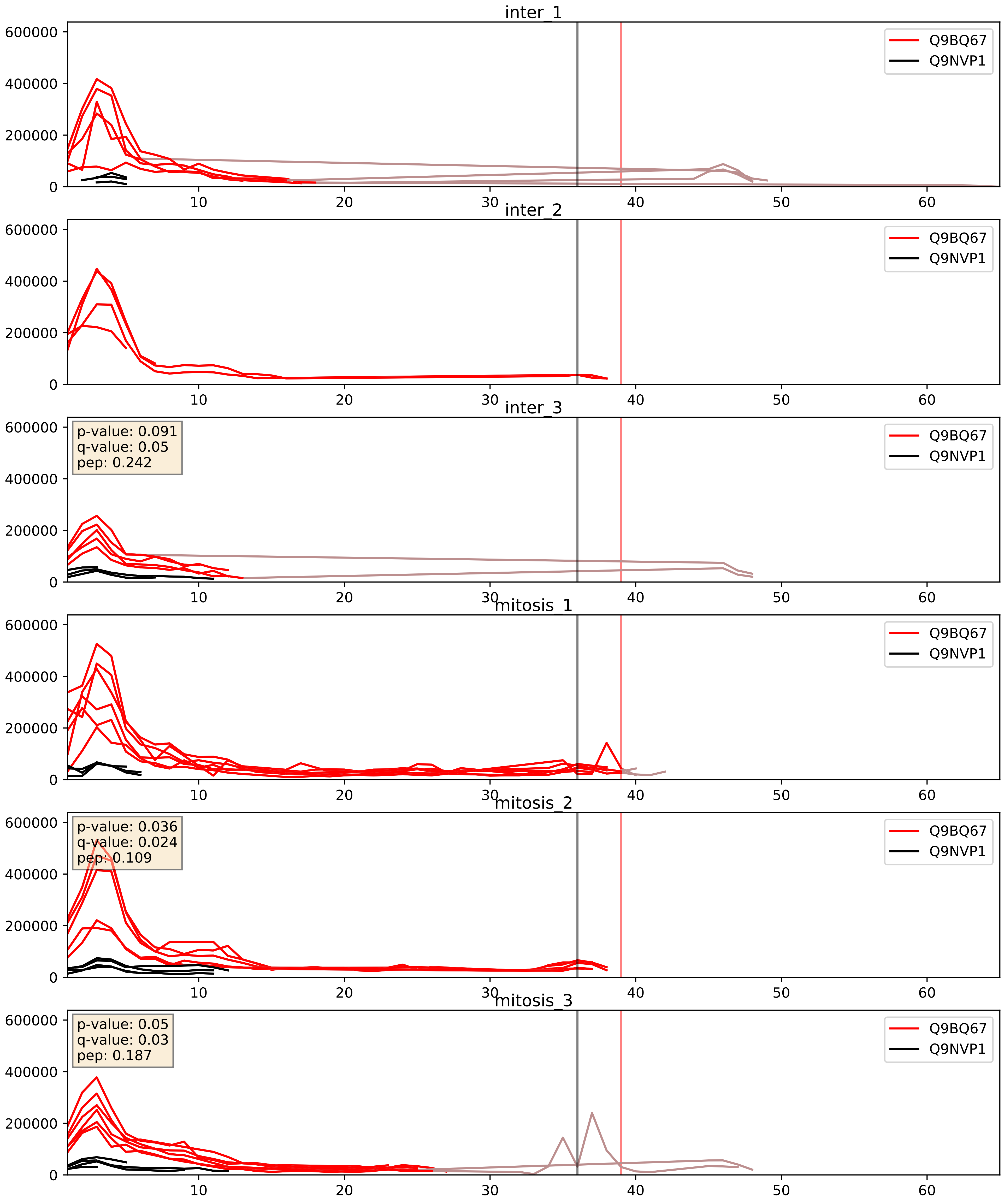


| level | condition_1 | condition_2 | pvalue | pvalue_adjusted |
| --- | --- | --- | --- | --- |
| interactor_ratio | mitosis | inter | 0.05392358354238998 | 0.38487308410443505 |
| interactor_abundance | mitosis | inter | 0.14019369867573772 | 0.3269656225748469 |
| complex_abundance | mitosis | inter | 0.22993119905718284 | 0.3831280826025919 |

Q9BQ67_Q9Y3T9
GRWD1_HUMAN vs NOC2L_HUMAN
p-value: 0.078 q-value: 0.046

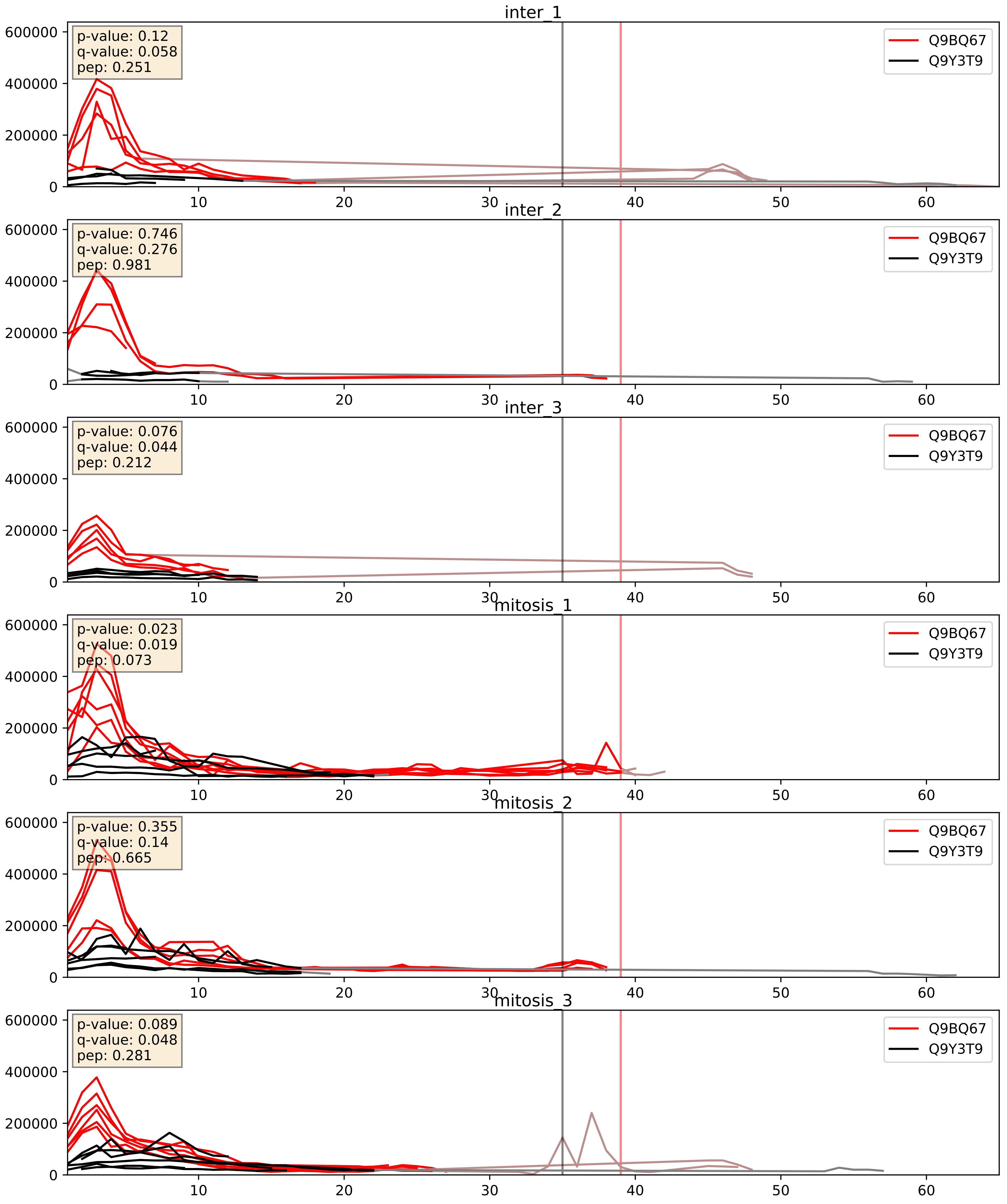


| level | condition_1 | condition_2 | pvalue | pvalue_adjusted |
|---|---|---|---|---|
| complex_abundance | mitosis | inter | 0.012364424967787958 | 0.08259410436140613 |
| interactor_ratio | mitosis | inter | 0.020920483835626235 | 0.21906942949212715 |
| interactor_abundance | mitosis | inter | 0.08943831949006091 | 0.25354926803607264 |
