## Supplementary Data 2 for "SECAT: Quantifying differential protein-protein interaction states by network-centric analysis": hela_string_000090_Q14137.pdf

Q14137_Q14137

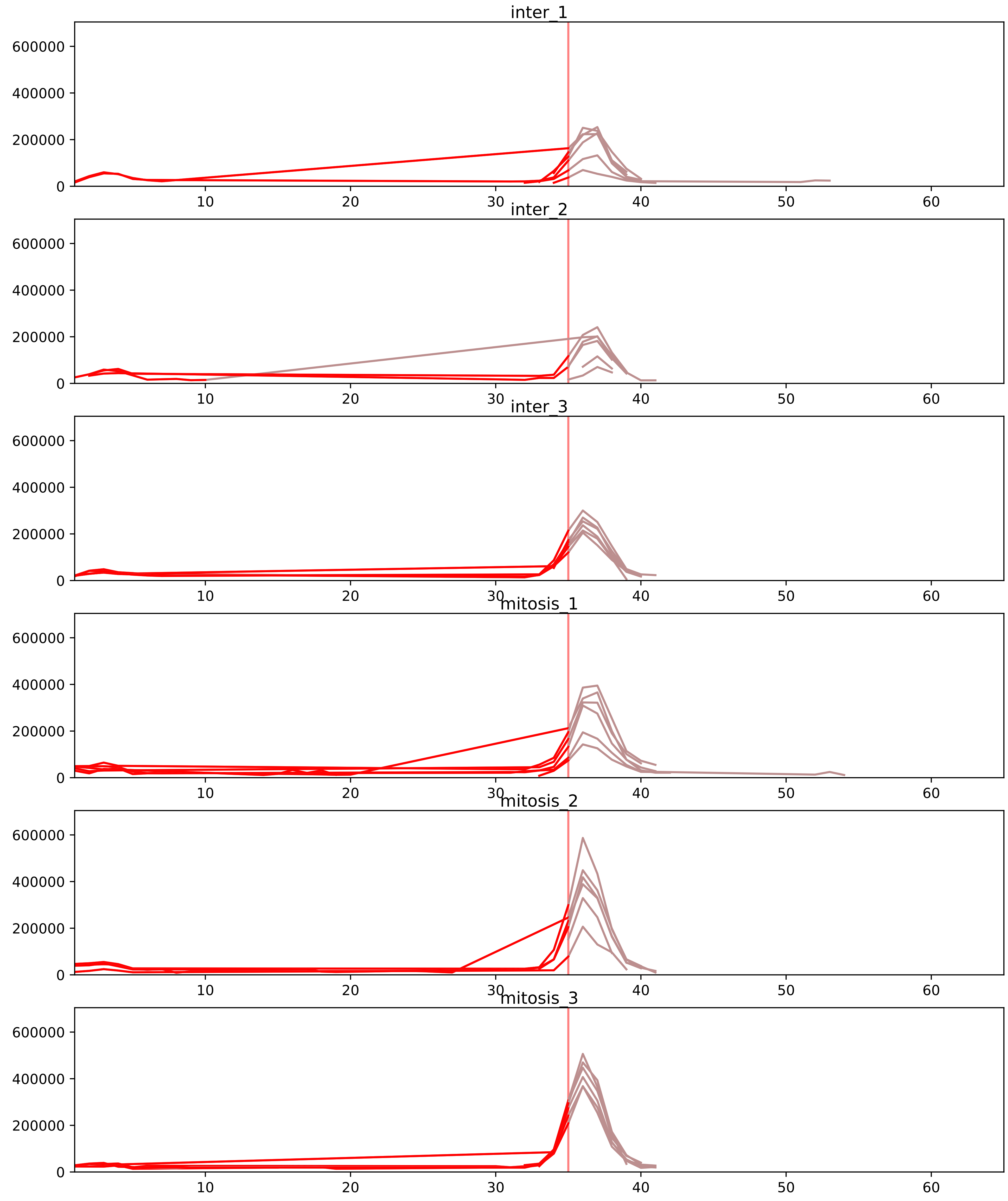


| level | condition_1 | condition_2 | pvalue | pvalue_adjusted |
|---|---|---|---|---|
| interactor_ratio | mitosis | inter | 6.093259807518079e-06 | 0.00014948797394444355 |
| monomer_abundance | mitosis | inter | 0.01039643685935092 | 0.18445730550018685 |
| total_abundance | mitosis | inter | 0.01235367864380042 | 0.1388357129703928 |
| complex_abundance | mitosis | inter | 0.05402885402602191 | 0.12944412943734415 |
| interactor_abundance | mitosis | inter | 0.06548677220757526 | 0.1721366583741978 |
| assembled_abundance | mitosis | inter | 0.9924774821916711 | 0.9967361283285159 |

O00541_Q14137
PESC_HUMAN vs BOP1_HUMAN
p-value: 0.006 q-value: 0.006

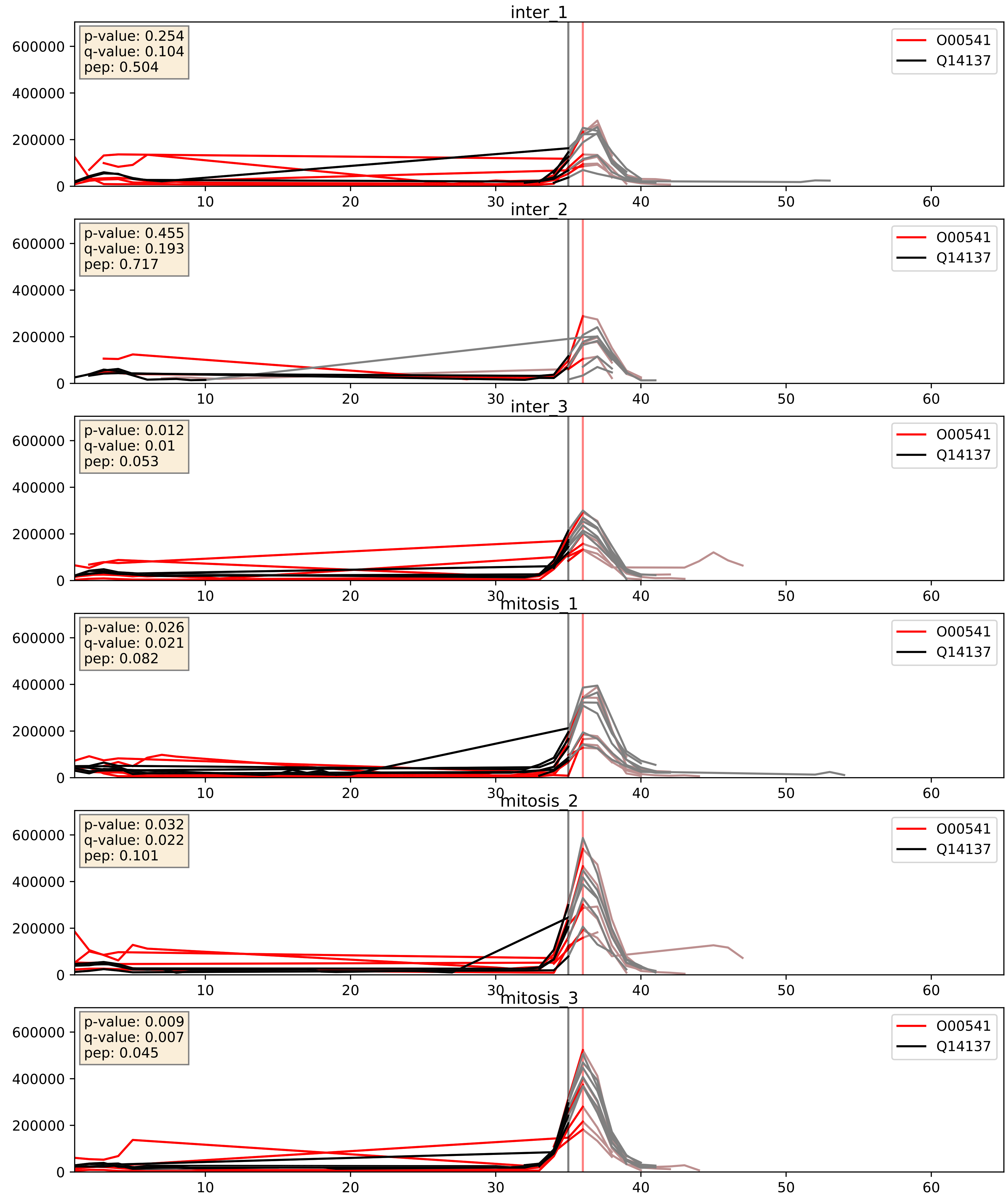


| level | condition_1 | condition_2 | pvalue | pvalue_adjusted |
|---|---|---|---|---|
| complex_abundance | mitosis | inter | 0.17765950619607593 | 0.3280339458667838 |
| interactor_ratio | mitosis | inter | 0.28437737580497385 | 0.7552808988180504 |
| interactor_abundance | mitosis | inter | 0.4628151344310173 | 0.6346840036413822 |

O76021_Q14137
RL1D1_HUMAN vs BOP1_HUMAN
p-value: 0.003 q-value: 0.003

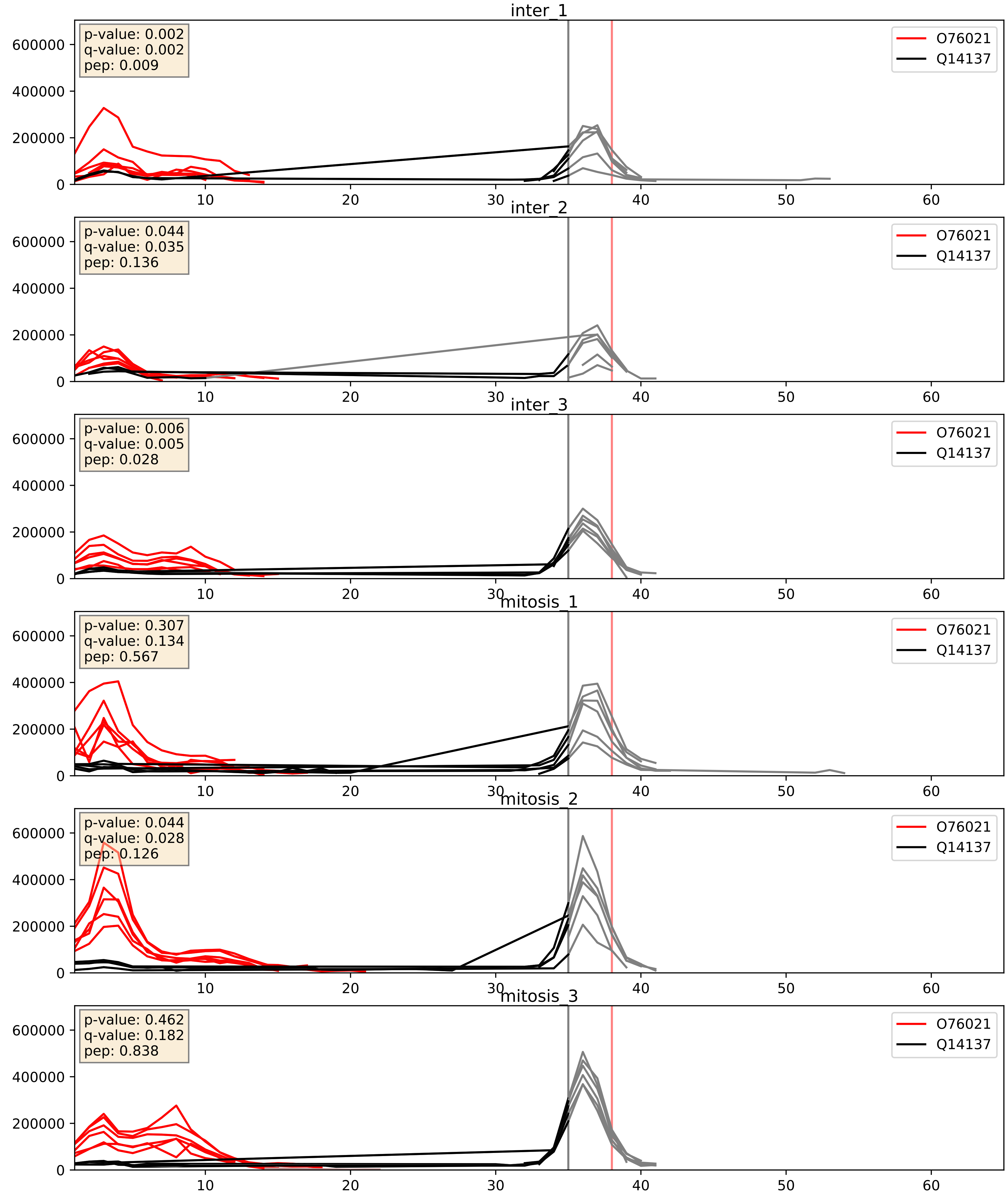


| level | condition_1 | condition_2 | pvalue | pvalue_adjusted |
|---|---|---|---|---|
| interactor_ratio | mitosis | inter | 0.014429314087919975 | 0.17619818629471468 |
| interactor_abundance | mitosis | inter | 0.04106316041394885 | 0.16296731794217187 |
| complex_abundance | mitosis | inter | 0.7782942865568719 | 0.8546759580406446 |

P46087_Q14137
NOP2_HUMAN vs BOP1_HUMAN
p-value: 0.044 q-value: 0.029

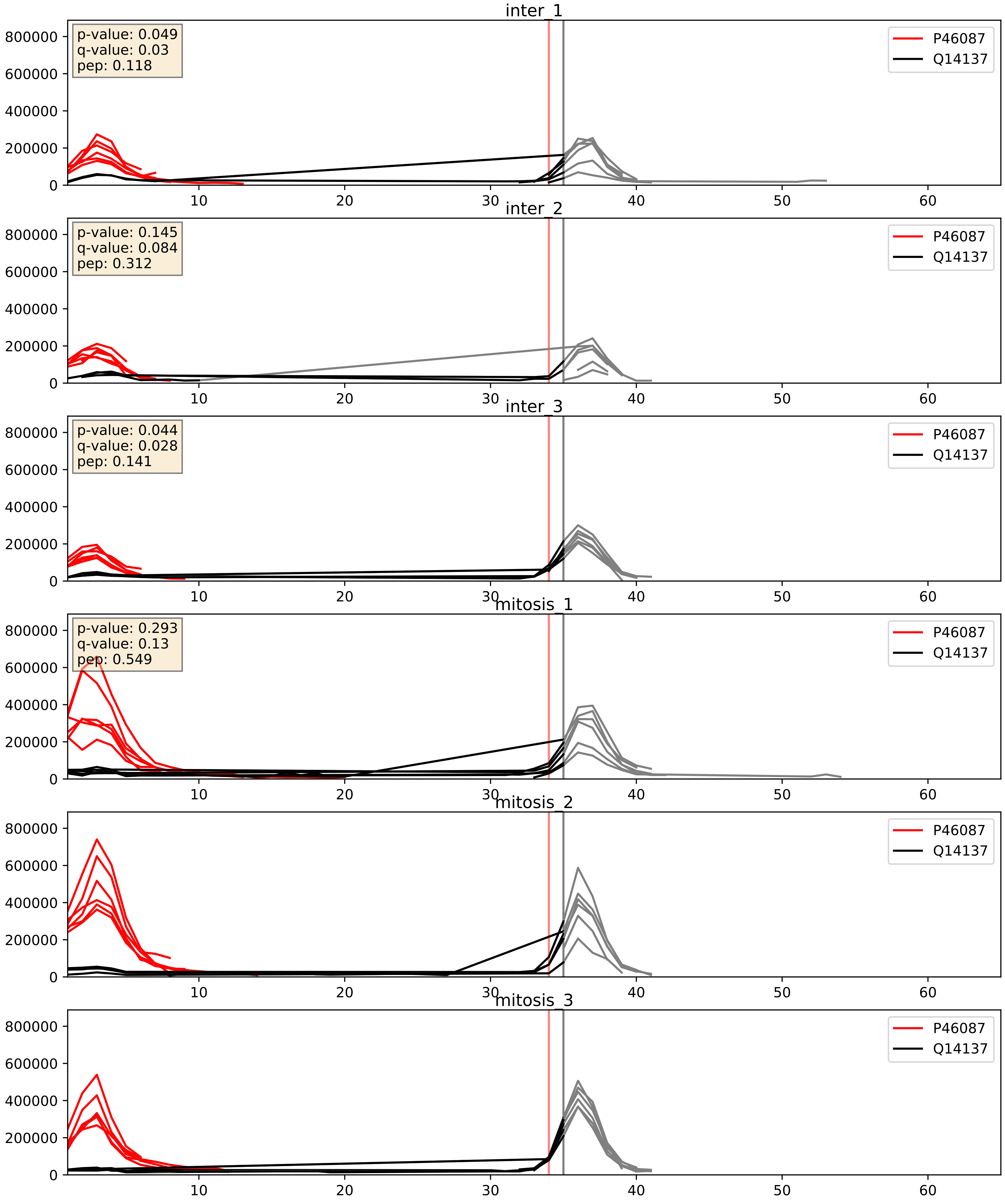


| level | condition_1 | condition_2 | pvalue | pvalue_adjusted |
|---|---|---|---|---|
| interactor_ratio | mitosis | inter | 0.00214770110748558 | 0.05734759220344108 |
| interactor_abundance | mitosis | inter | 0.017814919958046103 | 0.11522154502521695 |
| complex_abundance | mitosis | inter | 0.3752348769755155 | 0.5222928817512863 |

P56182_Q14137
RRP1_HUMAN vs BOP1_HUMAN
p-value: 0.004 q-value: 0.004

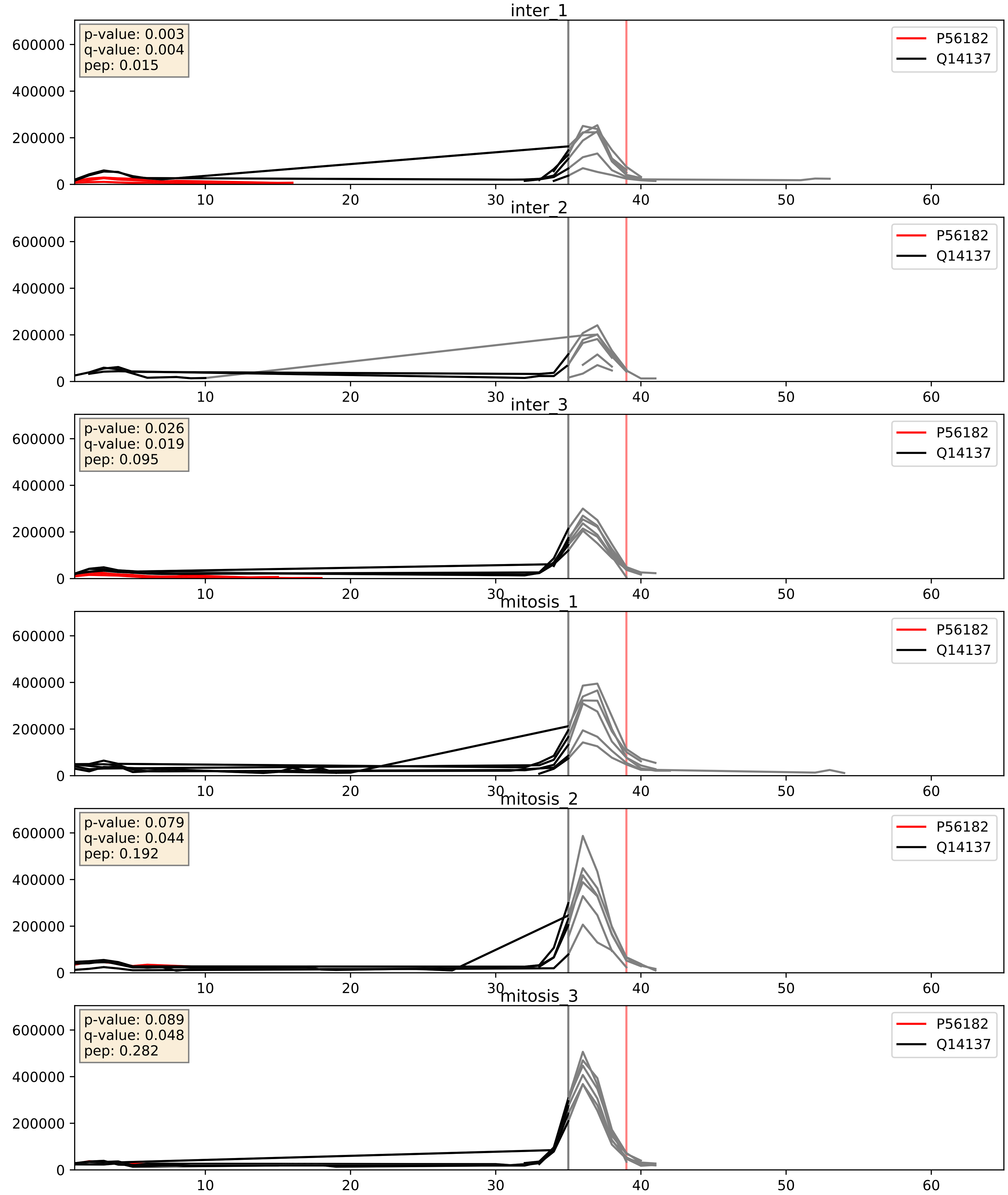


| level | condition_1 | condition_2 | pvalue | pvalue_adjusted |
|---|---|---|---|---|
| interactor_abundance | mitosis | inter | 0.02881223976524978 | 0.14041148442387597 |
| complex_abundance | mitosis | inter | 0.043717231909126686 | 0.14963536699613839 |
| interactor_ratio | mitosis | inter | 0.15446840222674124 | 0.6115862733861726 |

Q14137_Q14684
BOP1_HUMAN vs RRP1B_HUMAN
p-value: 0.068 q-value: 0.041

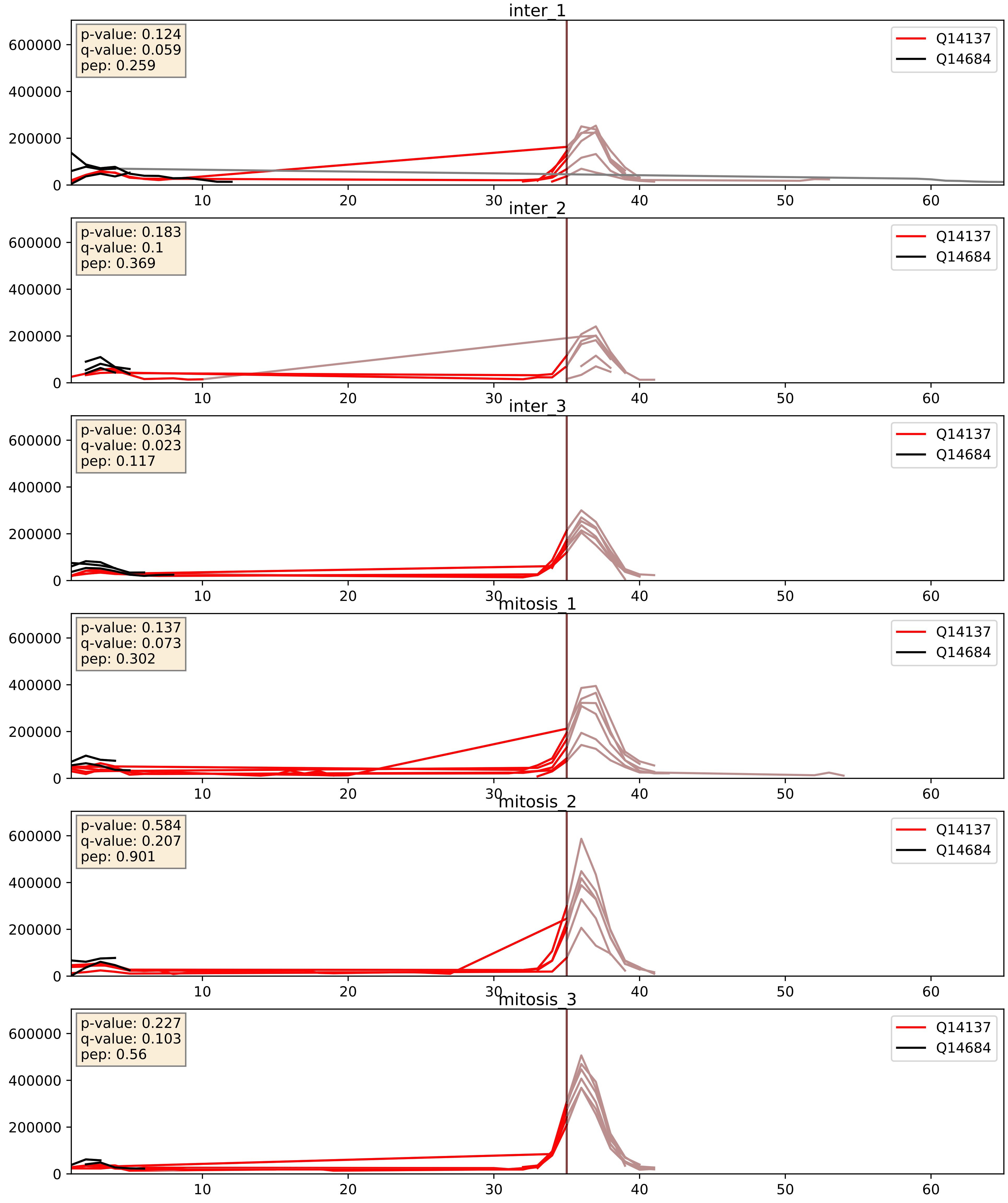


| level | condition_1 | condition_2 | pvalue | pvalue_adjusted |
|---|---|---|---|---|
| interactor_ratio | mitosis | inter | 0.012448259091787752 | 0.16096238342251235 |
| complex_abundance | mitosis | inter | 0.025482462219124032 | 0.11692128174565487 |
| interactor_abundance | mitosis | inter | 0.06447655496169065 | 0.20933787615098498 |

Q14137_Q15269
BOP1_HUMAN vs PWP2_HUMAN
p-value: 0.058 q-value: 0.037

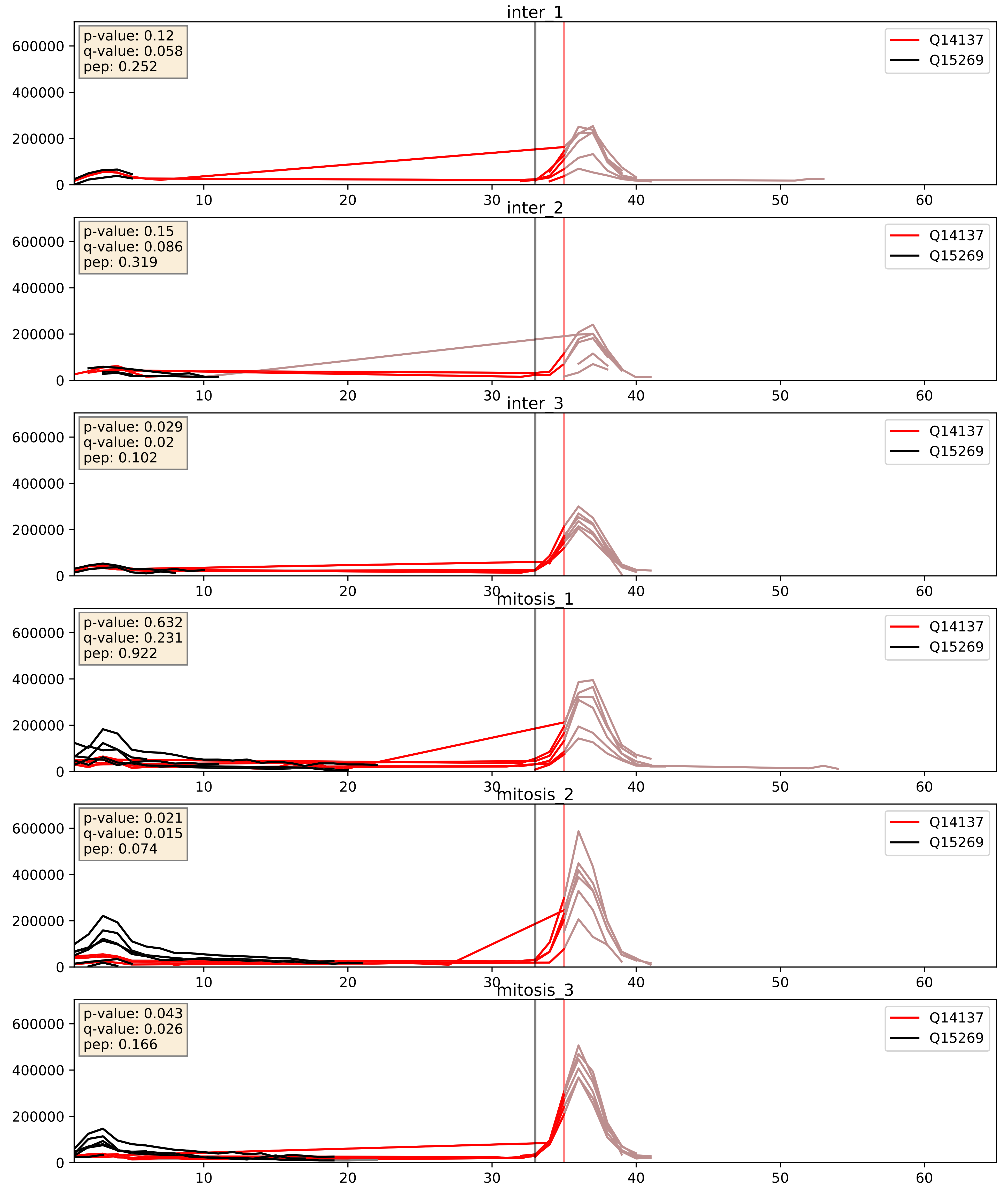


| level | condition_1 | condition_2 | pvalue | pvalue_adjusted |
|---|---|---|---|---|
| interactor_ratio | mitosis | inter | 0.0035194966342699525 | 0.07646419083591573 |
| interactor_abundance | mitosis | inter | 0.16567605704869157 | 0.3537339202733041 |
| complex_abundance | mitosis | inter | 0.7966276306979535 | 0.8676845042339333 |

Q14137_Q8IY81
BOP1_HUMAN vs SPB1_HUMAN
p-value: 0.041 q-value: 0.028

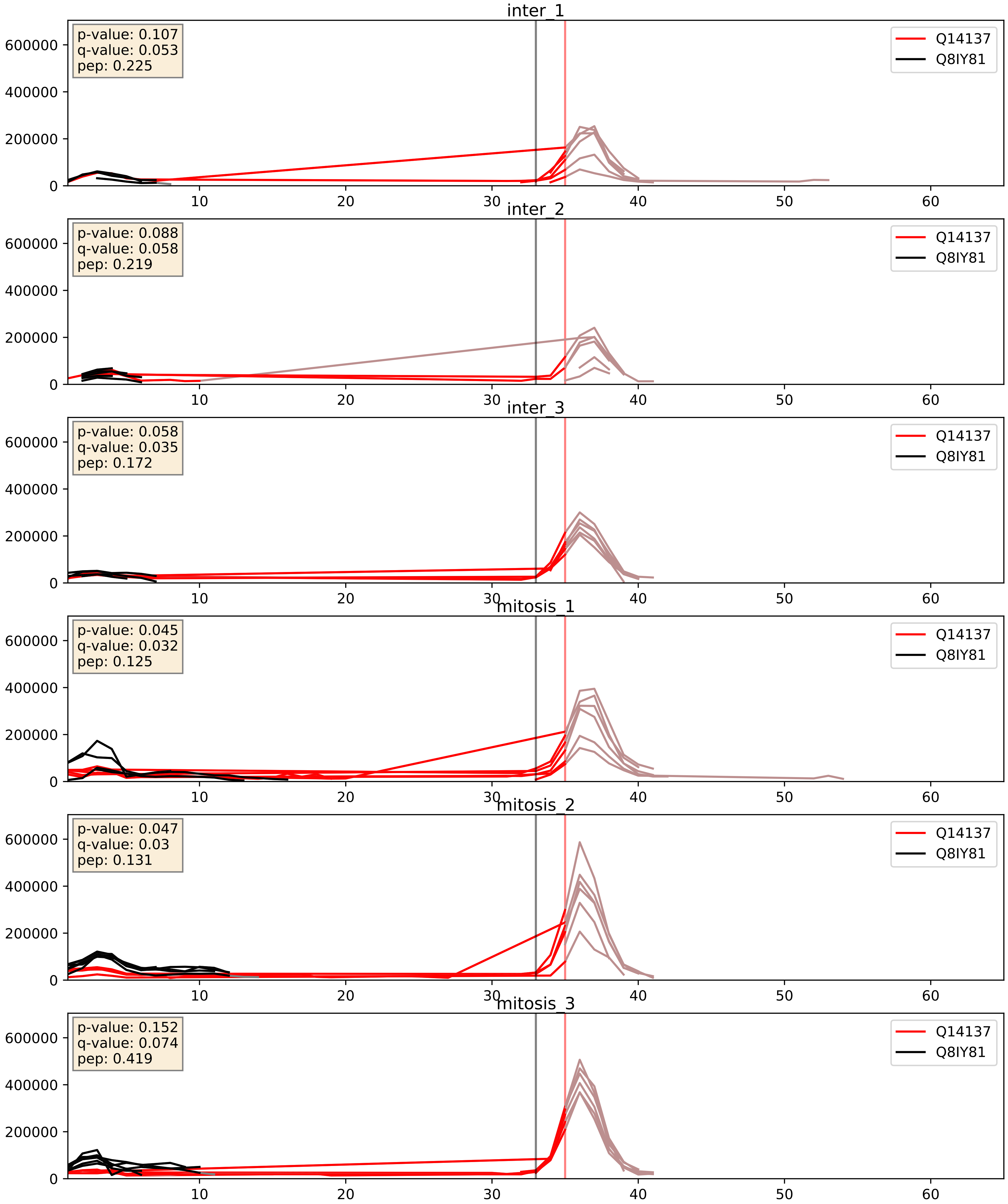


| level | condition_1 | condition_2 | pvalue | pvalue_adjusted |
|---|---|---|---|---|
| interactor_ratio | mitosis | inter | 0.06357934216723345 | 0.4189677974992443 |
| interactor_abundance | mitosis | inter | 0.11208372992542083 | 0.2886587657103679 |
| complex_abundance | mitosis | inter | 0.4002338471426604 | 0.5465861090525165 |

Q14137_Q8NI36
BOP1_HUMAN vs WDR36_HUMAN
p-value: 0.029 q-value: 0.021

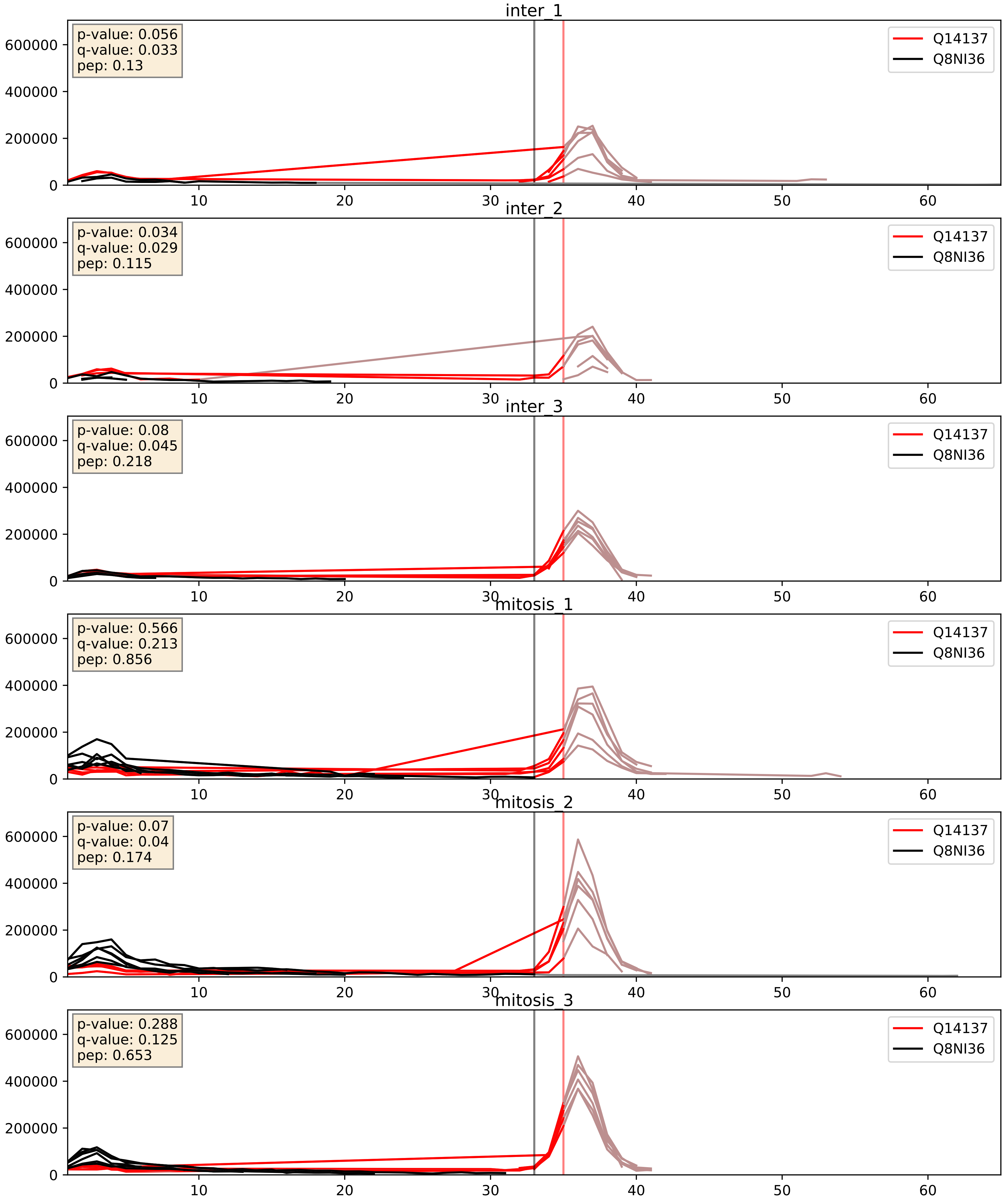


| level | condition_1 | condition_2 | pvalue | pvalue_adjusted |
|---|---|---|---|---|
| interactor_ratio | mitosis | inter | 0.0022534435788810043 | 0.05896690574521502 |
| interactor_abundance | mitosis | inter | 0.22206489816658428 | 0.4181424391346154 |
| complex_abundance | mitosis | inter | 0.5957970214043927 | 0.7204524977097283 |

Q14137_Q8TDN6
BOP1_HUMAN vs BRX1_HUMAN
p-value: 0.017 q-value: 0.014

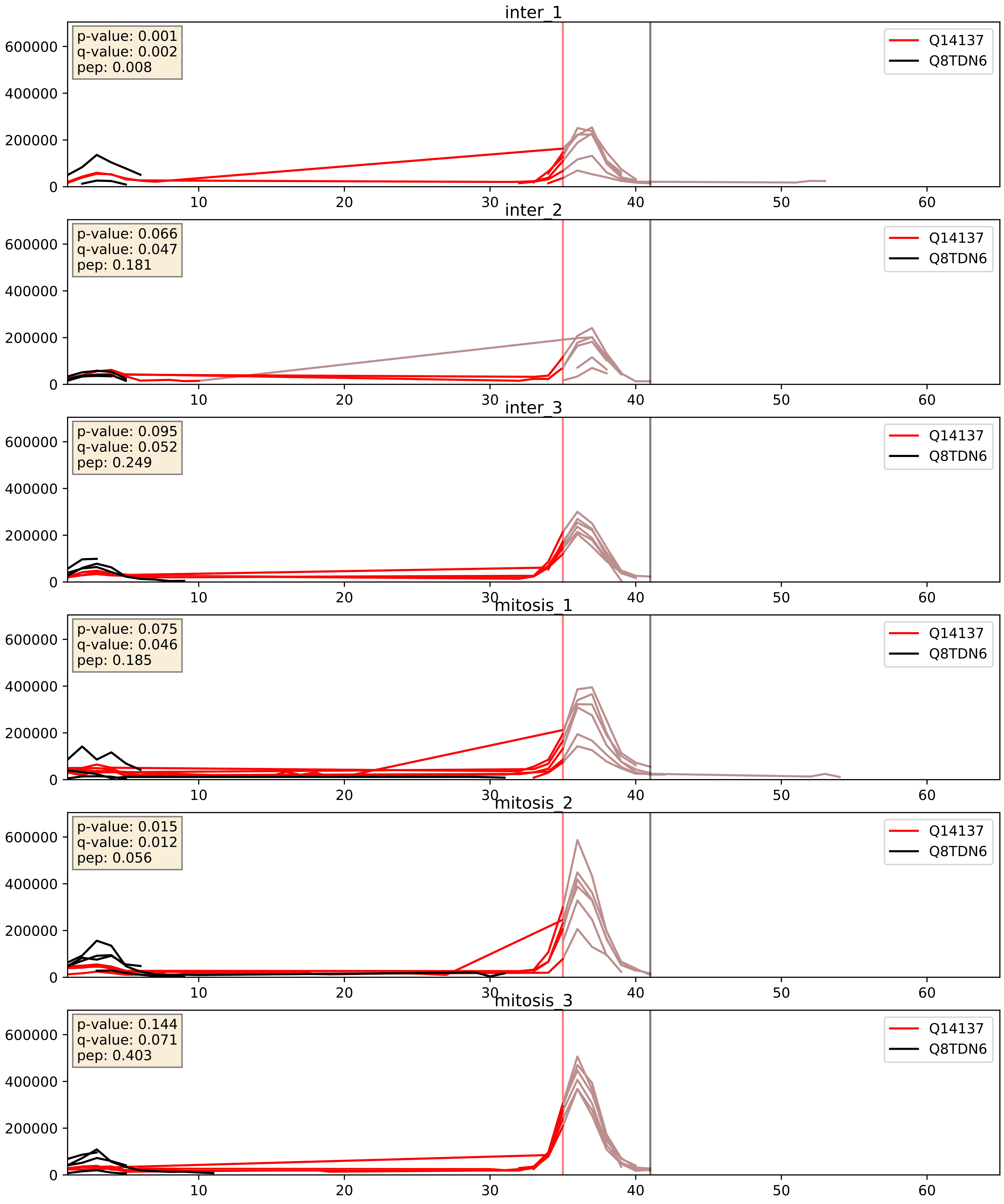


| level | condition_1 | condition_2 | pvalue | pvalue_adjusted |
|---|---|---|---|---|
| complex_abundance | mitosis | inter | 0.04192376036340907 | 0.14726323635770058 |
| interactor_abundance | mitosis | inter | 0.09173442235831389 | 0.2580077724288375 |
| interactor_ratio | mitosis | inter | 0.4069359185939671 | 0.8325457607945408 |

Q14137_Q8TED0
BOP1_HUMAN vs UTP15_HUMAN
p-value: 0.078 q-value: 0.047

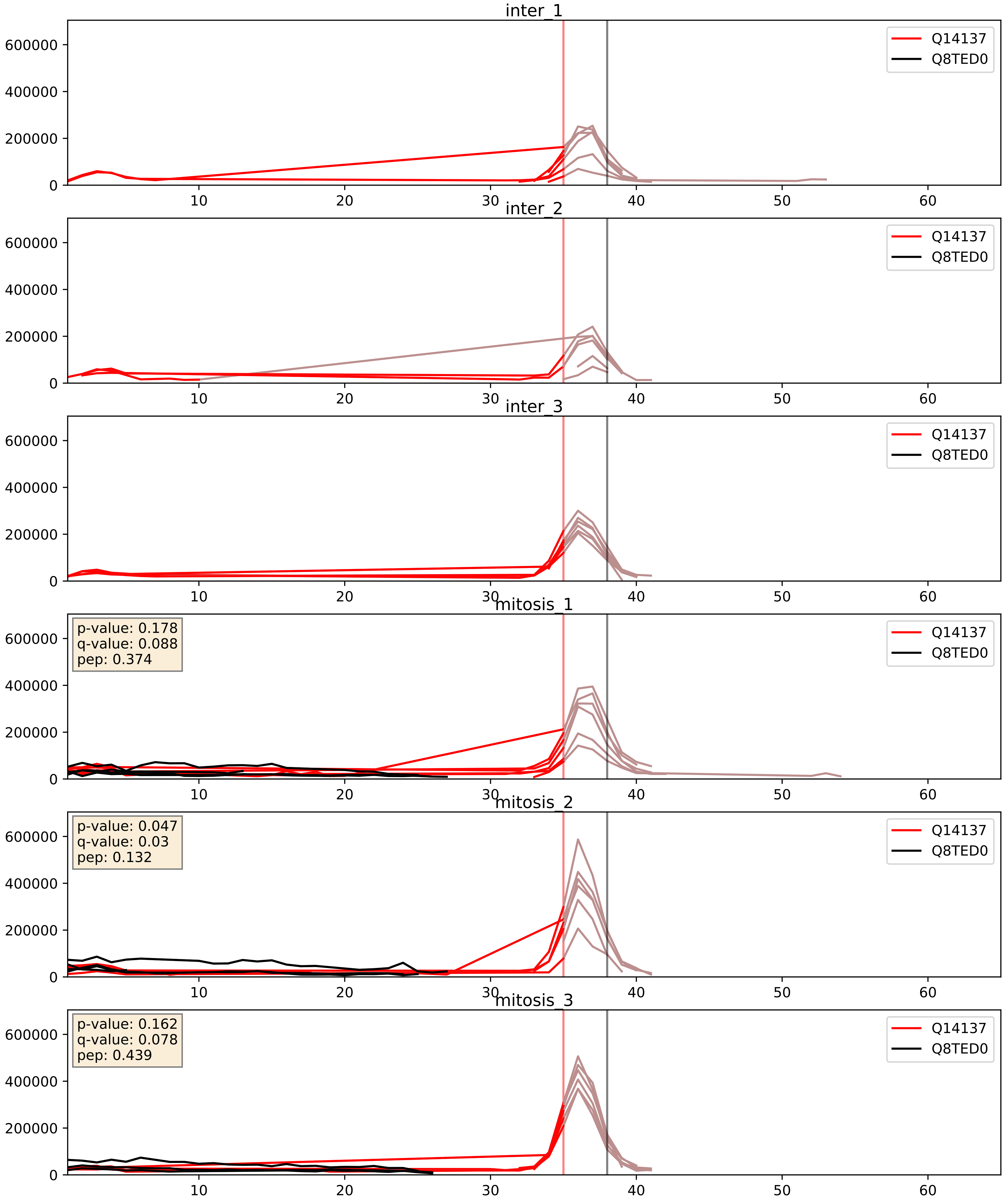


| level | condition_1 | condition_2 | pvalue | pvalue_adjusted |
|---|---|---|---|---|
| interactor_ratio | mitosis | inter | 0.01642019120281155 | 0.1887653035437297 |
| complex_abundance | mitosis | inter | 0.04358916305272642 | 0.14954839107468462 |
| interactor_abundance | mitosis | inter | 0.1276013680260762 | 0.3110987497303367 |

Q14137_Q969X6
BOP1_HUMAN vs UTP4_HUMAN
p-value: 0.055 q-value: 0.035

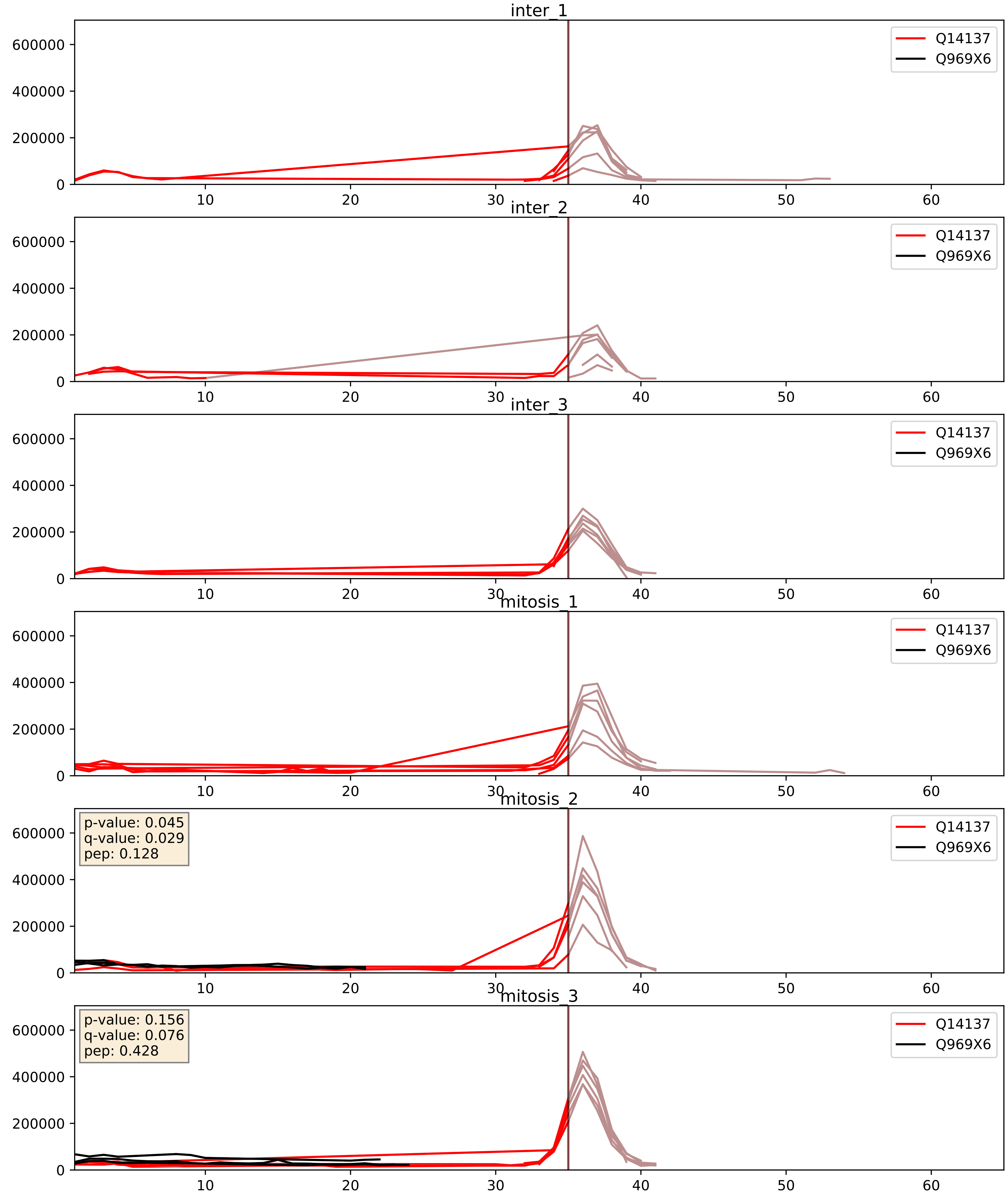


| level | condition_1 | condition_2 | pvalue | pvalue_adjusted |
|---|---|---|---|---|
| interactor_ratio | mitosis | inter | 0.2225872473581139 | 0.7003665360598988 |
| complex_abundance | mitosis | inter | 0.49247579418590043 | 0.6316441112123625 |
| interactor_abundance | mitosis | inter | 0.6875908701890414 | 0.7949860647871448 |

Q14137_Q9BV38
BOP1_HUMAN vs WDR18_HUMAN
p-value: 0.073 q-value: 0.044

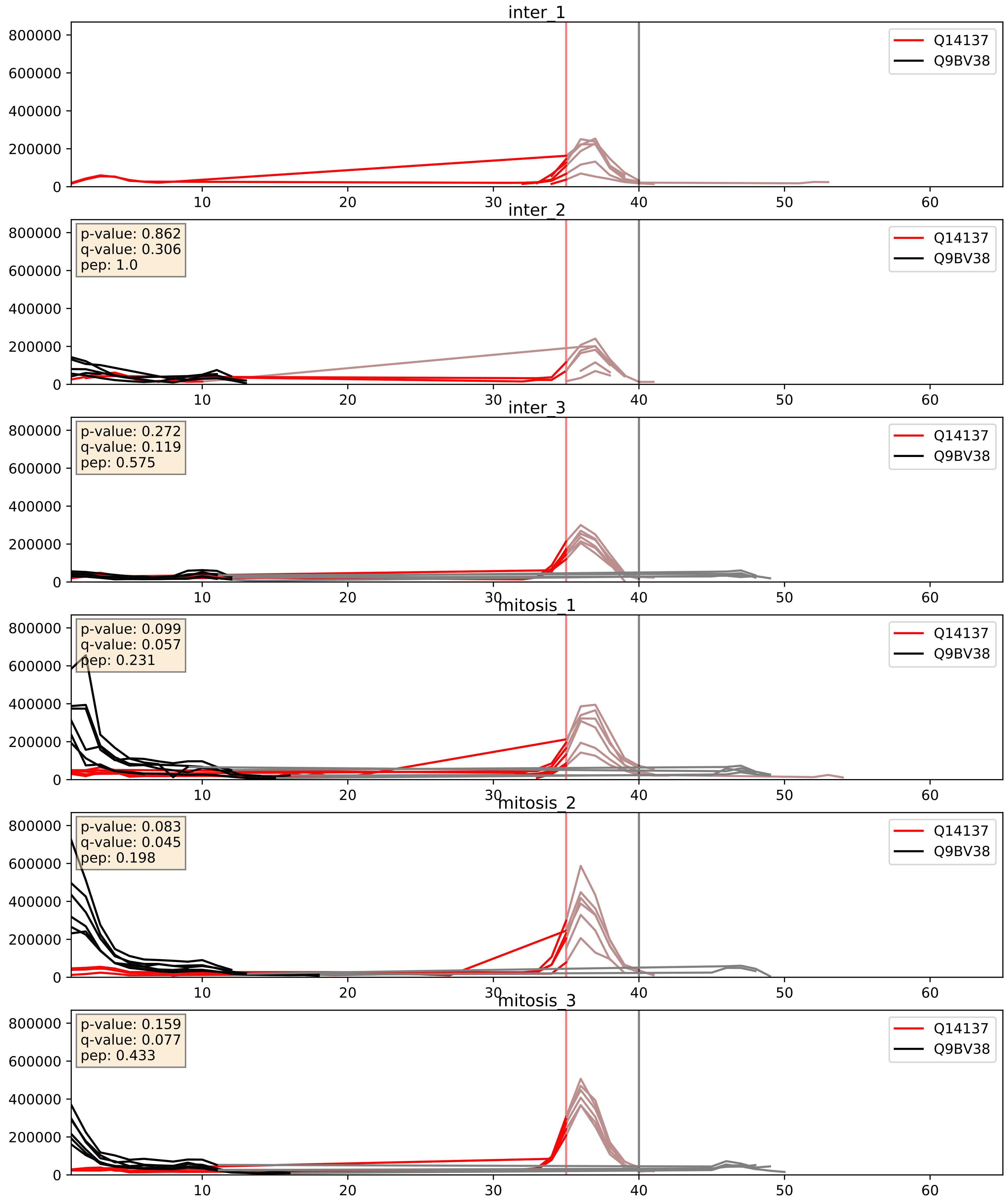


| level | condition_1 | condition_2 | pvalue | pvalue_adjusted |
|---|---|---|---|---|
| interactor_ratio | mitosis | inter | 0.001869302341360184 | 0.0524630427607973 |
| complex_abundance | mitosis | inter | 0.21507677809306397 | 0.3695903124243094 |
| interactor_abundance | mitosis | inter | 0.5302080199241265 | 0.689353305367806 |

Q14137_Q9BVP2
BOP1_HUMAN vs GNL3_HUMAN
p-value: 0.042 q-value: 0.028

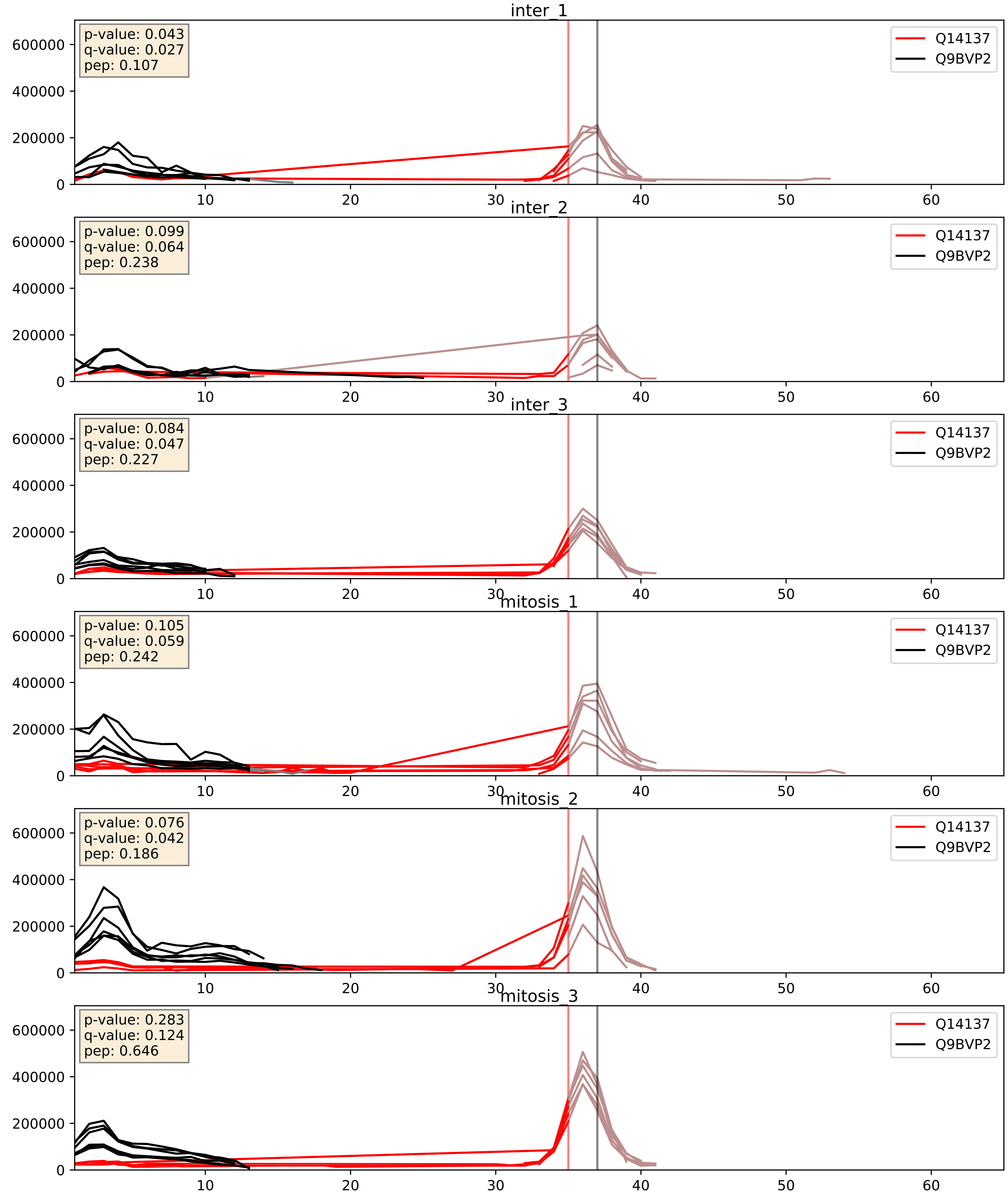


| level | condition_1 | condition_2 | pvalue | pvalue_adjusted |
|---|---|---|---|---|
| interactor_ratio | mitosis | inter | 0.011122406259462985 | 0.1518775136410363 |
| interactor_abundance | mitosis | inter | 0.09412782094612172 | 0.26266801867931605 |
| complex_abundance | mitosis | inter | 0.6045895512232708 | 0.7277860439419489 |

Q14137_Q9BZE4
BOP1_HUMAN vs NOG1_HUMAN
p-value: 0.053 q-value: 0.034

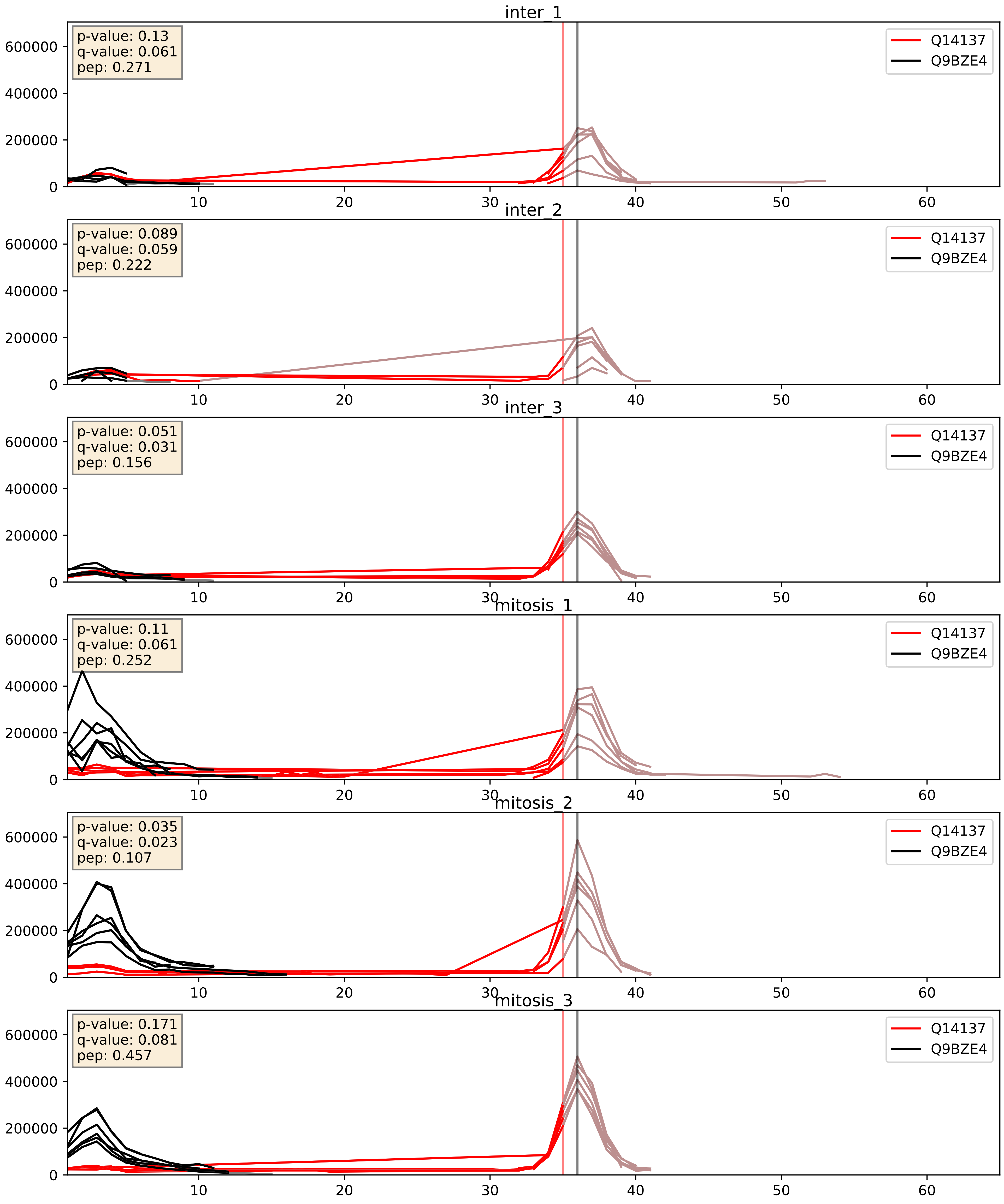


| level | condition_1 | condition_2 | pvalue | pvalue_adjusted |
|---|---|---|---|---|
| interactor_ratio | mitosis | inter | 0.0007612082988845985 | 0.03178508799244958 |
| complex_abundance | mitosis | inter | 0.0967292144756393 | 0.23055517679920082 |
| interactor_abundance | mitosis | inter | 0.09980342138358718 | 0.2711705935253959 |

Q14137_Q9GZL7
BOP1_HUMAN vs WDR12_HUMAN
p-value: 0.059 q-value: 0.037

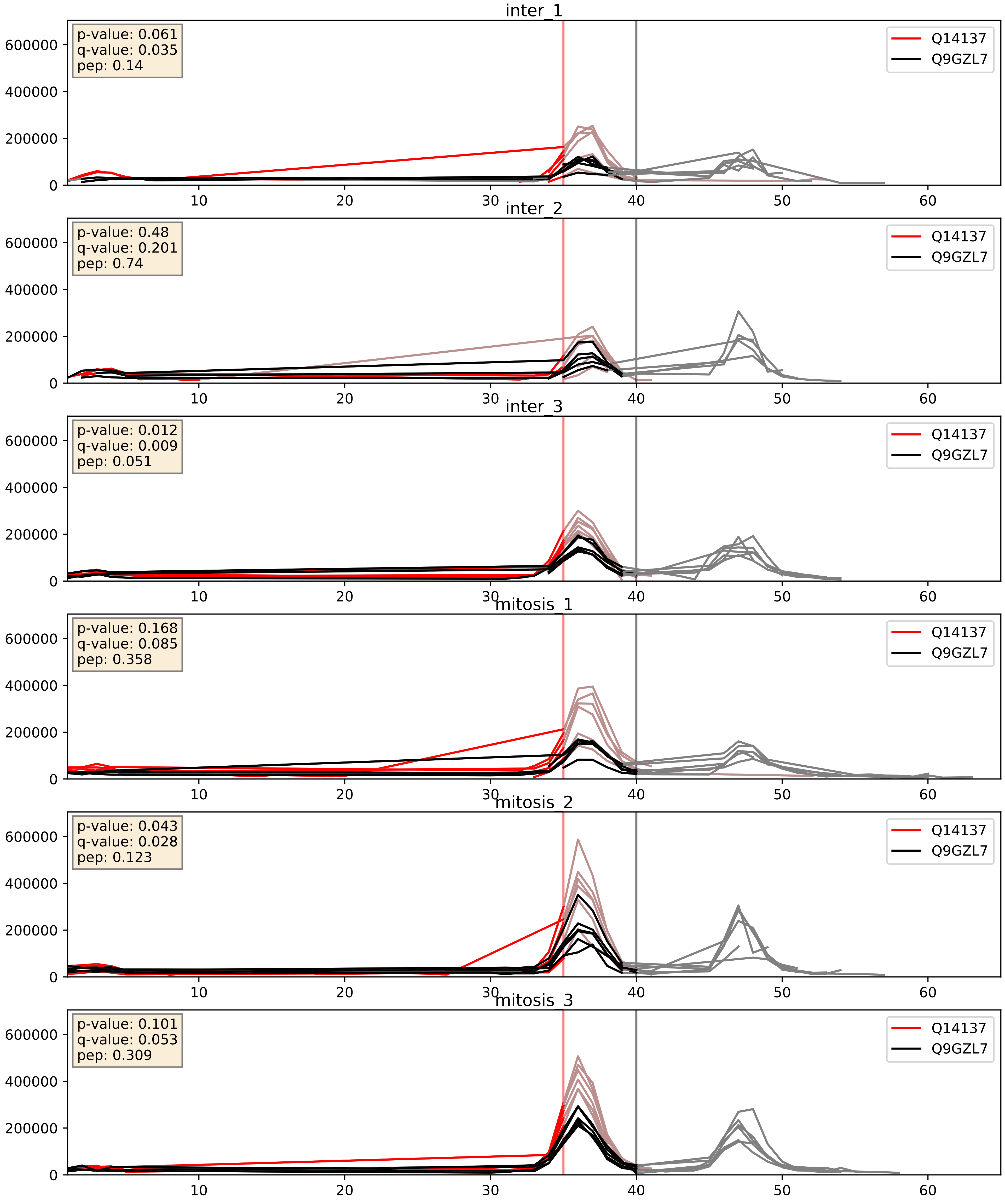


| level | condition_1 | condition_2 | pvalue | pvalue_adjusted |
|---|---|---|---|---|
| complex_abundance | mitosis | inter | 0.026301074213455553 | 0.11880590779270687 |
| interactor_abundance | mitosis | inter | 0.04259625139749957 | 0.16634302553038152 |
| interactor_ratio | mitosis | inter | 0.17705596082494385 | 0.6427476779735027 |

Q14137_Q9NXF1
BOP1_HUMAN vs TEX10_HUMAN
p-value: 0.049 q-value: 0.032

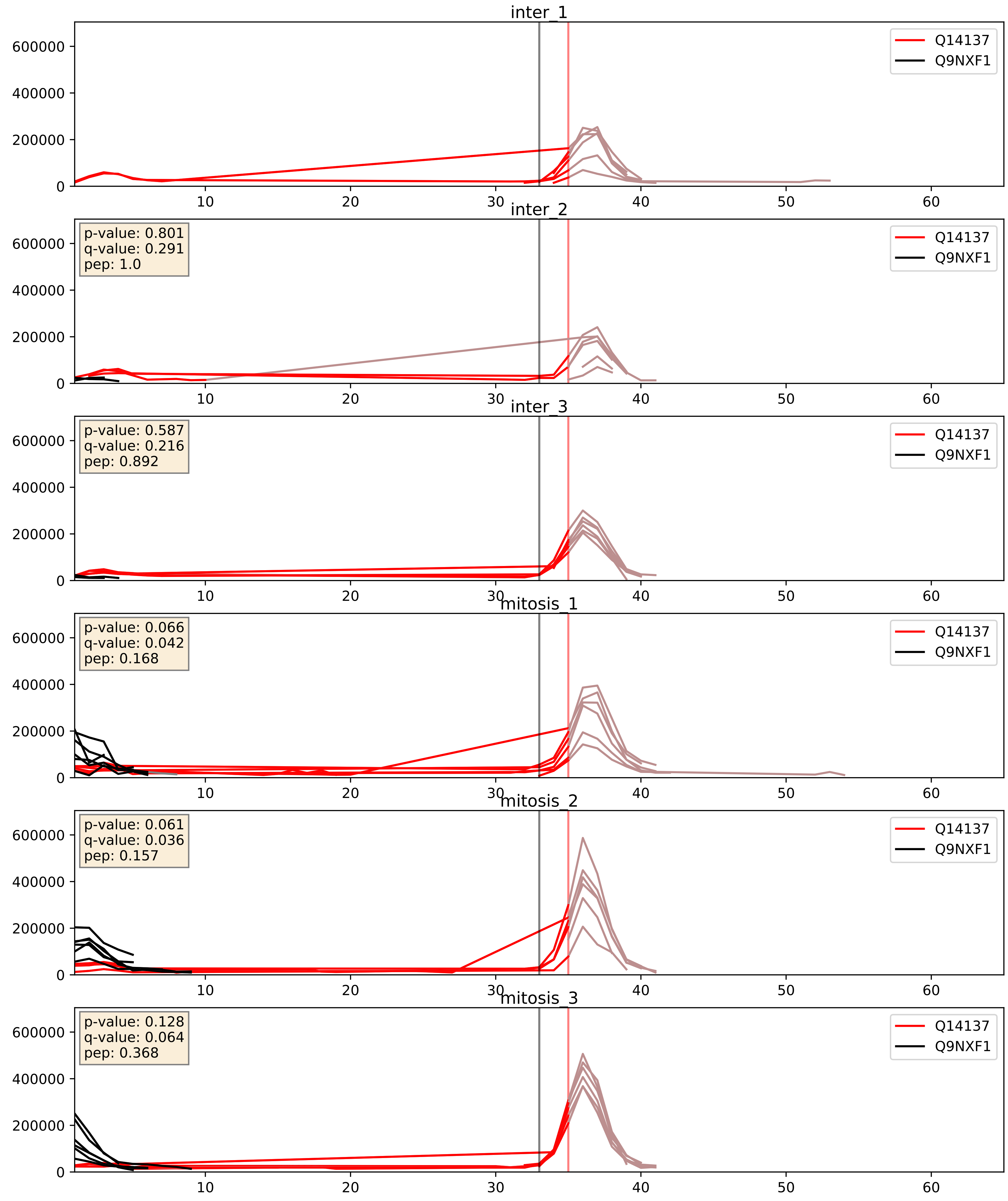


| level | condition_1 | condition_2 | pvalue | pvalue_adjusted |
|---|---|---|---|---|
| interactor_ratio | mitosis | inter | 0.01263103011635328 | 0.16210137600597313 |
| complex_abundance | mitosis | inter | 0.18912949473358198 | 0.34097482622566594 |
| interactor_abundance | mitosis | inter | 0.4858283549734151 | 0.655531323860724 |

Q14137_Q9UNX4
BOP1_HUMAN vs WDR3_HUMAN
p-value: 0.058 q-value: 0.037

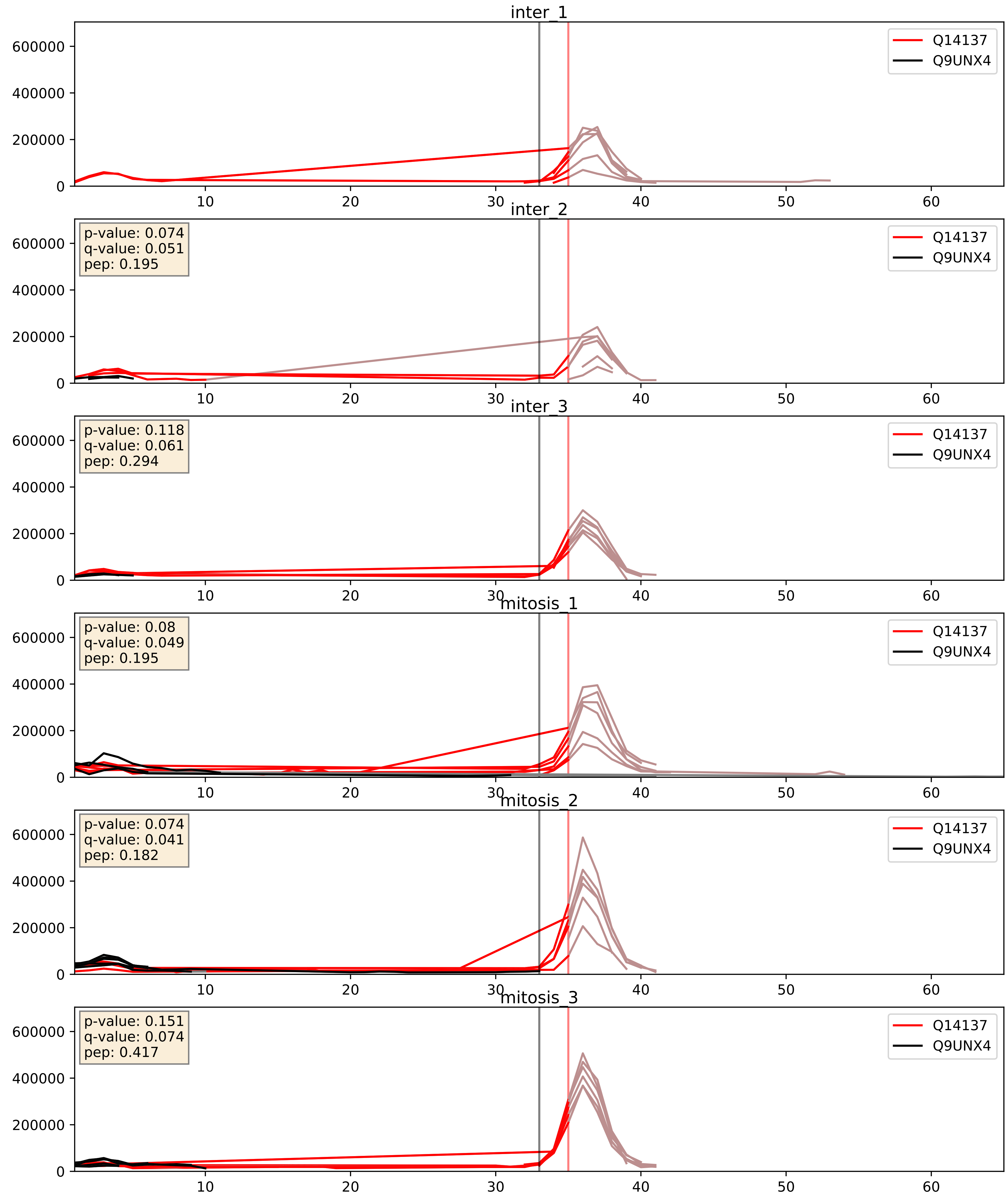


| level | condition_1 | condition_2 | pvalue | pvalue_adjusted |
|---|---|---|---|---|
| interactor_ratio | mitosis | inter | 0.025700362102627074 | 0.25374290611128925 |
| interactor_abundance | mitosis | inter | 0.5664835021564785 | 0.7138375943558745 |
| complex_abundance | mitosis | inter | 0.8968520804744315 | 0.9361137352430374 |

Q14137_Q9Y221
BOP1_HUMAN vs NIP7_HUMAN
p-value: 0.011 q-value: 0.01

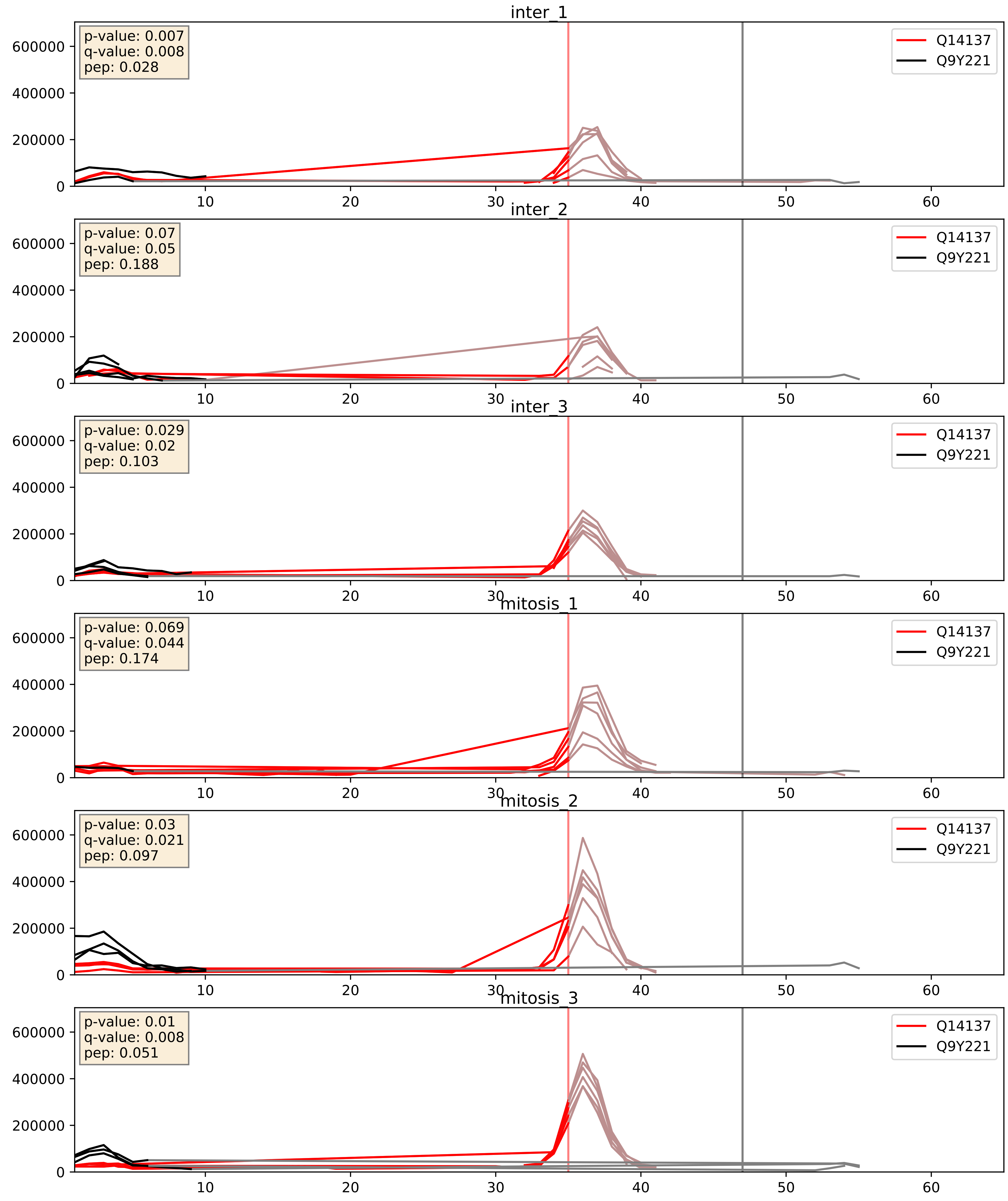


| level | condition_1 | condition_2 | pvalue | pvalue_adjusted |
|---|---|---|---|---|
| complex_abundance | mitosis | inter | 0.05642335124251802 | 0.16923051388786065 |
| interactor_abundance | mitosis | inter | 0.06916248570360795 | 0.21846157845862876 |
| interactor_ratio | mitosis | inter | 0.6364332158336101 | 0.9197408380221228 |
