## Supplementary Data 2 for "SECAT: Quantifying differential protein-protein interaction states by network-centric analysis": hela_string_000091_P31930.pdf

P31930_P31930

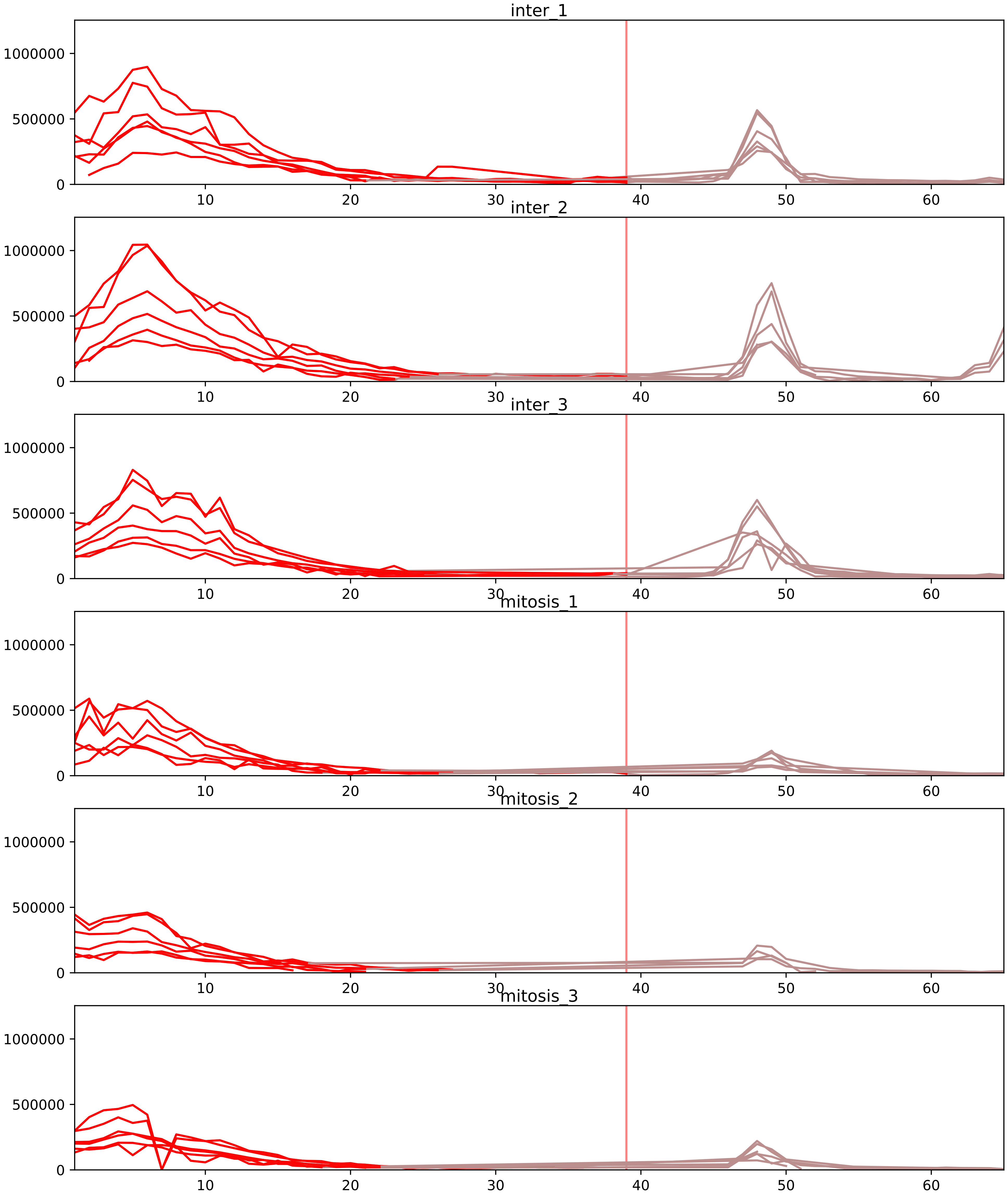


| level | condition_1 | condition_2 | pvalue | pvalue_adjusted |
|---|---|---|---|---|
| interactor_abundance | mitosis | inter | 6.163847080650688e-06 | 0.0003658541493031376 |
| complex_abundance | mitosis | inter | 3.4724527307517334e-05 | 0.0011409487543898552 |
| monomer_abundance | mitosis | inter | 0.0004065879488960143 | 0.055065561212150206 |
| total_abundance | mitosis | inter | 0.000526988700346485 | 0.027376525239428108 |
| assembled_abundance | mitosis | inter | 0.001524066442555335 | 0.036398750089803686 |
| interactor_ratio | mitosis | inter | 0.002561690706572467З | 0.0235675545004667 |

O00217_P31930
NDUS8_HUMAN vs QCR1_HUMAN
p-value: 0.022 q-value: 0.017

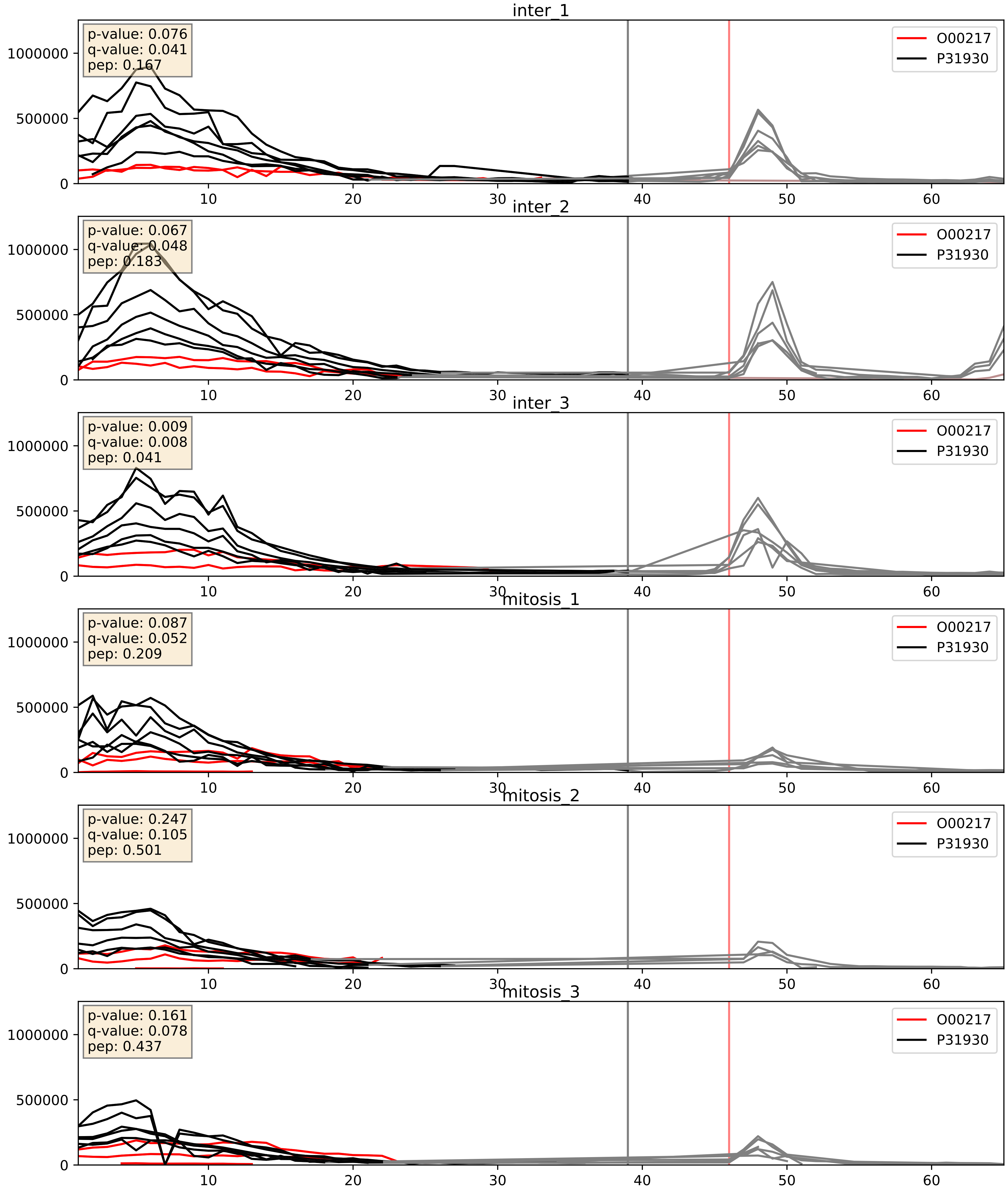


| level | condition_1 | condition_2 | pvalue | pvalue_adjusted |
|---|---|---|---|---|
| interactor_ratio | mitosis | inter | 0.0032267047125980487 | 0.07405554735675686 |
| complex_abundance | mitosis | inter | 0.012331335264185605 | 0.08253028136155495 |
| interactor_abundance | mitosis | inter | 0.46581523118537366 | 0.6378784800746758 |

O14949_P31930
QCR8_HUMAN vs QCR1_HUMAN
p-value: 0.023 q-value: 0.018

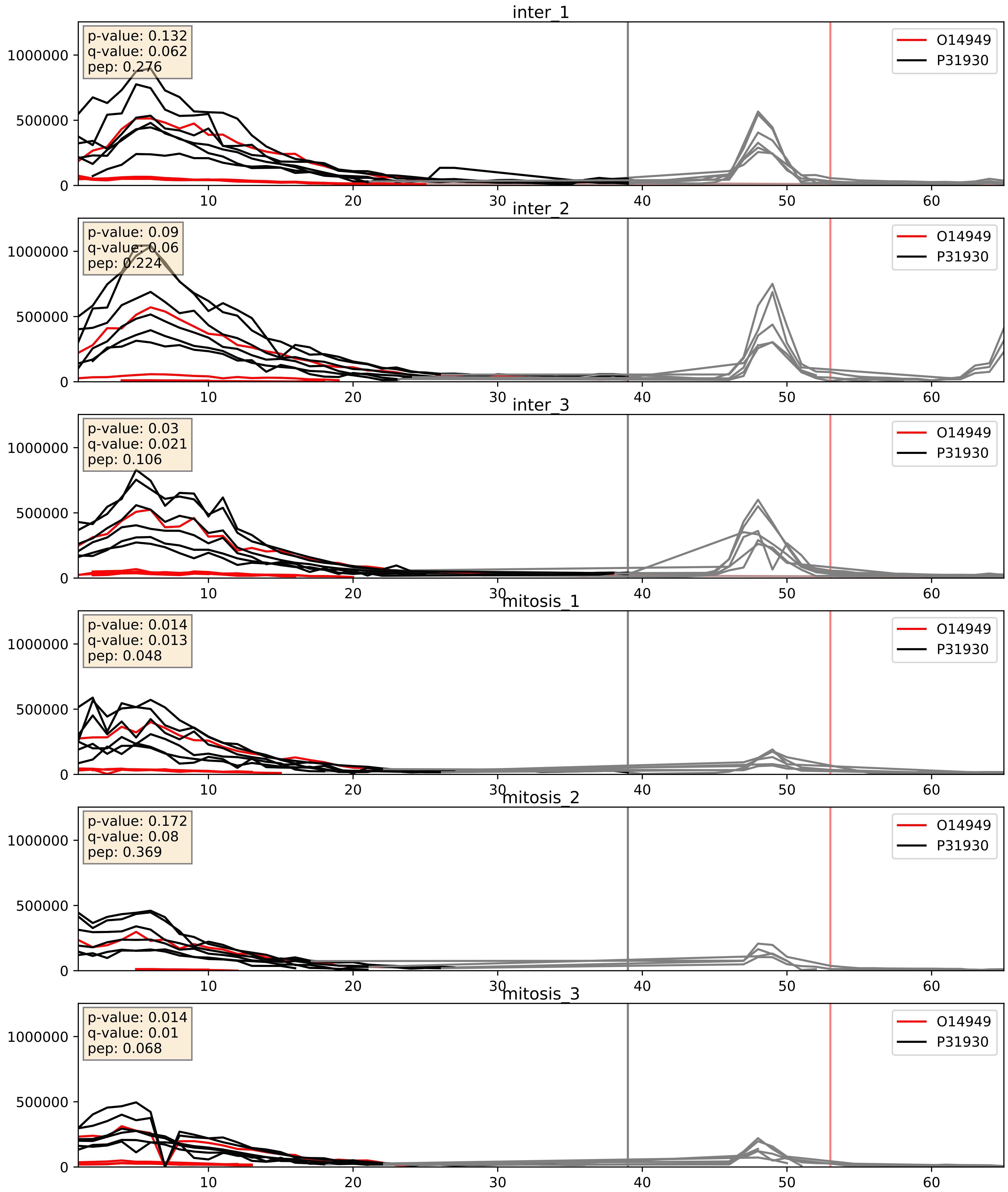


| level | condition_1 | condition_2 | pvalue | pvalue_adjusted |
|---|---|---|---|---|
| complex_abundance | mitosis | inter | 0.009157797517479936 | 0.07075397186708442 |
| interactor_abundance | mitosis | inter | 0.07395456278311857 | 0.22673988198390962 |
| interactor_ratio | mitosis | inter | 0.8834701325281622 | 0.9802747470617126 |

O75251_P31930
NDUS7_HUMAN vs QCR1_HUMAN
p-value: 0.008 q-value: 0.007

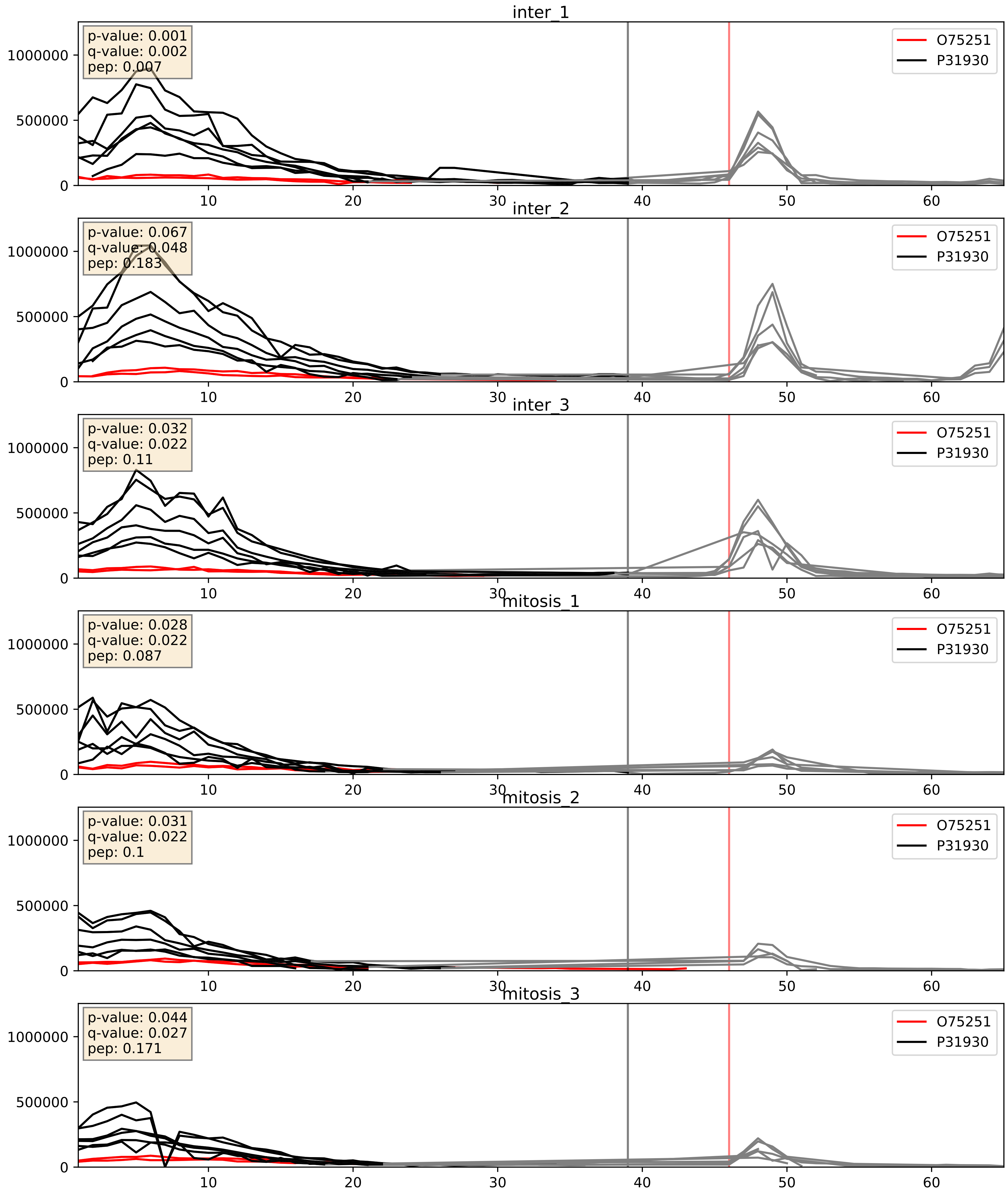


| level | condition_1 | condition_2 | pvalue | pvalue_adjusted |
|---|---|---|---|---|
| interactor_ratio | mitosis | inter | 0.0005090353089778639 | 0.0260918697296438 |
| complex_abundance | mitosis | inter | 0.0014943023962151634 | 0.026841427652794205 |
| interactor_abundance | mitosis | inter | 0.014149120616424192 | 0.10541033287779904 |

O75306_P31930
NDUS2_HUMAN vs QCR1_HUMAN
p-value: 0.012 q-value: 0.01

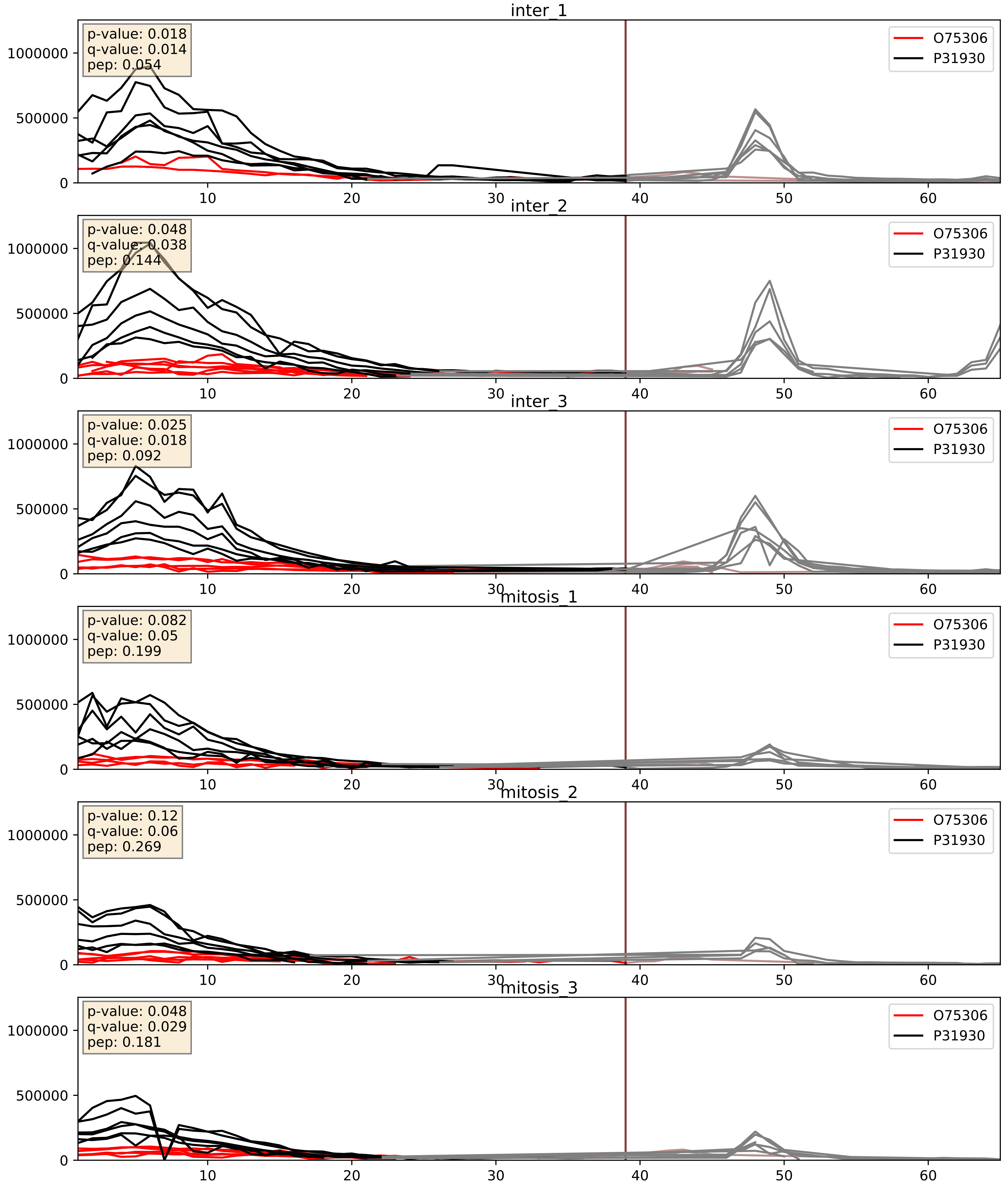


| level | condition_1 | condition_2 | pvalue | pvalue_adjusted |
|---|---|---|---|---|
| complex_abundance | mitosis | inter | 0.1026484368799358 | 0.23799312559378397 |
| interactor_ratio | mitosis | inter | 0.2393369386683702 | 0.7181554255891563 |
| interactor_abundance | mitosis | inter | 0.6391389840477945 | 0.759127196260458 |

P07919_P31930
QCR6_HUMAN vs QCR1_HUMAN
p-value: 0.006 q-value: 0.006

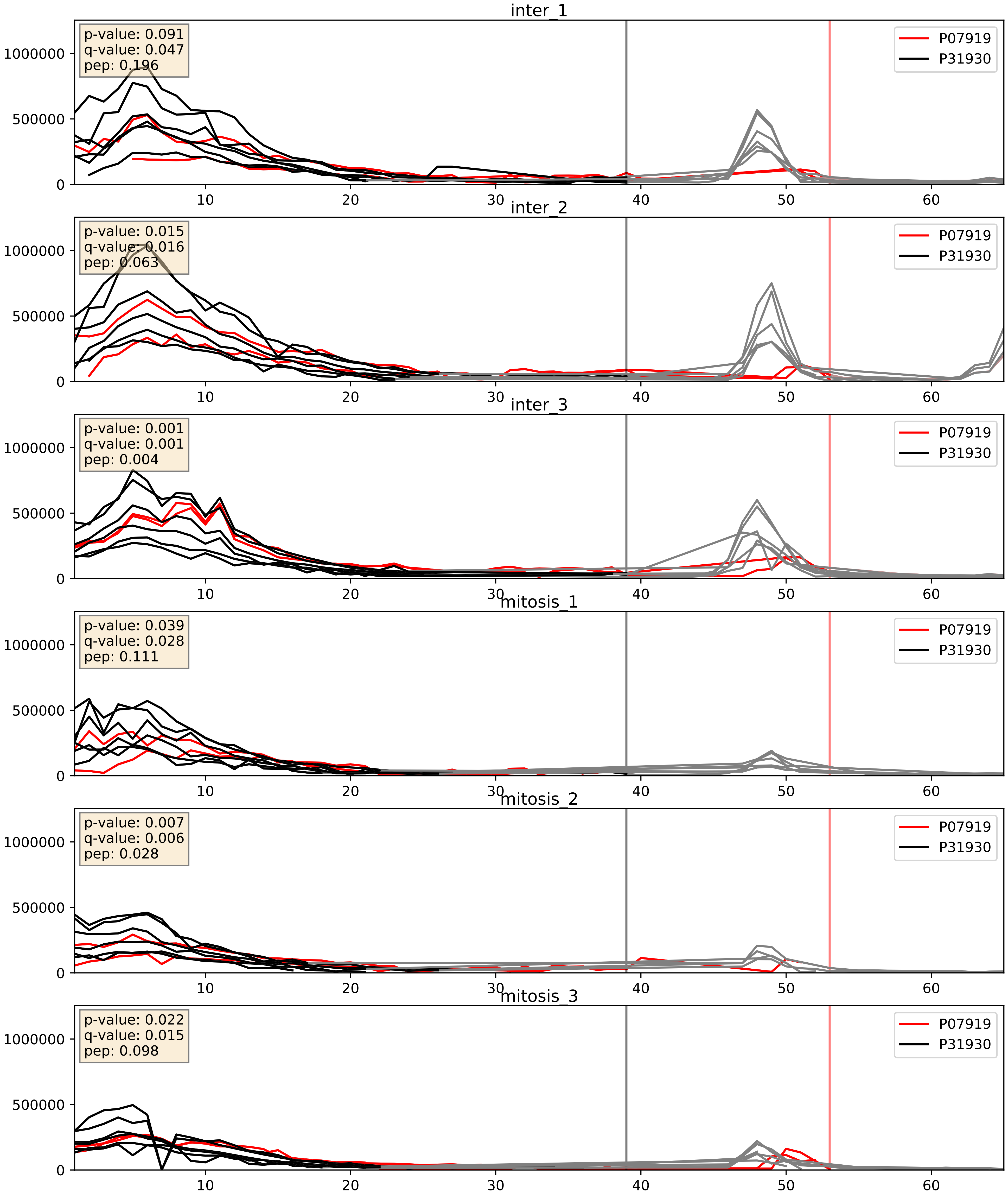


| level | condition_1 | condition_2 | pvalue | pvalue_adjusted |
|---|---|---|---|---|
| complex_abundance | mitosis | inter | 0.002305575437713905 | 0.03502731659907978 |
| interactor_abundance | mitosis | inter | 0.020711684860323237 | 0.12034491659677074 |
| interactor_ratio | mitosis | inter | 0.05720376836361693 | 0.3955284791539264 |

P08574_P31930
CY1_HUMAN vs QCR1_HUMAN
p-value: 0.008 q-value: 0.008

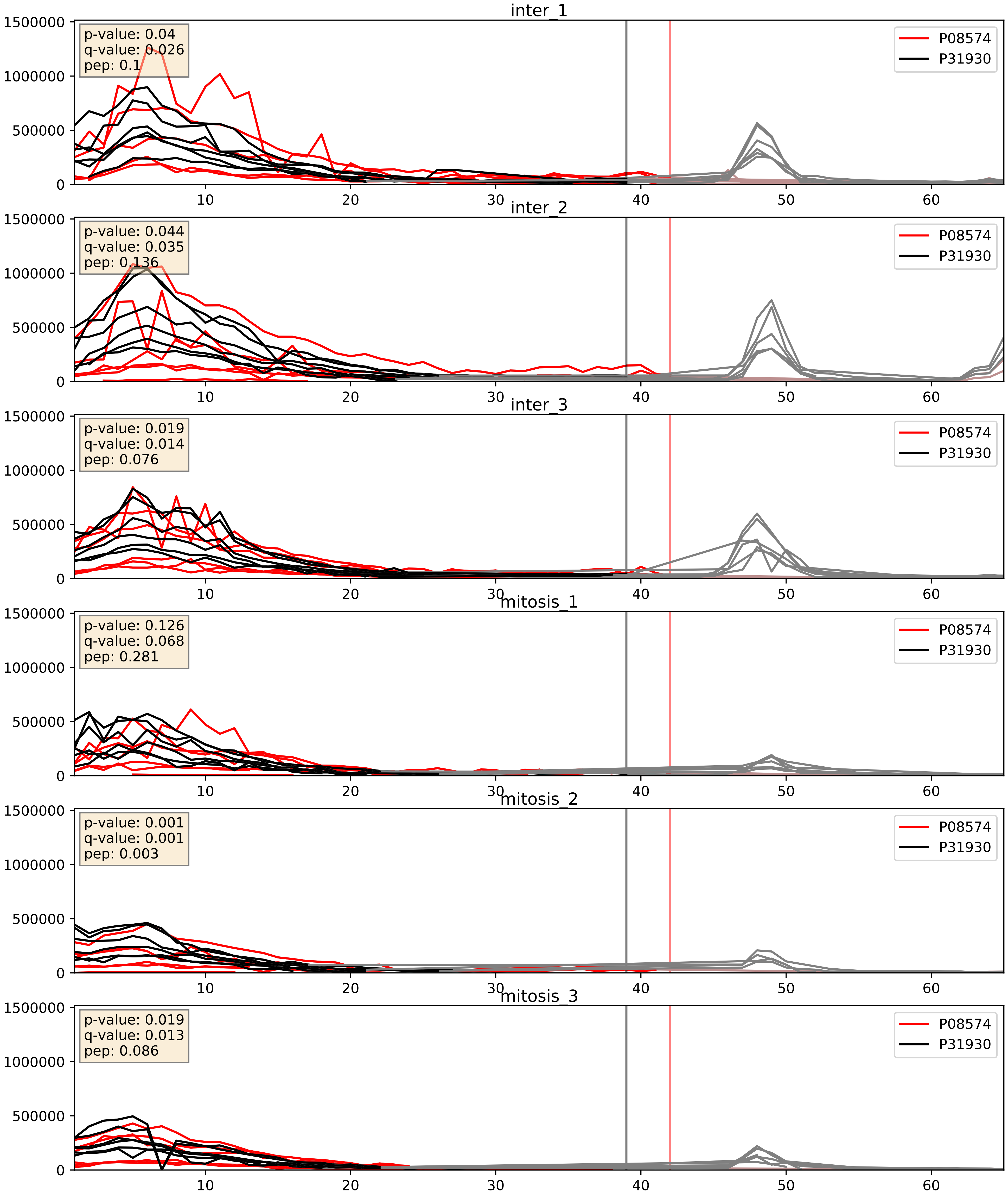


| level | condition_1 | condition_2 | pvalue | pvalue_adjusted |
|---|---|---|---|---|
| complex_abundance | mitosis | inter | 0.0009739141526850073 | 0.020686613267949534 |
| interactor_abundance | mitosis | inter | 0.006546882782702666 | 0.06901639977824485 |
| interactor_ratio | mitosis | inter | 0.8678606927151378 | 0.9786441219393465 |

P10606_P31930
COX5B_HUMAN vs QCR1_HUMAN
p-value: 0.005 q-value: 0.005

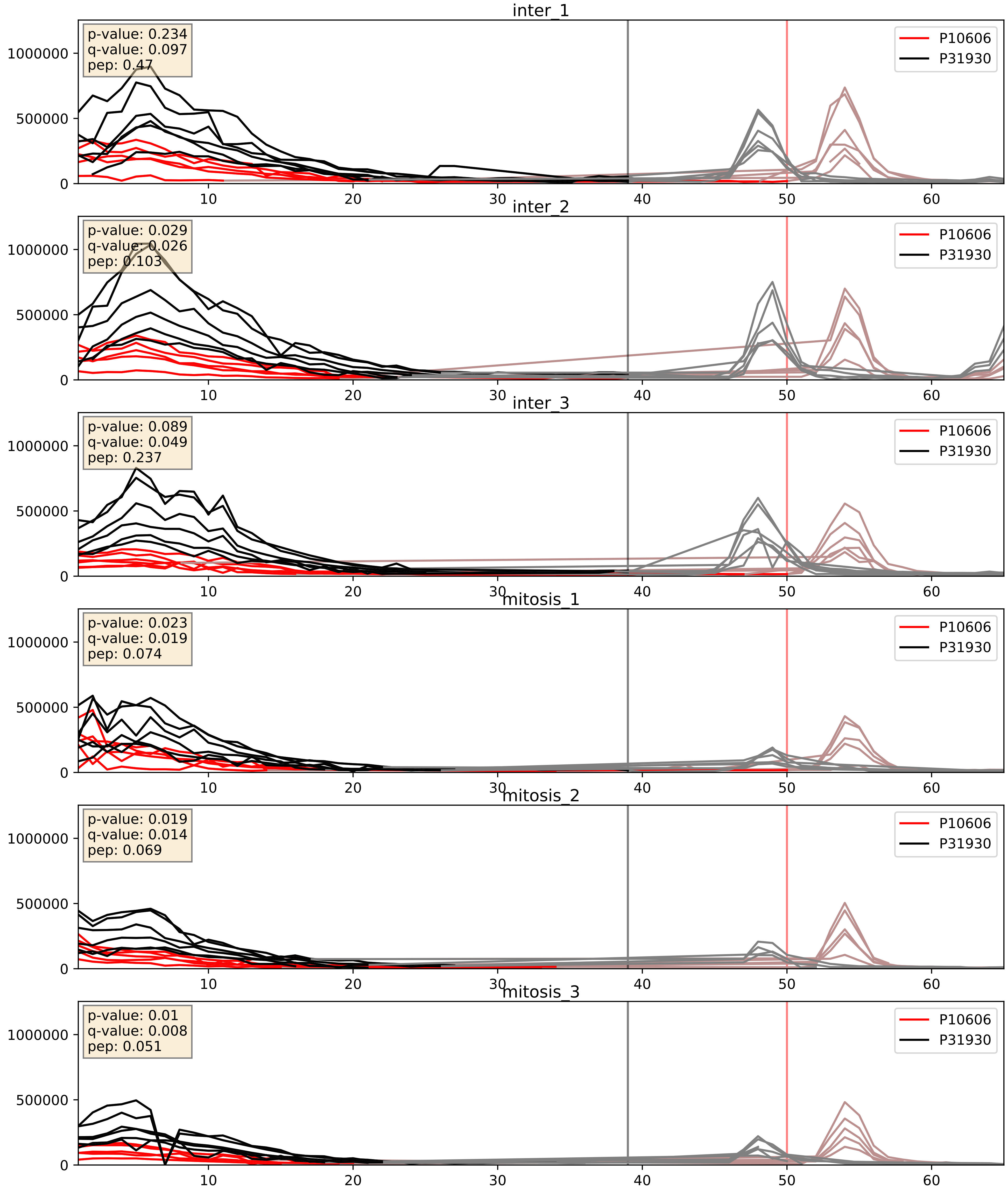


| level | condition_1 | condition_2 | pvalue | pvalue_adjusted |
|---|---|---|---|---|
| complex_abundance | mitosis | inter | 0.01146676888341548 | 0.07947817137007004 |
| interactor_abundance | mitosis | inter | 0.07861311074803888 | 0.2327263454965287 |
| interactor_ratio | mitosis | inter | 0.3161419540021466 | 0.7783421708721073 |

P14927_P31930
QCR7_HUMAN vs QCR1_HUMAN
p-value: 0.005 q-value: 0.005

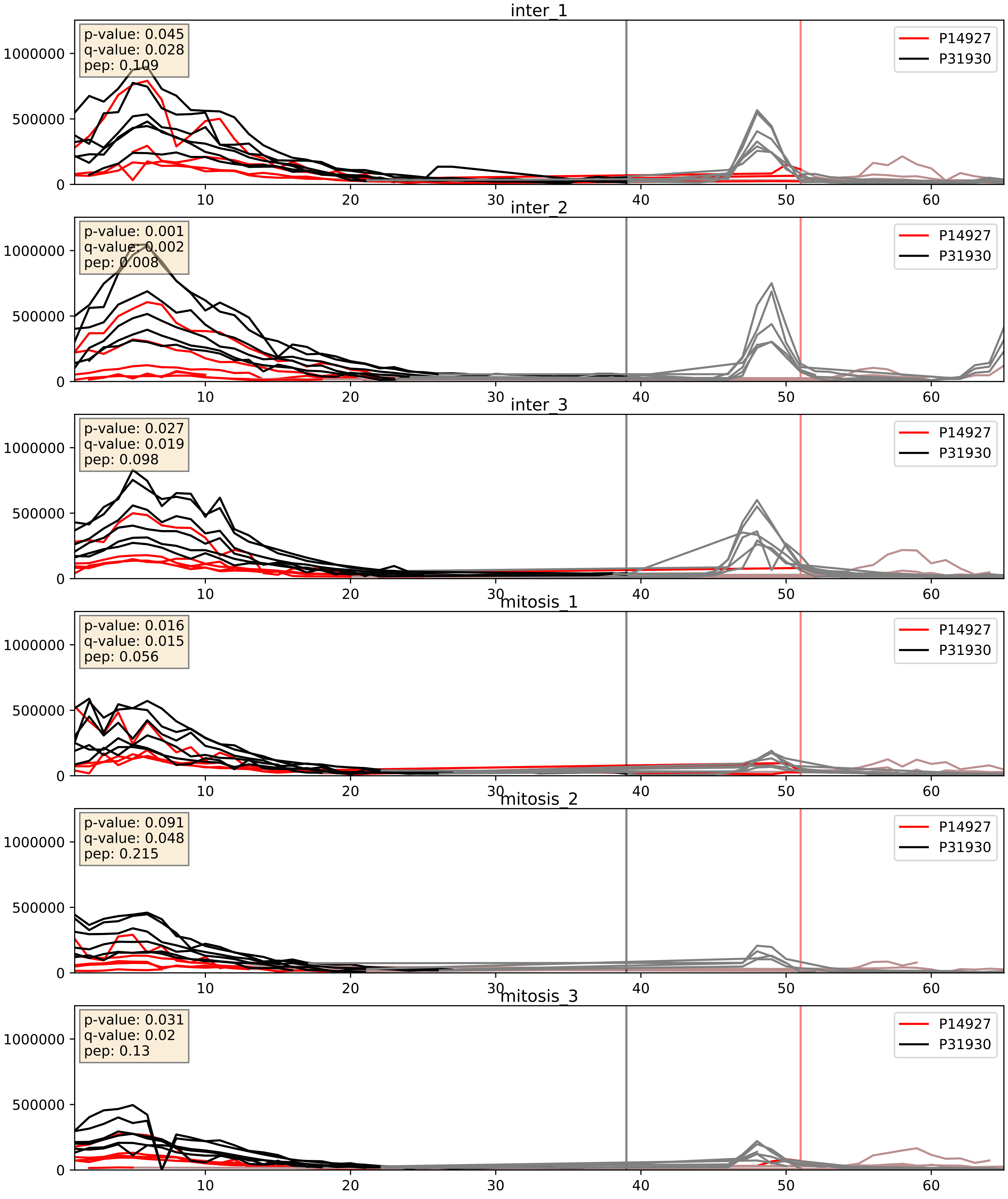


| level | condition_1 | condition_2 | pvalue | pvalue_adjusted |
|---|---|---|---|---|
| complex_abundance | mitosis | inter | 0.004145705864417472 | 0.04789101511391844 |
| interactor_abundance | mitosis | inter | 0.022350644269404238 | 0.12329937058940194 |
| interactor_ratio | mitosis | inter | 0.8696221698218296 | 0.9788064725083166 |

P20674_P31930
COX5A_HUMAN vs QCR1_HUMAN
p-value: 0.001 q-value: 0.002

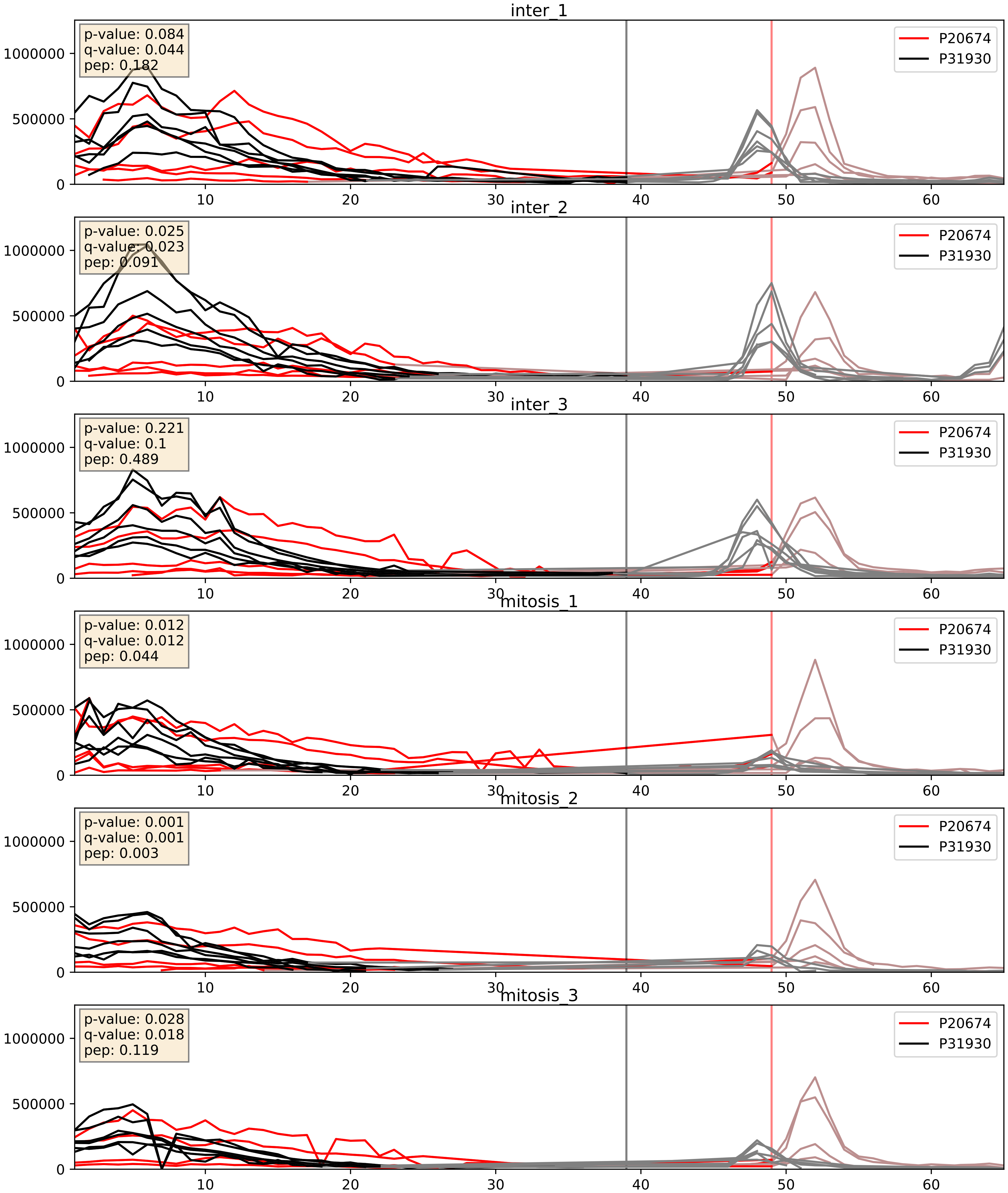


| level | condition_1 | condition_2 | pvalue | pvalue_adjusted |
|---|---|---|---|---|
| complex_abundance | mitosis | inter | 0.001073864635671266 | 0.021833569470040306 |
| interactor_abundance | mitosis | inter | 0.006684830843853993 | 0.07002298502274237 |
| interactor_ratio | mitosis | inter | 0.2152784393117739 | 0.6924580509898395 |

P22695_P31930
QCR2_HUMAN vs QCR1_HUMAN
p-value: 0.0 q-value: 0.0

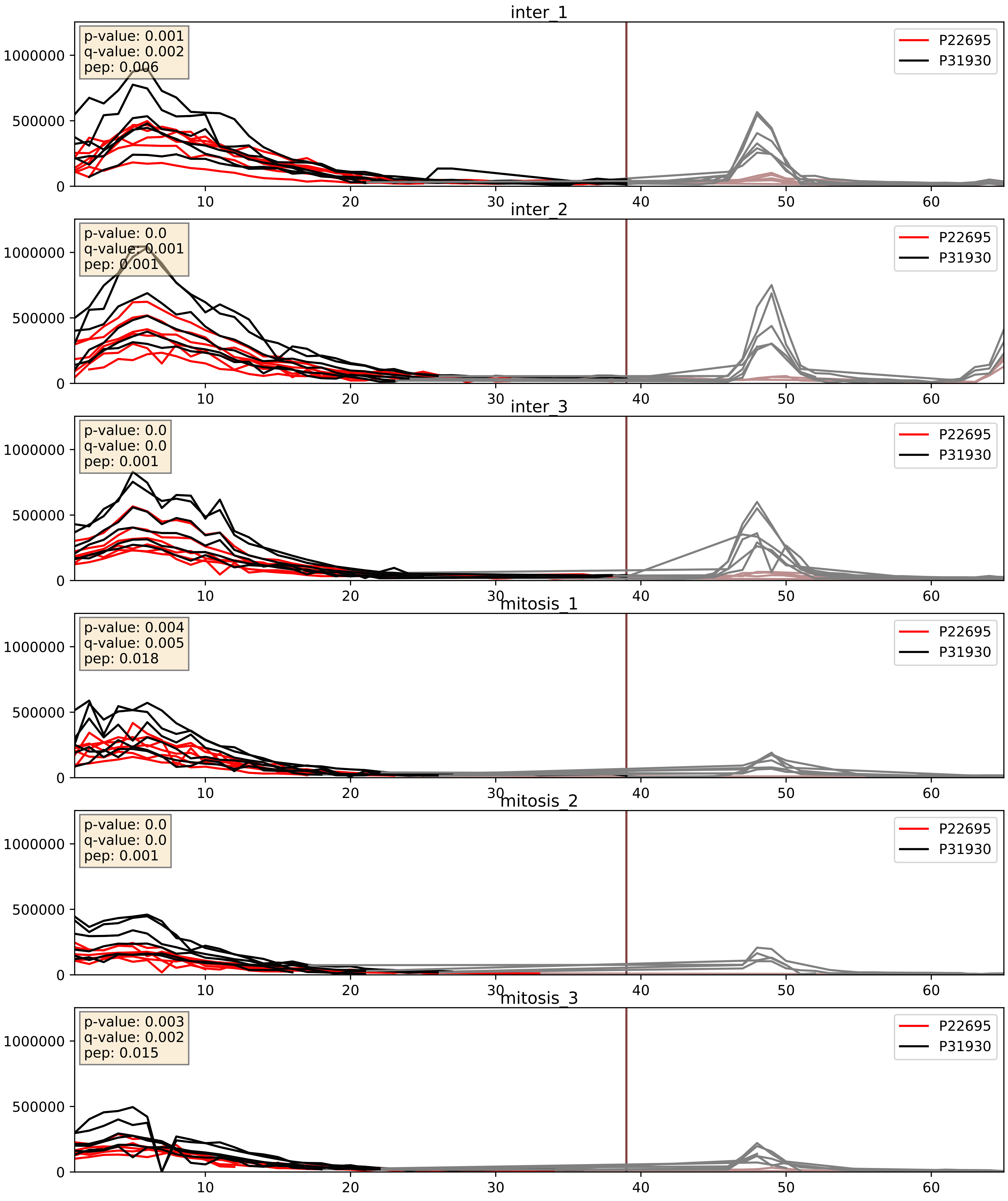


| level | condition_1 | condition_2 | pvalue | pvalue_adjusted |
|---|---|---|---|---|
| complex_abundance | mitosis | inter | 0.0008434285050207741 | 0.019252661341274204 |
| interactor_abundance | mitosis | inter | 0.0015444039397945596 | 0.028907984680718663 |
| interactor_ratio | mitosis | inter | 0.9461758960577978 | 0.9912117205440141 |

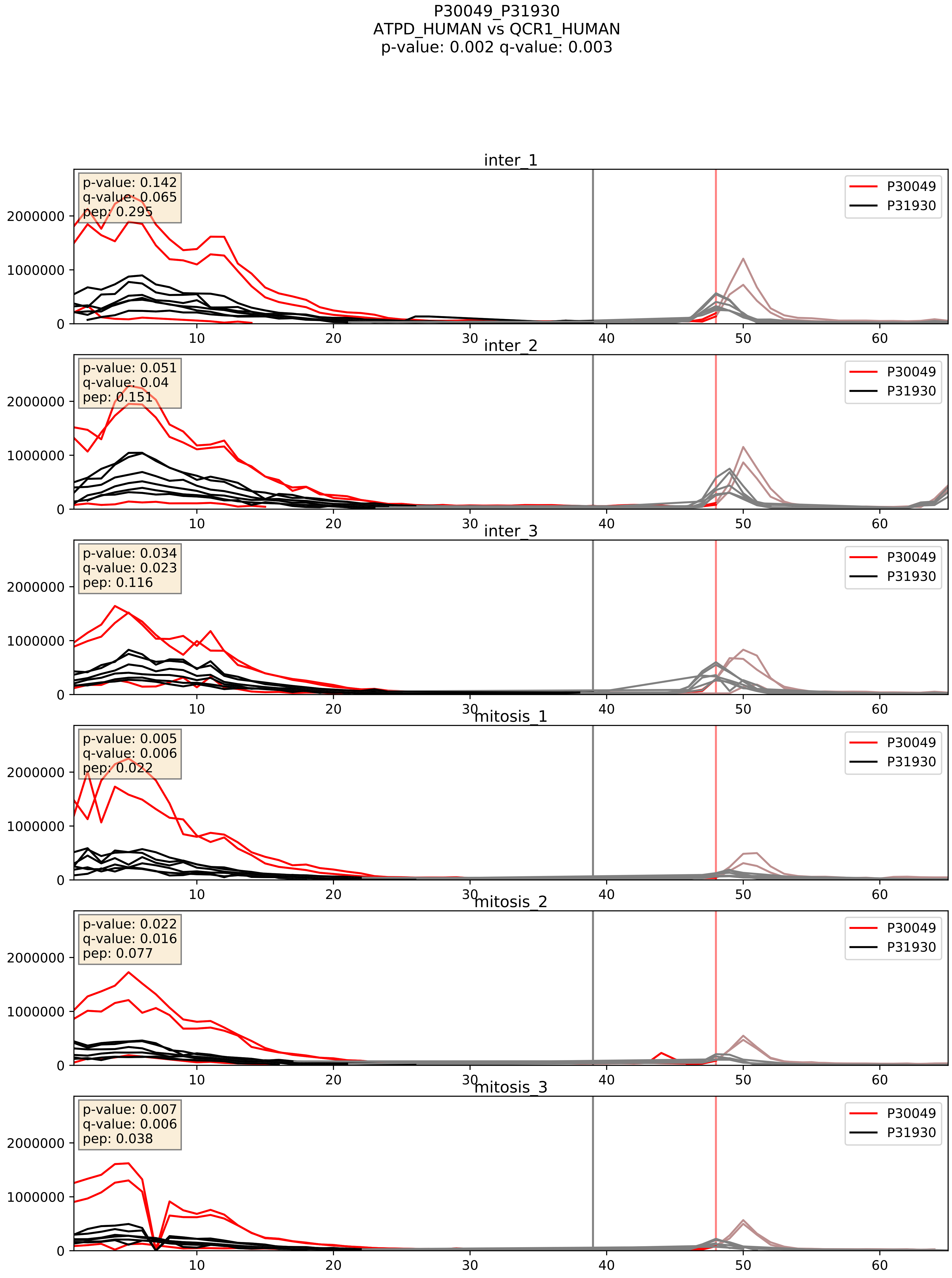


| level | condition_1 | condition_2 | pvalue | pvalue_adjusted |
|---|---|---|---|---|
| complex_abundance | mitosis | inter | 0.00031509670816226386 | 0.01165207110963347 |
| interactor_abundance | mitosis | inter | 0.0020066300443835885 | 0.03264187006238393 |
| interactor_ratio | mitosis | inter | 0.5719445498051304 | 0.8945917155252349 |

P31930_P36542
QCR1_HUMAN vs ATPG_HUMAN
p-value: 0.001 q-value: 0.024

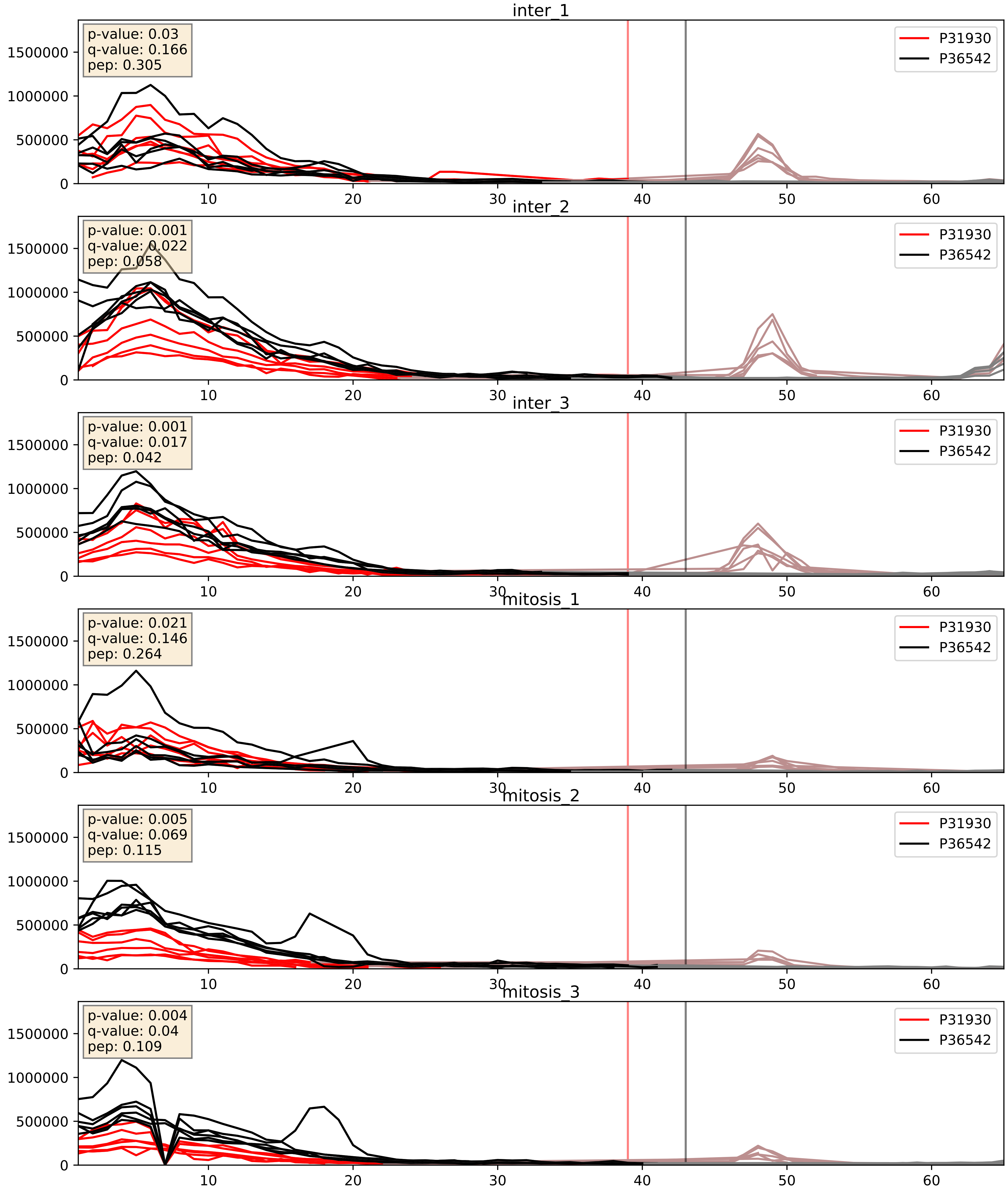


| level | condition_1 | condition_2 | pvalue | pvalue_adjusted |
|---|---|---|---|---|
| interactor_abundance | mitosis | inter | 0.0007825960512041081 | 0.020500371135067 71 |
| complex_abundance | mitosis | inter | 0.0067564519235486 | 0.060815171888092545 |
| interactor_ratio | mitosis | inter | 0.5286129521356591 | 0.8816356313046353 |

P31930_P45880
QCR1_HUMAN vs VDAC2_HUMAN
p-value: 0.0 q-value: 0.043

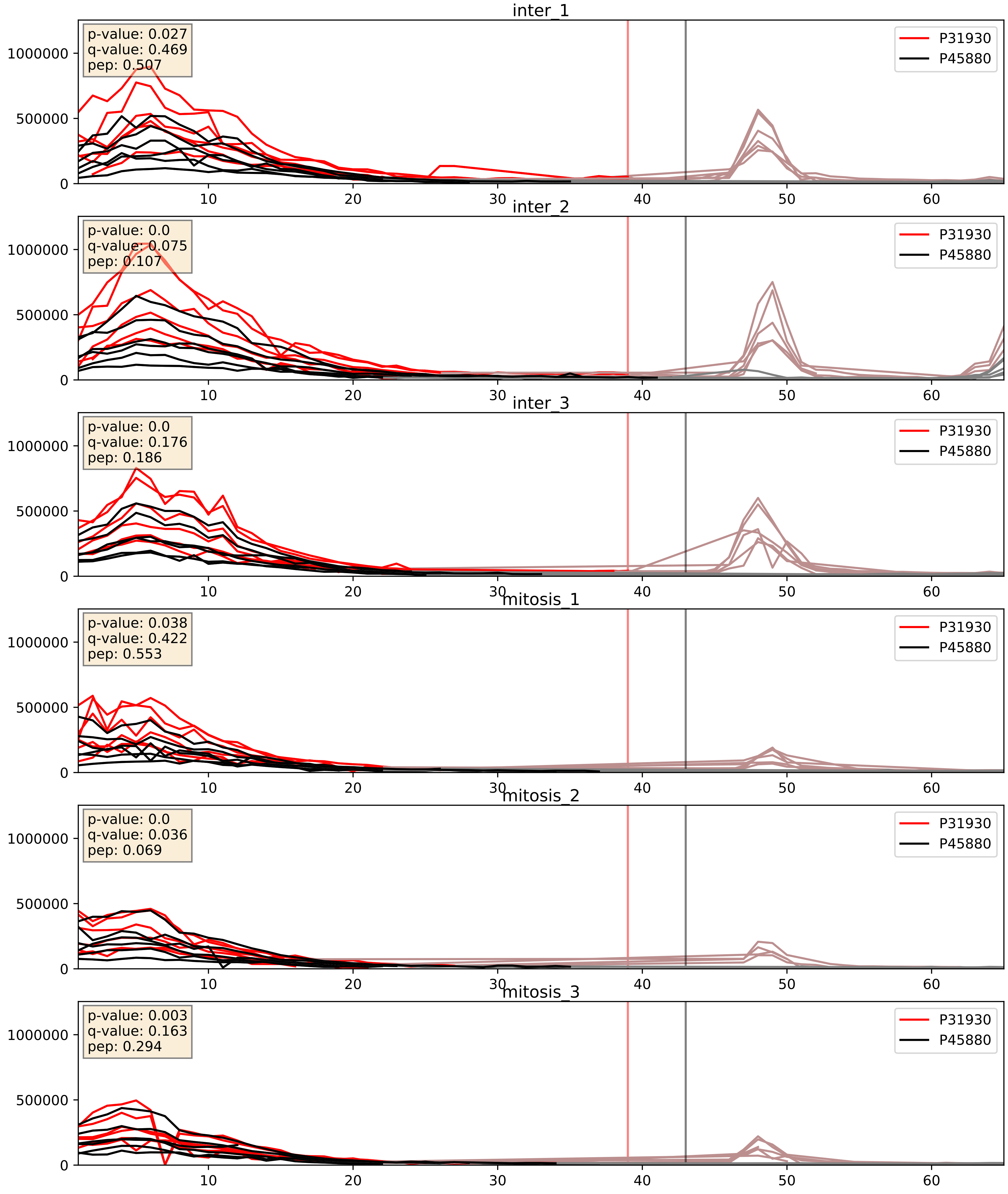


| level | condition_1 | condition_2 | pvalue | pvalue_adjusted |
|---|---|---|---|---|
| complex_abundance | mitosis | inter | 0.0006470922952819564 | 0.016393529992559892 |
| interactor_abundance | mitosis | inter | 0.0008808746421517374 | 0.021954049575556266 |
| interactor_ratio | mitosis | inter | 0.008824713236414494 | 0.13290071501637724 |

P31930_Q96EY1
QCR1_HUMAN vs DNJA3_HUMAN
p-value: 0.0 q-value: 0.025

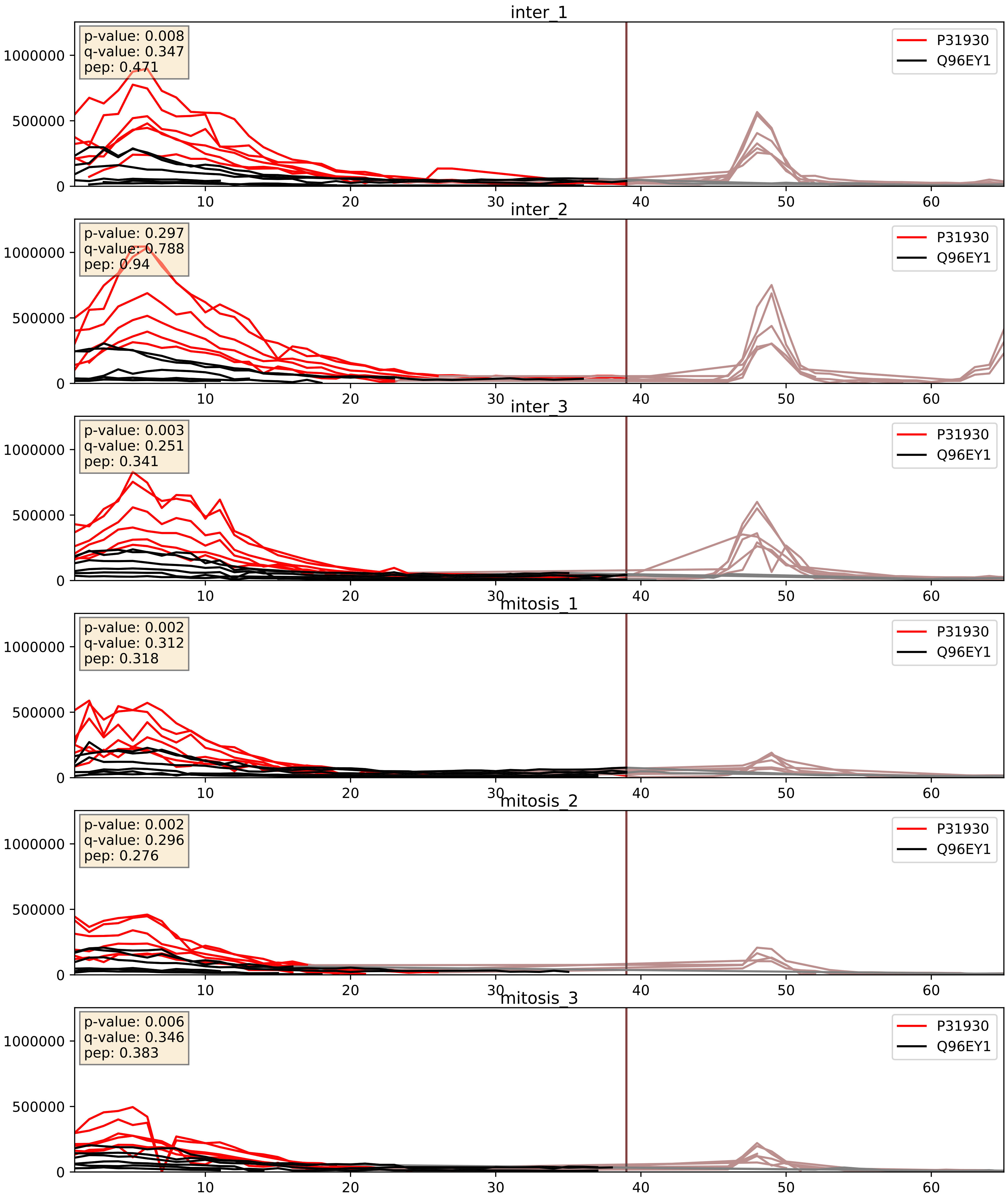


| level | condition_1 | condition_2 | pvalue | pvalue_adjusted |
|---|---|---|---|---|
| interactor_abundance | mitosis | inter | 0.0006576324138796844 | 0.018247434239254776 |
| complex_abundance | mitosis | inter | 0.005328077546970102 | 0.054883686885757 |
| interactor_ratio | mitosis | inter | 0.06633110039489377 | 0.4296400448702392 |
