## Supplementary Data 2 for "SECAT: Quantifying differential protein-protein interaction states by network-centric analysis": hela_string_000092_Q9BUQ8.pdf

Q9BUQ8_Q9BUQ8

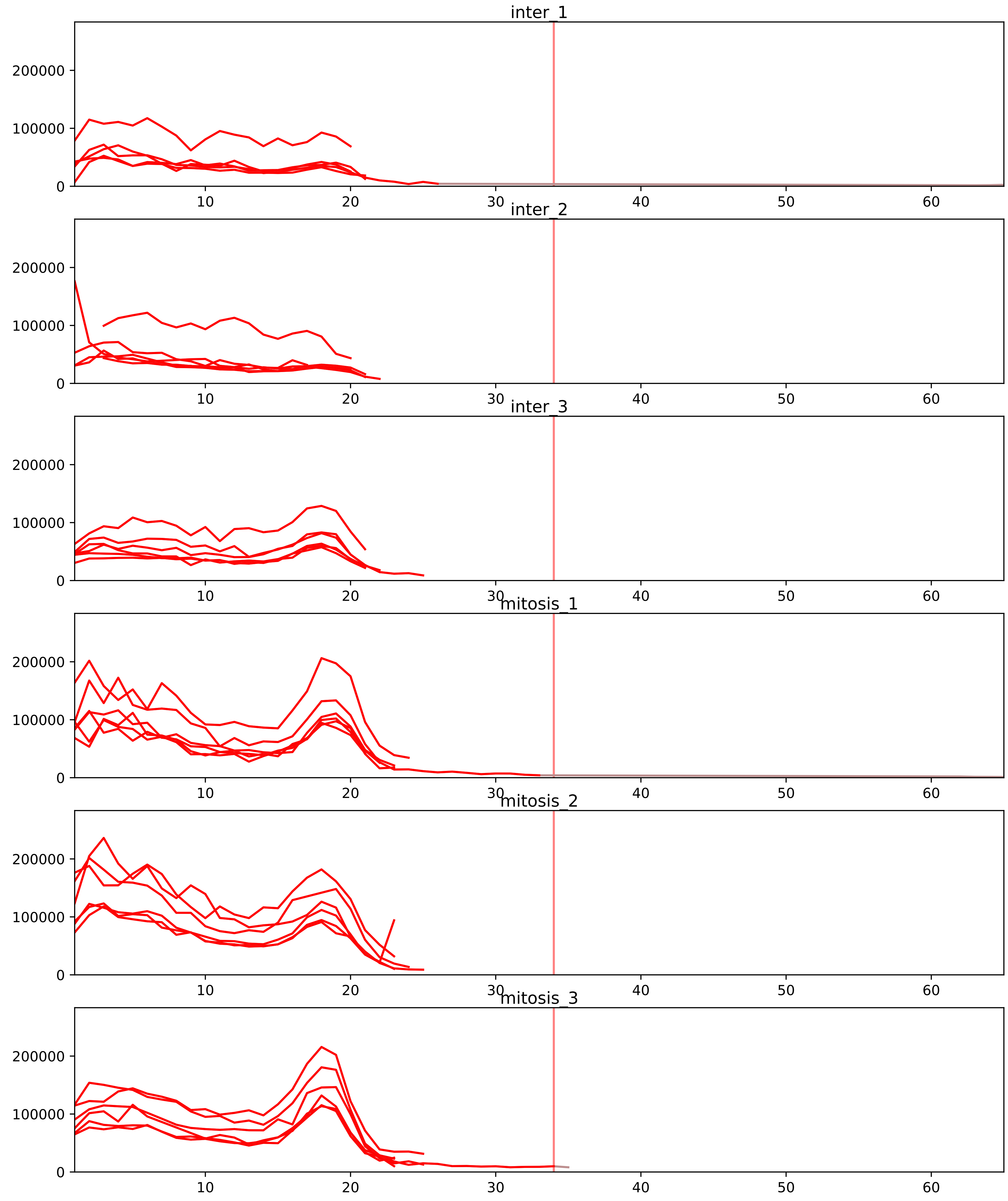


| level | condition_1 | condition_2 | pvalue | pvalue_adjusted |
|---|---|---|---|---|
| complex_abundance | mitosis | inter | 6.220078107625079e-06 | 0.000357654491188442 |
| interactor_abundance | mitosis | inter | 0.00037975711263337934 | 0.006022977368794052 |
| interactor_ratio | mitosis | inter | 0.004263968705078991 | 0.035662283715206106 |
| monomer_abundance | mitosis | inter | 0.3259380816140039 | 0.6718717495121003 |
| assembled_abundance | mitosis | inter | 0.7999813439998277 | 0.9176772141679141 |
| total_abundance | mitosis | inter | 0.8138573254228502 | 0.9272086367190218 |

O43143_Q9BUQ8
DHX15_HUMAN vs DDX23_HUMAN
p-value: 0.023 q-value: 0.017

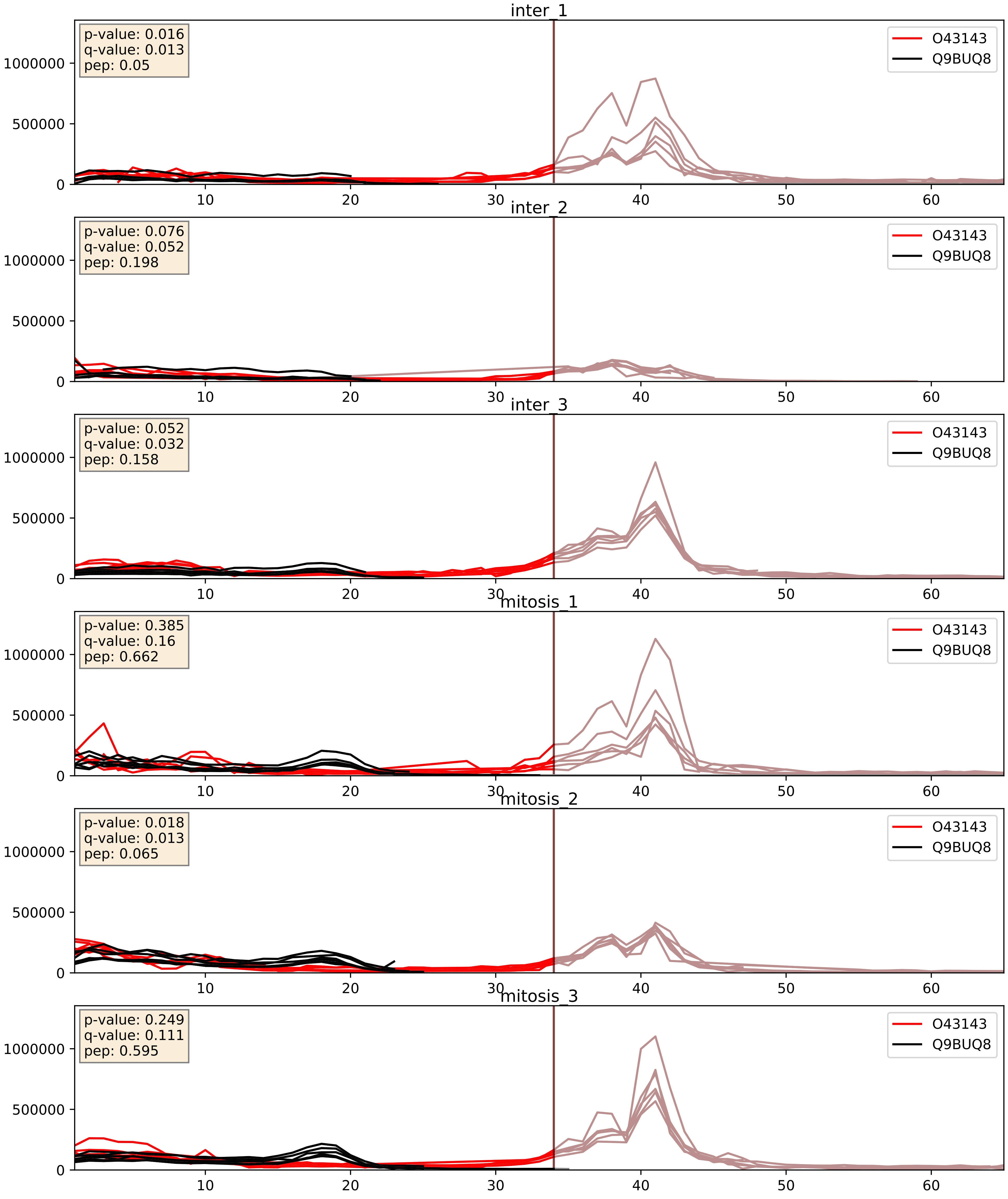


| level | condition_1 | condition_2 | pvalue | pvalue_adjusted |
|---|---|---|---|---|
| complex_abundance | mitosis | inter | 0.020349387064744866 | 0.10515559050023798 |
| interactor_abundance | mitosis | inter | 0.07495183037537691 | 0.22682965105647032 |
| interactor_ratio | mitosis | inter | 0.19225724787519052 | 0.6660896702330326 |

O43172_Q9BUQ8
PRP4_HUMAN vs DDX23_HUMAN
p-value: 0.005 q-value: 0.005

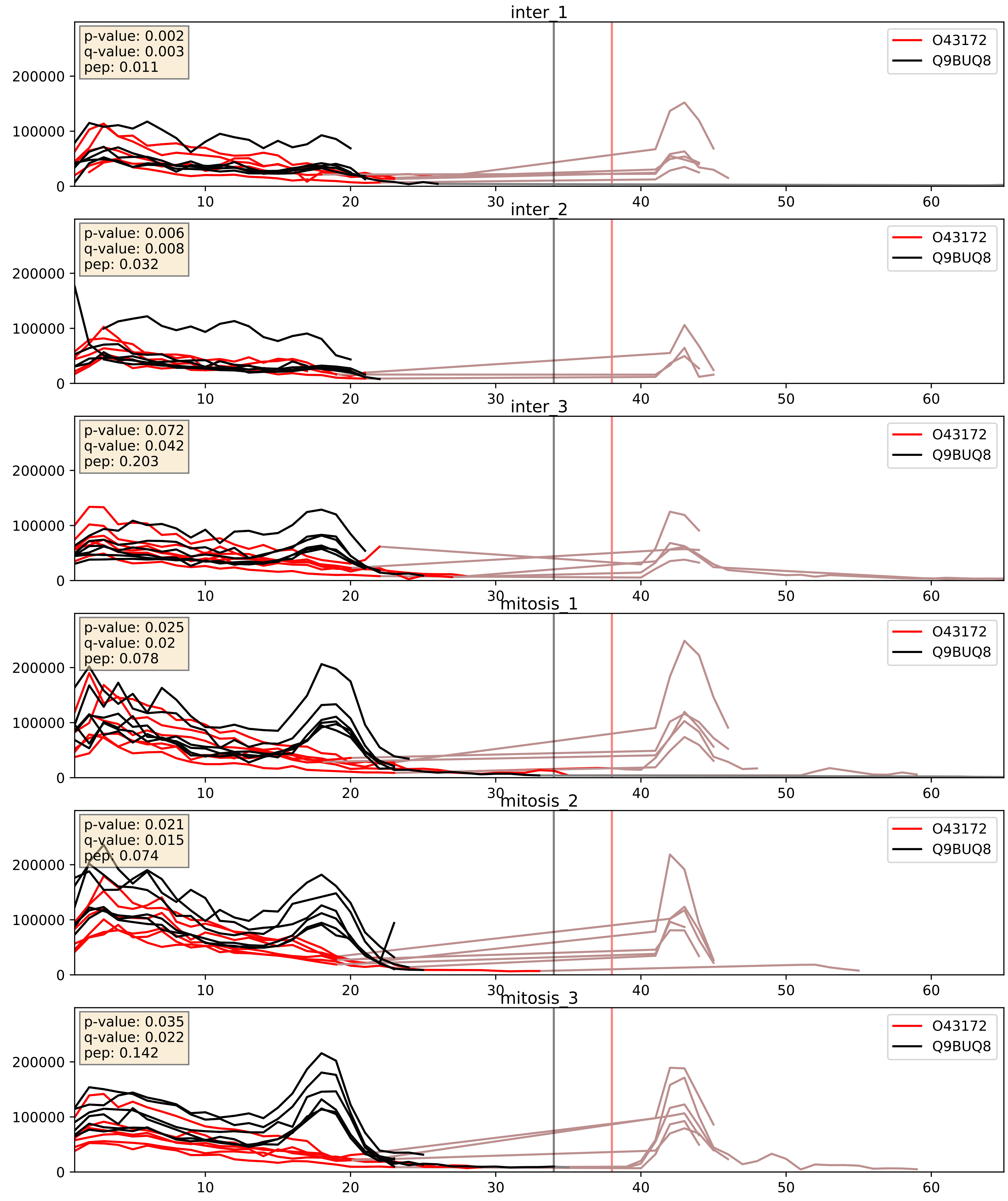


| level | condition_1 | condition_2 | pvalue | pvalue_adjusted |
|---|---|---|---|---|
| complex_abundance | mitosis | inter | 0.016765705806694113 | 0.09664272168707179 |
| interactor_abundance | mitosis | inter | 0.20701848199376688 | 0.40422974203168 |
| interactor_ratio | mitosis | inter | 0.2524563550030642 | 0.7251677296572153 |

O43290_Q9BUQ8
SNUT1_HUMAN vs DDX23_HUMAN
p-value: 0.069 q-value: 0.042

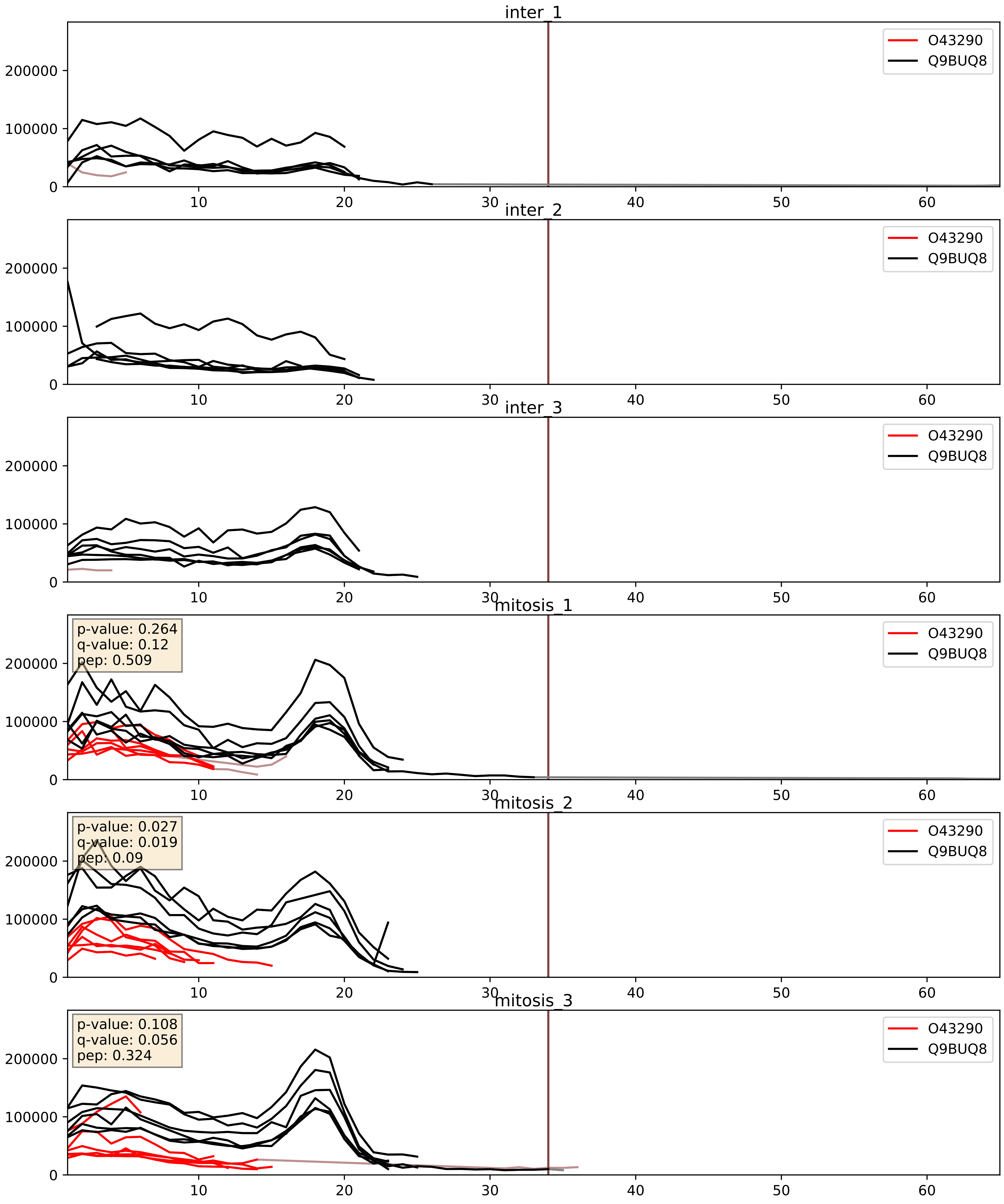


| level | condition_1 | condition_2 | pvalue | pvalue_adjusted |
|---|---|---|---|---|
| complex_abundance | mitosis | inter | 0.0002340547270430609 | 0.010221981956574496 |
| interactor_abundance | mitosis | inter | 0.0010647499919498099 | 0.024049720606162197 |
| interactor_ratio | mitosis | inter | 0.005120825250895749 | 0.09650938060012829 |

O94906_Q9BUQ8
PRP6_HUMAN vs DDX23_HUMAN
p-value: 0.008 q-value: 0.007

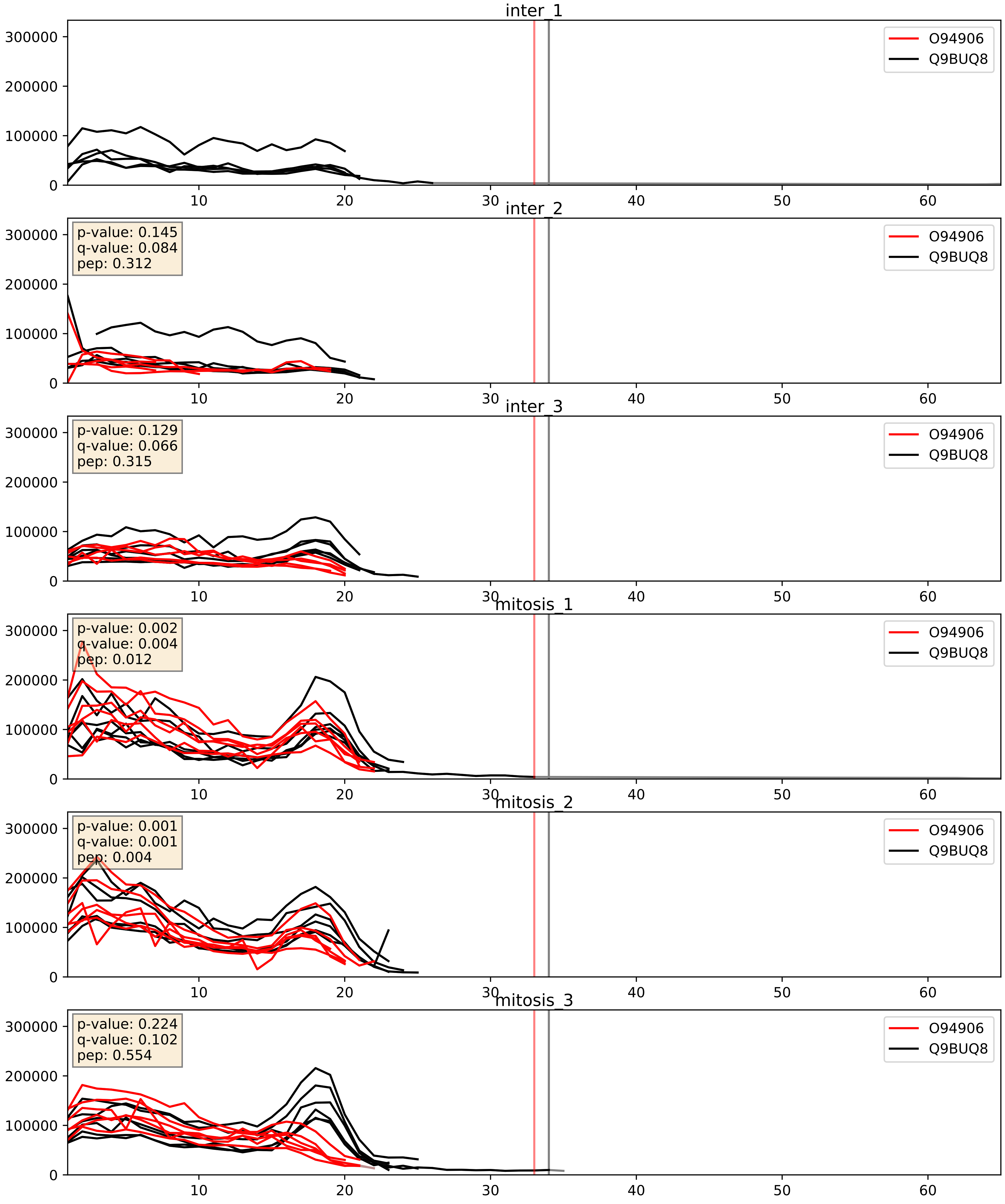


| level | condition_1 | condition_2 | pvalue | pvalue_adjusted |
|---|---|---|---|---|
| interactor_abundance | mitosis | inter | 0.04665310860590202 | 0.1724311786124876 |
| complex_abundance | mitosis | inter | 0.07150374961488644 | 0.19422056657737993 |
| interactor_ratio | mitosis | inter | 0.3936541743101139 | 0.8267123974716818 |

O95400_Q9BUQ8
CD2B2_HUMAN vs DDX23_HUMAN
p-value: 0.0 q-value: 0.0

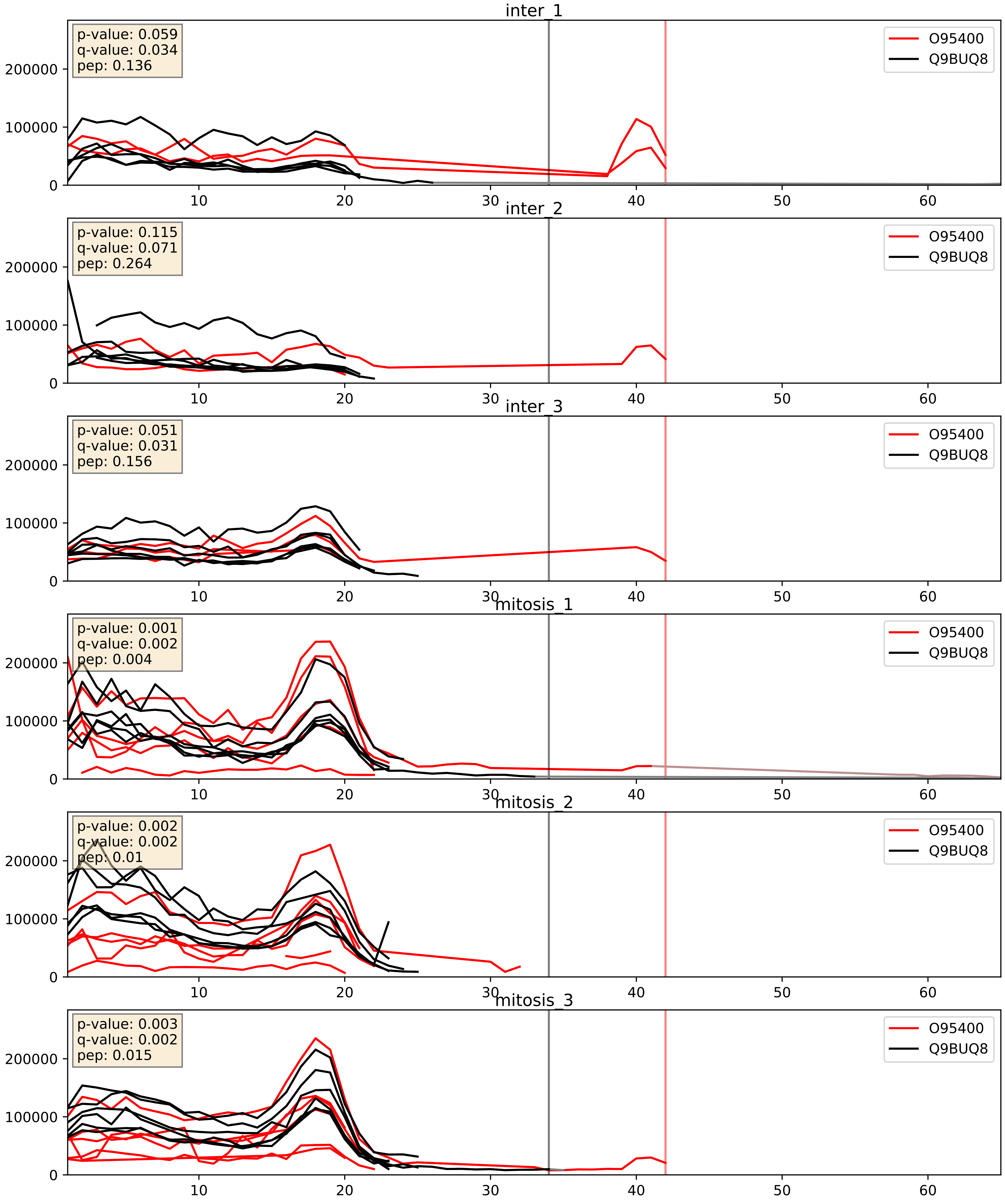


| level | condition_1 | condition_2 | pvalue | pvalue_adjusted |
|---|---|---|---|---|
| complex_abundance | mitosis | inter | 0.006814407597826003 | 0.061143950772946105 |
| interactor_abundance | mitosis | inter | 0.010469124642070678 | 0.08918423299129571 |
| interactor_ratio | mitosis | inter | 0.8759342926733549 | 0.9802747470617126 |

P55769_Q9BUQ8
NH2L1_HUMAN vs DDX23_HUMAN
p-value: 0.008 q-value: 0.007

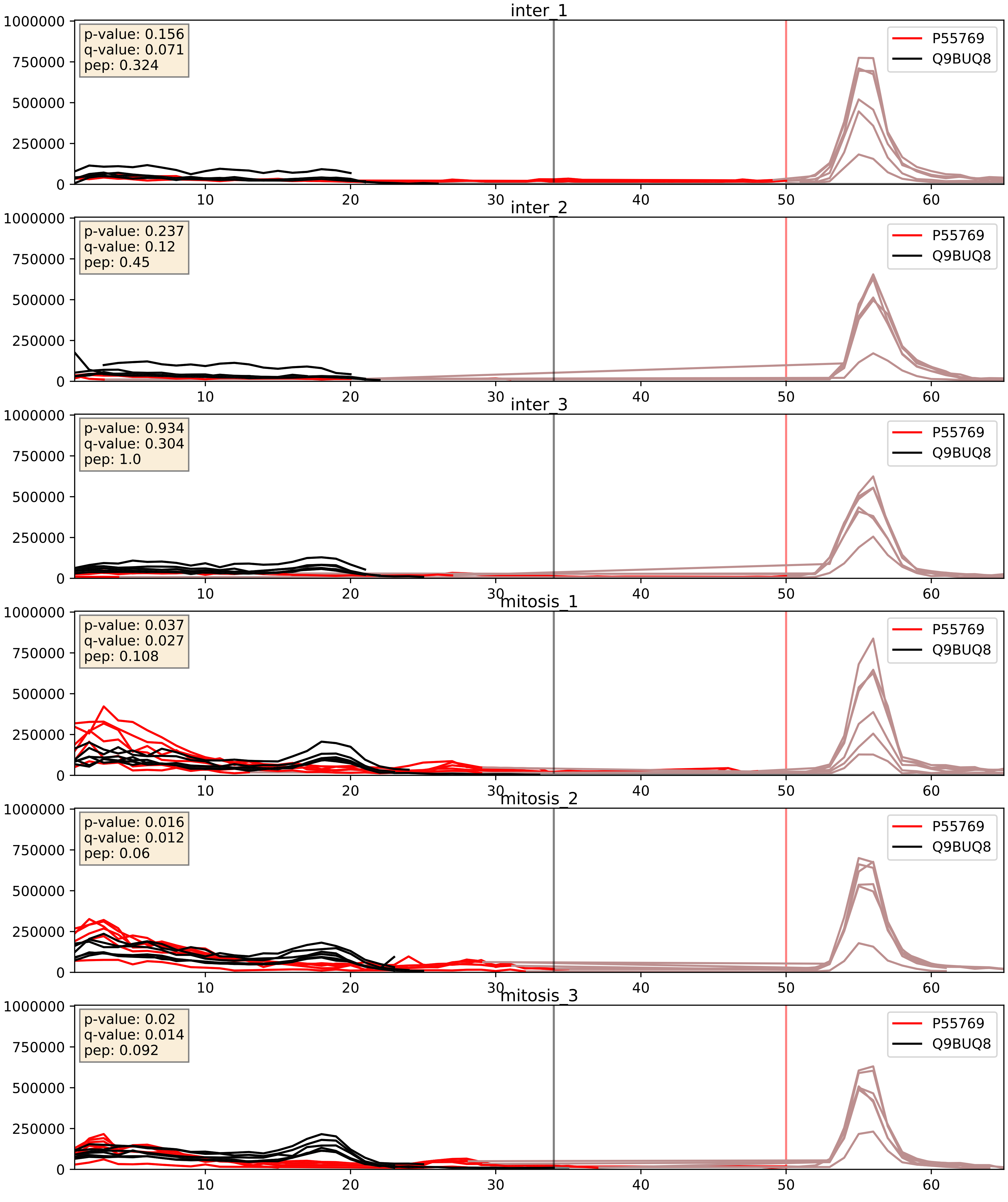


| level | condition_1 | condition_2 | pvalue | pvalue_adjusted |
|---|---|---|---|---|
| complex_abundance | mitosis | inter | 5.162211395155756e-05 | 0.005691857151652958 |
| interactor_abundance | mitosis | inter | 0.0010679120199147774 | 0.024049720606162197 |
| interactor_ratio | mitosis | inter | 0.1115601989567642 | 0.5532962393902757 |

Q15029_Q9BUQ8
U5S1_HUMAN vs DDX23_HUMAN
p-value: 0.081 q-value: 0.048

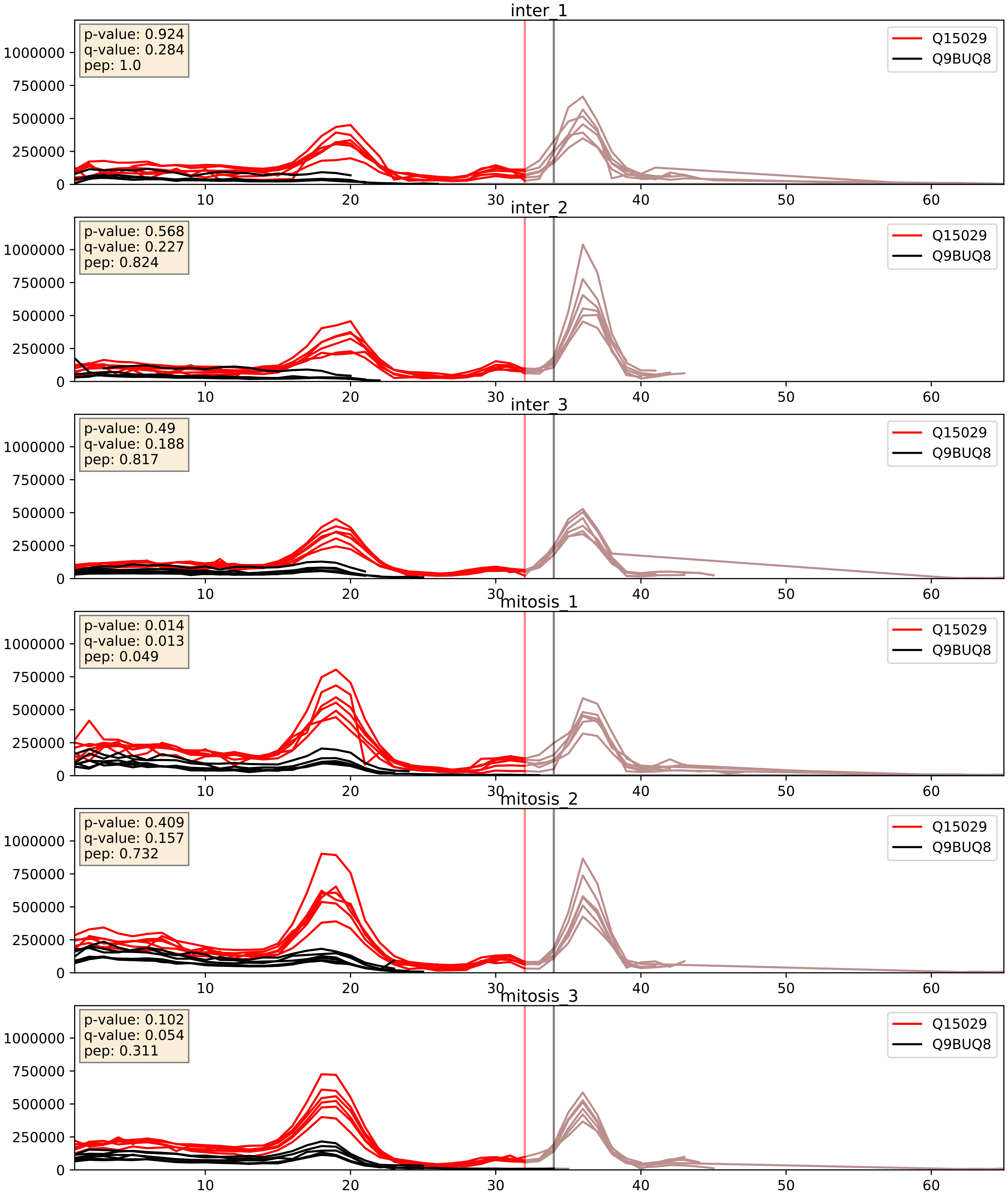


| level | condition_1 | condition_2 | pvalue | pvalue_adjusted |
|---|---|---|---|---|
| complex_abundance | mitosis | inter | 0.002609068629028324 | 0.03664546283589145 |
| interactor_abundance | mitosis | inter | 0.0037452506168265823 | 0.047601106577632585 |
| interactor_ratio | mitosis | inter | 0.15400658130323275 | 0.611454701278141 |

Q53GS9_Q9BUQ8
SNUT2_HUMAN vs DDX23_HUMAN
p-value: 0.024 q-value: 0.019

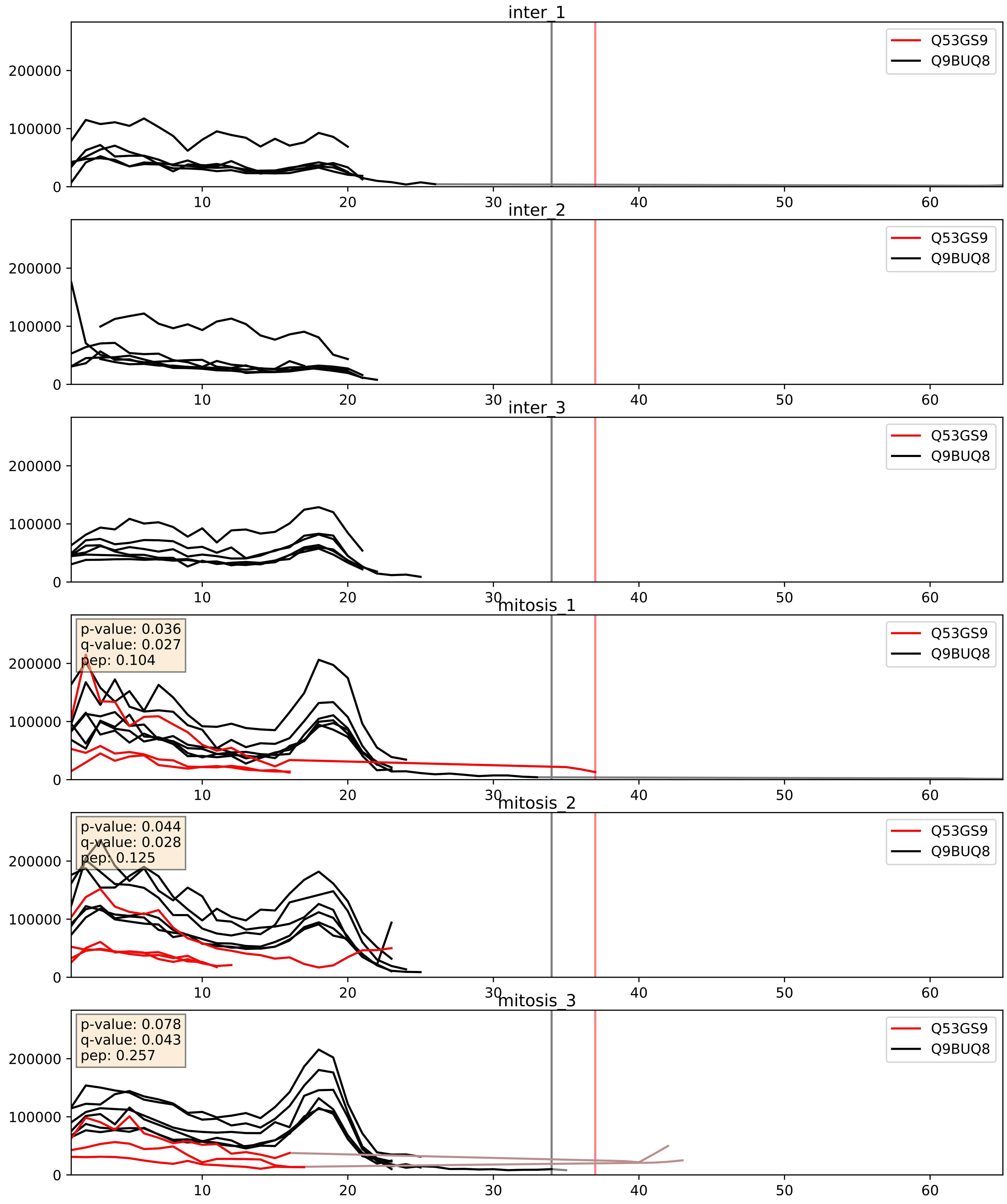


| level | condition_1 | condition_2 | pvalue | pvalue_adjusted |
|---|---|---|---|---|
| complex_abundance | mitosis | inter | 0.00014008468161613128 | 0.00812325774702682 |
| interactor_abundance | mitosis | inter | 0.0013451098335009928 | 0.027124005123129562 |
| interactor_ratio | mitosis | inter | 0.007320278961347636 | 0.11910596594407505 |

Q56P03_Q9BUQ8
EAPP_HUMAN vs DDX23_HUMAN
p-value: 0.0 q-value: 0.016

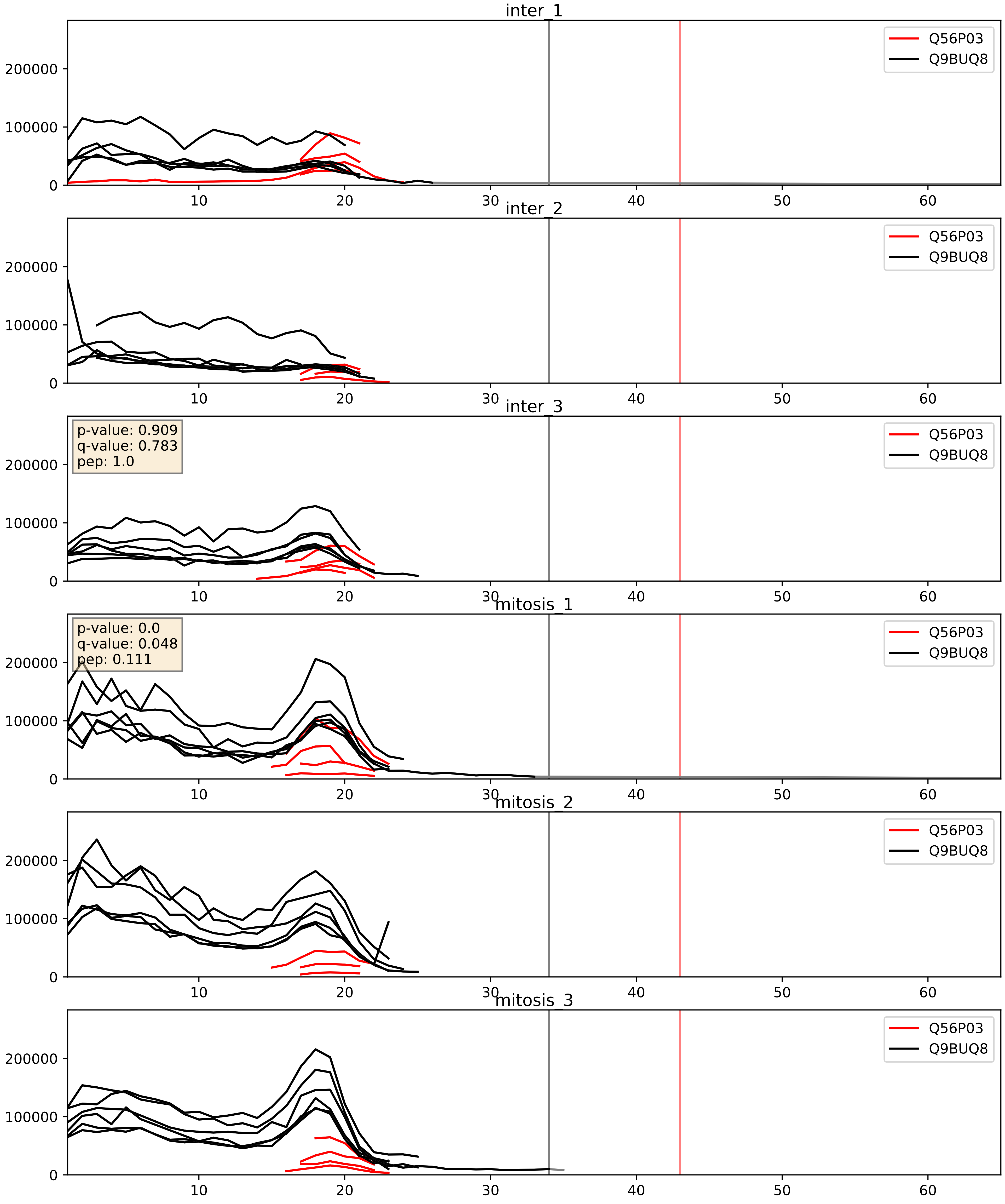


| level | condition_1 | condition_2 | pvalue | pvalue_adjusted |
|---|---|---|---|---|
| interactor_ratio | mitosis | inter | 0.036048508588732084 | 0.30857523351954663 |
| interactor_abundance | mitosis | inter | 0.1801513607733652 | 0.3727569853081958 |
| complex_abundance | mitosis | inter | 0.49805674536920475 | 0.6363232448299094 |

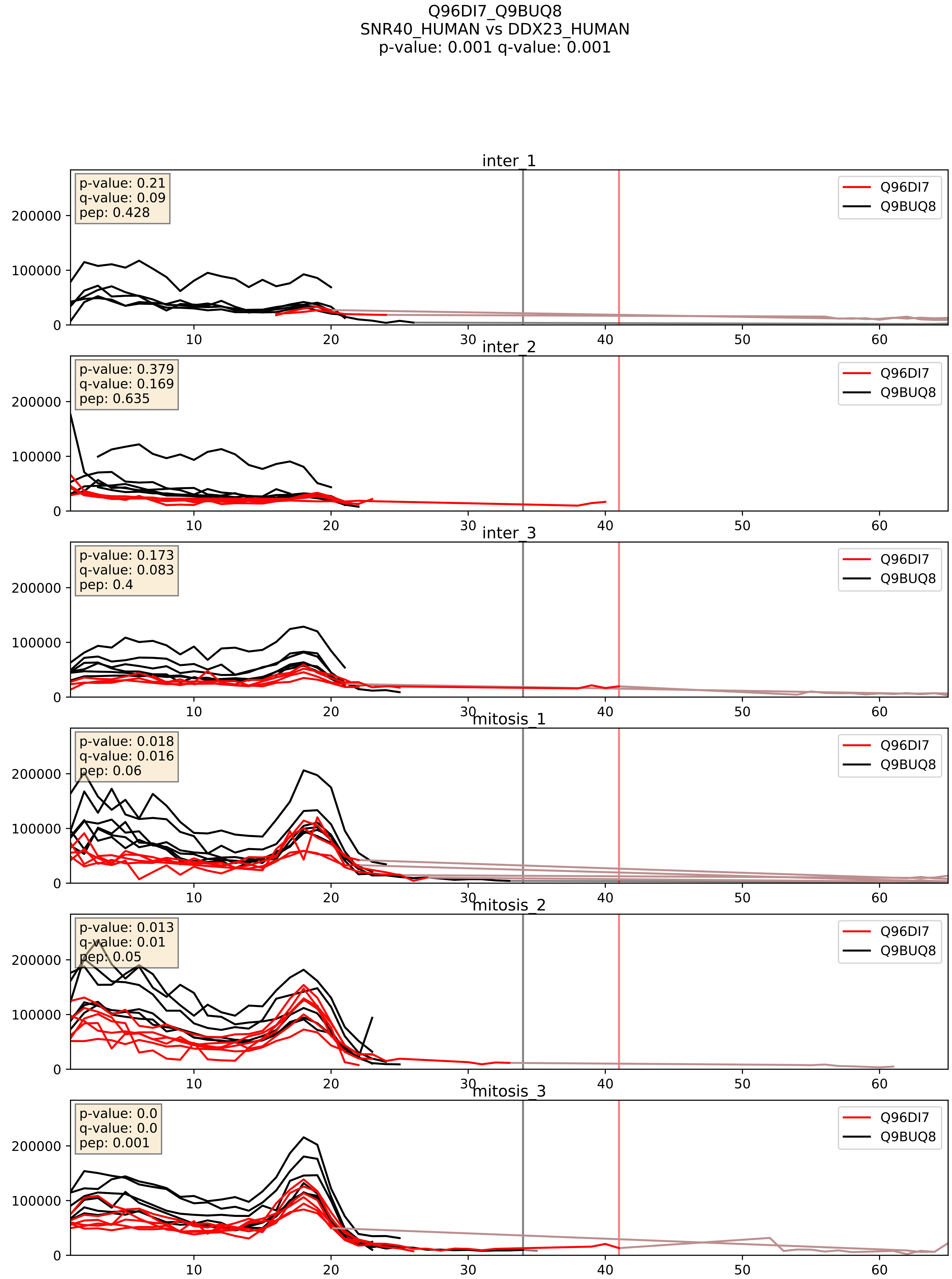


| level | condition_1 | condition_2 | pvalue | pvalue_adjusted |
|---|---|---|---|---|
| interactor_abundance | mitosis | inter | 0.015546485297274840 | 0.10925937121894304 |
| complex_abundance | mitosis | inter | 0.033901488710403466 | 0.13408245588106055 |
| interactor_ratio | mitosis | inter | 0.5458492076094994 | 0.8859730852936224 |

Q9BUQ8_Q9BZJ0
DDX23_HUMAN vs CRNL1_HUMAN
p-value: 0.036 q-value: 0.025

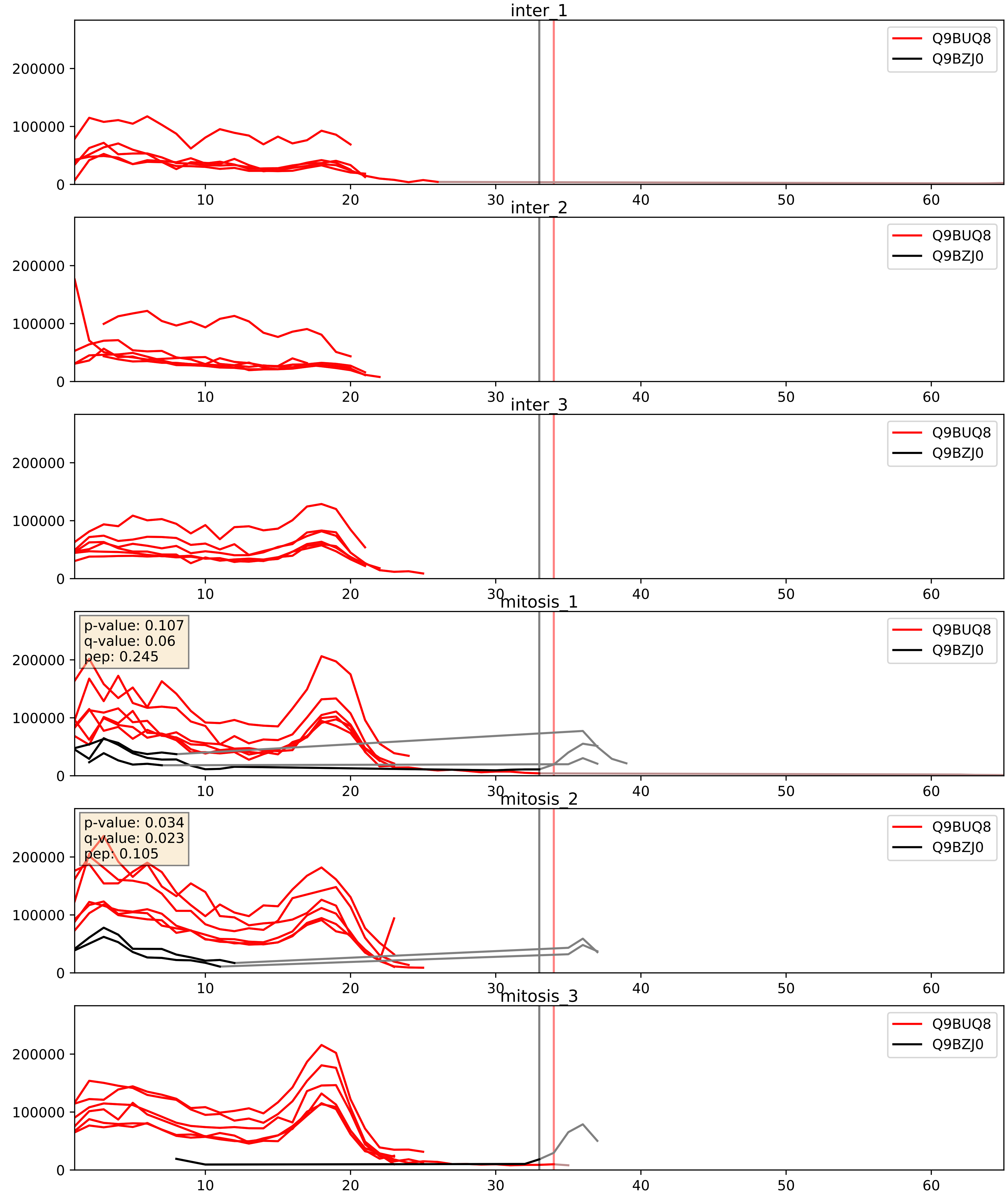


| level | condition_1 | condition_2 | pvalue | pvalue_adjusted |
|---|---|---|---|---|
| interactor_ratio | mitosis | inter | 0.01085137089615999 | 0.1495776729003696 |
| interactor_abundance | mitosis | inter | 0.04326640025801008 | 0.16685321787696922 |
| complex_abundance | mitosis | inter | 0.12438106921664024 | 0.26637526957579194 |
