## Supplementary Data 2 for "SECAT: Quantifying differential protein-protein interaction states by network-centric analysis": hela_string_000093_Q8WWY3.pdf

Q8WWY3_Q8WWY3

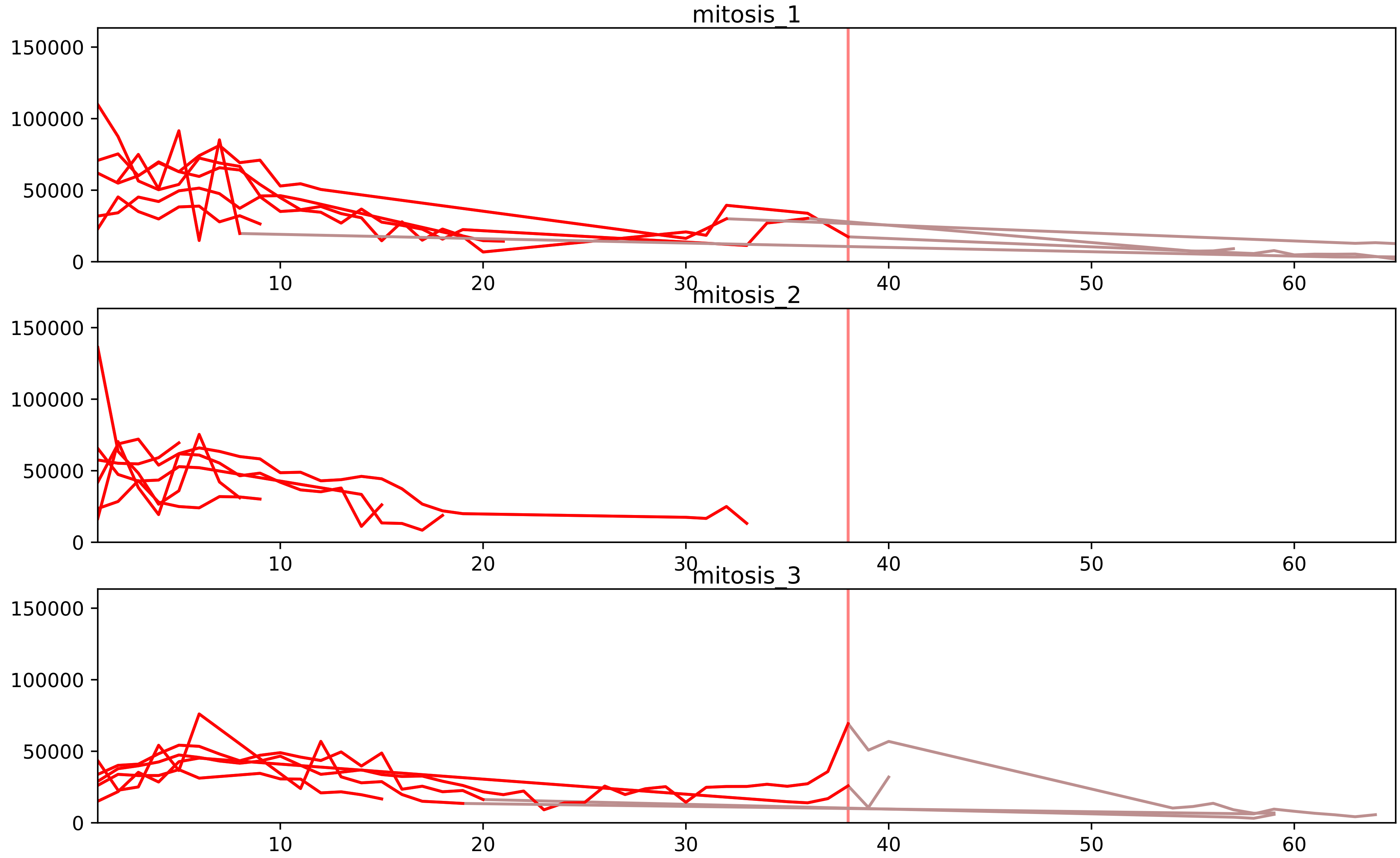


| level | condition_1 | condition_2 | pvalue | pvalue_adjusted |
|---|---|---|---|---|
| interactor_ratio | mitosis | inter | 6.290898600462644e-06 | 0.00015160452489447503 |
| complex_abundance | mitosis | inter | 3.331779060877442e-05 | 0.0011146315403662713 |
| interactor_abundance | mitosis | inter | 0.0003302293832642738 | 0.005513071612019375 |
| assembled_abundance | mitosis | inter | 0.0007677116292235458 | 0.024955959280523736 |
| total_abundance | mitosis | inter | 0.0008289058251877882 | 0.03196939057599265 |
| monomer_abundance | mitosis | inter | 0.11226351229754611 | 0.439428372316888 |

O43172_Q8WWY3
PRP4_HUMAN vs PRP31_HUMAN
p-value: 0.012 q-value: 0.01

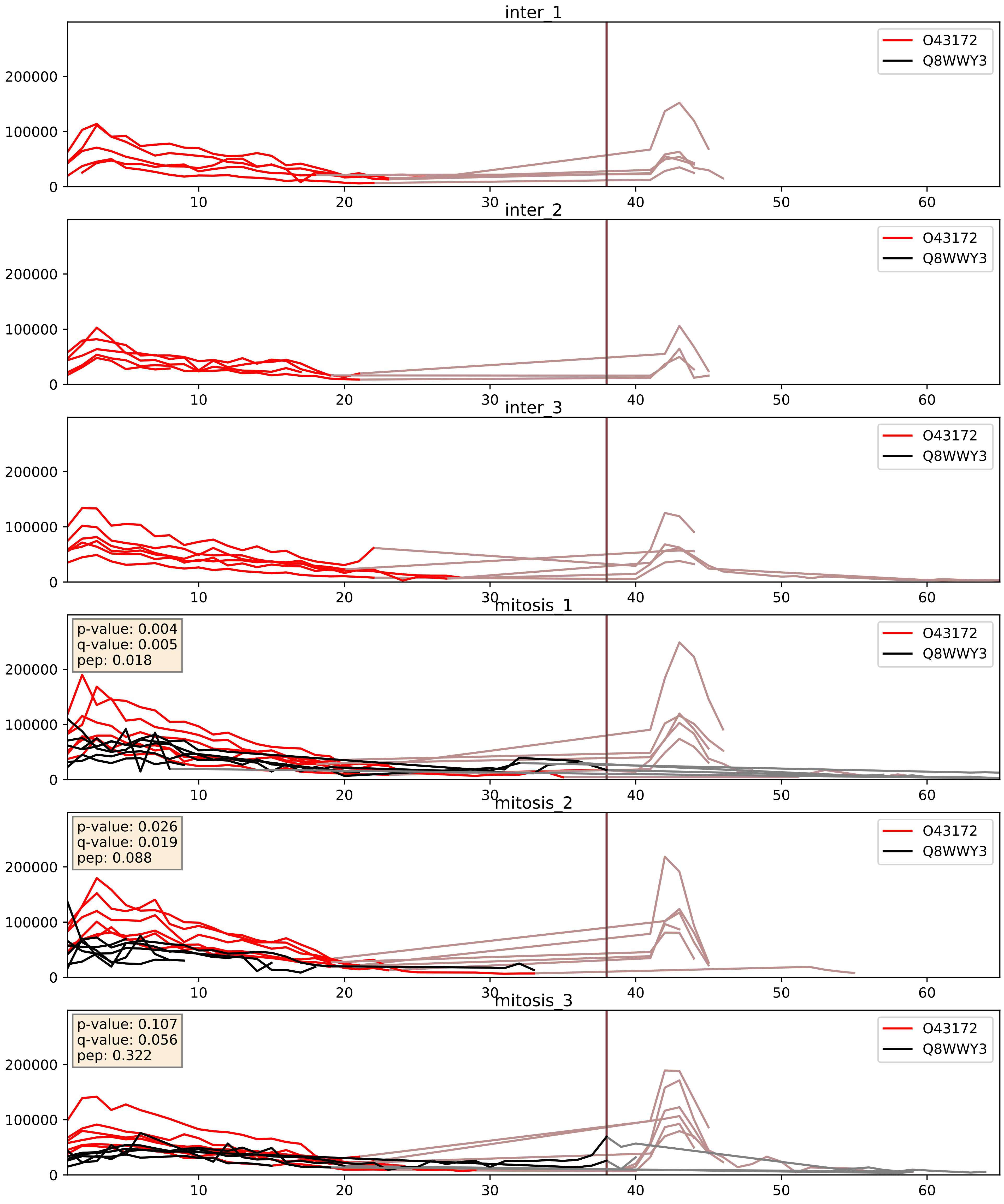


| level | condition_1 | condition_2 | pvalue | pvalue_adjusted |
|---|---|---|---|---|
| interactor_abundance | mitosis | inter | 7.464209076014017e-05 | 0.006870282762438709 |
| complex_abundance | mitosis | inter | 0.00042479332670986605 | 0.013091389064164169 |
| interactor_ratio | mitosis | inter | 0.0033003956960577104 | 0.07473911946628042 |

O43395_Q8WWY3
PRPF3_HUMAN vs PRP31_HUMAN
p-value: 0.036 q-value: 0.025

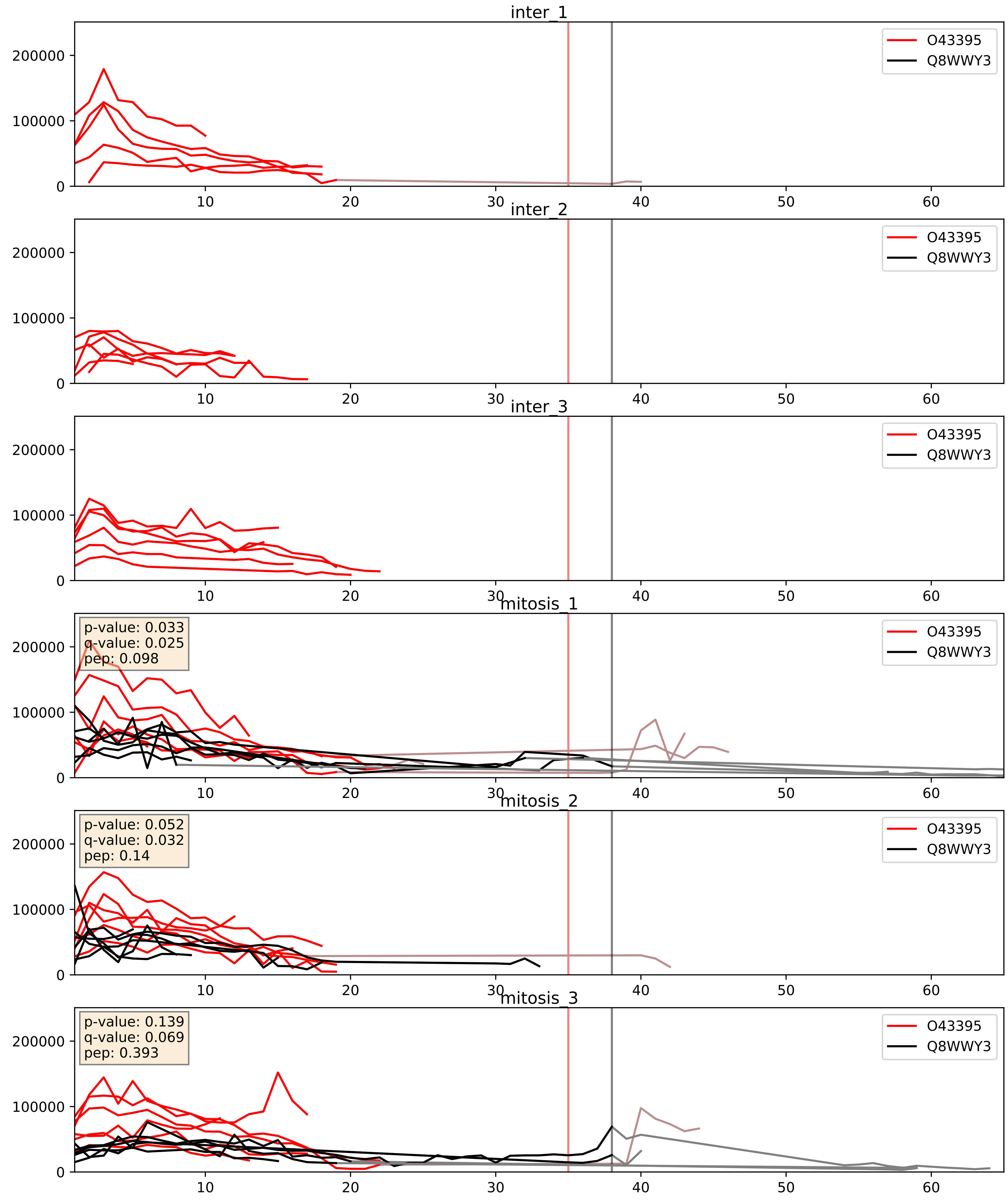


| level | condition_1 | condition_2 | pvalue | pvalue_adjusted |
|---|---|---|---|---|
| interactor_abundance | mitosis | inter | 0.0003488542111186024 | 0.01356427145975096 |
| complex_abundance | mitosis | inter | 0.0006558599407001601 | 0.01642175463112936 |
| interactor_ratio | mitosis | inter | 0.011383459471981348 | 0.15421453104903593 |

P55769_Q8WWY3
NH2L1_HUMAN vs PRP31_HUMAN
p-value: 0.032 q-value: 0.023

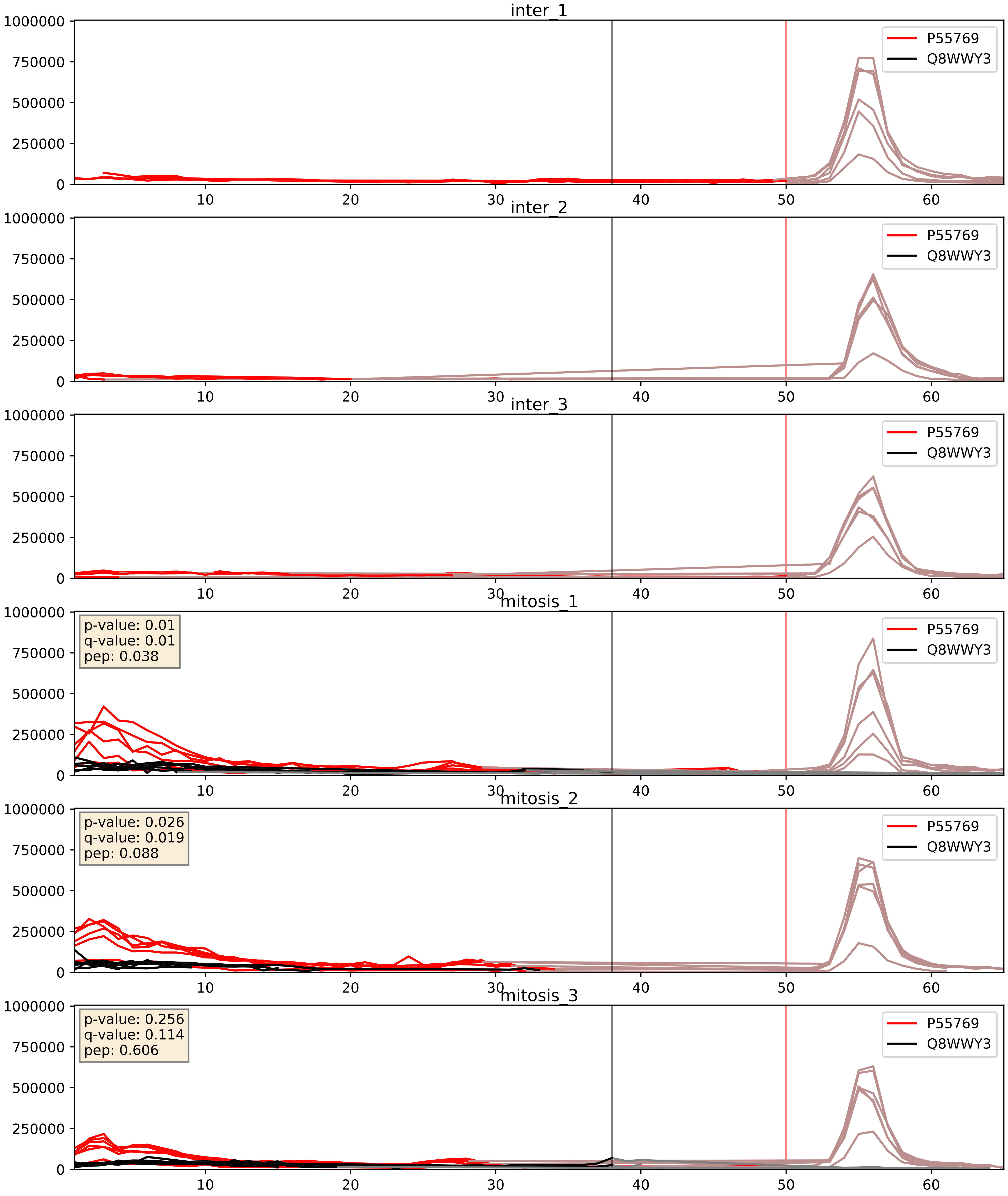


| level | condition_1 | condition_2 | pvalue | pvalue_adjusted |
|---|---|---|---|---|
| interactor_abundance | mitosis | inter | 6.0879231696132886e-05 | 0.006459477818931243 |
| interactor_ratio | mitosis | inter | 0.00020587750302125888 | 0.014683183137172115 |
| complex_abundance | mitosis | inter | 0.00030195135818513163 | 0.01165207110963347 |

P62312_Q8WWY3
LSM6_HUMAN vs PRP31_HUMAN
p-value: 0.085 q-value: 0.05

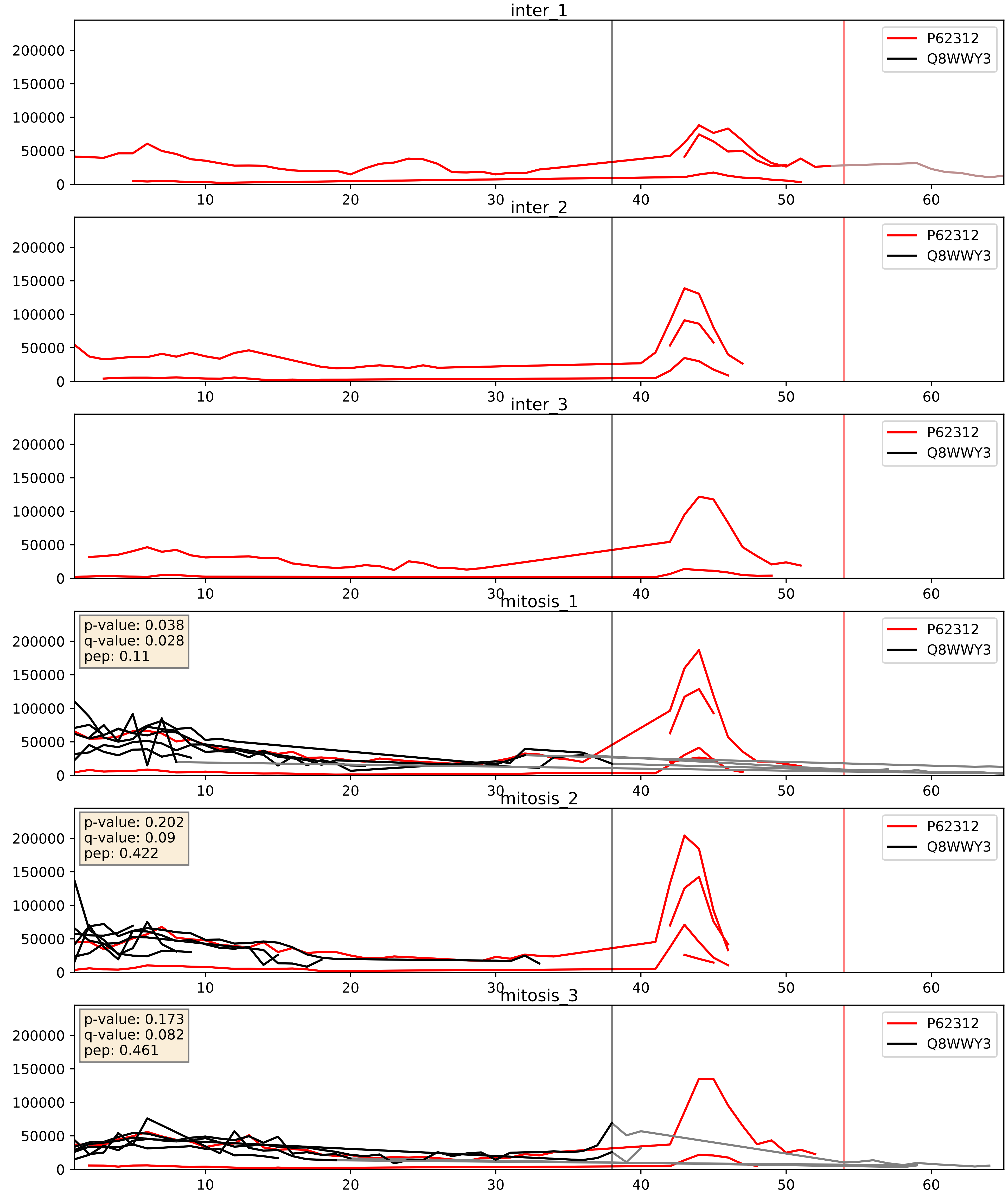


| level | condition_1 | condition_2 | pvalue | pvalue_adjusted |
|---|---|---|---|---|
| interactor_ratio | mitosis | inter | 0.0012760566415934561 | 0.0423373831474418 |
| complex_abundance | mitosis | inter | 0.0055591609825677984 | 0.05624098855659709 |
| interactor_abundance | mitosis | inter | 0.010224616160651864 | 0.0879625269700301 |

Q53GS9_Q8WWY3
SNUT2_HUMAN vs PRP31_HUMAN
p-value: 0.075 q-value: 0.045

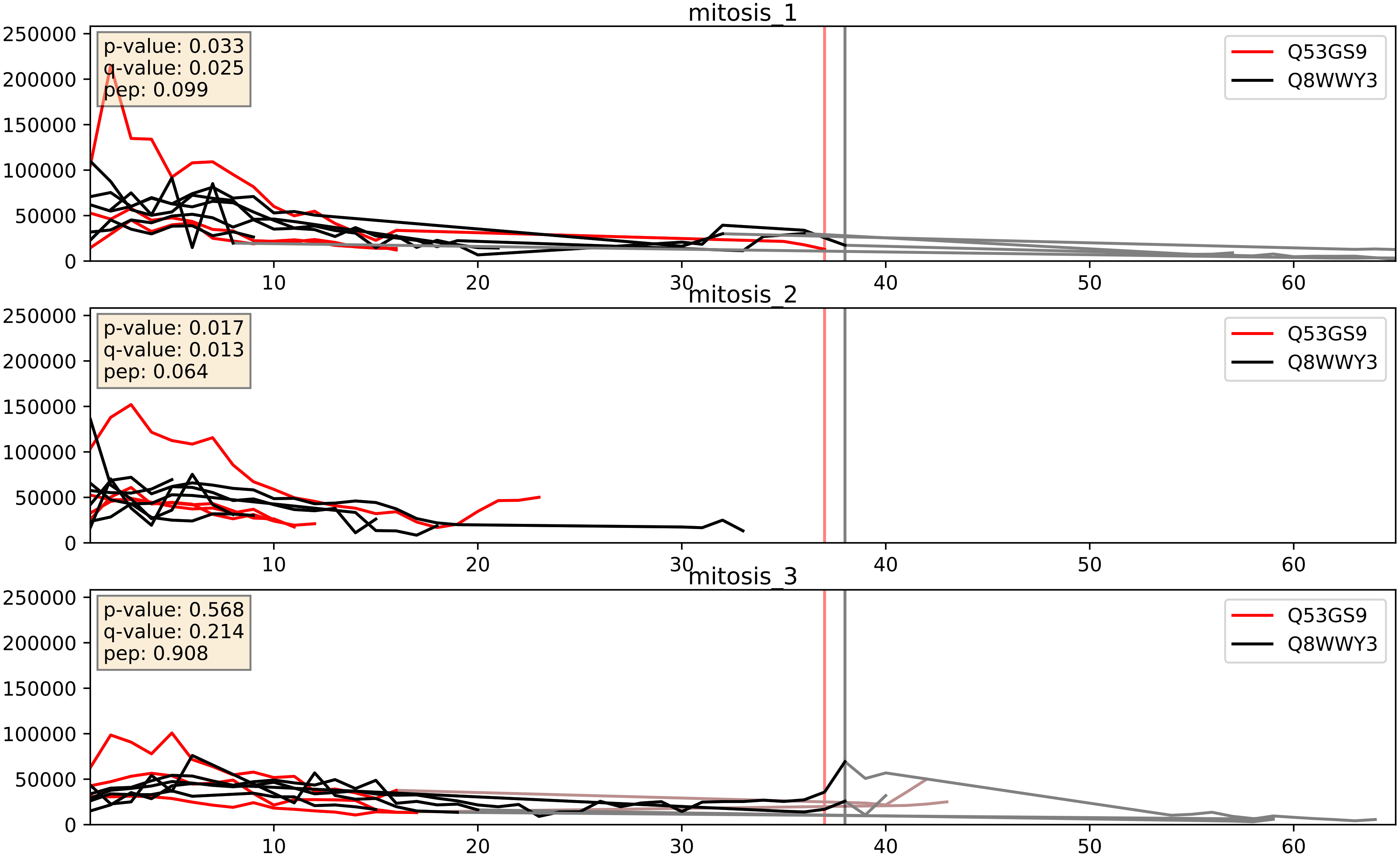


| level | condition_1 | condition_2 | pvalue | pvalue_adjusted |
|---|---|---|---|---|
| interactor_abundance | mitosis | inter | 0.0016940513754034087 | 0.030042528708242176 |
| complex_abundance | mitosis | inter | 0.0028637689480298383 | 0.03901194445107684 |
| interactor_ratio | mitosis | inter | 0.2851403101310909 | 0.755432081312949 |
