## Supplementary Data 2 for "SECAT: Quantifying differential protein-protein interaction states by network-centric analysis": hela_string_000094_Q13435.pdf

Q13435_Q13435

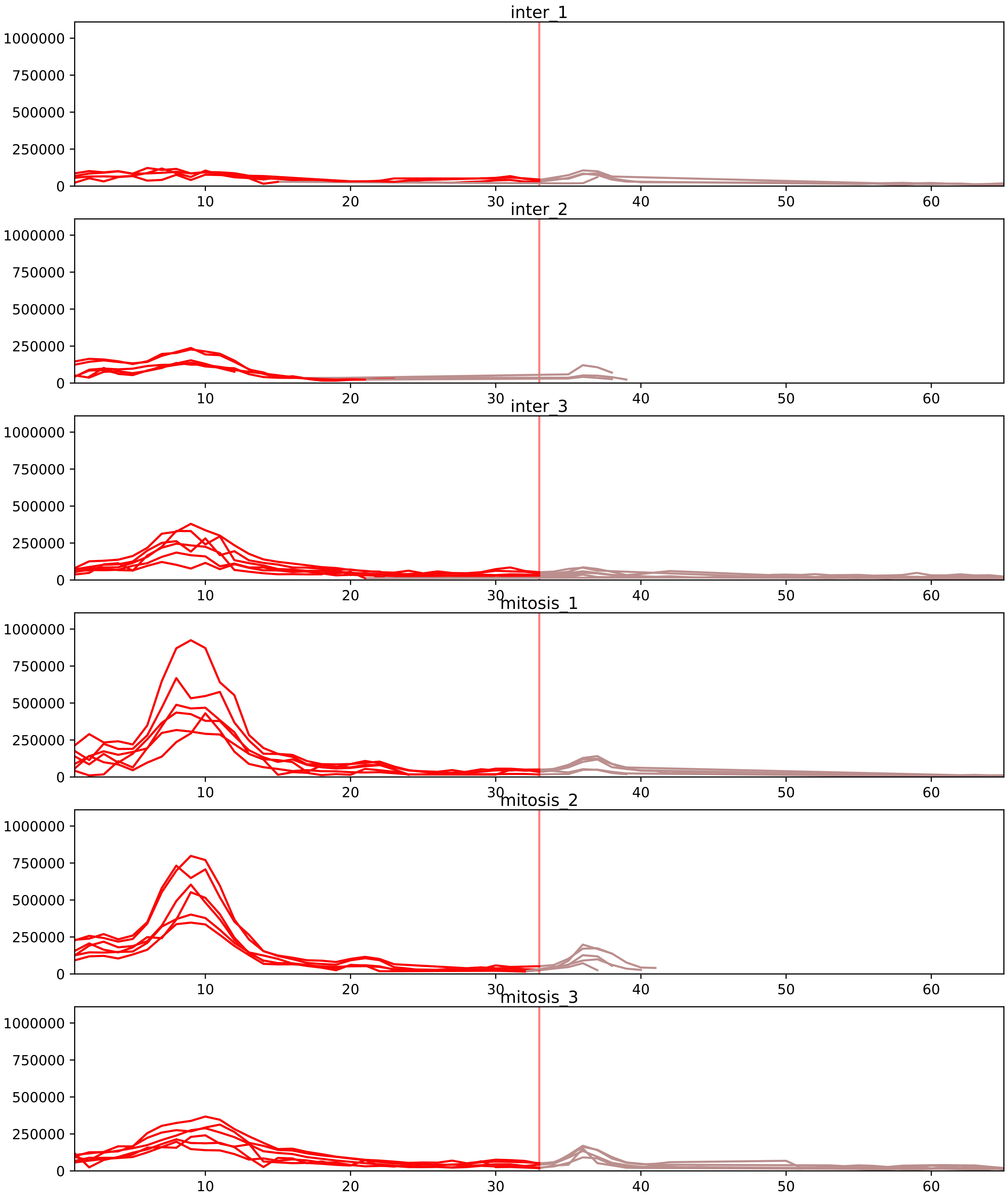


| level | condition_1 | condition_2 | pvalue | pvalue_adjusted |
|---|---|---|---|---|
| interactor_ratio | mitosis | inter | 6.3443197917796615e-06 | 0.00015160452489447503 |
| complex_abundance | mitosis | inter | 0.0024689594874622504 | 0.019037048720612647 |
| interactor_abundance | mitosis | inter | 0.006091618295058646 | 0.03889853441851164 |
| assembled_abundance | mitosis | inter | 0.06254226337237366 | 0.297242307517884 |
| total_abundance | mitosis | inter | 0.07307520506329829 | 0.33680096849796026 |
| monomer_abundance | mitosis | inter | 0.1803582493730838 | 0.5467791907178109 |

O75533_Q13435
SF3B1_HUMAN vs SF3B2_HUMAN
p-value: 0.001 q-value: 0.001

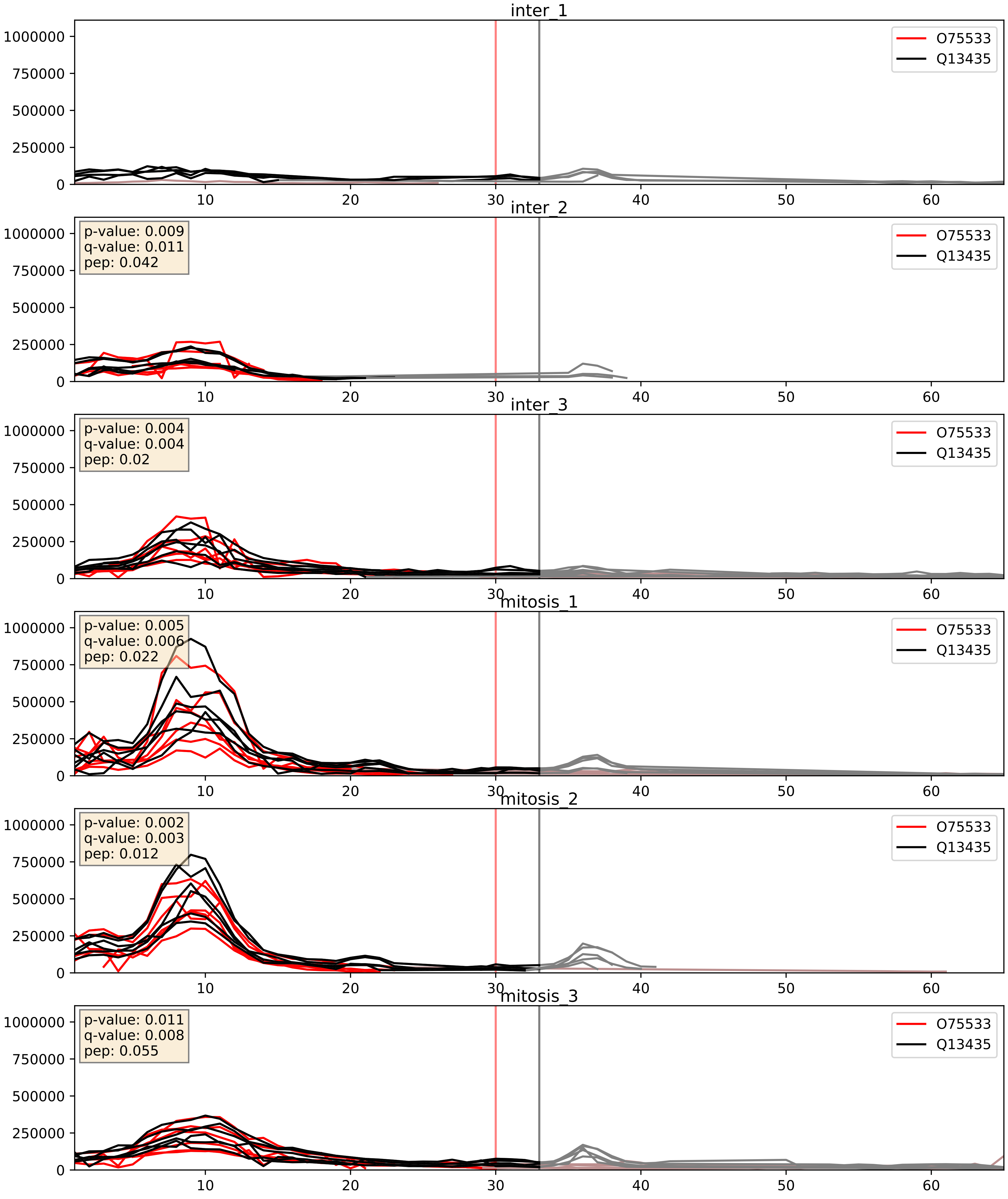


| level | condition_1 | condition_2 | pvalue | pvalue_adjusted |
|---|---|---|---|---|
| complex_abundance | mitosis | inter | 0.1915359666929163 | 0.34278650949014505 |
| interactor_abundance | mitosis | inter | 0.19730409548515387 | 0.3912230867423214 |
| interactor_ratio | mitosis | inter | 0.36652614850366505 | 0.8108720709709106 |

P07910_Q13435
HNRPC_HUMAN vs SF3B2_HUMAN
p-value: 0.002 q-value: 0.031

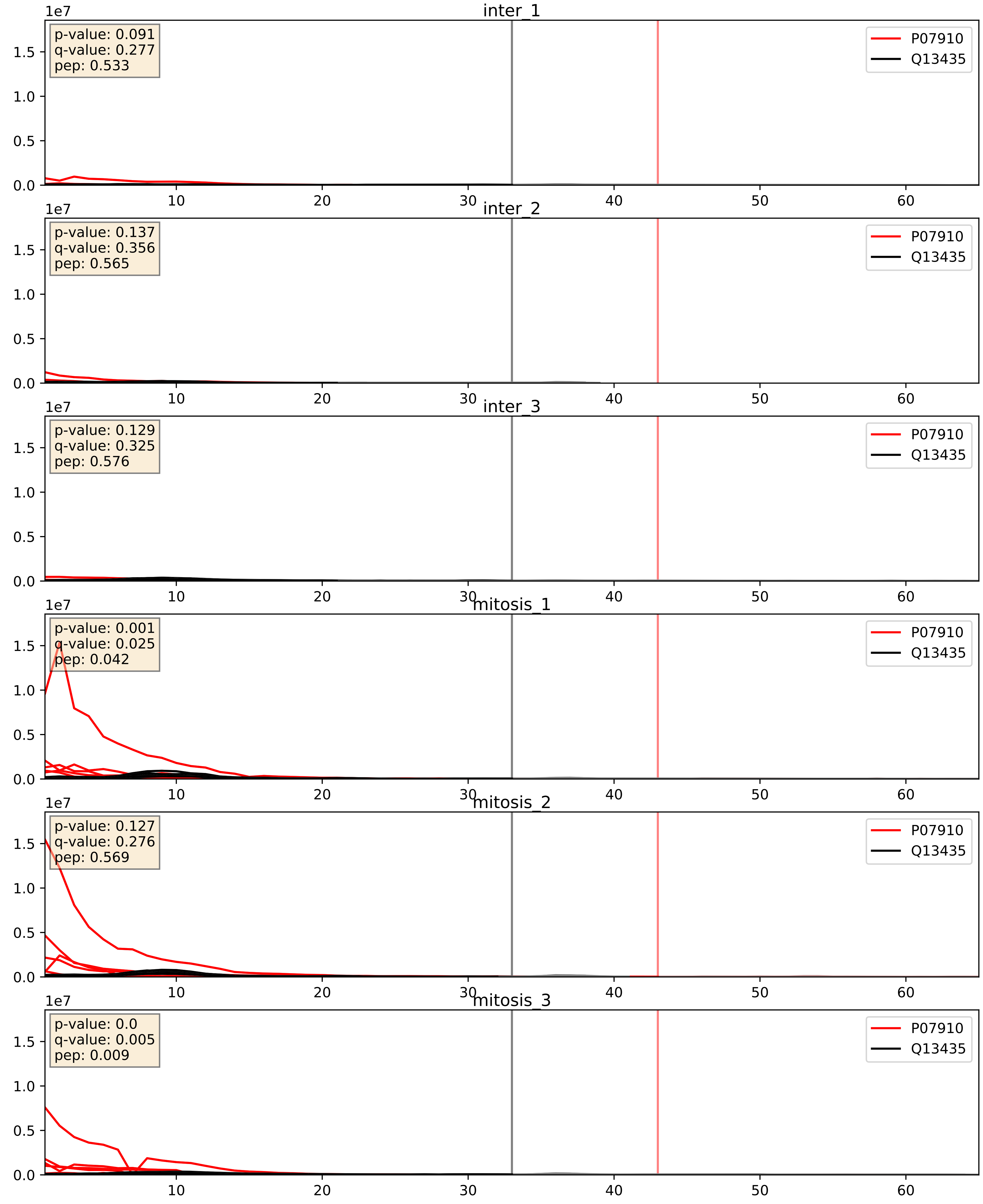


| level | condition_1 | condition_2 | pvalue | pvalue_adjusted |
|---|---|---|---|---|
| interactor_abundance | mitosis | inter | 9.628495089762232e-06 | 0.0033734327106189023 |
| complex_abundance | mitosis | inter | 3.2188890627744467e-05 | 0.005691857151652958 |
| interactor_ratio | mitosis | inter | 0.0016860726879665898 | 0.049768214513772446 |

P08579_Q13435
RU2B_HUMAN vs SF3B2_HUMAN
p-value: 0.059 q-value: 0.037

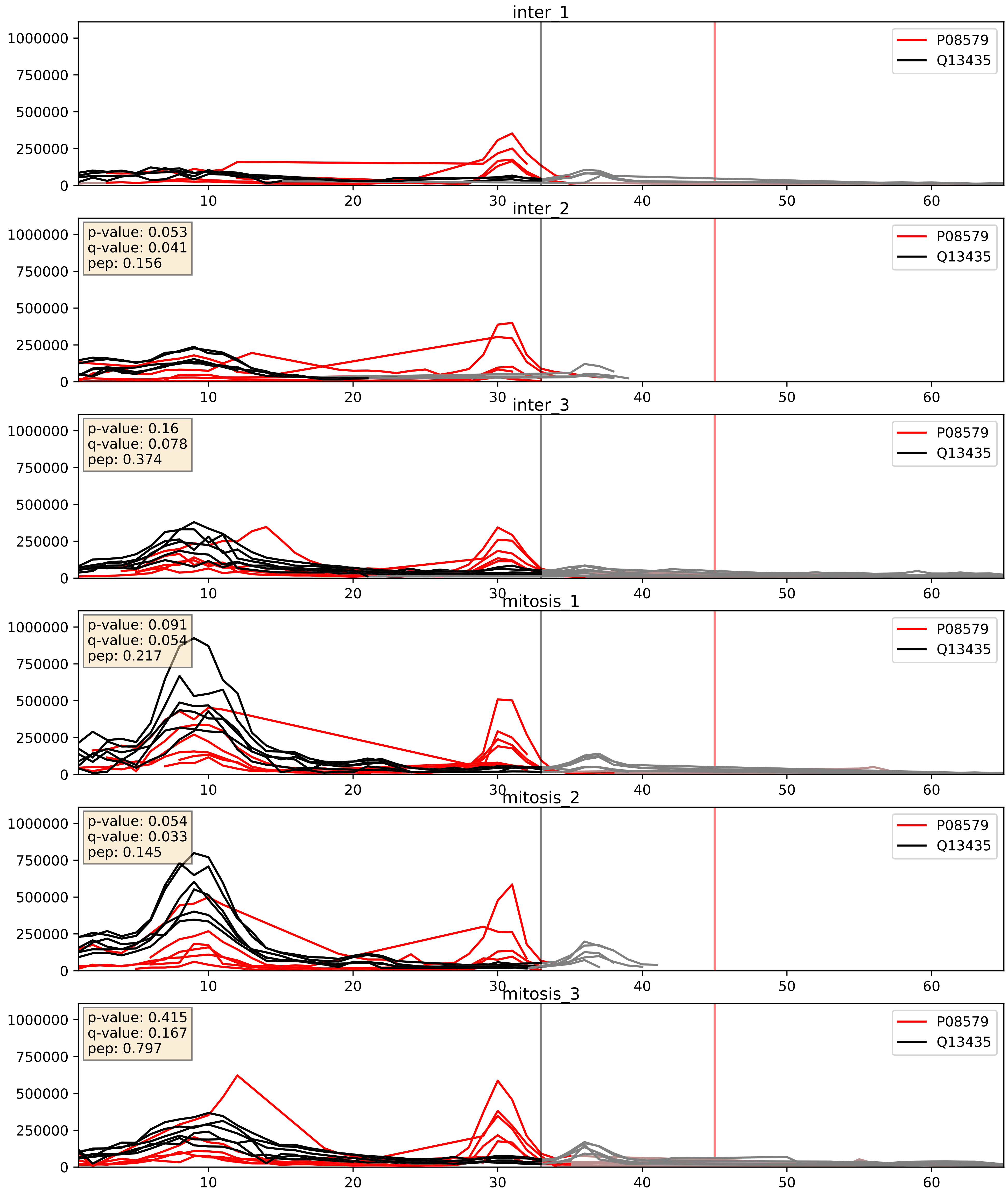


| level | condition_1 | condition_2 | pvalue | pvalue_adjusted |
|---|---|---|---|---|
| complex_abundance | mitosis | inter | 0.06554372227058855 | 0.18507877232212874 |
| interactor_abundance | mitosis | inter | 0.13904023715593933 | 0.3255326440824951 |
| interactor_ratio | mitosis | inter | 0.544800808174376 | 0.8859730852936224 |

P09661_Q13435
RU2A_HUMAN vs SF3B2_HUMAN
p-value: 0.03 q-value: 0.022

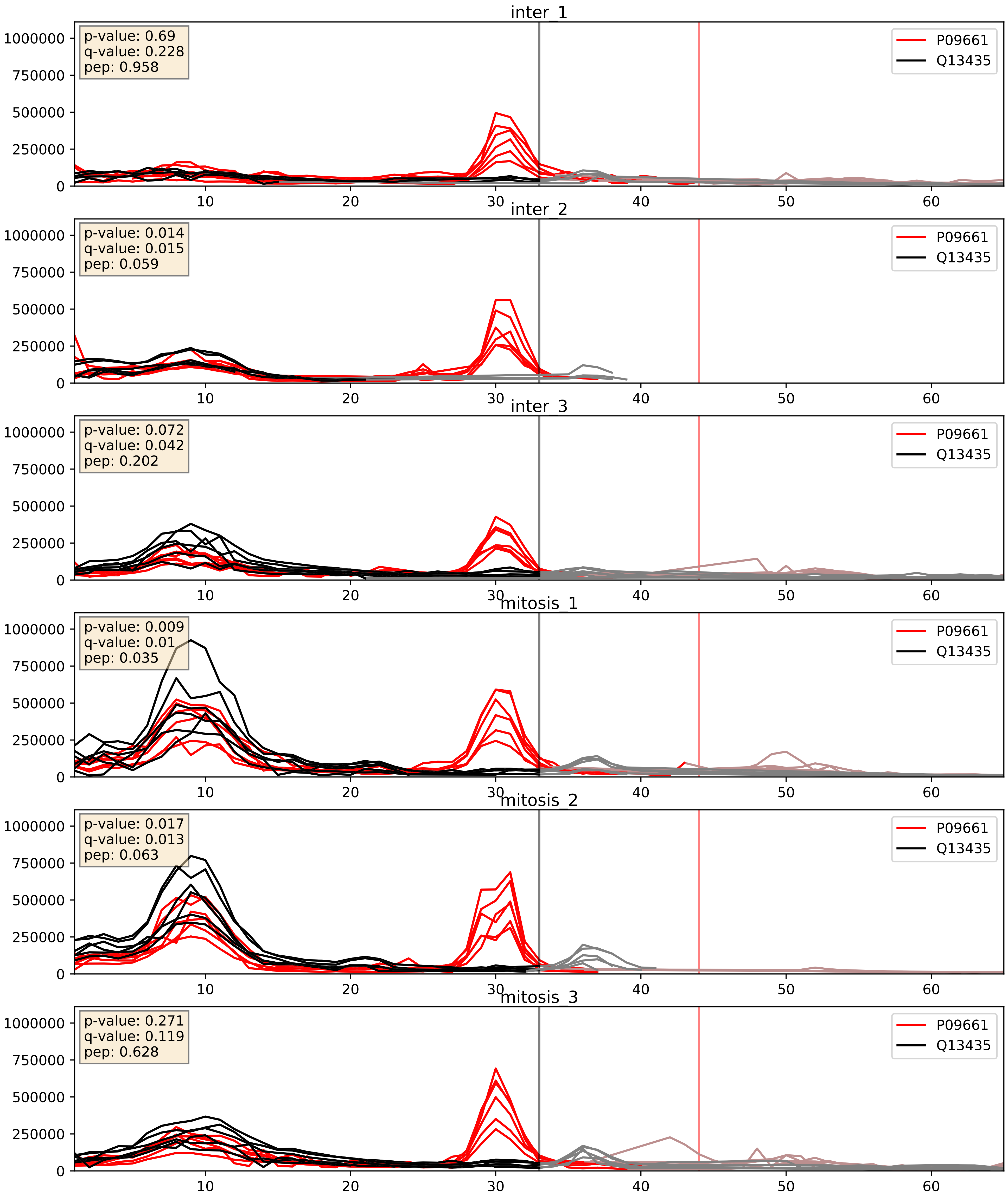


| level | condition_1 | condition_2 | pvalue | pvalue_adjusted |
|---|---|---|---|---|
| complex_abundance | mitosis | inter | 0.04454735172246902 | 0.15090040789249498 |
| interactor_abundance | mitosis | inter | 0.1323652811548473 | 0.3178814938442557 |
| interactor_ratio | mitosis | inter | 0.5154031872674406 | 0.8774122409401364 |

P62310_Q13435
LSM3_HUMAN vs SF3B2_HUMAN
p-value: 0.018 q-value: 0.014

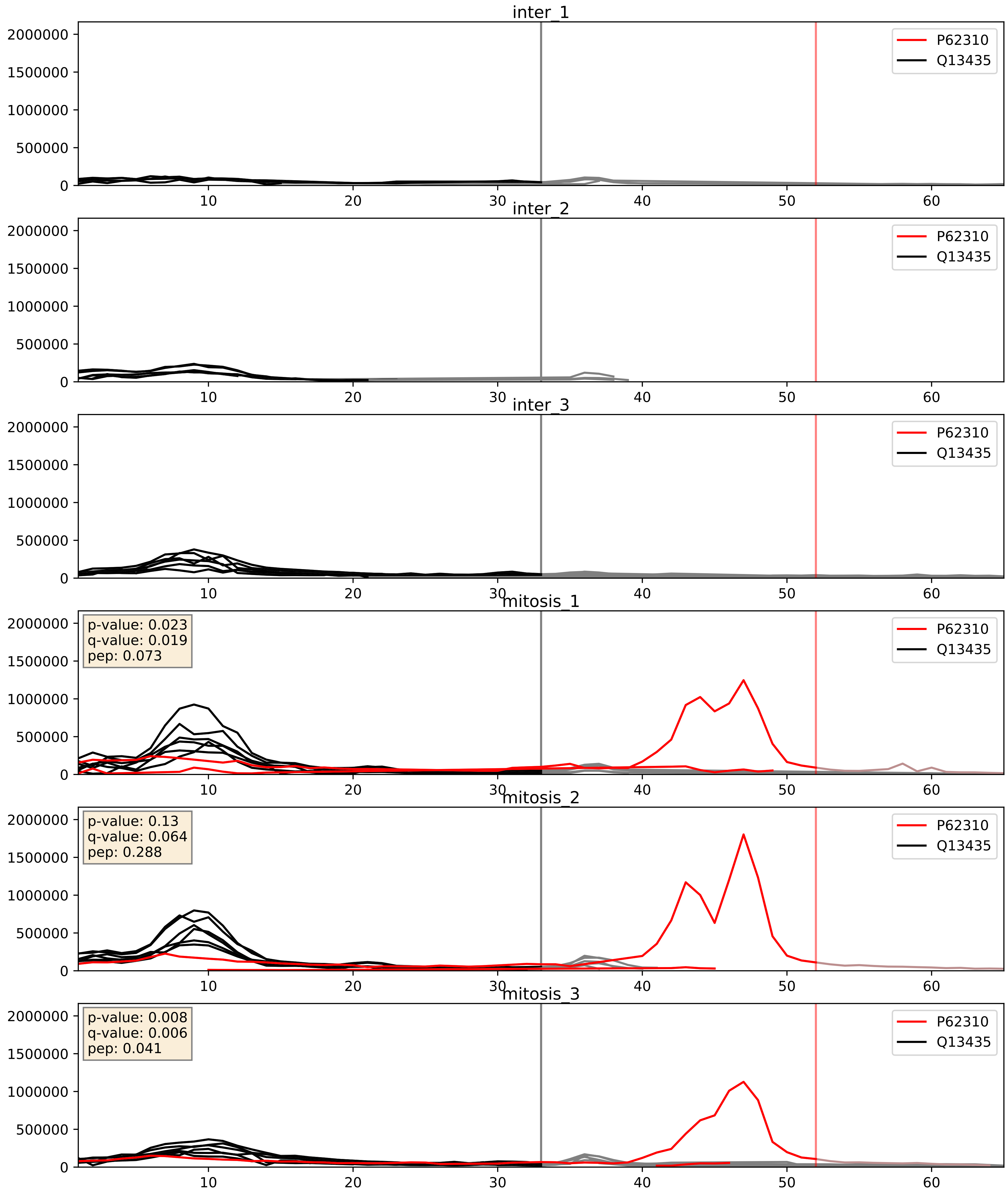


| level | condition_1 | condition_2 | pvalue | pvalue_adjusted |
|---|---|---|---|---|
| interactor_ratio | mitosis | inter | 4.374856816950016e-06 | 0.002520386835947235 |
| complex_abundance | mitosis | inter | 0.00047615701417451177 | 0.013816623868928207 |
| interactor_abundance | mitosis | inter | 0.011614432178085941 | 0.0950928163026453 |

Q07065_Q13435
CKAP4_HUMAN vs SF3B2_HUMAN
p-value: 0.0 q-value: 0.047

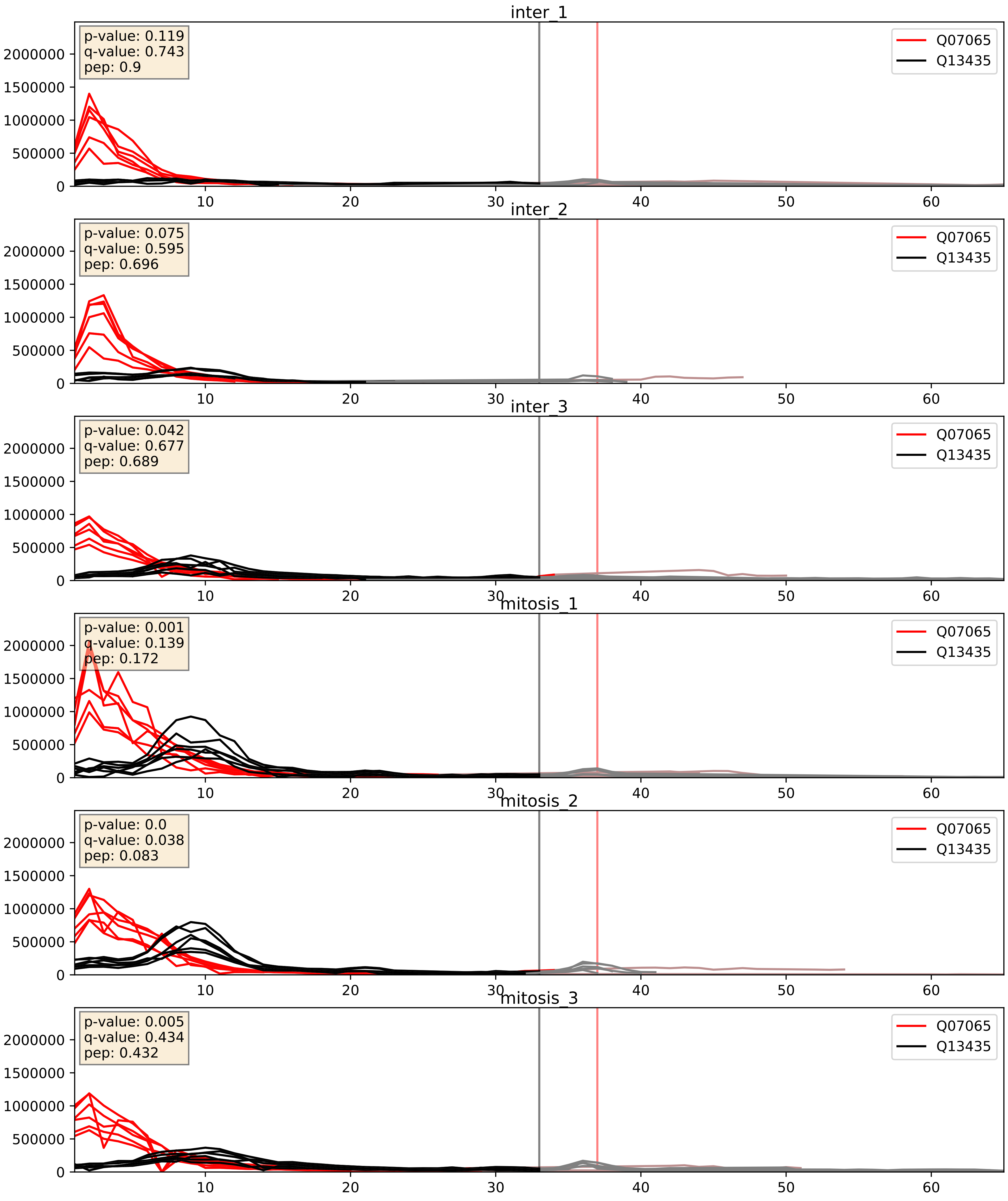


| level | condition_1 | condition_2 | pvalue | pvalue_adjusted |
|---|---|---|---|---|
| complex_abundance | mitosis | inter | 0.05729511254626756 | 0.17076816274235734 |
| interactor_ratio | mitosis | inter | 0.08634828062792886 | 0.4940783971758496 |
| interactor_abundance | mitosis | inter | 0.2900551488527625 | 0.48443083025939737 |

Q13435_Q15029
SF3B2_HUMAN vs U5S1_HUMAN
p-value: 0.029 q-value: 0.021

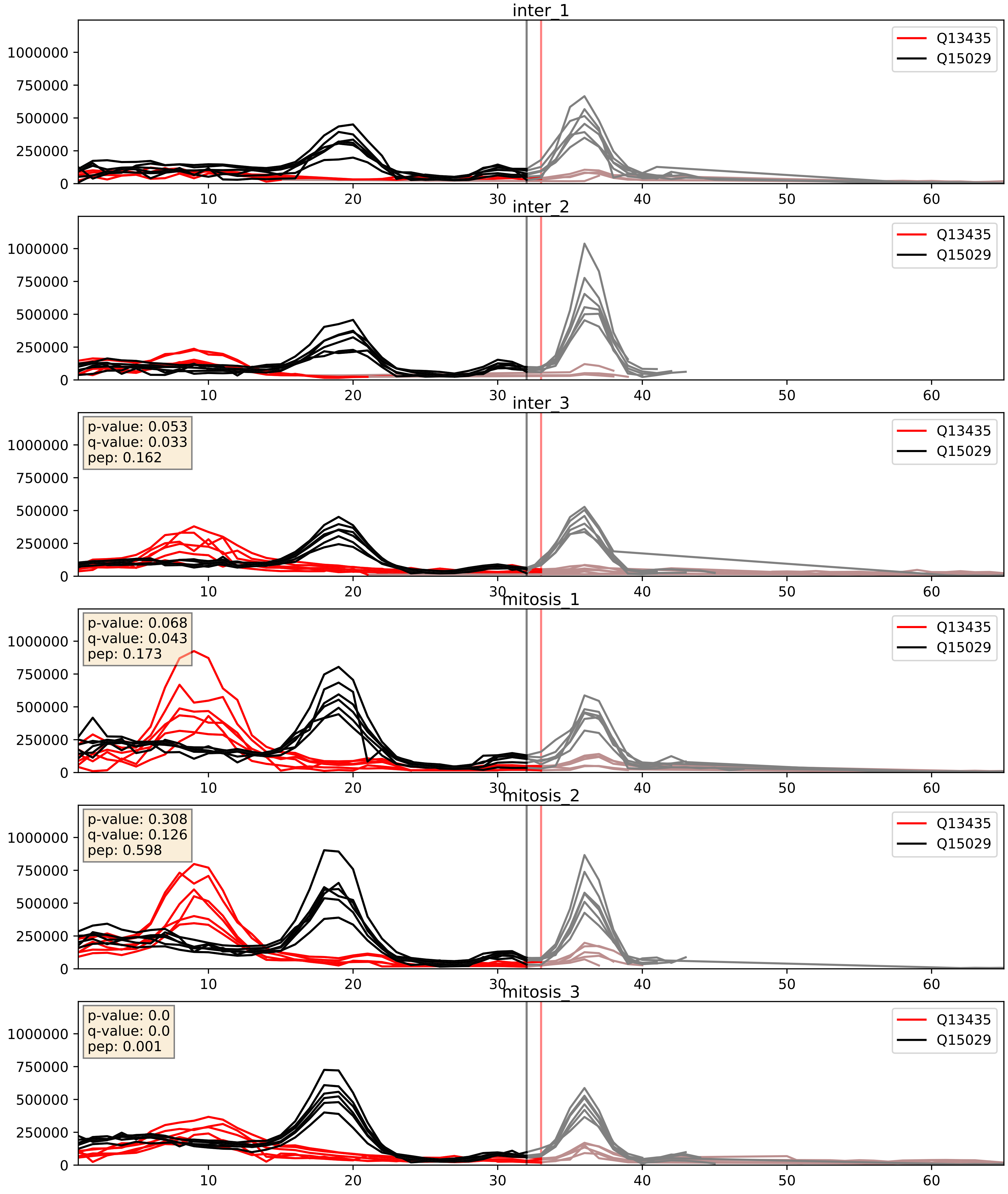


| level | condition_1 | condition_2 | pvalue | pvalue_adjusted |
|---|---|---|---|---|
| complex_abundance | mitosis | inter | 0.013064970538229406 | 0.08453223568196803 |
| interactor_abundance | mitosis | inter | 0.04372825972762542 | 0.16685321787696922 |
| interactor_ratio | mitosis | inter | 0.24309245013543793 | 0.7200661897289671 |

Q13435_Q15393
SF3B2_HUMAN vs SF3B3_HUMAN
p-value: 0.0 q-value: 0.0

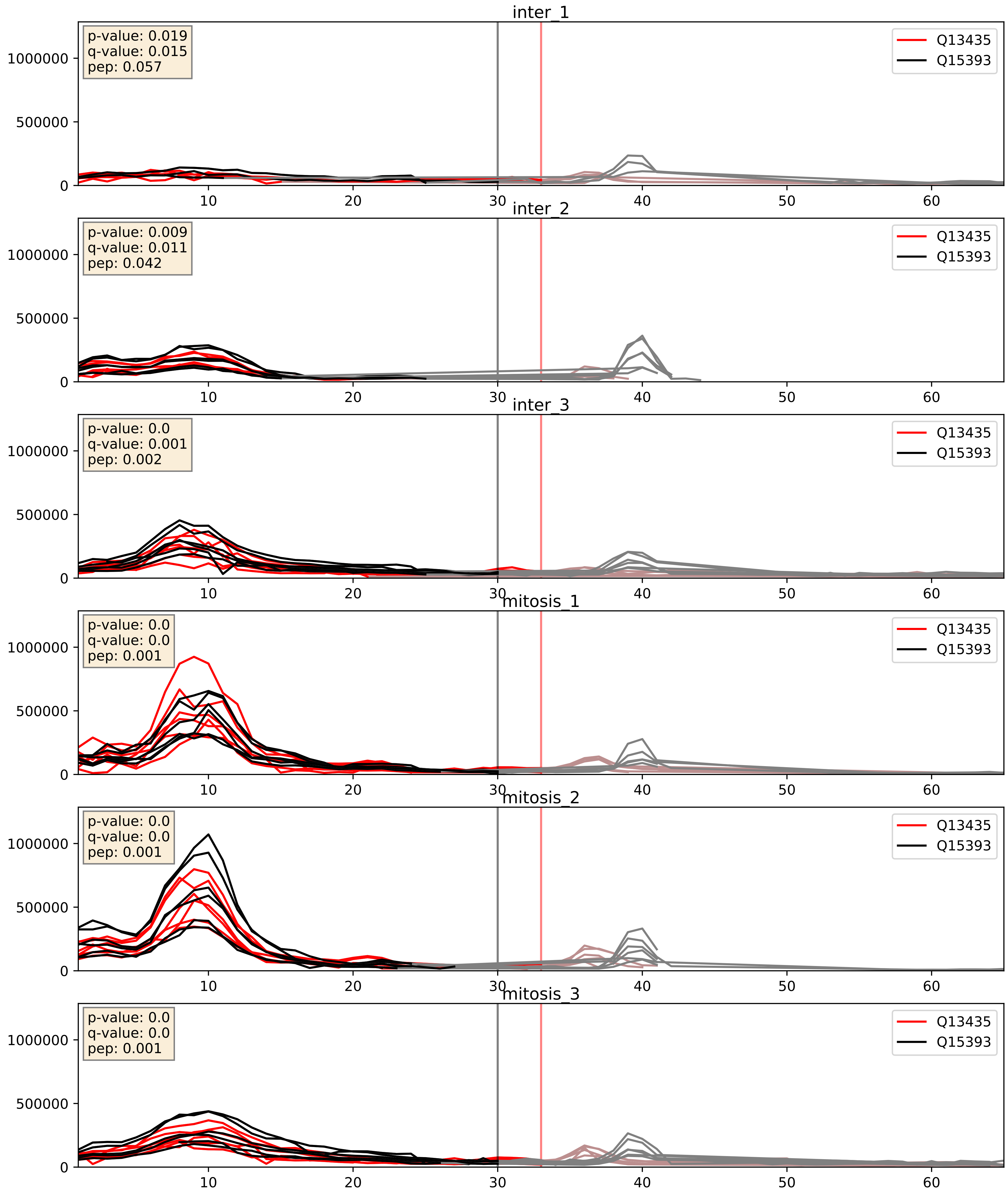


| level | condition_1 | condition_2 | pvalue | pvalue_adjusted |
|---|---|---|---|---|
| interactor_abundance | mitosis | inter | 0.0450455229676067 | 0.16891232828208008 |
| complex_abundance | mitosis | inter | 0.09093641383443817 | 0.22177085539110847 |
| interactor_ratio | mitosis | inter | 0.7846881966520124 | 0.95900240448312 |

Q13435_Q15459
SF3B2_HUMAN vs SF3A1_HUMAN
p-value: 0.024 q-value: 0.018

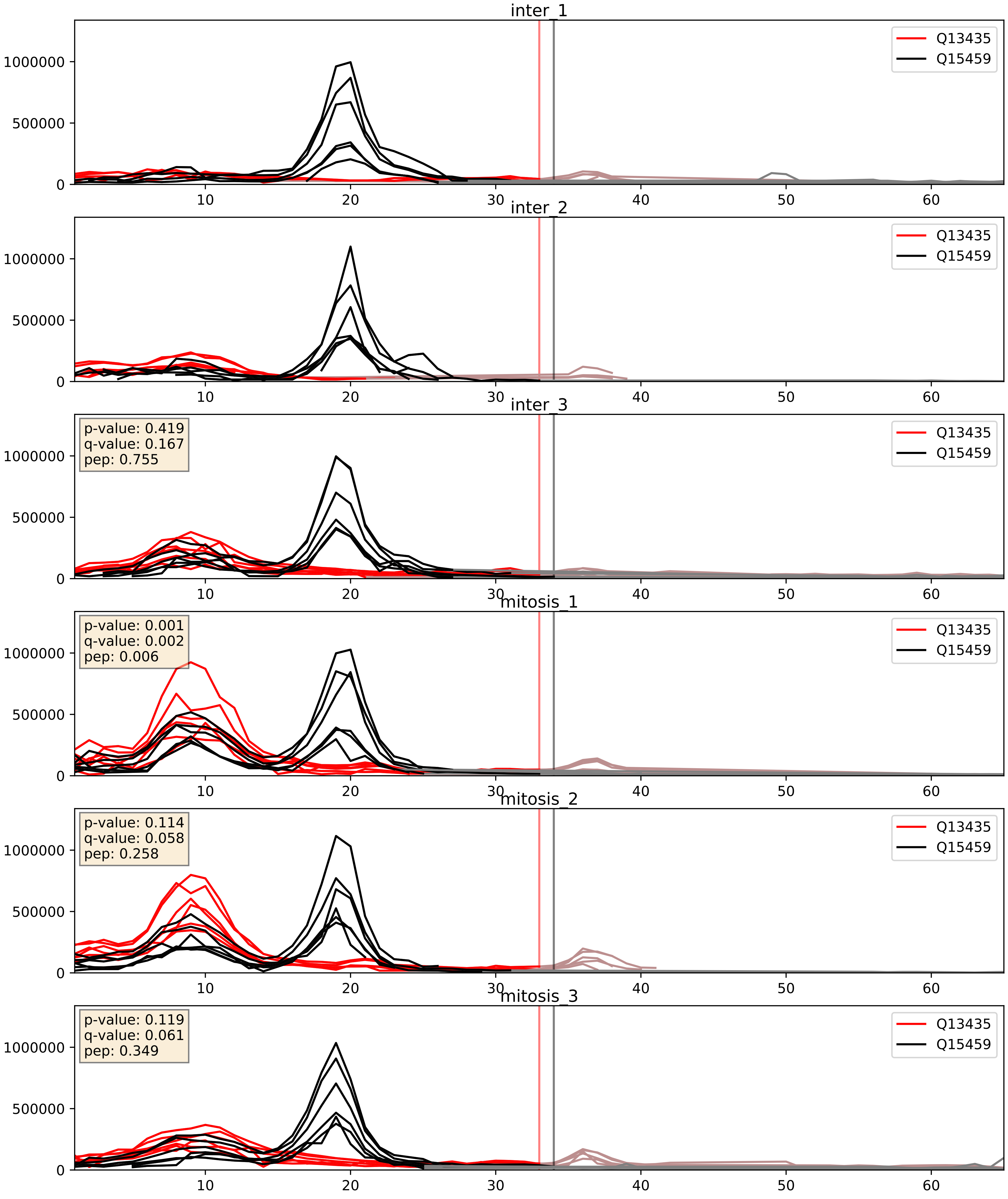


| level | condition_1 | condition_2 | pvalue | pvalue_adjusted |
|---|---|---|---|---|
| interactor_abundance | mitosis | inter | 0.04420415818007957 | 0.16761355216898388 |
| interactor_ratio | mitosis | inter | 0.05213934207934958 | 0.37764936302518703 |
| complex_abundance | mitosis | inter | 0.08983876847370324 | 0.22016028002716856 |

Q13435_Q7RTV0
SF3B2_HUMAN vs PHF5A_HUMAN
p-value: 0.047 q-value: 0.031

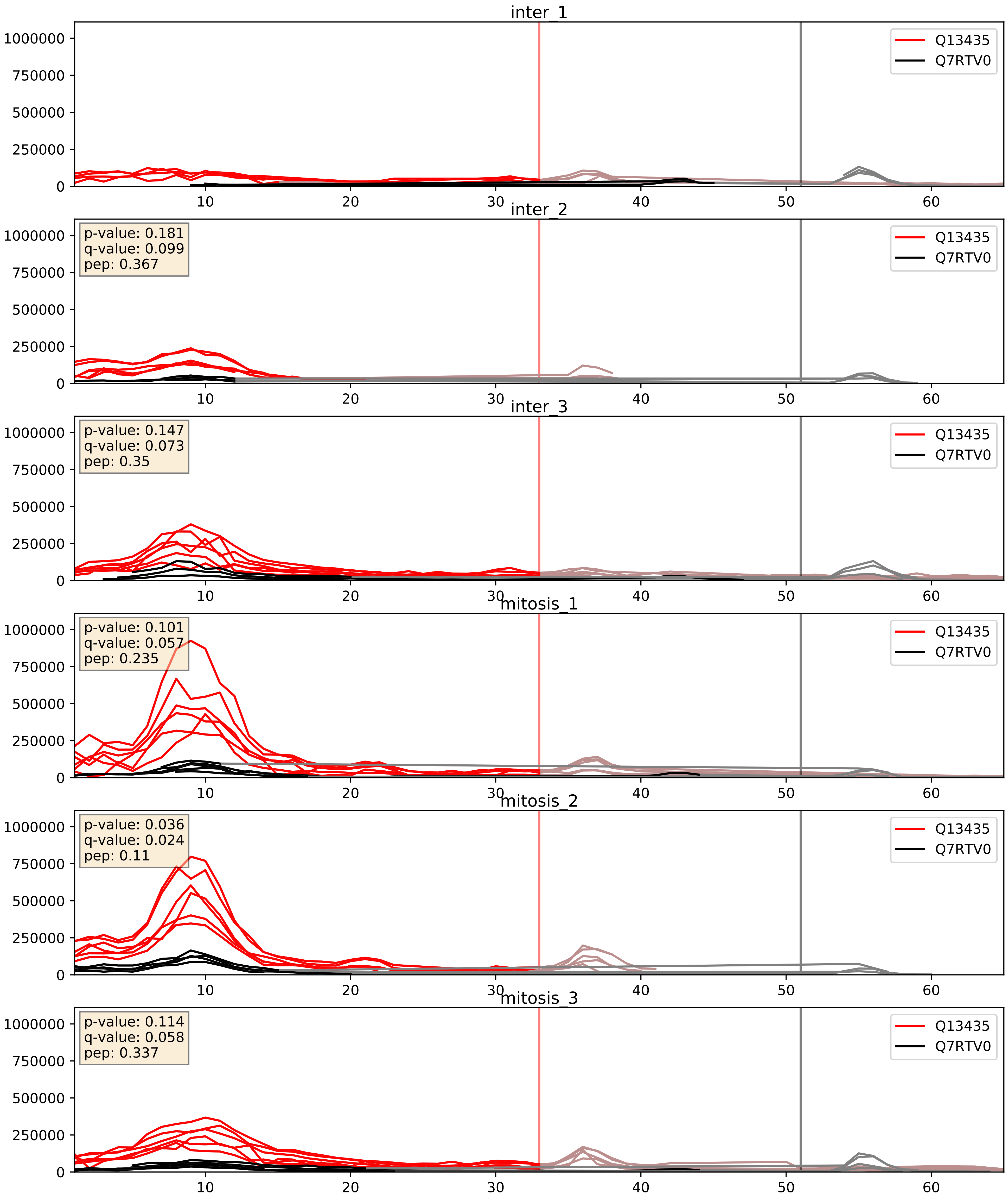


| level | condition_1 | condition_2 | pvalue | pvalue_adjusted |
|---|---|---|---|---|
| interactor_ratio | mitosis | inter | 0.06268509562303749 | 0.4159147709510605 |
| interactor_abundance | mitosis | inter | 0.09496314304923419 | 0.26396639210957773 |
| complex_abundance | mitosis | inter | 0.12404703822362097 | 0.2659260323551704 |

Q13435_Q9Y3B4
SF3B2_HUMAN vs SF3B6_HUMAN
p-value: 0.015 q-value: 0.012

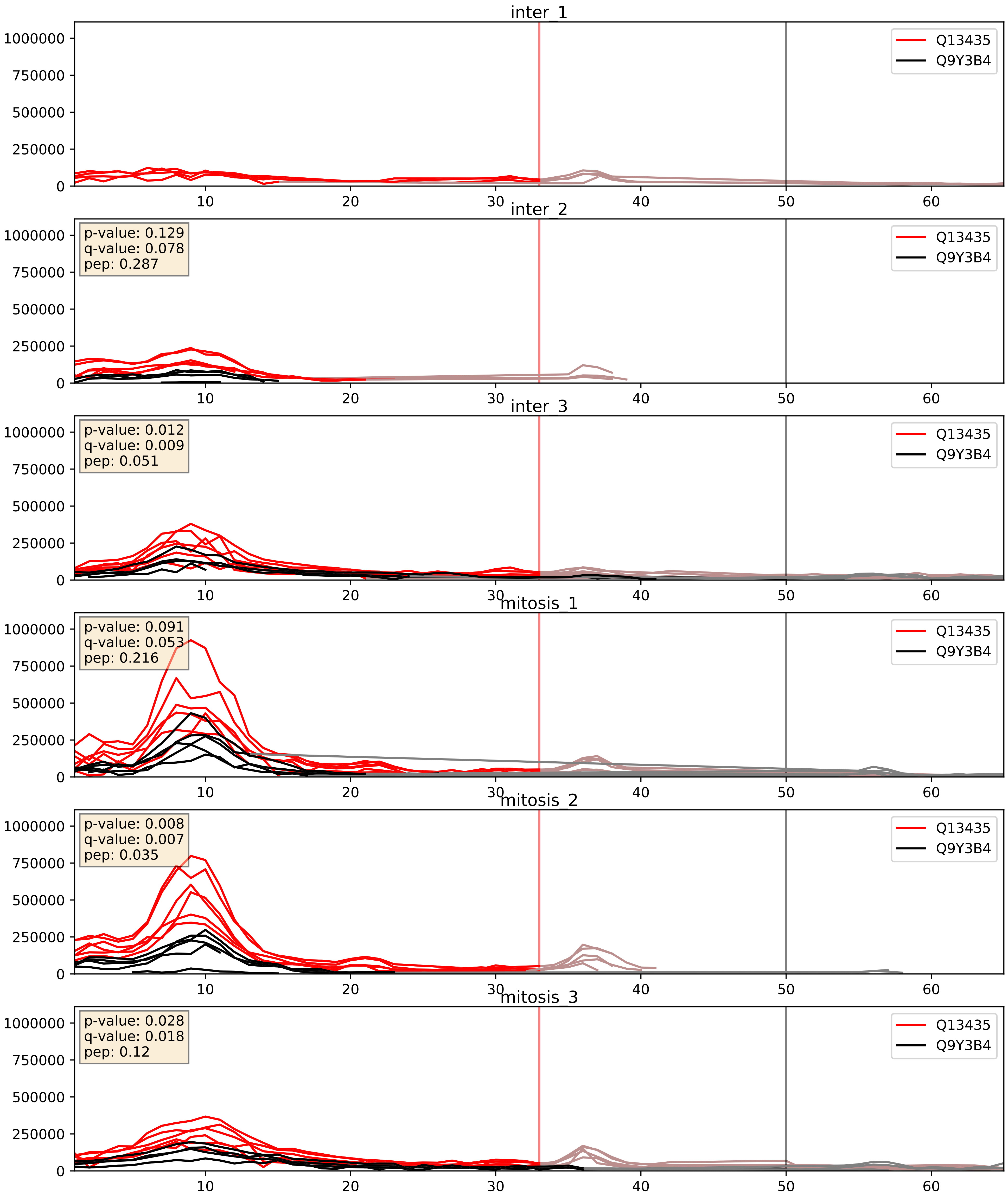


| level | condition_1 | condition_2 | pvalue | pvalue_adjusted |
|---|---|---|---|---|
| complex_abundance | mitosis | inter | 0.17263184262979864 | 0.32222602985413784 |
| interactor_abundance | mitosis | inter | 0.18112536448311178 | 0.3738686086268235 |
| interactor_ratio | mitosis | inter | 0.36928858964194816 | 0.8138388063636502 |
