## Supplementary Data 2 for "SECAT: Quantifying differential protein-protein interaction states by network-centric analysis": hela_string_000095_Q14103.pdf

# Q14103_Q14103

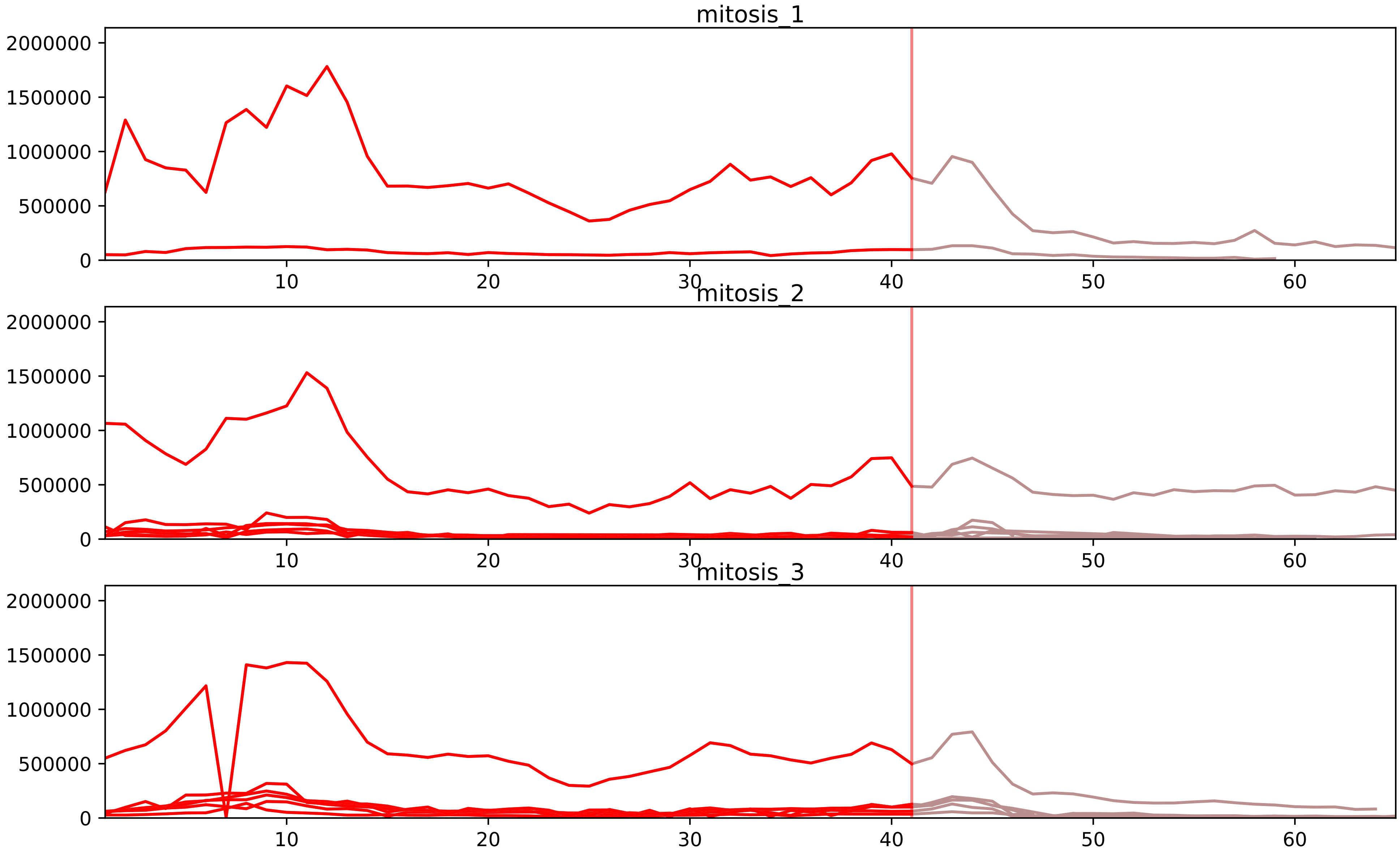


| level | condition_1 | condition_2 | pvalue | pvalue_adjusted |
|---|---|---|---|---|
| complex_abundance | mitosis | inter | 6.788620900151557e-06 | 0.0003785170441296625 |
| interactor_abundance | mitosis | inter | 0.0010317845493509612 | 0.011834086381599693 |
| assembled_abundance | mitosis | inter | 0.014082355765515184 | 0.13563684637526044 |
| total_abundance | mitosis | inter | 0.014854171900012666 | 0.15464742156025454 |
| monomer_abundance | mitosis | inter | 0.015150747154044283 | 0.21065310922095573 |
| interactor_ratio | mitosis | inter | 0.07284872971955096 | 0.26710158379363347 |

P09651_Q14103
ROA1_HUMAN vs HNRPD_HUMAN
p-value: 0.031 q-value: 0.023

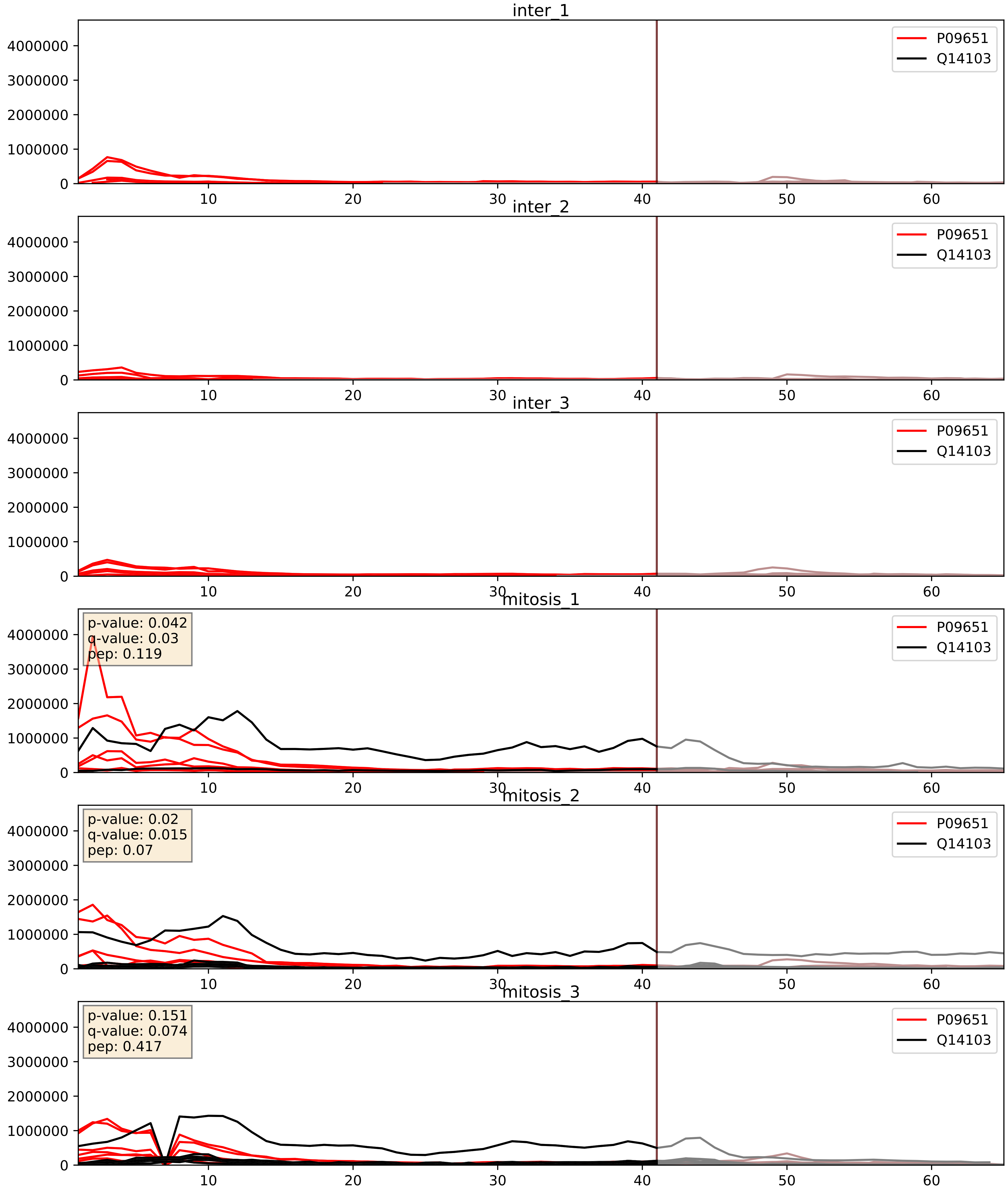


| level | condition_1 | condition_2 | pvalue | pvalue_adjusted |
|---|---|---|---|---|
| interactor_abundance | mitosis | inter | 6.959157312257744e-05 | 0.006732838333317079 |
| complex_abundance | mitosis | inter | 0.00029696456492007517 | 0.011617922212881656 |
| interactor_ratio | mitosis | inter | 0.26529417826222884 | 0.7404362309130512 |

P22626_Q14103
ROA2_HUMAN vs HNRPD_HUMAN
p-value: 0.036 q-value: 0.025

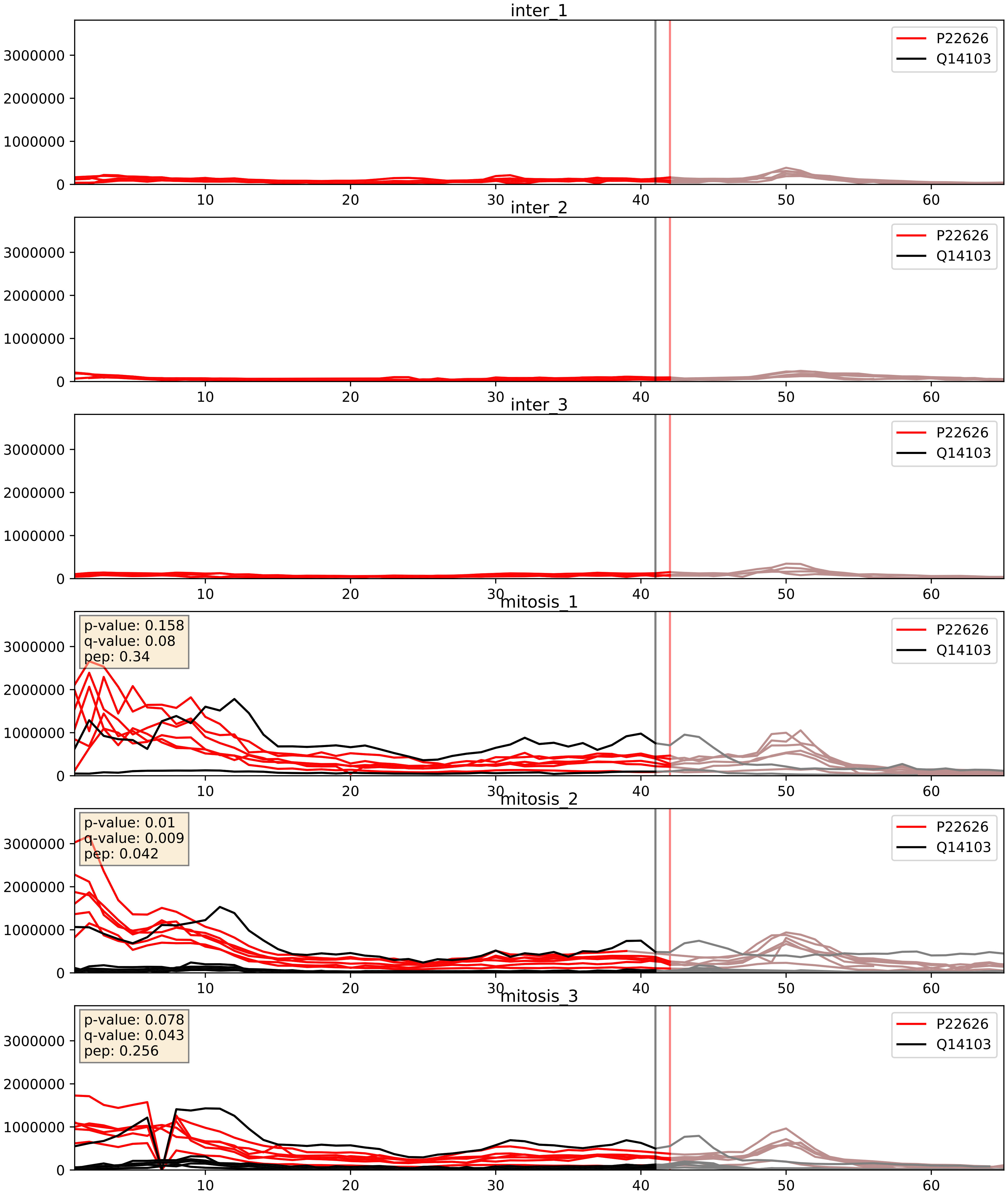


| level | condition_1 | condition_2 | pvalue | pvalue_adjusted |
|---|---|---|---|---|
| interactor_abundance | mitosis | inter | 2.1731713857435016e-05 | 0.004180302710553792 |
| complex_abundance | mitosis | inter | 0.0002607874150841398 | 0.010784252527150902 |
| interactor_ratio | mitosis | inter | 0.026748360525100004 | 0.2569763929235197 |

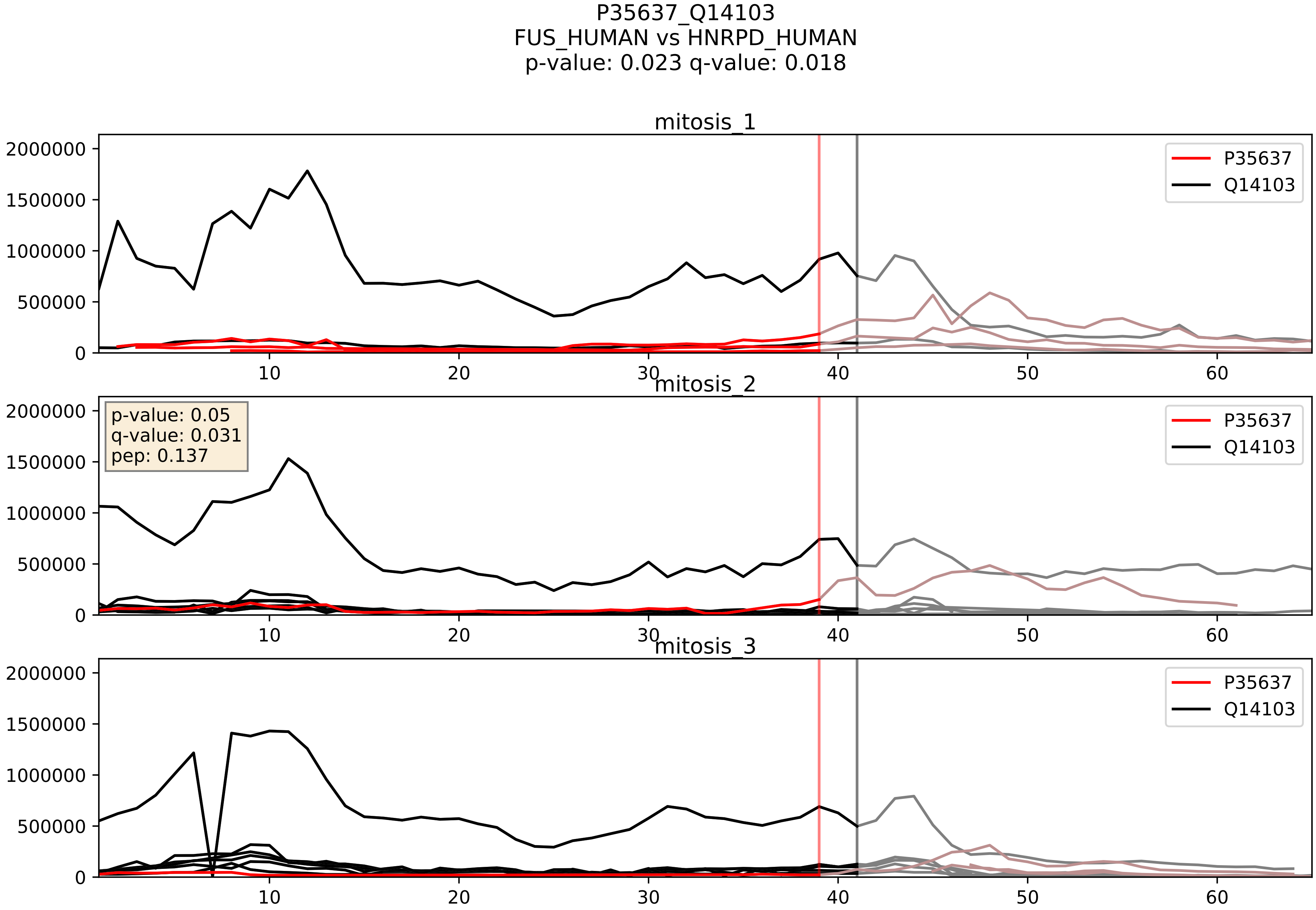


| level | condition_1 | condition_2 | pvalue | pvalue_adjusted |
|---|---|---|---|---|
| complex_abundance | mitosis | inter | 0.0004181087028958004 | 0.013091389064164169 |
| interactor_ratio | mitosis | inter | 0.112797983158615 | 0.553985317646331 |
| interactor_abundance | mitosis | inter | 0.34864704029911386 | 0.5474992964521033 |

P51991_Q14103
ROA3_HUMAN vs HNRPD_HUMAN
p-value: 0.037 q-value: 0.026

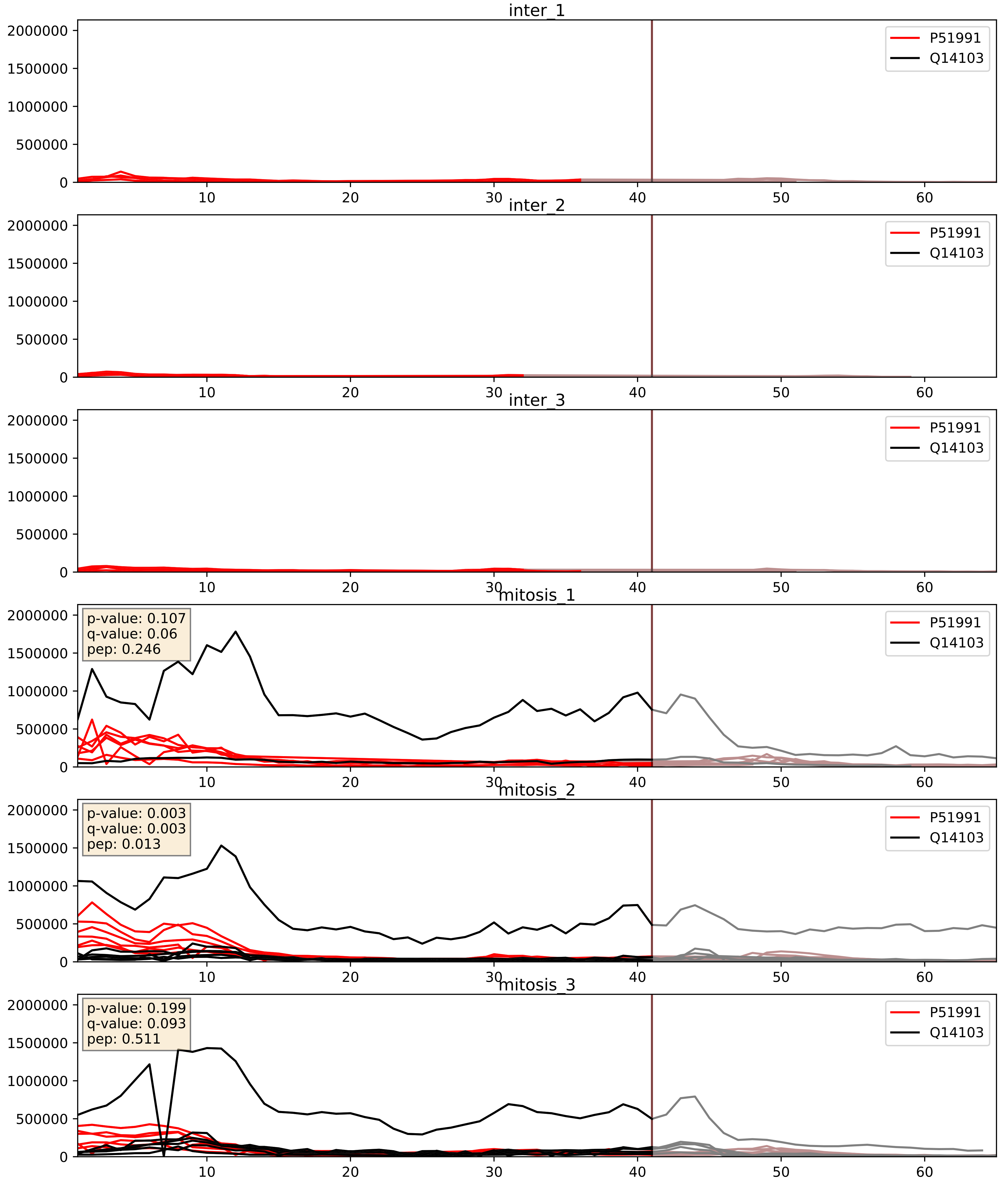


| level | condition_1 | condition_2 | pvalue | pvalue_adjusted |
|---|---|---|---|---|
| interactor_abundance | mitosis | inter | 1.150670630946923e-05 | 0.0033803141163598503 |
| complex_abundance | mitosis | inter | 0.0005845801120302033 | 0.015650423933737376 |
| interactor_ratio | mitosis | inter | 0.748127922946446 | 0.9491588884573259 |

Q13151_Q14103
ROA0_HUMAN vs HNRPD_HUMAN
p-value: 0.076 q-value: 0.046

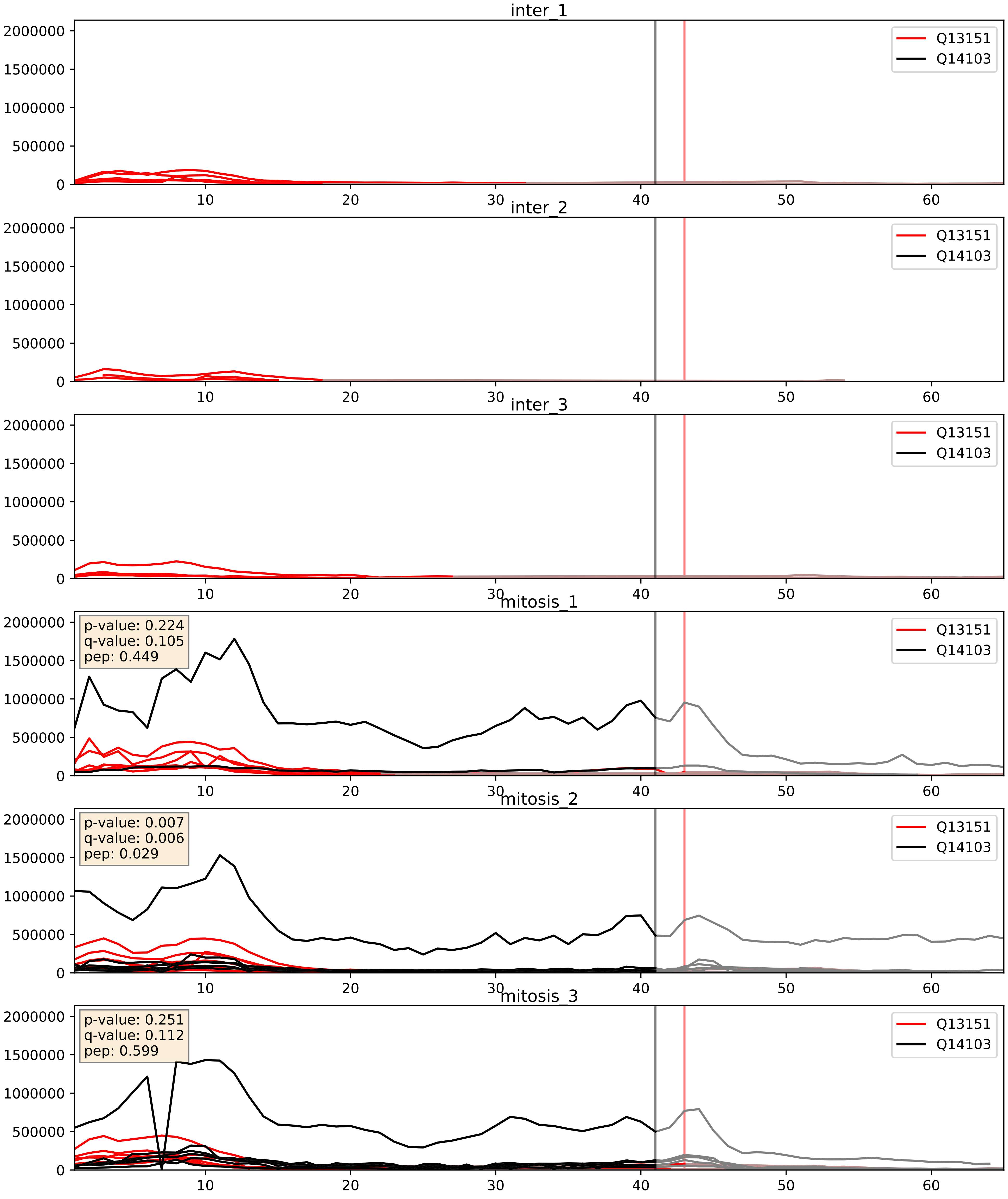


| level | condition_1 | condition_2 | pvalue | pvalue_adjusted |
|---|---|---|---|---|
| interactor_abundance | mitosis | inter | 2.9704194402271474e-05 | 0.0048431981730179775 |
| complex_abundance | mitosis | inter | 0.0009732578966867303 | 0.020686613267949534 |
| interactor_ratio | mitosis | inter | 0.7783913756132336 | 0.9574694891865612 |

Q14103_Q92973
HNRPD_HUMAN vs TNPO1_HUMAN
p-value: 0.032 q-value: 0.023

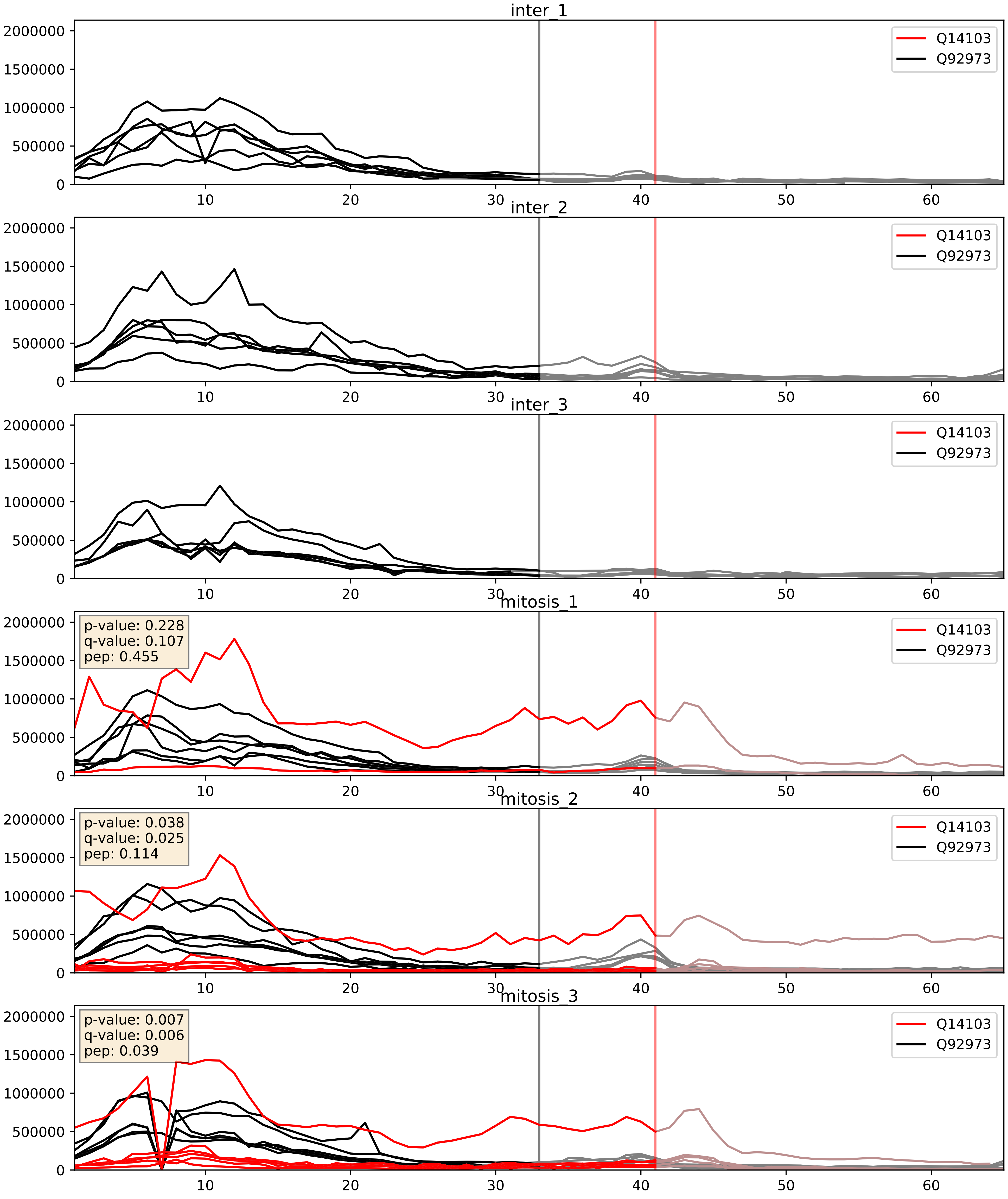


| level | condition_1 | condition_2 | pvalue | pvalue_adjusted |
|---|---|---|---|---|
| complex_abundance | mitosis | inter | 0.0010020915510917888 | 0.021075930411168824 |
| interactor_abundance | mitosis | inter | 0.010151249925242783 | 0.08768385404649669 |
| interactor_ratio | mitosis | inter | 0.05817972916925262 | 0.3974608792408638 |
