## Supplementary Data 2 for "SECAT: Quantifying differential protein-protein interaction states by network-centric analysis": hela_string_000096_Q96I24.pdf

Q96I24_Q96I24

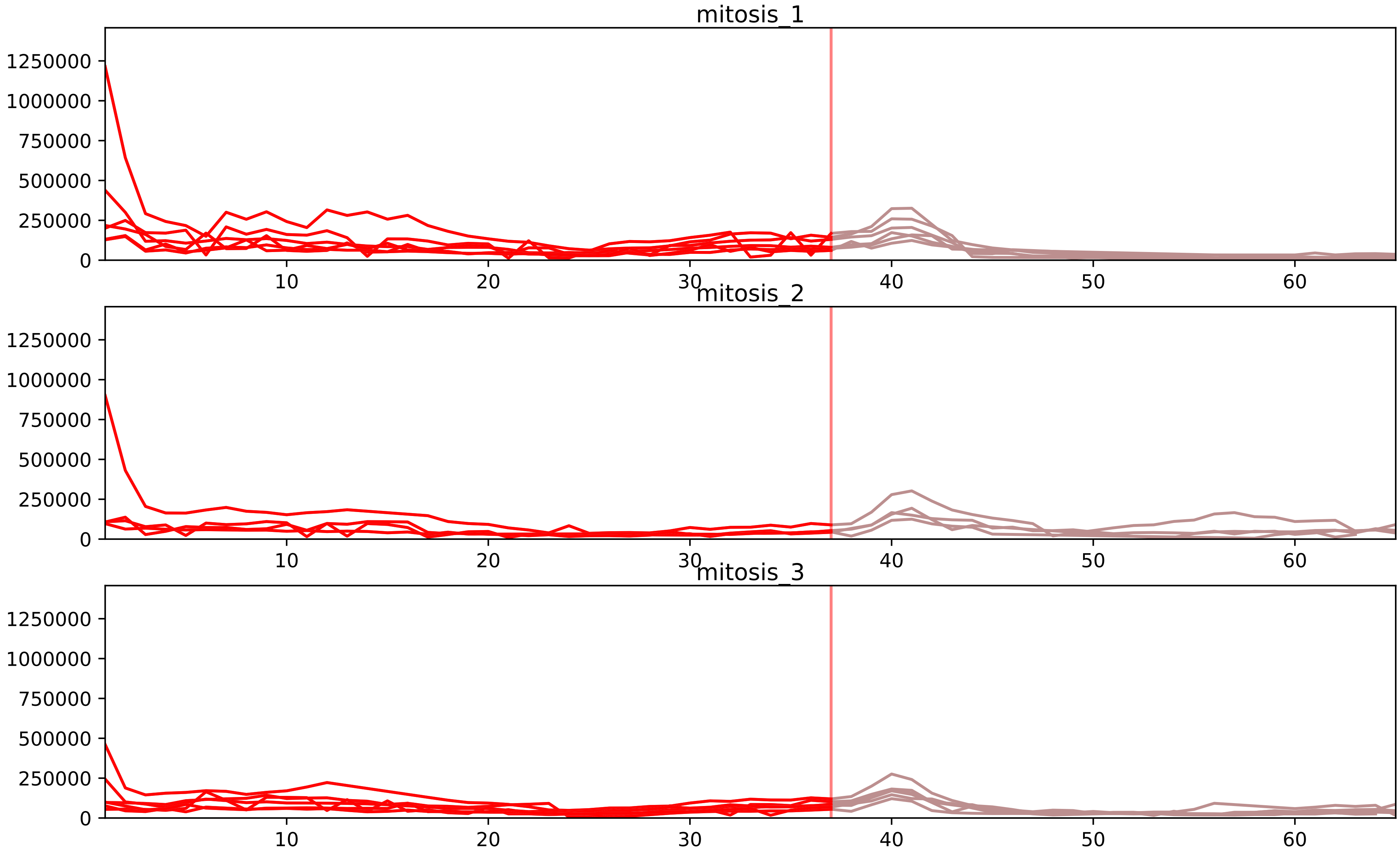


| level | condition_1 | condition_2 | pvalue | pvalue_adjusted |
|---|---|---|---|---|
| total_abundance | mitosis | inter | 7.043042412049715e-06 | 0.003984014324416122 |
| assembled_abundance | mitosis | inter | 4.4520238201982594e-05 | 0.007270487507555603 |
| monomer_abundance | mitosis | inter | 8.184919012349413e-05 | 0.03261473898341094 |
