## Supplementary Data 2 for "SECAT: Quantifying differential protein-protein interaction states by network-centric analysis": hela_string_000097_Q9BUJ2.pdf

Q9BUJ2_Q9BUJ2

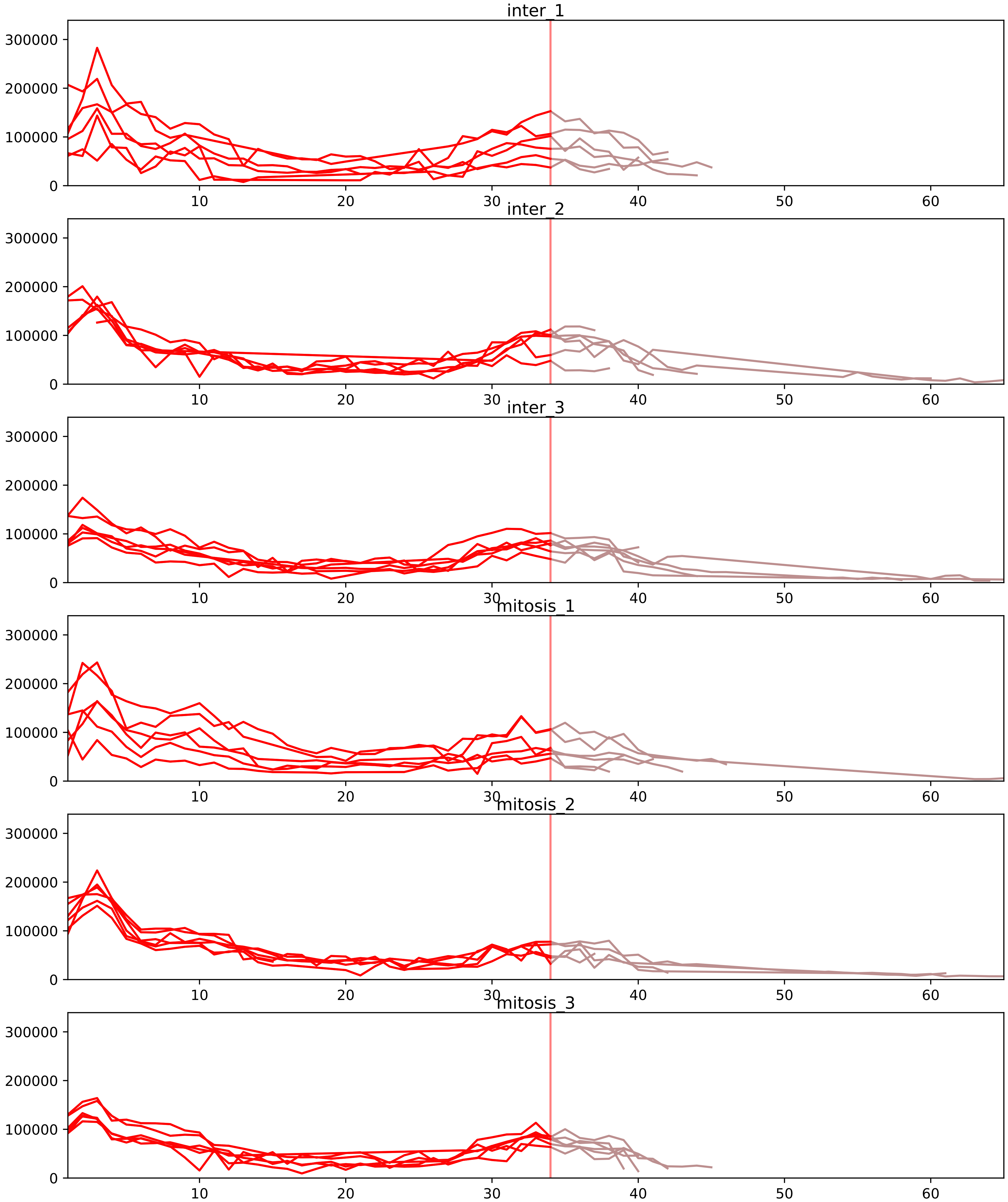


| level | condition_1 | condition_2 | pvalue | pvalue_adjusted |
|---|---|---|---|---|
| interactor_ratio | mitosis | inter | 7.232503685076959e-06 | 0.0001706129074428411 |
| complex_abundance | mitosis | inter | 8.807052603219957e-05 | 0.0020512628848005977 |
| interactor_abundance | mitosis | inter | 0.010798103352442946 | 0.05428554690845634 |
| monomer_abundance | mitosis | inter | 0.011972323632027077 | 0.1902553554136748 |
| total_abundance | mitosis | inter | 0.4061229479189579 | 0.7038902774003427 |
| assembled_abundance | mitosis | inter | 0.8009516216466347 | 0.918259745512588 |

O14979_Q9BUJ2
HNRDL_HUMAN vs HNRL1_HUMAN
p-value: 0.0 q-value: 0.041

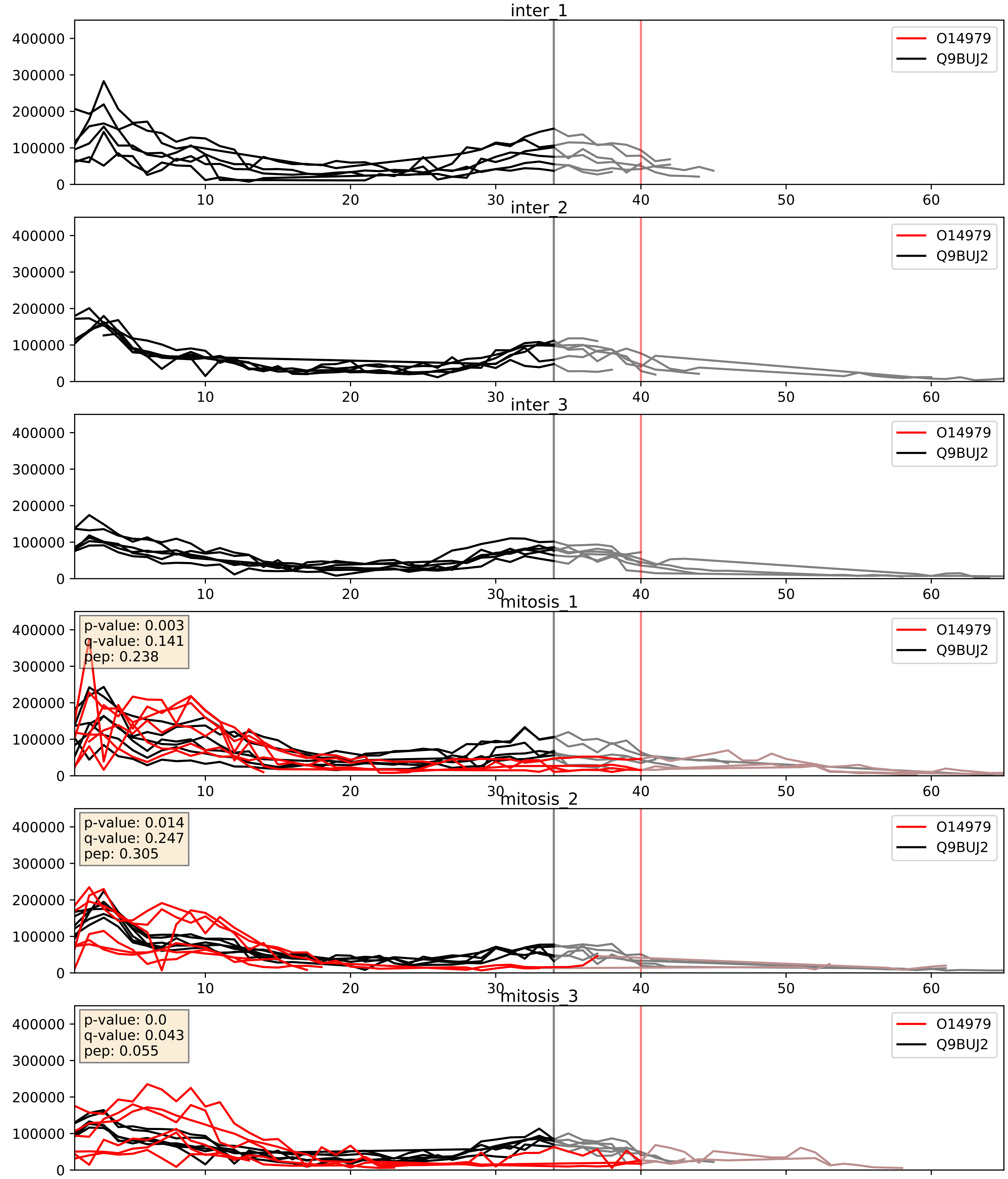


| level | condition_1 | condition_2 | pvalue | pvalue_adjusted |
|---|---|---|---|---|
| complex_abundance | mitosis | inter | 0.00016207257540637443 | 0.008563834848633118 |
| interactor_abundance | mitosis | inter | 0.0007200751823208013 | 0.019561481000604793 |
| interactor_ratio | mitosis | inter | 0.5211793406809895 | 0.8786895401167326 |

O43143_Q9BUJ2
DHX15_HUMAN vs HNRL1_HUMAN
p-value: 0.001 q-value: 0.042

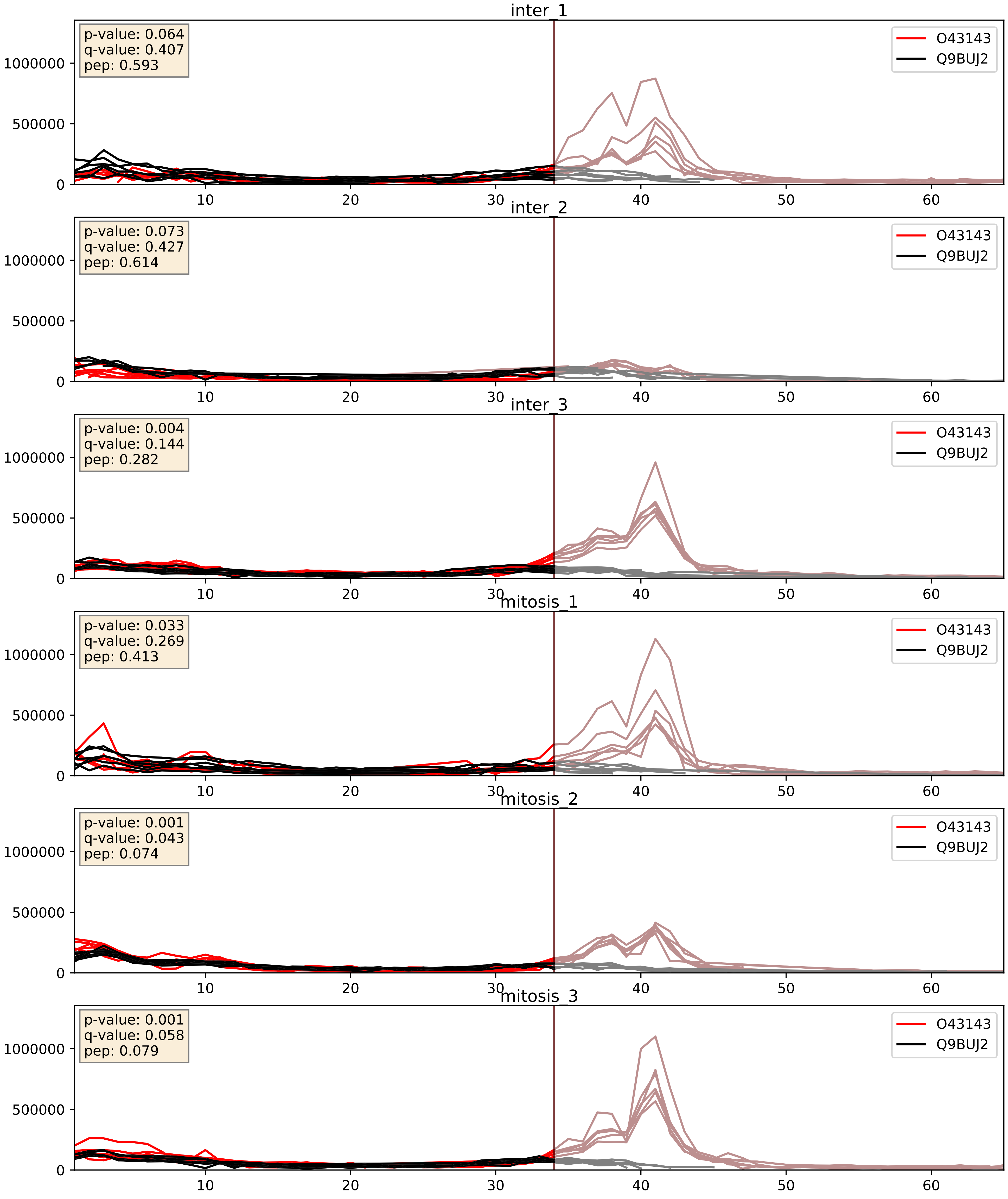


| level | condition_1 | condition_2 | pvalue | pvalue_adjusted |
|---|---|---|---|---|
| interactor_ratio | mitosis | inter | 0.19310834153893694 | 0.6660896702330326 |
| interactor_abundance | mitosis | inter | 0.31329514070062037 | 0.5103827965357903 |
| complex_abundance | mitosis | inter | 0.5050035747258289 | 0.6415134047602874 |

P31943_Q9BUJ2
HNRH1_HUMAN vs HNRL1_HUMAN
p-value: 0.001 q-value: 0.002

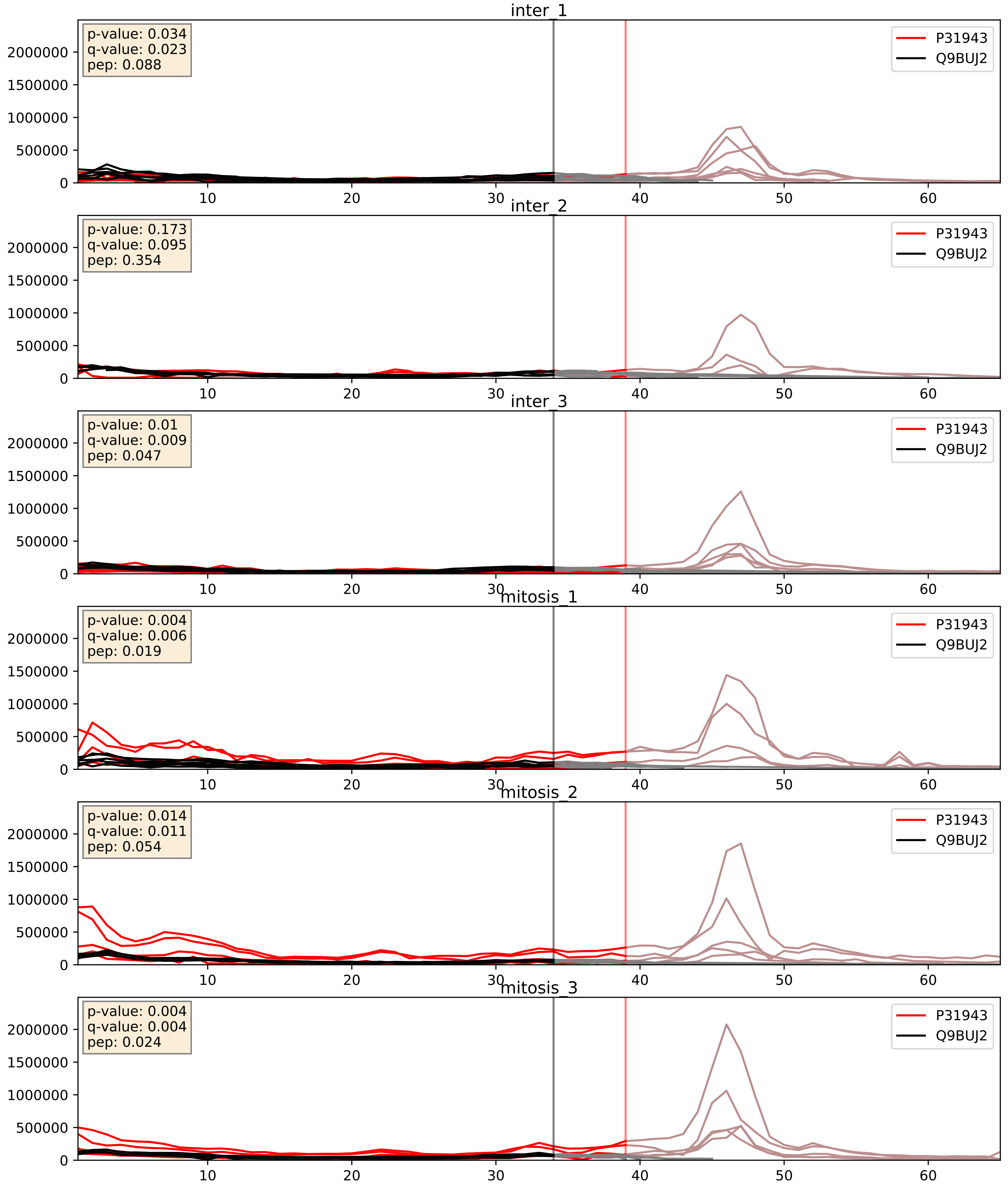


| level | condition_1 | condition_2 | pvalue | pvalue_adjusted |
|---|---|---|---|---|
| interactor_ratio | mitosis | inter | 0.001394553898793512 | 0.044212523606194314 |
| interactor_abundance | mitosis | inter | 0.0027782426171848577 | 0.03993577968615009 |
| complex_abundance | mitosis | inter | 0.007692182496874762 | 0.06519315066658214 |

P51991_Q9BUJ2
ROA3_HUMAN vs HNRL1_HUMAN
p-value: 0.003 q-value: 0.039

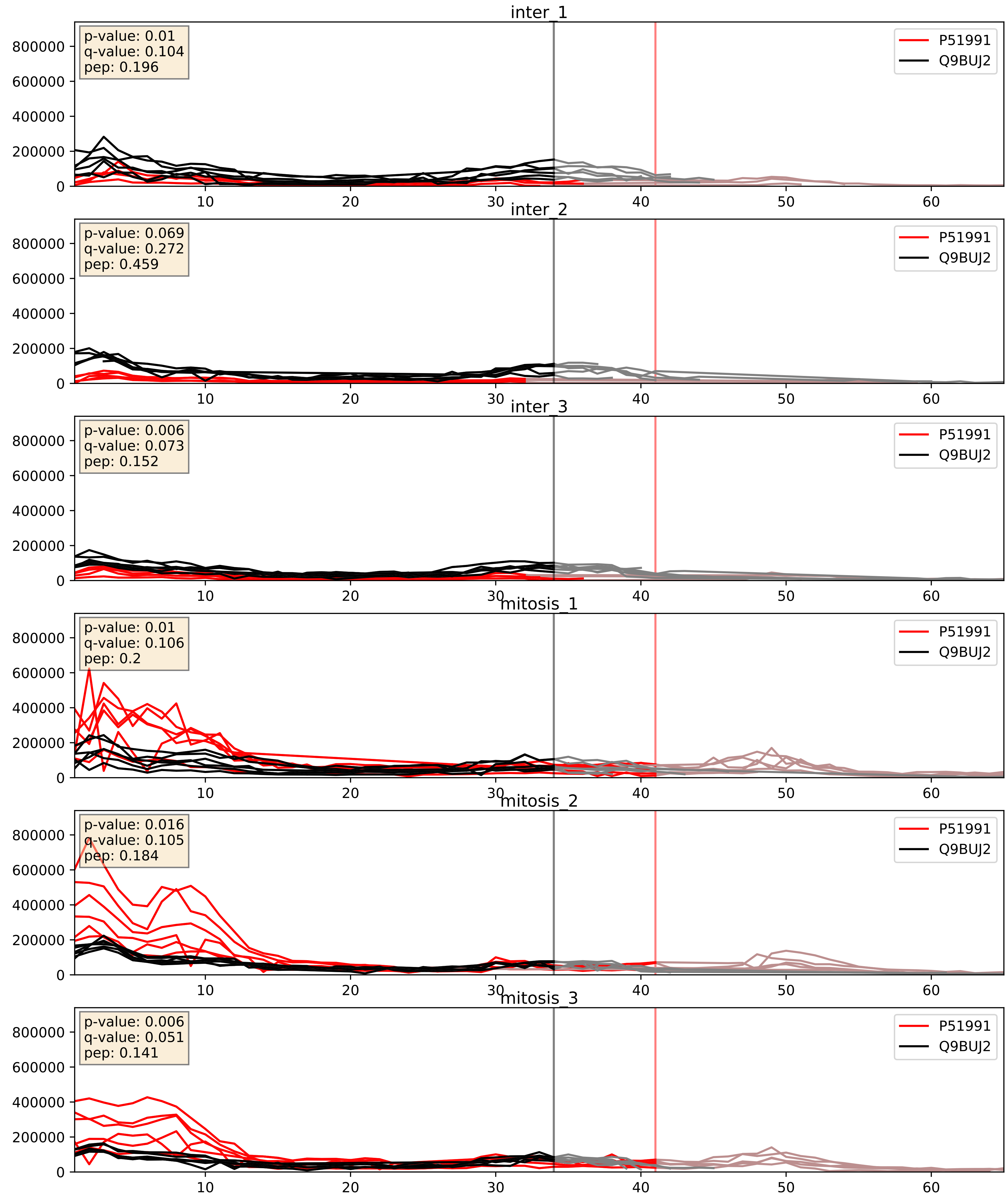


| level | condition_1 | condition_2 | pvalue | pvalue_adjusted |
|---|---|---|---|---|
| interactor_abundance | mitosis | inter | 1.1649447013156027e-05 | 0.0033803141163598503 |
| interactor_ratio | mitosis | inter | 7.24788381117057e-05 | 0.008863126489088583 |
| complex_abundance | mitosis | inter | 0.0001877343549764503 | 0.009111526782692203 |

P52597_Q9BUJ2
HNRPF_HUMAN vs HNRL1_HUMAN
p-value: 0.0 q-value: 0.0

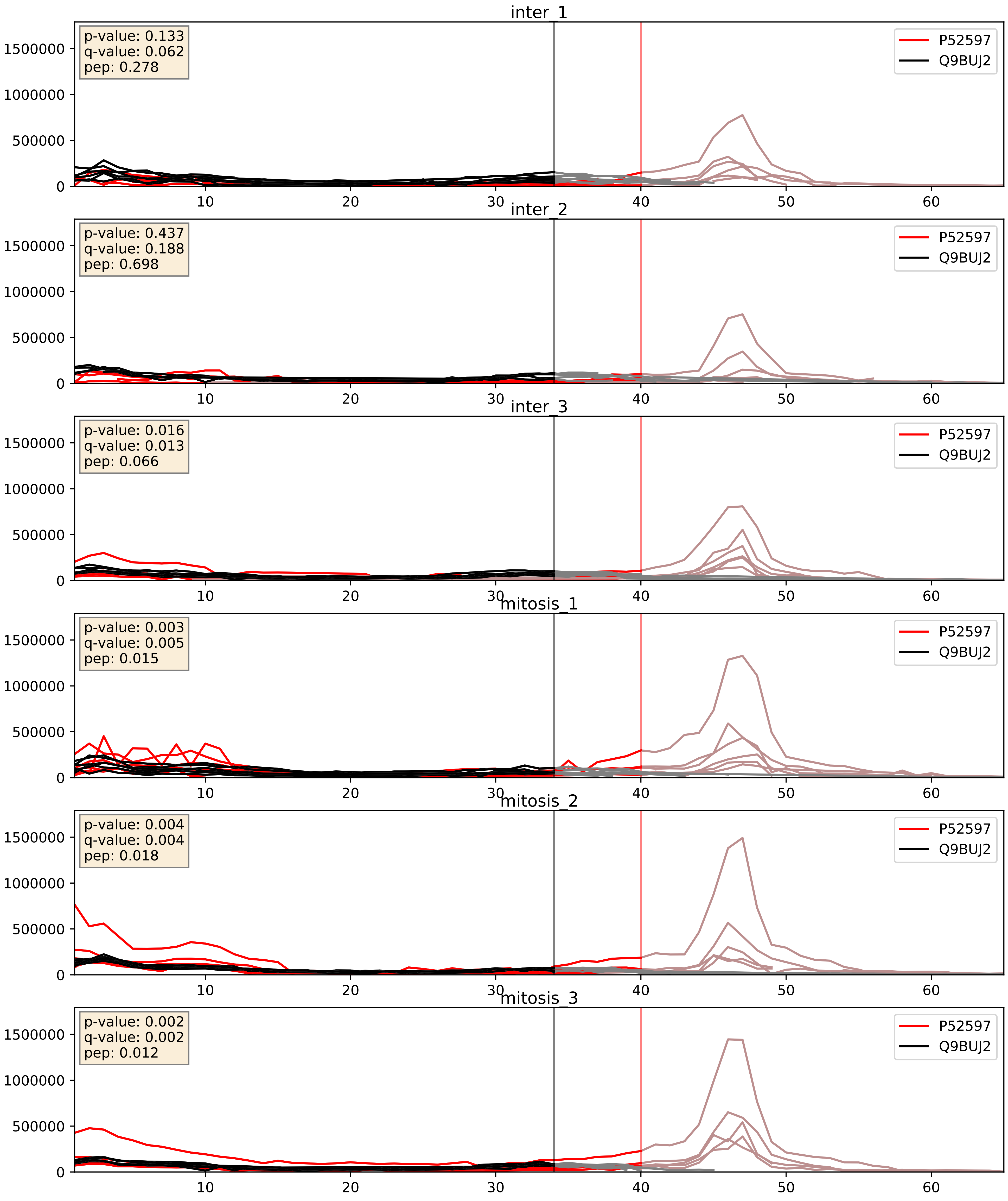


| level | condition_1 | condition_2 | pvalue | pvalue_adjusted |
|---|---|---|---|---|
| interactor_abundance | mitosis | inter | 0.005867622300035323 | 0.06493451439987377 |
| interactor_ratio | mitosis | inter | 0.00655810316750251 | 0.1124141137448166 |
| complex_abundance | mitosis | inter | 0.00962378637334349 | 0.07290231093435423 |

Q13151_Q9BUJ2
ROA0_HUMAN vs HNRL1_HUMAN
p-value: 0.002 q-value: 0.035

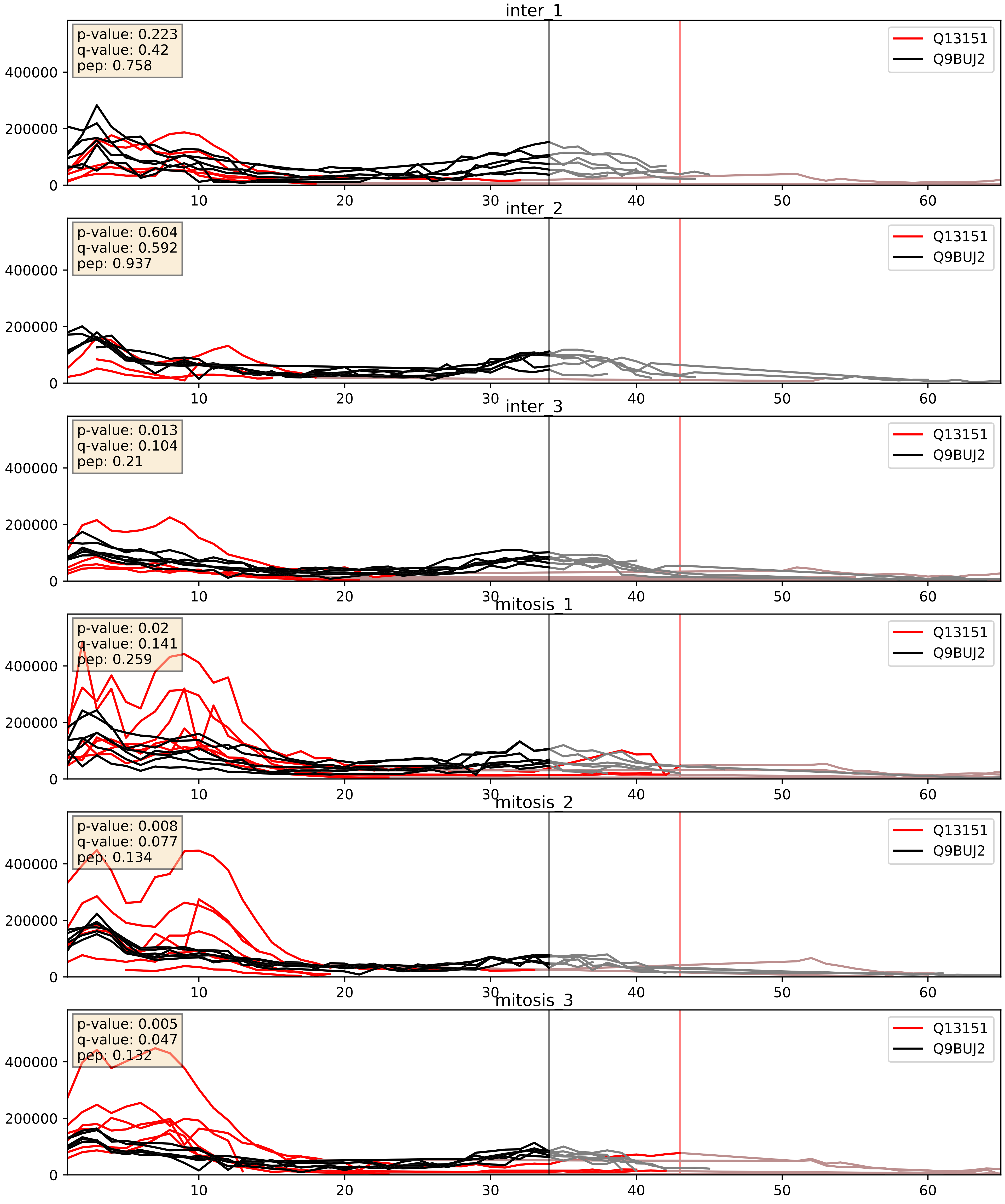


| level | condition_1 | condition_2 | pvalue | pvalue_adjusted |
|---|---|---|---|---|
| interactor_ratio | mitosis | inter | 0.002095252898243131 | 0.05622371413467461 |
| interactor_abundance | mitosis | inter | 0.008471091441998648 | 0.08017189393261118 |
| complex_abundance | mitosis | inter | 0.02812116174369059 | 0.1220004556068812 |
