## Supplementary Data 2 for "SECAT: Quantifying differential protein-protein interaction states by network-centric analysis": hela_string_000098_Q9Y3T9.pdf

Q9Y3T9_Q9Y3T9

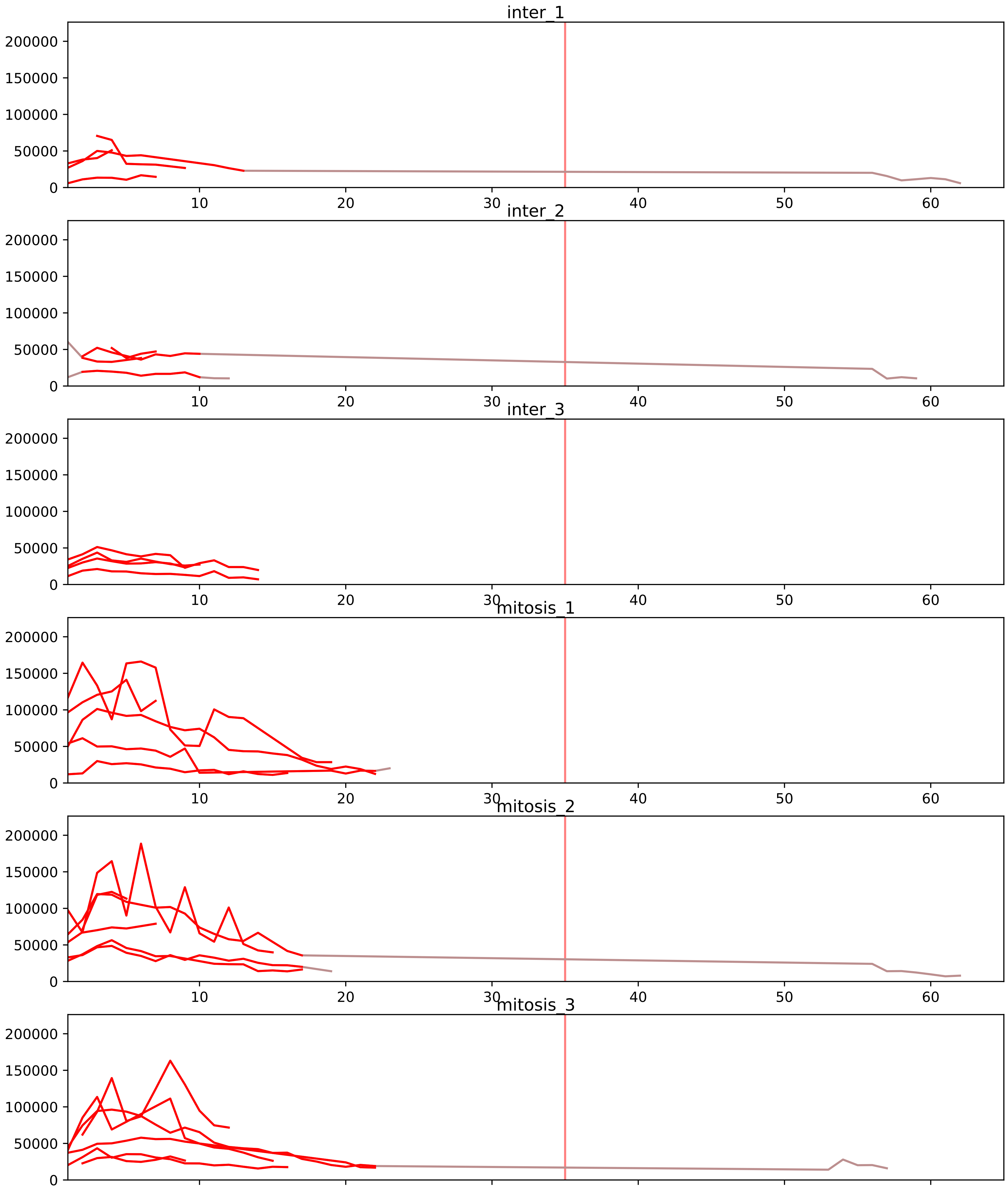


| level | condition_1 | condition_2 | pvalue | pvalue_adjusted |
|---|---|---|---|---|
| interactor_ratio | mitosis | inter | 9.448552445914398e-06 | 0.00021463378395657396 |
| interactor_abundance | mitosis | inter | 4.2180050899535e-05 | 0.0013616016430727086 |
| complex_abundance | mitosis | inter | 5.846893541774038e-05 | 0.0016300430480097317 |
| total_abundance | mitosis | inter | 0.41690155568050113 | 0.7105398385869994 |
| assembled_abundance | mitosis | inter | 0.44396959453332924 | 0.7129405392831953 |
| monomer_abundance | mitosis | inter | 0.9938496323593273 | 0.9981662859499865 |

P22087_Q9Y3T9
FBRL_HUMAN vs NOC2L_HUMAN
p-value: 0.029 q-value: 0.021

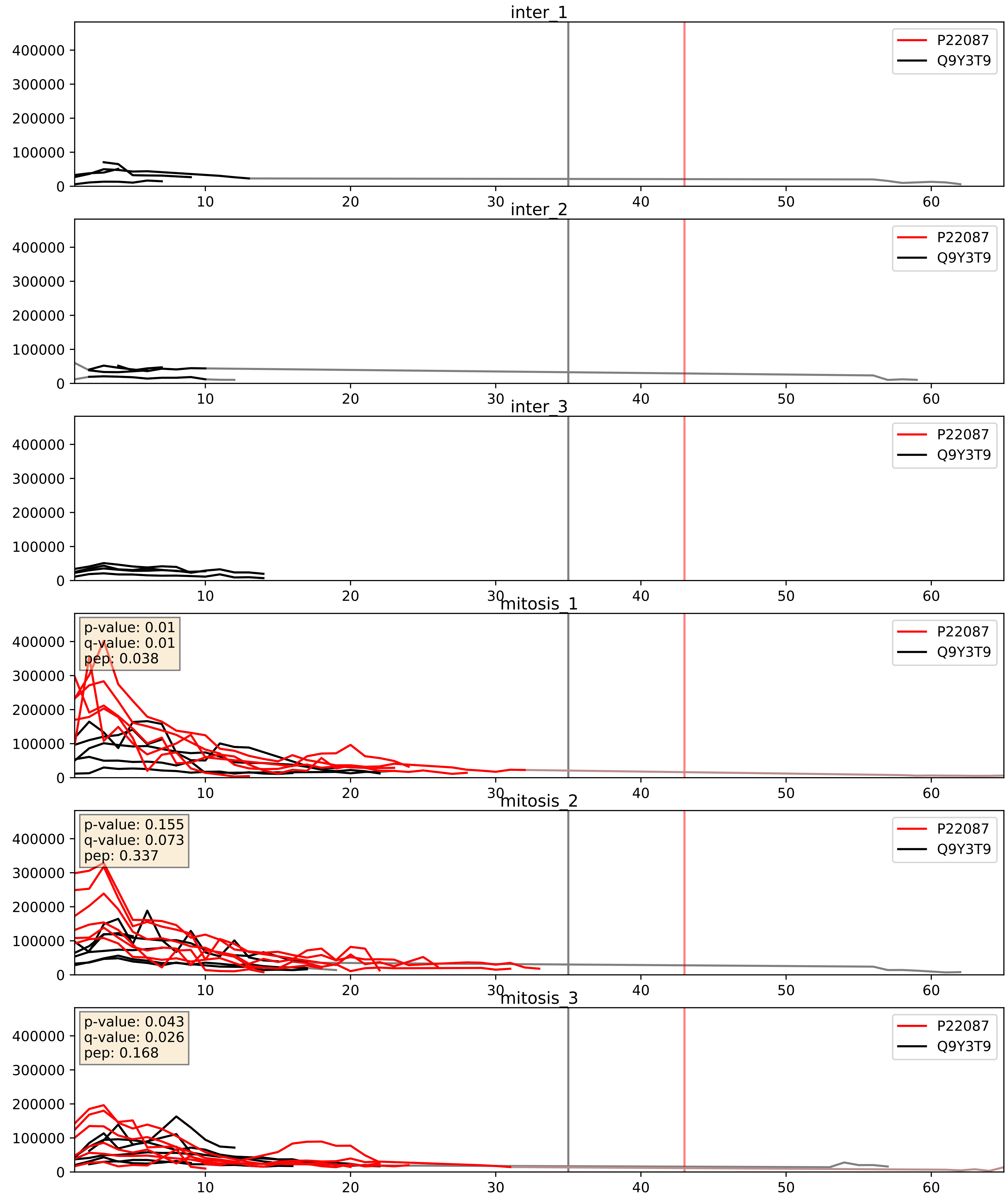


| level | condition_1 | condition_2 | pvalue | pvalue_adjusted |
|---|---|---|---|---|
| complex_abundance | mitosis | inter | 0.00035590071739036747 | 0.012039775462644836 |
| interactor_abundance | mitosis | inter | 0.00035742084283765416 | 0.01356427145975096 |
| interactor_ratio | mitosis | inter | 0.009198067791017207 | 0.13668839544866102 |

Q12788_Q9Y3T9
TBL3_HUMAN vs NOC2L_HUMAN
p-value: 0.028 q-value: 0.02

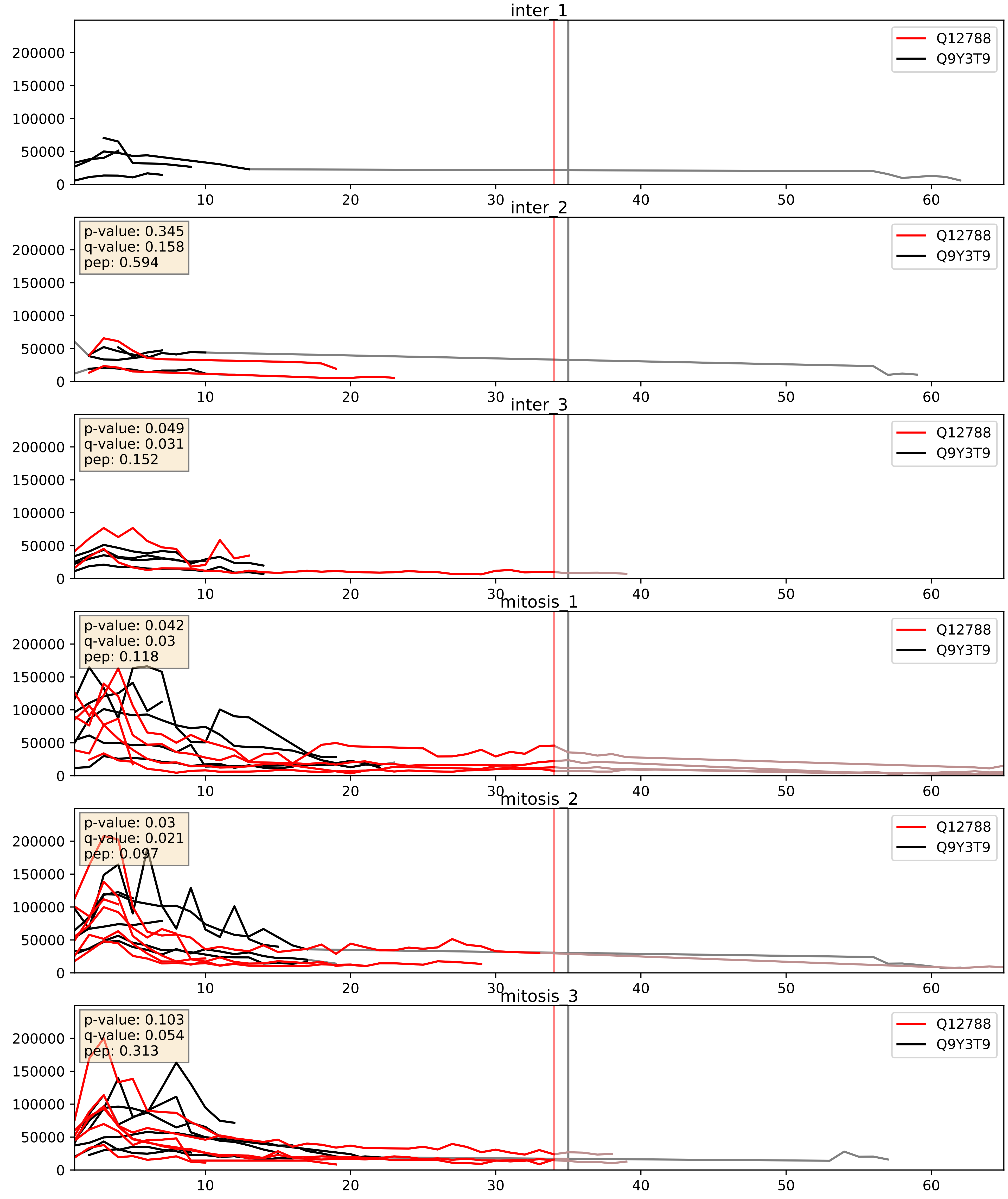


| level | condition_1 | condition_2 | pvalue | pvalue_adjusted |
|---|---|---|---|---|
| complex_abundance | mitosis | inter | 0.005487193845178426 | 0.05591711823181825 |
| interactor_abundance | mitosis | inter | 0.011367688817575375 | 0.09419885409336419 |
| interactor_ratio | mitosis | inter | 0.13663518010683853 | 0.58714715949525 |

Q9BQ67_Q9Y3T9
GRWD1_HUMAN vs NOC2L_HUMAN
p-value: 0.078 q-value: 0.046

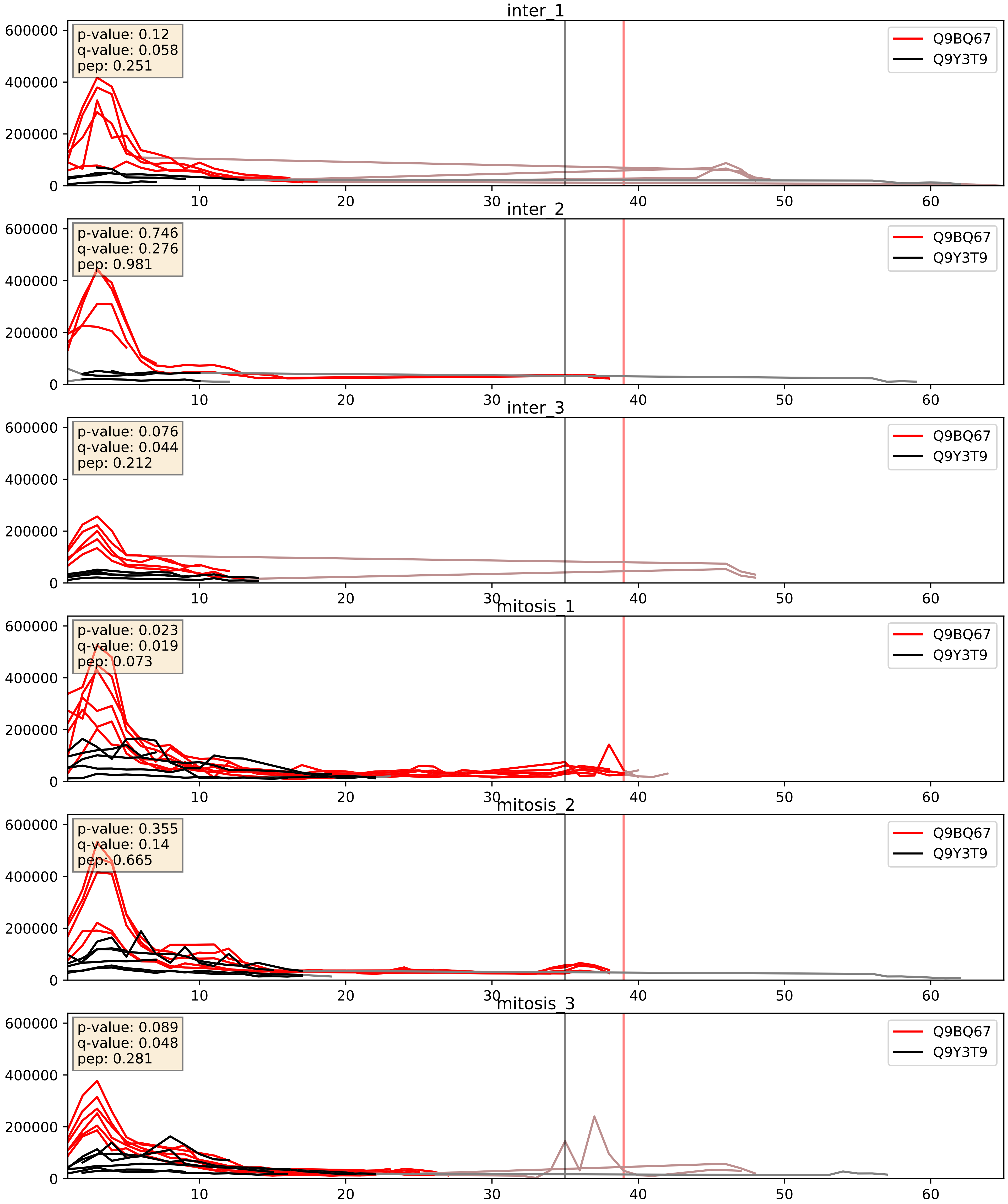


| level | condition_1 | condition_2 | pvalue | pvalue_adjusted |
|---|---|---|---|---|
| complex_abundance | mitosis | inter | 0.012364424967787958 | 0.08259410436140613 |
| interactor_ratio | mitosis | inter | 0.020920483835626235 | 0.21906942949212715 |
| interactor_abundance | mitosis | inter | 0.08943831949006091 | 0.25354926803607264 |

Q9BZE4_Q9Y3T9
NOG1_HUMAN vs NOC2L_HUMAN
p-value: 0.015 q-value: 0.012

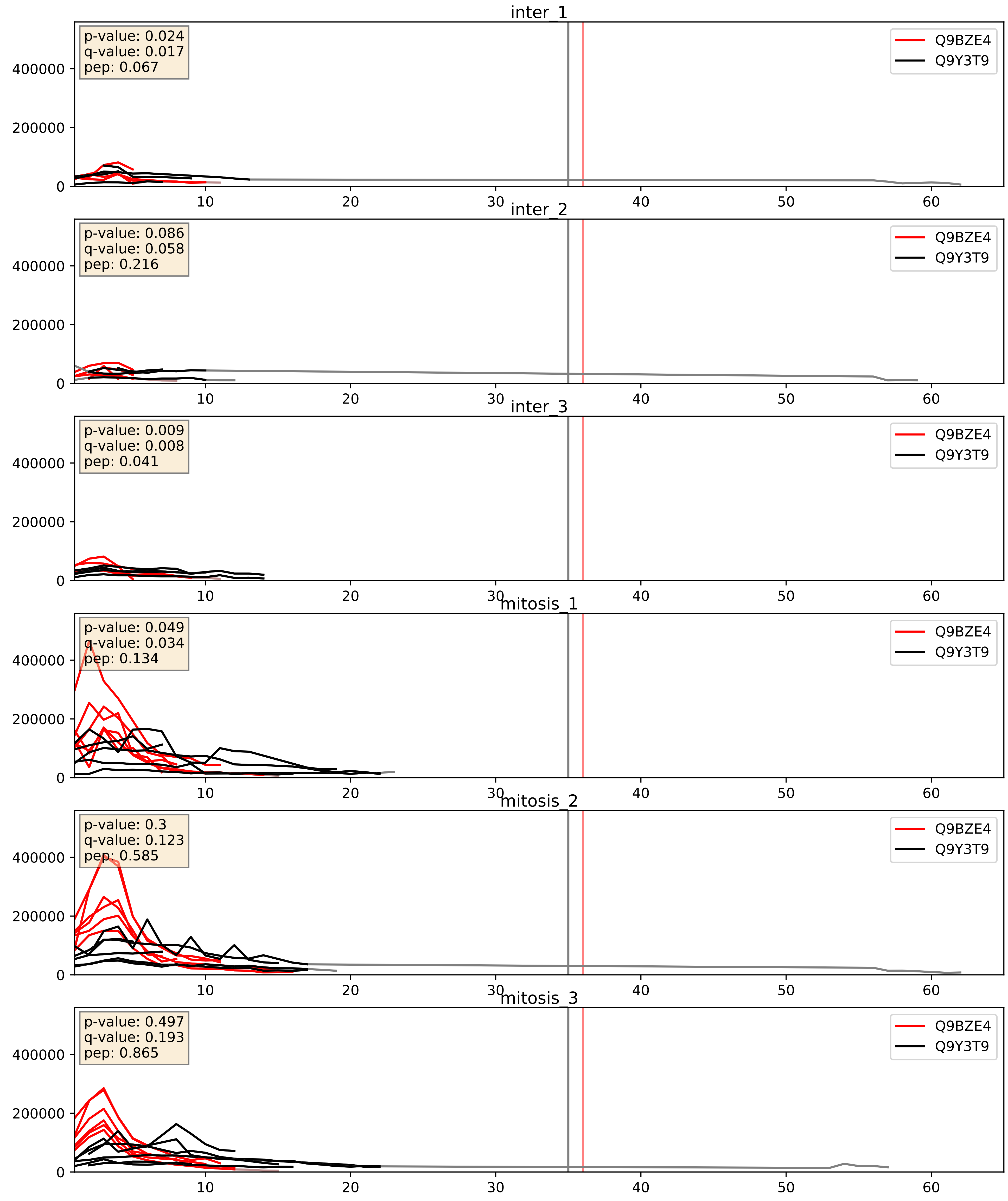


| level | condition_1 | condition_2 | pvalue | pvalue_adjusted |
|---|---|---|---|---|
| complex_abundance | mitosis | inter | 0.0007505306717870162 | 0.017895661700548352 |
| interactor_abundance | mitosis | inter | 0.0009449316519668341 | 0.022846219799594687 |
| interactor_ratio | mitosis | inter | 0.13624795726384295 | 0.5866546850118911 |

Q9NVP1_Q9Y3T9
DDX18_HUMAN vs NOC2L_HUMAN
p-value: 0.037 q-value: 0.026

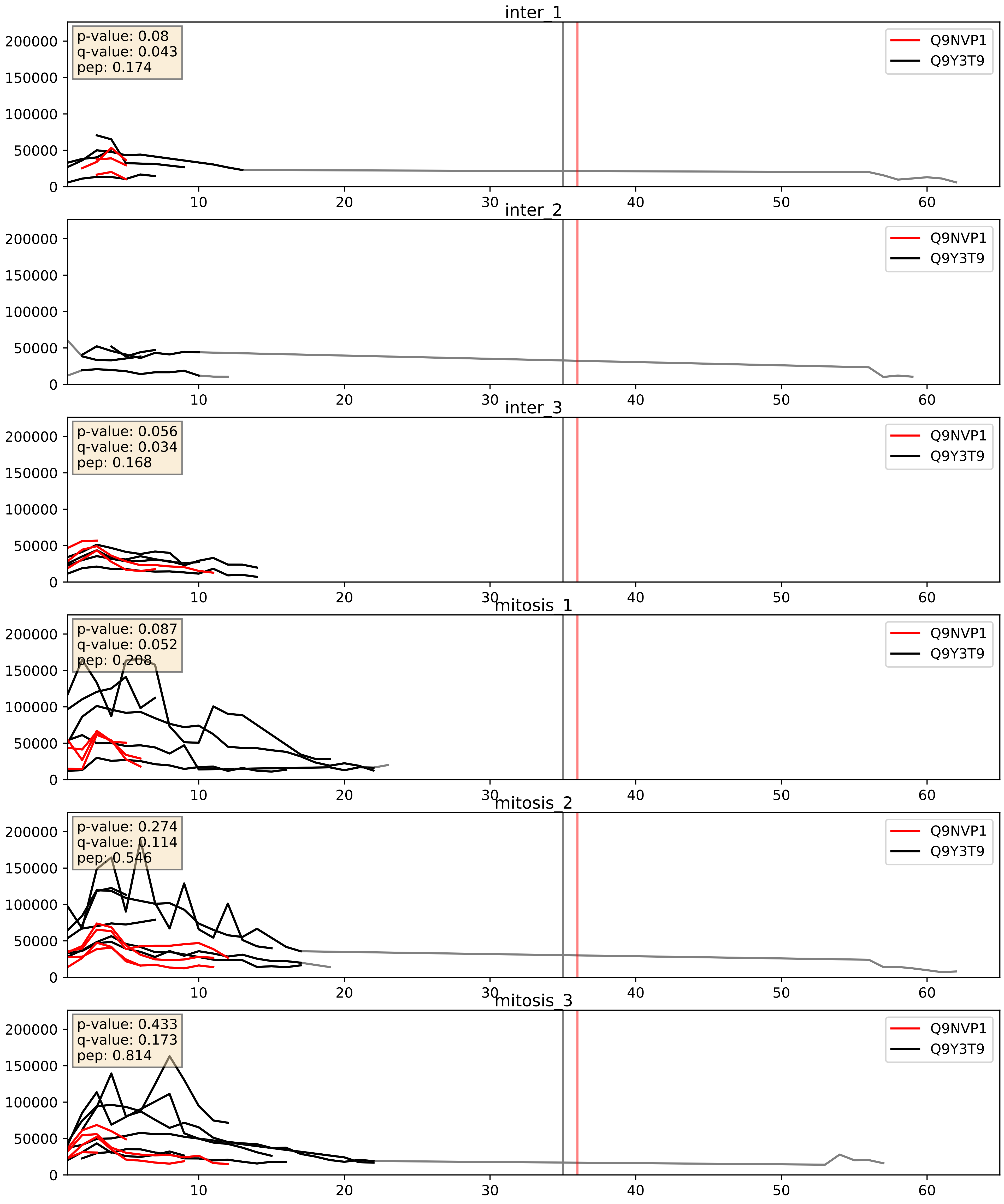


| level | condition_1 | condition_2 | pvalue | pvalue_adjusted |
|---|---|---|---|---|
| interactor_ratio | mitosis | inter | 0.0013447851463711195 | 0.04334578357589157 |
| complex_abundance | mitosis | inter | 0.07126313143207737 | 0.19408603406254607 |
| interactor_abundance | mitosis | inter | 0.7680100411481787 | 0.8624119050542318 |

Q9Y2X3_Q9Y3T9
NOP58_HUMAN vs NOC2L_HUMAN
p-value: 0.04 q-value: 0.027

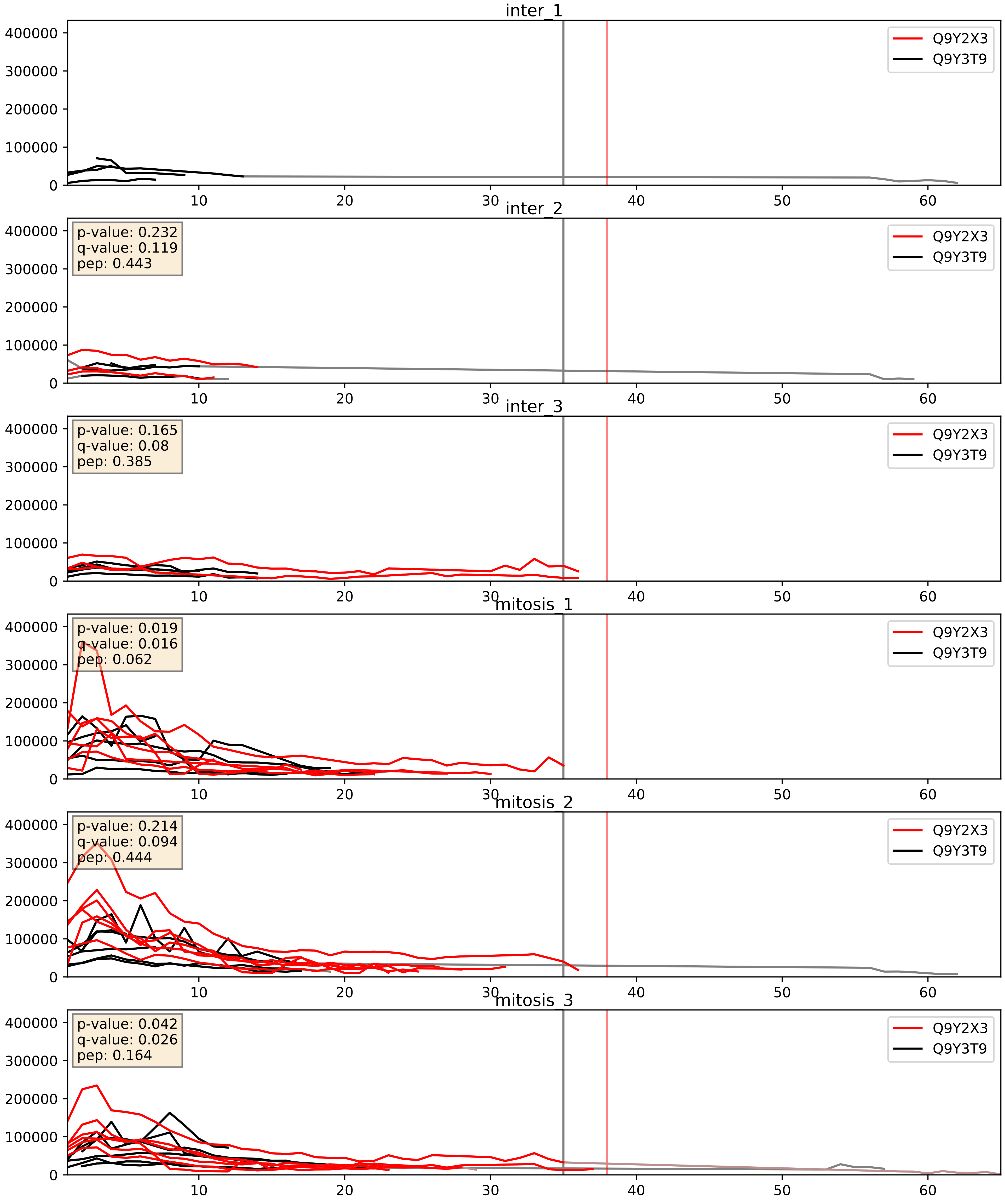


| level | condition_1 | condition_2 | pvalue | pvalue_adjusted |
|---|---|---|---|---|
| interactor_abundance | mitosis | inter | 0.0015817191617625384 | 0.029025269221018554 |
| complex_abundance | mitosis | inter | 0.0023478262388268484 | 0.035168510428347066 |
| interactor_ratio | mitosis | inter | 0.03146494532124448 | 0.28316826136323564 |
