## Supplementary Data 2 for "SECAT: Quantifying differential protein-protein interaction states by network-centric analysis": hela_string_000099_Q99733.pdf

# Q99733_Q99733

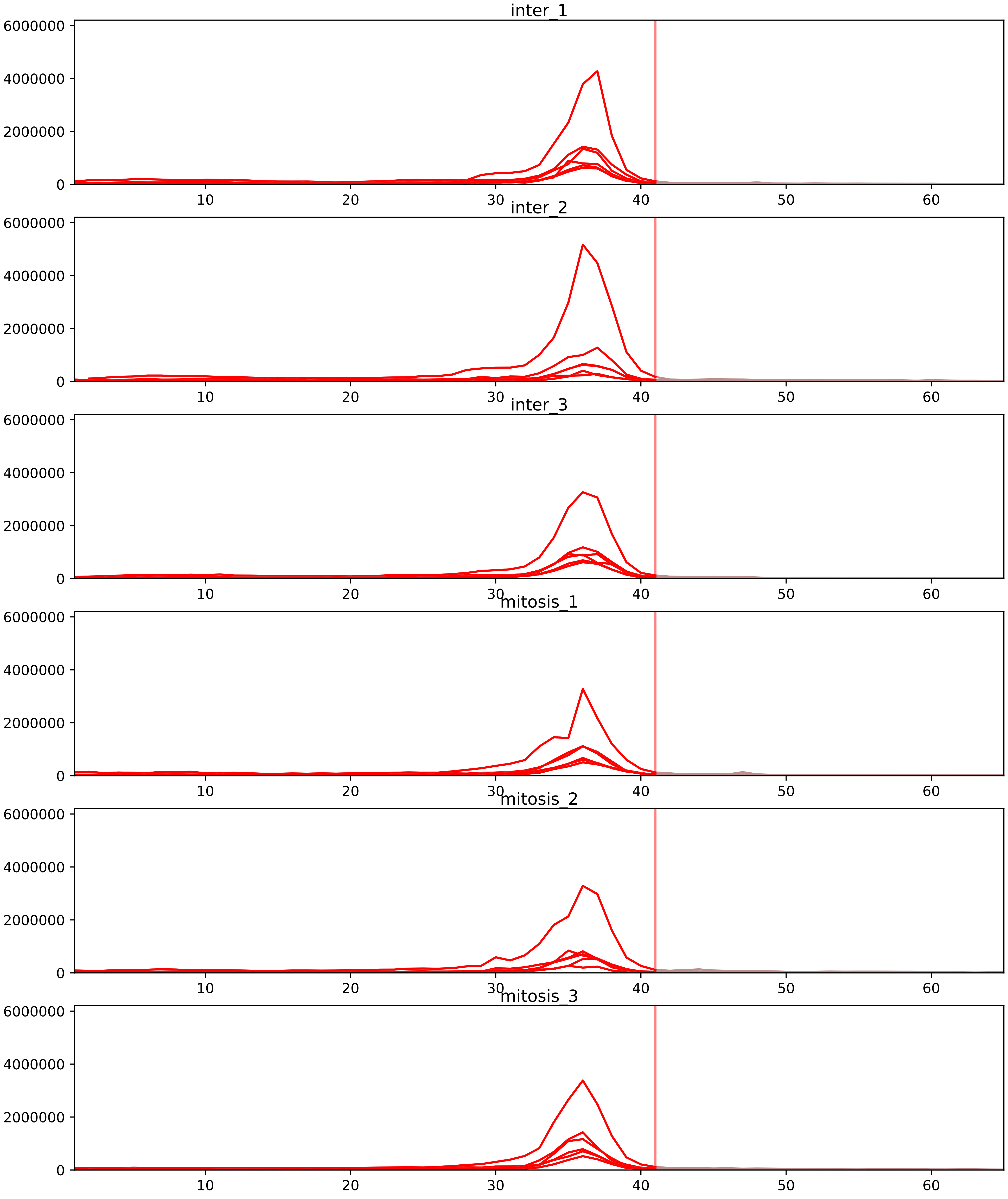


| level | condition_1 | condition_2 | pvalue | pvalue_adjusted |
|---|---|---|---|---|
| assembled_abundance | mitosis | inter | 9.48971827382153e-06 | 0.003651762585223375 |
| total_abundance | mitosis | inter | 9.630369152469451e-06 | 0.004123130635959455 |
| interactor_abundance | mitosis | inter | 0.0032588062997727352 | 0.026184295159745994 |
| complex_abundance | mitosis | inter | 0.003532865431671394 | 0.024025612923551223 |
| monomer_abundance | mitosis | inter | 0.01970447327428159 | 0.24279745128665506 |
| interactor_ratio | mitosis | inter | 0.08572761885861353 | 0.29762041264122435 |

P55209_Q99733
NP1L1_HUMAN vs NP1L4_HUMAN
p-value: 0.0 q-value: 0.008

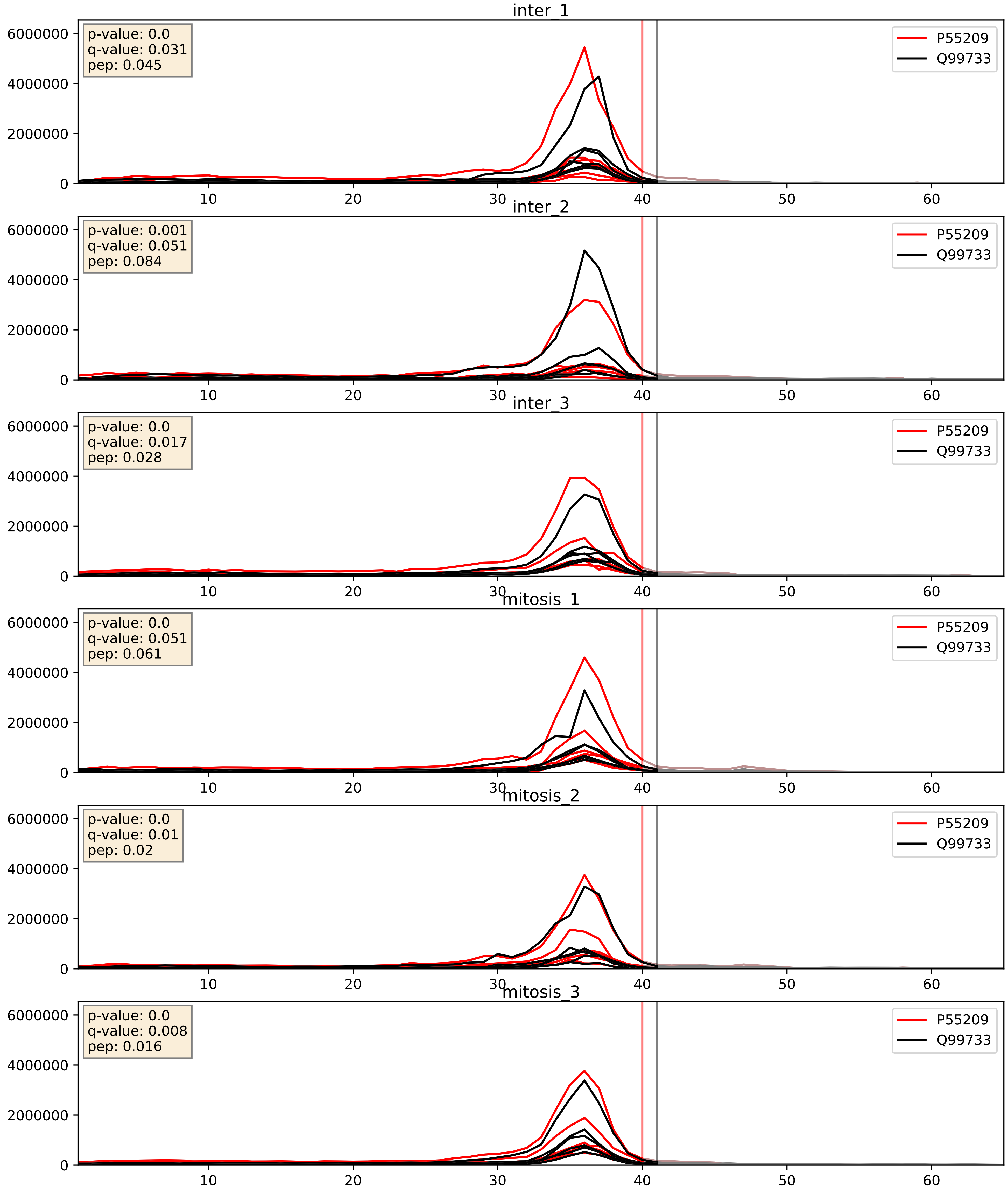


| level | condition_1 | condition_2 | pvalue | pvalue_adjusted |
|---|---|---|---|---|
| interactor_ratio | mitosis | inter | 0.03573226260715765 | 0.30712516692473574 |
| complex_abundance | mitosis | inter | 0.06776321895380114 | 0.18828567944900448 |
| interactor_abundance | mitosis | inter | 0.33853014625700684 | 0.5379085869158292 |

Q96AE4_Q99733
FUBP1_HUMAN vs NP1L4_HUMAN
p-value: 0.0 q-value: 0.047

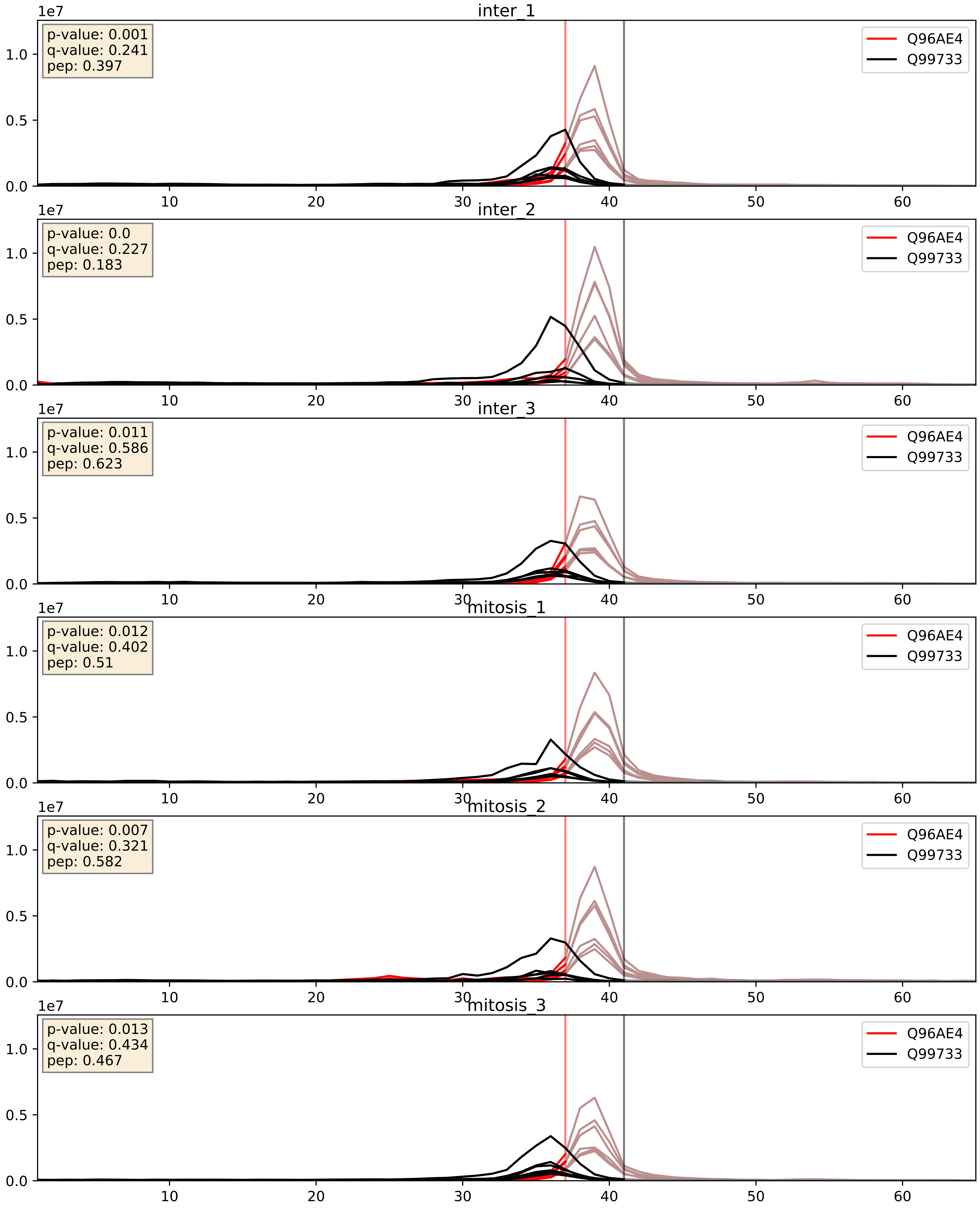


| level | condition_1 | condition_2 | pvalue | pvalue_adjusted |
|---|---|---|---|---|
| complex_abundance | mitosis | inter | 0.001295465539081554 | 0.024459508269436846 |
| interactor_abundance | mitosis | inter | 0.02767213951058815 | 0.13815894675452584 |
| interactor_ratio | mitosis | inter | 0.47205787855233544 | 0.8622324786853773 |
