## Supplementary Data 2 for "SECAT: Quantifying differential protein-protein interaction states by network-centric analysis": hela_string_000100_P42695.pdf

# P42695_P42695

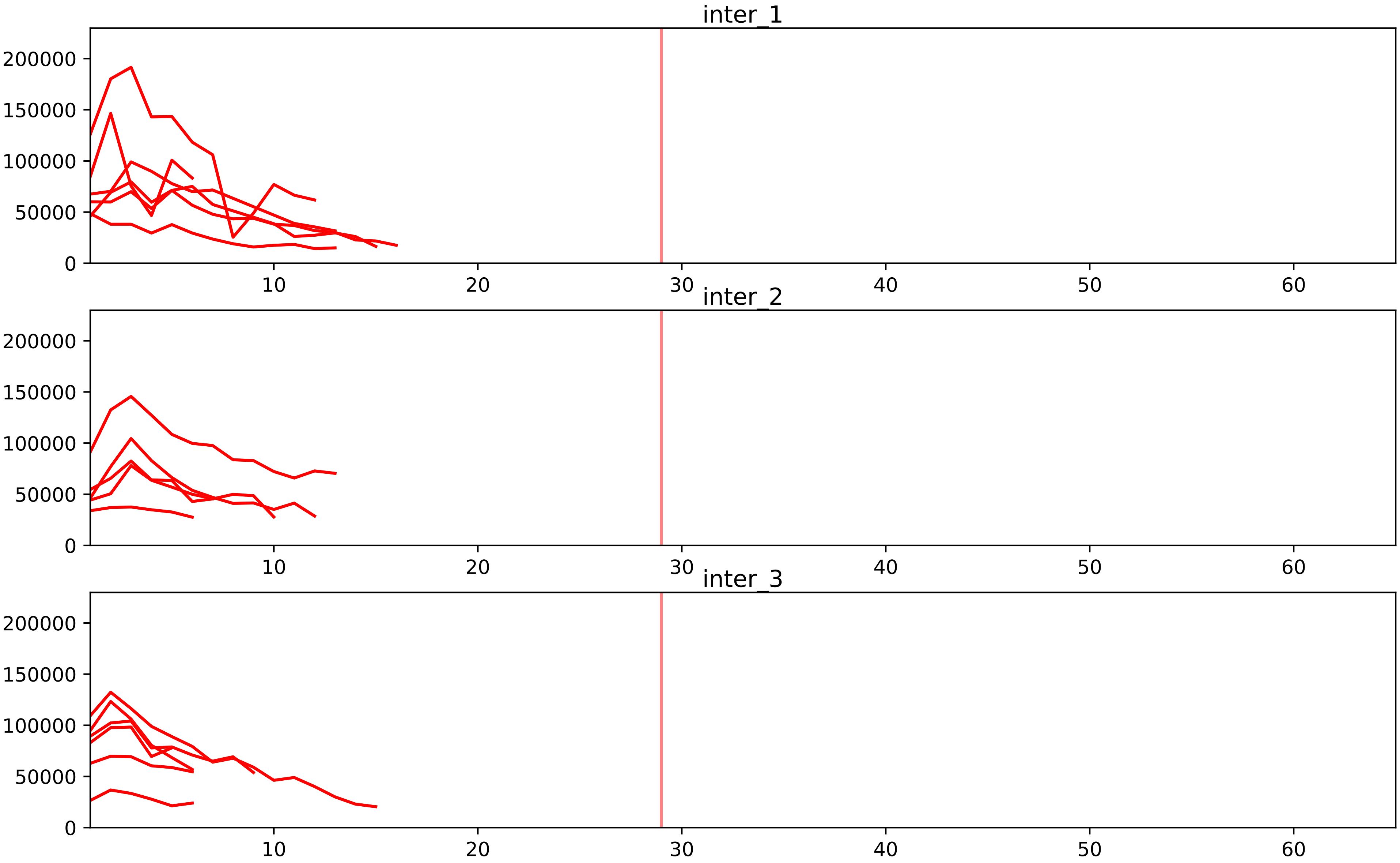


| level | condition_1 | condition_2 | pvalue | pvalue_adjusted |
|---|---|---|---|---|
| assembled_abundance | mitosis | inter | 1.0141617946572074e-05 | 0.003651762585223375 |
| total_abundance | mitosis | inter | 2.4052439204893496e-05 | 0.005915549590004797 |
| complex_abundance | mitosis | inter | 4.169943848123789e-05 | 0.0013004570644996223 |
| interactor_abundance | mitosis | inter | 4.6298705482385756e-05 | 0.0014198269681264966 |
| interactor_ratio | mitosis | inter | 0.041948379801130725 | 0.18465315510545582 |

P42695_Q6IBW4
CNDD3_HUMAN vs CNDH2_HUMAN
p-value: 0.023 q-value: 0.017

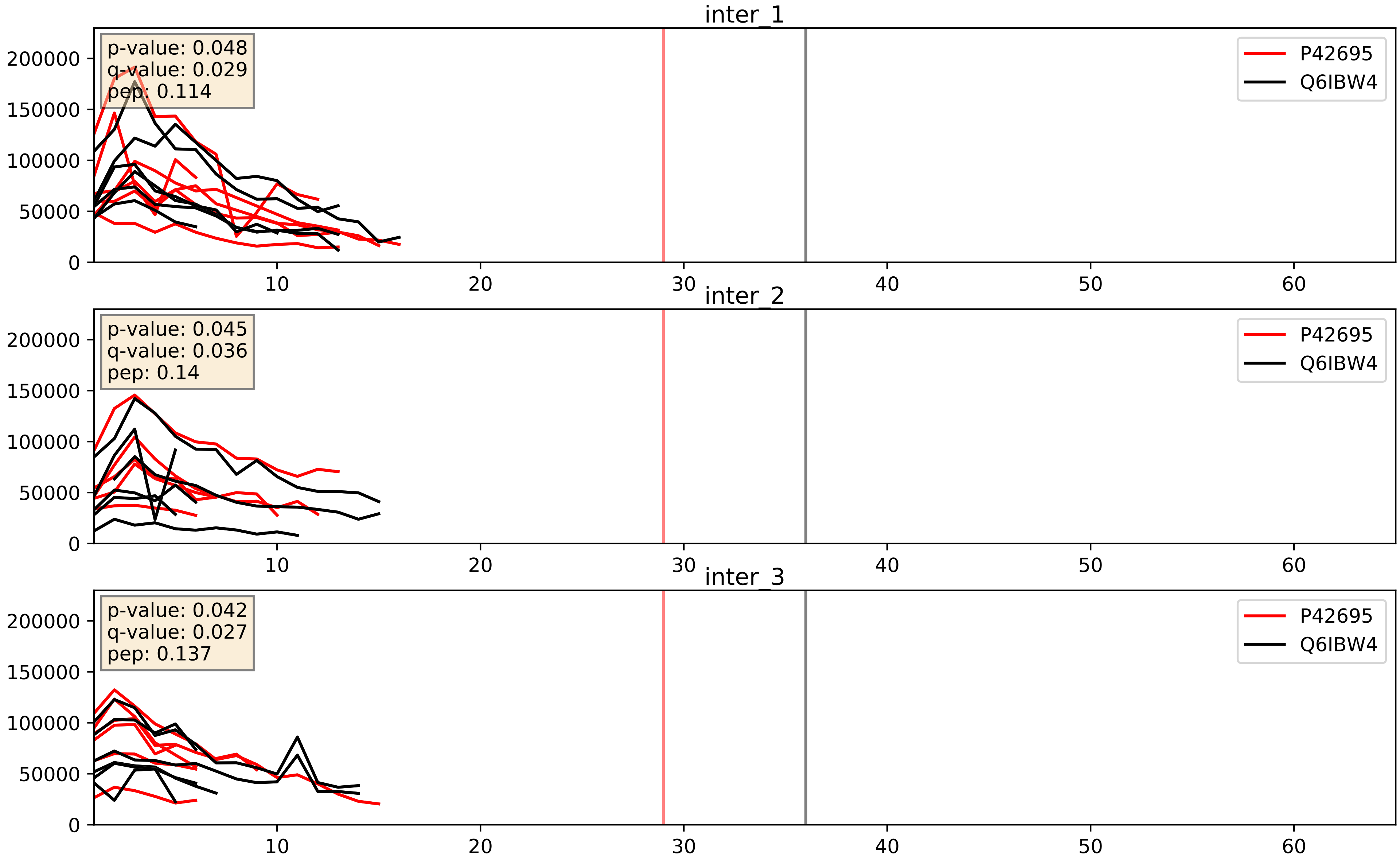


| level | condition_1 | condition_2 | pvalue | pvalue_adjusted |
|---|---|---|---|---|
| interactor_abundance | mitosis | inter | 0.0003511758884899916 | 0.01356427145975096 |
| complex_abundance | mitosis | inter | 0.0003576603614696945 | 0.012039775462644836 |
| interactor_ratio | mitosis | inter | 0.8058686900687333 | 0.9637099730355346 |

P42695_Q86XI2
CNDD3_HUMAN vs CNDG2_HUMAN
p-value: 0.002 q-value: 0.003

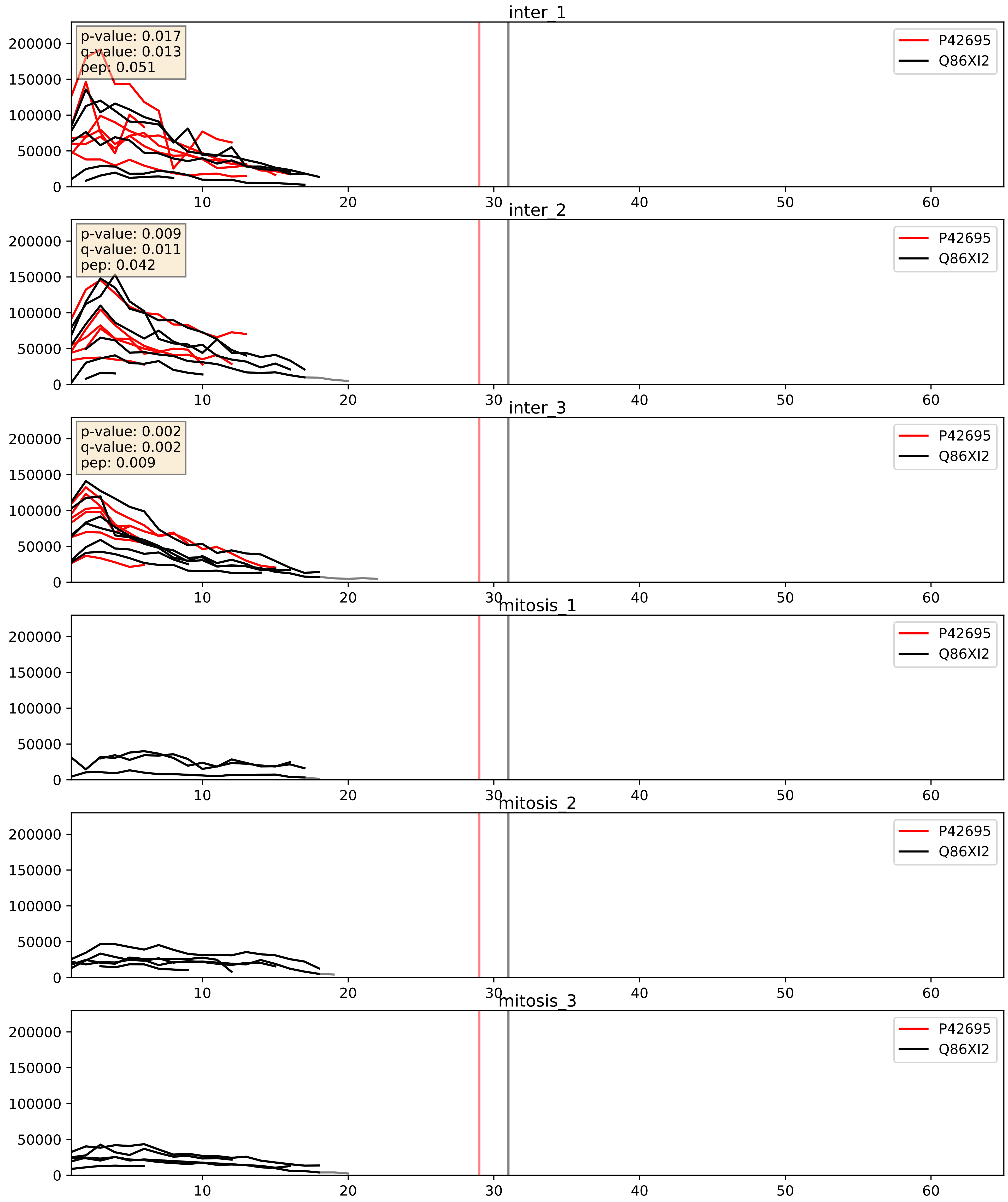


| level | condition_1 | condition_2 | pvalue | pvalue_adjusted |
|---|---|---|---|---|
| complex_abundance | mitosis | inter | 0.00030362639283545225 | 0.01165207110963347 |
| interactor_abundance | mitosis | inter | 0.0003437432204183121 | 0.01356427145975096 |
| interactor_ratio | mitosis | inter | 0.008622180363075611 | 0.13203195690147984 |
