## Supplementary Data 2 for "SECAT: Quantifying differential protein-protein interaction states by network-centric analysis": hela_string_000101_O60499.pdf

O60499_O60499

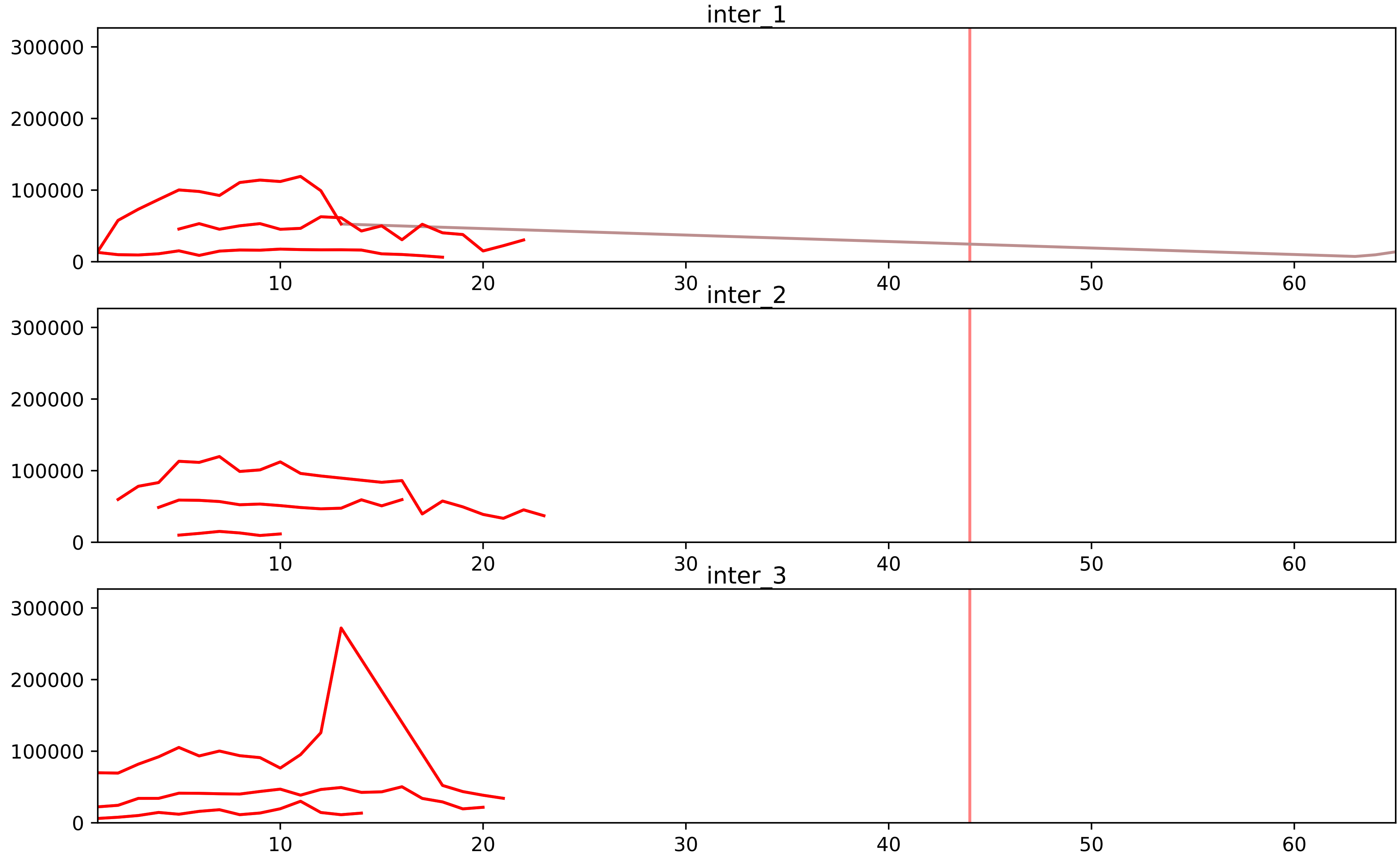


| level | condition_1 | condition_2 | pvalue | pvalue_adjusted |
|---|---|---|---|---|
| complex_abundance | mitosis | inter | 1.0204329890494878e-05 | 0.00048143505124386094 |
| interactor_abundance | mitosis | inter | 1.1580142313424275e-05 | 0.000579690055685502 |
| interactor_ratio | mitosis | inter | 0.00015476748069843209 | 0.002473765594533838 |
| total_abundance | mitosis | inter | 0.006952207965221883 | 0.09777262638382488 |
| assembled_abundance | mitosis | inter | 0.008267673658390871 | 0.09804934714714468 |
| monomer_abundance | mitosis | inter | 0.31503595841084214 | 0.6718670942432774 |

O14662_O60499
STX16_HUMAN vs STX10_HUMAN
p-value: 0.003 q-value: 0.003

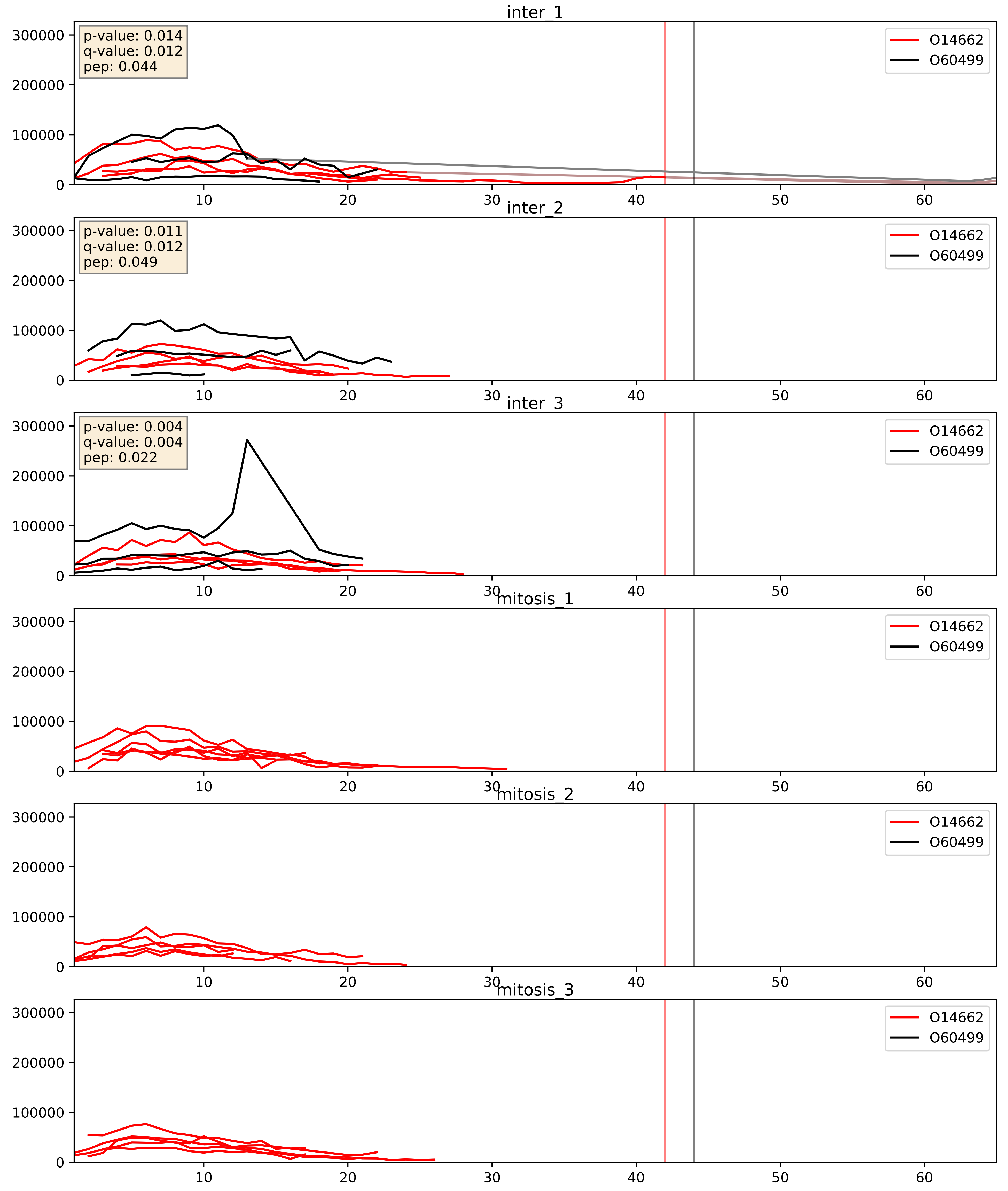


| level | condition_1 | condition_2 | pvalue | pvalue_adjusted |
|---|---|---|---|---|
| complex_abundance | mitosis | inter | 0.0004261048385601943 | 0.013091389064164169 |
| interactor_abundance | mitosis | inter | 0.0006985905693554247 | 0.019073880342257484 |
| interactor_ratio | mitosis | inter | 0.8536180534556025 | 0.9755634896635457 |

O60499_O95249
STX10_HUMAN vs GOSR1_HUMAN
p-value: 0.001 q-value: 0.032

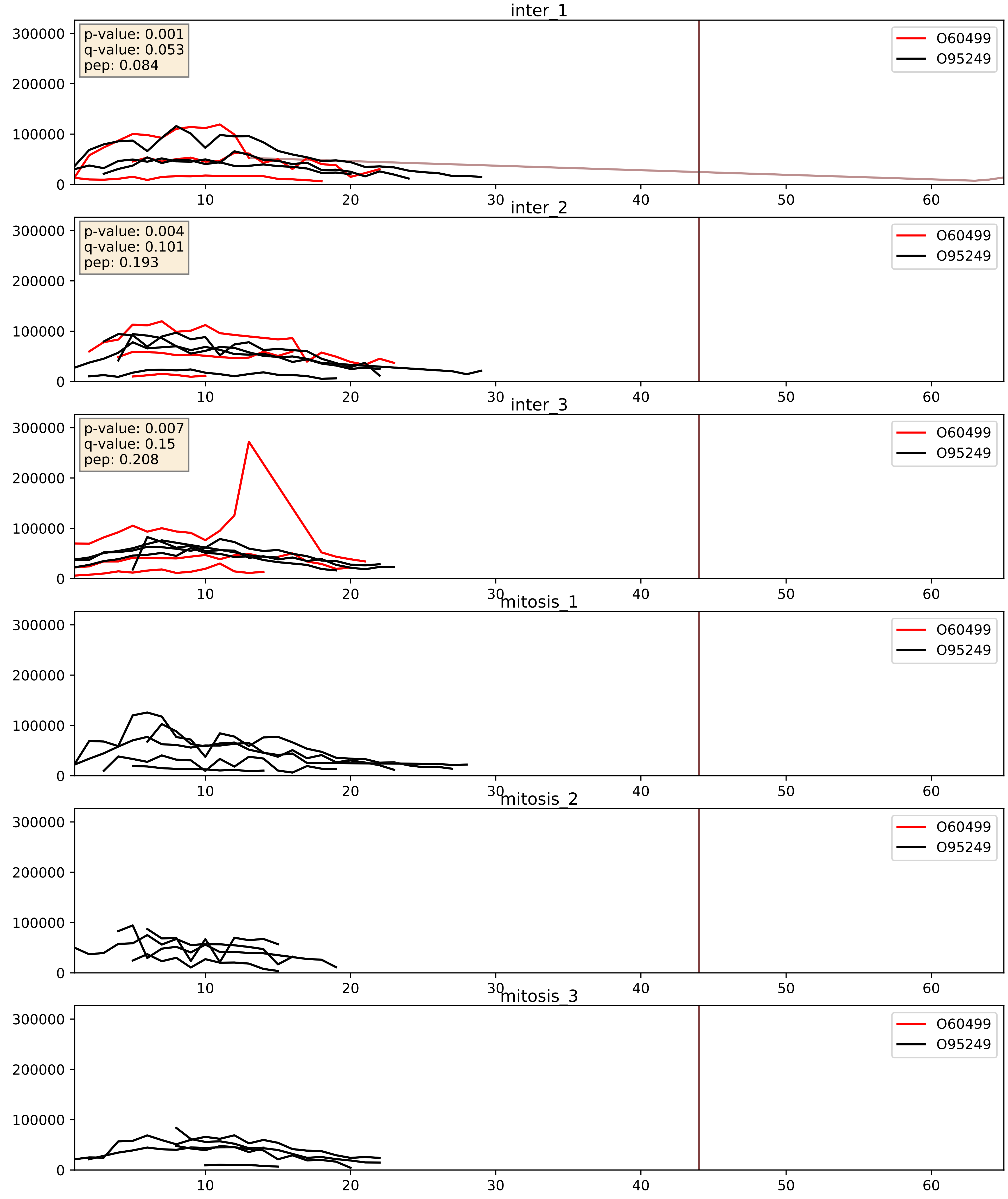


| level | condition_1 | condition_2 | pvalue | pvalue_adjusted |
|---|---|---|---|---|
| interactor_abundance | mitosis | inter | 0.000300864944976679 | 0.01300709055050693 |
| complex_abundance | mitosis | inter | 0.0003812214784971195 | 0.01249066060683217 |
| interactor_ratio | mitosis | inter | 0.5238543119343951 | 0.8797710241629237 |

O60499_Q12846
STX10_HUMAN vs STX4_HUMAN
p-value: 0.001 q-value: 0.039

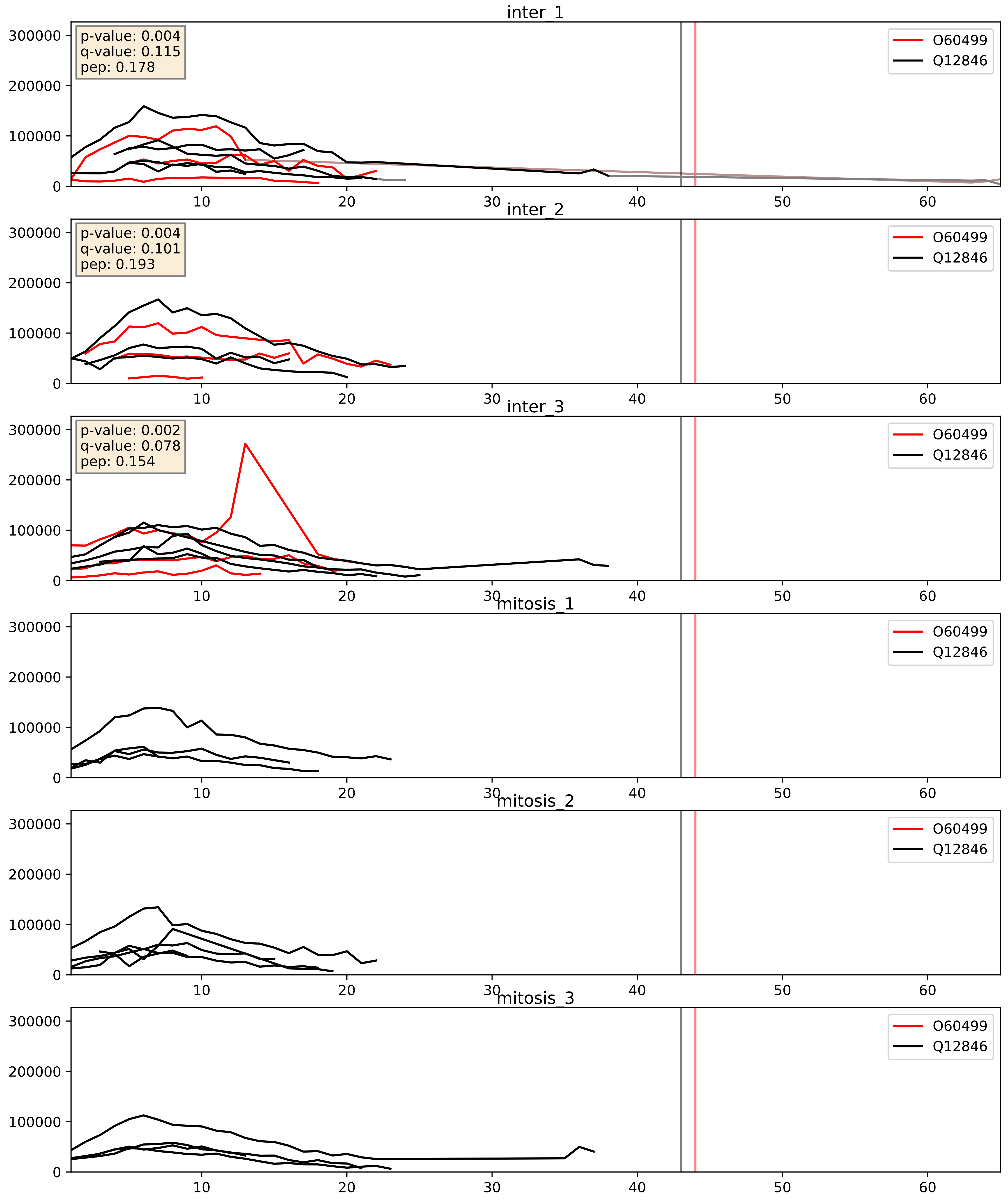


| level | condition_1 | condition_2 | pvalue | pvalue_adjusted |
|---|---|---|---|---|
| complex_abundance | mitosis | inter | 0.0002388077139889138 | 0.010310695784295044 |
| interactor_abundance | mitosis | inter | 0.00030071181320304993 | 0.01300709055050693 |
| interactor_ratio | mitosis | inter | 0.0009542417842175732 | 0.035669474554159065 |

O60499_Q86Y82
STX10_HUMAN vs STX12_HUMAN
p-value: 0.002 q-value: 0.03

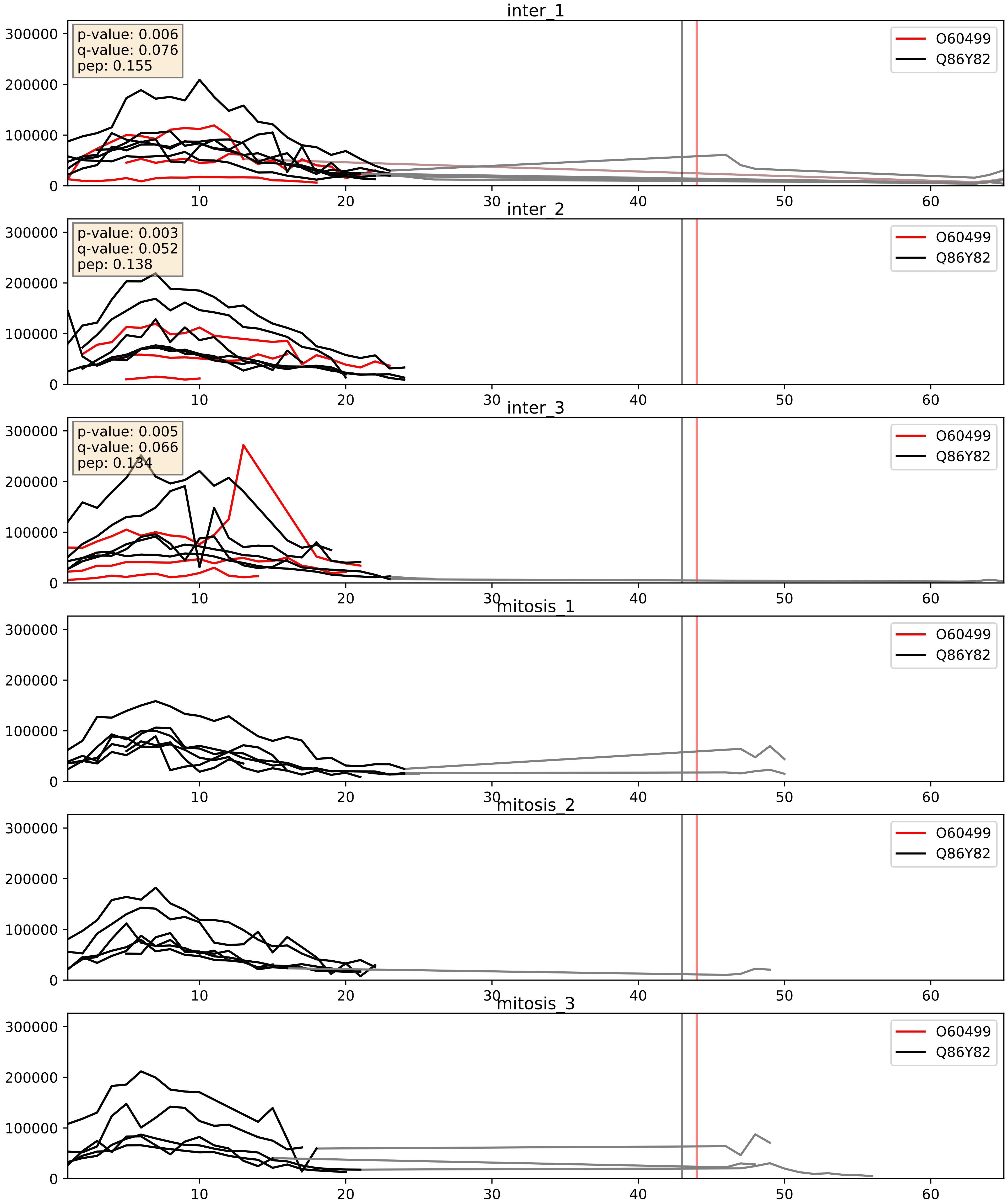


| level | condition_1 | condition_2 | pvalue | pvalue_adjusted |
|---|---|---|---|---|
| interactor_ratio | mitosis | inter | 4.00133417798734e-05 | 0.006715964816386594 |
| complex_abundance | mitosis | inter | 0.00013171704976947633 | 0.007884601021165855 |
| interactor_abundance | mitosis | inter | 0.00030071181320304993 | 0.01300709055050693 |

O60499_Q96AJ9
STX10_HUMAN vs VTI1A_HUMAN
p-value: 0.034 q-value: 0.024

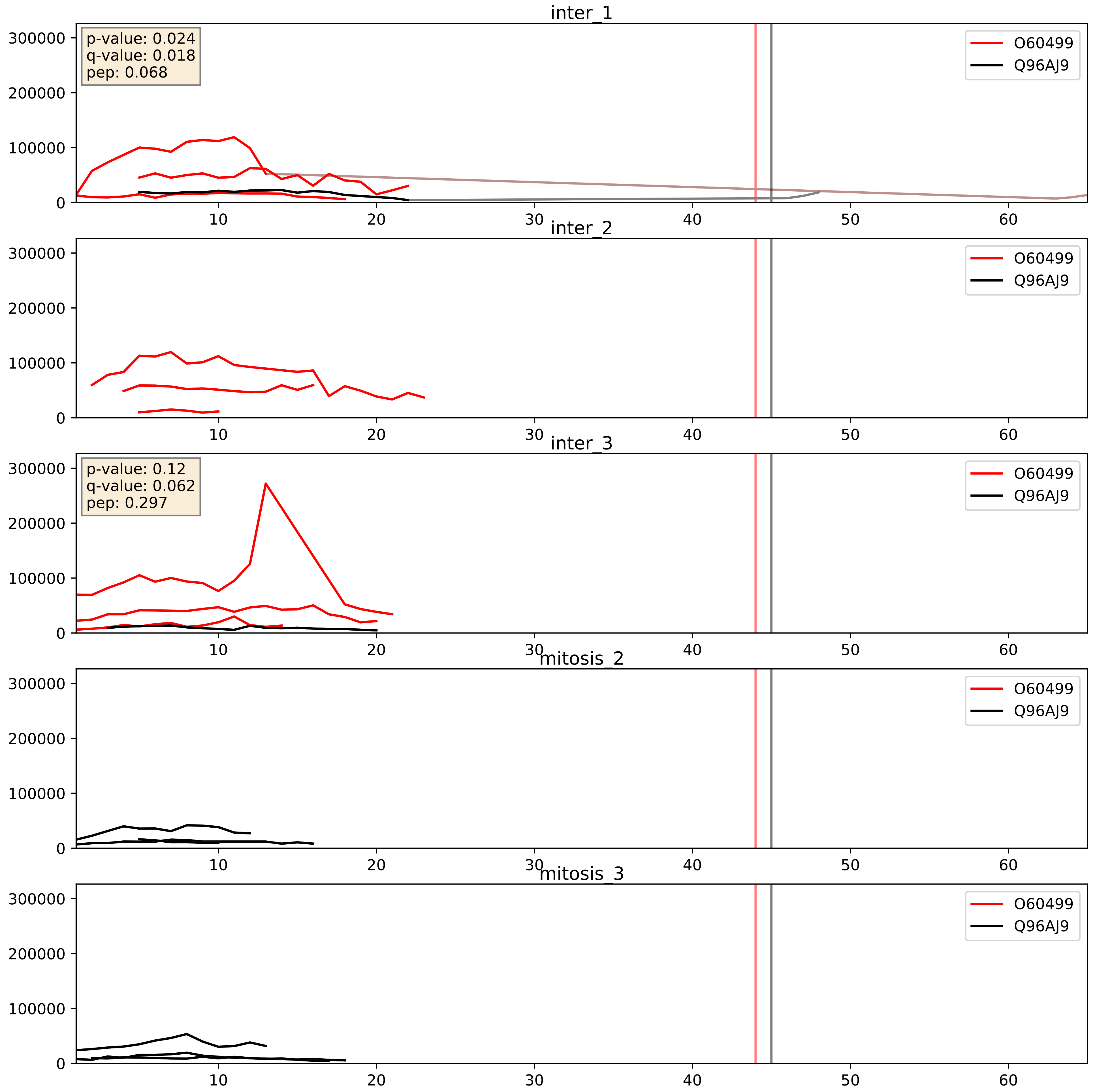


| level | condition_1 | condition_2 | pvalue | pvalue_adjusted |
|---|---|---|---|---|
| complex_abundance | mitosis | inter | 0.03150033222561792 | 0.12902786265749616 |
| interactor_abundance | mitosis | inter | 0.0331714951170254 | 0.15192509267080653 |
| interactor_ratio | mitosis | inter | 0.03777293625149884 | 0.3185579648402267 |

O60499_Q9UEU0
STX10_HUMAN vs VTI1B_HUMAN
p-value: 0.017 q-value: 0.014

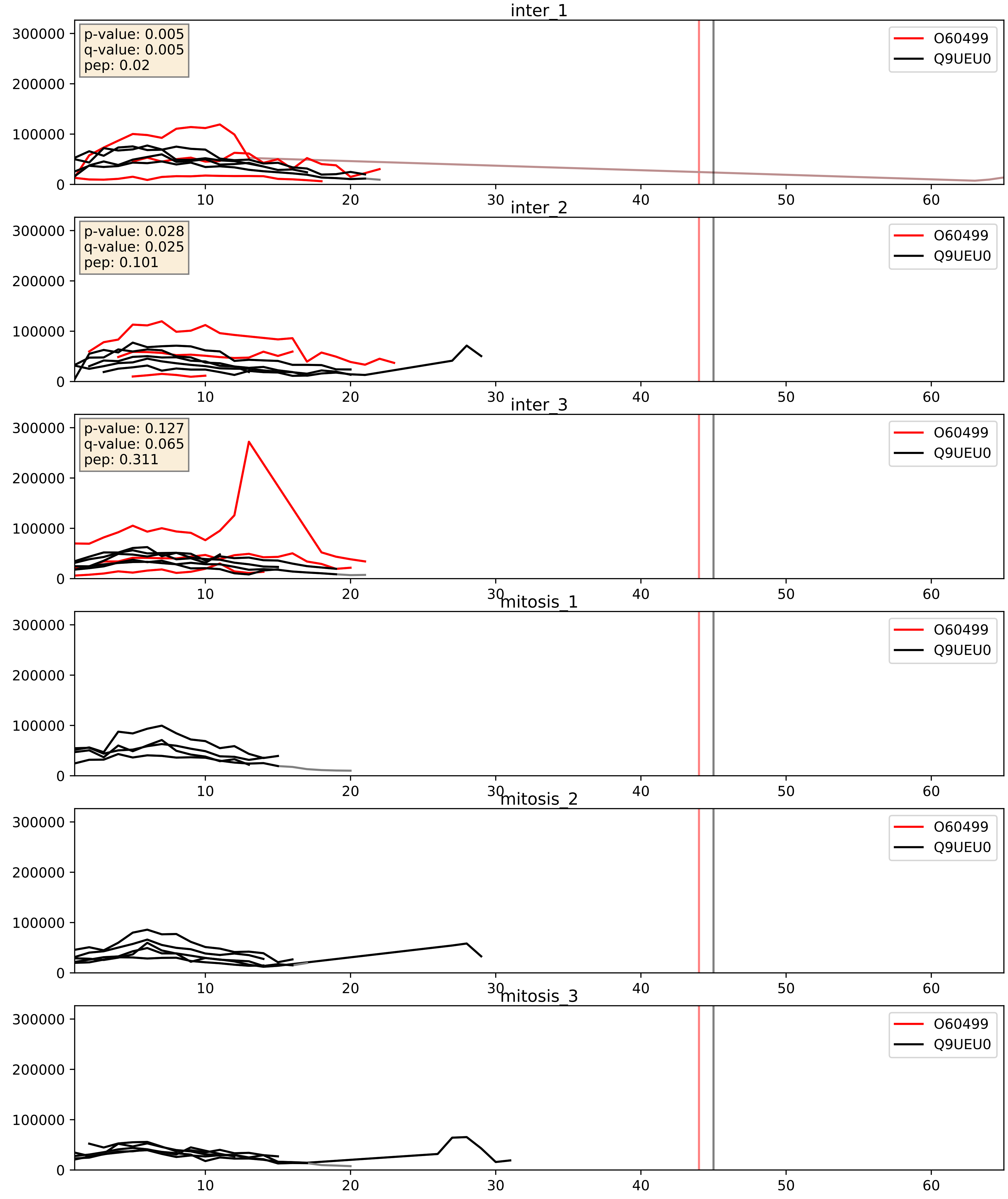


| level | condition_1 | condition_2 | pvalue | pvalue_adjusted |
|---|---|---|---|---|
| interactor_abundance | mitosis | inter | 0.0003261409774703485 | 0.013486795976551611 |
| complex_abundance | mitosis | inter | 0.00044380113923488167 | 0.01328299913234471 |
| interactor_ratio | mitosis | inter | 0.8886321278216783 | 0.9805281175683012 |
