## Supplementary Data 2 for "SECAT: Quantifying differential protein-protein interaction states by network-centric analysis": hela_string_000102_Q99595.pdf

Q99595_Q99595

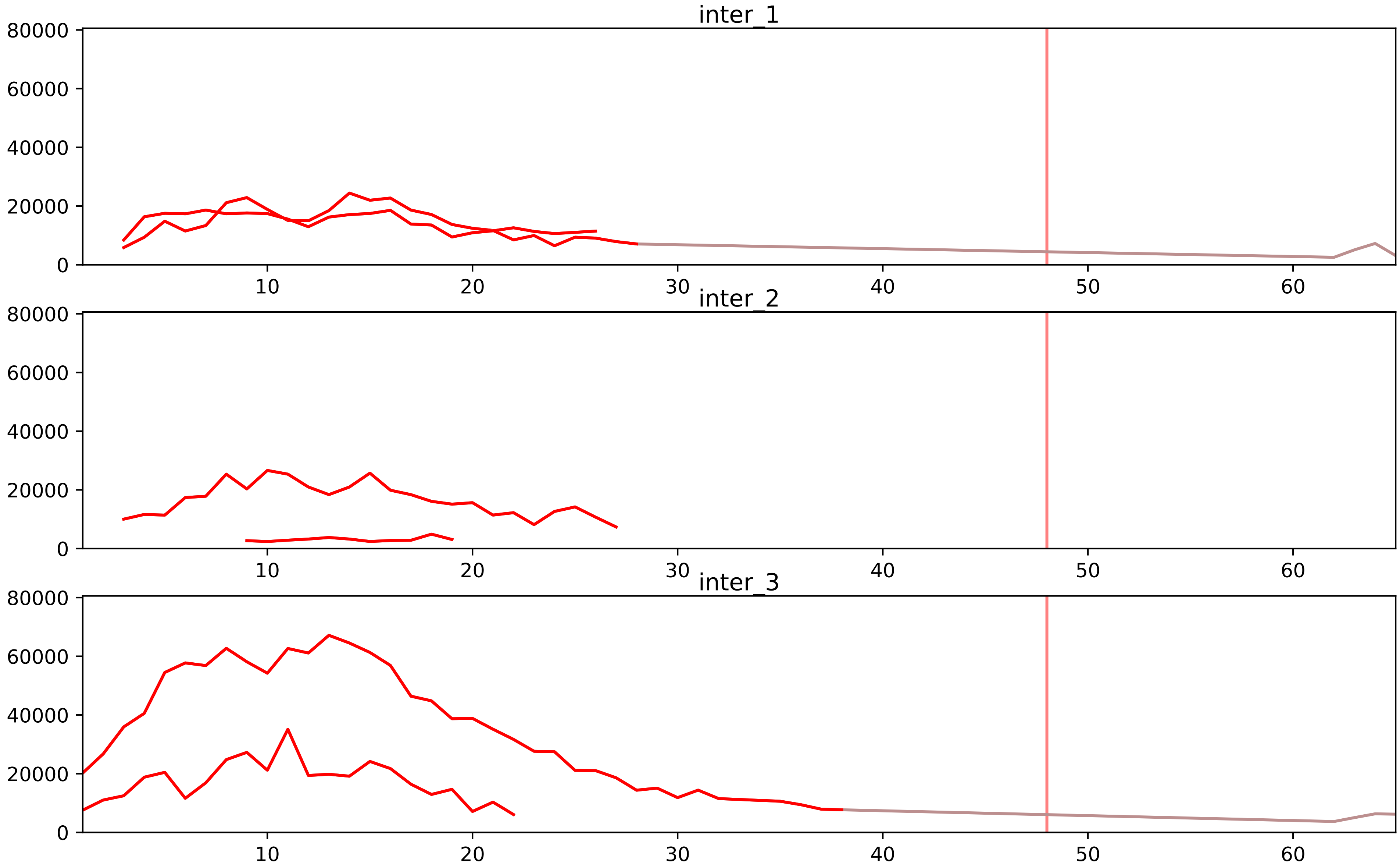


| level | condition_1 | condition_2 | pvalue | pvalue_adjusted |
|---|---|---|---|---|
| interactor_ratio | mitosis | inter | 1.0780151354088703e-05 | 0.00023898166857256882 |
| complex_abundance | mitosis | inter | 0.0001091471937437673 | 0.002323510983773901 |
| interactor_abundance | mitosis | inter | 0.0005489454749424384 | 0.007990604304242772 |
| assembled_abundance | mitosis | inter | 0.0012569822314976444 | 0.0335412168412003 |
| total_abundance | mitosis | inter | 0.0014698085542945035 | 0.04111426016435889 |
| monomer_abundance | mitosis | inter | 0.02915882130689906 | 0.2935880629314749 |

Q96DA6_Q99595
TIM14_HUMAN vs TI17A_HUMAN
p-value: 0.047 q-value: 0.031

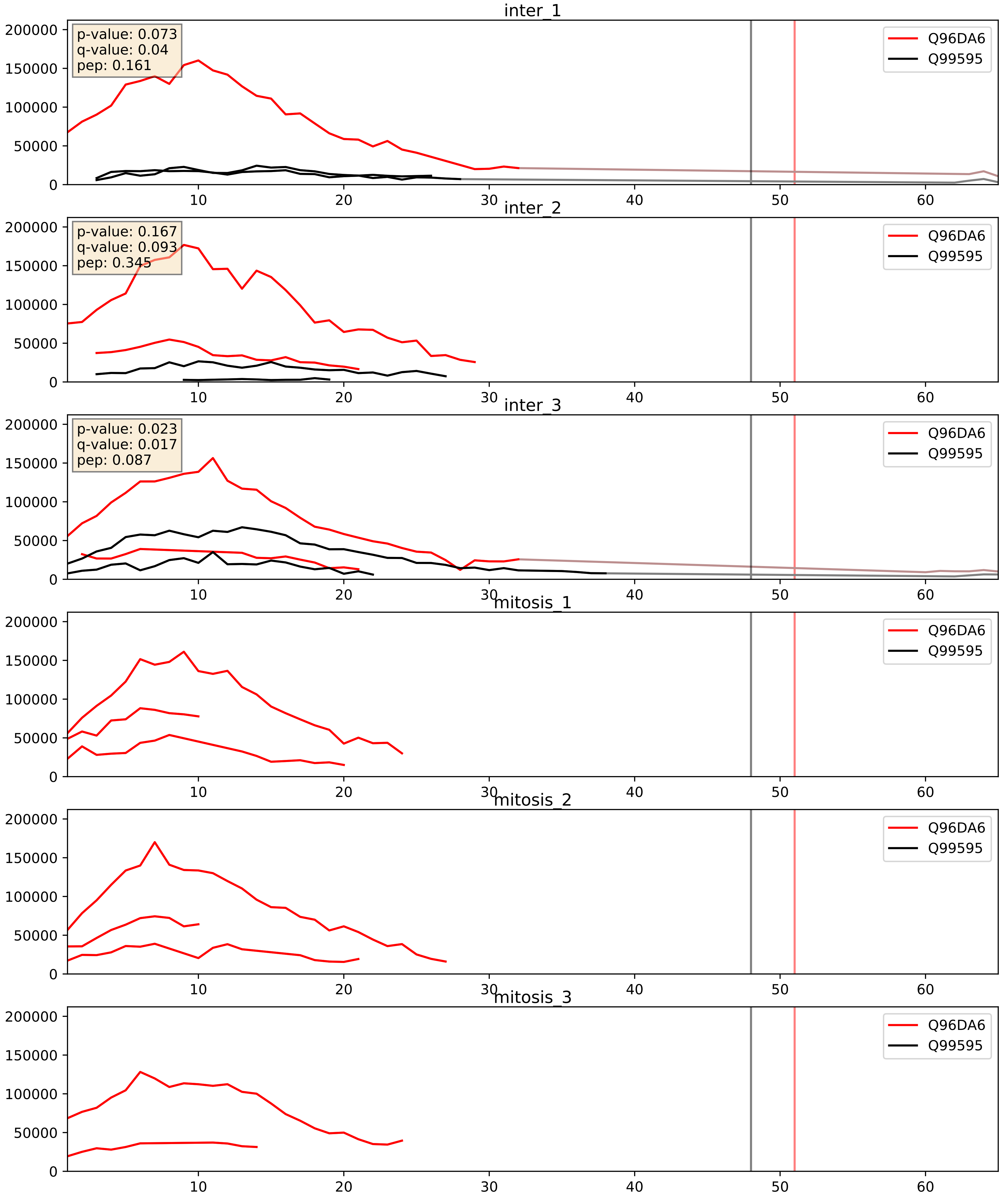


| level | condition_1 | condition_2 | pvalue | pvalue_adjusted |
|---|---|---|---|---|
| complex_abundance | mitosis | inter | 0.0005035020752987389 | 0.01422434905794457 |
| interactor_abundance | mitosis | inter | 0.0010282624686751918 | 0.023950821039073856 |
| interactor_ratio | mitosis | inter | 0.09826447396473732 | 0.5214779275500009 |

Q99595_Q9BVV7
TI17A_HUMAN vs TIM21_HUMAN
p-value: 0.08 q-value: 0.047

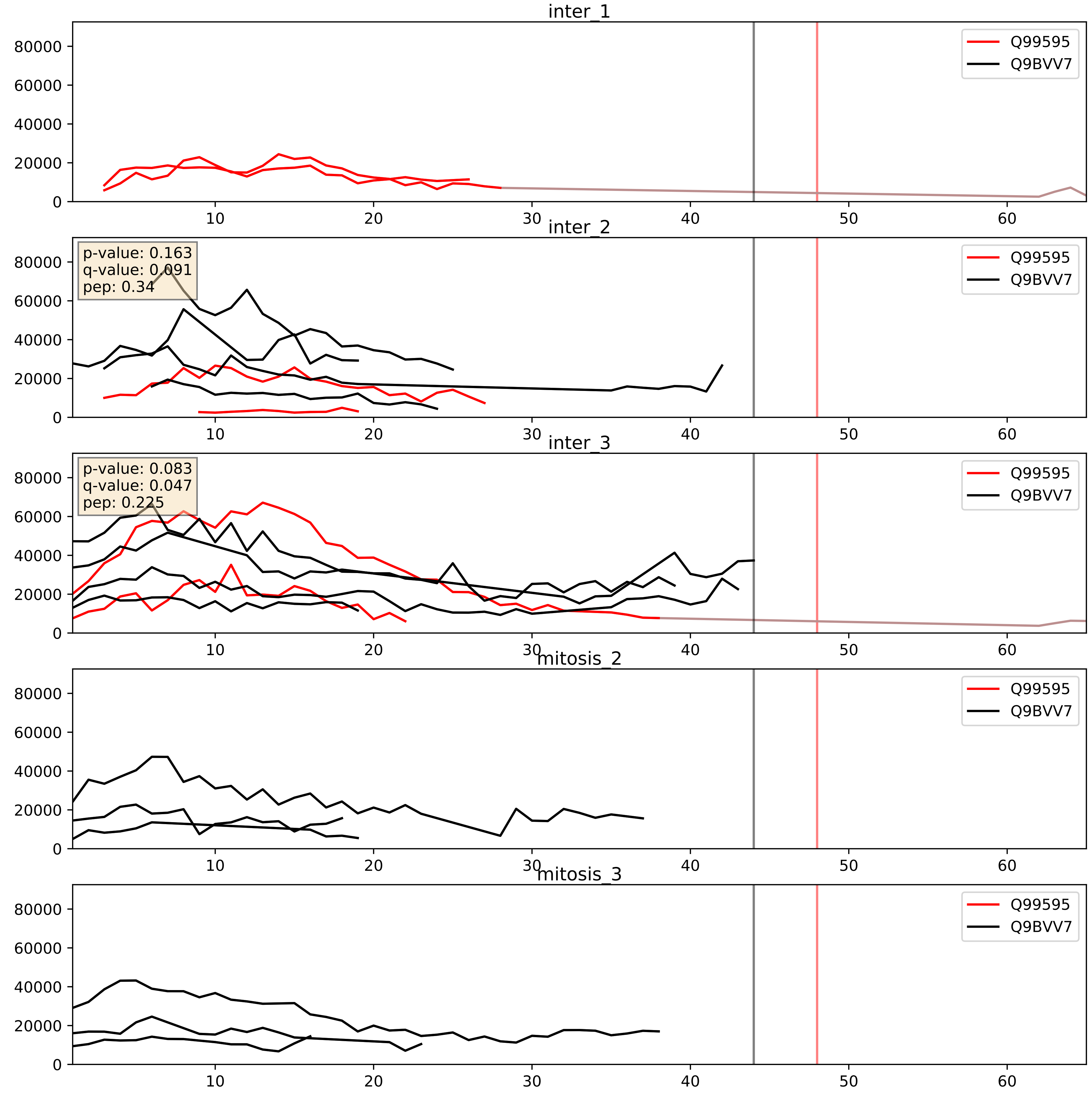


| level | condition_1 | condition_2 | pvalue | pvalue_adjusted |
|---|---|---|---|---|
| complex_abundance | mitosis | inter | 0.028759193802691027 | 0.1234203919435165 |
| interactor_abundance | mitosis | inter | 0.02898882659345003 | 0.14095106824193823 |
| interactor_ratio | mitosis | inter | 0.04818385724985605 | 0.3618015947883928 |

Q99595_Q9Y3D7
TI17A_HUMAN vs TIM16_HUMAN
p-value: 0.061 q-value: 0.038

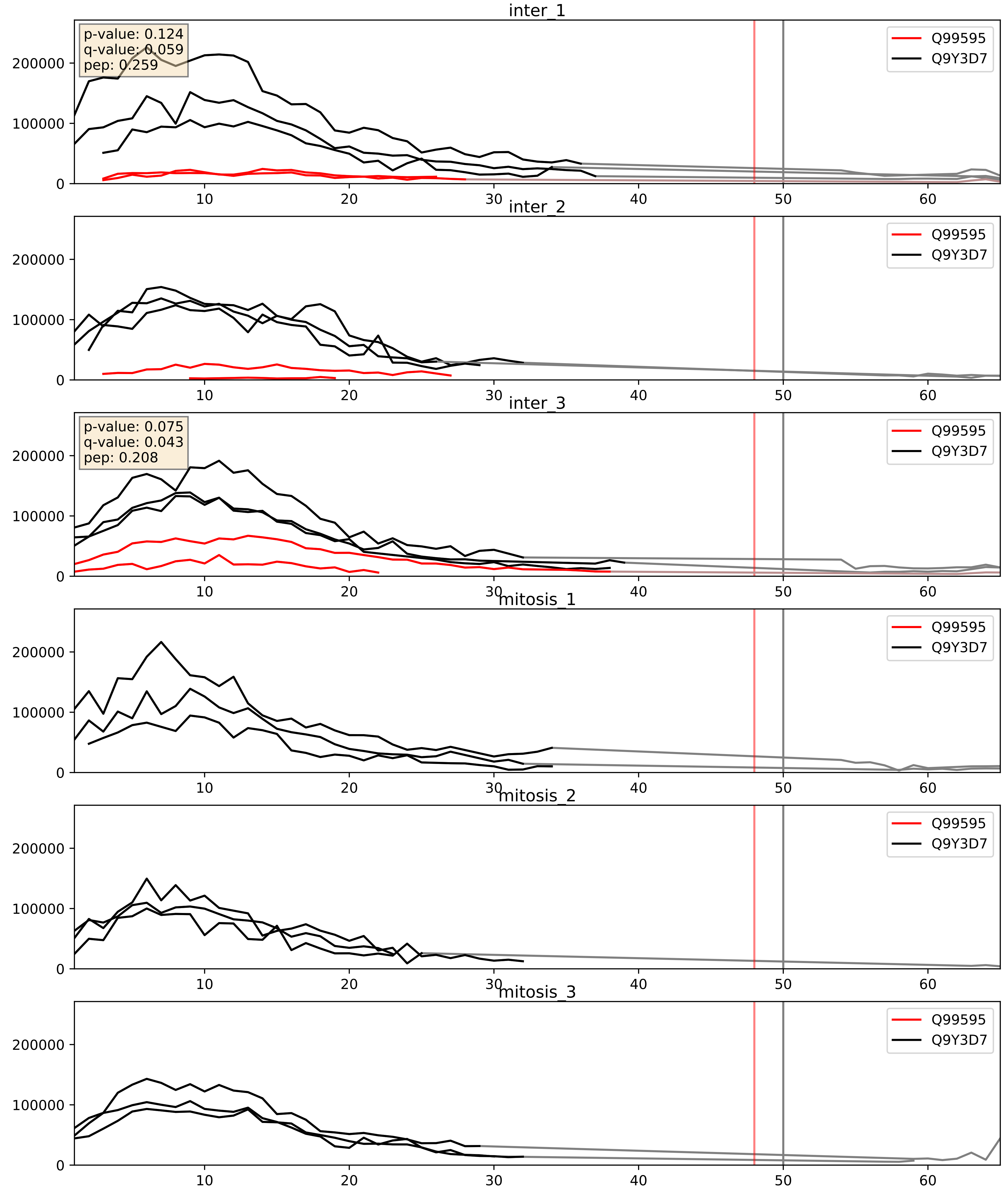


| level | condition_1 | condition_2 | pvalue | pvalue_adjusted |
|---|---|---|---|---|
| interactor_ratio | mitosis | inter | 1.1576148957150758e-05 | 0.003960471554059879 |
| complex_abundance | mitosis | inter | 0.00014027877323287218 | 0.00812325774702682 |
| interactor_abundance | mitosis | inter | 0.0013014868904229176 | 0.026652458808660705 |
