## Supplementary Data 2 for "SECAT: Quantifying differential protein-protein interaction states by network-centric analysis": hela_string_000103_O14734.pdf

O14734_O14734

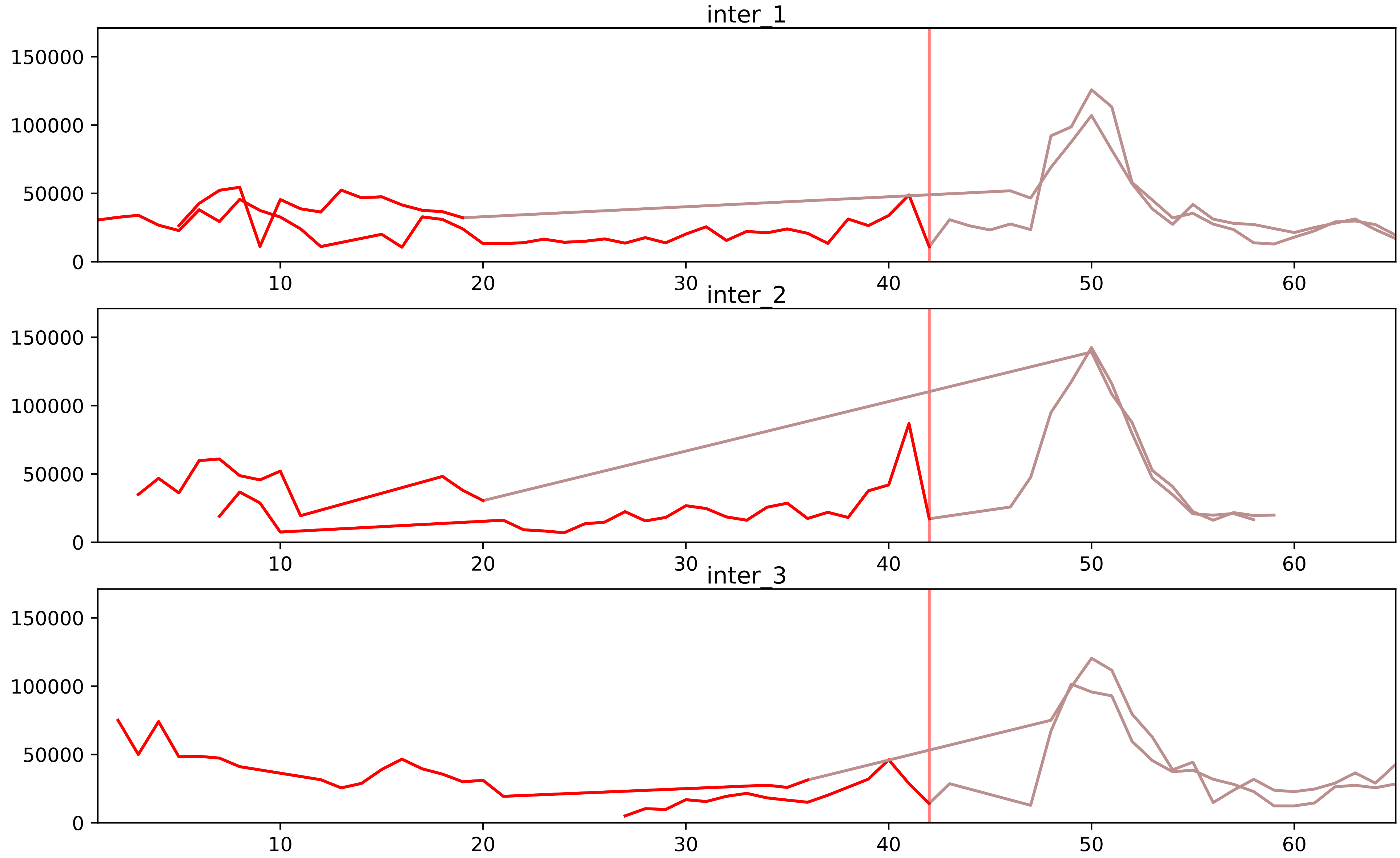


| level | condition_1 | condition_2 | pvalue | pvalue_adjusted |
|---|---|---|---|---|
| total_abundance | mitosis | inter | 1.0882506426934185e-05 | 0.0042617569399632255 |
| monomer_abundance | mitosis | inter | 5.055760073109308e-05 | 0.0267281636508792 |
| assembled_abundance | mitosis | inter | 7.696326290157016e-05 | 0.008577738896244046 |
