## Supplementary Data 2 for "SECAT: Quantifying differential protein-protein interaction states by network-centric analysis": hela_string_000104_Q9Y266.pdf

Q9Y266_Q9Y266

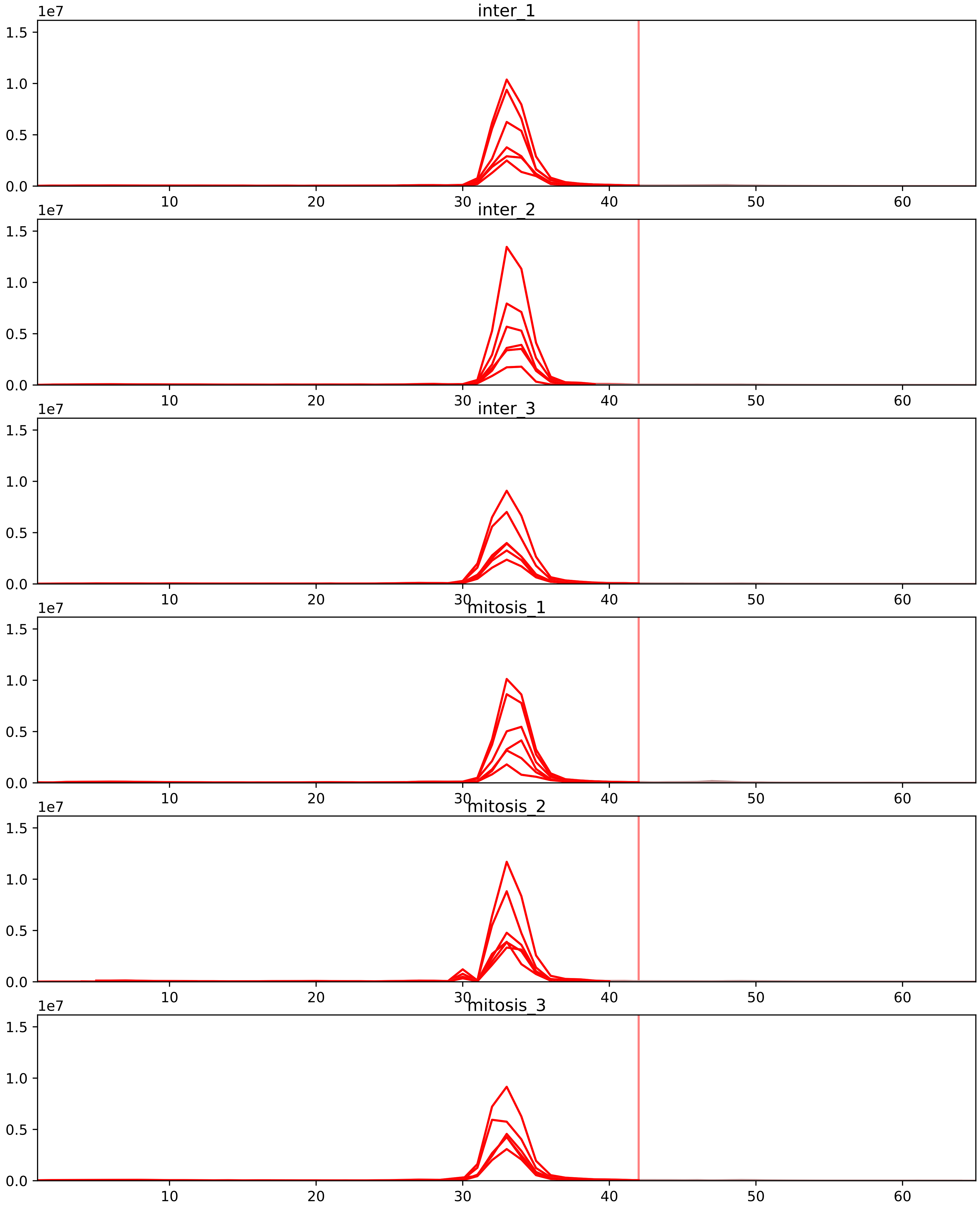


| level | condition_1 | condition_2 | pvalue | pvalue_adjusted |
|---|---|---|---|---|
| complex_abundance | mitosis | inter | 1.1936559716790173e-05 | 0.000549081746972348 |
| interactor_abundance | mitosis | inter | 1.4543245862946409e-05 | 0.0006689893096955349 |
| interactor_ratio | mitosis | inter | 0.0001559547874814811 | 0.002473765594533838 |
| assembled_abundance | mitosis | inter | 0.13088753279004076 | 0.4284507279651614 |
| total_abundance | mitosis | inter | 0.2765890298522224 | 0.6089270051792506 |
| monomer_abundance | mitosis | inter | 0.6147436587184092 | 0.8197254628726277 |

P53350_Q9Y266
PLK1_HUMAN vs NUDC_HUMAN
p-value: 0.073 q-value: 0.044

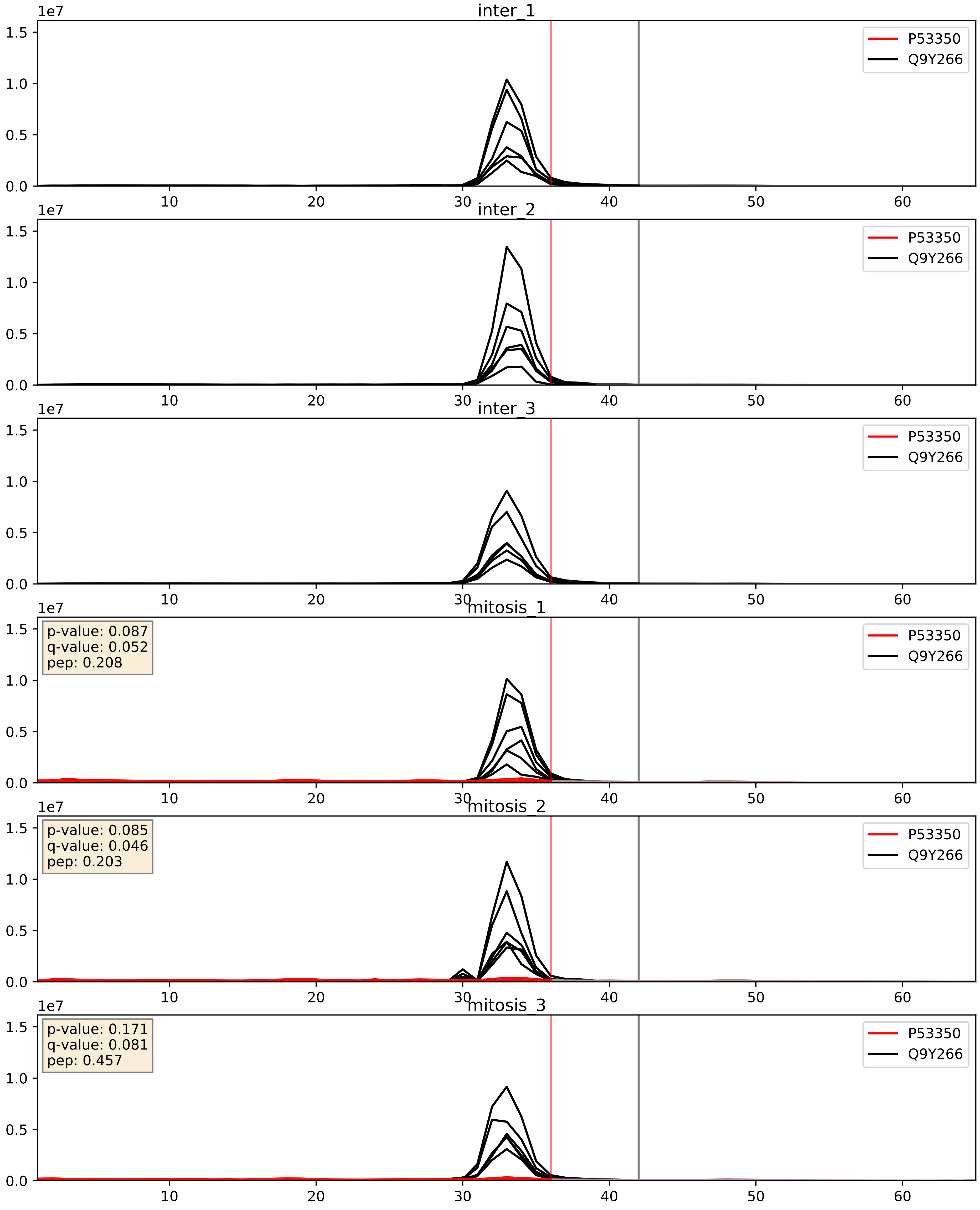


| level | condition_1 | condition_2 | pvalue | pvalue_adjusted |
|---|---|---|---|---|
| complex_abundance | mitosis | inter | 1.1936559716790173e-05 | 0.005691857151652958 |
| interactor_abundance | mitosis | inter | 1.8795469290468416e-05 | 0.004081879629435812 |
| interactor_ratio | mitosis | inter | 0.0001559547874814811 | 0.012836278661937291 |
