## Supplementary Data 2 for "SECAT: Quantifying differential protein-protein interaction states by network-centric analysis": hela_string_000105_Q9BVI4.pdf

Q9BVI4_Q9BVI4

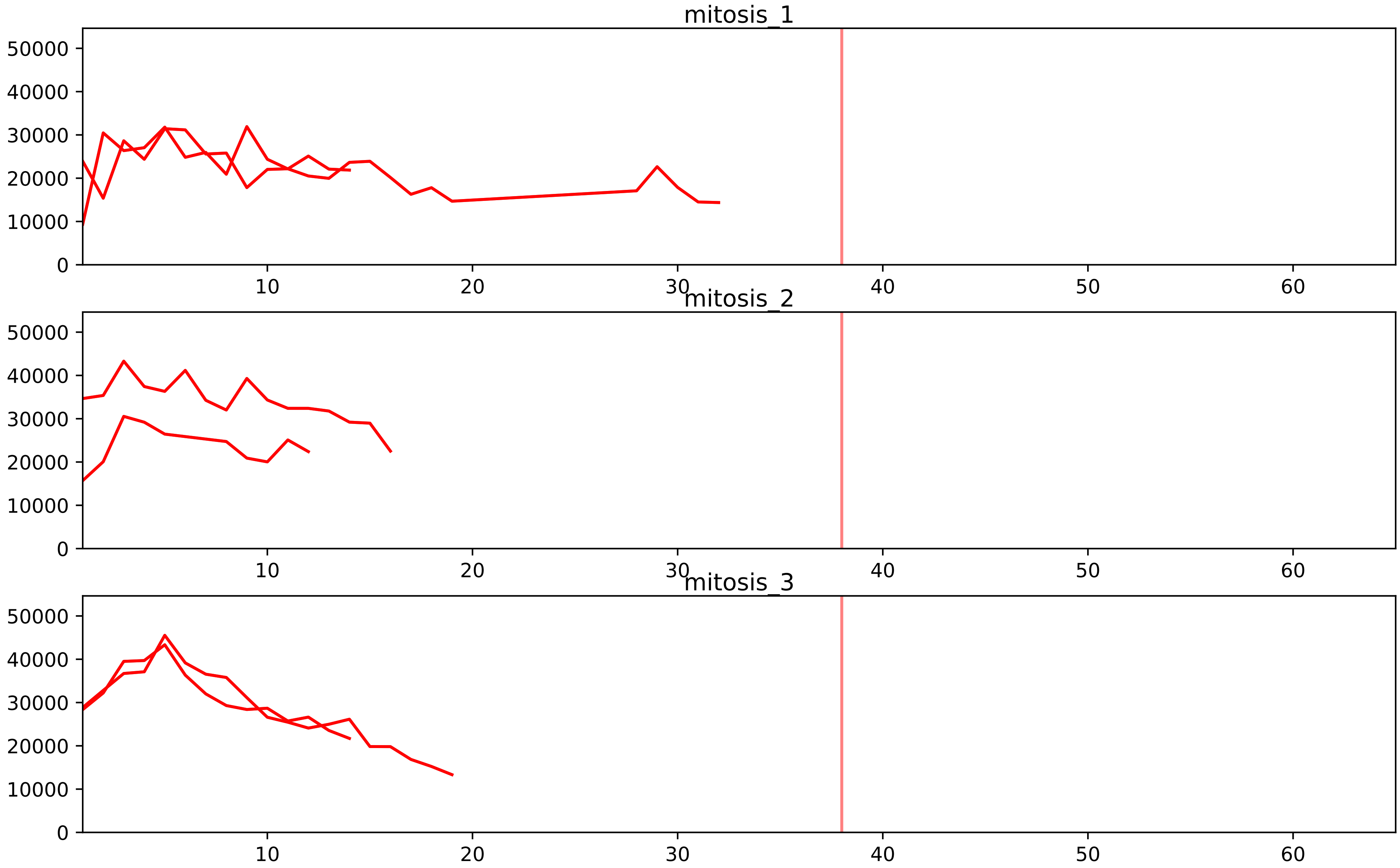


| level | condition_1 | condition_2 | pvalue | pvalue_adjusted |
|---|---|---|---|---|
| interactor_ratio | mitosis | inter | 1.265862488983855e-05 | 0.0002740219976153286 |
| assembled_abundance | mitosis | inter | 0.00012268879319921302 | 0.010835966810670115 |
| total_abundance | mitosis | inter | 0.0001737474892125411 | 0.016082699410564488 |
| complex_abundance | mitosis | inter | 0.0003100717640601649 | 0.005048956158147818 |
| interactor_abundance | mitosis | inter | 0.0037557272239833396 | 0.028793908717205604 |

Q13601_Q9BVI4
KRR1_HUMAN vs NOC4L_HUMAN
p-value: 0.076 q-value: 0.046

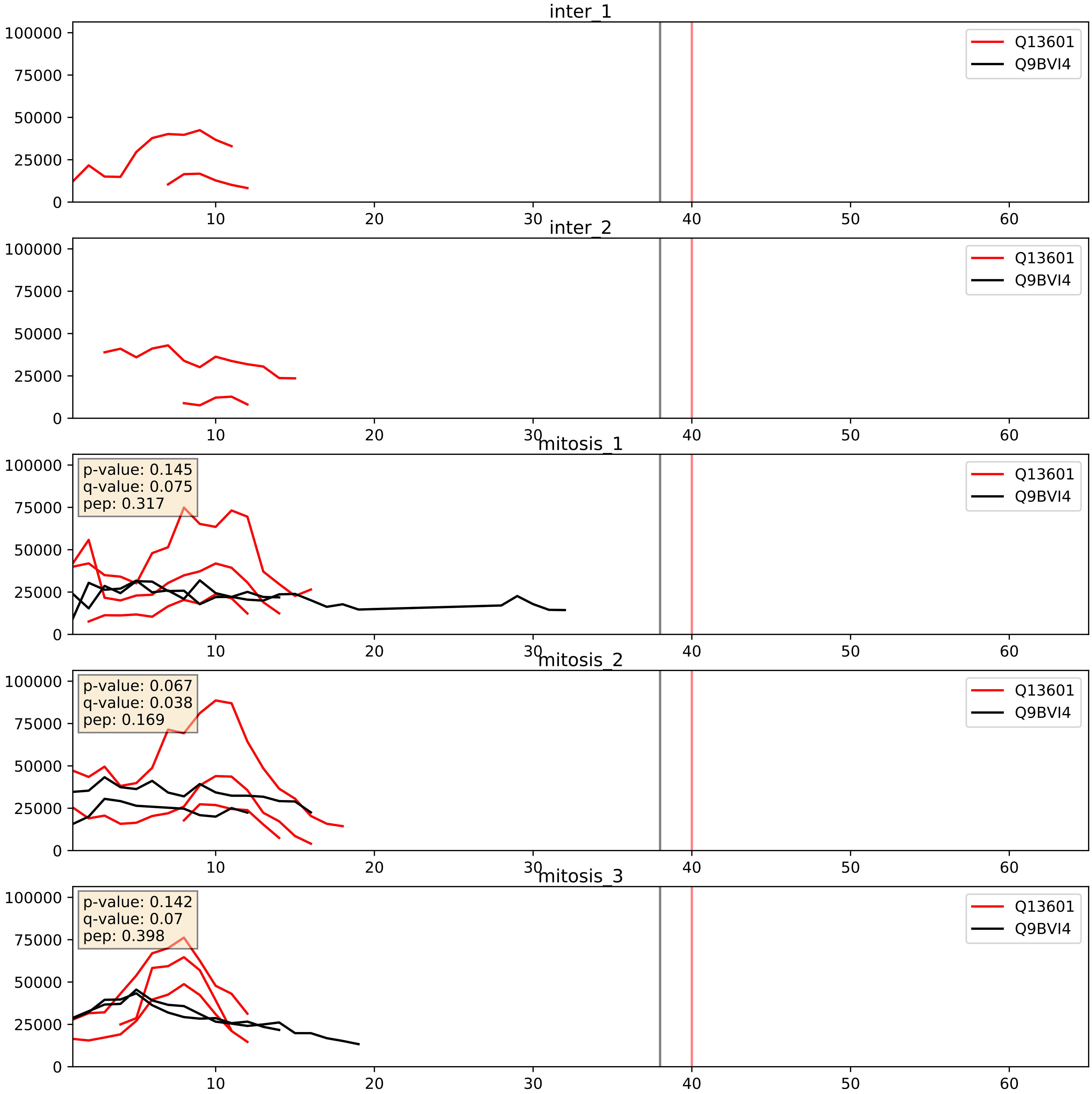


| level | condition_1 | condition_2 | pvalue | pvalue_adjusted |
|---|---|---|---|---|
| interactor_ratio | mitosis | inter | 0.004800392448507704 | 0.09212312397066487 |
| interactor_abundance | mitosis | inter | 0.007172766741519979 | 0.07326835716874823 |
| complex_abundance | mitosis | inter | 0.007934669910300927 | 0.0659424994487145 |

Q9BVI4_Q9Y2X3
NOC4L_HUMAN vs NOP58_HUMAN
p-value: 0.081 q-value: 0.048

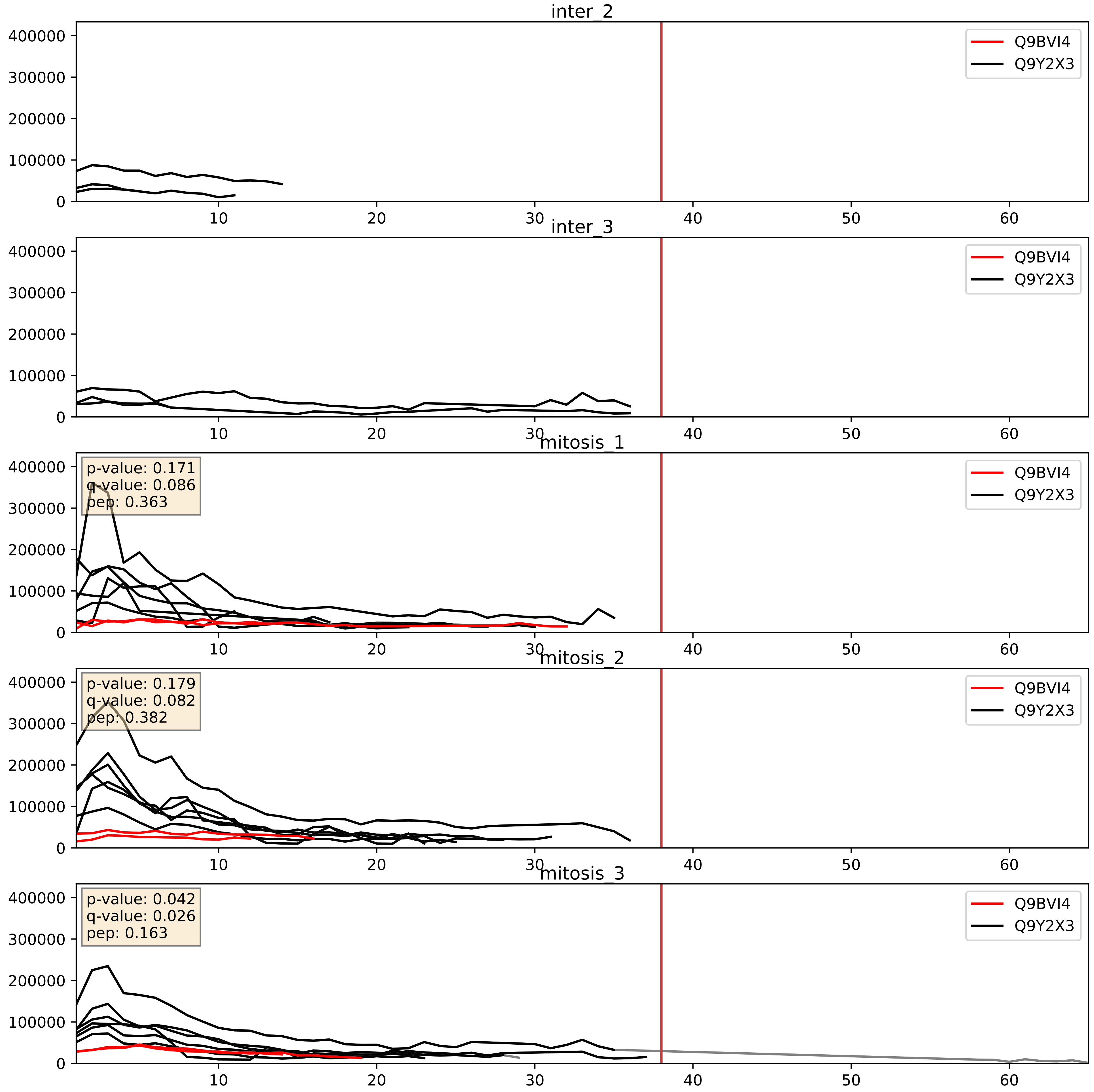


| level | condition_1 | condition_2 | pvalue | pvalue_adjusted |
|---|---|---|---|---|
| interactor_ratio | mitosis | inter | 0.00017598563399961114 | 0.01369488206396974 |
| complex_abundance | mitosis | inter | 0.0002596632330815969 | 0.010784252527150902 |
| interactor_abundance | mitosis | inter | 0.009419913204717079 | 0.08408839833172471 |
