## Supplementary Data 2 for "SECAT: Quantifying differential protein-protein interaction states by network-centric analysis": hela_string_000106_O95169.pdf

O95169_O95169

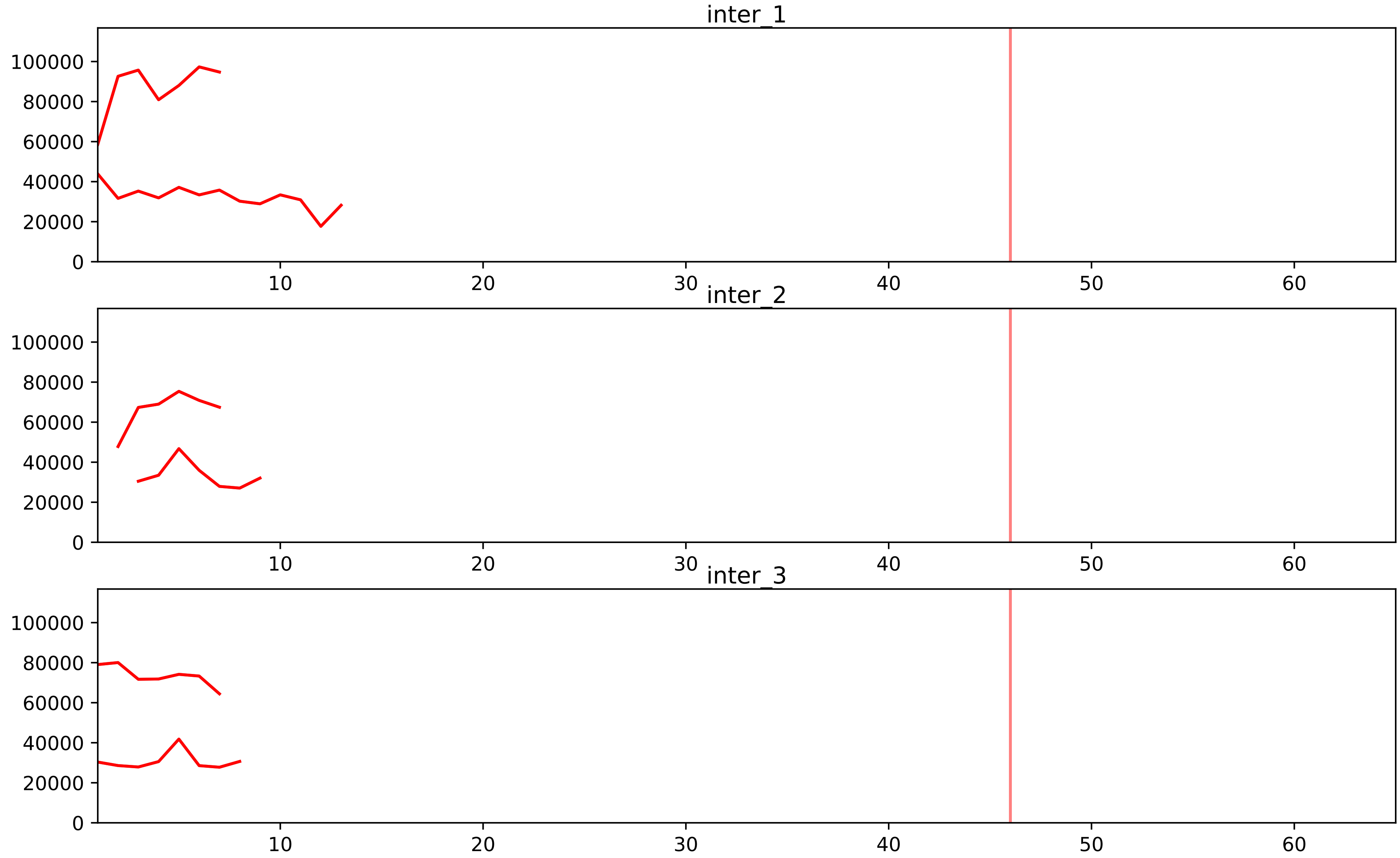


| level | condition_1 | condition_2 | pvalue | pvalue_adjusted |
|---|---|---|---|---|
| interactor_ratio | mitosis | inter | 1.3417239739498226e-05 | 0.0002870665246590318 |
| assembled_abundance | mitosis | inter | 0.00010362932801090962 | 0.010189410303009105 |
| total_abundance | mitosis | inter | 0.00020908857881109716 | 0.017741165912121593 |
| complex_abundance | mitosis | inter | 0.0011372662437390557 | 0.011530879234310282 |
| interactor_abundance | mitosis | inter | 0.0011734886263281338 | 0.013038452664356043 |

O95168_O95169
NDUB4_HUMAN vs NDUB8_HUMAN
p-value: 0.015 q-value: 0.012

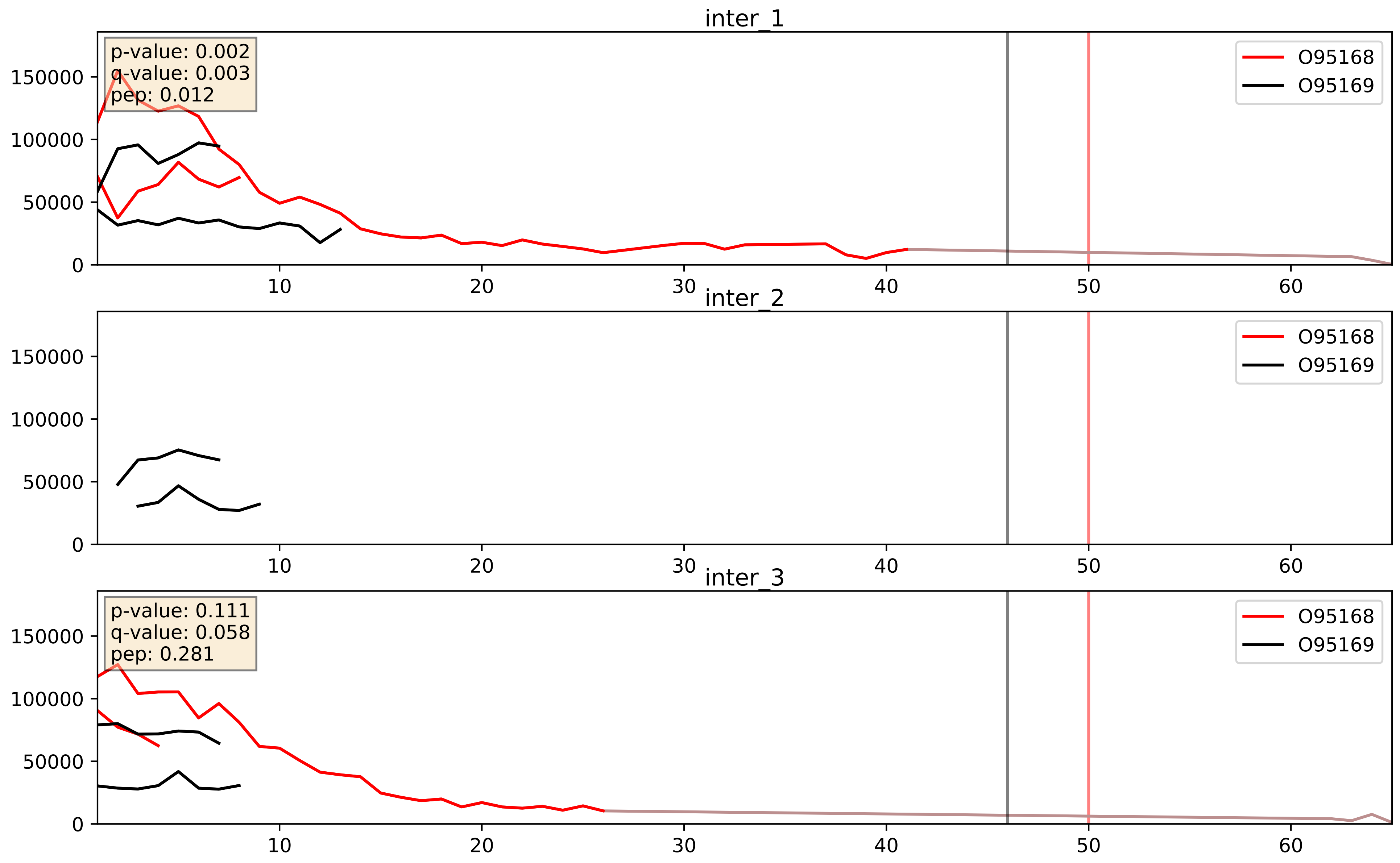


| level | condition_1 | condition_2 | pvalue | pvalue_adjusted |
|---|---|---|---|---|
| complex_abundance | mitosis | inter | 0.03489152424285779 | 0.13568043947020356 |
| interactor_abundance | mitosis | inter | 0.03825501507876028 | 0.15936046986653518 |
| interactor_ratio | mitosis | inter | 0.1232973270180264 | 0.5739294160480145 |

O95169_P03905
NDUB8_HUMAN vs NU4M_HUMAN
p-value: 0.051 q-value: 0.033

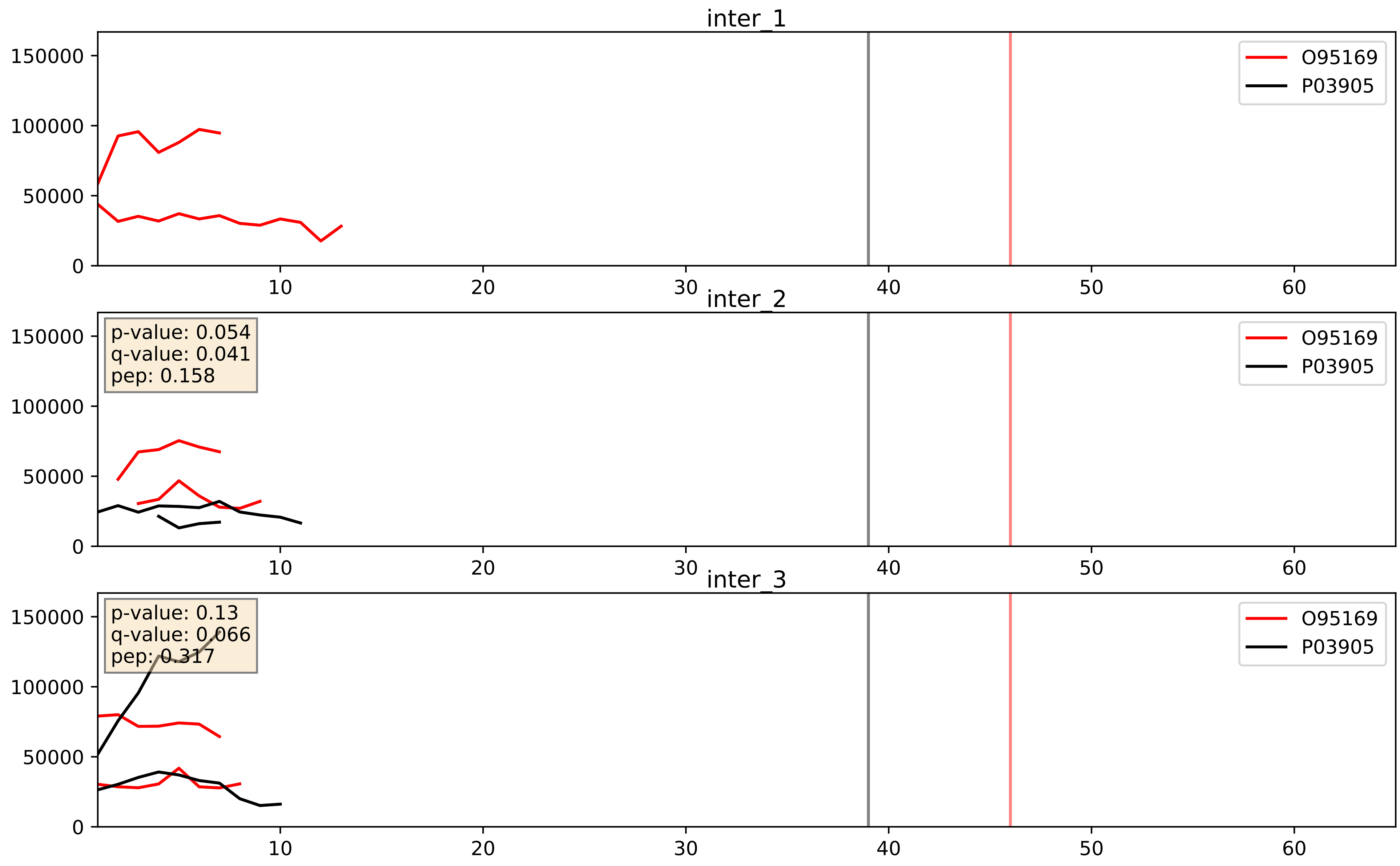


| level | condition_1 | condition_2 | pvalue | pvalue_adjusted |
|---|---|---|---|---|
| complex_abundance | mitosis | inter | 0.019441272961079645 | 0.1040108103417761 |
| interactor_abundance | mitosis | inter | 0.020443769332039115 | 0.12002651953515421 |
| interactor_ratio | mitosis | inter | 0.8216233008694807 | 0.9678127776858063 |

O95169_P17568
NDUB8_HUMAN vs NDUB7_HUMAN
p-value: 0.048 q-value: 0.032

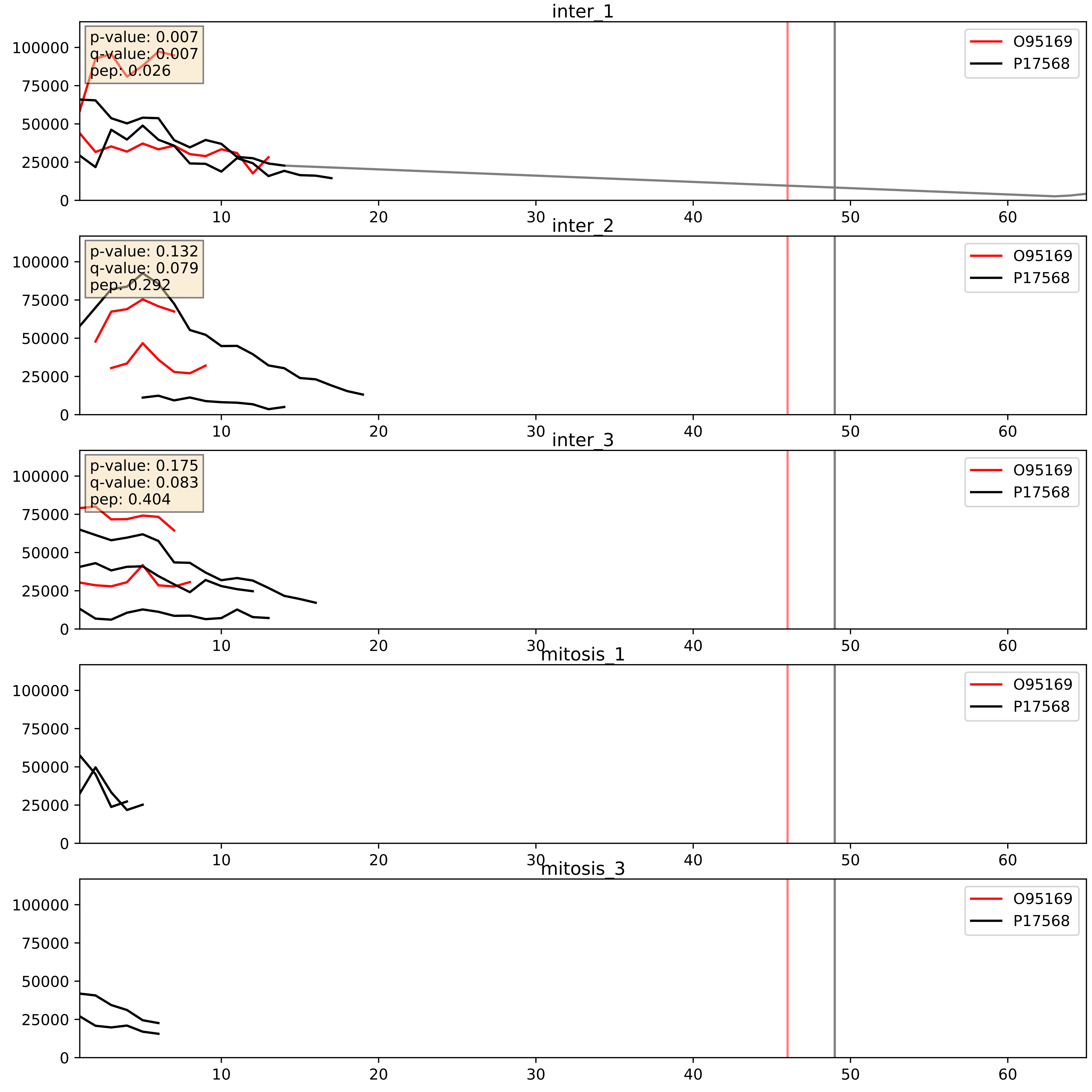


| level | condition_1 | condition_2 | pvalue | pvalue_adjusted |
|---|---|---|---|---|
| interactor_abundance | mitosis | inter | 0.0013596111197408754 | 0.027287857409101744 |
| complex_abundance | mitosis | inter | 0.0014852922757540659 | 0.02676653027464169 |
| interactor_ratio | mitosis | inter | 0.020004025436845558 | 0.21404307217424748 |

O95169_P47985
NDUB8_HUMAN vs UCRI_HUMAN
p-value: 0.07 q-value: 0.042

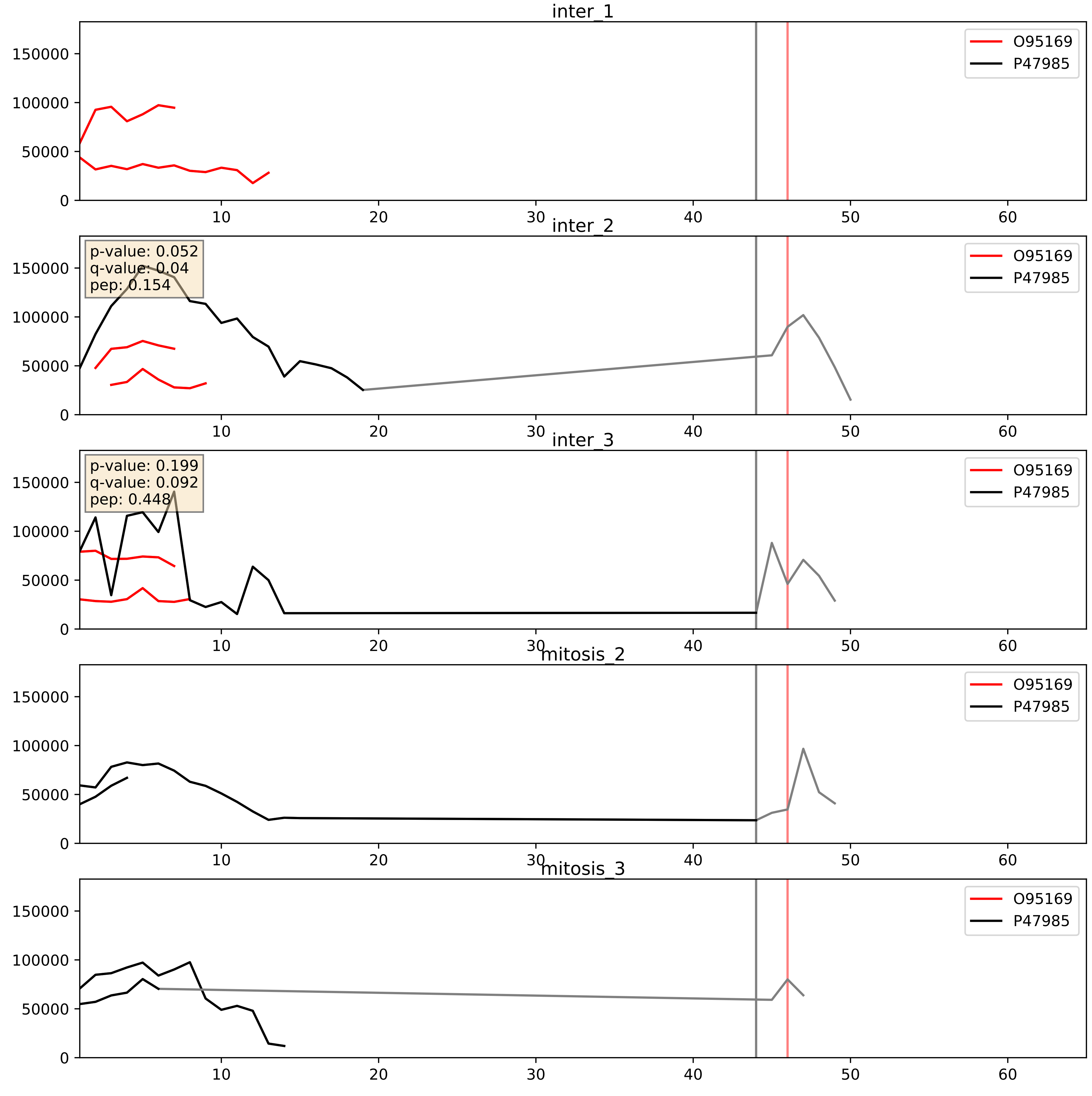


| level | condition_1 | condition_2 | pvalue | pvalue_adjusted |
|---|---|---|---|---|
| interactor_ratio | mitosis | inter | 0.011437192423823754 | 0.1546640871215345 |
| interactor_abundance | mitosis | inter | 0.020443769332039115 | 0.12002651953515421 |
| complex_abundance | mitosis | inter | 0.02559502909058073 | 0.11709002654681128 |

O95169_P49821
NDUB8_HUMAN vs NDUV1_HUMAN
p-value: 0.065 q-value: 0.04

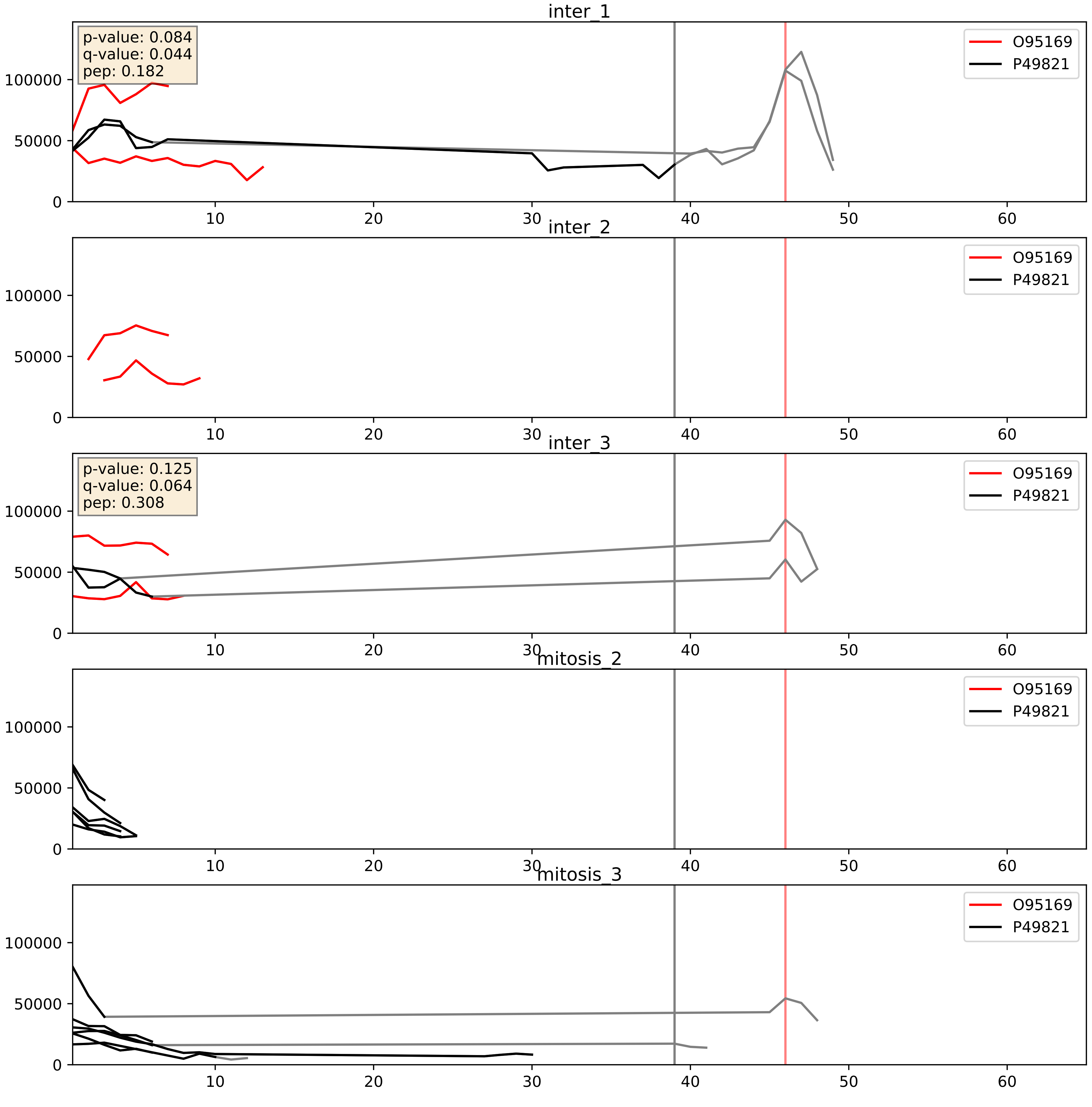


| level | condition_1 | condition_2 | pvalue | pvalue_adjusted |
|---|---|---|---|---|
| complex_abundance | mitosis | inter | 0.03022146279086701 | 0.12600863199699056 |
| interactor_abundance | mitosis | inter | 0.03071944481758559 | 0.1455735312929244 |
| interactor_ratio | mitosis | inter | 0.12623110897717676 | 0.5750062399685051 |

O95169_Q9P0J0
NDUB8_HUMAN vs NDUAD_HUMAN
p-value: 0.046 q-value: 0.031

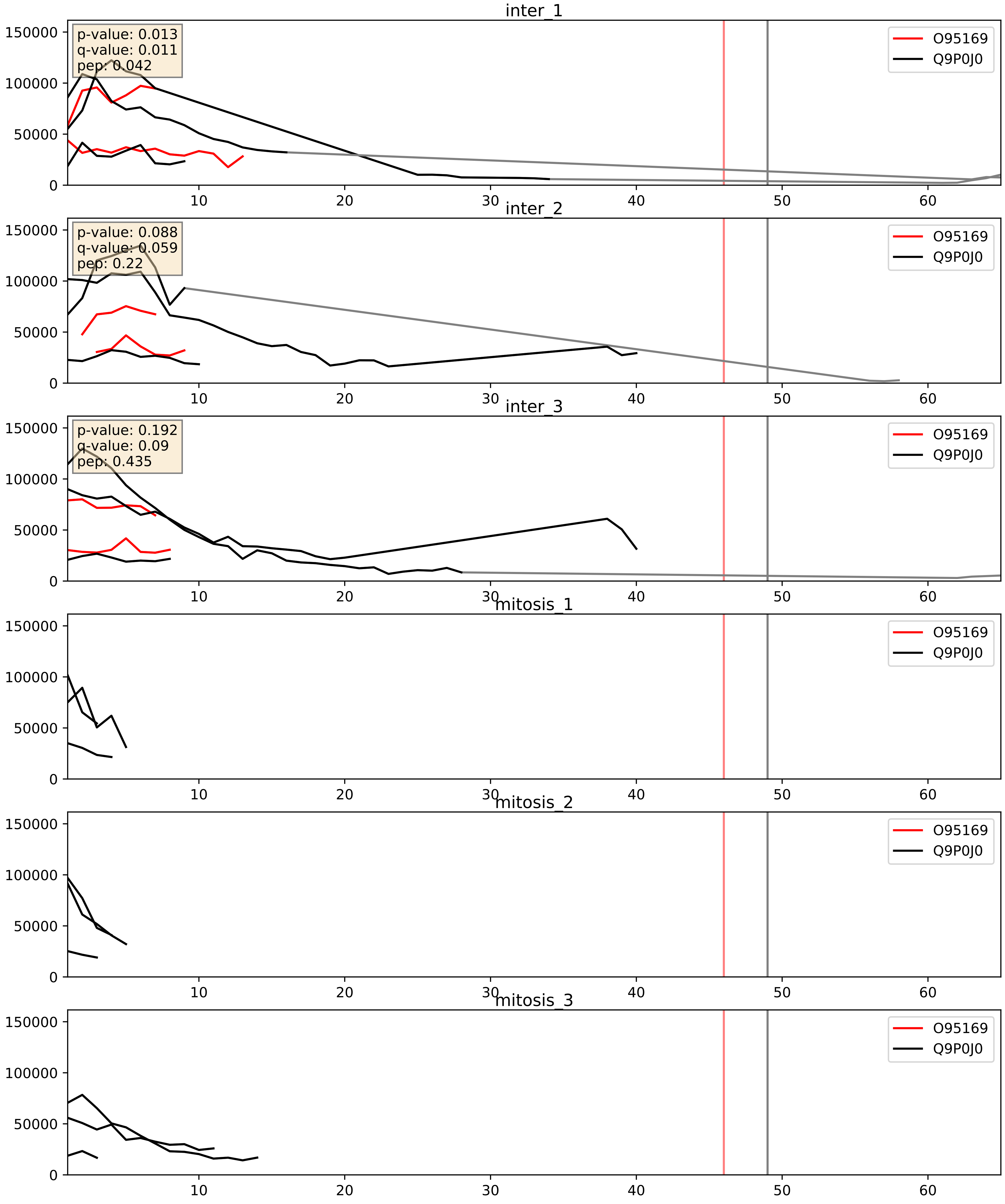


| level | condition_1 | condition_2 | pvalue | pvalue_adjusted |
|---|---|---|---|---|
| interactor_ratio | mitosis | inter | 0.000806772542671012 | 0.032604652001472285 |
| complex_abundance | mitosis | inter | 0.0009089003714998096 | 0.0197847007881119 |
| interactor_abundance | mitosis | inter | 0.0013596111197408754 | 0.02728785740910174 |

O95169_Q9Y6M9
NDUB8_HUMAN vs NDUB9_HUMAN
p-value: 0.076 q-value: 0.045

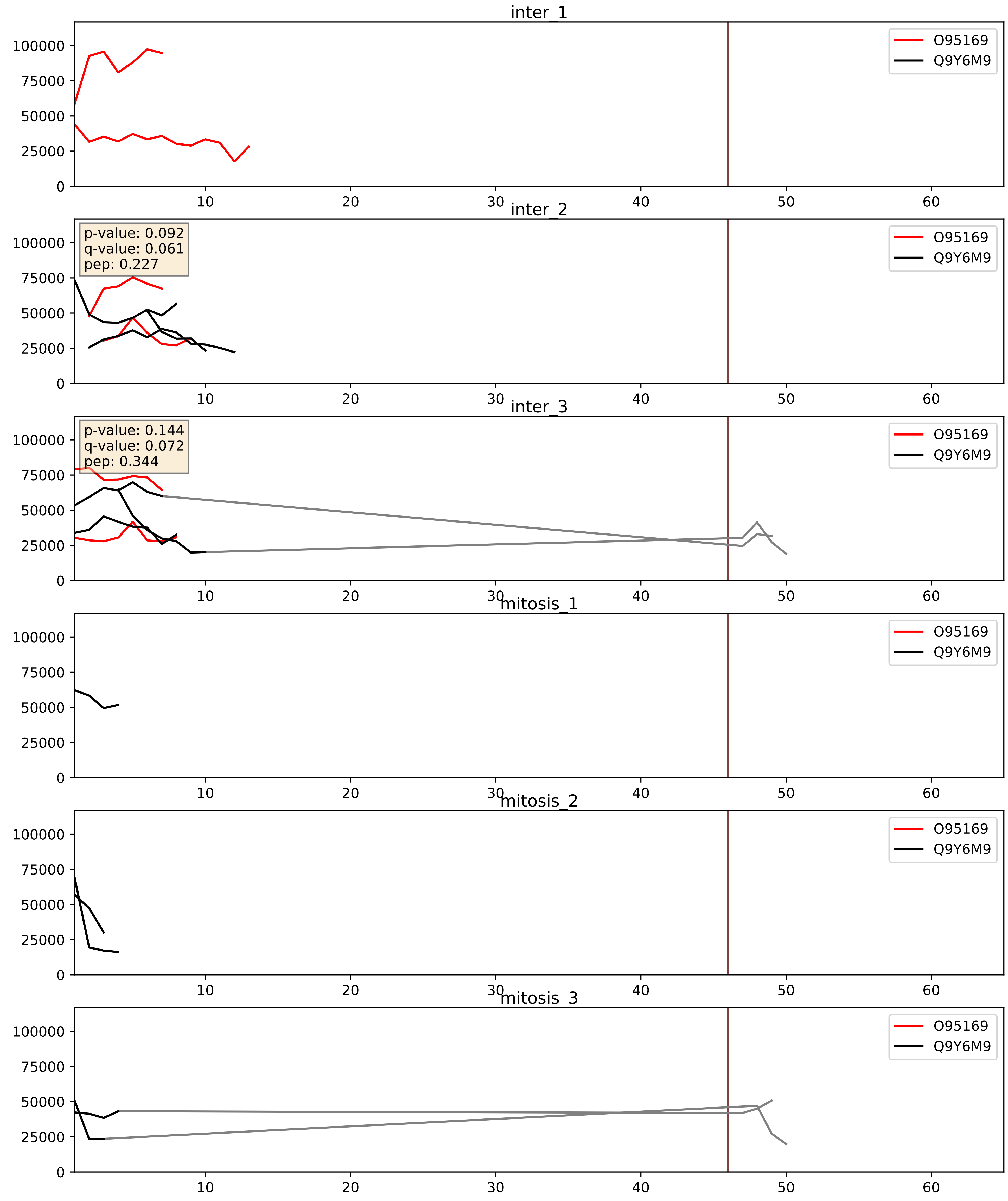


| level | condition_1 | condition_2 | pvalue | pvalue_adjusted |
|---|---|---|---|---|
| interactor_ratio | mitosis | inter | 0.016428755974308252 | 0.1887653035437297 |
| complex_abundance | mitosis | inter | 0.019400801532379355 | 0.1039241934400296 |
| interactor_abundance | mitosis | inter | 0.020443769332039115 | 0.12002651953515421 |
