## Supplementary Data 2 for "SECAT: Quantifying differential protein-protein interaction states by network-centric analysis": hela_string_000107_Q6IBW4.pdf

Q6IBW4_Q6IBW4

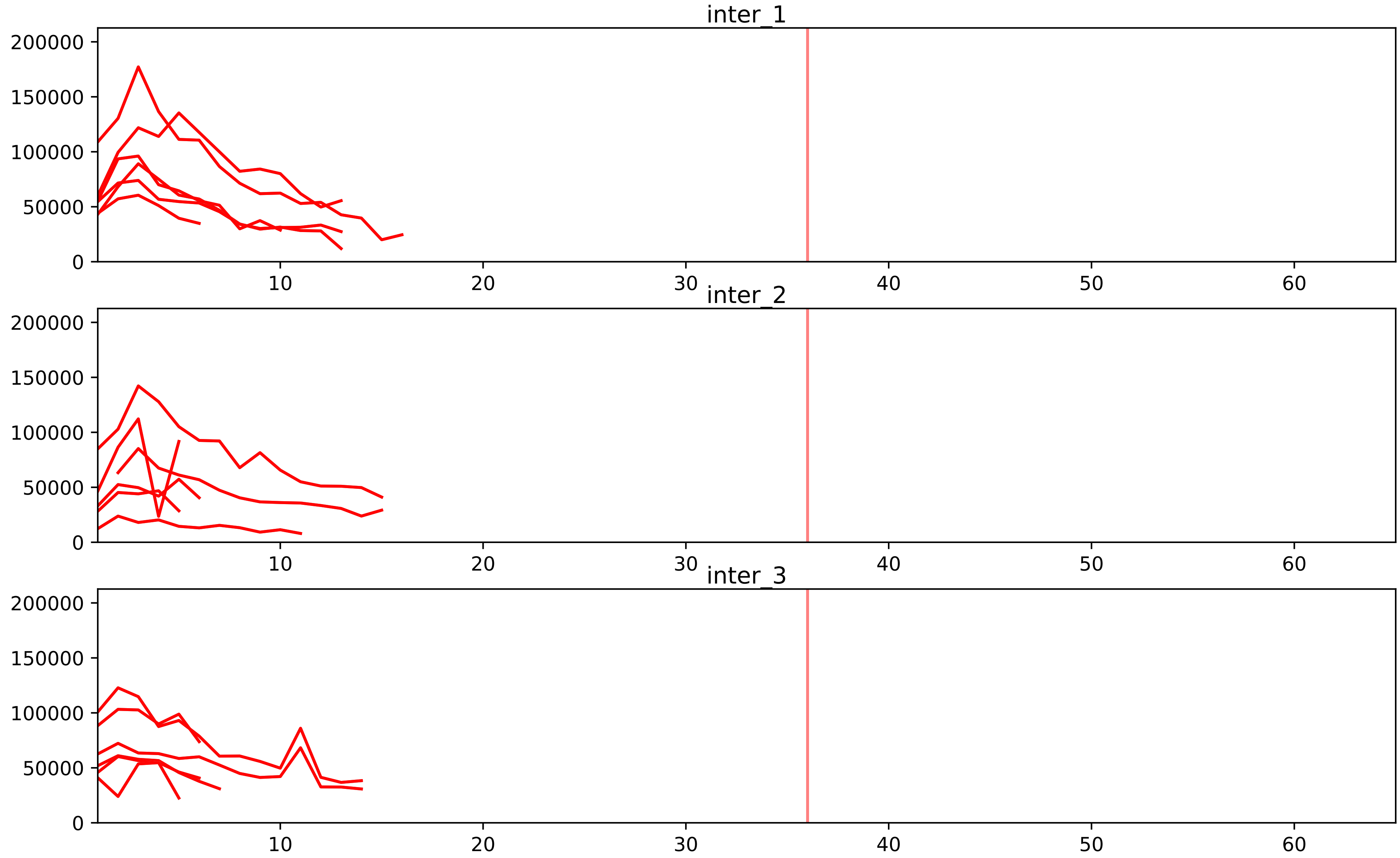


| level | condition_1 | condition_2 | pvalue | pvalue_adjusted |
|---|---|---|---|---|
| assembled_abundance | mitosis | inter | 1.3598888714635645e-05 | 0.004546885576657818 |
| total_abundance | mitosis | inter | 2.440120632294259e-05 | 0.005915549590004797 |
| complex_abundance | mitosis | inter | 4.35032354149368e-05 | 0.0013340992193913953 |
| interactor_abundance | mitosis | inter | 5.510497458340139e-05 | 0.0015598946651301318 |
| interactor_ratio | mitosis | inter | 0.40856902988493016 | 0.6993181534774618 |

P42695_Q6IBW4
CNDD3_HUMAN vs CNDH2_HUMAN
p-value: 0.023 q-value: 0.017

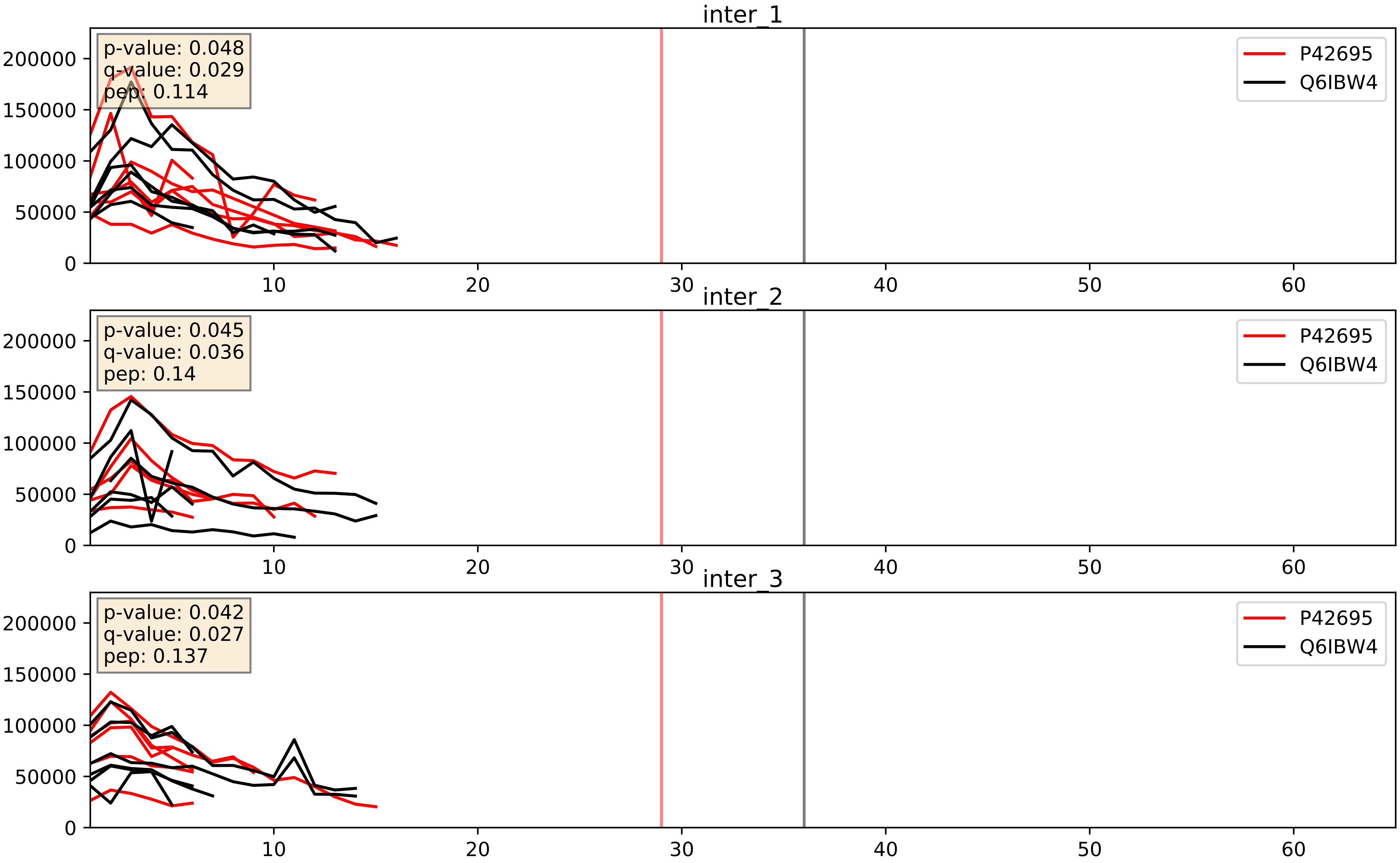


| level | condition_1 | condition_2 | pvalue | pvalue_adjusted |
|---|---|---|---|---|
| interactor_abundance | mitosis | inter | 0.0003511758884899916 | 0.01356427145975096 |
| complex_abundance | mitosis | inter | 0.0003576603614696945 | 0.012039775462644836 |
| interactor_ratio | mitosis | inter | 0.8058686900687333 | 0.9637099730355346 |

Q6IBW4_Q86XI2
CNDH2_HUMAN vs CNDG2_HUMAN
p-value: 0.003 q-value: 0.003

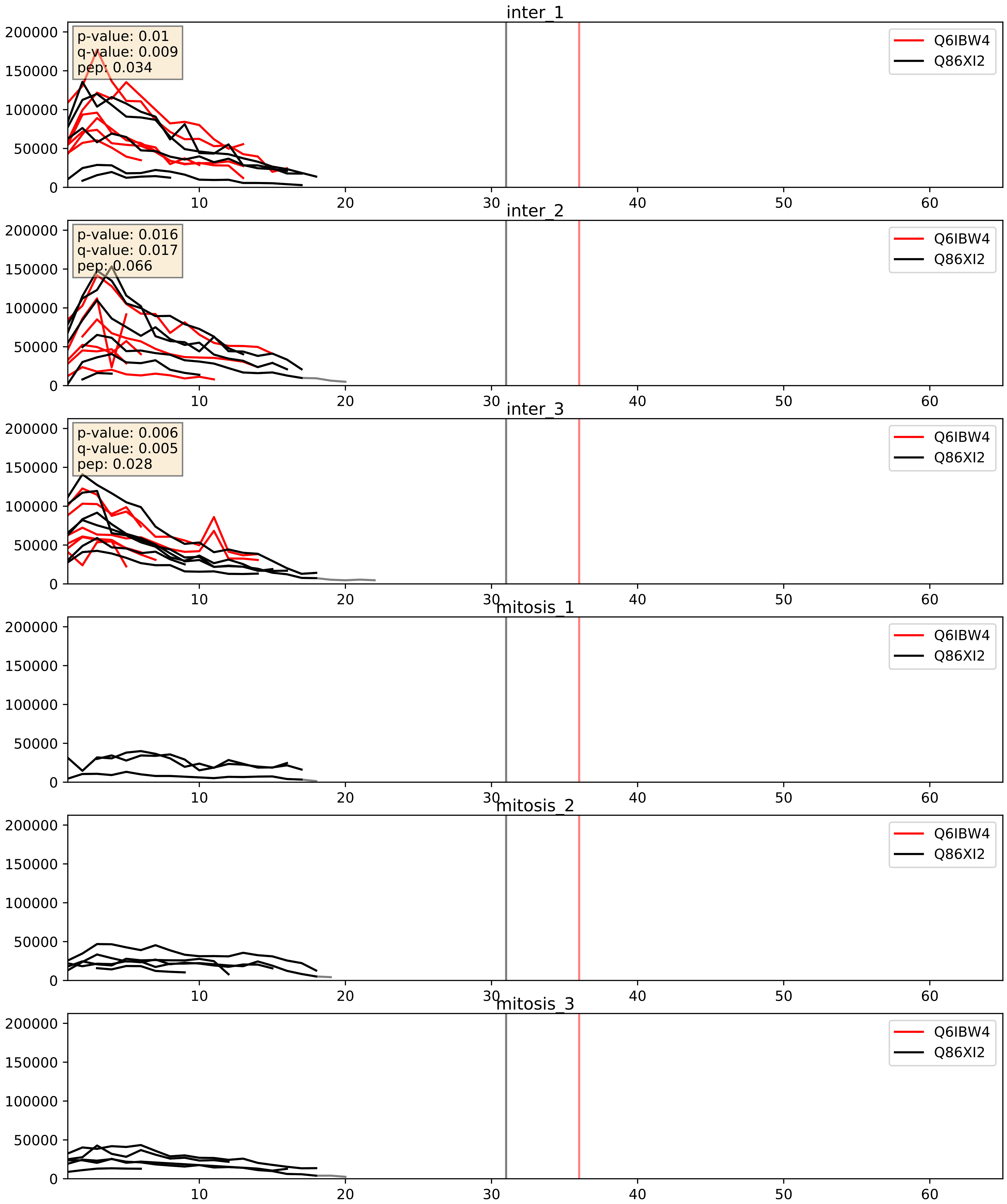


| level | condition_1 | condition_2 | pvalue | pvalue_adjusted |
|---|---|---|---|---|
| complex_abundance | mitosis | inter | 0.0003076550020109643 | 0.01165207110963347 |
| interactor_abundance | mitosis | inter | 0.0003831907621725412 | 0.013582248133320717 |
| interactor_ratio | mitosis | inter | 0.16977884276934044 | 0.6328499622031528 |
