## Supplementary Data 2 for "SECAT: Quantifying differential protein-protein interaction states by network-centric analysis": hela_string_000108_Q99623.pdf

Q99623_Q99623

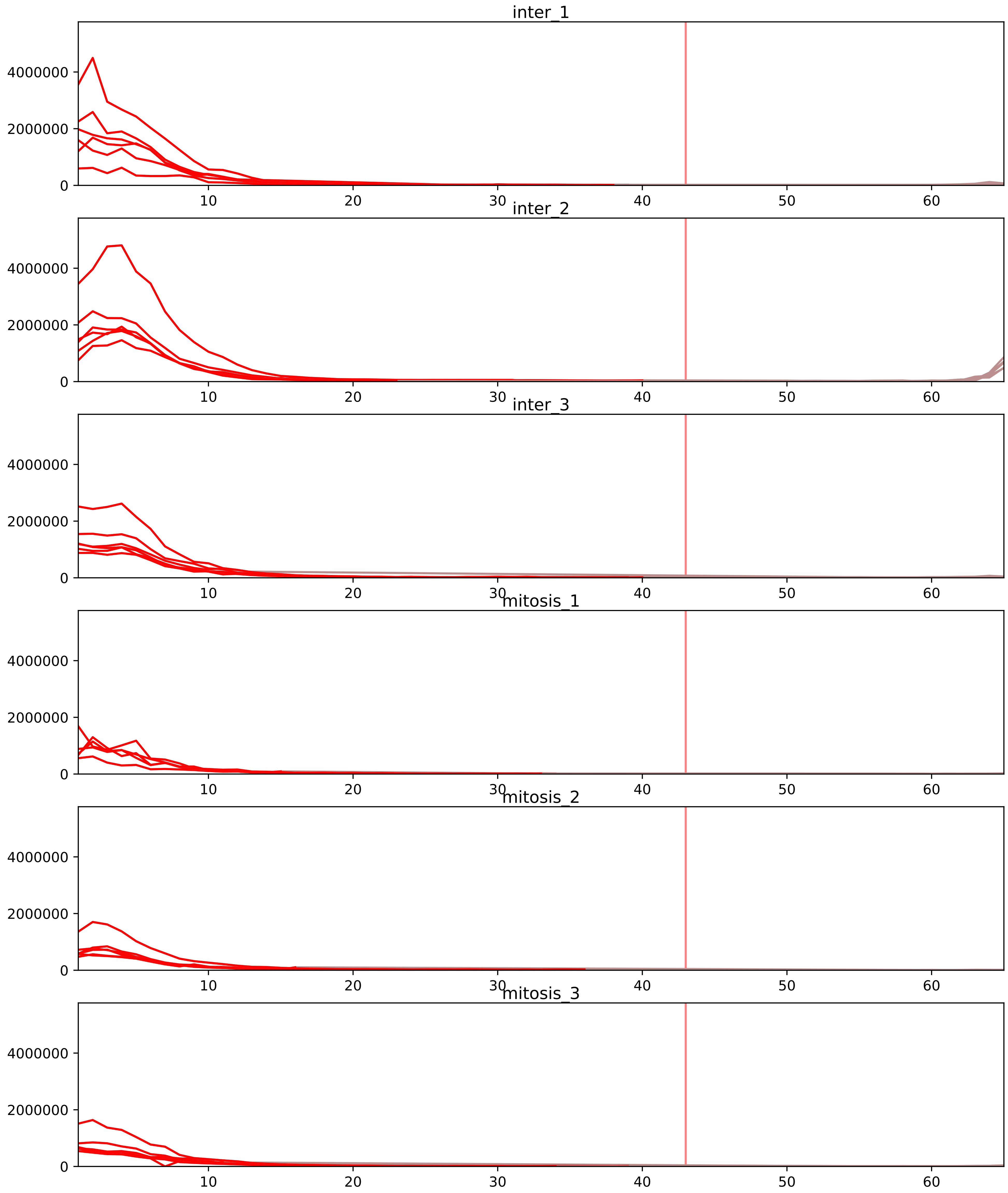


| level | condition_1 | condition_2 | pvalue | pvalue_adjusted |
|---|---|---|---|---|
| interactor_abundance | mitosis | inter | 1.3800862181799118e-05 | 0.0006682522740660626 |
| complex_abundance | mitosis | inter | 5.271522811972335e-05 | 0.0015705035793823847 |
| interactor_ratio | mitosis | inter | 0.0030488178026814084 | 0.026713451223494247 |
| monomer_abundance | mitosis | inter | 0.03952441445315753 | 0.3157957803087734 |
| total_abundance | mitosis | inter | 0.13466645567392124 | 0.4537305928761966 |
| assembled_abundance | mitosis | inter | 0.1422013191650812 | 0.4464415660709222 |

P35232_Q99623
PHB_HUMAN vs PHB2_HUMAN
p-value: 0.0 q-value: 0.0

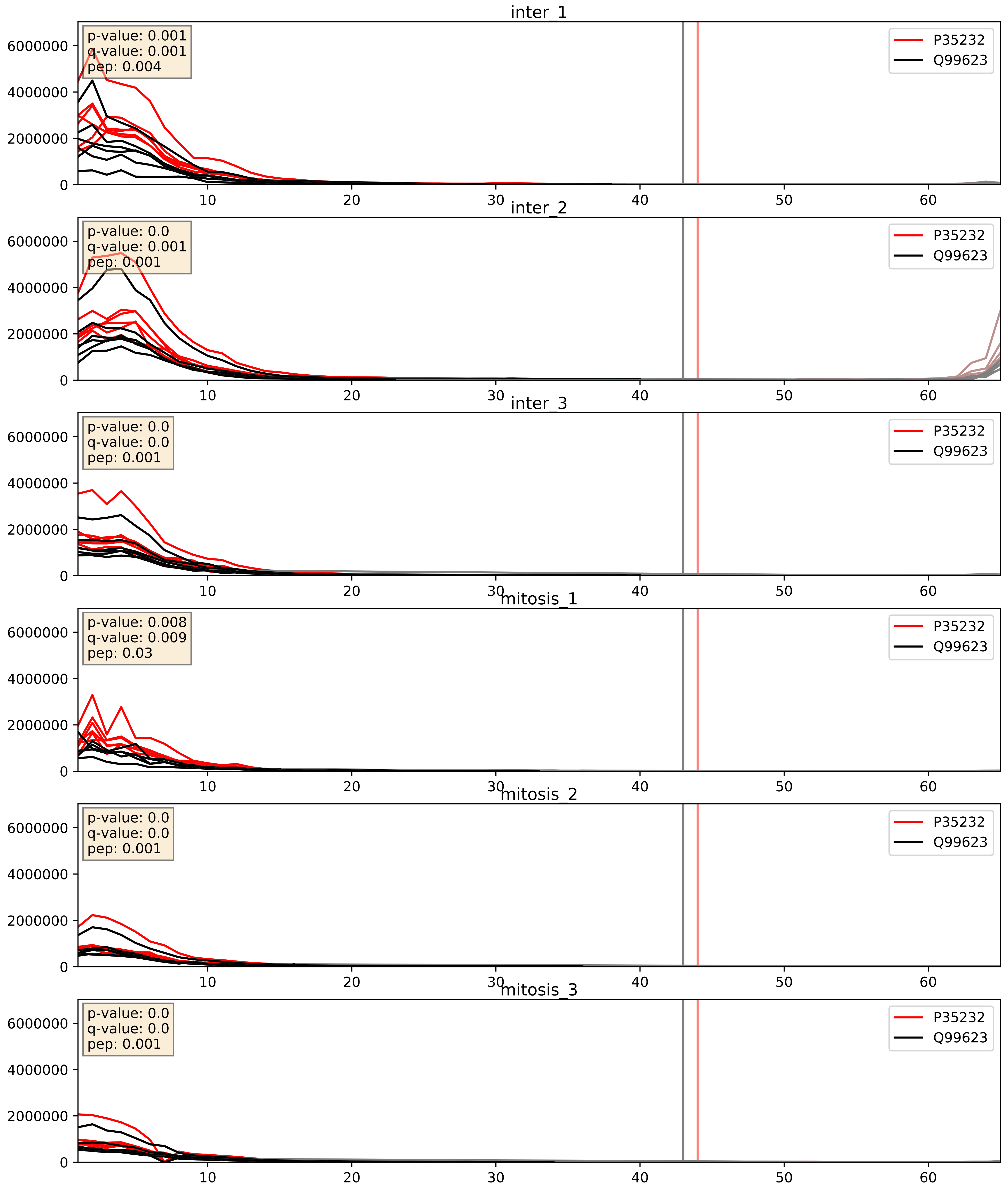


| level | condition_1 | condition_2 | pvalue | pvalue_adjusted |
|---|---|---|---|---|
| complex_abundance | mitosis | inter | 0.0005098332523952253 | 0.014318857255407222 |
| interactor_abundance | mitosis | inter | 0.0019578595900461063 | 0.032370554261591626 |
| interactor_ratio | mitosis | inter | 0.3720179690761096 | 0.8138388063636502 |

P39023_Q99623
RL3_HUMAN vs PHB2_HUMAN
p-value: 0.0 q-value: 0.042

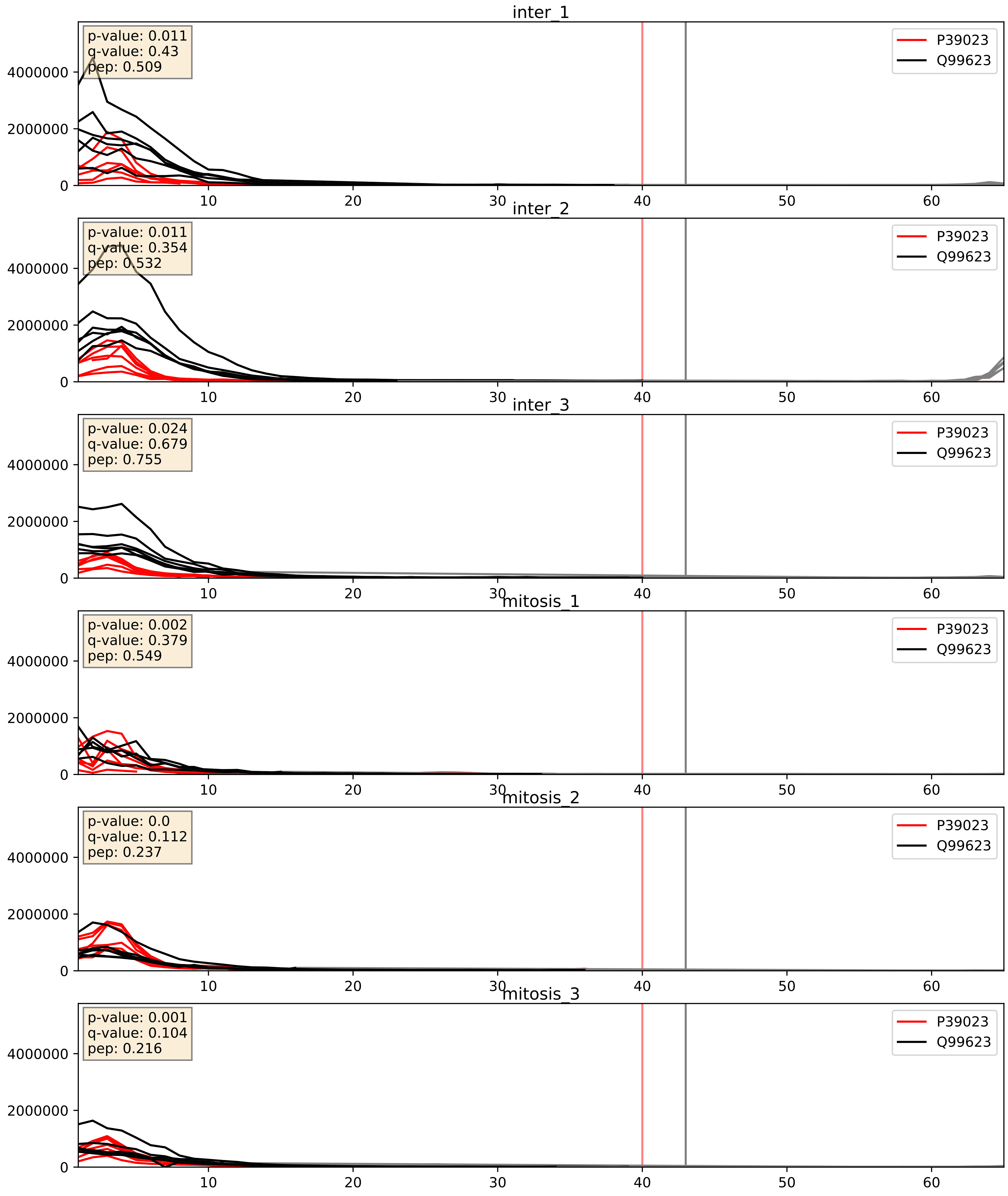


| level | condition_1 | condition_2 | pvalue | pvalue_adjusted |
|---|---|---|---|---|
| interactor_ratio | mitosis | inter | 0.0006263449140705945 | 0.029219626944512057 |
| complex_abundance | mitosis | inter | 0.0039012352647042063 | 0.0462092186673193 |
| interactor_abundance | mitosis | inter | 0.4577908838109454 | 0.629609570279835 |

Q99623_Q9Y4W6
PHB2_HUMAN vs AFG32_HUMAN
p-value: 0.012 q-value: 0.01

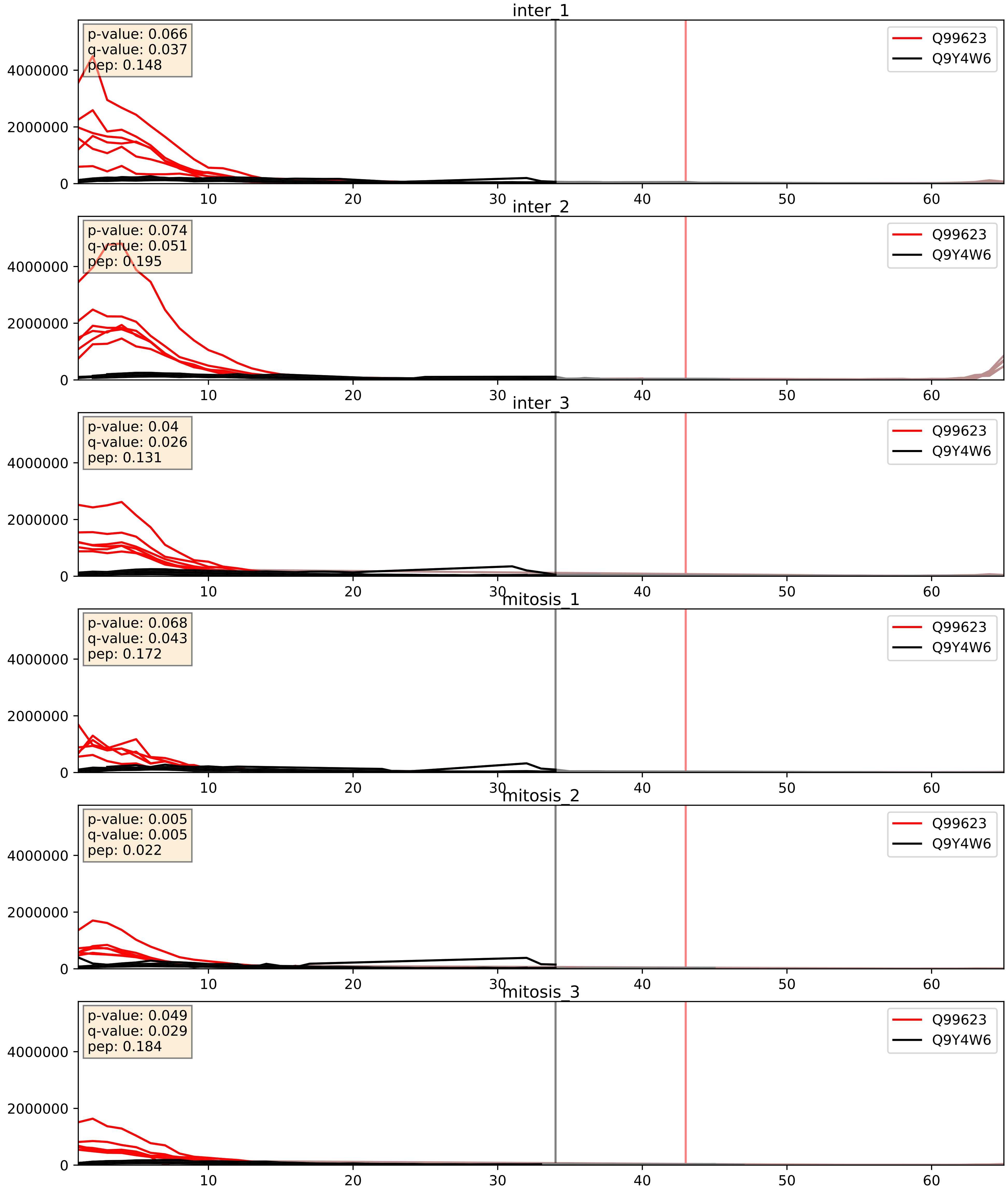


| level | condition_1 | condition_2 | pvalue | pvalue_adjusted |
|---|---|---|---|---|
| interactor_abundance | mitosis | inter | 0.00029573395803663804 | 0.01300709055050693 |
| complex_abundance | mitosis | inter | 0.0030999413960934785 | 0.040863210865152835 |
| interactor_ratio | mitosis | inter | 0.21911276598925933 | 0.6959574311198738 |
