## Supplementary Data 2 for "SECAT: Quantifying differential protein-protein interaction states by network-centric analysis": hela_string_000109_P32455.pdf

P32455_P32455

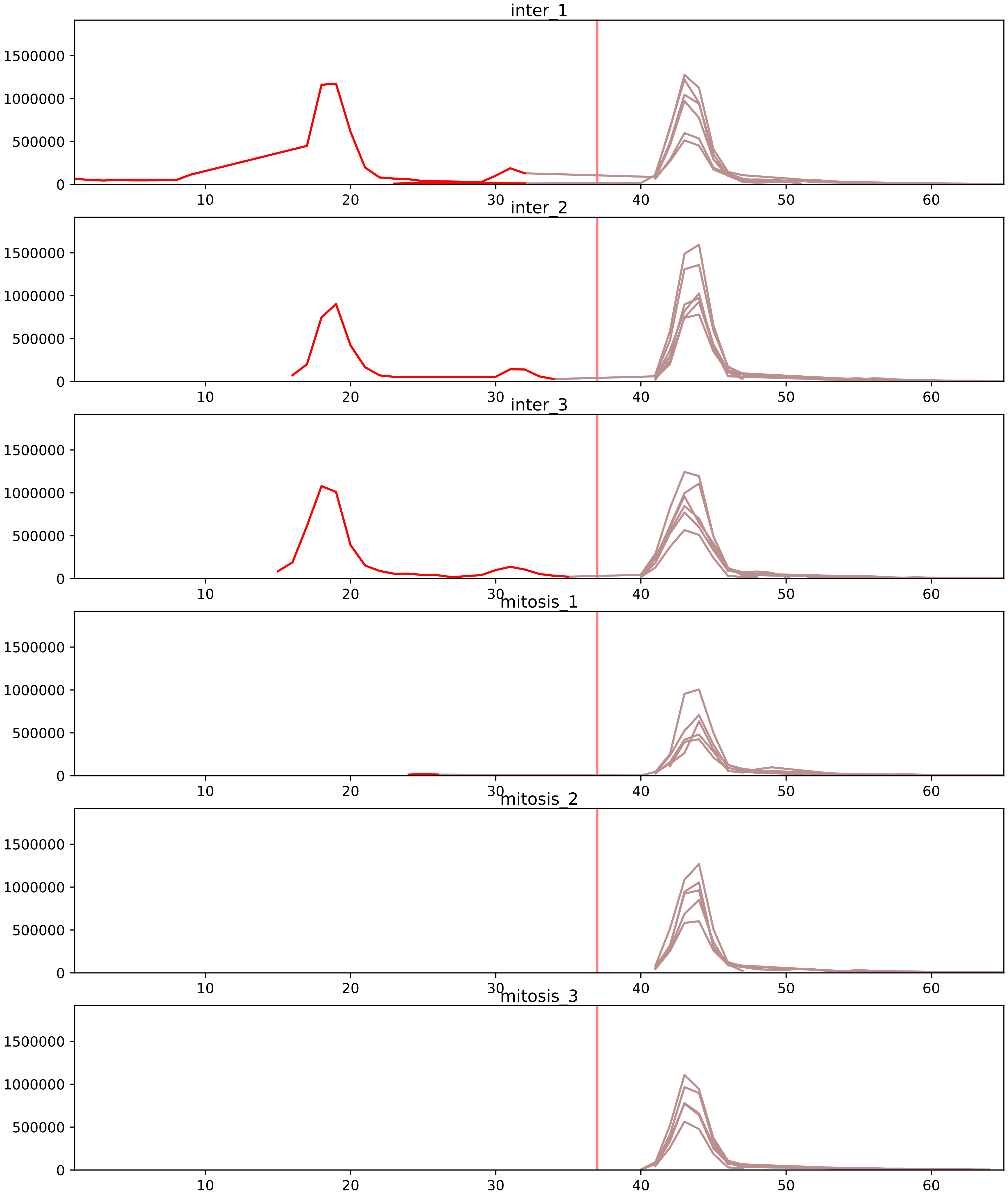


| level | condition_1 | condition_2 | pvalue | pvalue_adjusted |
|---|---|---|---|---|
| total_abundance | mitosis | inter | 1.4843795770896684e-05 | 0.00539784030497393 |
| monomer_abundance | mitosis | inter | 9.619825203400103e-05 | 0.03261473898341094 |
| assembled_abundance | mitosis | inter | 0.0018836554576943723 | 0.04067828623906237 |
