## Supplementary Data 2 for "SECAT: Quantifying differential protein-protein interaction states by network-centric analysis": hela_string_000110_Q00587.pdf

Q00587_Q00587

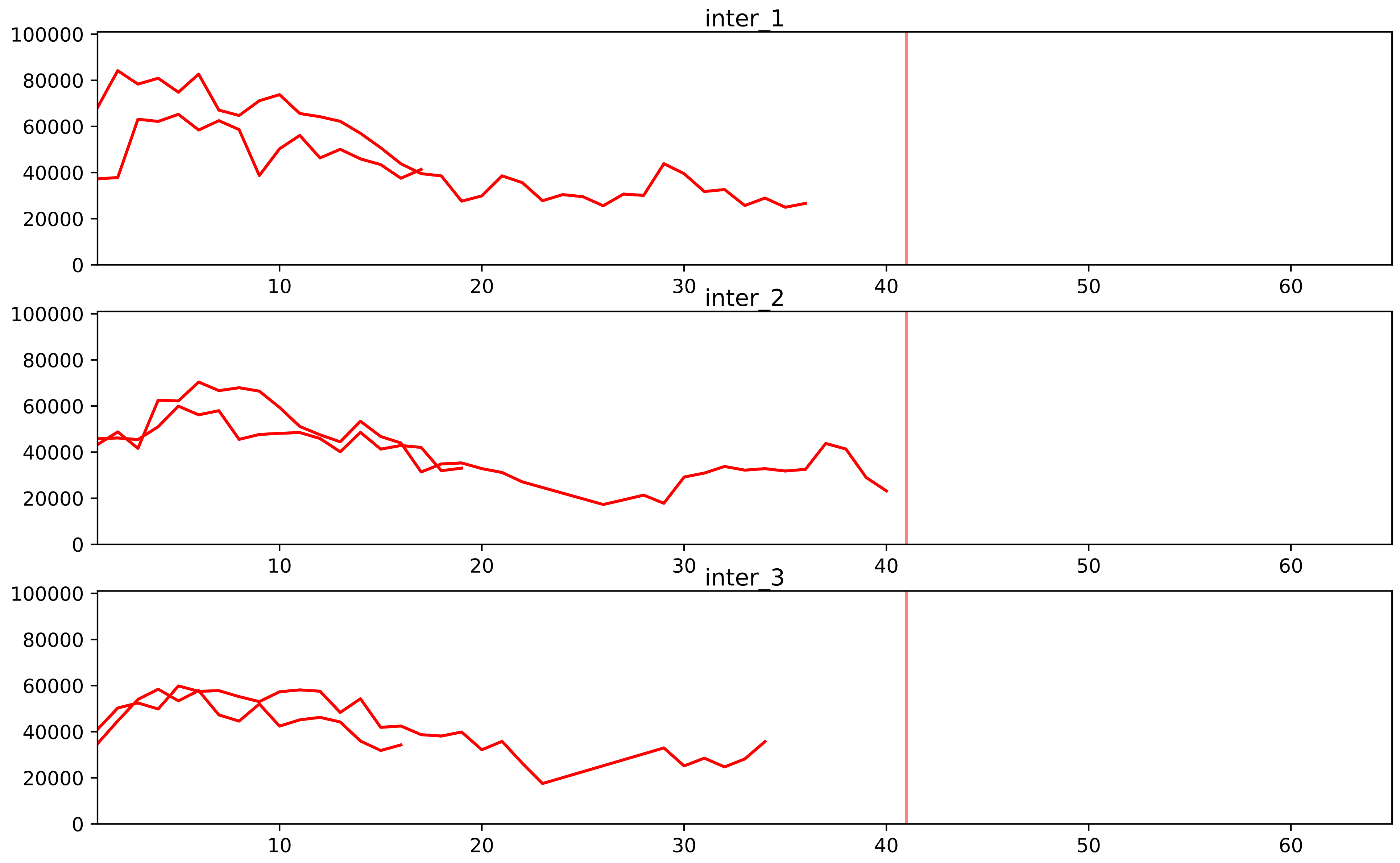


| level | condition_1 | condition_2 | pvalue | pvalue_adjusted |
|---|---|---|---|---|
| assembled_abundance | mitosis | inter | 1.5402529192725396e-05 | 0.004641305545861687 |
| total_abundance | mitosis | inter | 3.022339459721595e-05 | 0.006993968267928473 |
