## Supplementary Data 2 for "SECAT: Quantifying differential protein-protein interaction states by network-centric analysis": hela_string_000111_O00499.pdf

O00499_O00499

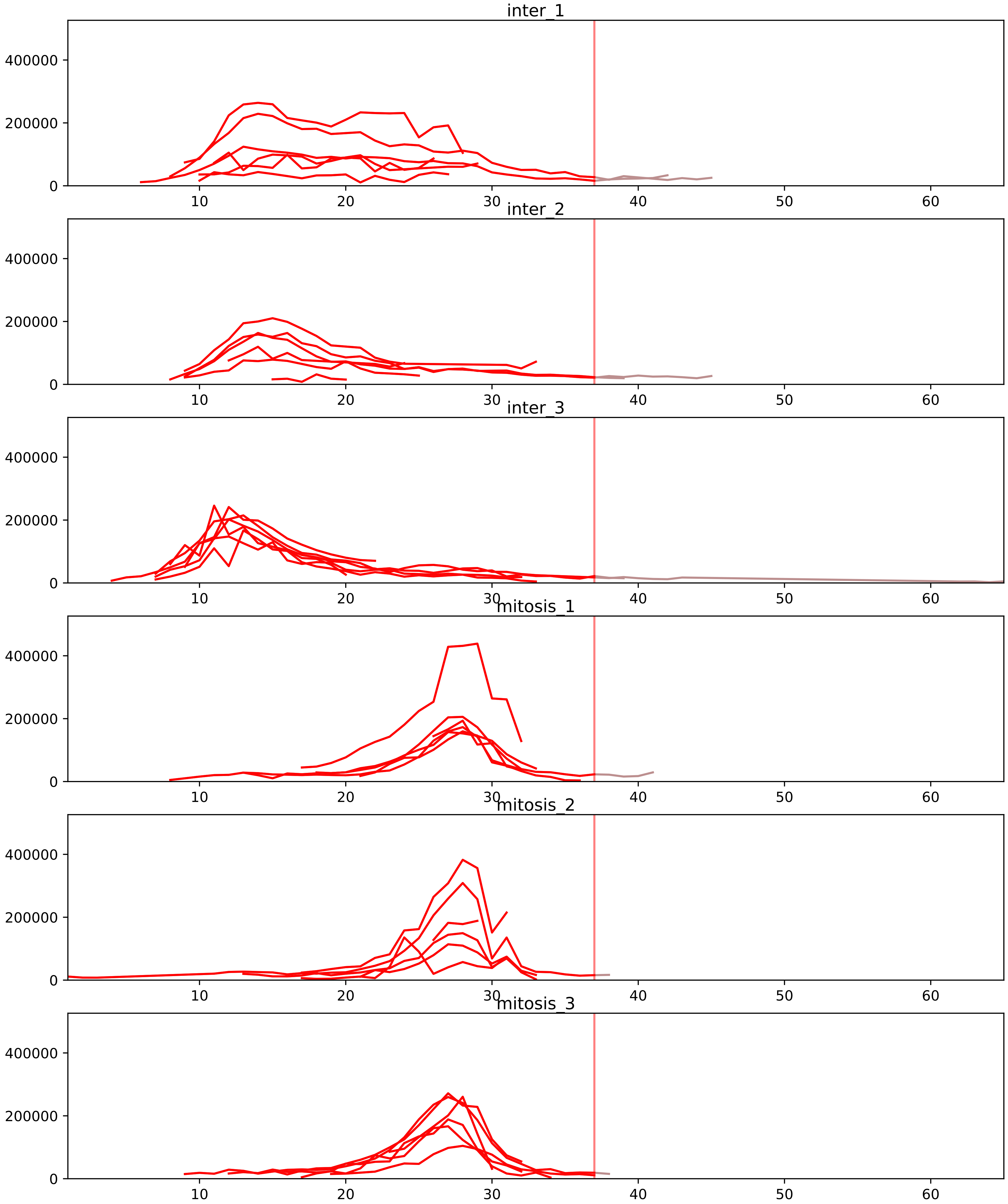


| level | condition_1 | condition_2 | pvalue | pvalue_adjusted |
|---|---|---|---|---|
| assembled_abundance | mitosis | inter | 1.5864321455626358e-05 | 0.004641305545861687 |
| total_abundance | mitosis | inter | 2.0639685508939232e-05 | 0.0058684532992606024 |
| monomer_abundance | mitosis | inter | 0.012476701646757392 | 0.19422543597998193 |
