## Supplementary Data 2 for "SECAT: Quantifying differential protein-protein interaction states by network-centric analysis": hela_string_000112_Q9NR30.pdf

Q9NR30_Q9NR30

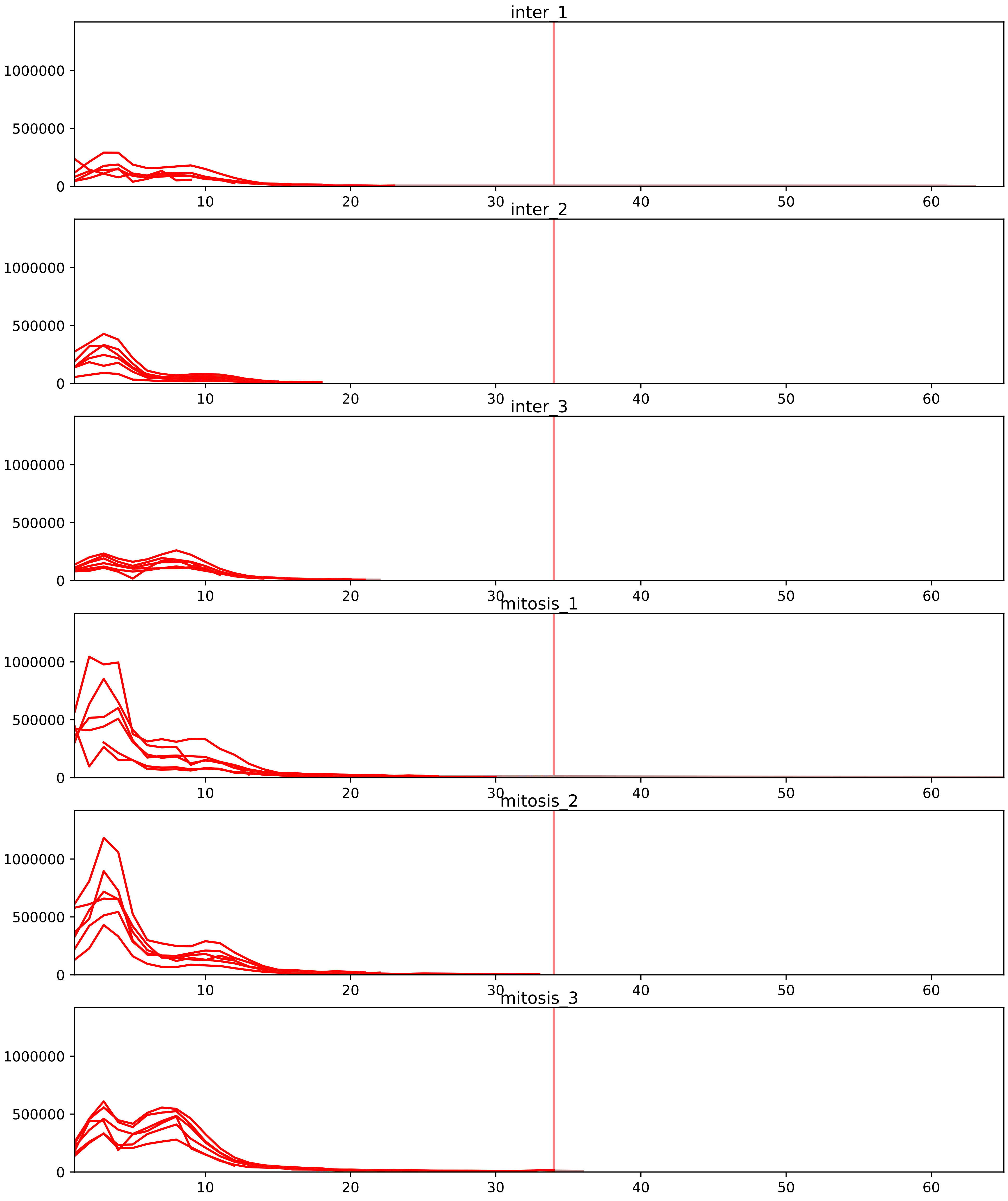


| level | condition_1 | condition_2 | pvalue | pvalue_adjusted |
|---|---|---|---|---|
| interactor_abundance | mitosis | inter | 1.62327608132201e-05 | 0.0007284946316176825 |
| complex_abundance | mitosis | inter | 5.291914234875427e-05 | 0.0015705035793823847 |
| interactor_ratio | mitosis | inter | 0.006073086035413971 | 0.04469791322064683 |
| monomer_abundance | mitosis | inter | 0.2733641310855585 | 0.6533402732944849 |
| total_abundance | mitosis | inter | 0.8051039559509369 | 0.9235287199690467 |
| assembled_abundance | mitosis | inter | 0.8269301507398982 | 0.9347770036244542 |

P55769_Q9NR30
NH2L1_HUMAN vs DDX21_HUMAN
p-value: 0.0 q-value: 0.025

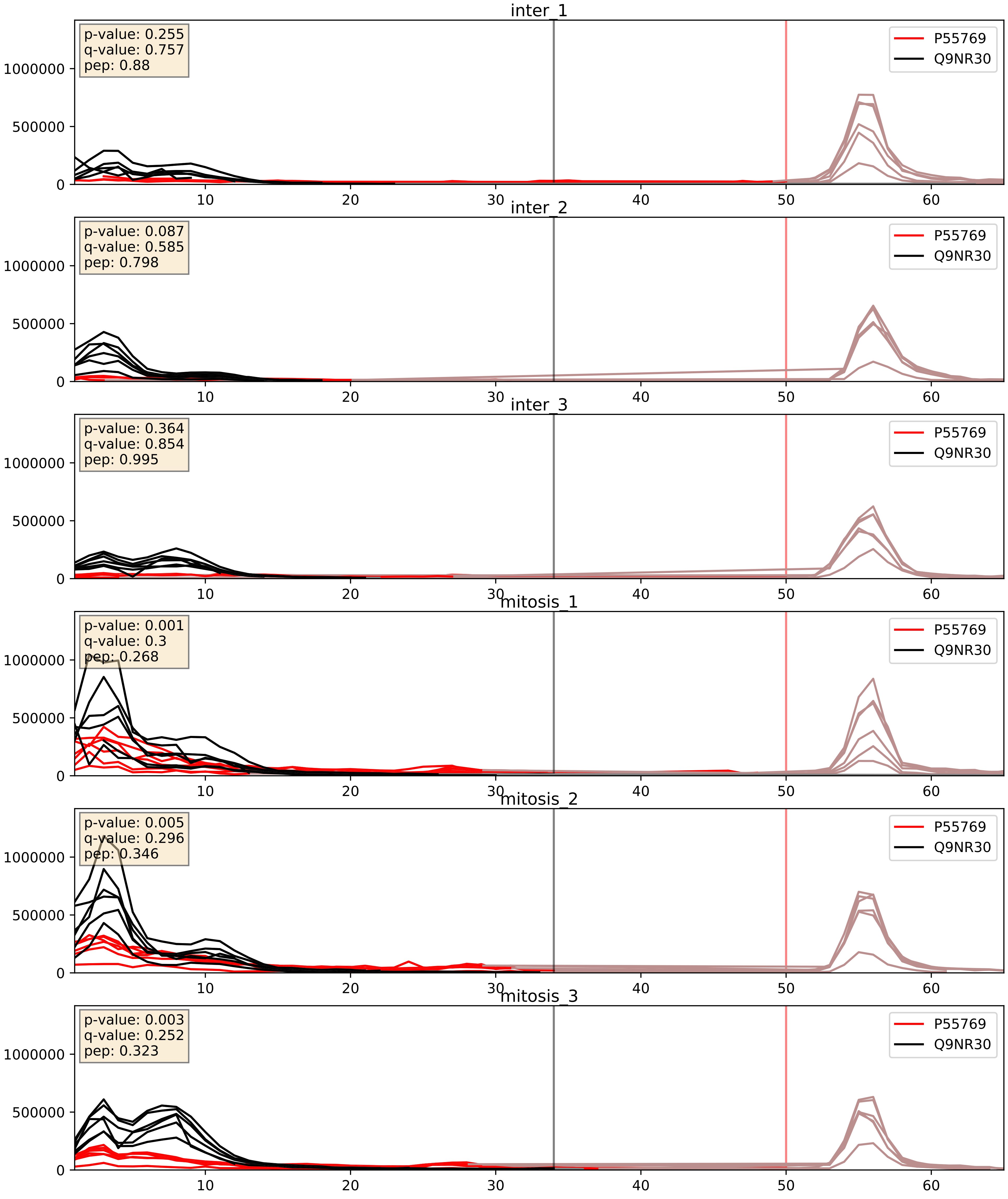


| level | condition_1 | condition_2 | pvalue | pvalue_adjusted |
|---|---|---|---|---|
| complex_abundance | mitosis | inter | 0.0002757050388916281 | 0.011185000629916286 |
| interactor_abundance | mitosis | inter | 0.0010777766016947538 | 0.024049720606162197 |
| interactor_ratio | mitosis | inter | 0.02503180125222349 | 0.24857565976685972 |

P62750_Q9NR30
RL23A_HUMAN vs DDX21_HUMAN
p-value: 0.0 q-value: 0.025

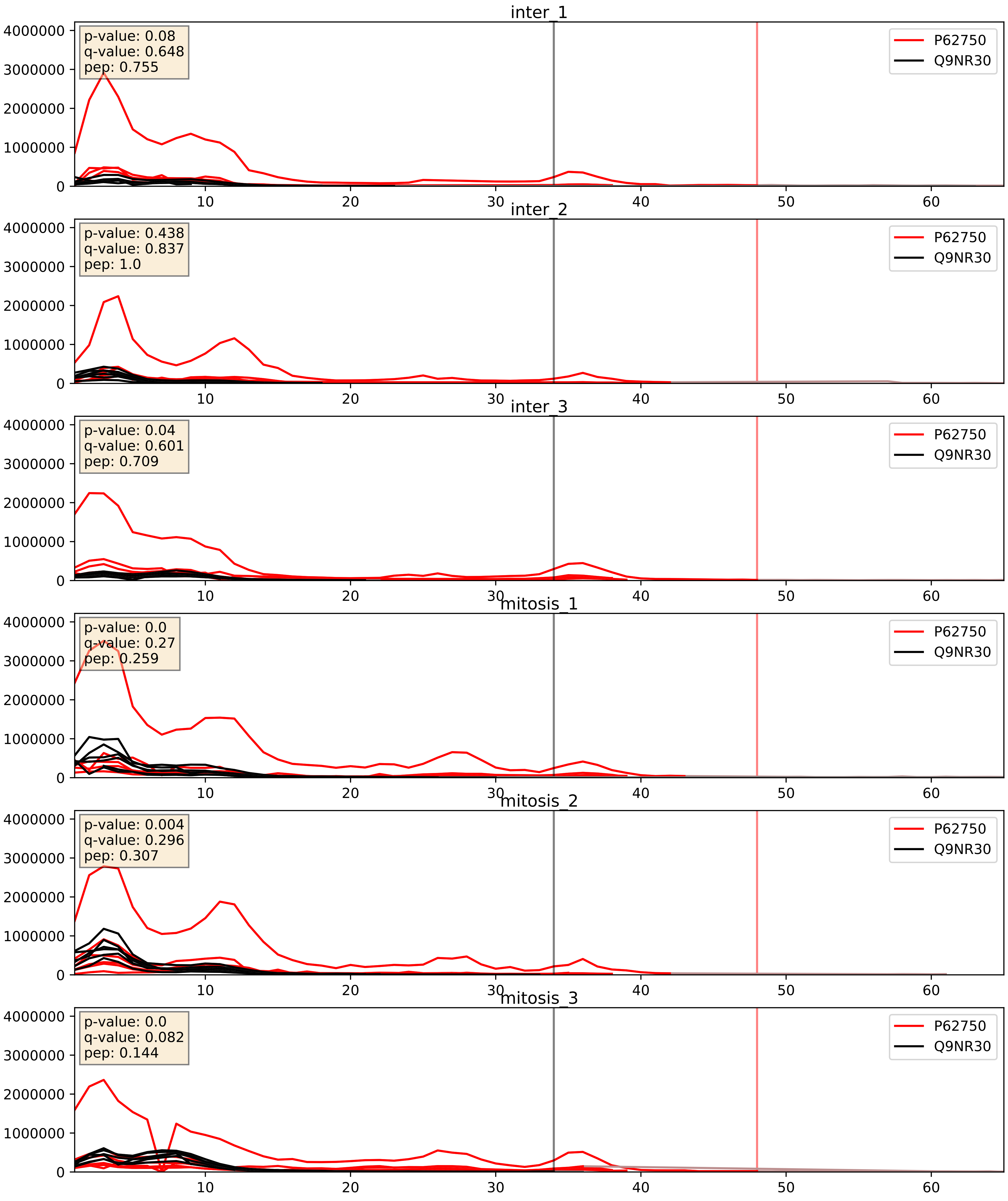


| level | condition_1 | condition_2 | pvalue | pvalue_adjusted |
|---|---|---|---|---|
| complex_abundance | mitosis | inter | 0.0017963512637637516 | 0.030065523984879133 |
| interactor_ratio | mitosis | inter | 0.02689083096680076 | 0.25724626346131724 |
| interactor_abundance | mitosis | inter | 0.07914995682557545 | 0.2337192500802537 |

Q99459_Q9NR30
CDC5L_HUMAN vs DDX21_HUMAN
p-value: 0.016 q-value: 0.013

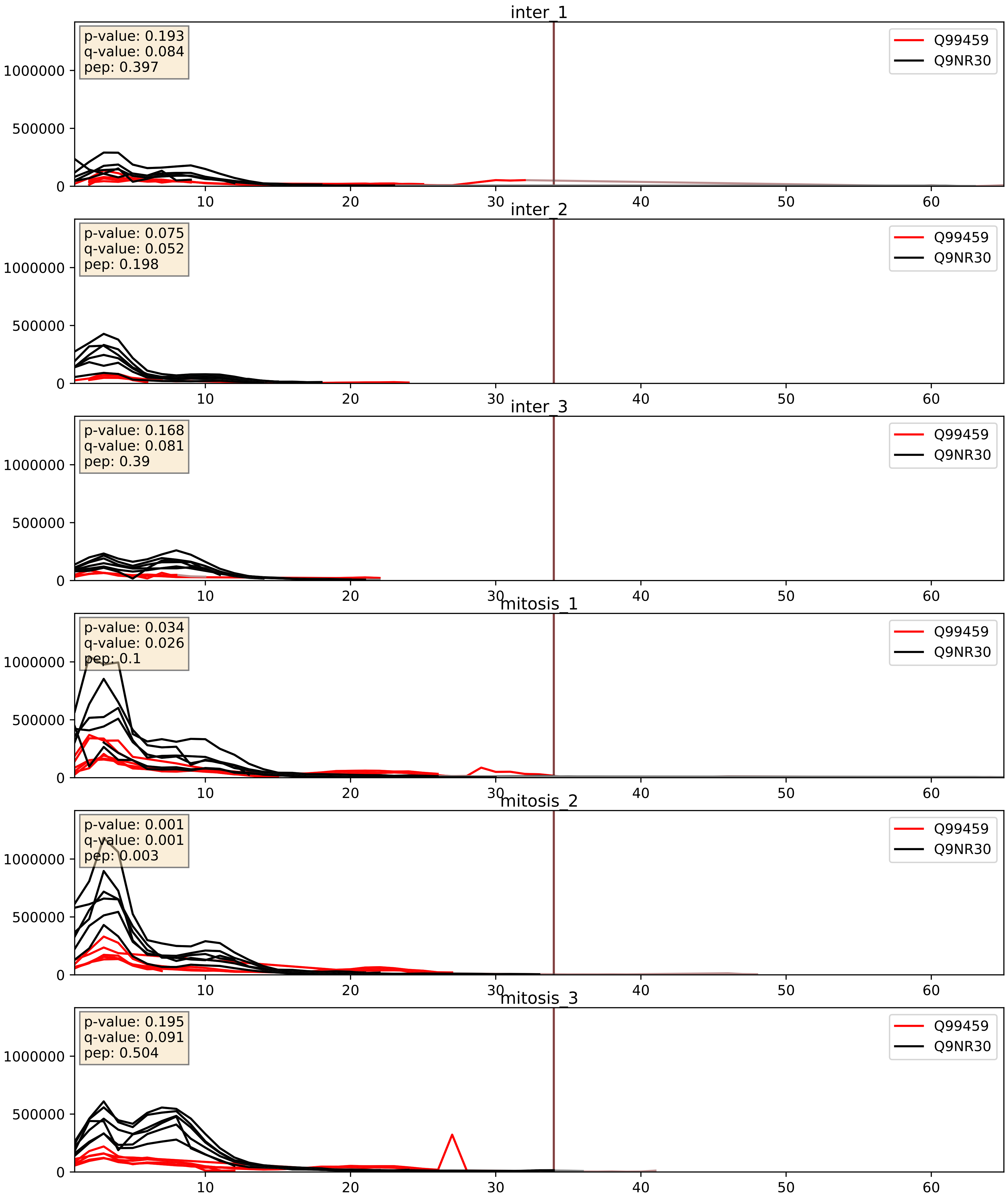


| level | condition_1 | condition_2 | pvalue | pvalue_adjusted |
|---|---|---|---|---|
| complex_abundance | mitosis | inter | 0.006365618502101007 | 0.05948545934570088 |
| interactor_abundance | mitosis | inter | 0.013030345214924086 | 0.10247106572324315 |
| interactor_ratio | mitosis | inter | 0.17752272019023063 | 0.6441689210802773 |
