## Supplementary Data 2 for "SECAT: Quantifying differential protein-protein interaction states by network-centric analysis": hela_string_000113_Q15050.pdf

Q15050_Q15050

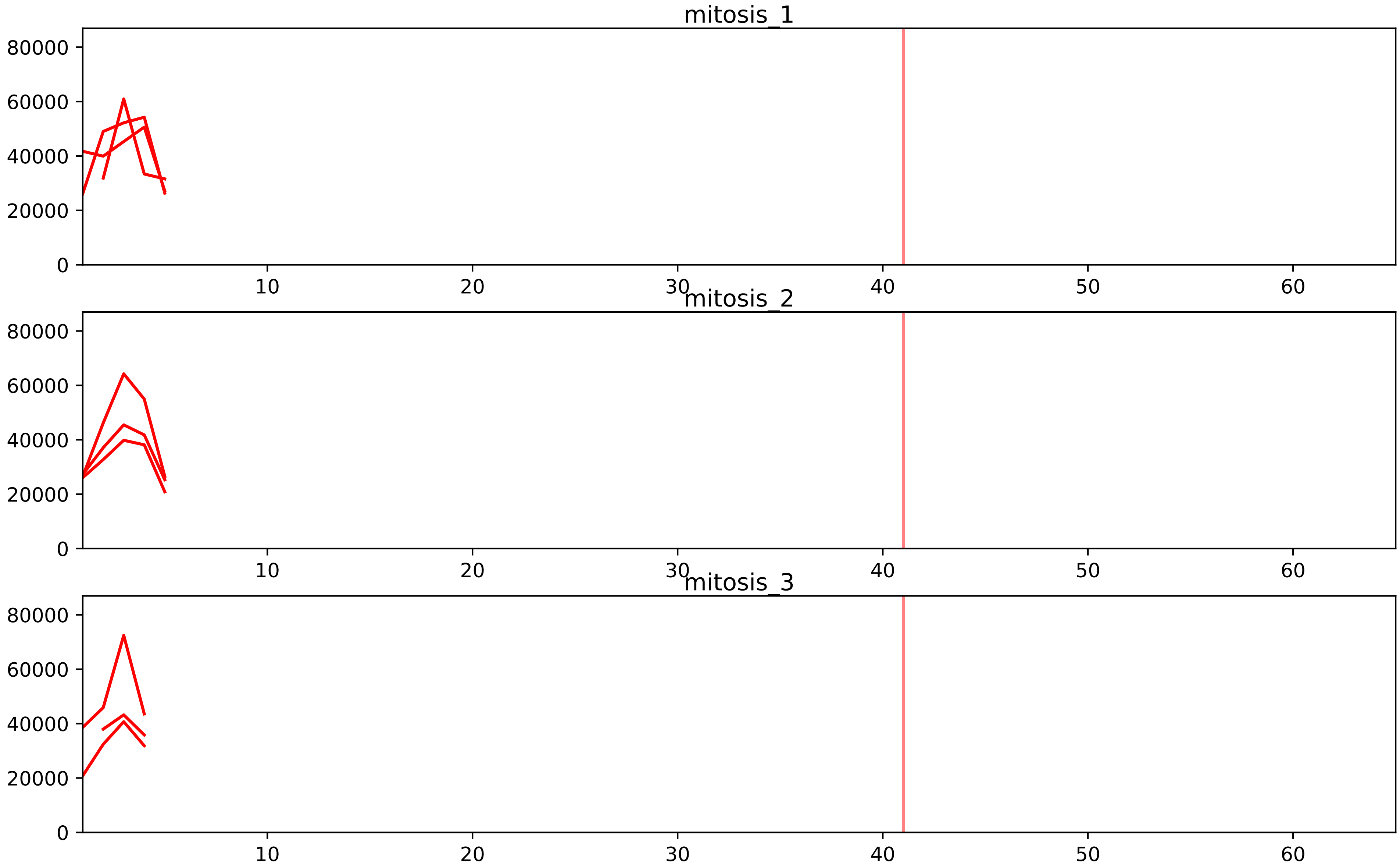


| level | condition_1 | condition_2 | pvalue | pvalue_adjusted |
|---|---|---|---|---|
| interactor_ratio | mitosis | inter | 1.7625106596891224e-05 | 0.000372760875152642 |
| assembled_abundance | mitosis | inter | 0.0014821568932065251 | 0.036157402973529304 |
| total_abundance | mitosis | inter | 0.008364931853731961 | 0.10935946807354902 |
| complex_abundance | mitosis | inter | 0.032385127433997365 | 0.09488636063464197 |
| interactor_abundance | mitosis | inter | 0.19101085592365186 | 0.3406202154081173 |

O75683_Q15050
SURF6_HUMAN vs RRS1_HUMAN
p-value: 0.026 q-value: 0.019

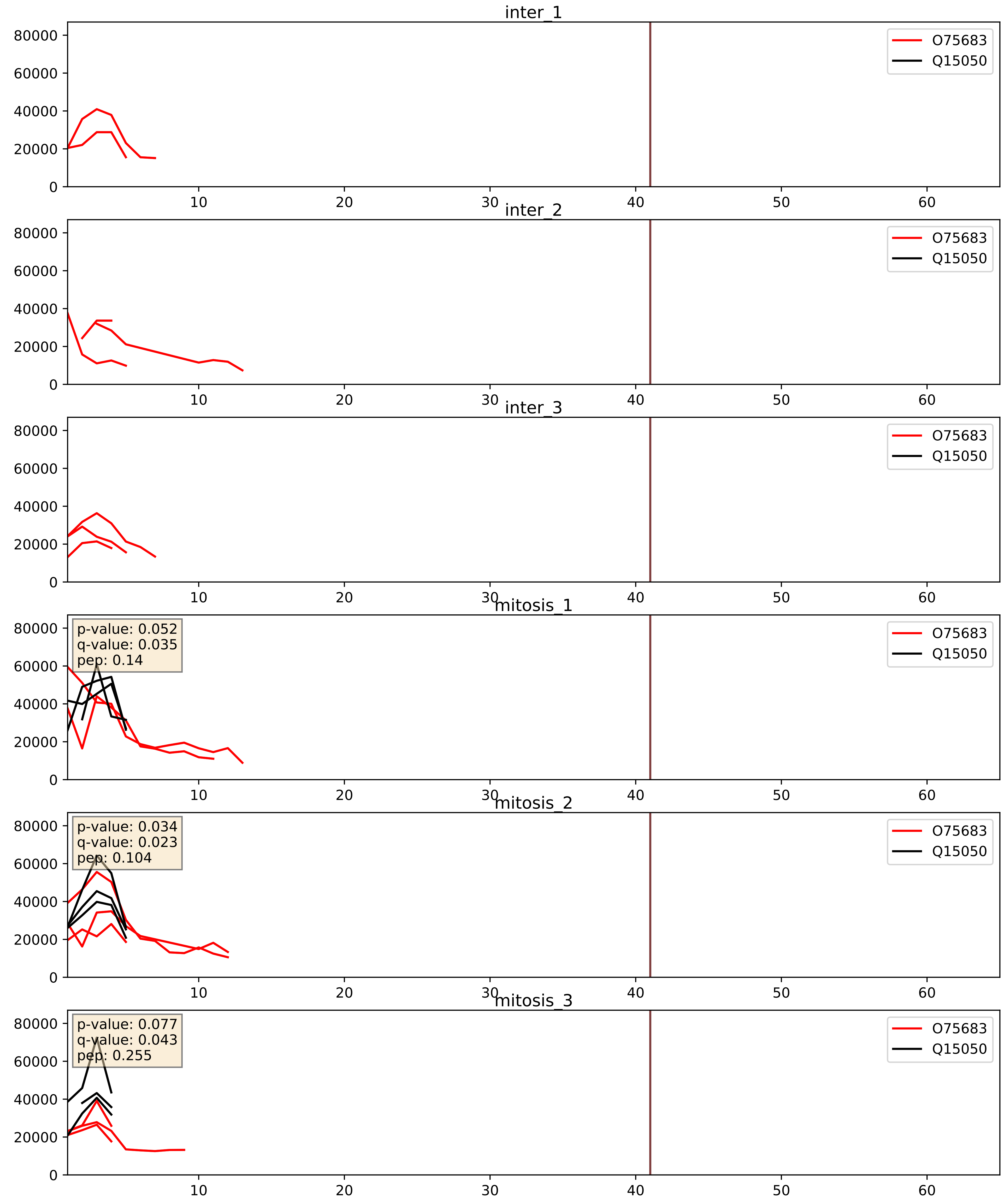


| level | condition_1 | condition_2 | pvalue | pvalue_adjusted |
|---|---|---|---|---|
| interactor_ratio | mitosis | inter | 0.08087148069121791 | 0.47583520079221503 |
| complex_abundance | mitosis | inter | 0.28444287840666316 | 0.43324395714609193 |
| interactor_abundance | mitosis | inter | 0.4127685785945814 | 0.5959853306518709 |

O76021_Q15050
RL1D1_HUMAN vs RRS1_HUMAN
p-value: 0.056 q-value: 0.036

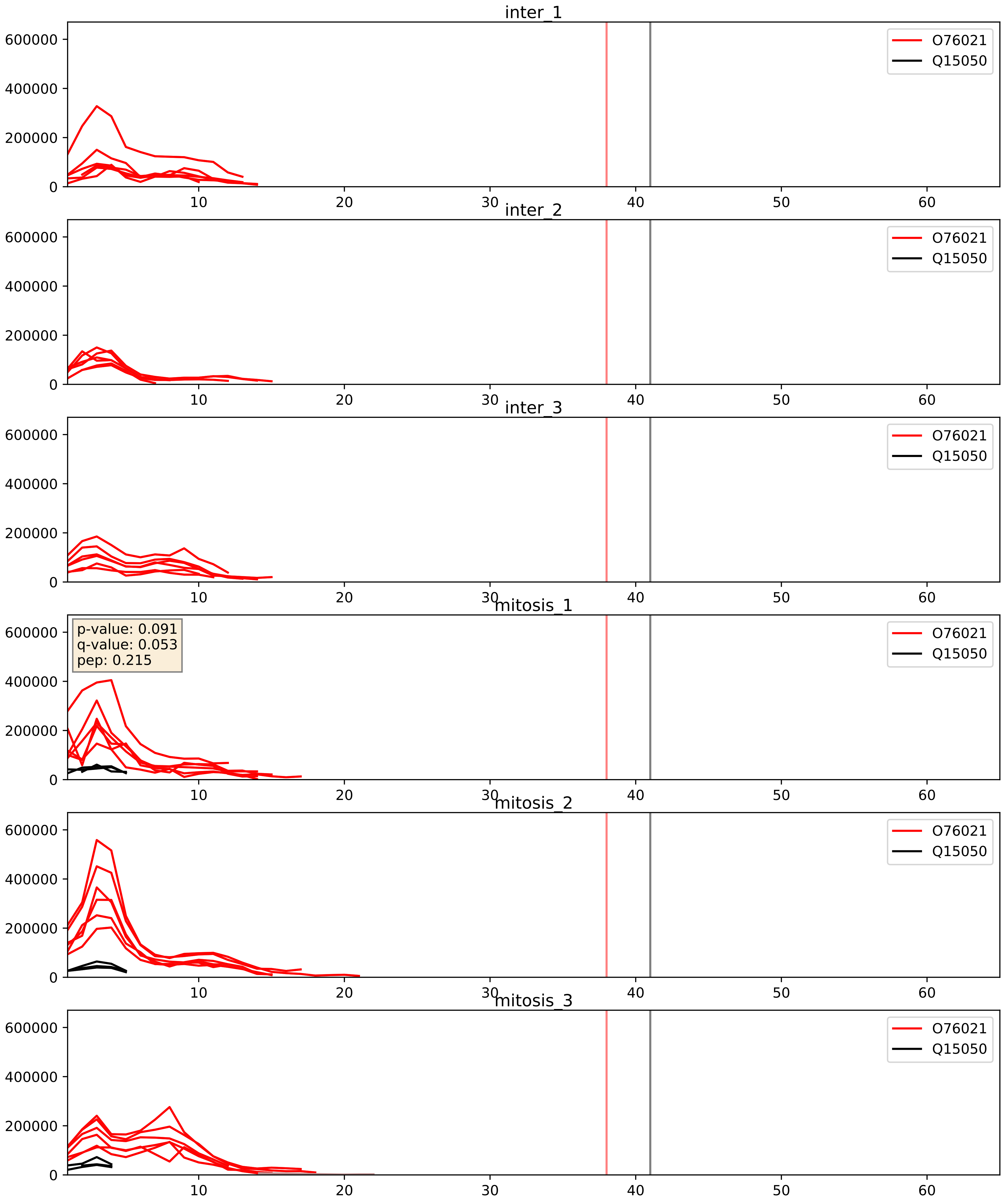


| level | condition_1 | condition_2 | pvalue | pvalue_adjusted |
|---|---|---|---|---|
| interactor_abundance | mitosis | inter | 0.002514210492437318 | 0.03746151752004081 |
| interactor_ratio | mitosis | inter | 0.003845030598611075 | 0.08075090146433855 |
| complex_abundance | mitosis | inter | 0.0081844105880903 | 0.06710589524334576 |

Q15050_Q15397
RRS1_HUMAN vs PUM3_HUMAN
p-value: 0.003 q-value: 0.044

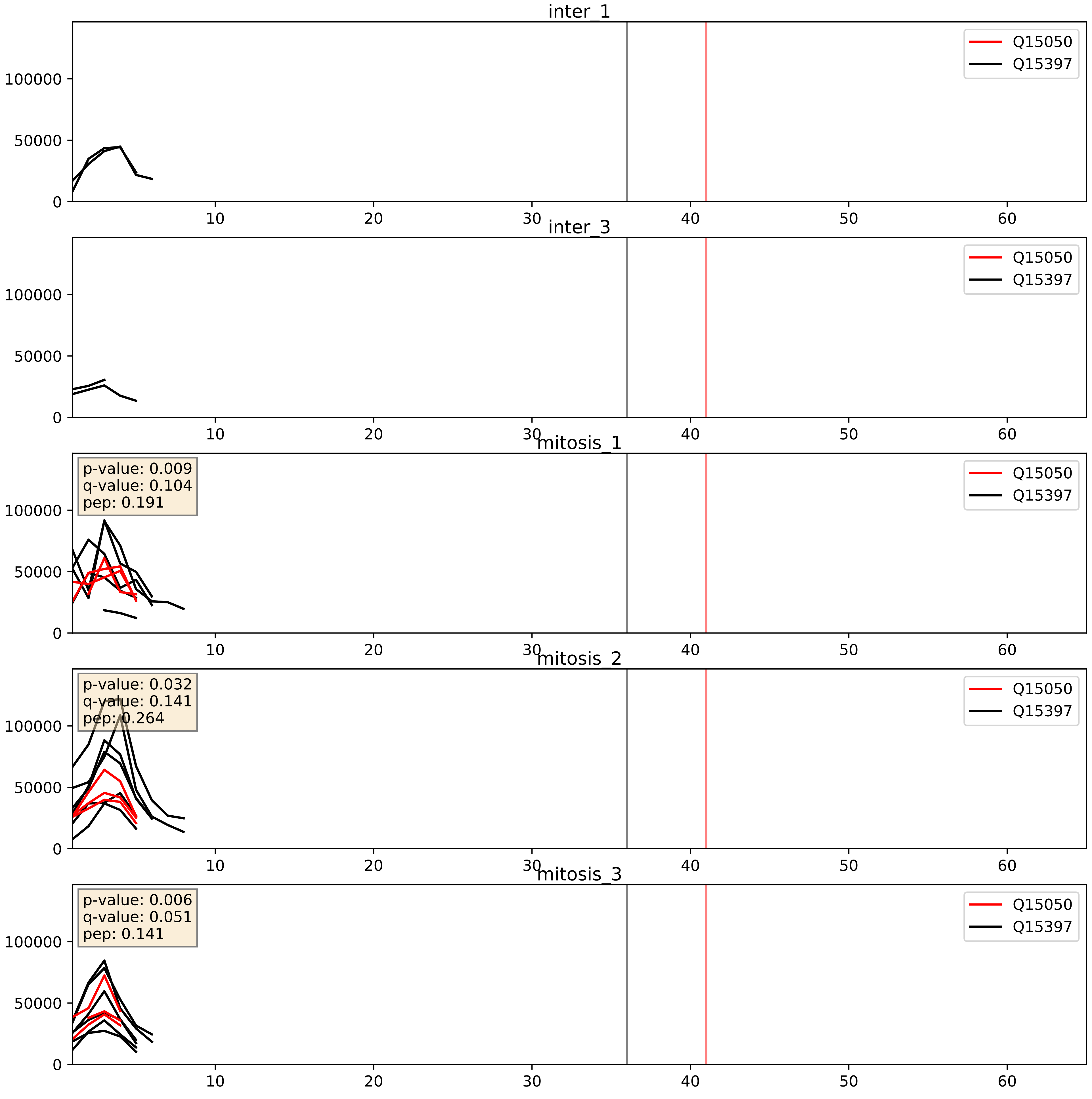


| level | condition_1 | condition_2 | pvalue | pvalue_adjusted |
|---|---|---|---|---|
| interactor_ratio | mitosis | inter | 0.05255410305415452 | 0.3793112328360562 |
| complex_abundance | mitosis | inter | 0.09861422506736761 | 0.23280136971226328 |
| interactor_abundance | mitosis | inter | 0.2162538244564617 | 0.4119120465837366 |

Q15050_Q8IY81
RRS1_HUMAN vs SPB1_HUMAN
p-value: 0.017 q-value: 0.014

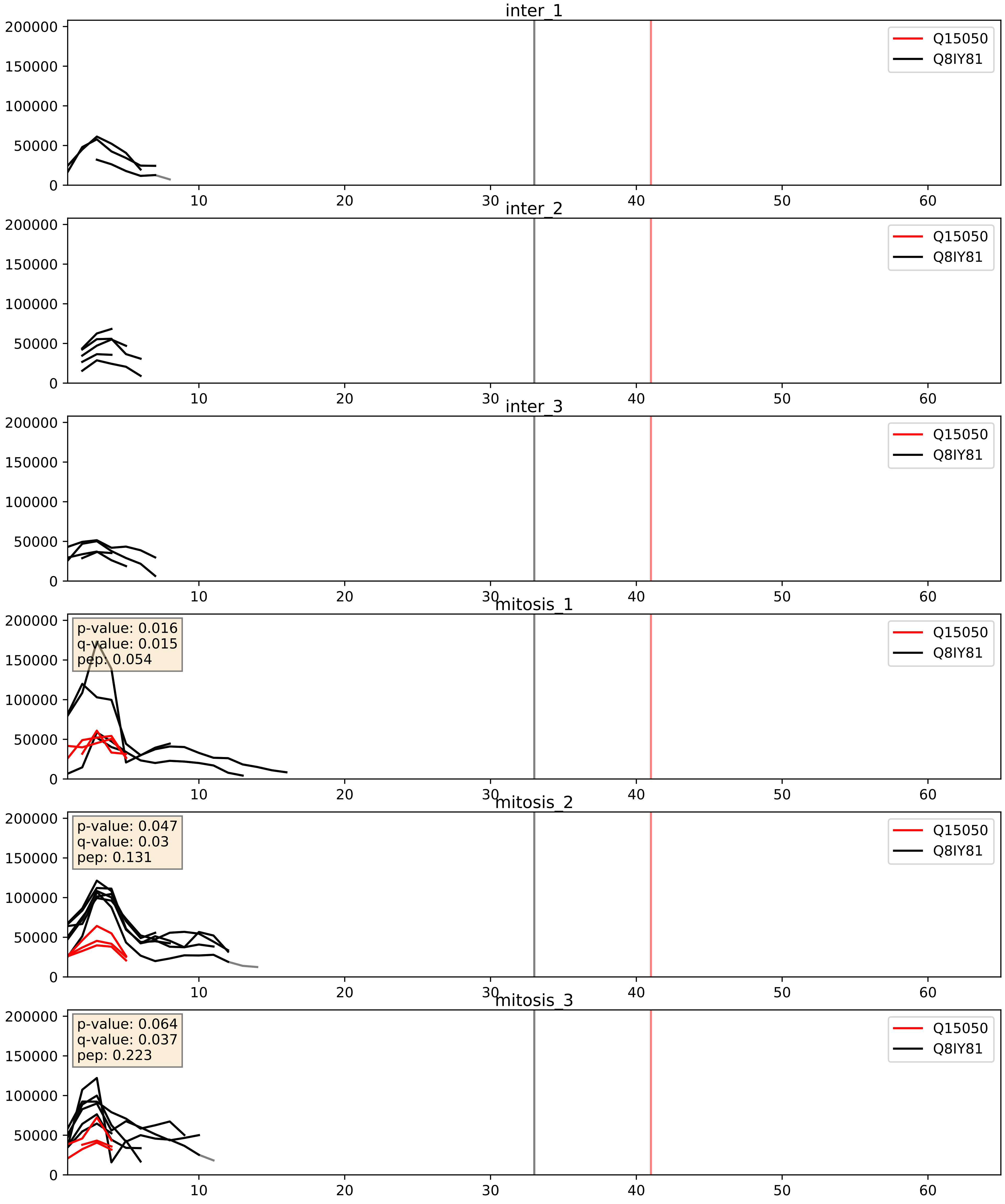


| level | condition_1 | condition_2 | pvalue | pvalue_adjusted |
|---|---|---|---|---|
| complex_abundance | mitosis | inter | 0.05036965452854774 | 0.15980547590484867 |
| interactor_ratio | mitosis | inter | 0.13938380188657049 | 0.5877140456747689 |
| interactor_abundance | mitosis | inter | 0.2162538244564617 | 0.4119120465837366 |

Q15050_Q9BYG3
RRS1_HUMAN vs MK67I_HUMAN
p-value: 0.033 q-value: 0.023

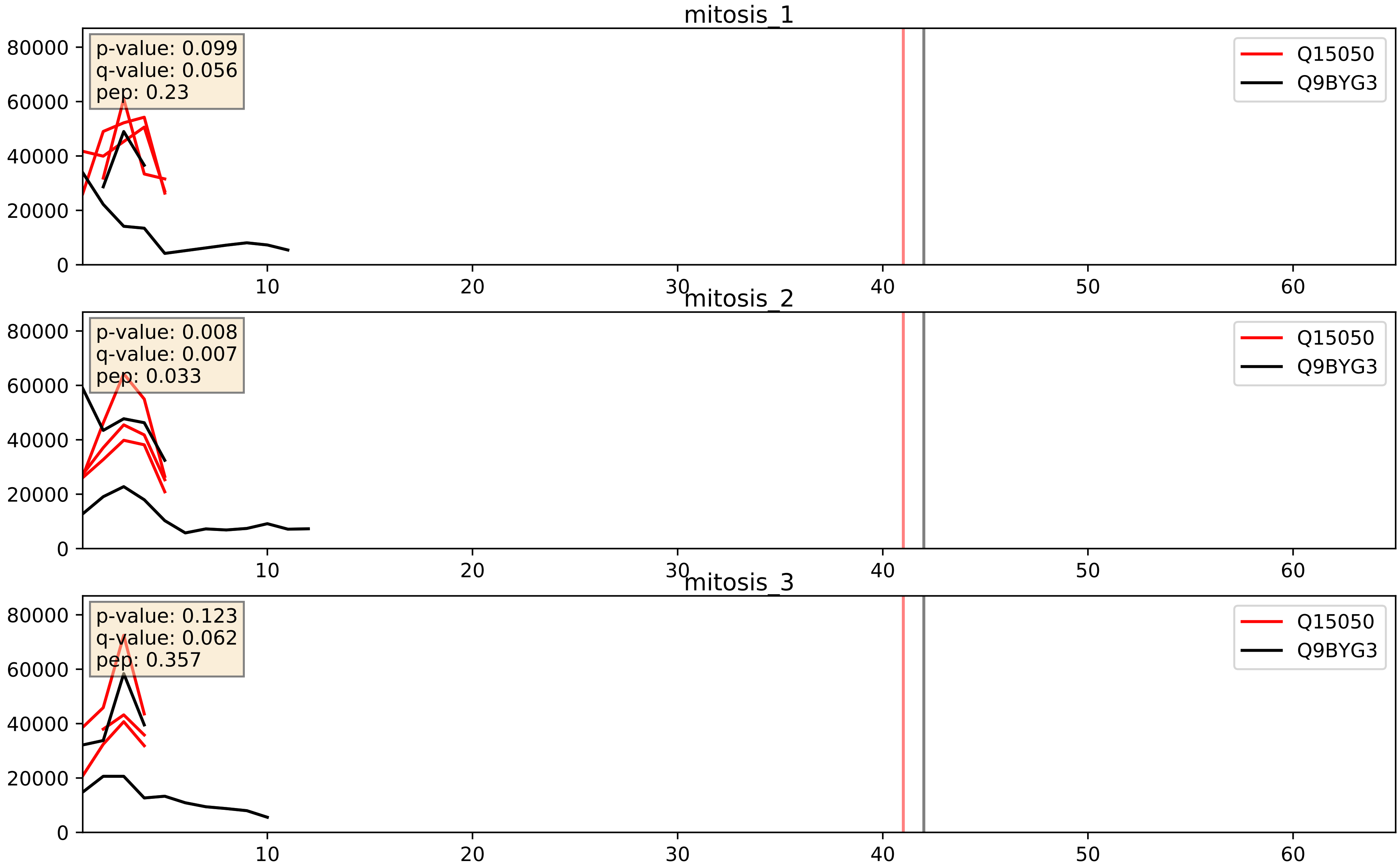


| level | condition_1 | condition_2 | pvalue | pvalue_adjusted |
|---|---|---|---|---|
| interactor_ratio | mitosis | inter | 0.12735748141619438 | 0.5750062399685051 |
| interactor_abundance | mitosis | inter | 0.2162538244564617 | 0.4119120465837366 |
| complex_abundance | mitosis | inter | 0.251694037814913 | 0.40201339026612704 |

Q15050_Q9BZE4
RRS1_HUMAN vs NOG1_HUMAN
p-value: 0.008 q-value: 0.007

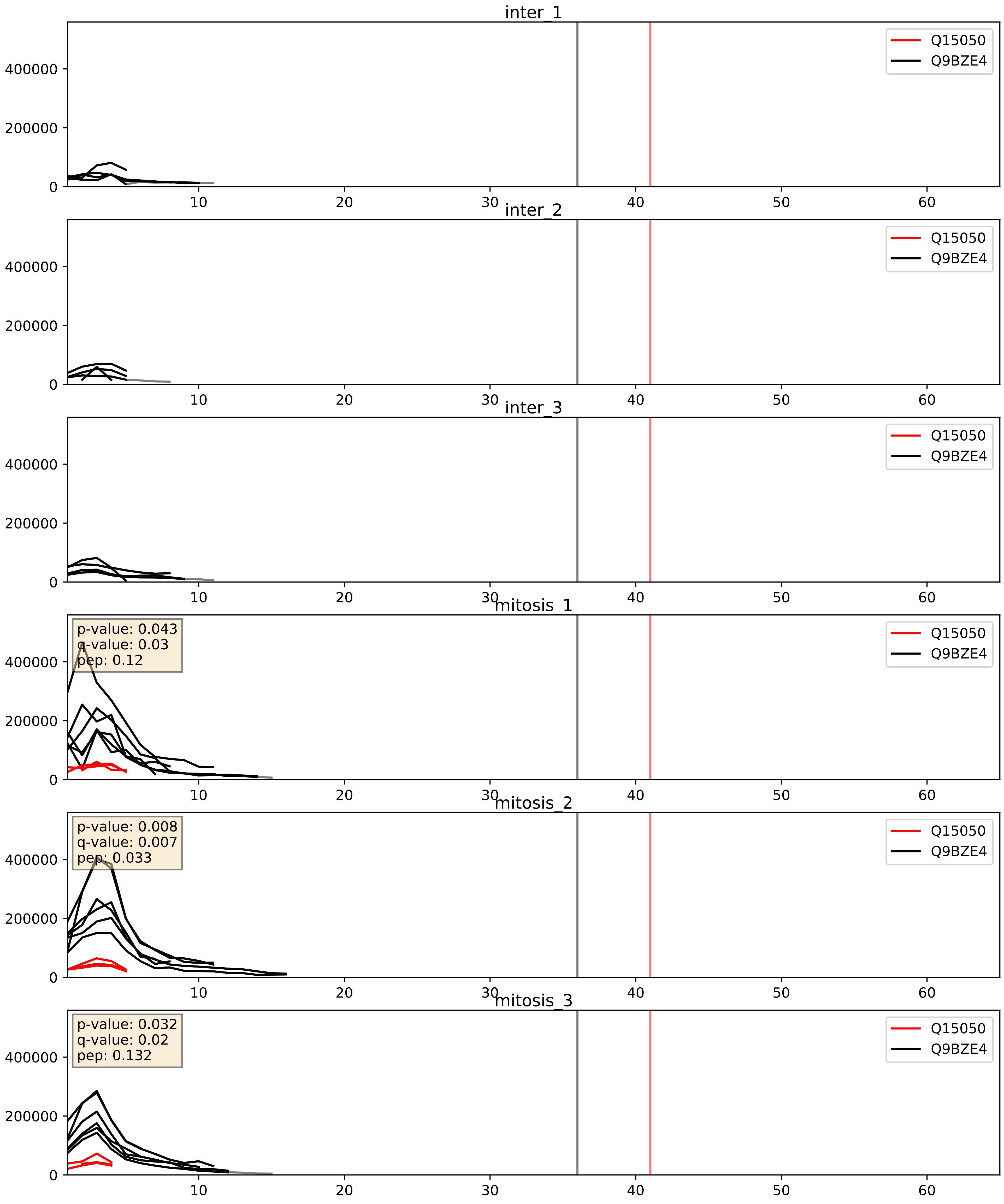


| level | condition_1 | condition_2 | pvalue | pvalue_adjusted |
|---|---|---|---|---|
| interactor_ratio | mitosis | inter | 0.0022417858763910544 | 0.05896690574521502 |
| complex_abundance | mitosis | inter | 0.005630838424790495 | 0.05650642076929266 |
| interactor_abundance | mitosis | inter | 0.2162538244564617 | 0.4119120465837366 |

Q15050_Q9H7B2
RRS1_HUMAN vs RPF2_HUMAN
p-value: 0.068 q-value: 0.042

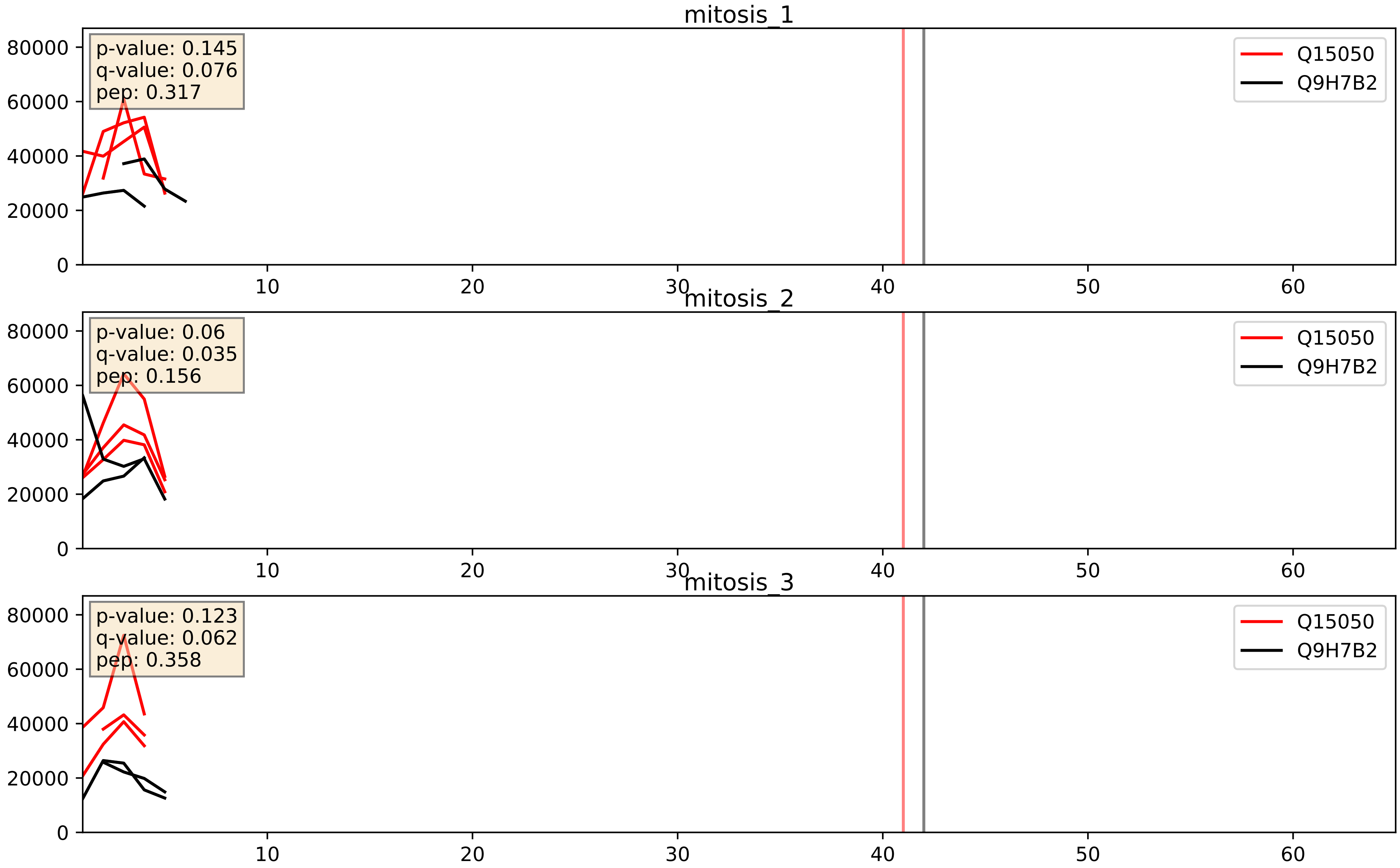


| level | condition_1 | condition_2 | pvalue | pvalue_adjusted |
|---|---|---|---|---|
| interactor_ratio | mitosis | inter | 0.05378980819845297 | 0.38466228753446735 |
| interactor_abundance | mitosis | inter | 0.2162538244564617 | 0.4119120465837366 |
| complex_abundance | mitosis | inter | 0.3048765855504356 | 0.4552142983275299 |

Q15050_Q9NY93
RRS1_HUMAN vs DDX56_HUMAN
p-value: 0.031 q-value: 0.023

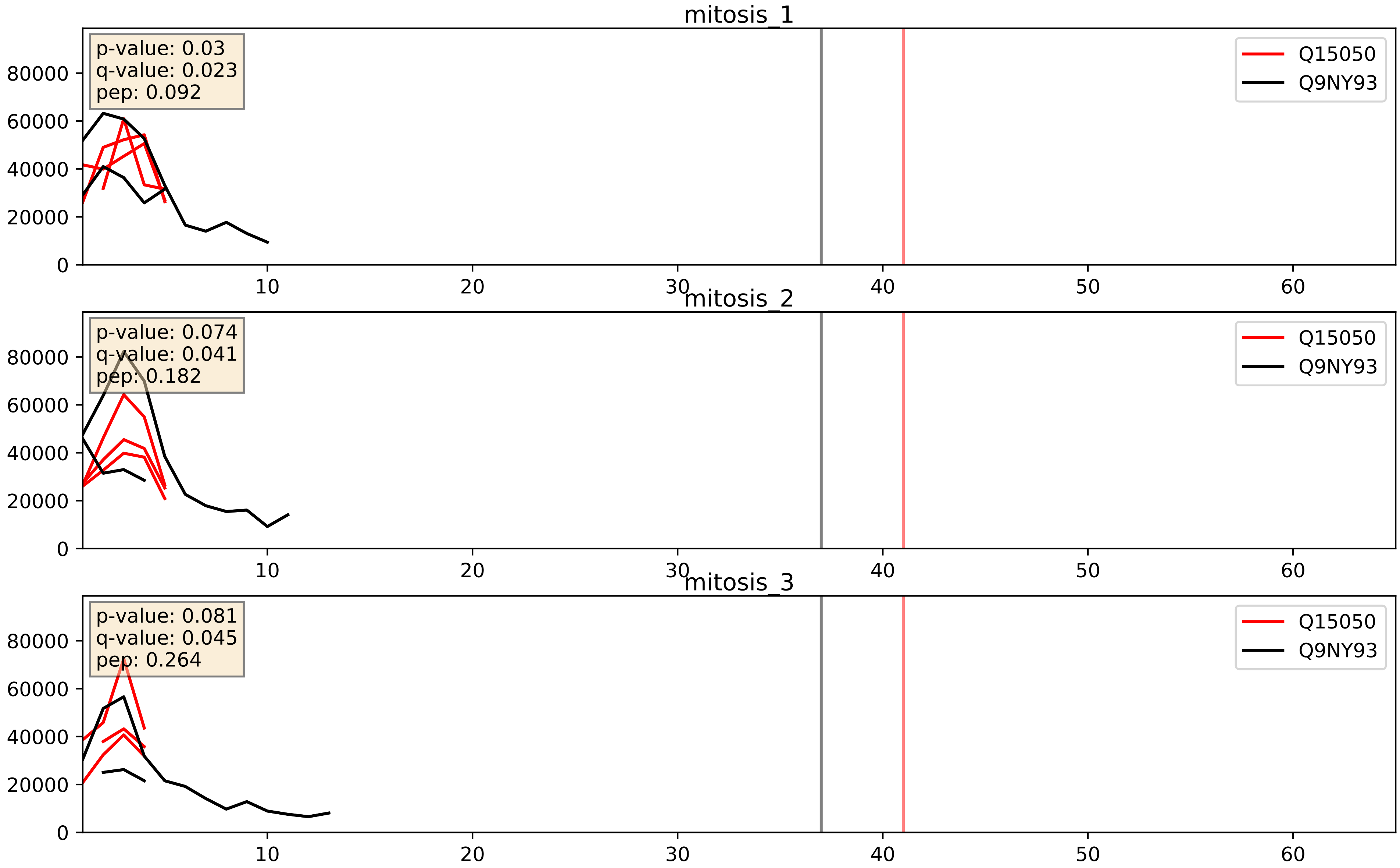


| level | condition_1 | condition_2 | pvalue | pvalue_adjusted |
|---|---|---|---|---|
| complex_abundance | mitosis | inter | 0.2143602303989282 | 0.36911920488133304 |
| interactor_abundance | mitosis | inter | 0.2162538244564617 | 0.4119120465837366 |
| interactor_ratio | mitosis | inter | 0.27768564921583294 | 0.7493660647186412 |
