## Supplementary Data 2 for "SECAT: Quantifying differential protein-protein interaction states by network-centric analysis": hela_string_000114_Q13309.pdf

Q13309_Q13309

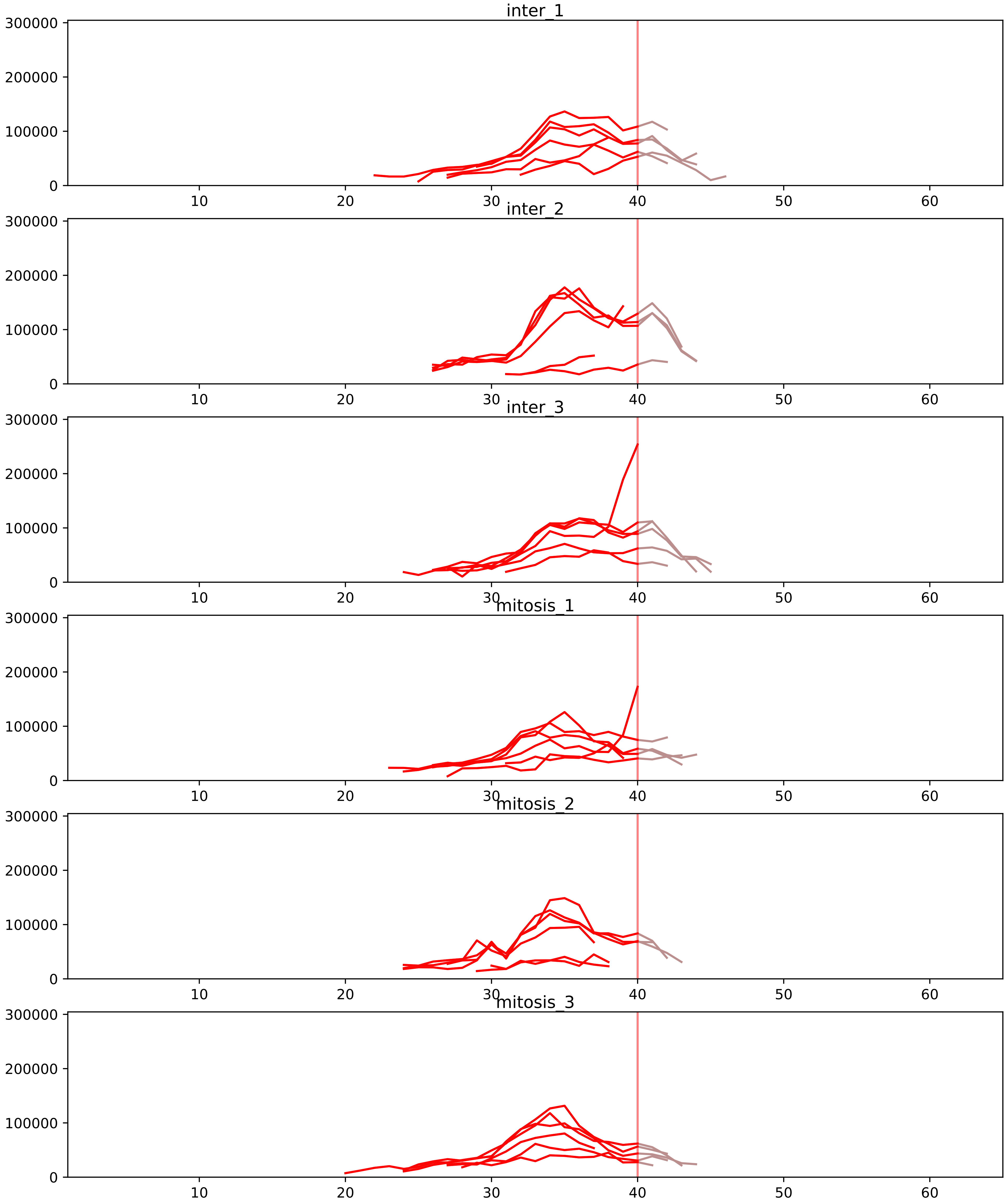


| level | condition_1 | condition_2 | pvalue | pvalue_adjusted |
|---|---|---|---|---|
| interactor_ratio | mitosis | inter | 1.793068691116623e-05 | 0.0003749143626880211 |
| complex_abundance | mitosis | inter | 0.0011405543590459082 | 0.011530879234310282 |
| interactor_abundance | mitosis | inter | 0.003541557463669617 | 0.027999199914880046 |
| monomer_abundance | mitosis | inter | 0.05149891284992204 | 0.3374840046923117 |
| total_abundance | mitosis | inter | 0.1280477064245759 | 0.44263875914078465 |
| assembled_abundance | mitosis | inter | 0.24991008879050655 | 0.5558292010635905 |

P20248_Q13309
CCNA2_HUMAN vs SKP2_HUMAN
p-value: 0.003 q-value: 0.003

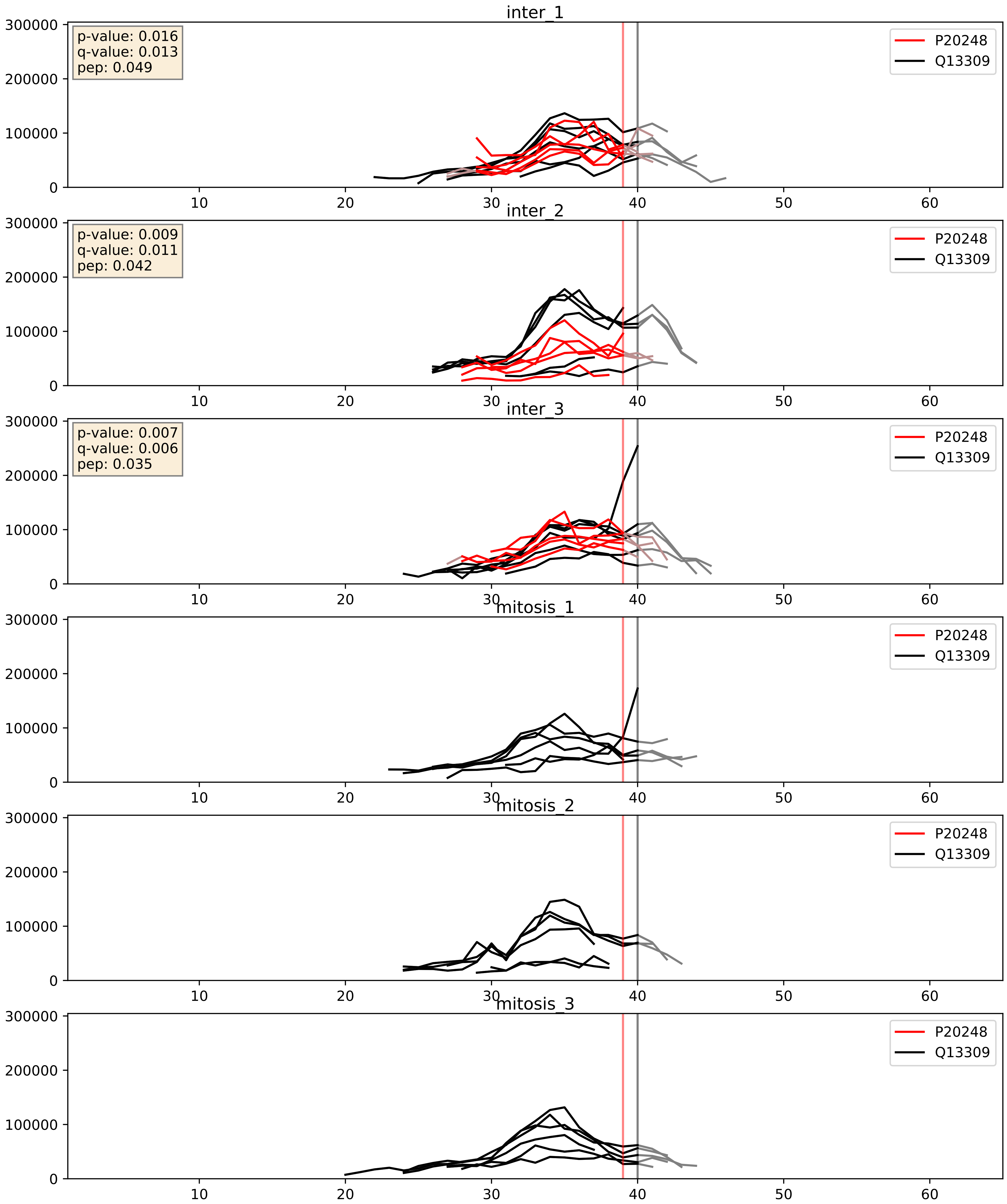


| level | condition_1 | condition_2 | pvalue | pvalue_adjusted |
|---|---|---|---|---|
| complex_abundance | mitosis | inter | 0.0002650544596844505 | 0.010908010456244694 |
| interactor_abundance | mitosis | inter | 0.0004058346268662555 | 0.013951583959739548 |
| interactor_ratio | mitosis | inter | 0.22591075557383503 | 0.7051563050171857 |

P24941_Q13309
CDK2_HUMAN vs SKP2_HUMAN
p-value: 0.0 q-value: 0.0

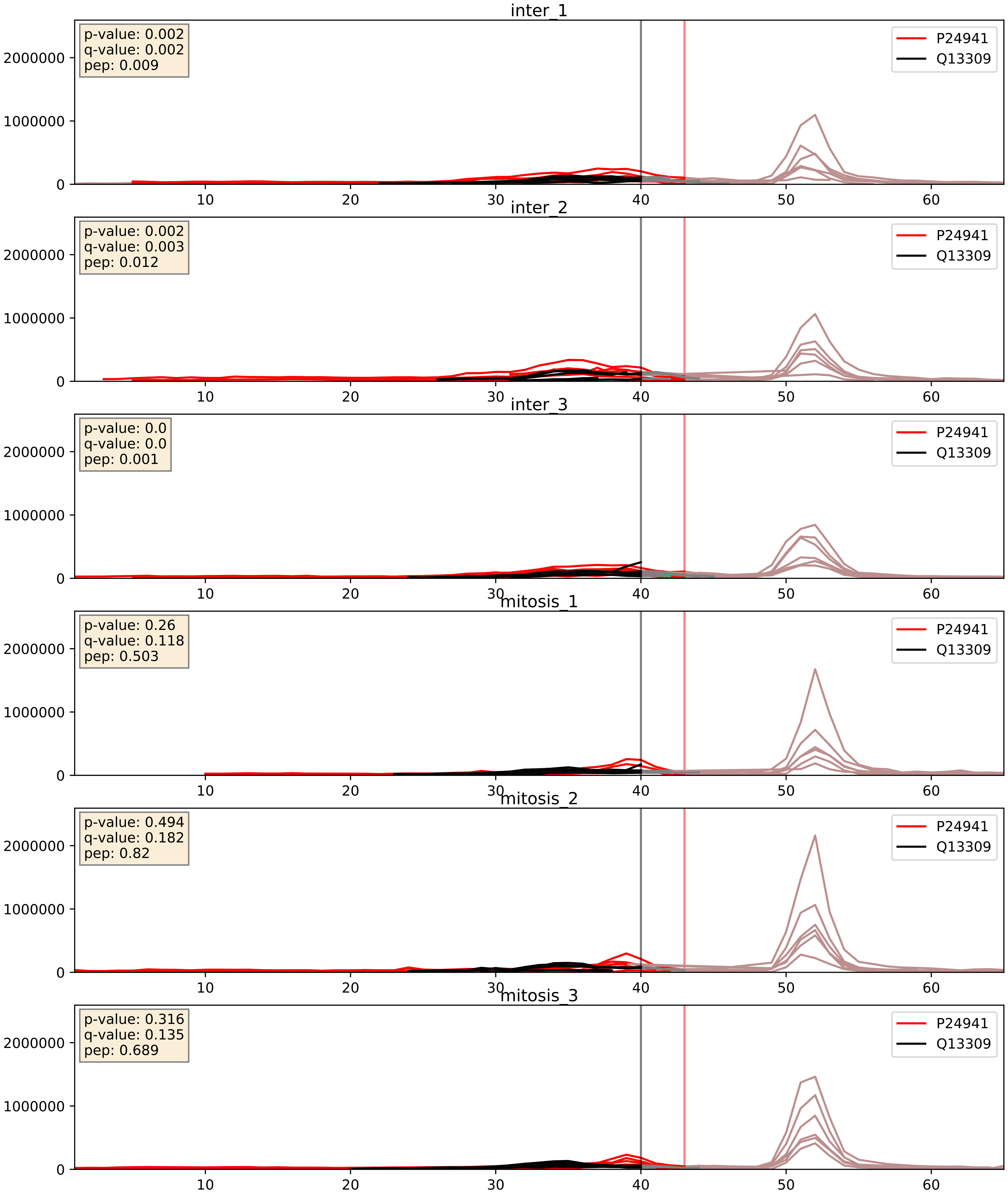


| level | condition_1 | condition_2 | pvalue | pvalue_adjusted |
|---|---|---|---|---|
| interactor_ratio | mitosis | inter | 4.400737078739184e-05 | 0.006796360849516482 |
| interactor_abundance | mitosis | inter | 0.0009100691980148597 | 0.022289534577989125 |
| complex_abundance | mitosis | inter | 0.004846689670913454 | 0.05231735634680853 |

P63208_Q13309
SKP1_HUMAN vs SKP2_HUMAN
p-value: 0.06 q-value: 0.038

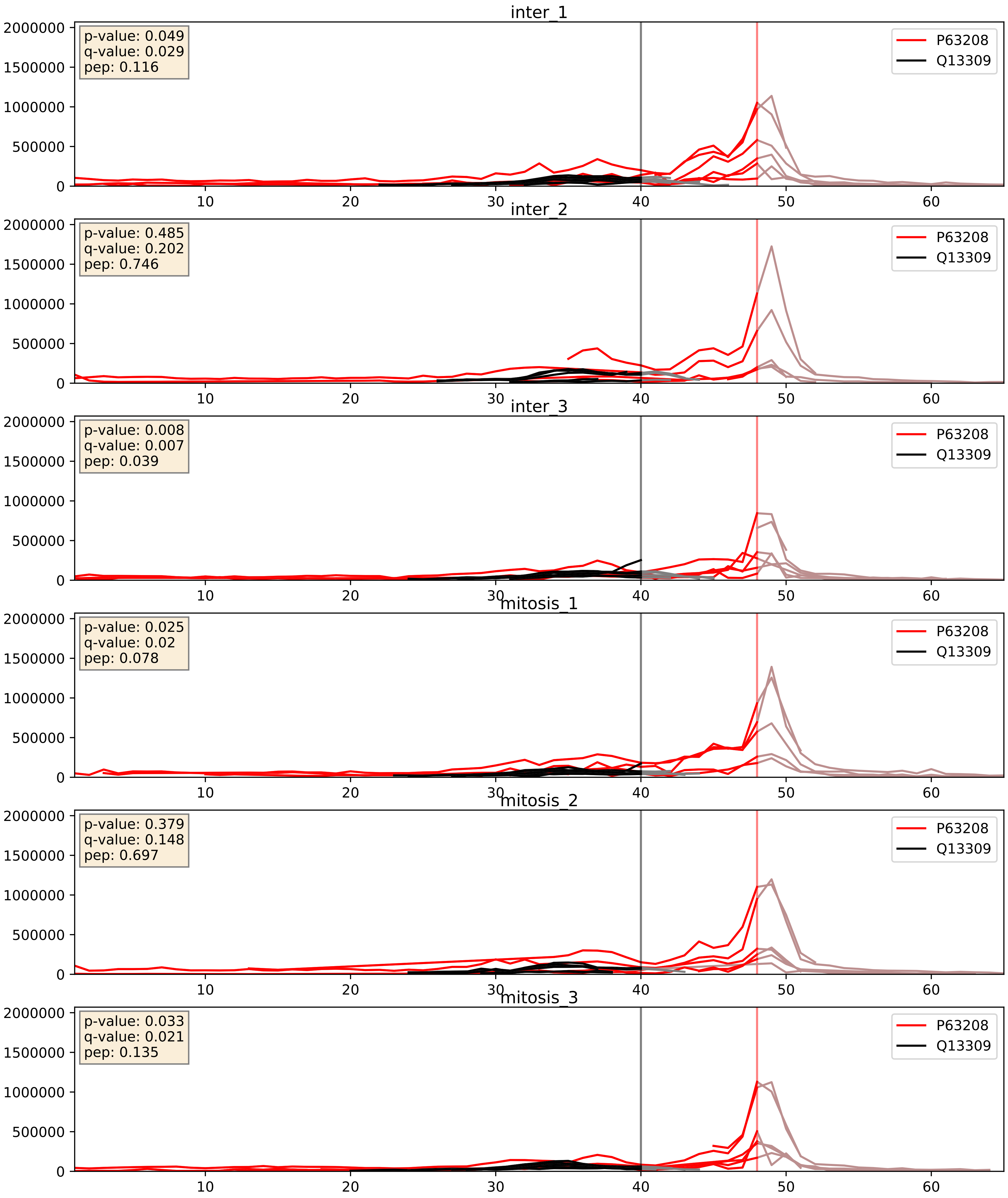


| level | condition_1 | condition_2 | pvalue | pvalue_adjusted |
|---|---|---|---|---|
| complex_abundance | mitosis | inter | 0.2581479959633784 | 0.4086070350307912 |
| interactor_ratio | mitosis | inter | 0.4569176661684025 | 0.8558457817071171 |
| interactor_abundance | mitosis | inter | 0.6841000850910031 | 0.7940602057914786 |

Q13309_Q13616
SKP2_HUMAN vs CUL1_HUMAN
p-value: 0.002 q-value: 0.003

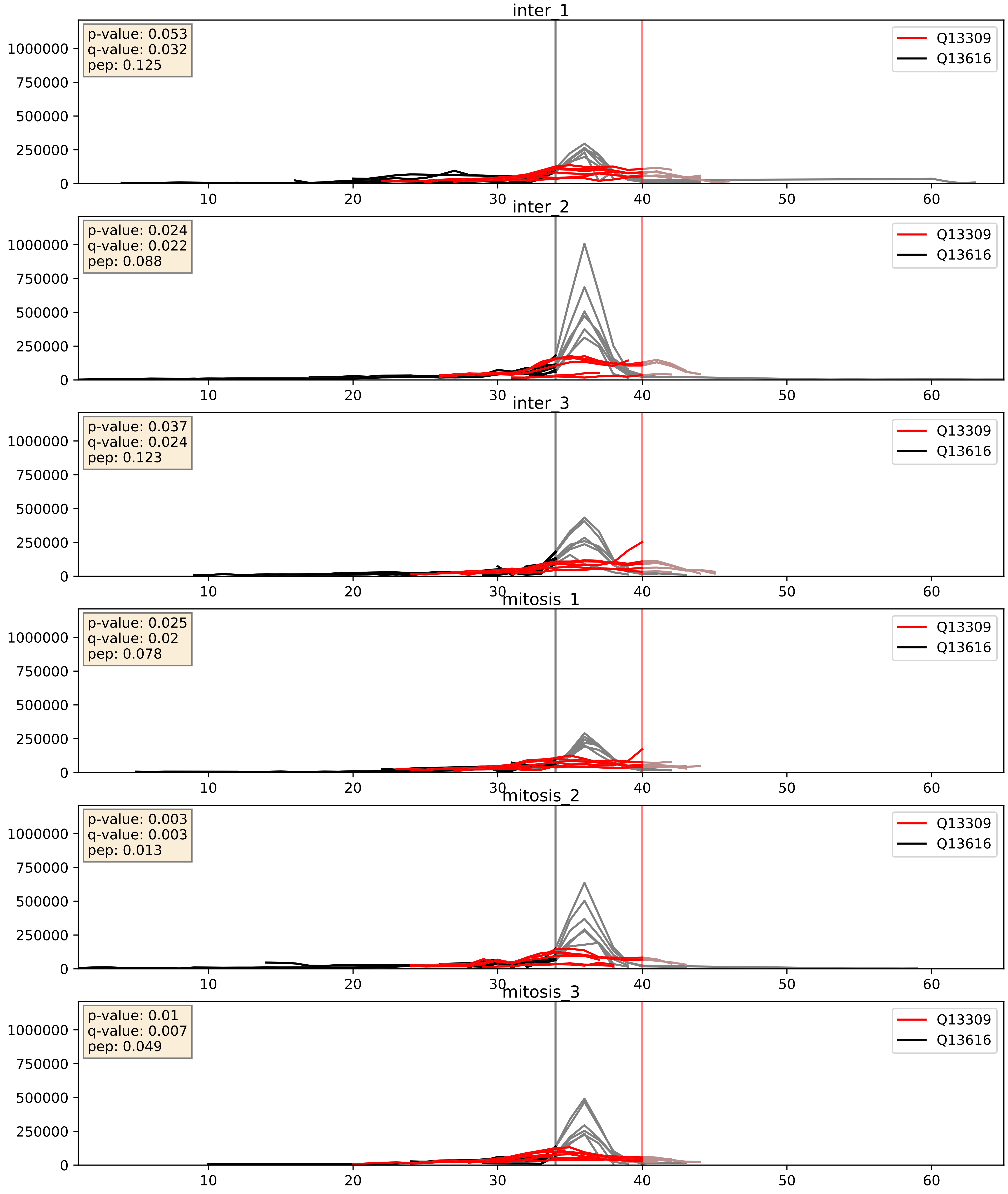


| level | condition_1 | condition_2 | pvalue | pvalue_adjusted |
|---|---|---|---|---|
| interactor_ratio | mitosis | inter | 0.003436823858222204 | 0.07572939655107606 |
| complex_abundance | mitosis | inter | 0.049301991929202635 | 0.15823961414097285 |
| interactor_abundance | mitosis | inter | 0.13276269888884848 | 0.31841791358713833 |
