## Supplementary Data 2 for "SECAT: Quantifying differential protein-protein interaction states by network-centric analysis": hela_string_000115_P06730.pdf

P06730_P06730

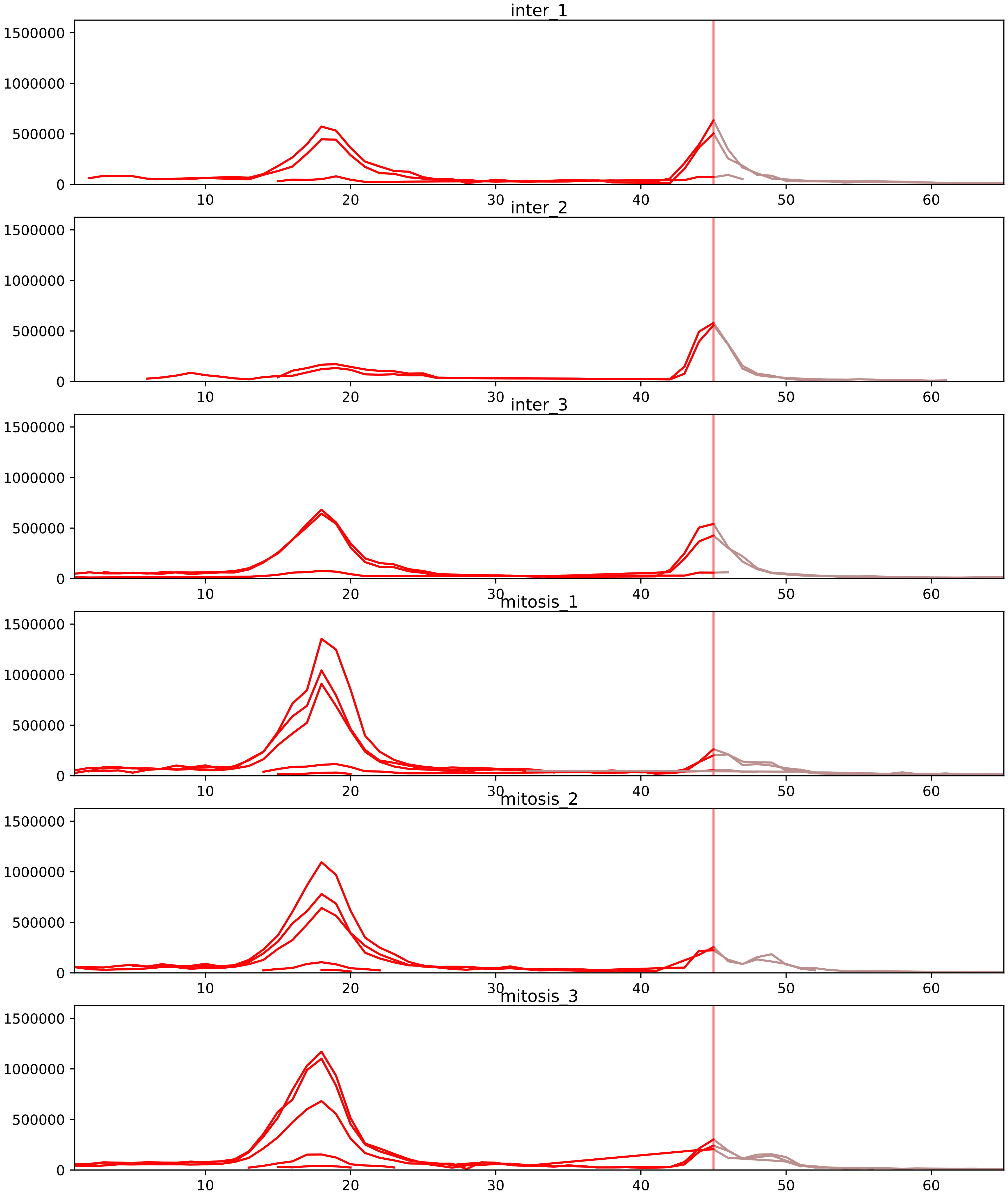


| level | condition_1 | condition_2 | pvalue | pvalue_adjusted |
|---|---|---|---|---|
| interactor_ratio | mitosis | inter | 1.9157690870760655e-05 | 0.00039606911463145627 |
| interactor_abundance | mitosis | inter | 0.007107693526799941 | 0.04205194884023117 |
| complex_abundance | mitosis | inter | 0.06809878848460033 | 0.15082718467073153 |
| assembled_abundance | mitosis | inter | 0.07195904034159487 | 0.31796498663492684 |
| total_abundance | mitosis | inter | 0.08816020280974954 | 0.36390714325112294 |
| monomer_abundance | mitosis | inter | 0.2749561090162295 | 0.6547210428856292 |

P06730_P11940
IF4E_HUMAN vs PABP1_HUMAN
p-value: 0.077 q-value: 0.046

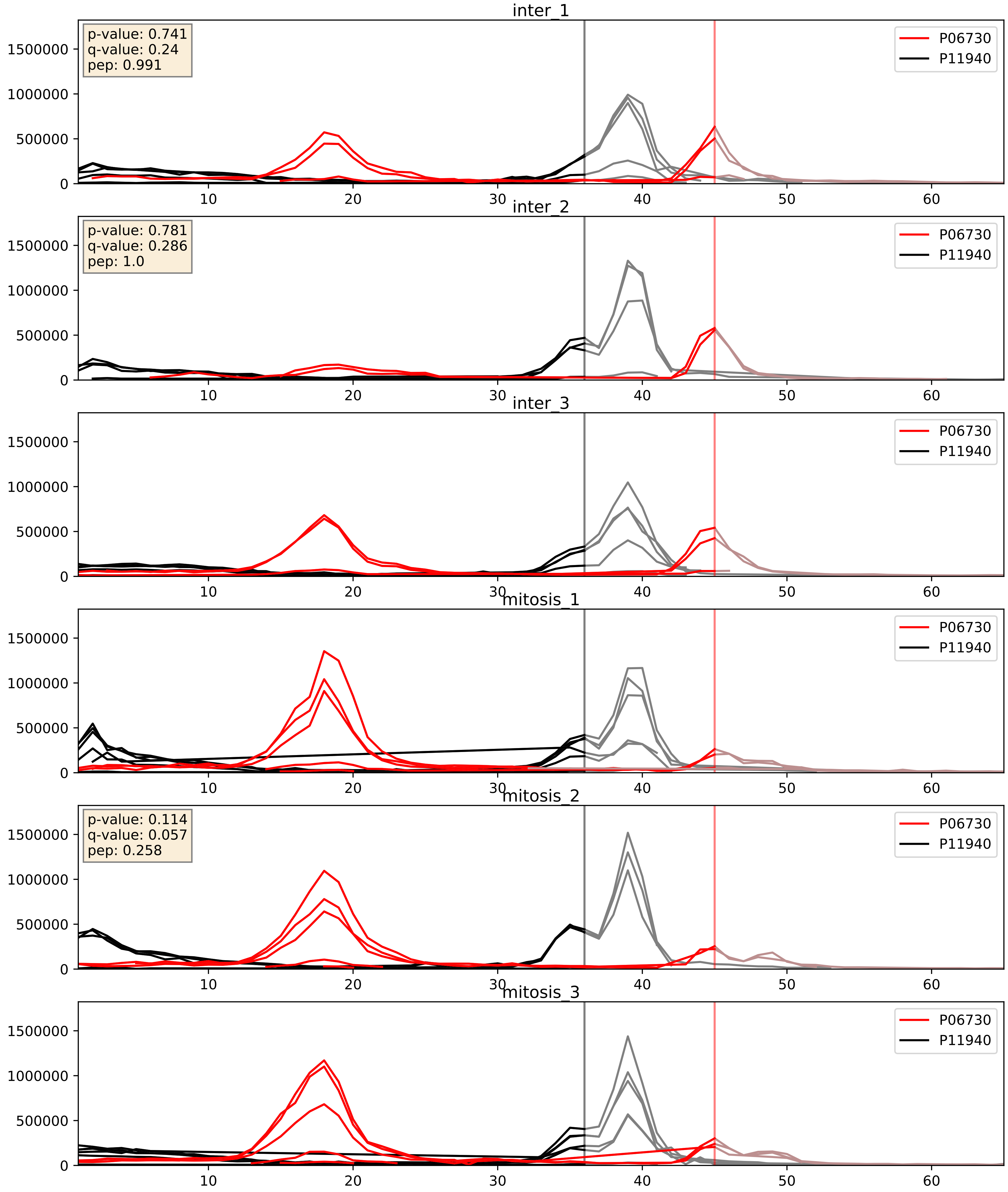


| level | condition_1 | condition_2 | pvalue | pvalue_adjusted |
|---|---|---|---|---|
| interactor_ratio | mitosis | inter | 0.005276695702534133 | 0.0974780445871482 |
| interactor_abundance | mitosis | inter | 0.008995173699102407 | 0.08274979781226932 |
| complex_abundance | mitosis | inter | 0.030746854054412083 | 0.12751204260889337 |

P06730_P52298
IF4E_HUMAN vs NCBP2_HUMAN
p-value: 0.078 q-value: 0.047

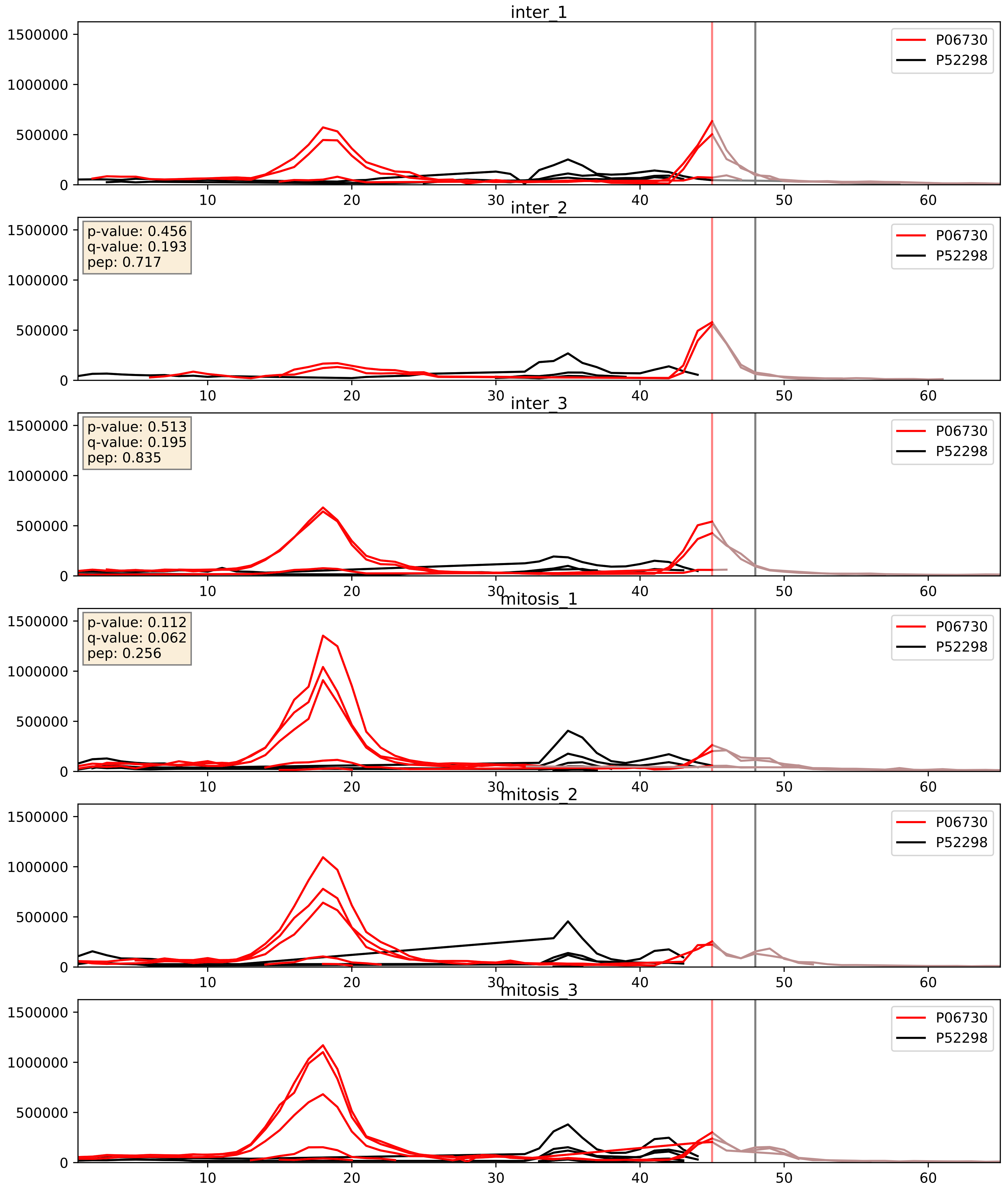


| level | condition_1 | condition_2 | pvalue | pvalue_adjusted |
|---|---|---|---|---|
| interactor_ratio | mitosis | inter | 0.11815934867104683 | 0.566001132973789 |
| complex_abundance | mitosis | inter | 0.651369168107556 | 0.7659441330091319 |
| interactor_abundance | mitosis | inter | 0.8431674357002676 | 0.9148738682117235 |

P06730_Q04637
IF4E_HUMAN vs IF4G1_HUMAN
p-value: 0.002 q-value: 0.002

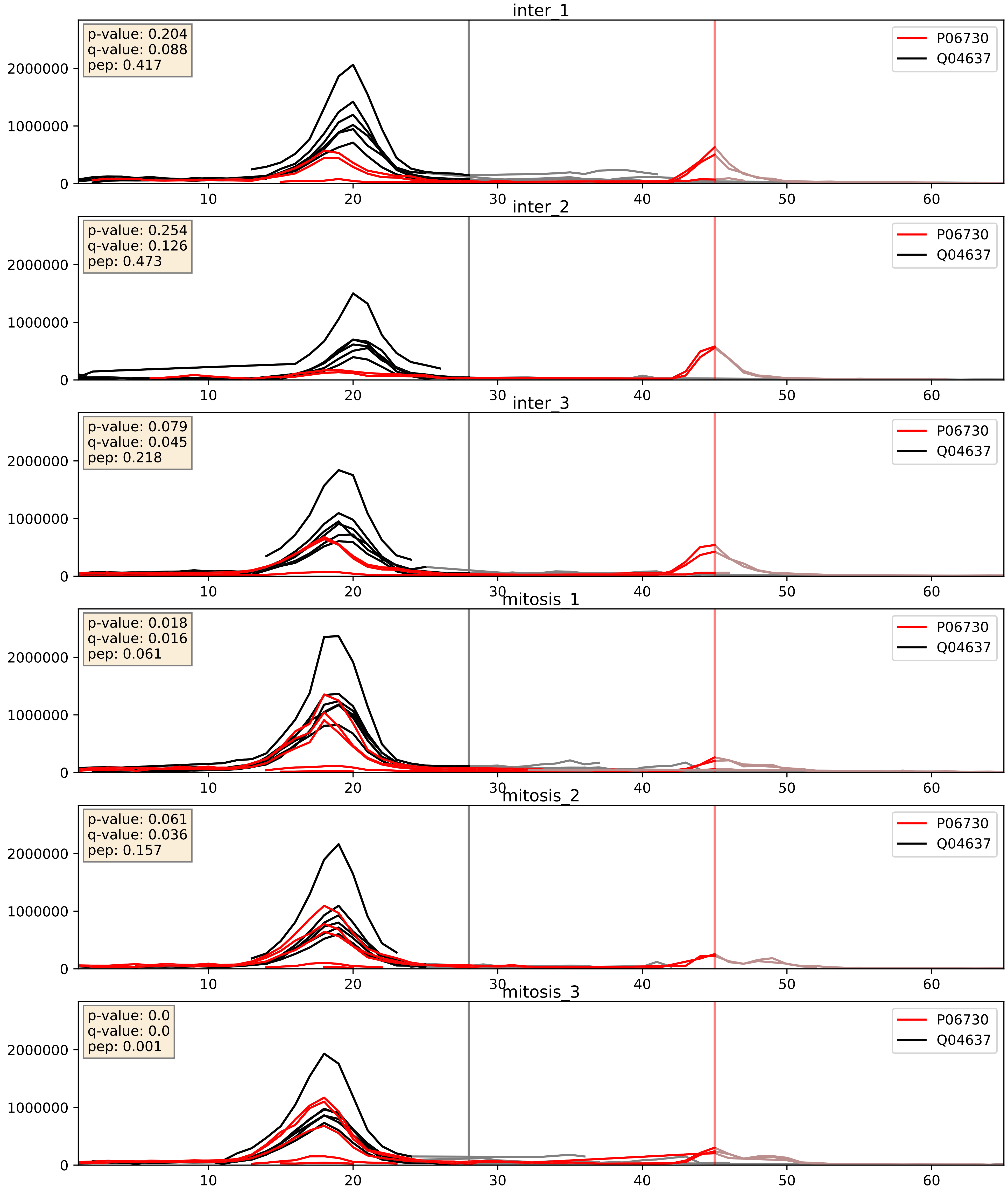


| level | condition_1 | condition_2 | pvalue | pvalue_adjusted |
|---|---|---|---|---|
| interactor_ratio | mitosis | inter | 8.260519479990175e-05 | 0.009820839826210542 |
| interactor_abundance | mitosis | inter | 0.003941007135026407 | 0.04939241738773945 |
| complex_abundance | mitosis | inter | 0.05576803101327905 | 0.16848105578480535 |
