## Supplementary Data 2 for "SECAT: Quantifying differential protein-protein interaction states by network-centric analysis": hela_string_000116_P62857.pdf

P62857_P62857

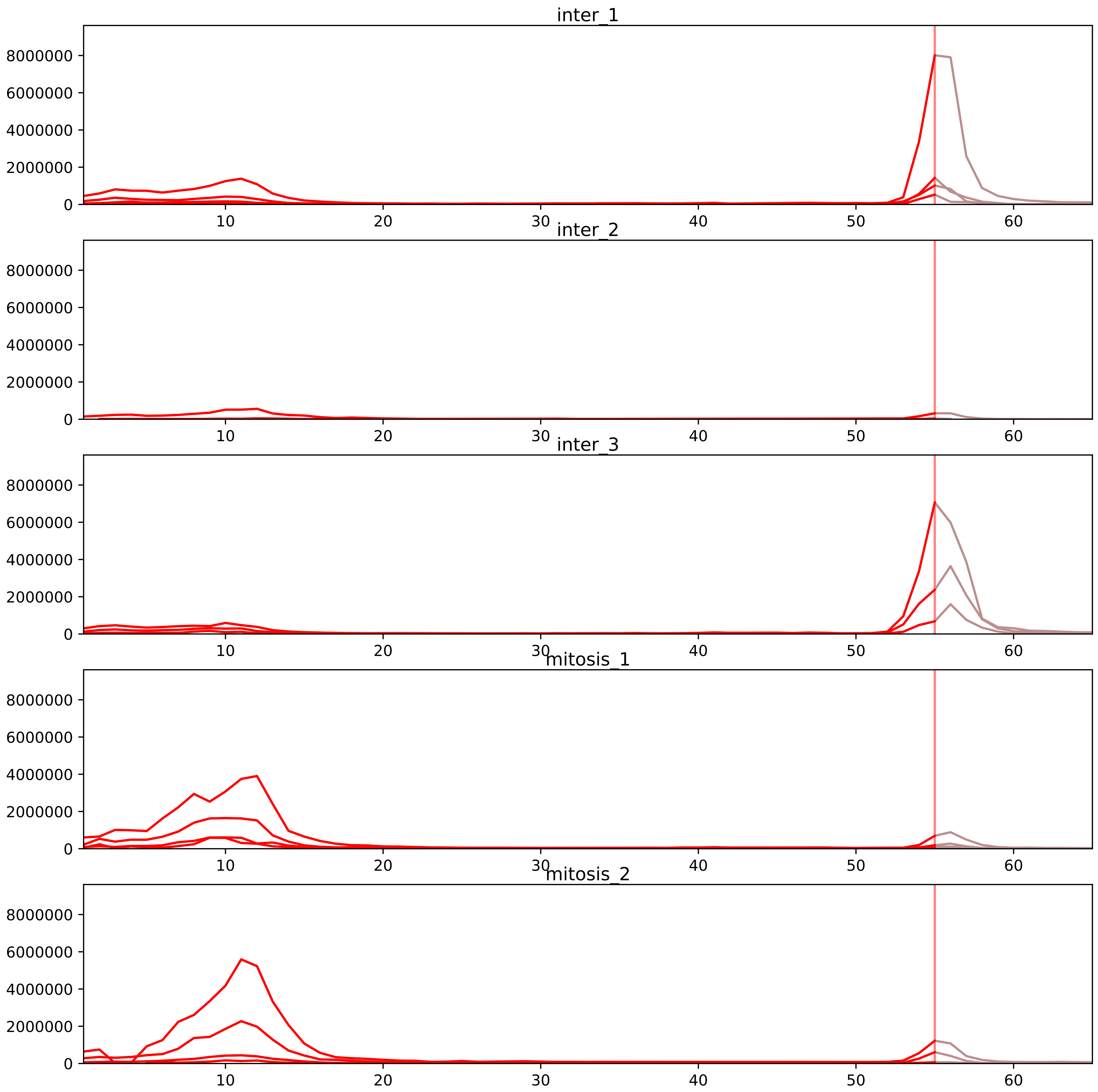


| level | condition_1 | condition_2 | pvalue | pvalue_adjusted |
|---|---|---|---|---|
| interactor_ratio | mitosis | inter | 1.9960504276149255e-05 | 0.0004080814207568292 |
| monomer_abundance | mitosis | inter | 0.3581387290098867 | 0.6771224734198139 |
| total_abundance | mitosis | inter | 0.8003699806125246 | 0.921568039209447 |
| complex_abundance | mitosis | inter | 0.9076838426298922 | 0.9340818067332224 |
| interactor_abundance | mitosis | inter | 0.9725941258707124 | 0.9849054439197087 |
| assembled_abundance | mitosis | inter | 0.9800051741031092 | 0.993589824556347 |

P08708_P62857
RS17_HUMAN vs RS28_HUMAN
p-value: 0.001 q-value: 0.001

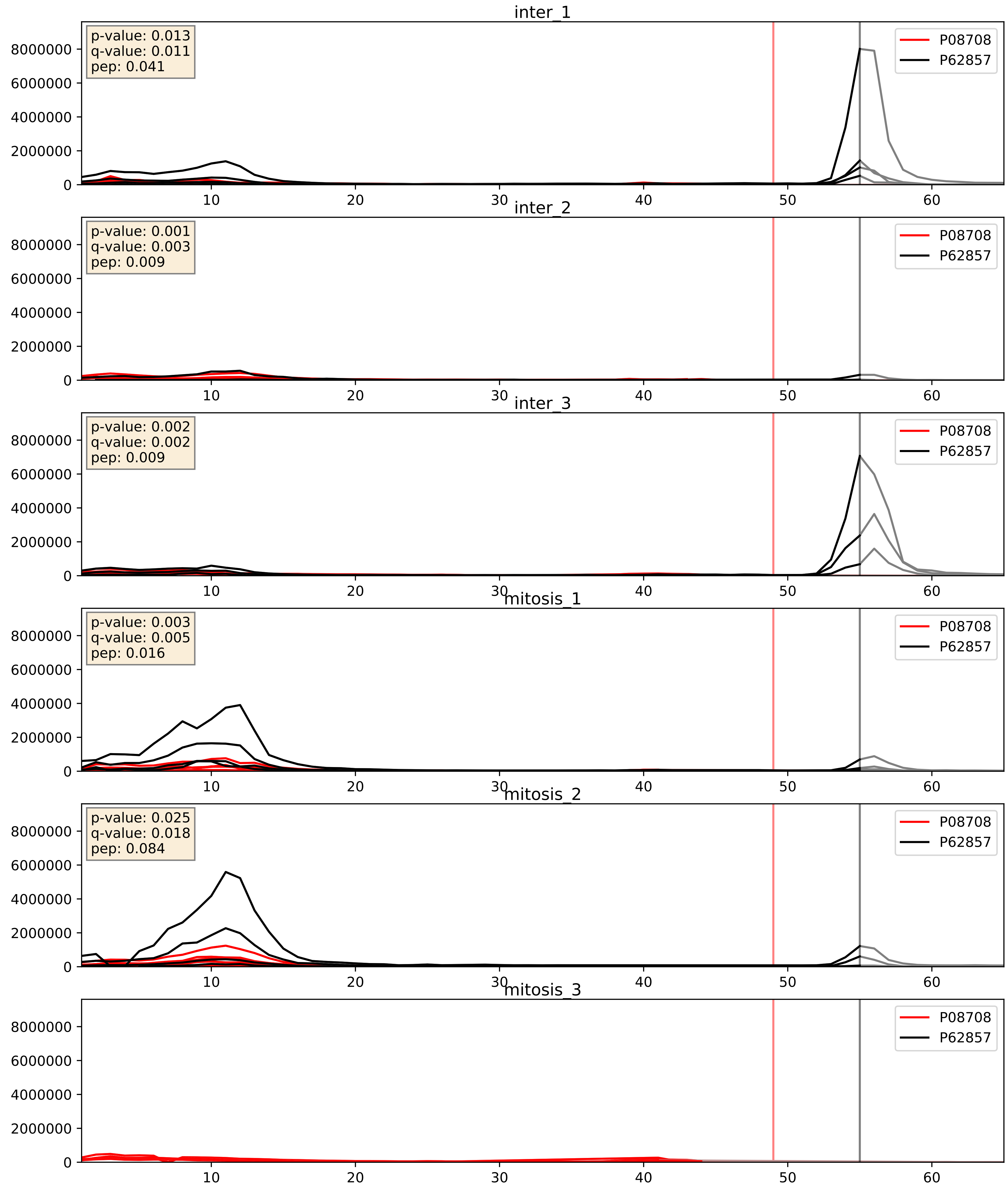


| level | condition_1 | condition_2 | pvalue | pvalue_adjusted |
|---|---|---|---|---|
| interactor_ratio | mitosis | inter | 0.150741423552149 | 0.606604569117897 |
| interactor_abundance | mitosis | inter | 0.49530593329685924 | 0.6615930075713685 |
| complex_abundance | mitosis | inter | 0.8344052552652697 | 0.8930607203023997 |

P08865_P62857
RSSA_HUMAN vs RS28_HUMAN
p-value: 0.004 q-value: 0.004

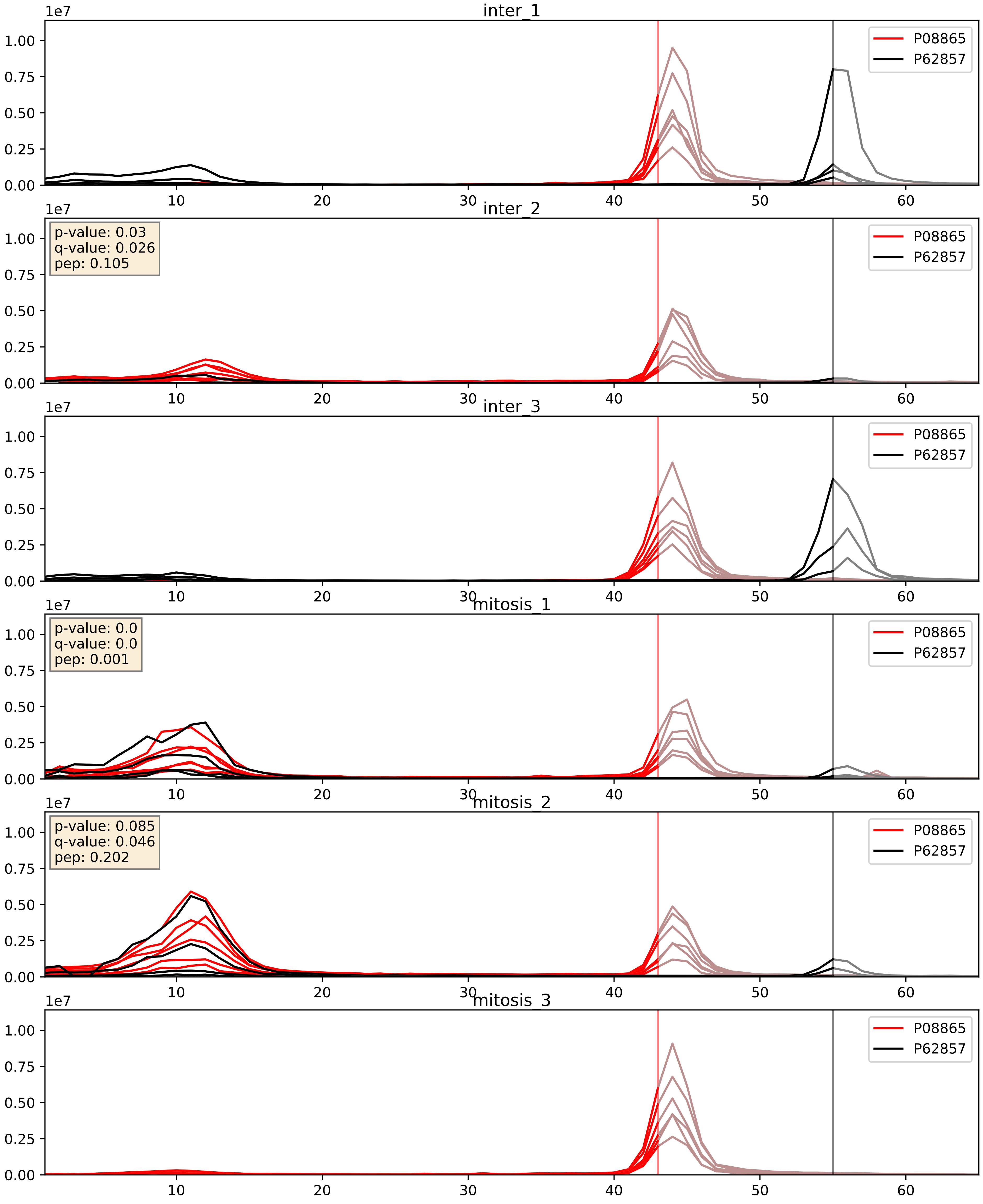


| level | condition_1 | condition_2 | pvalue | pvalue_adjusted |
|---|---|---|---|---|
| interactor_ratio | mitosis | inter | 0.6889717142374845 | 0.9349347226390419 |
| interactor_abundance | mitosis | inter | 0.9513685401036985 | 0.9696996354688533 |
| complex_abundance | mitosis | inter | 0.9655445972905117 | 0.979504829676082 |

P15880_P62857
RS2_HUMAN vs RS28_HUMAN
p-value: 0.002 q-value: 0.003

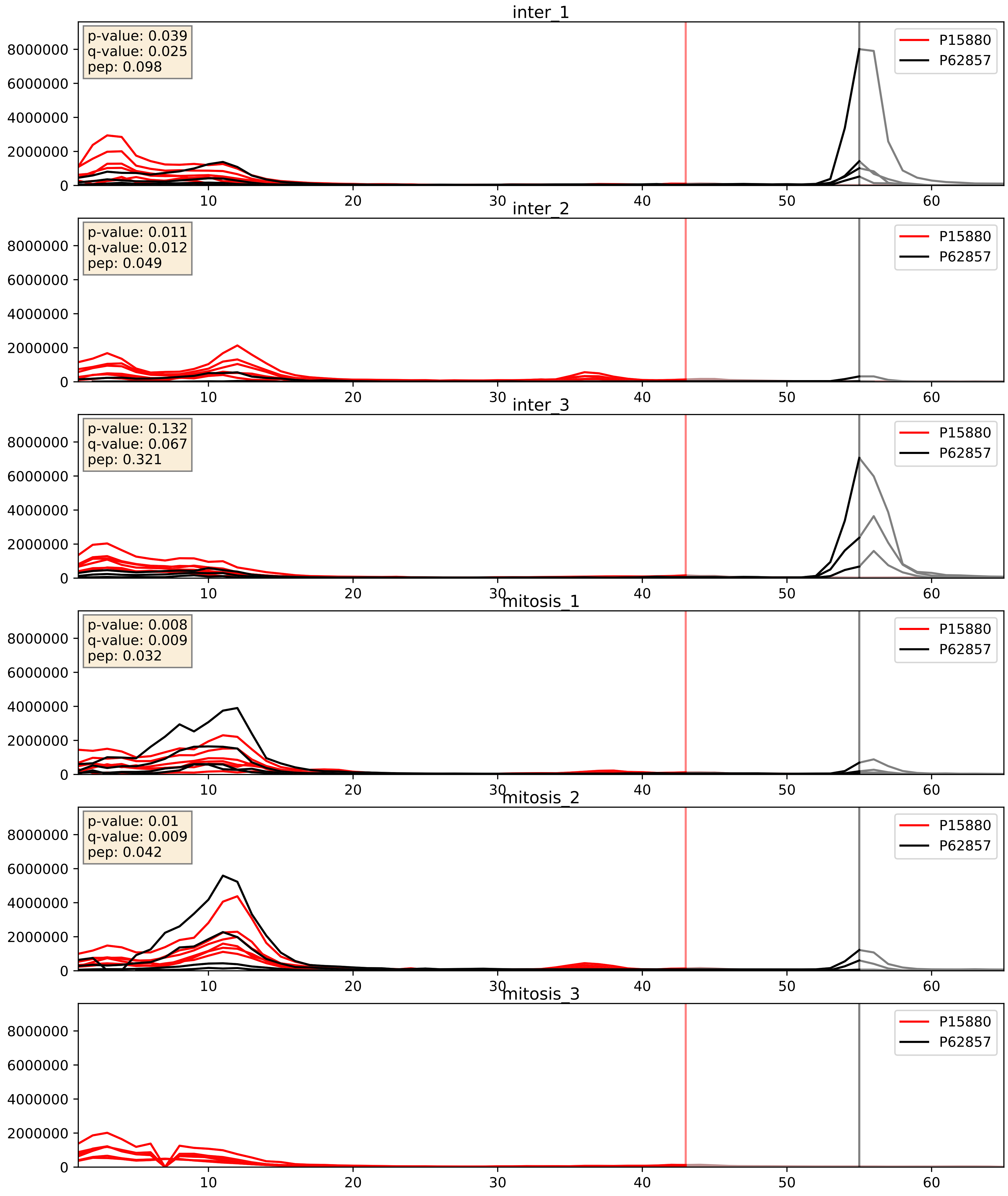


| level | condition_1 | condition_2 | pvalue | pvalue_adjusted |
|---|---|---|---|---|
| interactor_ratio | mitosis | inter | 0.016173375238959505 | 0.18789564671895648 |
| interactor_abundance | mitosis | inter | 0.3703384603774174 | 0.564526261389136 |
| complex_abundance | mitosis | inter | 0.6882105927713111 | 0.7910675488572768 |

P18124_P62857
RL7_HUMAN vs RS28_HUMAN
p-value: 0.028 q-value: 0.021

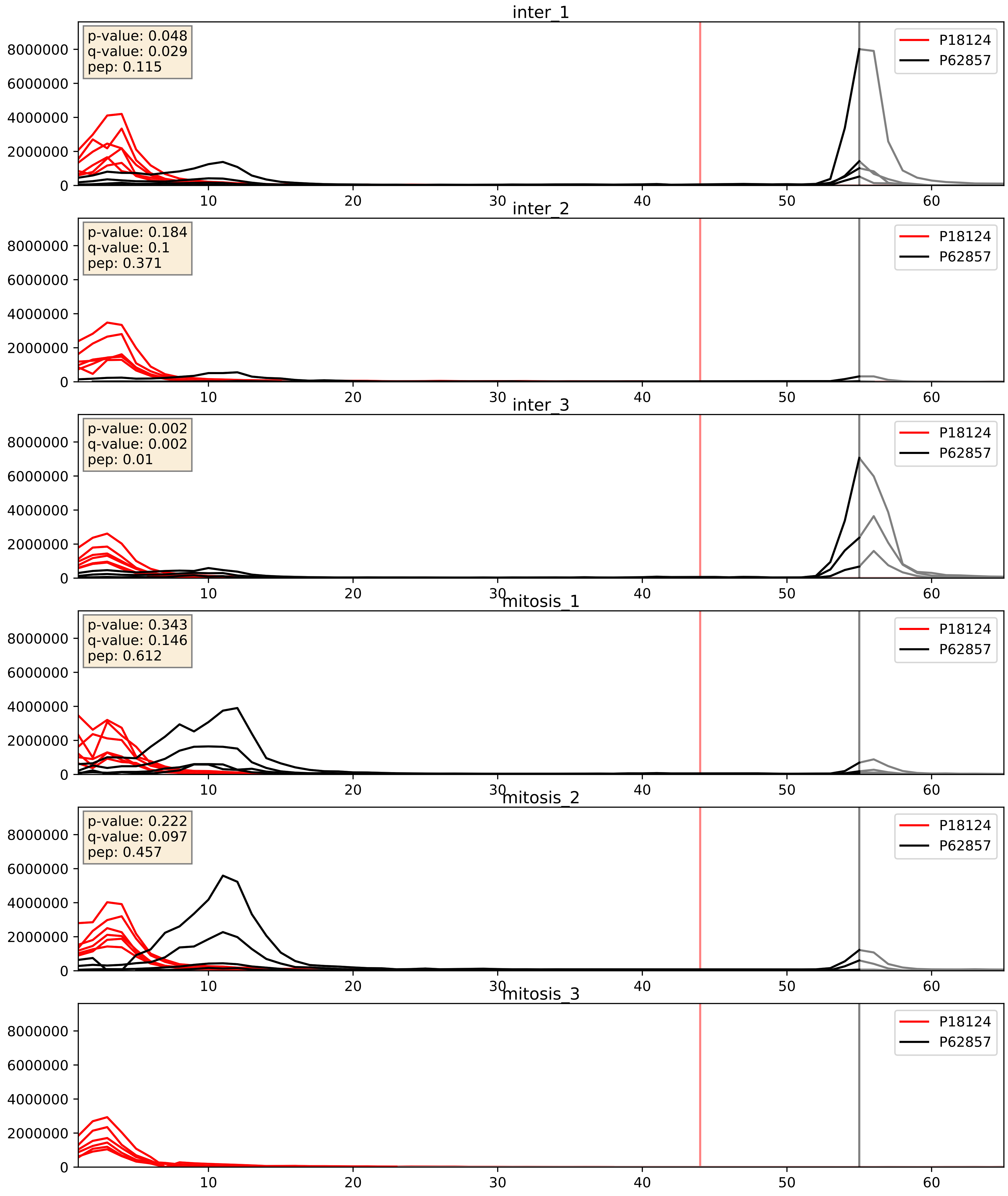


| level | condition_1 | condition_2 | pvalue | pvalue_adjusted |
|---|---|---|---|---|
| interactor_ratio | mitosis | inter | 0.022248222416681212 | 0.2287907703572271 |
| interactor_abundance | mitosis | inter | 0.3525214497021208 | 0.5503698117722354 |
| complex_abundance | mitosis | inter | 0.6932315326325629 | 0.7936588942398386 |

P23396_P62857
RS3_HUMAN vs RS28_HUMAN
p-value: 0.003 q-value: 0.003

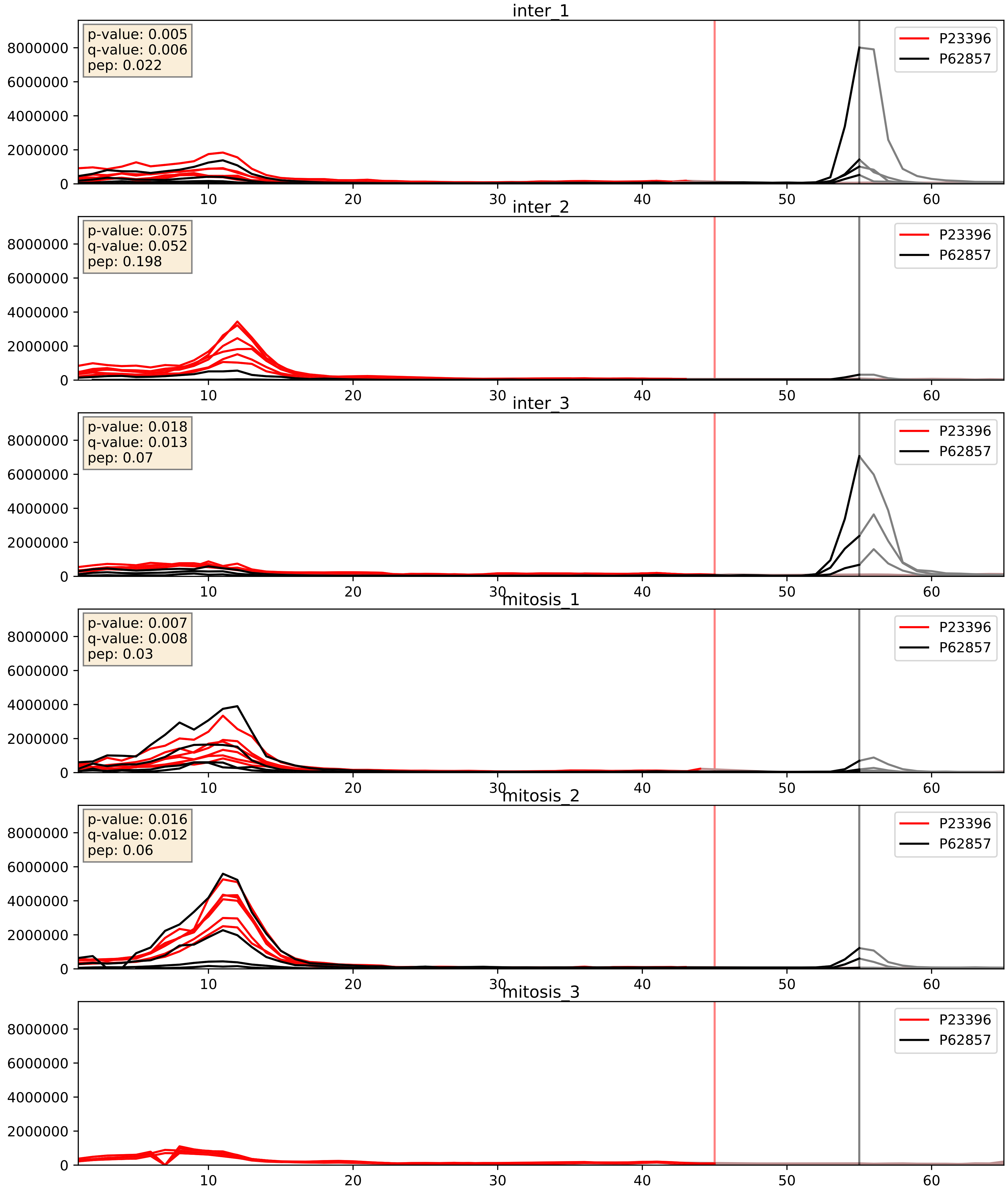


| level | condition_1 | condition_2 | pvalue | pvalue_adjusted |
|---|---|---|---|---|
| interactor_ratio | mitosis | inter | 0.09455592333659874 | 0.5100180868061028 |
| interactor_abundance | mitosis | inter | 0.4616393427835608 | 0.6333869717541278 |
| complex_abundance | mitosis | inter | 0.7397783081654679 | 0.8273454818260262 |

P25398_P62857
RS12_HUMAN vs RS28_HUMAN
p-value: 0.0 q-value: 0.0

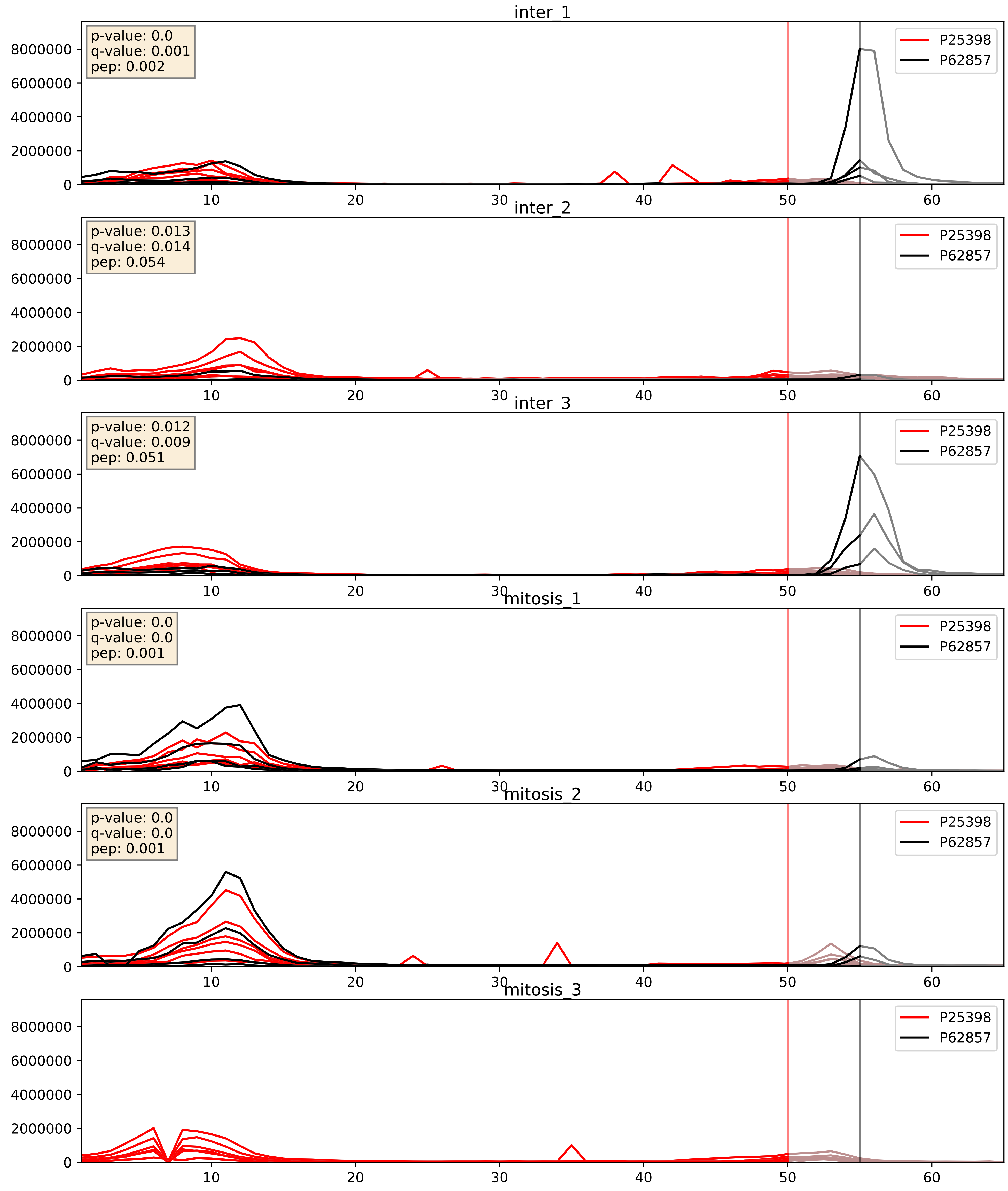


| level | condition_1 | condition_2 | pvalue | pvalue_adjusted |
|---|---|---|---|---|
| interactor_ratio | mitosis | inter | 0.04308720299269909 | 0.34205635861393335 |
| interactor_abundance | mitosis | inter | 0.39830631844871844 | 0.5885688295826003 |
| complex_abundance | mitosis | inter | 0.7140105674967991 | 0.8086472671317149 |

P26373_P62857
RL13_HUMAN vs RS28_HUMAN
p-value: 0.001 q-value: 0.001

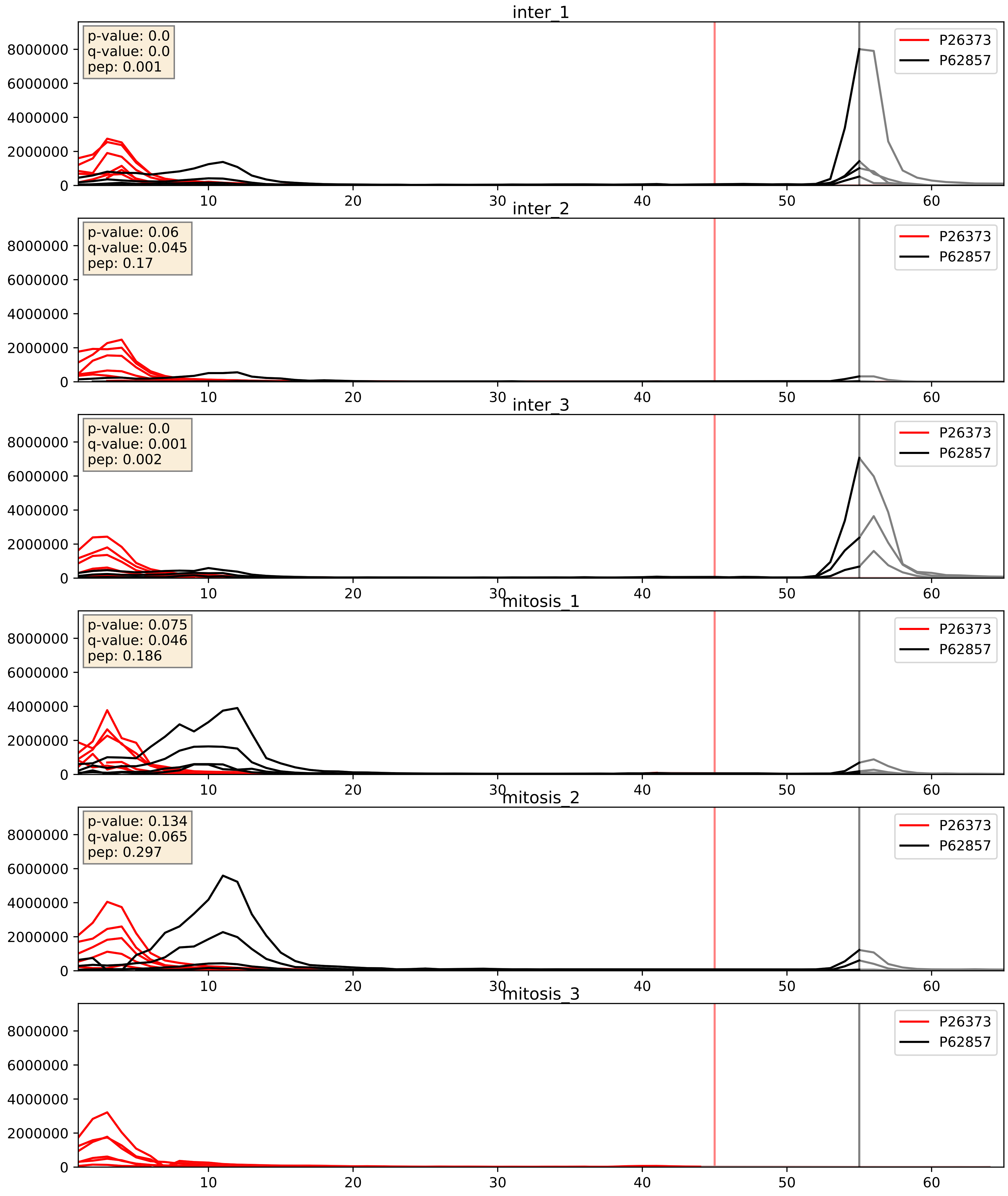


| level | condition_1 | condition_2 | pvalue | pvalue_adjusted |
|---|---|---|---|---|
| interactor_ratio | mitosis | inter | 0.022487011714898645 | 0.2302497850233641 |
| interactor_abundance | mitosis | inter | 0.3776830676469938 | 0.5708606348348745 |
| complex_abundance | mitosis | inter | 0.7187565919134282 | 0.8117900022138733 |

P30050_P62857
RL12_HUMAN vs RS28_HUMAN
p-value: 0.004 q-value: 0.004

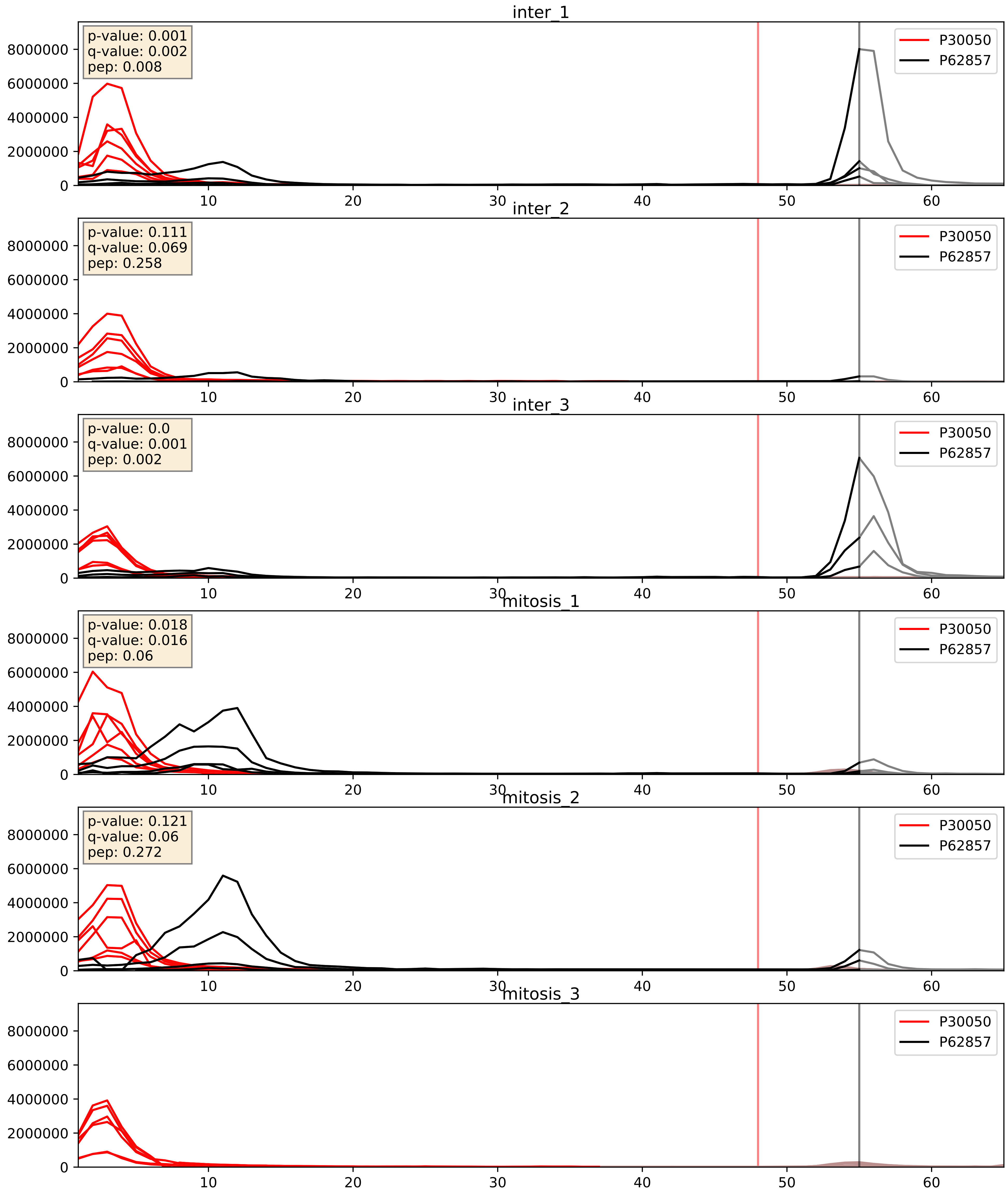


| level | condition_1 | condition_2 | pvalue | pvalue_adjusted |
|---|---|---|---|---|
| interactor_ratio | mitosis | inter | 0.014378853953145131 | 0.1758328426270319 |
| interactor_abundance | mitosis | inter | 0.35537611323055135 | 0.5531447457502536 |
| complex_abundance | mitosis | inter | 0.6787383751968641 | 0.785657028505082 |

P32969_P62857
RL9_HUMAN vs RS28_HUMAN
p-value: 0.05 q-value: 0.033

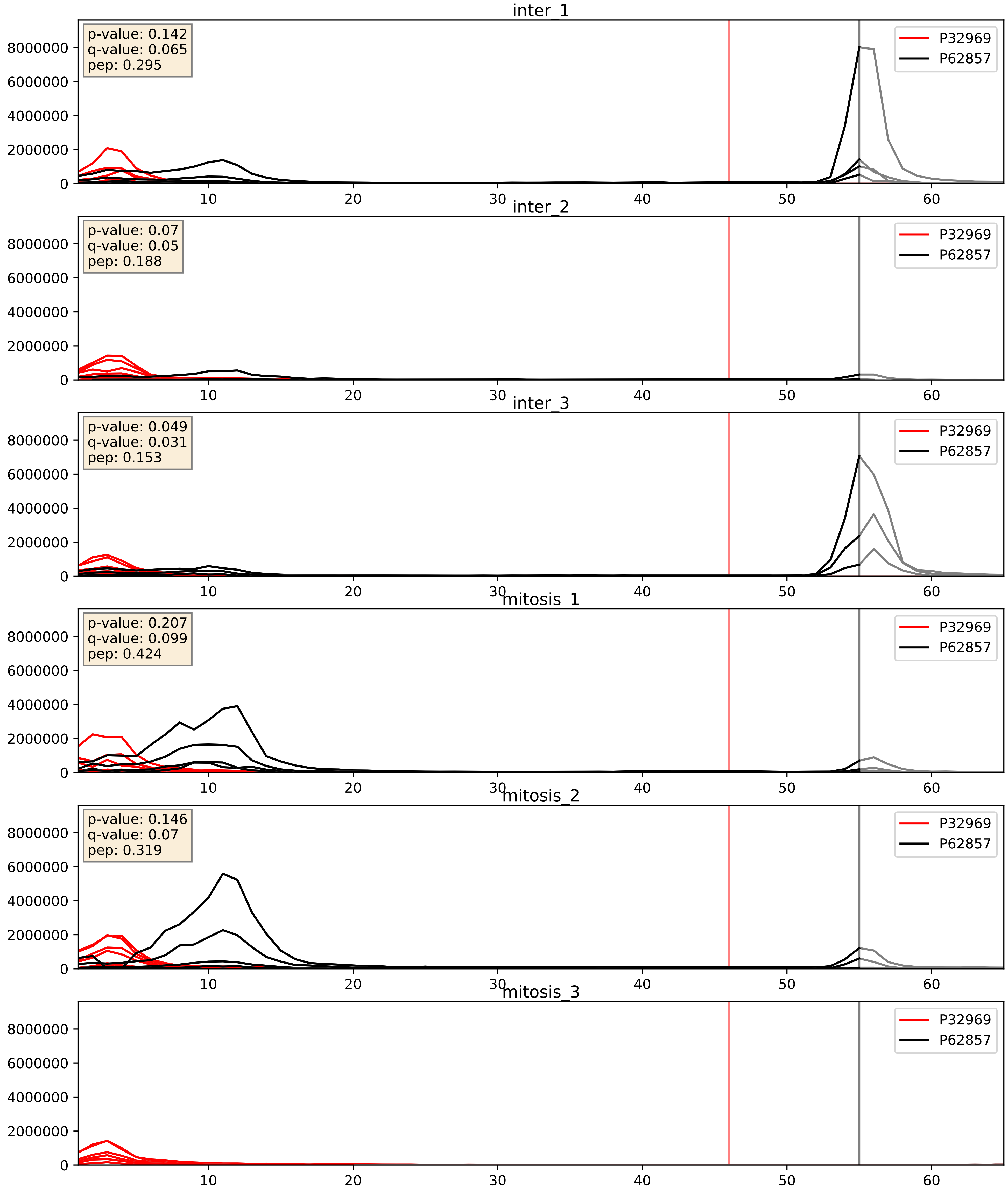


| level | condition_1 | condition_2 | pvalue | pvalue_adjusted |
|---|---|---|---|---|
| interactor_ratio | mitosis | inter | 0.07728850547621614 | 0.4662365094266457 |
| interactor_abundance | mitosis | inter | 0.42850403224410444 | 0.6085432627141492 |
| complex_abundance | mitosis | inter | 0.7857588294981157 | 0.8600045493547973 |

P35268_P62857
RL22_HUMAN vs RS28_HUMAN
p-value: 0.006 q-value: 0.006

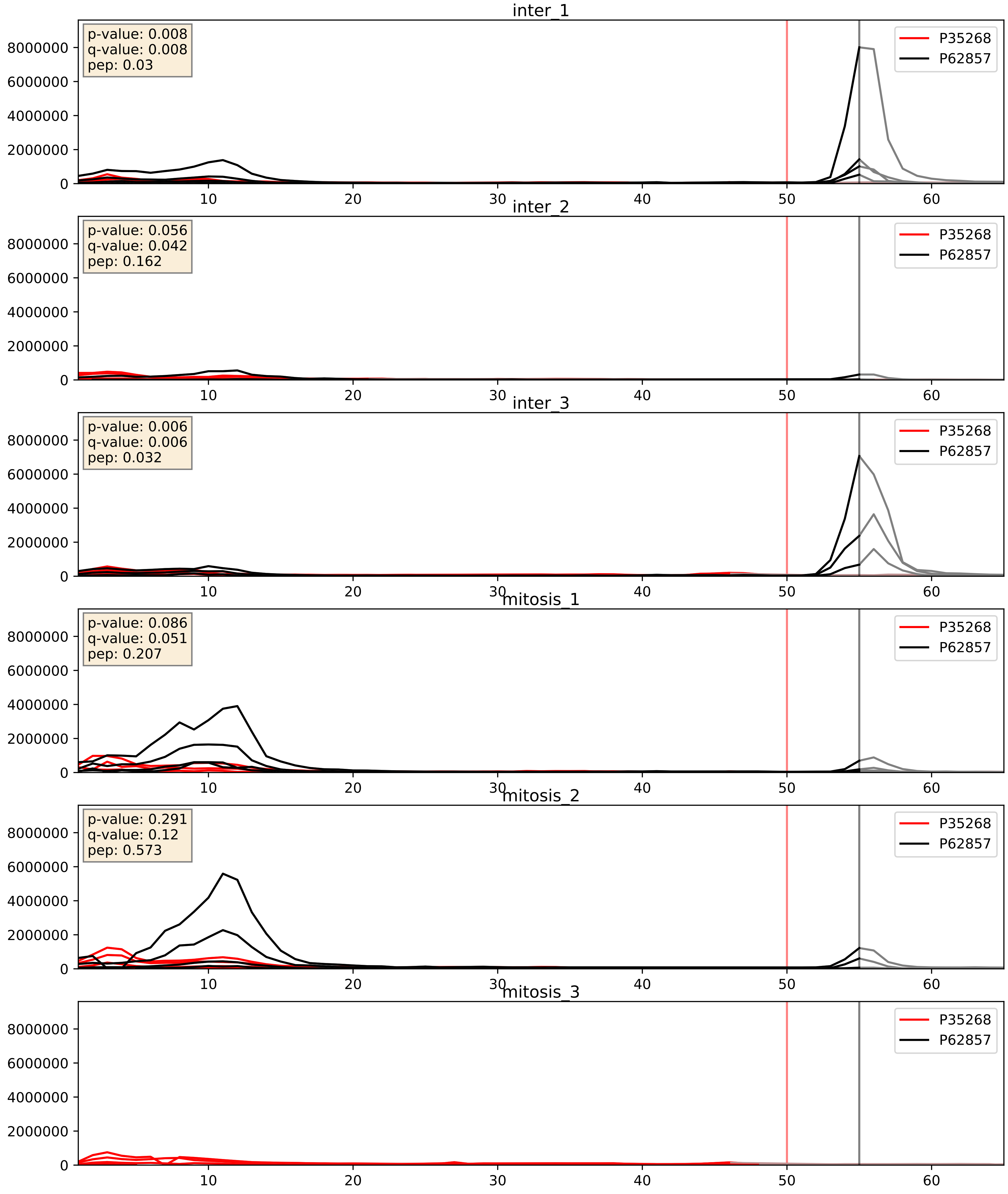


| level | condition_1 | condition_2 | pvalue | pvalue_adjusted |
|---|---|---|---|---|
| interactor_ratio | mitosis | inter | 0.1614099983853562 | 0.6192541702766019 |
| interactor_abundance | mitosis | inter | 0.5288098927802732 | 0.6880389589497613 |
| complex_abundance | mitosis | inter | 0.8513857011409774 | 0.9052993800369485 |

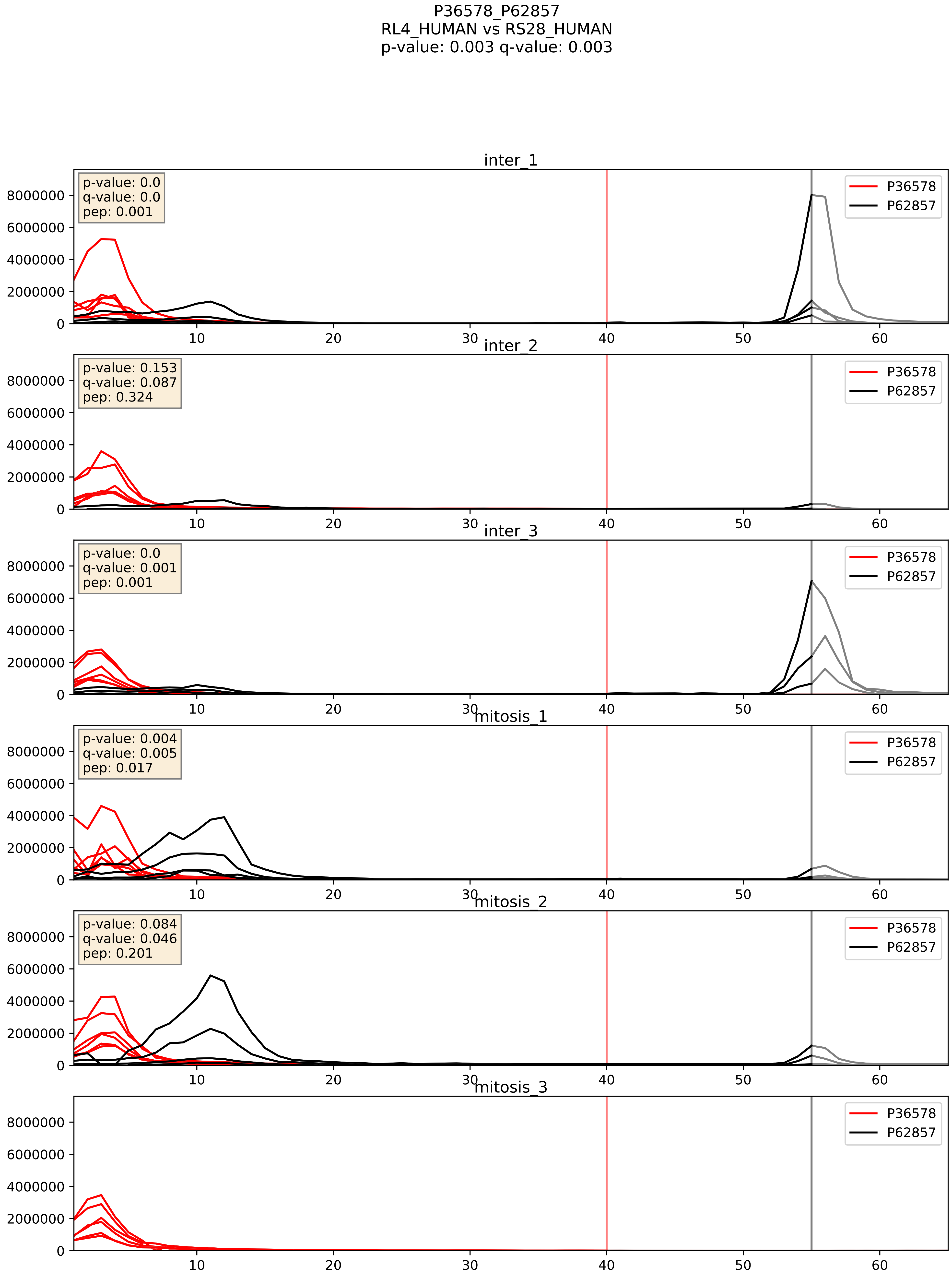


| level | condition_1 | condition_2 | pvalue | pvalue_adjusted |
|---|---|---|---|---|
| interactor_ratio | mitosis | inter | 0.018063266301216453 | 0.20237106227759052 |
| interactor_abundance | mitosis | inter | 0.35584761687354177 | 0.5536772880918873 |
| complex_abundance | mitosis | inter | 0.6983459735318287 | 0.7969562387832062 |

P37108_P62857
SRP14_HUMAN vs RS28_HUMAN
p-value: 0.084 q-value: 0.049

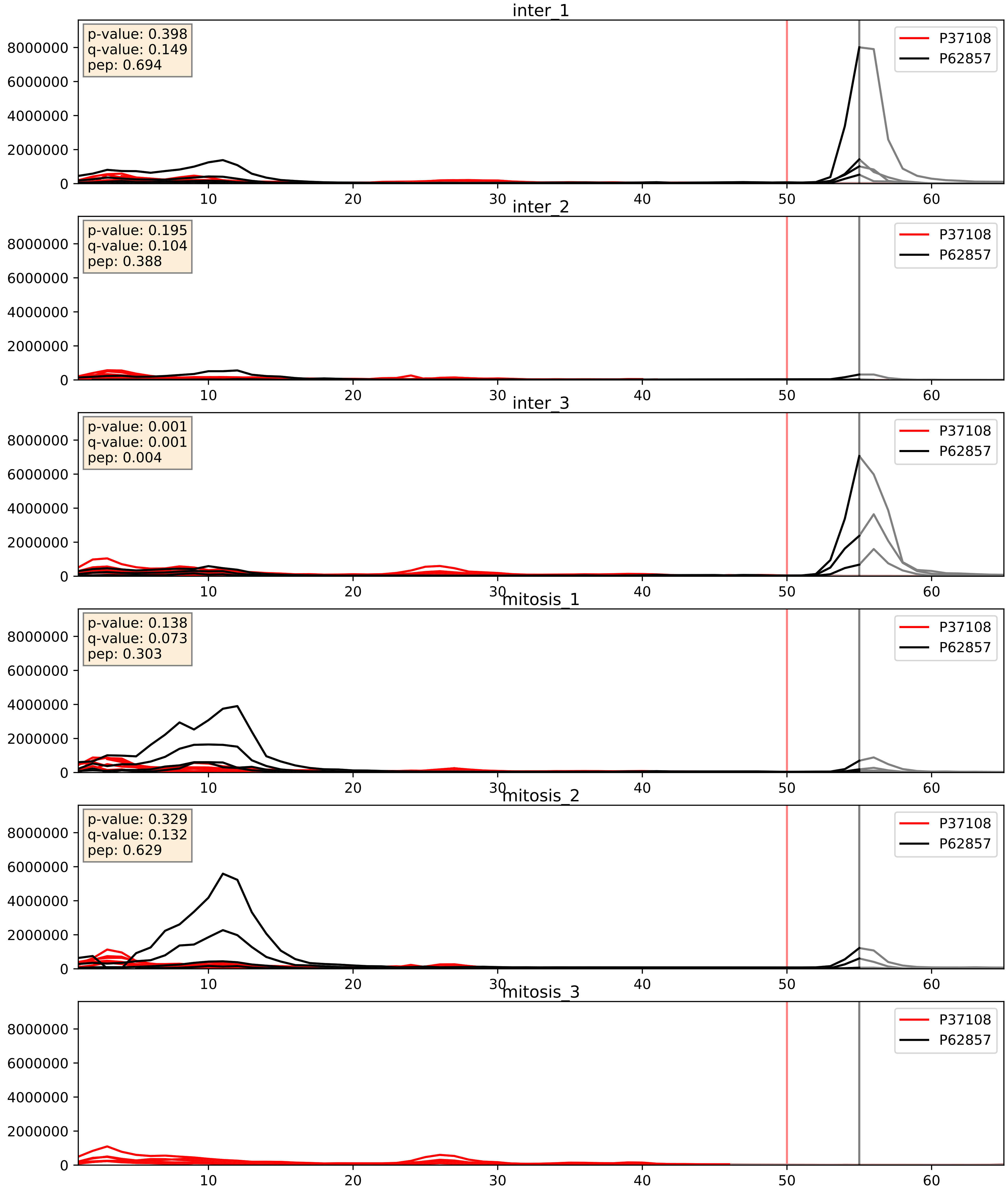


| level | condition_1 | condition_2 | pvalue | pvalue_adjusted |
|---|---|---|---|---|
| interactor_ratio | mitosis | inter | 0.021048944299145986 | 0.21980620419464833 |
| interactor_abundance | mitosis | inter | 0.4921294199526981 | 0.659384920364587 |
| complex_abundance | mitosis | inter | 0.8068295794400693 | 0.8747892590255849 |

P39019_P62857
RS19_HUMAN vs RS28_HUMAN
p-value: 0.0 q-value: 0.0

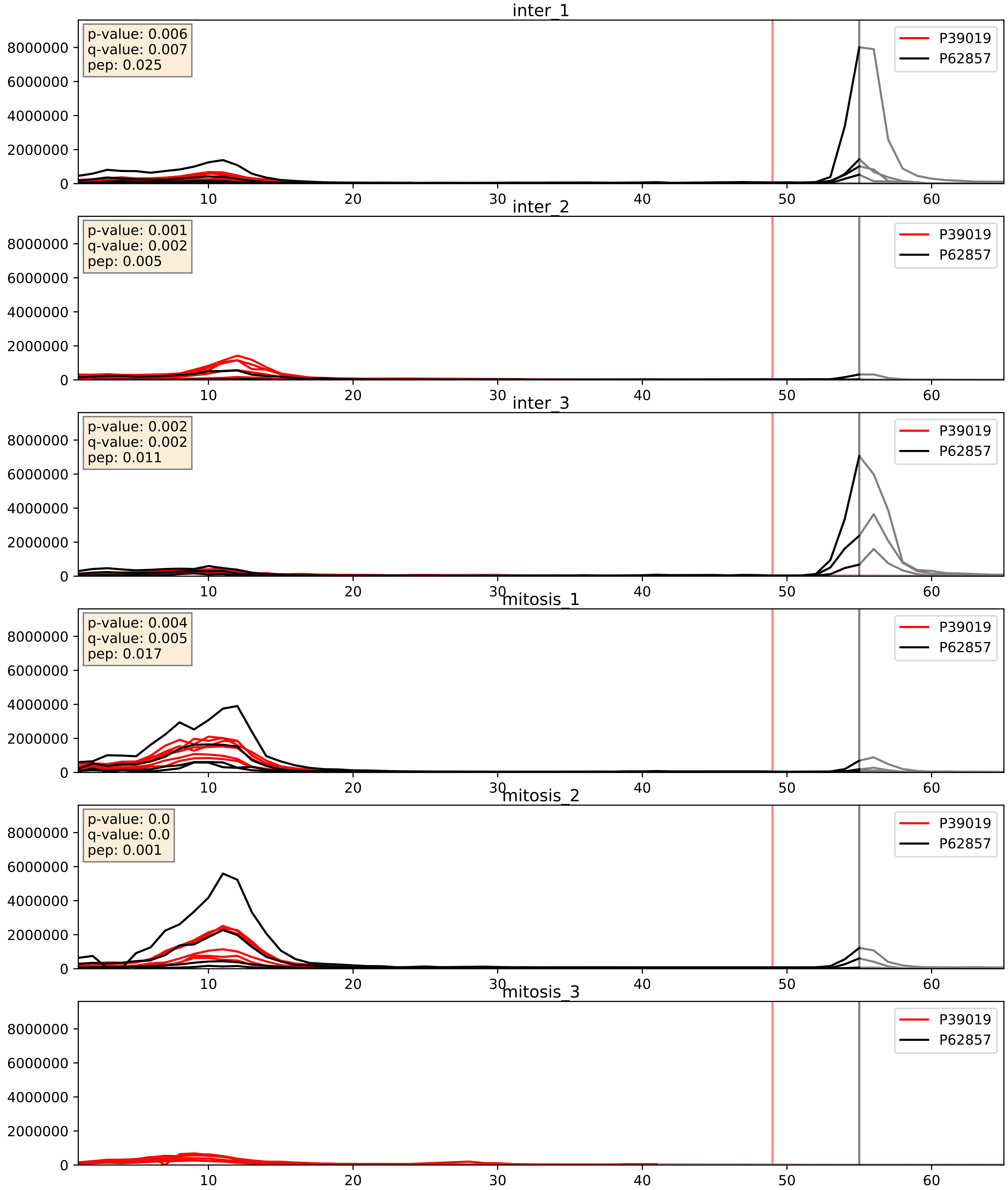


| level | condition_1 | condition_2 | pvalue | pvalue_adjusted |
|---|---|---|---|---|
| interactor_ratio | mitosis | inter | 0.27944122629842755 | 0.7507789882022703 |
| interactor_abundance | mitosis | inter | 0.7030564323942983 | 0.8052131470825787 |
| complex_abundance | mitosis | inter | 0.9125282532220204 | 0.9449668203271161 |

P42766_P62857
RL35_HUMAN vs RS28_HUMAN
p-value: 0.049 q-value: 0.032

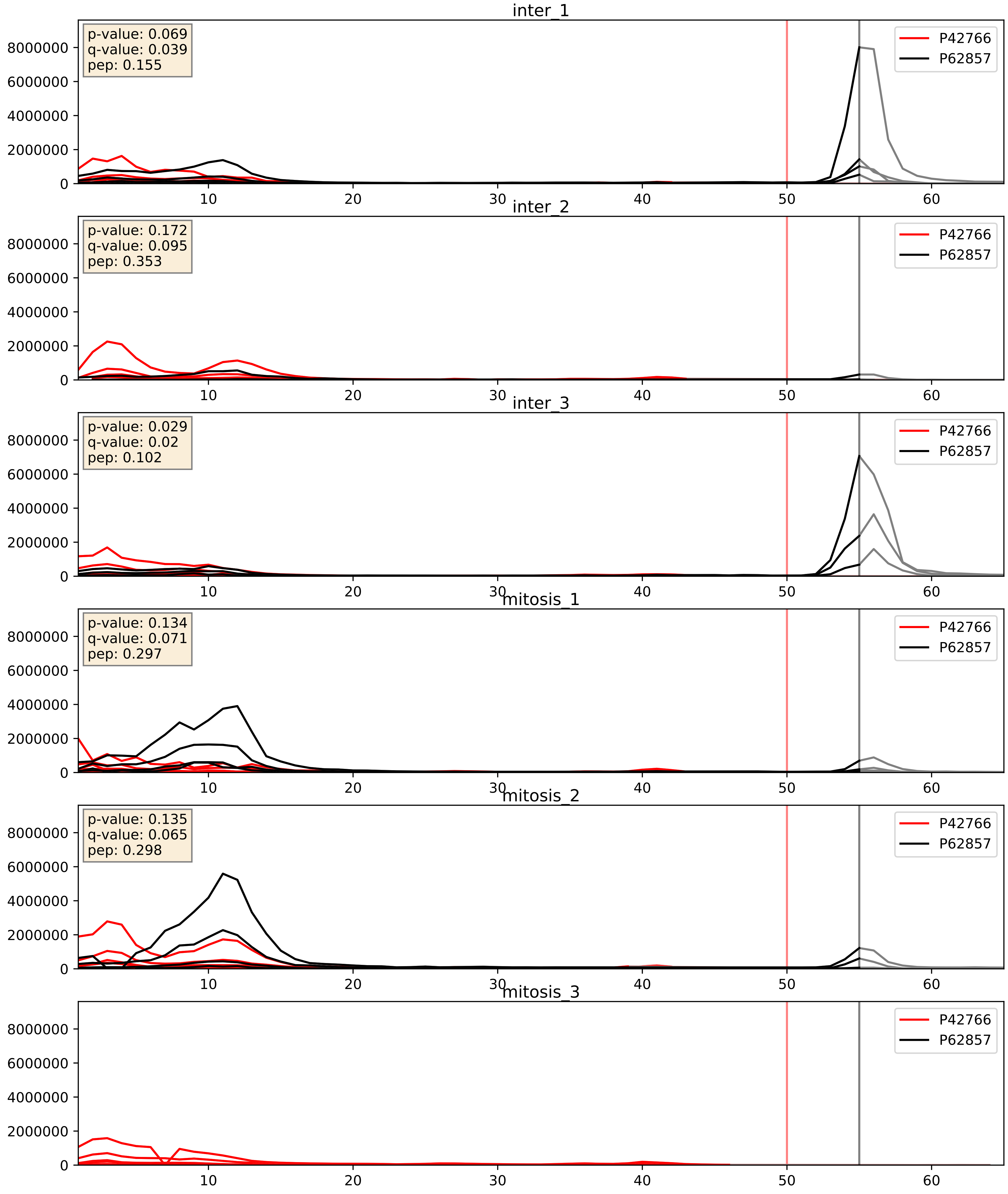


| level | condition_1 | condition_2 | pvalue | pvalue_adjusted |
|---|---|---|---|---|
| interactor_ratio | mitosis | inter | 0.06688928218297749 | 0.4304195561540654 |
| interactor_abundance | mitosis | inter | 0.3716053209992721 | 0.5654091211572675 |
| complex_abundance | mitosis | inter | 0.7414239158434656 | 0.828558226844317 |

P46776_P62857
RL27A_HUMAN vs RS28_HUMAN
p-value: 0.038 q-value: 0.026

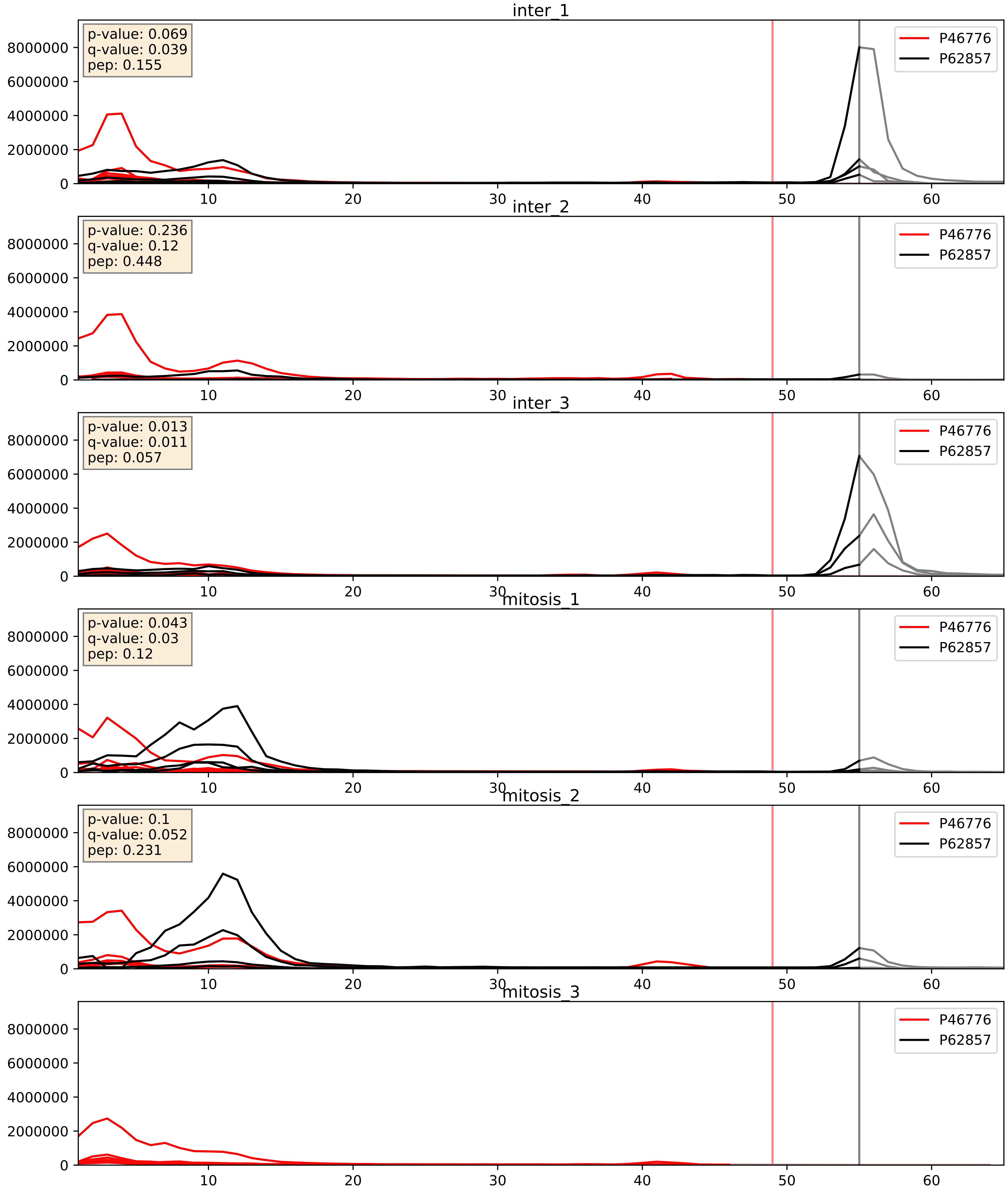


| level | condition_1 | condition_2 | pvalue | pvalue_adjusted |
|---|---|---|---|---|
| interactor_ratio | mitosis | inter | 0.03349952713335618 | 0.29380732813681243 |
| interactor_abundance | mitosis | inter | 0.36165743070370227 | 0.558806427224493 |
| complex_abundance | mitosis | inter | 0.7240717936785328 | 0.8162853356892191 |

P46779_P62857
RL28_HUMAN vs RS28_HUMAN
p-value: 0.062 q-value: 0.039

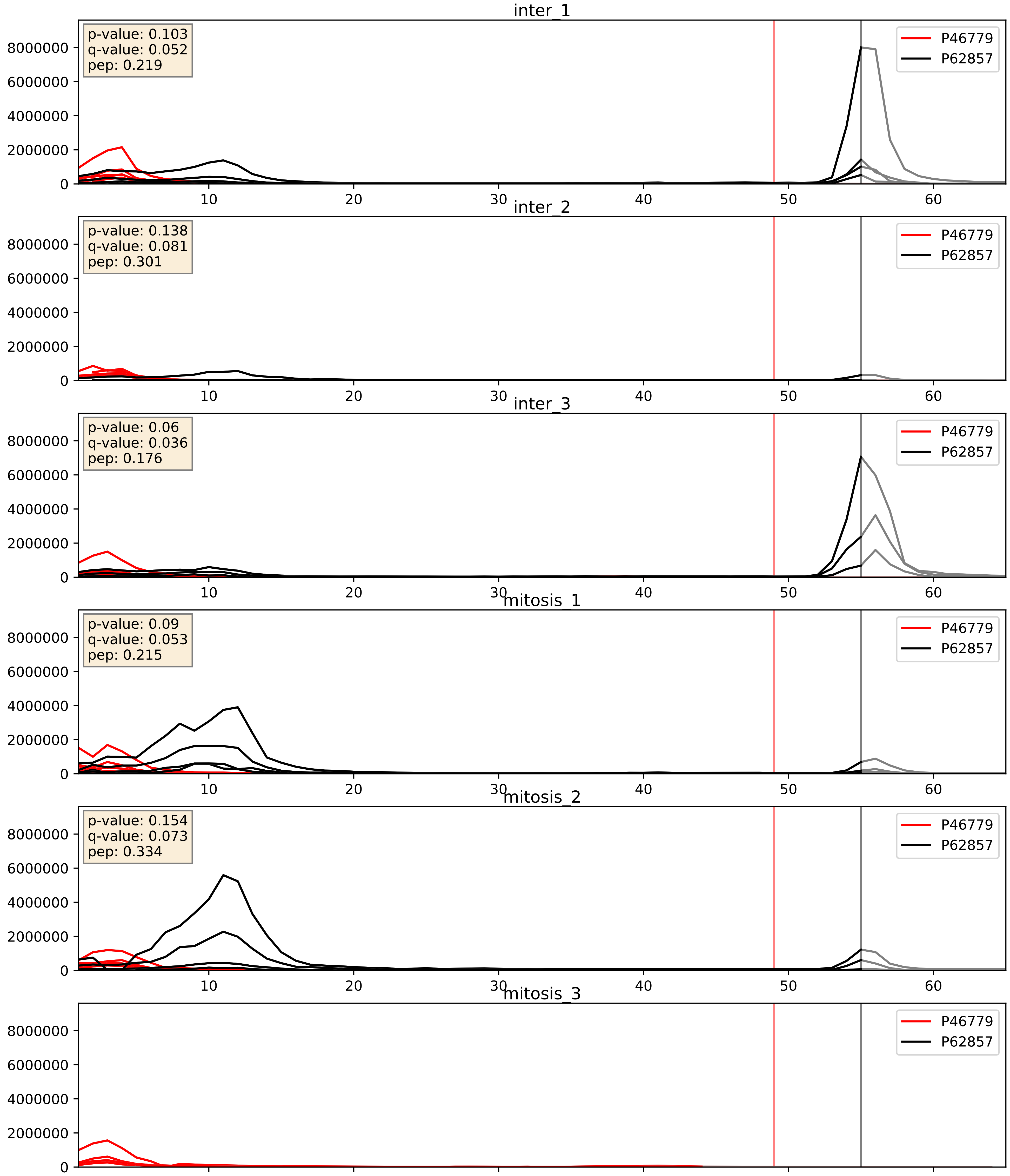


| level | condition_1 | condition_2 | pvalue | pvalue_adjusted |
|---|---|---|---|---|
| interactor_ratio | mitosis | inter | 0.10735105521769368 | 0.5447095629303248 |
| interactor_abundance | mitosis | inter | 0.2989205573614621 | 0.49344303591965044 |
| complex_abundance | mitosis | inter | 0.734872567853974 | 0.823472860430687 |

P46781_P62857
RS9_HUMAN vs RS28_HUMAN
p-value: 0.002 q-value: 0.003

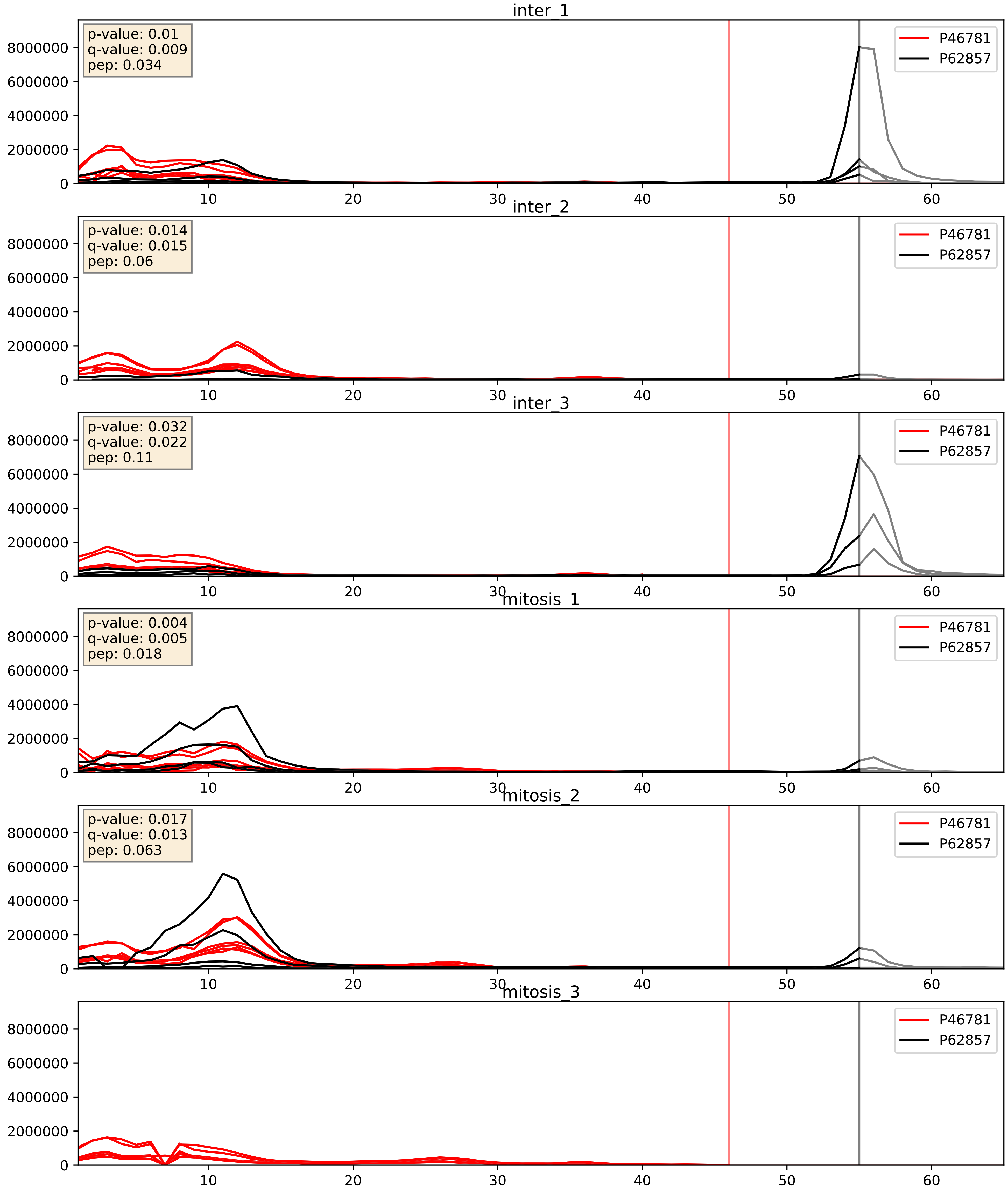


| level | condition_1 | condition_2 | pvalue | pvalue_adjusted |
|---|---|---|---|---|
| interactor_ratio | mitosis | inter | 0.027033678971341568 | 0.2576929754951936 |
| interactor_abundance | mitosis | inter | 0.363204567359319 | 0.5599839871390077 |
| complex_abundance | mitosis | inter | 0.6849765264031737 | 0.7893470281620353 |

P46782_P62857
RS5_HUMAN vs RS28_HUMAN
p-value: 0.002 q-value: 0.002

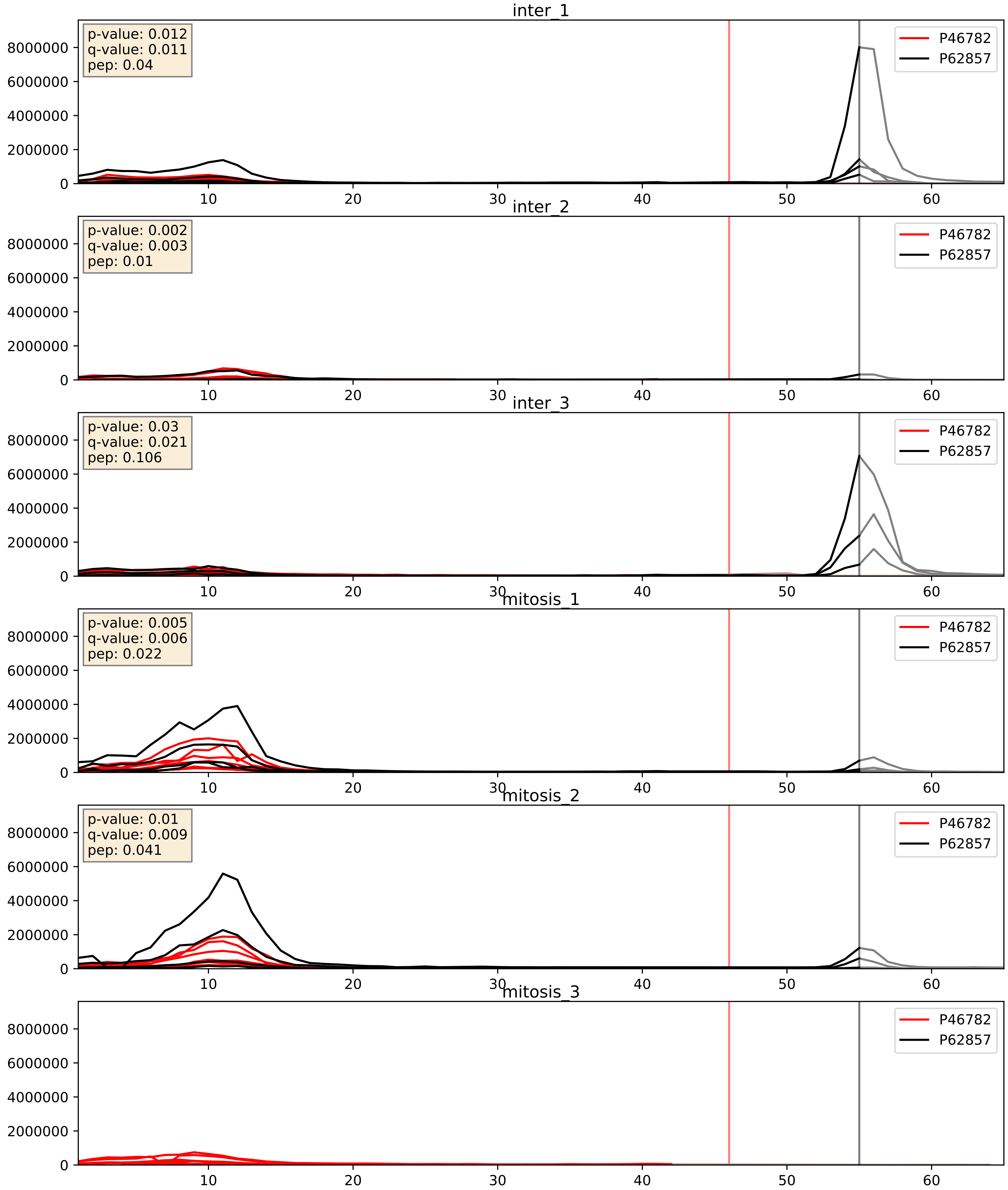


| level | condition_1 | condition_2 | pvalue | pvalue_adjusted |
|---|---|---|---|---|
| interactor_ratio | mitosis | inter | 0.12564163207144144 | 0.5750062399685051 |
| interactor_abundance | mitosis | inter | 0.5920816521270316 | 0.7318727714378903 |
| complex_abundance | mitosis | inter | 0.8607562996800598 | 0.9114391297948184 |

P46783_P62857
RS10_HUMAN vs RS28_HUMAN
p-value: 0.005 q-value: 0.005

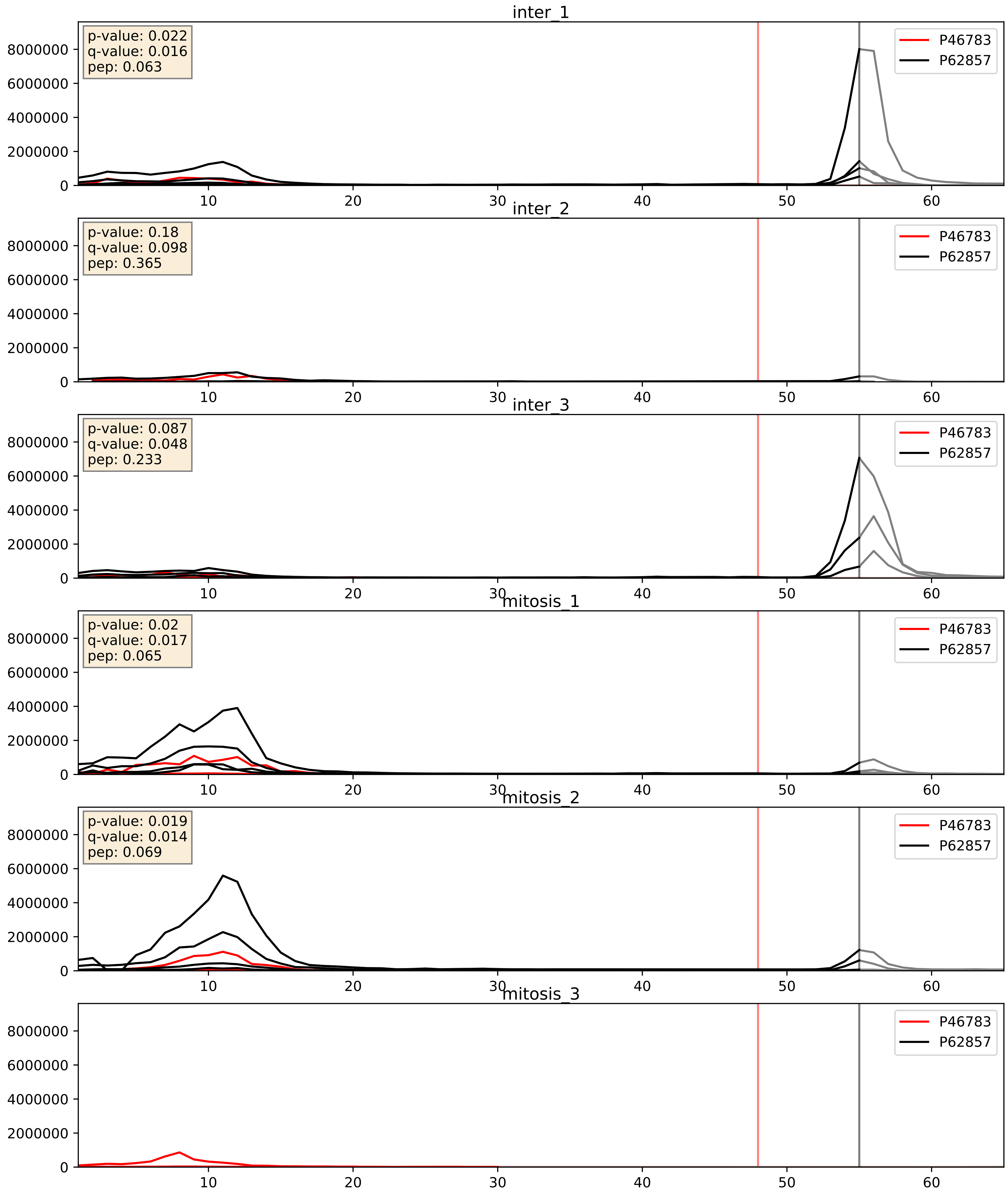


| level | condition_1 | condition_2 | pvalue | pvalue_adjusted |
|---|---|---|---|---|
| interactor_ratio | mitosis | inter | 0.4209555967180571 | 0.8388785723072948 |
| interactor_abundance | mitosis | inter | 0.4836565513371456 | 0.6544702818064209 |
| complex_abundance | mitosis | inter | 0.9125816934062856 | 0.9449668203271161 |

P47914_P62857
RL29_HUMAN vs RS28_HUMAN
p-value: 0.044 q-value: 0.029

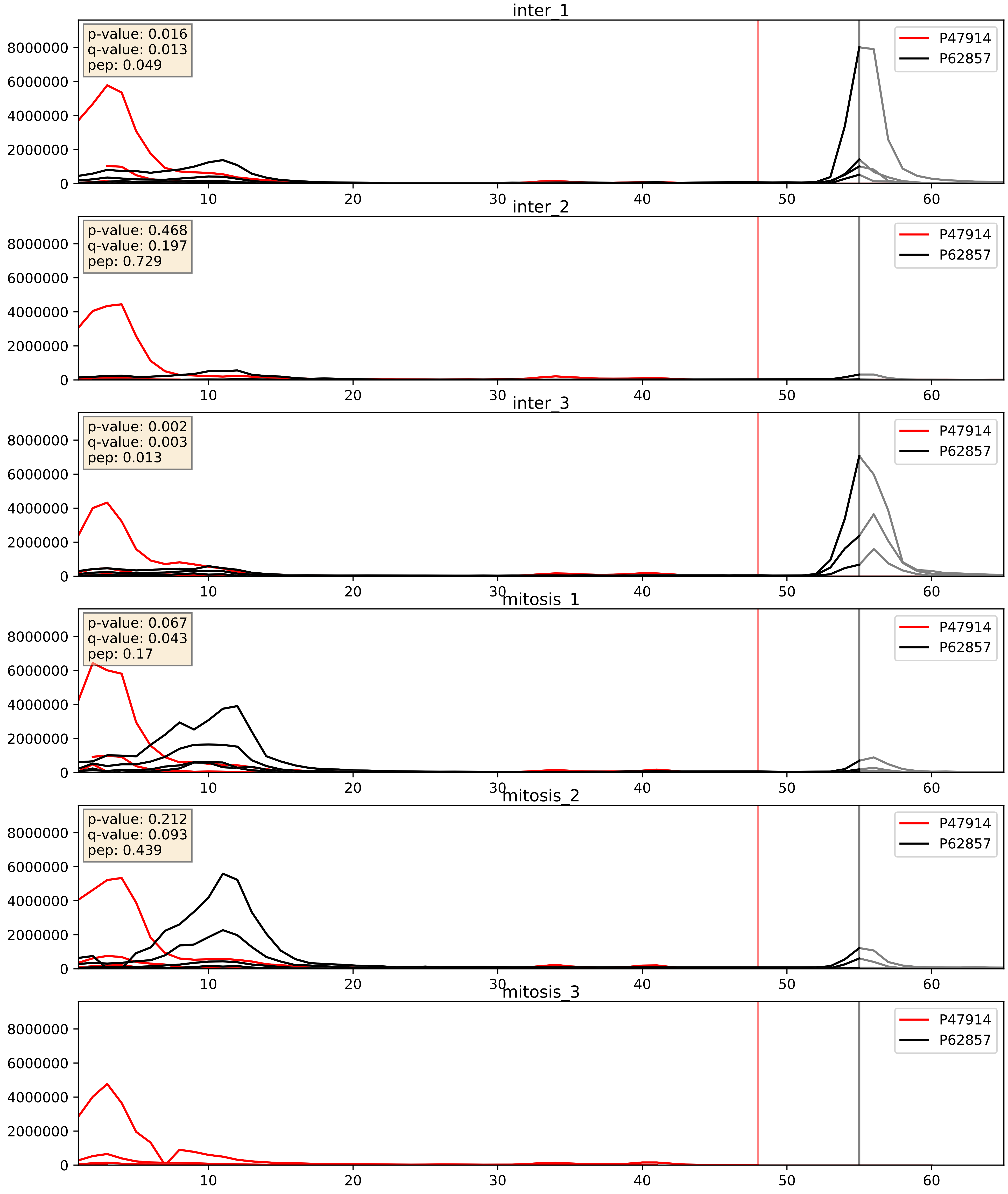


| level | condition_1 | condition_2 | pvalue | pvalue_adjusted |
|---|---|---|---|---|
| interactor_ratio | mitosis | inter | 0.03402844037520282 | 0.29692502508841606 |
| interactor_abundance | mitosis | inter | 0.5204014274281796 | 0.6820756727584164 |
| complex_abundance | mitosis | inter | 0.838718707828872 | 0.8963846703115416 |

P49458_P62857
SRP09_HUMAN vs RS28_HUMAN
p-value: 0.038 q-value: 0.026

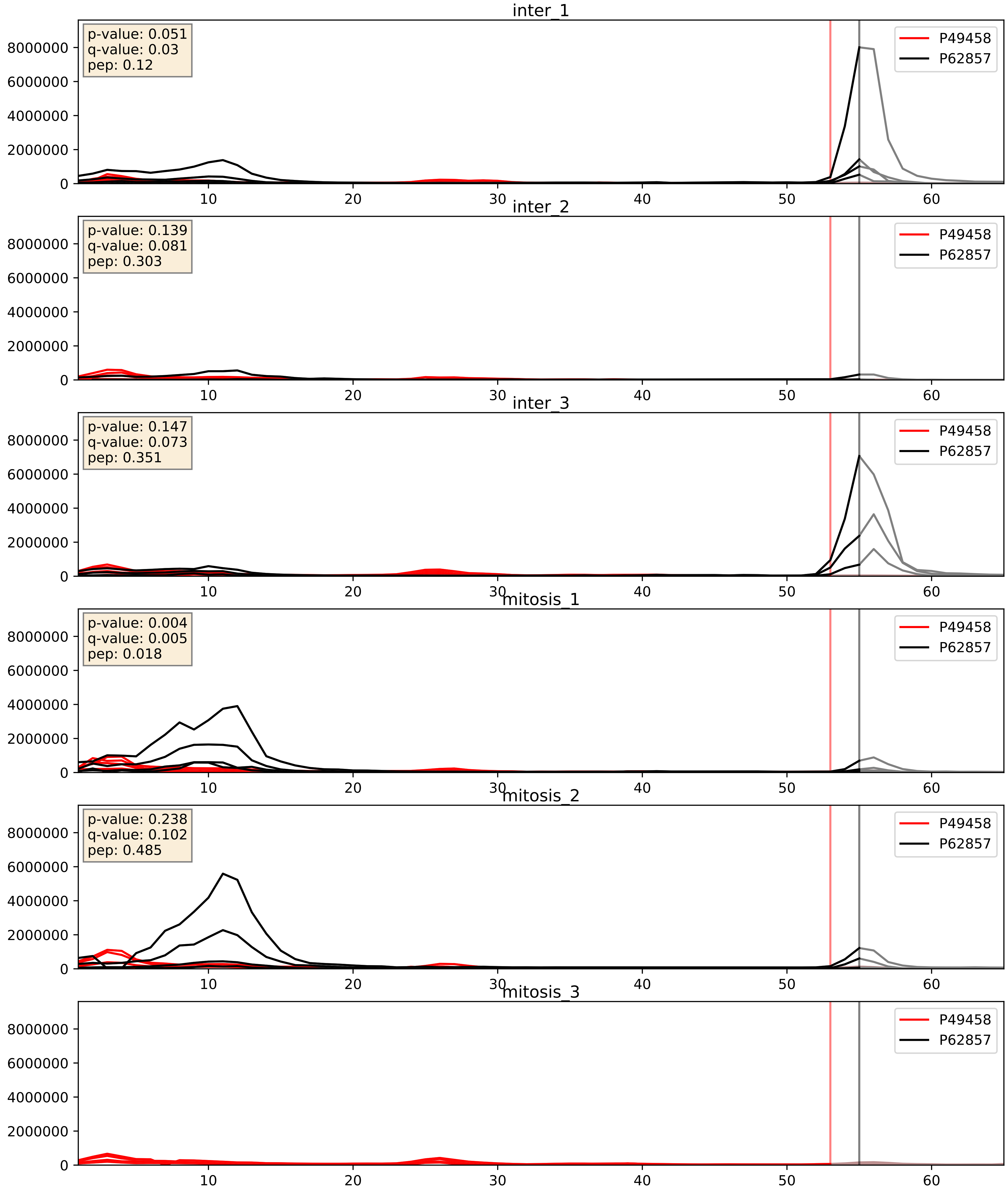


| level | condition_1 | condition_2 | pvalue | pvalue_adjusted |
|---|---|---|---|---|
| interactor_ratio | mitosis | inter | 0.06770194715755185 | 0.4323044818991264 |
| interactor_abundance | mitosis | inter | 0.4311755458807304 | 0.6094555272026176 |
| complex_abundance | mitosis | inter | 0.7847422341142323 | 0.8594413413533557 |

P50914_P62857
RL14_HUMAN vs RS28_HUMAN
p-value: 0.013 q-value: 0.011

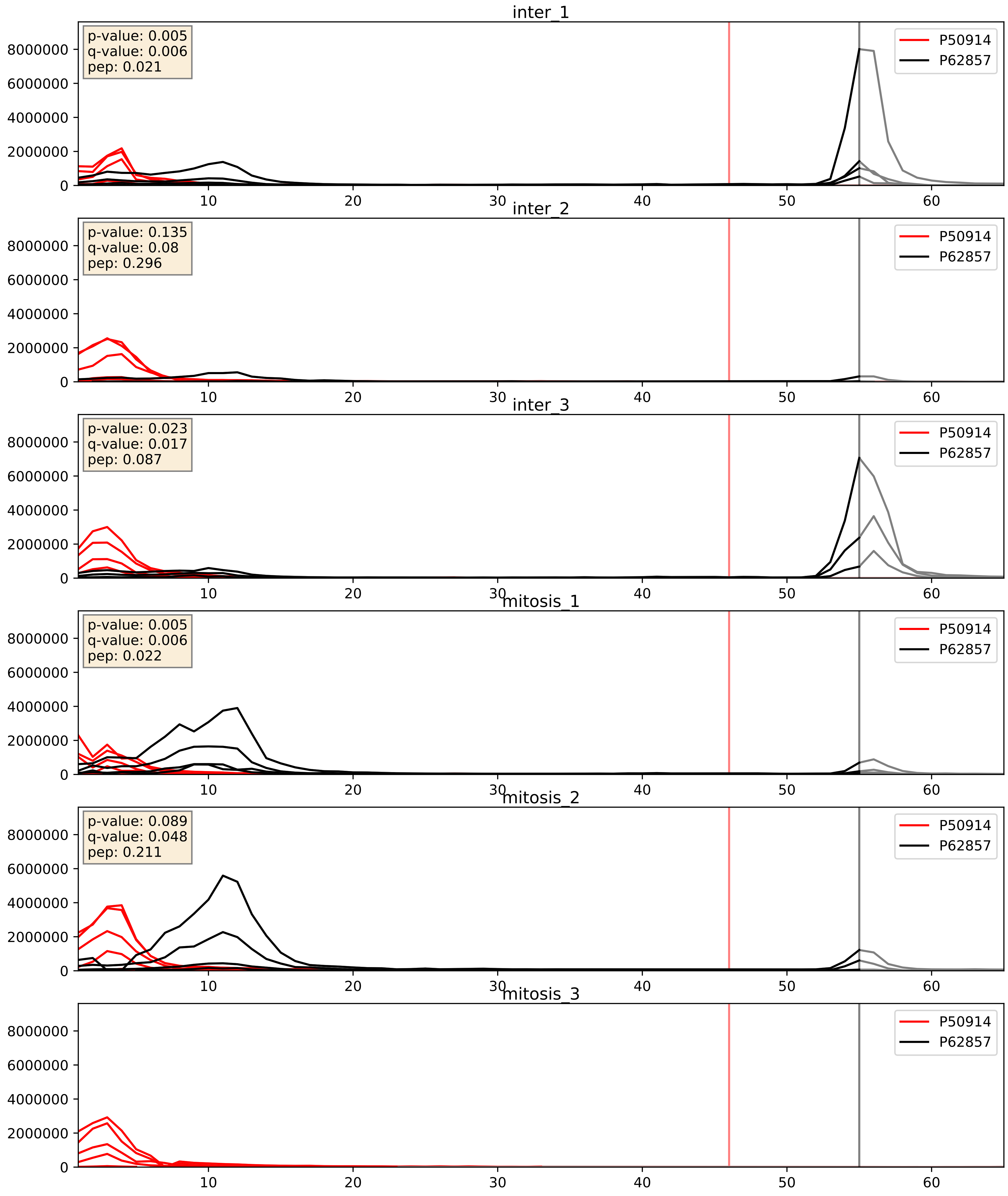


| level | condition_1 | condition_2 | pvalue | pvalue_adjusted |
|---|---|---|---|---|
| interactor_ratio | mitosis | inter | 0.047148653890849074 | 0.3590680403075339 |
| interactor_abundance | mitosis | inter | 0.33973900915626565 | 0.5386342939180133 |
| complex_abundance | mitosis | inter | 0.6997301324313905 | 0.7980932623068224 |

P60468_P62857
SC61B_HUMAN vs RS28_HUMAN
p-value: 0.017 q-value: 0.014

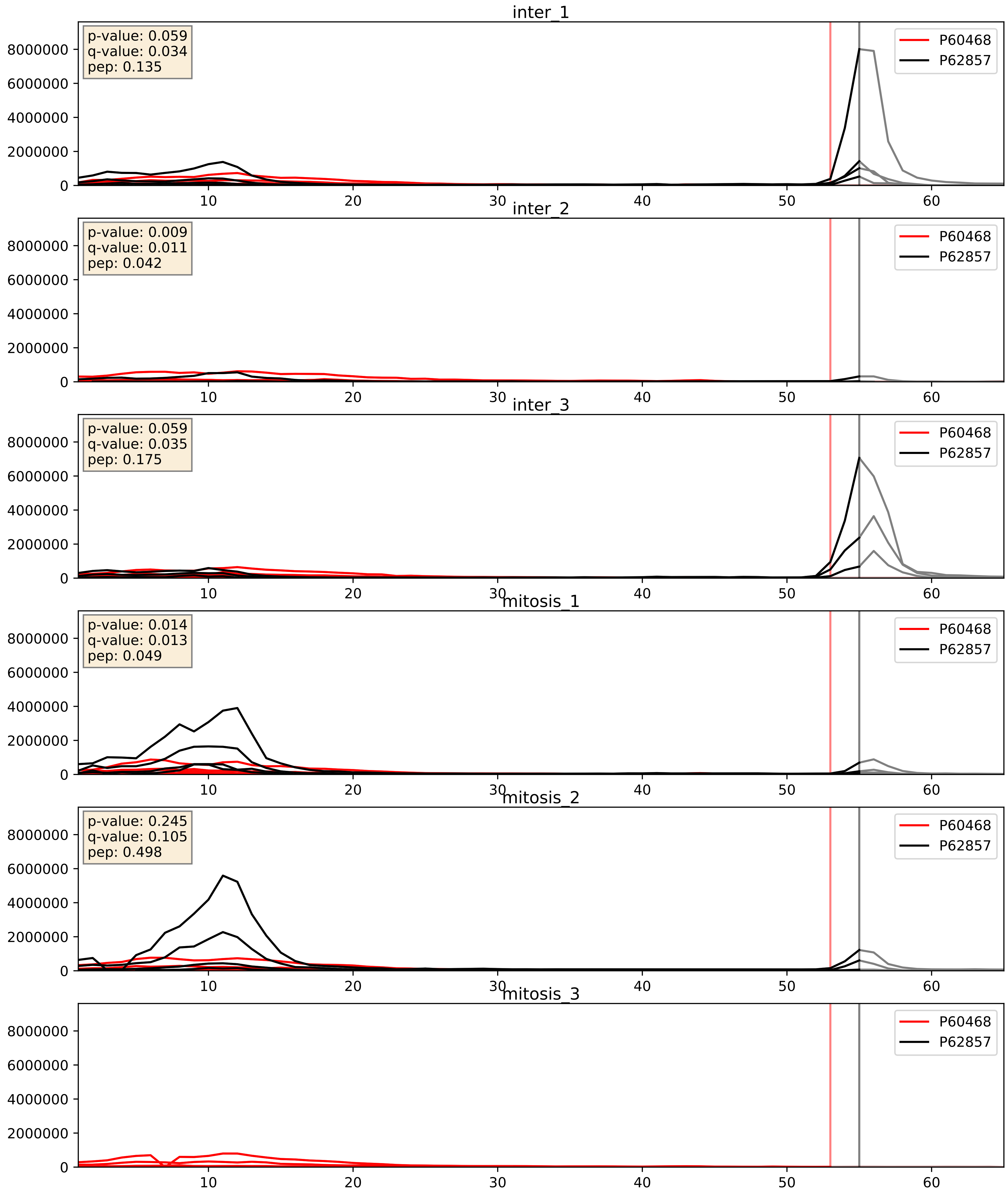


| level | condition_1 | condition_2 | pvalue | pvalue_adjusted |
|---|---|---|---|---|
| interactor_ratio | mitosis | inter | 0.04634661437352505 | 0.3574623087810929 |
| interactor_abundance | mitosis | inter | 0.2868998573476821 | 0.48088168766323847 |
| complex_abundance | mitosis | inter | 0.6886664134055517 | 0.7910675488572768 |

P60866_P62857
RS20_HUMAN vs RS28_HUMAN
p-value: 0.0 q-value: 0.0

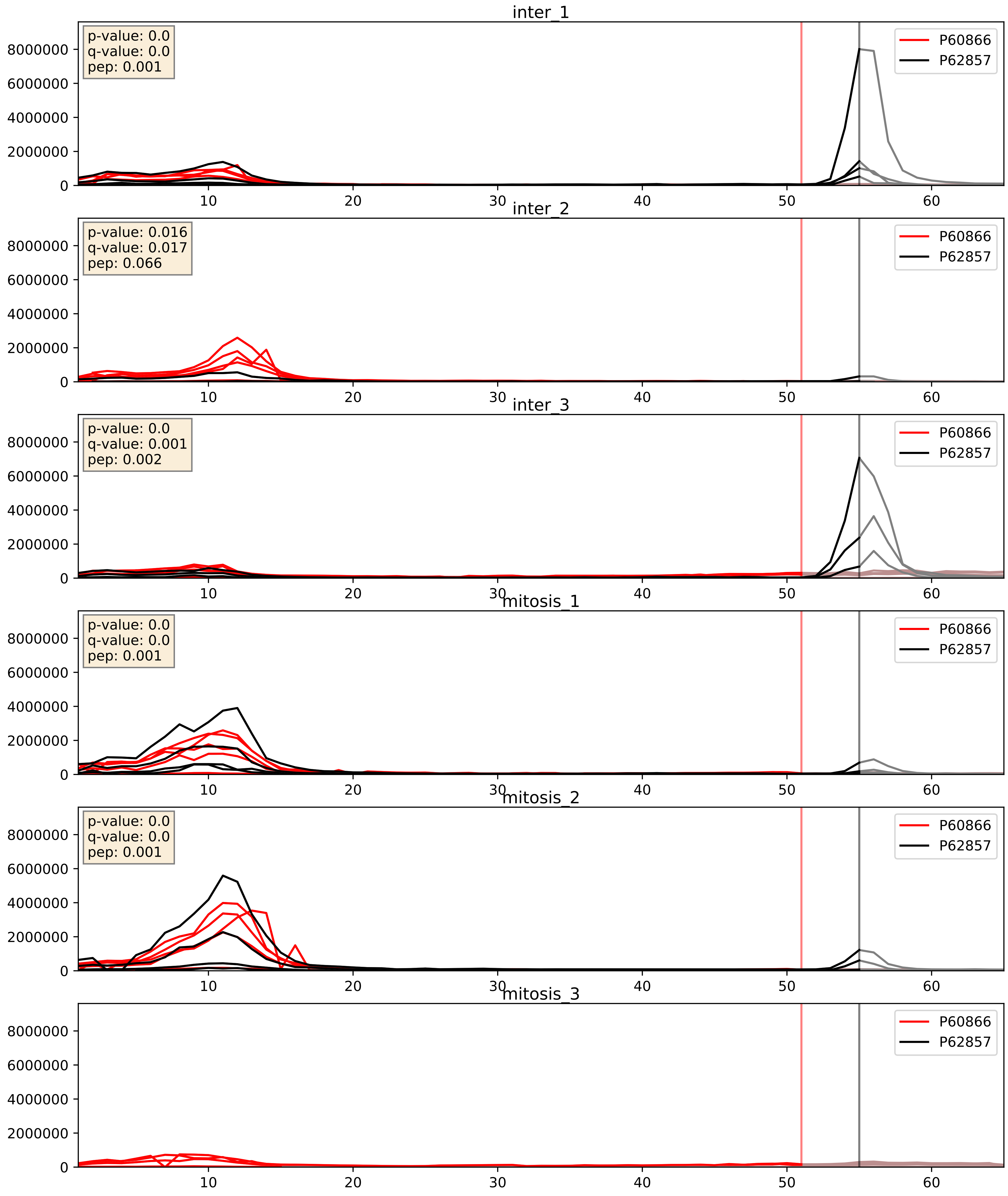


| level | condition_1 | condition_2 | pvalue | pvalue_adjusted |
|---|---|---|---|---|
| interactor_ratio | mitosis | inter | 0.10488091340820324 | 0.5400196384540948 |
| interactor_abundance | mitosis | inter | 0.540353731702065 | 0.6977436392201549 |
| complex_abundance | mitosis | inter | 0.8047672848235567 | 0.8737974444577015 |

P61247_P62857
RS3A_HUMAN vs RS28_HUMAN
p-value: 0.001 q-value: 0.002

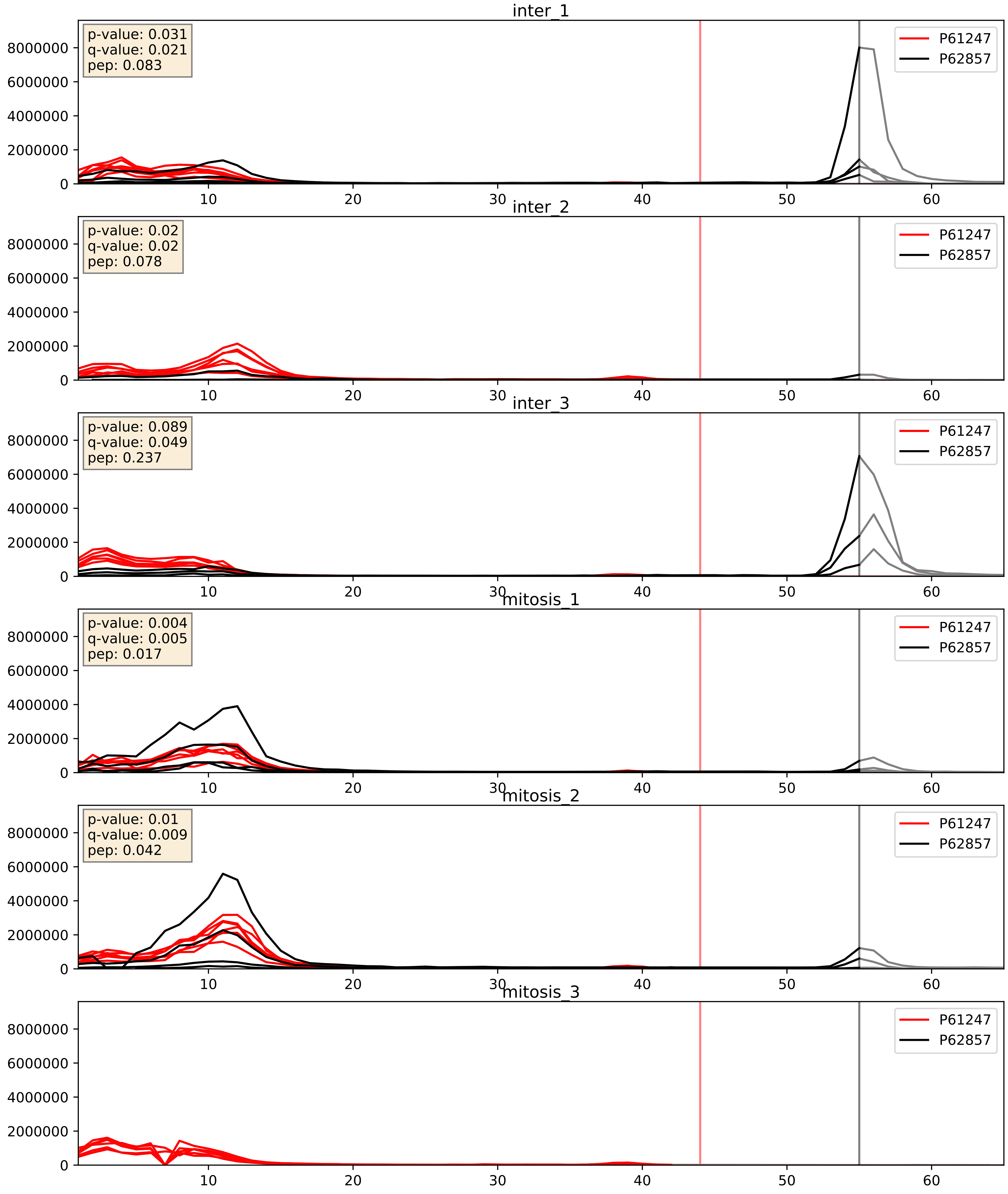


| level | condition_1 | condition_2 | pvalue | pvalue_adjusted |
|---|---|---|---|---|
| interactor_ratio | mitosis | inter | 0.030243439049970258 | 0.2765852972945998 |
| interactor_abundance | mitosis | inter | 0.39906612858990037 | 0.5887601601705548 |
| complex_abundance | mitosis | inter | 0.7141544174393843 | 0.8086472671317149 |

P62241_P62857
RS8_HUMAN vs RS28_HUMAN
p-value: 0.037 q-value: 0.025

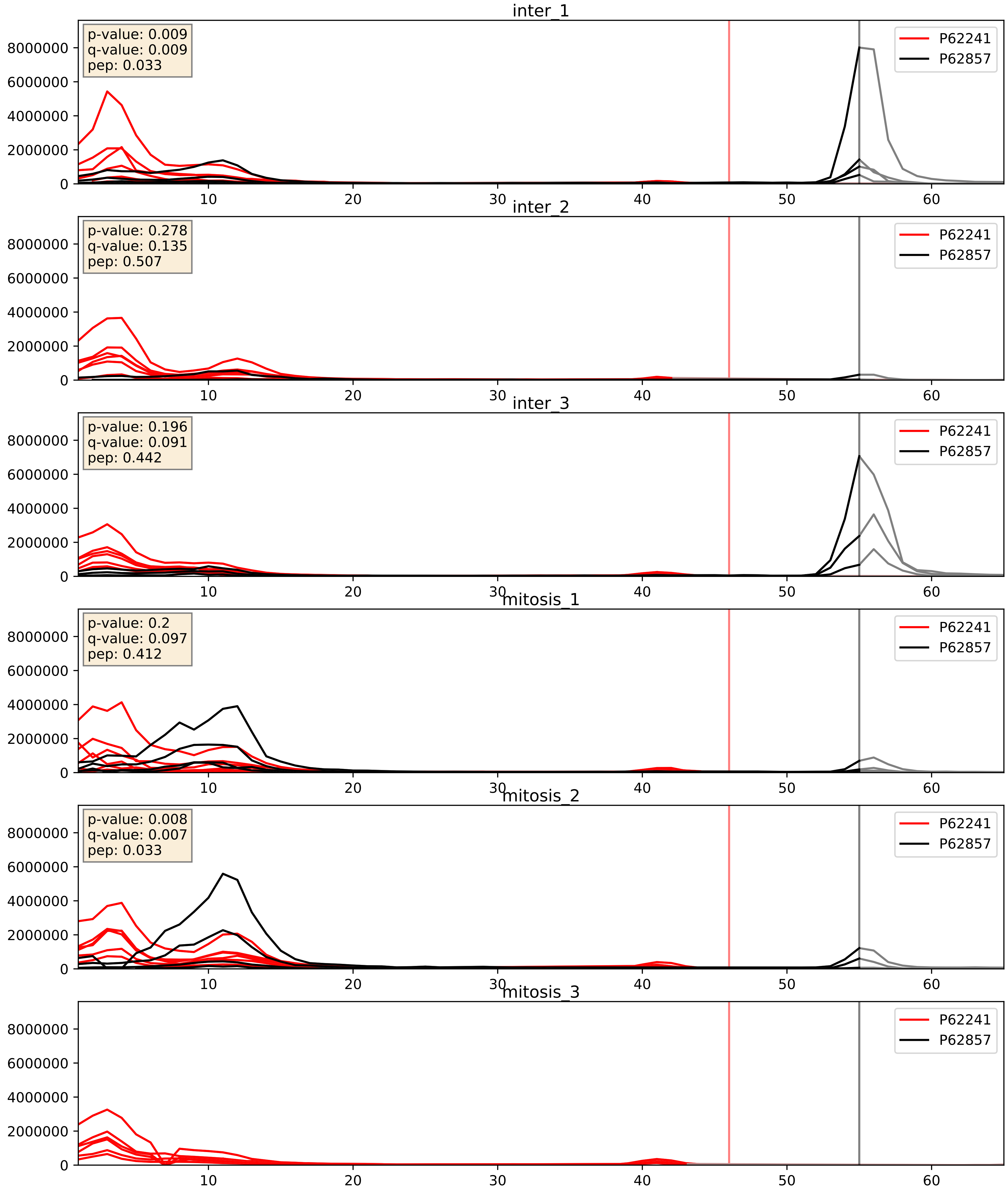


| level | condition_1 | condition_2 | pvalue | pvalue_adjusted |
|---|---|---|---|---|
| interactor_ratio | mitosis | inter | 0.013488651093478545 | 0.16880534116984847 |
| interactor_abundance | mitosis | inter | 0.34052031626807566 | 0.5389141513343335 |
| complex_abundance | mitosis | inter | 0.6665869465752479 | 0.7781241323720335 |

P62244_P62857
RS15A_HUMAN vs RS28_HUMAN
p-value: 0.076 q-value: 0.046

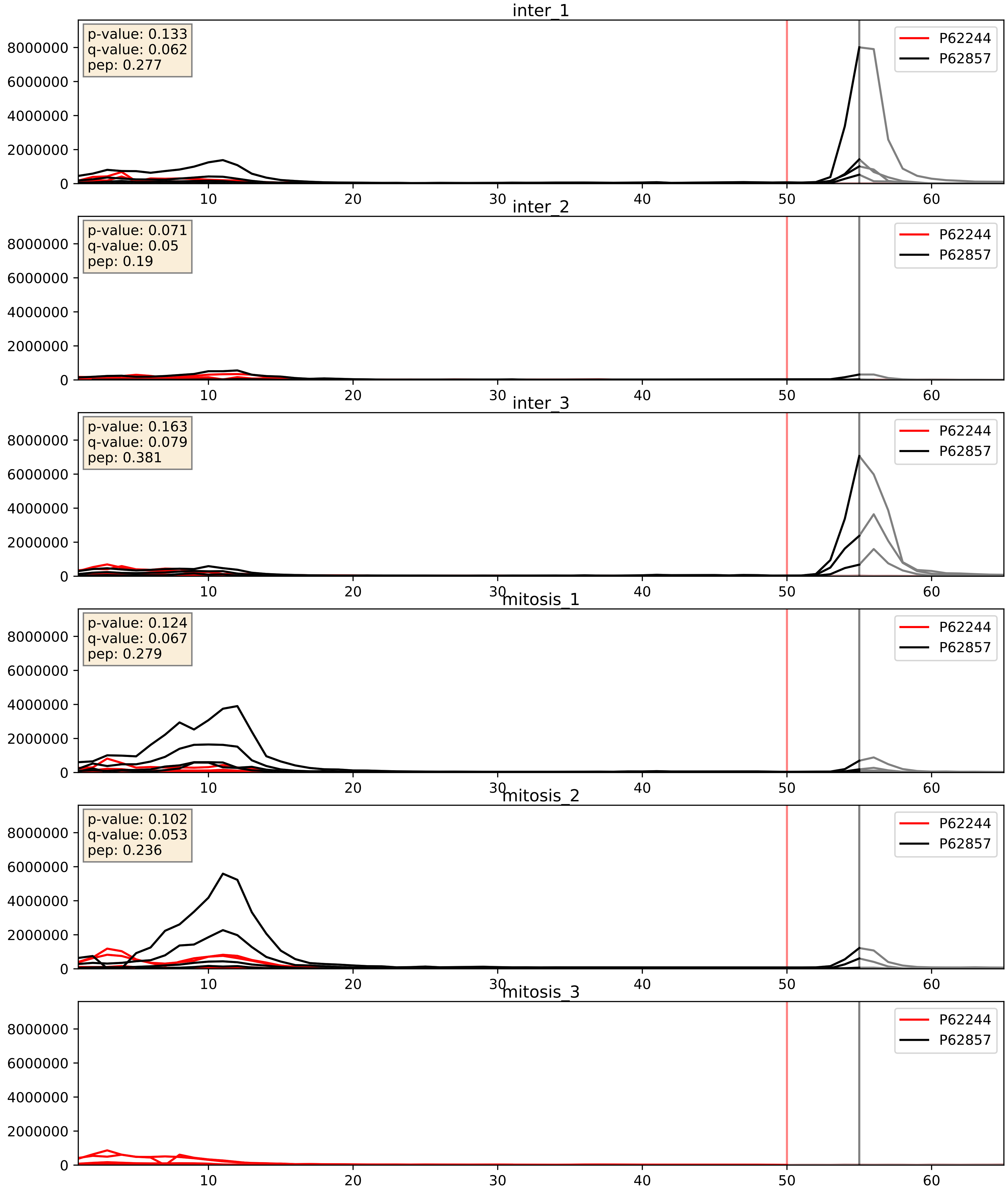


| level | condition_1 | condition_2 | pvalue | pvalue_adjusted |
|---|---|---|---|---|
| interactor_ratio | mitosis | inter | 0.1579546329737128 | 0.6144687163742579 |
| interactor_abundance | mitosis | inter | 0.49863842081742454 | 0.6640754386926728 |
| complex_abundance | mitosis | inter | 0.8528526052435929 | 0.9063213284773625 |

P62249_P62857
RS16_HUMAN vs RS28_HUMAN
p-value: 0.0 q-value: 0.001

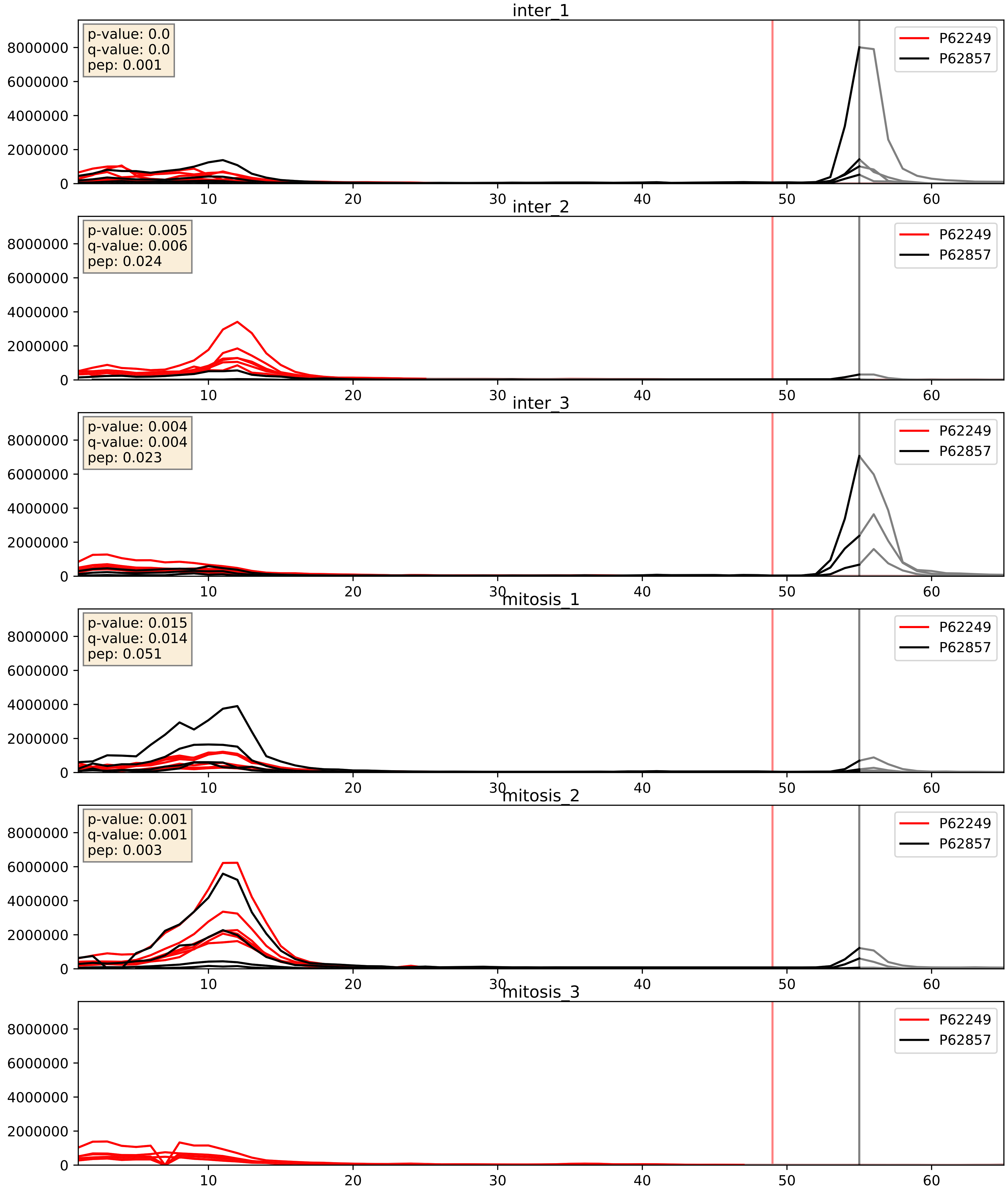


| level | condition_1 | condition_2 | pvalue | pvalue_adjusted |
|---|---|---|---|---|
| interactor_ratio | mitosis | inter | 0.11228759842177043 | 0.5536525308757383 |
| interactor_abundance | mitosis | inter | 0.4666711881887012 | 0.6386588740159269 |
| complex_abundance | mitosis | inter | 0.7583651834413061 | 0.8407883047708208 |

P62263_P62857
RS14_HUMAN vs RS28_HUMAN
p-value: 0.0 q-value: 0.0

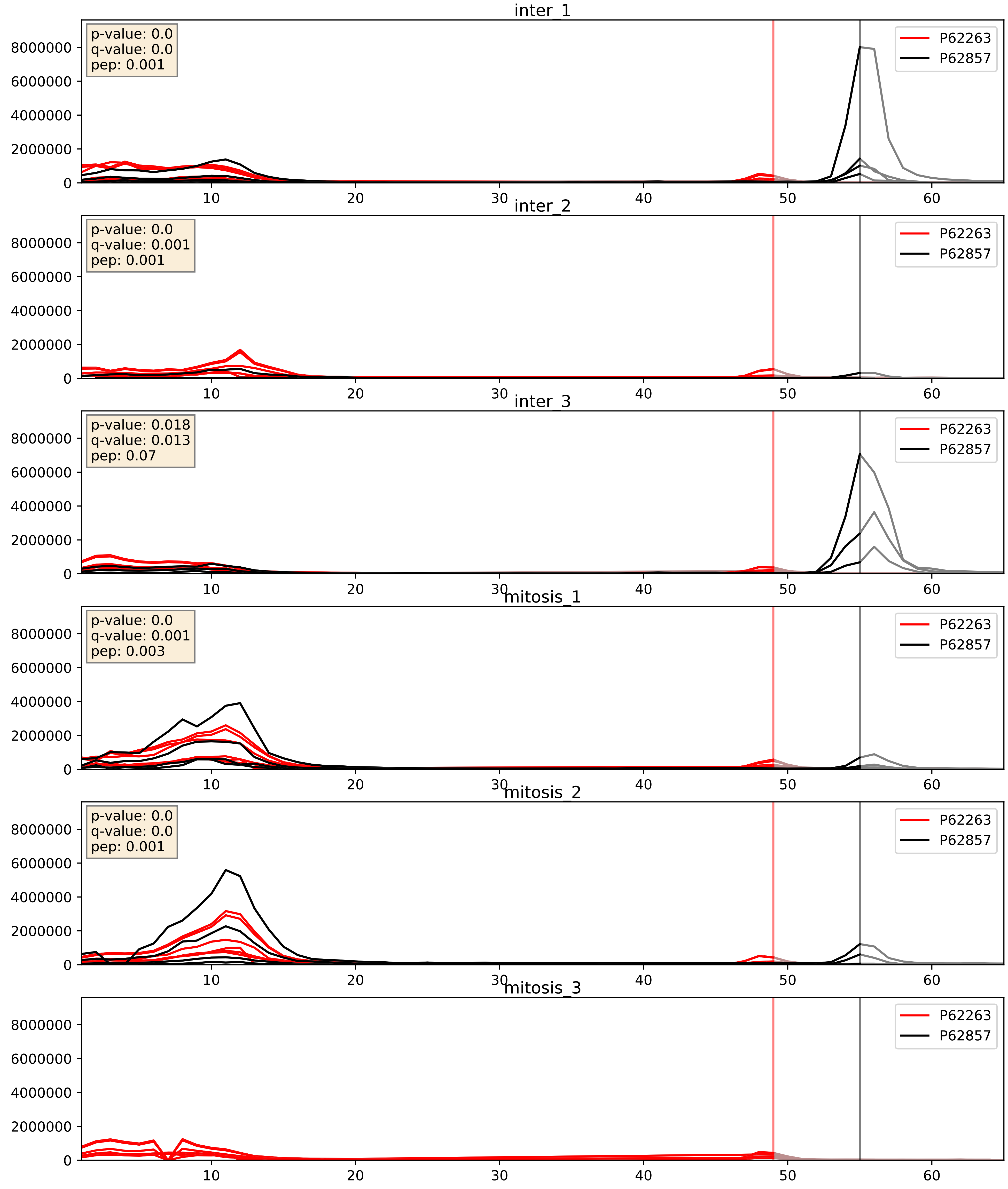


| level | condition_1 | condition_2 | pvalue | pvalue_adjusted |
|---|---|---|---|---|
| interactor_ratio | mitosis | inter | 0.03528881529567821 | 0.3050841933368649 |
| interactor_abundance | mitosis | inter | 0.5009443038875305 | 0.6659353717303512 |
| complex_abundance | mitosis | inter | 0.7816699189934629 | 0.8569537021752105 |

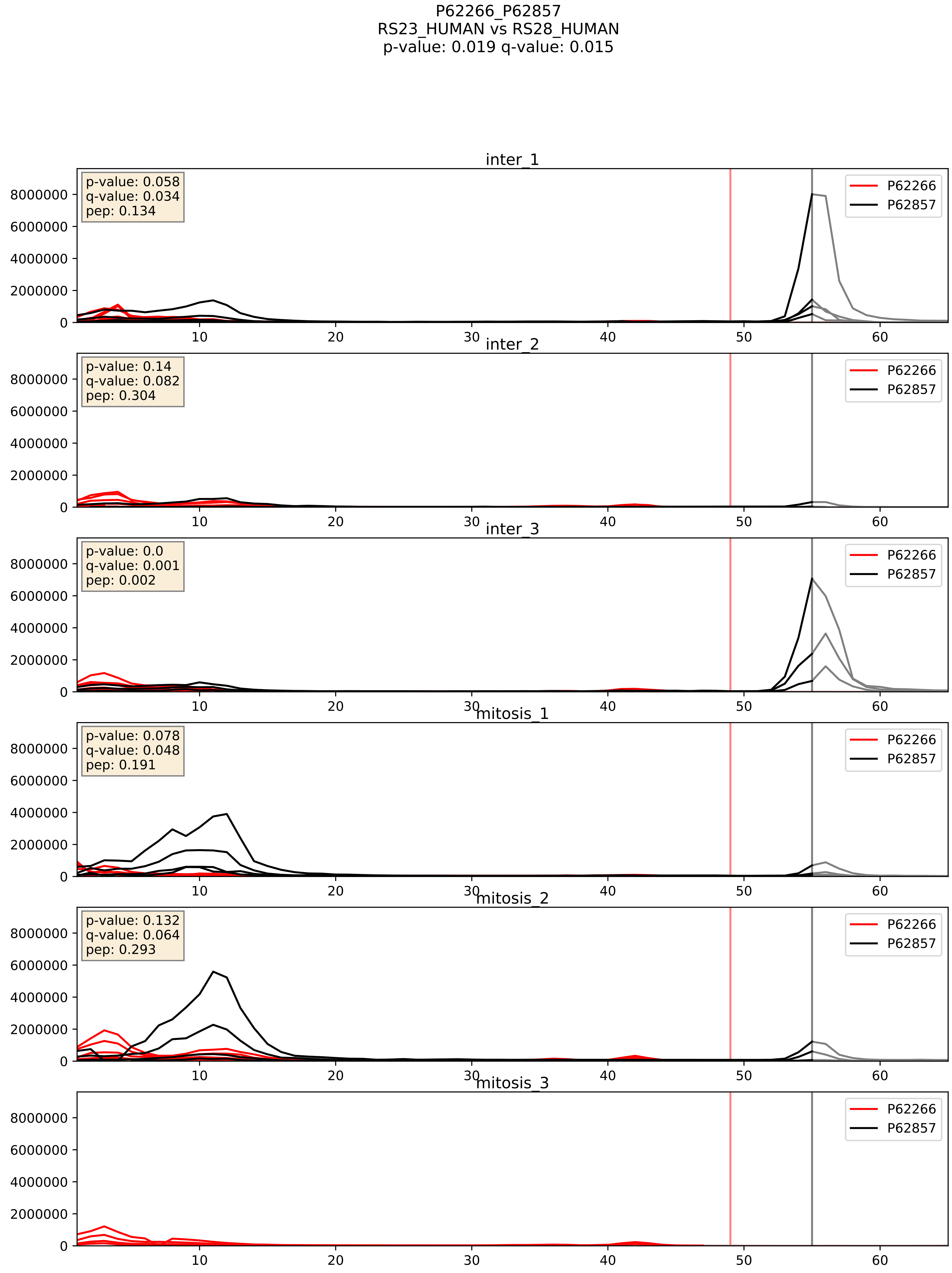


| level | condition_1 | condition_2 | pvalue | pvalue_adjusted |
|---|---|---|---|---|
| interactor_ratio | mitosis | inter | 0.06667980482252875 | 0.4304073446320155 |
| interactor_abundance | mitosis | inter | 0.33633952688648855 | 0.536138985130045 |
| complex_abundance | mitosis | inter | 0.7235006057102968 | 0.8159637924743268 |

P62269_P62857
RS18_HUMAN vs RS28_HUMAN
p-value: 0.0 q-value: 0.0

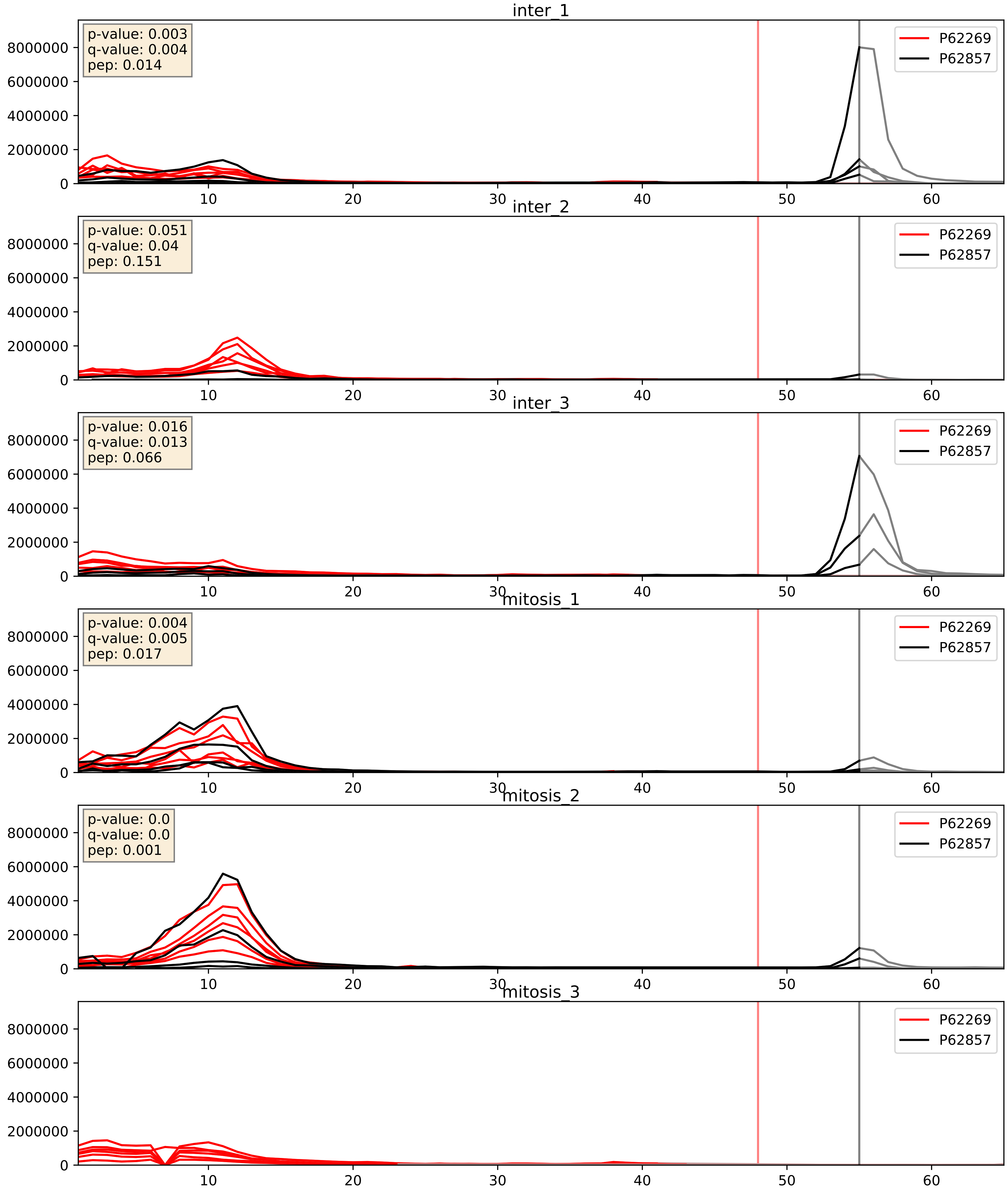


| level | condition_1 | condition_2 | pvalue | pvalue_adjusted |
|---|---|---|---|---|
| interactor_ratio | mitosis | inter | 0.0735062628030063 | 0.4539780732999524 |
| interactor_abundance | mitosis | inter | 0.5314249146780222 | 0.6900760554415658 |
| complex_abundance | mitosis | inter | 0.7942825341989727 | 0.8664532296091763 |

P62273_P62857
RS29_HUMAN vs RS28_HUMAN
p-value: 0.017 q-value: 0.014

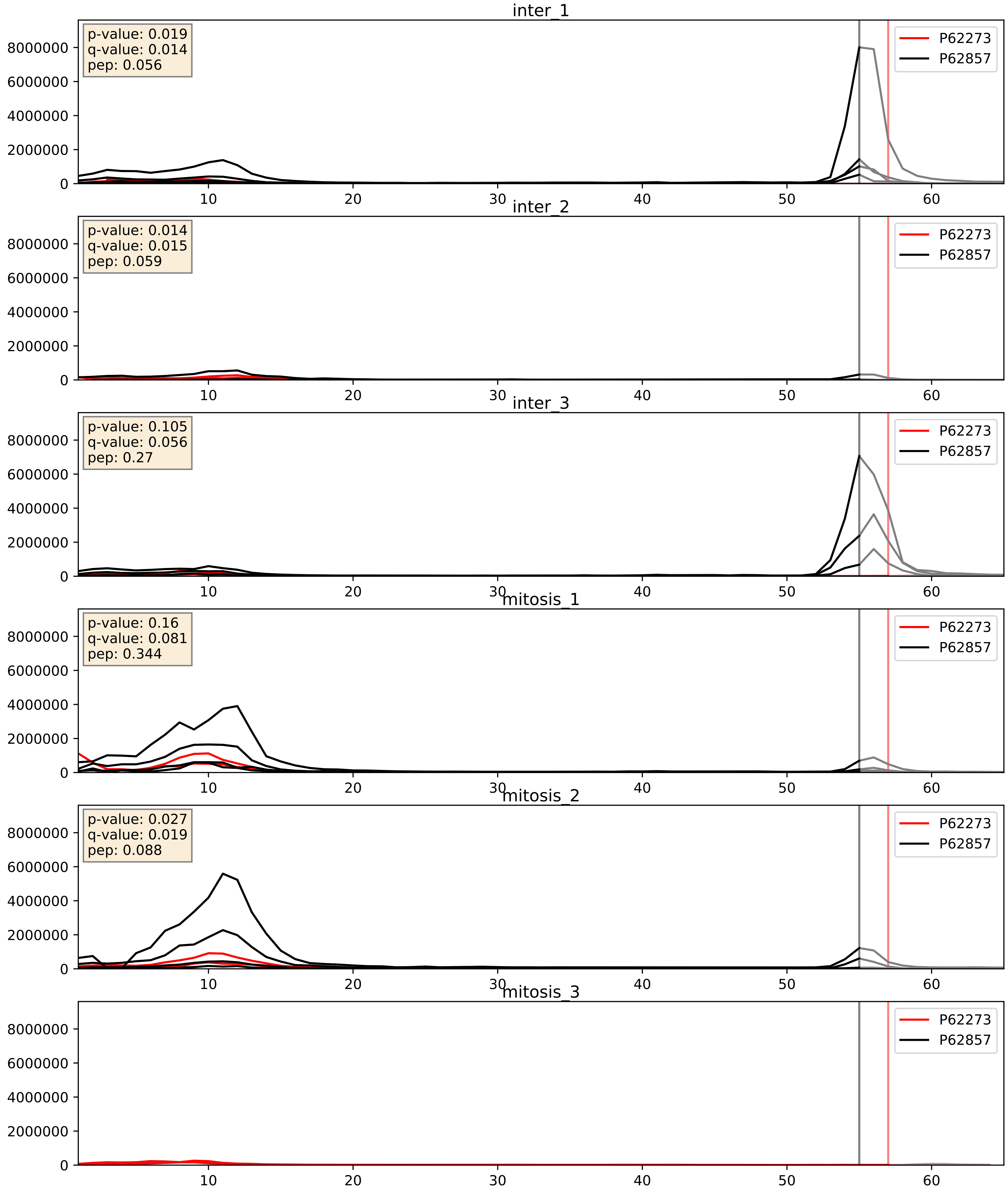


| level | condition_1 | condition_2 | pvalue | pvalue_adjusted |
|---|---|---|---|---|
| interactor_ratio | mitosis | inter | 0.7201148821644197 | 0.9442433206576832 |
| interactor_abundance | mitosis | inter | 0.7225706732953101 | 0.8232669989894655 |
| complex_abundance | mitosis | inter | 0.9285920119737423 | 0.955852456657777 |

P62277_P62857
RS13_HUMAN vs RS28_HUMAN
p-value: 0.014 q-value: 0.011

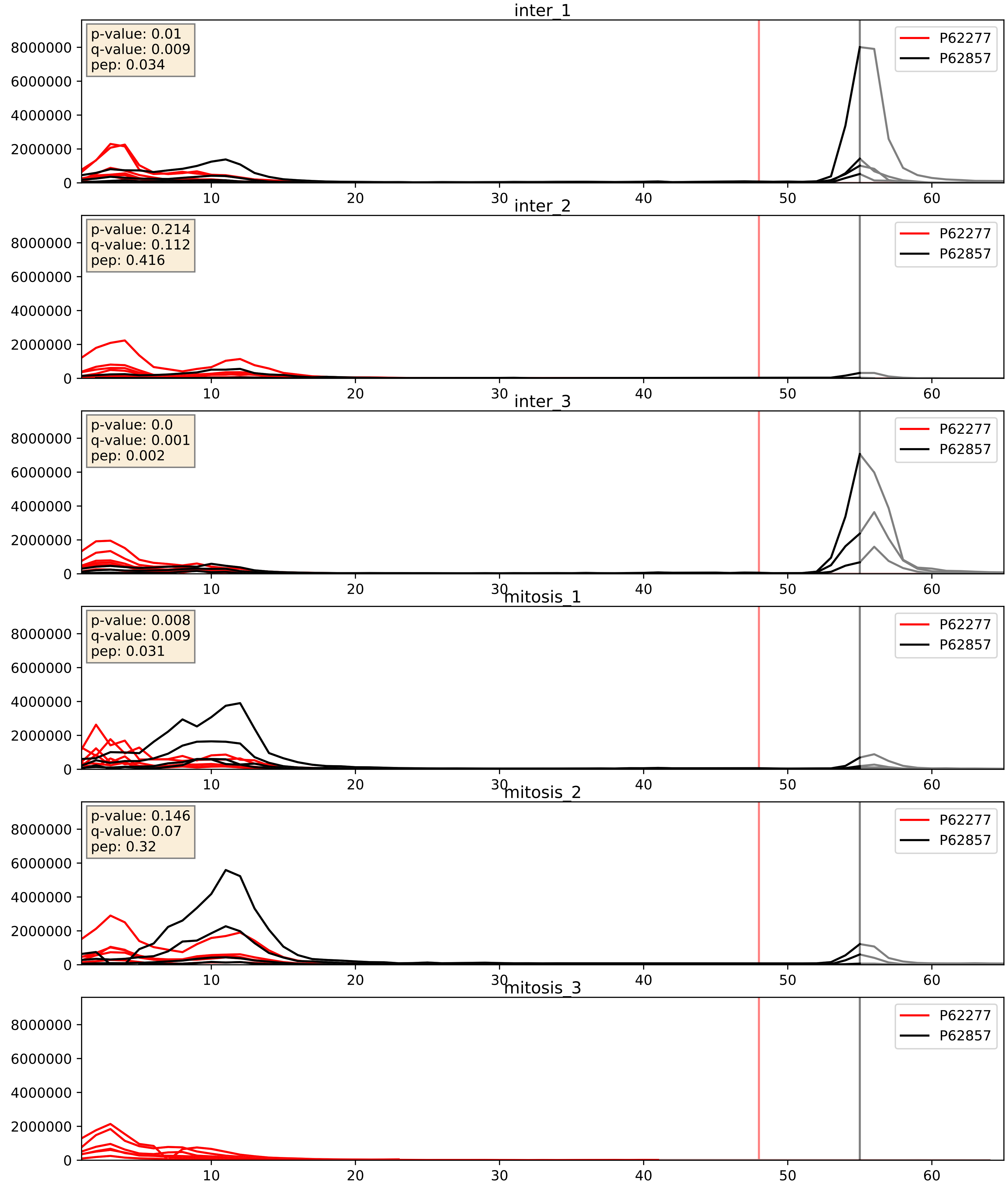


| level | condition_1 | condition_2 | pvalue | pvalue_adjusted |
|---|---|---|---|---|
| interactor_ratio | mitosis | inter | 0.03459494999630018 | 0.3009479389922048 |
| interactor_abundance | mitosis | inter | 0.37573907732032213 | 0.5689592255195396 |
| complex_abundance | mitosis | inter | 0.7279114996164175 | 0.8188884790007274 |

P62280_P62857
RS11_HUMAN vs RS28_HUMAN
p-value: 0.044 q-value: 0.029

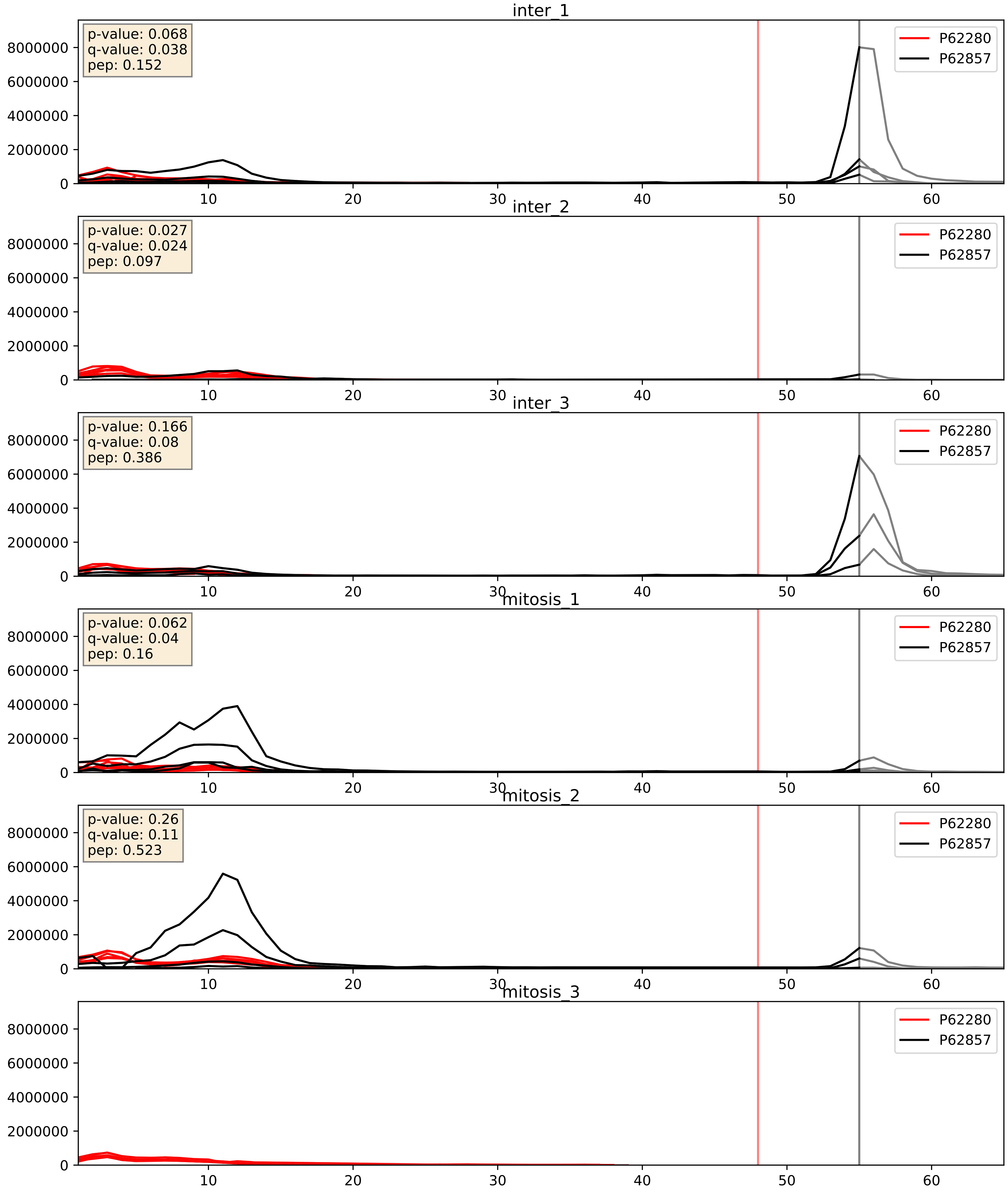


| level | condition_1 | condition_2 | pvalue | pvalue_adjusted |
|---|---|---|---|---|
| interactor_ratio | mitosis | inter | 0.1001842972215708 | 0.5276924928228902 |
| interactor_abundance | mitosis | inter | 0.39567221034491895 | 0.586815904862854 |
| complex_abundance | mitosis | inter | 0.767457501522378 | 0.847211428815272 |

P62424_P62857
RL7A_HUMAN vs RS28_HUMAN
p-value: 0.009 q-value: 0.008

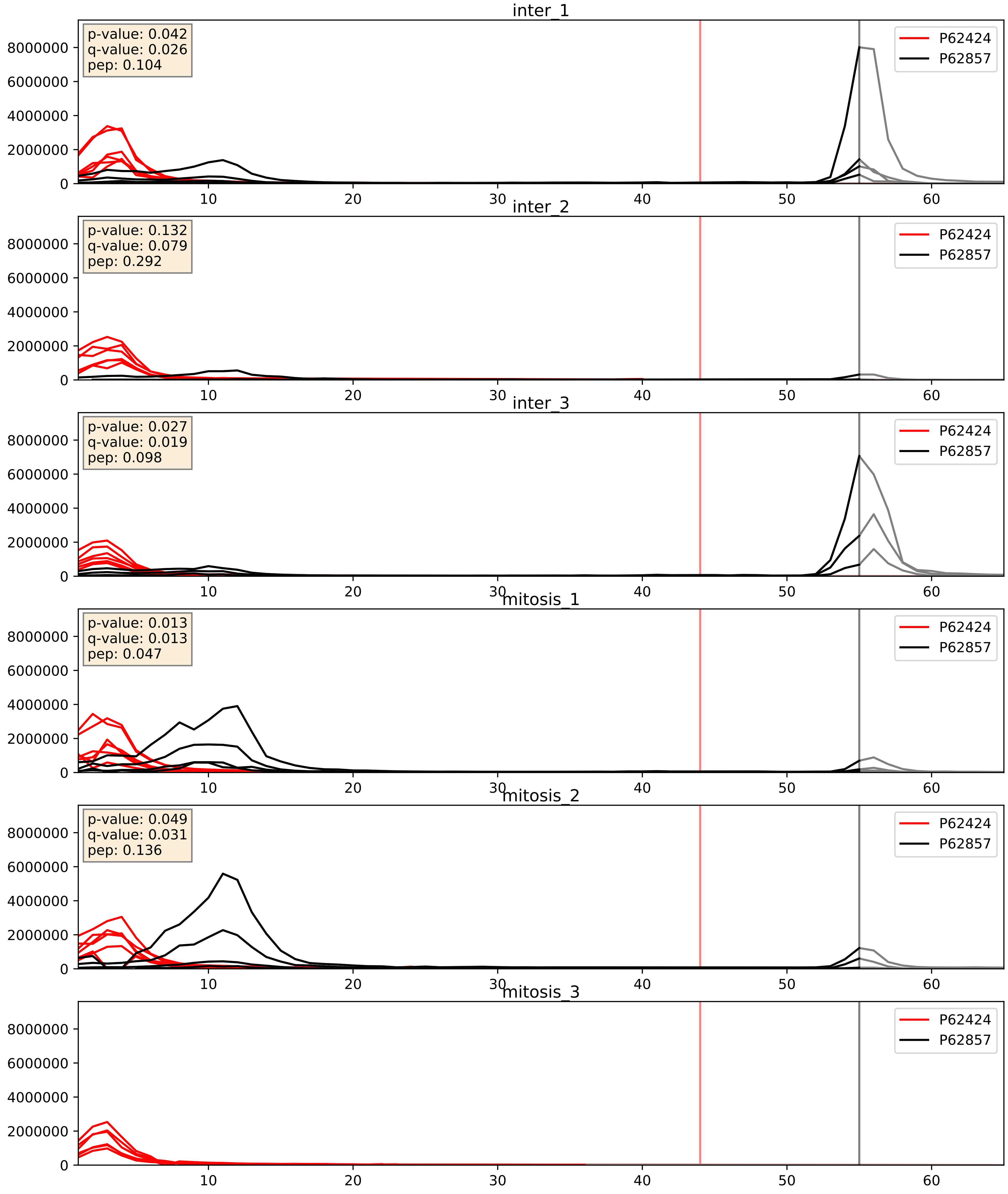


| level | condition_1 | condition_2 | pvalue | pvalue_adjusted |
|---|---|---|---|---|
| interactor_ratio | mitosis | inter | 0.01947093632770743 | 0.2112436184602986 |
| interactor_abundance | mitosis | inter | 0.37728511152723326 | 0.5704425602178075 |
| complex_abundance | mitosis | inter | 0.7251672959235822 | 0.8170611513420925 |

P62701_P62857
RS4X_HUMAN vs RS28_HUMAN
p-value: 0.006 q-value: 0.006

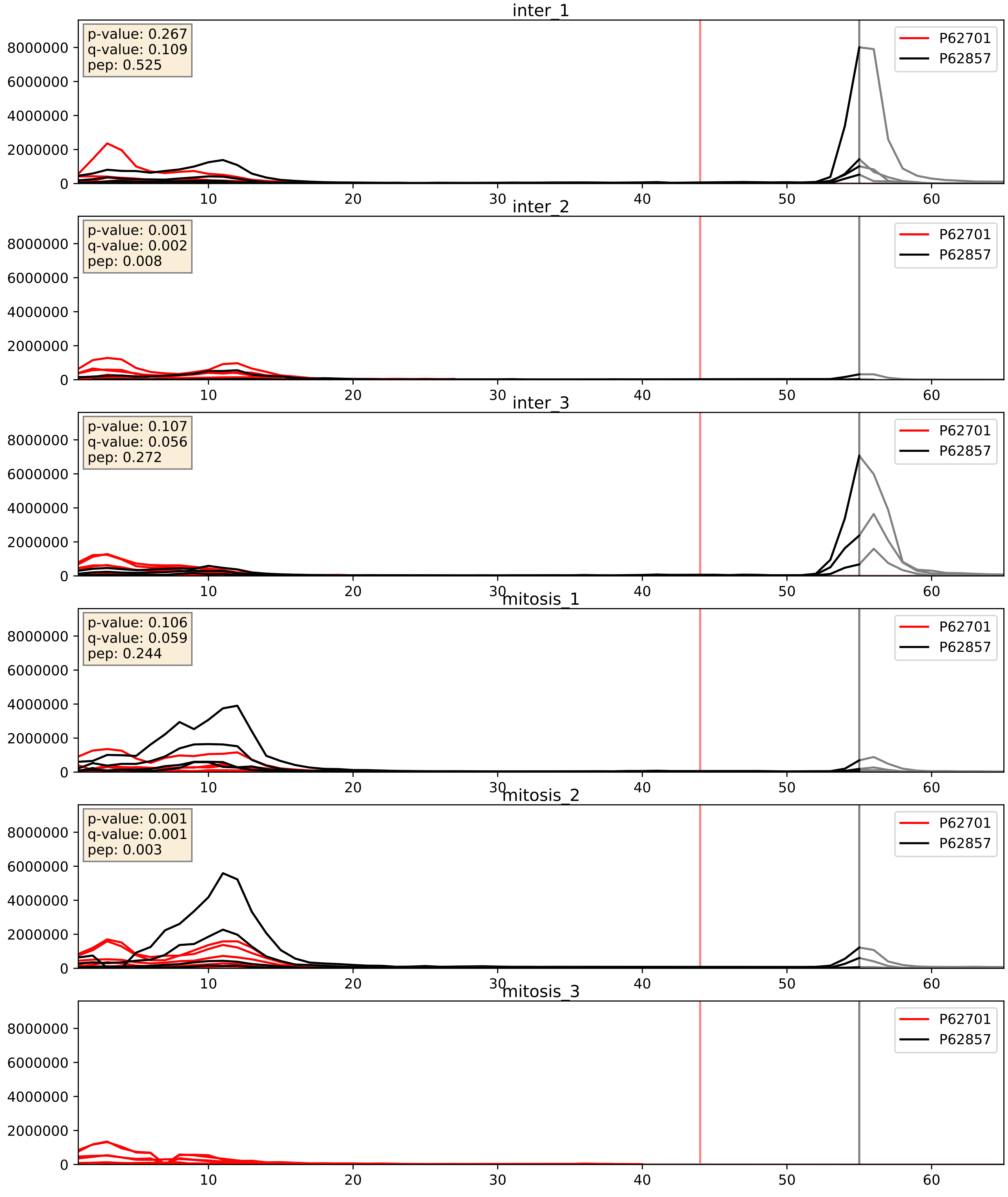


| level | condition_1 | condition_2 | pvalue | pvalue_adjusted |
|---|---|---|---|---|
| interactor_ratio | mitosis | inter | 0.060322096884244704 | 0.40594115513296747 |
| interactor_abundance | mitosis | inter | 0.3995524135705706 | 0.5889227137604967 |
| complex_abundance | mitosis | inter | 0.7475313540363294 | 0.833422024166816 |

P62750_P62857
RL23A_HUMAN vs RS28_HUMAN
p-value: 0.017 q-value: 0.014

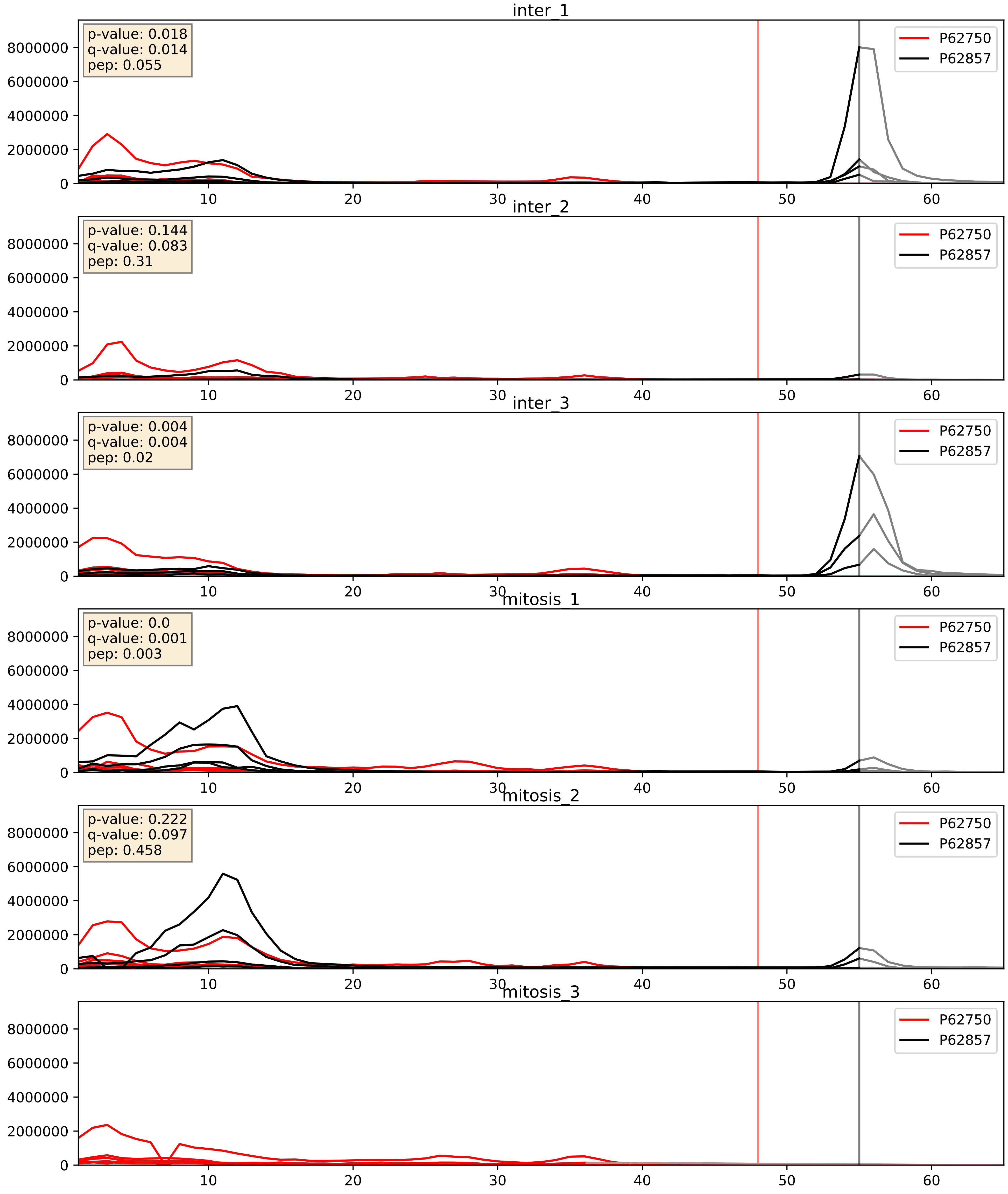


| level | condition_1 | condition_2 | pvalue | pvalue_adjusted |
|---|---|---|---|---|
| interactor_ratio | mitosis | inter | 0.03200609030832789 | 0.28568522736109153 |
| interactor_abundance | mitosis | inter | 0.464144924391648 | 0.6359981675672334 |
| complex_abundance | mitosis | inter | 0.7849922016599901 | 0.8596051229639906 |

P62753_P62857
RS6_HUMAN vs RS28_HUMAN
p-value: 0.003 q-value: 0.003

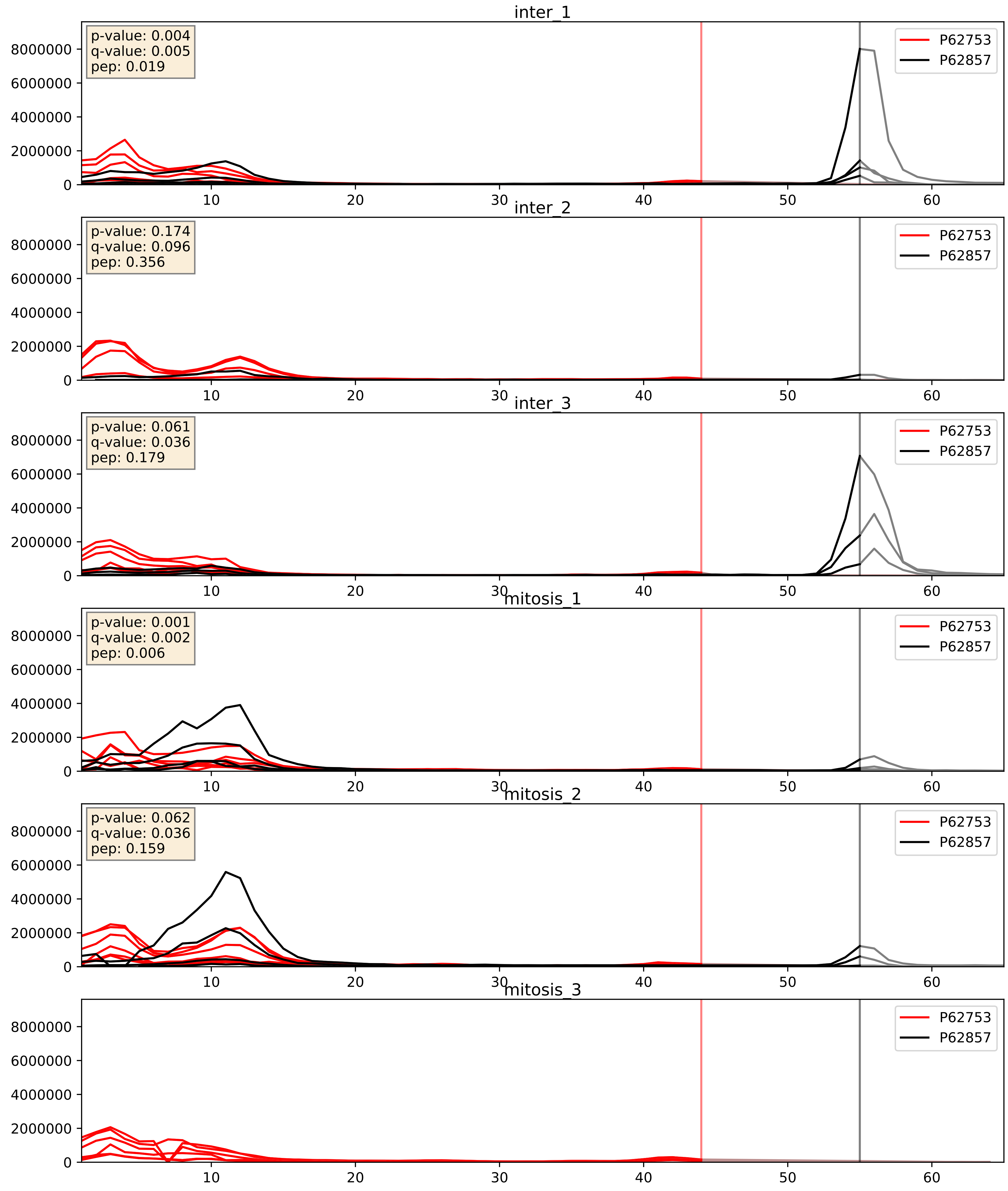


| level | condition_1 | condition_2 | pvalue | pvalue_adjusted |
|---|---|---|---|---|
| interactor_ratio | mitosis | inter | 0.02554058174948061 | 0.25245655863227945 |
| interactor_abundance | mitosis | inter | 0.3539193063864515 | 0.551880729149836 |
| complex_abundance | mitosis | inter | 0.6751256585054908 | 0.7832848518307132 |

P62829_P62857
RL23_HUMAN vs RS28_HUMAN
p-value: 0.023 q-value: 0.018

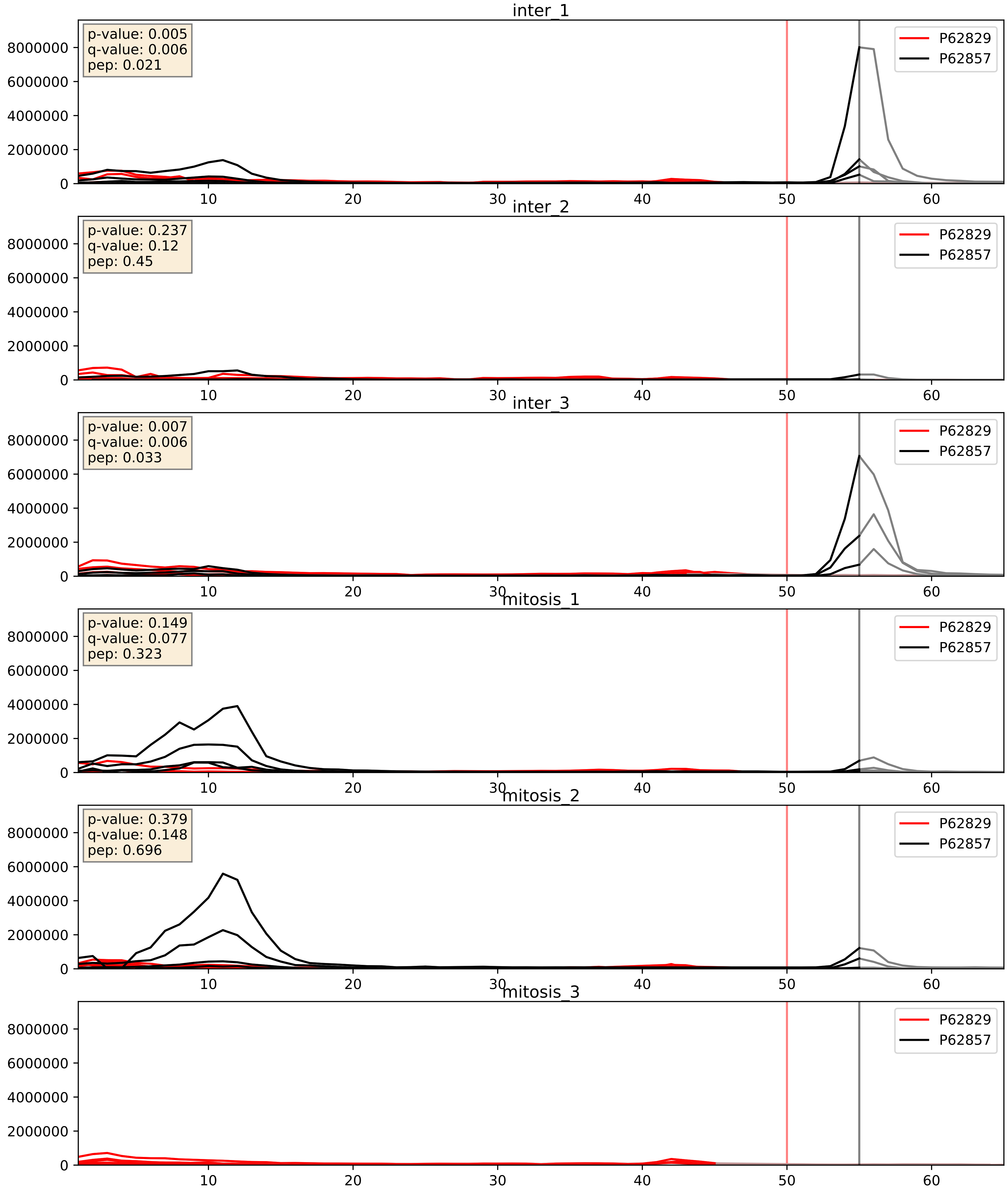


| level | condition_1 | condition_2 | pvalue | pvalue_adjusted |
|---|---|---|---|---|
| interactor_ratio | mitosis | inter | 0.047999143153140654 | 0.361488803041275 |
| interactor_abundance | mitosis | inter | 0.05463187832433133 | 0.18883459659046078 |
| complex_abundance | mitosis | inter | 0.4720766228631067 | 0.612826189218713 |

P62841_P62857
RS15_HUMAN vs RS28_HUMAN
p-value: 0.002 q-value: 0.002

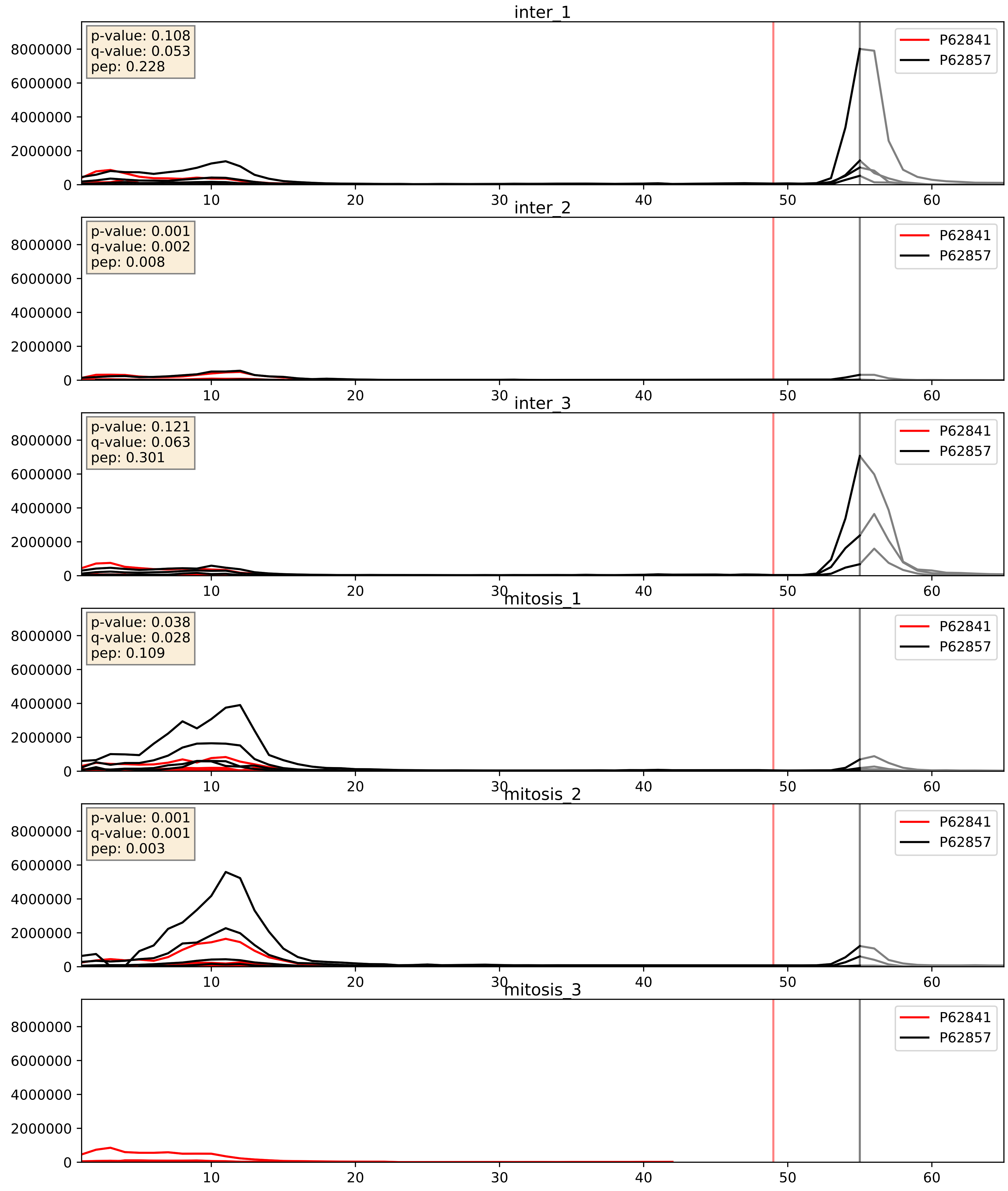


| level | condition_1 | condition_2 | pvalue | pvalue_adjusted |
|---|---|---|---|---|
| interactor_ratio | mitosis | inter | 0.25080013661969686 | 0.7250265613445431 |
| interactor_abundance | mitosis | inter | 0.5179160288491863 | 0.6806209682826628 |
| complex_abundance | mitosis | inter | 0.8859363437566812 | 0.9295067647735584 |

P62847_P62857
RS24_HUMAN vs RS28_HUMAN
p-value: 0.003 q-value: 0.003

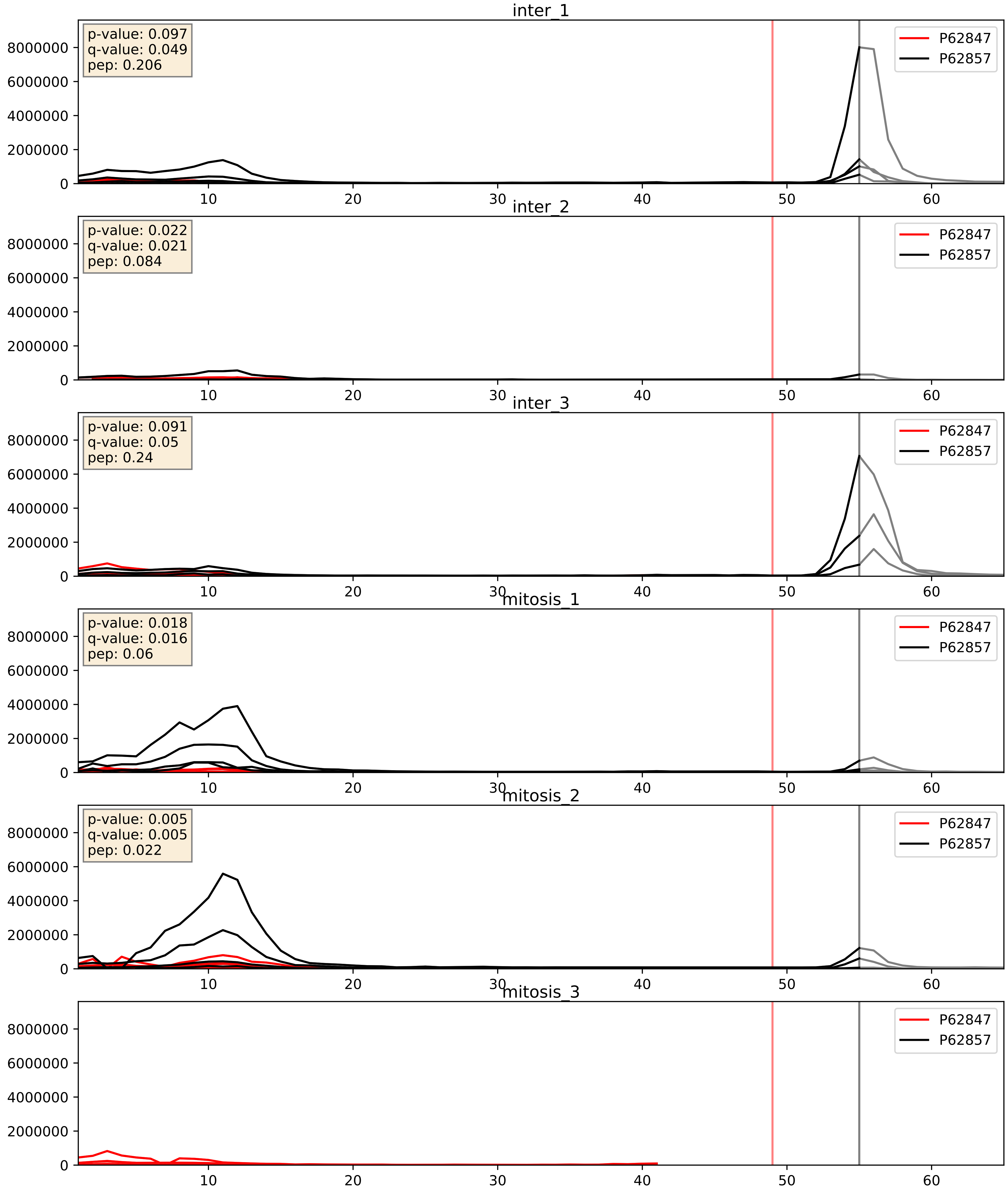


| level | condition_1 | condition_2 | pvalue | pvalue_adjusted |
|---|---|---|---|---|
| interactor_ratio | mitosis | inter | 0.18915969154861376 | 0.6603617290604135 |
| interactor_abundance | mitosis | inter | 0.505590943454336 | 0.6698434415677319 |
| complex_abundance | mitosis | inter | 0.8658429986439387 | 0.9152403146940128 |

P62851_P62857
RS25_HUMAN vs RS28_HUMAN
p-value: 0.0 q-value: 0.0

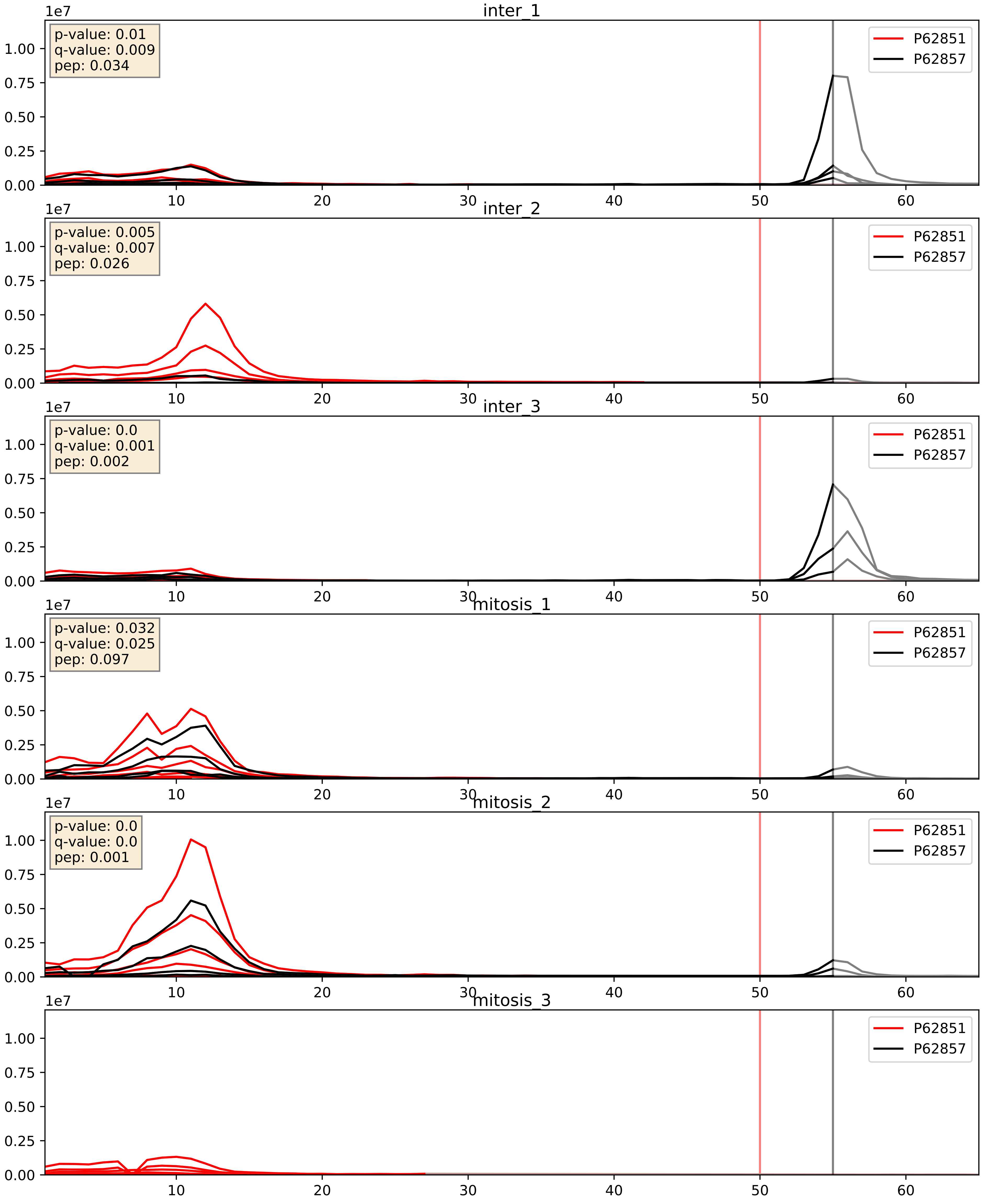


| level | condition_1 | condition_2 | pvalue | pvalue_adjusted |
|---|---|---|---|---|
| interactor_ratio | mitosis | inter | 0.3953553953753749 | 0.8288616665229511 |
| interactor_abundance | mitosis | inter | 0.72588490144551 | 0.8262732388794635 |
| complex_abundance | mitosis | inter | 0.9038489020000728 | 0.9400907170255921 |

P62857_P62861
RS28_HUMAN vs RS30_HUMAN
p-value: 0.018 q-value: 0.014

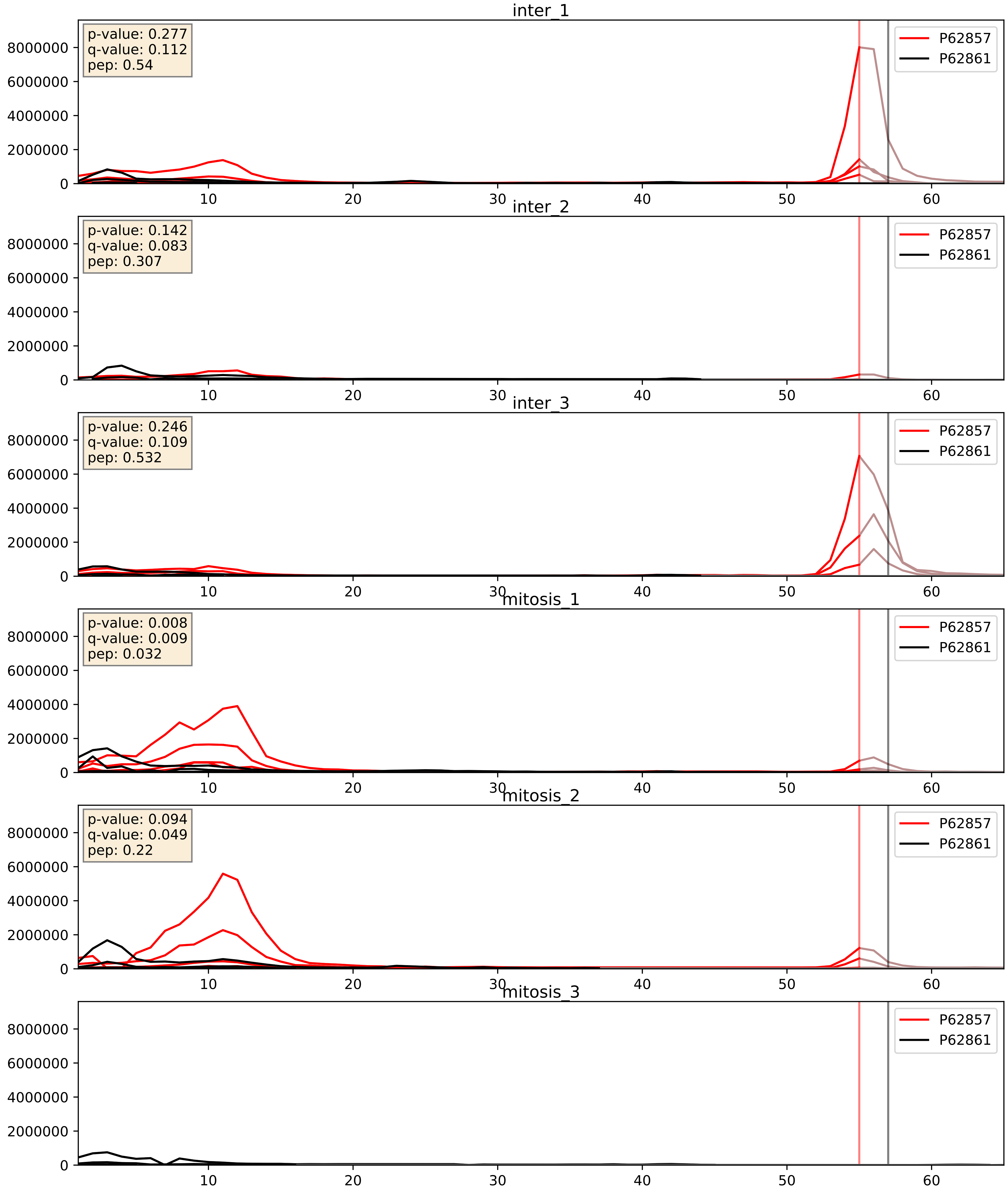


| level | condition_1 | condition_2 | pvalue | pvalue_adjusted |
|---|---|---|---|---|
| interactor_ratio | mitosis | inter | 0.3884484758569801 | 0.8251906958403389 |
| interactor_abundance | mitosis | inter | 0.8611740024278115 | 0.925645901403921 |
| complex_abundance | mitosis | inter | 0.9685242102500851 | 0.981480672397387 |

P62857_P62888
RS28_HUMAN vs RL30_HUMAN
p-value: 0.066 q-value: 0.041

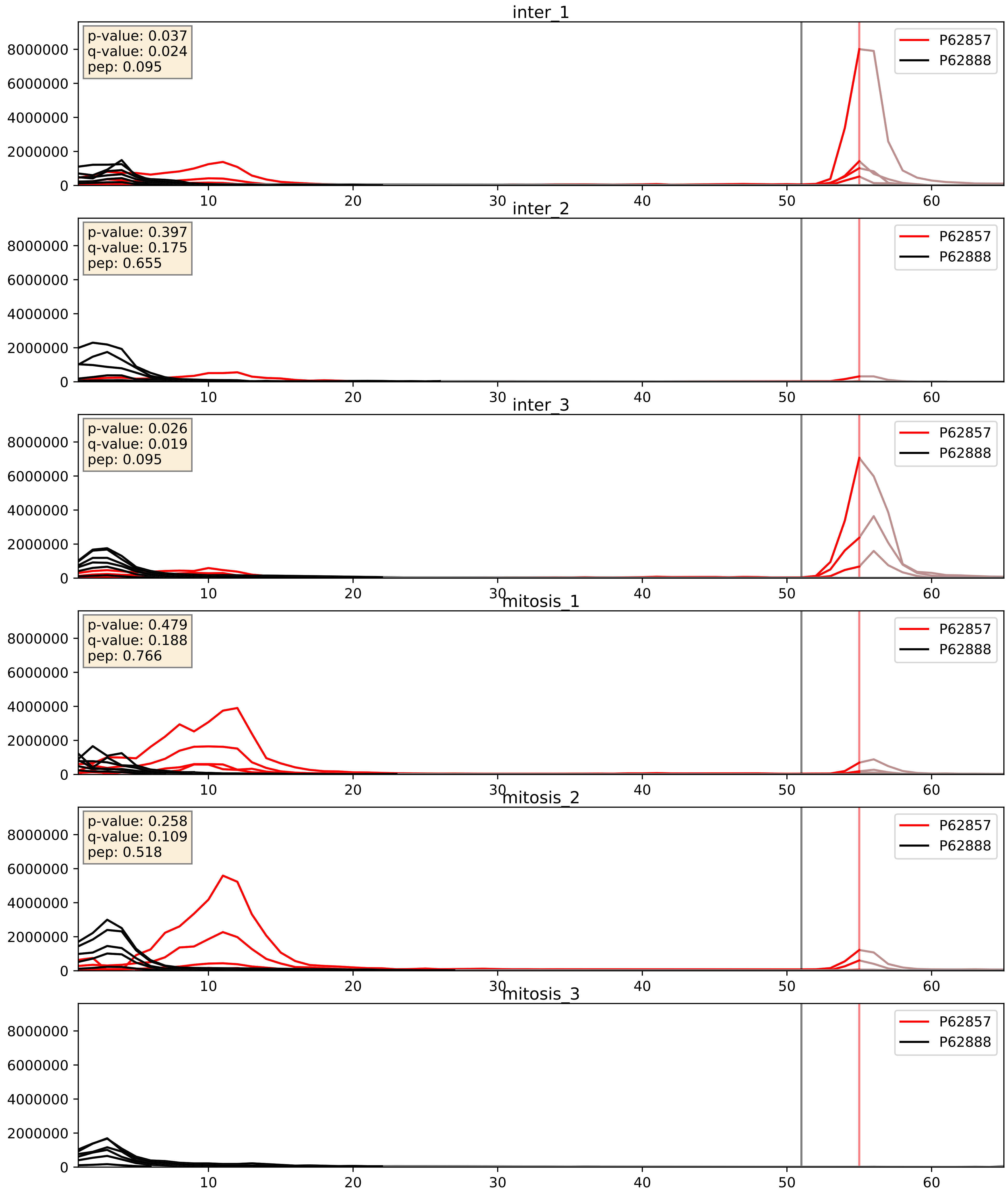


| level | condition_1 | condition_2 | pvalue | pvalue_adjusted |
|---|---|---|---|---|
| interactor_ratio | mitosis | inter | 0.07298171441413502 | 0.45138979435332066 |
| complex_abundance | mitosis | inter | 0.7002385789690855 | 0.7983540538059899 |
| interactor_abundance | mitosis | inter | 0.8559495452677266 | 0.9227678686579346 |

P62857_P62899
RS28_HUMAN vs RL31_HUMAN
p-value: 0.0 q-value: 0.0

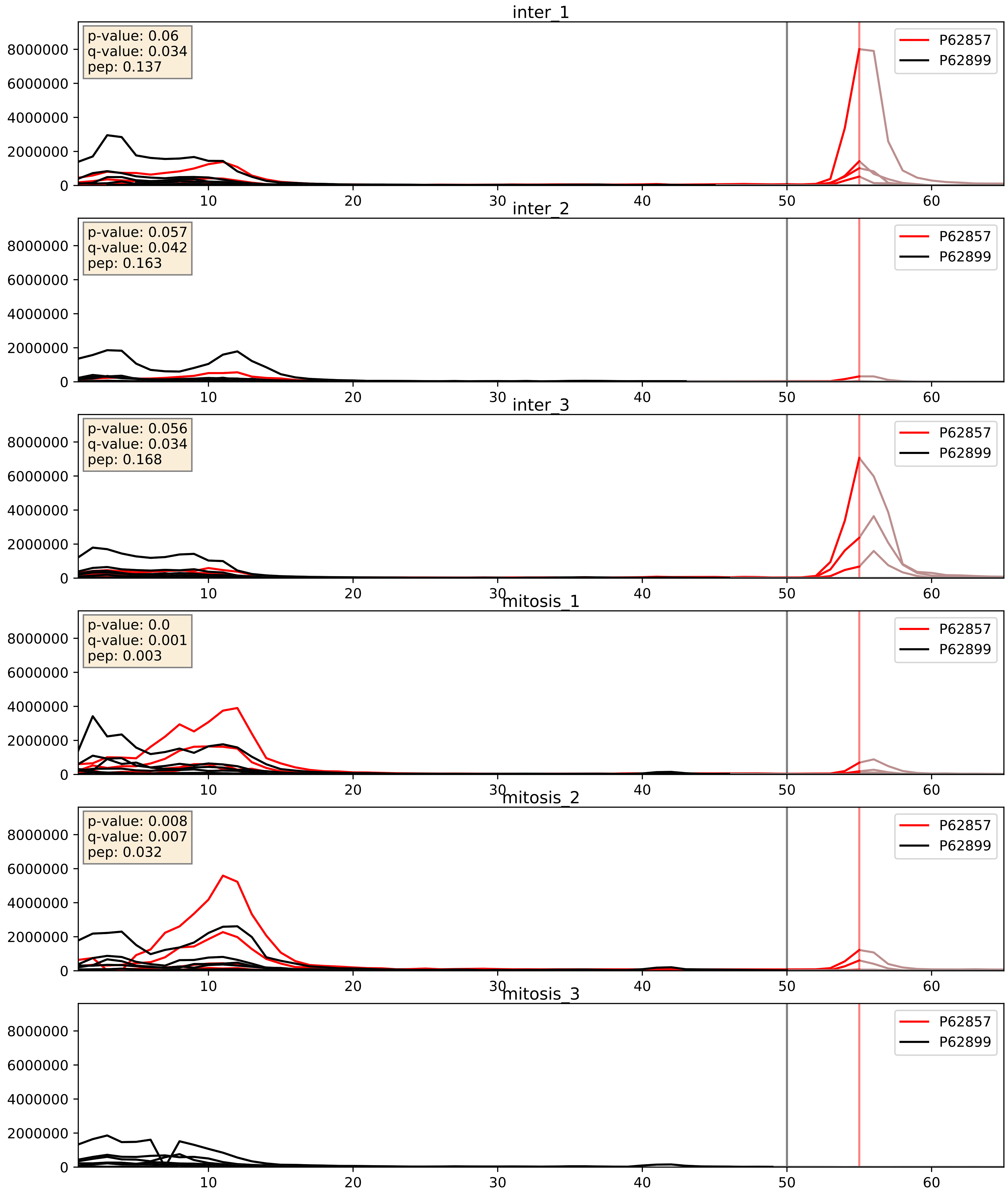


| level | condition_1 | condition_2 | pvalue | pvalue_adjusted |
|---|---|---|---|---|
| interactor_ratio | mitosis | inter | 0.02340619085301693 | 0.23710886828618336 |
| complex_abundance | mitosis | inter | 0.7655638411370577 | 0.8461234964665221 |
| interactor_abundance | mitosis | inter | 0.873382331824328 | 0.9298990881681792 |

P62857_P62913
RS28_HUMAN vs RL11_HUMAN
p-value: 0.048 q-value: 0.032

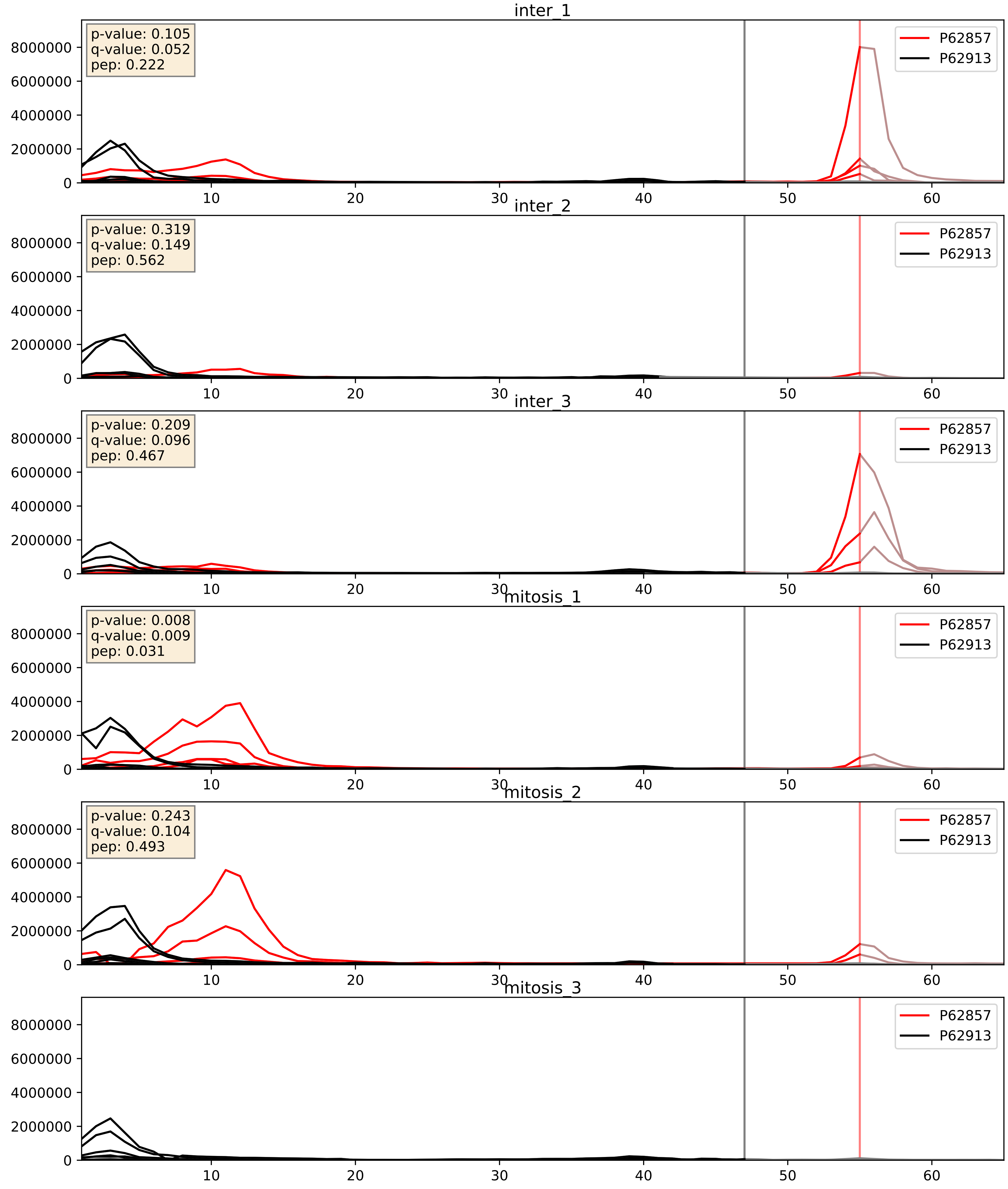


| level | condition_1 | condition_2 | pvalue | pvalue_adjusted |
|---|---|---|---|---|
| interactor_ratio | mitosis | inter | 0.06305963031352059 | 0.41679953365215705 |
| complex_abundance | mitosis | inter | 0.7572023722793146 | 0.8402453081035692 |
| interactor_abundance | mitosis | inter | 0.8740108272397997 | 0.9300183580300436 |

P62857_P62917
RS28_HUMAN vs RL8_HUMAN
p-value: 0.069 q-value: 0.042

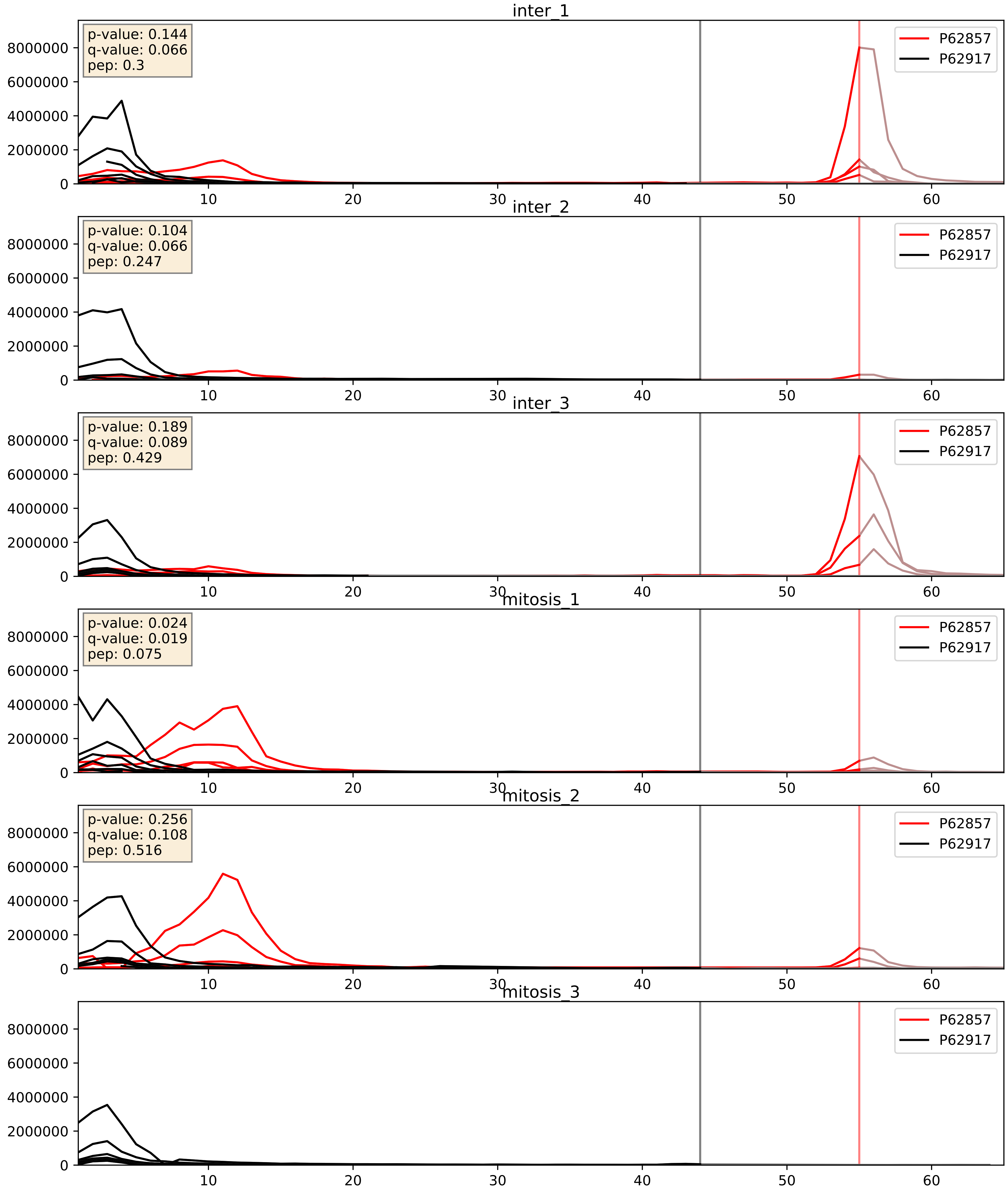


| level | condition_1 | condition_2 | pvalue | pvalue_adjusted |
|---|---|---|---|---|
| interactor_ratio | mitosis | inter | 0.009493181686084056 | 0.13913572462569762 |
| complex_abundance | mitosis | inter | 0.8053659260707171 | 0.8739772220037194 |
| interactor_abundance | mitosis | inter | 0.8485474914288413 | 0.9176963394353609 |

P62857_P63173
RS28_HUMAN vs RL38_HUMAN
p-value: 0.02 q-value: 0.016

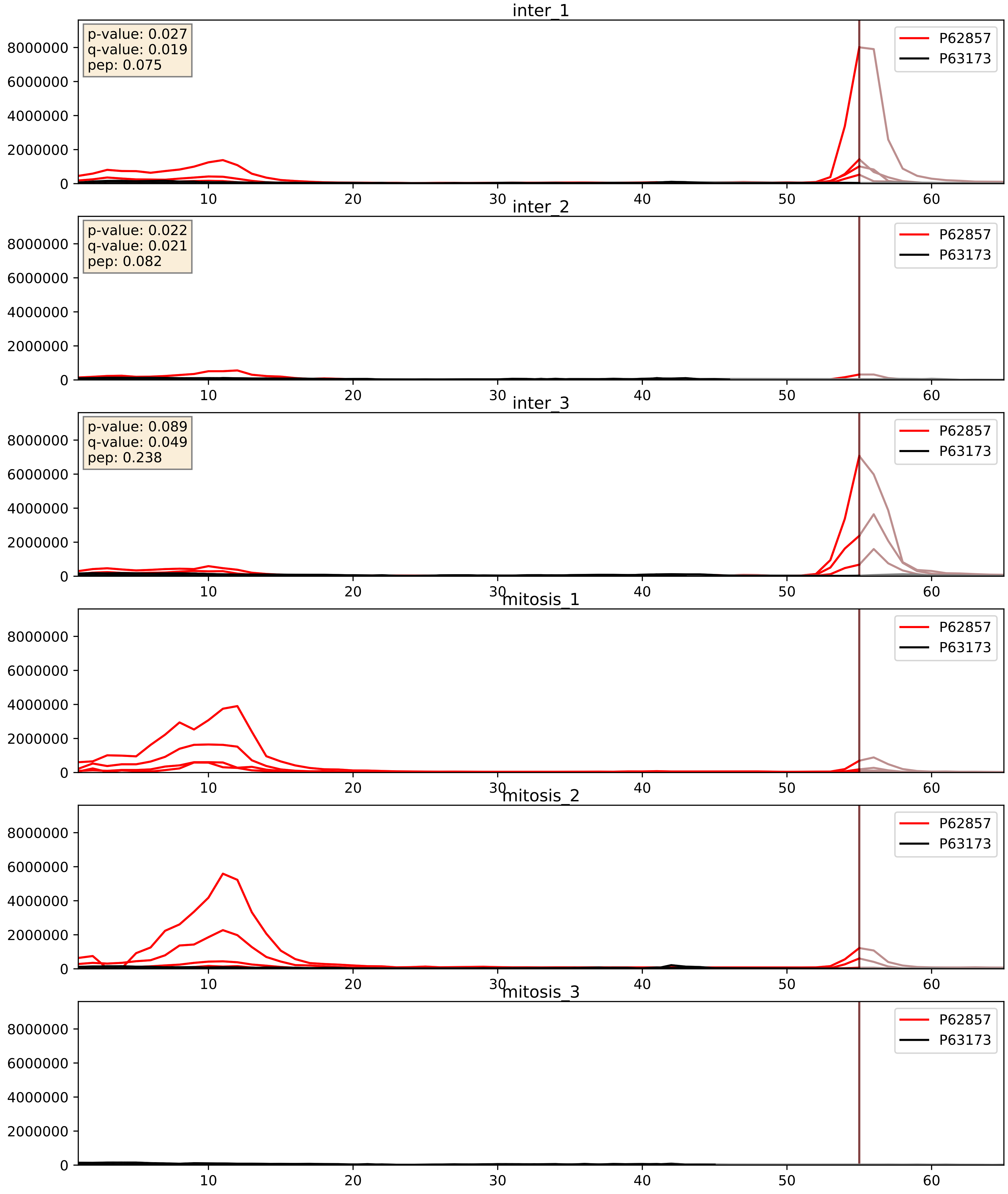


| level | condition_1 | condition_2 | pvalue | pvalue_adjusted |
|---|---|---|---|---|
| complex_abundance | mitosis | inter | 0.22291794216418093 | 0.3768494435238015 |
| interactor_abundance | mitosis | inter | 0.4485022630327217 | 0.6226032671235112 |
| interactor_ratio | mitosis | inter | 0.47604604588706995 | 0.8646587168966383 |

P62857_P63220
RS28_HUMAN vs RS21_HUMAN
p-value: 0.0 q-value: 0.0

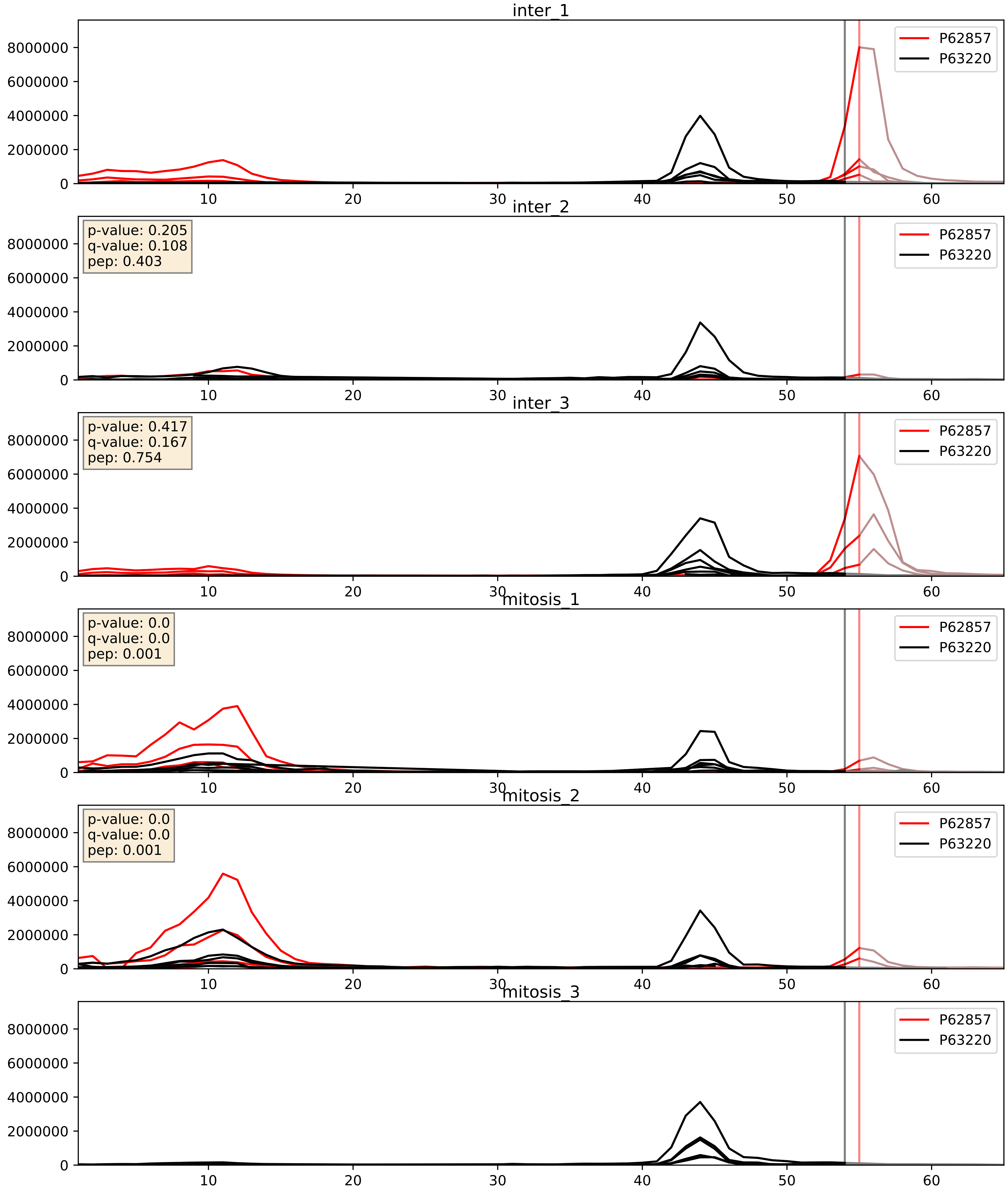


| level | condition_1 | condition_2 | pvalue | pvalue_adjusted |
|---|---|---|---|---|
| interactor_ratio | mitosis | inter | 0.2172203772291484 | 0.6935495819028386 |
| interactor_abundance | mitosis | inter | 0.7943294482148424 | 0.8808985116259767 |
| complex_abundance | mitosis | inter | 0.980794127518134 | 0.9887253383157429 |

P62857_P63244
RS28_HUMAN vs RACK1_HUMAN
p-value: 0.066 q-value: 0.04

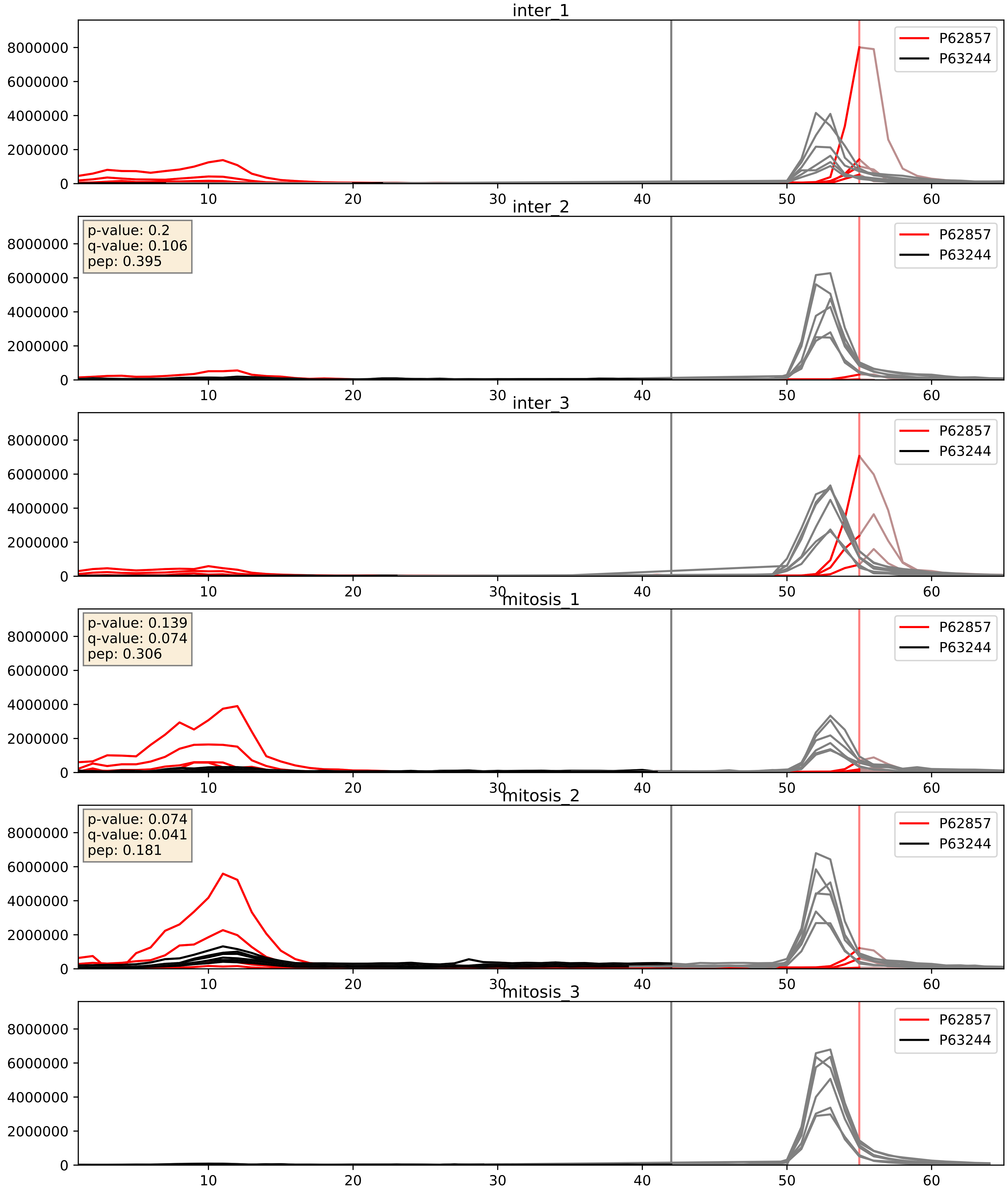


| level | condition_1 | condition_2 | pvalue | pvalue_adjusted |
|---|---|---|---|---|
| complex_abundance | mitosis | inter | 0.6744999243356734 | 0.7828962471193829 |
| interactor_abundance | mitosis | inter | 0.7424815232975018 | 0.8418626186405213 |
| interactor_ratio | mitosis | inter | 0.8669466625517663 | 0.9783867411263176 |

P62857_P83731
RS28_HUMAN vs RL24_HUMAN
p-value: 0.006 q-value: 0.006

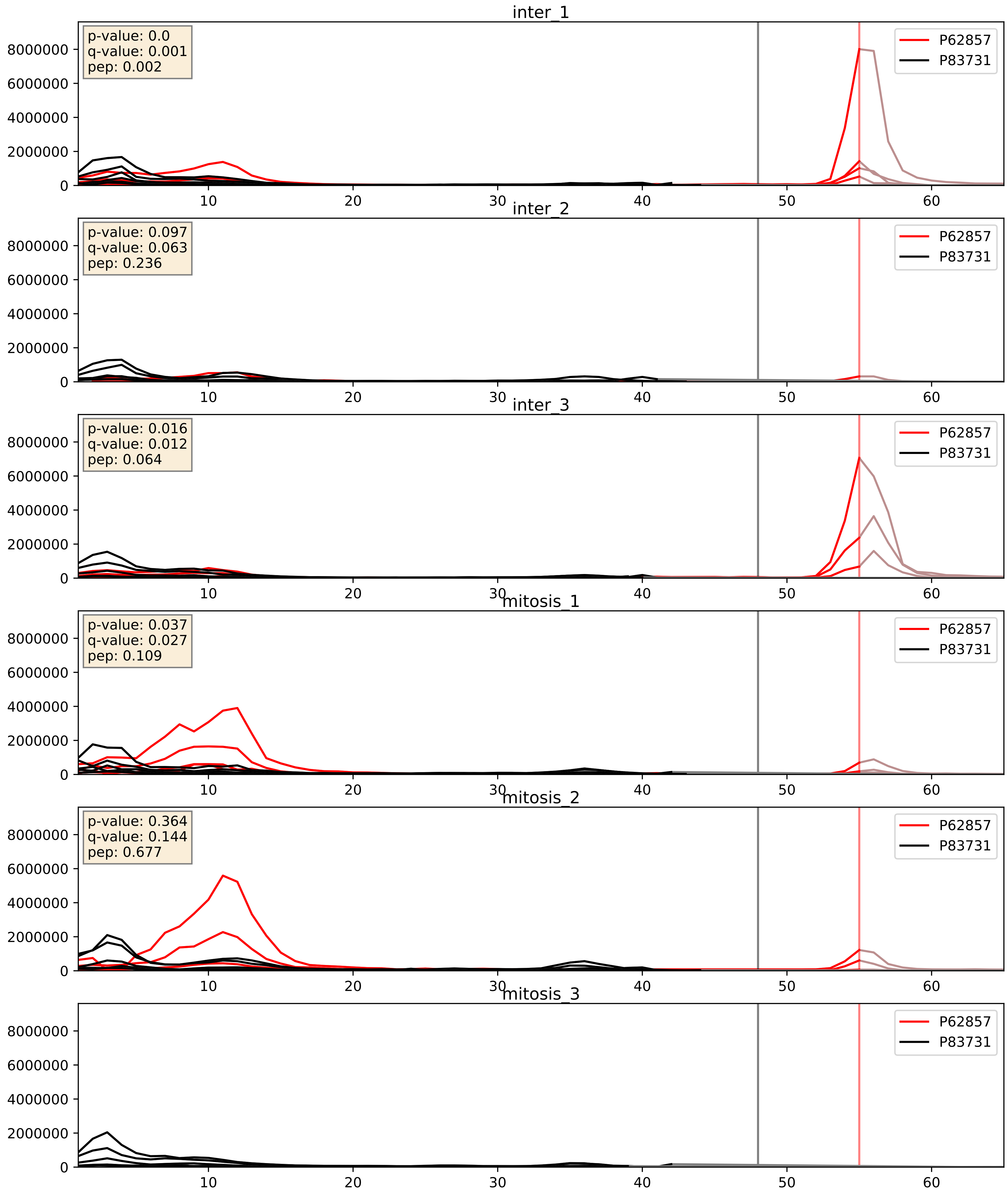


| level | condition_1 | condition_2 | pvalue | pvalue_adjusted |
|---|---|---|---|---|
| interactor_ratio | mitosis | inter | 0.02640800076918747 | 0.25543205403291264 |
| complex_abundance | mitosis | inter | 0.7582396111091129 | 0.8407883047708208 |
| interactor_abundance | mitosis | inter | 0.8724286860520415 | 0.9298990881681792 |

P62857_P84098
RS28_HUMAN vs RL19_HUMAN
p-value: 0.036 q-value: 0.025

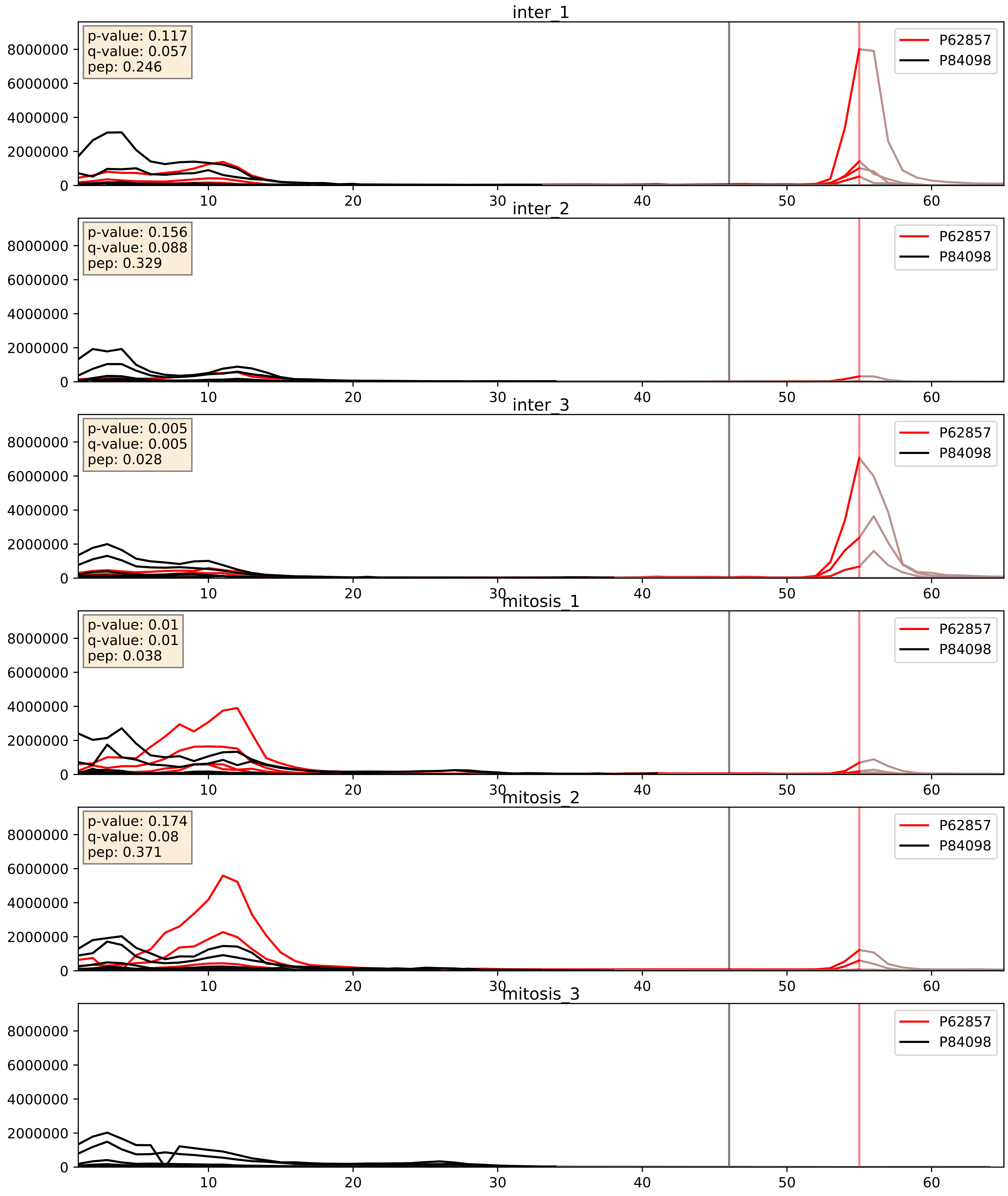


| level | condition_1 | condition_2 | pvalue | pvalue_adjusted |
|---|---|---|---|---|
| interactor_ratio | mitosis | inter | 0.02196422863494046 | 0.2273443737788275 |
| complex_abundance | mitosis | inter | 0.7298072285313559 | 0.8200511782919936 |
| interactor_abundance | mitosis | inter | 0.8648228845712524 | 0.9265766183027625 |

P62857_Q02878
RS28_HUMAN vs RL6_HUMAN
p-value: 0.04 q-value: 0.027

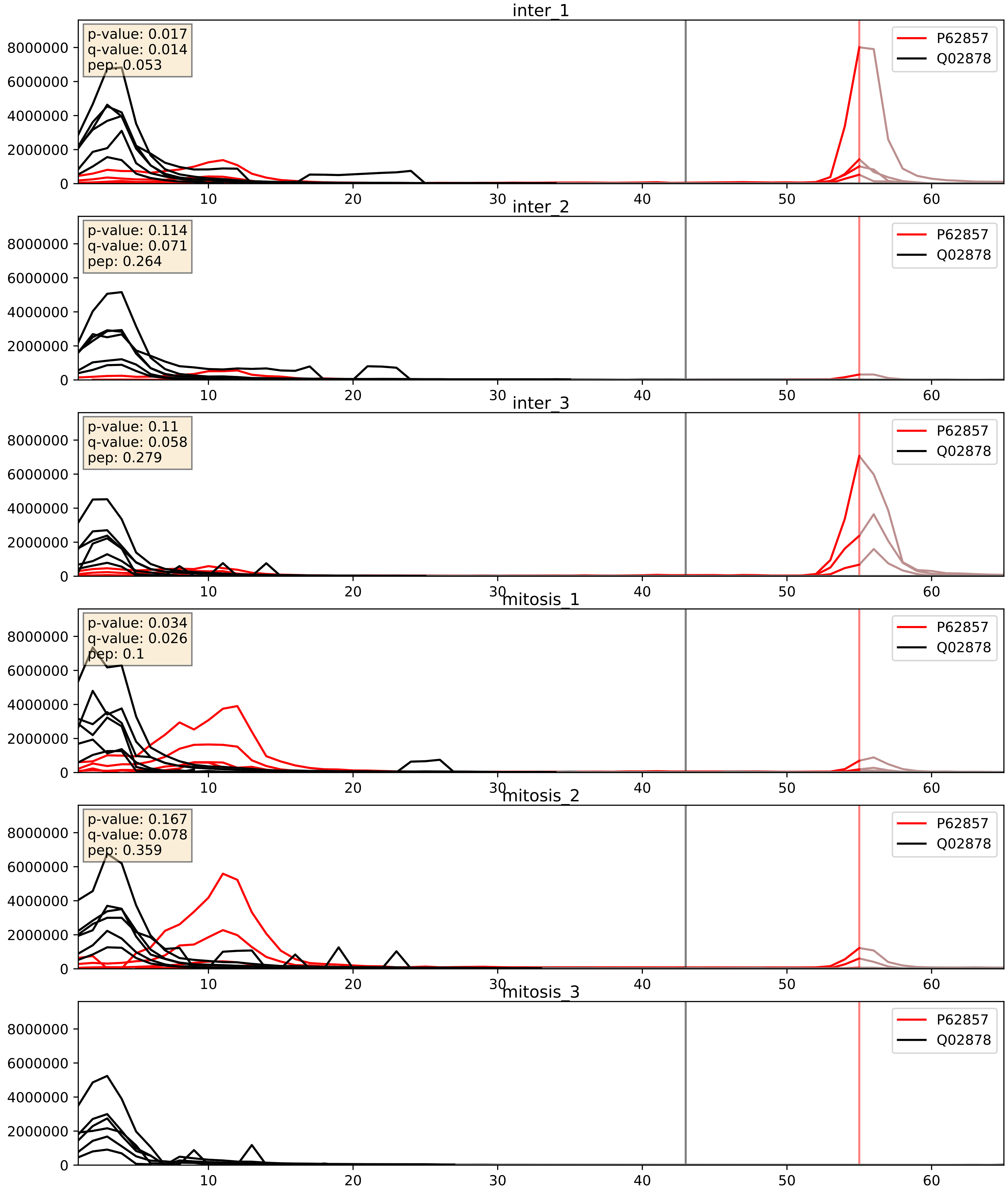


| level | condition_1 | condition_2 | pvalue | pvalue_adjusted |
|---|---|---|---|---|
| interactor_ratio | mitosis | inter | 0.021452797506055936 | 0.2228566460444407 |
| complex_abundance | mitosis | inter | 0.6621718243644792 | 0.7743833398247955 |
| interactor_abundance | mitosis | inter | 0.8627524151139239 | 0.925645901403921 |

P62857_Q07020
RS28_HUMAN vs RL18_HUMAN
p-value: 0.011 q-value: 0.01

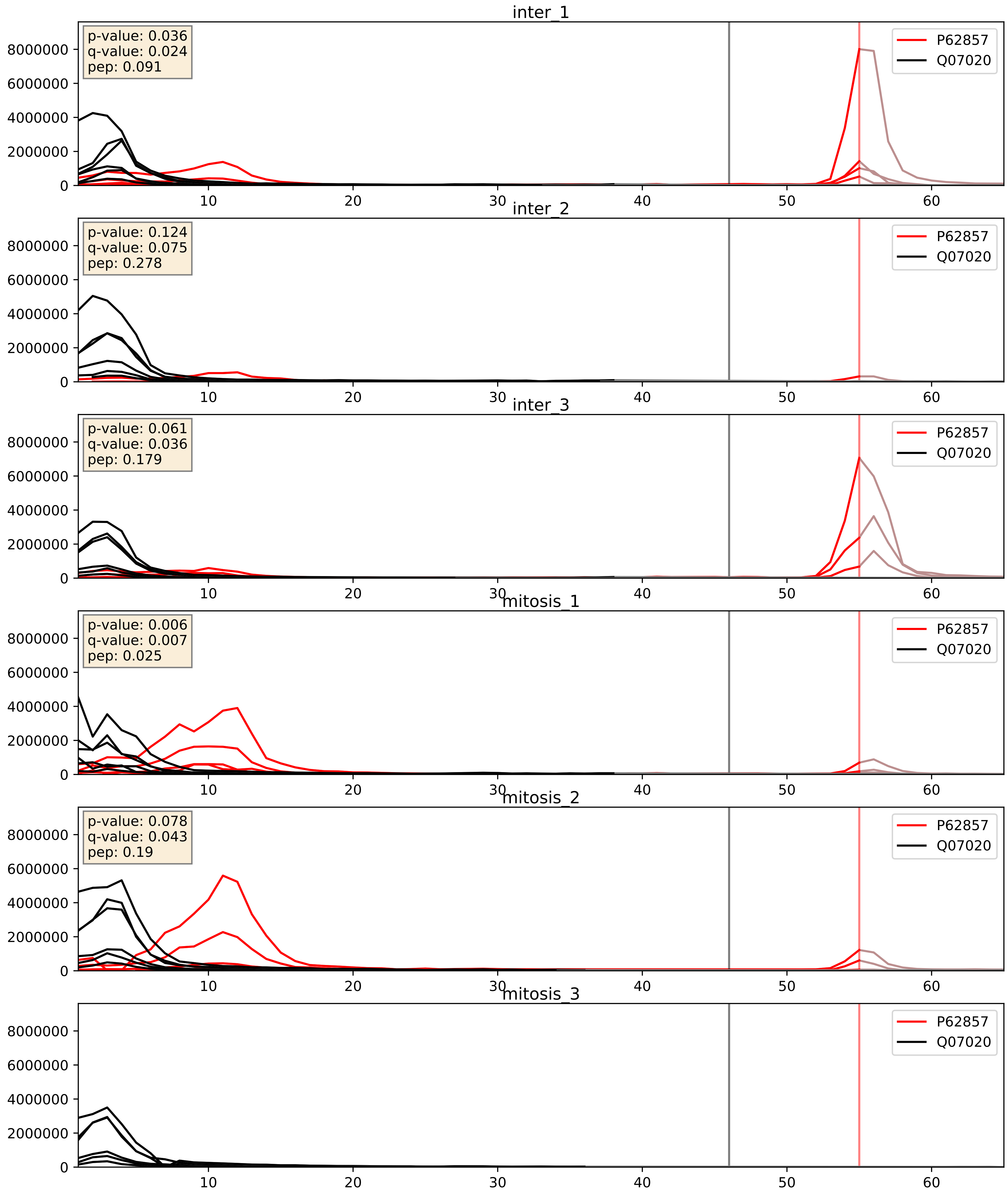


| level | condition_1 | condition_2 | pvalue | pvalue_adjusted |
|---|---|---|---|---|
| interactor_ratio | mitosis | inter | 0.031464048034990026 | 0.28316826136323564 |
| complex_abundance | mitosis | inter | 0.6582355845856114 | 0.7710204156642745 |
| interactor_abundance | mitosis | inter | 0.8696398385231697 | 0.9292949276272808 |

P62857_Q9Y3U8
RS28_HUMAN vs RL36_HUMAN
p-value: 0.003 q-value: 0.003

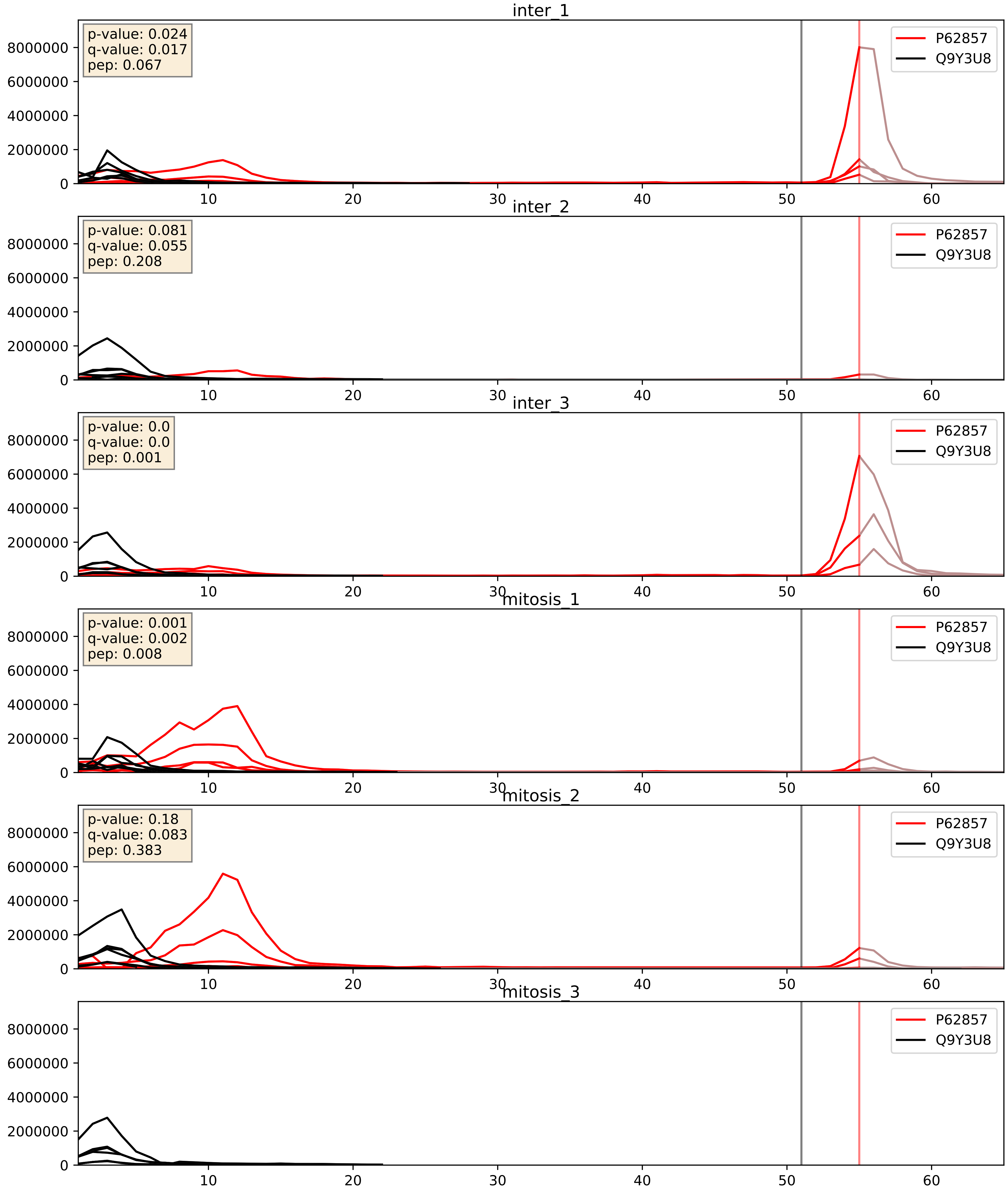


| level | condition_1 | condition_2 | pvalue | pvalue_adjusted |
|---|---|---|---|---|
| interactor_ratio | mitosis | inter | 0.060984749168564736 | 0.4087189096368508 |
| complex_abundance | mitosis | inter | 0.7644080768267483 | 0.8450643339321924 |
| interactor_abundance | mitosis | inter | 0.857751673400255 | 0.9236246712757007 |
