## Supplementary Data 2 for "SECAT: Quantifying differential protein-protein interaction states by network-centric analysis": hela_string_000117_Q14694.pdf

Q14694_Q14694

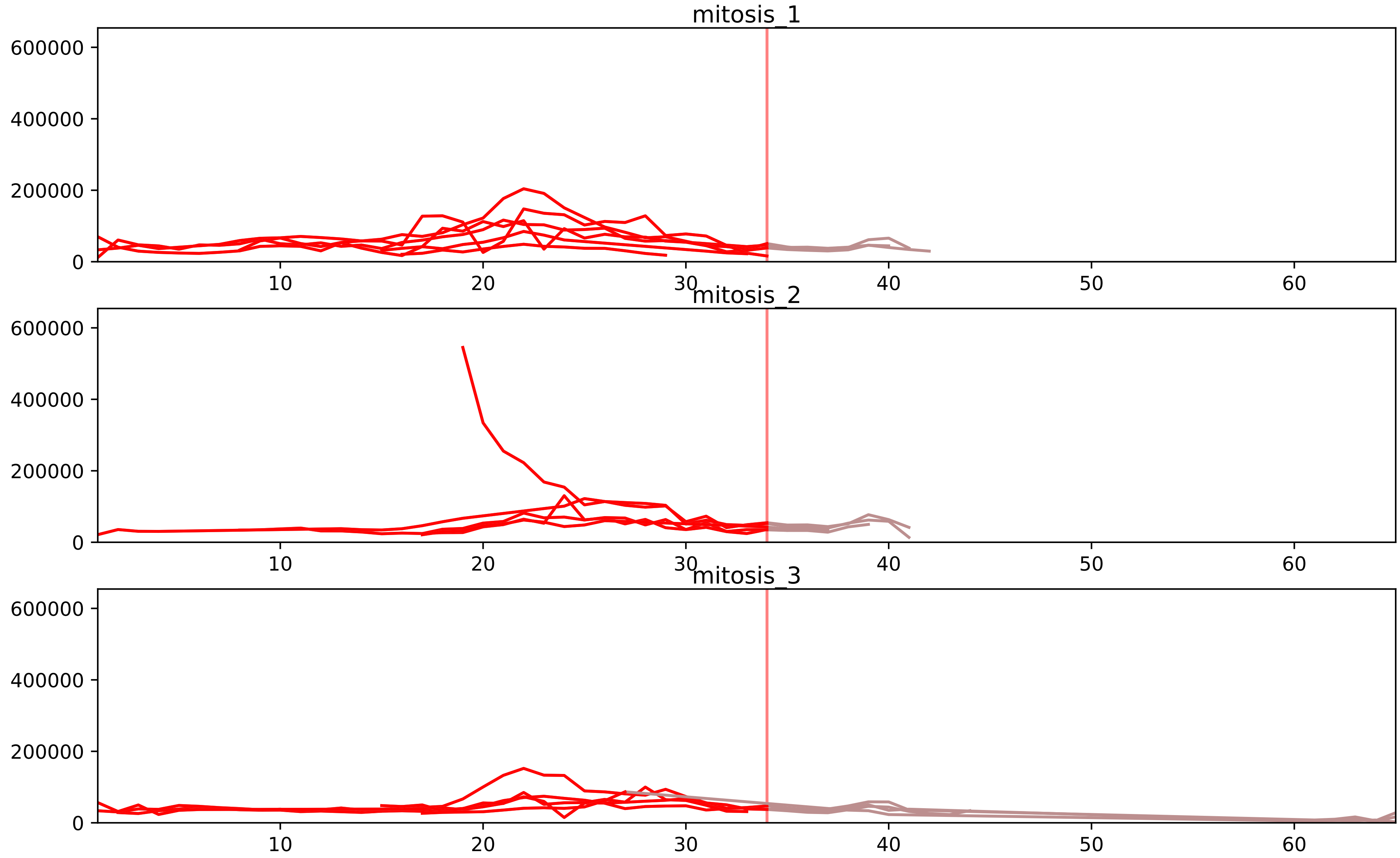


| level | condition_1 | condition_2 | pvalue | pvalue_adjusted |
|---|---|---|---|---|
| total_abundance | mitosis | inter | 2.0568065789596454e-05 | 0.0058684532992606024 |
| assembled_abundance | mitosis | inter | 3.068245787683866e-05 | 0.006314764940805662 |
| monomer_abundance | mitosis | inter | 0.0003668052181300632 | 0.05243686845257356 |
