## Supplementary Data 2 for "SECAT: Quantifying differential protein-protein interaction states by network-centric analysis": hela_string_000118_Q12788.pdf

Q12788_Q12788

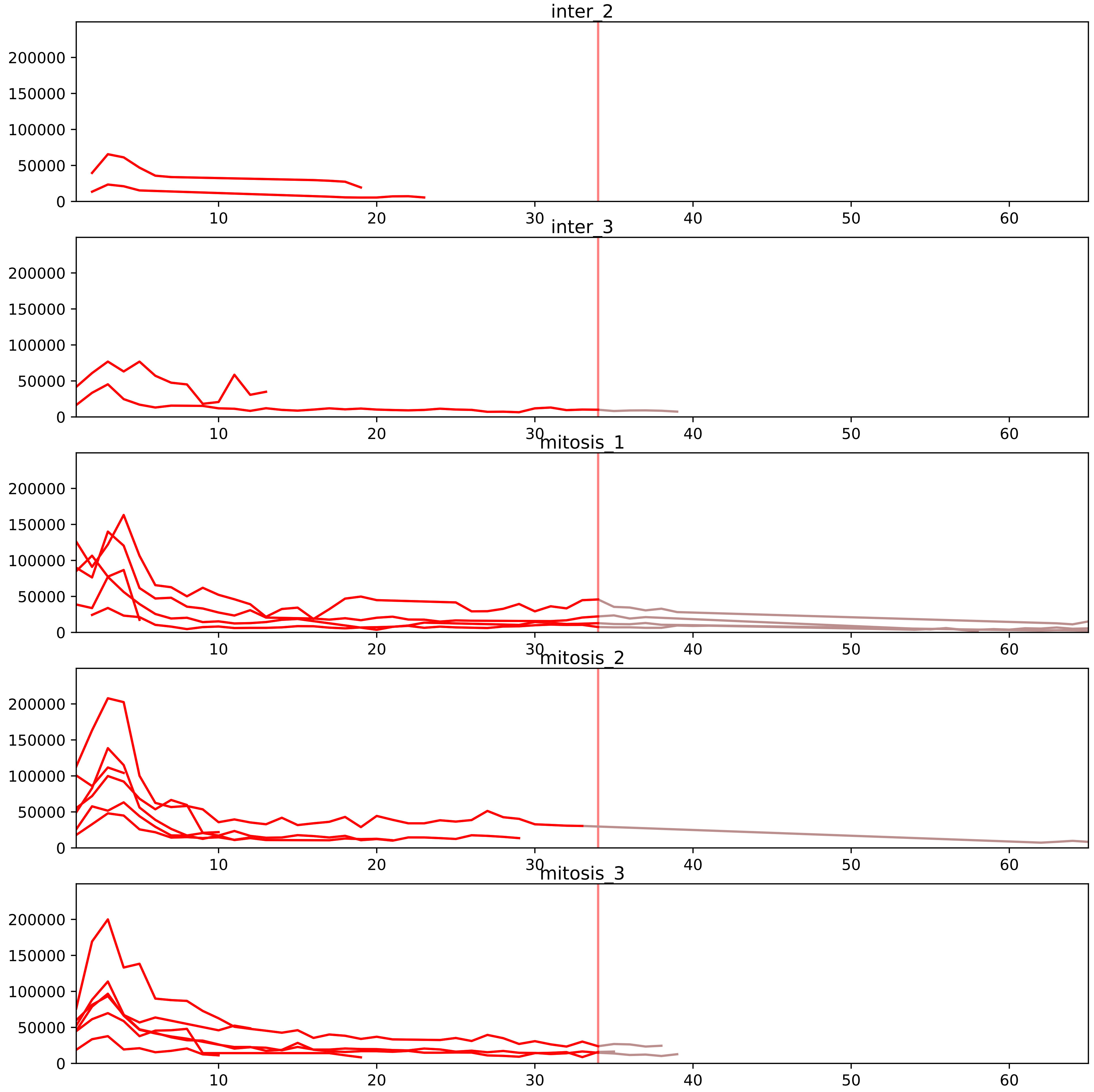


| level | condition_1 | condition_2 | pvalue | pvalue_adjusted |
|---|---|---|---|---|
| interactor_abundance | mitosis | inter | 2.1478673489699038e-05 | 0.0008896139540965403 |
| complex_abundance | mitosis | inter | 7.323605909769629e-05 | 0.0018871206097520157 |
| interactor_ratio | mitosis | inter | 0.00017076886820888872 | 0.0026184559792029605 |
| monomer_abundance | mitosis | inter | 0.04217350135692965 | 0.3157957803087734 |
| total_abundance | mitosis | inter | 0.07129872945302053 | 0.33263709944322245 |
| assembled_abundance | mitosis | inter | 0.08194534154929285 | 0.33692014801381864 |

O00567_Q12788
NOP56_HUMAN vs TBL3_HUMAN
p-value: 0.007 q-value: 0.007

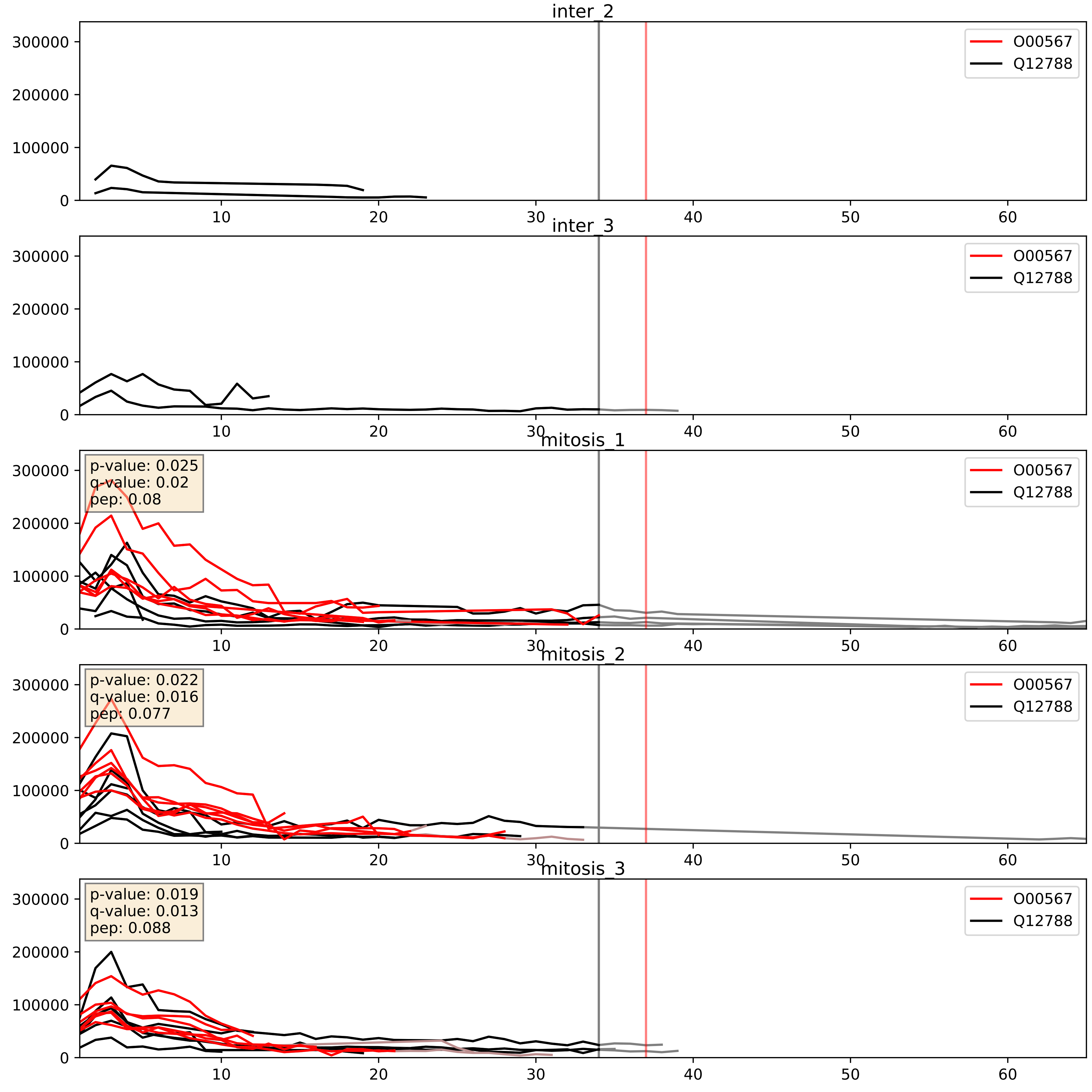


| level | condition_1 | condition_2 | pvalue | pvalue_adjusted |
|---|---|---|---|---|
| complex_abundance | mitosis | inter | 0.00030967416140721313 | 0.011652071109633347 |
| interactor_abundance | mitosis | inter | 0.0011111765577599547 | 0.024049720606162197 |
| interactor_ratio | mitosis | inter | 0.1581429879180654 | 0.6144687163742579 |

O60832_Q12788
DKC1_HUMAN vs TBL3_HUMAN
p-value: 0.005 q-value: 0.005

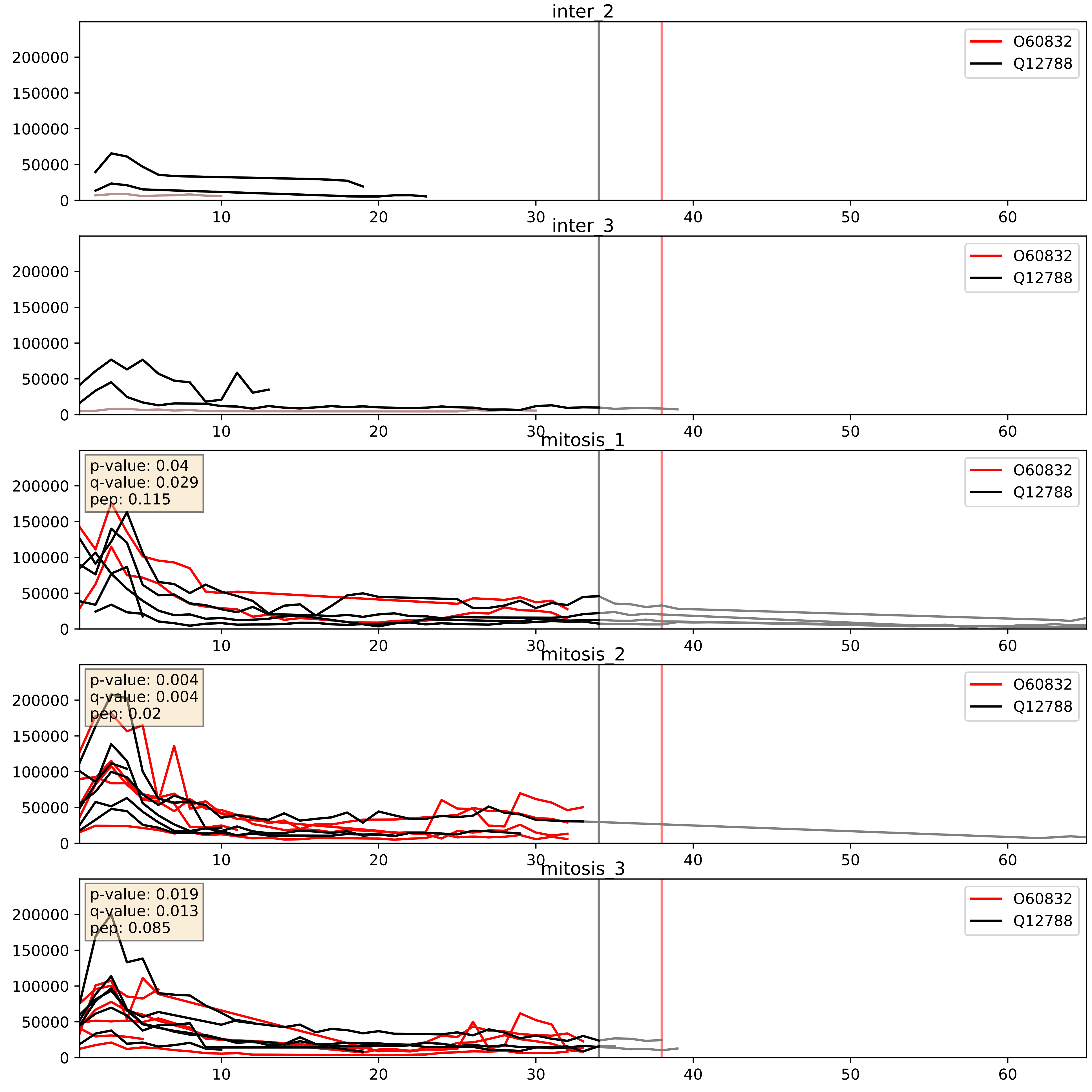


| level | condition_1 | condition_2 | pvalue | pvalue_adjusted |
|---|---|---|---|---|
| complex_abundance | mitosis | inter | 0.0037688012662694115 | 0.04531030735851989 |
| interactor_abundance | mitosis | inter | 0.04824150833940766 | 0.17504902777779313 |
| interactor_ratio | mitosis | inter | 0.4938582007919115 | 0.86984863222988 |

O76021_Q12788
RL1D1_HUMAN vs TBL3_HUMAN
p-value: 0.008 q-value: 0.007

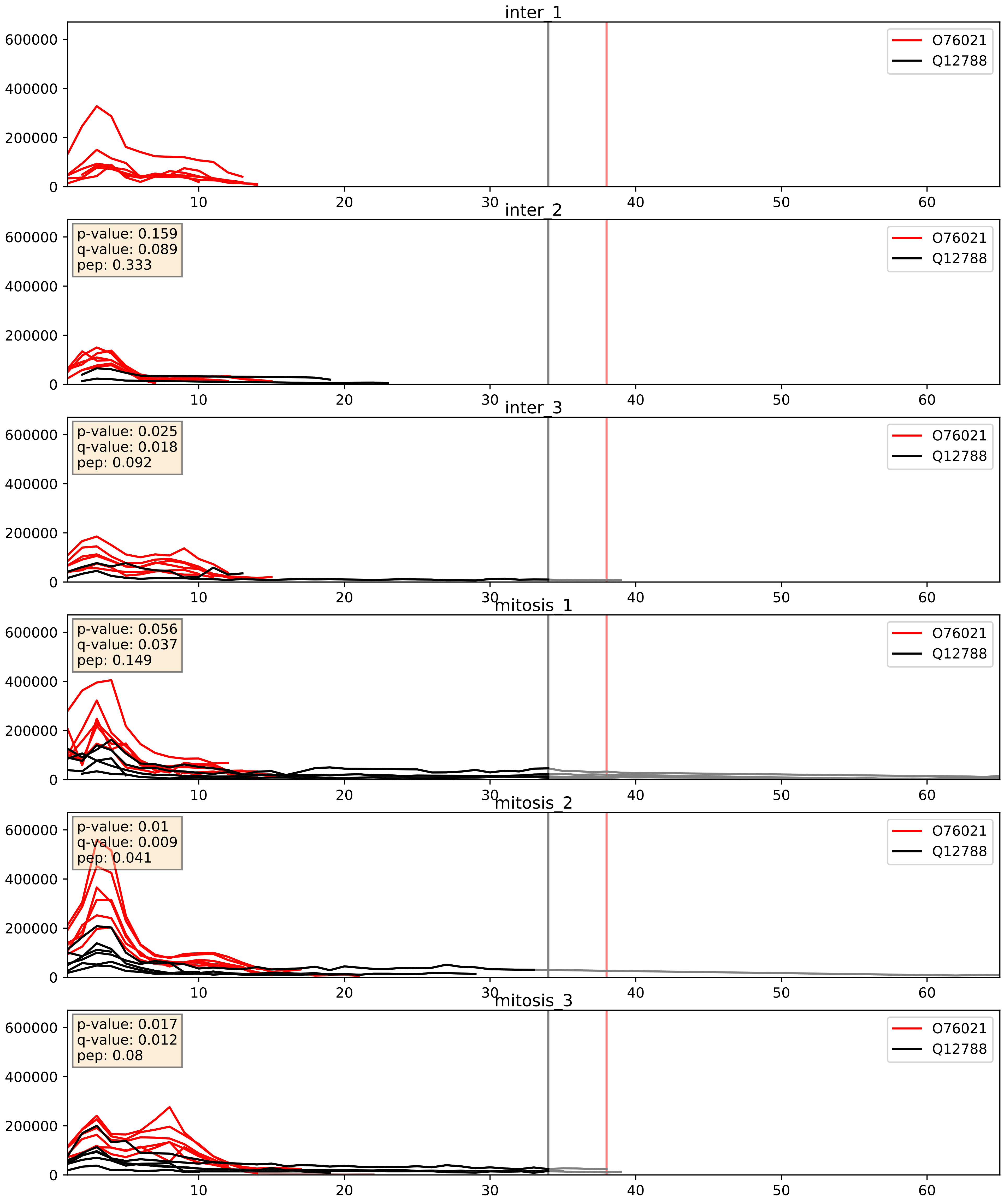


| level | condition_1 | condition_2 | pvalue | pvalue_adjusted |
|---|---|---|---|---|
| complex_abundance | mitosis | inter | 0.028907339226821576 | 0.12366158110024622 |
| interactor_abundance | mitosis | inter | 0.06155864533413795 | 0.2029091986005969 |
| interactor_ratio | mitosis | inter | 0.37222432746257683 | 0.8138388063636502 |

P22087_Q12788
FBRL_HUMAN vs TBL3_HUMAN
p-value: 0.009 q-value: 0.008

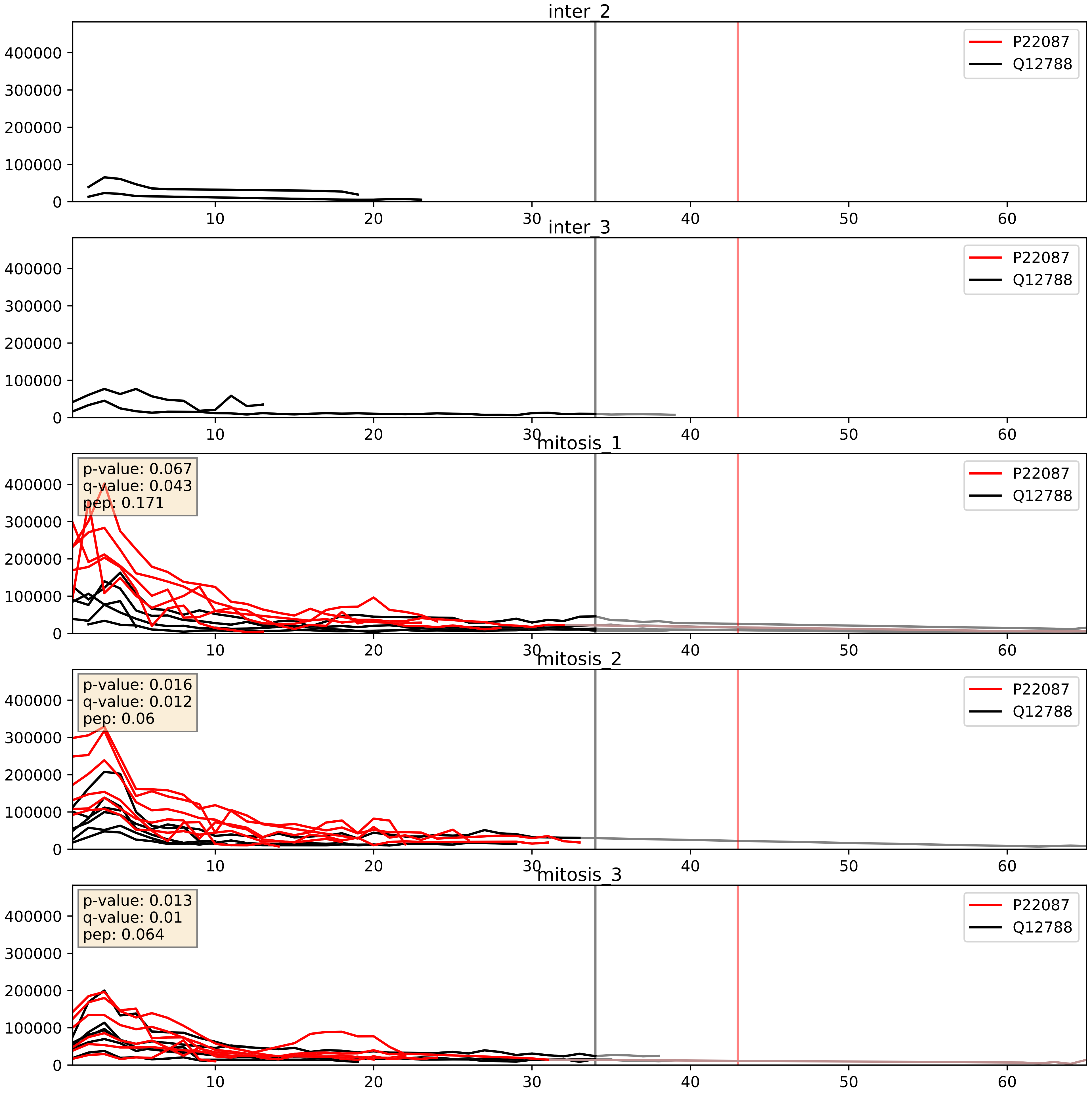


| level | condition_1 | condition_2 | pvalue | pvalue_adjusted |
|---|---|---|---|---|
| complex_abundance | mitosis | inter | 0.0001327659247700999 | 0.007892196639111494 |
| interactor_abundance | mitosis | inter | 0.0003671184791785453 | 0.01356427145975096 |
| interactor_ratio | mitosis | inter | 0.02640428302391608 | 0.25543205403291264 |

P42285_Q12788
MTREX_HUMAN vs TBL3_HUMAN
p-value: 0.029 q-value: 0.021

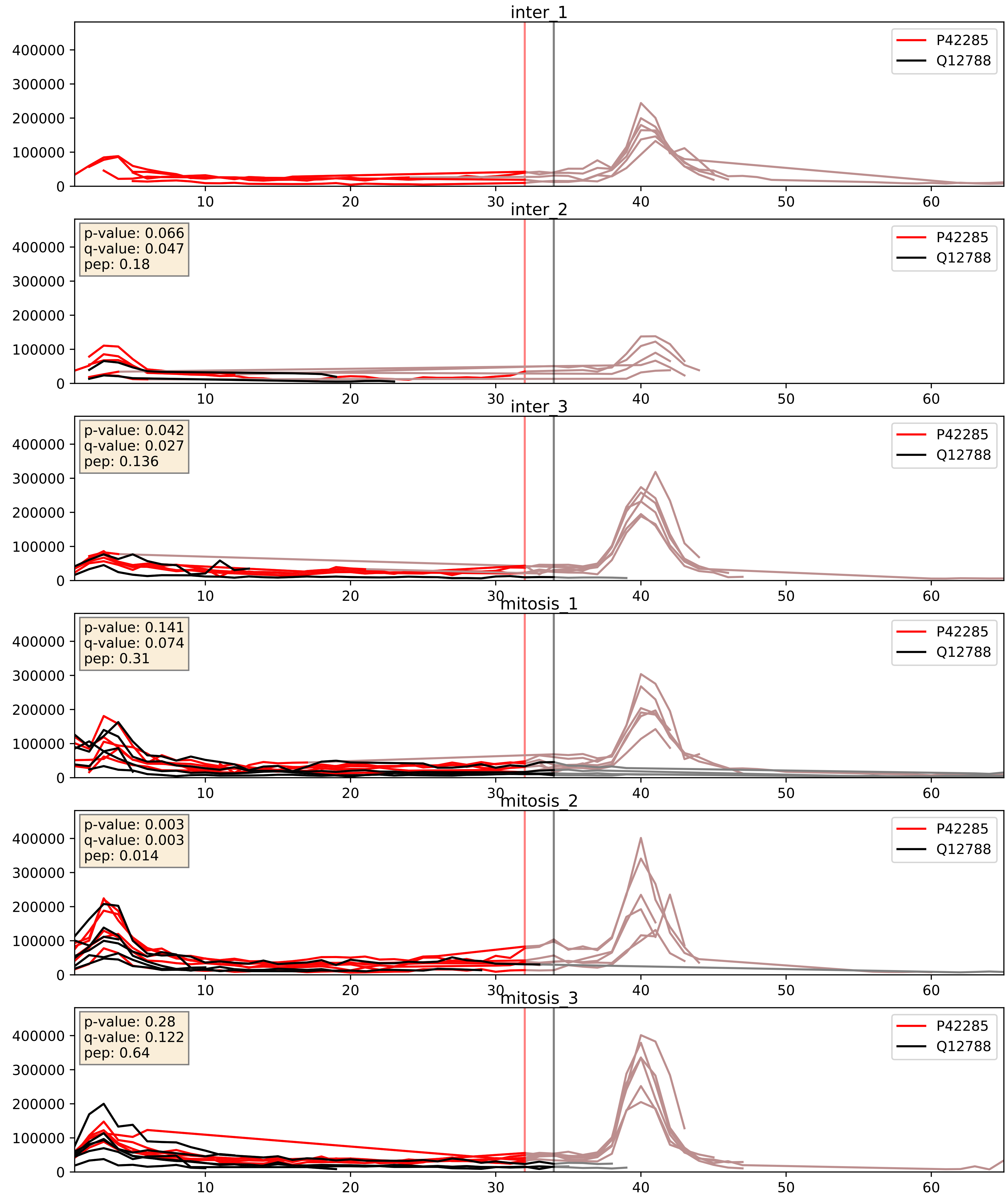


| level | condition_1 | condition_2 | pvalue | pvalue_adjusted |
|---|---|---|---|---|
| complex_abundance | mitosis | inter | 0.01601492374058697 | 0.09447811662262195 |
| interactor_abundance | mitosis | inter | 0.04149089666829527 | 0.1637021393260352 |
| interactor_ratio | mitosis | inter | 0.9340665467242752 | 0.9881103513214468 |

P55769_Q12788
NH2L1_HUMAN vs TBL3_HUMAN
p-value: 0.004 q-value: 0.004

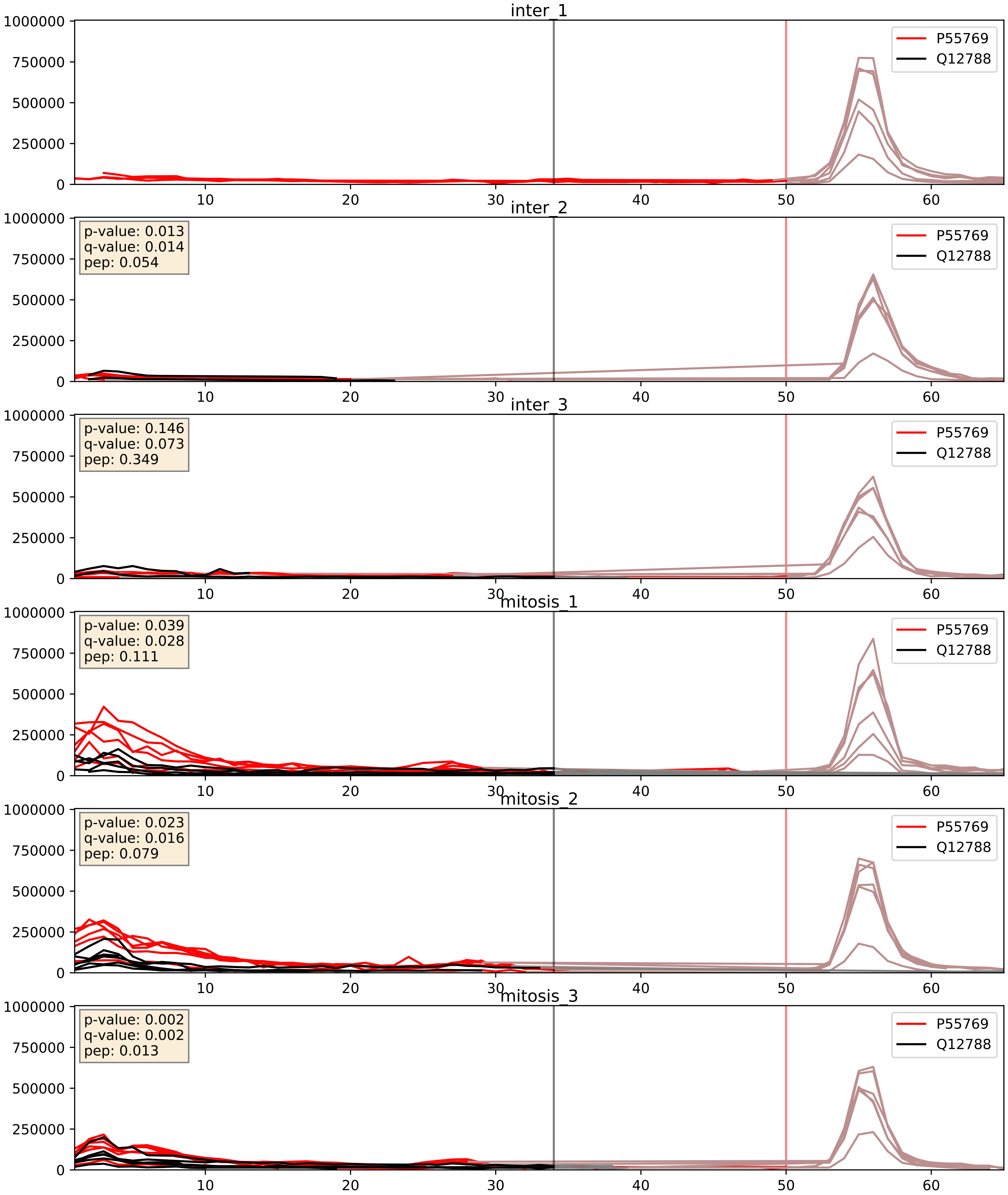


| level | condition_1 | condition_2 | pvalue | pvalue_adjusted |
|---|---|---|---|---|
| complex_abundance | mitosis | inter | 0.002265540023863082 | 0.03487953705803594 |
| interactor_abundance | mitosis | inter | 0.002465726822584136 | 0.037061411436434794 |
| interactor_ratio | mitosis | inter | 0.02210957812286063 | 0.22829672947127502 |

P62277_Q12788
RS13_HUMAN vs TBL3_HUMAN
p-value: 0.076 q-value: 0.046

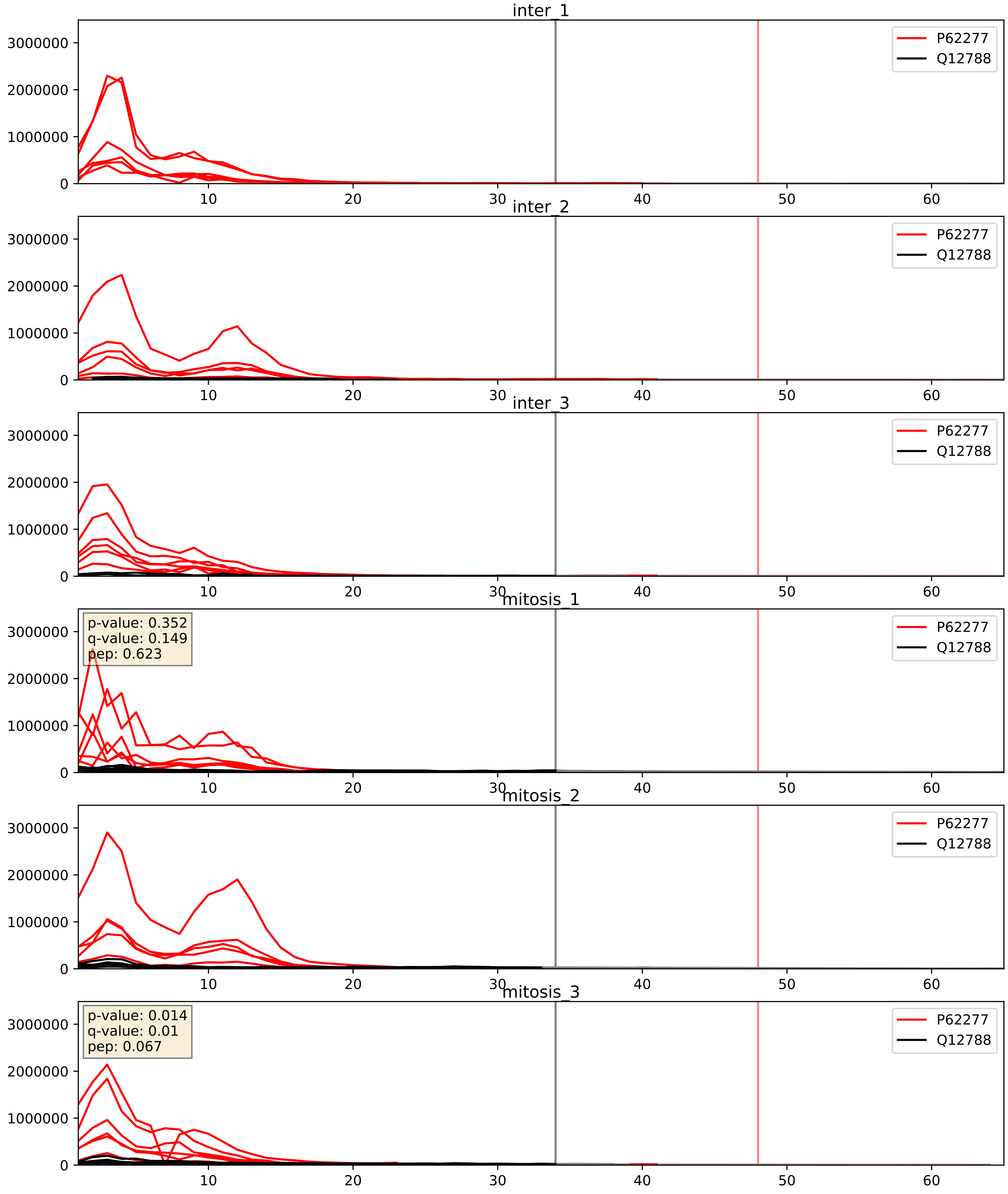


| level | condition_1 | condition_2 | pvalue | pvalue_adjusted |
|---|---|---|---|---|
| complex_abundance | mitosis | inter | 0.12337544066769168 | 0.26563809276703054 |
| interactor_abundance | mitosis | inter | 0.2502285495741613 | 0.44397479207271645 |
| interactor_ratio | mitosis | inter | 0.6398092576189784 | 0.9203357608237295 |

Q12788_Q15061
TBL3_HUMAN vs WDR43_HUMAN
p-value: 0.034 q-value: 0.024

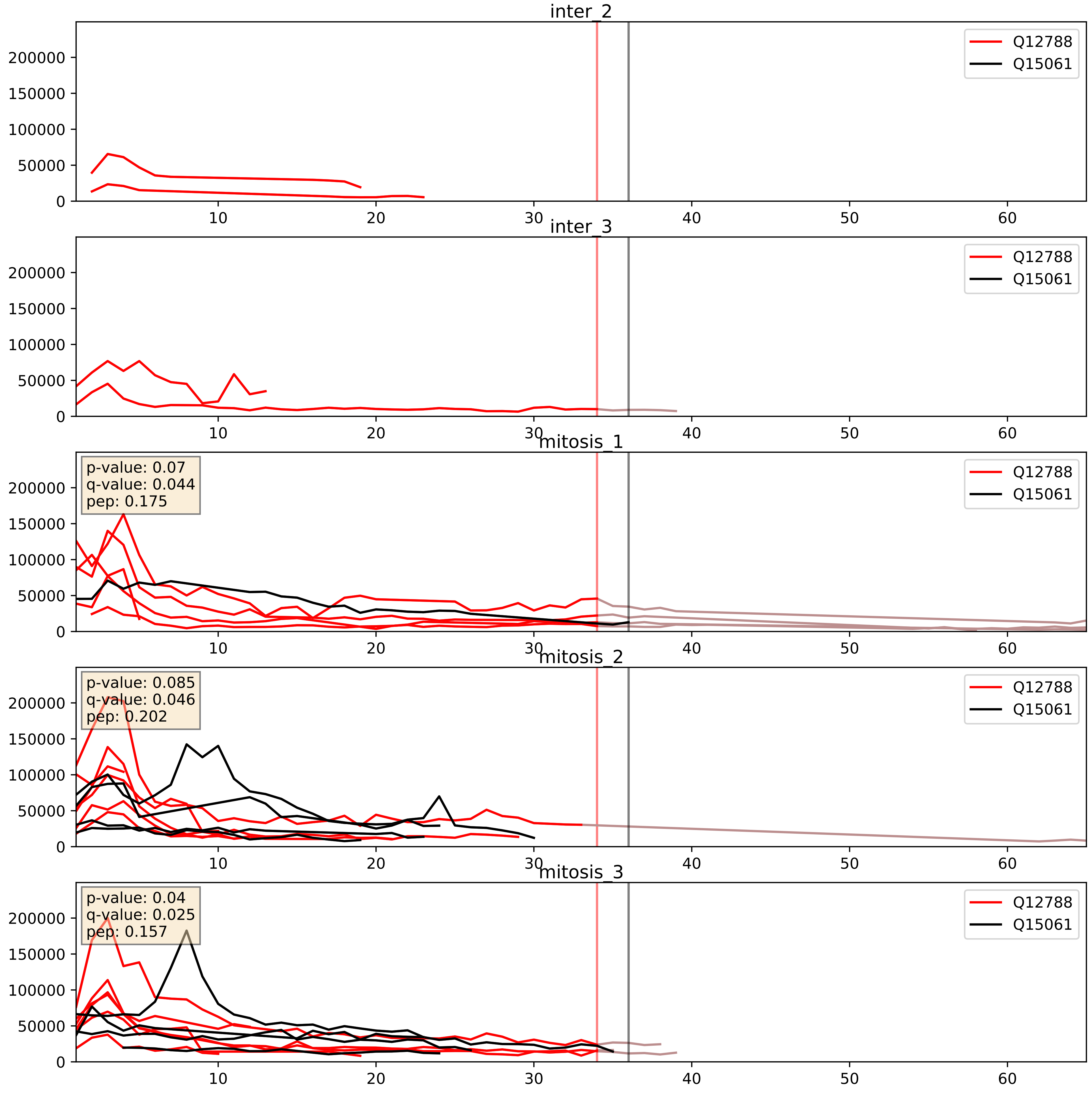


| level | condition_1 | condition_2 | pvalue | pvalue_adjusted |
|---|---|---|---|---|
| interactor_abundance | mitosis | inter | 0.0012685582694706194 | 0.026229127504030197 |
| complex_abundance | mitosis | inter | 0.00806041973070591 | 0.06653038413698939 |
| interactor_ratio | mitosis | inter | 0.23955938278422004 | 0.7181694669961206 |

Q12788_Q15269
TBL3_HUMAN vs PWP2_HUMAN
p-value: 0.015 q-value: 0.012

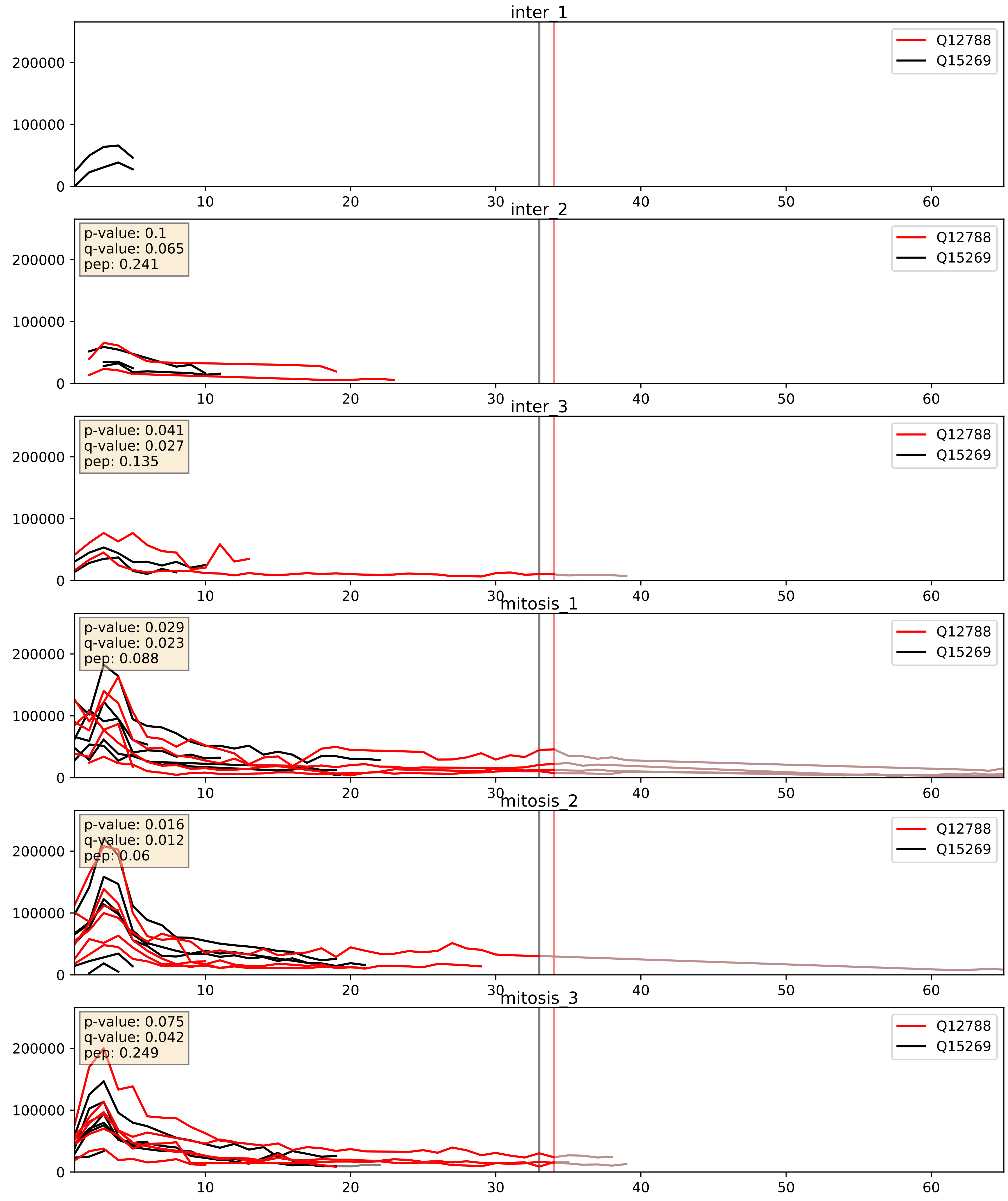


| level | condition_1 | condition_2 | pvalue | pvalue_adjusted |
|---|---|---|---|---|
| interactor_abundance | mitosis | inter | 0.0045441861105829885 | 0.05425137113889872 |
| complex_abundance | mitosis | inter | 0.005891801045557714 | 0.05764342671241339 |
| interactor_ratio | mitosis | inter | 0.6966472160731734 | 0.9356388300120577 |

Q12788_Q8IY81
TBL3_HUMAN vs SPB1_HUMAN
p-value: 0.049 q-value: 0.032

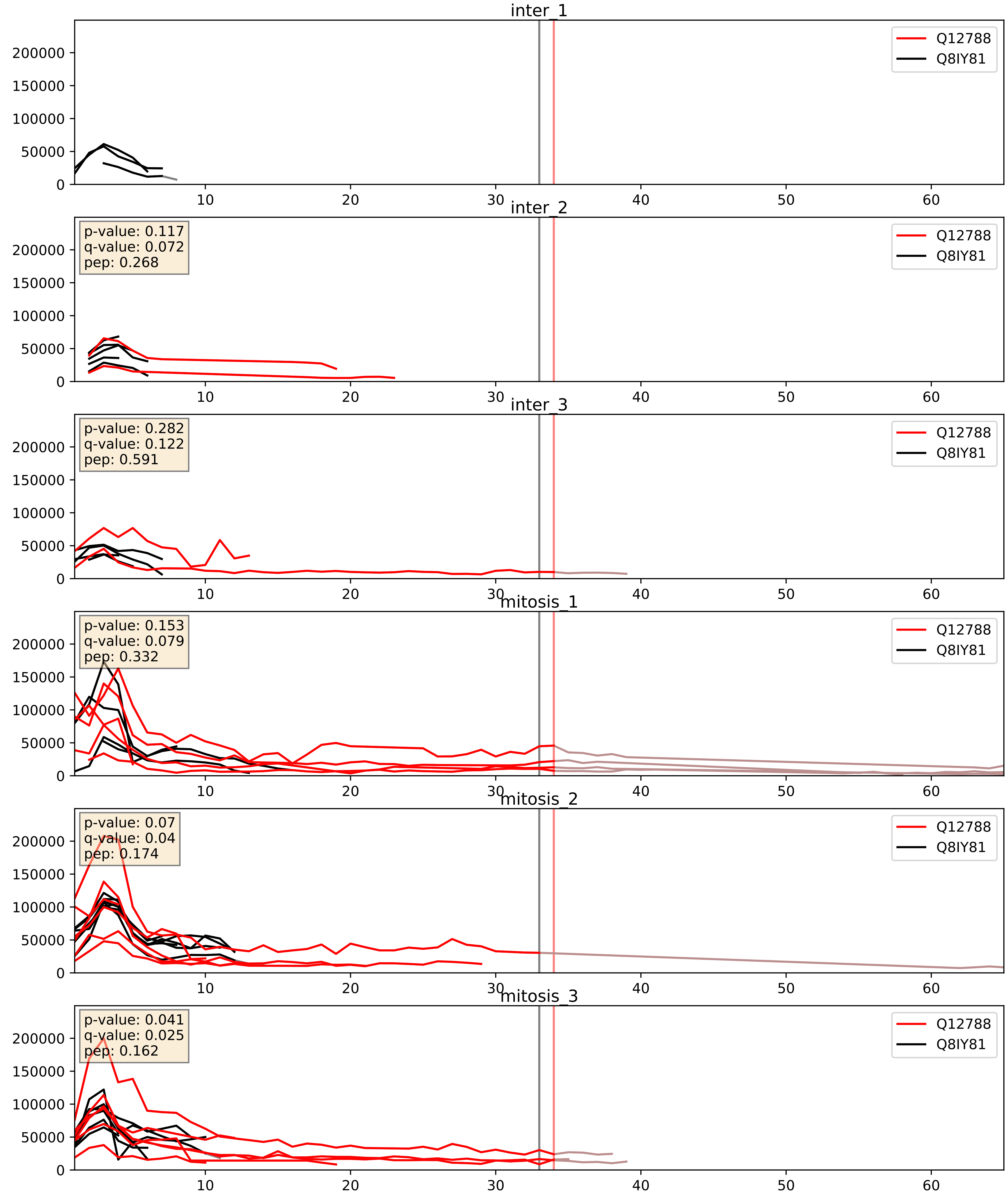


| level | condition_1 | condition_2 | pvalue | pvalue_adjusted |
|---|---|---|---|---|
| interactor_abundance | mitosis | inter | 0.007901002763394643 | 0.07711604472197223 |
| complex_abundance | mitosis | inter | 0.030978129908028652 | 0.127942946637983 |
| interactor_ratio | mitosis | inter | 0.1691647109442906 | 0.6320565072989578 |

Q12788_Q8NI36
TBL3_HUMAN vs WDR36_HUMAN
p-value: 0.0 q-value: 0.001

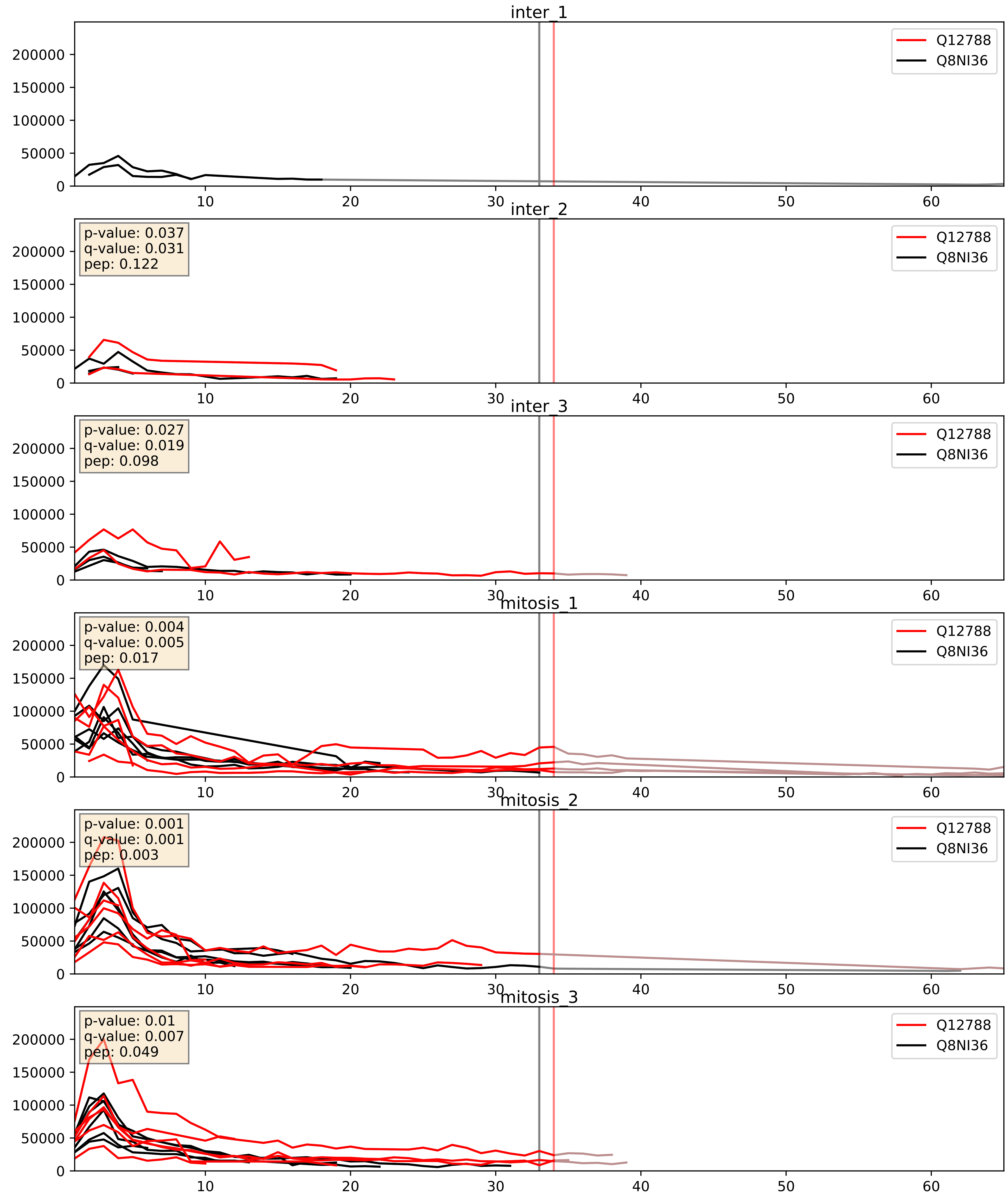


| level | condition_1 | condition_2 | pvalue | pvalue_adjusted |
|---|---|---|---|---|
| complex_abundance | mitosis | inter | 0.003360478542340517 | 0.04258563238370332 |
| interactor_abundance | mitosis | inter | 0.006662312924366897 | 0.06988896891247627 |
| interactor_ratio | mitosis | inter | 0.3561802533008551 | 0.8030936685700529 |

Q12788_Q9H0A0
TBL3_HUMAN vs NAT10_HUMAN
p-value: 0.028 q-value: 0.02

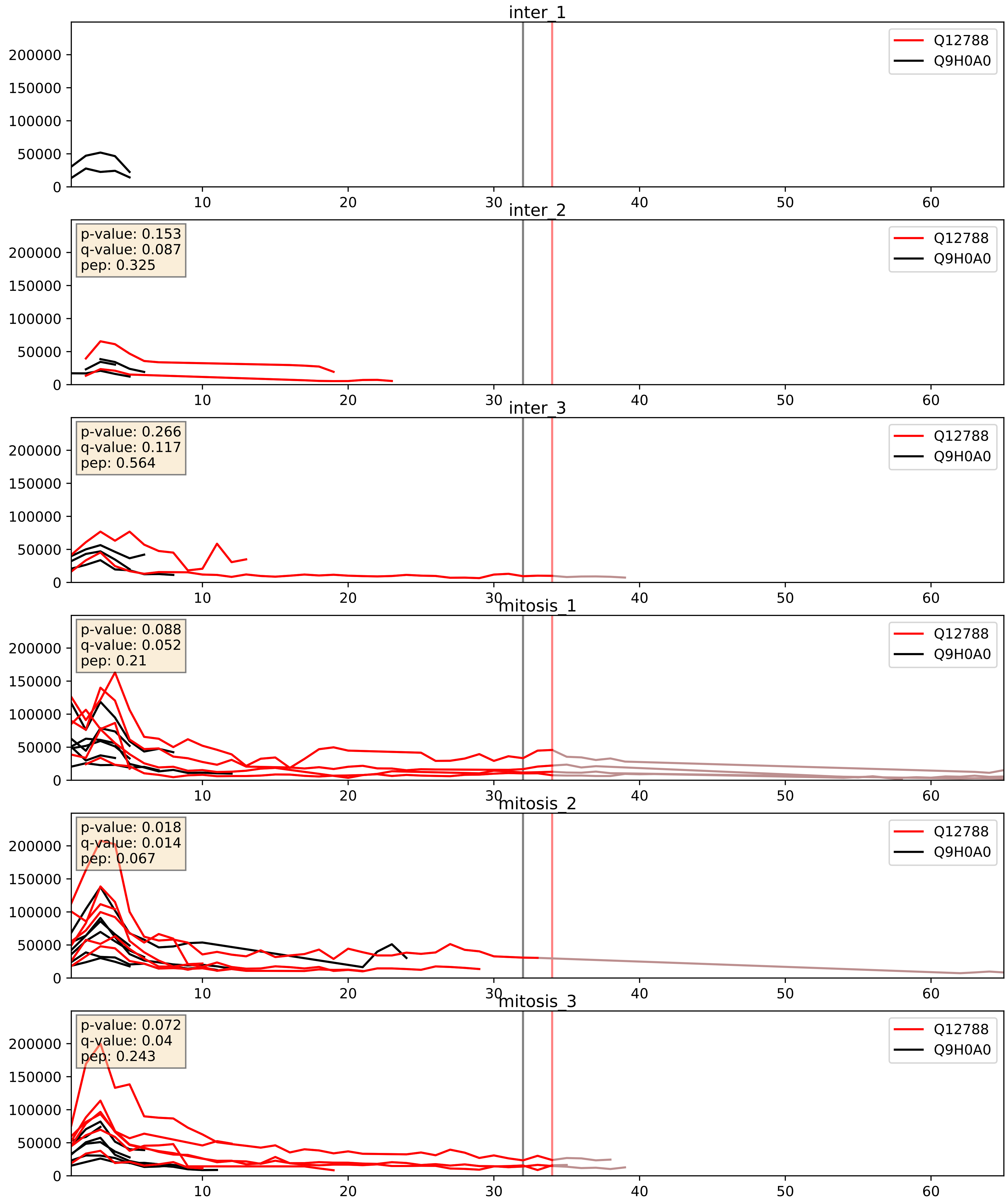


| level | condition_1 | condition_2 | pvalue | pvalue_adjusted |
|---|---|---|---|---|
| interactor_abundance | mitosis | inter | 0.005345390931153837 | 0.061666504542691174 |
| interactor_ratio | mitosis | inter | 0.04444676005266599 | 0.3493703085866124 |
| complex_abundance | mitosis | inter | 0.05040593282045461 | 0.15980547590484867 |

Q12788_Q9H0S4
TBL3_HUMAN vs DDX47_HUMAN
p-value: 0.041 q-value: 0.028

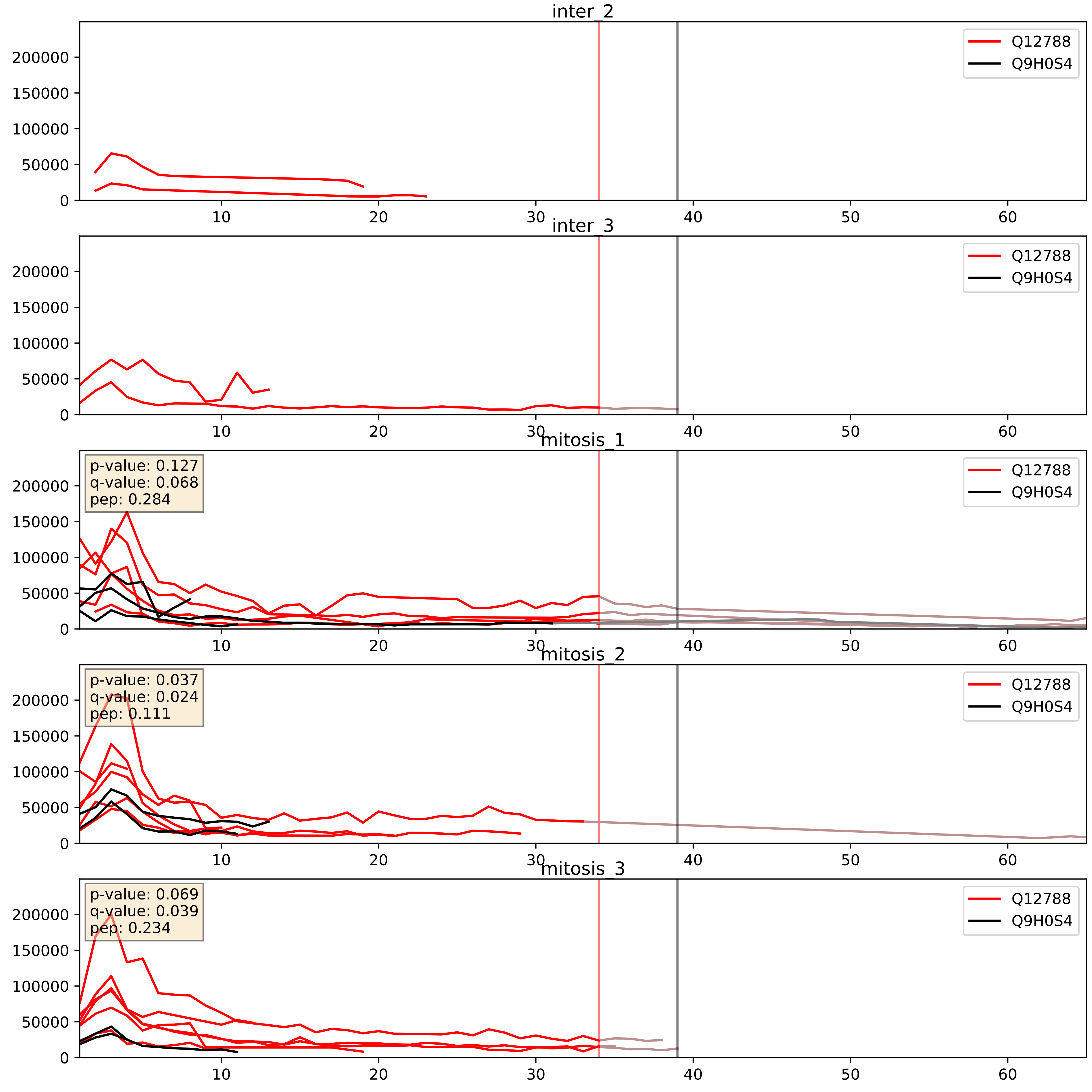


| level | condition_1 | condition_2 | pvalue | pvalue_adjusted |
|---|---|---|---|---|
| interactor_abundance | mitosis | inter | 0.0015075846392158566 | 0.02867761002597274 |
| interactor_ratio | mitosis | inter | 0.01308259178830462 | 0.16493001984594508 |
| complex_abundance | mitosis | inter | 0.02359161739620337 | 0.11181851877713225 |

Q12788_Q9H583
TBL3_HUMAN vs HEAT1_HUMAN
p-value: 0.023 q-value: 0.017

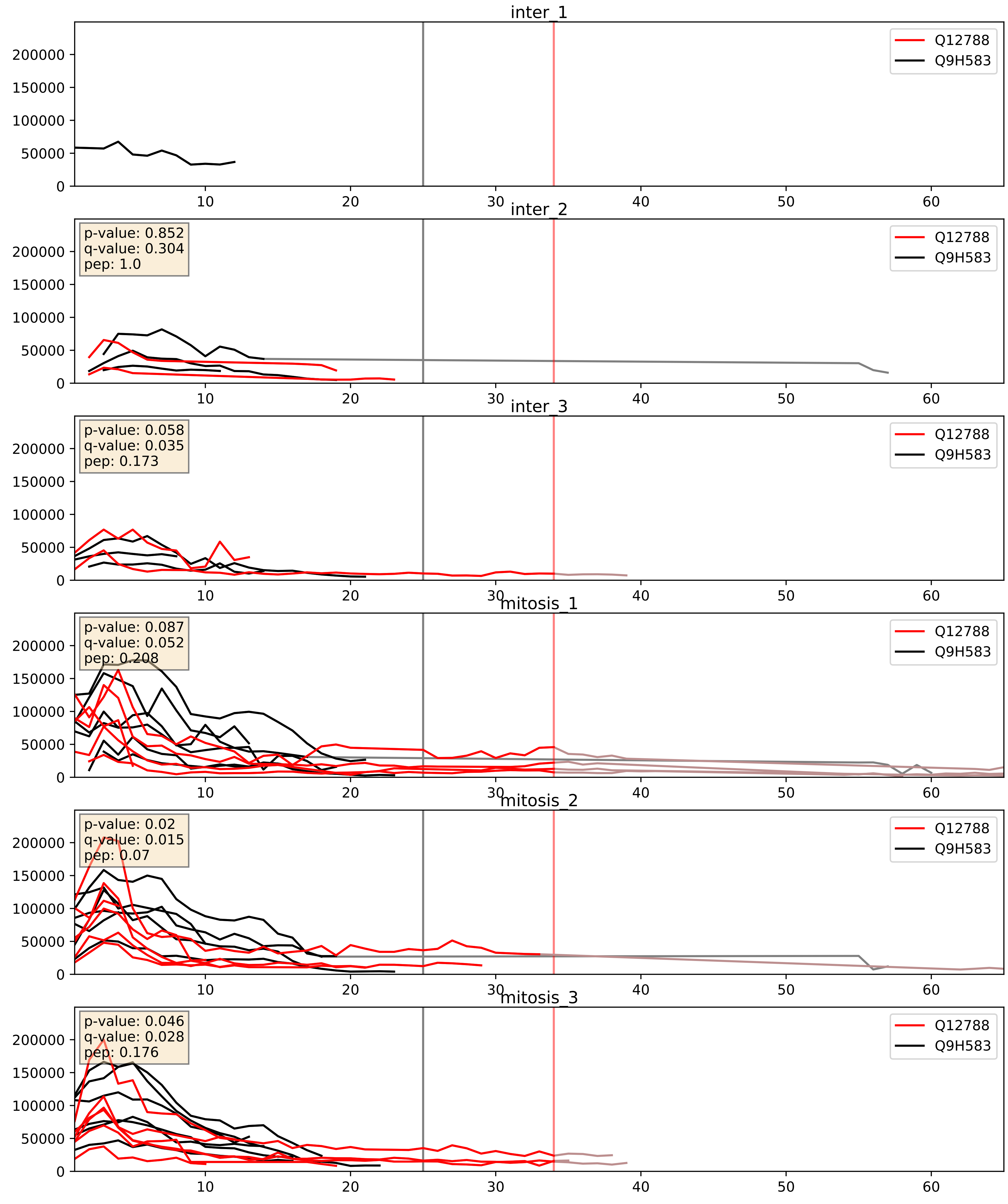


| level | condition_1 | condition_2 | pvalue | pvalue_adjusted |
|---|---|---|---|---|
| interactor_ratio | mitosis | inter | 0.0032358533534543603 | 0.07405554735675686 |
| complex_abundance | mitosis | inter | 0.004224521690450446 | 0.048345498563686945 |
| interactor_abundance | mitosis | inter | 0.009154209095090269 | 0.08324021891678159 |

Q12788_Q9NVP1
TBL3_HUMAN vs DDX18_HUMAN
p-value: 0.08 q-value: 0.047

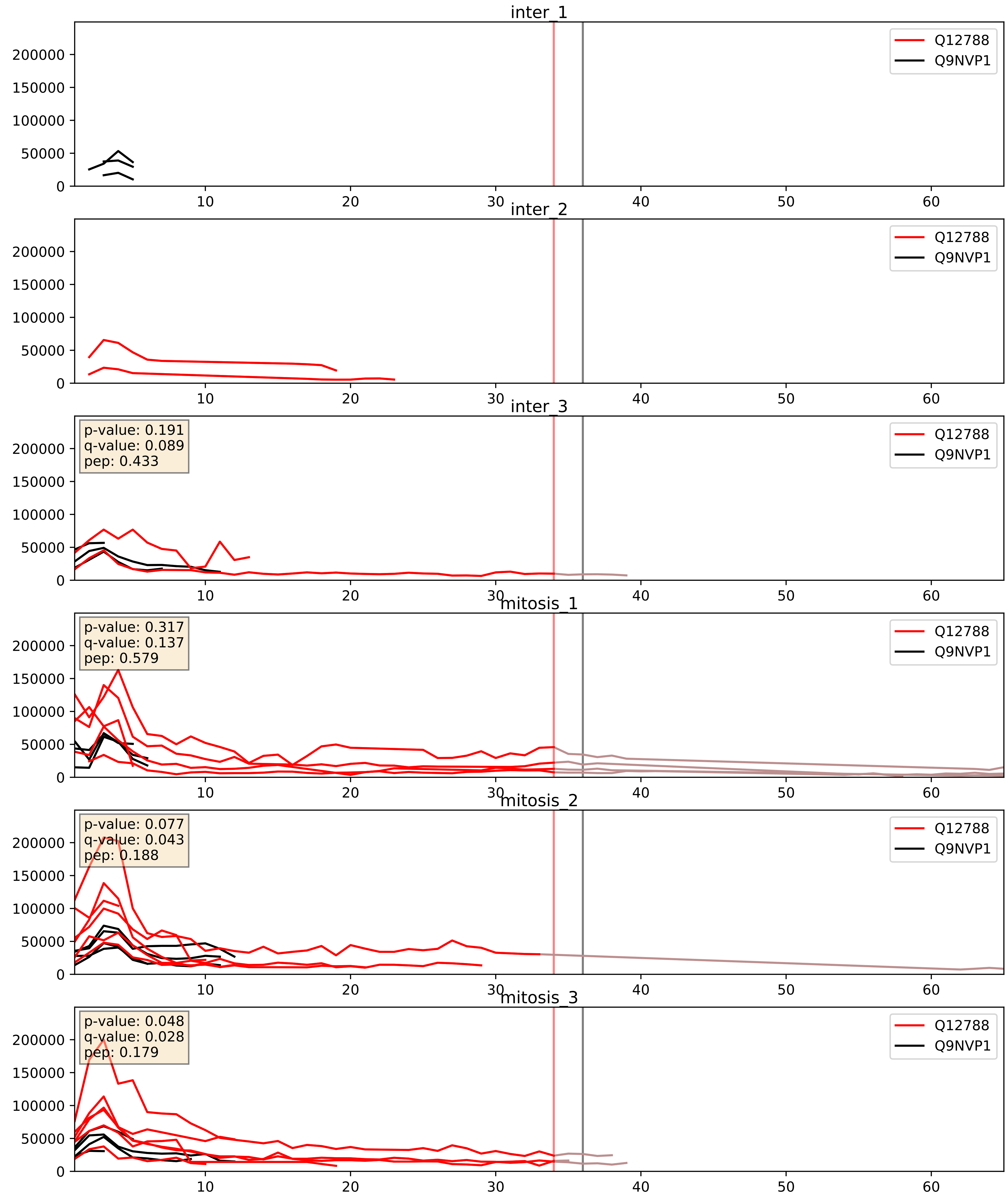


| level | condition_1 | condition_2 | pvalue | pvalue_adjusted |
|---|---|---|---|---|
| interactor_abundance | mitosis | inter | 0.007893035270042996 | 0.07711604472197223 |
| complex_abundance | mitosis | inter | 0.022357984284711807 | 0.10987189501272765 |
| interactor_ratio | mitosis | inter | 0.02782927283201316 | 0.2618295178868221 |

Q12788_Q9UNX4
TBL3_HUMAN vs WDR3_HUMAN
p-value: 0.052 q-value: 0.034

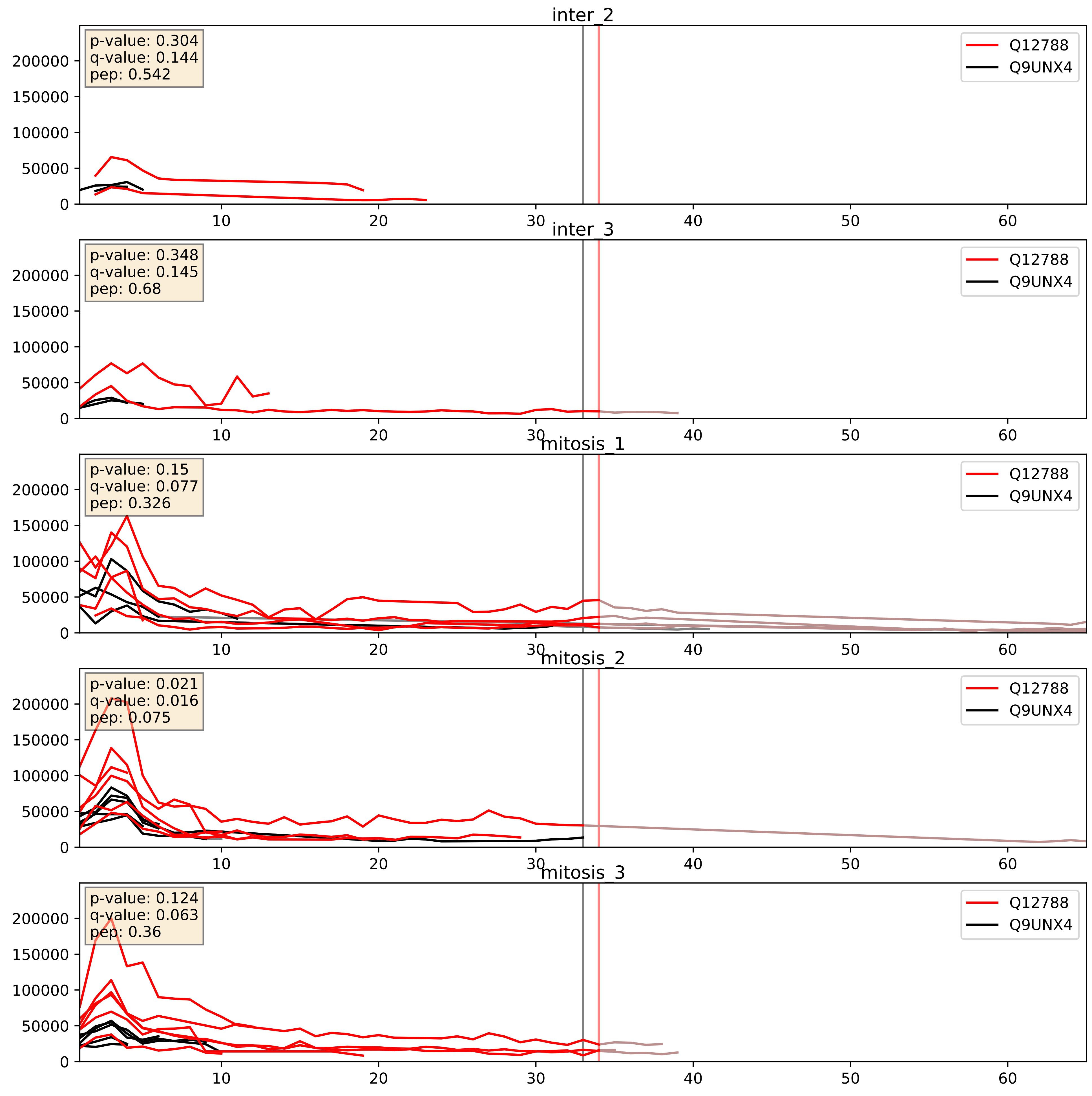


| level | condition_1 | condition_2 | pvalue | pvalue_adjusted |
|---|---|---|---|---|
| interactor_abundance | mitosis | inter | 0.005053479358839491 | 0.05906252380726447 |
| interactor_ratio | mitosis | inter | 0.006624735222091087 | 0.11281850825965009 |
| complex_abundance | mitosis | inter | 0.02123880532057976 | 0.10740063977781633 |

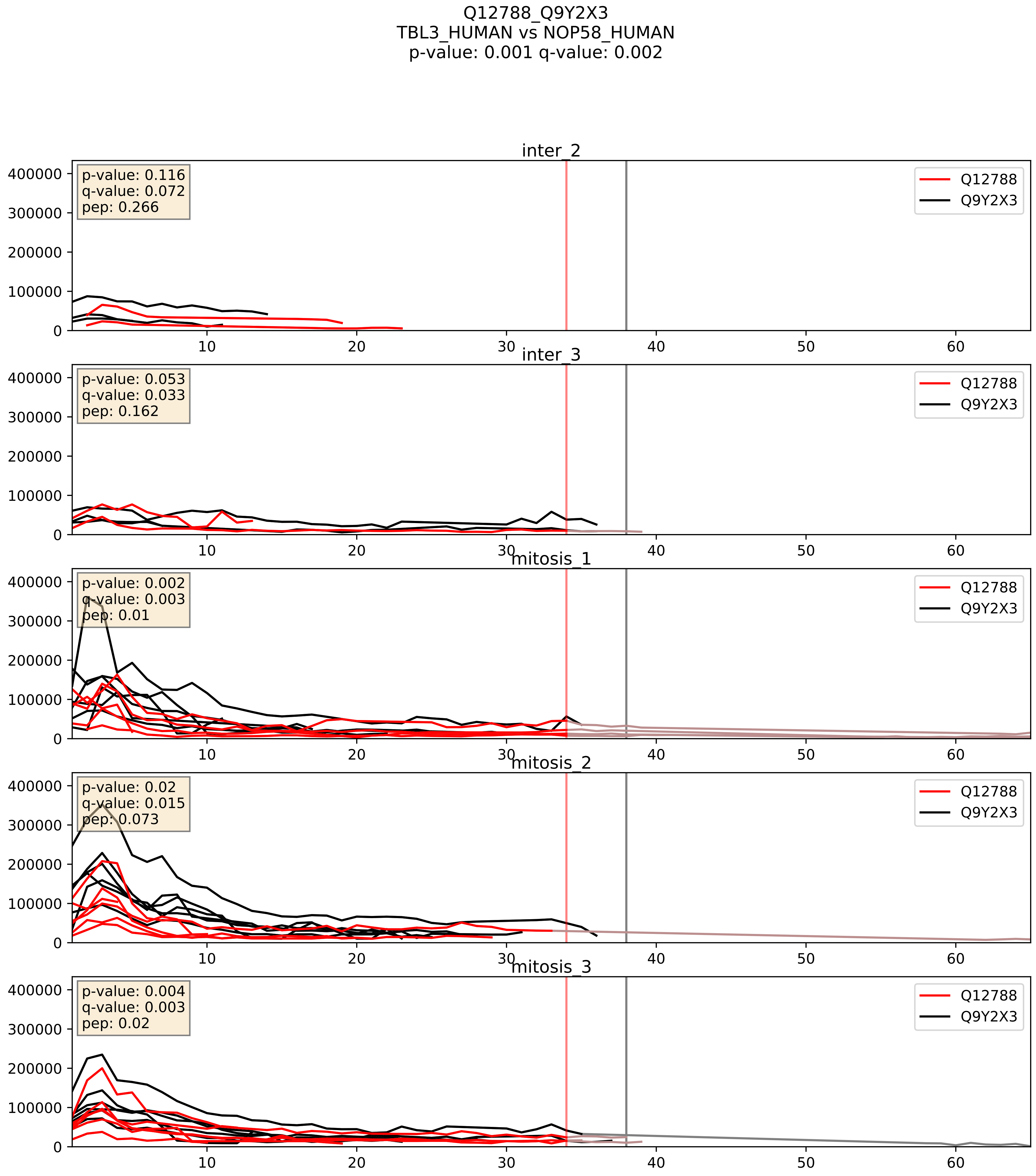


| level | condition_1 | condition_2 | pvalue | pvalue_adjusted |
|---|---|---|---|---|
| complex_abundance | mitosis | inter | 0.002118335372975017 | 0.033489340249041655 |
| interactor_abundance | mitosis | inter | 0.005521190842296777 | 0.06317594435380872 |
| interactor_ratio | mitosis | inter | 0.029449555755858125 | 0.27164676429972584 |

Q12788_Q9Y3T9
TBL3_HUMAN vs NOC2L_HUMAN
p-value: 0.028 q-value: 0.02

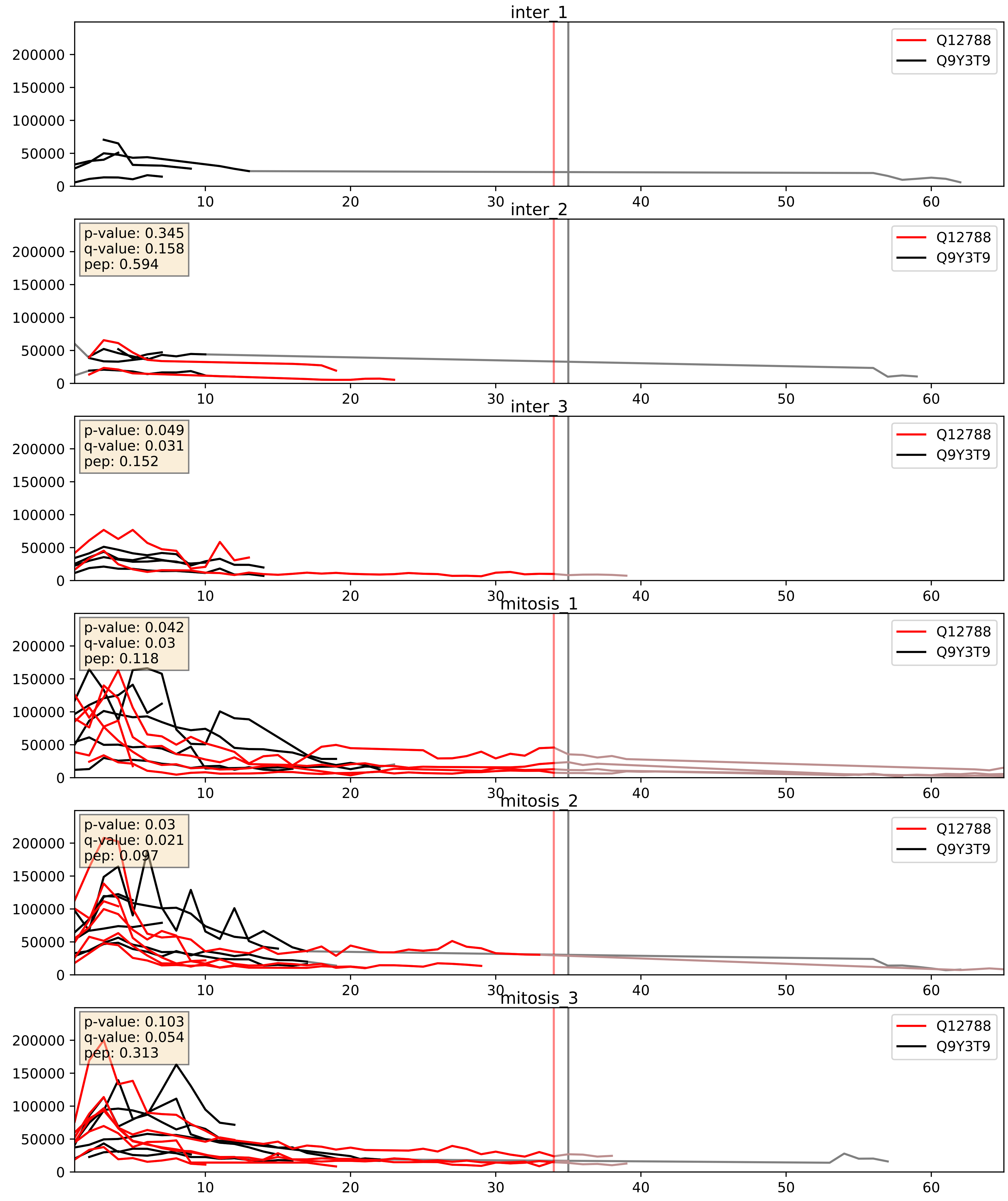


| level | condition_1 | condition_2 | pvalue | pvalue_adjusted |
|---|---|---|---|---|
| complex_abundance | mitosis | inter | 0.005487193845178426 | 0.05591711823181825 |
| interactor_abundance | mitosis | inter | 0.011367688817575375 | 0.09419885409336419 |
| interactor_ratio | mitosis | inter | 0.13663518010683853 | 0.58714715949525 |
