## Supplementary Data 2 for "SECAT: Quantifying differential protein-protein interaction states by network-centric analysis": hela_string_000119_O75643.pdf

# O75643_O75643

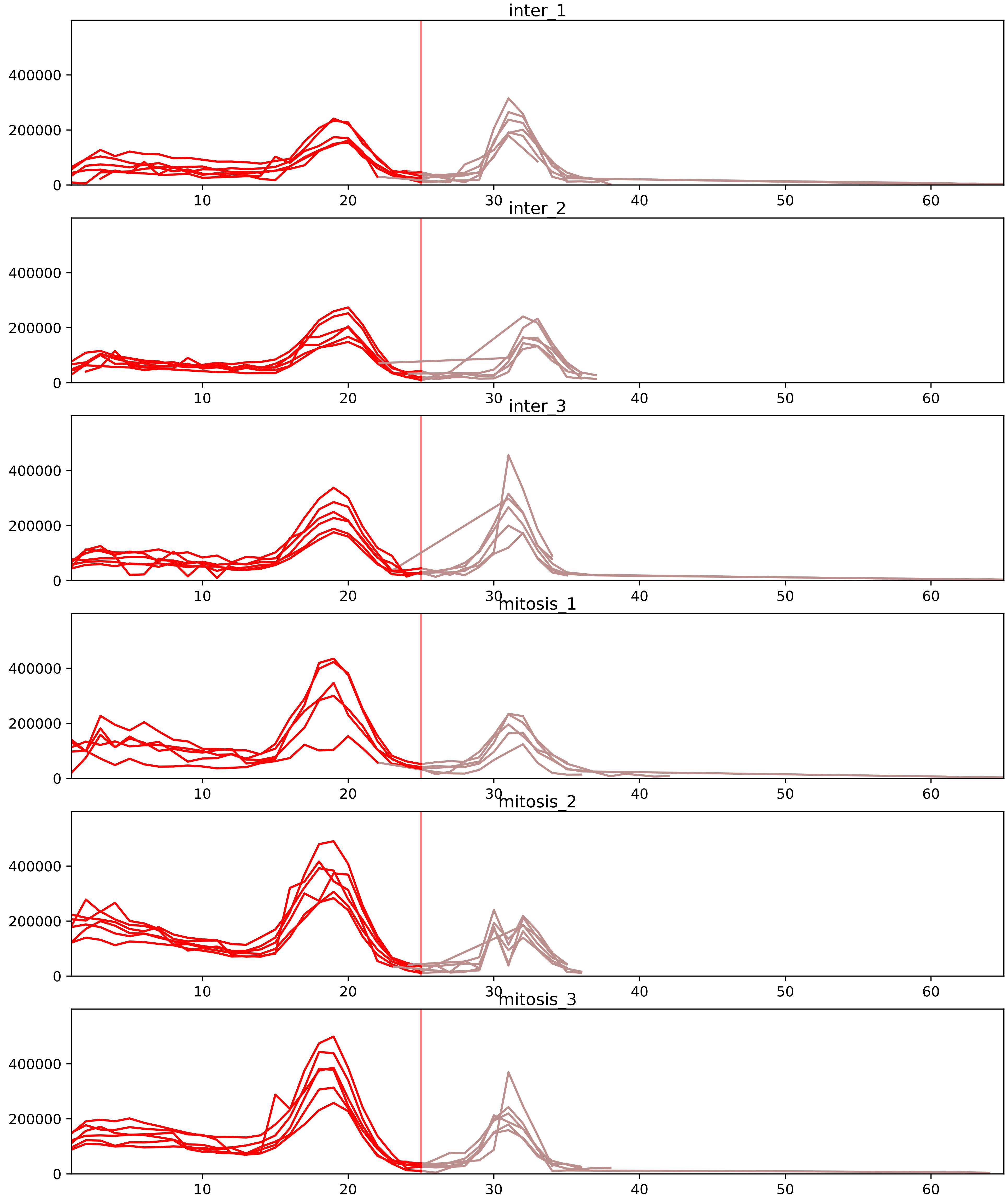


| level | condition_1 | condition_2 | pvalue | pvalue_adjusted |
|---|---|---|---|---|
| interactor_abundance | mitosis | inter | 2.2564549823031396e-05 | 0.0009025819929212558 |
| complex_abundance | mitosis | inter | 3.219926120733498e-05 | 0.0010971600115091918 |
| total_abundance | mitosis | inter | 0.0003314221524676441 | 0.023797906320223484 |
| assembled_abundance | mitosis | inter | 0.0004946368898010256 | 0.020328788531338803 |
| interactor_ratio | mitosis | inter | 0.03264918383318886 | 0.15170327841683715 |
| monomer_abundance | mitosis | inter | 0.9504802332368698 | 0.9904593966764303 |

O75643_Q15029
U520_HUMAN vs U5S1_HUMAN
p-value: 0.008 q-value: 0.007

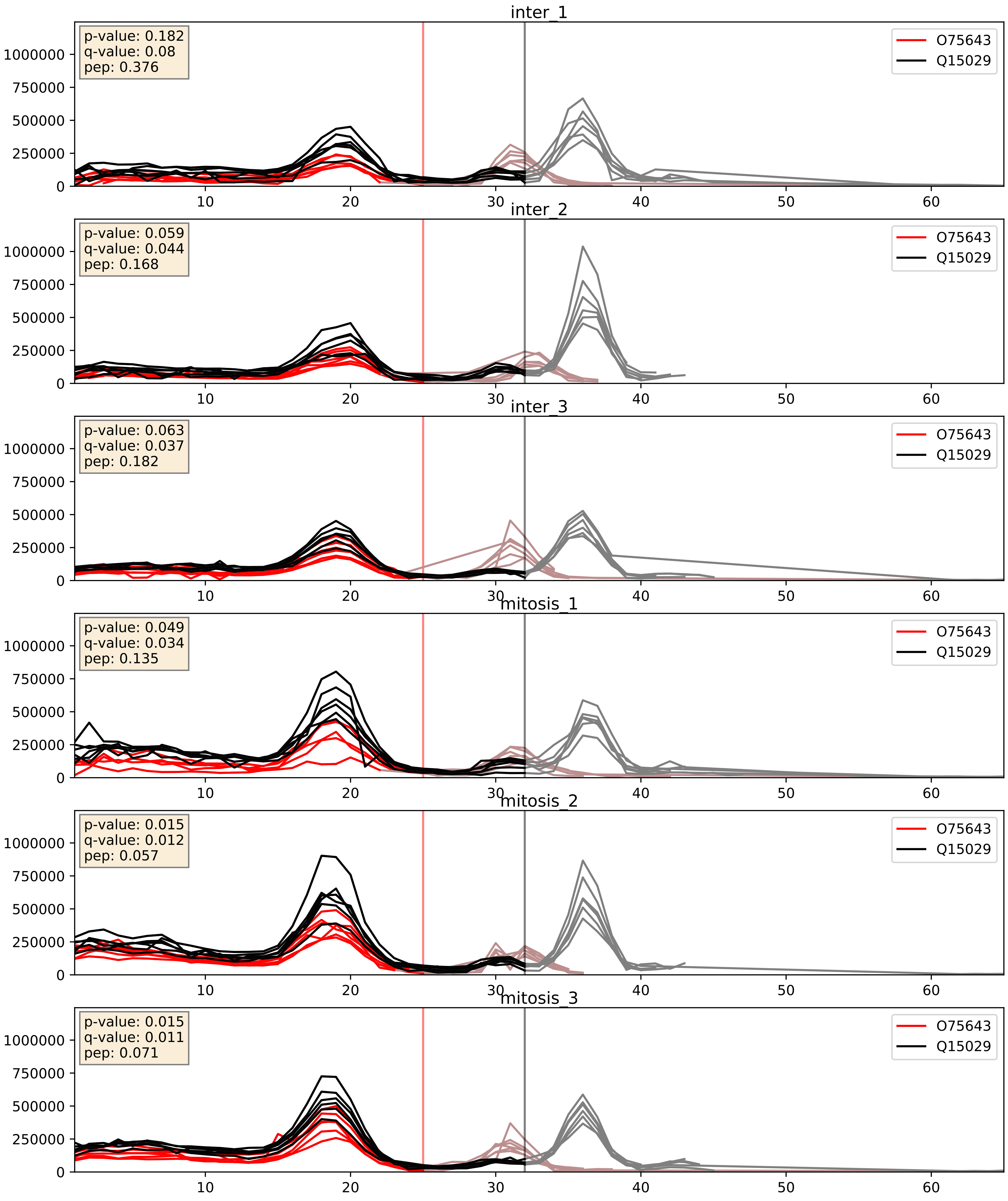


| level | condition_1 | condition_2 | pvalue | pvalue_adjusted |
|---|---|---|---|---|
| interactor_abundance | mitosis | inter | 0.00025670893364430084 | 0.012207935955528973 |
| complex_abundance | mitosis | inter | 0.0005272663919559008 | 0.014606473511788058 |
| interactor_ratio | mitosis | inter | 0.11708918598701334 | 0.5635744908591998 |

O75643_Q6P2Q9
U520_HUMAN vs PRP8_HUMAN
p-value: 0.005 q-value: 0.005

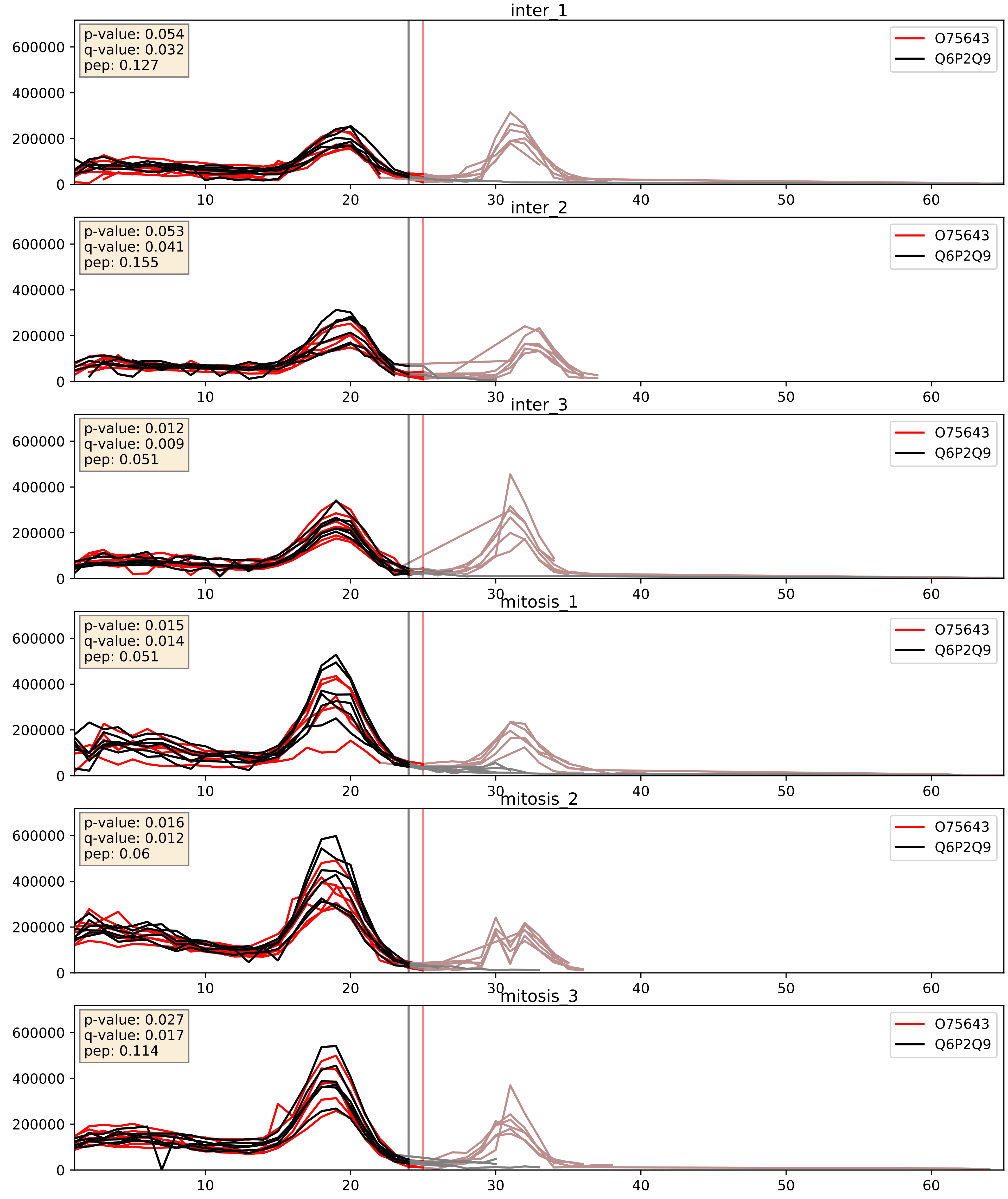


| level | condition_1 | condition_2 | pvalue | pvalue_adjusted |
|---|---|---|---|---|
| complex_abundance | mitosis | inter | 6.70384363368097e-05 | 0.005691857151652958 |
| interactor_abundance | mitosis | inter | 0.00020559566637153397 | 0.01080585011527384 |
| interactor_ratio | mitosis | inter | 0.6976944764935629 | 0.9359699707113329 |

O75643_Q9BZJ0
U520_HUMAN vs CRNL1_HUMAN
p-value: 0.034 q-value: 0.024

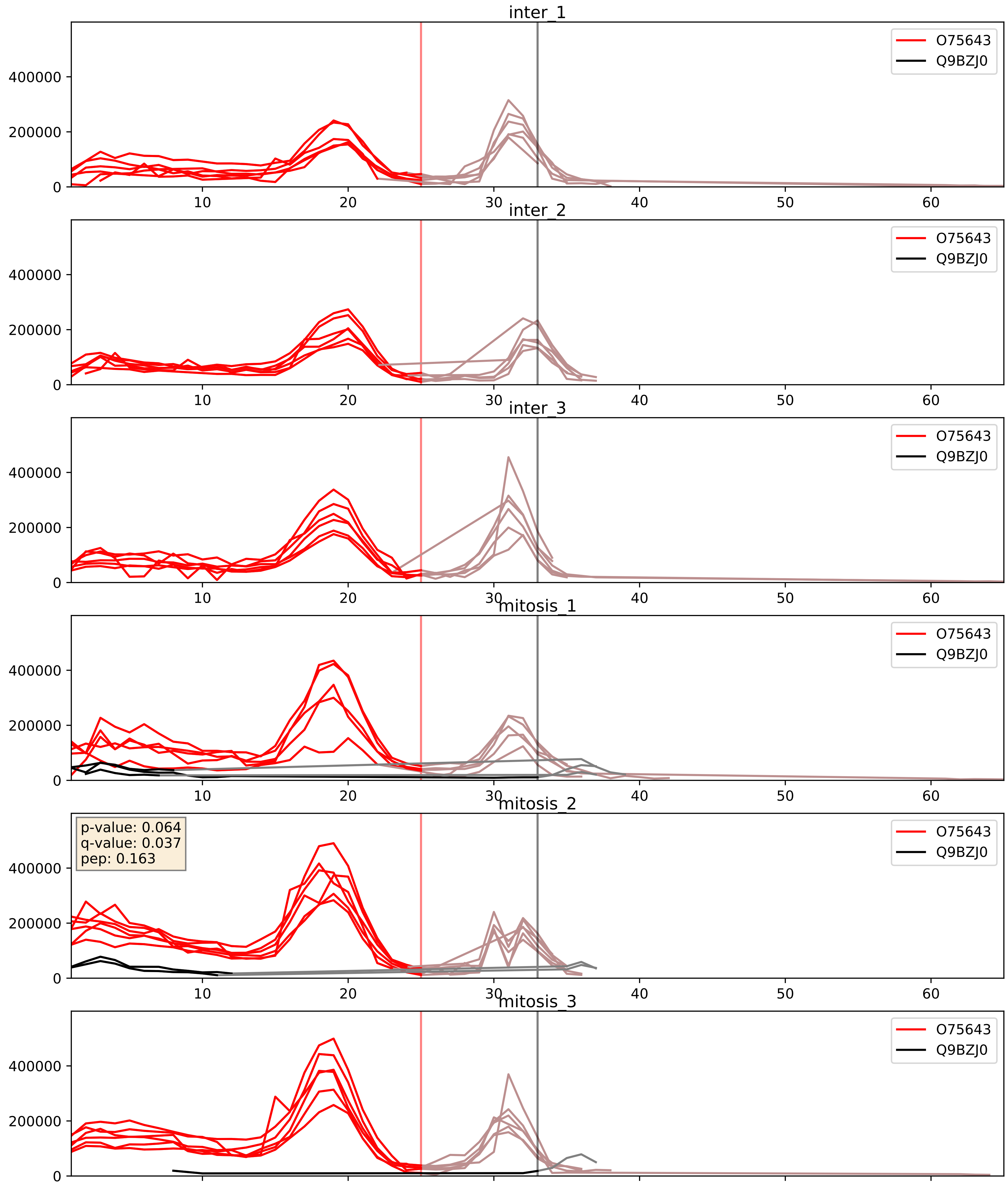


| level | condition_1 | condition_2 | pvalue | pvalue_adjusted |
|---|---|---|---|---|
| interactor_ratio | mitosis | inter | 0.004715096858563263 | 0.09159088101537749 |
| interactor_abundance | mitosis | inter | 0.030570403815191056 | 0.14519524113741764 |
| complex_abundance | mitosis | inter | 0.10340924979824517 | 0.23908191575103507 |
