## Supplementary Data 2 for "SECAT: Quantifying differential protein-protein interaction states by network-centric analysis": hela_string_000120_Q8NC56.pdf

Q8NC56_Q8NC56

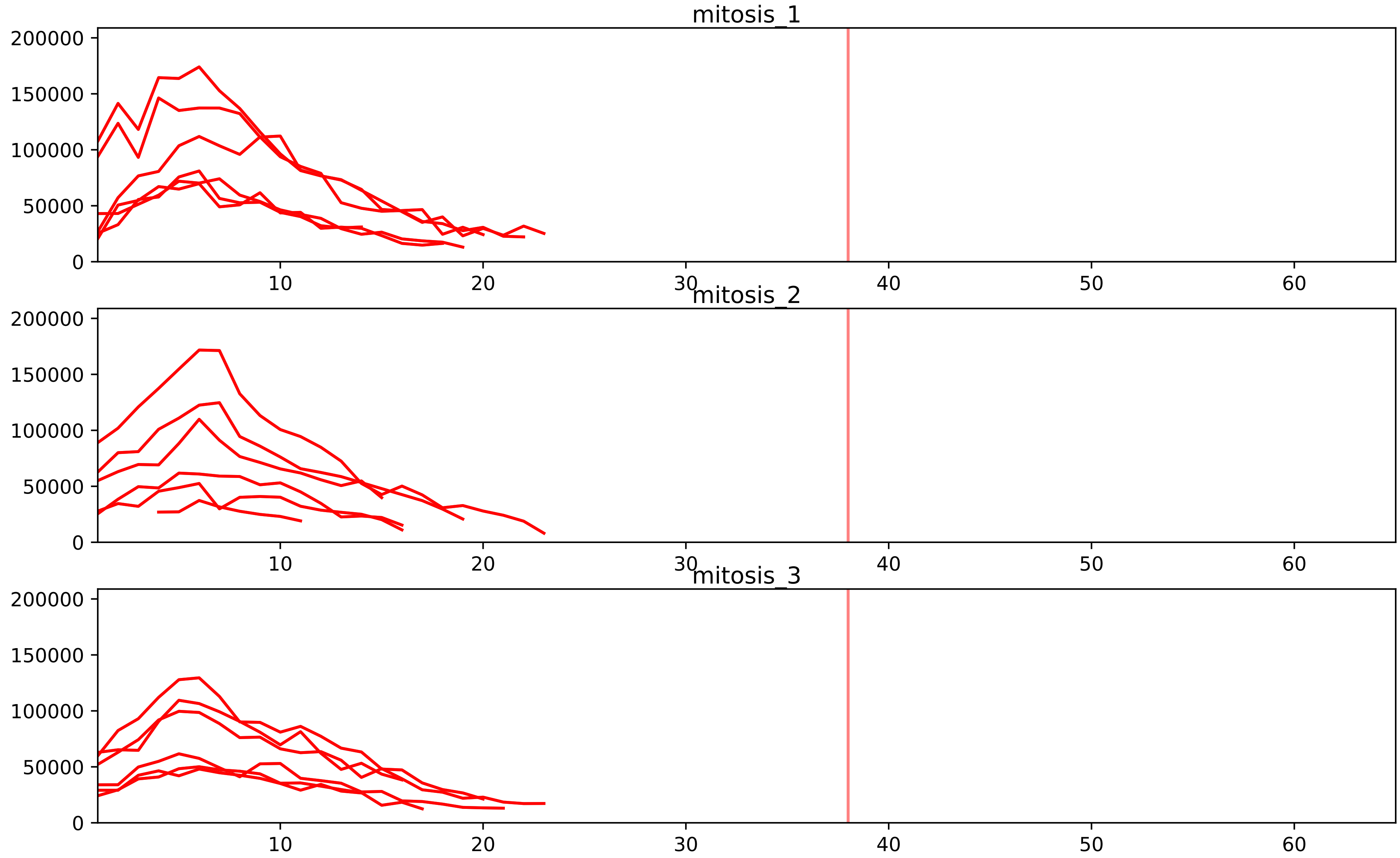


| level | condition_1 | condition_2 | pvalue | pvalue_adjusted |
|---|---|---|---|---|
| assembled_abundance | mitosis | inter | 2.272991455267281e-05 | 0.0059110405567256346 |
| total_abundance | mitosis | inter | 3.406740942241302e-05 | 0.007226549223729363 |
