## Supplementary Data 2 for "SECAT: Quantifying differential protein-protein interaction states by network-centric analysis": hela_string_000121_Q05519.pdf

Q05519_Q05519

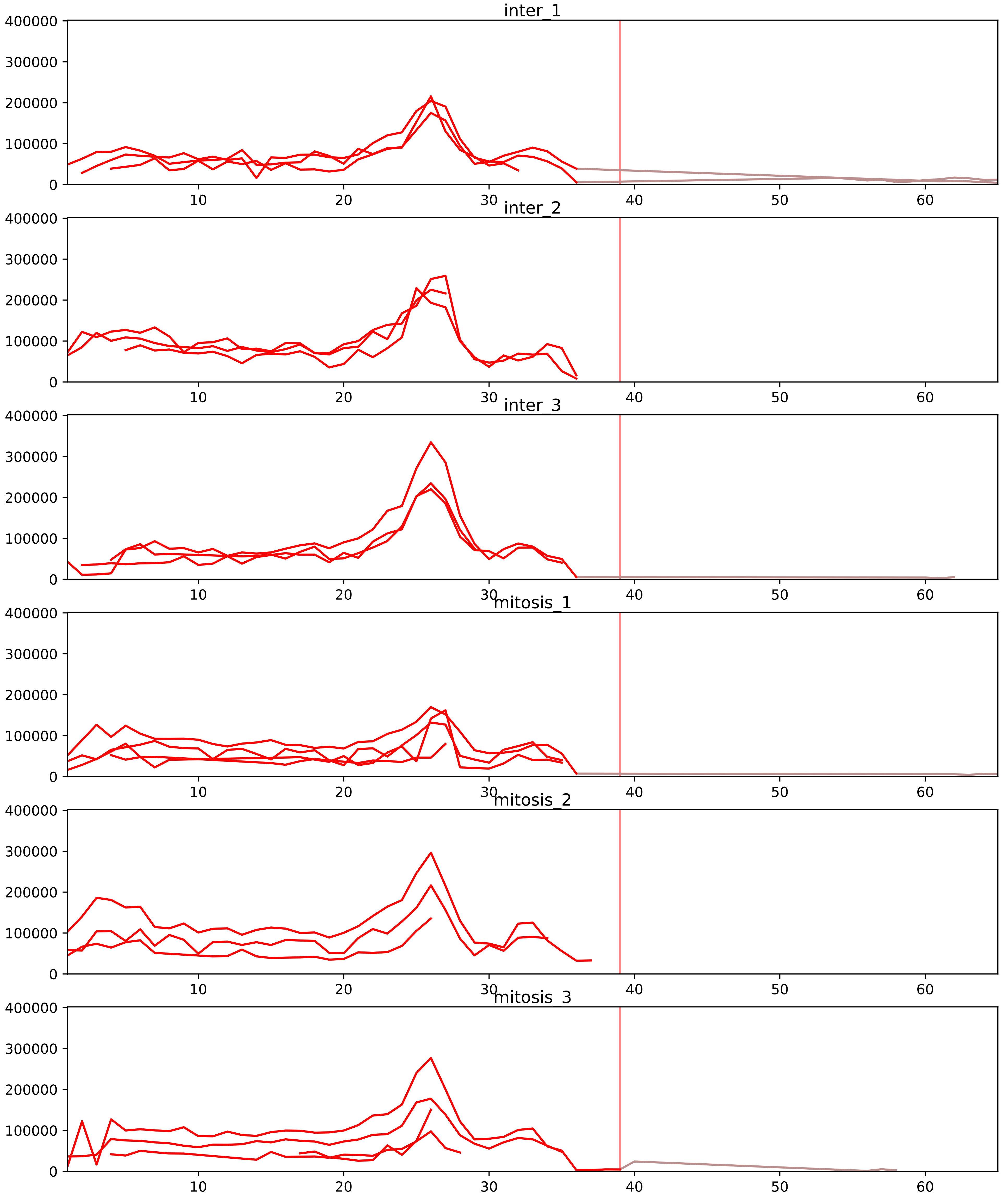


| level | condition_1 | condition_2 | pvalue | pvalue_adjusted |
|---|---|---|---|---|
| complex_abundance | mitosis | inter | 2.4413174450807402e-05 | 0.0008984048197897125 |
| interactor_abundance | mitosis | inter | 6.634639281590001e-05 | 0.0017952553350184708 |
| interactor_ratio | mitosis | inter | 0.05621811676799316 | 0.22801489072917164 |
| monomer_abundance | mitosis | inter | 0.36944368170317465 | 0.6798014065844532 |
| total_abundance | mitosis | inter | 0.6649613119785112 | 0.853619564229835 |
| assembled_abundance | mitosis | inter | 0.6741765672439284 | 0.861213929992511 |

Q05519_Q16629
SRS11_HUMAN vs SRSF7_HUMAN
p-value: 0.067 q-value: 0.041

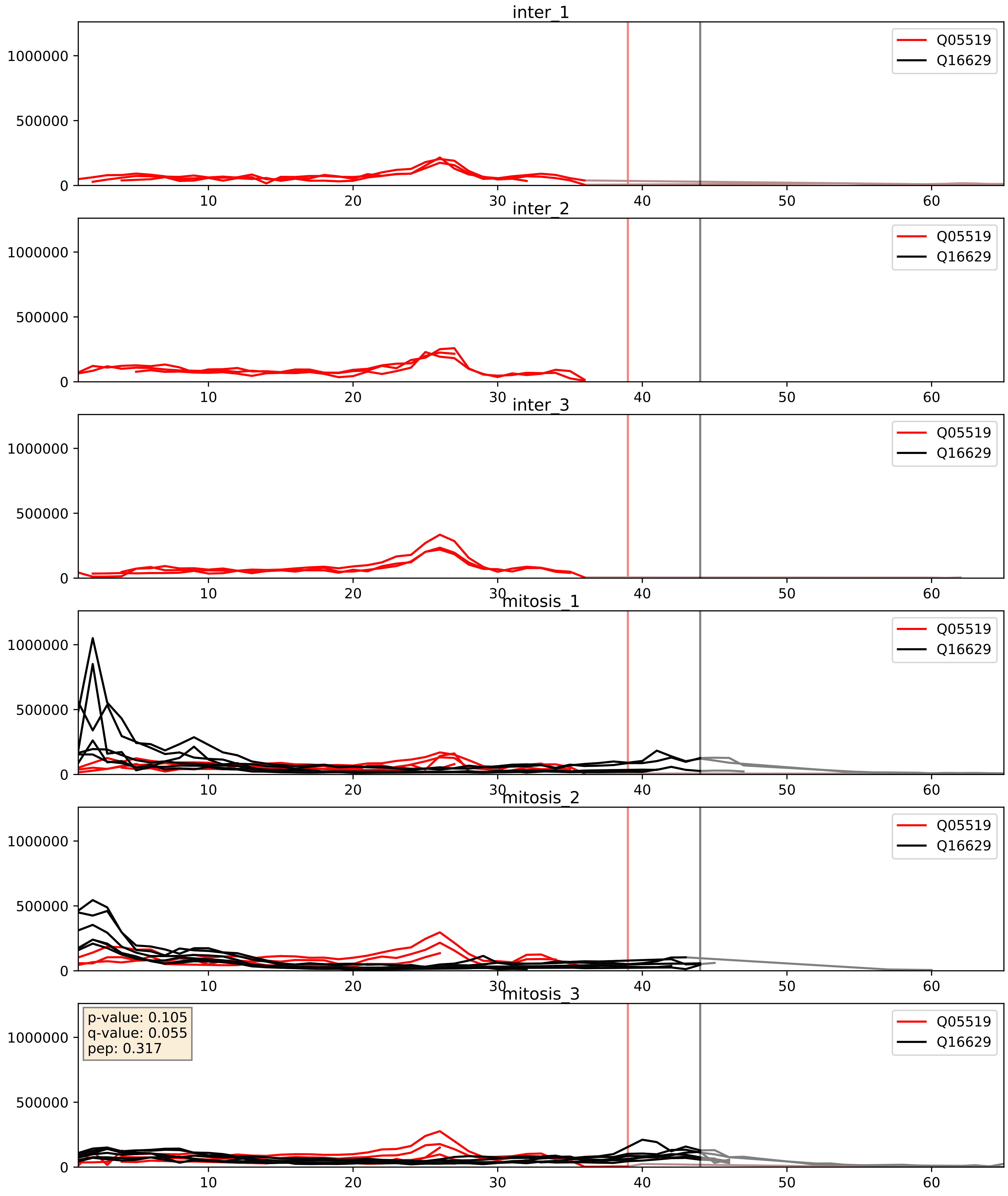


| level | condition_1 | condition_2 | pvalue | pvalue_adjusted |
|---|---|---|---|---|
| complex_abundance | mitosis | inter | 2.4413174450807402e-05 | 0.005691857151652958 |
| interactor_abundance | mitosis | inter | 6.634639281590001e-05 | 0.006642399093615253 |
| interactor_ratio | mitosis | inter | 0.05621811676799316 | 0.3926647223857623 |
