## Supplementary Data 2 for "SECAT: Quantifying differential protein-protein interaction states by network-centric analysis": hela_string_000122_Q86Y82.pdf

Q86Y82_Q86Y82

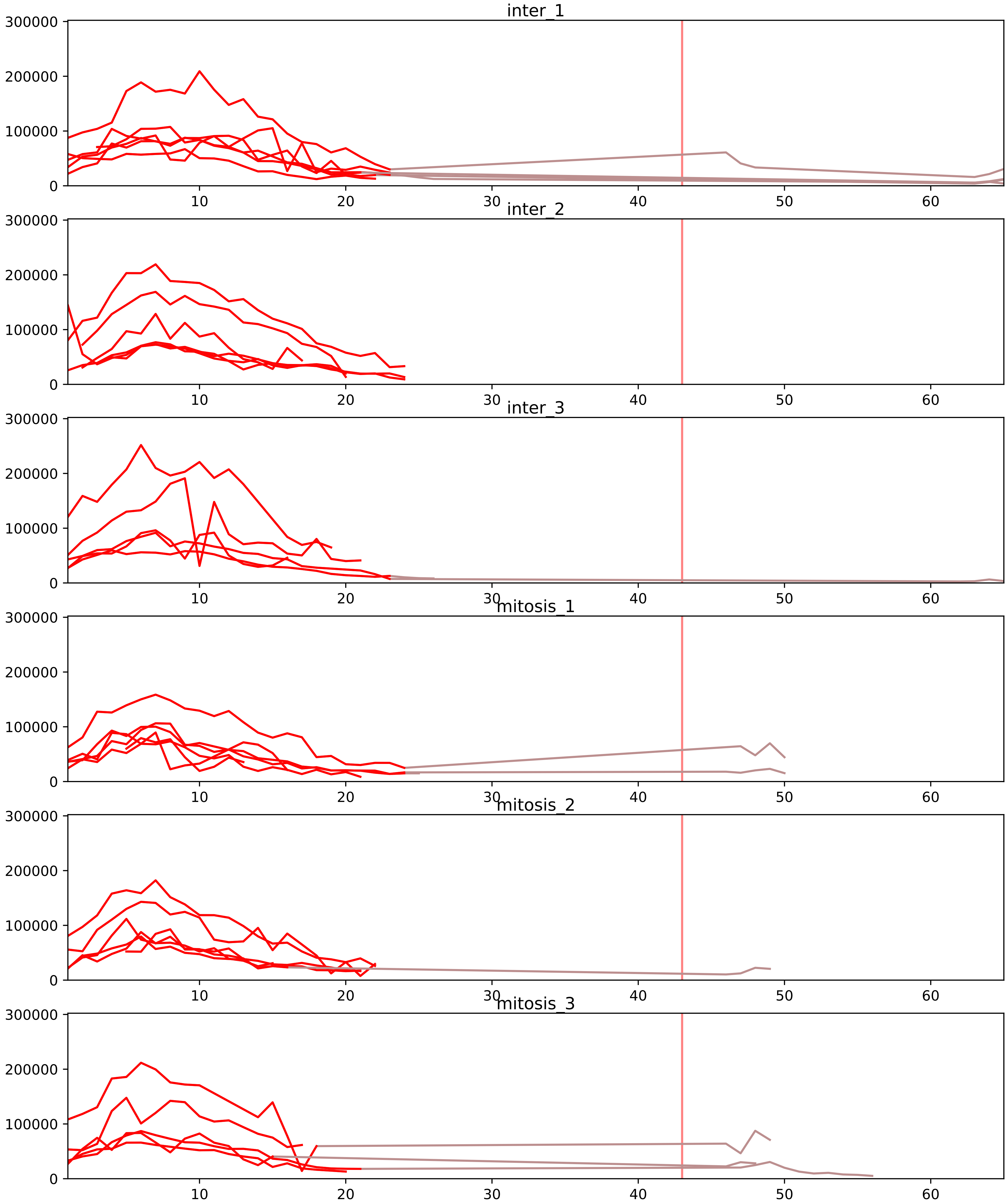


| level | condition_1 | condition_2 | pvalue | pvalue_adjusted |
|---|---|---|---|---|
| interactor_abundance | mitosis | inter | 2.702031308037437e-05 | 0.0009943475213577769 |
| complex_abundance | mitosis | inter | 0.0001863152109619952 | 0.0034628281633340523 |
| interactor_ratio | mitosis | inter | 0.0010508285535743539 | 0.011373673756334184 |
| monomer_abundance | mitosis | inter | 0.1251128138149701 | 0.46937521932615284 |
| assembled_abundance | mitosis | inter | 0.5119218545687104 | 0.7555040013311375 |
| total_abundance | mitosis | inter | 0.5373151100679913 | 0.7824745380966159 |

O00161_Q86Y82
SNP23_HUMAN vs STX12_HUMAN
p-value: 0.002 q-value: 0.002

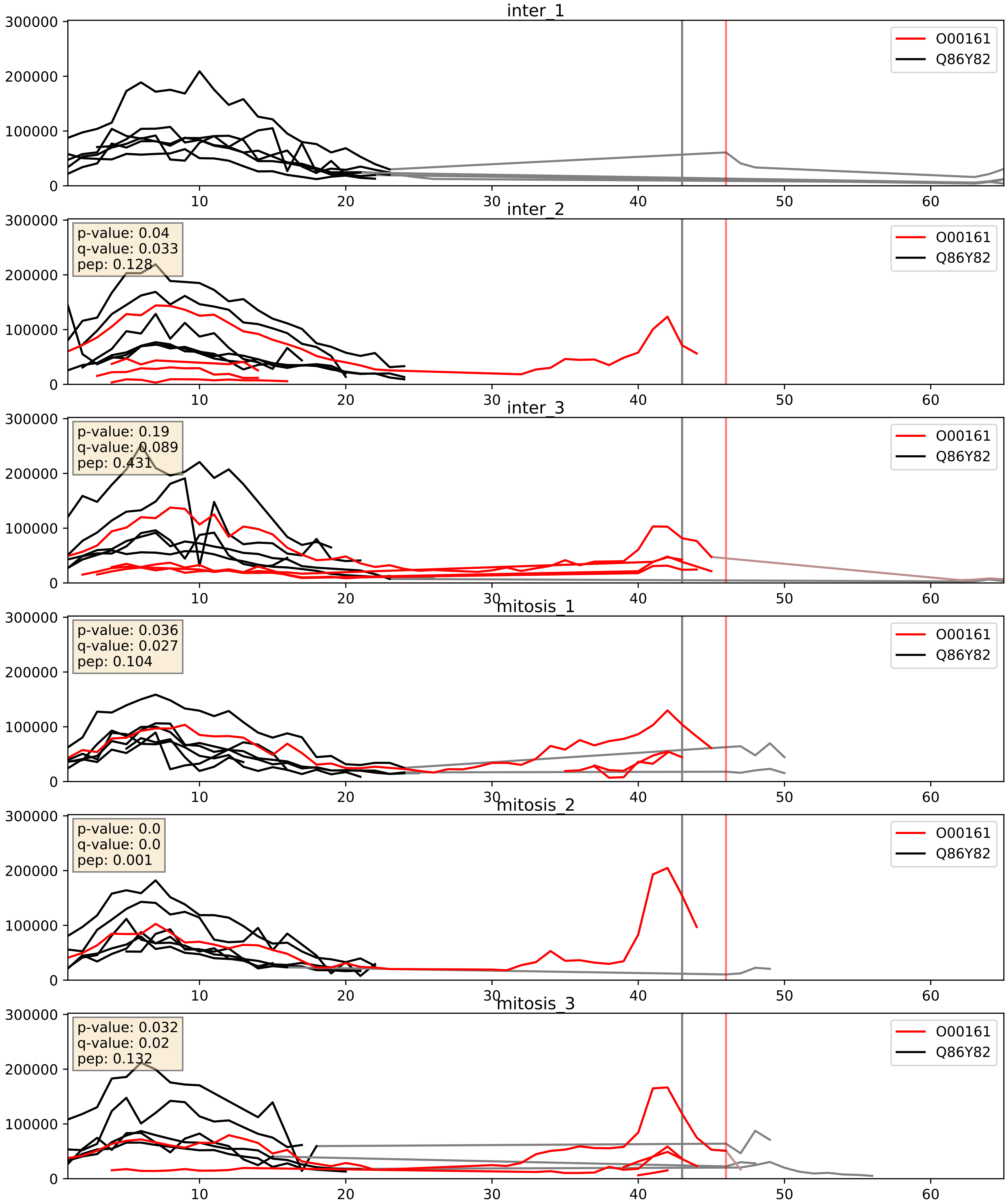


| level | condition_1 | condition_2 | pvalue | pvalue_adjusted |
|---|---|---|---|---|
| interactor_ratio | mitosis | inter | 0.05879801552416702 | 0.40040653372066 |
| interactor_abundance | mitosis | inter | 0.22731684012527184 | 0.4204477423233204 |
| complex_abundance | mitosis | inter | 0.8081554996004131 | 0.8757831468008022 |

O43752_Q86Y82
STX6_HUMAN vs STX12_HUMAN
p-value: 0.038 q-value: 0.026

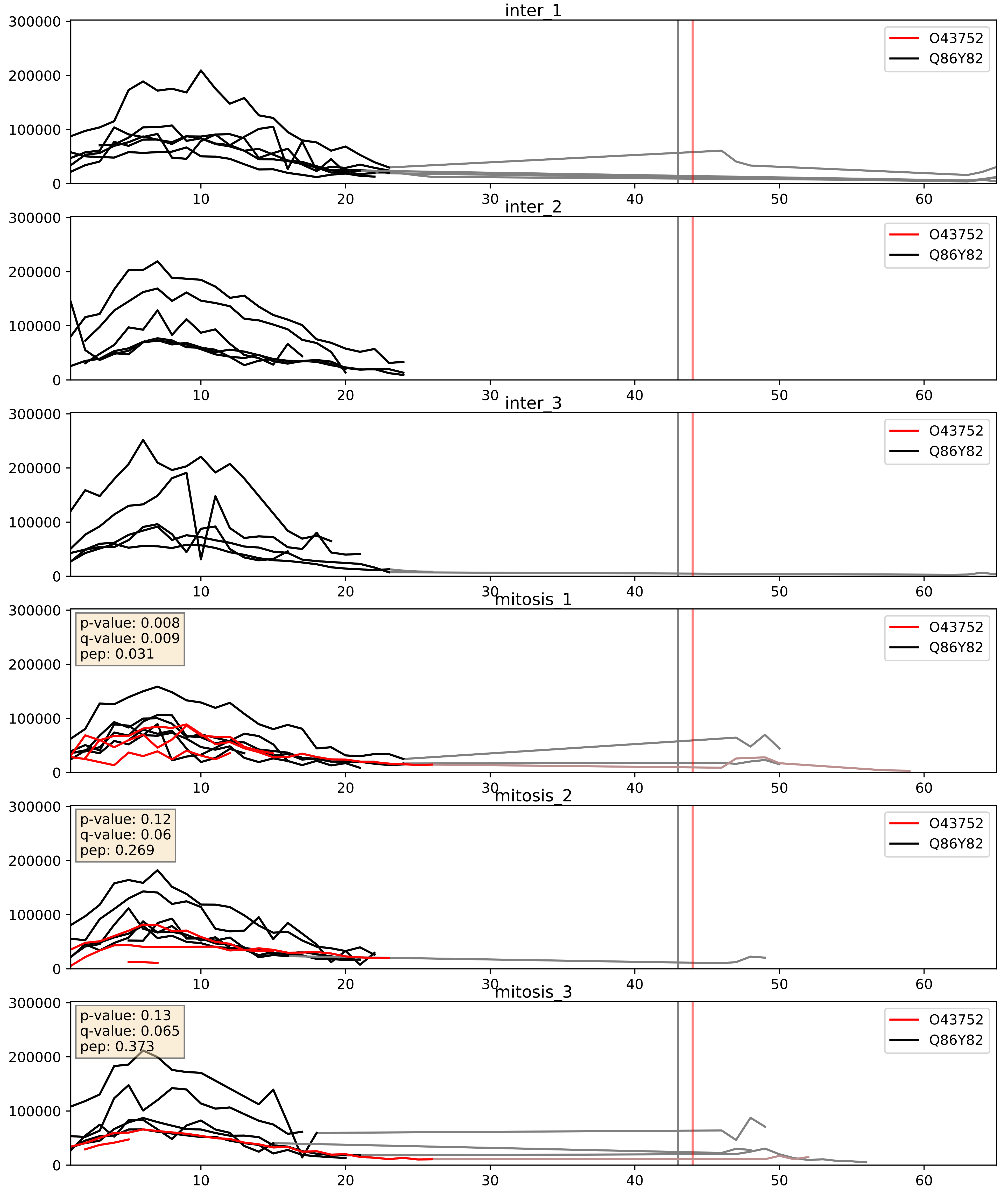


| level | condition_1 | condition_2 | pvalue | pvalue_adjusted |
|---|---|---|---|---|
| complex_abundance | mitosis | inter | 0.0019223394458935076 | 0.03128369896739244 |
| interactor_ratio | mitosis | inter | 0.02889933776935565 | 0.2680155268750643 |
| interactor_abundance | mitosis | inter | 0.08932656564997186 | 0.25331634983063084 |

O60499_Q86Y82
STX10_HUMAN vs STX12_HUMAN
p-value: 0.002 q-value: 0.03

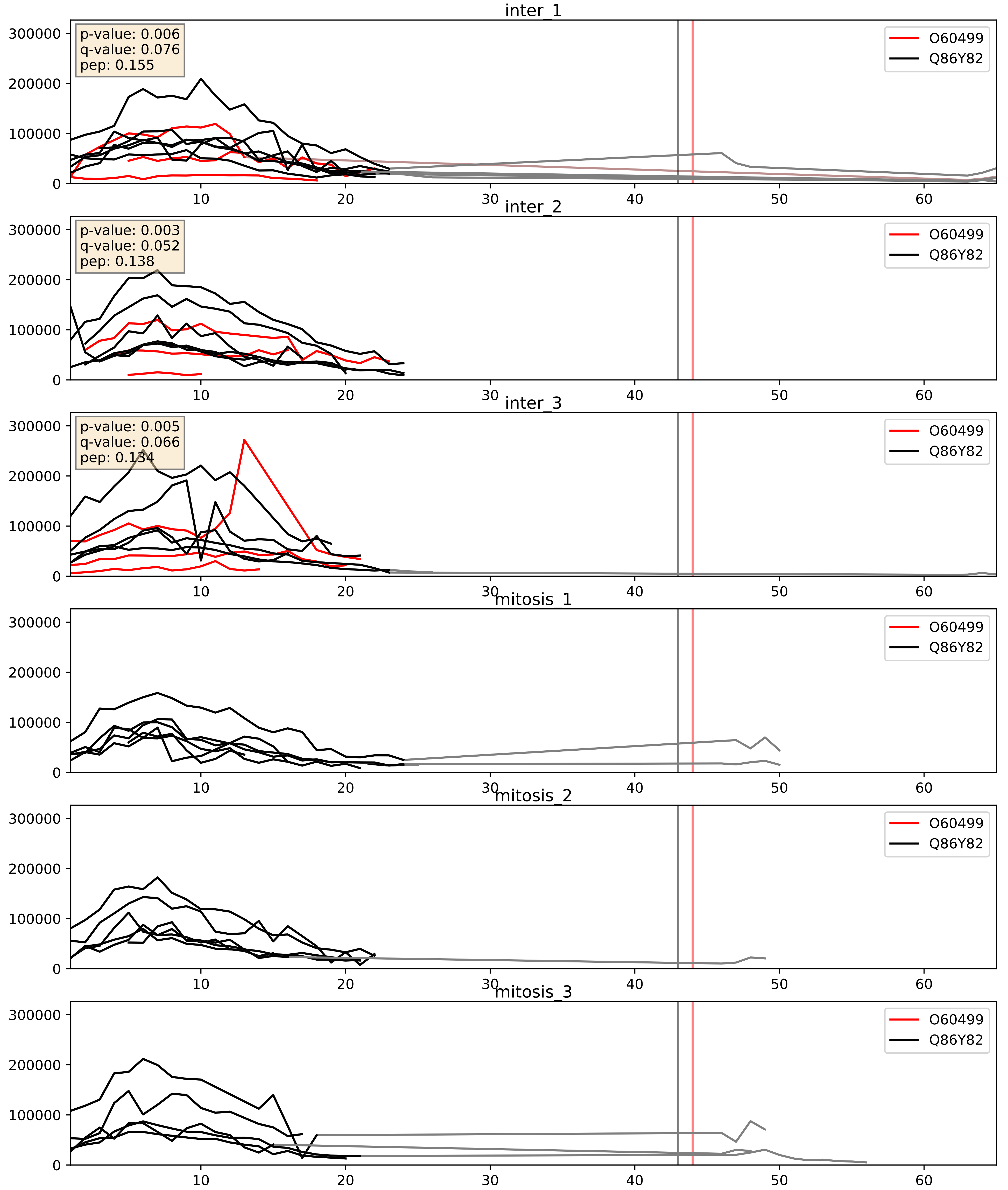


| level | condition_1 | condition_2 | pvalue | pvalue_adjusted |
|---|---|---|---|---|
| interactor_ratio | mitosis | inter | 4.00133417798734e-05 | 0.006715964816386594 |
| complex_abundance | mitosis | inter | 0.00013171704976947633 | 0.007884601021165855 |
| interactor_abundance | mitosis | inter | 0.00030071181320304993 | 0.01300709055050693 |

O95249_Q86Y82
GOSR1_HUMAN vs STX12_HUMAN
p-value: 0.002 q-value: 0.03

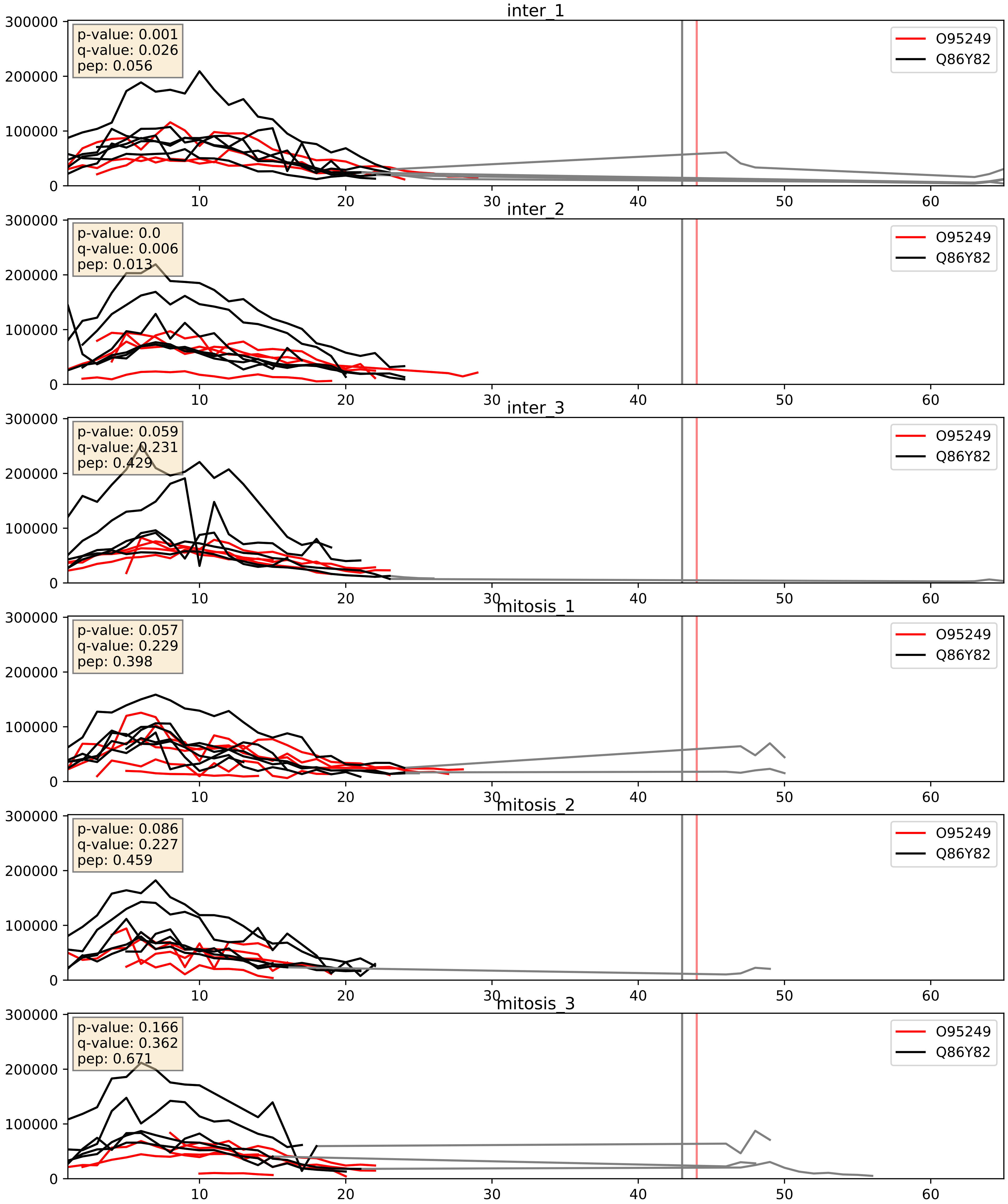


| level | condition_1 | condition_2 | pvalue | pvalue_adjusted |
|---|---|---|---|---|
| complex_abundance | mitosis | inter | 0.06880946579591136 | 0.18995481896965266 |
| interactor_abundance | mitosis | inter | 0.3152832548832075 | 0.512805168837875 |
| interactor_ratio | mitosis | inter | 0.7458582185341482 | 0.9491588884573259 |

P51809_Q86Y82
VAMP7_HUMAN vs STX12_HUMAN
p-value: 0.0 q-value: 0.0

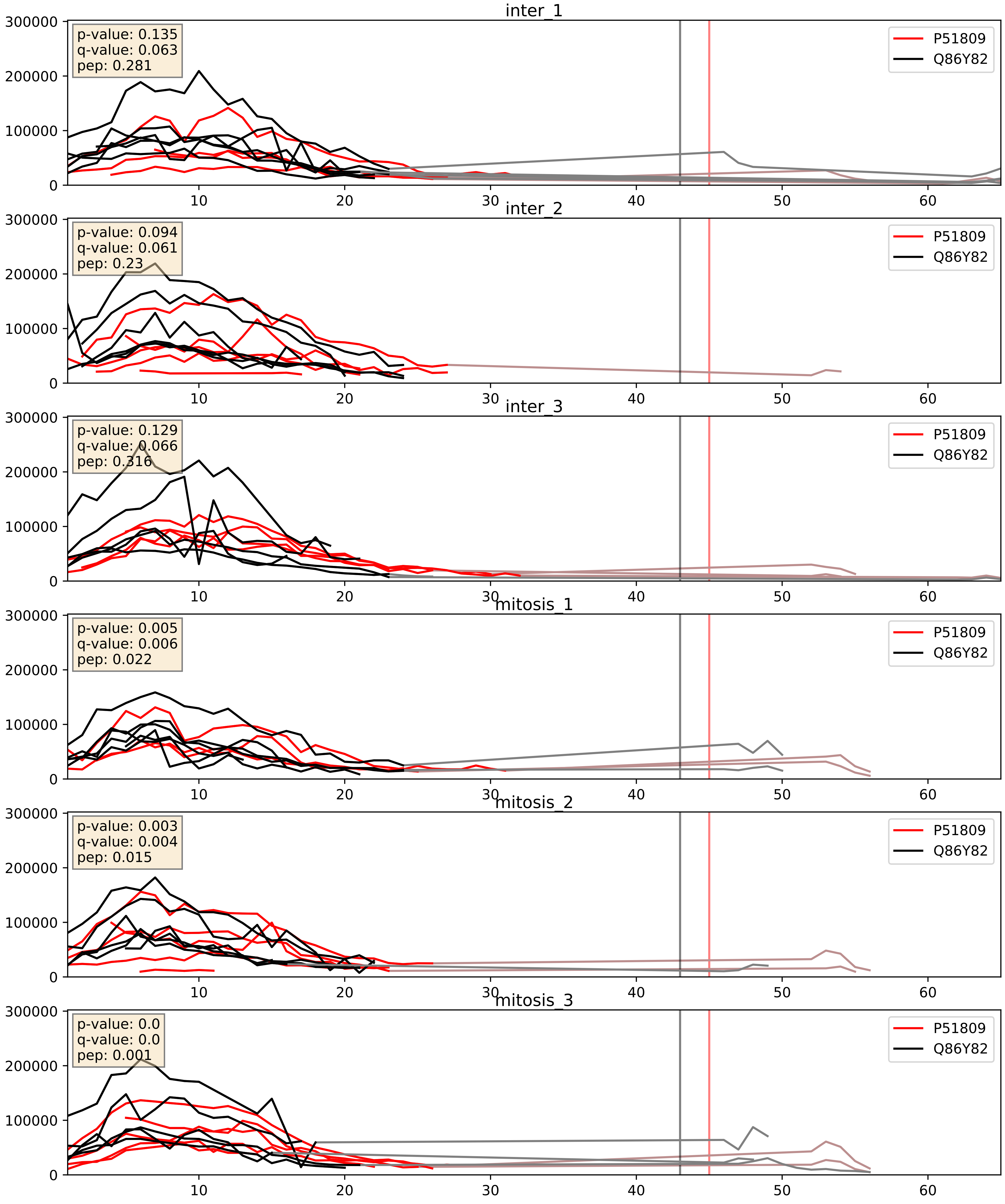


| level | condition_1 | condition_2 | pvalue | pvalue_adjusted |
|---|---|---|---|---|
| interactor_ratio | mitosis | inter | 0.015215532181280899 | 0.1808957714885618 |
| complex_abundance | mitosis | inter | 0.025114580703355222 | 0.11592191390630502 |
| interactor_abundance | mitosis | inter | 0.14657621085464068 | 0.33244573835524666 |

P63027_Q86Y82
VAMP2_HUMAN vs STX12_HUMAN
p-value: 0.0 q-value: 0.013

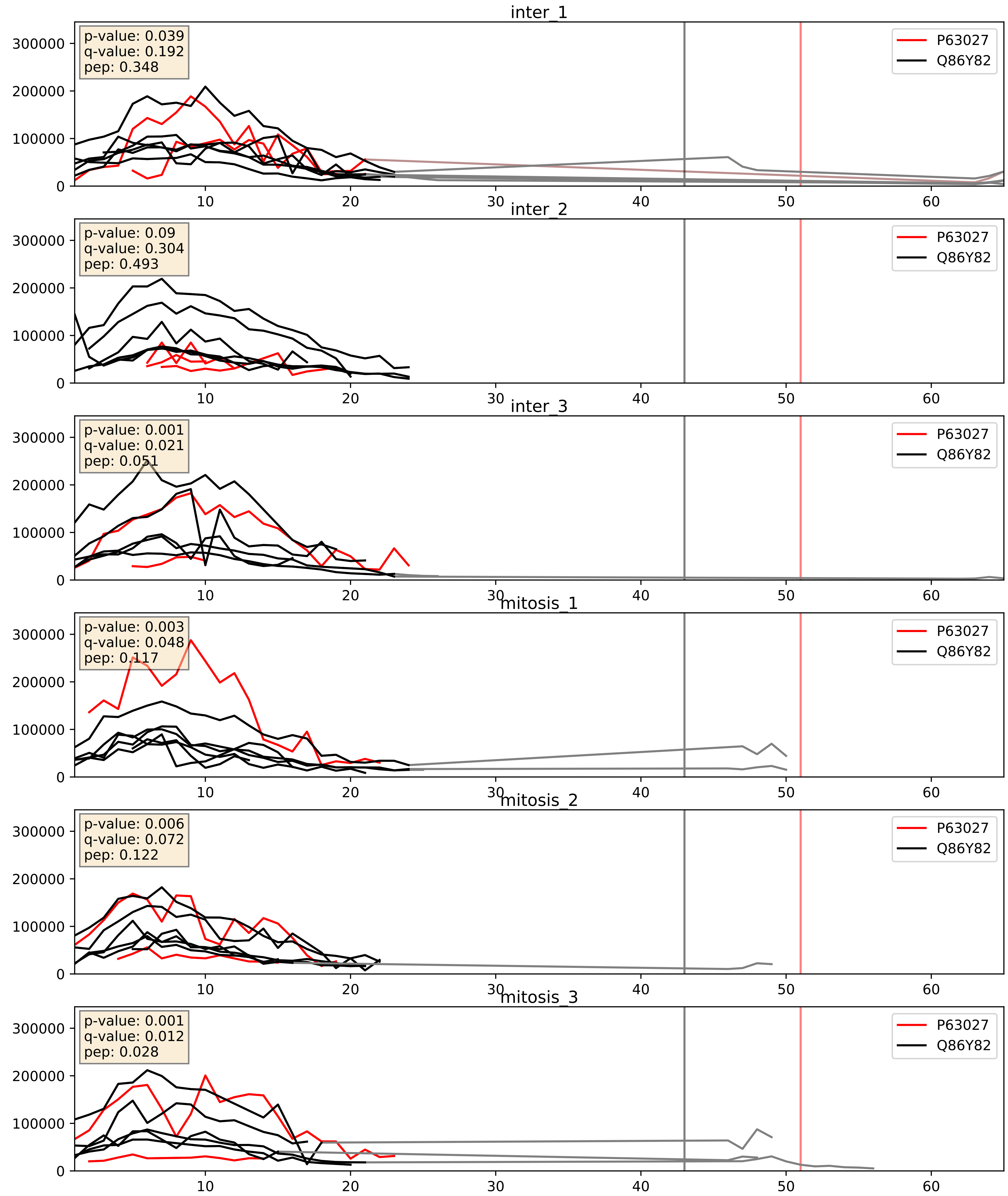


| level | condition_1 | condition_2 | pvalue | pvalue_adjusted |
|---|---|---|---|---|
| complex_abundance | mitosis | inter | 0.039178423829672036 | 0.14448905839702758 |
| interactor_abundance | mitosis | inter | 0.049998993505827004 | 0.17873935452490256 |
| interactor_ratio | mitosis | inter | 0.5476132549448776 | 0.8869573249438318 |

Q12846_Q86Y82
STX4_HUMAN vs STX12_HUMAN
p-value: 0.0 q-value: 0.003

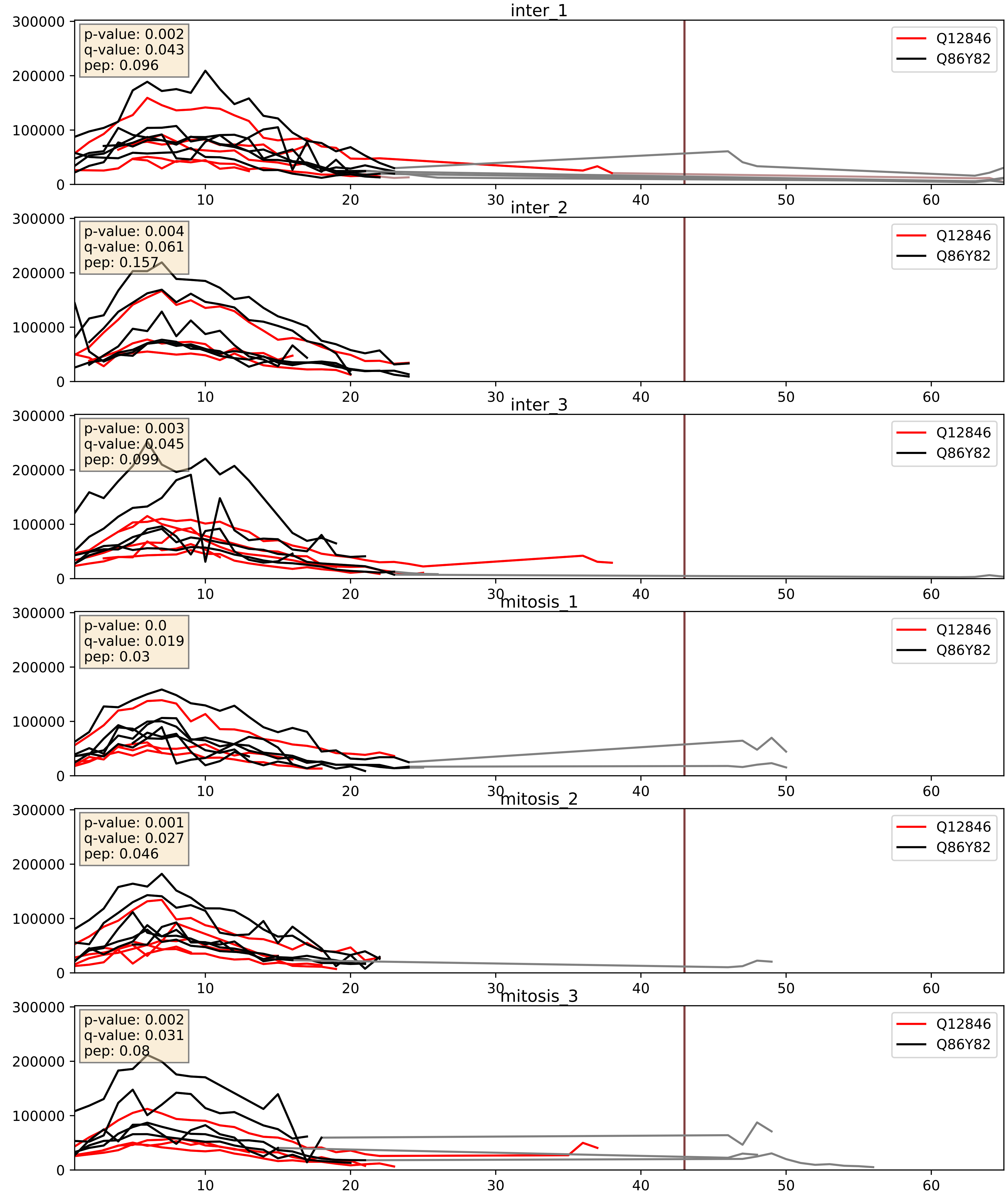


| level | condition_1 | condition_2 | pvalue | pvalue_adjusted |
|---|---|---|---|---|
| complex_abundance | mitosis | inter | 0.006030841312933395 | 0.05806974312565789 |
| interactor_abundance | mitosis | inter | 0.020897799627076514 | 0.12074597692053658 |
| interactor_ratio | mitosis | inter | 0.7196316839325936 | 0.9440685386150193 |

Q12981_Q86Y82
SEC20_HUMAN vs STX12_HUMAN
p-value: 0.0 q-value: 0.026

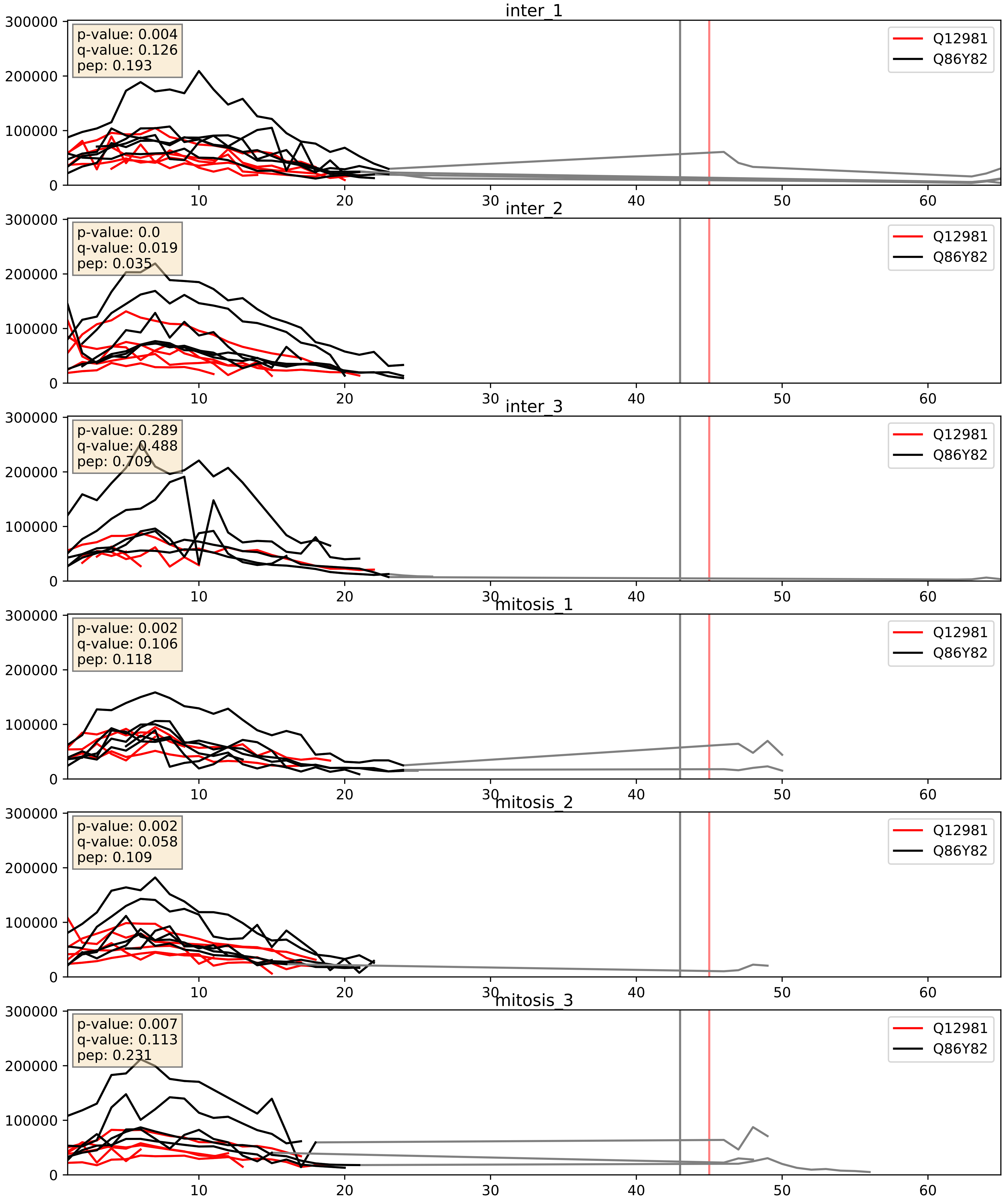


| level | condition_1 | condition_2 | pvalue | pvalue_adjusted |
|---|---|---|---|---|
| interactor_ratio | mitosis | inter | 0.23121834557969595 | 0.7093426594678715 |
| complex_abundance | mitosis | inter | 0.2457342747477258 | 0.3965847269684263 |
| interactor_abundance | mitosis | inter | 0.9117382207069966 | 0.9497966617076659 |

Q13190_Q86Y82
STX5_HUMAN vs STX12_HUMAN
p-value: 0.0 q-value: 0.003

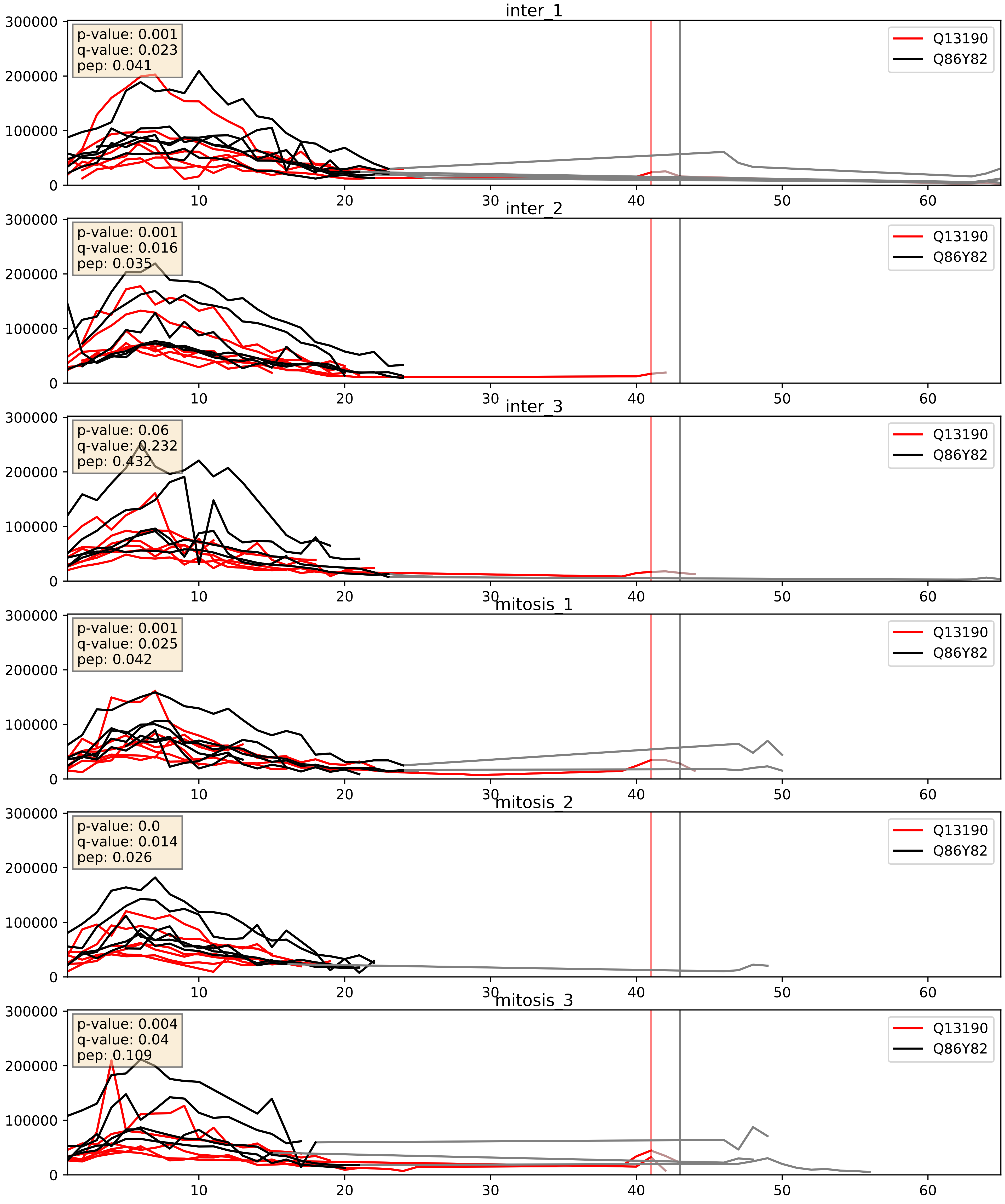


| level | condition_1 | condition_2 | pvalue | pvalue_adjusted |
|---|---|---|---|---|
| complex_abundance | mitosis | inter | 0.012170016918556072 | 0.08185897984306292 |
| interactor_abundance | mitosis | inter | 0.043034055988443856 | 0.1668005626295396 |
| interactor_ratio | mitosis | inter | 0.8963302871005671 | 0.9826571795057447 |

Q15836_Q86Y82
VAMP3_HUMAN vs STX12_HUMAN
p-value: 0.017 q-value: 0.014

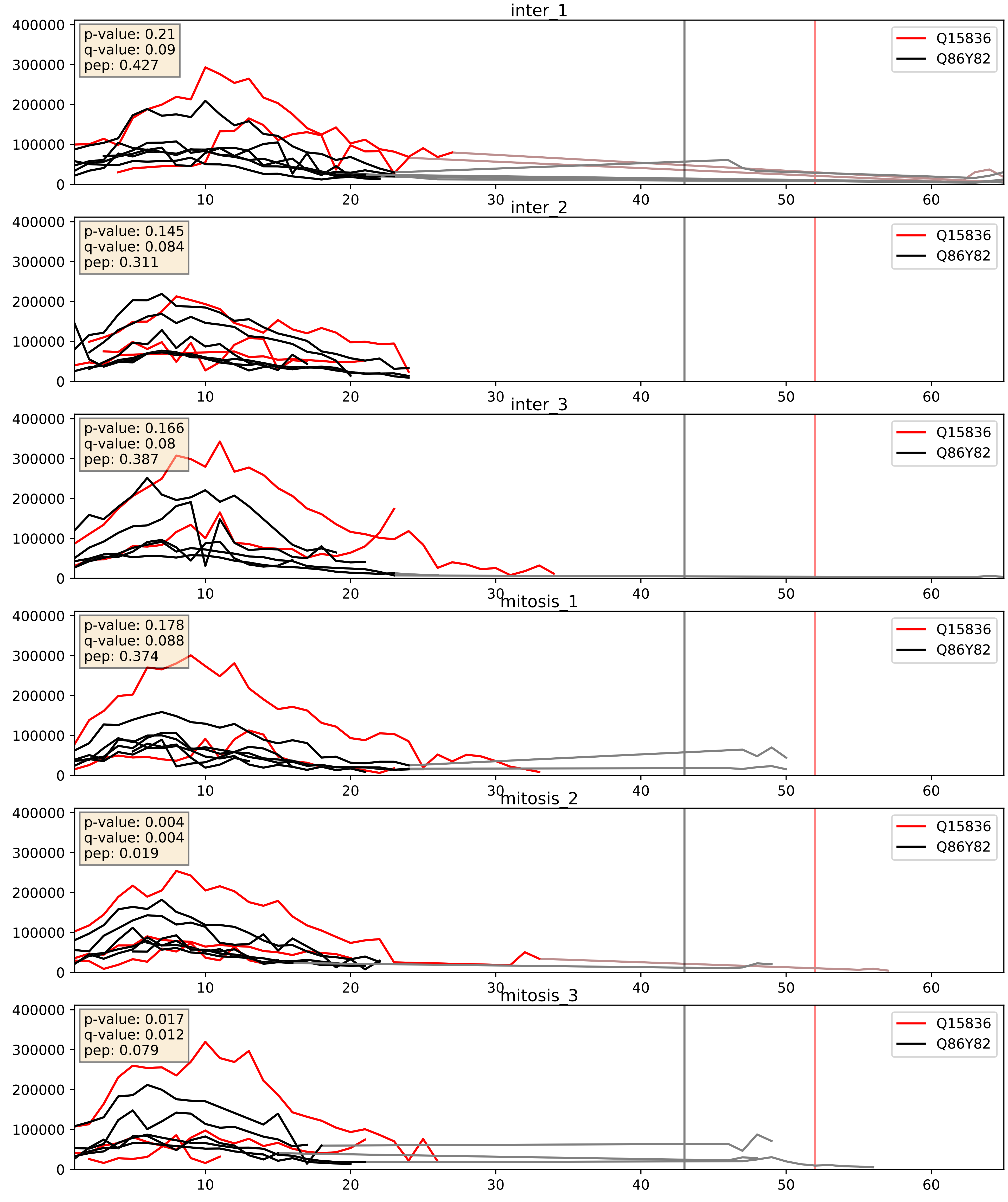


| level | condition_1 | condition_2 | pvalue | pvalue_adjusted |
|---|---|---|---|---|
| complex_abundance | mitosis | inter | 0.029129589590037797 | 0.12403912014070187 |
| interactor_ratio | mitosis | inter | 0.16206745945634524 | 0.6206299225379937 |
| interactor_abundance | mitosis | inter | 0.16837464658827767 | 0.35728482270591394 |

Q86Y82_Q9BV40
STX12_HUMAN vs VAMP8_HUMAN
p-value: 0.015 q-value: 0.013

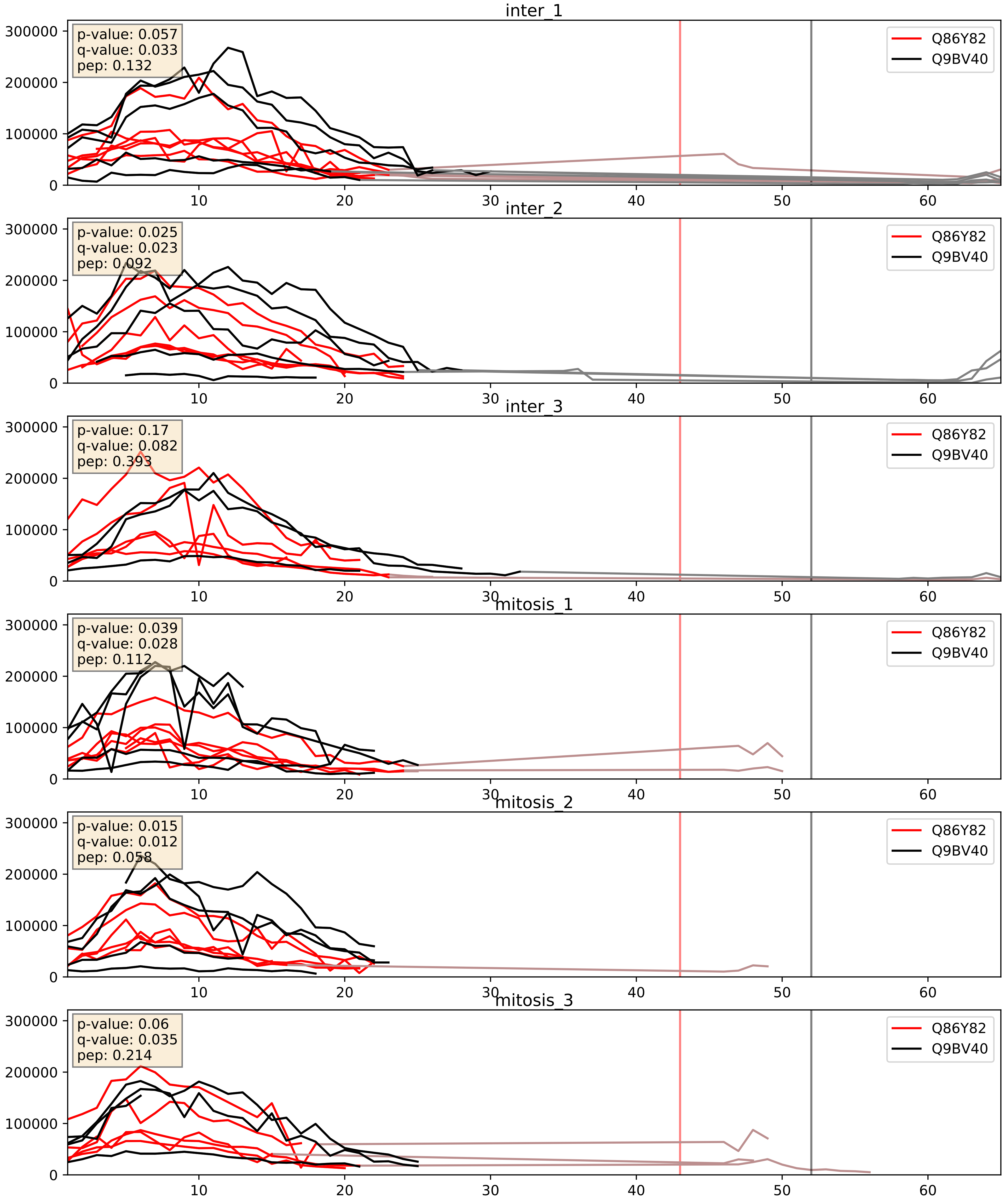


| level | condition_1 | condition_2 | pvalue | pvalue_adjusted |
|---|---|---|---|---|
| interactor_abundance | mitosis | inter | 0.005888490429492295 | 0.06496469125710559 |
| complex_abundance | mitosis | inter | 0.2846177949807808 | 0.43343325476525246 |
| interactor_ratio | mitosis | inter | 0.5733296496597162 | 0.8946257776299471 |

Q86Y82_Q9UNK0
STX12_HUMAN vs STX8_HUMAN
p-value: 0.001 q-value: 0.023

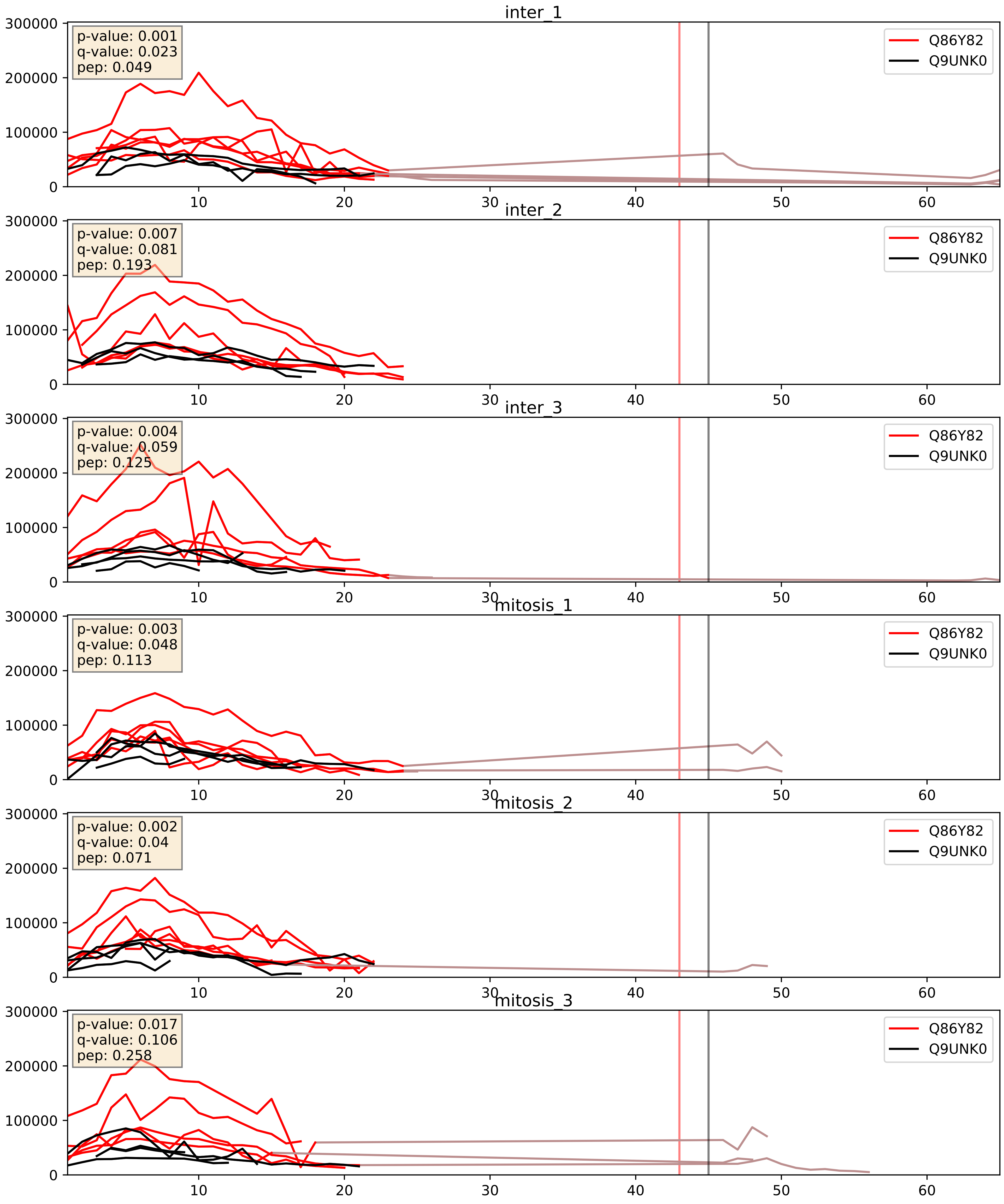


| level | condition_1 | condition_2 | pvalue | pvalue_adjusted |
|---|---|---|---|---|
| interactor_abundance | mitosis | inter | 0.0072230259778860864 | 0.07346242905026662 |
| complex_abundance | mitosis | inter | 0.009166606869228577 | 0.07075397186708442 |
| interactor_ratio | mitosis | inter | 0.884813405019416 | 0.9802747470617126 |
