## Supplementary Data 2 for "SECAT: Quantifying differential protein-protein interaction states by network-centric analysis": hela_string_000123_Q6FIF0.pdf

Q6FIF0_Q6FIF0

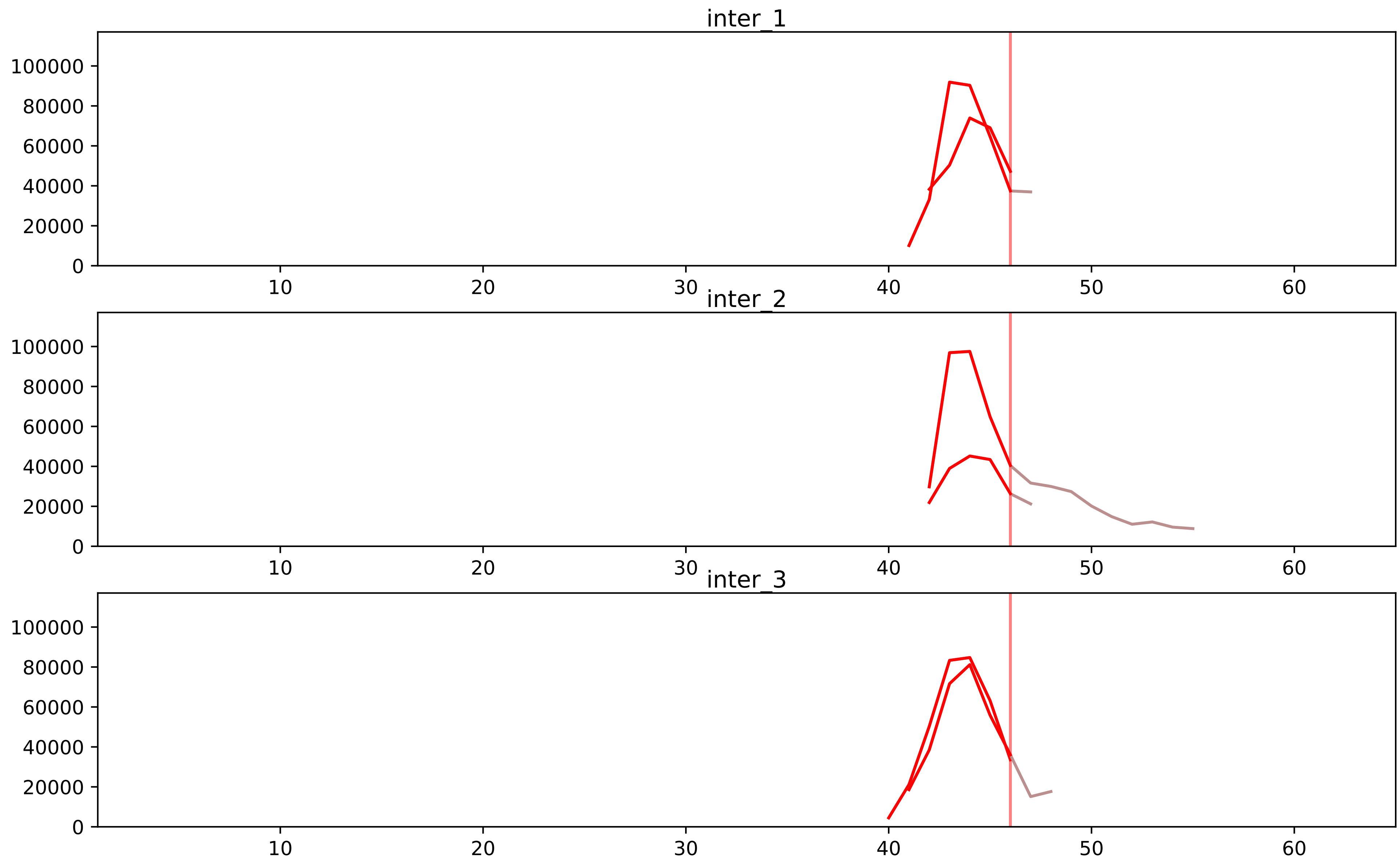


| level | condition_1 | condition_2 | pvalue | pvalue_adjusted |
|---|---|---|---|---|
| assembled_abundance | mitosis | inter | 2.7998623105398545e-05 | 0.006314764940805662 |
| total_abundance | mitosis | inter | 0.00010636816002230238 | 0.01273261737241483 |
| monomer_abundance | mitosis | inter | 0.011567709238707757 | 0.1902553554136748 |
