## Supplementary Data 2 for "SECAT: Quantifying differential protein-protein interaction states by network-centric analysis": hela_string_000124_Q8TED0.pdf

Q8TED0_Q8TED0

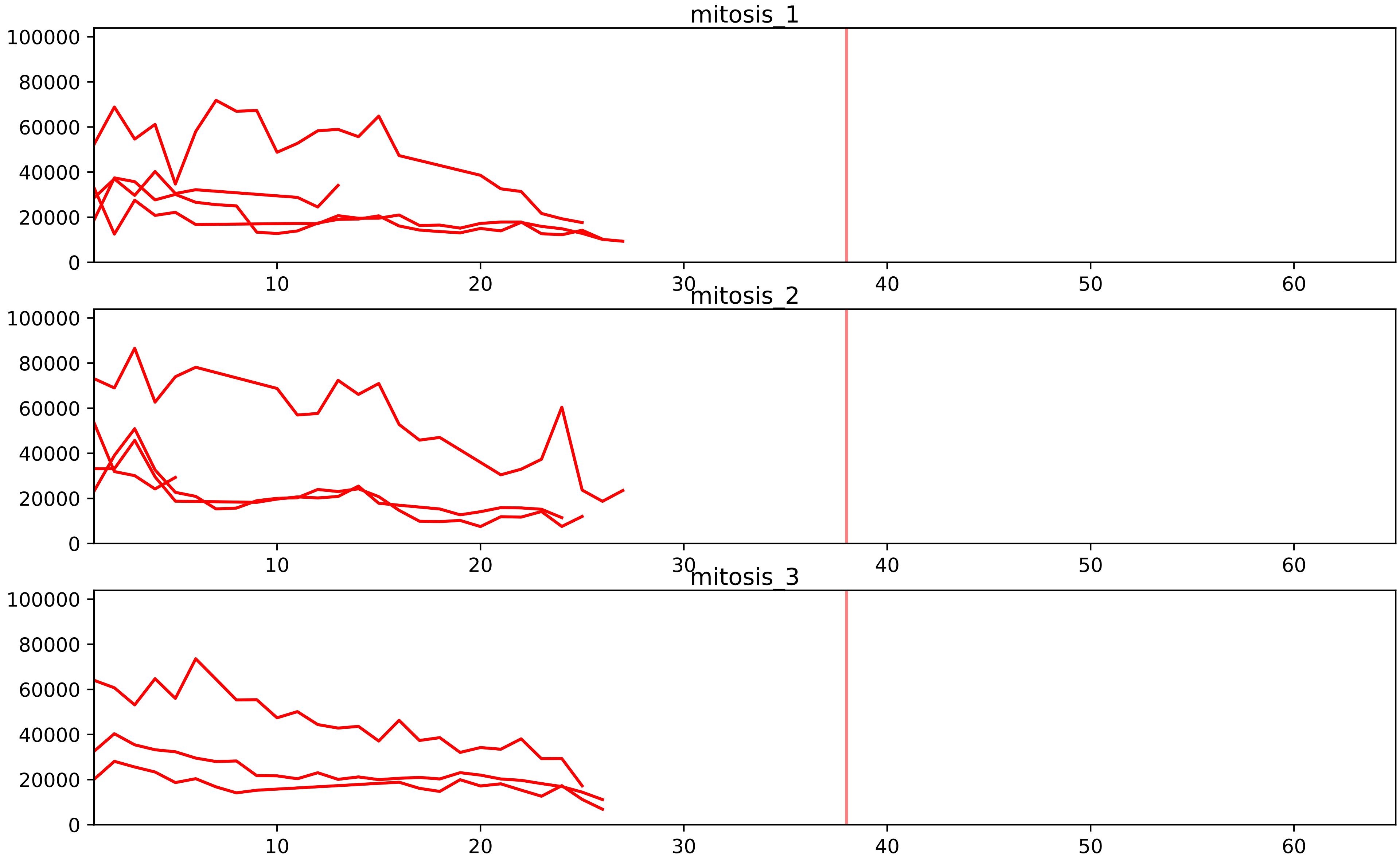


| level | condition_1 | condition_2 | pvalue | pvalue_adjusted |
|---|---|---|---|---|
| assembled_abundance | mitosis | inter | 2.838950175144202e-05 | 0.006314764940805662 |
| total_abundance | mitosis | inter | 5.311650963812582e-05 | 0.010015412983988834 |
| interactor_ratio | mitosis | inter | 6.241541236454692e-05 | 0.0011484435875076633 |
| complex_abundance | mitosis | inter | 0.0003760376695577583 | 0.005765910933218961 |
| interactor_abundance | mitosis | inter | 0.0005563094833760093 | 0.007990604304242772 |

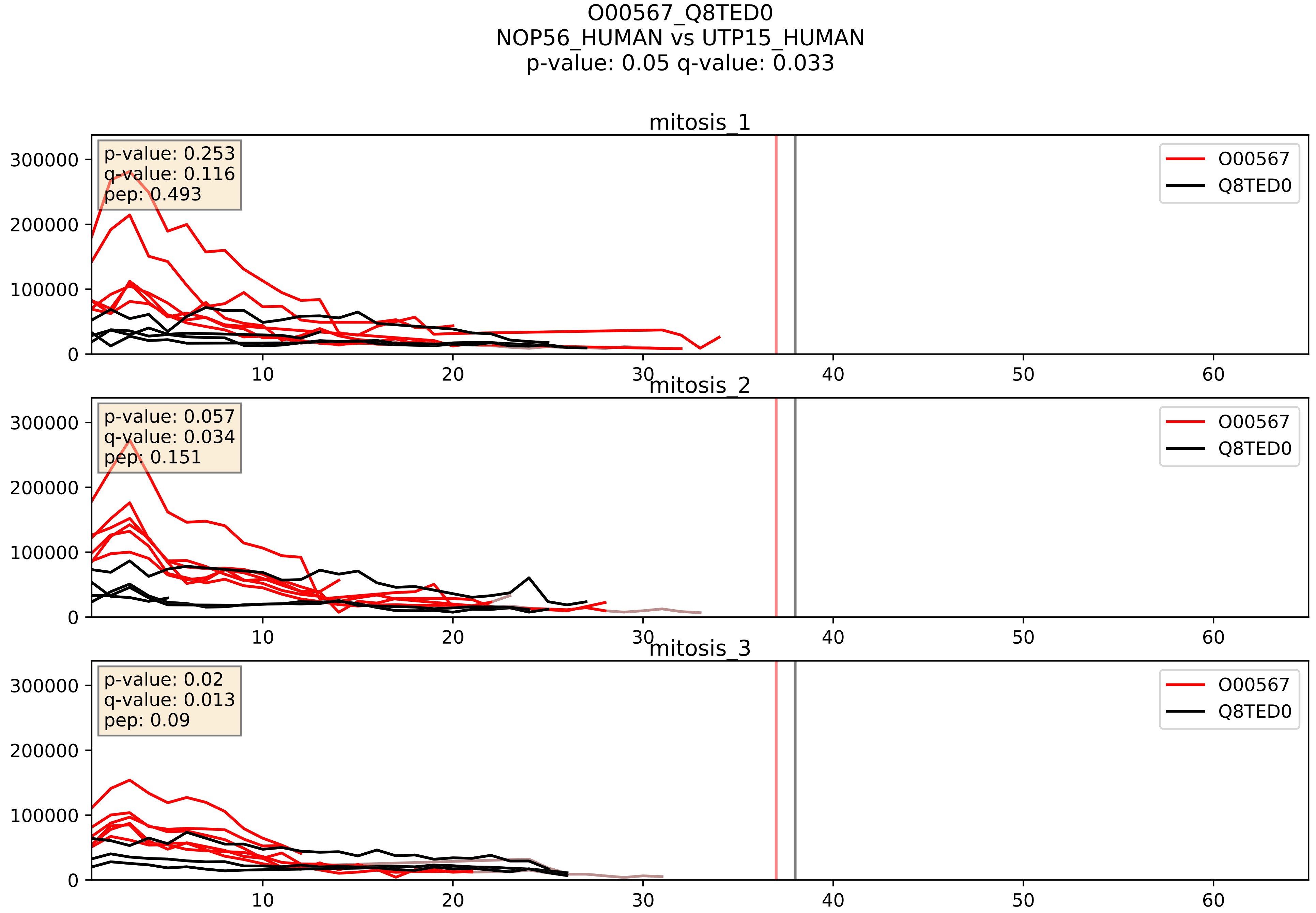


| level | condition_1 | condition_2 | pvalue | pvalue_adjusted |
|---|---|---|---|---|
| complex_abundance | mitosis | inter | 0.0008768577861510543 | 0.019513211095311817 |
| interactor_abundance | mitosis | inter | 0.0010560524122632741 | 0.024049720606162197 |
| interactor_ratio | mitosis | inter | 0.0075917466514874325 | 0.12238295920288592 |

O76021_Q8TED0
RL1D1_HUMAN vs UTP15_HUMAN
p-value: 0.066 q-value: 0.041

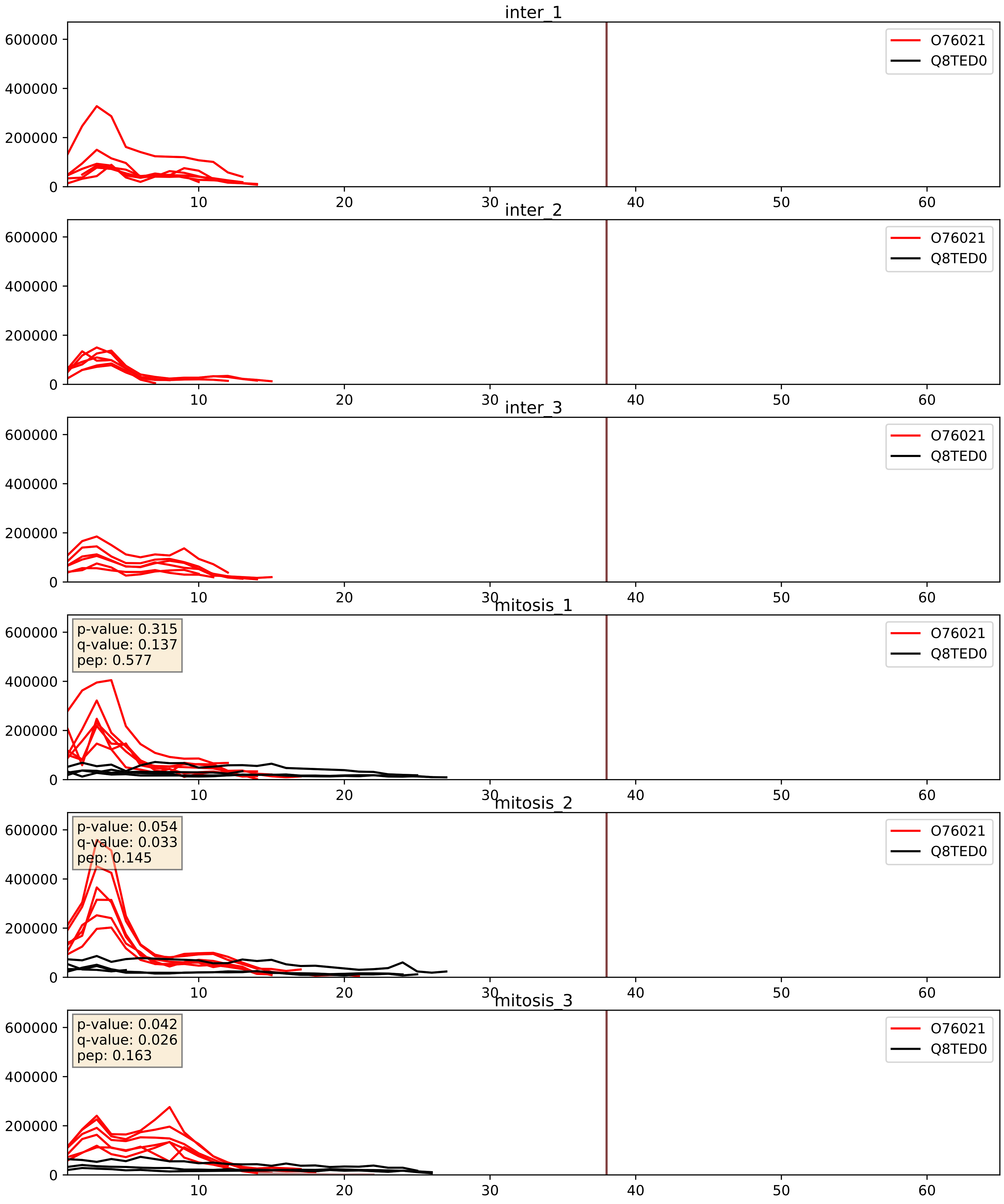


| level | condition_1 | condition_2 | pvalue | pvalue_adjusted |
|---|---|---|---|---|
| interactor_abundance | mitosis | inter | 8.109392026765088e-05 | 0.007119630332419645 |
| complex_abundance | mitosis | inter | 0.00016019902877882495 | 0.008563834848633118 |
| interactor_ratio | mitosis | inter | 0.0006707240247119572 | 0.029848225645041686 |

P56182_Q8TED0
RRP1_HUMAN vs UTP15_HUMAN
p-value: 0.058 q-value: 0.036

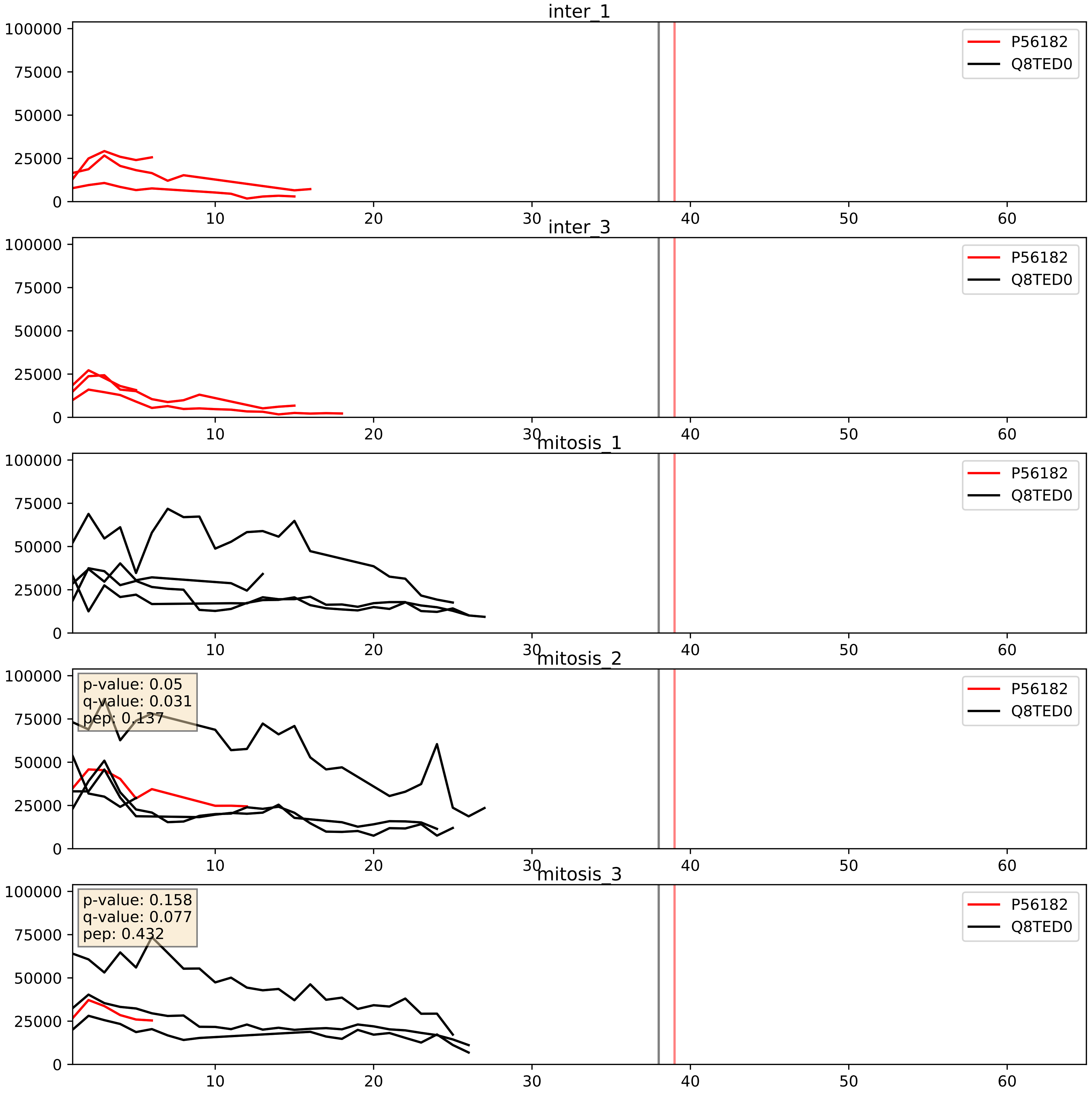


| level | condition_1 | condition_2 | pvalue | pvalue_adjusted |
|---|---|---|---|---|
| interactor_ratio | mitosis | inter | 0.3790689182104837 | 0.8176766193507878 |
| complex_abundance | mitosis | inter | 0.44437232470299726 | 0.5873794555794607 |
| interactor_abundance | mitosis | inter | 0.5376123657004697 | 0.6952129088897984 |

Q14137_Q8TED0
BOP1_HUMAN vs UTP15_HUMAN
p-value: 0.078 q-value: 0.047

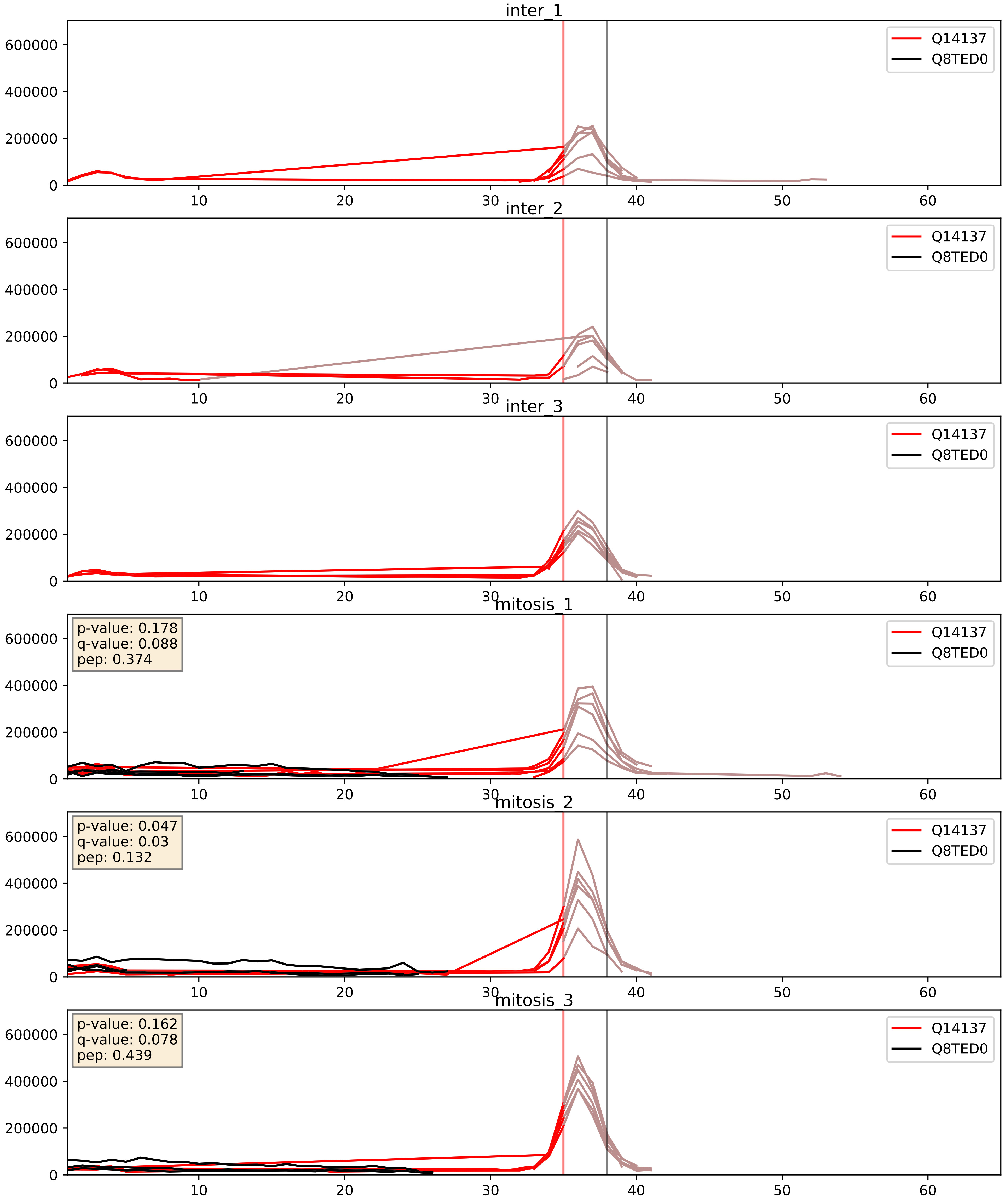


| level | condition_1 | condition_2 | pvalue | pvalue_adjusted |
|---|---|---|---|---|
| interactor_ratio | mitosis | inter | 0.01642019120281155 | 0.1887653035437297 |
| complex_abundance | mitosis | inter | 0.04358916305272642 | 0.14954839107468462 |
| interactor_abundance | mitosis | inter | 0.1276013680260762 | 0.3110987497303367 |

Q15061_Q8TED0
WDR43_HUMAN vs UTP15_HUMAN
p-value: 0.014 q-value: 0.012

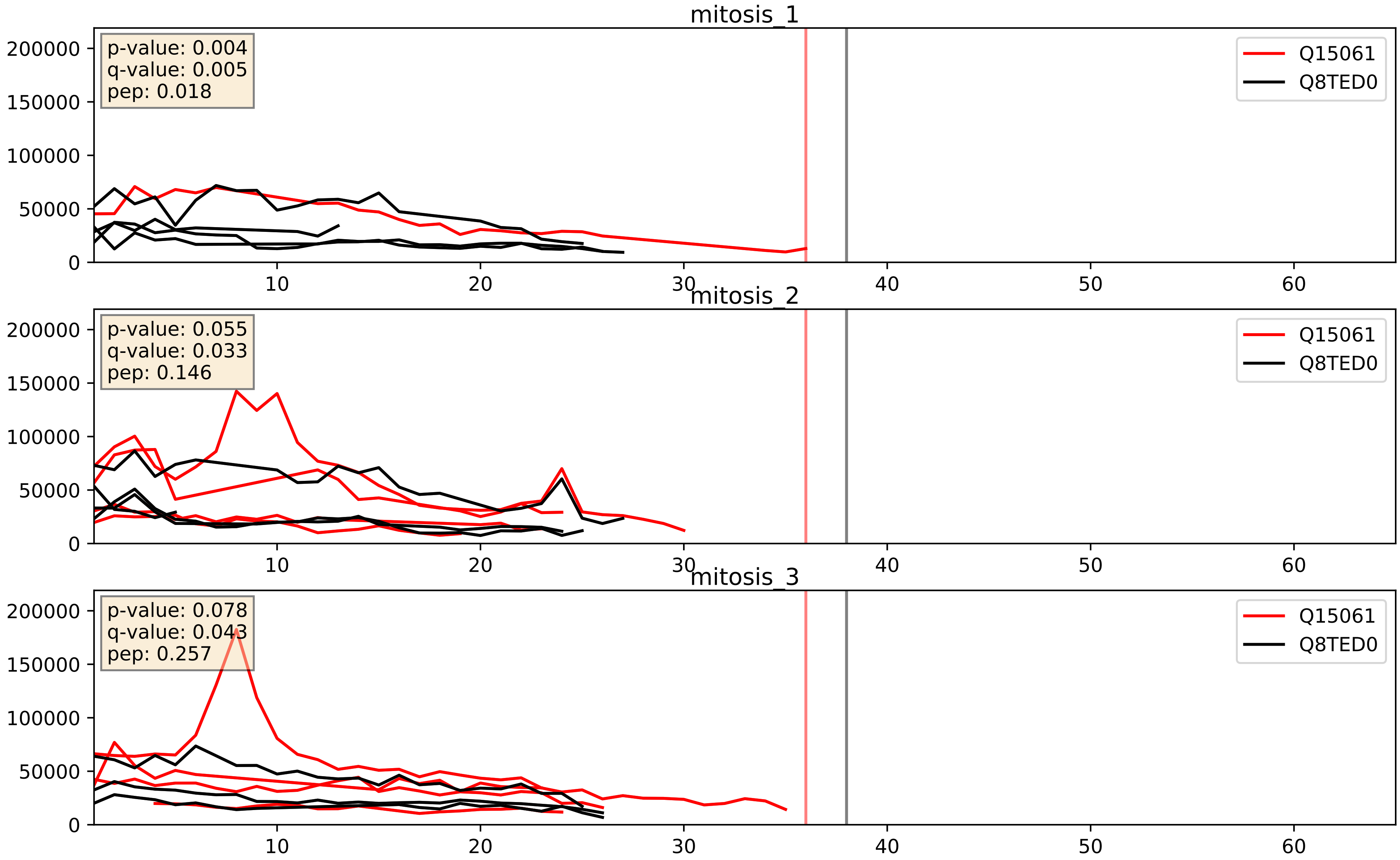


| level | condition_1 | condition_2 | pvalue | pvalue_adjusted |
|---|---|---|---|---|
| complex_abundance | mitosis | inter | 0.00607593990941489 | 0.05817678481497925 |
| interactor_abundance | mitosis | inter | 0.04157442246858351 | 0.1638853586604075 |
| interactor_ratio | mitosis | inter | 0.9786487798482965 | 0.996103871046542 |

Q8NI36_Q8TED0
WDR36_HUMAN vs UTP15_HUMAN
p-value: 0.034 q-value: 0.024

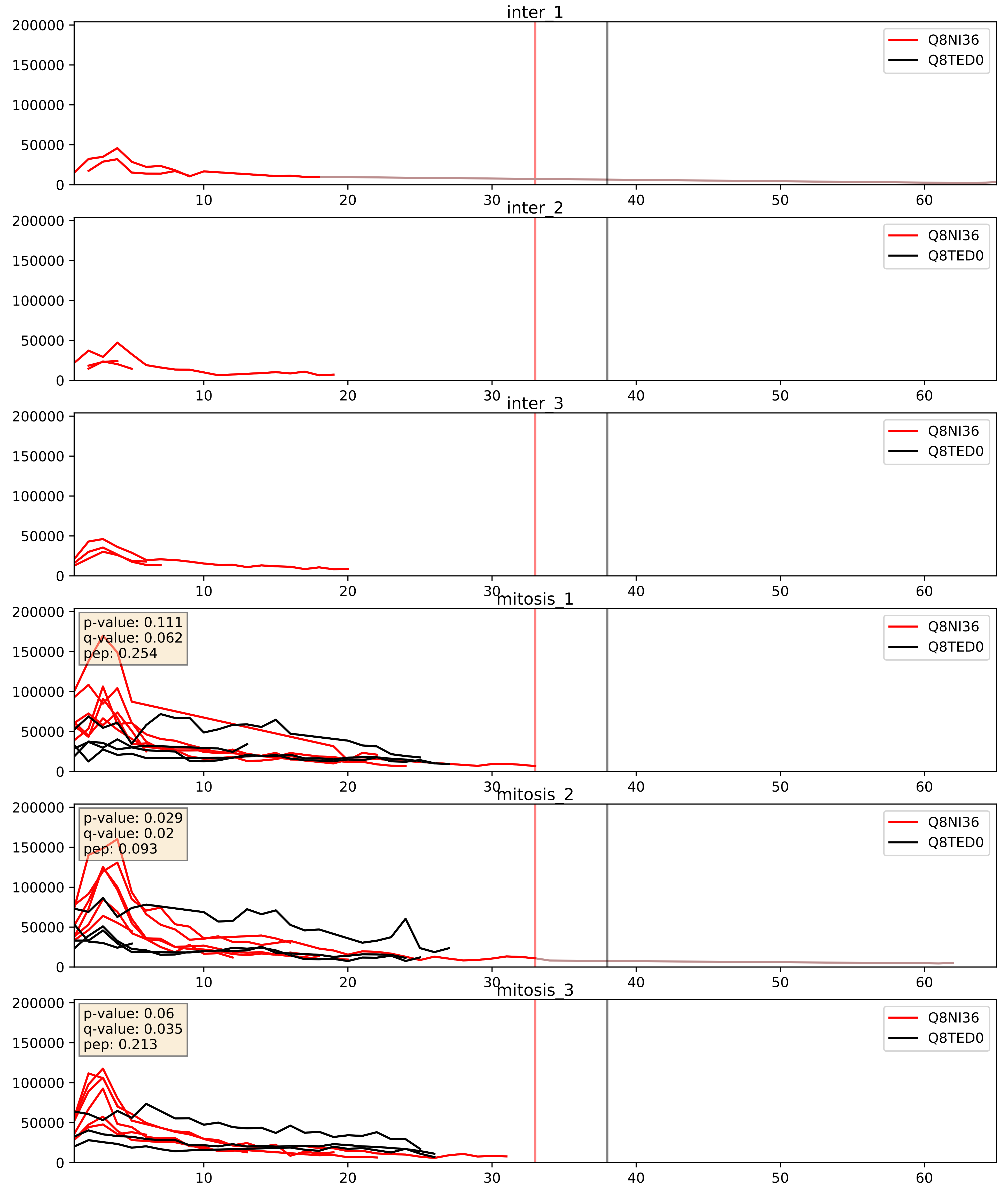


| level | condition_1 | condition_2 | pvalue | pvalue_adjusted |
|---|---|---|---|---|
| interactor_abundance | mitosis | inter | 0.0005649580704027154 | 0.017415659477552636 |
| complex_abundance | mitosis | inter | 0.0006846248474034801 | 0.016840197395901695 |
| interactor_ratio | mitosis | inter | 0.15977555659771123 | 0.616090239021059 |

Q8TED0_Q969X6
UTP15_HUMAN vs UTP4_HUMAN
p-value: 0.071 q-value: 0.043

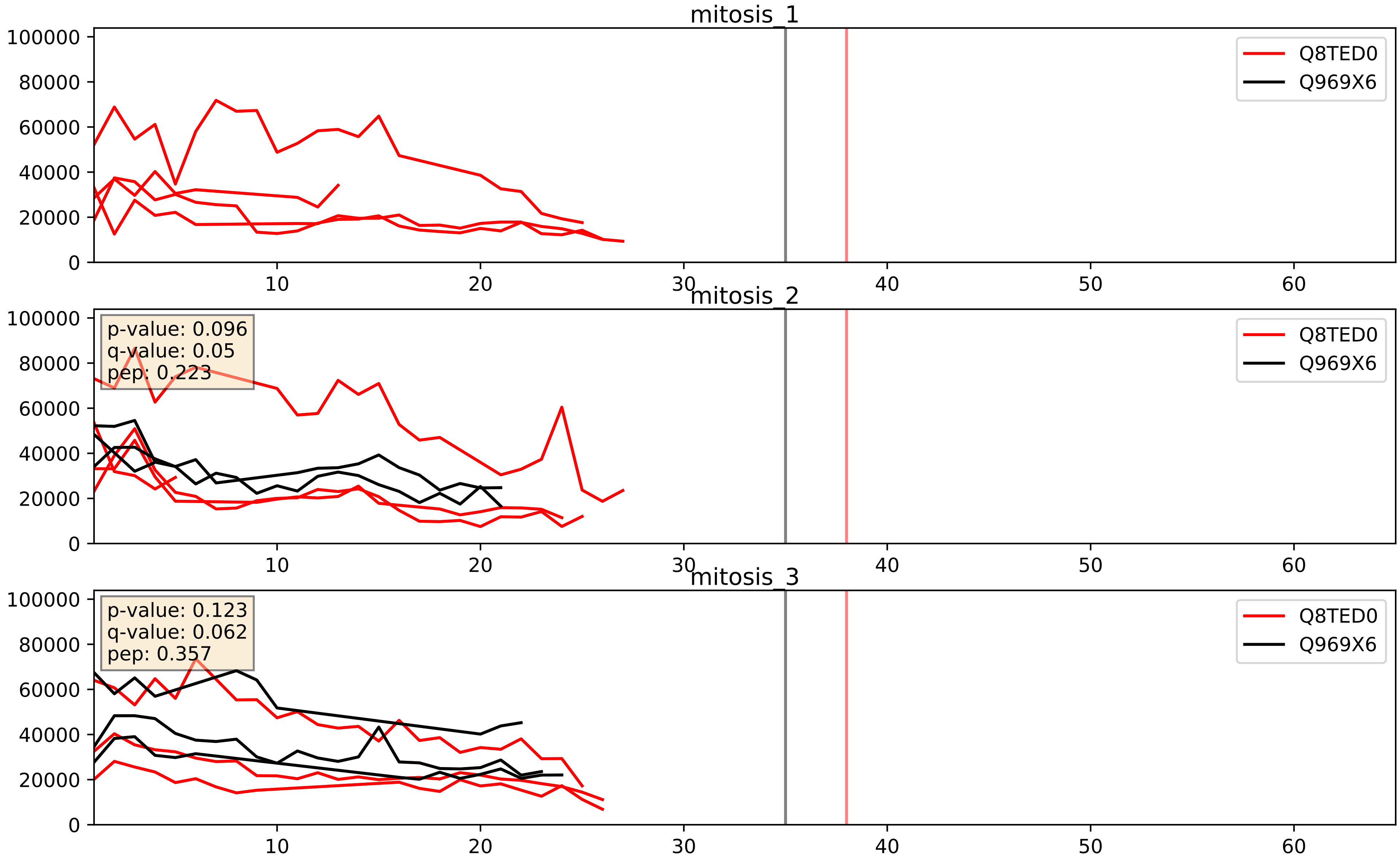


| level | condition_1 | condition_2 | pvalue | pvalue_adjusted |
|---|---|---|---|---|
| interactor_ratio | mitosis | inter | 0.13900618306294427 | 0.5876014454413842 |
| interactor_abundance | mitosis | inter | 0.22392082766632176 | 0.4190762663832913 |
| complex_abundance | mitosis | inter | 0.23996634234660613 | 0.3916523538998532 |

Q8TED0_Q9H583
UTP15_HUMAN vs HEAT1_HUMAN
p-value: 0.048 q-value: 0.032

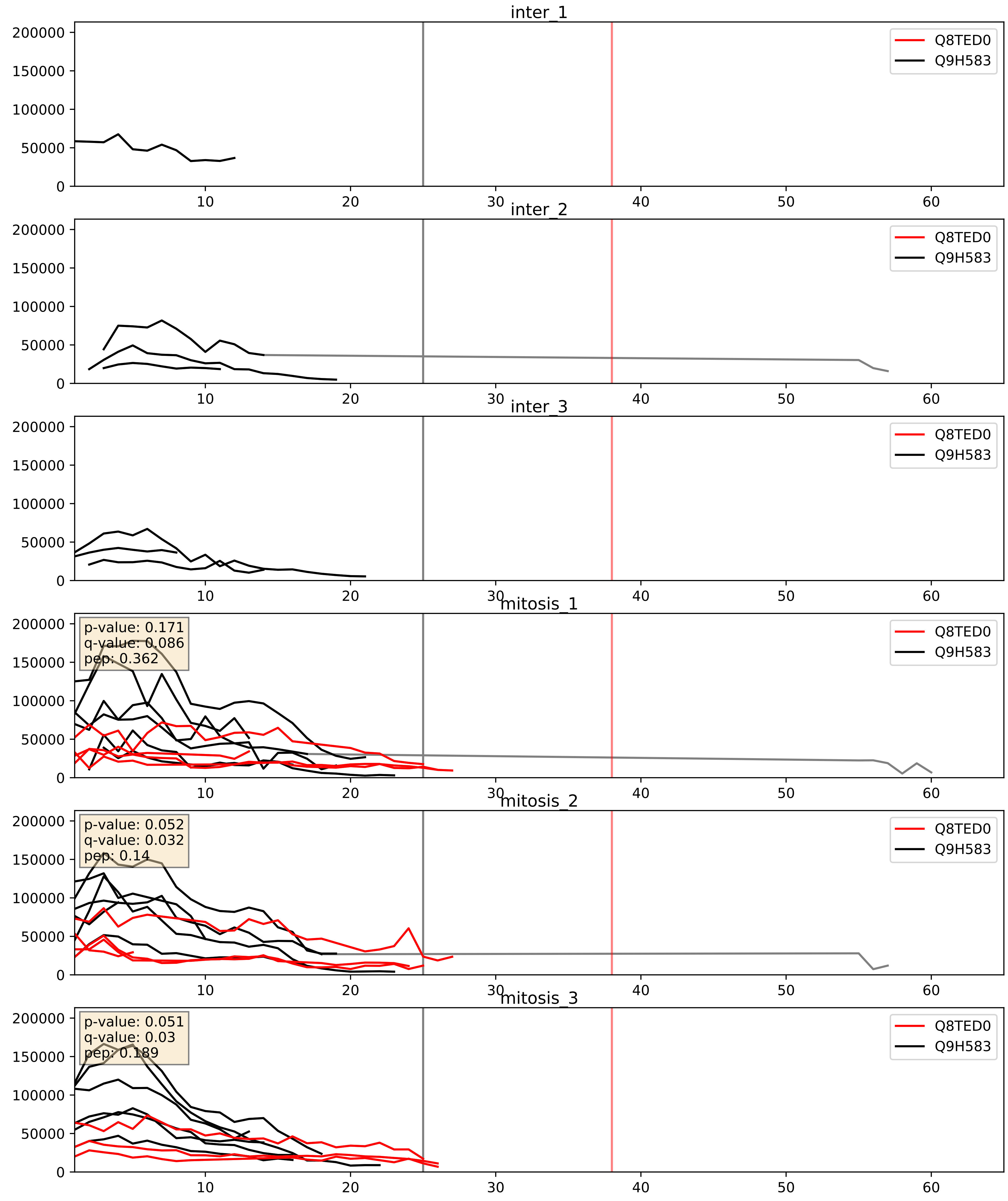


| level | condition_1 | condition_2 | pvalue | pvalue_adjusted |
|---|---|---|---|---|
| complex_abundance | mitosis | inter | 0.00031488057374661957 | 0.01165207110963347 |
| interactor_abundance | mitosis | inter | 0.0015696169880338438 | 0.029025269221018554 |
| interactor_ratio | mitosis | inter | 0.003125922786166029 | 0.0729098066746082 |

Q8TED0_Q9H8H0
UTP15_HUMAN vs NOL11_HUMAN
p-value: 0.045 q-value: 0.03

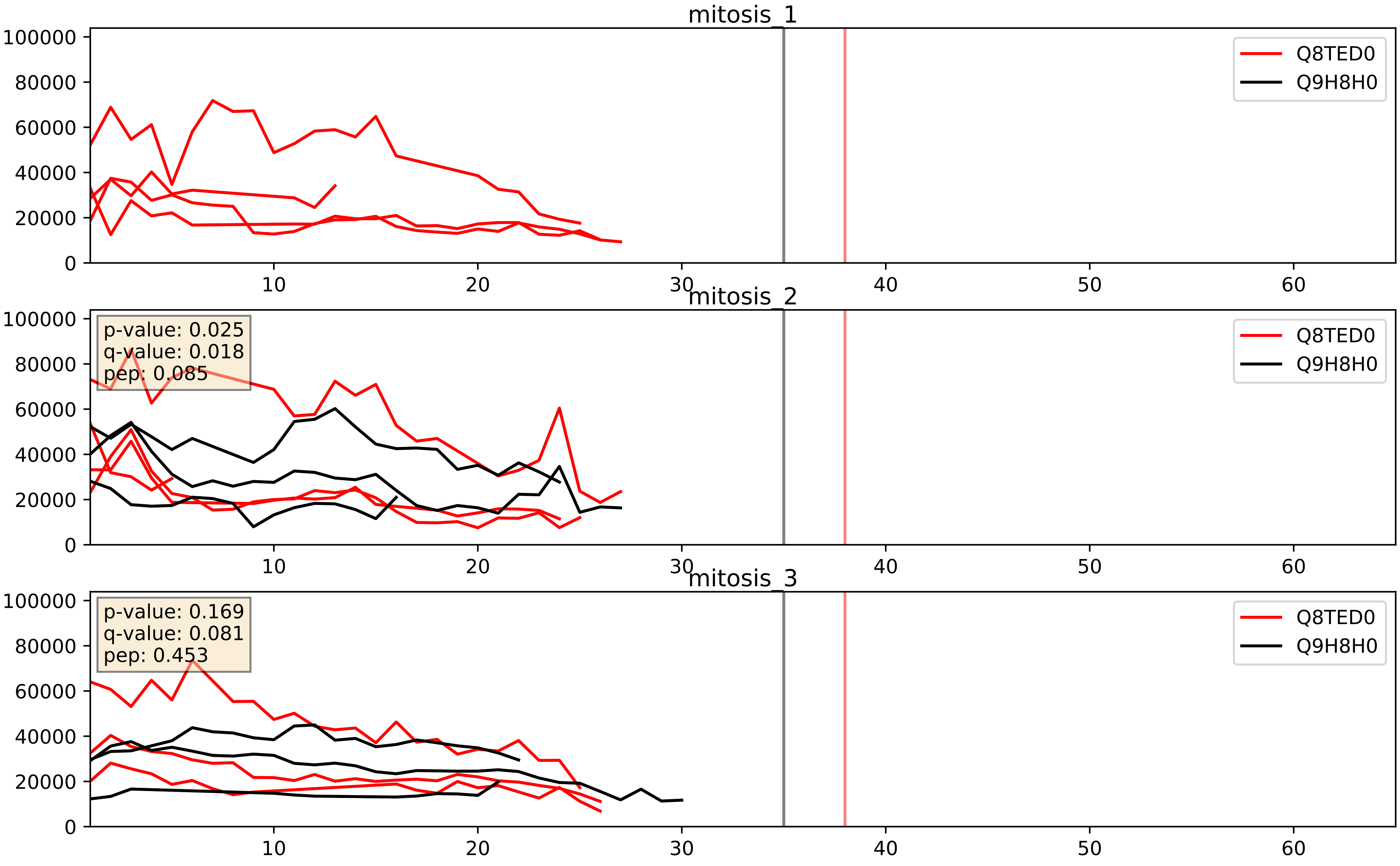


| level | condition_1 | condition_2 | pvalue | pvalue_adjusted |
| --- | --- | --- | --- | --- |
| interactor_abundance | mitosis | inter | 0.21184370799976018 | 0.40715039907095185 |
| complex_abundance | mitosis | inter | 0.2184326512322685 | 0.37304877445088935 |
| interactor_ratio | mitosis | inter | 0.24474365756638608 | 0.7204195520862359 |
