## Supplementary Data 2 for "SECAT: Quantifying differential protein-protein interaction states by network-centric analysis": hela_string_000125_Q969G3.pdf

Q969G3_Q969G3

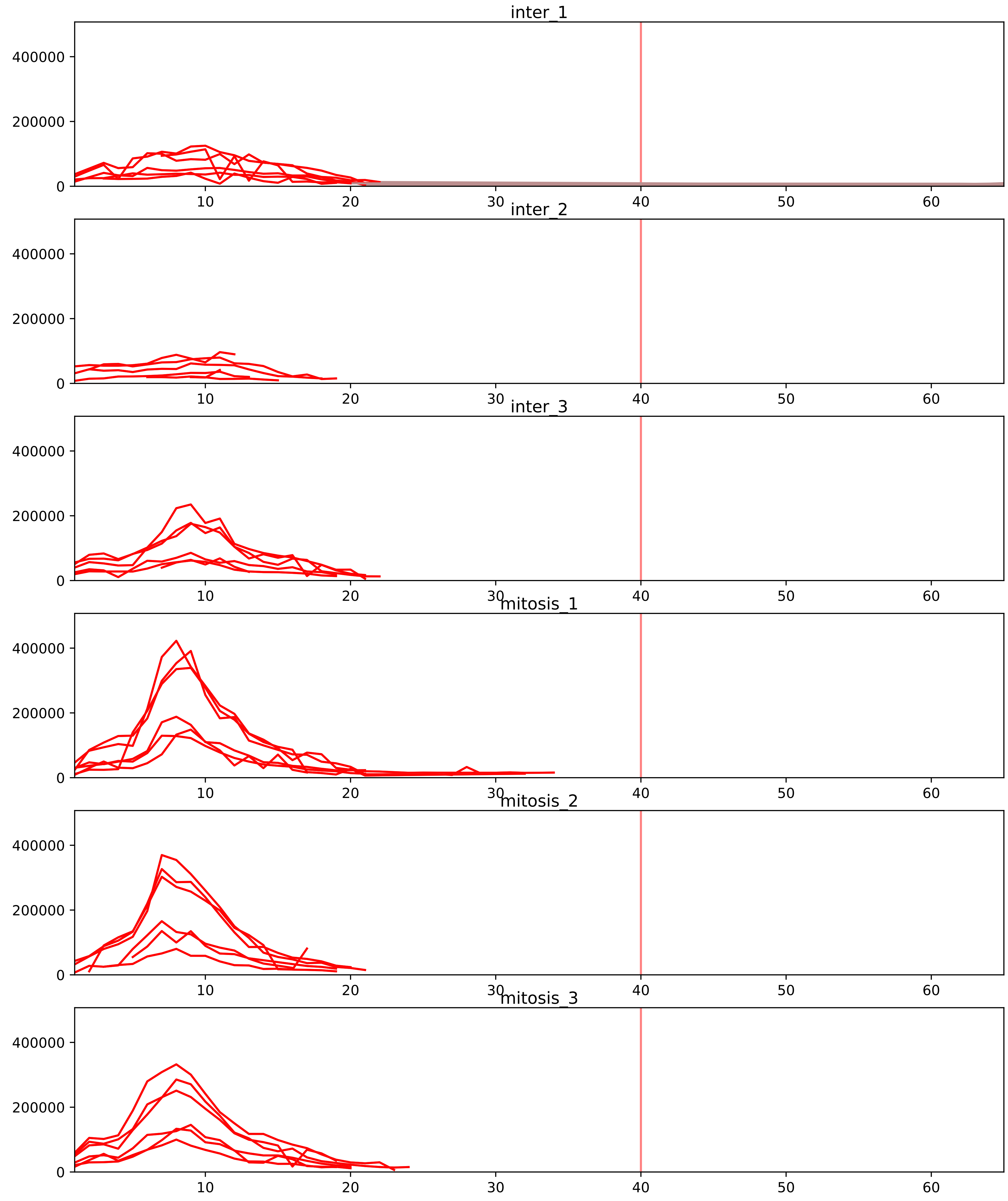


| level | condition_1 | condition_2 | pvalue | pvalue_adjusted |
|---|---|---|---|---|
| interactor_ratio | mitosis | inter | 2.8696775001420228e-05 | 0.0005739355000284046 |
| interactor_abundance | mitosis | inter | 0.0002473218663972253 | 0.0044181770307853835 |
| complex_abundance | mitosis | inter | 0.0003052602911709884 | 0.005014990497809095 |
| total_abundance | mitosis | inter | 0.04890788317167642 | 0.277074883094003 |
| assembled_abundance | mitosis | inter | 0.054730419603357054 | 0.2745906689853316 |
| monomer_abundance | mitosis | inter | 0.31344646763528394 | 0.6718670942432774 |

O14497_Q969G3
ARI1A_HUMAN vs SMCE1_HUMAN
p-value: 0.063 q-value: 0.039

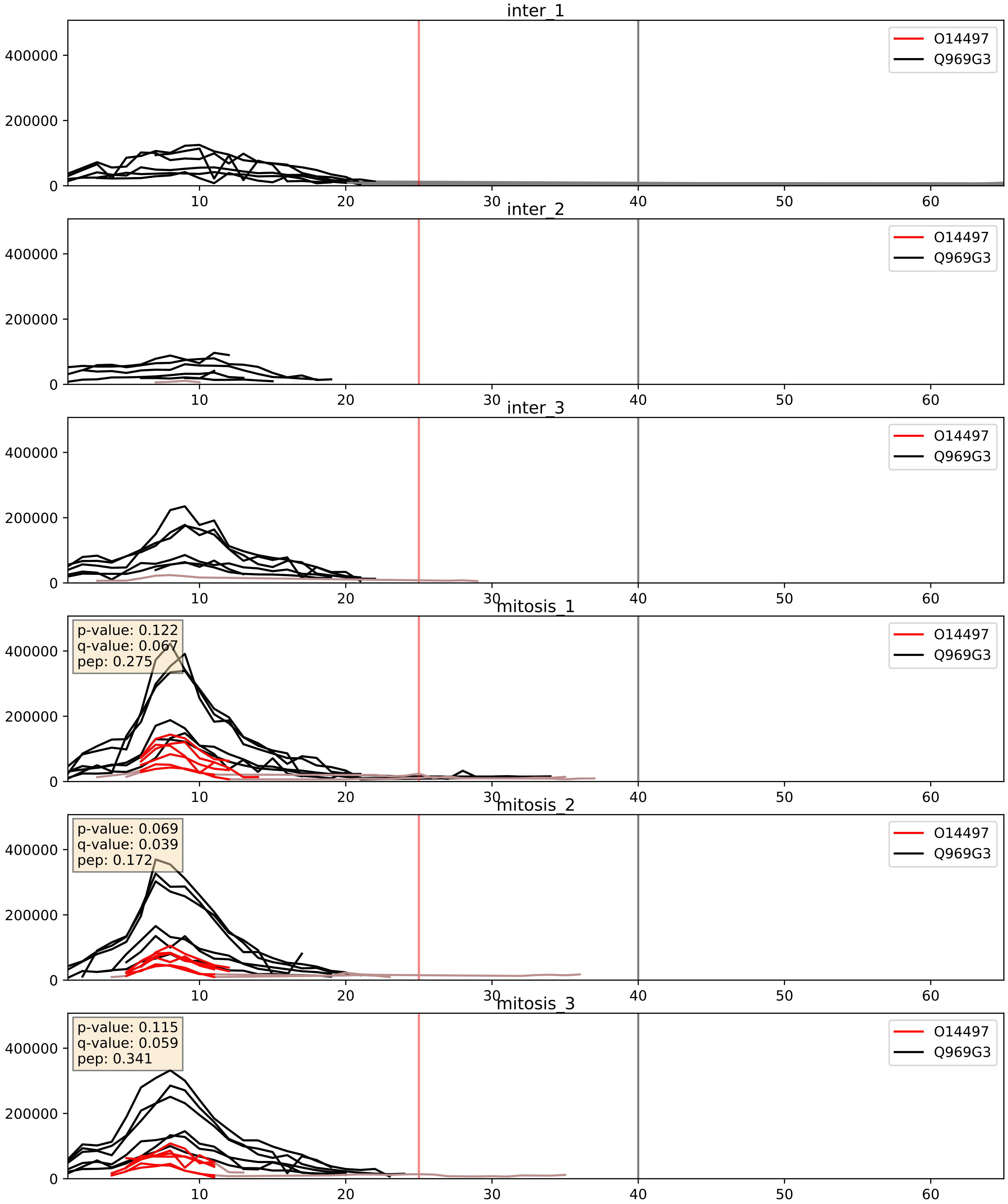


| level | condition_1 | condition_2 | pvalue | pvalue_adjusted |
|---|---|---|---|---|
| interactor_ratio | mitosis | inter | 2.3917376242737105e-05 | 0.004993481478971454 |
| complex_abundance | mitosis | inter | 0.0008870588476678302 | 0.019570164268135633 |
| interactor_abundance | mitosis | inter | 0.012640483206095423 | 0.10027749130318384 |

O96019_Q969G3
ACL6A_HUMAN vs SMCE1_HUMAN
p-value: 0.001 q-value: 0.001

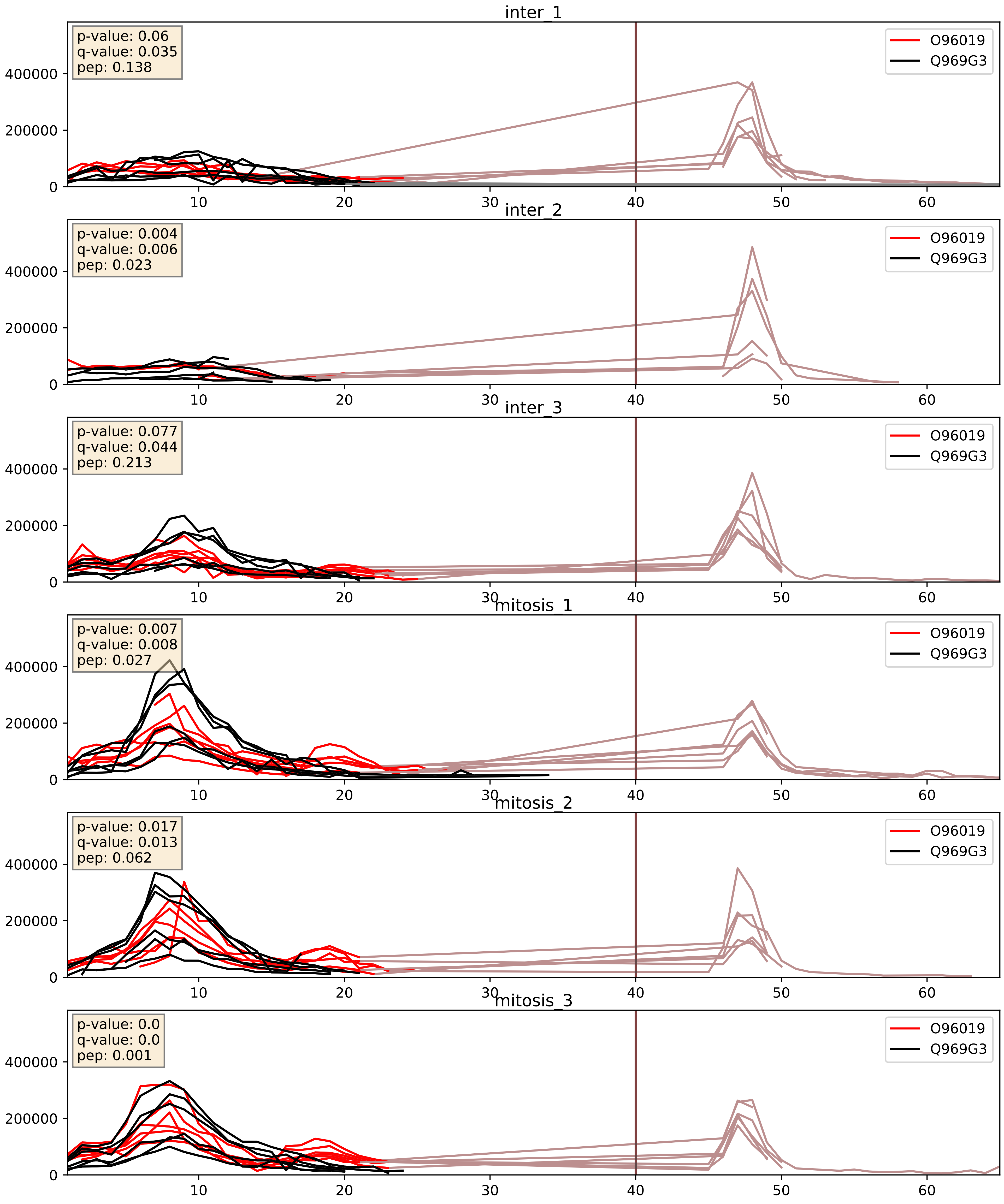


| level | condition_1 | condition_2 | pvalue | pvalue_adjusted |
|---|---|---|---|---|
| complex_abundance | mitosis | inter | 0.031796261929511016 | 0.12971126164945707 |
| interactor_abundance | mitosis | inter | 0.037529994025157706 | 0.15821558673004185 |
| interactor_ratio | mitosis | inter | 0.9284585062171995 | 0.987346851953133 |

P51531_Q969G3
SMCA2_HUMAN vs SMCE1_HUMAN
p-value: 0.047 q-value: 0.031

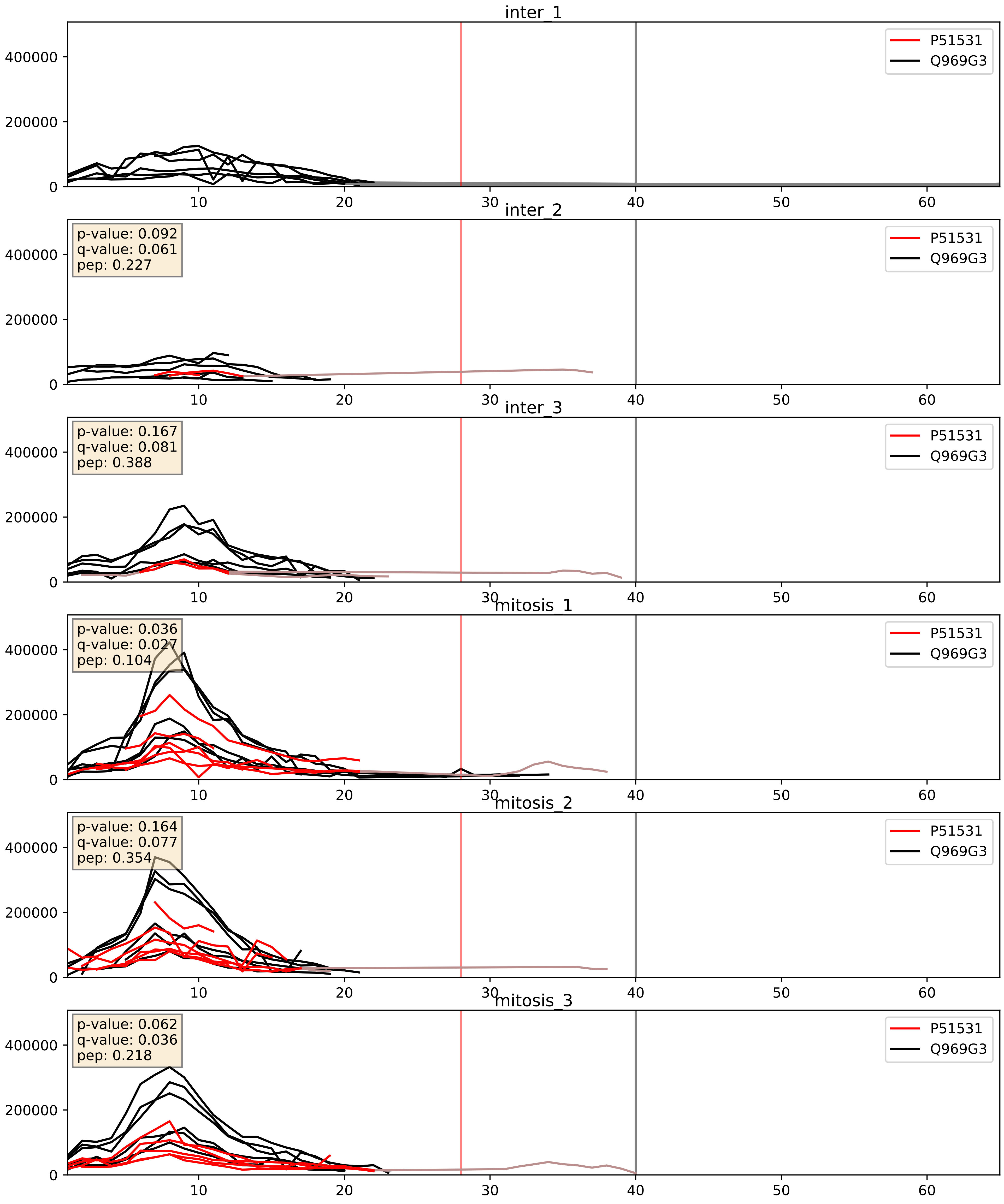


| level | condition_1 | condition_2 | pvalue | pvalue_adjusted |
|---|---|---|---|---|
| complex_abundance | mitosis | inter | 0.008183943251571598 | 0.06710589524334576 |
| interactor_abundance | mitosis | inter | 0.013657409124963755 | 0.10336642096347458 |
| interactor_ratio | mitosis | inter | 0.4403853634924024 | 0.8463612688491312 |

Q8TAQ2_Q969G3
SMRC2_HUMAN vs SMCE1_HUMAN
p-value: 0.001 q-value: 0.001

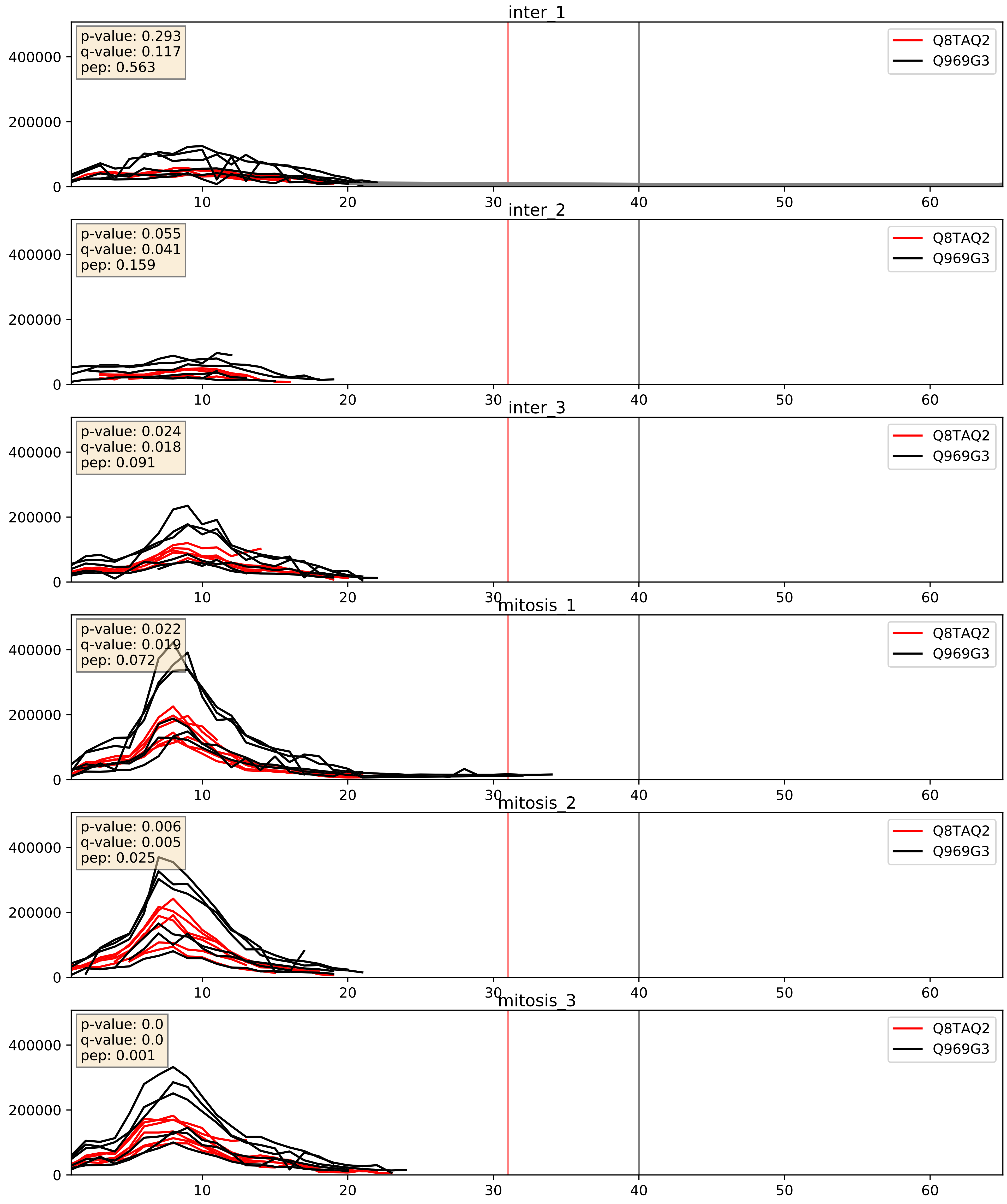


| level | condition_1 | condition_2 | pvalue | pvalue_adjusted |
|---|---|---|---|---|
| interactor_abundance | mitosis | inter | 0.018061703010538034 | 0.11569520685146502 |
| complex_abundance | mitosis | inter | 0.02664451025294988 | 0.11941204595039318 |
| interactor_ratio | mitosis | inter | 0.16673623168381013 | 0.6292809151959985 |

Q92785_Q969G3
REQU_HUMAN vs SMCE1_HUMAN
p-value: 0.0 q-value: 0.013

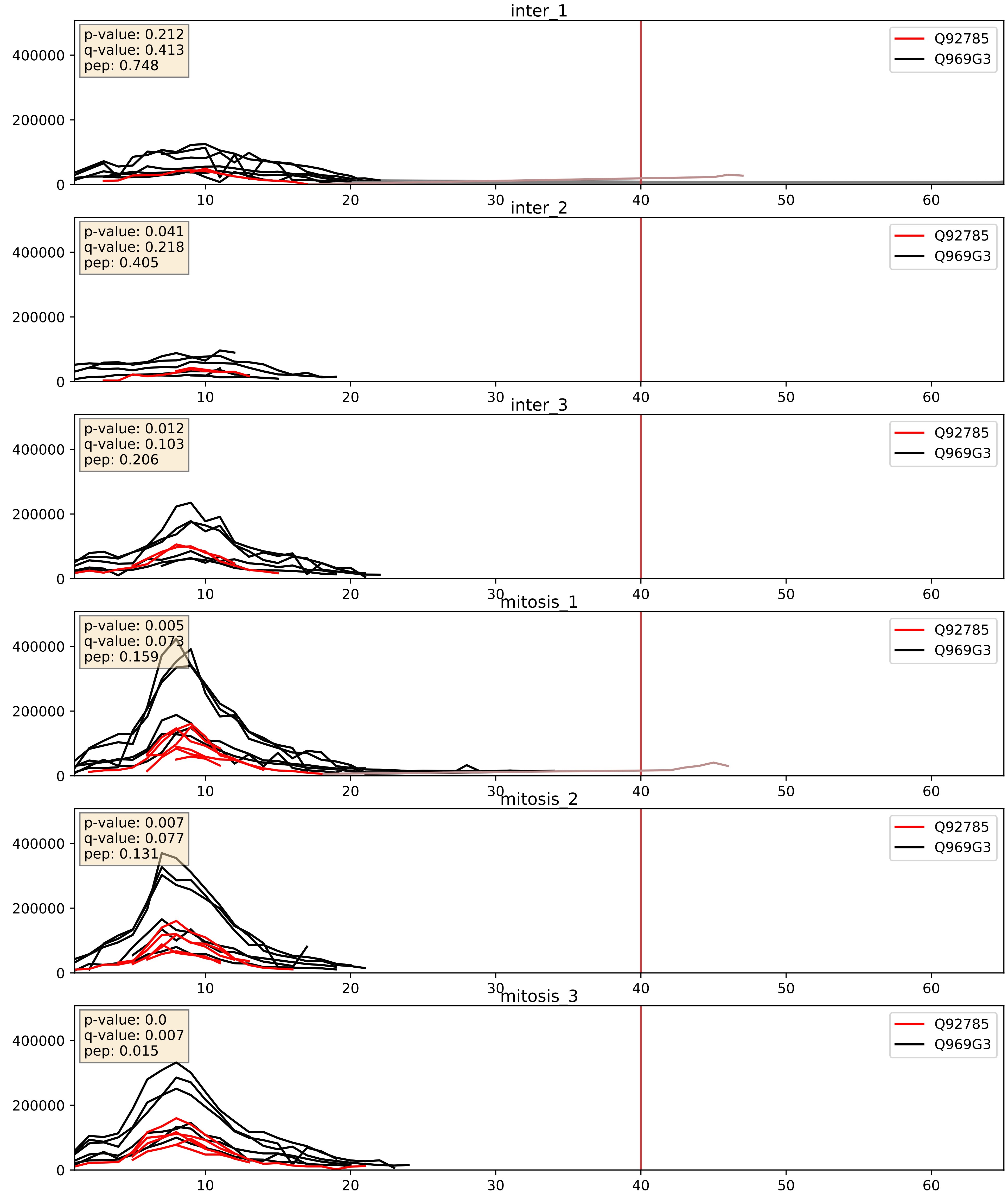


| level | condition_1 | condition_2 | pvalue | pvalue_adjusted |
|---|---|---|---|---|
| interactor_abundance | mitosis | inter | 0.0034125104988964945 | 0.04511548235131213 |
| complex_abundance | mitosis | inter | 0.009769144173742679 | 0.07329383942917299 |
| interactor_ratio | mitosis | inter | 0.14847644305248364 | 0.60240057967345 |

Q92922_Q969G3
SMRC1_HUMAN vs SMCE1_HUMAN
p-value: 0.0 q-value: 0.0

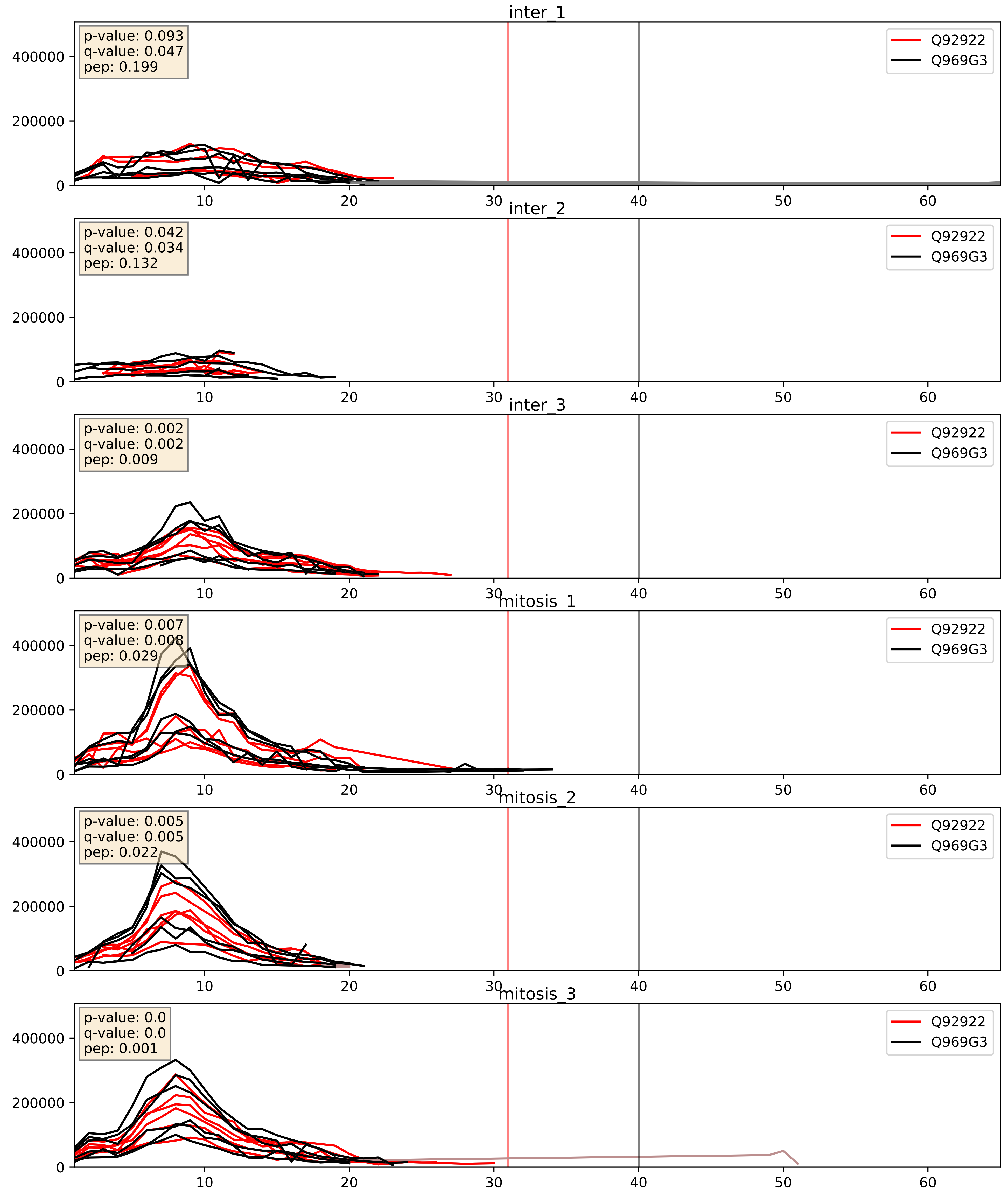


| level | condition_1 | condition_2 | pvalue | pvalue_adjusted |
|---|---|---|---|---|
| complex_abundance | mitosis | inter | 0.05577982617922177 | 0.16848105578480535 |
| interactor_abundance | mitosis | inter | 0.07736330381217478 | 0.23055134583284403 |
| interactor_ratio | mitosis | inter | 0.21333639791135564 | 0.6904192954236995 |

Q92925_Q969G3
SMRD2_HUMAN vs SMCE1_HUMAN
p-value: 0.017 q-value: 0.014

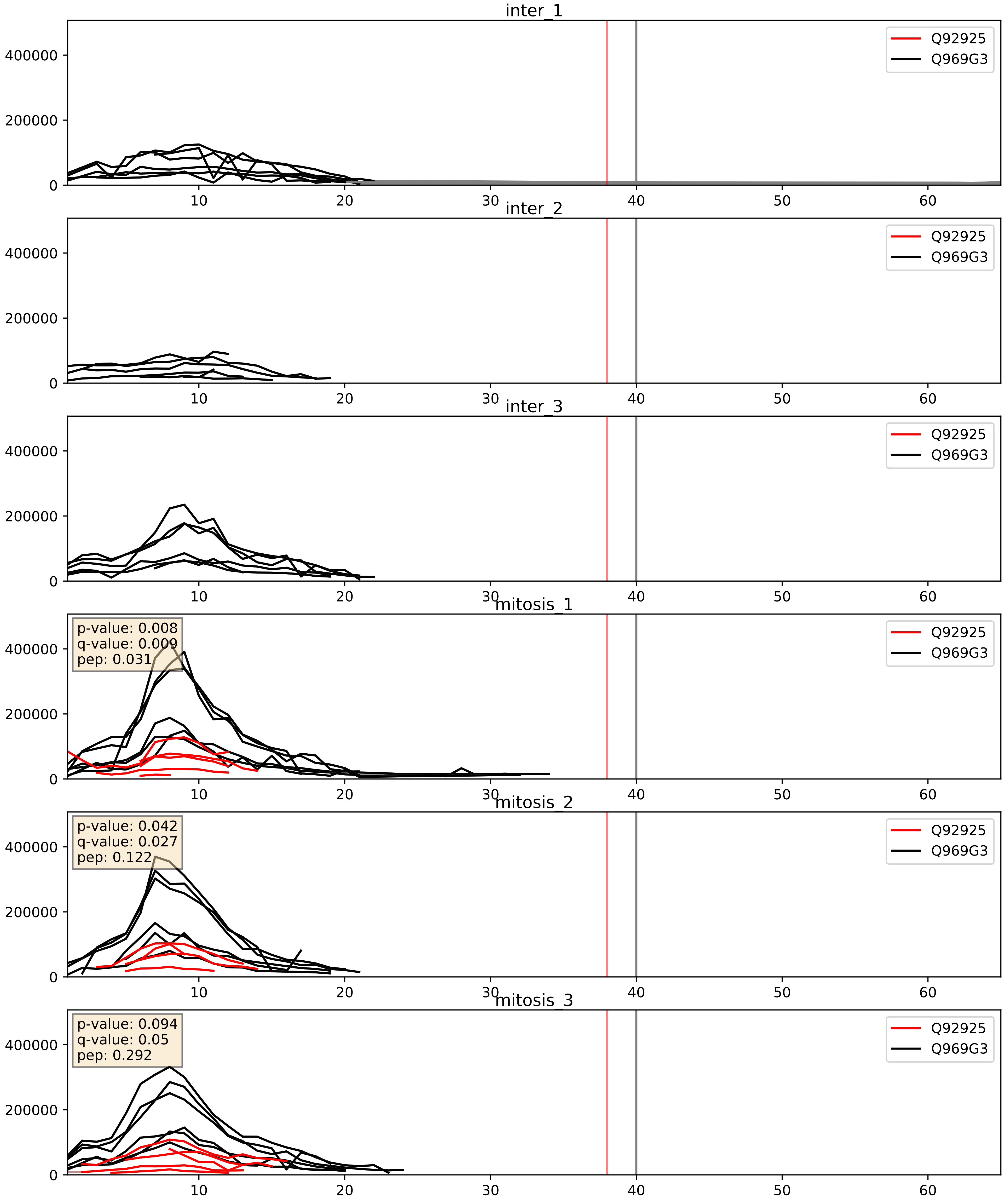


| level | condition_1 | condition_2 | pvalue | pvalue_adjusted |
|---|---|---|---|---|
| interactor_ratio | mitosis | inter | 3.4231436616441946e-06 | 0.002520386835947235 |
| complex_abundance | mitosis | inter | 0.0003362431142253742 | 0.011893558089955385 |
| interactor_abundance | mitosis | inter | 0.004123135752406863 | 0.050966125690401085 |

Q969G3_Q96GM5
SMCE1_HUMAN vs SMRD1_HUMAN
p-value: 0.0 q-value: 0.0

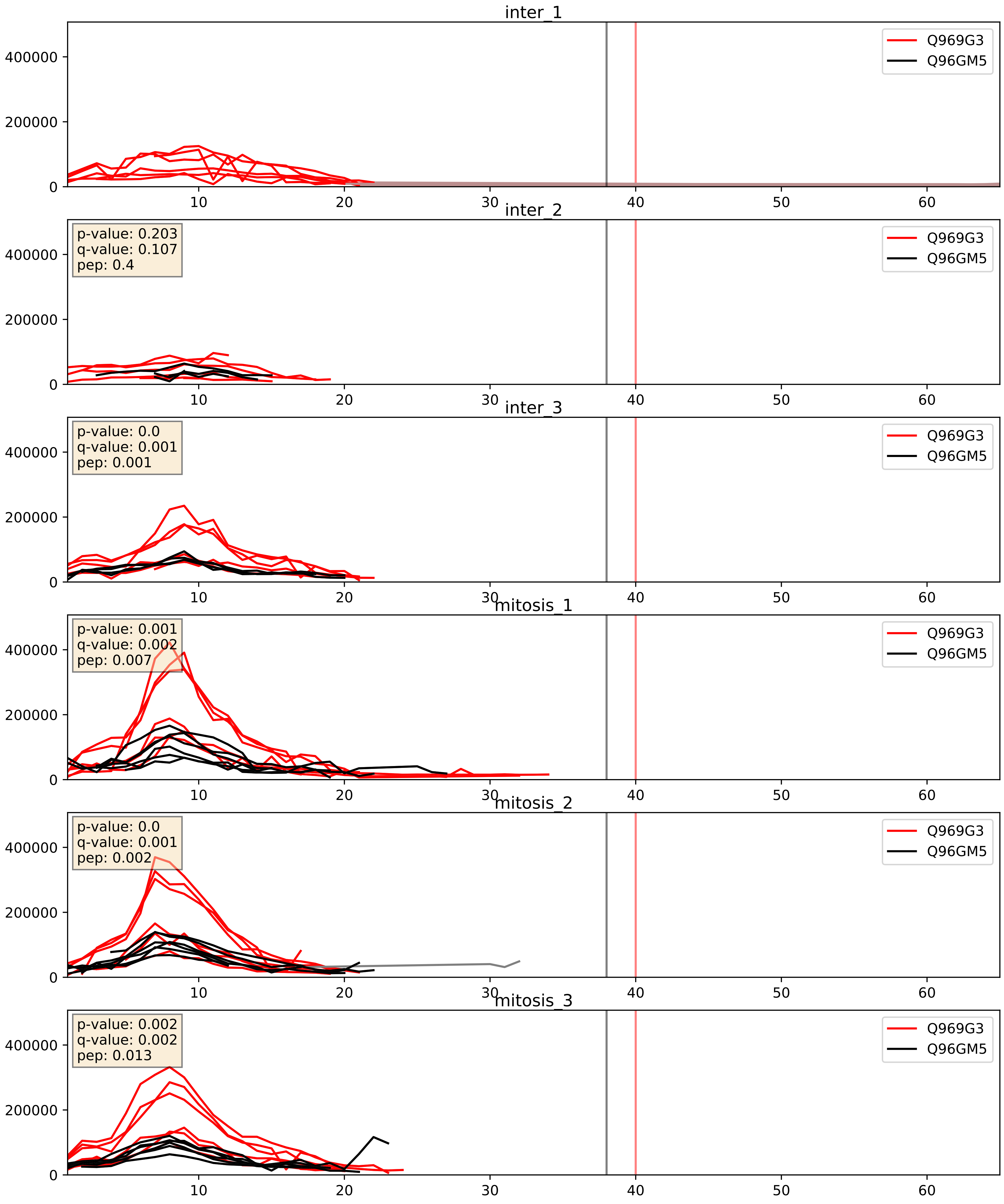


| level | condition_1 | condition_2 | pvalue | pvalue_adjusted |
|---|---|---|---|---|
| complex_abundance | mitosis | inter | 0.05012708341796544 | 0.1593936976440506 |
| interactor_abundance | mitosis | inter | 0.10625982548010601 | 0.27884246048734135 |
| interactor_ratio | mitosis | inter | 0.6739310697288592 | 0.9319765113016878 |
