## Supplementary Data 2 for "SECAT: Quantifying differential protein-protein interaction states by network-centric analysis": hela_string_000126_Q13247.pdf

Q13247_Q13247

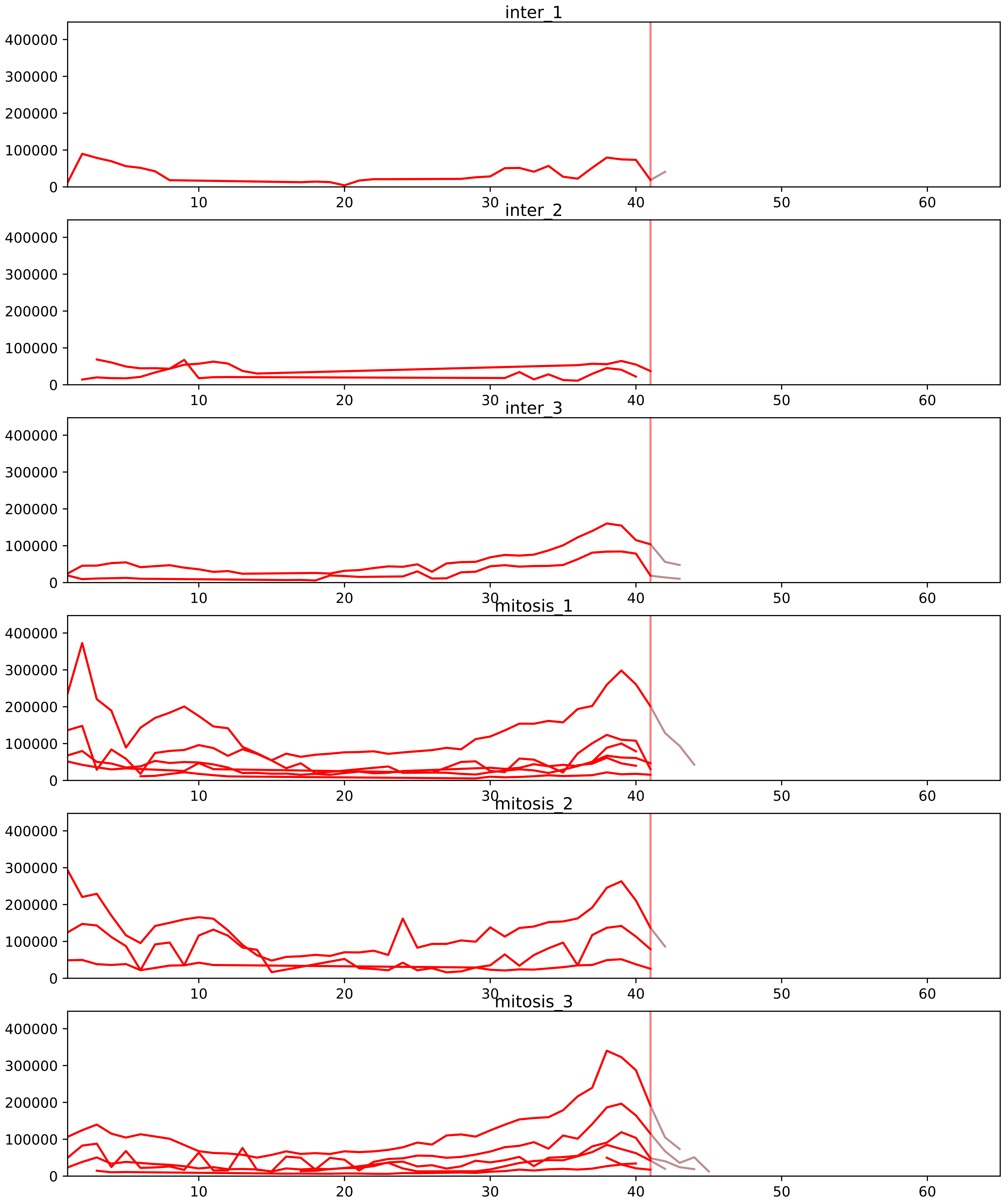


| level | condition_1 | condition_2 | pvalue | pvalue_adjusted |
|---|---|---|---|---|
| complex_abundance | mitosis | inter | 3.0509655457769694e-05 | 0.001059203132873514 |
| interactor_abundance | mitosis | inter | 0.0008894069348561209 | 0.010406300872833717 |
| total_abundance | mitosis | inter | 0.026075455590452712 | 0.2072098874208553 |
| assembled_abundance | mitosis | inter | 0.027110412932334814 | 0.19931285478519598 |
| monomer_abundance | mitosis | inter | 0.06296091407949753 | 0.3605569711864924 |
| interactor_ratio | mitosis | inter | 0.20485556576140154 | 0.5097668221526757 |

O43390_Q13247
HNRPR_HUMAN vs SRSF6_HUMAN
p-value: 0.0 q-value: 0.005

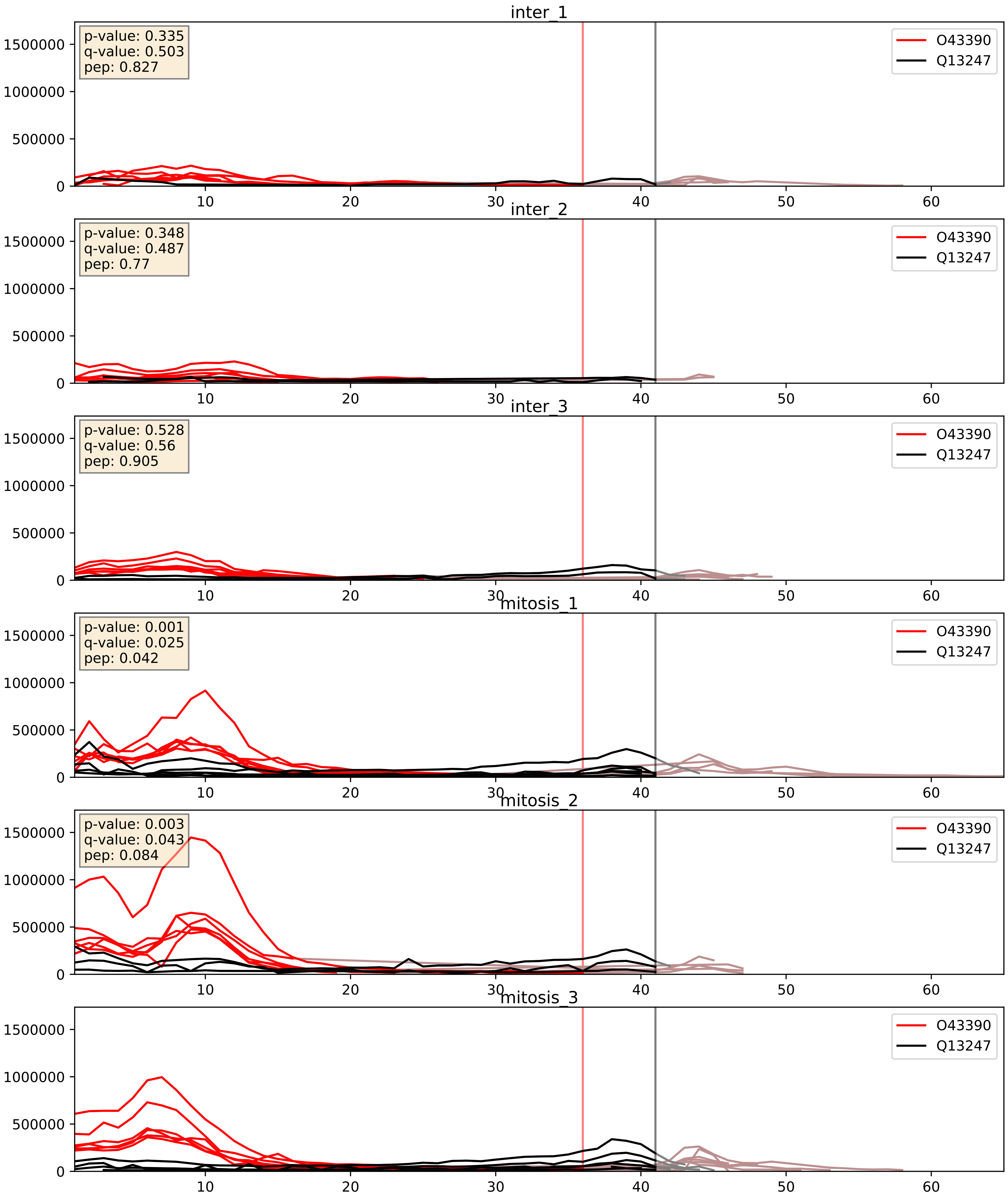


| level | condition_1 | condition_2 | pvalue | pvalue_adjusted |
|---|---|---|---|---|
| interactor_abundance | mitosis | inter | 0.0005707867388178207 | 0.017415659477552636 |
| complex_abundance | mitosis | inter | 0.0006004542403235107 | 0.01581504091436693 |
| interactor_ratio | mitosis | inter | 0.9652626032384787 | 0.9937294965826962 |

P11940_Q13247
PABP1_HUMAN vs SRSF6_HUMAN
p-value: 0.0 q-value: 0.047

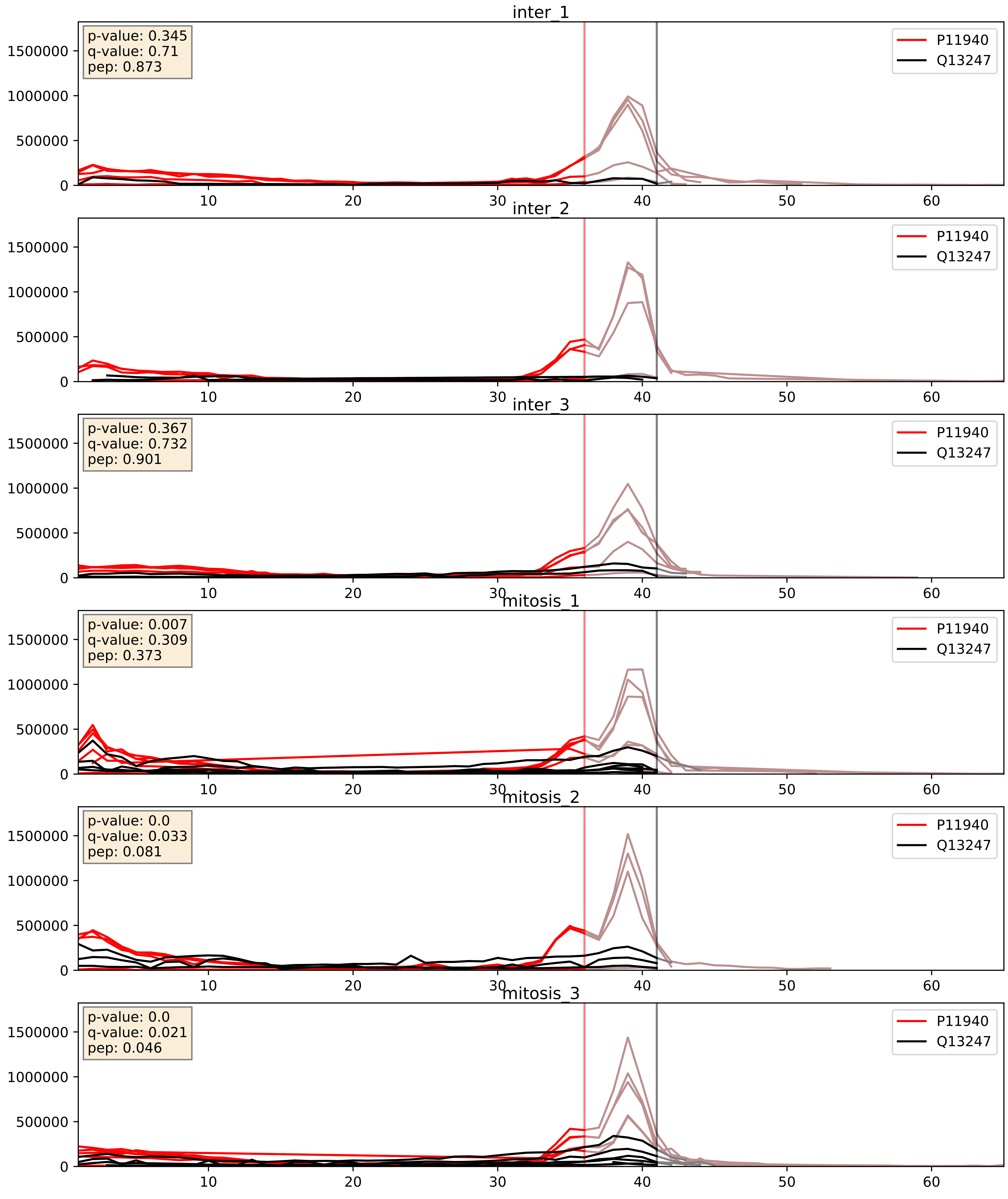


| level | condition_1 | condition_2 | pvalue | pvalue_adjusted |
|---|---|---|---|---|
| complex_abundance | mitosis | inter | 0.007836804827711203 | 0.0657361191517291 |
| interactor_abundance | mitosis | inter | 0.008345387686835815 | 0.07950642025521934 |
| interactor_ratio | mitosis | inter | 0.04176762891071437 | 0.3395355208696249 |

P17844_Q13247
DDX5_HUMAN vs SRSF6_HUMAN
p-value: 0.0 q-value: 0.005

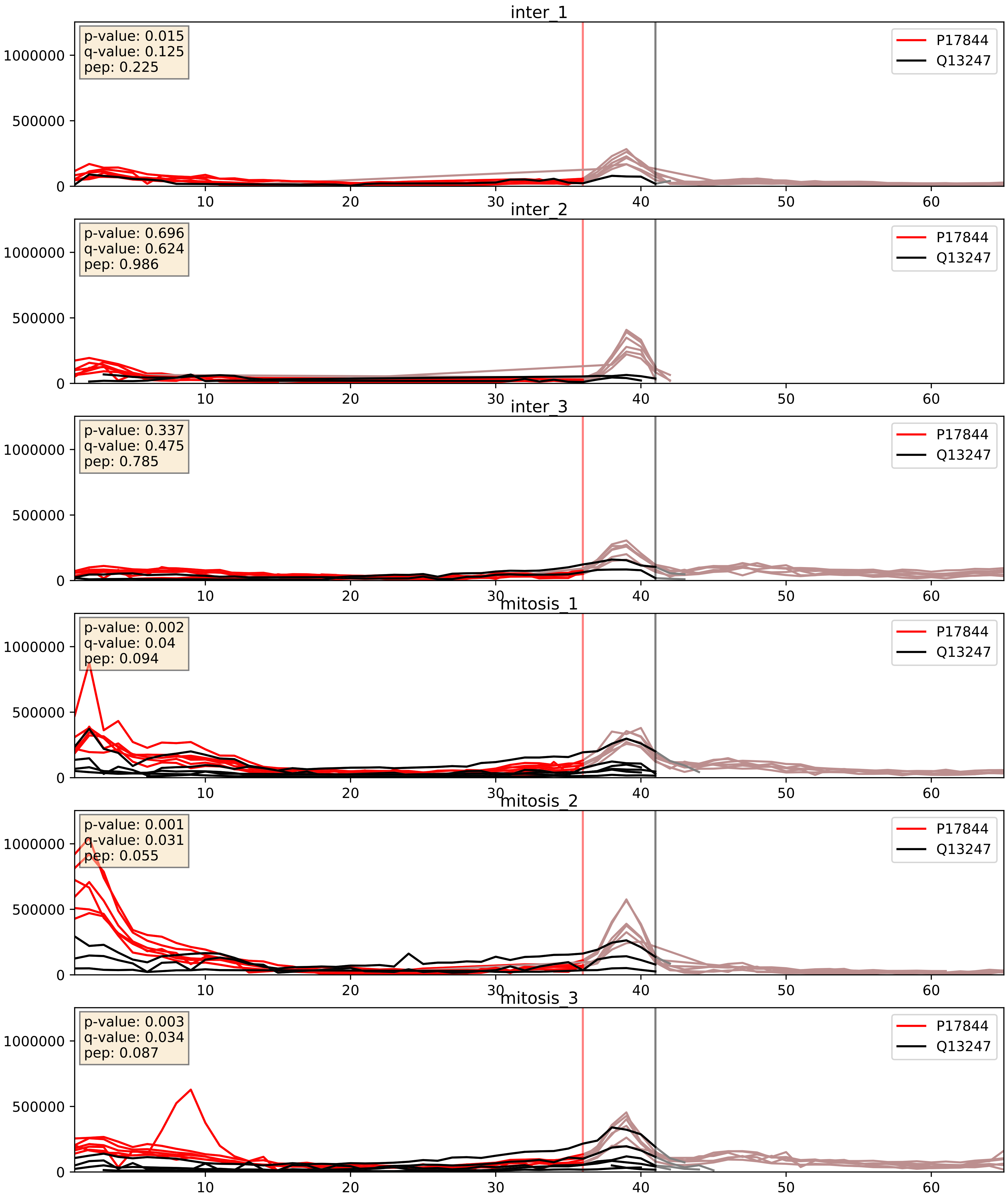


| level | condition_1 | condition_2 | pvalue | pvalue_adjusted |
|---|---|---|---|---|
| complex_abundance | mitosis | inter | 0.002718775066662734 | 0.03753663640424678 |
| interactor_abundance | mitosis | inter | 0.003078864127685547 | 0.042536422819834926 |
| interactor_ratio | mitosis | inter | 0.6545547922536914 | 0.9236711212811735 |

P26368_Q13247
U2AF2_HUMAN vs SRSF6_HUMAN
p-value: 0.001 q-value: 0.001

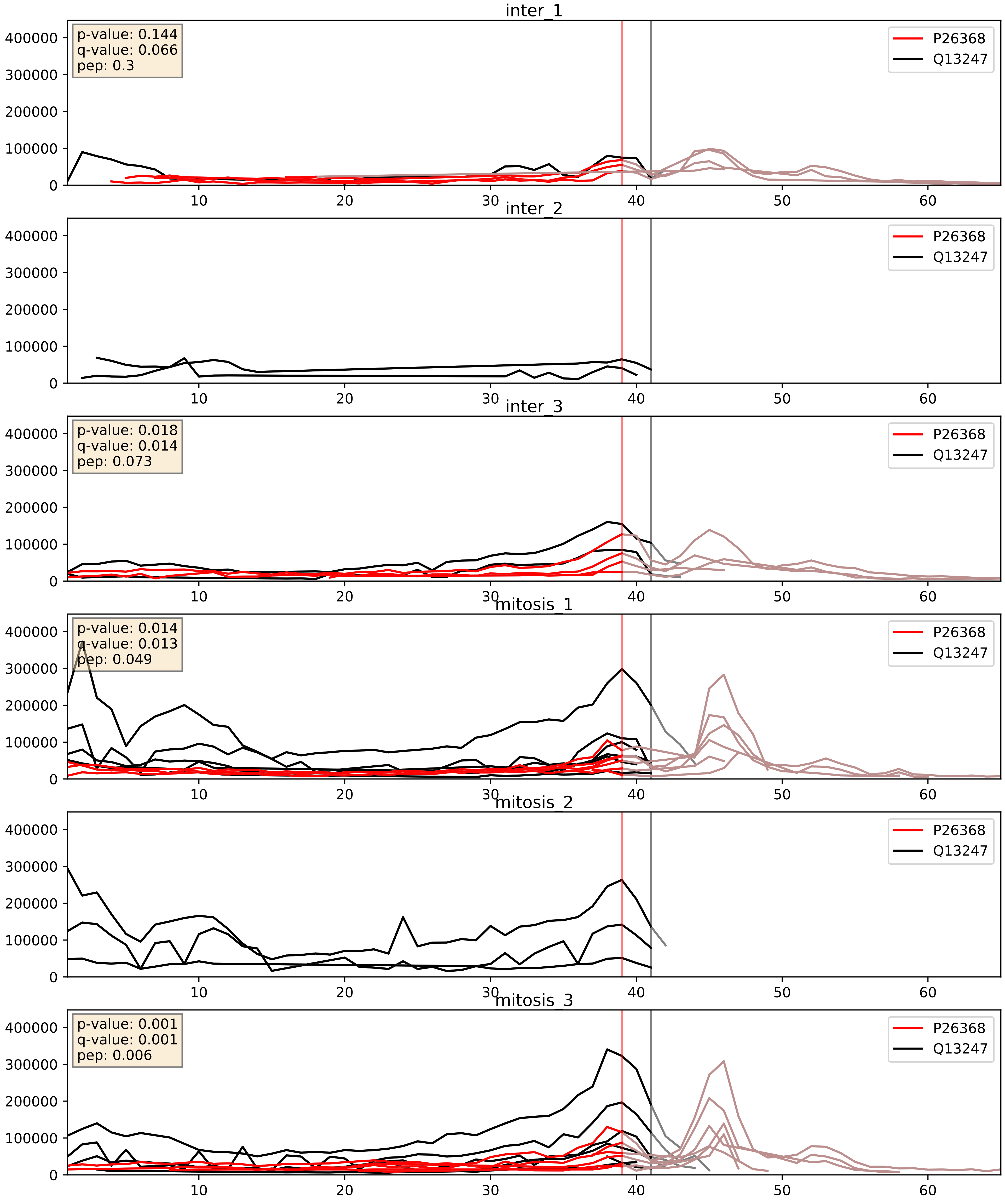


| level | condition_1 | condition_2 | pvalue | pvalue_adjusted |
|---|---|---|---|---|
| interactor_ratio | mitosis | inter | 0.09507039565668088 | 0.512147631731396 |
| complex_abundance | mitosis | inter | 0.7508450084124039 | 0.835375407525308 |
| interactor_abundance | mitosis | inter | 0.8487972712124756 | 0.9177345764277872 |

P26599_Q13247
PTBP1_HUMAN vs SRSF6_HUMAN
p-value: 0.002 q-value: 0.034

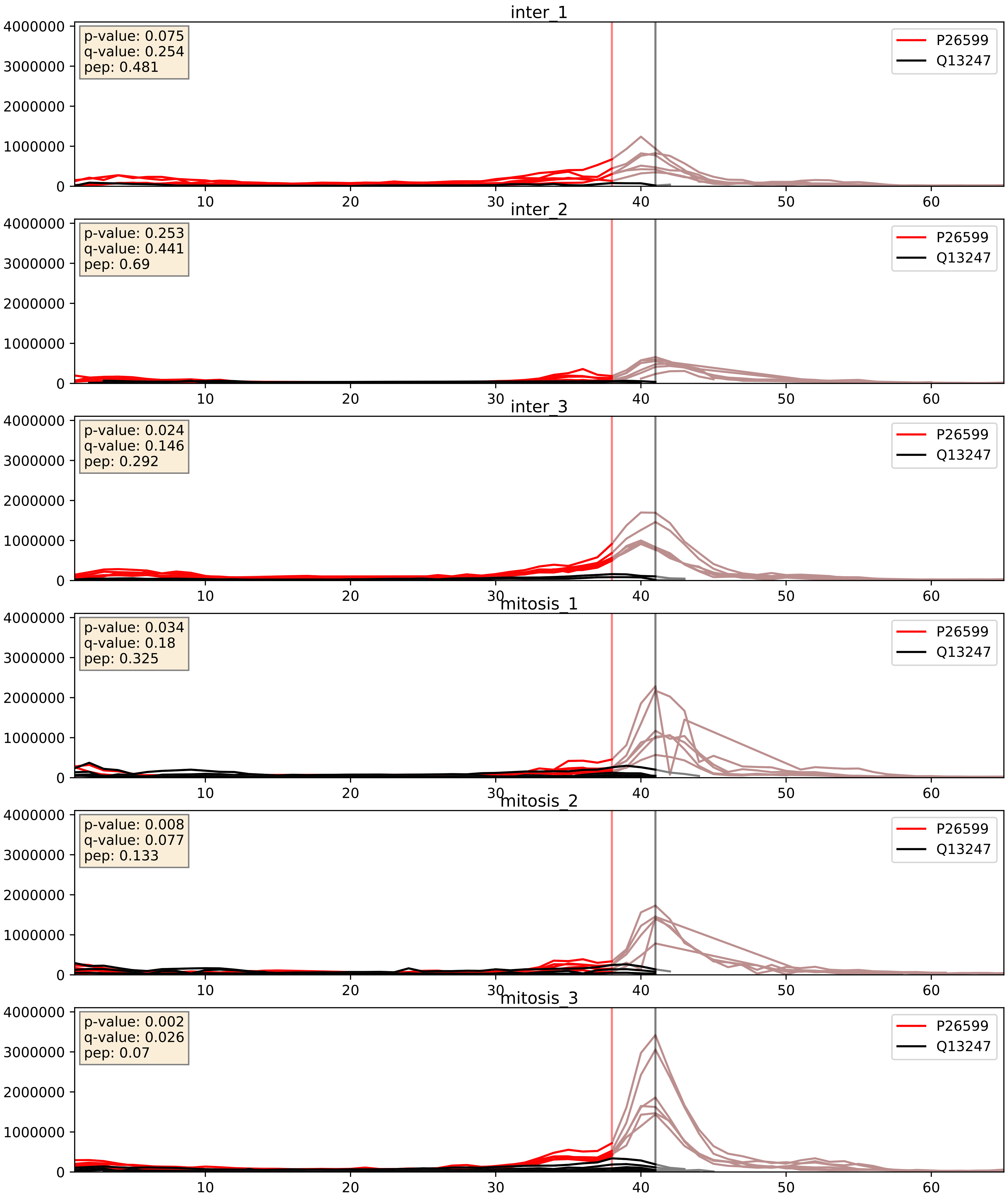


| level | condition_1 | condition_2 | pvalue | pvalue_adjusted |
|---|---|---|---|---|
| interactor_ratio | mitosis | inter | 0.019132910025978646 | 0.20874187291953322 |
| complex_abundance | mitosis | inter | 0.4046989653308828 | 0.5499639852726396 |
| interactor_abundance | mitosis | inter | 0.5028274627208574 | 0.6675252917013864 |

P31943_Q13247
HNRH1_HUMAN vs SRSF6_HUMAN
p-value: 0.005 q-value: 0.005

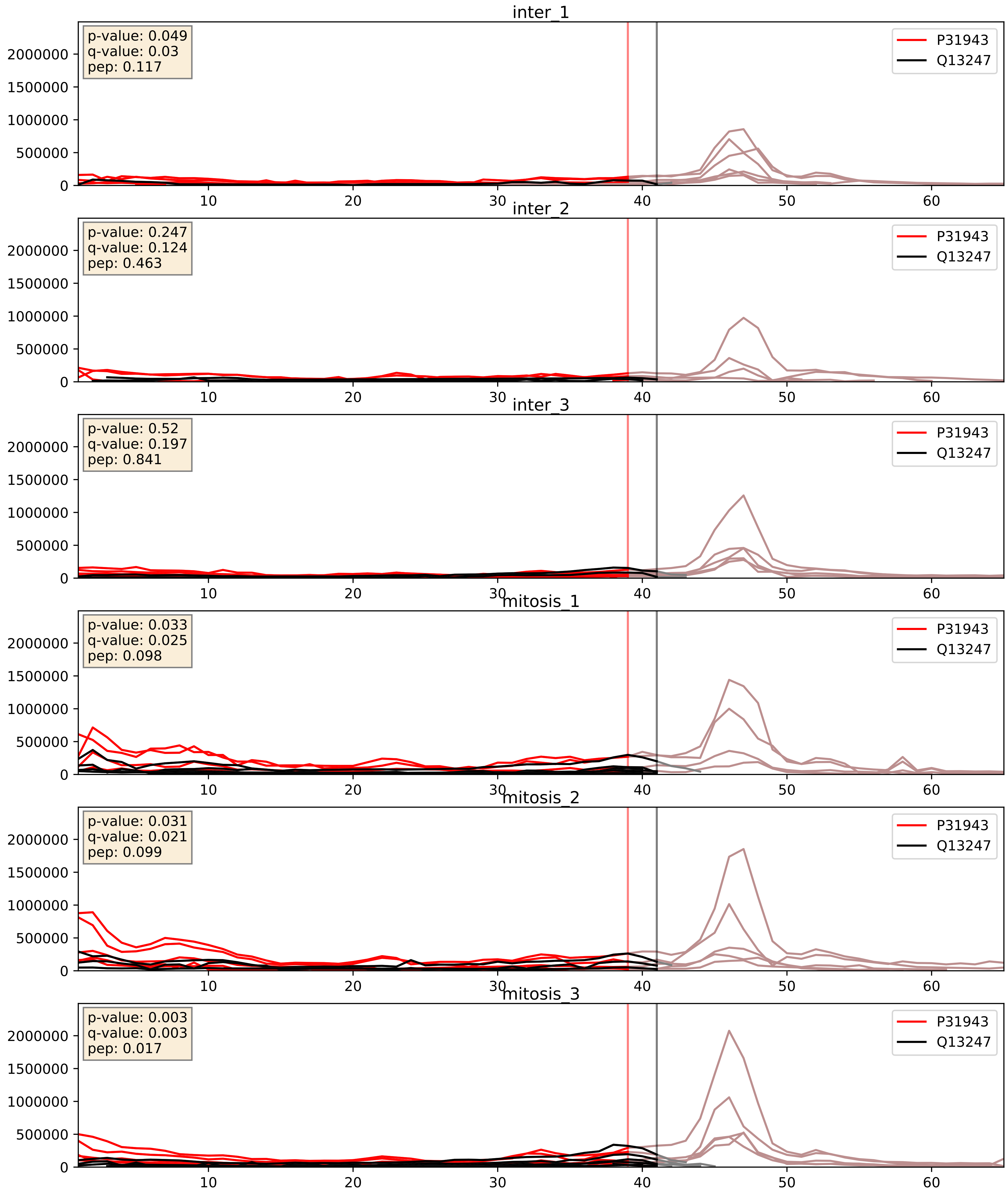


| level | condition_1 | condition_2 | pvalue | pvalue_adjusted |
| --- | --- | --- | --- | --- |
| complex_abundance | mitosis | inter | 0.004753245691634782 | 0.05150352293720726 |
| interactor_abundance | mitosis | inter | 0.0056571357548407175 | 0.06348577161904892 |
| interactor_ratio | mitosis | inter | 0.9922029355017021 | 0.9981863229331772 |

P52597_Q13247
HNRPF_HUMAN vs SRSF6_HUMAN
p-value: 0.002 q-value: 0.03

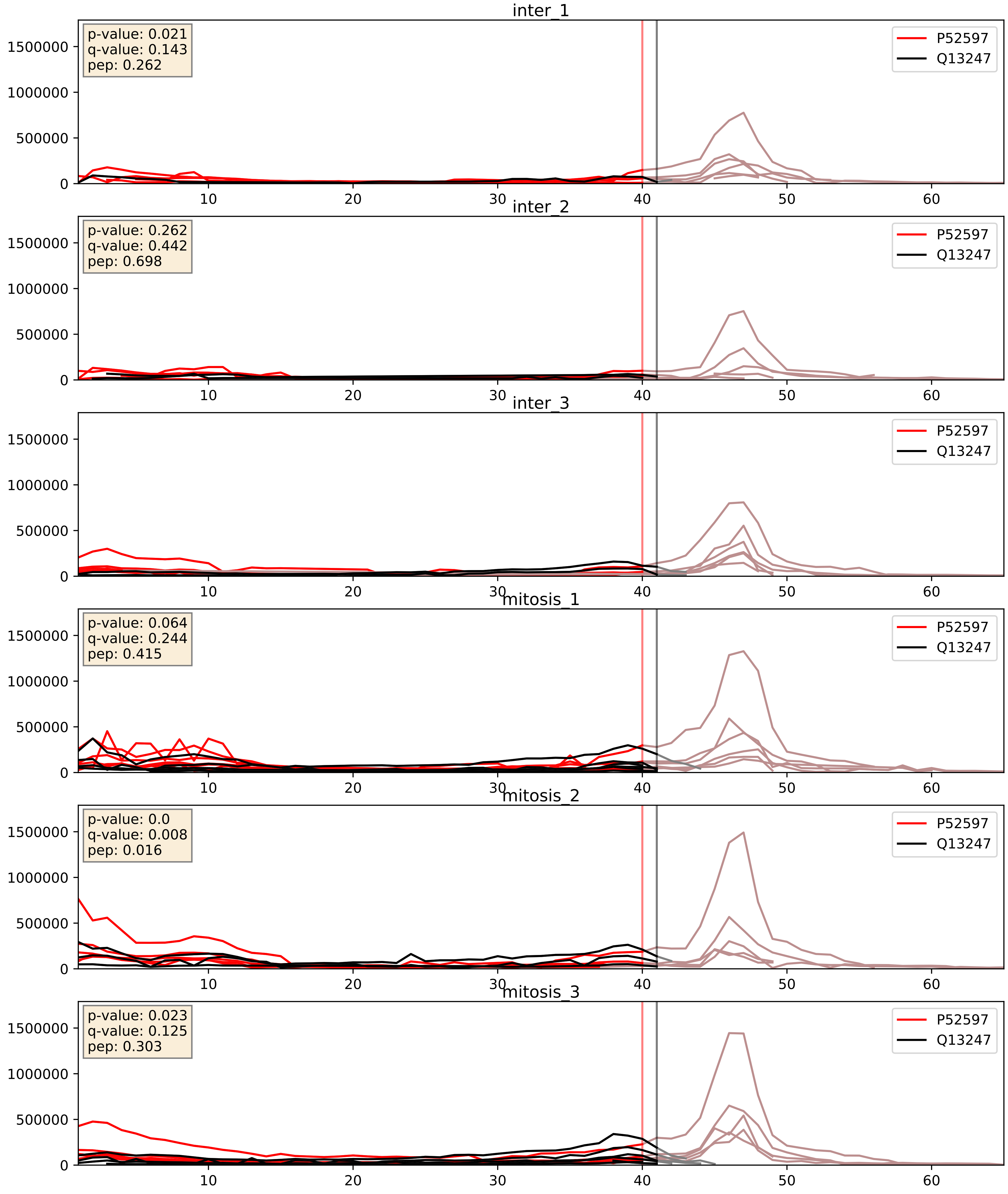


| level | condition_1 | condition_2 | pvalue | pvalue_adjusted |
|---|---|---|---|---|
| interactor_abundance | mitosis | inter | 0.005847440986472472 | 0.06479494478214157 |
| complex_abundance | mitosis | inter | 0.010125200125526758 | 0.07439632023563009 |
| interactor_ratio | mitosis | inter | 0.2895904474858344 | 0.7585355662419652 |

P62995_Q13247
TRA2B_HUMAN vs SRSF6_HUMAN
p-value: 0.007 q-value: 0.006

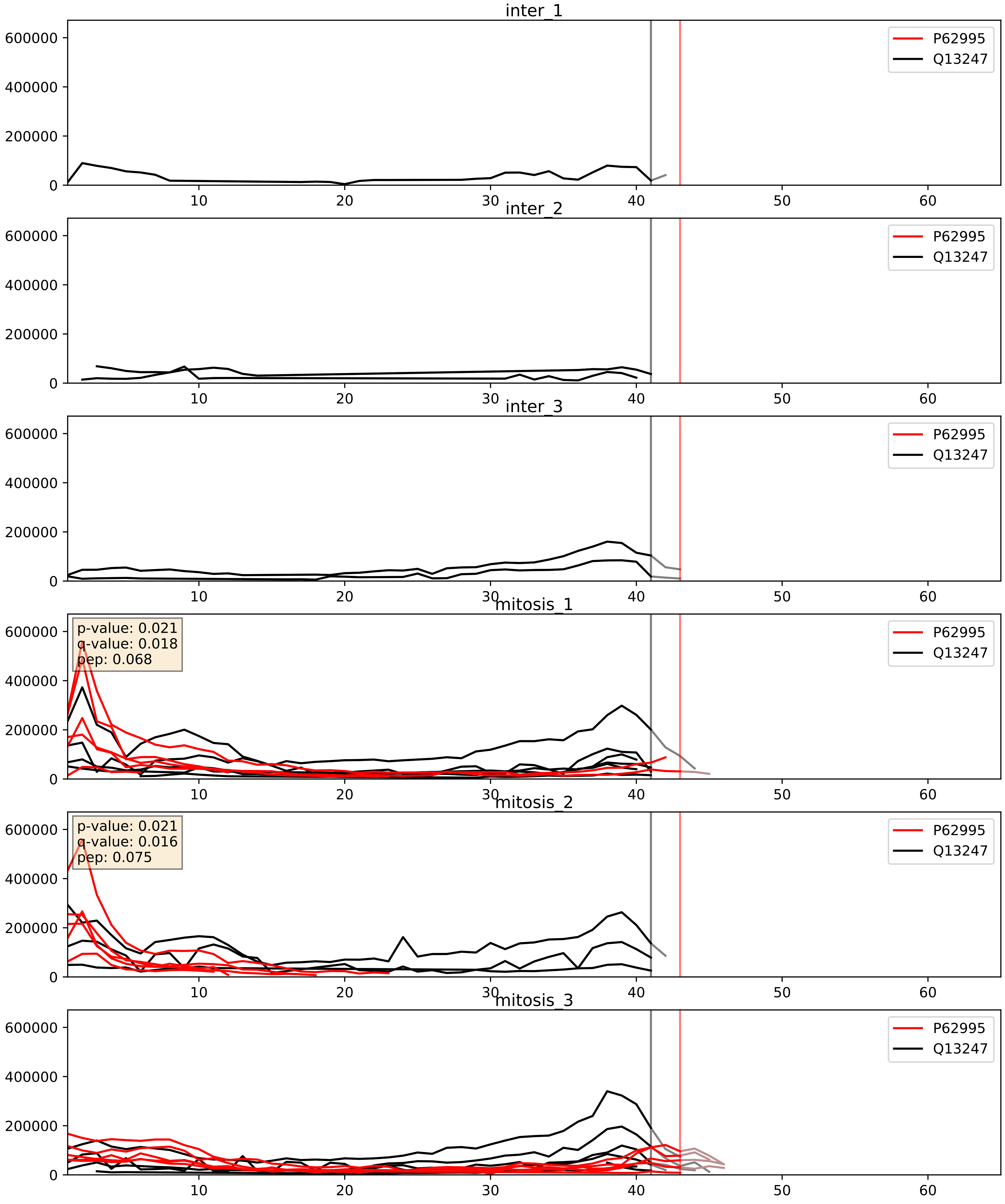


| level | condition_1 | condition_2 | pvalue | pvalue_adjusted |
|---|---|---|---|---|
| interactor_abundance | mitosis | inter | 0.00038990687589259026 | 0.01362698060740691 |
| complex_abundance | mitosis | inter | 0.002674131184473713 | 0.037220427543243874 |
| interactor_ratio | mitosis | inter | 0.9430800845236742 | 0.9912117205440141 |

Q07955_Q13247
SRSF1_HUMAN vs SRSF6_HUMAN
p-value: 0.0 q-value: 0.0

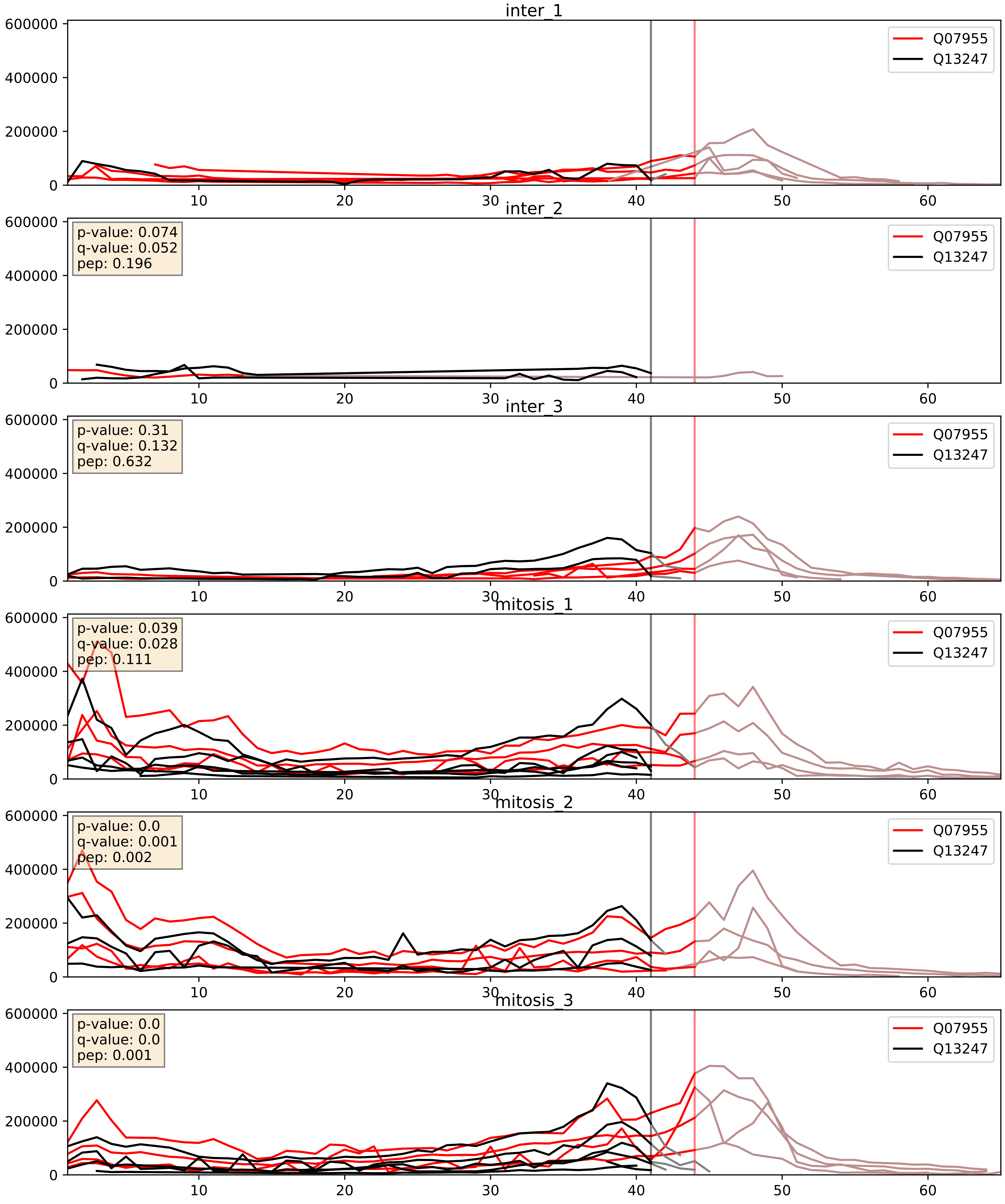


| level | condition_1 | condition_2 | pvalue | pvalue_adjusted |
|---|---|---|---|---|
| complex_abundance | mitosis | inter | 0.0021933030847482732 | 0.034197949736694384 |
| interactor_abundance | mitosis | inter | 0.009132210010912856 | 0.08324021891678159 |
| interactor_ratio | mitosis | inter | 0.6314956834780092 | 0.9183953086181231 |

Q13247_Q16629
SRSF6_HUMAN vs SRSF7_HUMAN
p-value: 0.001 q-value: 0.002

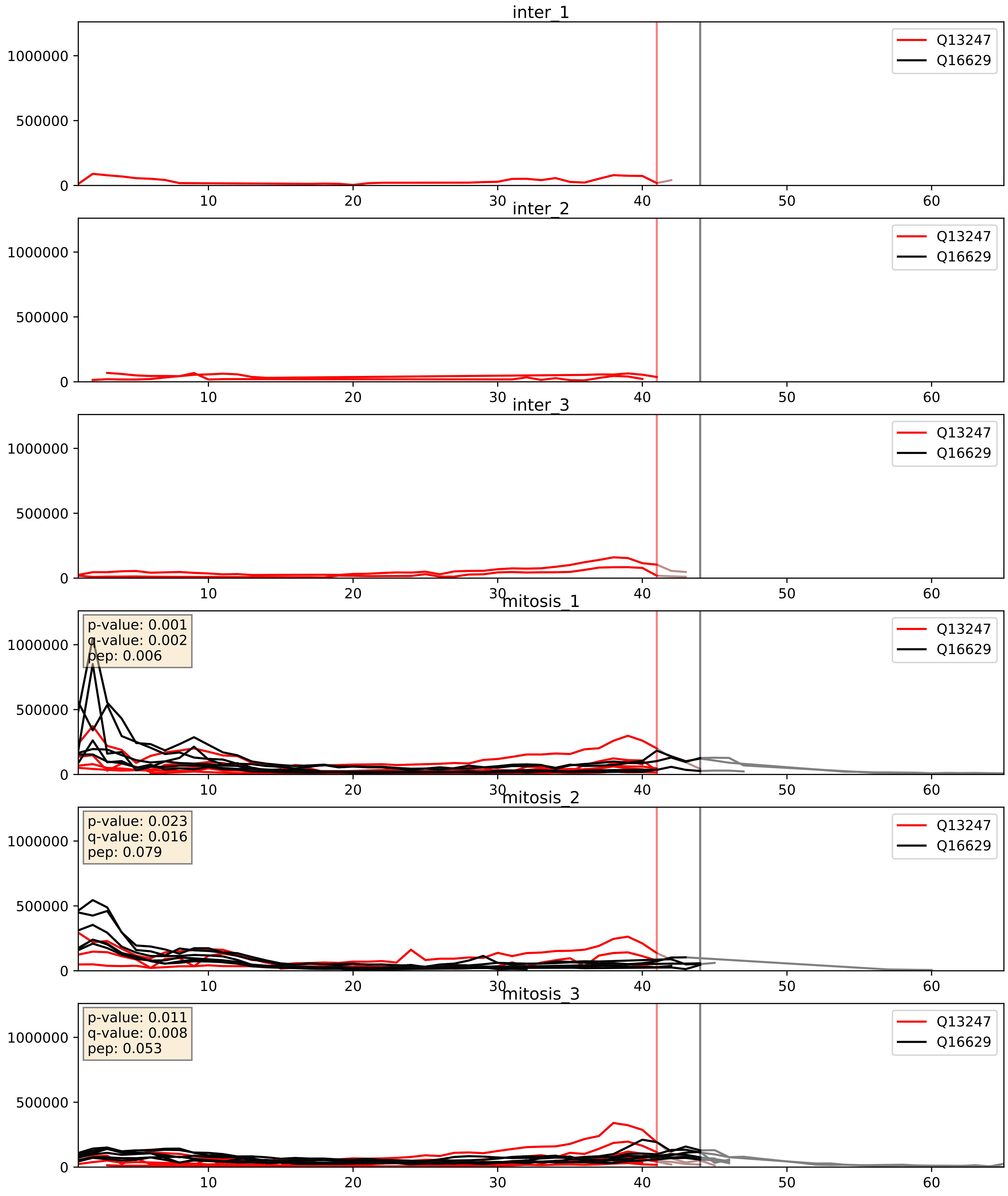


| level | condition_1 | condition_2 | pvalue | pvalue_adjusted |
|---|---|---|---|---|
| complex_abundance | mitosis | inter | 0.00033327015539205967 | 0.011886635542316795 |
| interactor_abundance | mitosis | inter | 0.00279726046188116 | 0.040175418714266324 |
| interactor_ratio | mitosis | inter | 0.11156417163406727 | 0.5532962393902757 |
