## Supplementary Data 2 for "SECAT: Quantifying differential protein-protein interaction states by network-centric analysis": hela_string_000127_O75400.pdf

# O75400_O75400

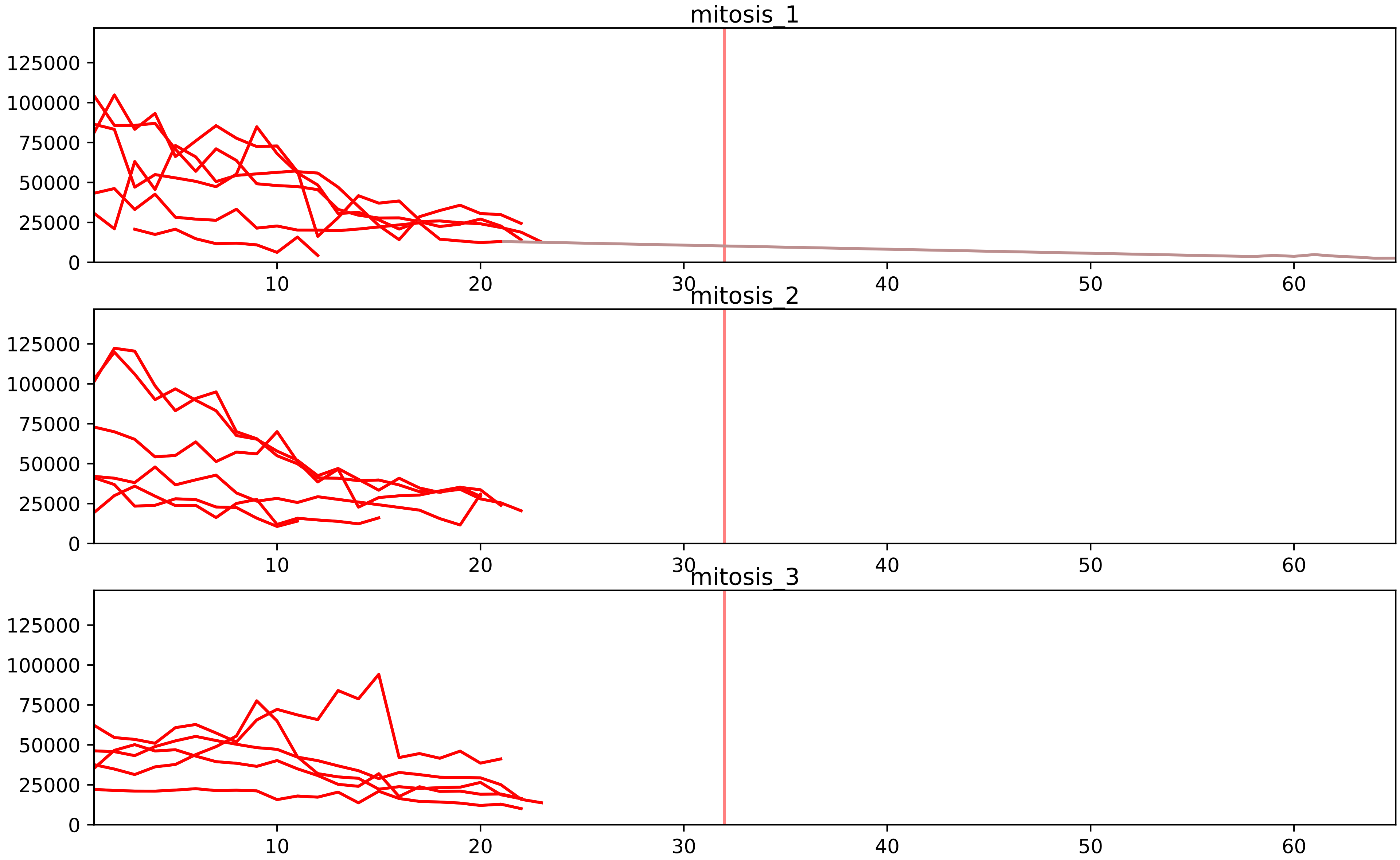


| level | condition_1 | condition_2 | pvalue | pvalue_adjusted |
|---|---|---|---|---|
| assembled_abundance | mitosis | inter | 3.1788048094639004e-05 | 0.006314764940805662 |
| total_abundance | mitosis | inter | 5.996022035115571e-05 | 0.010526120062335645 |
