## Supplementary Data 2 for "SECAT: Quantifying differential protein-protein interaction states by network-centric analysis": hela_string_000128_Q96HD1.pdf

Q96HD1_Q96HD1

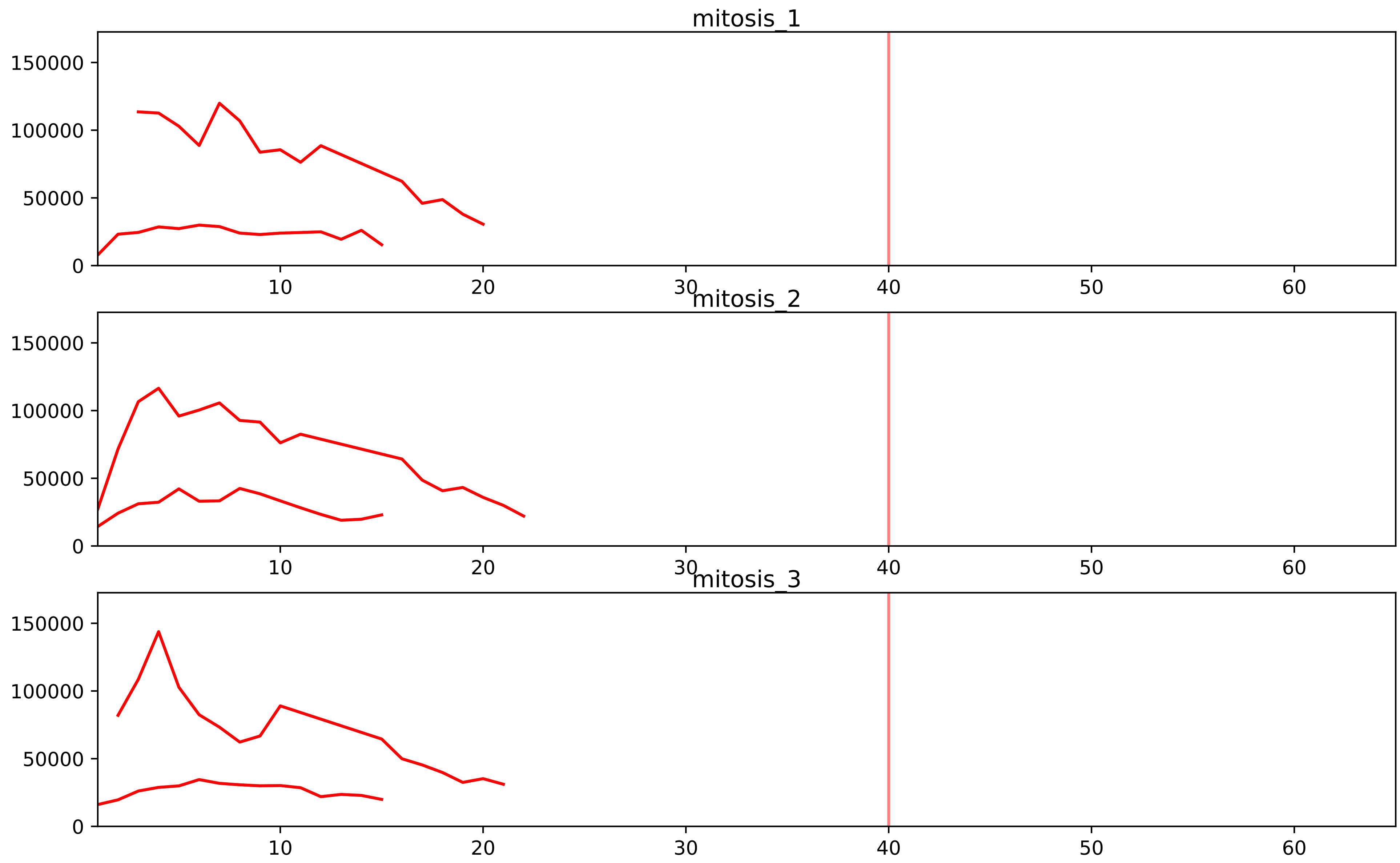


| level | condition_1 | condition_2 | pvalue | pvalue_adjusted |
|---|---|---|---|---|
| assembled_abundance | mitosis | inter | 3.283924537218159e-05 | 0.006314764940805662 |
| total_abundance | mitosis | inter | 7.319517014119239e-05 | 0.011644894099650326 |
