## Supplementary Data 2 for "SECAT: Quantifying differential protein-protein interaction states by network-centric analysis": hela_string_000129_P22695.pdf

P22695_P22695

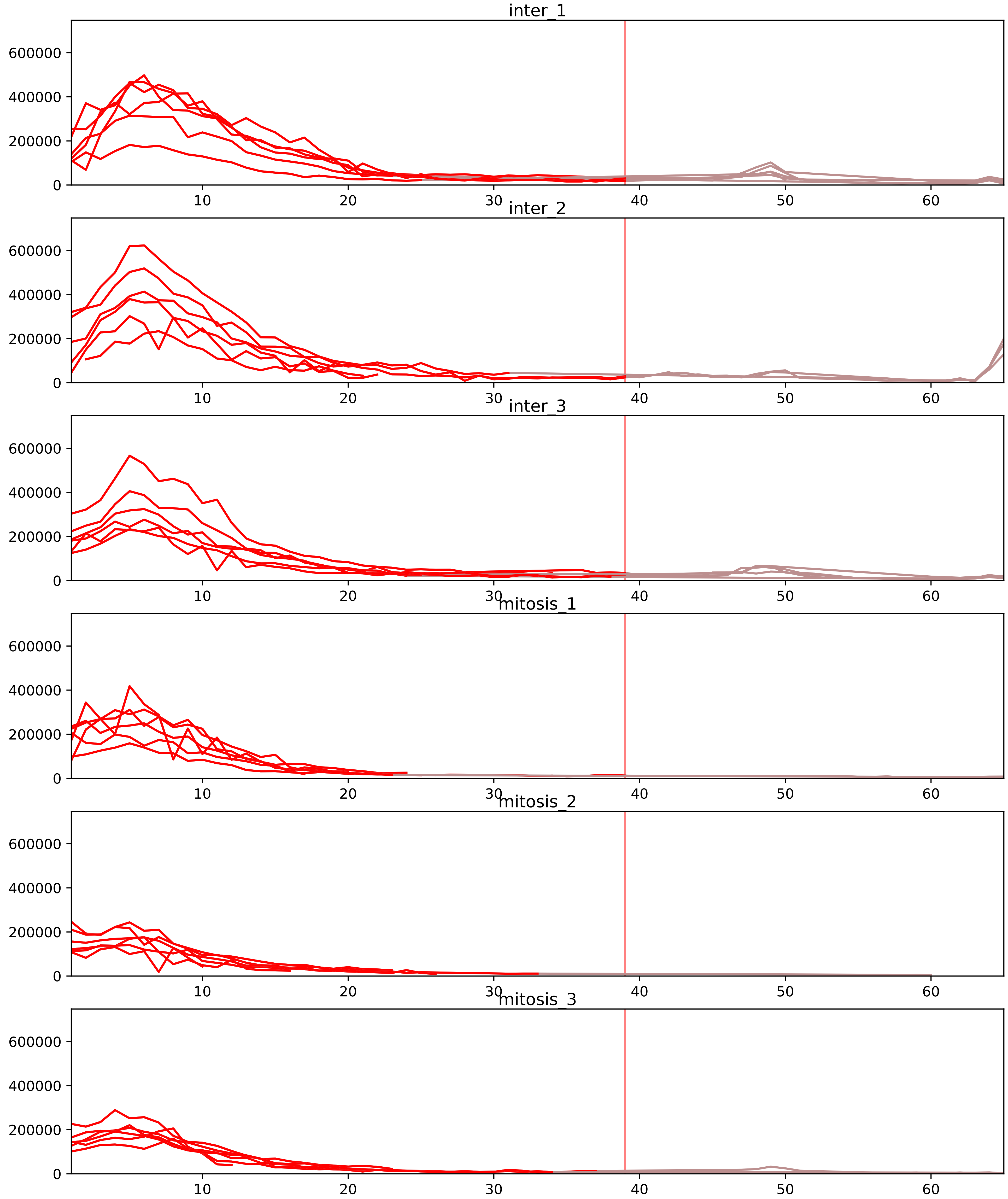


| level | condition_1 | condition_2 | pvalue | pvalue_adjusted |
|---|---|---|---|---|
| interactor_abundance | mitosis | inter | 3.296528016439947e-05 | 0.0011457923708108099 |
| complex_abundance | mitosis | inter | 9.917385386845972e-05 | 0.0022253645258288523 |
| monomer_abundance | mitosis | inter | 0.0001351598950842242 | 0.038277056377303 |
| interactor_ratio | mitosis | inter | 0.00016304411325931083 | 0.0025641125504028374 |
| total_abundance | mitosis | inter | 0.003149461656622661 | 0.06263245817916394 |
| assembled_abundance | mitosis | inter | 0.003913296826482518 | 0.06212029154760582 |

O14949_P22695
QCR8_HUMAN vs QCR2_HUMAN
p-value: 0.009 q-value: 0.008

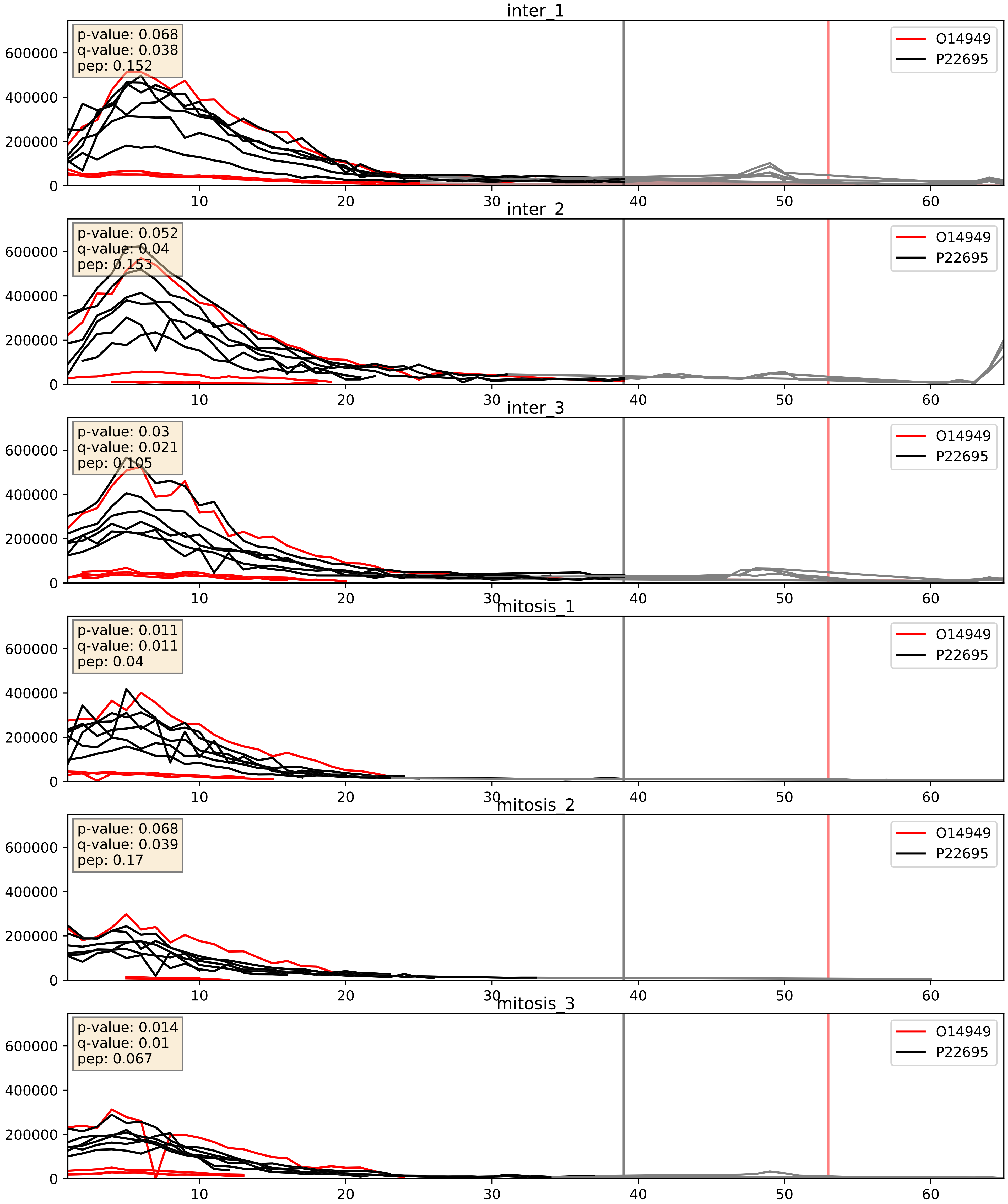


| level | condition_1 | condition_2 | pvalue | pvalue_adjusted |
|---|---|---|---|---|
| complex_abundance | mitosis | inter | 0.015810889343351366 | 0.09374757296036364 |
| interactor_abundance | mitosis | inter | 0.07395456278311857 | 0.22673988198390962 |
| interactor_ratio | mitosis | inter | 0.88612935614183 | 0.9802747470617126 |

O43674_P22695
NDUB5_HUMAN vs QCR2_HUMAN
p-value: 0.009 q-value: 0.008

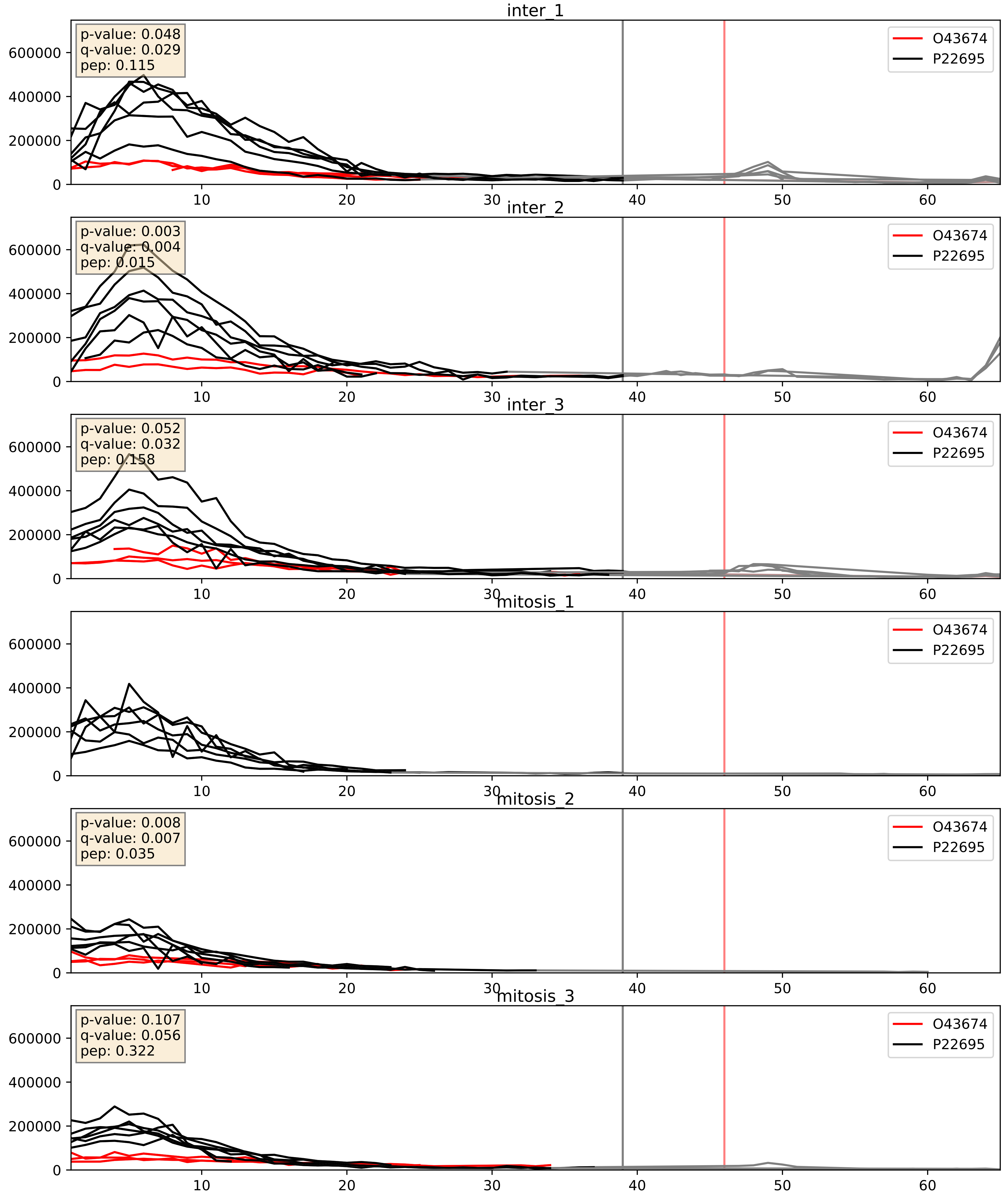


| level | condition_1 | condition_2 | pvalue | pvalue_adjusted |
|---|---|---|---|---|
| complex_abundance | mitosis | inter | 0.09864491867022618 | 0.23280962332978664 |
| interactor_abundance | mitosis | inter | 0.1092864182144783 | 0.2841594926196597 |
| interactor_ratio | mitosis | inter | 0.2016678292884779 | 0.6766624597719332 |

O75439_P22695
MPPB_HUMAN vs QCR2_HUMAN
p-value: 0.08 q-value: 0.047

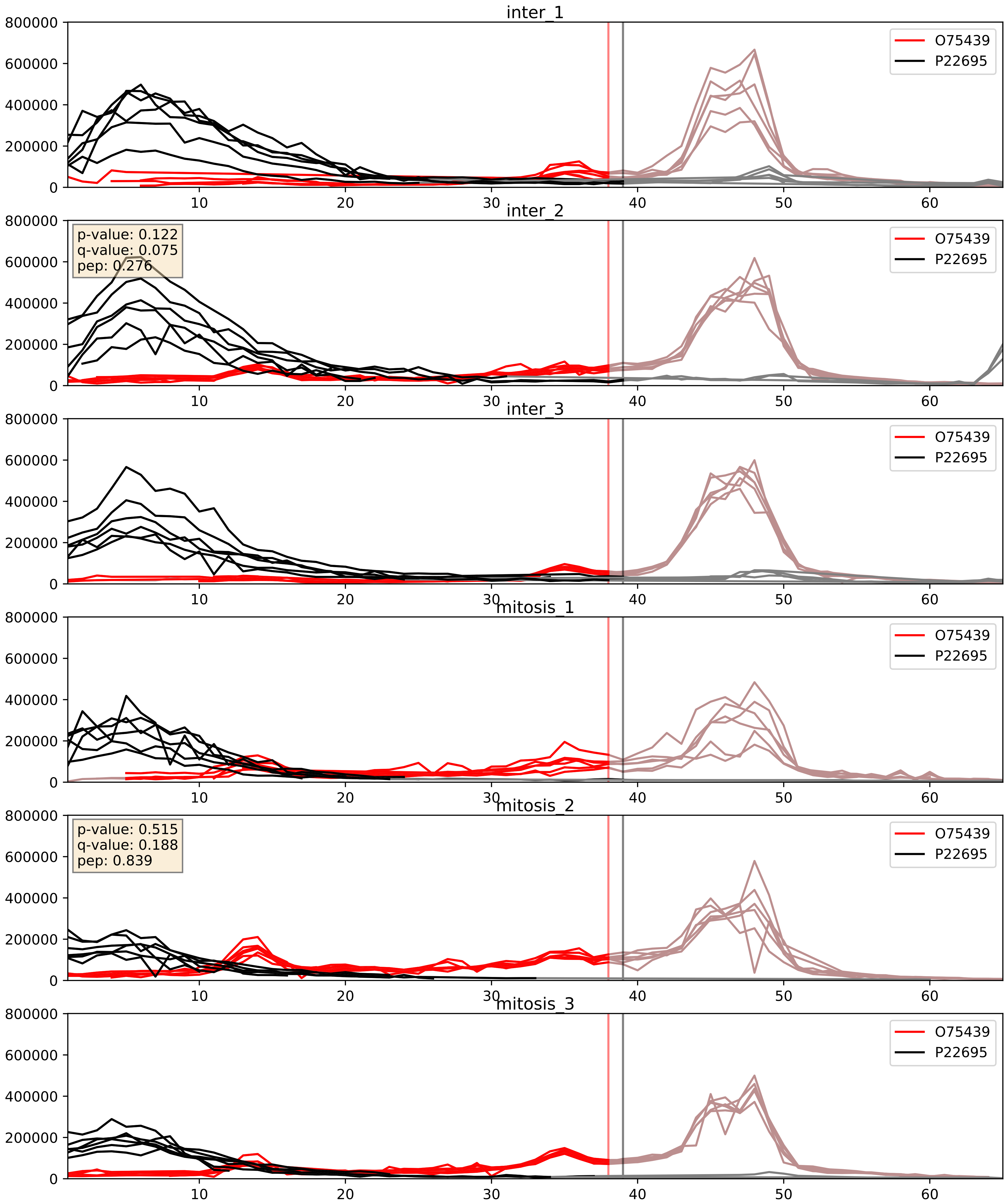


| level | condition_1 | condition_2 | pvalue | pvalue_adjusted |
|---|---|---|---|---|
| interactor_ratio | mitosis | inter | 0.0006300723229304282 | 0.029219626944512057 |
| interactor_abundance | mitosis | inter | 0.19941681144583612 | 0.393611418559031 |
| complex_abundance | mitosis | inter | 0.29806595099498284 | 0.4477299056847811 |

O75489_P22695
NDUS3_HUMAN vs QCR2_HUMAN
p-value: 0.004 q-value: 0.004

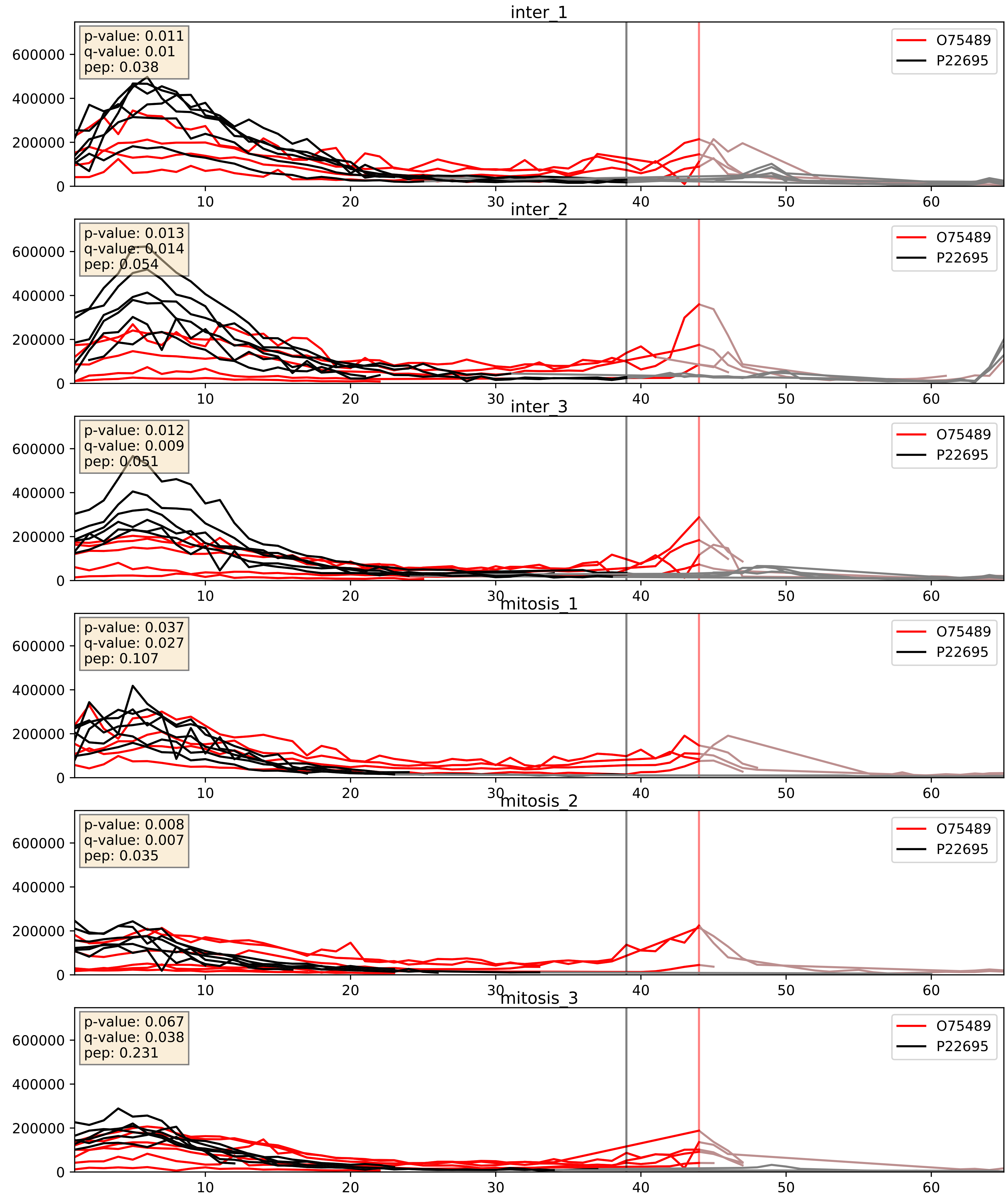


| level | condition_1 | condition_2 | pvalue | pvalue_adjusted |
|---|---|---|---|---|
| interactor_ratio | mitosis | inter | 0.0036794568894891852 | 0.0787300451287945 |
| complex_abundance | mitosis | inter | 0.007507361267008286 | 0.06444768685385507 |
| interactor_abundance | mitosis | inter | 0.06657853699399839 | 0.21422397339471275 |

P00403_P22695
COX2_HUMAN vs QCR2_HUMAN
p-value: 0.01 q-value: 0.009

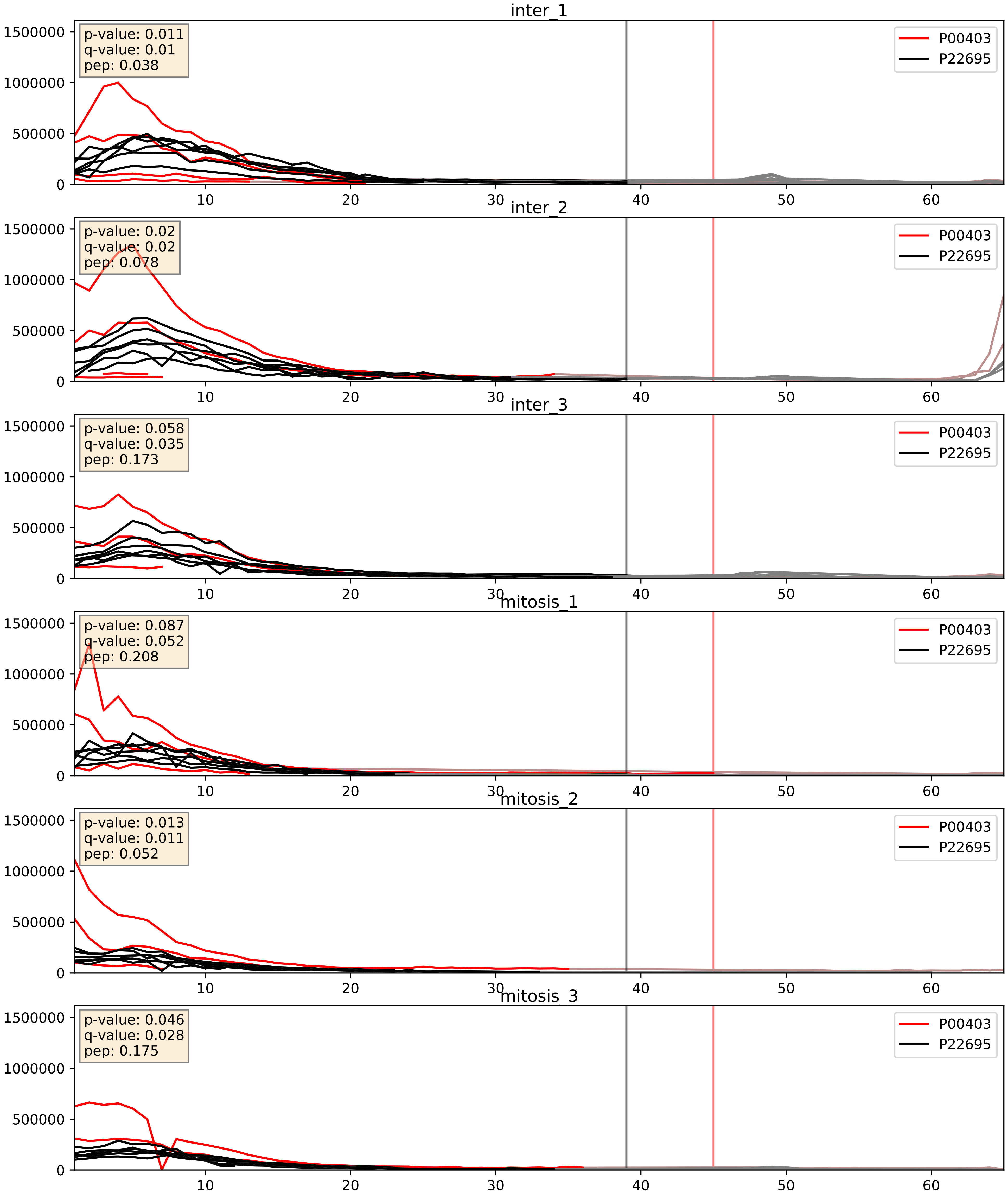


| level | condition_1 | condition_2 | pvalue | pvalue_adjusted |
|---|---|---|---|---|
| complex_abundance | mitosis | inter | 0.007639655089387588 | 0.06507009707975896 |
| interactor_abundance | mitosis | inter | 0.04799835291854166 | 0.17504902777779313 |
| interactor_ratio | mitosis | inter | 0.4897202497476749 | 0.8692763386835093 |

P07919_P22695
QCR6_HUMAN vs QCR2_HUMAN
p-value: 0.021 q-value: 0.016

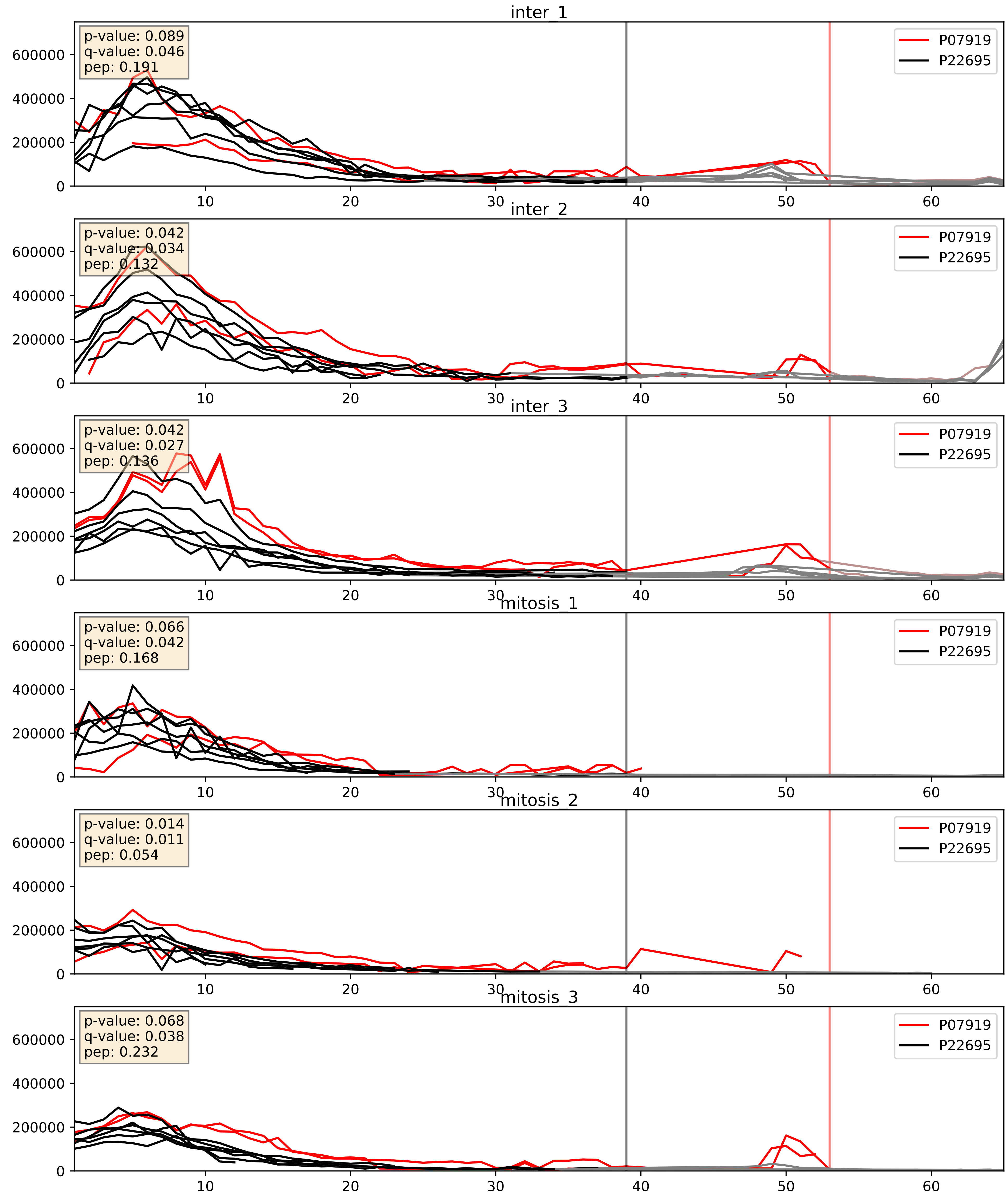


| level | condition_1 | condition_2 | pvalue | pvalue_adjusted |
|---|---|---|---|---|
| complex_abundance | mitosis | inter | 0.002608285993433769 | 0.03664546283589145 |
| interactor_abundance | mitosis | inter | 0.021463020495520537 | 0.1218501618383326 |
| interactor_ratio | mitosis | inter | 0.08011628646589991 | 0.47352742543809706 |

P08574_P22695
CY1_HUMAN vs QCR2_HUMAN
p-value: 0.003 q-value: 0.004

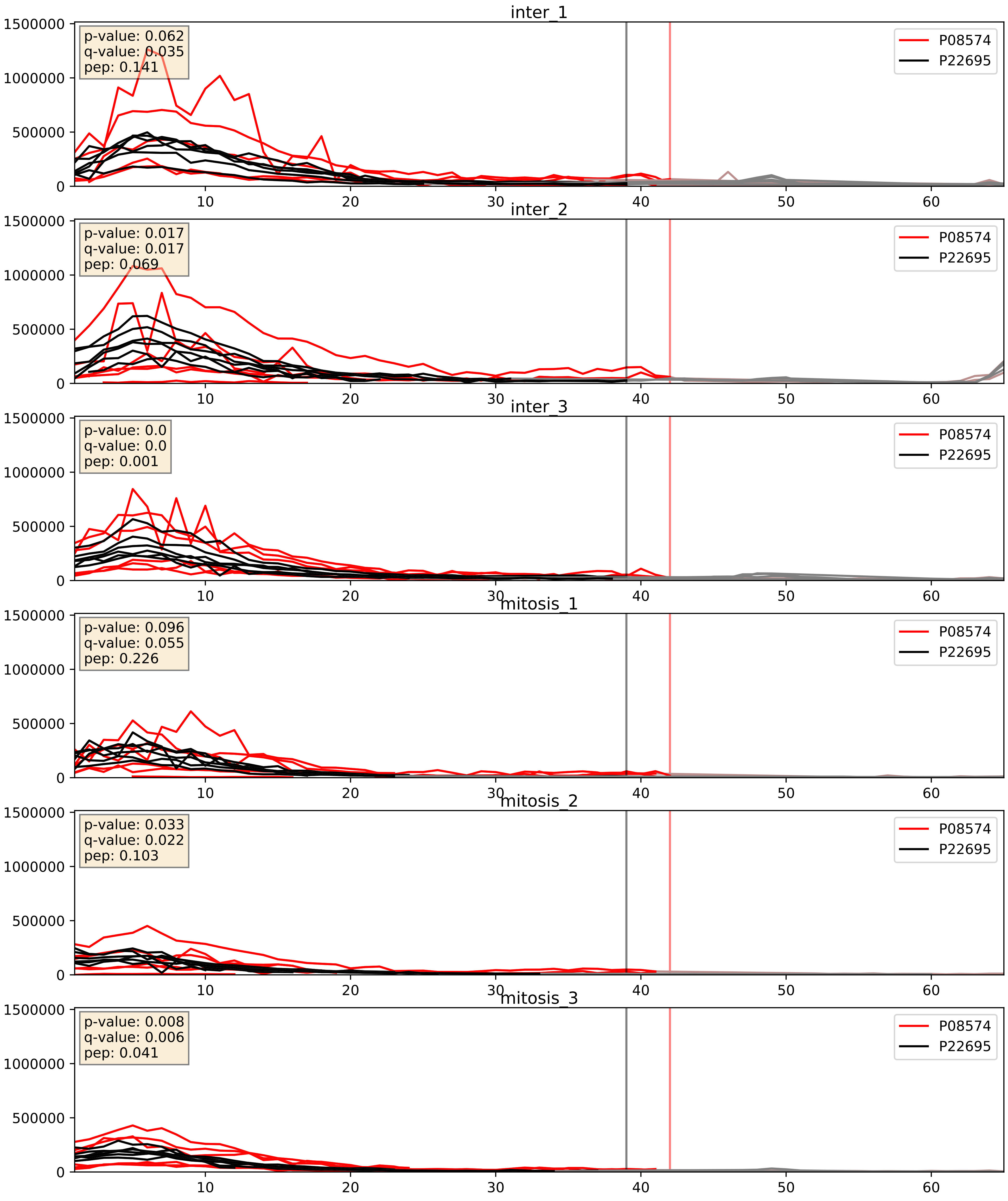


| level | condition_1 | condition_2 | pvalue | pvalue_adjusted |
|---|---|---|---|---|
| complex_abundance | mitosis | inter | 0.002527814595984375 | 0.036409709098919725 |
| interactor_abundance | mitosis | inter | 0.007419881684355014 | 0.07459761261938556 |
| interactor_ratio | mitosis | inter | 0.8436266256088374 | 0.9755634896635457 |

P10606_P22695
COX5B_HUMAN vs QCR2_HUMAN
p-value: 0.001 q-value: 0.002

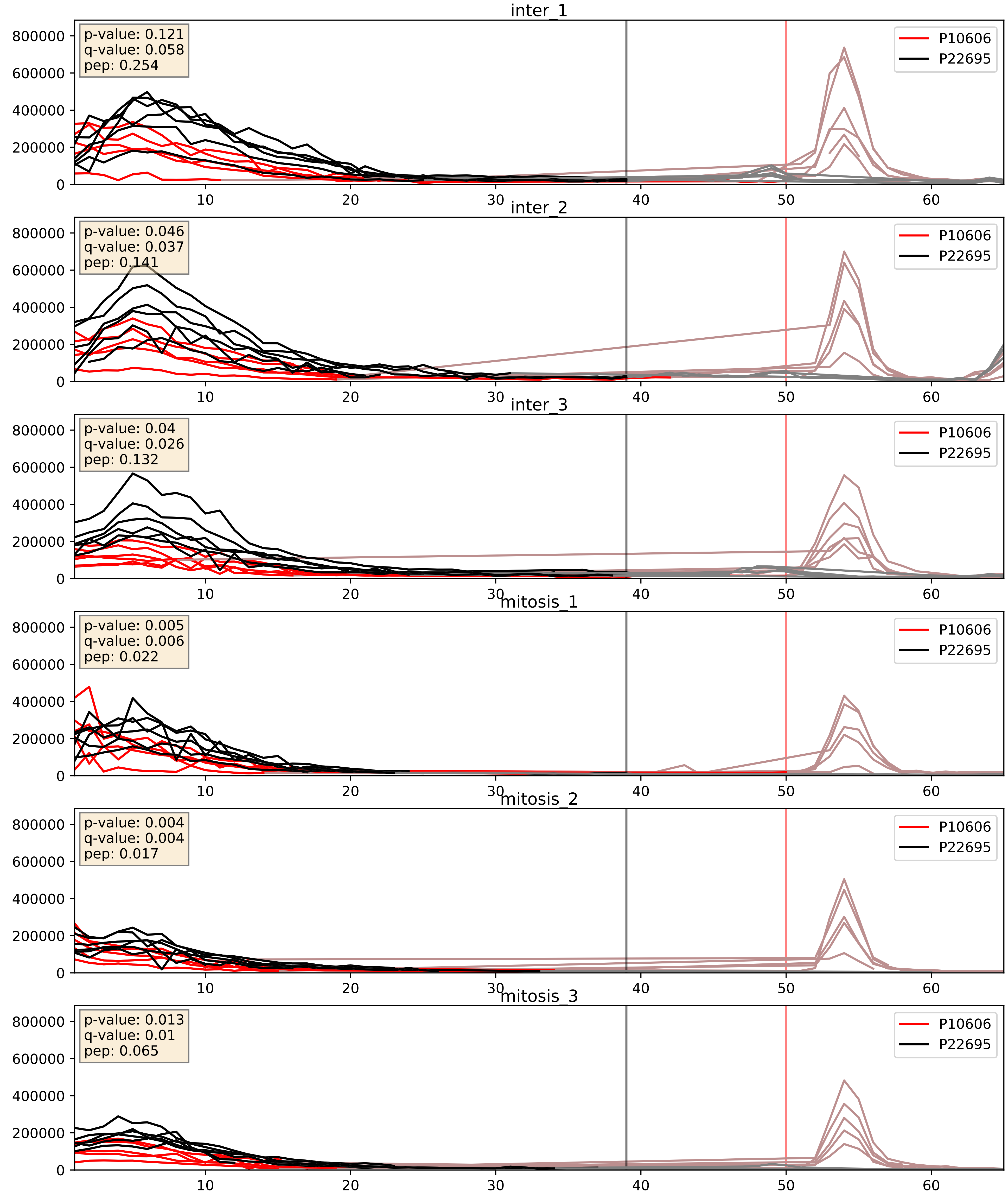


| level | condition_1 | condition_2 | pvalue | pvalue_adjusted |
|---|---|---|---|---|
| complex_abundance | mitosis | inter | 0.01770352106384553 | 0.09969877651744588 |
| interactor_abundance | mitosis | inter | 0.07823589389834215 | 0.23202668713788538 |
| interactor_ratio | mitosis | inter | 0.19324221330416416 | 0.6660896702330326 |

P14927_P22695
QCR7_HUMAN vs QCR2_HUMAN
p-value: 0.002 q-value: 0.003

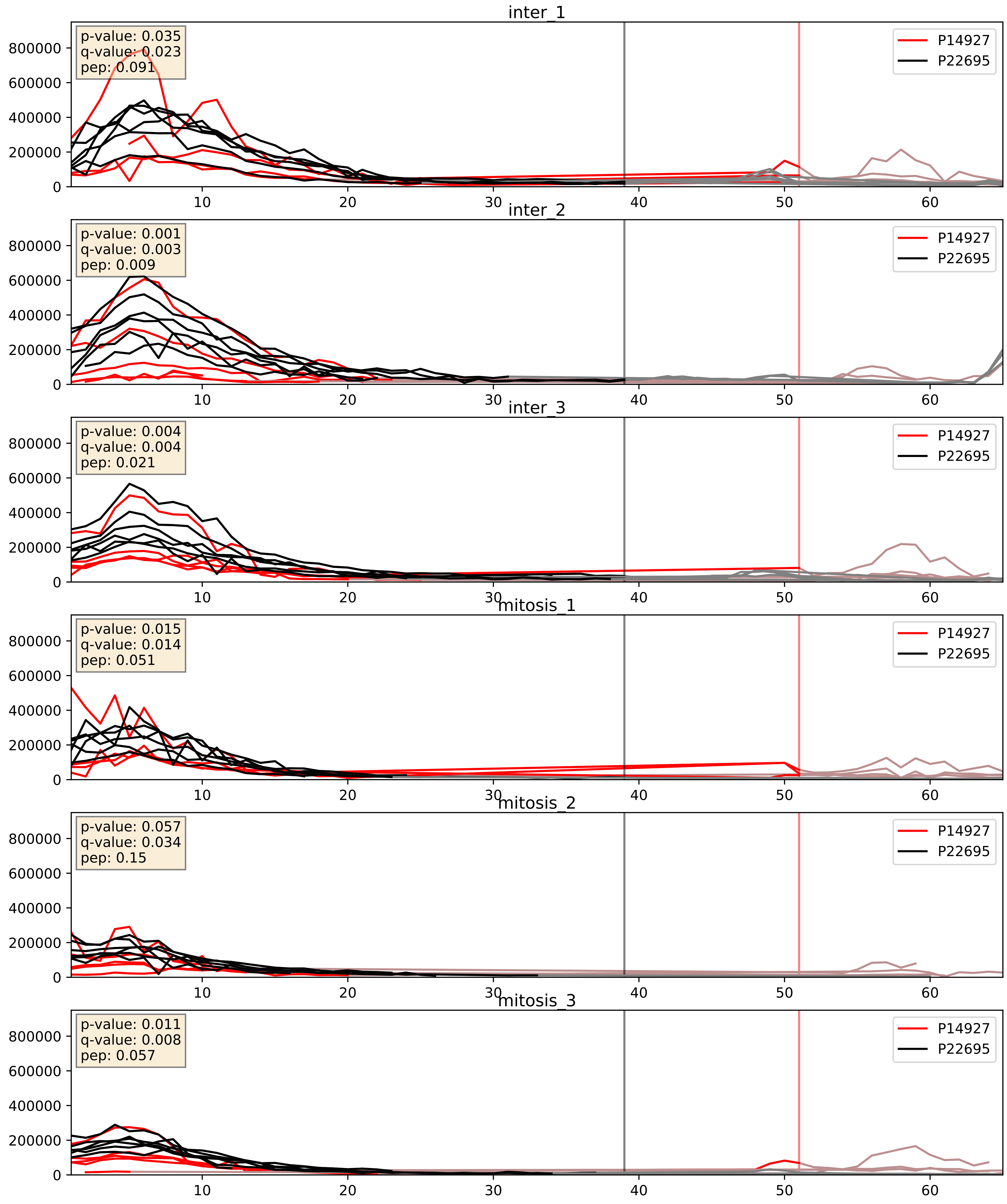


| level | condition_1 | condition_2 | pvalue | pvalue_adjusted |
|---|---|---|---|---|
| complex_abundance | mitosis | inter | 0.007887164881313349 | 0.06591342526929356 |
| interactor_abundance | mitosis | inter | 0.022350644269404238 | 0.12329937058940194 |
| interactor_ratio | mitosis | inter | 0.8166003486958785 | 0.9666192201555632 |

P20674_P22695
COX5A_HUMAN vs QCR2_HUMAN
p-value: 0.0 q-value: 0.001

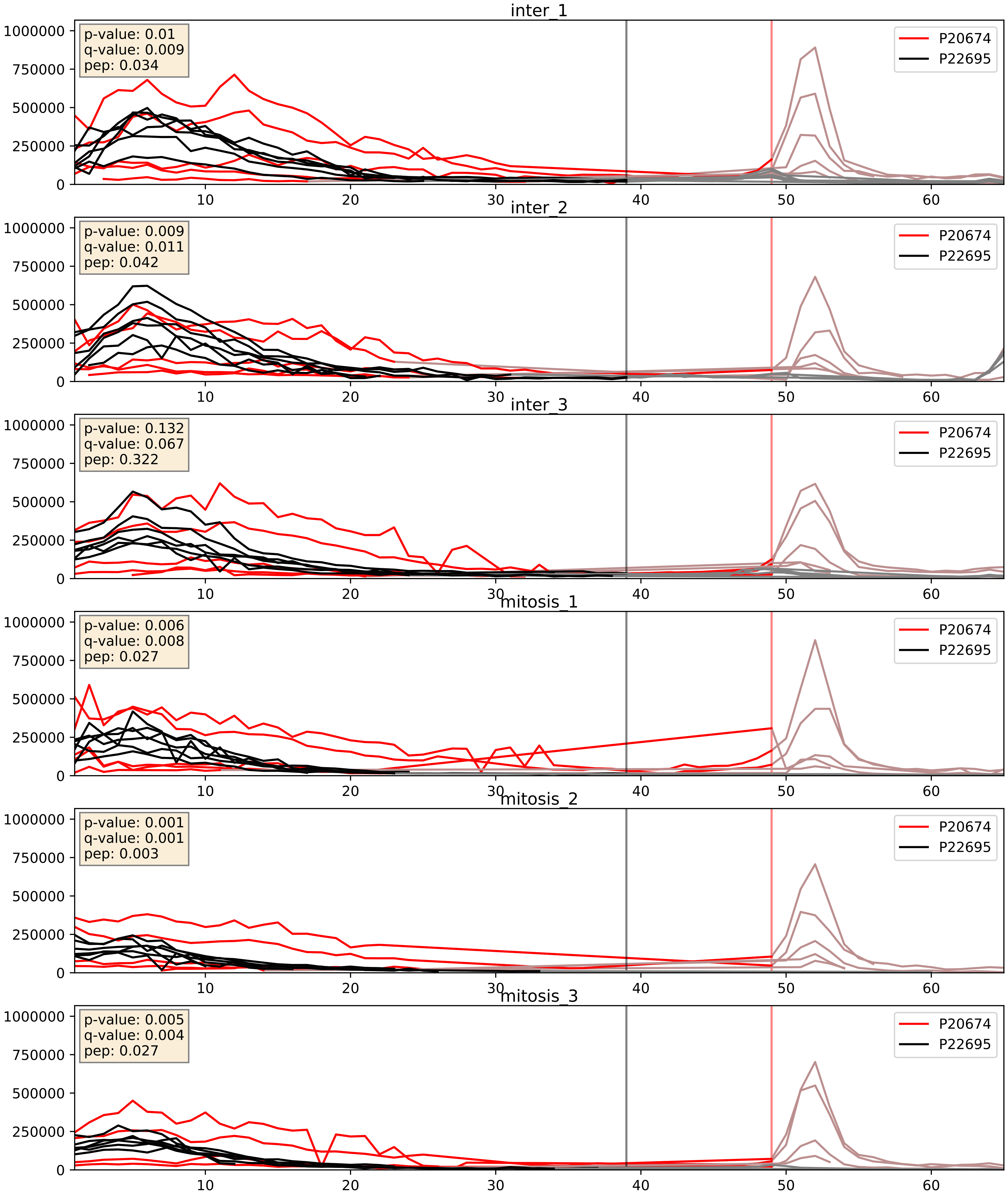


| level | condition_1 | condition_2 | pvalue | pvalue_adjusted |
|---|---|---|---|---|
| complex_abundance | mitosis | inter | 0.0021306539940250053 | 0.033489340249041655 |
| interactor_abundance | mitosis | inter | 0.005856311363255288 | 0.064851261668131 |
| interactor_ratio | mitosis | inter | 0.12059051910908962 | 0.5703368335321749 |

P21912_P22695
SDHB_HUMAN vs QCR2_HUMAN
p-value: 0.05 q-value: 0.033

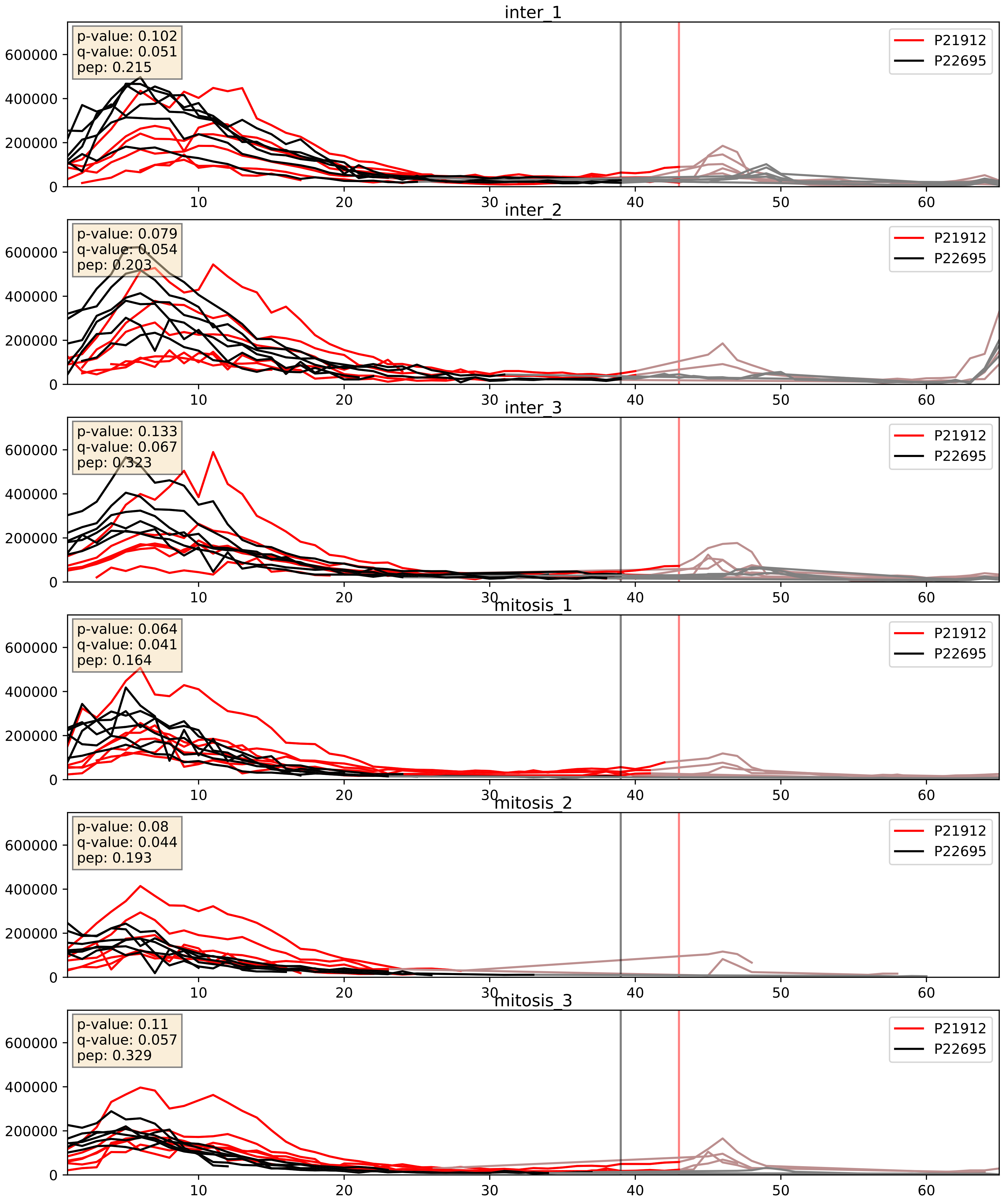


| level | condition_1 | condition_2 | pvalue | pvalue_adjusted |
|---|---|---|---|---|
| interactor_ratio | mitosis | inter | 0.00020876624523135206 | 0.014683183137172115 |
| complex_abundance | mitosis | inter | 0.0045406701215263525 | 0.05028219435998134 |
| interactor_abundance | mitosis | inter | 0.018796059017627726 | 0.11808618106318929 |

P22695_P24539
QCR2_HUMAN vs AT5F1_HUMAN
p-value: 0.001 q-value: 0.015

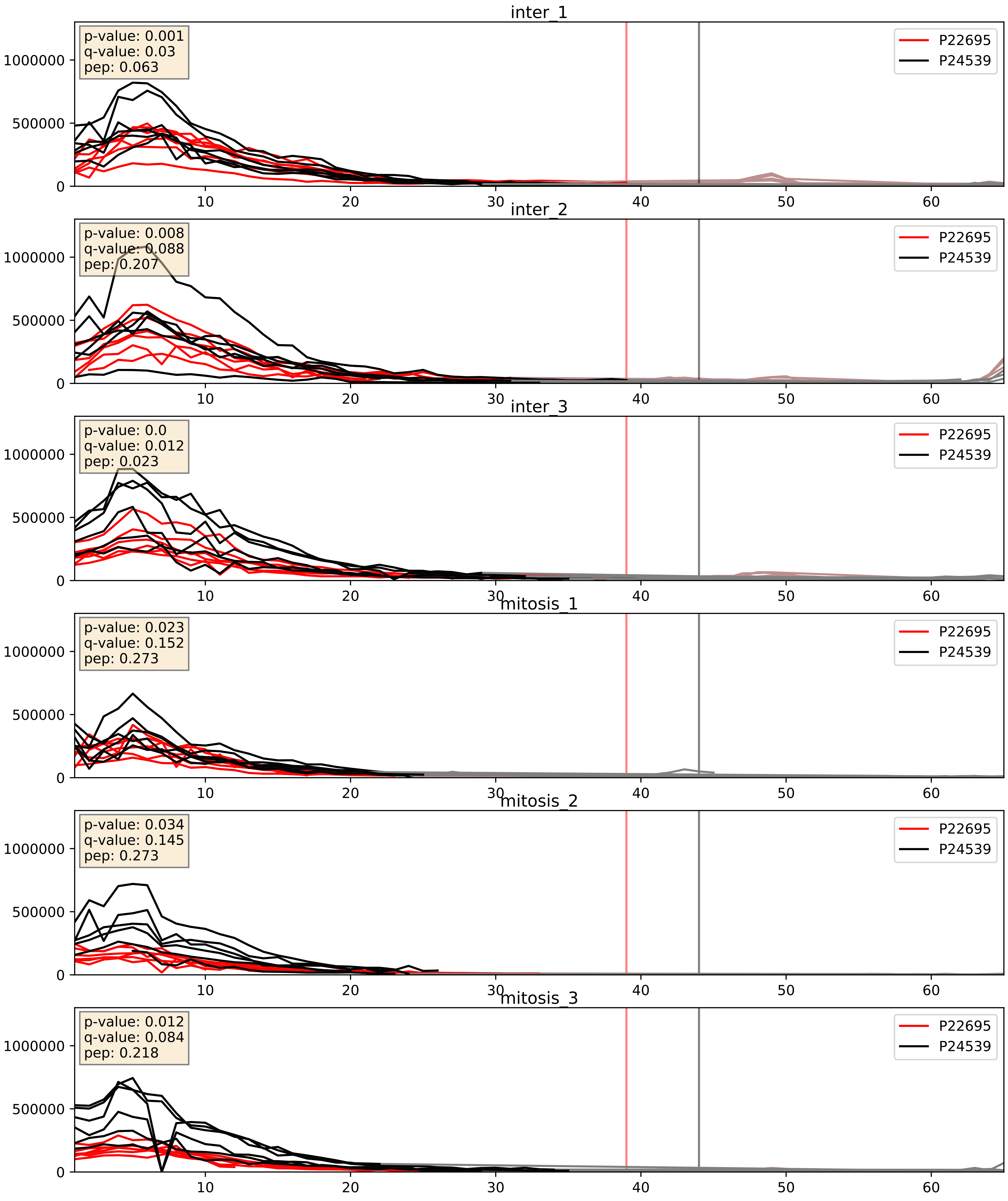


| level | condition_1 | condition_2 | pvalue | pvalue_adjusted |
|---|---|---|---|---|
| interactor_abundance | mitosis | inter | 0.0014834042890461618 | 0.028438836985968968 |
| complex_abundance | mitosis | inter | 0.003551064505513013 | 0.043600374082146835 |
| interactor_ratio | mitosis | inter | 0.3178441158985159 | 0.7783421708721073 |

P22695_P28331
QCR2_HUMAN vs NDUS1_HUMAN
p-value: 0.043 q-value: 0.029

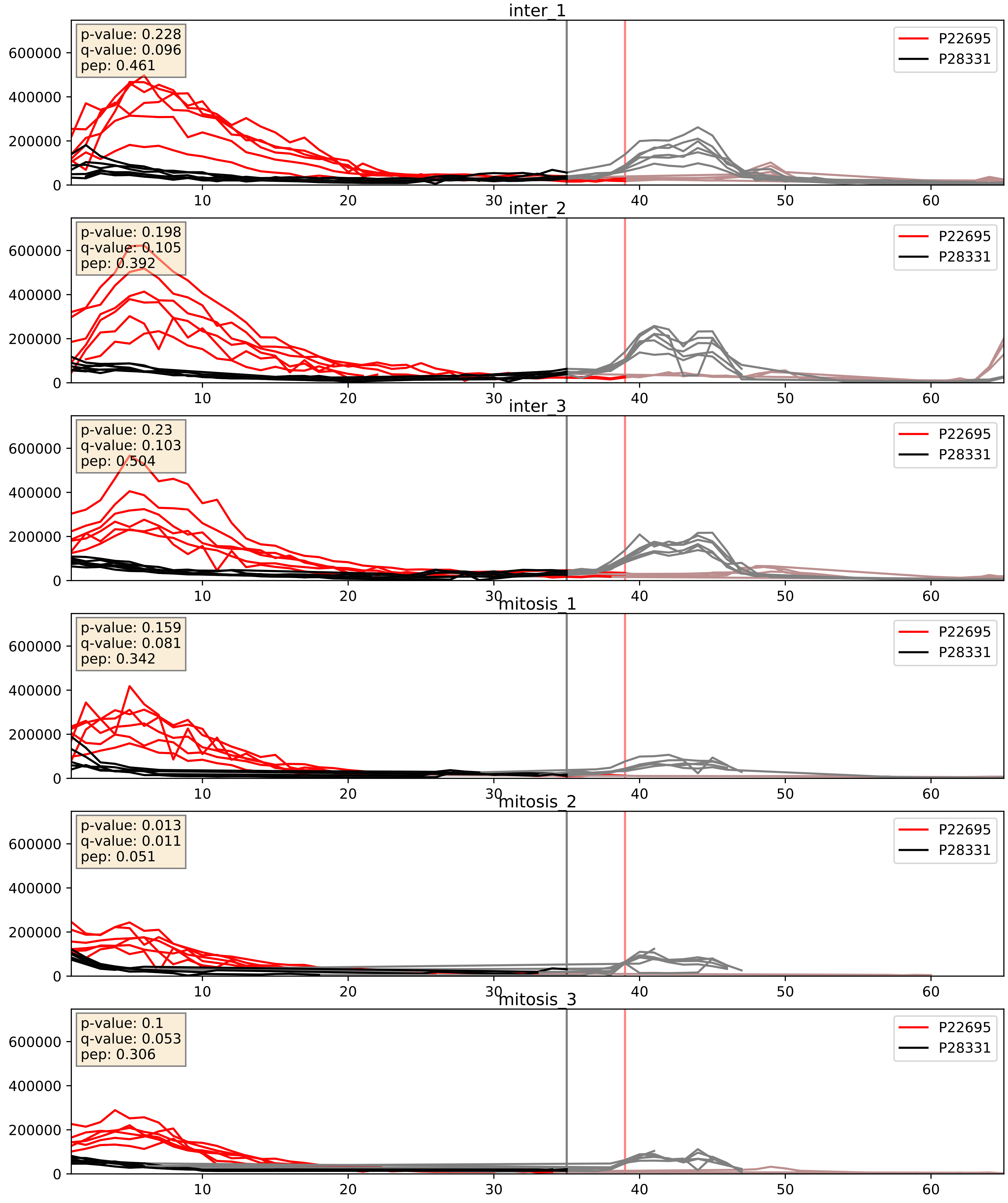


| level | condition_1 | condition_2 | pvalue | pvalue_adjusted |
|---|---|---|---|---|
| complex_abundance | mitosis | inter | 0.0016073371358469538 | 0.02813661734734136 |
| interactor_abundance | mitosis | inter | 0.0017502652133548574 | 0.030576061686362406 |
| interactor_ratio | mitosis | inter | 0.2970697251002717 | 0.7646944825997081 |

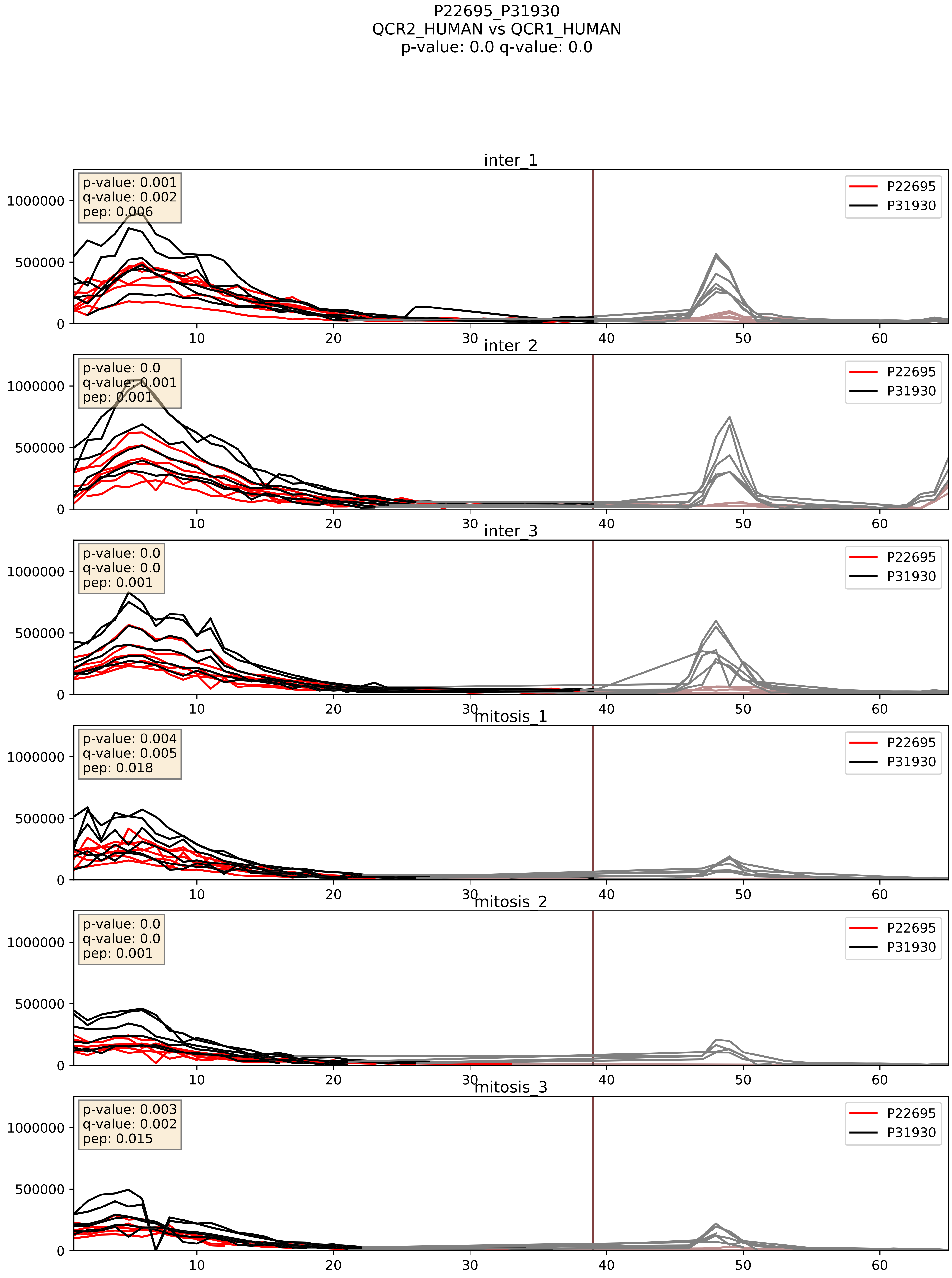


| level | condition_1 | condition_2 | pvalue | pvalue_adjusted |
|---|---|---|---|---|
| complex_abundance | mitosis | inter | 0.0008434285050207741 | 0.019252661341274204 |
| interactor_abundance | mitosis | inter | 0.0015444039397945596 | 0.028907984680718663 |
| interactor_ratio | mitosis | inter | 0.9461758960577978 | 0.9912117205440141 |

P22695_P36542
QCR2_HUMAN vs ATPG_HUMAN
p-value: 0.0 q-value: 0.0

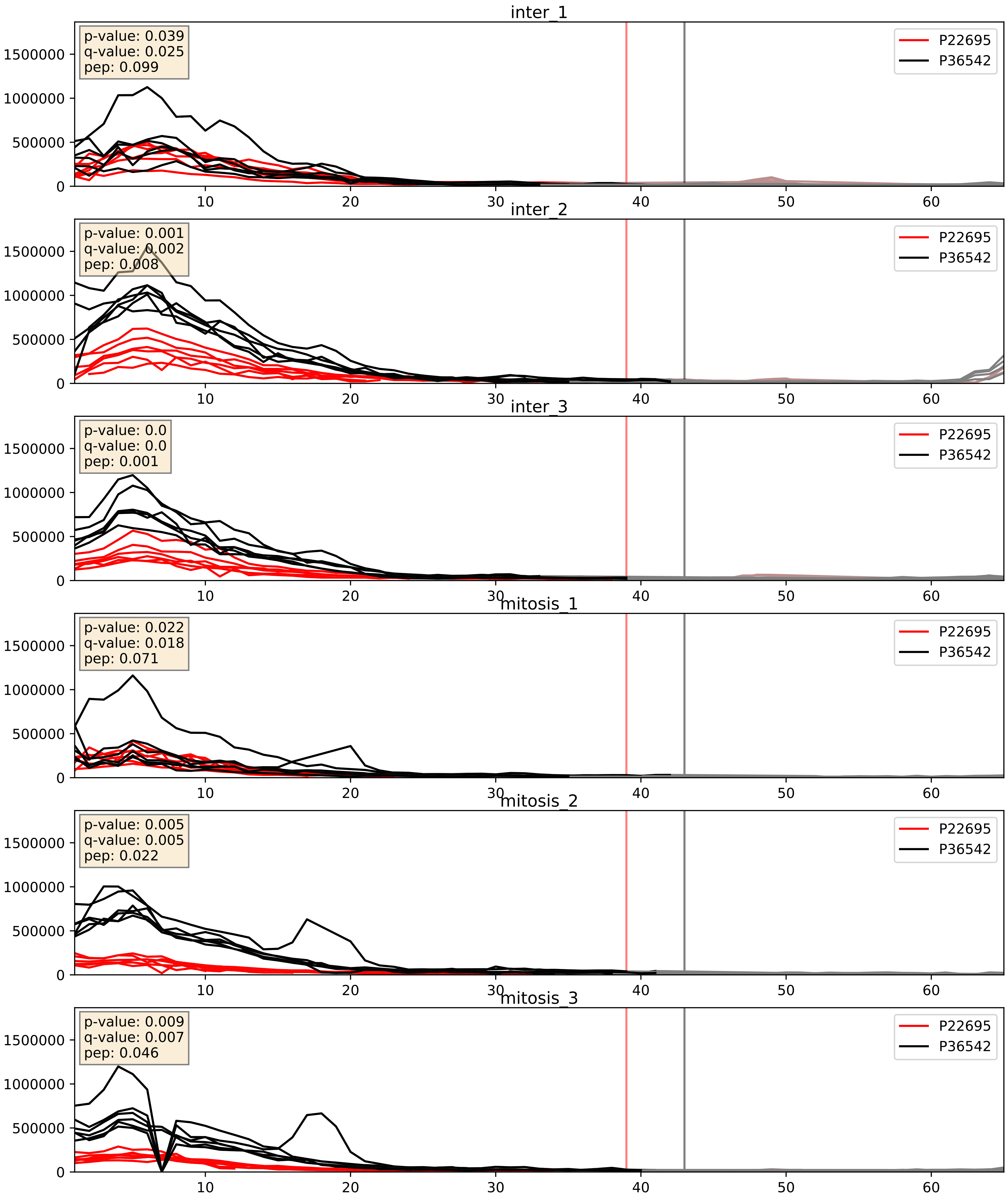


| level | condition_1 | condition_2 | pvalue | pvalue_adjusted |
|---|---|---|---|---|
| interactor_abundance | mitosis | inter | 0.0015349905250669595 | 0.028907984680718663 |
| complex_abundance | mitosis | inter | 0.004837624358669671 | 0.0522854349876419 |
| interactor_ratio | mitosis | inter | 0.5886014574668961 | 0.8994900937741748 |

P22695_P45880
QCR2_HUMAN vs VDAC2_HUMAN
p-value: 0.0 q-value: 0.008

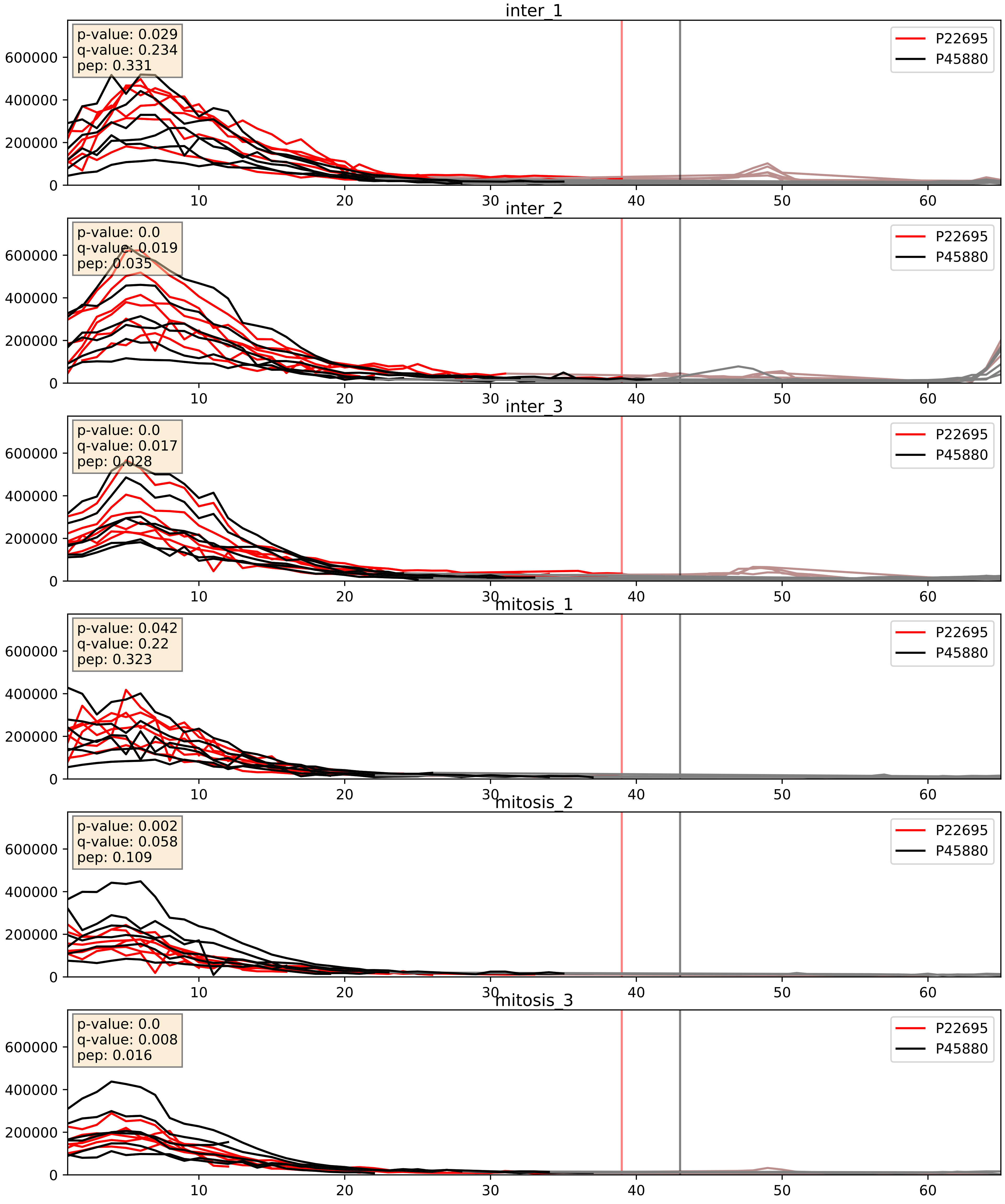


| level | condition_1 | condition_2 | pvalue | pvalue_adjusted |
|---|---|---|---|---|
| complex_abundance | mitosis | inter | 0.0010980792513486063 | 0.022013017310407657 |
| interactor_abundance | mitosis | inter | 0.001583504757735475 | 0.029025269221018554 |
| interactor_ratio | mitosis | inter | 0.006983683590651673 | 0.11653432577764955 |

P22695_P47985
QCR2_HUMAN vs UCRI_HUMAN
p-value: 0.02 q-value: 0.016

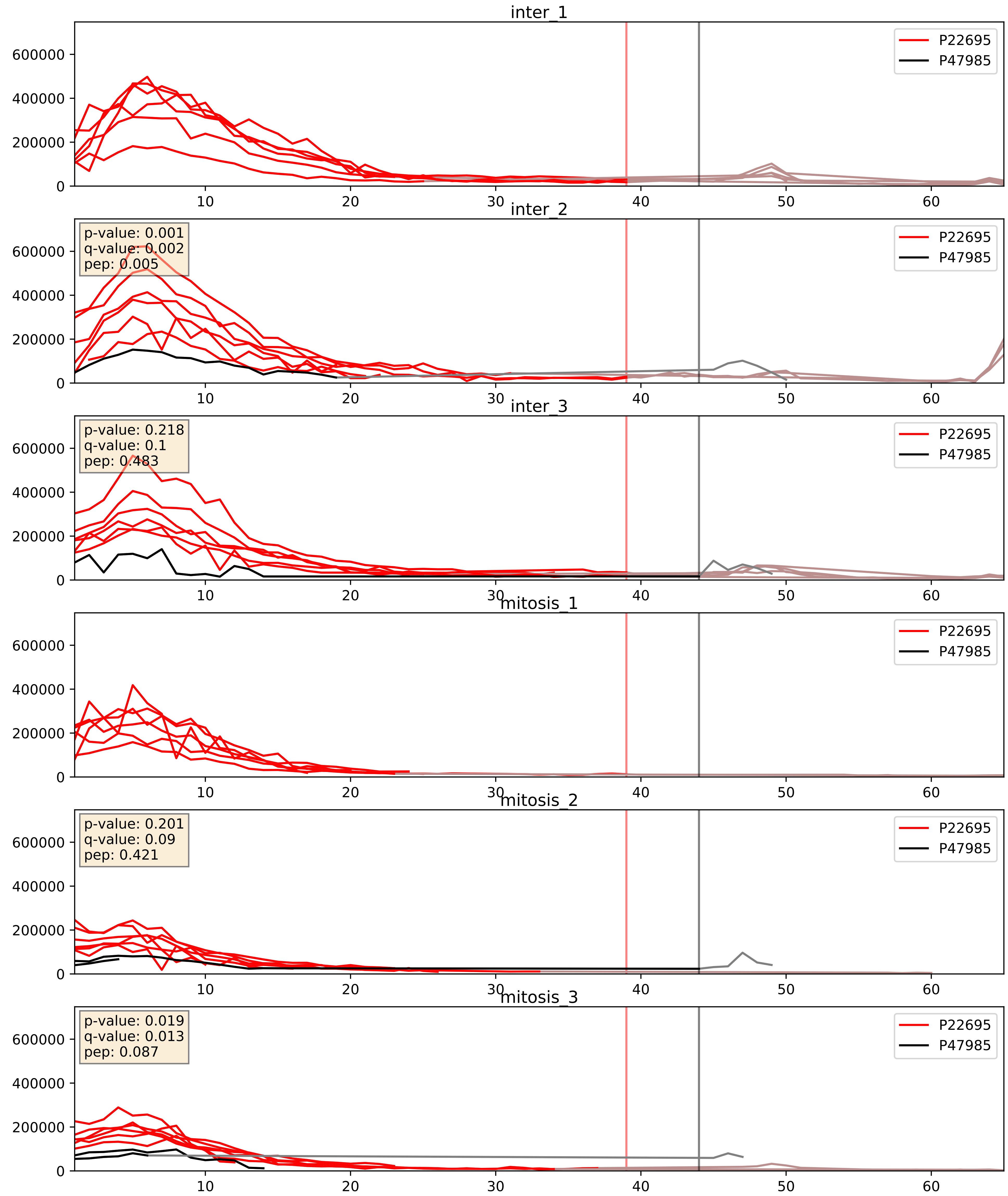


| level | condition_1 | condition_2 | pvalue | pvalue_adjusted |
|---|---|---|---|---|
| interactor_ratio | mitosis | inter | 0.5644555660667254 | 0.8918137840744521 |
| interactor_abundance | mitosis | inter | 0.605302039936046 | 0.7407270138459692 |
| complex_abundance | mitosis | inter | 0.635045503173613 | 0.751553920526217 |

P22695_P48047
QCR2_HUMAN vs ATPO_HUMAN
p-value: 0.002 q-value: 0.003

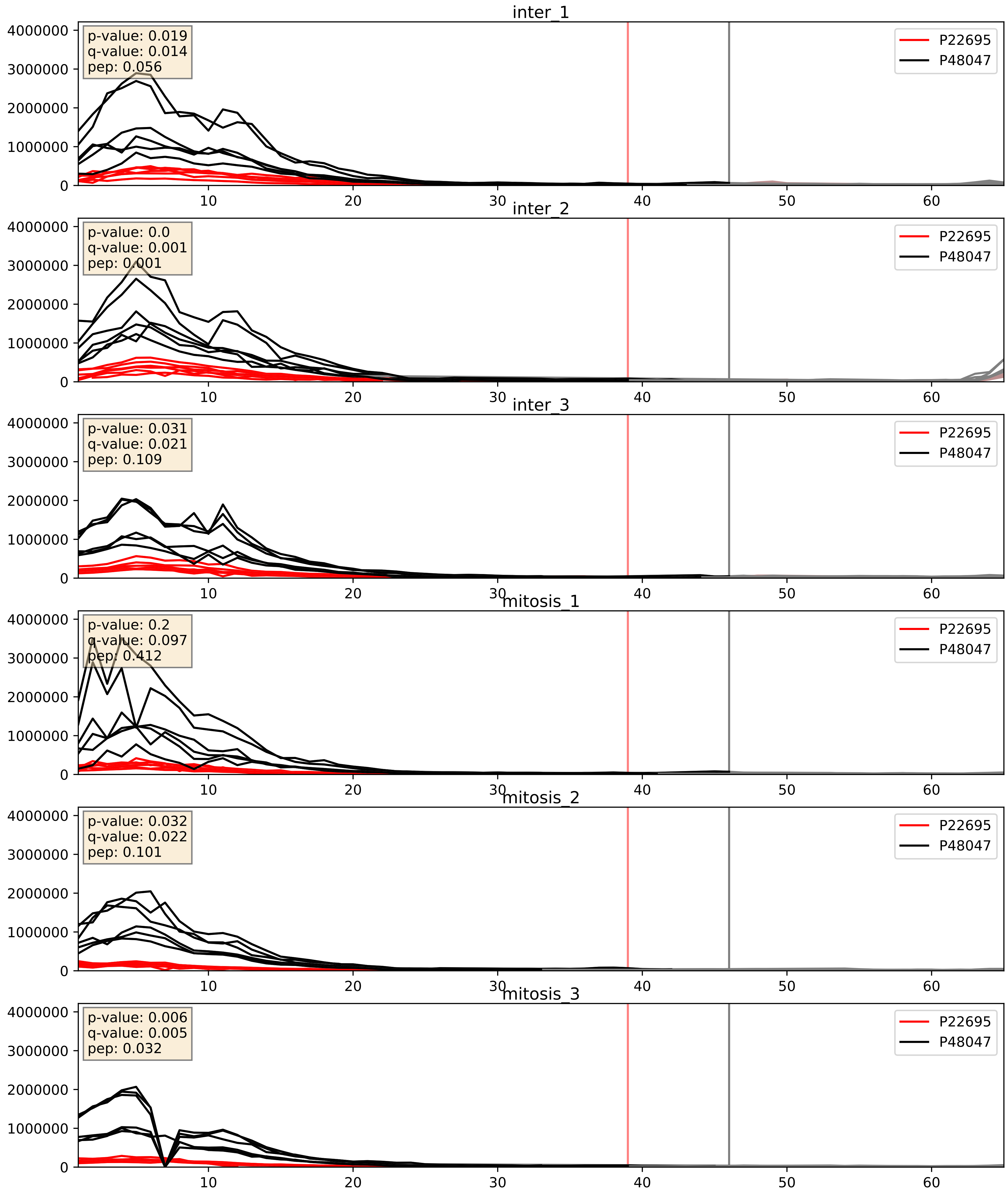


| level | condition_1 | condition_2 | pvalue | pvalue_adjusted |
|---|---|---|---|---|
| interactor_abundance | mitosis | inter | 0.0016984570860933794 | 0.030042528708242176 |
| complex_abundance | mitosis | inter | 0.009987333949595958 | 0.07371585975104354 |
| interactor_ratio | mitosis | inter | 0.631777354213924 | 0.9183953086181231 |

P22695_Q16795
QCR2_HUMAN vs NDUA9_HUMAN
p-value: 0.059 q-value: 0.037

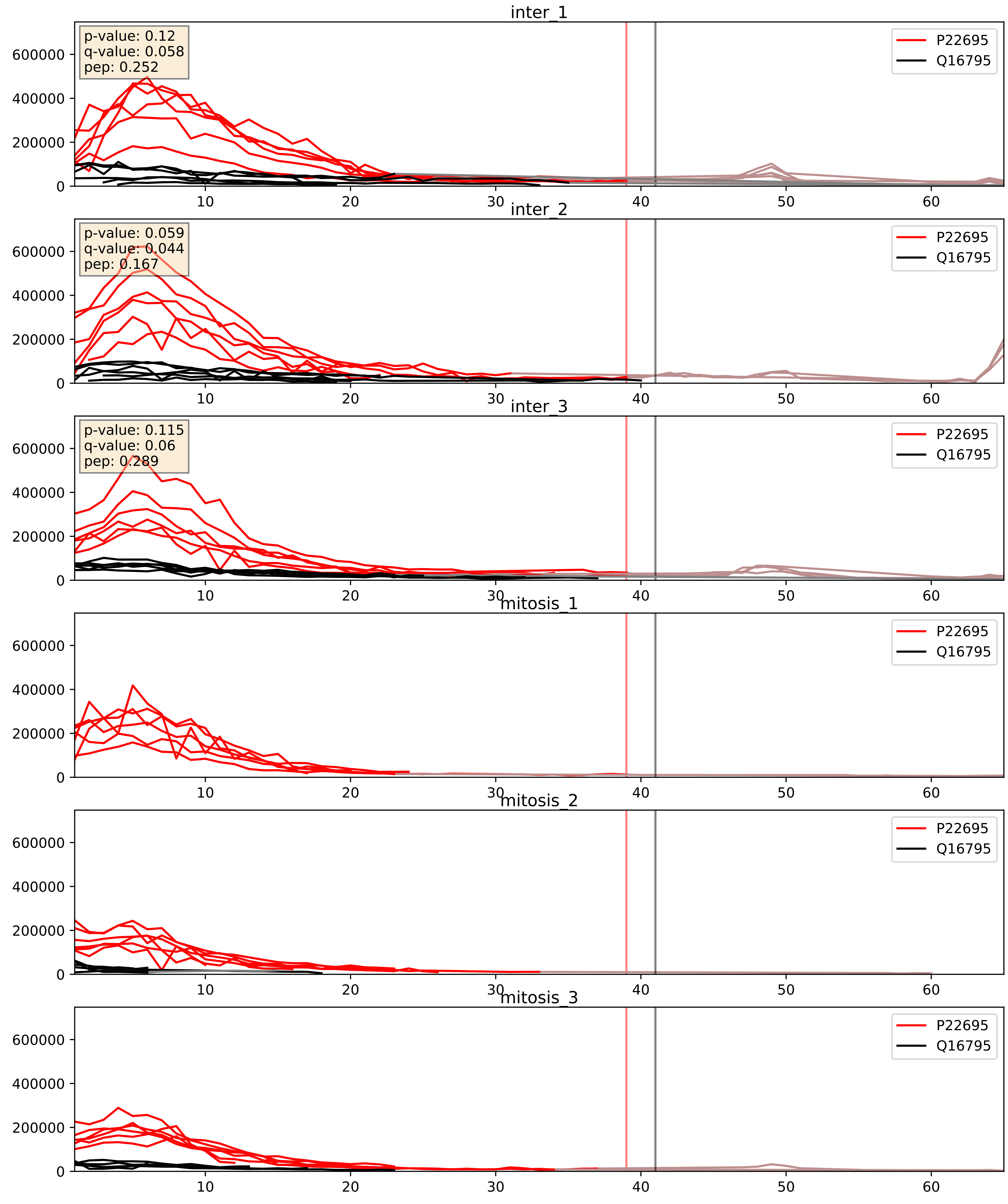


| level | condition_1 | condition_2 | pvalue | pvalue_adjusted |
|---|---|---|---|---|
| complex_abundance | mitosis | inter | 0.010152476802326144 | 0.07446889582511722 |
| interactor_abundance | mitosis | inter | 0.05984572023708979 | 0.19991389862614187 |
| interactor_ratio | mitosis | inter | 0.7776608823156874 | 0.9571917866015084 |
