## Supplementary Data 2 for "SECAT: Quantifying differential protein-protein interaction states by network-centric analysis": hela_string_000130_Q9Y5X2.pdf

Q9Y5X2_Q9Y5X2

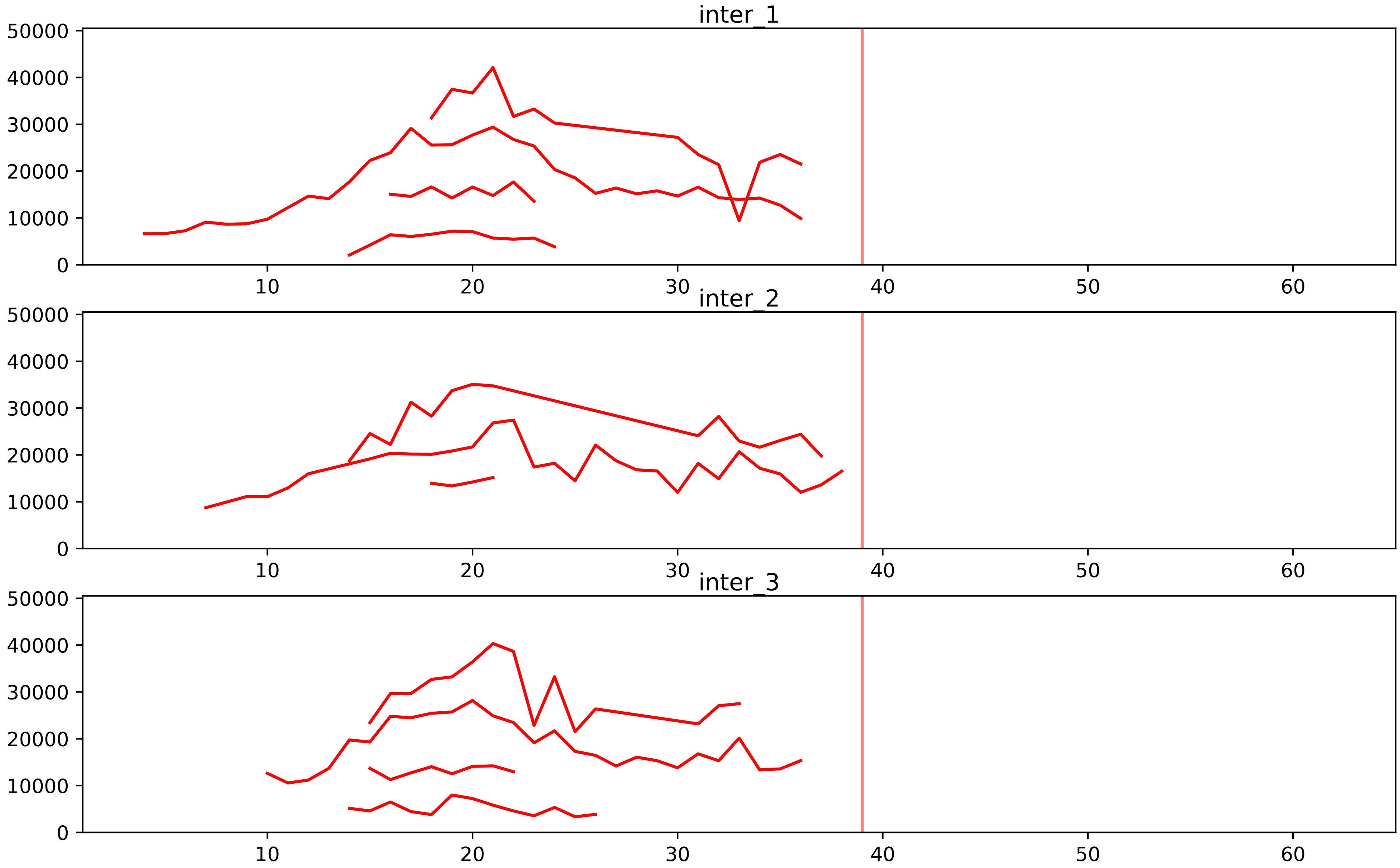


| level | condition_1 | condition_2 | pvalue | pvalue_adjusted |
|---|---|---|---|---|
| assembled_abundance | mitosis | inter | 3.3725512394817676e-05 | 0.006314764940805662 |
| total_abundance | mitosis | inter | 0.00010378565287769052 | 0.01273261737241483 |
