## Supplementary Data 2 for "SECAT: Quantifying differential protein-protein interaction states by network-centric analysis": hela_string_000131_Q9H7B2.pdf

Q9H7B2_Q9H7B2

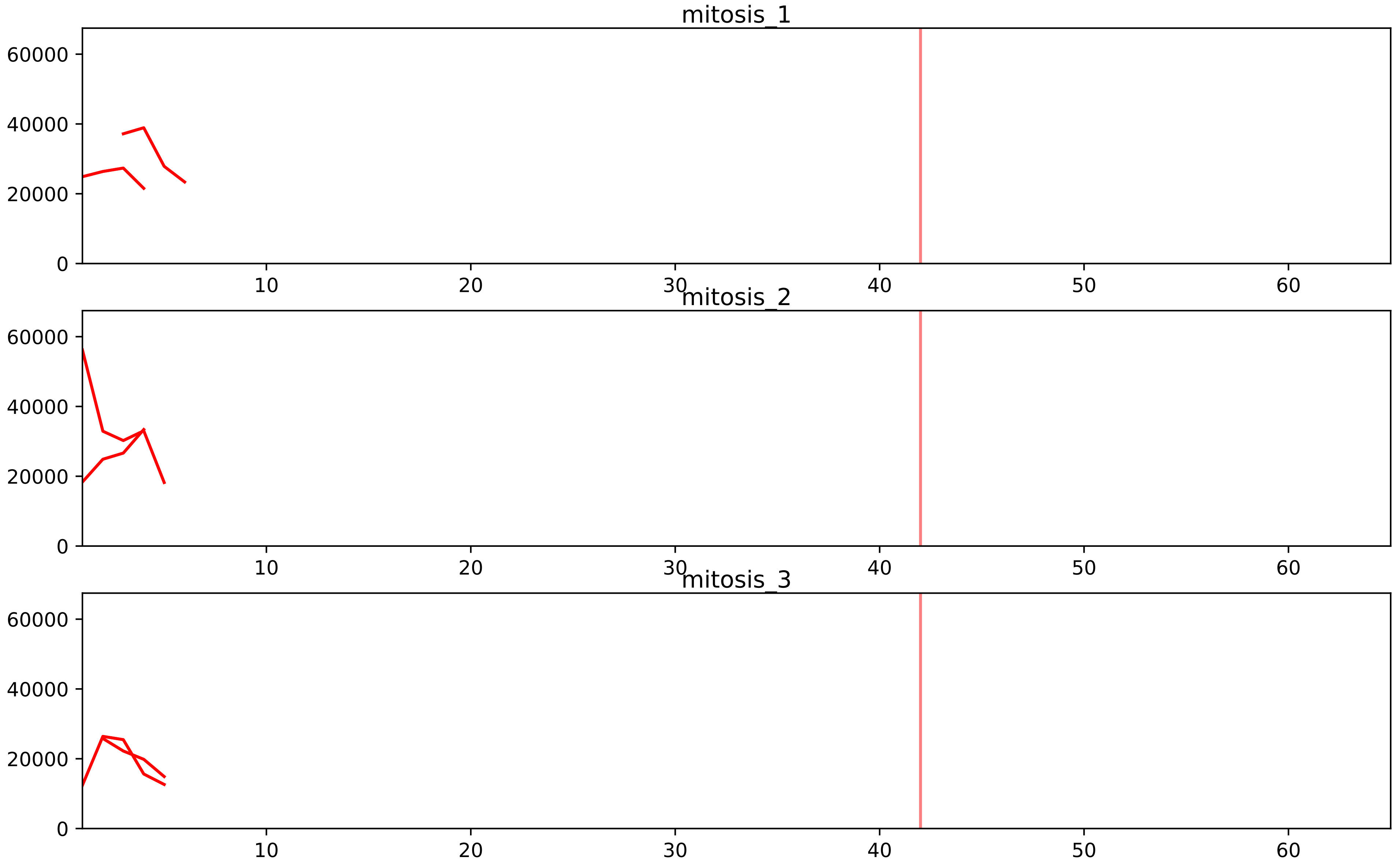


| level | condition_1 | condition_2 | pvalue | pvalue_adjusted |
|---|---|---|---|---|
| interactor_ratio | mitosis | inter | 3.6087533929128946e-05 | 0.0007088834744667758 |
| assembled_abundance | mitosis | inter | 0.003931482757690892 | 0.06212029154760582 |
| total_abundance | mitosis | inter | 0.02652769235754605 | 0.20745388908182325 |
| complex_abundance | mitosis | inter | 0.07259893153191785 | 0.15662330533046523 |
| interactor_abundance | mitosis | inter | 0.590389249501613 | 0.7320190155545606 |

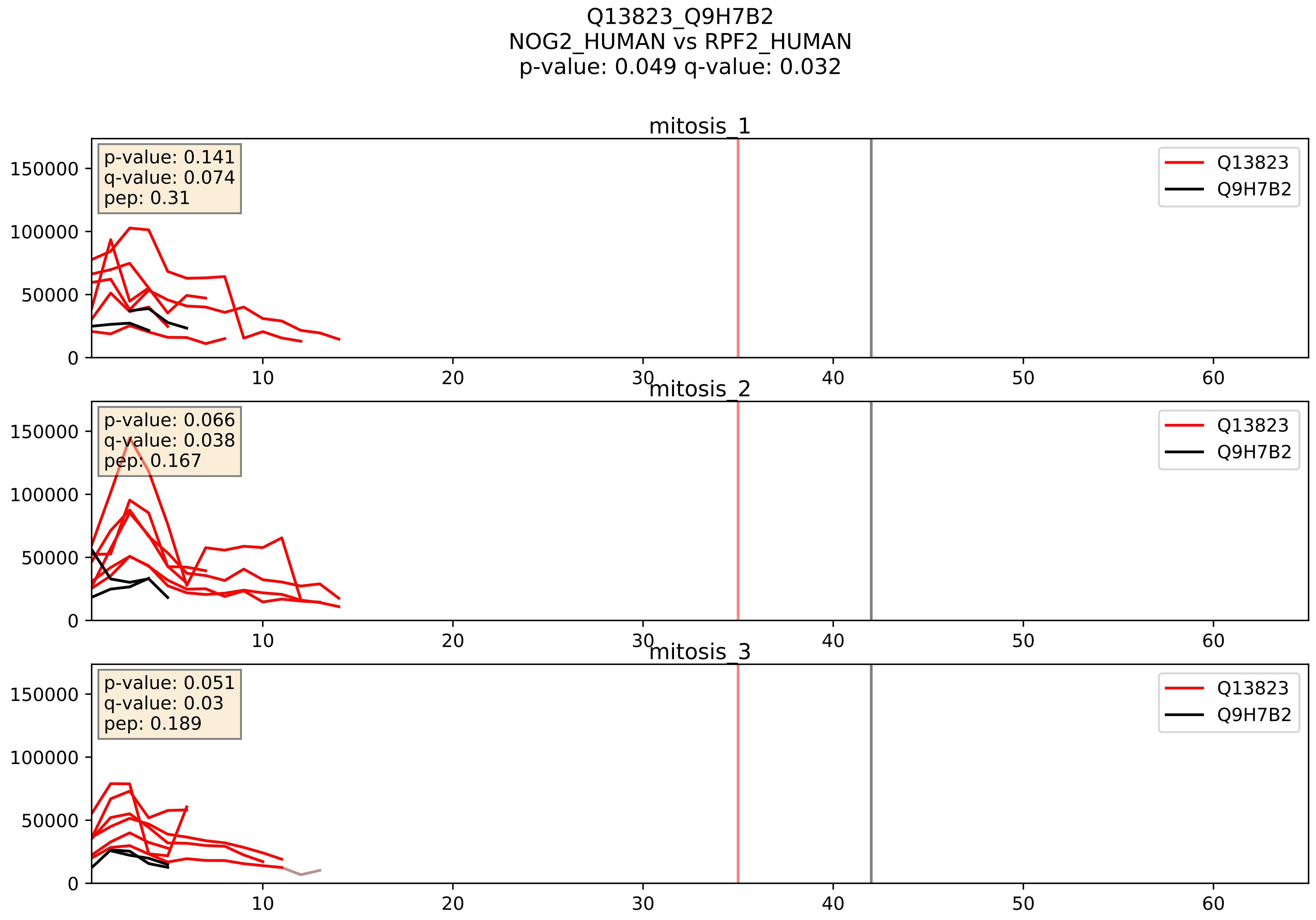


| level | condition_1 | condition_2 | pvalue | pvalue_adjusted |
|---|---|---|---|---|
| interactor_ratio | mitosis | inter | 0.0008412384979360982 | 0.03318433890476037 |
| interactor_abundance | mitosis | inter | 0.03007008855576401 | 0.14418523491160162 |
| complex_abundance | mitosis | inter | 0.08703996497906602 | 0.21596002904950873 |

Q15050_Q9H7B2
RRS1_HUMAN vs RPF2_HUMAN
p-value: 0.068 q-value: 0.042

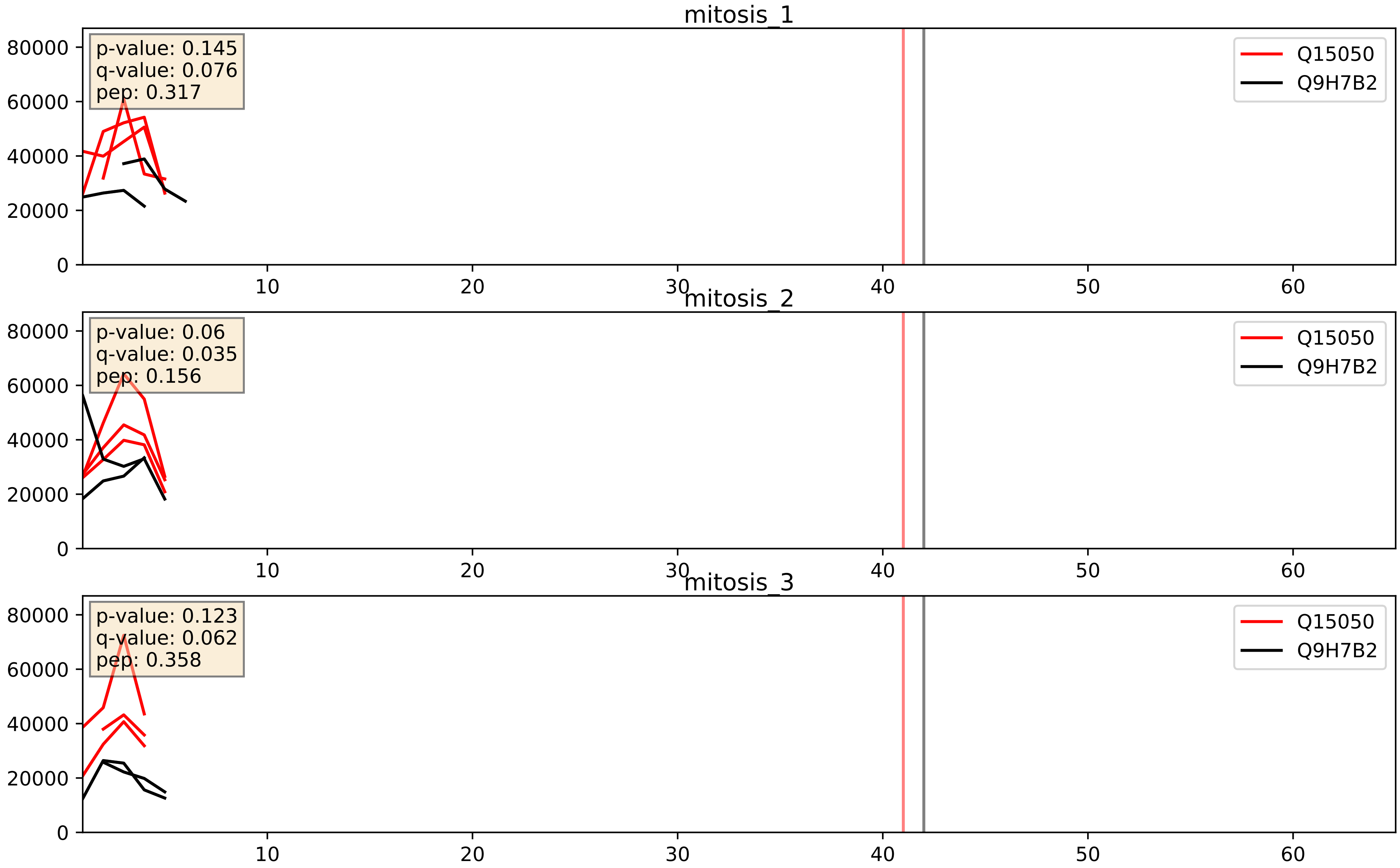


| level | condition_1 | condition_2 | pvalue | pvalue_adjusted |
|---|---|---|---|---|
| interactor_ratio | mitosis | inter | 0.05378980819845297 | 0.38466228753446735 |
| interactor_abundance | mitosis | inter | 0.2162538244564617 | 0.4119120465837366 |
| complex_abundance | mitosis | inter | 0.3048765855504356 | 0.4552142983275299 |

Q15397_Q9H7B2
PUM3_HUMAN vs RPF2_HUMAN
p-value: 0.066 q-value: 0.041

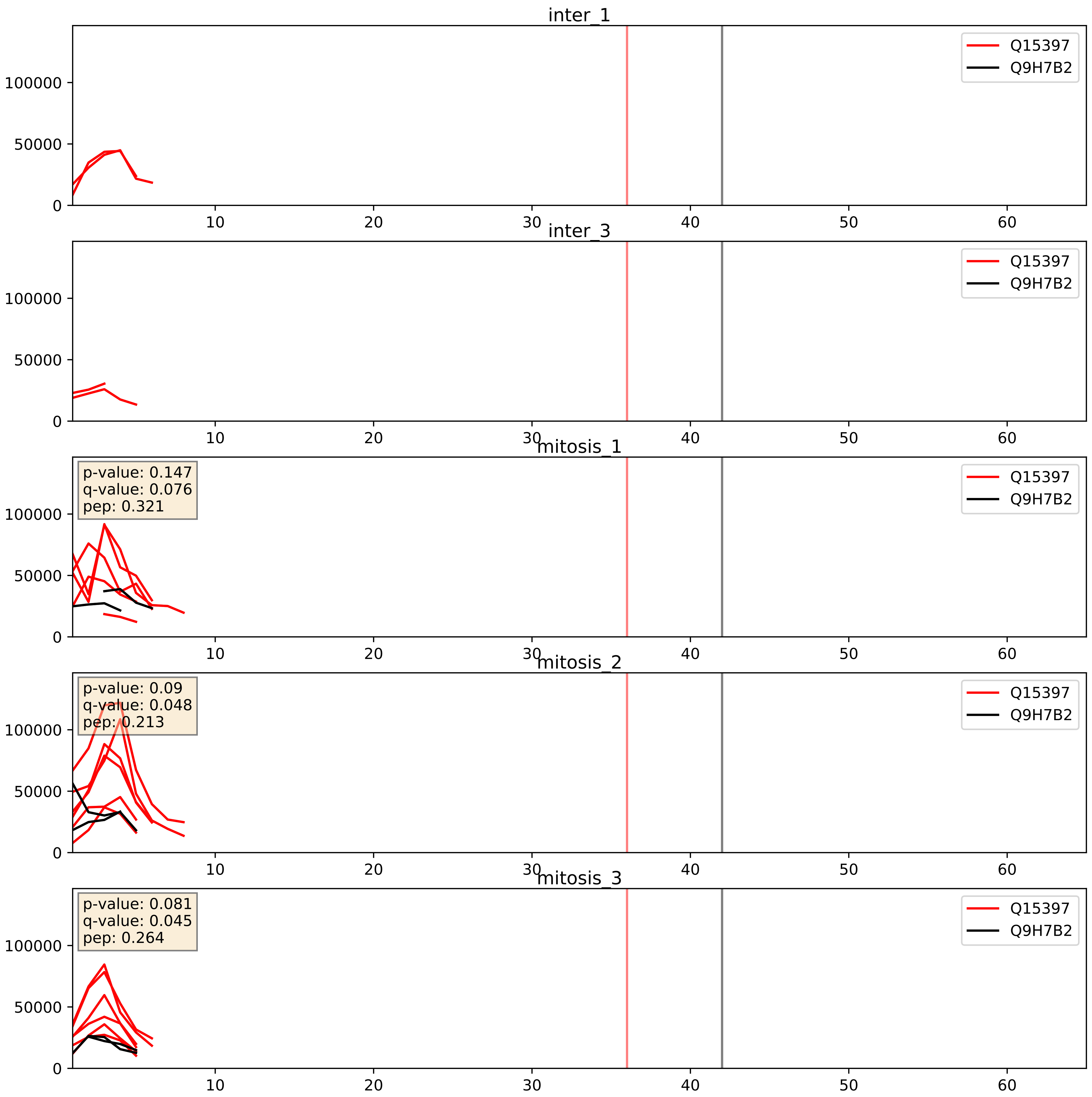


| level | condition_1 | condition_2 | pvalue | pvalue_adjusted |
|---|---|---|---|---|
| interactor_ratio | mitosis | inter | 0.00133235664863674 | 0.04334578357589157 |
| interactor_abundance | mitosis | inter | 0.037615662043017056 | 0.15827677864782586 |
| complex_abundance | mitosis | inter | 0.10346993290830749 | 0.23908191575103507 |

Q8IY81_Q9H7B2
SPB1_HUMAN vs RPF2_HUMAN
p-value: 0.059 q-value: 0.037

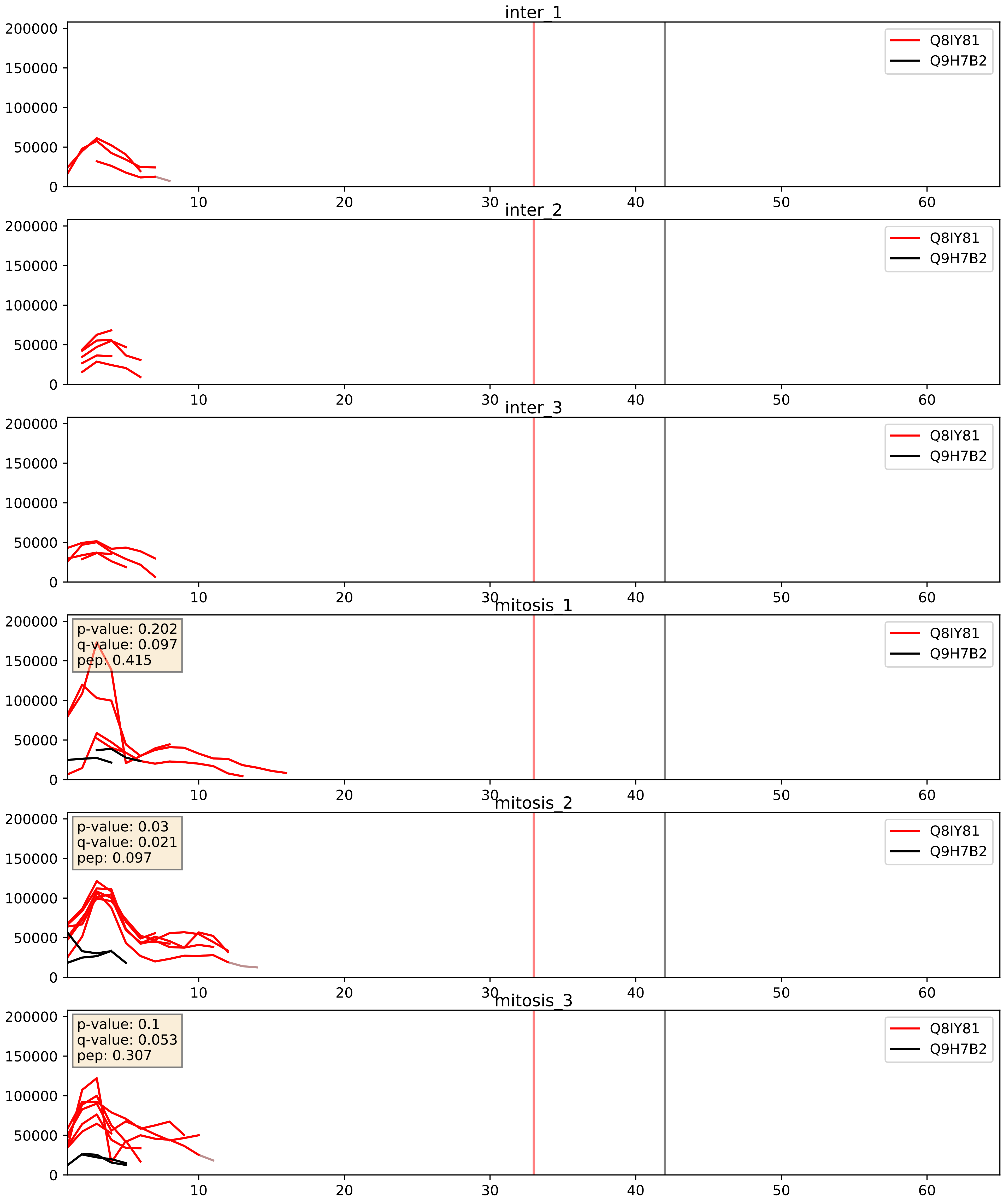


| level | condition_1 | condition_2 | pvalue | pvalue_adjusted |
|---|---|---|---|---|
| interactor_ratio | mitosis | inter | 0.006296065976282134 | 0.10931911715410765 |
| interactor_abundance | mitosis | inter | 0.00774173526098488 | 0.07639107070205253 |
| complex_abundance | mitosis | inter | 0.03990807958361974 | 0.1449976963395235 |

Q8TDN6_Q9H7B2
BRX1_HUMAN vs RPF2_HUMAN
p-value: 0.075 q-value: 0.045

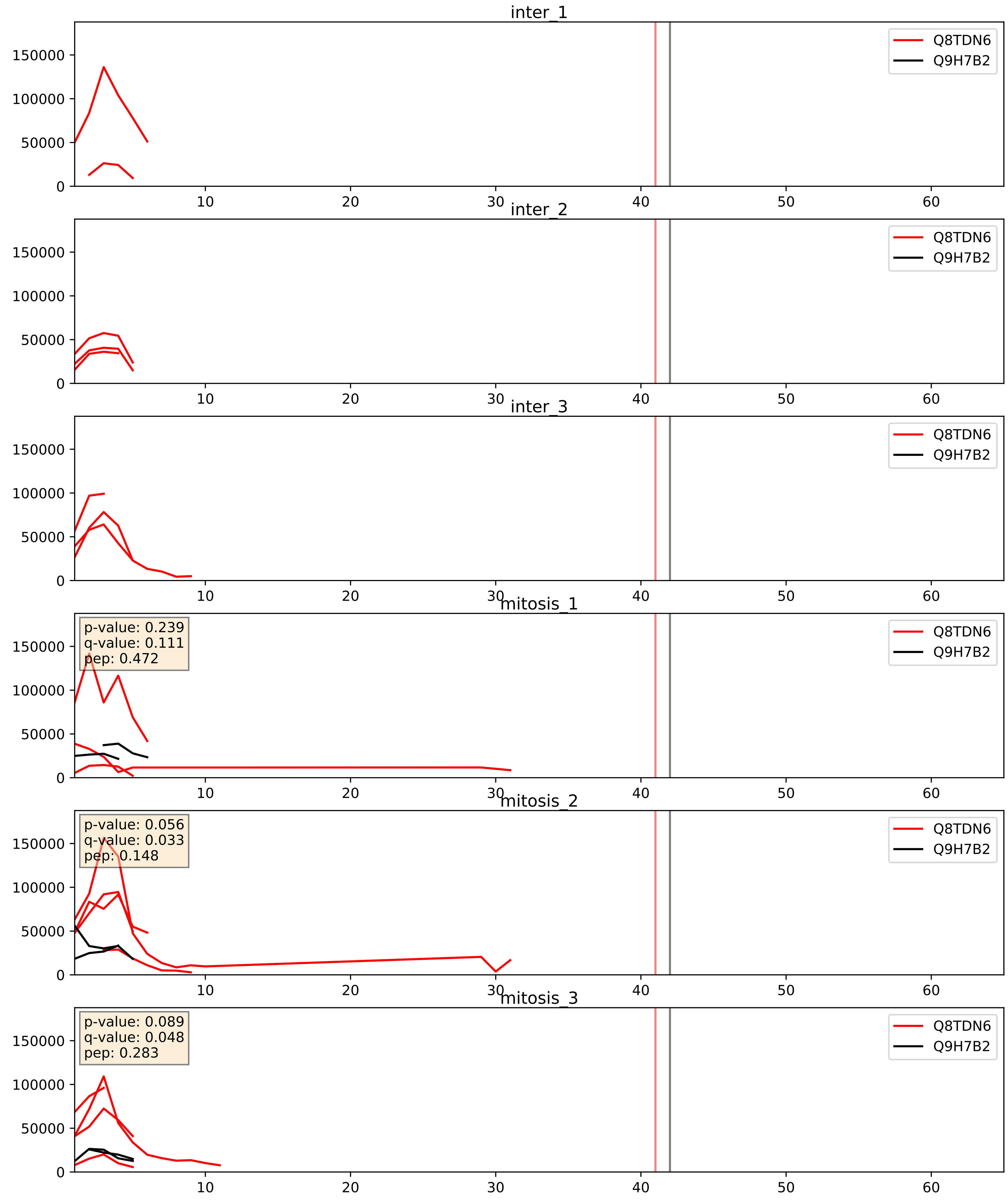


| level | condition_1 | condition_2 | pvalue | pvalue_adjusted |
|---|---|---|---|---|
| interactor_abundance | mitosis | inter | 0.015110876907849336 | 0.10761156932711341 |
| interactor_ratio | mitosis | inter | 0.039906808552959576 | 0.3303697110380406 |
| complex_abundance | mitosis | inter | 0.05121810455375765 | 0.161227792796871 |

Q9BYG3_Q9H7B2
MK67I_HUMAN vs RPF2_HUMAN
p-value: 0.036 q-value: 0.025

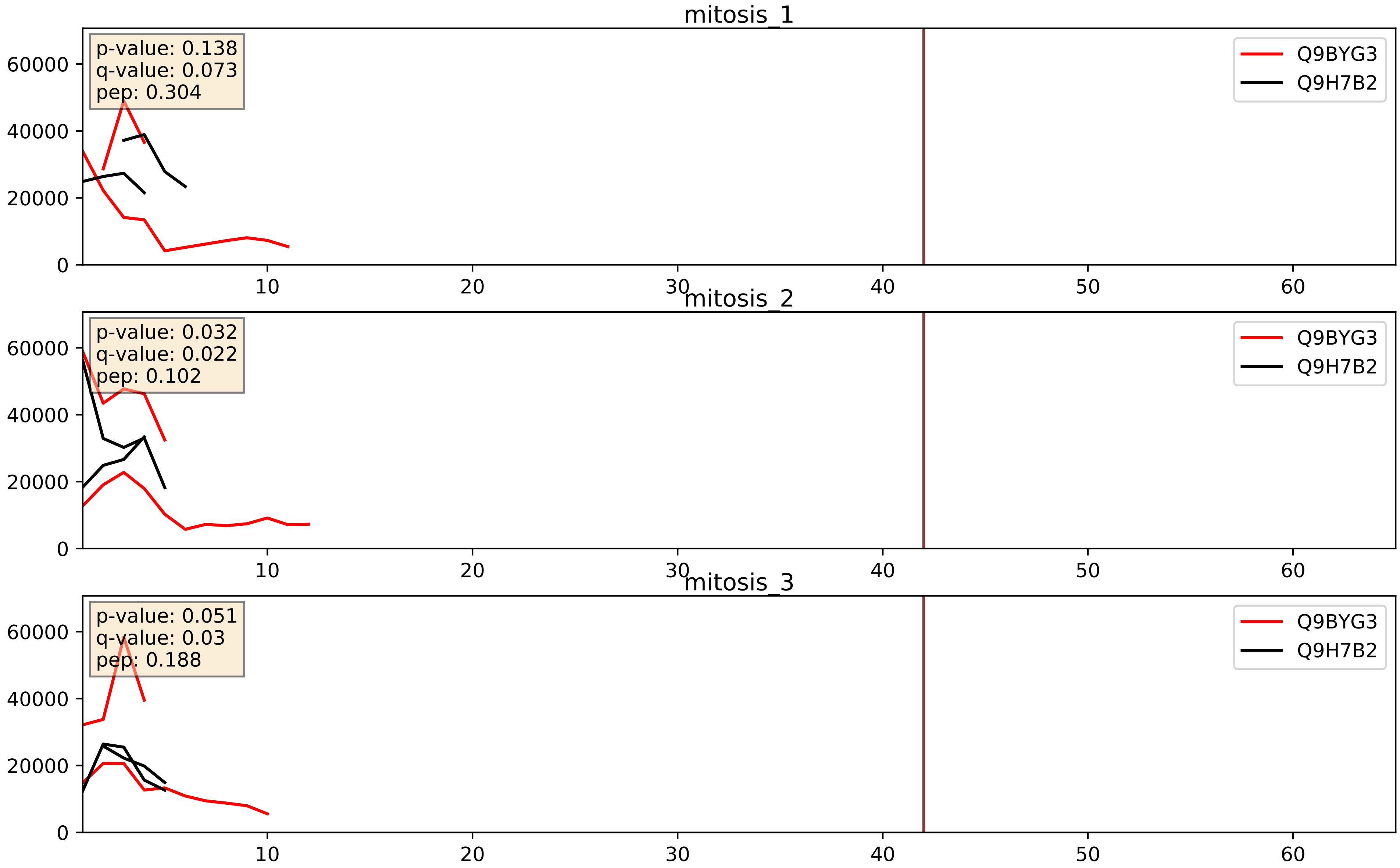


| level | condition_1 | condition_2 | pvalue | pvalue_adjusted |
|---|---|---|---|---|
| interactor_ratio | mitosis | inter | 0.15944016446652348 | 0.616090239021059 |
| interactor_abundance | mitosis | inter | 0.3187687602568554 | 0.5167917779921747 |
| complex_abundance | mitosis | inter | 0.3970451119903028 | 0.5437591915331307 |

Q9H7B2_Q9NVP1
RPF2_HUMAN vs DDX18_HUMAN
p-value: 0.038 q-value: 0.026

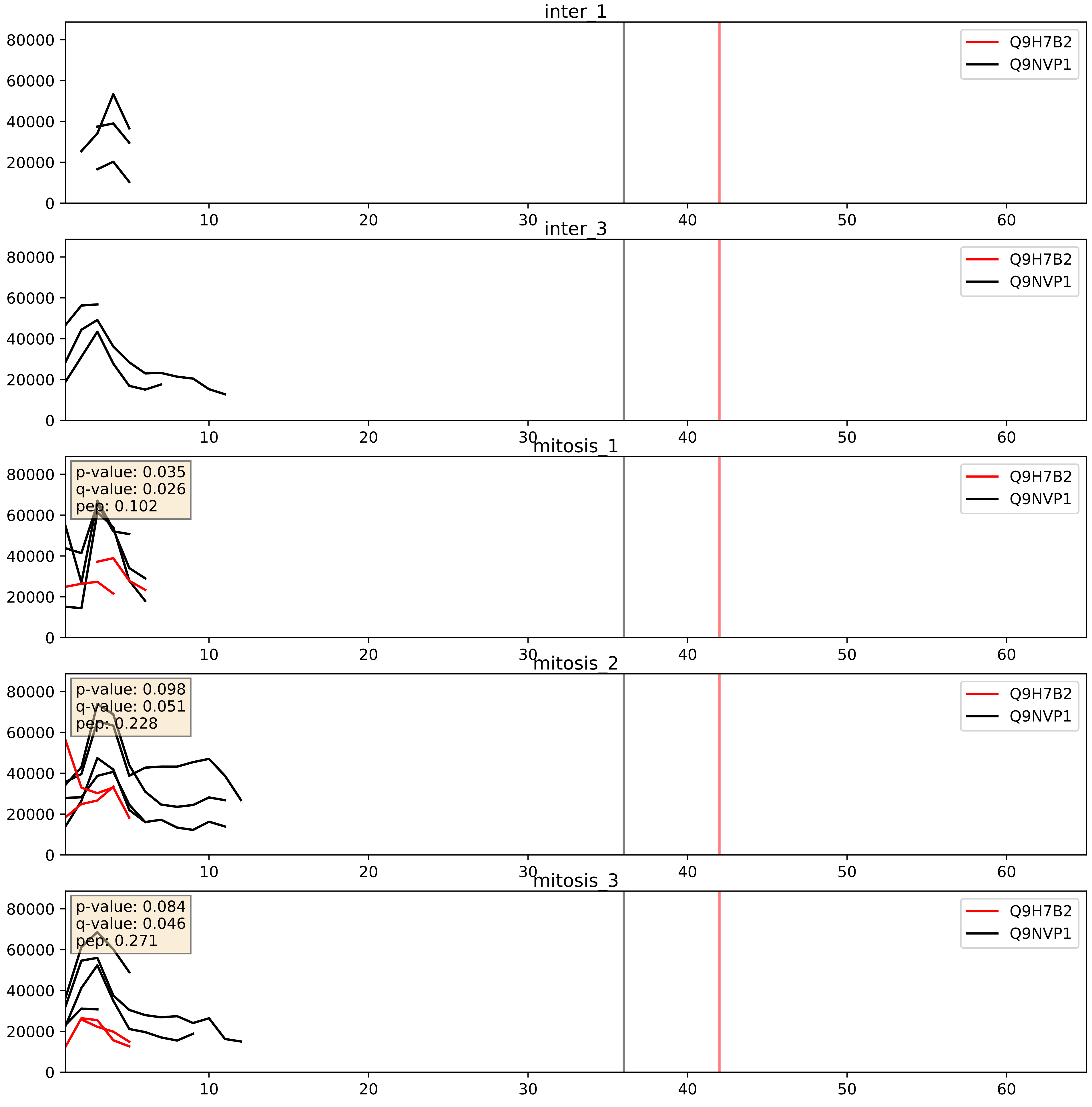


| level | condition_1 | condition_2 | pvalue | pvalue_adjusted |
|---|---|---|---|---|
| interactor_ratio | mitosis | inter | 0.009946672546552762 | 0.14263182888175546 |
| complex_abundance | mitosis | inter | 0.17688335514007095 | 0.3273476593888062 |
| interactor_abundance | mitosis | inter | 0.4740076787521983 | 0.6453802656463842 |

Q9H7B2_Q9NW13
RPF2_HUMAN vs RBM28_HUMAN
p-value: 0.004 q-value: 0.047

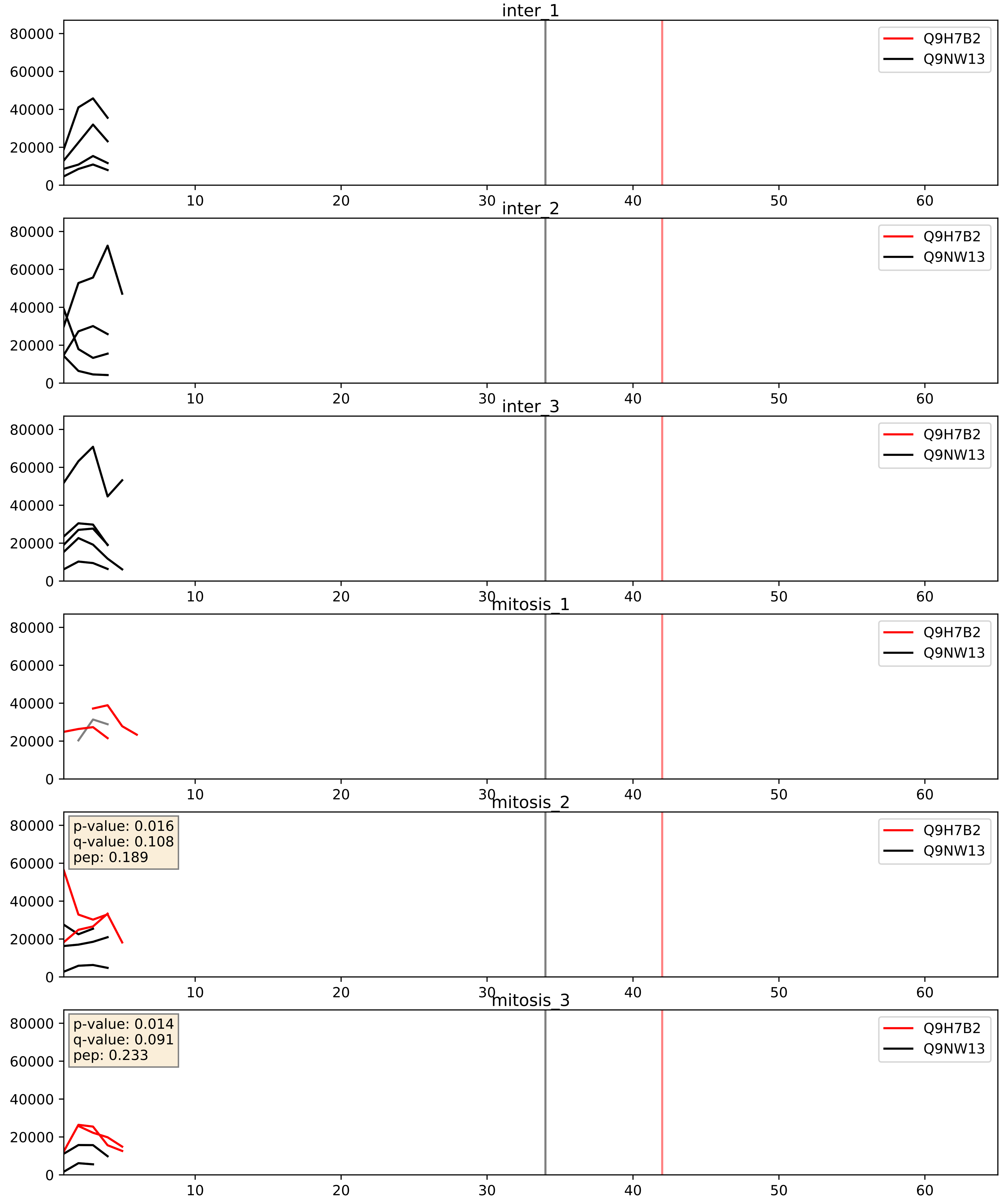


| level | condition_1 | condition_2 | pvalue | pvalue_adjusted |
|---|---|---|---|---|
| complex_abundance | mitosis | inter | 0.4624647684711027 | 0.6037280571941613 |
| interactor_abundance | mitosis | inter | 0.5682420339490499 | 0.7156832025681775 |
| interactor_ratio | mitosis | inter | 0.90372503512248 | 0.9850867567360791 |

Q9H7B2_Q9Y221
RPF2_HUMAN vs NIP7_HUMAN
p-value: 0.037 q-value: 0.026

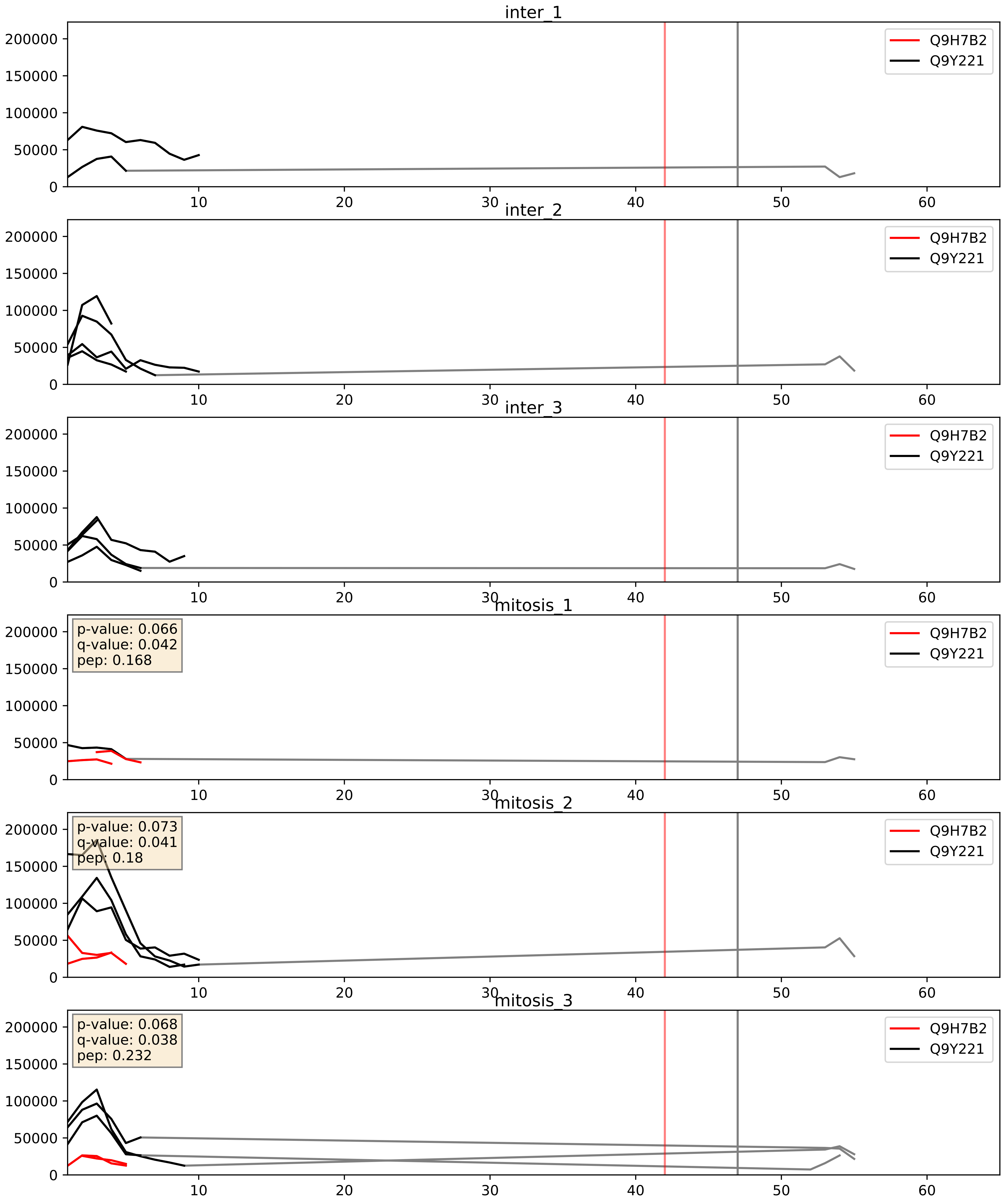


| level | condition_1 | condition_2 | pvalue | pvalue_adjusted |
|---|---|---|---|---|
| complex_abundance | mitosis | inter | 0.06296674817008943 | 0.18087092762951862 |
| interactor_ratio | mitosis | inter | 0.272602302630665 | 0.7468898440591442 |
| interactor_abundance | mitosis | inter | 0.48294198041389746 | 0.6544702818064209 |
