## Supplementary Data 2 for "SECAT: Quantifying differential protein-protein interaction states by network-centric analysis": hela_string_000132_Q86XI2.pdf

Q86XI2_Q86XI2

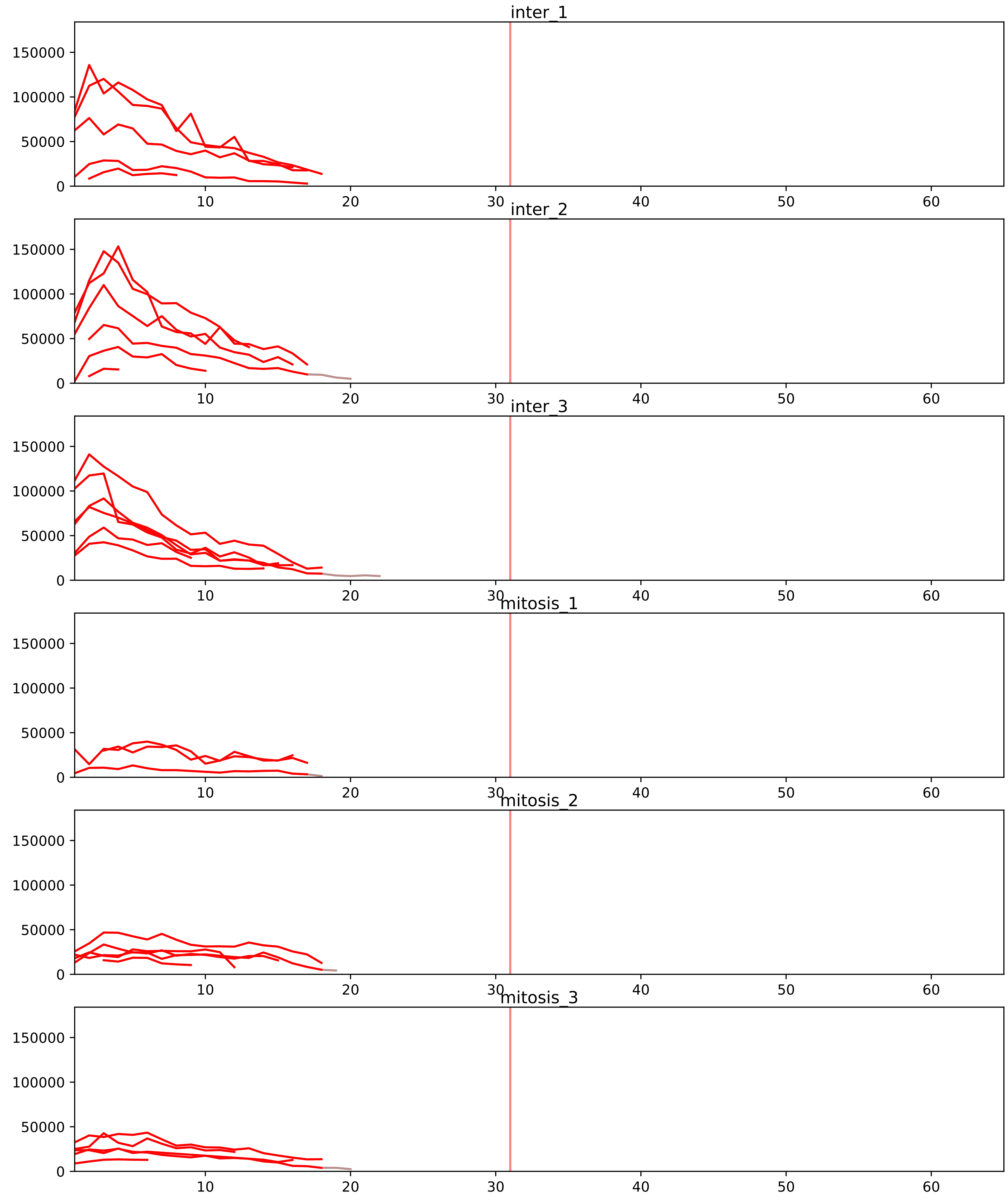


| level | condition_1 | condition_2 | pvalue | pvalue_adjusted |
|---|---|---|---|---|
| interactor_abundance | mitosis | inter | 3.688274774467412e-05 | 0.001219258698154584 |
| complex_abundance | mitosis | inter | 3.766620731323839e-05 | 0.0011949279561441145 |
| total_abundance | mitosis | inter | 0.0022095124419999027 | 0.050217088581346 |
| assembled_abundance | mitosis | inter | 0.002509298501915236 | 0.04841532160005554 |
| interactor_ratio | mitosis | inter | 0.02441686728190865 | 0.12106603474500952 |

P42695_Q86XI2
CNDD3_HUMAN vs CNDG2_HUMAN
p-value: 0.002 q-value: 0.003

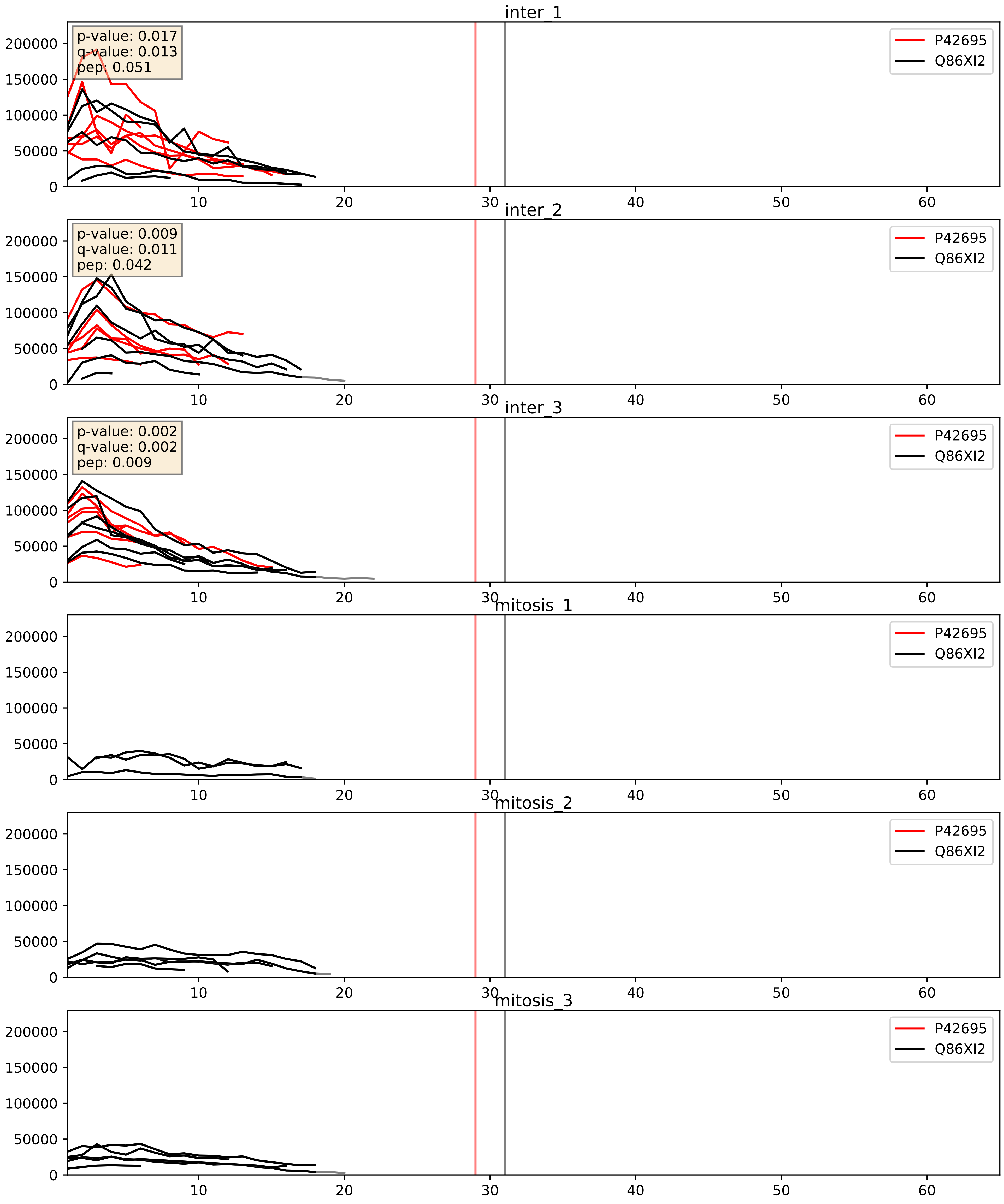


| level | condition_1 | condition_2 | pvalue | pvalue_adjusted |
|---|---|---|---|---|
| complex_abundance | mitosis | inter | 0.00030362639283545225 | 0.011652071109633347 |
| interactor_abundance | mitosis | inter | 0.0003437432204183121 | 0.013564271459750096 |
| interactor_ratio | mitosis | inter | 0.008622180363075611 | 0.13203195690147984 |

Q6IBW4_Q86XI2
CNDH2_HUMAN vs CNDG2_HUMAN
p-value: 0.003 q-value: 0.003

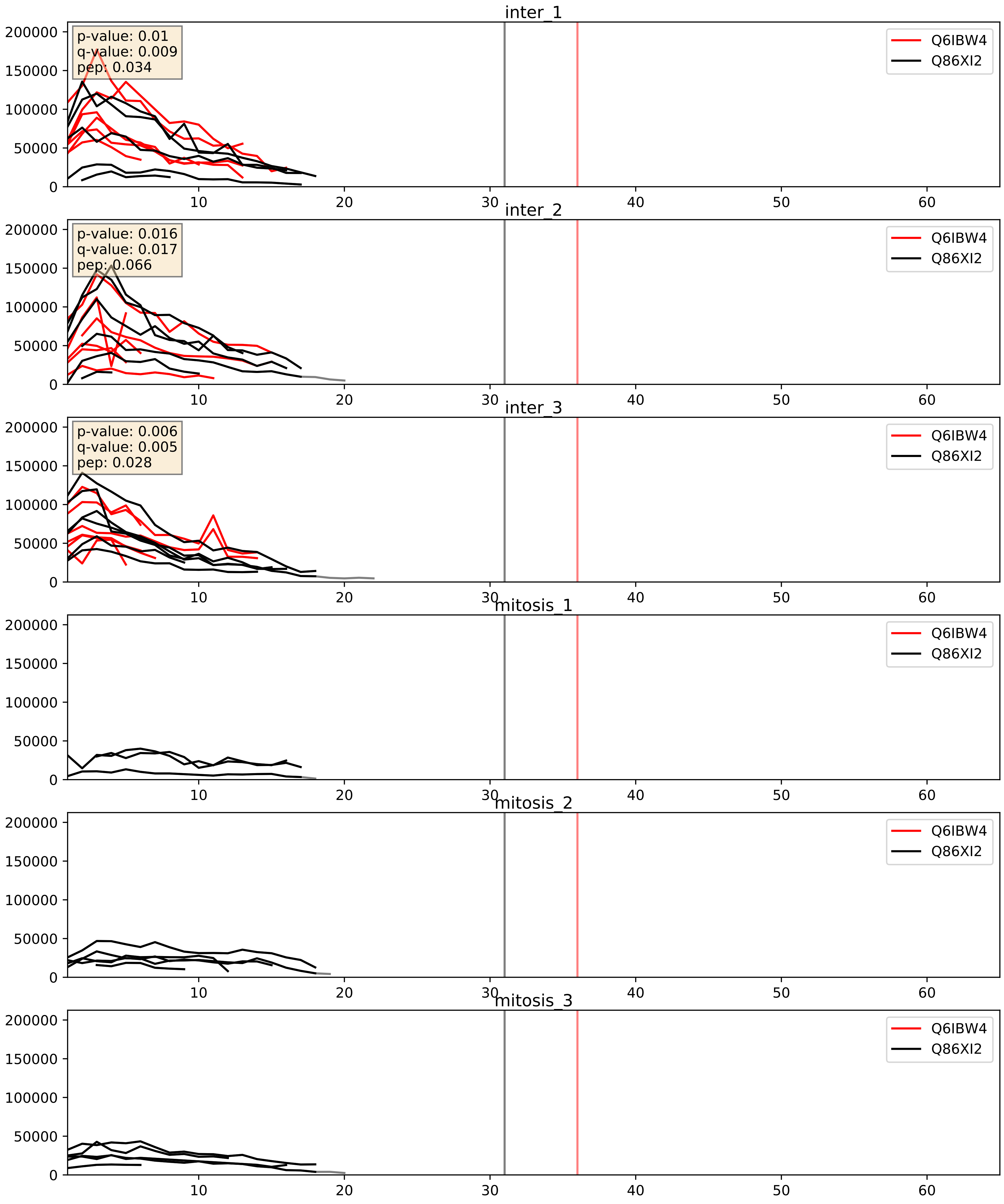


| level | condition_1 | condition_2 | pvalue | pvalue_adjusted |
|---|---|---|---|---|
| complex_abundance | mitosis | inter | 0.0003076550020109643 | 0.01165207110963347 |
| interactor_abundance | mitosis | inter | 0.0003831907621725412 | 0.013582248133320717 |
| interactor_ratio | mitosis | inter | 0.16977884276934044 | 0.6328499622031528 |
