## Supplementary Data 2 for "SECAT: Quantifying differential protein-protein interaction states by network-centric analysis": hela_string_000133_Q08499.pdf

Q08499_Q08499

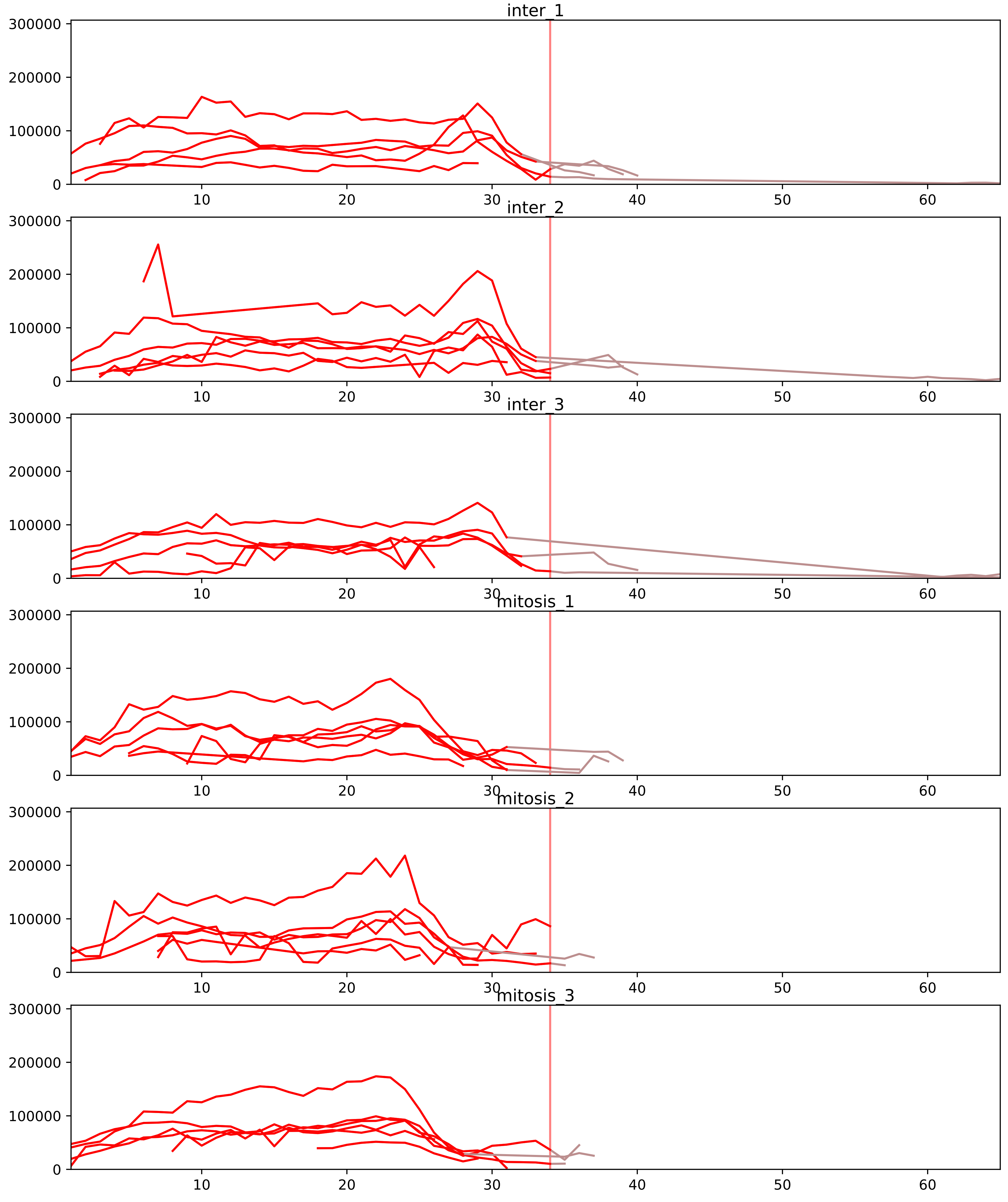


| level | condition_1 | condition_2 | pvalue | pvalue_adjusted |
|---|---|---|---|---|
| assembled_abundance | mitosis | inter | 3.717459969599214e-05 | 0.006692857737574585 |
| total_abundance | mitosis | inter | 8.063625309227403e-05 | 0.012439974681599002 |
| monomer_abundance | mitosis | inter | 0.087450153294486 | 0.3862064922124963 |
