## Supplementary Data 2 for "SECAT: Quantifying differential protein-protein interaction states by network-centric analysis": hela_string_000134_Q9BVC5.pdf

Q9BVC5_Q9BVC5

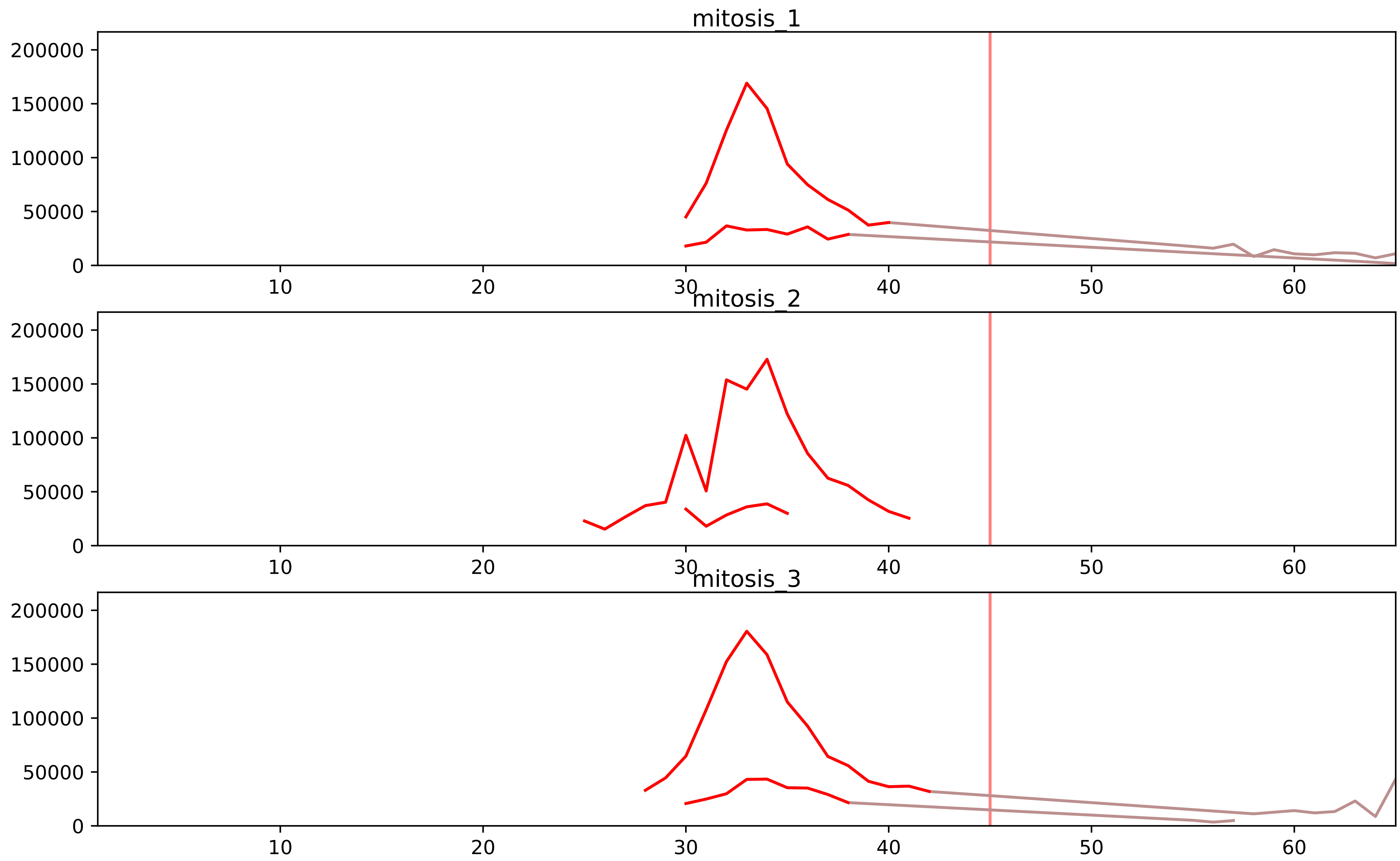


| level | condition_1 | condition_2 | pvalue | pvalue_adjusted |
|---|---|---|---|---|
| assembled_abundance | mitosis | inter | 4.140300046956673e-05 | 0.007178053525853403 |
| total_abundance | mitosis | inter | 6.850974437623276e-05 | 0.011251068019980676 |
| complex_abundance | mitosis | inter | 0.00040108613453436\93 | 0.005999987703603573 |
| interactor_ratio | mitosis | inter | 0.001101580008624925 | 0.01178434427831315 |
| interactor_abundance | mitosis | inter | 0.00561516501975561 | 0.037165120994065916 |
| monomer_abundance | mitosis | inter | 0.06882587040778249 | 0.37224850260048076 |

Q92499_Q9BVC5
DDX1_HUMAN vs ASHWN_HUMAN
p-value: 0.027 q-value: 0.02

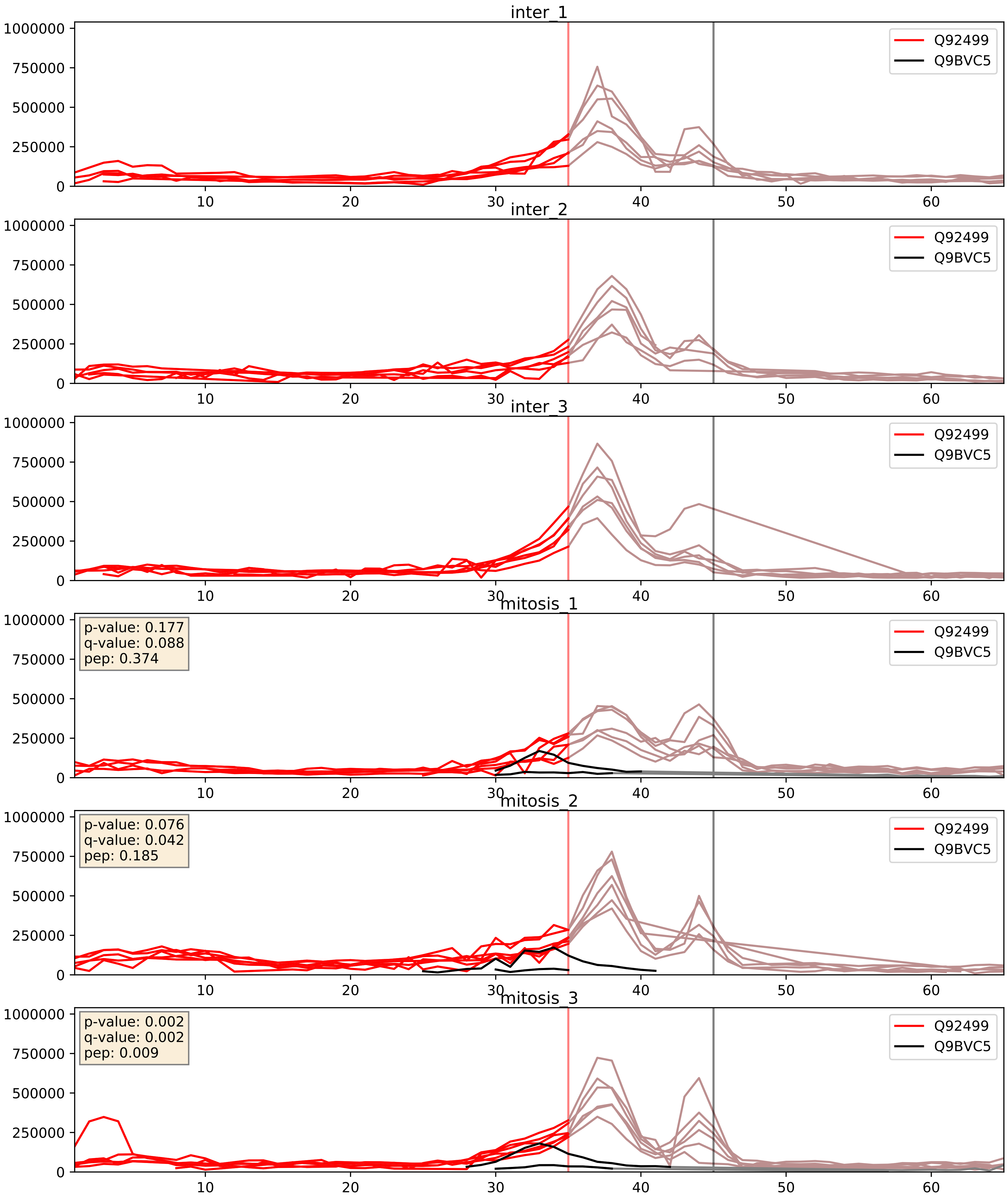


| level | condition_1 | condition_2 | pvalue | pvalue_adjusted |
|---|---|---|---|---|
| interactor_abundance | mitosis | inter | 0.0003457704548032222 | 0.01356427145975096 |
| complex_abundance | mitosis | inter | 0.0004010861345343693 | 0.01272633620489383 |
| interactor_ratio | mitosis | inter | 0.001101580008624925 | 0.0388046291104088 |
