## Supplementary Data 2 for "SECAT: Quantifying differential protein-protein interaction states by network-centric analysis": hela_string_000135_Q12805.pdf

# Q12805_Q12805

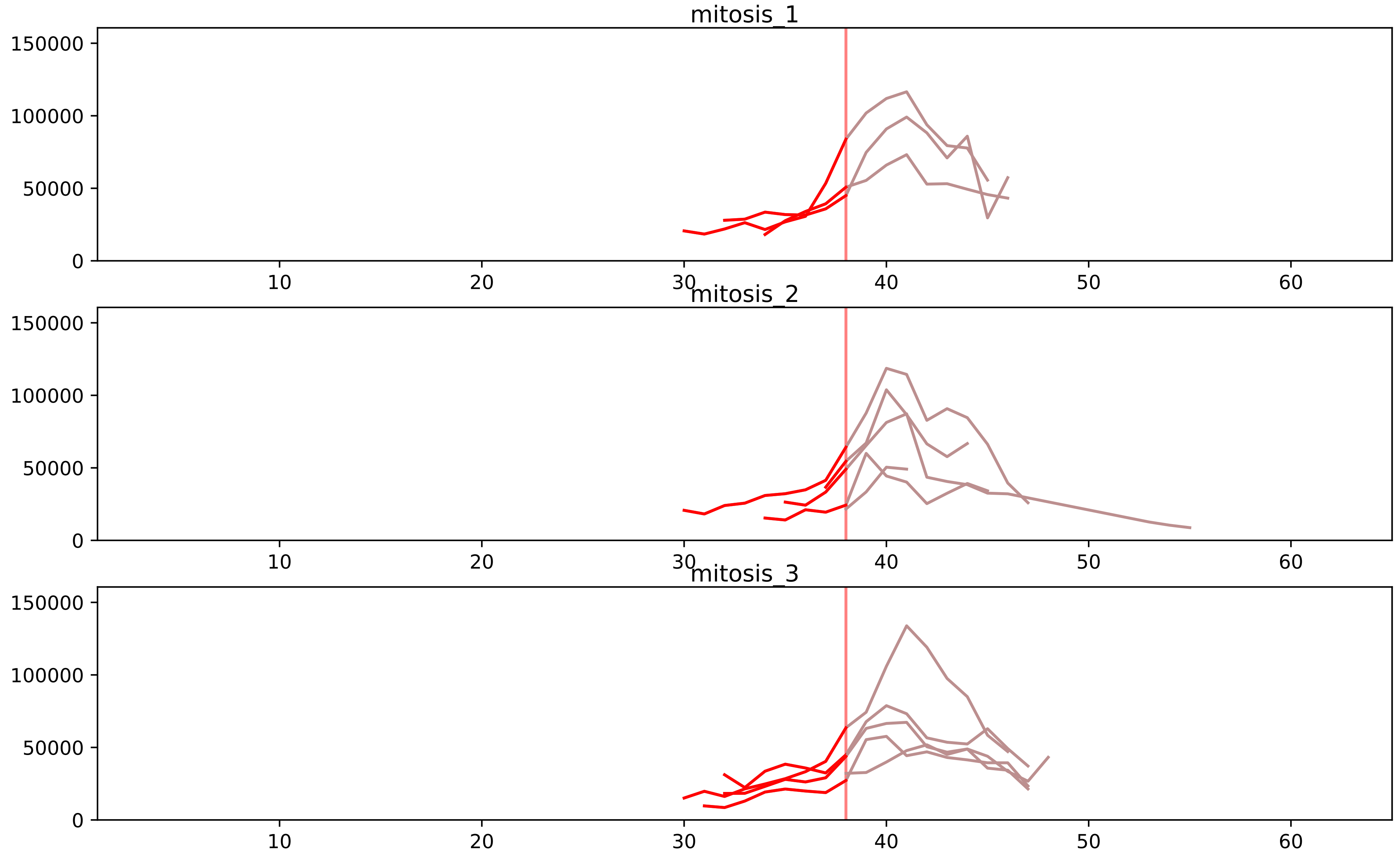


| level | condition_1 | condition_2 | pvalue | pvalue_adjusted |
|---|---|---|---|---|
| total_abundance | mitosis | inter | 4.368075235351529e-05 | 0.008895148409269854 |
| monomer_abundance | mitosis | inter | 0.00016378798225056255 | 0.038277056377303 |
| assembled_abundance | mitosis | inter | 0.0012890205392981858 | 0.0335412168412003 |
