## Supplementary Data 2 for "SECAT: Quantifying differential protein-protein interaction states by network-centric analysis": hela_string_000136_Q92973.pdf

Q92973_Q92973

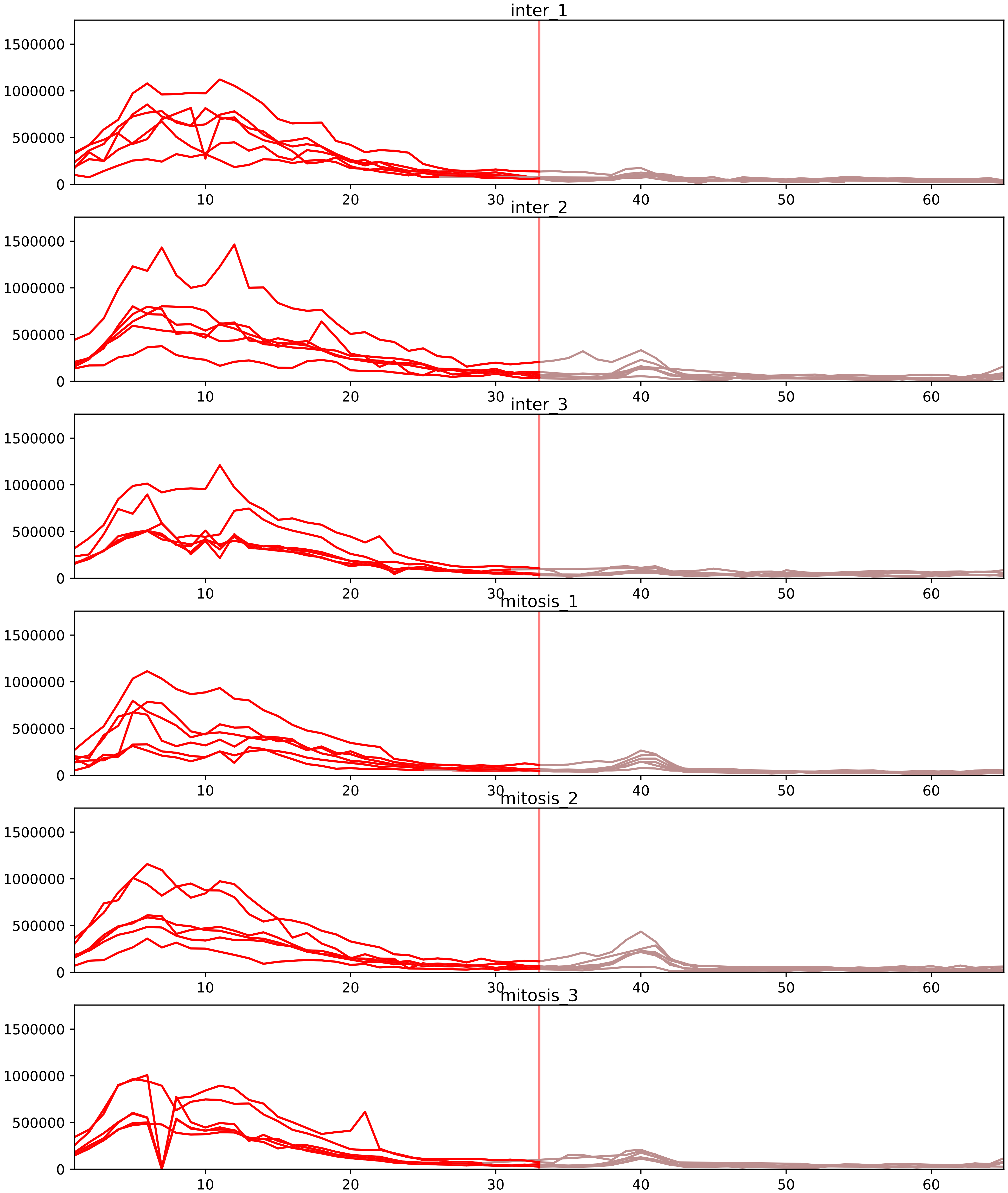


| level | condition_1 | condition_2 | pvalue | pvalue_adjusted |
|---|---|---|---|---|
| interactor_abundance | mitosis | inter | 4.4549836631914046e-05 | 0.0014009190465561038 |
| complex_abundance | mitosis | inter | 0.0010020915510917888 | 0.010782739497128019 |
| interactor_ratio | mitosis | inter | 0.05817972916925262 | 0.23271891667701047 |
| assembled_abundance | mitosis | inter | 0.1764151834170807 | 0.4852547347909627 |
| total_abundance | mitosis | inter | 0.19953737885128914 | 0.536986746555105 |
| monomer_abundance | mitosis | inter | 0.7421413698658288 | 0.8954341477630327 |

Q14103_Q92973
HNRPD_HUMAN vs TNPO1_HUMAN
p-value: 0.032 q-value: 0.023

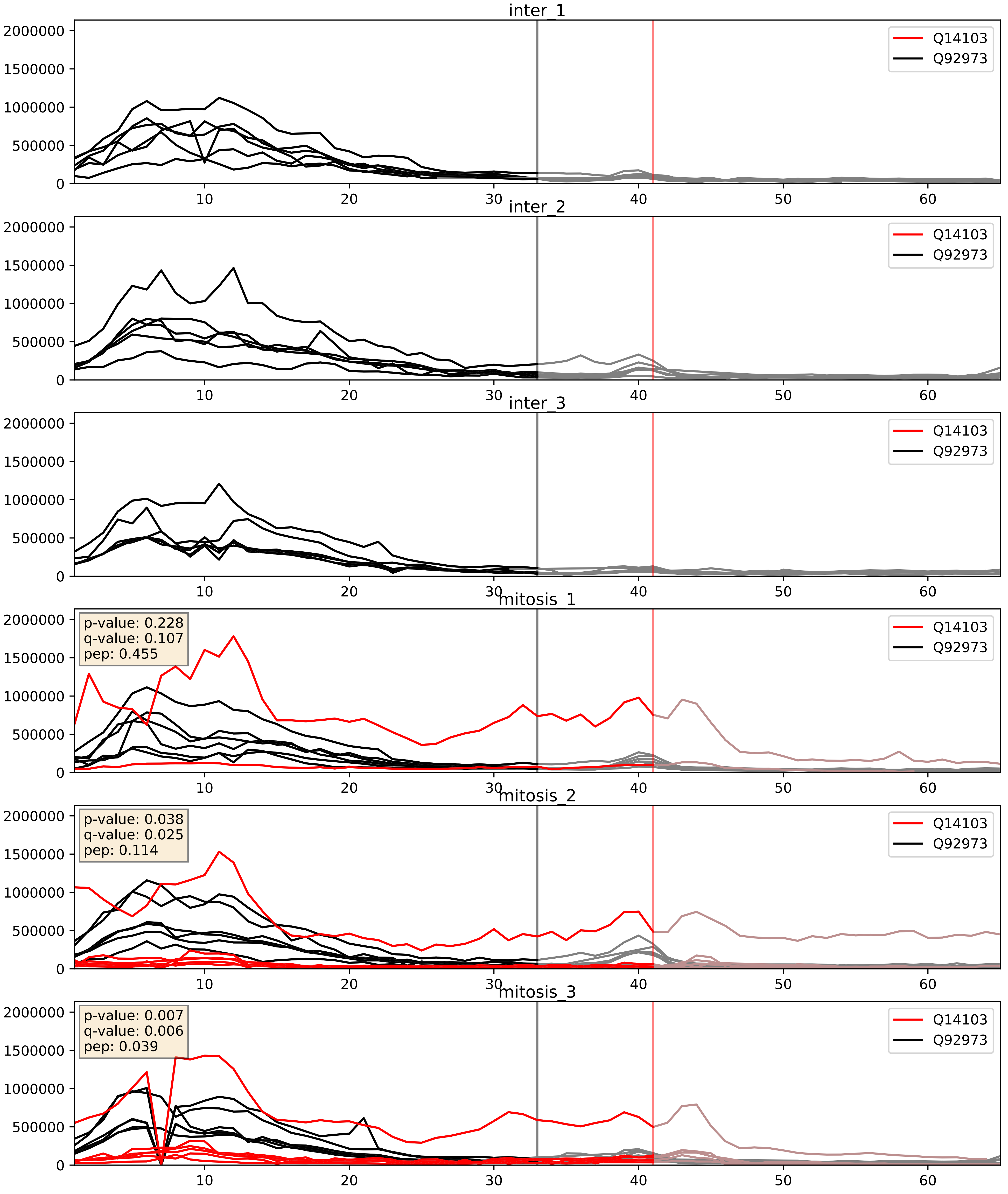


| level | condition_1 | condition_2 | pvalue | pvalue_adjusted |
|---|---|---|---|---|
| complex_abundance | mitosis | inter | 0.0010020915510917888 | 0.021075930411168824 |
| interactor_abundance | mitosis | inter | 0.010151249925242783 | 0.08768385404649669 |
| interactor_ratio | mitosis | inter | 0.05817972916925262 | 0.3974608792408638 |
