## Supplementary Data 2 for "SECAT: Quantifying differential protein-protein interaction states by network-centric analysis": hela_string_000137_Q15005.pdf

Q15005_Q15005

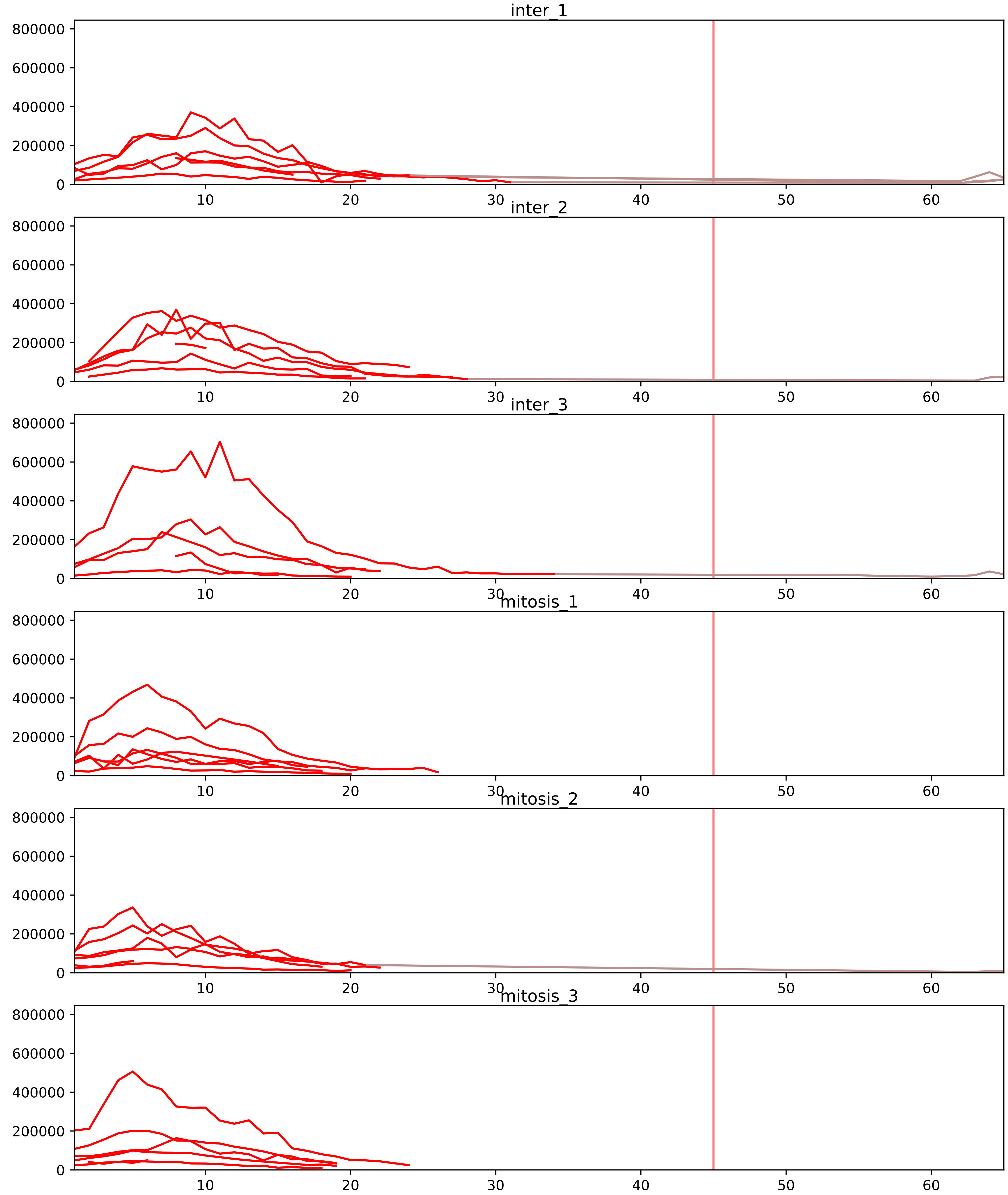


| level | condition_1 | condition_2 | pvalue | pvalue_adjusted |
|---|---|---|---|---|
| interactor_abundance | mitosis | inter | 4.492077377544029e-05 | 0.0014009190465561038 |
| complex_abundance | mitosis | inter | 0.0009821960069712062 | 0.010746183439302845 |
| interactor_ratio | mitosis | inter | 0.014547209687236523 | 0.0815319611305561 |
| monomer_abundance | mitosis | inter | 0.07355451091463493 | 0.37224850260048076 |
| assembled_abundance | mitosis | inter | 0.39424255664451646 | 0.6679266230185432 |
| total_abundance | mitosis | inter | 0.4118091866511959 | 0.7078010147538231 |

O00264_Q15005
PGRC1_HUMAN vs SPCS2_HUMAN
p-value: 0.0 q-value: 0.023

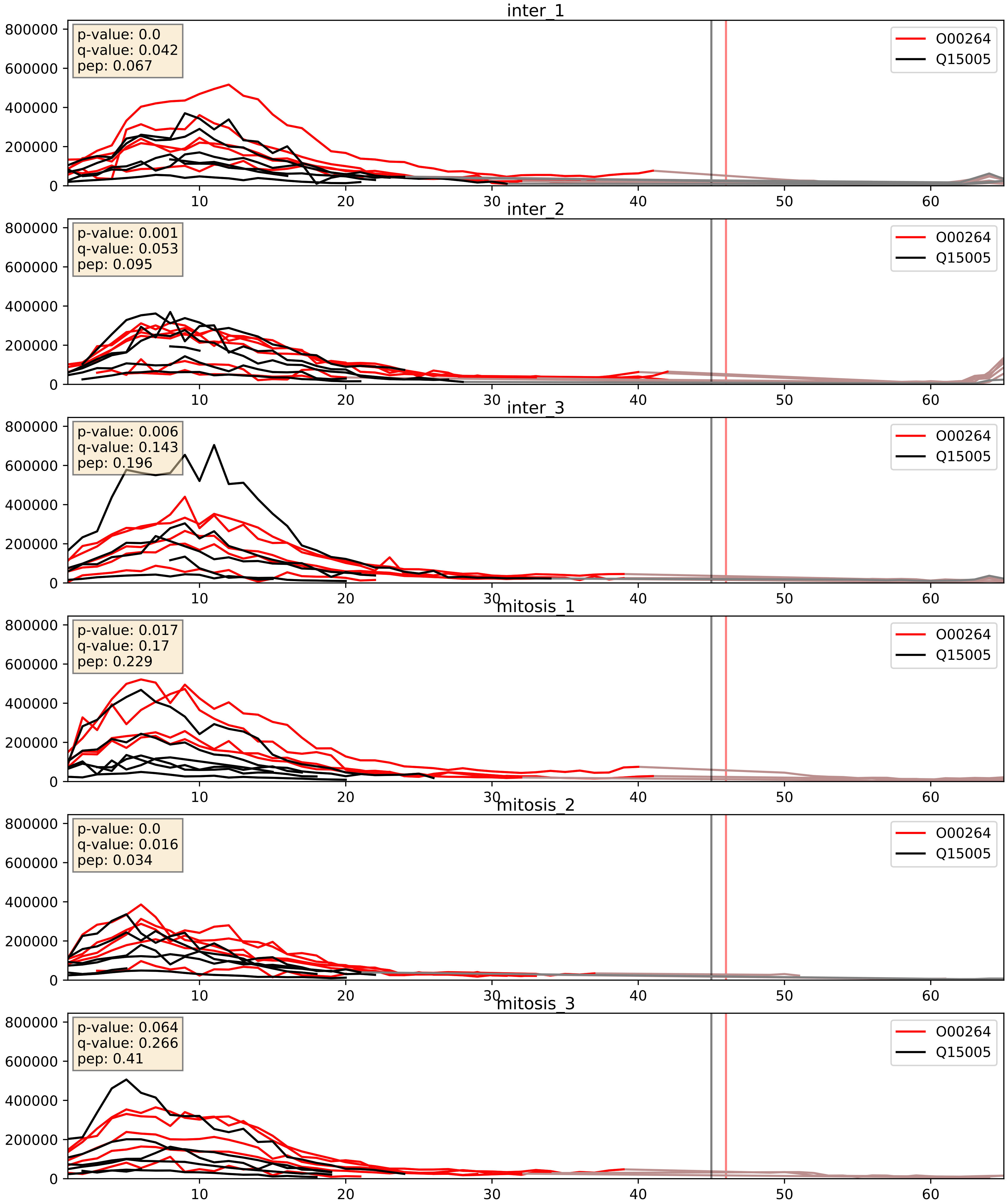


| level | condition_1 | condition_2 | pvalue | pvalue_adjusted |
|---|---|---|---|---|
| complex_abundance | mitosis | inter | 0.018133561615901368 | 0.10072893408962733 |
| interactor_abundance | mitosis | inter | 0.11344496820395018 | 0.2903539924729597 |
| interactor_ratio | mitosis | inter | 0.181404298452545 | 0.650805027139055 |

P43307_Q15005
SSRA_HUMAN vs SPCS2_HUMAN
p-value: 0.002 q-value: 0.035

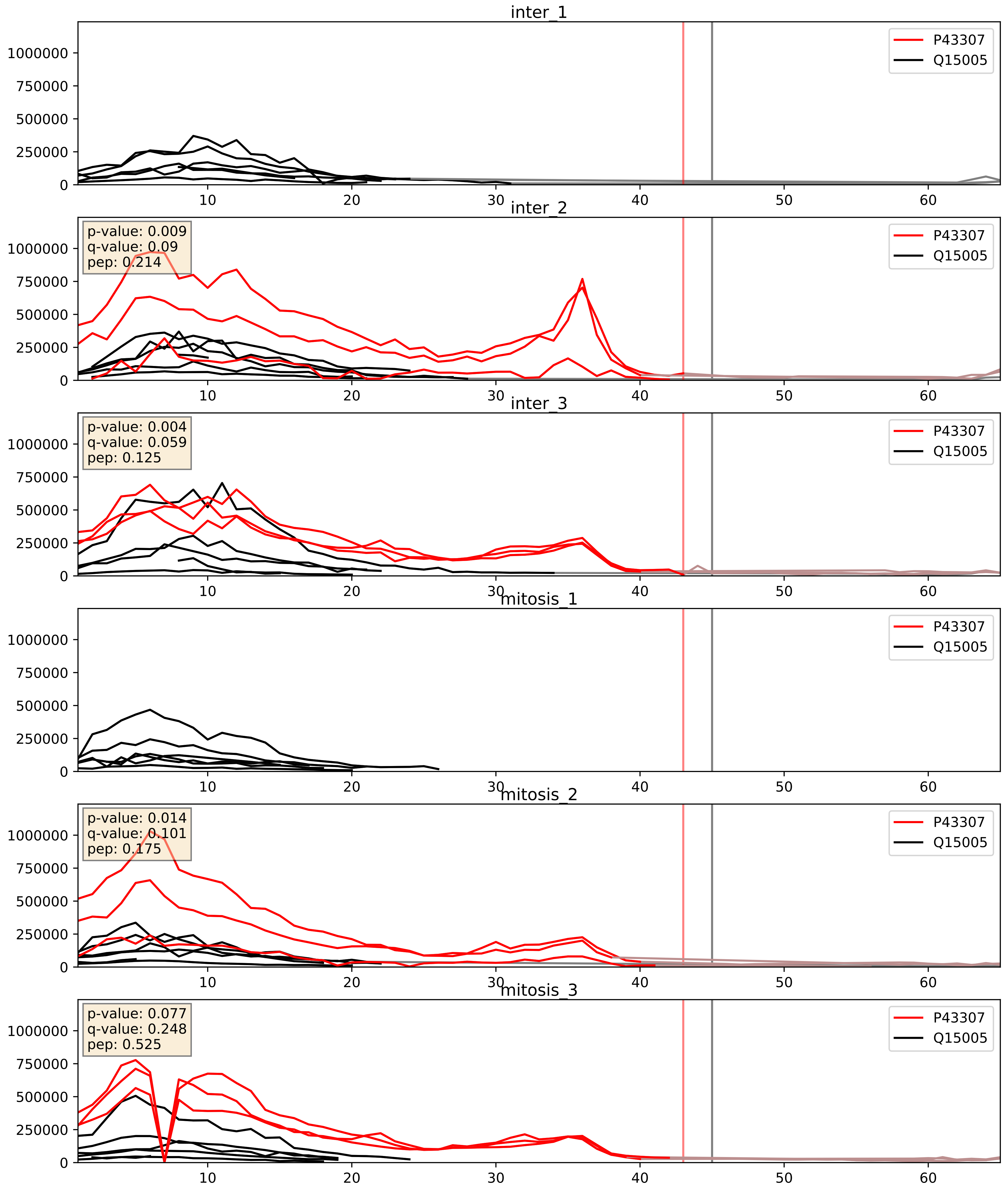


| level | condition_1 | condition_2 | pvalue | pvalue_adjusted |
|---|---|---|---|---|
| complex_abundance | mitosis | inter | 0.7854985630691979 | 0.8599395958399199 |
| interactor_abundance | mitosis | inter | 0.8296810608791333 | 0.9061652160925526 |
| interactor_ratio | mitosis | inter | 0.8801373268065507 | 0.9802747470617126 |

P51571_Q15005
SSRD_HUMAN vs SPCS2_HUMAN
p-value: 0.0 q-value: 0.024

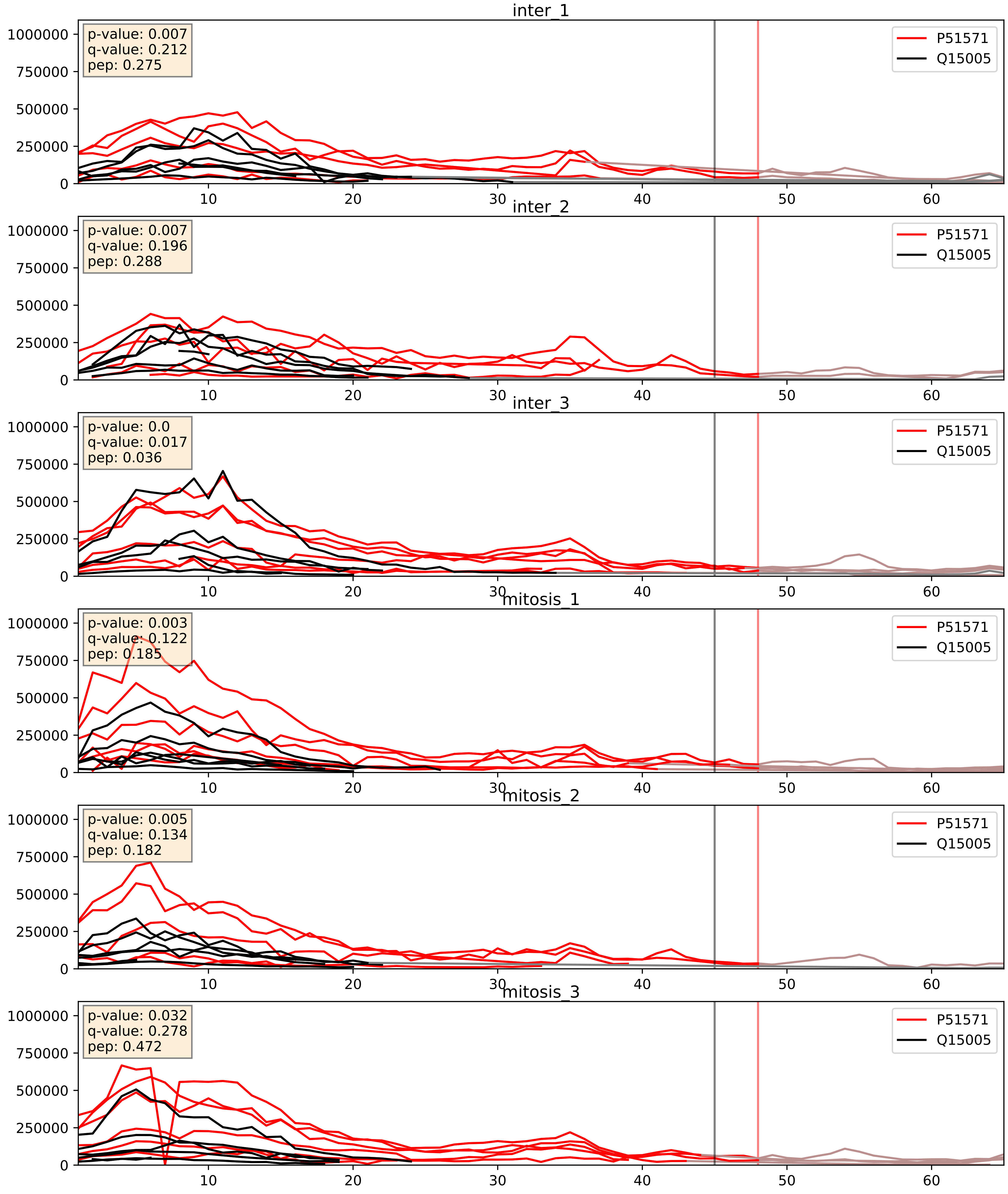


| level | condition_1 | condition_2 | pvalue | pvalue_adjusted |
|---|---|---|---|---|
| complex_abundance | mitosis | inter | 0.05251164624444799 | 0.1635138929983539 |
| interactor_ratio | mitosis | inter | 0.17805931906580372 | 0.6446022882213848 |
| interactor_abundance | mitosis | inter | 0.3488760277590967 | 0.5477081701270734 |

P61009_Q15005
SPCS3_HUMAN vs SPCS2_HUMAN
p-value: 0.0 q-value: 0.0

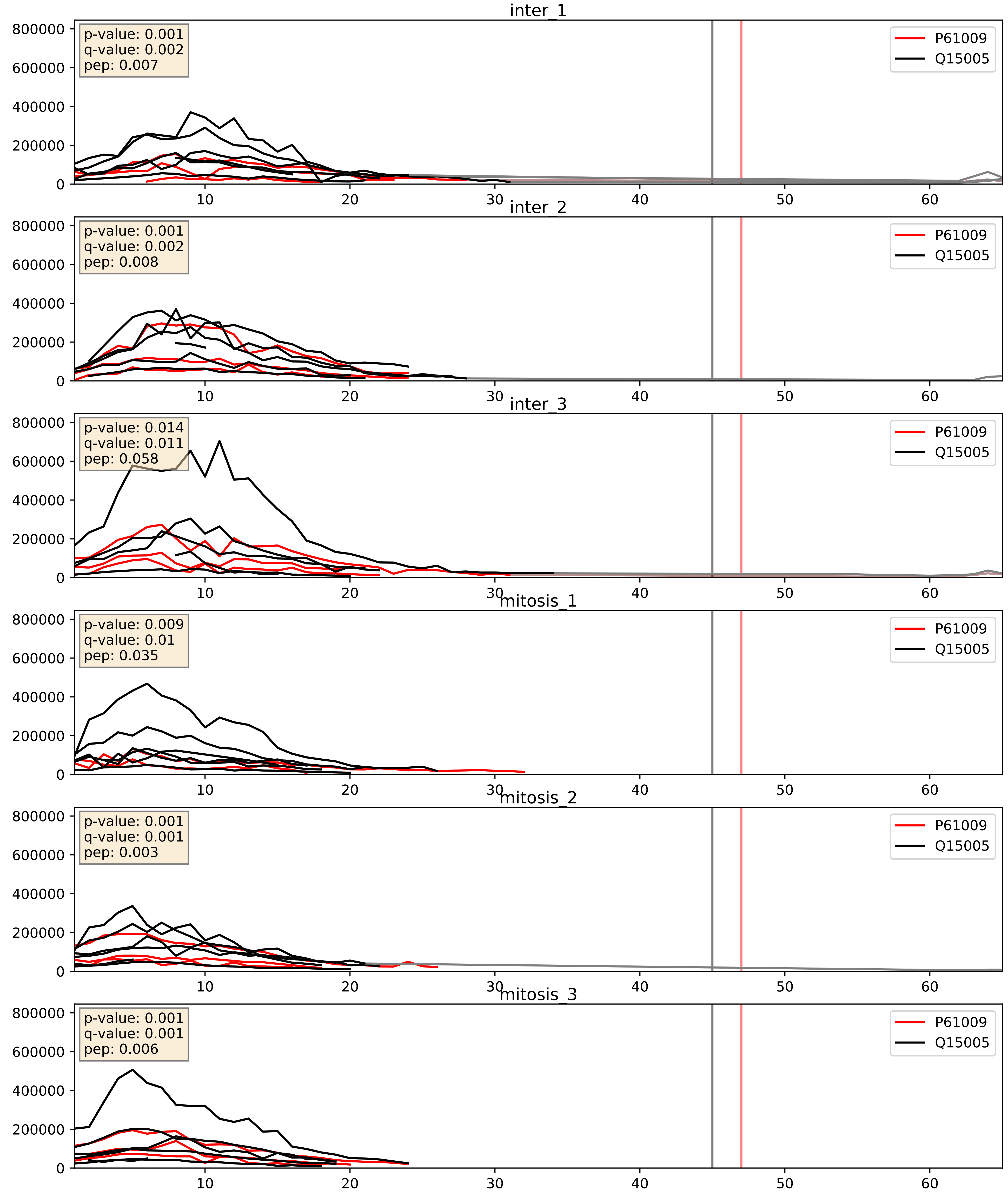


| level | condition_1 | condition_2 | pvalue | pvalue_adjusted |
|---|---|---|---|---|
| complex_abundance | mitosis | inter | 0.030881393666621796 | 0.12776448998853676 |
| interactor_abundance | mitosis | inter | 0.09221145992737646 | 0.2587423824685193 |
| interactor_ratio | mitosis | inter | 0.5895851499264092 | 0.8994900937741748 |

P61619_Q15005
S61A1_HUMAN vs SPCS2_HUMAN
p-value: 0.003 q-value: 0.047

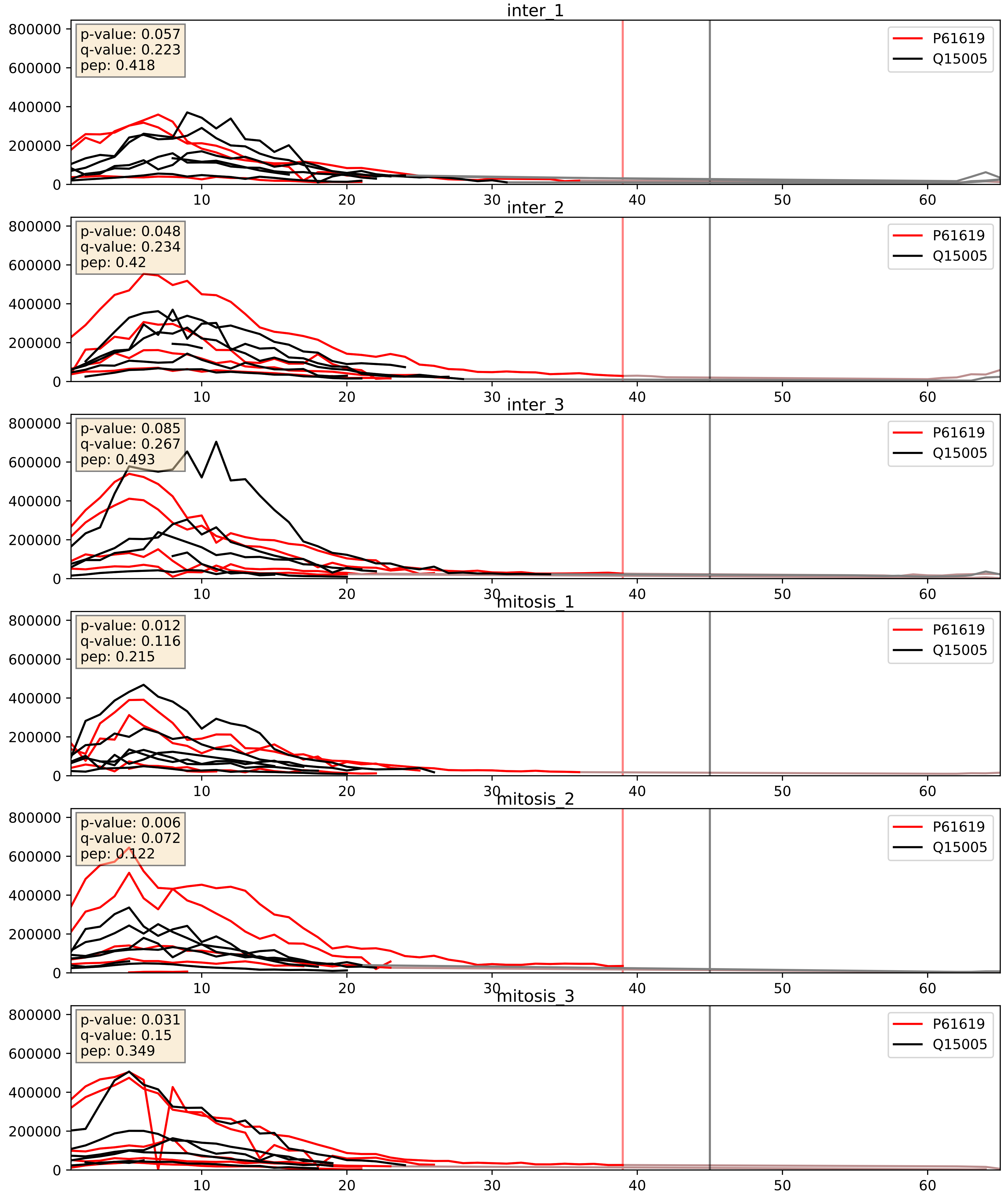


| level | condition_1 | condition_2 | pvalue | pvalue_adjusted |
|---|---|---|---|---|
| complex_abundance | mitosis | inter | 0.10990587953465765 | 0.24809977025756053 |
| interactor_ratio | mitosis | inter | 0.23775869015793552 | 0.7166151429234675 |
| interactor_abundance | mitosis | inter | 0.5550883311164501 | 0.7073926030008653 |

P67812_Q15005
SC11A_HUMAN vs SPCS2_HUMAN
p-value: 0.0 q-value: 0.0

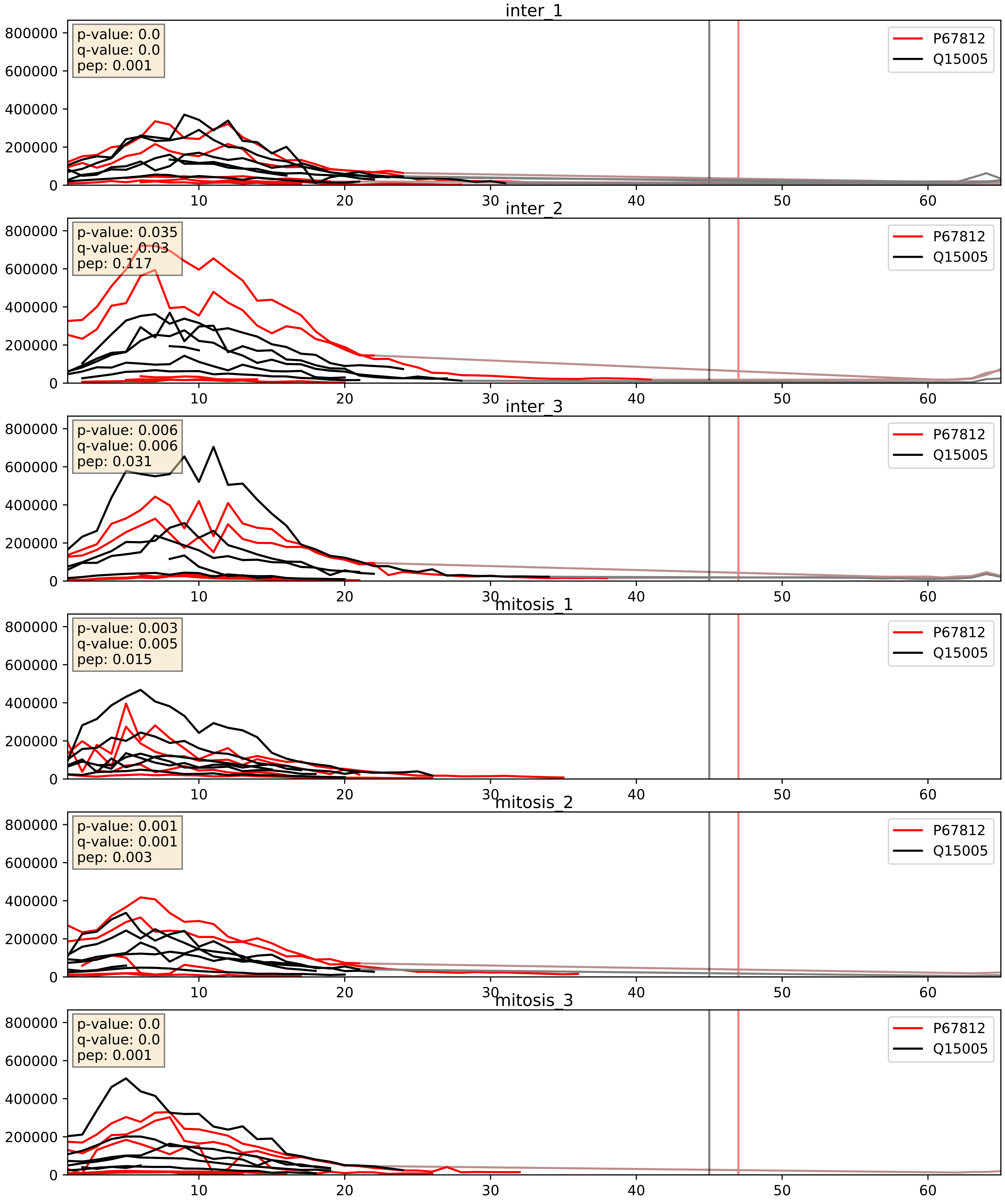


| level | condition_1 | condition_2 | pvalue | pvalue_adjusted |
|---|---|---|---|---|
| interactor_ratio | mitosis | inter | 0.046353173218109014 | 0.3574623087810929 |
| complex_abundance | mitosis | inter | 0.23967432288111756 | 0.39152904653861953 |
| interactor_abundance | mitosis | inter | 0.7658787207549315 | 0.8606343369553001 |

Q15005_Q9UNL2
SPCS2_HUMAN vs SSRG_HUMAN
p-value: 0.002 q-value: 0.003

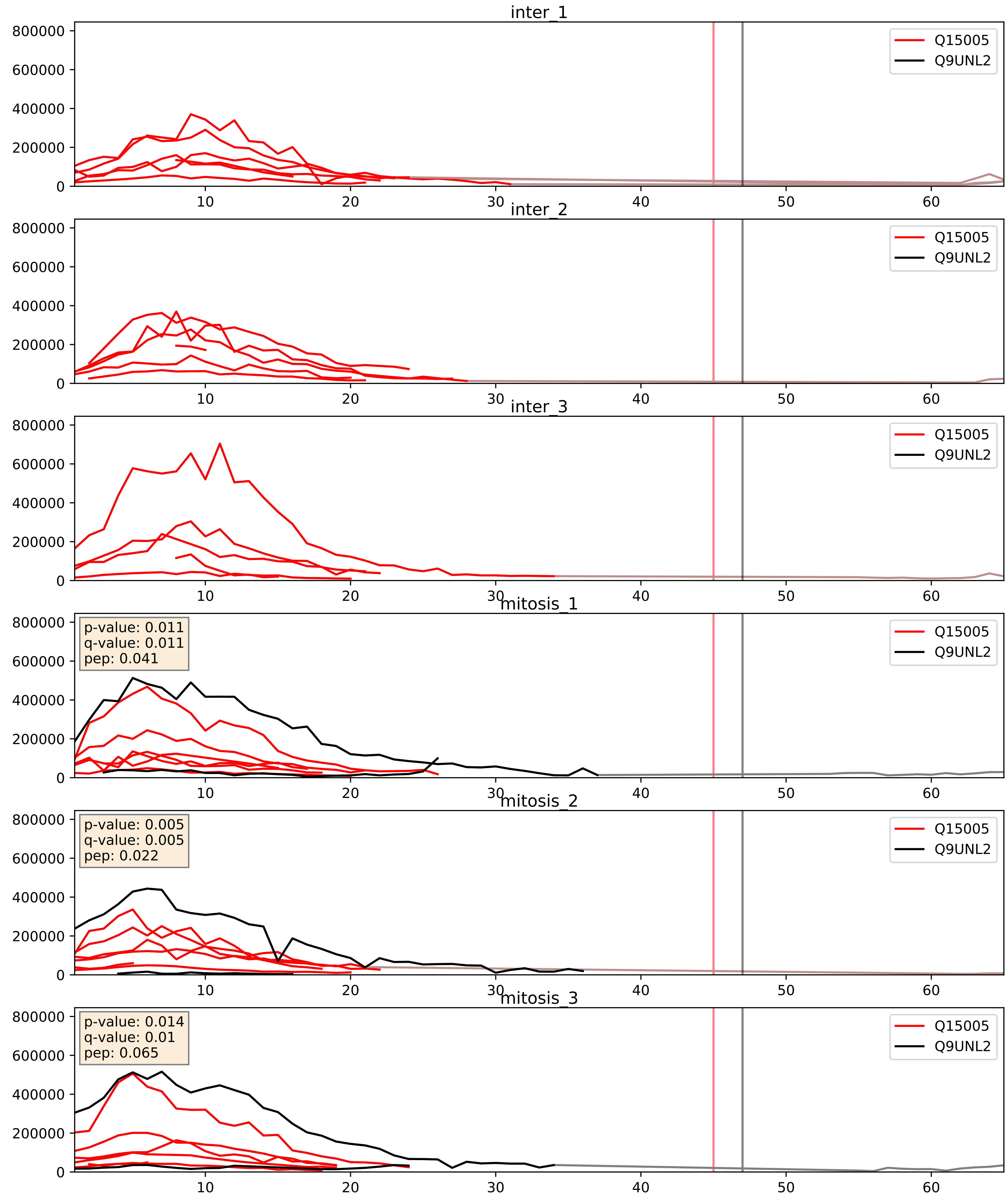


| level | condition_1 | condition_2 | pvalue | pvalue_adjusted |
|---|---|---|---|---|
| interactor_abundance | mitosis | inter | 4.11755258236053e-05 | 0.005825826463637378 |
| complex_abundance | mitosis | inter | 0.00023041379763452546 | 0.010219389159334393 |
| interactor_ratio | mitosis | inter | 0.0006764667961609915 | 0.029848225645041686 |
