## Supplementary Data 2 for "SECAT: Quantifying differential protein-protein interaction states by network-centric analysis": hela_string_000138_P08574.pdf

P08574_P08574

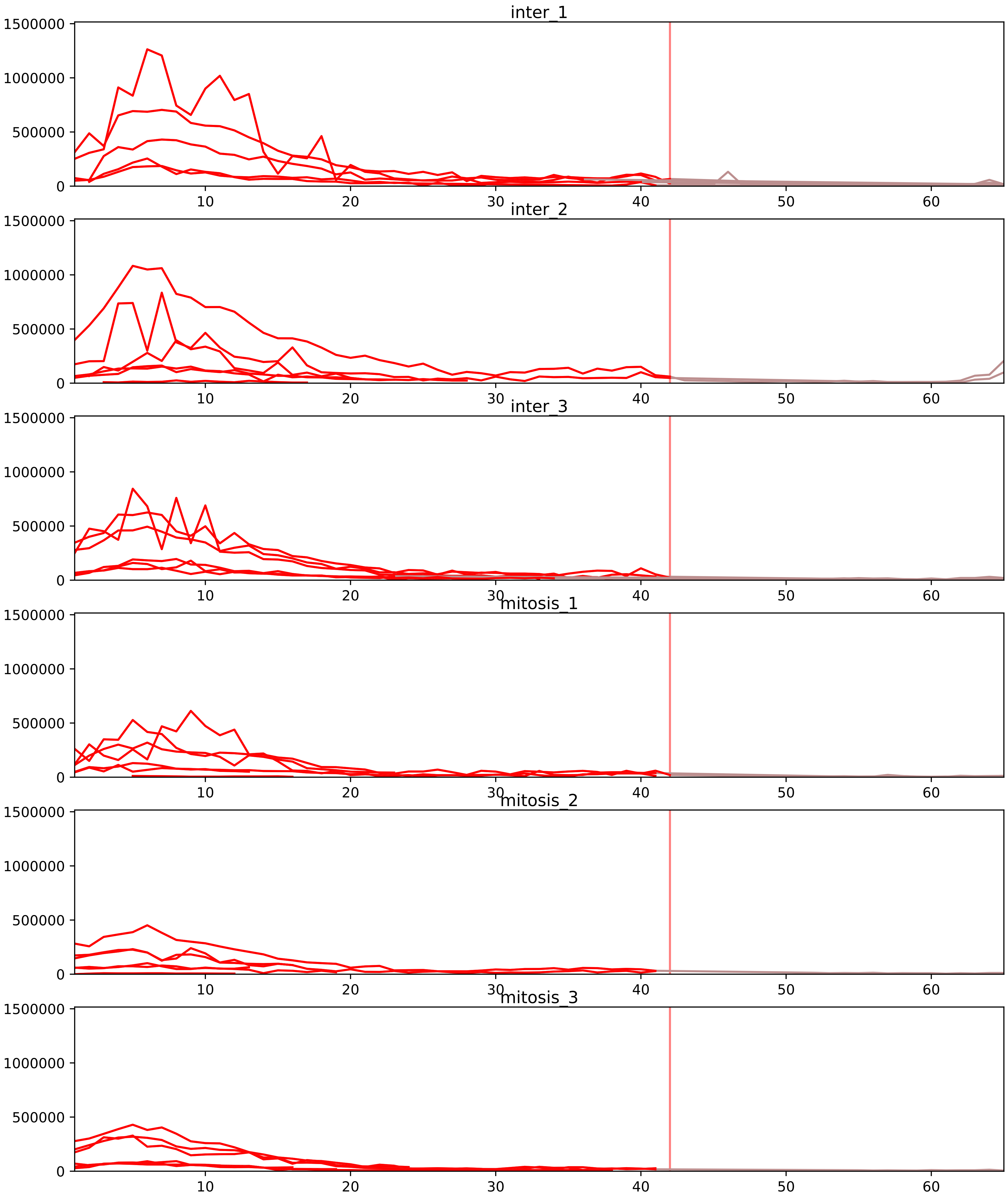


| level | condition_1 | condition_2 | pvalue | pvalue_adjusted |
|---|---|---|---|---|
| interactor_ratio | mitosis | inter | 4.8347682984173804e-05 | 0.0009266639238633313 |
| complex_abundance | mitosis | inter | 0.00018580810352361167 | 0.0034628281633340523 |
| interactor_abundance | mitosis | inter | 0.0002923832638481655 | 0.005027899116641351 |
| monomer_abundance | mitosis | inter | 0.041939757229383826 | 0.3157957803087734 |
| total_abundance | mitosis | inter | 0.05633960073634679 | 0.29753621094267796 |
| assembled_abundance | mitosis | inter | 0.05834219483251838 | 0.28656853516371306 |

O00217_P08574
NDUS8_HUMAN vs CY1_HUMAN
p-value: 0.008 q-value: 0.008

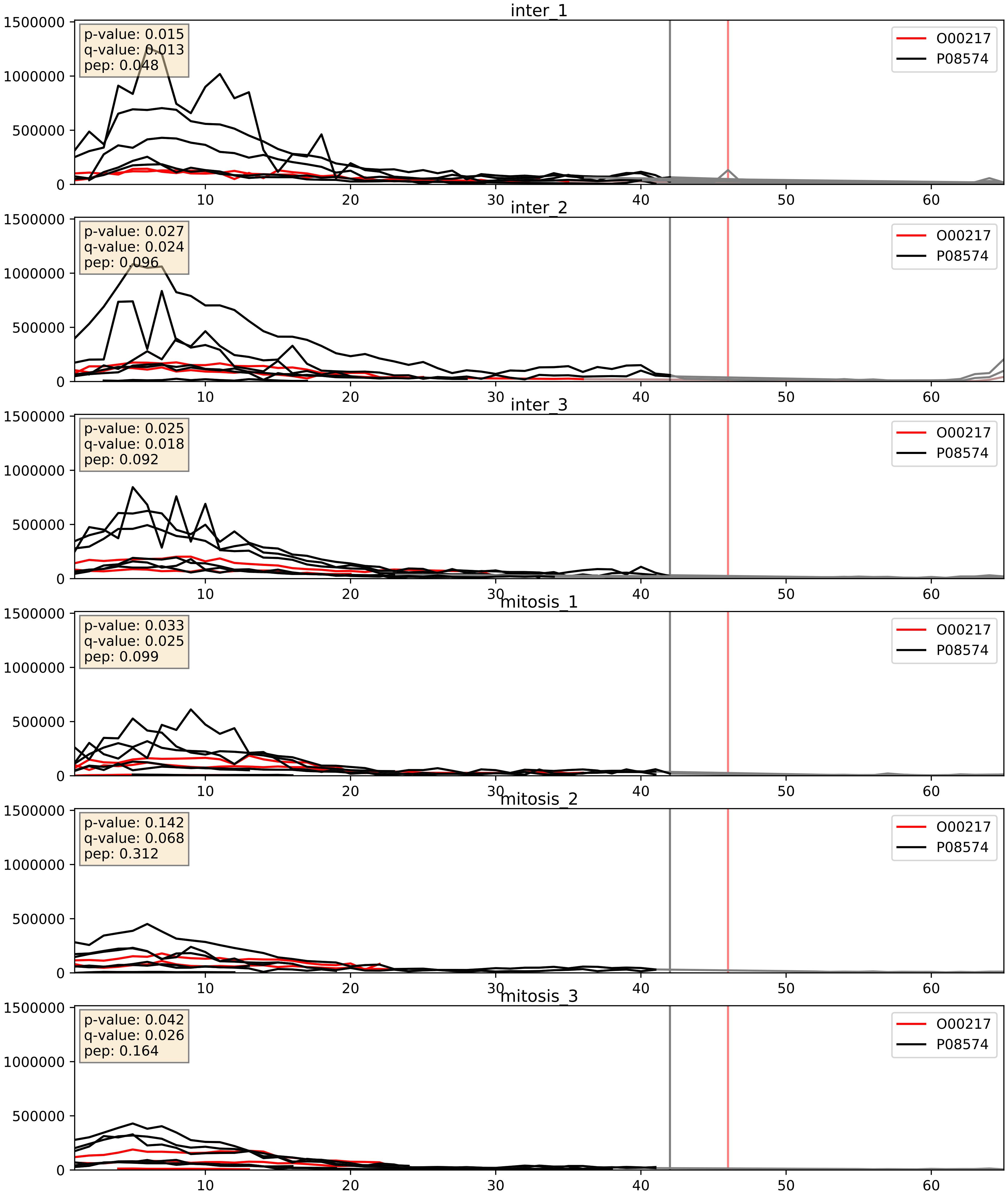


| level | condition_1 | condition_2 | pvalue | pvalue_adjusted |
|---|---|---|---|---|
| interactor_ratio | mitosis | inter | 0.0032999834690576797 | 0.07473911946628042 |
| complex_abundance | mitosis | inter | 0.03926113441956983 | 0.14448905839702758 |
| interactor_abundance | mitosis | inter | 0.4865513439277089 | 0.6558960893652855 |

O14949_P08574
QCR8_HUMAN vs CY1_HUMAN
p-value: 0.046 q-value: 0.03

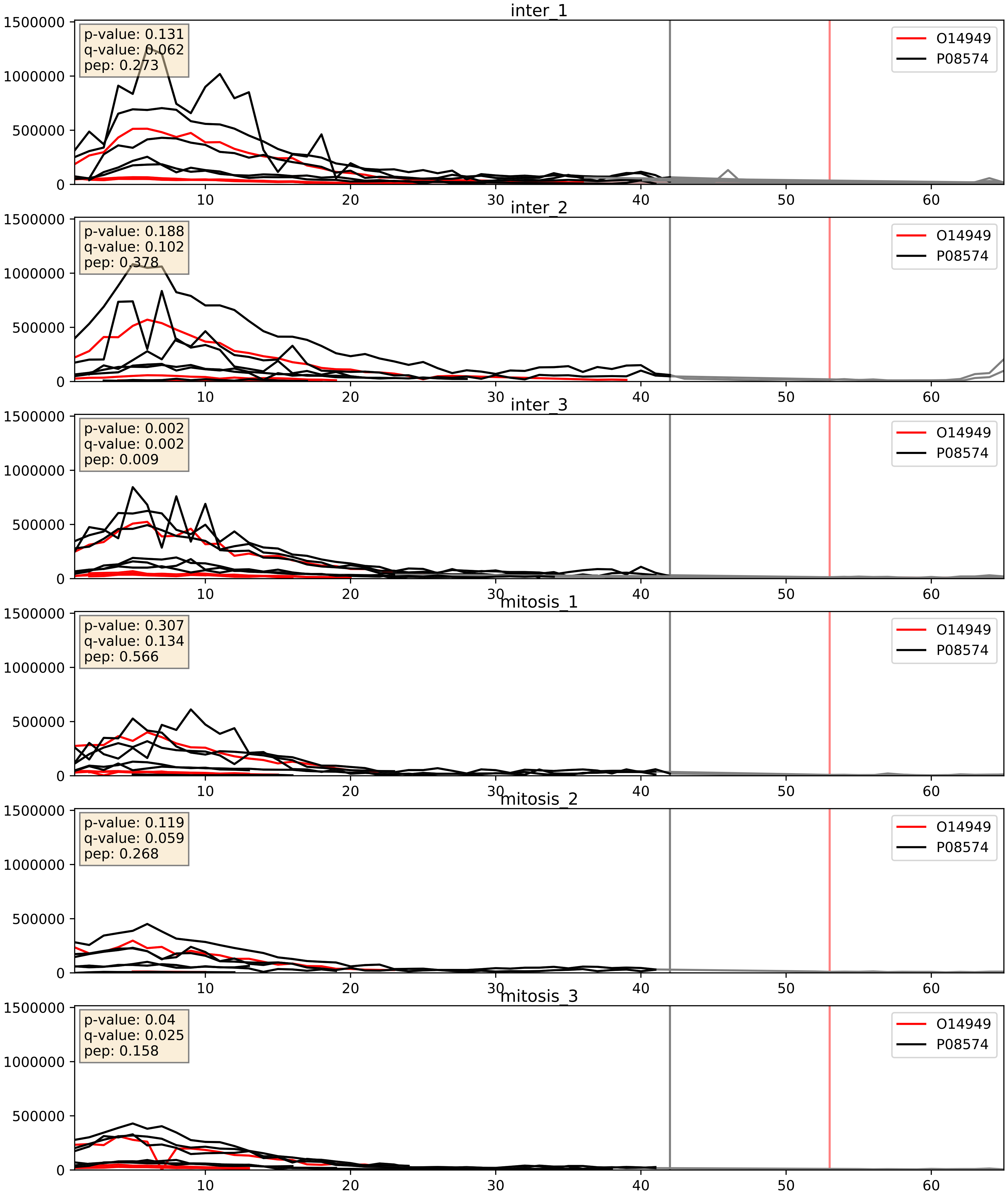


| level | condition_1 | condition_2 | pvalue | pvalue_adjusted |
|---|---|---|---|---|
| complex_abundance | mitosis | inter | 0.029938526405177388 | 0.12573011743528603 |
| interactor_abundance | mitosis | inter | 0.07386499194203666 | 0.22673988198390962 |
| interactor_ratio | mitosis | inter | 0.7469313480450761 | 0.9491588884573259 |

O75251_P08574
NDUS7_HUMAN vs CY1_HUMAN
p-value: 0.007 q-value: 0.006

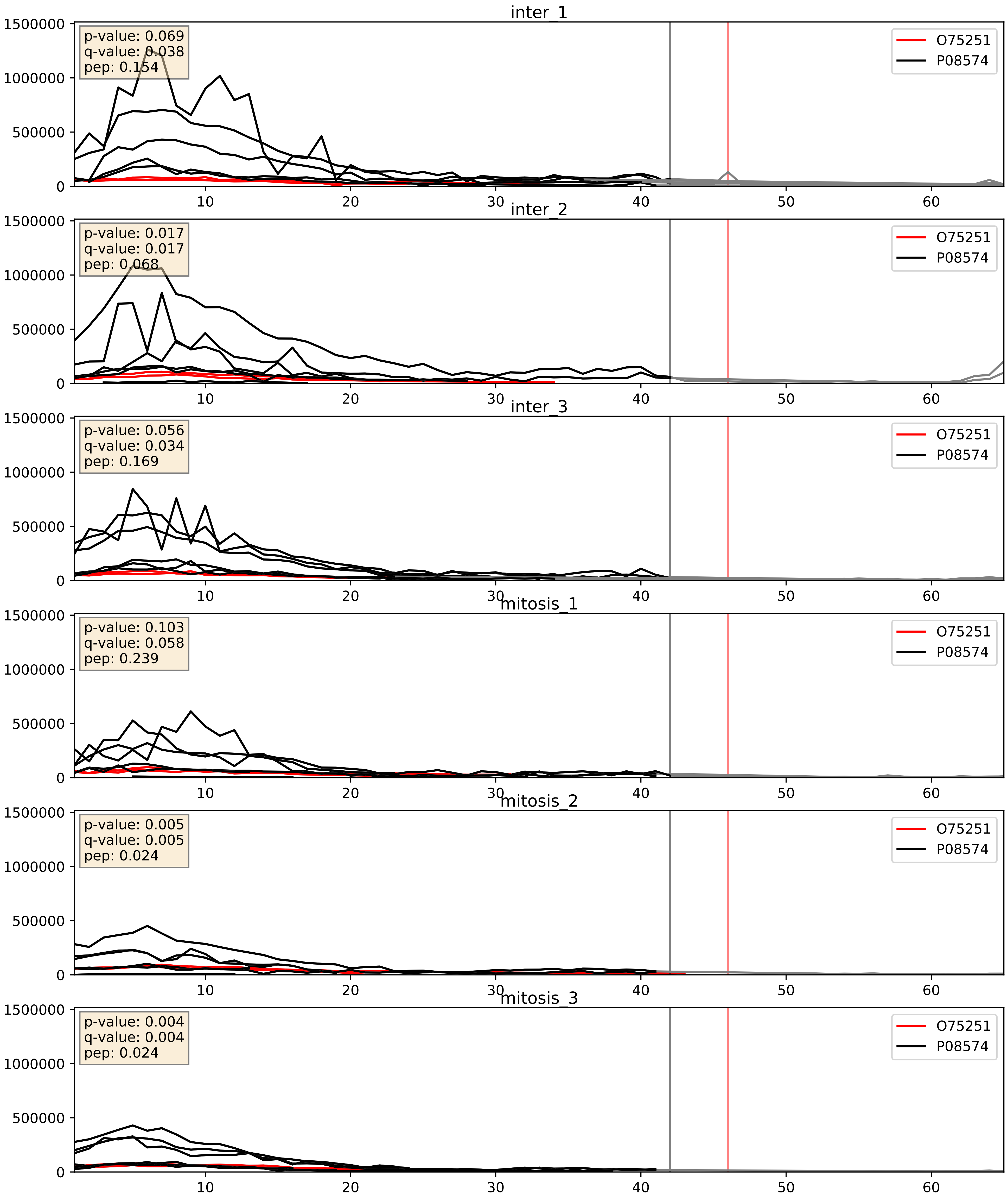


| level | condition_1 | condition_2 | pvalue | pvalue_adjusted |
|---|---|---|---|---|
| complex_abundance | mitosis | inter | 0.006505441299260121 | 0.05955783692156859 |
| interactor_ratio | mitosis | inter | 0.012832792382496886 | 0.16355562016423936 |
| interactor_abundance | mitosis | inter | 0.016601804854909166 | 0.11277132084292767 |

O75306_P08574
NDUS2_HUMAN vs CY1_HUMAN
p-value: 0.04 q-value: 0.027

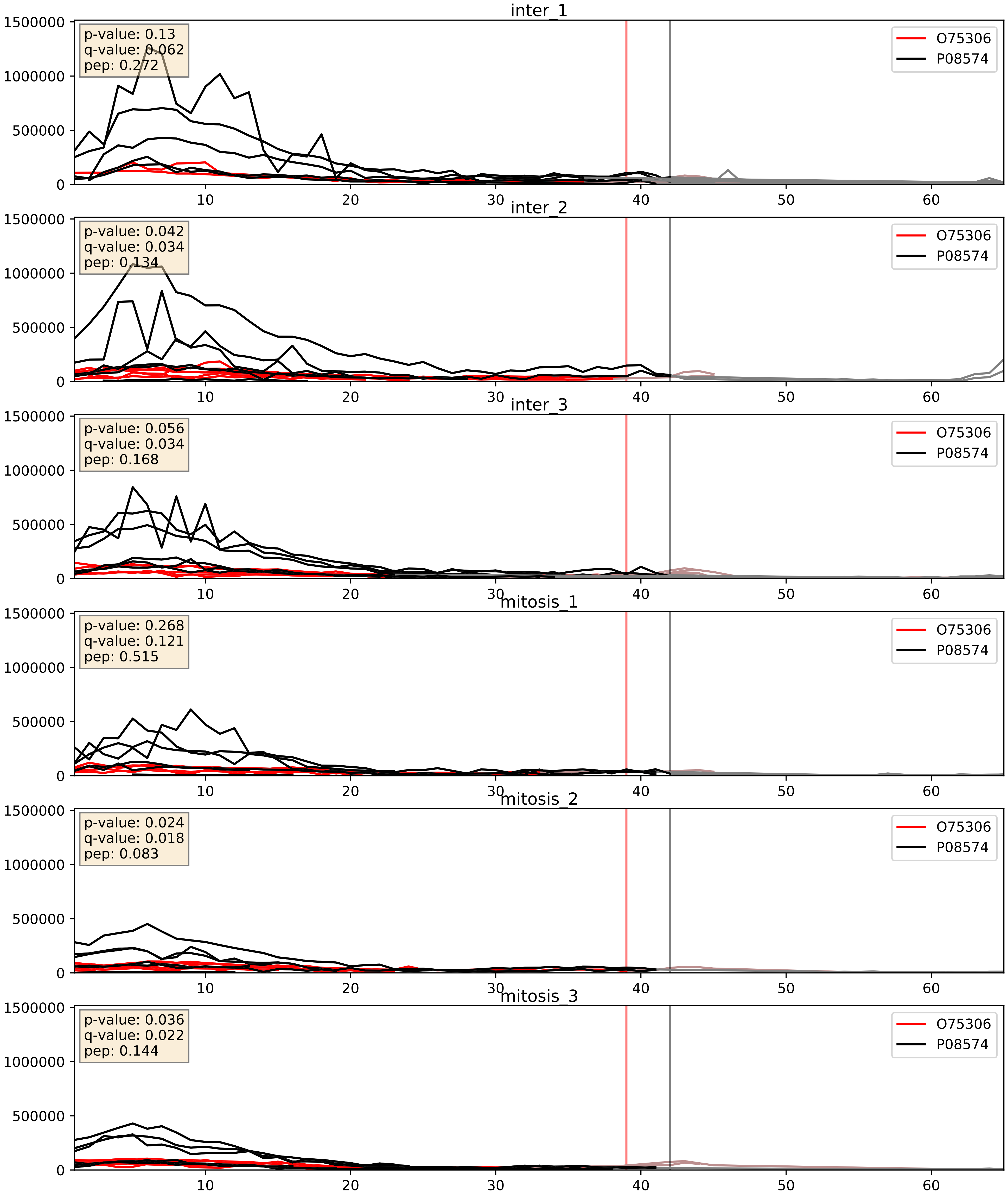


| level | condition_1 | condition_2 | pvalue | pvalue_adjusted |
|---|---|---|---|---|
| complex_abundance | mitosis | inter | 0.04490609809020621 | 0.15163558171683042 |
| interactor_ratio | mitosis | inter | 0.3299753539706618 | 0.7875542793987639 |
| interactor_abundance | mitosis | inter | 0.6718484200519785 | 0.7850689339492098 |

O75489_P08574
NDUS3_HUMAN vs CY1_HUMAN
p-value: 0.0 q-value: 0.0

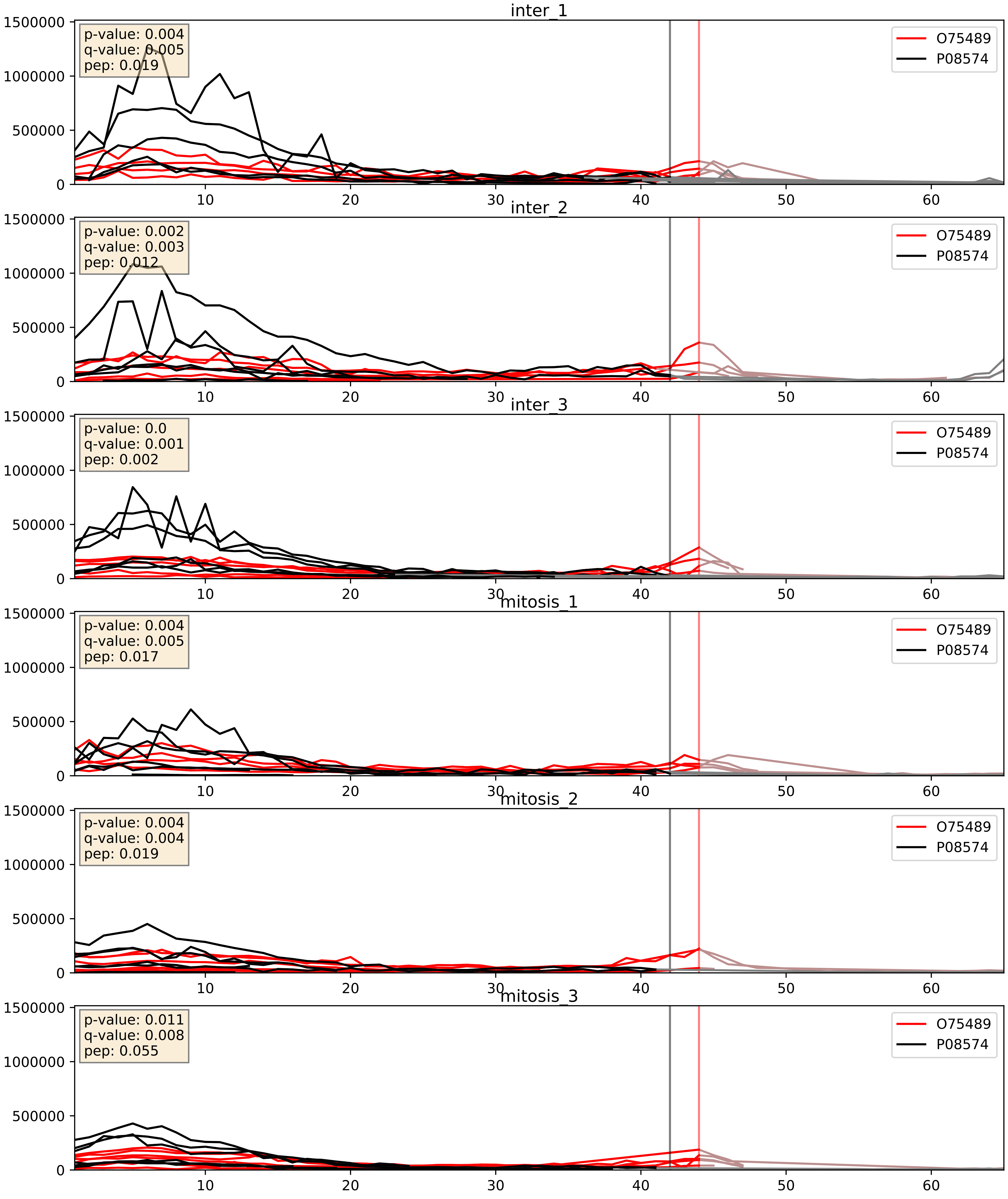


| level | condition_1 | condition_2 | pvalue | pvalue_adjusted |
|---|---|---|---|---|
| interactor_ratio | mitosis | inter | 0.0025061399092815868 | 0.06403748544313548 |
| complex_abundance | mitosis | inter | 0.015294714772104251 | 0.09174685245214603 |
| interactor_abundance | mitosis | inter | 0.07448847062620823 | 0.22673988198390962 |

O75947_P08574
ATP5H_HUMAN vs CY1_HUMAN
p-value: 0.0 q-value: 0.015

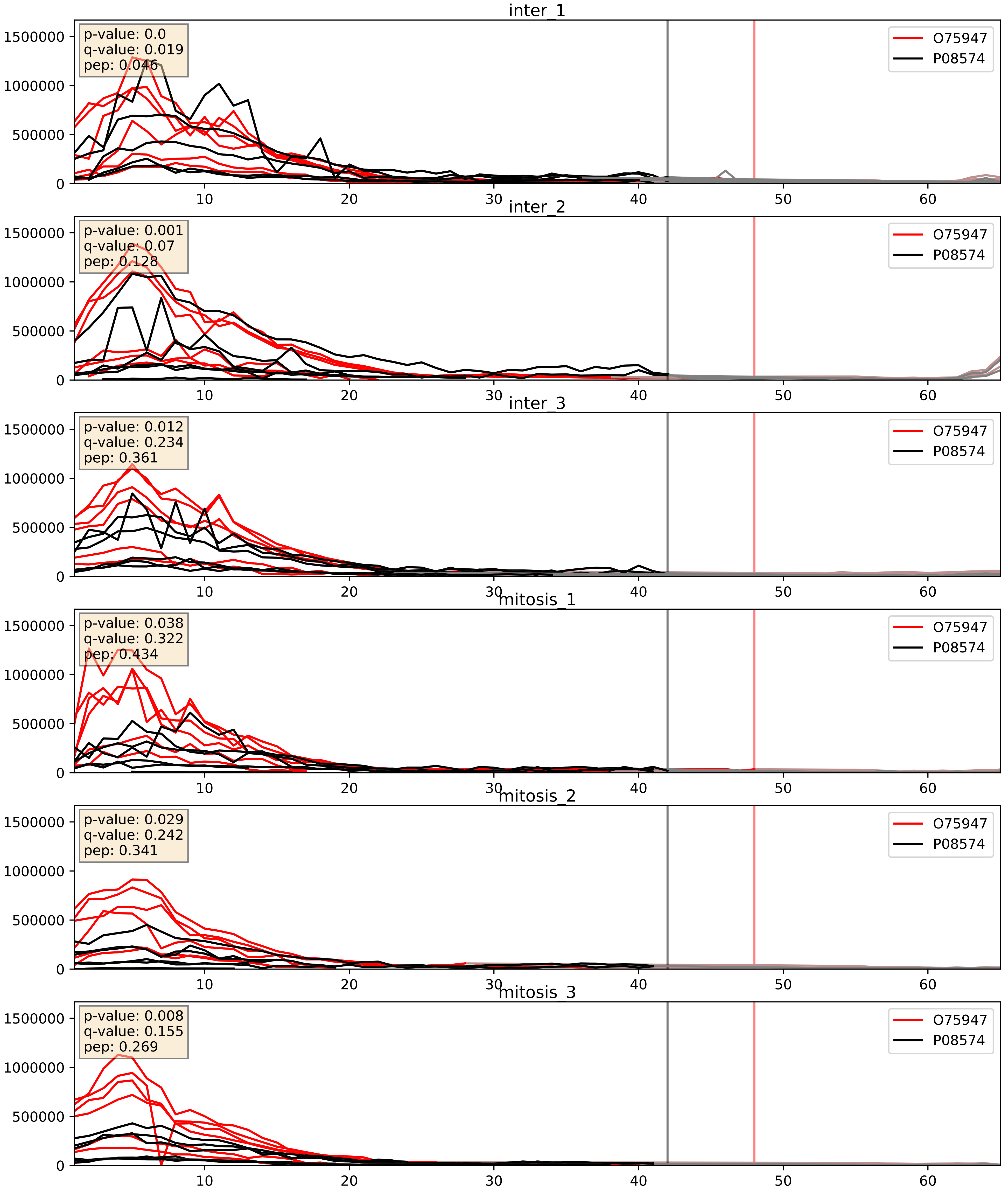


| level | condition_1 | condition_2 | pvalue | pvalue_adjusted |
|---|---|---|---|---|
| complex_abundance | mitosis | inter | 0.005159719217035711 | 0.0539125642450646 |
| interactor_abundance | mitosis | inter | 0.009009320082318547 | 0.08277270927005473 |
| interactor_ratio | mitosis | inter | 0.08702802799589962 | 0.49442537105387907 |

O75964_P08574
ATP5L_HUMAN vs CY1_HUMAN
p-value: 0.0 q-value: 0.016

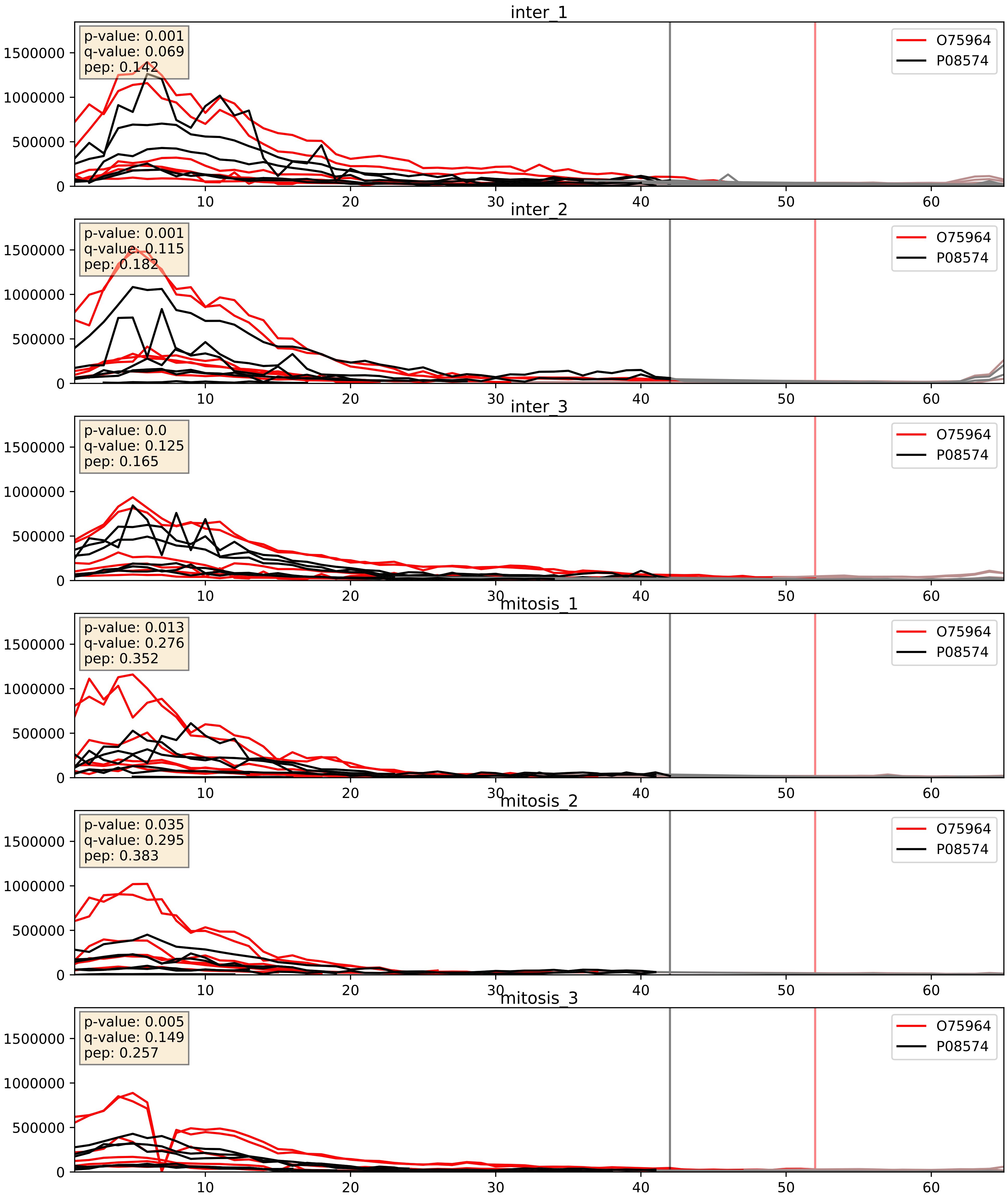


| level | condition_1 | condition_2 | pvalue | pvalue_adjusted |
|---|---|---|---|---|
| complex_abundance | mitosis | inter | 0.006965260773570615 | 0.061721151368286195 |
| interactor_abundance | mitosis | inter | 0.008230706235449504 | 0.07899680460559061 |
| interactor_ratio | mitosis | inter | 0.016771506213886653 | 0.1916743567301332 |

O96000_P08574
NDUBA_HUMAN vs CY1_HUMAN
p-value: 0.002 q-value: 0.03

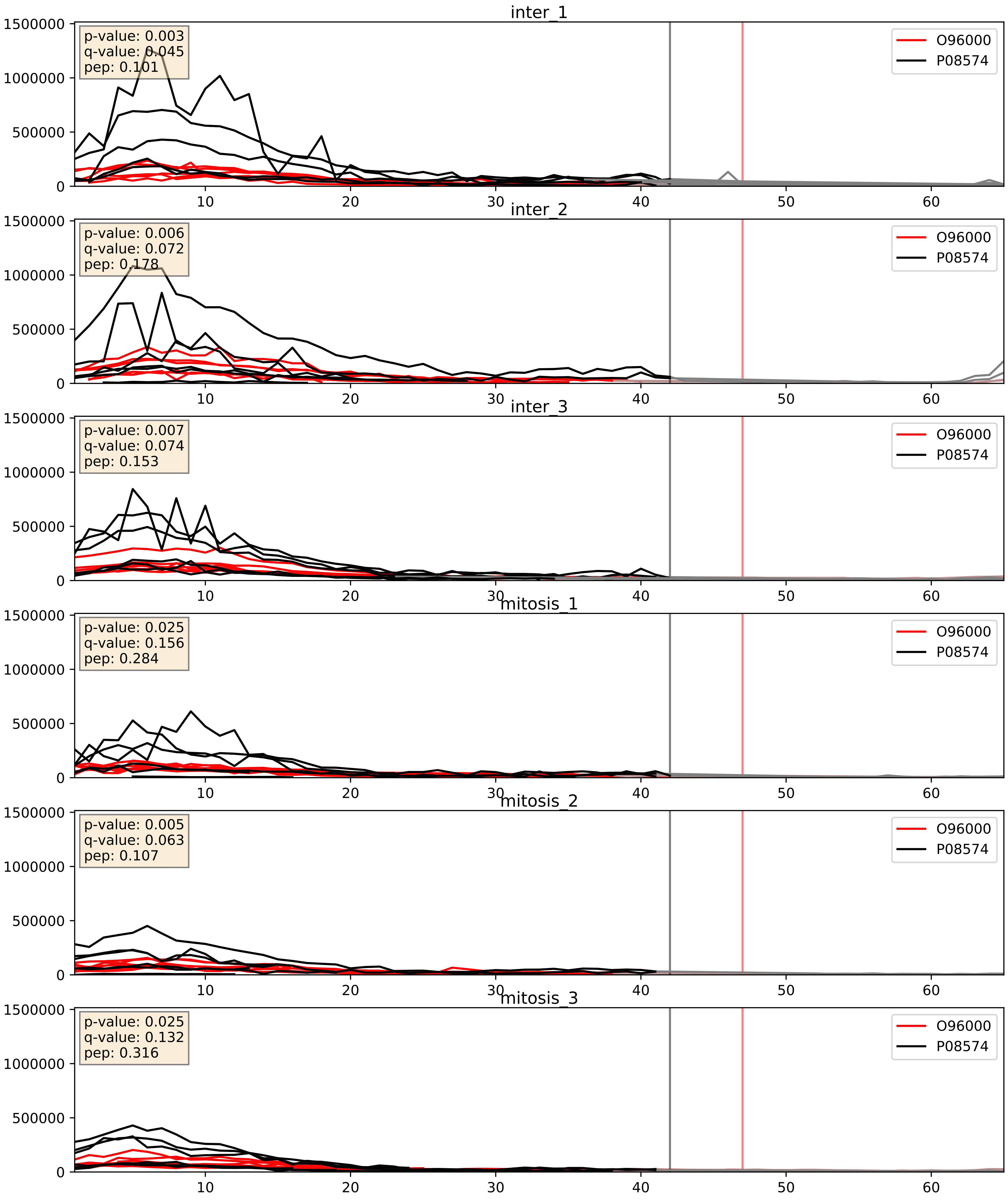


| level | condition_1 | condition_2 | pvalue | pvalue_adjusted |
|---|---|---|---|---|
| complex_abundance | mitosis | inter | 0.002334679939336067 | 0.035168510428347066 |
| interactor_abundance | mitosis | inter | 0.0030692858094850456 | 0.042465703978082026 |
| interactor_ratio | mitosis | inter | 0.3283581682329763 | 0.786600701916436 |

O96008_P08574
TOM40_HUMAN vs CY1_HUMAN
p-value: 0.015 q-value: 0.012

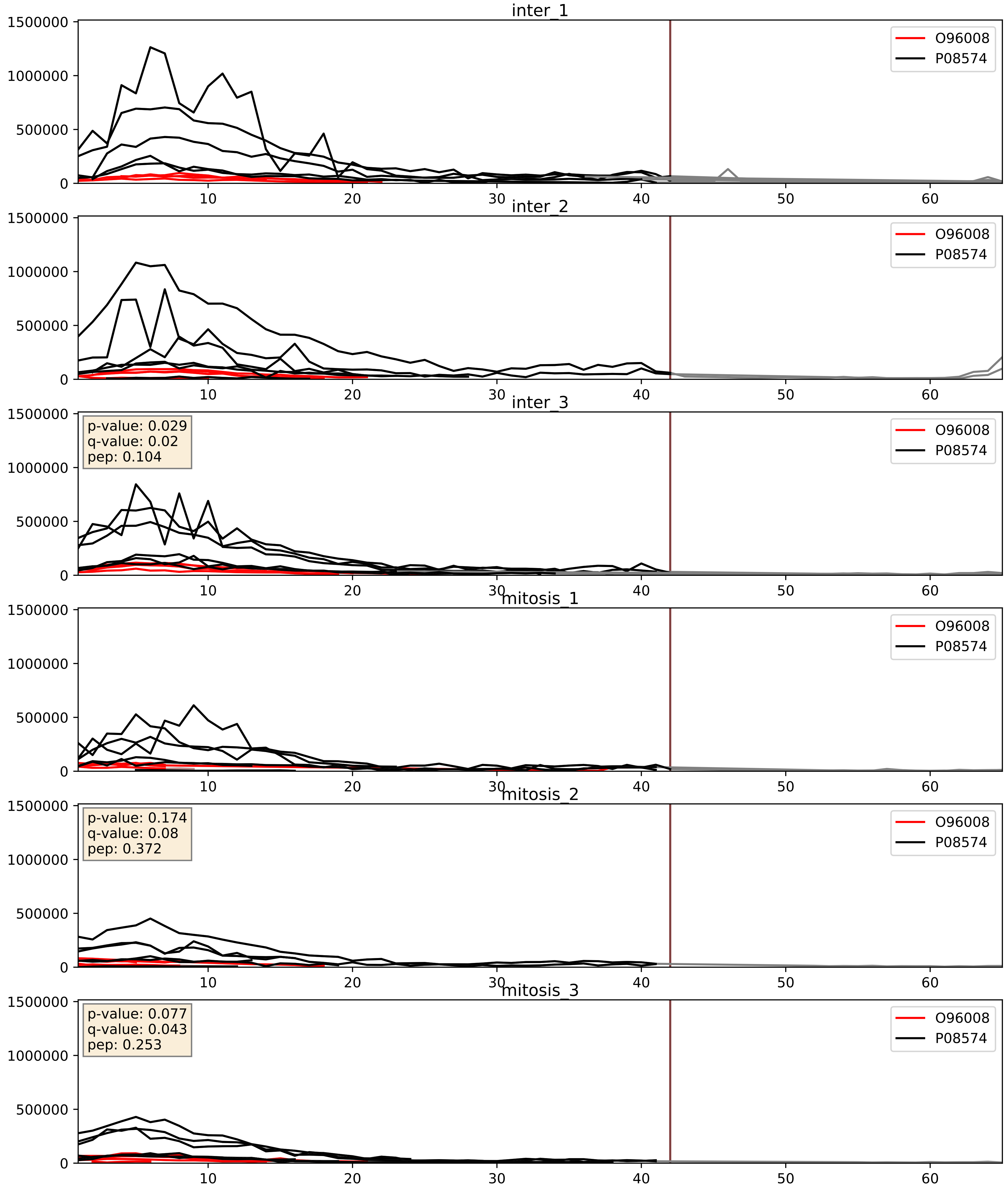


| level | condition_1 | condition_2 | pvalue | pvalue_adjusted |
|---|---|---|---|---|
| complex_abundance | mitosis | inter | 0.004378871638023938 | 0.04940908555989897 |
| interactor_abundance | mitosis | inter | 0.007789930253278921 | 0.07668982515016397 |
| interactor_ratio | mitosis | inter | 0.08281278739547326 | 0.4835453343146324 |

P06576_P08574
ATPB_HUMAN vs CY1_HUMAN
p-value: 0.056 q-value: 0.036

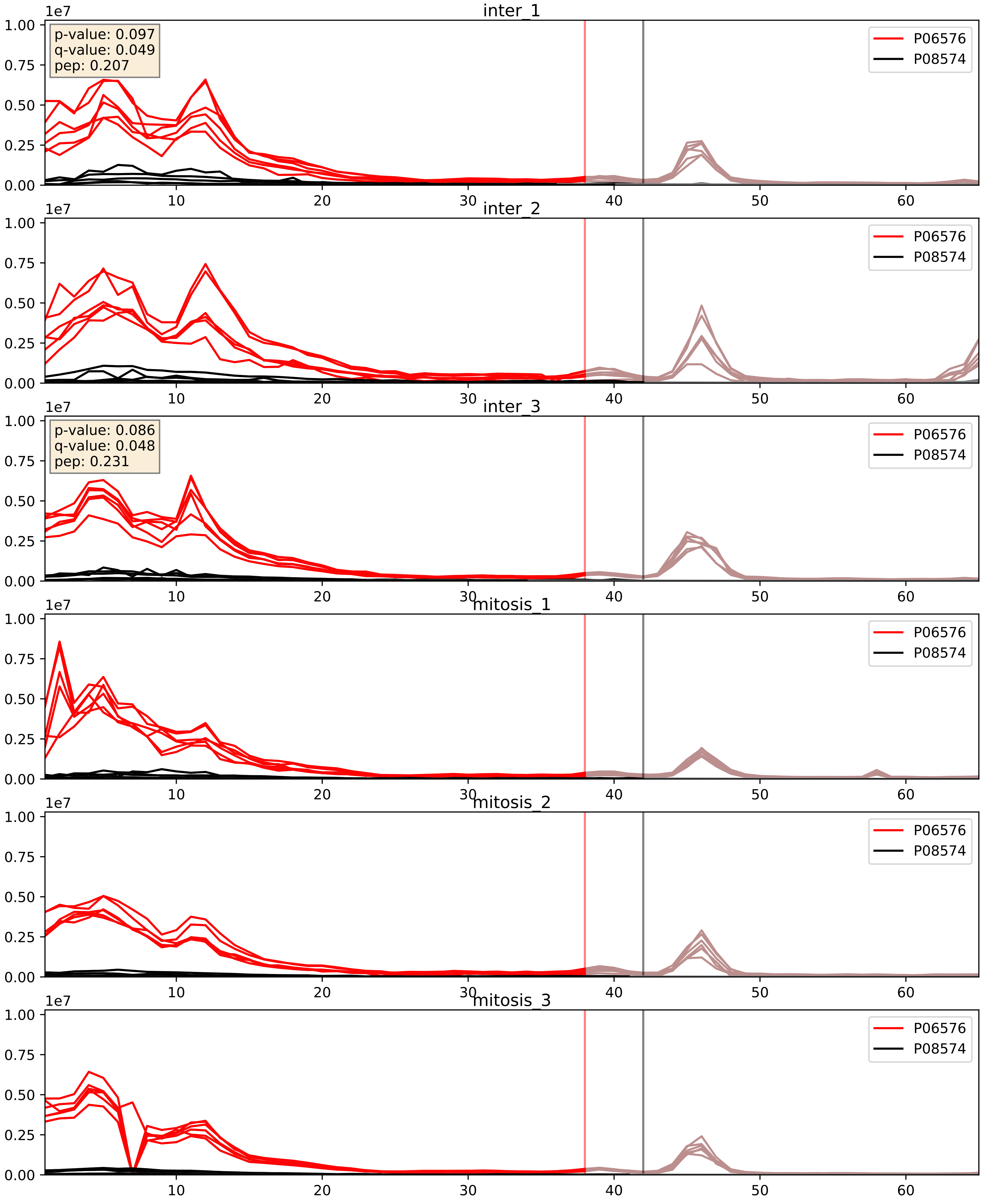


| level | condition_1 | condition_2 | pvalue | pvalue_adjusted |
|---|---|---|---|---|
| interactor_ratio | mitosis | inter | 0.006374976371333102 | 0.1102420156335583 |
| complex_abundance | mitosis | inter | 0.013557427971984582 | 0.0865877814270891 |
| interactor_abundance | mitosis | inter | 0.10278097277061035 | 0.2754555813764636 |

P07919_P08574
QCR6_HUMAN vs CY1_HUMAN
p-value: 0.0 q-value: 0.0

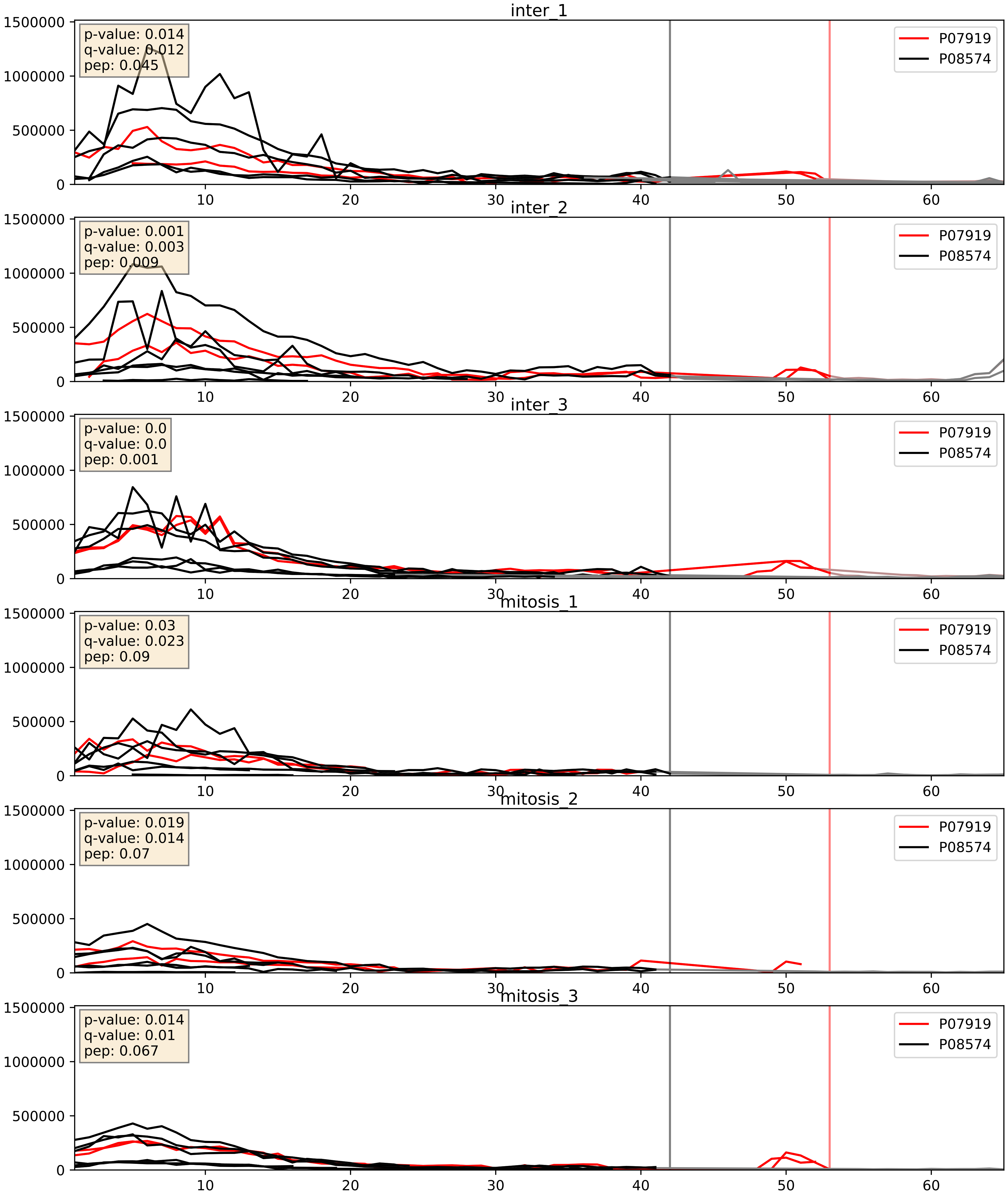


| level | condition_1 | condition_2 | pvalue | pvalue_adjusted |
|---|---|---|---|---|
| complex_abundance | mitosis | inter | 0.0032280338005122537 | 0.04165459892439789 |
| interactor_abundance | mitosis | inter | 0.016612815348068178 | 0.11277132084292767 |
| interactor_ratio | mitosis | inter | 0.12425944554959535 | 0.5739294160480145 |

P08574_P13073
CY1_HUMAN vs COX41_HUMAN
p-value: 0.001 q-value: 0.002

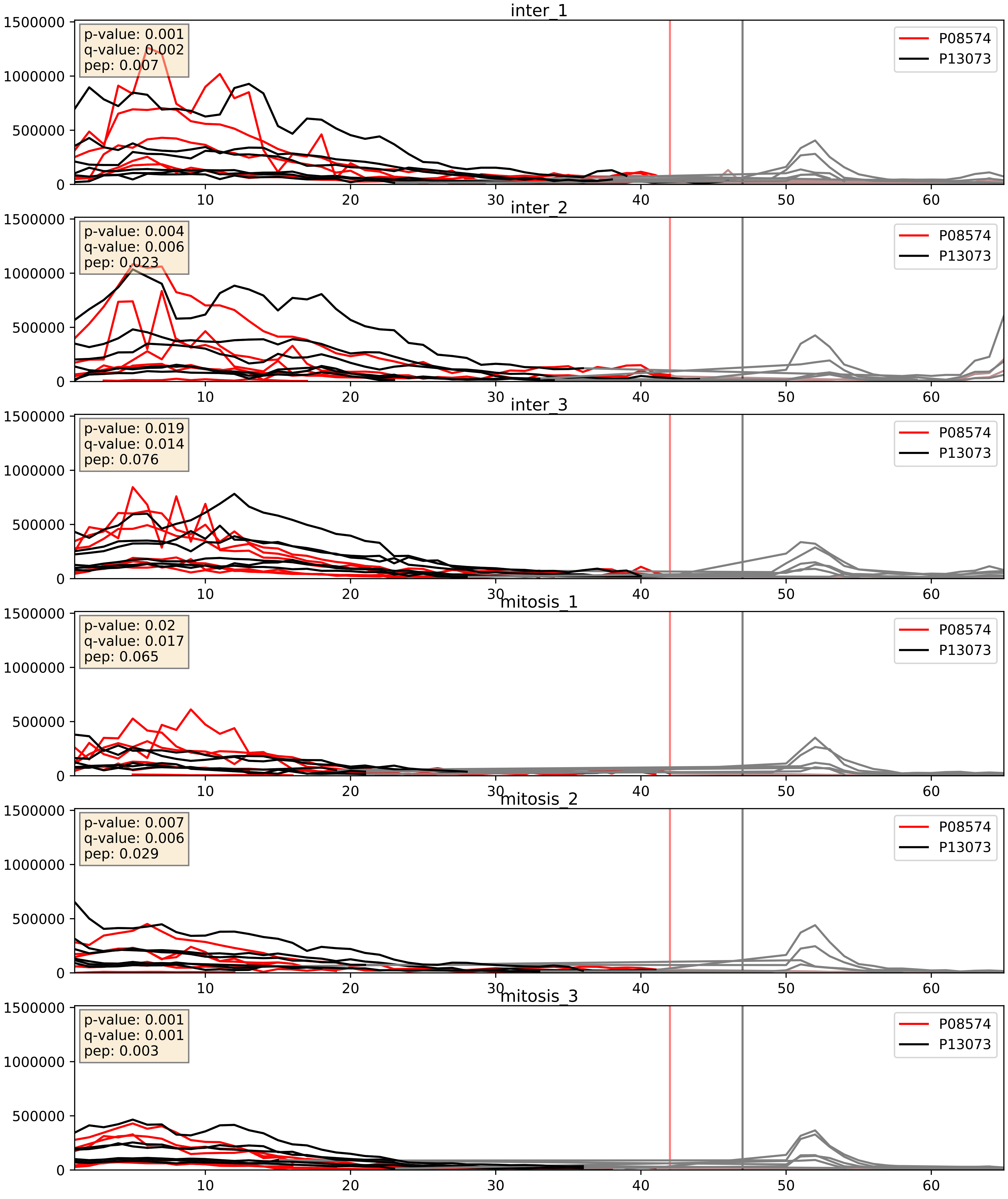


| level | condition_1 | condition_2 | pvalue | pvalue_adjusted |
|---|---|---|---|---|
| complex_abundance | mitosis | inter | 0.0024352323345376956 | 0.035755726901616935 |
| interactor_abundance | mitosis | inter | 0.006017826453889495 | 0.06595014057915963 |
| interactor_ratio | mitosis | inter | 0.4518645391568424 | 0.852724968073759 |

P08574_P14927
CY1_HUMAN vs QCR7_HUMAN
p-value: 0.01 q-value: 0.009

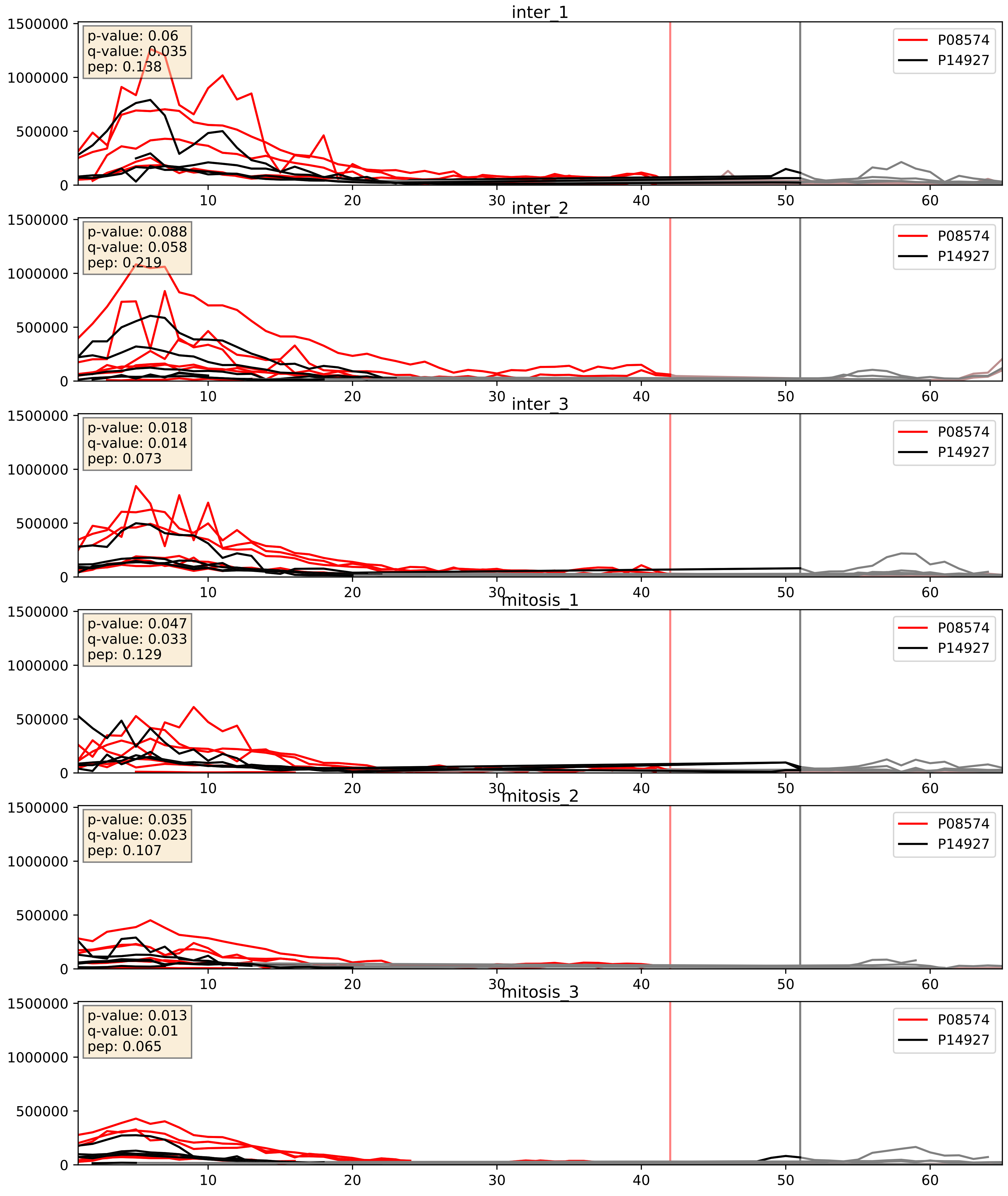


| level | condition_1 | condition_2 | pvalue | pvalue_adjusted |
|---|---|---|---|---|
| interactor_abundance | mitosis | inter | 0.008246700169002542 | 0.07899680460559061 |
| complex_abundance | mitosis | inter | 0.012168739002753967 | 0.08185897984306292 |
| interactor_ratio | mitosis | inter | 0.7939244280867633 | 0.9608508478447312 |

P08574_P20674
CY1_HUMAN vs COX5A_HUMAN
p-value: 0.002 q-value: 0.002

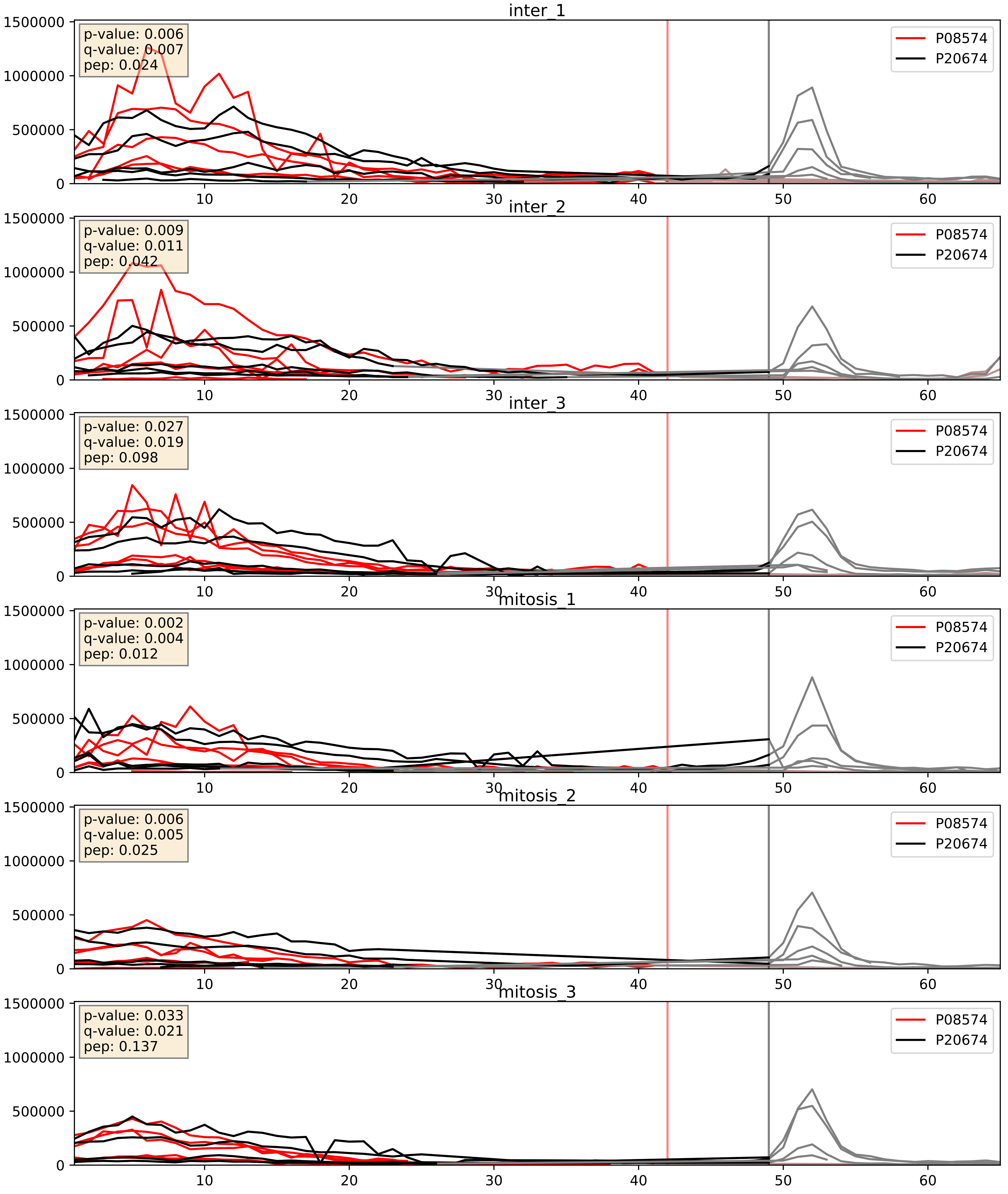


| level | condition_1 | condition_2 | pvalue | pvalue_adjusted |
|---|---|---|---|---|
| complex_abundance | mitosis | inter | 0.006782287808849994 | 0.06098359626444952 |
| interactor_abundance | mitosis | inter | 0.0071594169240471565 | 0.07321936543589445 |
| interactor_ratio | mitosis | inter | 0.022972280560885346 | 0.23382012080996262 |

P08574_P21796
CY1_HUMAN vs VDAC1_HUMAN
p-value: 0.0 q-value: 0.013

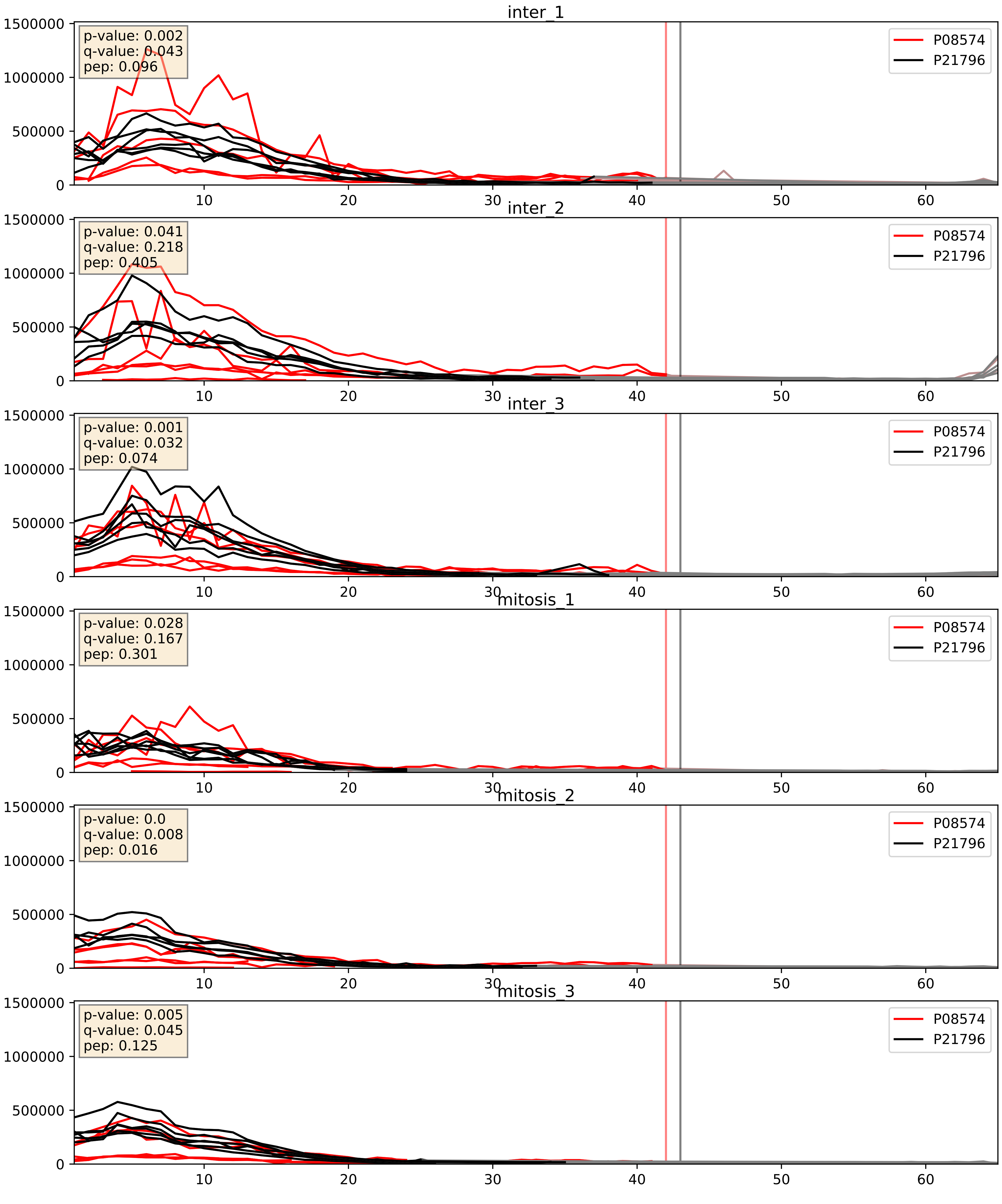


| level | condition_1 | condition_2 | pvalue | pvalue_adjusted |
|---|---|---|---|---|
| complex_abundance | mitosis | inter | 0.0013086979891824854 | 0.024459508269436846 |
| interactor_abundance | mitosis | inter | 0.005367004948843058 | 0.06171739355782327 |
| interactor_ratio | mitosis | inter | 0.5295368897566973 | 0.8821330249831557 |

P08574_P22695
CY1_HUMAN vs QCR2_HUMAN
p-value: 0.003 q-value: 0.004

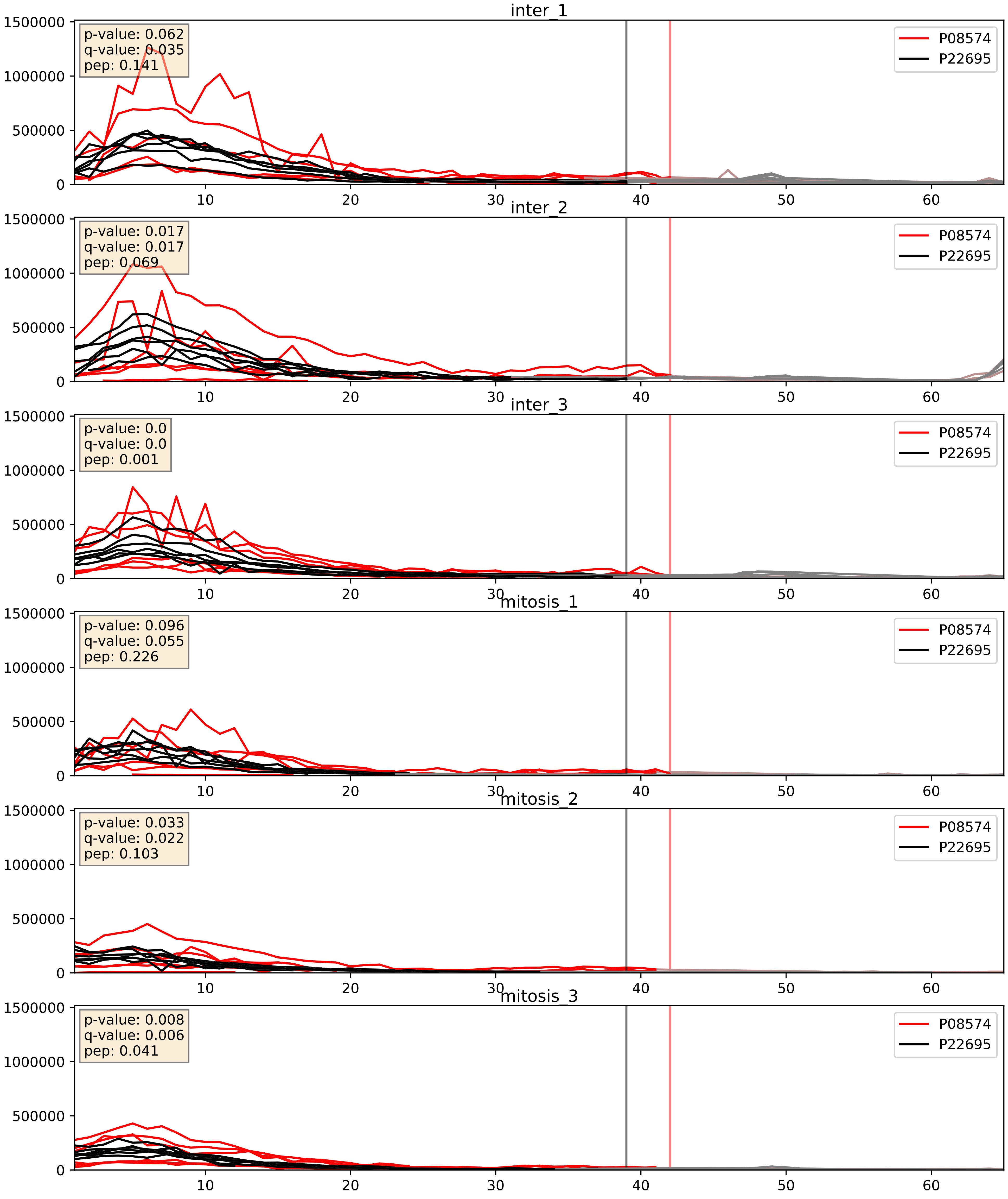


| level | condition_1 | condition_2 | pvalue | pvalue_adjusted |
|---|---|---|---|---|
| complex_abundance | mitosis | inter | 0.002527814595984375 | 0.036409709098919725 |
| interactor_abundance | mitosis | inter | 0.007419881684355014 | 0.07459761261938556 |
| interactor_ratio | mitosis | inter | 0.8436266256088374 | 0.9755634896635457 |

P08574_P31930
CY1_HUMAN vs QCR1_HUMAN
p-value: 0.008 q-value: 0.008

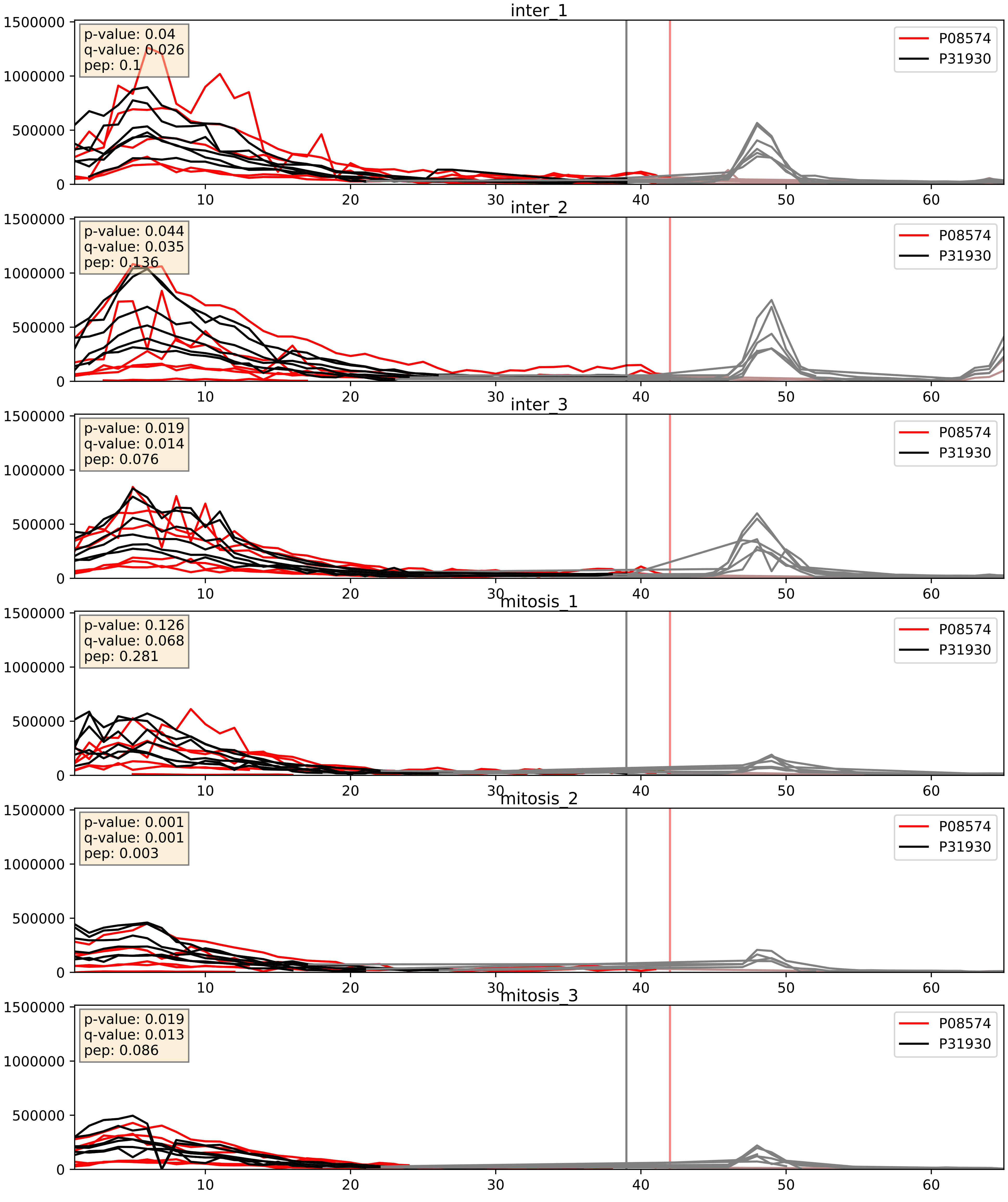


| level | condition_1 | condition_2 | pvalue | pvalue_adjusted |
|---|---|---|---|---|
| complex_abundance | mitosis | inter | 0.0009739141526850073 | 0.020686613267949534 |
| interactor_abundance | mitosis | inter | 0.006546882782702666 | 0.06901639977824485 |
| interactor_ratio | mitosis | inter | 0.8678606927151378 | 0.9786441219393465 |

P08574_Q3ZCQ8
CY1_HUMAN vs TIM50_HUMAN
p-value: 0.001 q-value: 0.027

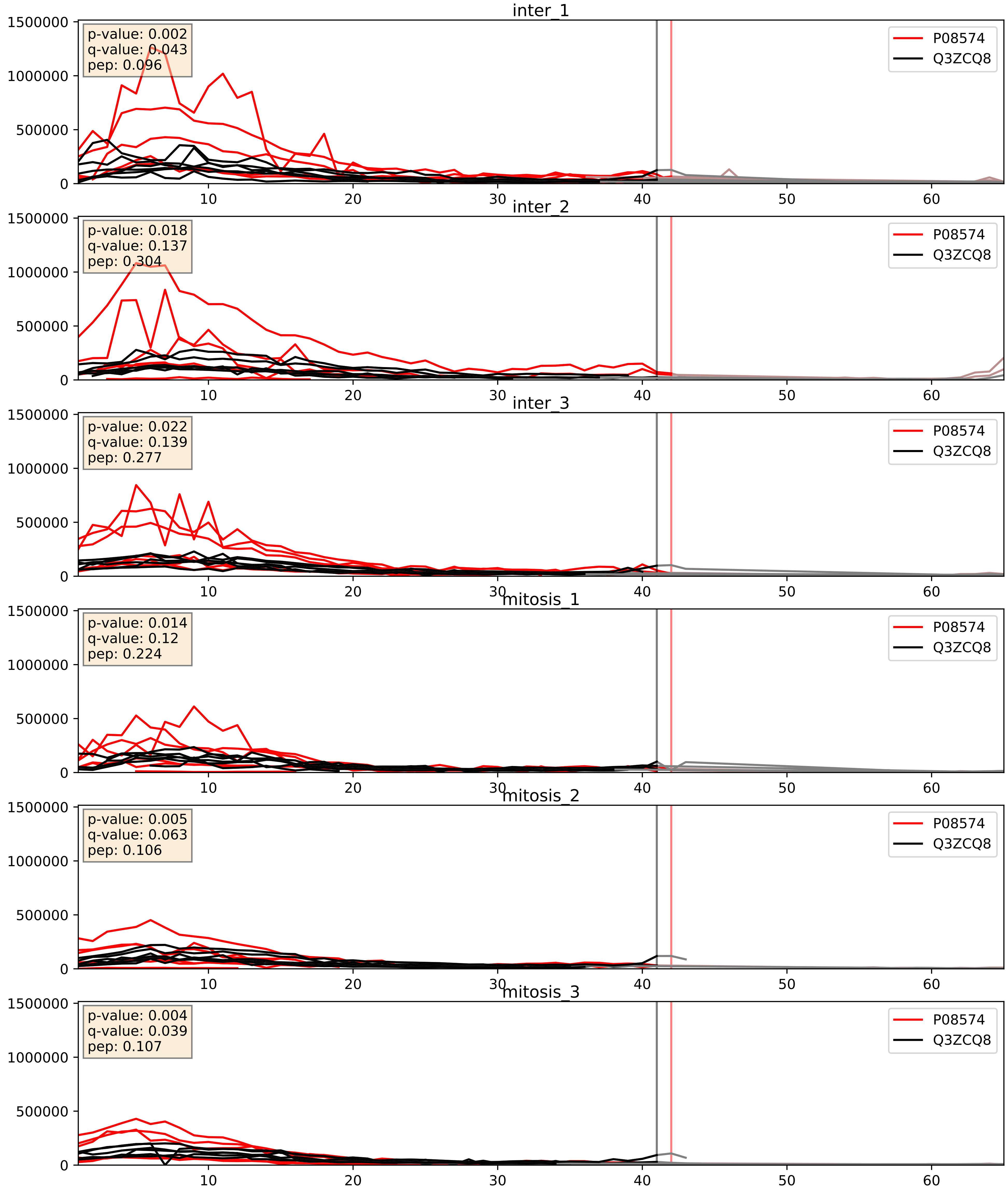


| level | condition_1 | condition_2 | pvalue | pvalue_adjusted |
|---|---|---|---|---|
| interactor_ratio | mitosis | inter | 0.0013400977657042504 | 0.04334578357589157 |
| interactor_abundance | mitosis | inter | 0.006249087821090977 | 0.06732812051420864 |
| complex_abundance | mitosis | inter | 0.012935955732803869 | 0.08411248670667204 |
