## Supplementary Data 2 for "SECAT: Quantifying differential protein-protein interaction states by network-centric analysis": hela_string_000139_P28331.pdf

P28331_P28331

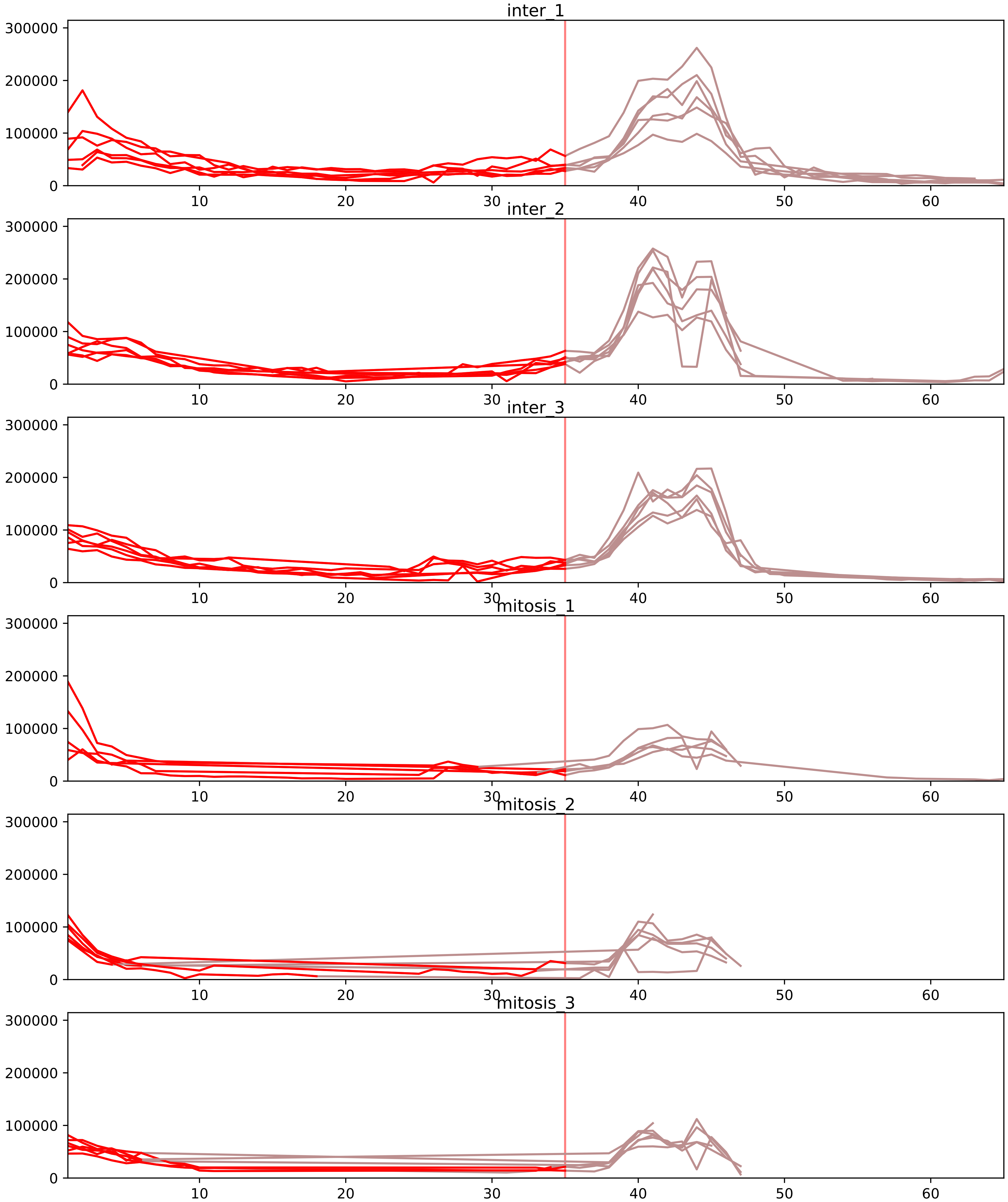


| level | condition_1 | condition_2 | pvalue | pvalue_adjusted |
|---|---|---|---|---|
| interactor_abundance | mitosis | inter | 5.004229607214721e-05 | 0.0014685477164666158 |
| monomer_abundance | mitosis | inter | 5.2108248385234455e-05 | 0.0267281636508792 |
| complex_abundance | mitosis | inter | 0.0001090848339676391 | 0.002323510983773901 |
| total_abundance | mitosis | inter | 0.00013134892997051346 | 0.013931195884997584 |
| assembled_abundance | mitosis | inter | 0.0008358164509817024 | 0.025739847414772032 |
| interactor_ratio | mitosis | inter | 0.007233229368594498 | 0.04966097775452939 |

O43678_P28331
NDUA2_HUMAN vs NDUS1_HUMAN
p-value: 0.085 q-value: 0.05

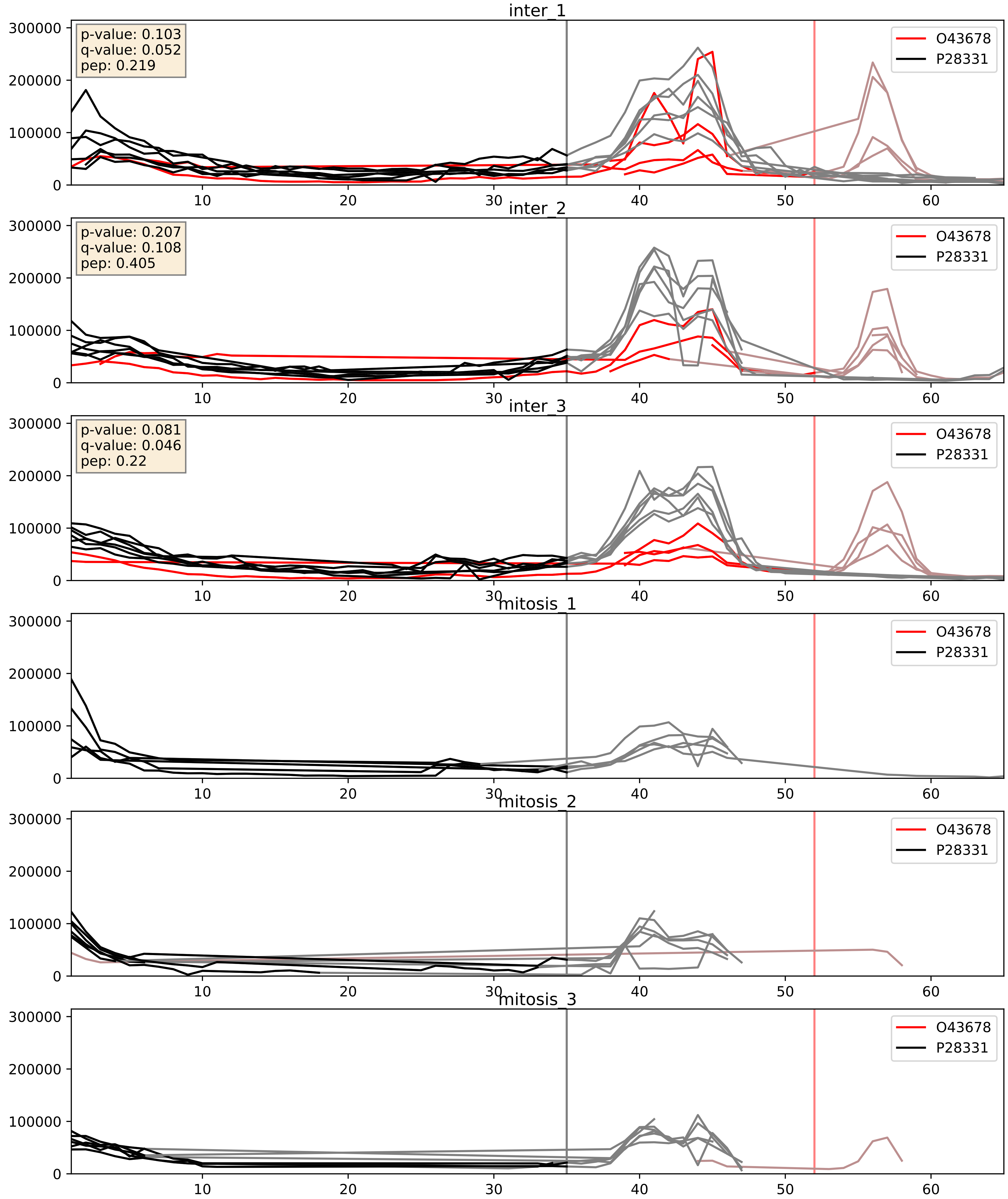


| level | condition_1 | condition_2 | pvalue | pvalue_adjusted |
|---|---|---|---|---|
| complex_abundance | mitosis | inter | 0.000623955510183607 | 0.01603921671823326 |
| interactor_ratio | mitosis | inter | 0.0019183710724063662 | 0.05317578467954594 |
| interactor_abundance | mitosis | inter | 0.01084111579398431 | 0.09135080923250061 |

O75251_P28331
NDUS7_HUMAN vs NDUS1_HUMAN
p-value: 0.042 q-value: 0.029

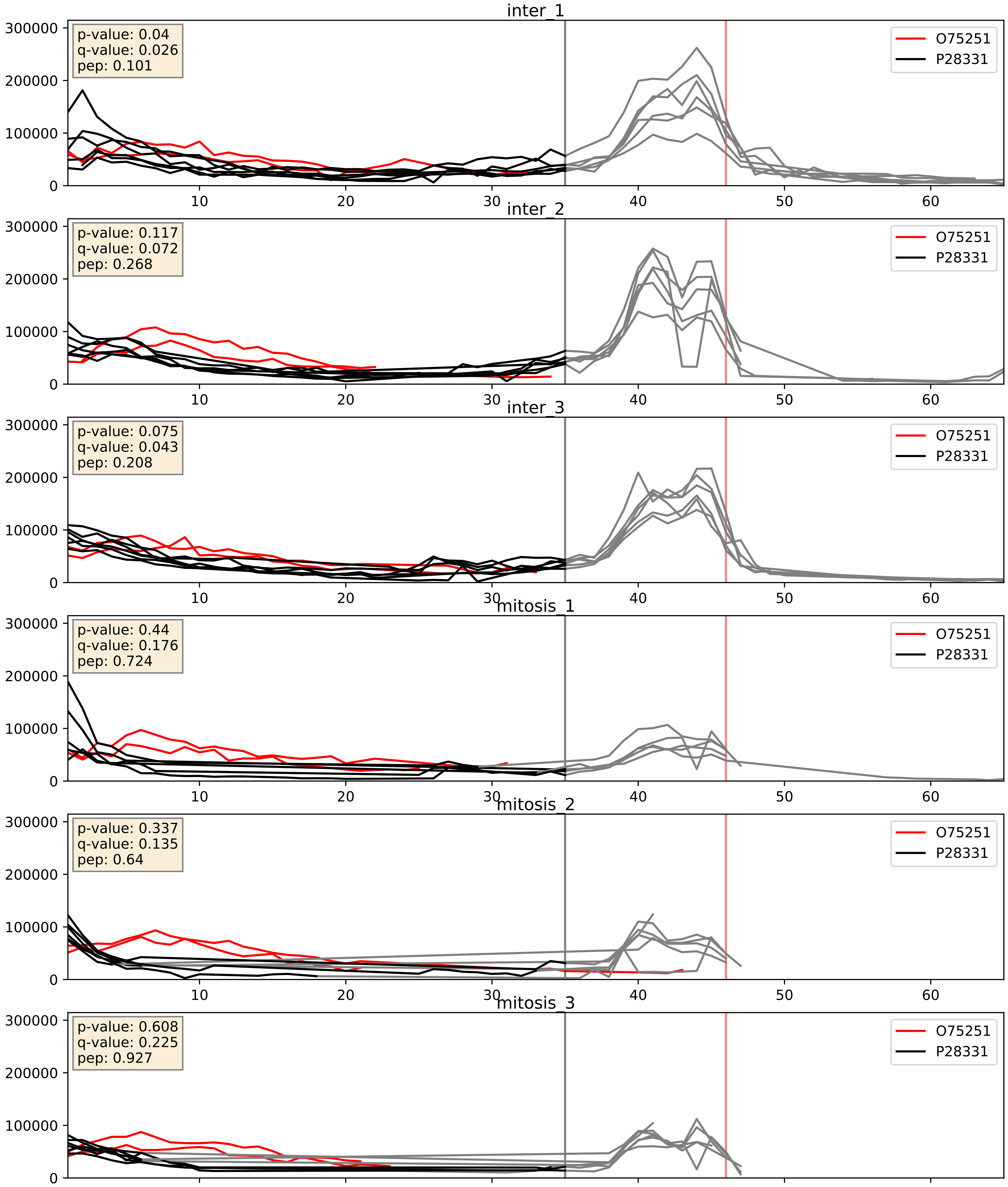


| level | condition_1 | condition_2 | pvalue | pvalue_adjusted |
|---|---|---|---|---|
| interactor_ratio | mitosis | inter | 0.00290870224888575 | 0.07007095913365735 |
| complex_abundance | mitosis | inter | 0.0036819100356348367 | 0.04464185538956686 |
| interactor_abundance | mitosis | inter | 0.019724483203473128 | 0.1190514126751731 |

O75306_P28331
NDUS2_HUMAN vs NDUS1_HUMAN
p-value: 0.045 q-value: 0.03

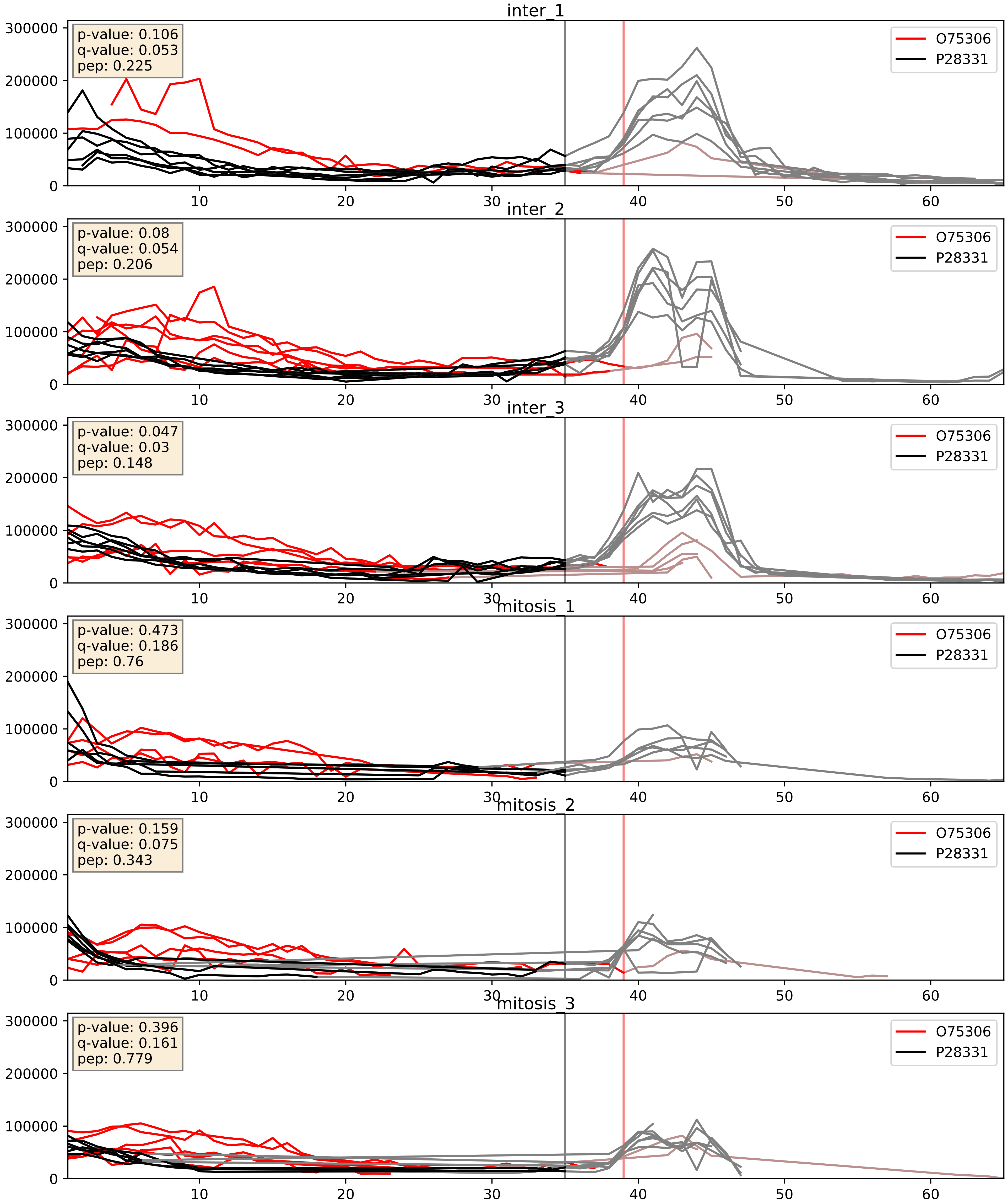


| level | condition_1 | condition_2 | pvalue | pvalue_adjusted |
|---|---|---|---|---|
| complex_abundance | mitosis | inter | 0.031008736876247462 | 0.12798205769560186 |
| interactor_abundance | mitosis | inter | 0.35172706552440497 | 0.5499224790849478 |
| interactor_ratio | mitosis | inter | 0.41093453794863094 | 0.8340755186867097 |

O75380_P28331
NDUS6_HUMAN vs NDUS1_HUMAN
p-value: 0.031 q-value: 0.022

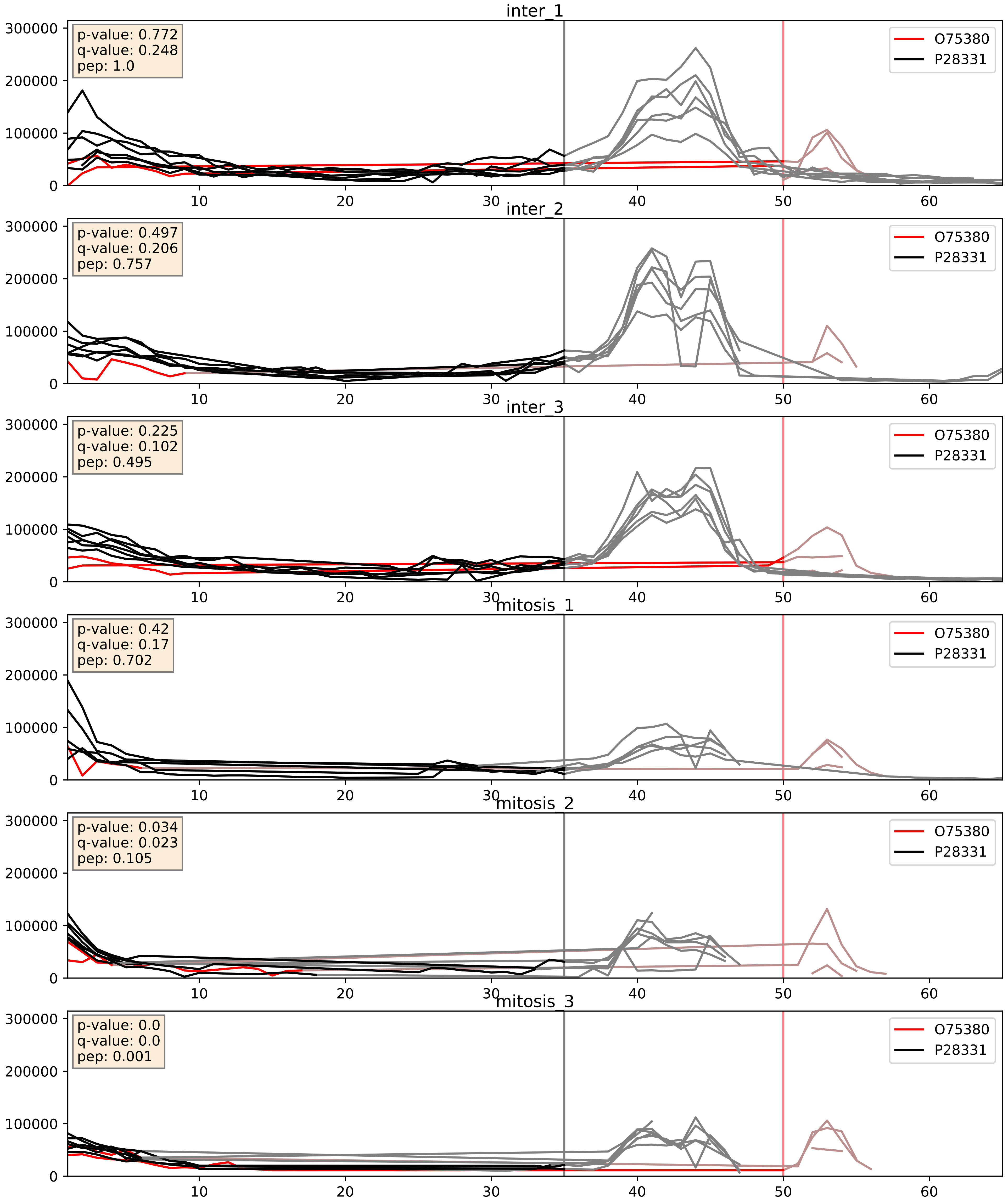


| level | condition_1 | condition_2 | pvalue | pvalue_adjusted |
|---|---|---|---|---|
| complex_abundance | mitosis | inter | 0.005545258000284267 | 0.05624098855659709 |
| interactor_abundance | mitosis | inter | 0.0333518832106805 | 0.15234371413203046 |
| interactor_ratio | mitosis | inter | 0.15165301561531105 | 0.6067576723168497 |

O75489_P28331
NDUS3_HUMAN vs NDUS1_HUMAN
p-value: 0.07 q-value: 0.043

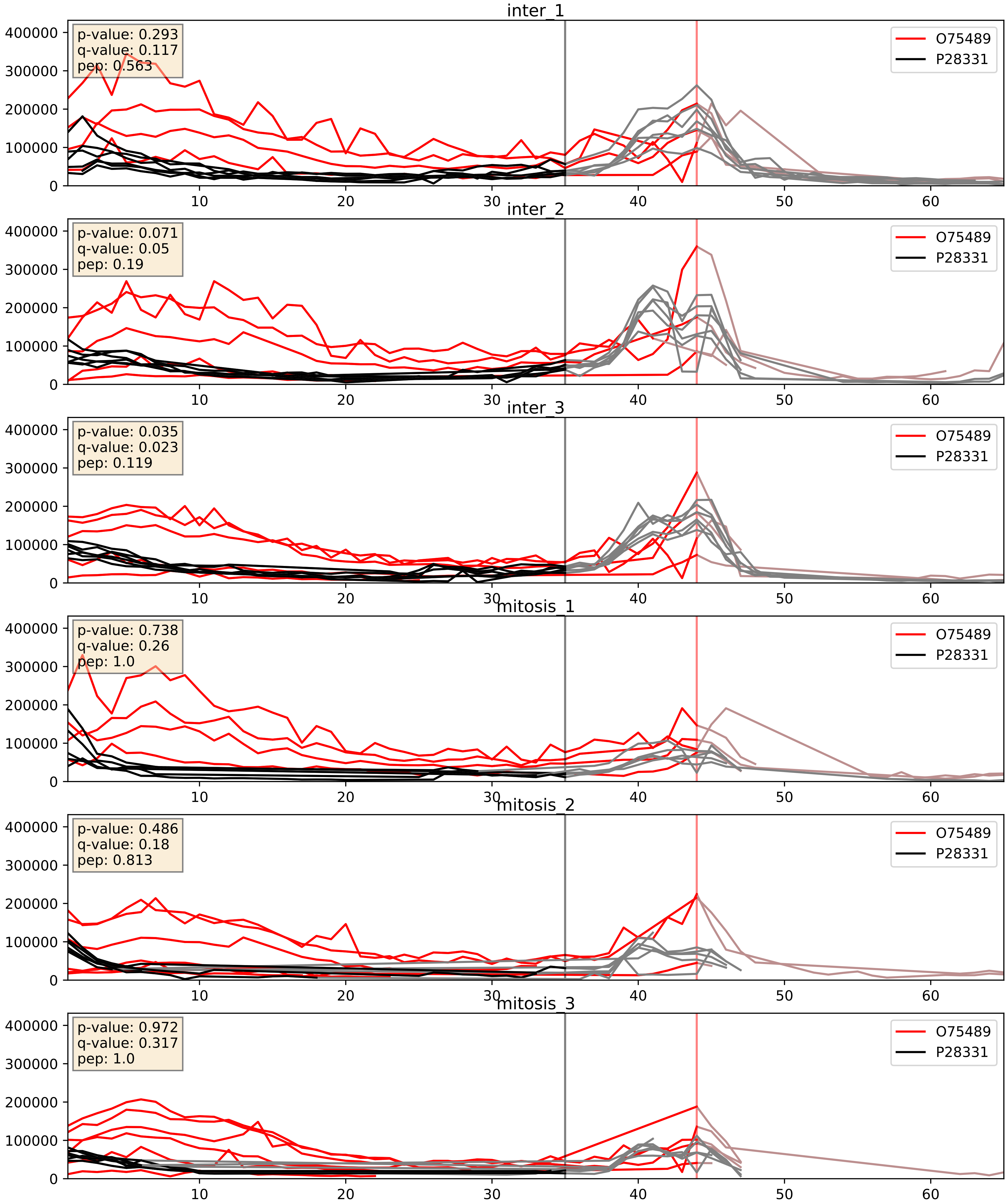


| level | condition_1 | condition_2 | pvalue | pvalue_adjusted |
|---|---|---|---|---|
| complex_abundance | mitosis | inter | 0.010772707718540754 | 0.07684531505892406 |
| interactor_ratio | mitosis | inter | 0.035771470960743165 | 0.30712516692473574 |
| interactor_abundance | mitosis | inter | 0.04205030485200938 | 0.16496361573473892 |

O95168_P28331
NDUB4_HUMAN vs NDUS1_HUMAN
p-value: 0.016 q-value: 0.013

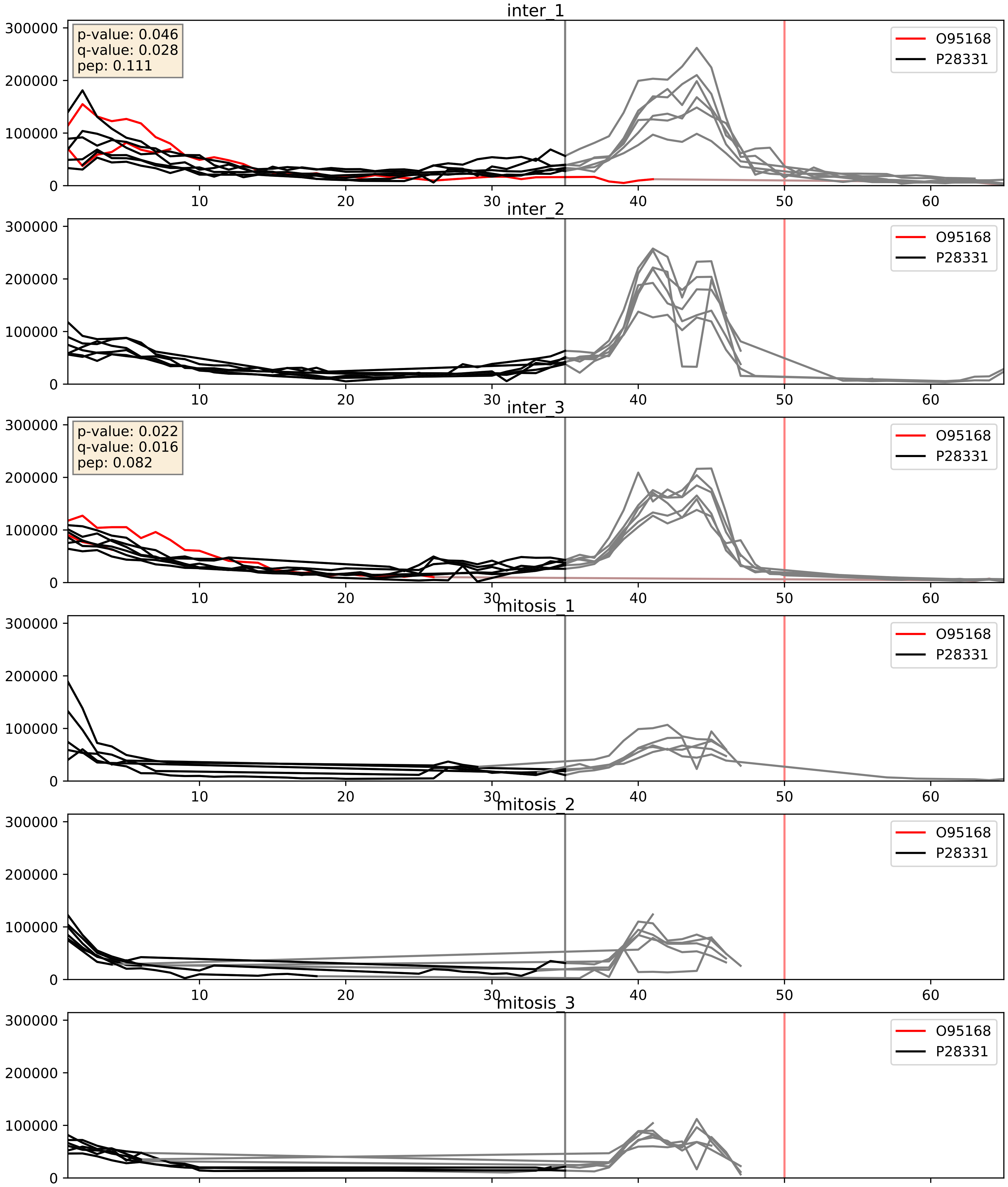


| level | condition_1 | condition_2 | pvalue | pvalue_adjusted |
|---|---|---|---|---|
| complex_abundance | mitosis | inter | 0.04009458528165575 | 0.14508340529050803 |
| interactor_abundance | mitosis | inter | 0.04013928536710259 | 0.16166968180878097 |
| interactor_ratio | mitosis | inter | 0.04525637880516615 | 0.35346222862429033 |

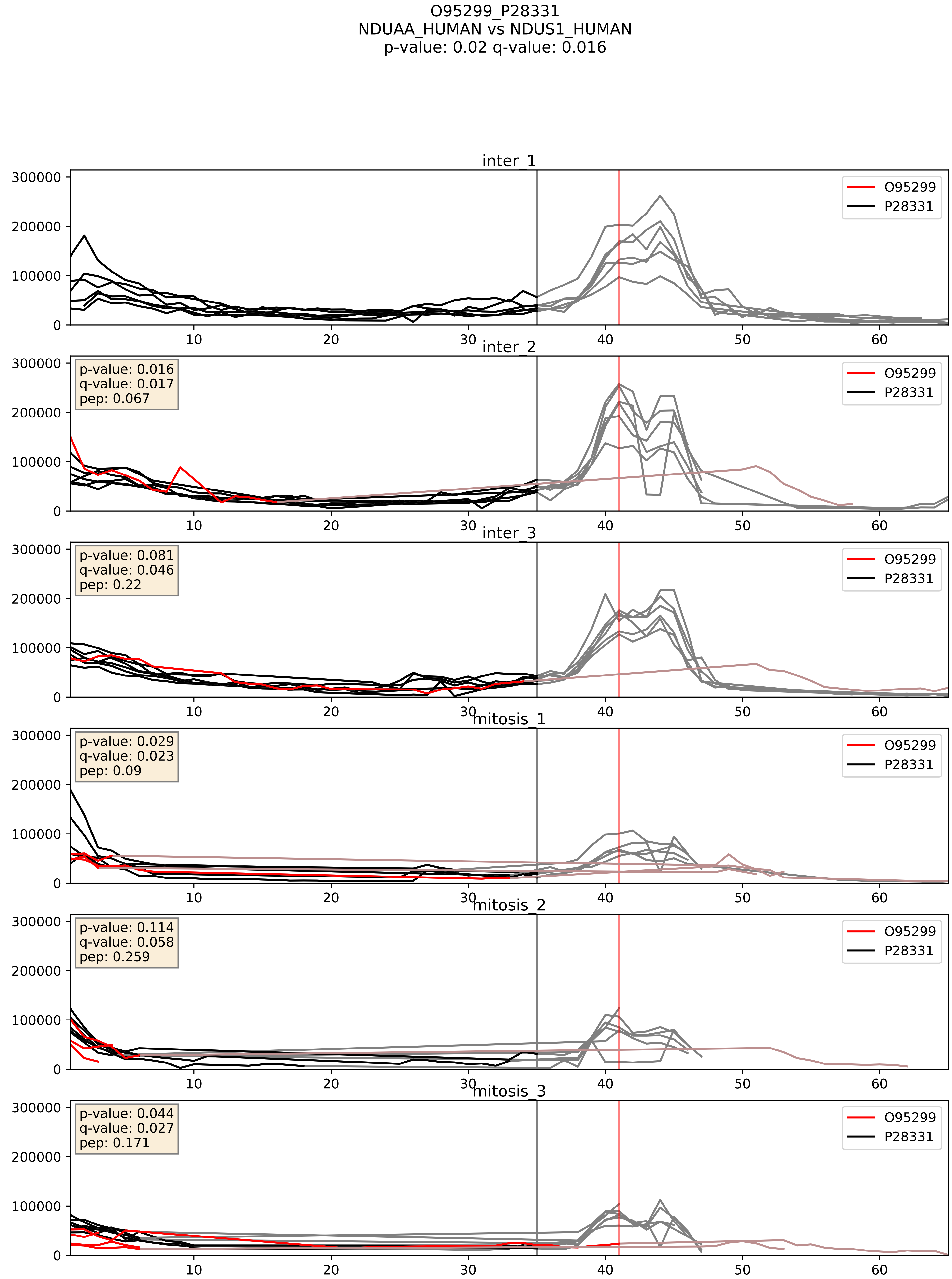


| level | condition_1 | condition_2 | pvalue | pvalue_adjusted |
|---|---|---|---|---|
| interactor_abundance | mitosis | inter | 0.3612415873555567 | 0.5585671943214533 |
| complex_abundance | mitosis | inter | 0.3876565439187527 | 0.5341822305126406 |
| interactor_ratio | mitosis | inter | 0.5404421061348641 | 0.8859730852936224 |

P17568_P28331
NDUB7_HUMAN vs NDUS1_HUMAN
p-value: 0.019 q-value: 0.015

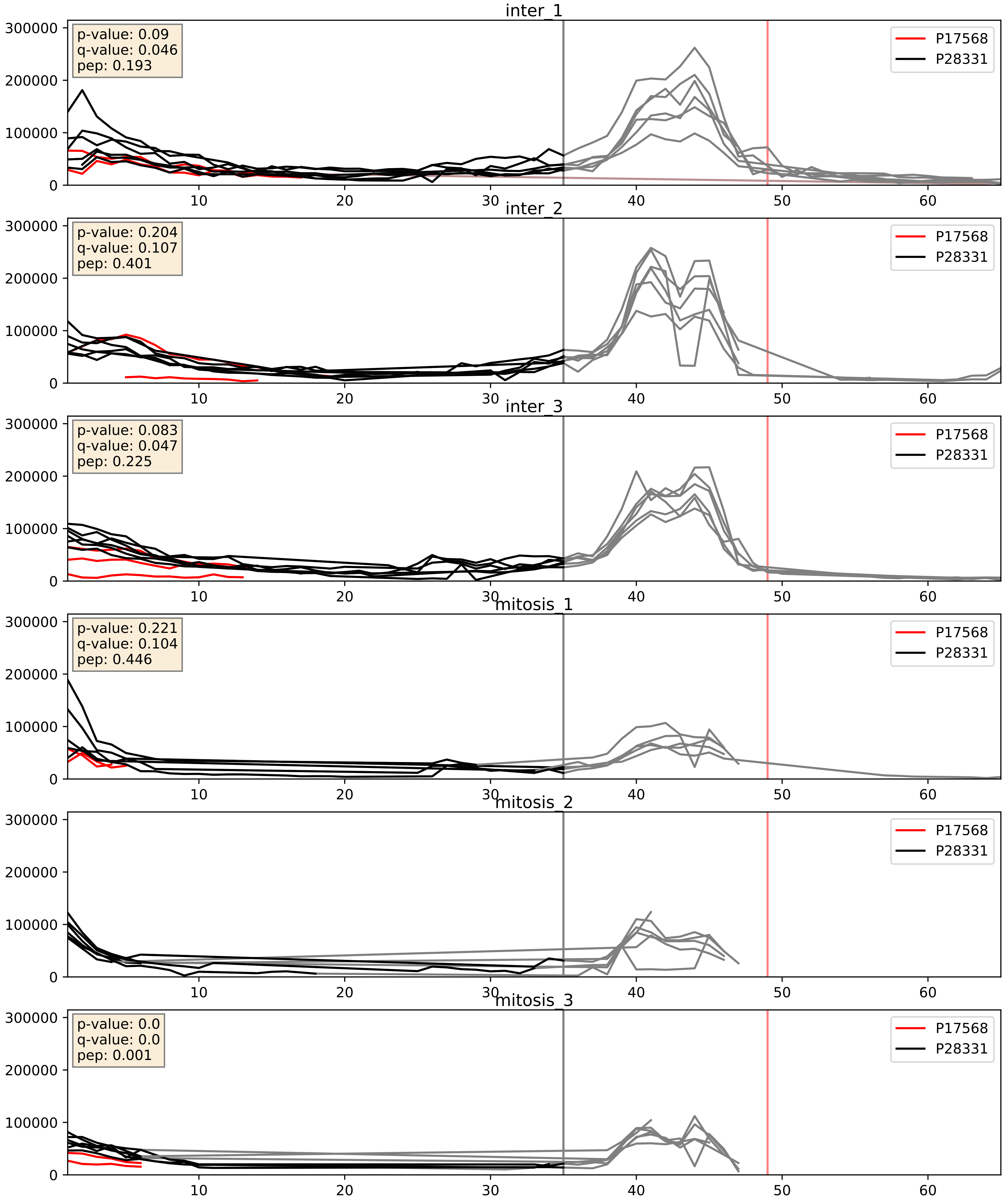


| level | condition_1 | condition_2 | pvalue | pvalue_adjusted |
|---|---|---|---|---|
| interactor_abundance | mitosis | inter | 0.02339687029671626 | 0.12556564873974368 |
| complex_abundance | mitosis | inter | 0.03639353842322165 | 0.1388274014718259 |
| interactor_ratio | mitosis | inter | 0.3453894832905555 | 0.797415151237597 |

P19404_P28331
NDUV2_HUMAN vs NDUS1_HUMAN
p-value: 0.001 q-value: 0.002

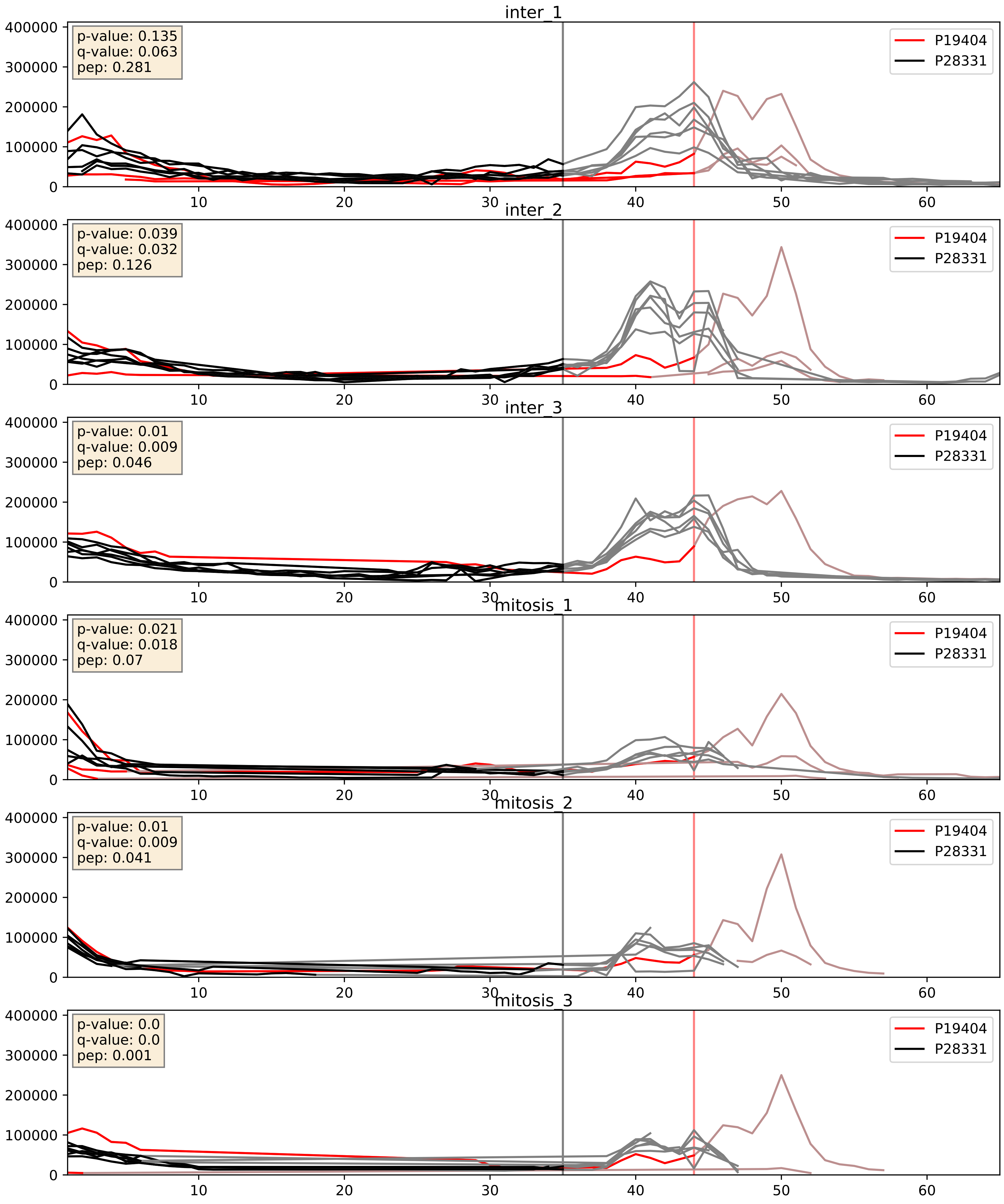


| level | condition_1 | condition_2 | pvalue | pvalue_adjusted |
|---|---|---|---|---|
| complex_abundance | mitosis | inter | 0.054772837546031 | 0.16691188657672673 |
| interactor_abundance | mitosis | inter | 0.10621351871074543 | 0.27876367320680084 |
| interactor_ratio | mitosis | inter | 0.8285729716024981 | 0.9701897807578924 |

P22695_P28331
QCR2_HUMAN vs NDUS1_HUMAN
p-value: 0.043 q-value: 0.029

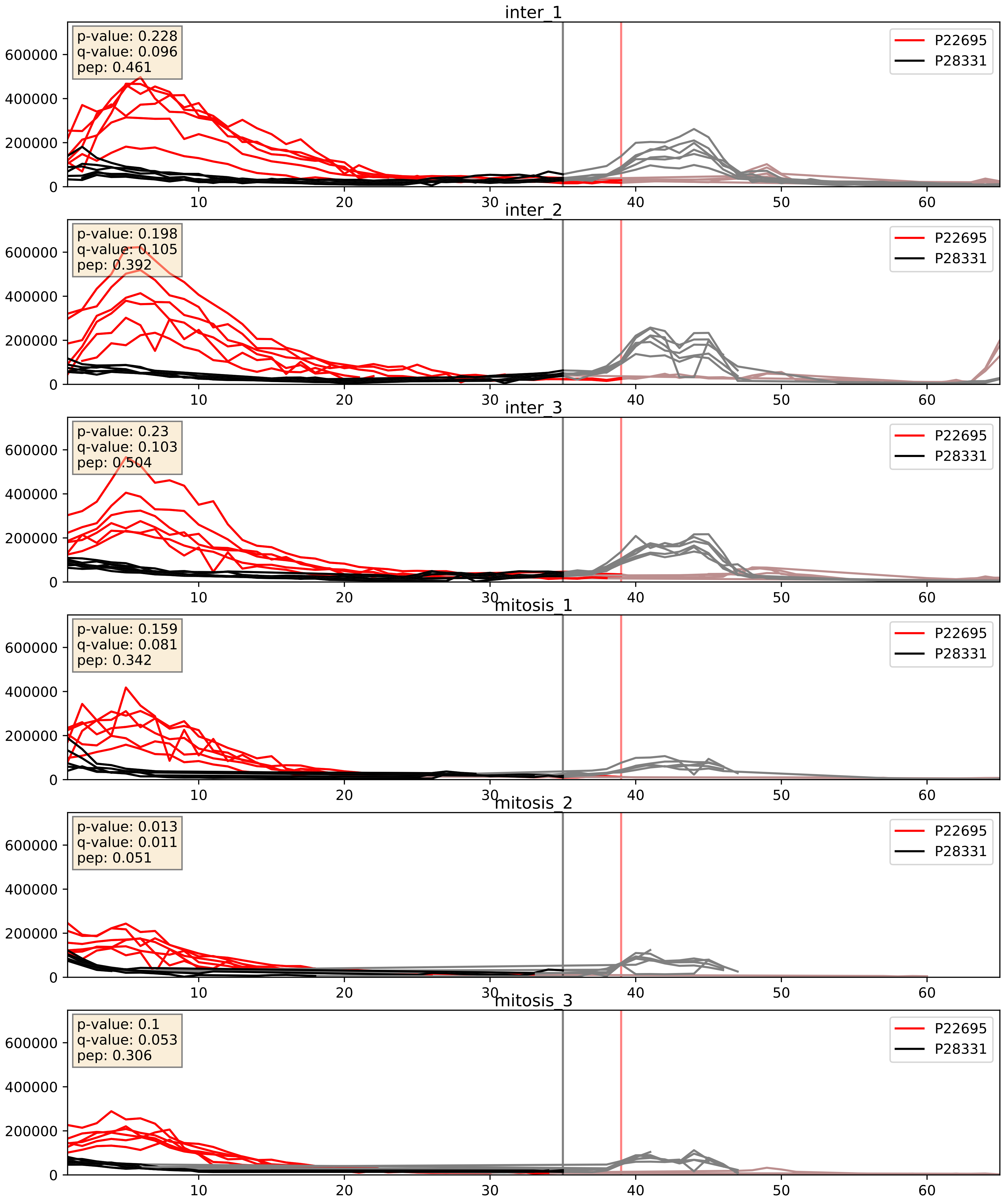


| level | condition_1 | condition_2 | pvalue | pvalue_adjusted |
|---|---|---|---|---|
| complex_abundance | mitosis | inter | 0.0016073371358469538 | 0.02813661734734136 |
| interactor_abundance | mitosis | inter | 0.0017502652133548574 | 0.030576061686362406 |
| interactor_ratio | mitosis | inter | 0.2970697251002717 | 0.7646944825997081 |

P28331_P47985
NDUS1_HUMAN vs UCRI_HUMAN
p-value: 0.075 q-value: 0.045

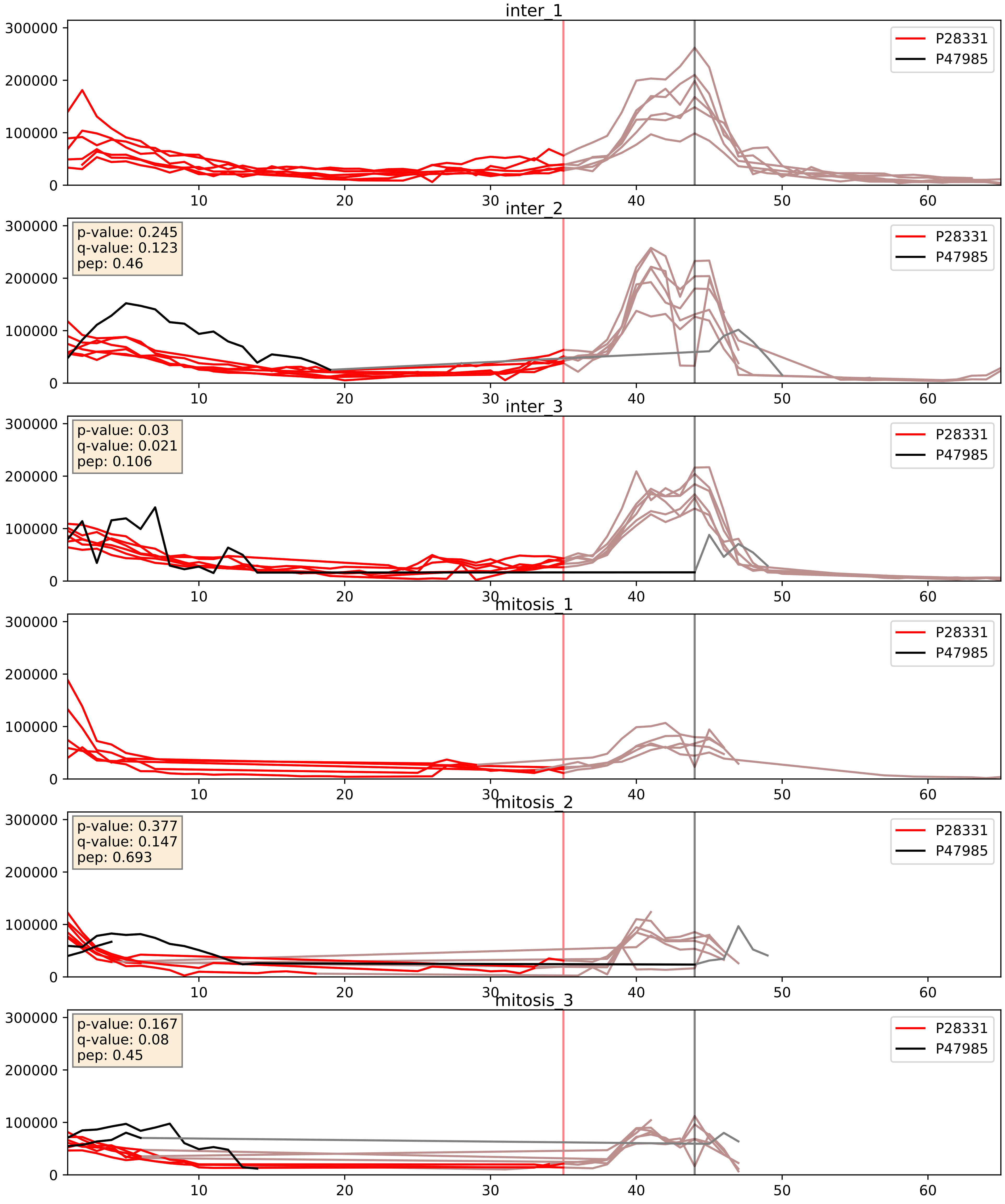


| level | condition_1 | condition_2 | pvalue | pvalue_adjusted |
|---|---|---|---|---|
| interactor_ratio | mitosis | inter | 0.10823198767629048 | 0.5471896468804465 |
| interactor_abundance | mitosis | inter | 0.26395317448967154 | 0.4561146740425073 |
| complex_abundance | mitosis | inter | 0.38920707846937663 | 0.5351988099113034 |

P28331_P49821
NDUS1_HUMAN vs NDUV1_HUMAN
p-value: 0.023 q-value: 0.018

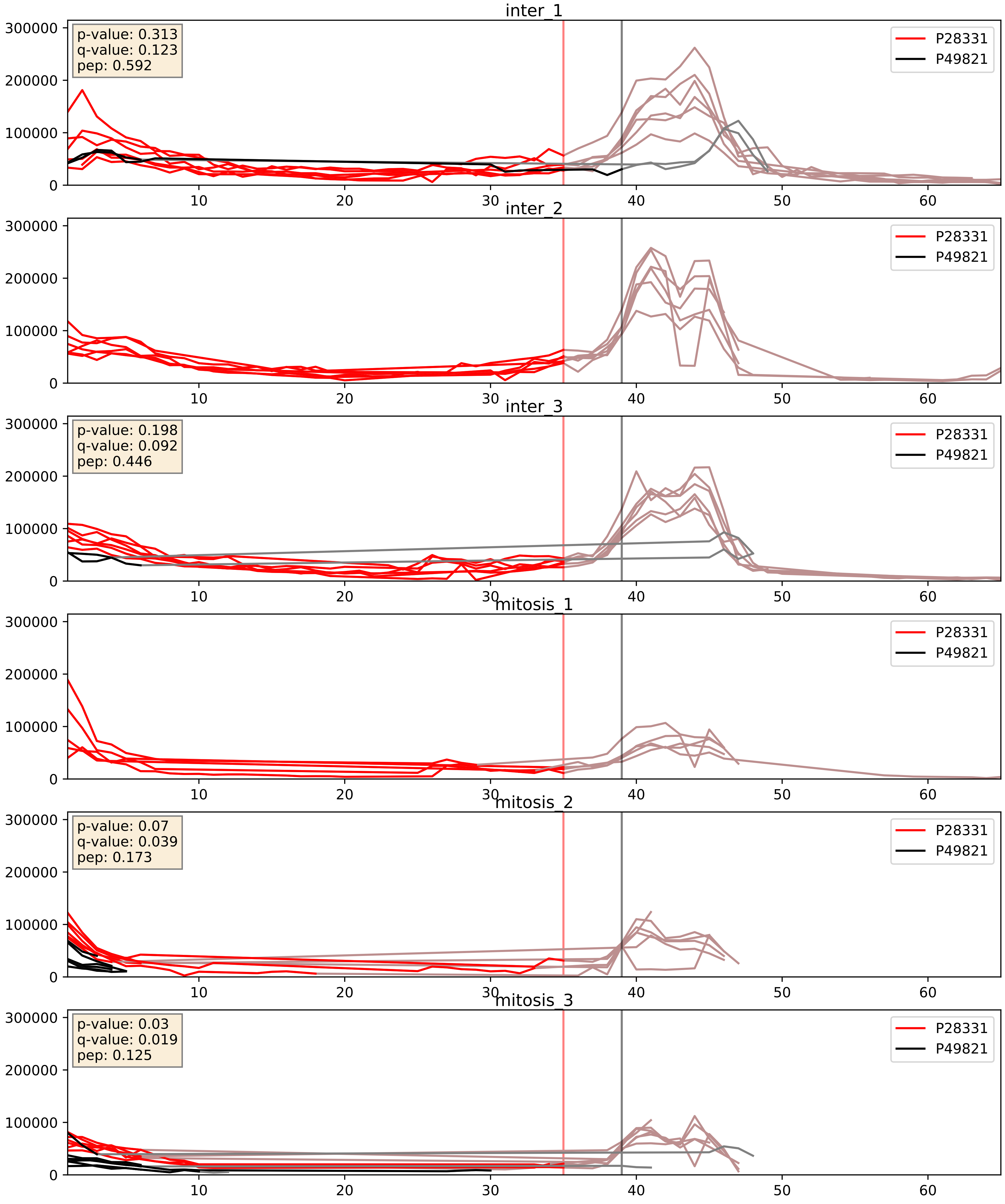


| level | condition_1 | condition_2 | pvalue | pvalue_adjusted |
|---|---|---|---|---|
| complex_abundance | mitosis | inter | 0.2504835493077138 | 0.40084860386502713 |
| interactor_abundance | mitosis | inter | 0.3146347995210318 | 0.5119805881380158 |
| interactor_ratio | mitosis | inter | 0.6478161205831032 | 0.9217731067492805 |

P28331_P56556
NDUS1_HUMAN vs NDUA6_HUMAN
p-value: 0.006 q-value: 0.006

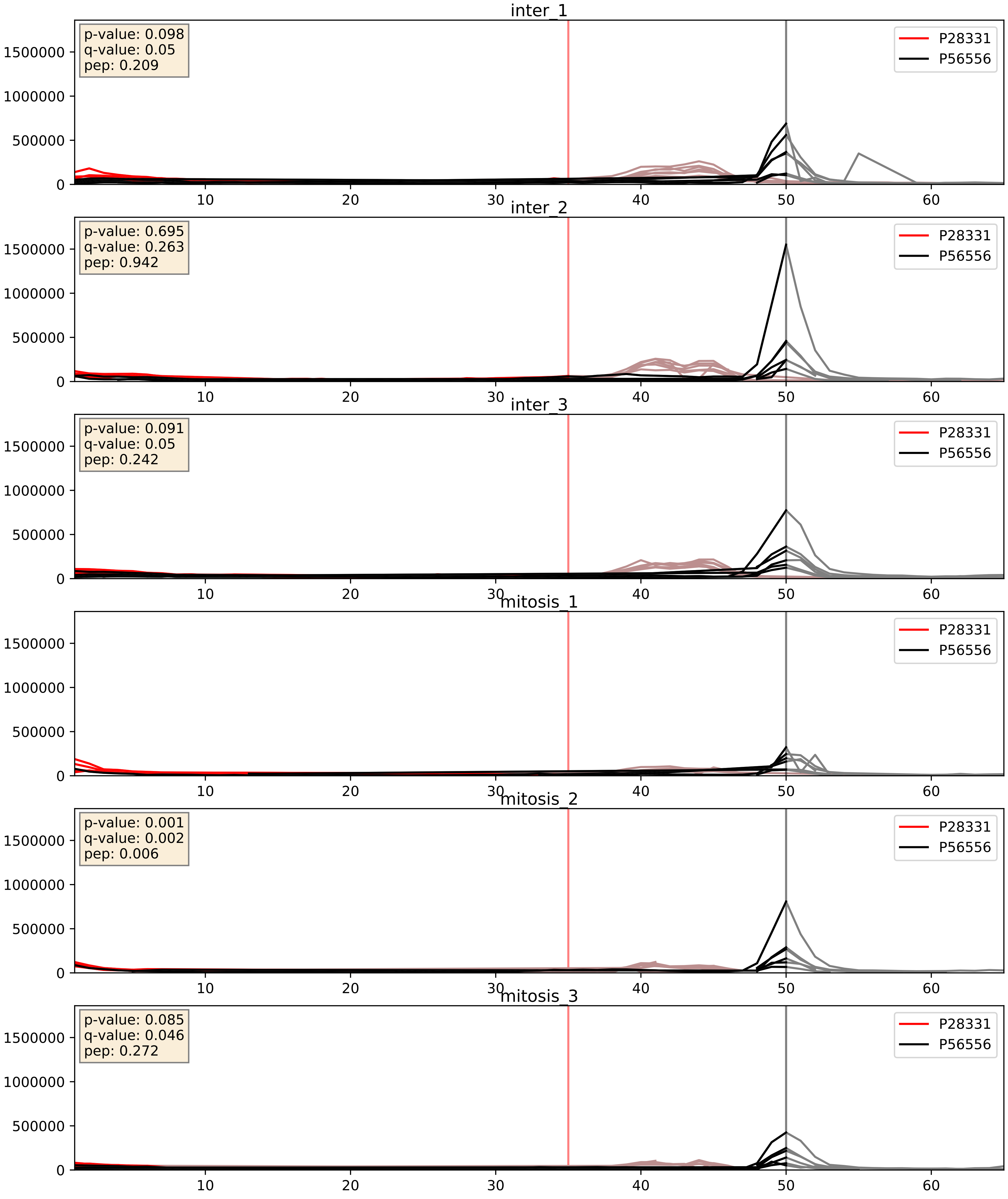


| level | condition_1 | condition_2 | pvalue | pvalue_adjusted |
|---|---|---|---|---|
| interactor_abundance | mitosis | inter | 0.002578412972883447 | 0.03806046117740333 |
| complex_abundance | mitosis | inter | 0.003507660020266425 | 0.04332694051007301 |
| interactor_ratio | mitosis | inter | 0.4030351428112066 | 0.8312490856002908 |

P28331_Q16795
NDUS1_HUMAN vs NDUA9_HUMAN
p-value: 0.032 q-value: 0.023

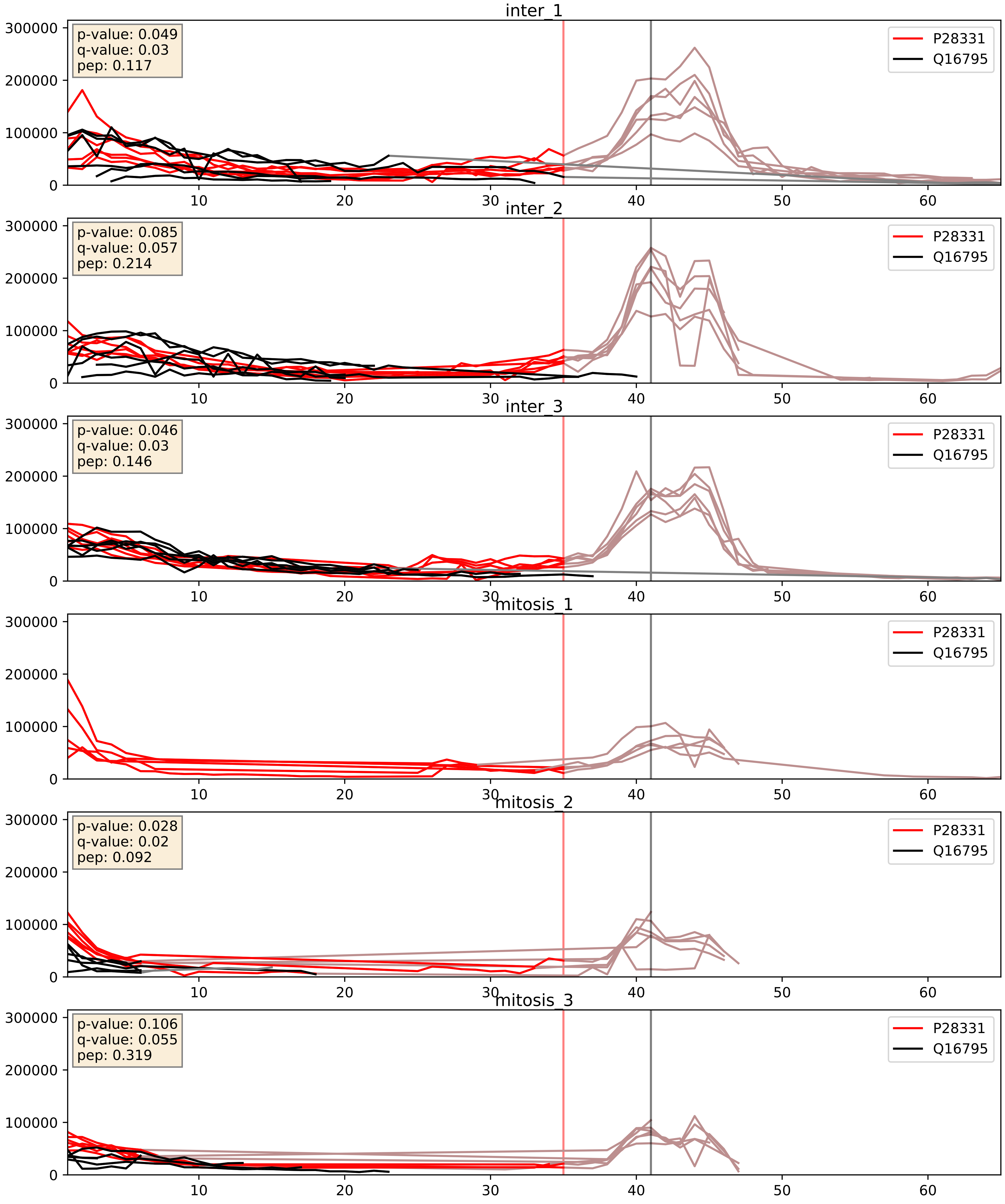


| level | condition_1 | condition_2 | pvalue | pvalue_adjusted |
|---|---|---|---|---|
| complex_abundance | mitosis | inter | 0.0004818900669497129 | 0.013833744180327739 |
| interactor_abundance | mitosis | inter | 0.005524204947292736 | 0.06317594435380872 |
| interactor_ratio | mitosis | inter | 0.15636129142680918 | 0.6134063495020562 |

P28331_Q9P0J0
NDUS1_HUMAN vs NDUAD_HUMAN
p-value: 0.01 q-value: 0.009

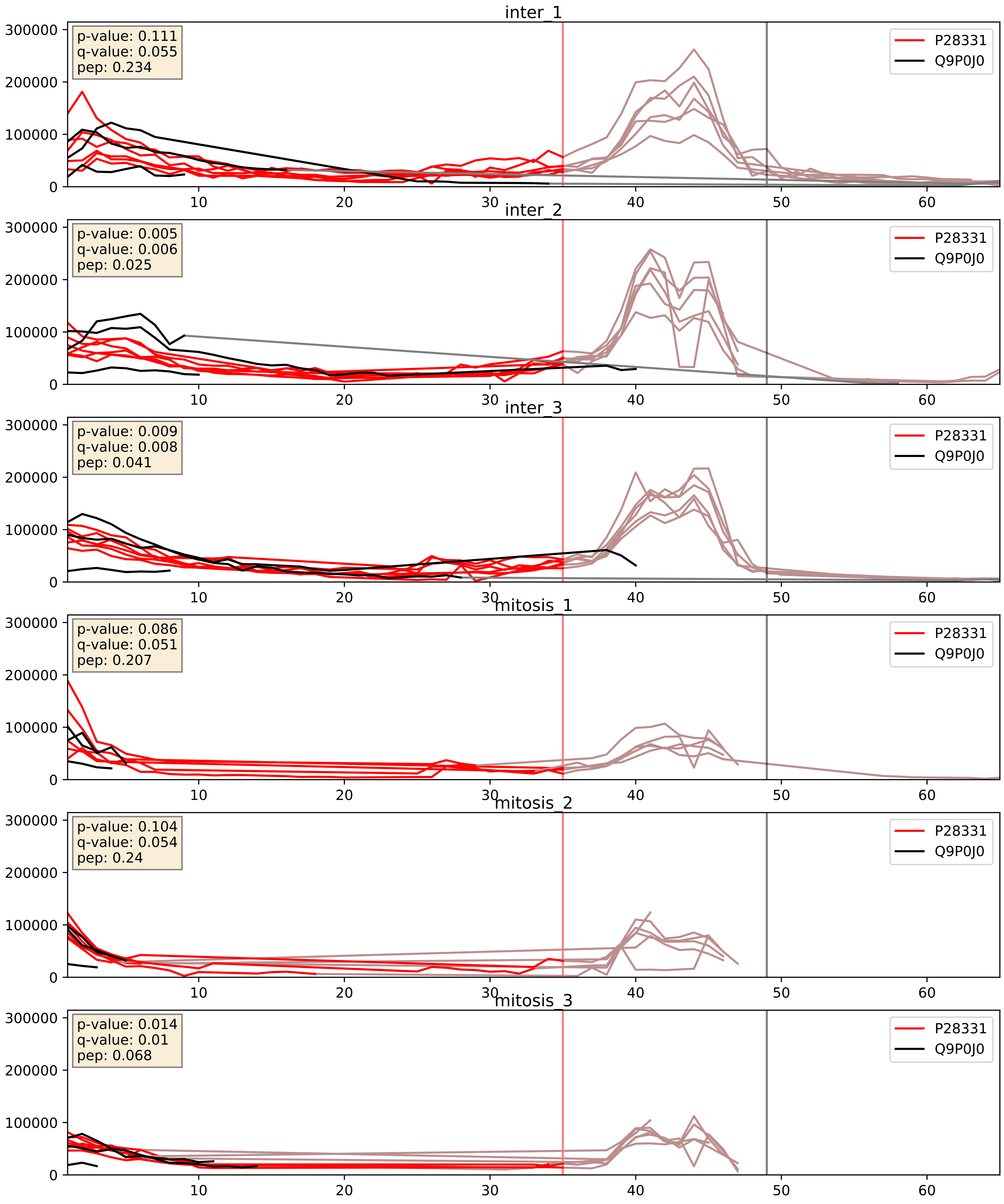


| level | condition_1 | condition_2 | pvalue | pvalue_adjusted |
|---|---|---|---|---|
| interactor_abundance | mitosis | inter | 0.0013232095430149022 | 0.026872298192663256 |
| complex_abundance | mitosis | inter | 0.0016453851528469353 | 0.02839616312171324 |
| interactor_ratio | mitosis | inter | 0.9164984746886587 | 0.9855669004419936 |
