## Supplementary Data 2 for "SECAT: Quantifying differential protein-protein interaction states by network-centric analysis": hela_string_000140_Q9UJX4.pdf

Q9UJX4_Q9UJX4

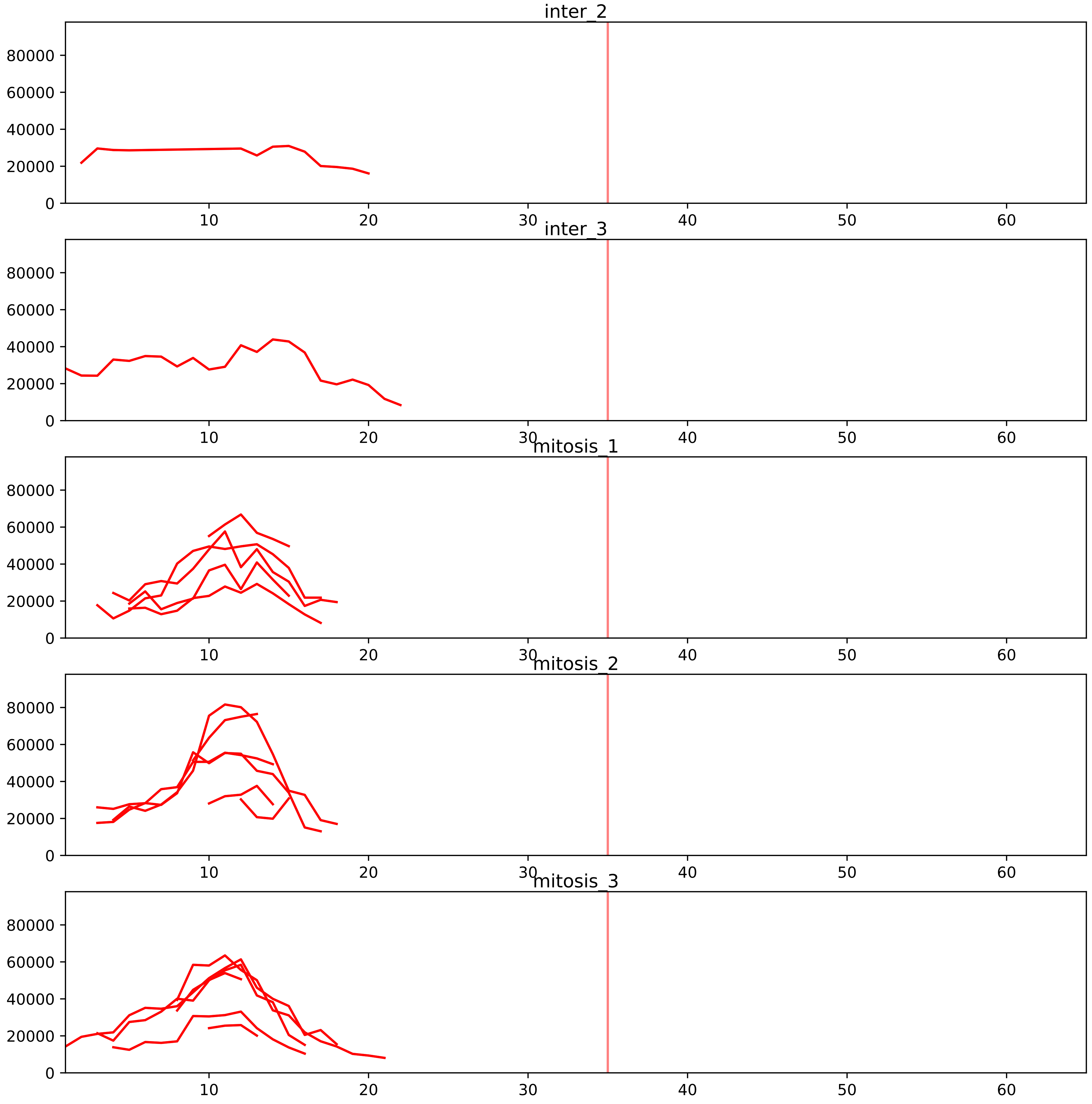


| level | condition_1 | condition_2 | pvalue | pvalue_adjusted |
|---|---|---|---|---|
| interactor_abundance | mitosis | inter | 5.083628512250066e-05 | 0.0014685477164666158 |
| assembled_abundance | mitosis | inter | 0.0023478789505937607 | 0.04701901122388186 |
| total_abundance | mitosis | inter | 0.002446662734107678 | 0.0530040850184774 |
| complex_abundance | mitosis | inter | 0.0036972044291633706 | 0.02417534628390644 |
| interactor_ratio | mitosis | inter | 0.08419686109995445 | 0.29452894377170374 |

P30260_Q9UJX4
CDC27_HUMAN vs APC5_HUMAN
p-value: 0.008 q-value: 0.008

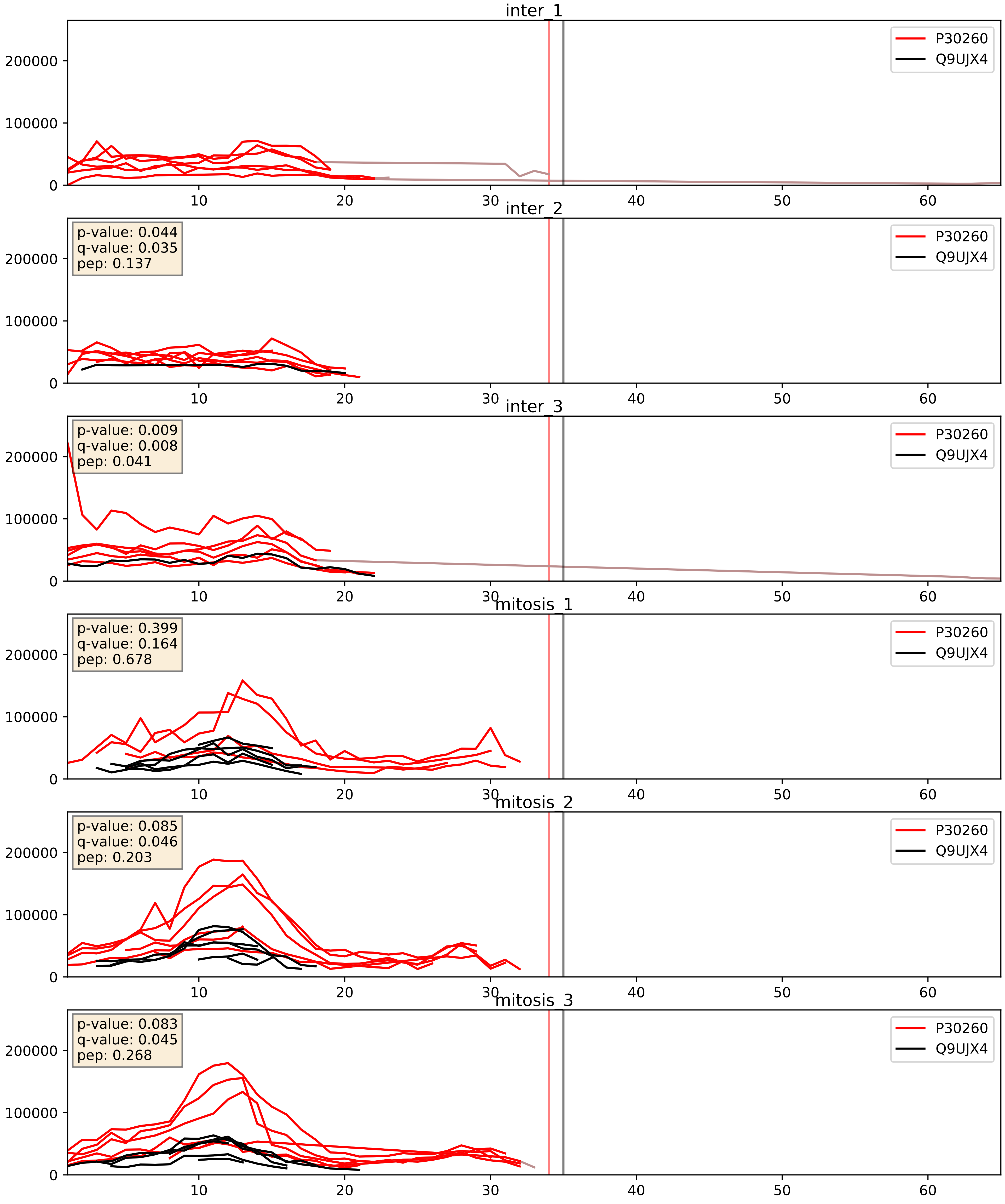


| level | condition_1 | condition_2 | pvalue | pvalue_adjusted |
|---|---|---|---|---|
| complex_abundance | mitosis | inter | 0.09885406543393127 | 0.23317464869508175 |
| interactor_abundance | mitosis | inter | 0.2935639836172767 | 0.488027781996528 |
| interactor_ratio | mitosis | inter | 0.9077478785320257 | 0.9852158746906624 |

Q13042_Q9UJX4
CDC16_HUMAN vs APC5_HUMAN
p-value: 0.0 q-value: 0.0

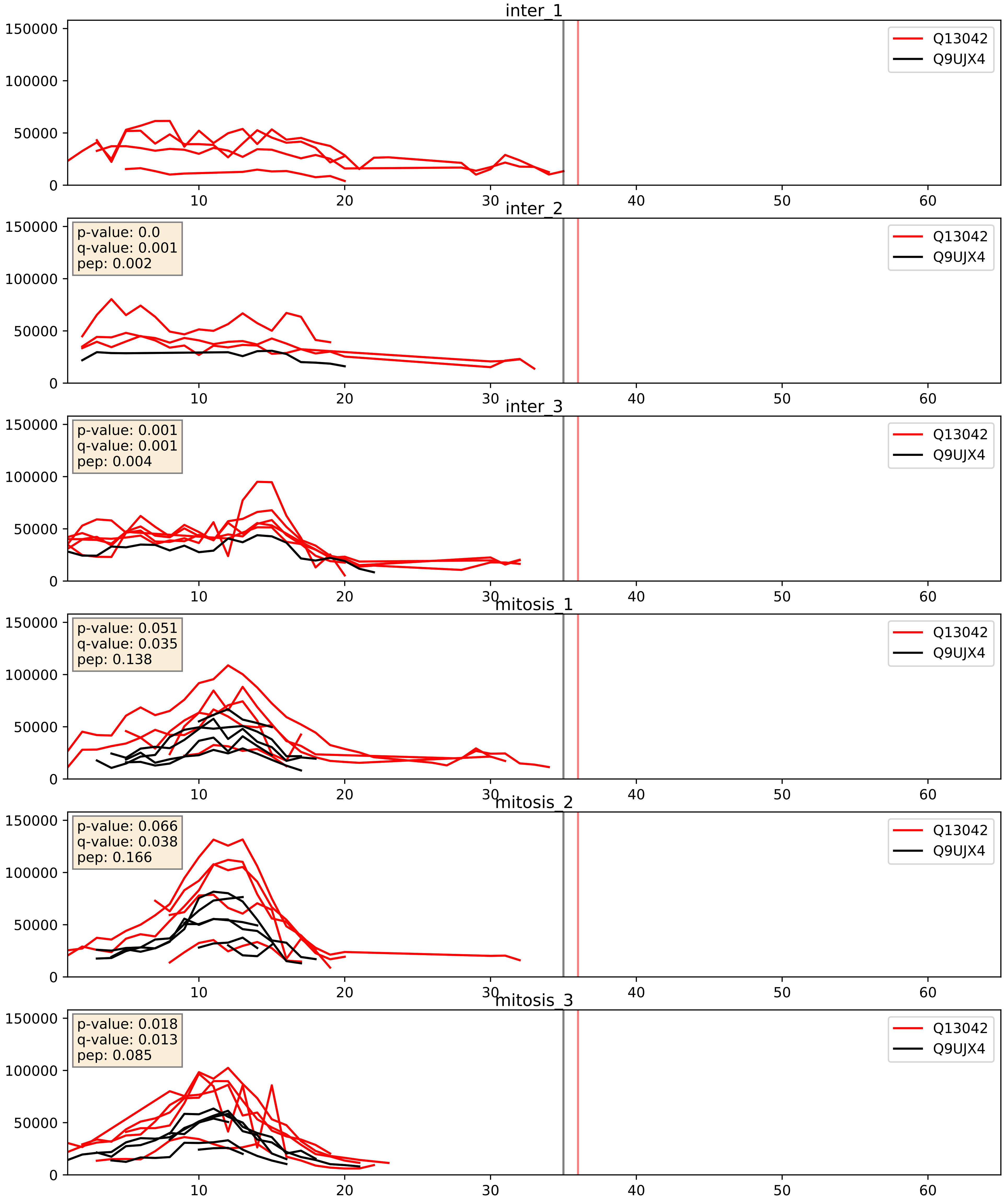


| level | condition_1 | condition_2 | pvalue | pvalue_adjusted |
|---|---|---|---|---|
| complex_abundance | mitosis | inter | 0.10709149167598454 | 0.2441286734344681 |
| interactor_abundance | mitosis | inter | 0.36757488444518643 | 0.5632087317955554 |
| interactor_ratio | mitosis | inter | 0.7020302830553122 | 0.9366476772081006 |

Q8NHZ8_Q9UJX4
CDC26_HUMAN vs APC5_HUMAN
p-value: 0.026 q-value: 0.02

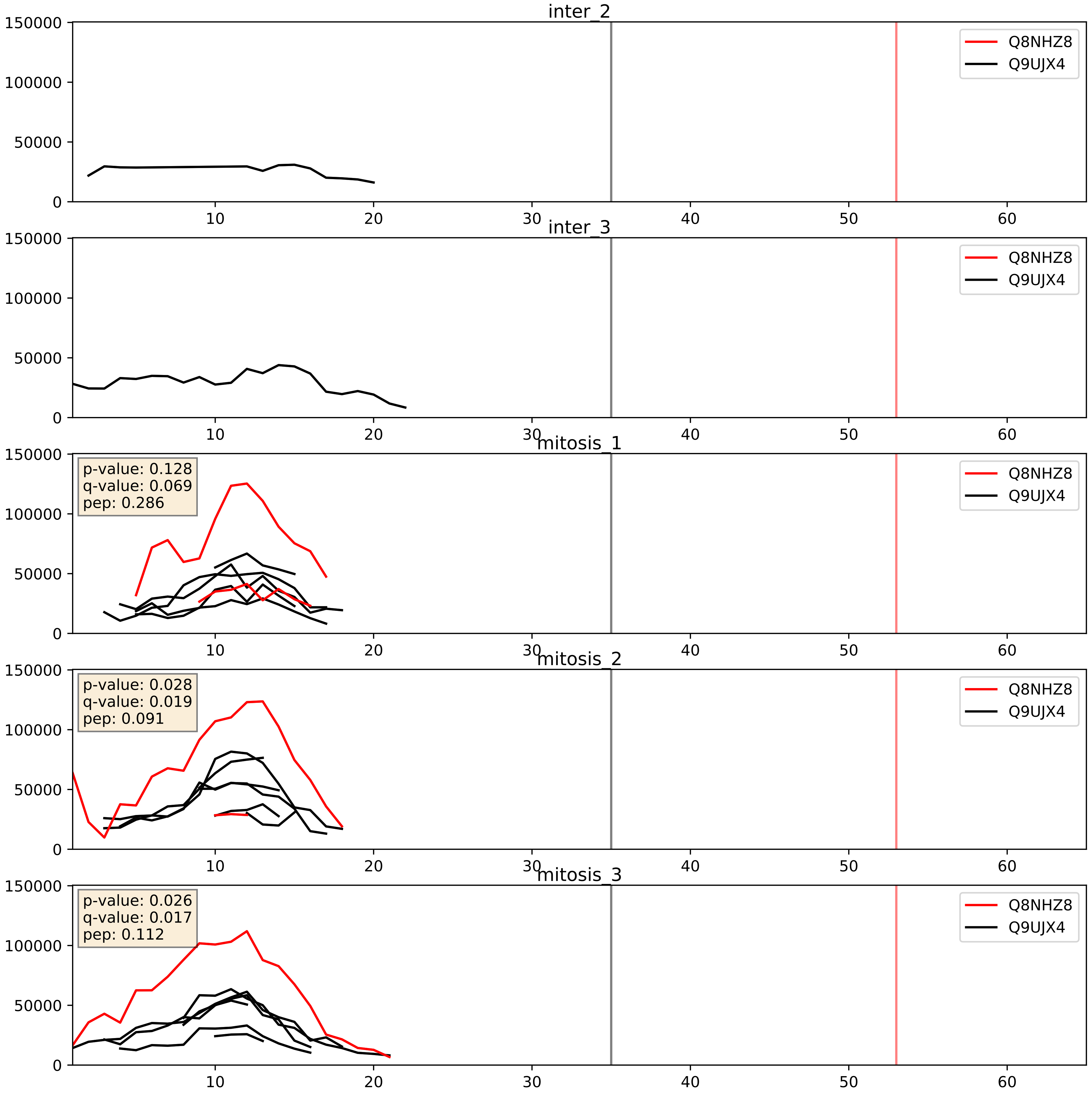


| level | condition_1 | condition_2 | pvalue | pvalue_adjusted |
|---|---|---|---|---|
| complex_abundance | mitosis | inter | 0.004557678589736247 | 0.0503402951330868 |
| interactor_abundance | mitosis | inter | 0.02065482954399308 | 0.12021882713294628 |
| interactor_ratio | mitosis | inter | 0.07564477353850585 | 0.4628166812754875 |

Q96DE5_Q9UJX4
APC16_HUMAN vs APC5_HUMAN
p-value: 0.06 q-value: 0.038

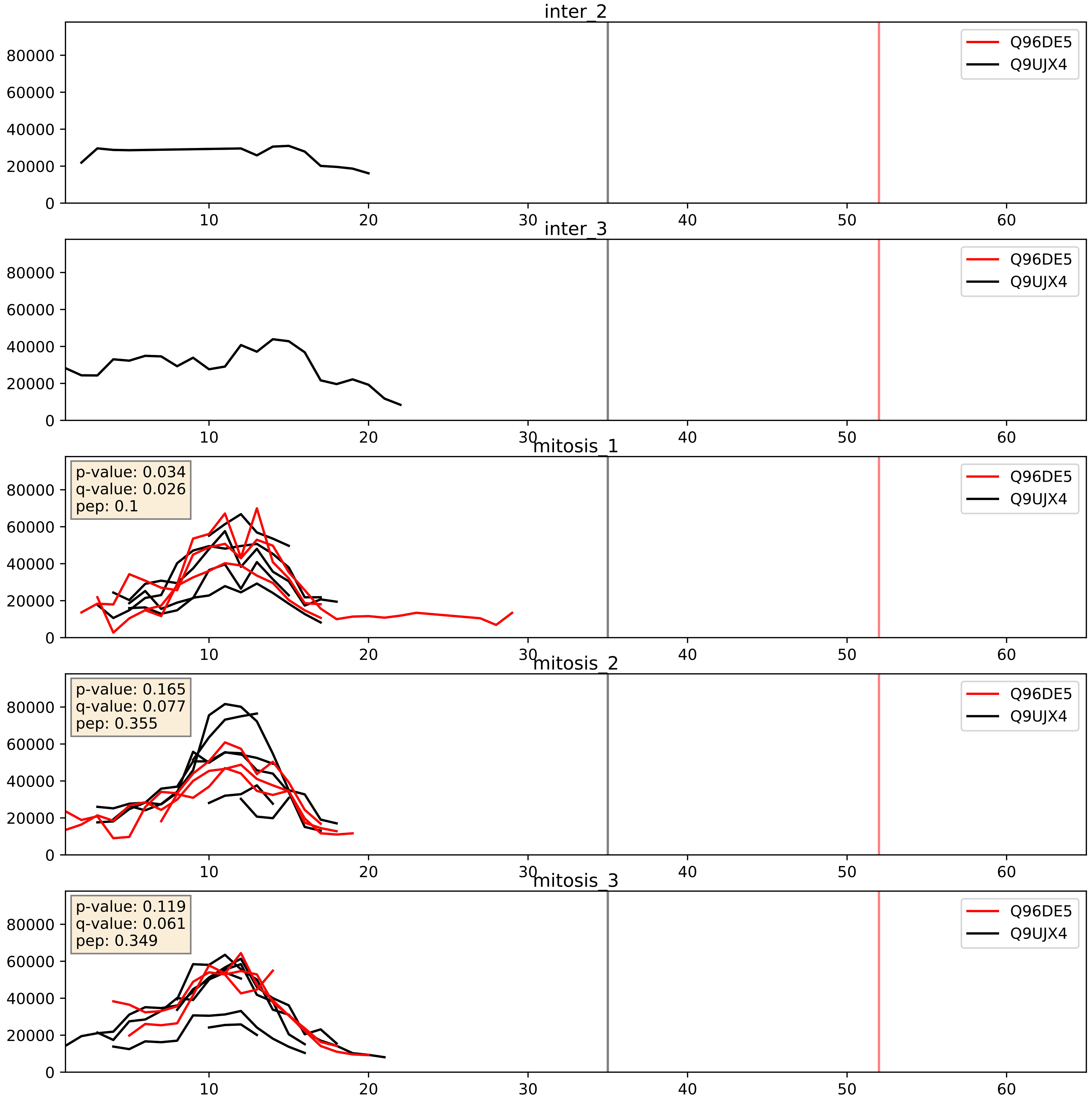


| level | condition_1 | condition_2 | pvalue | pvalue_adjusted |
|---|---|---|---|---|
| complex_abundance | mitosis | inter | 0.004058881031412856 | 0.04732787841836958 |
| interactor_abundance | mitosis | inter | 0.007466750033025538 | 0.07484236567060727 |
| interactor_ratio | mitosis | inter | 0.3148006520149434 | 0.7783421708721073 |

Q9H1A4_Q9UJX4
APC1_HUMAN vs APC5_HUMAN
p-value: 0.032 q-value: 0.023

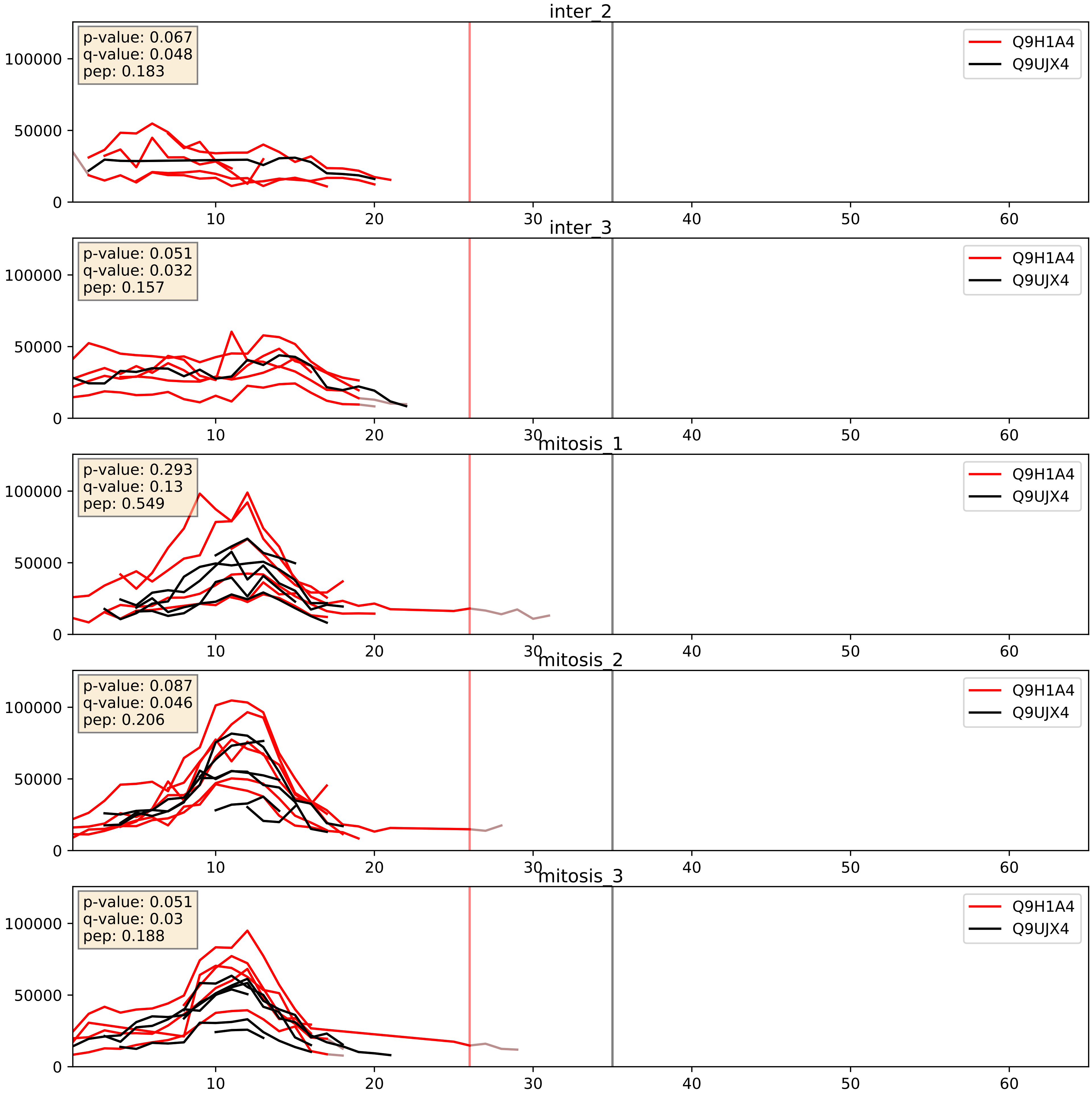


| level | condition_1 | condition_2 | pvalue | pvalue_adjusted |
|---|---|---|---|---|
| complex_abundance | mitosis | inter | 0.05869870036719124 | 0.17326237073901965 |
| interactor_abundance | mitosis | inter | 0.3361434946847083 | 0.5359263018254987 |
| interactor_ratio | mitosis | inter | 0.49268986640519663 | 0.86984863222988 |

Q9UJX2_Q9UJX4
CDC23_HUMAN vs APC5_HUMAN
p-value: 0.002 q-value: 0.002

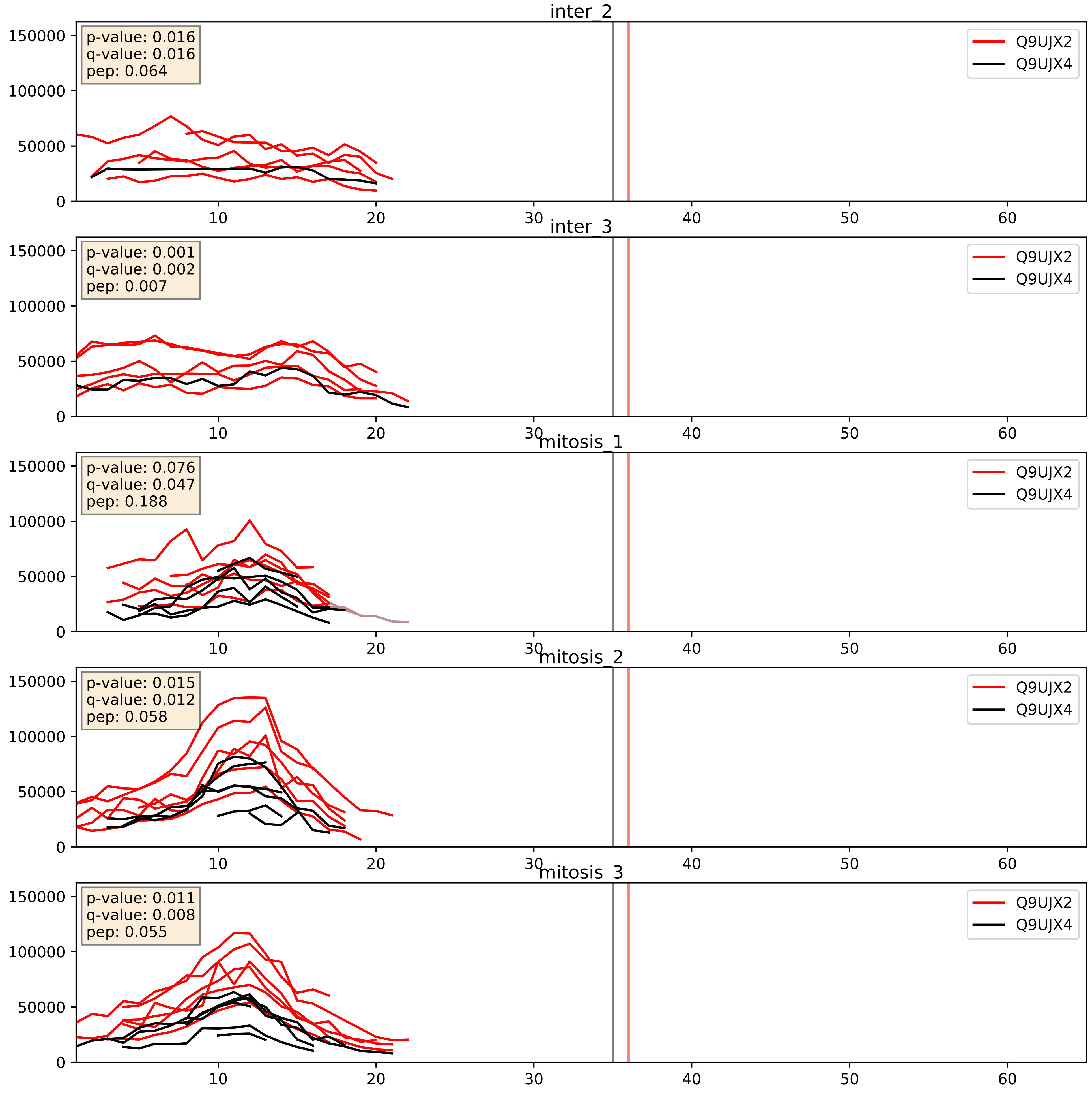


| level | condition_1 | condition_2 | pvalue | pvalue_adjusted |
|---|---|---|---|---|
| complex_abundance | mitosis | inter | 0.12139220629116379 | 0.26373535173917817 |
| interactor_abundance | mitosis | inter | 0.41298963126896665 | 0.596254215493735 |
| interactor_ratio | mitosis | inter | 0.7032938827213633 | 0.9369954297423922 |

Q9UJX3_Q9UJX4
APC7_HUMAN vs APC5_HUMAN
p-value: 0.008 q-value: 0.007

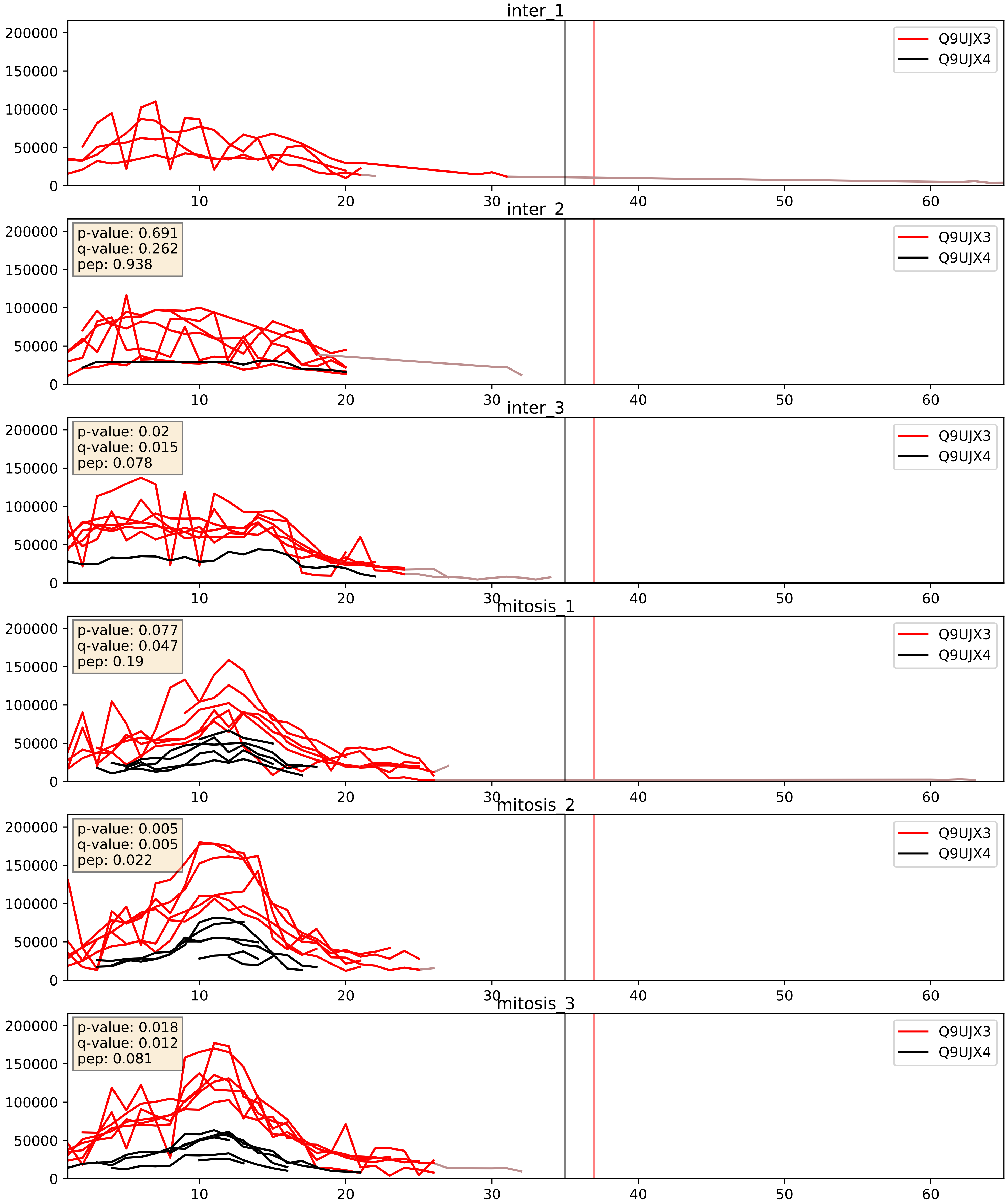


| level | condition_1 | condition_2 | pvalue | pvalue_adjusted |
|---|---|---|---|---|
| complex_abundance | mitosis | inter | 0.09492098790435125 | 0.22791687418267786 |
| interactor_abundance | mitosis | inter | 0.2616462882407935 | 0.45356262198080044 |
| interactor_ratio | mitosis | inter | 0.8161411056759111 | 0.9666192201555632 |

Q9UJX4_Q9UJX5
APC5_HUMAN vs APC4_HUMAN
p-value: 0.0 q-value: 0.001

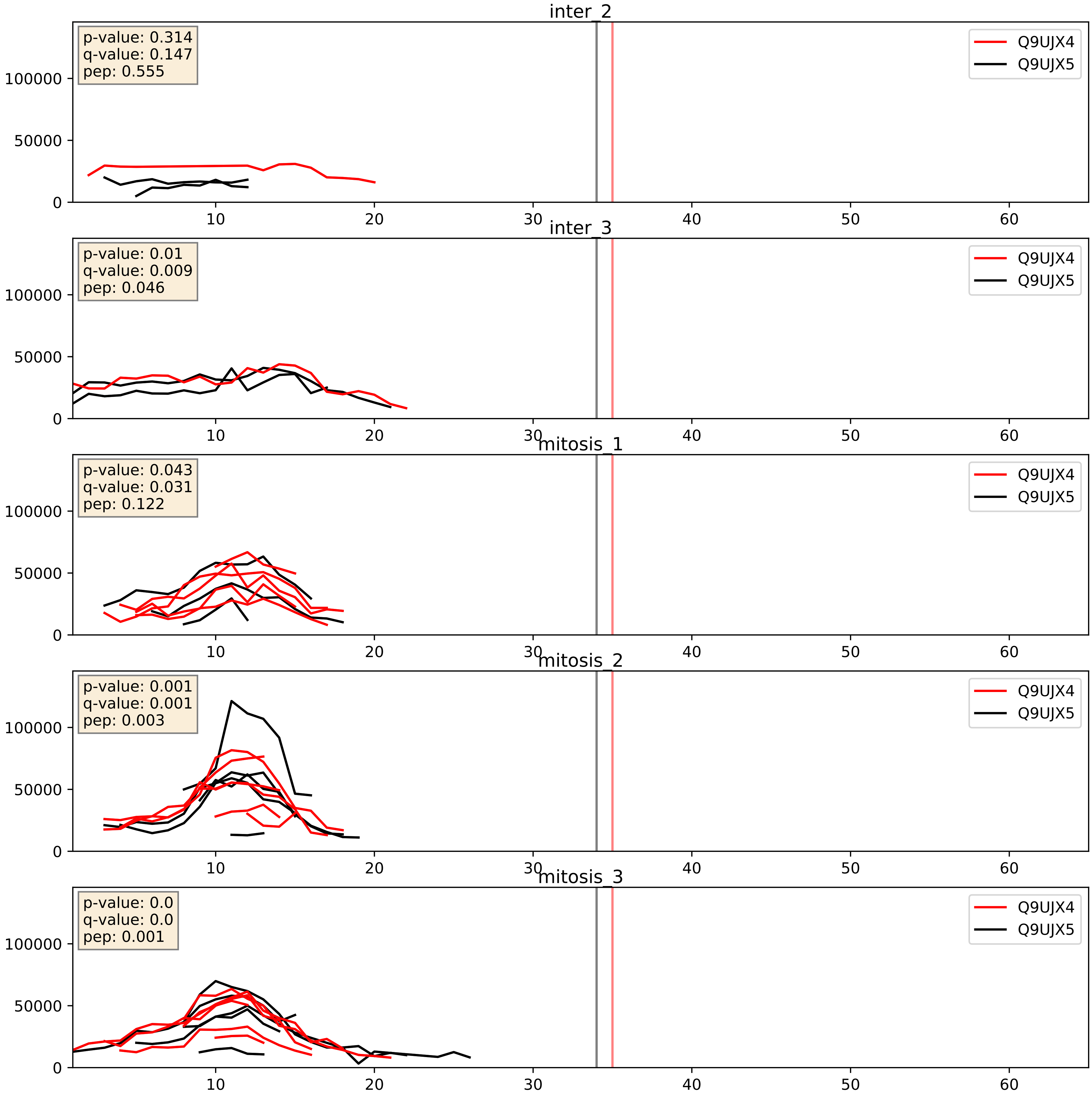


| level | condition_1 | condition_2 | pvalue | pvalue_adjusted |
|---|---|---|---|---|
| interactor_abundance | mitosis | inter | 0.0020000297974639616 | 0.03264187006238393 |
| complex_abundance | mitosis | inter | 0.01246613611220189 | 0.08296588784734041 |
| interactor_ratio | mitosis | inter | 0.05641809227912415 | 0.3926647223857623 |

Q9UJX4_Q9UJX6
APC5_HUMAN vs ANC2_HUMAN
p-value: 0.001 q-value: 0.002

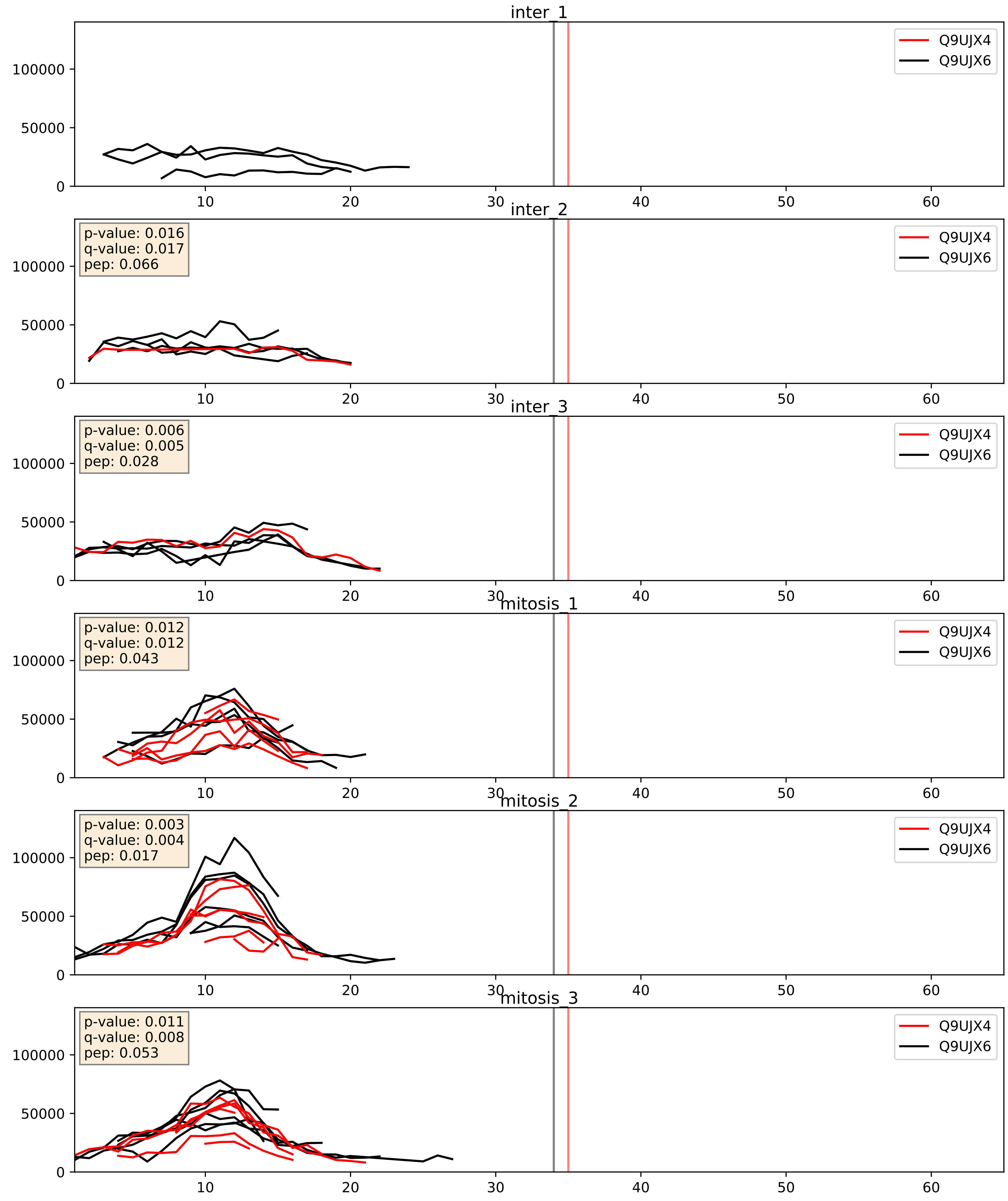


| level | condition_1 | condition_2 | pvalue | pvalue_adjusted |
|---|---|---|---|---|
| interactor_abundance | mitosis | inter | 0.0043377971196649975 | 0.052557032334511505 |
| complex_abundance | mitosis | inter | 0.06738313915890881 | 0.1874552067599153 |
| interactor_ratio | mitosis | inter | 0.42946970889461056 | 0.8432094269126428 |

Q9UJX4_Q9UM13
APC5_HUMAN vs APC10_HUMAN
p-value: 0.02 q-value: 0.016

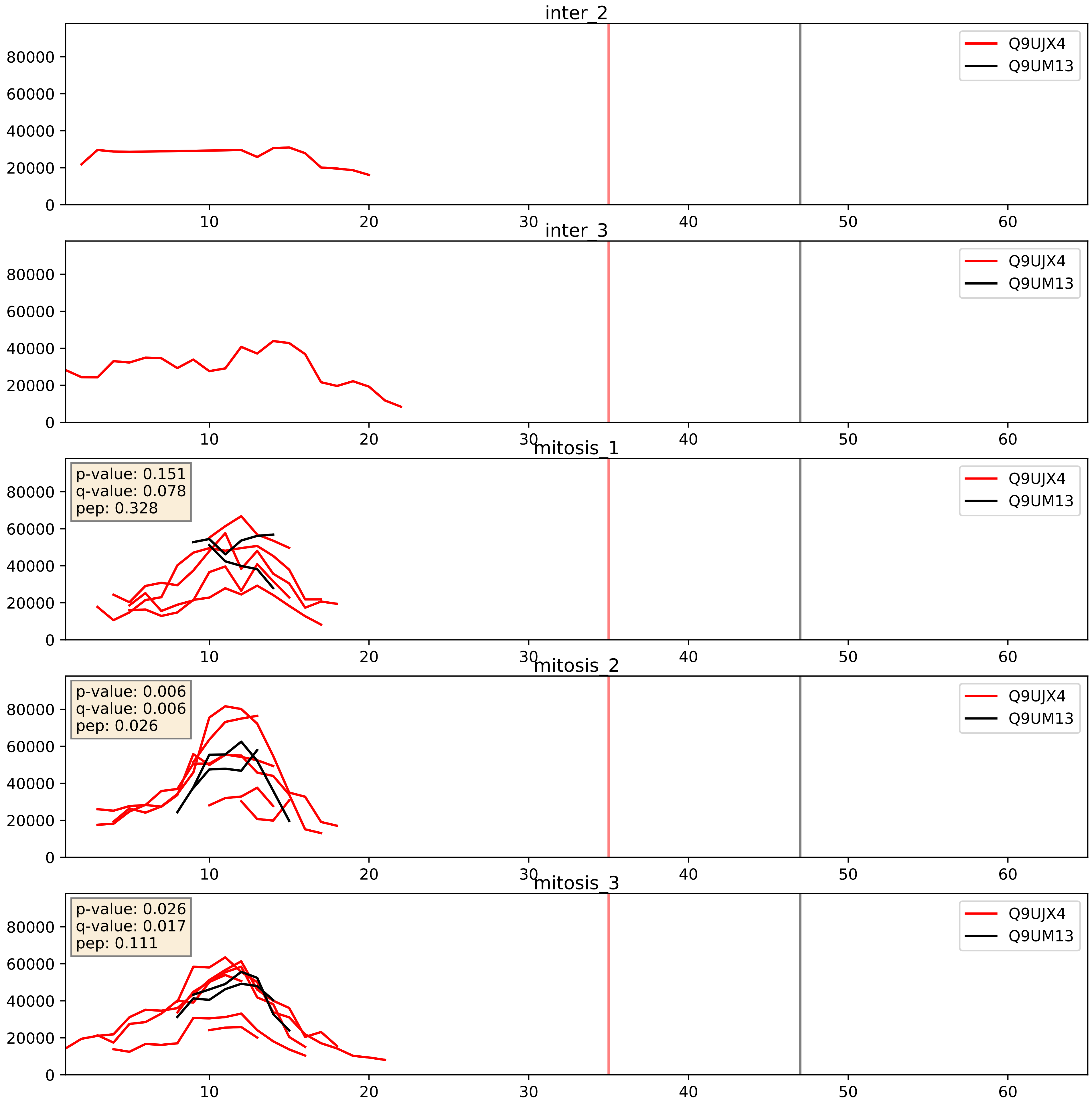


| level | condition_1 | condition_2 | pvalue | pvalue_adjusted |
|---|---|---|---|---|
| interactor_ratio | mitosis | inter | 0.004423717277729796 | 0.08831792861907116 |
| interactor_abundance | mitosis | inter | 0.005757966398448644 | 0.0644290096349286 |
| complex_abundance | mitosis | inter | 0.01019927853196216 | 0.07468419523832 |
