## Supplementary Data 2 for "SECAT: Quantifying differential protein-protein interaction states by network-centric analysis": hela_string_000141_Q15287.pdf

Q15287_Q15287

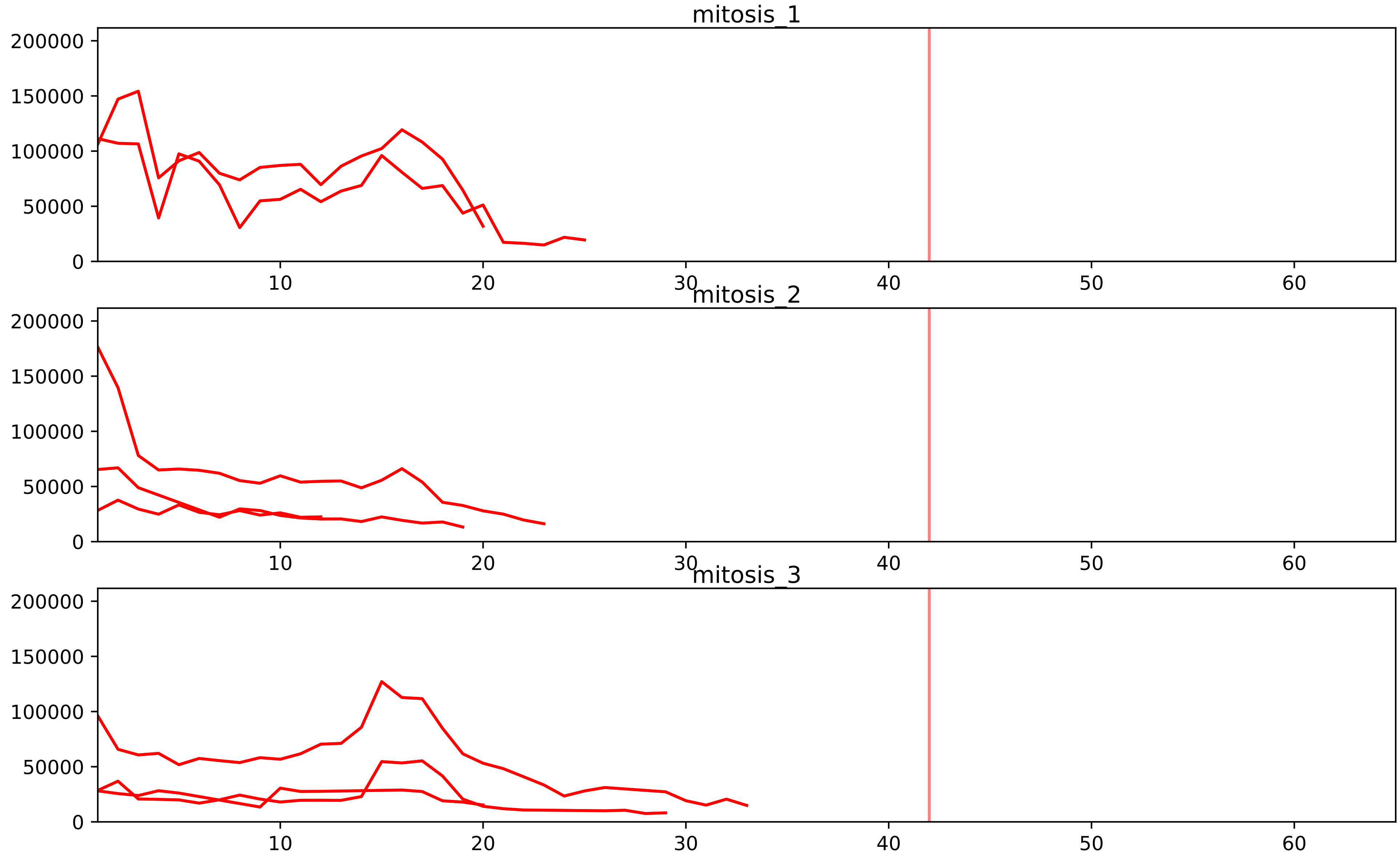


| level | condition_1 | condition_2 | pvalue | pvalue_adjusted |
|---|---|---|---|---|
| total_abundance | mitosis | inter | 5.089588603973985e-05 | 0.009965805993396753 |
| assembled_abundance | mitosis | inter | 5.427474961411258e-05 | 0.007270487507555603 |
