## Supplementary Data 2 for "SECAT: Quantifying differential protein-protein interaction states by network-centric analysis": hela_string_000142_P49790.pdf

P49790_P49790

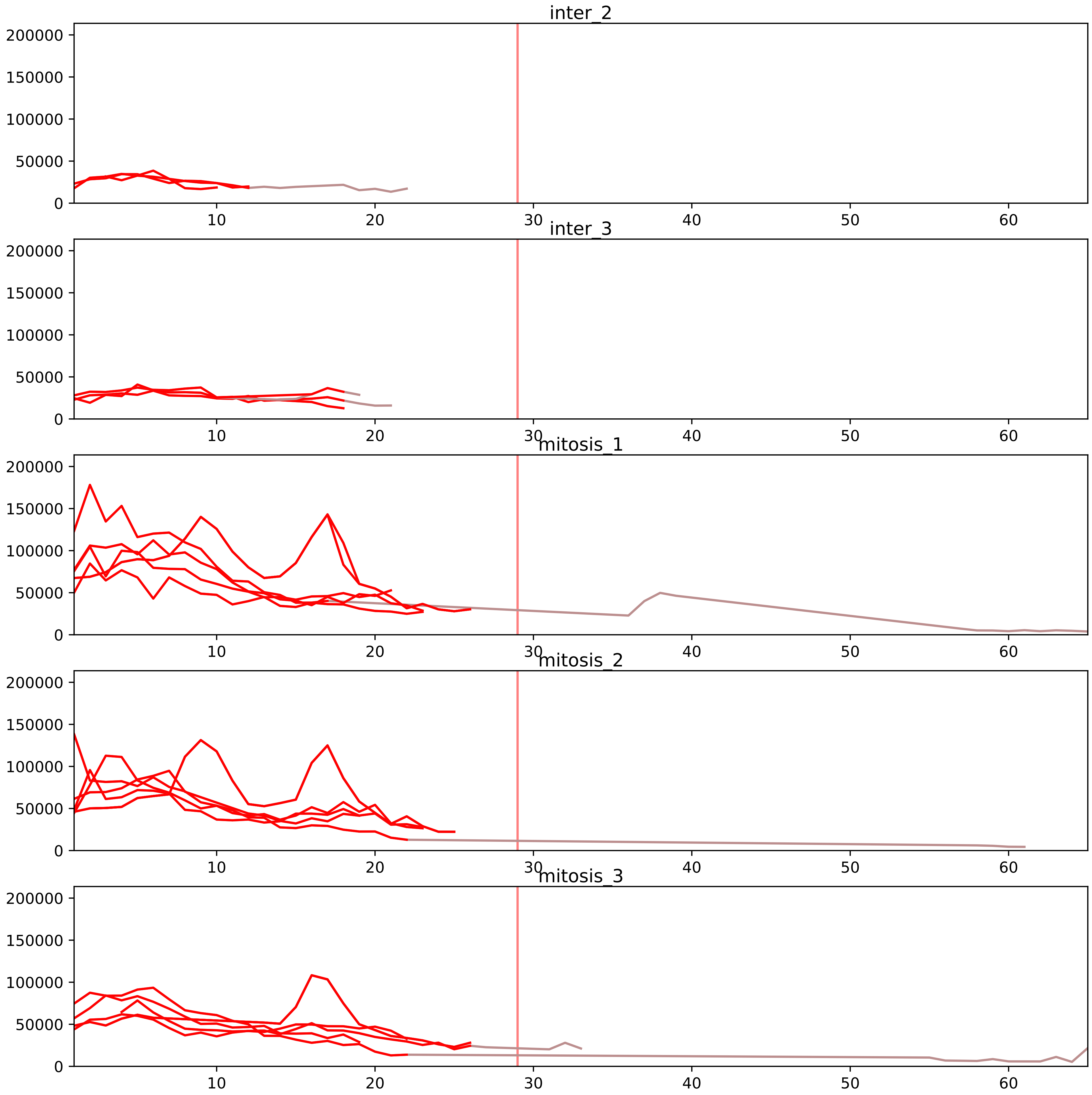


| level | condition_1 | condition_2 | pvalue | pvalue_adjusted |
|---|---|---|---|---|
| interactor_abundance | mitosis | inter | 5.100888742476215e-05 | 0.0014685477164666158 |
| complex_abundance | mitosis | inter | 0.0010140383355446271 | 0.01078514761504112 |
| monomer_abundance | mitosis | inter | 0.00674195451928637 | 0.15959869910773627 |
| total_abundance | mitosis | inter | 0.033761687191675584 | 0.2310311433694902 |
| assembled_abundance | mitosis | inter | 0.054552651938748684 | 0.2745906689853316 |
| interactor_ratio | mitosis | inter | 0.5431471676582453 | 0.7950935929783239 |

P46060_P49790
RAGP1_HUMAN vs NU153_HUMAN
p-value: 0.056 q-value: 0.035

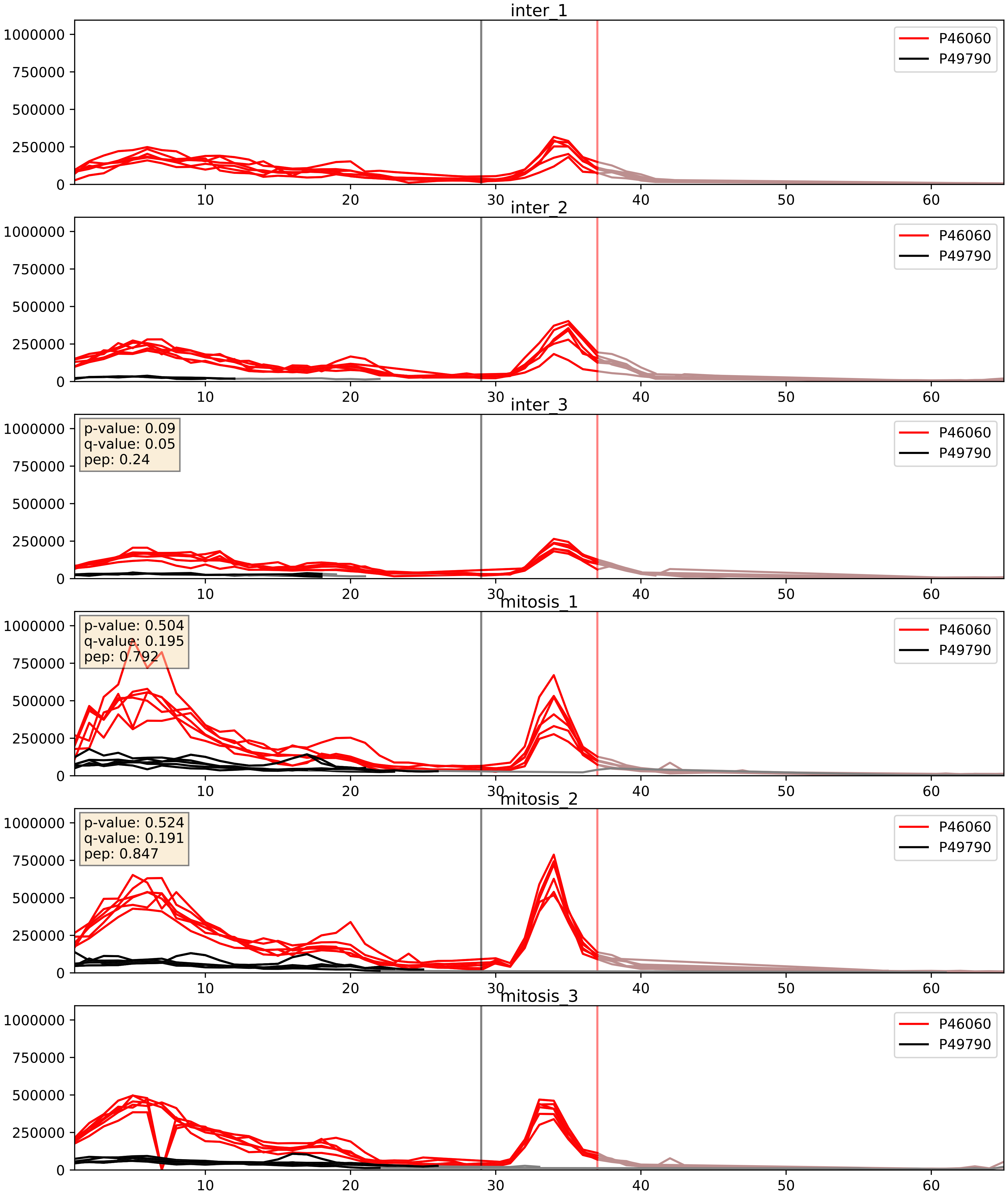


| level | condition_1 | condition_2 | pvalue | pvalue_adjusted |
|---|---|---|---|---|
| complex_abundance | mitosis | inter | 0.02027125341735831 | 0.10510110796643679 |
| interactor_abundance | mitosis | inter | 0.10382630907907112 | 0.27639658084803254 |
| interactor_ratio | mitosis | inter | 0.992216422503107 | 0.9981863229331772 |

P49790_P52948
NU153_HUMAN vs NUP98_HUMAN
p-value: 0.045 q-value: 0.03

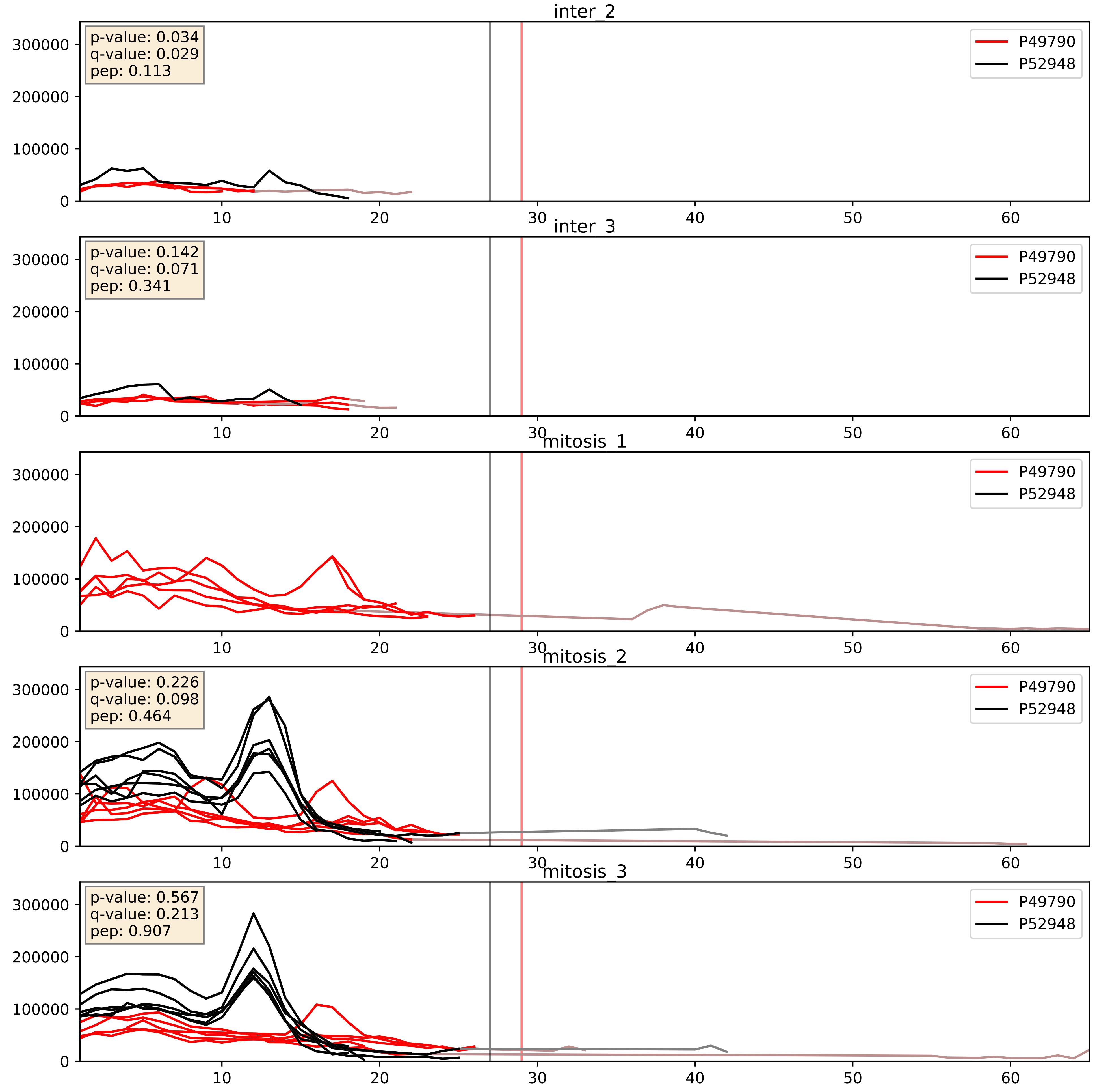


| level | condition_1 | condition_2 | pvalue | pvalue_adjusted |
|---|---|---|---|---|
| interactor_ratio | mitosis | inter | 0.15689149288799156 | 0.6136850697404735 |
| complex_abundance | mitosis | inter | 0.23611602546823163 | 0.38845416722667625 |
| interactor_abundance | mitosis | inter | 0.2573903637600927 | 0.4503344943865904 |

P49790_P57740
NU153_HUMAN vs NU107_HUMAN
p-value: 0.028 q-value: 0.02

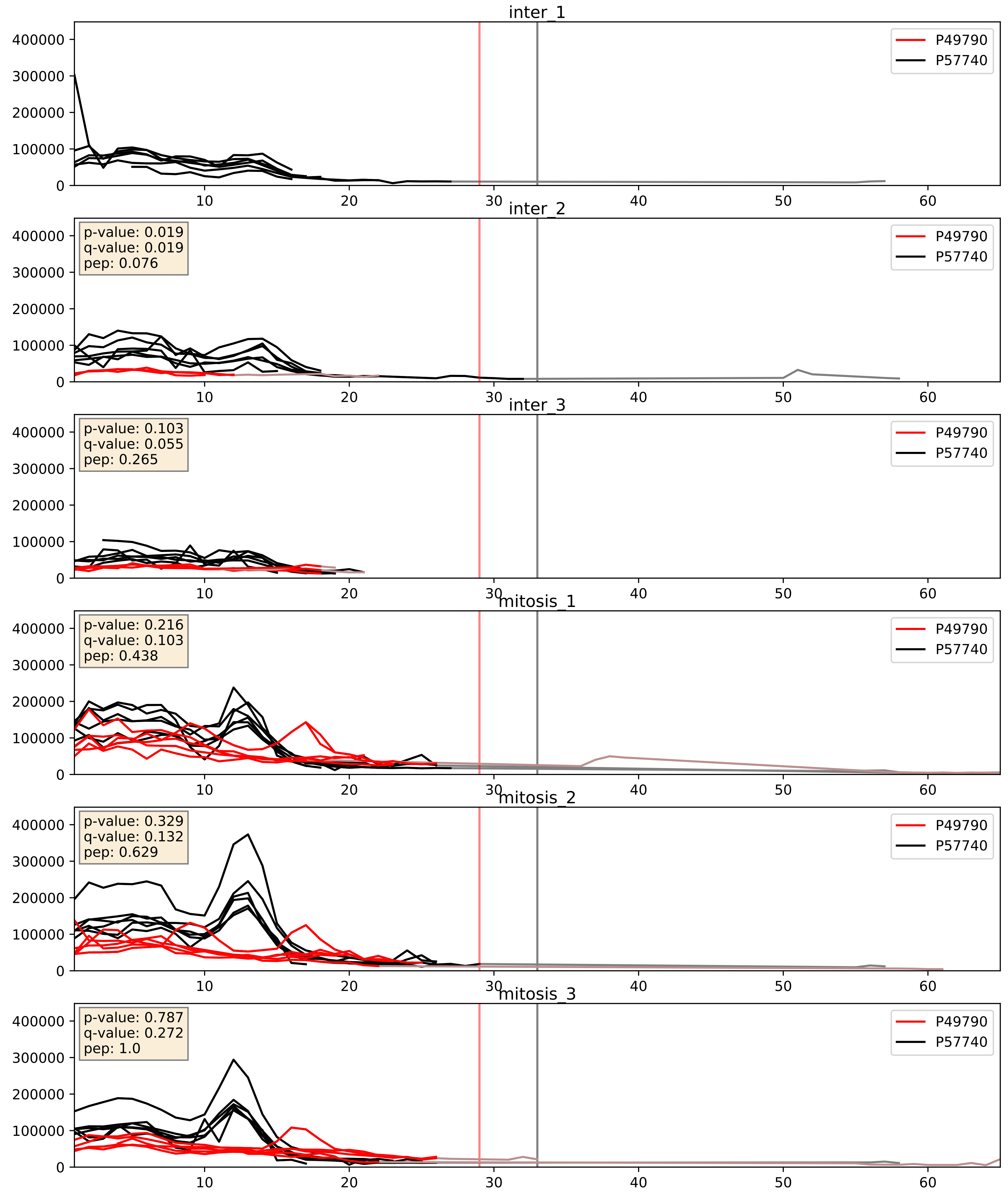


| level | condition_1 | condition_2 | pvalue | pvalue_adjusted |
|---|---|---|---|---|
| interactor_abundance | mitosis | inter | 0.0013903875369213287 | 0.02753913185123736 |
| complex_abundance | mitosis | inter | 0.009235045771373991 | 0.07106059027160733 |
| interactor_ratio | mitosis | inter | 0.5433837440418895 | 0.8859730852936224 |

P49790_Q7Z3B4
NU153_HUMAN vs NUP54_HUMAN
p-value: 0.036 q-value: 0.025

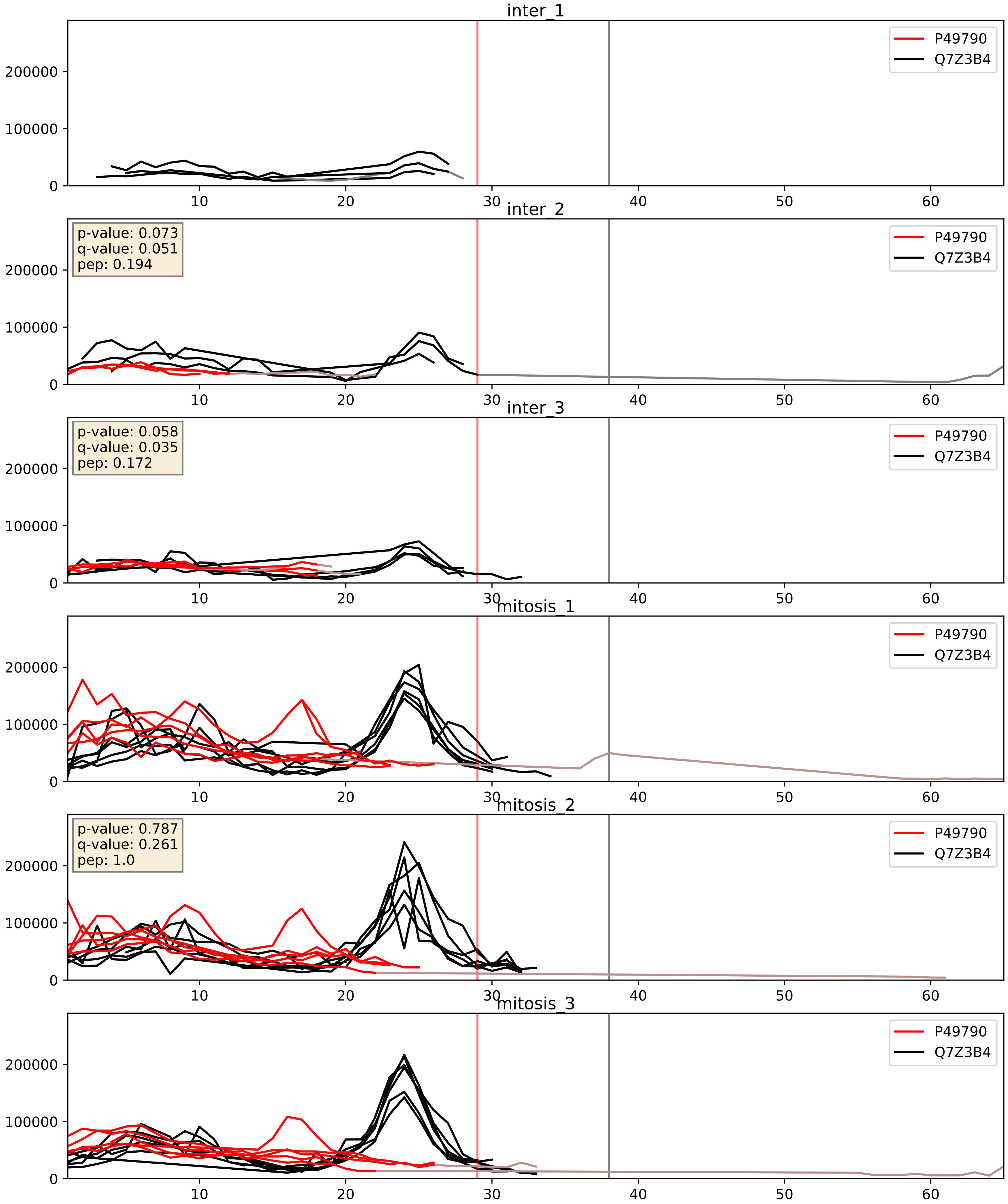


| level | condition_1 | condition_2 | pvalue | pvalue_adjusted |
|---|---|---|---|---|
| complex_abundance | mitosis | inter | 0.0010513693196996844 | 0.021612845224852777 |
| interactor_abundance | mitosis | inter | 0.0013903875369213287 | 0.02753913185123736 |
| interactor_ratio | mitosis | inter | 0.4651985580743366 | 0.8591965497319327 |

P49790_Q8TEM1
NU153_HUMAN vs PO210_HUMAN
p-value: 0.036 q-value: 0.025

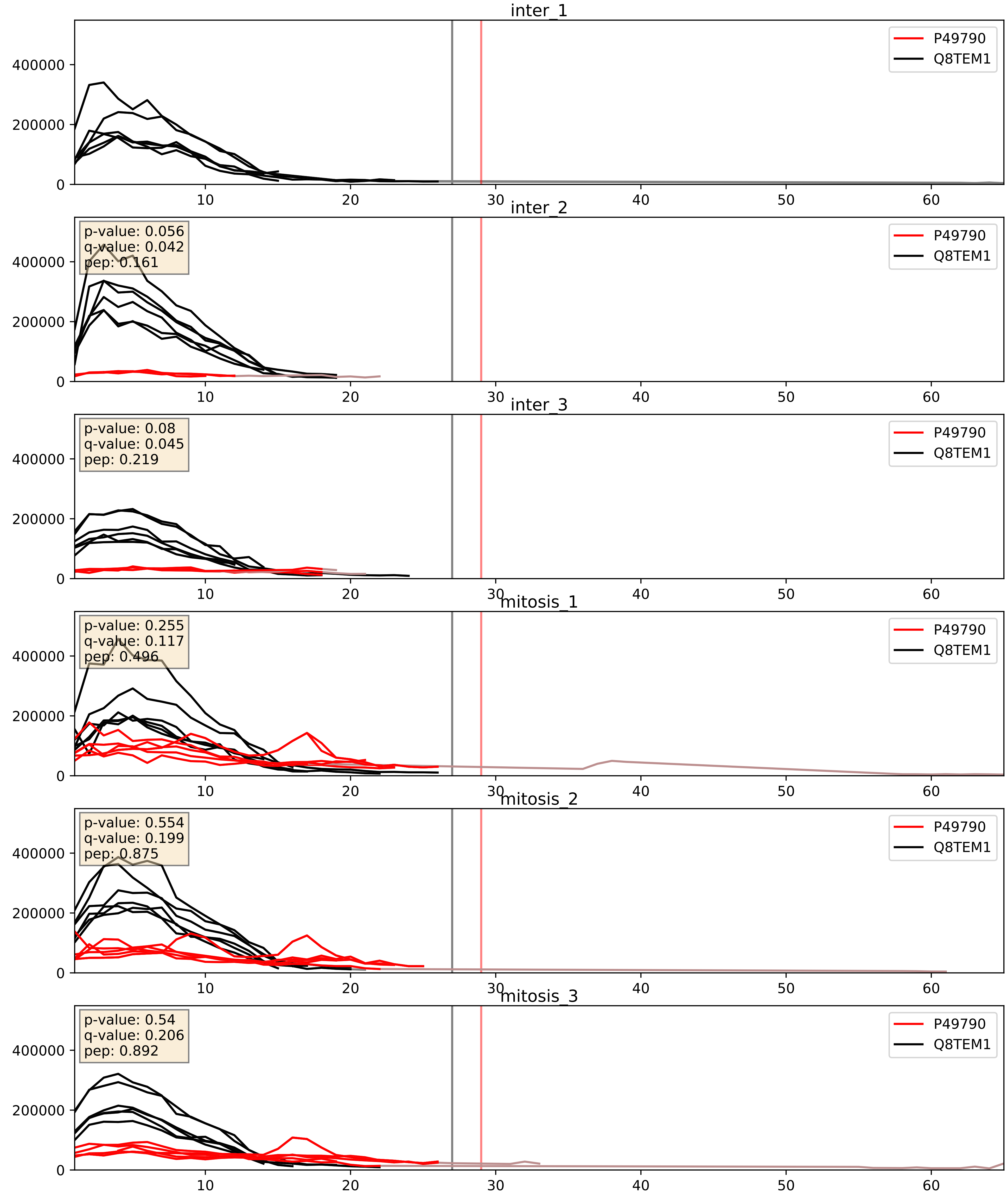


| level | condition_1 | condition_2 | pvalue | pvalue_adjusted |
|---|---|---|---|---|
| interactor_abundance | mitosis | inter | 0.0022806847342712756 | 0.03523946087610491 |
| complex_abundance | mitosis | inter | 0.052461510990486494 | 0.16341722491941935 |
| interactor_ratio | mitosis | inter | 0.40064090751427167 | 0.8305378551148518 |

P49790_Q92621
NU153_HUMAN vs NU205_HUMAN
p-value: 0.06 q-value: 0.037

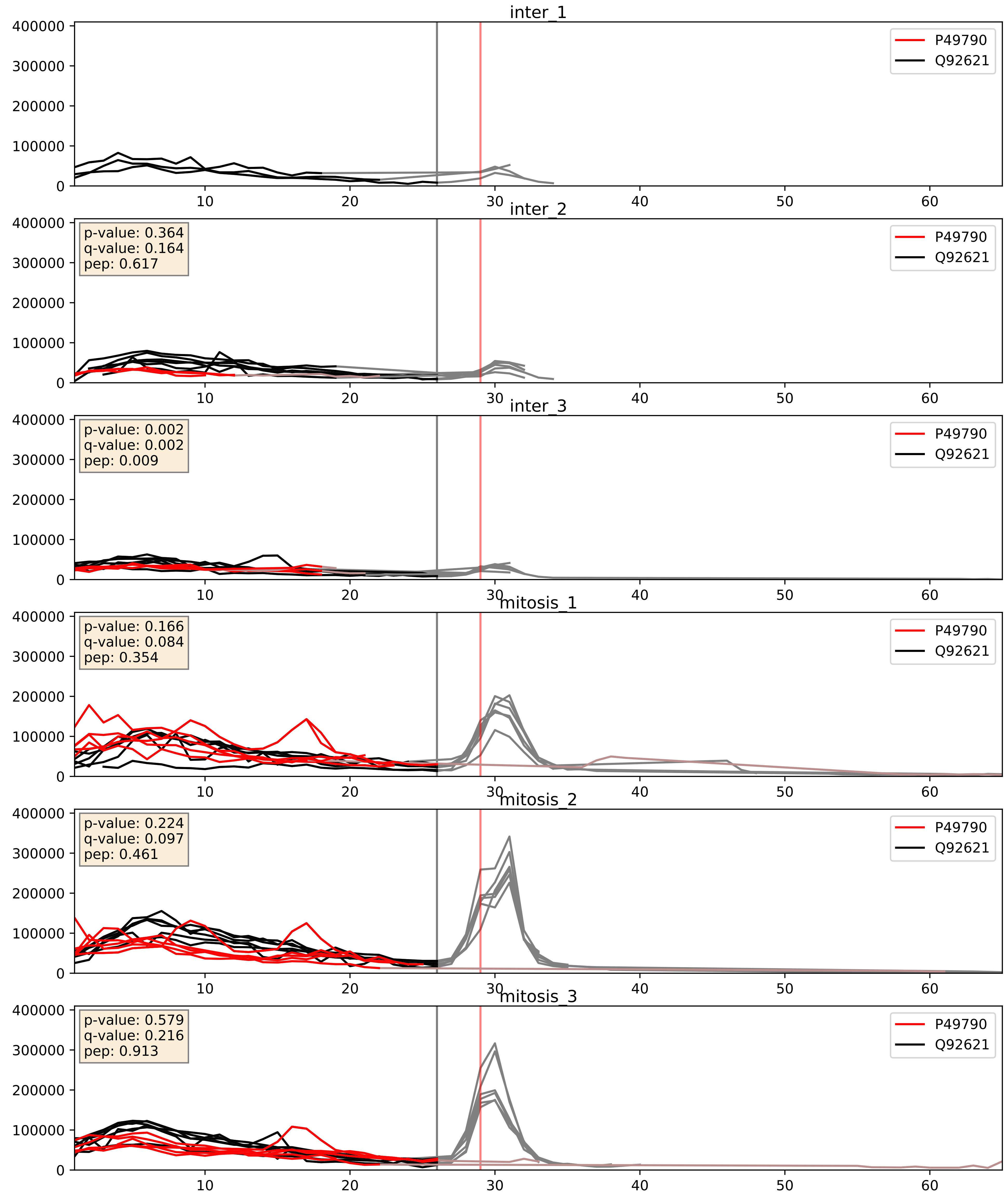


| level | condition_1 | condition_2 | pvalue | pvalue_adjusted |
|---|---|---|---|---|
| interactor_abundance | mitosis | inter | 0.0014533751808639753 | 0.028146813457456173 |
| complex_abundance | mitosis | inter | 0.004500543036216297 | 0.04990239428757967 |
| interactor_ratio | mitosis | inter | 0.31798500928032697 | 0.7783676521131252 |
