## Supplementary Data 2 for "SECAT: Quantifying differential protein-protein interaction states by network-centric analysis": hela_string_000143_P42285.pdf

P42285_P42285

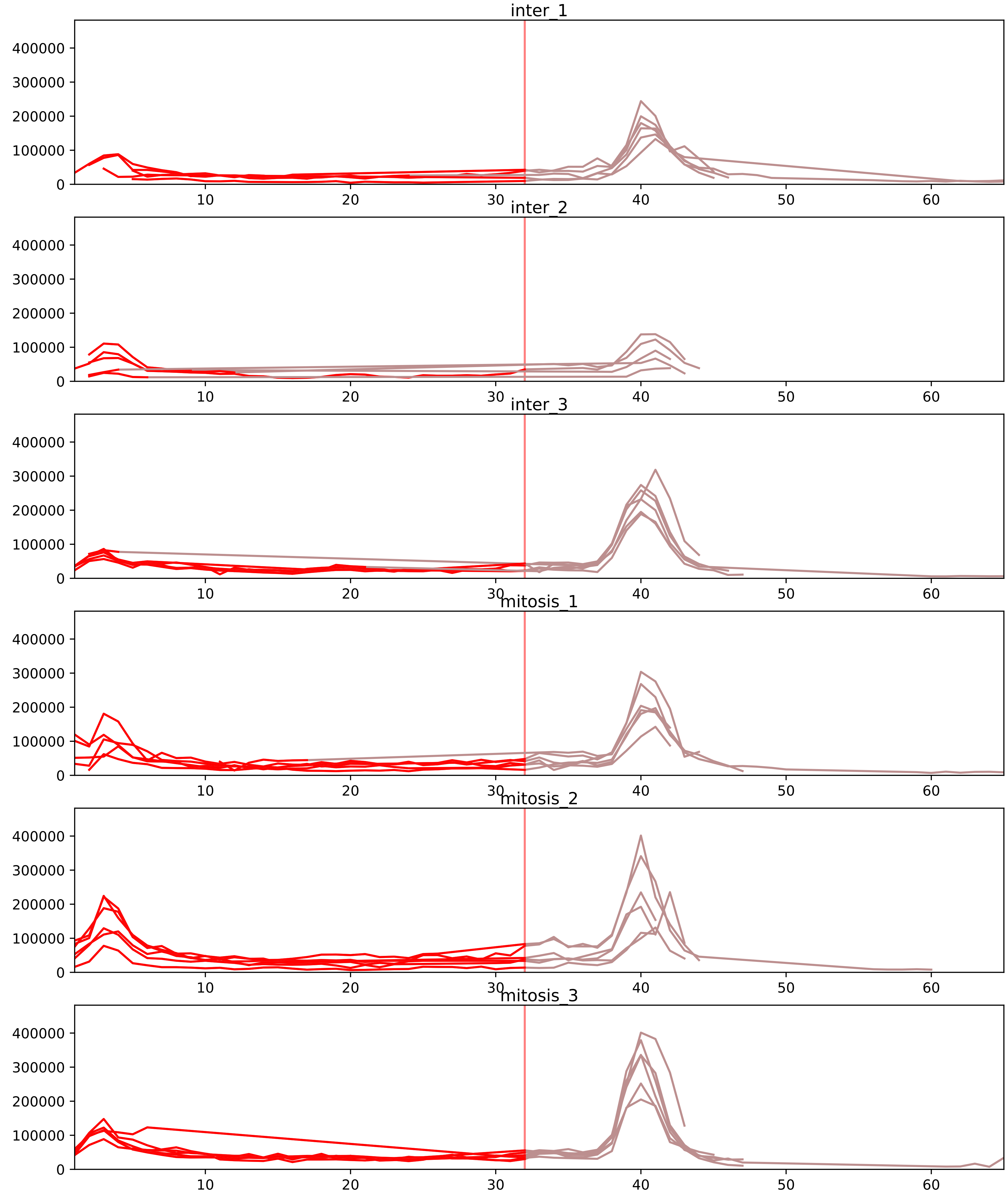


| level | condition_1 | condition_2 | pvalue | pvalue_adjusted |
|---|---|---|---|---|
| interactor_abundance | mitosis | inter | 5.1079920572751854e-05 | 0.0014685477164666158 |
| complex_abundance | mitosis | inter | 6.713374659379714e-05 | 0.0018436730407848769 |
| interactor_ratio | mitosis | inter | 0.0022352643329447168 | 0.020772153397062013 |
| assembled_abundance | mitosis | inter | 0.009599870665656396 | 0.10724819710247635 |
| total_abundance | mitosis | inter | 0.08424961145432264 | 0.35547800715590927 |
| monomer_abundance | mitosis | inter | 0.20184168010612893 | 0.5769564403452059 |

O00567_P42285
NOP56_HUMAN vs MTREX_HUMAN
p-value: 0.031 q-value: 0.023

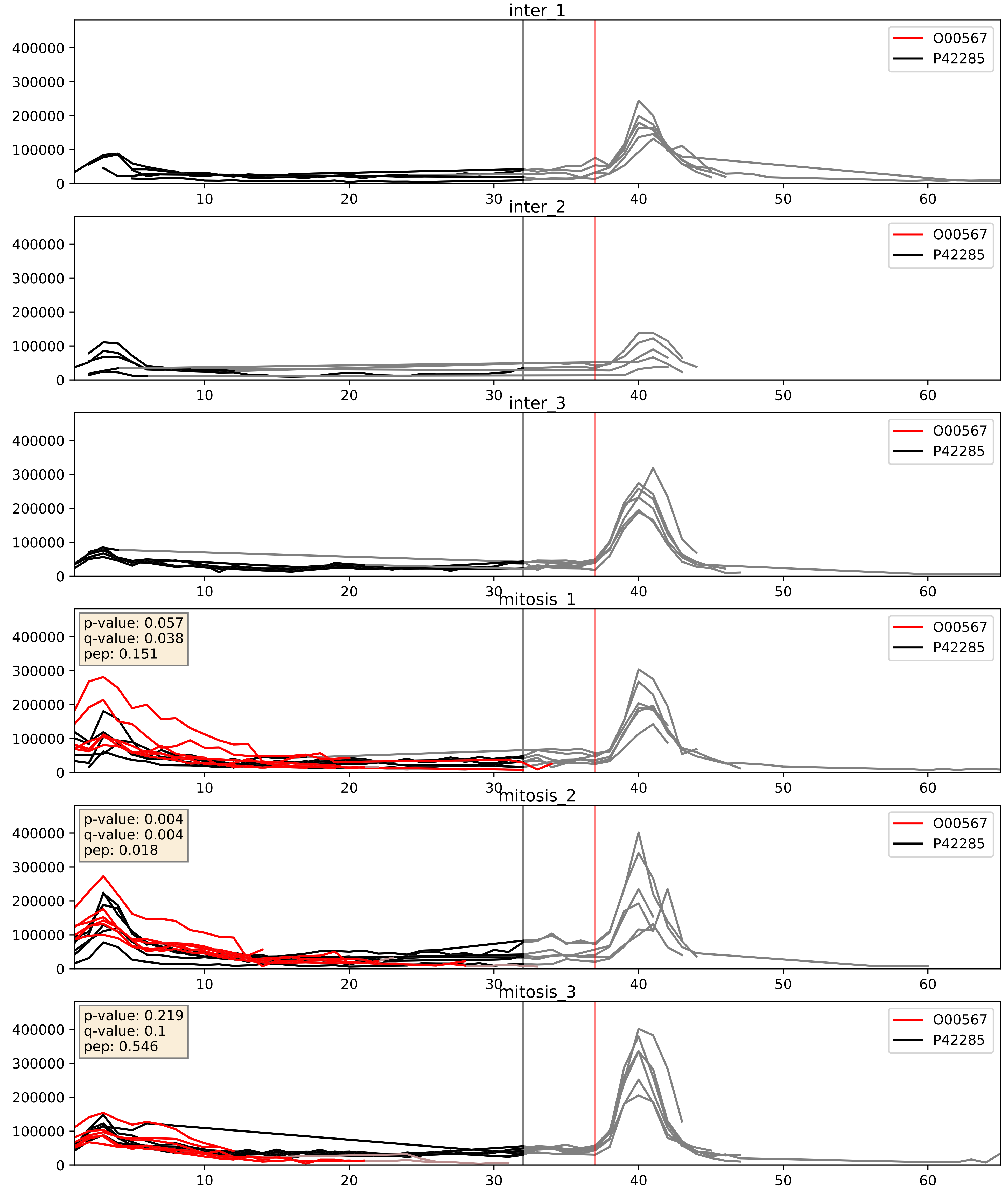


| level | condition_1 | condition_2 | pvalue | pvalue_adjusted |
|---|---|---|---|---|
| complex_abundance | mitosis | inter | 0.0005777242975888954 | 0.015600378509025063 |
| interactor_abundance | mitosis | inter | 0.0010892229270805026 | 0.024049720606162197 |
| interactor_ratio | mitosis | inter | 0.2336592312420024 | 0.7130563349132052 |

P17844_P42285
DDX5_HUMAN vs MTREX_HUMAN
p-value: 0.047 q-value: 0.031

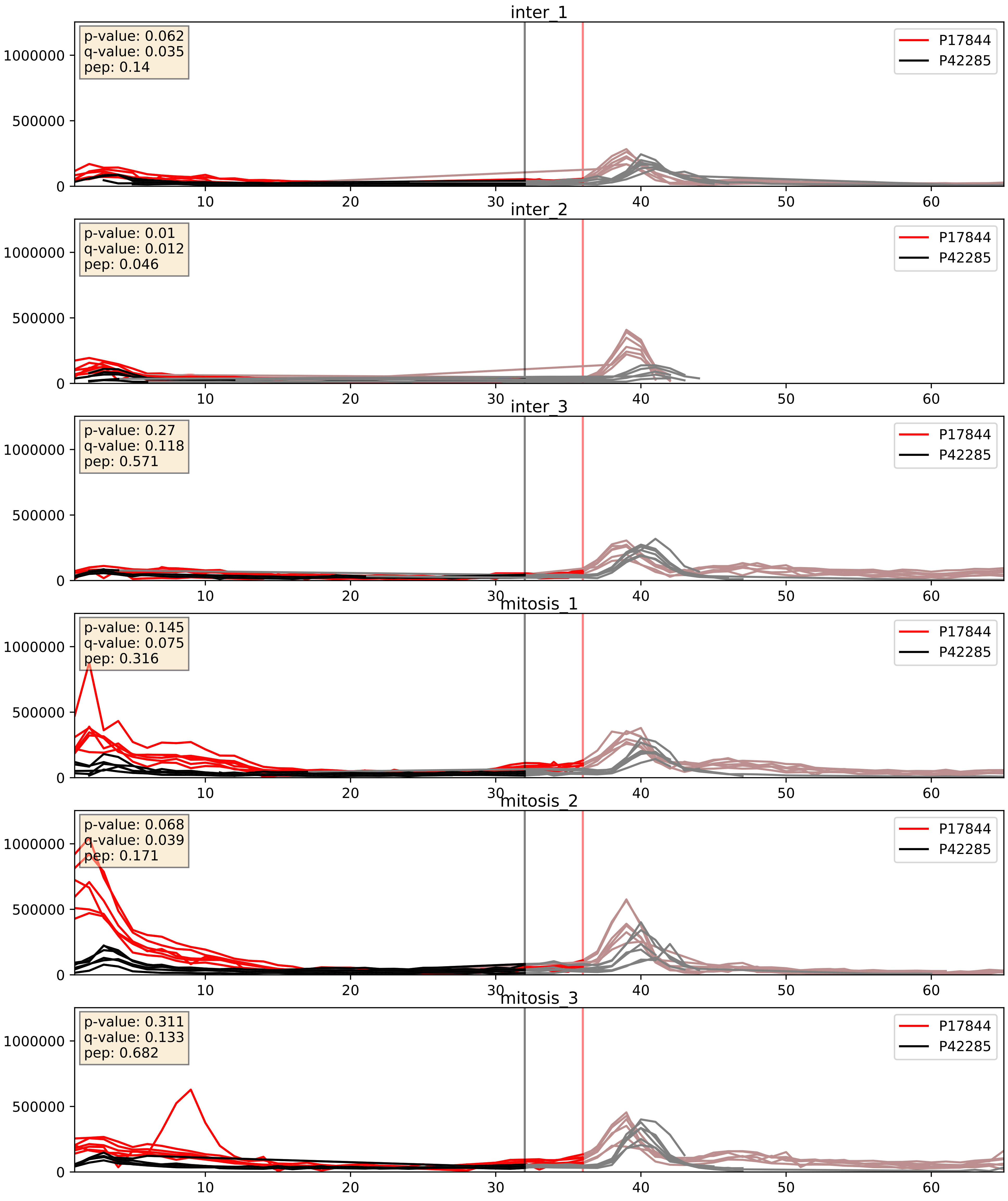


| level | condition_1 | condition_2 | pvalue | pvalue_adjusted |
|---|---|---|---|---|
| interactor_abundance | mitosis | inter | 0.0019402894813686398 | 0.032281589816356766 |
| complex_abundance | mitosis | inter | 0.0026967538514306786 | 0.03746658311311581 |
| interactor_ratio | mitosis | inter | 0.012520475632328332 | 0.1615861374955141 |

P42285_Q01780
MTREX_HUMAN vs EXOSX_HUMAN
p-value: 0.035 q-value: 0.025

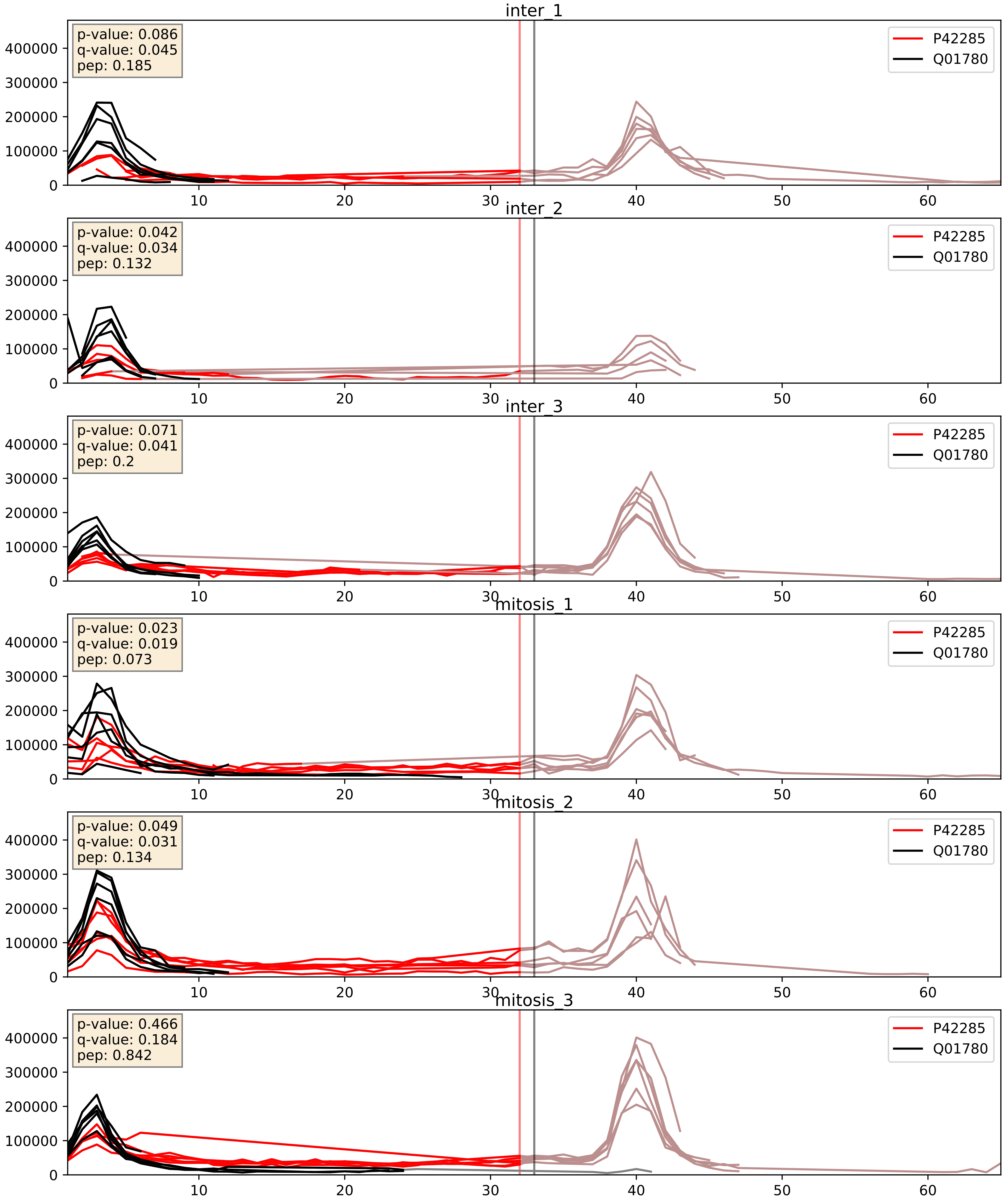


| level | condition_1 | condition_2 | pvalue | pvalue_adjusted |
|---|---|---|---|---|
| interactor_abundance | mitosis | inter | 0.0027241793218210184 | 0.03935691982242687 |
| interactor_ratio | mitosis | inter | 0.013227482029593517 | 0.16610126135854406 |
| complex_abundance | mitosis | inter | 0.04923814278230084 | 0.15816515545495635 |

P42285_Q06265
MTREX_HUMAN vs EXOS9_HUMAN
p-value: 0.042 q-value: 0.029

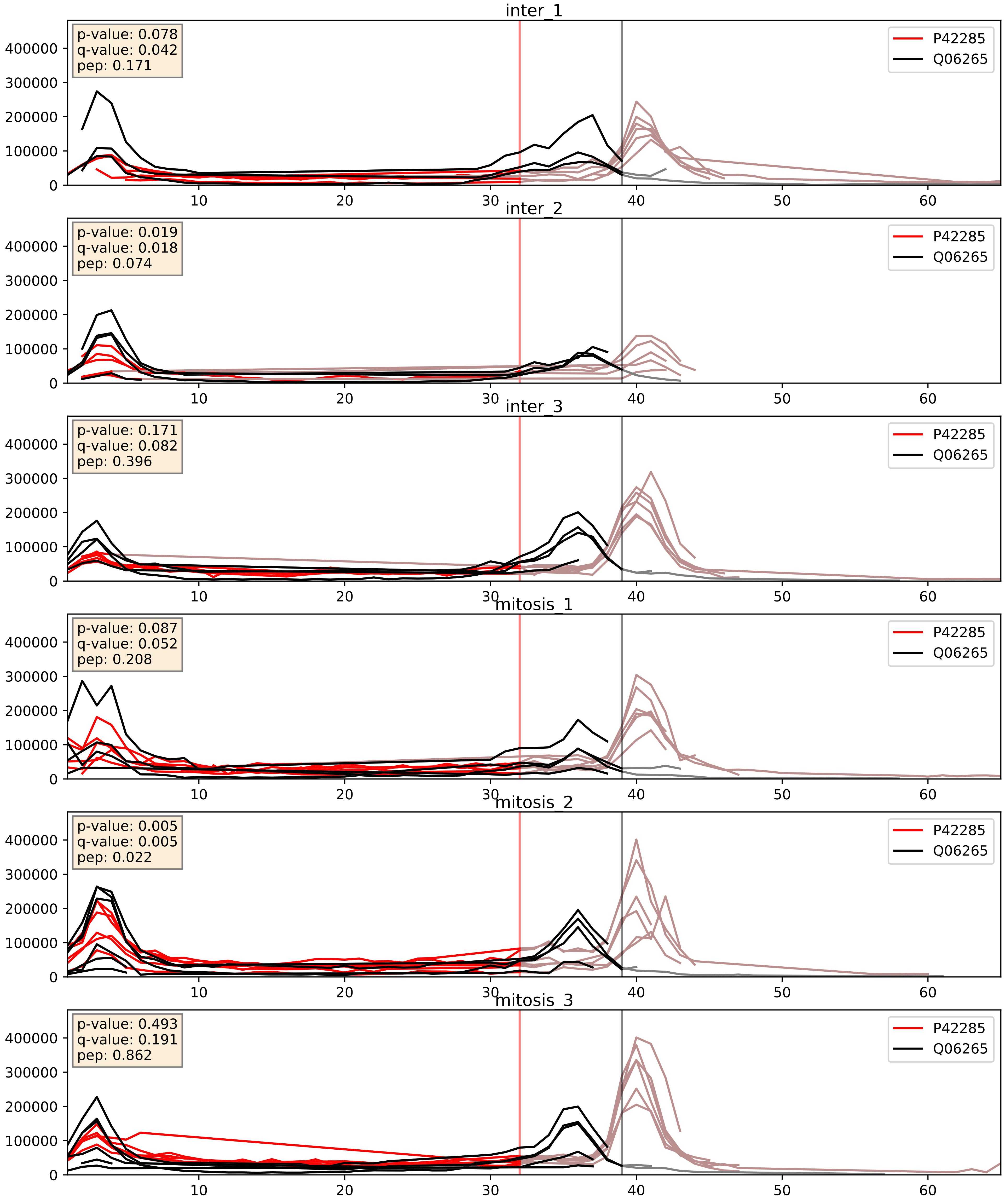


| level | condition_1 | condition_2 | pvalue | pvalue_adjusted |
|---|---|---|---|---|
| complex_abundance | mitosis | inter | 0.003363070968619561 | 0.04258563238370332 |
| interactor_abundance | mitosis | inter | 0.004752087633392944 | 0.056379584396179634 |
| interactor_ratio | mitosis | inter | 0.016199414401704425 | 0.18789564671895648 |

P42285_Q12788
MTREX_HUMAN vs TBL3_HUMAN
p-value: 0.029 q-value: 0.021

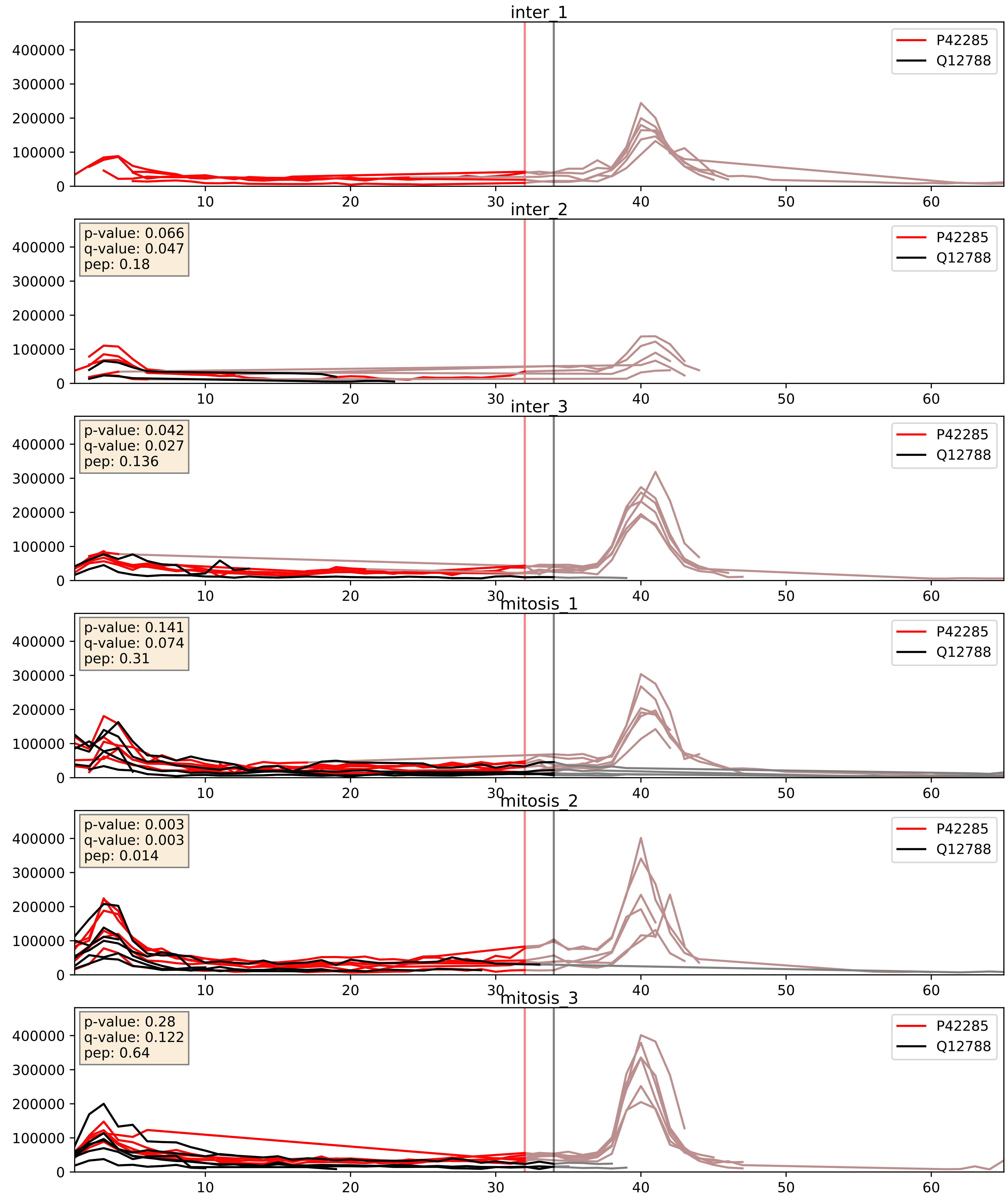


| level | condition_1 | condition_2 | pvalue | pvalue_adjusted |
|---|---|---|---|---|
| complex_abundance | mitosis | inter | 0.01601492374058697 | 0.09447811662262195 |
| interactor_abundance | mitosis | inter | 0.04149089666829527 | 0.1637021393260352 |
| interactor_ratio | mitosis | inter | 0.9340665467242752 | 0.9881103513214468 |

P42285_Q15024
MTREX_HUMAN vs EXOS7_HUMAN
p-value: 0.072 q-value: 0.044

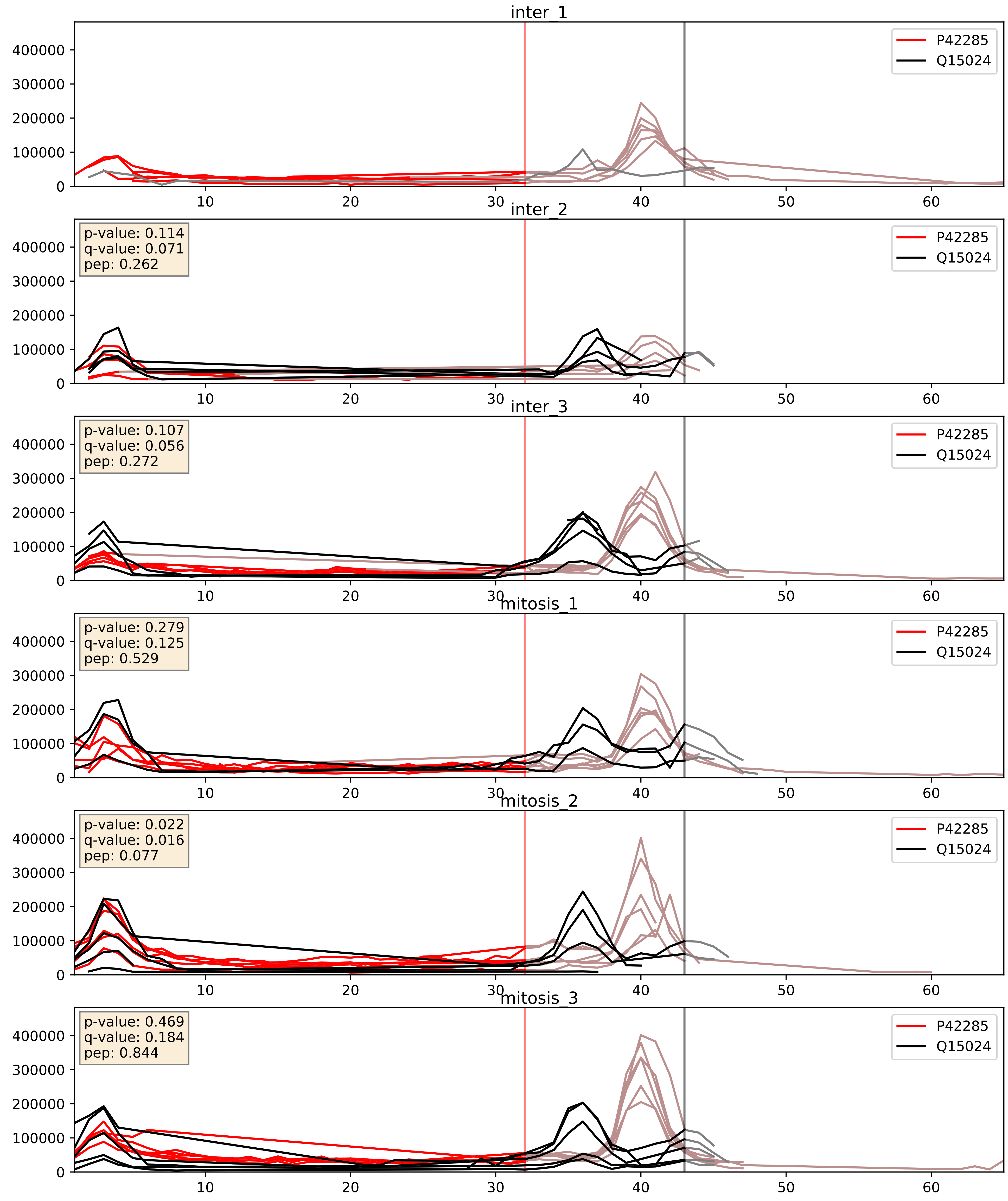


| level | condition_1 | condition_2 | pvalue | pvalue_adjusted |
|---|---|---|---|---|
| interactor_abundance | mitosis | inter | 0.04029663812757574 | 0.16188554627742371 |
| interactor_ratio | mitosis | inter | 0.04221950426886838 | 0.3399226239910554 |
| complex_abundance | mitosis | inter | 0.11976153335577917 | 0.261720379250822 |

P42285_Q15269
MTREX_HUMAN vs PWP2_HUMAN
p-value: 0.08 q-value: 0.048

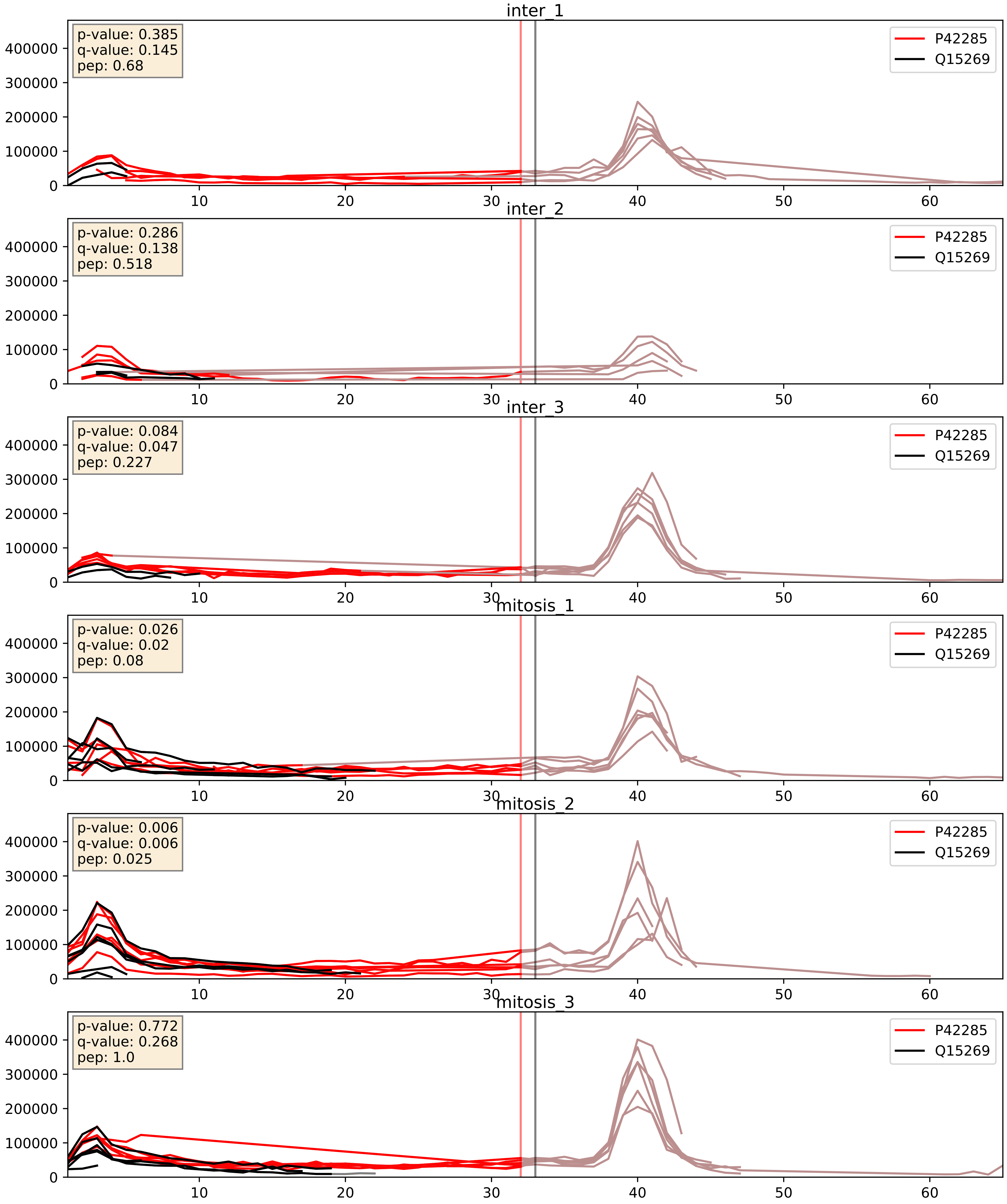


| level | condition_1 | condition_2 | pvalue | pvalue_adjusted |
|---|---|---|---|---|
| interactor_abundance | mitosis | inter | 0.0023540448297948925 | 0.036079899271341594 |
| complex_abundance | mitosis | inter | 0.005088813183140754 | 0.053382648097653 |
| interactor_ratio | mitosis | inter | 0.5603810643672665 | 0.8918094730984788 |

P42285_Q5RKV6
MTREX_HUMAN vs EXOS6_HUMAN
p-value: 0.028 q-value: 0.02

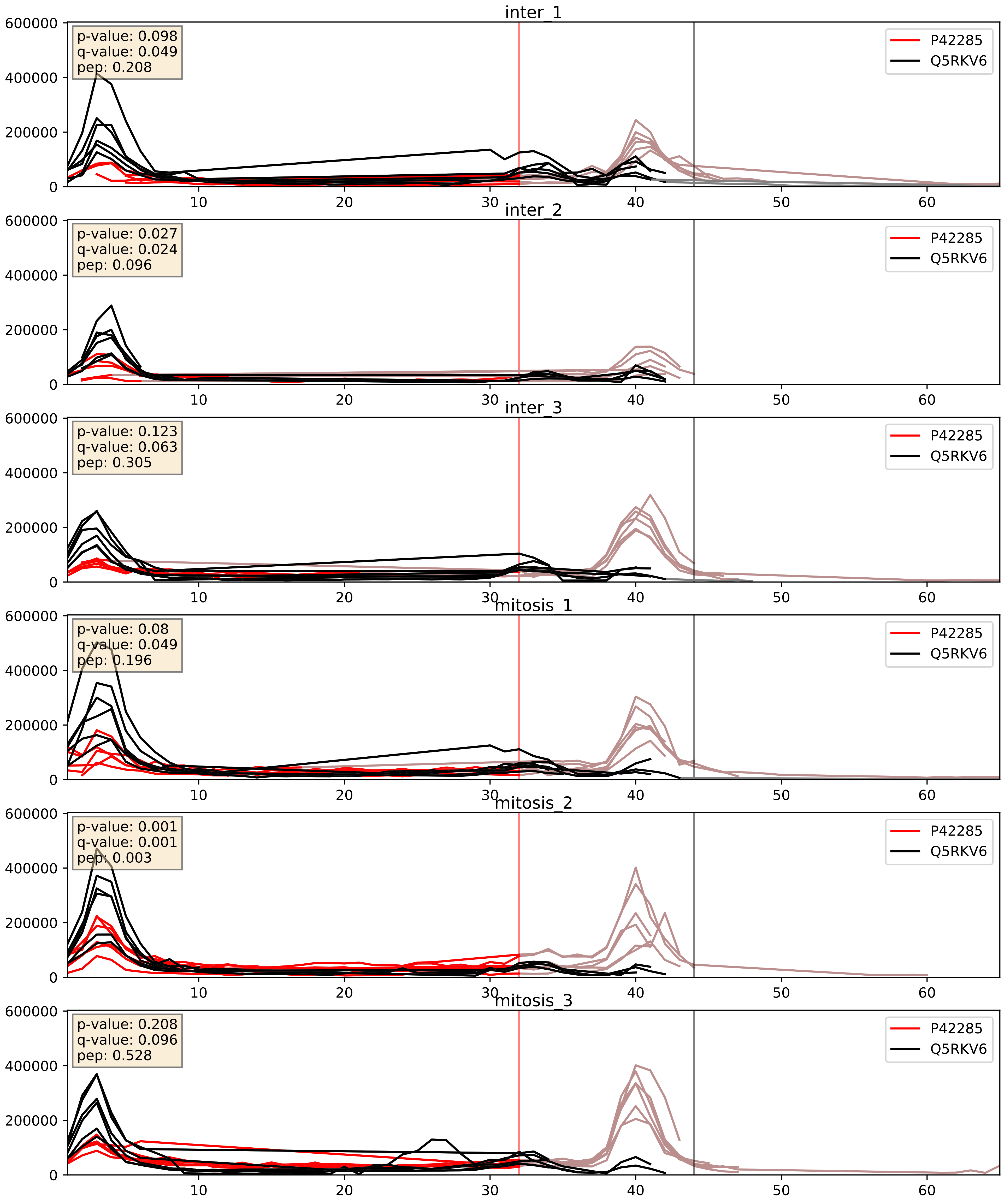


| level | condition_1 | condition_2 | pvalue | pvalue_adjusted |
|---|---|---|---|---|
| interactor_abundance | mitosis | inter | 0.006473415695686758 | 0.0687067394609051 |
| complex_abundance | mitosis | inter | 0.013240175400162837 | 0.08527908308908494 |
| interactor_ratio | mitosis | inter | 0.2156967647918259 | 0.6924580509898395 |

P42285_Q8NI36
MTREX_HUMAN vs WDR36_HUMAN
p-value: 0.043 q-value: 0.029

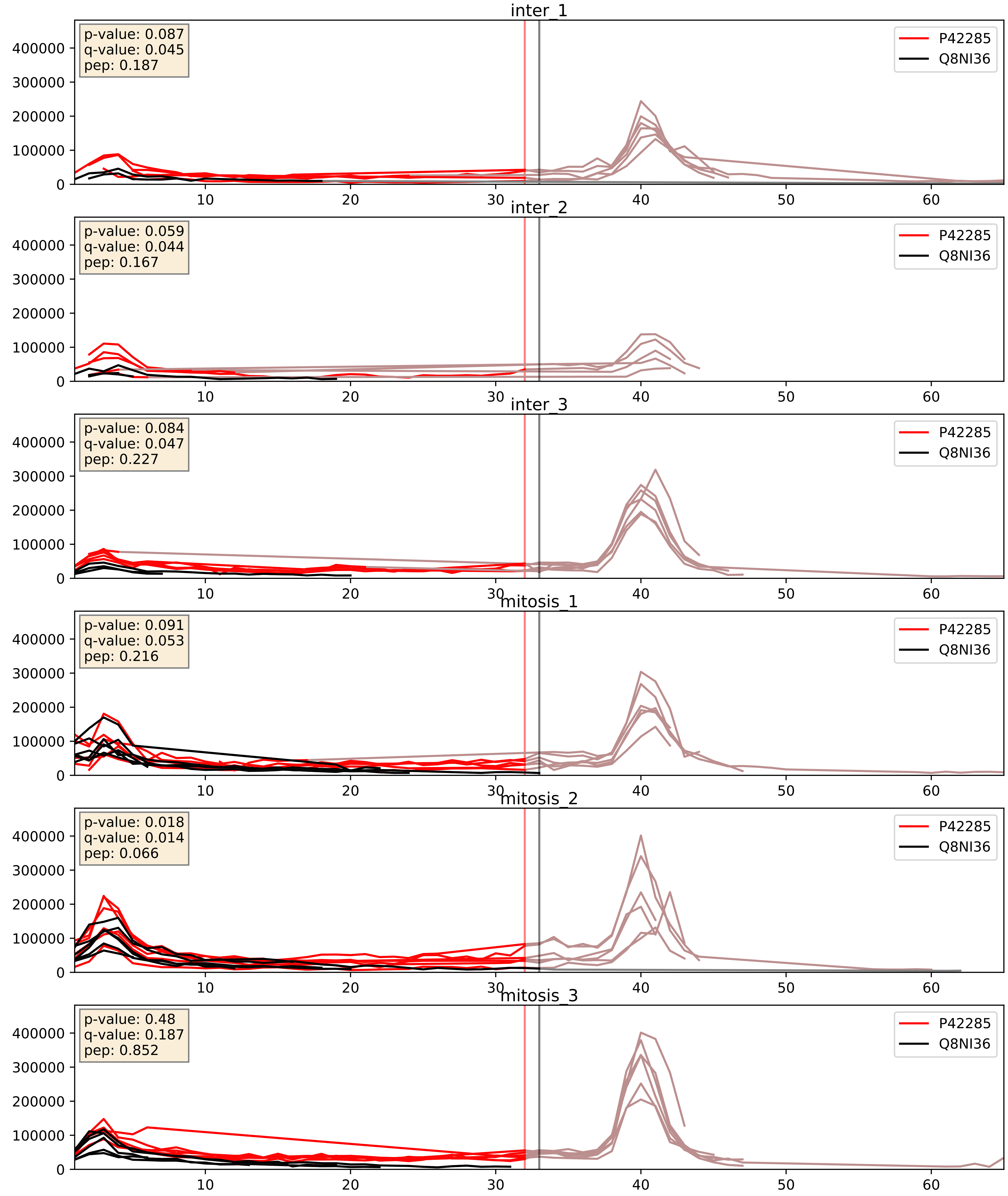


| level | condition_1 | condition_2 | pvalue | pvalue_adjusted |
|---|---|---|---|---|
| complex_abundance | mitosis | inter | 6.003680961344212e-05 | 0.005691857151652958 |
| interactor_abundance | mitosis | inter | 0.0011713522879652522 | 0.0247880731396355 |
| interactor_ratio | mitosis | inter | 0.5585467090945081 | 0.8918094730984788 |

P42285_Q96B26
MTREX_HUMAN vs EXOS8_HUMAN
p-value: 0.023 q-value: 0.018

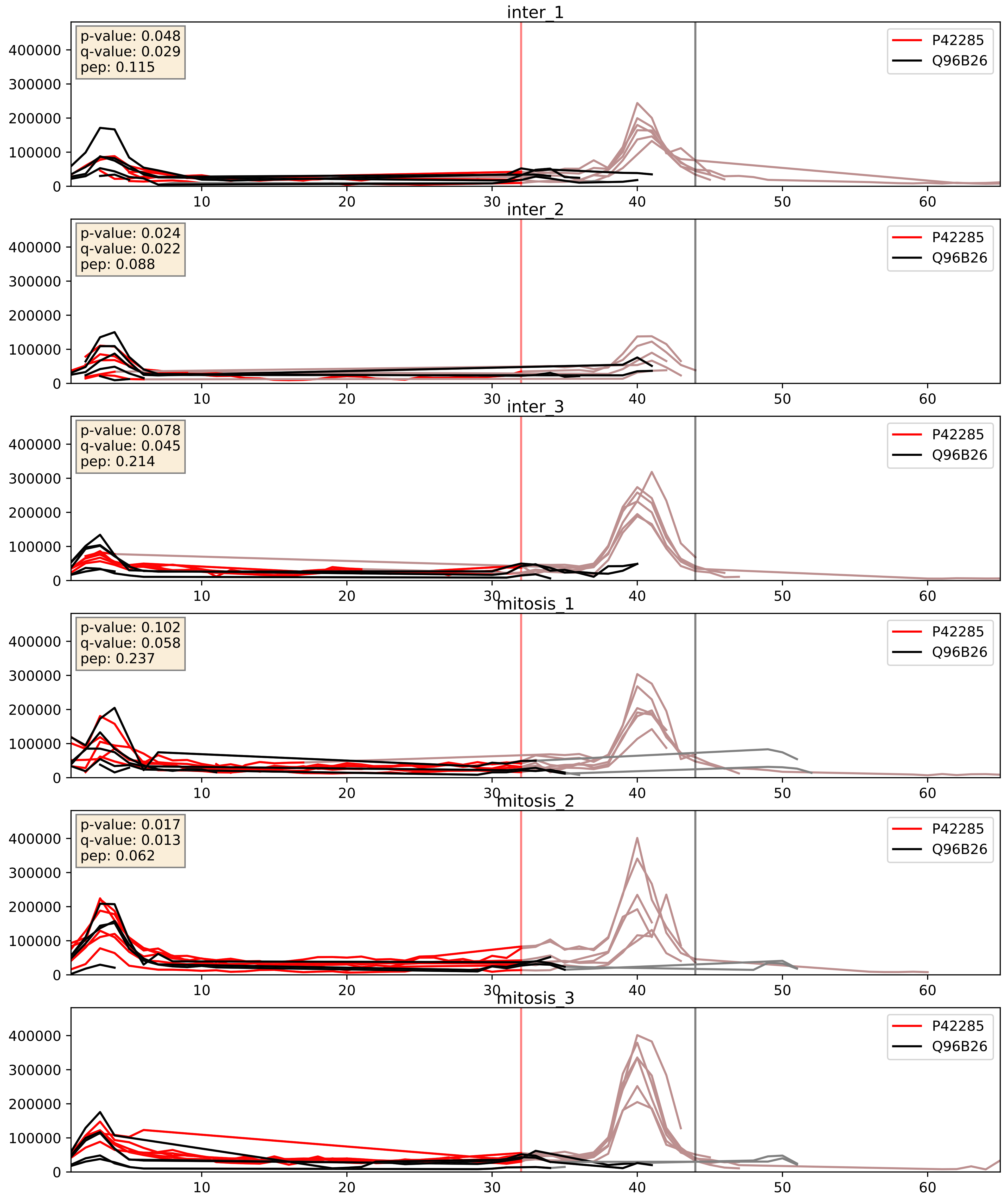


| level | condition_1 | condition_2 | pvalue | pvalue_adjusted |
|---|---|---|---|---|
| interactor_ratio | mitosis | inter | 0.008075244075749367 | 0.1269414661815779 |
| interactor_abundance | mitosis | inter | 0.024618987127700397 | 0.12960549188998488 |
| complex_abundance | mitosis | inter | 0.14610027982522583 | 0.29192772999624955 |

P42285_Q9H583
MTREX_HUMAN vs HEAT1_HUMAN
p-value: 0.032 q-value: 0.023

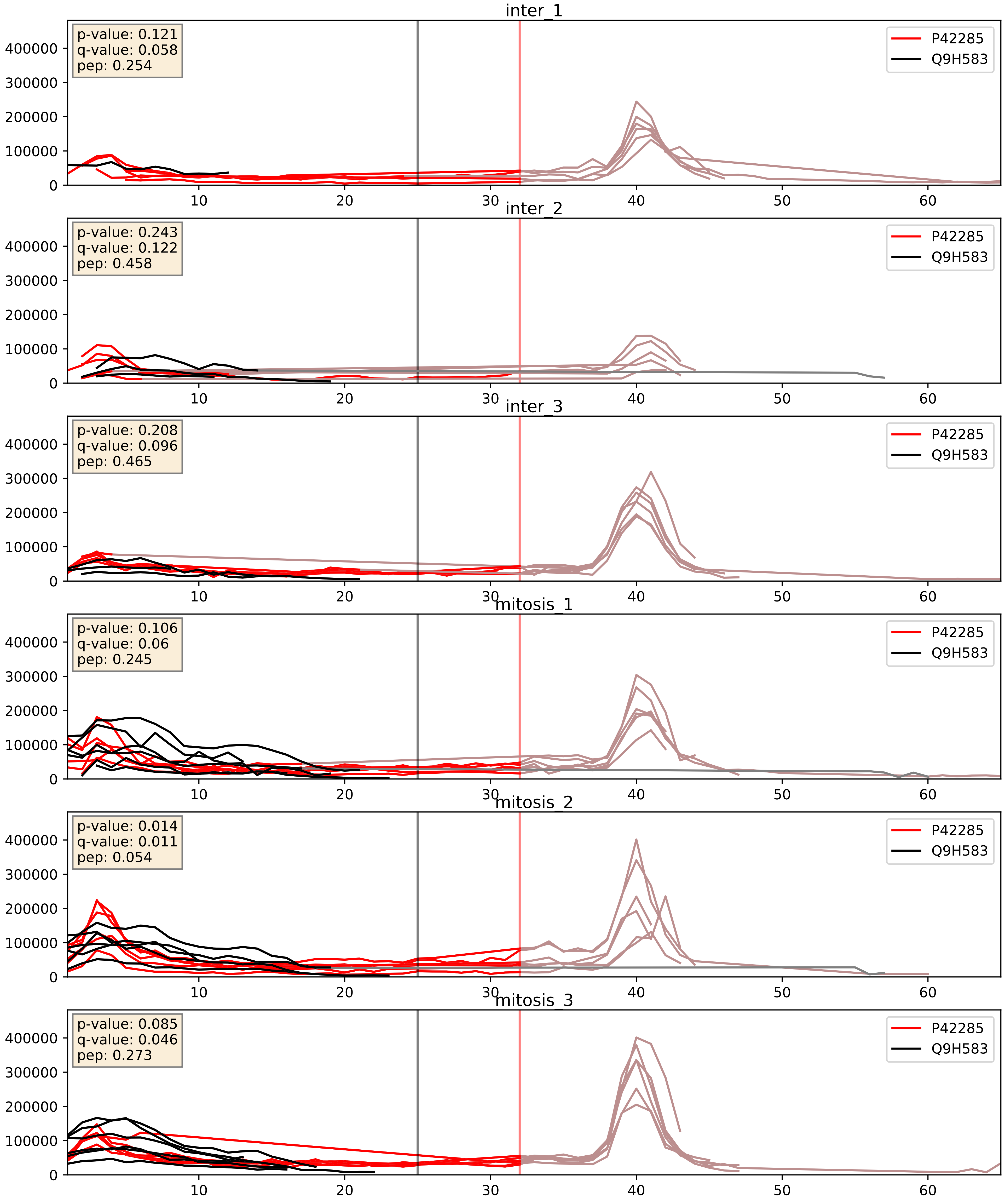


| level | condition_1 | condition_2 | pvalue | pvalue_adjusted |
|---|---|---|---|---|
| complex_abundance | mitosis | inter | 0.00024714820281548363 | 0.010525316498012636 |
| interactor_abundance | mitosis | inter | 0.00721018643363647 | 0.07343152394042615 |
| interactor_ratio | mitosis | inter | 0.11491151406247362 | 0.559317550599762 |

P42285_Q9NQT4
MTREX_HUMAN vs EXOS5_HUMAN
p-value: 0.079 q-value: 0.047

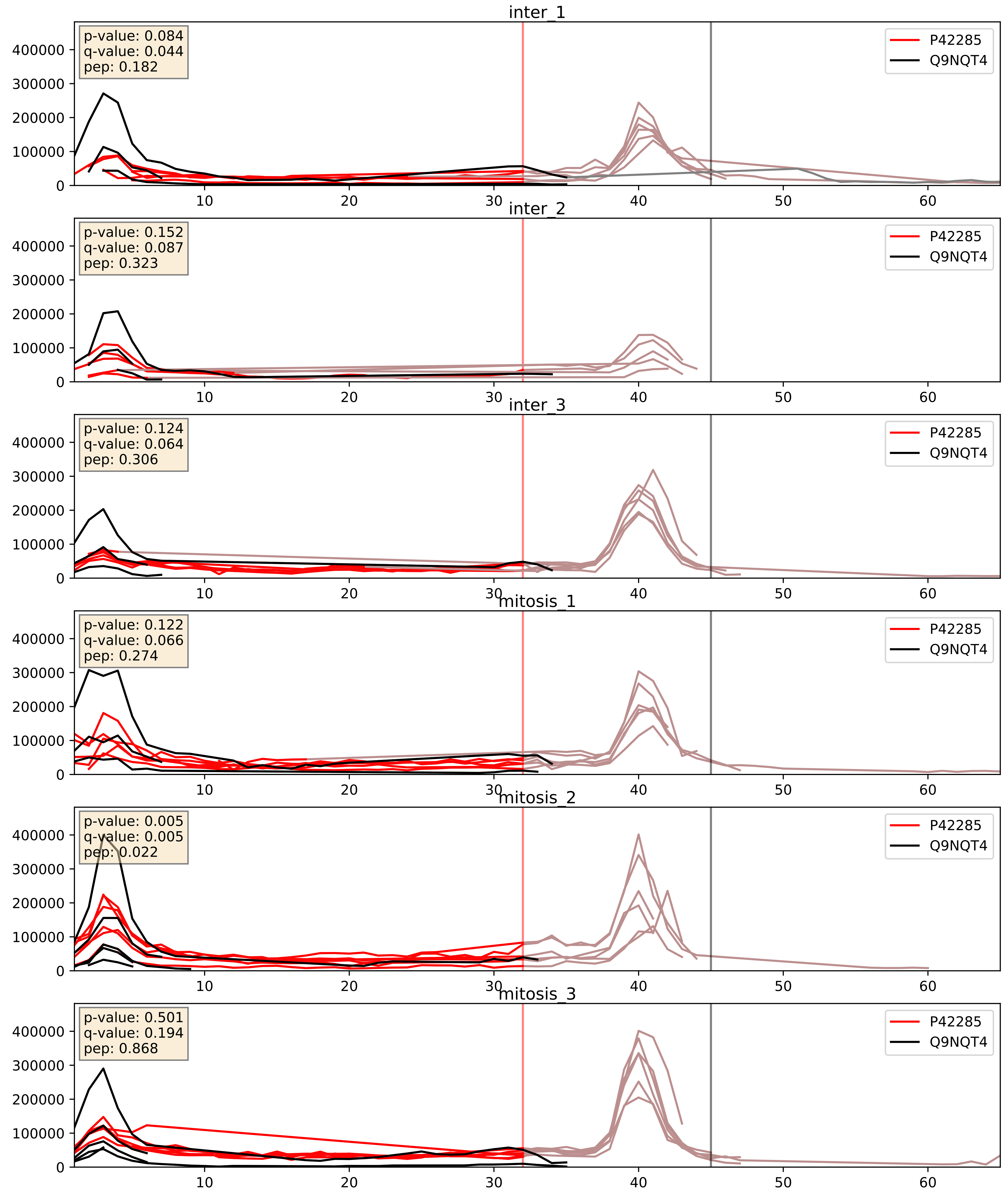


| level | condition_1 | condition_2 | pvalue | pvalue_adjusted |
|---|---|---|---|---|
| interactor_abundance | mitosis | inter | 0.000947405279672421 | 0.022846219799594687 |
| interactor_ratio | mitosis | inter | 0.0027077888480365724 | 0.06718455808461757 |
| complex_abundance | mitosis | inter | 0.056036993646254994 | 0.16874258962594654 |

P42285_Q9Y2X3
MTREX_HUMAN vs NOP58_HUMAN
p-value: 0.061 q-value: 0.038

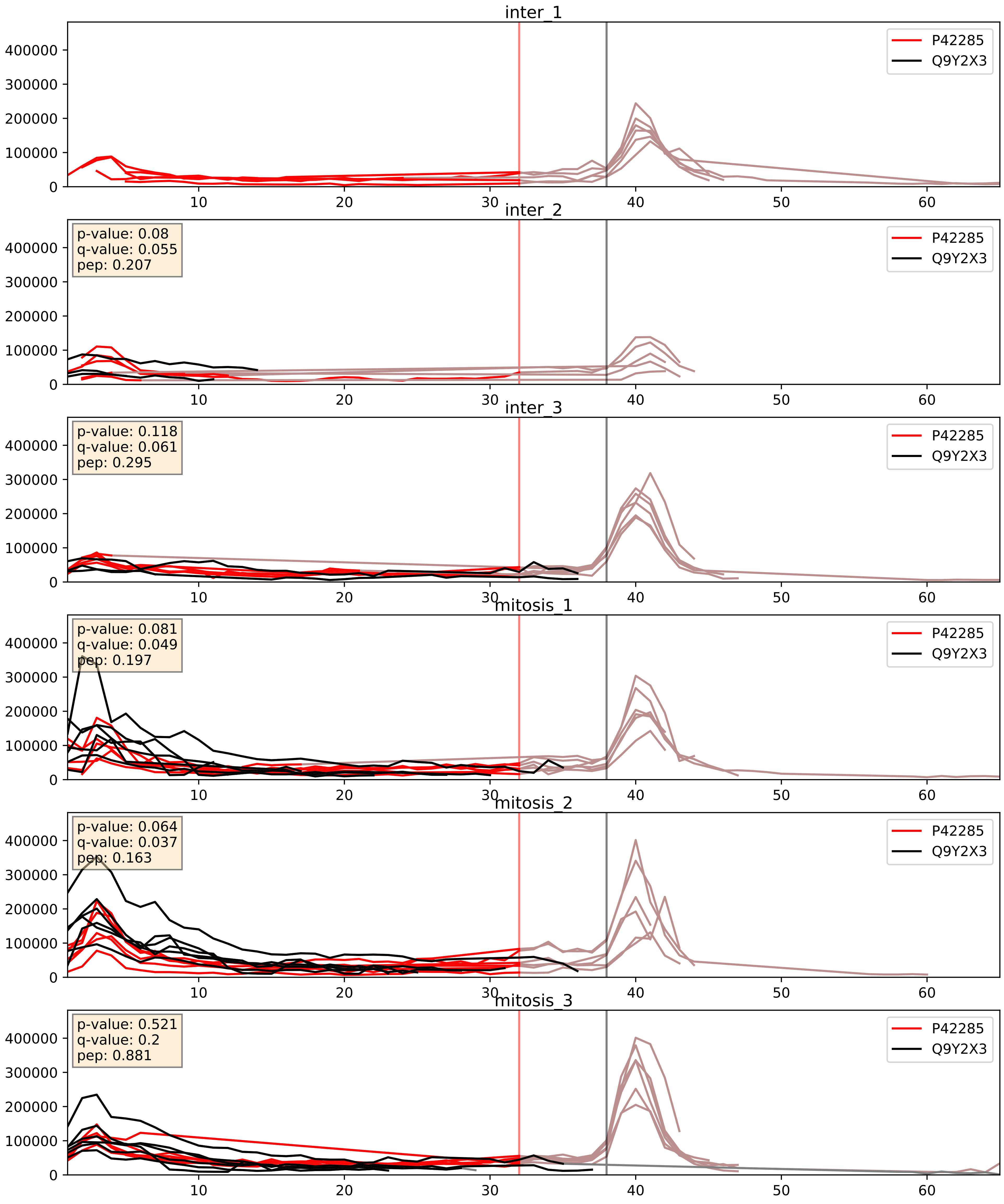


| level | condition_1 | condition_2 | pvalue | pvalue_adjusted |
|---|---|---|---|---|
| complex_abundance | mitosis | inter | 0.010746925639457542 | 0.07672534067869605 |
| interactor_ratio | mitosis | inter | 0.03074091262697153 | 0.2793441741898899 |
| interactor_abundance | mitosis | inter | 0.04261221300826687 | 0.16636566759504254 |

P42285_Q9Y3B2
MTREX_HUMAN vs EXOS1_HUMAN
p-value: 0.071 q-value: 0.043

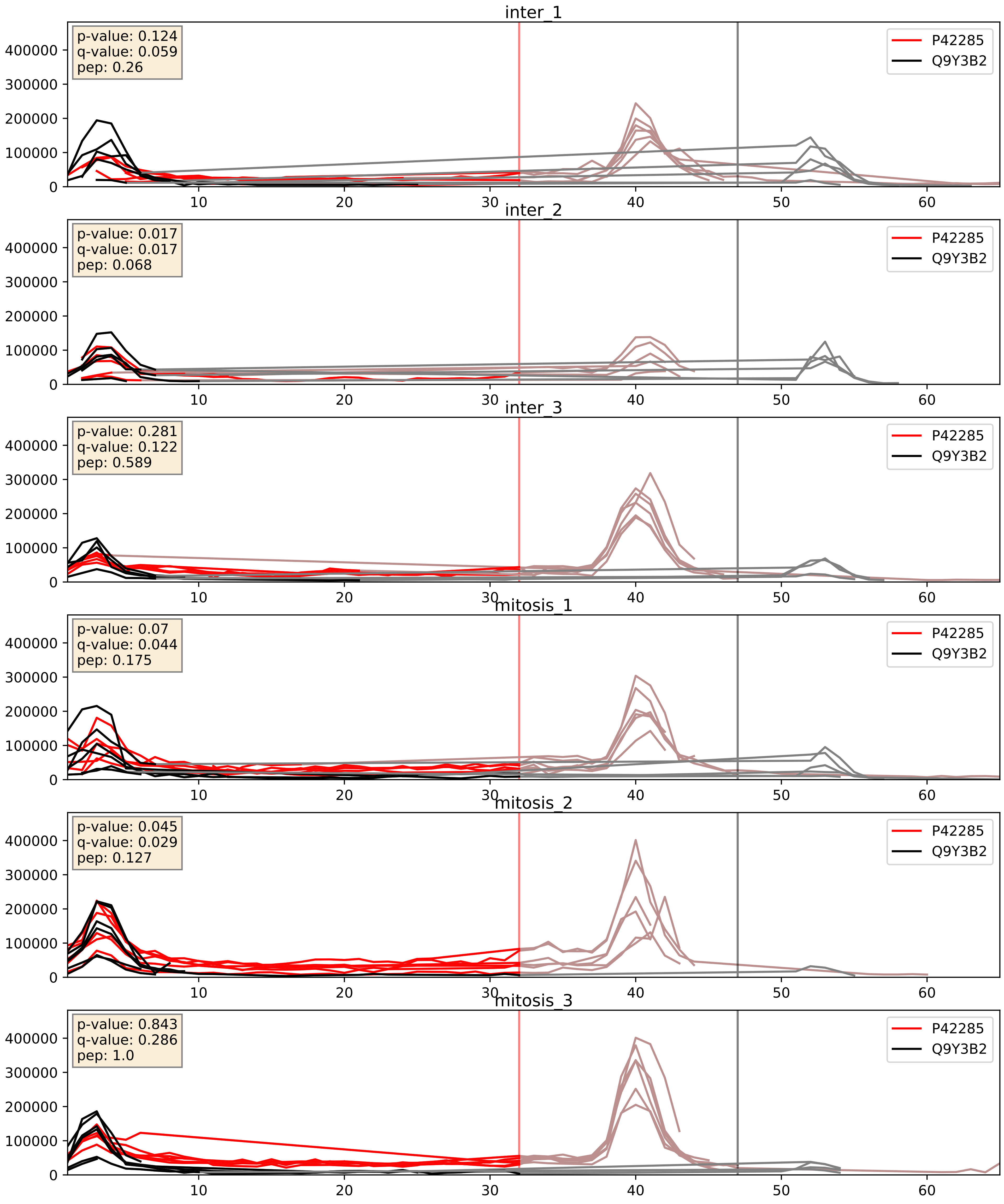


| level | condition_1 | condition_2 | pvalue | pvalue_adjusted |
|---|---|---|---|---|
| interactor_abundance | mitosis | inter | 0.0041414877724340535 | 0.05108961712397815 |
| complex_abundance | mitosis | inter | 0.05170294906287137 | 0.16205684510369056 |
| interactor_ratio | mitosis | inter | 0.11756875363783922 | 0.5644694535170984 |
