## Supplementary Data 2 for "SECAT: Quantifying differential protein-protein interaction states by network-centric analysis": hela_string_000144_P00966.pdf

P00966_P00966

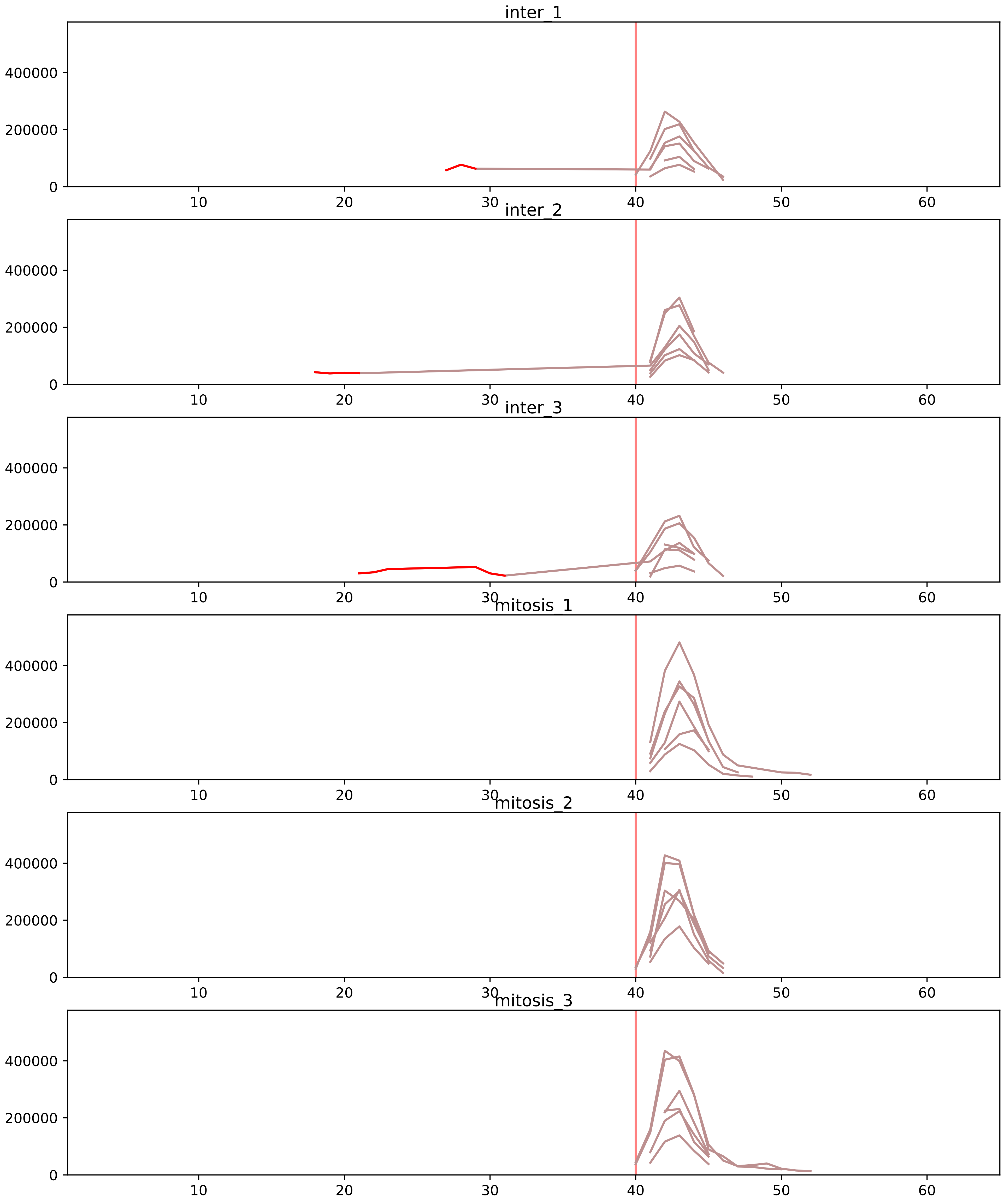


| level | condition_1 | condition_2 | pvalue | pvalue_adjusted |
|---|---|---|---|---|
| assembled_abundance | mitosis | inter | 5.2635990561479474e-05 | 0.007270487507555603 |
| monomer_abundance | mitosis | inter | 0.0052511025451213444 | 0.14513761660427227 |
| total_abundance | mitosis | inter | 0.013269545946618087 | 0.14349118205454814 |
