## Supplementary Data 2 for "SECAT: Quantifying differential protein-protein interaction states by network-centric analysis": hela_string_000145_P09651.pdf

P09651_P09651

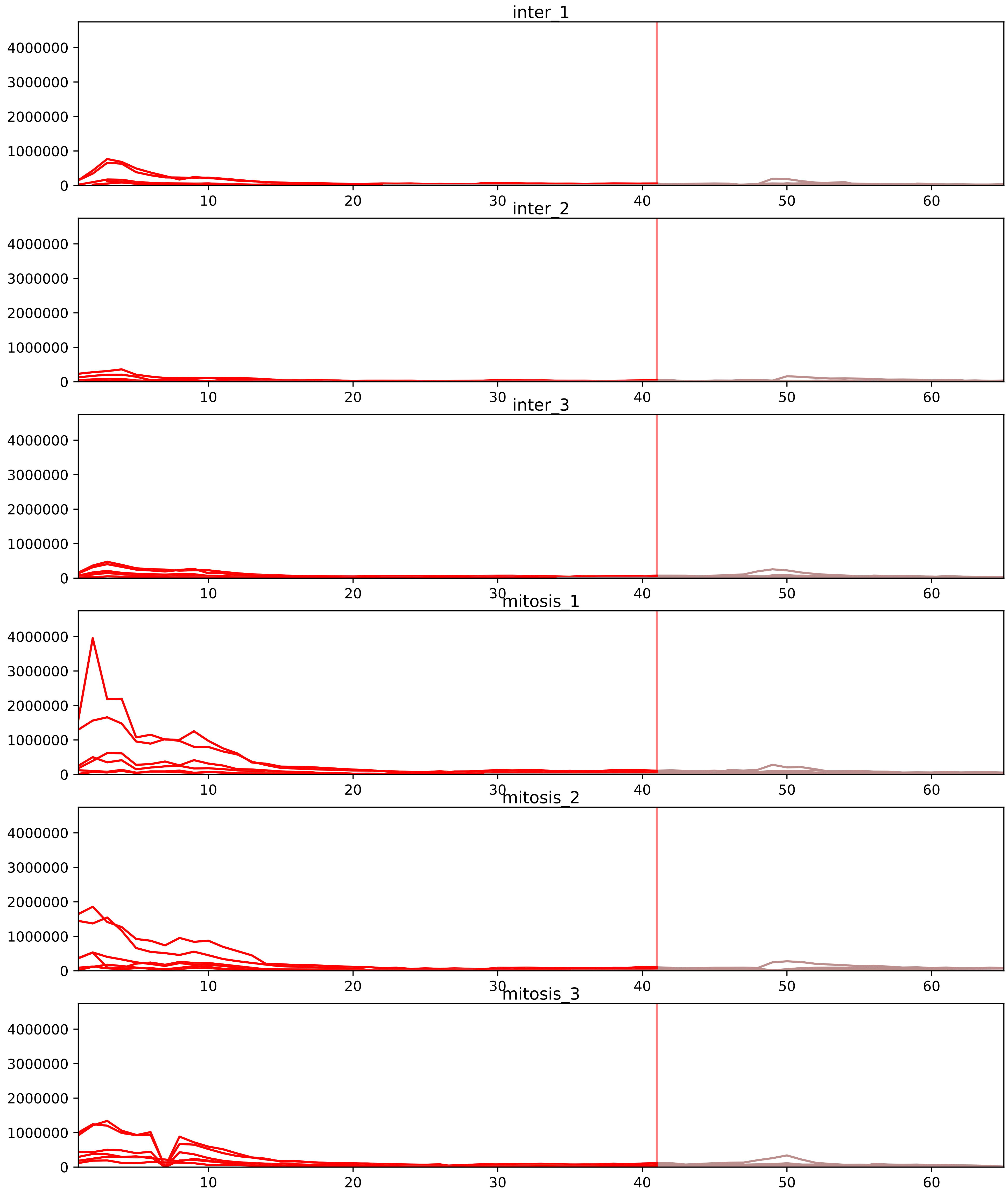


| level | condition_1 | condition_2 | pvalue | pvalue_adjusted |
|---|---|---|---|---|
| interactor_ratio | mitosis | inter | 5.3193194956259236e-05 | 0.0010090255538094536 |
| complex_abundance | mitosis | inter | 7.406505121087693e-05 | 0.0018871206097520157 |
| interactor_abundance | mitosis | inter | 0.00030630180763079393 | 0.005170599321473953 |
| assembled_abundance | mitosis | inter | 0.010579055044115337 | 0.11480207137326143 |
| total_abundance | mitosis | inter | 0.01600571108612868 | 0.16008855626617113 |
| monomer_abundance | mitosis | inter | 0.09576377873186462 | 0.4074222334948335 |

O43390_P09651
HNRPR_HUMAN vs ROA1_HUMAN
p-value: 0.065 q-value: 0.04

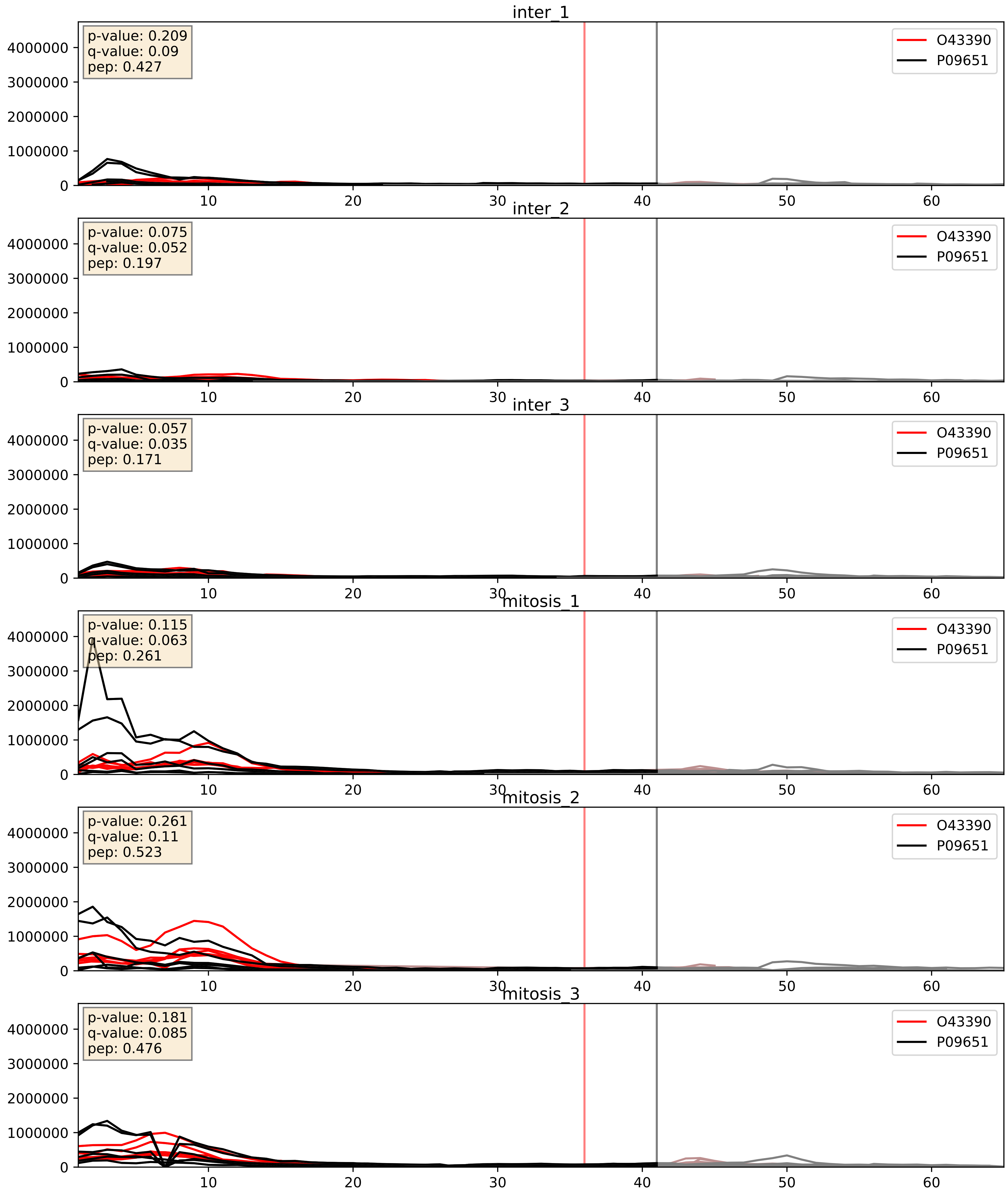


| level | condition_1 | condition_2 | pvalue | pvalue_adjusted |
|---|---|---|---|---|
| interactor_abundance | mitosis | inter | 0.00010796061997593195 | 0.007876481428251637 |
| complex_abundance | mitosis | inter | 0.0009535473068718876 | 0.020405912367058396 |
| interactor_ratio | mitosis | inter | 0.3502980208802063 | 0.8007410362093738 |

P05198_P09651
IF2A_HUMAN vs ROA1_HUMAN
p-value: 0.0 q-value: 0.038

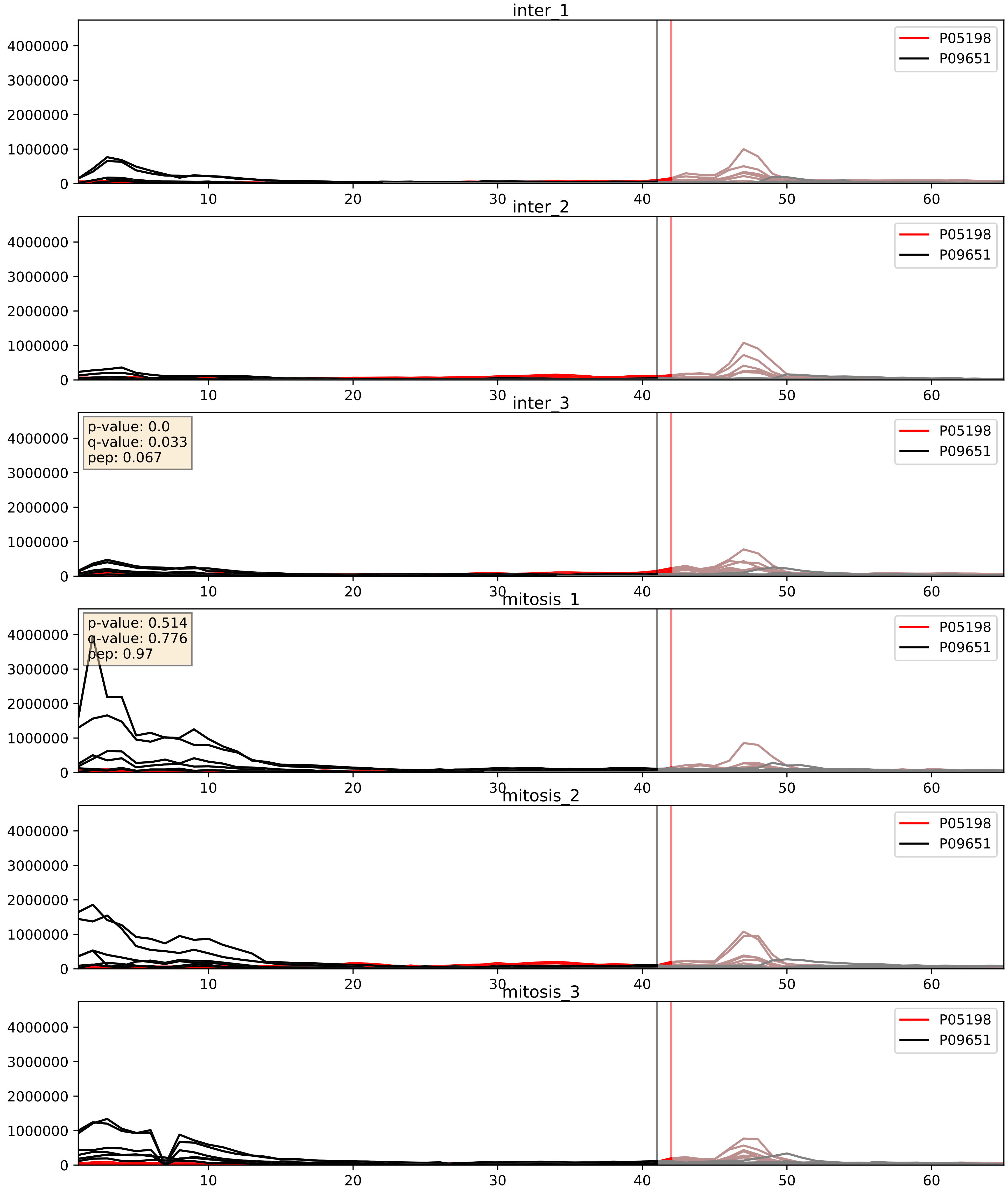


| level | condition_1 | condition_2 | pvalue | pvalue_adjusted |
| --- | --- | --- | --- | --- |
| interactor_ratio | mitosis | inter | 0.06223260567916239 | 0.41456117090554867 |
| complex_abundance | mitosis | inter | 0.07026639337343314 | 0.19241213284599734 |
| interactor_abundance | mitosis | inter | 0.47373227991024996 | 0.6453132266123074 |

P05387_P09651
RLA2_HUMAN vs ROA1_HUMAN
p-value: 0.0 q-value: 0.023

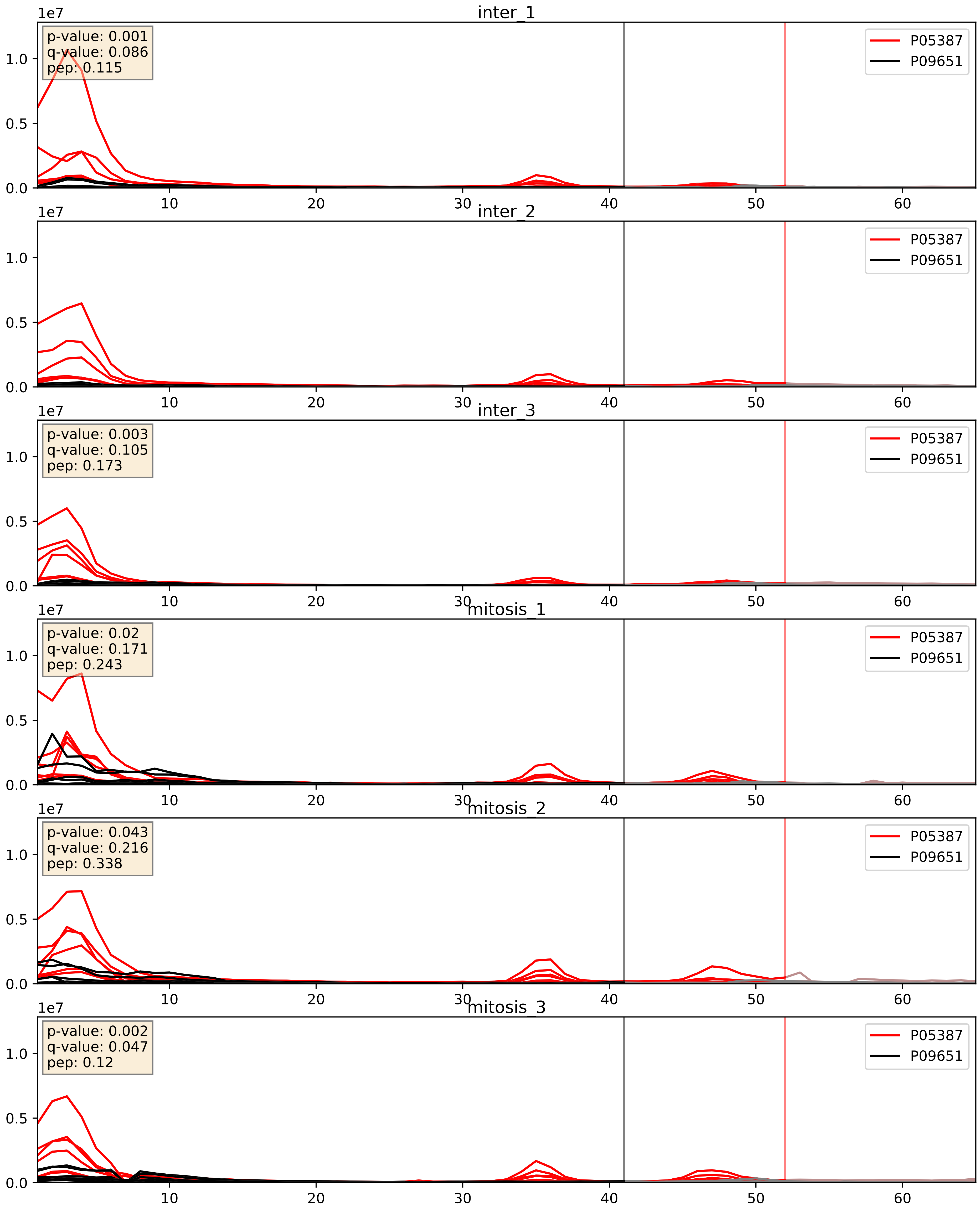


| level | condition_1 | condition_2 | pvalue | pvalue_adjusted |
|---|---|---|---|---|
| interactor_ratio | mitosis | inter | 0.008674913163209456 | 0.13211821468820312 |
| complex_abundance | mitosis | inter | 0.04199765259200707 | 0.14726323635770058 |
| interactor_abundance | mitosis | inter | 0.9422825537275921 | 0.9643061342468991 |

P07910_P09651
HNRPC_HUMAN vs ROA1_HUMAN
p-value: 0.004 q-value: 0.004

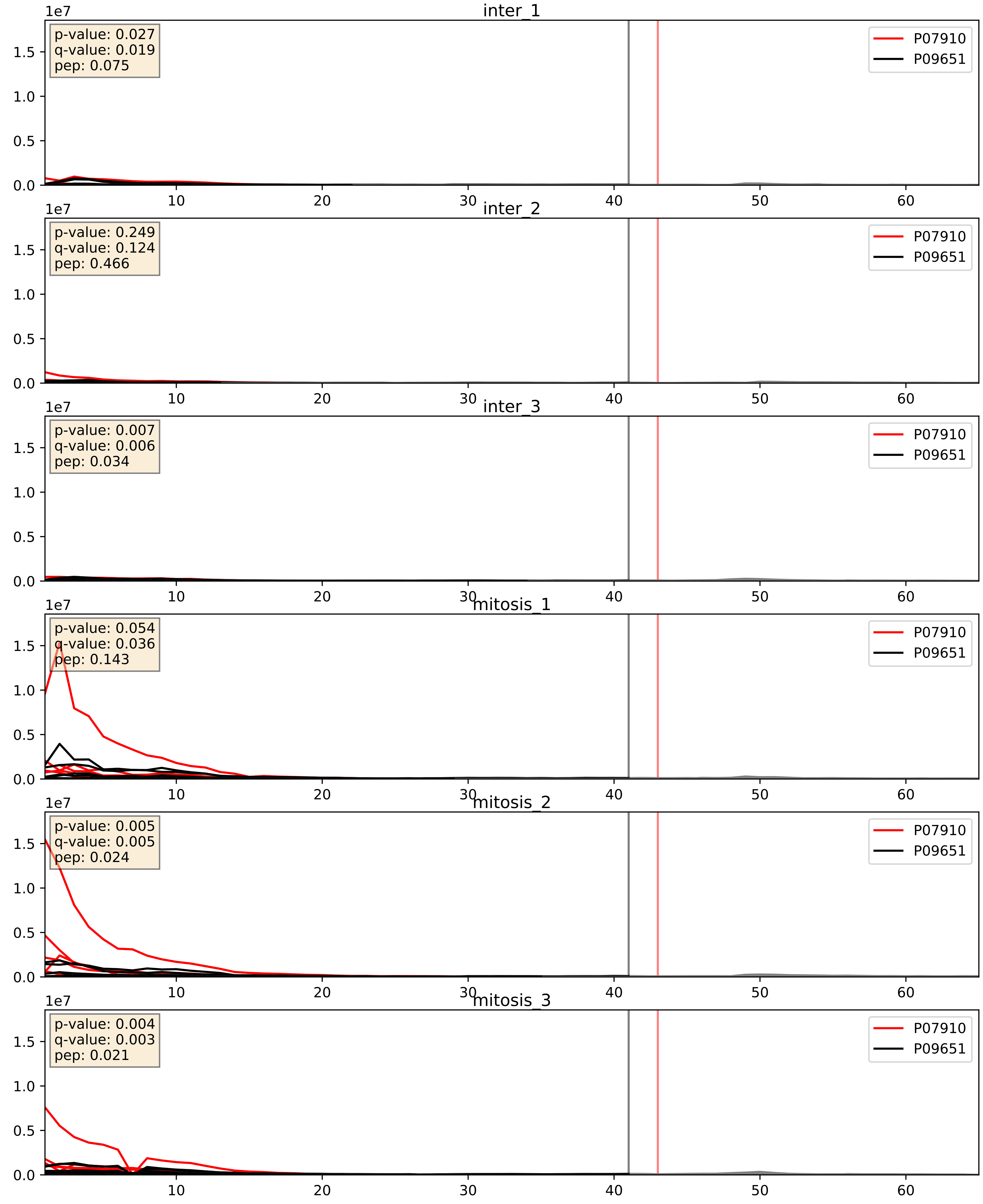


| level | condition_1 | condition_2 | pvalue | pvalue_adjusted |
|---|---|---|---|---|
| interactor_abundance | mitosis | inter | 9.357211976477008e-06 | 0.0033734327106189023 |
| complex_abundance | mitosis | inter | 7.435901811379124e-05 | 0.005893640694944935 |
| interactor_ratio | mitosis | inter | 0.02654415193625456 | 0.25614196664208316 |

P08708_P09651
RS17_HUMAN vs ROA1_HUMAN
p-value: 0.0 q-value: 0.038

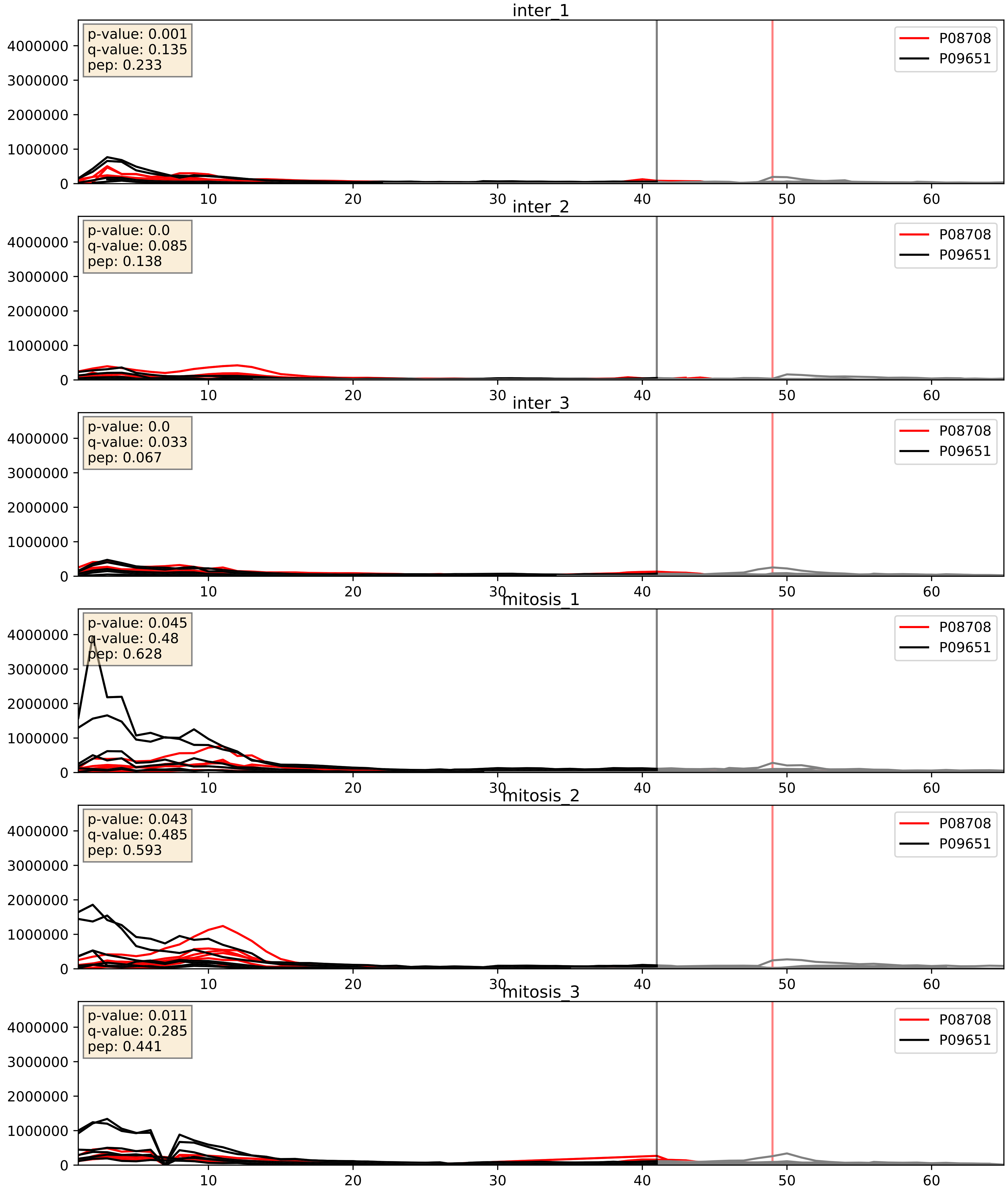


| level | condition_1 | condition_2 | pvalue | pvalue_adjusted |
|---|---|---|---|---|
| complex_abundance | mitosis | inter | 0.005620107195053823 | 0.05646492674842808 |
| interactor_ratio | mitosis | inter | 0.055141866994355686 | 0.3890548630198597 |
| interactor_abundance | mitosis | inter | 0.12458602588785918 | 0.30782403856258467 |

P09651_P14866
ROA1_HUMAN vs HNRPL_HUMAN
p-value: 0.005 q-value: 0.005

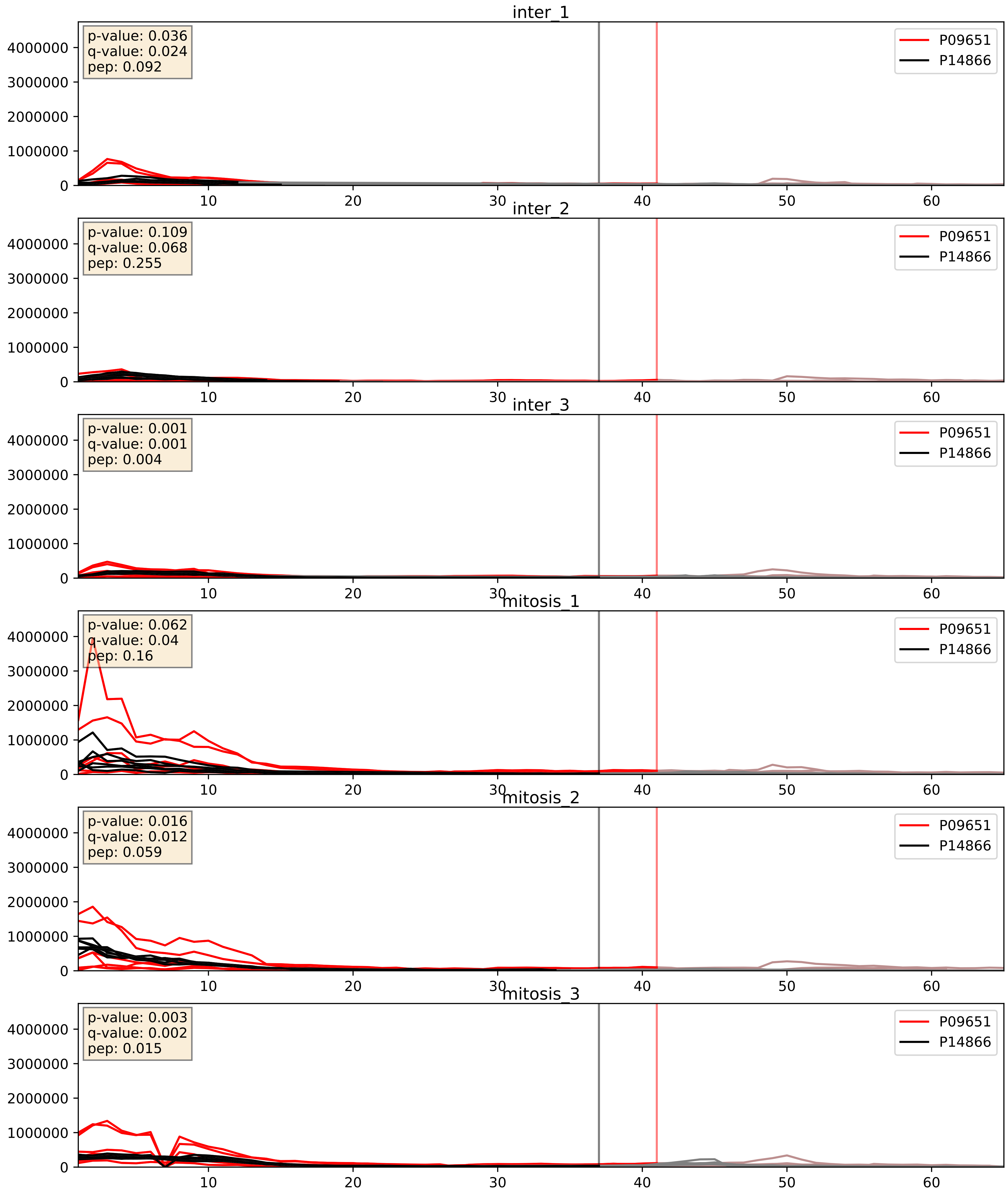


| level | condition_1 | condition_2 | pvalue | pvalue_adjusted |
|---|---|---|---|---|
| complex_abundance | mitosis | inter | 0.0022461160486105898 | 0.03476808928771546 |
| interactor_abundance | mitosis | inter | 0.010916100555075968 | 0.0916545568920552 |
| interactor_ratio | mitosis | inter | 0.343200993452667 | 0.7955051459395694 |

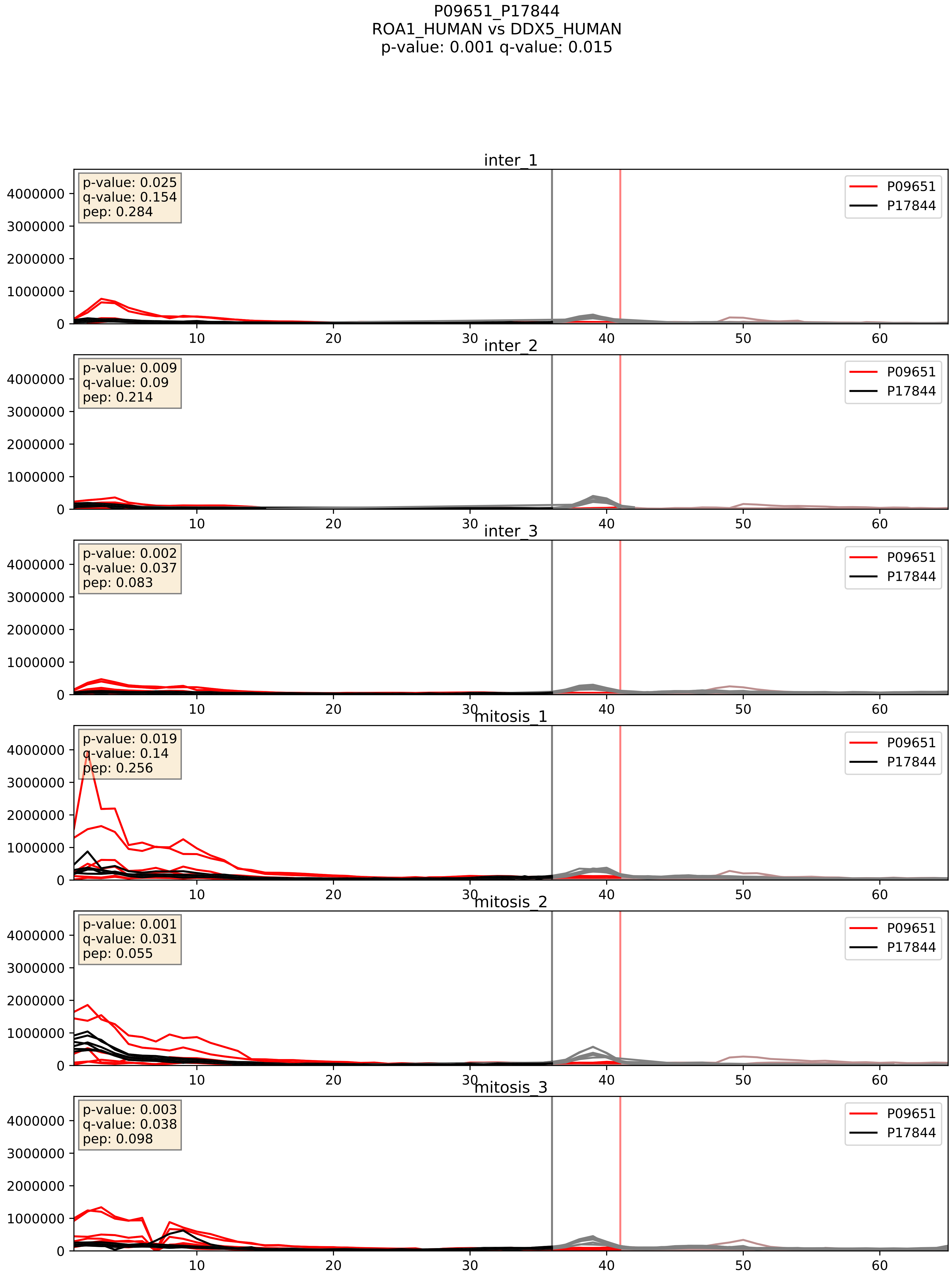


| level | condition_1 | condition_2 | pvalue | pvalue_adjusted |
|---|---|---|---|---|
| complex_abundance | mitosis | inter | 0.005592113315961333 | 0.056382202573179045 |
| interactor_abundance | mitosis | inter | 0.017017205188648715 | 0.11334462047152971 |
| interactor_ratio | mitosis | inter | 0.1872358107783095 | 0.657442464198634 |

P09651_P22626
ROA1_HUMAN vs ROA2_HUMAN
p-value: 0.0 q-value: 0.0

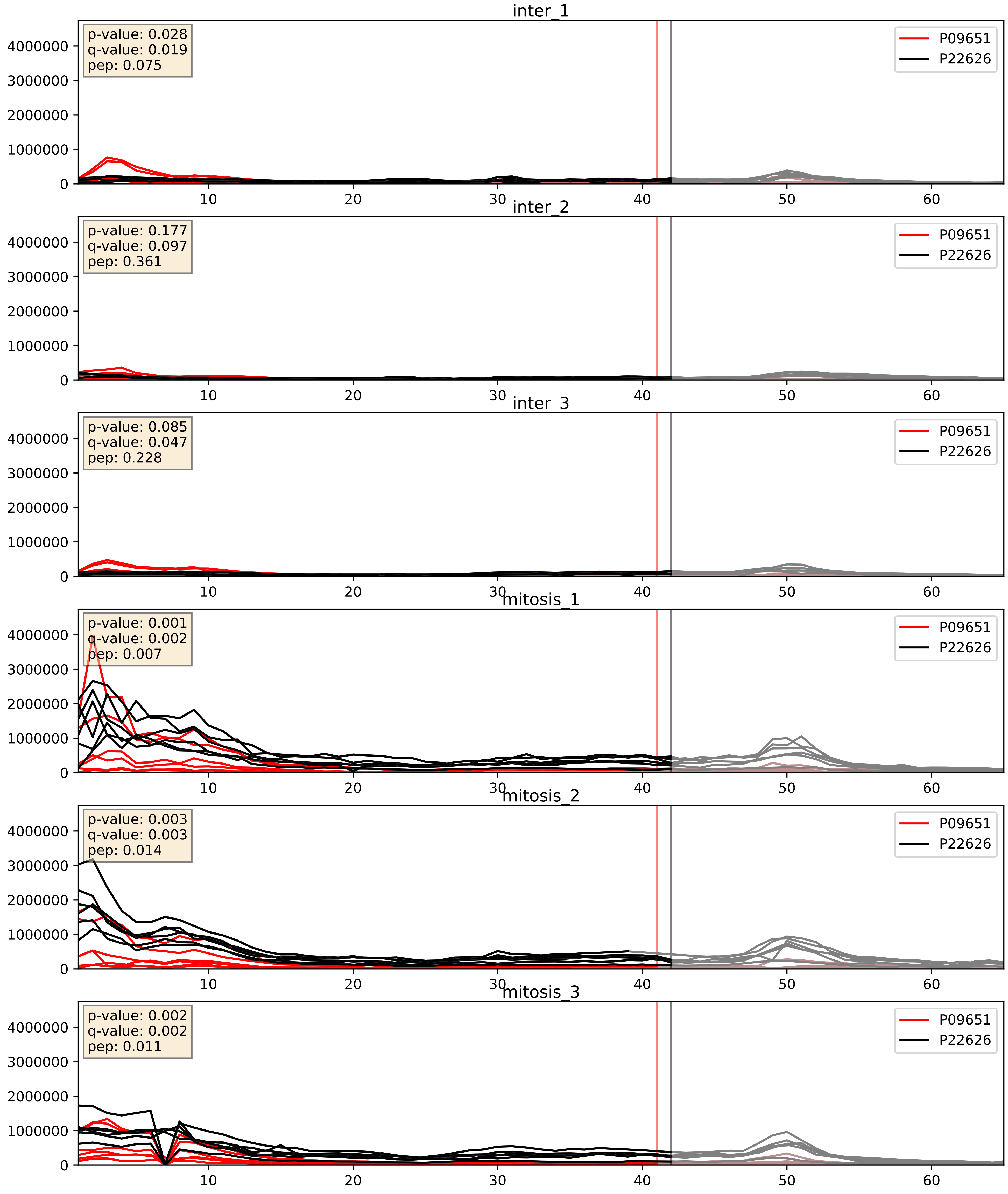


| level | condition_1 | condition_2 | pvalue | pvalue_adjusted |
|---|---|---|---|---|
| complex_abundance | mitosis | inter | 0.0022885483088337477 | 0.03502731659907978 |
| interactor_abundance | mitosis | inter | 0.016856414137678265 | 0.11277132084292767 |
| interactor_ratio | mitosis | inter | 0.066322140233217 | 0.4296400448702392 |

P09651_P31943
ROA1_HUMAN vs HNRH1_HUMAN
p-value: 0.0 q-value: 0.001

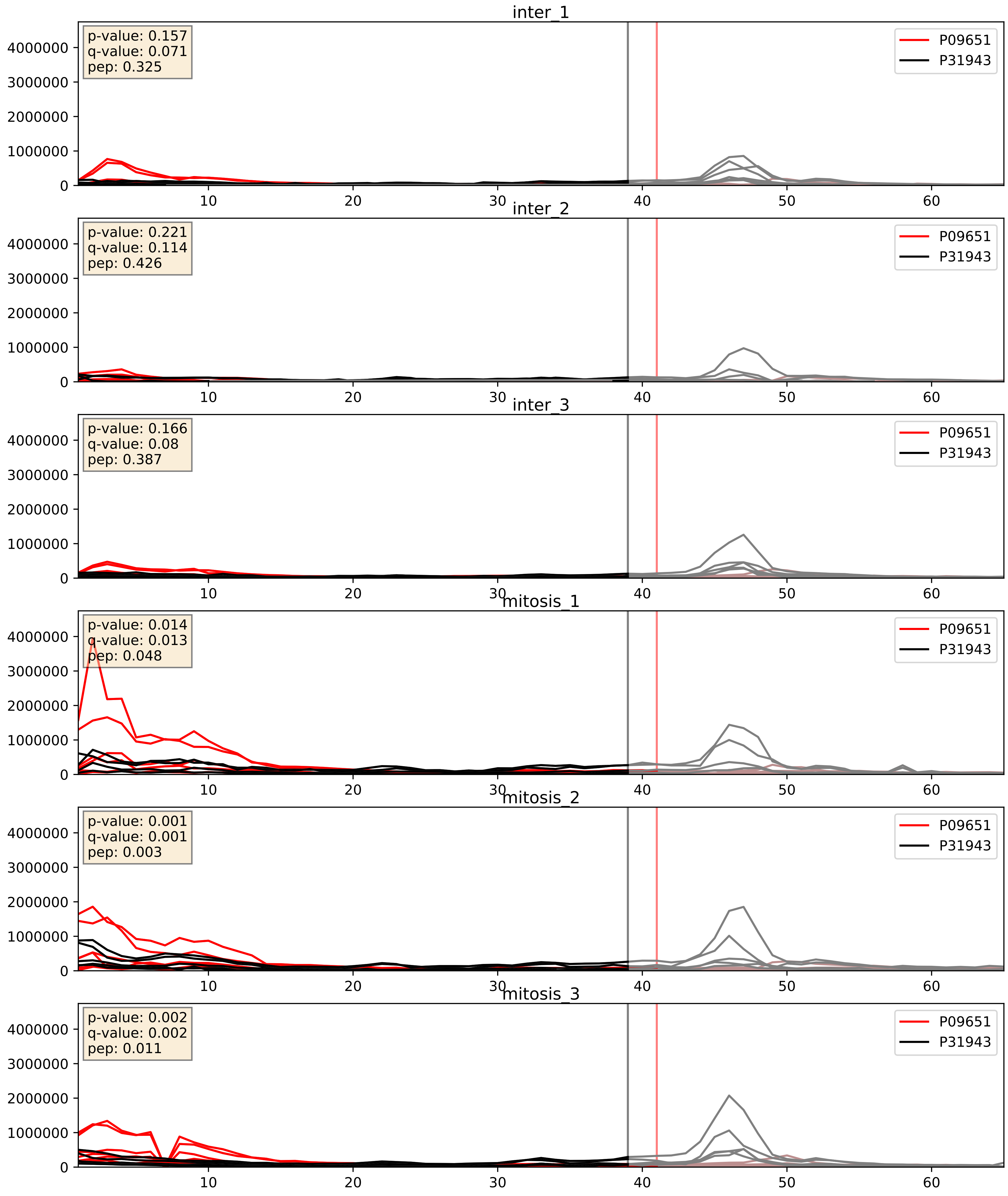


| level | condition_1 | condition_2 | pvalue | pvalue_adjusted |
|---|---|---|---|---|
| complex_abundance | mitosis | inter | 0.007967826015523555 | 0.06596188655017565 |
| interactor_abundance | mitosis | inter | 0.016854435383293762 | 0.11277132084292767 |
| interactor_ratio | mitosis | inter | 0.2314283726697915 | 0.7093426594678715 |

P09651_P38159
ROA1_HUMAN vs RBMX_HUMAN
p-value: 0.004 q-value: 0.004

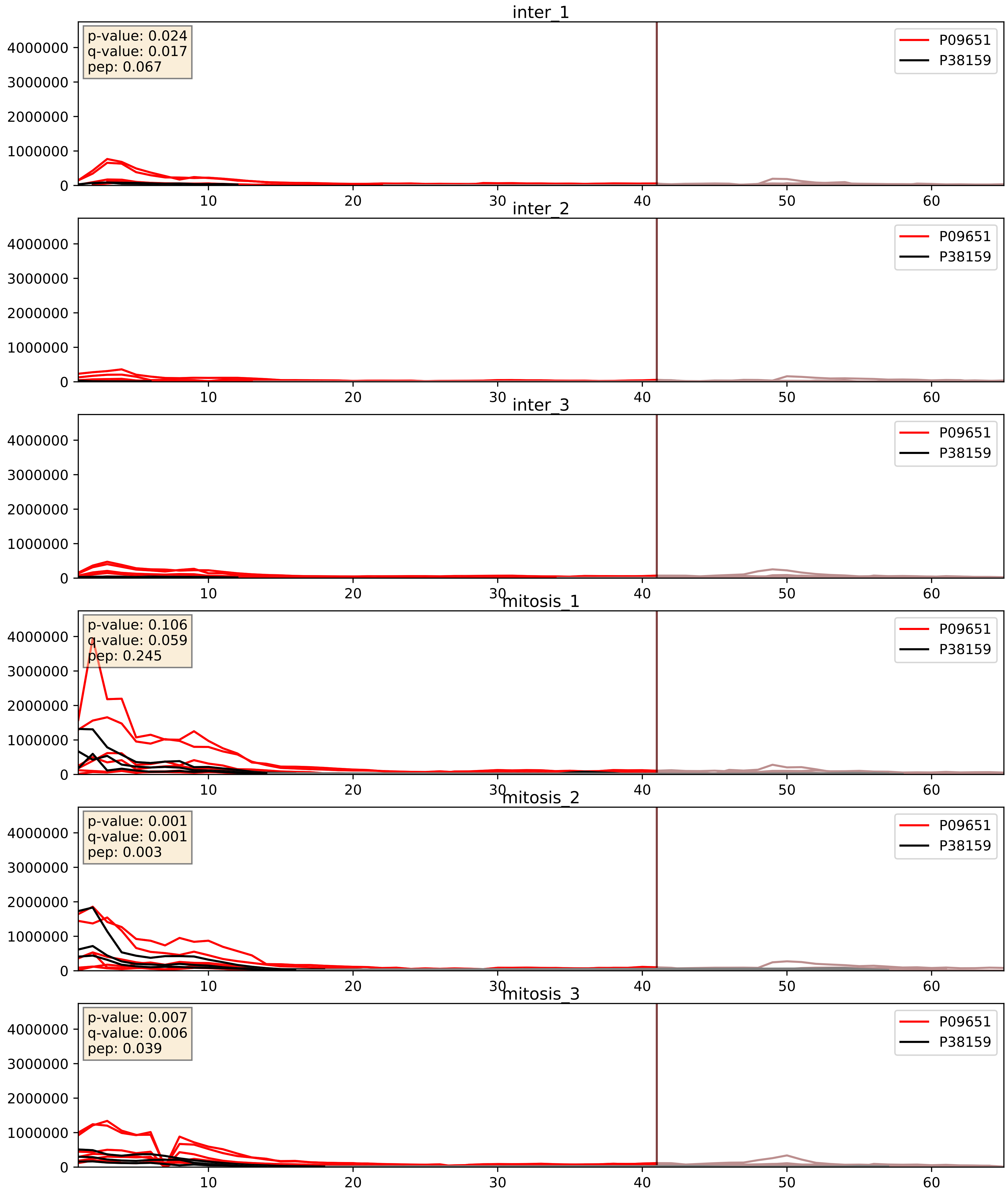


| level | condition_1 | condition_2 | pvalue | pvalue_adjusted |
|---|---|---|---|---|
| complex_abundance | mitosis | inter | 0.003021295973201694 | 0.04033974863104599 |
| interactor_abundance | mitosis | inter | 0.013856881209314718 | 0.10436859054266079 |
| interactor_ratio | mitosis | inter | 0.2375025223286475 | 0.7164022336563056 |

P09651_P46776
ROA1_HUMAN vs RL27A_HUMAN
p-value: 0.0 q-value: 0.023

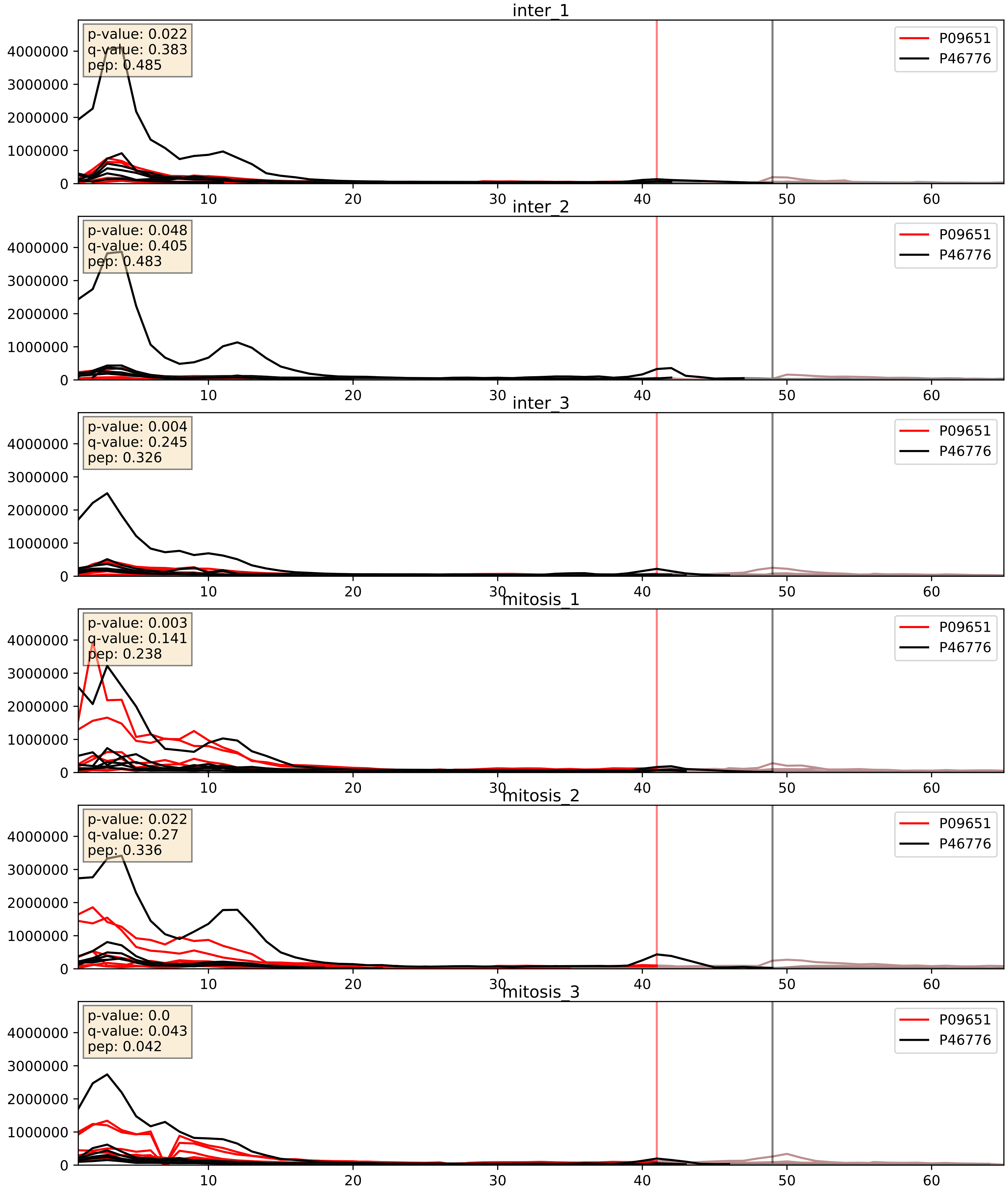


| level | condition_1 | condition_2 | pvalue | pvalue_adjusted |
|---|---|---|---|---|
| interactor_ratio | mitosis | inter | 0.012839880461491688 | 0.16355562016423936 |
| interactor_abundance | mitosis | inter | 0.017291258615792112 | 0.11372506627059584 |
| complex_abundance | mitosis | inter | 0.030487060445578192 | 0.1267456228334868 |

P09651_P46781
ROA1_HUMAN vs RS9_HUMAN
p-value: 0.0 q-value: 0.041

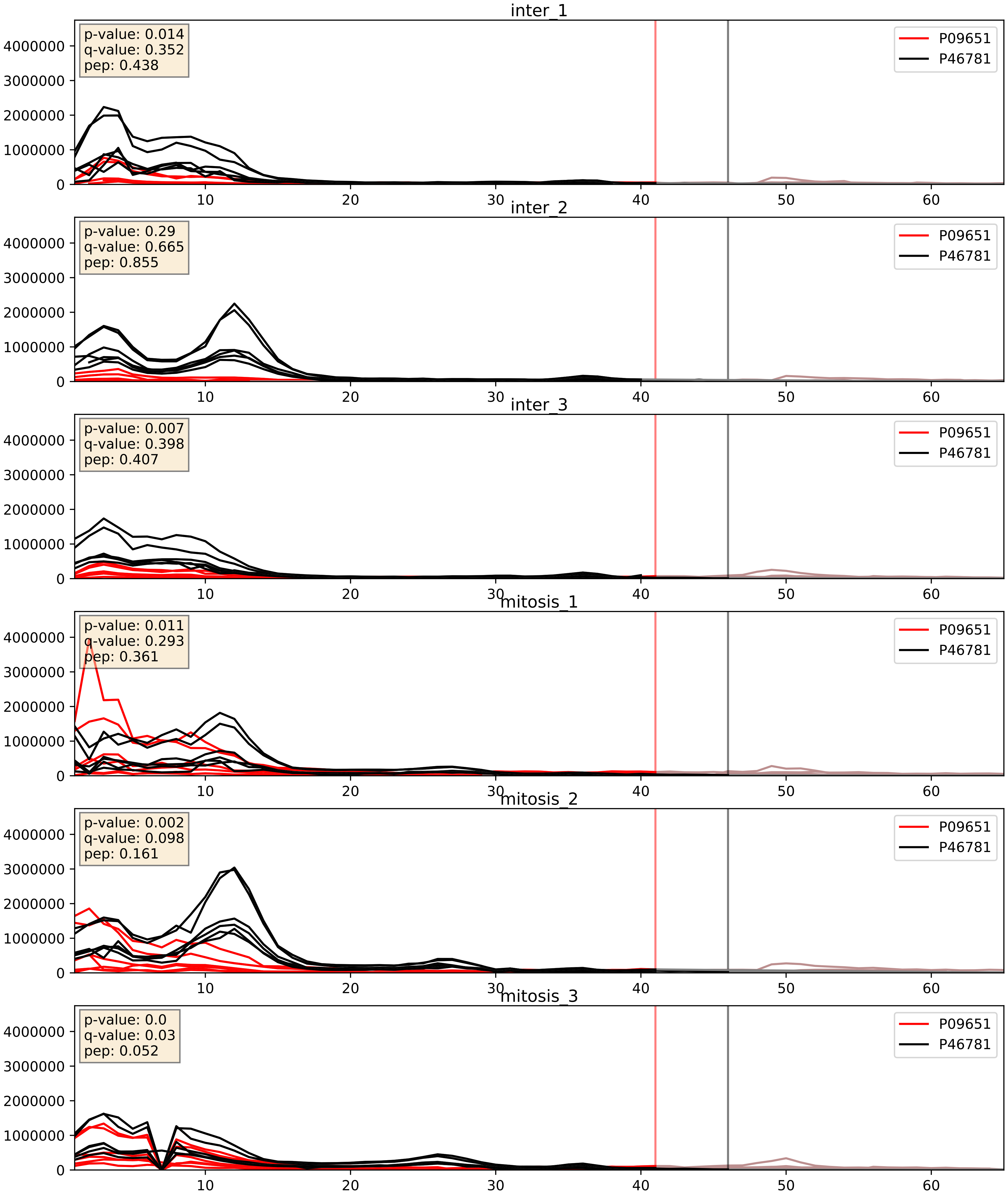


| level | condition_1 | condition_2 | pvalue | pvalue_adjusted |
|---|---|---|---|---|
| complex_abundance | mitosis | inter | 0.008305813174941331 | 0.06742444735376033 |
| interactor_abundance | mitosis | inter | 0.016856414137678265 | 0.11277132084292767 |
| interactor_ratio | mitosis | inter | 0.03745460002517187 | 0.31712302296287953 |

P09651_P51991
ROA1_HUMAN vs ROA3_HUMAN
p-value: 0.0 q-value: 0.0

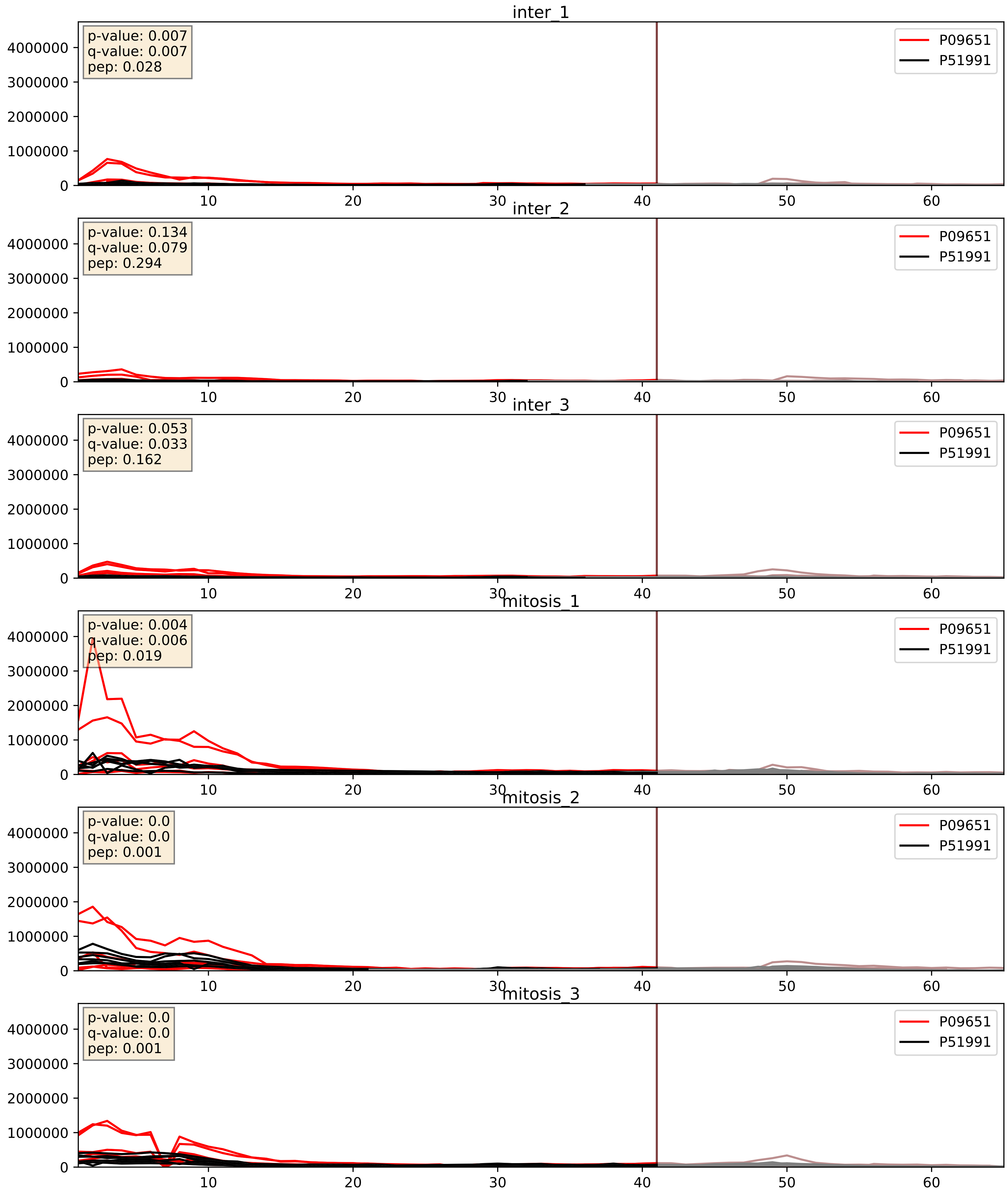


| level | condition_1 | condition_2 | pvalue | pvalue_adjusted |
|---|---|---|---|---|
| complex_abundance | mitosis | inter | 0.0011795042359659484 | 0.023096652182529428 |
| interactor_abundance | mitosis | inter | 0.016104784849936434 | 0.11126469597696198 |
| interactor_ratio | mitosis | inter | 0.19038795235618836 | 0.6627604950548505 |

P09651_P52272
ROA1_HUMAN vs HNRPM_HUMAN
p-value: 0.0 q-value: 0.001

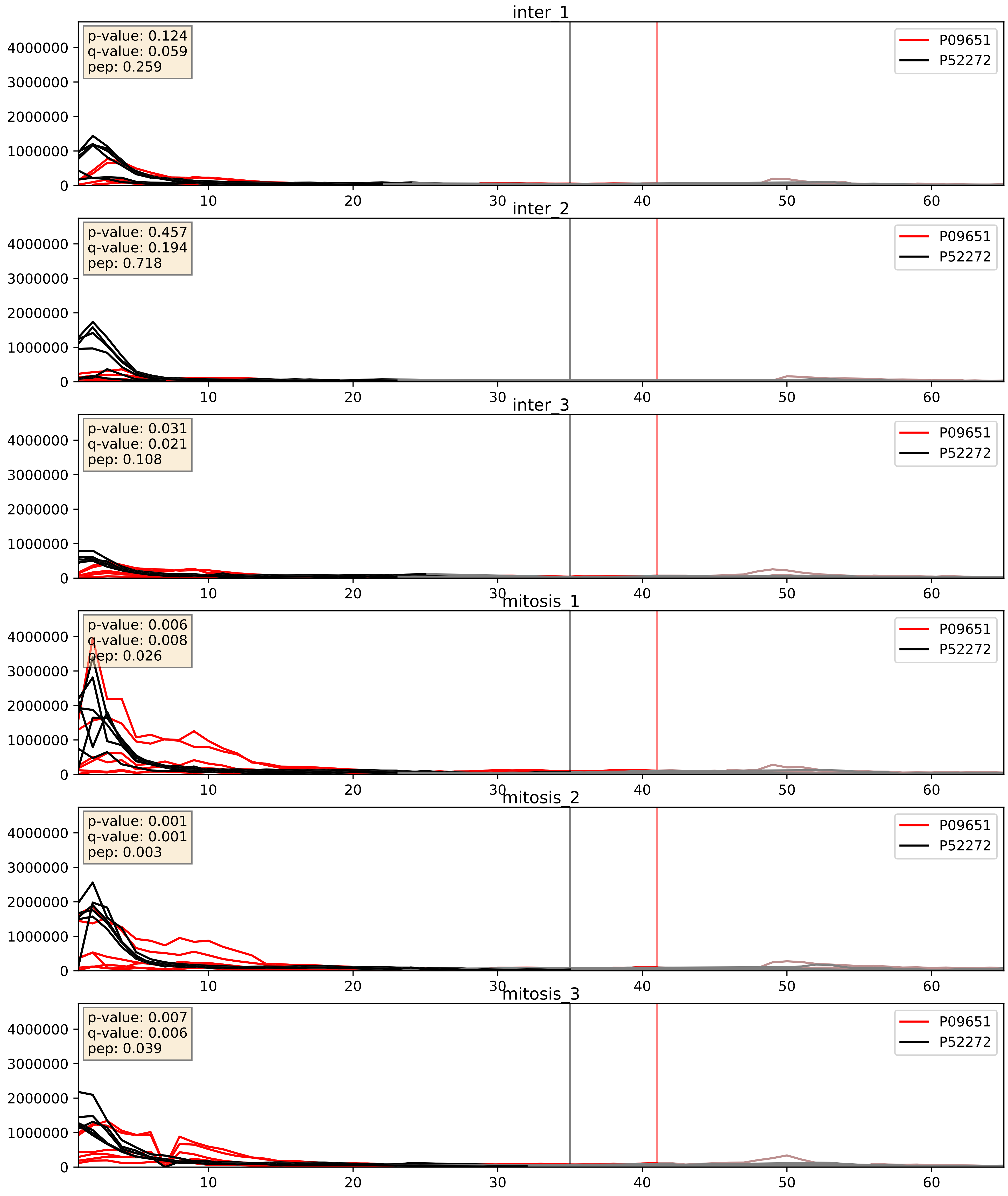


| level | condition_1 | condition_2 | pvalue | pvalue_adjusted |
|---|---|---|---|---|
| interactor_abundance | mitosis | inter | 0.01476418936640577 | 0.1063369465514795 |
| complex_abundance | mitosis | inter | 0.016581499584325202 | 0.09584596180627945 |
| interactor_ratio | mitosis | inter | 0.04323656308648082 | 0.34205635861393335 |

P09651_P52597
ROA1_HUMAN vs HNRPF_HUMAN
p-value: 0.001 q-value: 0.001

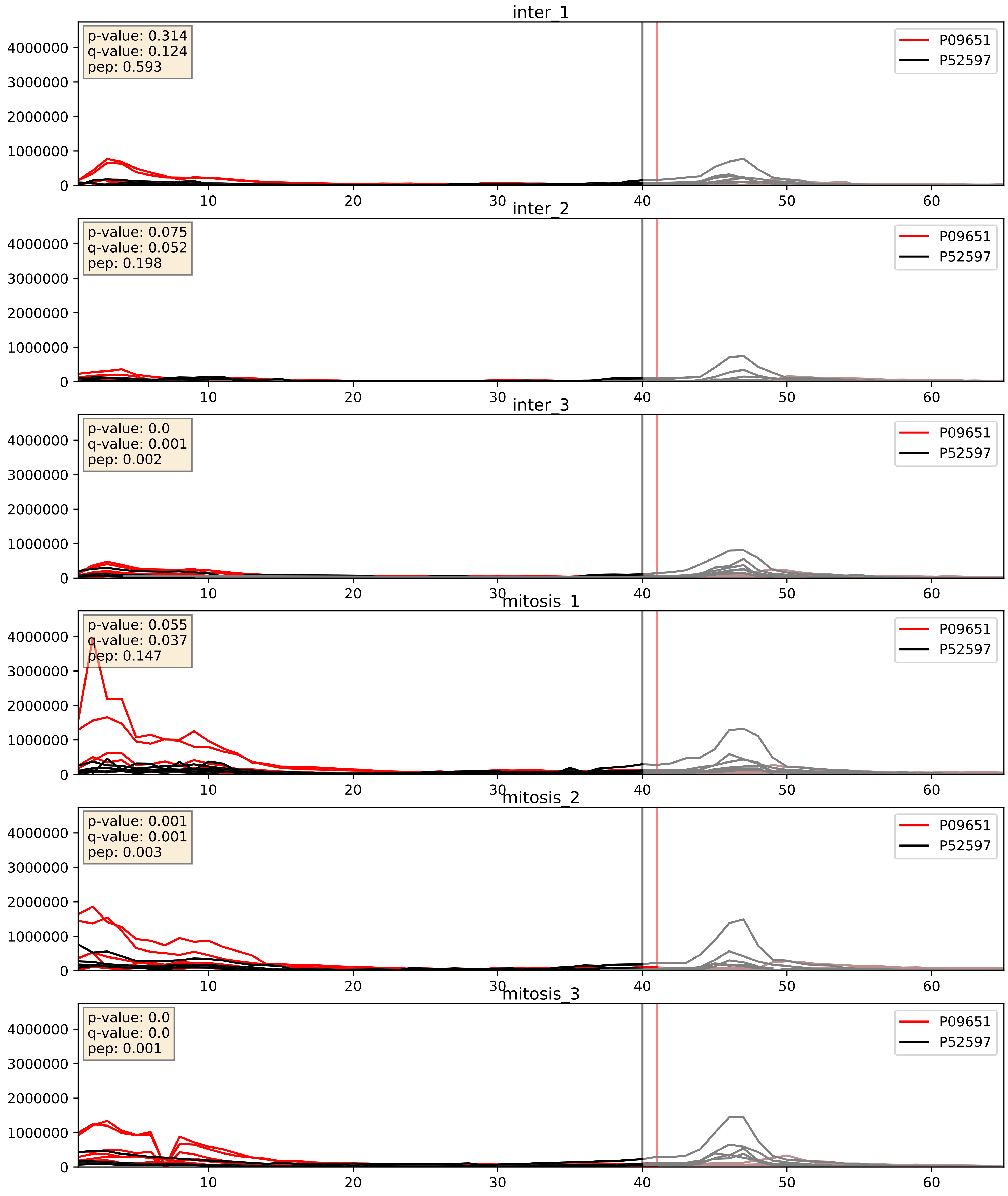


| level | condition_1 | condition_2 | pvalue | pvalue_adjusted |
|---|---|---|---|---|
| complex_abundance | mitosis | inter | 0.006743082334176311 | 0.06075872082163076 |
| interactor_abundance | mitosis | inter | 0.017050532289726136 | 0.11334462047152971 |
| interactor_ratio | mitosis | inter | 0.11810419448591807 | 0.566001132973789 |

P09651_P55769
ROA1_HUMAN vs NH2L1_HUMAN
p-value: 0.0 q-value: 0.003

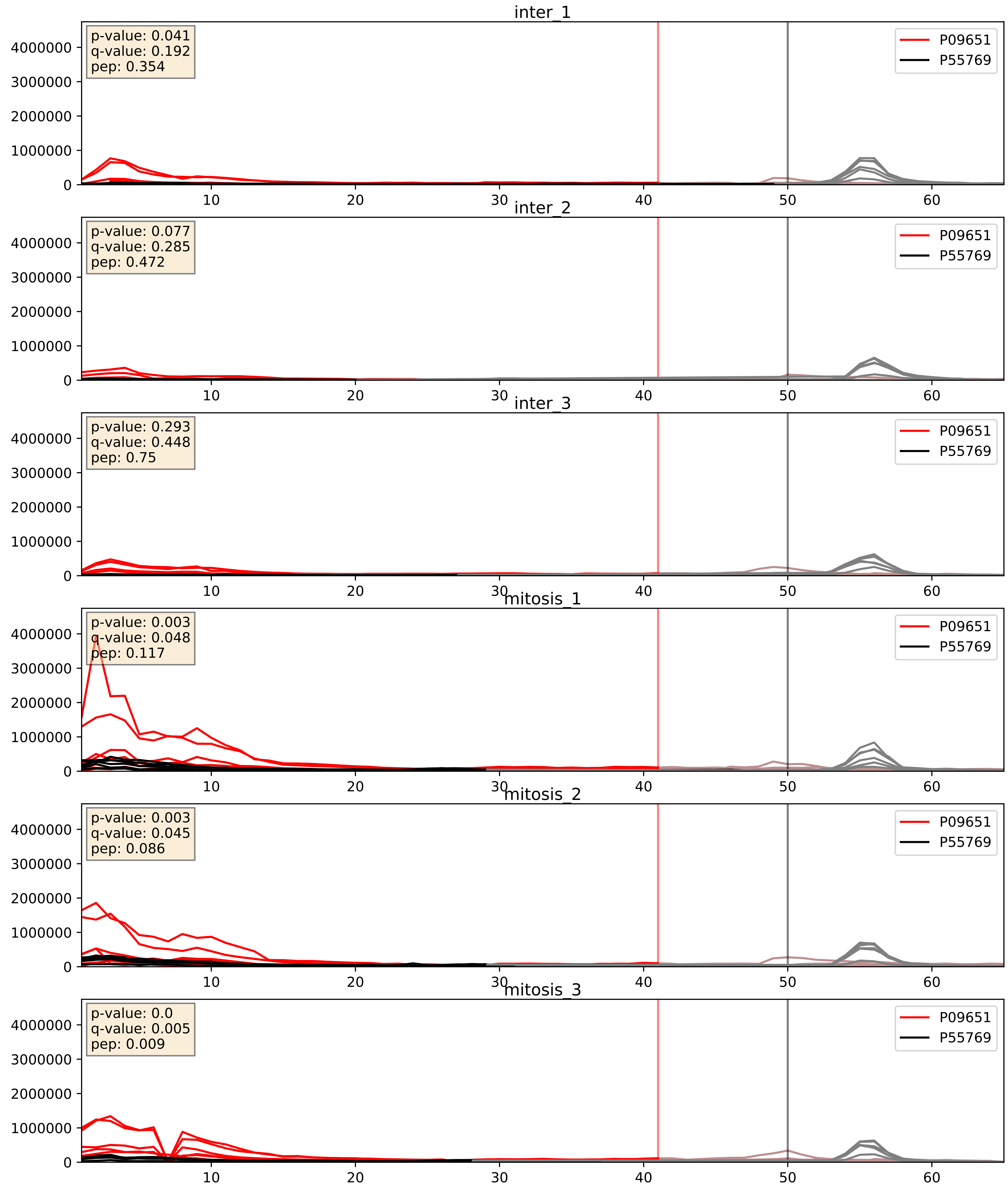


| level | condition_1 | condition_2 | pvalue | pvalue_adjusted |
|---|---|---|---|---|
| complex_abundance | mitosis | inter | 0.004746622165762423 | 0.05149694010003338 |
| interactor_abundance | mitosis | inter | 0.01925771606241603 | 0.11885079271397347 |
| interactor_ratio | mitosis | inter | 0.7812992714023176 | 0.9589625145419588 |

P09651_P61978
ROA1_HUMAN vs HNRPK_HUMAN
p-value: 0.0 q-value: 0.0

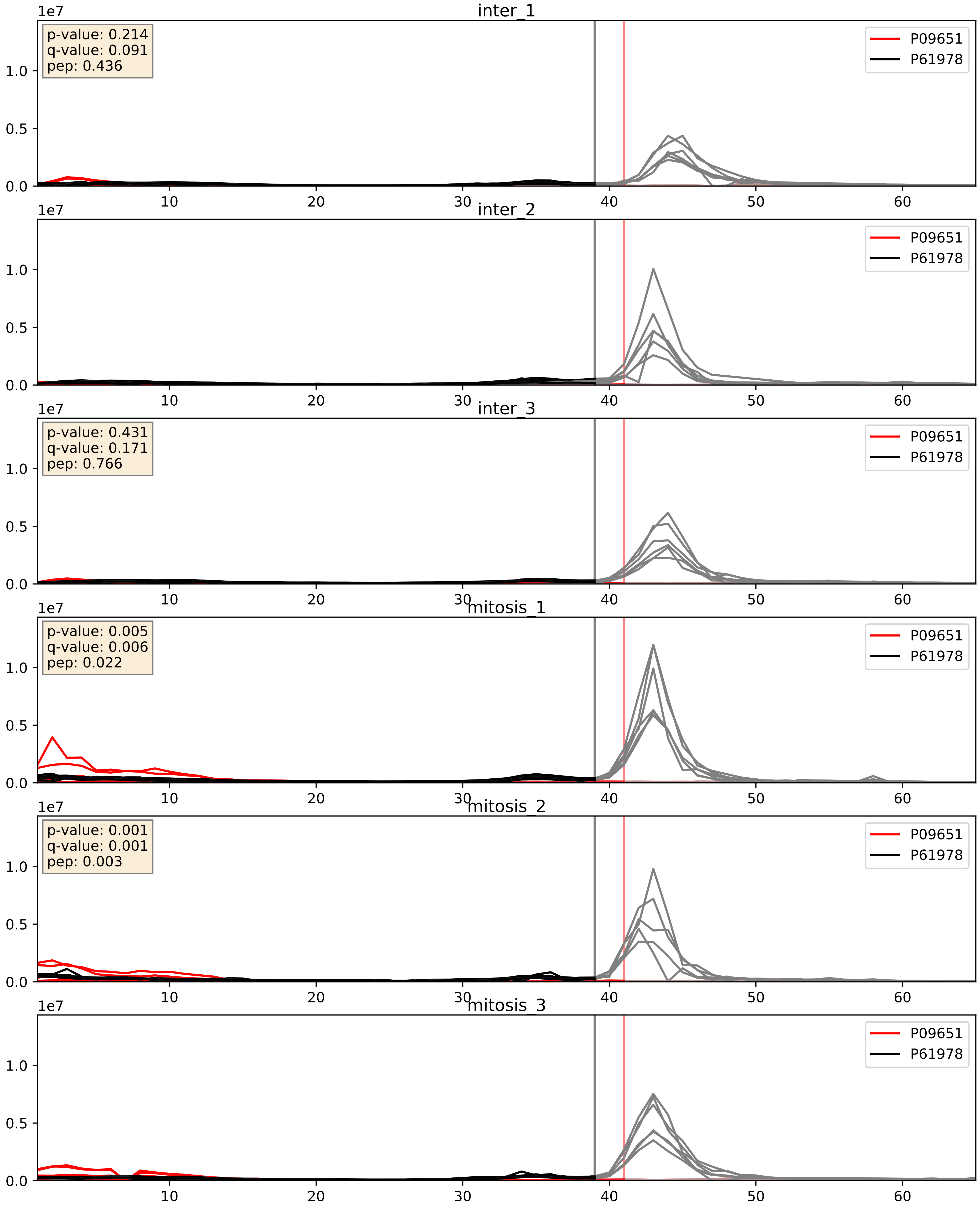


| level | condition_1 | condition_2 | pvalue | pvalue_adjusted |
|---|---|---|---|---|
| interactor_abundance | mitosis | inter | 0.016854435383293762 | 0.11277132084292767 |
| interactor_ratio | mitosis | inter | 0.021159436558454855 | 0.2203464439663912 |
| complex_abundance | mitosis | inter | 0.2471044668148845 | 0.3977727299820493 |

P09651_P62750
ROA1_HUMAN vs RL23A_HUMAN
p-value: 0.0 q-value: 0.046

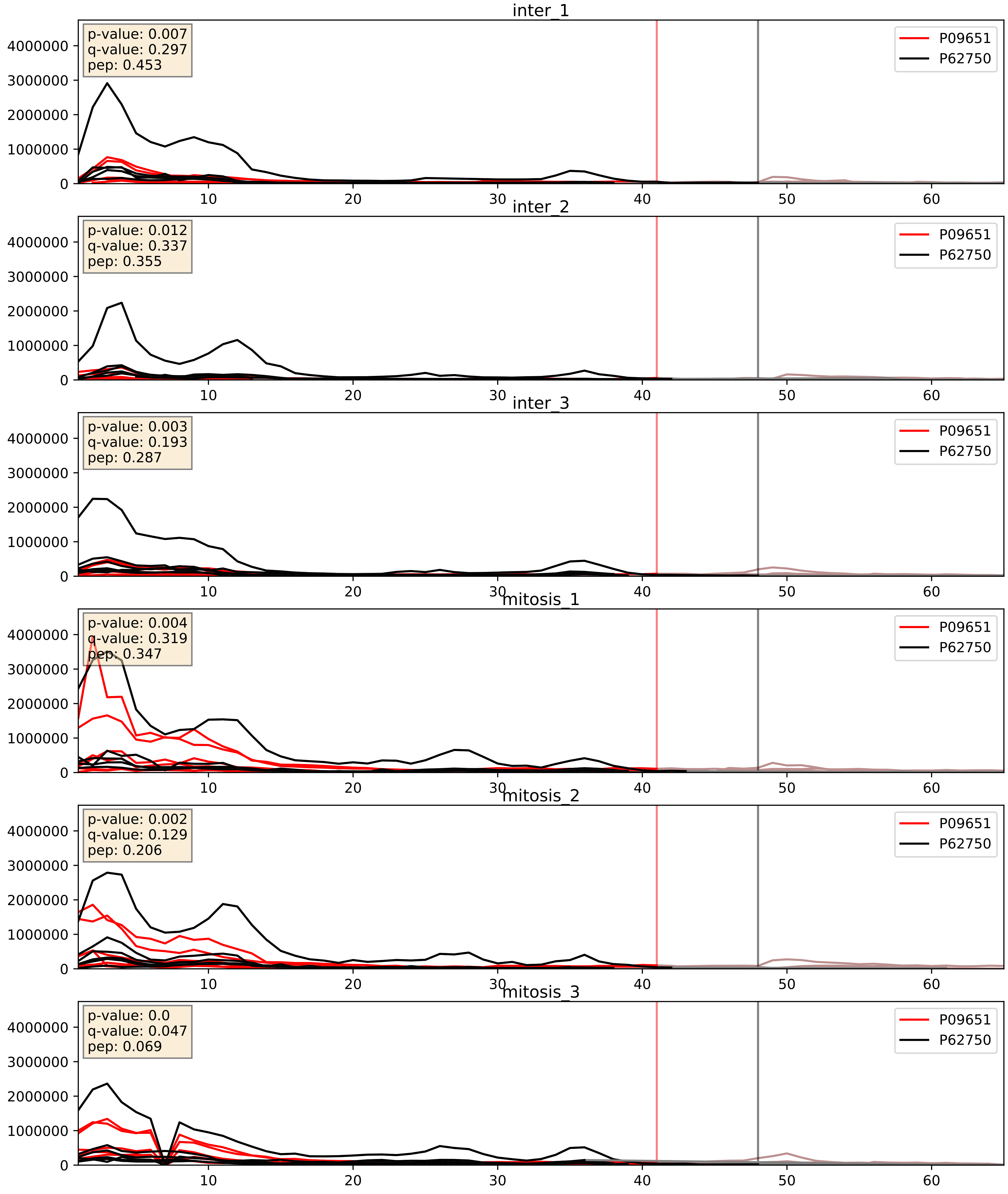


| level | condition_1 | condition_2 | pvalue | pvalue_adjusted |
|---|---|---|---|---|
| interactor_ratio | mitosis | inter | 0.004701192208311226 | 0.09159088101537749 |
| interactor_abundance | mitosis | inter | 0.015952360904200196 | 0.11088021938056777 |
| complex_abundance | mitosis | inter | 0.030385197940464886 | 0.12650646613345304 |

P09651_P62913
ROA1_HUMAN vs RL11_HUMAN
p-value: 0.0 q-value: 0.049

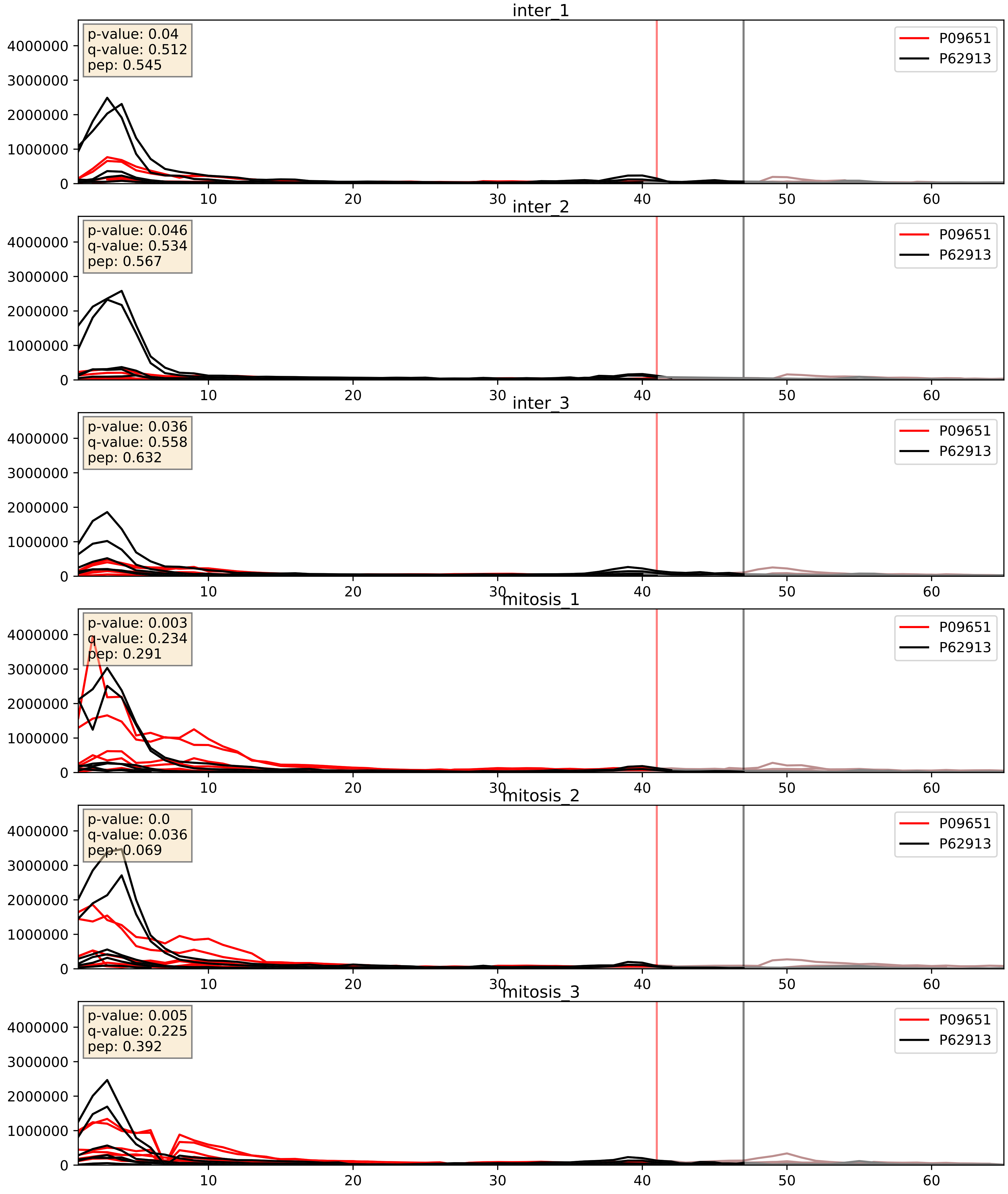


| level | condition_1 | condition_2 | pvalue | pvalue_adjusted |
|---|---|---|---|---|
| complex_abundance | mitosis | inter | 0.008793072340019222 | 0.06968788570668541 |
| interactor_abundance | mitosis | inter | 0.016856414137678265 | 0.11277132084292767 |
| interactor_ratio | mitosis | inter | 0.03158501771911432 | 0.28348436581583686 |

P09651_P62995
ROA1_HUMAN vs TRA2B_HUMAN
p-value: 0.006 q-value: 0.005

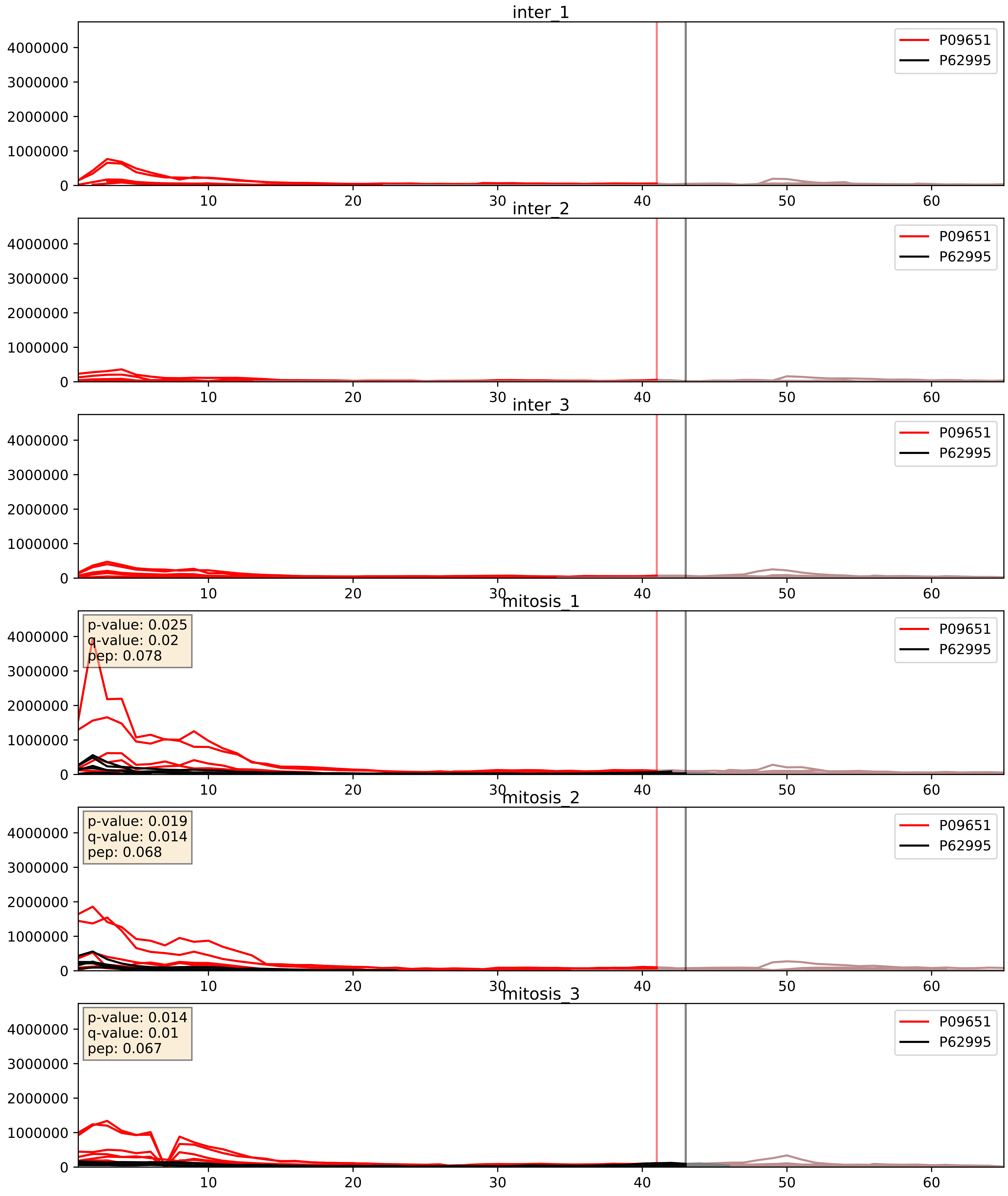


| level | condition_1 | condition_2 | pvalue | pvalue_adjusted |
|---|---|---|---|---|
| interactor_ratio | mitosis | inter | 9.380515584716844e-06 | 0.0035800160365075866 |
| interactor_abundance | mitosis | inter | 9.497380795475017e-05 | 0.007548664551888891 |
| complex_abundance | mitosis | inter | 0.0001646369536294834 | 0.008645965172198638 |

P09651_P67809
ROA1_HUMAN vs YBOX1_HUMAN
p-value: 0.001 q-value: 0.021

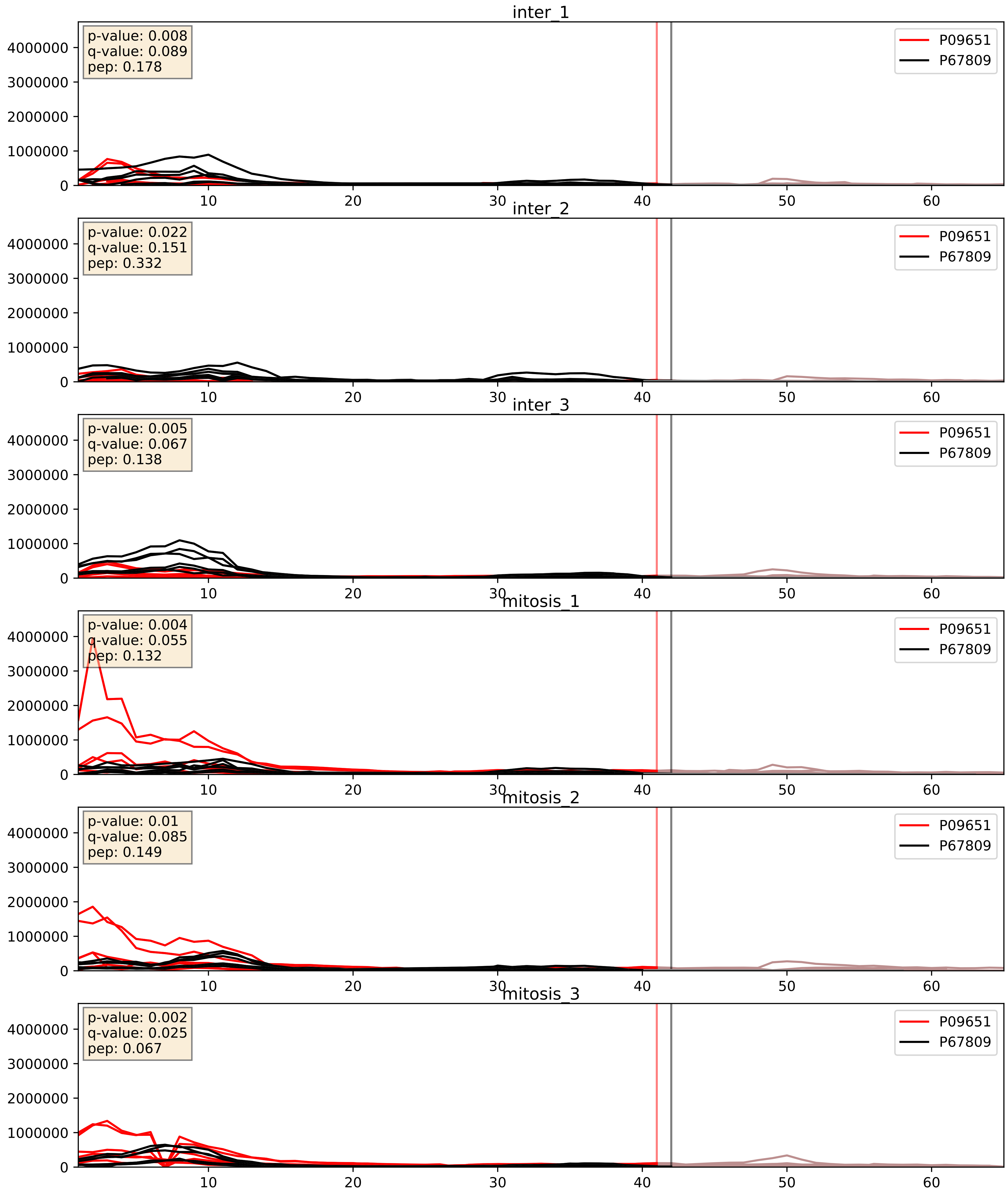


| level | condition_1 | condition_2 | pvalue | pvalue_adjusted |
|---|---|---|---|---|
| interactor_ratio | mitosis | inter | 0.003328707930510469 | 0.07518137172867972 |
| interactor_abundance | mitosis | inter | 0.016856414137678265 | 0.11277132084292767 |
| complex_abundance | mitosis | inter | 0.16292822492460565 | 0.3115178926411937 |

P09651_P84103
ROA1_HUMAN vs SRSF3_HUMAN
p-value: 0.051 q-value: 0.033

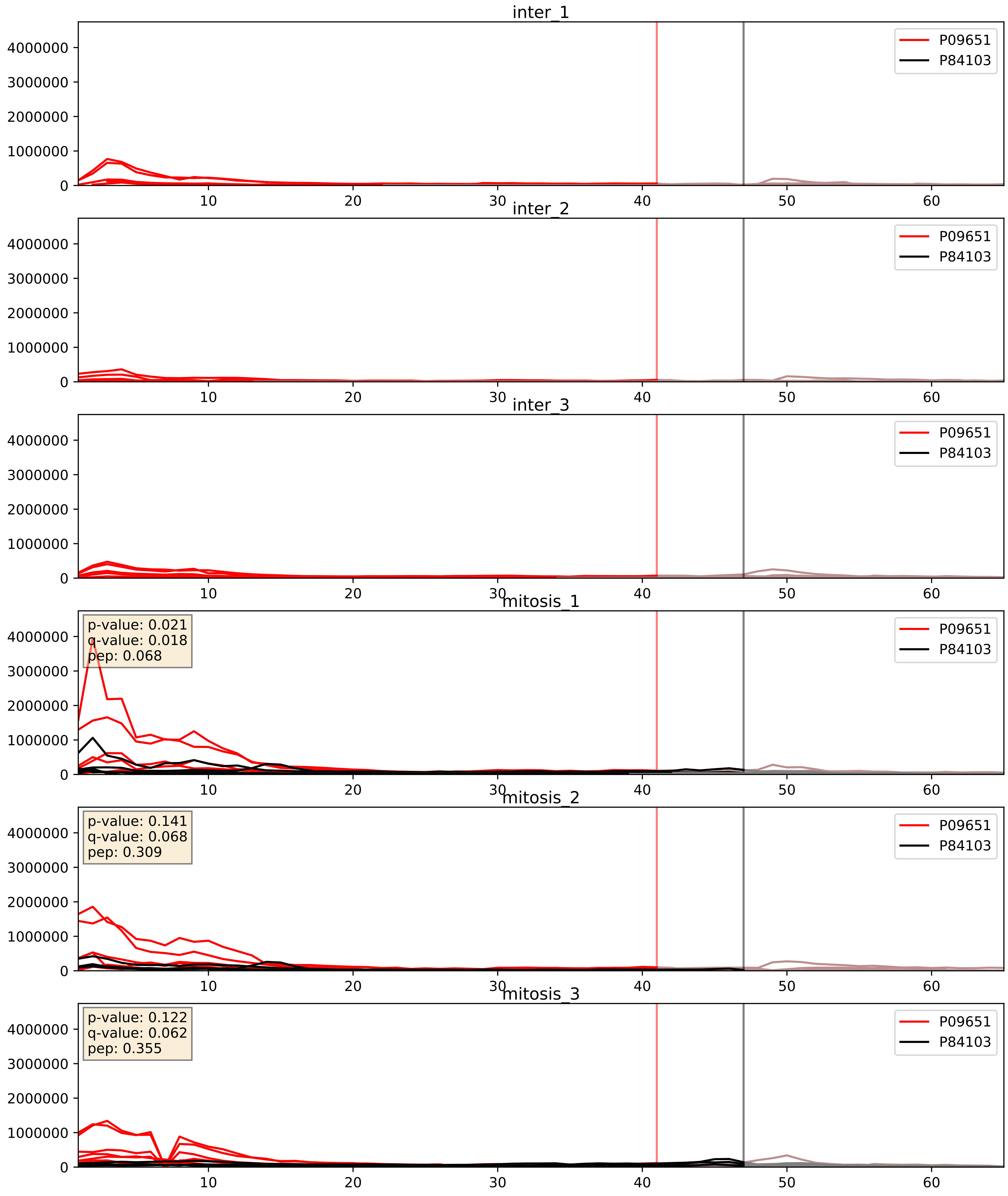


| level | condition_1 | condition_2 | pvalue | pvalue_adjusted |
|---|---|---|---|---|
| interactor_abundance | mitosis | inter | 7.226077128975986e-05 | 0.006834830963981706 |
| complex_abundance | mitosis | inter | 0.0002291506843386366 | 0.010219389159334393 |
| interactor_ratio | mitosis | inter | 0.004188942000443196 | 0.08517183734867877 |

P09651_Q02878
ROA1_HUMAN vs RL6_HUMAN
p-value: 0.0 q-value: 0.023

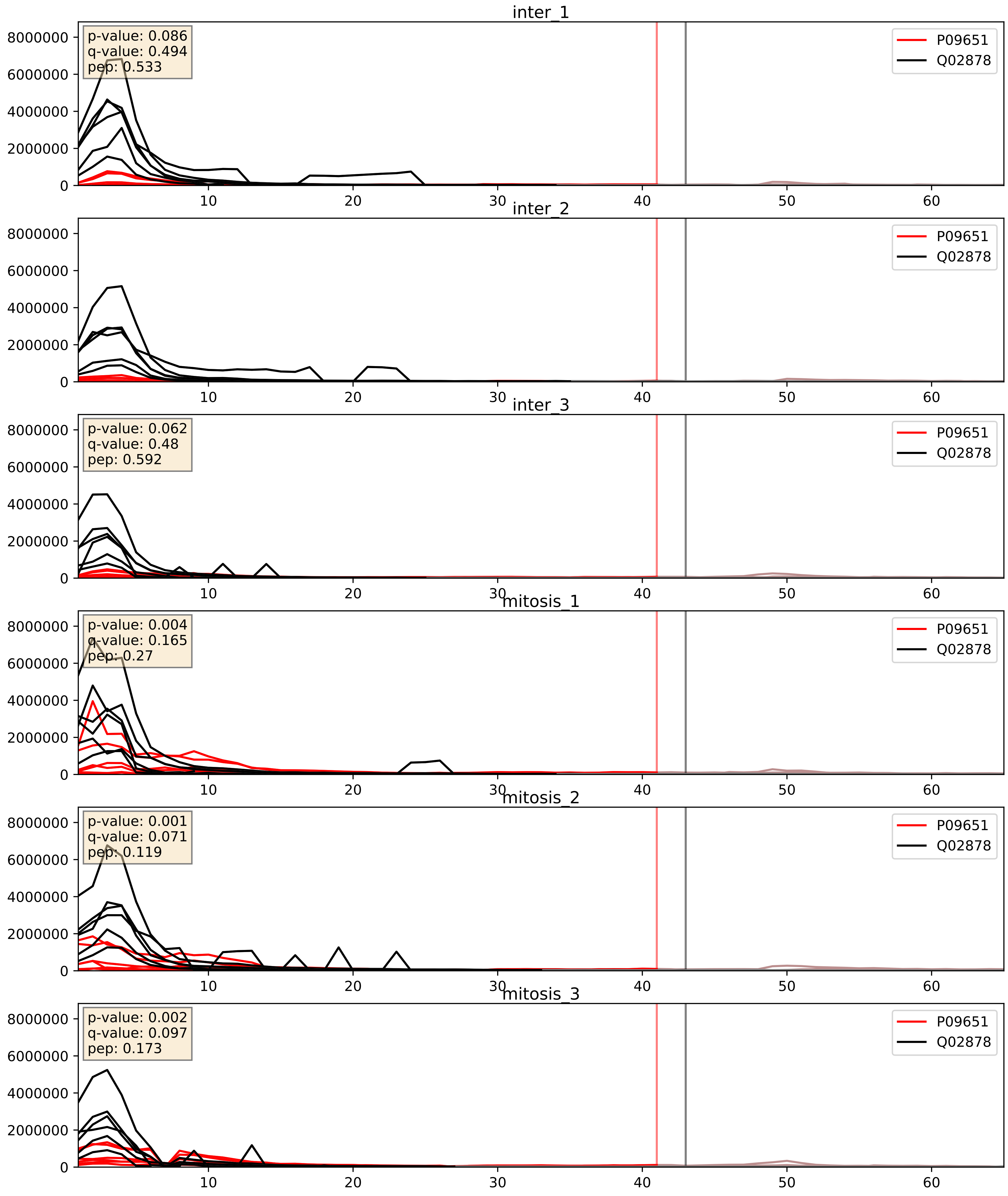


| level | condition_1 | condition_2 | pvalue | pvalue_adjusted |
|---|---|---|---|---|
| interactor_abundance | mitosis | inter | 0.01399567717616296 | 0.10490630177579242 |
| interactor_ratio | mitosis | inter | 0.014778673204475712 | 0.17767618346953945 |
| complex_abundance | mitosis | inter | 0.05322209695940319 | 0.16470757410429912 |

P09651_Q07955
ROA1_HUMAN vs SRSF1_HUMAN
p-value: 0.0 q-value: 0.0

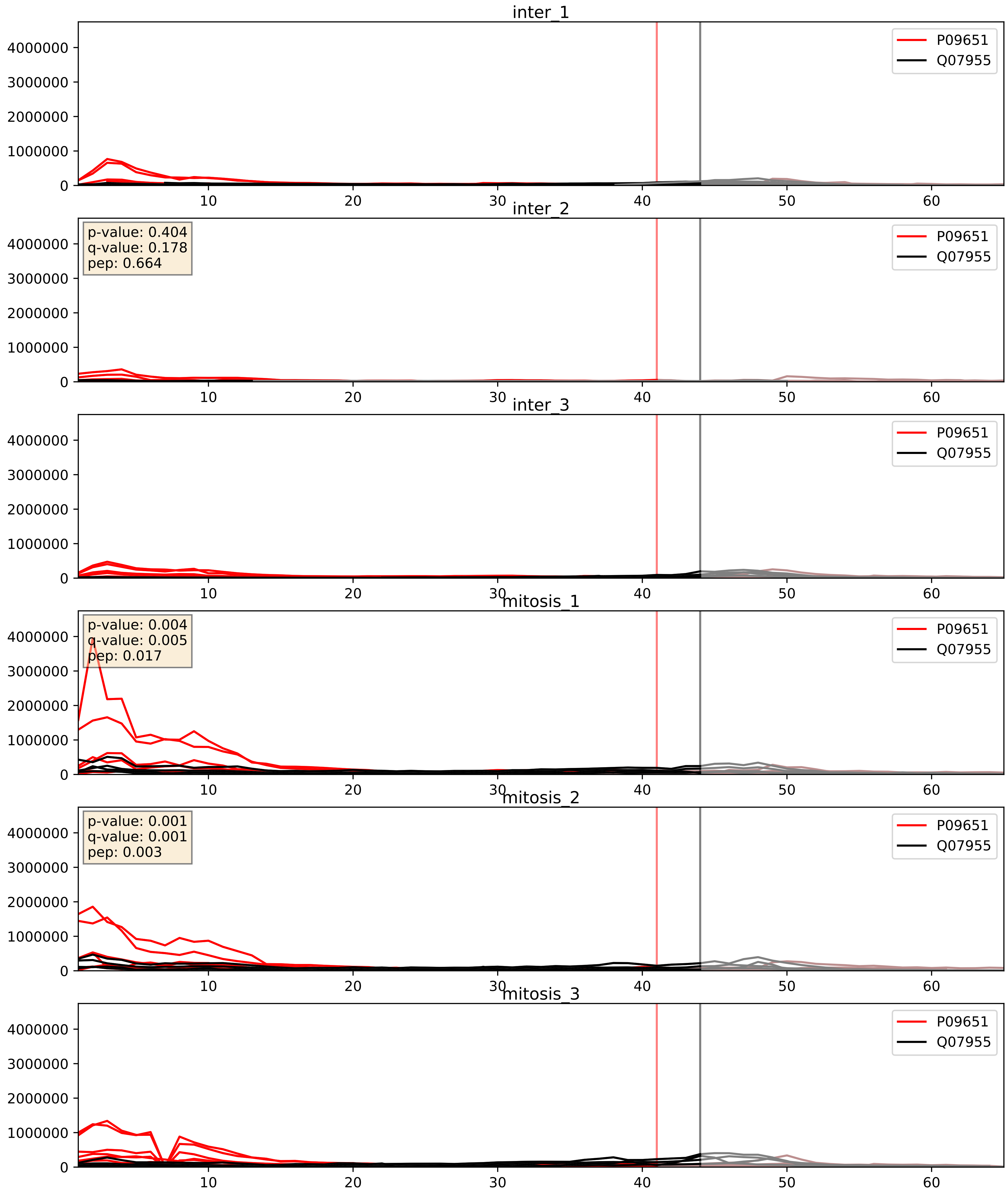


| level | condition_1 | condition_2 | pvalue | pvalue_adjusted |
|---|---|---|---|---|
| complex_abundance | mitosis | inter | 0.008509151419512374 | 0.0686506467021922 |
| interactor_abundance | mitosis | inter | 0.016490670357221612 | 0.11276964141946393 |
| interactor_ratio | mitosis | inter | 0.9147749642922947 | 0.9855669004419936 |

P09651_Q08211
ROA1_HUMAN vs DHX9_HUMAN
p-value: 0.003 q-value: 0.004

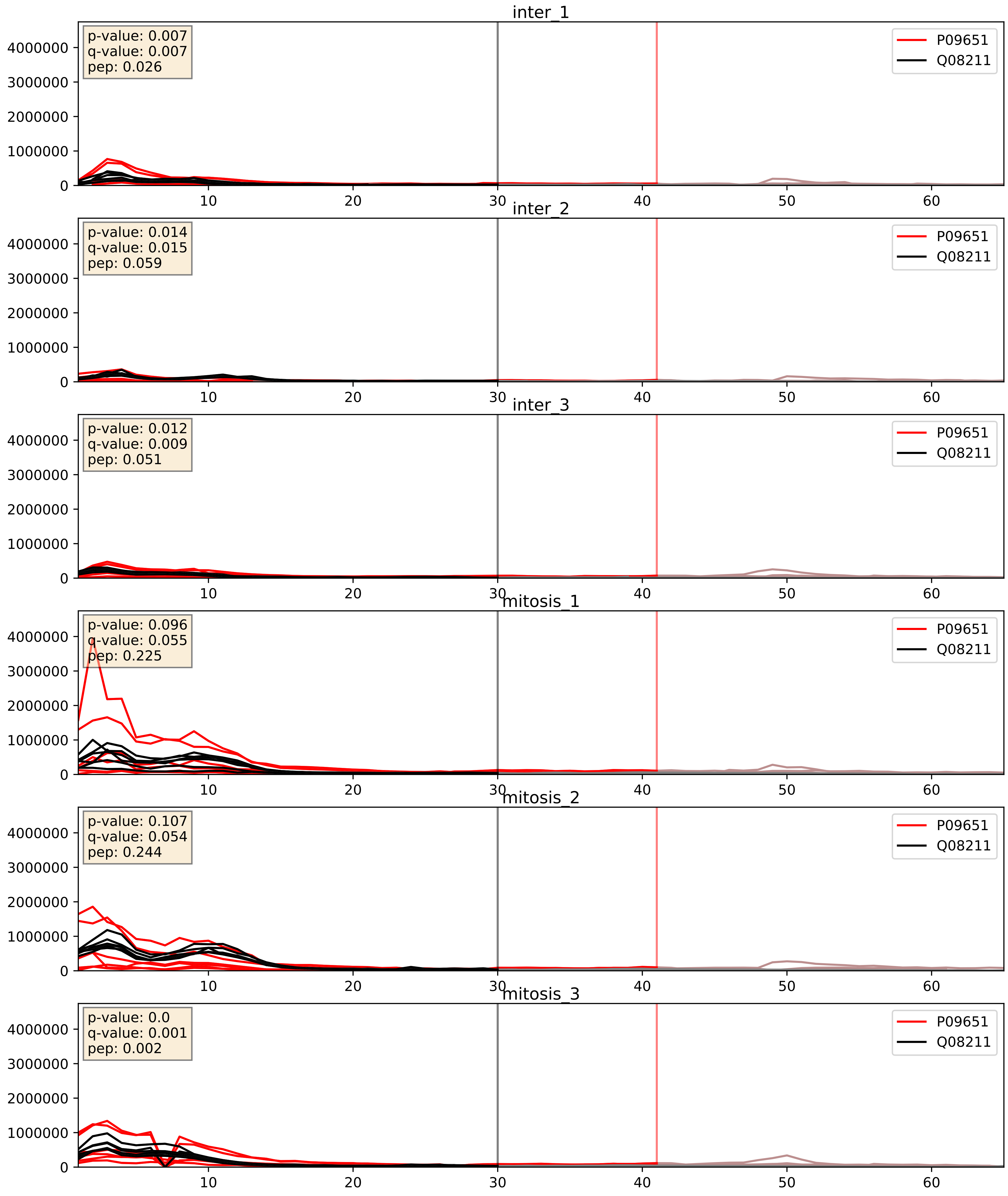


| level | condition_1 | condition_2 | pvalue | pvalue_adjusted |
|---|---|---|---|---|
| complex_abundance | mitosis | inter | 0.002467831850261896 | 0.03614494596242307 |
| interactor_abundance | mitosis | inter | 0.013517041923941197 | 0.10304923015527873 |
| interactor_ratio | mitosis | inter | 0.27952484058418636 | 0.7507789882022703 |

P09651_Q12905
ROA1_HUMAN vs ILF2_HUMAN
p-value: 0.0 q-value: 0.0

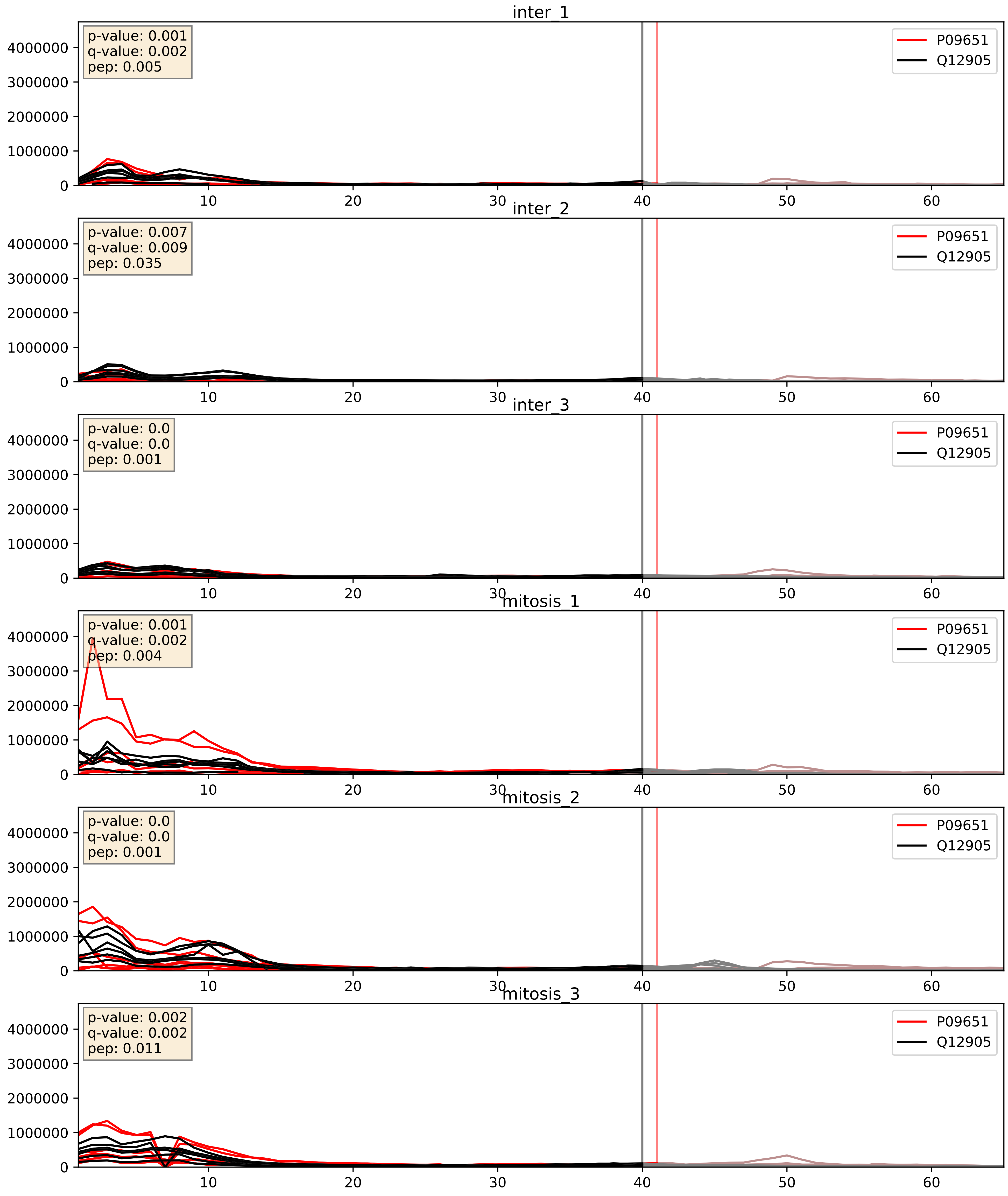


| level | condition_1 | condition_2 | pvalue | pvalue_adjusted |
|---|---|---|---|---|
| complex_abundance | mitosis | inter | 0.005618403472428057 | 0.05646492674842808 |
| interactor_abundance | mitosis | inter | 0.017050532289726136 | 0.11334462047152971 |
| interactor_ratio | mitosis | inter | 0.07777248885697595 | 0.4669171100029366 |

P09651_Q12906
ROA1_HUMAN vs ILF3_HUMAN
p-value: 0.001 q-value: 0.024

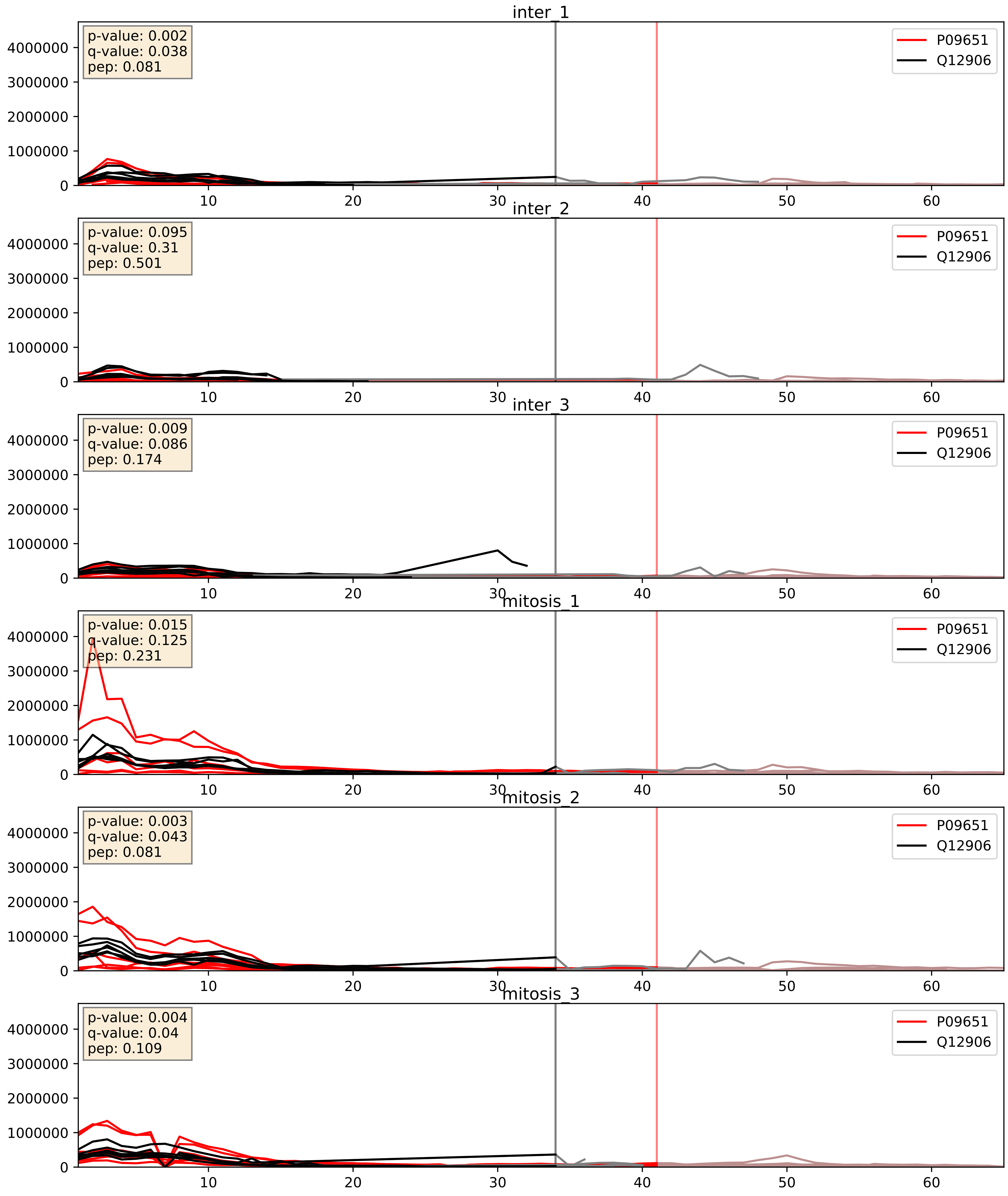


| level | condition_1 | condition_2 | pvalue | pvalue_adjusted |
|---|---|---|---|---|
| interactor_abundance | mitosis | inter | 0.013249945143764147 | 0.10304923015527873 |
| complex_abundance | mitosis | inter | 0.013672198236740039 | 0.08692368307384146 |
| interactor_ratio | mitosis | inter | 0.01455608166892149 | 0.17686306829592754 |

P09651_Q13151
ROA1_HUMAN vs ROA0_HUMAN
p-value: 0.019 q-value: 0.015

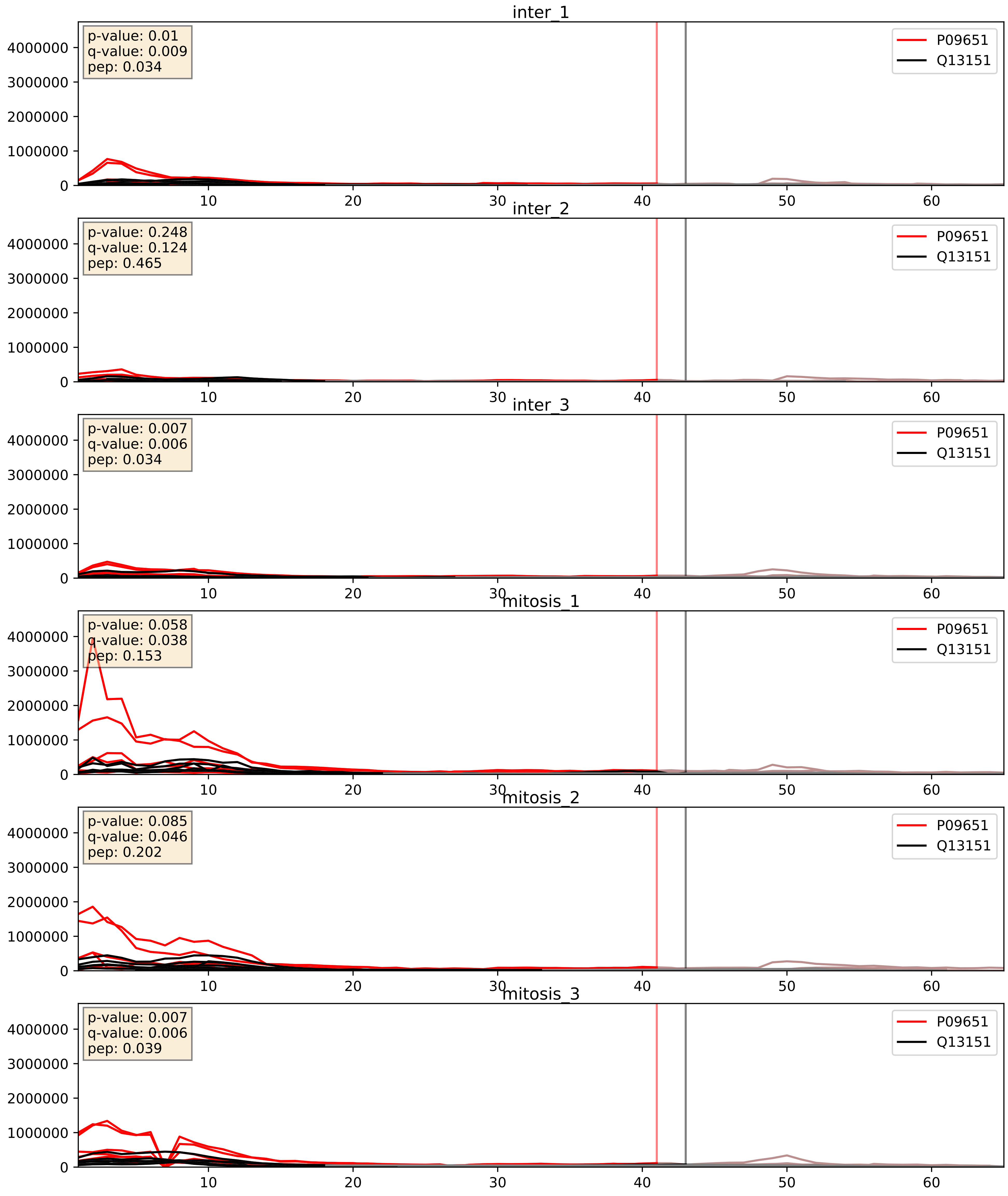


| level | condition_1 | condition_2 | pvalue | pvalue_adjusted |
|---|---|---|---|---|
| complex_abundance | mitosis | inter | 0.0063288454482083225 | 0.05946752693376865 |
| interactor_abundance | mitosis | inter | 0.014789027585019647 | 0.10635646274438217 |
| interactor_ratio | mitosis | inter | 0.3390820858283831 | 0.7924528012231136 |

P09651_Q14103
ROA1_HUMAN vs HNRPD_HUMAN
p-value: 0.031 q-value: 0.023

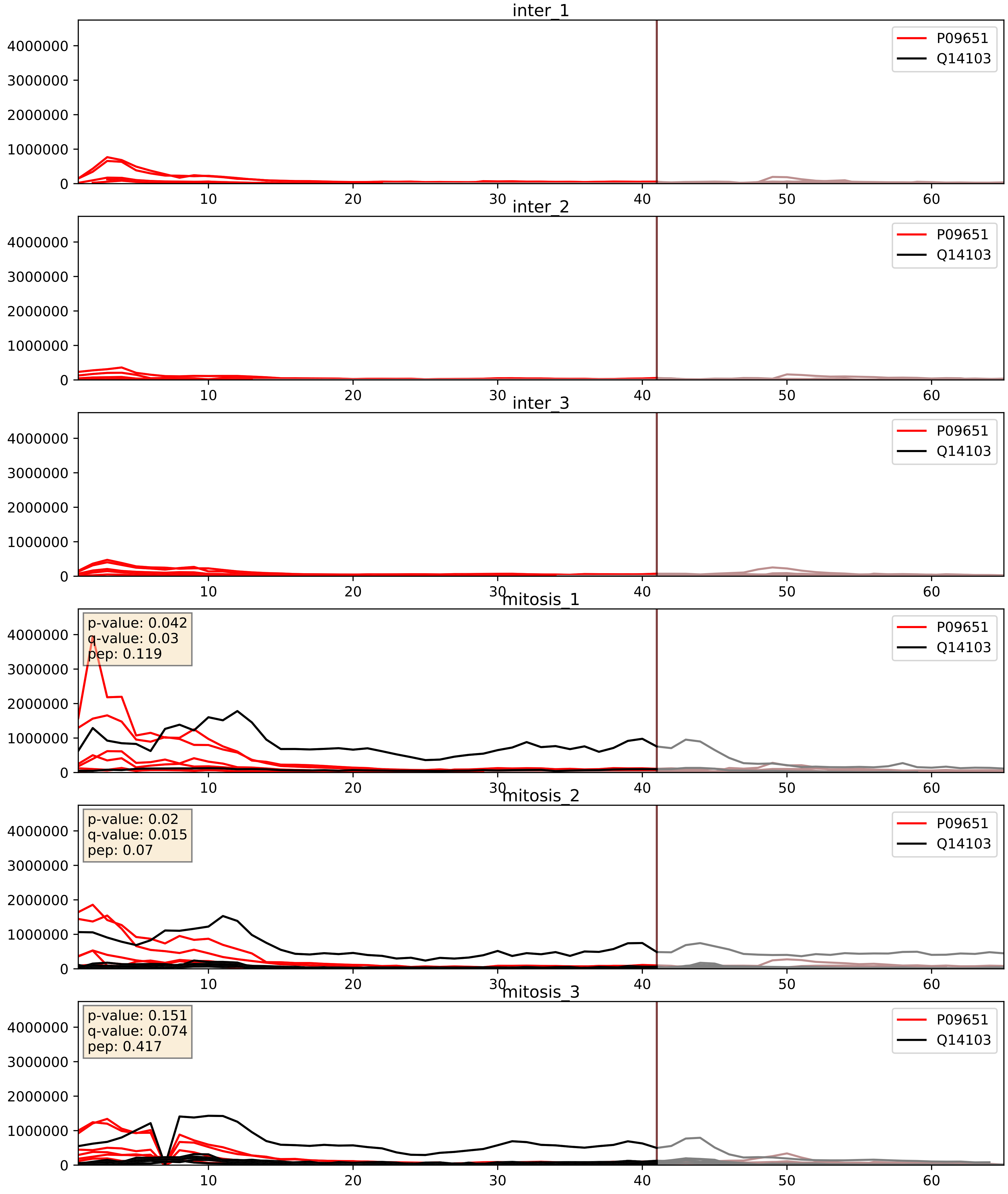


| level | condition_1 | condition_2 | pvalue | pvalue_adjusted |
|---|---|---|---|---|
| interactor_abundance | mitosis | inter | 6.959157312257744e-05 | 0.006732838333317079 |
| complex_abundance | mitosis | inter | 0.00029696456492007517 | 0.011617922212881656 |
| interactor_ratio | mitosis | inter | 0.26529417826222884 | 0.7404362309130512 |

P09651_Q15366
ROA1_HUMAN vs PCBP2_HUMAN
p-value: 0.078 q-value: 0.046

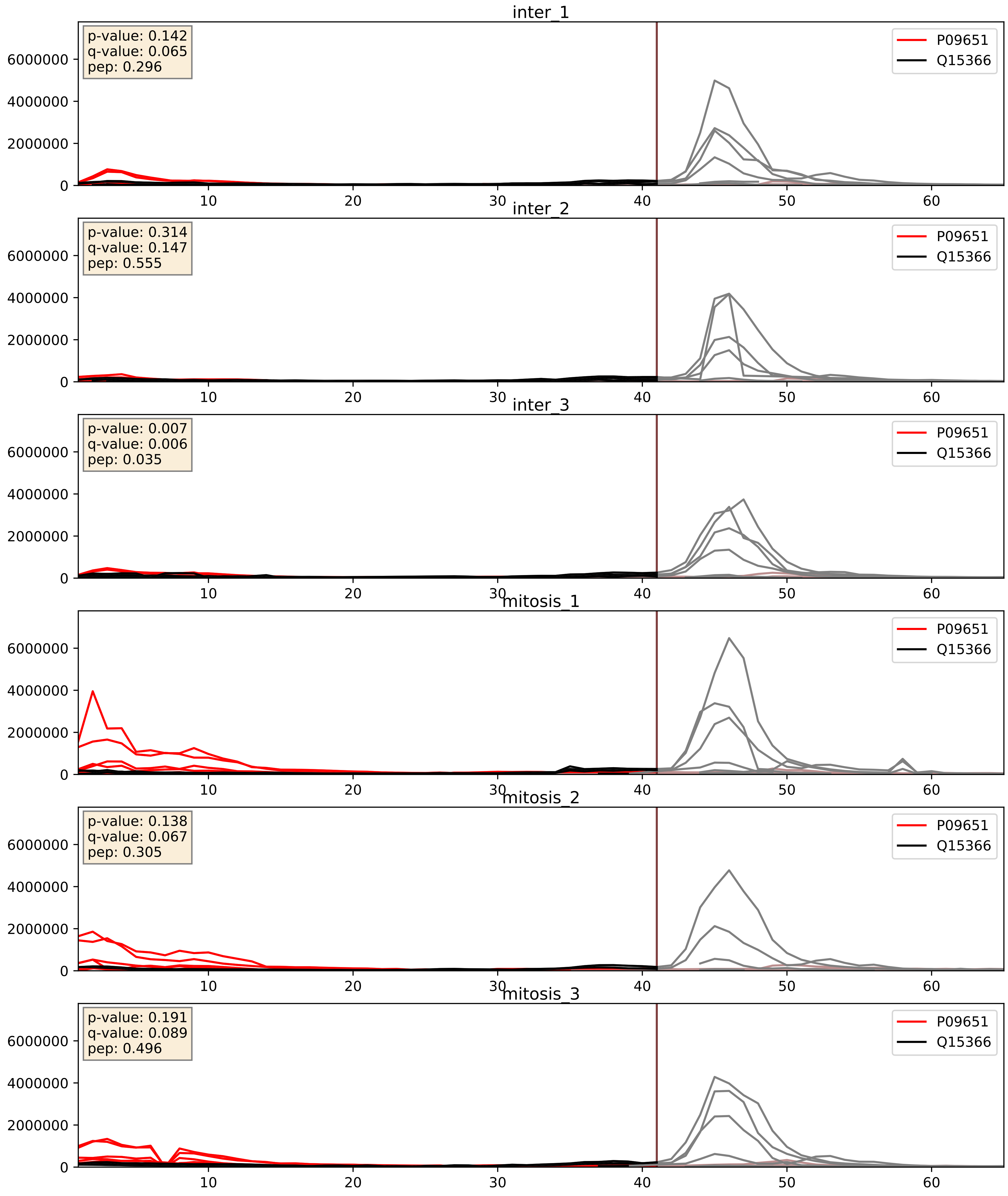


| level | condition_1 | condition_2 | pvalue | pvalue_adjusted |
|---|---|---|---|---|
| interactor_ratio | mitosis | inter | 0.011566658837338225 | 0.15494522140242886 |
| interactor_abundance | mitosis | inter | 0.016856414137678265 | 0.11277132084292767 |
| complex_abundance | mitosis | inter | 0.261542523081216 | 0.4114893083547594 |

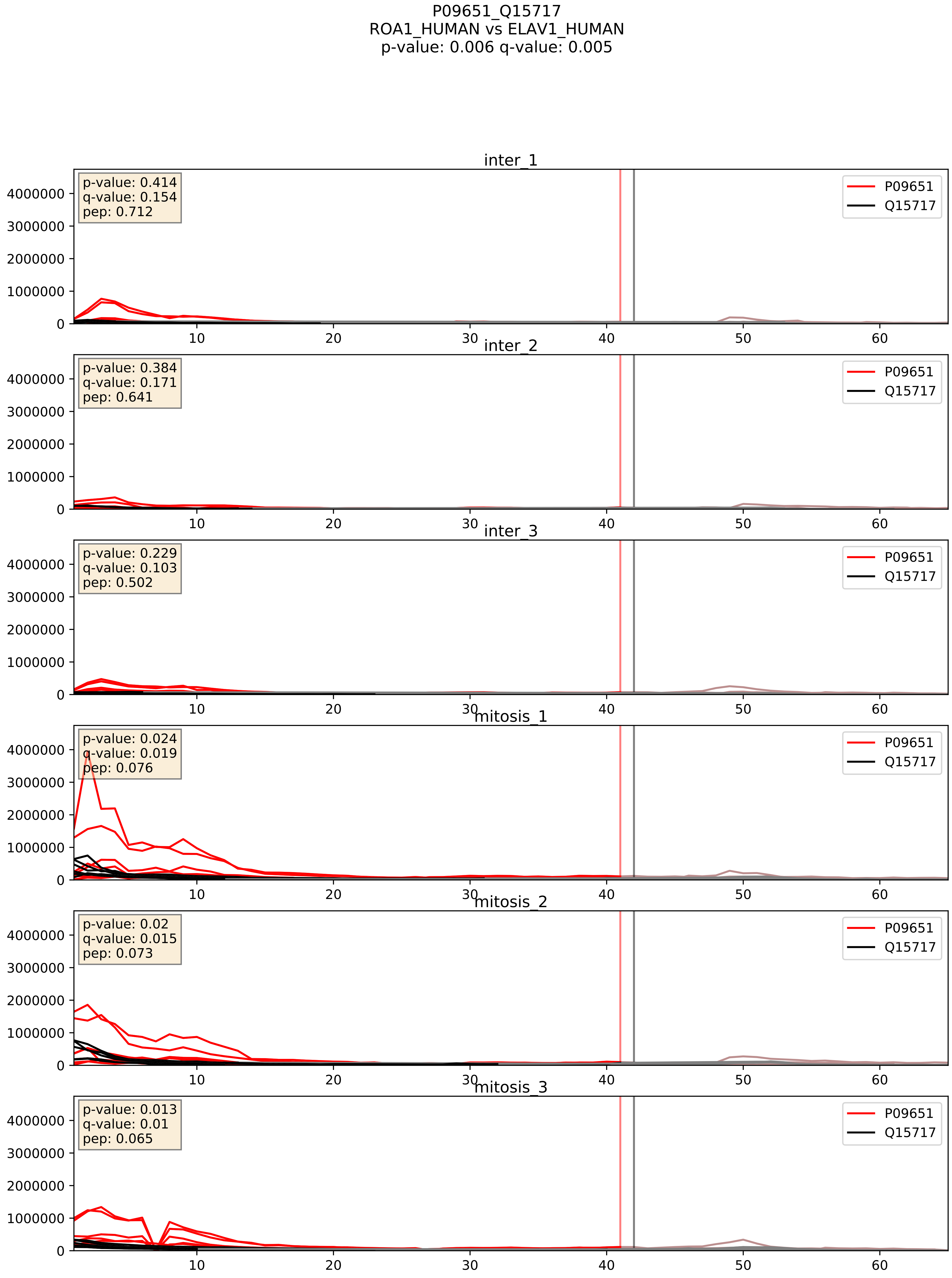


| level | condition_1 | condition_2 | pvalue | pvalue_adjusted |
|---|---|---|---|---|
| complex_abundance | mitosis | inter | 0.0009095527070255489 | 0.0197847007881119 |
| interactor_abundance | mitosis | inter | 0.012699265612423818 | 0.10046738783950822 |
| interactor_ratio | mitosis | inter | 0.4221604689601121 | 0.8388785723072948 |

P09651_Q16629
ROA1_HUMAN vs SRSF7_HUMAN
p-value: 0.008 q-value: 0.007

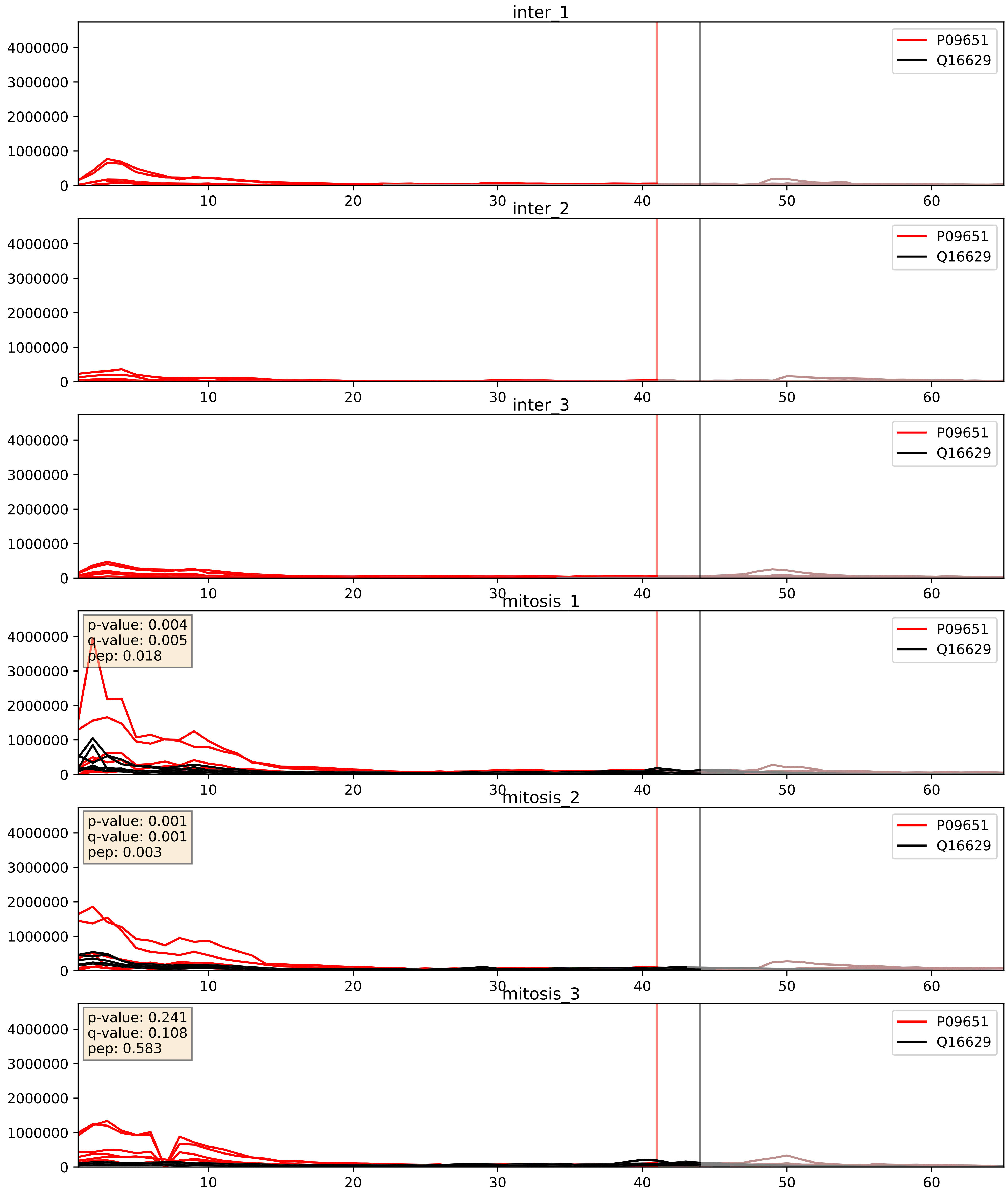


| level | condition_1 | condition_2 | pvalue | pvalue_adjusted |
|---|---|---|---|---|
| complex_abundance | mitosis | inter | 5.081957409361551e-05 | 0.005691857151652958 |
| interactor_abundance | mitosis | inter | 6.959157312257744e-05 | 0.006732838333317079 |
| interactor_ratio | mitosis | inter | 0.00173672861406567 | 0.05073855609693561 |

P09651_Q9UMS4
ROA1_HUMAN vs PRP19_HUMAN
p-value: 0.02 q-value: 0.016

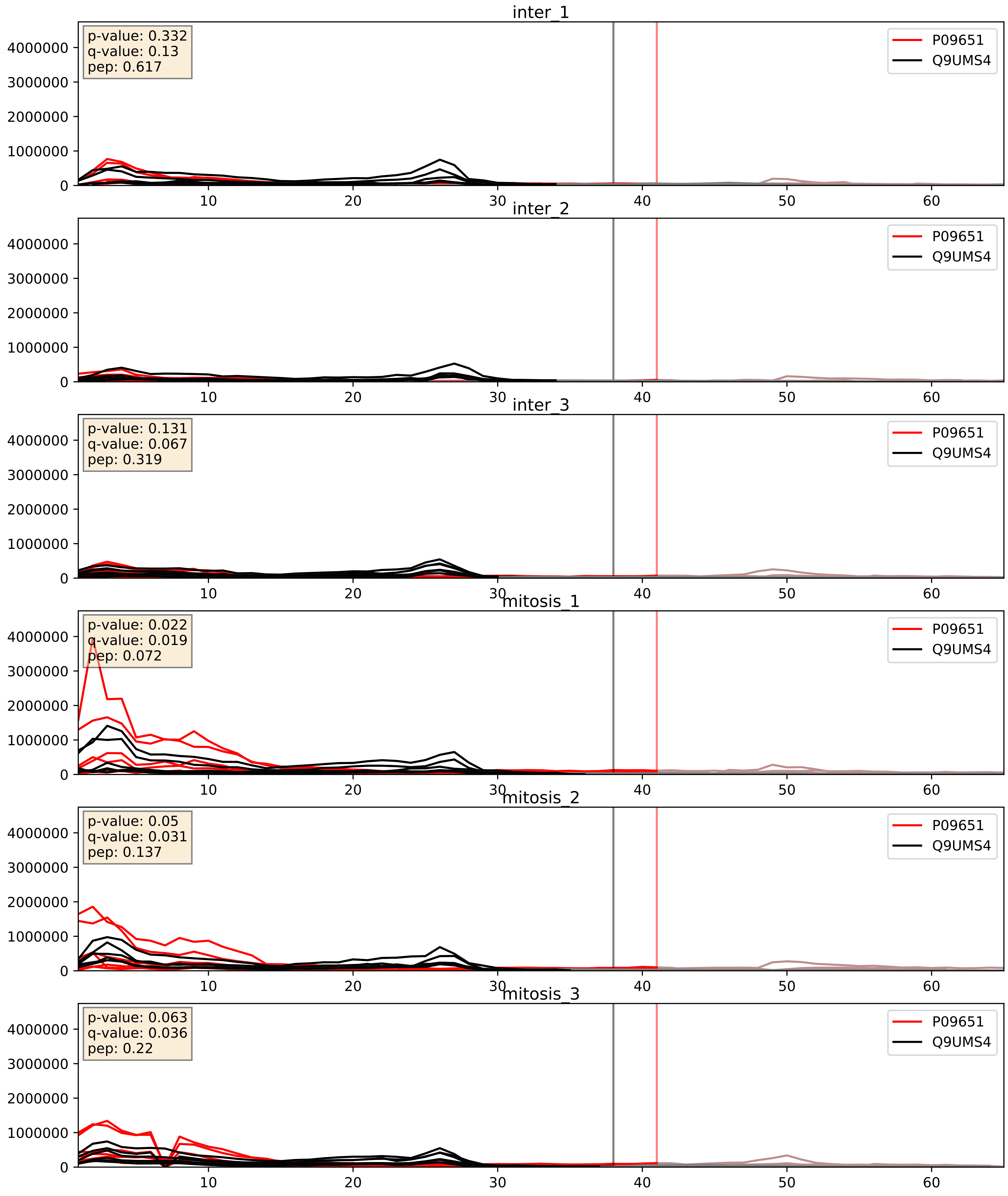


| level | condition_1 | condition_2 | pvalue | pvalue_adjusted |
|---|---|---|---|---|
| interactor_ratio | mitosis | inter | 0.0062861241323112415 | 0.10931911715410765 |
| interactor_abundance | mitosis | inter | 0.015540296295012546 | 0.10925937121894304 |
| complex_abundance | mitosis | inter | 0.025085574239161013 | 0.11592191390630502 |
