## Supplementary Data 2 for "SECAT: Quantifying differential protein-protein interaction states by network-centric analysis": hela_string_000146_Q9UL03.pdf

Q9UL03_Q9UL03

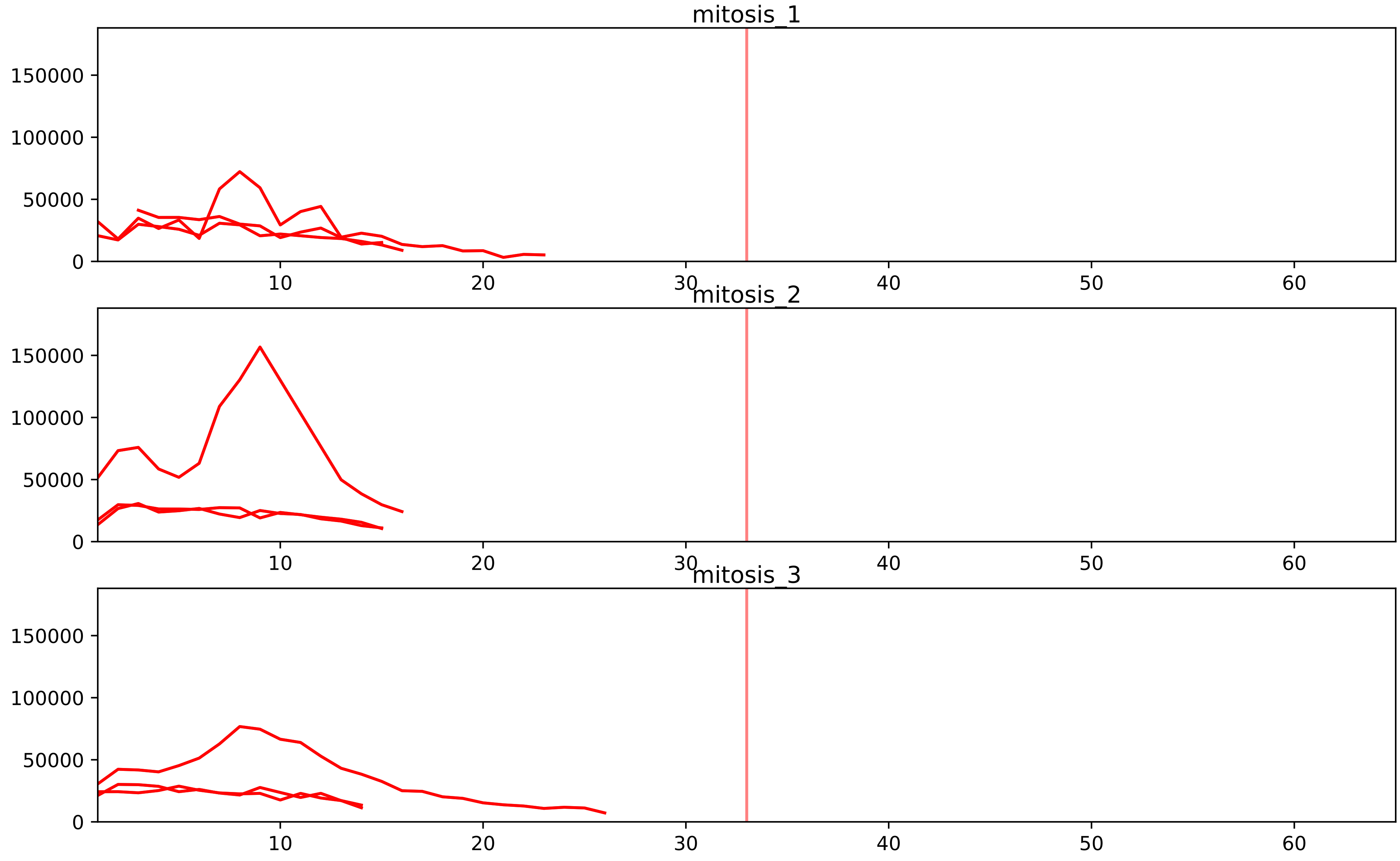


| level | condition_1 | condition_2 | pvalue | pvalue_adjusted |
|---|---|---|---|---|
| assembled_abundance | mitosis | inter | 5.4374426560626974e-05 | 0.007270487507555603 |
| total_abundance | mitosis | inter | 0.00012855998062251173 | 0.013925507688281004 |
| interactor_abundance | mitosis | inter | 0.007977928386047226 | 0.045306753797305234 |
| complex_abundance | mitosis | inter | 0.06612431021463074 | 0.14729870556285782 |
| interactor_ratio | mitosis | inter | 0.1392816302572313 | 0.40031290158651084 |

P52434_Q9UL03
RPAB3_HUMAN vs INT6_HUMAN
p-value: 0.079 q-value: 0.047

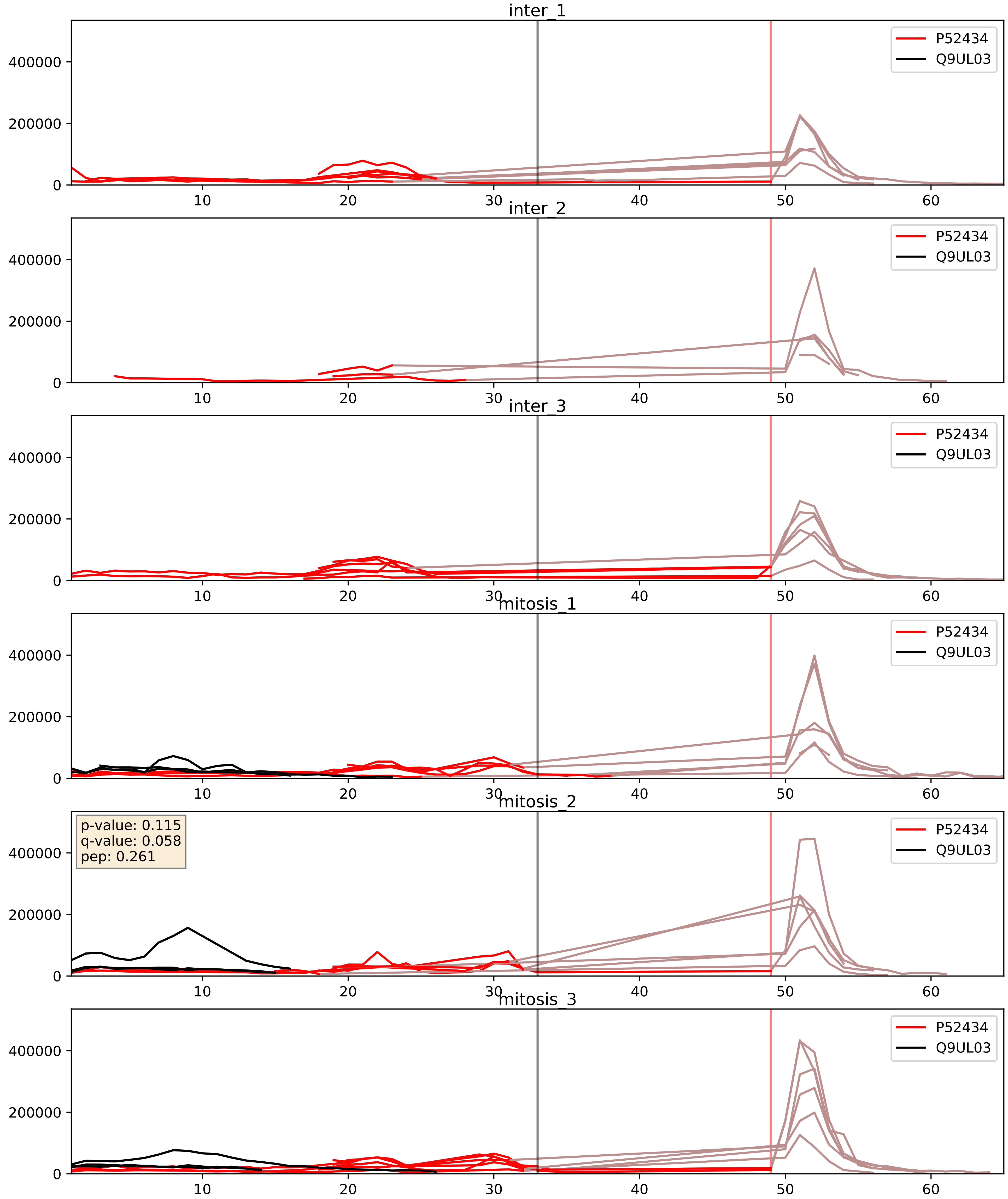


| level | condition_1 | condition_2 | pvalue | pvalue_adjusted |
|---|---|---|---|---|
| complex_abundance | mitosis | inter | 0.06612431021463074 | 0.18558167063516037 |
| interactor_ratio | mitosis | inter | 0.1392816302572313 | 0.5876051035001971 |
| interactor_abundance | mitosis | inter | 0.27796856819288246 | 0.4714038521507823 |
