## Supplementary Data 2 for "SECAT: Quantifying differential protein-protein interaction states by network-centric analysis": hela_string_000147_Q9Y5J1.pdf

Q9Y5J1_Q9Y5J1

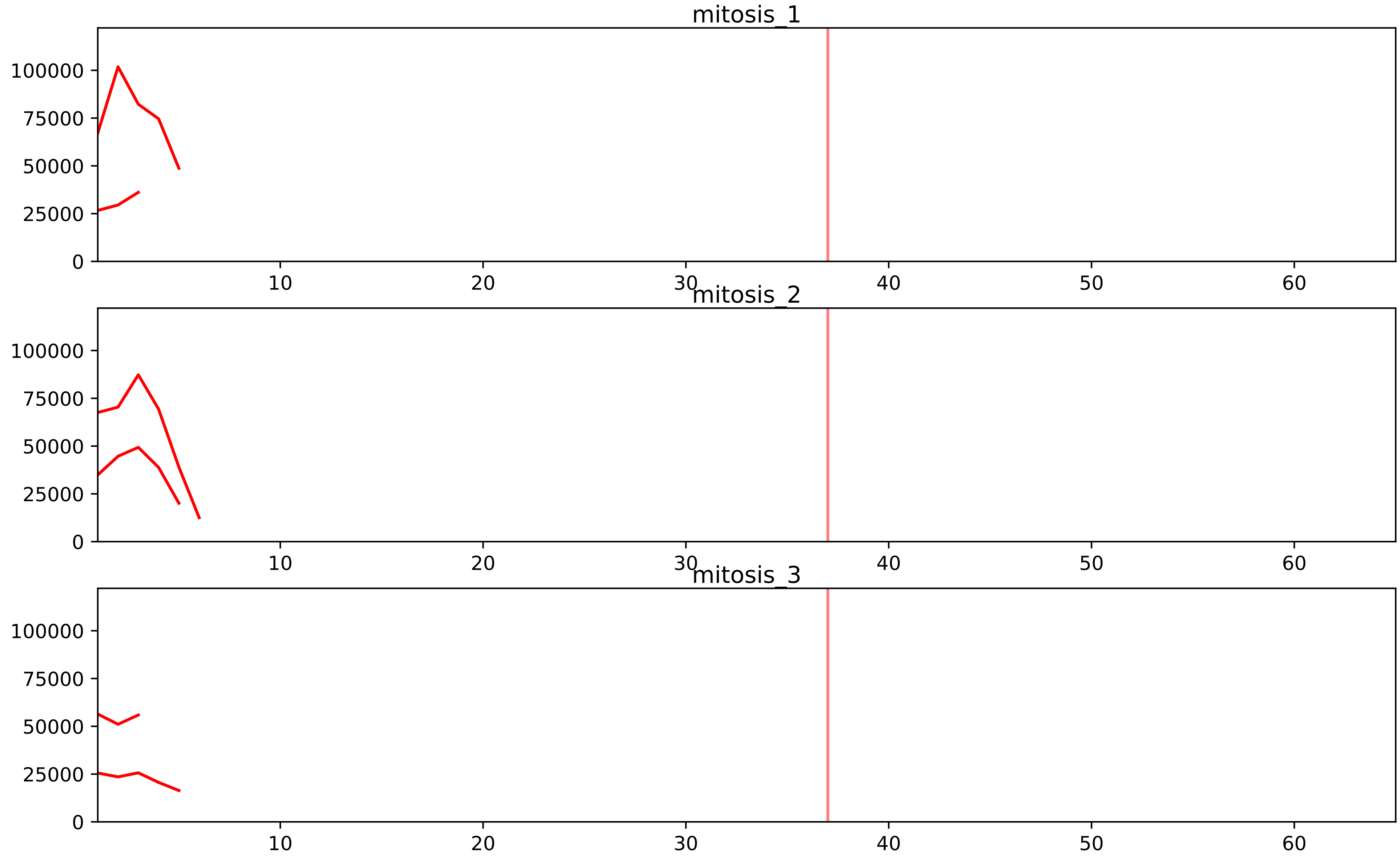


| level | condition_1 | condition_2 | pvalue | pvalue_adjusted |
|---|---|---|---|---|
| interactor_ratio | mitosis | inter | 5.456615007259685e-05 | 0.001024507307485492 |
| assembled_abundance | mitosis | inter | 0.003029389630227466 | 0.053918527981348933 |
| complex_abundance | mitosis | inter | 0.004422316426769359 | 0.02689873750764849 |
| total_abundance | mitosis | inter | 0.016260732958888646 | 0.16200272307965183 |
| interactor_abundance | mitosis | inter | 0.11303148731605983 | 0.246132235178146 |

O76021_Q9Y5J1
RL1D1_HUMAN vs UTP18_HUMAN
p-value: 0.068 q-value: 0.042

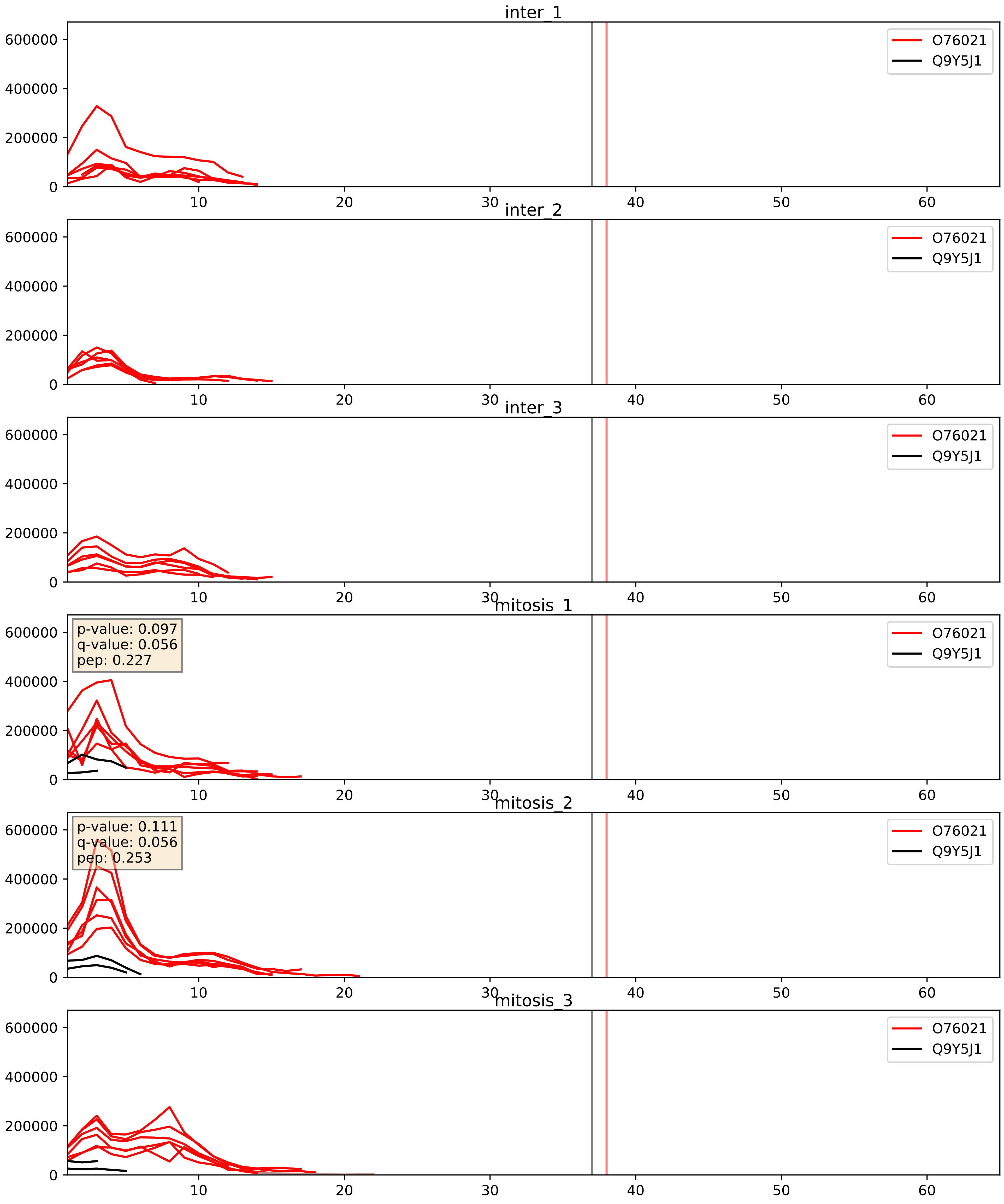


| level | condition_1 | condition_2 | pvalue | pvalue_adjusted |
|---|---|---|---|---|
| interactor_ratio | mitosis | inter | 0.0008389471802542103 | 0.03318433890476037 |
| interactor_abundance | mitosis | inter | 0.0012303243090360165 | 0.02571813451855507 |
| complex_abundance | mitosis | inter | 0.0035472690192235807 | 0.043600374082146835 |

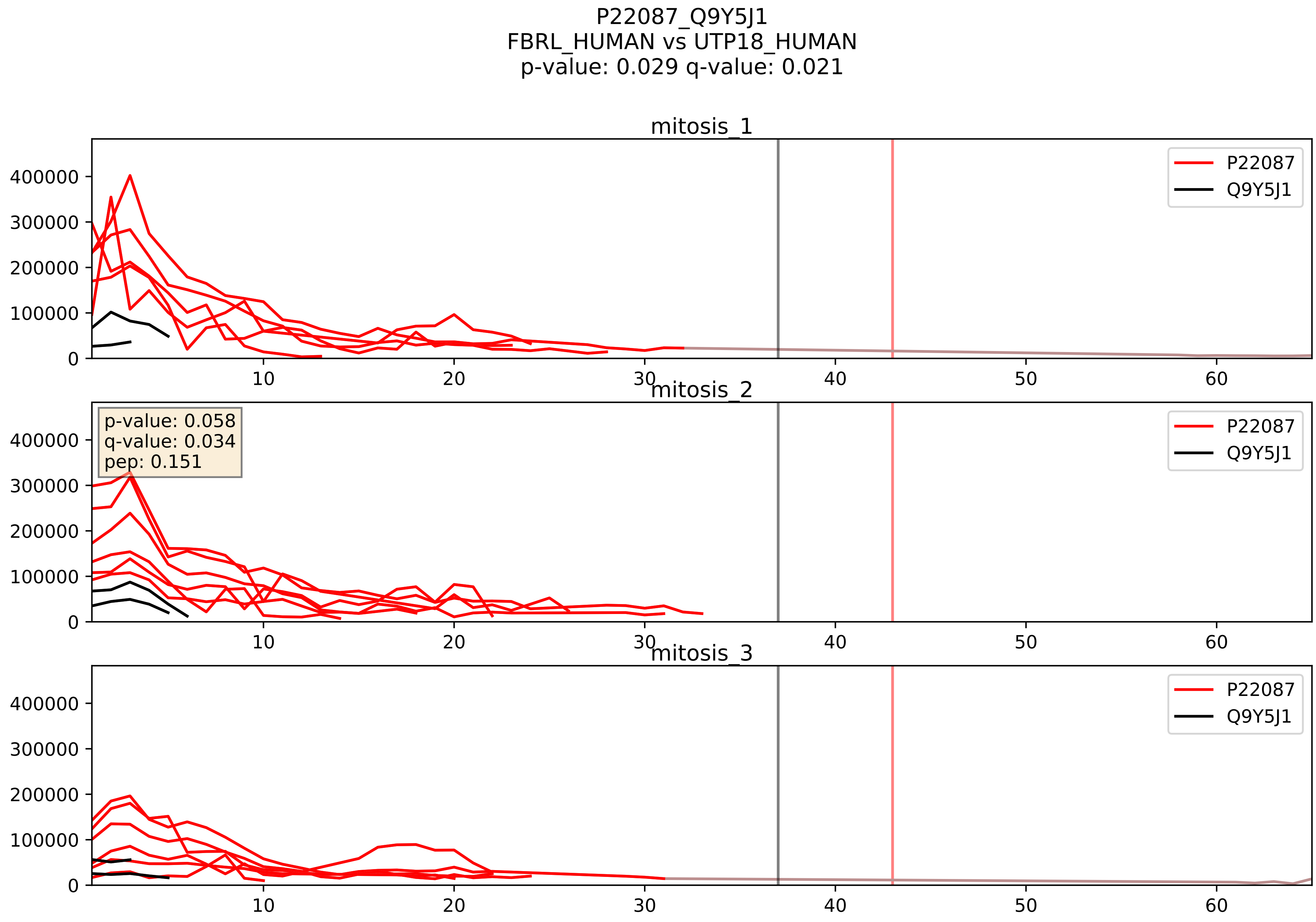


| level | condition_1 | condition_2 | pvalue | pvalue_adjusted |
|---|---|---|---|---|
| interactor_ratio | mitosis | inter | 0.00023104780740332214 | 0.015451322120097167 |
| interactor_abundance | mitosis | inter | 0.0020671551707800175 | 0.03307448273248028 |
| complex_abundance | mitosis | inter | 0.0066943970868762175 | 0.06051112889510077 |

Q8IY81_Q9Y5J1
SPB1_HUMAN vs UTP18_HUMAN
p-value: 0.076 q-value: 0.046

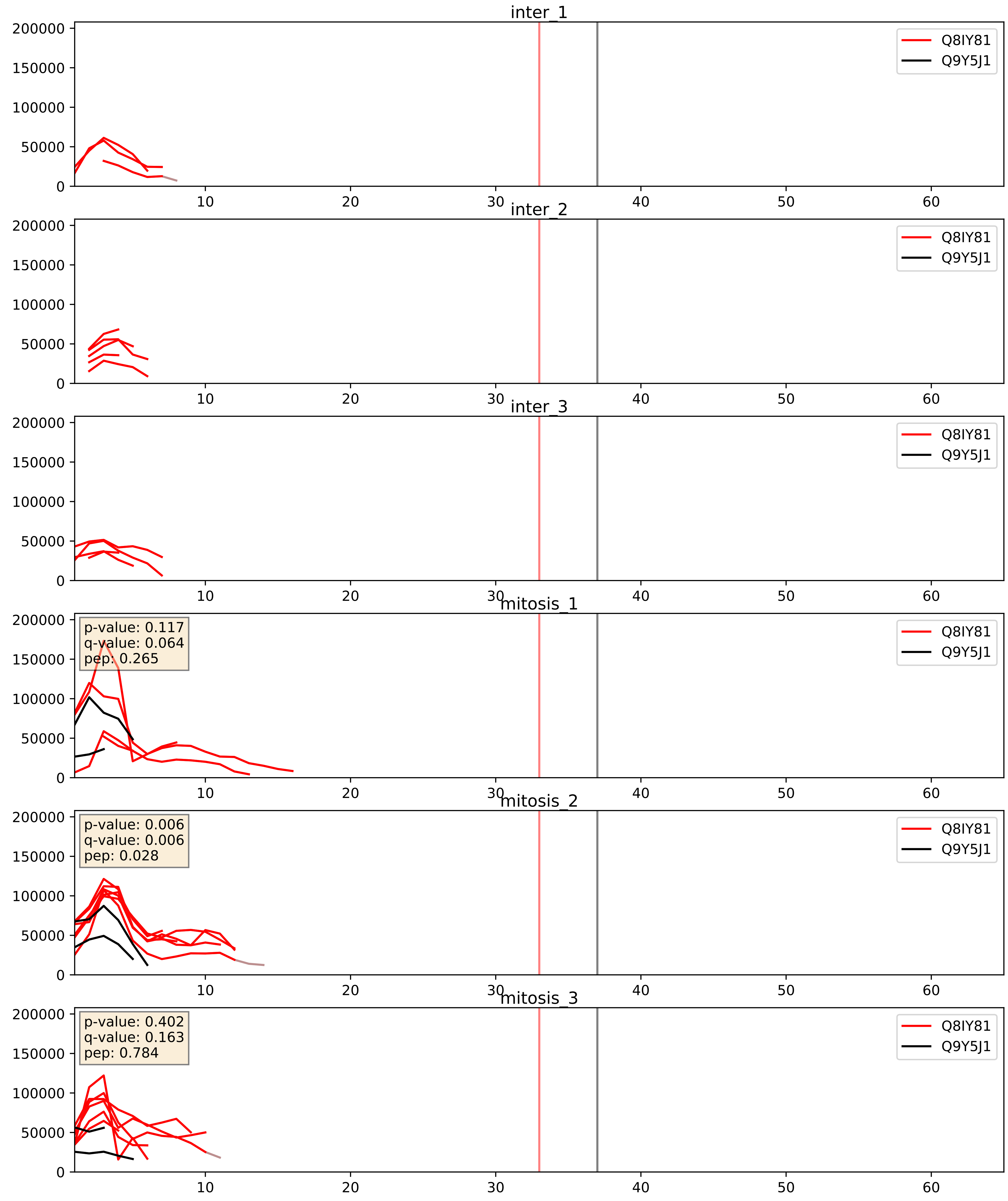


| level | condition_1 | condition_2 | pvalue | pvalue_adjusted |
|---|---|---|---|---|
| interactor_abundance | mitosis | inter | 0.008355526262714321 | 0.07951451340615297 |
| complex_abundance | mitosis | inter | 0.023105806231831845 | 0.11091884921279084 |
| interactor_ratio | mitosis | inter | 0.0241747810665698 | 0.24345426579980883 |

Q9H0A0_Q9Y5J1
NAT10_HUMAN vs UTP18_HUMAN
p-value: 0.064 q-value: 0.04

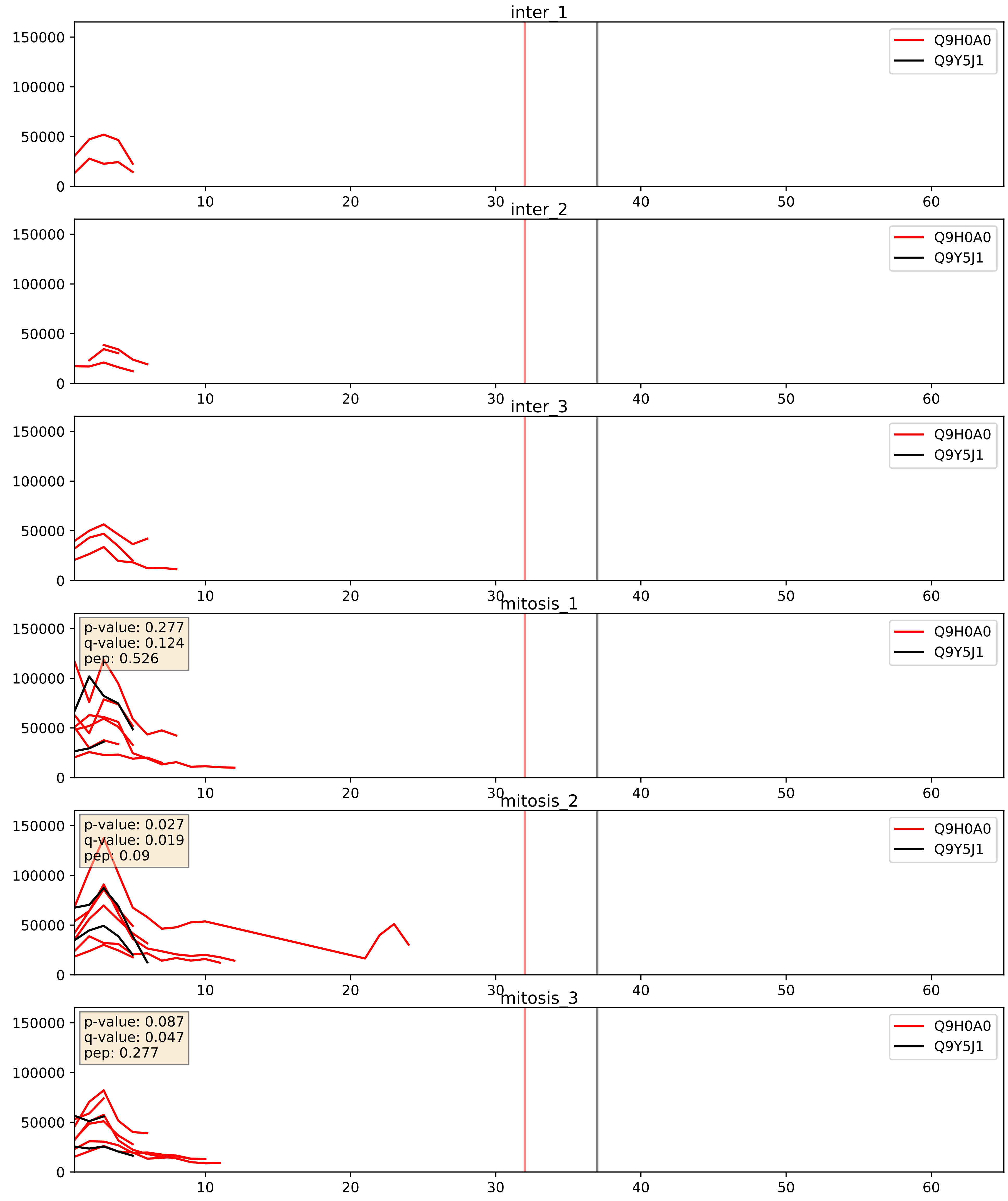


| level | condition_1 | condition_2 | pvalue | pvalue_adjusted |
|---|---|---|---|---|
| interactor_ratio | mitosis | inter | 0.0038335460165161953 | 0.08075090146433855 |
| interactor_abundance | mitosis | inter | 0.03832306629411007 | 0.15936046986653518 |
| complex_abundance | mitosis | inter | 0.06582643627069286 | 0.18541437791284332 |

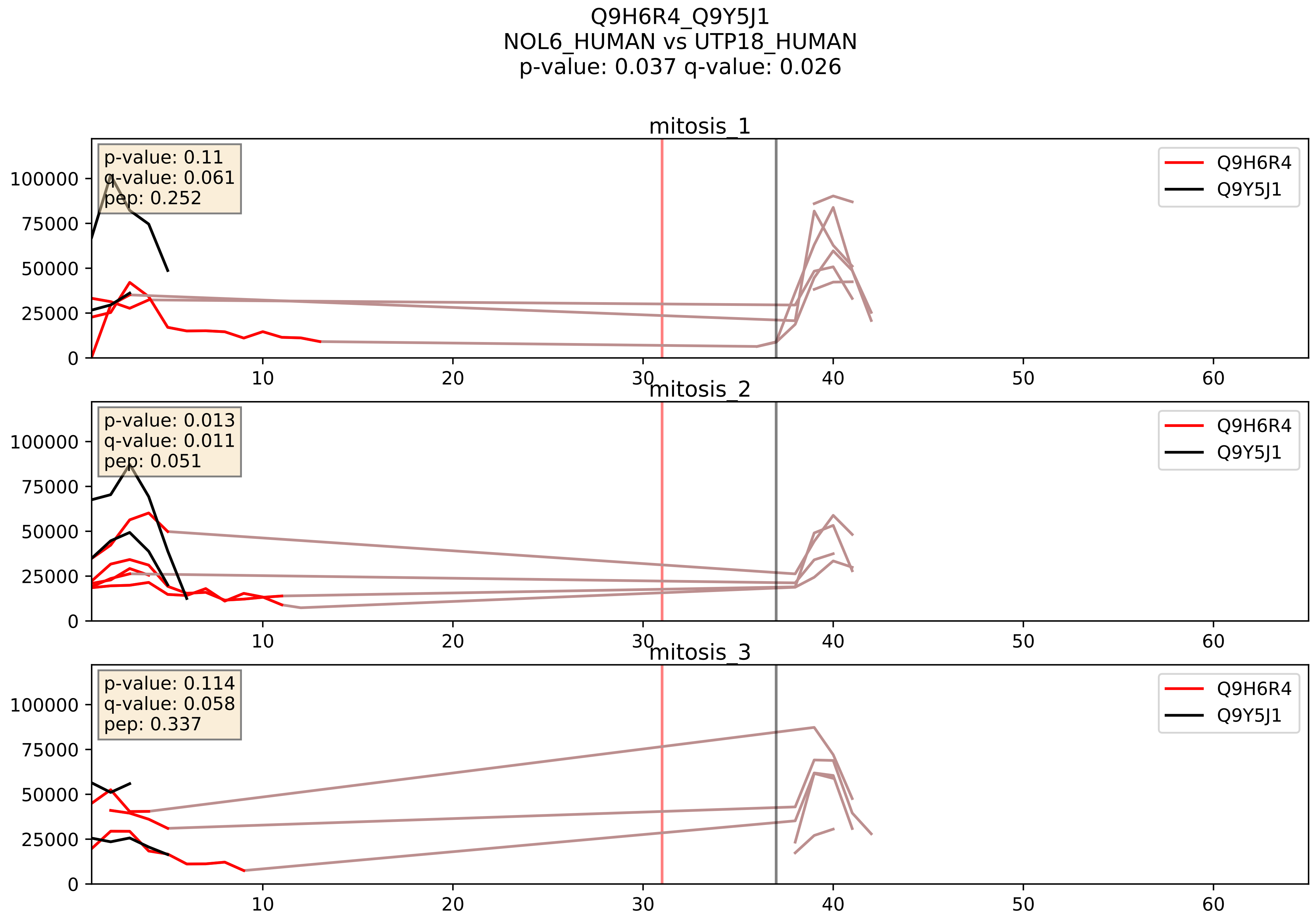


| level | condition_1 | condition_2 | pvalue | pvalue_adjusted |
|---|---|---|---|---|
| interactor_ratio | mitosis | inter | 0.2463865098672941 | 0.7216070053502578 |
| complex_abundance | mitosis | inter | 0.30028671272589114 | 0.4500094994631702 |
| interactor_abundance | mitosis | inter | 0.6347233985504436 | 0.7573495751476084 |

Q9NXF1_Q9Y5J1
TEX10_HUMAN vs UTP18_HUMAN
p-value: 0.036 q-value: 0.025

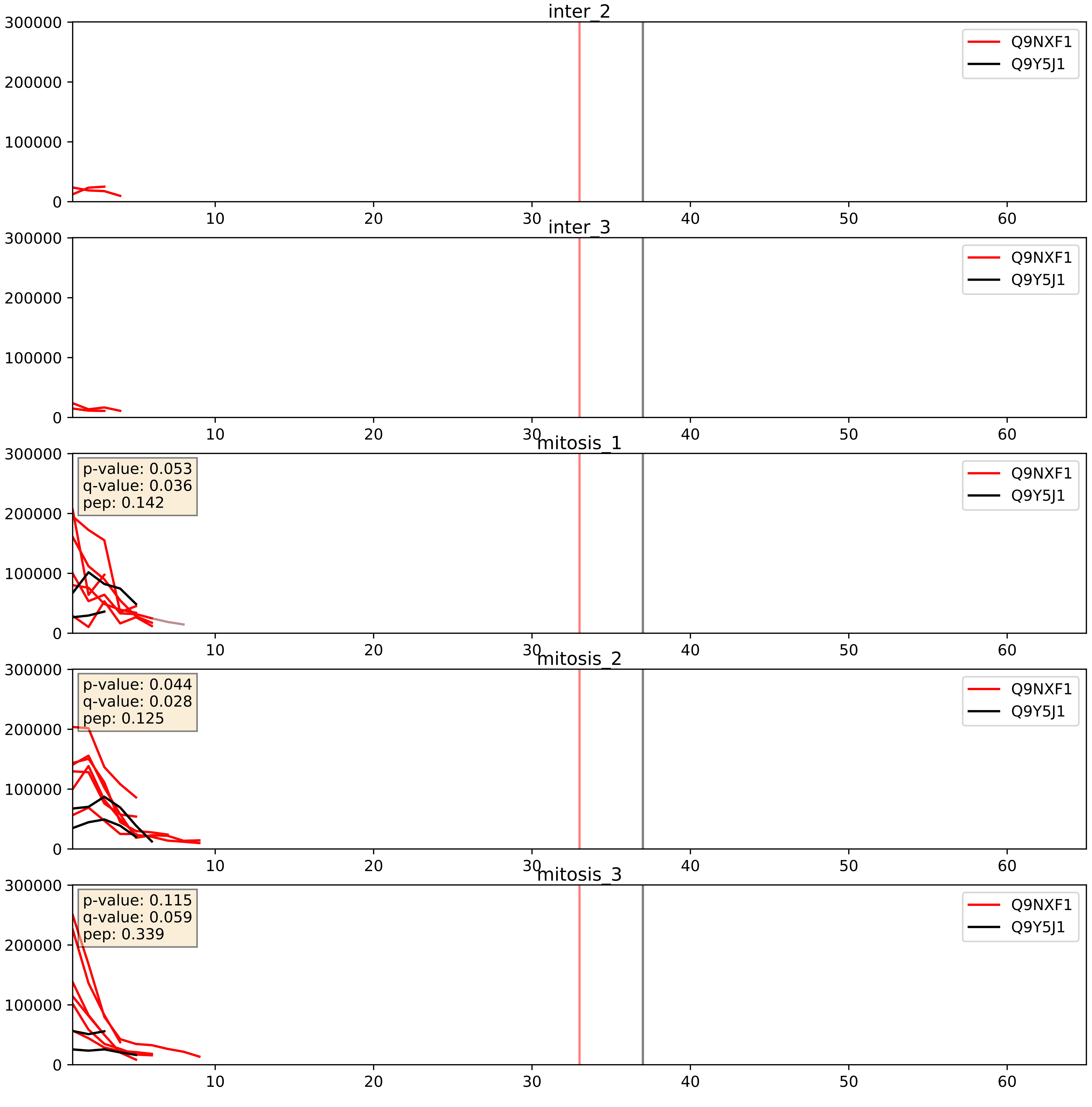


| level | condition_1 | condition_2 | pvalue | pvalue_adjusted |
|---|---|---|---|---|
| interactor_ratio | mitosis | inter | 0.0019222246008891858 | 0.05317578467954594 |
| interactor_abundance | mitosis | inter | 0.0022253892890756657 | 0.03466841143289118 |
| complex_abundance | mitosis | inter | 0.011308378166680228 | 0.07877001668010823 |

Q9Y221_Q9Y5J1
NIP7_HUMAN vs UTP18_HUMAN
p-value: 0.016 q-value: 0.013

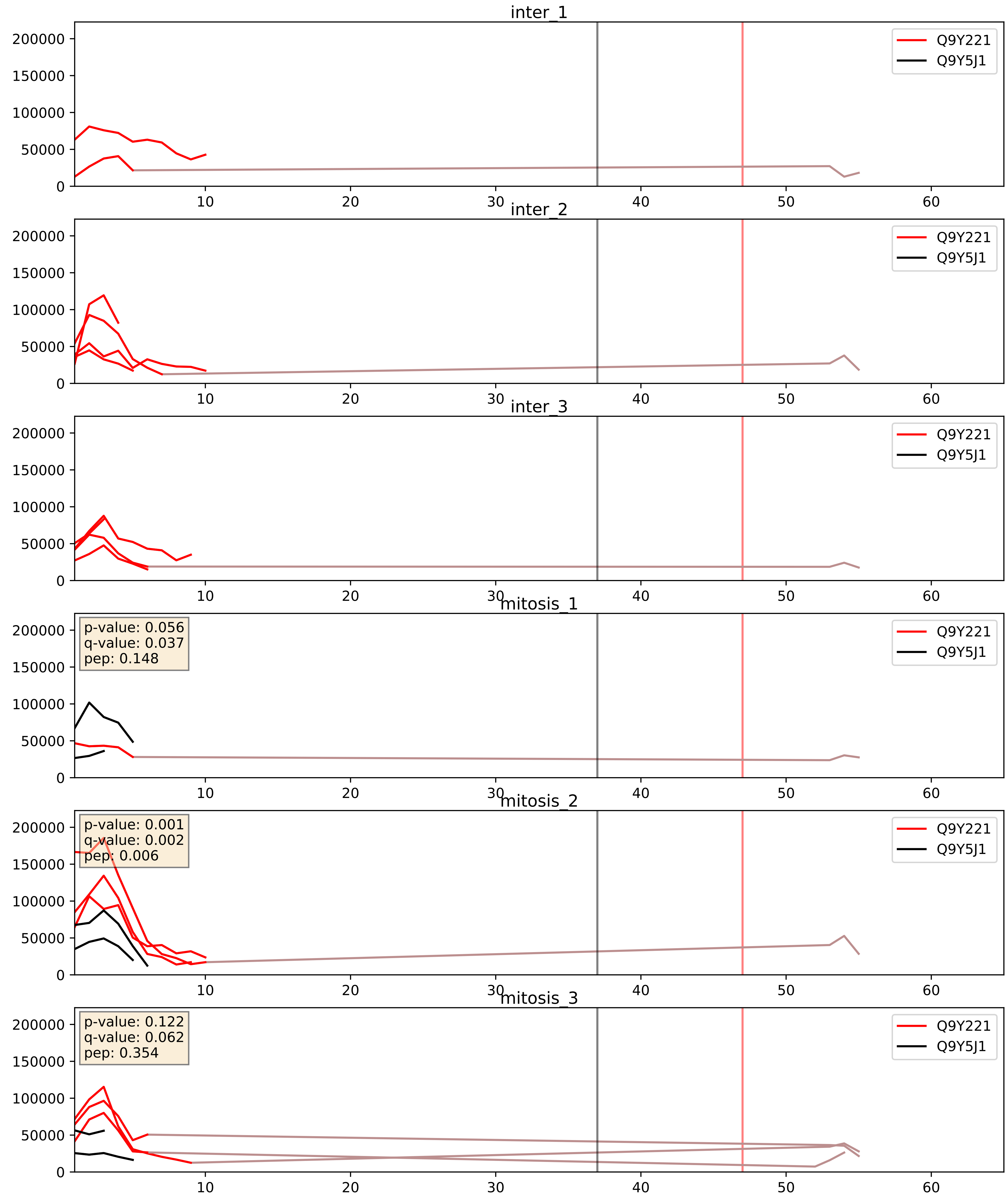


| level | condition_1 | condition_2 | pvalue | pvalue_adjusted |
|---|---|---|---|---|
| complex_abundance | mitosis | inter | 0.041655460807756714 | 0.14685780251828562 |
| interactor_abundance | mitosis | inter | 0.11599014326606938 | 0.2944907686067192 |
| interactor_ratio | mitosis | inter | 0.4397443002310046 | 0.8459188043830358 |
