## Supplementary Data 2 for "SECAT: Quantifying differential protein-protein interaction states by network-centric analysis": hela_string_000148_Q7Z3B4.pdf

Q7Z3B4_Q7Z3B4

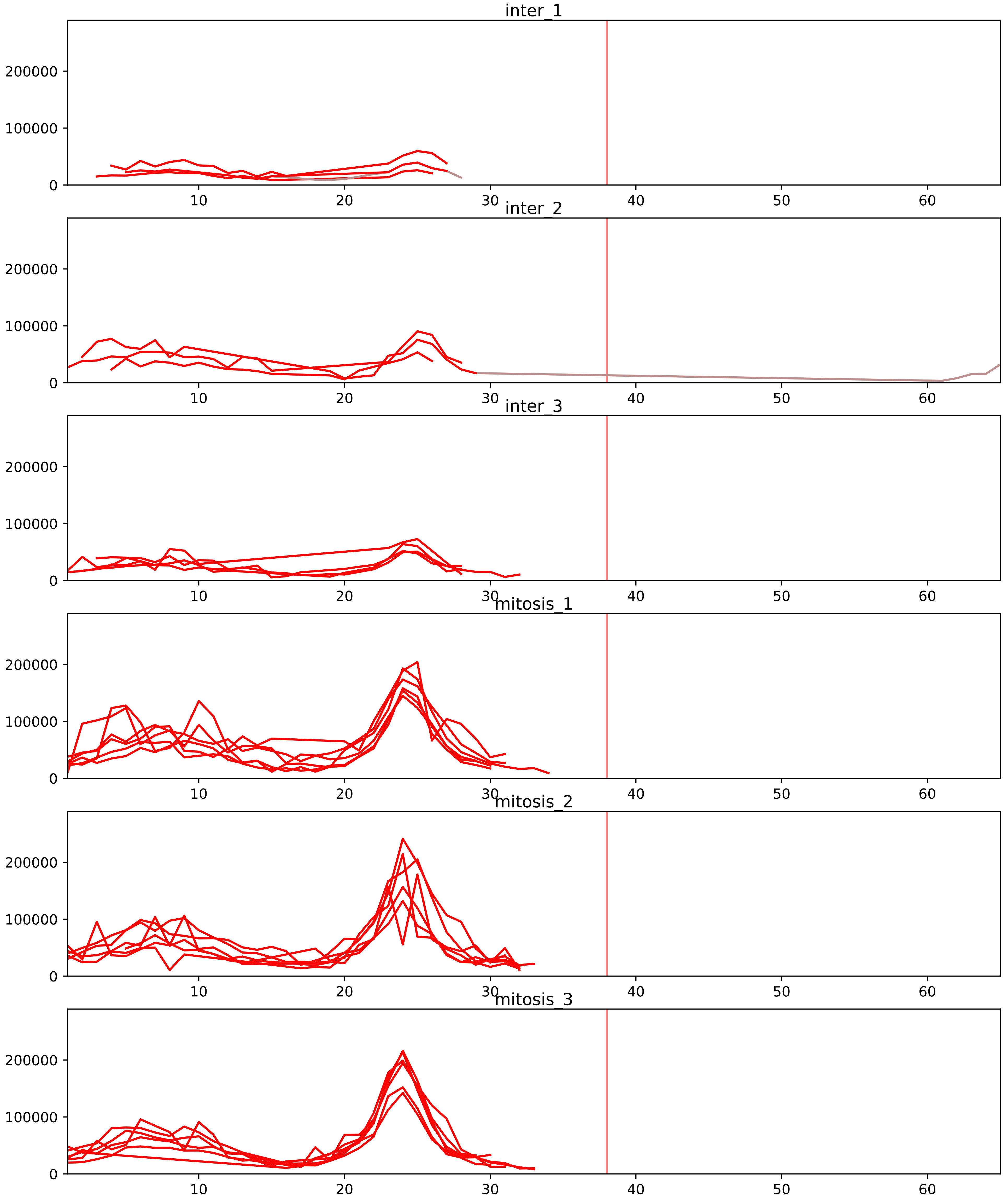


| level | condition_1 | condition_2 | pvalue | pvalue_adjusted |
|---|---|---|---|---|
| complex_abundance | mitosis | inter | 5.49591353032237e-05 | 0.0015824001673152665 |
| interactor_abundance | mitosis | inter | 5.853431294026908e-05 | 0.001614131618466031 |
| total_abundance | mitosis | inter | 0.0064173103795235 | 0.09361182562221816 |
| assembled_abundance | mitosis | inter | 0.00990125270237469 | 0.10905356211721394 |
| interactor_ratio | mitosis | inter | 0.2463684571144744 | 0.5529941201573977 |
| monomer_abundance | mitosis | inter | 0.5340163572597816 | 0.7727103819717502 |

P37198_Q7Z3B4
NUP62_HUMAN vs NUP54_HUMAN
p-value: 0.036 q-value: 0.025

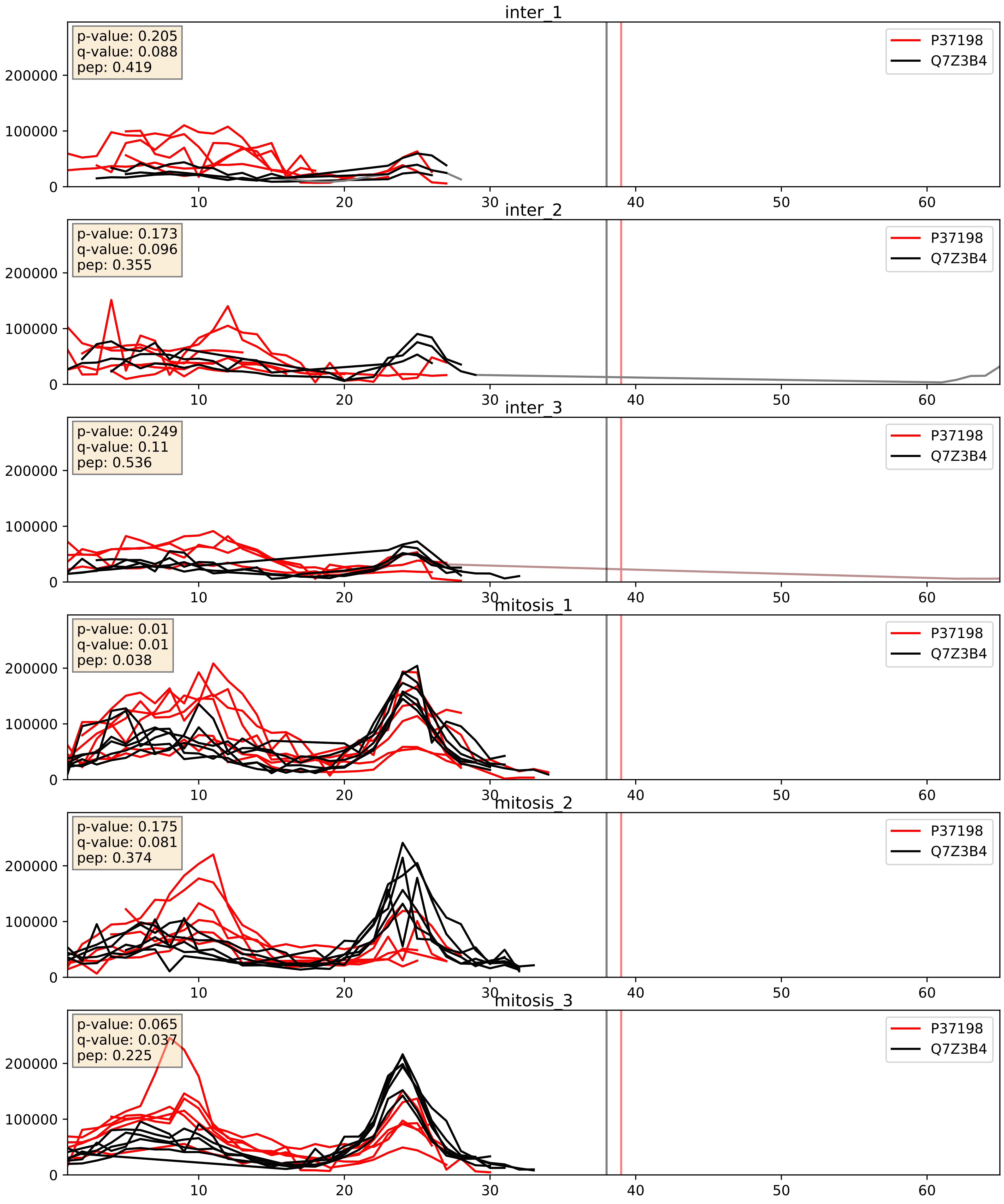


| level | condition_1 | condition_2 | pvalue | pvalue_adjusted |
|---|---|---|---|---|
| complex_abundance | mitosis | inter | 0.0018287581812309788 | 0.03051495132814265 |
| interactor_abundance | mitosis | inter | 0.04419861880302638 | 0.16761355216898388 |
| interactor_ratio | mitosis | inter | 0.4268597951580075 | 0.842479306395587 |

P49790_Q7Z3B4
NU153_HUMAN vs NUP54_HUMAN
p-value: 0.036 q-value: 0.025

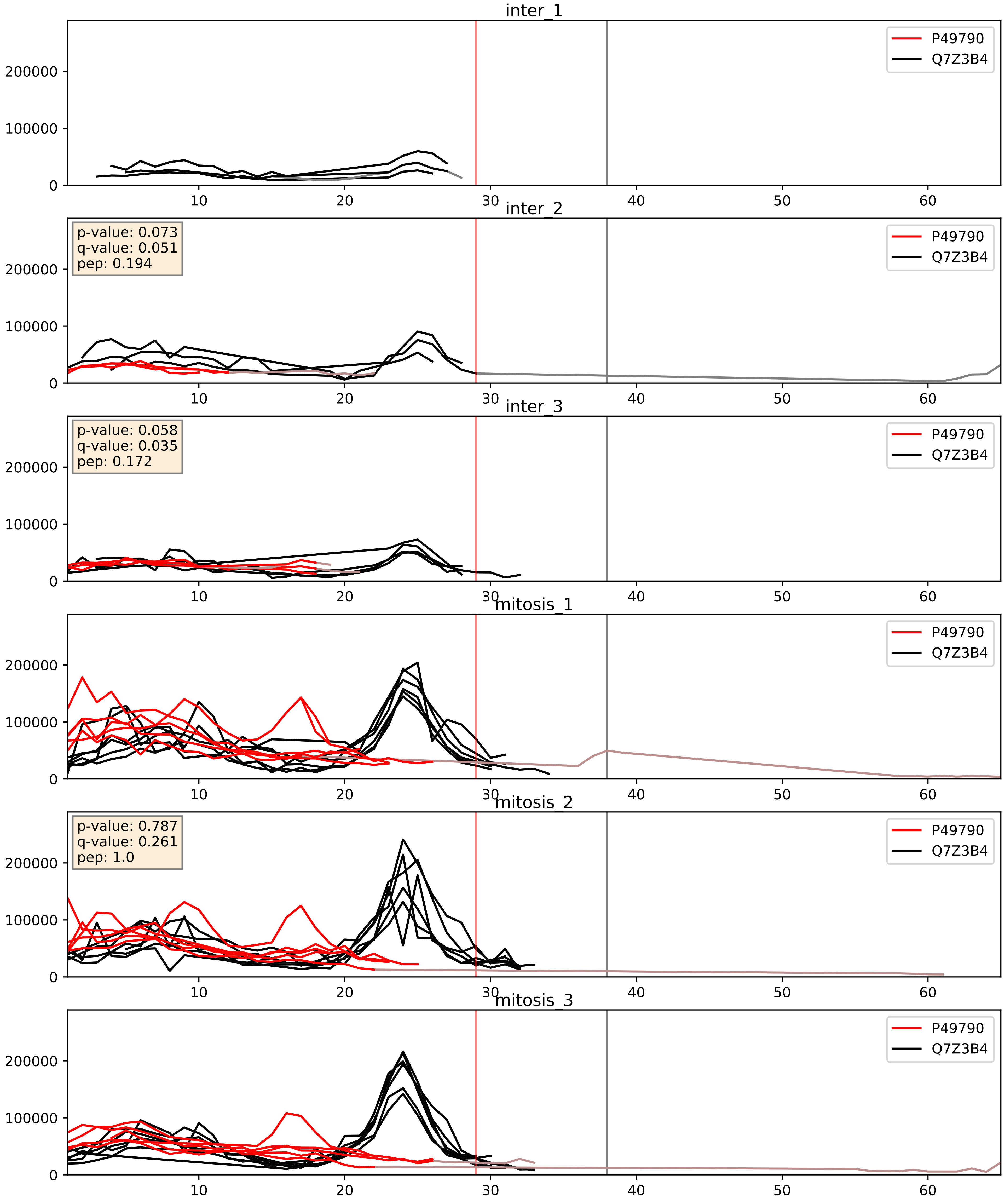


| level | condition_1 | condition_2 | pvalue | pvalue_adjusted |
|---|---|---|---|---|
| complex_abundance | mitosis | inter | 0.0010513693196996844 | 0.021612845224852777 |
| interactor_abundance | mitosis | inter | 0.0013903875369213287 | 0.02753913185123736 |
| interactor_ratio | mitosis | inter | 0.4651985580743366 | 0.8591965497319327 |

Q5SRE5_Q7Z3B4
NU188_HUMAN vs NUP54_HUMAN
p-value: 0.048 q-value: 0.032

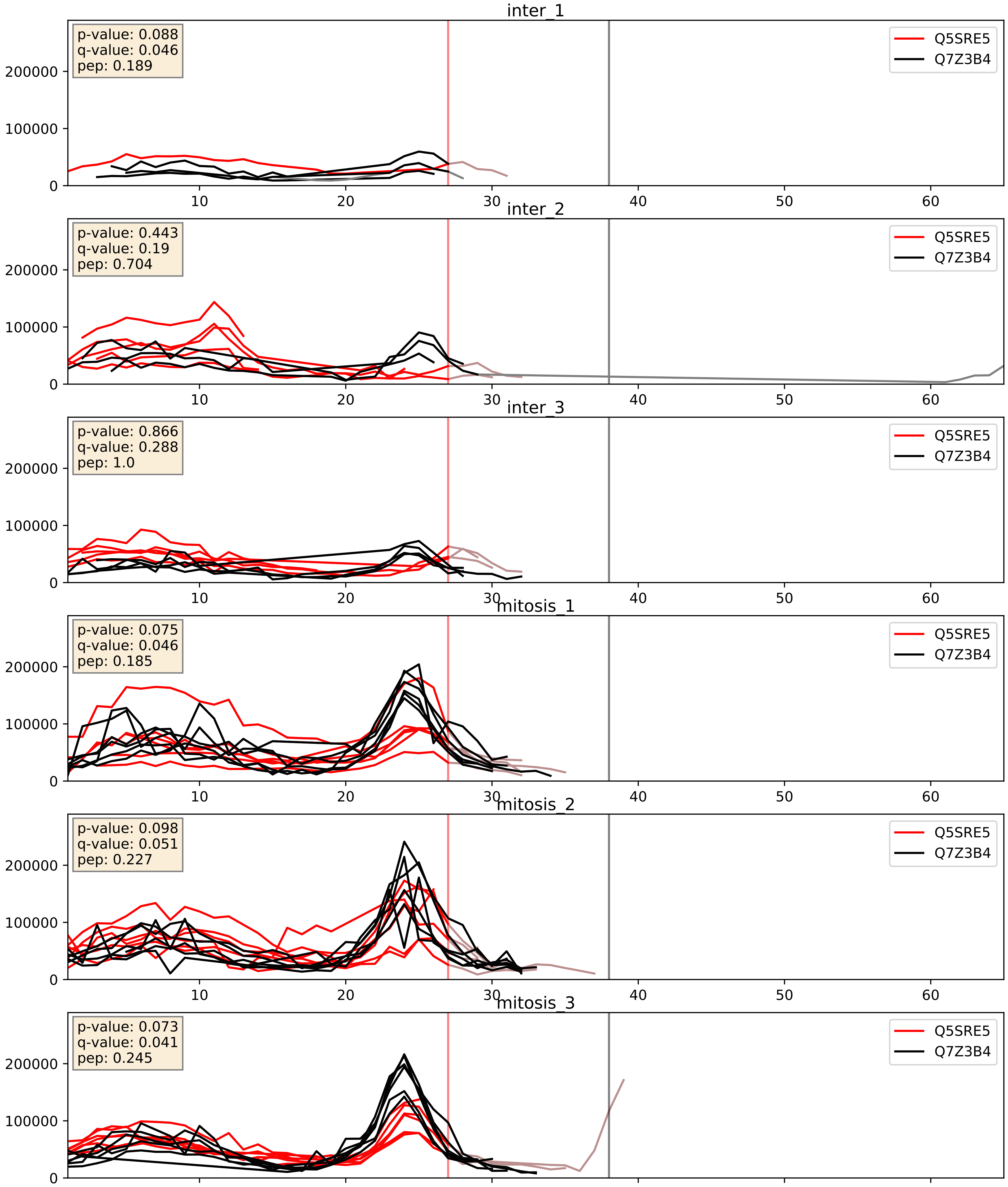


| level | condition_1 | condition_2 | pvalue | pvalue_adjusted |
|---|---|---|---|---|
| complex_abundance | mitosis | inter | 0.01634921709991212 | 0.09565912397487884 |
| interactor_abundance | mitosis | inter | 0.07916693523454389 | 0.2337192500802537 |
| interactor_ratio | mitosis | inter | 0.40641780524135535 | 0.8325457607945408 |

Q7Z3B4_Q9BVL2
NUP54_HUMAN vs NUP58_HUMAN
p-value: 0.008 q-value: 0.008

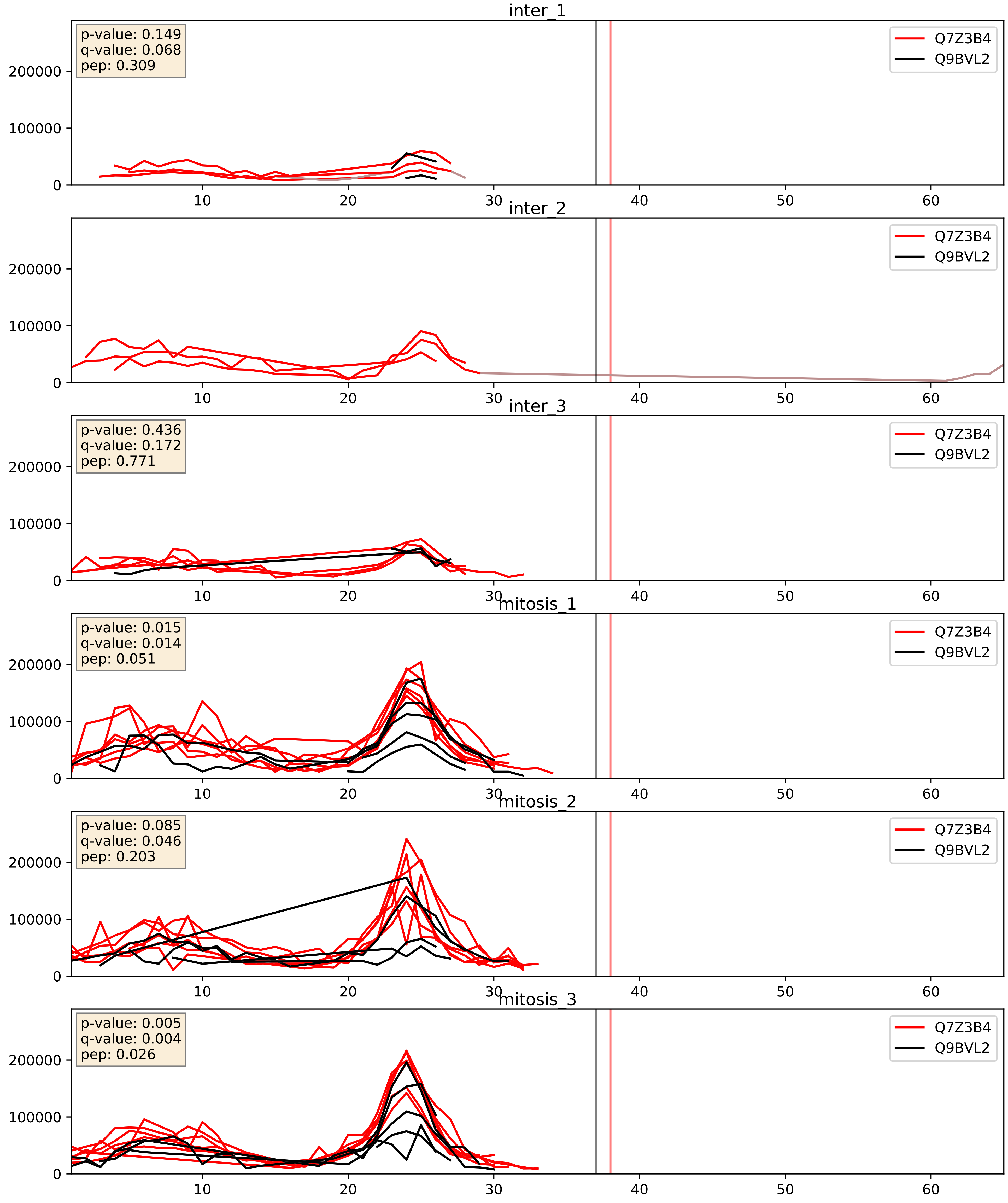


| level | condition_1 | condition_2 | pvalue | pvalue_adjusted |
|---|---|---|---|---|
| interactor_abundance | mitosis | inter | 0.0018625829583547879 | 0.03172877636520793 |
| complex_abundance | mitosis | inter | 0.001908787386402109 | 0.03112232386209915 |
| interactor_ratio | mitosis | inter | 0.07282740231921711 | 0.4507610729229924 |
