## Supplementary Data 2 for "SECAT: Quantifying differential protein-protein interaction states by network-centric analysis": hela_string_000149_Q6PJT7.pdf

Q6PJT7_Q6PJT7

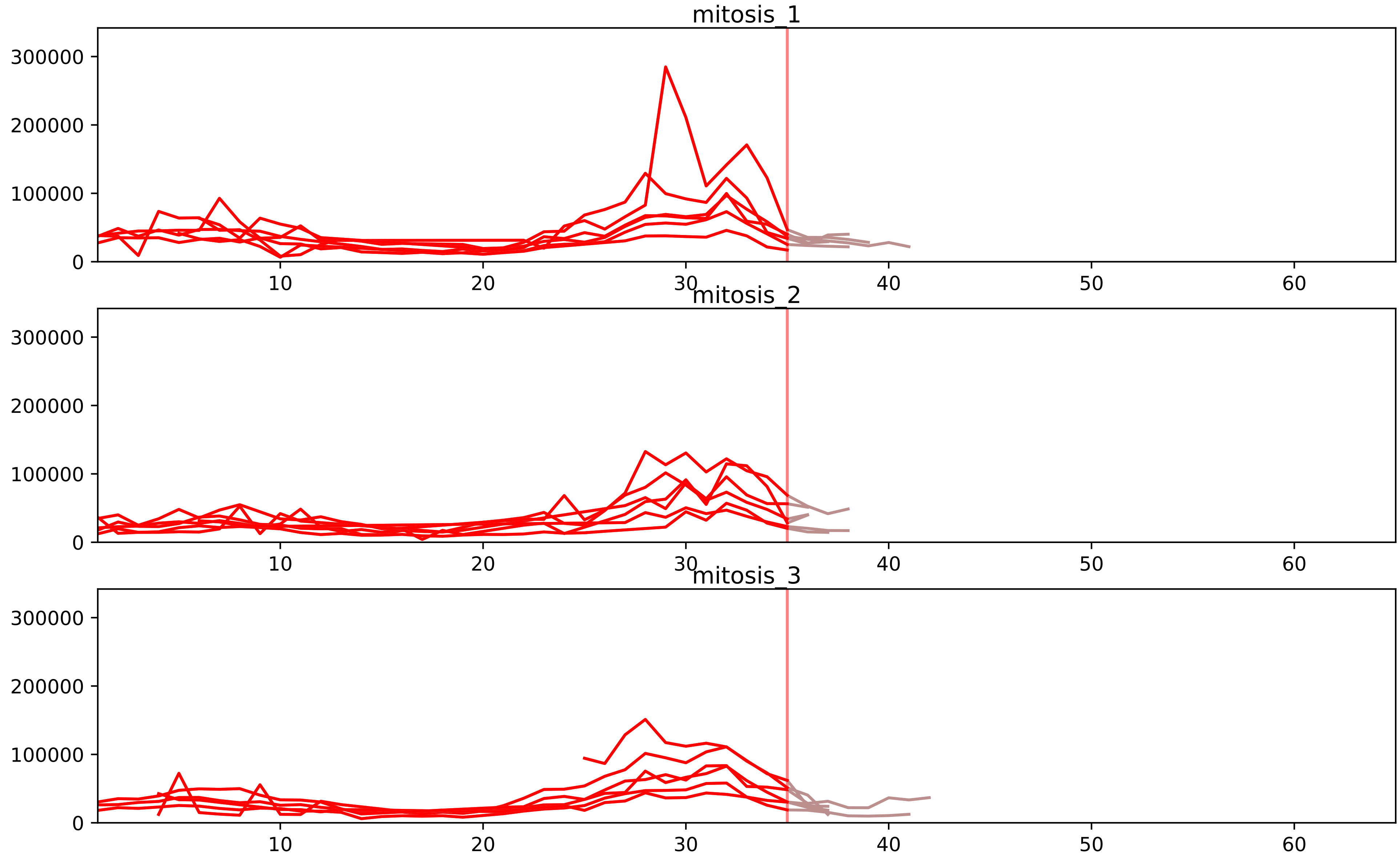


| level | condition_1 | condition_2 | pvalue | pvalue_adjusted |
|---|---|---|---|---|
| assembled_abundance | mitosis | inter | 5.5071825797152436e-05 | 0.007270487507555603 |
| total_abundance | mitosis | inter | 5.8363723241318766e-05 | 0.010526120062335645 |
| monomer_abundance | mitosis | inter | 0.001506105097076222 | 0.08447898271593504 |
