## Supplementary Data 2 for "SECAT: Quantifying differential protein-protein interaction states by network-centric analysis": hela_string_000150_P62906.pdf

P62906_P62906

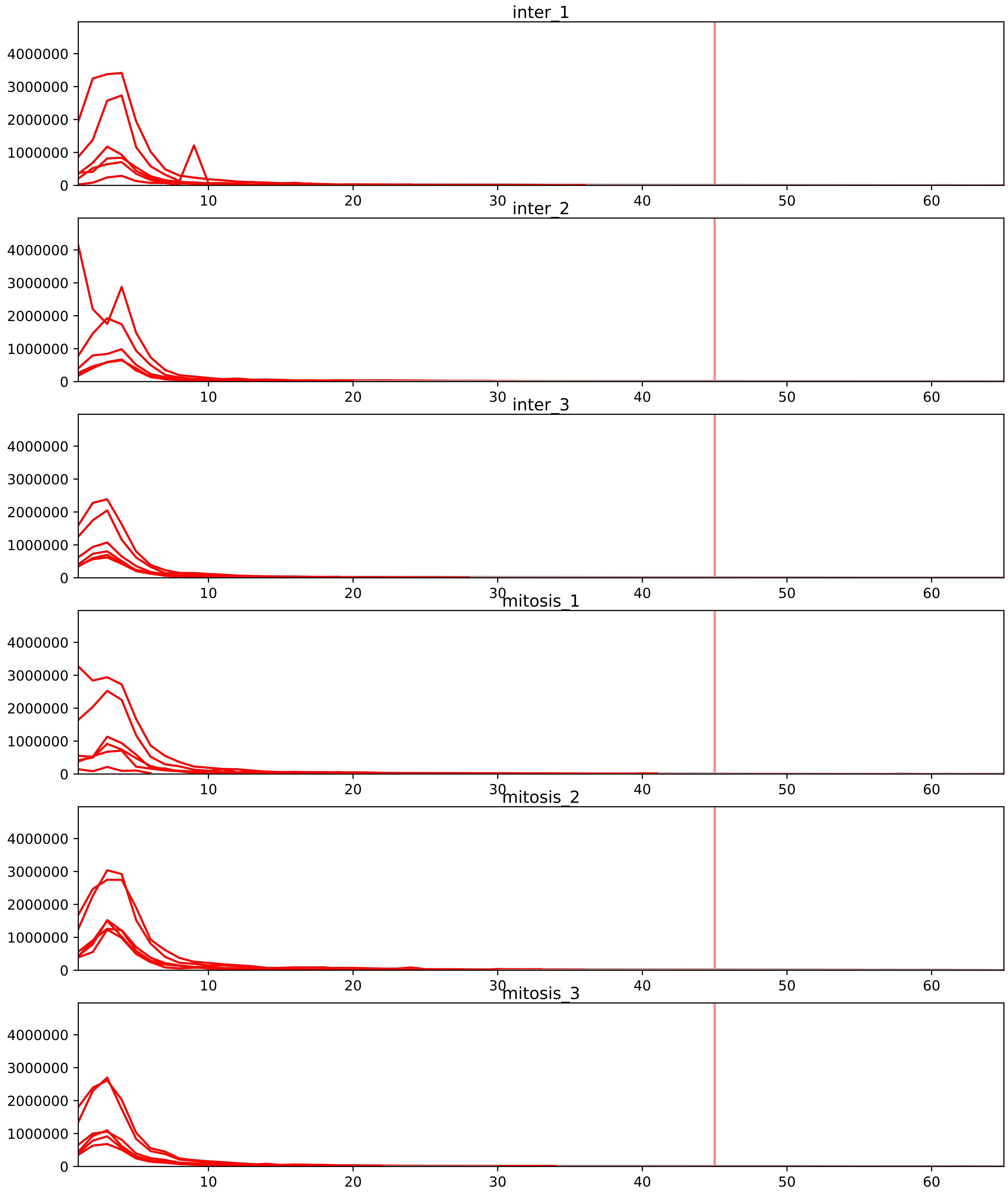


| level | condition_1 | condition_2 | pvalue | pvalue_adjusted |
|---|---|---|---|---|
| assembled_abundance | mitosis | inter | 5.636304290272102e-05 | 0.007270487507555603 |
| total_abundance | mitosis | inter | 9.195117402595856e-05 | 0.01273261737241483 |
| interactor_ratio | mitosis | inter | 0.05817790521507821 | 0.23271891667701047 |
| complex_abundance | mitosis | inter | 0.14717291875183972 | 0.2552808017084434 |
| monomer_abundance | mitosis | inter | 0.17852112920841234 | 0.5465948364534886 |
| interactor_abundance | mitosis | inter | 0.763724438028244 | 0.8610618664043928 |

P05387_P62906
RLA2_HUMAN vs RL10A_HUMAN
p-value: 0.0 q-value: 0.0

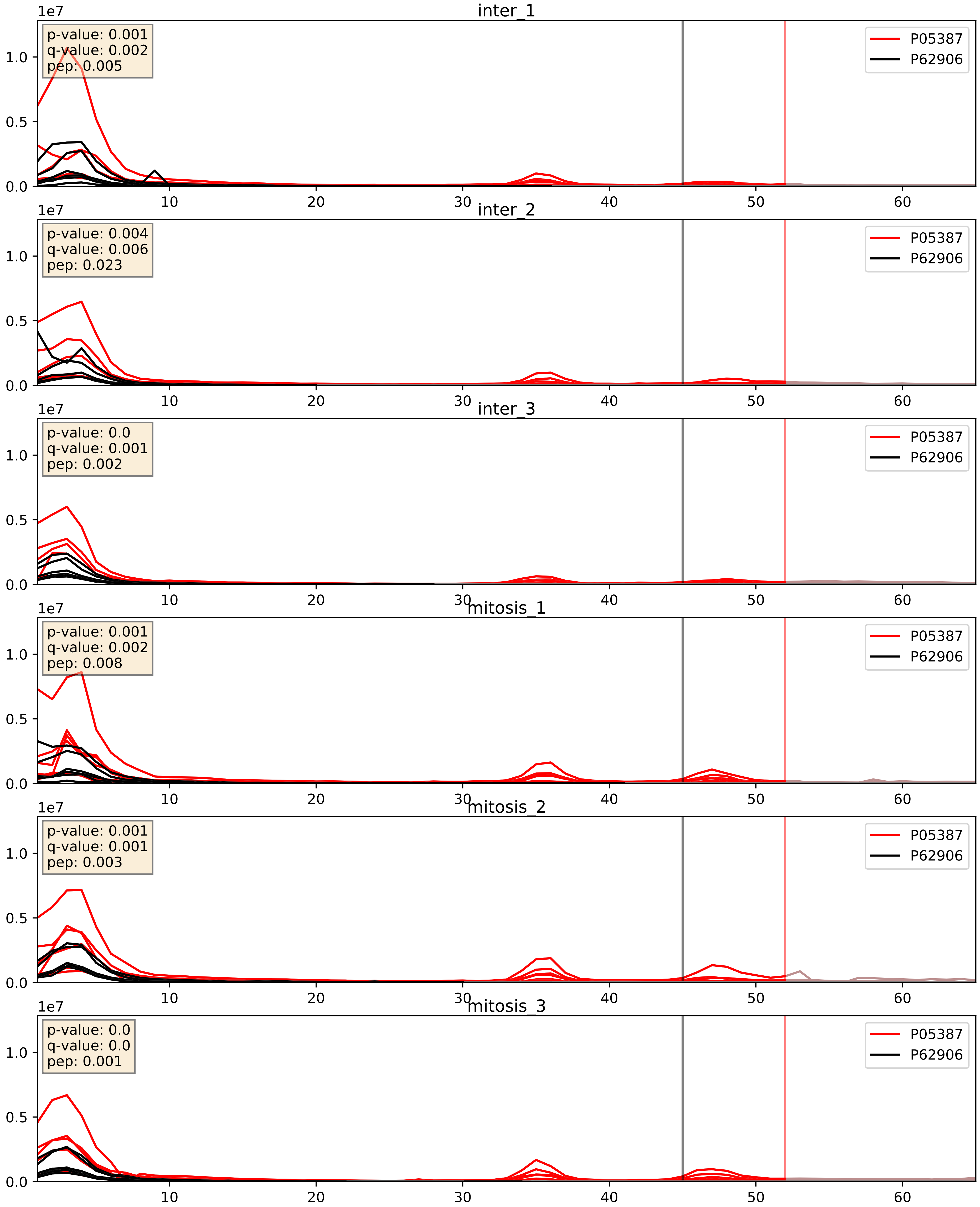


| level | condition_1 | condition_2 | pvalue | pvalue_adjusted |
|---|---|---|---|---|
| interactor_ratio | mitosis | inter | 0.8066492807203018 | 0.964095391868208 |
| interactor_abundance | mitosis | inter | 0.8990108830106276 | 0.9441584502781086 |
| complex_abundance | mitosis | inter | 0.9974958362940012 | 0.9988999610731659 |

P05388_P62906
RLA0_HUMAN vs RL10A_HUMAN
p-value: 0.0 q-value: 0.0

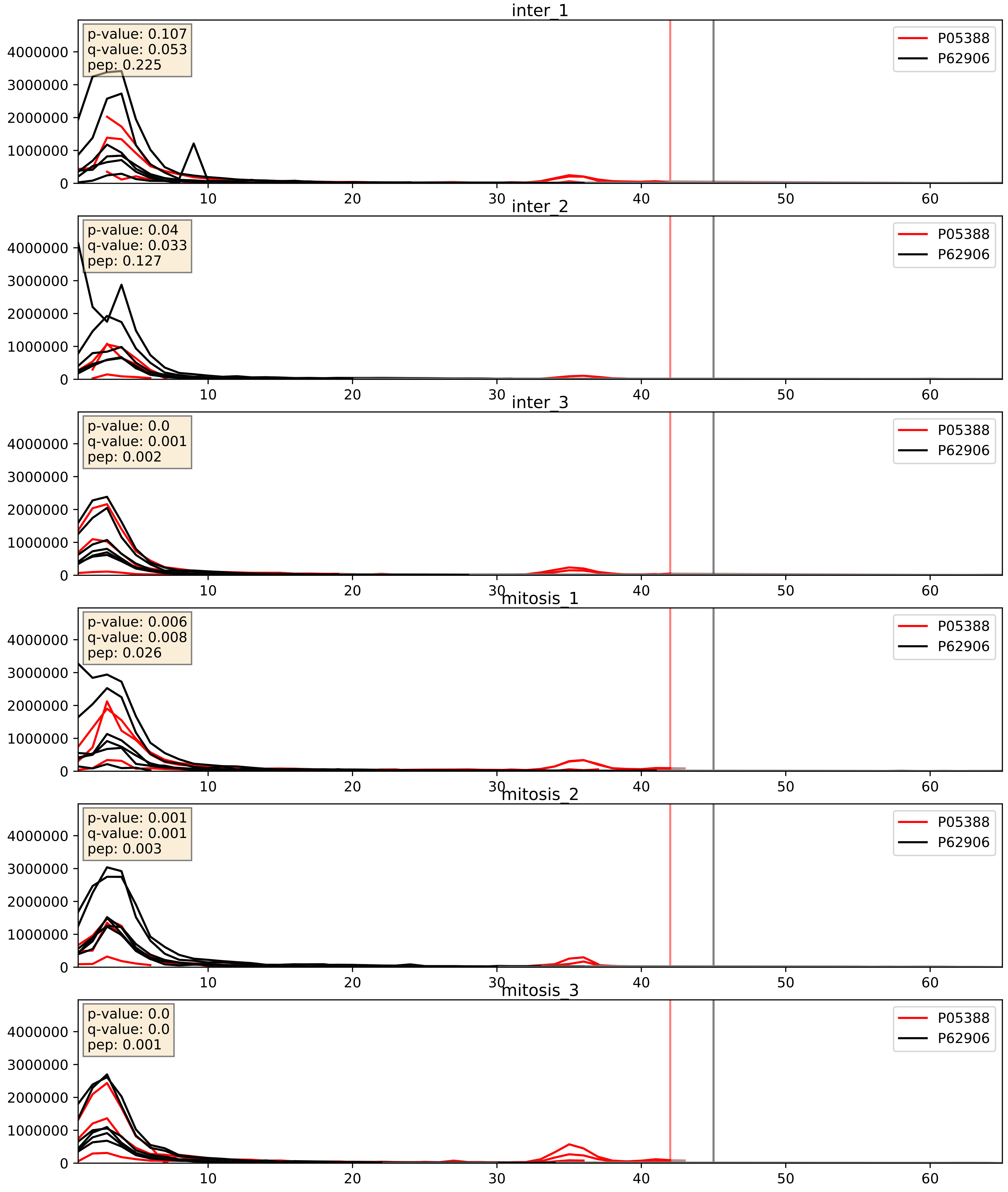


| level | condition_1 | condition_2 | pvalue | pvalue_adjusted |
|---|---|---|---|---|
| interactor_ratio | mitosis | inter | 0.45576100709392253 | 0.8549372939008518 |
| interactor_abundance | mitosis | inter | 0.6307422906206419 | 0.756956599441332 |
| complex_abundance | mitosis | inter | 0.754792958706123 | 0.838441179149288 |

P08708_P62906
RS17_HUMAN vs RL10A_HUMAN
p-value: 0.0 q-value: 0.001

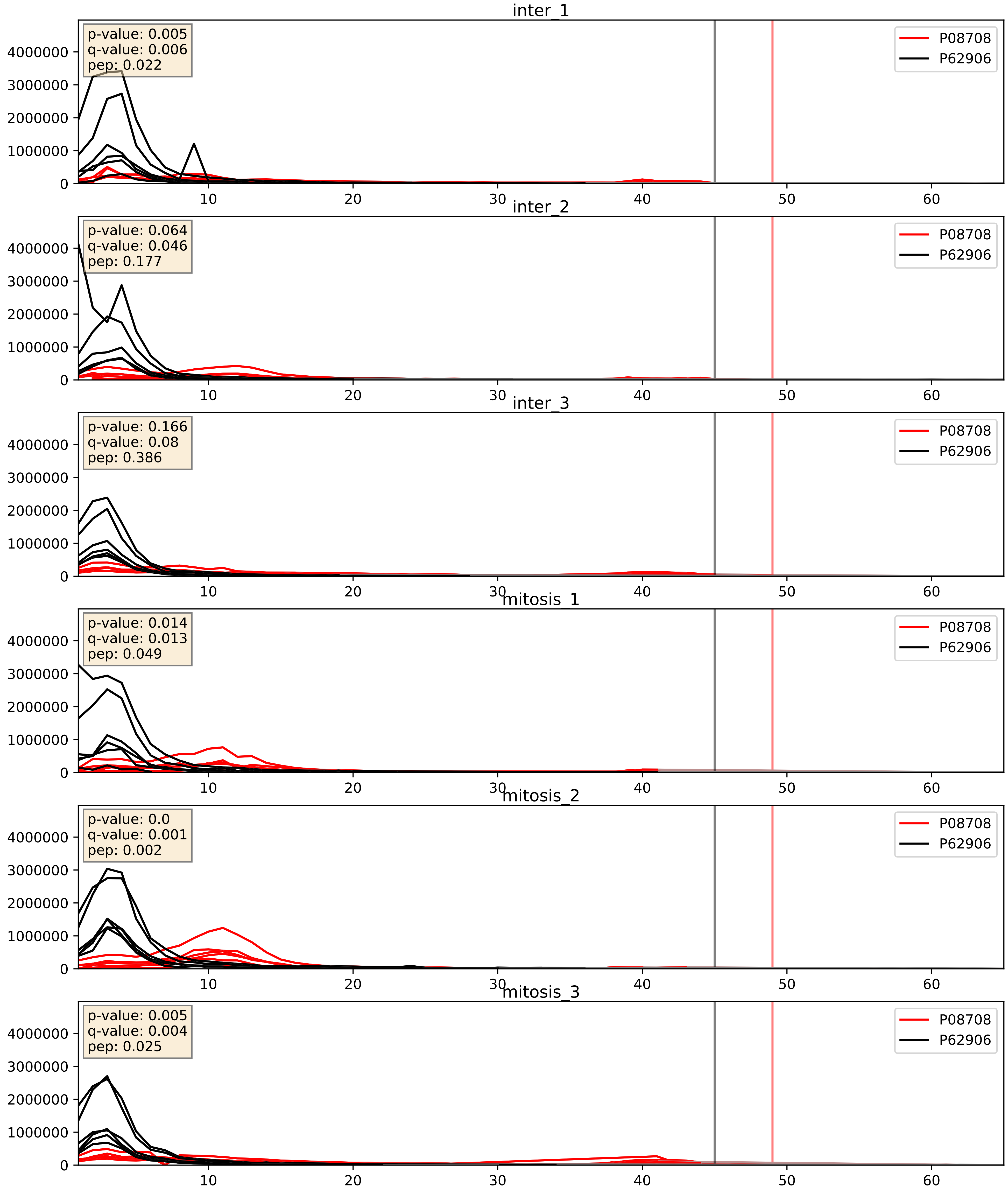


| level | condition_1 | condition_2 | pvalue | pvalue_adjusted |
|---|---|---|---|---|
| interactor_abundance | mitosis | inter | 0.14210869486476208 | 0.32880142136715784 |
| complex_abundance | mitosis | inter | 0.17022657991370388 | 0.318988512272615 |
| interactor_ratio | mitosis | inter | 0.23586128772487683 | 0.7155827200888129 |

P11940_P62906
PABP1_HUMAN vs RL10A_HUMAN
p-value: 0.034 q-value: 0.024

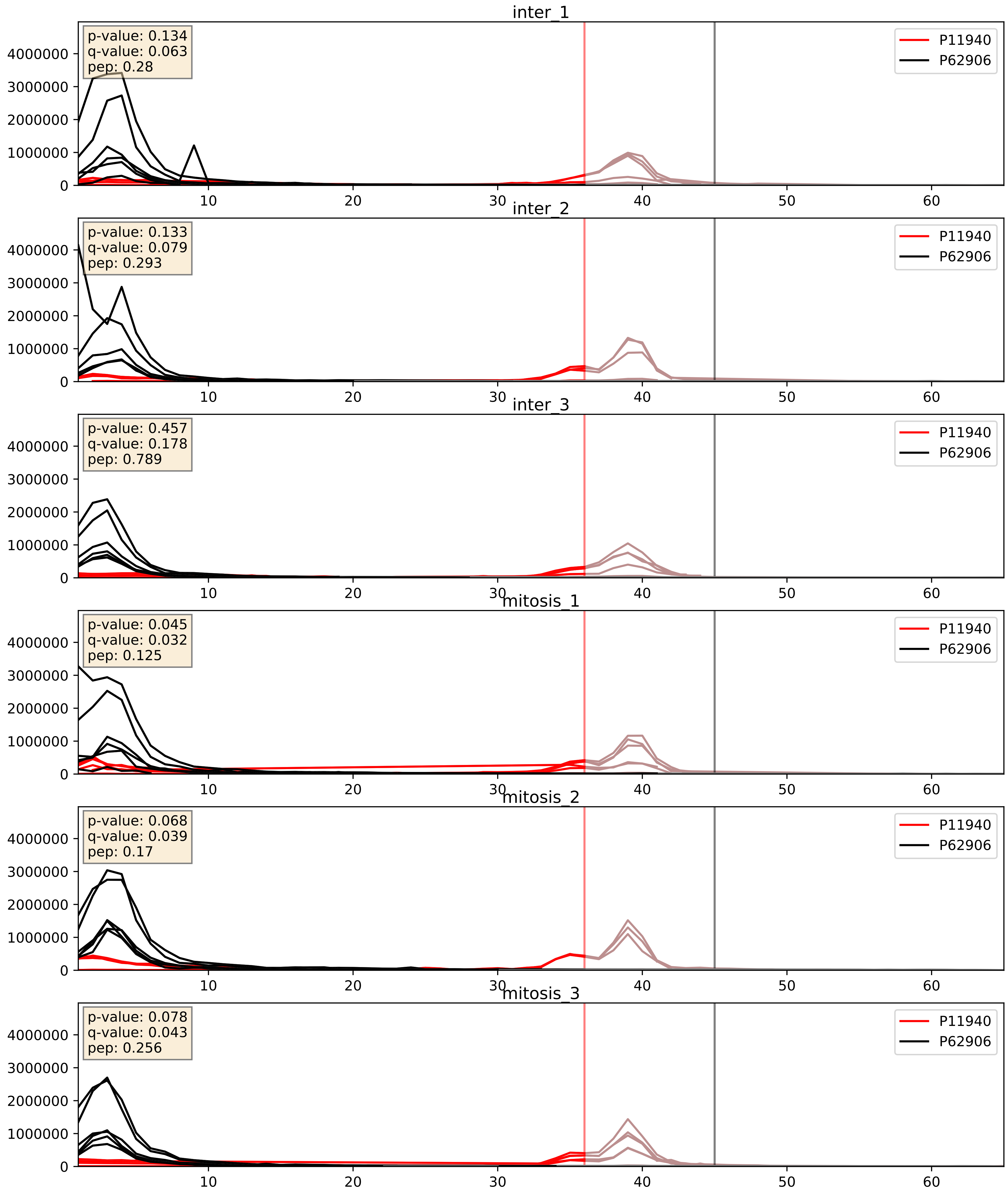


| level | condition_1 | condition_2 | pvalue | pvalue_adjusted |
|---|---|---|---|---|
| interactor_ratio | mitosis | inter | 0.13717642330363286 | 0.5873558393508659 |
| interactor_abundance | mitosis | inter | 0.22174568684515863 | 0.41786308847431103 |
| complex_abundance | mitosis | inter | 0.38602269073306805 | 0.5326732558823649 |

P15880_P62906
RS2_HUMAN vs RL10A_HUMAN
p-value: 0.029 q-value: 0.021

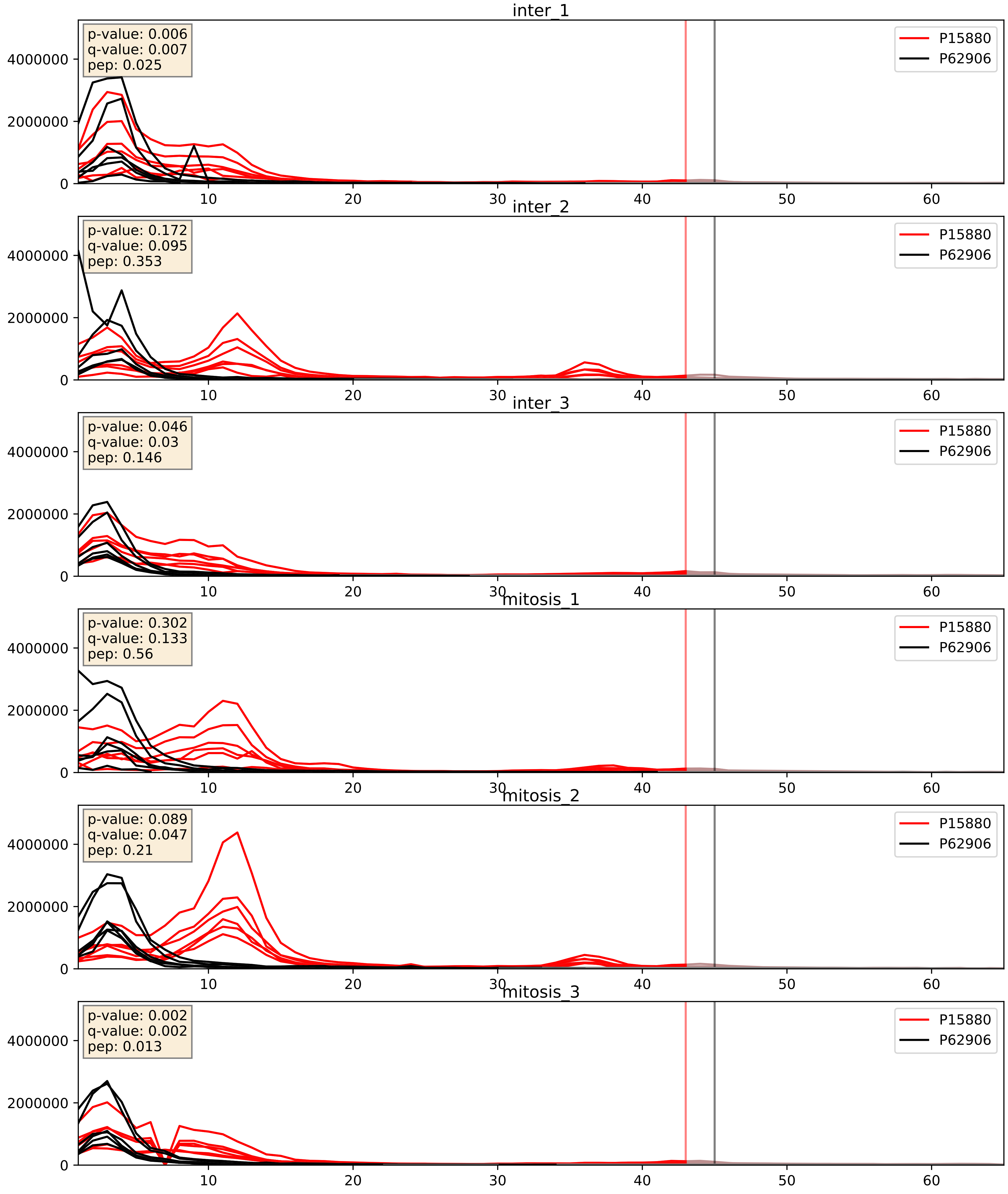


| level | condition_1 | condition_2 | pvalue | pvalue_adjusted |
|---|---|---|---|---|
| complex_abundance | mitosis | inter | 0.6286961384222463 | 0.7467238719154194 |
| interactor_abundance | mitosis | inter | 0.6728980422640023 | 0.7859834184117398 |
| interactor_ratio | mitosis | inter | 0.8621794287181117 | 0.9774309046518314 |

P18077_P62906
RL35A_HUMAN vs RL10A_HUMAN
p-value: 0.0 q-value: 0.0

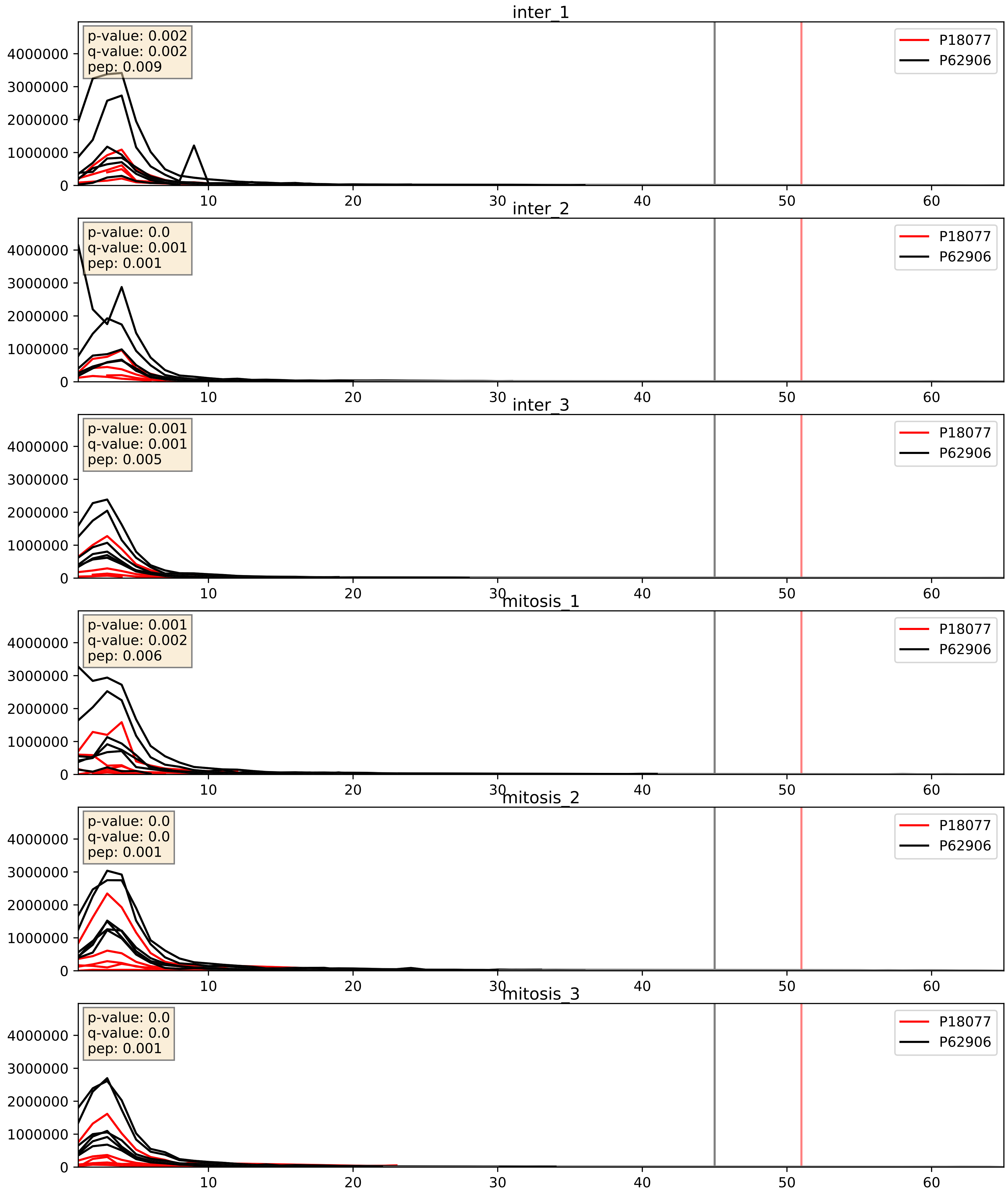


| level | condition_1 | condition_2 | pvalue | pvalue_adjusted |
|---|---|---|---|---|
| interactor_ratio | mitosis | inter | 0.8040153823785711 | 0.9634160836222222 |
| interactor_abundance | mitosis | inter | 0.8878397503920612 | 0.9361222224987428 |
| complex_abundance | mitosis | inter | 0.9760422511586349 | 0.9857075068317154 |

P18124_P62906
RL7_HUMAN vs RL10A_HUMAN
p-value: 0.0 q-value: 0.0

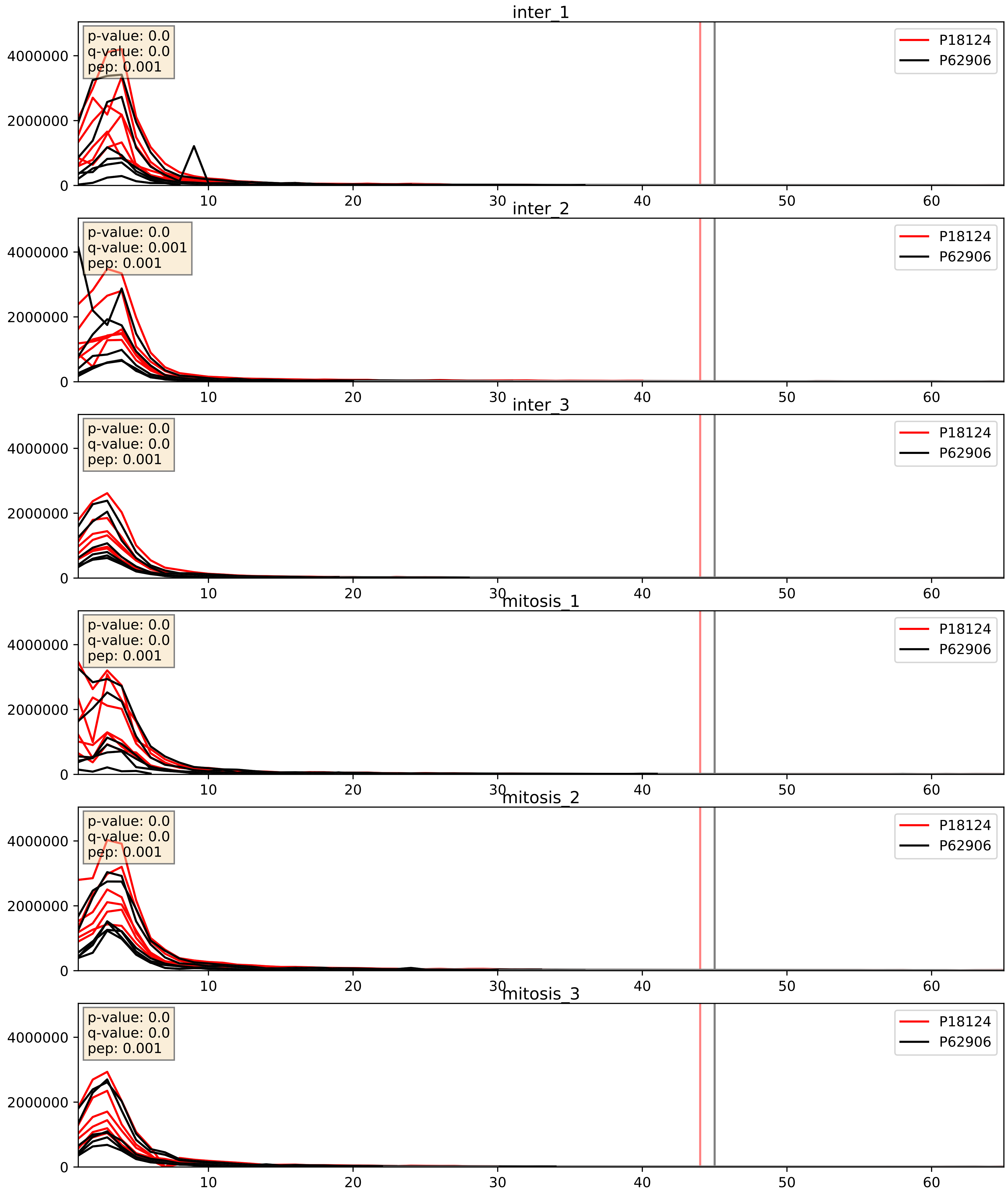


| level | condition_1 | condition_2 | pvalue | pvalue_adjusted |
|---|---|---|---|---|
| interactor_ratio | mitosis | inter | 0.3706101988477591 | 0.8138388063636502 |
| interactor_abundance | mitosis | inter | 0.5309783321918246 | 0.6897601522971422 |
| complex_abundance | mitosis | inter | 0.6421187593417945 | 0.7578845538368789 |

P22087_P62906
FBRL_HUMAN vs RL10A_HUMAN
p-value: 0.0 q-value: 0.017

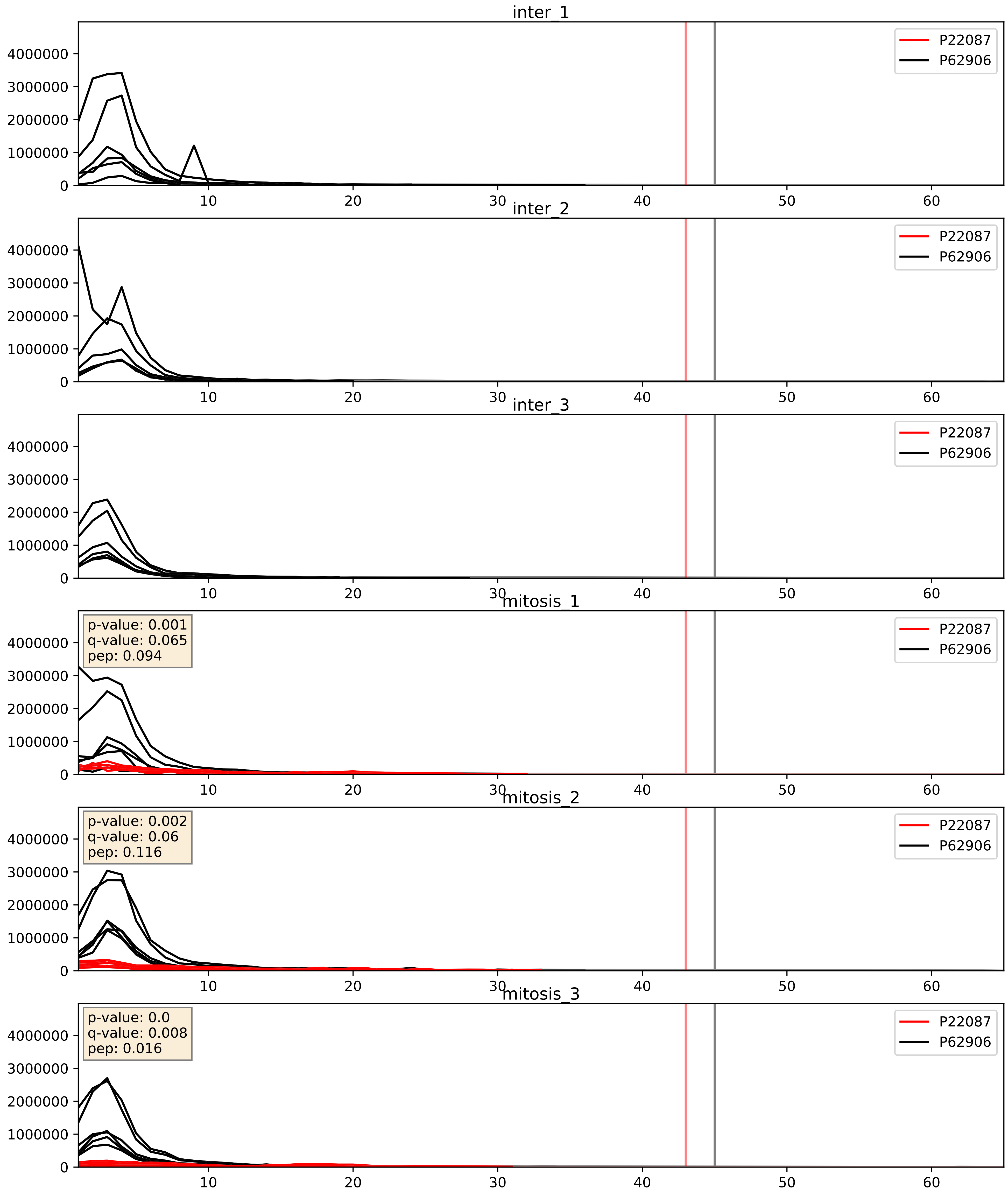


| level | condition_1 | condition_2 | pvalue | pvalue_adjusted |
|---|---|---|---|---|
| complex_abundance | mitosis | inter | 2.264226142142095e-05 | 0.005691857151652958 |
| interactor_abundance | mitosis | inter | 0.0003841155059274424 | 0.013586895581565731 |
| interactor_ratio | mitosis | inter | 0.0014007263931960893 | 0.04424434658951485 |

P23396_P62906
RS3_HUMAN vs RL10A_HUMAN
p-value: 0.012 q-value: 0.01

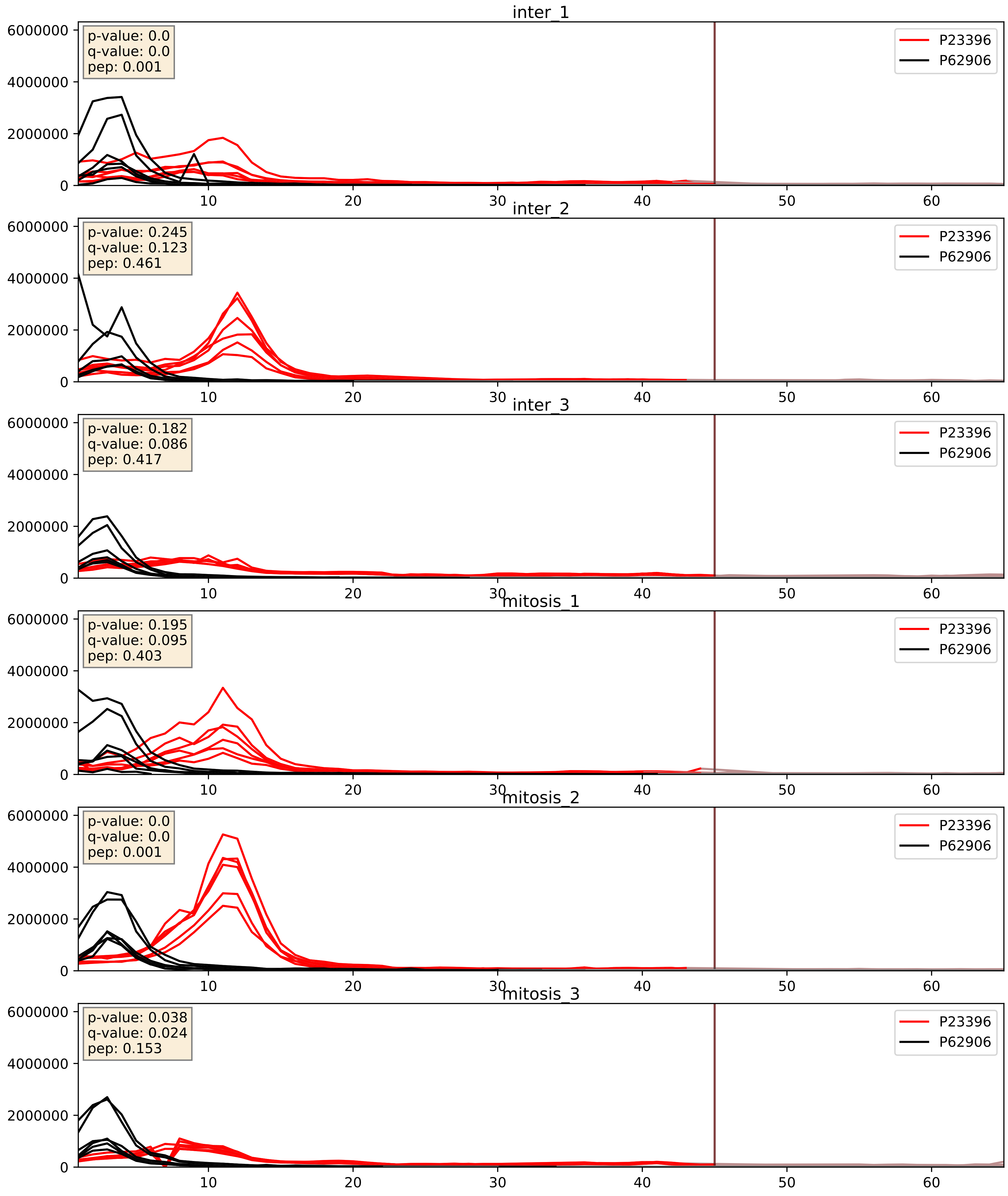


| level | condition_1 | condition_2 | pvalue | pvalue_adjusted |
|---|---|---|---|---|
| interactor_abundance | mitosis | inter | 0.682167401823537 | 0.7935330223039571 |
| interactor_ratio | mitosis | inter | 0.6876412068319663 | 0.9349347226390419 |
| complex_abundance | mitosis | inter | 0.692083815388447 | 0.7929571389337764 |

P25398_P62906
RS12_HUMAN vs RL10A_HUMAN
p-value: 0.003 q-value: 0.003

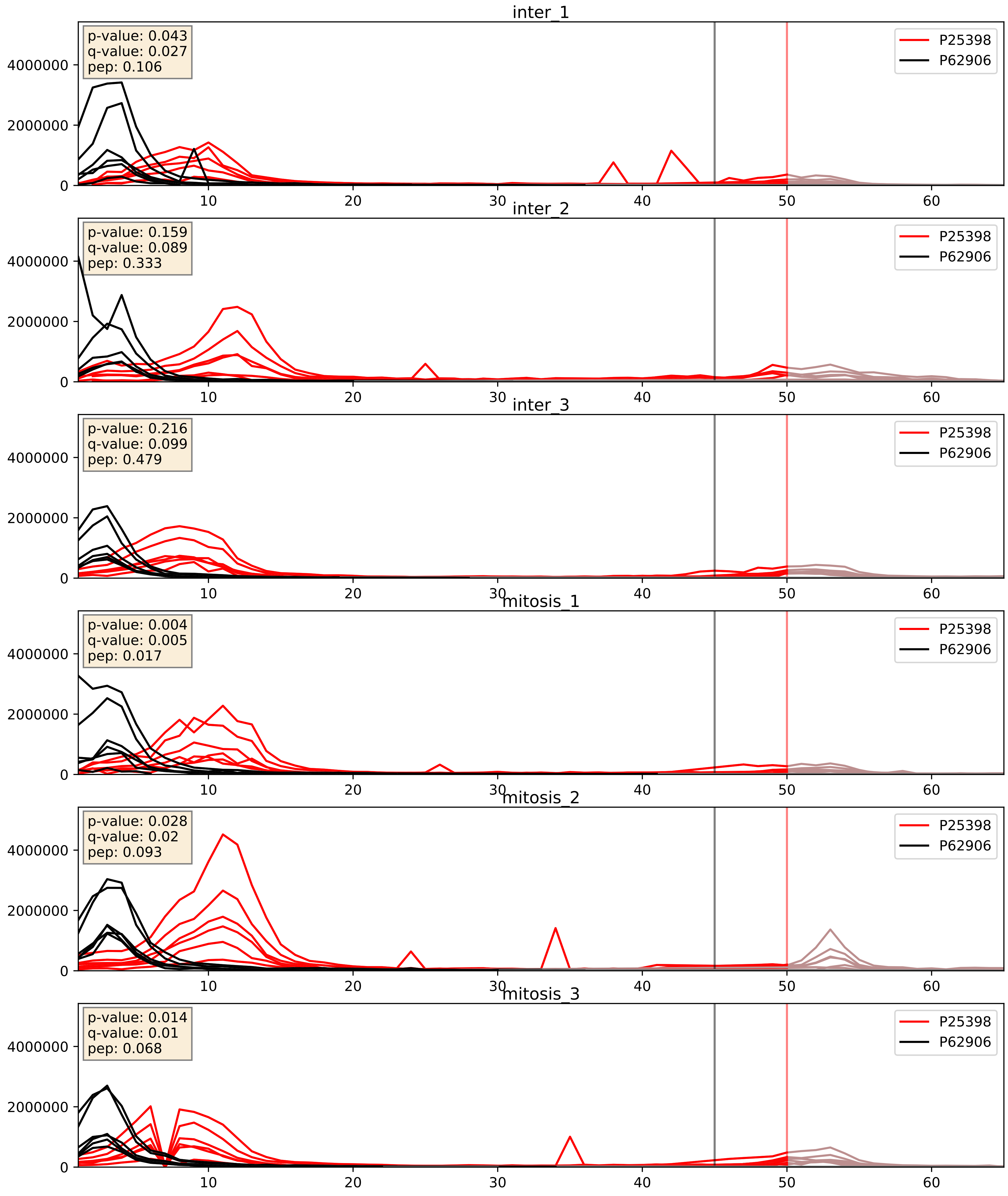


| level | condition_1 | condition_2 | pvalue | pvalue_adjusted |
|---|---|---|---|---|
| complex_abundance | mitosis | inter | 0.27333273715839607 | 0.42234722113987533 |
| interactor_abundance | mitosis | inter | 0.4583125795804698 | 0.630225812242381 |
| interactor_ratio | mitosis | inter | 0.5736148263269321 | 0.8947053413554188 |

P26373_P62906
RL13_HUMAN vs RL10A_HUMAN
p-value: 0.0 q-value: 0.0

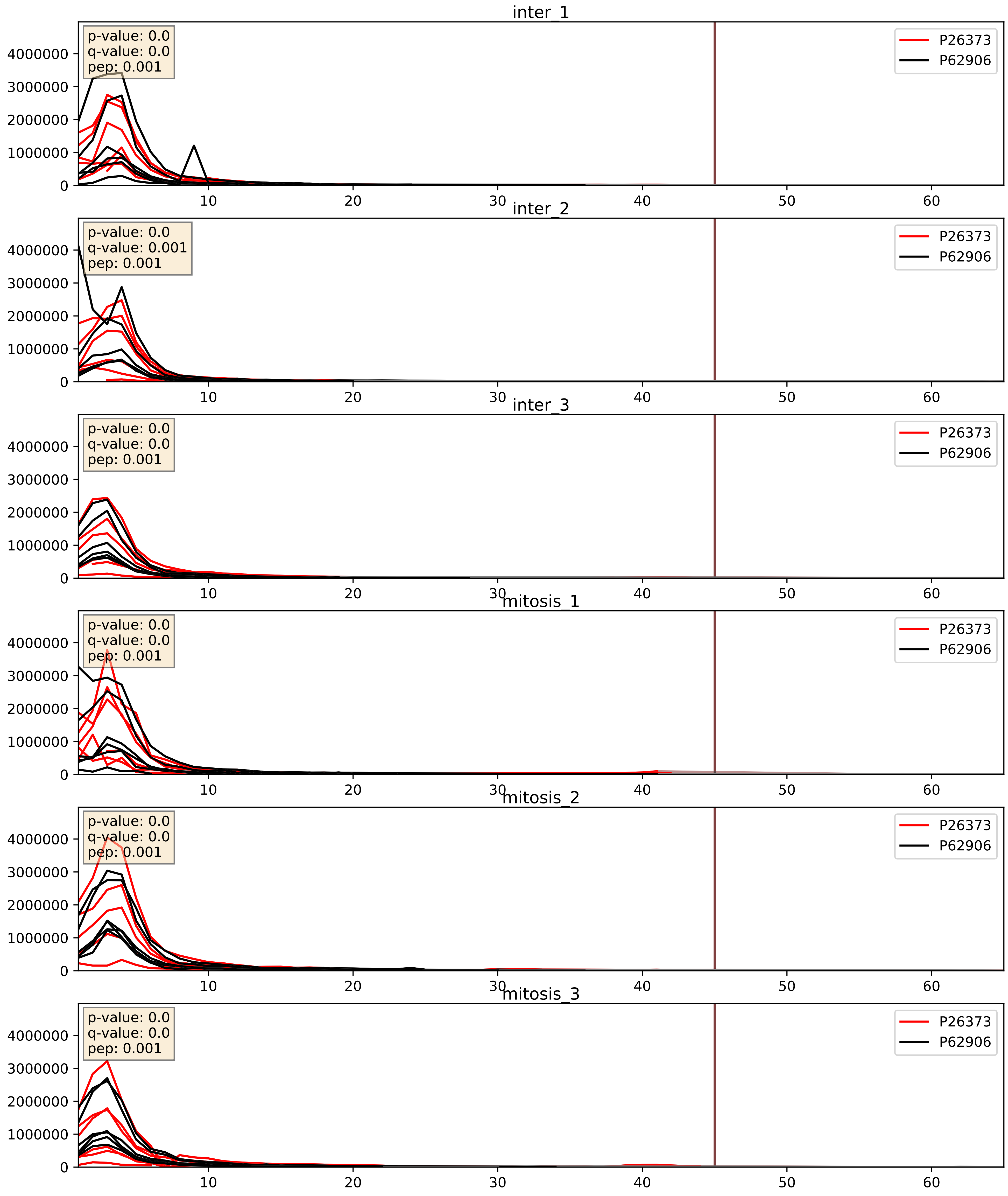


| level | condition_1 | condition_2 | pvalue | pvalue_adjusted |
|---|---|---|---|---|
| interactor_ratio | mitosis | inter | 0.3797048213571721 | 0.818376527351592 |
| interactor_abundance | mitosis | inter | 0.6748858480619343 | 0.7874832952969395 |
| complex_abundance | mitosis | inter | 0.9050933038607721 | 0.9409276999086967 |

P27635_P62906
RL10_HUMAN vs RL10A_HUMAN
p-value: 0.039 q-value: 0.027

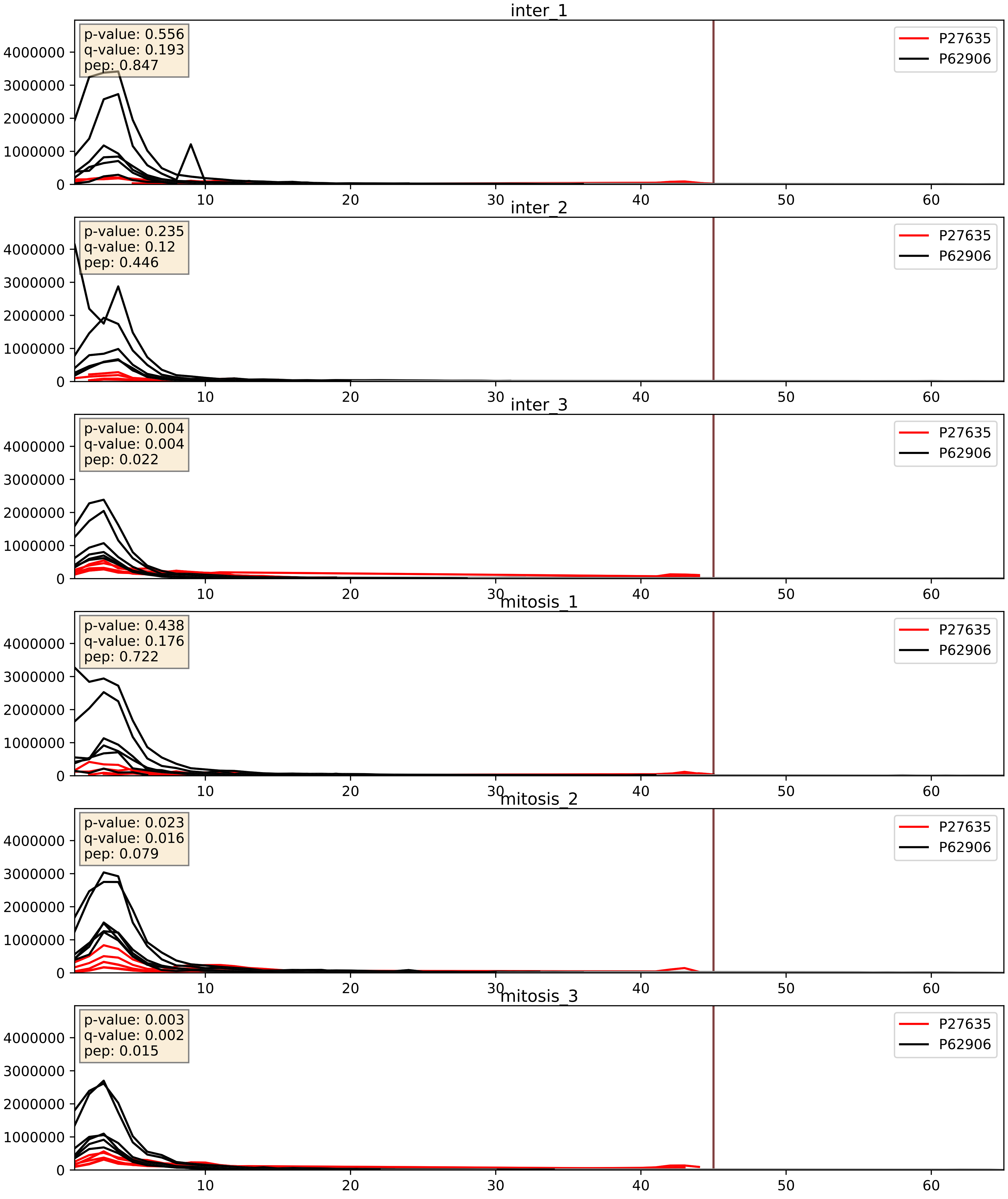


| level | condition_1 | condition_2 | pvalue | pvalue_adjusted |
|---|---|---|---|---|
| interactor_abundance | mitosis | inter | 0.5971657517650126 | 0.7352375167362112 |
| interactor_ratio | mitosis | inter | 0.6098938709015818 | 0.9070298390929 |
| complex_abundance | mitosis | inter | 0.609974219950882 | 0.7320262570289169 |

P30050_P62906
RL12_HUMAN vs RL10A_HUMAN
p-value: 0.0 q-value: 0.0

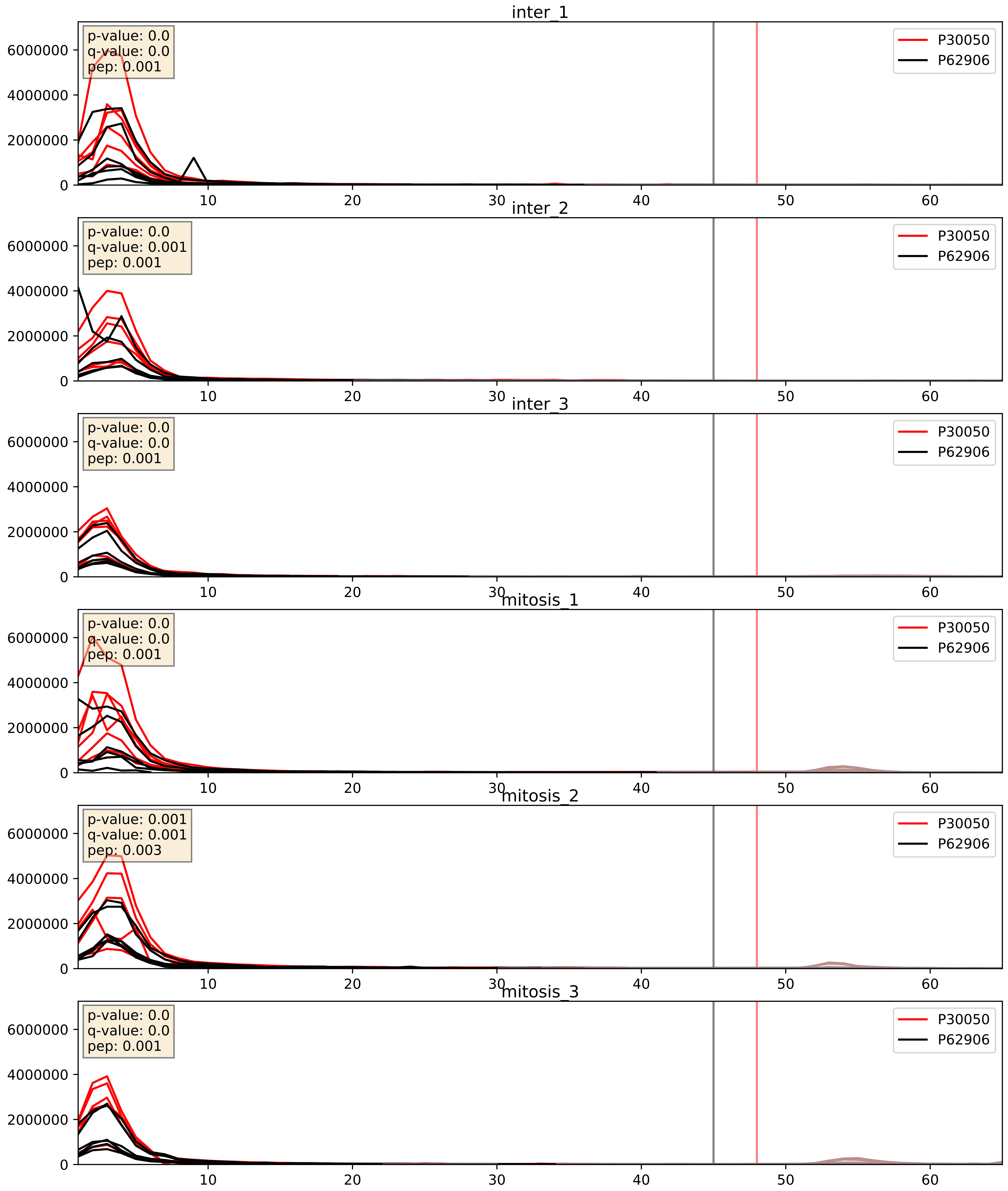


| level | condition_1 | condition_2 | pvalue | pvalue_adjusted |
|---|---|---|---|---|
| interactor_ratio | mitosis | inter | 0.9061399990575636 | 0.9852158746906624 |
| complex_abundance | mitosis | inter | 0.9211561127964304 | 0.95104285677692 |
| interactor_abundance | mitosis | inter | 0.9578403210458195 | 0.9739977605312682 |

P32969_P62906
RL9_HUMAN vs RL10A_HUMAN
p-value: 0.0 q-value: 0.0

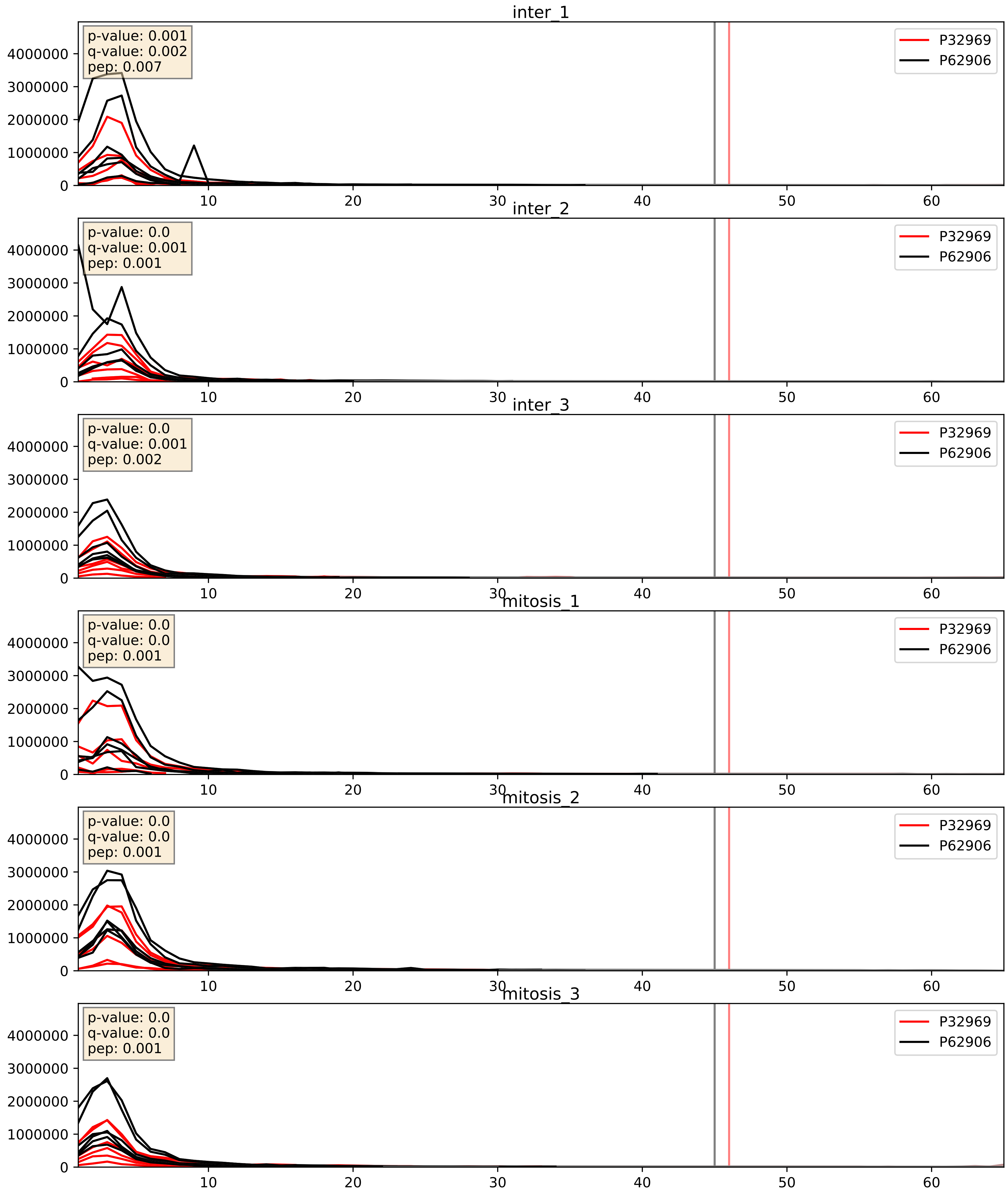


| level | condition_1 | condition_2 | pvalue | pvalue_adjusted |
|---|---|---|---|---|
| interactor_abundance | mitosis | inter | 0.2565315458198631 | 0.4494289873553066 |
| interactor_ratio | mitosis | inter | 0.34070518223703217 | 0.7929177515417035 |
| complex_abundance | mitosis | inter | 0.5660081471975618 | 0.6964251458978193 |

P35268_P62906
RL22_HUMAN vs RL10A_HUMAN
p-value: 0.001 q-value: 0.001

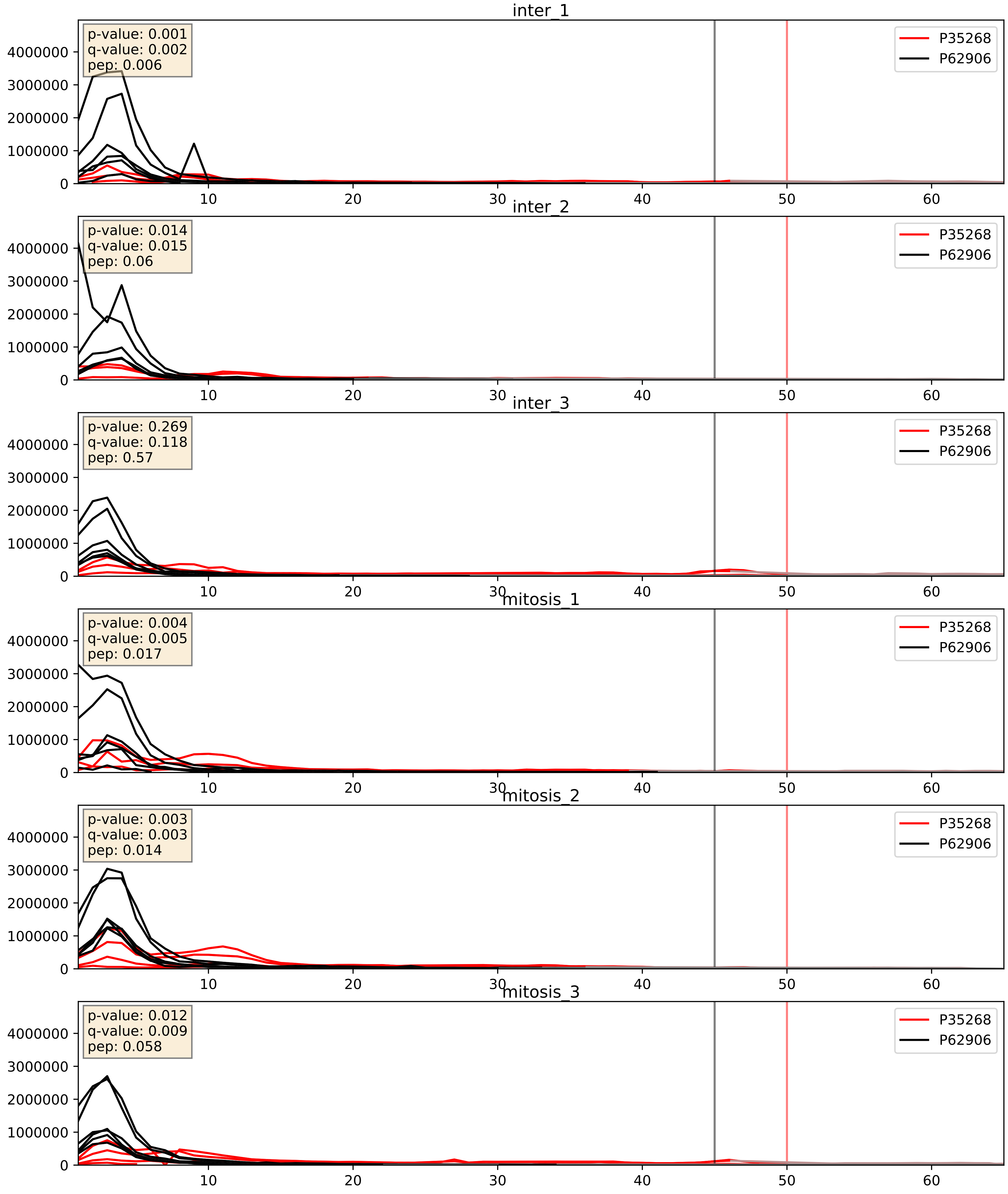


| level | condition_1 | condition_2 | pvalue | pvalue_adjusted |
|---|---|---|---|---|
| complex_abundance | mitosis | inter | 0.002410662275627882 | 0.03561197063475765 |
| interactor_abundance | mitosis | inter | 0.007905967289639496 | 0.07711604472197223 |
| interactor_ratio | mitosis | inter | 0.07929887789520691 | 0.4717153542619675 |

P36578_P62906
RL4_HUMAN vs RL10A_HUMAN
p-value: 0.0 q-value: 0.0

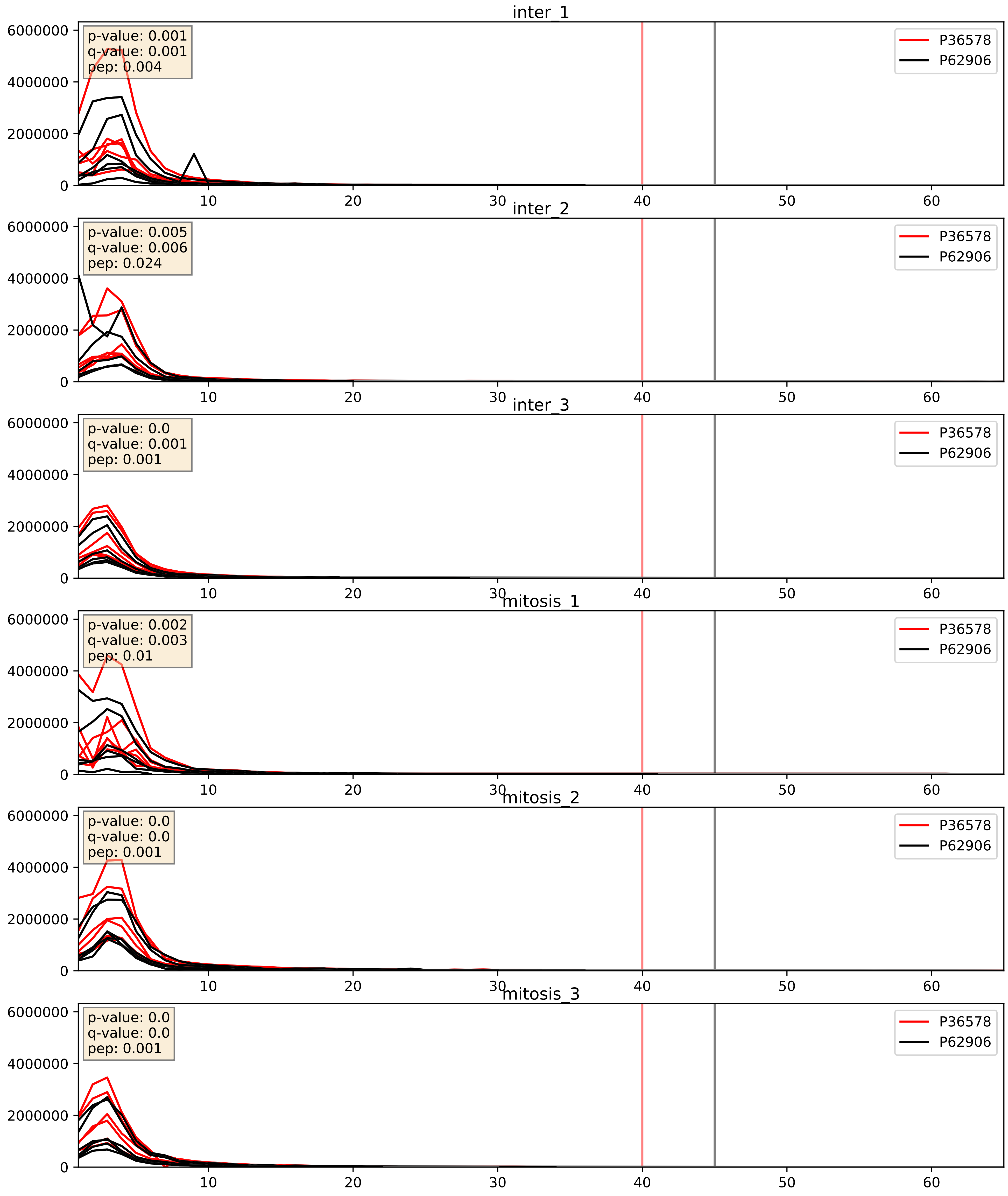


| level | condition_1 | condition_2 | pvalue | pvalue_adjusted |
|---|---|---|---|---|
| interactor_ratio | mitosis | inter | 0.8606670225775136 | 0.9774309046518314 |
| complex_abundance | mitosis | inter | 0.9154934693729797 | 0.9466808526011967 |
| interactor_abundance | mitosis | inter | 0.9799863548434697 | 0.988299151444404 |

P39019_P62906
RS19_HUMAN vs RL10A_HUMAN
p-value: 0.006 q-value: 0.006

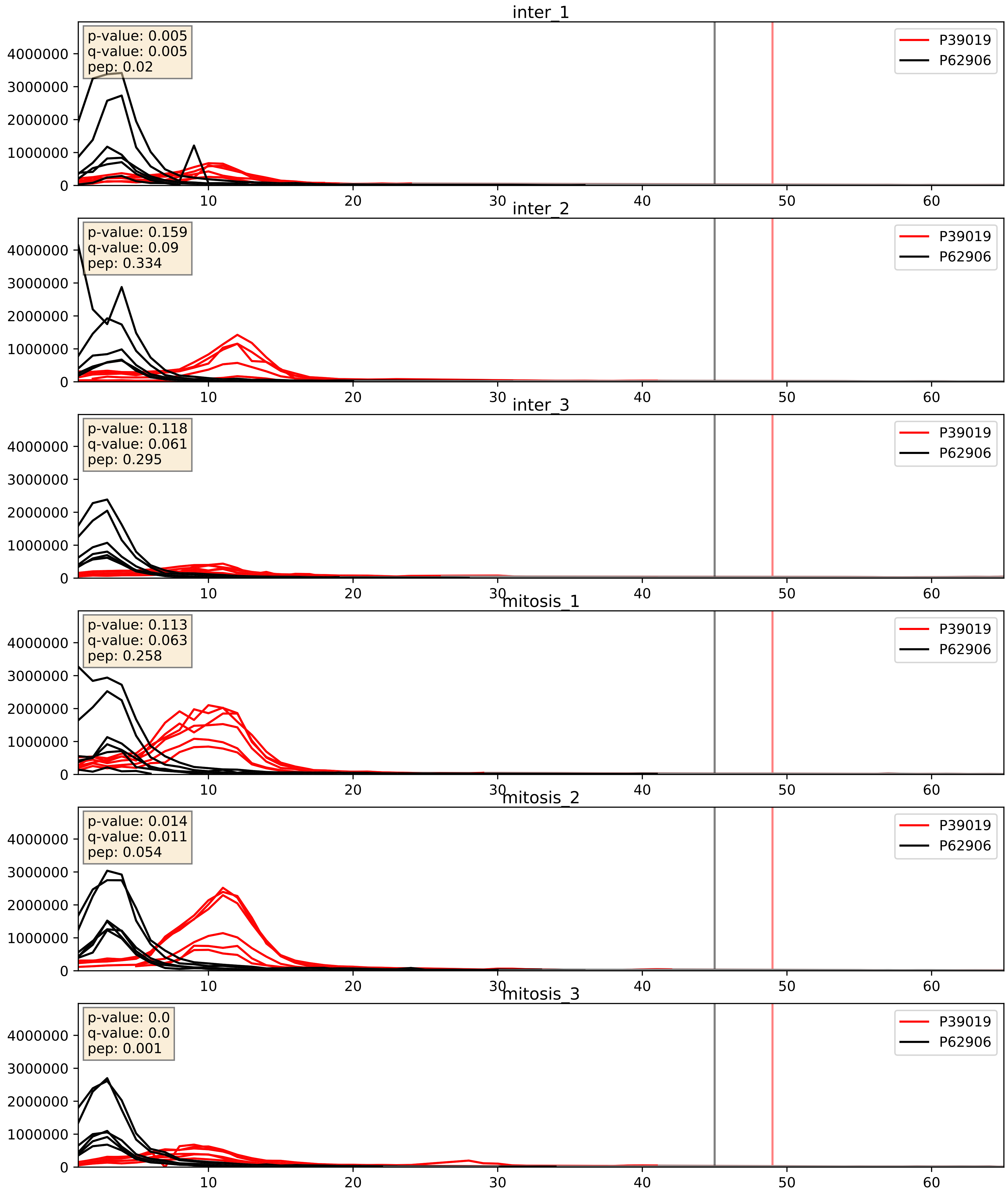


| level | condition_1 | condition_2 | pvalue | pvalue_adjusted |
|---|---|---|---|---|
| interactor_abundance | mitosis | inter | 0.22979336338511047 | 0.42246864822439595 |
| complex_abundance | mitosis | inter | 0.24080805297036958 | 0.3920344110738615 |
| interactor_ratio | mitosis | inter | 0.2441605334522016 | 0.7200661897289671 |

P39023_P62906
RL3_HUMAN vs RL10A_HUMAN
p-value: 0.0 q-value: 0.0

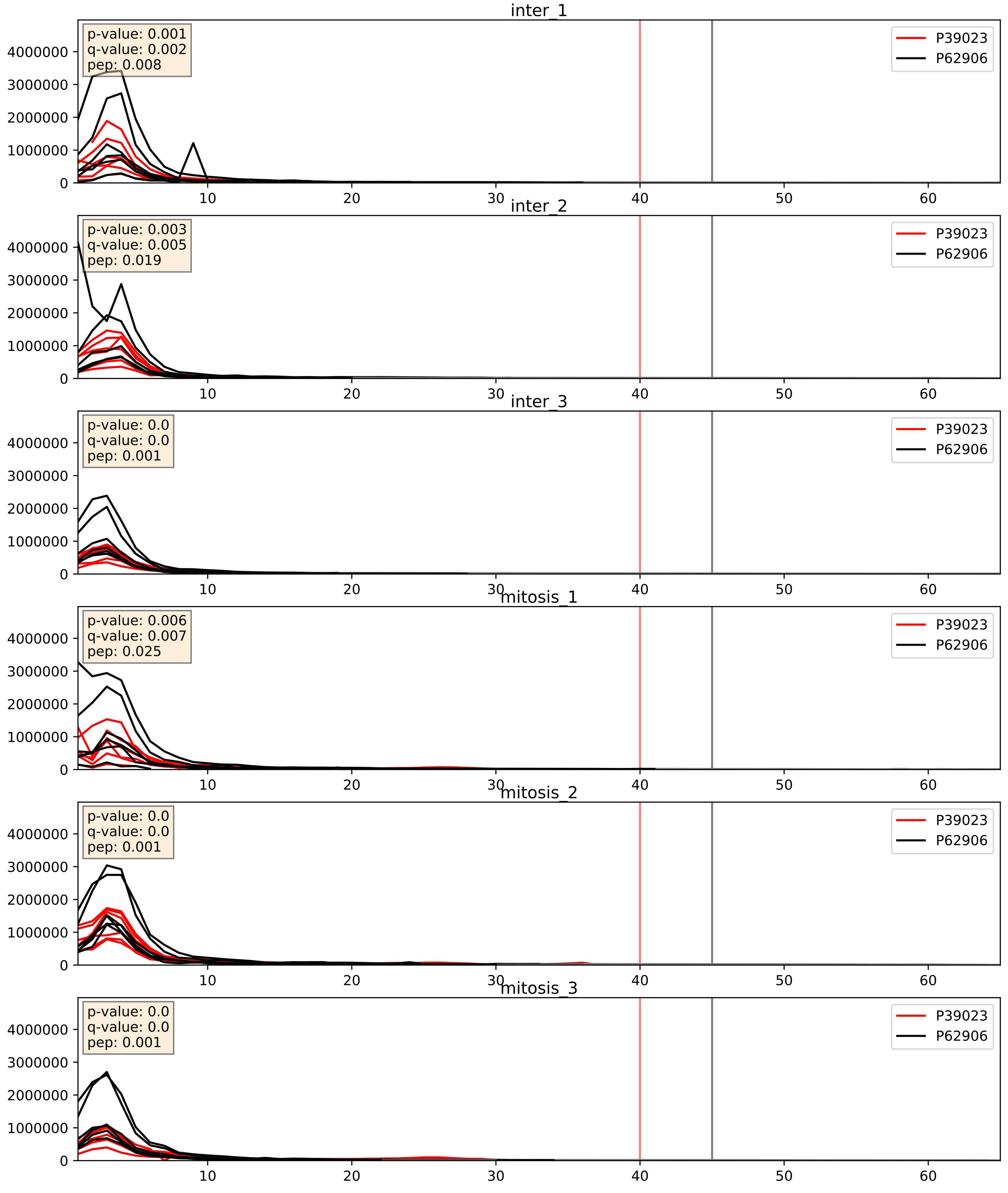


| level | condition_1 | condition_2 | pvalue | pvalue_adjusted |
|---|---|---|---|---|
| interactor_abundance | mitosis | inter | 0.7088955621271644 | 0.8107075500078192 |
| interactor_ratio | mitosis | inter | 0.7248875341586889 | 0.9456877650408758 |
| complex_abundance | mitosis | inter | 0.8123022164271102 | 0.8779427995727352 |

P40429_P62906
RL13A_HUMAN vs RL10A_HUMAN
p-value: 0.0 q-value: 0.0

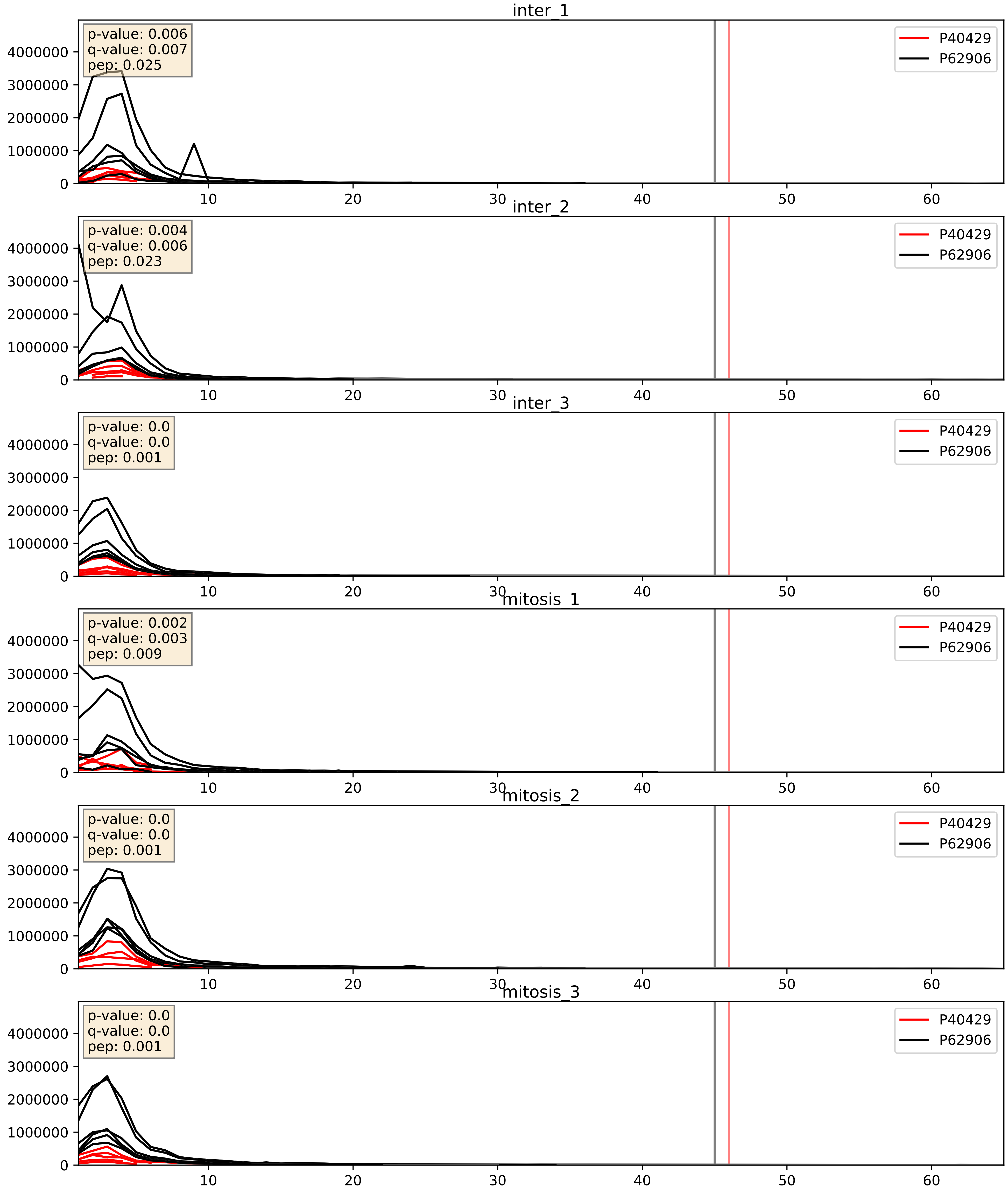


| level | condition_1 | condition_2 | pvalue | pvalue_adjusted |
|---|---|---|---|---|
| complex_abundance | mitosis | inter | 0.682738364743998 | 0.7883772294898992 |
| interactor_ratio | mitosis | inter | 0.8622549694005426 | 0.9774309046518314 |
| interactor_abundance | mitosis | inter | 0.9364707510987881 | 0.962096690999235 |

P42677_P62906
RS27_HUMAN vs RL10A_HUMAN
p-value: 0.004 q-value: 0.004

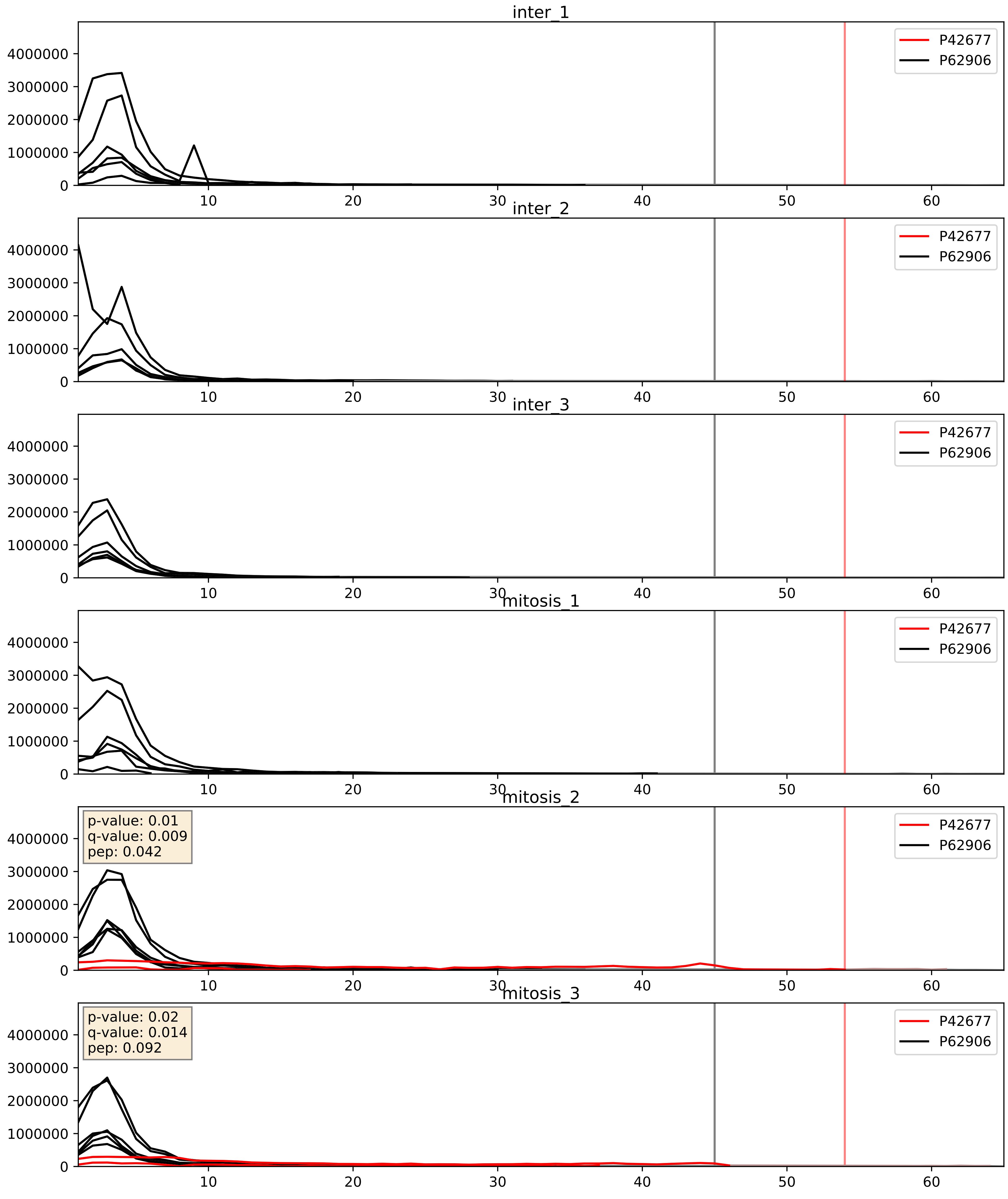


| level | condition_1 | condition_2 | pvalue | pvalue_adjusted |
|---|---|---|---|---|
| interactor_ratio | mitosis | inter | 0.1268648284343642 | 0.5750062399685051 |
| complex_abundance | mitosis | inter | 0.14258082961888732 | 0.28819951692442236 |
| interactor_abundance | mitosis | inter | 0.1575610663577279 | 0.34350000523081714 |

P42766_P62906
RL35_HUMAN vs RL10A_HUMAN
p-value: 0.0 q-value: 0.0

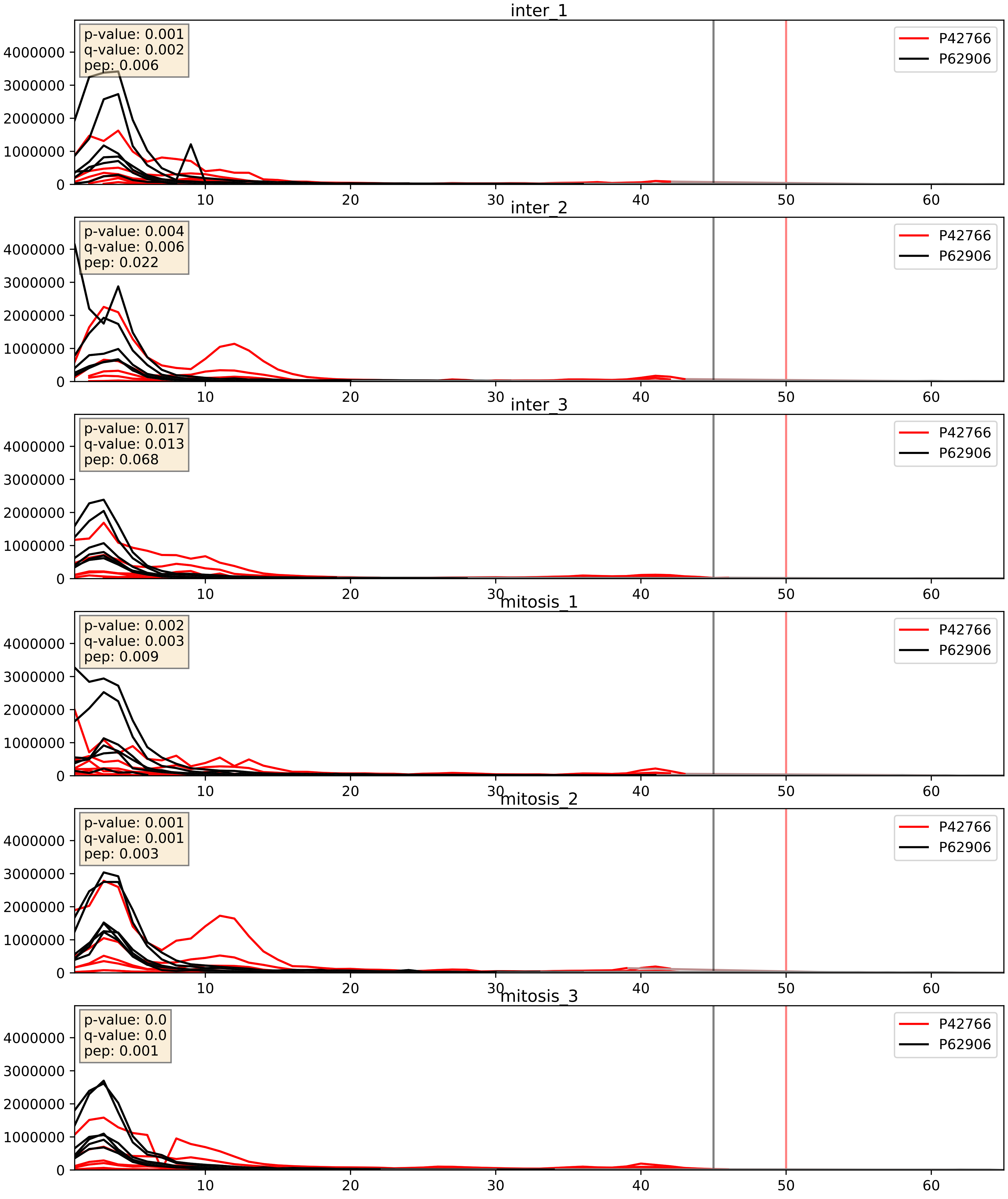


| level | condition_1 | condition_2 | pvalue | pvalue_adjusted |
|---|---|---|---|---|
| interactor_ratio | mitosis | inter | 0.8790124618306582 | 0.9802747470617126 |
| interactor_abundance | mitosis | inter | 0.9320899201060098 | 0.9614037493803402 |
| complex_abundance | mitosis | inter | 0.9607459875559152 | 0.9769524416106717 |

P46776_P62906
RL27A_HUMAN vs RL10A_HUMAN
p-value: 0.0 q-value: 0.0

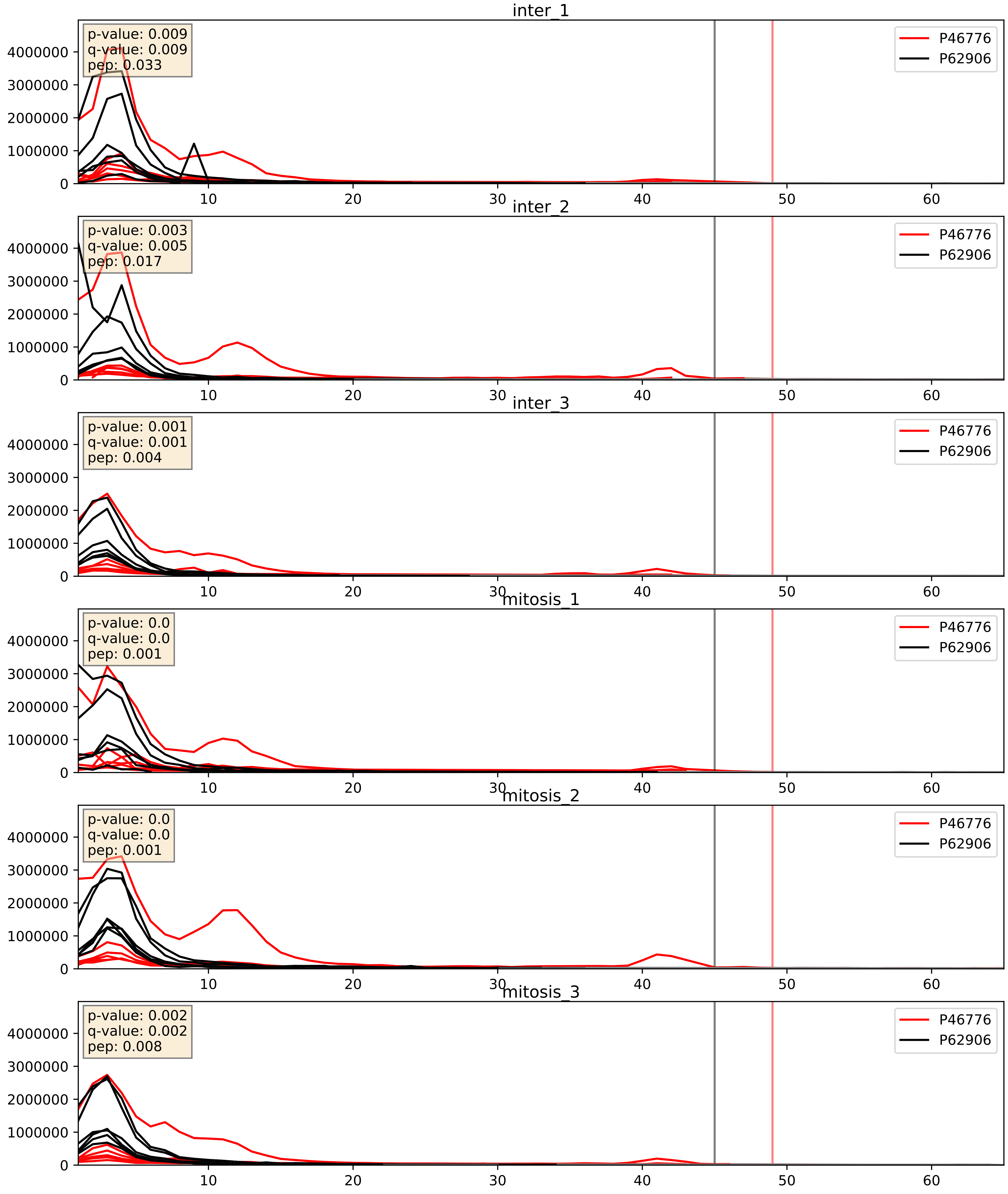


| level | condition_1 | condition_2 | pvalue | pvalue_adjusted |
|---|---|---|---|---|
| complex_abundance | mitosis | inter | 0.879871418393896 | 0.9250429060982253 |
| interactor_abundance | mitosis | inter | 0.9036357089158716 | 0.9461832168094921 |
| interactor_ratio | mitosis | inter | 0.9258503749074931 | 0.9871879493798719 |

P46777_P62906
RL5_HUMAN vs RL10A_HUMAN
p-value: 0.0 q-value: 0.0

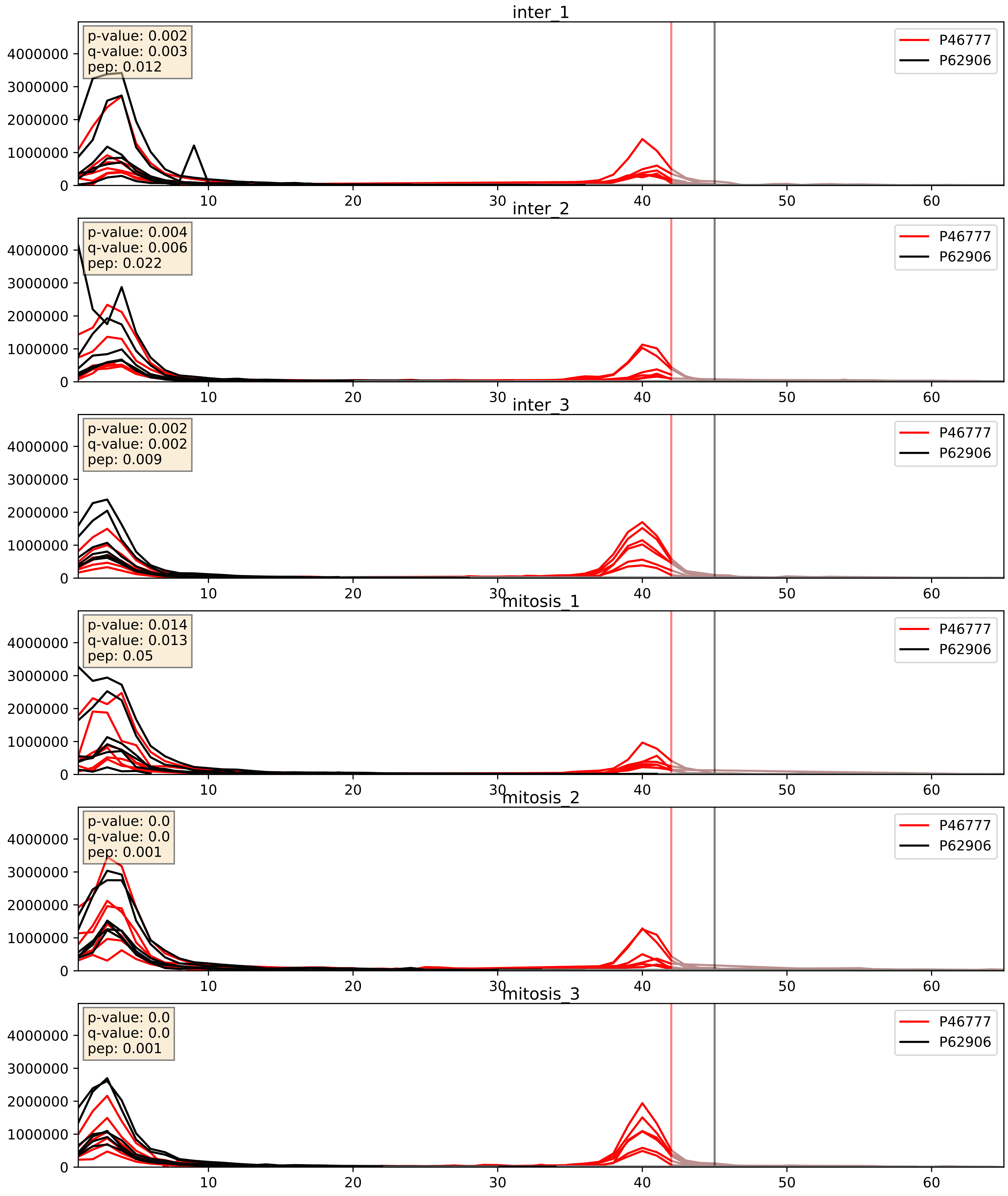


| level | condition_1 | condition_2 | pvalue | pvalue_adjusted |
|---|---|---|---|---|
| interactor_abundance | mitosis | inter | 0.11888633325337002 | 0.2983485818378327 |
| complex_abundance | mitosis | inter | 0.12340967201102081 | 0.26563809276703054 |
| interactor_ratio | mitosis | inter | 0.23031405748017306 | 0.7084040000108809 |

P46778_P62906
RL21_HUMAN vs RL10A_HUMAN
p-value: 0.0 q-value: 0.0

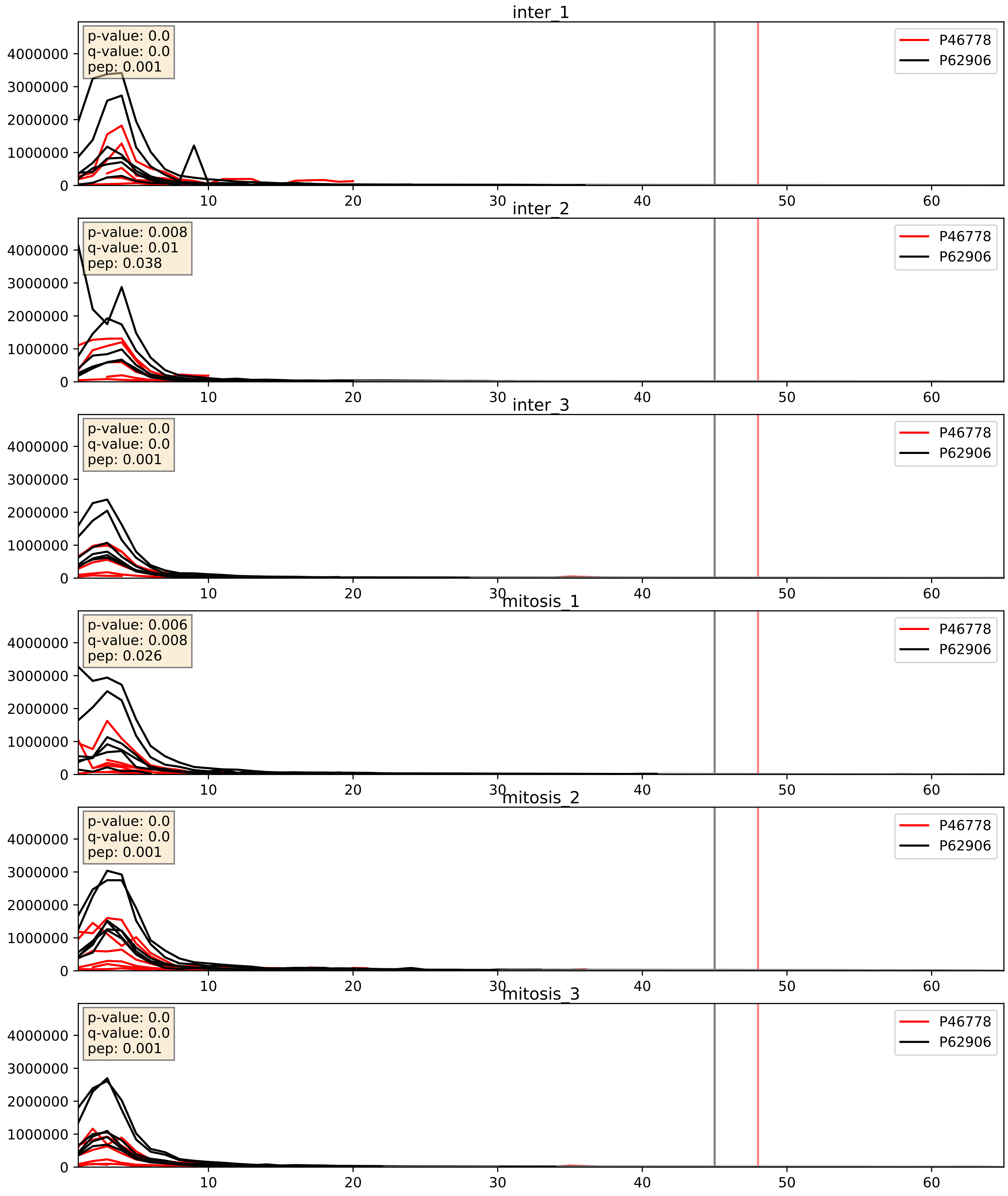


| level | condition_1 | condition_2 | pvalue | pvalue_adjusted |
|---|---|---|---|---|
| complex_abundance | mitosis | inter | 0.3954663126624109 | 0.5419775274399995 |
| interactor_abundance | mitosis | inter | 0.5008063444764381 | 0.6659291057040283 |
| interactor_ratio | mitosis | inter | 0.6988492197292088 | 0.9359699707113329 |

P46779_P62906
RL28_HUMAN vs RL10A_HUMAN
p-value: 0.0 q-value: 0.0

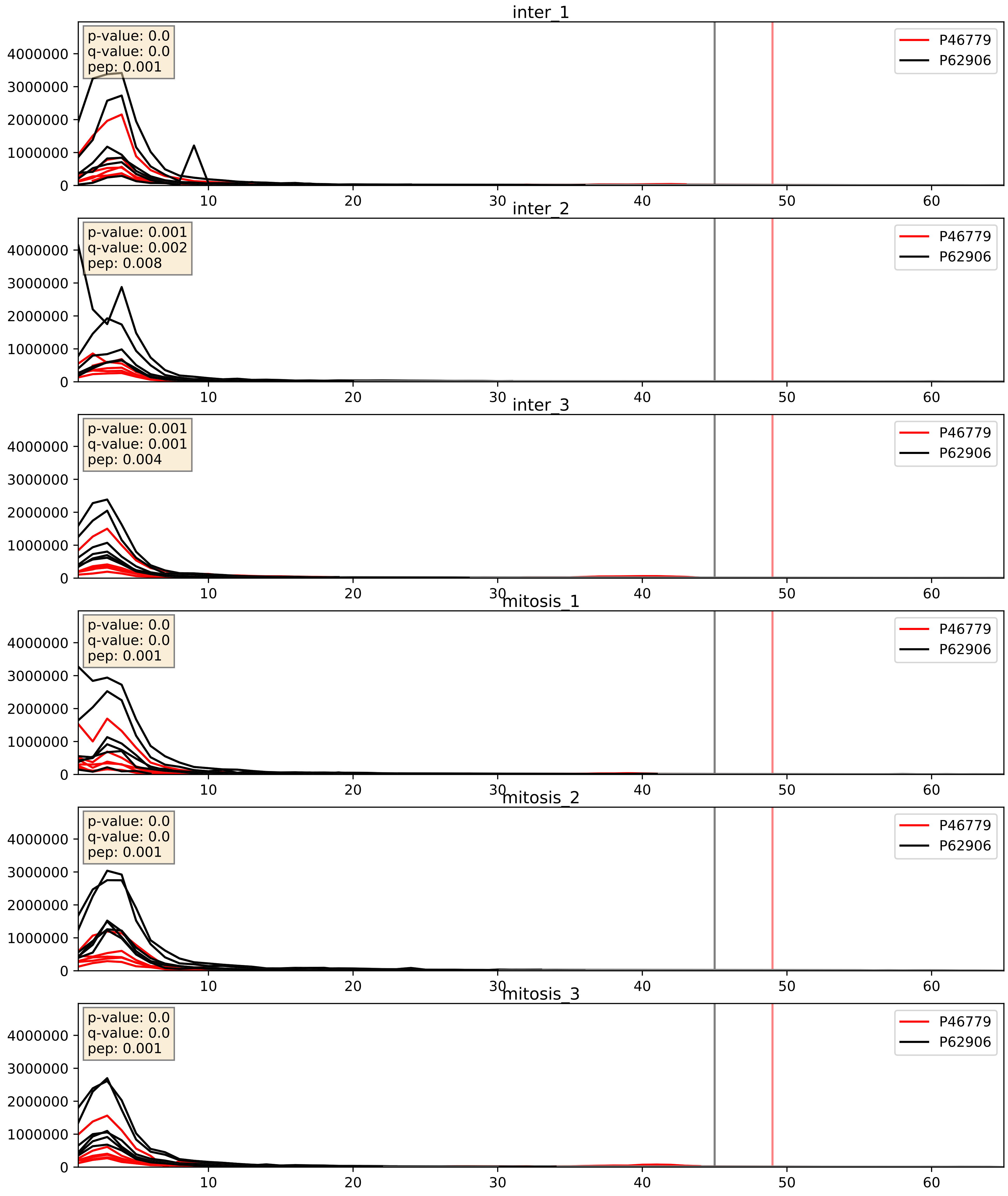


| level | condition_1 | condition_2 | pvalue | pvalue_adjusted |
|---|---|---|---|---|
| interactor_abundance | mitosis | inter | 0.18776556728119148 | 0.3800752453282745 |
| interactor_ratio | mitosis | inter | 0.31238772314116064 | 0.7777940684278926 |
| complex_abundance | mitosis | inter | 0.3634457332175016 | 0.5112728802533795 |

P46781_P62906
RS9_HUMAN vs RL10A_HUMAN
p-value: 0.01 q-value: 0.009

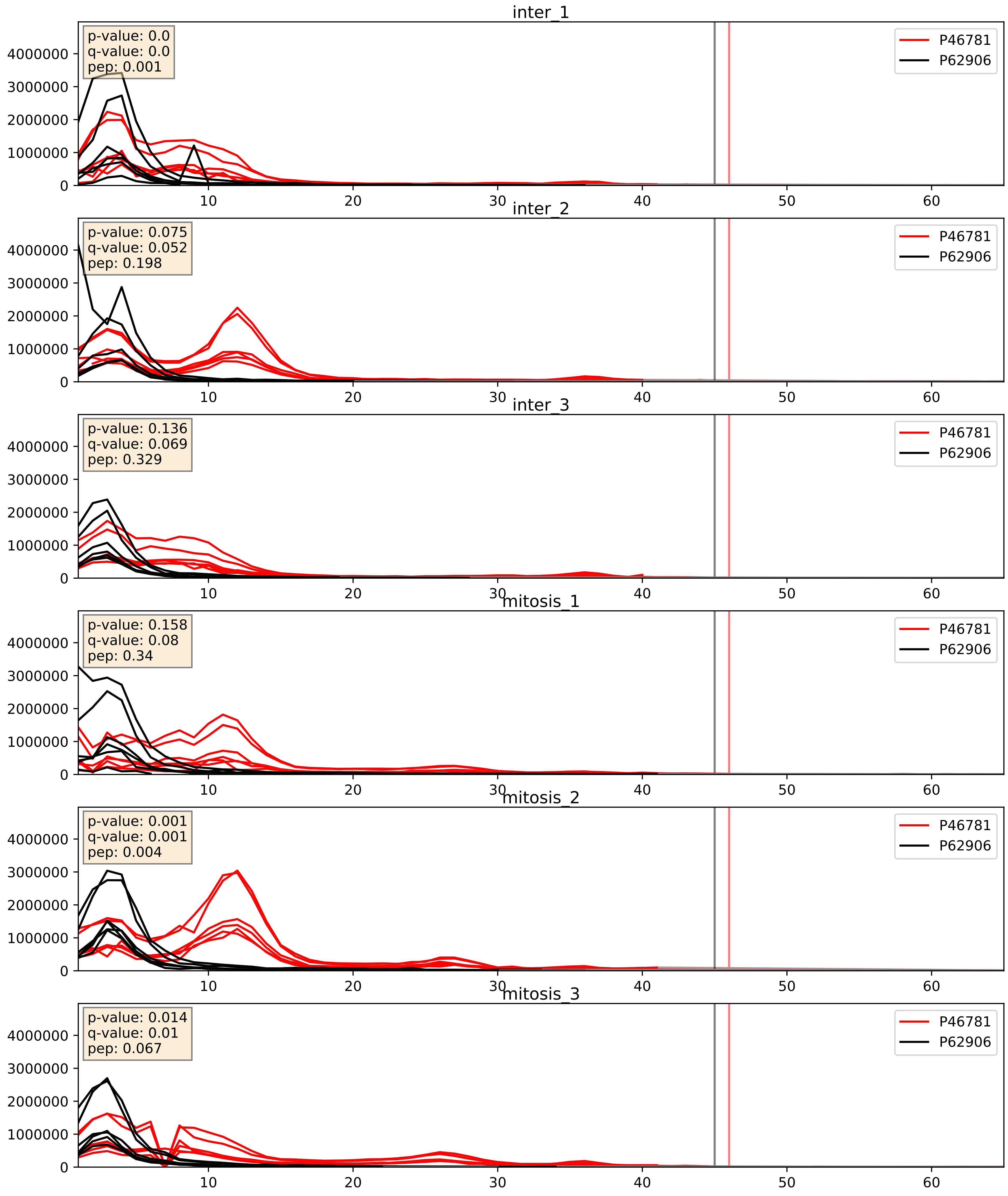


| level | condition_1 | condition_2 | pvalue | pvalue_adjusted |
|---|---|---|---|---|
| complex_abundance | mitosis | inter | 0.7687664473142581 | 0.8479114532933965 |
| interactor_abundance | mitosis | inter | 0.8509196088867155 | 0.9192745443390276 |
| interactor_ratio | mitosis | inter | 0.9839801381406212 | 0.9976654700477962 |

P46782_P62906
RS5_HUMAN vs RL10A_HUMAN
p-value: 0.005 q-value: 0.005

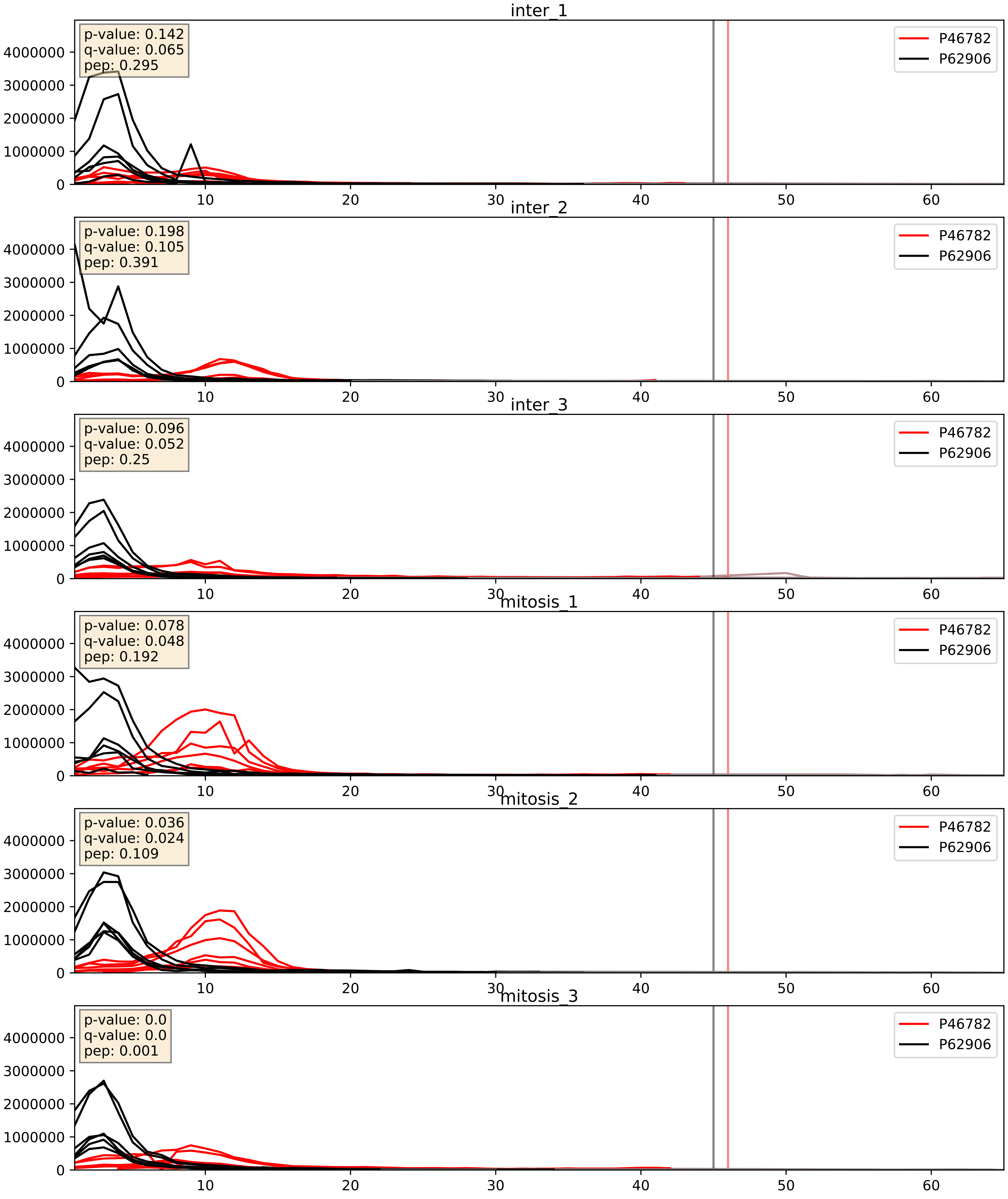


| level | condition_1 | condition_2 | pvalue | pvalue_adjusted |
|---|---|---|---|---|
| interactor_abundance | mitosis | inter | 0.12326945614198187 | 0.30568259135663056 |
| interactor_ratio | mitosis | inter | 0.15183118389050143 | 0.6067576723168497 |
| complex_abundance | mitosis | inter | 0.1571196543127301 | 0.3038735293531336 |

P46783_P62906
RS10_HUMAN vs RL10A_HUMAN
p-value: 0.068 q-value: 0.042

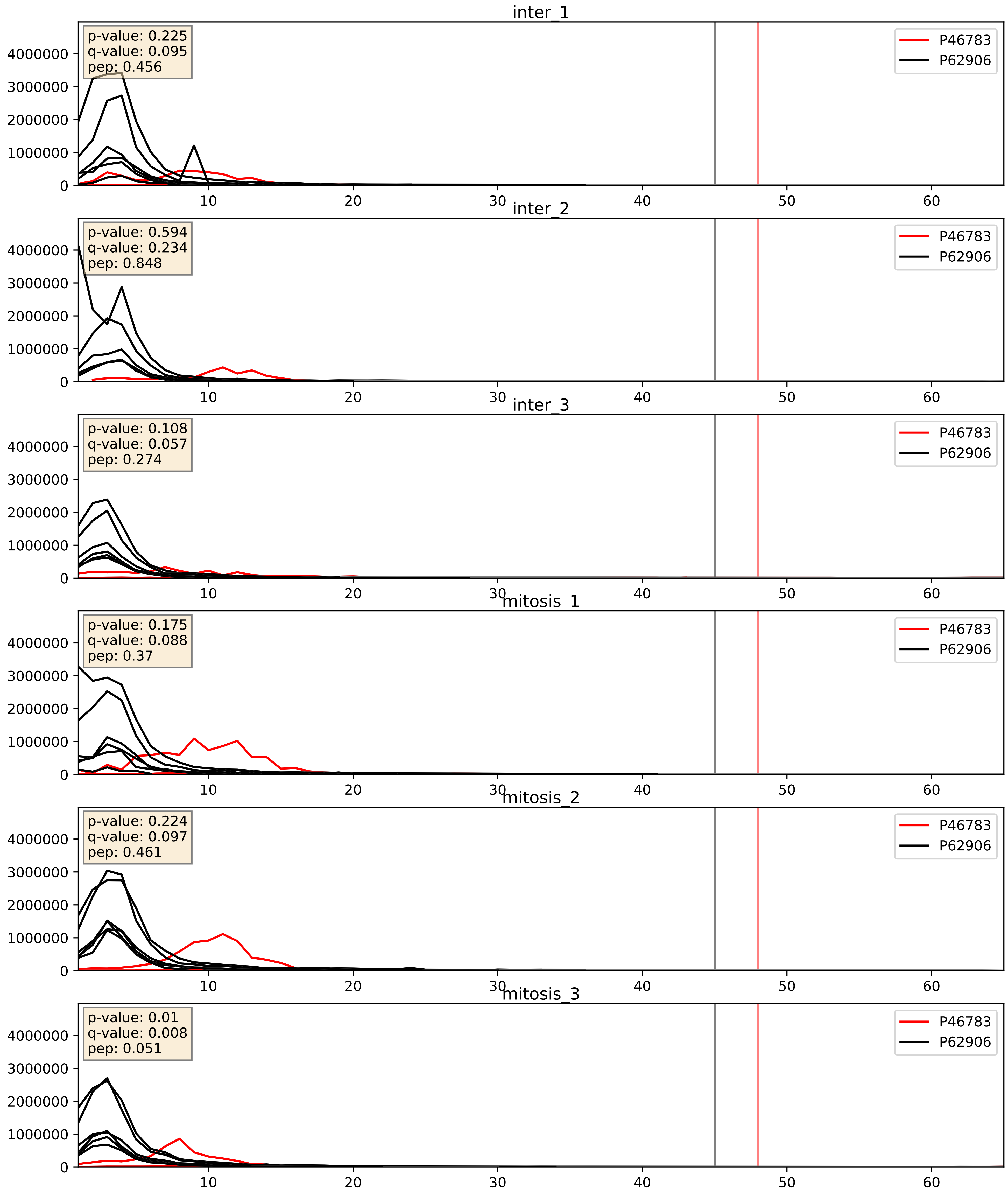


| level | condition_1 | condition_2 | pvalue | pvalue_adjusted |
|---|---|---|---|---|
| complex_abundance | mitosis | inter | 0.8343474390746579 | 0.8930607203023997 |
| interactor_ratio | mitosis | inter | 0.8509116146425202 | 0.9755634896635457 |
| interactor_abundance | mitosis | inter | 0.9498385396348883 | 0.9691041583035597 |

P47914_P62906
RL29_HUMAN vs RL10A_HUMAN
p-value: 0.0 q-value: 0.0

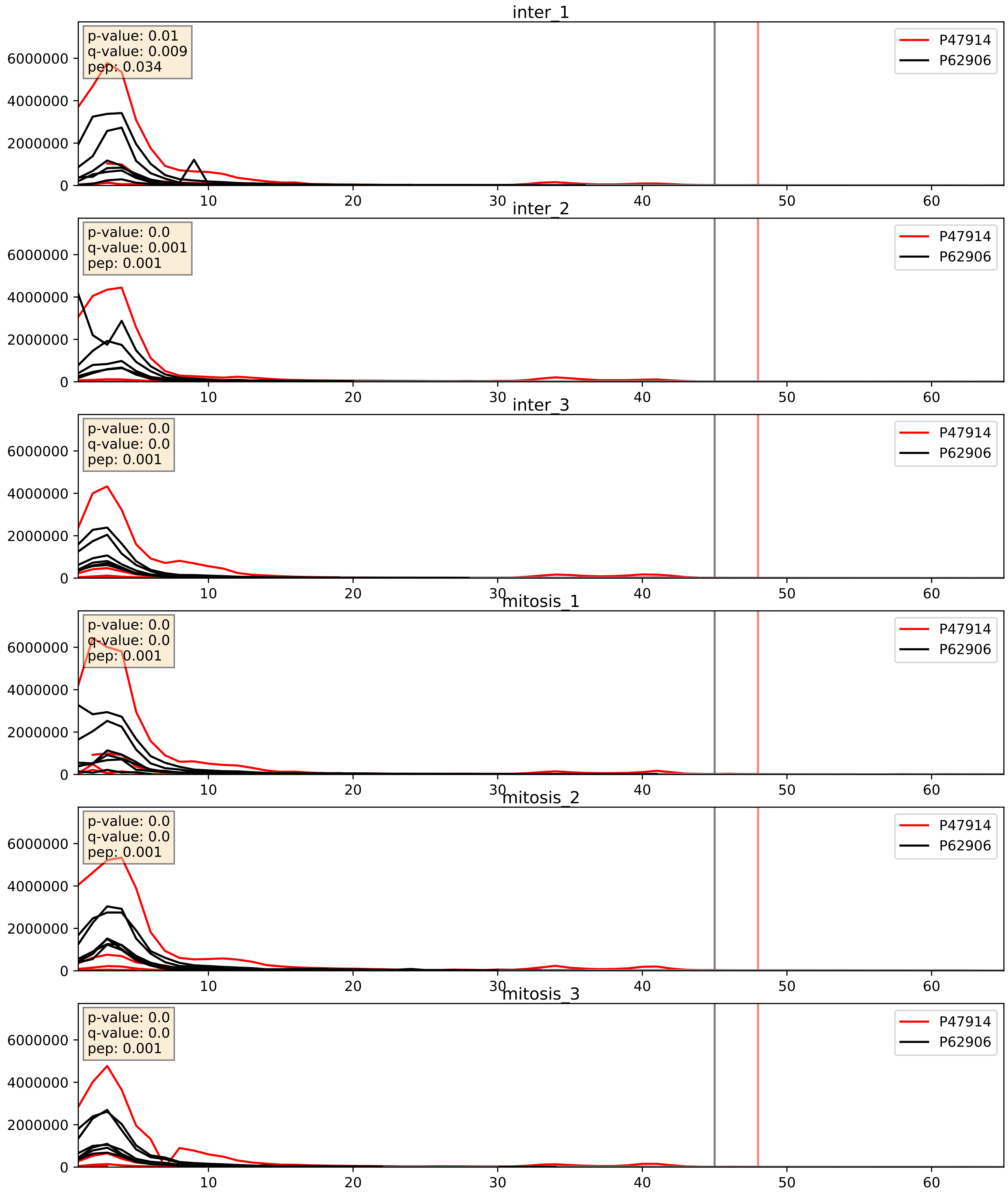


| level | condition_1 | condition_2 | pvalue | pvalue_adjusted |
|---|---|---|---|---|
| interactor_ratio | mitosis | inter | 0.35043002572570564 | 0.8007410362093738 |
| interactor_abundance | mitosis | inter | 0.4056879197306024 | 0.5920136290887666 |
| complex_abundance | mitosis | inter | 0.49897977625367135 | 0.6370399577165258 |

P49207_P62906
RL34_HUMAN vs RL10A_HUMAN
p-value: 0.0 q-value: 0.0

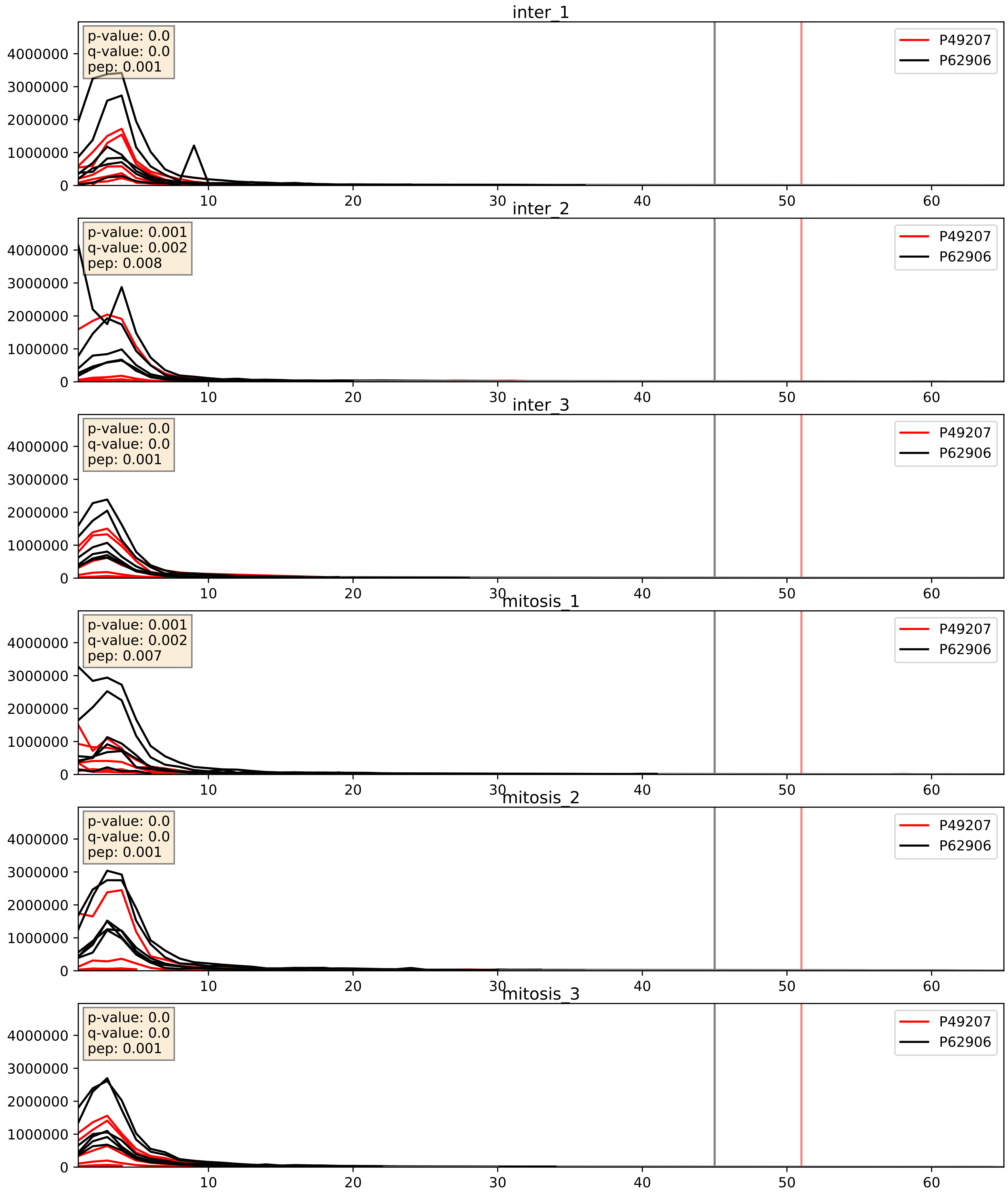


| level | condition_1 | condition_2 | pvalue | pvalue_adjusted |
|---|---|---|---|---|
| interactor_ratio | mitosis | inter | 0.6851051419636982 | 0.9349347226390419 |
| interactor_abundance | mitosis | inter | 0.6867637966708919 | 0.7949860647871448 |
| complex_abundance | mitosis | inter | 0.695652315046499 | 0.794722782533779 |

P50914_P62906
RL14_HUMAN vs RL10A_HUMAN
p-value: 0.0 q-value: 0.0

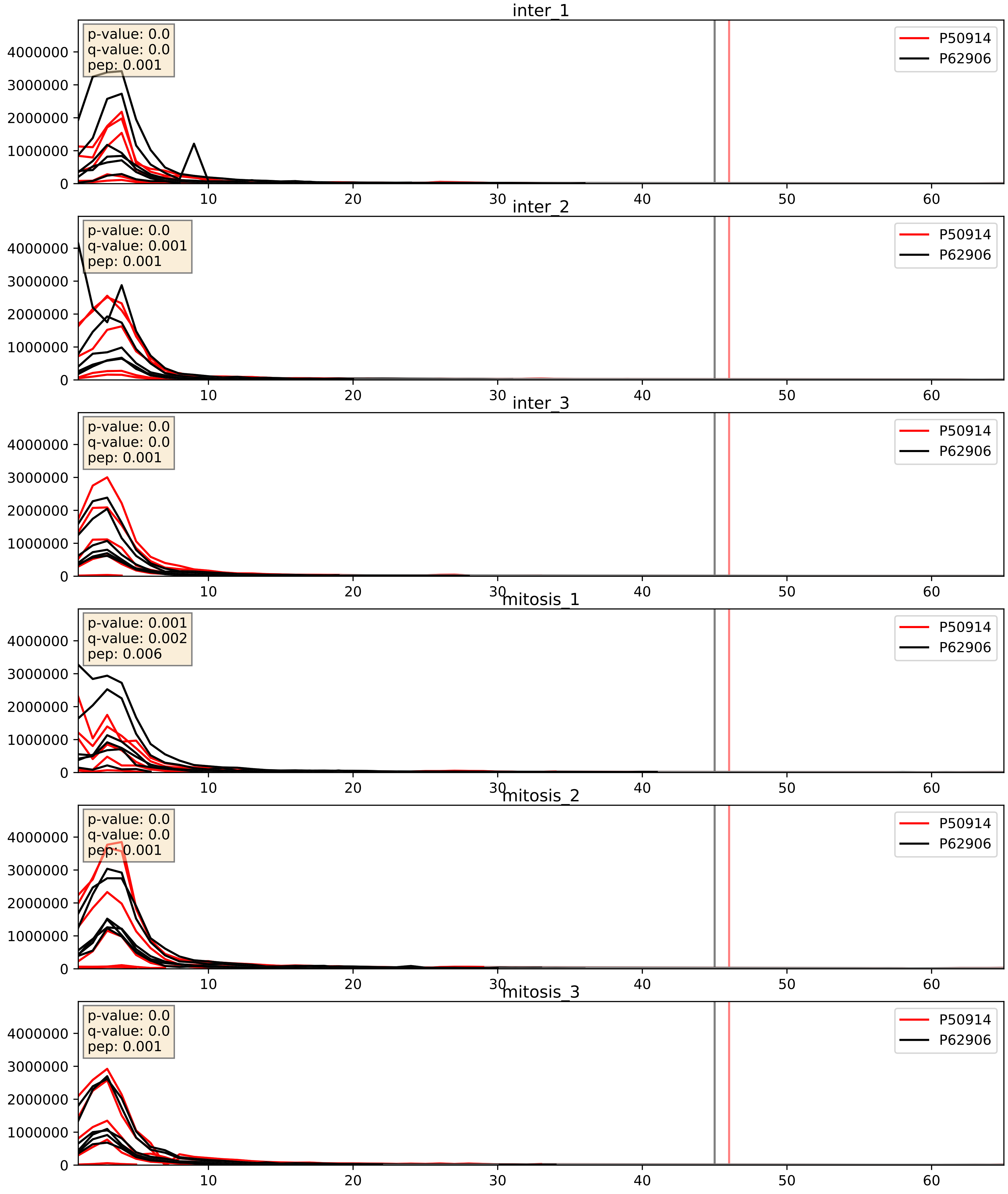


| level | condition_1 | condition_2 | pvalue | pvalue_adjusted |
|---|---|---|---|---|
| complex_abundance | mitosis | inter | 0.7503723469859708 | 0.835375407525308 |
| interactor_abundance | mitosis | inter | 0.8580026131539118 | 0.9237786766363731 |
| interactor_ratio | mitosis | inter | 0.9417251208705126 | 0.99110227089768 |

P55769_P62906
NH2L1_HUMAN vs RL10A_HUMAN
p-value: 0.0 q-value: 0.016

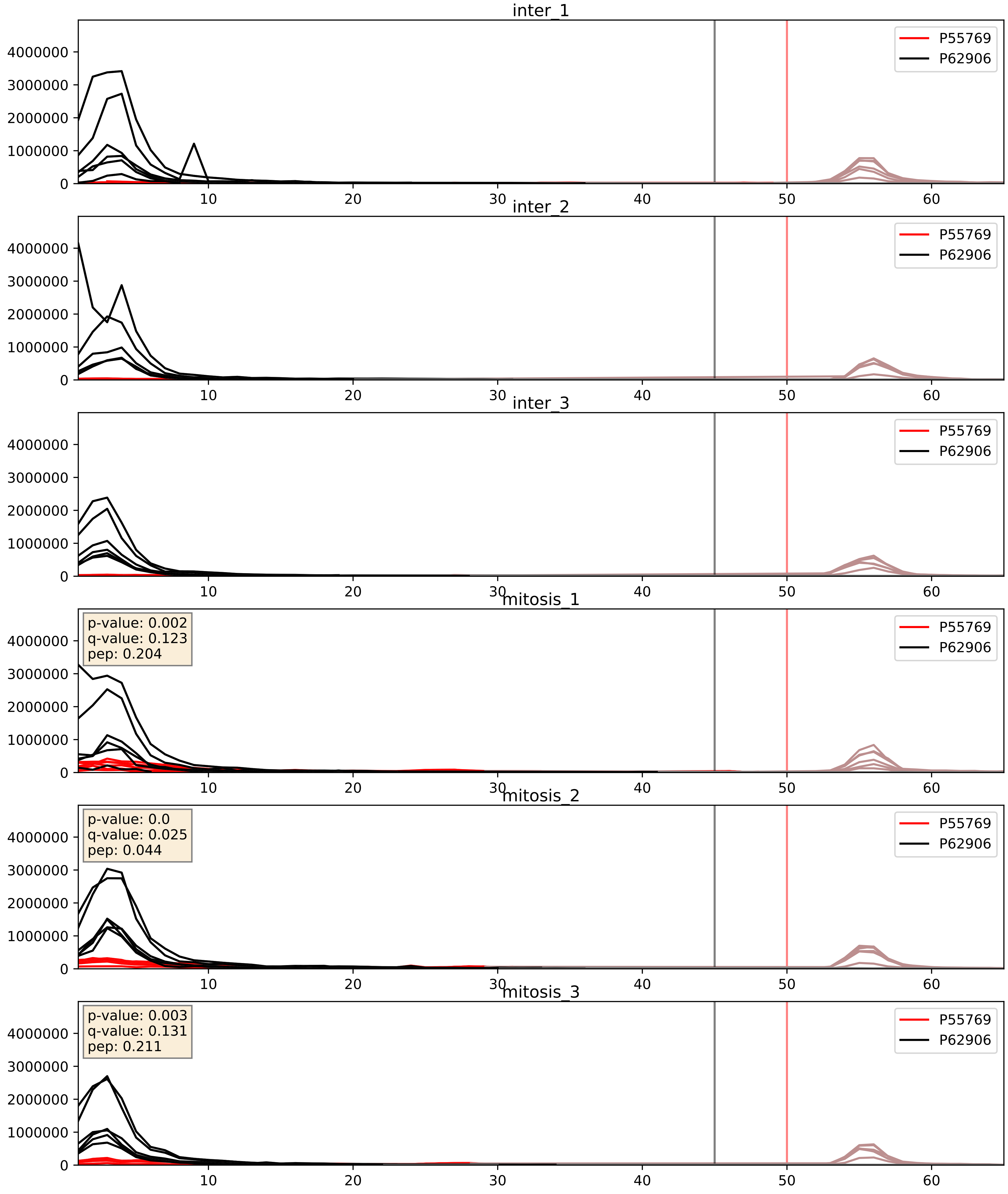


| level | condition_1 | condition_2 | pvalue | pvalue_adjusted |
|---|---|---|---|---|
| interactor_ratio | mitosis | inter | 0.002581226673103492 | 0.0651778770553566 |
| interactor_abundance | mitosis | inter | 0.002912682752214205 | 0.040906586314936176 |
| complex_abundance | mitosis | inter | 0.0072955578923150815 | 0.06327386507688666 |

P56537_P62906
IF6_HUMAN vs RL10A_HUMAN
p-value: 0.0 q-value: 0.014

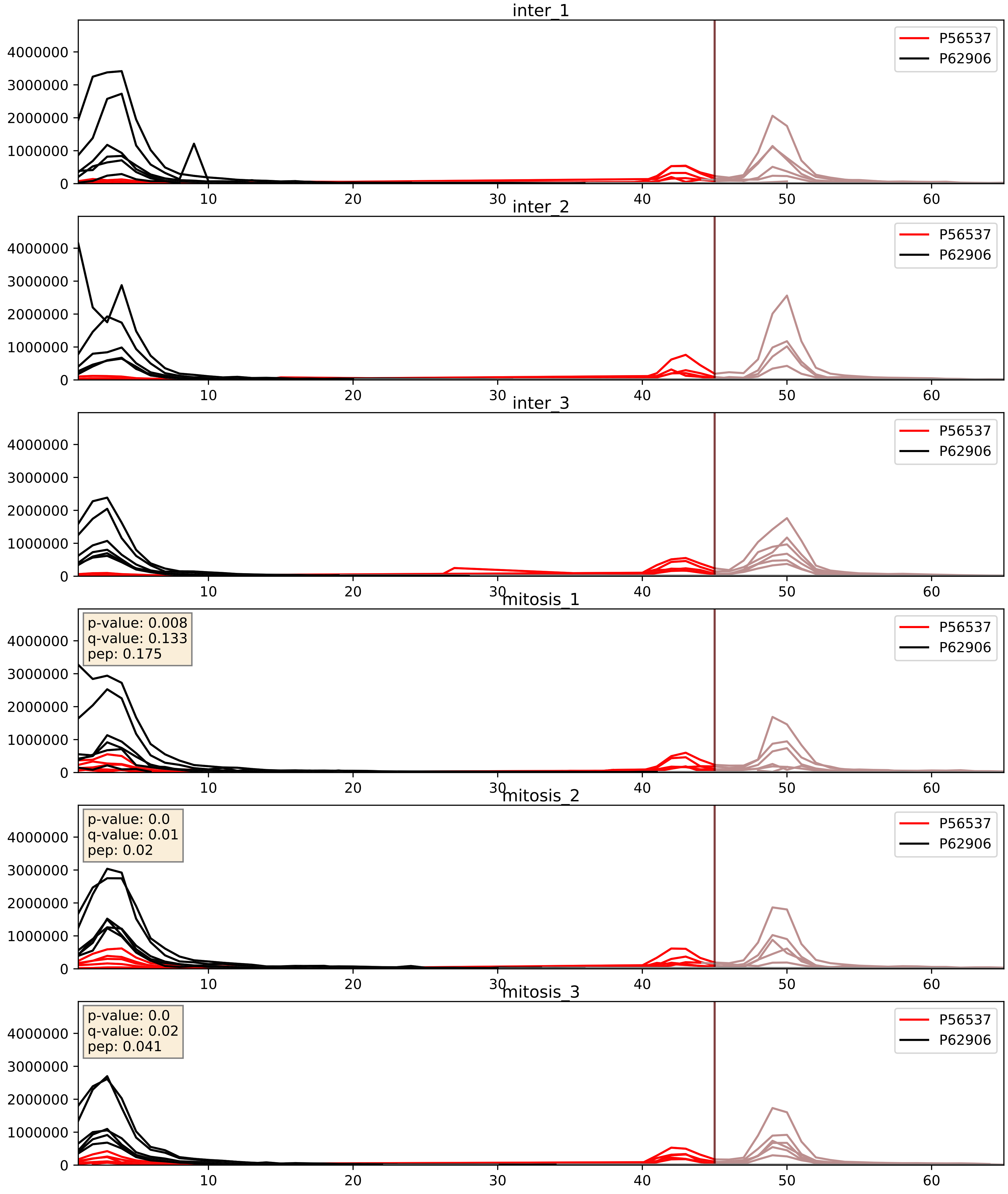


| level | condition_1 | condition_2 | pvalue | pvalue_adjusted |
|---|---|---|---|---|
| interactor_abundance | mitosis | inter | 0.004768879721580431 | 0.05653962661596744 |
| interactor_ratio | mitosis | inter | 0.007006627641535918 | 0.1166862502170184 |
| complex_abundance | mitosis | inter | 0.011403424595661052 | 0.07923158647634627 |

P60866_P62906
RS20_HUMAN vs RL10A_HUMAN
p-value: 0.024 q-value: 0.018

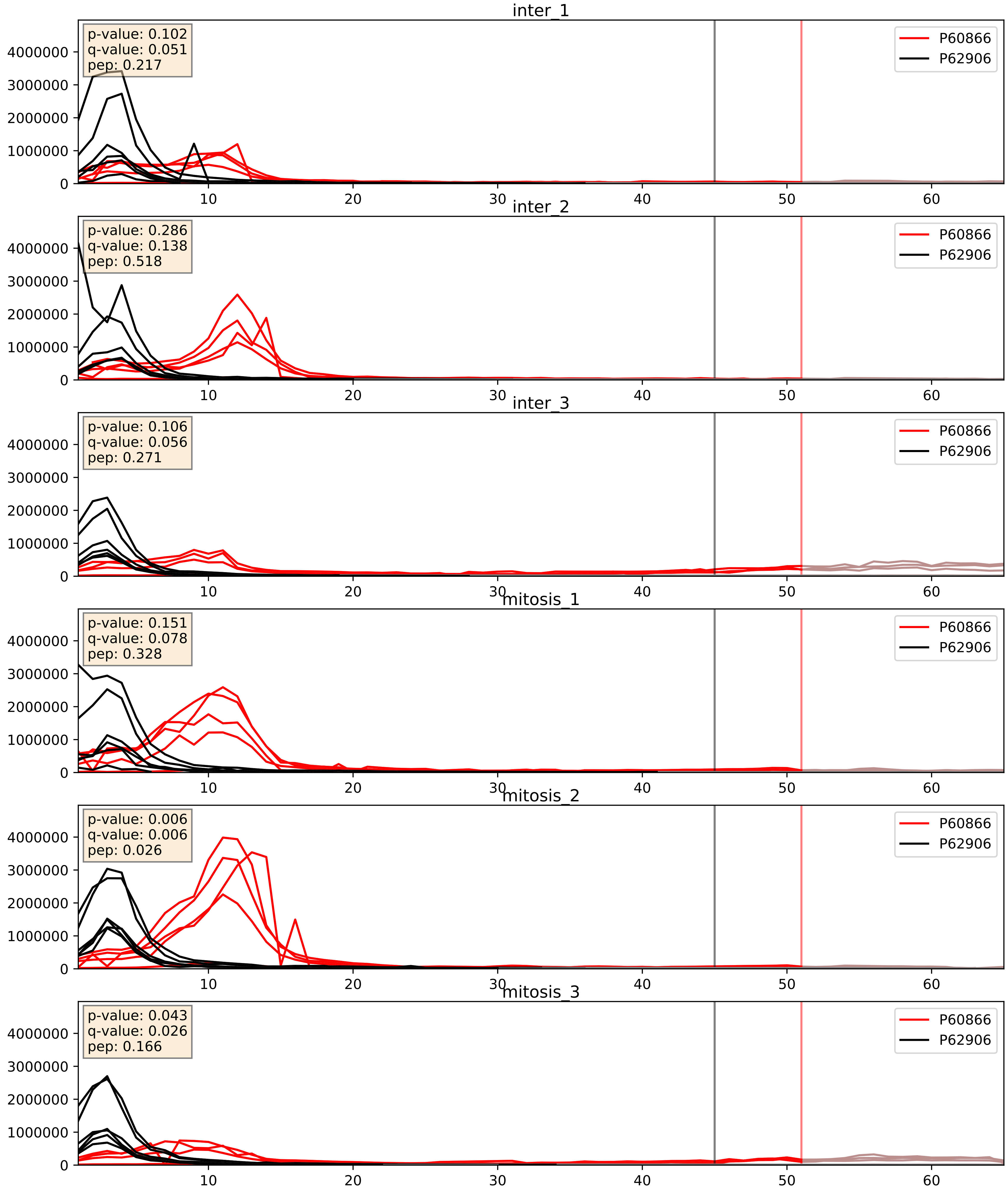


| level | condition_1 | condition_2 | pvalue | pvalue_adjusted |
|---|---|---|---|---|
| interactor_abundance | mitosis | inter | 0.4375548439362783 | 0.615777963681799 |
| complex_abundance | mitosis | inter | 0.45026267370462403 | 0.5925055321924031 |
| interactor_ratio | mitosis | inter | 0.4556813592651873 | 0.8549372939008518 |

P61247_P62906
RS3A_HUMAN vs RL10A_HUMAN
p-value: 0.007 q-value: 0.007

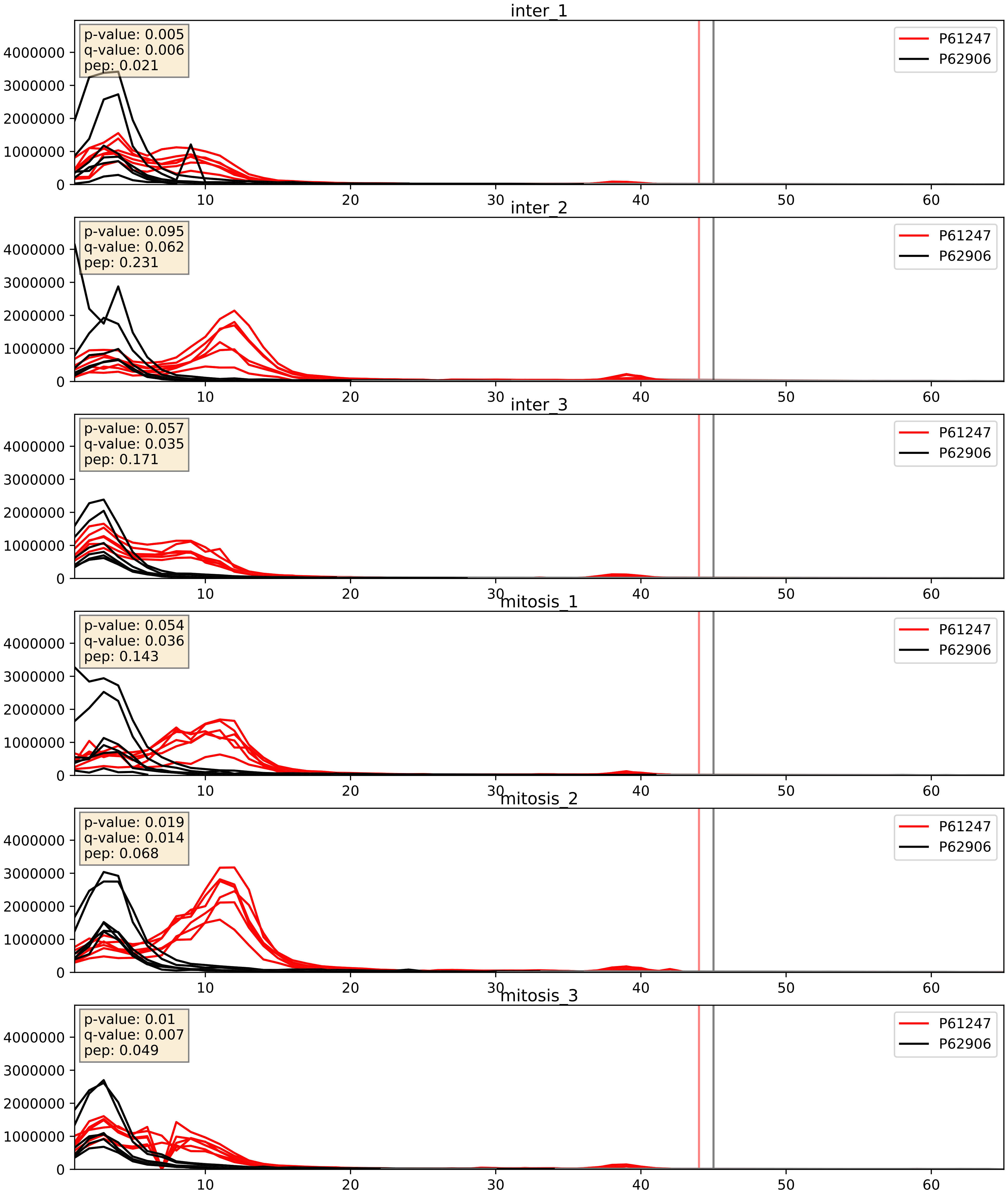


| level | condition_1 | condition_2 | pvalue | pvalue_adjusted |
|---|---|---|---|---|
| complex_abundance | mitosis | inter | 0.27212783148209513 | 0.42138463051496644 |
| interactor_abundance | mitosis | inter | 0.4668271168037885 | 0.6386588740159269 |
| interactor_ratio | mitosis | inter | 0.6186423062950968 | 0.9118004871703665 |

P61254_P62906
RL26_HUMAN vs RL10A_HUMAN
p-value: 0.009 q-value: 0.008

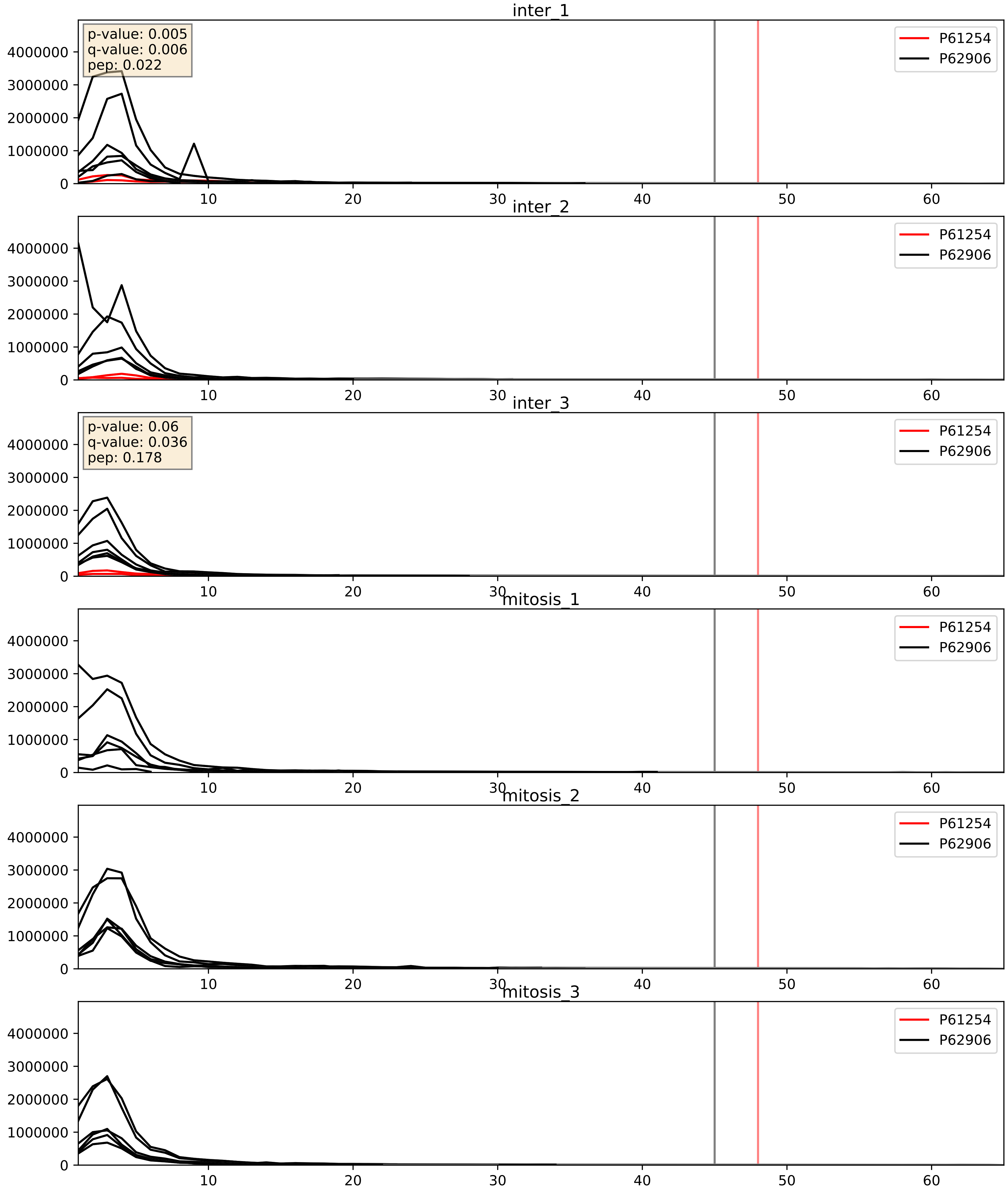


| level | condition_1 | condition_2 | pvalue | pvalue_adjusted |
|---|---|---|---|---|
| interactor_ratio | mitosis | inter | 6.337284687624959e-07 | 0.0013561789231517412 |
| complex_abundance | mitosis | inter | 6.359477139163931e-05 | 0.005691857151652958 |
| interactor_abundance | mitosis | inter | 0.0006144309768949645 | 0.017415659477552636 |

P61313_P62906
RL15_HUMAN vs RL10A_HUMAN
p-value: 0.0 q-value: 0.0

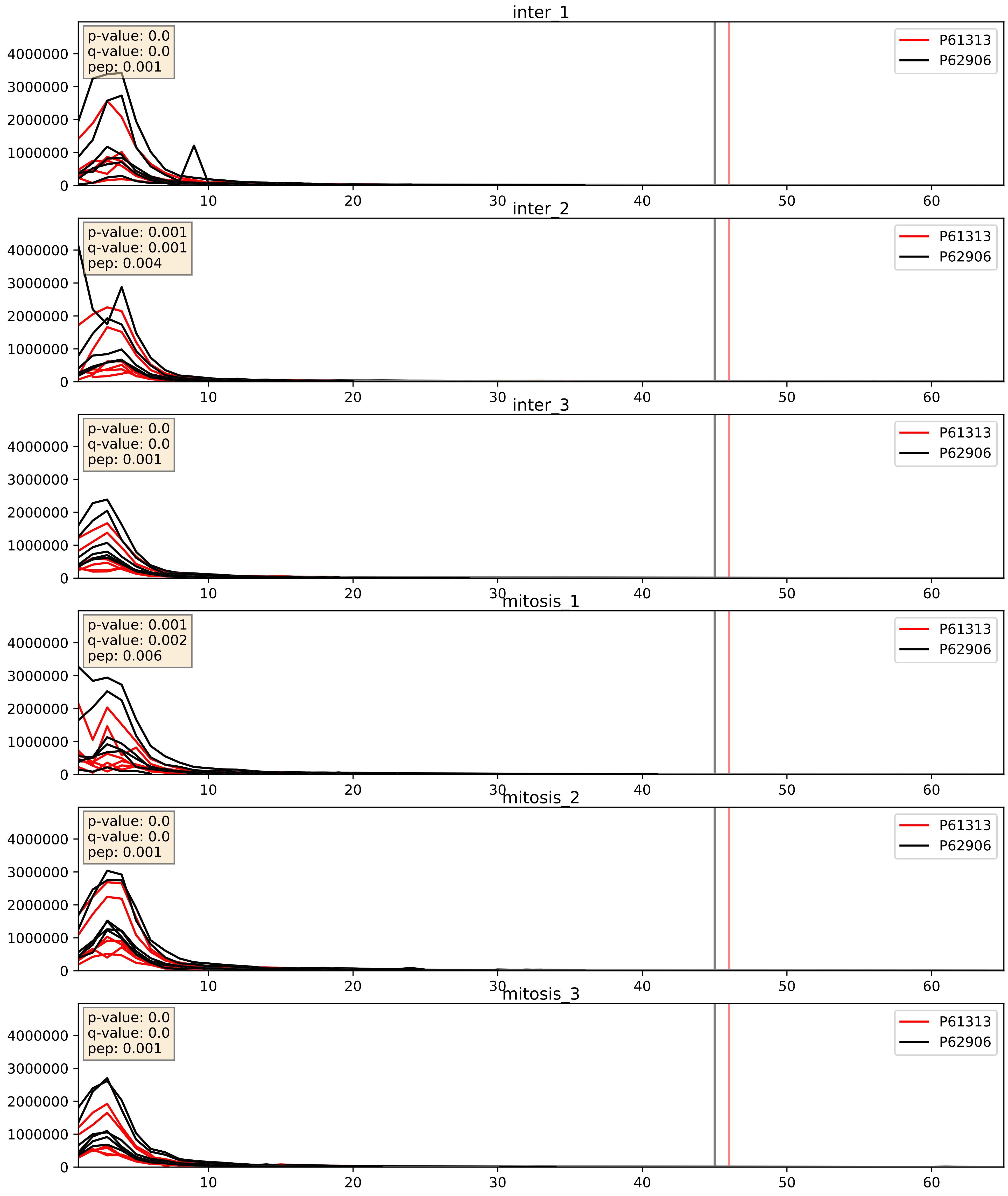


| level | condition_1 | condition_2 | pvalue | pvalue_adjusted |
|---|---|---|---|---|
| interactor_ratio | mitosis | inter | 0.7728456202580132 | 0.9554532798105997 |
| interactor_abundance | mitosis | inter | 0.8908468698415253 | 0.9385414407191947 |
| complex_abundance | mitosis | inter | 0.9981289348112699 | 0.9992744262328542 |

P61353_P62906
RL27_HUMAN vs RL10A_HUMAN
p-value: 0.0 q-value: 0.0

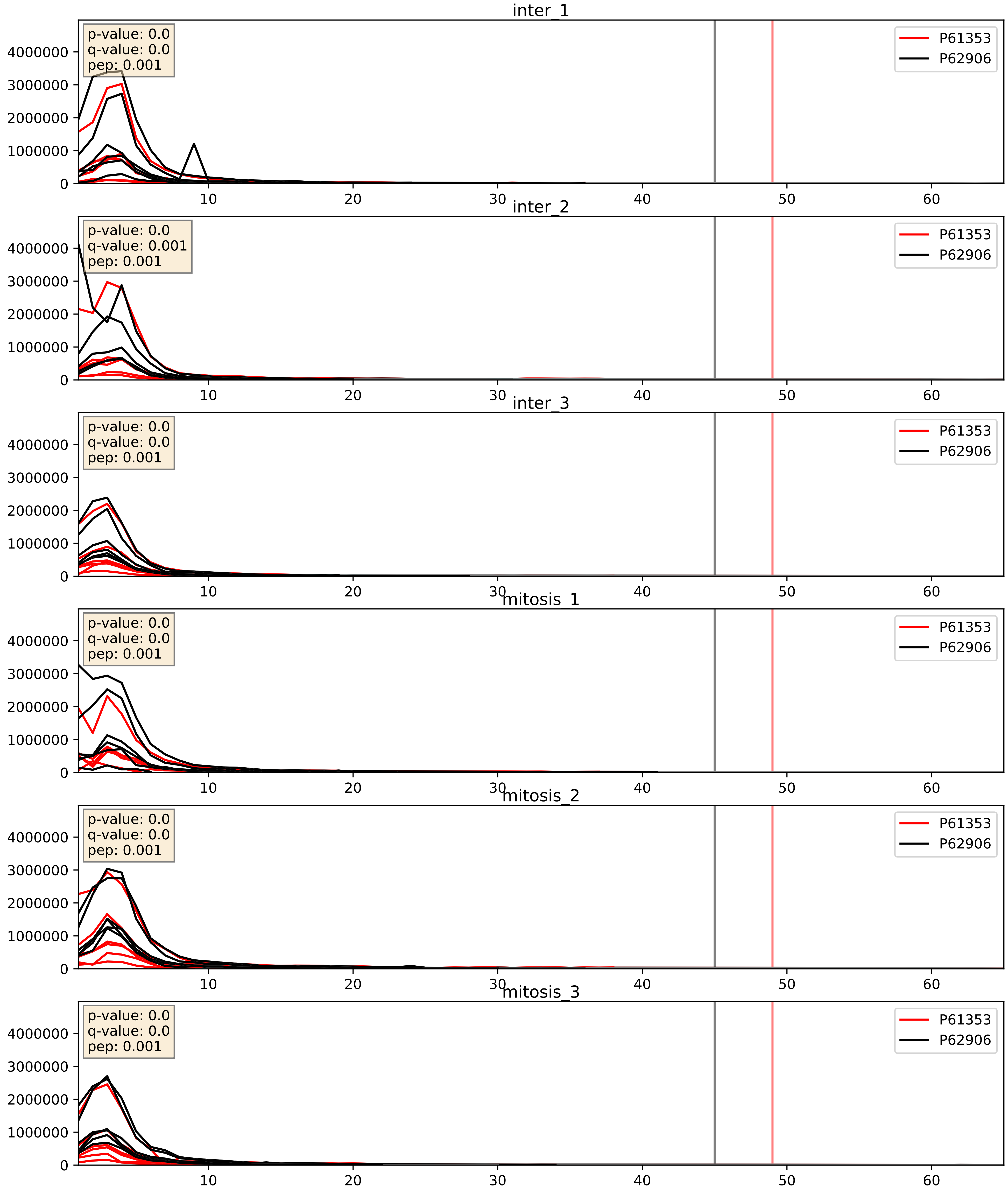


| level | condition_1 | condition_2 | pvalue | pvalue_adjusted |
|---|---|---|---|---|
| interactor_abundance | mitosis | inter | 0.6045661252362317 | 0.7402823513317188 |
| complex_abundance | mitosis | inter | 0.6582496726255278 | 0.7710204156642745 |
| interactor_ratio | mitosis | inter | 0.7860664790677256 | 0.9590548832411249 |

P61513_P62906
RL37A_HUMAN vs RL10A_HUMAN
p-value: 0.0 q-value: 0.0

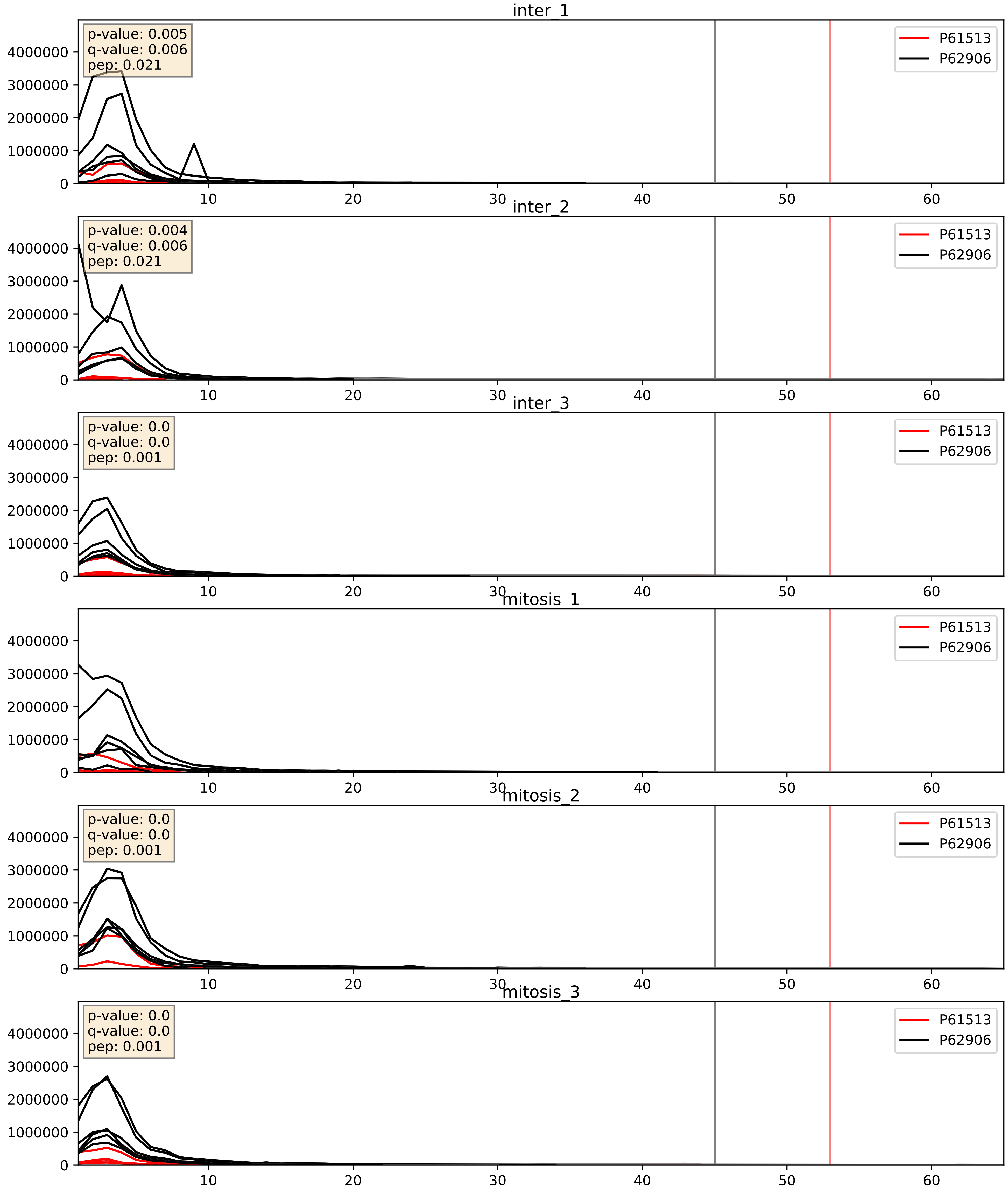


| level | condition_1 | condition_2 | pvalue | pvalue_adjusted |
|---|---|---|---|---|
| interactor_ratio | mitosis | inter | 0.041225439884434244 | 0.33704848654322556 |
| interactor_abundance | mitosis | inter | 0.04383858707081264 | 0.16685321787696922 |
| complex_abundance | mitosis | inter | 0.13696140910630644 | 0.2820982332168934 |

P62081_P62906
RS7_HUMAN vs RL10A_HUMAN
p-value: 0.001 q-value: 0.001

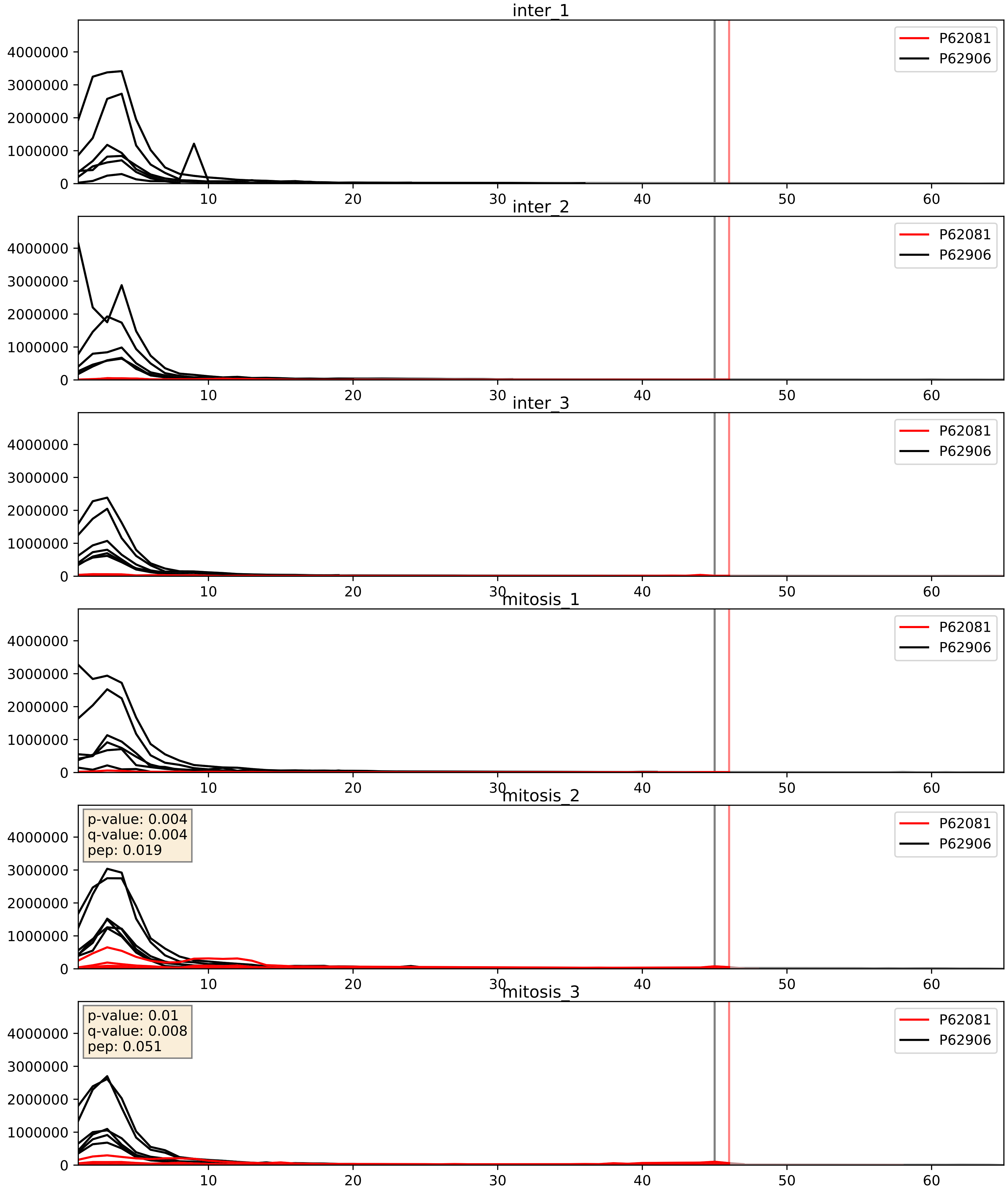


| level | condition_1 | condition_2 | pvalue | pvalue_adjusted |
|---|---|---|---|---|
| complex_abundance | mitosis | inter | 0.24315407181622054 | 0.39415524335368746 |
| interactor_abundance | mitosis | inter | 0.2568833674363821 | 0.4496312494132361 |
| interactor_ratio | mitosis | inter | 0.6641442086188893 | 0.9283996733722243 |

P62241_P62906
RS8_HUMAN vs RL10A_HUMAN
p-value: 0.0 q-value: 0.0

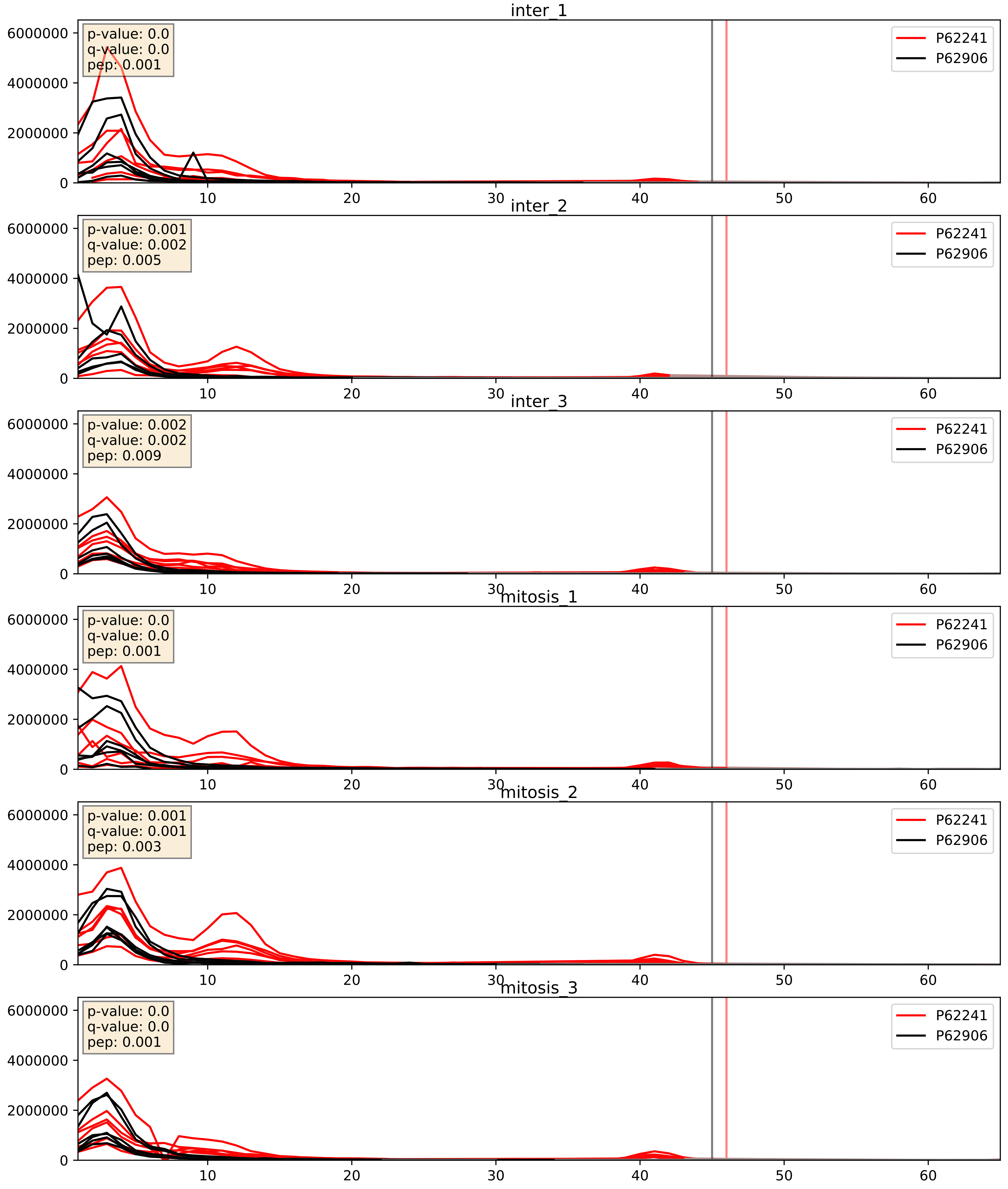


| level | condition_1 | condition_2 | pvalue | pvalue_adjusted |
|---|---|---|---|---|
| interactor_abundance | mitosis | inter | 0.27007566395238486 | 0.4623500366987127 |
| interactor_ratio | mitosis | inter | 0.4194588021263685 | 0.8388785723072948 |
| complex_abundance | mitosis | inter | 0.5828381553164882 | 0.7111239938454823 |

P62244_P62906
RS15A_HUMAN vs RL10A_HUMAN
p-value: 0.0 q-value: 0.0

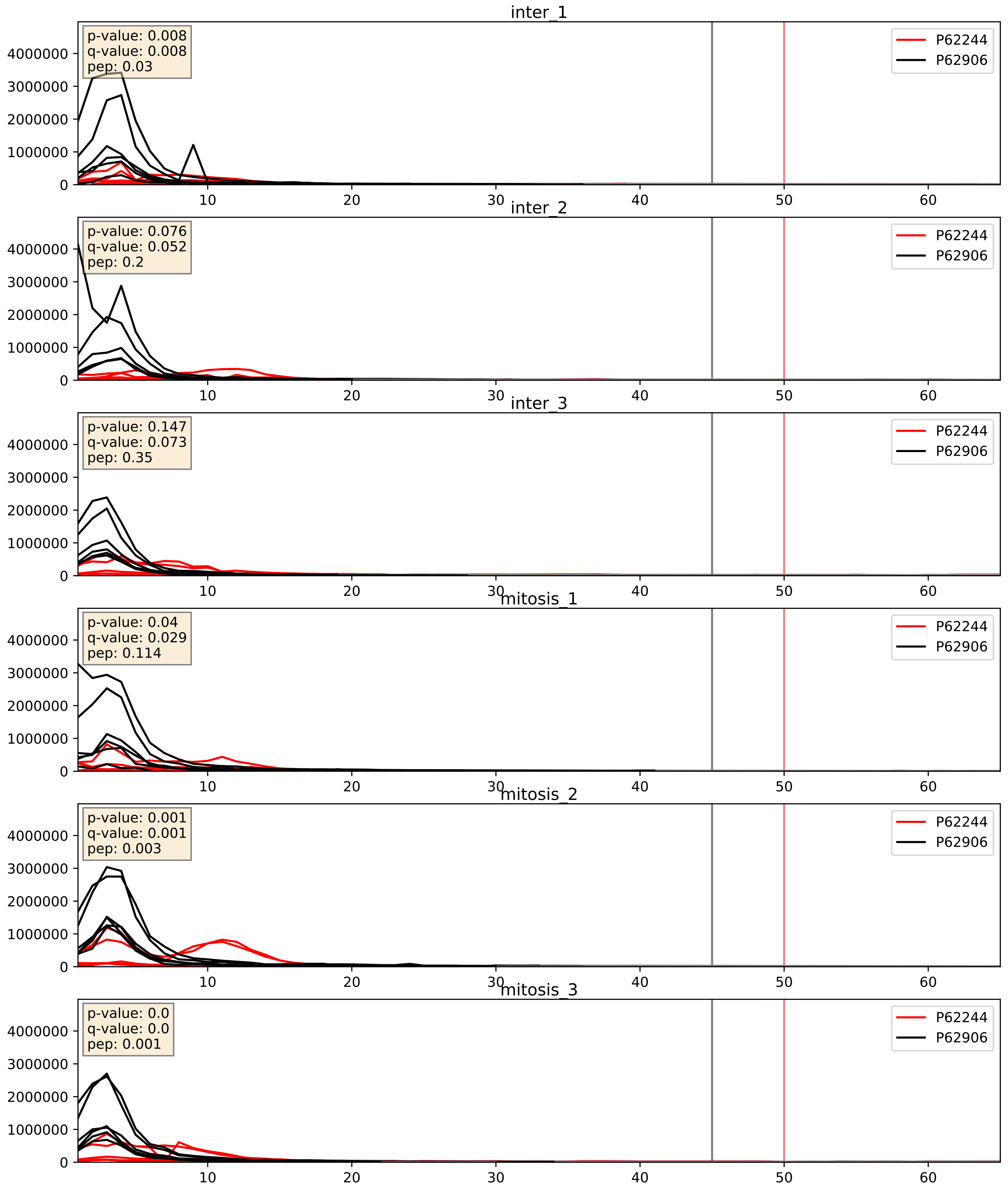


| level | condition_1 | condition_2 | pvalue | pvalue_adjusted |
|---|---|---|---|---|
| interactor_abundance | mitosis | inter | 0.15968425167992217 | 0.346006124384289 |
| interactor_ratio | mitosis | inter | 0.2374762429377228 | 0.7164022336563056 |
| complex_abundance | mitosis | inter | 0.265520456527718 | 0.4154852422081414 |

P62249_P62906
RS16_HUMAN vs RL10A_HUMAN
p-value: 0.004 q-value: 0.004

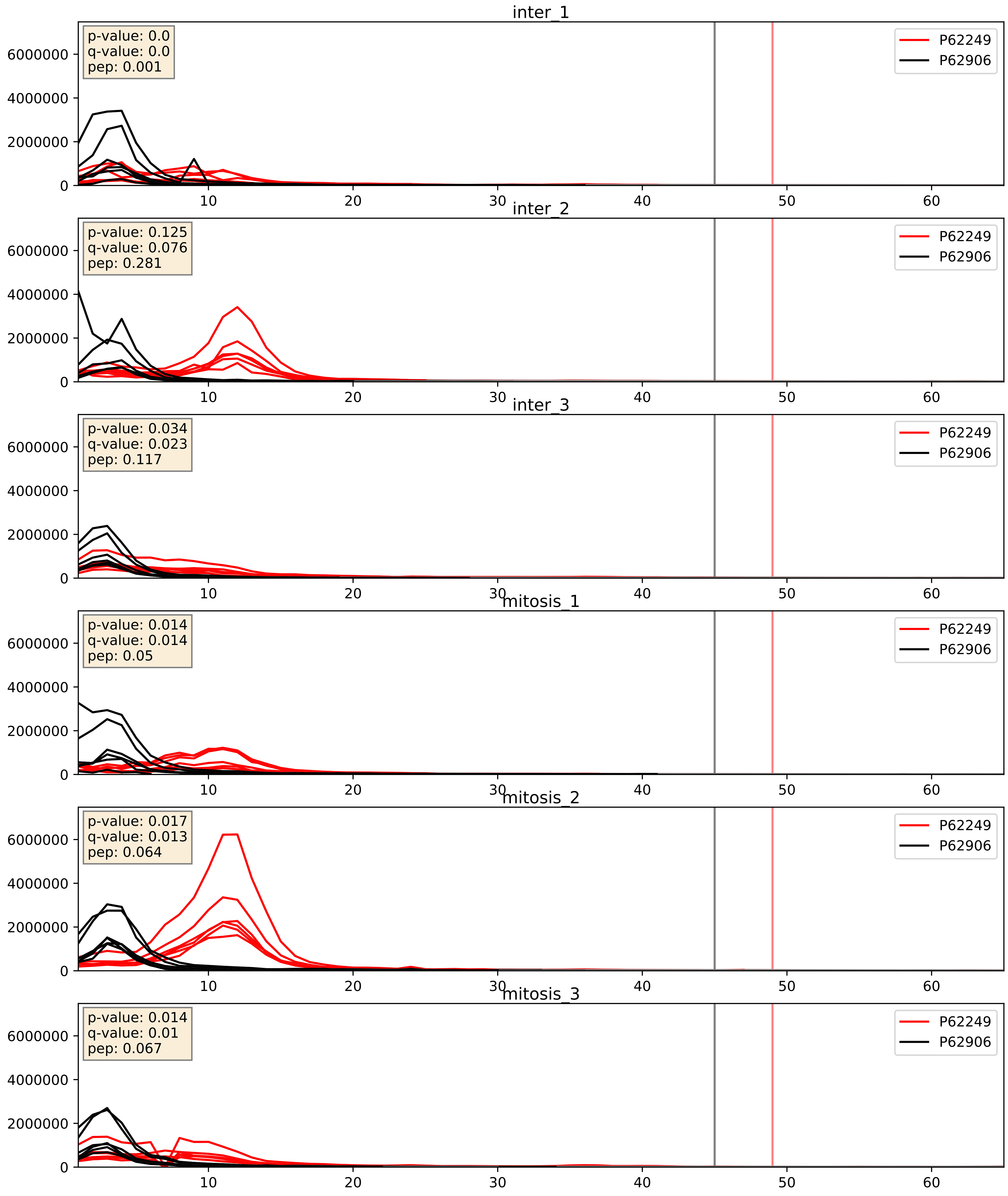


| level | condition_1 | condition_2 | pvalue | pvalue_adjusted |
|---|---|---|---|---|
| complex_abundance | mitosis | inter | 0.5521638177376935 | 0.6854005626210349 |
| interactor_abundance | mitosis | inter | 0.5655454997176717 | 0.7130754864609322 |
| interactor_ratio | mitosis | inter | 0.5993406054916213 | 0.9019028605875125 |

P62263_P62906
RS14_HUMAN vs RL10A_HUMAN
p-value: 0.0 q-value: 0.0

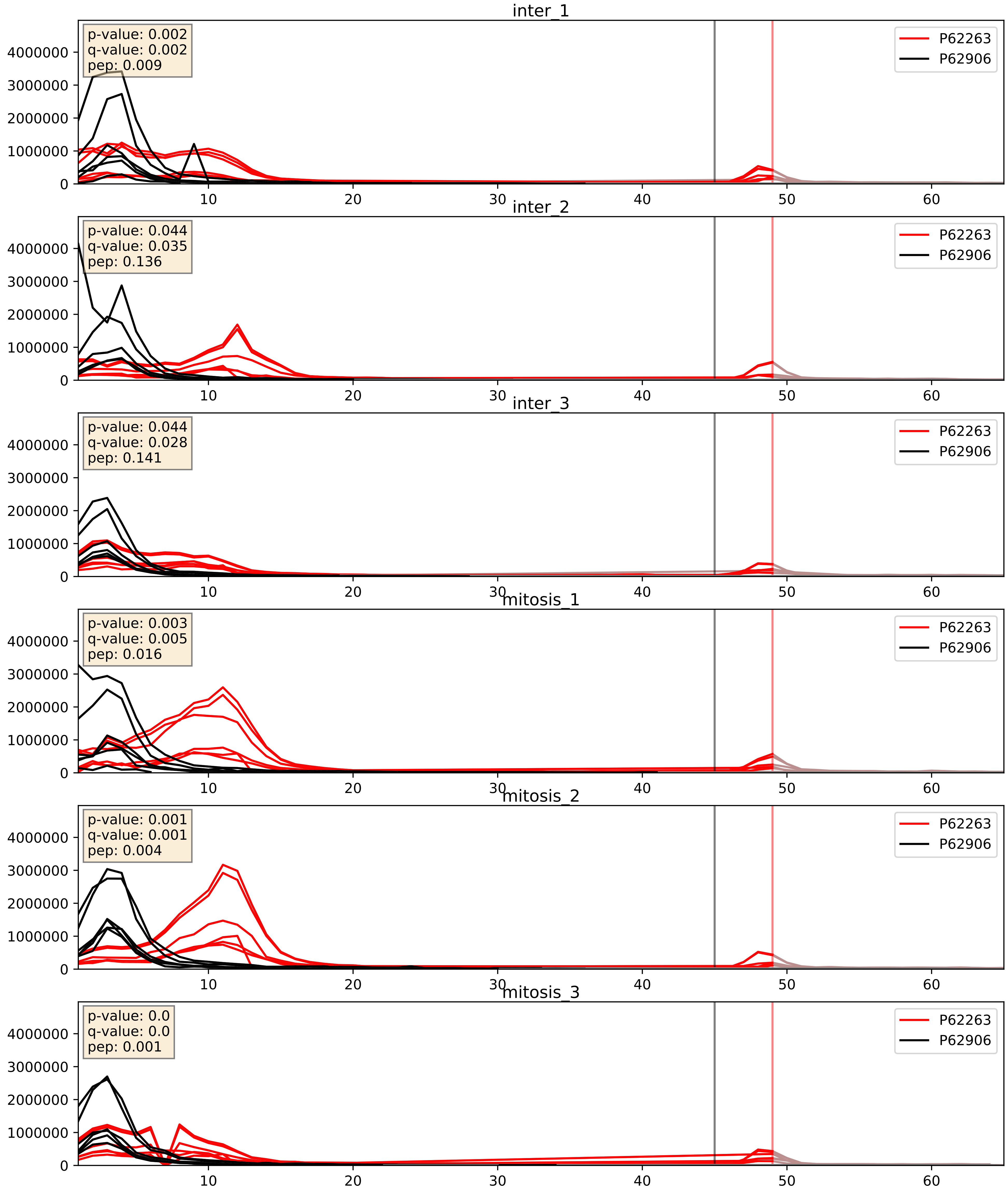


| level | condition_1 | condition_2 | pvalue | pvalue_adjusted |
|---|---|---|---|---|
| interactor_ratio | mitosis | inter | 0.2869397444633757 | 0.7573864362030516 |
| interactor_abundance | mitosis | inter | 0.3678292541008239 | 0.5632087317955554 |
| complex_abundance | mitosis | inter | 0.48730362747079103 | 0.6268889466711709 |

P62266_P62906
RS23_HUMAN vs RL10A_HUMAN
p-value: 0.0 q-value: 0.0

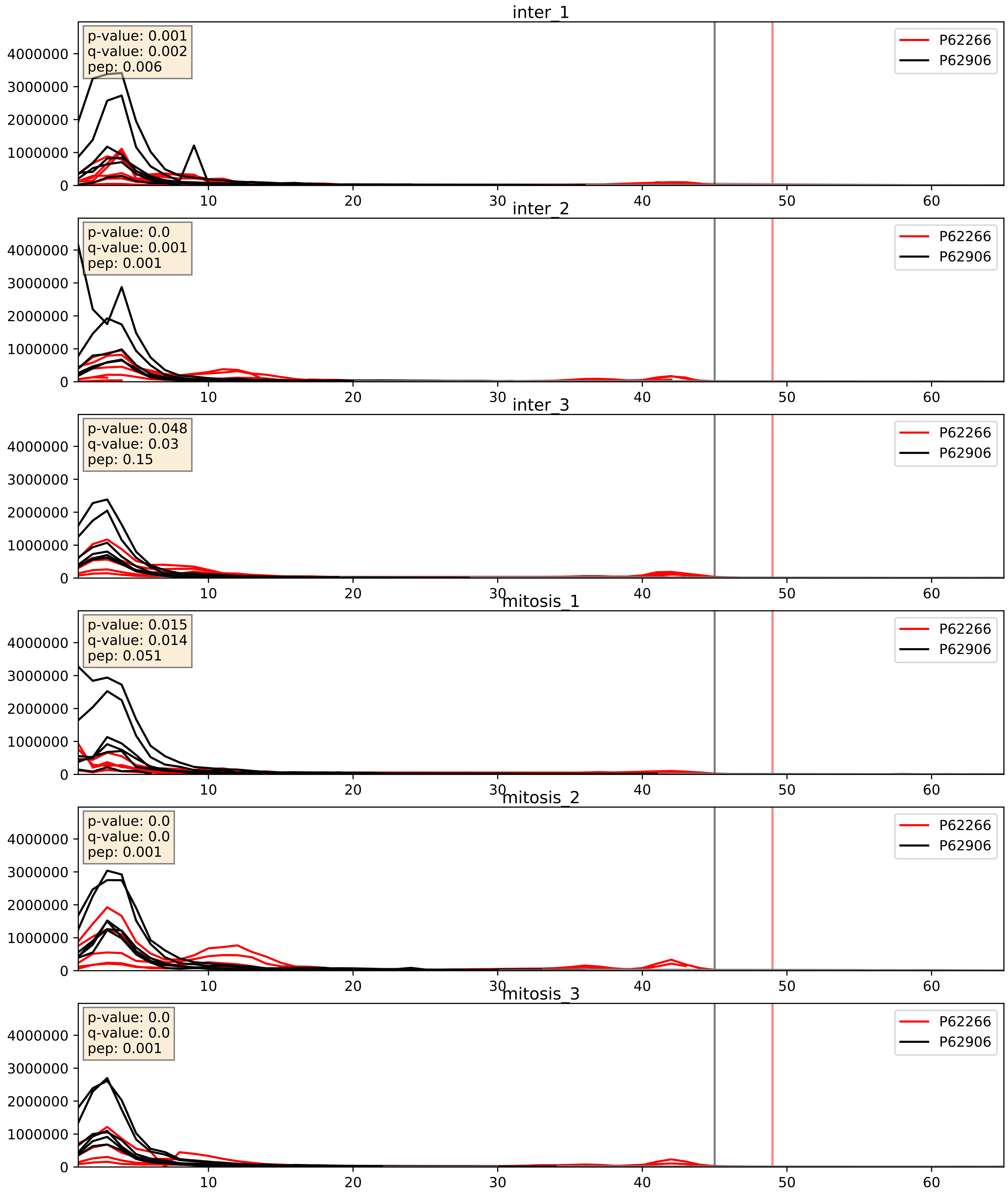


| level | condition_1 | condition_2 | pvalue | pvalue_adjusted |
|---|---|---|---|---|
| complex_abundance | mitosis | inter | 0.25222135852974487 | 0.4025009002637241 |
| interactor_abundance | mitosis | inter | 0.30581655170252814 | 0.5014922763551036 |
| interactor_ratio | mitosis | inter | 0.38264570299788037 | 0.8203229067498934 |

P62269_P62906
RS18_HUMAN vs RL10A_HUMAN
p-value: 0.006 q-value: 0.006

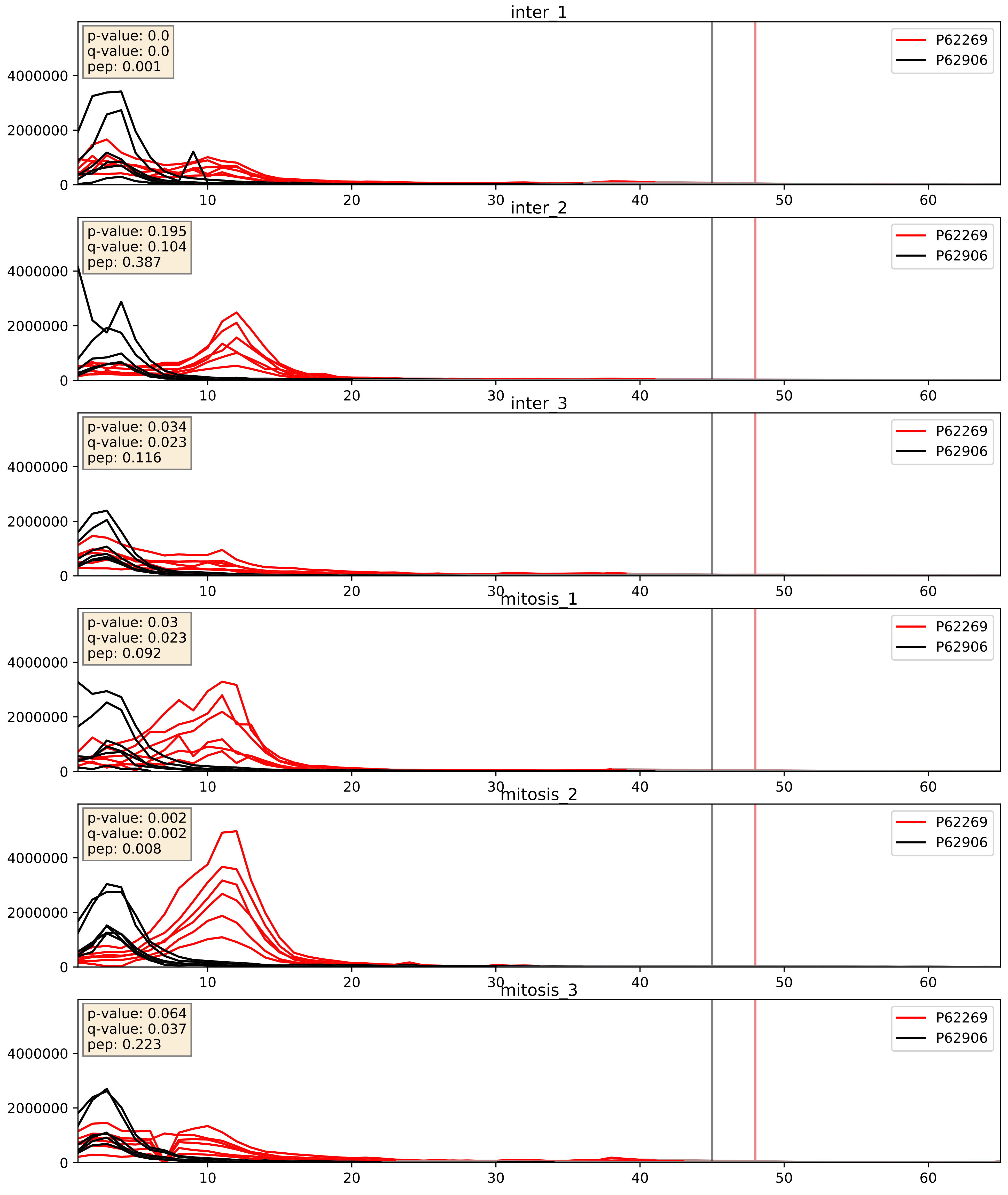


| level | condition_1 | condition_2 | pvalue | pvalue_adjusted |
|---|---|---|---|---|
| complex_abundance | mitosis | inter | 0.006023236527422844 | 0.05806974312565789 |
| interactor_abundance | mitosis | inter | 0.01739271702981435 | 0.11421684524373674 |
| interactor_ratio | mitosis | inter | 0.0760206439167499 | 0.46316861816154964 |

P62273_P62906
RS29_HUMAN vs RL10A_HUMAN
p-value: 0.015 q-value: 0.013

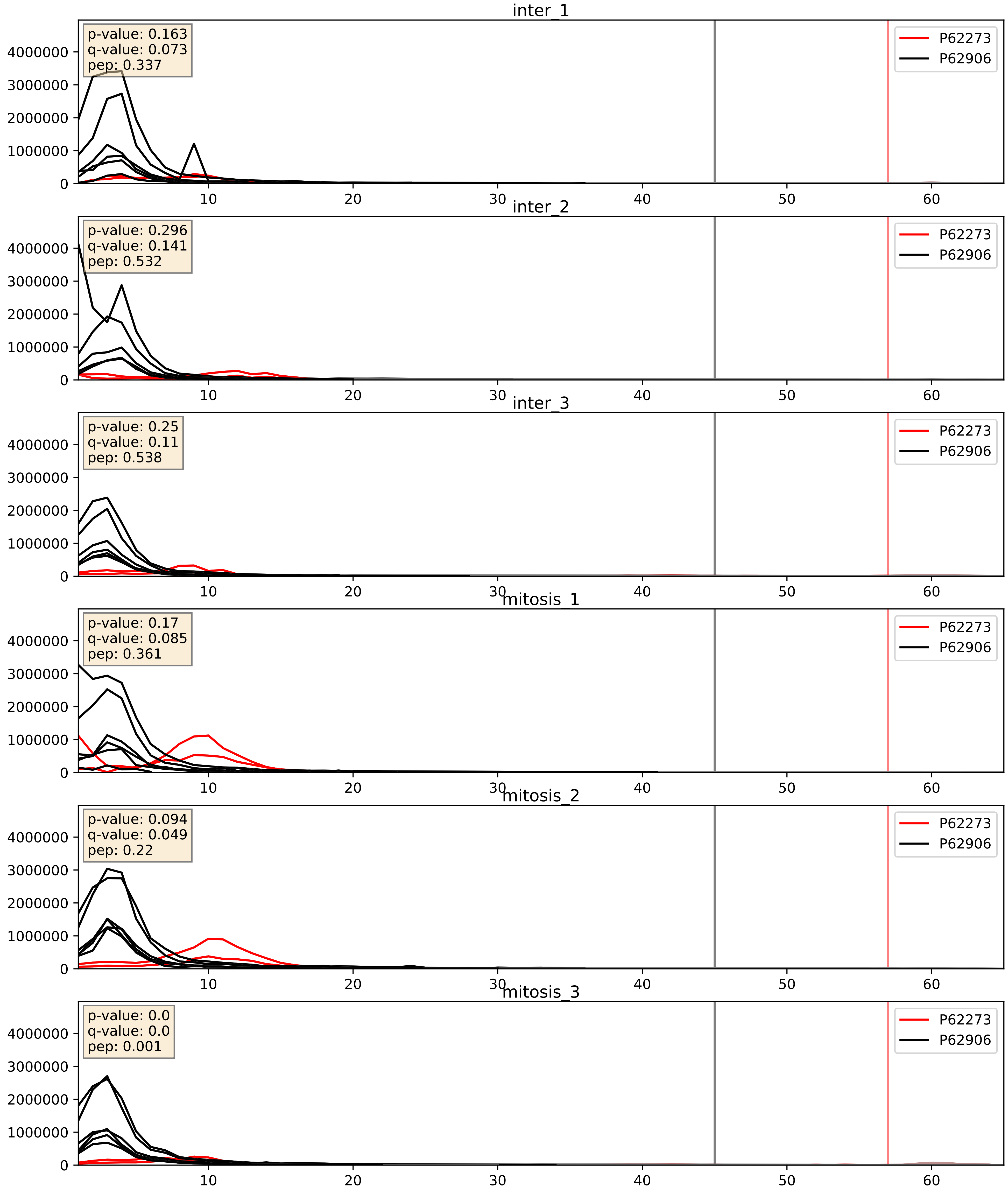


| level | condition_1 | condition_2 | pvalue | pvalue_adjusted |
|---|---|---|---|---|
| interactor_ratio | mitosis | inter | 0.23525888402246042 | 0.7151335395000927 |
| interactor_abundance | mitosis | inter | 0.2388653609573371 | 0.43041521730234833 |
| complex_abundance | mitosis | inter | 0.33126820553385883 | 0.4796440864969269 |

P62277_P62906
RS13_HUMAN vs RL10A_HUMAN
p-value: 0.0 q-value: 0.0

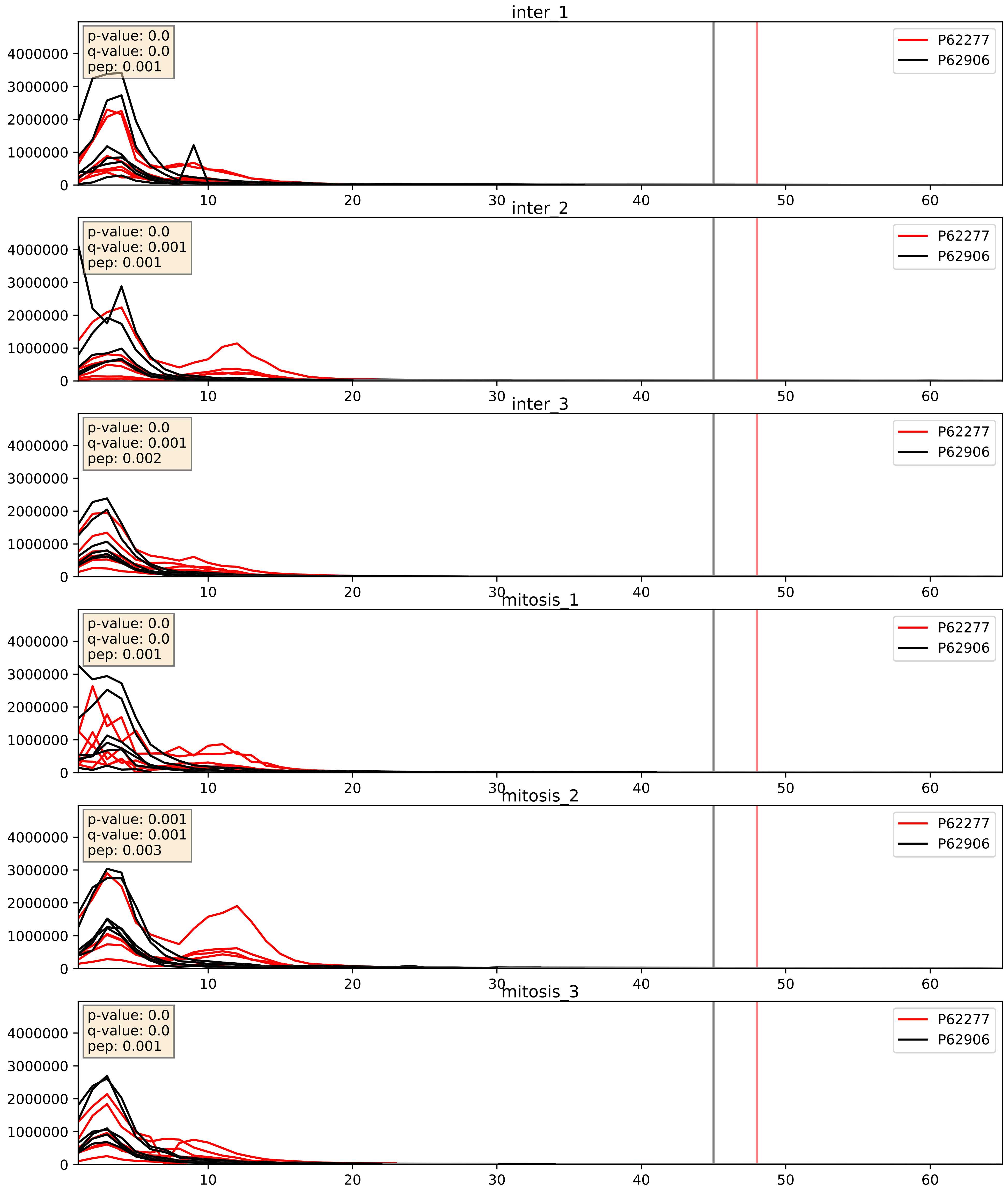


| level | condition_1 | condition_2 | pvalue | pvalue_adjusted |
|---|---|---|---|---|
| interactor_abundance | mitosis | inter | 0.37805473060280287 | 0.5712530439470419 |
| interactor_ratio | mitosis | inter | 0.4838194083928914 | 0.8676177308124418 |
| complex_abundance | mitosis | inter | 0.7384919210152213 | 0.8265547651530196 |

P62280_P62906
RS11_HUMAN vs RL10A_HUMAN
p-value: 0.0 q-value: 0.0

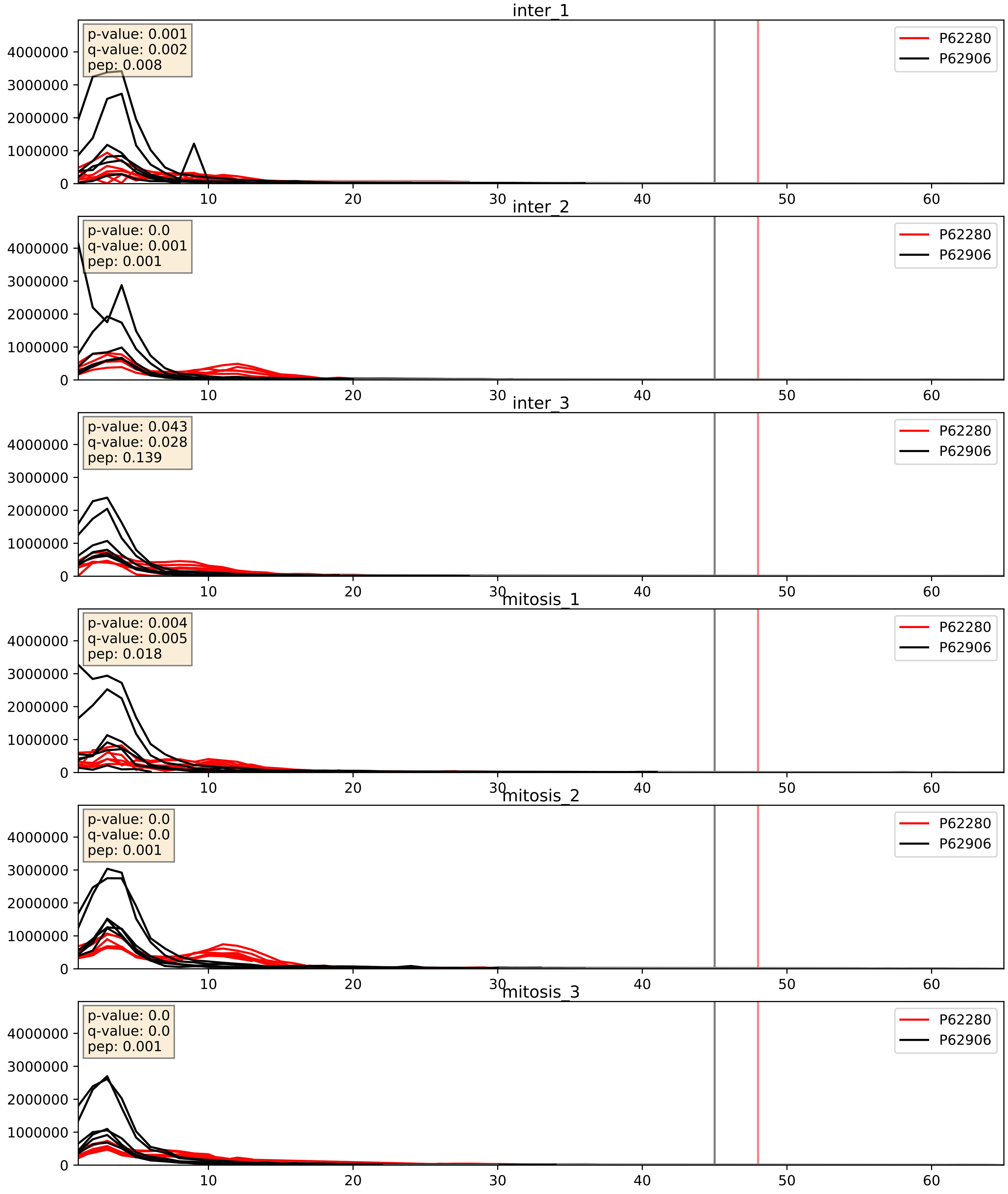


| level | condition_1 | condition_2 | pvalue | pvalue_adjusted |
|---|---|---|---|---|
| interactor_abundance | mitosis | inter | 0.8773389974984234 | 0.9318222989746887 |
| interactor_ratio | mitosis | inter | 0.8838713720605 | 0.9802747470617126 |
| complex_abundance | mitosis | inter | 0.926140892502378 | 0.9545774882384535 |

P62424_P62906
RL7A_HUMAN vs RL10A_HUMAN
p-value: 0.0 q-value: 0.0

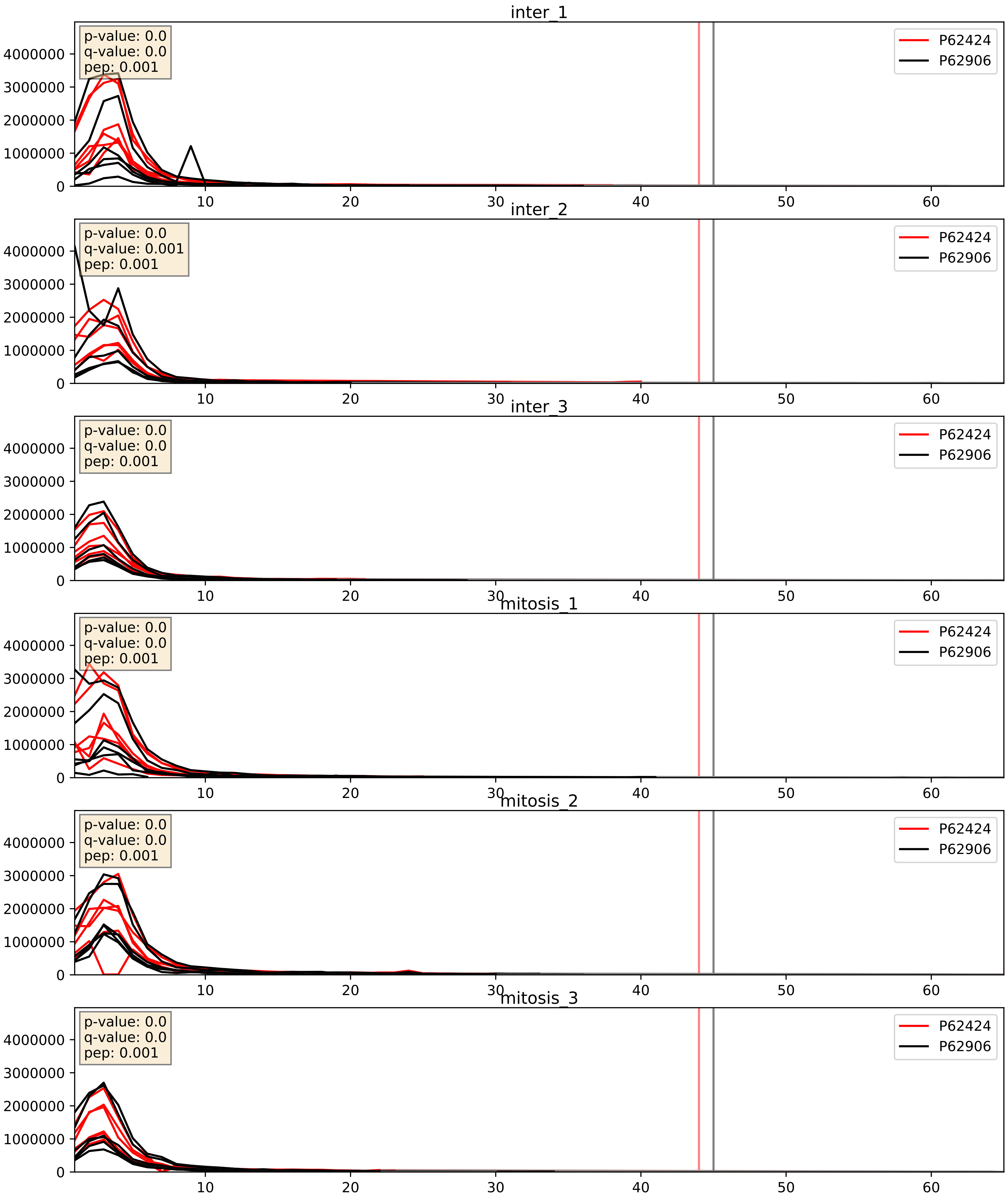


| level | condition_1 | condition_2 | pvalue | pvalue_adjusted |
|---|---|---|---|---|
| complex_abundance | mitosis | inter | 0.9080660755316258 | 0.9424158106875262 |
| interactor_abundance | mitosis | inter | 0.9353774579133864 | 0.962096690999235 |
| interactor_ratio | mitosis | inter | 0.9782000303795223 | 0.9959018321340639 |

P62701_P62906
RS4X_HUMAN vs RL10A_HUMAN
p-value: 0.003 q-value: 0.004

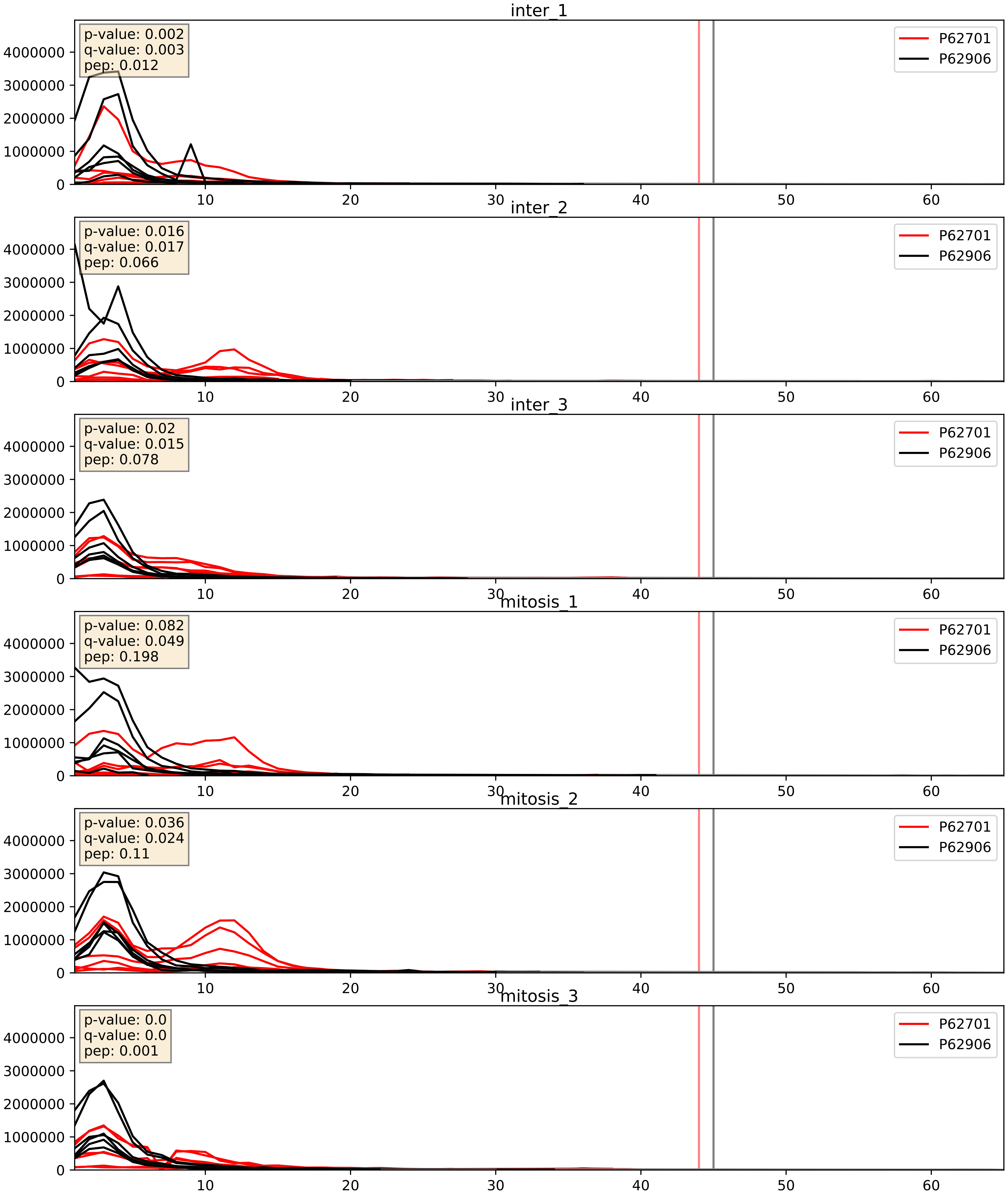


| level | condition_1 | condition_2 | pvalue | pvalue_adjusted |
|---|---|---|---|---|
| complex_abundance | mitosis | inter | 0.5666967139733629 | 0.6965980671643014 |
| interactor_abundance | mitosis | inter | 0.6493034798304969 | 0.7658336599860907 |
| interactor_ratio | mitosis | inter | 0.7300321436679927 | 0.9475669293562753 |

P62750_P62906
RL23A_HUMAN vs RL10A_HUMAN
p-value: 0.0 q-value: 0.0

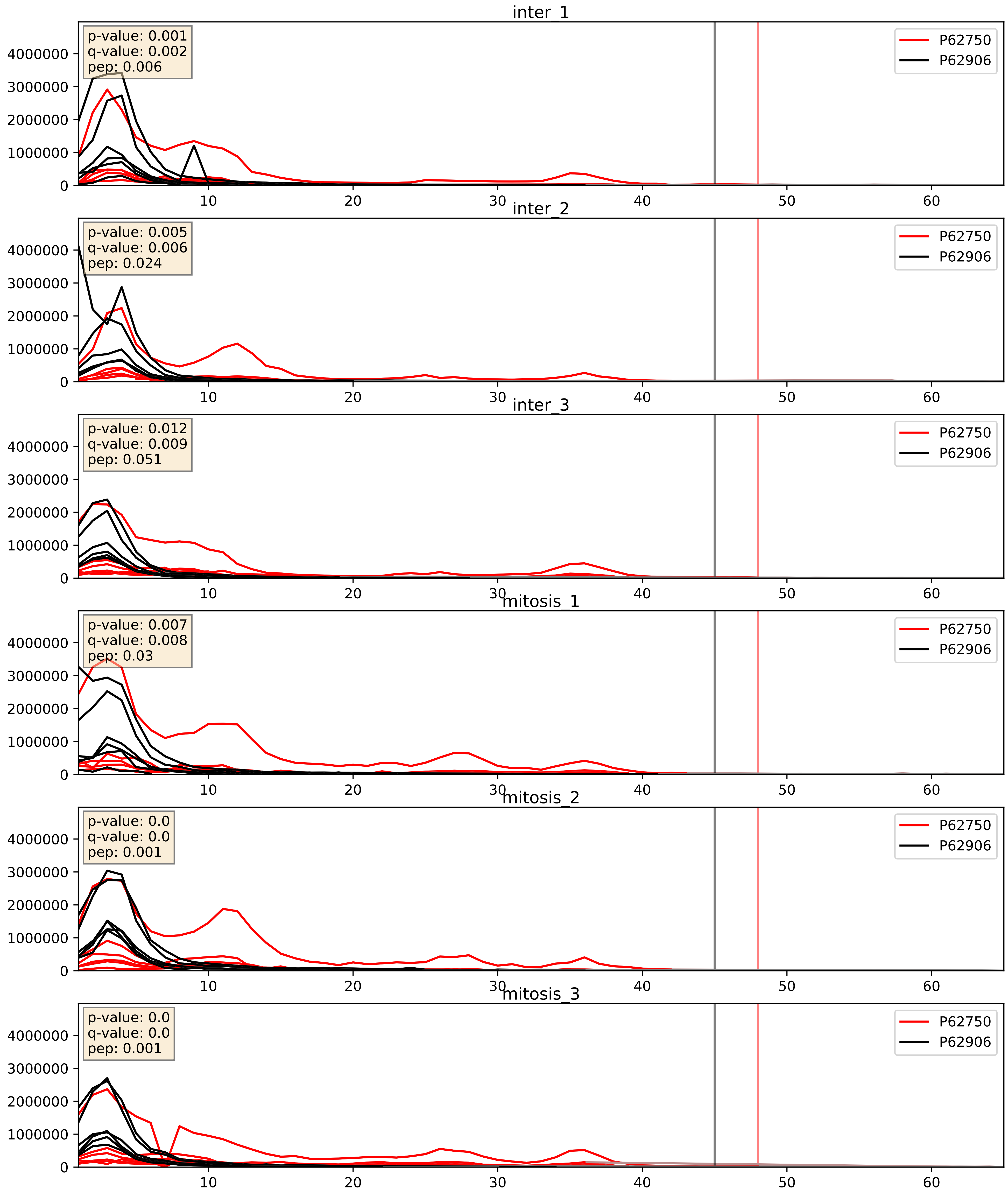


| level | condition_1 | condition_2 | pvalue | pvalue_adjusted |
|---|---|---|---|---|
| interactor_abundance | mitosis | inter | 0.11203685396590637 | 0.2886587657103679 |
| interactor_ratio | mitosis | inter | 0.14961768501258352 | 0.604401785610059 |
| complex_abundance | mitosis | inter | 0.2687045139844621 | 0.41824971083391327 |

P62753_P62906
RS6_HUMAN vs RL10A_HUMAN
p-value: 0.0 q-value: 0.0

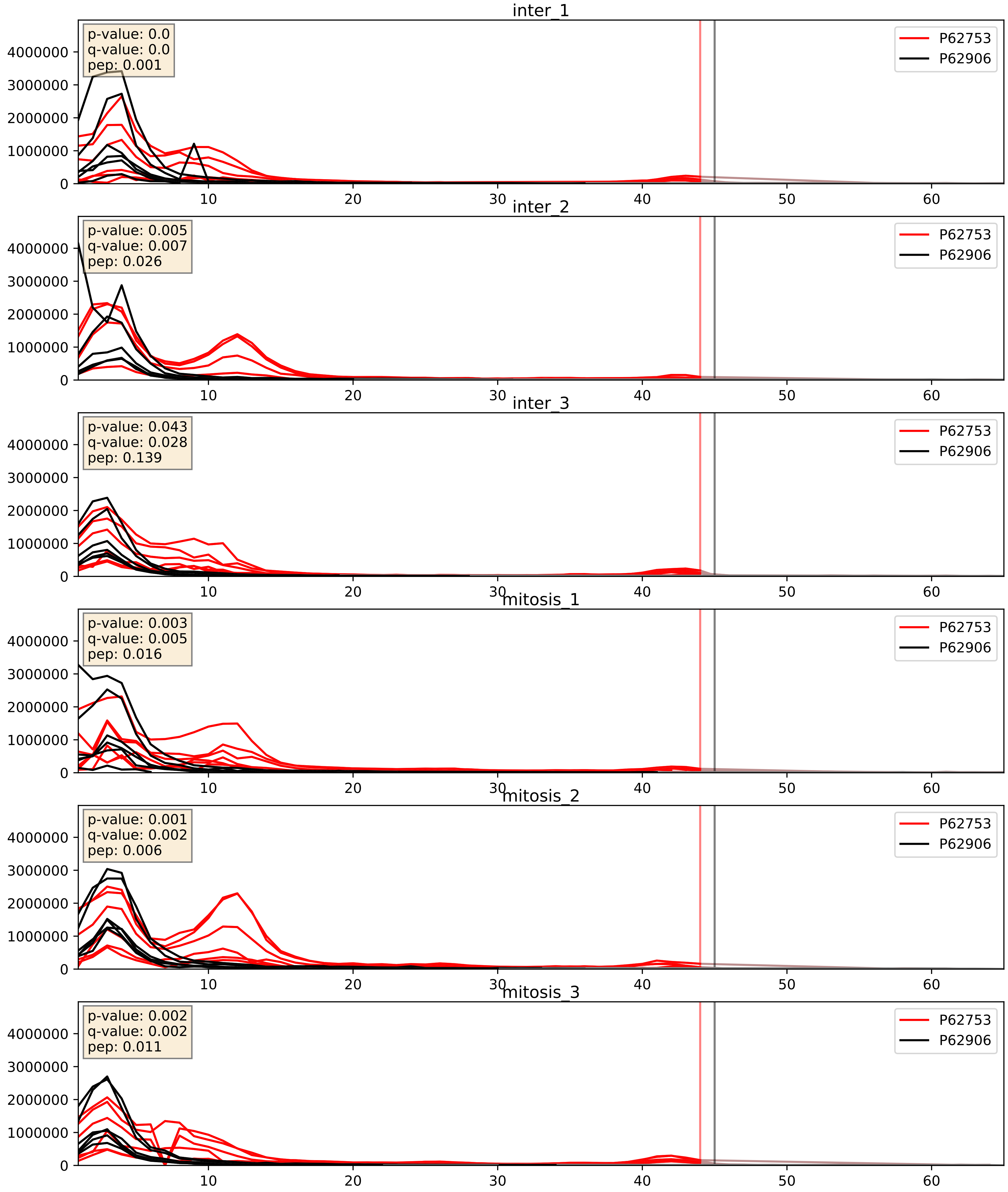


| level | condition_1 | condition_2 | pvalue | pvalue_adjusted |
|---|---|---|---|---|
| complex_abundance | mitosis | inter | 0.6526200707848244 | 0.7666293132864137 |
| interactor_abundance | mitosis | inter | 0.809410272369899 | 0.8902161032360704 |
| interactor_ratio | mitosis | inter | 0.9748231545083021 | 0.9957326006317586 |

P62829_P62906
RL23_HUMAN vs RL10A_HUMAN
p-value: 0.0 q-value: 0.0

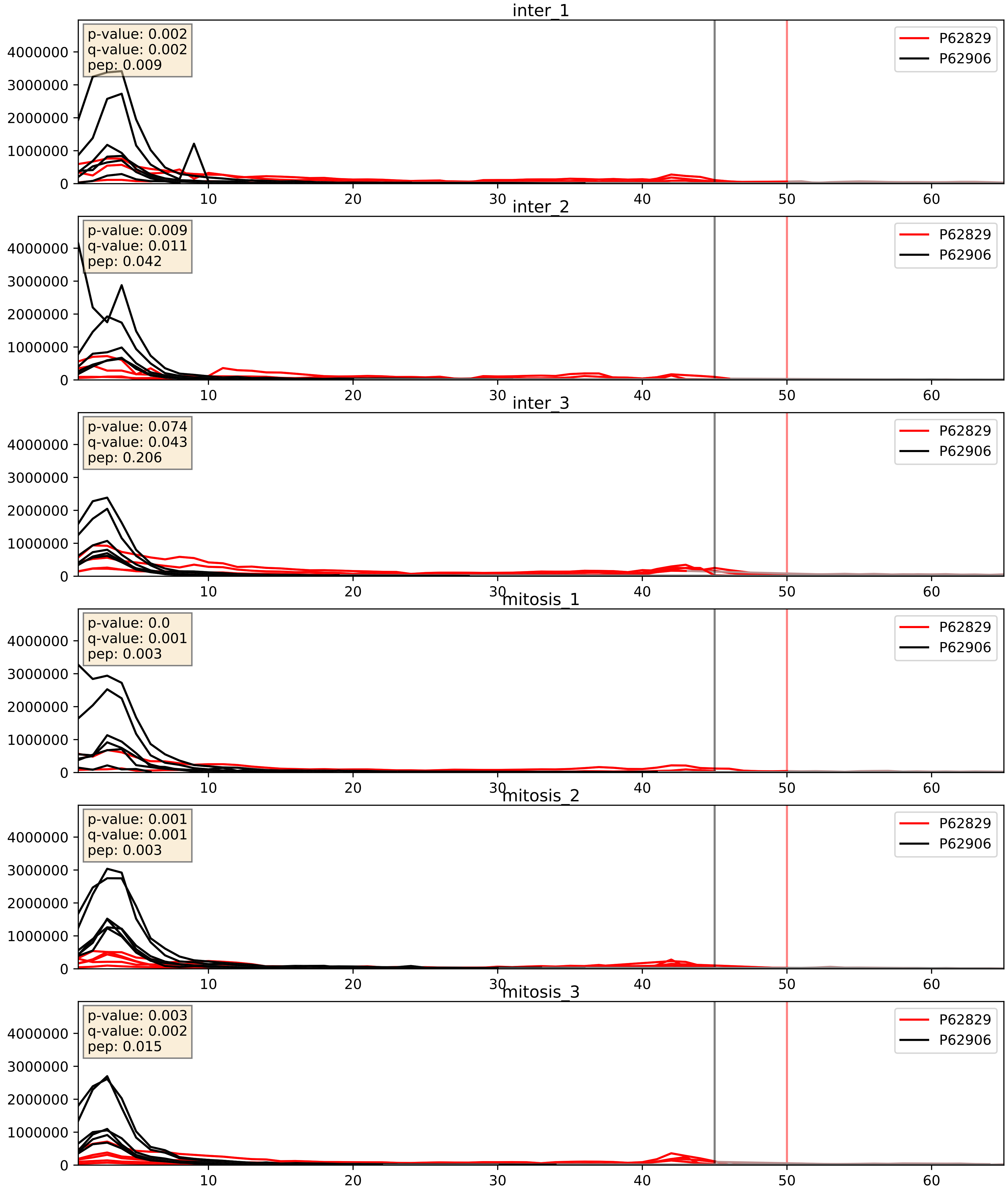


| level | condition_1 | condition_2 | pvalue | pvalue_adjusted |
|---|---|---|---|---|
| interactor_abundance | mitosis | inter | 0.002632665877349012 | 0.03838967005823521 |
| complex_abundance | mitosis | inter | 0.003632835025813253 | 0.04428054295684183 |
| interactor_ratio | mitosis | inter | 0.005822434720010065 | 0.10426786862612165 |

P62841_P62906
RS15_HUMAN vs RL10A_HUMAN
p-value: 0.004 q-value: 0.004

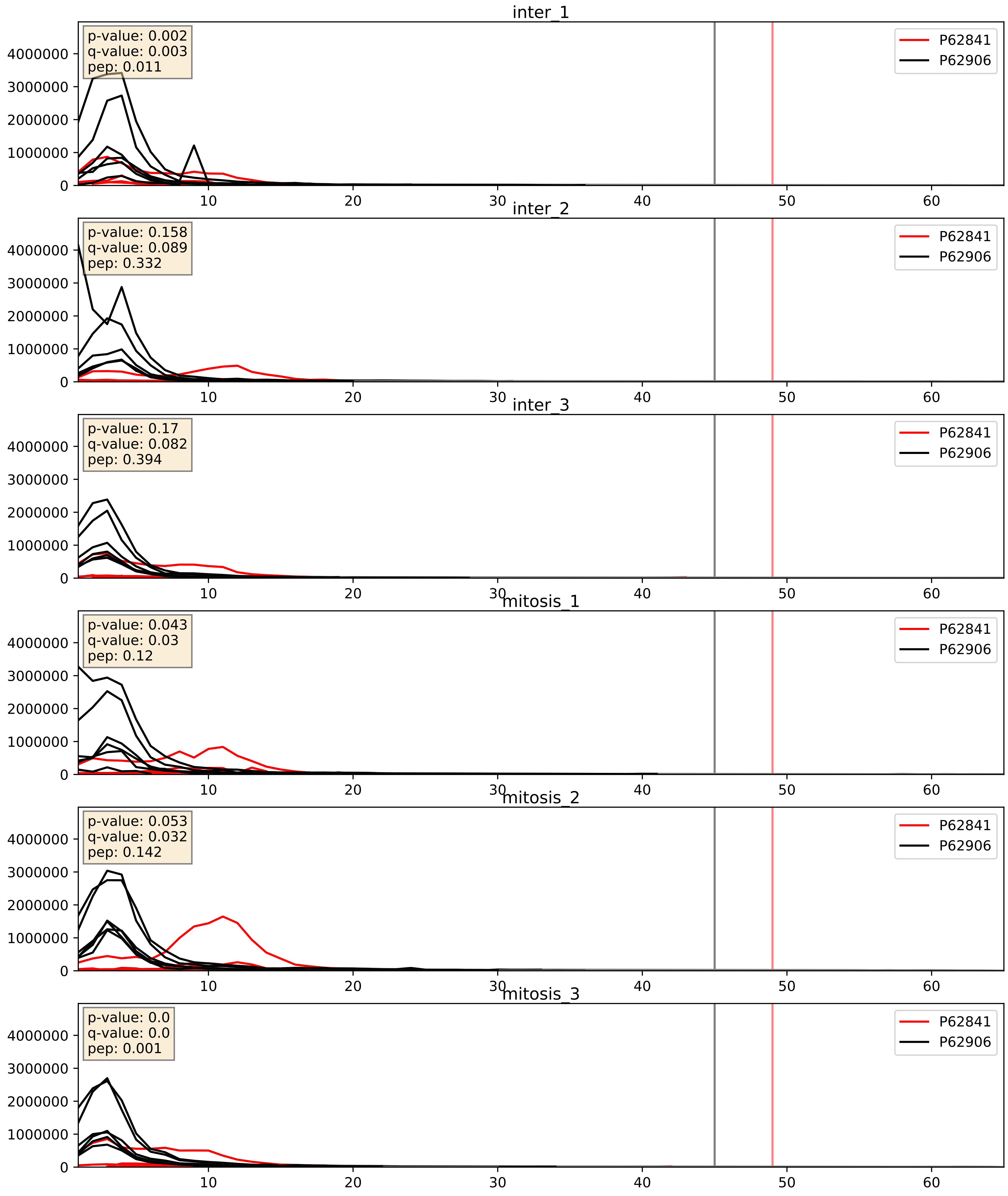


| level | condition_1 | condition_2 | pvalue | pvalue_adjusted |
|---|---|---|---|---|
| complex_abundance | mitosis | inter | 0.8793768098635149 | 0.9248765678210293 |
| interactor_abundance | mitosis | inter | 0.910108678282144 | 0.9488496785938922 |
| interactor_ratio | mitosis | inter | 0.9489389144637204 | 0.9912365517130919 |

P62847_P62906
RS24_HUMAN vs RL10A_HUMAN
p-value: 0.015 q-value: 0.012

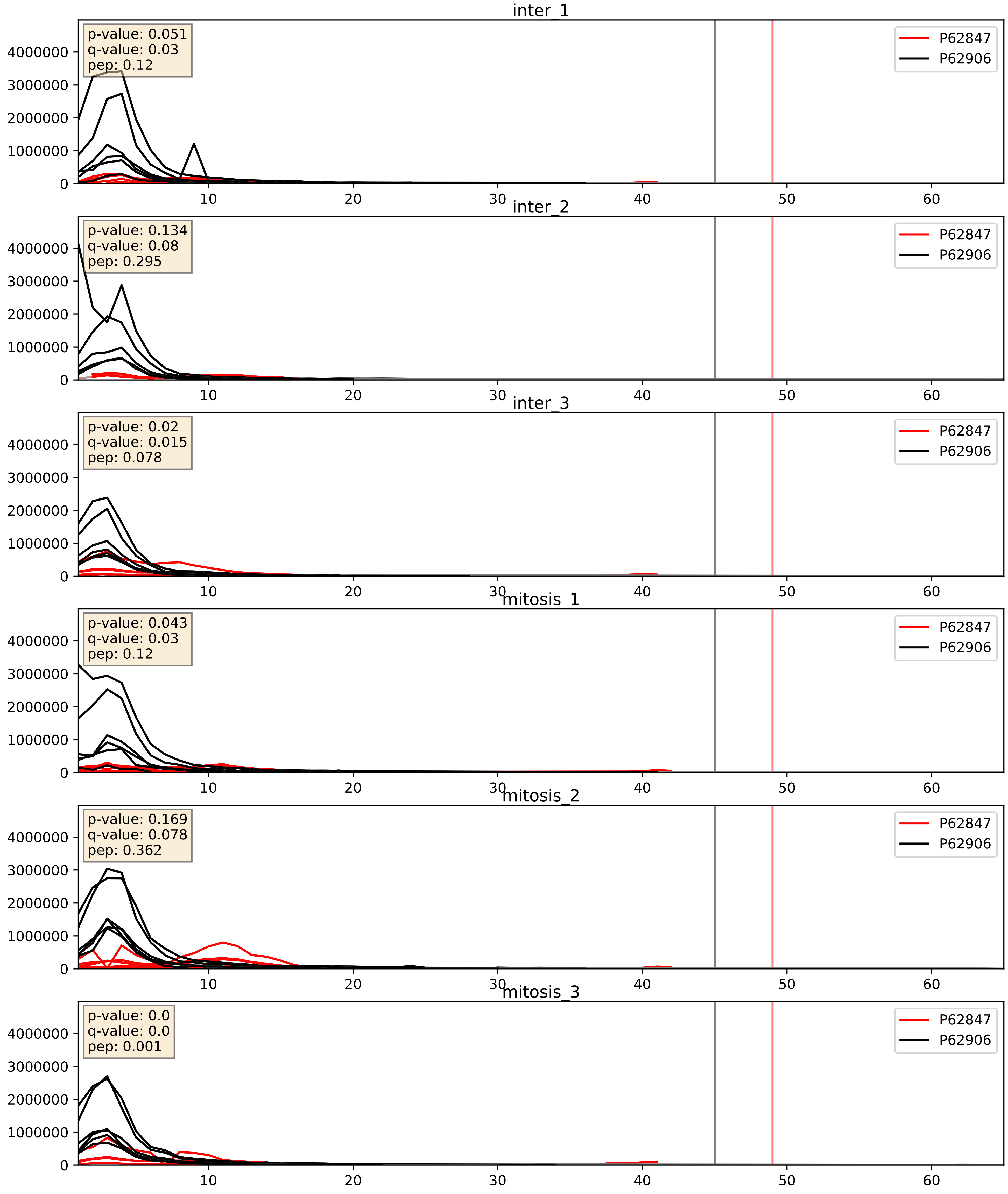


| level | condition_1 | condition_2 | pvalue | pvalue_adjusted |
|---|---|---|---|---|
| interactor_abundance | mitosis | inter | 0.11450727790006568 | 0.29180776981975654 |
| complex_abundance | mitosis | inter | 0.1508998891028792 | 0.29694029255027643 |
| interactor_ratio | mitosis | inter | 0.4246766463091138 | 0.841246386050888 |

P62851_P62906
RS25_HUMAN vs RL10A_HUMAN
p-value: 0.001 q-value: 0.002

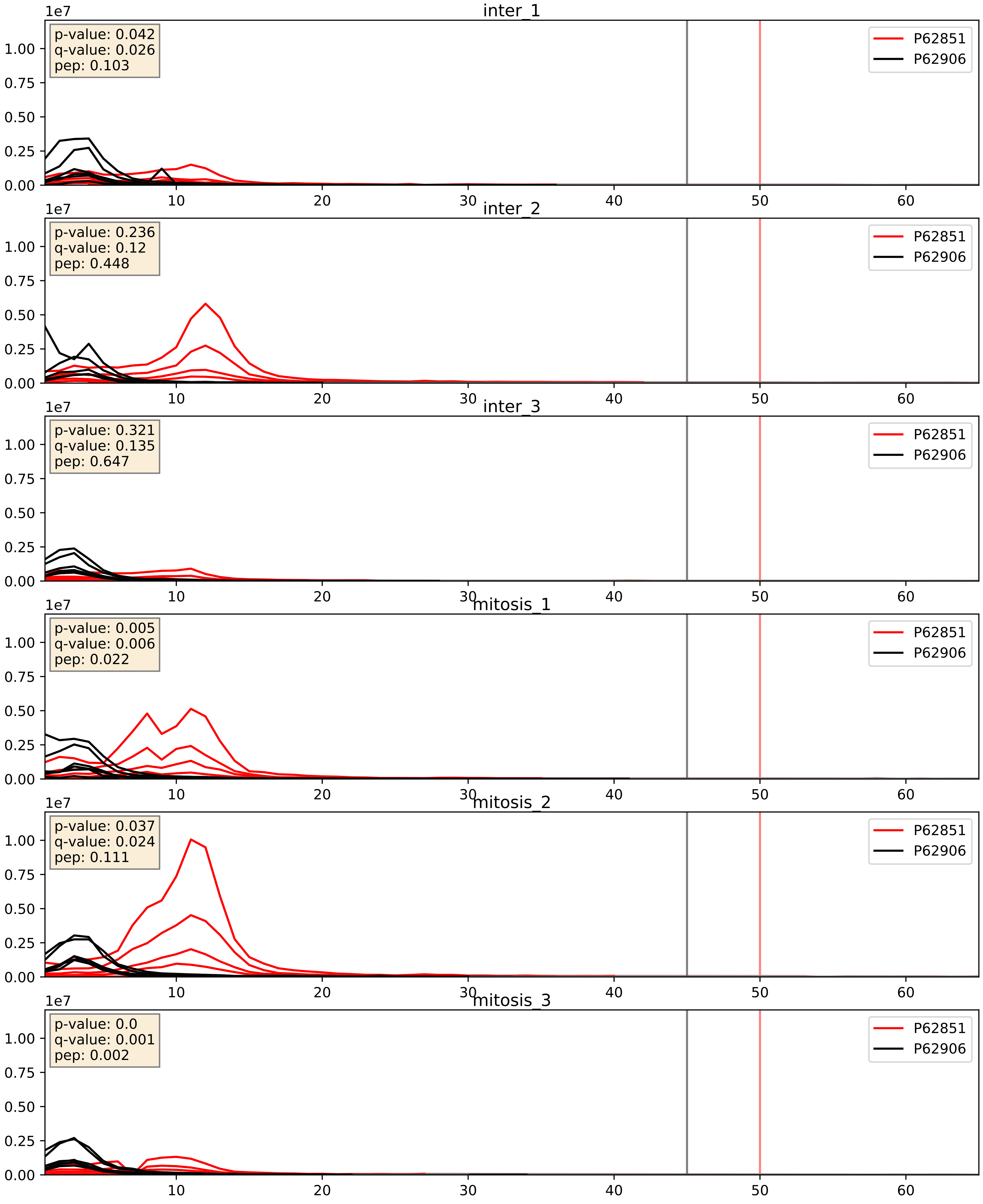


| level | condition_1 | condition_2 | pvalue | pvalue_adjusted |
|---|---|---|---|---|
| interactor_abundance | mitosis | inter | 0.4166905106069957 | 0.5982637417587343 |
| interactor_ratio | mitosis | inter | 0.4215129924159497 | 0.8388785723072948 |
| complex_abundance | mitosis | inter | 0.42229277997771897 | 0.5656864910932369 |

P62861_P62906
RS30_HUMAN vs RL10A_HUMAN
p-value: 0.007 q-value: 0.007

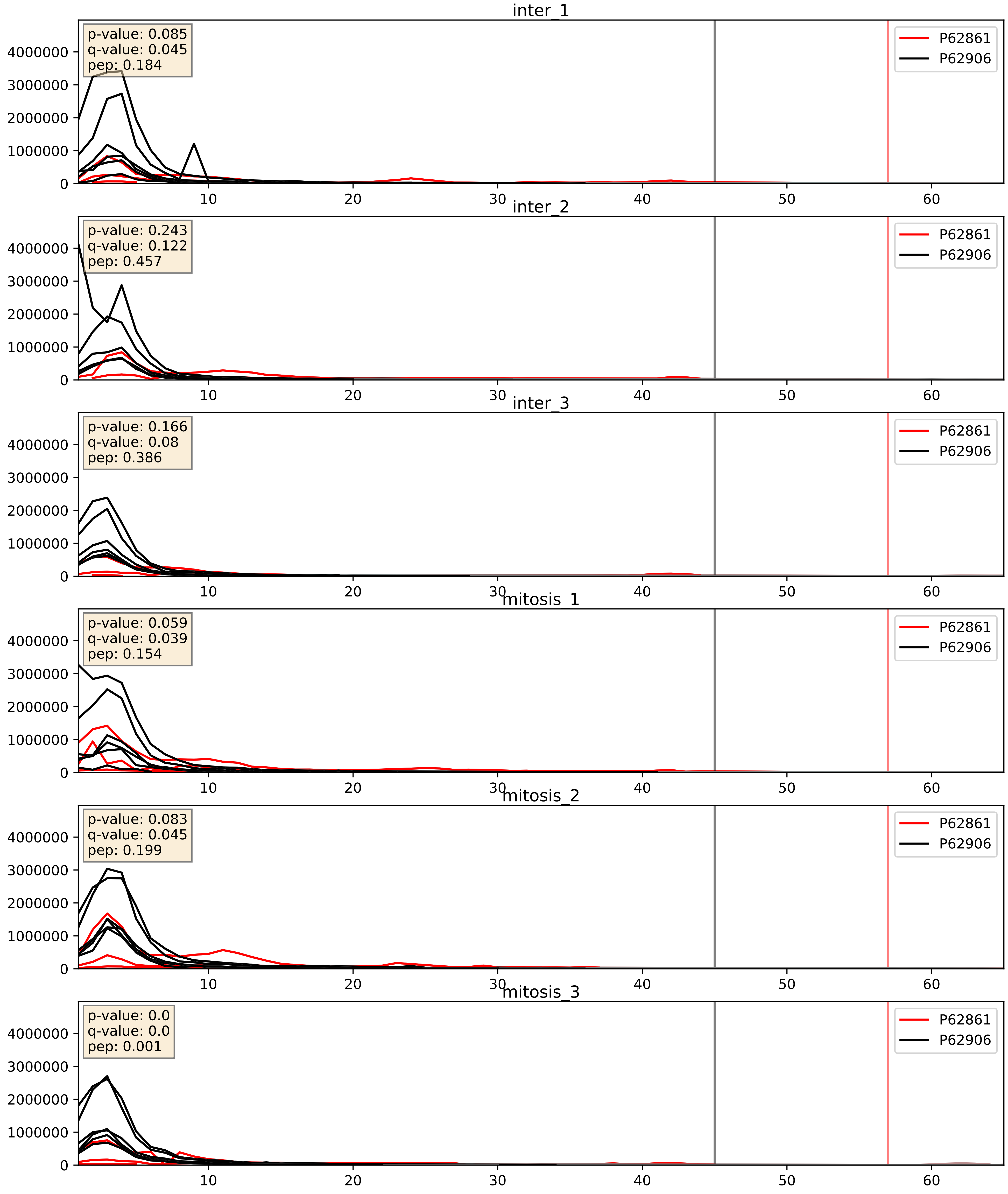


| level | condition_1 | condition_2 | pvalue | pvalue_adjusted |
|---|---|---|---|---|
| interactor_ratio | mitosis | inter | 0.35383159128615865 | 0.8027884343608738 |
| interactor_abundance | mitosis | inter | 0.4070932163799528 | 0.5931309092132087 |
| complex_abundance | mitosis | inter | 0.5009401777326098 | 0.6382942240168897 |

P62888_P62906
RL30_HUMAN vs RL10A_HUMAN
p-value: 0.0 q-value: 0.0

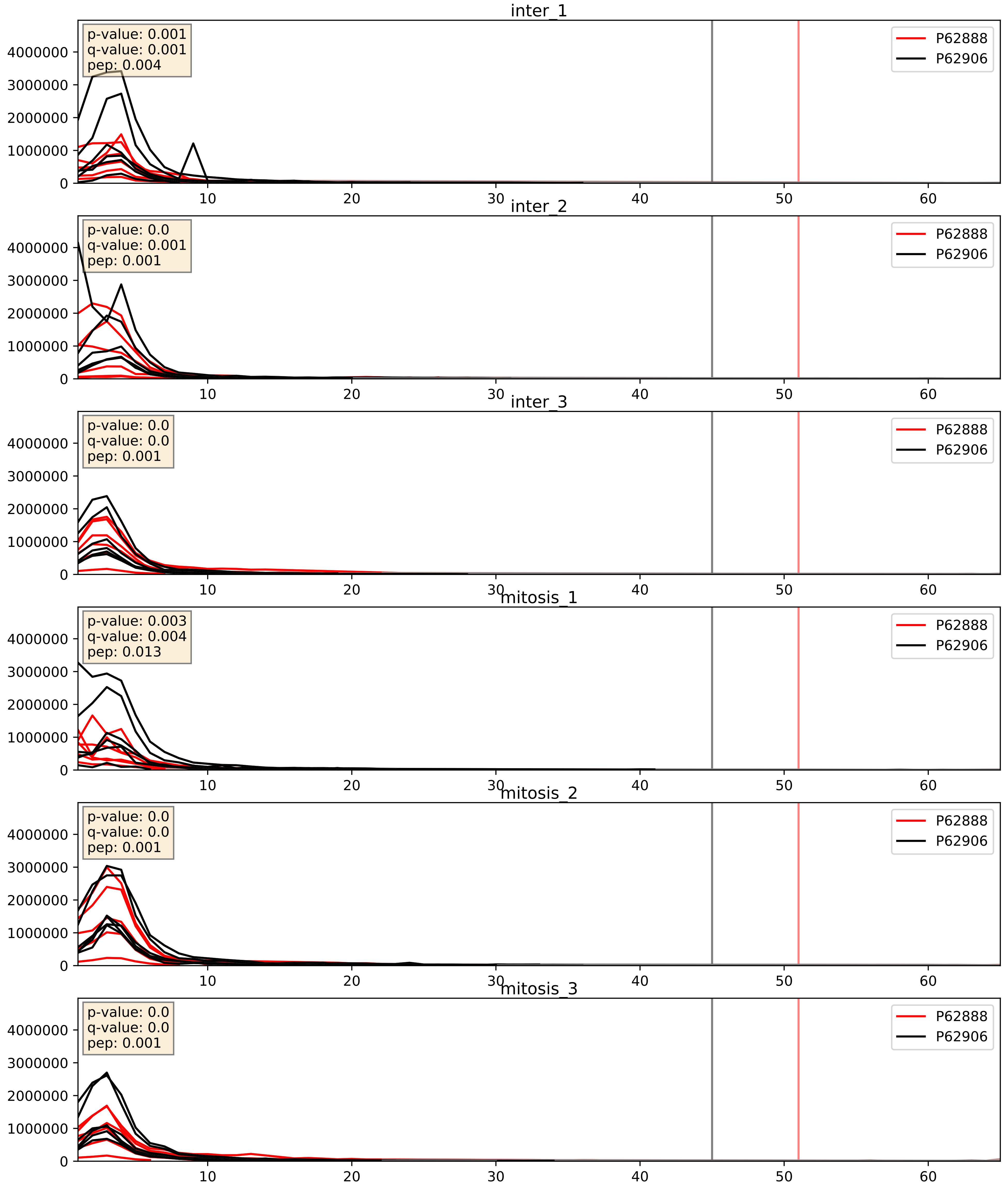


| level | condition_1 | condition_2 | pvalue | pvalue_adjusted |
|---|---|---|---|---|
| complex_abundance | mitosis | inter | 0.5022943572357844 | 0.6394603058701613 |
| interactor_abundance | mitosis | inter | 0.6205548267872167 | 0.750049035840717 |
| interactor_ratio | mitosis | inter | 0.7335675867402336 | 0.9488272200810516 |

P62899_P62906
RL31_HUMAN vs RL10A_HUMAN
p-value: 0.0 q-value: 0.0

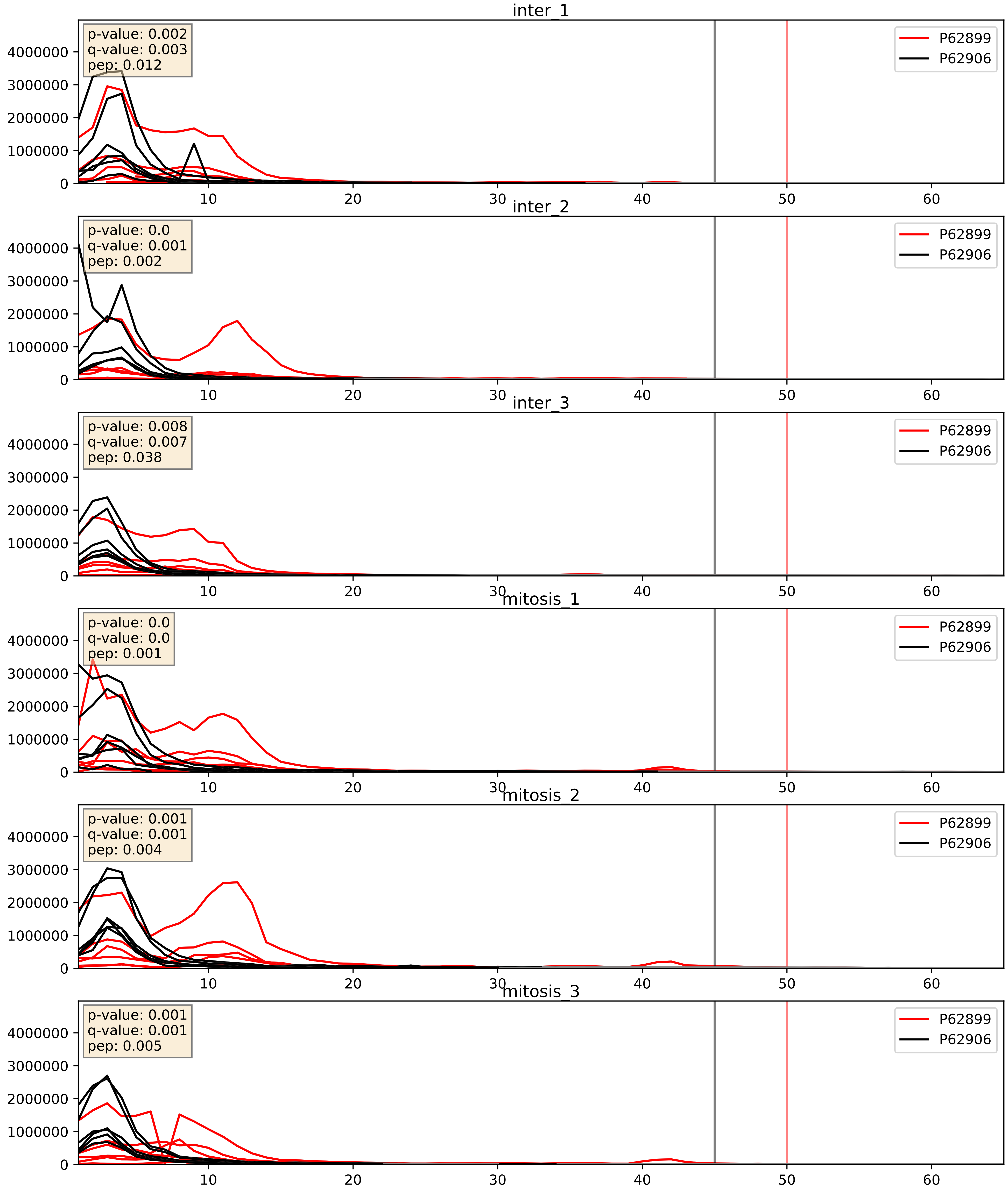


| level | condition_1 | condition_2 | pvalue | pvalue_adjusted |
|---|---|---|---|---|
| interactor_ratio | mitosis | inter | 0.1721568565331396 | 0.6357474943587899 |
| interactor_abundance | mitosis | inter | 0.274722386704829 | 0.46755423780703664 |
| complex_abundance | mitosis | inter | 0.500220180010493 | 0.6376596784646046 |

P62906_P62910
RL10A_HUMAN vs RL32_HUMAN
p-value: 0.0 q-value: 0.0

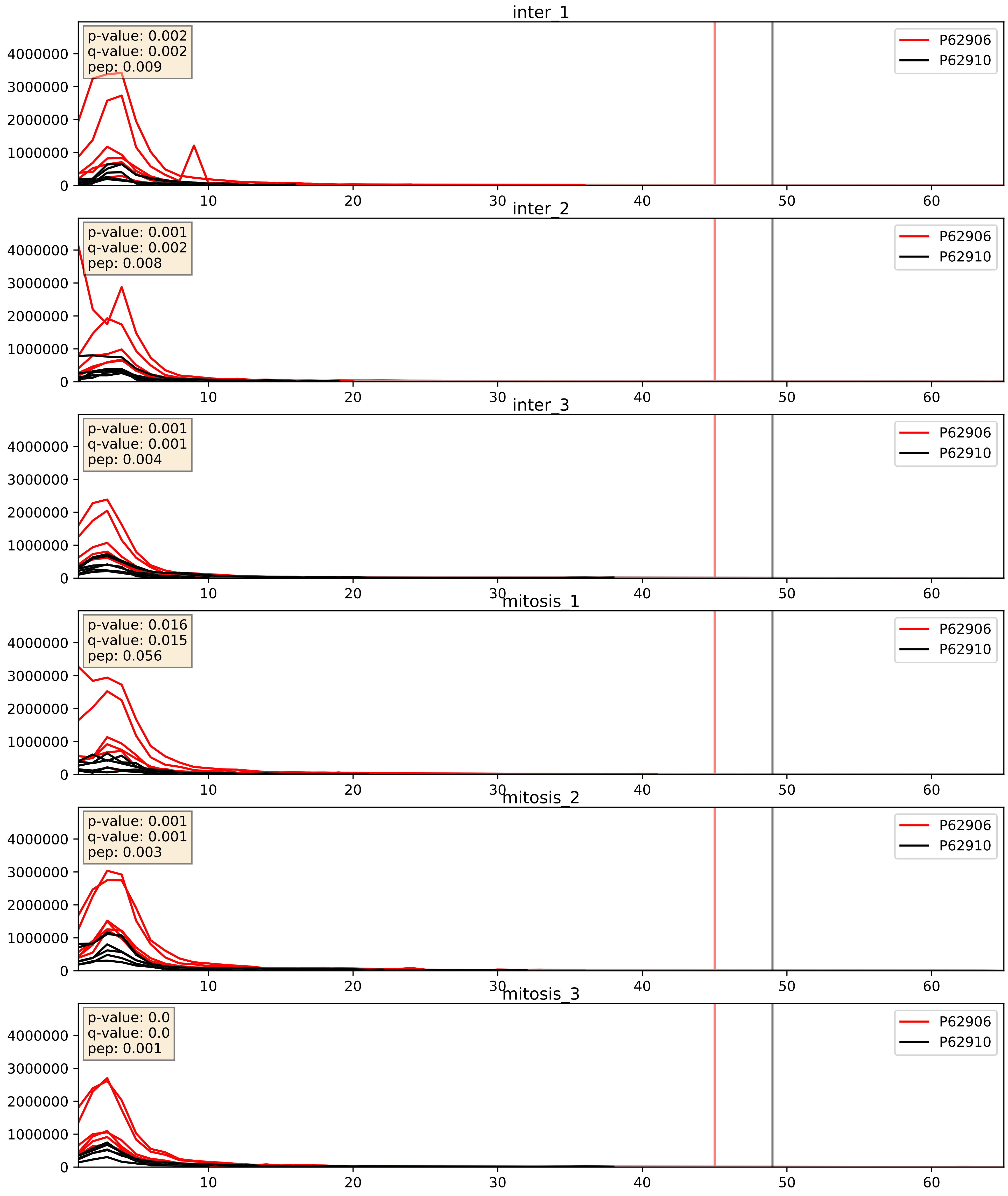


| level | condition_1 | condition_2 | pvalue | pvalue_adjusted |
|---|---|---|---|---|
| complex_abundance | mitosis | inter | 0.583351338076487 | 0.7113229991360013 |
| interactor_ratio | mitosis | inter | 0.6971383712402925 | 0.9356388300120577 |
| interactor_abundance | mitosis | inter | 0.8989071510304912 | 0.9441584502781086 |

P62906_P62913
RL10A_HUMAN vs RL11_HUMAN
p-value: 0.0 q-value: 0.0

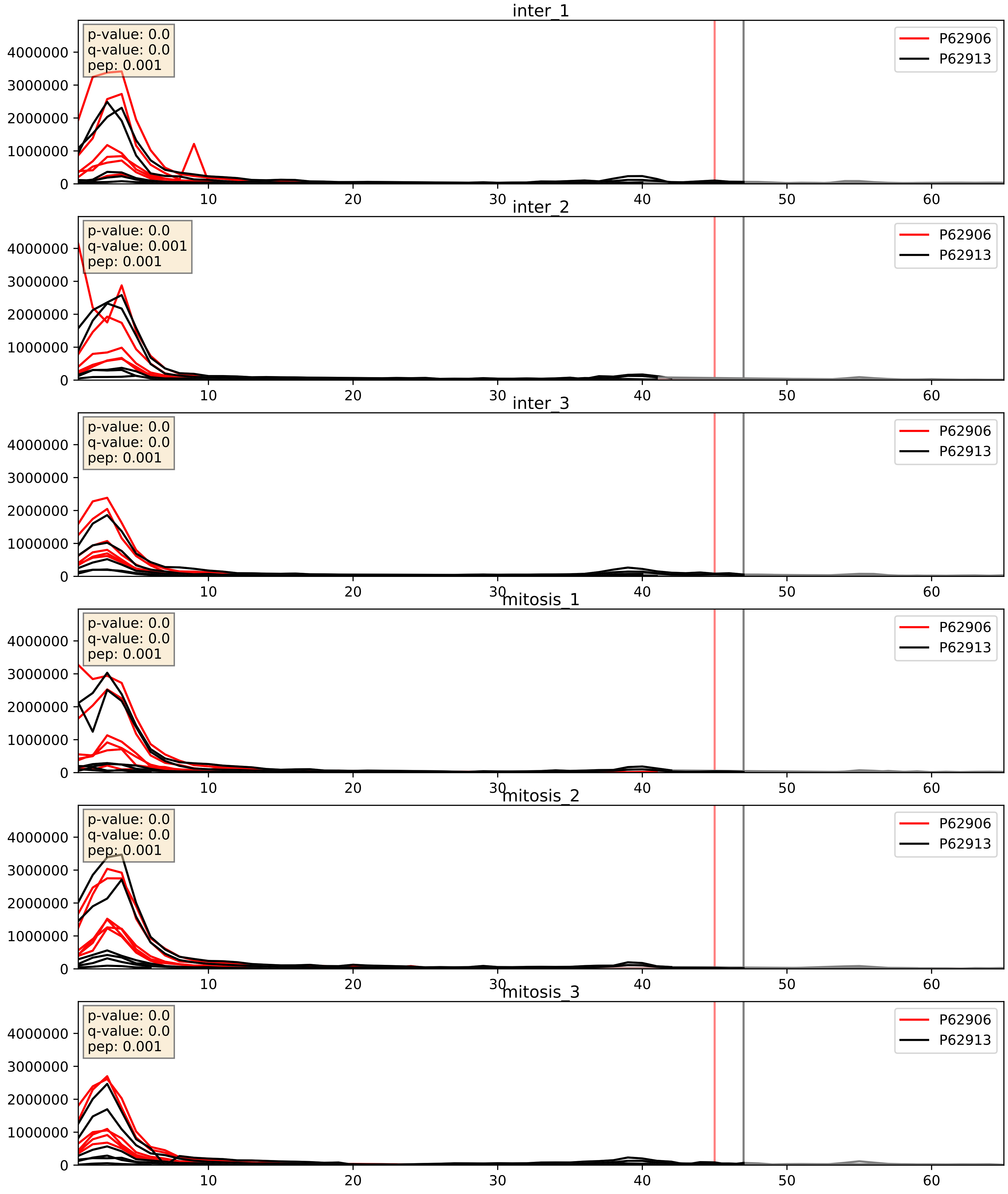


| level | condition_1 | condition_2 | pvalue | pvalue_adjusted |
|---|---|---|---|---|
| interactor_ratio | mitosis | inter | 0.19875952904116928 | 0.6733870464282405 |
| complex_abundance | mitosis | inter | 0.3508579851414222 | 0.4982323080309512 |
| interactor_abundance | mitosis | inter | 0.8633053801236219 | 0.925645901403921 |

P62906_P62917
RL10A_HUMAN vs RL8_HUMAN
p-value: 0.0 q-value: 0.0

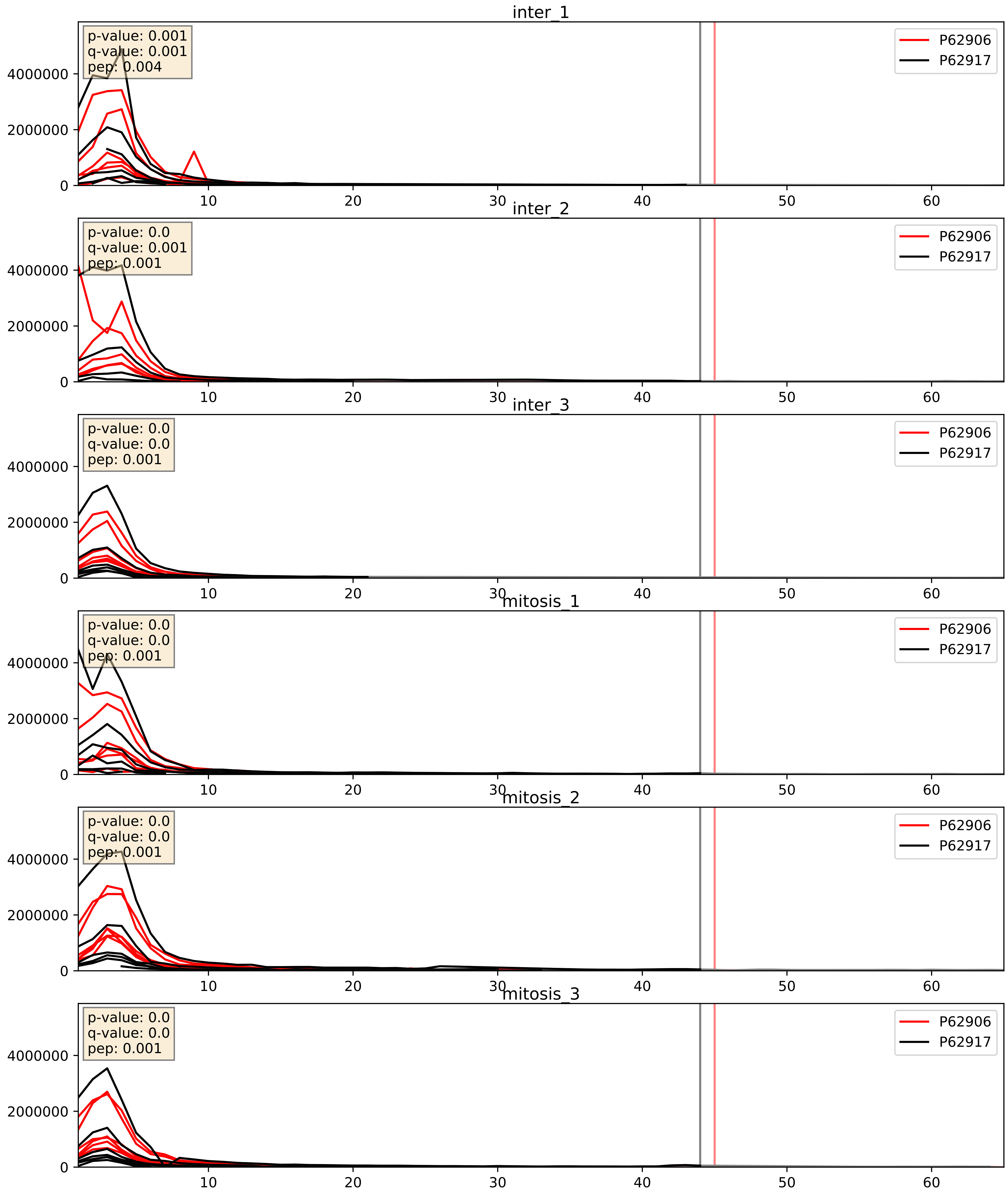


| level | condition_1 | condition_2 | pvalue | pvalue_adjusted |
|---|---|---|---|---|
| interactor_ratio | mitosis | inter | 0.44082434990540736 | 0.8466845385837265 |
| complex_abundance | mitosis | inter | 0.6443985492092159 | 0.7592638102176033 |
| interactor_abundance | mitosis | inter | 0.9099459448901732 | 0.9488496785938922 |

P62906_P63173
RL10A_HUMAN vs RL38_HUMAN
p-value: 0.024 q-value: 0.018

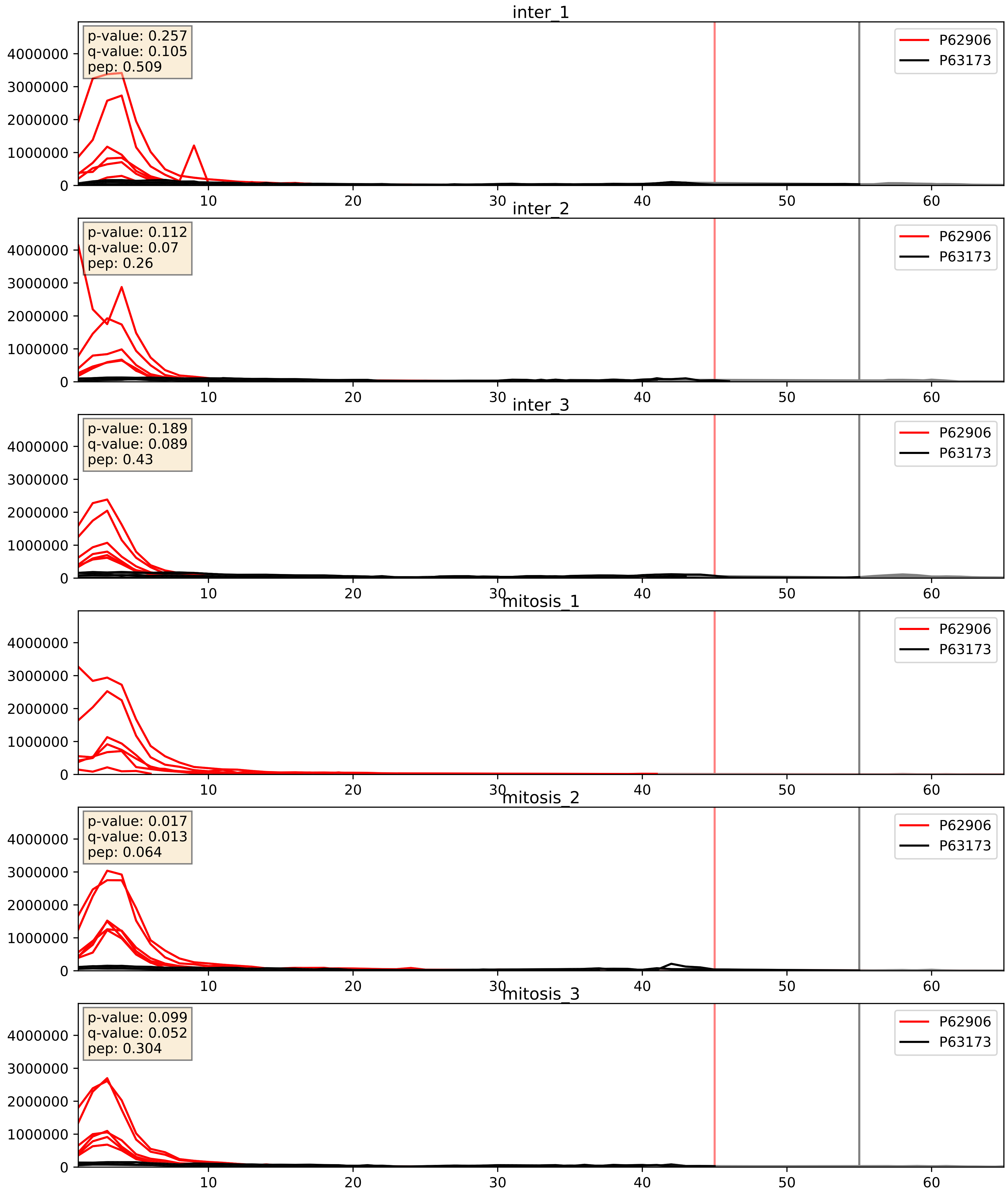


| level | condition_1 | condition_2 | pvalue | pvalue_adjusted |
|---|---|---|---|---|
| complex_abundance | mitosis | inter | 0.3144766160051418 | 0.46324554001101603 |
| interactor_abundance | mitosis | inter | 0.36132286911833944 | 0.5585919739304652 |
| interactor_ratio | mitosis | inter | 0.5746365467042631 | 0.8953201382942287 |

P62906_P83731
RL10A_HUMAN vs RL24_HUMAN
p-value: 0.0 q-value: 0.0

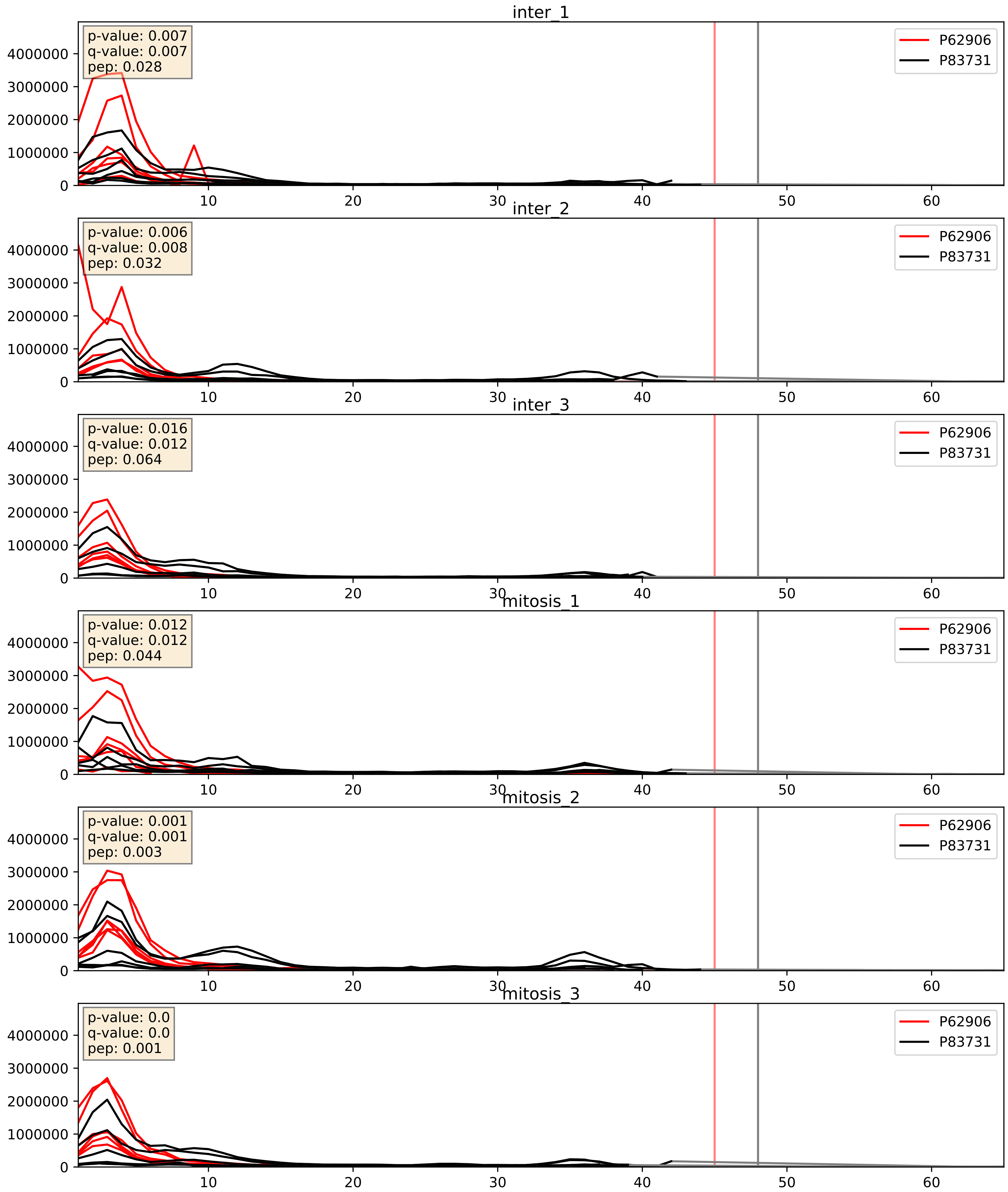


| level | condition_1 | condition_2 | pvalue | pvalue_adjusted |
|---|---|---|---|---|
| interactor_ratio | mitosis | inter | 0.3389996732831241 | 0.7924528012231136 |
| complex_abundance | mitosis | inter | 0.5621337516981424 | 0.6937521503079728 |
| interactor_abundance | mitosis | inter | 0.8633053801236219 | 0.925645901403921 |

P62906_P84098
RL10A_HUMAN vs RL19_HUMAN
p-value: 0.0 q-value: 0.0

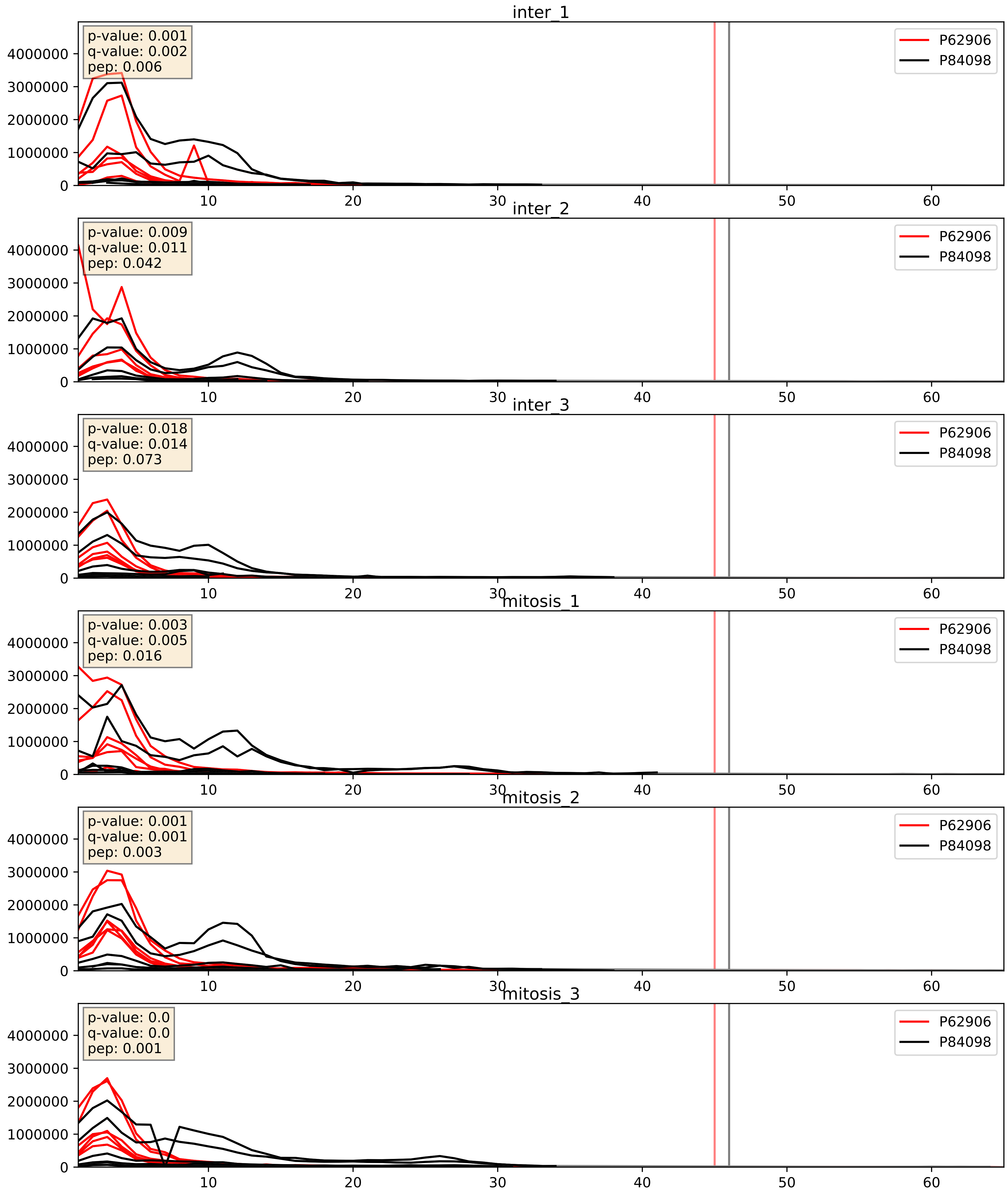


| level | condition_1 | condition_2 | pvalue | pvalue_adjusted |
|---|---|---|---|---|
| interactor_ratio | mitosis | inter | 0.5291257527906434 | 0.8821330249831557 |
| complex_abundance | mitosis | inter | 0.7386943549735082 | 0.8265651867415987 |
| interactor_abundance | mitosis | inter | 0.8638783411956787 | 0.9259702730572263 |

P62906_Q02543
RL10A_HUMAN vs RL18A_HUMAN
p-value: 0.0 q-value: 0.0

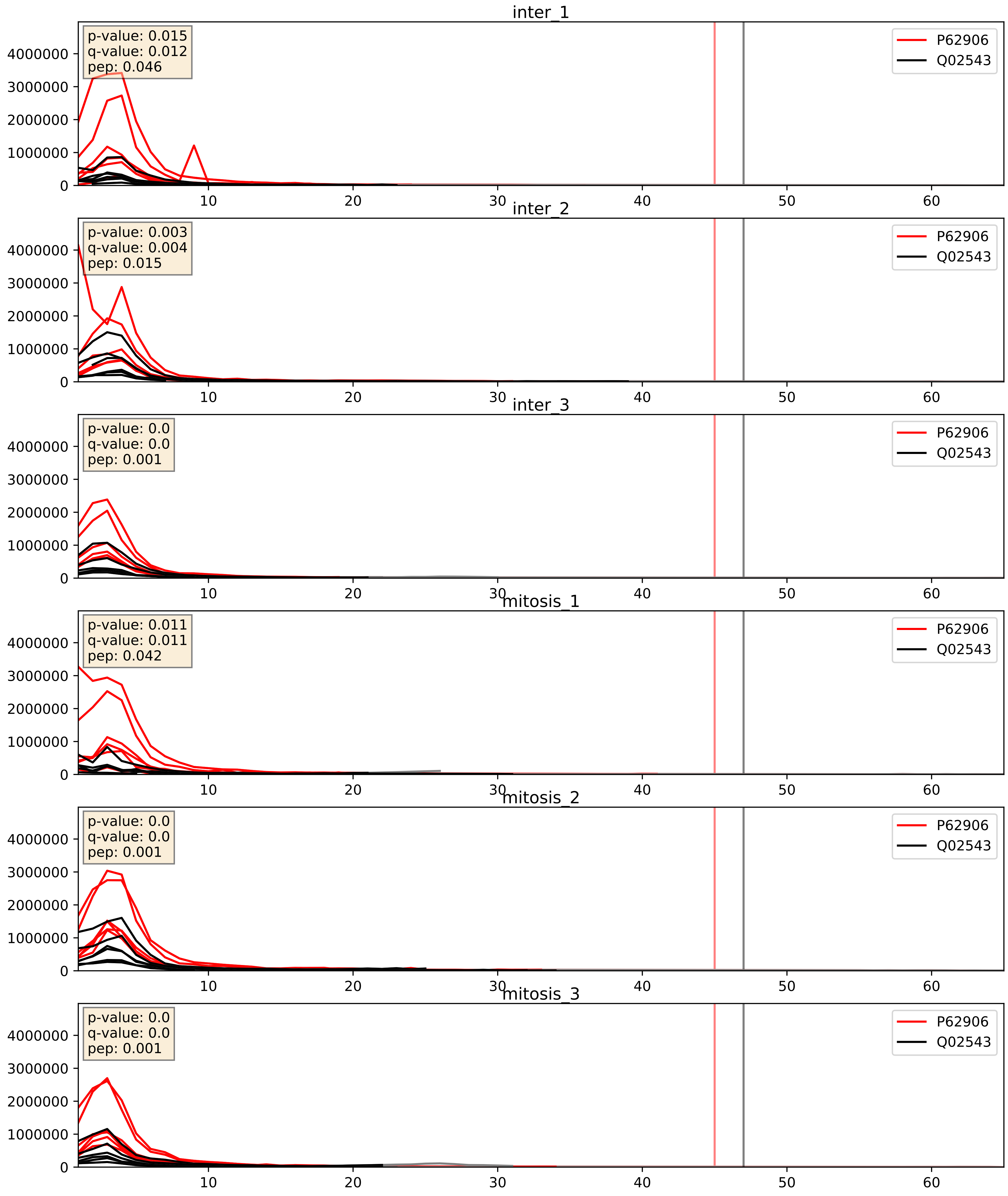


| level | condition_1 | condition_2 | pvalue | pvalue_adjusted |
|---|---|---|---|---|
| complex_abundance | mitosis | inter | 0.5649764777781392 | 0.6960562247813574 |
| interactor_ratio | mitosis | inter | 0.7367543462984597 | 0.9491588884573259 |
| interactor_abundance | mitosis | inter | 0.8383435441530803 | 0.911819840761536 |

P62906_Q02878
RL10A_HUMAN vs RL6_HUMAN
p-value: 0.0 q-value: 0.0

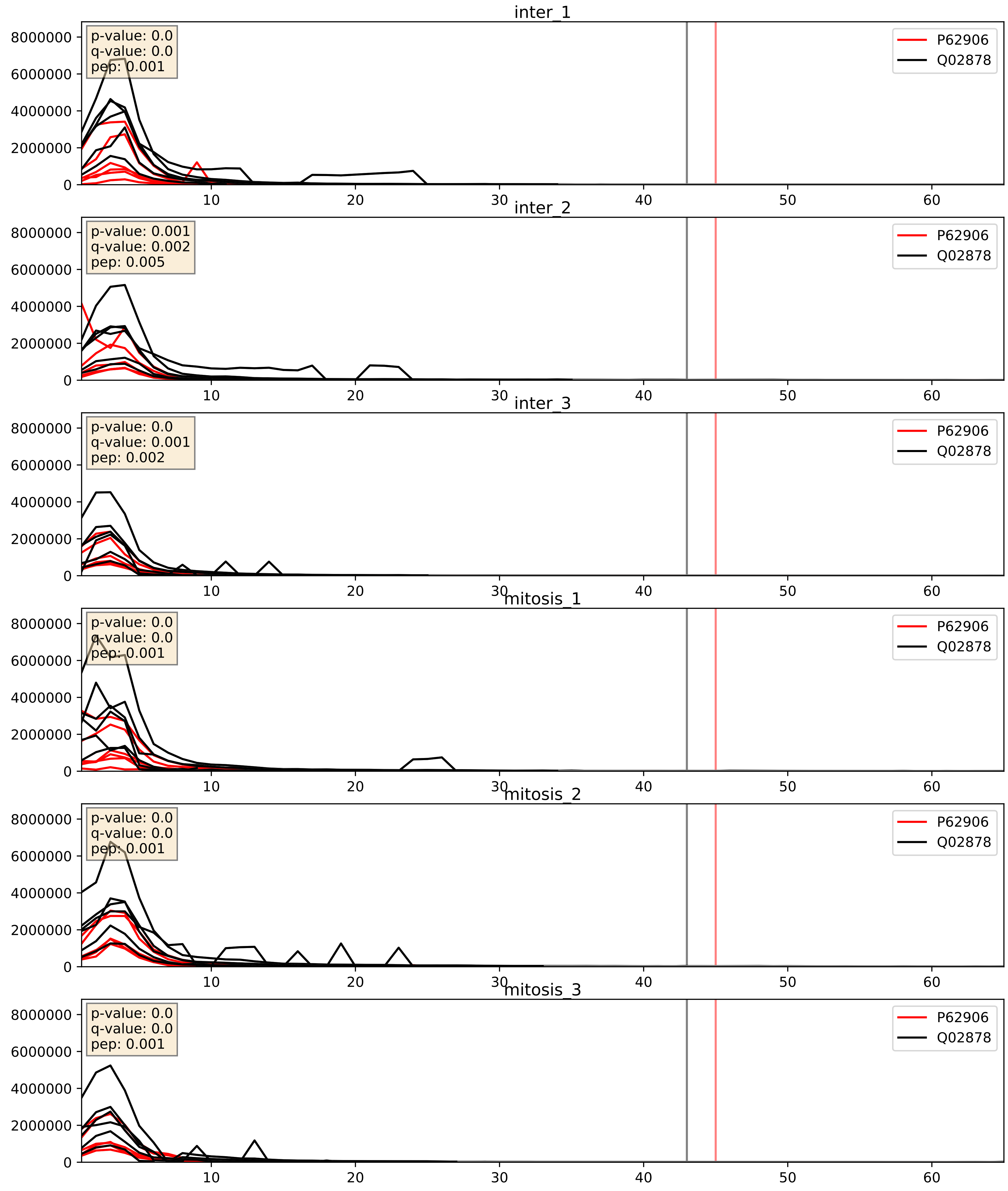


| level | condition_1 | condition_2 | pvalue | pvalue_adjusted |
|---|---|---|---|---|
| interactor_ratio | mitosis | inter | 0.4226622222730265 | 0.8391460036711063 |
| complex_abundance | mitosis | inter | 0.6190009750645107 | 0.7394705093209963 |
| interactor_abundance | mitosis | inter | 0.8582434073944609 | 0.9239217213477439 |

P62906_Q07020
RL10A_HUMAN vs RL18_HUMAN
p-value: 0.0 q-value: 0.0

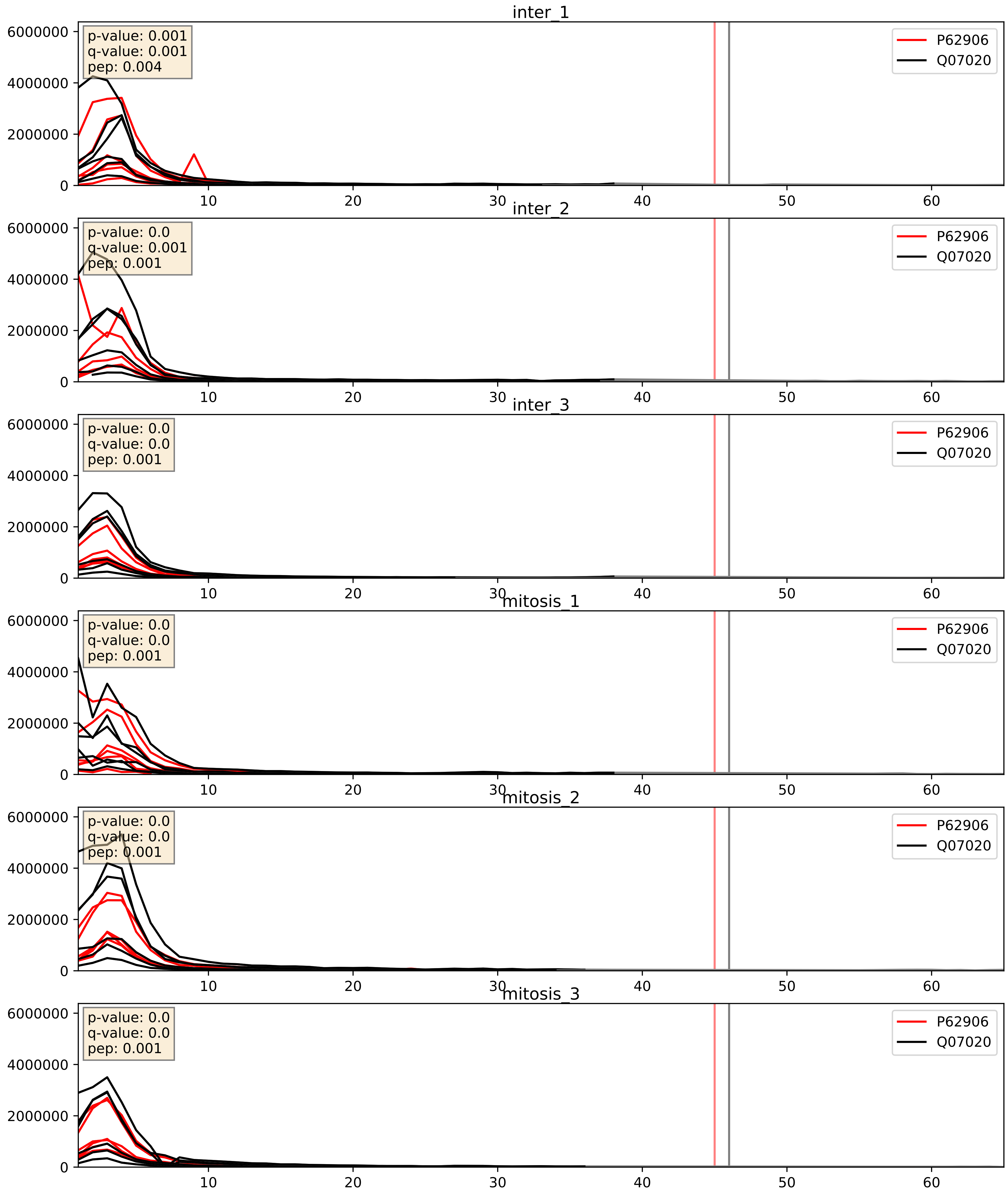


| level | condition_1 | condition_2 | pvalue | pvalue_adjusted |
|---|---|---|---|---|
| complex_abundance | mitosis | inter | 0.47411656520159695 | 0.6148216631003893 |
| interactor_ratio | mitosis | inter | 0.7326361374205206 | 0.9485774717892954 |
| interactor_abundance | mitosis | inter | 0.856311843263932 | 0.9227678686579346 |

P62906_Q12905
RL10A_HUMAN vs ILF2_HUMAN
p-value: 0.0 q-value: 0.041

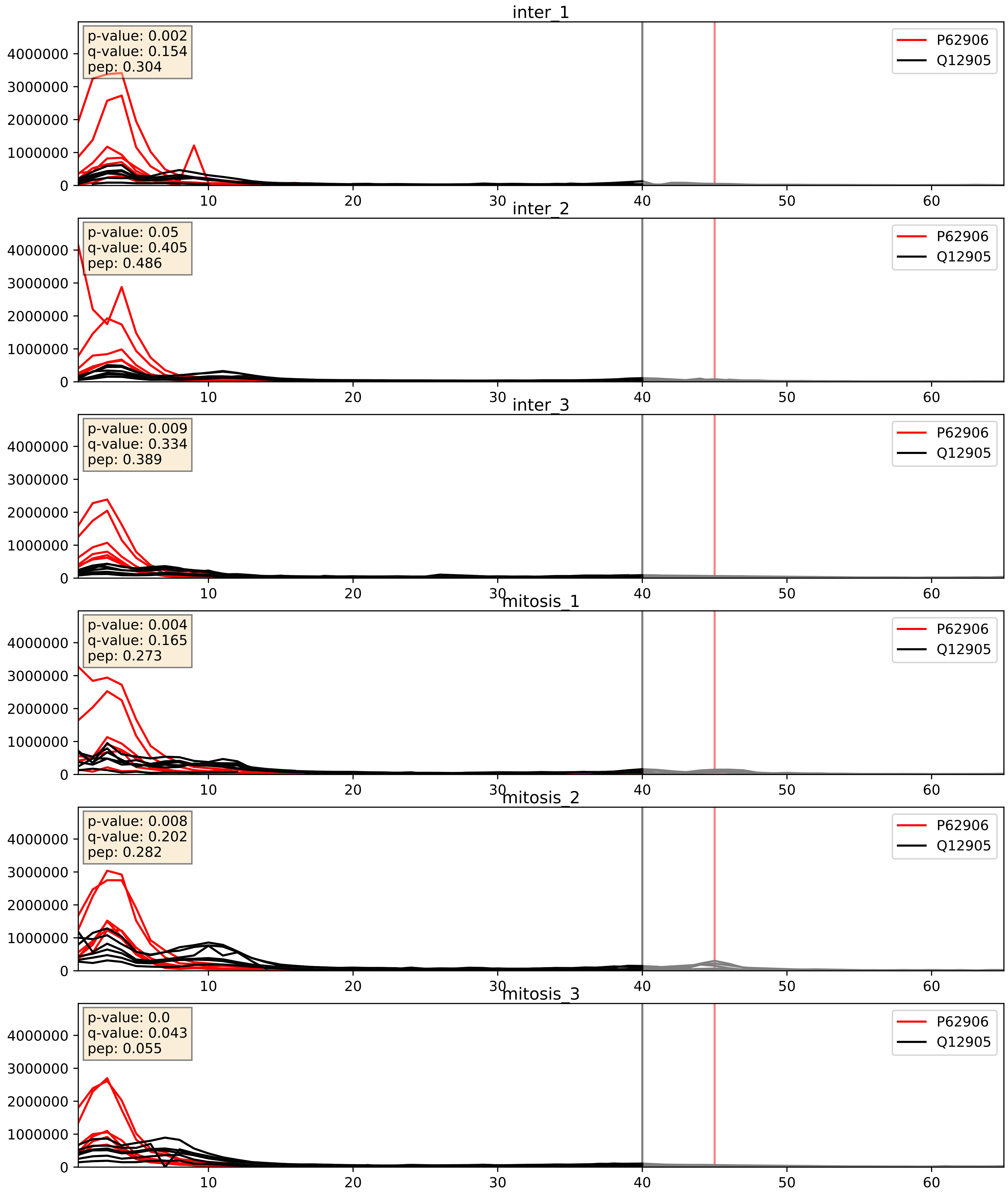


| level | condition_1 | condition_2 | pvalue | pvalue_adjusted |
|---|---|---|---|---|
| interactor_ratio | mitosis | inter | 0.019821751658541914 | 0.213852931597635 |
| complex_abundance | mitosis | inter | 0.05391602986386887 | 0.16559785275734393 |
| interactor_abundance | mitosis | inter | 0.8597142149458314 | 0.9247491429927516 |

P62906_Q969Q0
RL10A_HUMAN vs RL36L_HUMAN
p-value: 0.004 q-value: 0.004

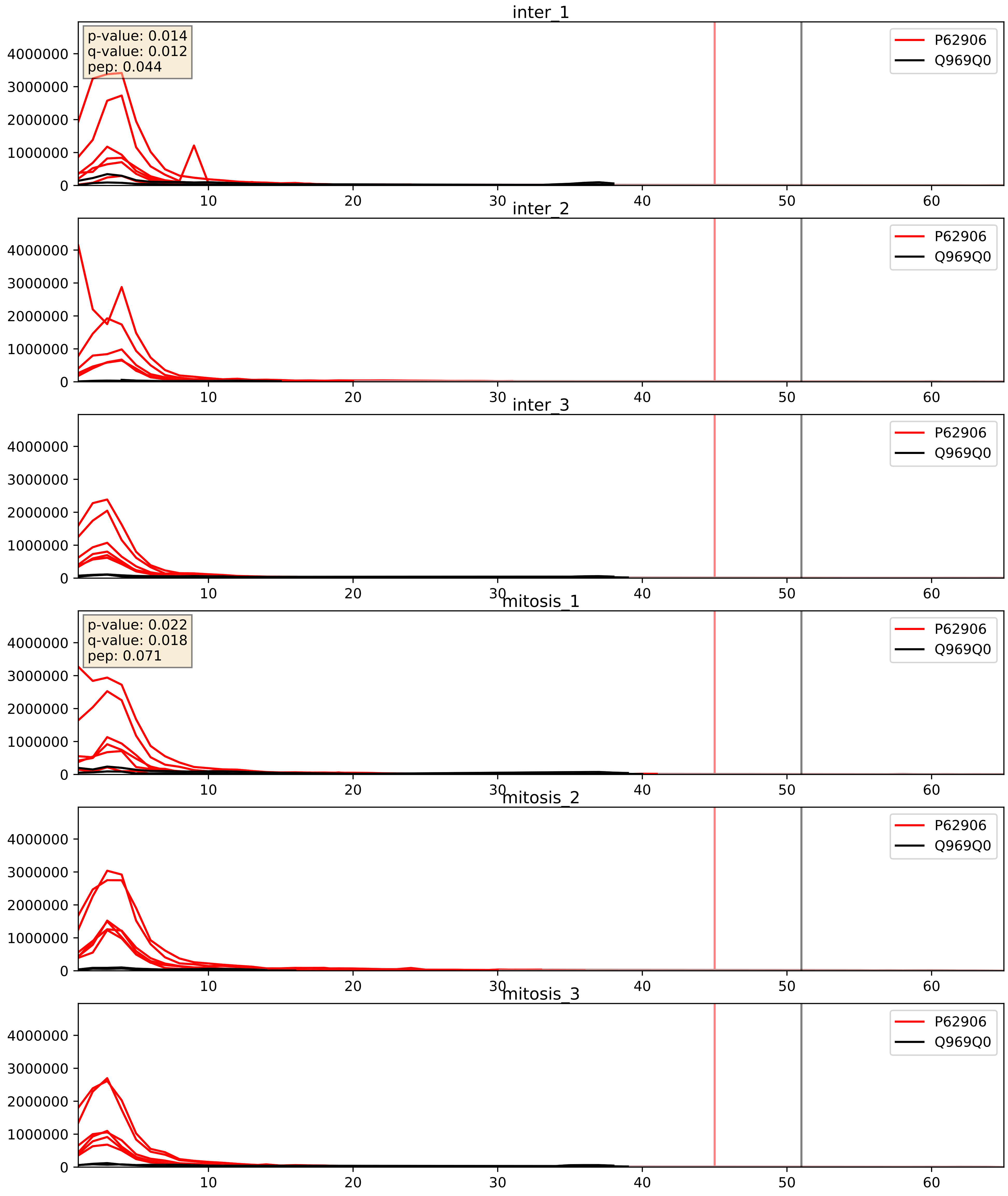


| level | condition_1 | condition_2 | pvalue | pvalue_adjusted |
|---|---|---|---|---|
| interactor_abundance | mitosis | inter | 0.7846633807796286 | 0.8753135517656377 |
| complex_abundance | mitosis | inter | 0.7945528824923207 | 0.8666136928694865 |
| interactor_ratio | mitosis | inter | 0.8680742436766177 | 0.9786759416121144 |

P62906_Q9BZE4
RL10A_HUMAN vs NOG1_HUMAN
p-value: 0.001 q-value: 0.041

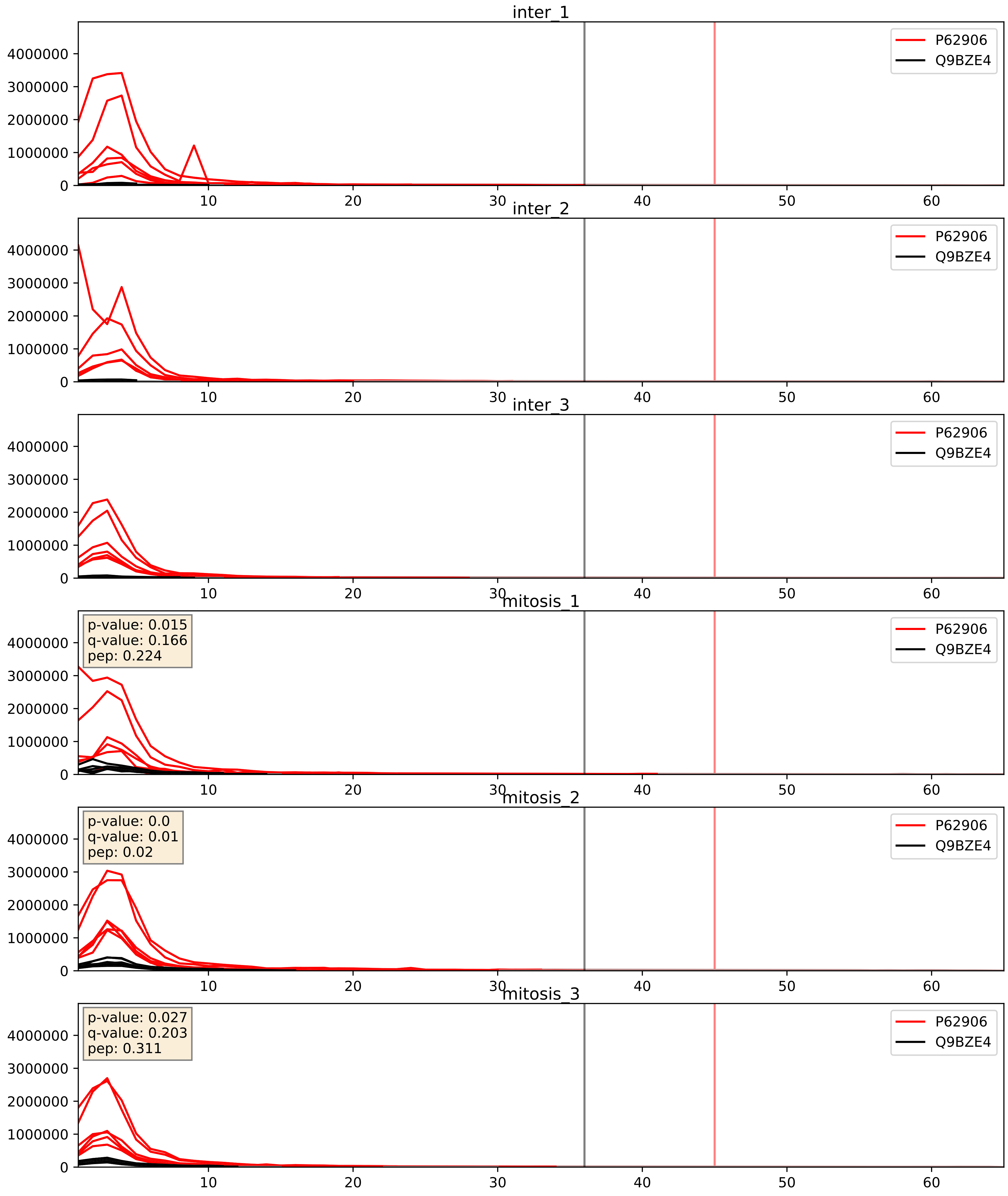


| level | condition_1 | condition_2 | pvalue | pvalue_adjusted |
|---|---|---|---|---|
| complex_abundance | mitosis | inter | 0.0017417866920582635 | 0.0295241467010272 |
| interactor_ratio | mitosis | inter | 0.002923484829602876 | 0.07007095913365735 |
| interactor_abundance | mitosis | inter | 0.40955067394198835 | 0.594379996833825 |

P62906_Q9Y3U8
RL10A_HUMAN vs RL36_HUMAN
p-value: 0.0 q-value: 0.0

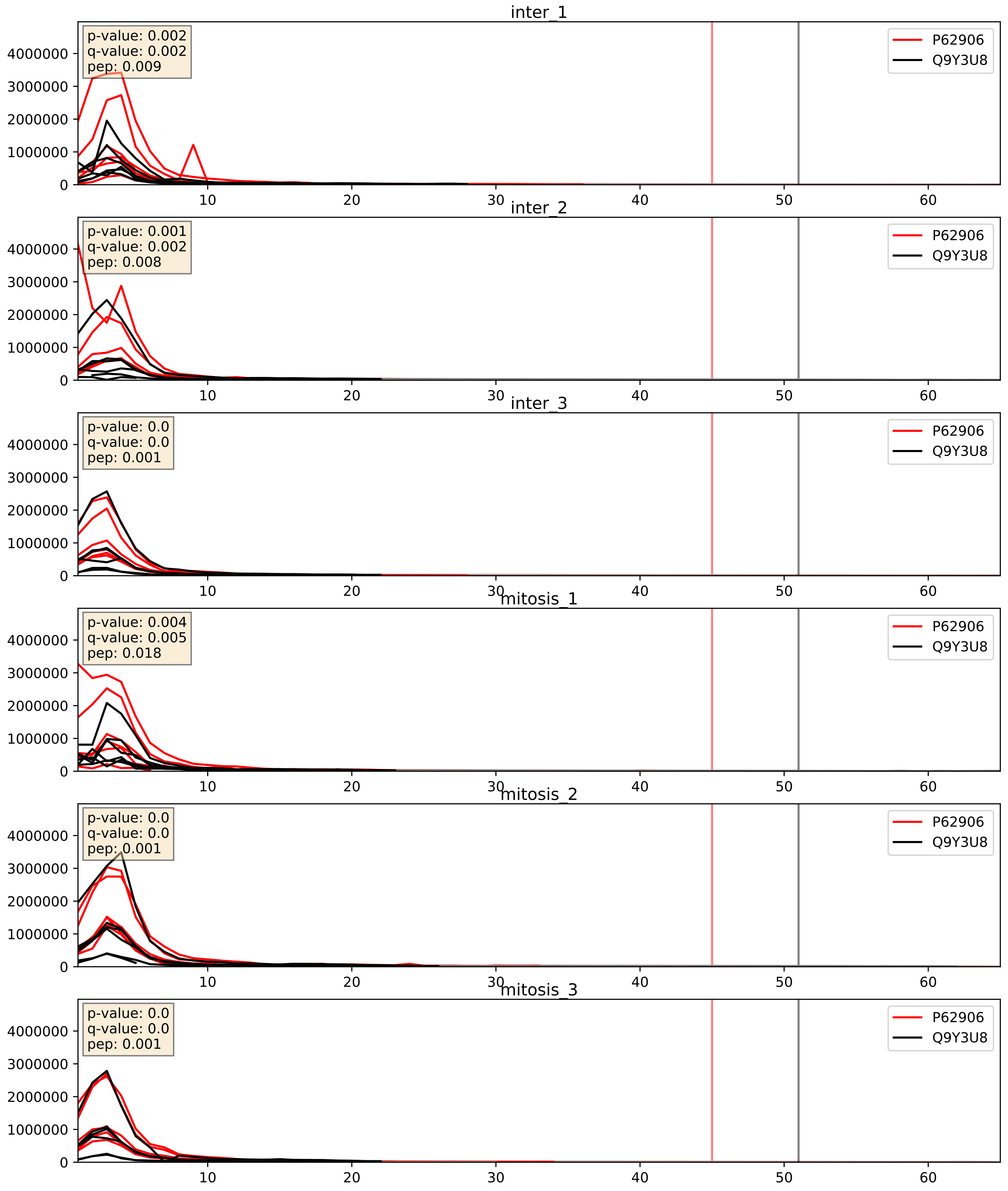


| level | condition_1 | condition_2 | pvalue | pvalue_adjusted |
|---|---|---|---|---|
| interactor_ratio | mitosis | inter | 0.6073009346985436 | 0.9055803577343394 |
| complex_abundance | mitosis | inter | 0.6227270576890505 | 0.7418989024103371 |
| interactor_abundance | mitosis | inter | 0.8620061177901677 | 0.925645901403921 |
