## Supplementary Data 2 for "SECAT: Quantifying differential protein-protein interaction states by network-centric analysis": hela_string_000151_Q12834.pdf

Q12834_Q12834

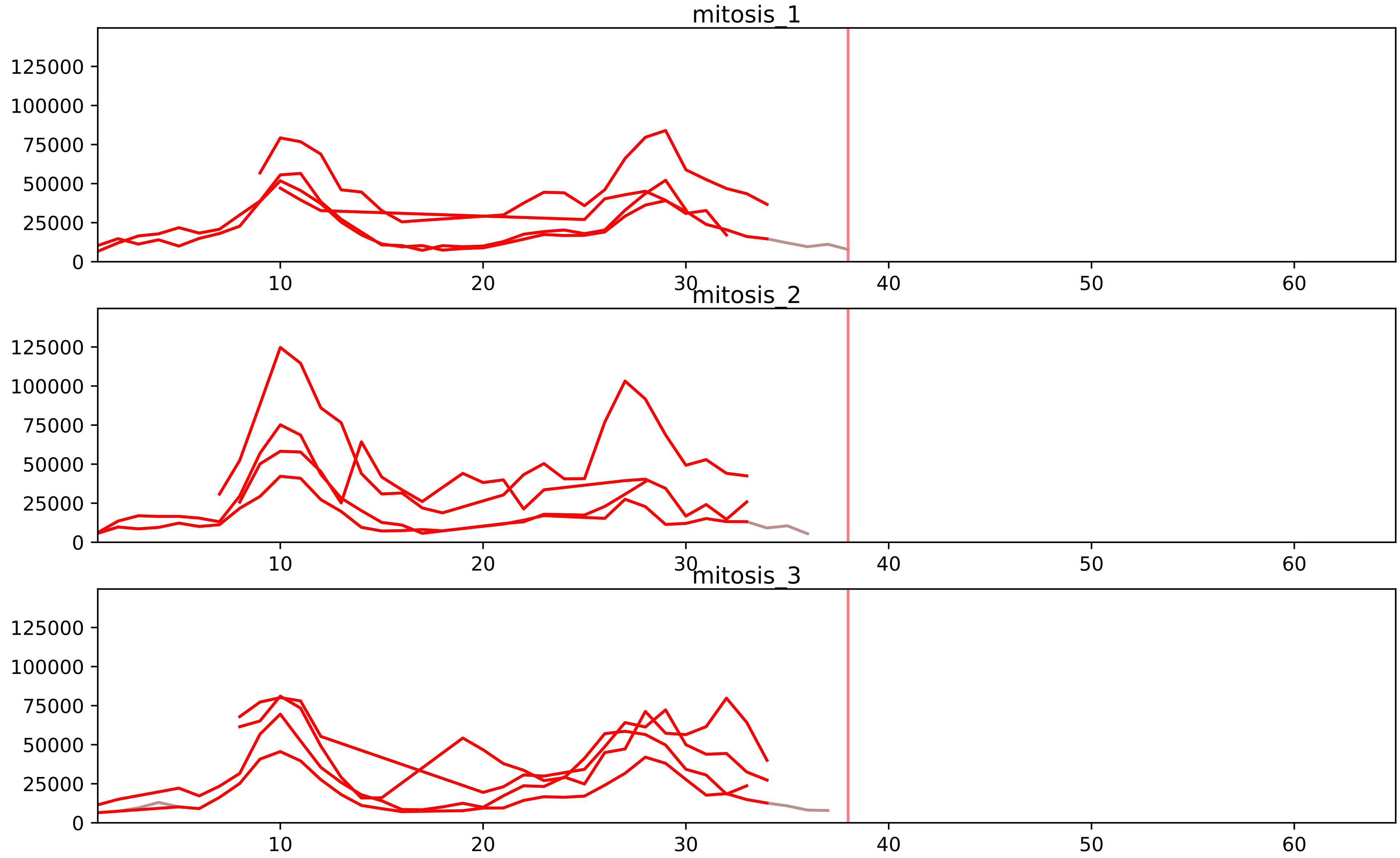


| level | condition_1 | condition_2 | pvalue | pvalue_adjusted |
|---|---|---|---|---|
| assembled_abundance | mitosis | inter | 5.6432793748147563e-05 | 0.007270487507555603 |
| total_abundance | mitosis | inter | 9.549500824992085e-05 | 0.01273261737241483 |
| interactor_ratio | mitosis | inter | 0.0005374916663425212 | 0.006682328824798913 |
| complex_abundance | mitosis | inter | 0.0009106509474901972 | 0.010356234336273524 |
| interactor_abundance | mitosis | inter | 0.002579417553320174 | 0.022600610943376766 |

Q12834_Q13042
CDC20_HUMAN vs CDC16_HUMAN
p-value: 0.048 q-value: 0.032

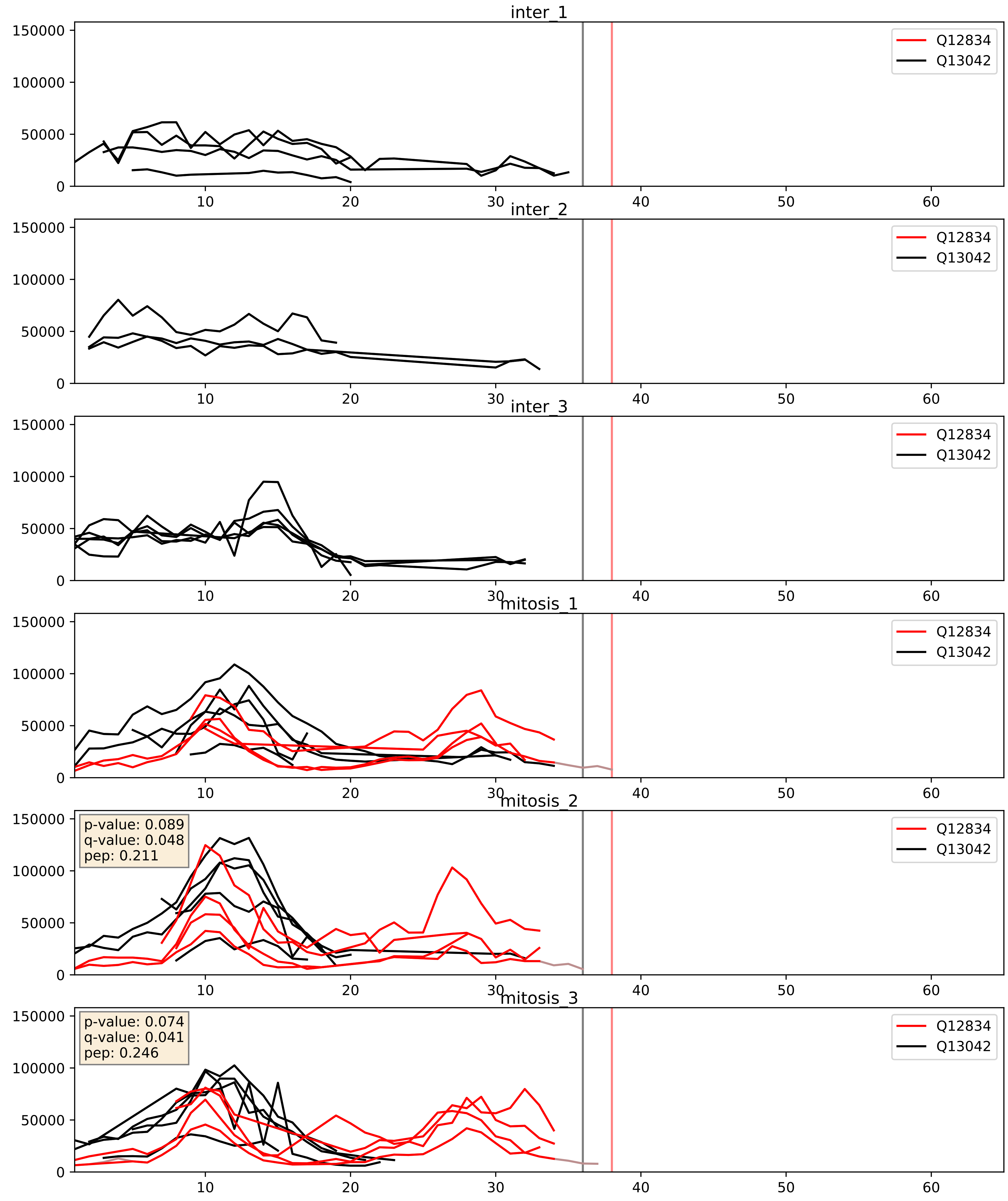


| level | condition_1 | condition_2 | pvalue | pvalue_adjusted |
|---|---|---|---|---|
| interactor_ratio | mitosis | inter | 0.0005374916663425212 | 0.02690601557831568 |
| complex_abundance | mitosis | inter | 0.0009106509474901972 | 0.0197847007881119 |
| interactor_abundance | mitosis | inter | 0.002579417553320174 | 0.03806046117740333 |
