## Supplementary Data 2 for "SECAT: Quantifying differential protein-protein interaction states by network-centric analysis": hela_string_000152_Q9UFC0.pdf

### Q9UFC0_Q9UFC0

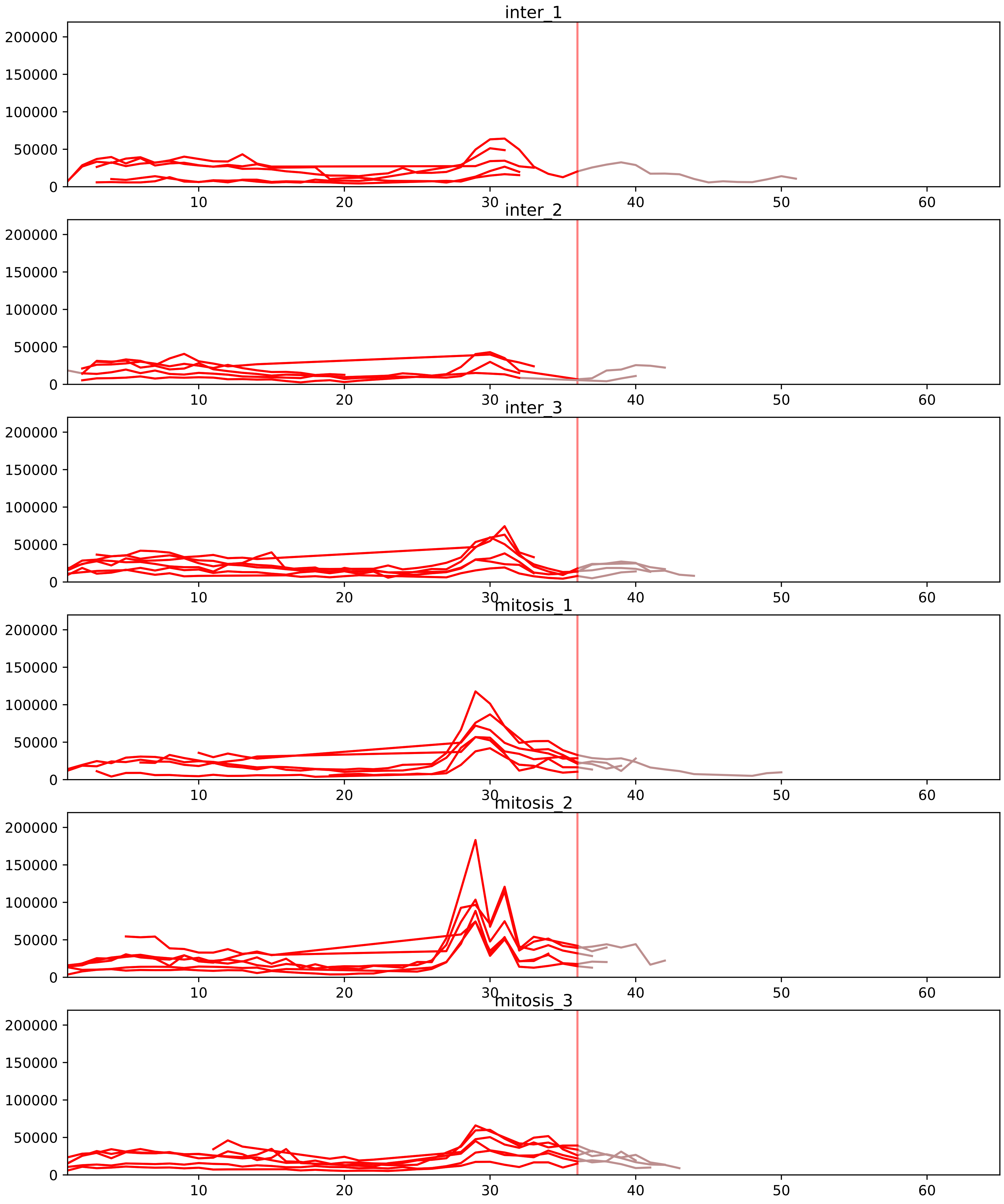


| level | condition_1 | condition_2 | pvalue | pvalue_adjusted |
|---|---|---|---|---|
| interactor_abundance | mitosis | inter | 6.741999743748337e-05 | 0.00179786659833289 |
| complex_abundance | mitosis | inter | 0.00011742609215818978 | 0.0024276855007985304 |
| assembled_abundance | mitosis | inter | 0.00682871900552158 | 0.08854635364223412 |
| total_abundance | mitosis | inter | 0.008026376549363584 | 0.1064121953458594 |
| monomer_abundance | mitosis | inter | 0.03282181639001813 | 0.30153627455136073 |
| interactor_ratio | mitosis | inter | 0.5079779625380085 | 0.7790392652192673 |

O43913_Q9UFC0
ORC5_HUMAN vs LRWD1_HUMAN
p-value: 0.014 q-value: 0.011

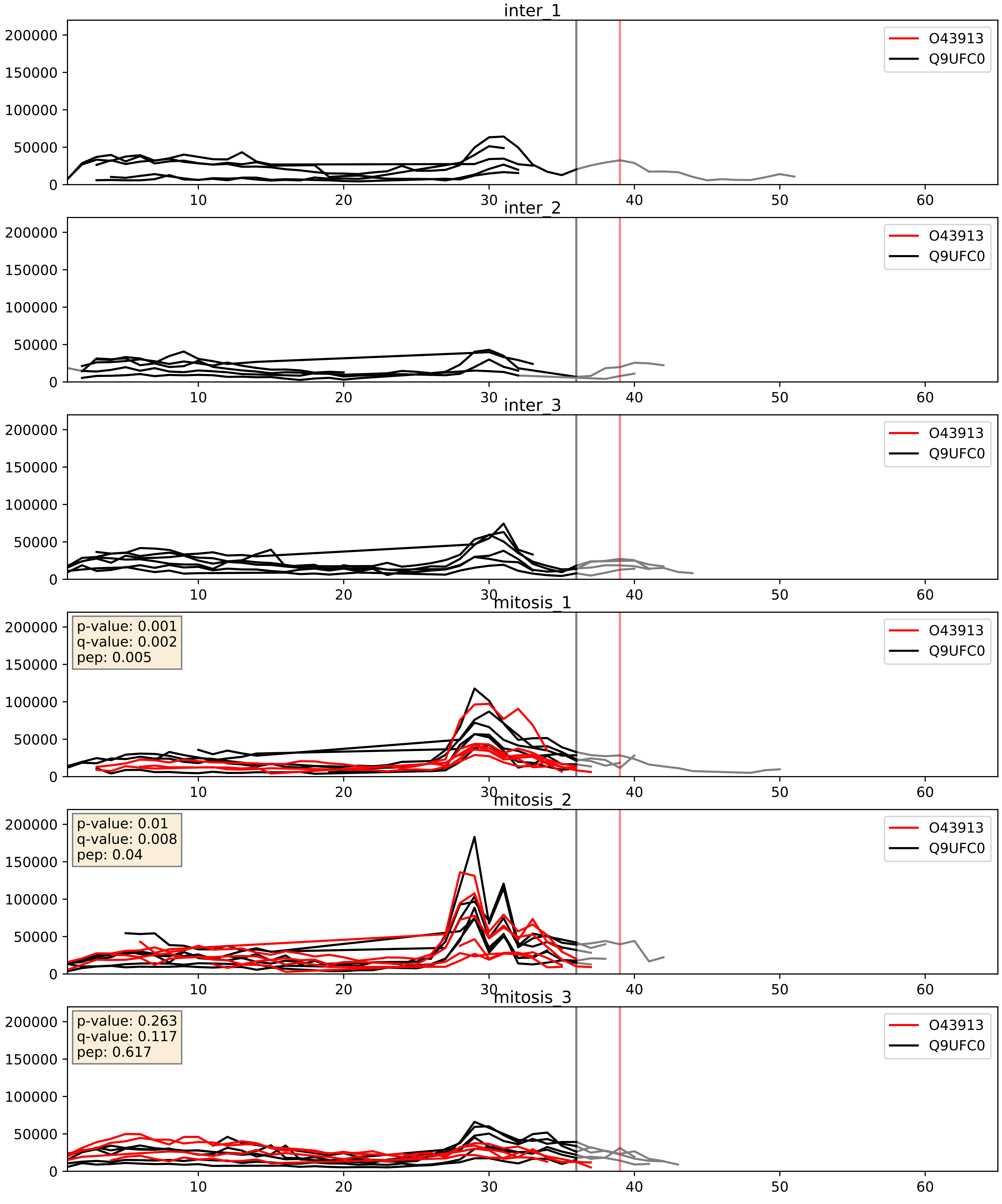


| level | condition_1 | condition_2 | pvalue | pvalue_adjusted |
|---|---|---|---|---|
| complex_abundance | mitosis | inter | 0.0005544121160053182 | 0.015065929247636585 |
| interactor_abundance | mitosis | inter | 0.0028274540106219656 | 0.04037198720754633 |
| interactor_ratio | mitosis | inter | 0.29660021331883807 | 0.7646944825997081 |

Q13416_Q9UFC0
ORC2_HUMAN vs LRWD1_HUMAN
p-value: 0.043 q-value: 0.029

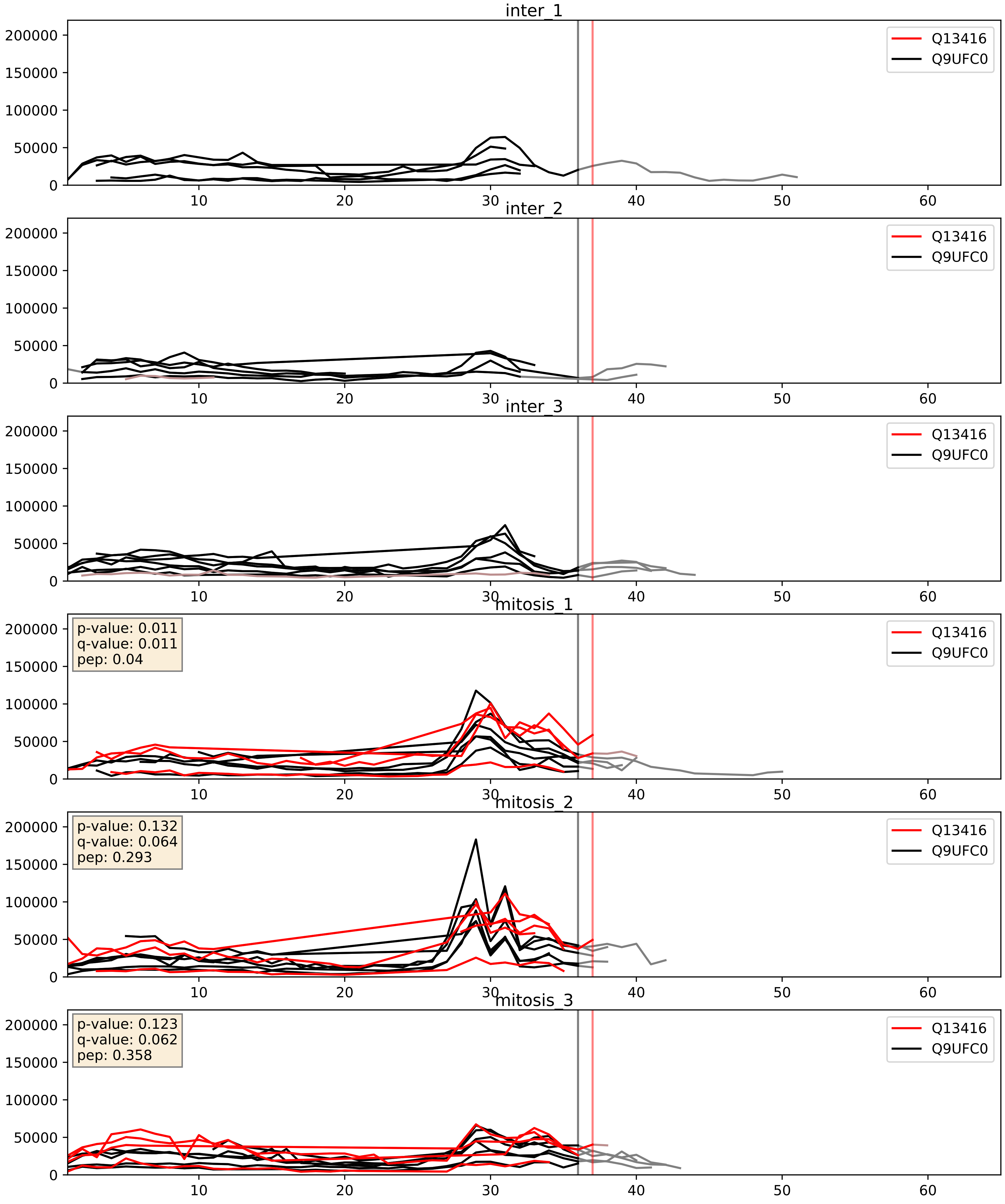


| level | condition_1 | condition_2 | pvalue | pvalue_adjusted |
|---|---|---|---|---|
| complex_abundance | mitosis | inter | 0.0004014148102010904 | 0.01272633620489383 |
| interactor_abundance | mitosis | inter | 0.0014603150123462959 | 0.028218917238081776 |
| interactor_ratio | mitosis | inter | 0.326104446487663 | 0.7849334771084482 |

Q9UBD5_Q9UFC0
ORC3_HUMAN vs LRWD1_HUMAN
p-value: 0.003 q-value: 0.003

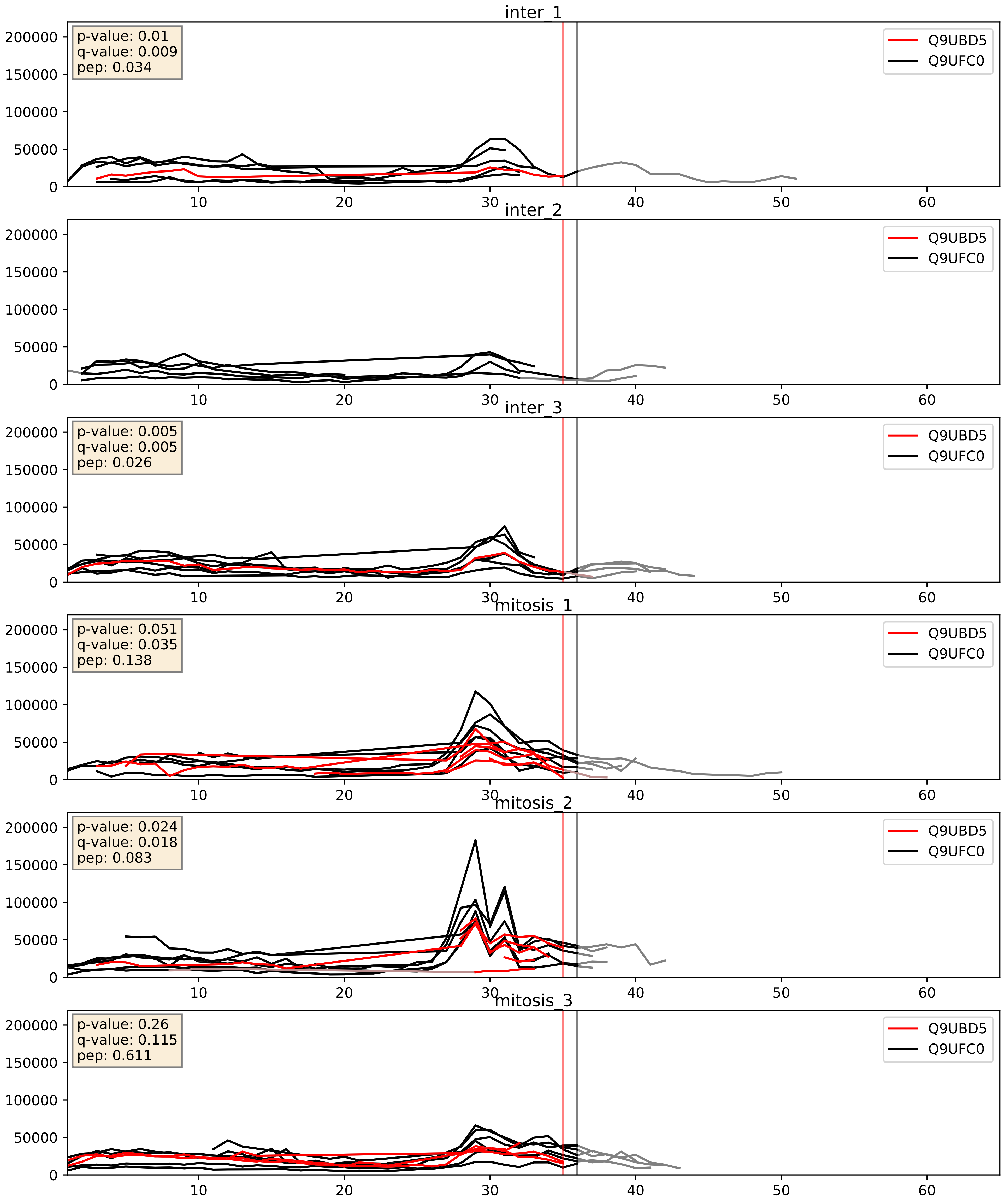


| level | condition_1 | condition_2 | pvalue | pvalue_adjusted |
|---|---|---|---|---|
| interactor_abundance | mitosis | inter | 0.033303639821968566 | 0.15216394815908776 |
| complex_abundance | mitosis | inter | 0.15281576553483447 | 0.29930365505571416 |
| interactor_ratio | mitosis | inter | 0.8238855440719605 | 0.9688776284181868 |
