## Supplementary Data 2 for "SECAT: Quantifying differential protein-protein interaction states by network-centric analysis": hela_string_000153_P02649.pdf

P02649_P02649

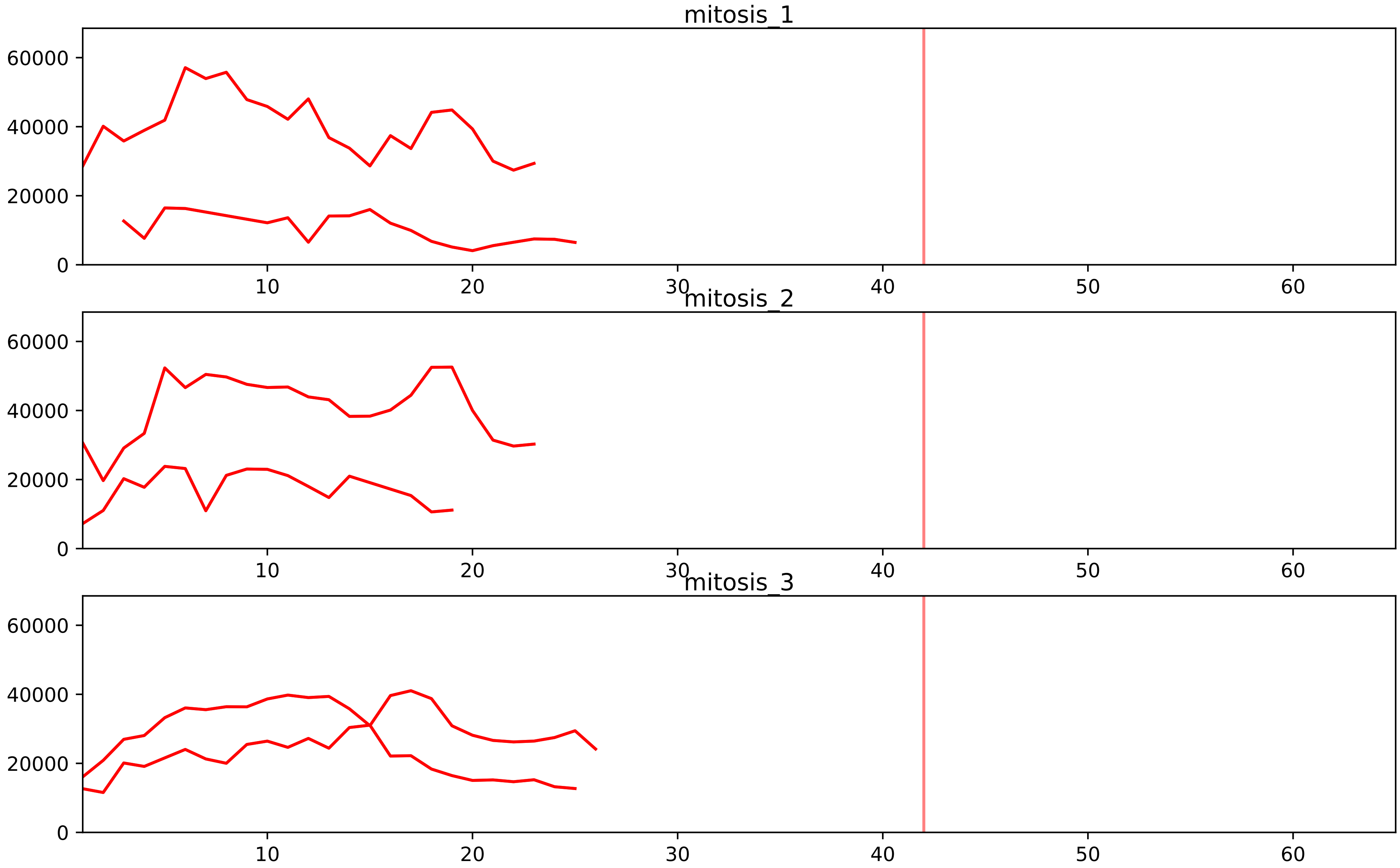


| level | condition_1 | condition_2 | pvalue | pvalue_adjusted |
|---|---|---|---|---|
| assembled_abundance | mitosis | inter | 6.89446540468017e-05 | 0.00849289277876523 |
| total_abundance | mitosis | inter | 0.00012647186935907918 | 0.013925507688281004 |
