## Supplementary Data 2 for "SECAT: Quantifying differential protein-protein interaction states by network-centric analysis": hela_string_000154_Q9NXF1.pdf

Q9NXF1_Q9NXF1

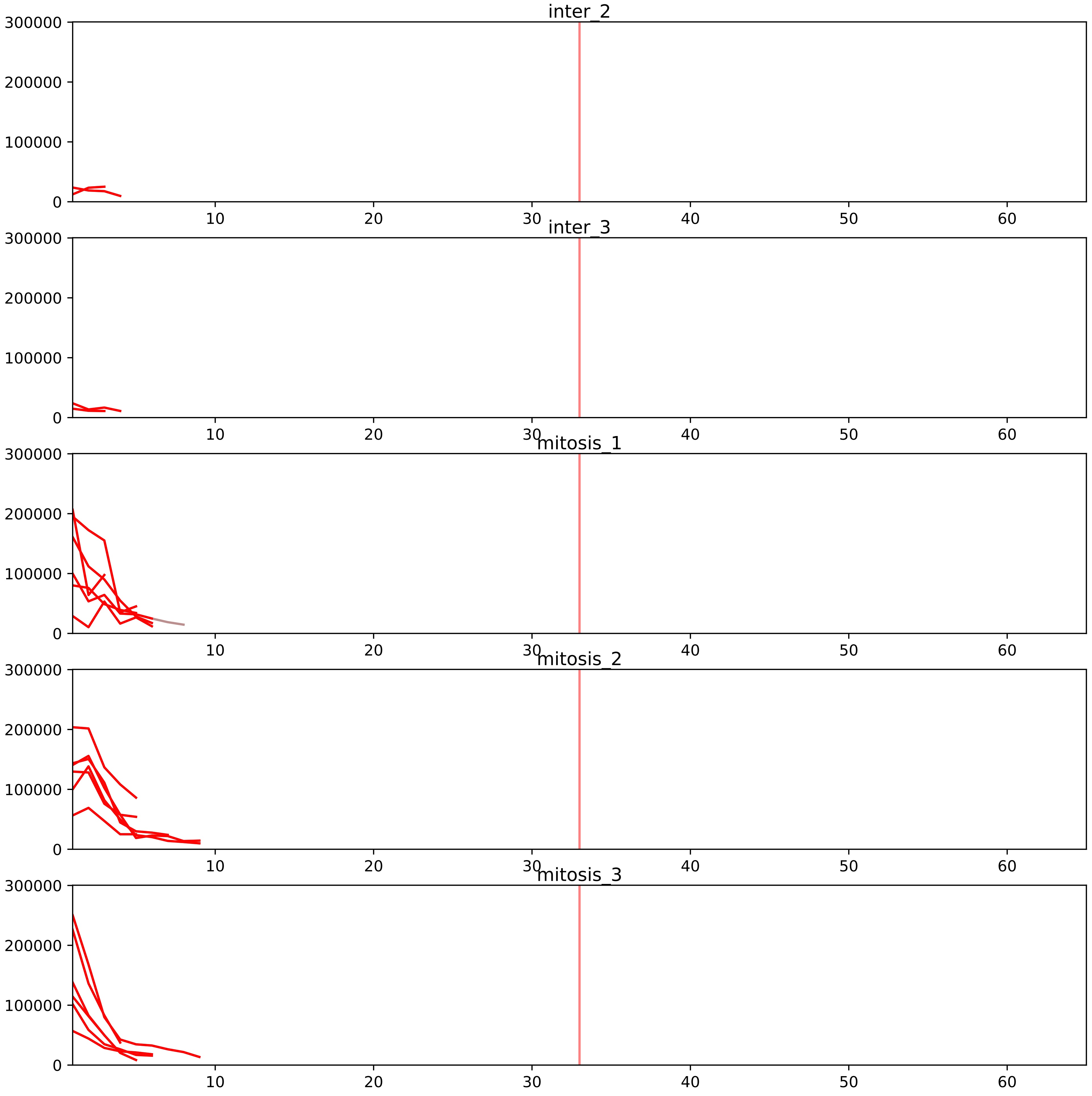


| level | condition_1 | condition_2 | pvalue | pvalue_adjusted |
|---|---|---|---|---|
| interactor_ratio | mitosis | inter | 7.109986776871762e-05 | 0.0012825858499454943 |
| interactor_abundance | mitosis | inter | 0.0001458194249685149 | 0.002948436724638104 |
| assembled_abundance | mitosis | inter | 0.0006795400989689796 | 0.024360801173422066 |
| total_abundance | mitosis | inter | 0.0014004994377979888 | 0.04051103771494069 |
| complex_abundance | mitosis | inter | 0.006062615814497111 | 0.03261758215986749 |

### Q14137_Q9NXF1
BOP1_HUMAN vs TEX10_HUMAN
p-value: 0.049 q-value: 0.032

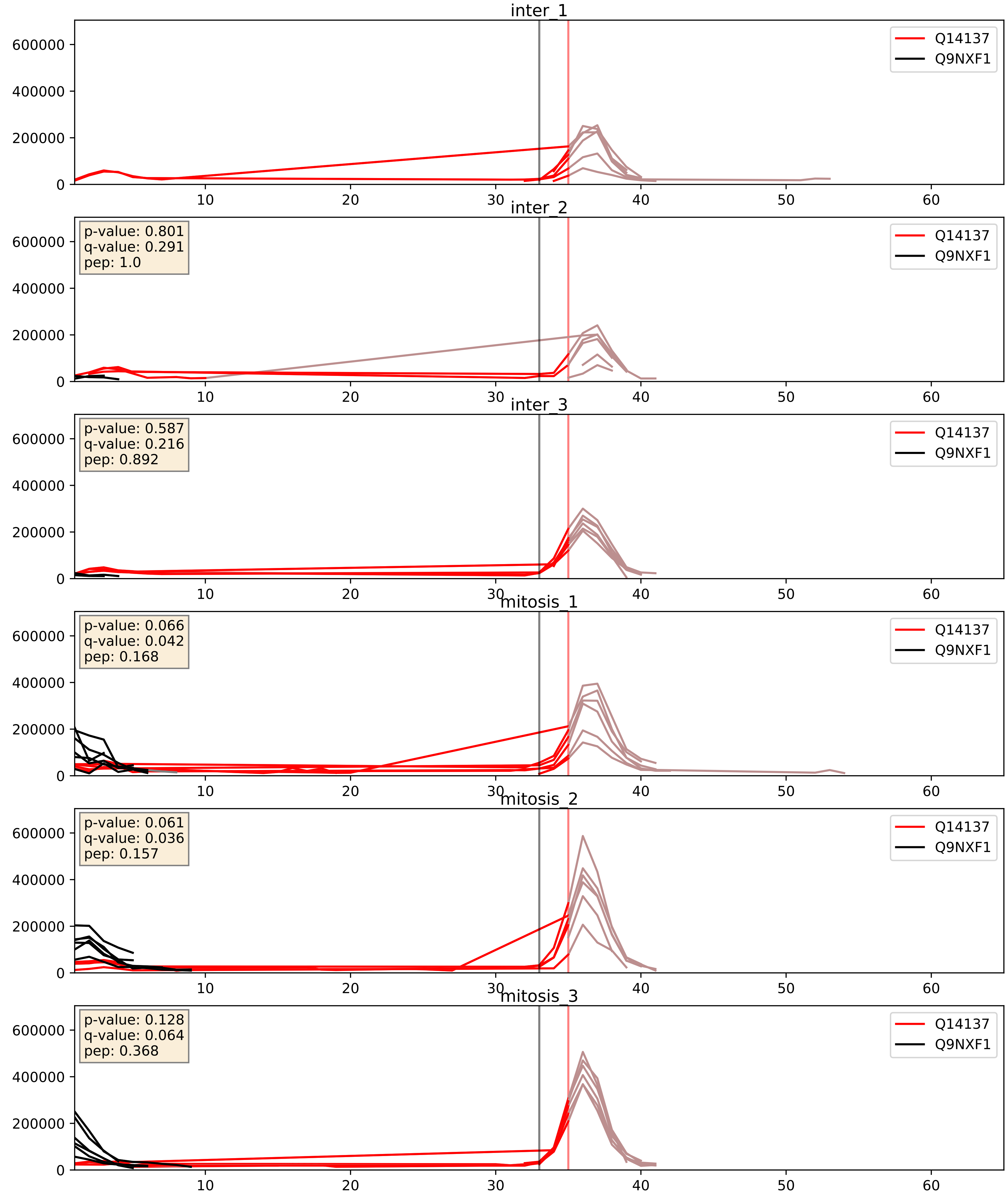


| level | condition_1 | condition_2 | pvalue | pvalue_adjusted |
|---|---|---|---|---|
| interactor_ratio | mitosis | inter | 0.01263103011635328 | 0.16210137600597313 |
| complex_abundance | mitosis | inter | 0.18912949473358198 | 0.34097482622566594 |
| interactor_abundance | mitosis | inter | 0.4858283549734151 | 0.655531323860724 |

Q5SY16_Q9NXF1
NOL9_HUMAN vs TEX10_HUMAN
p-value: 0.065 q-value: 0.04

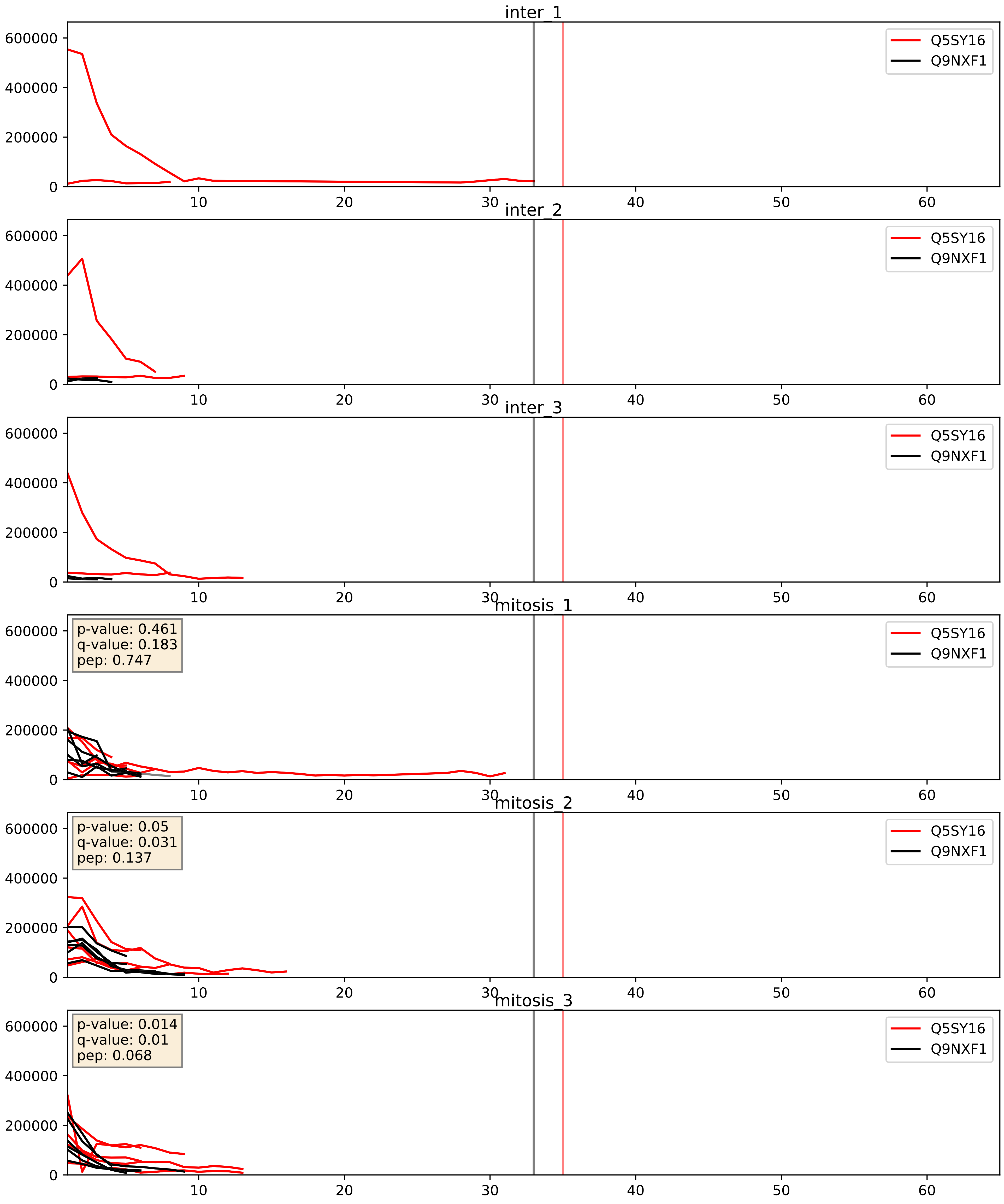


| level | condition_1 | condition_2 | pvalue | pvalue_adjusted |
|---|---|---|---|---|
| complex_abundance | mitosis | inter | 0.029768884749929257 | 0.12558977499230872 |
| interactor_abundance | mitosis | inter | 0.21855537322261576 | 0.413901326279998 |
| interactor_ratio | mitosis | inter | 0.5707862700213564 | 0.8945917155252349 |

Q9BV38_Q9NXF1
WDR18_HUMAN vs TEX10_HUMAN
p-value: 0.005 q-value: 0.005

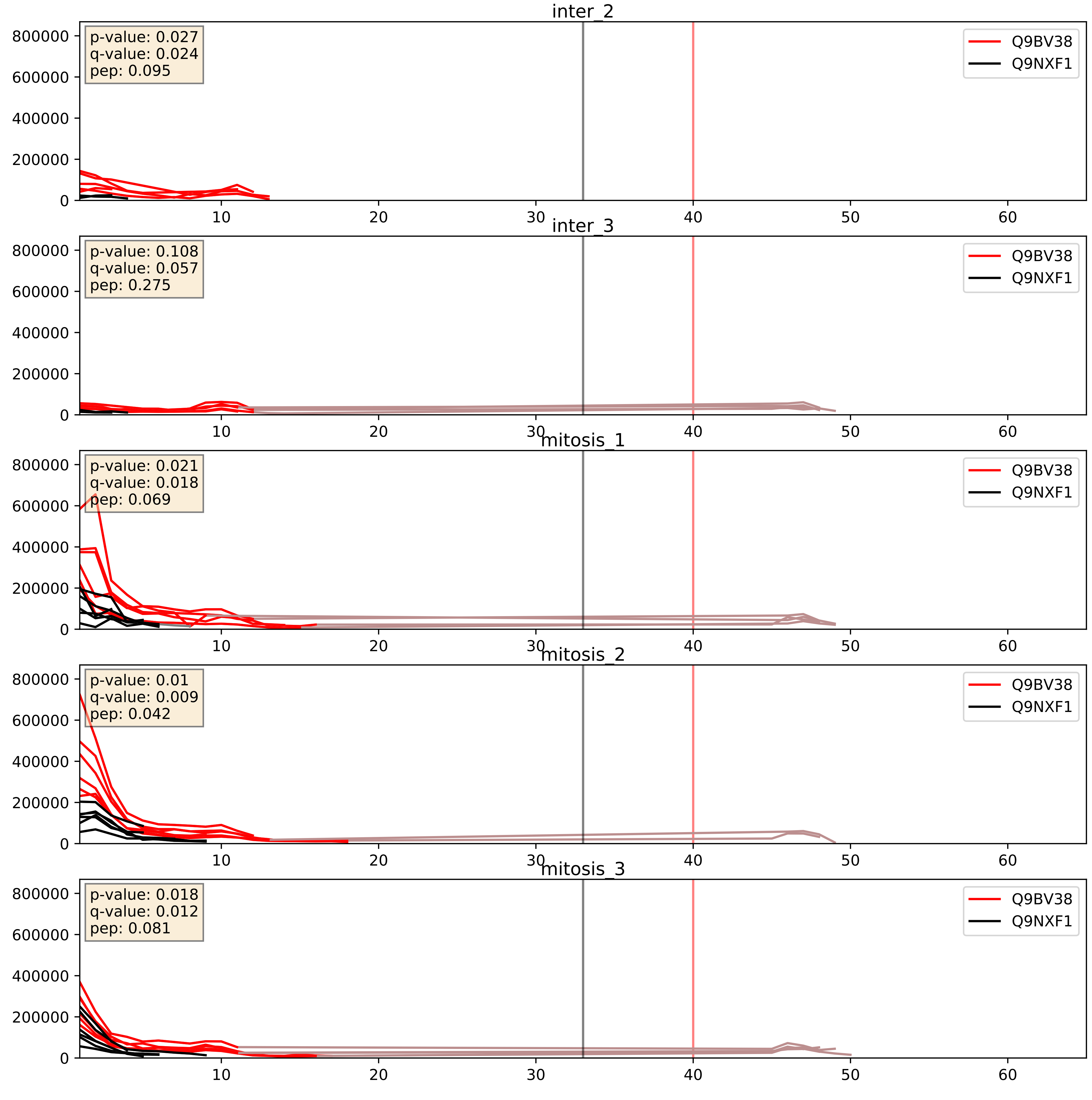


| level | condition_1 | condition_2 | pvalue | pvalue_adjusted |
|---|---|---|---|---|
| complex_abundance | mitosis | inter | 0.0064532229320805835 | 0.05955783692156859 |
| interactor_abundance | mitosis | inter | 0.015148920743567303 | 0.10774803619853436 |
| interactor_ratio | mitosis | inter | 0.10491269377910745 | 0.5400196384540948 |

Q9GZL7_Q9NXF1
WDR12_HUMAN vs TEX10_HUMAN
p-value: 0.06 q-value: 0.037

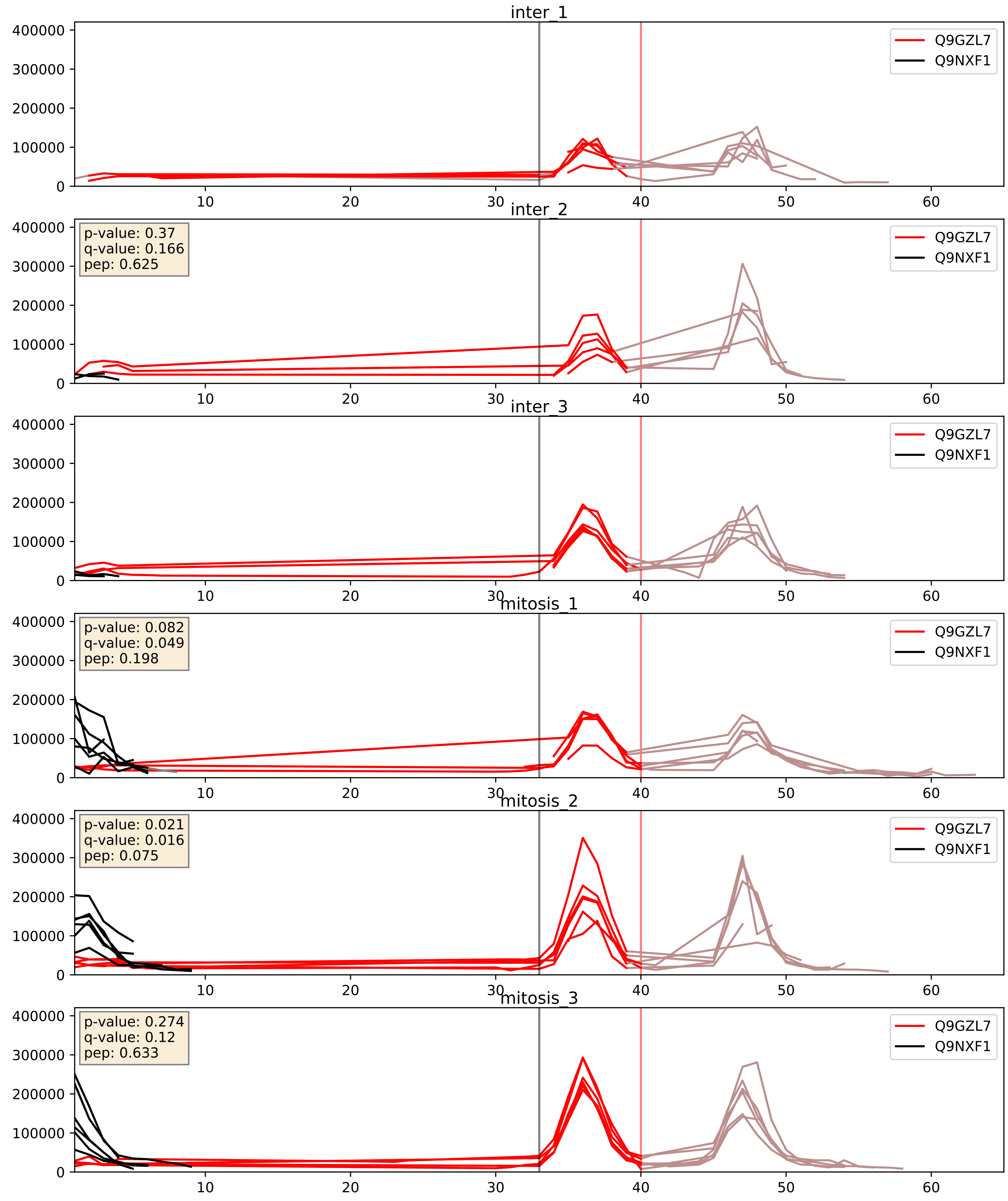


| level | condition_1 | condition_2 | pvalue | pvalue_adjusted |
|---|---|---|---|---|
| interactor_ratio | mitosis | inter | 0.002940307259930933 | 0.07007095913365735 |
| interactor_abundance | mitosis | inter | 0.1368111058888913 | 0.32360609311684 |
| complex_abundance | mitosis | inter | 0.14649069181537597 | 0.2923014462256658 |

Q9H4L4_Q9NXF1
SENP3_HUMAN vs TEX10_HUMAN
p-value: 0.01 q-value: 0.009

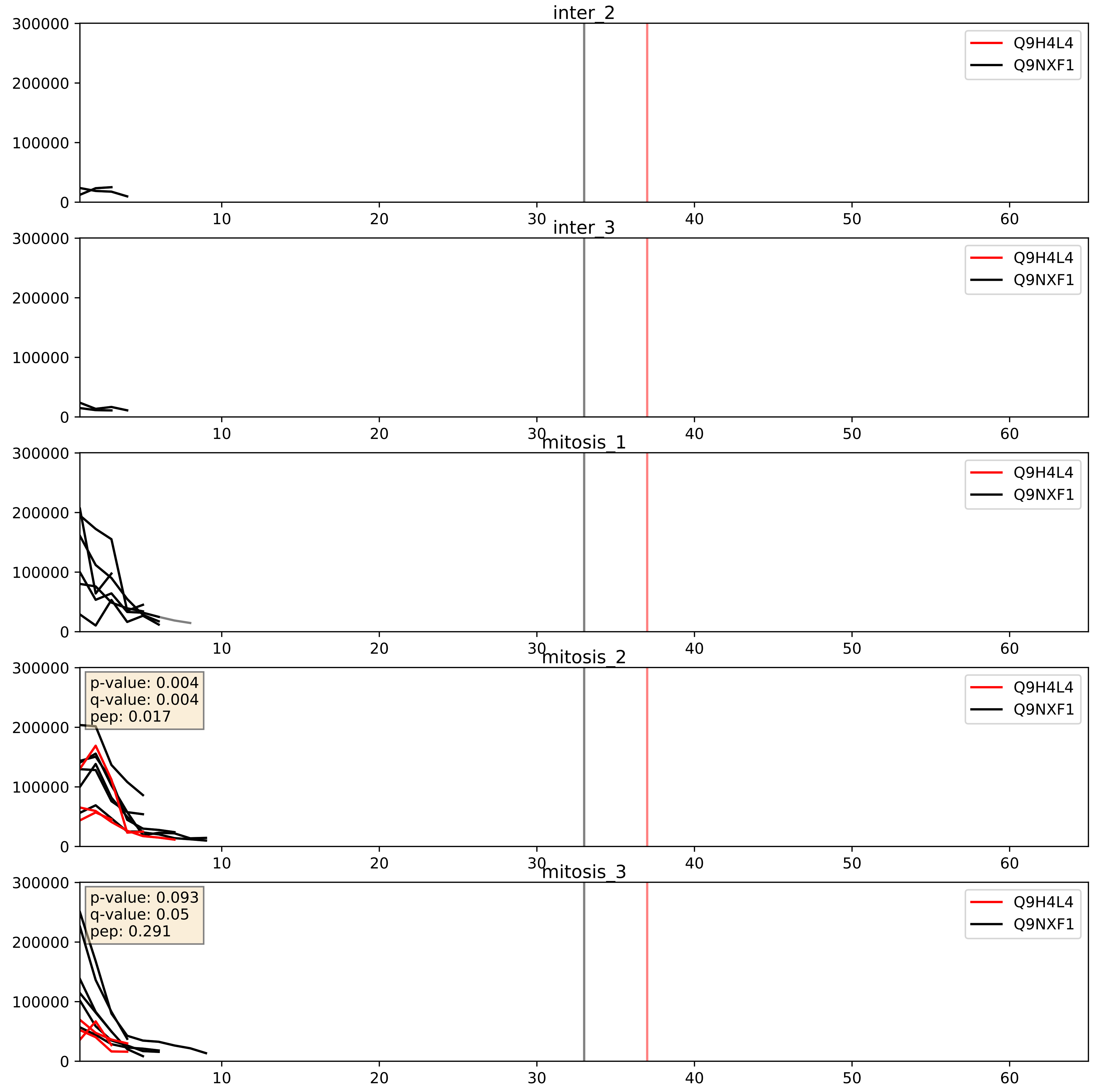


| level | condition_1 | condition_2 | pvalue | pvalue_adjusted |
|---|---|---|---|---|
| interactor_ratio | mitosis | inter | 0.12097775337355715 | 0.5703368335321749 |
| complex_abundance | mitosis | inter | 0.35005400115369945 | 0.49766853510640546 |
| interactor_abundance | mitosis | inter | 0.6644427812888789 | 0.779127425730521 |

Q9NXF1_Q9Y4W2
TEX10_HUMAN vs LAS1L_HUMAN
p-value: 0.069 q-value: 0.042

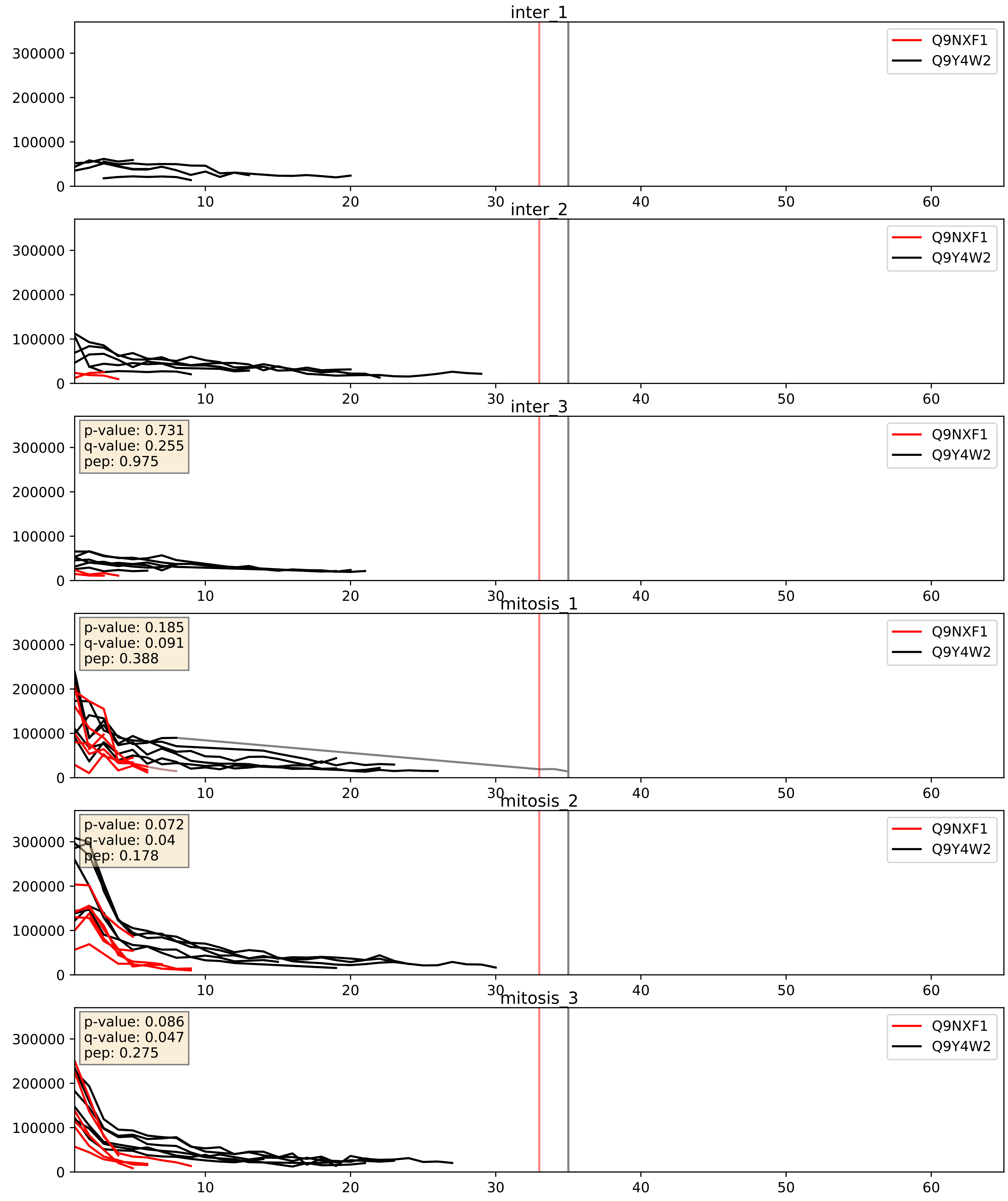


| level | condition_1 | condition_2 | pvalue | pvalue_adjusted |
|---|---|---|---|---|
| interactor_abundance | mitosis | inter | 0.0019024202547968964 | 0.03196215383917848 |
| complex_abundance | mitosis | inter | 0.009889564976144676 | 0.07371585975104354 |
| interactor_ratio | mitosis | inter | 0.690715792753423 | 0.9349347226390419 |

Q9NXF1_Q9Y5J1
TEX10_HUMAN vs UTP18_HUMAN
p-value: 0.036 q-value: 0.025

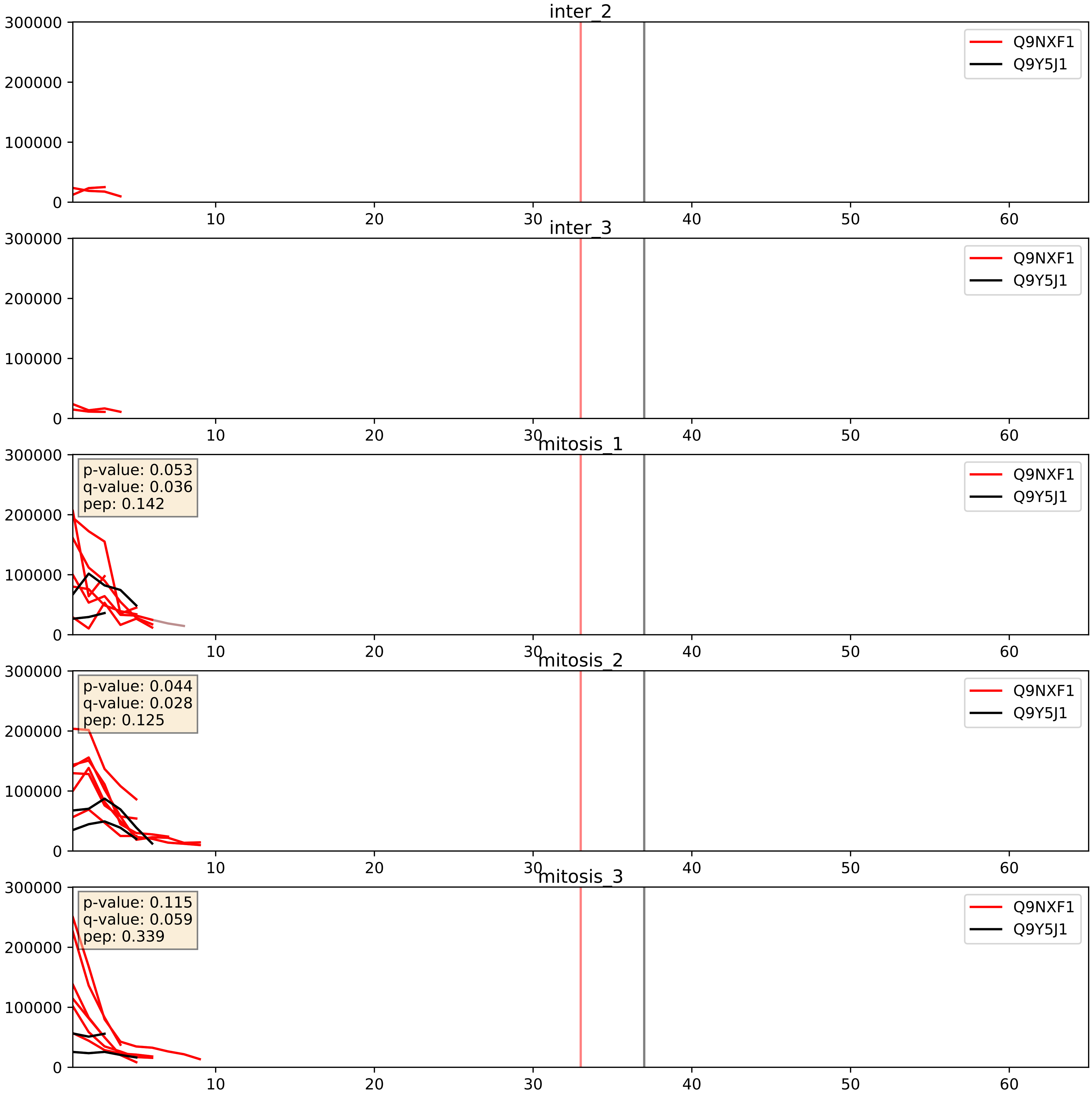


| level | condition_1 | condition_2 | pvalue | pvalue_adjusted |
| --- | --- | --- | --- | --- |
| interactor_ratio | mitosis | inter | 0.0019222246008891858 | 0.05317578467954594 |
| interactor_abundance | mitosis | inter | 0.0022253892890756657 | 0.03466841143289118 |
| complex_abundance | mitosis | inter | 0.011308378166680228 | 0.07877001668010823 |
