## Supplementary Data 2 for "SECAT: Quantifying differential protein-protein interaction states by network-centric analysis": hela_string_000155_P20248.pdf

P20248_P20248

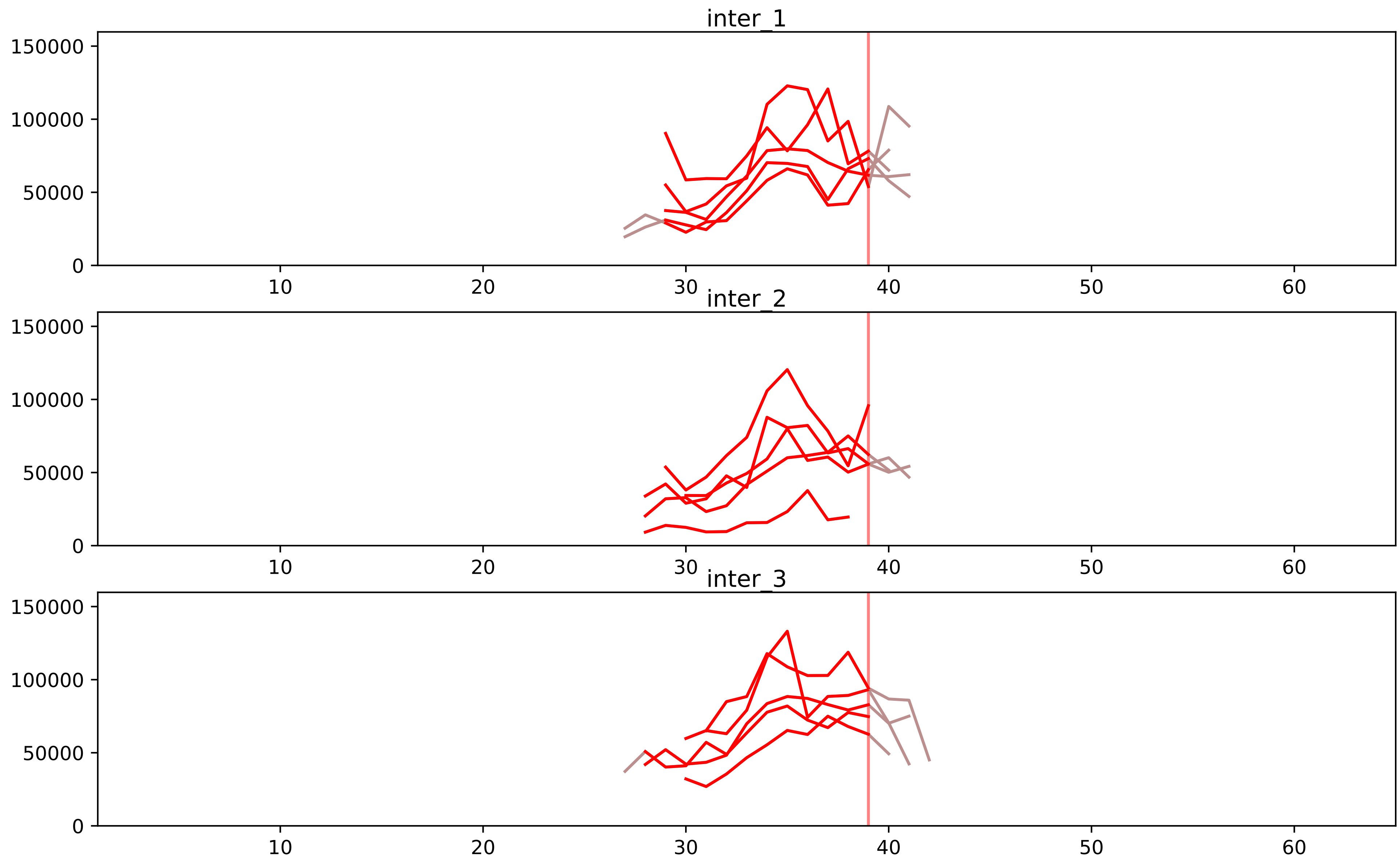


| level | condition_1 | condition_2 | pvalue | pvalue_adjusted |
|---|---|---|---|---|
| complex_abundance | mitosis | inter | 7.177079227126583e-05 | 0.0018865465397018446 |
| interactor_abundance | mitosis | inter | 0.00016653337796498116 | 0.003225488583742793 |
| assembled_abundance | mitosis | inter | 0.0006883819956320218 | 0.024360801173422066 |
| total_abundance | mitosis | inter | 0.0008853631535674589 | 0.032841764788842195 |
| monomer_abundance | mitosis | inter | 0.001754110319167246 | 0.09123294054534994 |
| interactor_ratio | mitosis | inter | 0.023170476536805575 | 0.11712548579044577 |

P11802_P20248
CDK4_HUMAN vs CCNA2_HUMAN
p-value: 0.083 q-value: 0.049

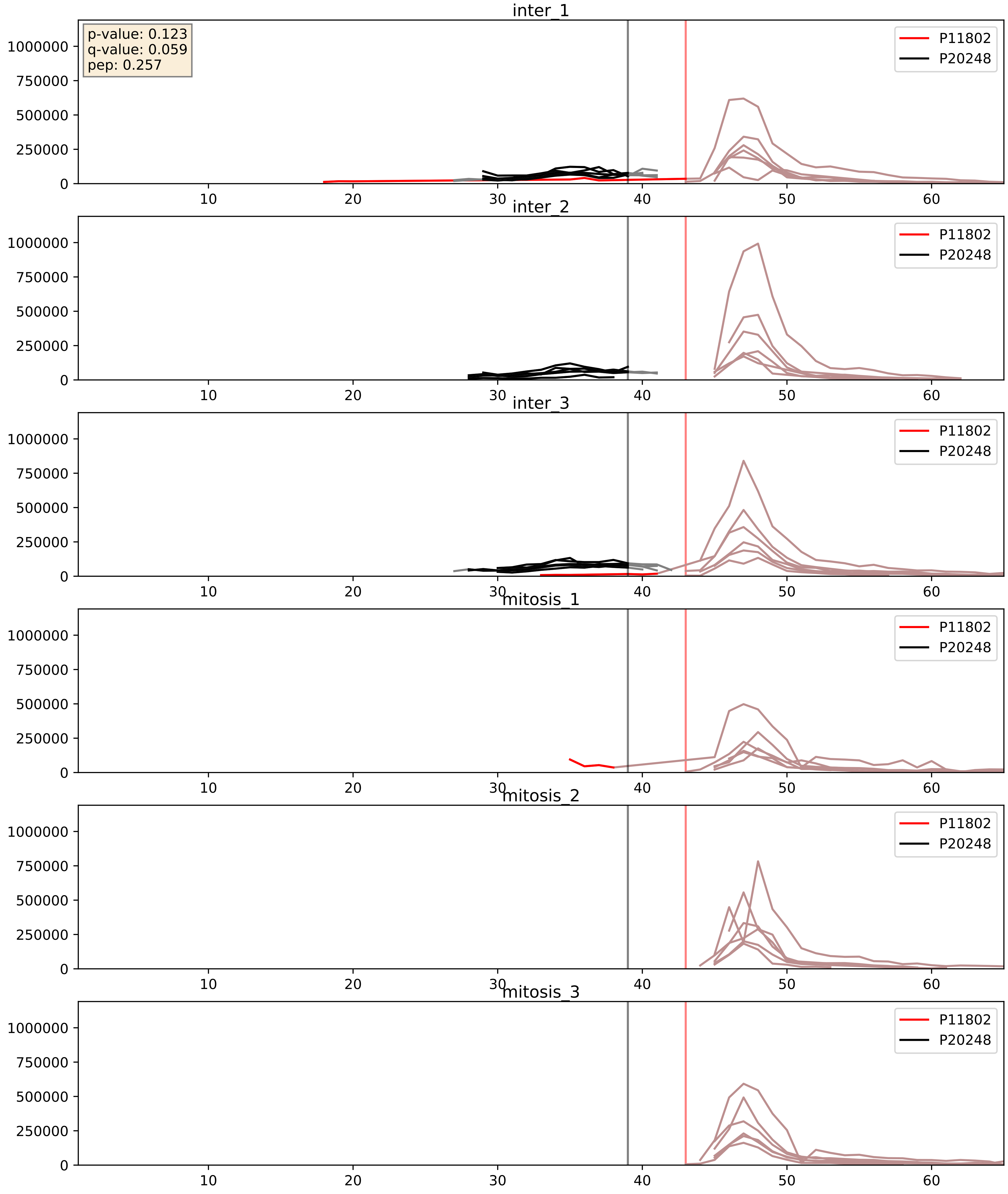


| level | condition_1 | condition_2 | pvalue | pvalue_adjusted |
|---|---|---|---|---|
| complex_abundance | mitosis | inter | 0.022550980606221196 | 0.10990318464300756 |
| interactor_abundance | mitosis | inter | 0.02832171390921066 | 0.13974940313044457 |
| interactor_ratio | mitosis | inter | 0.05061179134837742 | 0.372196678644425 |

P20248_P24941
CCNA2_HUMAN vs CDK2_HUMAN
p-value: 0.008 q-value: 0.007

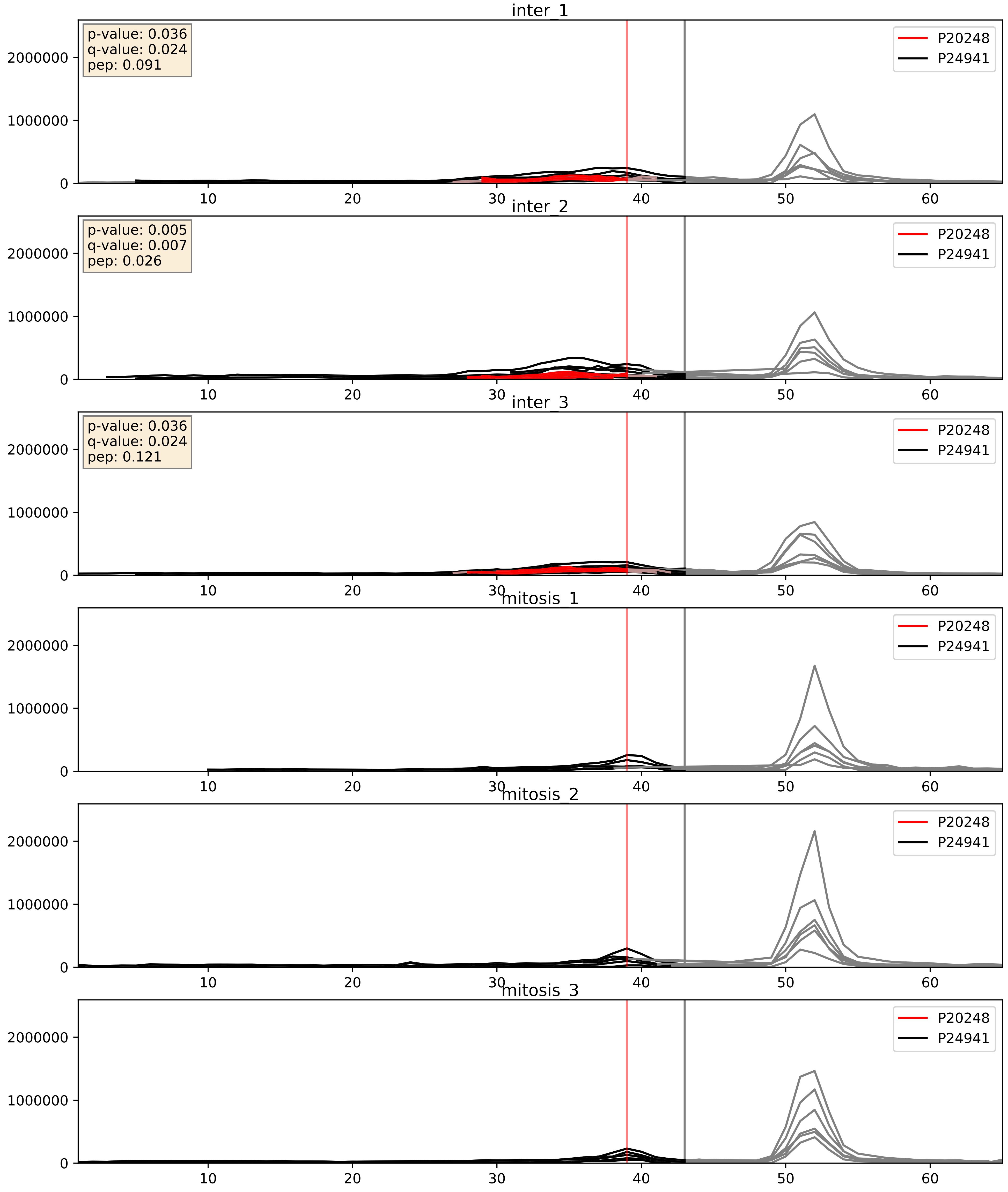


| level | condition_1 | condition_2 | pvalue | pvalue_adjusted |
|---|---|---|---|---|
| complex_abundance | mitosis | inter | 0.00017174227842104968 | 0.008803077265174762 |
| interactor_abundance | mitosis | inter | 0.0004058346268662555 | 0.013951583959739548 |
| interactor_ratio | mitosis | inter | 0.01412332765988096 | 0.17445264757370998 |

P20248_Q13309
CCNA2_HUMAN vs SKP2_HUMAN
p-value: 0.003 q-value: 0.003

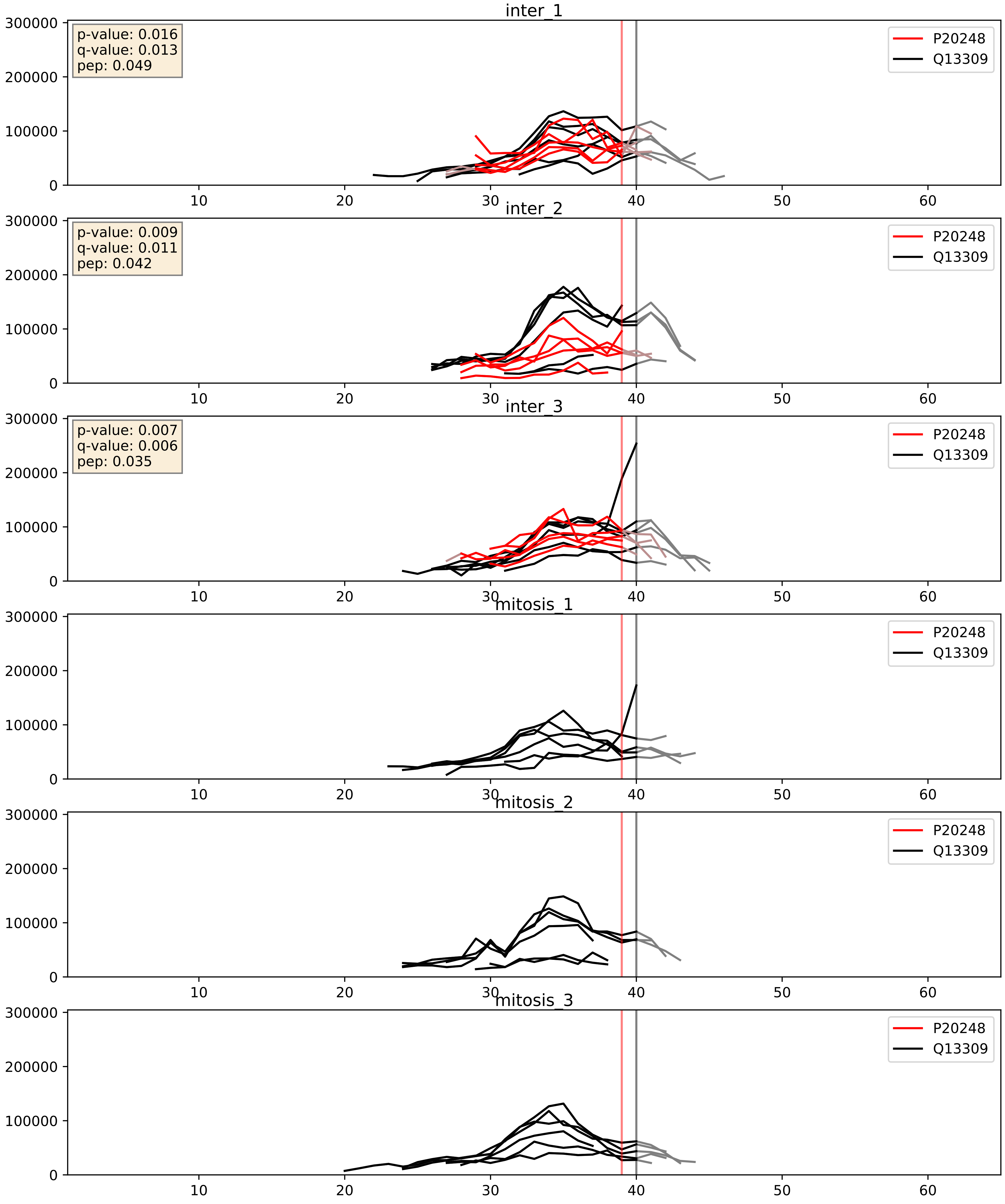


| level | condition_1 | condition_2 | pvalue | pvalue_adjusted |
|---|---|---|---|---|
| complex_abundance | mitosis | inter | 0.0002650544596844505 | 0.010908010456244694 |
| interactor_abundance | mitosis | inter | 0.0004058346268662555 | 0.013951583959739548 |
| interactor_ratio | mitosis | inter | 0.22591075557383503 | 0.7051563050171857 |
