## Supplementary Data 2 for "SECAT: Quantifying differential protein-protein interaction states by network-centric analysis": hela_string_000156_Q9P0J0.pdf

Q9P0J0_Q9P0J0

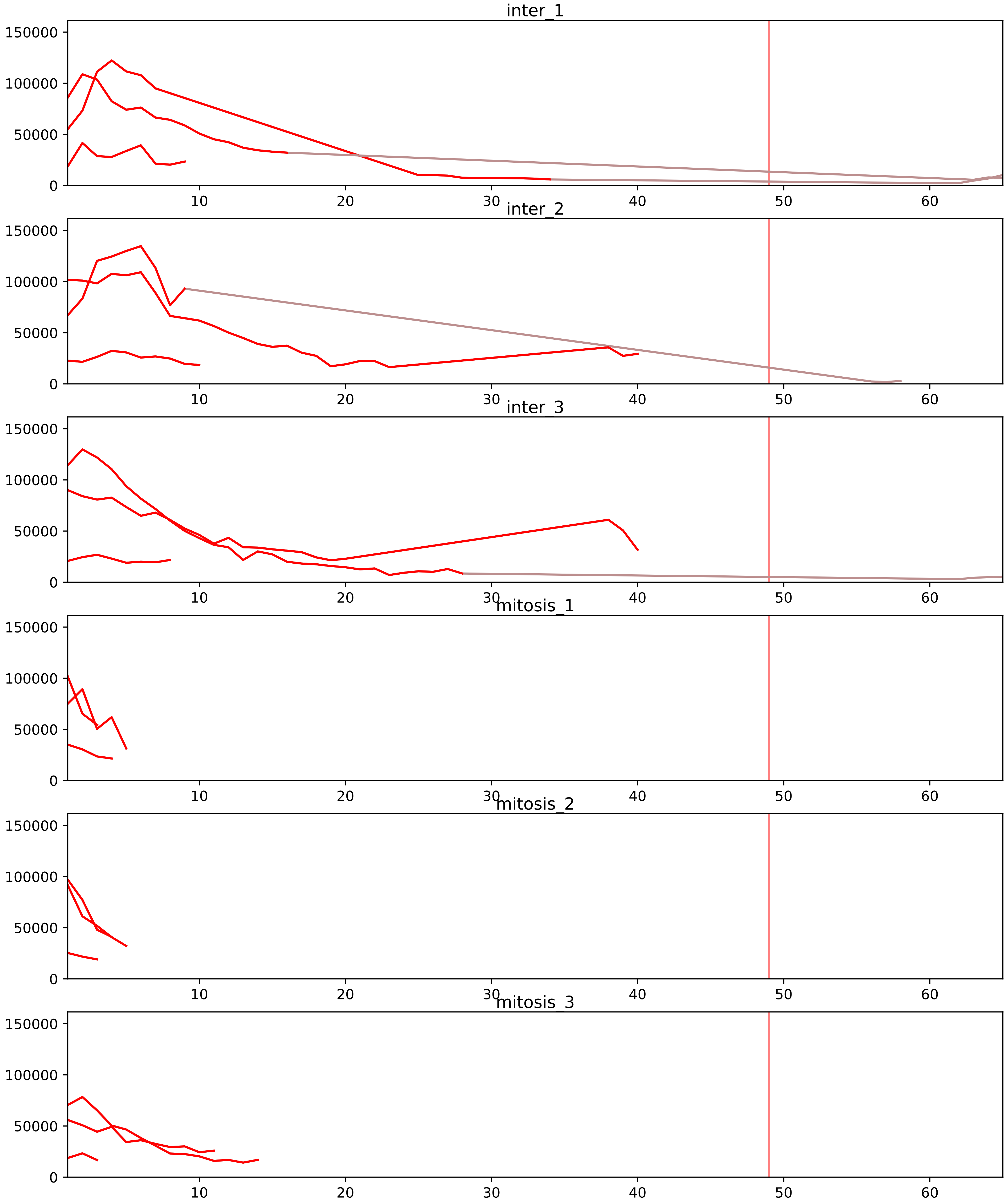


| level | condition_1 | condition_2 | pvalue | pvalue_adjusted |
|---|---|---|---|---|
| interactor_abundance | mitosis | inter | 7.213334625195255e-05 | 0.0018479549585806118 |
| complex_abundance | mitosis | inter | 0.00042691891111223214 | 0.006234371400369105 |
| interactor_ratio | mitosis | inter | 0.006694620170762521 | 0.04761458131729075 |
| monomer_abundance | mitosis | inter | 0.05616507758026643 | 0.3478638875131441 |
| assembled_abundance | mitosis | inter | 0.7333802655497785 | 0.8914445658370587 |
| total_abundance | mitosis | inter | 0.7496668818411386 | 0.896680762009549 |

O00217_Q9P0J0
NDUS8_HUMAN vs NDUAD_HUMAN
p-value: 0.07 q-value: 0.043

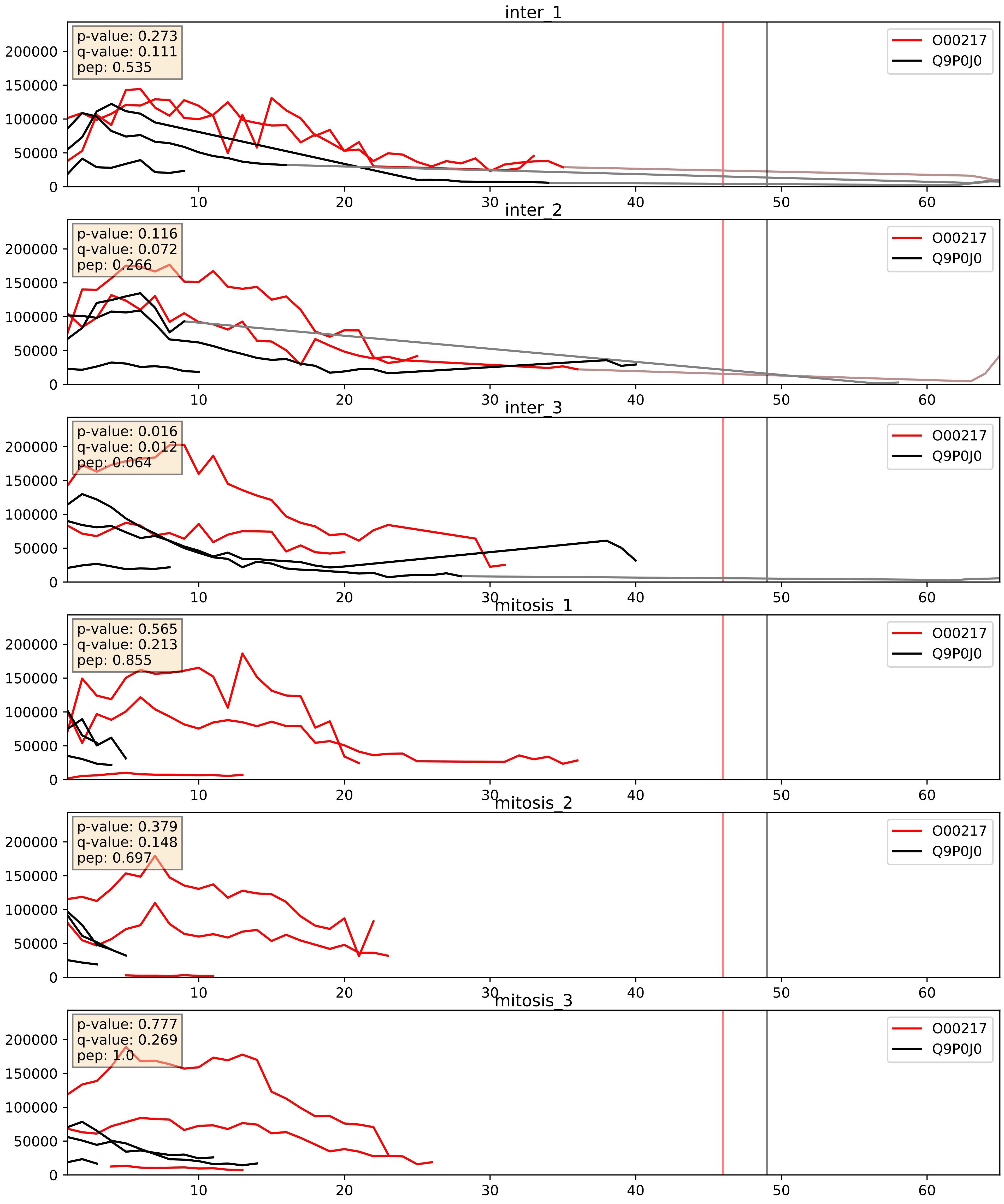


| level | condition_1 | condition_2 | pvalue | pvalue_adjusted |
|---|---|---|---|---|
| complex_abundance | mitosis | inter | 0.01305971628033173 | 0.08453223568196803 |
| interactor_abundance | mitosis | inter | 0.05594699255729353 | 0.19207567336447973 |
| interactor_ratio | mitosis | inter | 0.3419898548484073 | 0.7944187673005065 |

O75251_Q9P0J0
NDUS7_HUMAN vs NDUAD_HUMAN
p-value: 0.059 q-value: 0.037

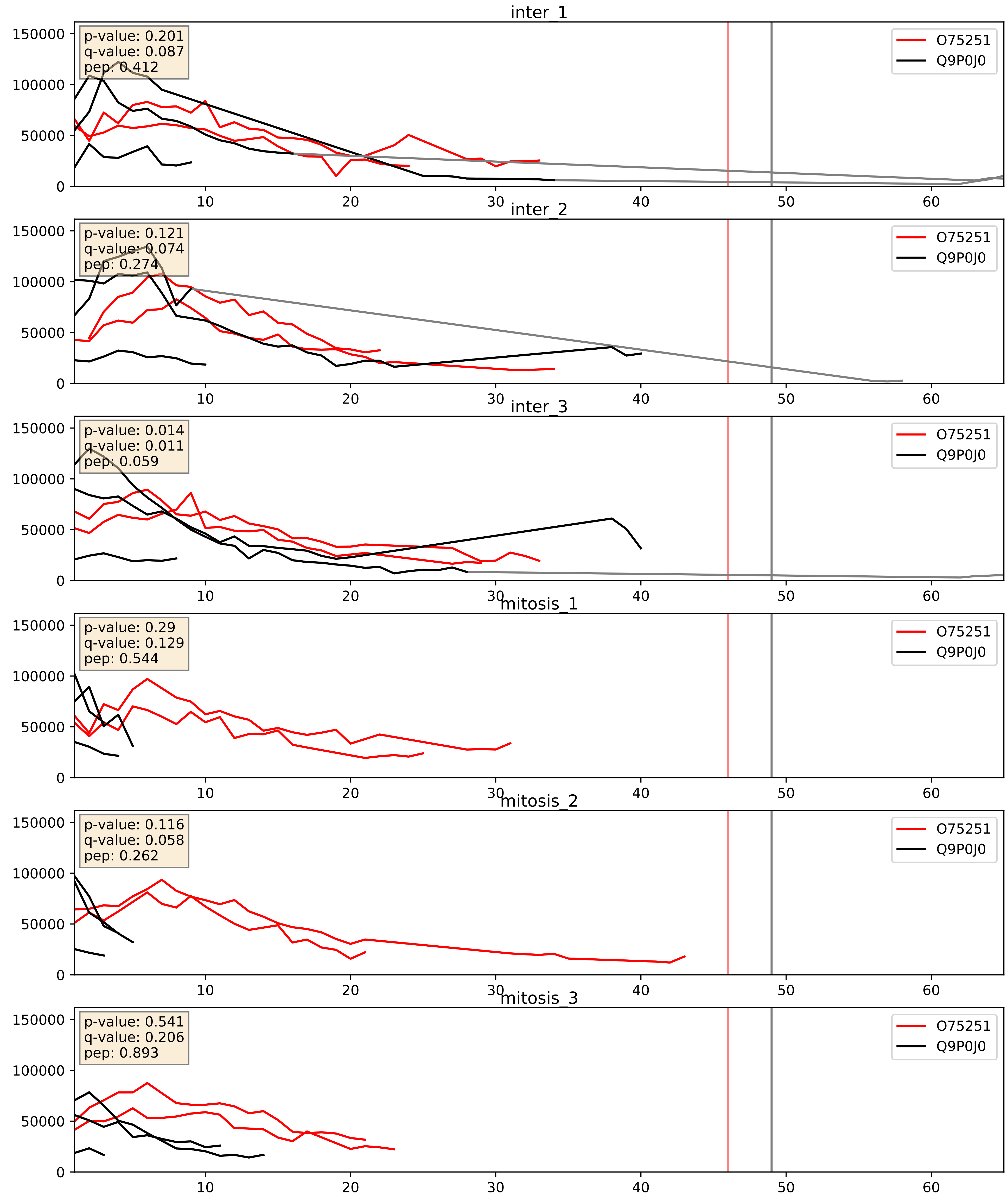


| level | condition_1 | condition_2 | pvalue | pvalue_adjusted |
|---|---|---|---|---|
| complex_abundance | mitosis | inter | 0.00560161555307883 | 0.05641156368747622 |
| interactor_abundance | mitosis | inter | 0.023219329977434956 | 0.1254449714090221 |
| interactor_ratio | mitosis | inter | 0.087277751184758 | 0.4946978435260244 |

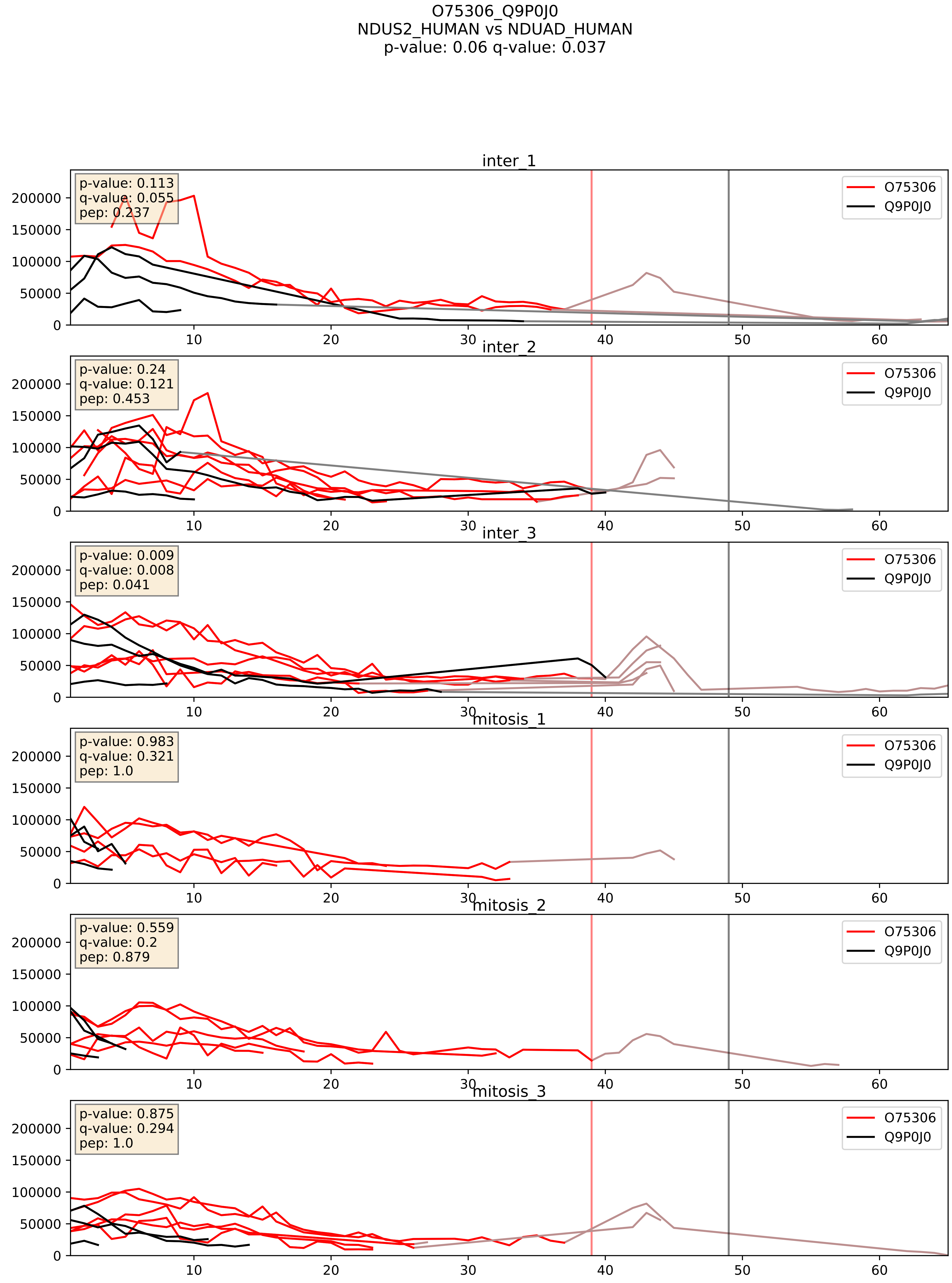


| level | condition_1 | condition_2 | pvalue | pvalue_adjusted |
|---|---|---|---|---|
| complex_abundance | mitosis | inter | 0.03899334396121441 | 0.14411521647412365 |
| interactor_abundance | mitosis | inter | 0.19132788109129 | 0.38330984314428235 |
| interactor_ratio | mitosis | inter | 0.8352367458594616 | 0.9728867369194887 |

O75380_Q9P0J0
NDUS6_HUMAN vs NDUAD_HUMAN
p-value: 0.0 q-value: 0.0

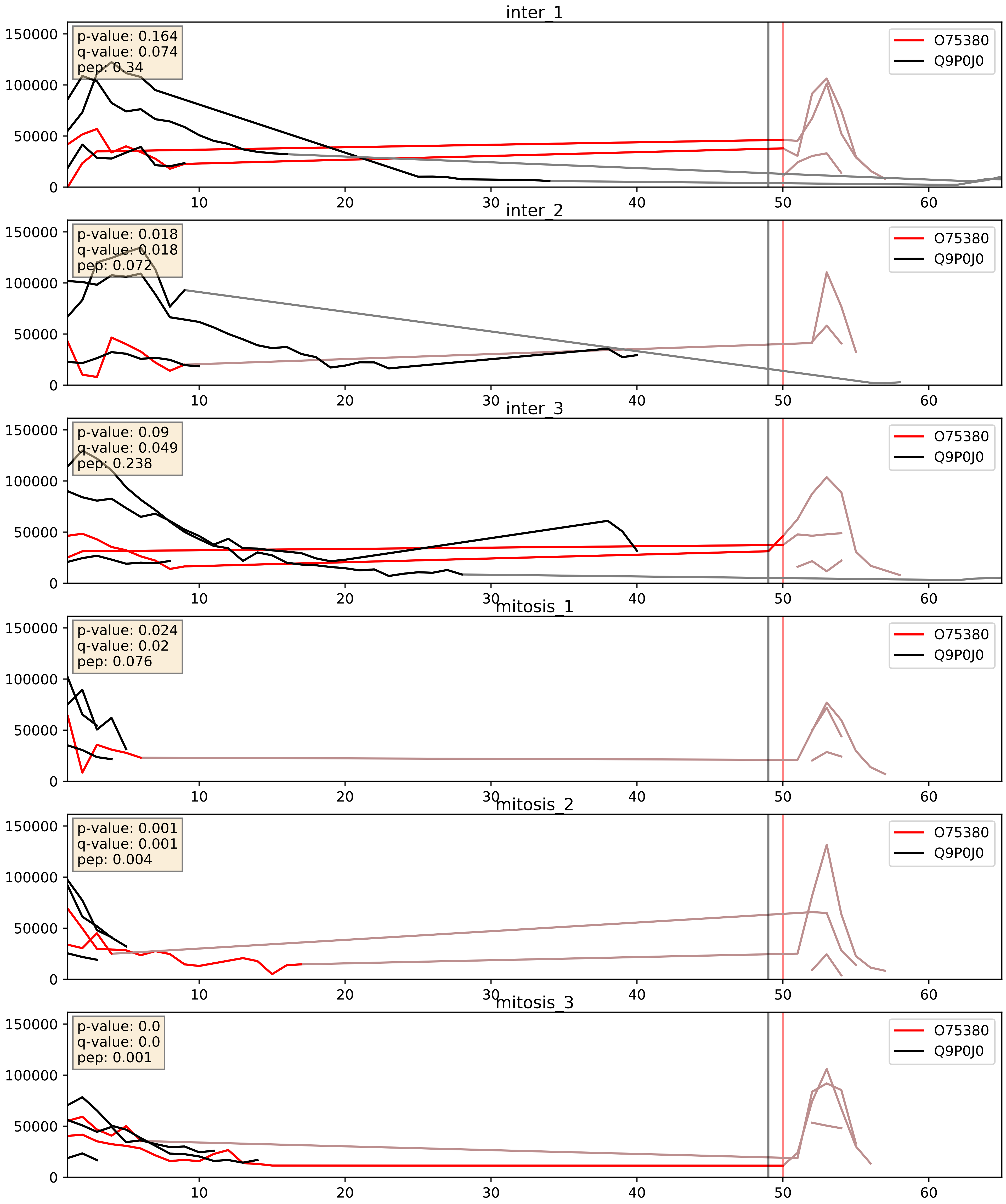


| level | condition_1 | condition_2 | pvalue | pvalue_adjusted |
|---|---|---|---|---|
| complex_abundance | mitosis | inter | 0.011510431423233639 | 0.07958747413802902 |
| interactor_ratio | mitosis | inter | 0.03836879199944975 | 0.3214392061698628 |
| interactor_abundance | mitosis | inter | 0.04518880429095441 | 0.1691739185351278 |

O95168_Q9P0J0
NDUB4_HUMAN vs NDUAD_HUMAN
p-value: 0.004 q-value: 0.004

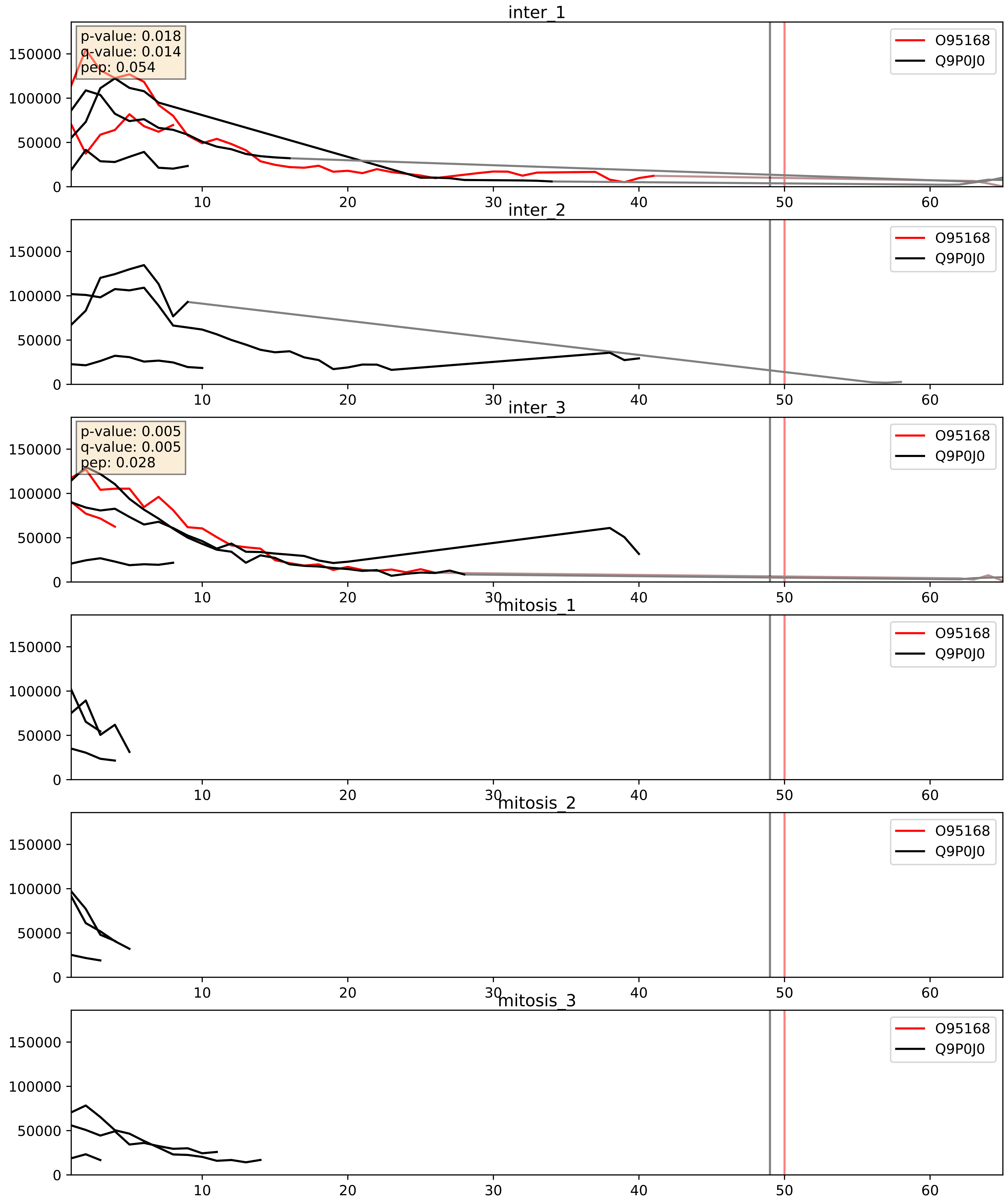


| level | condition_1 | condition_2 | pvalue | pvalue_adjusted |
|---|---|---|---|---|
| interactor_ratio | mitosis | inter | 0.03262410456690737 | 0.28849414782306515 |
| complex_abundance | mitosis | inter | 0.03587008157578323 | 0.1376167508298101 |
| interactor_abundance | mitosis | inter | 0.03857806571915839 | 0.1597234546824647 |

O95169_Q9P0J0
NDUB8_HUMAN vs NDUAD_HUMAN
p-value: 0.046 q-value: 0.031

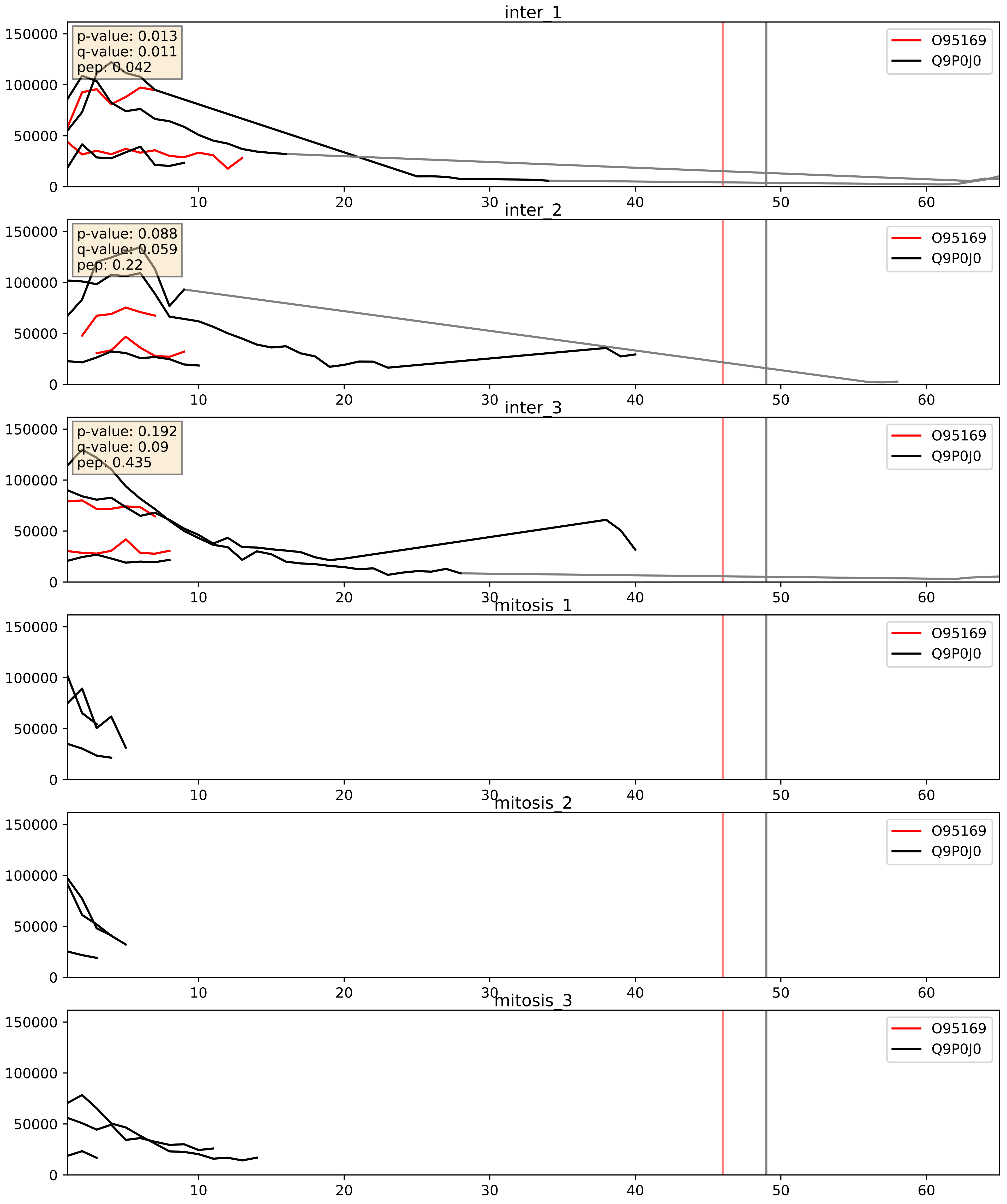


| level | condition_1 | condition_2 | pvalue | pvalue_adjusted |
|---|---|---|---|---|
| interactor_ratio | mitosis | inter | 0.000806772542671012 | 0.032604652001472285 |
| complex_abundance | mitosis | inter | 0.0009089003714998096 | 0.0197847007881119 |
| interactor_abundance | mitosis | inter | 0.0013596111197408754 | 0.02728785740910174 |

O95299_Q9P0J0
NDUAA_HUMAN vs NDUAD_HUMAN
p-value: 0.007 q-value: 0.006

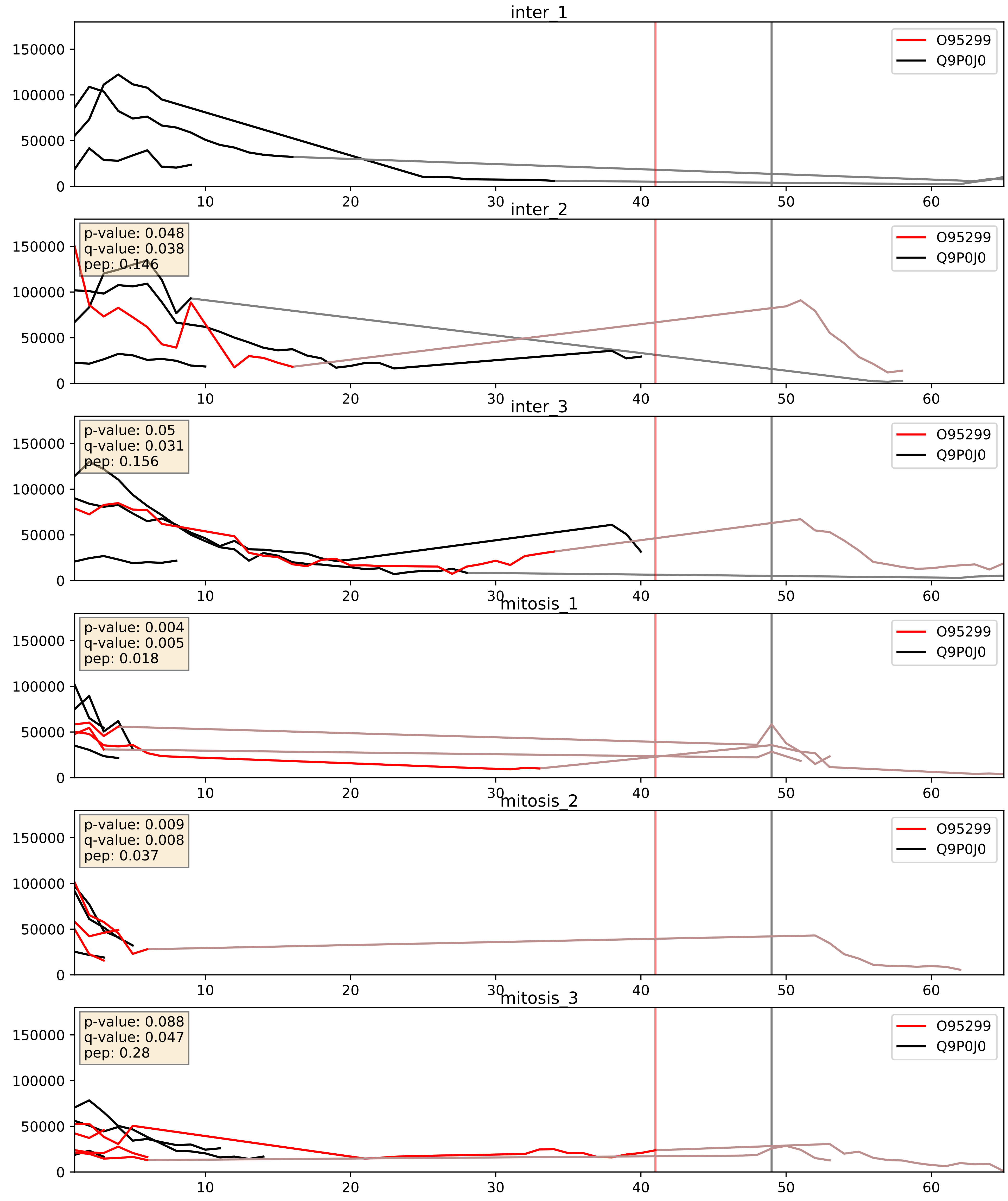


| level | condition_1 | condition_2 | pvalue | pvalue_adjusted |
|---|---|---|---|---|
| complex_abundance | mitosis | inter | 0.2647762425660163 | 0.41468590131852623 |
| interactor_ratio | mitosis | inter | 0.2839044596985366 | 0.7544930689287406 |
| interactor_abundance | mitosis | inter | 0.3232823409943367 | 0.5222300130046277 |

P10606_Q9P0J0
COX5B_HUMAN vs NDUAD_HUMAN
p-value: 0.0 q-value: 0.017

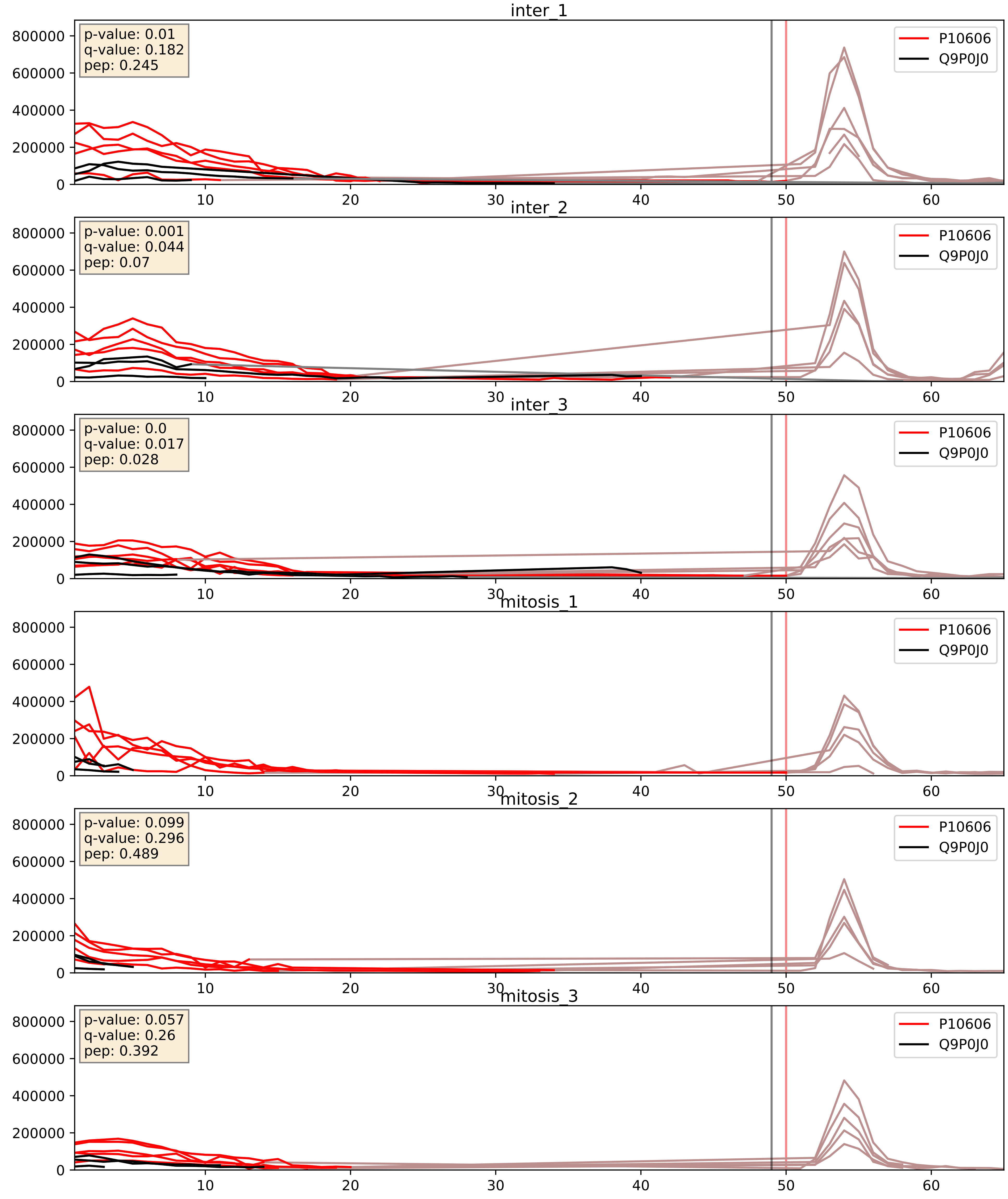


| level | condition_1 | condition_2 | pvalue | pvalue_adjusted |
|---|---|---|---|---|
| complex_abundance | mitosis | inter | 0.007931969085931577 | 0.0659424994487145 |
| interactor_abundance | mitosis | inter | 0.019670182719559937 | 0.1190514126751731 |
| interactor_ratio | mitosis | inter | 0.7523020484092622 | 0.9503276416012604 |

P17568_Q9P0J0
NDUB7_HUMAN vs NDUAD_HUMAN
p-value: 0.0 q-value: 0.0

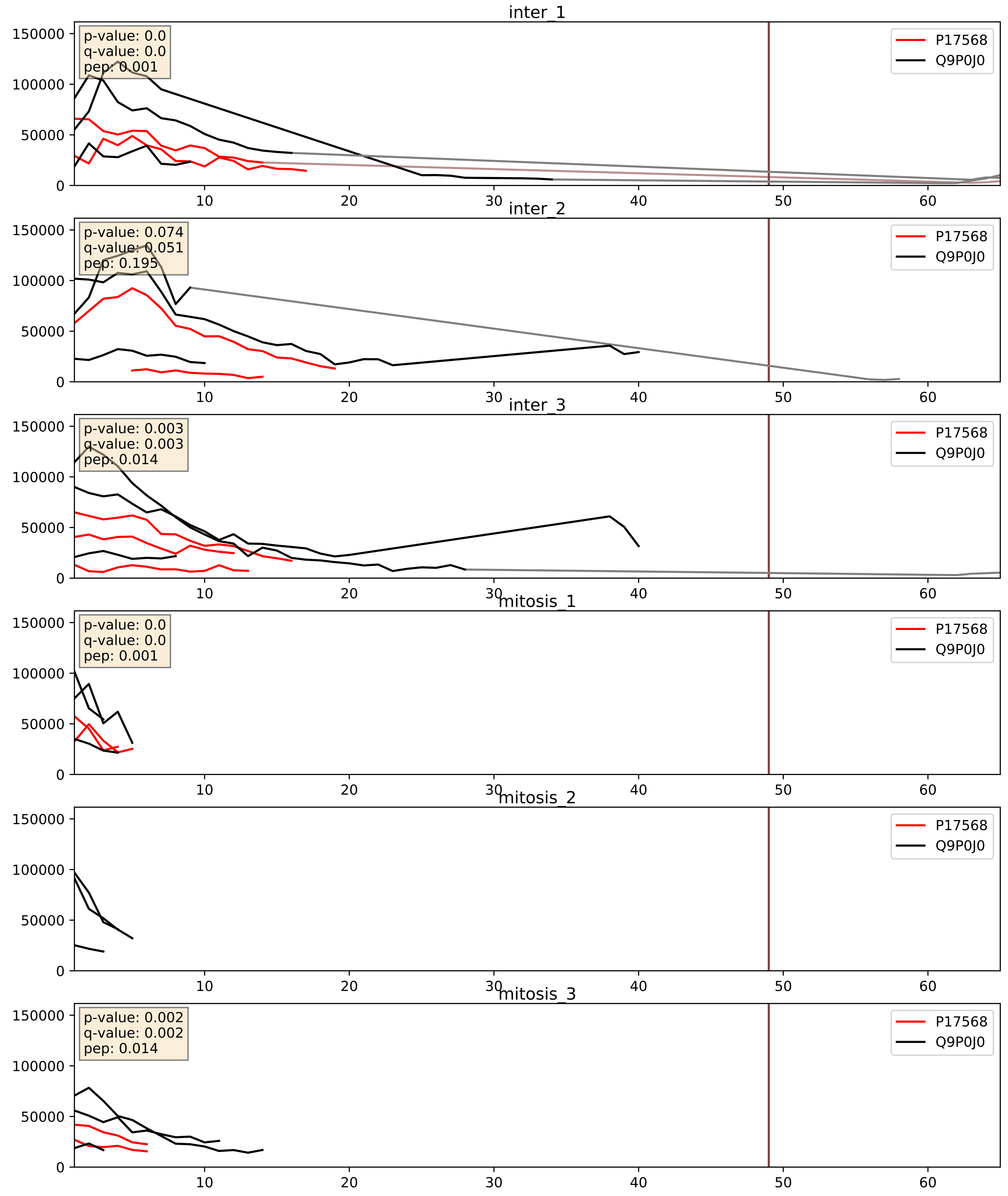


| level | condition_1 | condition_2 | pvalue | pvalue_adjusted |
|---|---|---|---|---|
| interactor_abundance | mitosis | inter | 0.023596815251976218 | 0.12607602513772825 |
| complex_abundance | mitosis | inter | 0.027154627244607016 | 0.12025018583230009 |
| interactor_ratio | mitosis | inter | 0.1461391374999046 | 0.5999765069540447 |

P19404_Q9P0J0
NDUV2_HUMAN vs NDUAD_HUMAN
p-value: 0.052 q-value: 0.034

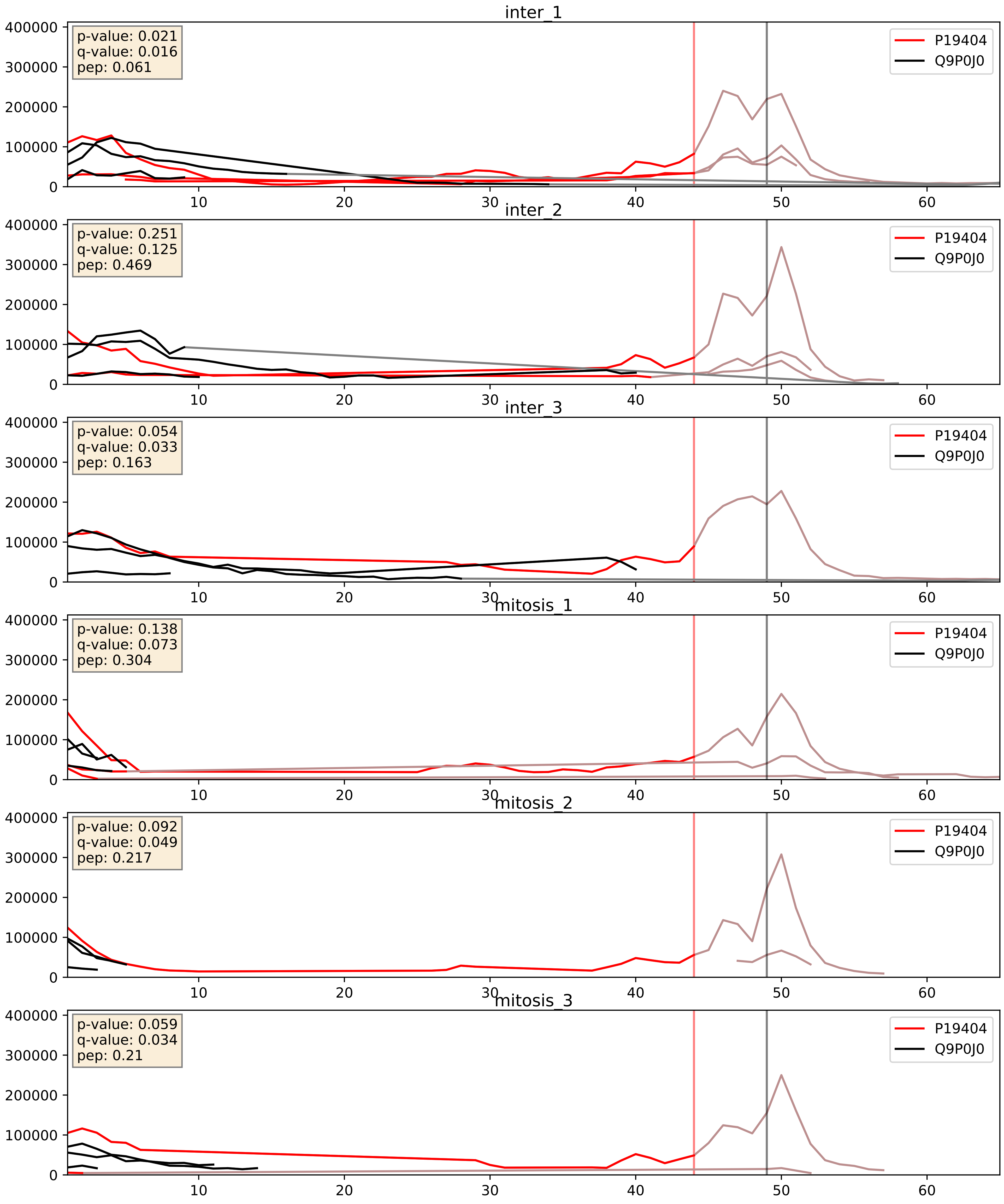


| level | condition_1 | condition_2 | pvalue | pvalue_adjusted |
|---|---|---|---|---|
| complex_abundance | mitosis | inter | 0.021119423117817886 | 0.10728917619496801 |
| interactor_abundance | mitosis | inter | 0.06298937153801733 | 0.2057063105514646 |
| interactor_ratio | mitosis | inter | 0.41989765797674794 | 0.8388785723072948 |

P28331_Q9P0J0
NDUS1_HUMAN vs NDUAD_HUMAN
p-value: 0.01 q-value: 0.009

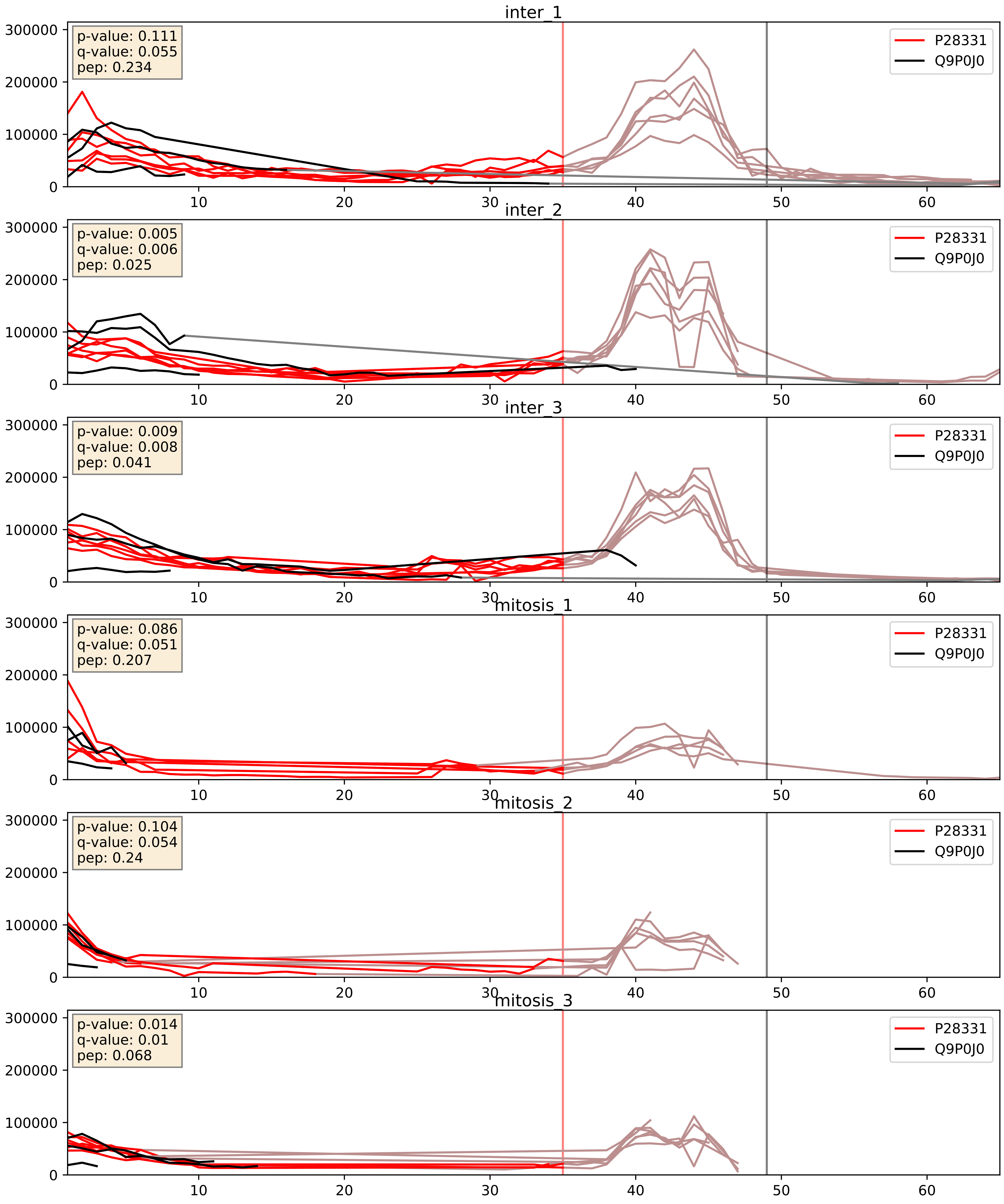


| level | condition_1 | condition_2 | pvalue | pvalue_adjusted |
|---|---|---|---|---|
| interactor_abundance | mitosis | inter | 0.0013232095430149022 | 0.026872298192663256 |
| complex_abundance | mitosis | inter | 0.0016453851528469353 | 0.02839616312171324 |
| interactor_ratio | mitosis | inter | 0.9164984746886587 | 0.9855669004419936 |

P47985_Q9P0J0
UCRI_HUMAN vs NDUAD_HUMAN
p-value: 0.03 q-value: 0.022

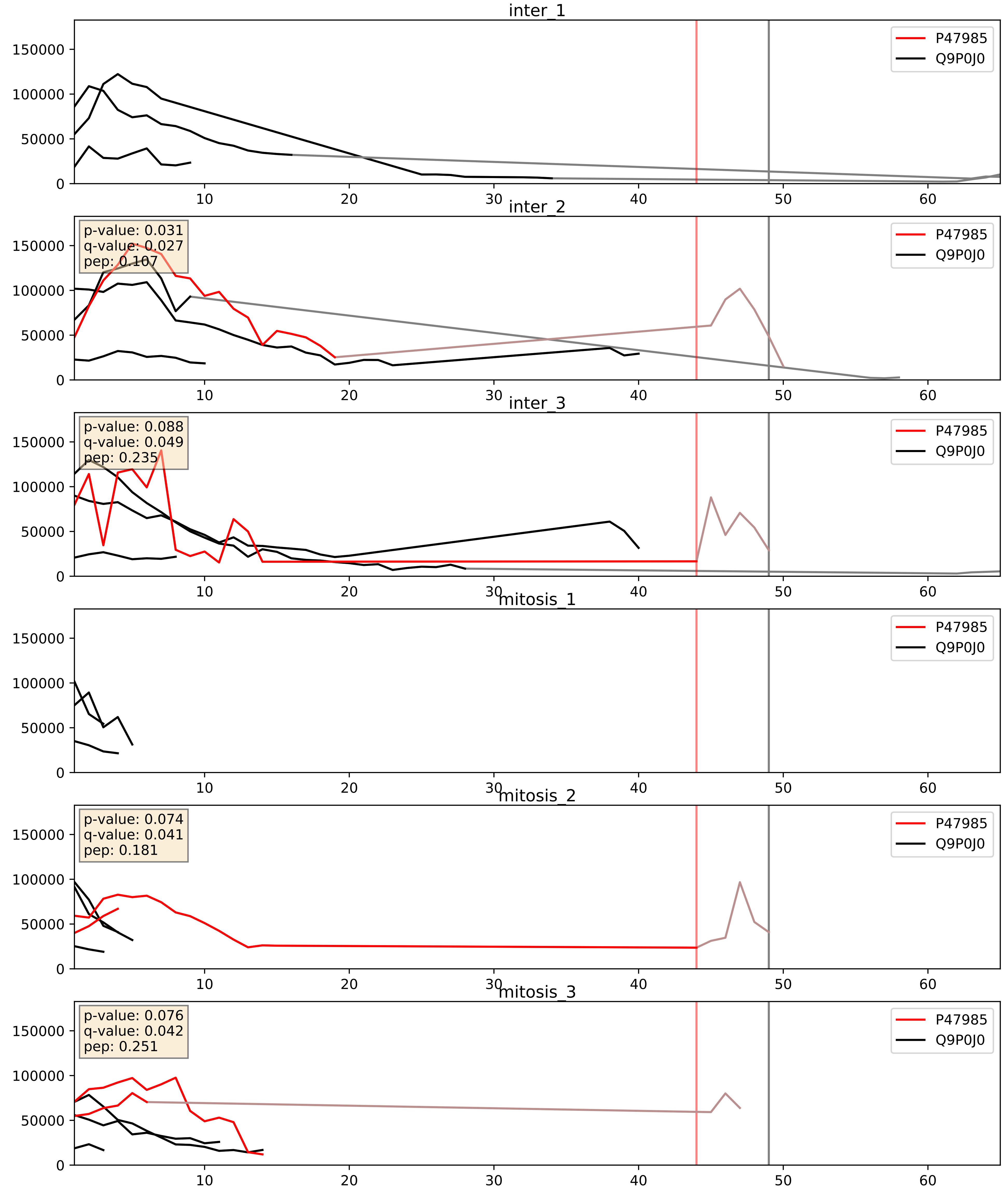


| level | condition_1 | condition_2 | pvalue | pvalue_adjusted |
|---|---|---|---|---|
| interactor_ratio | mitosis | inter | 0.06197779484707071 | 0.4131852989804714 |
| complex_abundance | mitosis | inter | 0.2715453067673854 | 0.4207870792774836 |
| interactor_abundance | mitosis | inter | 0.6173108539886365 | 0.747718522088818 |

P49821_Q9P0J0
NDUV1_HUMAN vs NDUAD_HUMAN
p-value: 0.007 q-value: 0.007

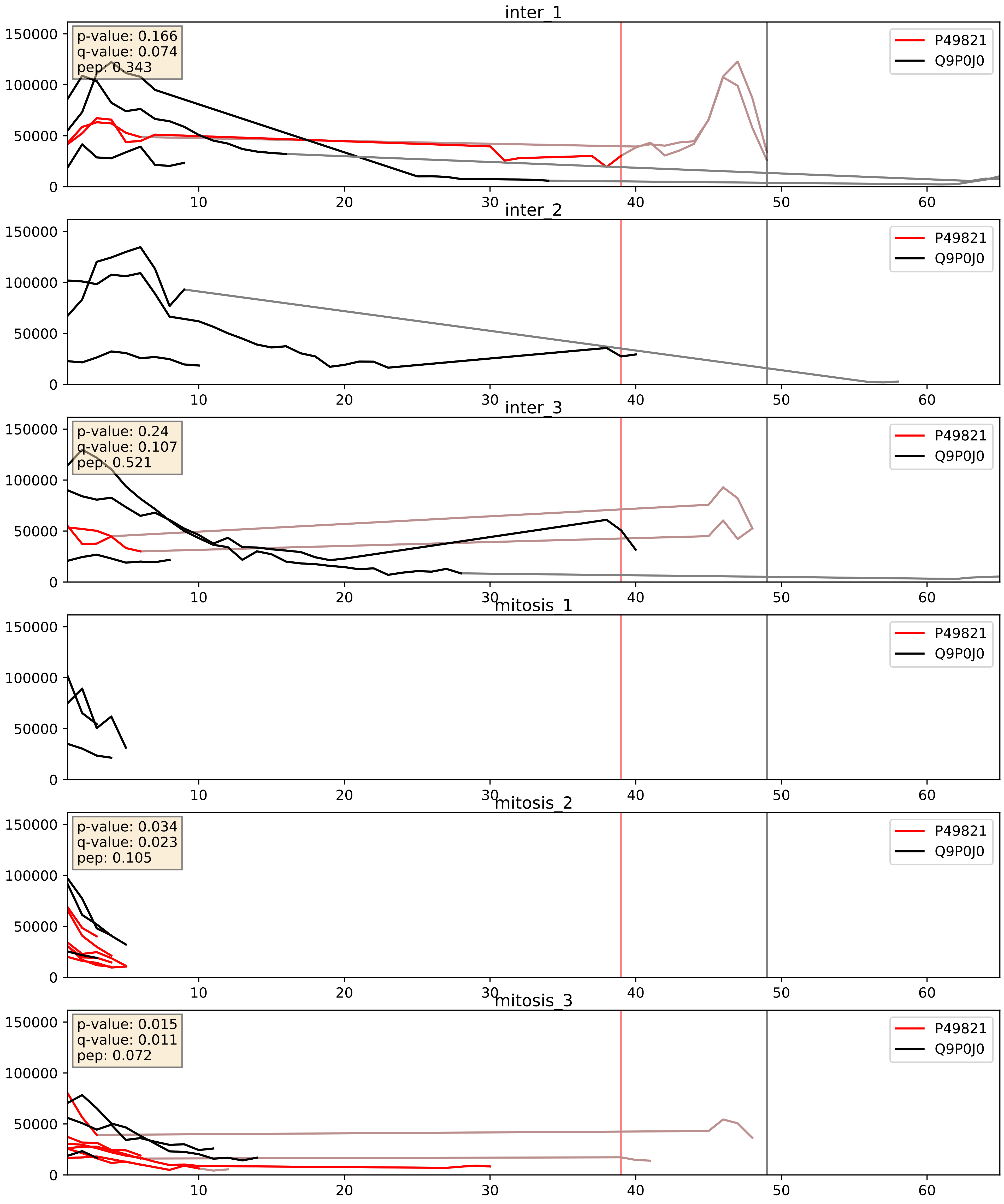


| level | condition_1 | condition_2 | pvalue | pvalue_adjusted |
|---|---|---|---|---|
| interactor_abundance | mitosis | inter | 0.17522303609267548 | 0.36556165648983 |
| complex_abundance | mitosis | inter | 0.18476430623631043 | 0.3364154646219575 |
| interactor_ratio | mitosis | inter | 0.3071287050006226 | 0.7739981394021083 |

P56556_Q9P0J0
NDUA6_HUMAN vs NDUAD_HUMAN
p-value: 0.048 q-value: 0.031

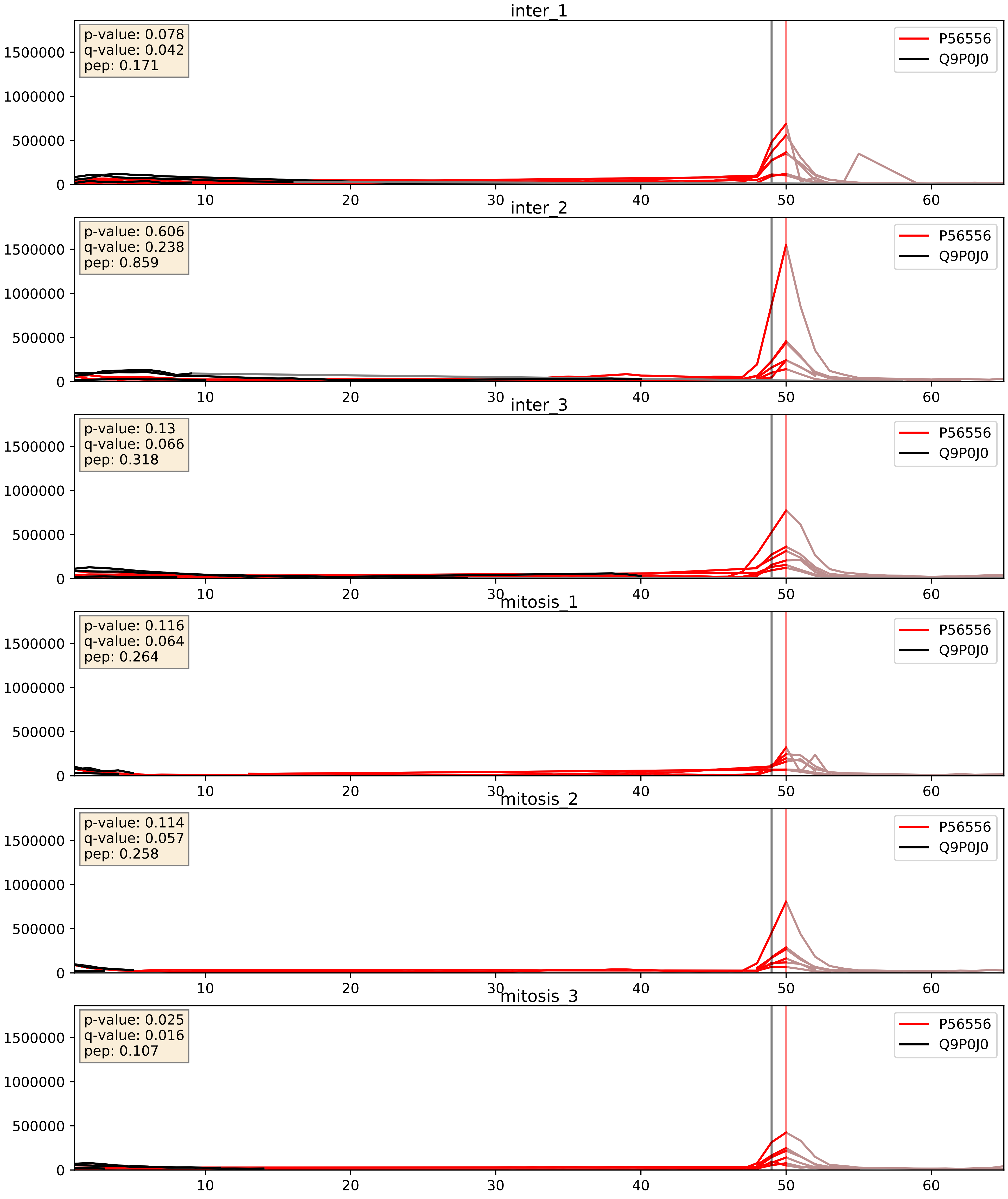


| level | condition_1 | condition_2 | pvalue | pvalue_adjusted |
|---|---|---|---|---|
| interactor_abundance | mitosis | inter | 0.0016940579623344308 | 0.030042528708242176 |
| complex_abundance | mitosis | inter | 0.0018394931343961804 | 0.030581181485790792 |
| interactor_ratio | mitosis | inter | 0.10850493173860024 | 0.5473201035252906 |

Q16718_Q9P0J0
NDUA5_HUMAN vs NDUAD_HUMAN
p-value: 0.019 q-value: 0.015

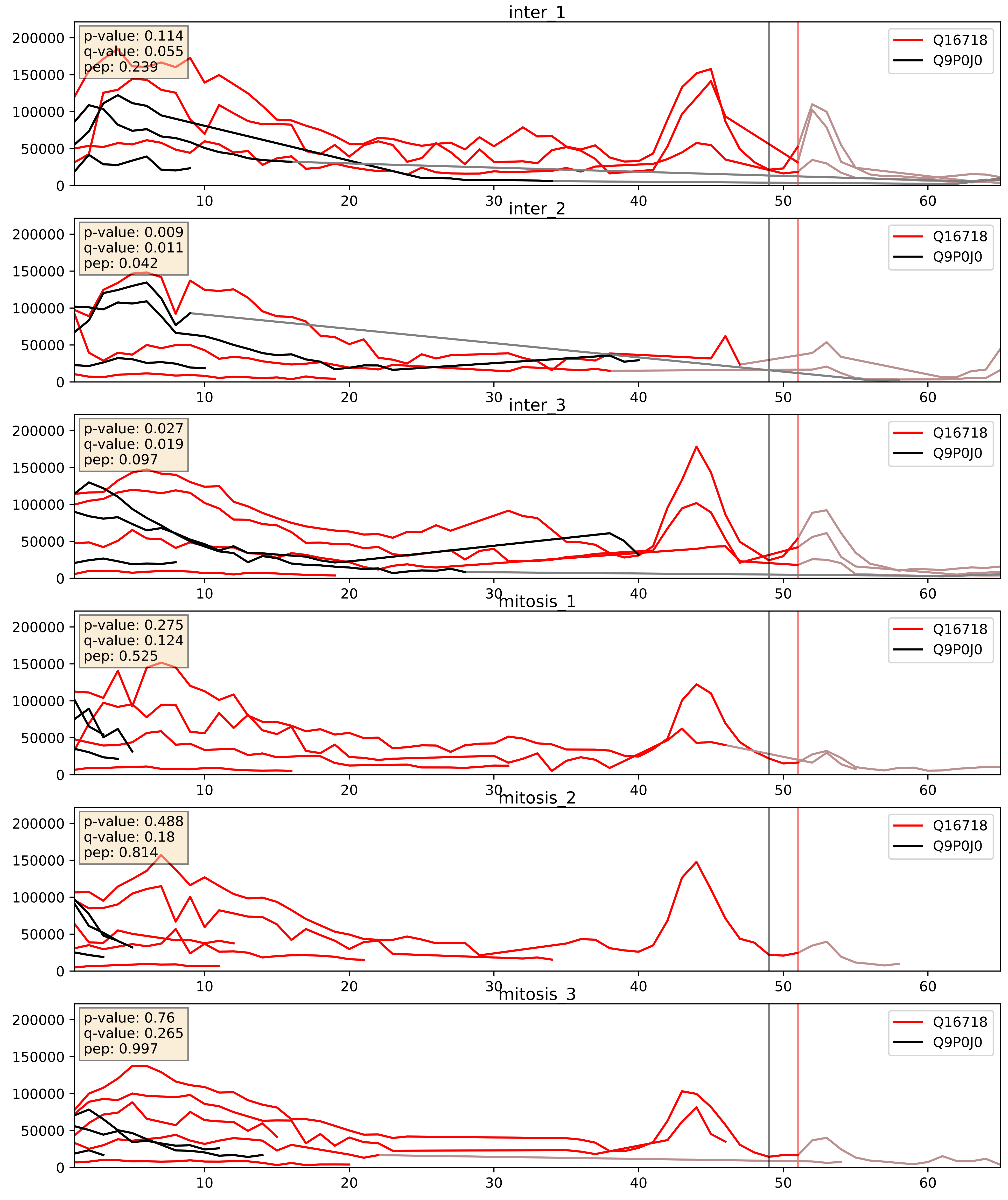


| level | condition_1 | condition_2 | pvalue | pvalue_adjusted |
|---|---|---|---|---|
| complex_abundance | mitosis | inter | 0.012017132690947261 | 0.08151082078804164 |
| interactor_abundance | mitosis | inter | 0.07531430280610488 | 0.22772533805025 |
| interactor_ratio | mitosis | inter | 0.9073367726401307 | 0.9852158746906624 |

Q16795_Q9P0J0
NDUA9_HUMAN vs NDUAD_HUMAN
p-value: 0.006 q-value: 0.006

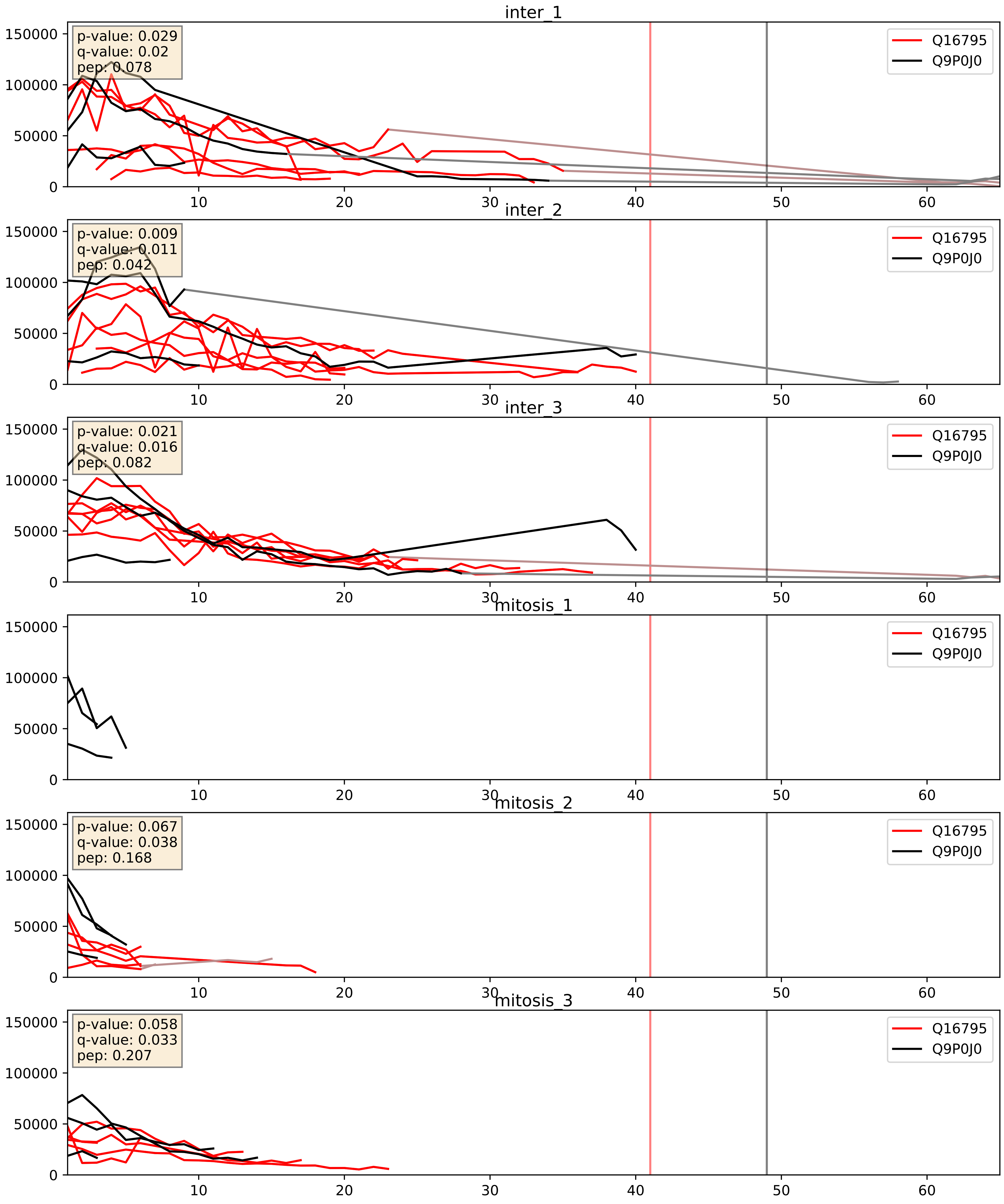


| level | condition_1 | condition_2 | pvalue | pvalue_adjusted |
|---|---|---|---|---|
| interactor_abundance | mitosis | inter | 5.119480063602888e-05 | 0.006283598333582311 |
| complex_abundance | mitosis | inter | 0.0004070925892423535 | 0.012858717948024156 |
| interactor_ratio | mitosis | inter | 0.040201029565406006 | 0.33216294698829674 |

Q9P0J0_Q9Y6M9
NDUAD_HUMAN vs NDUB9_HUMAN
p-value: 0.025 q-value: 0.019

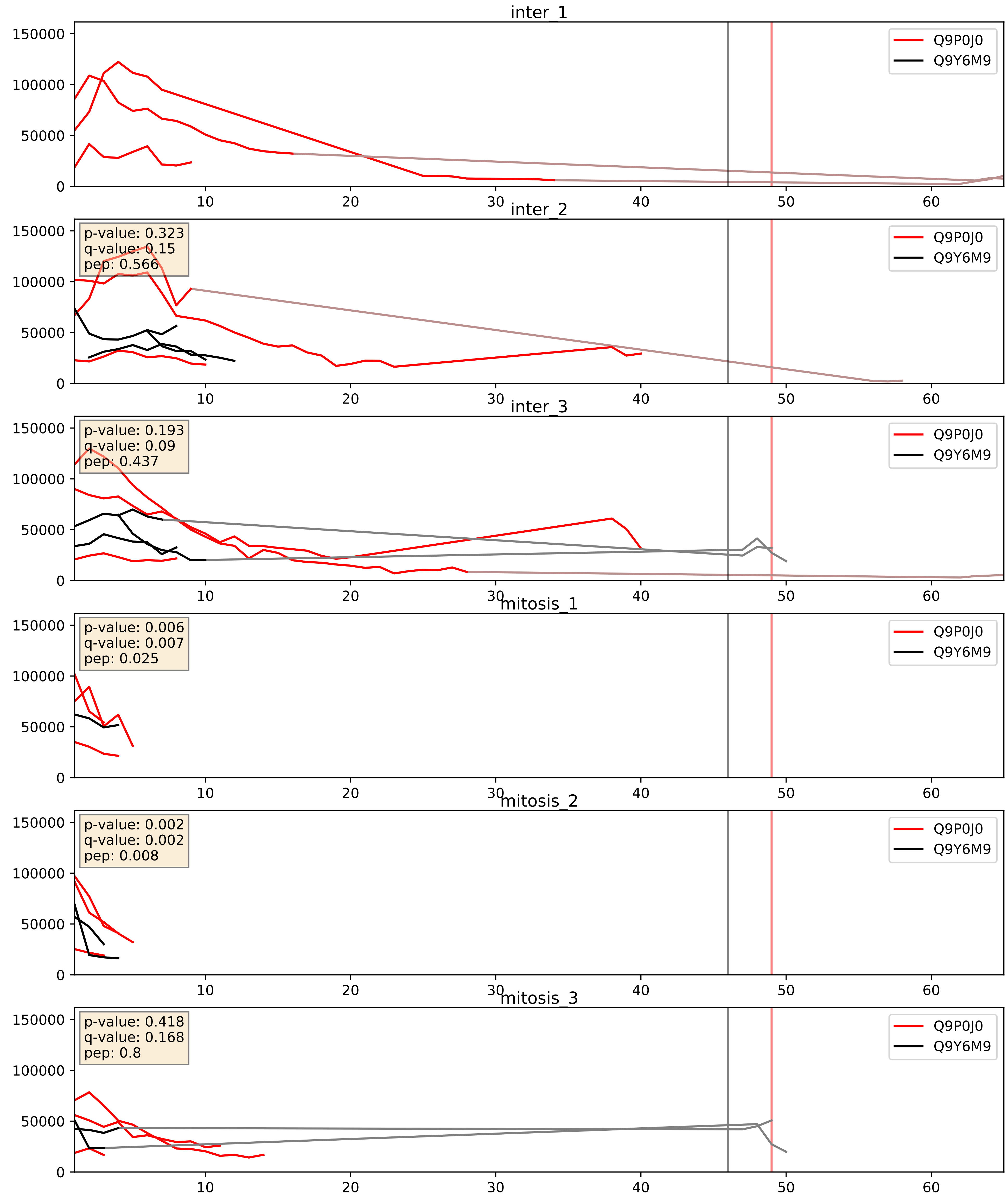


| level | condition_1 | condition_2 | pvalue | pvalue_adjusted |
|---|---|---|---|---|
| complex_abundance | mitosis | inter | 0.1330143407775346 | 0.2769649129301134 |
| interactor_abundance | mitosis | inter | 0.2394794531305443 | 0.43084155502258503 |
| interactor_ratio | mitosis | inter | 0.33306329554243636 | 0.7901935887782929 |
