## Supplementary Data 2 for "SECAT: Quantifying differential protein-protein interaction states by network-centric analysis": hela_string_000157_Q6P1K2.pdf

Q6P1K2_Q6P1K2

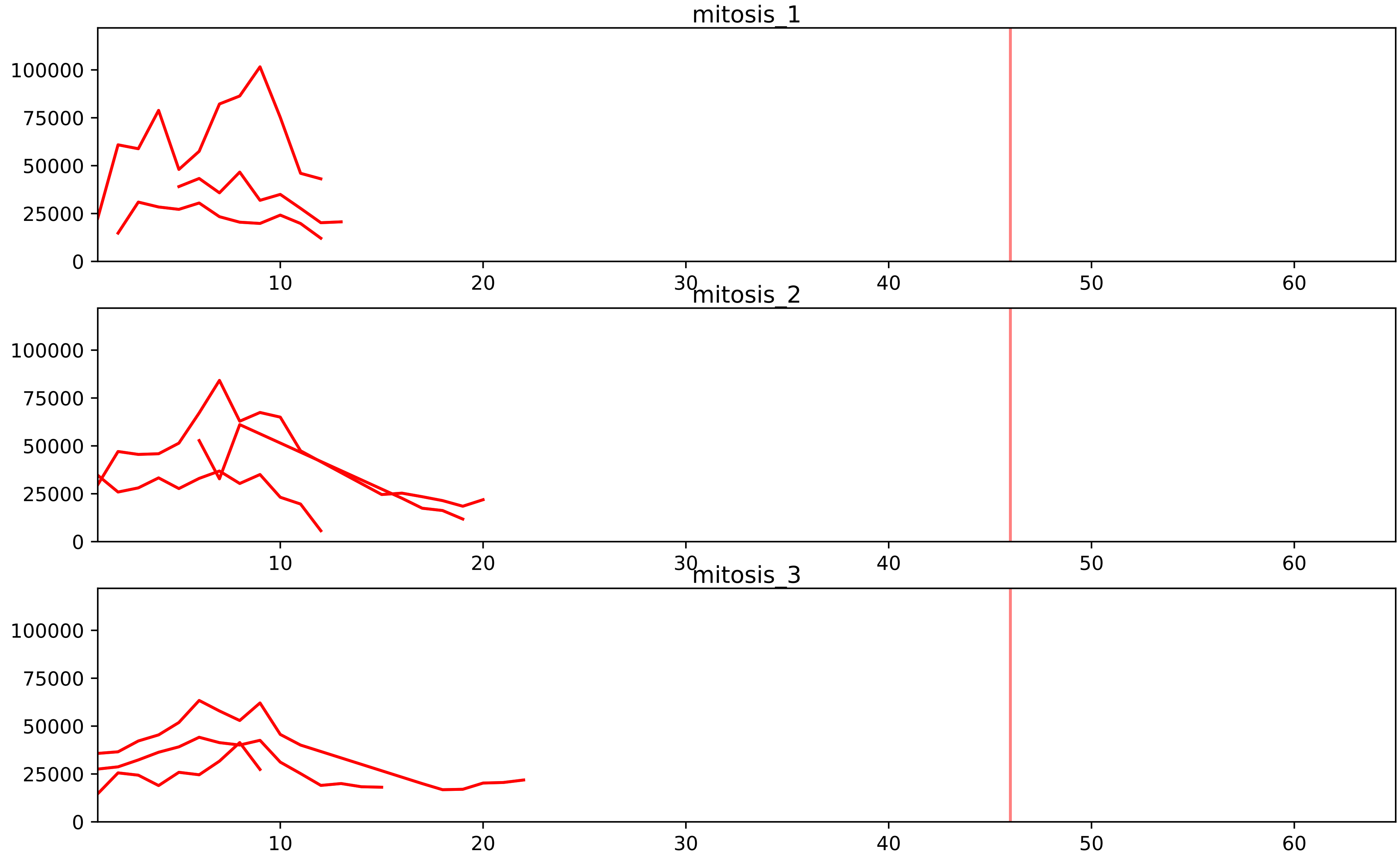


| level | condition_1 | condition_2 | pvalue | pvalue_adjusted |
|---|---|---|---|---|
| assembled_abundance | mitosis | inter | 7.270860409274554e-05 | 0.008577738896244046 |
| total_abundance | mitosis | inter | 0.0001560968480970854 | 0.015536779207822805 |
| complex_abundance | mitosis | inter | 0.0020388752880554344 | 0.017052411500099996 |
| interactor_abundance | mitosis | inter | 0.0036594031158886666 | 0.028607714909554285 |
| interactor_ratio | mitosis | inter | 0.06365921099324959 | 0.2481630259058882 |

O95229_Q6P1K2
ZWINT_HUMAN vs PMF1_HUMAN
p-value: 0.013 q-value: 0.011

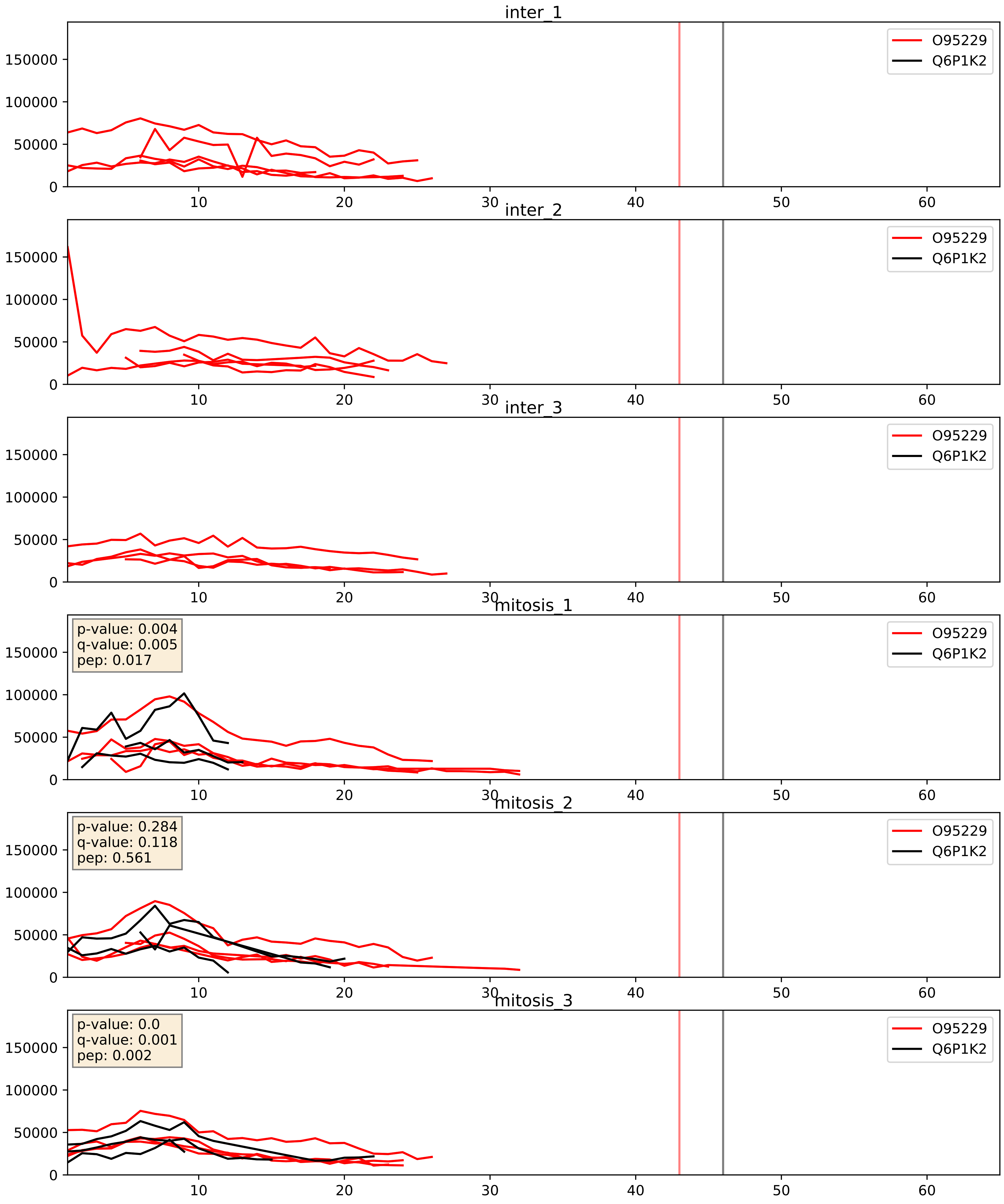


| level | condition_1 | condition_2 | pvalue | pvalue_adjusted |
|---|---|---|---|---|
| interactor_abundance | mitosis | inter | 0.0026424981971553425 | 0.03840370894337815 |
| complex_abundance | mitosis | inter | 0.0040116918026668474 | 0.04692687308892084 |
| interactor_ratio | mitosis | inter | 0.017457081123981564 | 0.19792399261096977 |

Q6P1K2_Q96IY1
PMF1_HUMAN vs NSL1_HUMAN
p-value: 0.003 q-value: 0.003

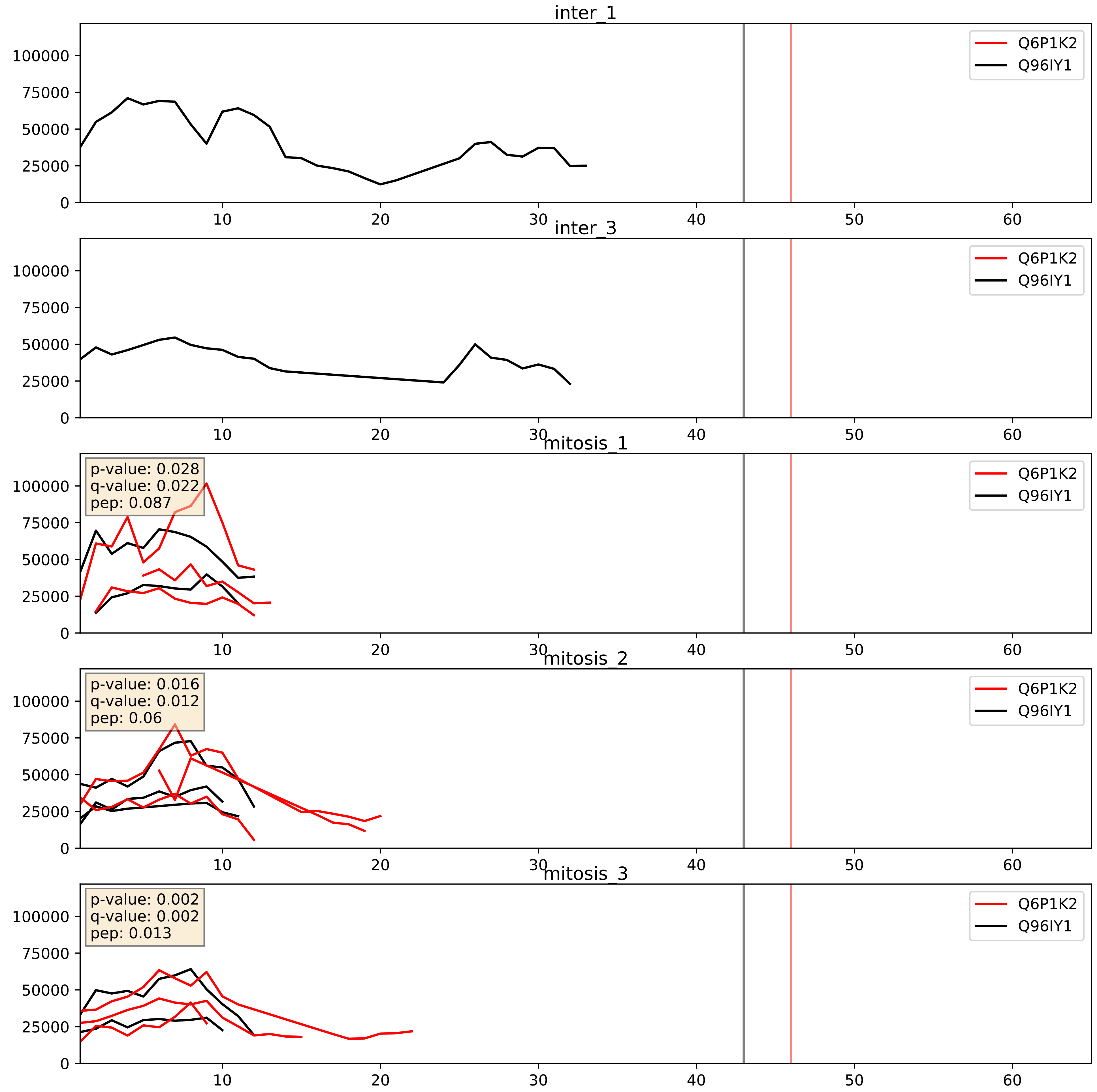


| level | condition_1 | condition_2 | pvalue | pvalue_adjusted |
|---|---|---|---|---|
| interactor_abundance | mitosis | inter | 0.016645545806478236 | 0.11277132084292767 |
| complex_abundance | mitosis | inter | 0.018508480590145496 | 0.10154500794221556 |
| interactor_ratio | mitosis | inter | 0.1596239527784914 | 0.616090239021059 |

Q6P1K2_Q9H081
PMF1_HUMAN vs MIS12_HUMAN
p-value: 0.008 q-value: 0.007

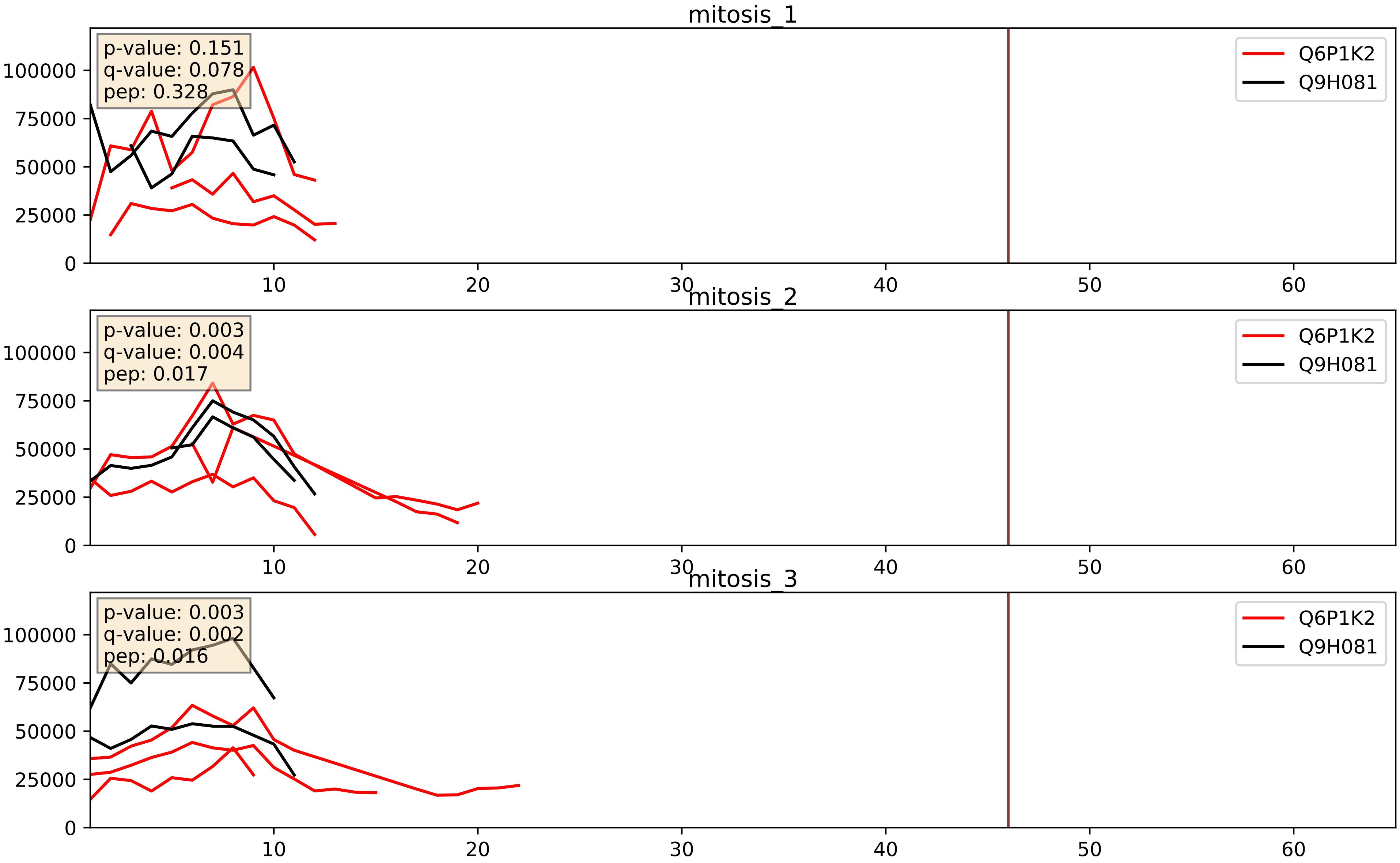


| level | condition_1 | condition_2 | pvalue | pvalue_adjusted |
|---|---|---|---|---|
| complex_abundance | mitosis | inter | 0.009136809477122337 | 0.07075397186708442 |
| interactor_abundance | mitosis | inter | 0.015727061238973217 | 0.10998663742288459 |
| interactor_ratio | mitosis | inter | 0.9234496167064464 | 0.9871879493798719 |
