## Supplementary Data 2 for "SECAT: Quantifying differential protein-protein interaction states by network-centric analysis": hela_string_000158_P35268.pdf

P35268_P35268

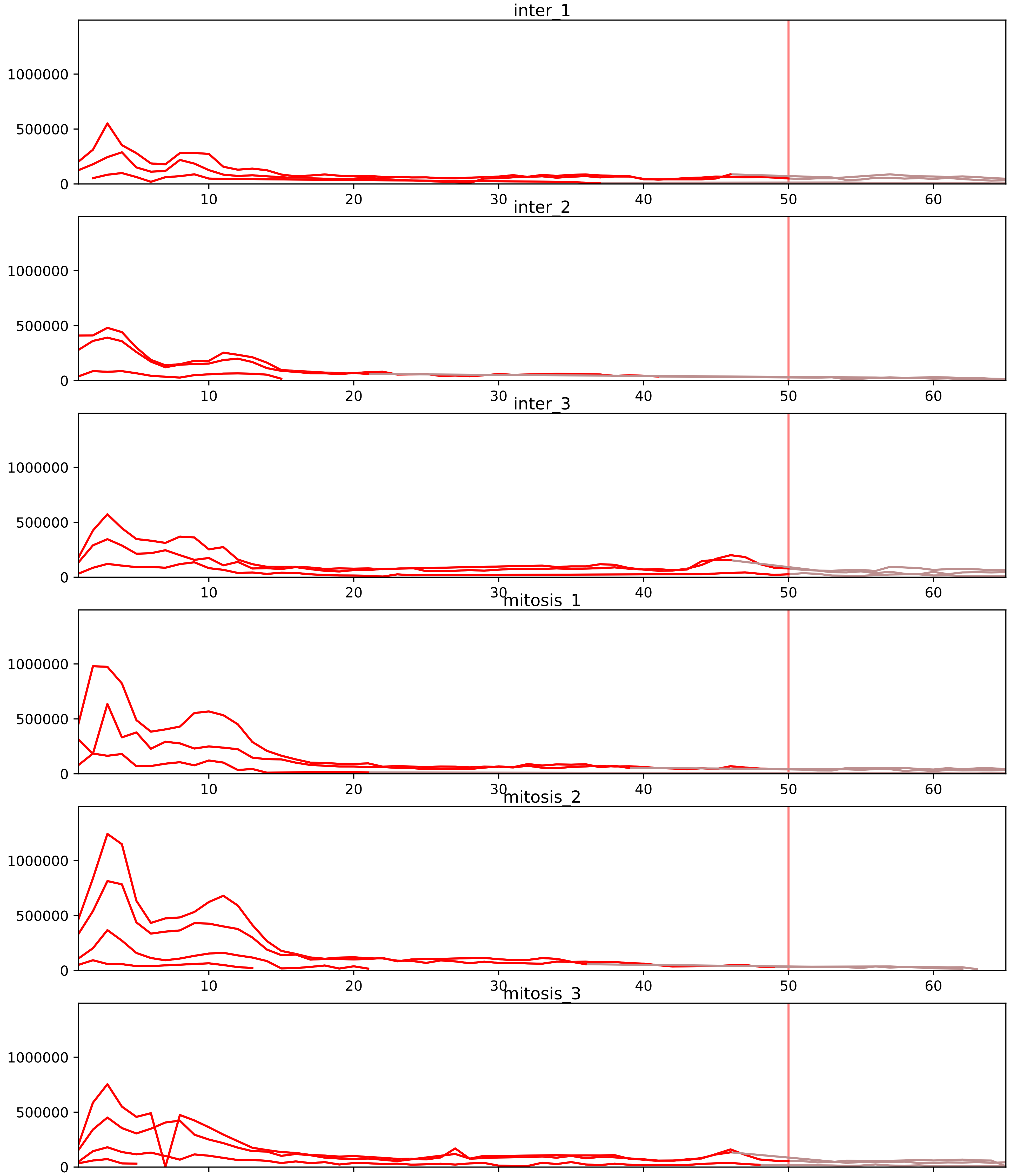


| level | condition_1 | condition_2 | pvalue | pvalue_adjusted |
|---|---|---|---|---|
| interactor_ratio | mitosis | inter | 7.277145615235256e-05 | 0.0012960110367126292 |
| interactor_abundance | mitosis | inter | 0.0011221410120851667 | 0.012589874769736017 |
| complex_abundance | mitosis | inter | 0.0040381331655963155 | 0.025017390655546194 |
| assembled_abundance | mitosis | inter | 0.4297742846282324 | 0.6978734506663661 |
| total_abundance | mitosis | inter | 0.49088155552536317 | 0.756776782072043 |
| monomer_abundance | mitosis | inter | 0.4969184658172587 | 0.7532862551906346 |

P04843_P35268
RPN1_HUMAN vs RL22_HUMAN
p-value: 0.012 q-value: 0.011

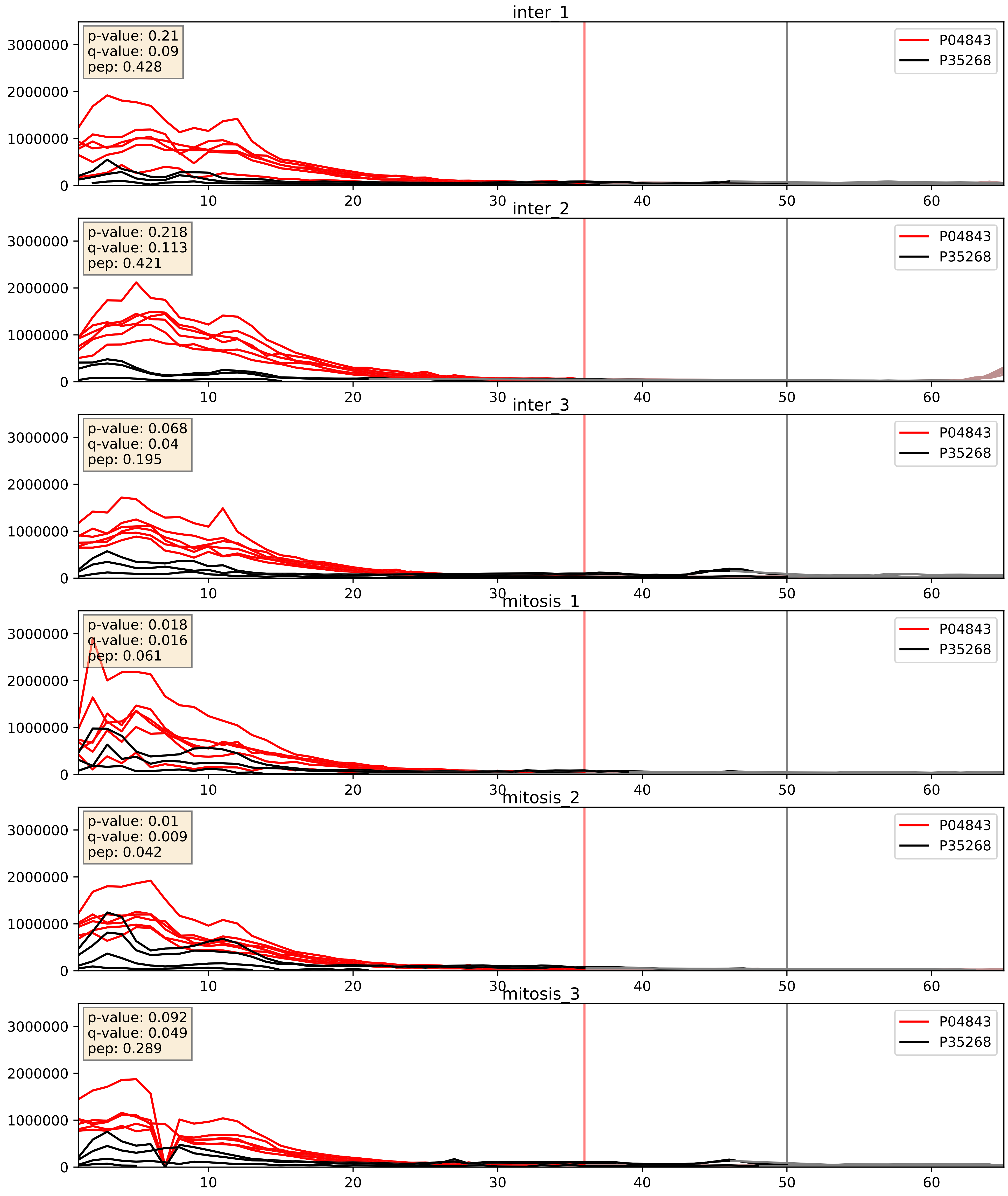


| level | condition_1 | condition_2 | pvalue | pvalue_adjusted |
|---|---|---|---|---|
| interactor_ratio | mitosis | inter | 0.013024268071311108 | 0.1649226844532886 |
| interactor_abundance | mitosis | inter | 0.055512062499012145 | 0.19106684961461357 |
| complex_abundance | mitosis | inter | 0.670943411025327 | 0.7808668386644185 |

P05387_P35268
RLA2_HUMAN vs RL22_HUMAN
p-value: 0.001 q-value: 0.002

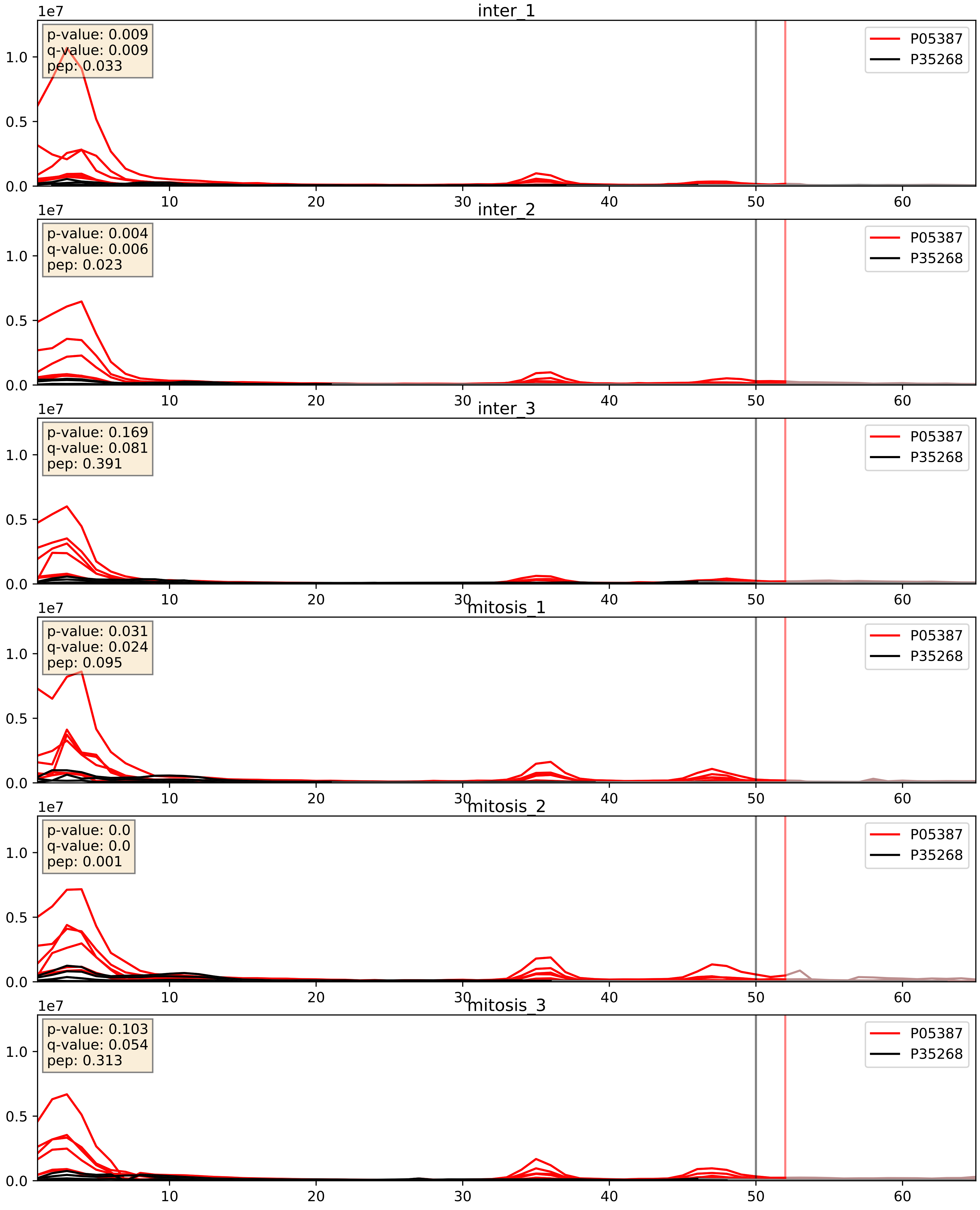


| level | condition_1 | condition_2 | pvalue | pvalue_adjusted |
|---|---|---|---|---|
| interactor_ratio | mitosis | inter | 0.13522811787324118 | 0.5858060166978465 |
| complex_abundance | mitosis | inter | 0.18501349370583747 | 0.33653113177262406 |
| interactor_abundance | mitosis | inter | 0.7355817141095475 | 0.83558880933949 |

P05388_P35268
RLA0_HUMAN vs RL22_HUMAN
p-value: 0.009 q-value: 0.008

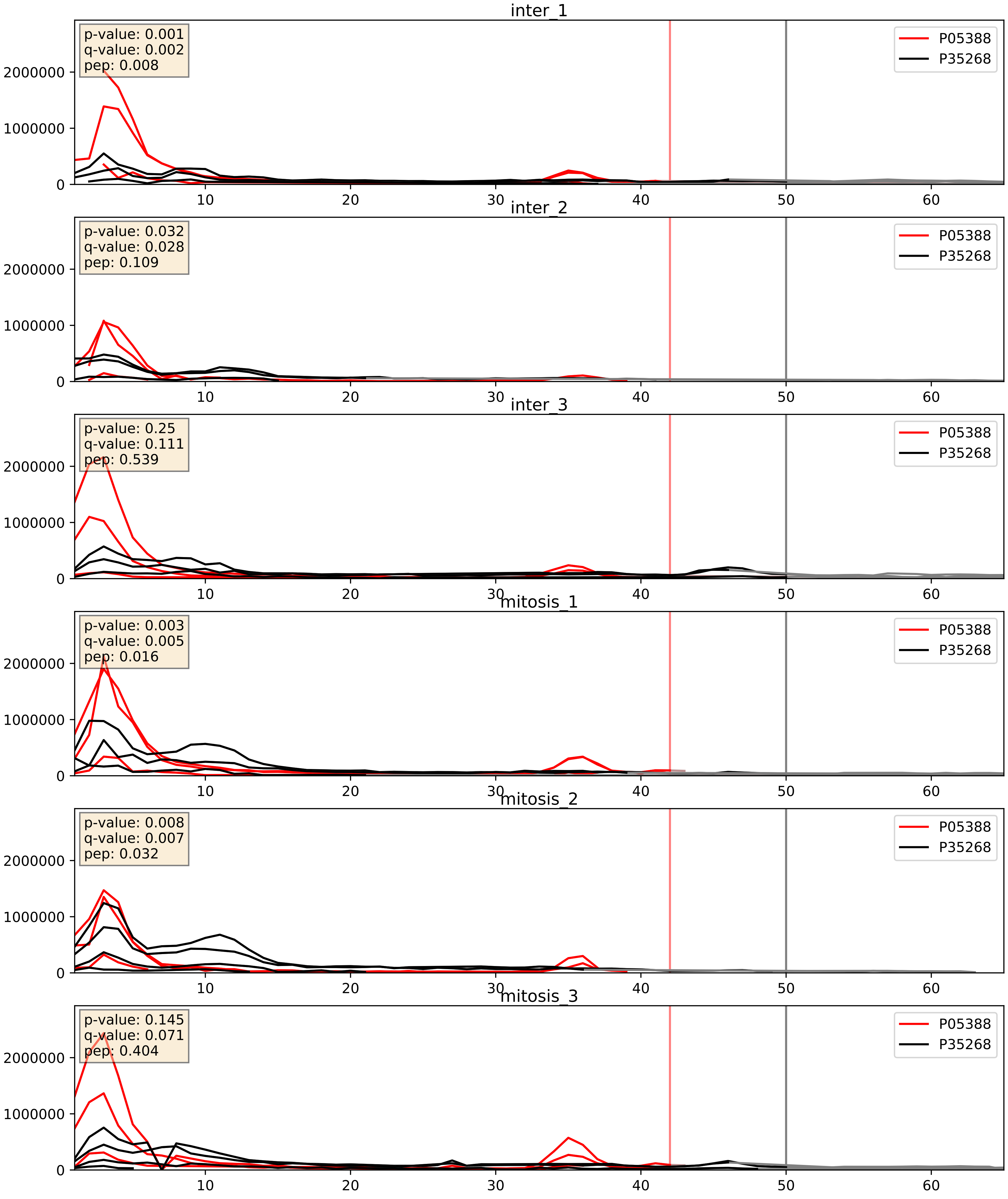


| level | condition_1 | condition_2 | pvalue | pvalue_adjusted |
|---|---|---|---|---|
| complex_abundance | mitosis | inter | 0.11338851248912898 | 0.2531244338436389 |
| interactor_ratio | mitosis | inter | 0.5305546372675891 | 0.8821330249831557 |
| interactor_abundance | mitosis | inter | 0.5648057672917774 | 0.712615132731611 |

P08708_P35268
RS17_HUMAN vs RL22_HUMAN
p-value: 0.0 q-value: 0.0

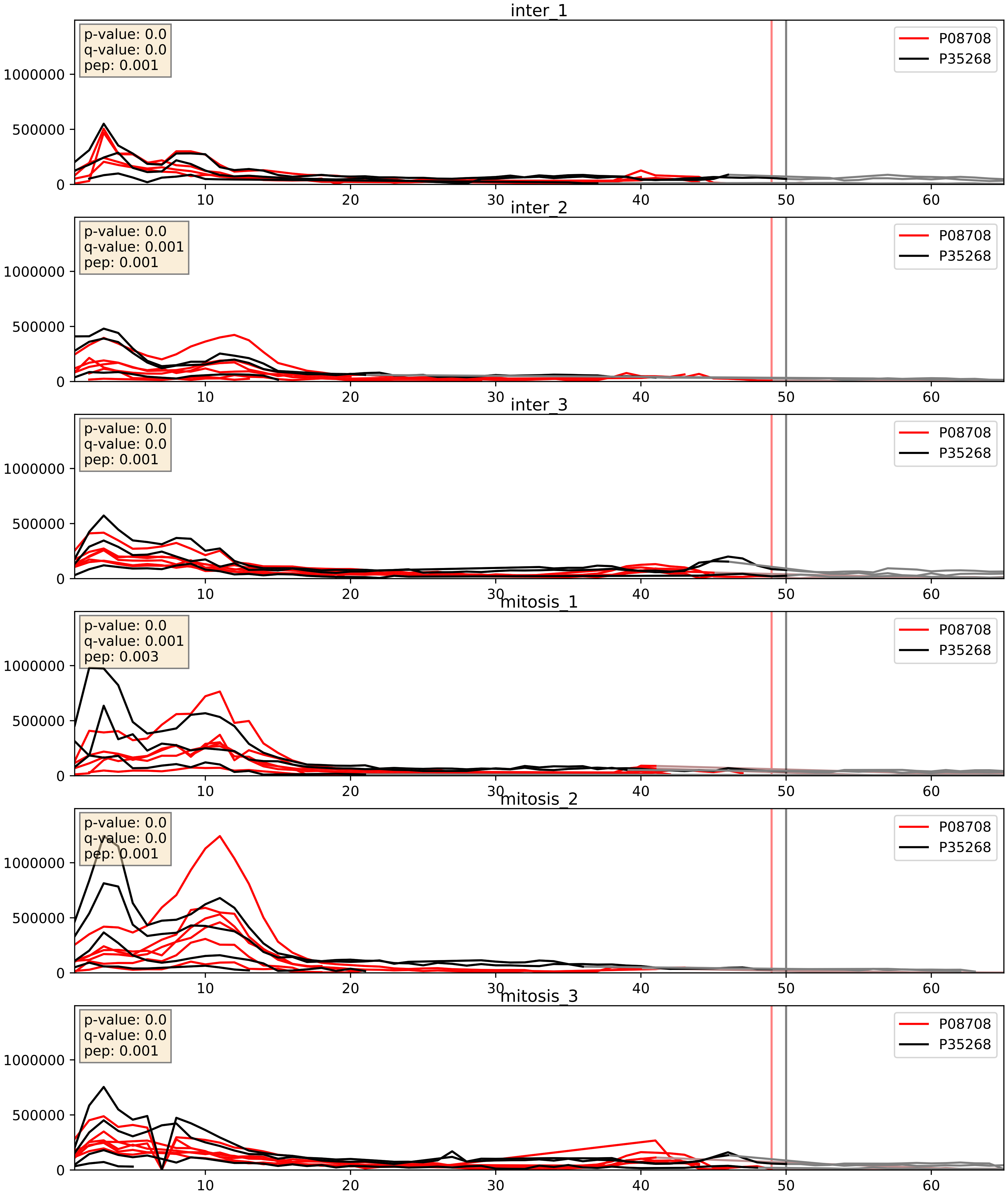


| level | condition_1 | condition_2 | pvalue | pvalue_adjusted |
|---|---|---|---|---|
| complex_abundance | mitosis | inter | 0.014045970322417972 | 0.08823558742482049 |
| interactor_abundance | mitosis | inter | 0.026849293089140424 | 0.13605729000585648 |
| interactor_ratio | mitosis | inter | 0.31289369186171695 | 0.7783421708721073 |

P08865_P35268
RSSA_HUMAN vs RL22_HUMAN
p-value: 0.028 q-value: 0.021

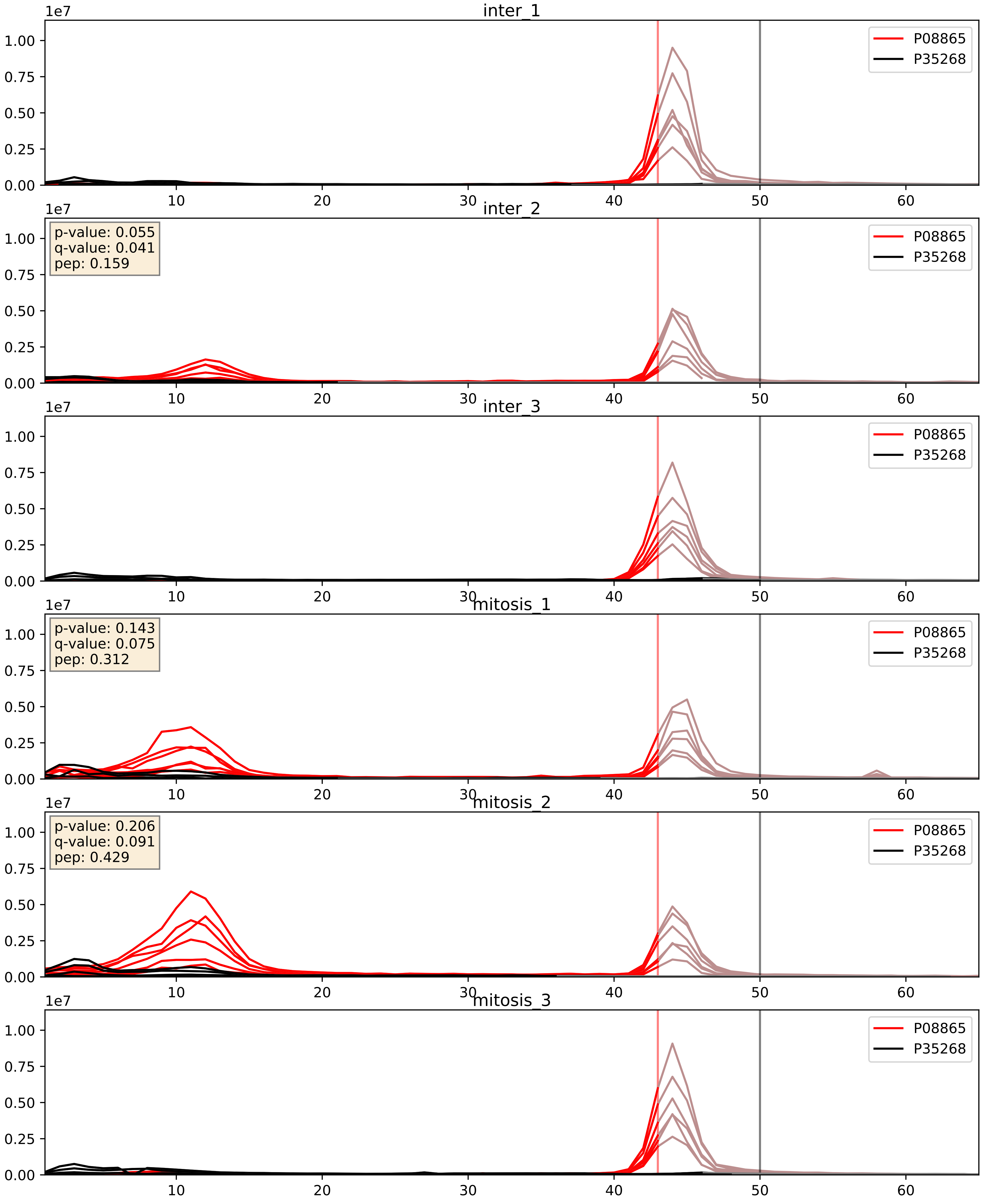


| level | condition_1 | condition_2 | pvalue | pvalue_adjusted |
|---|---|---|---|---|
| complex_abundance | mitosis | inter | 0.12504258102224622 | 0.2668650533019743 |
| interactor_abundance | mitosis | inter | 0.24392062888572197 | 0.4368321372728773 |
| interactor_ratio | mitosis | inter | 0.46692624378010217 | 0.85989947136842 |

P15880_P35268
RS2_HUMAN vs RL22_HUMAN
p-value: 0.004 q-value: 0.004

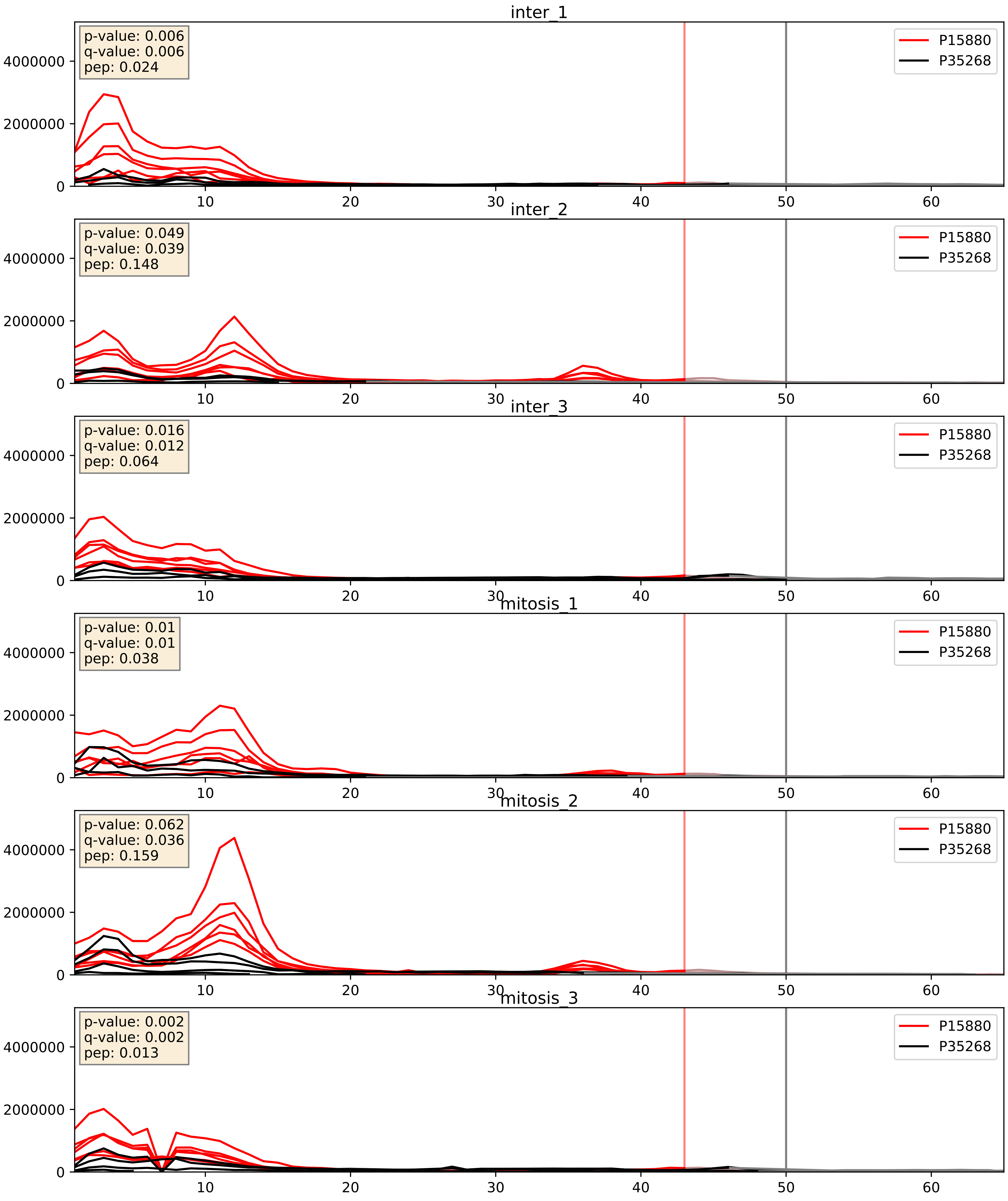


| level | condition_1 | condition_2 | pvalue | pvalue_adjusted |
|---|---|---|---|---|
| interactor_ratio | mitosis | inter | 0.00442621394597915 | 0.08831792861907116 |
| complex_abundance | mitosis | inter | 0.12181597272358244 | 0.2641110761636074 |
| interactor_abundance | mitosis | inter | 0.3099770845913117 | 0.5063273428985764 |

P18124_P35268
RL7_HUMAN vs RL22_HUMAN
p-value: 0.008 q-value: 0.007

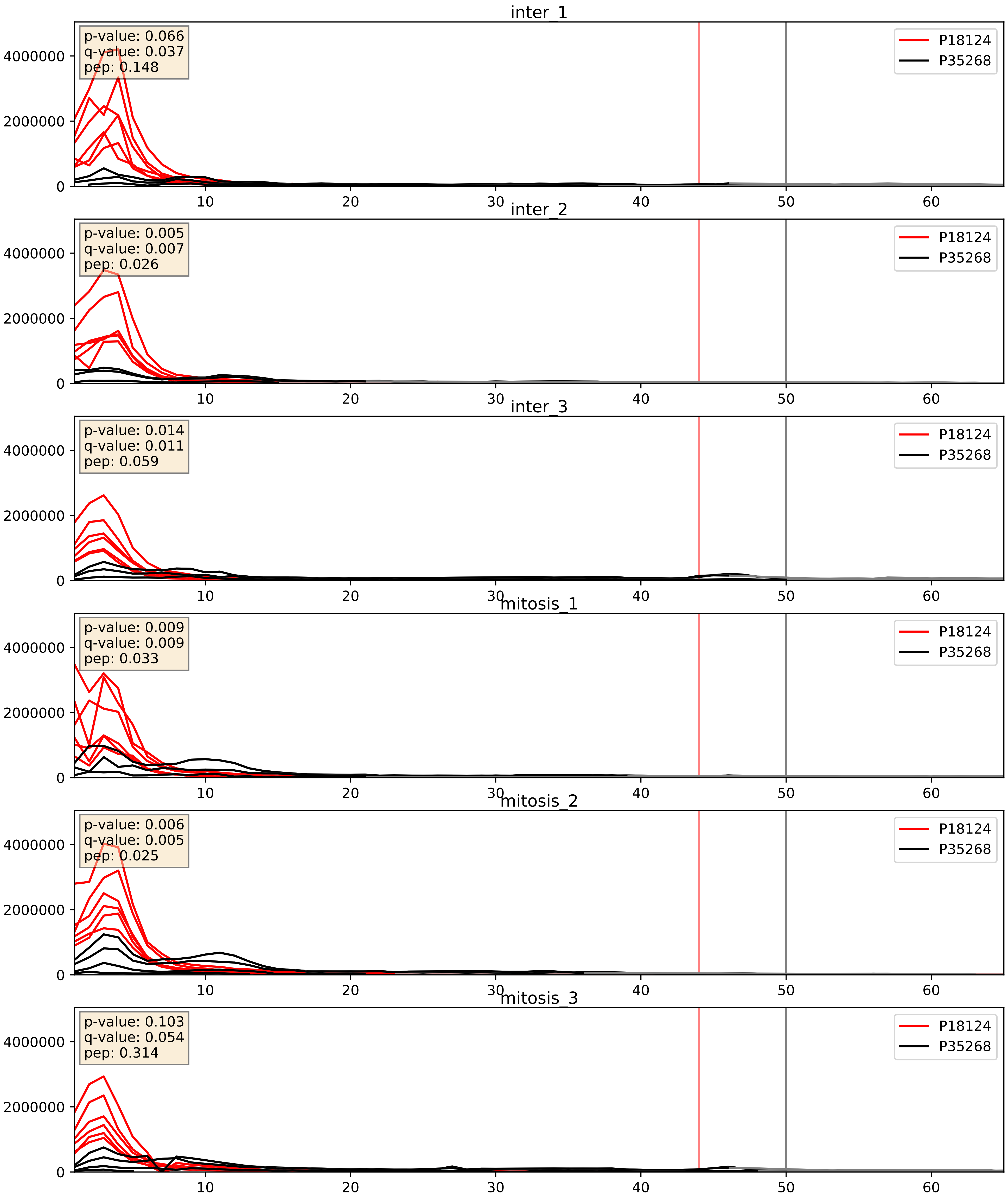


| level | condition_1 | condition_2 | pvalue | pvalue_adjusted |
|---|---|---|---|---|
| complex_abundance | mitosis | inter | 0.022078182232444676 | 0.10952717692621856 |
| interactor_ratio | mitosis | inter | 0.08794018198868897 | 0.4946978435260244 |
| interactor_abundance | mitosis | inter | 0.49463356866583236 | 0.6611591736070463 |

P23396_P35268
RS3_HUMAN vs RL22_HUMAN
p-value: 0.001 q-value: 0.001

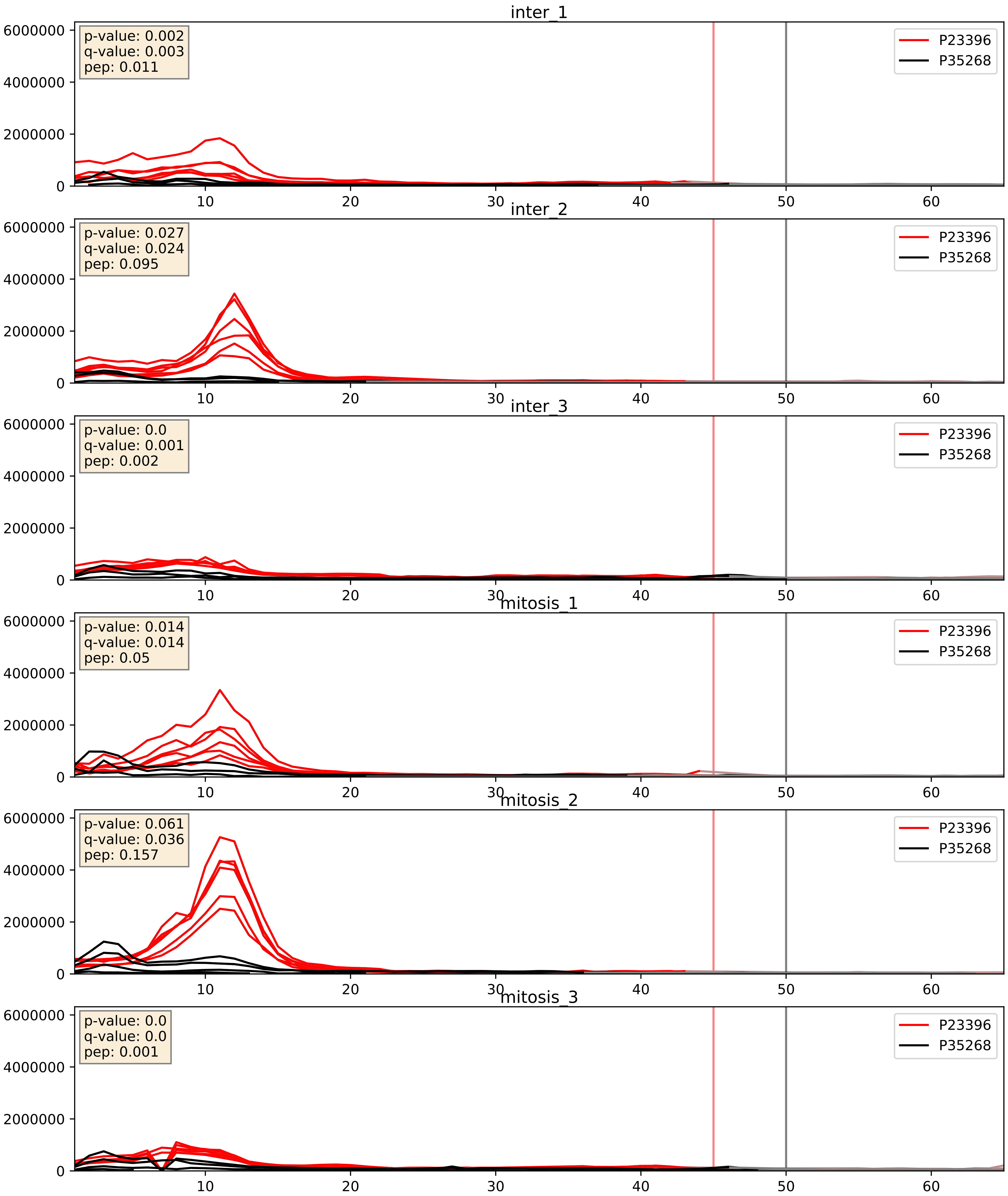


| level | condition_1 | condition_2 | pvalue | pvalue_adjusted |
|---|---|---|---|---|
| complex_abundance | mitosis | inter | 0.23649540112164197 | 0.38870969615590656 |
| interactor_ratio | mitosis | inter | 0.4153128789603051 | 0.8362922239238325 |
| interactor_abundance | mitosis | inter | 0.7907562442052162 | 0.8798171768891467 |

P25398_P35268
RS12_HUMAN vs RL22_HUMAN
p-value: 0.0 q-value: 0.0

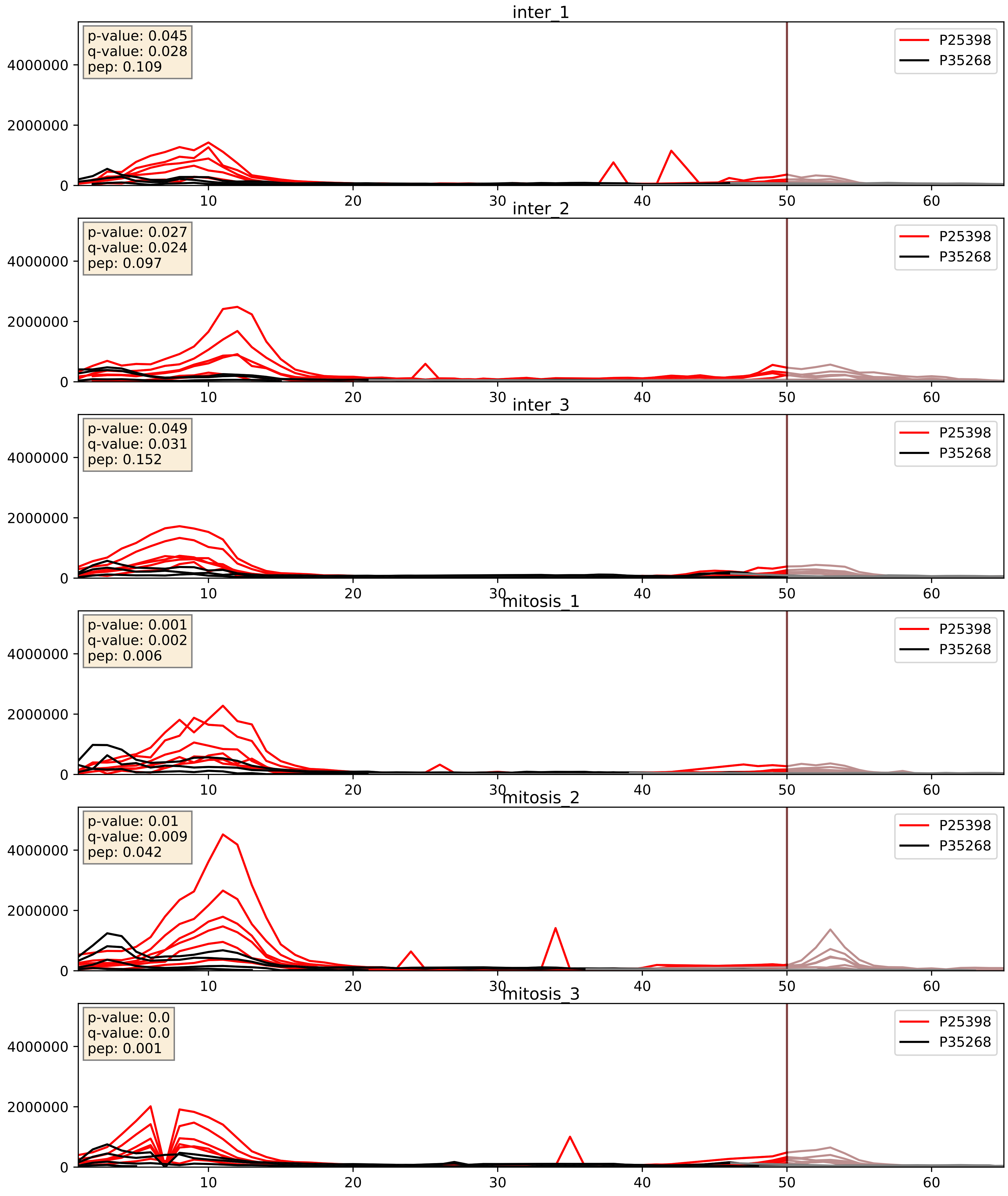


| level | condition_1 | condition_2 | pvalue | pvalue_adjusted |
|---|---|---|---|---|
| interactor_ratio | mitosis | inter | 0.06630166015377828 | 0.4296400448702392 |
| complex_abundance | mitosis | inter | 0.19027991889227136 | 0.34191584029671307 |
| interactor_abundance | mitosis | inter | 0.6487998653481982 | 0.7652923863002035 |

P26373_P35268
RL13_HUMAN vs RL22_HUMAN
p-value: 0.0 q-value: 0.0

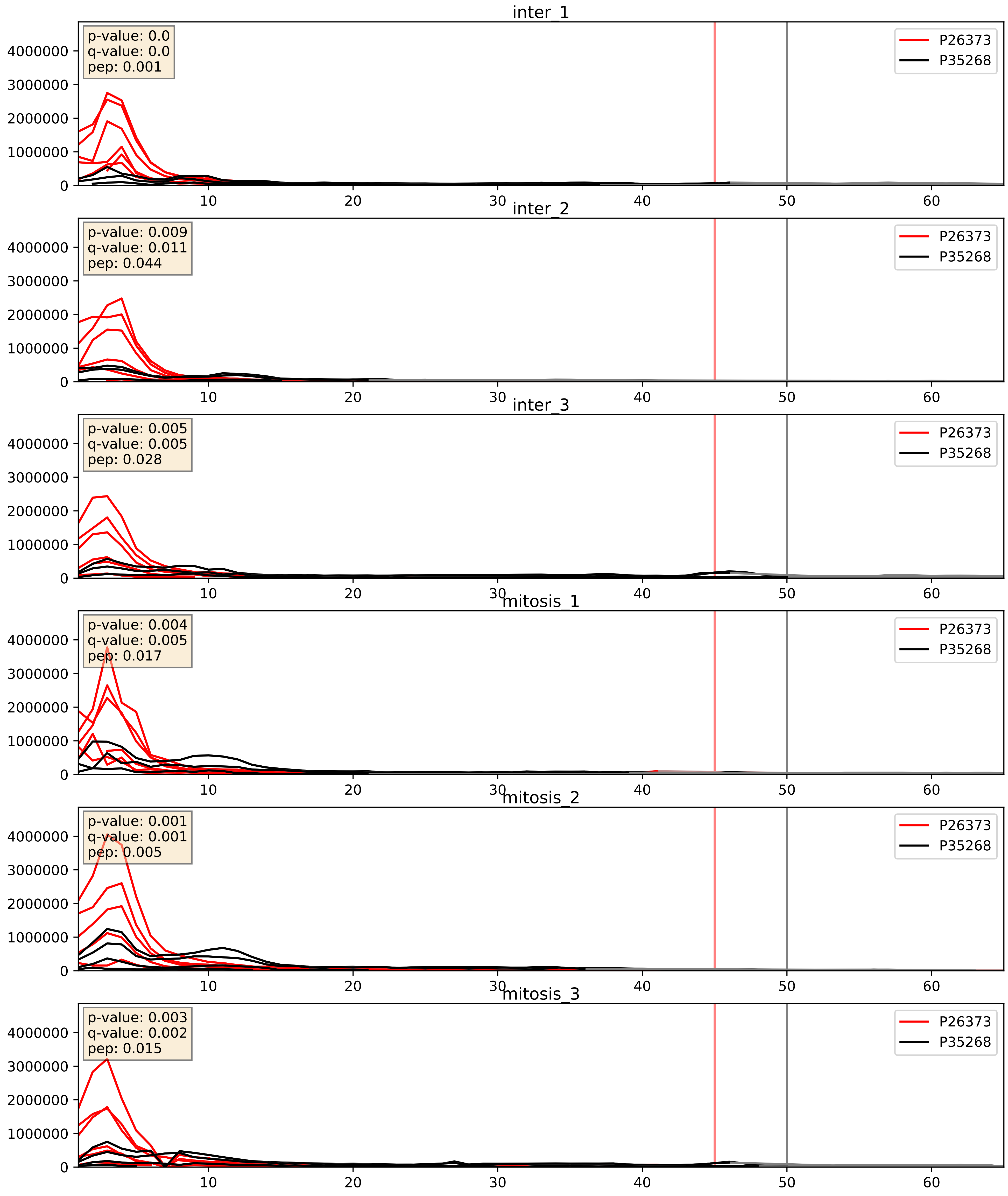


| level | condition_1 | condition_2 | pvalue | pvalue_adjusted |
|---|---|---|---|---|
| complex_abundance | mitosis | inter | 0.02374693448489318 | 0.11228870016928881 |
| interactor_ratio | mitosis | inter | 0.07802395073353222 | 0.46705246033499004 |
| interactor_abundance | mitosis | inter | 0.6159278988347678 | 0.7471505673469989 |

P27635_P35268
RL10_HUMAN vs RL22_HUMAN
p-value: 0.002 q-value: 0.003

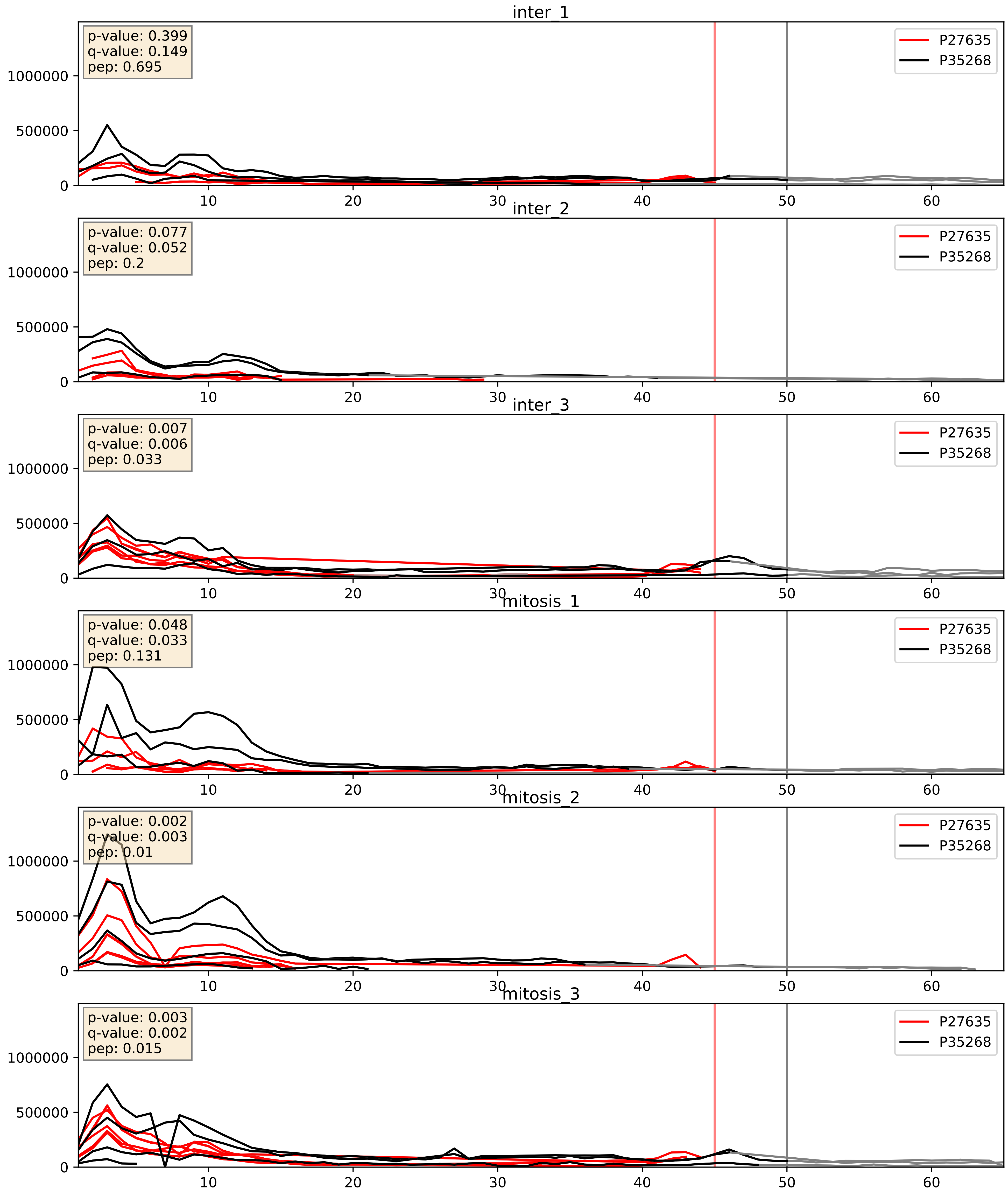


| level | condition_1 | condition_2 | pvalue | pvalue_adjusted |
|---|---|---|---|---|
| complex_abundance | mitosis | inter | 0.2540733040006283 | 0.40468649902523524 |
| interactor_ratio | mitosis | inter | 0.4988641649519024 | 0.8702488381316982 |
| interactor_abundance | mitosis | inter | 0.6249837242695993 | 0.7532958585150921 |

P30050_P35268
RL12_HUMAN vs RL22_HUMAN
p-value: 0.0 q-value: 0.0

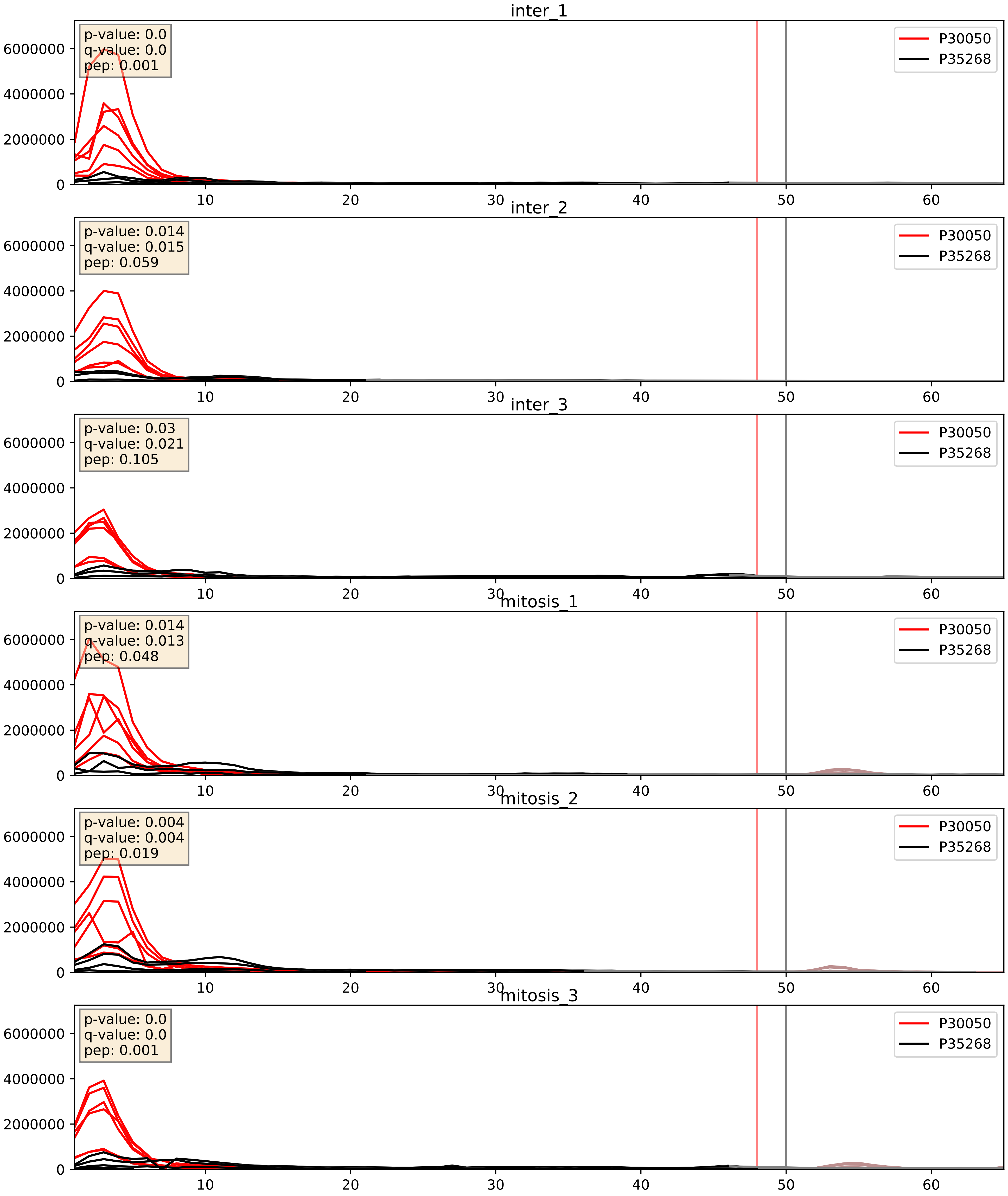


| level | condition_1 | condition_2 | pvalue | pvalue_adjusted |
|---|---|---|---|---|
| complex_abundance | mitosis | inter | 0.09727542503607842 | 0.2310352286309006 |
| interactor_ratio | mitosis | inter | 0.17662594485135244 | 0.6420034343641515 |
| interactor_abundance | mitosis | inter | 0.8832806096565684 | 0.9352897103736053 |

P32969_P35268
RL9_HUMAN vs RL22_HUMAN
p-value: 0.0 q-value: 0.0

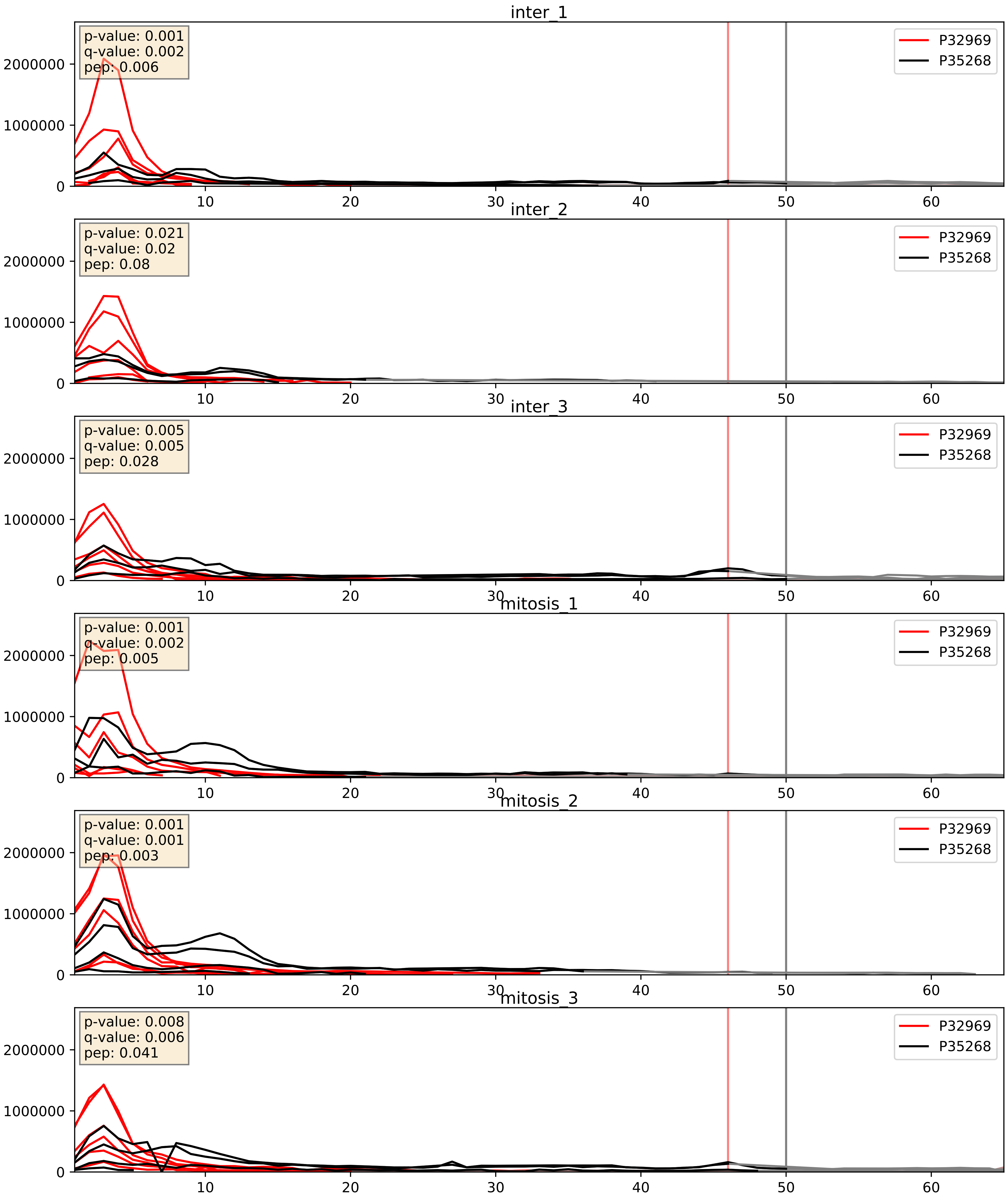


| level | condition_1 | condition_2 | pvalue | pvalue_adjusted |
|---|---|---|---|---|
| complex_abundance | mitosis | inter | 0.00556496697516796 | 0.05624098855659709 |
| interactor_ratio | mitosis | inter | 0.11613059889012095 | 0.5618454386254771 |
| interactor_abundance | mitosis | inter | 0.2103491843605969 | 0.40652154270556234 |

P35268_P36578
RL22_HUMAN vs RL4_HUMAN
p-value: 0.002 q-value: 0.002

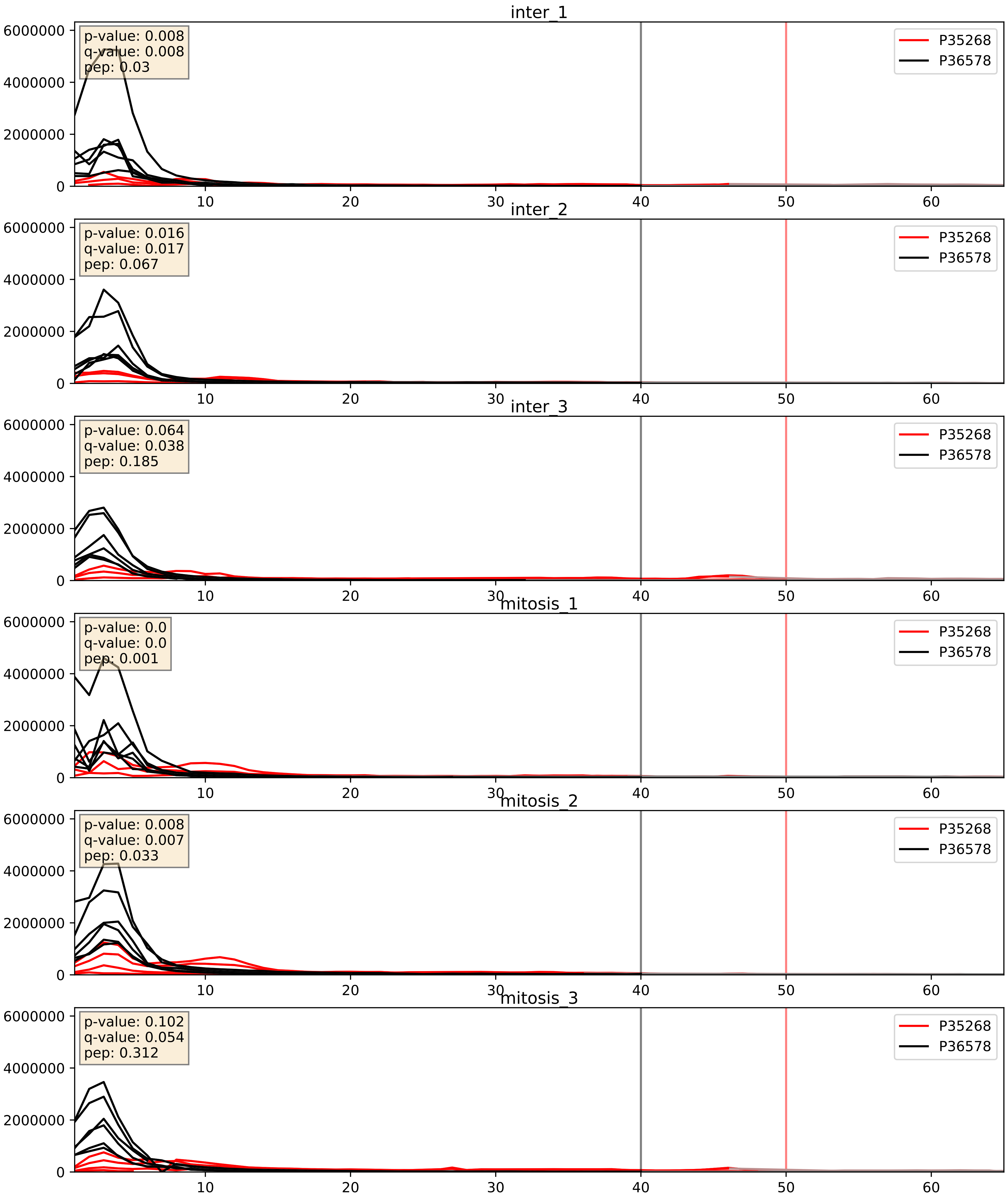


| level | condition_1 | condition_2 | pvalue | pvalue_adjusted |
|---|---|---|---|---|
| interactor_abundance | mitosis | inter | 0.008711956748677724 | 0.0807312115073317 |
| interactor_ratio | mitosis | inter | 0.015289267825559236 | 0.18126888169915106 |
| complex_abundance | mitosis | inter | 0.0481004310752405 | 0.15661456447472755 |

P35268_P37108
RL22_HUMAN vs SRP14_HUMAN
p-value: 0.0 q-value: 0.0

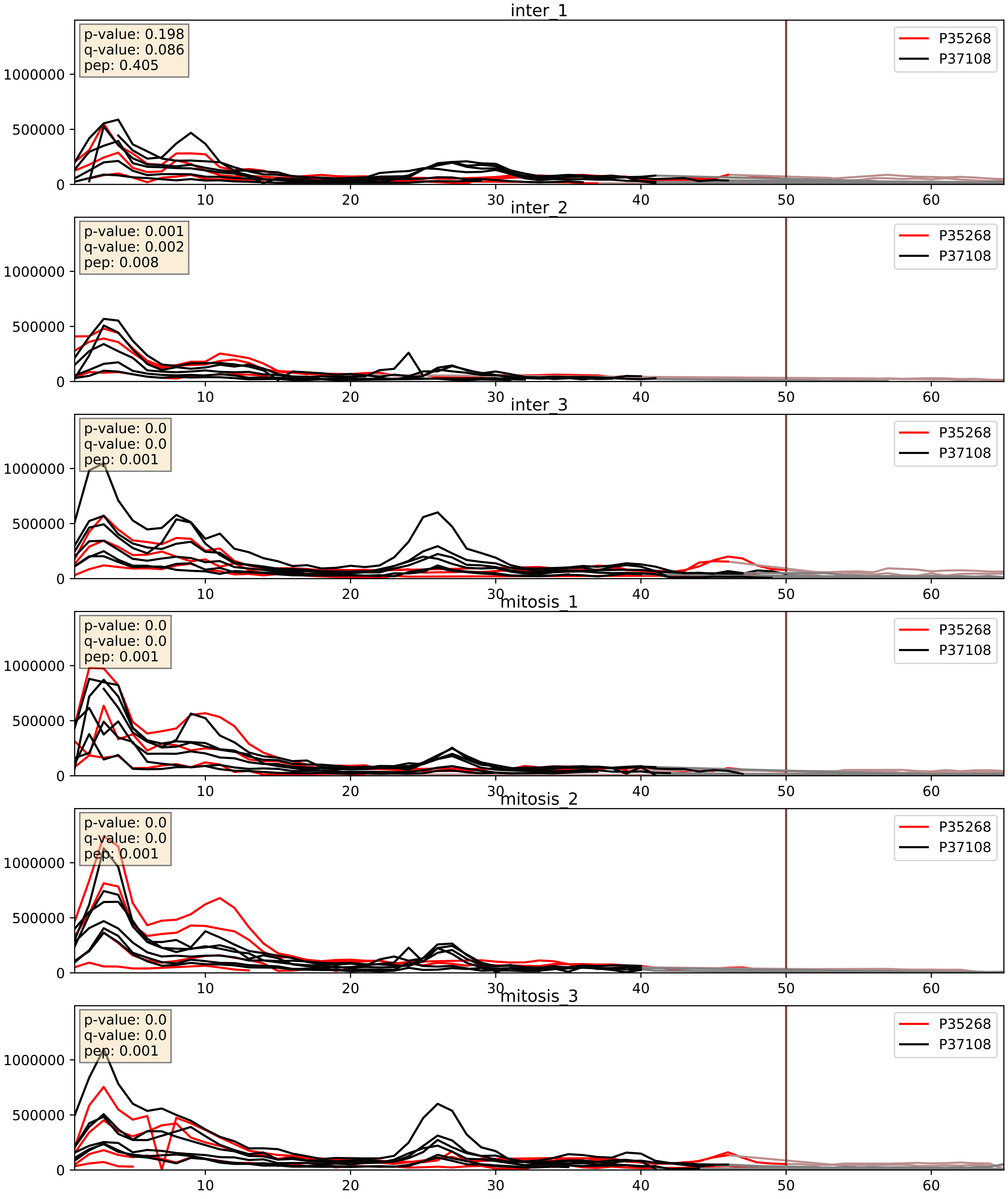


| level | condition_1 | condition_2 | pvalue | pvalue_adjusted |
|---|---|---|---|---|
| interactor_abundance | mitosis | inter | 0.04296378245255331 | 0.16669366057920354 |
| complex_abundance | mitosis | inter | 0.3098570997574628 | 0.45904493130205165 |
| interactor_ratio | mitosis | inter | 0.9321348367212055 | 0.9878882449502933 |

P35268_P39019
RL22_HUMAN vs RS19_HUMAN
p-value: 0.004 q-value: 0.004

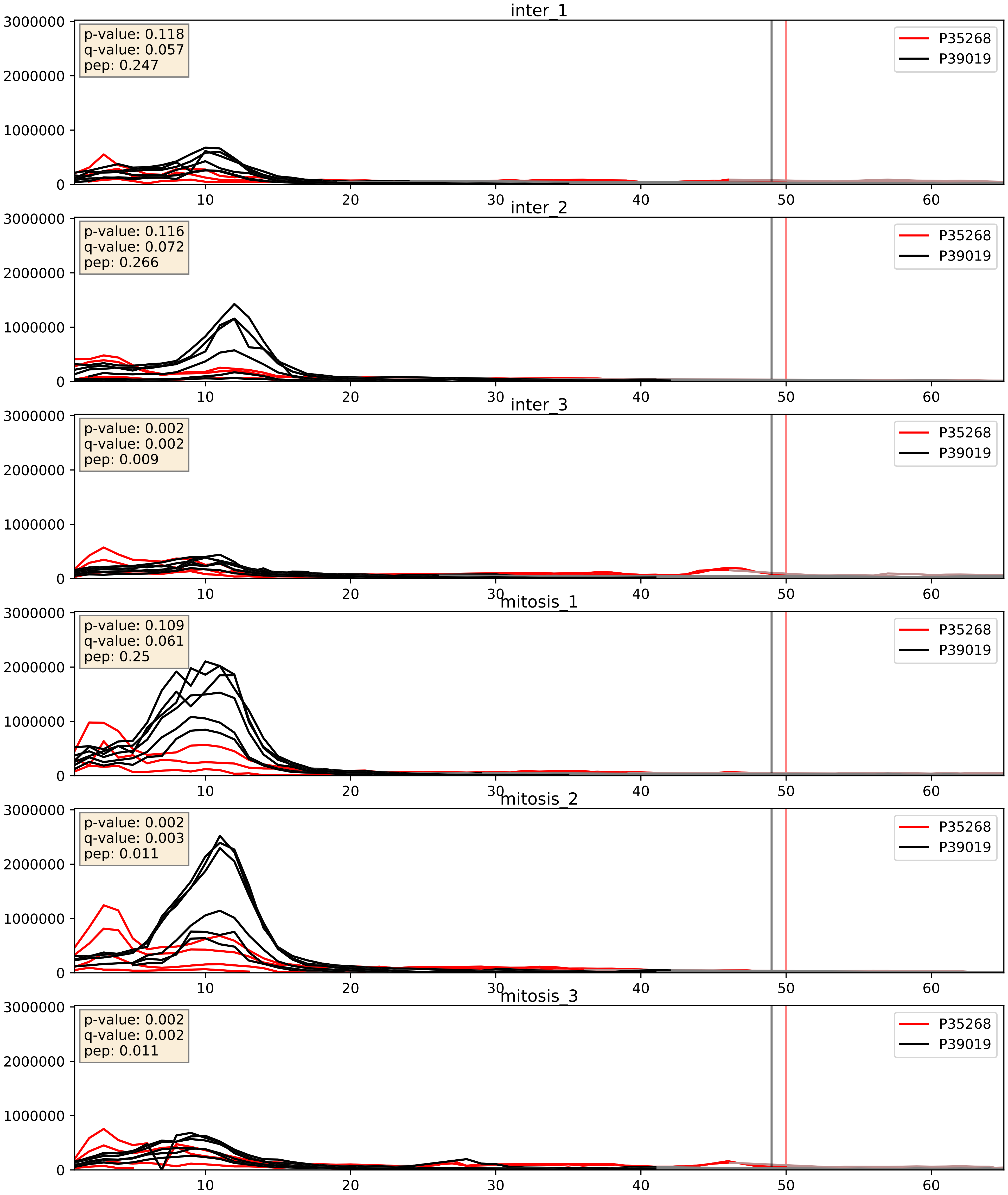


| level | condition_1 | condition_2 | pvalue | pvalue_adjusted |
| --- | --- | --- | --- | --- |
| interactor_abundance | mitosis | inter | 0.01853194053537669 | 0.11724568439233146 |
| complex_abundance | mitosis | inter | 0.04889073402824224 | 0.15795119509349145 |
| interactor_ratio | mitosis | inter | 0.7480489295514303 | 0.9491588884573259 |

P35268_P39023
RL22_HUMAN vs RL3_HUMAN
p-value: 0.0 q-value: 0.0

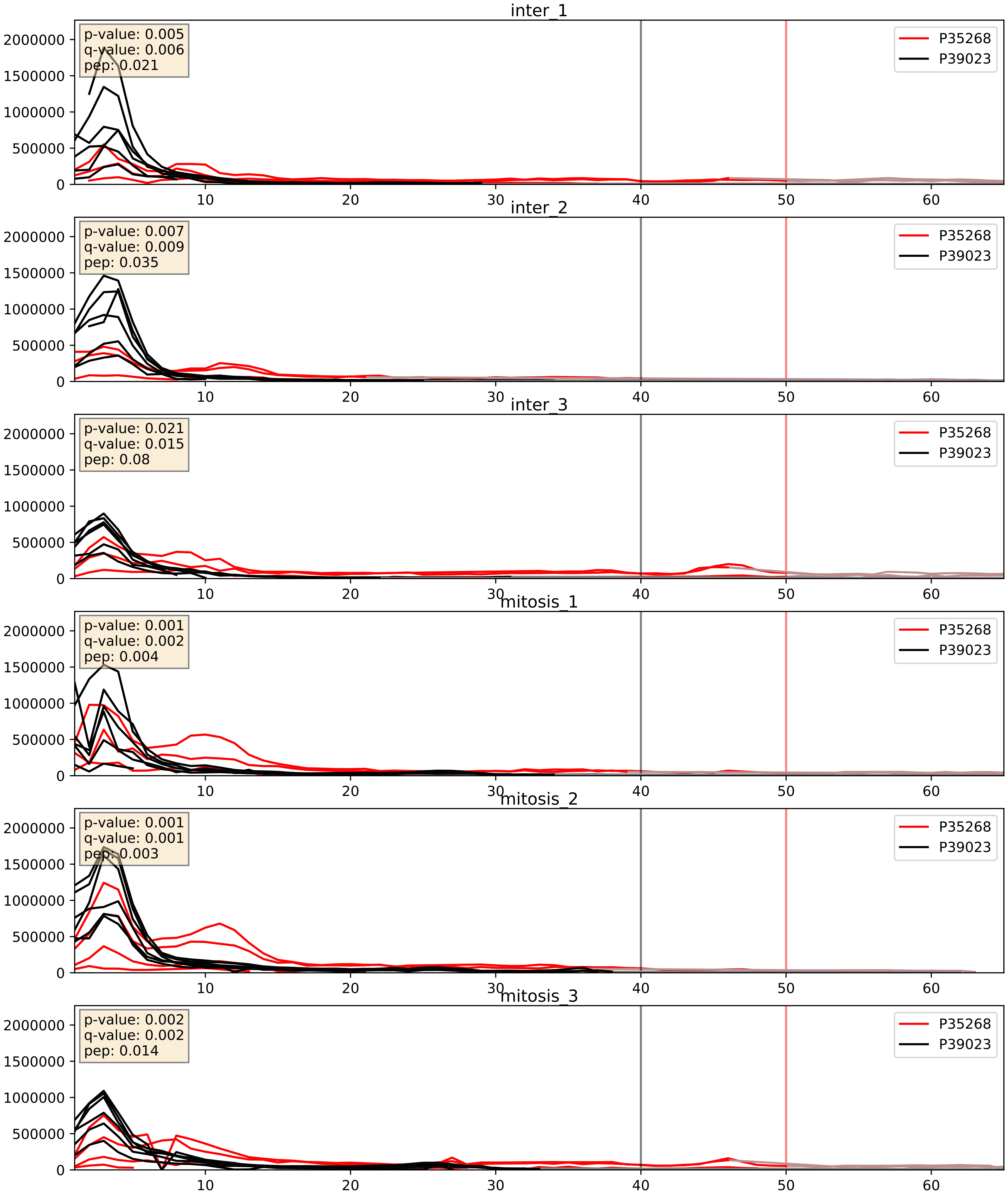


| level | condition_1 | condition_2 | pvalue | pvalue_adjusted |
|---|---|---|---|---|
| interactor_abundance | mitosis | inter | 0.00742054522726715 | 0.07459761261938556 |
| complex_abundance | mitosis | inter | 0.030772500800209473 | 0.12751204260889337 |
| interactor_ratio | mitosis | inter | 0.061718037558987436 | 0.41209547699292703 |

P35268_P41252
RL22_HUMAN vs SYIC_HUMAN
p-value: 0.0 q-value: 0.043

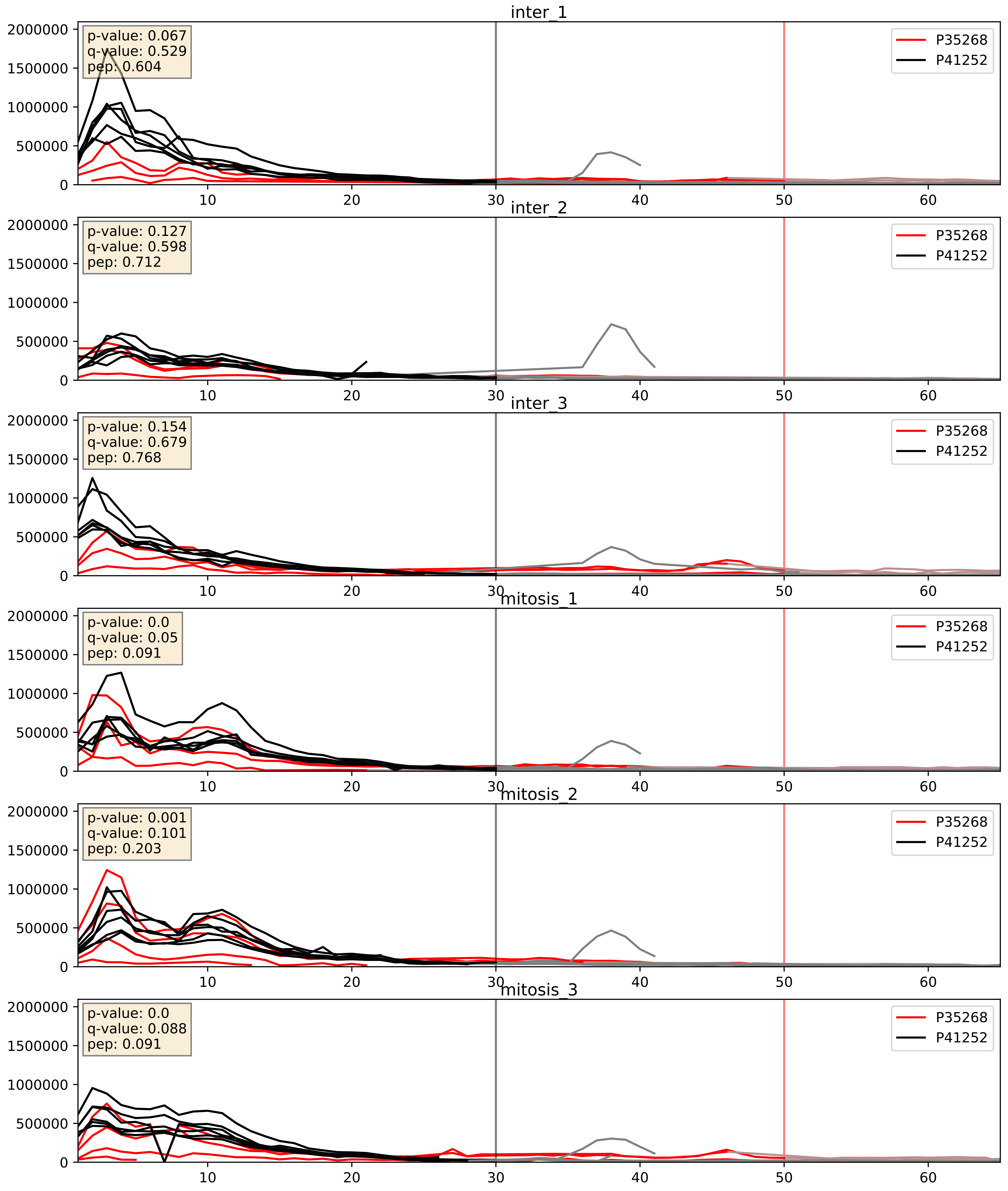


| level | condition_1 | condition_2 | pvalue | pvalue_adjusted |
|---|---|---|---|---|
| interactor_abundance | mitosis | inter | 0.009605436360937513 | 0.08507272274138435 |
| complex_abundance | mitosis | inter | 0.03983389750077914 | 0.1449976963395235 |
| interactor_ratio | mitosis | inter | 0.16802923527714494 | 0.6302368772162471 |

P35268_P42677
RL22_HUMAN vs RS27_HUMAN
p-value: 0.0 q-value: 0.0

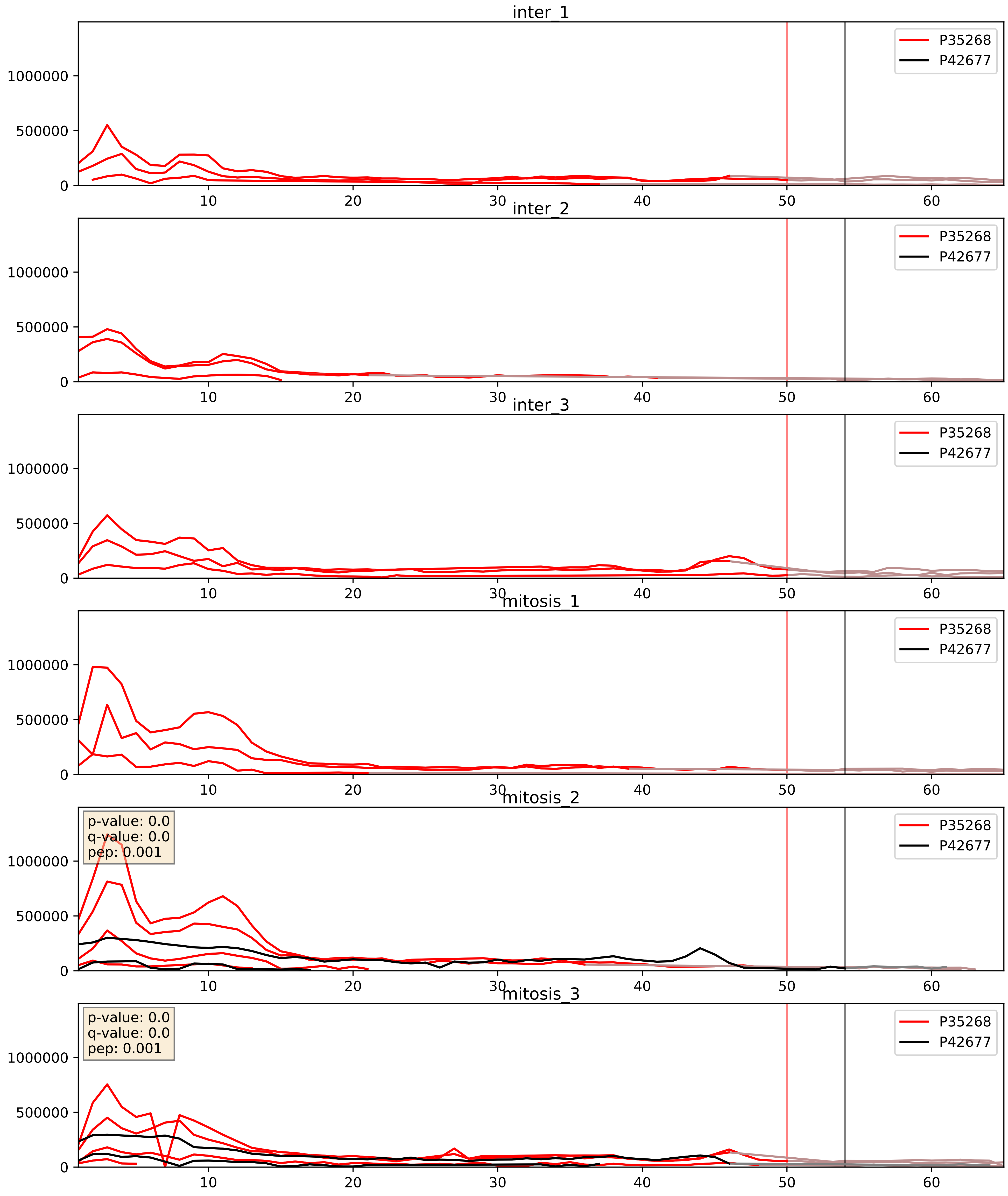


| level | condition_1 | condition_2 | pvalue | pvalue_adjusted |
|---|---|---|---|---|
| interactor_ratio | mitosis | inter | 0.13274569025832464 | 0.5815266676618521 |
| interactor_abundance | mitosis | inter | 0.14148540361303055 | 0.3287704445636205 |
| complex_abundance | mitosis | inter | 0.1456401137377613 | 0.29155229851491943 |

P35268_P42766
RL22_HUMAN vs RL35_HUMAN
p-value: 0.0 q-value: 0.0

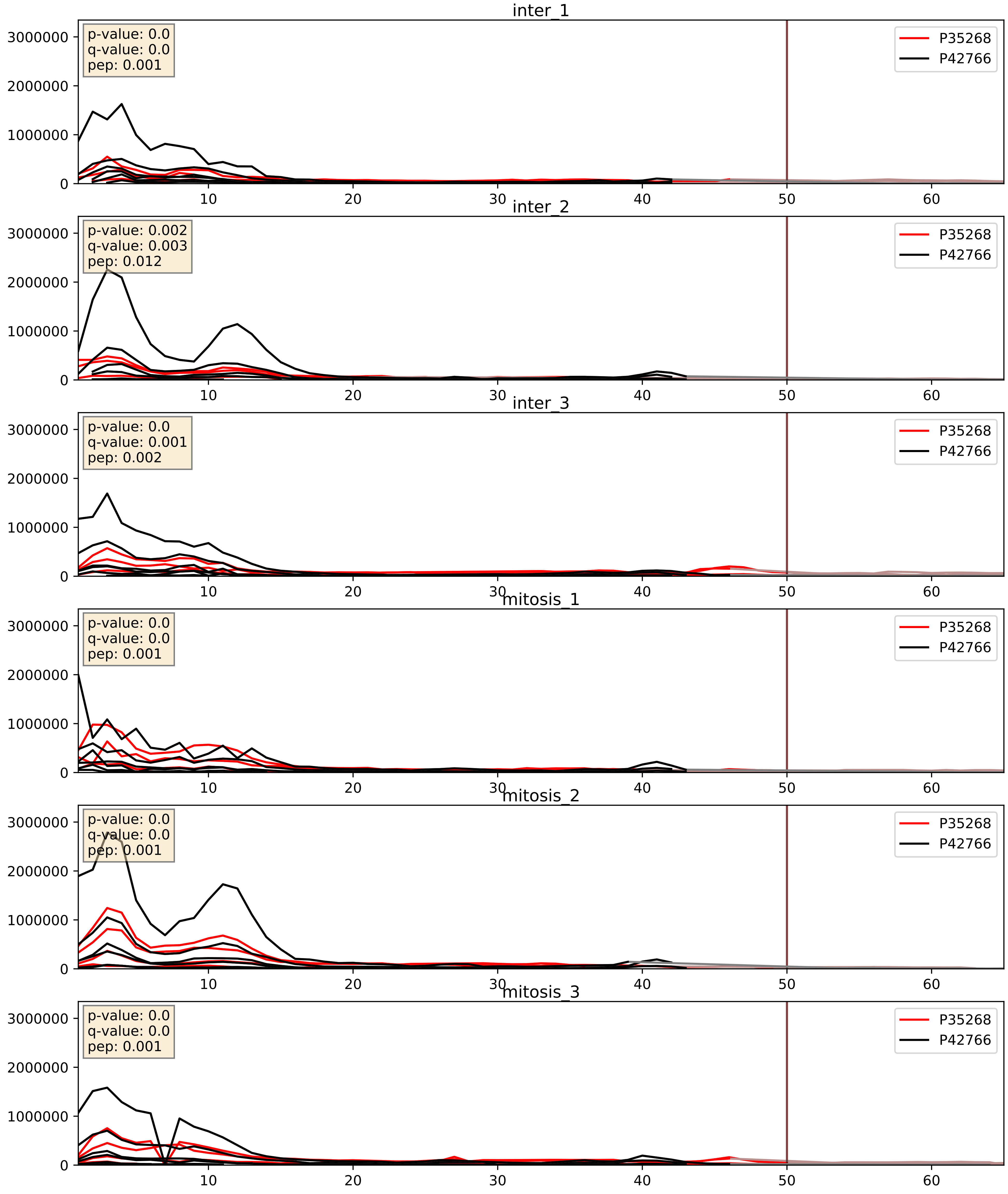


| level | condition_1 | condition_2 | pvalue | pvalue_adjusted |
|---|---|---|---|---|
| interactor_abundance | mitosis | inter | 0.028762389464174098 | 0.14032832933219166 |
| interactor_ratio | mitosis | inter | 0.048117240695319266 | 0.3616185955679832 |
| complex_abundance | mitosis | inter | 0.09155070953784378 | 0.22288639049953404 |

P35268_P46776
RL22_HUMAN vs RL27A_HUMAN
p-value: 0.0 q-value: 0.0

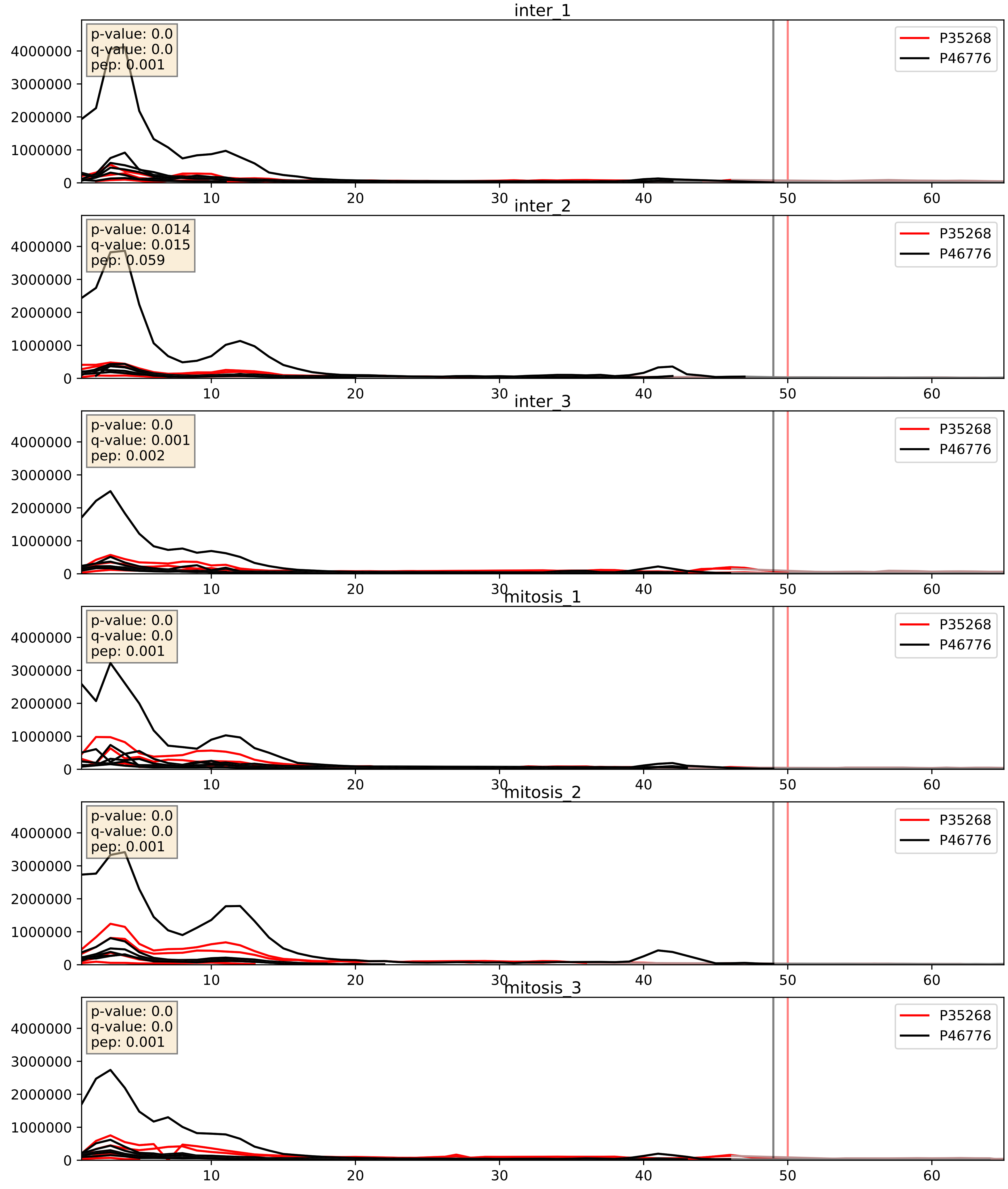


| level | condition_1 | condition_2 | pvalue | pvalue_adjusted |
|---|---|---|---|---|
| complex_abundance | mitosis | inter | 0.01413726847873977 | 0.08846811134929532 |
| interactor_abundance | mitosis | inter | 0.03125050448751661 | 0.1469400265933217 |
| interactor_ratio | mitosis | inter | 0.10926655659212704 | 0.5488977256036429 |

P35268_P46777
RL22_HUMAN vs RL5_HUMAN
p-value: 0.054 q-value: 0.035

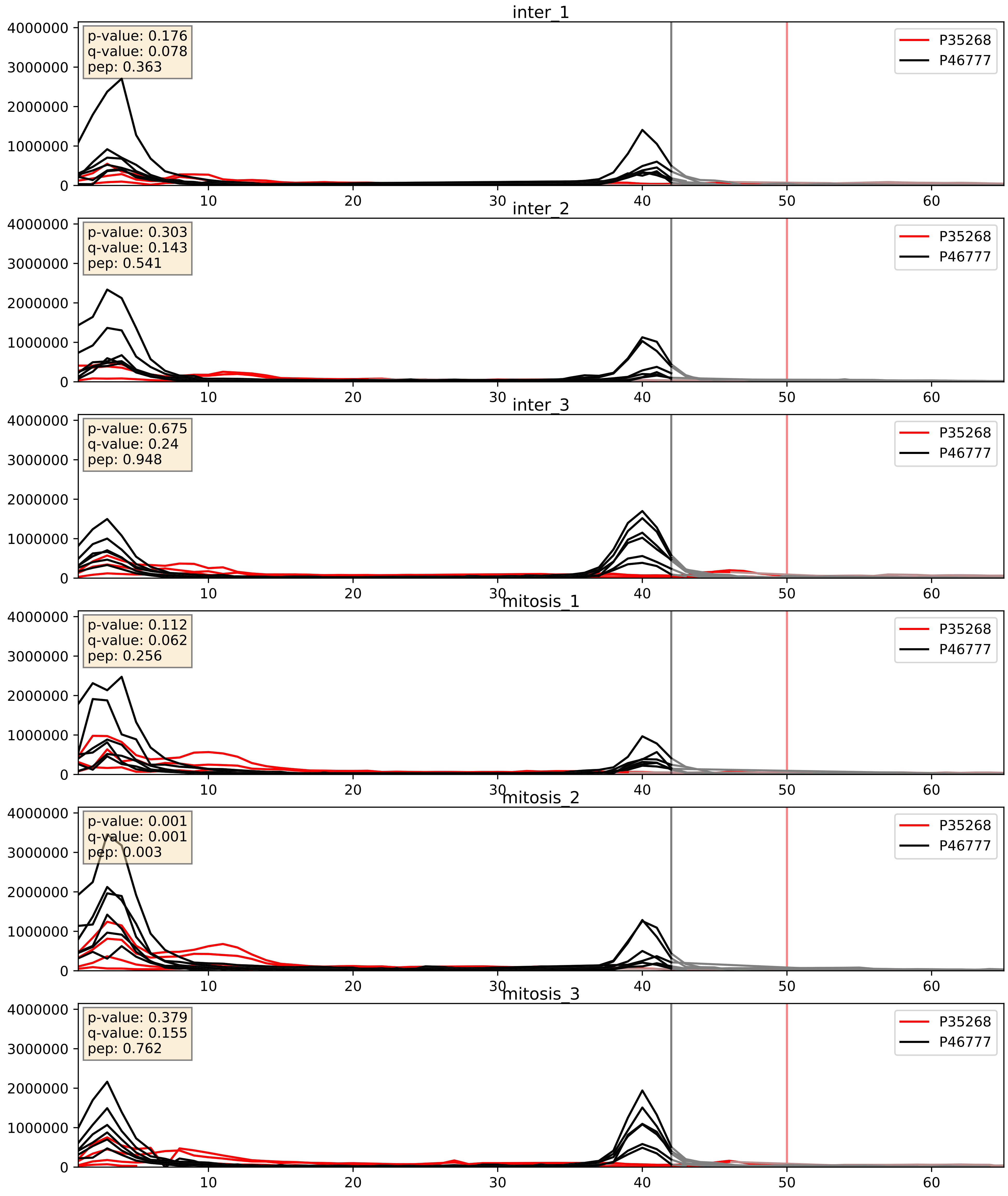


| level | condition_1 | condition_2 | pvalue | pvalue_adjusted |
|---|---|---|---|---|
| interactor_abundance | mitosis | inter | 0.010296112912537297 | 0.08839992631025002 |
| interactor_ratio | mitosis | inter | 0.07631250674185508 | 0.4639941947788648 |
| complex_abundance | mitosis | inter | 0.16429960921075804 | 0.313300212707527 |

P35268_P46778
RL22_HUMAN vs RL21_HUMAN
p-value: 0.0 q-value: 0.0

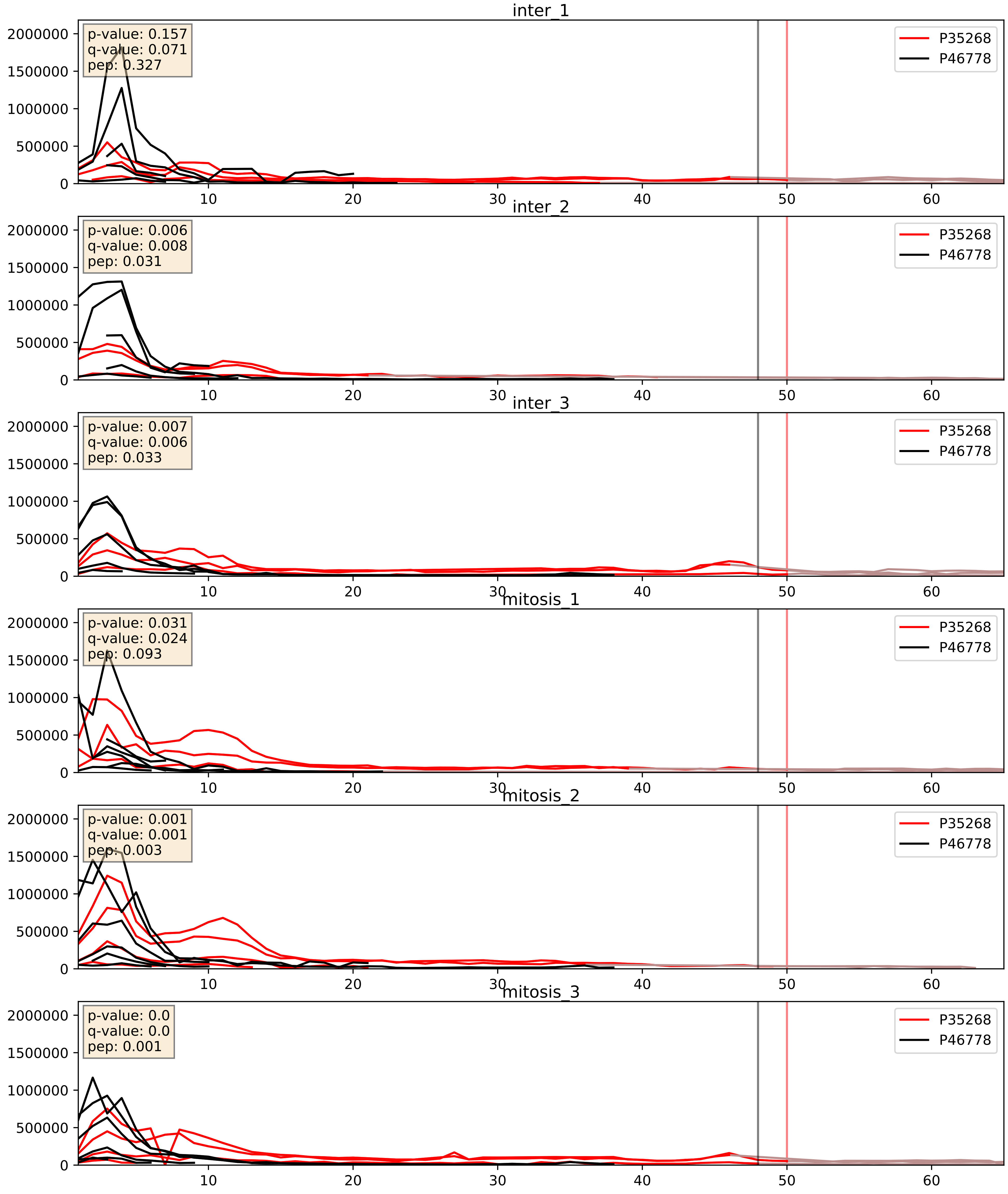


| level | condition_1 | condition_2 | pvalue | pvalue_adjusted |
|---|---|---|---|---|
| interactor_ratio | mitosis | inter | 0.007854201071049864 | 0.12516692290607134 |
| interactor_abundance | mitosis | inter | 0.030086876453833896 | 0.14418523491160162 |
| complex_abundance | mitosis | inter | 0.2995877636523552 | 0.4492766742929503 |

P35268_P46779
RL22_HUMAN vs RL28_HUMAN
p-value: 0.0 q-value: 0.0

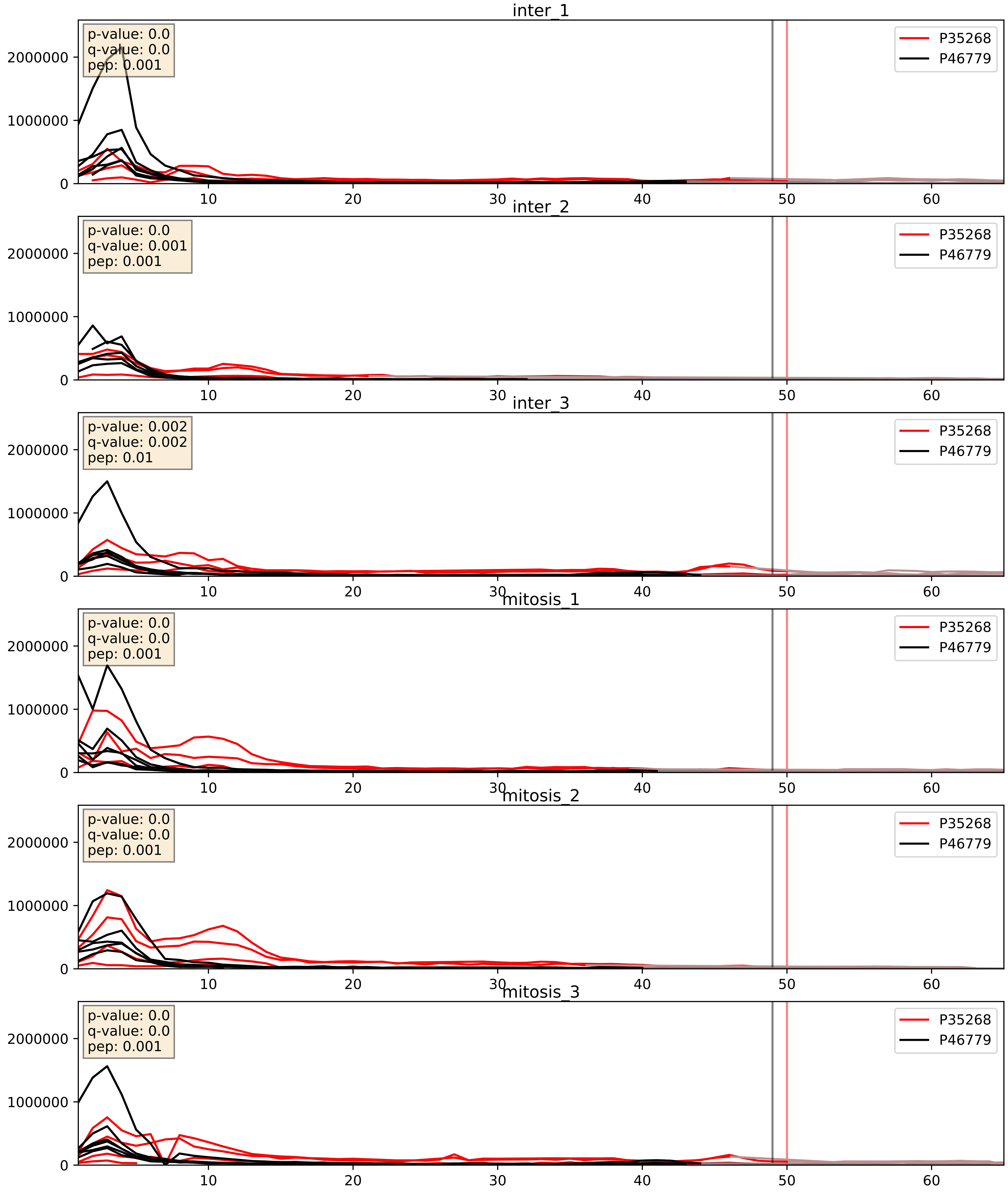


| level | condition_1 | condition_2 | pvalue | pvalue_adjusted |
|---|---|---|---|---|
| interactor_ratio | mitosis | inter | 0.015677349611296285 | 0.18459162678500163 |
| interactor_abundance | mitosis | inter | 0.016318721607803916 | 0.1120194522556548 |
| complex_abundance | mitosis | inter | 0.1489732054740355 | 0.29488459277225637 |

P35268_P46781
RL22_HUMAN vs RS9_HUMAN
p-value: 0.0 q-value: 0.0

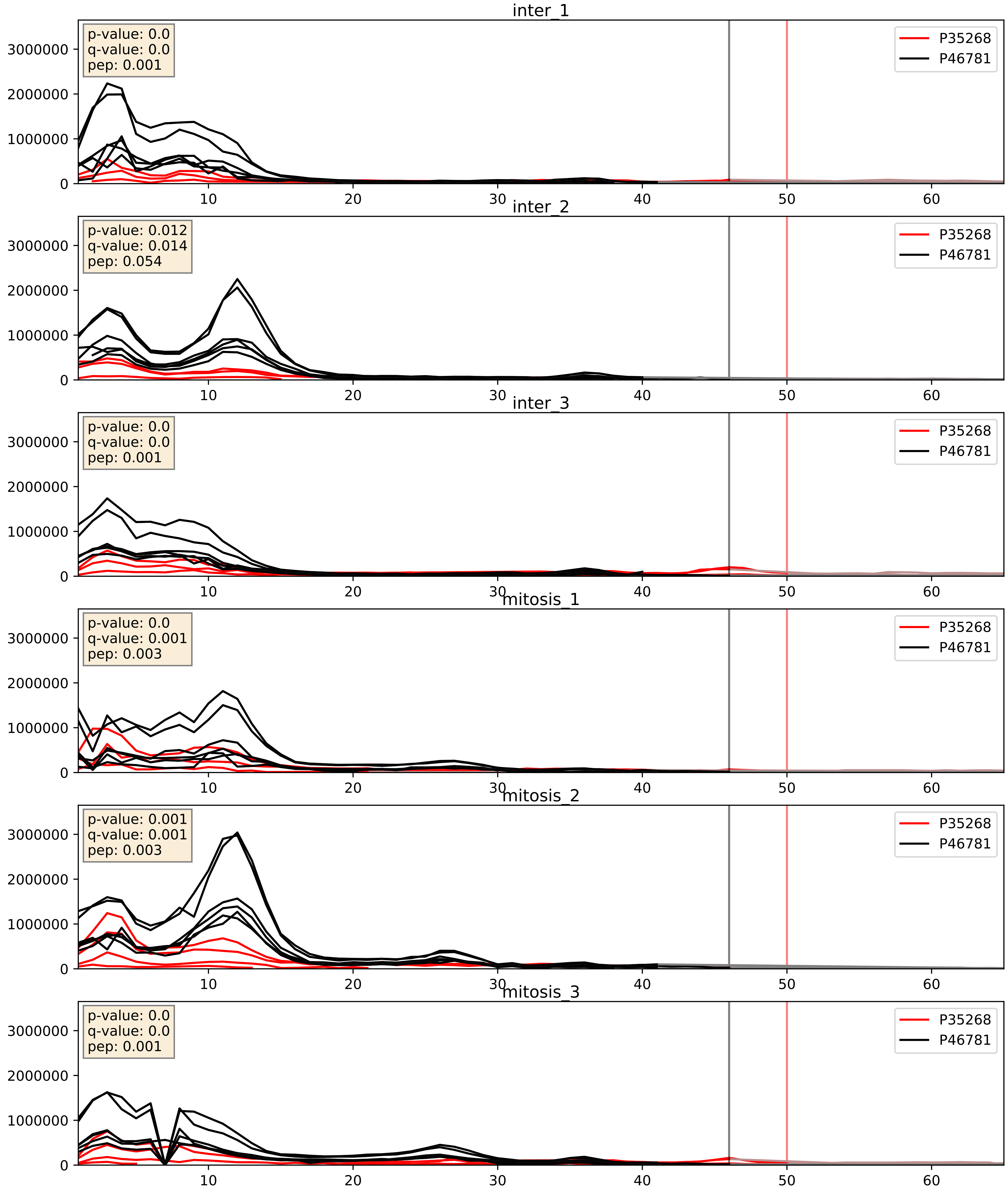


| level | condition_1 | condition_2 | pvalue | pvalue_adjusted |
|---|---|---|---|---|
| interactor_abundance | mitosis | inter | 0.02184175207454888 | 0.12300355115667001 |
| interactor_ratio | mitosis | inter | 0.030507036392276504 | 0.27810461290509786 |
| complex_abundance | mitosis | inter | 0.09721297481016311 | 0.23097403067083264 |

P35268_P46782
RL22_HUMAN vs RS5_HUMAN
p-value: 0.001 q-value: 0.001

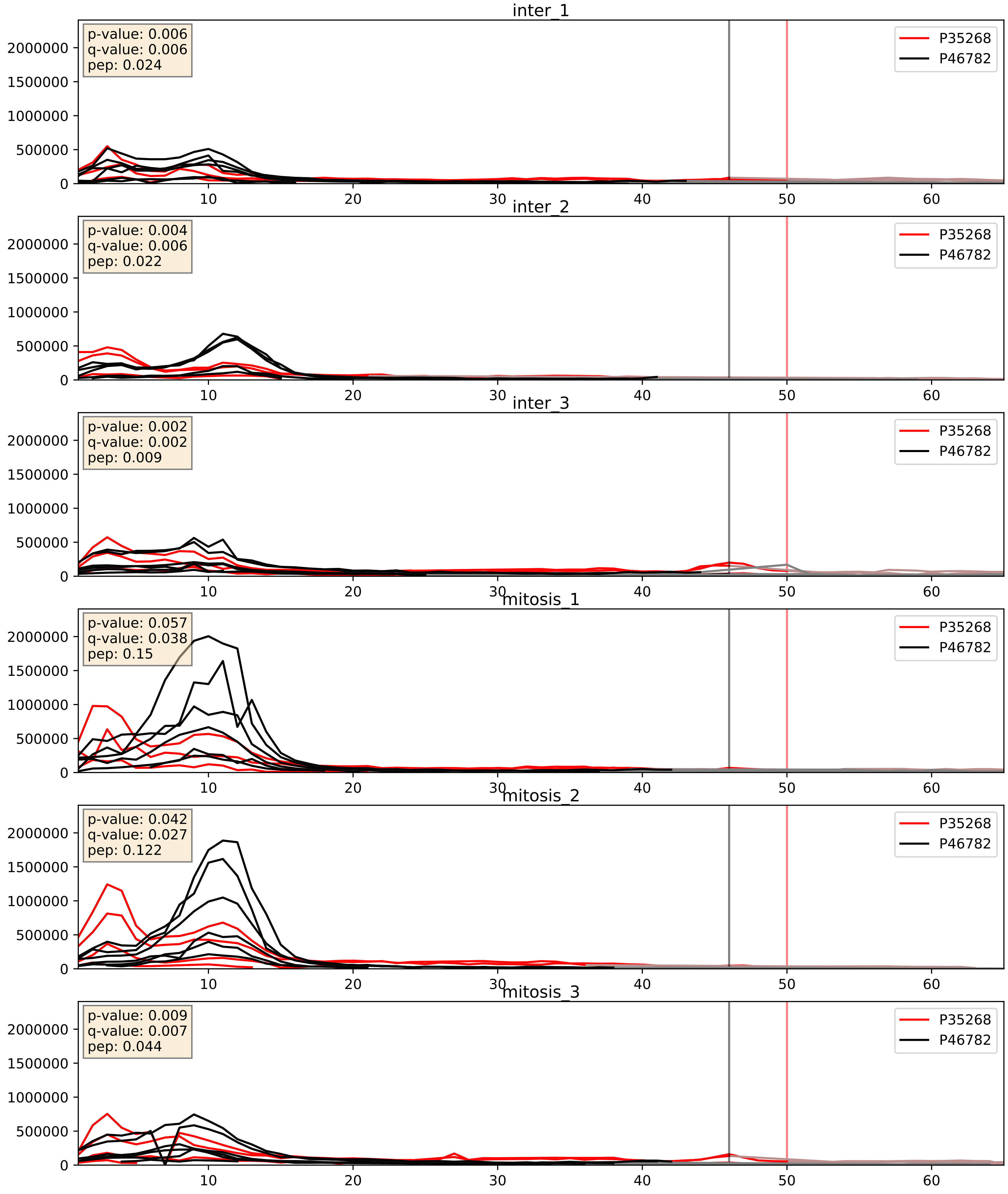


| level | condition_1 | condition_2 | pvalue | pvalue_adjusted |
|---|---|---|---|---|
| complex_abundance | mitosis | inter | 0.016041114677862726 | 0.09450236864590841 |
| interactor_abundance | mitosis | inter | 0.02337378226101933 | 0.12556564873974368 |
| interactor_ratio | mitosis | inter | 0.9399372146826102 | 0.9903819002564184 |

P35268_P46783
RL22_HUMAN vs RS10_HUMAN
p-value: 0.008 q-value: 0.008

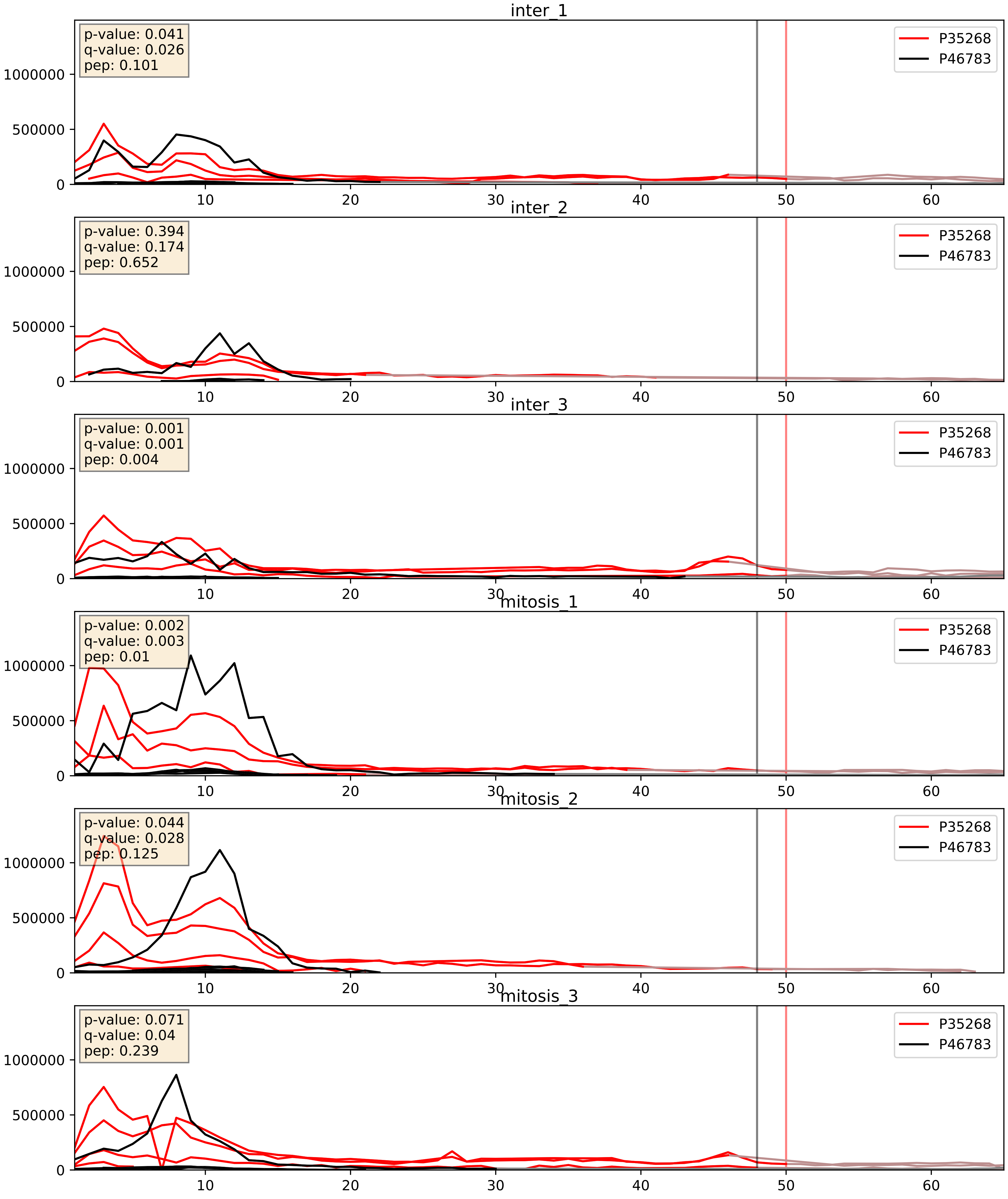


| level | condition_1 | condition_2 | pvalue | pvalue_adjusted |
|---|---|---|---|---|
| interactor_abundance | mitosis | inter | 0.02214083462429393 | 0.12329937058940194 |
| complex_abundance | mitosis | inter | 0.07857209659196283 | 0.2043232668723216 |
| interactor_ratio | mitosis | inter | 0.22593708830381132 | 0.7051563050171857 |

P35268_P47914
RL22_HUMAN vs RL29_HUMAN
p-value: 0.0 q-value: 0.0

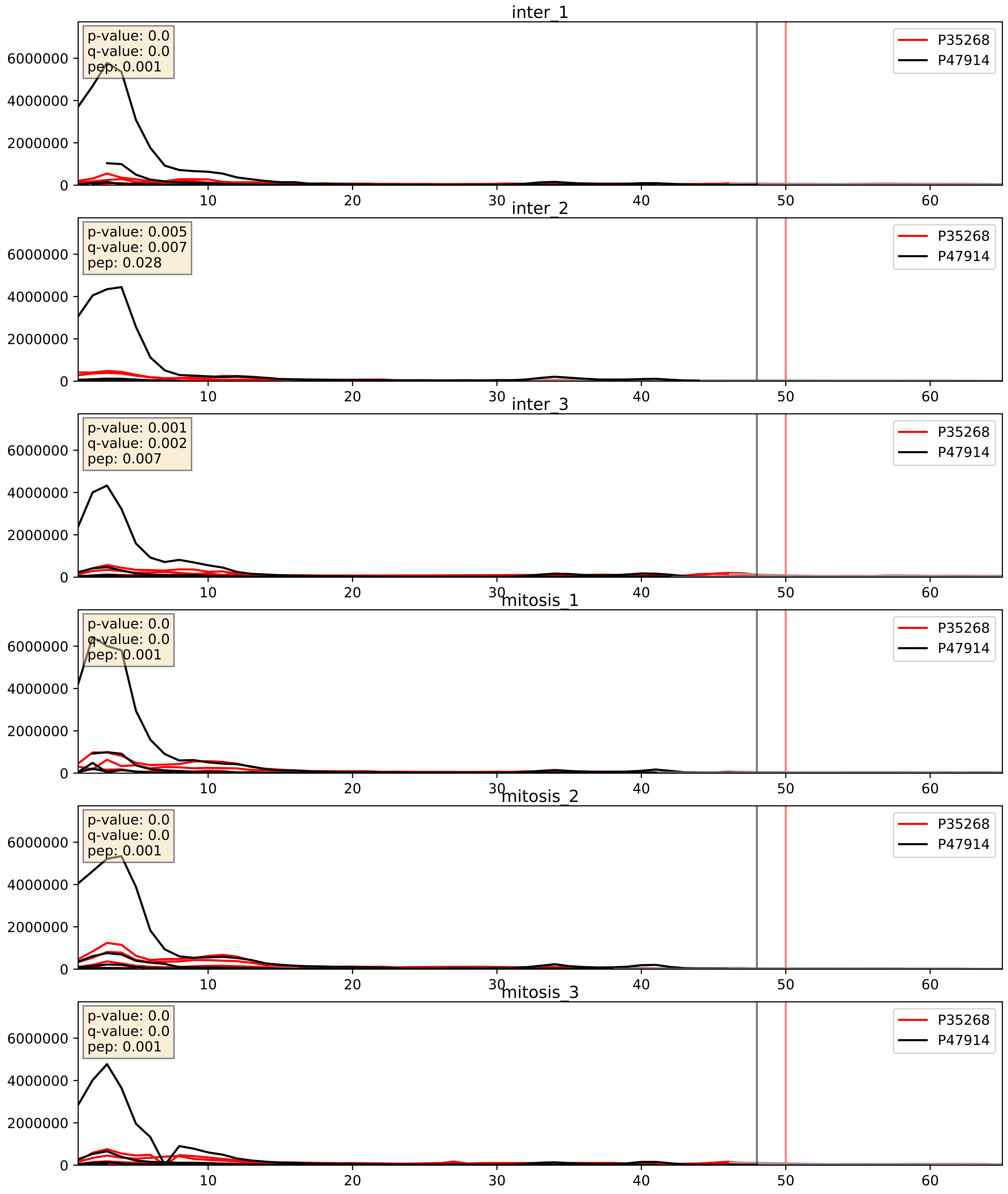


| level | condition_1 | condition_2 | pvalue | pvalue_adjusted |
|---|---|---|---|---|
| interactor_abundance | mitosis | inter | 0.02824331336117877 | 0.13974940313044457 |
| complex_abundance | mitosis | inter | 0.12521034869759495 | 0.26706591785818723 |
| interactor_ratio | mitosis | inter | 0.7637254962784351 | 0.9536594850393604 |

P35268_P49207
RL22_HUMAN vs RL34_HUMAN
p-value: 0.0 q-value: 0.0

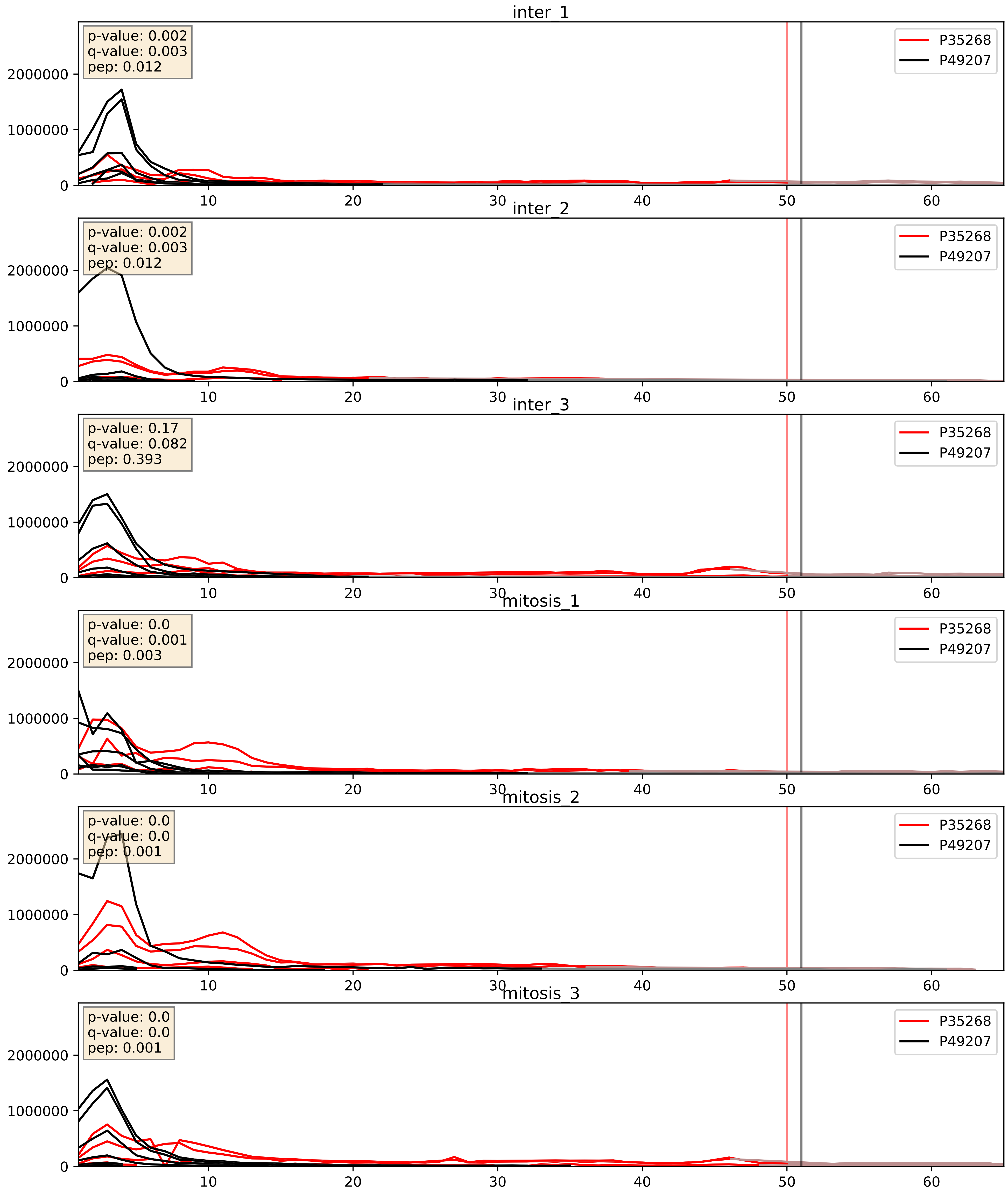


| level | condition_1 | condition_2 | pvalue | pvalue_adjusted |
|---|---|---|---|---|
| interactor_abundance | mitosis | inter | 0.006138385291398371 | 0.06671148765013551 |
| interactor_ratio | mitosis | inter | 0.3171486556067397 | 0.7783421708721073 |
| complex_abundance | mitosis | inter | 0.6716378467965999 | 0.7812501655902833 |

P35268_P49458
RL22_HUMAN vs SRP09_HUMAN
p-value: 0.0 q-value: 0.0

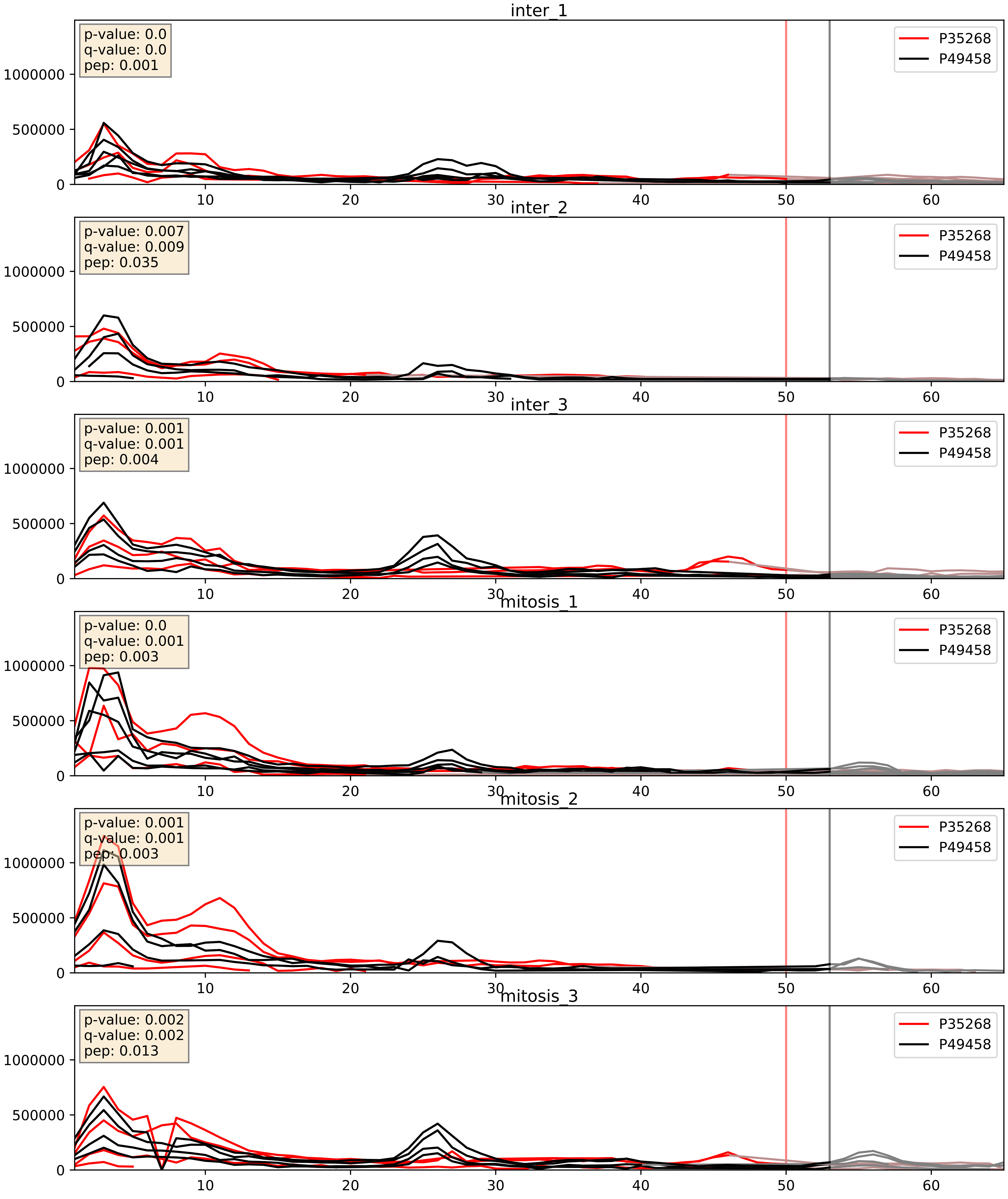


| level | condition_1 | condition_2 | pvalue | pvalue_adjusted |
|---|---|---|---|---|
| interactor_abundance | mitosis | inter | 0.038878589075571256 | 0.16041561241136493 |
| complex_abundance | mitosis | inter | 0.12285425111174318 | 0.26509513221994496 |
| interactor_ratio | mitosis | inter | 0.17006382700324327 | 0.6328516675108425 |

P35268_P50914
RL22_HUMAN vs RL14_HUMAN
p-value: 0.0 q-value: 0.0

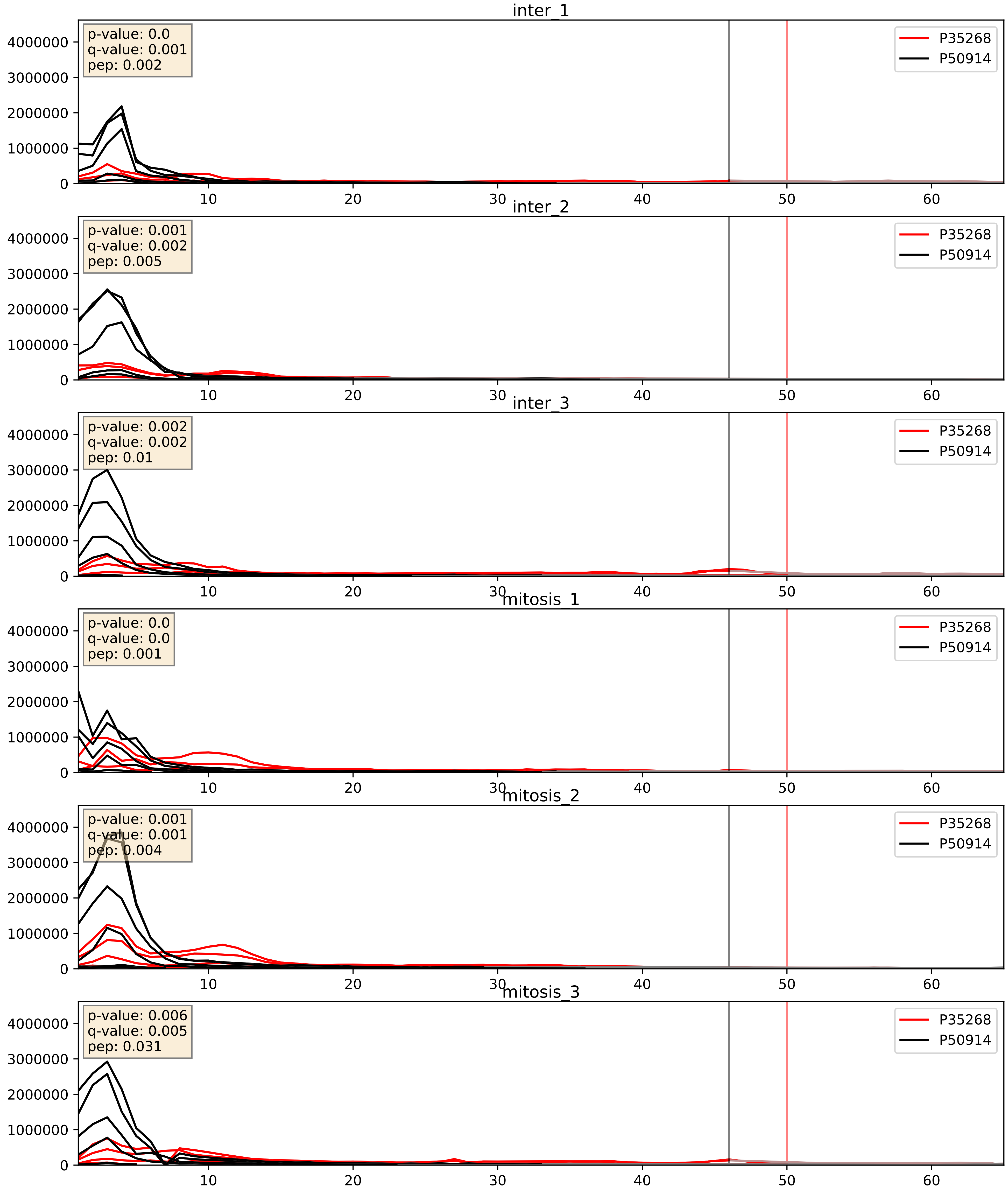


| level | condition_1 | condition_2 | pvalue | pvalue_adjusted |
|---|---|---|---|---|
| interactor_abundance | mitosis | inter | 0.009511085383422742 | 0.08467487351232311 |
| interactor_ratio | mitosis | inter | 0.05672354220694349 | 0.39385842361407786 |
| complex_abundance | mitosis | inter | 0.2915404434727278 | 0.4406049075082185 |

P35268_P55769
RL22_HUMAN vs NH2L1_HUMAN
p-value: 0.0 q-value: 0.016

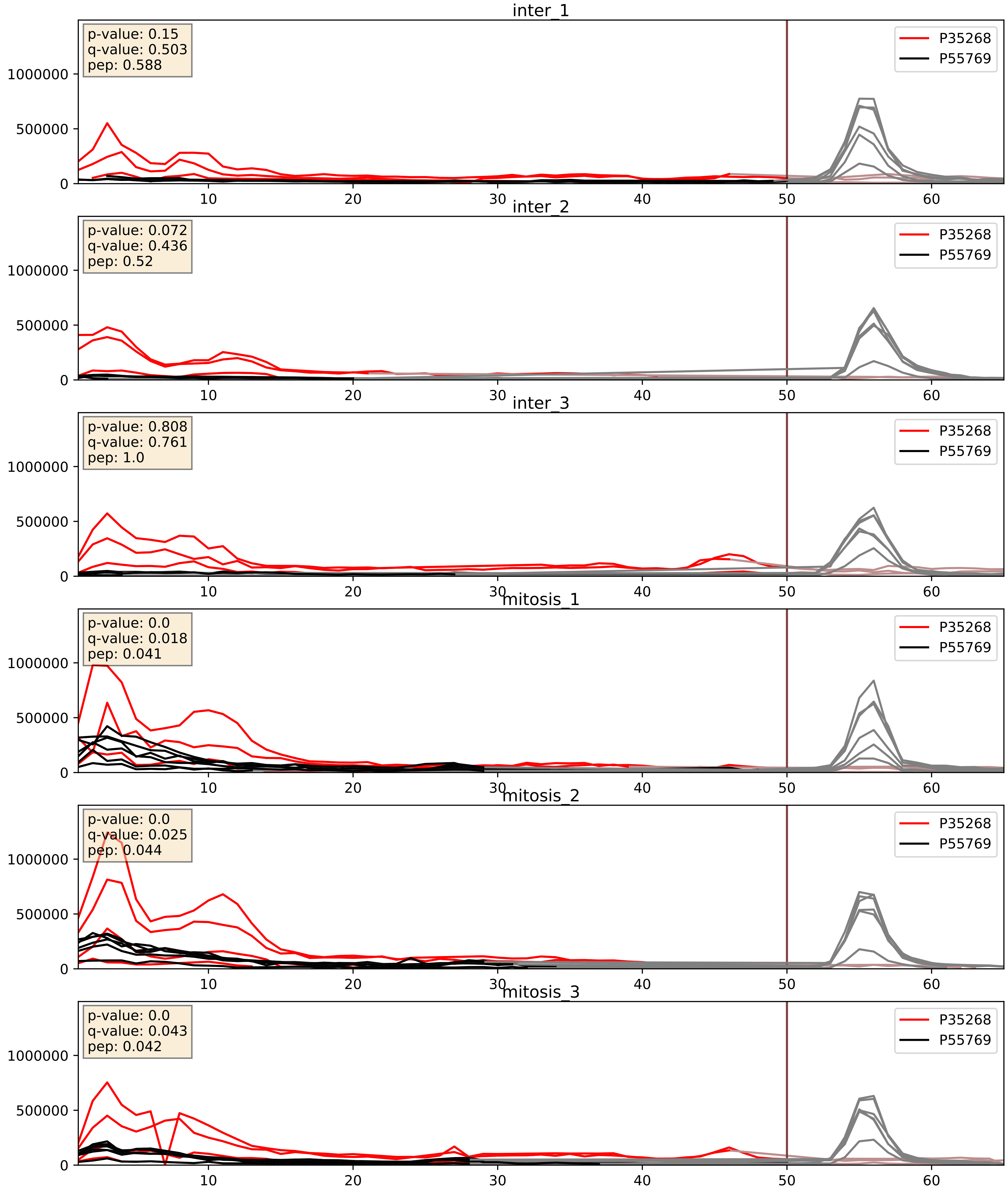


| level | condition_1 | condition_2 | pvalue | pvalue_adjusted |
|---|---|---|---|---|
| complex_abundance | mitosis | inter | 0.0010492460264906755 | 0.021612845224852777 |
| interactor_abundance | mitosis | inter | 0.014141208892344399 | 0.10541033287779904 |
| interactor_ratio | mitosis | inter | 0.024818760043489395 | 0.24732082185363125 |

P35268_P60468
RL22_HUMAN vs SC61B_HUMAN
p-value: 0.0 q-value: 0.0

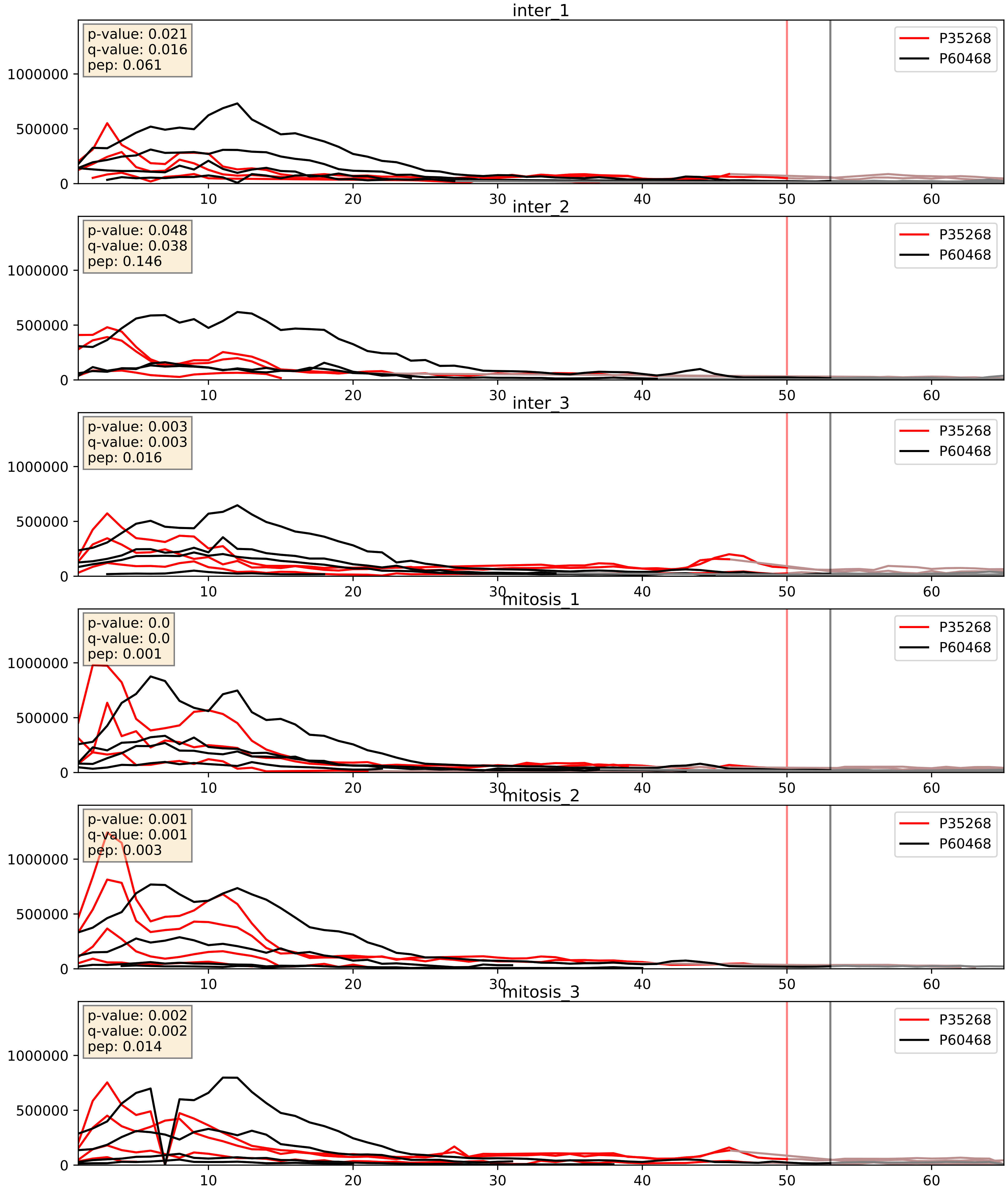


| level | condition_1 | condition_2 | pvalue | pvalue_adjusted |
|---|---|---|---|---|
| interactor_abundance | mitosis | inter | 0.040438240341609484 | 0.16224576392040177 |
| interactor_ratio | mitosis | inter | 0.06822206804345823 | 0.43247946911820456 |
| complex_abundance | mitosis | inter | 0.731253314838157 | 0.8212448668347709 |

P35268_P60866
RL22_HUMAN vs RS20_HUMAN
p-value: 0.003 q-value: 0.003

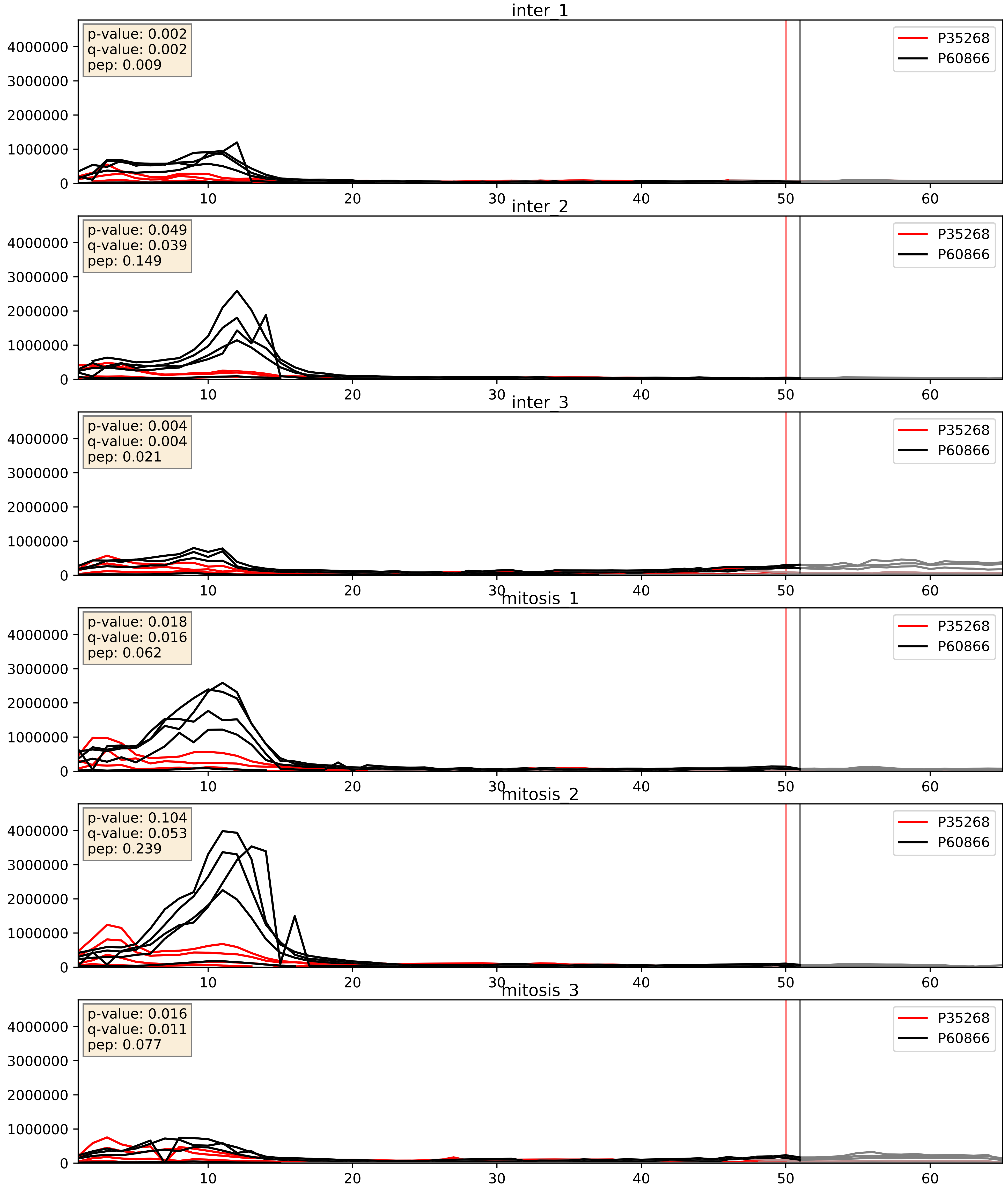


| level | condition_1 | condition_2 | pvalue | pvalue_adjusted |
|---|---|---|---|---|
| interactor_abundance | mitosis | inter | 0.040267692761700224 | 0.16188554627742371 |
| complex_abundance | mitosis | inter | 0.0688142399094501 | 0.18995481896965266 |
| interactor_ratio | mitosis | inter | 0.8194967016902116 | 0.9672664180415323 |

P35268_P61247
RL22_HUMAN vs RS3A_HUMAN
p-value: 0.0 q-value: 0.0

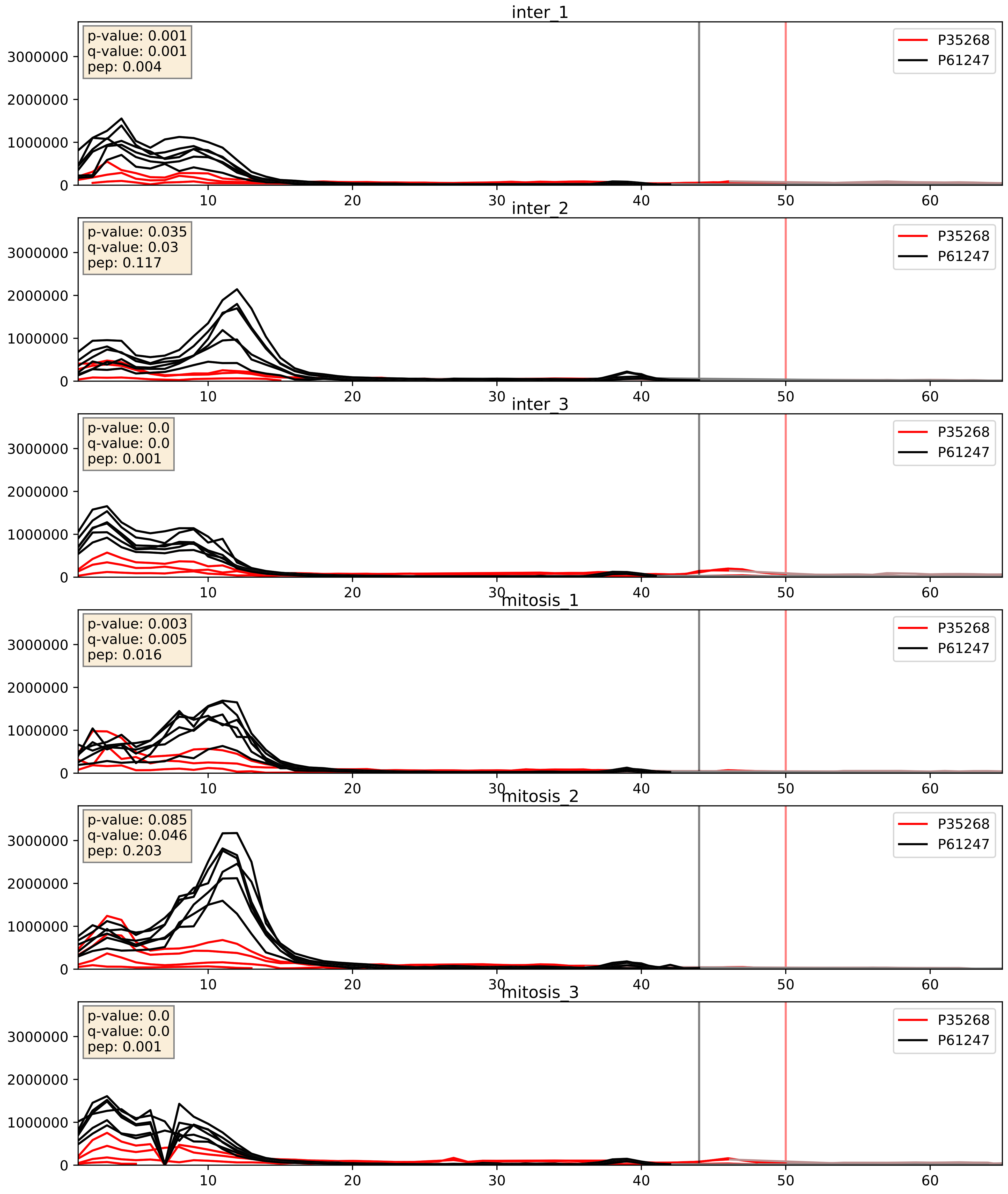


| level | condition_1 | condition_2 | pvalue | pvalue_adjusted |
|---|---|---|---|---|
| interactor_ratio | mitosis | inter | 0.0038237859648561343 | 0.08075090146433855 |
| interactor_abundance | mitosis | inter | 0.011597435274916442 | 0.0950928163026453 |
| complex_abundance | mitosis | inter | 0.10038583194691802 | 0.2350018038308818 |

P35268_P61254
RL22_HUMAN vs RL26_HUMAN
p-value: 0.002 q-value: 0.003

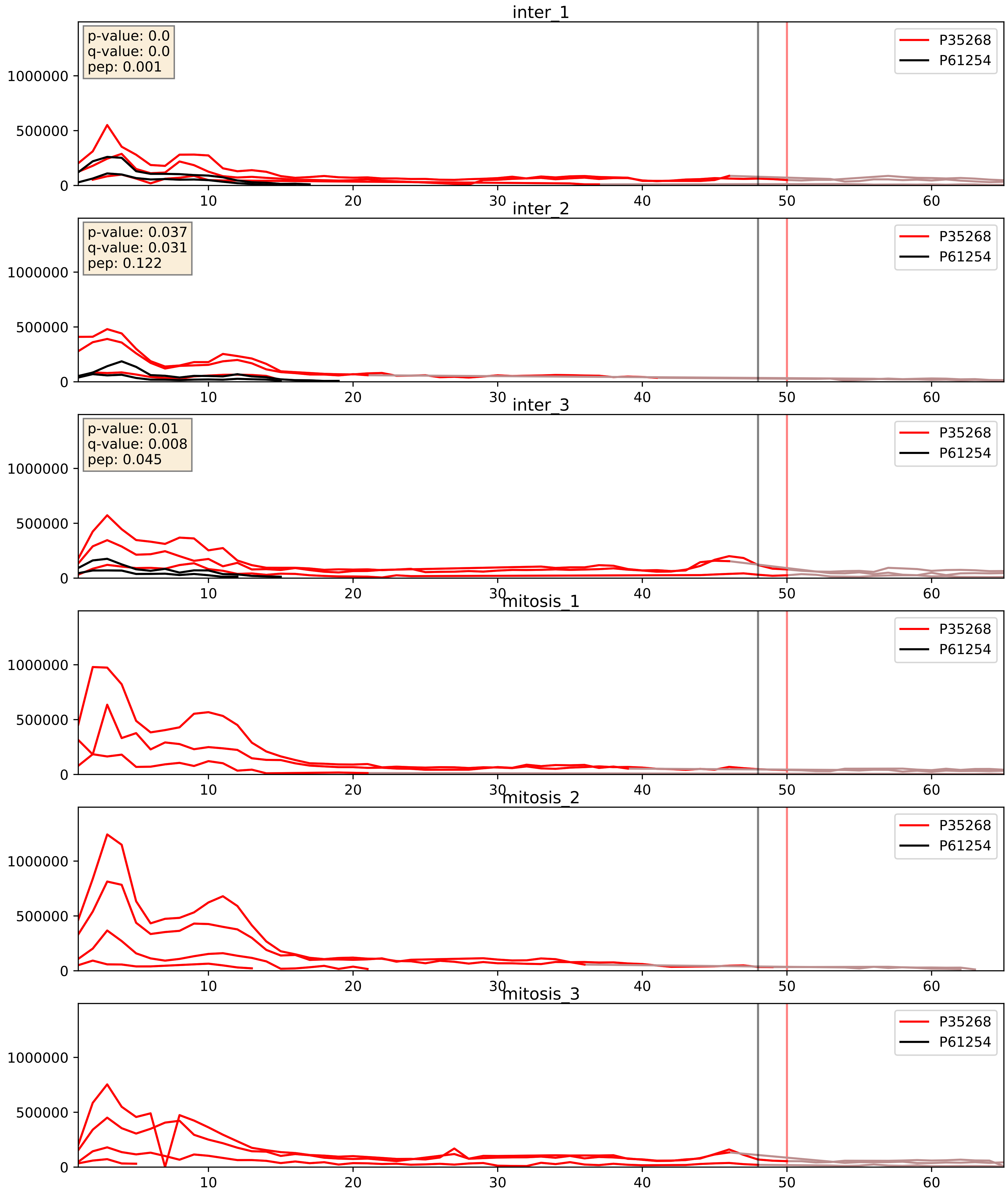


| level | condition_1 | condition_2 | pvalue | pvalue_adjusted |
|---|---|---|---|---|
| interactor_abundance | mitosis | inter | 7.421754572657264e-05 | 0.006870282762438709 |
| complex_abundance | mitosis | inter | 8.3817361471577e-05 | 0.006185143225833613 |
| interactor_ratio | mitosis | inter | 0.0008875457249814373 | 0.034218313408302205 |

P35268_P61313
RL22_HUMAN vs RL15_HUMAN
p-value: 0.0 q-value: 0.0

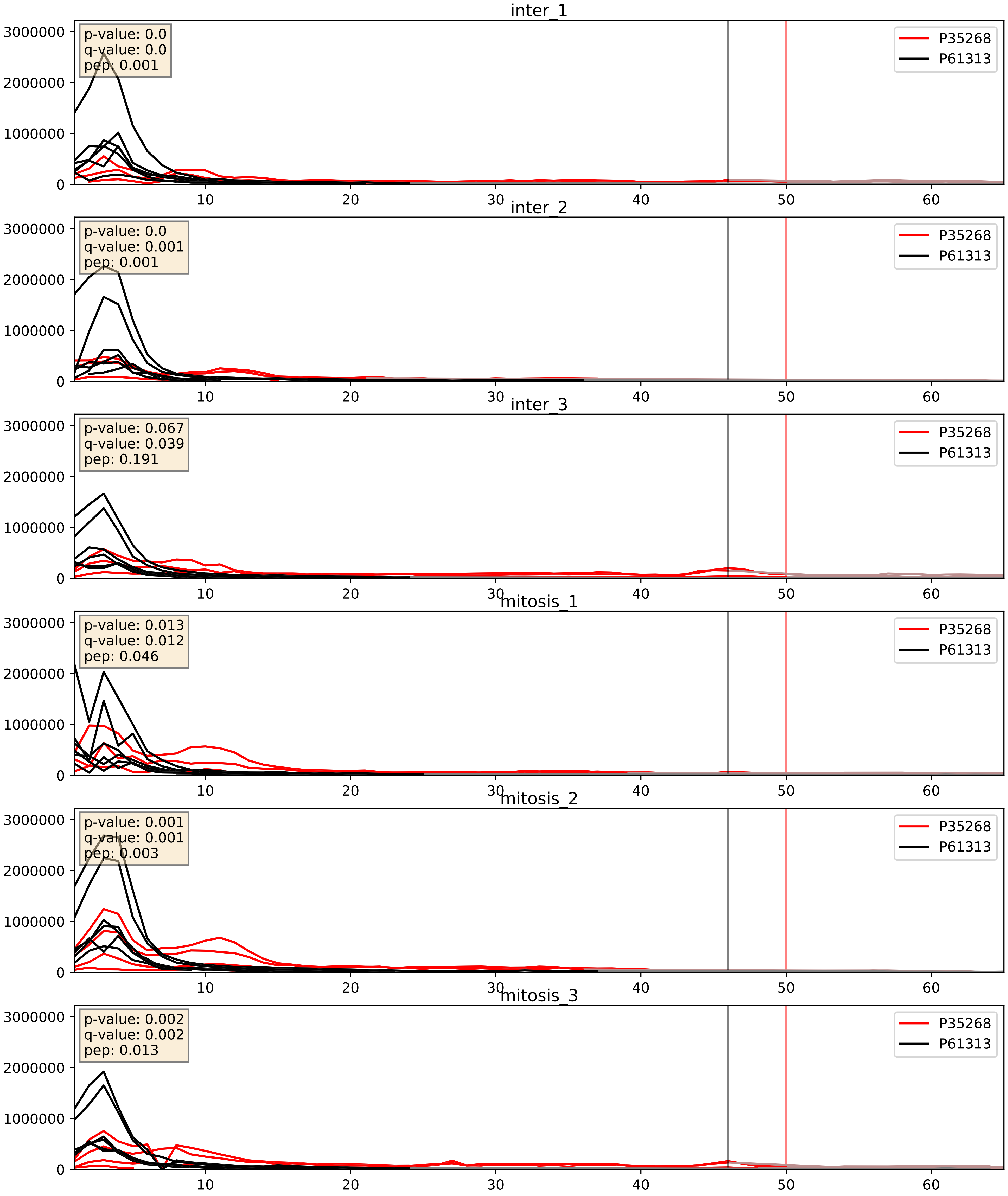


| level | condition_1 | condition_2 | pvalue | pvalue_adjusted |
|---|---|---|---|---|
| interactor_abundance | mitosis | inter | 0.025158206349733474 | 0.13151404357478994 |
| complex_abundance | mitosis | inter | 0.05792275015579216 | 0.17180136567345144 |
| interactor_ratio | mitosis | inter | 0.10549026842768912 | 0.5403929968527941 |

P35268_P61353
RL22_HUMAN vs RL27_HUMAN
p-value: 0.0 q-value: 0.0

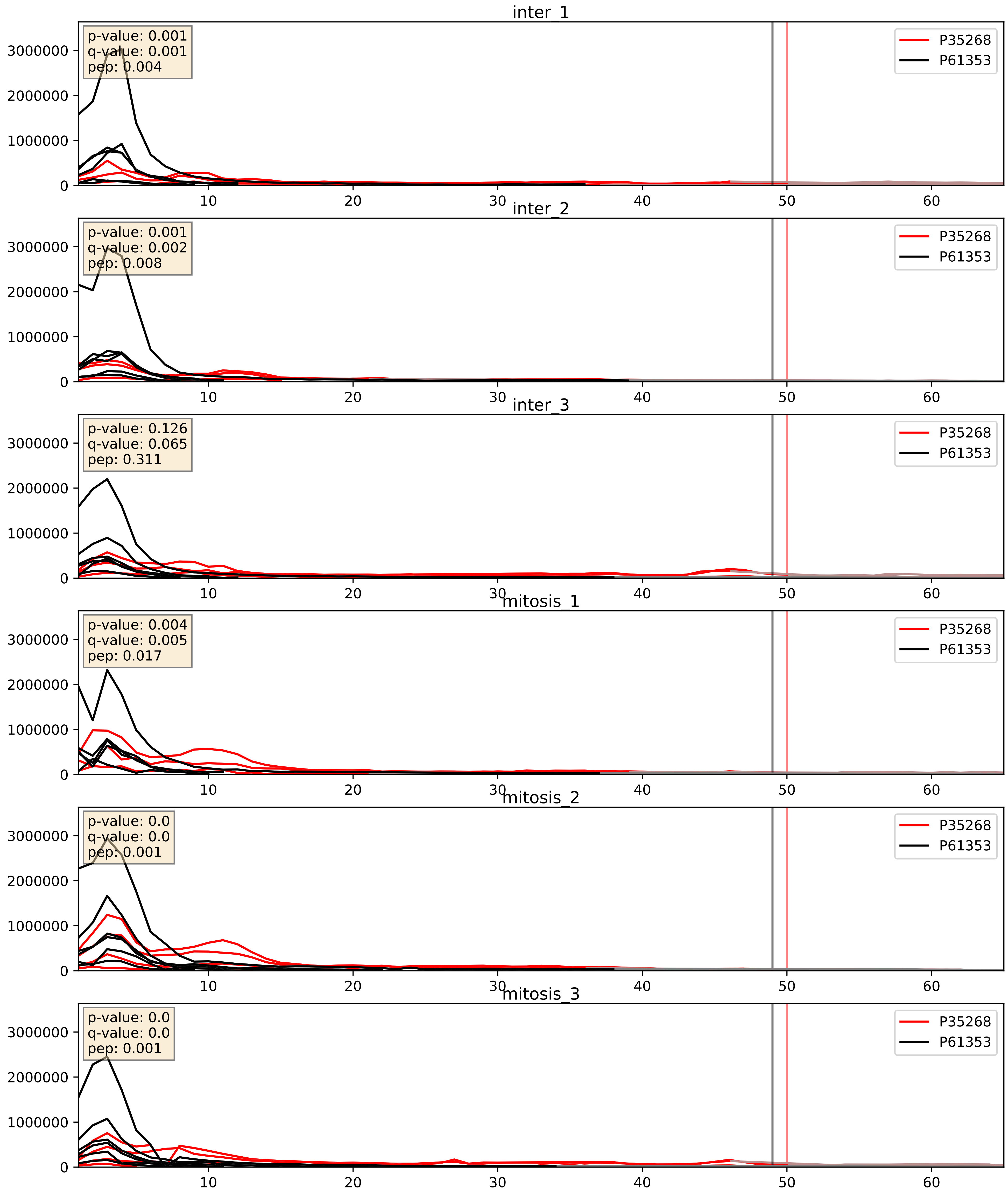


| level | condition_1 | condition_2 | pvalue | pvalue_adjusted |
|---|---|---|---|---|
| interactor_abundance | mitosis | inter | 0.011611818406732543 | 0.0950928163026453 |
| interactor_ratio | mitosis | inter | 0.028100497074638374 | 0.2625985316145245 |
| complex_abundance | mitosis | inter | 0.07576798894609839 | 0.1999303284151055 |

P35268_P61619
RL22_HUMAN vs S61A1_HUMAN
p-value: 0.001 q-value: 0.001

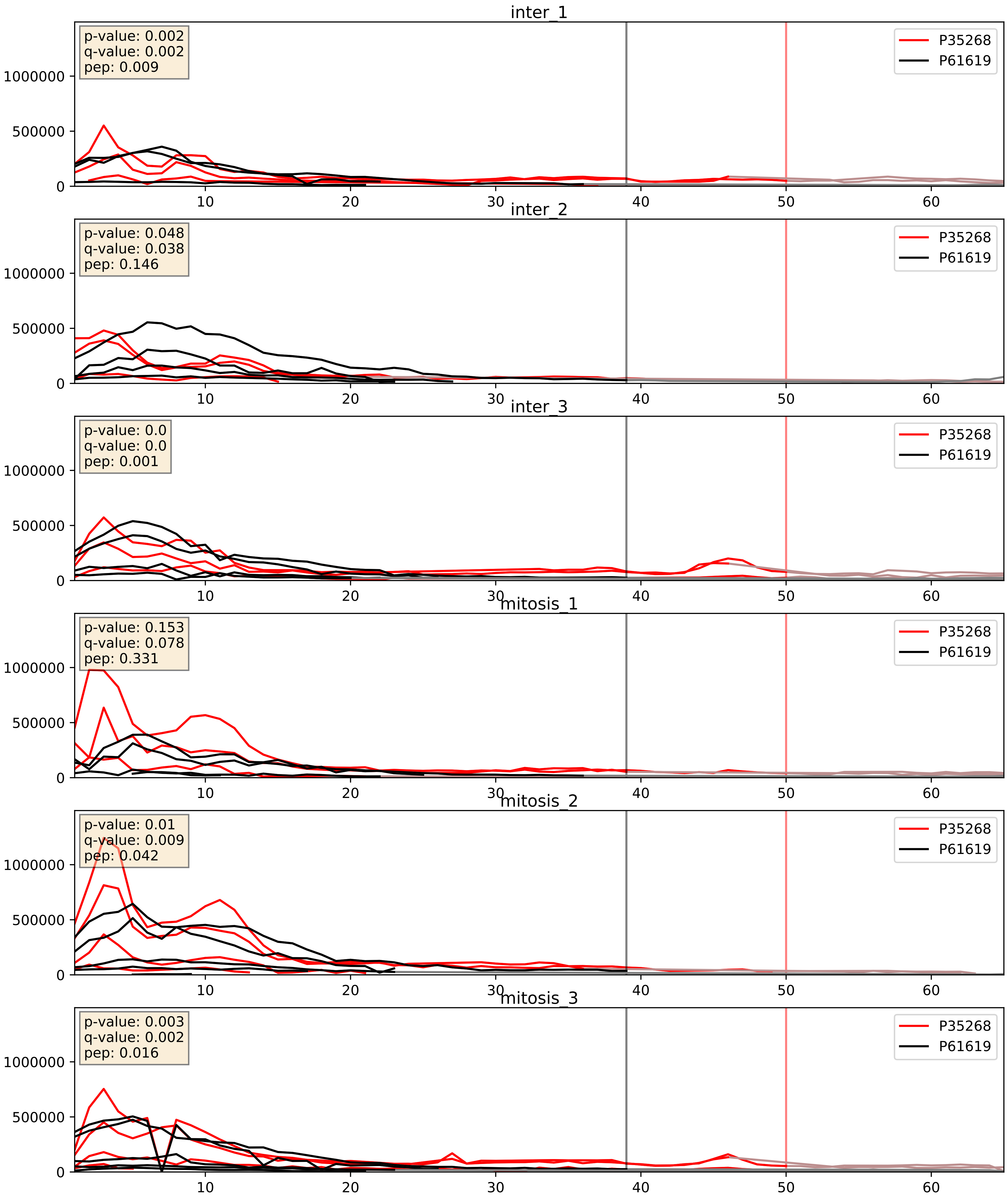


| level | condition_1 | condition_2 | pvalue | pvalue_adjusted |
|---|---|---|---|---|
| interactor_abundance | mitosis | inter | 0.01380959642827535 | 0.10419580910183958 |
| interactor_ratio | mitosis | inter | 0.02745093431290878 | 0.2593598208813456 |
| complex_abundance | mitosis | inter | 0.38008220322385666 | 0.5272247058169199 |

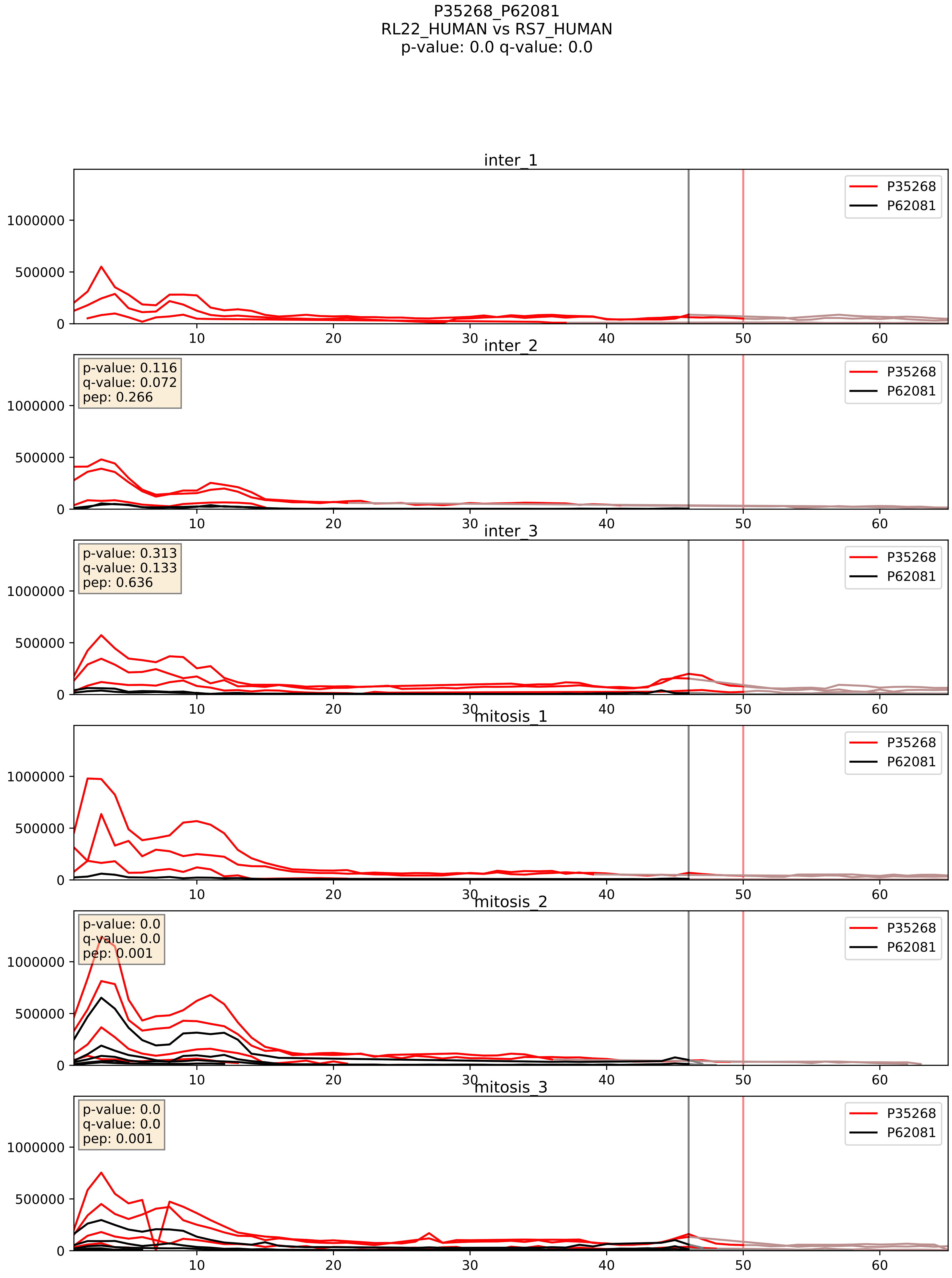


| level | condition_1 | condition_2 | pvalue | pvalue_adjusted |
|---|---|---|---|---|
| complex_abundance | mitosis | inter | 0.18050301115866102 | 0.331212384891348 |
| interactor_abundance | mitosis | inter | 0.23102492158232235 | 0.42318727300216413 |
| interactor_ratio | mitosis | inter | 0.5936263395102741 | 0.900166778779087 |

P35268_P62241
RL22_HUMAN vs RS8_HUMAN
p-value: 0.0 q-value: 0.0

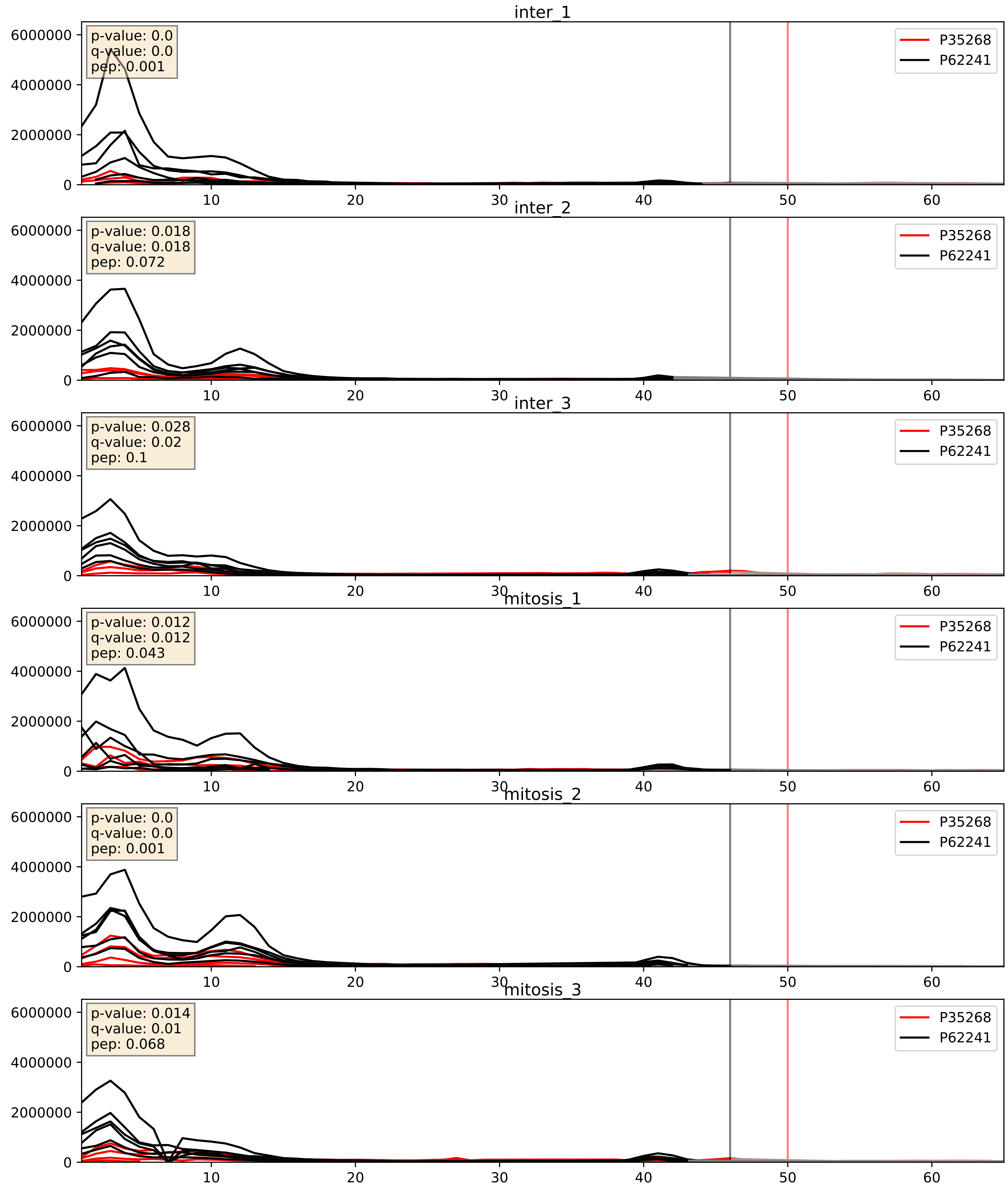


| level | condition_1 | condition_2 | pvalue | pvalue_adjusted |
|---|---|---|---|---|
| interactor_abundance | mitosis | inter | 0.02948973046273838 | 0.14263569727070707 |
| interactor_ratio | mitosis | inter | 0.031169809491203276 | 0.2814489127053798 |
| complex_abundance | mitosis | inter | 0.0659799883513077 | 0.18548068974948895 |

P35268_P62244
RL22_HUMAN vs RS15A_HUMAN
p-value: 0.0 q-value: 0.0

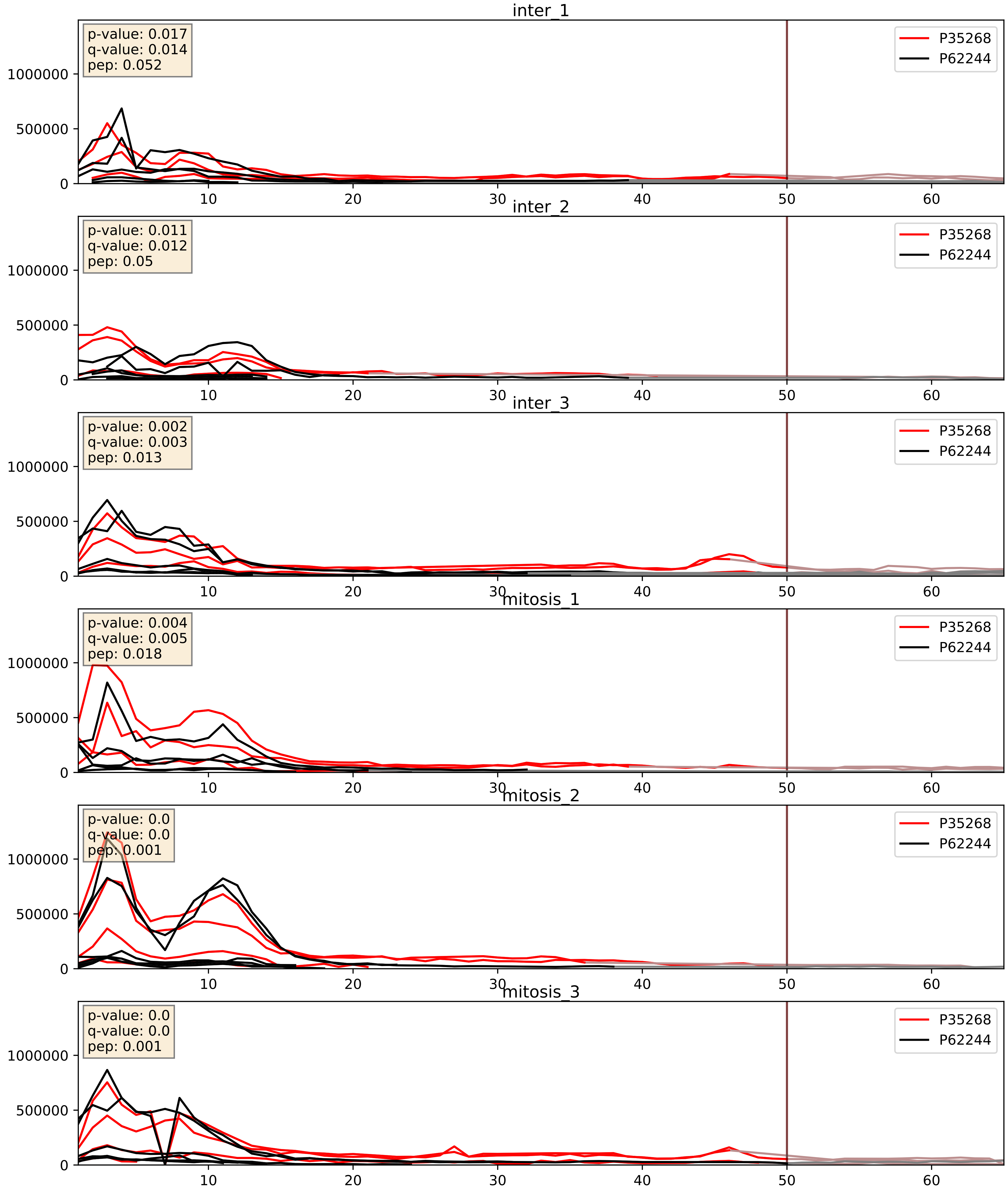


| level | condition_1 | condition_2 | pvalue | pvalue_adjusted |
|---|---|---|---|---|
| interactor_abundance | mitosis | inter | 0.04420014043586256 | 0.16761355216898388 |
| complex_abundance | mitosis | inter | 0.04500677214141851 | 0.15174465694825814 |
| interactor_ratio | mitosis | inter | 0.3571763419778247 | 0.8036815332643721 |

P35268_P62249
RL22_HUMAN vs RS16_HUMAN
p-value: 0.0 q-value: 0.0

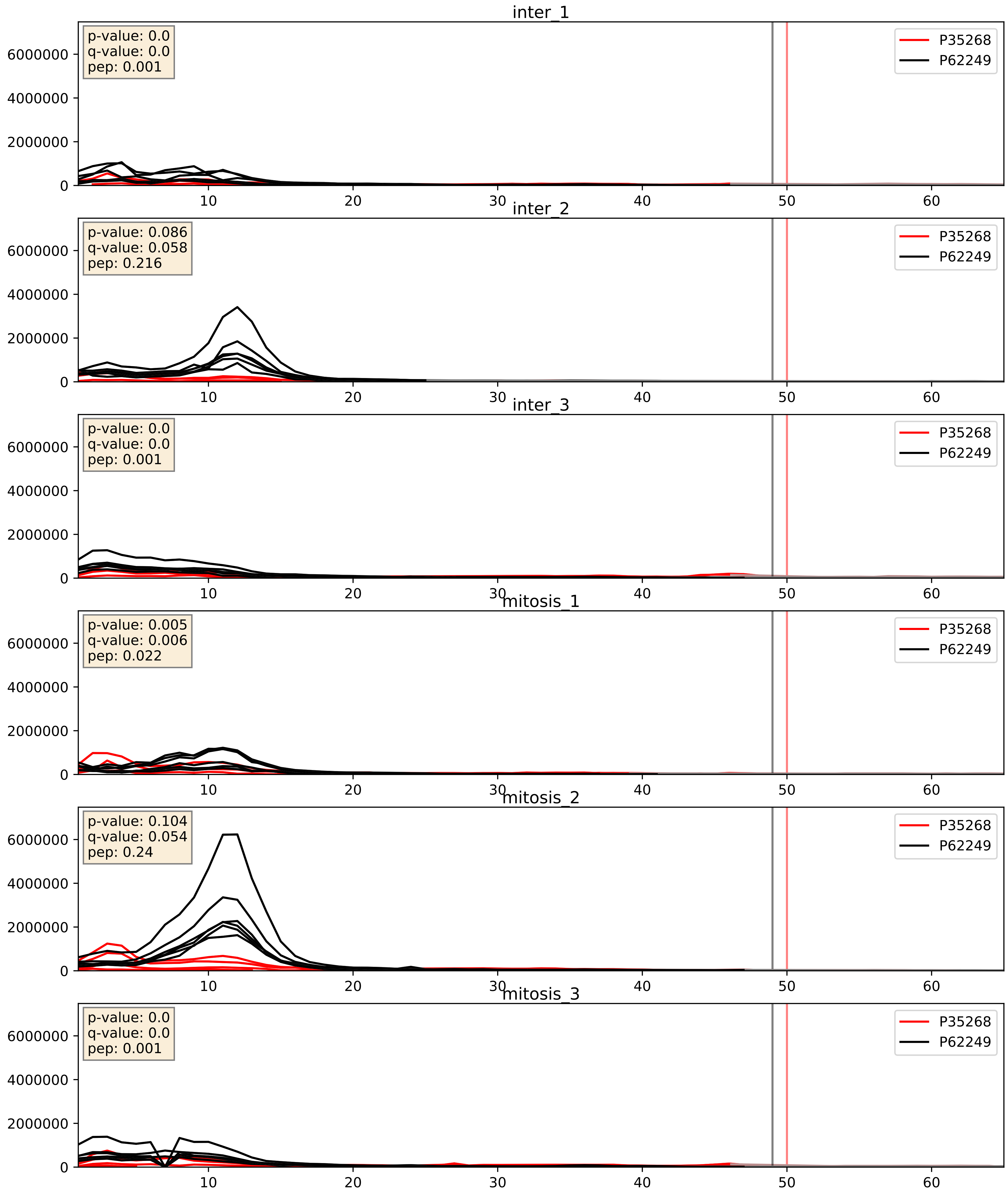


| level | condition_1 | condition_2 | pvalue | pvalue_adjusted |
|---|---|---|---|---|
| interactor_abundance | mitosis | inter | 0.038088099441202075 | 0.15907711297191632 |
| complex_abundance | mitosis | inter | 0.18505668917307902 | 0.33653818978575667 |
| interactor_ratio | mitosis | inter | 0.6078779283847071 | 0.9055803577343394 |

P35268_P62263
RL22_HUMAN vs RS14_HUMAN
p-value: 0.002 q-value: 0.002

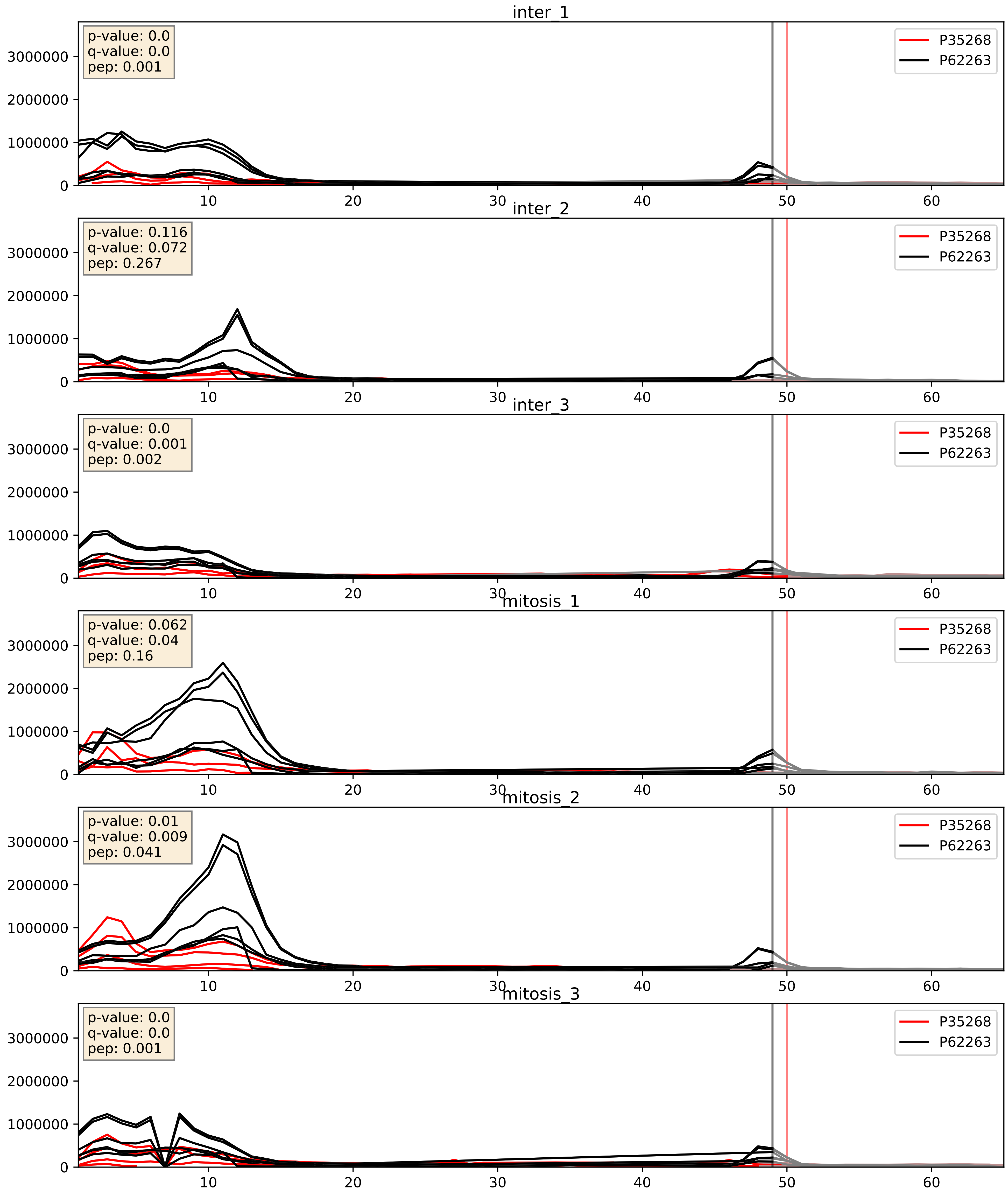


| level | condition_1 | condition_2 | pvalue | pvalue_adjusted |
|---|---|---|---|---|
| interactor_abundance | mitosis | inter | 0.03086131760129867 | 0.14599219600282765 |
| complex_abundance | mitosis | inter | 0.036271497578612456 | 0.138485289595416 |
| interactor_ratio | mitosis | inter | 0.7897932738716726 | 0.9601804323734581 |

P35268_P62266
RL22_HUMAN vs RS23_HUMAN
p-value: 0.0 q-value: 0.0

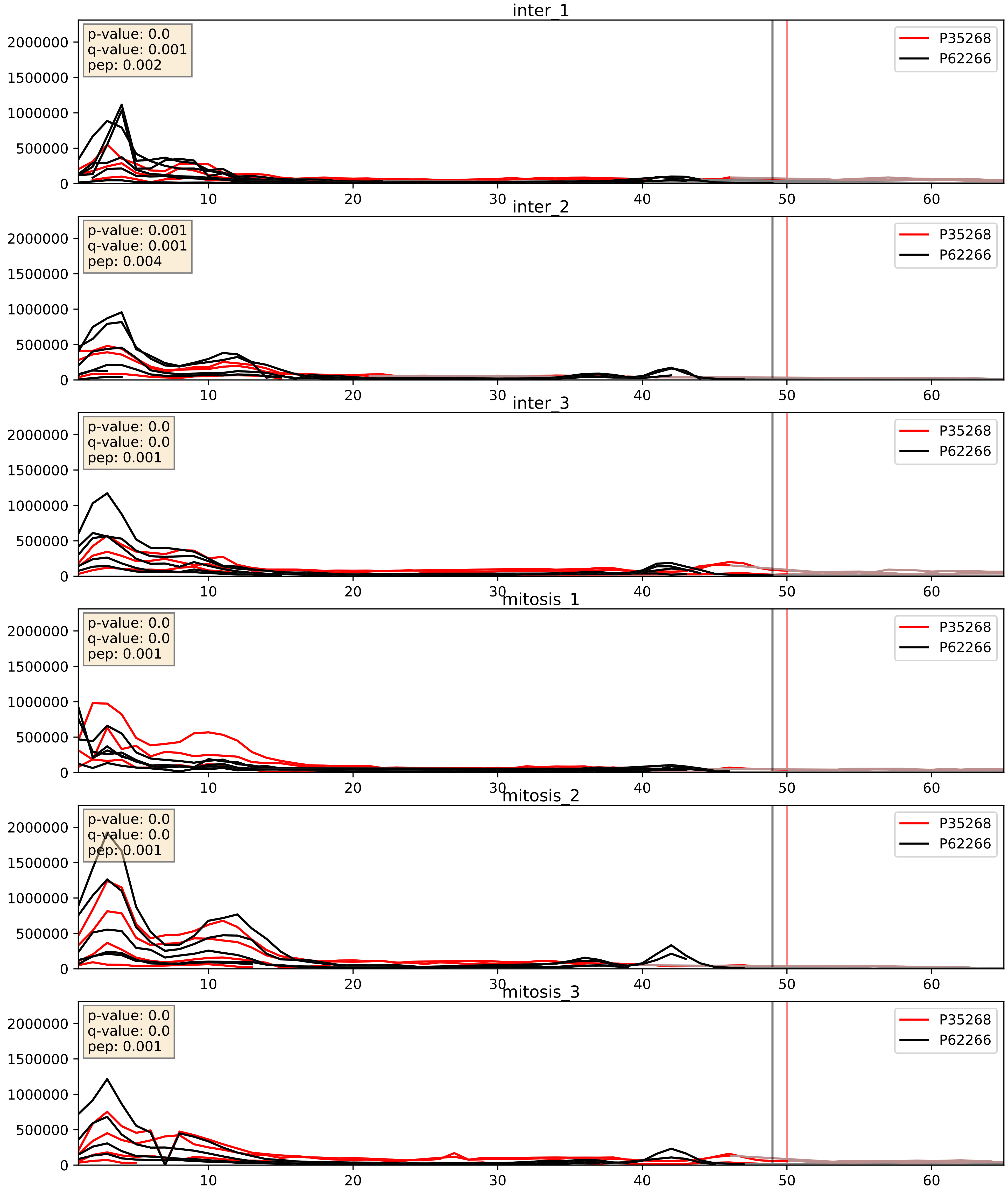


| level | condition_1 | condition_2 | pvalue | pvalue_adjusted |
|---|---|---|---|---|
| interactor_abundance | mitosis | inter | 0.041036812040372614 | 0.16296731794217187 |
| interactor_ratio | mitosis | inter | 0.11315441833776613 | 0.553985317646331 |
| complex_abundance | mitosis | inter | 0.8352185170231932 | 0.8934604481027909 |

P35268_P62269
RL22_HUMAN vs RS18_HUMAN
p-value: 0.0 q-value: 0.0

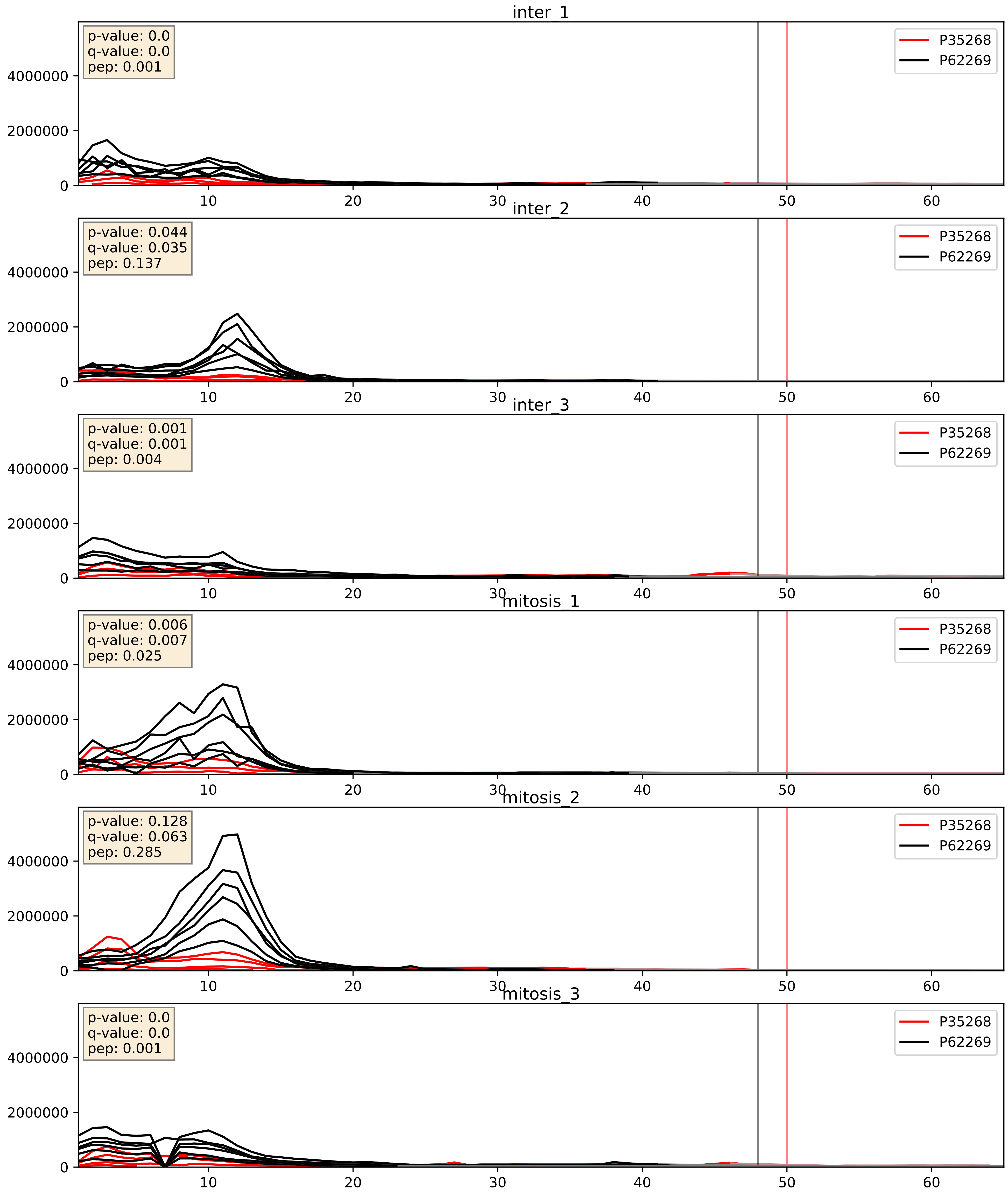


| level | condition_1 | condition_2 | pvalue | pvalue_adjusted |
|---|---|---|---|---|
| complex_abundance | mitosis | inter | 0.012824079295520931 | 0.08405635484106207 |
| interactor_abundance | mitosis | inter | 0.03731151707436548 | 0.15802829520754763 |
| interactor_ratio | mitosis | inter | 0.4899321816095601 | 0.8692763386835093 |

P35268_P62273
RL22_HUMAN vs RS29_HUMAN
p-value: 0.0 q-value: 0.0

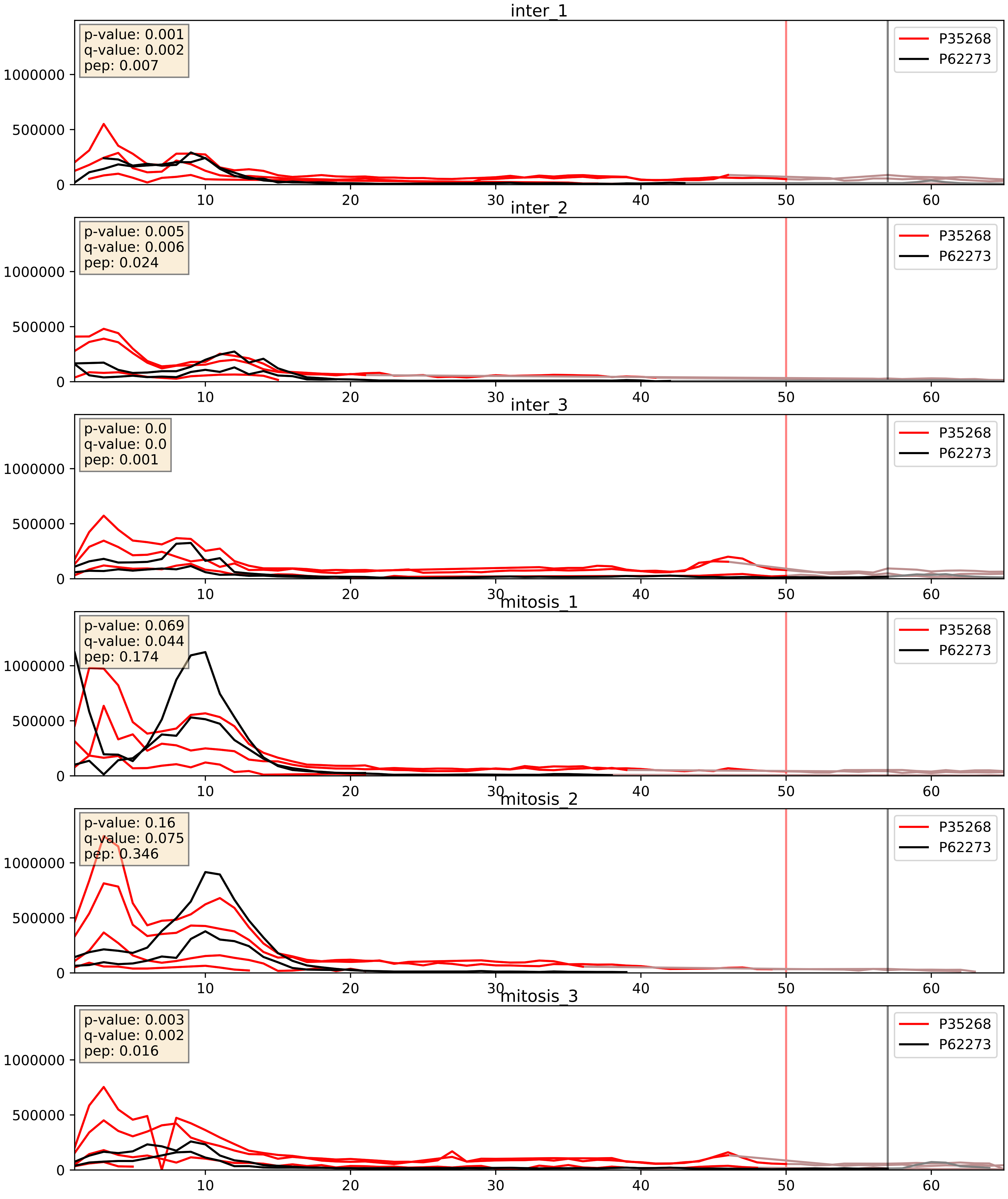


| level | condition_1 | condition_2 | pvalue | pvalue_adjusted |
|---|---|---|---|---|
| complex_abundance | mitosis | inter | 0.010338740912422375 | 0.07506222748092777 |
| interactor_abundance | mitosis | inter | 0.0528881924296407 | 0.18437097422020948 |
| interactor_ratio | mitosis | inter | 0.7578080963147068 | 0.953204216368304 |

P35268_P62277
RL22_HUMAN vs RS13_HUMAN
p-value: 0.0 q-value: 0.0

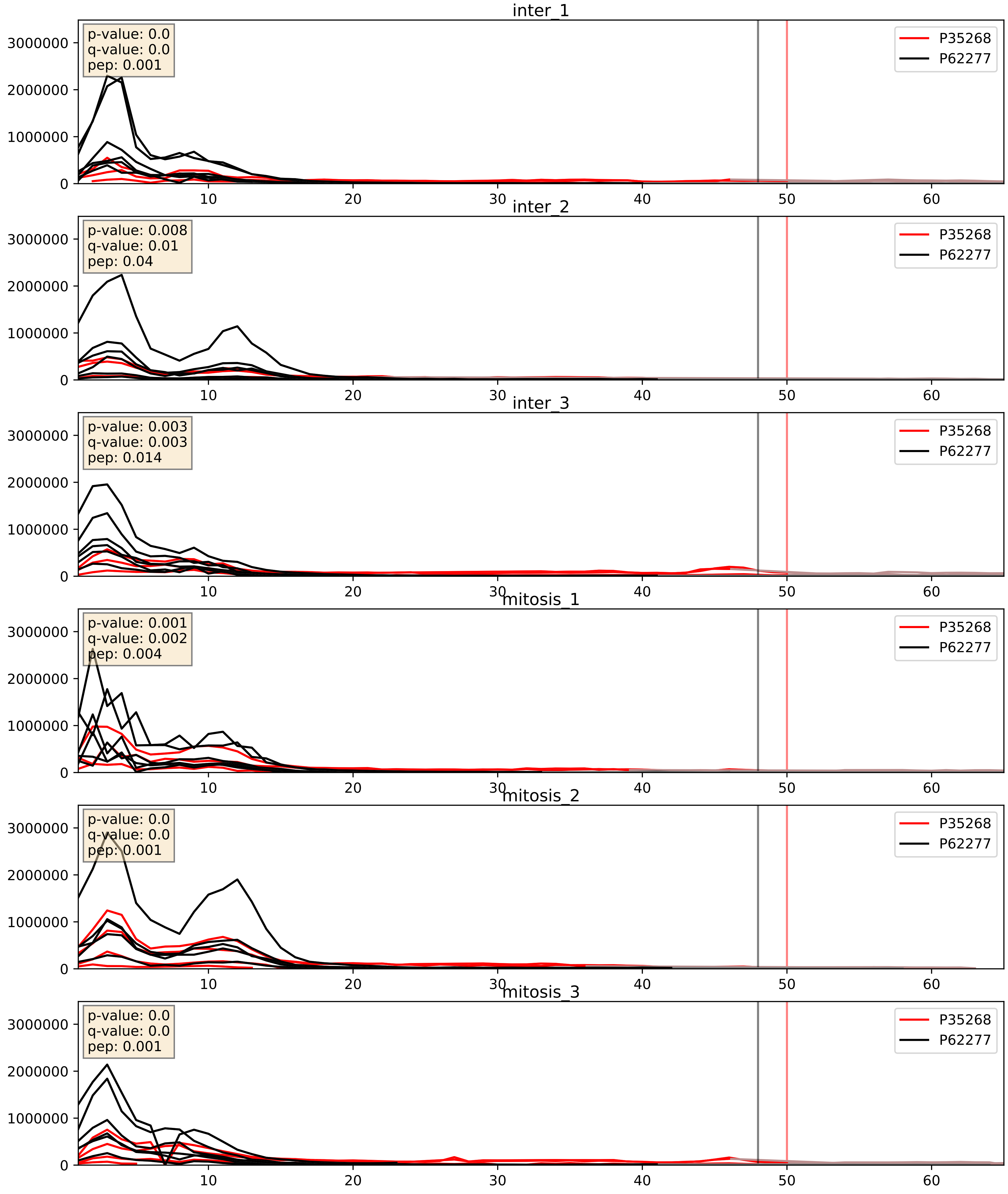


| level | condition_1 | condition_2 | pvalue | pvalue_adjusted |
|---|---|---|---|---|
| interactor_abundance | mitosis | inter | 0.009254031771362048 | 0.08329601678534083 |
| interactor_ratio | mitosis | inter | 0.01562191419072123 | 0.1843379164928052 |
| complex_abundance | mitosis | inter | 0.02366227599398235 | 0.11209135722661258 |

P35268_P62280
RL22_HUMAN vs RS11_HUMAN
p-value: 0.0 q-value: 0.0

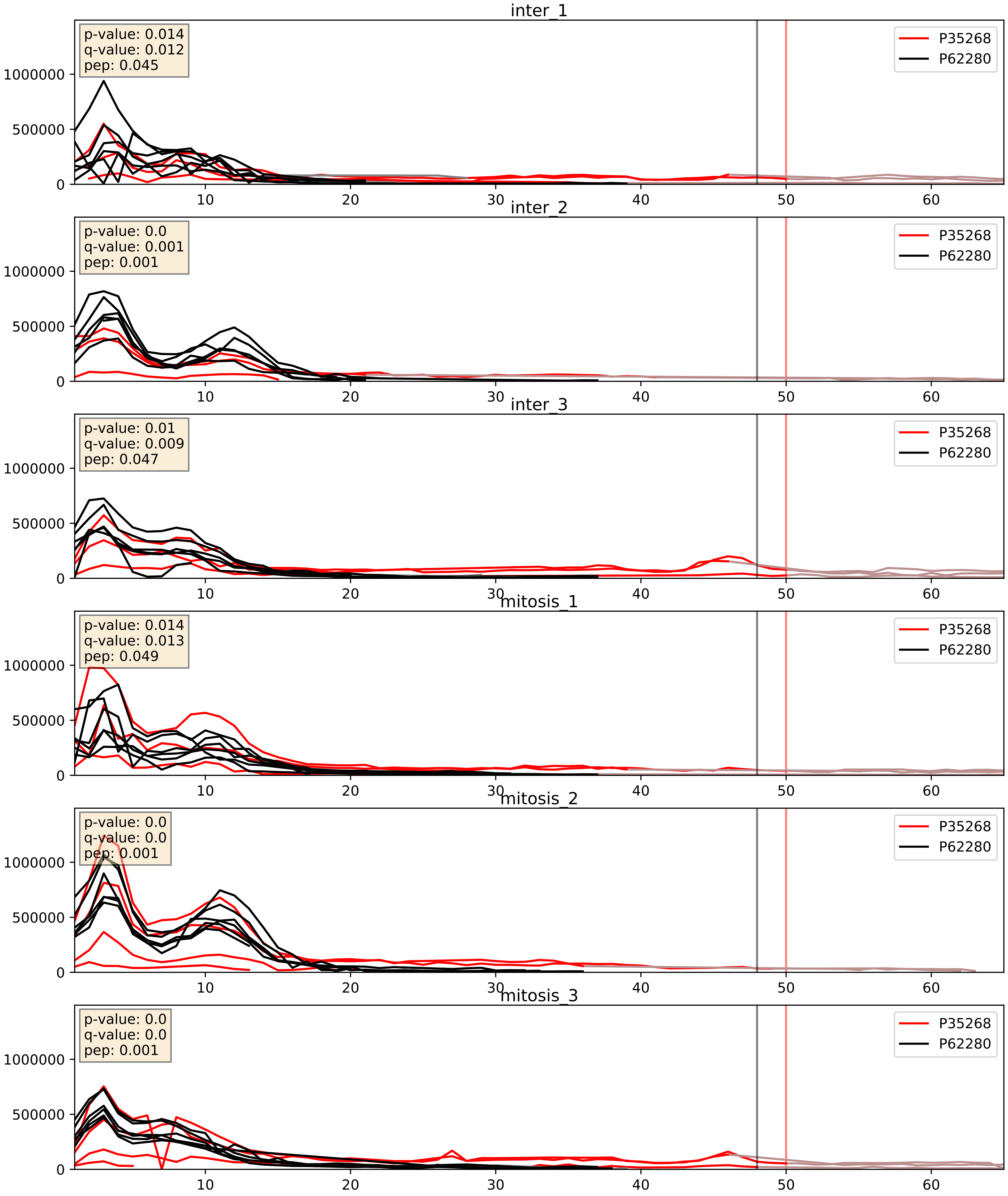


| level | condition_1 | condition_2 | pvalue | pvalue_adjusted |
|---|---|---|---|---|
| interactor_abundance | mitosis | inter | 0.009548600918579717 | 0.08483861064632973 |
| interactor_ratio | mitosis | inter | 0.014211394396282188 | 0.17478381613818322 |
| complex_abundance | mitosis | inter | 0.07688034796933005 | 0.20137569725136634 |

P35268_P62424
RL22_HUMAN vs RL7A_HUMAN
p-value: 0.001 q-value: 0.001

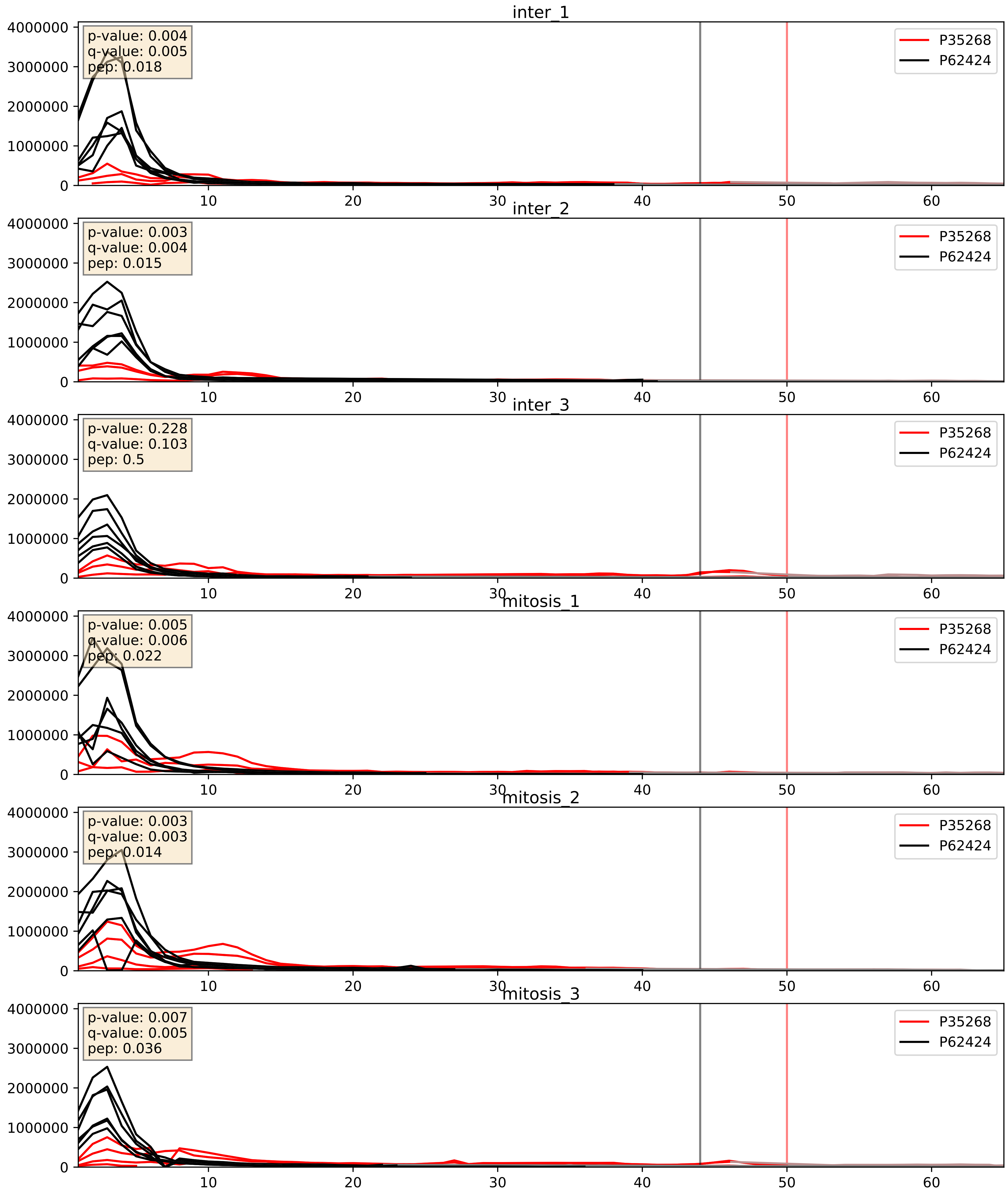


| level | condition_1 | condition_2 | pvalue | pvalue_adjusted |
|---|---|---|---|---|
| interactor_abundance | mitosis | inter | 0.0024700391617390243 | 0.037061411436434794 |
| complex_abundance | mitosis | inter | 0.0490055574884795 | 0.15809873188702586 |
| interactor_ratio | mitosis | inter | 0.14567085073811542 | 0.5994295998710799 |

P35268_P62701
RL22_HUMAN vs RS4X_HUMAN
p-value: 0.0 q-value: 0.001

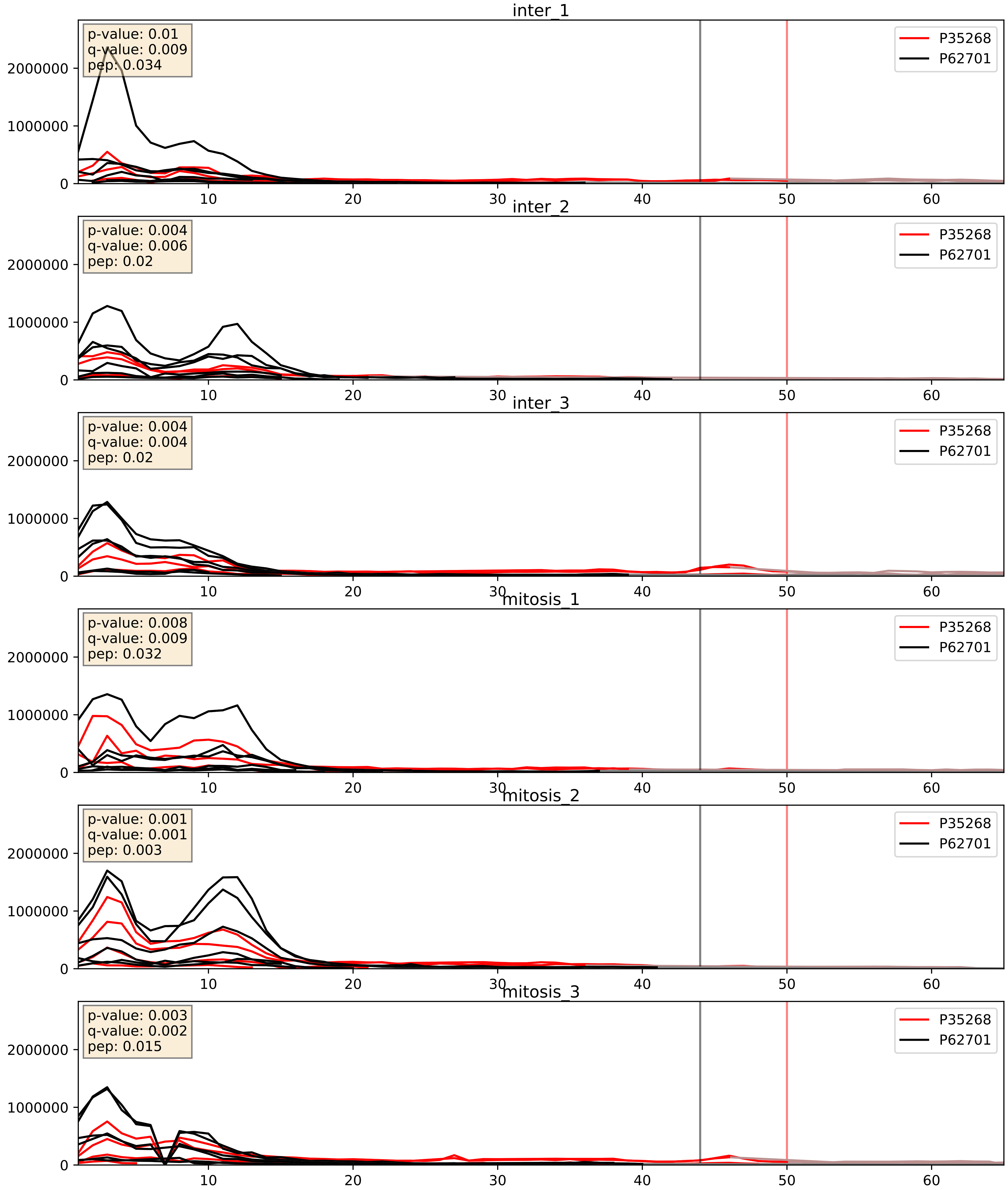


| level | condition_1 | condition_2 | pvalue | pvalue_adjusted |
|---|---|---|---|---|
| interactor_ratio | mitosis | inter | 0.005797322266886906 | 0.10426786862612165 |
| interactor_abundance | mitosis | inter | 0.012887295150753023 | 0.10167976967449956 |
| complex_abundance | mitosis | inter | 0.10072924412189838 | 0.2354669386569353 |

P35268_P62750
RL22_HUMAN vs RL23A_HUMAN
p-value: 0.0 q-value: 0.0

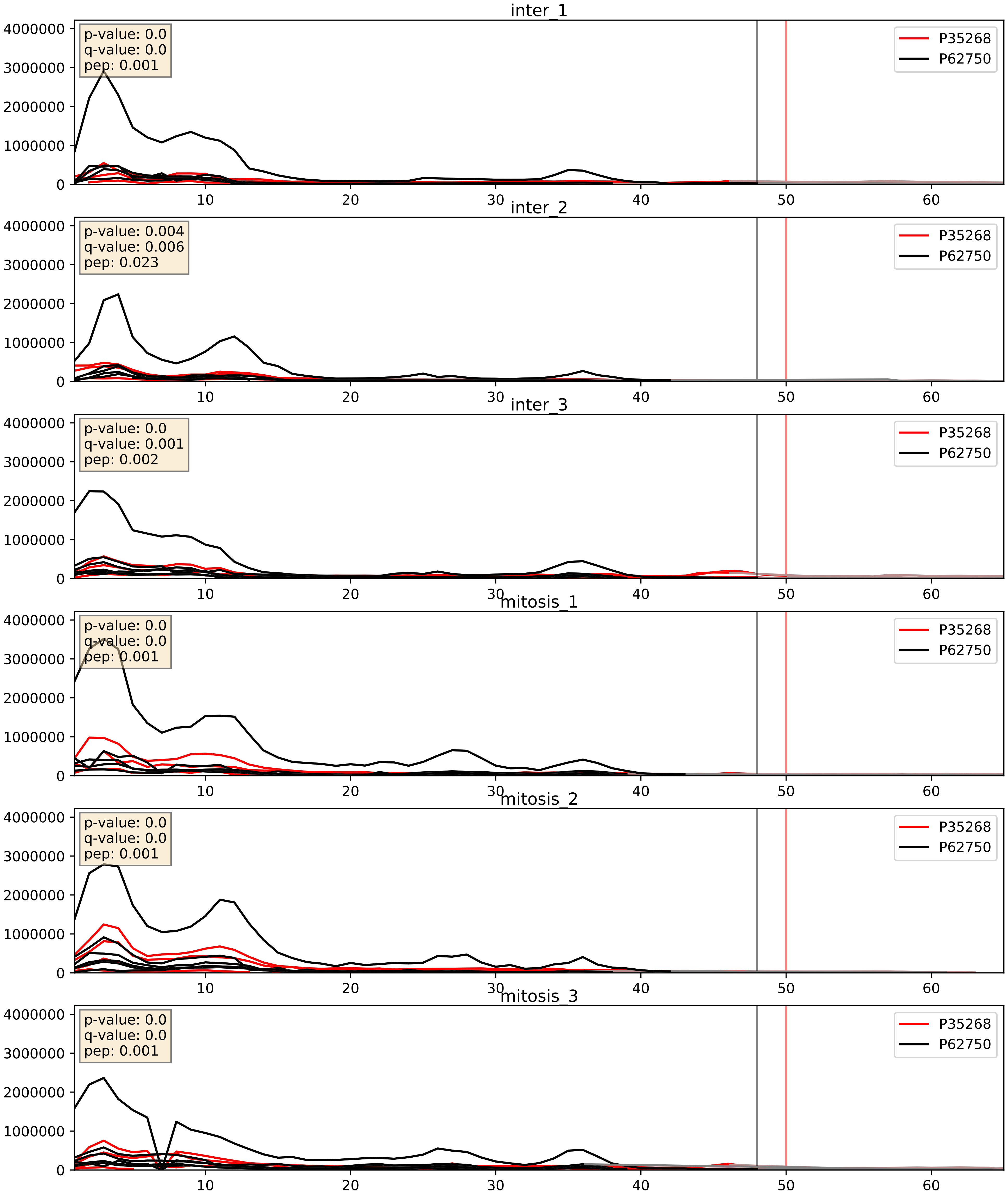


| level | condition_1 | condition_2 | pvalue | pvalue_adjusted |
|---|---|---|---|---|
| interactor_abundance | mitosis | inter | 0.019724728854510 78 | 0.1190514126751731 |
| complex_abundance | mitosis | inter | 0.03944285833610676 | 0.14457363202042897 |
| interactor_ratio | mitosis | inter | 0.662989350983256 | 0.9280766712046886 |

P35268_P62753
RL22_HUMAN vs RS6_HUMAN
p-value: 0.0 q-value: 0.0

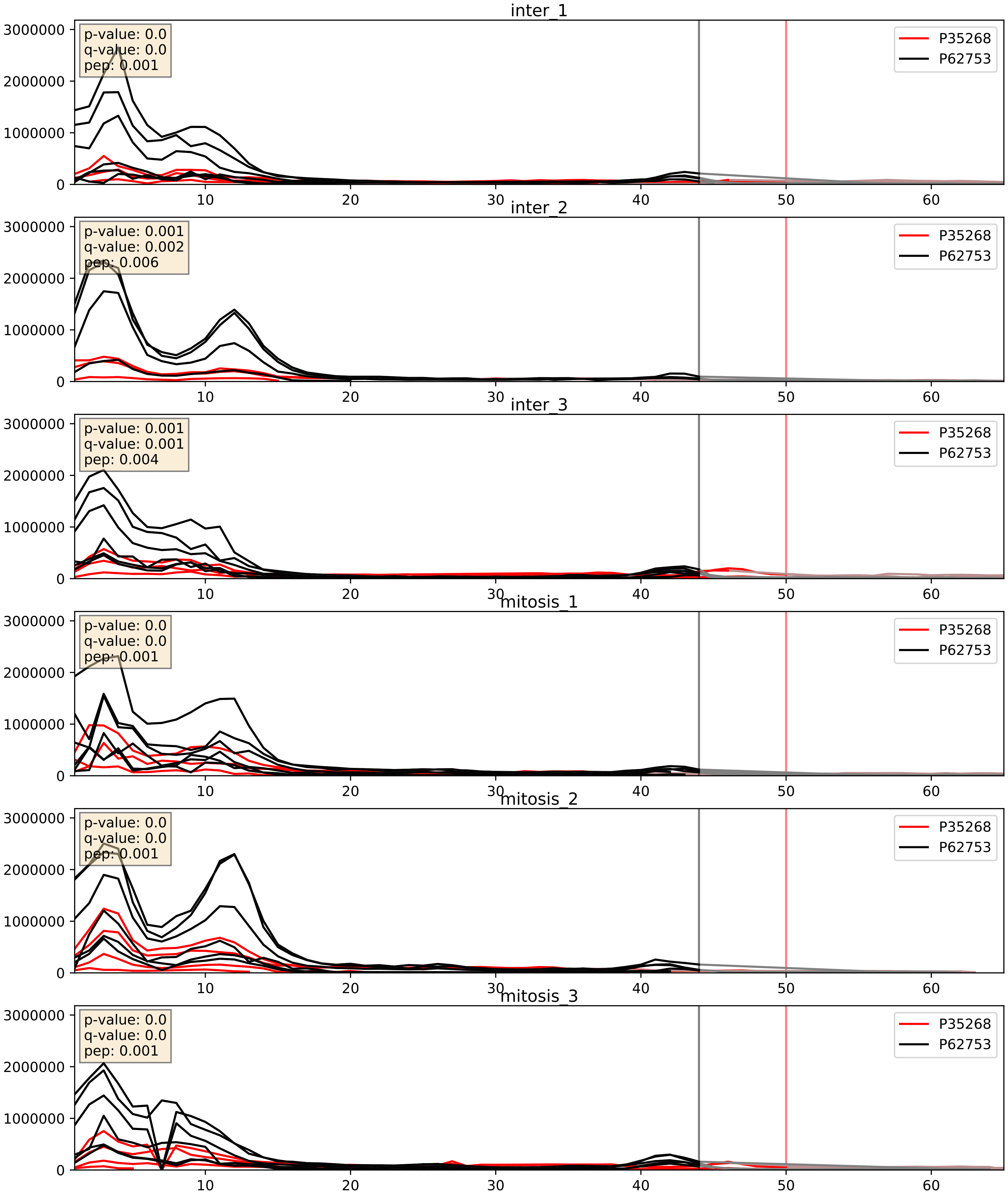


| level | condition_1 | condition_2 | pvalue | pvalue_adjusted |
|---|---|---|---|---|
| interactor_ratio | mitosis | inter | 0.004037990744645896 | 0.08271268336775048 |
| interactor_abundance | mitosis | inter | 0.011278934868275112 | 0.09369013340362442 |
| complex_abundance | mitosis | inter | 0.1164345432842469 | 0.25734048296234274 |

P35268_P62829
RL22_HUMAN vs RL23_HUMAN
p-value: 0.0 q-value: 0.0

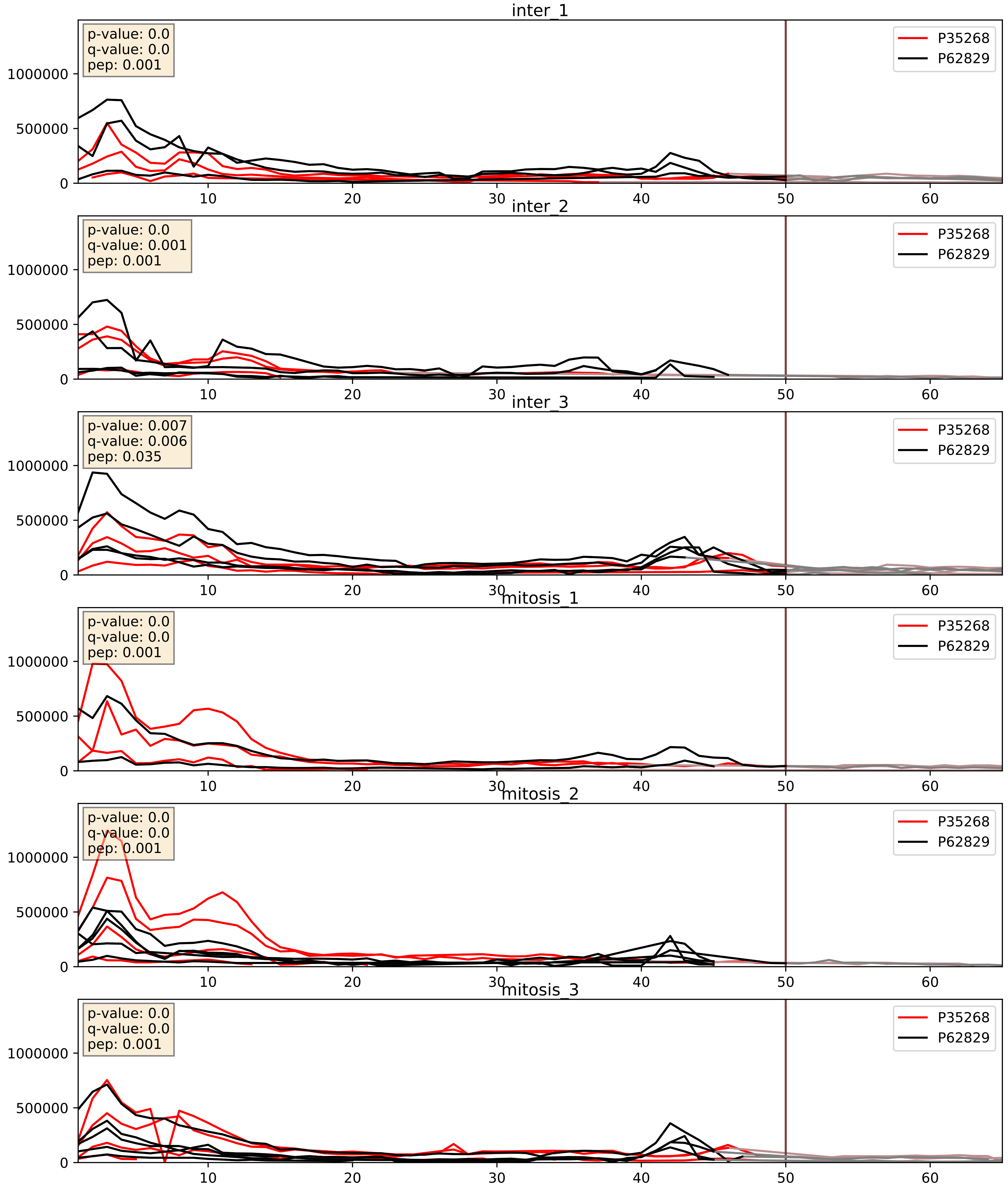


| level | condition_1 | condition_2 | pvalue | pvalue_adjusted |
|---|---|---|---|---|
| interactor_ratio | mitosis | inter | 0.0012700844385174963 | 0.0423373831474418 |
| interactor_abundance | mitosis | inter | 0.04055078168160862 | 0.1623548602406781 |
| complex_abundance | mitosis | inter | 0.07592611735990688 | 0.20016247754875358 |

P35268_P62841
RL22_HUMAN vs RS15_HUMAN
p-value: 0.001 q-value: 0.002

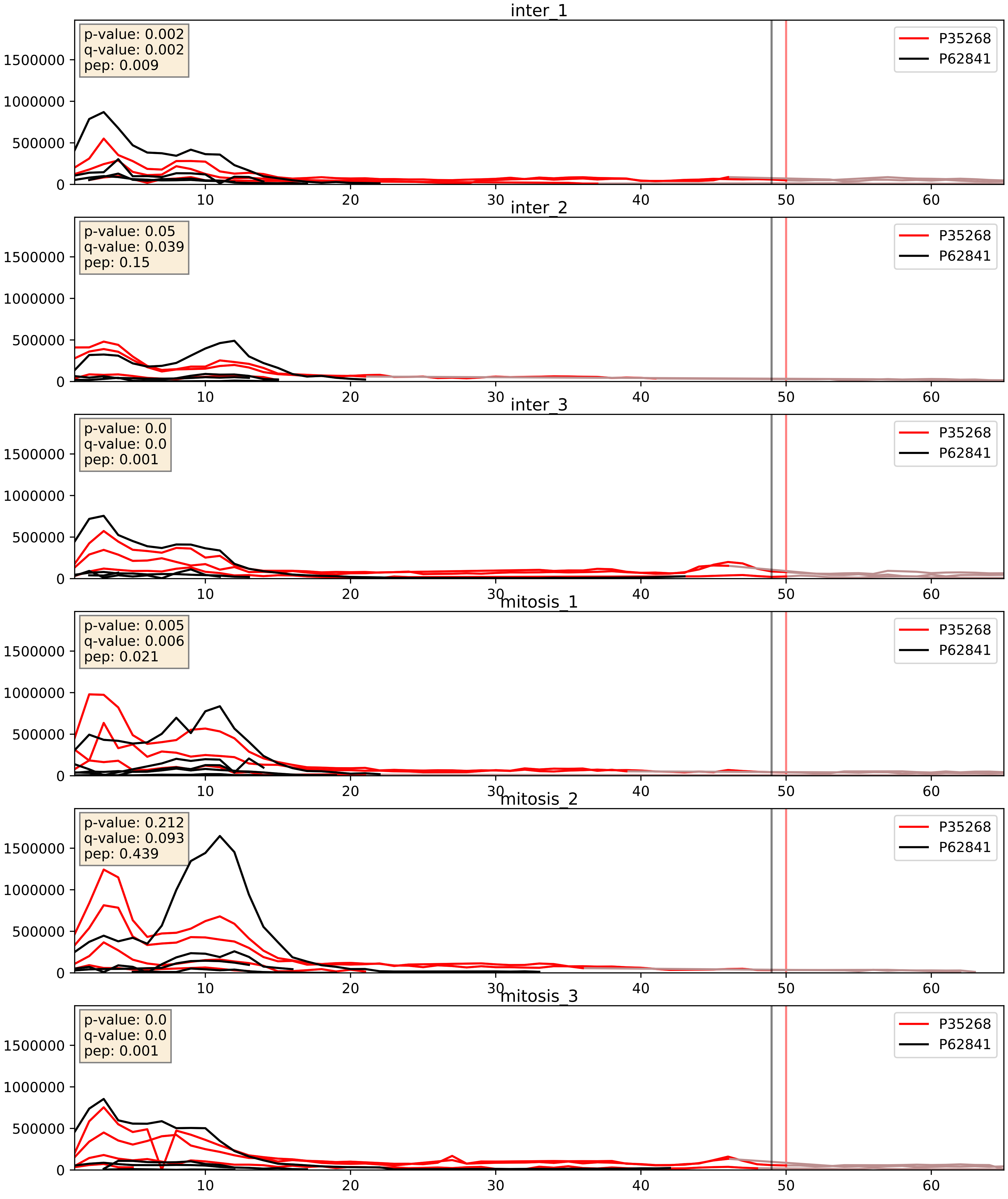


| level | condition_1 | condition_2 | pvalue | pvalue_adjusted |
|---|---|---|---|---|
| interactor_abundance | mitosis | inter | 0.022709228038218003 | 0.12357977877123083 |
| interactor_ratio | mitosis | inter | 0.2949706965551553 | 0.7639785665694794 |
| complex_abundance | mitosis | inter | 0.32166084814511153 | 0.4703118017536005 |

P35268_P62847
RL22_HUMAN vs RS24_HUMAN
p-value: 0.0 q-value: 0.0

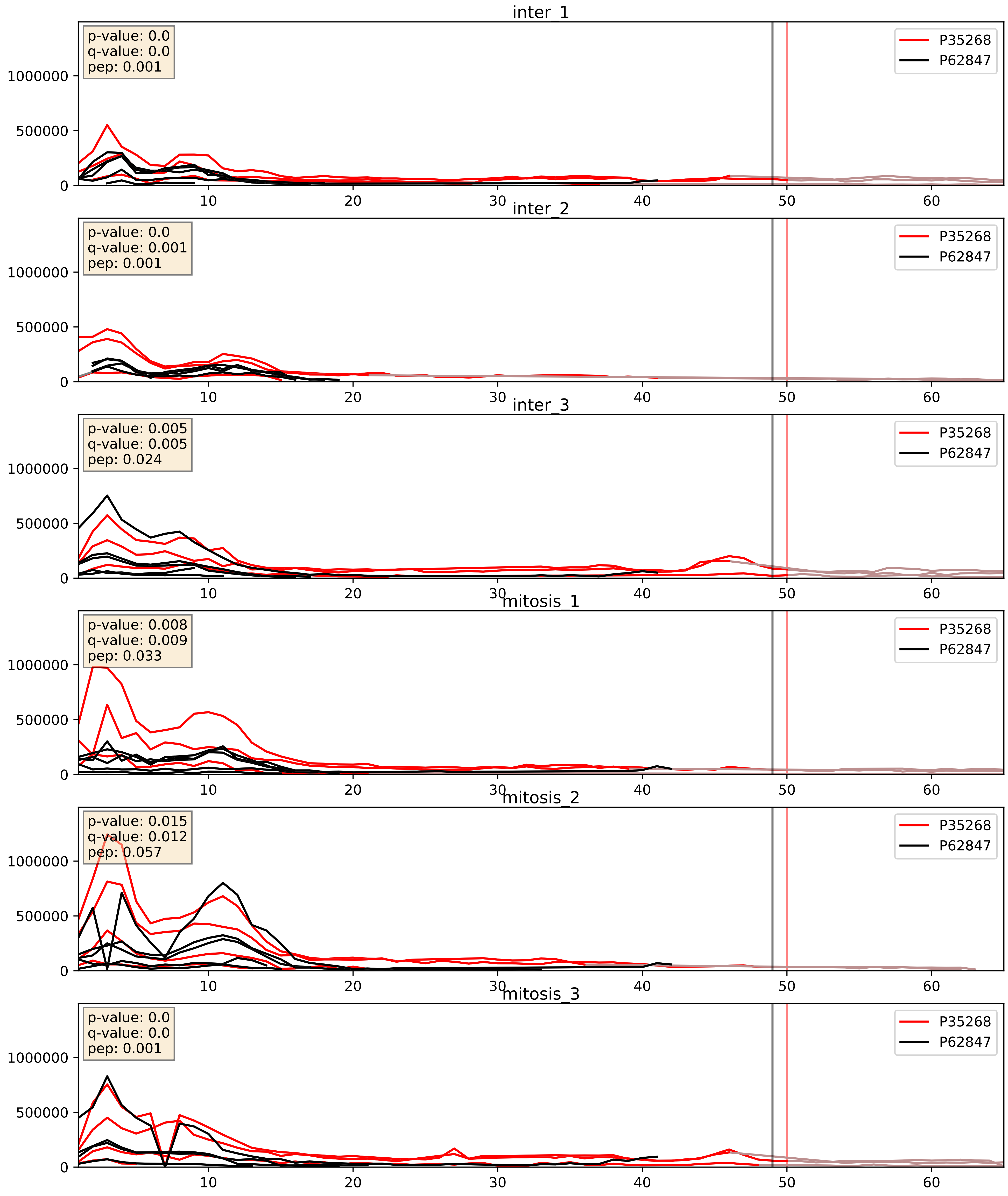


| level | condition_1 | condition_2 | pvalue | pvalue_adjusted |
|---|---|---|---|---|
| interactor_abundance | mitosis | inter | 0.017192532646947804 | 0.11357768828956229 |
| complex_abundance | mitosis | inter | 0.030172898921410098 | 0.12592882241212602 |
| interactor_ratio | mitosis | inter | 0.06577883822127165 | 0.42949416870639606 |

P35268_P62851
RL22_HUMAN vs RS25_HUMAN
p-value: 0.002 q-value: 0.003

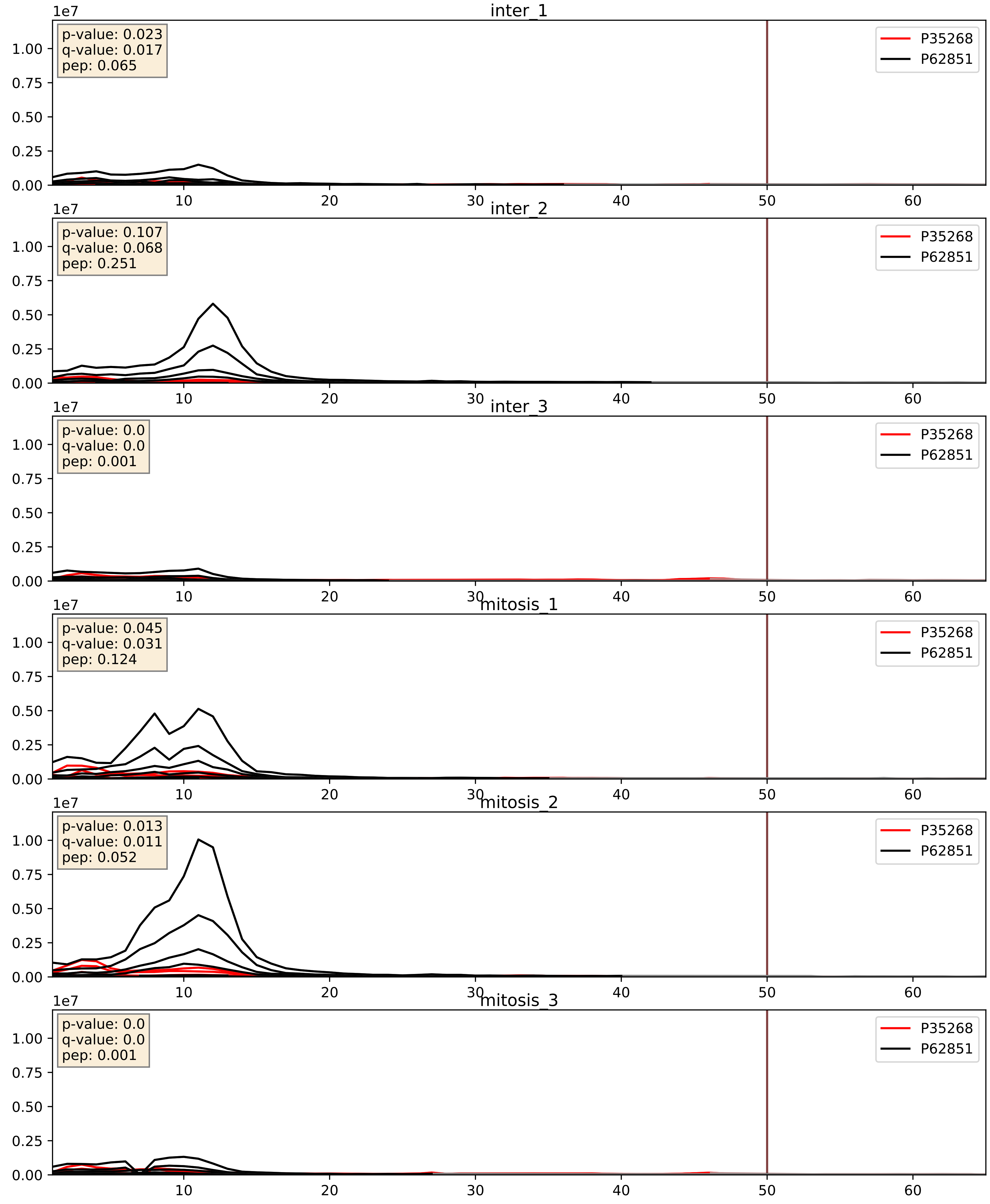


| level | condition_1 | condition_2 | pvalue | pvalue_adjusted |
|---|---|---|---|---|
| interactor_abundance | mitosis | inter | 0.012261362050313774 | 0.09846874754825552 |
| complex_abundance | mitosis | inter | 0.1859713933316283 | 0.3371987136027829 |
| interactor_ratio | mitosis | inter | 0.7822294100425186 | 0.95900240448312 |

P35268_P62857
RL22_HUMAN vs RS28_HUMAN
p-value: 0.006 q-value: 0.006

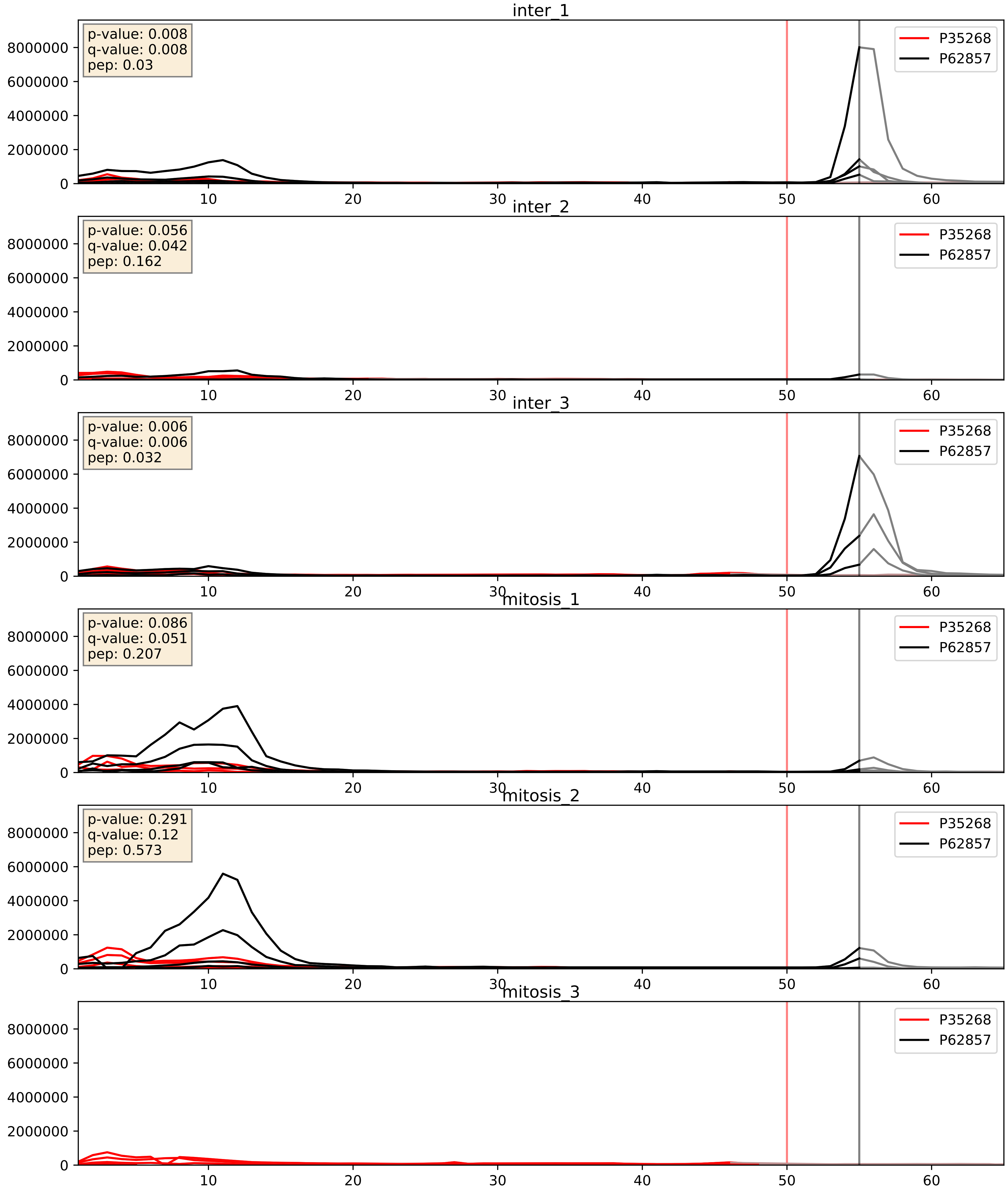


| level | condition_1 | condition_2 | pvalue | pvalue_adjusted |
|---|---|---|---|---|
| interactor_ratio | mitosis | inter | 0.1614099983853562 | 0.6192541702766019 |
| interactor_abundance | mitosis | inter | 0.5288098927802732 | 0.6880389589497613 |
| complex_abundance | mitosis | inter | 0.8513857011409774 | 0.9052993800369485 |

P35268_P62861
RL22_HUMAN vs RS30_HUMAN
p-value: 0.0 q-value: 0.0

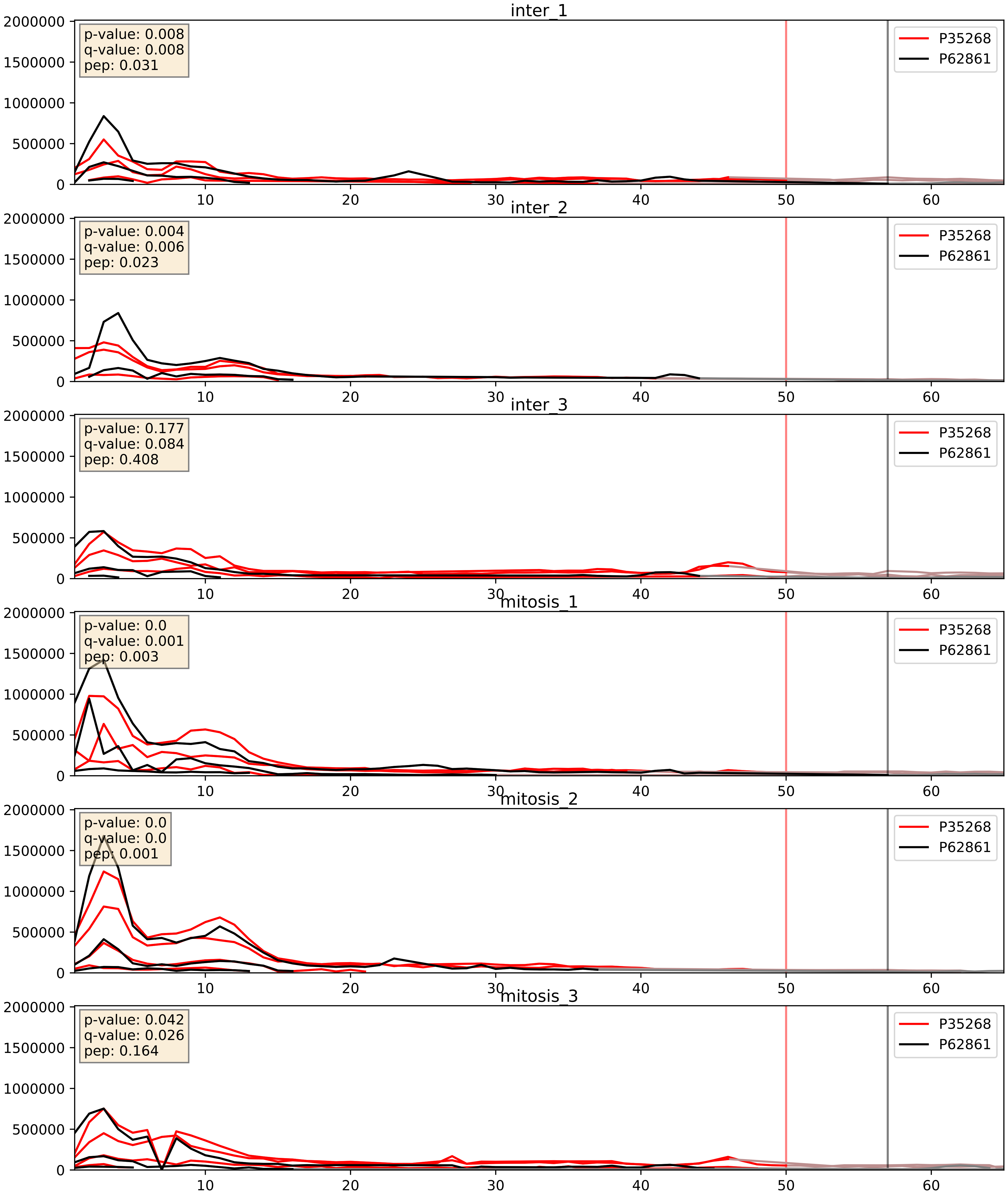


| level | condition_1 | condition_2 | pvalue | pvalue_adjusted |
|---|---|---|---|---|
| interactor_abundance | mitosis | inter | 0.0016605937970136154 | 0.02983144365673987 |
| complex_abundance | mitosis | inter | 0.06919427347047818 | 0.19057367468059627 |
| interactor_ratio | mitosis | inter | 0.7190769993585142 | 0.9438671263908177 |

P35268_P62888
RL22_HUMAN vs RL30_HUMAN
p-value: 0.0 q-value: 0.0

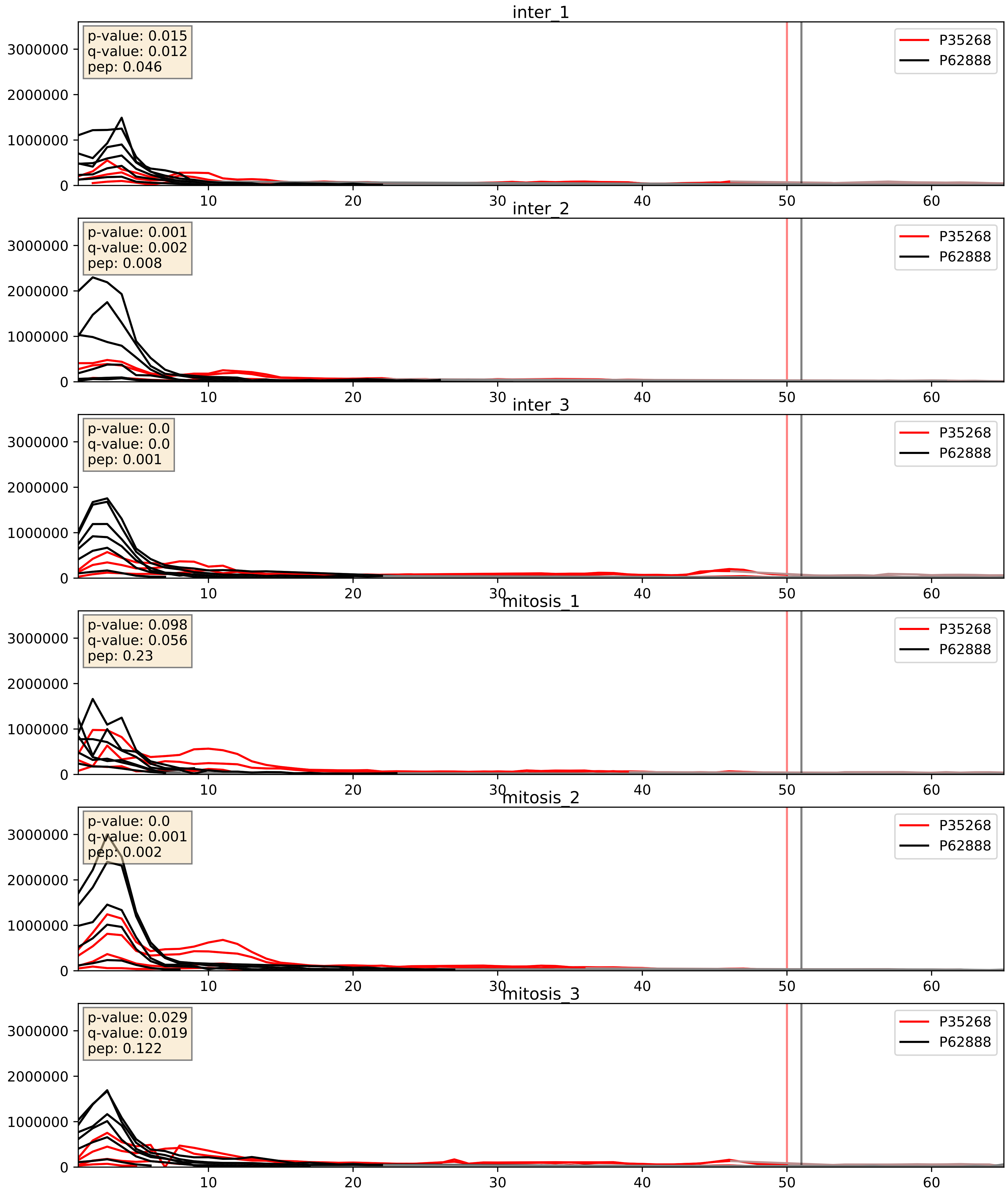


| level | condition_1 | condition_2 | pvalue | pvalue_adjusted |
|---|---|---|---|---|
| interactor_abundance | mitosis | inter | 0.016092022966764406 | 0.11124909769145716 |
| interactor_ratio | mitosis | inter | 0.0392069099918715 | 0.32615272063208944 |
| complex_abundance | mitosis | inter | 0.47328120964854536 | 0.614061811580691 |

P35268_P62899
RL22_HUMAN vs RL31_HUMAN
p-value: 0.0 q-value: 0.0

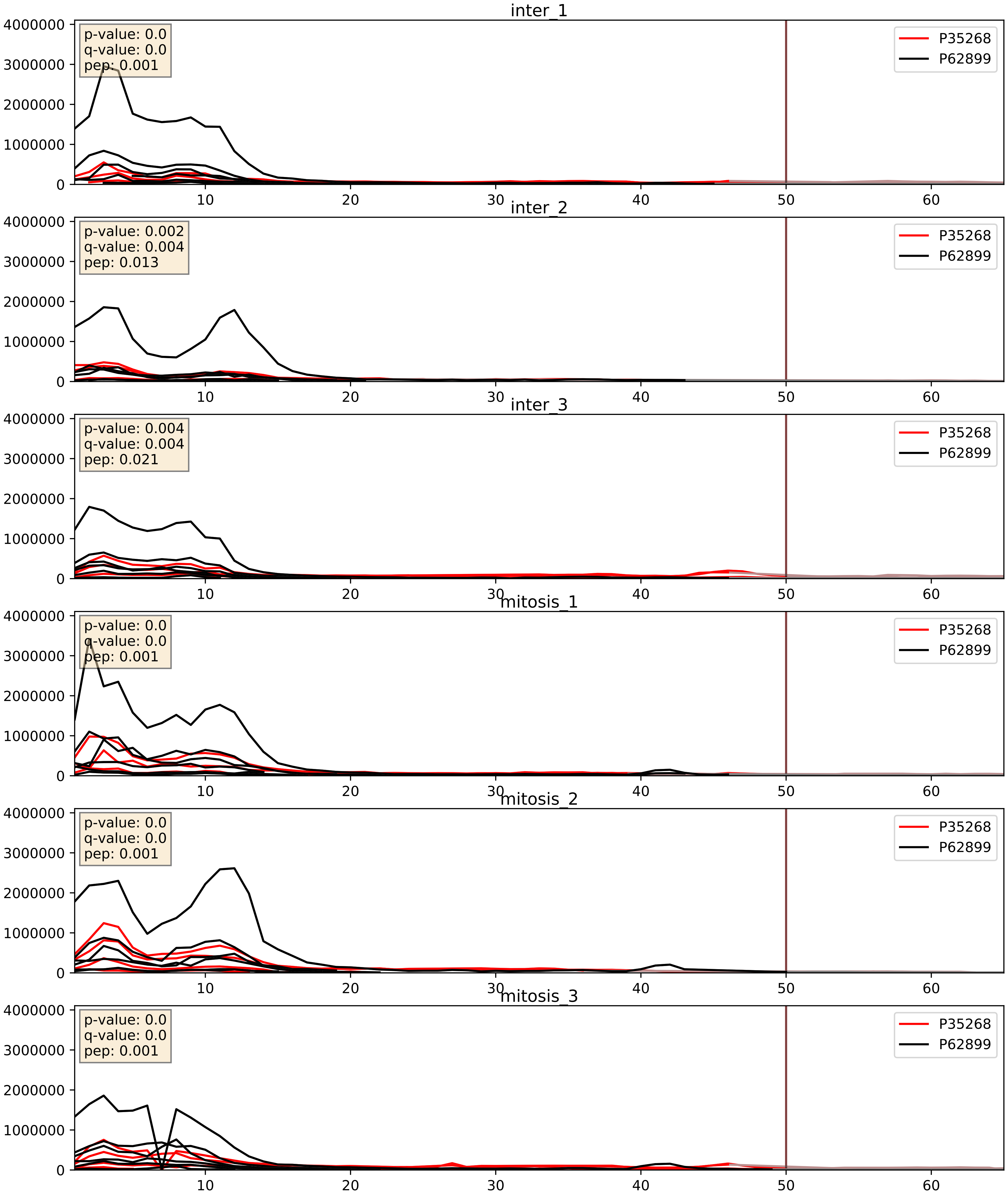


| level | condition_1 | condition_2 | pvalue | pvalue_adjusted |
|---|---|---|---|---|
| complex_abundance | mitosis | inter | 0.025366657595246995 | 0.1166784465423505 |
| interactor_abundance | mitosis | inter | 0.02905216201711013 | 0.14119387763757685 |
| interactor_ratio | mitosis | inter | 0.3845114796713773 | 0.8212585015521048 |

P35268_P62906
RL22_HUMAN vs RL10A_HUMAN
p-value: 0.001 q-value: 0.001

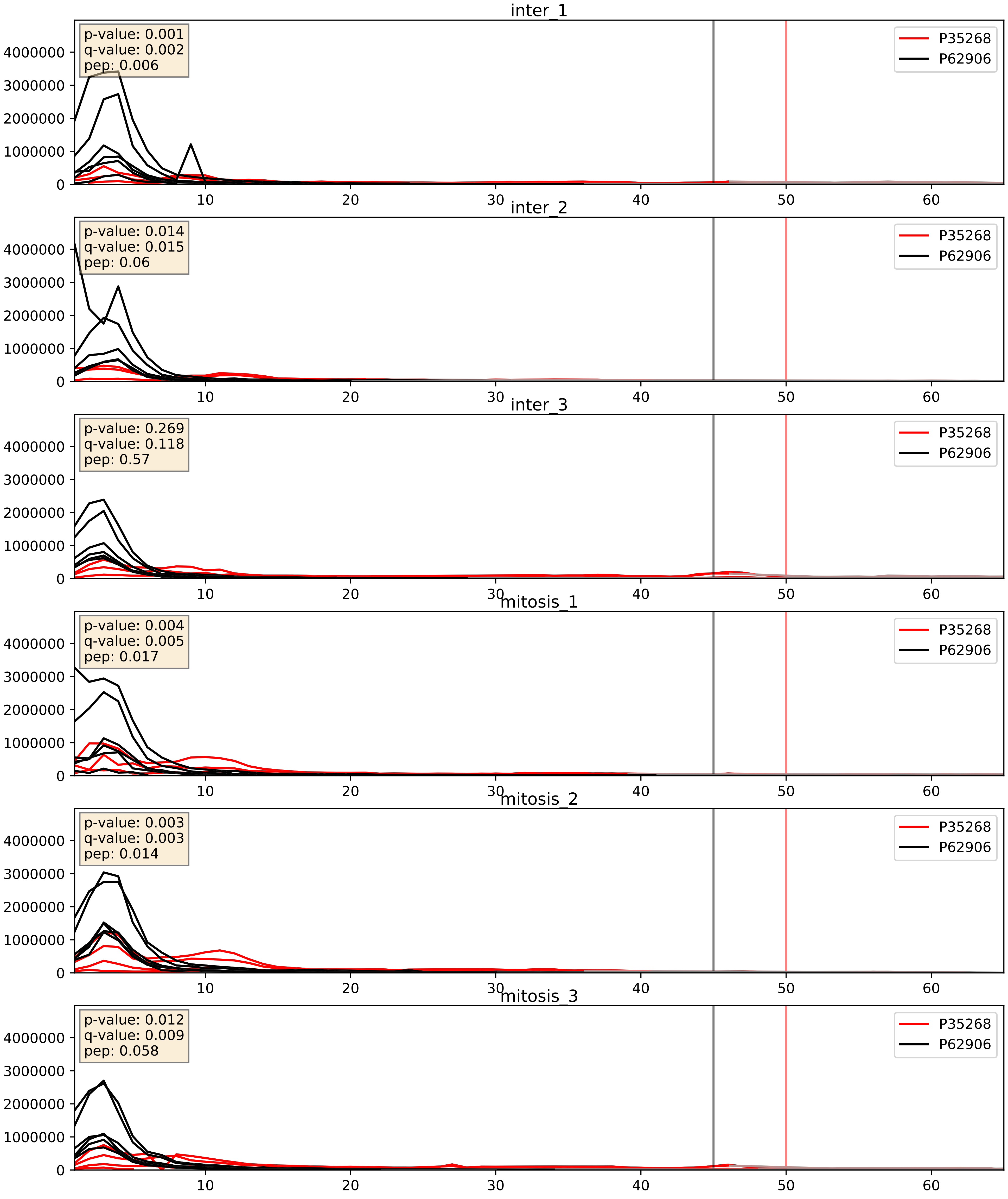


| level | condition_1 | condition_2 | pvalue | pvalue_adjusted |
|---|---|---|---|---|
| complex_abundance | mitosis | inter | 0.002410662275627882 | 0.03561197063475765 |
| interactor_abundance | mitosis | inter | 0.007905967289639496 | 0.07711604472197223 |
| interactor_ratio | mitosis | inter | 0.07929887789520691 | 0.4717153542619675 |

P35268_P62910
RL22_HUMAN vs RL32_HUMAN
p-value: 0.002 q-value: 0.002

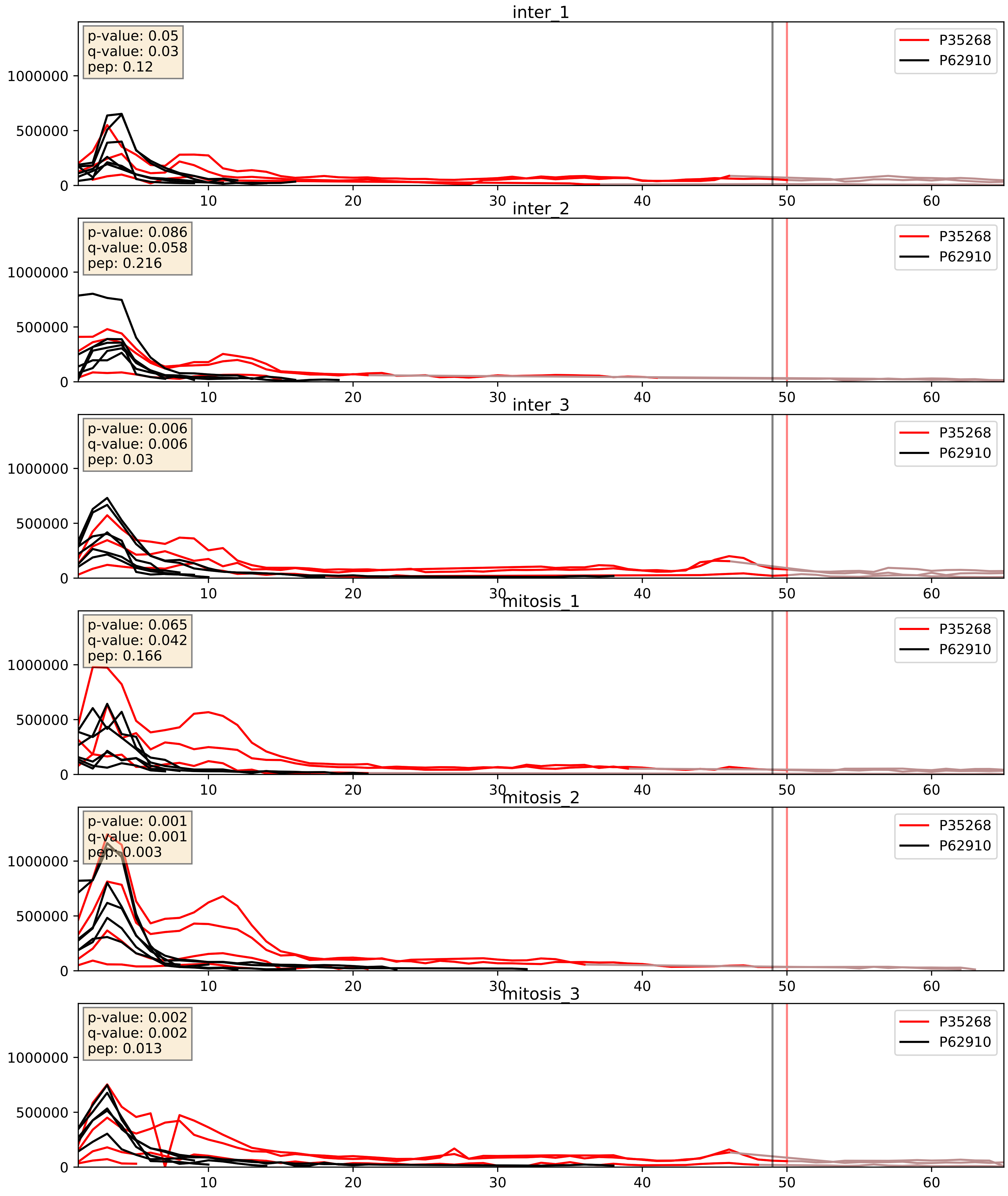


| level | condition_1 | condition_2 | pvalue | pvalue_adjusted |
| --- | --- | --- | --- | --- |
| interactor_abundance | mitosis | inter | 0.028525255352382952 | 0.13992953983177217 |
| interactor_ratio | mitosis | inter | 0.037414321803036776 | 0.31709563825148 |
| complex_abundance | mitosis | inter | 0.14429077945890056 | 0.29021207390906123 |

P35268_P62913
RL22_HUMAN vs RL11_HUMAN
p-value: 0.0 q-value: 0.0

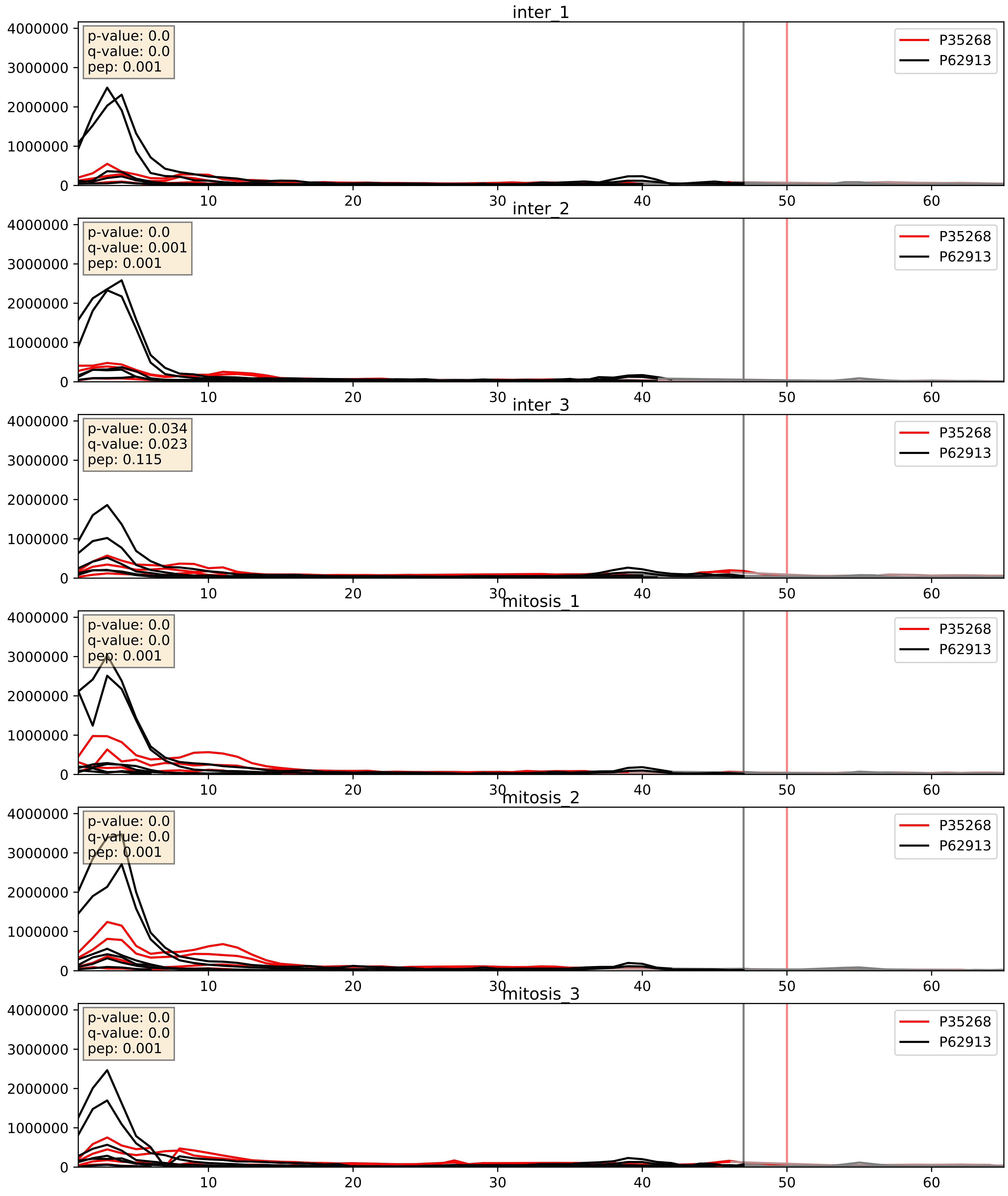


| level | condition_1 | condition_2 | pvalue | pvalue_adjusted |
|---|---|---|---|---|
| complex_abundance | mitosis | inter | 0.021106617186278903 | 0.10728779282336545 |
| interactor_abundance | mitosis | inter | 0.029151255470338998 | 0.14151303993549846 |
| interactor_ratio | mitosis | inter | 0.06691699243304988 | 0.4304195561540654 |

P35268_P62917
RL22_HUMAN vs RL8_HUMAN
p-value: 0.0 q-value: 0.0

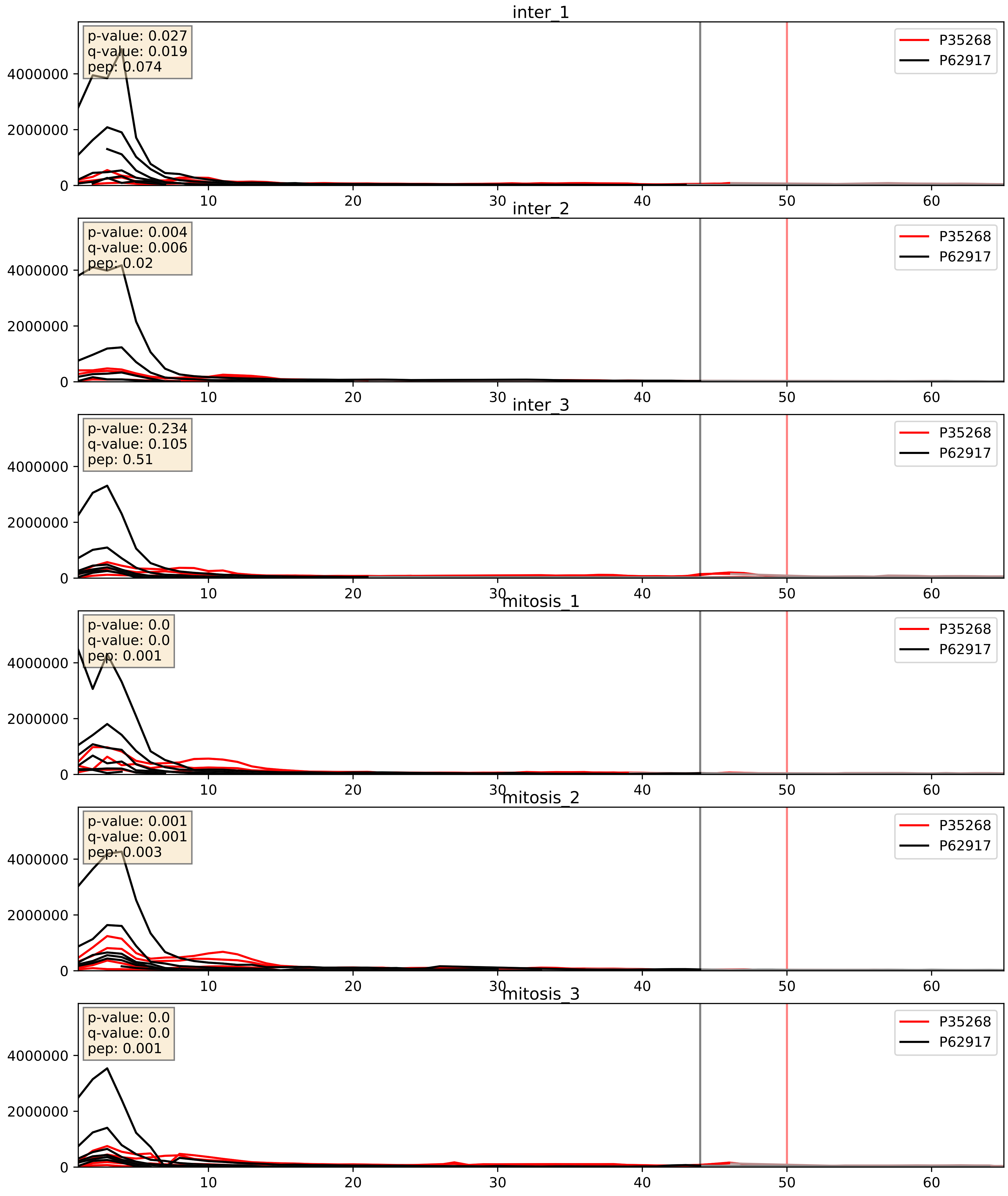


| level | condition_1 | condition_2 | pvalue | pvalue_adjusted |
|---|---|---|---|---|
| interactor_abundance | mitosis | inter | 0.003309871911744124 | 0.0443342273453124 |
| complex_abundance | mitosis | inter | 0.035557759061584884 | 0.13685900070466125 |
| interactor_ratio | mitosis | inter | 0.44745344795344283 | 0.8518291011066611 |

P35268_P63173
RL22_HUMAN vs RL38_HUMAN
p-value: 0.0 q-value: 0.0

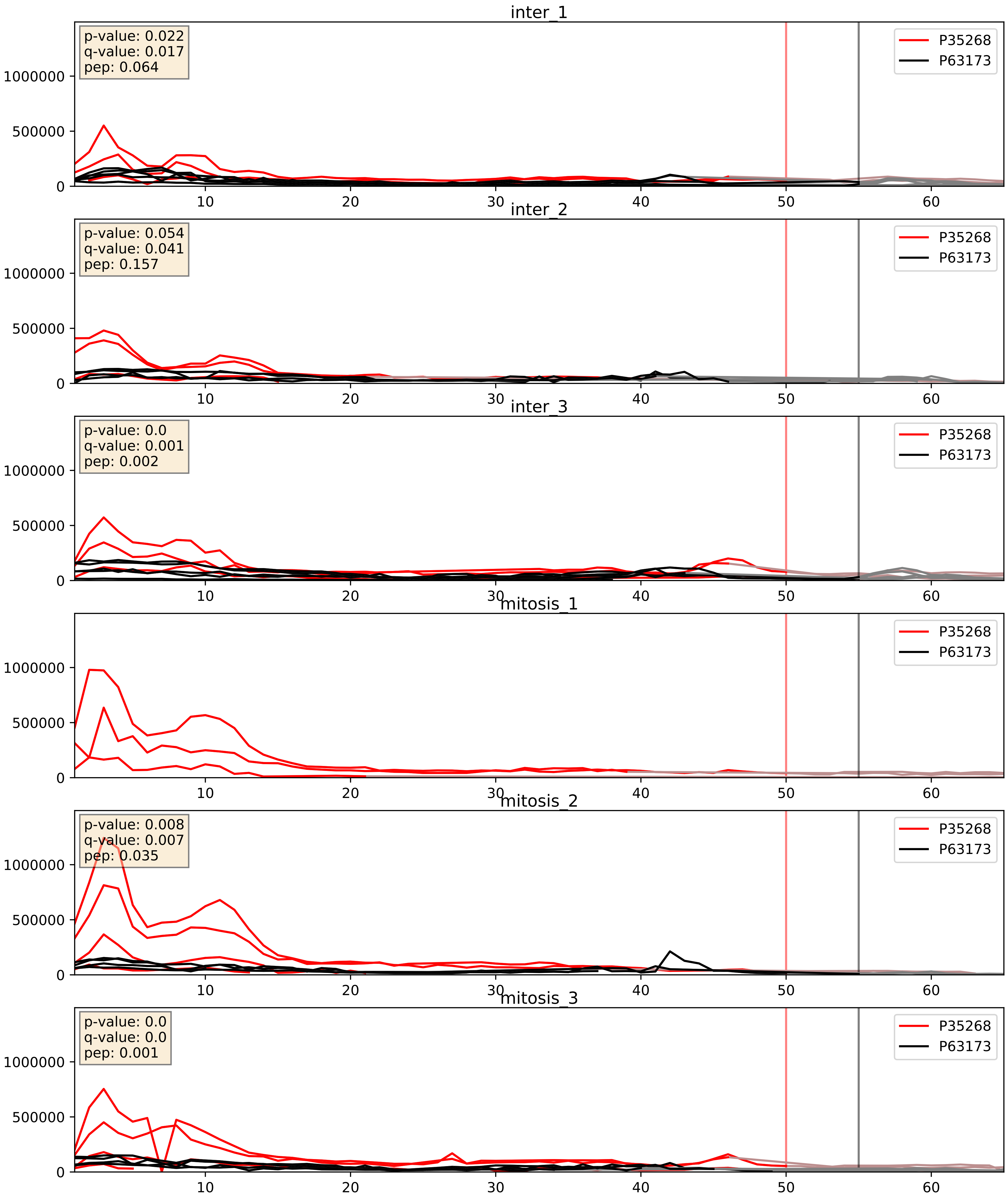


| level | condition_1 | condition_2 | pvalue | pvalue_adjusted |
|---|---|---|---|---|
| interactor_ratio | mitosis | inter | 0.20698097825597533 | 0.68434035298229 |
| complex_abundance | mitosis | inter | 0.40452105946112066 | 0.5498095060316279 |
| interactor_abundance | mitosis | inter | 0.5970590235913023 | 0.7351846908789775 |

P35268_P83731
RL22_HUMAN vs RL24_HUMAN
p-value: 0.0 q-value: 0.0

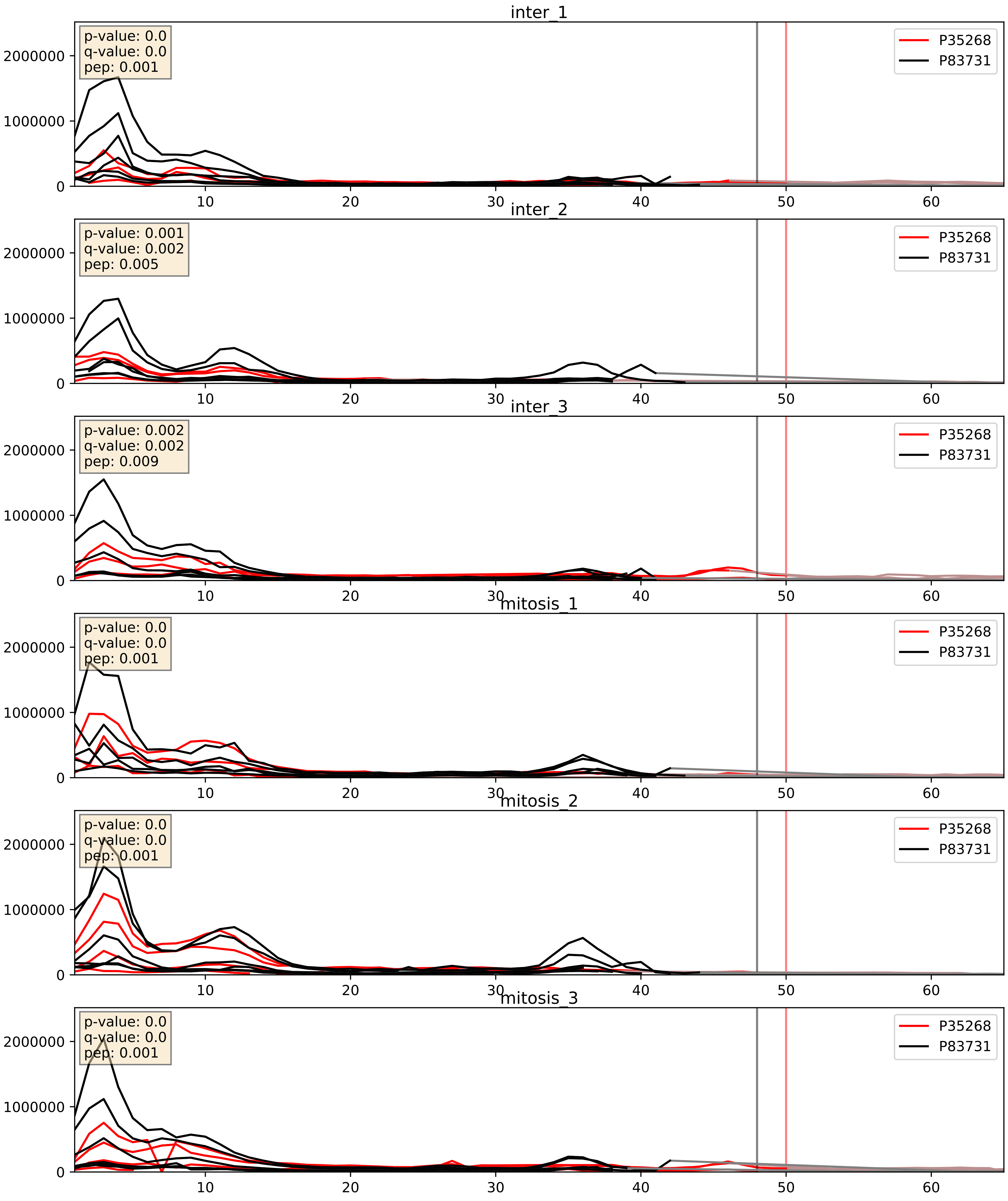


| level | condition_1 | condition_2 | pvalue | pvalue_adjusted |
|---|---|---|---|---|
| interactor_abundance | mitosis | inter | 0.009074387999507256 | 0.08316569729741126 |
| complex_abundance | mitosis | inter | 0.029296992400353123 | 0.12439595979515017 |
| interactor_ratio | mitosis | inter | 0.06023229715106691 | 0.40594115513296747 |

P35268_P84098
RL22_HUMAN vs RL19_HUMAN
p-value: 0.0 q-value: 0.0

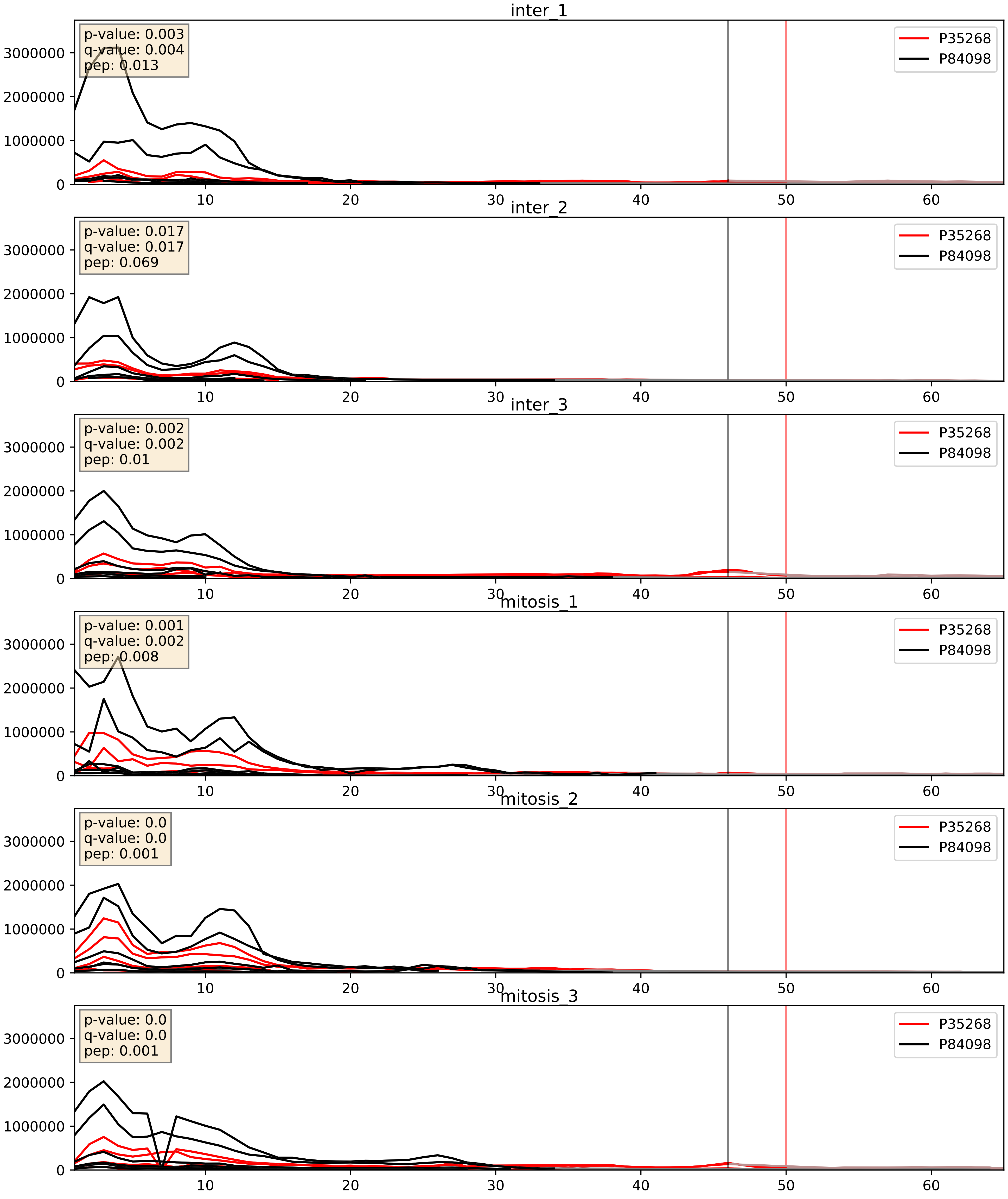


| level | condition_1 | condition_2 | pvalue | pvalue_adjusted |
|---|---|---|---|---|
| interactor_abundance | mitosis | inter | 0.014663451232014672 | 0.10605571190893752 |
| complex_abundance | mitosis | inter | 0.047356497544444036 | 0.1554235610965092 |
| interactor_ratio | mitosis | inter | 0.2382830624234638 | 0.7169430630386117 |

P35268_Q02543
RL22_HUMAN vs RL18A_HUMAN
p-value: 0.0 q-value: 0.0

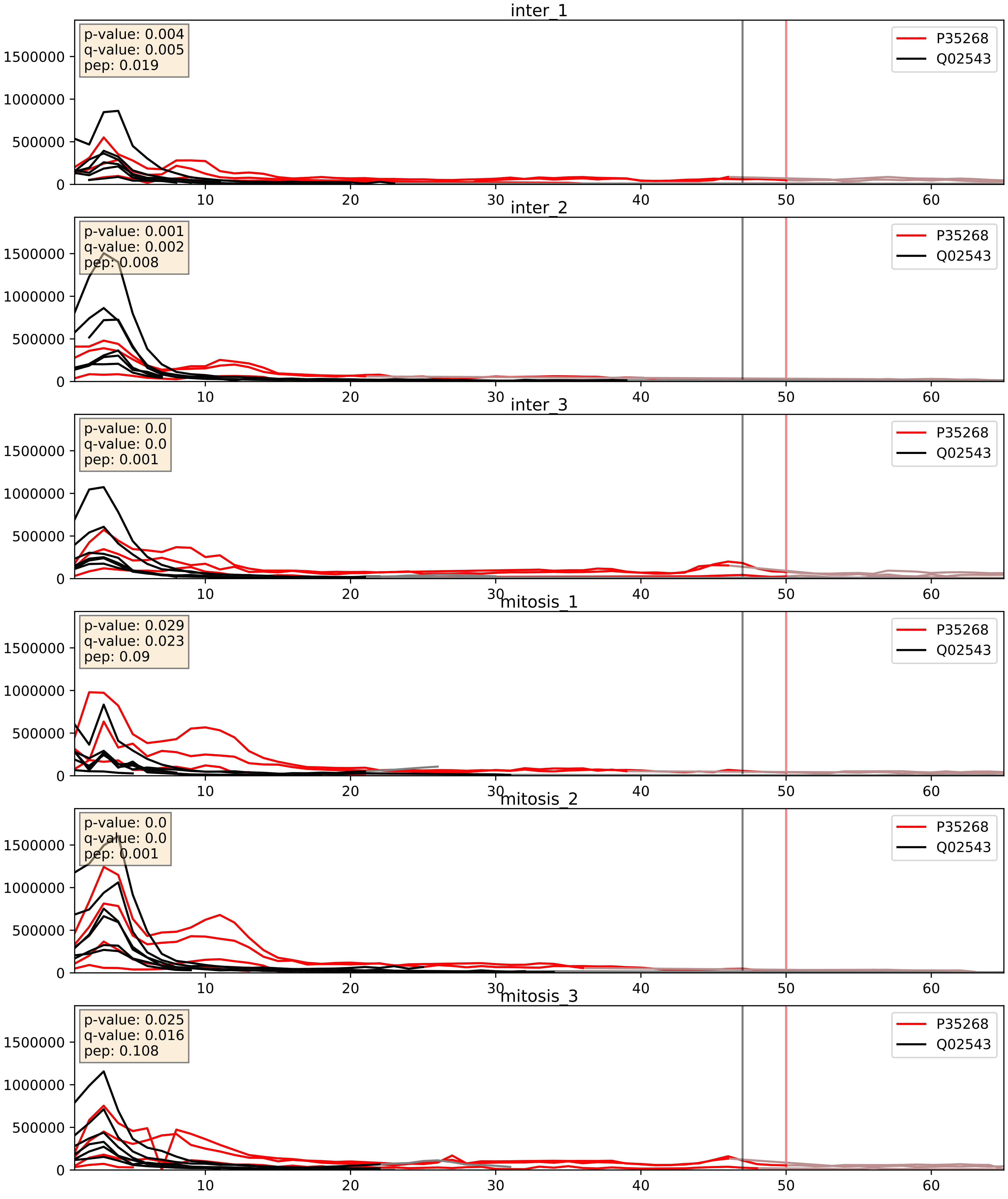


| level | condition_1 | condition_2 | pvalue | pvalue_adjusted |
|---|---|---|---|---|
| interactor_abundance | mitosis | inter | 0.027865956968170714 | 0.1386817393299659 |
| interactor_ratio | mitosis | inter | 0.11653798616523614 | 0.5623280359909346 |
| complex_abundance | mitosis | inter | 0.6385525788767992 | 0.7548667819387027 |

P35268_Q02878
RL22_HUMAN vs RL6_HUMAN
p-value: 0.003 q-value: 0.003

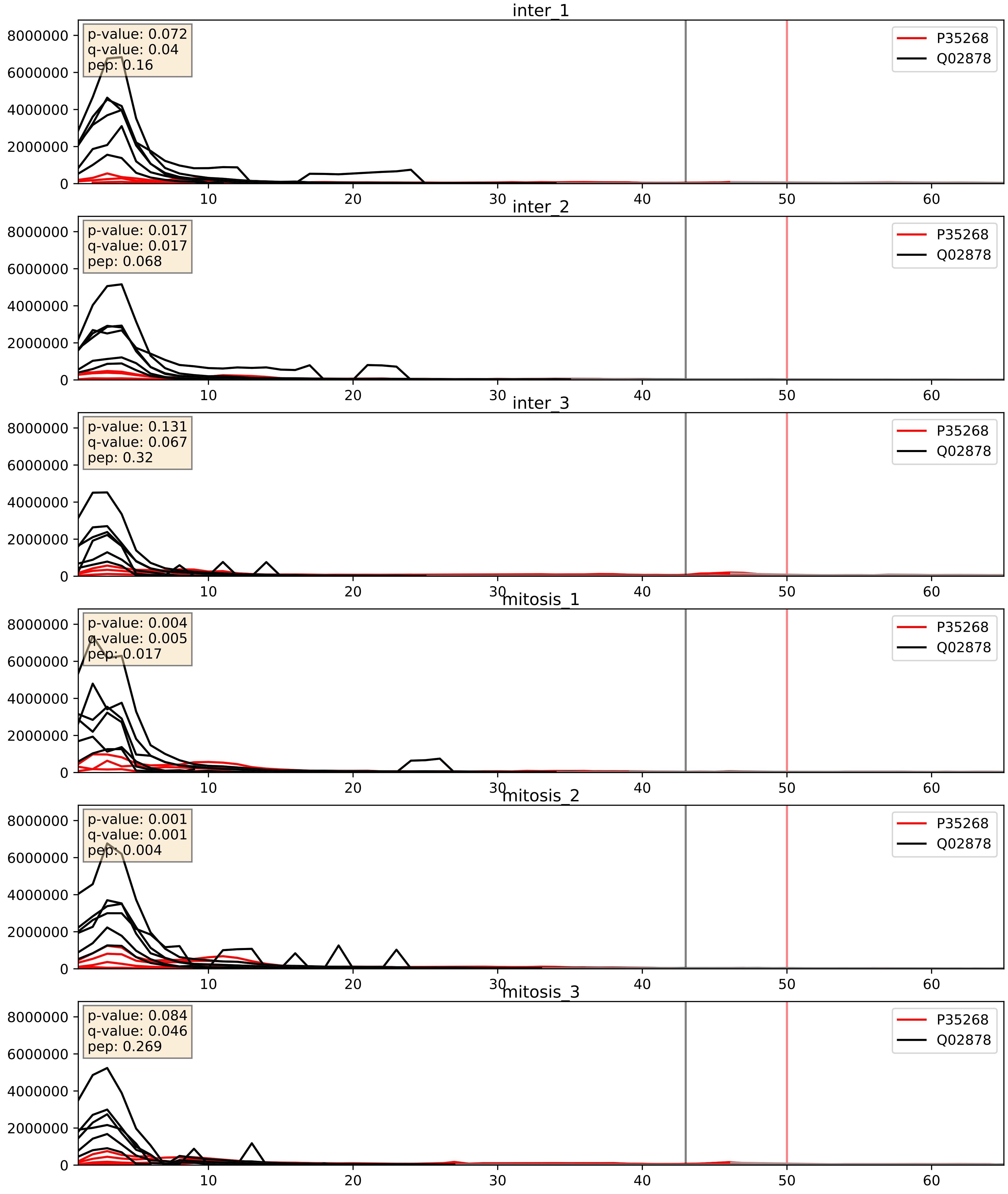


| level | condition_1 | condition_2 | pvalue | pvalue_adjusted |
|---|---|---|---|---|
| interactor_abundance | mitosis | inter | 0.003167903640315041 | 0.043387608257754805 |
| interactor_ratio | mitosis | inter | 0.11231869868494299 | 0.5536525308757383 |
| complex_abundance | mitosis | inter | 0.2662217964442123 | 0.4159260043005032 |

P35268_Q07020
RL22_HUMAN vs RL18_HUMAN
p-value: 0.0 q-value: 0.0

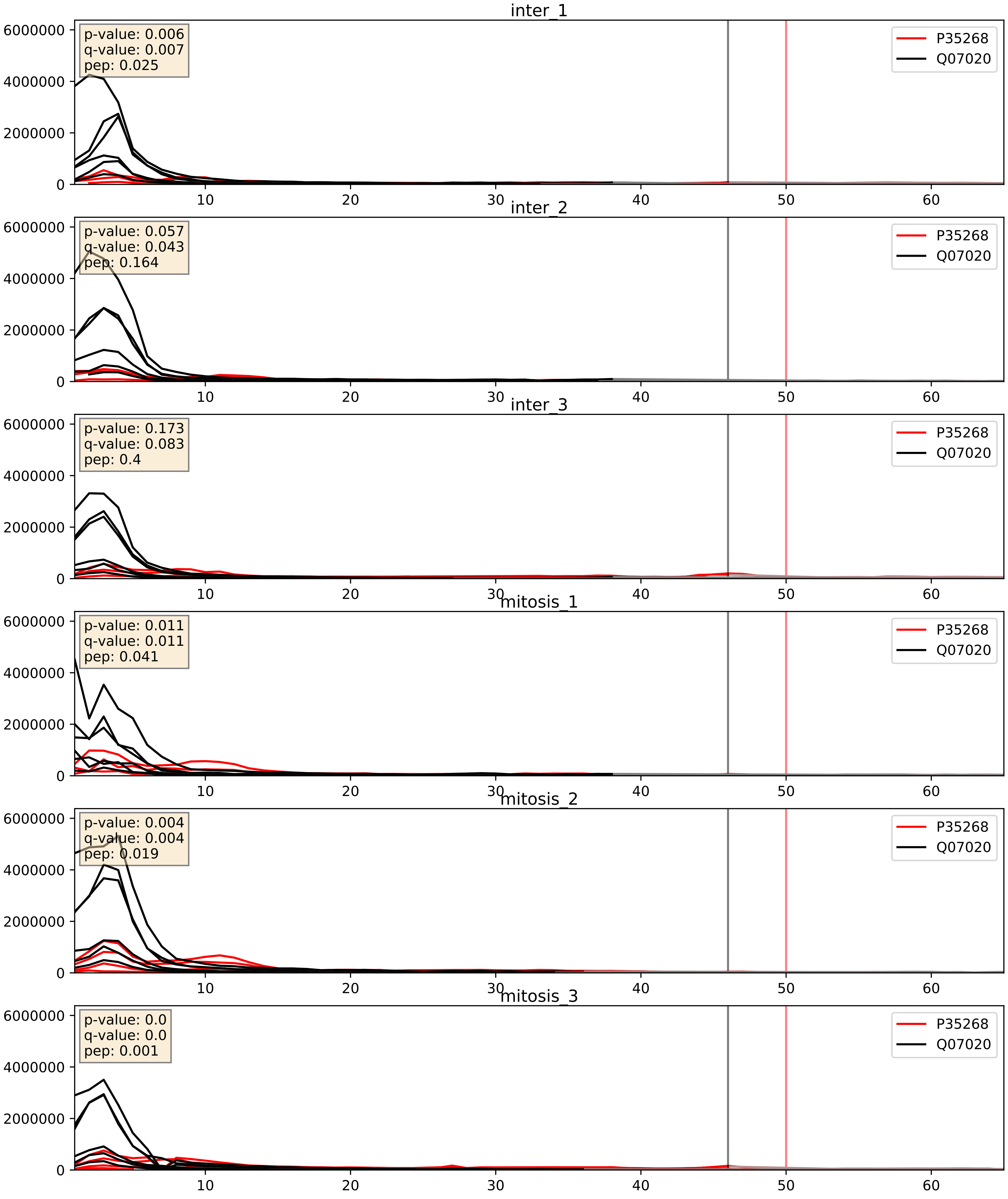


| level | condition_1 | condition_2 | pvalue | pvalue_adjusted |
|---|---|---|---|---|
| interactor_abundance | mitosis | inter | 0.00962037331935281 | 0.08507272274138435 |
| interactor_ratio | mitosis | inter | 0.027027410023691214 | 0.2576929754951936 |
| complex_abundance | mitosis | inter | 0.32938146471987556 | 0.47776871861026654 |

P35268_Q969Q0
RL22_HUMAN vs RL36L_HUMAN
p-value: 0.037 q-value: 0.025

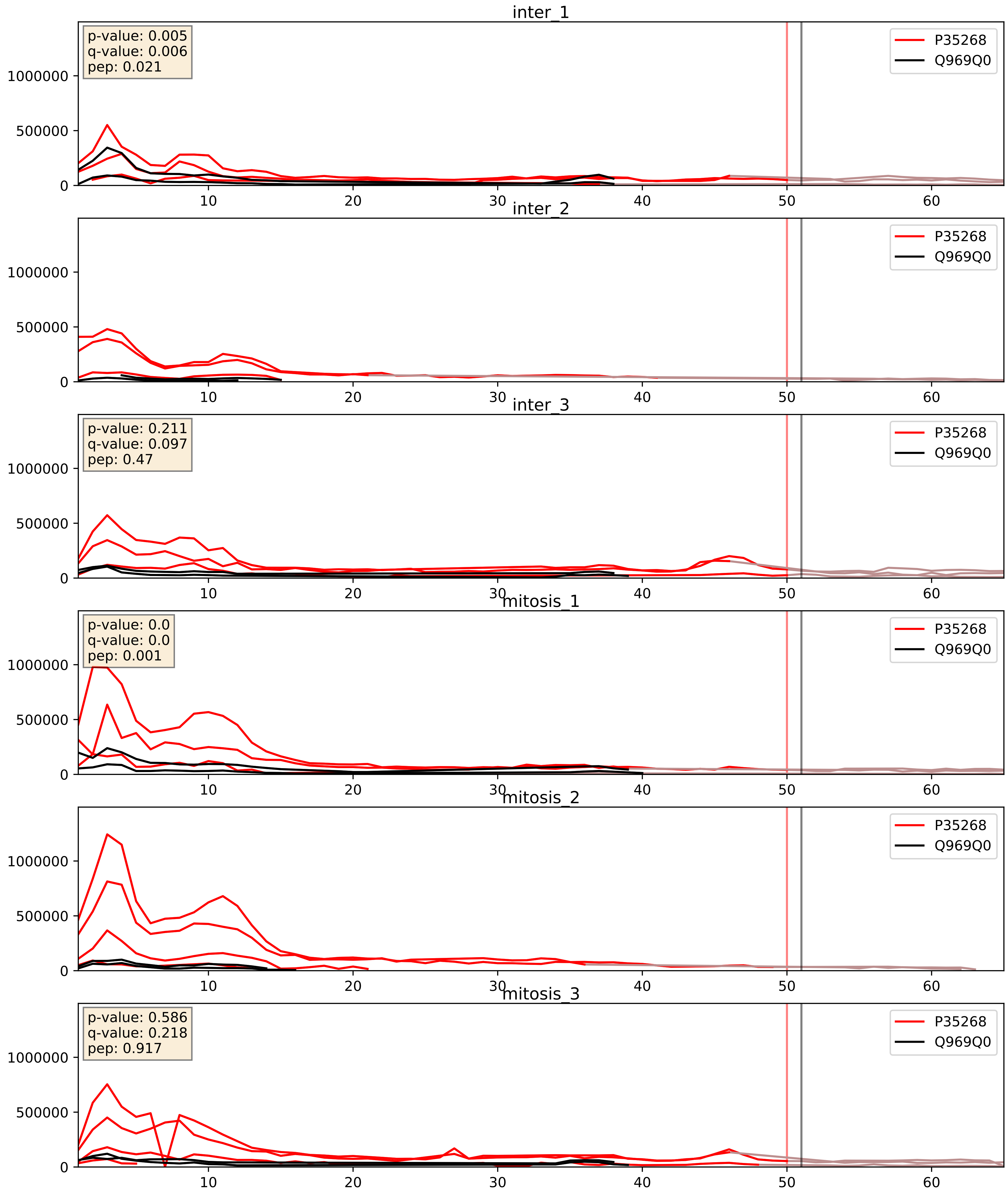


| level | condition_1 | condition_2 | pvalue | pvalue_adjusted |
|---|---|---|---|---|
| interactor_abundance | mitosis | inter | 0.017508386855220 89 | 0.11449308843051126 |
| complex_abundance | mitosis | inter | 0.23249327497235406 | 0.3849983251608077 |
| interactor_ratio | mitosis | inter | 0.25119018667533316 | 0.7251677296572153 |

P35268_Q9UHB9
RL22_HUMAN vs SRP68_HUMAN
p-value: 0.057 q-value: 0.036

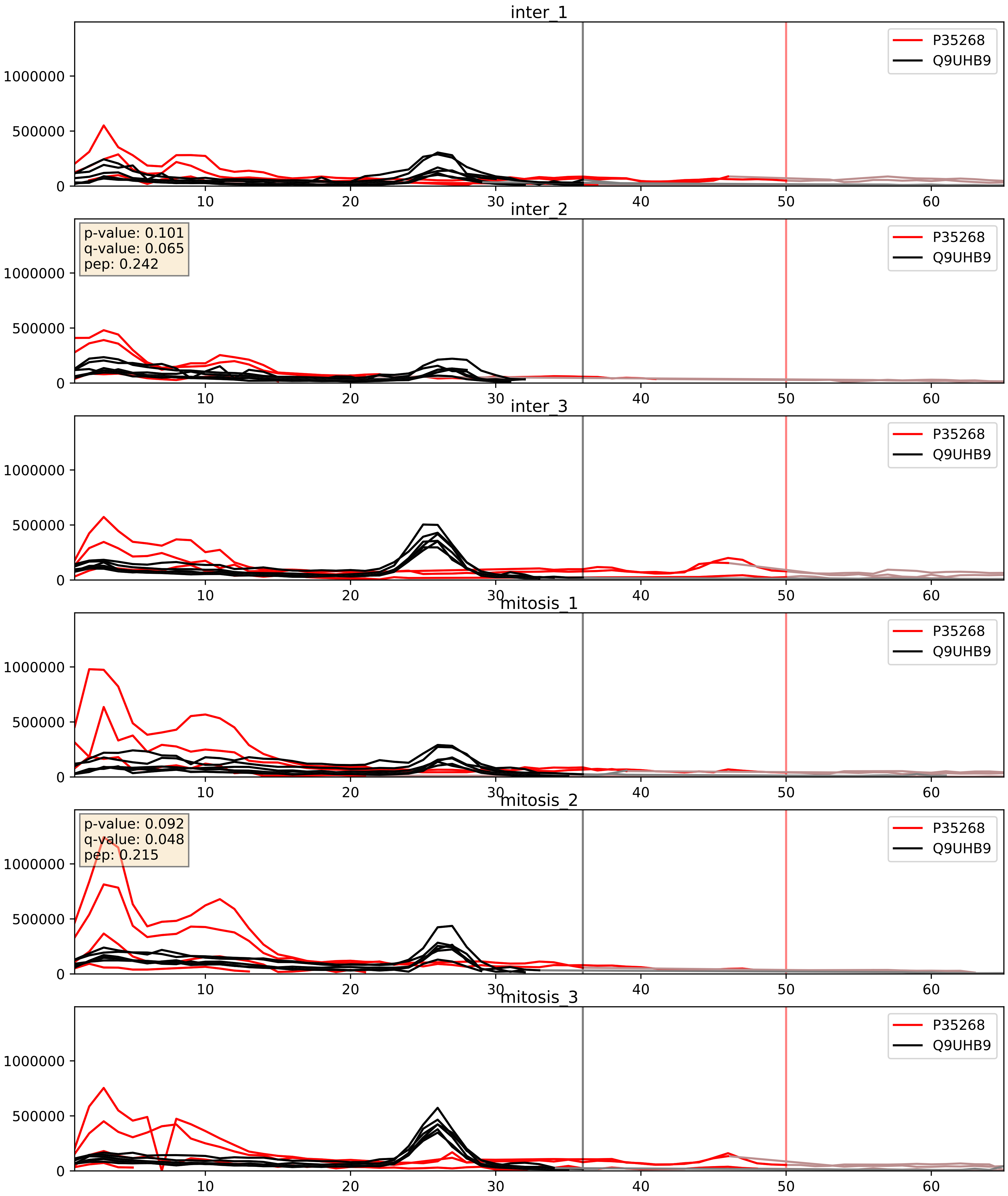


| level | condition_1 | condition_2 | pvalue | pvalue_adjusted |
|---|---|---|---|---|
| interactor_abundance | mitosis | inter | 0.0074254387770958654 | 0.07460299992011808 |
| interactor_ratio | mitosis | inter | 0.01795502474286222 | 0.20196453587240554 |
| complex_abundance | mitosis | inter | 0.053482115042832075 | 0.16500954966991963 |

P35268_Q9Y3U8
RL22_HUMAN vs RL36_HUMAN
p-value: 0.0 q-value: 0.0

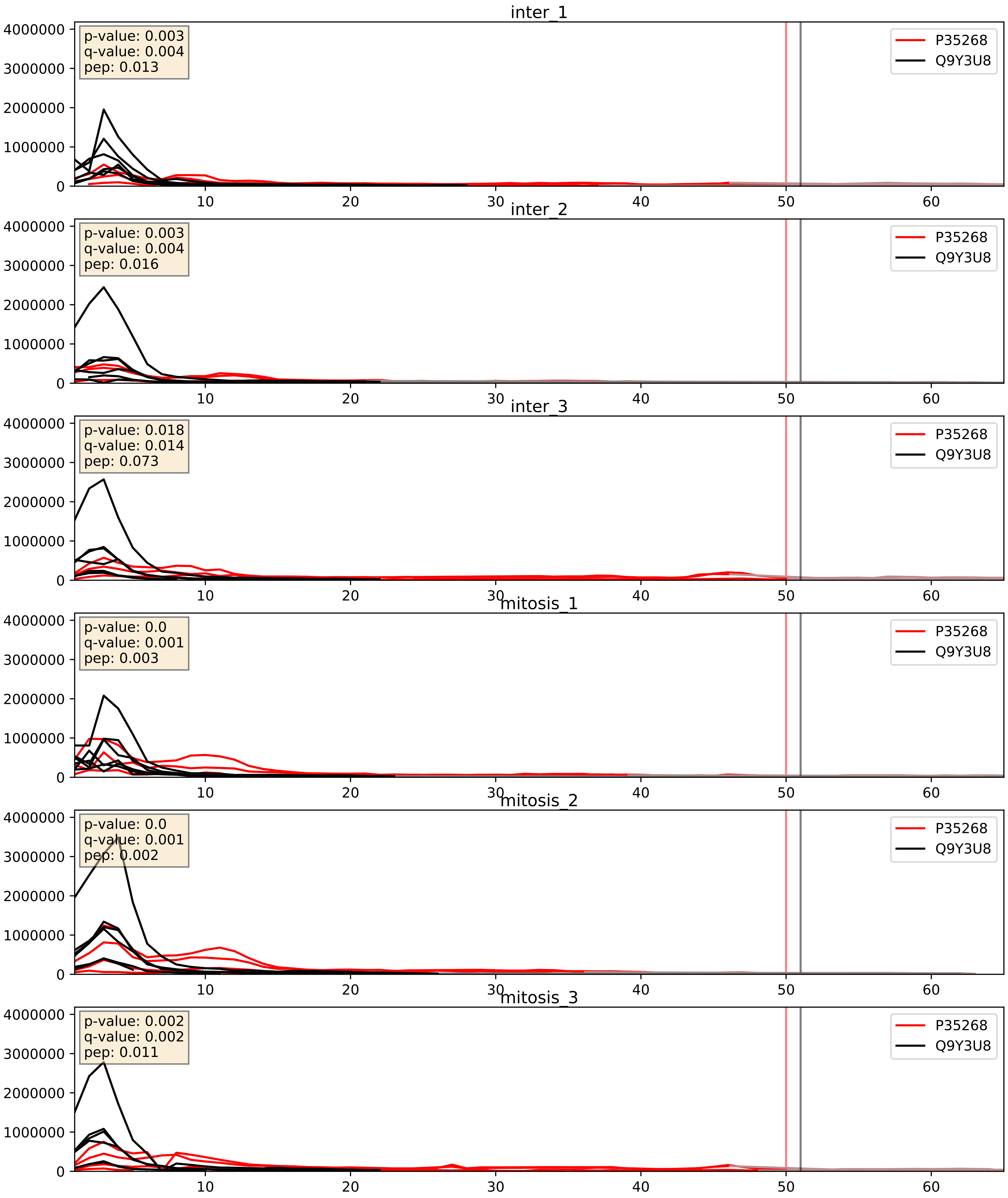


| level | condition_1 | condition_2 | pvalue | pvalue_adjusted |
|---|---|---|---|---|
| interactor_abundance | mitosis | inter | 0.01806386027534848 | 0.11569520685146502 |
| interactor_ratio | mitosis | inter | 0.026976802148841173 | 0.2576929754951936 |
| complex_abundance | mitosis | inter | 0.04219350760323655 | 0.1474505946364627 |
