## Supplementary Data 2 for "SECAT: Quantifying differential protein-protein interaction states by network-centric analysis": hela_string_000159_Q9P2I0.pdf

Q9P2I0_Q9P2I0

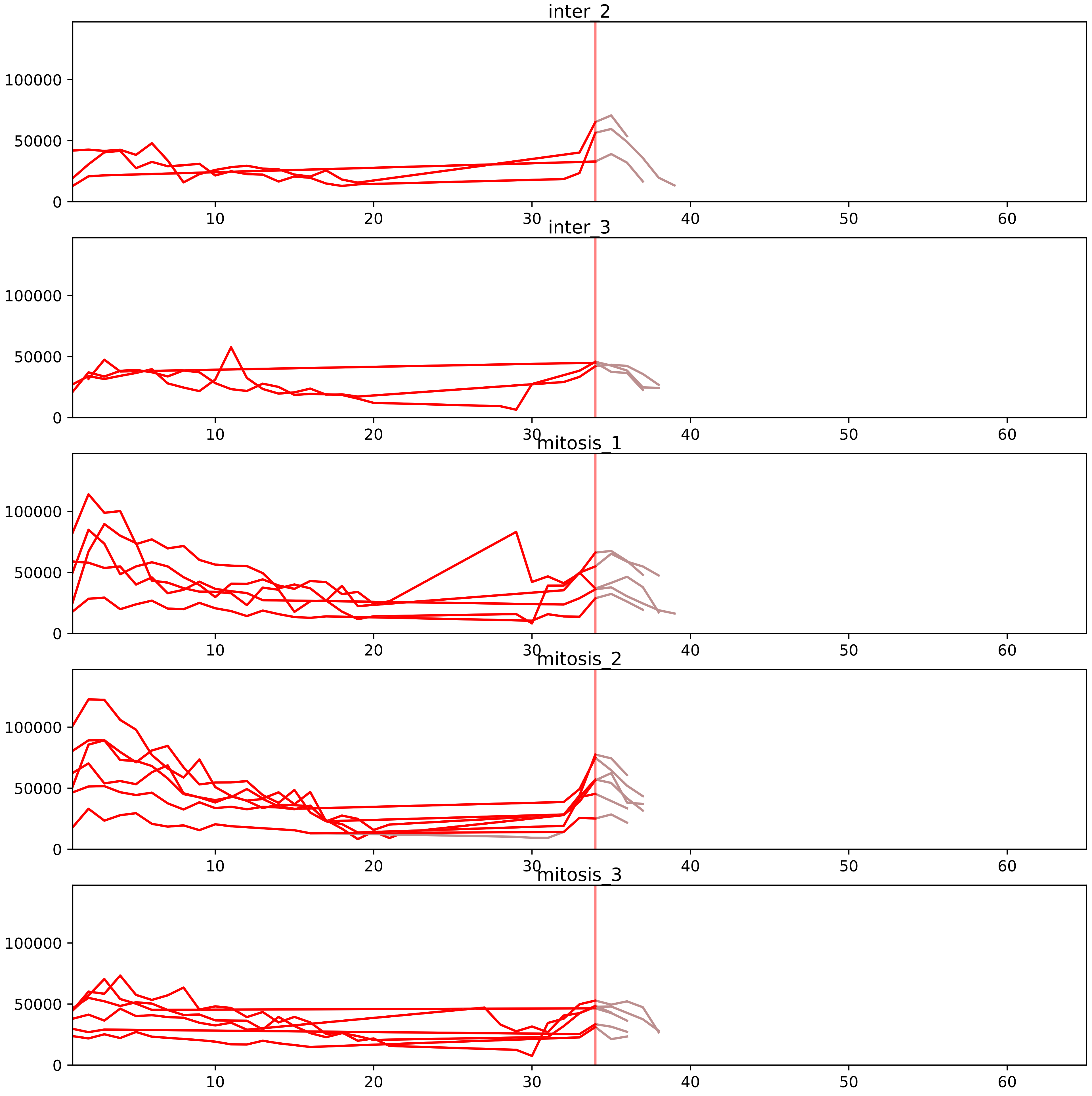


| level | condition_1 | condition_2 | pvalue | pvalue_adjusted |
|---|---|---|---|---|
| interactor_ratio | mitosis | inter | 7.325279772723556e-05 | 0.0012960110367126292 |
| interactor_abundance | mitosis | inter | 0.0008935845314715909 | 0.010406300872833717 |
| complex_abundance | mitosis | inter | 0.0025332213215515844 | 0.019346245430430554 |
| assembled_abundance | mitosis | inter | 0.026776913045488782 | 0.1989024030297464 |
| total_abundance | mitosis | inter | 0.031039636689276603 | 0.2245119859421039 |
| monomer_abundance | mitosis | inter | 0.12883698753176645 | 0.475444759619952 |

O95639_Q9P2I0
CPSF4_HUMAN vs CPSF2_HUMAN
p-value: 0.066 q-value: 0.041

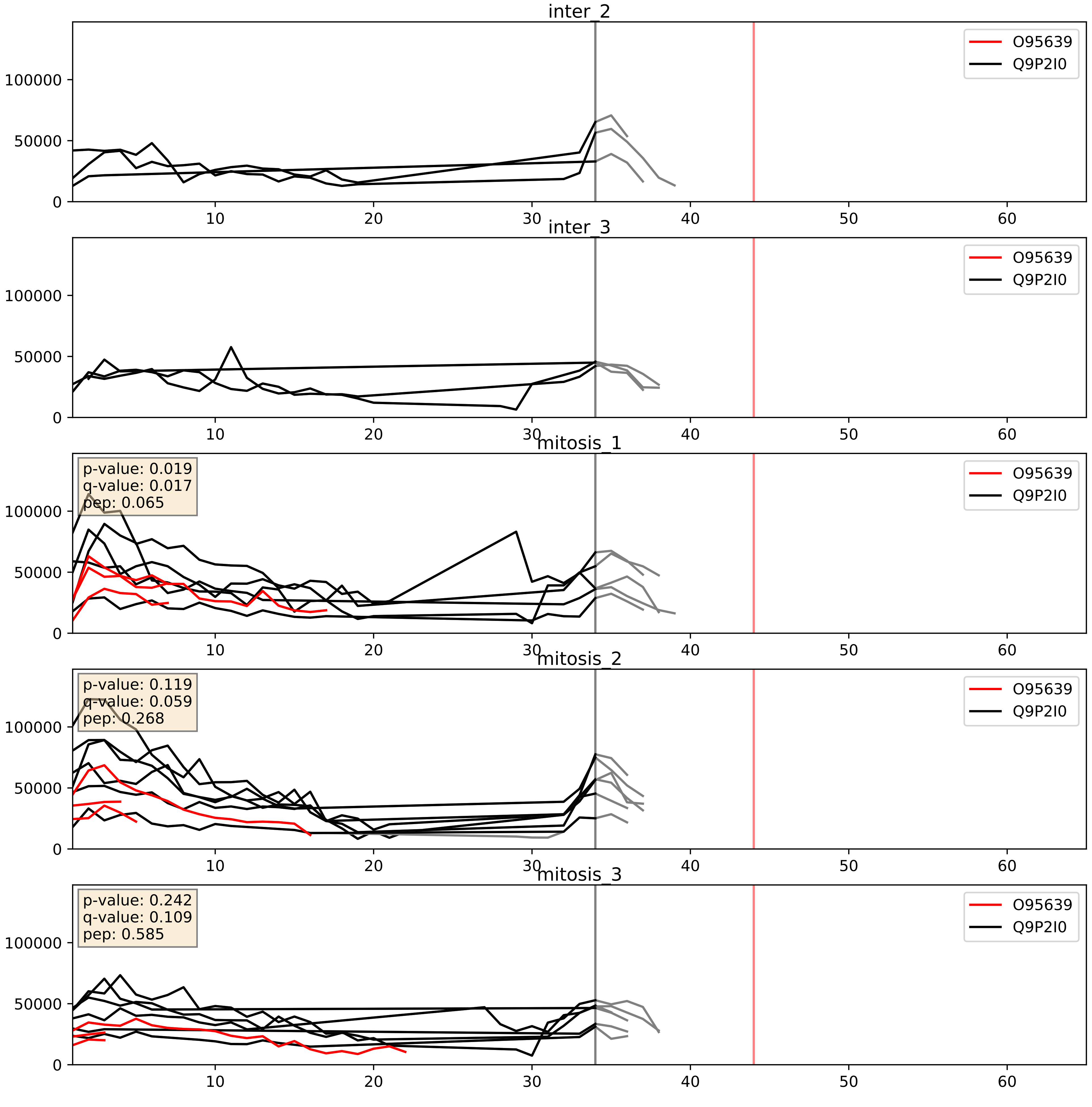


| level | condition_1 | condition_2 | pvalue | pvalue_adjusted |
|---|---|---|---|---|
| interactor_ratio | mitosis | inter | 0.0008914350338845084 | 0.034218313408302205 |
| complex_abundance | mitosis | inter | 0.007493000089970082 | 0.06444768685385507 |
| interactor_abundance | mitosis | inter | 0.09930756660188488 | 0.27089635758831565 |

Q10570_Q9P2I0
CPSF1_HUMAN vs CPSF2_HUMAN
p-value: 0.01 q-value: 0.009

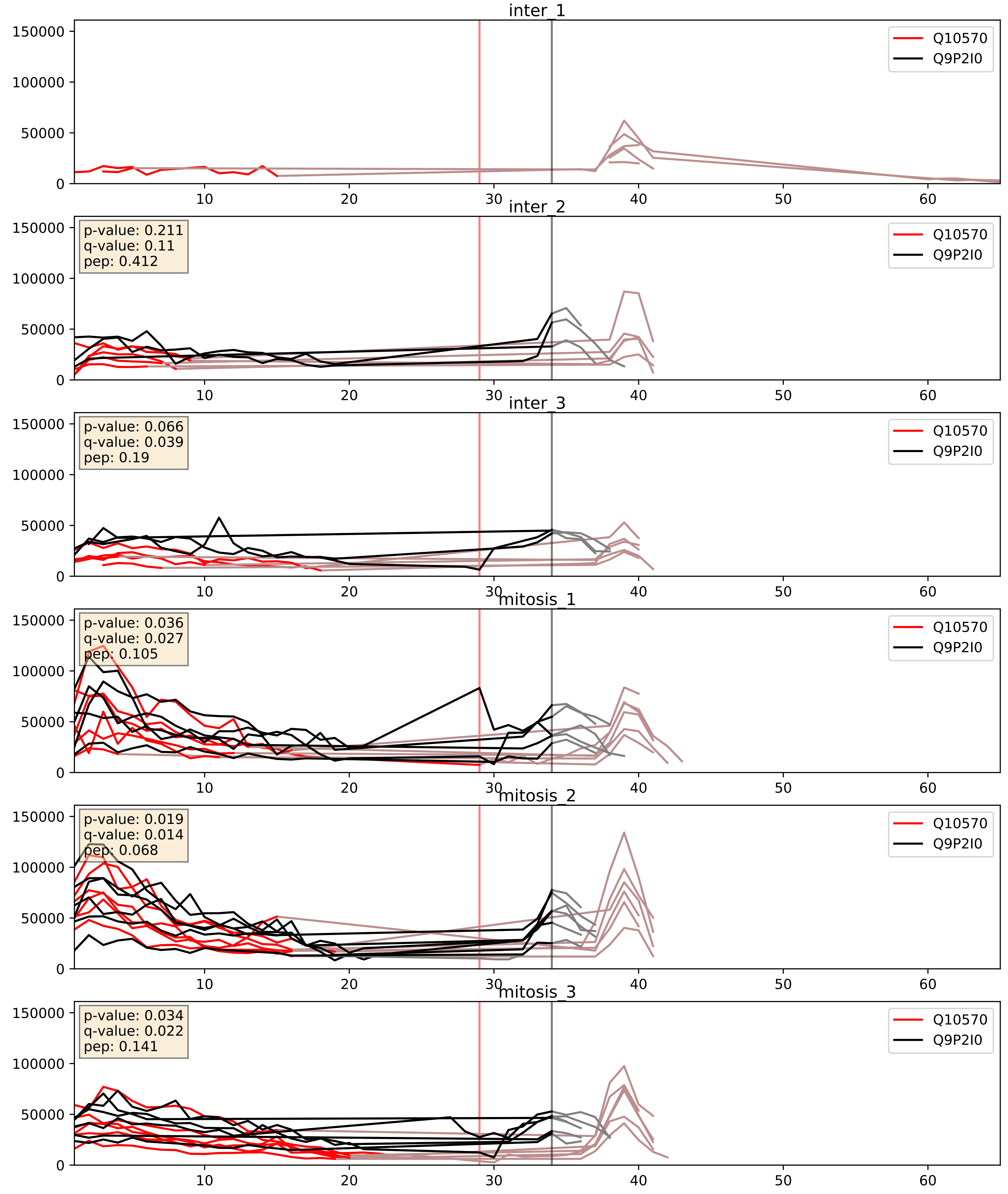


| level | condition_1 | condition_2 | pvalue | pvalue_adjusted |
|---|---|---|---|---|
| interactor_abundance | mitosis | inter | 0.011060306054893729 | 0.09263817987269111 |
| complex_abundance | mitosis | inter | 0.012379466810243465 | 0.08259410436140613 |
| interactor_ratio | mitosis | inter | 0.3884011215695155 | 0.8251906958403389 |

Q6UN15_Q9P2I0
FIP1_HUMAN vs CPSF2_HUMAN
p-value: 0.007 q-value: 0.006

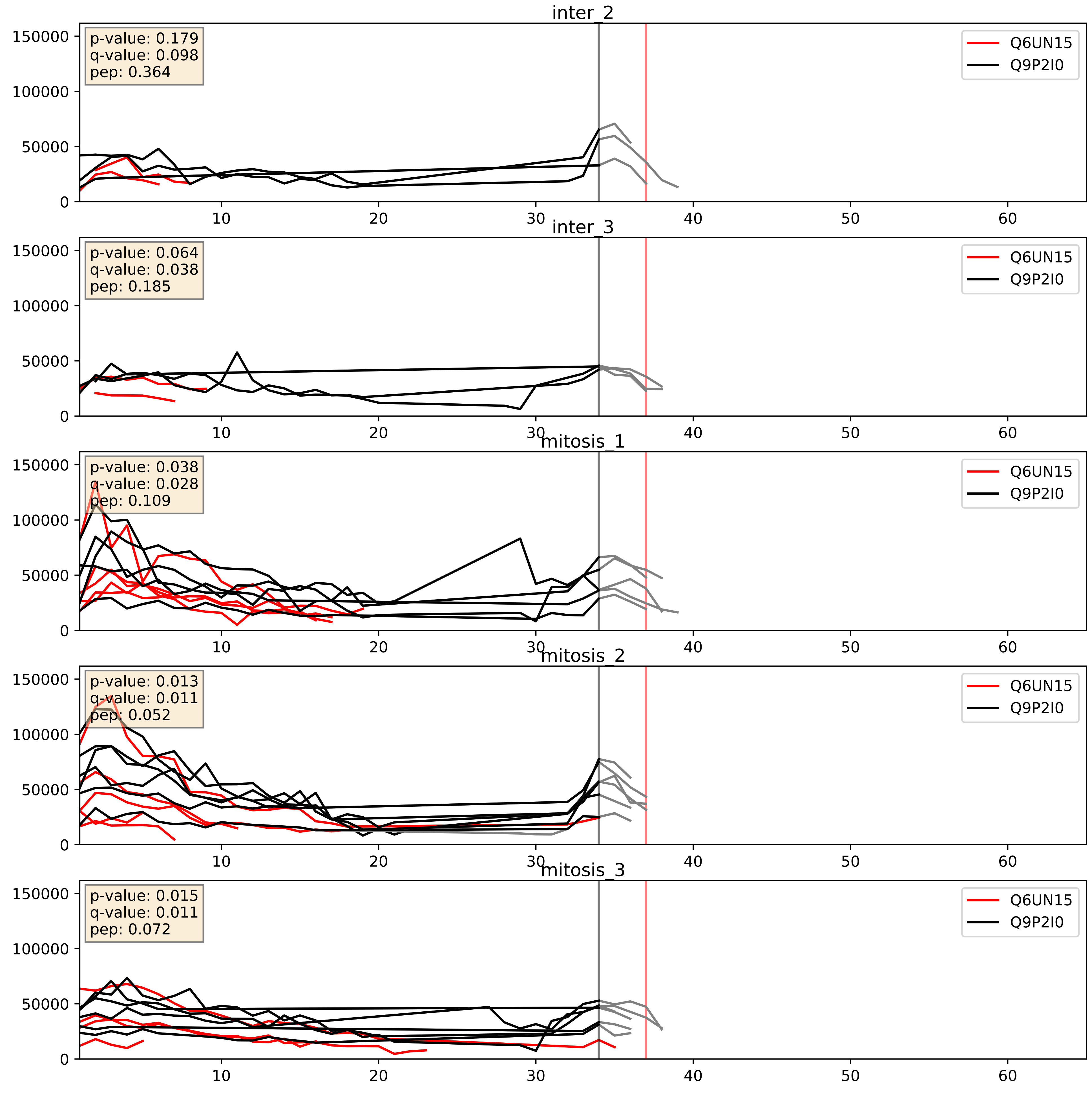


| level | condition_1 | condition_2 | pvalue | pvalue_adjusted |
|---|---|---|---|---|
| complex_abundance | mitosis | inter | 0.007845649398103807 | 0.0657361191517291 |
| interactor_abundance | mitosis | inter | 0.013852354432628004 | 0.10436859054266079 |
| interactor_ratio | mitosis | inter | 0.015100829020357177 | 0.18053505085790145 |

Q92797_Q9P2I0
SYMPK_HUMAN vs CPSF2_HUMAN
p-value: 0.017 q-value: 0.014

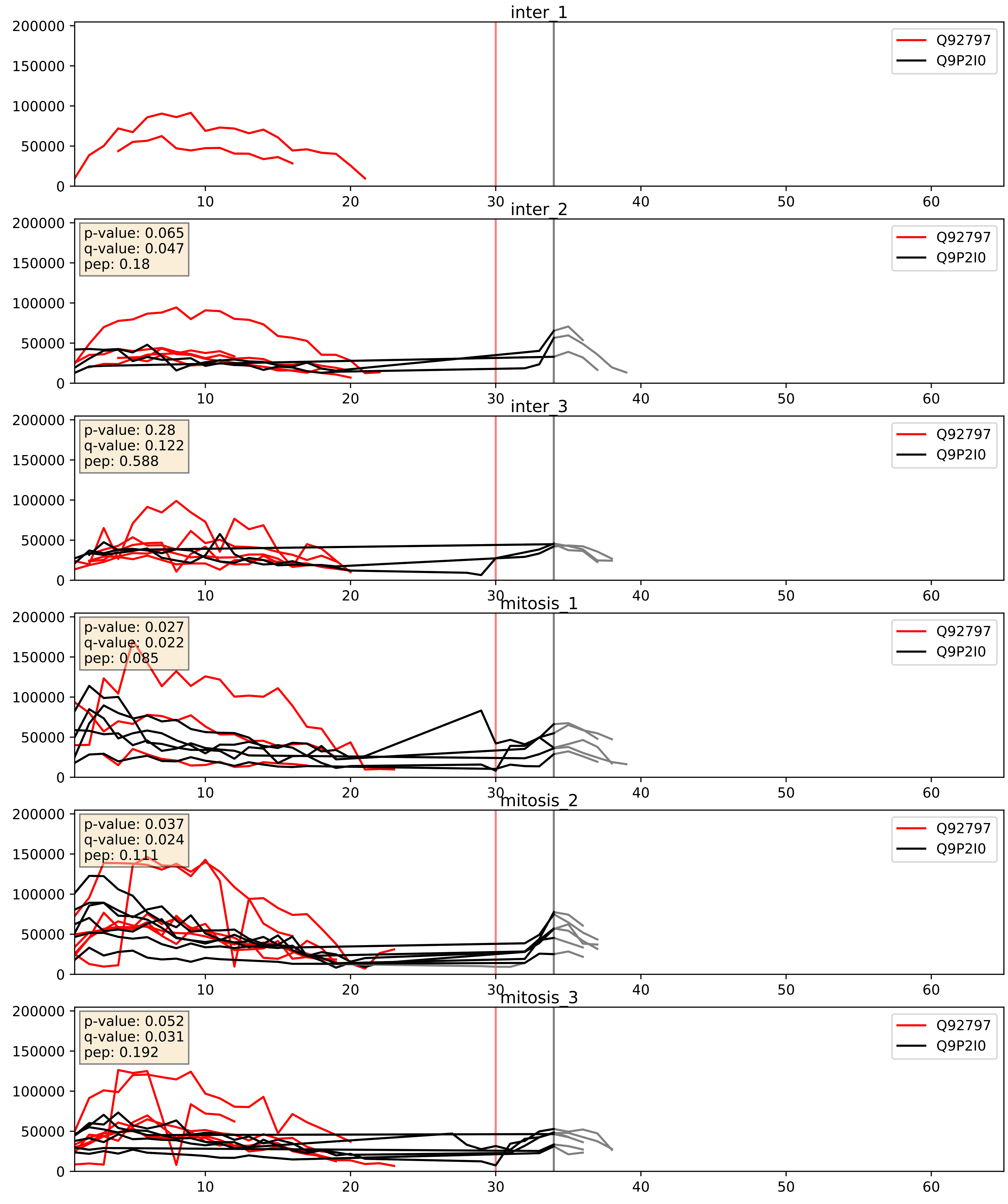


| level | condition_1 | condition_2 | pvalue | pvalue_adjusted |
|---|---|---|---|---|
| complex_abundance | mitosis | inter | 0.10377243798855093 | 0.23930282036152908 |
| interactor_abundance | mitosis | inter | 0.2466500302835656 | 0.44008843339808684 |
| interactor_ratio | mitosis | inter | 0.390491770134291 | 0.825249705827432 |

Q99459_Q9P2I0
CDC5L_HUMAN vs CPSF2_HUMAN
p-value: 0.017 q-value: 0.014

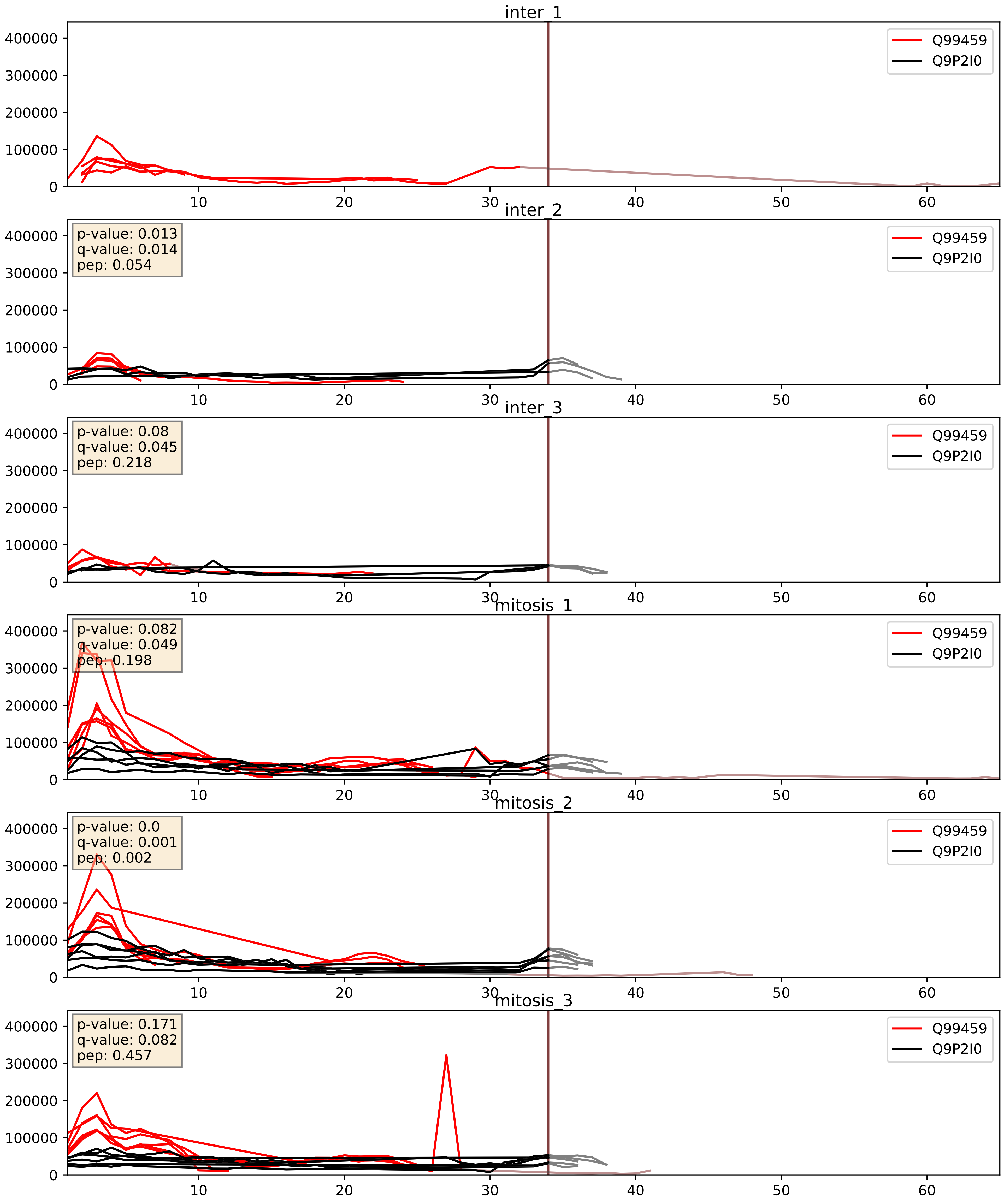


| level | condition_1 | condition_2 | pvalue | pvalue_adjusted |
|---|---|---|---|---|
| interactor_abundance | mitosis | inter | 0.005440530079986679 5 | 0.06238571665731544 |
| complex_abundance | mitosis | inter | 0.009246753476483927 | 0.07106059027160733 |
| interactor_ratio | mitosis | inter | 0.011940403429490735 | 0.15773125517969241 |

Q9P2I0_Q9UKF6
CPSF2_HUMAN vs CPSF3_HUMAN
p-value: 0.003 q-value: 0.003

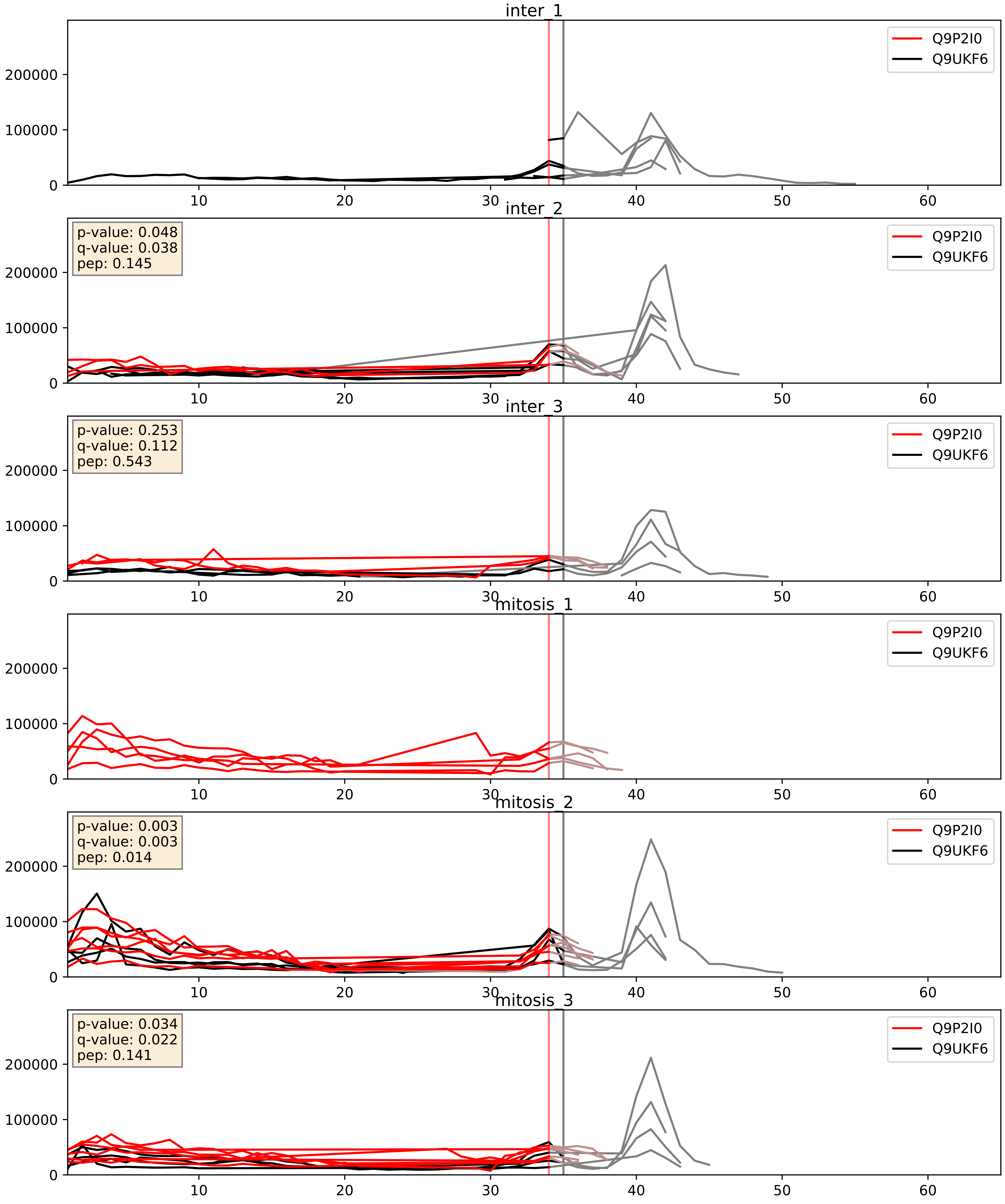


| level | condition_1 | condition_2 | pvalue | pvalue_adjusted |
|---|---|---|---|---|
| interactor_ratio | mitosis | inter | 0.08662265700536775 | 0.4940835073702731 |
| interactor_abundance | mitosis | inter | 0.6446062749996755 | 0.7620785462326113 |
| complex_abundance | mitosis | inter | 0.7906373023012381 | 0.8634671226969377 |
