## Supplementary Data 2 for "SECAT: Quantifying differential protein-protein interaction states by network-centric analysis": hela_string_000160_Q96DE5.pdf

Q96DE5_Q96DE5

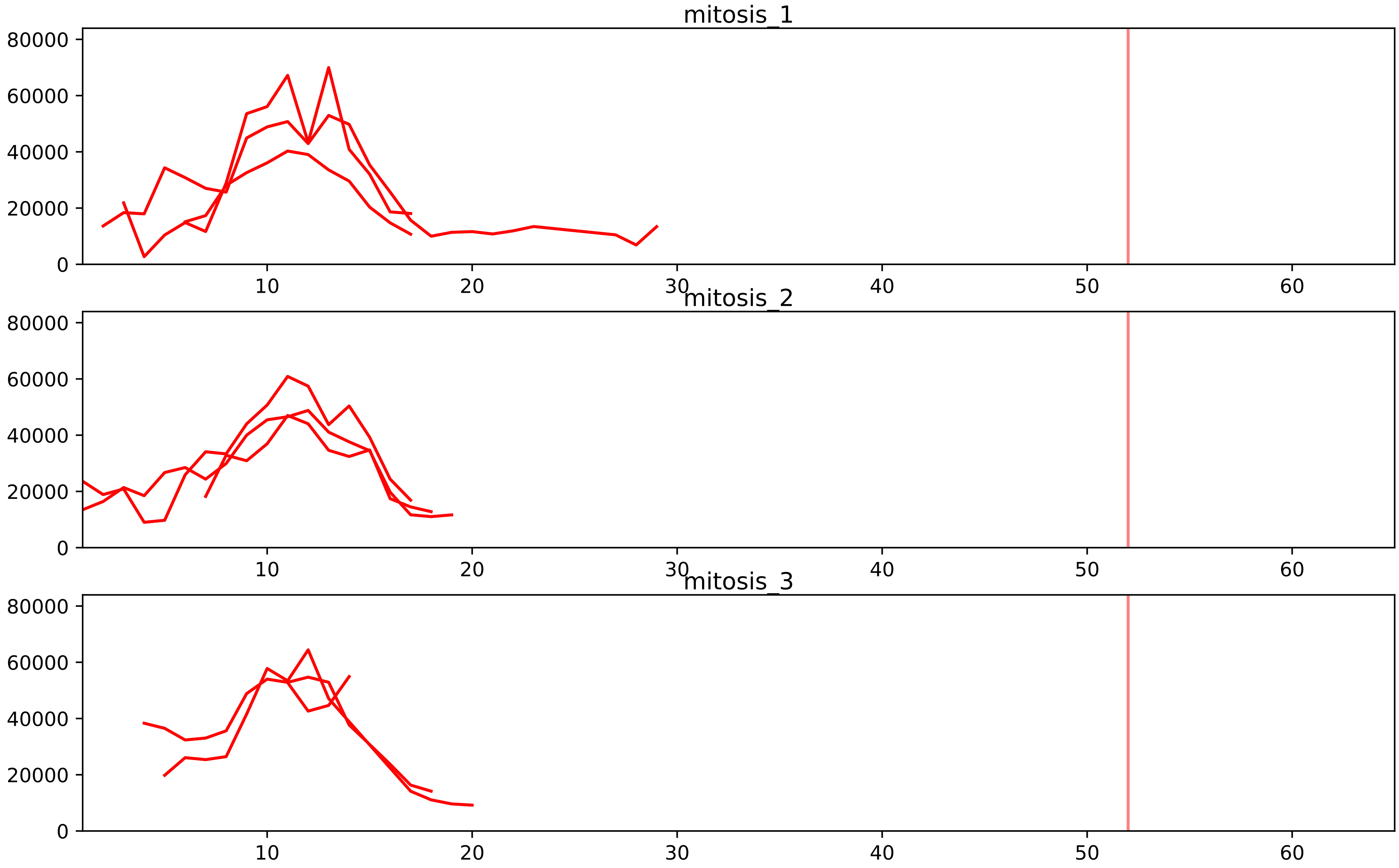


| level | condition_1 | condition_2 | pvalue | pvalue_adjusted |
|---|---|---|---|---|
| complex_abundance | mitosis | inter | 7.585234076493022e-05 | 0.0018871206097520157 |
| interactor_ratio | mitosis | inter | 9.247584039818141e-05 | 0.0015755143178949424 |
| assembled_abundance | mitosis | inter | 0.00024286823855681412 | 0.016114995197788636 |
| interactor_abundance | mitosis | inter | 0.0006676688547327848 | 0.008902251396437131 |
| total_abundance | mitosis | inter | 0.0007811465341362471 | 0.03143259708120346 |

Q13042_Q96DE5
CDC16_HUMAN vs APC16_HUMAN
p-value: 0.016 q-value: 0.013

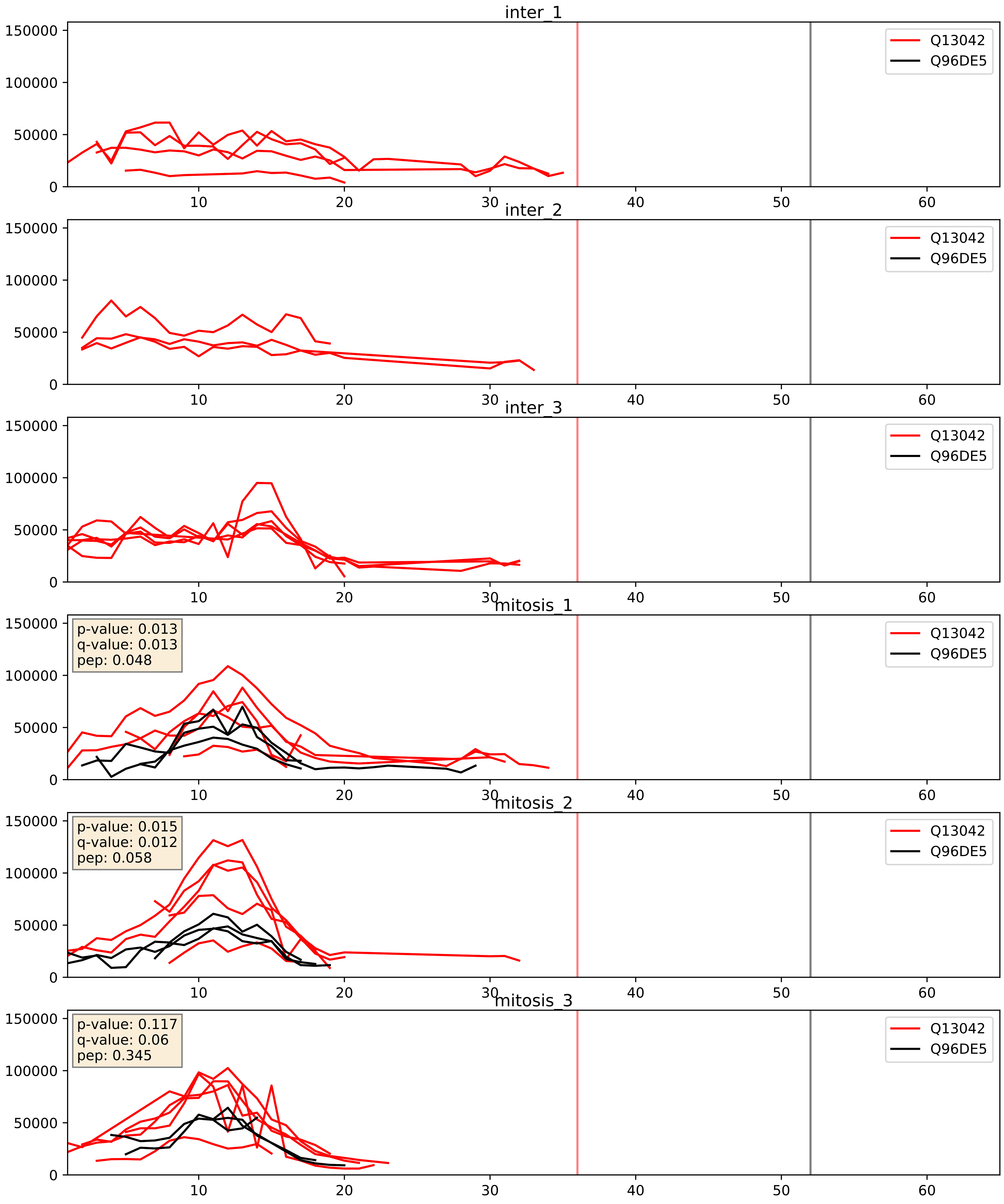


| level | condition_1 | condition_2 | pvalue | pvalue_adjusted |
|---|---|---|---|---|
| interactor_ratio | mitosis | inter | 0.00015295883421606864 | 0.012836278661937291 |
| interactor_abundance | mitosis | inter | 0.00040216400779502346 | 0.013883593575284832 |
| complex_abundance | mitosis | inter | 0.0014684336072848944 | 0.0265185478446386 |

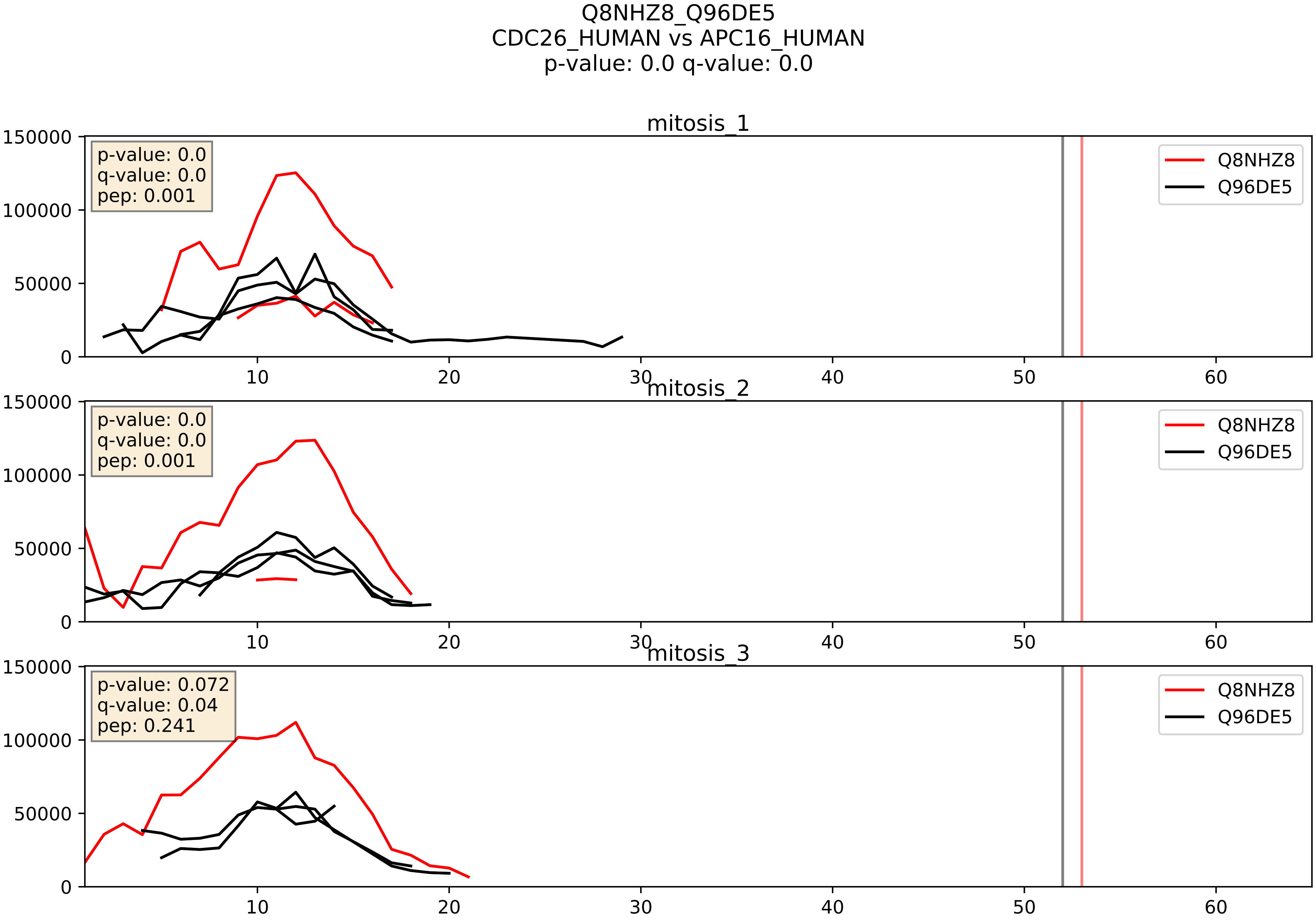


| level | condition_1 | condition_2 | pvalue | pvalue_adjusted |
|---|---|---|---|---|
| complex_abundance | mitosis | inter | 0.010729792027556192 | 0.07672534067869605 |
| interactor_ratio | mitosis | inter | 0.022311319891476004 | 0.2287907703572271 |
| interactor_abundance | mitosis | inter | 0.023574696254356885 | 0.12607602513772825 |

Q96DE5_Q9H1A4
APC16_HUMAN vs APC1_HUMAN
p-value: 0.015 q-value: 0.012

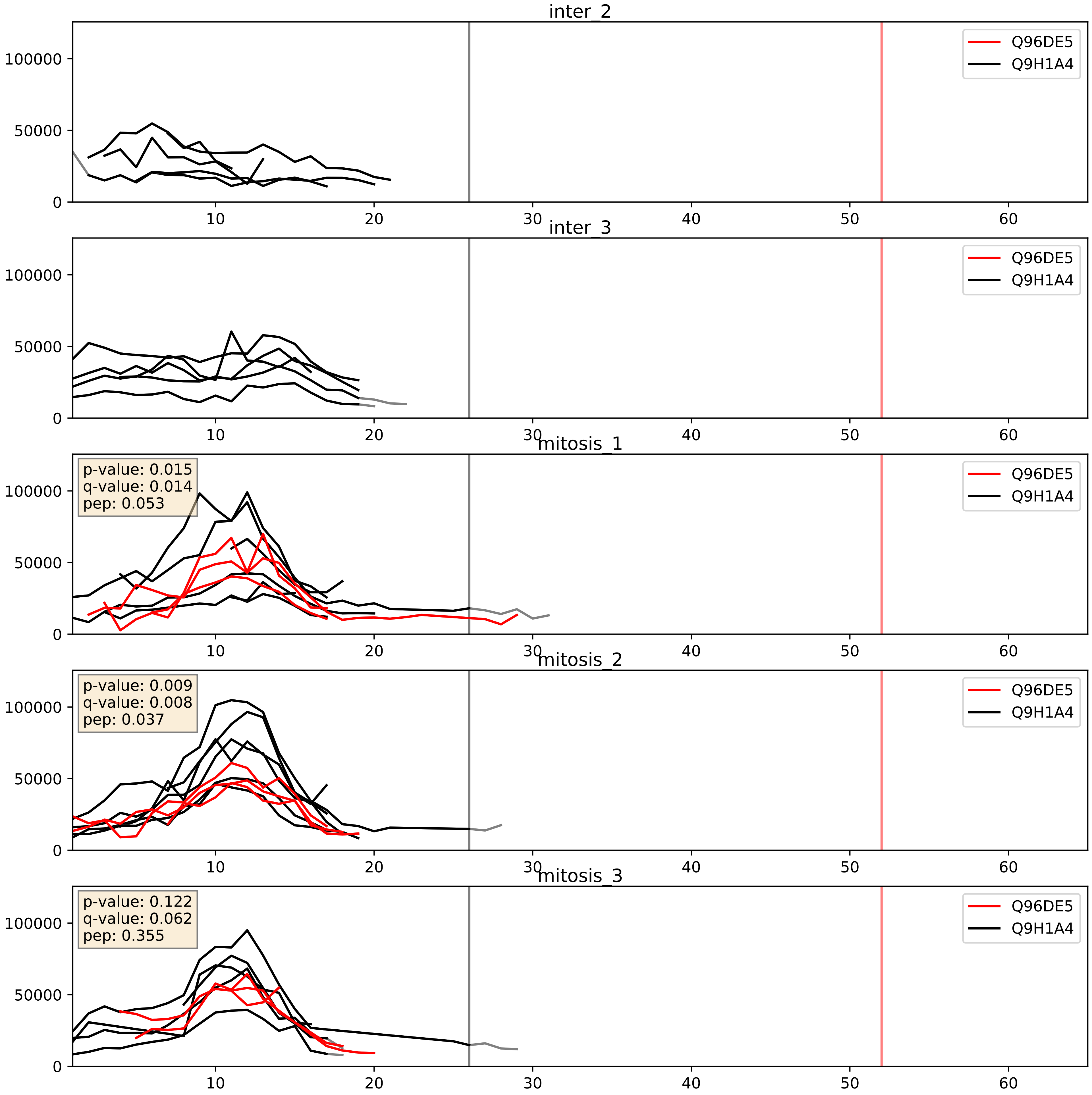


| level | condition_1 | condition_2 | pvalue | pvalue_adjusted |
|---|---|---|---|---|
| complex_abundance | mitosis | inter | 0.00184701762011143 | 0.030581181485790792 |
| interactor_abundance | mitosis | inter | 0.007821690016800924 | 0.07673772669778328 |
| interactor_ratio | mitosis | inter | 0.15083241465874134 | 0.606604569117897 |

Q96DE5_Q9UJX2
APC16_HUMAN vs CDC23_HUMAN
p-value: 0.008 q-value: 0.008

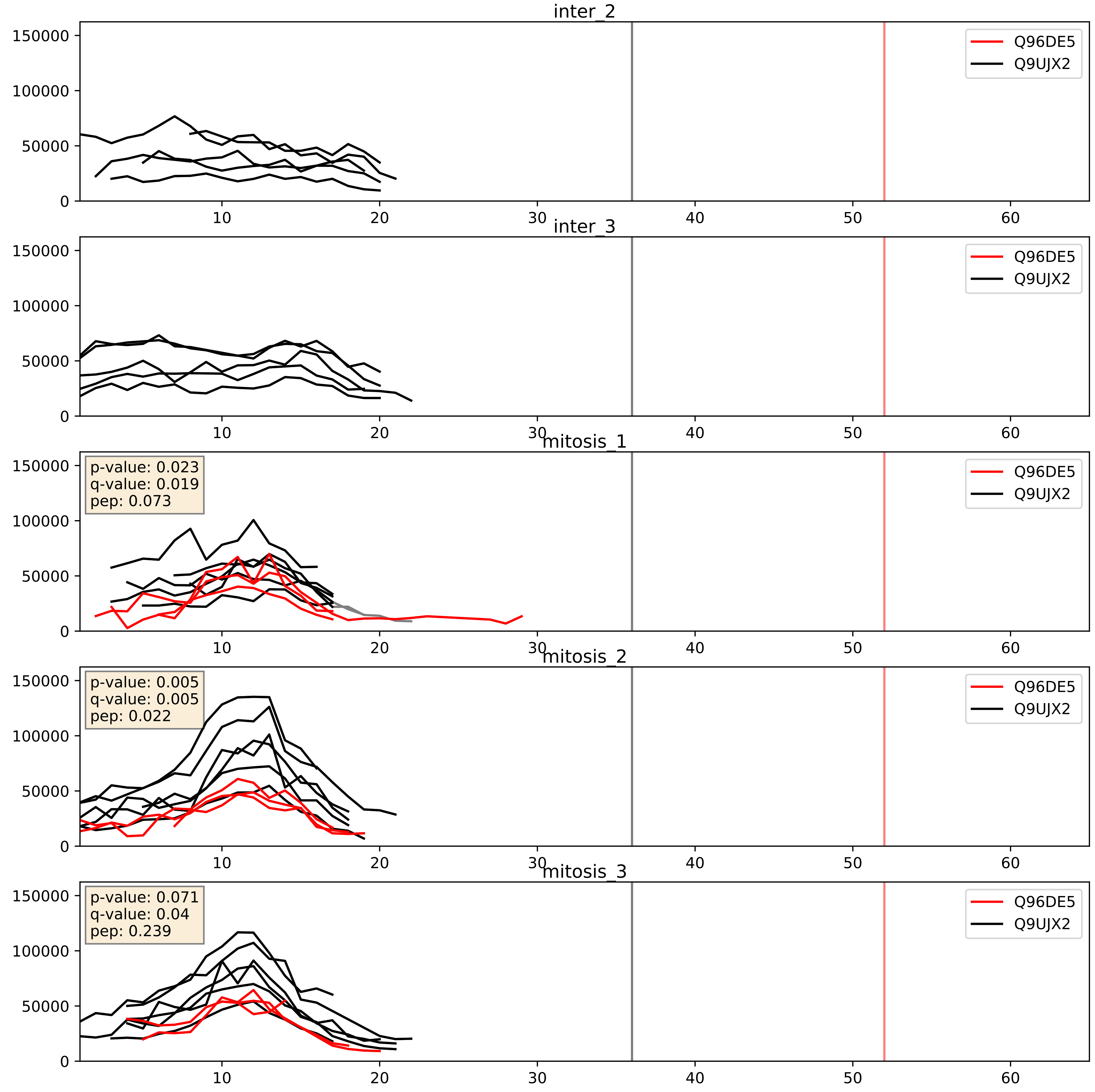


| level | condition_1 | condition_2 | pvalue | pvalue_adjusted |
|---|---|---|---|---|
| complex_abundance | mitosis | inter | 0.0006185536298757055 | 0.015948250216072407 |
| interactor_abundance | mitosis | inter | 0.006150388407561276 | 0.06671148765013551 |
| interactor_ratio | mitosis | inter | 0.020934438472495327 | 0.21906942949212715 |

Q96DE5_Q9UJX3
APC16_HUMAN vs APC7_HUMAN
p-value: 0.027 q-value: 0.02

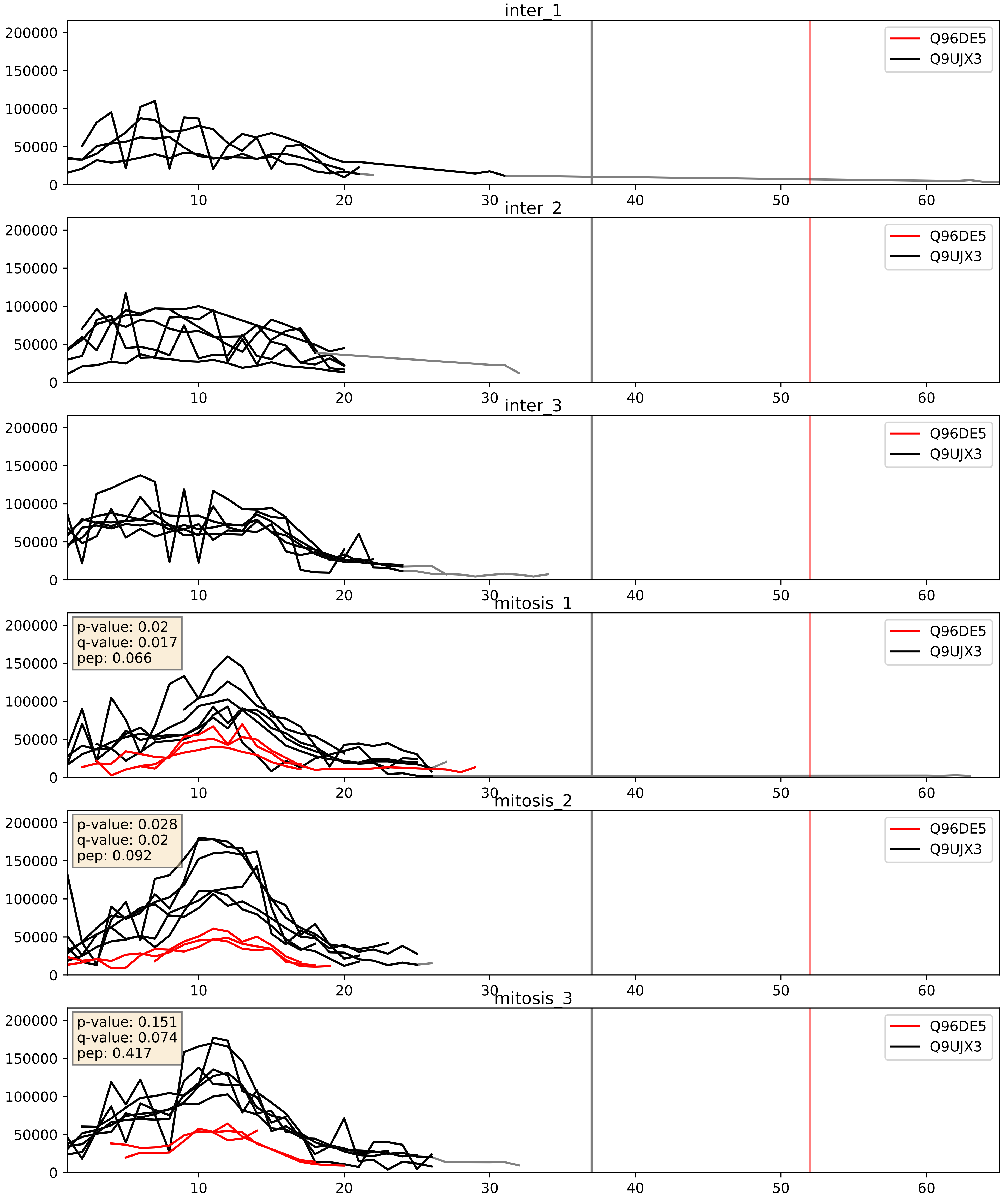


| level | condition_1 | condition_2 | pvalue | pvalue_adjusted |
|---|---|---|---|---|
| complex_abundance | mitosis | inter | 0.0003539069915483773 | 0.012039775462644836 |
| interactor_ratio | mitosis | inter | 0.0013063250170481464 | 0.0430082390228159 |
| interactor_abundance | mitosis | inter | 0.007099124223563132 | 0.0729076224999405 |

Q96DE5_Q9UJX4
APC16_HUMAN vs APC5_HUMAN
p-value: 0.06 q-value: 0.038

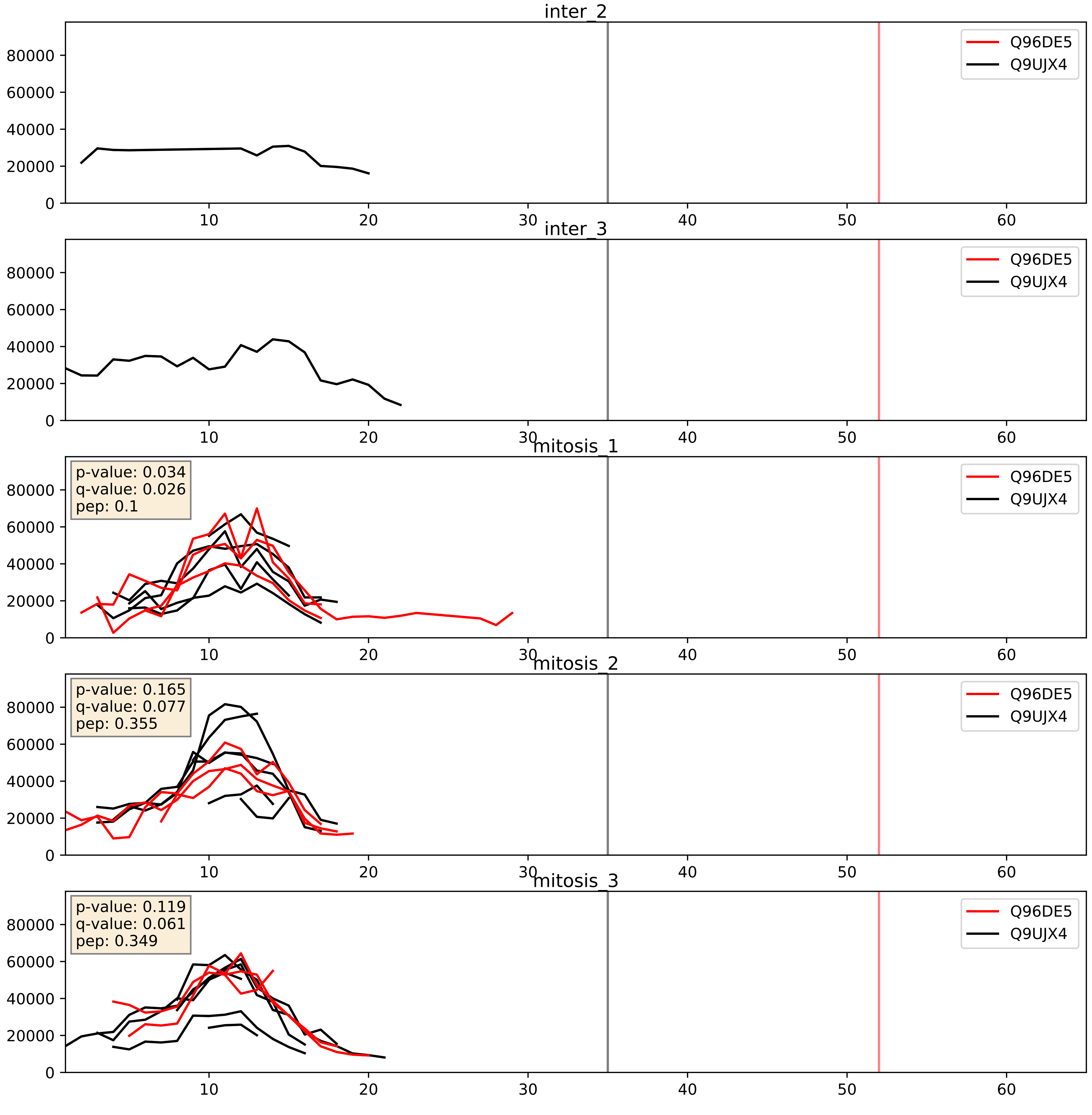


| level | condition_1 | condition_2 | pvalue | pvalue_adjusted |
|---|---|---|---|---|
| complex_abundance | mitosis | inter | 0.004058881031412856 | 0.04732787841836958 |
| interactor_abundance | mitosis | inter | 0.007466750033025538 | 0.07484236567060727 |
| interactor_ratio | mitosis | inter | 0.3148006520149434 | 0.7783421708721073 |

Q96DE5_Q9UJX5
APC16_HUMAN vs APC4_HUMAN
p-value: 0.024 q-value: 0.018

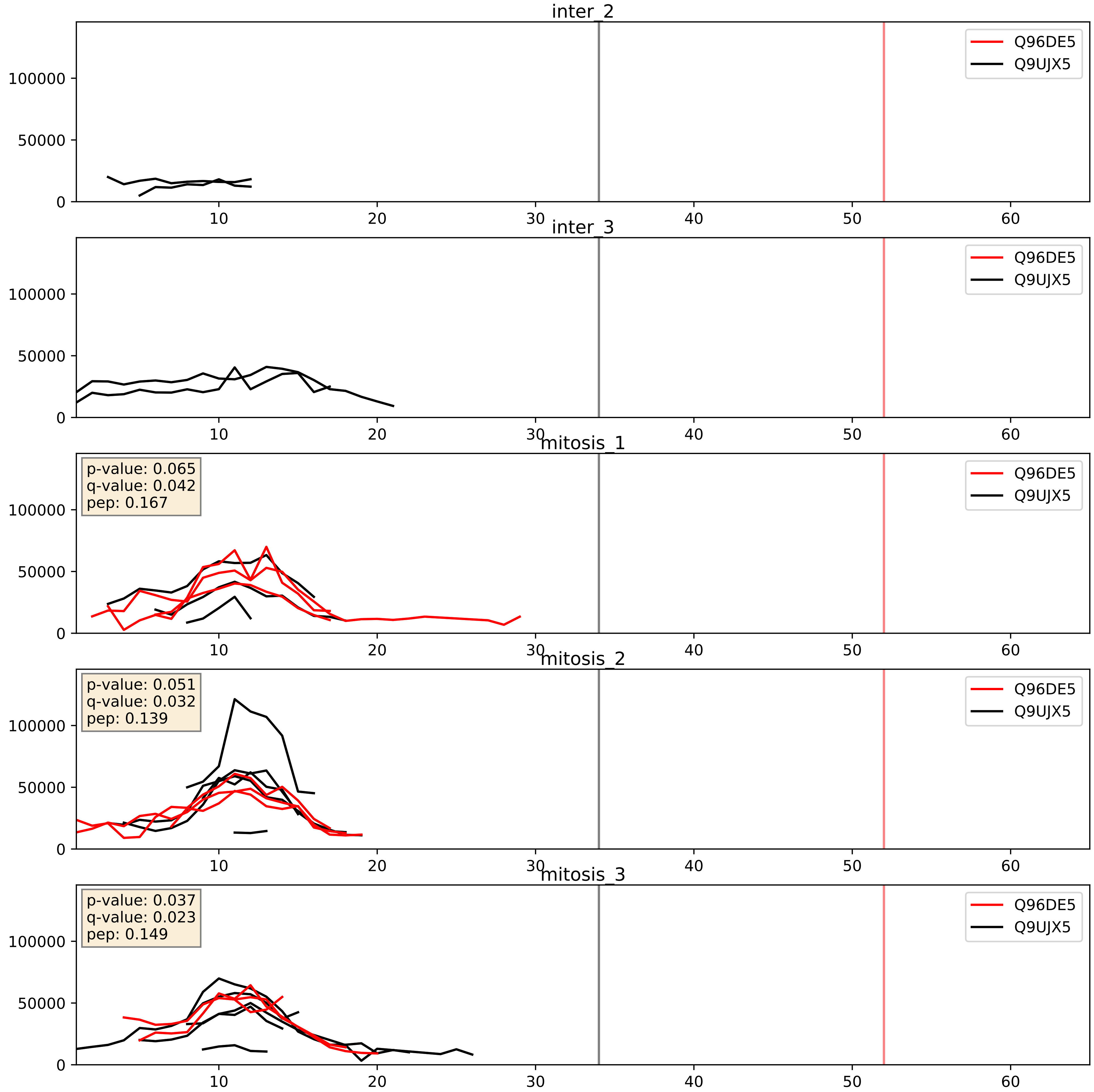


| level | condition_1 | condition_2 | pvalue | pvalue_adjusted |
|---|---|---|---|---|
| complex_abundance | mitosis | inter | 0.004254208343929297 | 0.04855469789871305 |
| interactor_abundance | mitosis | inter | 0.007377307691146444 | 0.07429382804260419 |
| interactor_ratio | mitosis | inter | 0.9008182788935496 | 0.9837485002527325 |

Q96DE5_Q9UJX6
APC16_HUMAN vs ANC2_HUMAN
p-value: 0.032 q-value: 0.023

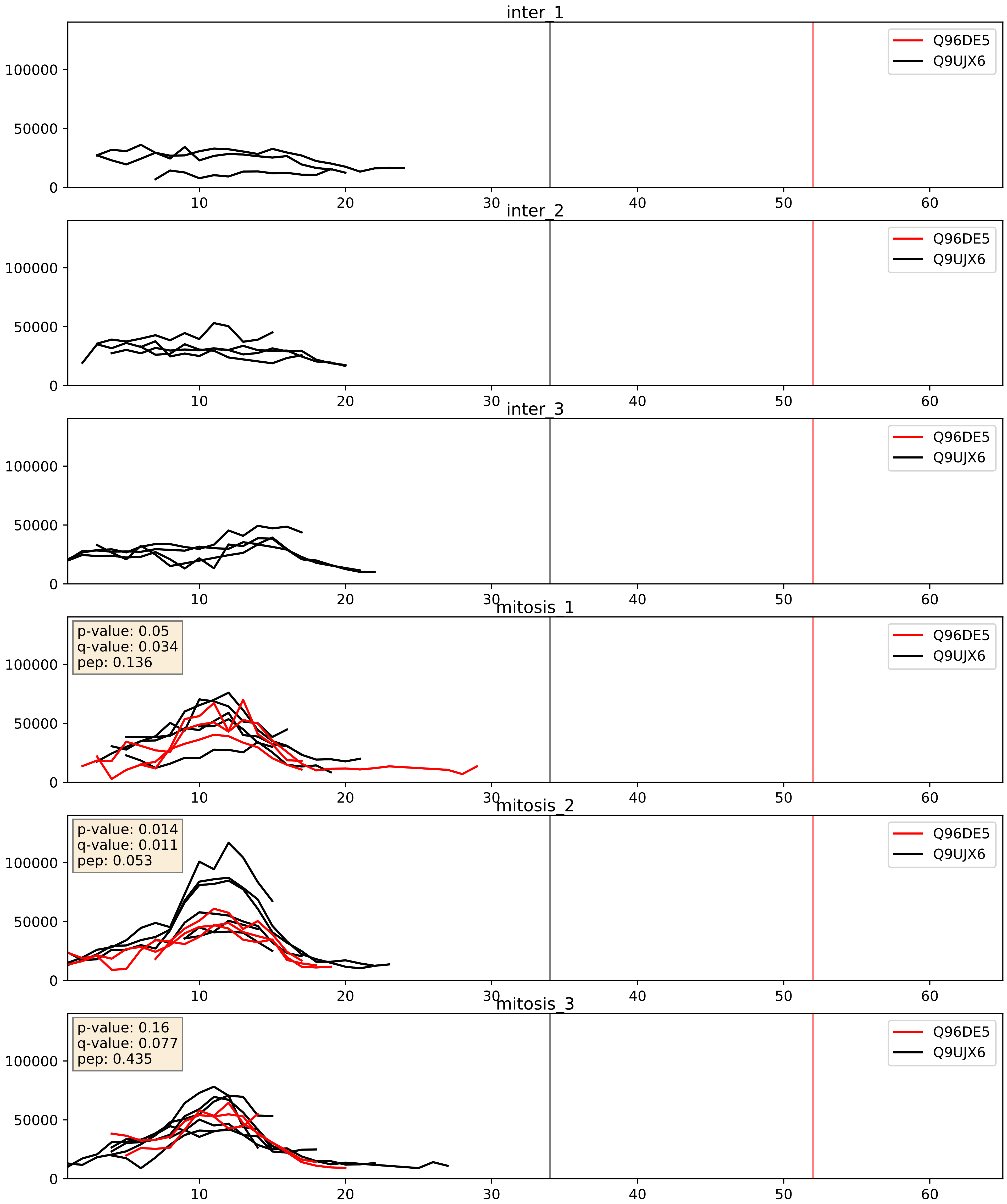


| level | condition_1 | condition_2 | pvalue | pvalue_adjusted |
|---|---|---|---|---|
| complex_abundance | mitosis | inter | 0.0015347885258439784 | 0.027256825272249905 |
| interactor_abundance | mitosis | inter | 0.006556558742981825 | 0.06907586811067622 |
| interactor_ratio | mitosis | inter | 0.03794917131731285 | 0.3197292386576752 |

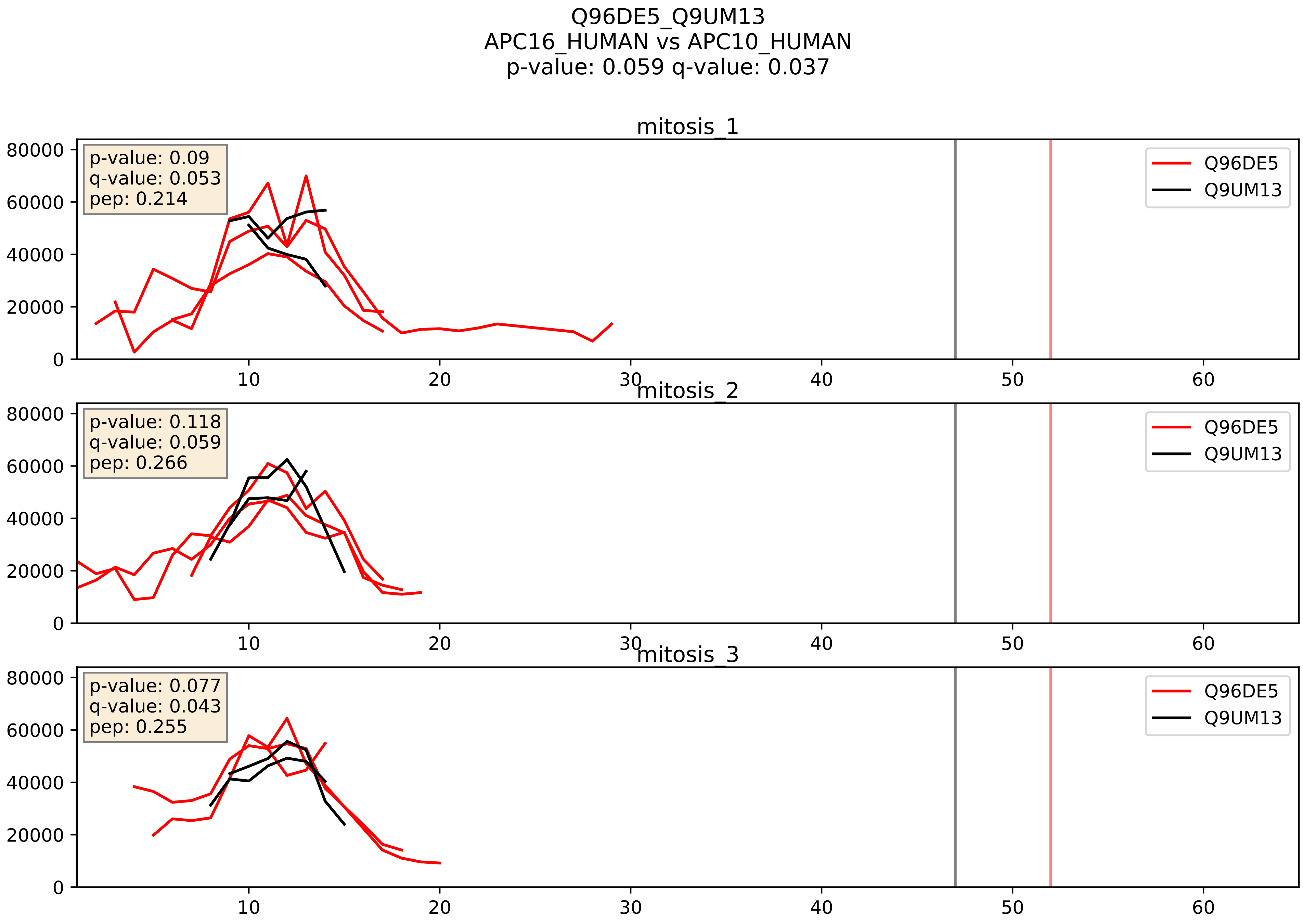


| level | condition_1 | condition_2 | pvalue | pvalue_adjusted |
|---|---|---|---|---|
| interactor_abundance | mitosis | inter | 0.0208935484573403 | 0.12074597692053658 |
| interactor_ratio | mitosis | inter | 0.02177167608911305 | 0.22562414930122 |
| complex_abundance | mitosis | inter | 0.022584098003647766 | 0.10996580142845556 |
