## Supplementary Data 2 for "SECAT: Quantifying differential protein-protein interaction states by network-centric analysis": hela_string_000161_Q99460.pdf

Q99460_Q99460

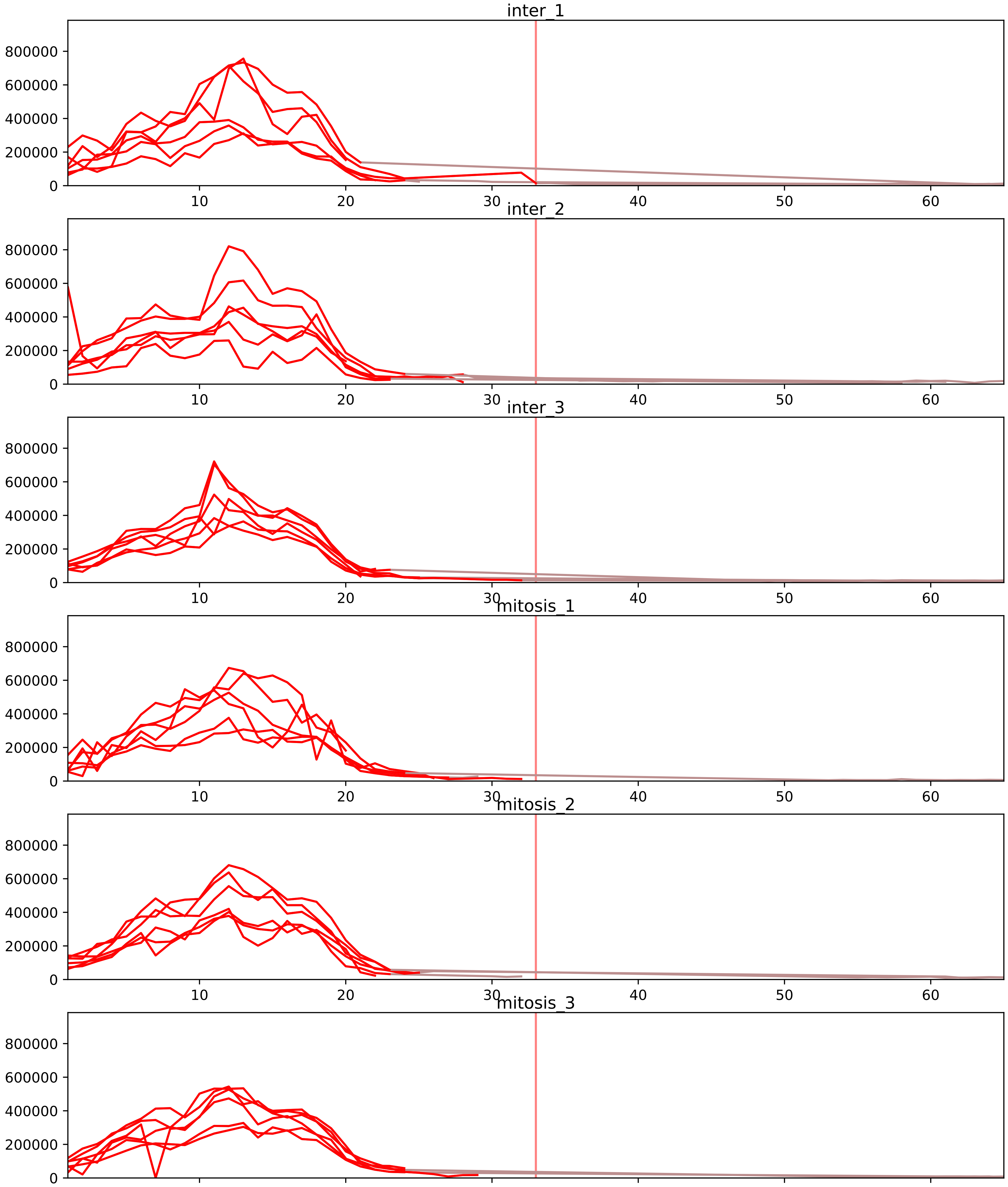


| level | condition_1 | condition_2 | pvalue | pvalue_adjusted |
|---|---|---|---|---|
| assembled_abundance | mitosis | inter | 7.631215210284456e-05 | 0.008577738896244046 |
| total_abundance | mitosis | inter | 9.50780375667225e-05 | 0.01273261737241483 |
| complex_abundance | mitosis | inter | 0.048578228555214126 | 0.12078910883999187 |
| interactor_abundance | mitosis | inter | 0.05823312694810028 | 0.15992381132015598 |
| monomer_abundance | mitosis | inter | 0.08057793451390323 | 0.3776302652748292 |
| interactor_ratio | mitosis | inter | 0.1171524487852447 | 0.3622865643106727 |

O00231_Q99460
PSD11_HUMAN vs PSMD1_HUMAN
p-value: 0.0 q-value: 0.0

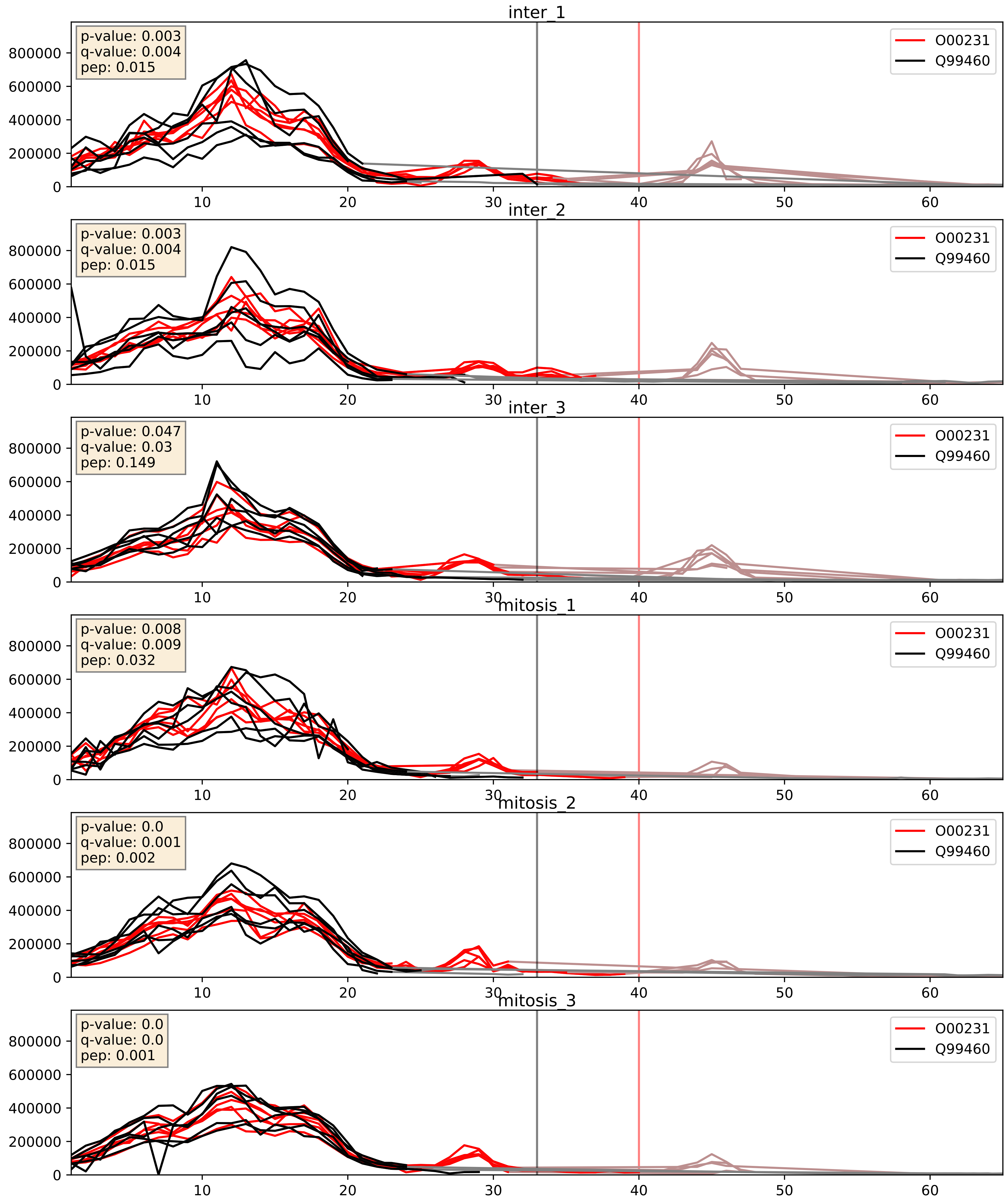


| level | condition_1 | condition_2 | pvalue | pvalue_adjusted |
|---|---|---|---|---|
| complex_abundance | mitosis | inter | 0.17711497210979785 | 0.32757488878789803 |
| interactor_abundance | mitosis | inter | 0.236588161925004 | 0.4277450936487804 |
| interactor_ratio | mitosis | inter | 0.5376608733934739 | 0.8851272942373031 |

O00232_Q99460
PSD12_HUMAN vs PSMD1_HUMAN
p-value: 0.005 q-value: 0.005

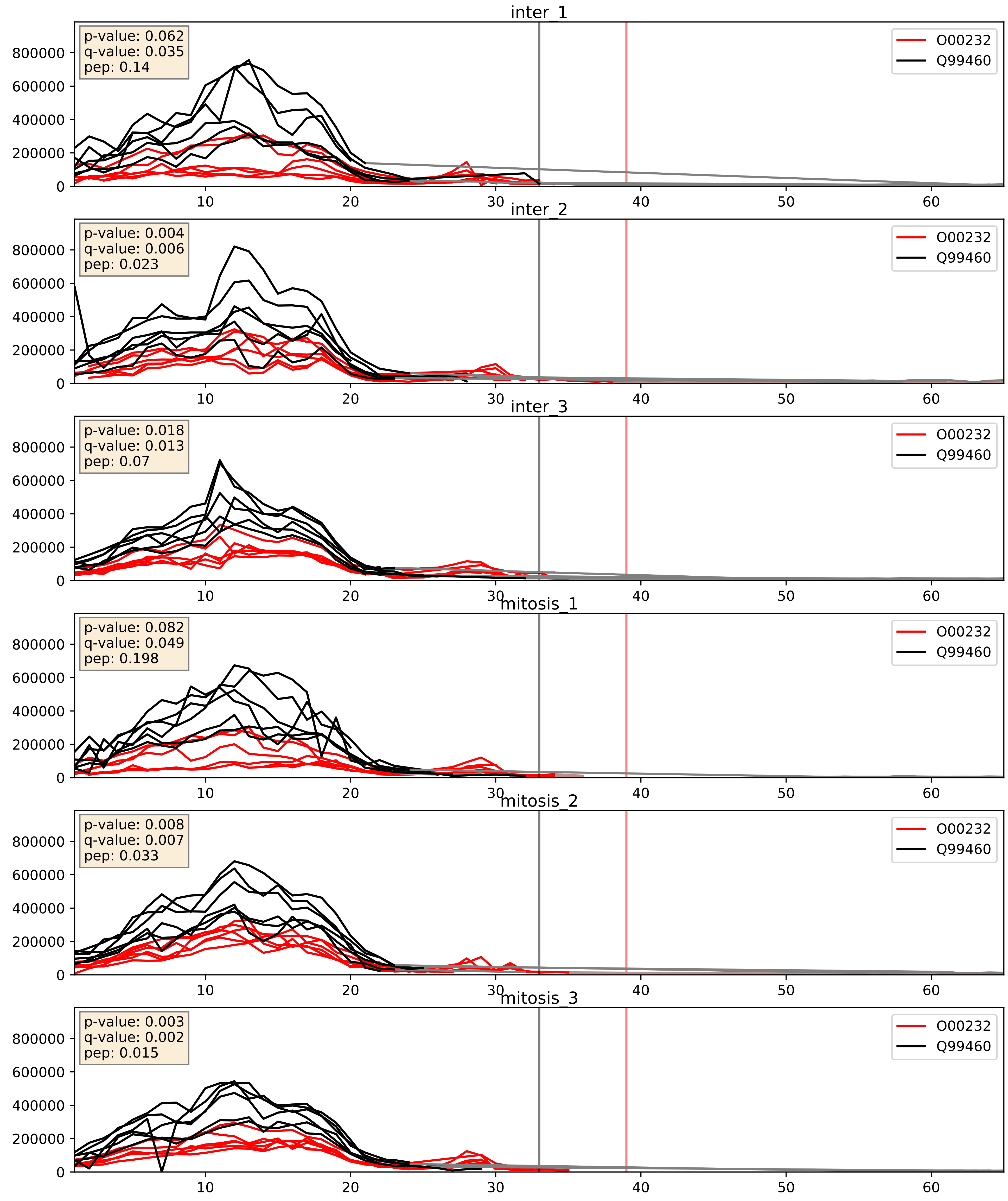


| level | condition_1 | condition_2 | pvalue | pvalue_adjusted |
|---|---|---|---|---|
| complex_abundance | mitosis | inter | 0.0351412030867749 | 0.13586662078716946 |
| interactor_abundance | mitosis | inter | 0.0737069172692737 | 0.22673988198390962 |
| interactor_ratio | mitosis | inter | 0.5412146966764987 | 0.8859730852936224 |

O00487_Q99460
PSDE_HUMAN vs PSMD1_HUMAN
p-value: 0.008 q-value: 0.007

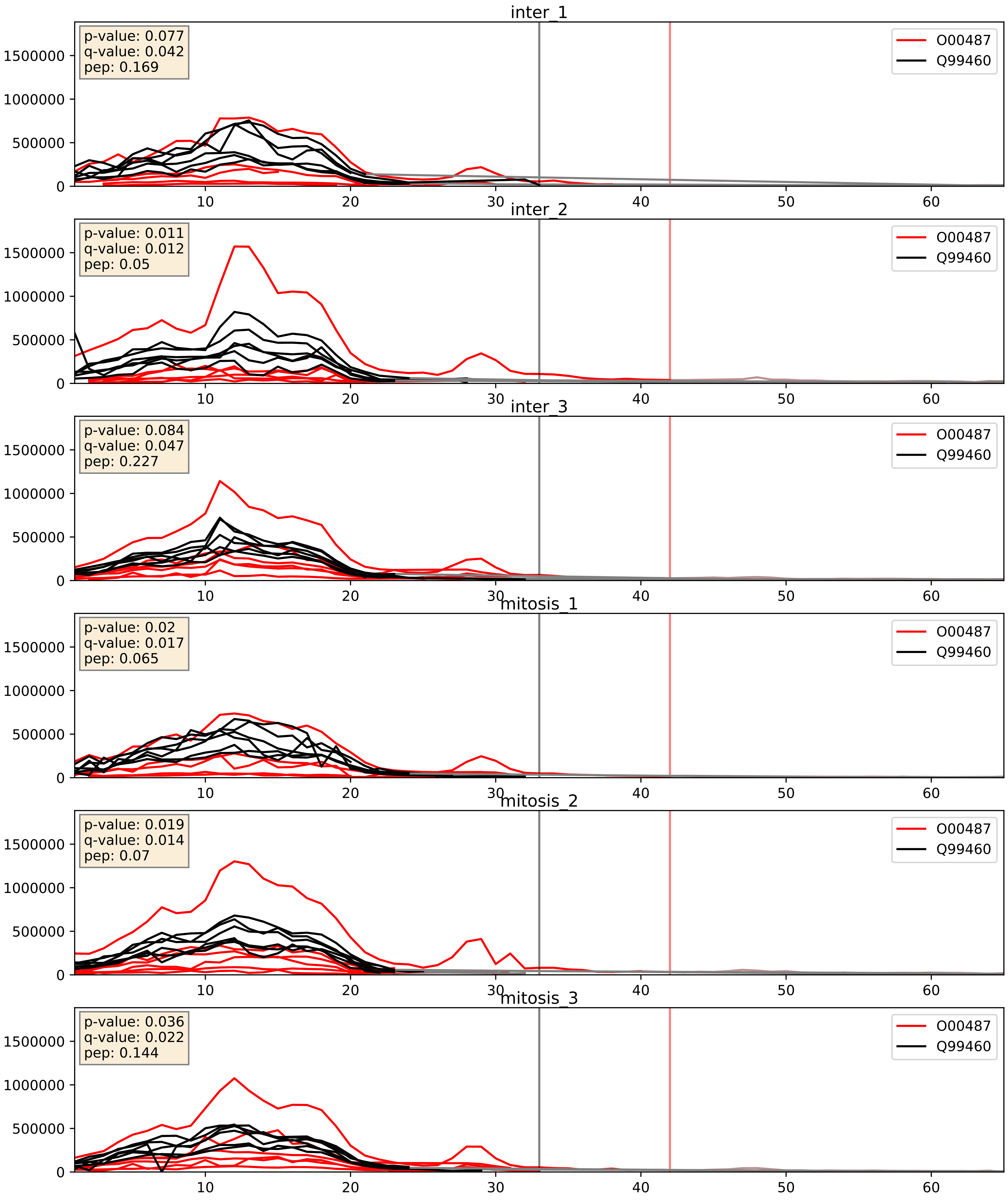


| level | condition_1 | condition_2 | pvalue | pvalue_adjusted |
|---|---|---|---|---|
| interactor_ratio | mitosis | inter | 0.5865777126785633 | 0.8994900937741748 |
| complex_abundance | mitosis | inter | 0.6205599806927463 | 0.7403475170355274 |
| interactor_abundance | mitosis | inter | 0.9781164380428207 | 0.9870601720480916 |

O43242_Q99460
PSMD3_HUMAN vs PSMD1_HUMAN
p-value: 0.0 q-value: 0.001

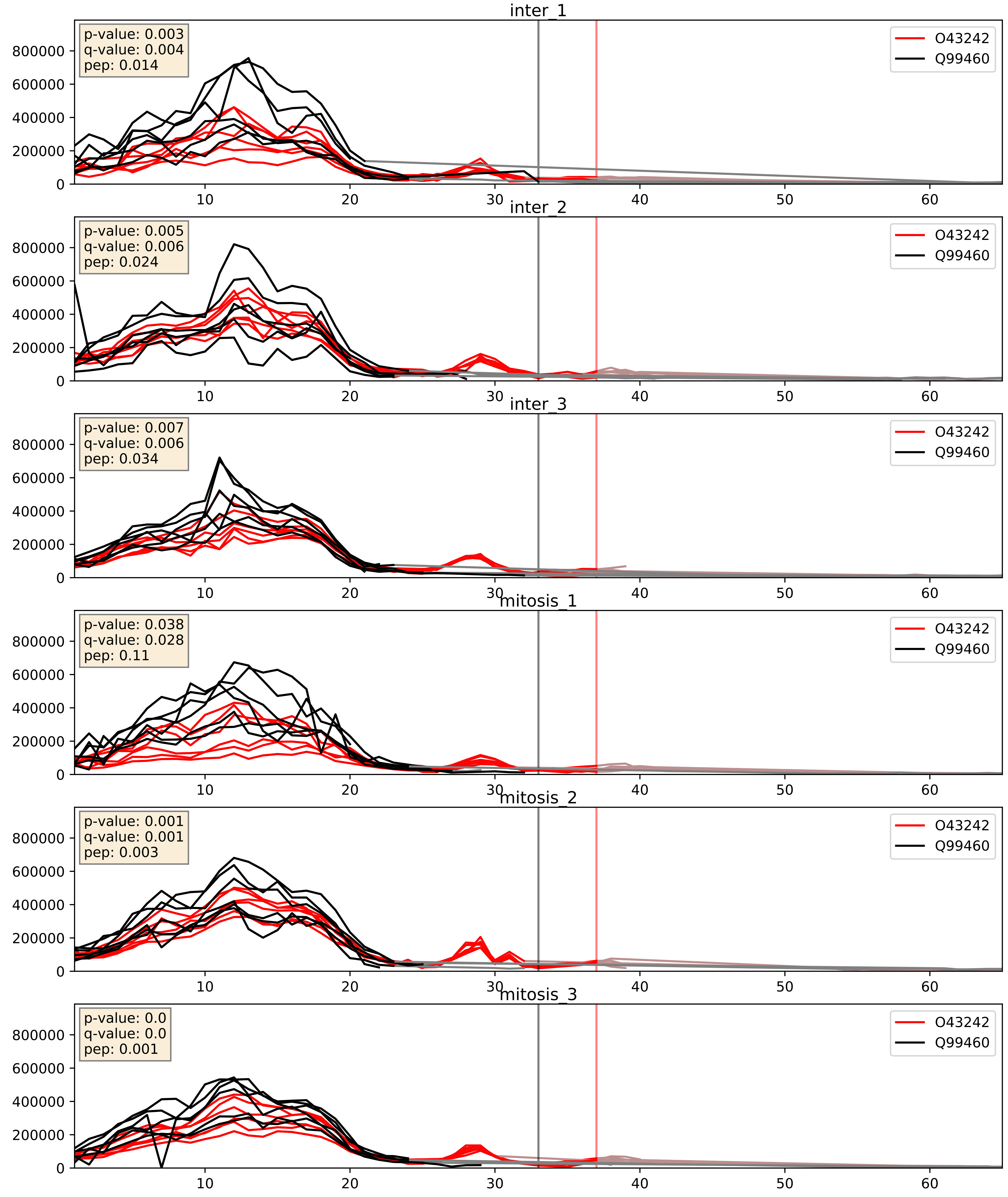


| level | condition_1 | condition_2 | pvalue | pvalue_adjusted |
|---|---|---|---|---|
| complex_abundance | mitosis | inter | 0.05842393052094995 | 0.1727491693469194 |
| interactor_abundance | mitosis | inter | 0.07120012317659774 | 0.22221305735359212 |
| interactor_ratio | mitosis | inter | 0.44608214983954037 | 0.850695399873193 |

O75832_Q99460
PSD10_HUMAN vs PSMD1_HUMAN
p-value: 0.027 q-value: 0.02

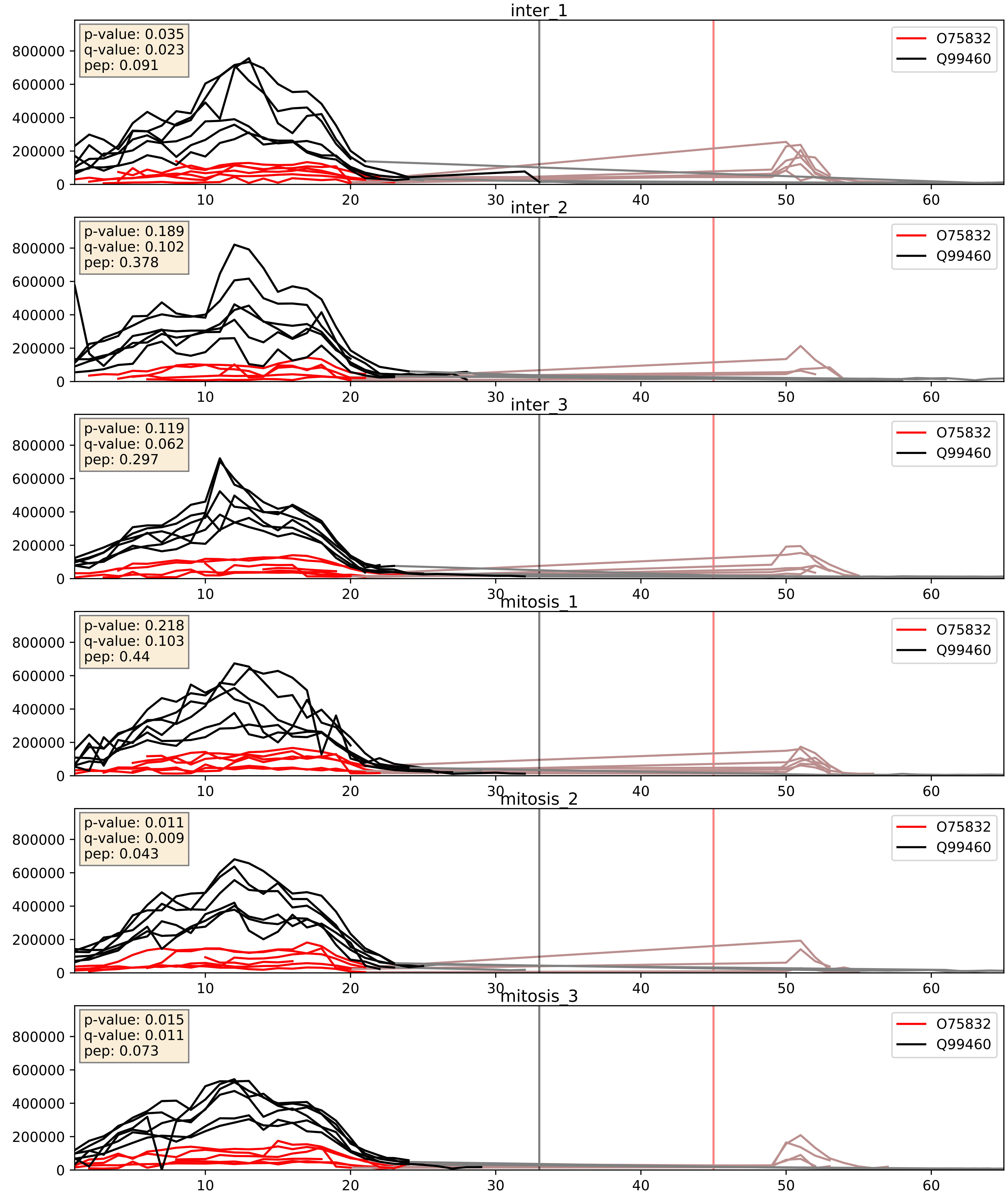


| level | condition_1 | condition_2 | pvalue | pvalue_adjusted |
|---|---|---|---|---|
| interactor_ratio | mitosis | inter | 0.005093770058038552 | 0.0962531384035541 |
| interactor_abundance | mitosis | inter | 0.14228828561323173 | 0.32880142136715784 |
| complex_abundance | mitosis | inter | 0.9632782096674886 | 0.9783278484761004 |

P17980_Q99460
PRS6A_HUMAN vs PSMD1_HUMAN
p-value: 0.0 q-value: 0.0

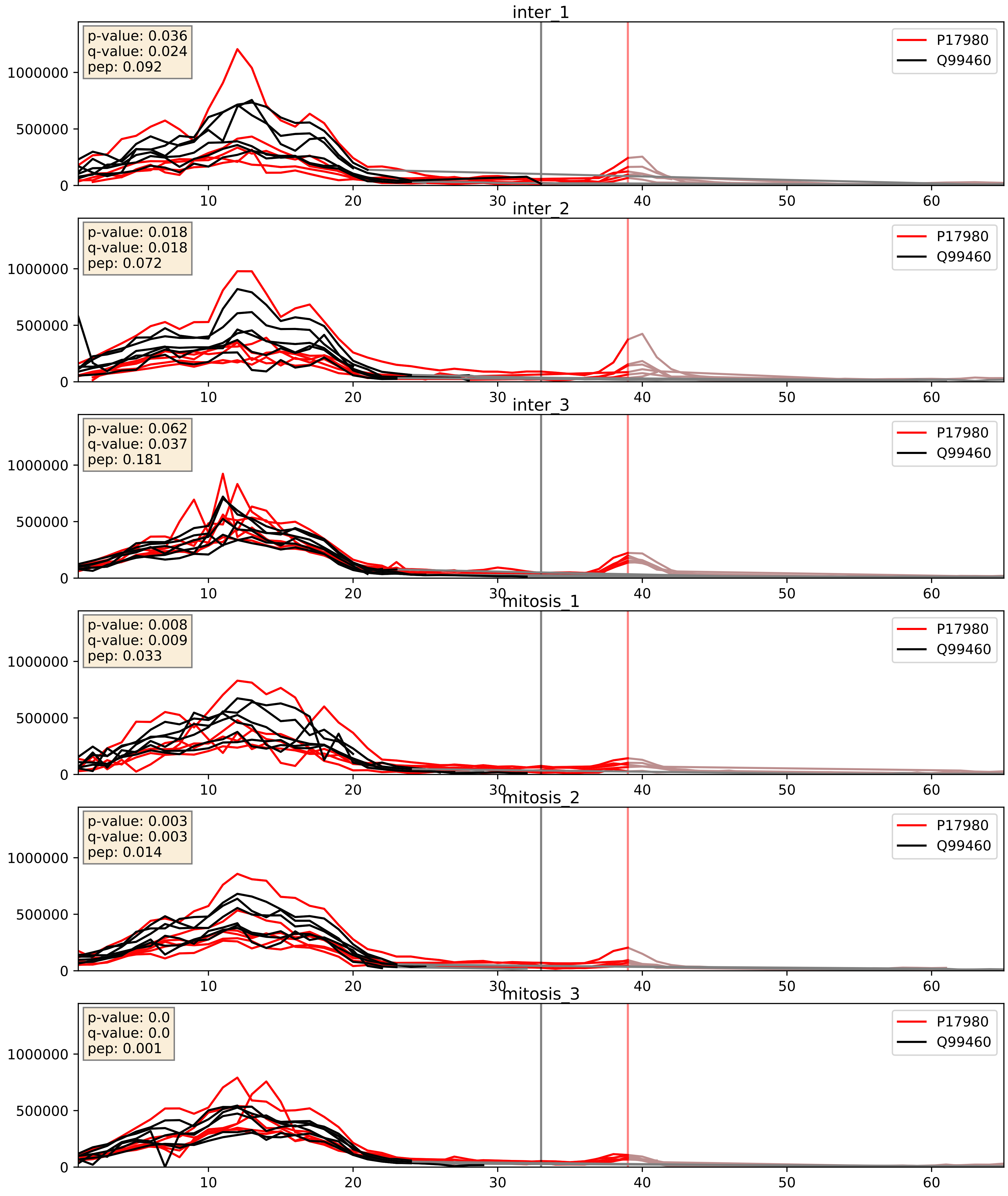


| level | condition_1 | condition_2 | pvalue | pvalue_adjusted |
|---|---|---|---|---|
| complex_abundance | mitosis | inter | 0.15953125042280933 | 0.30721878596608504 |
| interactor_abundance | mitosis | inter | 0.30490085909502707 | 0.5005660440838956 |
| interactor_ratio | mitosis | inter | 0.6228250544333103 | 0.9137335840337968 |

P35998_Q99460
PRS7_HUMAN vs PSMD1_HUMAN
p-value: 0.002 q-value: 0.002

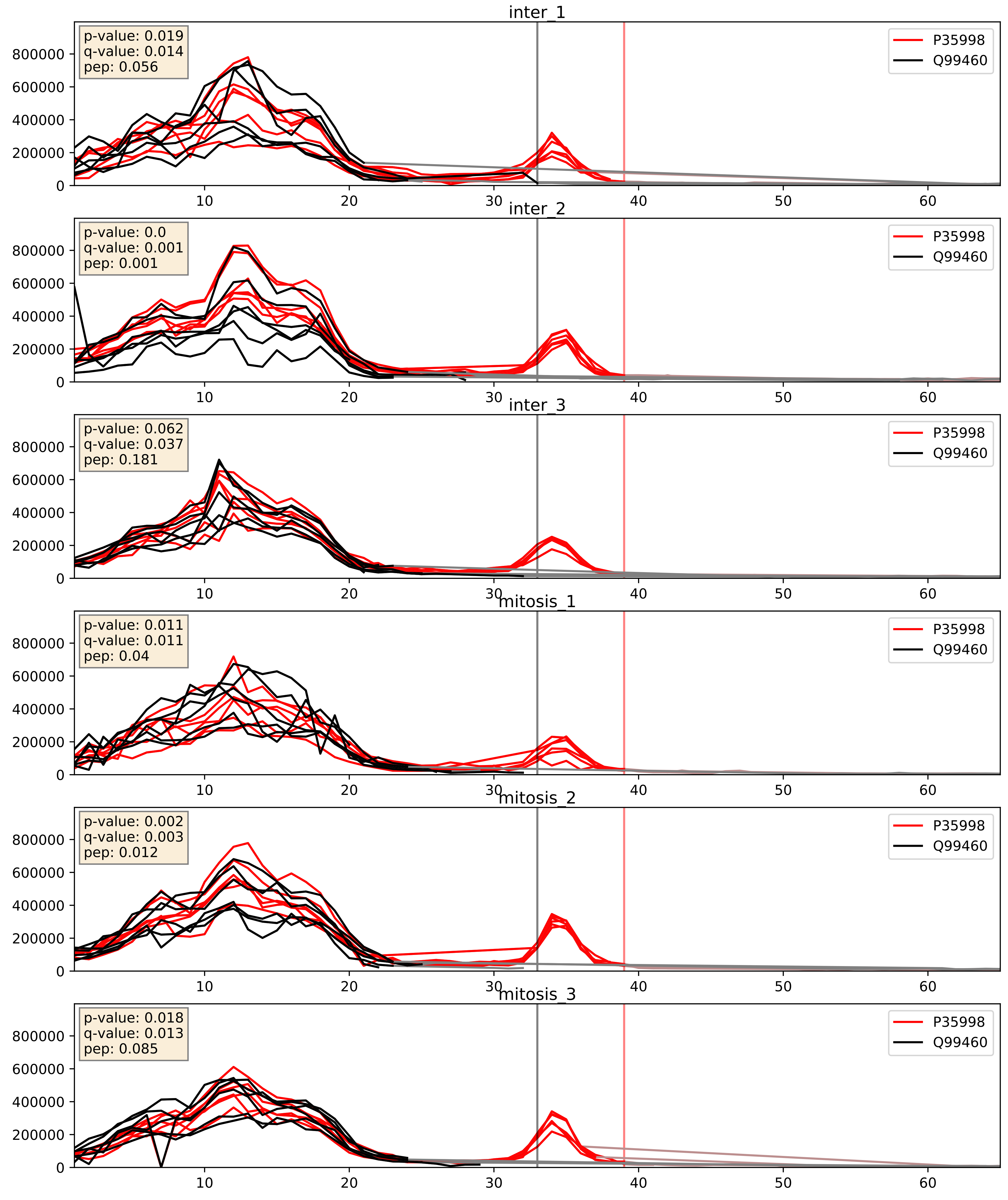


| level | condition_1 | condition_2 | pvalue | pvalue_adjusted |
|---|---|---|---|---|
| interactor_abundance | mitosis | inter | 0.01304944055912961 | 0.10257411495514183 |
| complex_abundance | mitosis | inter | 0.014514587463278805 | 0.08966829497179525 |
| interactor_ratio | mitosis | inter | 0.08722662806863365 | 0.4946978435260244 |

P43686_Q99460
PRS6B_HUMAN vs PSMD1_HUMAN
p-value: 0.0 q-value: 0.0

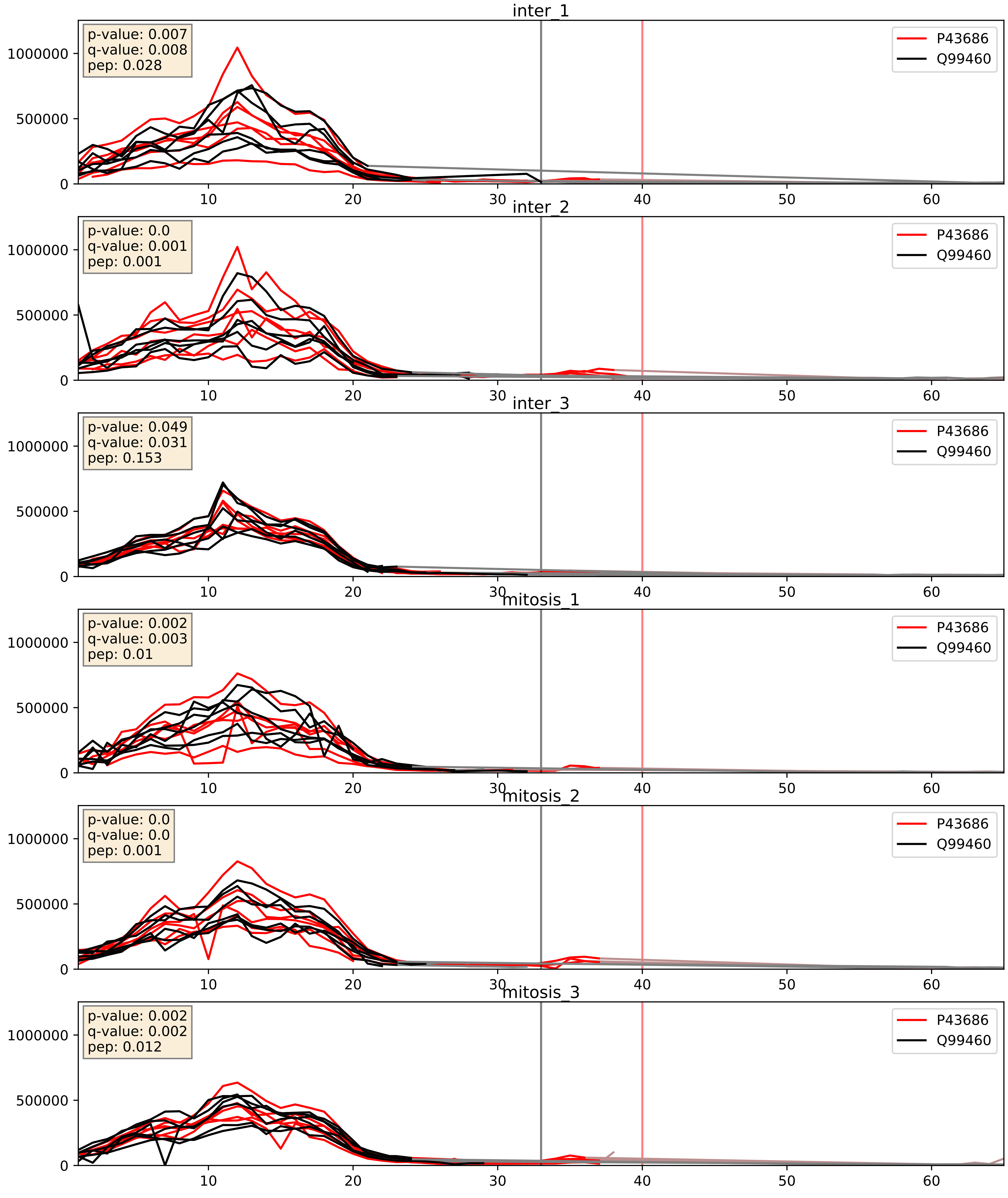


| level | condition_1 | condition_2 | pvalue | pvalue_adjusted |
|---|---|---|---|---|
| complex_abundance | mitosis | inter | 0.07151673199204457 | 0.19422056657737993 |
| interactor_abundance | mitosis | inter | 0.09323066004468038 | 0.2607169062340621 |
| interactor_ratio | mitosis | inter | 0.36170794814998475 | 0.8057976545208556 |

P48556_Q99460
PSMD8_HUMAN vs PSMD1_HUMAN
p-value: 0.0 q-value: 0.0

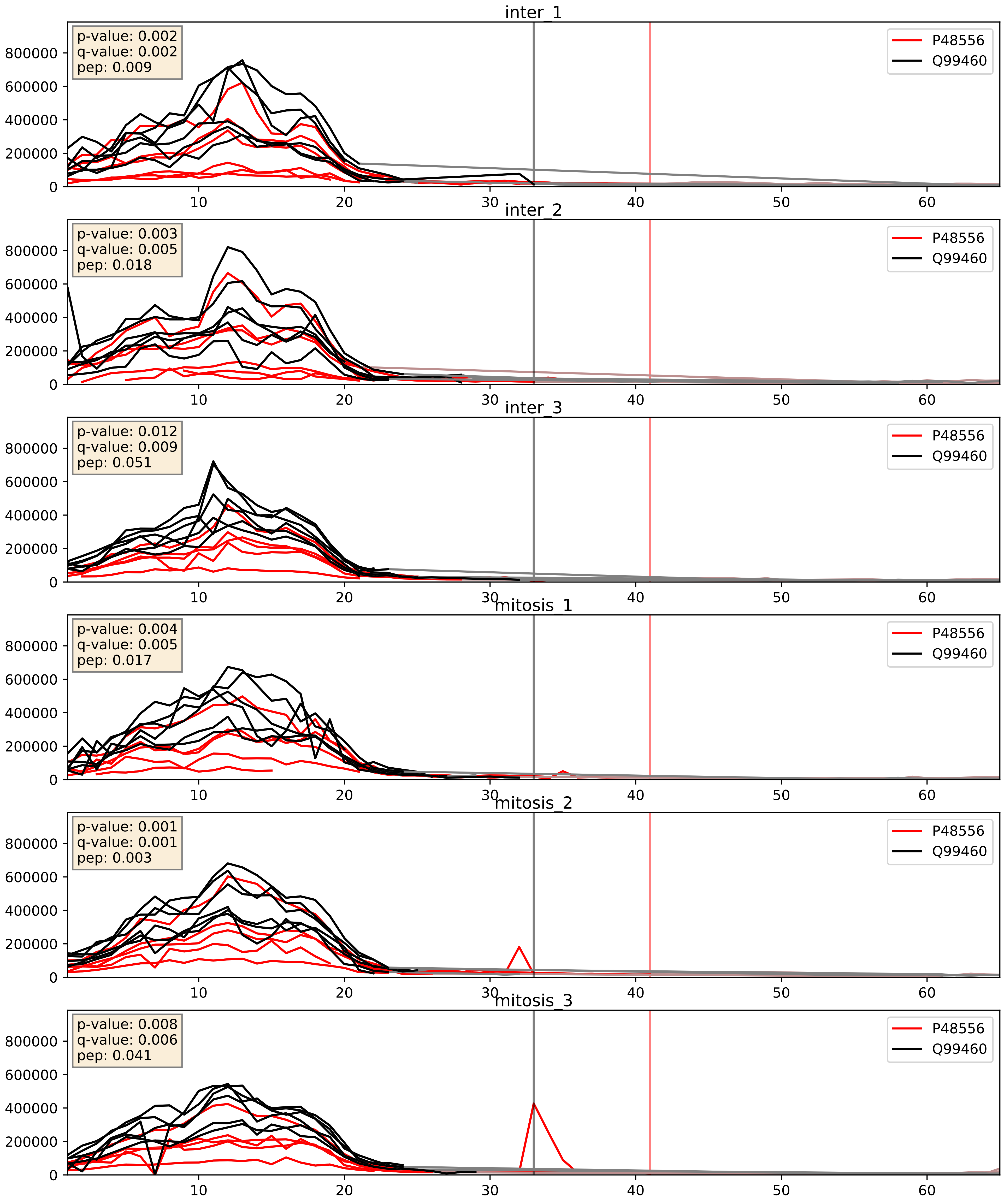


| level | condition_1 | condition_2 | pvalue | pvalue_adjusted |
|---|---|---|---|---|
| complex_abundance | mitosis | inter | 0.13001931130528305 | 0.2736575620293147 |
| interactor_abundance | mitosis | inter | 0.25191684013000043 | 0.4452169198952831 |
| interactor_ratio | mitosis | inter | 0.6387573375507691 | 0.9197898119332029 |

P51665_Q99460
PSMD7_HUMAN vs PSMD1_HUMAN
p-value: 0.001 q-value: 0.001

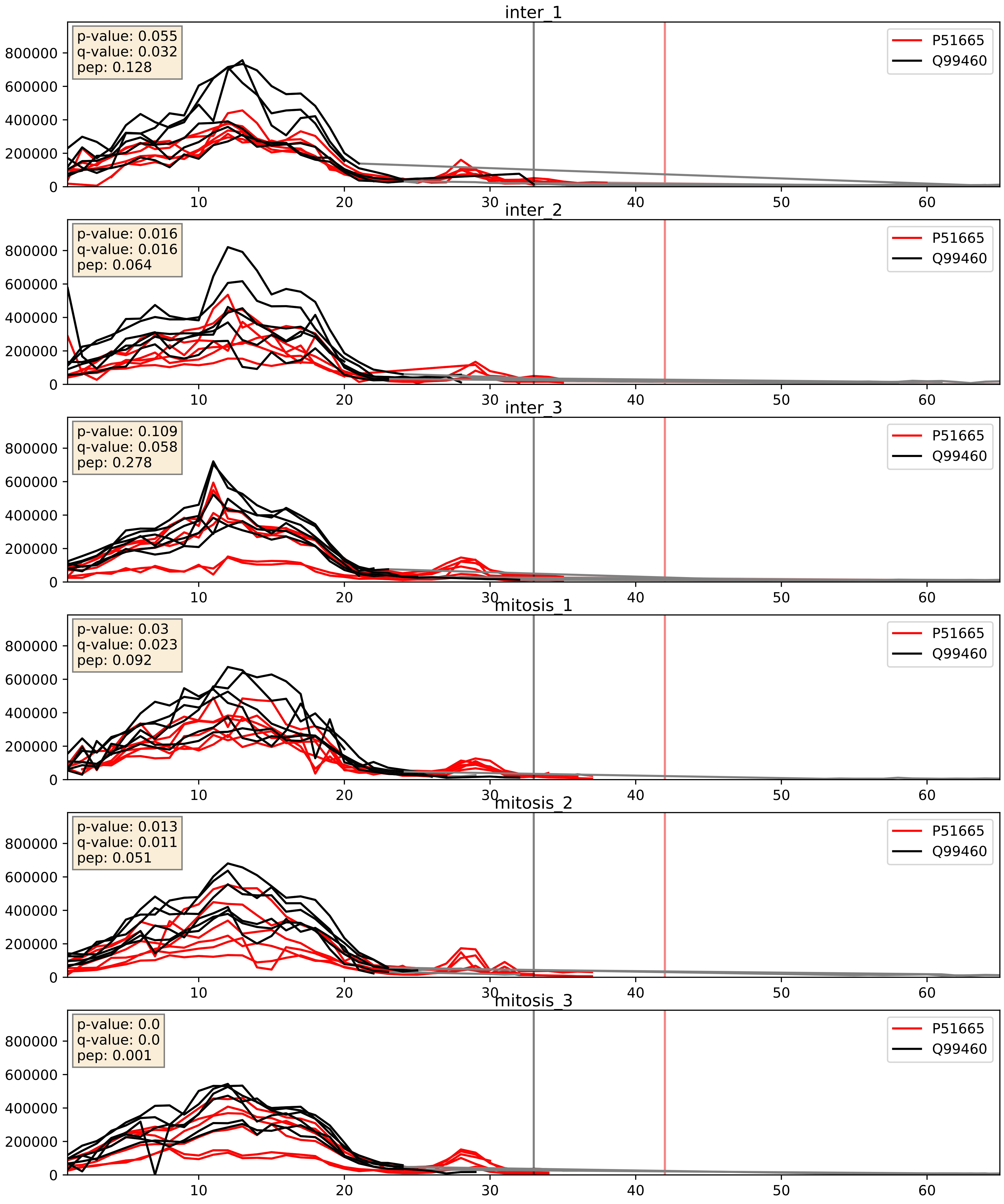


| level | condition_1 | condition_2 | pvalue | pvalue_adjusted |
|---|---|---|---|---|
| complex_abundance | mitosis | inter | 0.2851060339559172 | 0.4339451725929323 |
| interactor_ratio | mitosis | inter | 0.43915369311084773 | 0.8458946023917319 |
| interactor_abundance | mitosis | inter | 0.5821632514487826 | 0.7256432383893312 |

P55036_Q99460
PSMD4_HUMAN vs PSMD1_HUMAN
p-value: 0.0 q-value: 0.0

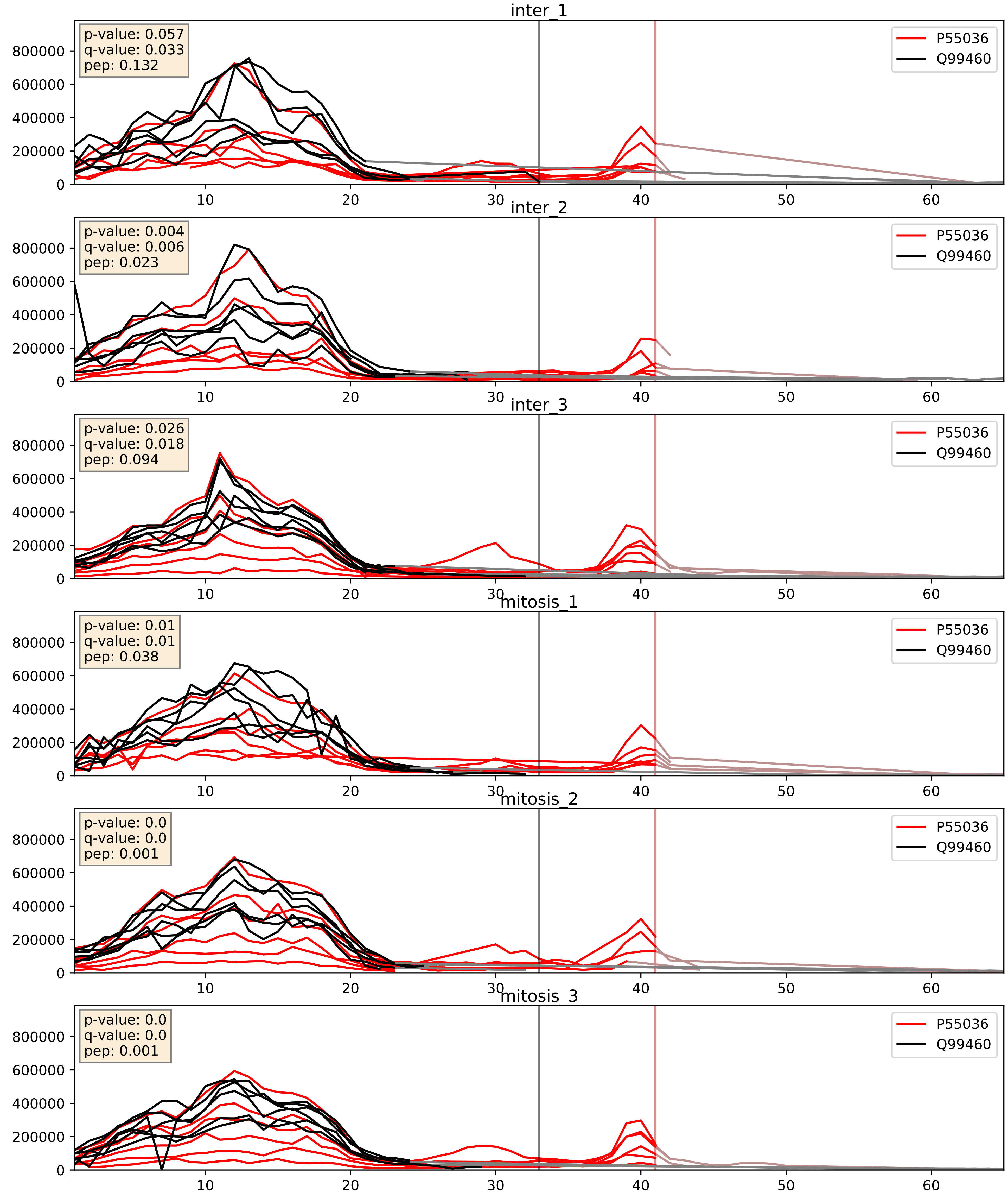


| level | condition_1 | condition_2 | pvalue | pvalue_adjusted |
|---|---|---|---|---|
| complex_abundance | mitosis | inter | 0.06122960017095163 | 0.1778907646484952 |
| interactor_abundance | mitosis | inter | 0.09075757280178176 | 0.2562285036884076 |
| interactor_ratio | mitosis | inter | 0.9583610828580089 | 0.9929524032138645 |

P62191_Q99460
PRS4_HUMAN vs PSMD1_HUMAN
p-value: 0.0 q-value: 0.0

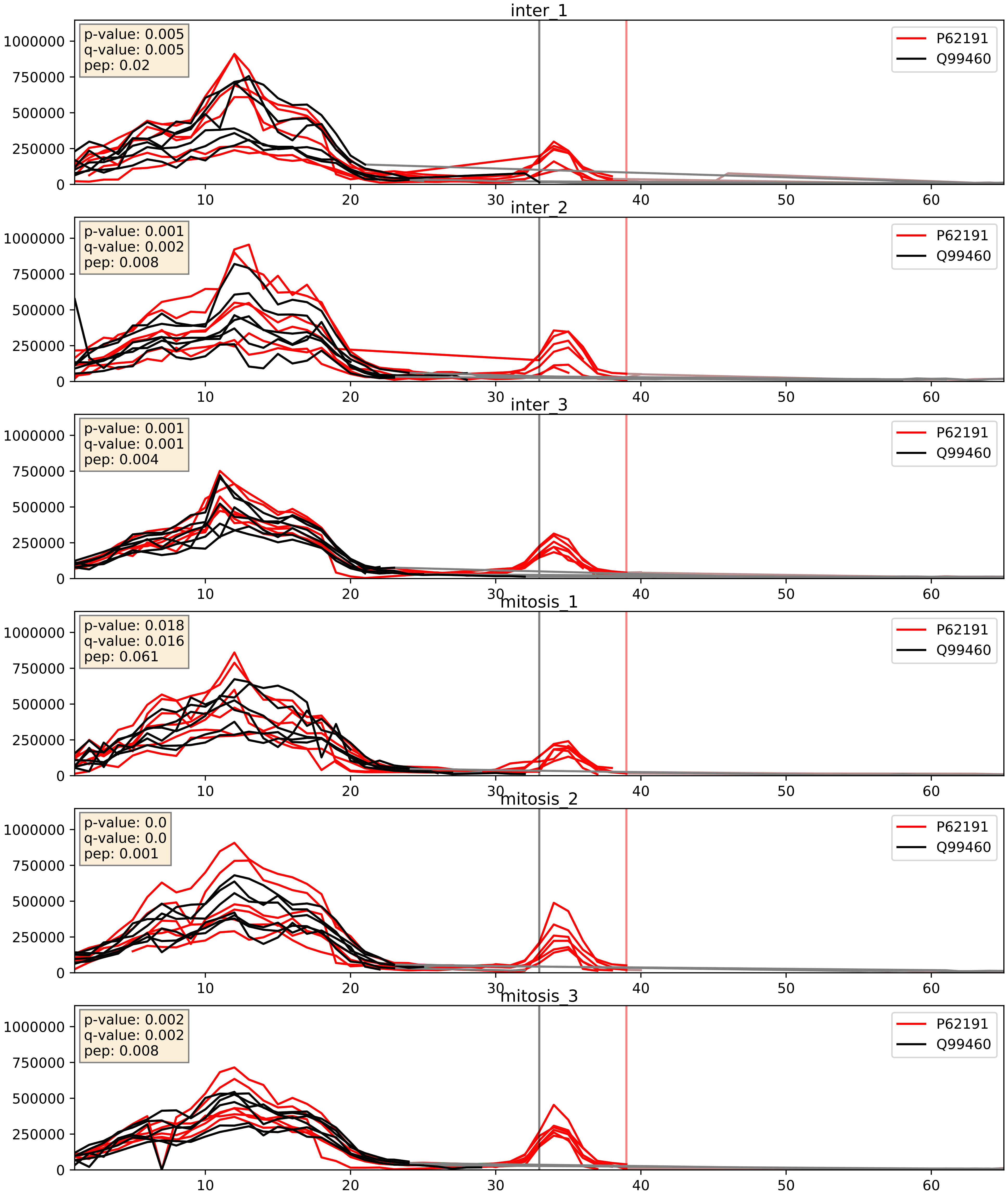


| level | condition_1 | condition_2 | pvalue | pvalue_adjusted |
|---|---|---|---|---|
| complex_abundance | mitosis | inter | 0.04969710710928881 | 0.15897087091075196 |
| interactor_abundance | mitosis | inter | 0.05109970805464057 | 0.18133868228631386 |
| interactor_ratio | mitosis | inter | 0.192536356203767 | 0.6660896702330326 |

P62195_Q99460
PRS8_HUMAN vs PSMD1_HUMAN
p-value: 0.0 q-value: 0.0

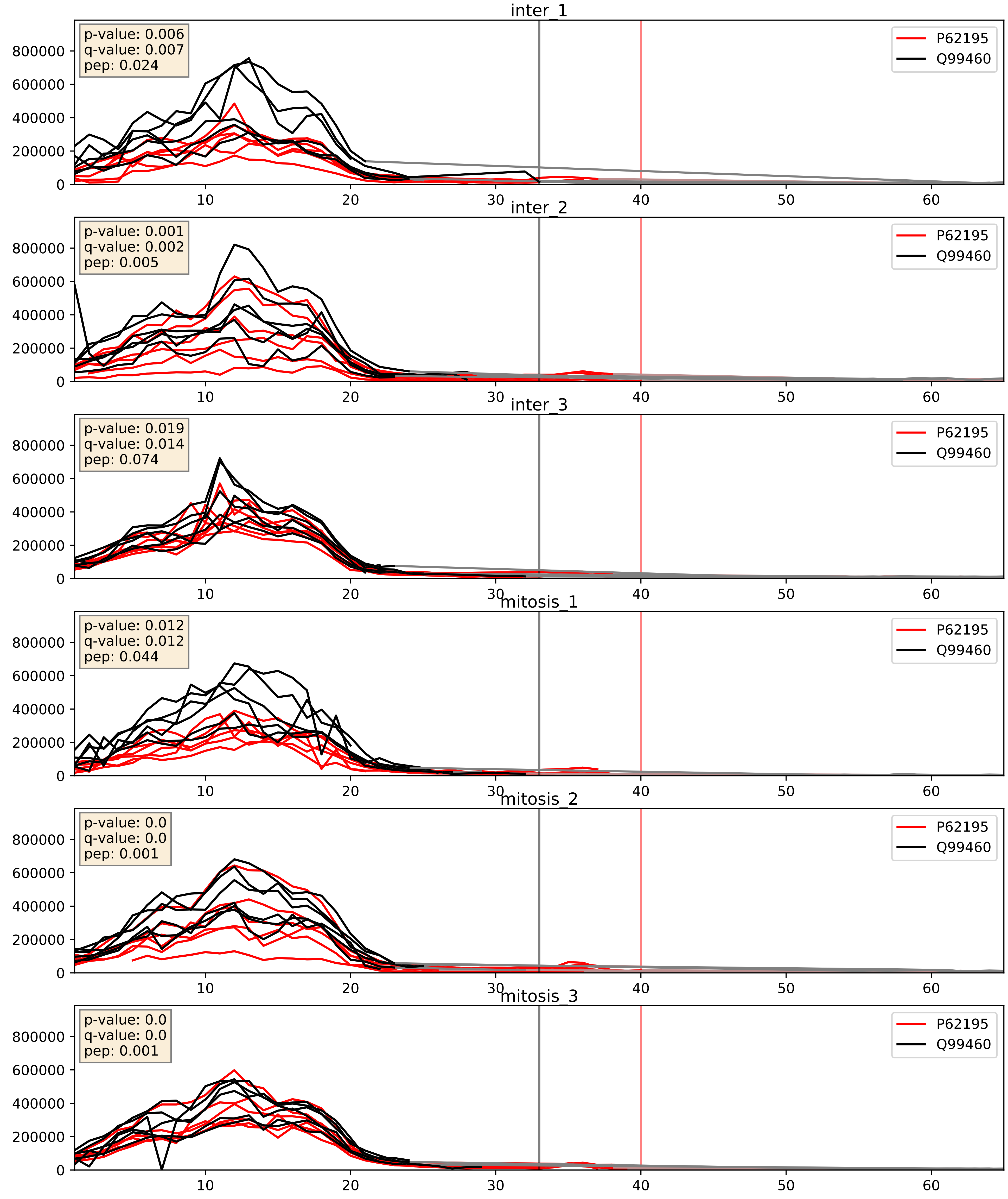


| level | condition_1 | condition_2 | pvalue | pvalue_adjusted |
|---|---|---|---|---|
| complex_abundance | mitosis | inter | 0.155618604964646 | 0.3025426132448931 |
| interactor_abundance | mitosis | inter | 0.35261932206785884 | 0.550455256113955 |
| interactor_ratio | mitosis | inter | 0.8470955036222981 | 0.9755634896635457 |

Q13200_Q99460
PSMD2_HUMAN vs PSMD1_HUMAN
p-value: 0.0 q-value: 0.0

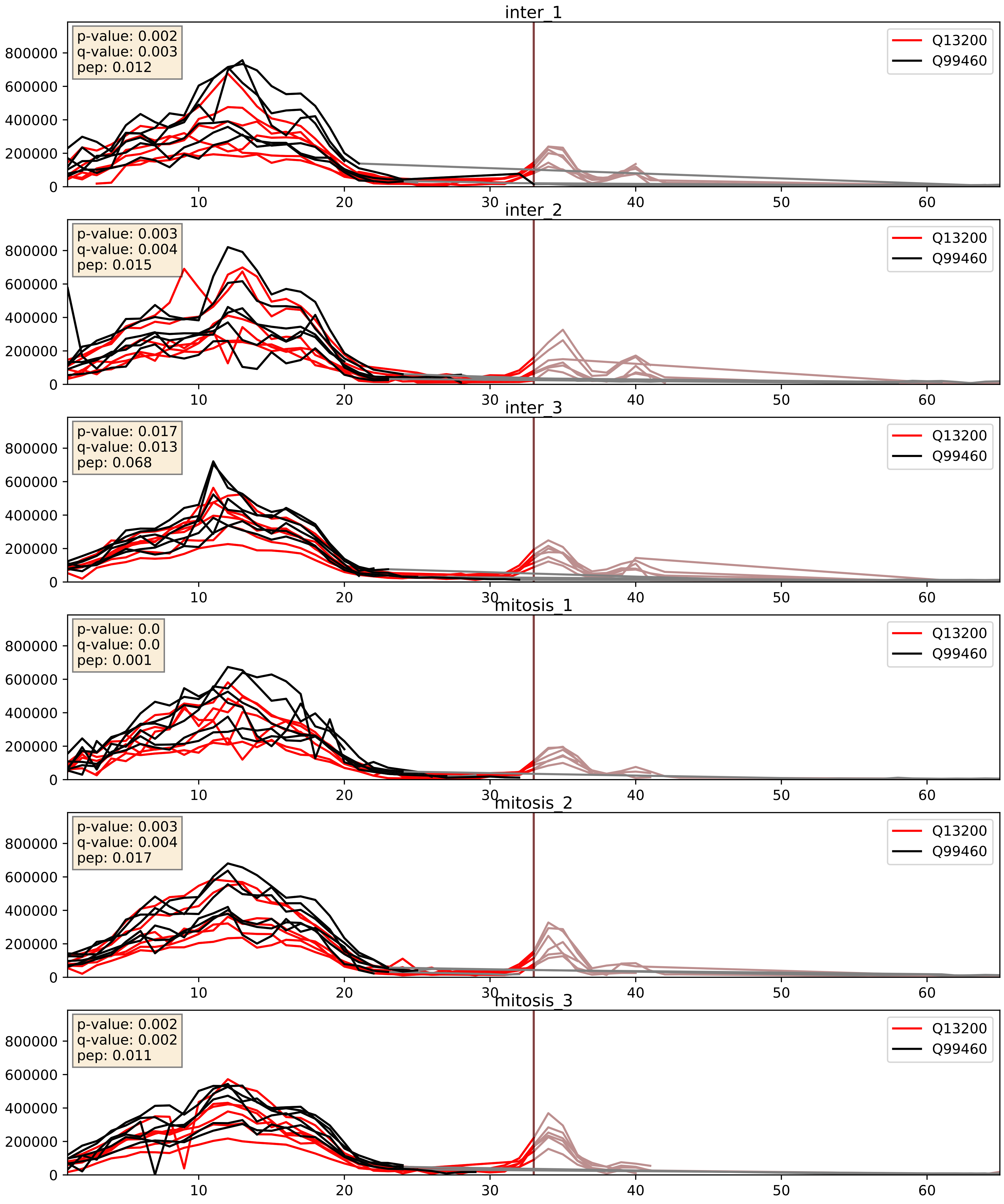


| level | condition_1 | condition_2 | pvalue | pvalue_adjusted |
|---|---|---|---|---|
| interactor_abundance | mitosis | inter | 0.014107588654821486 | 0.10527143000494996 |
| complex_abundance | mitosis | inter | 0.033440033765030824 | 0.13332402842508798 |
| interactor_ratio | mitosis | inter | 0.1808357904961016 | 0.6497321739897209 |

Q15008_Q99460
PSMD6_HUMAN vs PSMD1_HUMAN
p-value: 0.0 q-value: 0.0

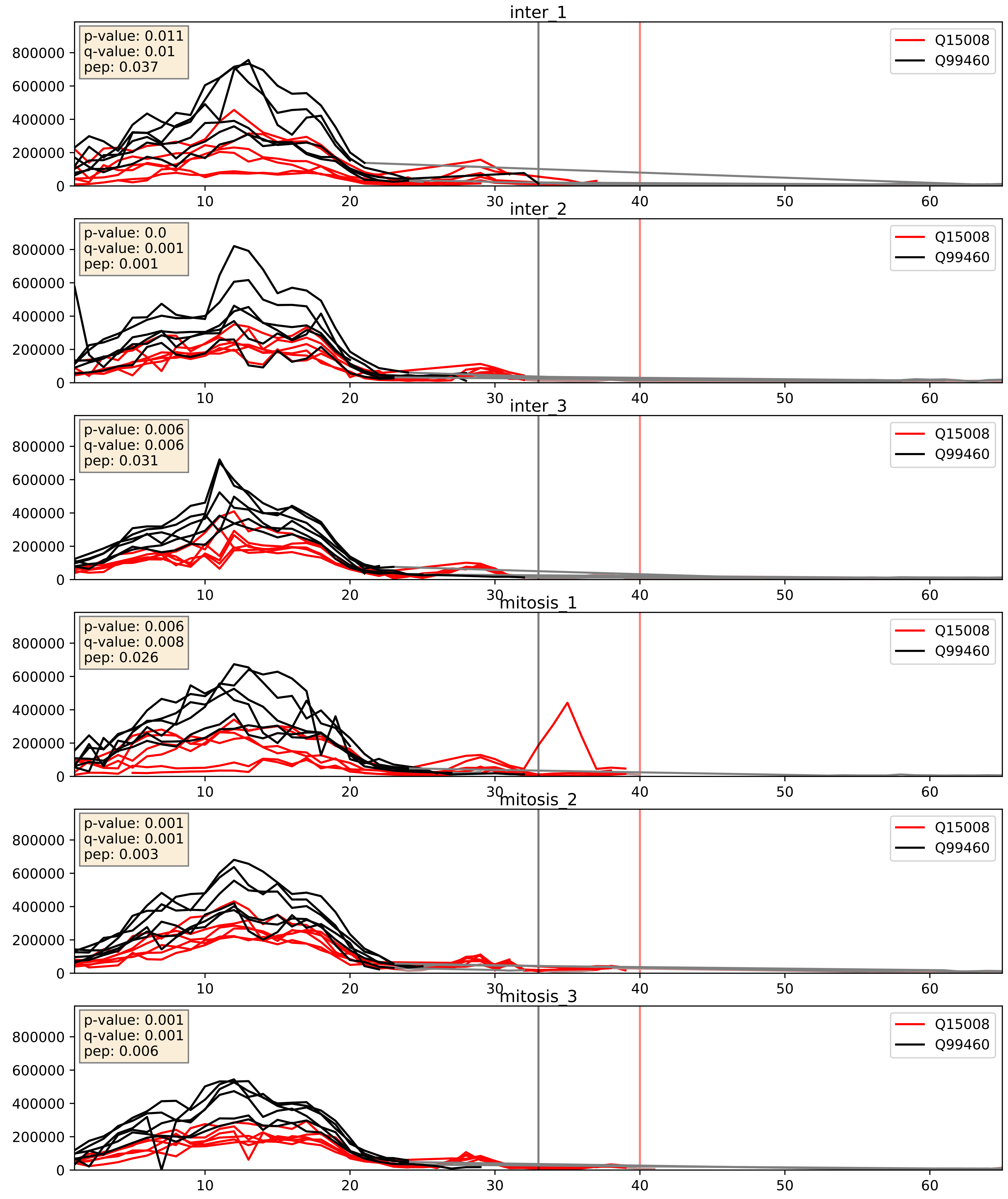


| level | condition_1 | condition_2 | pvalue | pvalue_adjusted |
|---|---|---|---|---|
| complex_abundance | mitosis | inter | 0.20859761961075882 | 0.3624838862907218 |
| interactor_ratio | mitosis | inter | 0.28686526563161785 | 0.7573864362030516 |
| interactor_abundance | mitosis | inter | 0.5470663407264278 | 0.7033573667862878 |

Q16186_Q99460
ADRM1_HUMAN vs PSMD1_HUMAN
p-value: 0.008 q-value: 0.008

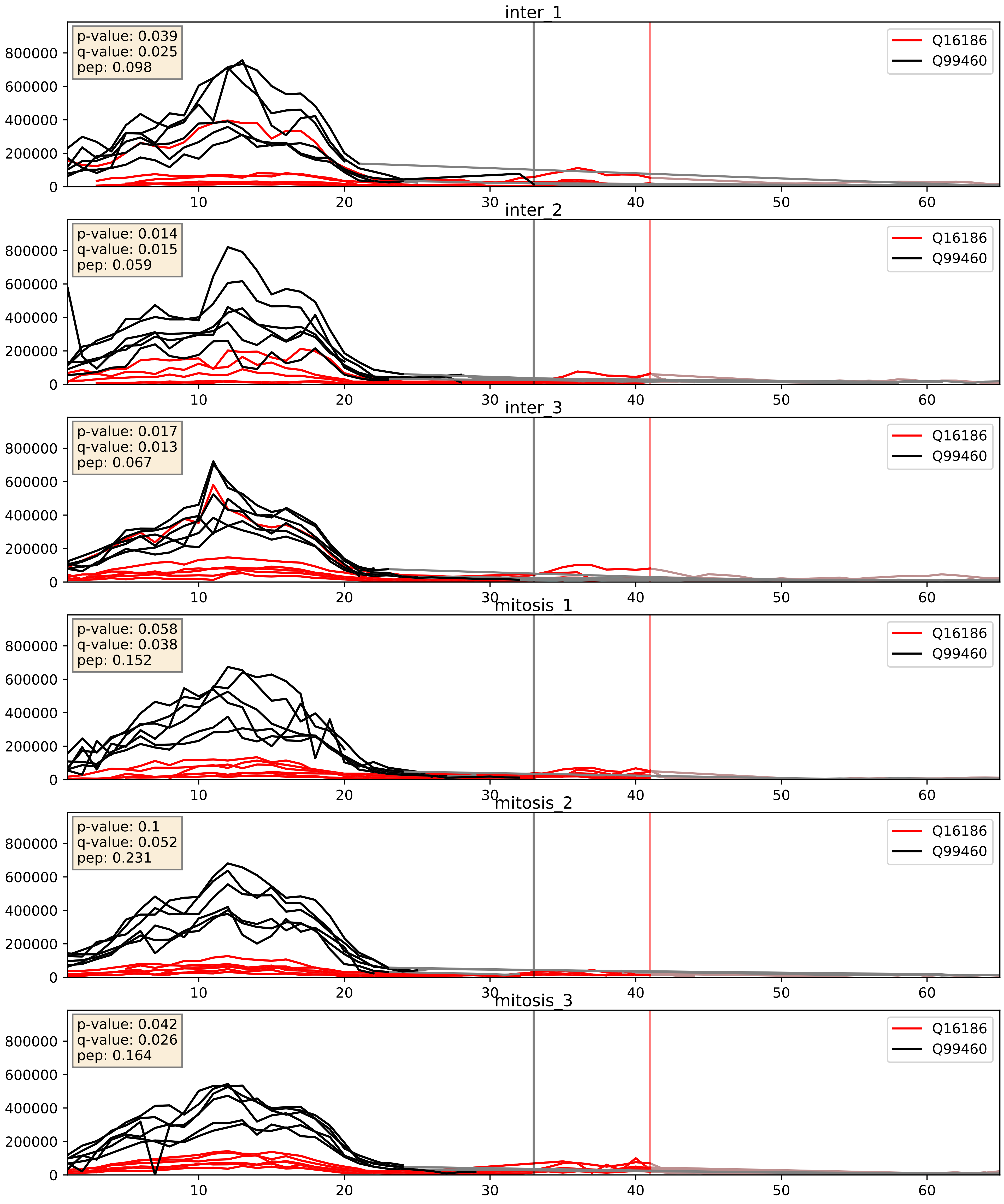


| level | condition_1 | condition_2 | pvalue | pvalue_adjusted |
|---|---|---|---|---|
| complex_abundance | mitosis | inter | 0.13697732494780485 | 0.2820982332168934 |
| interactor_abundance | mitosis | inter | 0.1602820083722657 | 0.34690380590339004 |
| interactor_ratio | mitosis | inter | 0.20519962474487793 | 0.6802900030271708 |

Q16401_Q99460
PSMD5_HUMAN vs PSMD1_HUMAN
p-value: 0.008 q-value: 0.007

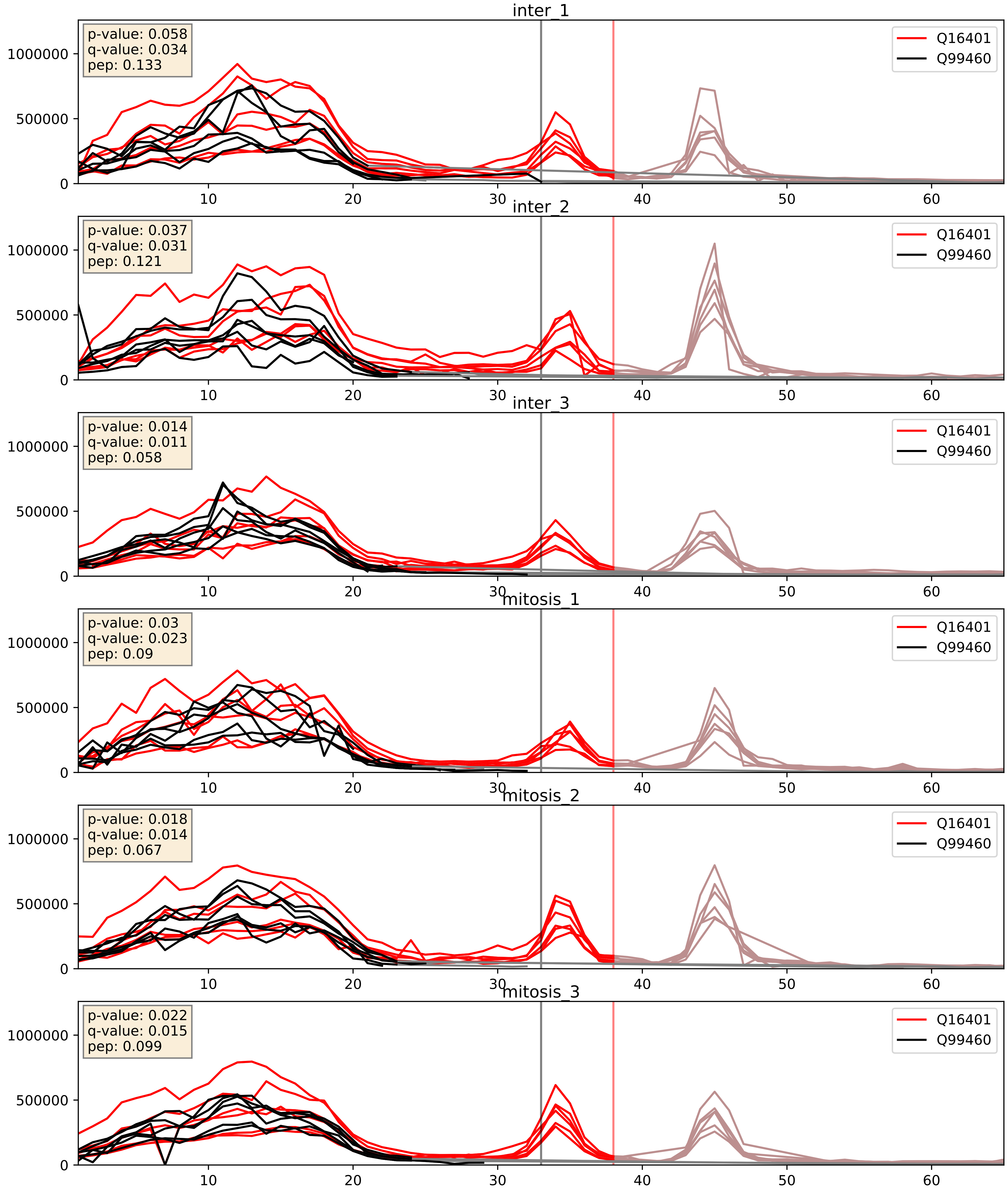


| level | condition_1 | condition_2 | pvalue | pvalue_adjusted |
|---|---|---|---|---|
| complex_abundance | mitosis | inter | 0.11306395964955188 | 0.2527501304171717 |
| interactor_abundance | mitosis | inter | 0.12445916533298877 | 0.30759938075657106 |
| interactor_ratio | mitosis | inter | 0.1788476262884211 | 0.6459428707889276 |

Q99460_Q9UNM6
PSMD1_HUMAN vs PSD13_HUMAN
p-value: 0.0 q-value: 0.001

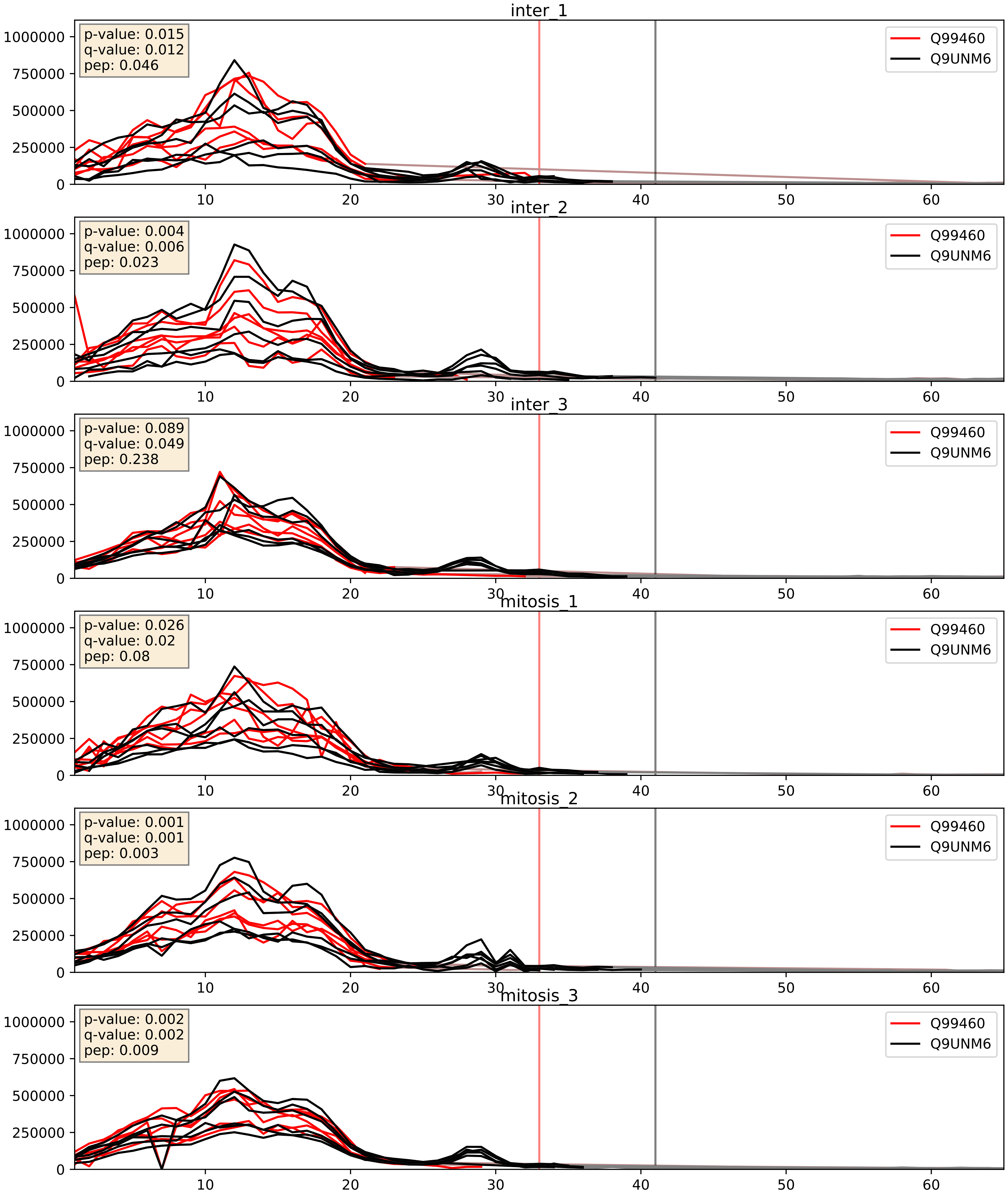


| level | condition_1 | condition_2 | pvalue | pvalue_adjusted |
|---|---|---|---|---|
| complex_abundance | mitosis | inter | 0.03974887232900887 | 0.14497245297670044 |
| interactor_ratio | mitosis | inter | 0.06960555103464508 | 0.4371412449424519 |
| interactor_abundance | mitosis | inter | 0.10334451592191912 | 0.2756657235420093 |

Q99460_Q9Y5K5
PSMD1_HUMAN vs UCHL5_HUMAN
p-value: 0.027 q-value: 0.02

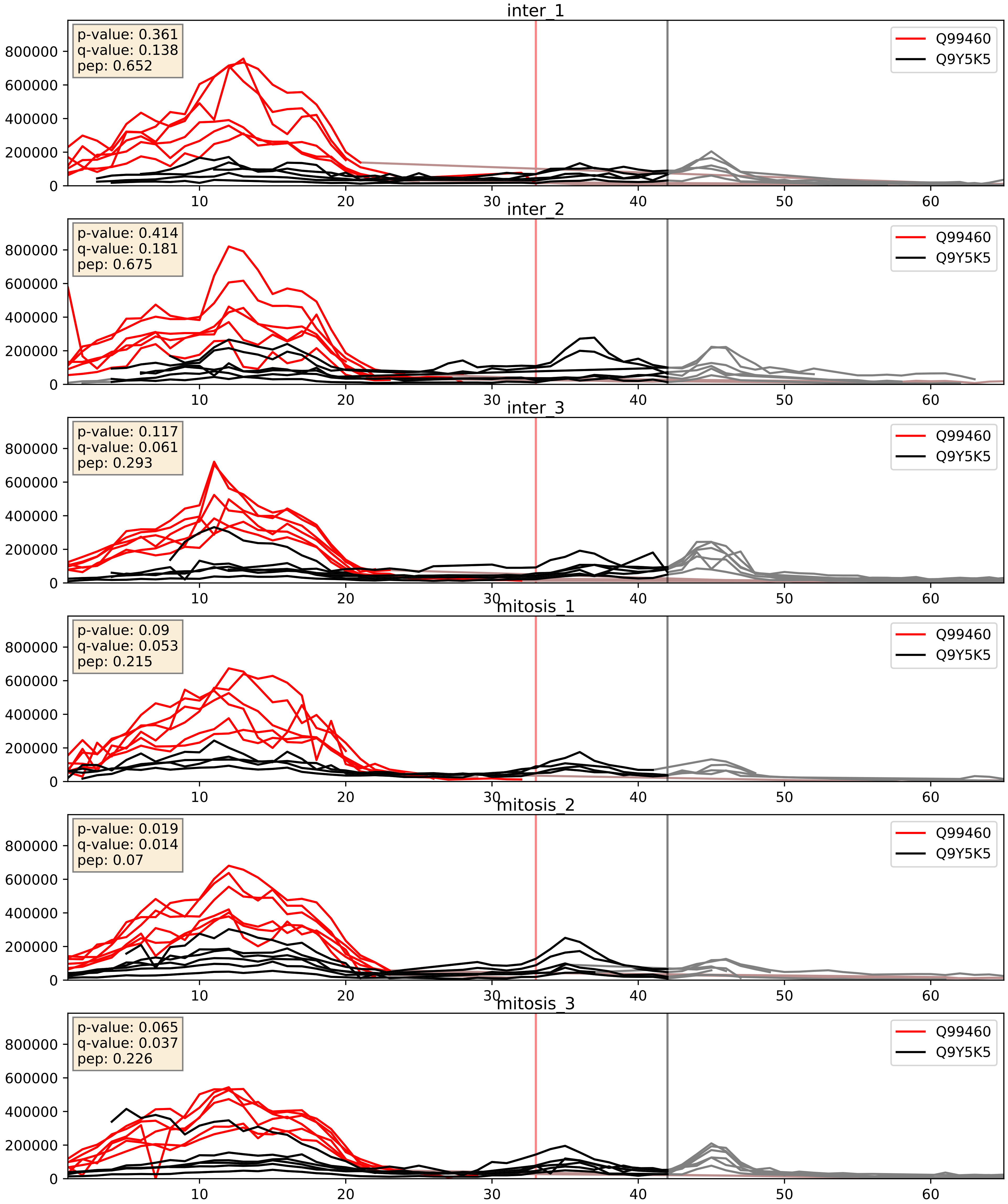


| level | condition_1 | condition_2 | pvalue | pvalue_adjusted |
|---|---|---|---|---|
| interactor_abundance | mitosis | inter | 0.2092841509656856 | 0.40652154270556234 |
| interactor_ratio | mitosis | inter | 0.6442916302536423 | 0.9210314554060085 |
| complex_abundance | mitosis | inter | 0.9364100694291304 | 0.9611057606429837 |
