## Supplementary Data 2 for "SECAT: Quantifying differential protein-protein interaction states by network-centric analysis": hela_string_000162_Q13895.pdf

Q13895_Q13895

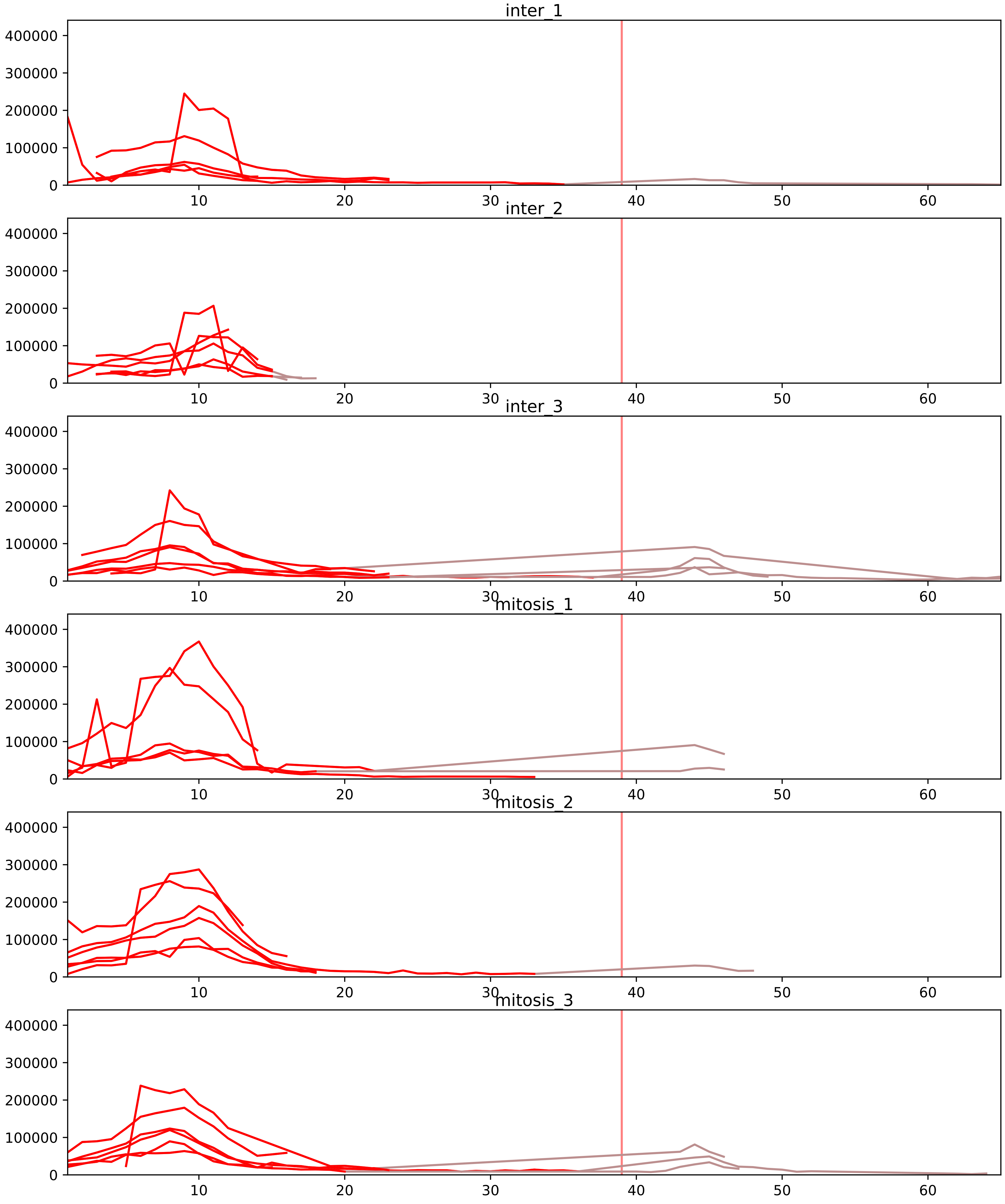


| level | condition_1 | condition_2 | pvalue | pvalue_adjusted |
|---|---|---|---|---|
| interactor_abundance | mitosis | inter | 7.917145803562673e-05 | 0.0019423397704740424 |
| interactor_ratio | mitosis | inter | 0.0006333039841803019 | 0.007616204777070297 |
| complex_abundance | mitosis | inter | 0.0014719597192170684 | 0.01361007981587641 |
| assembled_abundance | mitosis | inter | 0.5425127719535686 | 0.7750511655270553 |
| total_abundance | mitosis | inter | 0.5442071899704043 | 0.7867027001784113 |
| monomer_abundance | mitosis | inter | 0.7162853628552125 | 0.880298677943354 |

O00567_Q13895
NOP56_HUMAN vs BYST_HUMAN
p-value: 0.067 q-value: 0.041

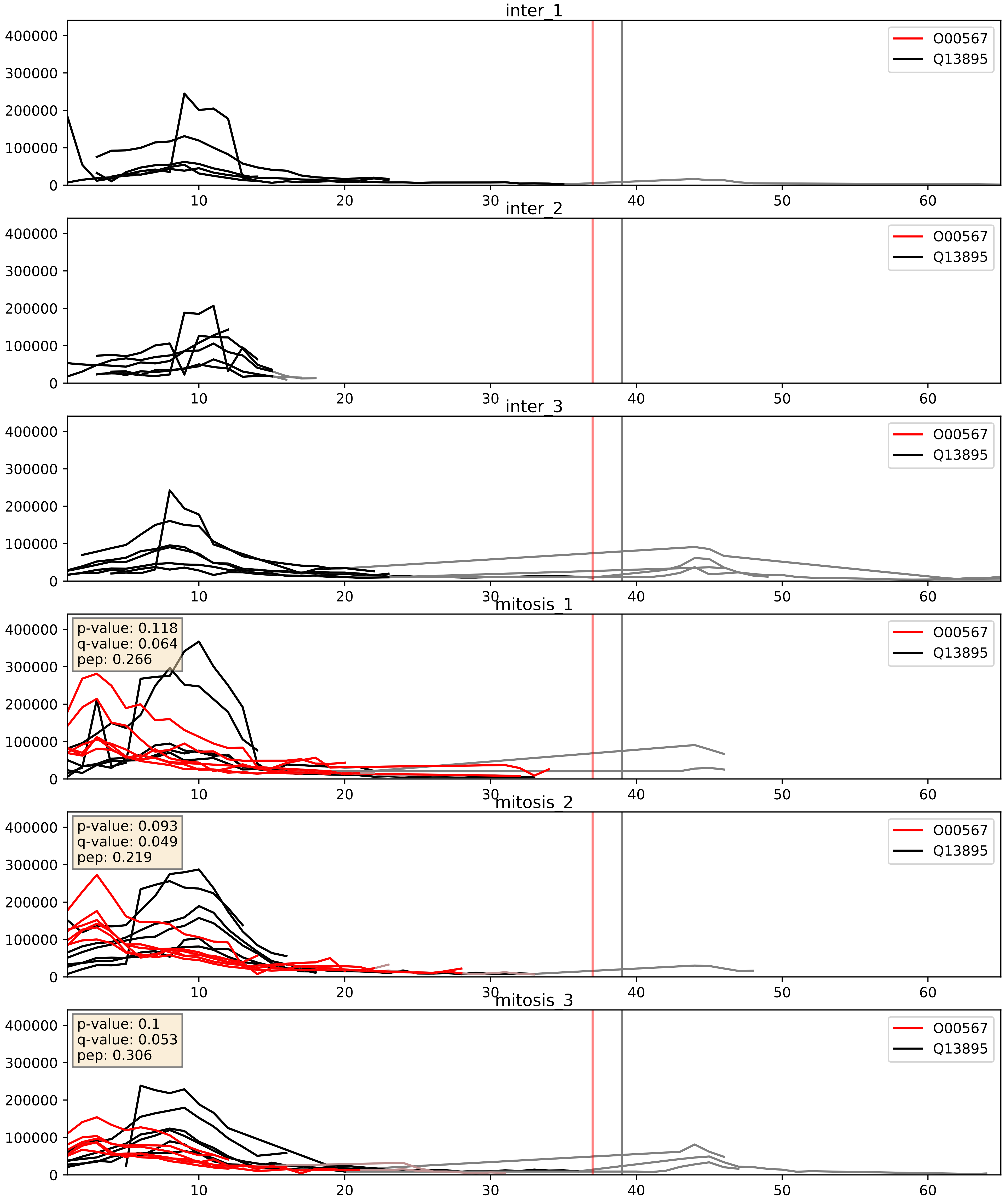


| level | condition_1 | condition_2 | pvalue | pvalue_adjusted |
|---|---|---|---|---|
| complex_abundance | mitosis | inter | 0.0003517935453936726 | 0.012039775462644836 |
| interactor_abundance | mitosis | inter | 0.0011111765577599547 | 0.024049720606162197 |
| interactor_ratio | mitosis | inter | 0.04276010684486895 | 0.3405601671696144 |

P08708_Q13895
RS17_HUMAN vs BYST_HUMAN
p-value: 0.02 q-value: 0.016

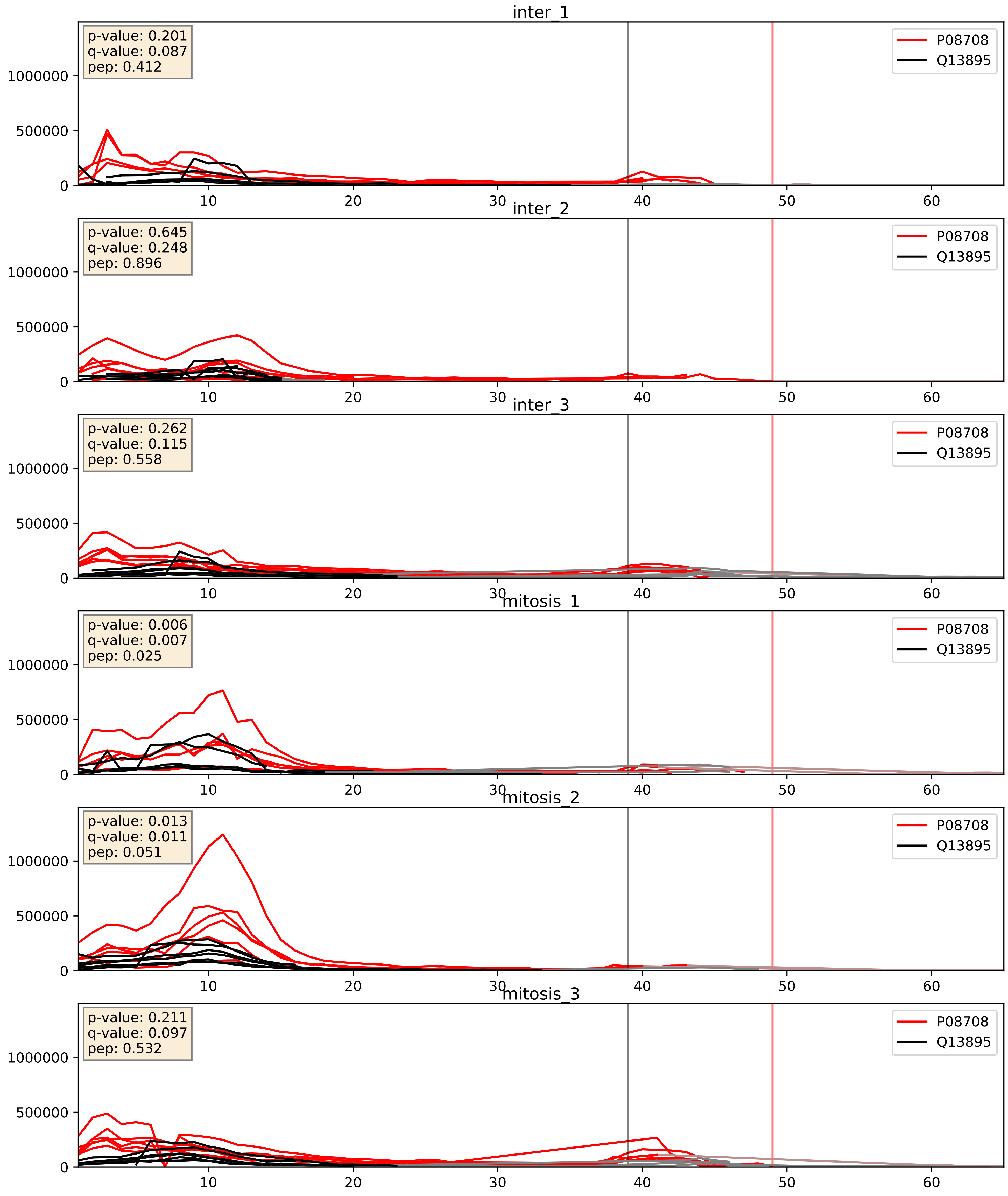


| level | condition_1 | condition_2 | pvalue | pvalue_adjusted |
|---|---|---|---|---|
| complex_abundance | mitosis | inter | 0.024597888856258736 | 0.11443365685302977 |
| interactor_abundance | mitosis | inter | 0.06107836245986043 | 0.20222183298452015 |
| interactor_ratio | mitosis | inter | 0.2700856757800291 | 0.7461847438669881 |

P22087_Q13895
FBRL_HUMAN vs BYST_HUMAN
p-value: 0.062 q-value: 0.039

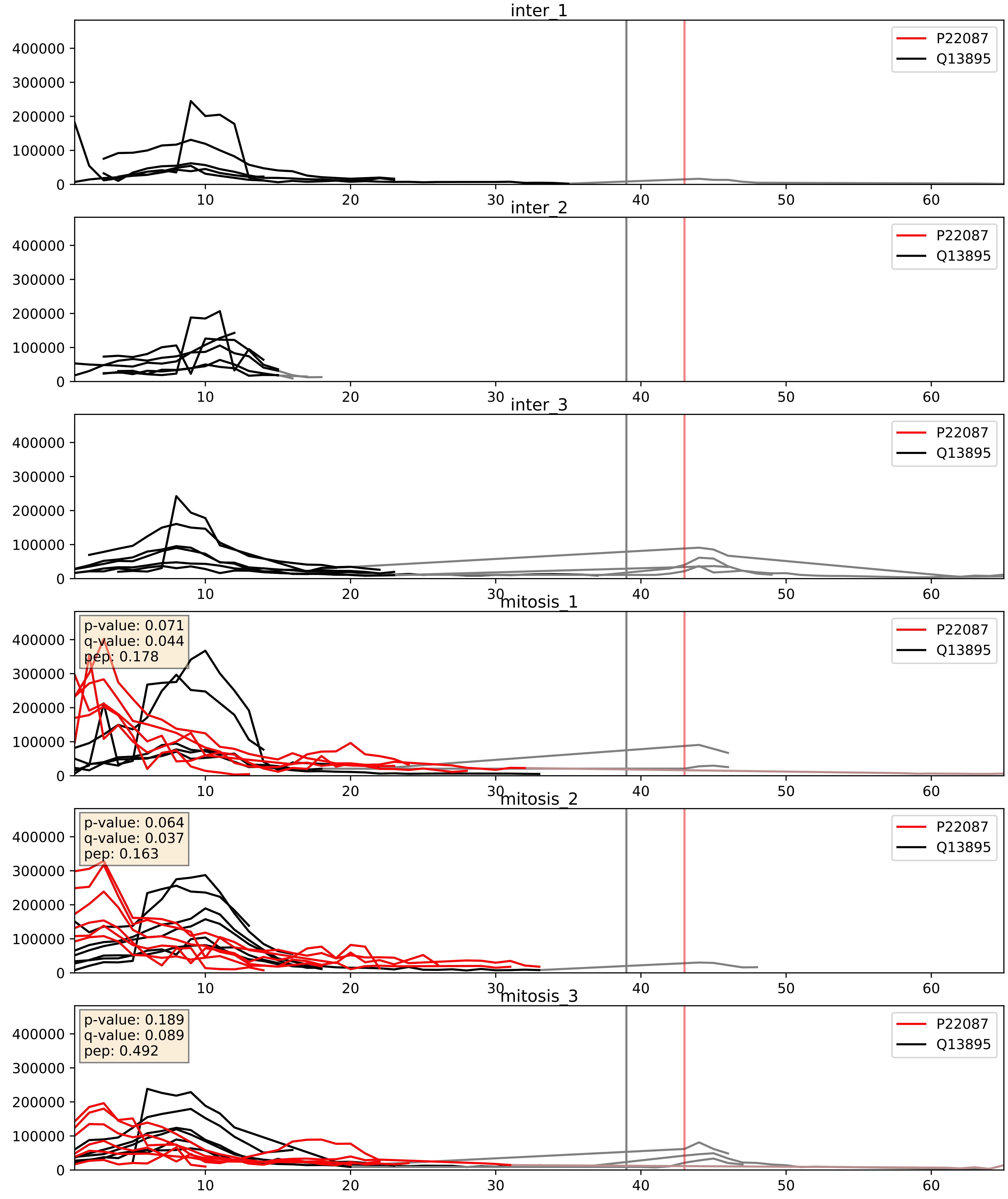


| level | condition_1 | condition_2 | pvalue | pvalue_adjusted |
|---|---|---|---|---|
| complex_abundance | mitosis | inter | 0.0001916121831474646 | 0.009214608358102793 |
| interactor_abundance | mitosis | inter | 0.00035333092874419484 | 0.01356427145975096 |
| interactor_ratio | mitosis | inter | 0.26434880442933373 | 0.7392439614227693 |

P39019_Q13895
RS19_HUMAN vs BYST_HUMAN
p-value: 0.058 q-value: 0.037

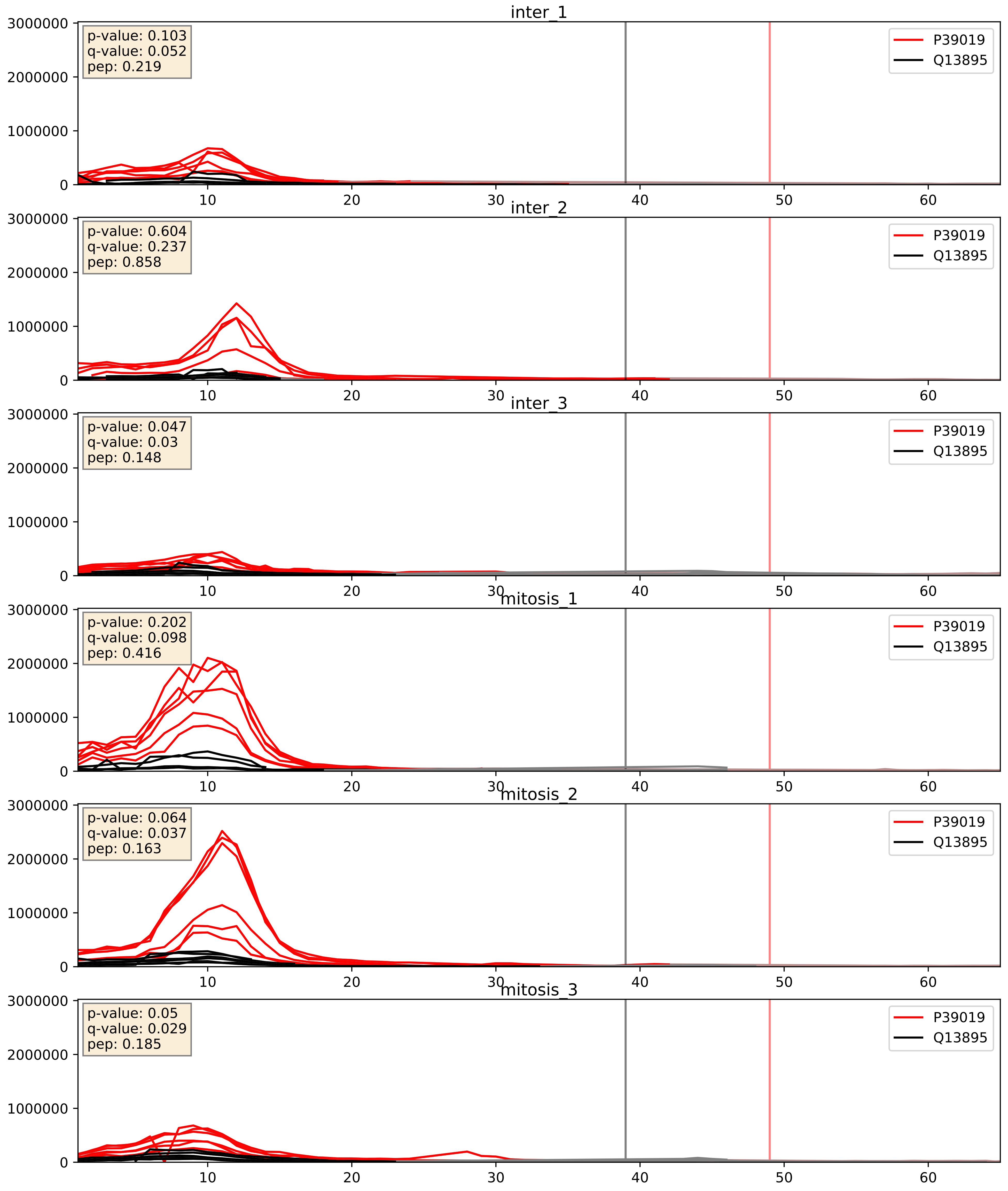


| level | condition_1 | condition_2 | pvalue | pvalue_adjusted |
|---|---|---|---|---|
| complex_abundance | mitosis | inter | 0.09008825005263829 | 0.22062267011438283 |
| interactor_abundance | mitosis | inter | 0.18936325296252154 | 0.38117564852655716 |
| interactor_ratio | mitosis | inter | 0.5172299413079594 | 0.8775869903454201 |

P46781_Q13895
RS9_HUMAN vs BYST_HUMAN
p-value: 0.006 q-value: 0.006

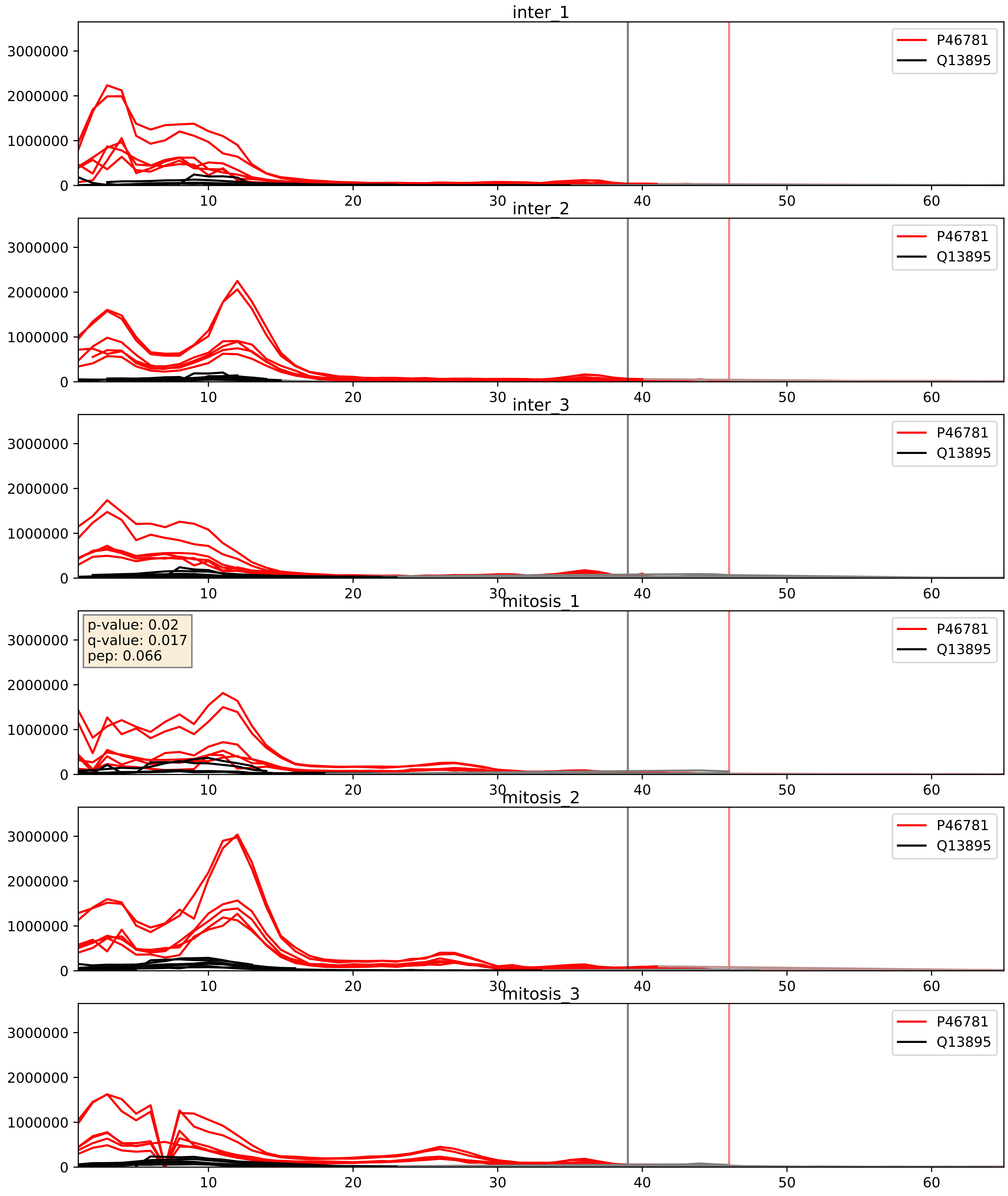


| level | condition_1 | condition_2 | pvalue | pvalue_adjusted |
|---|---|---|---|---|
| complex_abundance | mitosis | inter | 0.02600807583701099 | 0.11810151502907984 |
| interactor_ratio | mitosis | inter | 0.16702284467572145 | 0.6293444626080091 |
| interactor_abundance | mitosis | inter | 0.6354134395563686 | 0.7573495751476084 |

P46782_Q13895
RS5_HUMAN vs BYST_HUMAN
p-value: 0.022 q-value: 0.017

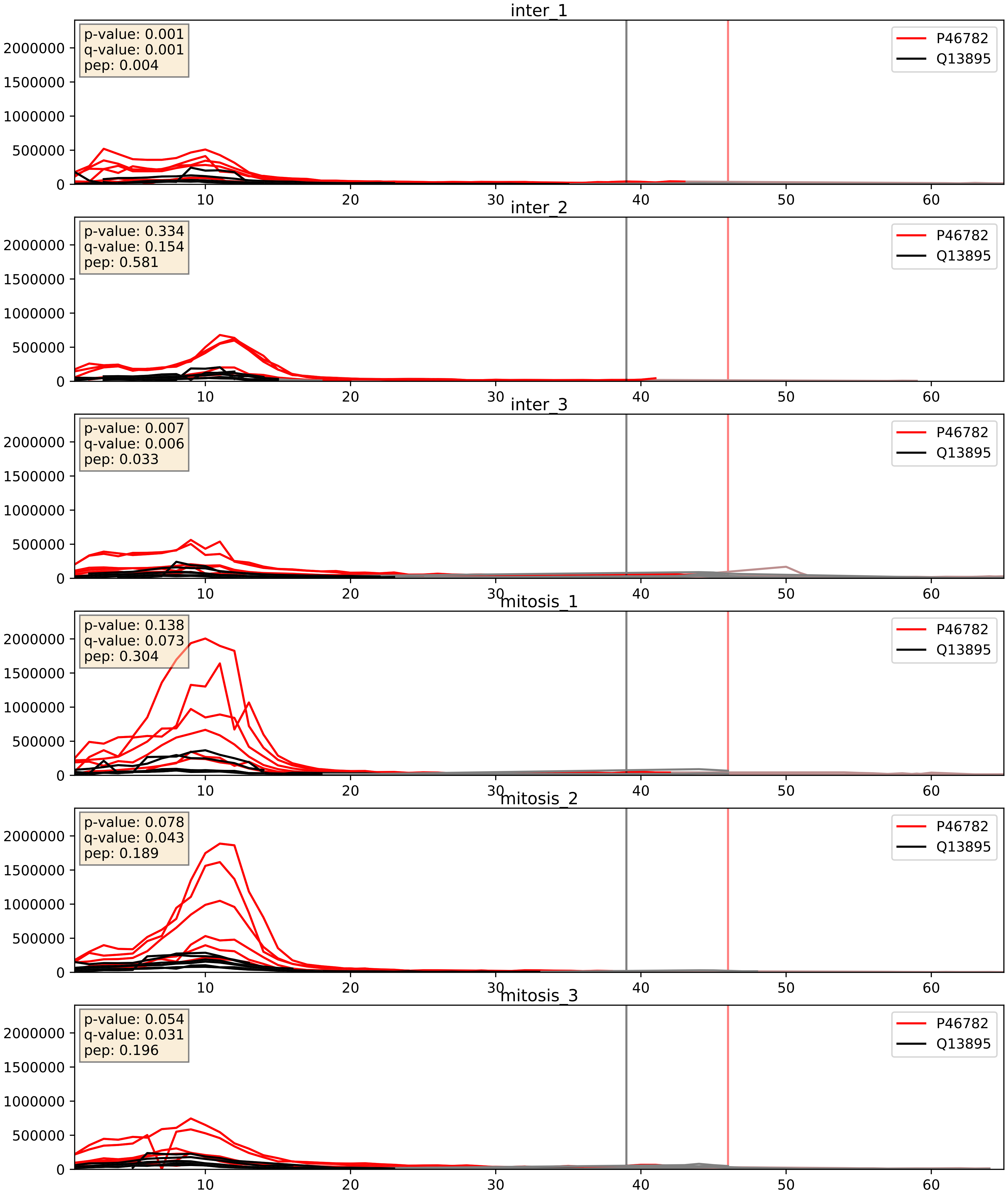


| level | condition_1 | condition_2 | pvalue | pvalue_adjusted |
|---|---|---|---|---|
| complex_abundance | mitosis | inter | 0.038644163281217184 | 0.14351151309640742 |
| interactor_abundance | mitosis | inter | 0.09712380843407231 | 0.2678414304754056 |
| interactor_ratio | mitosis | inter | 0.6376222947740969 | 0.9197898119332029 |

P46783_Q13895
RS10_HUMAN vs BYST_HUMAN
p-value: 0.002 q-value: 0.031

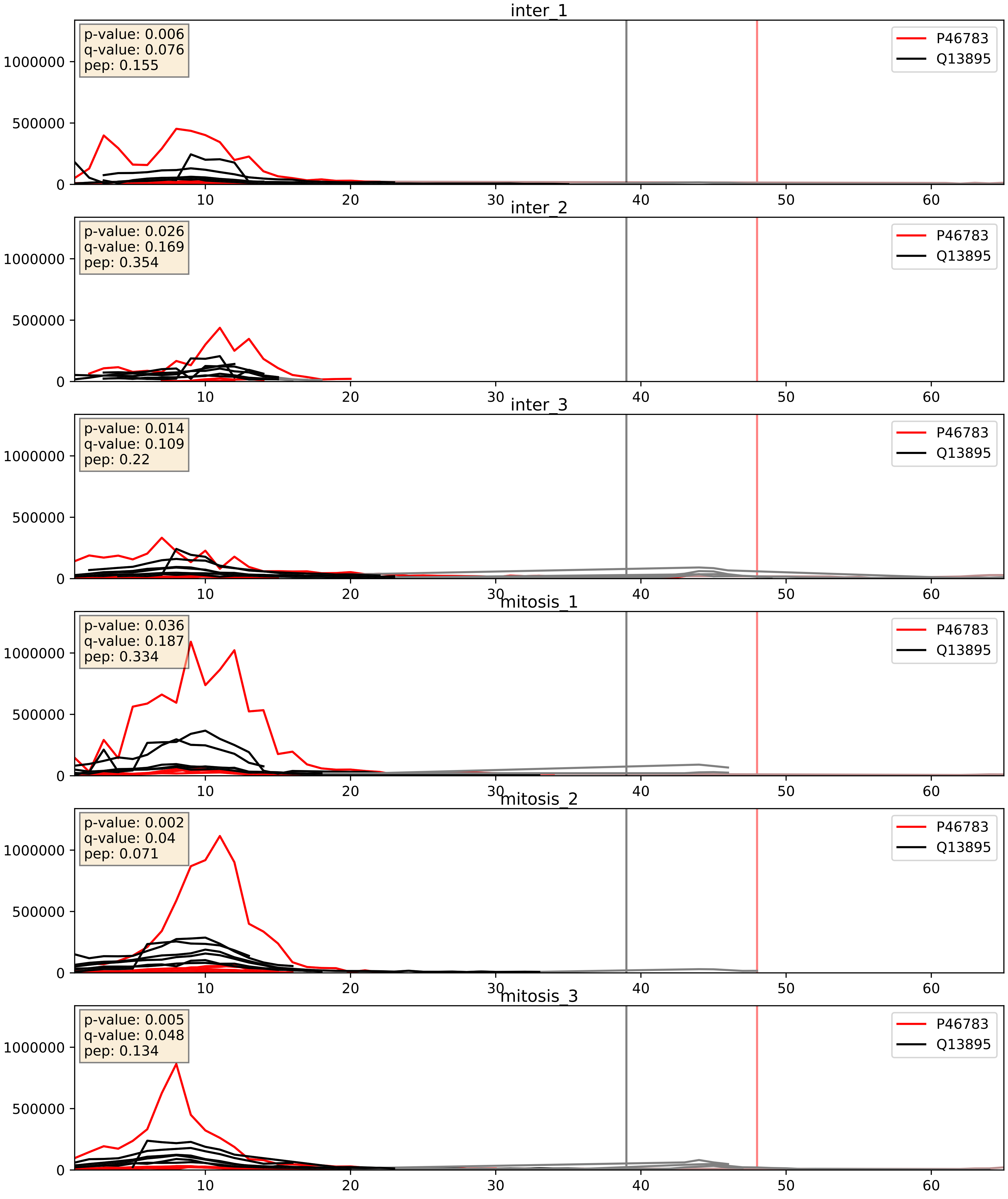


| level | condition_1 | condition_2 | pvalue | pvalue_adjusted |
|---|---|---|---|---|
| interactor_ratio | mitosis | inter | 0.16695138719958383 | 0.6293444626080091 |
| complex_abundance | mitosis | inter | 0.21397827155768678 | 0.36869041959214954 |
| interactor_abundance | mitosis | inter | 0.9360045143841014 | 0.962096690999235 |

P55769_Q13895
NH2L1_HUMAN vs BYST_HUMAN
p-value: 0.062 q-value: 0.039

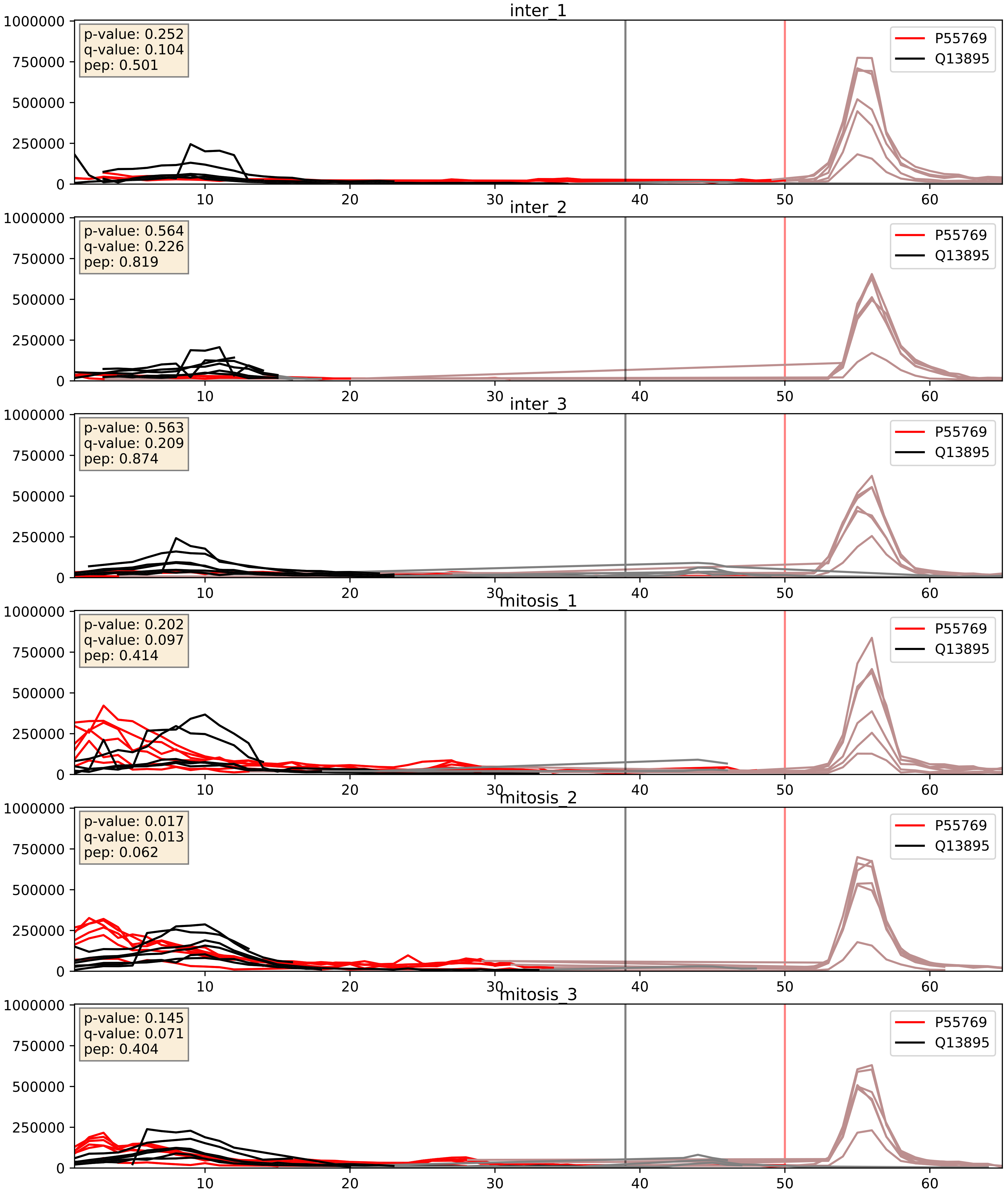


| level | condition_1 | condition_2 | pvalue | pvalue_adjusted |
|---|---|---|---|---|
| complex_abundance | mitosis | inter | 0.0017384148809047146 | 0.0295241467010272 |
| interactor_abundance | mitosis | inter | 0.00183168678013043 | 0.03145283618438612 |
| interactor_ratio | mitosis | inter | 0.0033897716918089944 | 0.07563355707045767 |

P60866_Q13895
RS20_HUMAN vs BYST_HUMAN
p-value: 0.022 q-value: 0.017

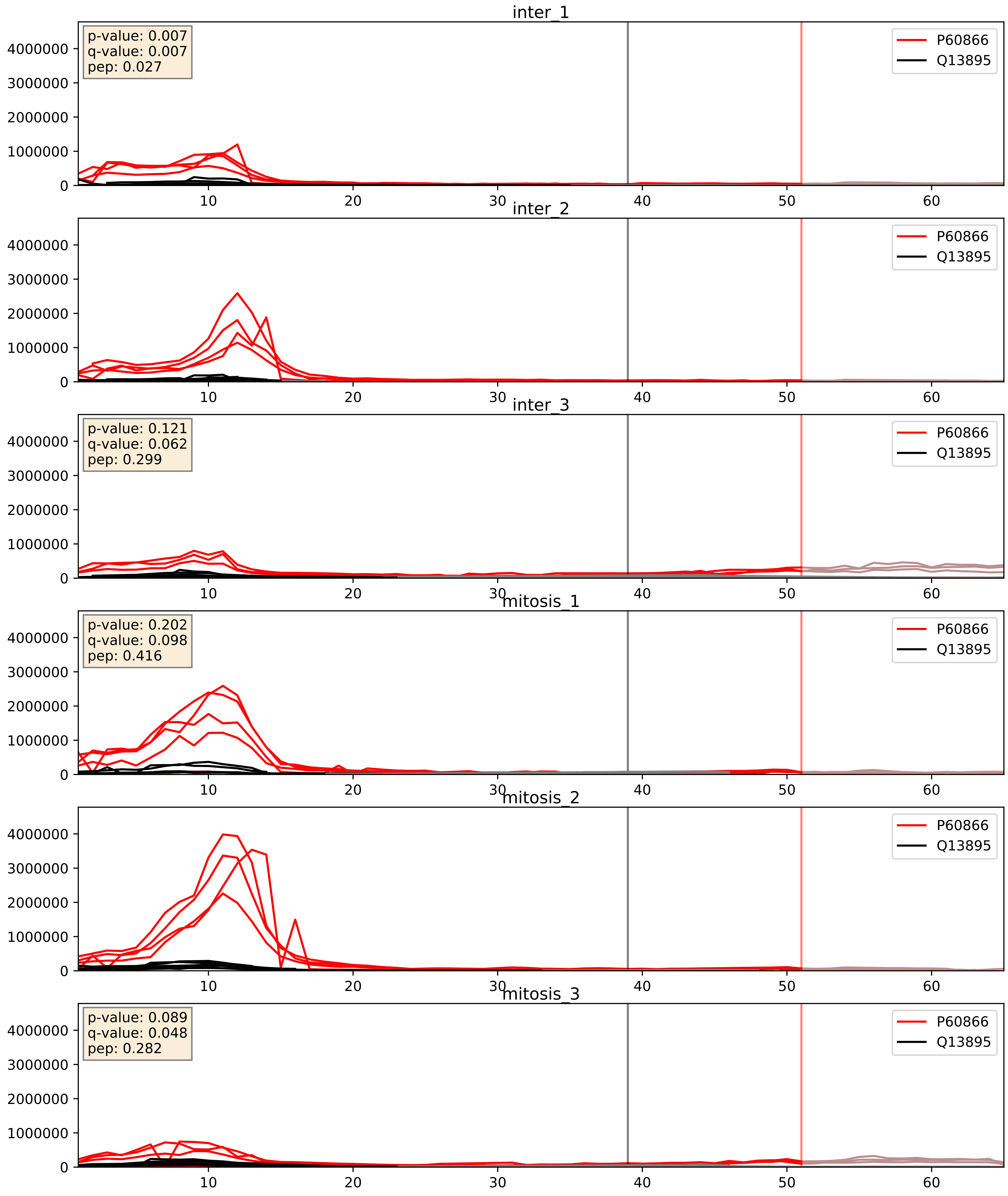


| level | condition_1 | condition_2 | pvalue | pvalue_adjusted |
|---|---|---|---|---|
| complex_abundance | mitosis | inter | 0.1522440225643645 | 0.29878471786517213 |
| interactor_abundance | mitosis | inter | 0.36941461392897135 | 0.5638896199509011 |
| interactor_ratio | mitosis | inter | 0.9130200995845674 | 0.9852905071335931 |

P62244_Q13895
RS15A_HUMAN vs BYST_HUMAN
p-value: 0.037 q-value: 0.026

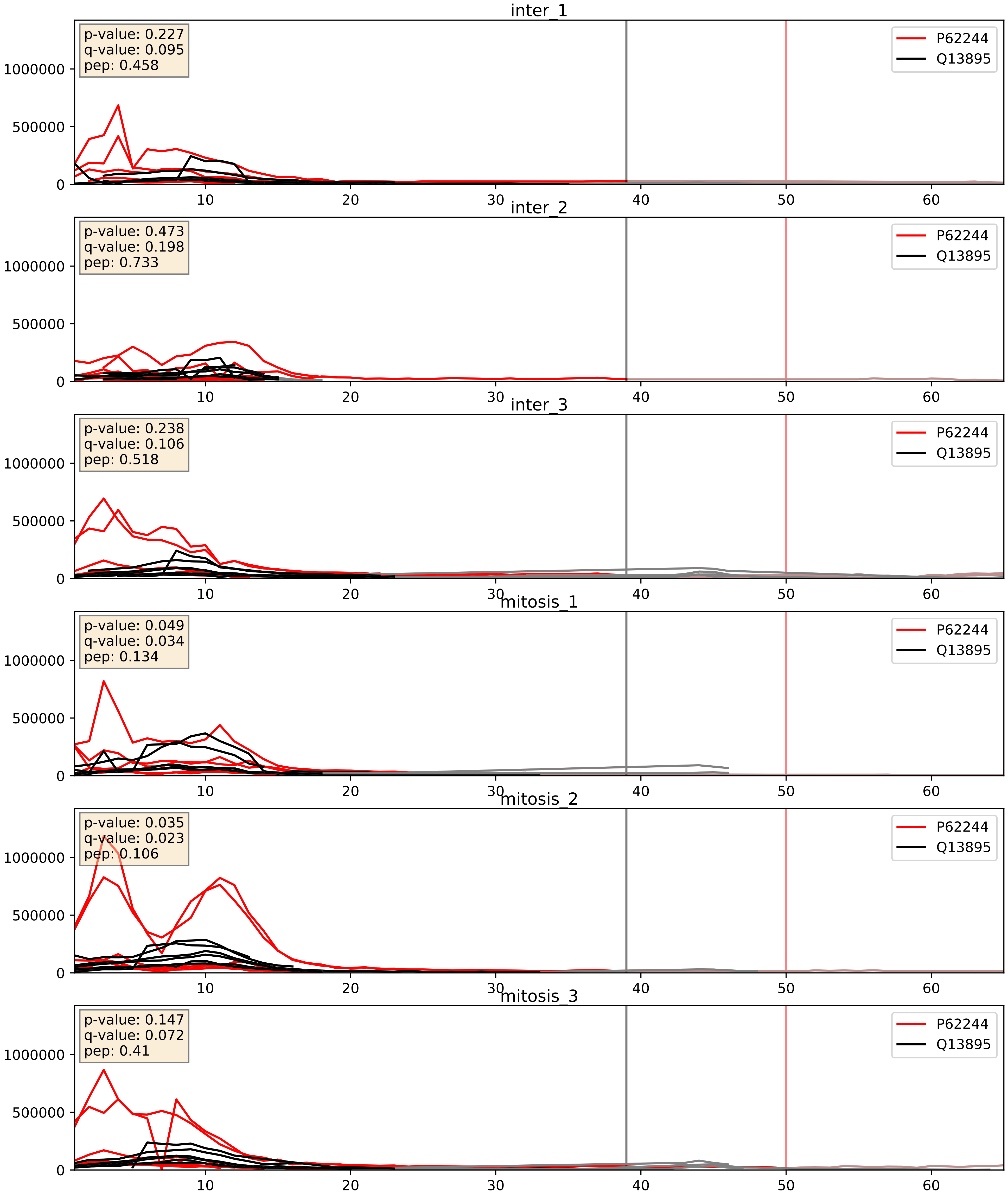


| level | condition_1 | condition_2 | pvalue | pvalue_adjusted |
|---|---|---|---|---|
| complex_abundance | mitosis | inter | 0.03424571255974177 | 0.13450656280738457 |
| interactor_abundance | mitosis | inter | 0.16638865974763825 | 0.35469727990033206 |
| interactor_ratio | mitosis | inter | 0.4595673835610124 | 0.8573956600848013 |

P62263_Q13895
RS14_HUMAN vs BYST_HUMAN
p-value: 0.037 q-value: 0.026

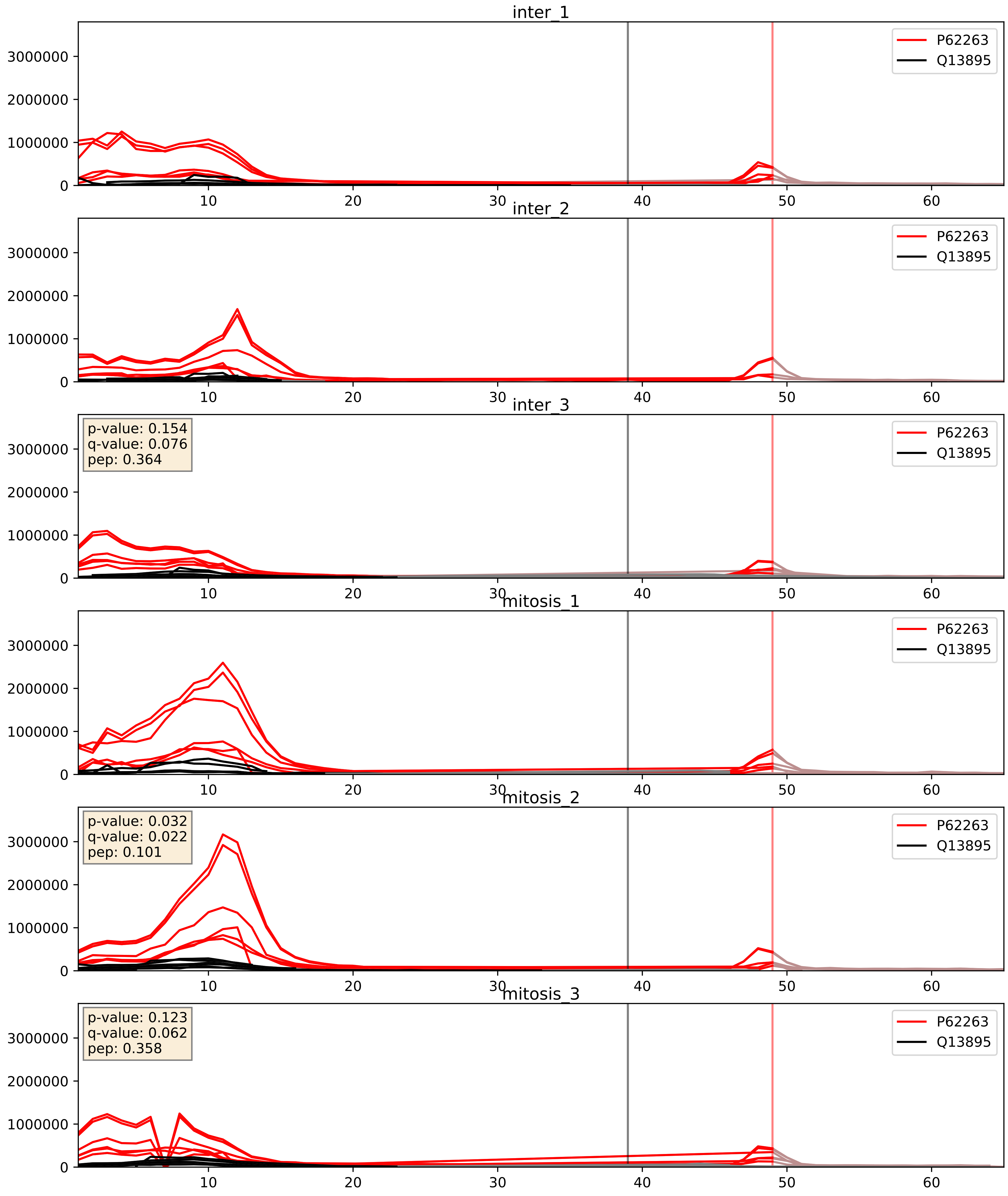


| level | condition_1 | condition_2 | pvalue | pvalue_adjusted |
|---|---|---|---|---|
| complex_abundance | mitosis | inter | 0.10024371386956872 | 0.2350018038308818 |
| interactor_abundance | mitosis | inter | 0.3205600376550994 | 0.5189095919681639 |
| interactor_ratio | mitosis | inter | 0.8706385543477446 | 0.9789394489973328 |

P62266_Q13895
RS23_HUMAN vs BYST_HUMAN
p-value: 0.059 q-value: 0.037

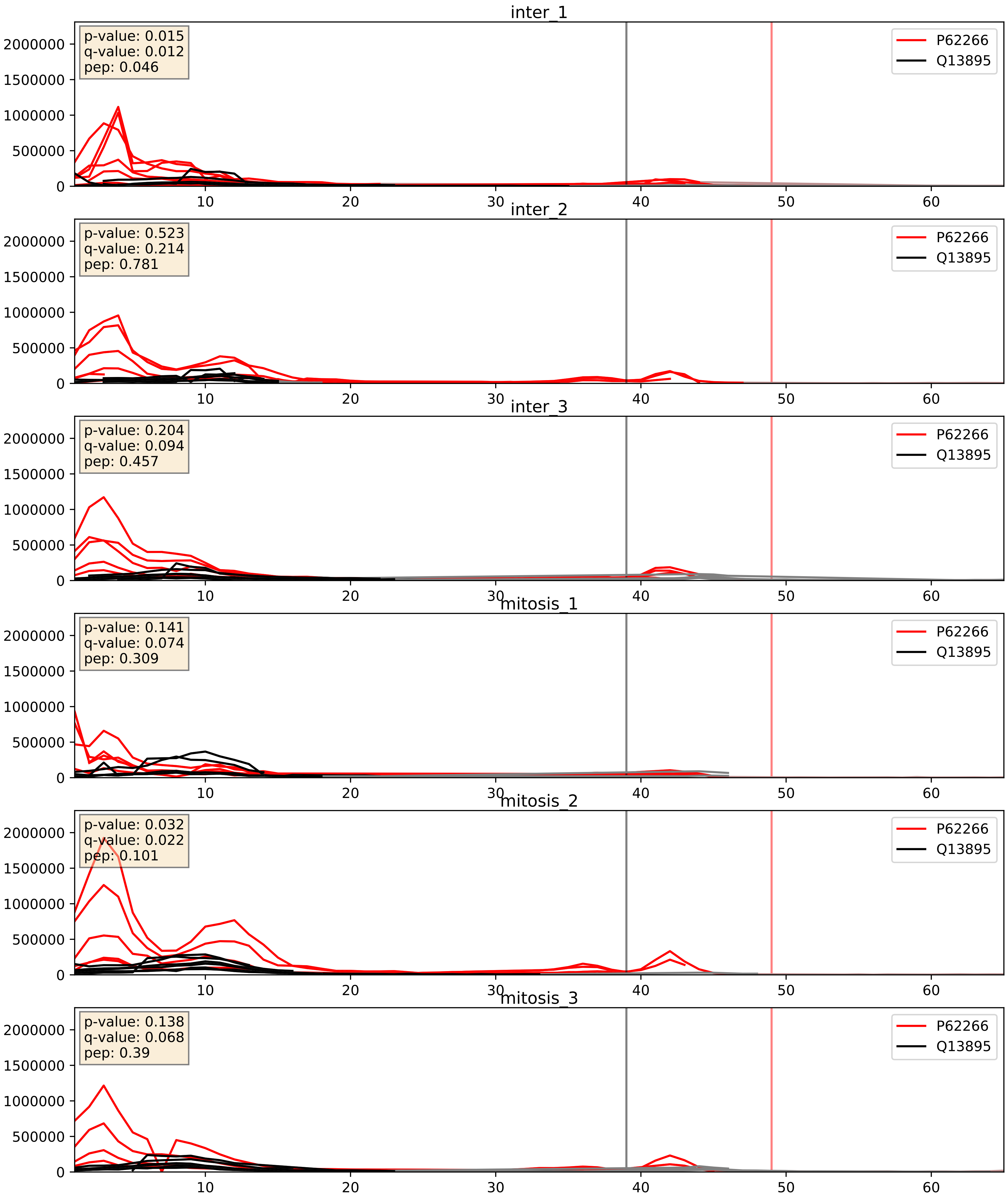


| level | condition_1 | condition_2 | pvalue | pvalue_adjusted |
|---|---|---|---|---|
| interactor_ratio | mitosis | inter | 0.05694156515797698 | 0.39422508341841167 |
| interactor_abundance | mitosis | inter | 0.29449099698533354 | 0.48914925171135015 |
| complex_abundance | mitosis | inter | 0.8442030634725546 | 0.9010446662500083 |

P62277_Q13895
RS13_HUMAN vs BYST_HUMAN
p-value: 0.032 q-value: 0.023

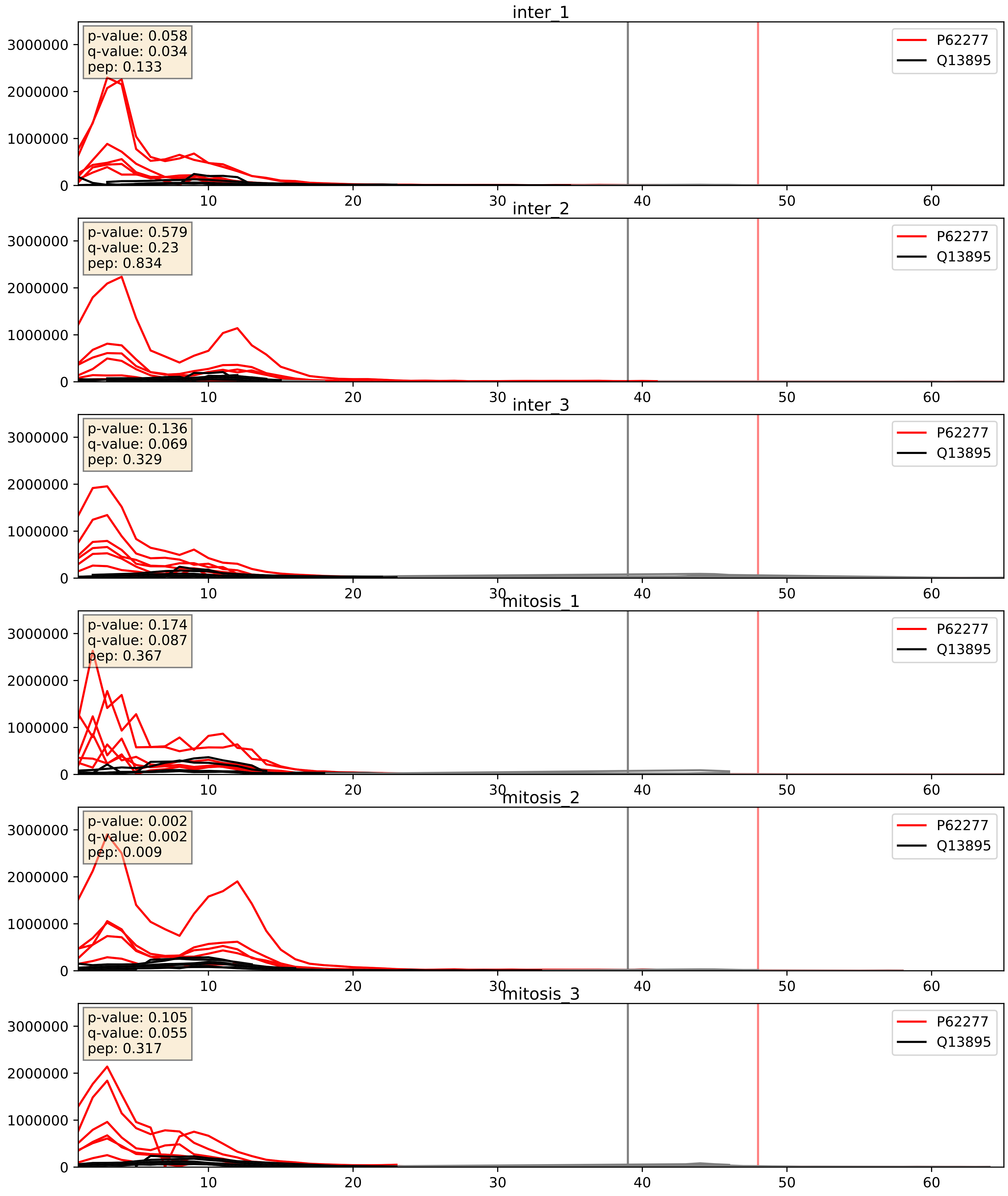


| level | condition_1 | condition_2 | pvalue | pvalue_adjusted |
|---|---|---|---|---|
| complex_abundance | mitosis | inter | 0.004224583285705355 | 0.048345498563686945 |
| interactor_ratio | mitosis | inter | 0.11133177333308898 | 0.5531333750848836 |
| interactor_abundance | mitosis | inter | 0.38055853451301636 | 0.5734168377805703 |

P62280_Q13895
RS11_HUMAN vs BYST_HUMAN
p-value: 0.069 q-value: 0.042

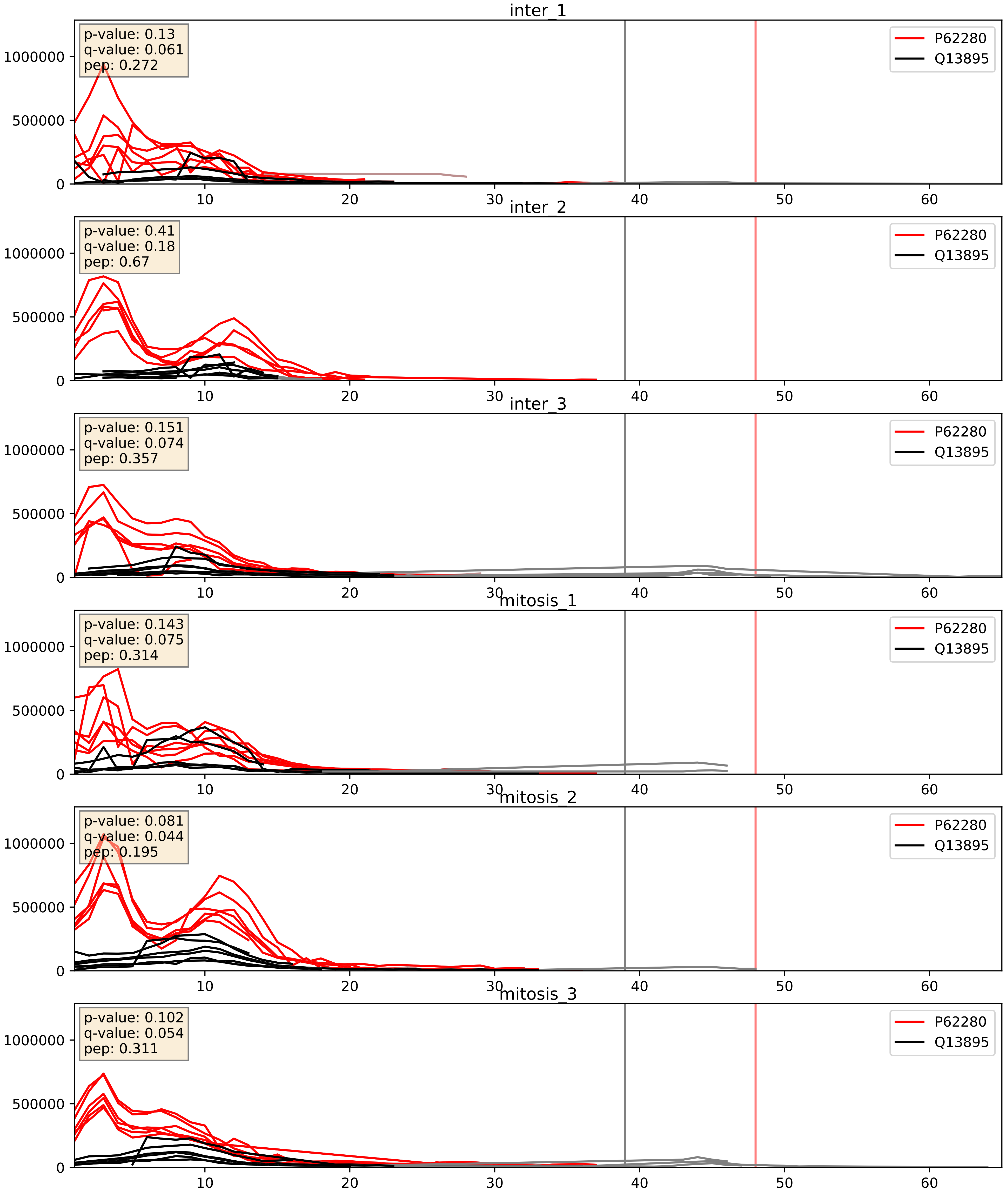


| level | condition_1 | condition_2 | pvalue | pvalue_adjusted |
|---|---|---|---|---|
| interactor_ratio | mitosis | inter | 0.046302272936529054 | 0.3574623087810929 |
| complex_abundance | mitosis | inter | 0.06569924217266986 | 0.1852389700257095 |
| interactor_abundance | mitosis | inter | 0.7476568989001916 | 0.8460497130789398 |

P62701_Q13895
RS4X_HUMAN vs BYST_HUMAN
p-value: 0.046 q-value: 0.031

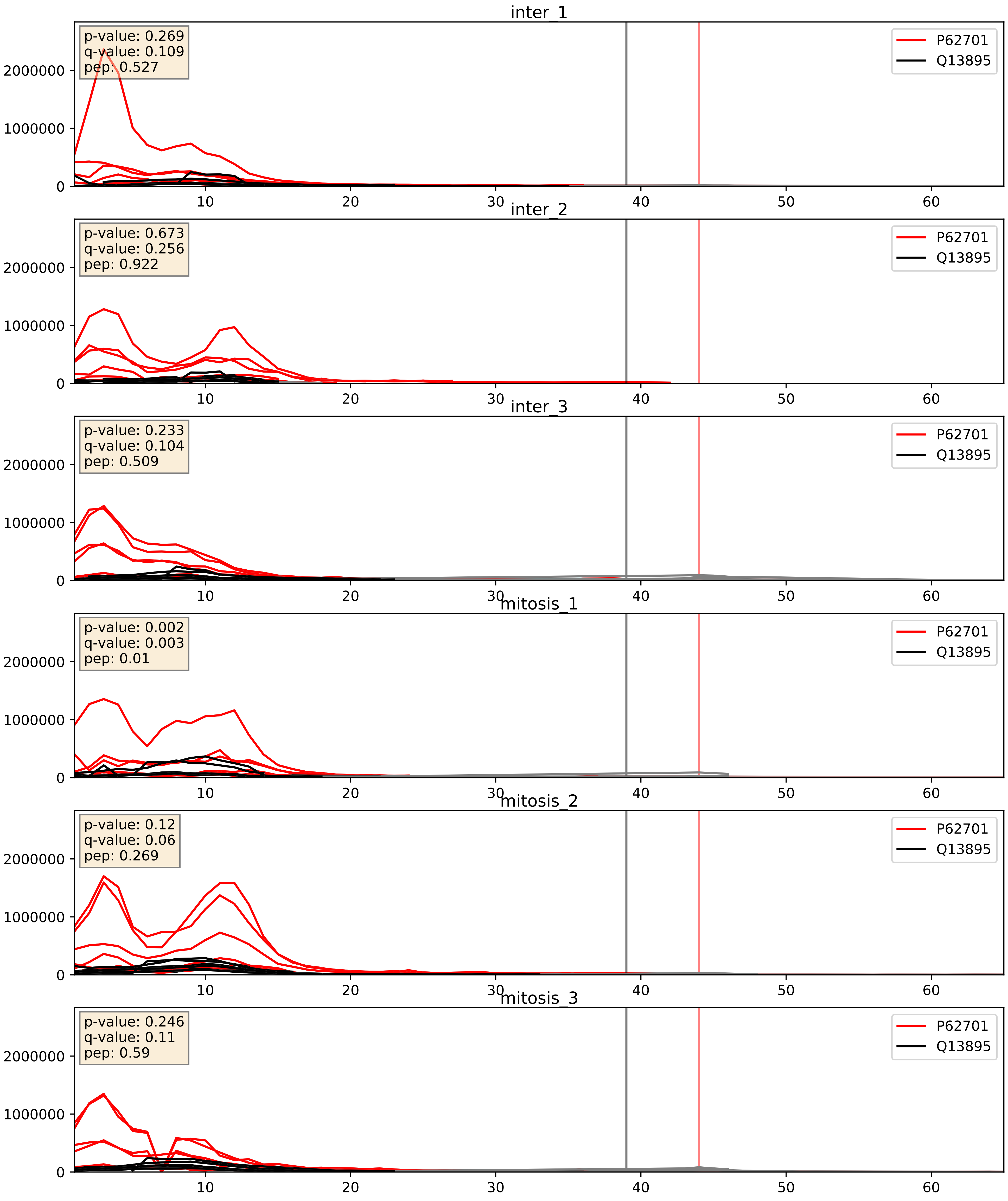


| level | condition_1 | condition_2 | pvalue | pvalue_adjusted |
|---|---|---|---|---|
| complex_abundance | mitosis | inter | 0.06331226394236993 | 0.18143722107354757 |
| interactor_ratio | mitosis | inter | 0.08394014387778045 | 0.4858108960883037 |
| interactor_abundance | mitosis | inter | 0.5450471717041256 | 0.7017528449706423 |

P62841_Q13895
RS15_HUMAN vs BYST_HUMAN
p-value: 0.026 q-value: 0.019

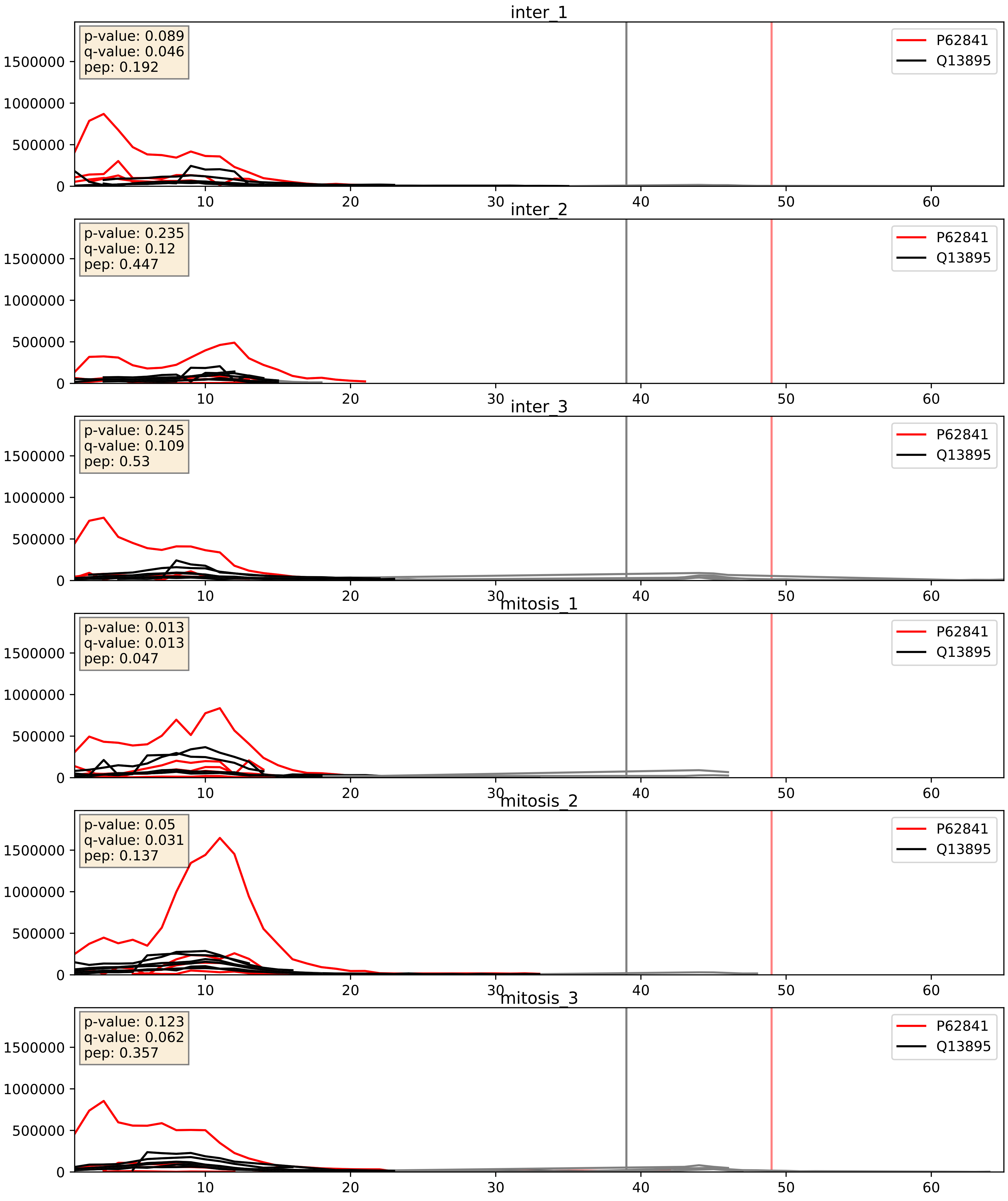


| level | condition_1 | condition_2 | pvalue | pvalue_adjusted |
|---|---|---|---|---|
| interactor_ratio | mitosis | inter | 0.22970951878082022 | 0.7075666146681794 |
| complex_abundance | mitosis | inter | 0.46789122187826215 | 0.60885374521723 |
| interactor_abundance | mitosis | inter | 0.9192413058805448 | 0.9527406319455572 |

P62851_Q13895
RS25_HUMAN vs BYST_HUMAN
p-value: 0.008 q-value: 0.008

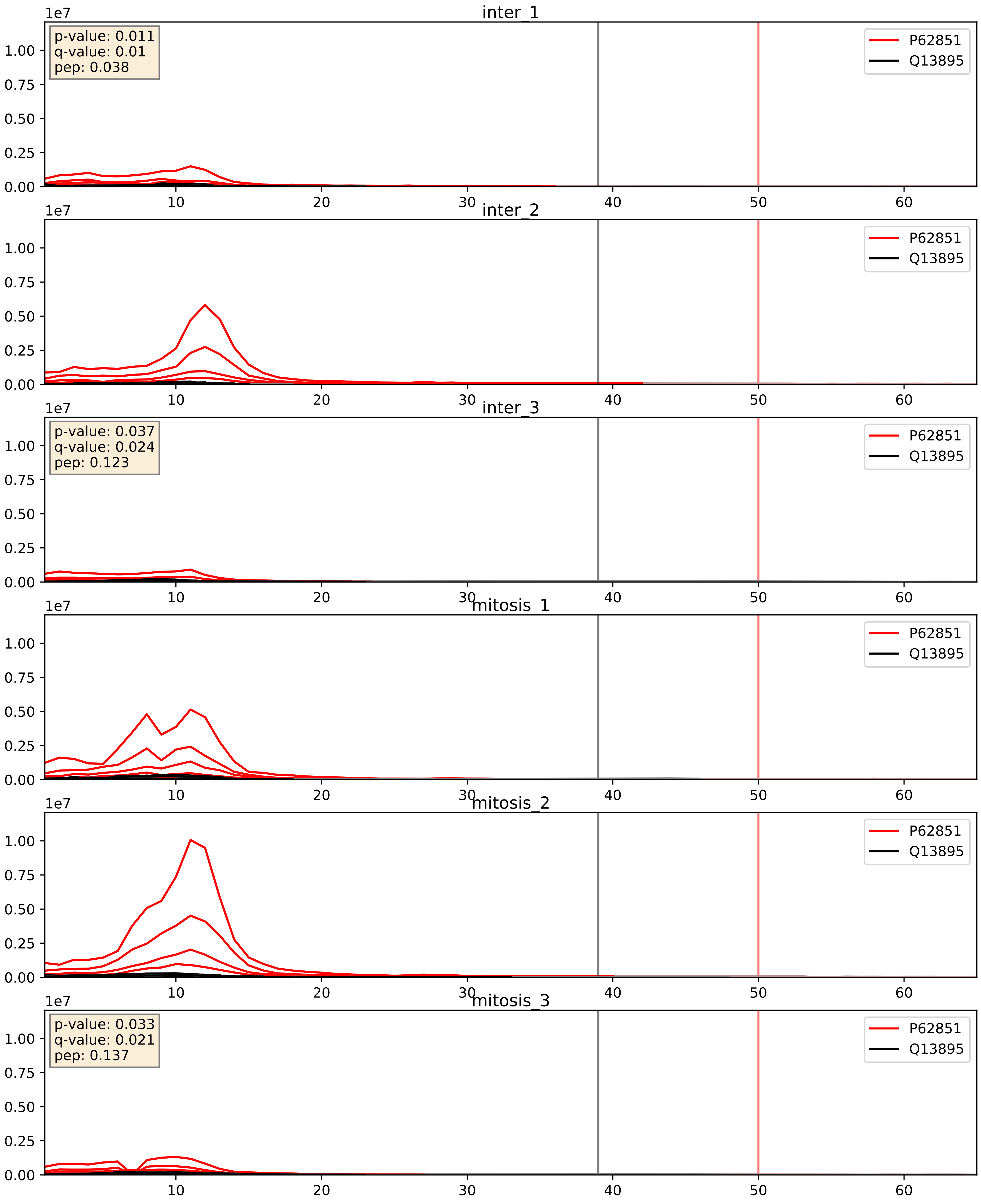


| level | condition_1 | condition_2 | pvalue | pvalue_adjusted |
|---|---|---|---|---|
| complex_abundance | mitosis | inter | 0.21396849490253195 | 0.36869041959214954 |
| interactor_abundance | mitosis | inter | 0.3681688256809099 | 0.5636278543912346 |
| interactor_ratio | mitosis | inter | 0.6580304779064501 | 0.925532925708346 |

Q13601_Q13895
KRR1_HUMAN vs BYST_HUMAN
p-value: 0.028 q-value: 0.02

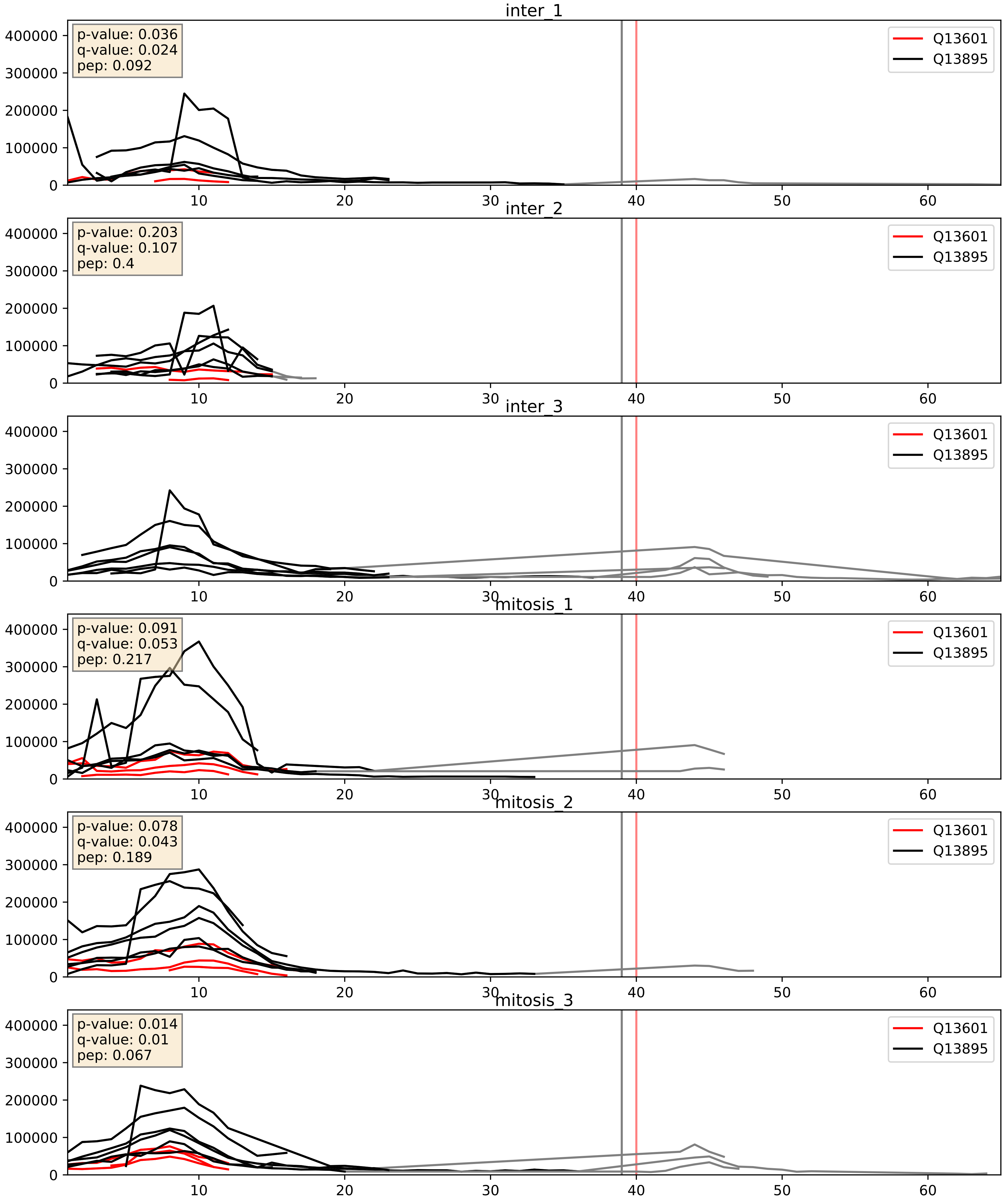


| level | condition_1 | condition_2 | pvalue | pvalue_adjusted |
|---|---|---|---|---|
| interactor_ratio | mitosis | inter | 0.16816133499555003 | 0.6302368772162471 |
| complex_abundance | mitosis | inter | 0.20104289081844154 | 0.3538817901307546 |
| interactor_abundance | mitosis | inter | 0.2635837344619755 | 0.45562988713383146 |

Q13895_Q15061
BYST_HUMAN vs WDR43_HUMAN
p-value: 0.076 q-value: 0.046

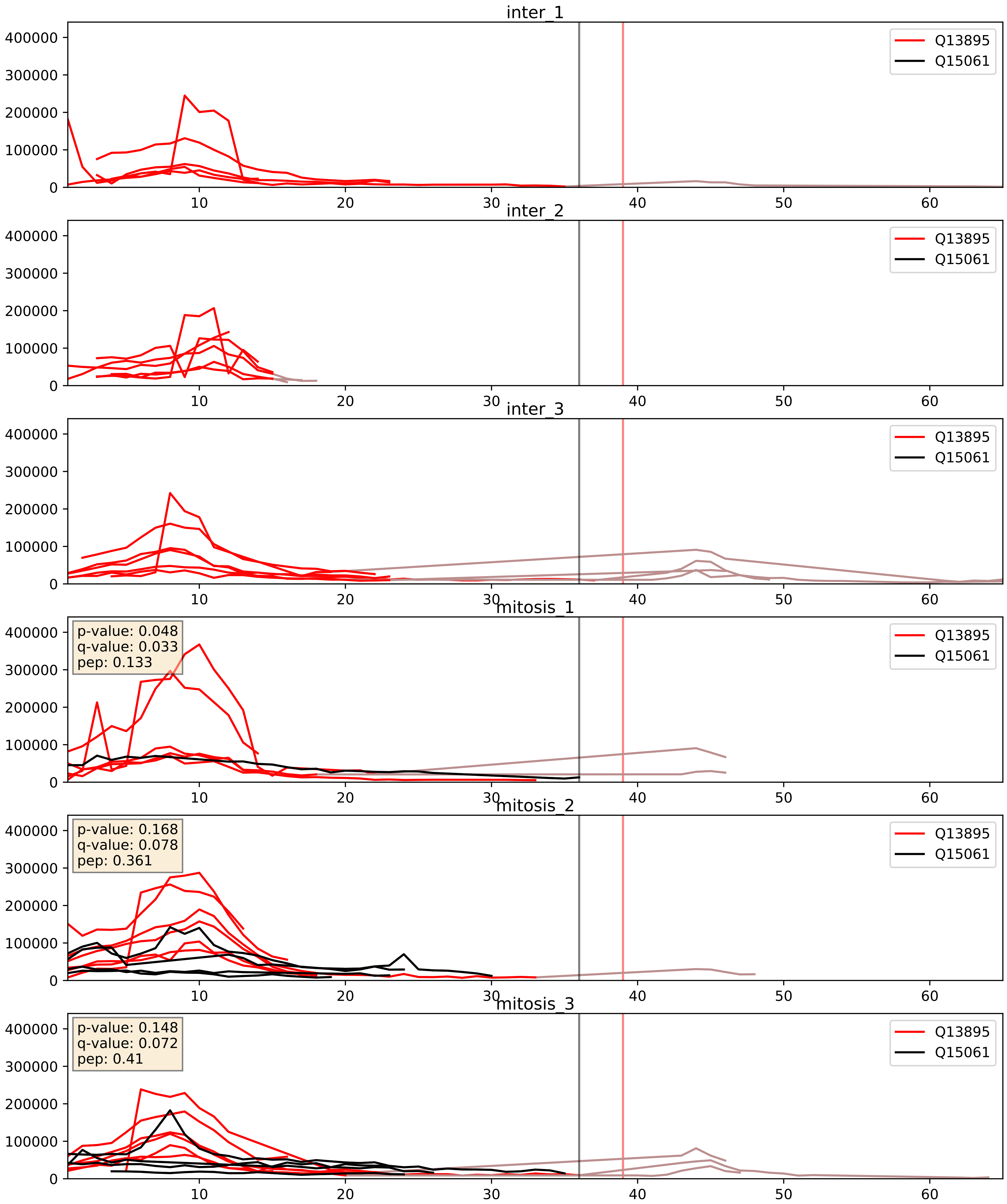


| level | condition_1 | condition_2 | pvalue | pvalue_adjusted |
| --- | --- | --- | --- | --- |
| interactor_abundance | mitosis | inter | 0.0006377390807793674 | 0.01786921941561828 |
| complex_abundance | mitosis | inter | 0.004882908275355024 | 0.052397293069493756 |
| interactor_ratio | mitosis | inter | 0.022909515726422306 | 0.2336681581804622 |

Q13895_Q2NL82
BYST_HUMAN vs TSR1_HUMAN
p-value: 0.08 q-value: 0.048

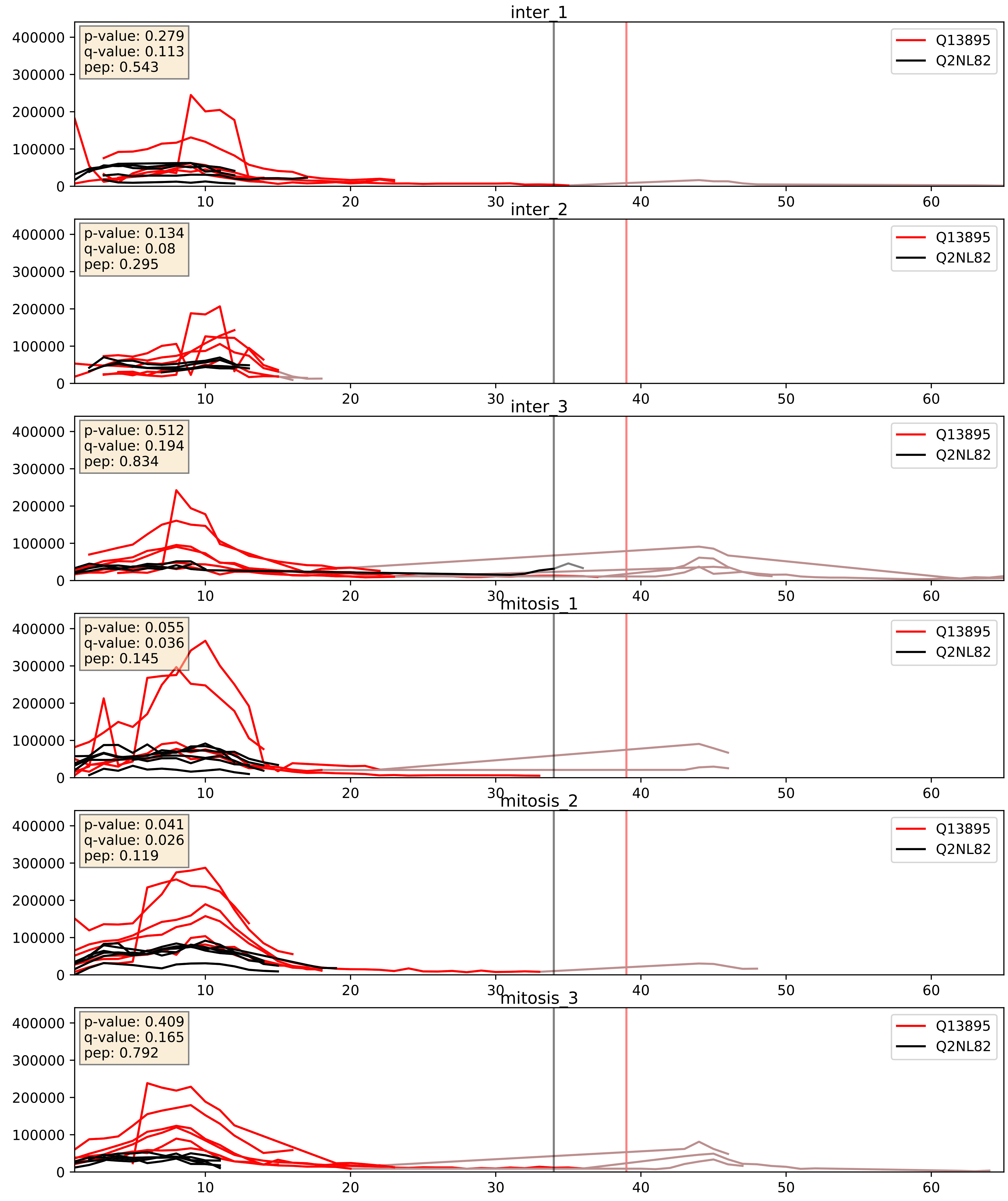


| level | condition_1 | condition_2 | pvalue | pvalue_adjusted |
|---|---|---|---|---|
| interactor_ratio | mitosis | inter | 0.009503652718623835 | 0.13913572462569762 |
| interactor_abundance | mitosis | inter | 0.0794137153007989 | 0.23432657806785198 |
| complex_abundance | mitosis | inter | 0.42223084636112357 | 0.5656864910932369 |

Q13895_Q96GA3
BYST_HUMAN vs LTV1_HUMAN
p-value: 0.001 q-value: 0.001

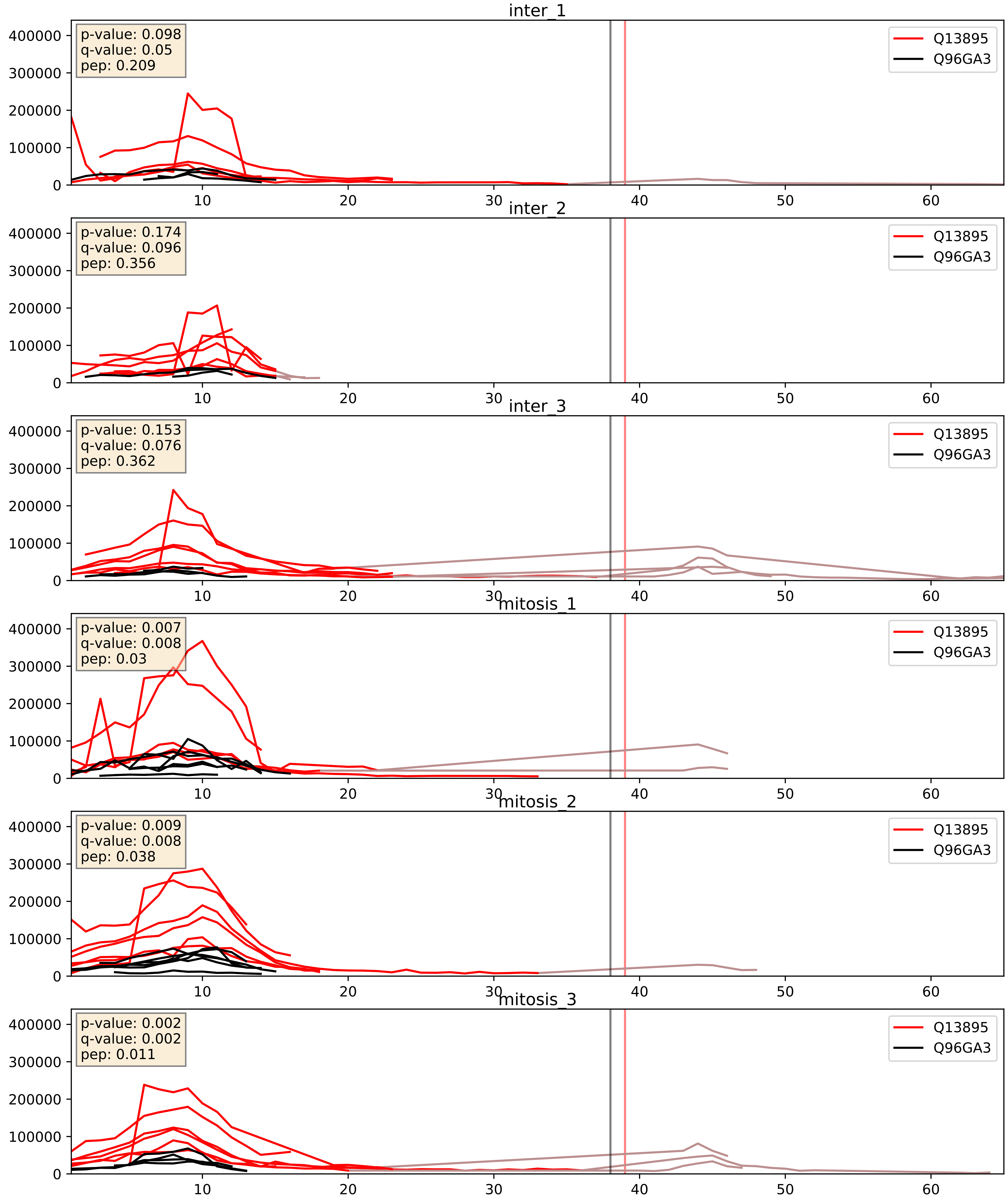


| level | condition_1 | condition_2 | pvalue | pvalue_adjusted |
|---|---|---|---|---|
| interactor_abundance | mitosis | inter | 0.033531541587113765 | 0.1528788260909155 |
| interactor_ratio | mitosis | inter | 0.04805774040431904 | 0.361488803041275 |
| complex_abundance | mitosis | inter | 0.2870108843701452 | 0.4354969066593979 |
