## Supplementary Data 2 for "SECAT: Quantifying differential protein-protein interaction states by network-centric analysis": hela_string_000163_O75934.pdf

O75934_O75934

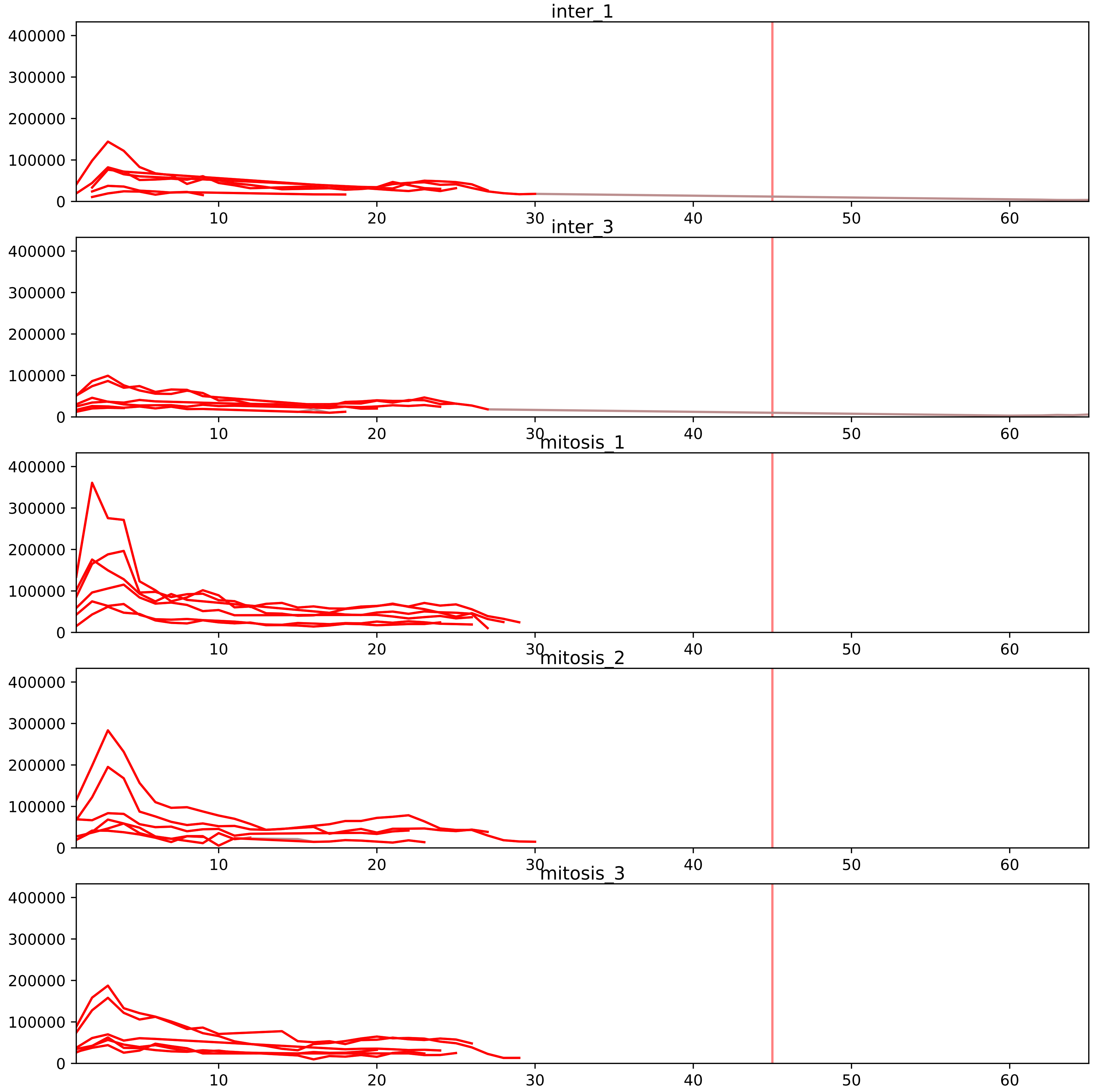


| level | condition_1 | condition_2 | pvalue | pvalue_adjusted |
|---|---|---|---|---|
| interactor_ratio | mitosis | inter | 8.151639571244083e-05 | 0.0014284777915322963 |
| assembled_abundance | mitosis | inter | 0.0013184327918779868 | 0.0335412168412003 |
| total_abundance | mitosis | inter | 0.0014875504308782847 | 0.04138316526558113 |
| complex_abundance | mitosis | inter | 0.0016745199648677198 | 0.01495687735610002 |
| interactor_abundance | mitosis | inter | 0.00256169278826435 | 0.022552702059360785 |
| monomer_abundance | mitosis | inter | 0.030501027006072094 | 0.2978982517444012 |

O43395_O75934
PRPF3_HUMAN vs SPF27_HUMAN
p-value: 0.017 q-value: 0.014

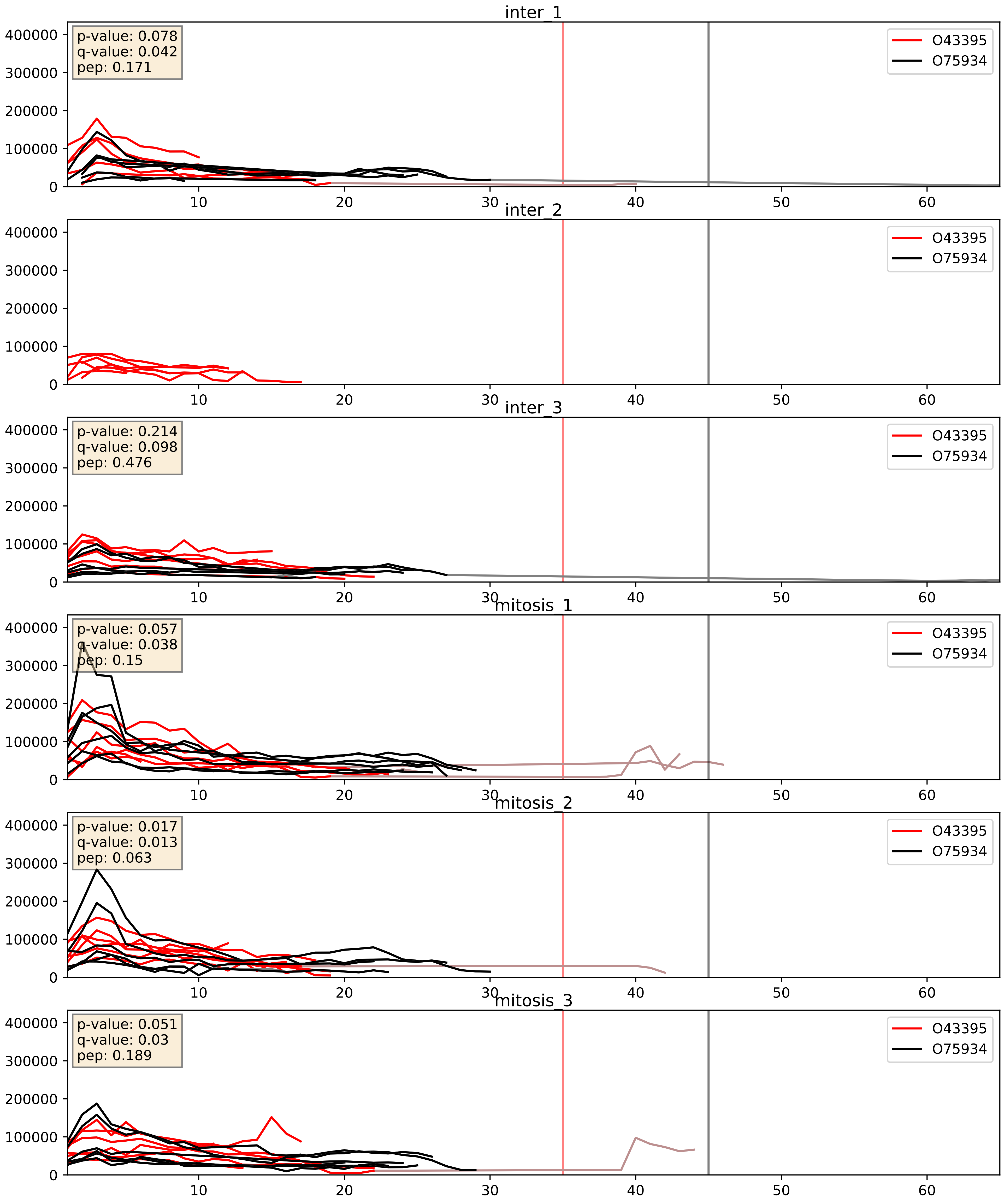


| level | condition_1 | condition_2 | pvalue | pvalue_adjusted |
|---|---|---|---|---|
| complex_abundance | mitosis | inter | 0.1851475550568205 | 0.336615734724054 |
| interactor_abundance | mitosis | inter | 0.29173386071592 | 0.4862448032696506 |
| interactor_ratio | mitosis | inter | 0.3146050758170687 | 0.7783421708721073 |

O43660_O75934
PLRG1_HUMAN vs SPF27_HUMAN
p-value: 0.064 q-value: 0.04

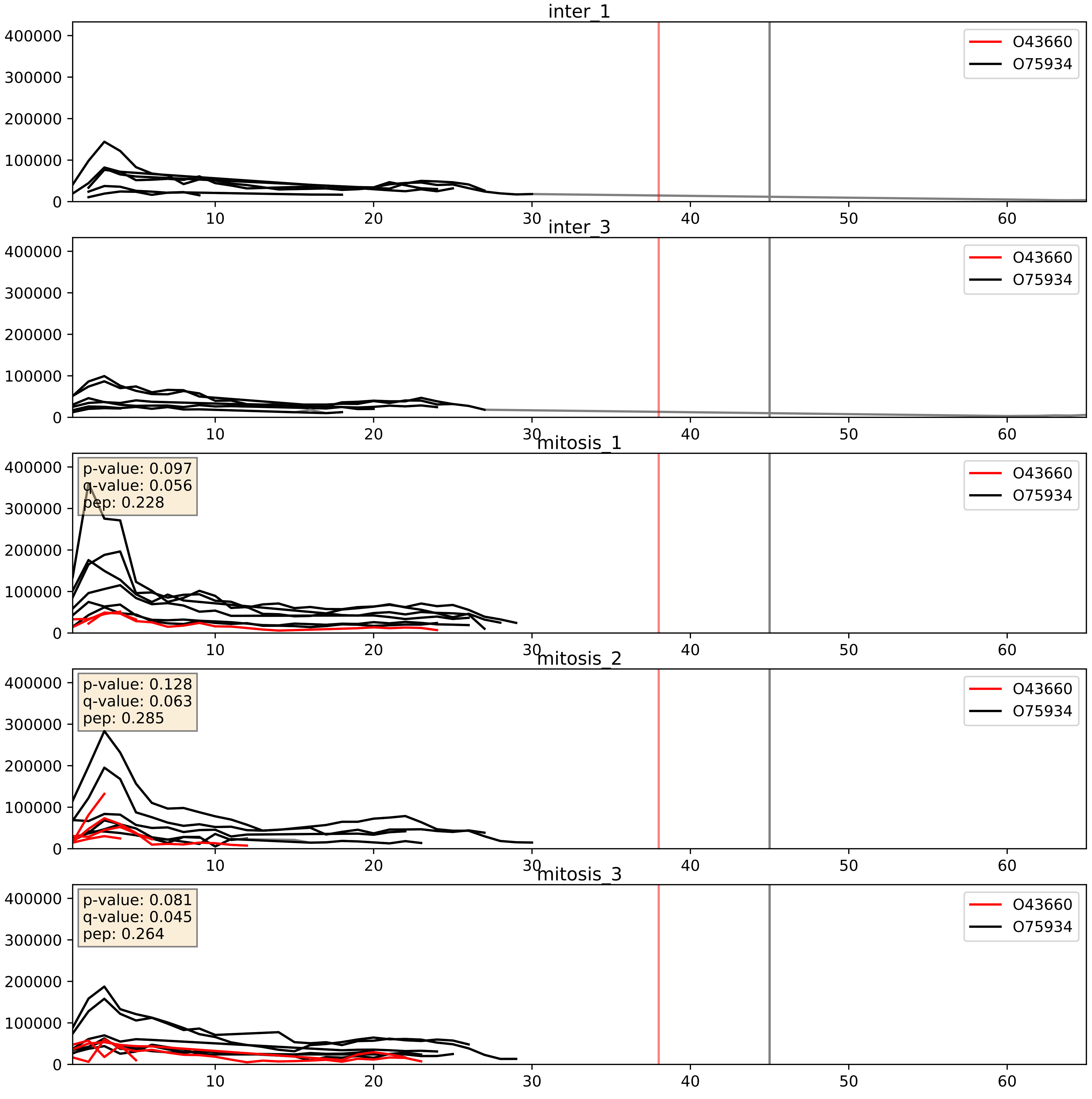


| level | condition_1 | condition_2 | pvalue | pvalue_adjusted |
|---|---|---|---|---|
| interactor_ratio | mitosis | inter | 0.0017960205665454114 | 0.051461731073100085 |
| complex_abundance | mitosis | inter | 0.0030660571959431095 | 0.040627630955530986 |
| interactor_abundance | mitosis | inter | 0.03668661671701946 | 0.15802829520754763 |

O75934_P62995
SPF27_HUMAN vs TRA2B_HUMAN
p-value: 0.004 q-value: 0.047

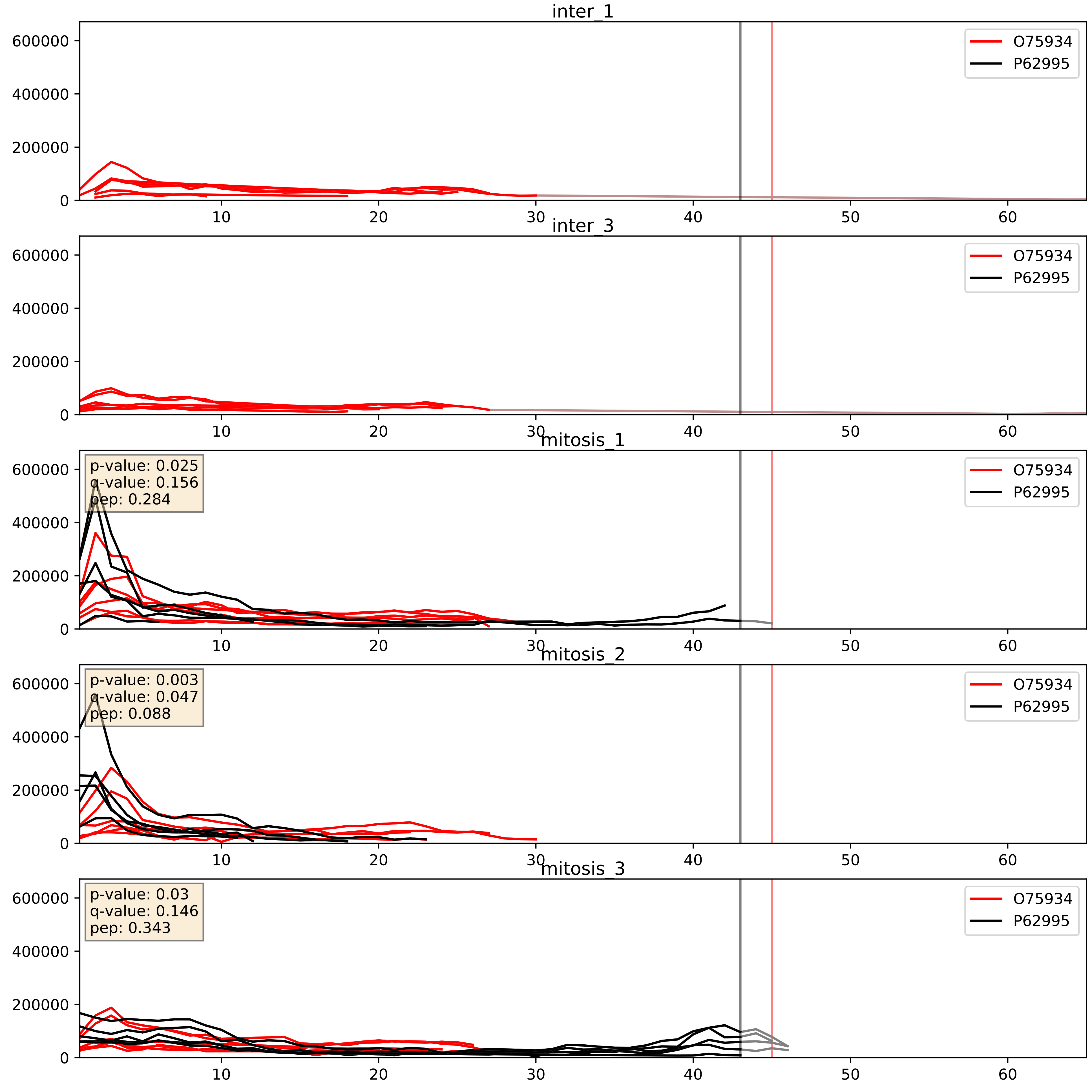


| level | condition_1 | condition_2 | pvalue | pvalue_adjusted |
|---|---|---|---|---|
| complex_abundance | mitosis | inter | 0.000556330786787743 | 0.01507022637627557 |
| interactor_abundance | mitosis | inter | 0.0007609849117950717 | 0.020324589219862133 |
| interactor_ratio | mitosis | inter | 0.23601861432310012 | 0.7155827200888129 |

O75934_Q6UN15
SPF27_HUMAN vs FIP1_HUMAN
p-value: 0.0 q-value: 0.033

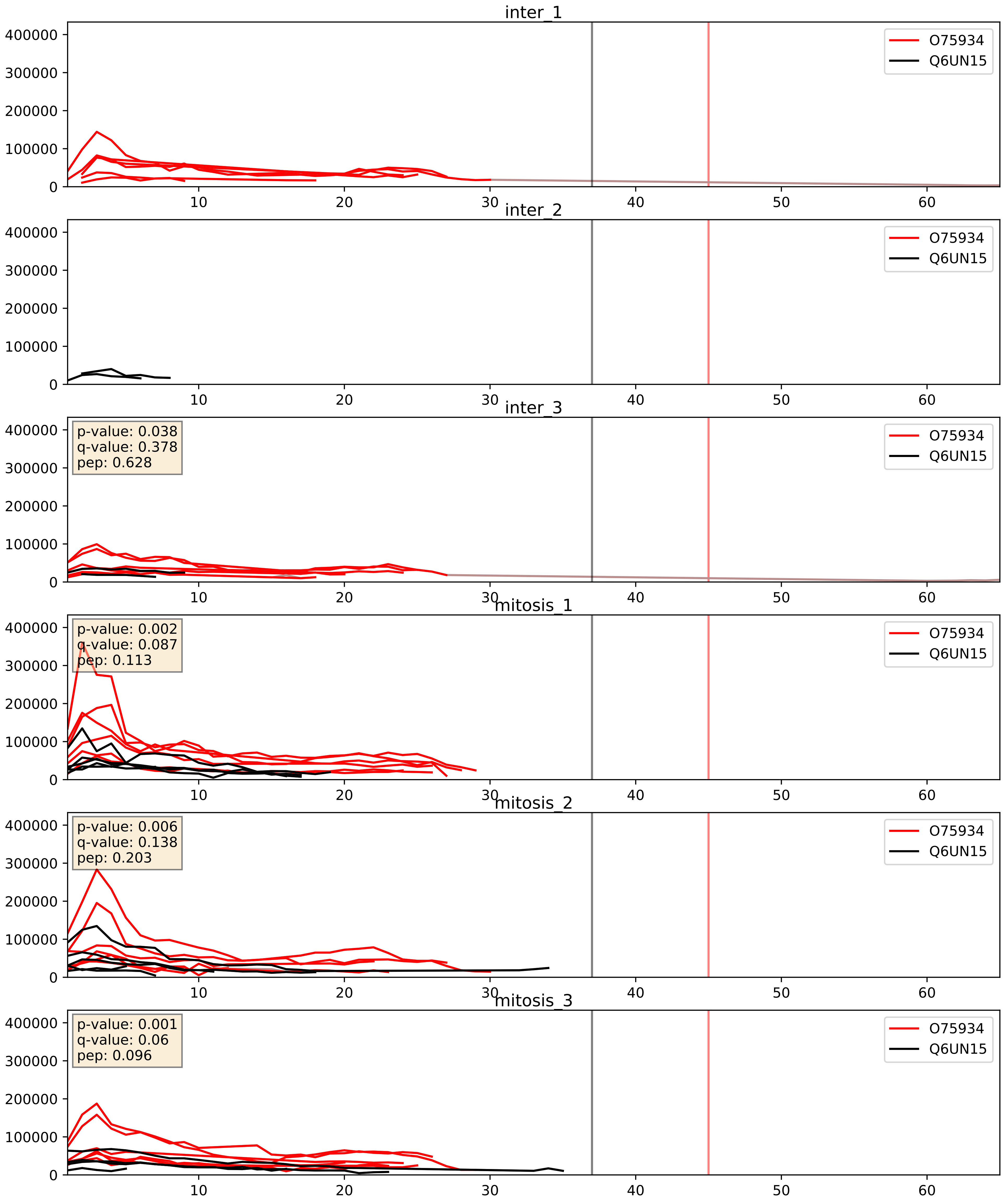


| level | condition_1 | condition_2 | pvalue | pvalue_adjusted |
|---|---|---|---|---|
| complex_abundance | mitosis | inter | 0.007783987211497894 | 0.06557020805367884 |
| interactor_abundance | mitosis | inter | 0.014772120850023046 | 0.10634933093035936 |
| interactor_ratio | mitosis | inter | 0.08024151017262679 | 0.47370160488116225 |

O75934_Q99459
SPF27_HUMAN vs CDC5L_HUMAN
p-value: 0.004 q-value: 0.004

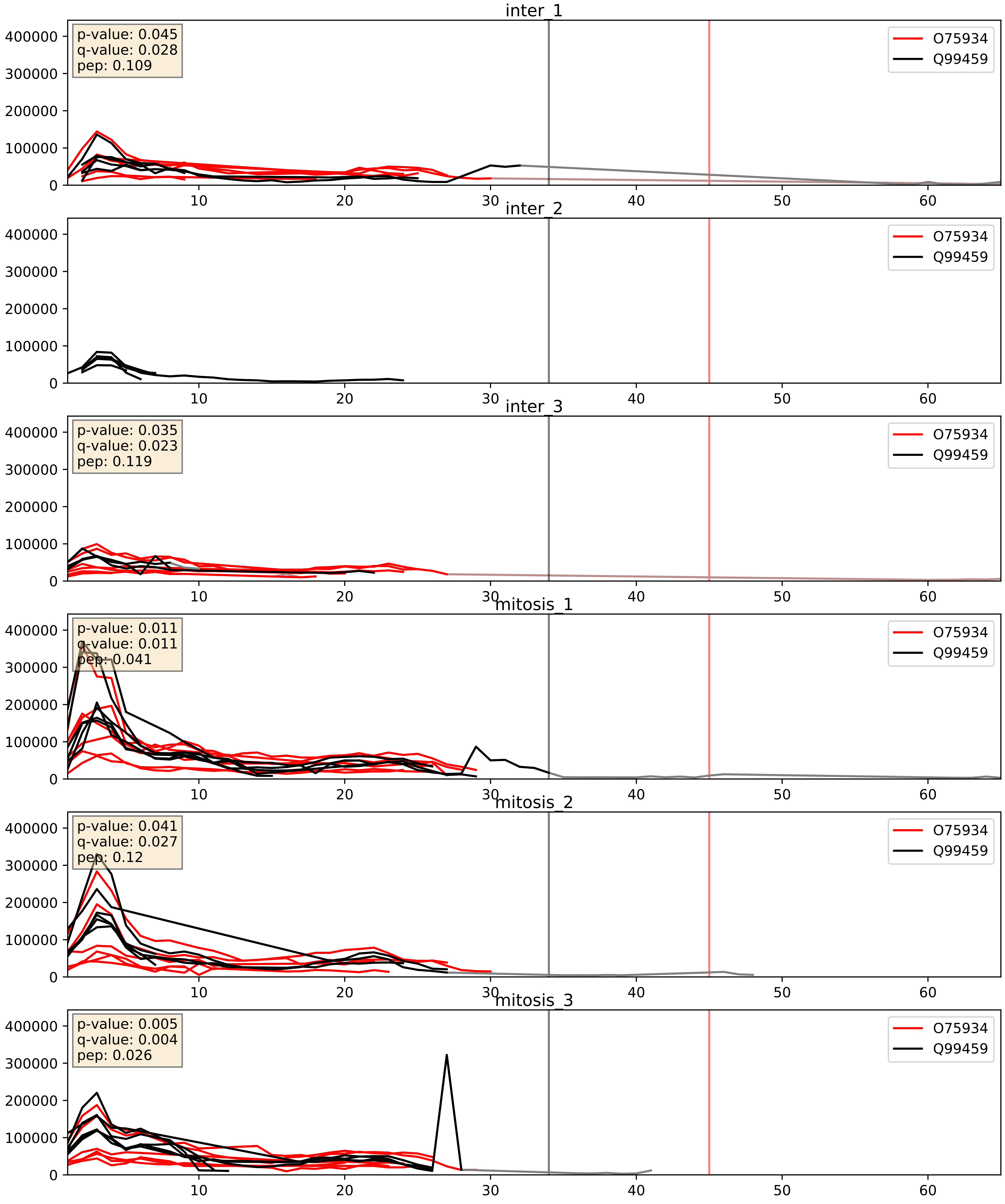


| level | condition_1 | condition_2 | pvalue | pvalue_adjusted |
|---|---|---|---|---|
| complex_abundance | mitosis | inter | 0.03282547885145826 | 0.13204233974082835 |
| interactor_abundance | mitosis | inter | 0.10154606889418748 | 0.27370850762629345 |
| interactor_ratio | mitosis | inter | 0.14845822739515527 | 0.60240057967345 |

O75934_Q9BZJ0
SPF27_HUMAN vs CRNL1_HUMAN
p-value: 0.022 q-value: 0.017

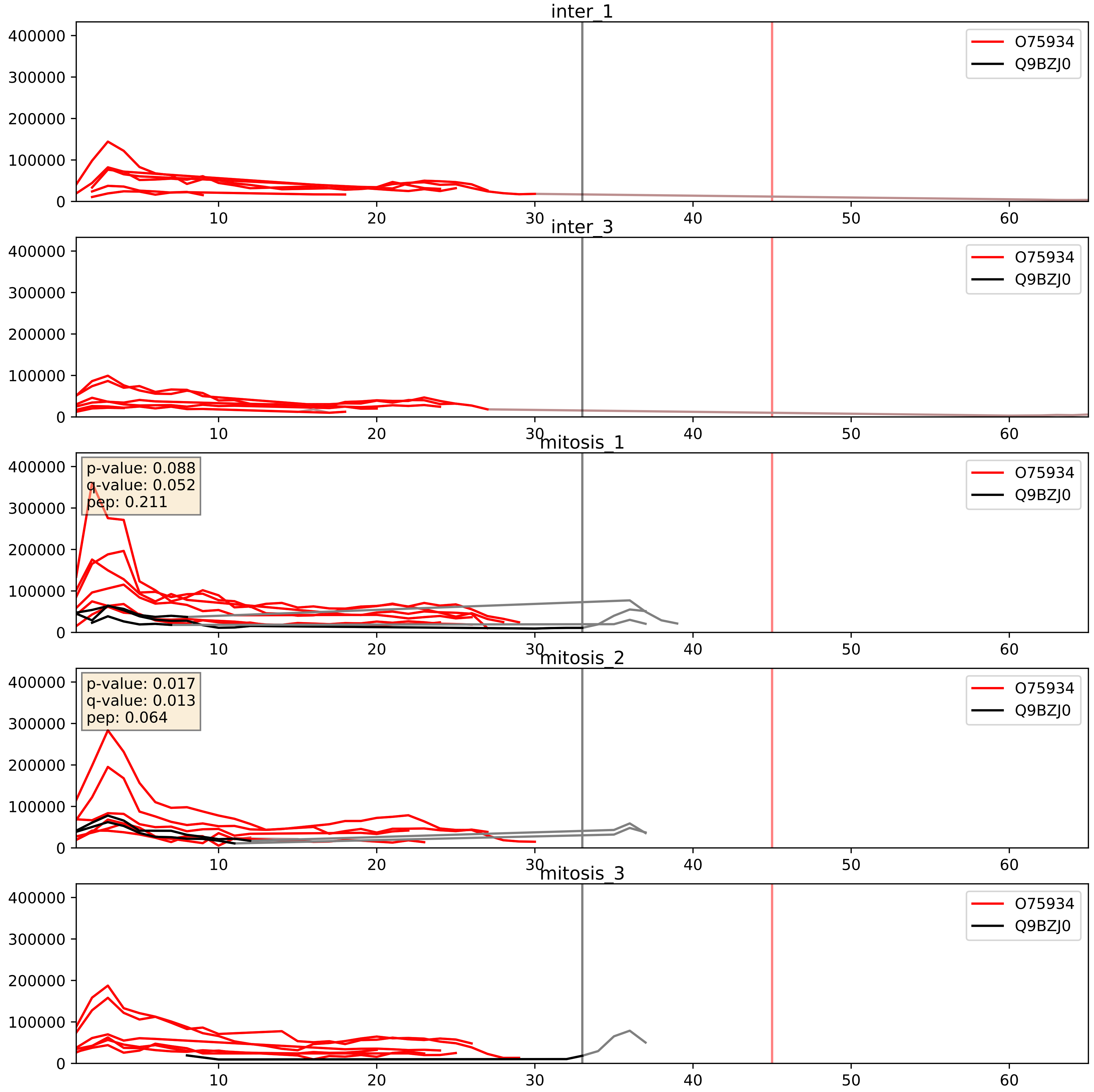


| level | condition_1 | condition_2 | pvalue | pvalue_adjusted |
|---|---|---|---|---|
| interactor_ratio | mitosis | inter | 0.014619333770107596 | 0.17725424514464733 |
| interactor_abundance | mitosis | inter | 0.12392143906875966 | 0.30662451753969727 |
| complex_abundance | mitosis | inter | 0.24775694395957115 | 0.39823025501804077 |

O75934_Q9UNP9
SPF27_HUMAN vs PPIE_HUMAN
p-value: 0.04 q-value: 0.027

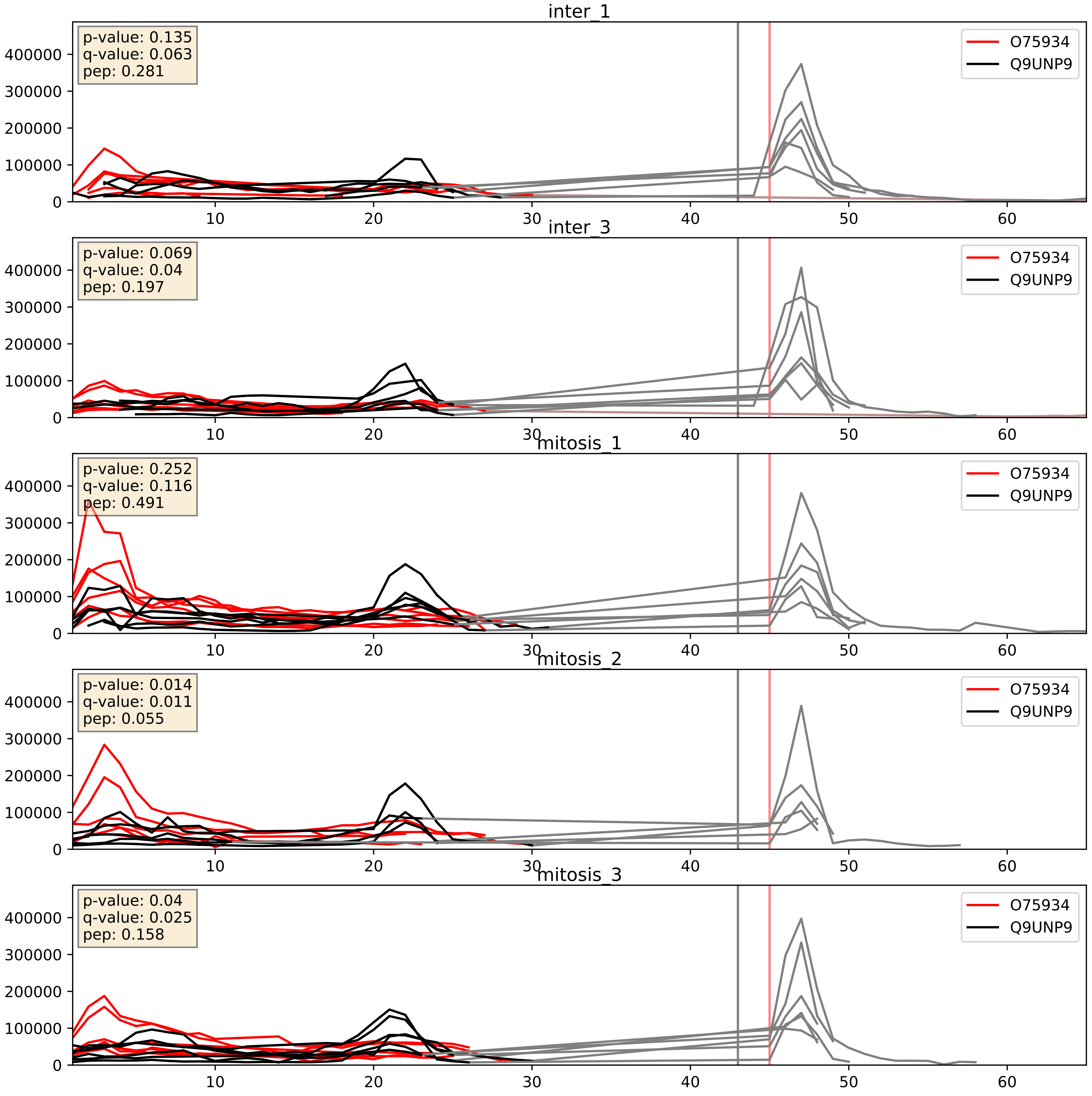


| level | condition_1 | condition_2 | pvalue | pvalue_adjusted |
|---|---|---|---|---|
| interactor_ratio | mitosis | inter | 0.01851041841948883 | 0.2047641739035816 |
| interactor_abundance | mitosis | inter | 0.13122128086562018 | 0.31618695685002357 |
| complex_abundance | mitosis | inter | 0.18995792099545952 | 0.34153325009895685 |
