## Supplementary Data 2 for "SECAT: Quantifying differential protein-protein interaction states by network-centric analysis": hela_string_000164_Q8NHZ8.pdf

Q8NHZ8_Q8NHZ8

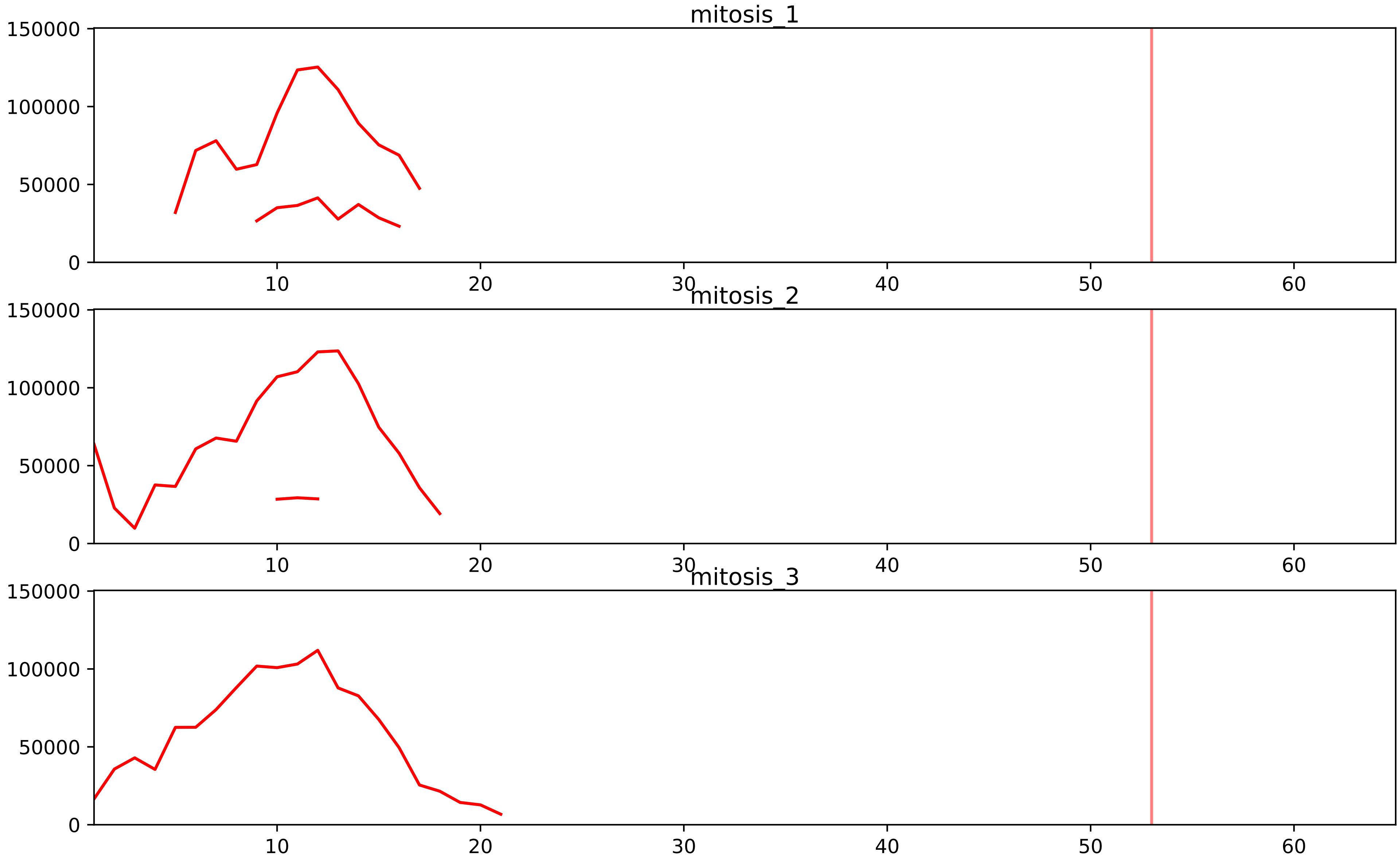


| level | condition_1 | condition_2 | pvalue | pvalue_adjusted |
|---|---|---|---|---|
| complex_abundance | mitosis | inter | 8.211499831629189e-05 | 0.001988047327657593 |
| total_abundance | mitosis | inter | 0.003978999975876761 | 0.07208928426045762 |
| assembled_abundance | mitosis | inter | 0.003994827615650287 | 0.06220699778667183 |
| interactor_abundance | mitosis | inter | 0.00467381417075401 | 0.03259526752575256 |
| interactor_ratio | mitosis | inter | 0.017797944183518773 | 0.09519830609789111 |

Q8NHZ8_Q96DE5
CDC26_HUMAN vs APC16_HUMAN
p-value: 0.0 q-value: 0.0

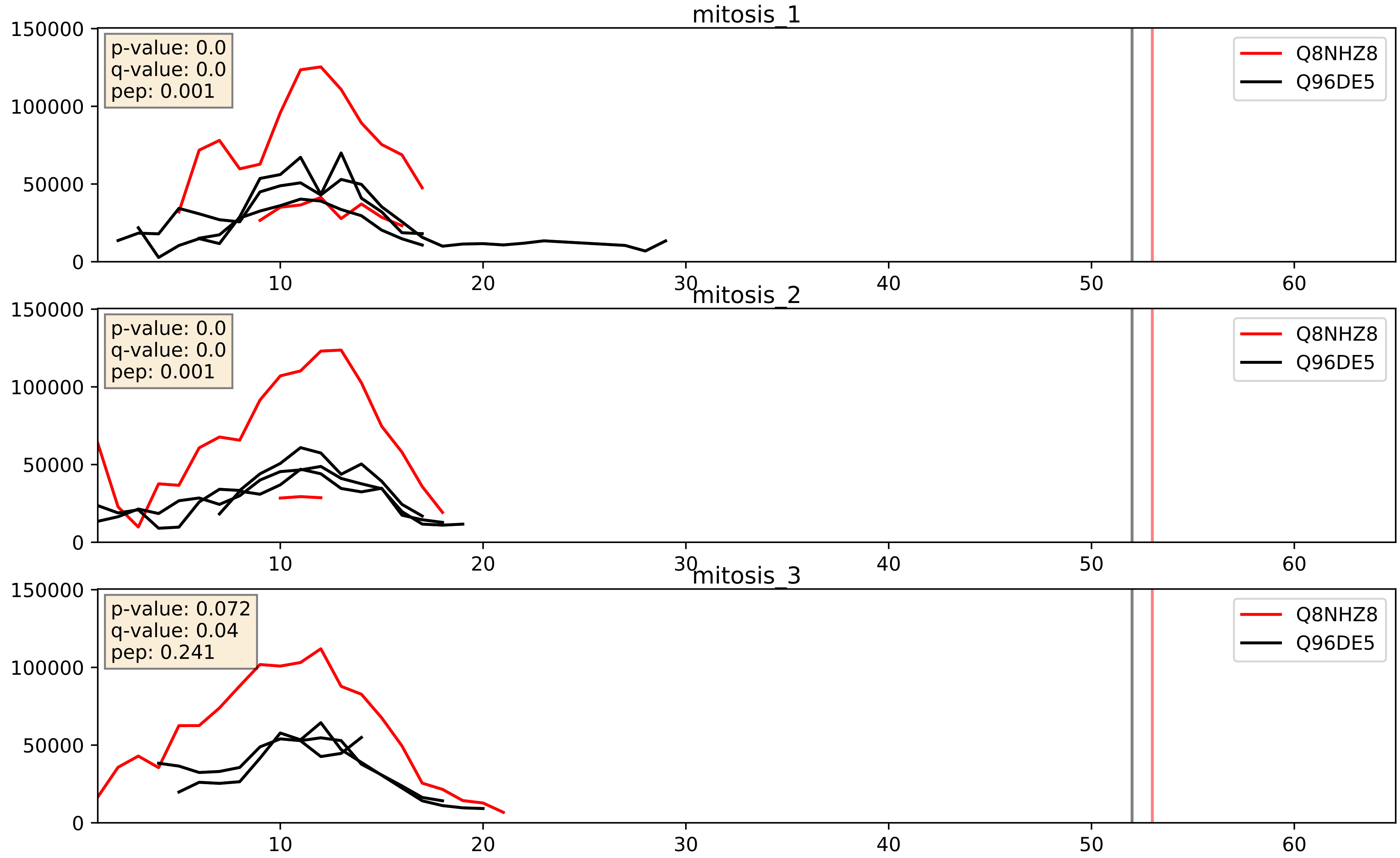


| level | condition_1 | condition_2 | pvalue | pvalue_adjusted |
|---|---|---|---|---|
| complex_abundance | mitosis | inter | 0.010729792027556192 | 0.07672534067869605 |
| interactor_ratio | mitosis | inter | 0.022311319891476004 | 0.2287907703572271 |
| interactor_abundance | mitosis | inter | 0.023574696254356885 | 0.12607602513772825 |

Q8NHZ8_Q9H1A4
CDC26_HUMAN vs APC1_HUMAN
p-value: 0.039 q-value: 0.027

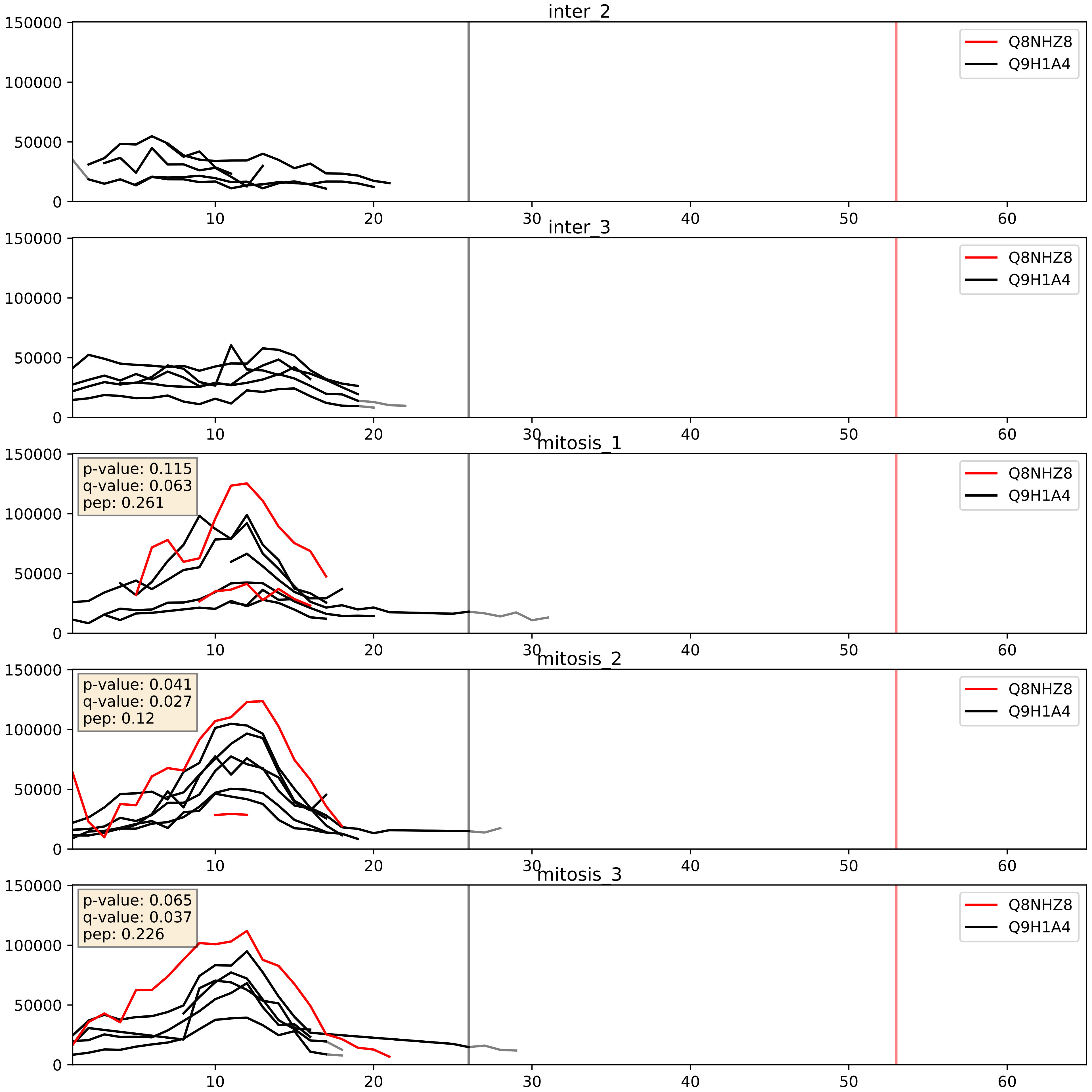


| level | condition_1 | condition_2 | pvalue | pvalue_adjusted |
|---|---|---|---|---|
| complex_abundance | mitosis | inter | 0.0013512642250712022 | 0.025090719667265705 |
| interactor_abundance | mitosis | inter | 0.02056550834926251 | 0.12021882713294628 |
| interactor_ratio | mitosis | inter | 0.07821558900815949 | 0.46754569965771314 |

Q8NHZ8_Q9UJX2
CDC26_HUMAN vs CDC23_HUMAN
p-value: 0.004 q-value: 0.004

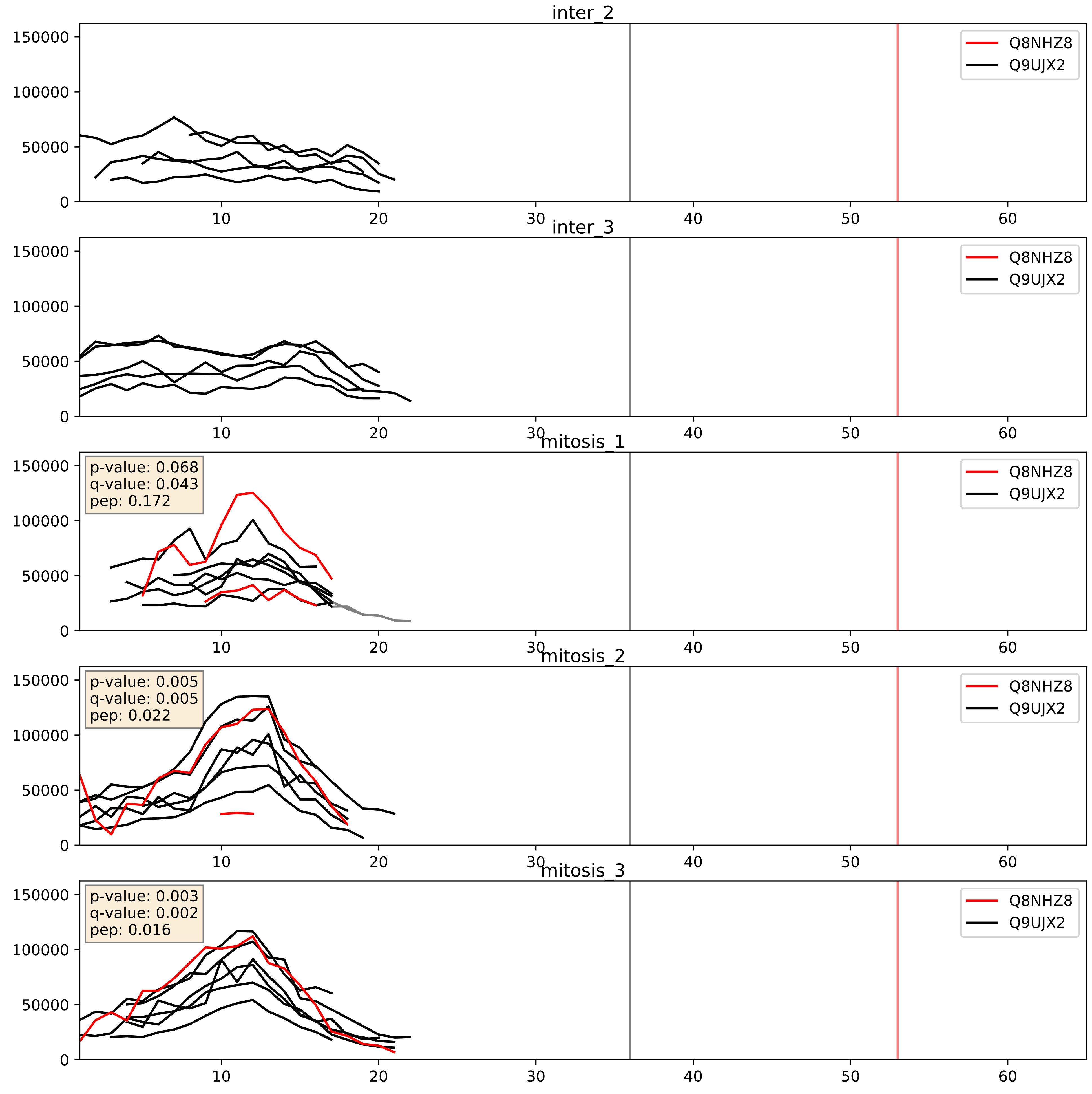


| level | condition_1 | condition_2 | pvalue | pvalue_adjusted |
|---|---|---|---|---|
| complex_abundance | mitosis | inter | 0.00046975875910792607 | 0.013771010198506326 |
| interactor_abundance | mitosis | inter | 0.01883033144290343 | 0.11808618106318929 |
| interactor_ratio | mitosis | inter | 0.019561537689939432 | 0.21168996539302345 |

Q8NHZ8_Q9UJX3
CDC26_HUMAN vs APC7_HUMAN
p-value: 0.02 q-value: 0.016

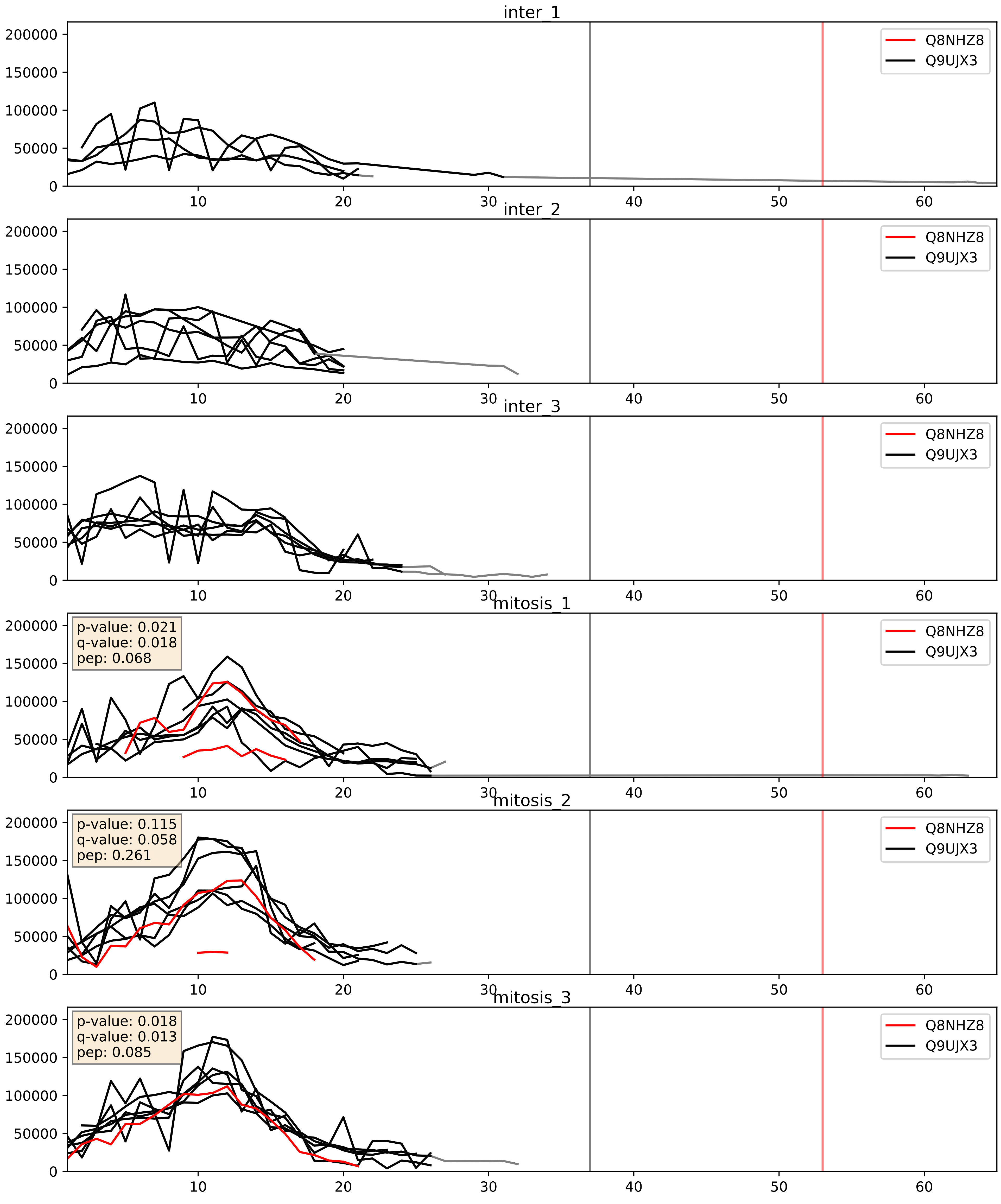


| level | condition_1 | condition_2 | pvalue | pvalue_adjusted |
|---|---|---|---|---|
| complex_abundance | mitosis | inter | 0.0002094786952569282 | 0.009840811891188612 |
| interactor_ratio | mitosis | inter | 0.0068537415293100255 | 0.11503534802136042 |
| interactor_abundance | mitosis | inter | 0.01883033144290343 | 0.11808618106318929 |

Q8NHZ8_Q9UJX4
CDC26_HUMAN vs APC5_HUMAN
p-value: 0.026 q-value: 0.02

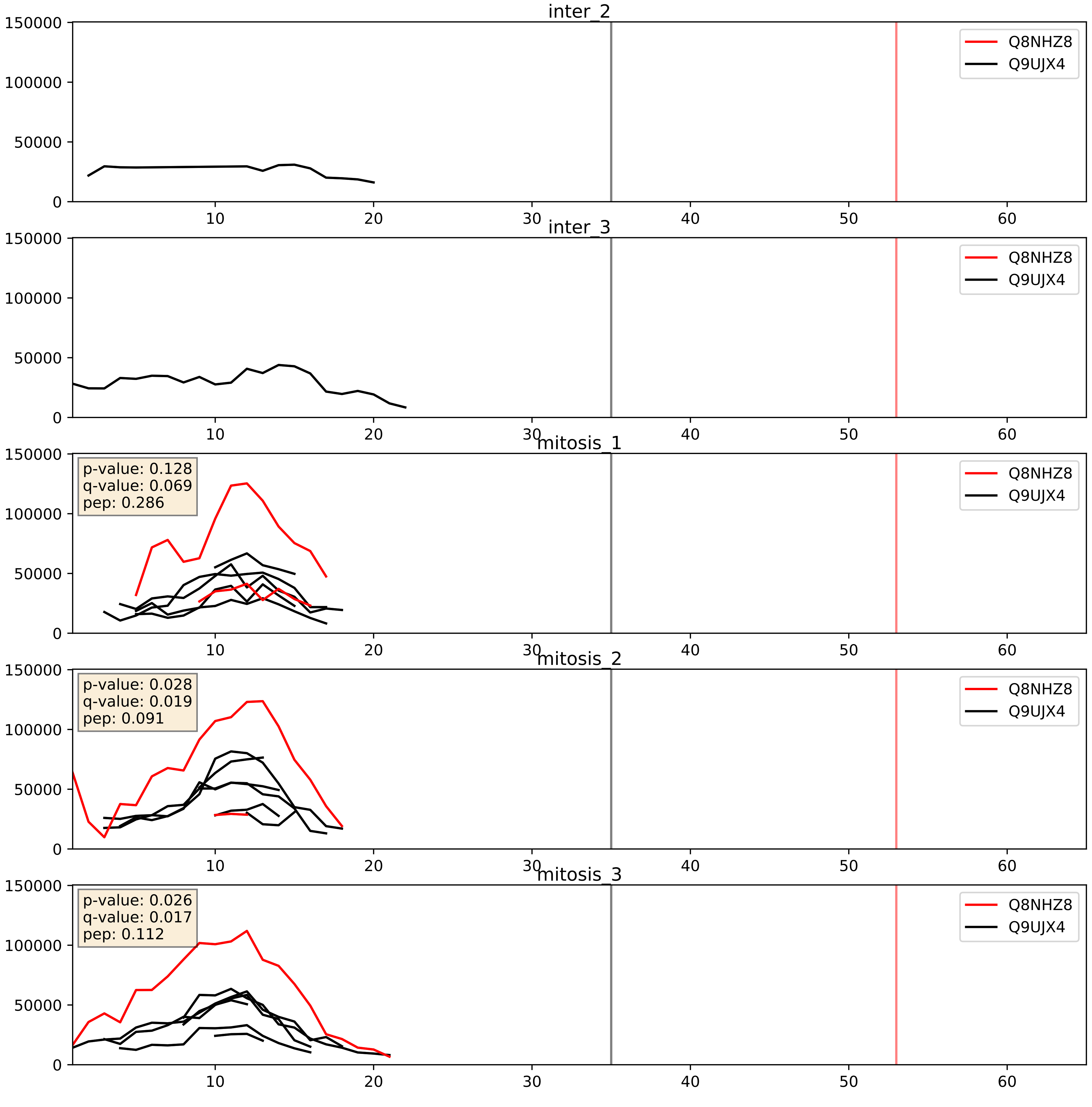


| level | condition_1 | condition_2 | pvalue | pvalue_adjusted |
|---|---|---|---|---|
| complex_abundance | mitosis | inter | 0.004557678589736247 | 0.0503402951330868 |
| interactor_abundance | mitosis | inter | 0.02065482954399308 | 0.12021882713294628 |
| interactor_ratio | mitosis | inter | 0.07564477353850585 | 0.4628166812754875 |

Q8NHZ8_Q9UJX5
CDC26_HUMAN vs APC4_HUMAN
p-value: 0.035 q-value: 0.025

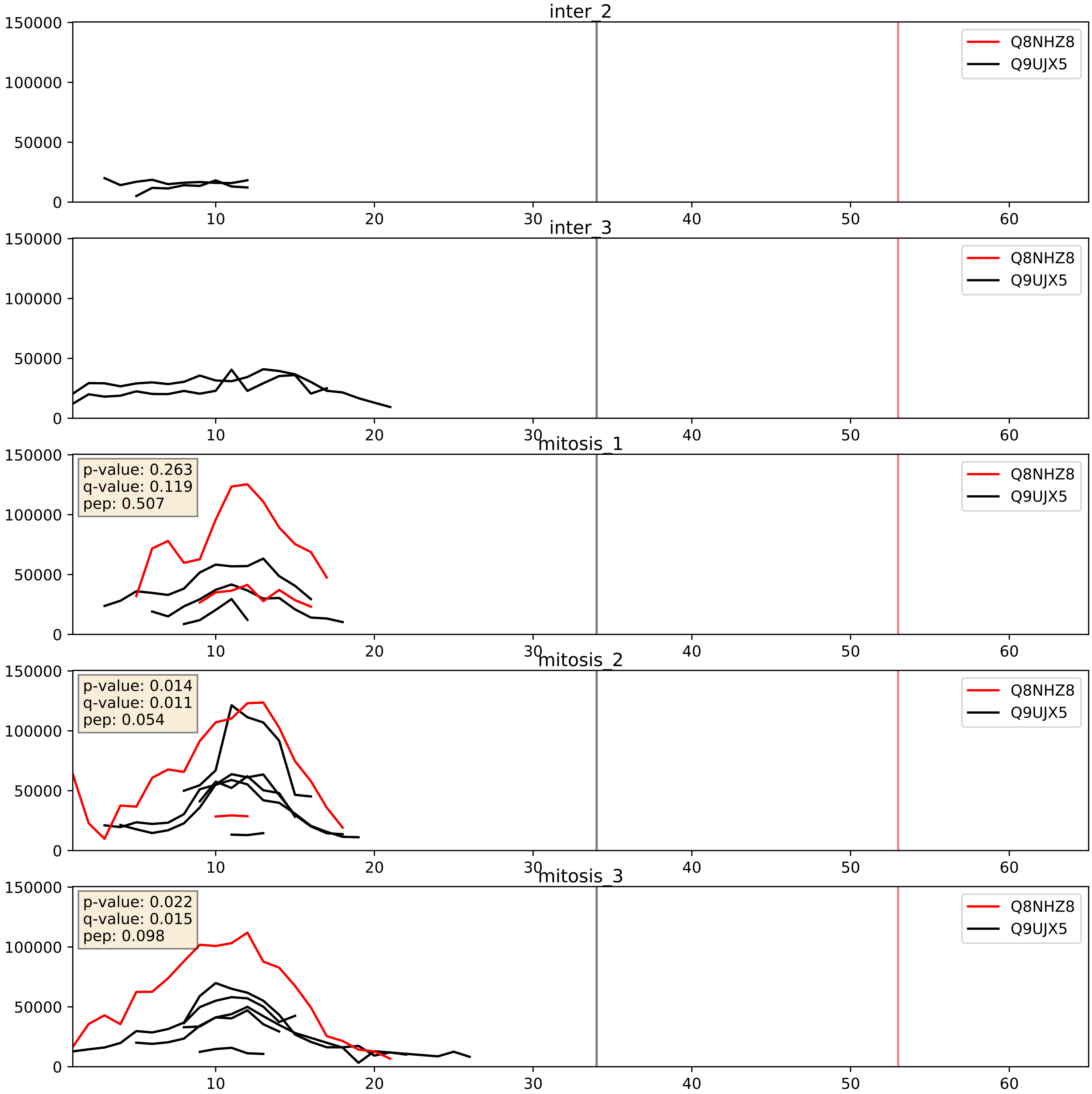


| level | condition_1 | condition_2 | pvalue | pvalue_adjusted |
|---|---|---|---|---|
| complex_abundance | mitosis | inter | 0.004857379598774094 | 0.05236671204723708 |
| interactor_abundance | mitosis | inter | 0.02065482954399308 | 0.12021882713294628 |
| interactor_ratio | mitosis | inter | 0.22523680336602545 | 0.7044307770599847 |

Q8NHZ8_Q9UJX6
CDC26_HUMAN vs ANC2_HUMAN
p-value: 0.028 q-value: 0.021

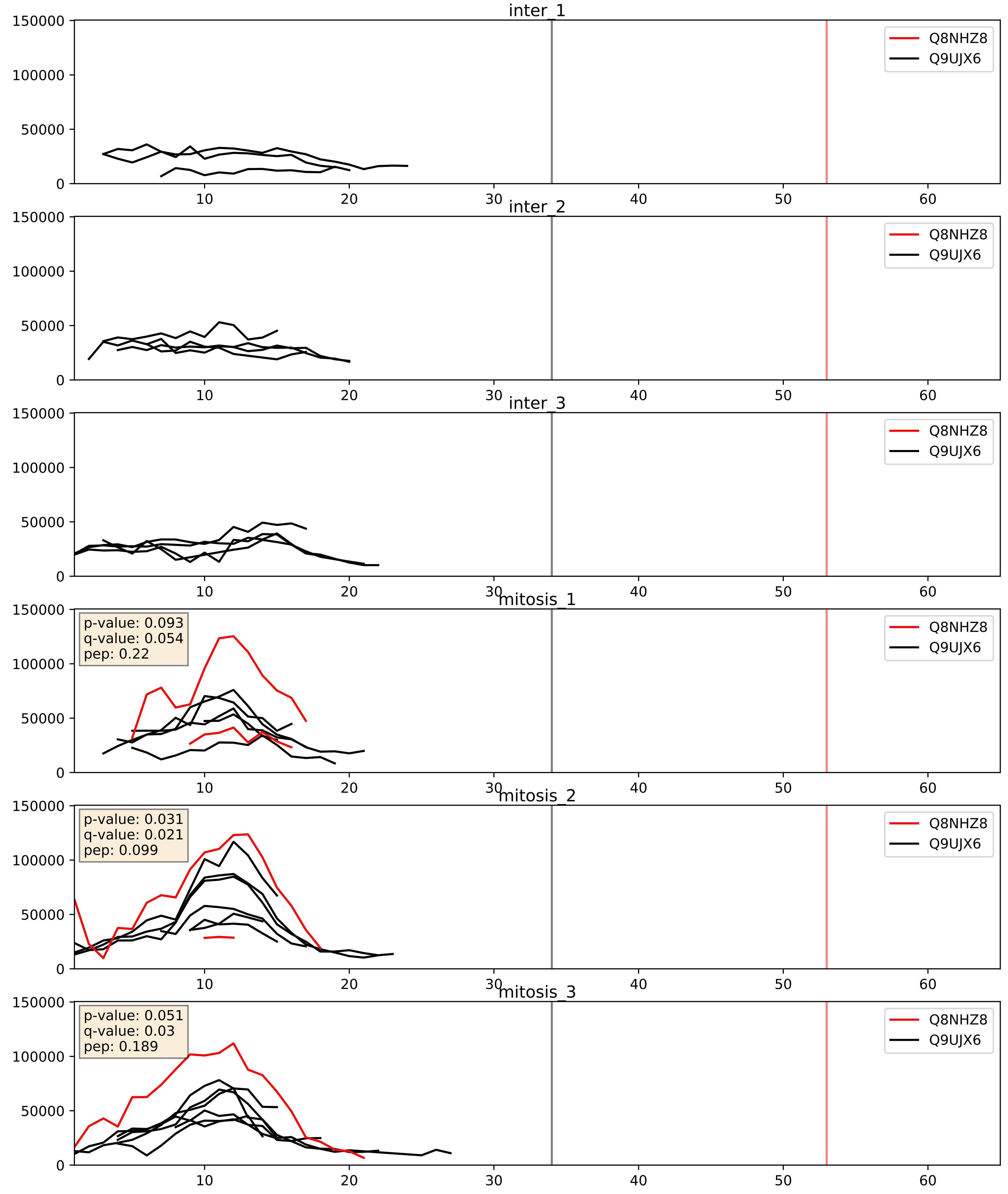


| level | condition_1 | condition_2 | pvalue | pvalue_adjusted |
|---|---|---|---|---|
| complex_abundance | mitosis | inter | 0.0013657306472519182 | 0.02519537573378539 |
| interactor_abundance | mitosis | inter | 0.01883033144290343 | 0.11808618106318929 |
| interactor_ratio | mitosis | inter | 0.029866466192683988 | 0.27401602423298493 |

Q8NHZ8_Q9UM13
CDC26_HUMAN vs APC10_HUMAN
p-value: 0.068 q-value: 0.042

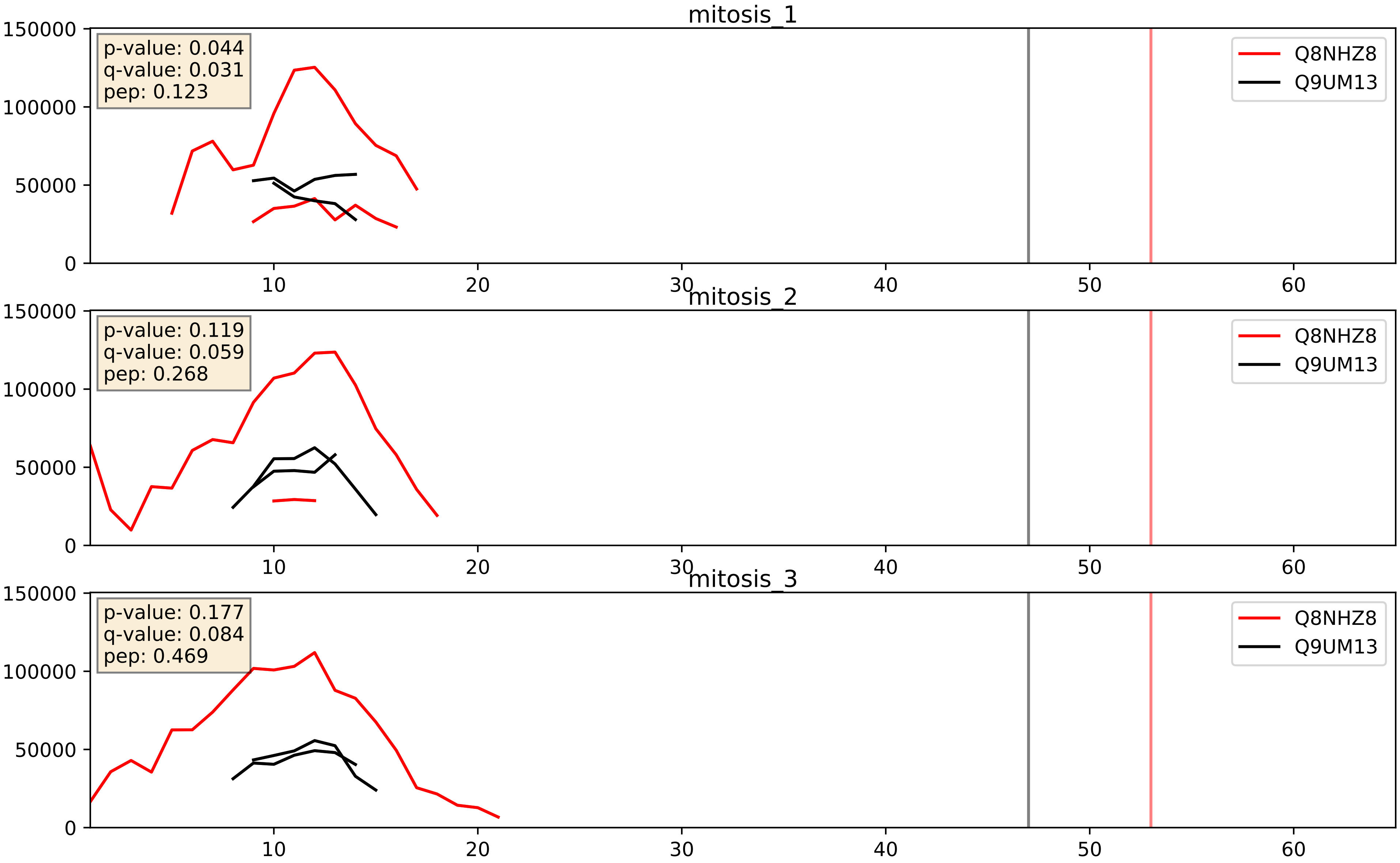


| level | condition_1 | condition_2 | pvalue | pvalue_adjusted |
|---|---|---|---|---|
| complex_abundance | mitosis | inter | 0.03319519889018759 | 0.1329671981750144 |
| interactor_abundance | mitosis | inter | 0.05065623838503444 | 0.18014848382878887 |
| interactor_ratio | mitosis | inter | 0.6435608944490359 | 0.9204479960708014 |
