## Supplementary Data 2 for "SECAT: Quantifying differential protein-protein interaction states by network-centric analysis": hela_string_000165_Q9Y5Y0.pdf

# Q9Y5Y0_Q9Y5Y0

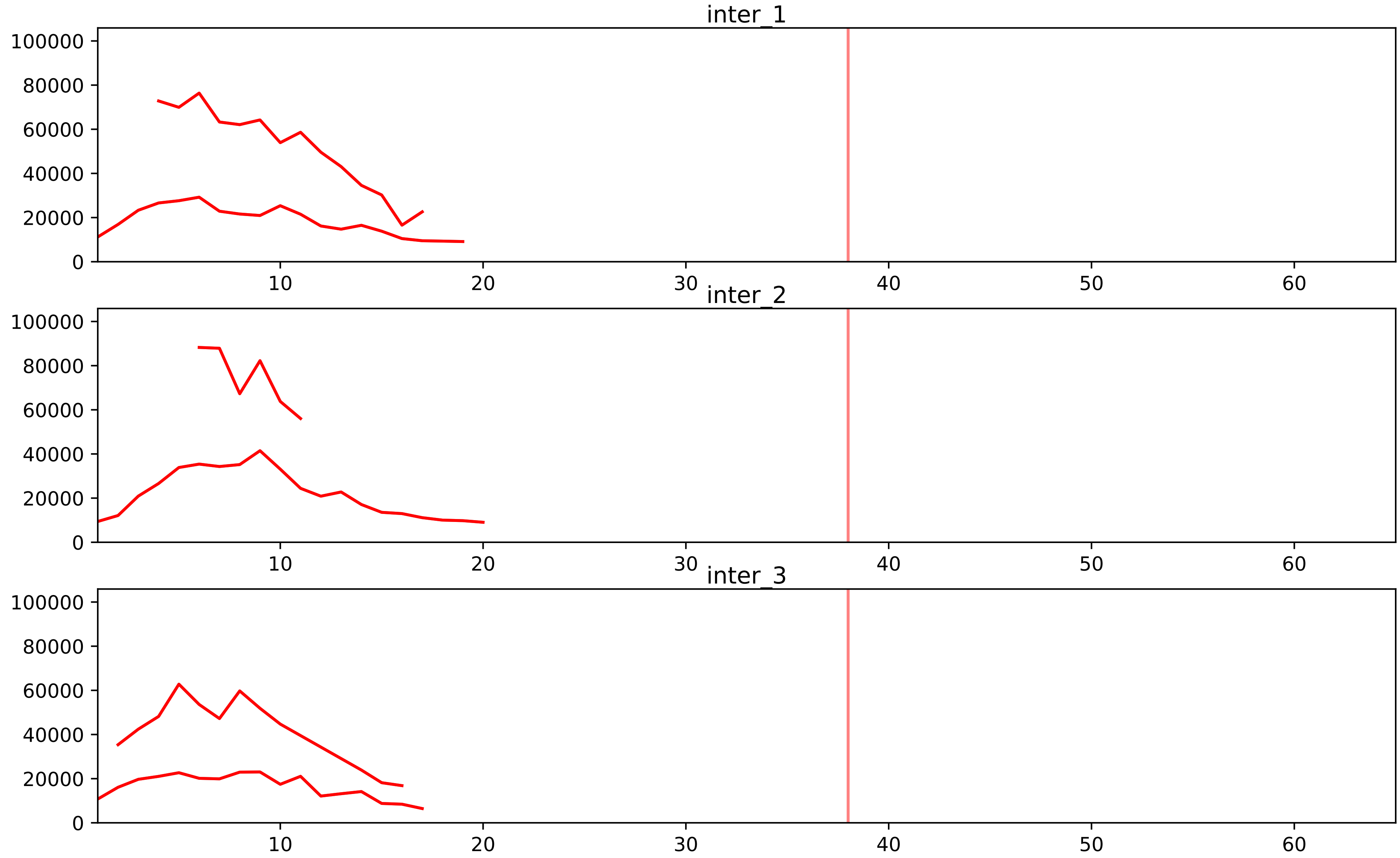


| level | condition_1 | condition_2 | pvalue | pvalue_adjusted |
|---|---|---|---|---|
| assembled_abundance | mitosis | inter | 8.454221036712893e-05 | 0.00899413833473933 |
| total_abundance | mitosis | inter | 0.00020576714506739365 | 0.017741165912121593 |
