## Supplementary Data 2 for "SECAT: Quantifying differential protein-protein interaction states by network-centric analysis": hela_string_000166_O43683.pdf

O43683_O43683

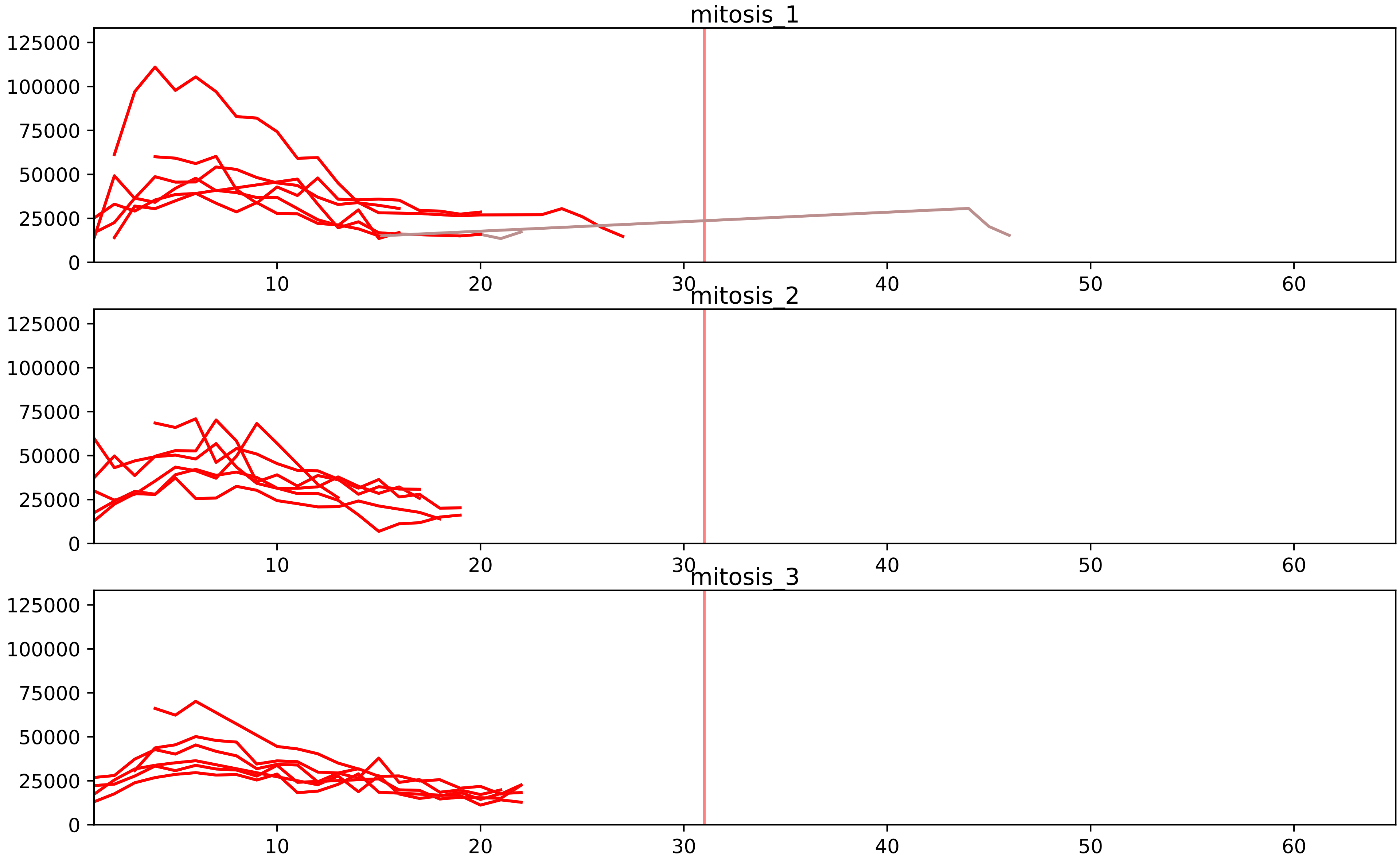


| level | condition_1 | condition_2 | pvalue | pvalue_adjusted |
|---|---|---|---|---|
| complex_abundance | mitosis | inter | 8.505514897849286e-05 | 0.0020324866768886606 |
| assembled_abundance | mitosis | inter | 0.0002595233590003358 | 0.016114995197788636 |
| total_abundance | mitosis | inter | 0.0005784952422312104 | 0.027952761634398485 |
| interactor_abundance | mitosis | inter | 0.0007631112715670559 | 0.009617292737557416 |
| interactor_ratio | mitosis | inter | 0.003468809370486694 | 0.02968655461253729 |

O43264_O43683
ZW10_HUMAN vs BUB1_HUMAN
p-value: 0.063 q-value: 0.039

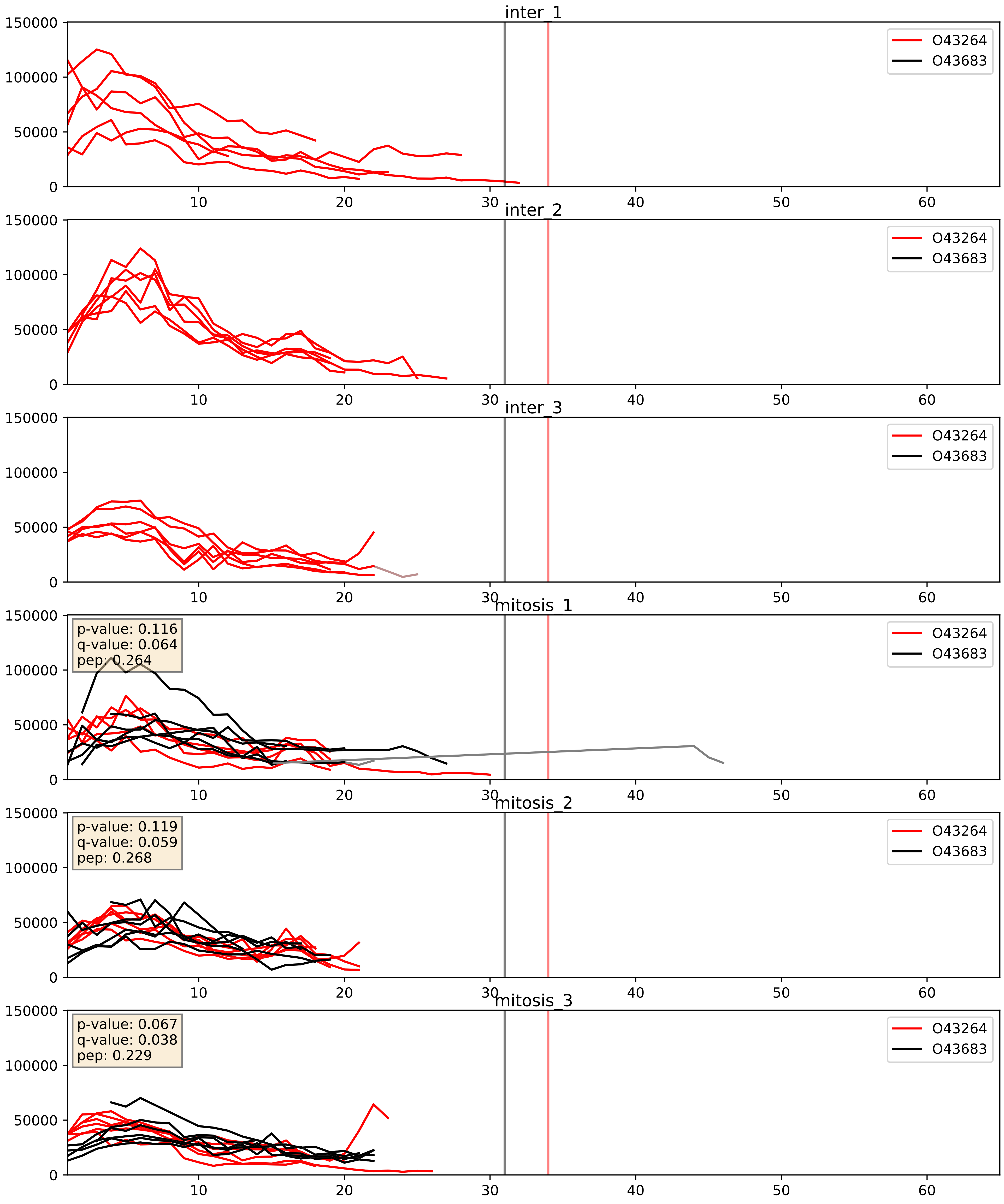


| level | condition_1 | condition_2 | pvalue | pvalue_adjusted |
|---|---|---|---|---|
| interactor_abundance | mitosis | inter | 0.0030695021355890283 | 0.042465703978082026 |
| complex_abundance | mitosis | inter | 0.003717228522607822 | 0.044942762928704735 |
| interactor_ratio | mitosis | inter | 0.129758286904364 | 0.5779454961622694 |

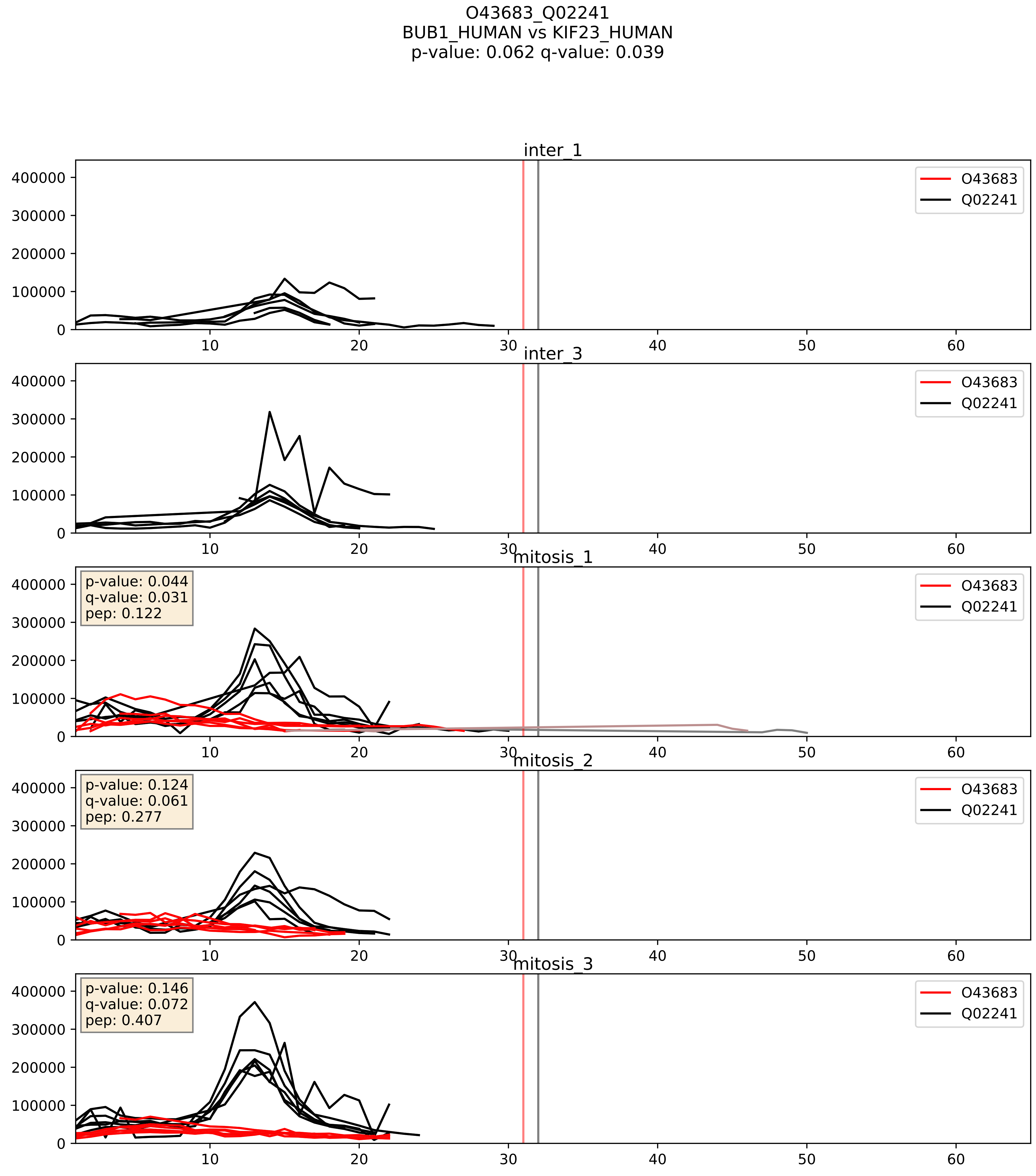


| level | condition_1 | condition_2 | pvalue | pvalue_adjusted |
|---|---|---|---|---|
| complex_abundance | mitosis | inter | 0.0007854526368926495 | 0.018471083988464504 |
| interactor_abundance | mitosis | inter | 0.004465595680965726 | 0.05368749863632952 |
| interactor_ratio | mitosis | inter | 0.01852309206901418 | 0.2047641739035816 |

O43683_Q9BPX3
BUB1_HUMAN vs CND3_HUMAN
p-value: 0.075 q-value: 0.045

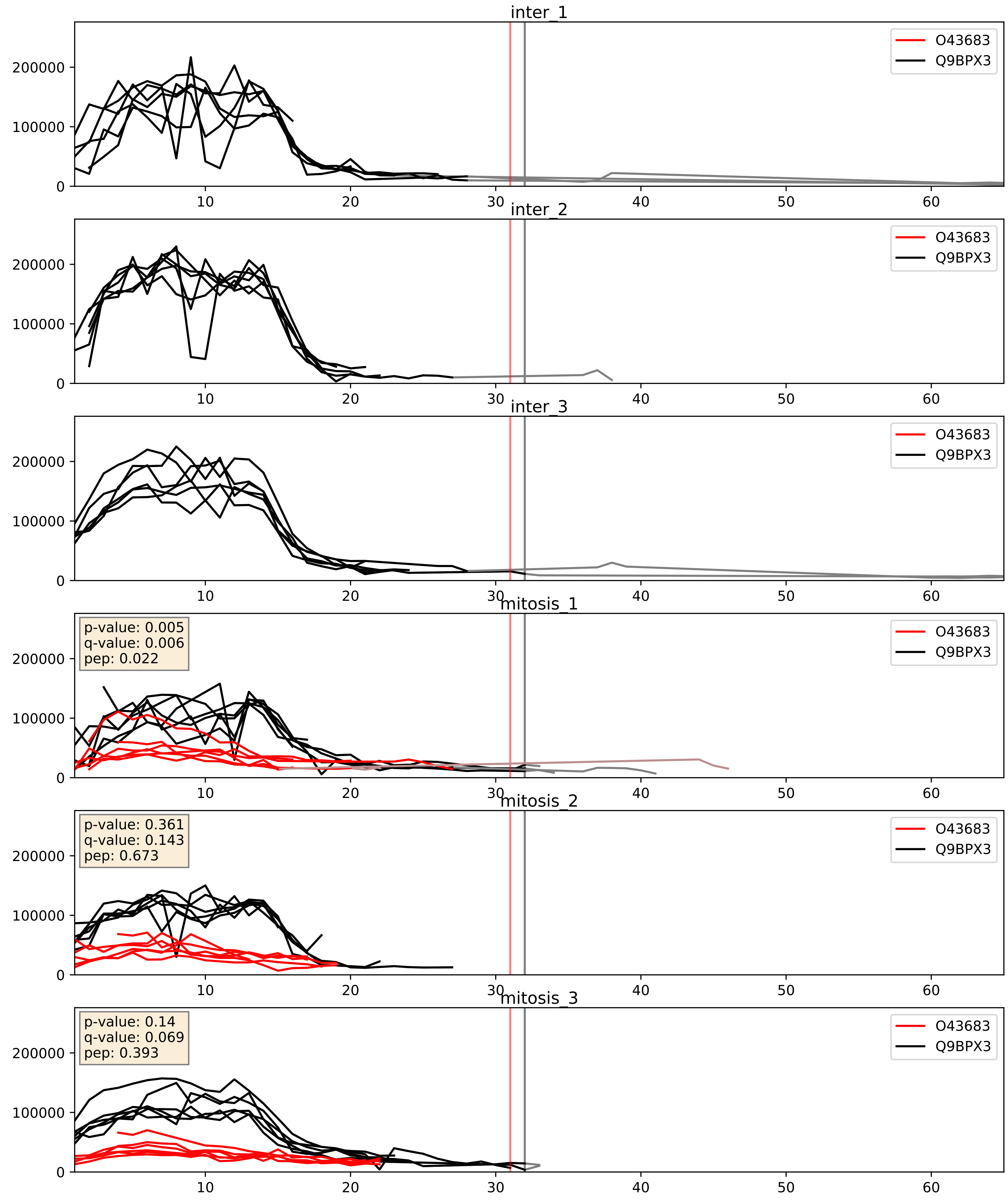


| level | condition_1 | condition_2 | pvalue | pvalue_adjusted |
|---|---|---|---|---|
| complex_abundance | mitosis | inter | 0.0004260739905454476 | 0.013091389064164169 |
| interactor_ratio | mitosis | inter | 0.0017705498104541042 | 0.05137595382199028 |
| interactor_abundance | mitosis | inter | 0.004465595680965726 | 0.05368749863632952 |
