## Supplementary Data 2 for "SECAT: Quantifying differential protein-protein interaction states by network-centric analysis": hela_string_000167_Q01780.pdf

Q01780_Q01780

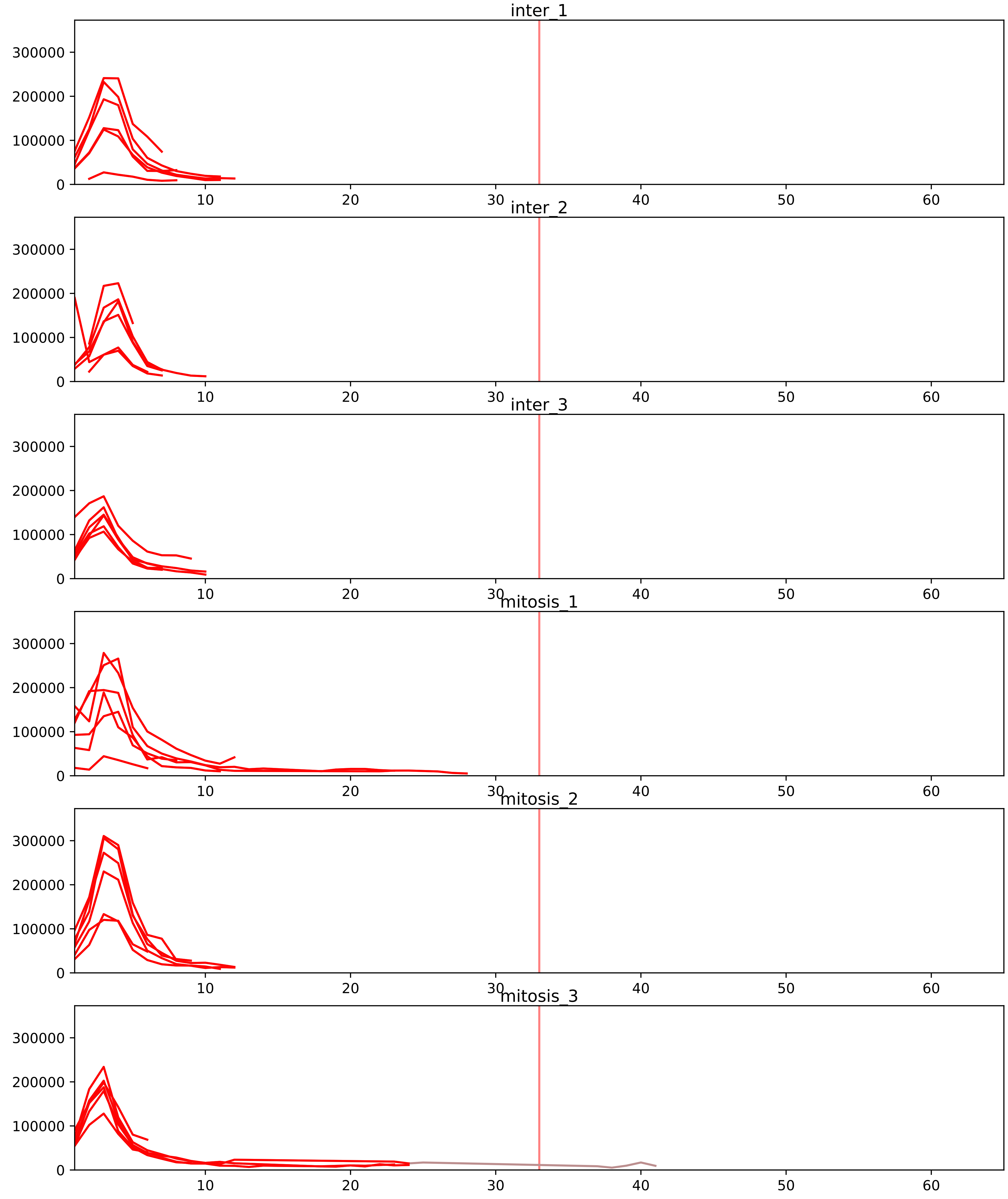


| level | condition_1 | condition_2 | pvalue | pvalue_adjusted |
|---|---|---|---|---|
| interactor_ratio | mitosis | inter | 8.610473425871309e-05 | 0.0014806795423928233 |
| assembled_abundance | mitosis | inter | 0.20276505791053548 | 0.5105665605590192 |
| total_abundance | mitosis | inter | 0.20879685197146403 | 0.5462665357872817 |
| complex_abundance | mitosis | inter | 0.23561227728540912 | 0.35740032168602864 |
| interactor_abundance | mitosis | inter | 0.4792571927753546 | 0.6498402613903113 |

P22087_Q01780
FBRL_HUMAN vs EXOSX_HUMAN
p-value: 0.002 q-value: 0.034

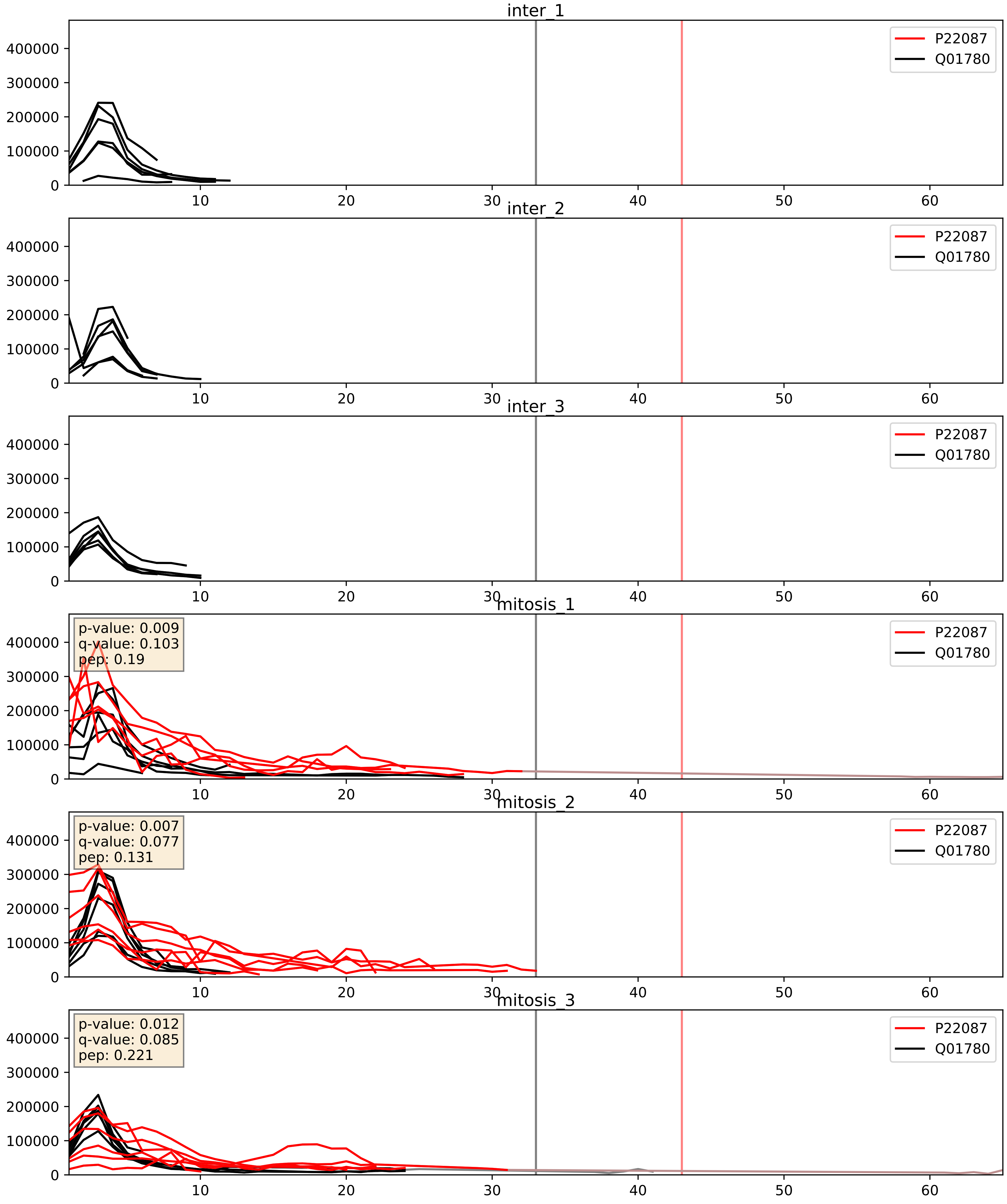


| level | condition_1 | condition_2 | pvalue | pvalue_adjusted |
|---|---|---|---|---|
| interactor_ratio | mitosis | inter | 3.1797532631946304e-06 | 0.002520386835947235 |
| interactor_abundance | mitosis | inter | 0.0004135262530475819 | 0.014019253191624928 |
| complex_abundance | mitosis | inter | 0.0006473146188650986 | 0.016393529992559892 |

P39023_Q01780
RL3_HUMAN vs EXOSX_HUMAN
p-value: 0.0 q-value: 0.013

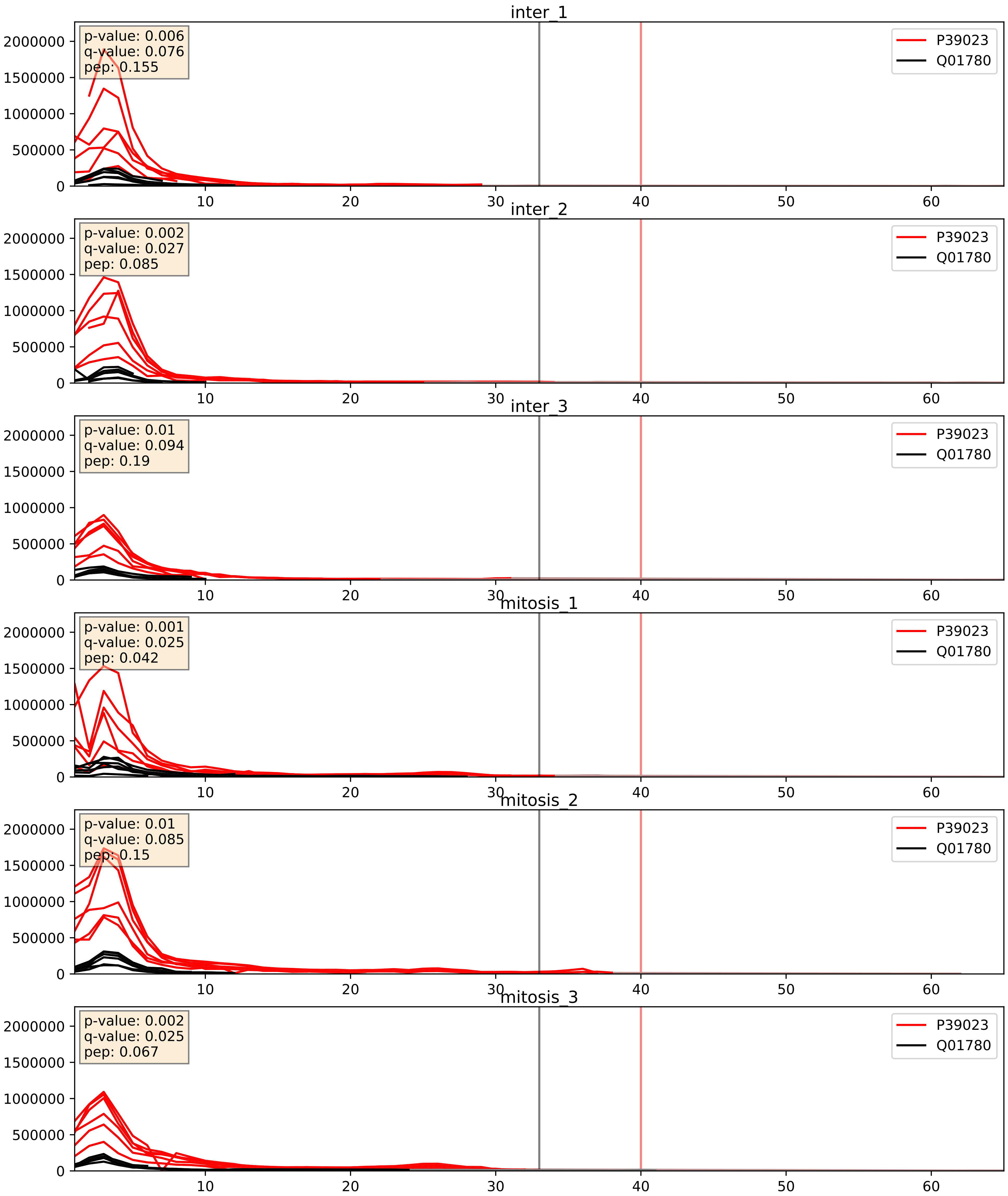


| level | condition_1 | condition_2 | pvalue | pvalue_adjusted |
|---|---|---|---|---|
| complex_abundance | mitosis | inter | 0.49119075386434813 | 0.6302792464516294 |
| interactor_abundance | mitosis | inter | 0.5762045835029546 | 0.7214673056074441 |
| interactor_ratio | mitosis | inter | 0.8879598402127304 | 0.9804177989881011 |

P42285_Q01780
MTREX_HUMAN vs EXOSX_HUMAN
p-value: 0.035 q-value: 0.025

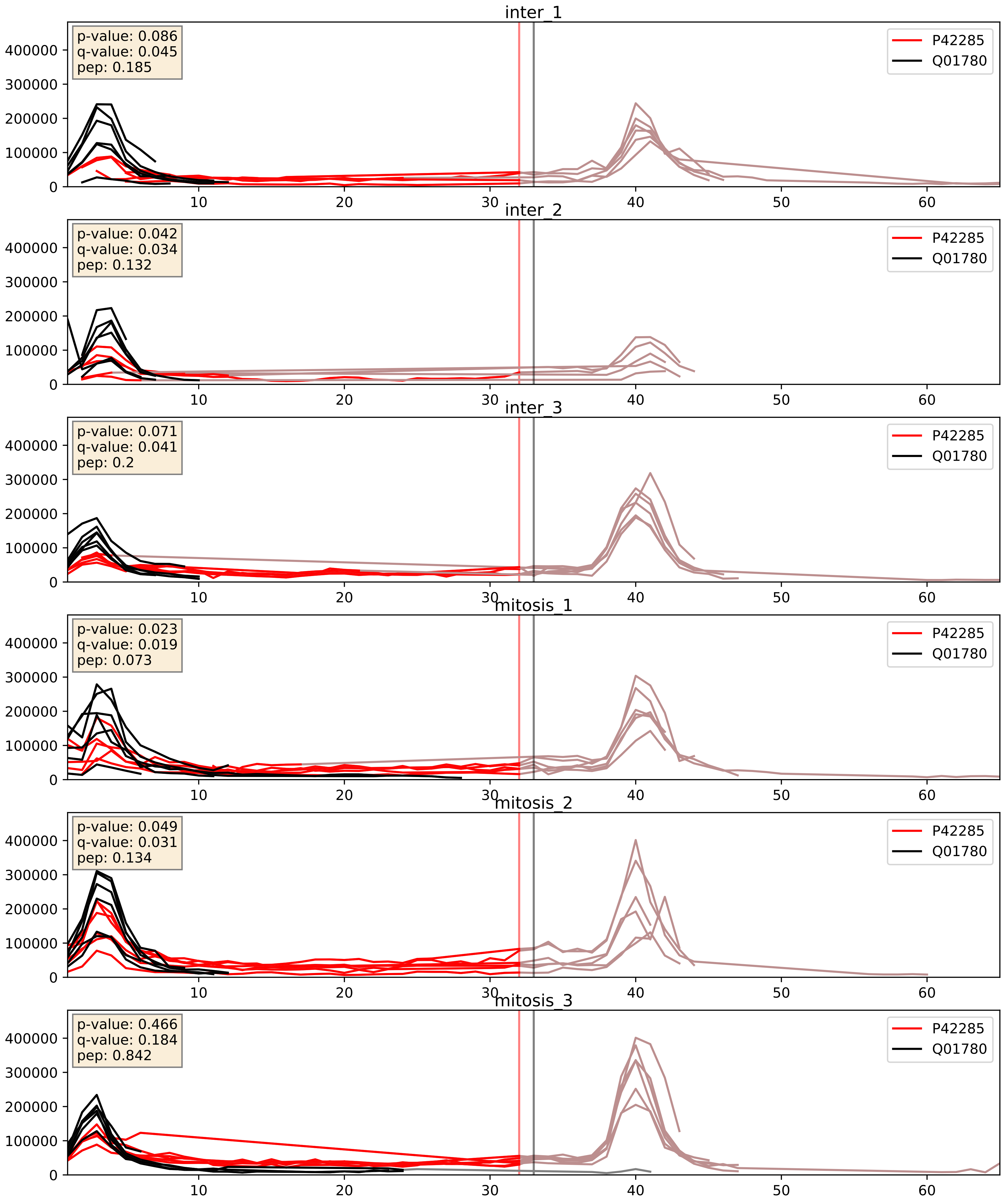


| level | condition_1 | condition_2 | pvalue | pvalue_adjusted |
|---|---|---|---|---|
| interactor_abundance | mitosis | inter | 0.0027241793218210184 | 0.03935691982242687 |
| interactor_ratio | mitosis | inter | 0.013227482029593517 | 0.16610126135854406 |
| complex_abundance | mitosis | inter | 0.04923814278230084 | 0.15816515545495635 |

P62910_Q01780
RL32_HUMAN vs EXOSX_HUMAN
p-value: 0.002 q-value: 0.035

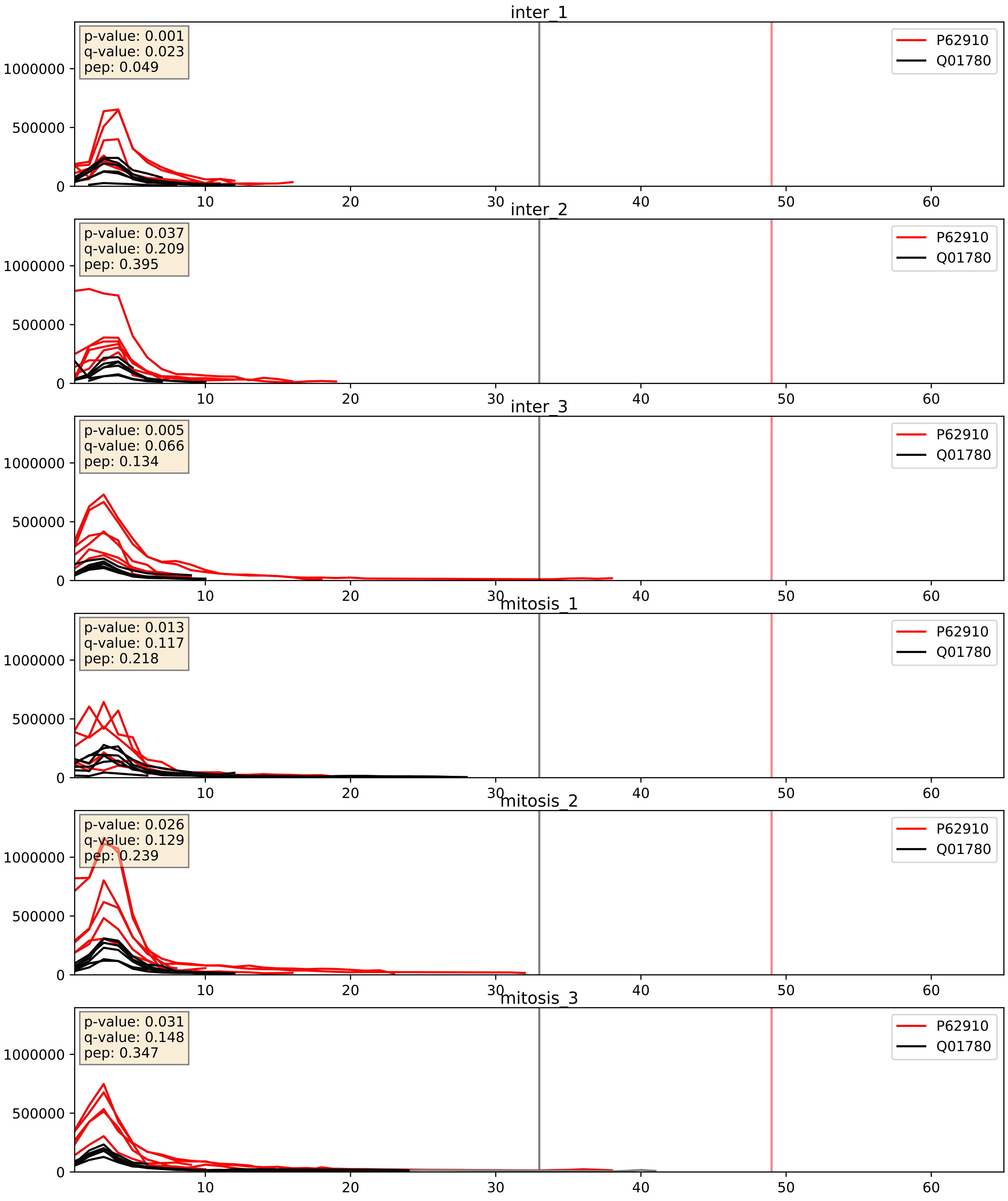


| level | condition_1 | condition_2 | pvalue | pvalue_adjusted |
|---|---|---|---|---|
| complex_abundance | mitosis | inter | 0.16923709094277398 | 0.31852144506036695 |
| interactor_abundance | mitosis | inter | 0.541072814936031 | 0.6983818320158905 |
| interactor_ratio | mitosis | inter | 0.9223312305539277 | 0.9869241197086243 |

P62913_Q01780
RL11_HUMAN vs EXOSX_HUMAN
p-value: 0.001 q-value: 0.049

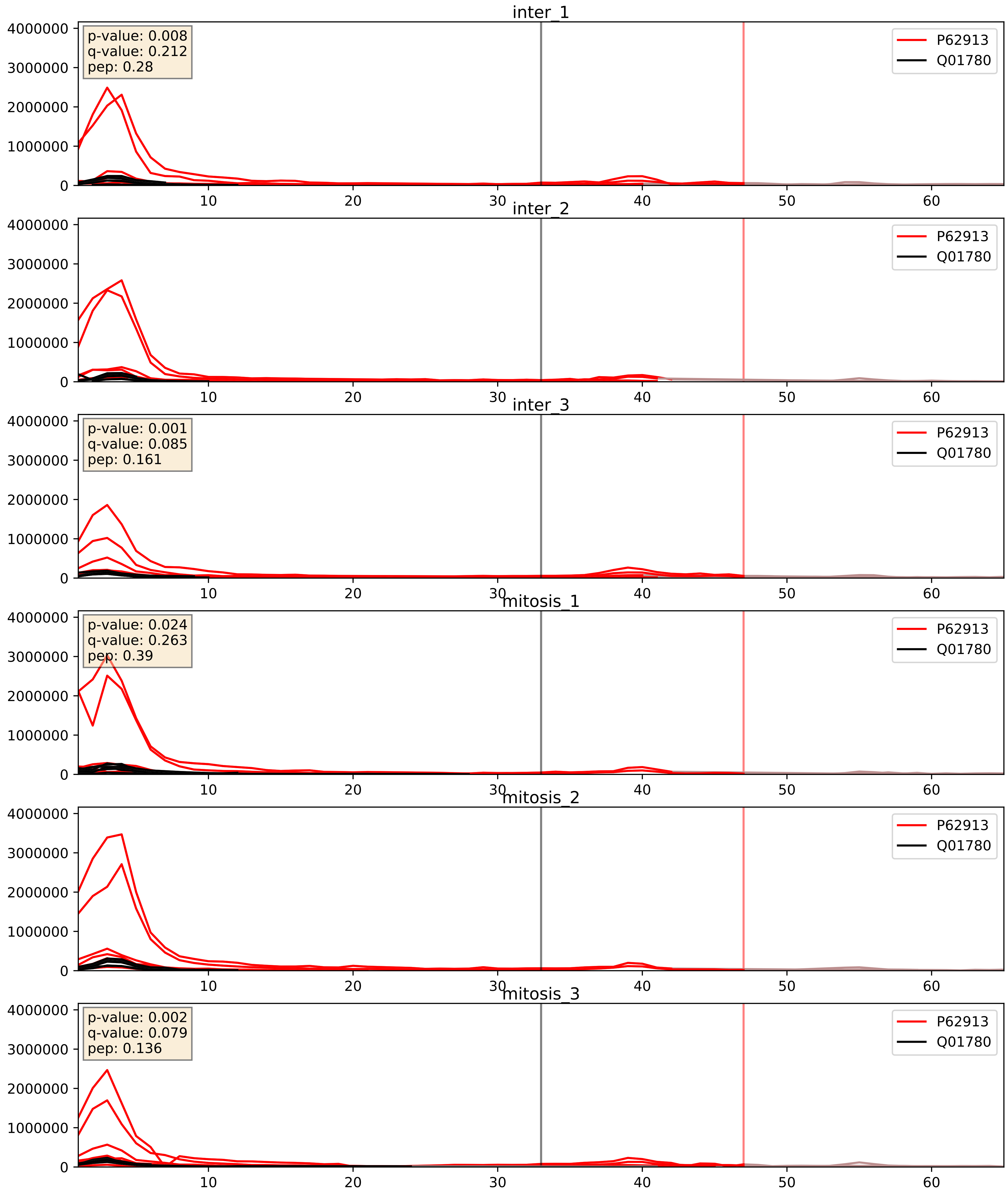


| level | condition_1 | condition_2 | pvalue | pvalue_adjusted |
|---|---|---|---|---|
| interactor_abundance | mitosis | inter | 0.055363585780035654 | 0.1907859477766124 |
| complex_abundance | mitosis | inter | 0.22477365578611566 | 0.37880969172162987 |
| interactor_ratio | mitosis | inter | 0.5780179766142616 | 0.8965922889176856 |

Q01780_Q02543
EXOSX_HUMAN vs RL18A_HUMAN
p-value: 0.004 q-value: 0.047

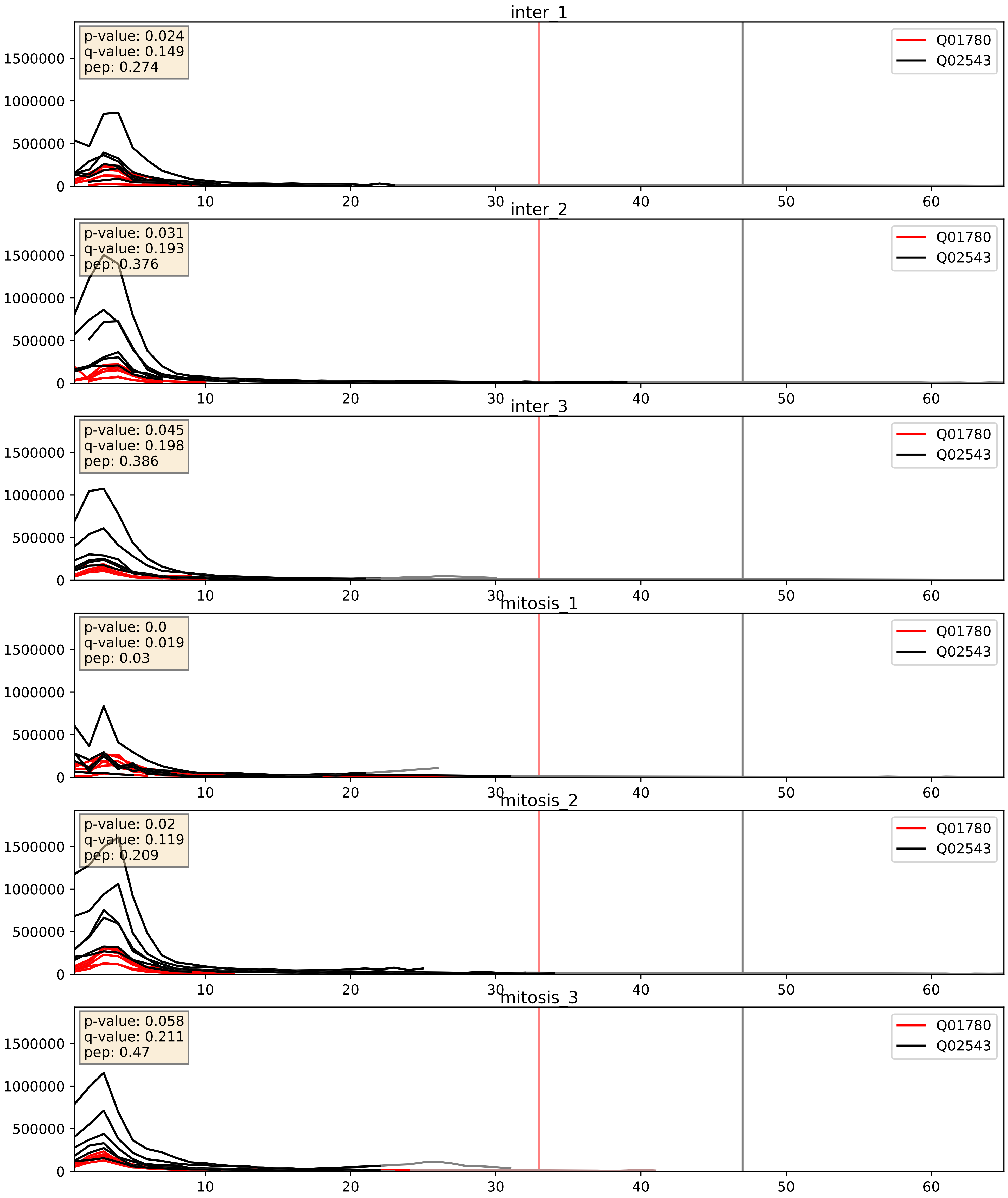


| level | condition_1 | condition_2 | pvalue | pvalue_adjusted |
|---|---|---|---|---|
| interactor_ratio | mitosis | inter | 0.6339514786449288 | 0.9189880875868909 |
| interactor_abundance | mitosis | inter | 0.645971301645025 | 0.7631923812070815 |
| complex_abundance | mitosis | inter | 0.8348141463903724 | 0.893145763220603 |

Q01780_Q06265
EXOSX_HUMAN vs EXOS9_HUMAN
p-value: 0.0 q-value: 0.001

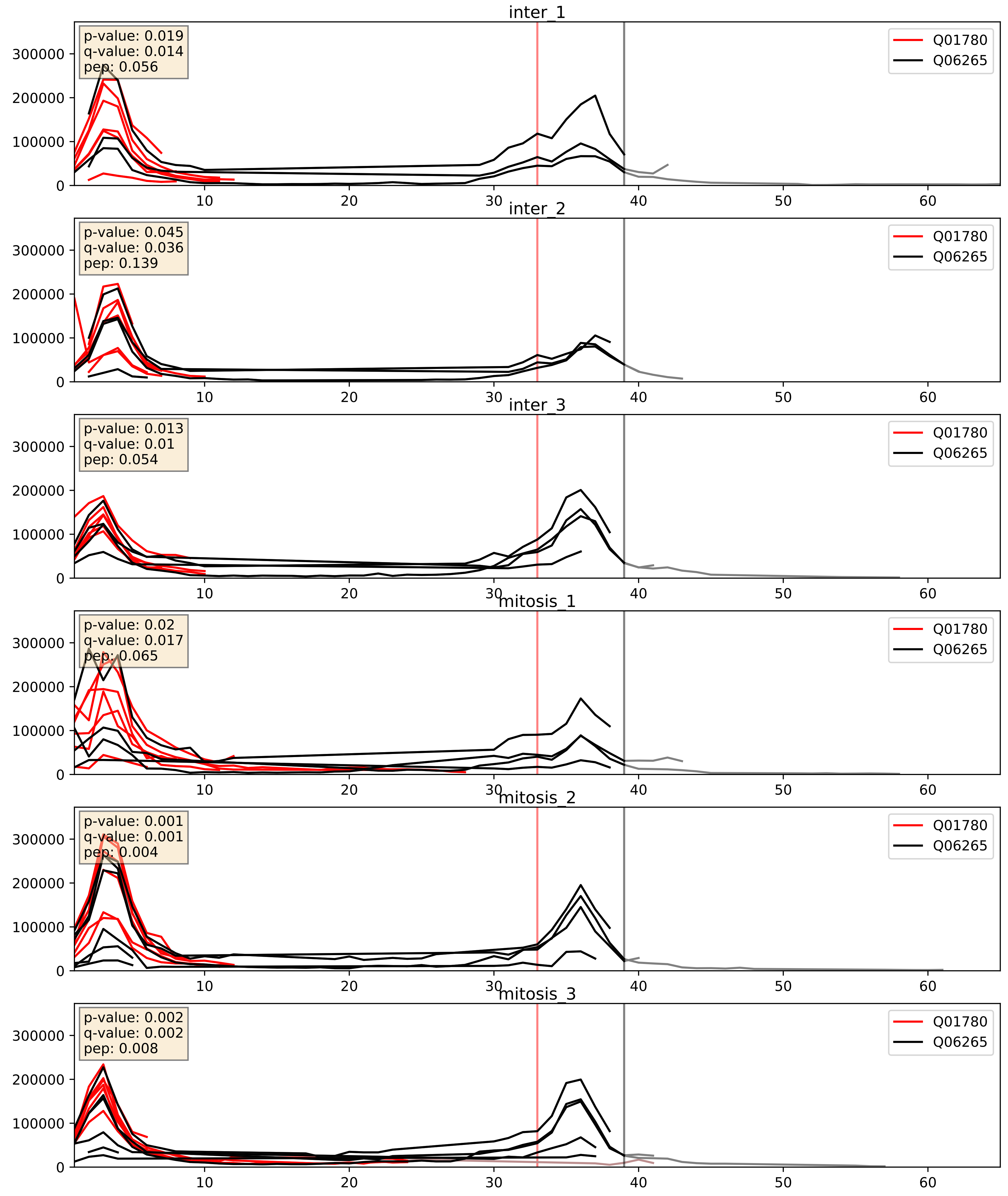


| level | condition_1 | condition_2 | pvalue | pvalue_adjusted |
|---|---|---|---|---|
| interactor_abundance | mitosis | inter | 0.6000759074560541 | 0.7373600537525452 |
| complex_abundance | mitosis | inter | 0.6006931169597226 | 0.7238081476879541 |
| interactor_ratio | mitosis | inter | 0.6751052260343209 | 0.9322491938895787 |

Q01780_Q13868
EXOSX_HUMAN vs EXOS2_HUMAN
p-value: 0.033 q-value: 0.023

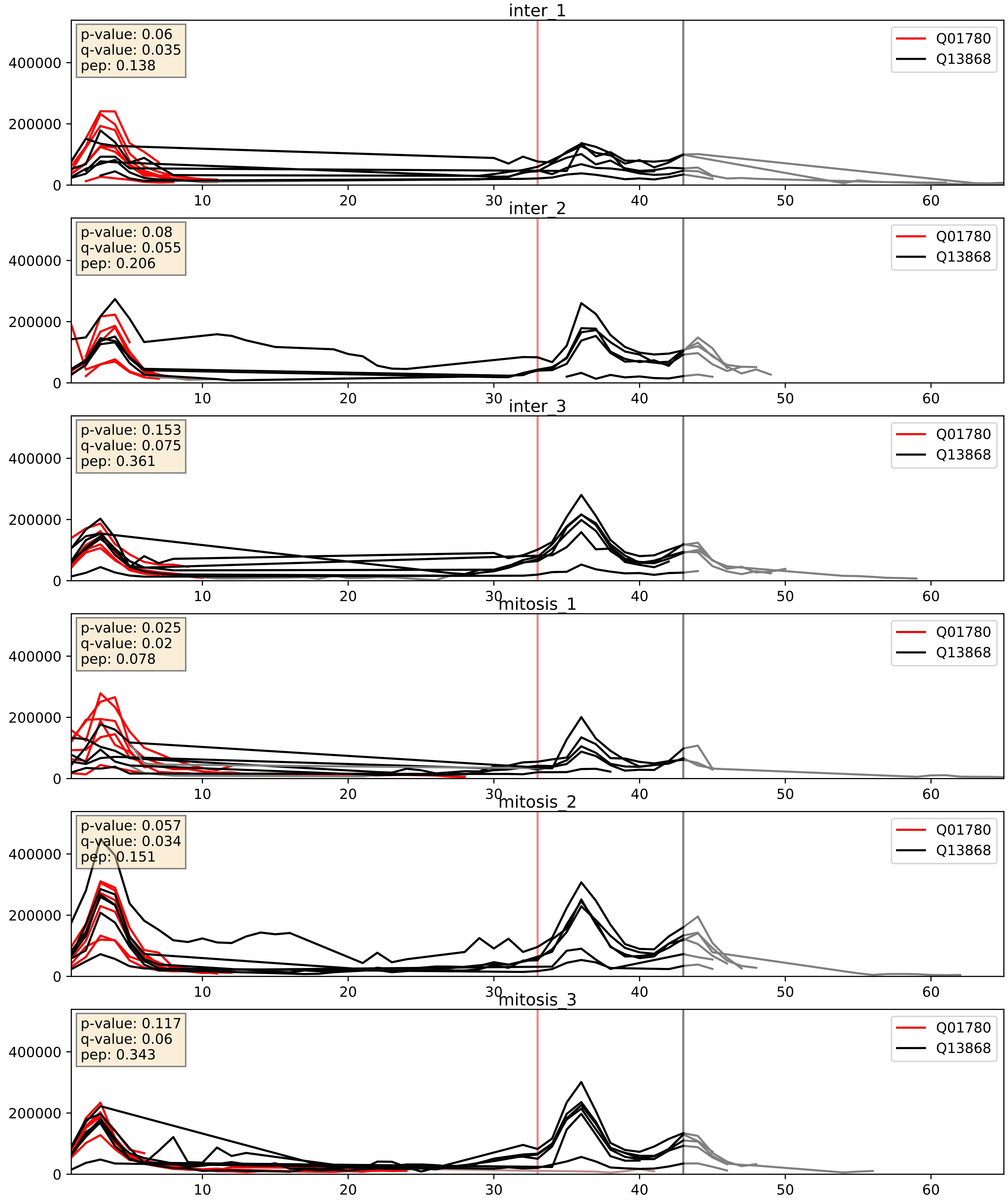


| level | condition_1 | condition_2 | pvalue | pvalue_adjusted |
|---|---|---|---|---|
| interactor_abundance | mitosis | inter | 0.7882514588834479 | 0.8777718860468732 |
| interactor_ratio | mitosis | inter | 0.847225951786114 | 0.9755634896635457 |
| complex_abundance | mitosis | inter | 0.9892400014275067 | 0.993417927289941 |

Q01780_Q15024
EXOSX_HUMAN vs EXOS7_HUMAN
p-value: 0.002 q-value: 0.002

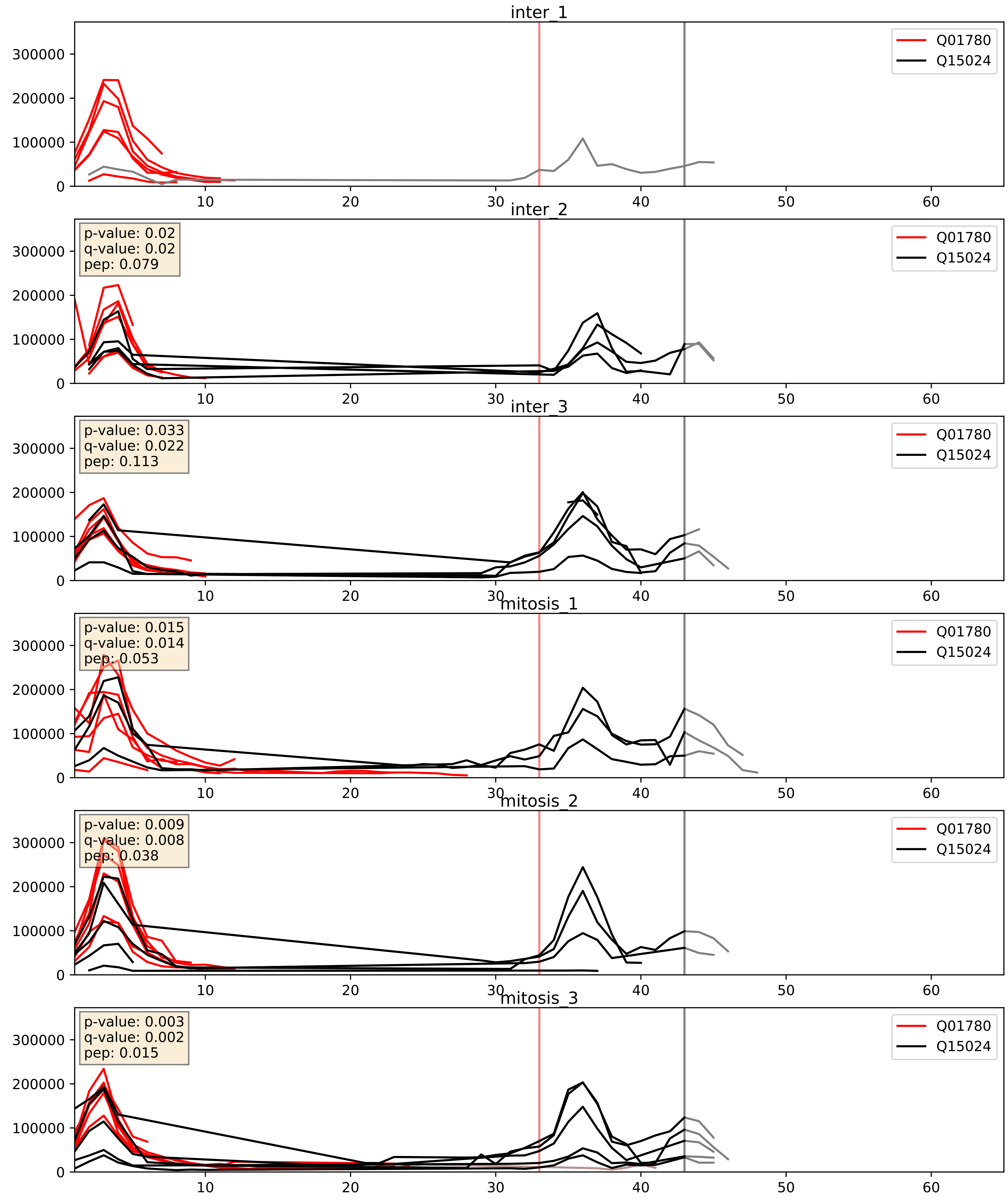


| level | condition_1 | condition_2 | pvalue | pvalue_adjusted |
|---|---|---|---|---|
| interactor_ratio | mitosis | inter | 0.21299696318226766 | 0.6904192954236995 |
| interactor_abundance | mitosis | inter | 0.24839210422378386 | 0.442025080683381 |
| complex_abundance | mitosis | inter | 0.29212097381937857 | 0.4412113289145512 |

Q01780_Q5RKV6
EXOSX_HUMAN vs EXOS6_HUMAN
p-value: 0.0 q-value: 0.0

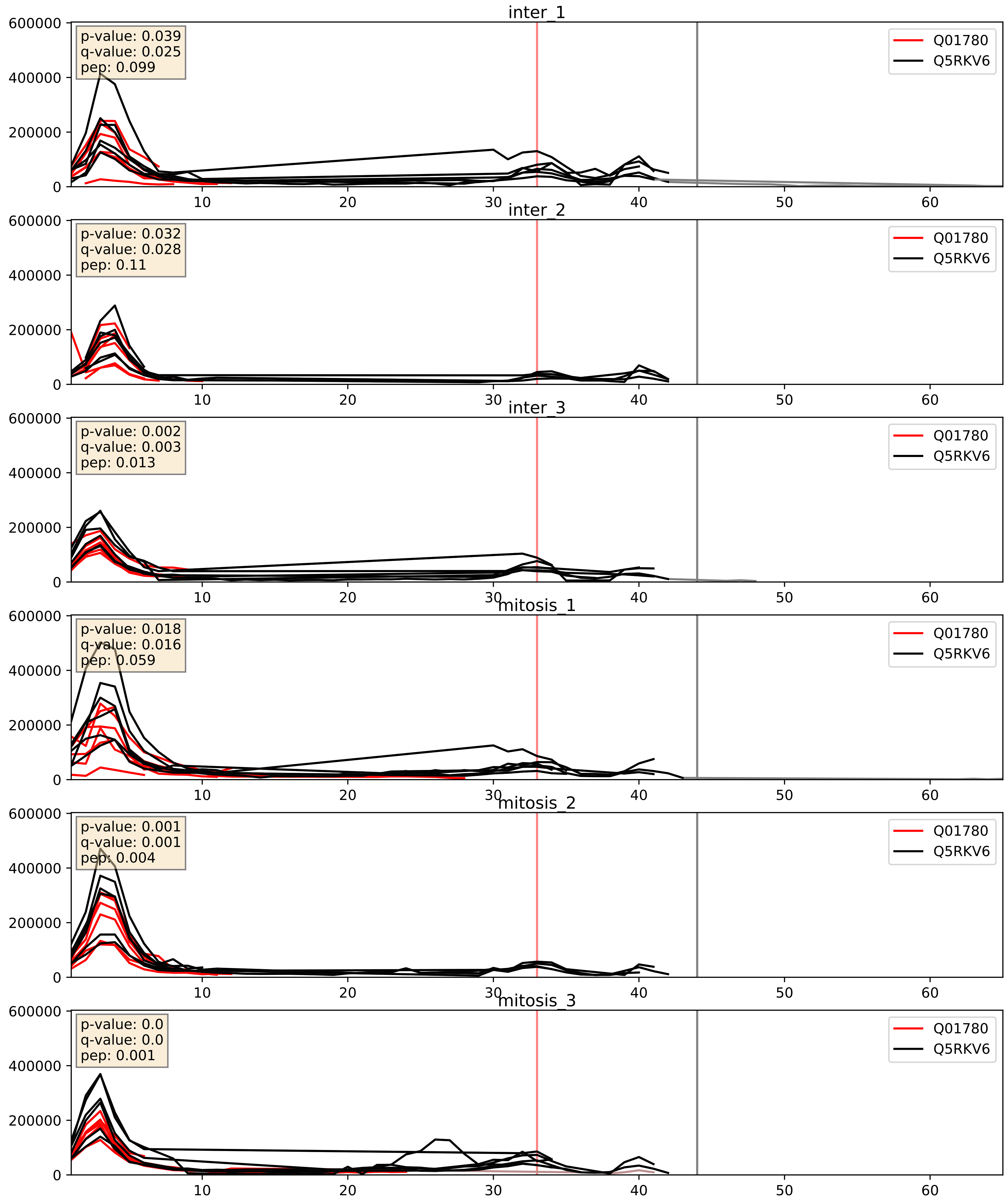


| level | condition_1 | condition_2 | pvalue | pvalue_adjusted |
|---|---|---|---|---|
| interactor_ratio | mitosis | inter | 0.022680955502768215 | 0.2316813593122863 |
| complex_abundance | mitosis | inter | 0.27897301299568367 | 0.4274224075967518 |
| interactor_abundance | mitosis | inter | 0.6000759074560541 | 0.7373600537525452 |

Q01780_Q96B26
EXOSX_HUMAN vs EXOS8_HUMAN
p-value: 0.0 q-value: 0.001

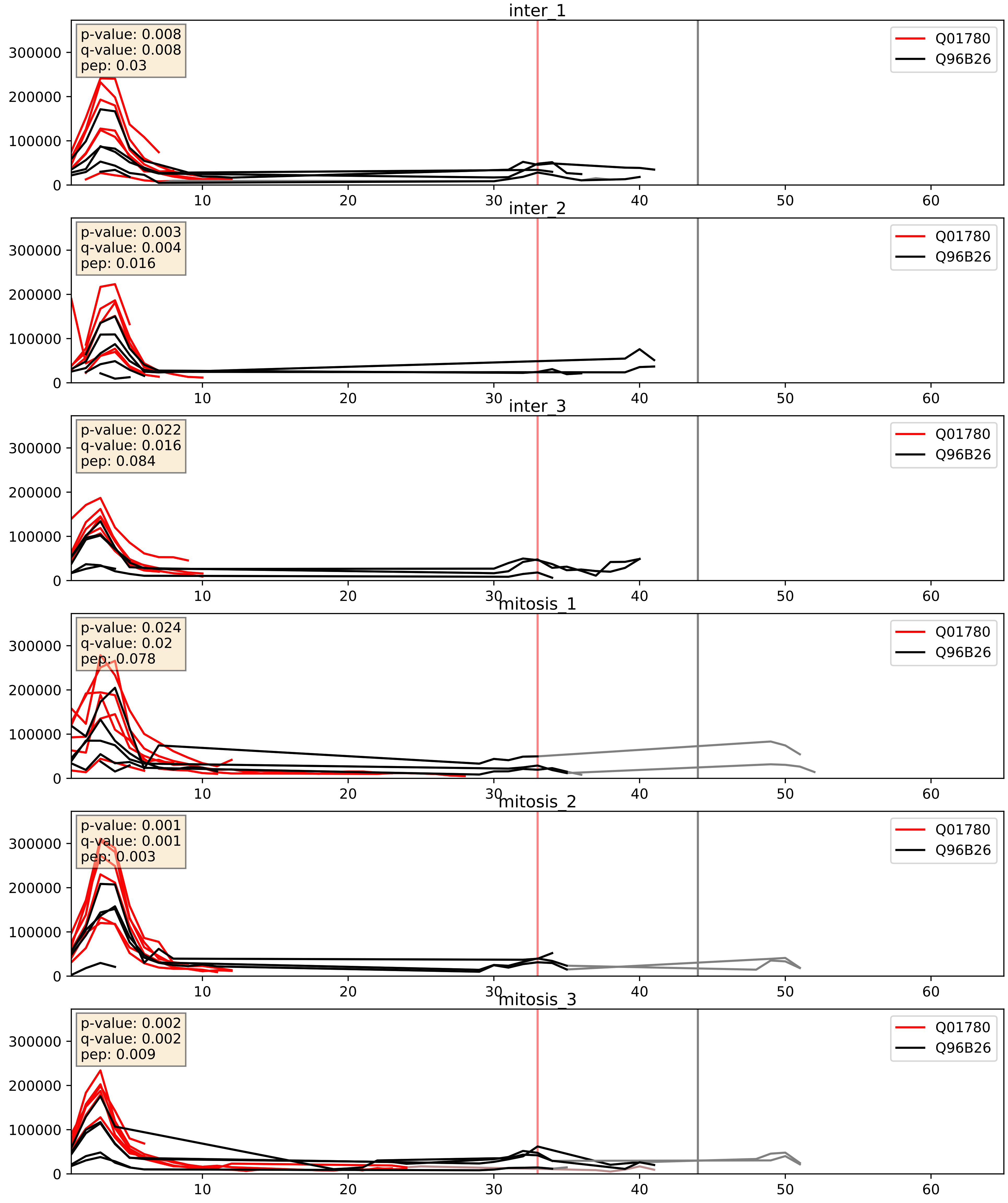


| level | condition_1 | condition_2 | pvalue | pvalue_adjusted |
|---|---|---|---|---|
| interactor_ratio | mitosis | inter | 0.050810198545751736 | 0.37301483666521 |
| interactor_abundance | mitosis | inter | 0.7200907402724626 | 0.8209331739671412 |
| complex_abundance | mitosis | inter | 0.9245214130727728 | 0.9533674612580333 |

Q01780_Q99459
EXOSX_HUMAN vs CDC5L_HUMAN
p-value: 0.007 q-value: 0.006

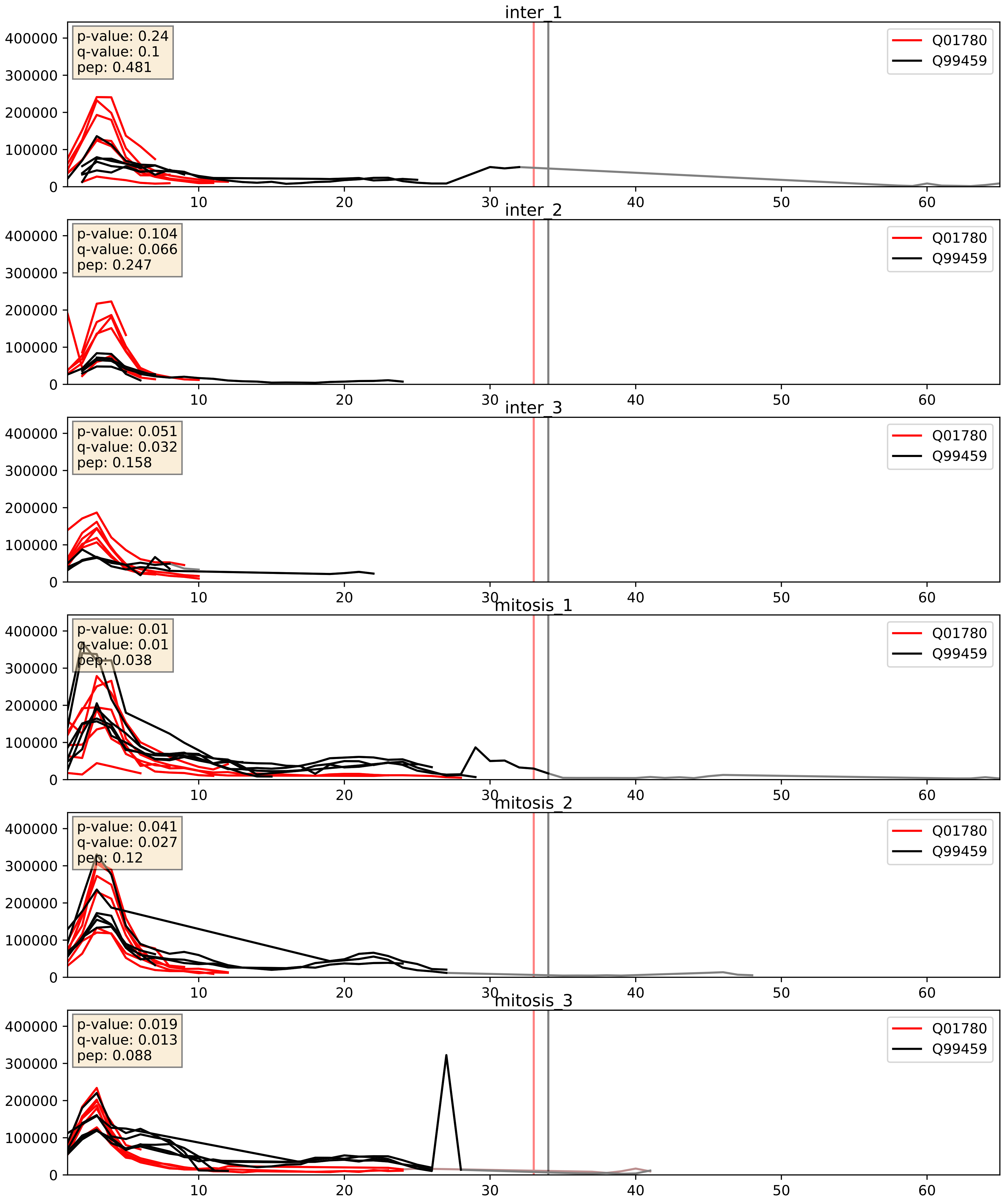


| level | condition_1 | condition_2 | pvalue | pvalue_adjusted |
|---|---|---|---|---|
| interactor_ratio | mitosis | inter | 0.010011115757435327 | 0.14263182888175546 |
| complex_abundance | mitosis | inter | 0.06311567682197815 | 0.1809950397306978 |
| interactor_abundance | mitosis | inter | 0.5783388797850517 | 0.722976387142759 |

Q01780_Q99547
EXOSX_HUMAN vs MPH6_HUMAN
p-value: 0.024 q-value: 0.018

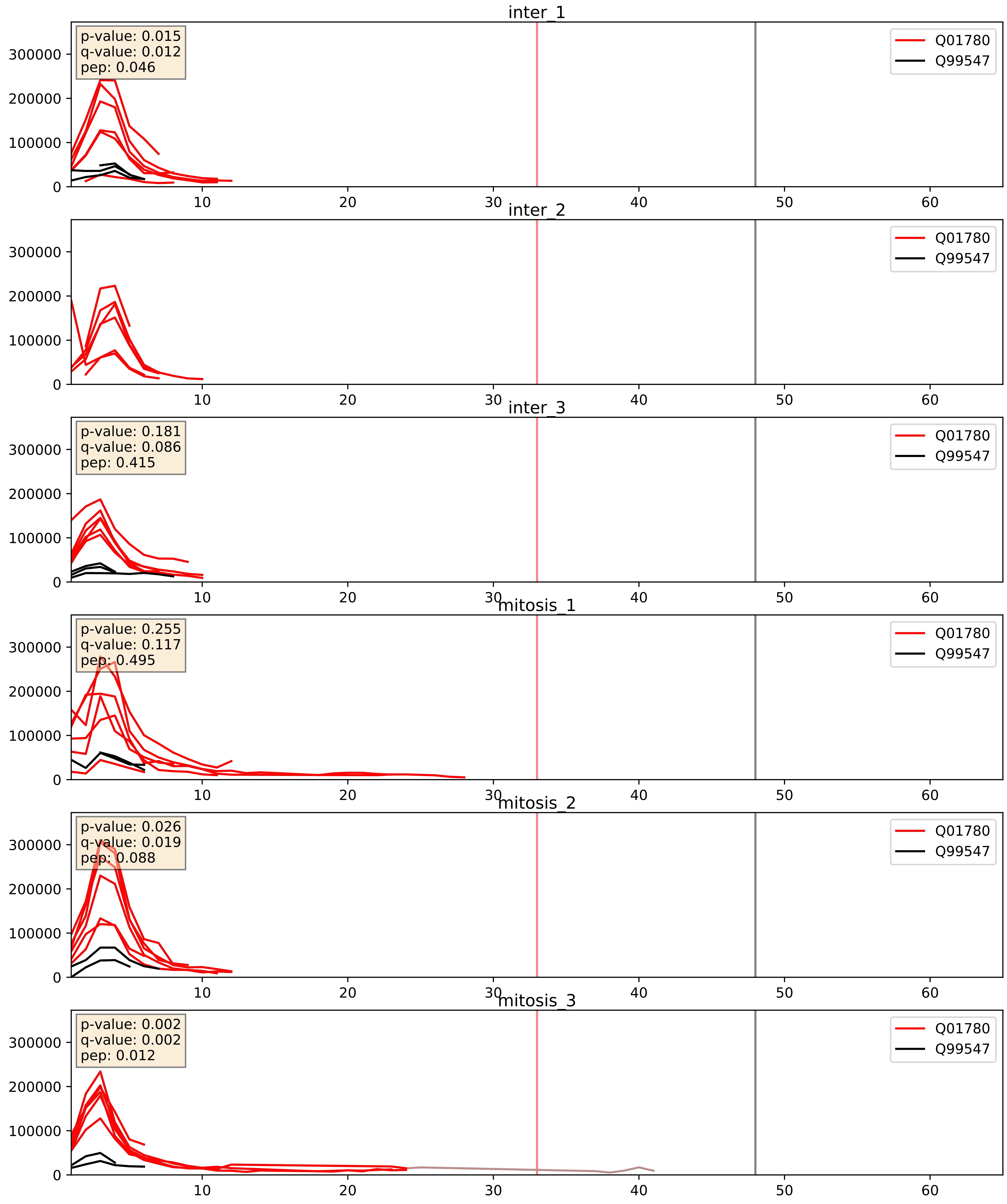


| level | condition_1 | condition_2 | pvalue | pvalue_adjusted |
|---|---|---|---|---|
| interactor_ratio | mitosis | inter | 0.027135379938690897 | 0.2577730310940899 |
| interactor_abundance | mitosis | inter | 0.3953301978901166 | 0.5864355764560087 |
| complex_abundance | mitosis | inter | 0.9932760166578614 | 0.9963021681030341 |

Q01780_Q9NPD3
EXOSX_HUMAN vs EXOS4_HUMAN
p-value: 0.004 q-value: 0.004

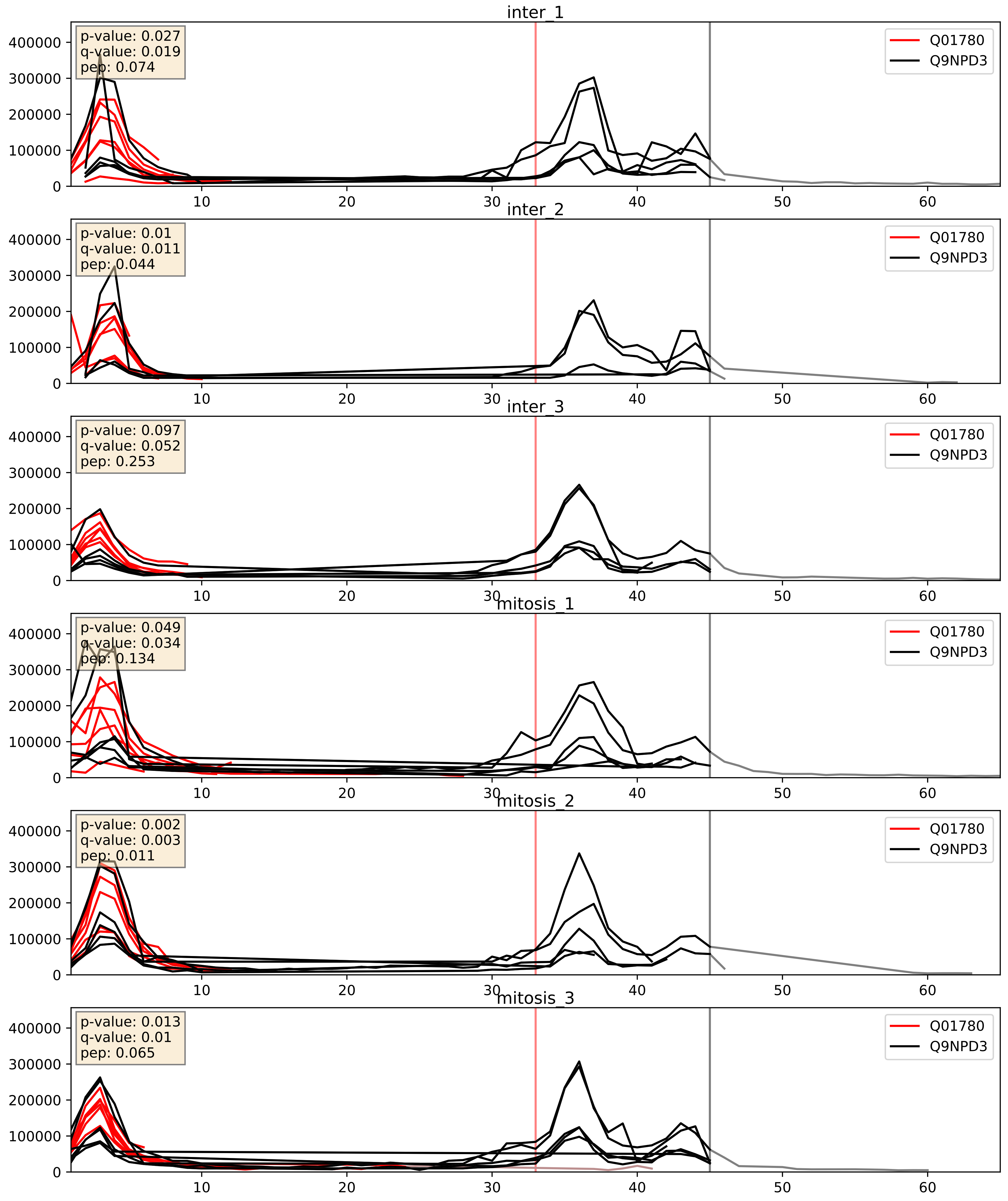


| level | condition_1 | condition_2 | pvalue | pvalue_adjusted |
|---|---|---|---|---|
| interactor_ratio | mitosis | inter | 0.2155538972480964 | 0.6924580509898395 |
| complex_abundance | mitosis | inter | 0.3792458463426227 | 0.5264911522369202 |
| interactor_abundance | mitosis | inter | 0.5840759329389282 | 0.7273863367945335 |

Q01780_Q9NQT4
EXOSX_HUMAN vs EXOS5_HUMAN
p-value: 0.001 q-value: 0.001

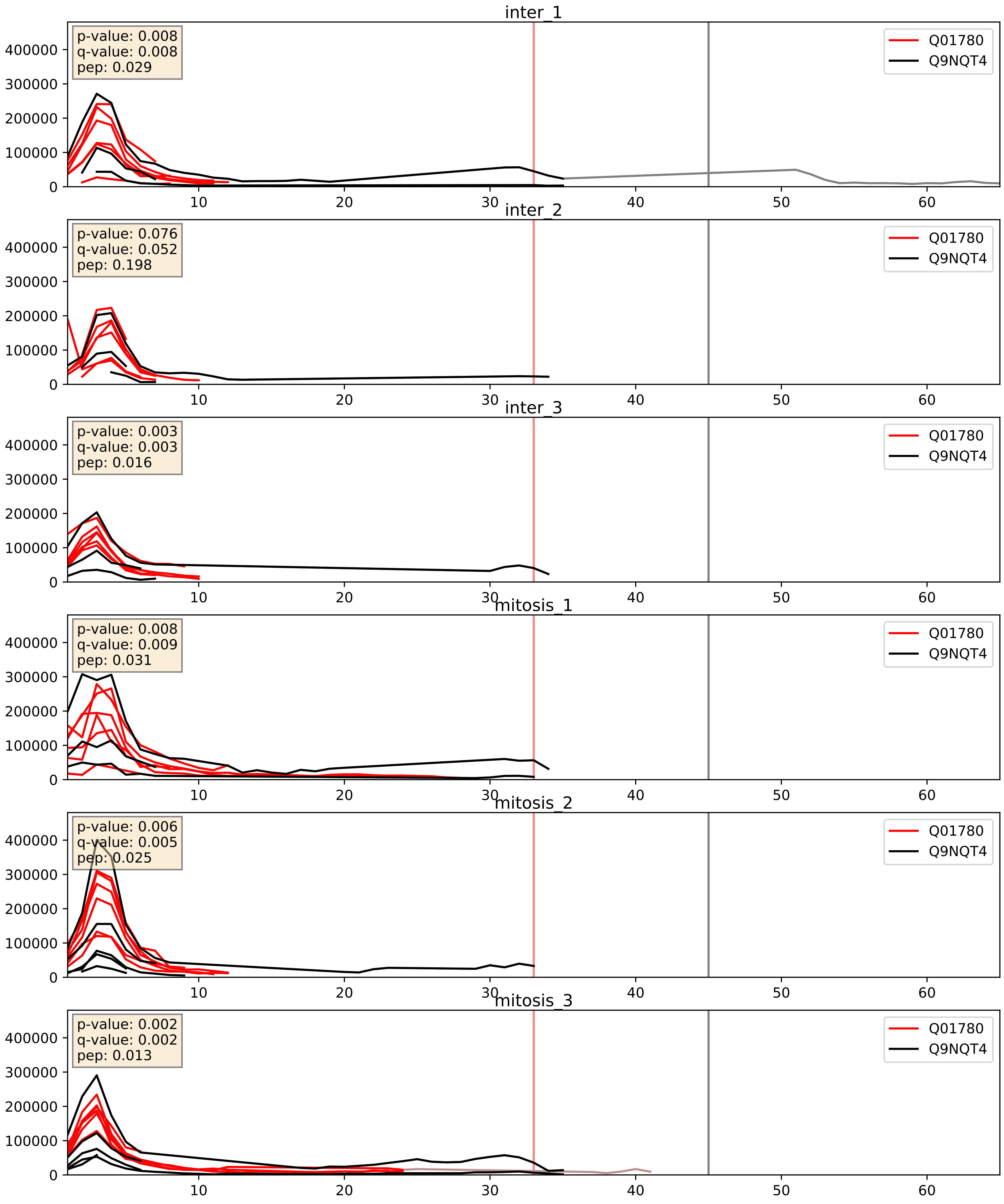


| level | condition_1 | condition_2 | pvalue | pvalue_adjusted |
|---|---|---|---|---|
| interactor_ratio | mitosis | inter | 0.06093952455390902 | 0.4087189096368508 |
| interactor_abundance | mitosis | inter | 0.6540838256807727 | 0.7700687586129621 |
| complex_abundance | mitosis | inter | 0.9002166656103112 | 0.9382201112485165 |

Q01780_Q9NQT5
EXOSX_HUMAN vs EXOS3_HUMAN
p-value: 0.017 q-value: 0.014

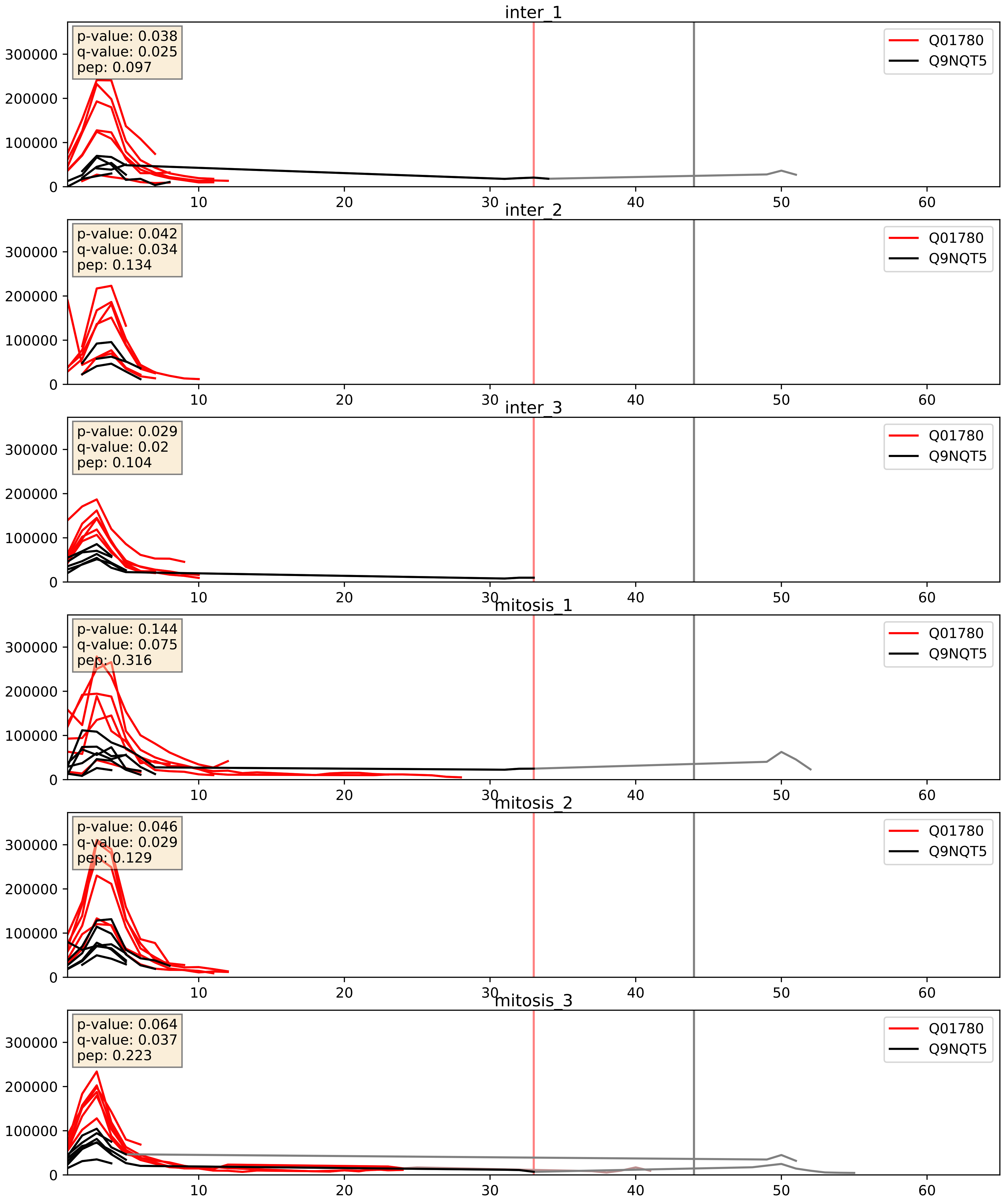


| level | condition_1 | condition_2 | pvalue | pvalue_adjusted |
|---|---|---|---|---|
| interactor_ratio | mitosis | inter | 0.04797918077346781 | 0.361488803041275 |
| complex_abundance | mitosis | inter | 0.7425116415401979 | 0.82917542145925 |
| interactor_abundance | mitosis | inter | 0.7589305936139555 | 0.8552456399862375 |

Q01780_Q9Y3B2
EXOSX_HUMAN vs EXOS1_HUMAN
p-value: 0.001 q-value: 0.001

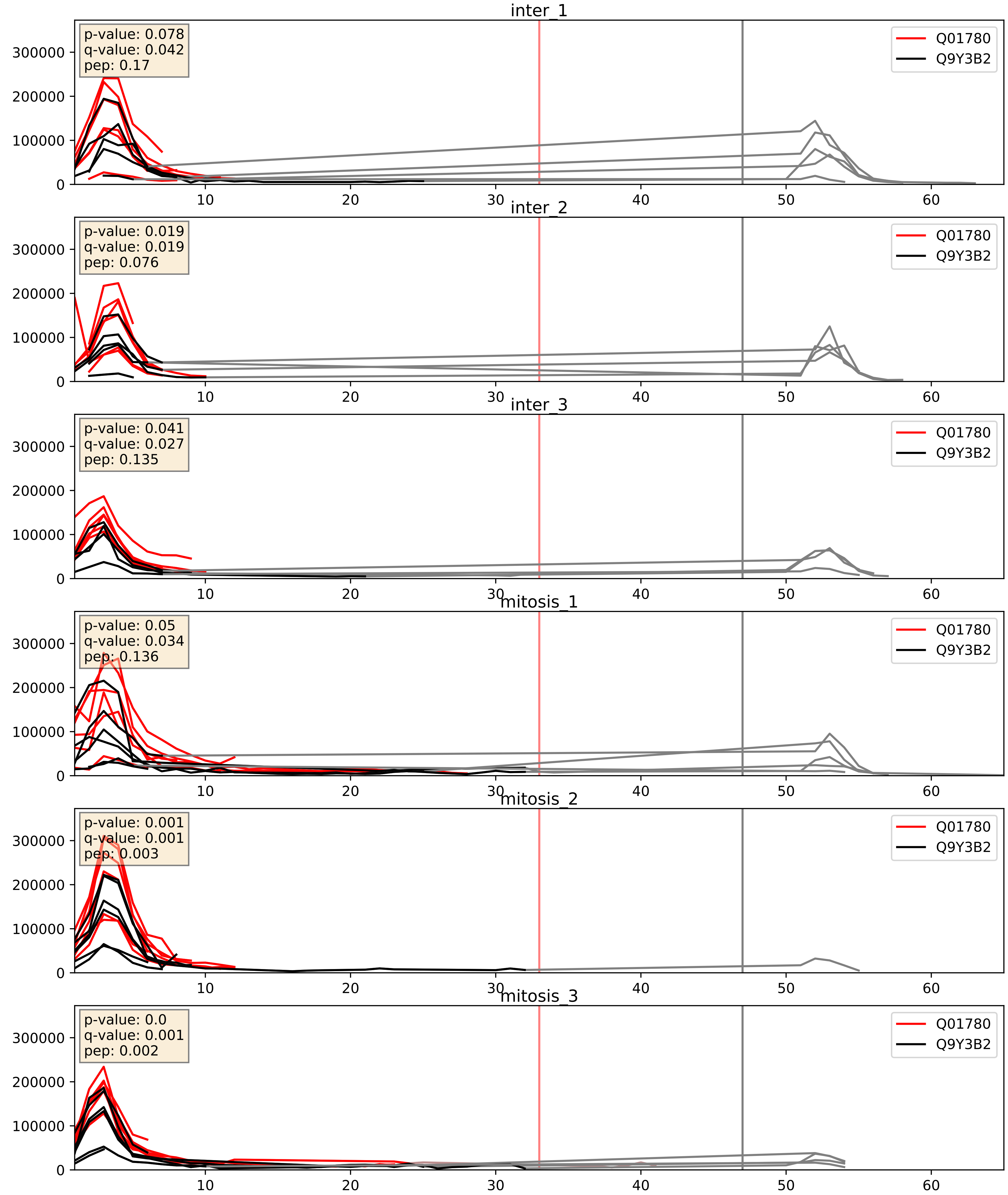


| level | condition_1 | condition_2 | pvalue | pvalue_adjusted |
|---|---|---|---|---|
| complex_abundance | mitosis | inter | 0.6132648655847321 | 0.7343051125199759 |
| interactor_abundance | mitosis | inter | 0.6257397034774399 | 0.7539881562171855 |
| interactor_ratio | mitosis | inter | 0.8867597462384897 | 0.9802747470617126 |
