## Supplementary Data 2 for "SECAT: Quantifying differential protein-protein interaction states by network-centric analysis": hela_string_000168_Q8NBN7.pdf

### Q8NBN7_Q8NBN7

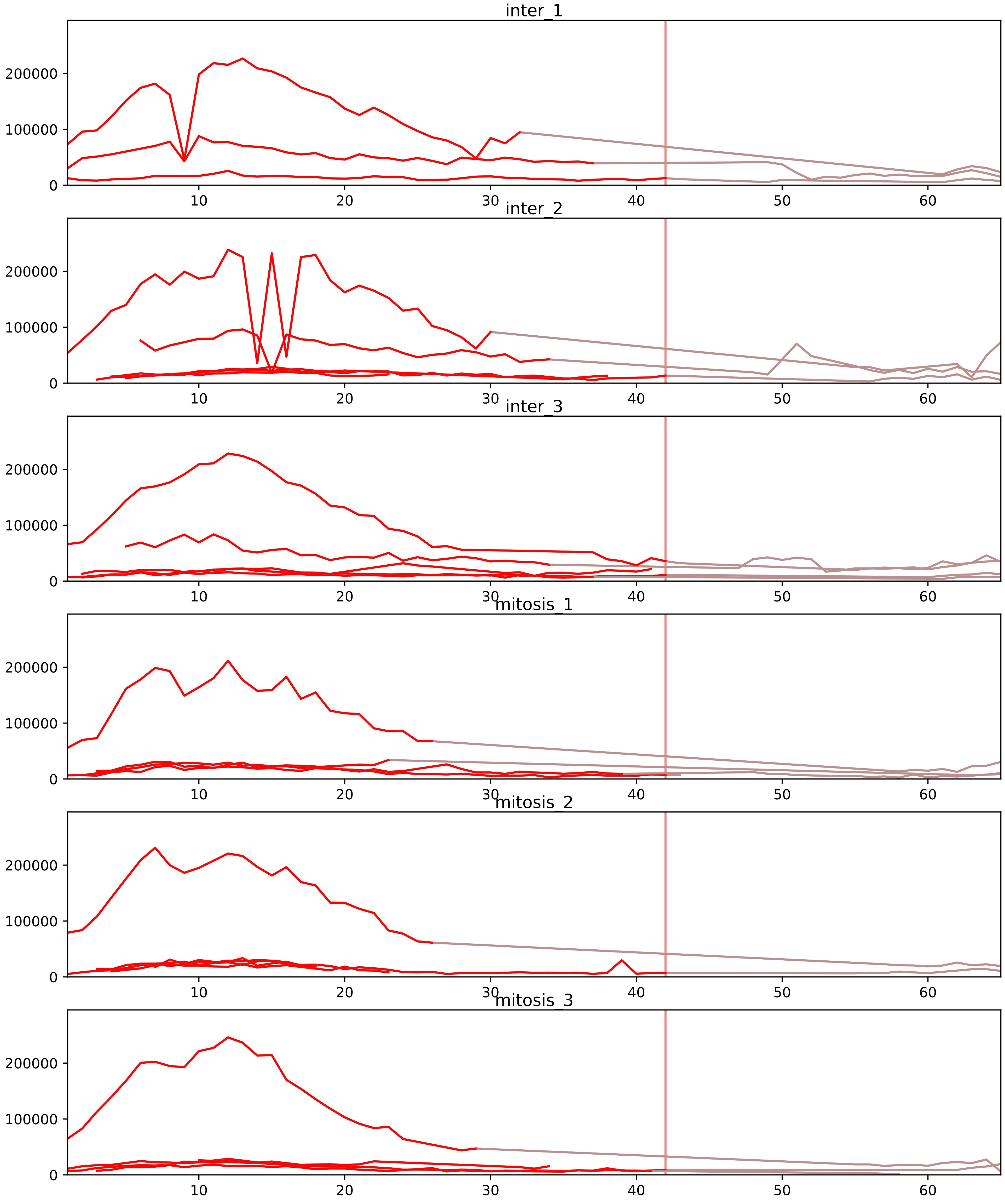


| level | condition_1 | condition_2 | pvalue | pvalue_adjusted |
|---|---|---|---|---|
| assembled_abundance | mitosis | inter | 8.688005042735817e-05 | 0.009037455912232524 |
| total_abundance | mitosis | inter | 0.00010055673757923784 | 0.01273261737241483 |
| monomer_abundance | mitosis | inter | 0.012700941349417837 | 0.19546941175259344 |
