## Supplementary Data 2 for "SECAT: Quantifying differential protein-protein interaction states by network-centric analysis": hela_string_000169_Q8IUX1.pdf

### Q8IUX1_Q8IUX1

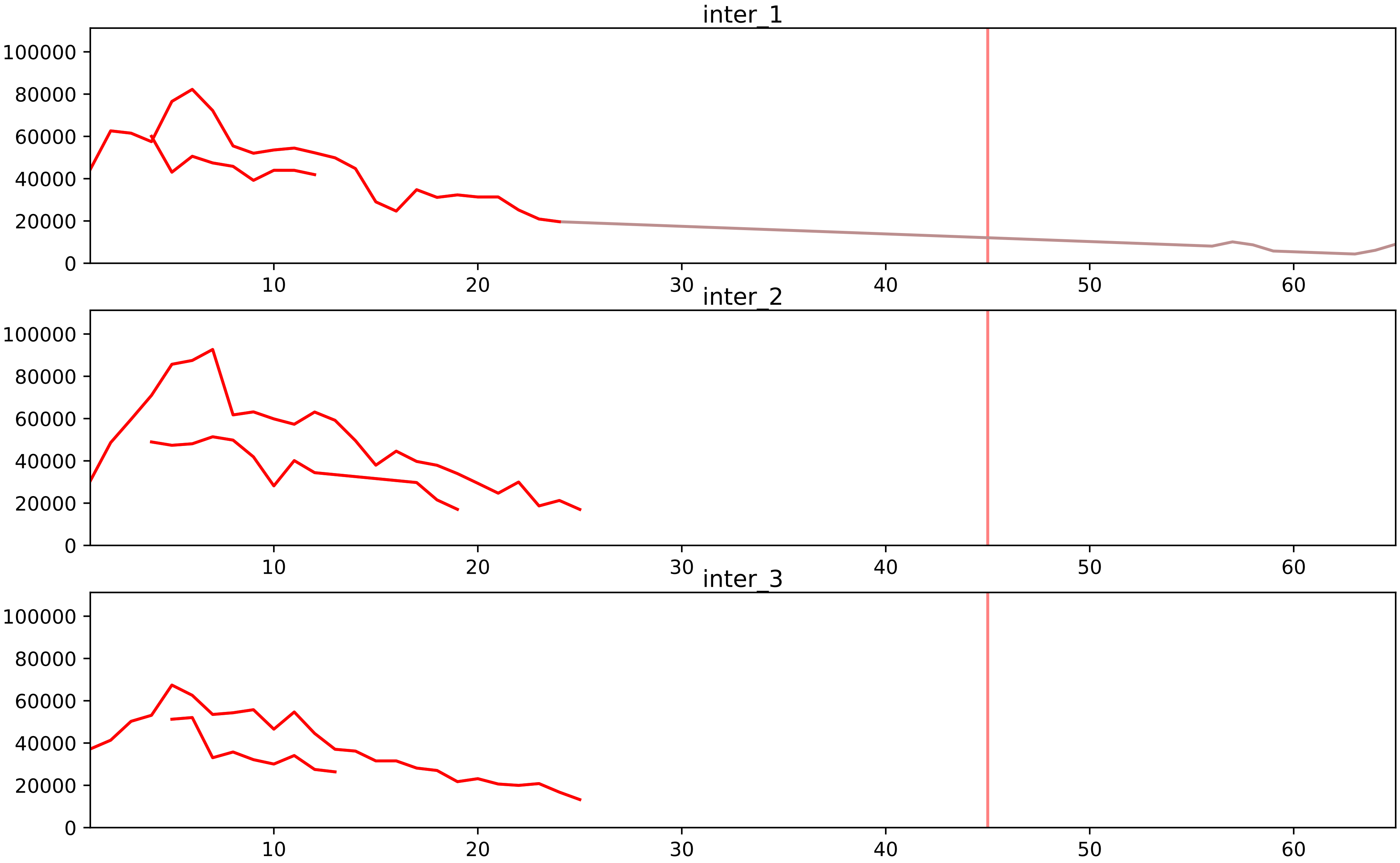


| level | condition_1 | condition_2 | pvalue | pvalue_adjusted |
|---|---|---|---|---|
| complex_abundance | mitosis | inter | 8.927825687788868e-05 | 0.0020533999081914396 |
| interactor_abundance | mitosis | inter | 0.00011843429865209775 | 0.002594275113331665 |
| interactor_ratio | mitosis | inter | 0.0006253207513218211 | 0.007569672252843098 |
| total_abundance | mitosis | inter | 0.001949185254732357 | 0.04766014179201654 |
| assembled_abundance | mitosis | inter | 0.0022114515826176756 | 0.04542643971557477 |
| monomer_abundance | mitosis | inter | 0.30810126715730446 | 0.6718670942432774 |

Q8IUX1_Q9BQ95
T126B_HUMAN vs ECSIT_HUMAN
p-value: 0.028 q-value: 0.021

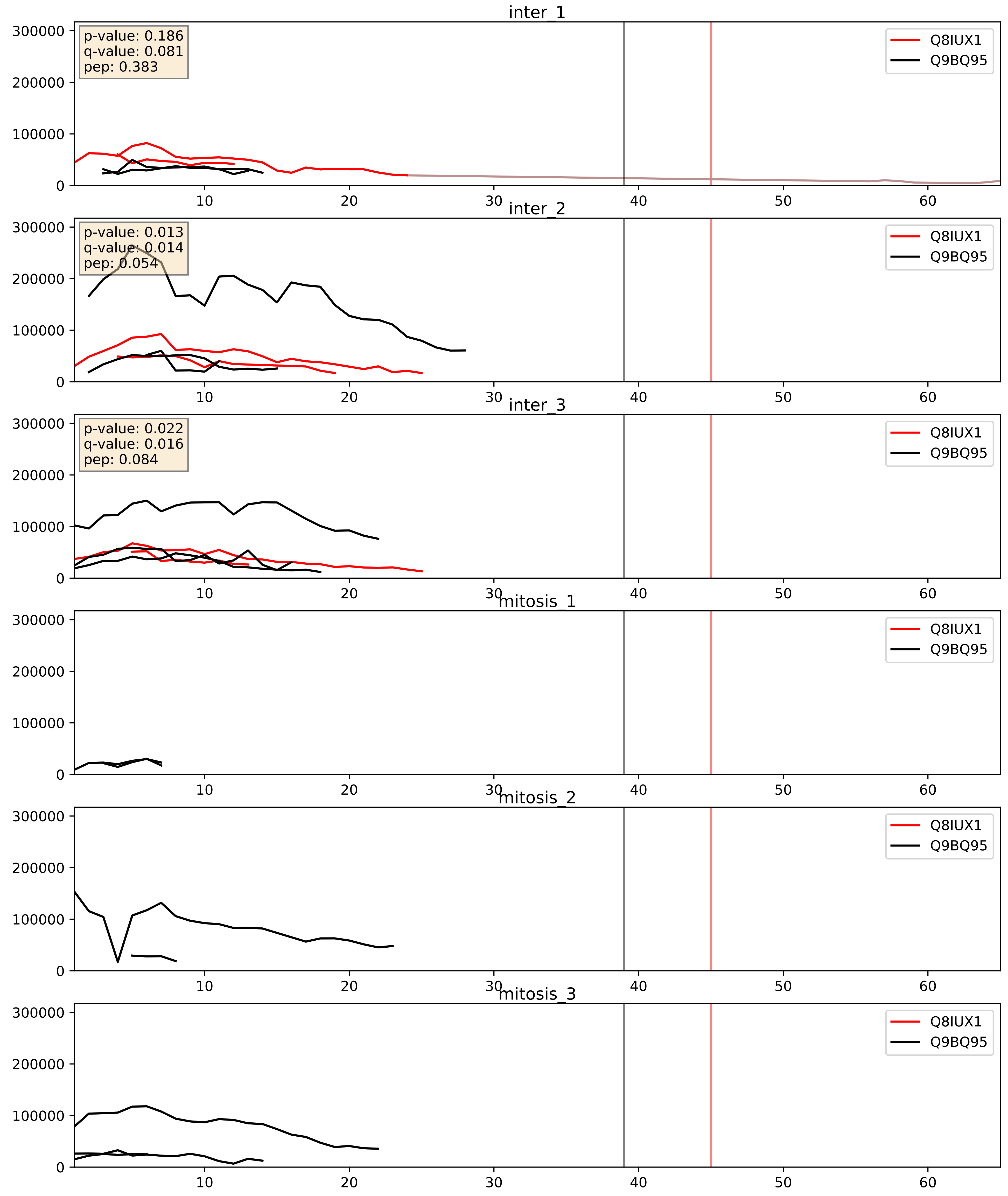


| level | condition_1 | condition_2 | pvalue | pvalue_adjusted |
|---|---|---|---|---|
| interactor_abundance | mitosis | inter | 0.0007479532056803059 | 0.020070468465904132 |
| complex_abundance | mitosis | inter | 0.0019629509291587523 | 0.03176344036597149 |
| interactor_ratio | mitosis | inter | 0.10054039444180876 | 0.5283153937519233 |

Q8IUX1_Q9H845
T126B_HUMAN vs ACAD9_HUMAN
p-value: 0.035 q-value: 0.025

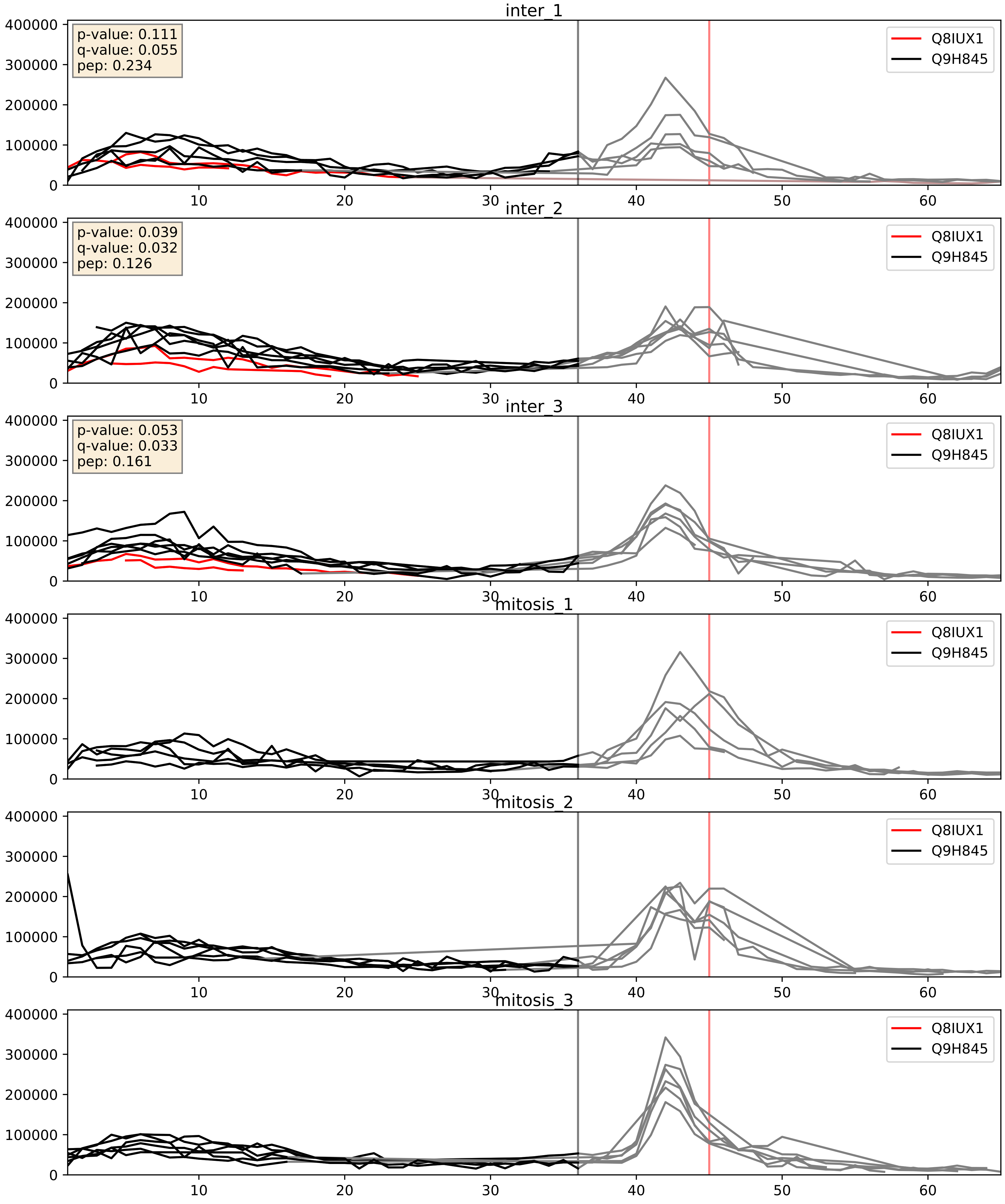


| level | condition_1 | condition_2 | pvalue | pvalue_adjusted |
|---|---|---|---|---|
| interactor_ratio | mitosis | inter | 6.375874247003068e-05 | 0.008269315690052464 |
| complex_abundance | mitosis | inter | 0.00017489733536780794 | 0.008807651911797798 |
| interactor_abundance | mitosis | inter | 0.0006537977201236383 | 0.018170482091747868 |
