## Supplementary Data 2 for "SECAT: Quantifying differential protein-protein interaction states by network-centric analysis": hela_string_000170_O75410.pdf

# O75410_O75410

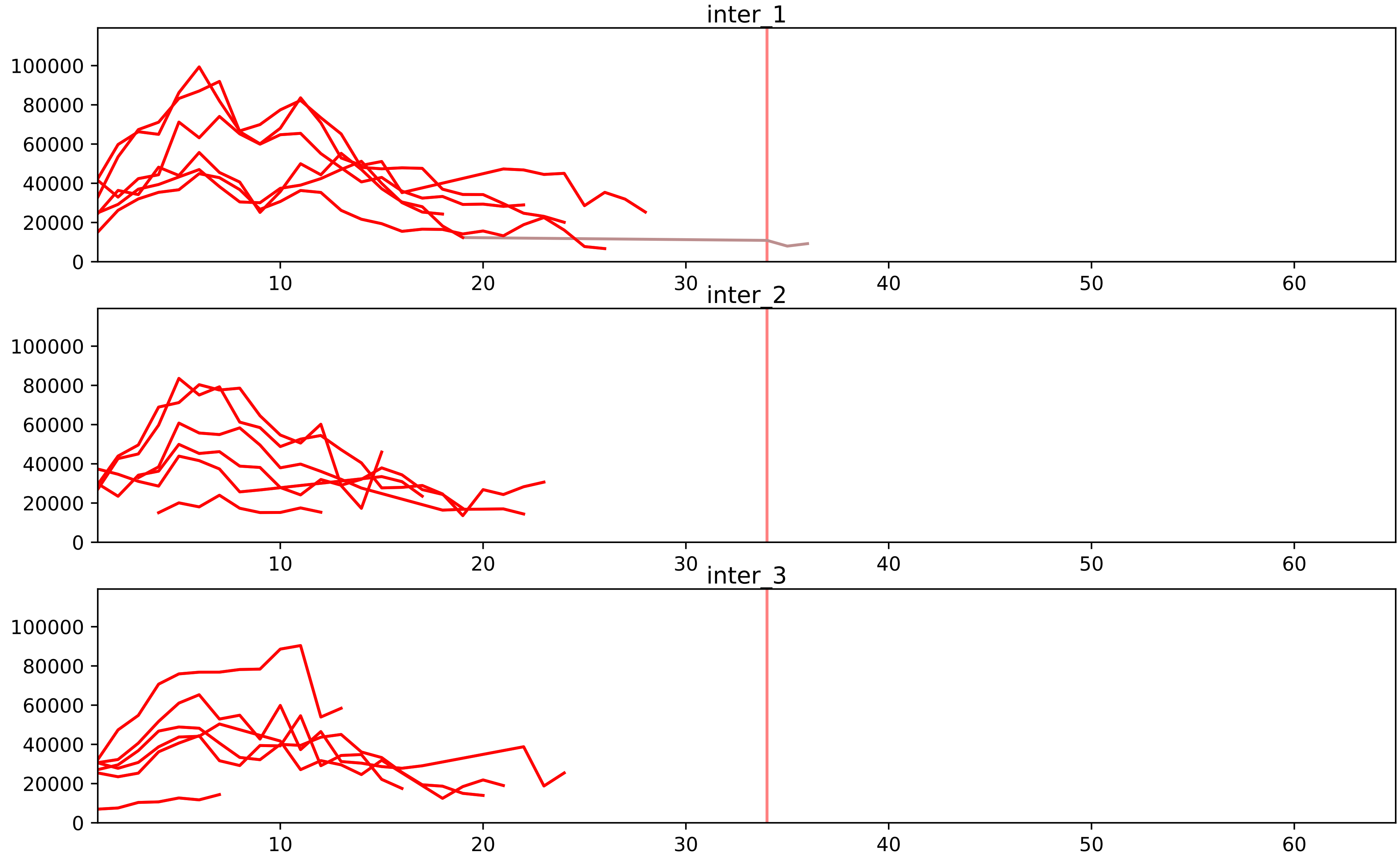


| level | condition_1 | condition_2 | pvalue | pvalue_adjusted |
|---|---|---|---|---|
| assembled_abundance | mitosis | inter | 9.154771819471233e-05 | 0.0093159754102054 |
| total_abundance | mitosis | inter | 0.00015869426808226003 | 0.015536779207822805 |
