## Supplementary Data 2 for "SECAT: Quantifying differential protein-protein interaction states by network-centric analysis": hela_string_000171_O14497.pdf

O14497_O14497

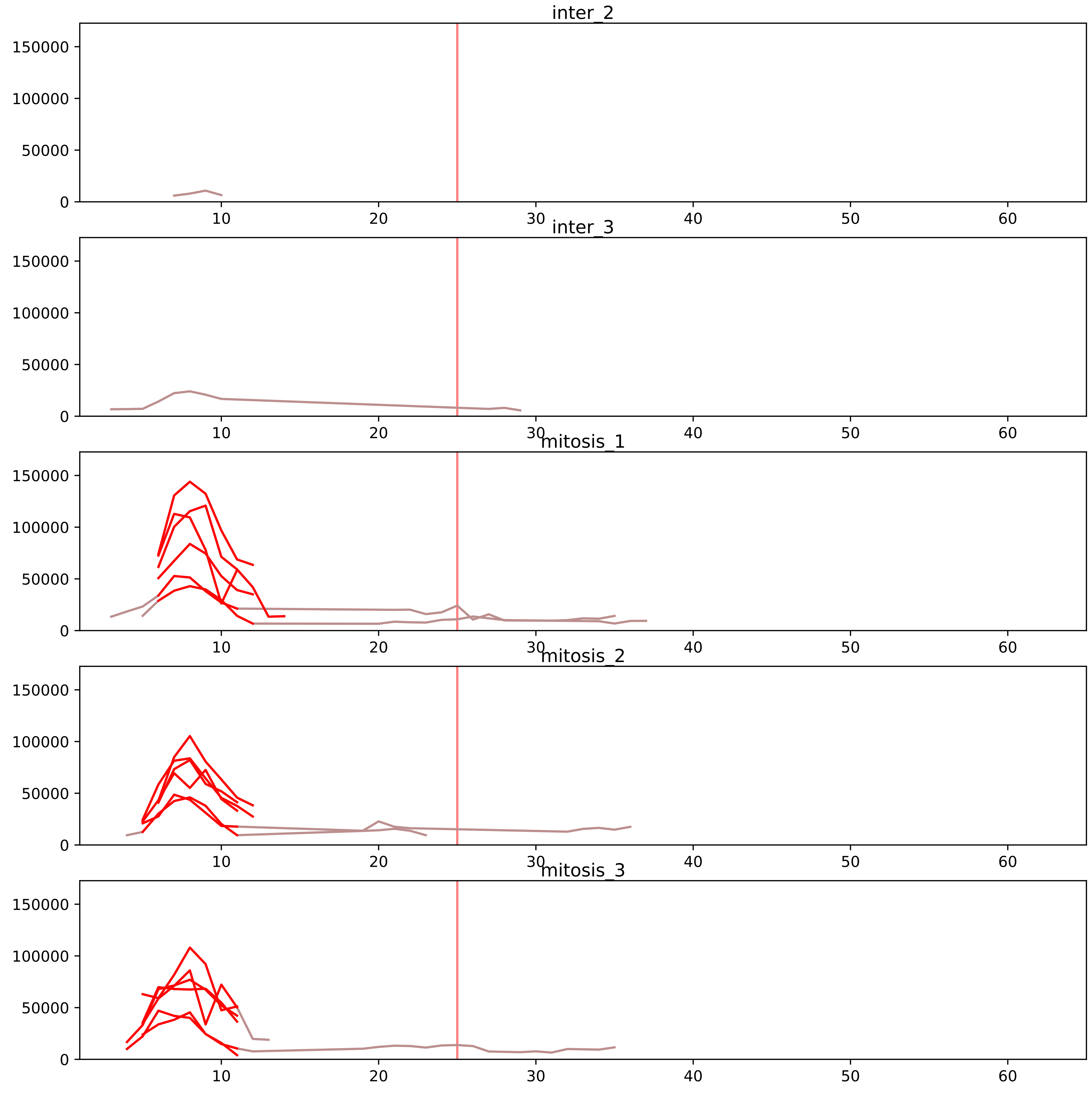


| level | condition_1 | condition_2 | pvalue | pvalue_adjusted |
|---|---|---|---|---|
| interactor_ratio | mitosis | inter | 0.00010160503024946163 | 0.001715167483110178 |
| complex_abundance | mitosis | inter | 0.00016544487608947752 | 0.003138335793862254 |
| assembled_abundance | mitosis | inter | 0.00025744902309385396 | 0.016114995197788636 |
| total_abundance | mitosis | inter | 0.0005247260370651838 | 0.027376525239428108 |
| interactor_abundance | mitosis | inter | 0.0020044105134013686 | 0.019153095904280958 |

O14497_P51531
ARI1A_HUMAN vs SMCA2_HUMAN
p-value: 0.037 q-value: 0.026

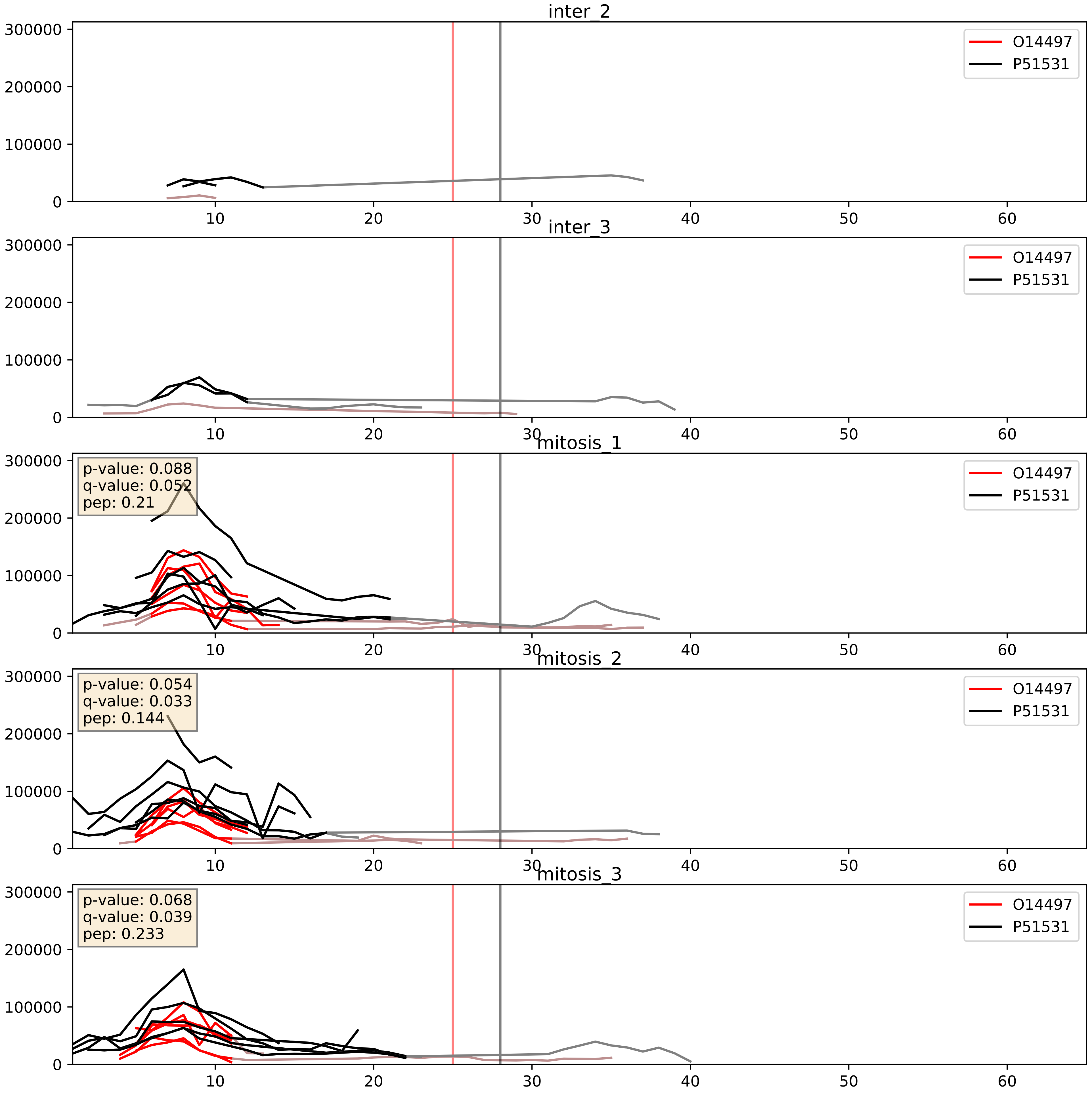


| level | condition_1 | condition_2 | pvalue | pvalue_adjusted |
|---|---|---|---|---|
| complex_abundance | mitosis | inter | 0.01207594557861336 | 0.08173689978386886 |
| interactor_abundance | mitosis | inter | 0.012640483206095423 | 0.10027749130318384 |
| interactor_ratio | mitosis | inter | 0.3232480159403212 | 0.7829618581024819 |

O14497_Q8TAQ2
ARI1A_HUMAN vs SMRC2_HUMAN
p-value: 0.012 q-value: 0.01

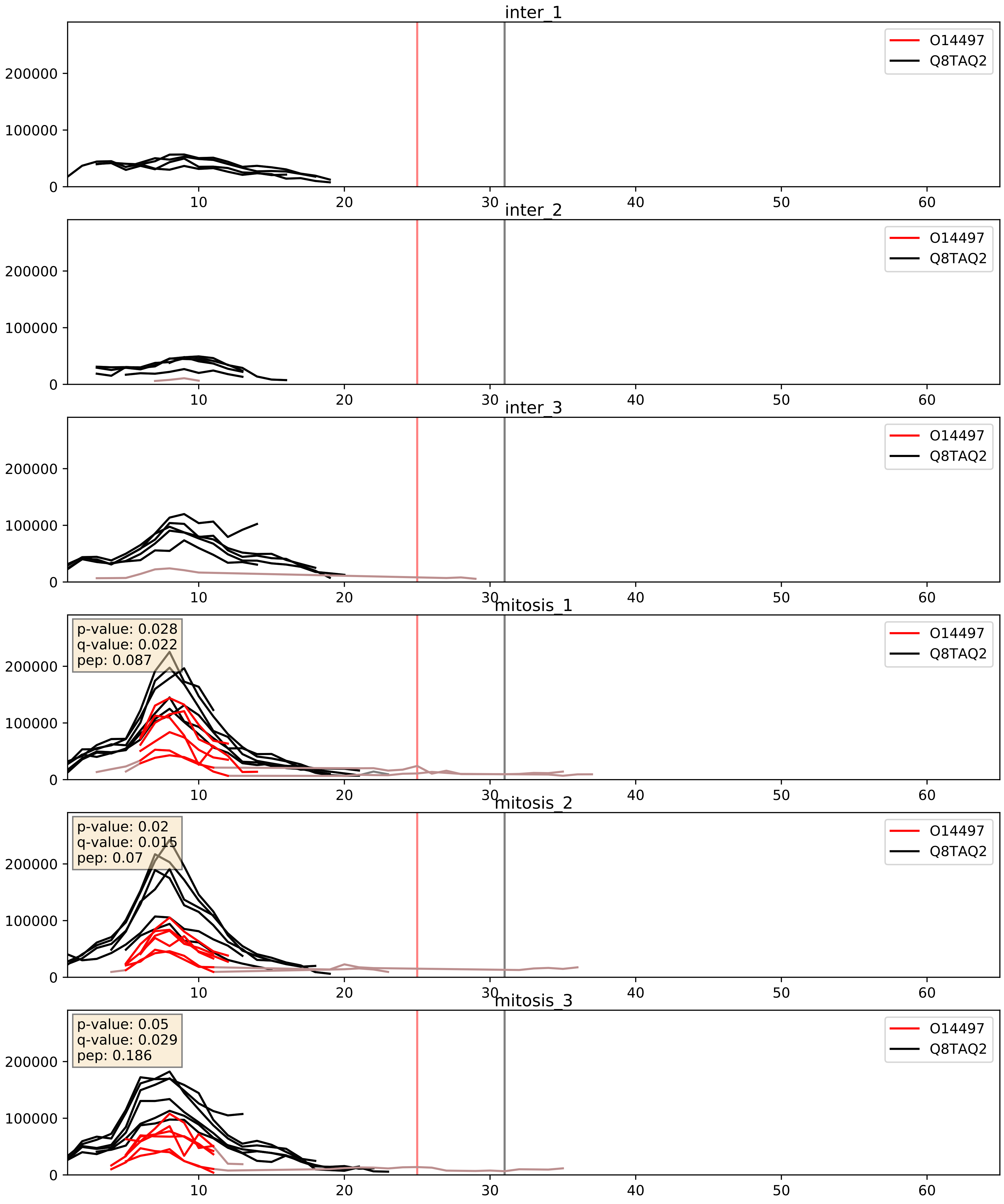


| level | condition_1 | condition_2 | pvalue | pvalue_adjusted |
|---|---|---|---|---|
| complex_abundance | mitosis | inter | 0.0012216884898636504 | 0.02350034488366932 |
| interactor_ratio | mitosis | inter | 0.007977797967995308 | 0.1259990886723217 |
| interactor_abundance | mitosis | inter | 0.012640483206095423 | 0.10027749130318384 |

O14497_Q92922
ARI1A_HUMAN vs SMRC1_HUMAN
p-value: 0.023 q-value: 0.018

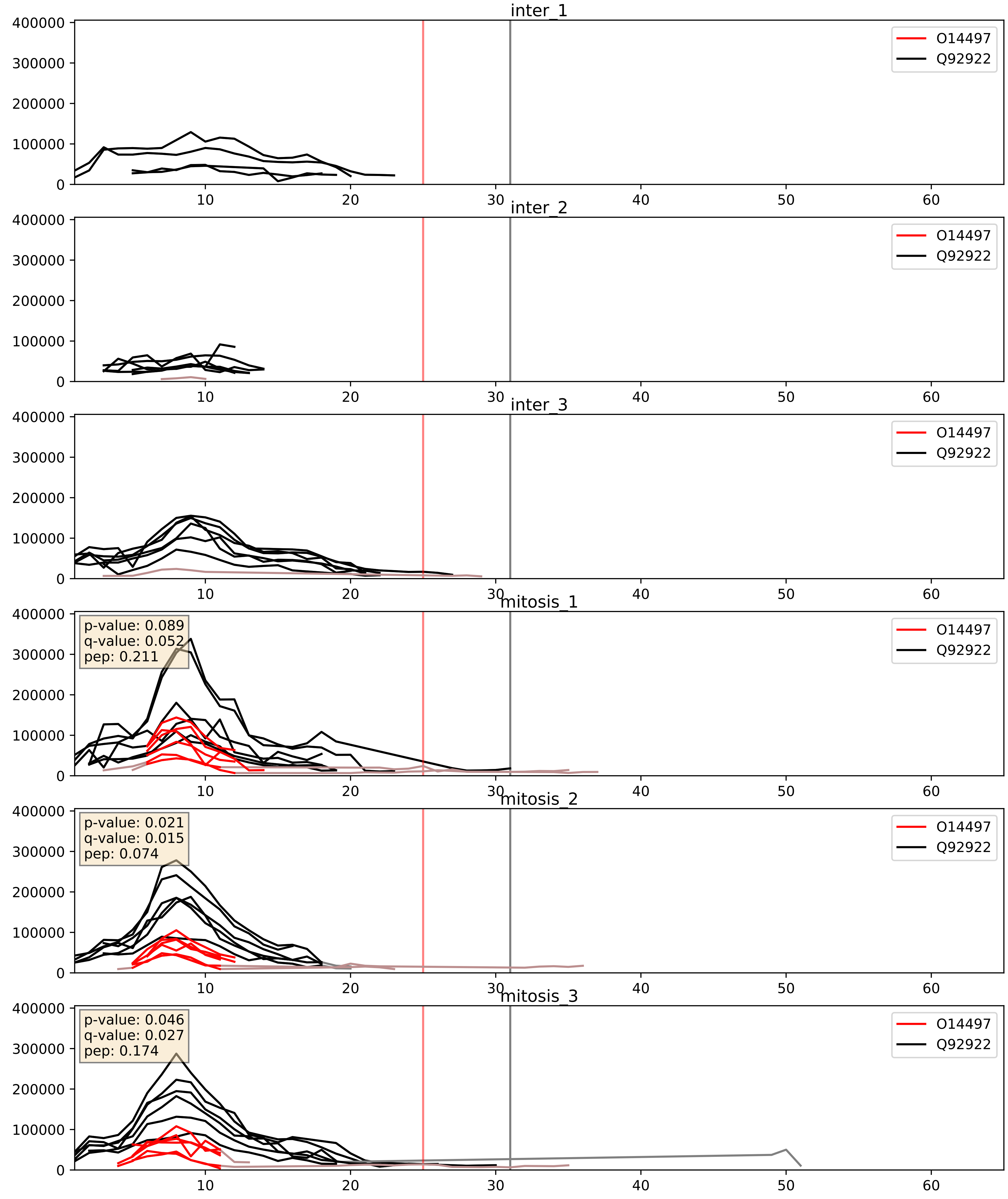


| level | condition_1 | condition_2 | pvalue | pvalue_adjusted |
|---|---|---|---|---|
| interactor_ratio | mitosis | inter | 0.0002786983396124844 | 0.017163005662466667 |
| complex_abundance | mitosis | inter | 0.0013036206631568974 | 0.024459508269436846 |
| interactor_abundance | mitosis | inter | 0.012640483206095423 | 0.10027749130318384 |

O14497_Q92925
ARI1A_HUMAN vs SMRD2_HUMAN
p-value: 0.035 q-value: 0.025

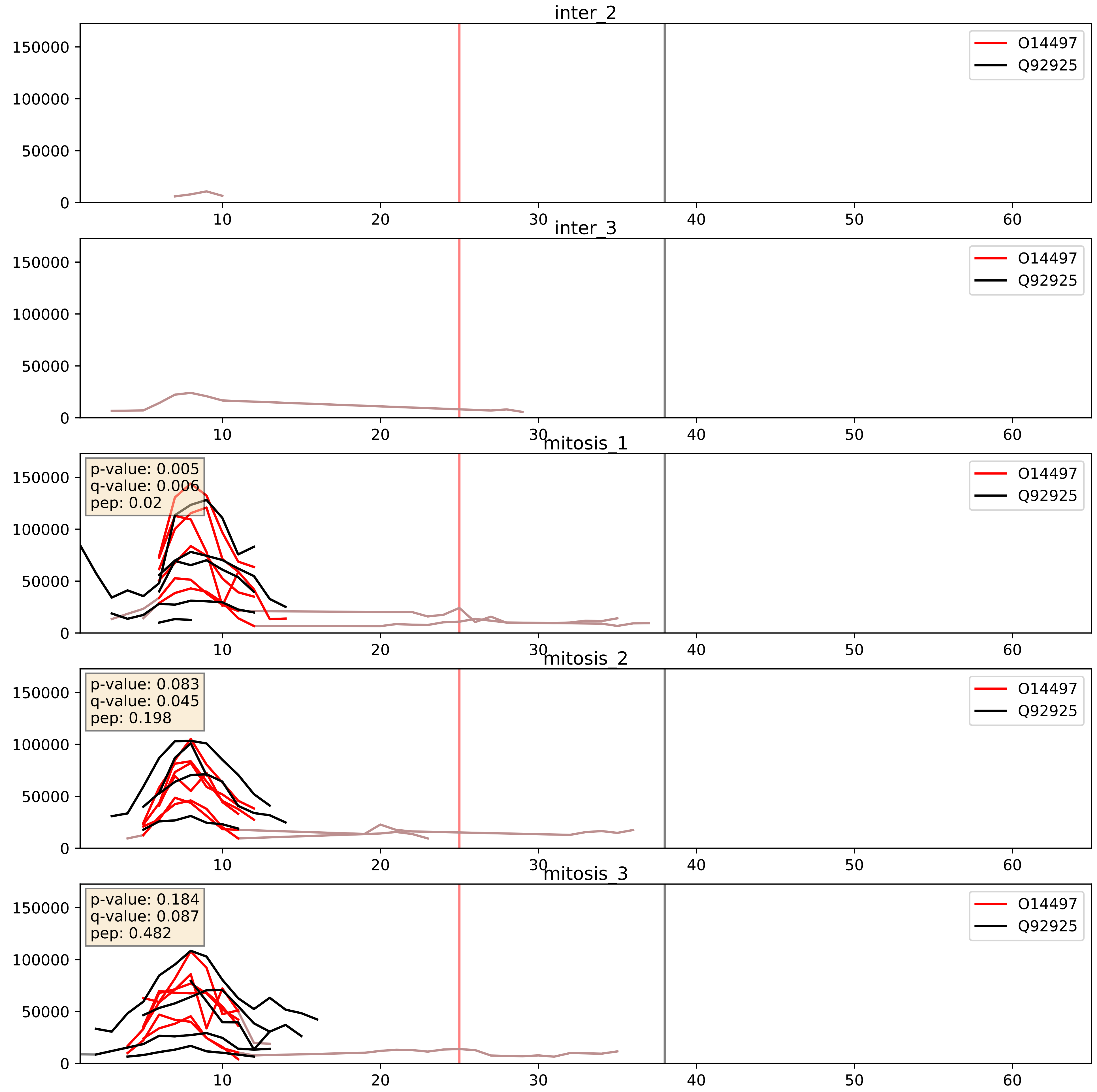


| level | condition_1 | condition_2 | pvalue | pvalue_adjusted |
|---|---|---|---|---|
| complex_abundance | mitosis | inter | 0.009992203841417977 | 0.07371585975104354 |
| interactor_abundance | mitosis | inter | 0.012640483206095423 | 0.10027749130318384 |
| interactor_ratio | mitosis | inter | 0.8861547475939666 | 0.9802747470617126 |

O14497_Q969G3
ARI1A_HUMAN vs SMCE1_HUMAN
p-value: 0.063 q-value: 0.039

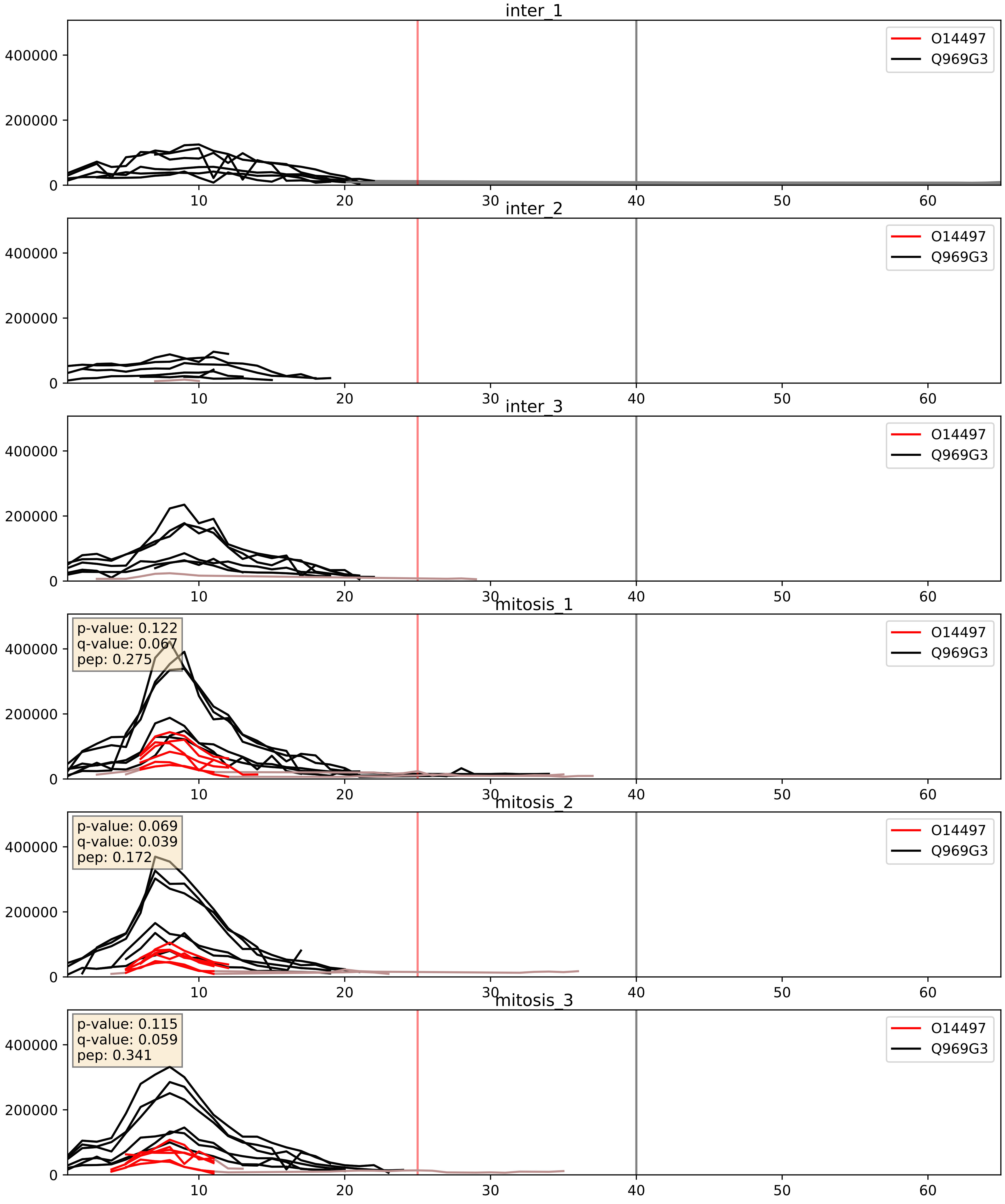


| level | condition_1 | condition_2 | pvalue | pvalue_adjusted |
|---|---|---|---|---|
| interactor_ratio | mitosis | inter | 2.3917376242737105e-05 | 0.004993481478971454 |
| complex_abundance | mitosis | inter | 0.0008870588476678302 | 0.019570164268135633 |
| interactor_abundance | mitosis | inter | 0.012640483206095423 | 0.10027749130318384 |

O14497_Q96GM5
ARI1A_HUMAN vs SMRD1_HUMAN
p-value: 0.03 q-value: 0.022

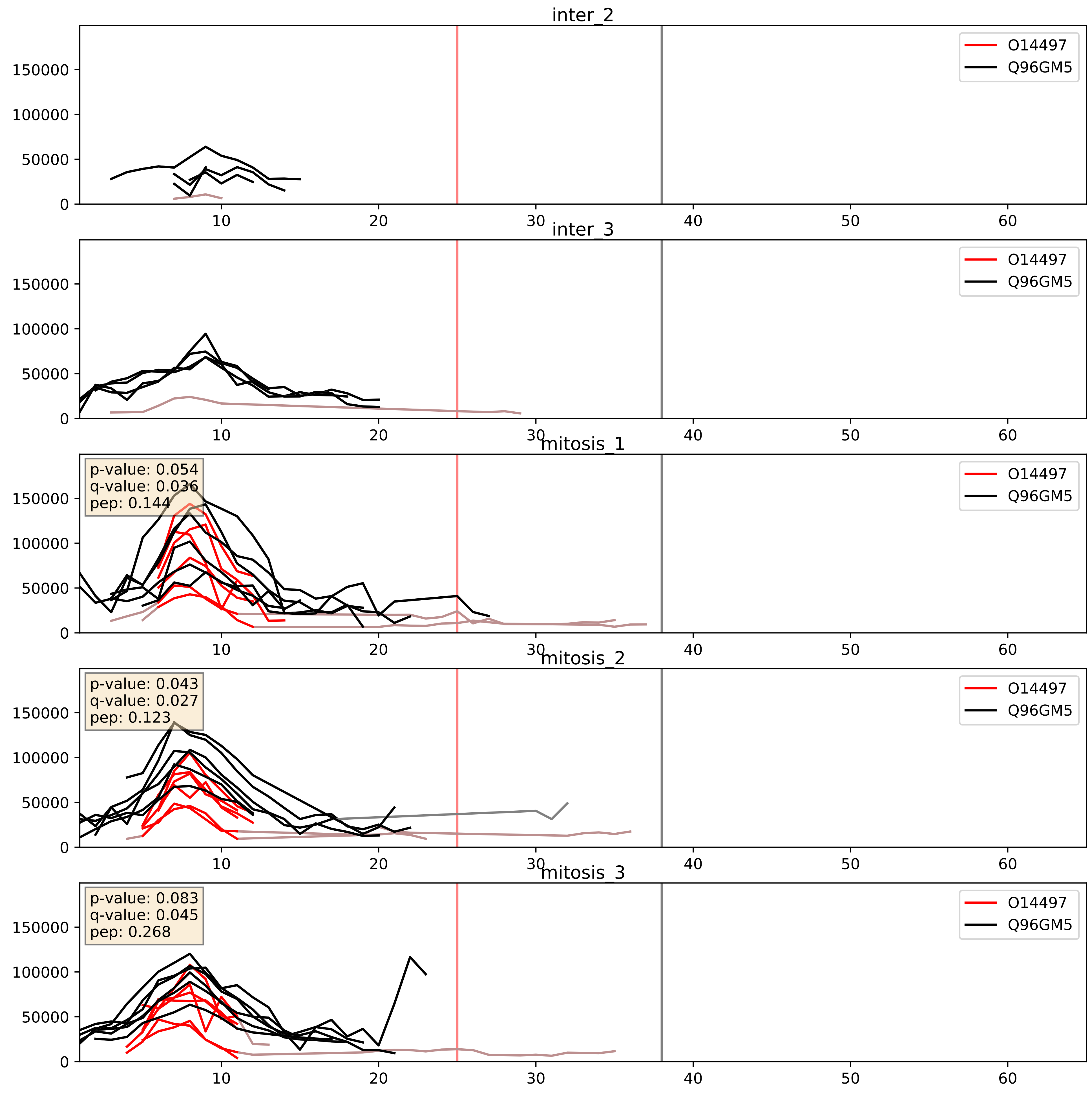


| level | condition_1 | condition_2 | pvalue | pvalue_adjusted |
|---|---|---|---|---|
| interactor_ratio | mitosis | inter | 0.0008196289026357029 | 0.03277774637652267 |
| complex_abundance | mitosis | inter | 0.00445543122107188 | 0.049788645279664145 |
| interactor_abundance | mitosis | inter | 0.012640483206095423 | 0.10027749130318384 |
