## Supplementary Data 2 for "SECAT: Quantifying differential protein-protein interaction states by network-centric analysis": hela_string_000172_Q8WXH0.pdf

Q8WXH0_Q8WXH0

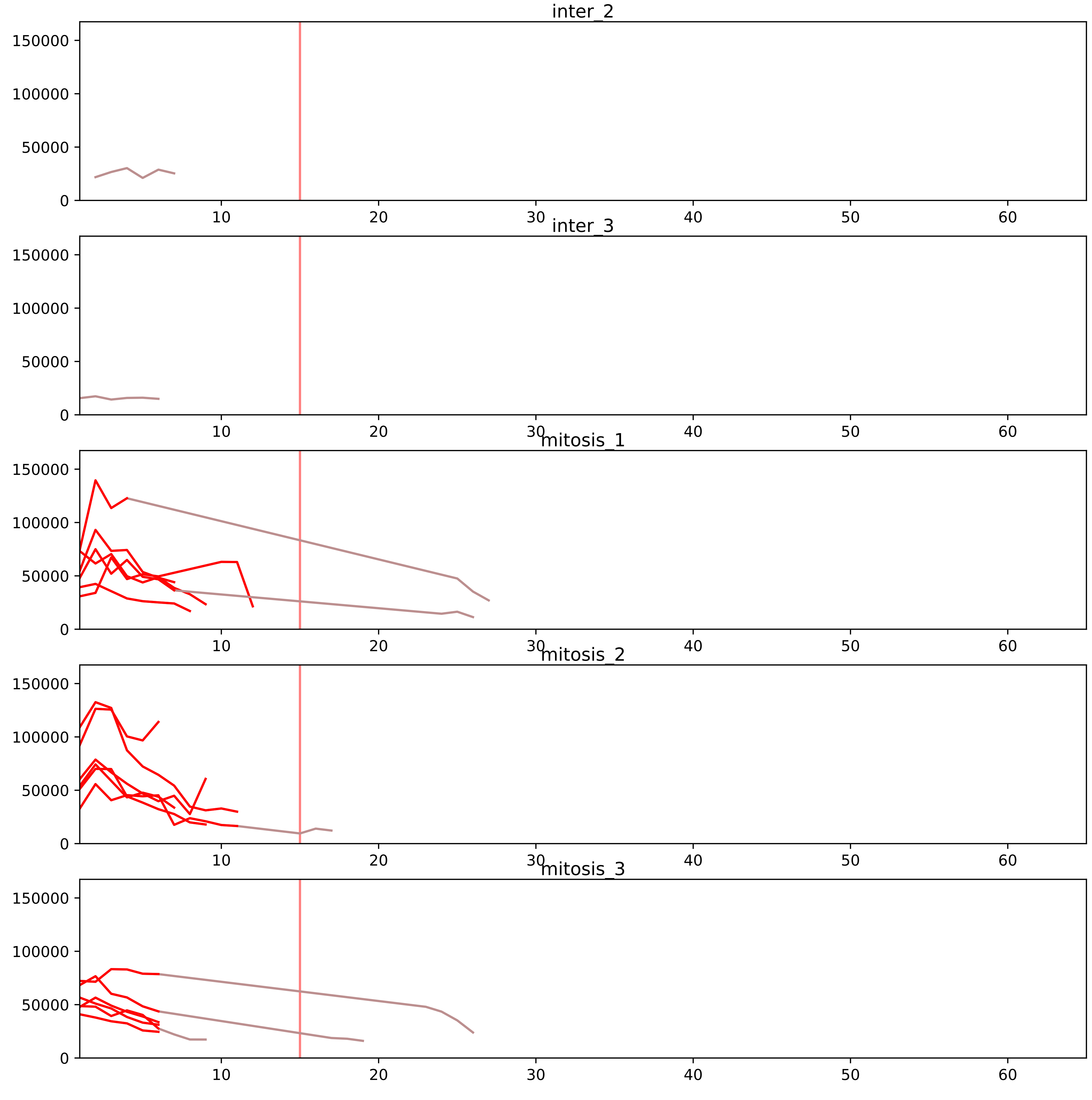


| level | condition_1 | condition_2 | pvalue | pvalue_adjusted |
|---|---|---|---|---|
| interactor_ratio | mitosis | inter | 0.00010545256052661315 | 0.0017480424447654794 |
| complex_abundance | mitosis | inter | 0.005050833438409132 | 0.028772549618182052 |
| interactor_abundance | mitosis | inter | 0.006139185516523765 | 0.03889853441851164 |
| total_abundance | mitosis | inter | 0.07040151318023956 | 0.3321724778504167 |
| assembled_abundance | mitosis | inter | 0.07385293961148502 | 0.3209575457643533 |
| monomer_abundance | mitosis | inter | 0.10749662792133326 | 0.42861511211420705 |

O94901_Q8WXH0
SUN1_HUMAN vs SYNE2_HUMAN
p-value: 0.065 q-value: 0.04

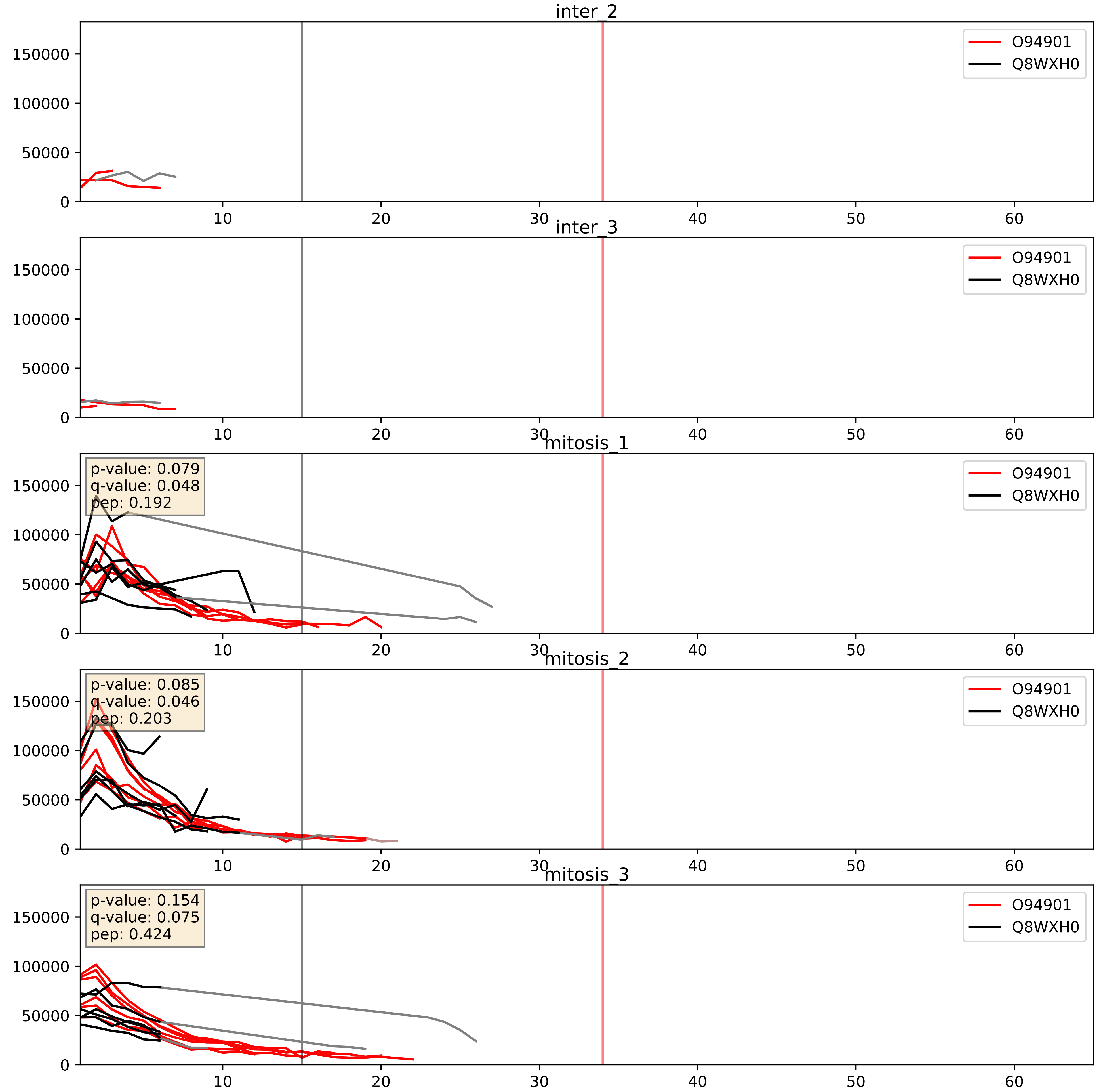


| level | condition_1 | condition_2 | pvalue | pvalue_adjusted |
|---|---|---|---|---|
| interactor_ratio | mitosis | inter | 0.00010545256052661315 | 0.010720128702277293 |
| interactor_abundance | mitosis | inter | 0.0041831623258390205 | 0.05137427476209758 |
| complex_abundance | mitosis | inter | 0.005050833438409132 | 0.05313654189341448 |
