## Supplementary Data 2 for "SECAT: Quantifying differential protein-protein interaction states by network-centric analysis": hela_string_000173_O94901.pdf

# O94901_O94901

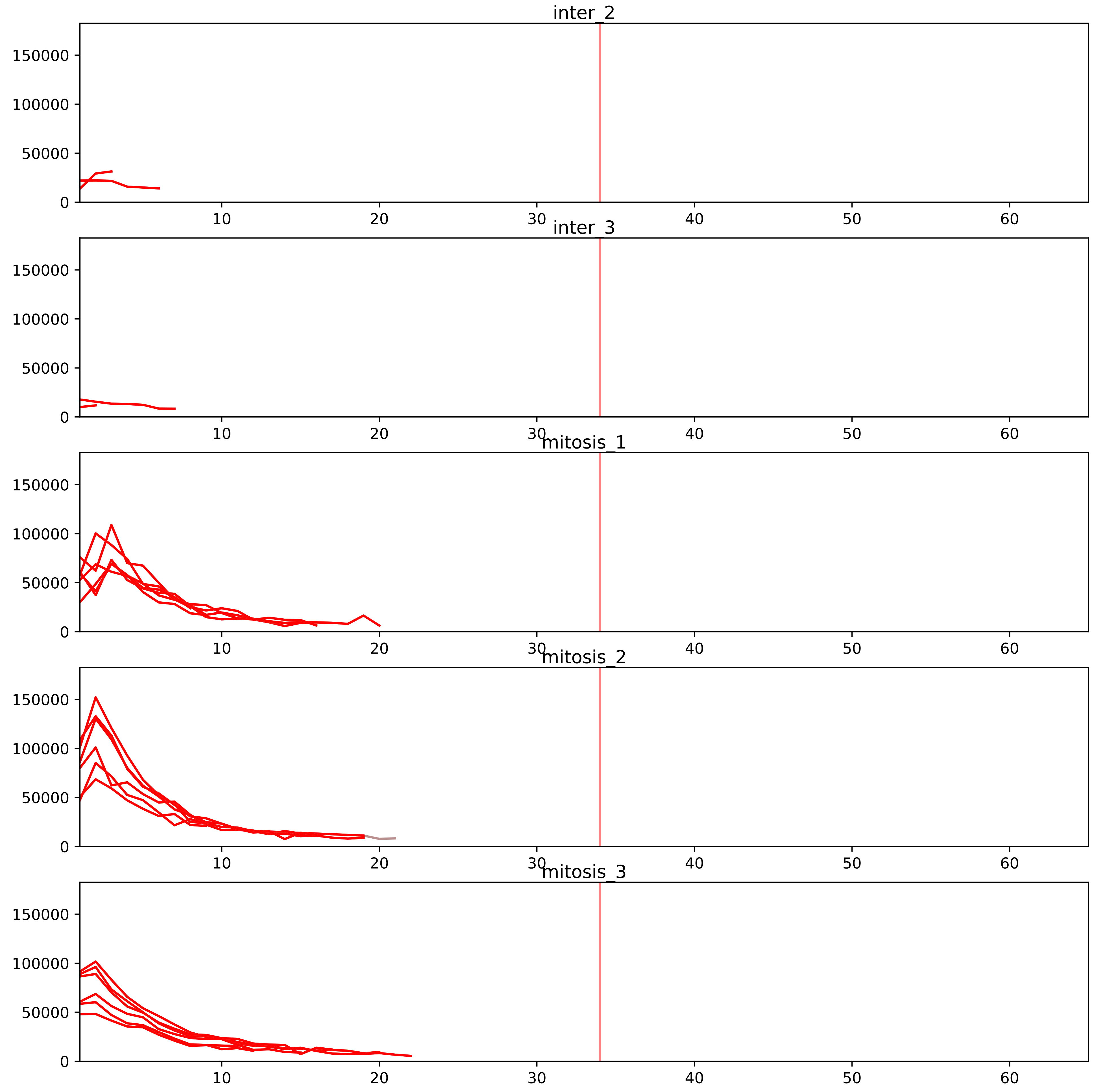


| level | condition_1 | condition_2 | pvalue | pvalue_adjusted |
|---|---|---|---|---|
| interactor_ratio | mitosis | inter | 0.00010545256052661315 | 0.0017480424447654794 |
| assembled_abundance | mitosis | inter | 0.00031017316045731016 | 0.017041946440071516 |
| total_abundance | mitosis | inter | 0.0004357526962895693 | 0.027376525239428108 |
| interactor_abundance | mitosis | inter | 0.0041831623258390205 | 0.03078807471817519 |
| complex_abundance | mitosis | inter | 0.005050833438409132 | 0.028772549618182052 |

O94901_Q8WXH0
SUN1_HUMAN vs SYNE2_HUMAN
p-value: 0.065 q-value: 0.04

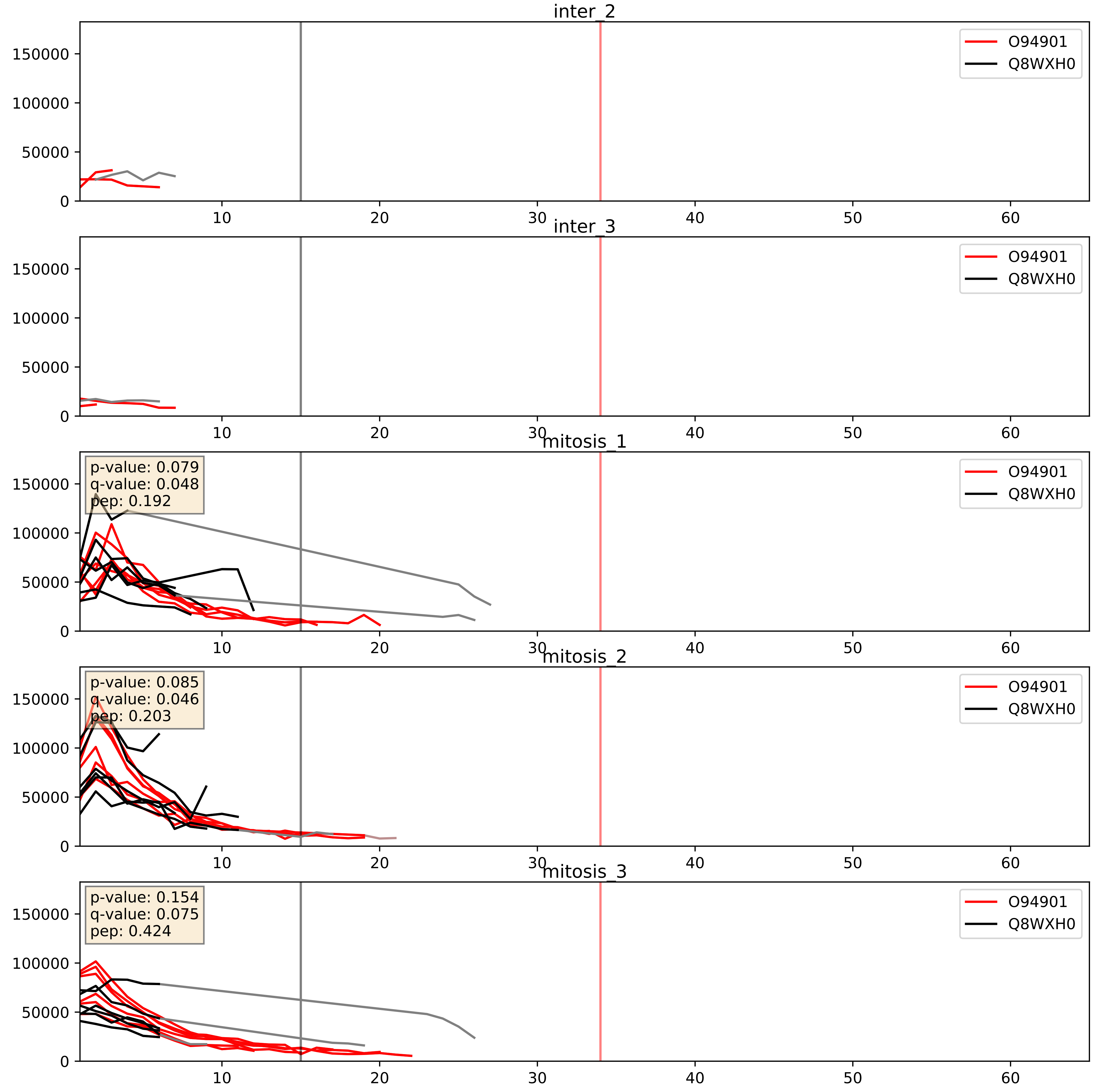


| level | condition_1 | condition_2 | pvalue | pvalue_adjusted |
|---|---|---|---|---|
| interactor_ratio | mitosis | inter | 0.00010545256052661315 | 0.010720128702277293 |
| interactor_abundance | mitosis | inter | 0.0041831623258390205 | 0.05137427476209758 |
| complex_abundance | mitosis | inter | 0.005050833438409132 | 0.05313654189341448 |
