## Supplementary Data 2 for "SECAT: Quantifying differential protein-protein interaction states by network-centric analysis": hela_string_000174_O43264.pdf

O43264_O43264

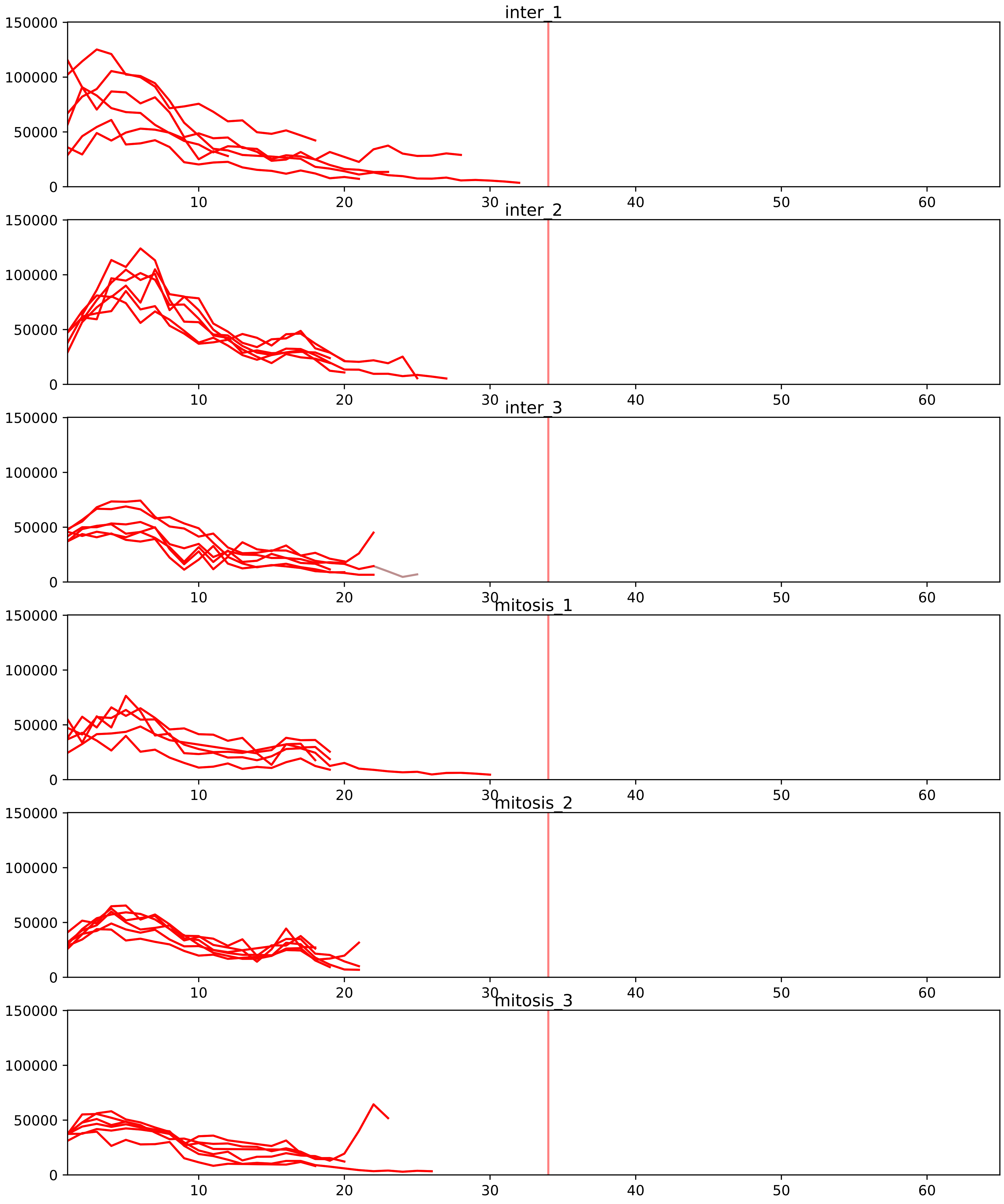


| level | condition_1 | condition_2 | pvalue | pvalue_adjusted |
|---|---|---|---|---|
| complex_abundance | mitosis | inter | 0.00010986166064583118 | 0.002323510983773901 |
| interactor_abundance | mitosis | inter | 0.0006474358524657071 | 0.008695488821437234 |
| assembled_abundance | mitosis | inter | 0.100455167017685 | 0.37055211726539283 |
| total_abundance | mitosis | inter | 0.11938219914963244 | 0.4272828770688029 |
| interactor_ratio | mitosis | inter | 0.5393775977445514 | 0.7939201847375204 |

O43264_O43683
ZW10_HUMAN vs BUB1_HUMAN
p-value: 0.063 q-value: 0.039

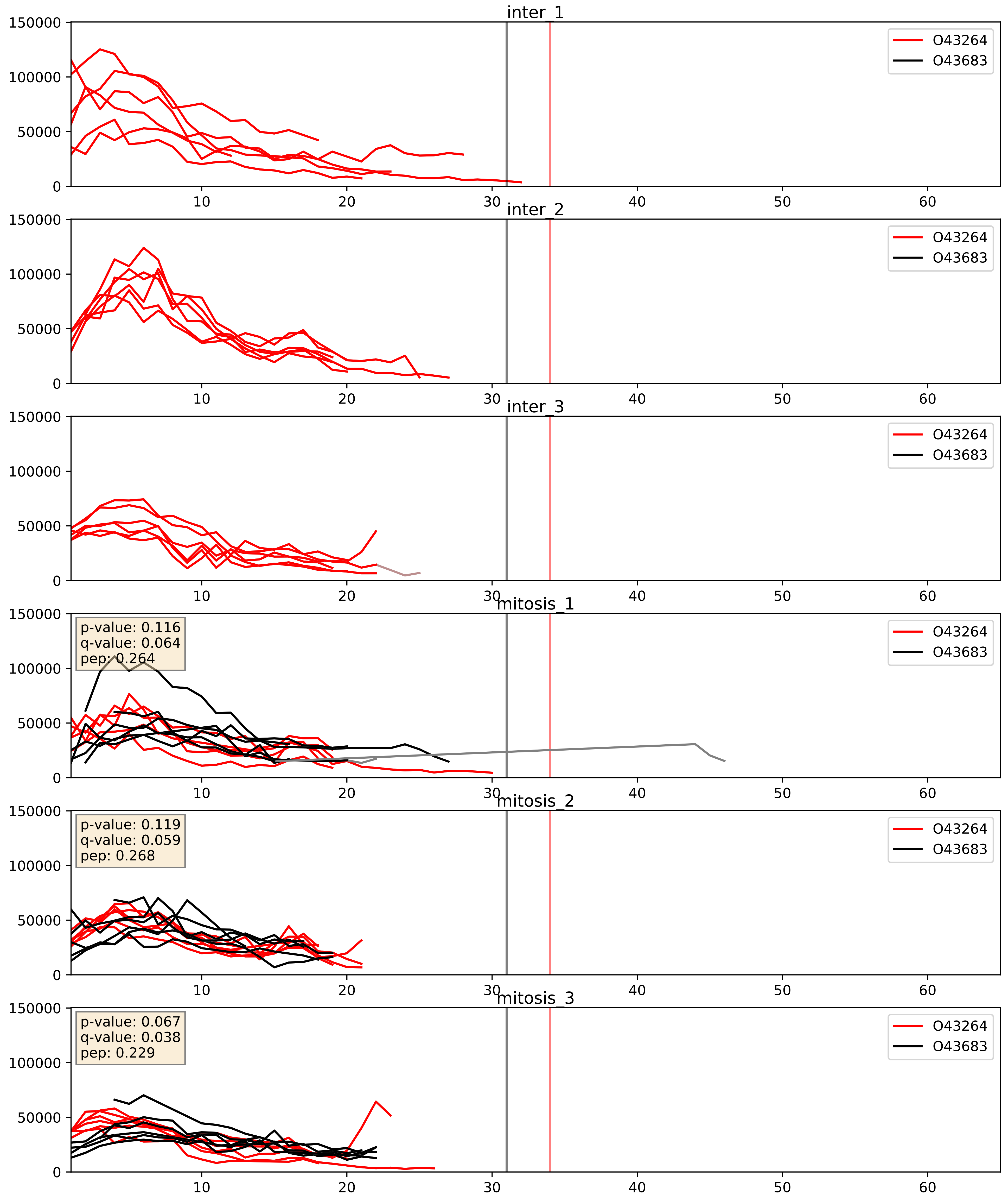


| level | condition_1 | condition_2 | pvalue | pvalue_adjusted |
|---|---|---|---|---|
| interactor_abundance | mitosis | inter | 0.0030695021355890283 | 0.042465703978082026 |
| complex_abundance | mitosis | inter | 0.003717228522607822 | 0.044942762928704735 |
| interactor_ratio | mitosis | inter | 0.129758286904364 | 0.5779454961622694 |

O43264_P50748
ZW10_HUMAN vs KNTC1_HUMAN
p-value: 0.004 q-value: 0.004

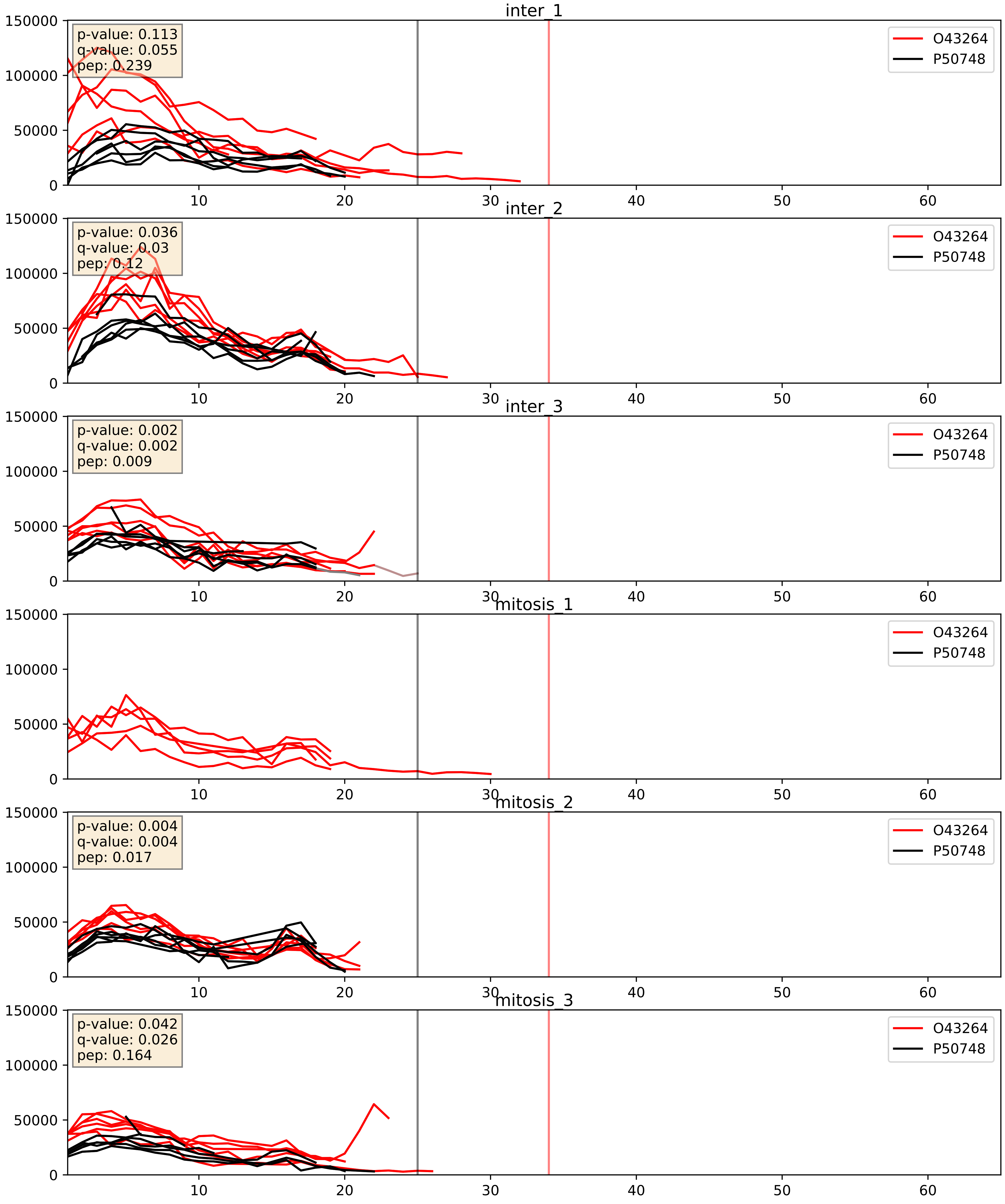


| level | condition_1 | condition_2 | pvalue | pvalue_adjusted |
|---|---|---|---|---|
| complex_abundance | mitosis | inter | 0.0577072812196418 | 0.1715081355118302 |
| interactor_abundance | mitosis | inter | 0.0669529080932372 | 0.21513396894823963 |
| interactor_ratio | mitosis | inter | 0.3125349074293607 | 0.7777940684278926 |

O43264_Q9H900
ZW10_HUMAN vs ZWILC_HUMAN
p-value: 0.062 q-value: 0.039

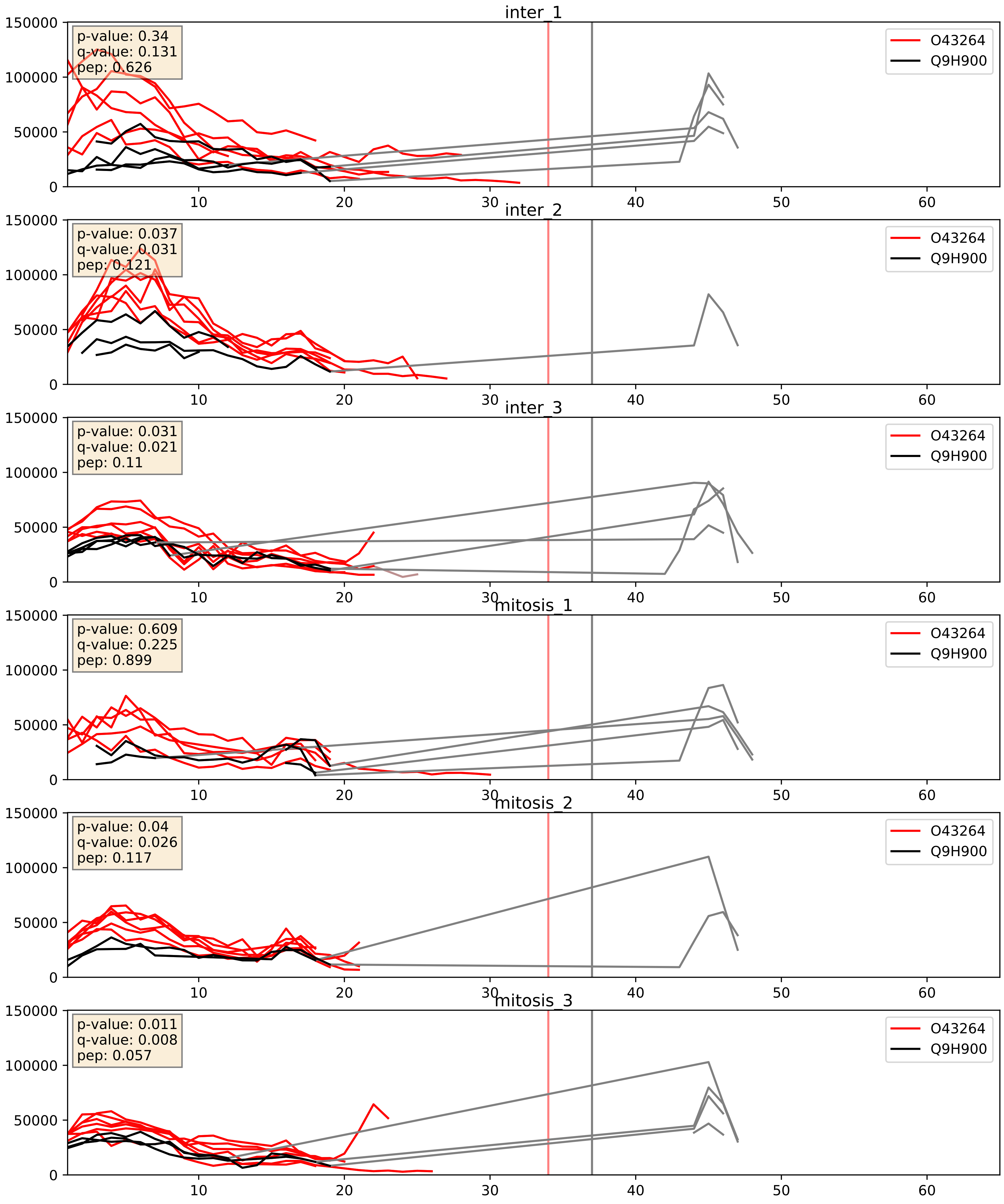


| level | condition_1 | condition_2 | pvalue | pvalue_adjusted |
|---|---|---|---|---|
| complex_abundance | mitosis | inter | 0.019634368166323 | 0.1042142626599879 |
| interactor_abundance | mitosis | inter | 0.040989623738849554 | 0.16289284085633804 |
| interactor_ratio | mitosis | inter | 0.9336404298130966 | 0.9881103513214468 |

O43264_Q9NZ43
ZW10_HUMAN vs USE1_HUMAN
p-value: 0.028 q-value: 0.021

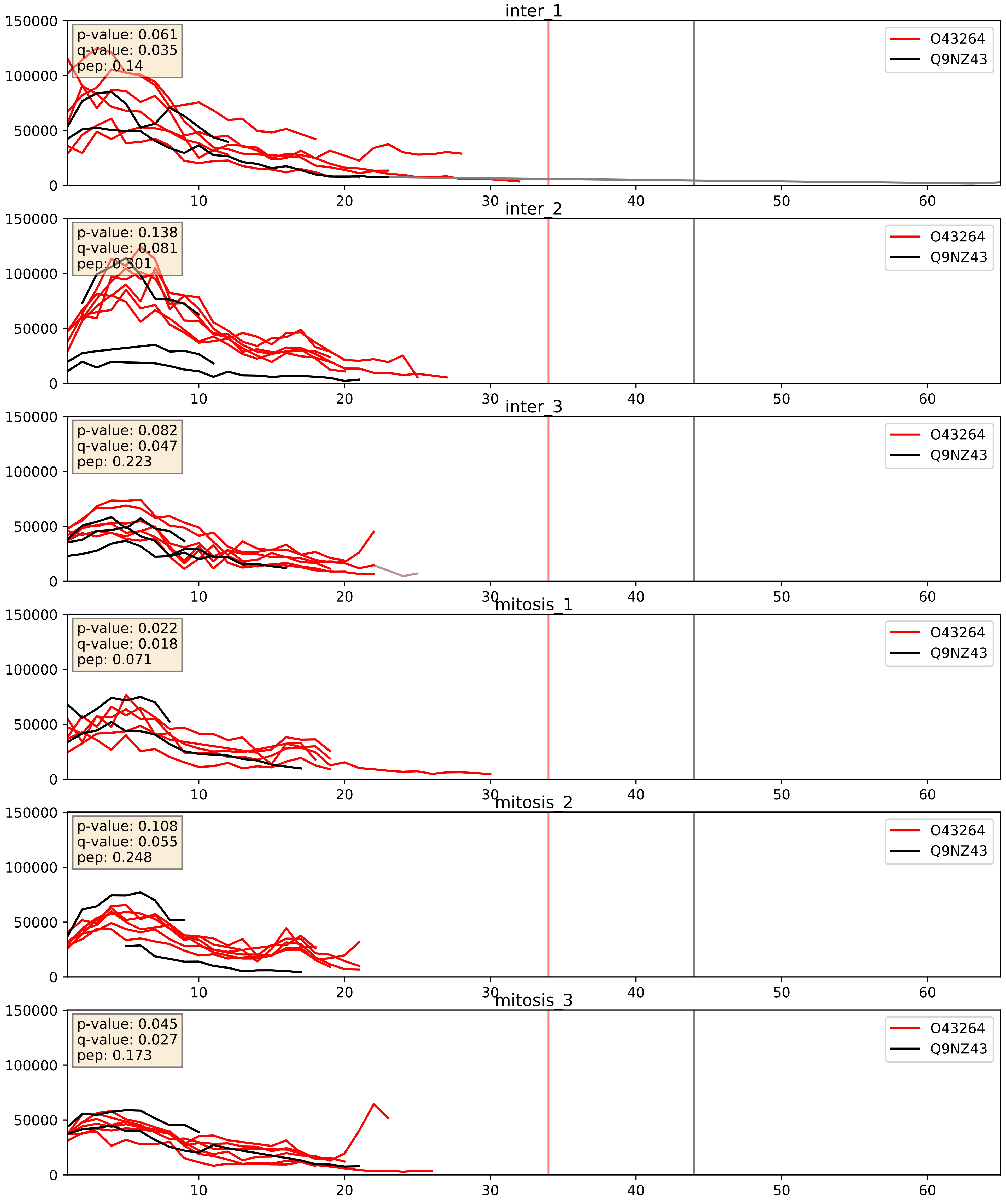


| level | condition_1 | condition_2 | pvalue | pvalue_adjusted |
|---|---|---|---|---|
| complex_abundance | mitosis | inter | 0.028788205534191227 | 0.1234203919435165 |
| interactor_abundance | mitosis | inter | 0.06536552993494153 | 0.21178233771502633 |
| interactor_ratio | mitosis | inter | 0.9133486468125236 | 0.9852905071335931 |
