## Supplementary Data 2 for "SECAT: Quantifying differential protein-protein interaction states by network-centric analysis": hela_string_000175_Q92934.pdf

Q92934_Q92934

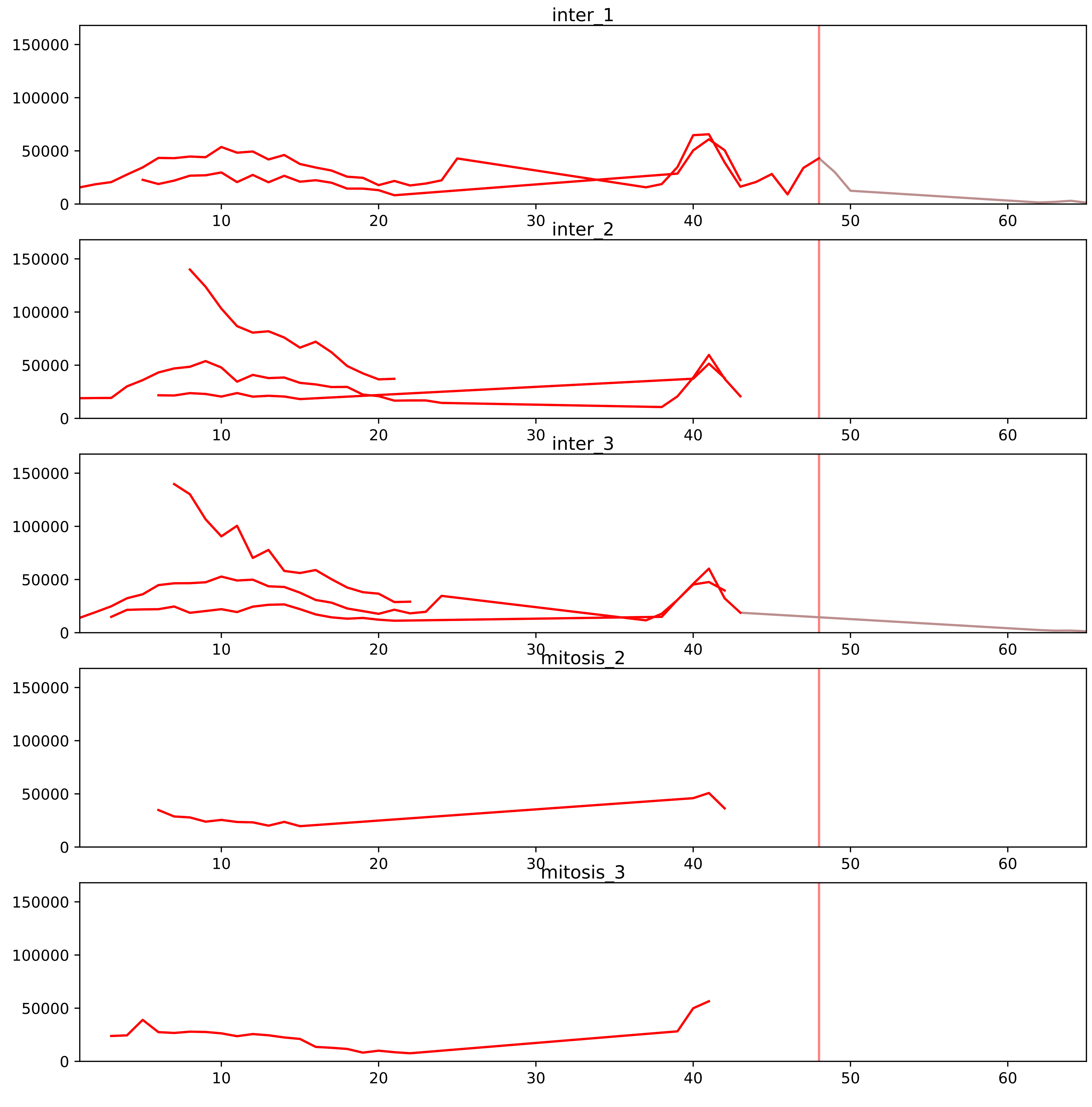


| level | condition_1 | condition_2 | pvalue | pvalue_adjusted |
|---|---|---|---|---|
| interactor_abundance | mitosis | inter | 0.00011567887126131087 | 0.002564447266515807 |
| complex_abundance | mitosis | inter | 0.0037071984496871123 | 0.02417534628390644 |
| monomer_abundance | mitosis | inter | 0.06919444779405431 | 0.37224850260048076 |
| total_abundance | mitosis | inter | 0.08150118968800386 | 0.35304392251041206 |
| assembled_abundance | mitosis | inter | 0.09742256884277352 | 0.36525609379817825 |
| interactor_ratio | mitosis | inter | 0.31884214779669445 | 0.6353858406274948 |

P32970_Q92934
CD70_HUMAN vs BAD_HUMAN
p-value: 0.0 q-value: 0.026

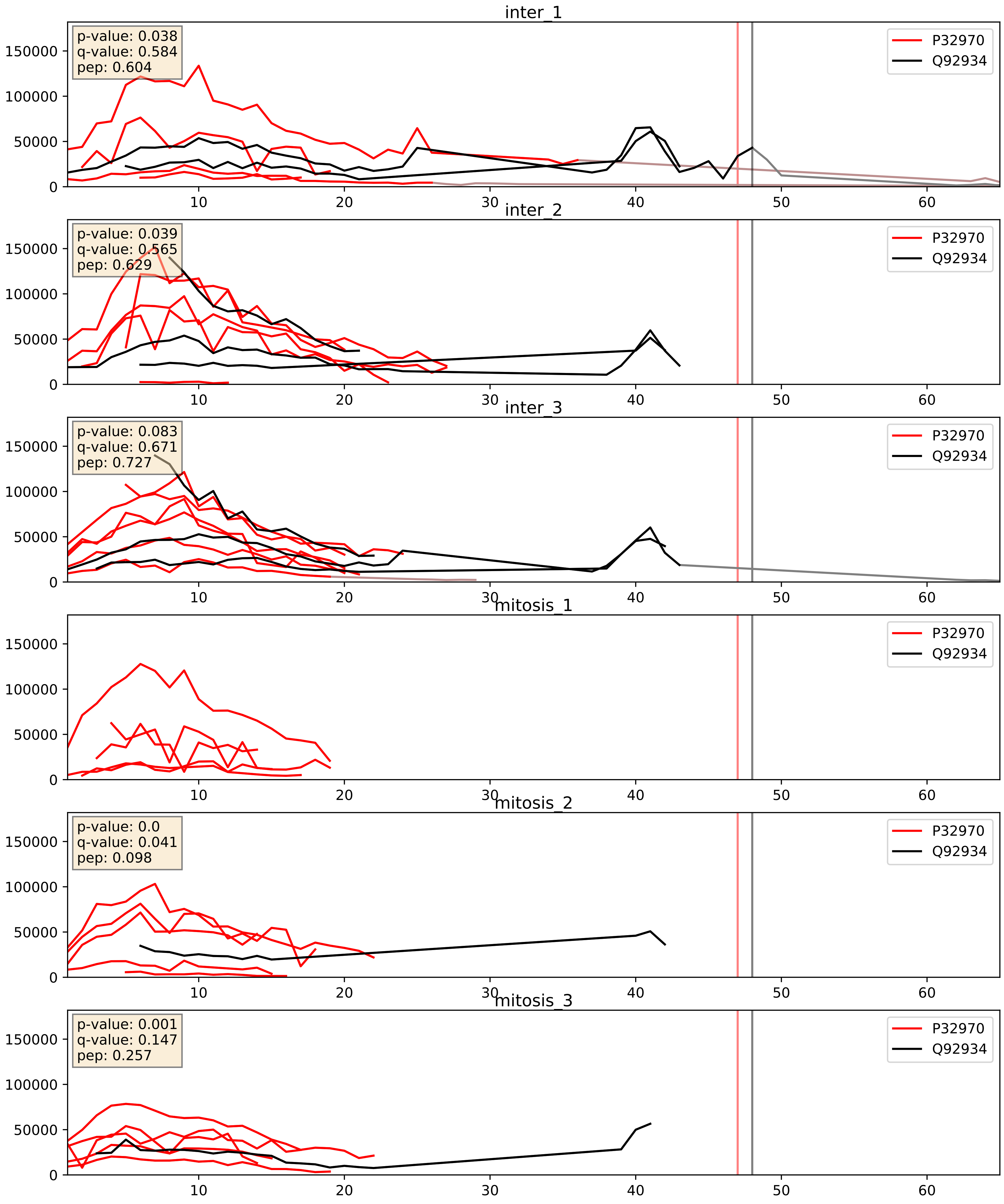


| level | condition_1 | condition_2 | pvalue | pvalue_adjusted |
|---|---|---|---|---|
| complex_abundance | mitosis | inter | 0.006892257384617678 | 0.06152004505977823 |
| interactor_abundance | mitosis | inter | 0.06633094922798757 | 0.21373722017375255 |
| interactor_ratio | mitosis | inter | 0.4175461342637969 | 0.8386191715856643 |

# Q07817_Q92934
### B2CL1_HUMAN vs BAD_HUMAN
p-value: 0.001 q-value: 0.002

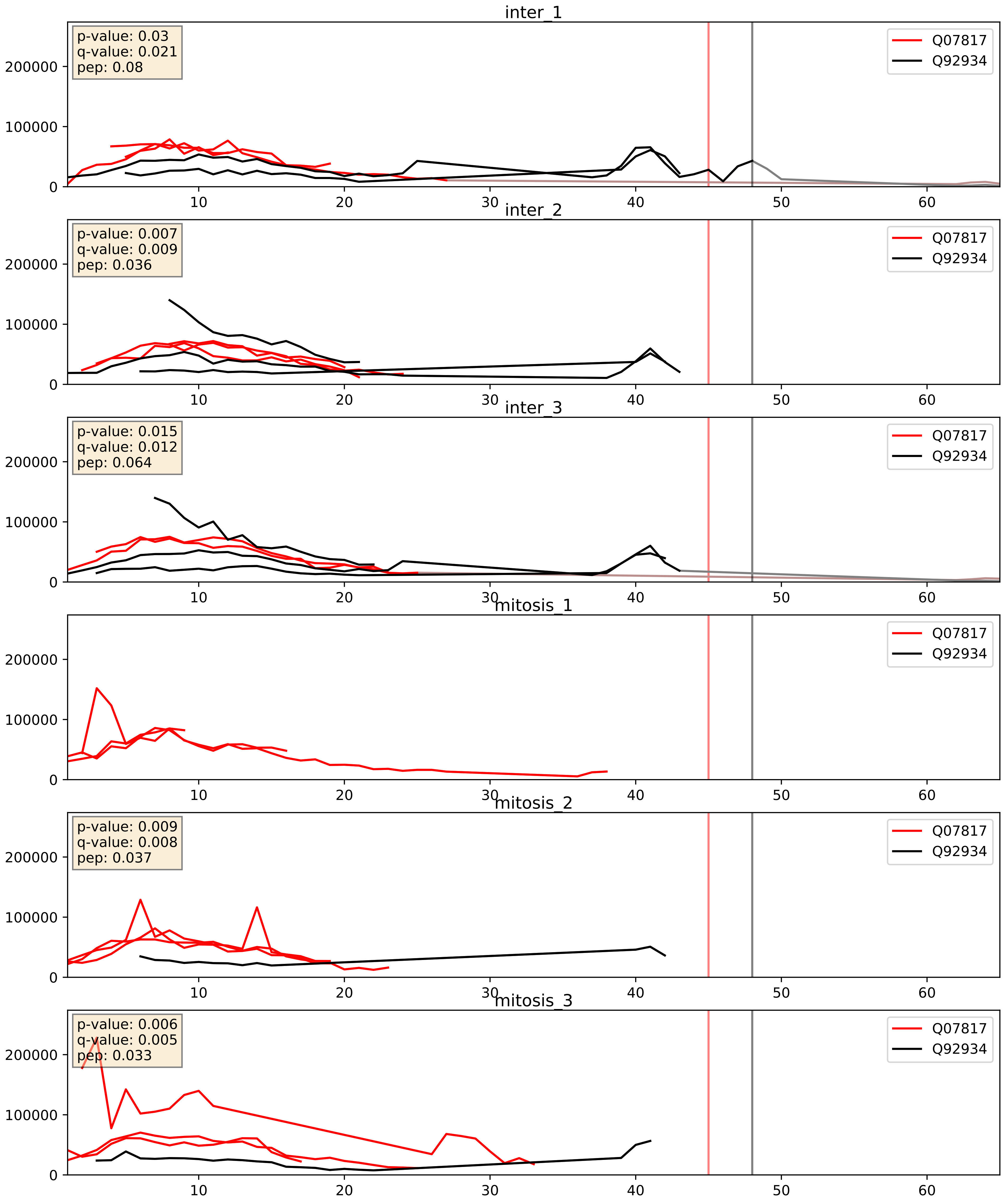


| level | condition_1 | condition_2 | pvalue | pvalue_adjusted |
|---|---|---|---|---|
| complex_abundance | mitosis | inter | 0.01300757610066597 | 0.08439667486957497 |
| interactor_ratio | mitosis | inter | 0.22987302419388697 | 0.7078104629854937 |
| interactor_abundance | mitosis | inter | 0.24693017430353273 | 0.44030845883252073 |
