## Supplementary Data 2 for "SECAT: Quantifying differential protein-protein interaction states by network-centric analysis": hela_string_000176_P00403.pdf

P00403_P00403

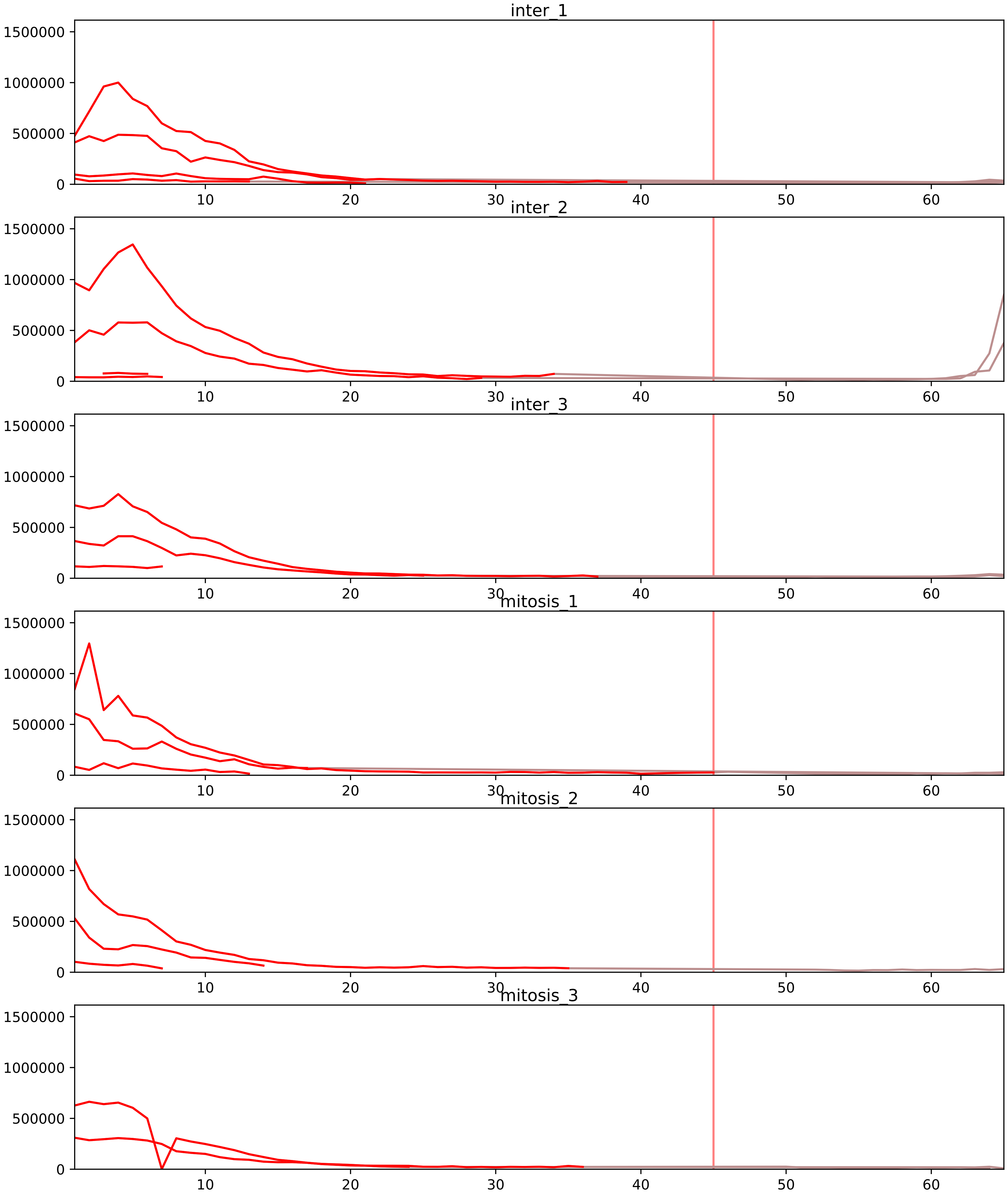


| level | condition_1 | condition_2 | pvalue | pvalue_adjusted |
|---|---|---|---|---|
| complex_abundance | mitosis | inter | 0.00012284706541505395 | 0.0024864499543178153 |
| interactor_abundance | mitosis | inter | 0.0034385631267166675 | 0.027508505013733334 |
| interactor_ratio | mitosis | inter | 0.012454196257759623 | 0.072748320997707 |
| total_abundance | mitosis | inter | 0.20002799203276148 | 0.5370308313401743 |
| assembled_abundance | mitosis | inter | 0.21274985875095648 | 0.5207127377522183 |
| monomer_abundance | mitosis | inter | 0.24442481692395726 | 0.6185083011948871 |

P00403_P03905
COX2_HUMAN vs NU4M_HUMAN
p-value: 0.004 q-value: 0.004

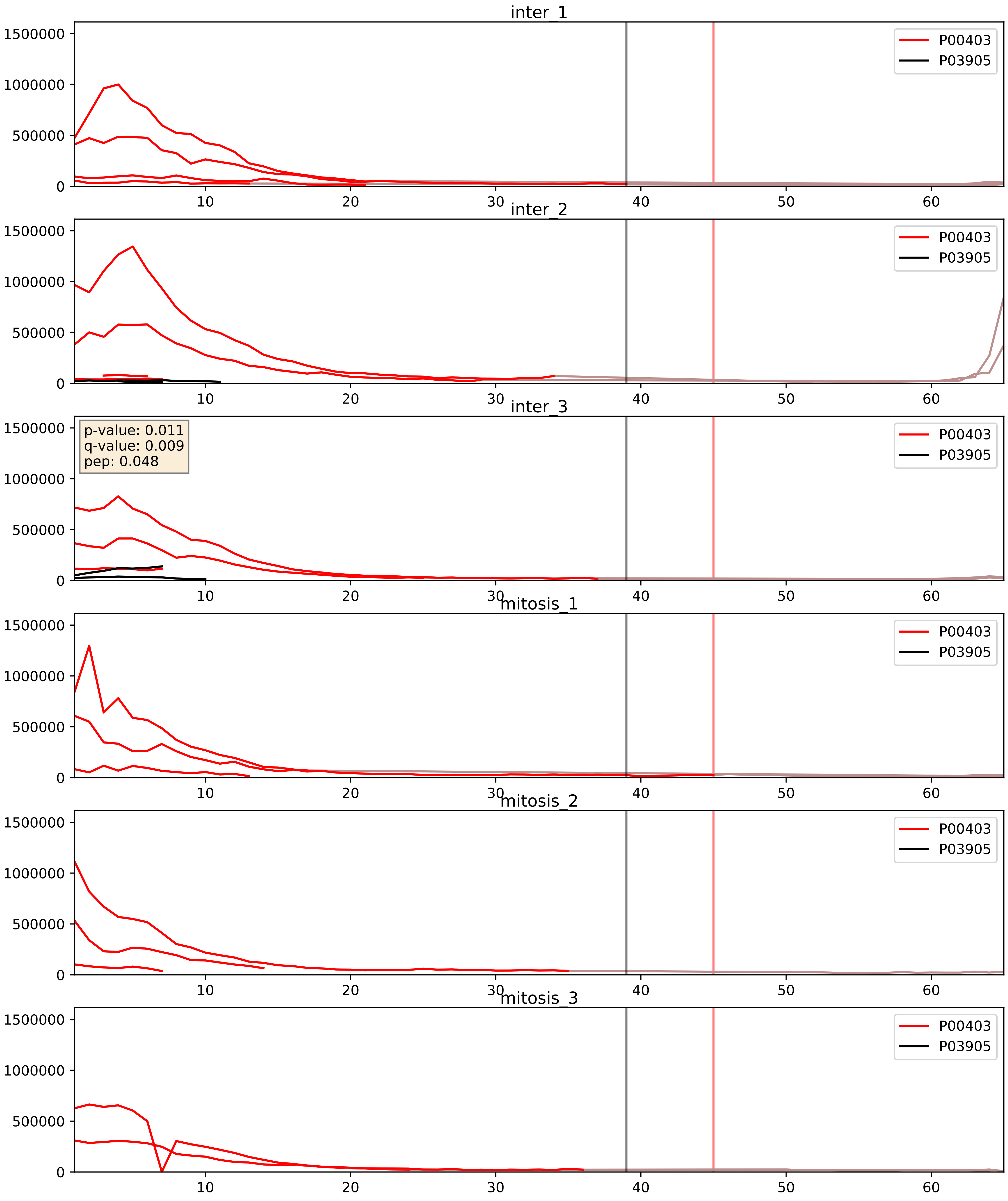


| level | condition_1 | condition_2 | pvalue | pvalue_adjusted |
|---|---|---|---|---|
| complex_abundance | mitosis | inter | 0.04223477259844842 | 0.1474505946364627 |
| interactor_abundance | mitosis | inter | 0.05371649766592087 | 0.18649838022436502 |
| interactor_ratio | mitosis | inter | 0.08312475013297722 | 0.4841037832929192 |

P00403_P09669
COX2_HUMAN vs COX6C_HUMAN
p-value: 0.004 q-value: 0.004

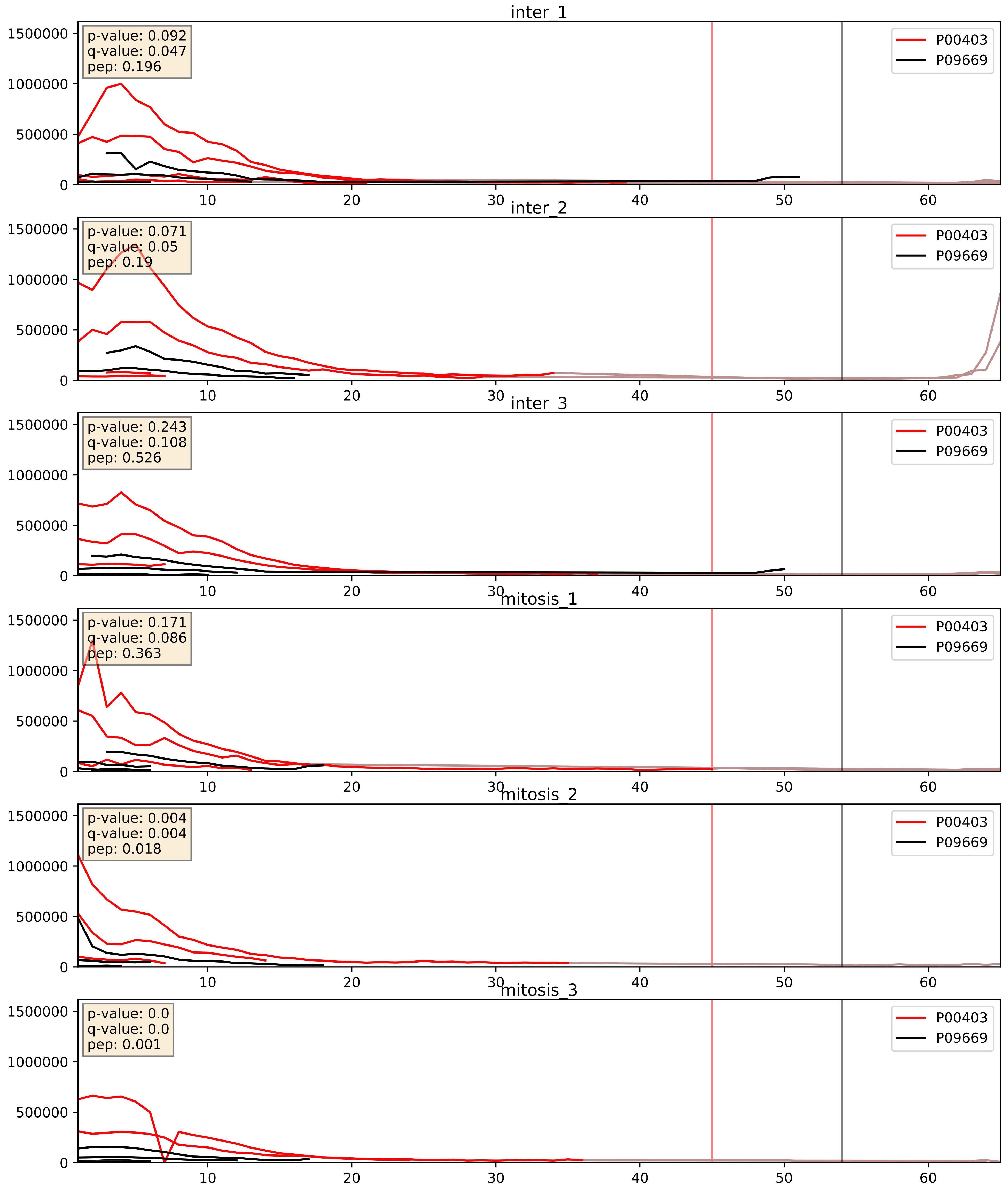


| level | condition_1 | condition_2 | pvalue | pvalue_adjusted |
|---|---|---|---|---|
| complex_abundance | mitosis | inter | 0.017954812729542805 | 0.10024201781571572 |
| interactor_abundance | mitosis | inter | 0.06299755760638603 | 0.2057063105514646 |
| interactor_ratio | mitosis | inter | 0.6520979495953283 | 0.9220833993116596 |

P00403_P10606
COX2_HUMAN vs COX5B_HUMAN
p-value: 0.001 q-value: 0.001

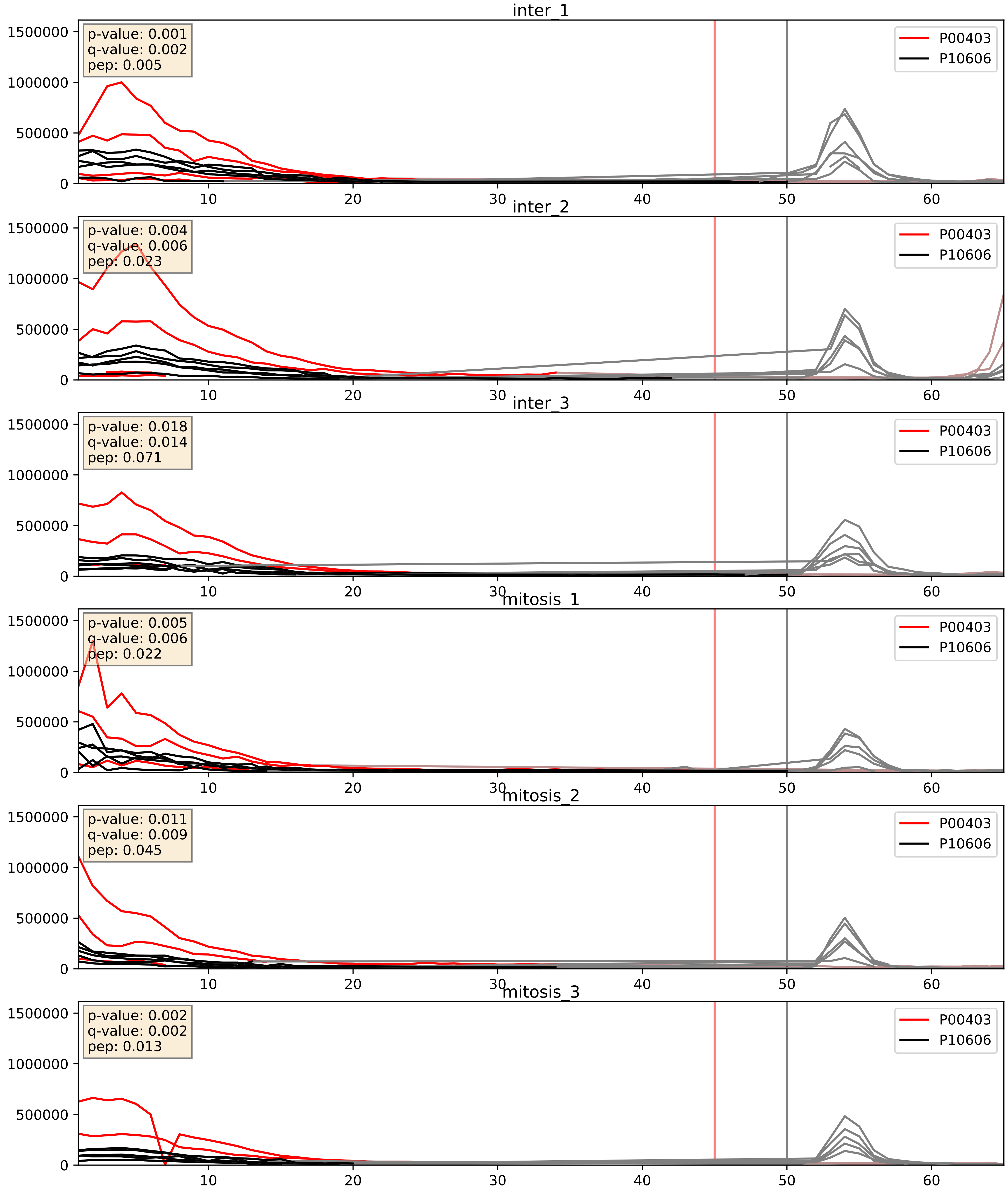


| level | condition_1 | condition_2 | pvalue | pvalue_adjusted |
|---|---|---|---|---|
| interactor_abundance | mitosis | inter | 0.05120559943433709 | 0.18161775695418983 |
| complex_abundance | mitosis | inter | 0.0542976469532346 | 0.16621801977276188 |
| interactor_ratio | mitosis | inter | 0.6344114624593611 | 0.9191879009228388 |

P00403_P13073
COX2_HUMAN vs COX41_HUMAN
p-value: 0.0 q-value: 0.0

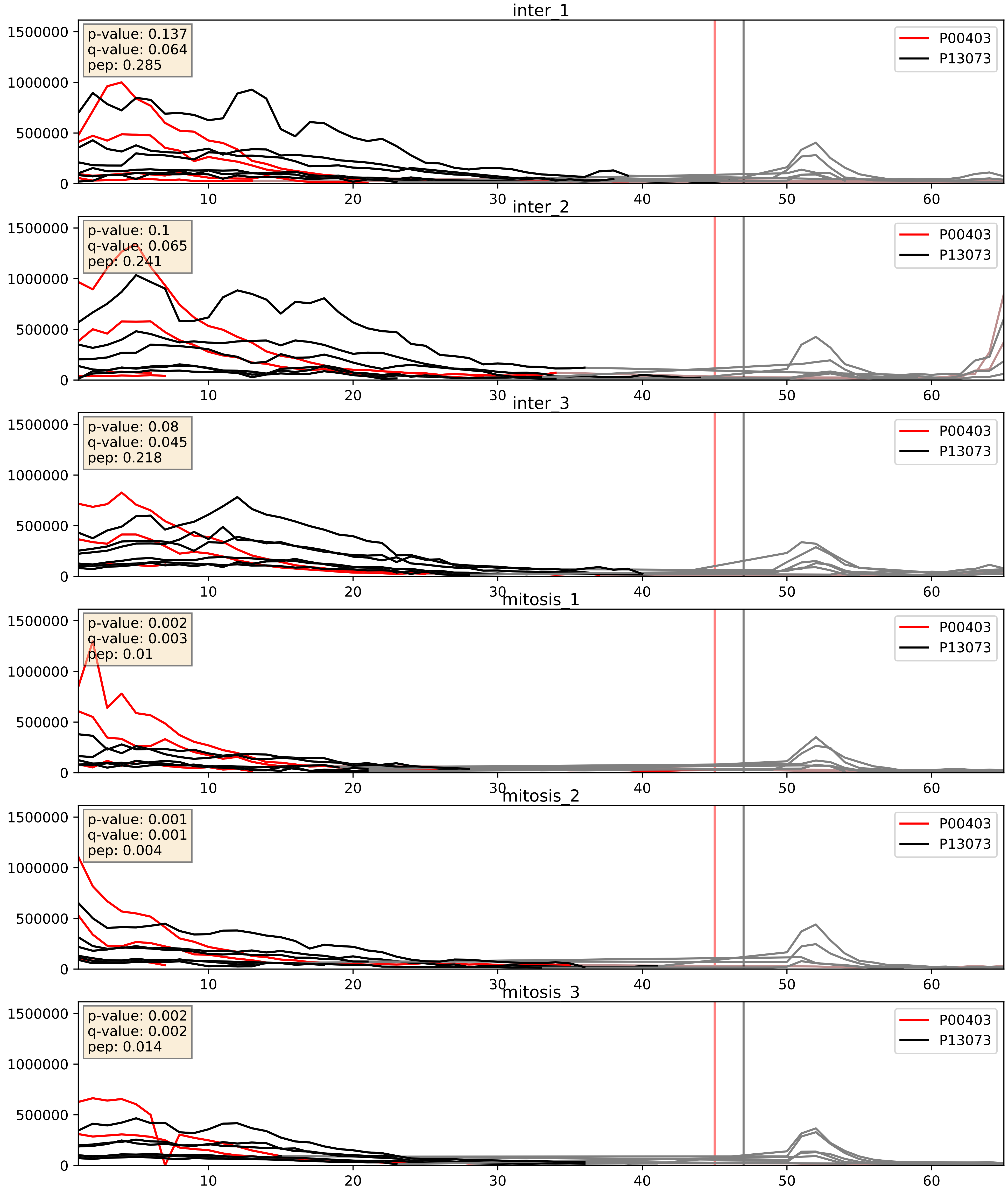


| level | condition_1 | condition_2 | pvalue | pvalue_adjusted |
|---|---|---|---|---|
| complex_abundance | mitosis | inter | 0.00033309103855573122 | 0.011886635542316795 |
| interactor_abundance | mitosis | inter | 0.04934755549120187 | 0.17741078328630325 |
| interactor_ratio | mitosis | inter | 0.37642293376682834 | 0.8148464407854535 |

P00403_P15954
COX2_HUMAN vs COX7C_HUMAN
p-value: 0.019 q-value: 0.015

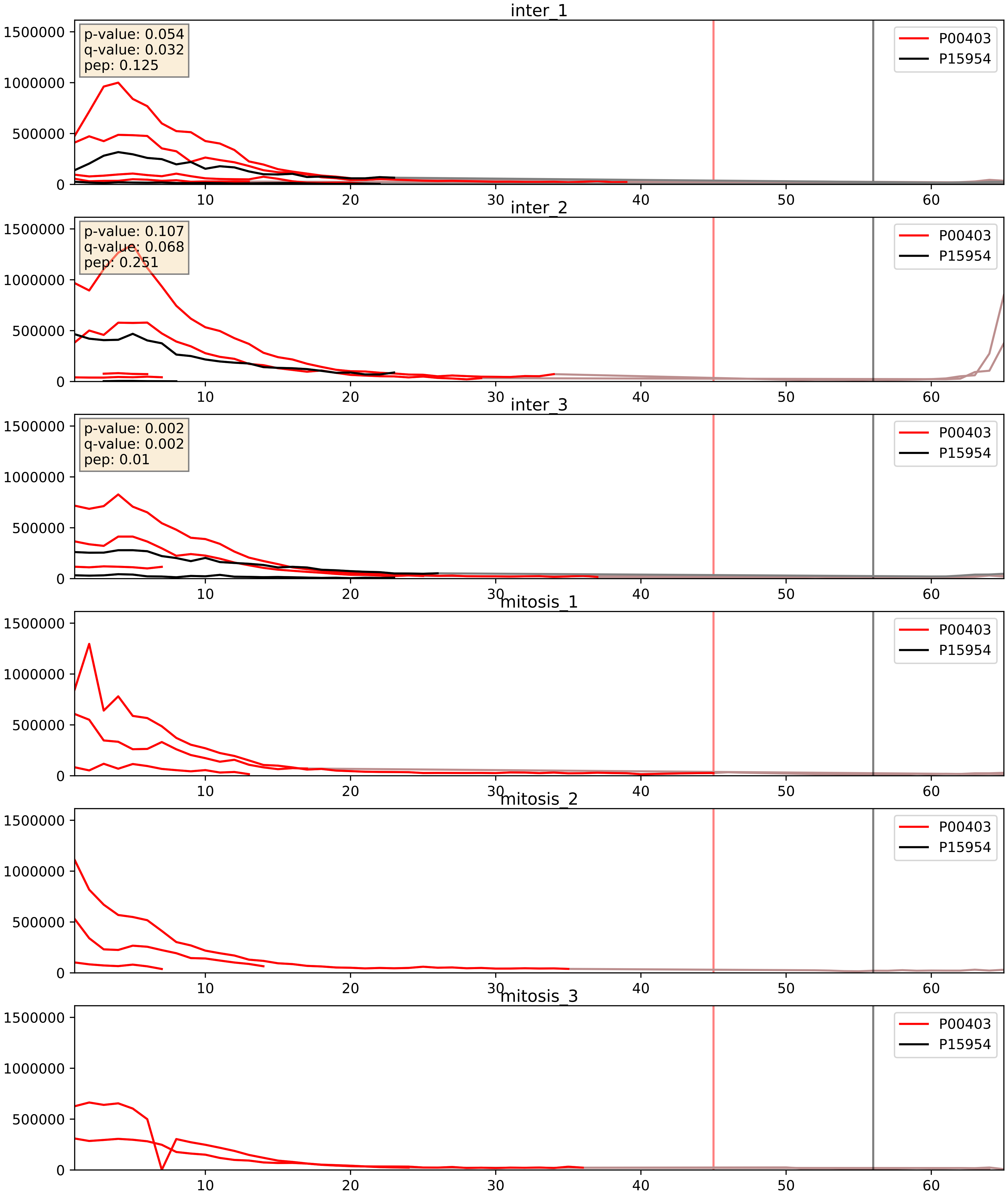


| level | condition_1 | condition_2 | pvalue | pvalue_adjusted |
|---|---|---|---|---|
| interactor_abundance | mitosis | inter | 4.4946683692739026e-05 | 0.006058954526139308 |
| complex_abundance | mitosis | inter | 5.2983607009485405e-05 | 0.005691857151652958 |
| interactor_ratio | mitosis | inter | 5.4319379841399415e-05 | 0.007596351369213947 |

P00403_P20674
COX2_HUMAN vs COX5A_HUMAN
p-value: 0.003 q-value: 0.004

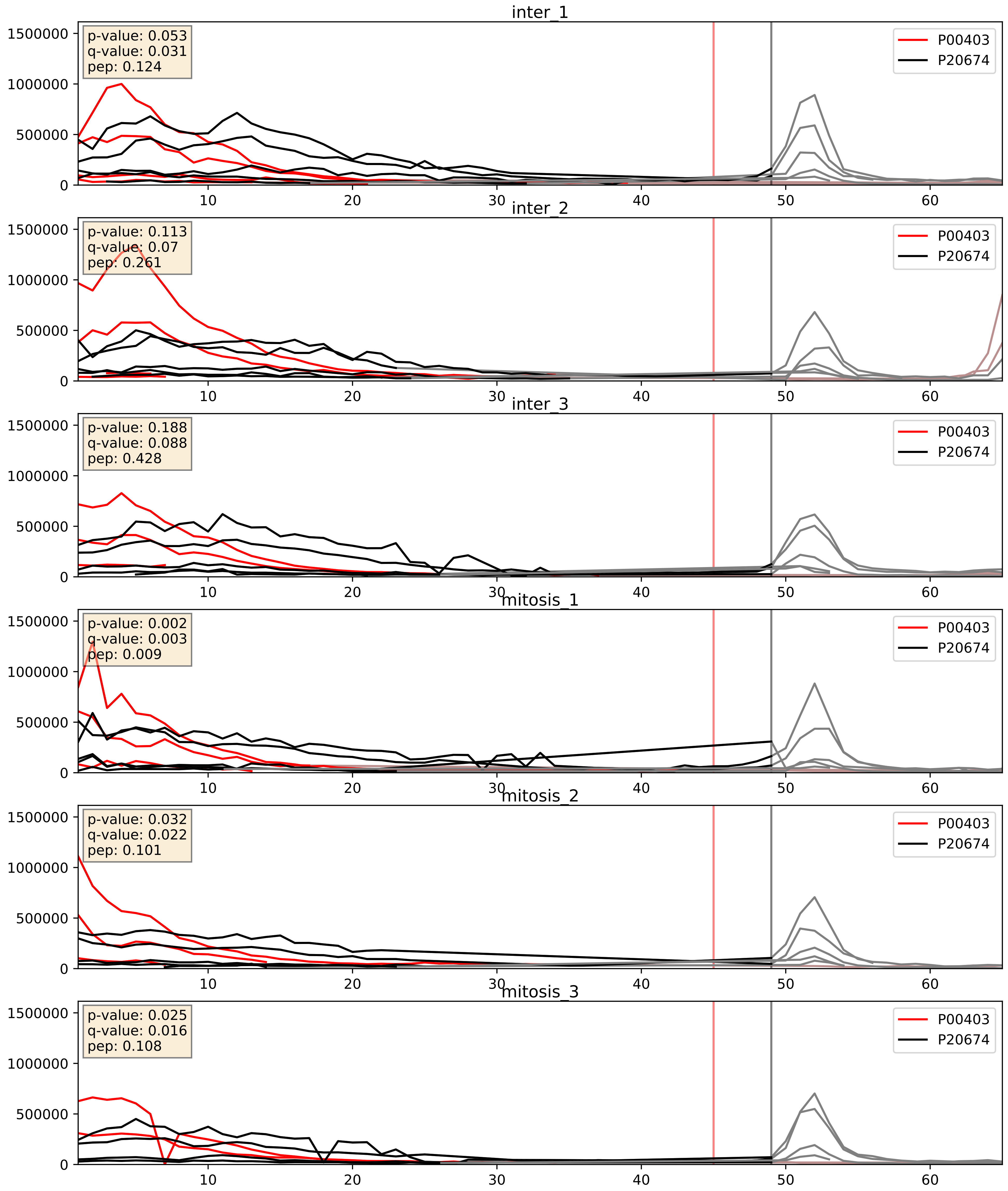


| level | condition_1 | condition_2 | pvalue | pvalue_adjusted |
|---|---|---|---|---|
| complex_abundance | mitosis | inter | 0.01723840389210783 | 0.0979818972884748 |
| interactor_abundance | mitosis | inter | 0.05043231763306499 | 0.17968809112967168 |
| interactor_ratio | mitosis | inter | 0.8992468989604984 | 0.9830552478042482 |

P00403_P22695
COX2_HUMAN vs QCR2_HUMAN
p-value: 0.01 q-value: 0.009

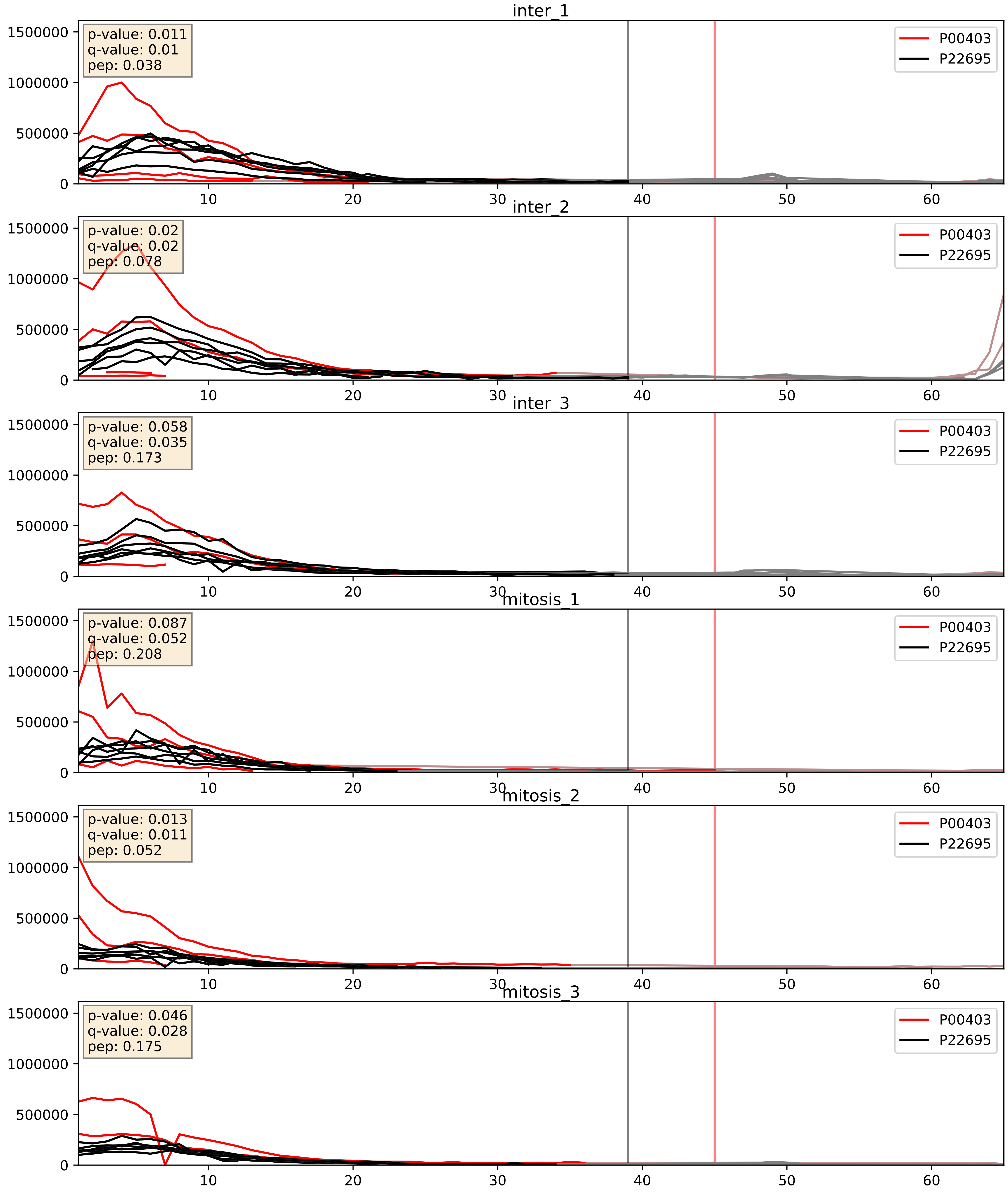


| level | condition_1 | condition_2 | pvalue | pvalue_adjusted |
|---|---|---|---|---|
| complex_abundance | mitosis | inter | 0.007639655089387588 | 0.06507009707975896 |
| interactor_abundance | mitosis | inter | 0.04799835291854166 | 0.17504902777779313 |
| interactor_ratio | mitosis | inter | 0.4897202497476749 | 0.8692763386835093 |

P00403_P35232
COX2_HUMAN vs PHB_HUMAN
p-value: 0.0 q-value: 0.022

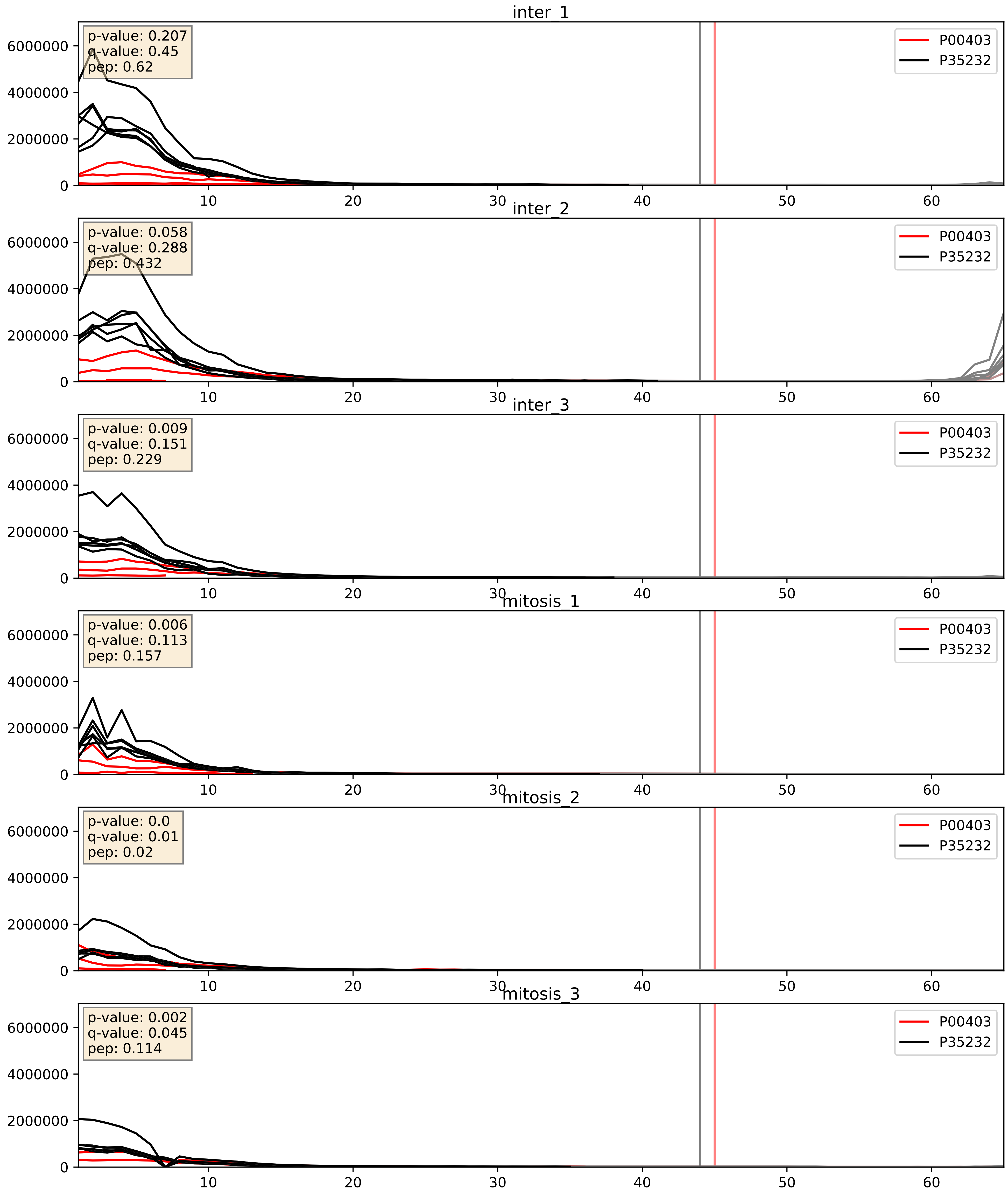


| level | condition_1 | condition_2 | pvalue | pvalue_adjusted |
|---|---|---|---|---|
| complex_abundance | mitosis | inter | 0.005505336892302833 | 0.055958215865660764 |
| interactor_ratio | mitosis | inter | 0.019625730779465705 | 0.21211648418210408 |
| interactor_abundance | mitosis | inter | 0.05021540360985647 | 0.17917626298473172 |

P00403_P47985
COX2_HUMAN vs UCRI_HUMAN
p-value: 0.014 q-value: 0.012

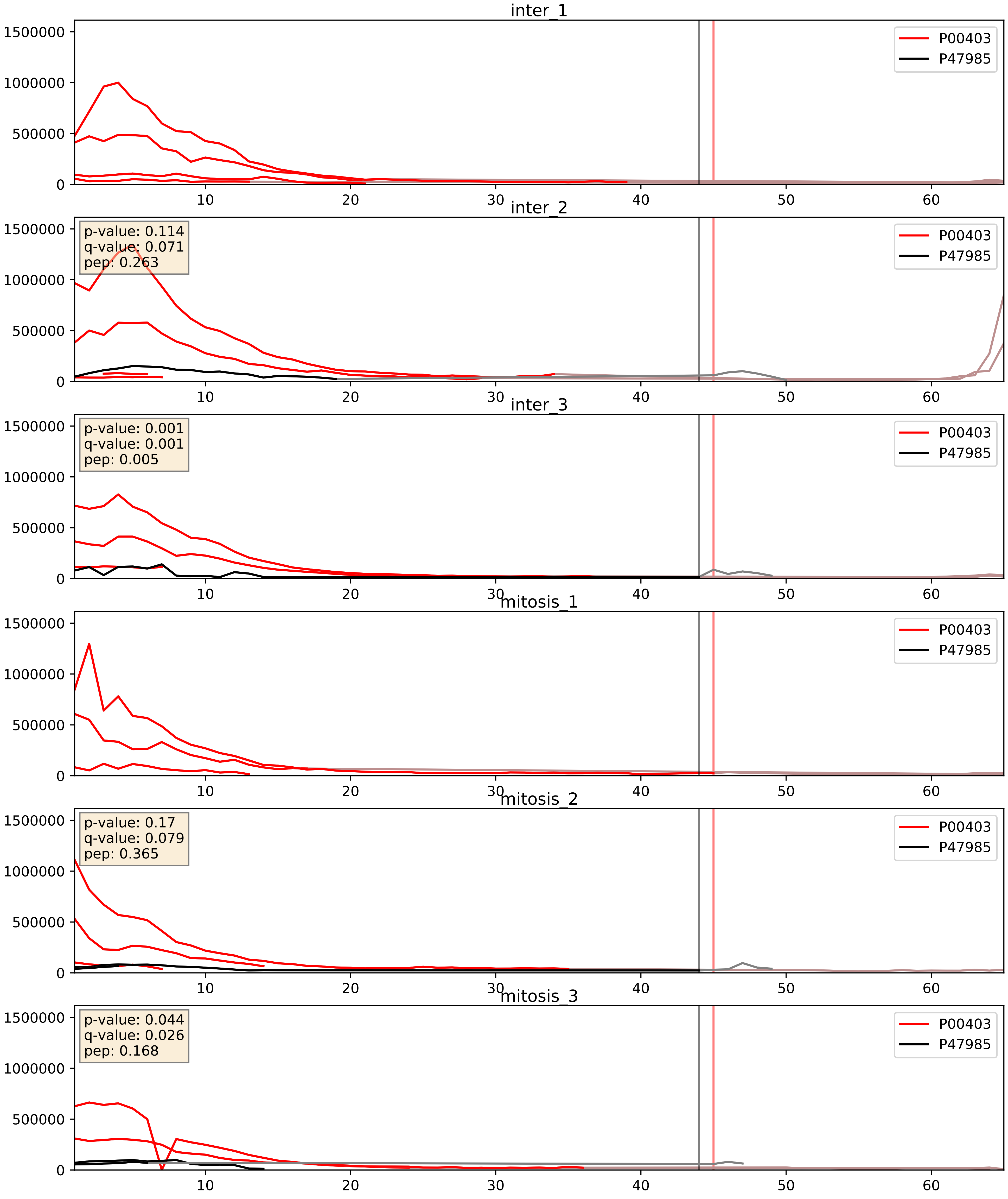


| level | condition_1 | condition_2 | pvalue | pvalue_adjusted |
|---|---|---|---|---|
| interactor_ratio | mitosis | inter | 0.5438799953751101 | 0.8859730852936224 |
| interactor_abundance | mitosis | inter | 0.5887080245027201 | 0.7299160906348905 |
| complex_abundance | mitosis | inter | 0.6227007082141295 | 0.7418989024103371 |
