## Supplementary Data 2 for "SECAT: Quantifying differential protein-protein interaction states by network-centric analysis": hela_string_000177_Q15061.pdf

# Q15061_Q15061

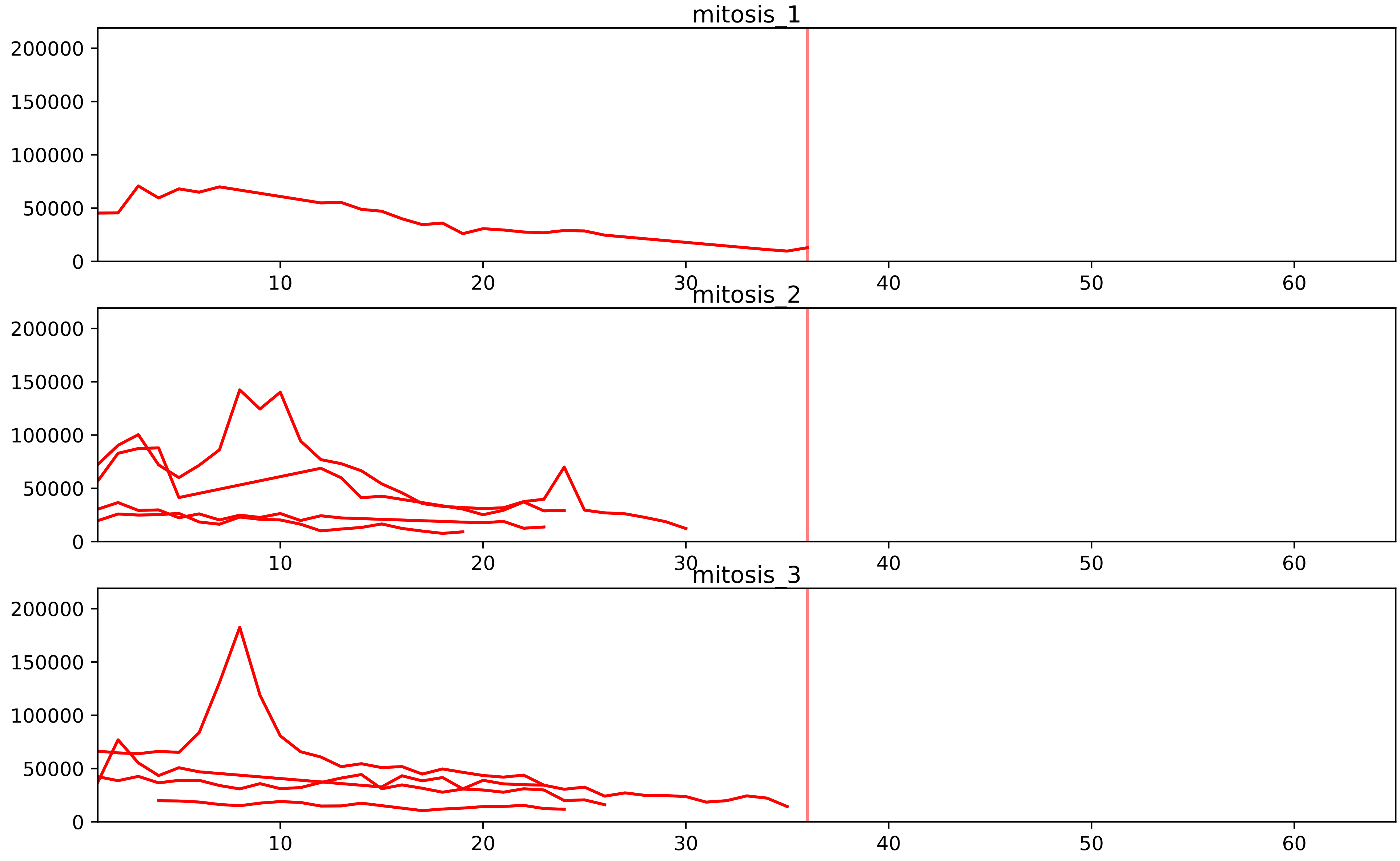


| level | condition_1 | condition_2 | pvalue | pvalue_adjusted |
|---|---|---|---|---|
| complex_abundance | mitosis | inter | 0.00012297116621897891 | 0.0024864499543178153 |
| interactor_ratio | mitosis | inter | 0.004890518584794627 | 0.03946734296500927 |
| interactor_abundance | mitosis | inter | 0.01397982660020302 | 0.06367049738706326 |
| assembled_abundance | mitosis | inter | 0.03307265972761888 | 0.21948082391689583 |
| total_abundance | mitosis | inter | 0.03428134198213041 | 0.2315066896680853 |
| monomer_abundance | mitosis | inter | 0.4590891064696274 | 0.7303363506601787 |

O00567_Q15061
NOP56_HUMAN vs WDR43_HUMAN
p-value: 0.021 q-value: 0.017

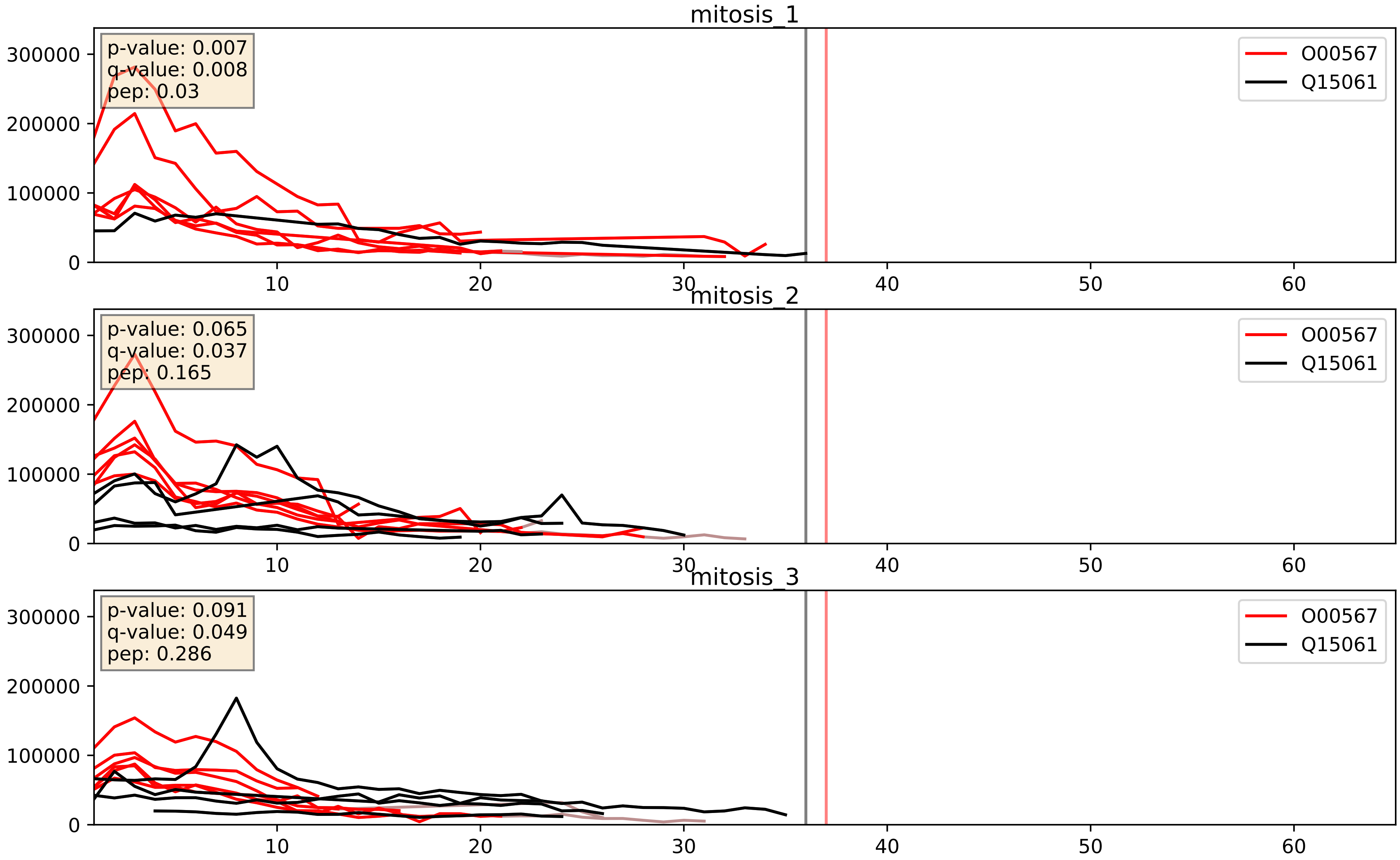


| level | condition_1 | condition_2 | pvalue | pvalue_adjusted |
|---|---|---|---|---|
| interactor_abundance | mitosis | inter | 0.0005738807154731015 | 0.017415659477552636 |
| complex_abundance | mitosis | inter | 0.0012557787786977876 | 0.02410194247904274 |
| interactor_ratio | mitosis | inter | 0.15614177644181174 | 0.6132288760320385 |

P22087_Q15061
FBRL_HUMAN vs WDR43_HUMAN
p-value: 0.073 q-value: 0.044

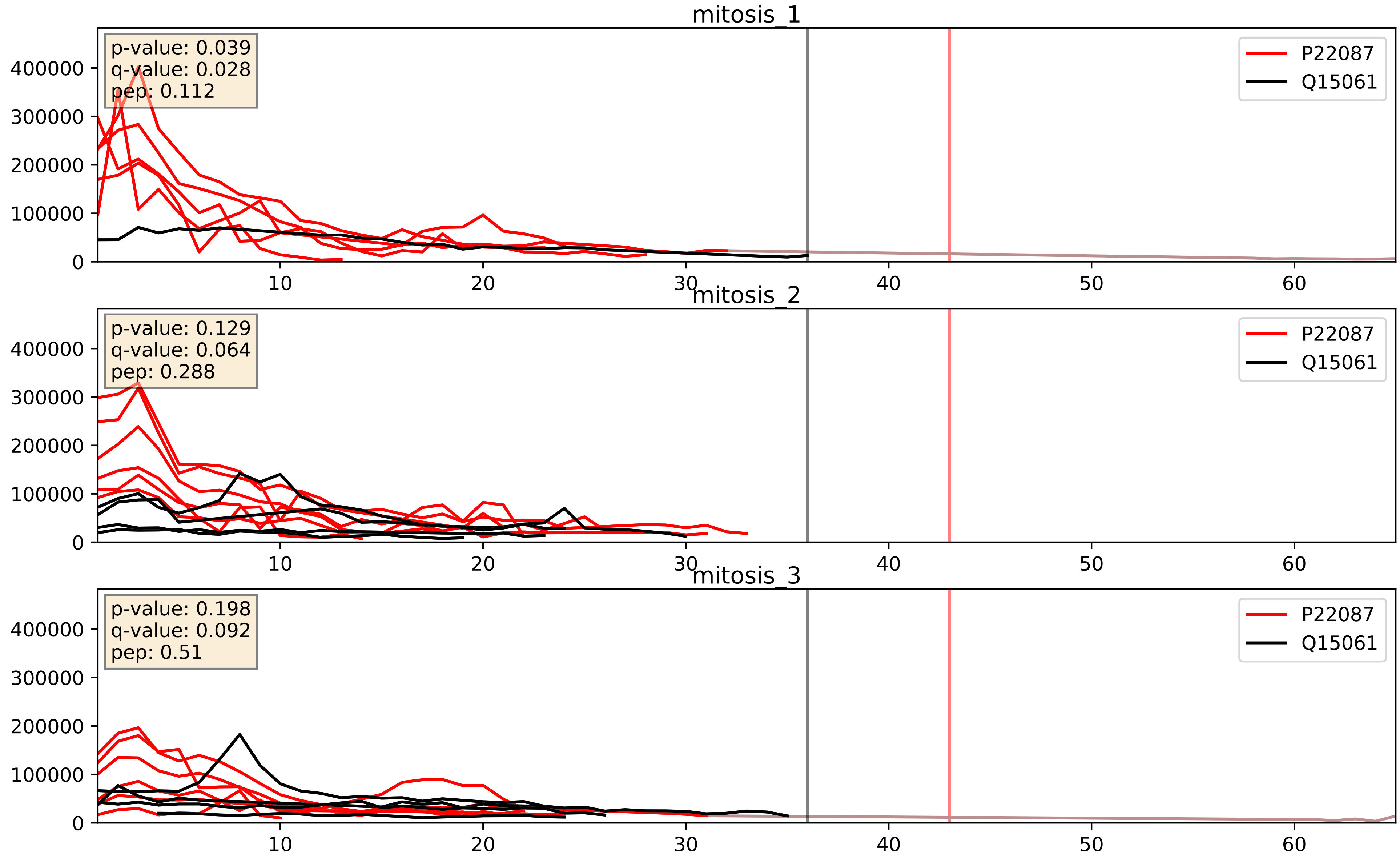


| level | condition_1 | condition_2 | pvalue | pvalue_adjusted |
|---|---|---|---|---|
| interactor_abundance | mitosis | inter | 0.00022145143300215012 | 0.011215116209912415 |
| complex_abundance | mitosis | inter | 0.000804026877839027 | 0.018753324453139156 |
| interactor_ratio | mitosis | inter | 0.05603485152930944 | 0.3926647223857623 |

P55769_Q15061
NH2L1_HUMAN vs WDR43_HUMAN
p-value: 0.036 q-value: 0.025

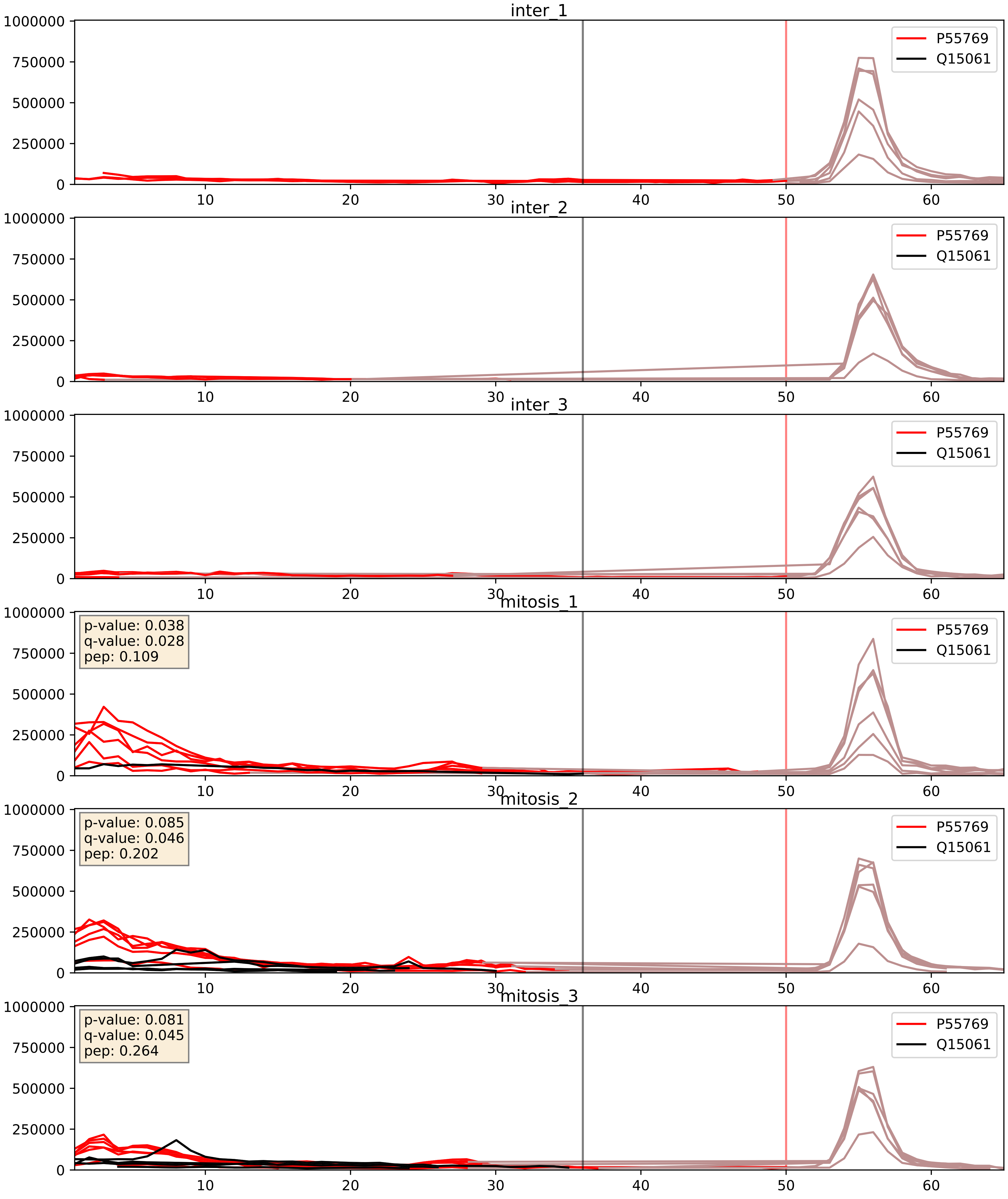


| level | condition_1 | condition_2 | pvalue | pvalue_adjusted |
|---|---|---|---|---|
| interactor_abundance | mitosis | inter | 5.175159842494777e-05 | 0.006283598333582311 |
| complex_abundance | mitosis | inter | 0.0008534383700319466 | 0.01942934161562091 |
| interactor_ratio | mitosis | inter | 0.020867400541259323 | 0.21890312332497525 |

Q12788_Q15061
TBL3_HUMAN vs WDR43_HUMAN
p-value: 0.034 q-value: 0.024

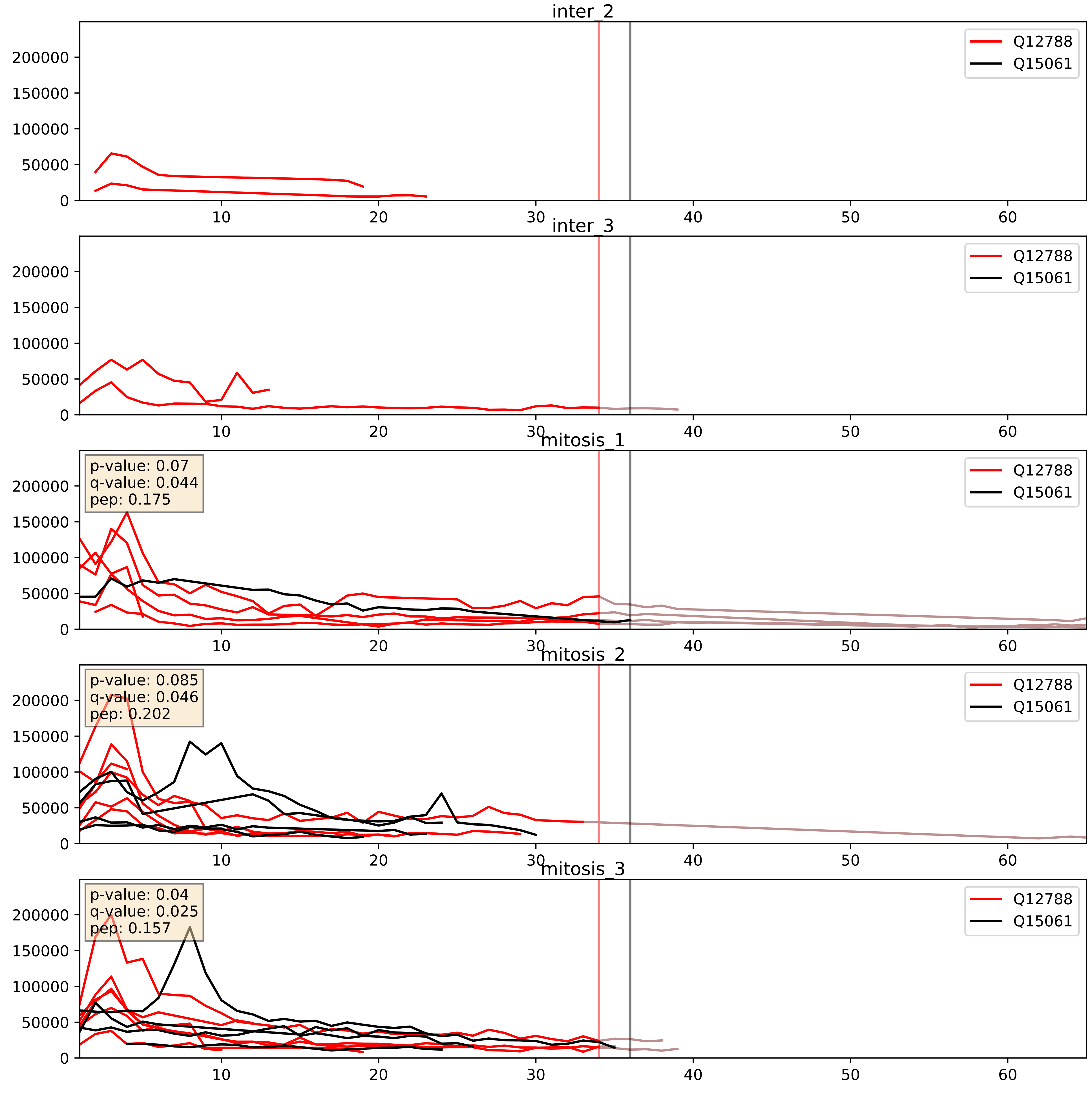


| level | condition_1 | condition_2 | pvalue | pvalue_adjusted |
|---|---|---|---|---|
| interactor_abundance | mitosis | inter | 0.0012685582694706194 | 0.026229127504030197 |
| complex_abundance | mitosis | inter | 0.00806041973070591 | 0.06653038413698939 |
| interactor_ratio | mitosis | inter | 0.23955938278422004 | 0.7181694669961206 |

Q13895_Q15061
BYST_HUMAN vs WDR43_HUMAN
p-value: 0.076 q-value: 0.046

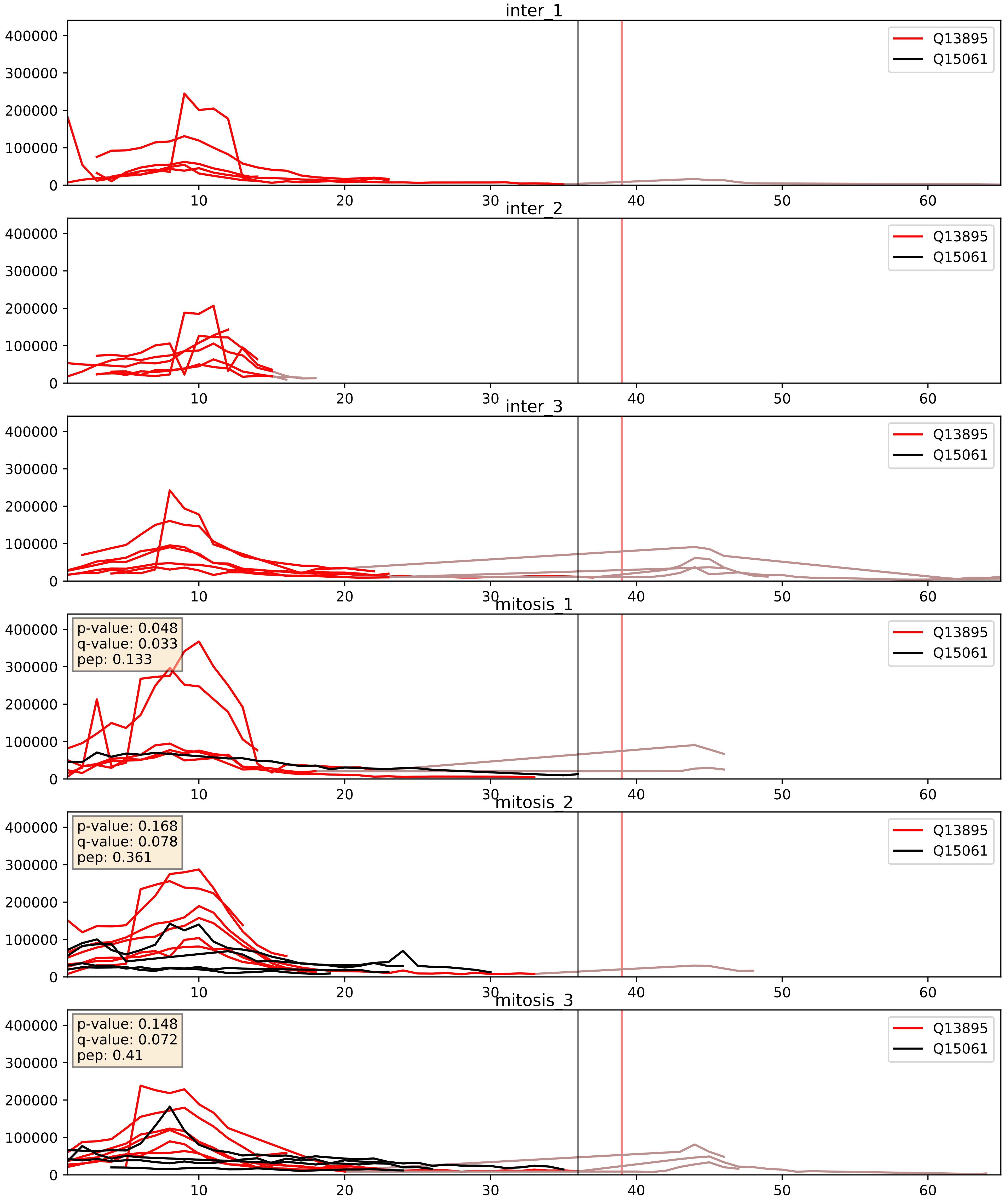


| level | condition_1 | condition_2 | pvalue | pvalue_adjusted |
|---|---|---|---|---|
| interactor_abundance | mitosis | inter | 0.0006377390807793674 | 0.01786921941561828 |
| complex_abundance | mitosis | inter | 0.004882908275355024 | 0.052397293069493756 |
| interactor_ratio | mitosis | inter | 0.022909515726422306 | 0.2336681581804622 |

Q15061_Q8NI36
WDR43_HUMAN vs WDR36_HUMAN
p-value: 0.08 q-value: 0.048

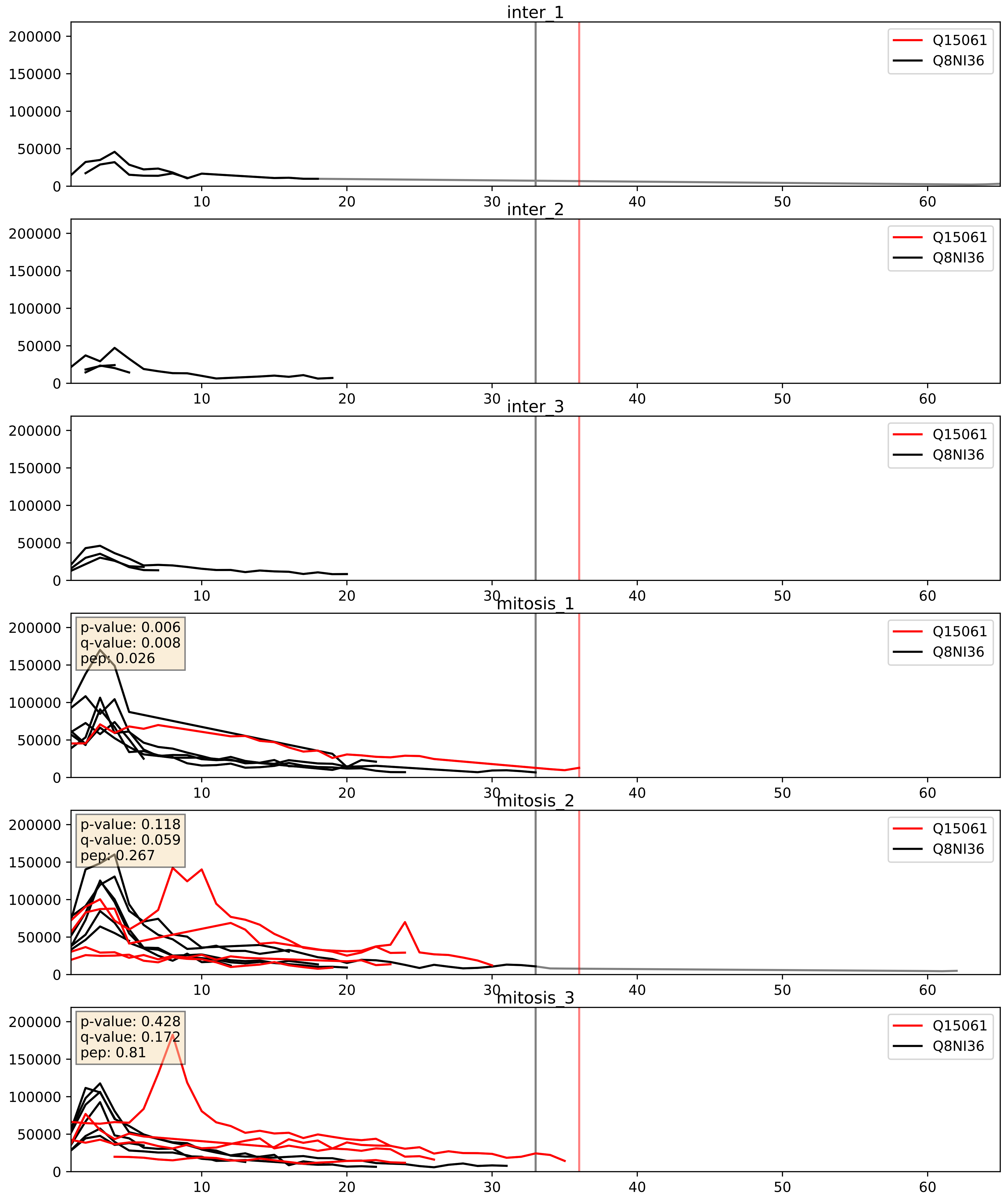


| level | condition_1 | condition_2 | pvalue | pvalue_adjusted |
|---|---|---|---|---|
| complex_abundance | mitosis | inter | 0.005860437094623565 | 0.05756820427608883 |
| interactor_abundance | mitosis | inter | 0.045200483218485674 | 0.16918064554011253 |
| interactor_ratio | mitosis | inter | 0.7727111896398332 | 0.9554532798105997 |

Q15061_Q8TED0
WDR43_HUMAN vs UTP15_HUMAN
p-value: 0.014 q-value: 0.012

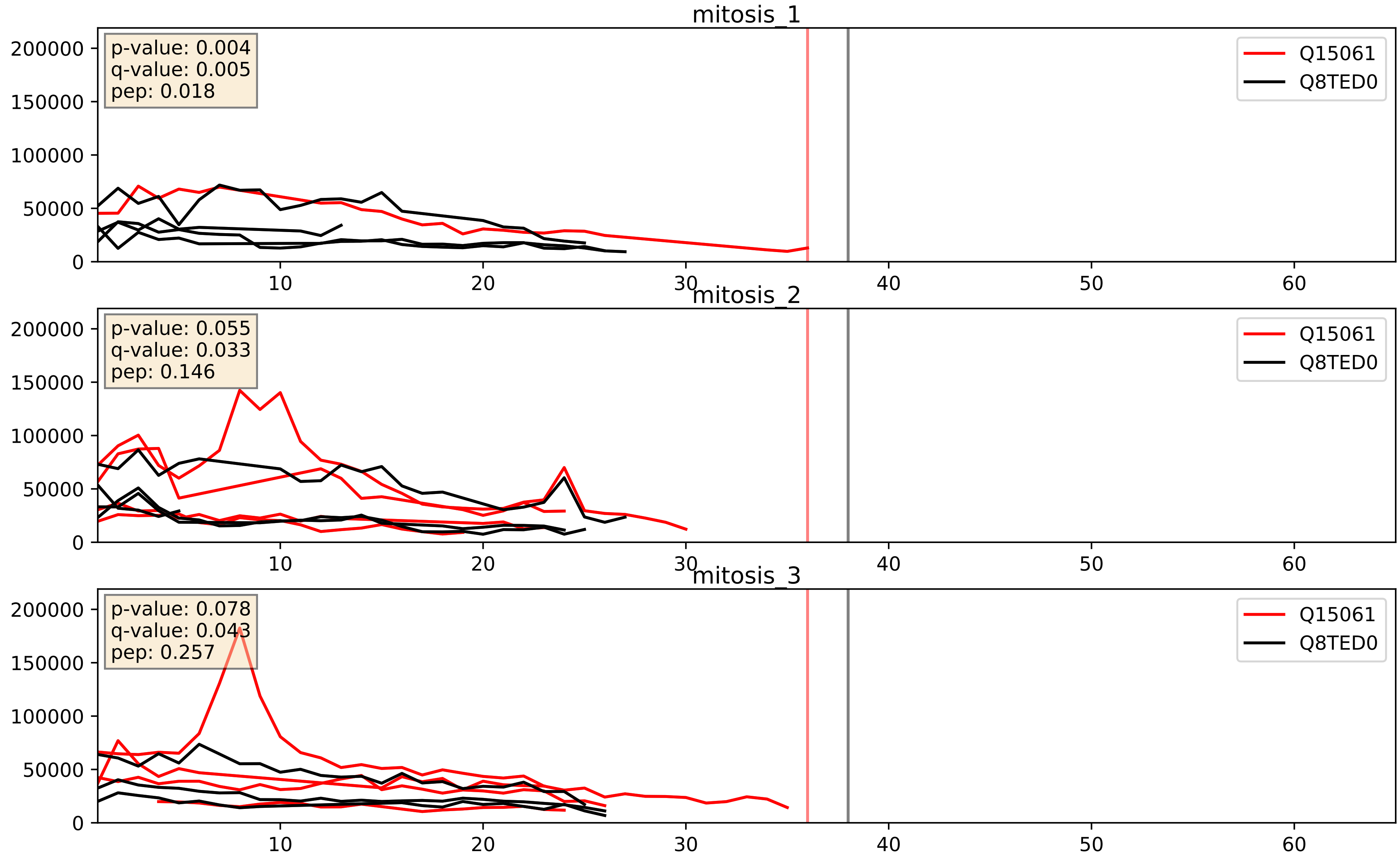


| level | condition_1 | condition_2 | pvalue | pvalue_adjusted |
|---|---|---|---|---|
| complex_abundance | mitosis | inter | 0.00607593990941489 | 0.05817678481497925 |
| interactor_abundance | mitosis | inter | 0.04157442246858351 | 0.1638853586604075 |
| interactor_ratio | mitosis | inter | 0.9786487798482965 | 0.996103871046542 |

Q15061_Q9H583
WDR43_HUMAN vs HEAT1_HUMAN
p-value: 0.034 q-value: 0.024

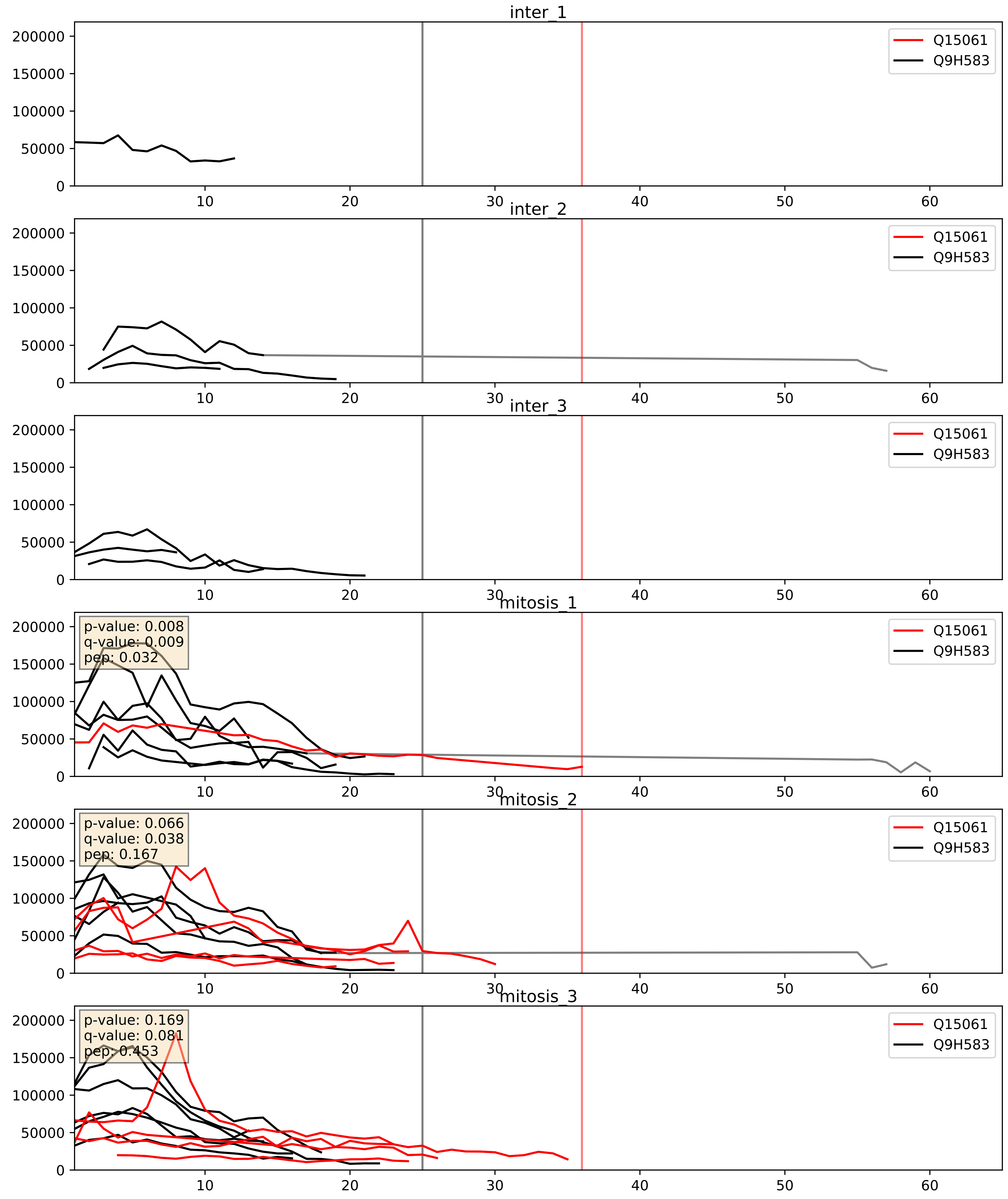


| level | condition_1 | condition_2 | pvalue | pvalue_adjusted |
|---|---|---|---|---|
| complex_abundance | mitosis | inter | 0.005287778237647574 | 0.054740033961712564 |
| interactor_ratio | mitosis | inter | 0.03948312491726528 | 0.32783954136208876 |
| interactor_abundance | mitosis | inter | 0.04327794470388063 | 0.16685321787696922 |
