## Supplementary Data 2 for "SECAT: Quantifying differential protein-protein interaction states by network-centric analysis": hela_string_000178_Q9BZJ0.pdf

Q9BZJ0_Q9BZJ0

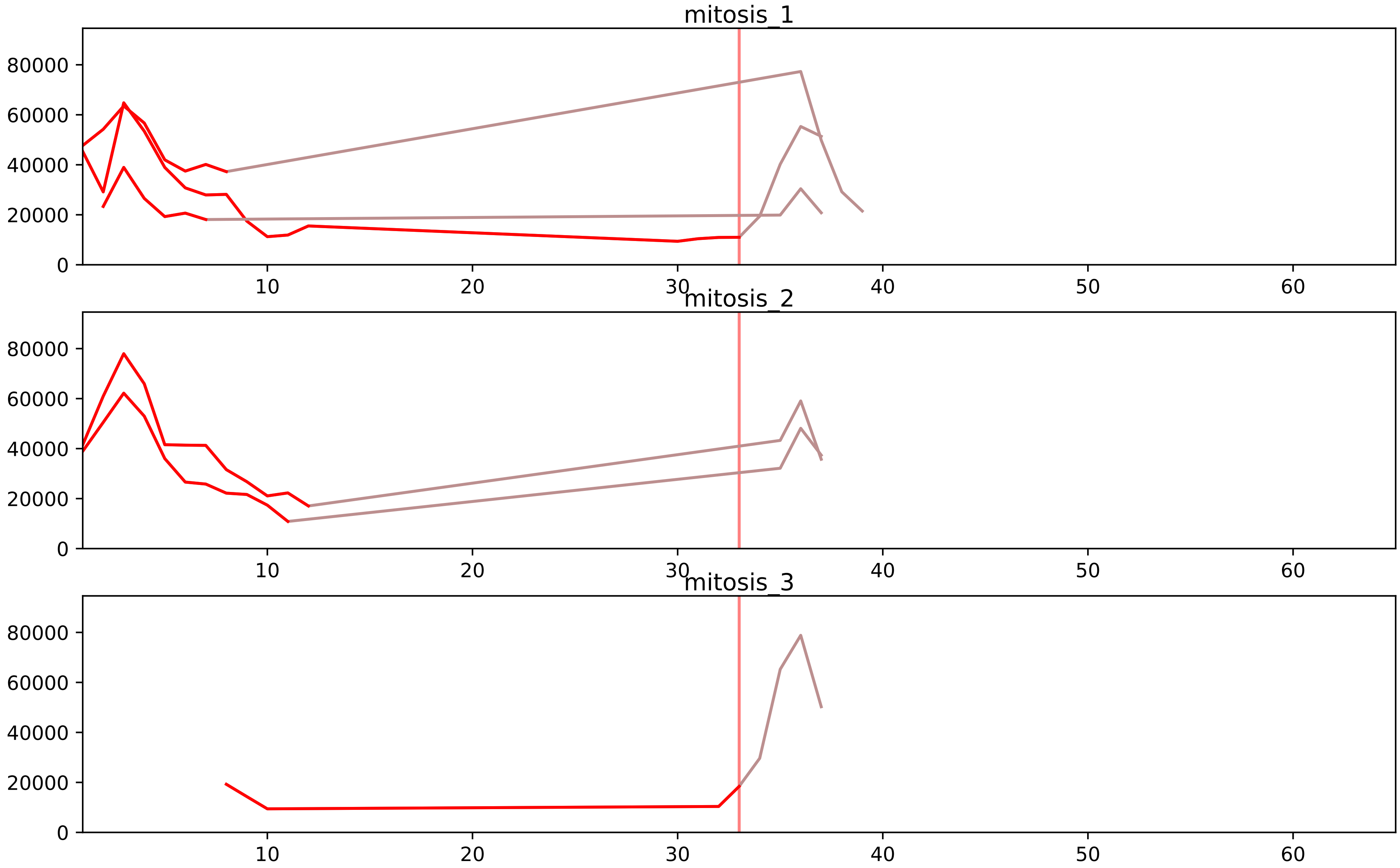


| level | condition_1 | condition_2 | pvalue | pvalue_adjusted |
|---|---|---|---|---|
| interactor_ratio | mitosis | inter | 0.00012431512287014085 | 0.0020242462485049483 |
| monomer_abundance | mitosis | inter | 0.038149562253736655 | 0.3150273663293861 |
| total_abundance | mitosis | inter | 0.1264767308116015 | 0.44100490543840143 |
| assembled_abundance | mitosis | inter | 0.17643741905879481 | 0.4852547347909627 |
| complex_abundance | mitosis | inter | 0.25131479355341696 | 0.3741255826361547 |
| interactor_abundance | mitosis | inter | 0.8480637760093213 | 0.9159366994483951 |

O43290_Q9BZJ0
SNUT1_HUMAN vs CRNL1_HUMAN
p-value: 0.047 q-value: 0.031

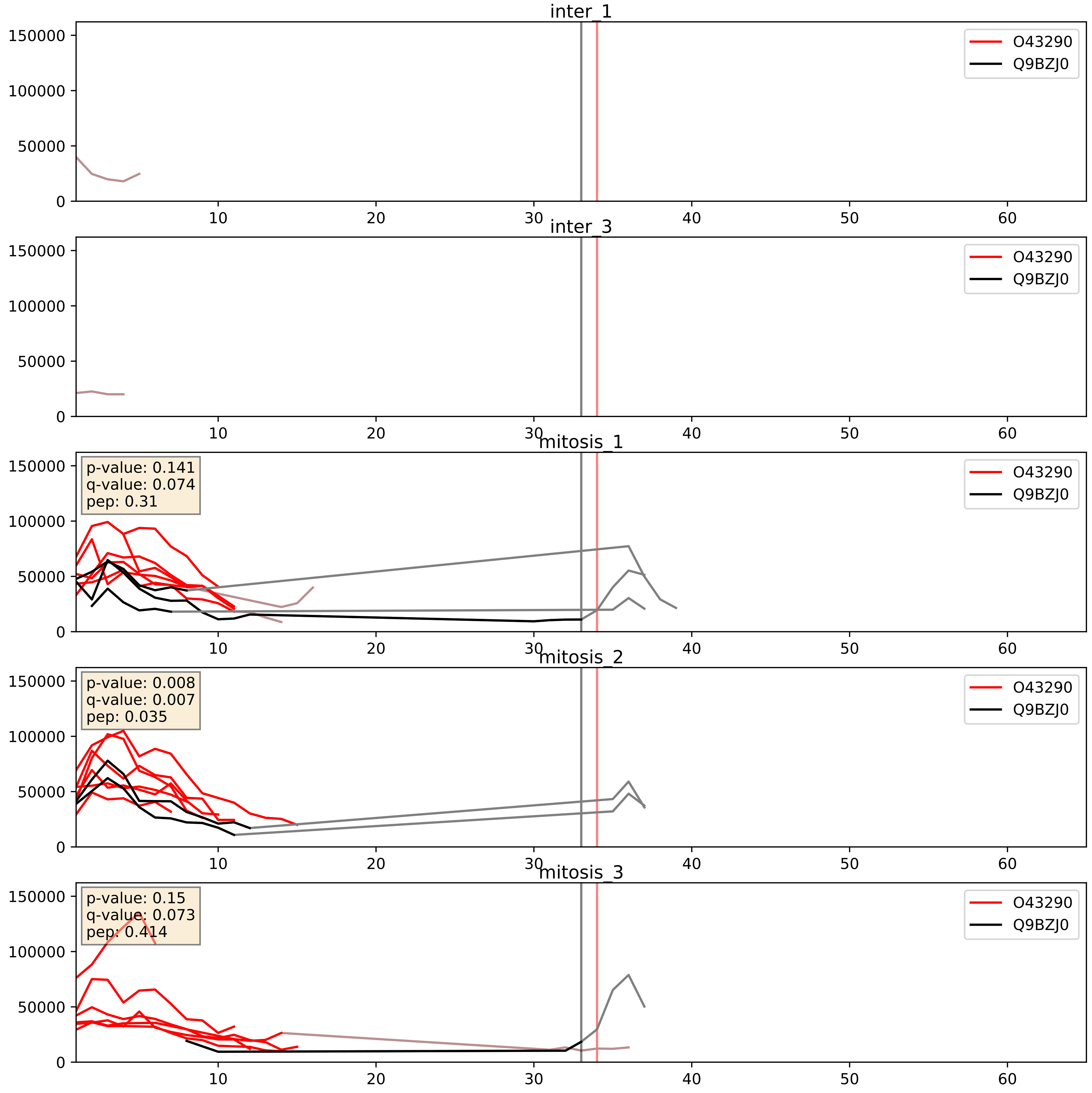


| level | condition_1 | condition_2 | pvalue | pvalue_adjusted |
|---|---|---|---|---|
| interactor_ratio | mitosis | inter | 0.0521009646673782 | 0.37764936302518703 |
| interactor_abundance | mitosis | inter | 0.2312435840999291 | 0.42318727300216413 |
| complex_abundance | mitosis | inter | 0.3821085485176027 | 0.5292636206004335 |

O43395_Q9BZJ0
PRPF3_HUMAN vs CRNL1_HUMAN
p-value: 0.022 q-value: 0.017

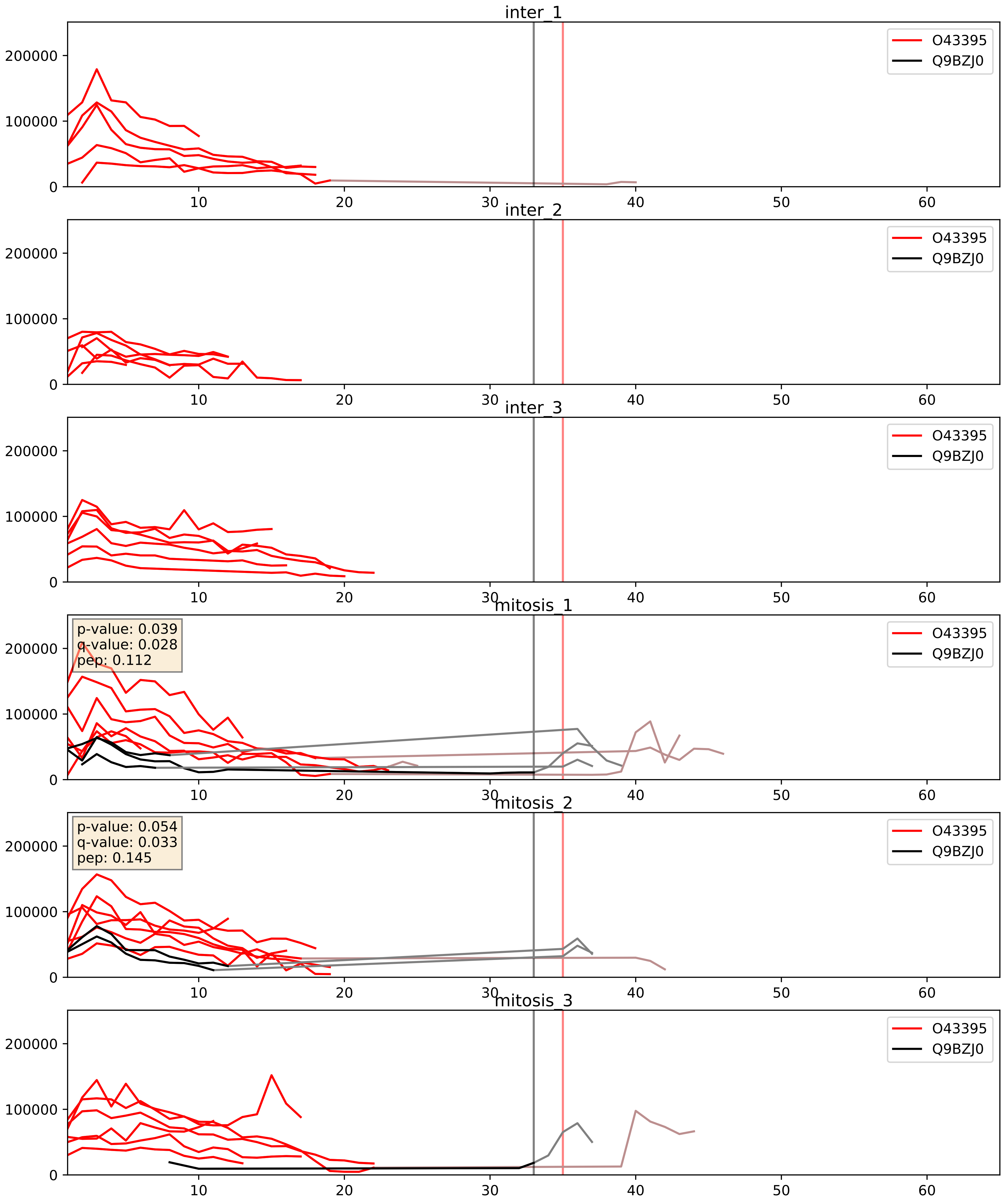


| level | condition_1 | condition_2 | pvalue | pvalue_adjusted |
|---|---|---|---|---|
| interactor_ratio | mitosis | inter | 0.0010612977904857902 | 0.038414698994273265 |
| interactor_abundance | mitosis | inter | 0.050901463188763756 | 0.18087028845820577 |
| complex_abundance | mitosis | inter | 0.16968315397865294 | 0.31873042673431795 |

O43660_Q9BZJ0
PLRG1_HUMAN vs CRNL1_HUMAN
p-value: 0.046 q-value: 0.031

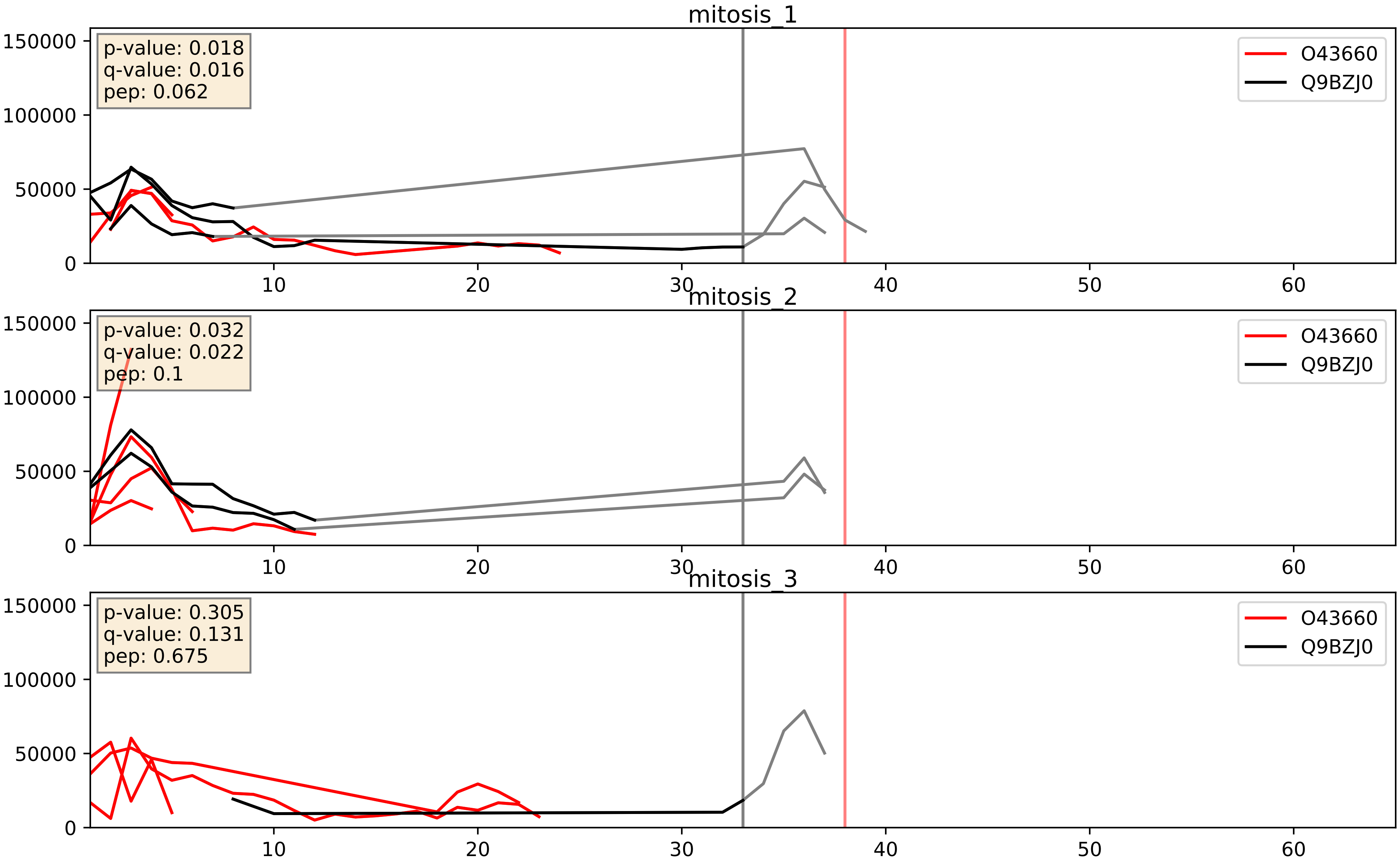


| level | condition_1 | condition_2 | pvalue | pvalue_adjusted |
|---|---|---|---|---|
| interactor_ratio | mitosis | inter | 0.5714409020468911 | 0.8945917155252349 |
| complex_abundance | mitosis | inter | 0.7395285783854642 | 0.8272358106284508 |
| interactor_abundance | mitosis | inter | 0.8186812854359015 | 0.8975871666130408 |

O75643_Q9BZJ0
U520_HUMAN vs CRNL1_HUMAN
p-value: 0.034 q-value: 0.024

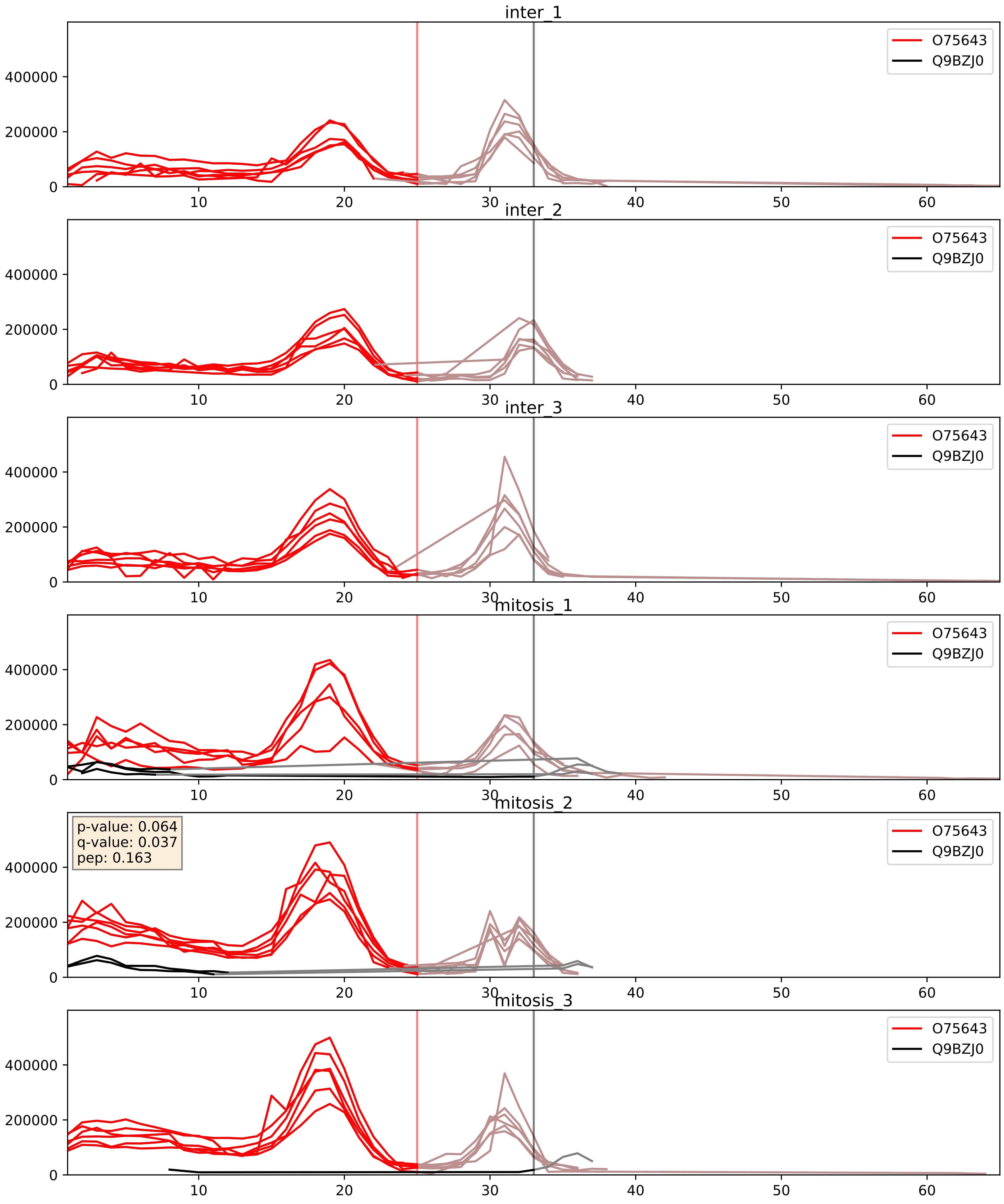


| level | condition_1 | condition_2 | pvalue | pvalue_adjusted |
|---|---|---|---|---|
| interactor_ratio | mitosis | inter | 0.004715096858563263 | 0.09159088101537749 |
| interactor_abundance | mitosis | inter | 0.030570403815191056 | 0.14519524113741764 |
| complex_abundance | mitosis | inter | 0.10340924979824517 | 0.23908191575103507 |

O75934_Q9BZJ0
SPF27_HUMAN vs CRNL1_HUMAN
p-value: 0.022 q-value: 0.017

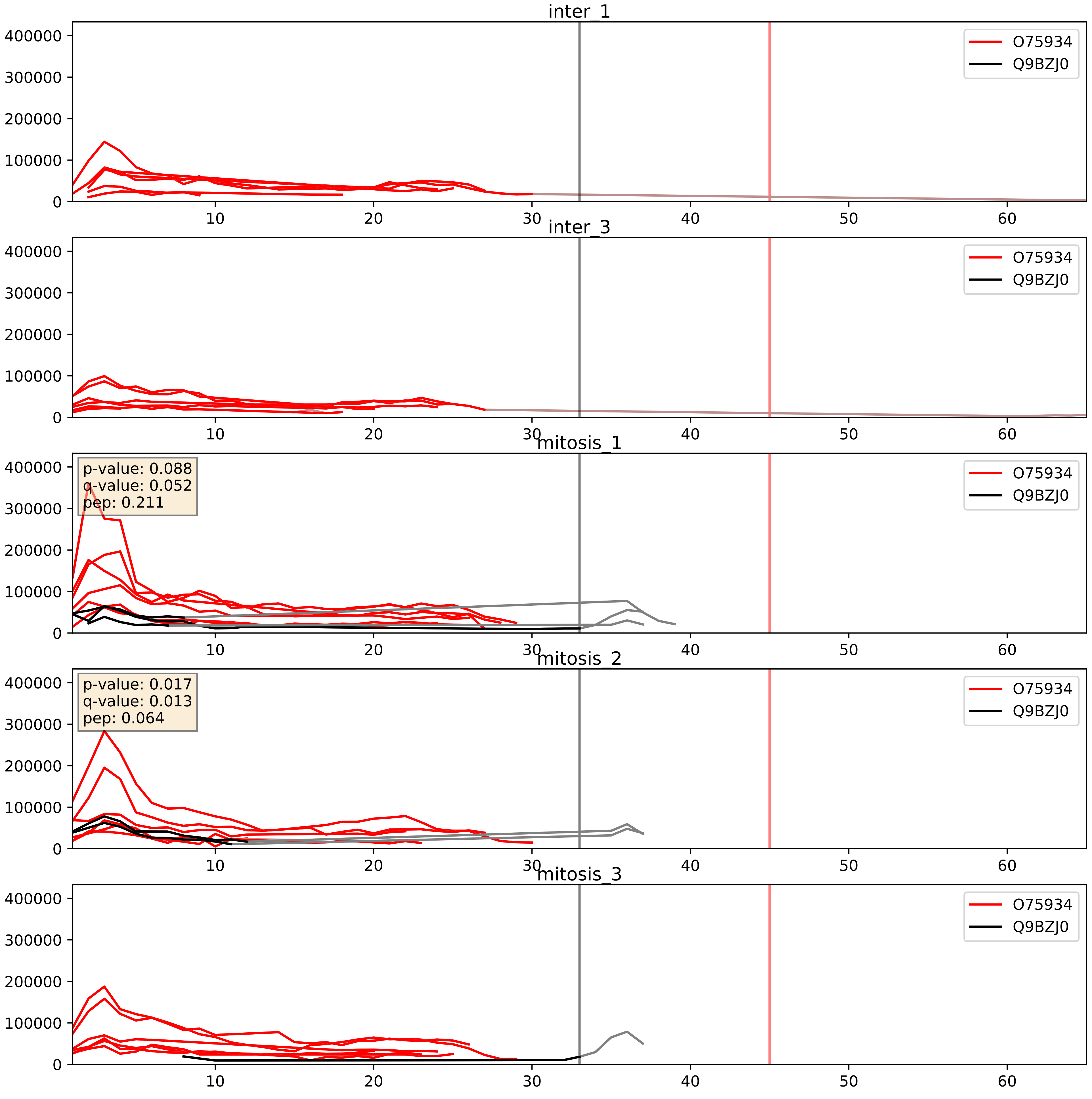


| level | condition_1 | condition_2 | pvalue | pvalue_adjusted |
|---|---|---|---|---|
| interactor_ratio | mitosis | inter | 0.014619333770107596 | 0.17725424514464733 |
| interactor_abundance | mitosis | inter | 0.12392143906875966 | 0.30662451753969727 |
| complex_abundance | mitosis | inter | 0.24775694395957115 | 0.39823025501804077 |

Q96DI7_Q9BZJ0
SNR40_HUMAN vs CRNL1_HUMAN
p-value: 0.051 q-value: 0.033

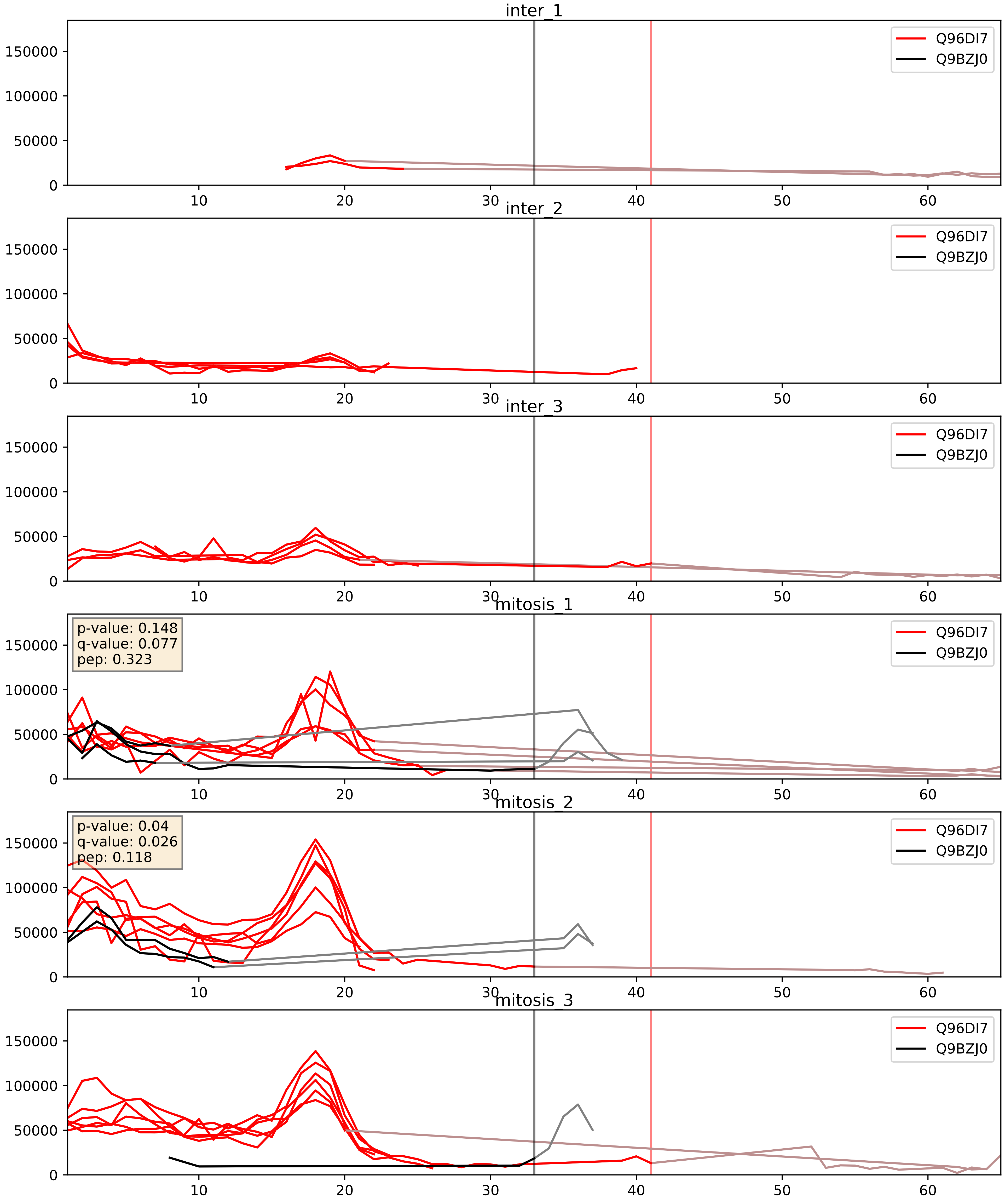


| level | condition_1 | condition_2 | pvalue | pvalue_adjusted |
|---|---|---|---|---|
| interactor_ratio | mitosis | inter | 0.06629577513681317 | 0.4296400448702392 |
| interactor_abundance | mitosis | inter | 0.08693438568941618 | 0.249591930740031 |
| complex_abundance | mitosis | inter | 0.2317976423310166 | 0.384588332584036 |

Q99459_Q9BZJ0
CDC5L_HUMAN vs CRNL1_HUMAN
p-value: 0.019 q-value: 0.015

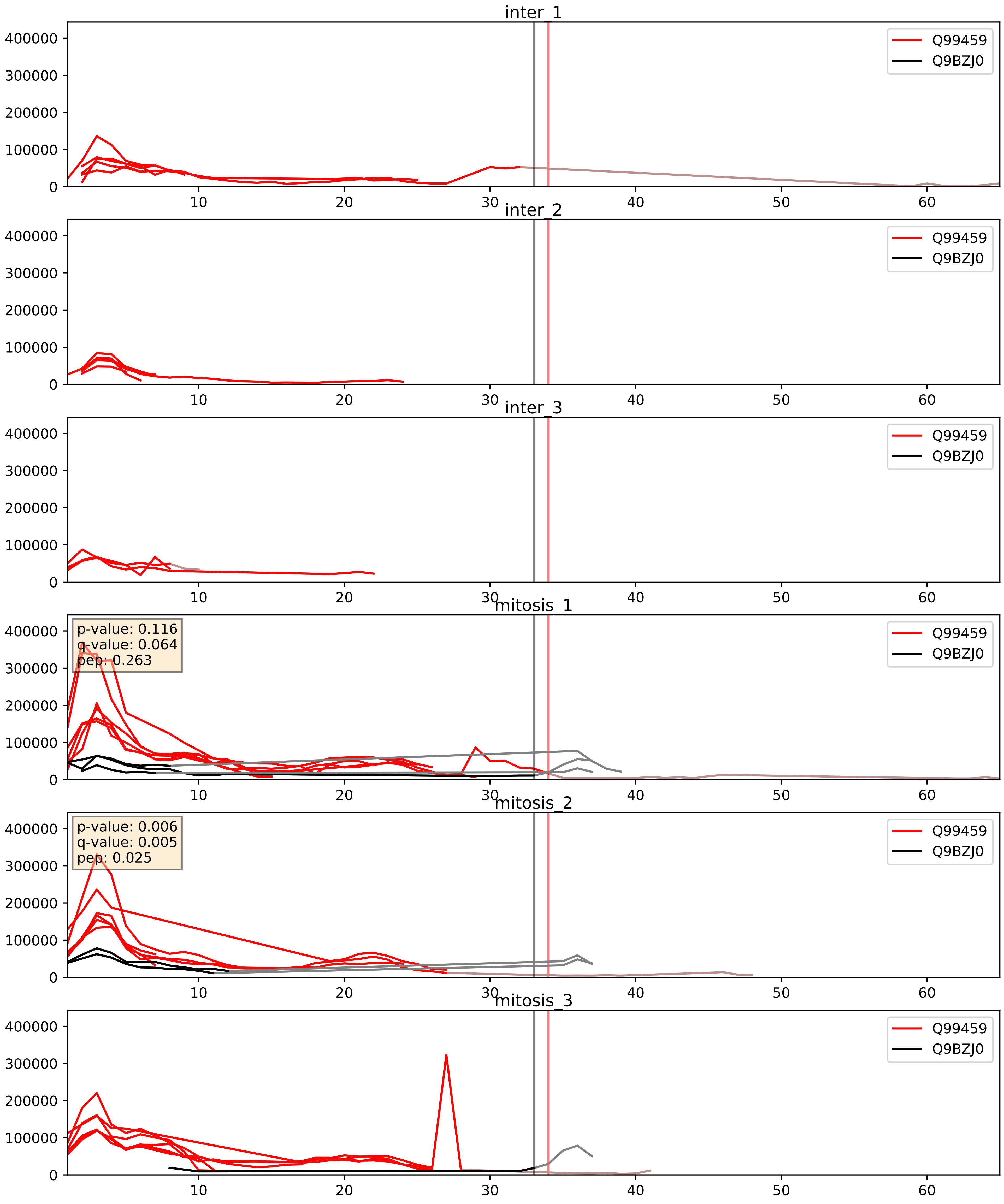


| level | condition_1 | condition_2 | pvalue | pvalue_adjusted |
|---|---|---|---|---|
| interactor_ratio | mitosis | inter | 0.022207834723738464 | 0.22875940461516397 |
| interactor_abundance | mitosis | inter | 0.14787363404624873 | 0.33422995072666123 |
| complex_abundance | mitosis | inter | 0.26937271491850834 | 0.41886111529562786 |

Q9BUQ8_Q9BZJ0
DDX23_HUMAN vs CRNL1_HUMAN
p-value: 0.036 q-value: 0.025

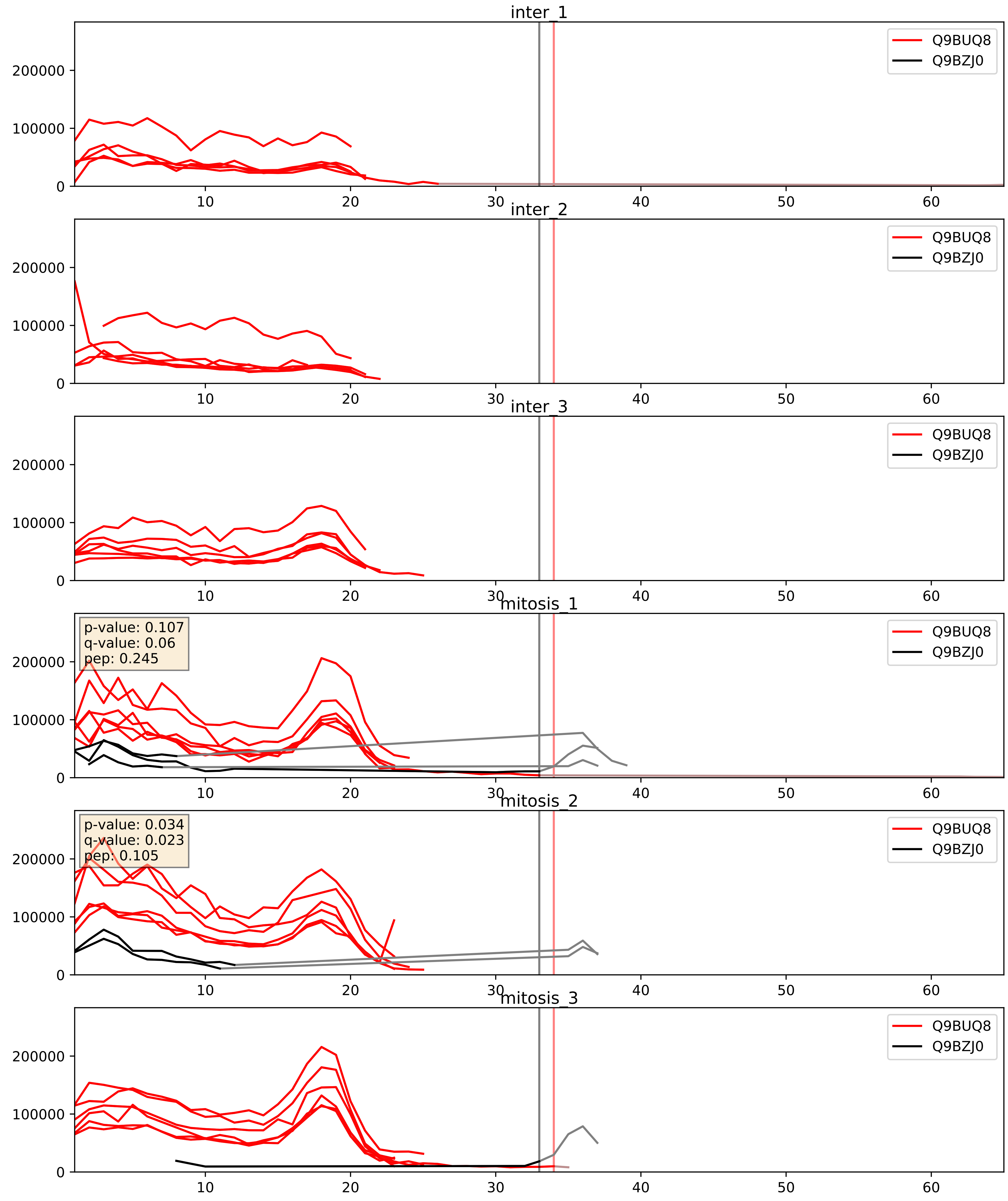


| level | condition_1 | condition_2 | pvalue | pvalue_adjusted |
|---|---|---|---|---|
| interactor_ratio | mitosis | inter | 0.01085137089615999 | 0.1495776729003696 |
| interactor_abundance | mitosis | inter | 0.04326640025801008 | 0.16685321787696922 |
| complex_abundance | mitosis | inter | 0.12438106921664024 | 0.26637526957579194 |

Q9BZJ0_Q9HCS7
CRNL1_HUMAN vs SYF1_HUMAN
p-value: 0.029 q-value: 0.021

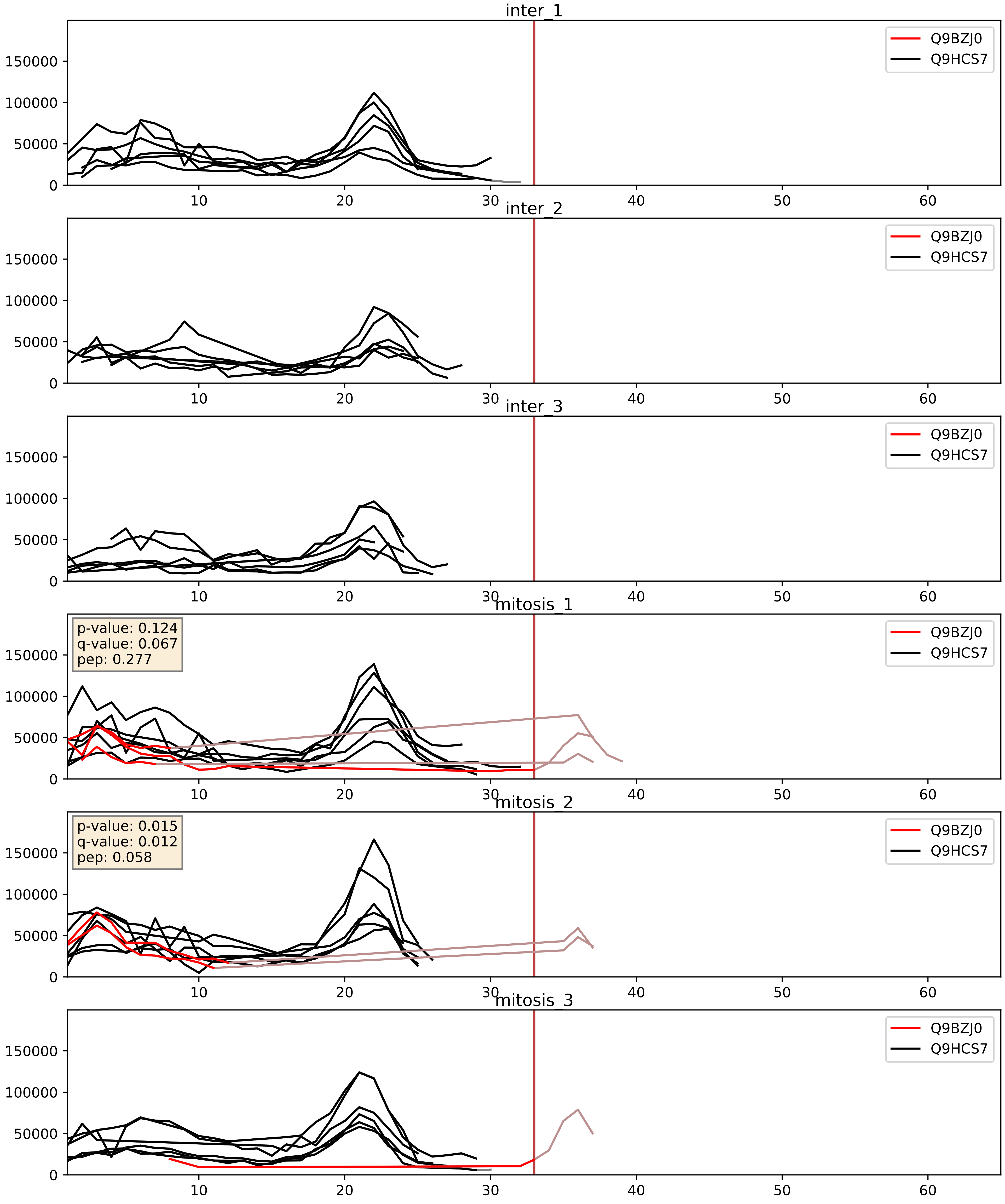


| level | condition_1 | condition_2 | pvalue | pvalue_adjusted |
|---|---|---|---|---|
| interactor_ratio | mitosis | inter | 0.0005171009168808326 | 0.02634752290773766 |
| complex_abundance | mitosis | inter | 0.3146115756662077 | 0.4632848938074553 |
| interactor_abundance | mitosis | inter | 0.6827514158457508 | 0.7939617062409492 |

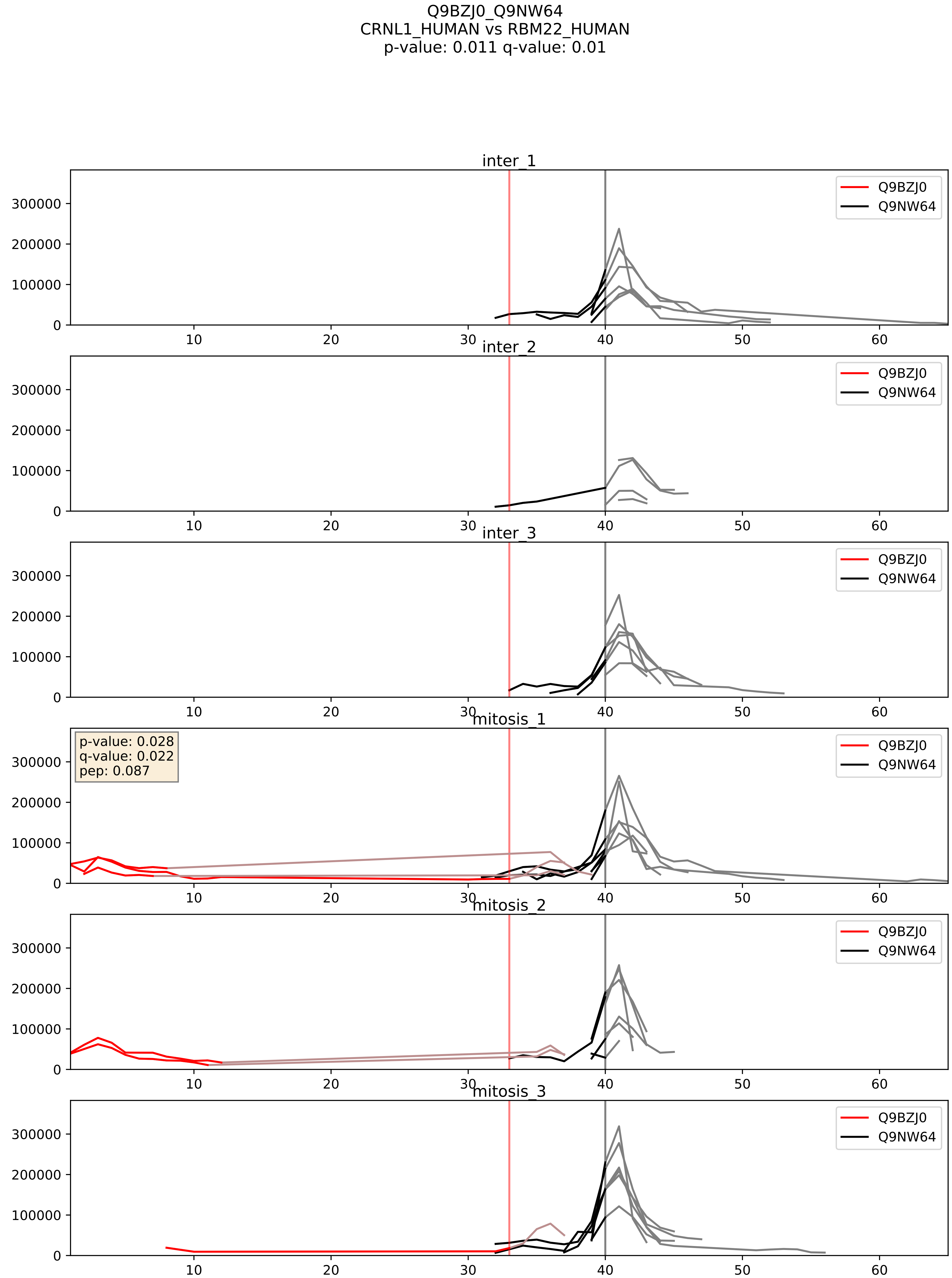


| level | condition_1 | condition_2 | pvalue | pvalue_adjusted |
|---|---|---|---|---|
| interactor_abundance | mitosis | inter | 0.6275856834215694 | 0.7553873584804905 |
| complex_abundance | mitosis | inter | 0.6333029739162097 | 0.7502177493388812 |
| interactor_ratio | mitosis | inter | 0.6446968165854603 | 0.9211491821017427 |
