## Supplementary Data 2 for "SECAT: Quantifying differential protein-protein interaction states by network-centric analysis": hela_string_000179_Q9Y3D7.pdf

Q9Y3D7_Q9Y3D7

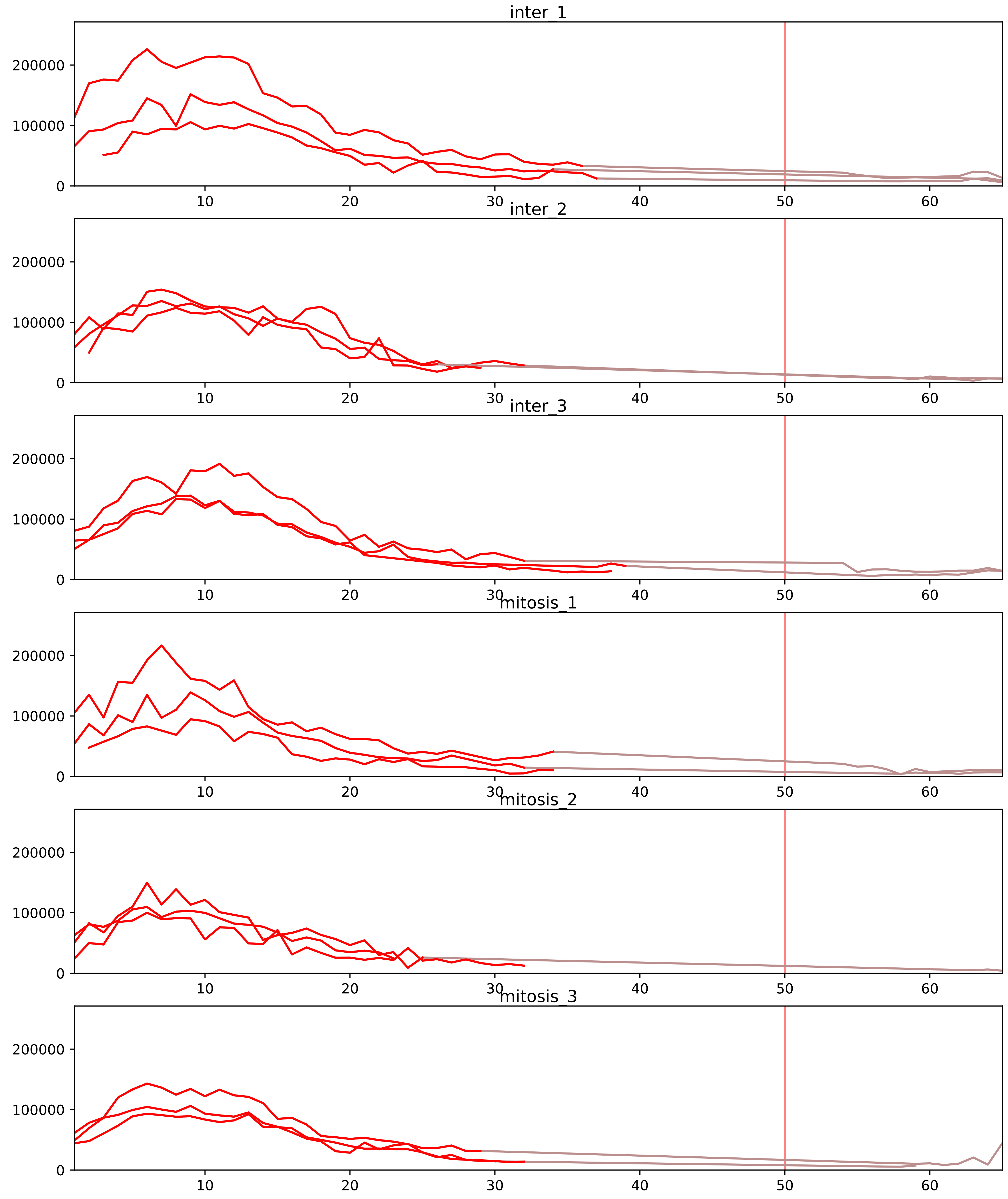


| level | condition_1 | condition_2 | pvalue | pvalue_adjusted |
|---|---|---|---|---|
| interactor_ratio | mitosis | inter | 0.00012830766676026397 | 0.0020709307617446113 |
| interactor_abundance | mitosis | inter | 0.00077933443857357 | 0.009703686176212032 |
| complex_abundance | mitosis | inter | 0.0019193187692909401 | 0.016349752479145045 |
| monomer_abundance | mitosis | inter | 0.0076385876624995444 | 0.16598262670786754 |
| total_abundance | mitosis | inter | 0.13550506007030969 | 0.4539835324486893 |
| assembled_abundance | mitosis | inter | 0.14333809923673363 | 0.4479076385361483 |

Q3ZCQ8_Q9Y3D7
TIM50_HUMAN vs TIM16_HUMAN
p-value: 0.012 q-value: 0.011

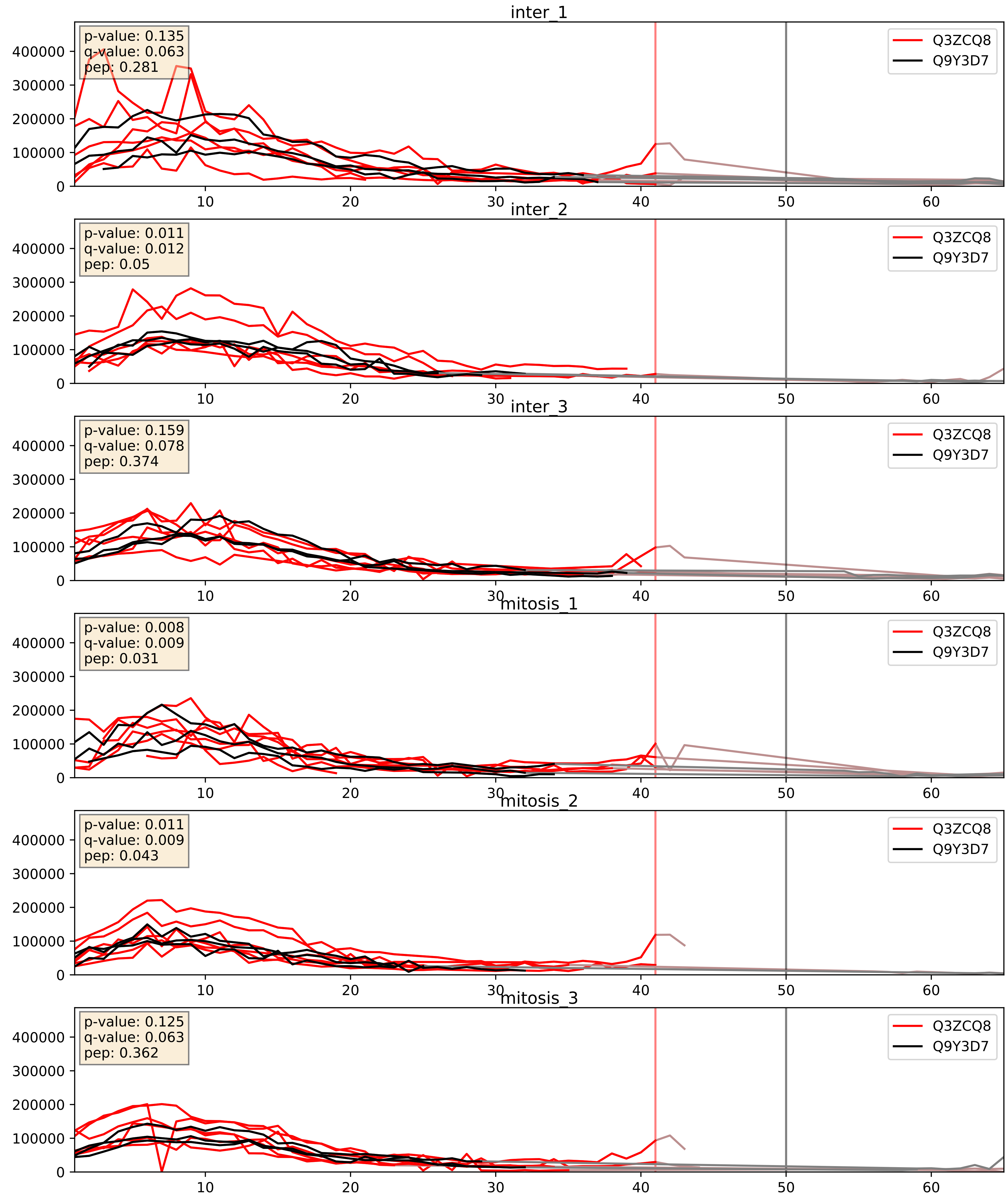


| level | condition_1 | condition_2 | pvalue | pvalue_adjusted |
|---|---|---|---|---|
| complex_abundance | mitosis | inter | 0.029895582448325072 | 0.12569066098117024 |
| interactor_abundance | mitosis | inter | 0.04221745465433784 | 0.16546767941443769 |
| interactor_ratio | mitosis | inter | 0.07698076388416783 | 0.46536348583259013 |

Q96DA6_Q9Y3D7
TIM14_HUMAN vs TIM16_HUMAN
p-value: 0.0 q-value: 0.0

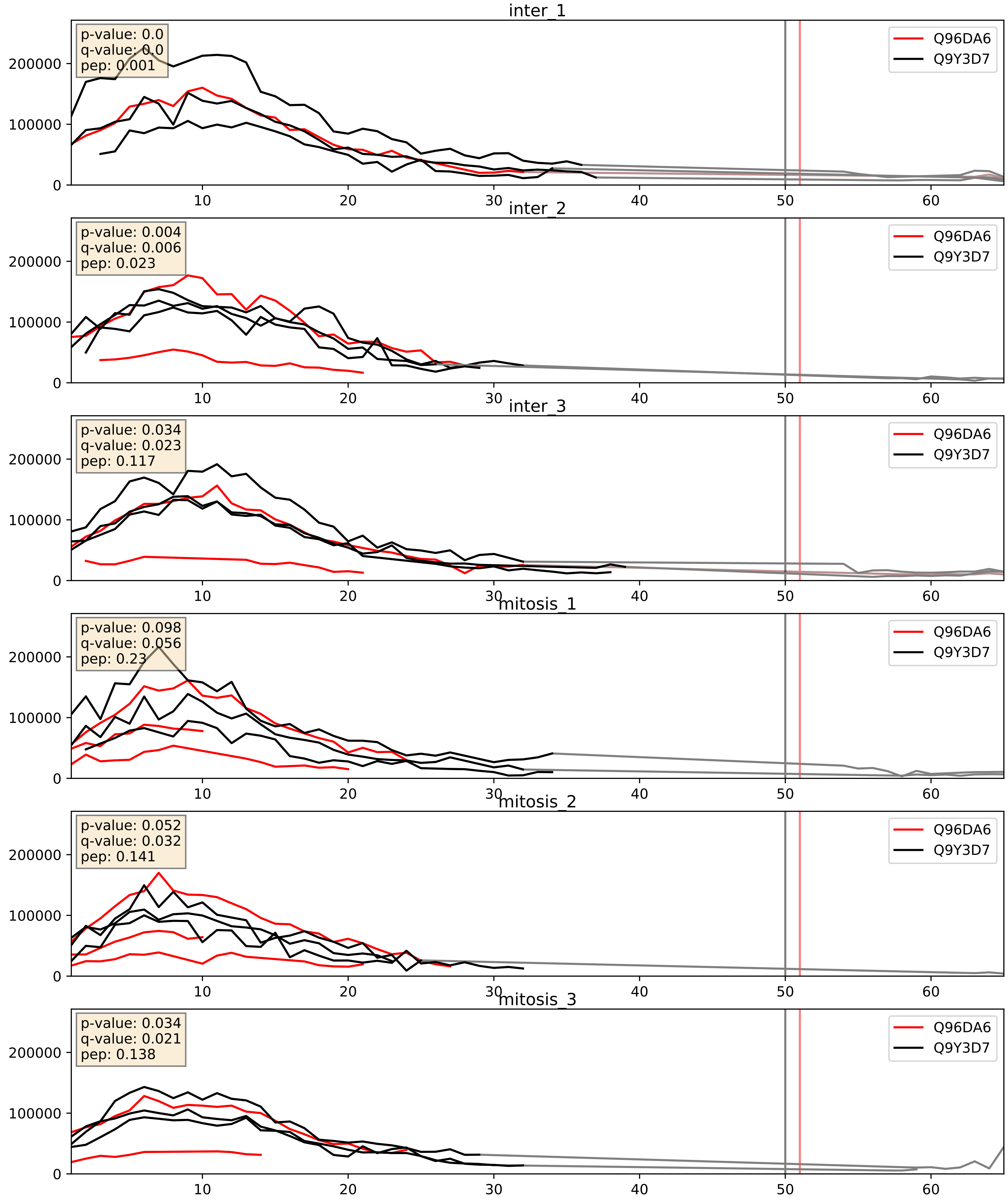


| level | condition_1 | condition_2 | pvalue | pvalue_adjusted |
| --- | --- | --- | --- | --- |
| interactor_ratio | mitosis | inter | 0.17866562431481217 | 0.6458520878947602 |
| complex_abundance | mitosis | inter | 0.2047236080758951 | 0.35822446548030706 |
| interactor_abundance | mitosis | inter | 0.8935549575136748 | 0.940642001637712 |

Q99595_Q9Y3D7
TI17A_HUMAN vs TIM16_HUMAN
p-value: 0.061 q-value: 0.038

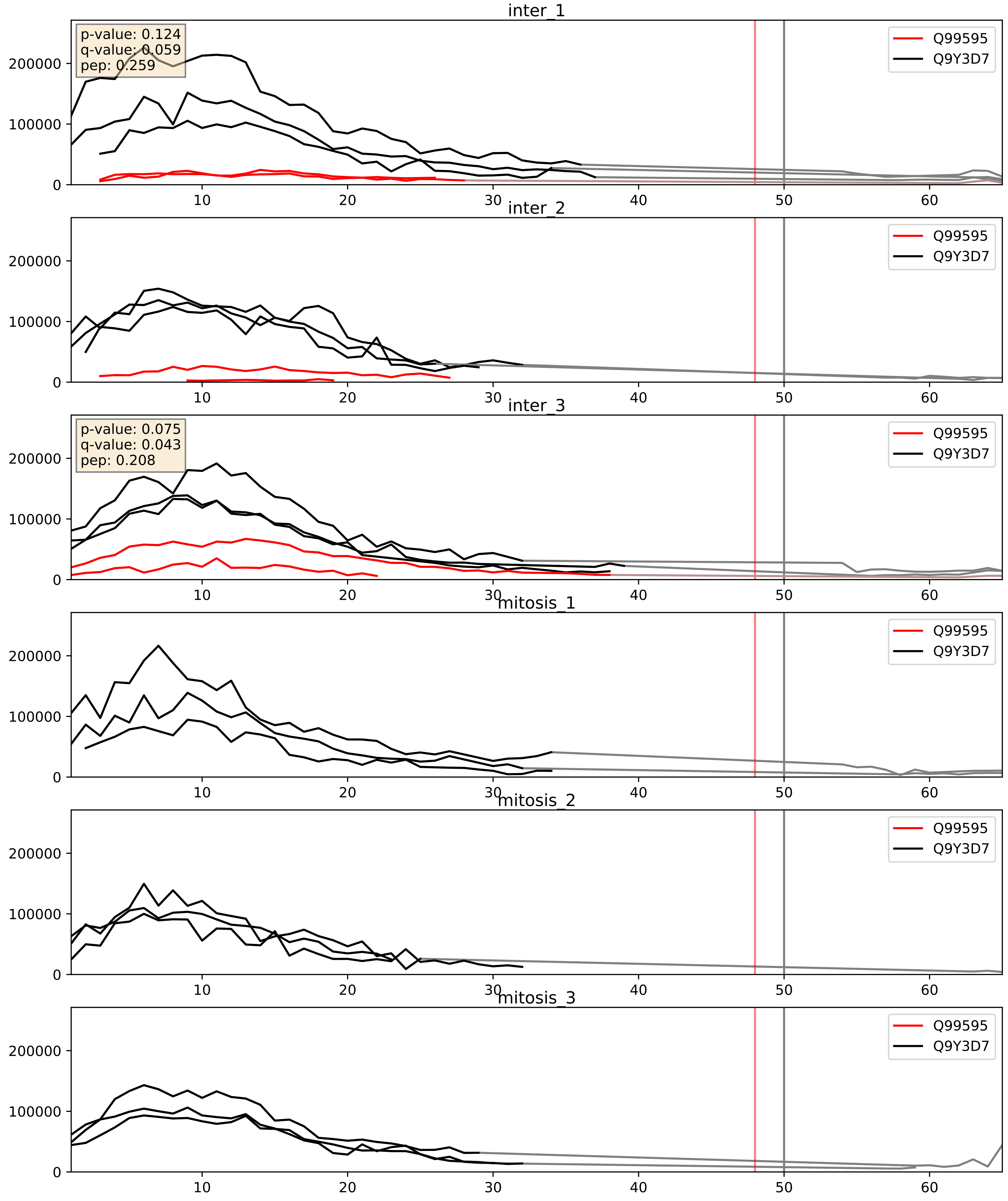


| level | condition_1 | condition_2 | pvalue | pvalue_adjusted |
|---|---|---|---|---|
| interactor_ratio | mitosis | inter | 1.1576148957150758e-05 | 0.003960471554059879 |
| complex_abundance | mitosis | inter | 0.00014027877323287218 | 0.00812325774702682 |
| interactor_abundance | mitosis | inter | 0.0013014868904229176 | 0.026652458808660705 |

Q9BVV7_Q9Y3D7
TIM21_HUMAN vs TIM16_HUMAN
p-value: 0.008 q-value: 0.008

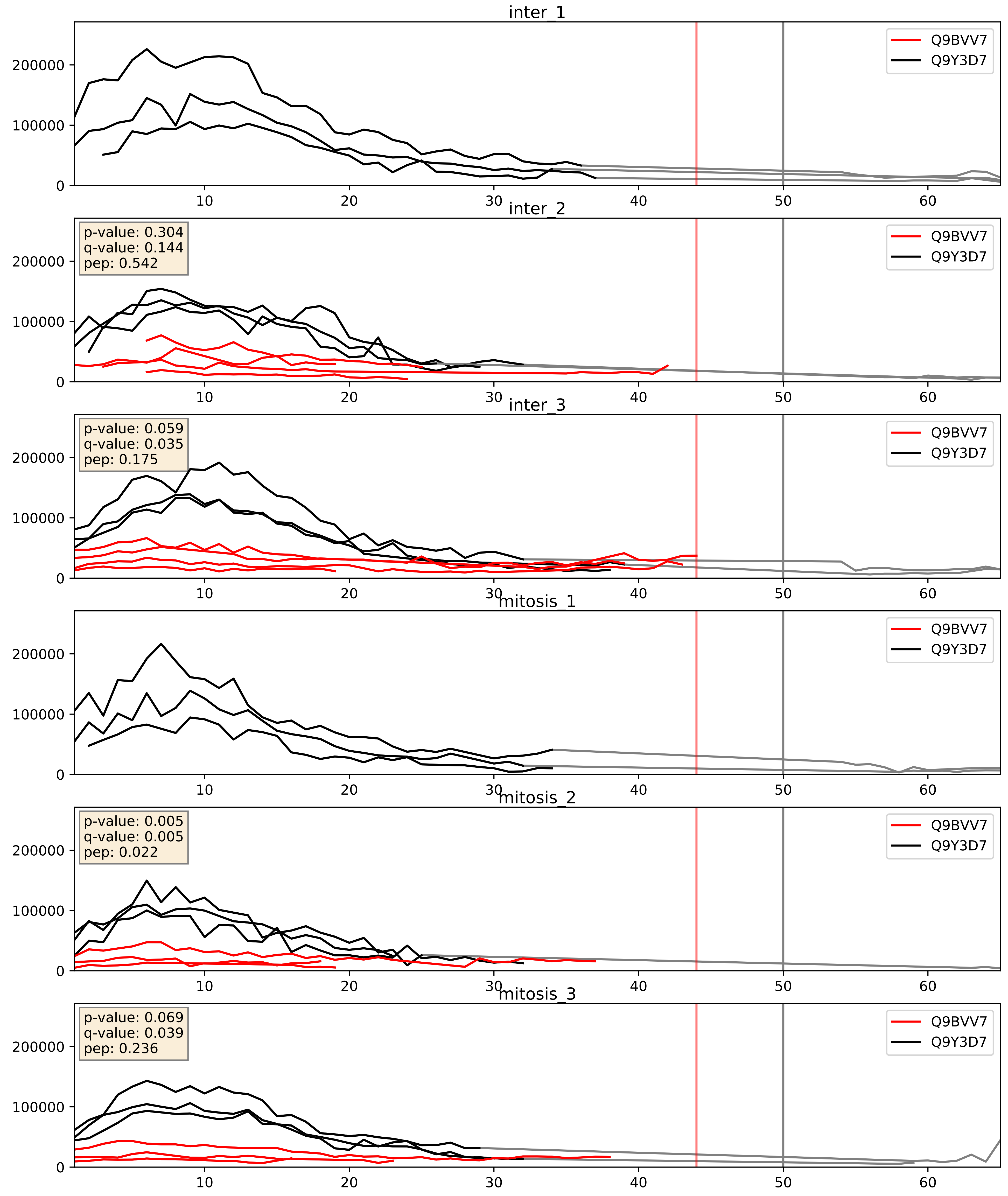


| level | condition_1 | condition_2 | pvalue | pvalue_adjusted |
|---|---|---|---|---|
| interactor_abundance | mitosis | inter | 0.13698015422537446 | 0.323712785188194 |
| complex_abundance | mitosis | inter | 0.4550999296256824 | 0.5969913112594304 |
| interactor_ratio | mitosis | inter | 0.572852652118993 | 0.8946257776299471 |

Q9Y3D7_Q9Y584
TIM16_HUMAN vs TIM22_HUMAN
p-value: 0.0 q-value: 0.003

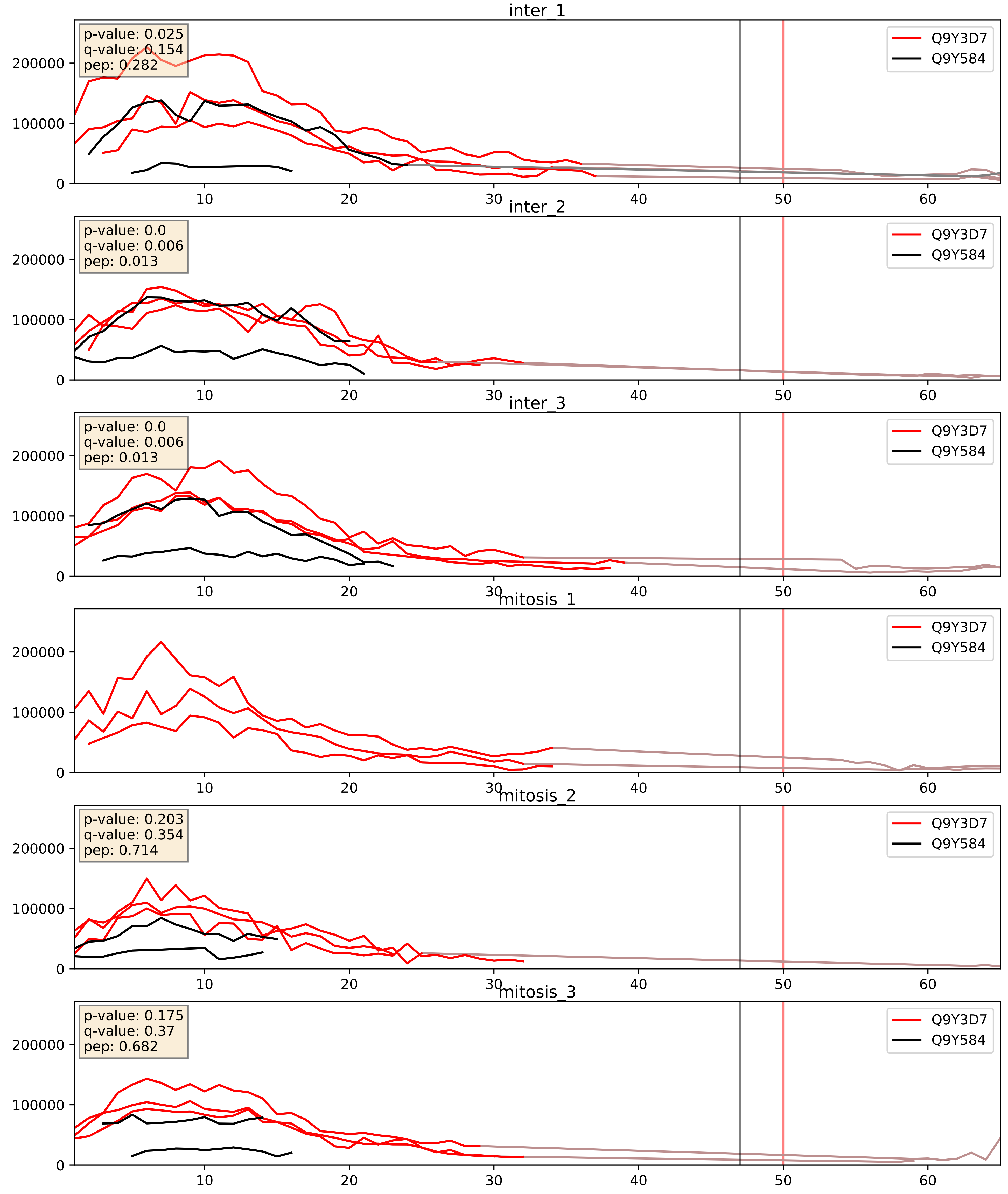


| level | condition_1 | condition_2 | pvalue | pvalue_adjusted |
|---|---|---|---|---|
| complex_abundance | mitosis | inter | 0.05995804731561505 | 0.17540700103269474 |
| interactor_abundance | mitosis | inter | 0.09551648789820179 | 0.2649882146843647 |
| interactor_ratio | mitosis | inter | 0.2850030135171922 | 0.7553021039341069 |
