## Supplementary Data 2 for "SECAT: Quantifying differential protein-protein interaction states by network-centric analysis": hela_string_000180_Q9BWJ5.pdf

### Q9BWJ5_Q9BWJ5

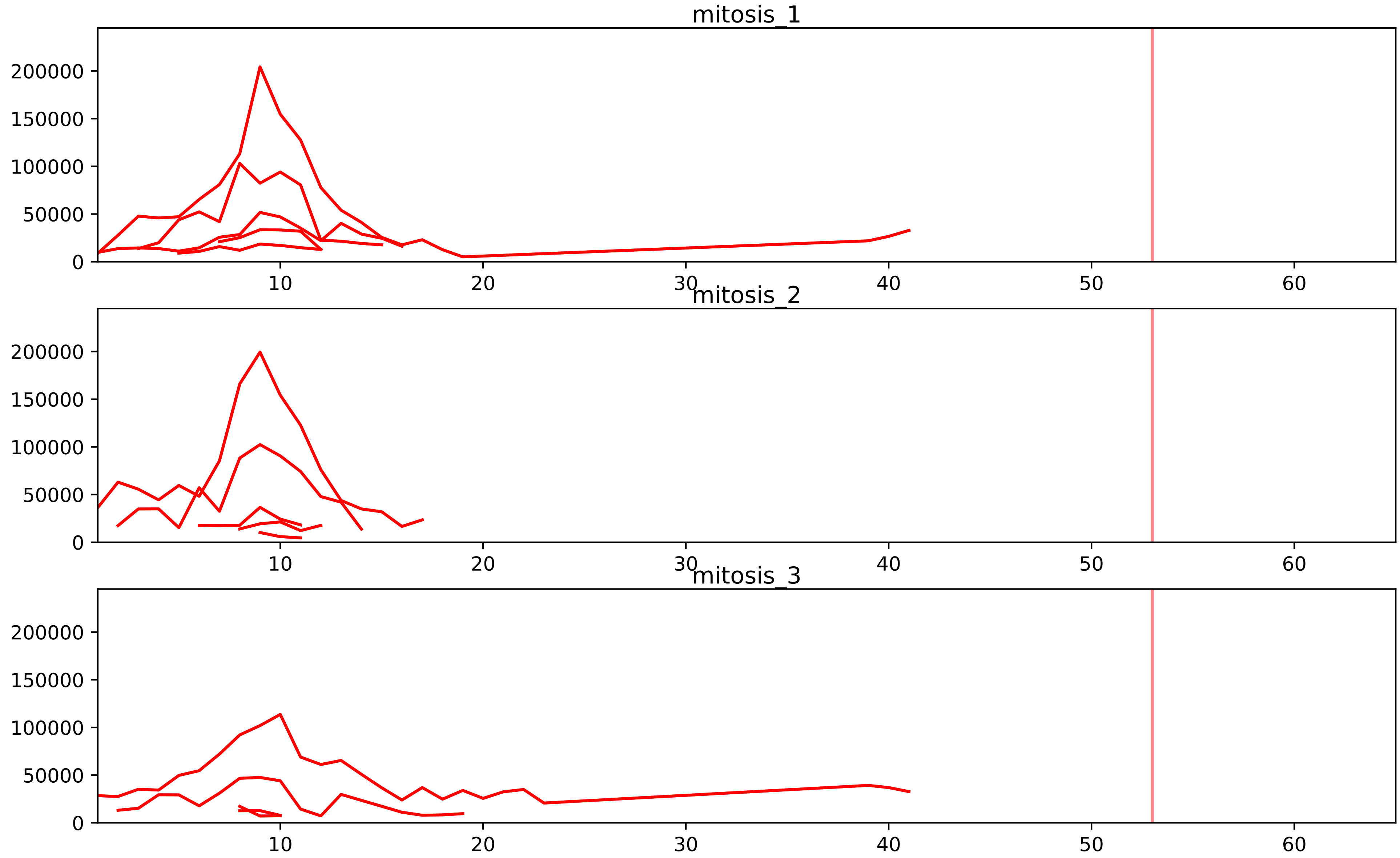


| level | condition_1 | condition_2 | pvalue | pvalue_adjusted |
|---|---|---|---|---|
| complex_abundance | mitosis | inter | 0.00013640255715634733 | 0.0027139113990035054 |
| assembled_abundance | mitosis | inter | 0.0008623173149009361 | 0.02604198291000827 |
| total_abundance | mitosis | inter | 0.0014696640216107972 | 0.04111426016435889 |
| interactor_abundance | mitosis | inter | 0.0038870443400276333 | 0.029192496267962633 |
| interactor_ratio | mitosis | inter | 0.005111212383093453 | 0.040889699064747626 |

Q7RTV0_Q9BWJ5
PHF5A_HUMAN vs SF3B5_HUMAN
p-value: 0.0 q-value: 0.001

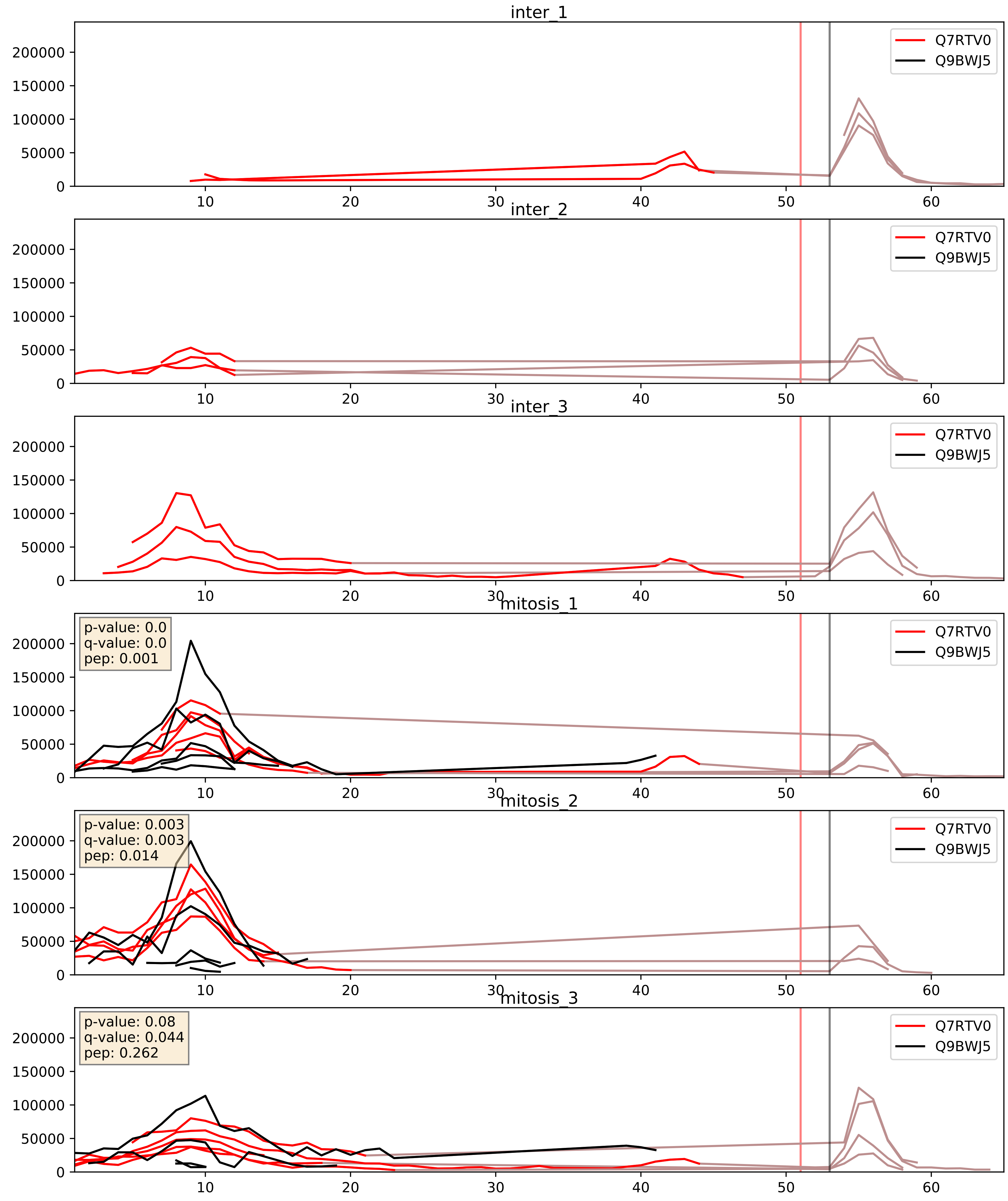


| level | condition_1 | condition_2 | pvalue | pvalue_adjusted |
|---|---|---|---|---|
| interactor_abundance | mitosis | inter | 0.000736287308134052 | 0.019819557728388316 |
| complex_abundance | mitosis | inter | 0.0017983117149834248 | 0.030065523984879133 |
| interactor_ratio | mitosis | inter | 0.33243641433023874 | 0.7893553151713977 |

Q9BWJ5_Q9Y3B4
SF3B5_HUMAN vs SF3B6_HUMAN
p-value: 0.0 q-value: 0.0

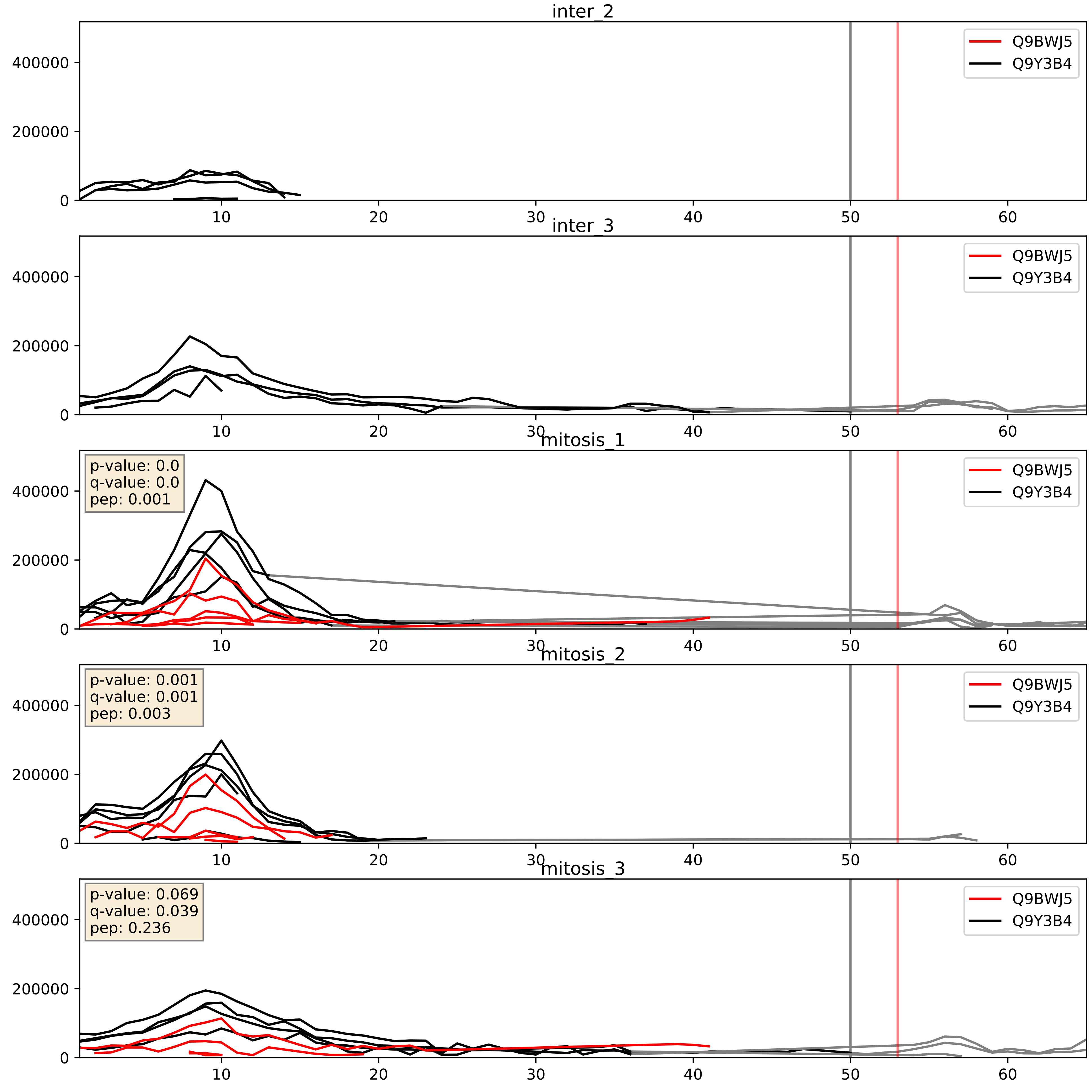


| level | condition_1 | condition_2 | pvalue | pvalue_adjusted |
|---|---|---|---|---|
| complex_abundance | mitosis | inter | 0.0003578301856744203 | 0.012039775462644836 |
| interactor_ratio | mitosis | inter | 0.001829233271177215 | 0.0519767486809081 |
| interactor_abundance | mitosis | inter | 0.00949433585822683 | 0.08456973459565209 |
