## Supplementary Data 2 for "SECAT: Quantifying differential protein-protein interaction states by network-centric analysis": hela_string_000181_Q969M3.pdf

Q969M3_Q969M3

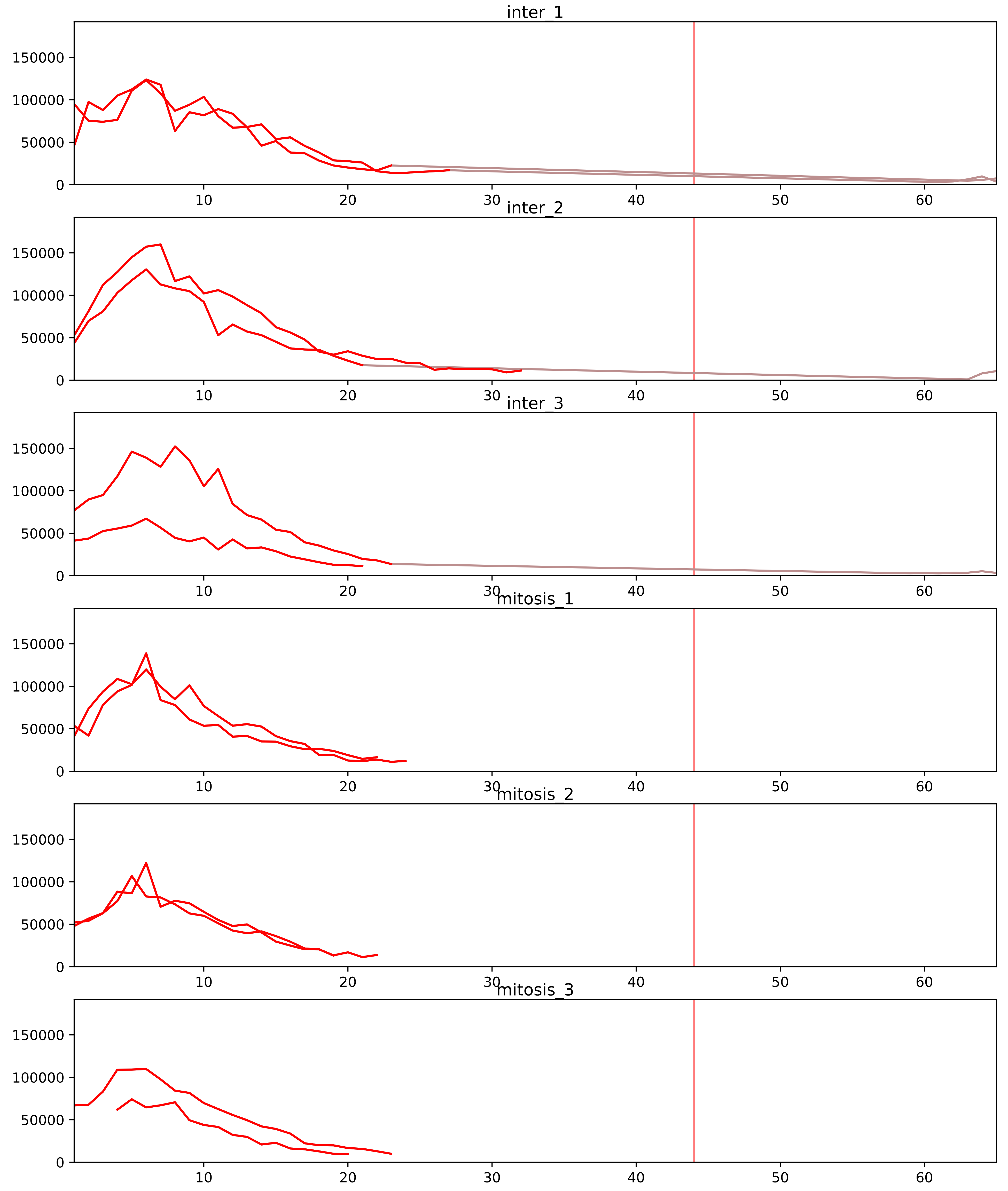


| level | condition_1 | condition_2 | pvalue | pvalue_adjusted |
|---|---|---|---|---|
| complex_abundance | mitosis | inter | 0.0001371990402873377 | 0.0027139113990035054 |
| interactor_abundance | mitosis | inter | 0.0008562581094817792 | 0.01035560027477911 |
| interactor_ratio | mitosis | inter | 0.010210390044533127 | 0.06411985556976435 |
| monomer_abundance | mitosis | inter | 0.050123803978309074 | 0.33495561770373317 |
| assembled_abundance | mitosis | inter | 0.32356291959440475 | 0.6139432617030436 |
| total_abundance | mitosis | inter | 0.36983303211457913 | 0.6787382719882201 |

O95070_Q969M3
YIF1A_HUMAN vs YIPF5_HUMAN
p-value: 0.0 q-value: 0.0

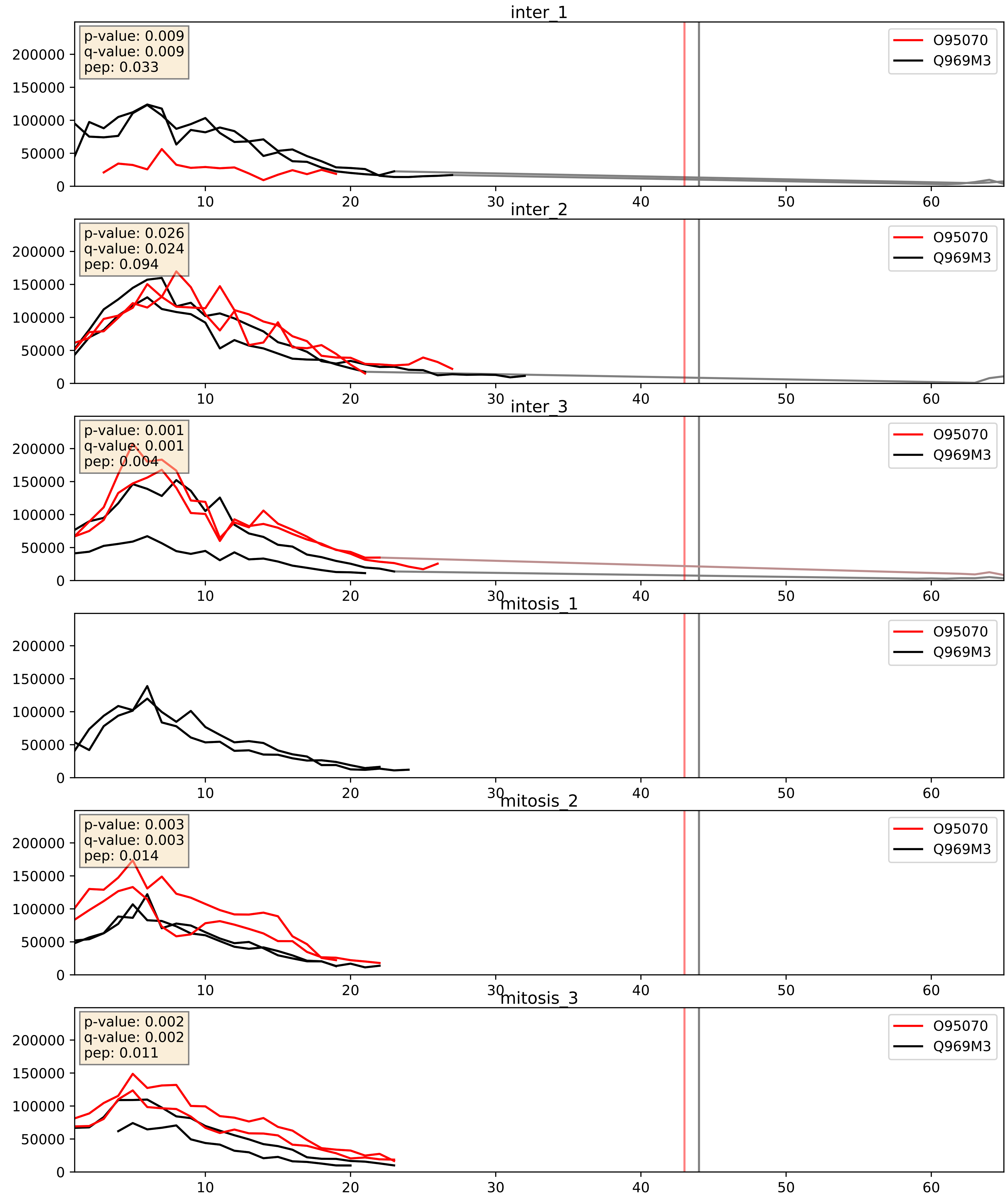


| level | condition_1 | condition_2 | pvalue | pvalue_adjusted |
|---|---|---|---|---|
| complex_abundance | mitosis | inter | 0.3164321592635194 | 0.46461882893325146 |
| interactor_ratio | mitosis | inter | 0.3845331862407519 | 0.8212585015521048 |
| interactor_abundance | mitosis | inter | 0.6062865688545522 | 0.7415077910265706 |

Q5BJH7_Q969M3
YIF1B_HUMAN vs YIPF5_HUMAN
p-value: 0.001 q-value: 0.046

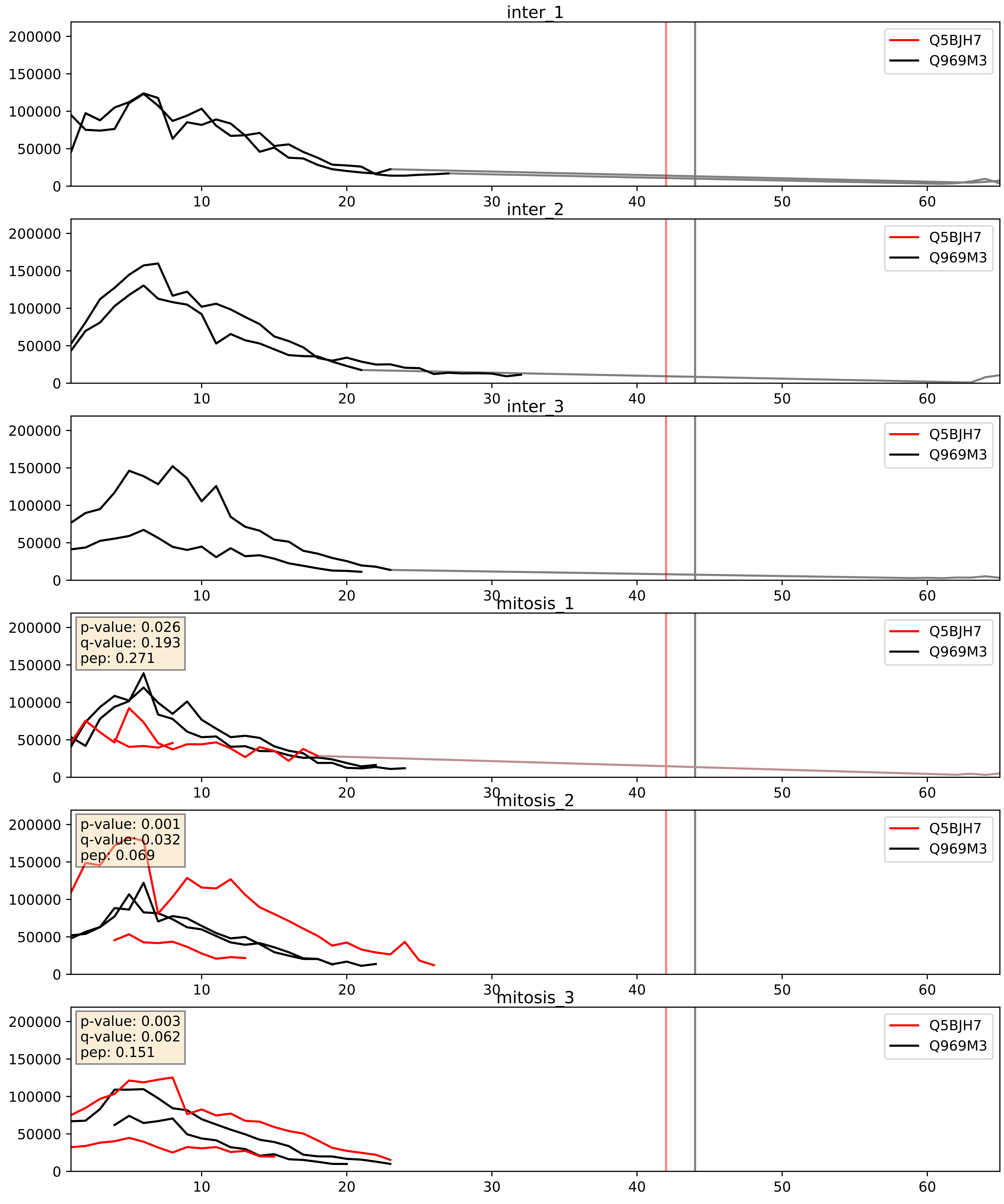


| level | condition_1 | condition_2 | pvalue | pvalue_adjusted |
|---|---|---|---|---|
| complex_abundance | mitosis | inter | 0.00022211895660948554 | 0.010113501428602107 |
| interactor_abundance | mitosis | inter | 0.0015399695522842847 | 0.028907984680718663 |
| interactor_ratio | mitosis | inter | 0.35140728464106313 | 0.8012909846903304 |

Q969M3_Q96EC8
YIPF5_HUMAN vs YIPF6_HUMAN
p-value: 0.0 q-value: 0.008

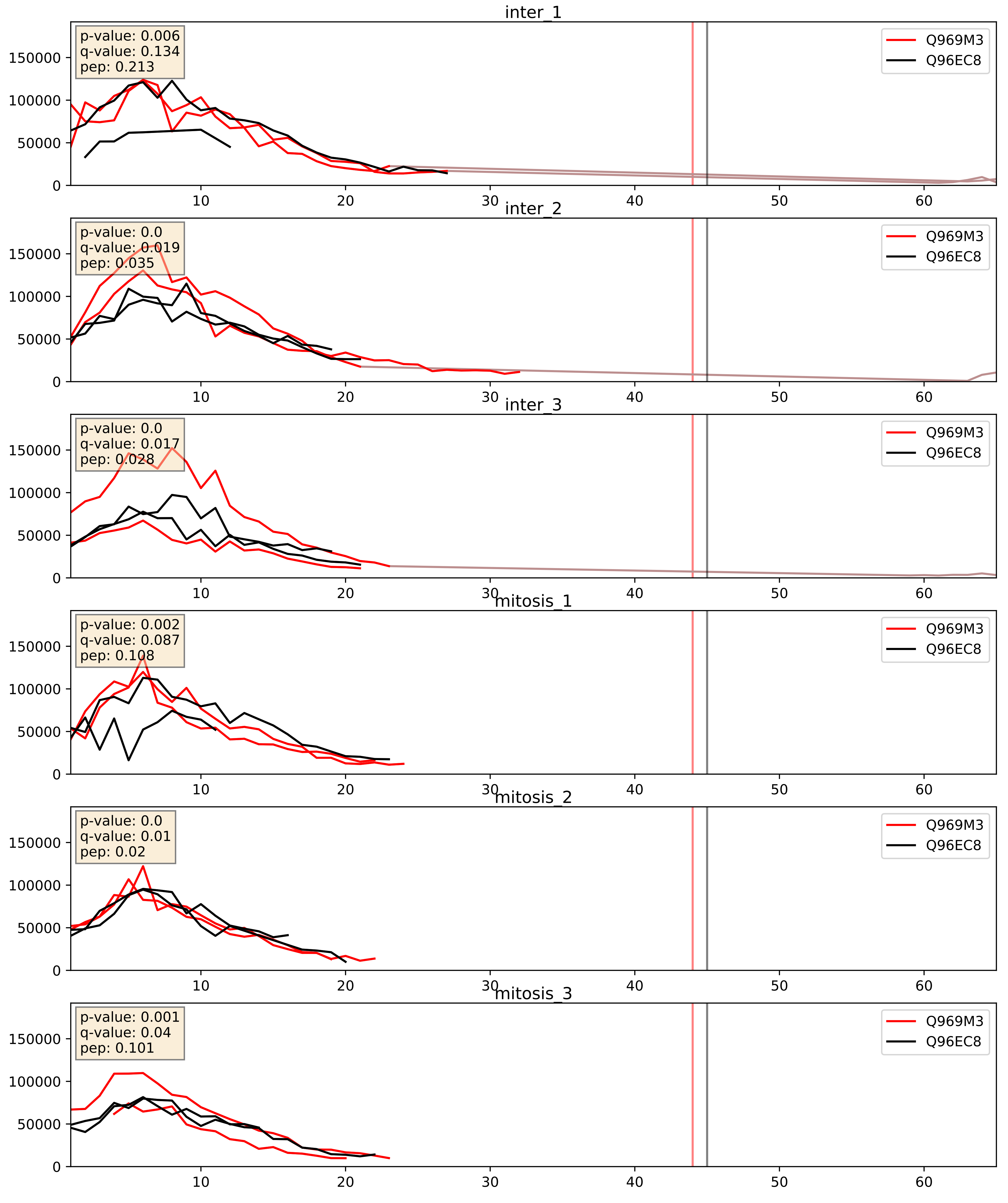


| level | condition_1 | condition_2 | pvalue | pvalue_adjusted |
|---|---|---|---|---|
| complex_abundance | mitosis | inter | 0.04065834692844474 | 0.14525686548726502 |
| interactor_abundance | mitosis | inter | 0.049629357480397145 | 0.1778264127384678 |
| interactor_ratio | mitosis | inter | 0.2650964039288194 | 0.7404362309130512 |

Q969M3_Q96N66
YIPF5_HUMAN vs MBOA7_HUMAN
p-value: 0.0 q-value: 0.047

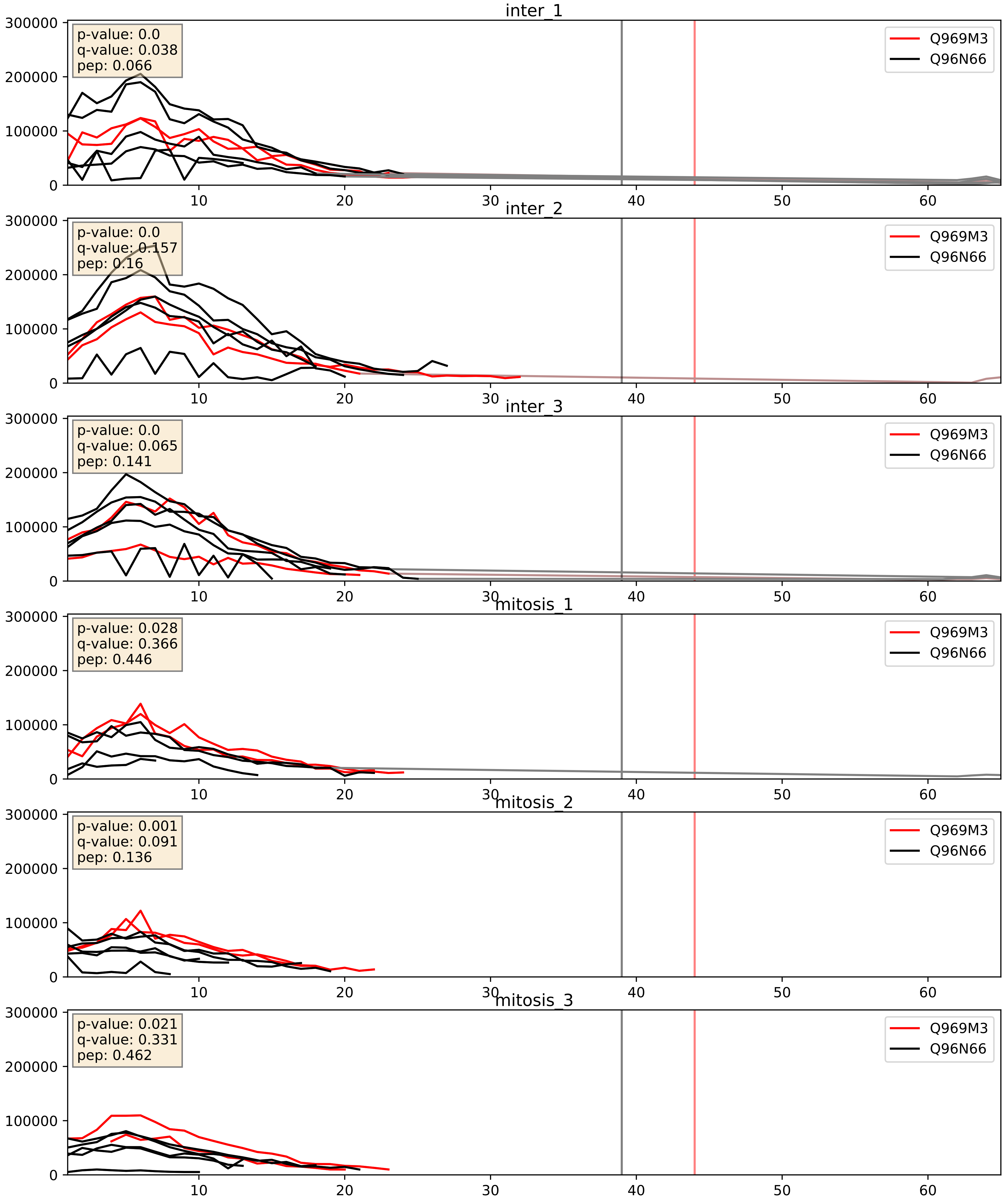


| level | condition_1 | condition_2 | pvalue | pvalue_adjusted |
|---|---|---|---|---|
| interactor_ratio | mitosis | inter | 0.0003149130268567279 | 0.01878749154928596 |
| complex_abundance | mitosis | inter | 0.004936834350413667 | 0.052626777135169354 |
| interactor_abundance | mitosis | inter | 0.04686004299581258 | 0.17297195689700548 |

Q969M3_Q9BUN8
YIPF5_HUMAN vs DERL1_HUMAN
p-value: 0.0 q-value: 0.04

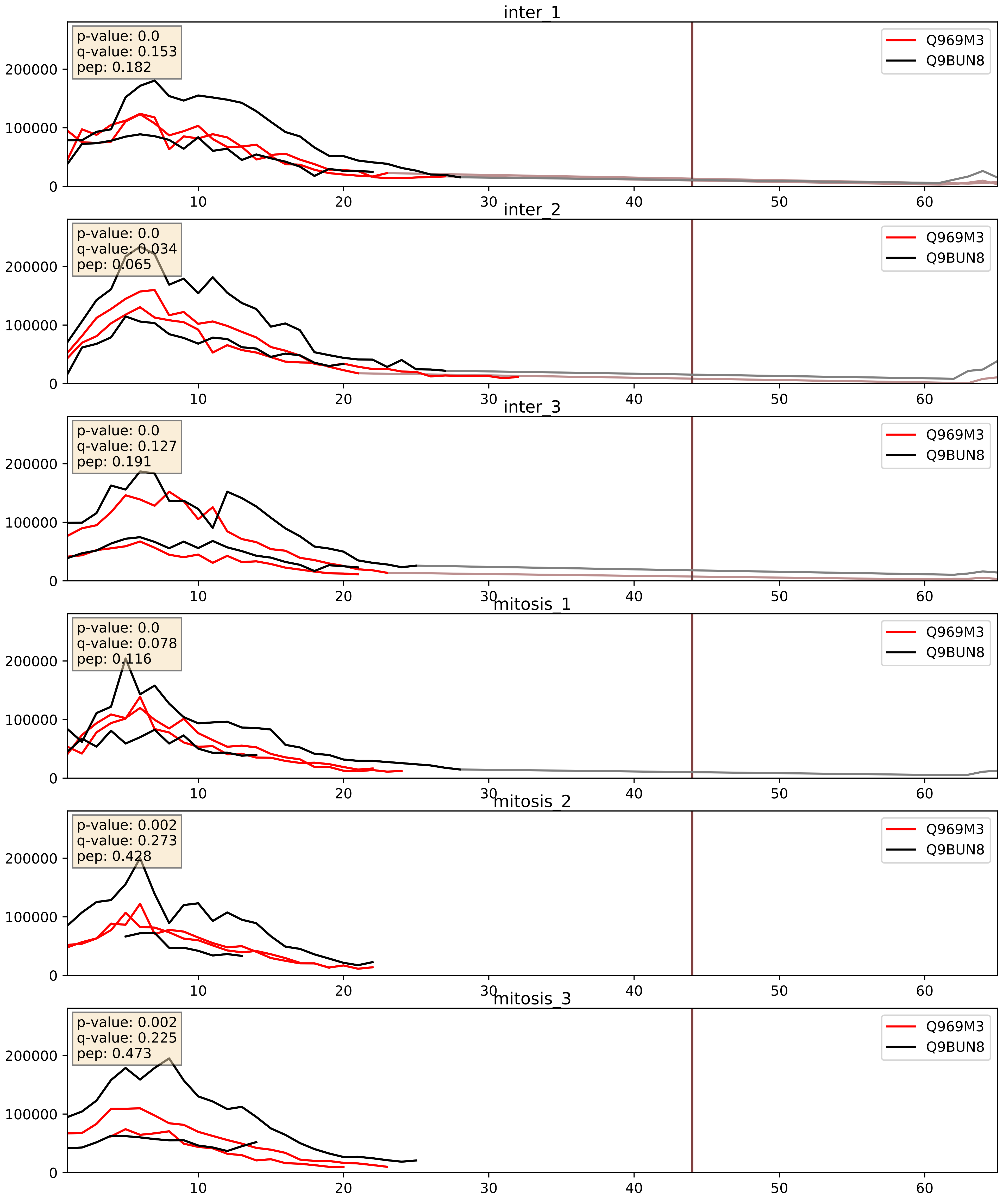


| level | condition_1 | condition_2 | pvalue | pvalue_adjusted |
|---|---|---|---|---|
| complex_abundance | mitosis | inter | 0.028324778373801218 | 0.12245459741400931 |
| interactor_abundance | mitosis | inter | 0.04597569190886313 | 0.17107234198646748 |
| interactor_ratio | mitosis | inter | 0.6139748691712491 | 0.90762799337104 |

Q969M3_Q9GZM5
YIPF5_HUMAN vs YIPF3_HUMAN
p-value: 0.0 q-value: 0.008

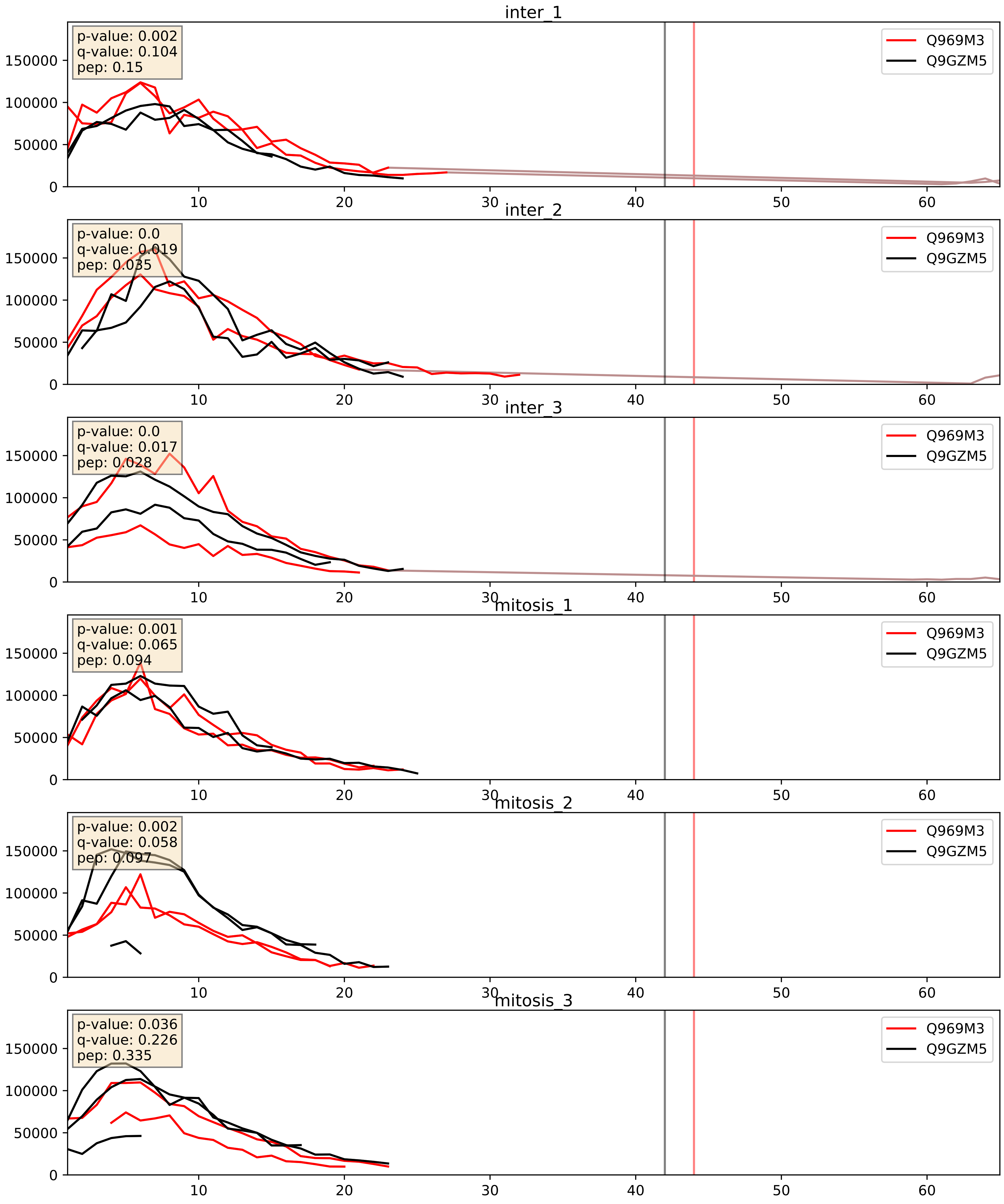


| level | condition_1 | condition_2 | pvalue | pvalue_adjusted |
|---|---|---|---|---|
| interactor_abundance | mitosis | inter | 0.049295148137983115 | 0.17730702850330524 |
| complex_abundance | mitosis | inter | 0.0643997844610902 | 0.18320443834726888 |
| interactor_ratio | mitosis | inter | 0.14692912049294174 | 0.6009141287241191 |

Q969M3_Q9UKR5
YIPF5_HUMAN vs ERG28_HUMAN
p-value: 0.0 q-value: 0.04

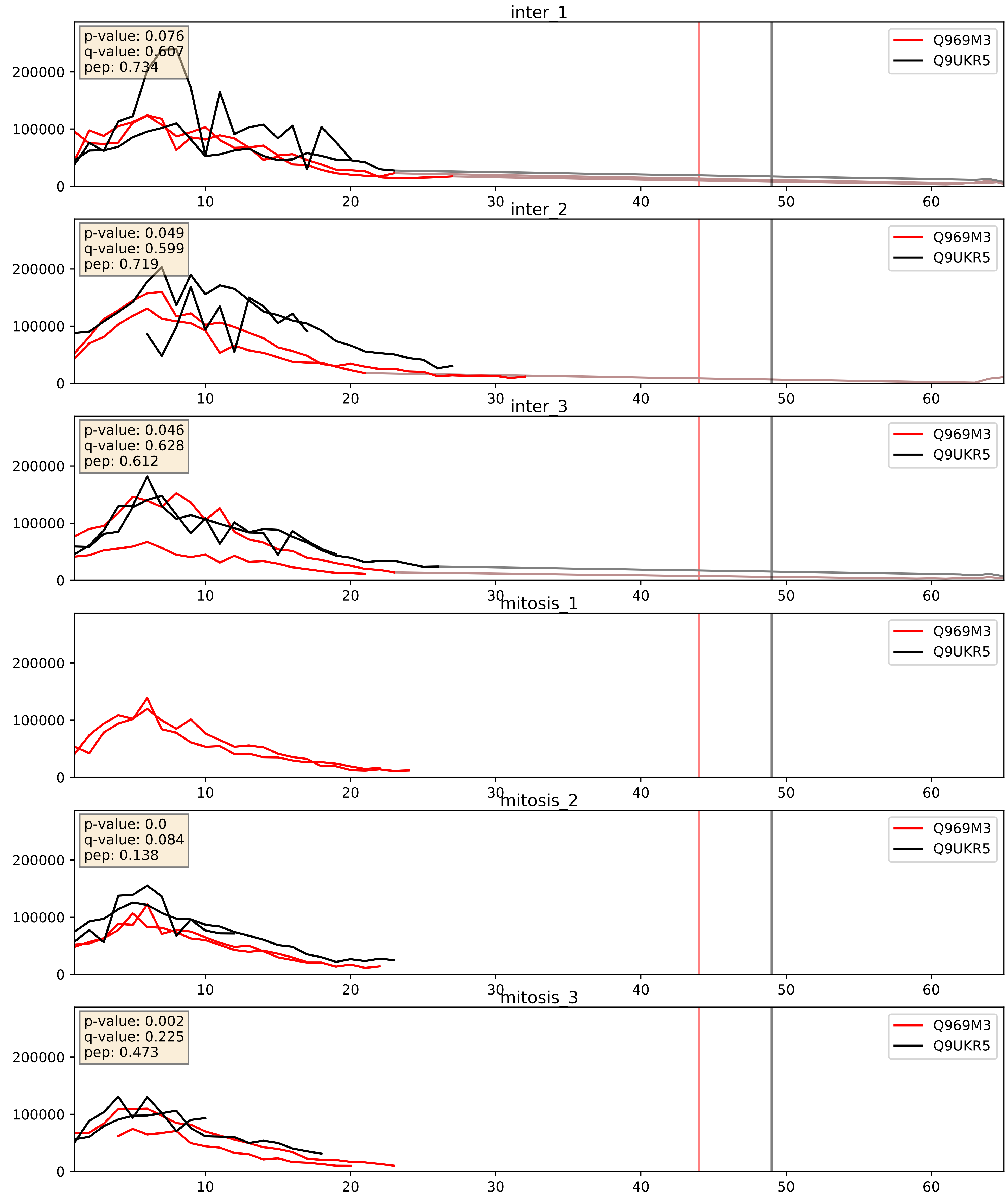


| level | condition_1 | condition_2 | pvalue | pvalue_adjusted |
|---|---|---|---|---|
| interactor_abundance | mitosis | inter | 0.09732700525946608 | 0.268228964913403 |
| complex_abundance | mitosis | inter | 0.10494668608291362 | 0.2407782452076496 |
| interactor_ratio | mitosis | inter | 0.47747330577073394 | 0.865462474268604 |
