## Supplementary Data 2 for "SECAT: Quantifying differential protein-protein interaction states by network-centric analysis": hela_string_000182_O43678.pdf

O43678_O43678

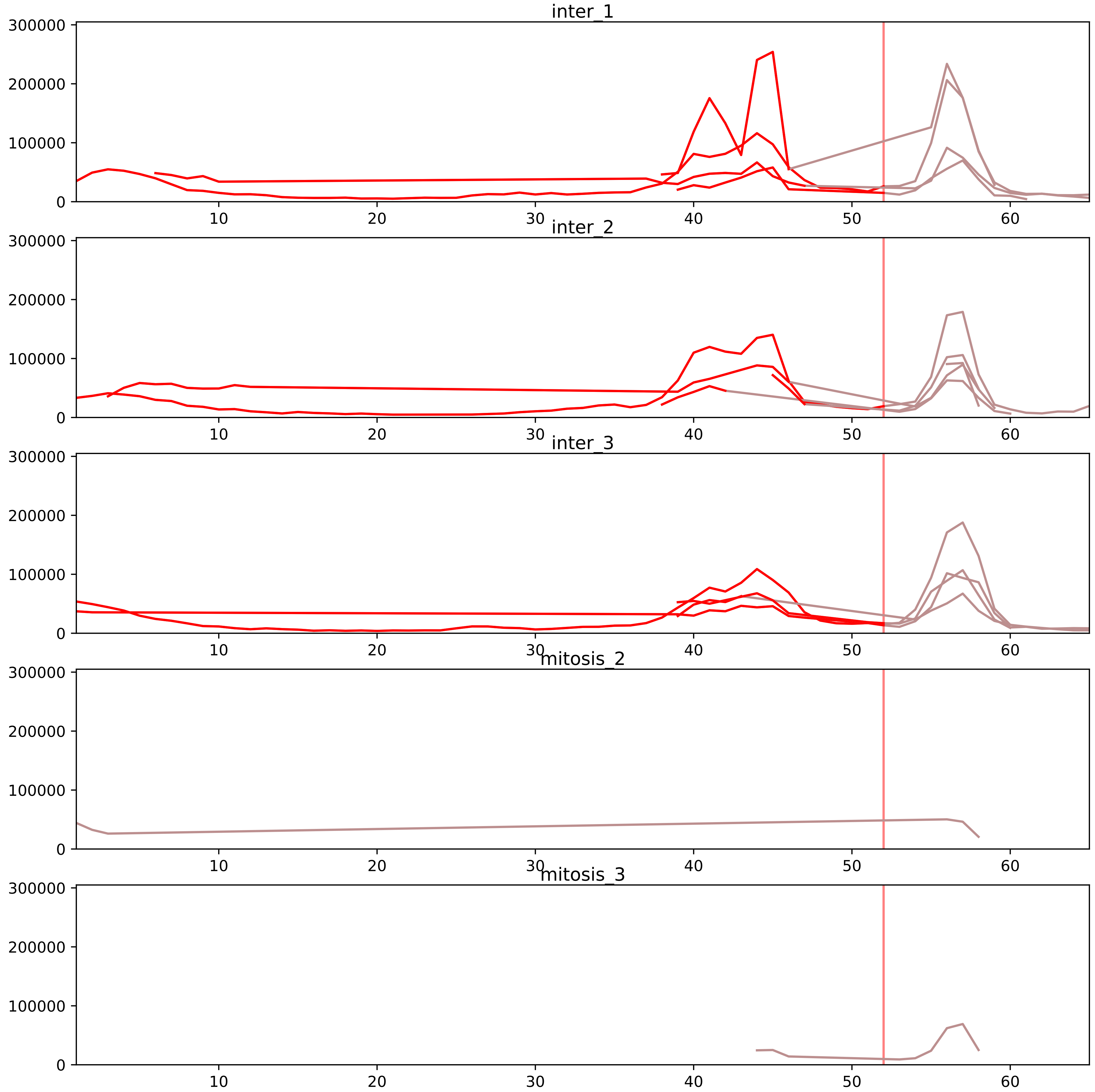


| level | condition_1 | condition_2 | pvalue | pvalue_adjusted |
|---|---|---|---|---|
| complex_abundance | mitosis | inter | 0.00013938989471800356 | 0.0027139113990035054 |
| interactor_abundance | mitosis | inter | 0.002989801196775818 | 0.024892462452794143 |
| total_abundance | mitosis | inter | 0.0032004717971851194 | 0.06315349581189707 |
| monomer_abundance | mitosis | inter | 0.0038057792847432247 | 0.12994017843623296 |
| assembled_abundance | mitosis | inter | 0.0043409721673620776 | 0.06554867972716737 |
| interactor_ratio | mitosis | inter | 0.006255648776225347 | 0.045316510819900145 |

O43678_P28331
NDUA2_HUMAN vs NDUS1_HUMAN
p-value: 0.085 q-value: 0.05

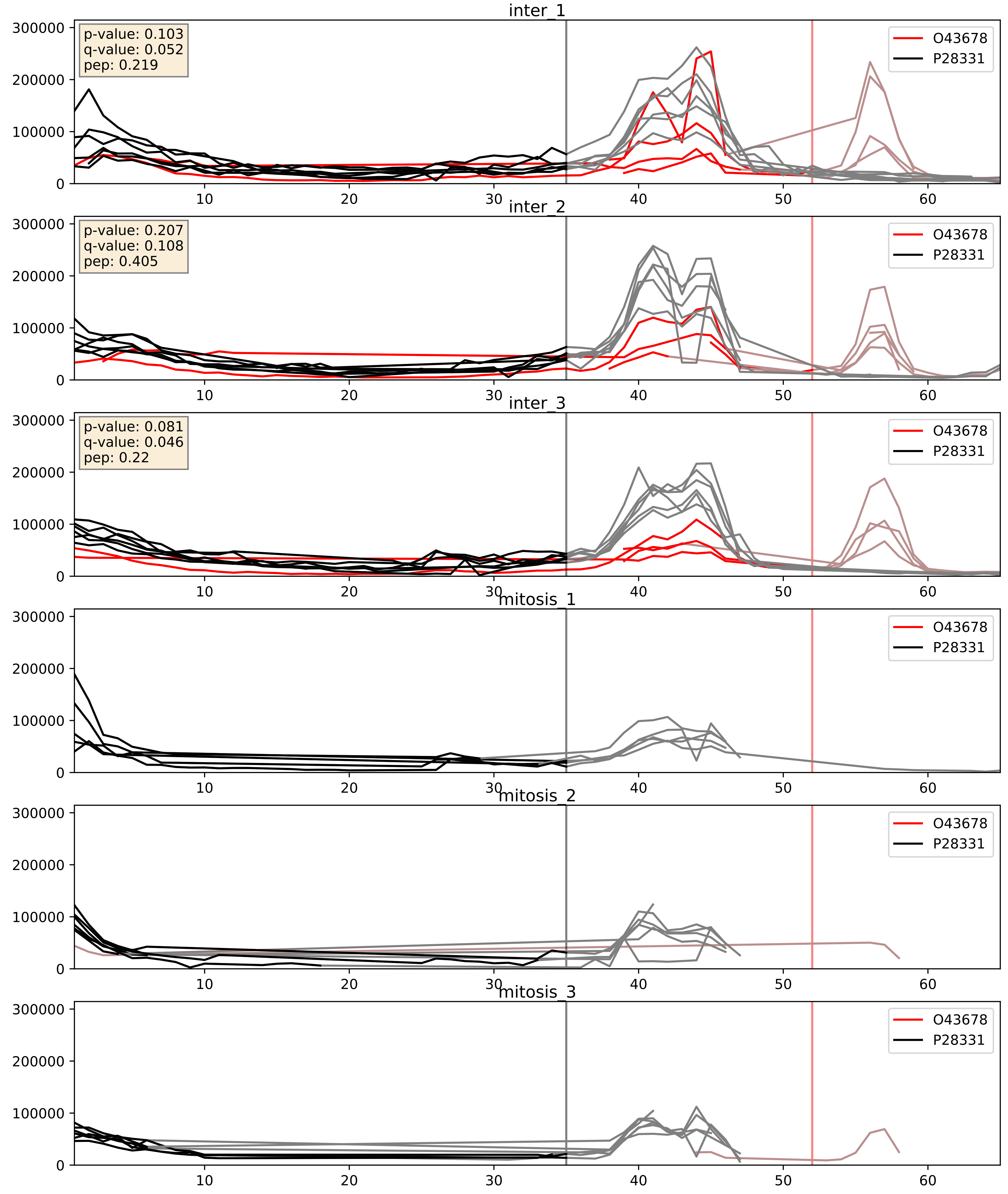


| level | condition_1 | condition_2 | pvalue | pvalue_adjusted |
|---|---|---|---|---|
| complex_abundance | mitosis | inter | 0.000623955510183607 | 0.01603921671823326 |
| interactor_ratio | mitosis | inter | 0.0019183710724063662 | 0.05317578467954594 |
| interactor_abundance | mitosis | inter | 0.01084111579398431 | 0.09135080923250061 |

O43678_P56556
NDUA2_HUMAN vs NDUA6_HUMAN
p-value: 0.063 q-value: 0.039

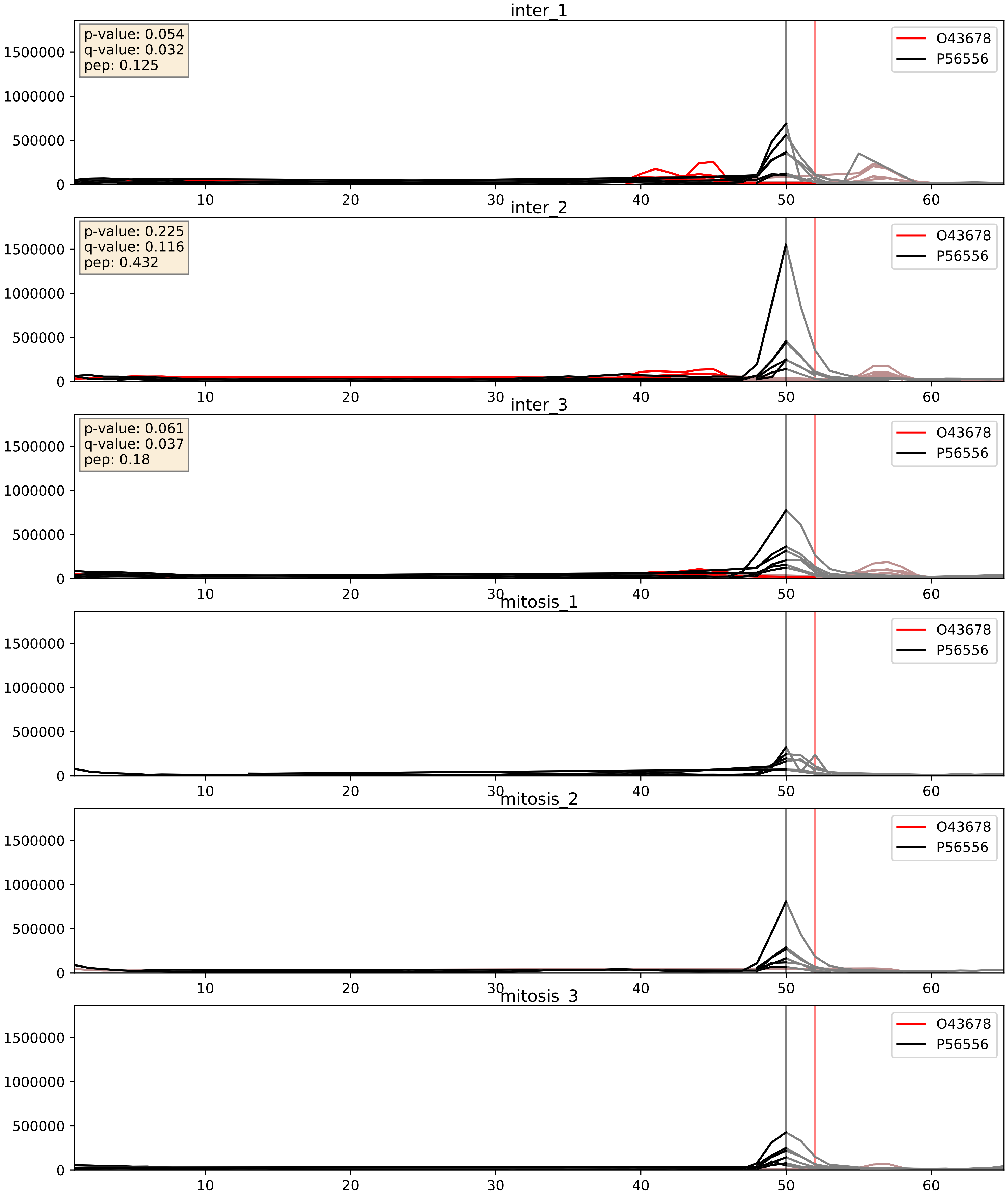


| level | condition_1 | condition_2 | pvalue | pvalue_adjusted |
|---|---|---|---|---|
| complex_abundance | mitosis | inter | 0.0010427131847453482 | 0.02161168247317235 |
| interactor_abundance | mitosis | inter | 0.005906515623201406 | 0.06511239373419707 |
| interactor_ratio | mitosis | inter | 0.15173525543561164 | 0.6067576723168497 |
