## Supplementary Data 2 for "SECAT: Quantifying differential protein-protein interaction states by network-centric analysis": hela_string_000183_P13073.pdf

P13073_P13073

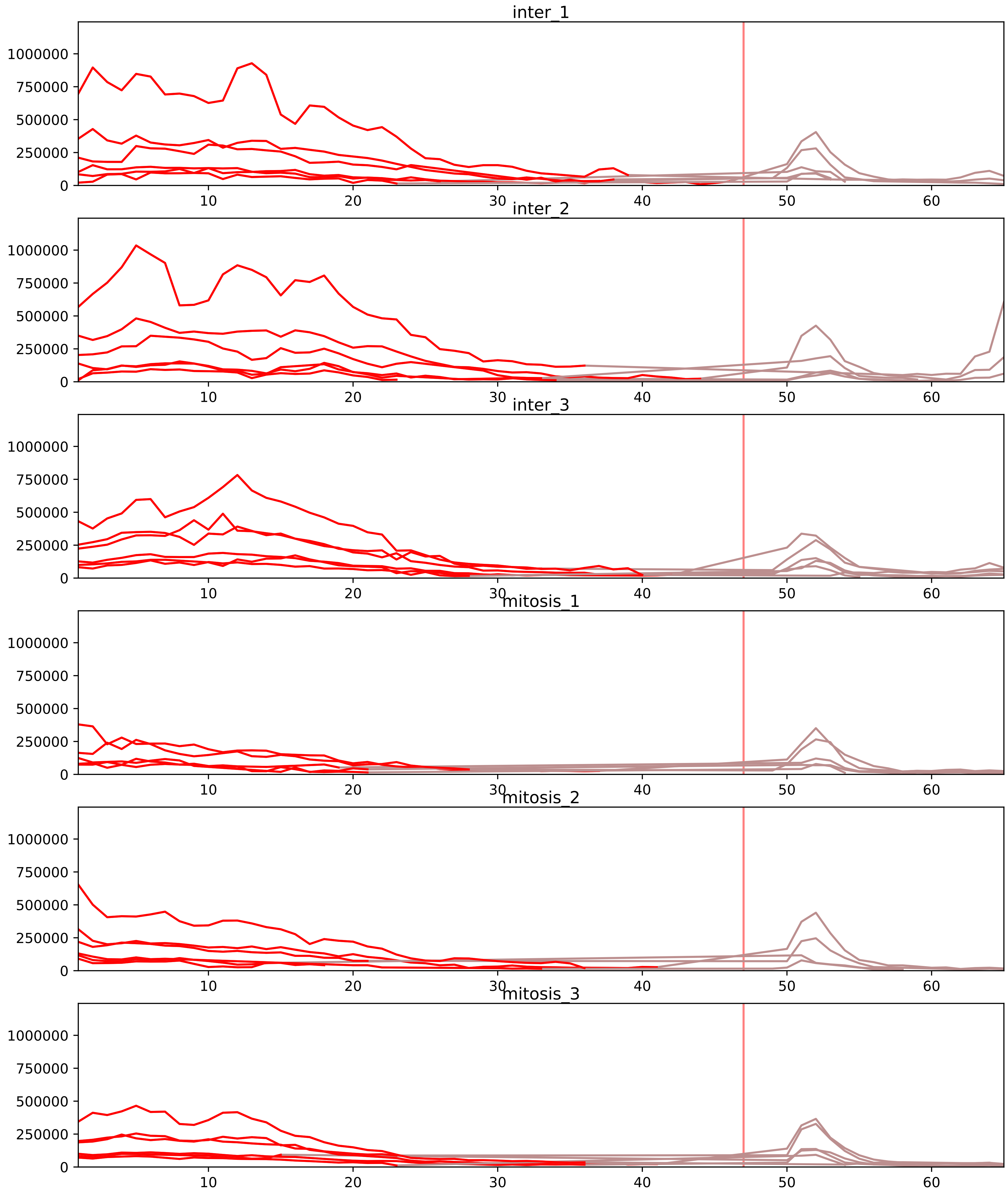


| level | condition_1 | condition_2 | pvalue | pvalue_adjusted |
|---|---|---|---|---|
| complex_abundance | mitosis | inter | 0.00014012042549202882 | 0.0027139113990035054 |
| interactor_abundance | mitosis | inter | 0.004613273559283442 | 0.03239856240107455 |
| monomer_abundance | mitosis | inter | 0.025423502609103428 | 0.2792215738541247 |
| interactor_ratio | mitosis | inter | 0.026194557911263436 | 0.12784611818759872 |
| assembled_abundance | mitosis | inter | 0.03710162689246421 | 0.23094775995162894 |
| total_abundance | mitosis | inter | 0.039705923103559006 | 0.25173456353700985 |

O00483_P13073
NDUA4_HUMAN vs COX41_HUMAN
p-value: 0.0 q-value: 0.007

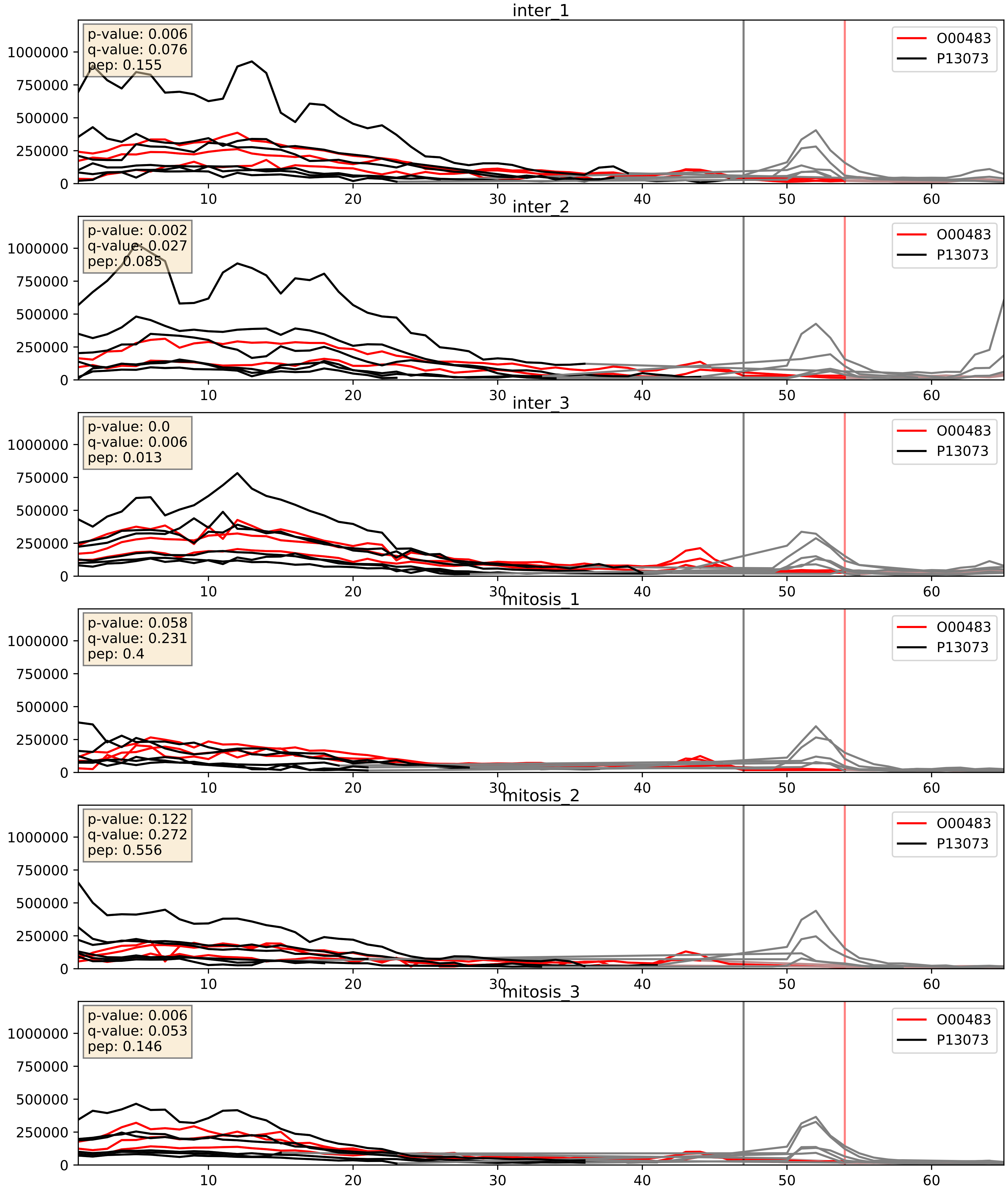


| level | condition_1 | condition_2 | pvalue | pvalue_adjusted |
|---|---|---|---|---|
| complex_abundance | mitosis | inter | 0.04986678092573984 | 0.15897087091075196 |
| interactor_ratio | mitosis | inter | 0.15446428442401383 | 0.6115862733861726 |
| interactor_abundance | mitosis | inter | 0.5965173439203346 | 0.7348091040377125 |

O75251_P13073
NDUS7_HUMAN vs COX41_HUMAN
p-value: 0.0 q-value: 0.023

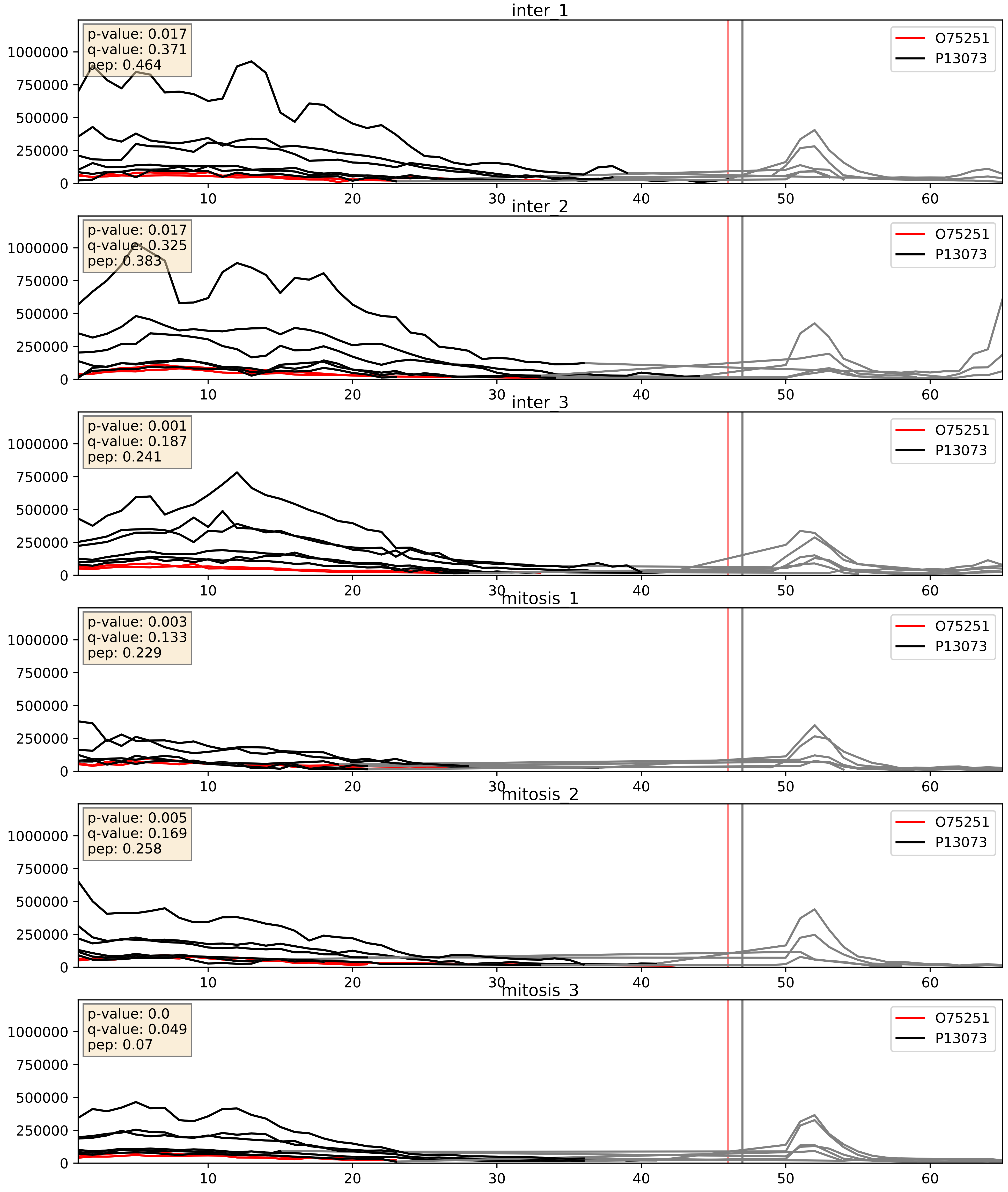


| level | condition_1 | condition_2 | pvalue | pvalue_adjusted |
|---|---|---|---|---|
| complex_abundance | mitosis | inter | 0.01217365436217513 | 0.08185897984306292 |
| interactor_abundance | mitosis | inter | 0.016601804854909166 | 0.11277132084292767 |
| interactor_ratio | mitosis | inter | 0.05378202562470084 | 0.38466228753446735 |

O75489_P13073
NDUS3_HUMAN vs COX41_HUMAN
p-value: 0.001 q-value: 0.032

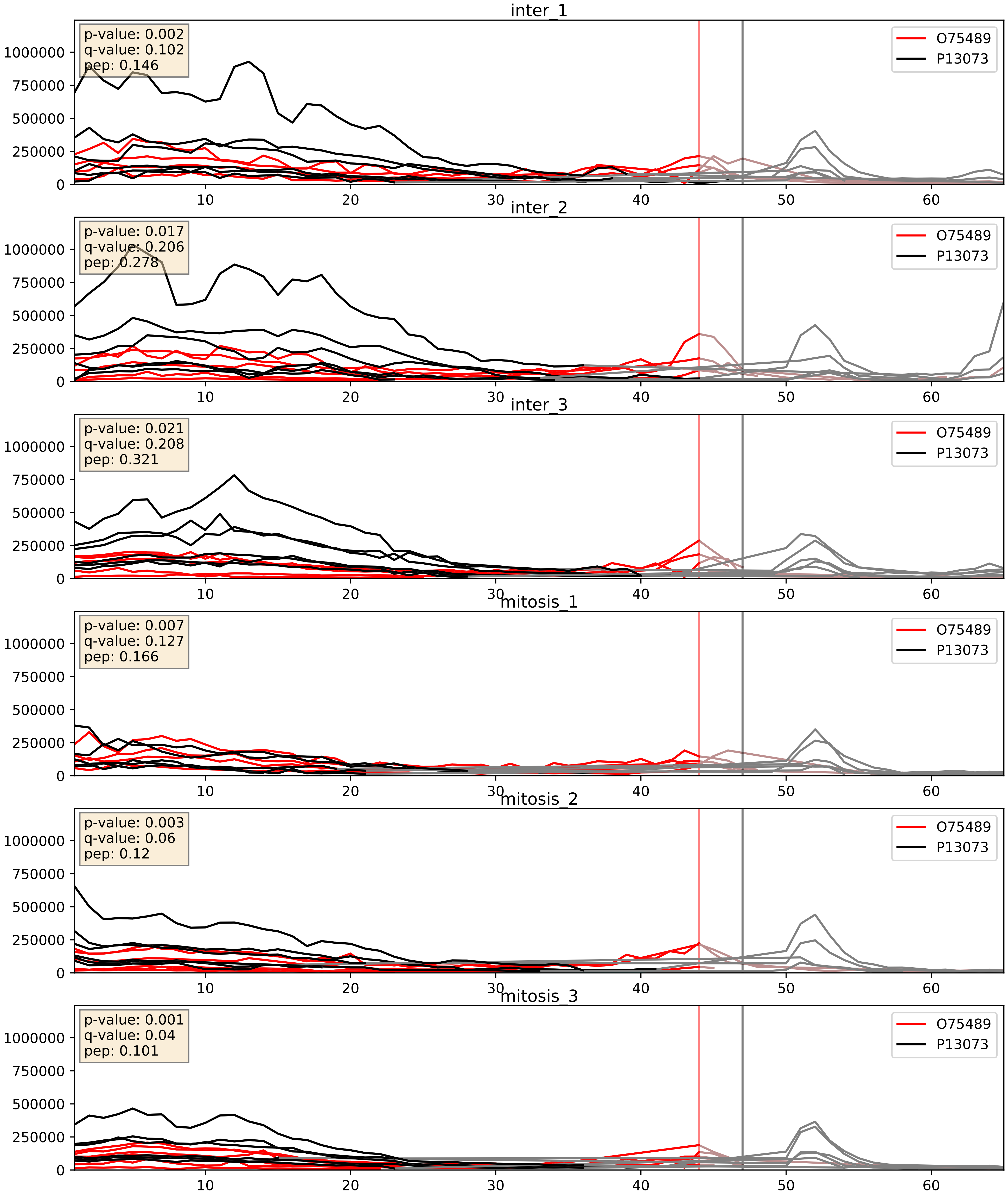


| level | condition_1 | condition_2 | pvalue | pvalue_adjusted |
|---|---|---|---|---|
| complex_abundance | mitosis | inter | 0.005711097468885759 | 0.056713450503088285 |
| interactor_abundance | mitosis | inter | 0.046783256688479116 | 0.17280029223446872 |
| interactor_ratio | mitosis | inter | 0.1298311132519205 | 0.5779454961622694 |

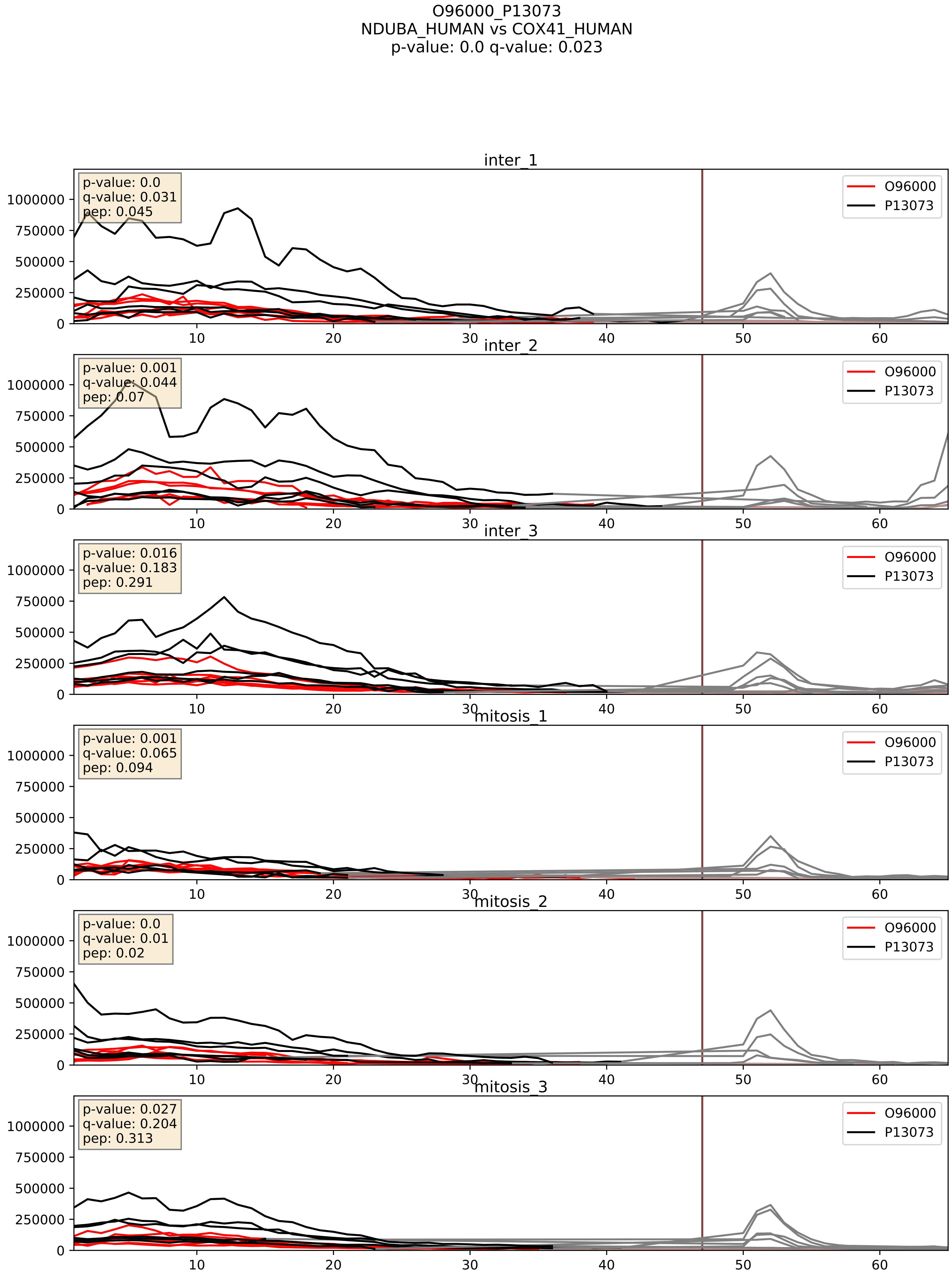


| level | condition_1 | condition_2 | pvalue | pvalue_adjusted |
|---|---|---|---|---|
| interactor_abundance | mitosis | inter | 0.0026153349281260135 | 0.038268832452578935 |
| complex_abundance | mitosis | inter | 0.00346532263349165 | 0.04302508960916016 |
| interactor_ratio | mitosis | inter | 0.25486933343791507 | 0.7264298204376385 |

P00403_P13073
COX2_HUMAN vs COX41_HUMAN
p-value: 0.0 q-value: 0.0

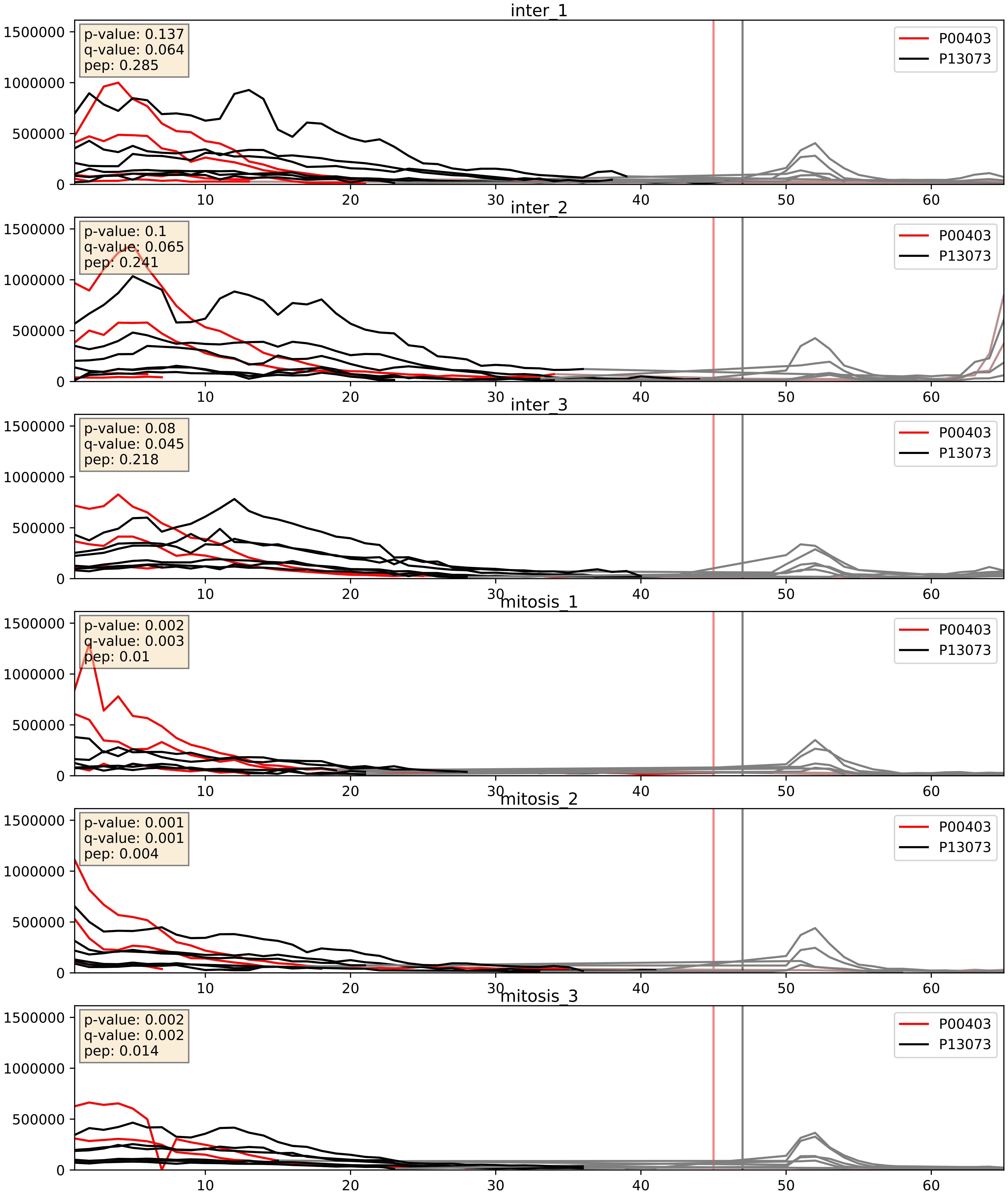


| level | condition_1 | condition_2 | pvalue | pvalue_adjusted |
|---|---|---|---|---|
| complex_abundance | mitosis | inter | 0.0003330910385573122 | 0.011886635542316795 |
| interactor_abundance | mitosis | inter | 0.04934755549120187 | 0.17741078328630325 |
| interactor_ratio | mitosis | inter | 0.37642293376682834 | 0.8148464407854535 |

P07919_P13073
QCR6_HUMAN vs COX41_HUMAN
p-value: 0.001 q-value: 0.046

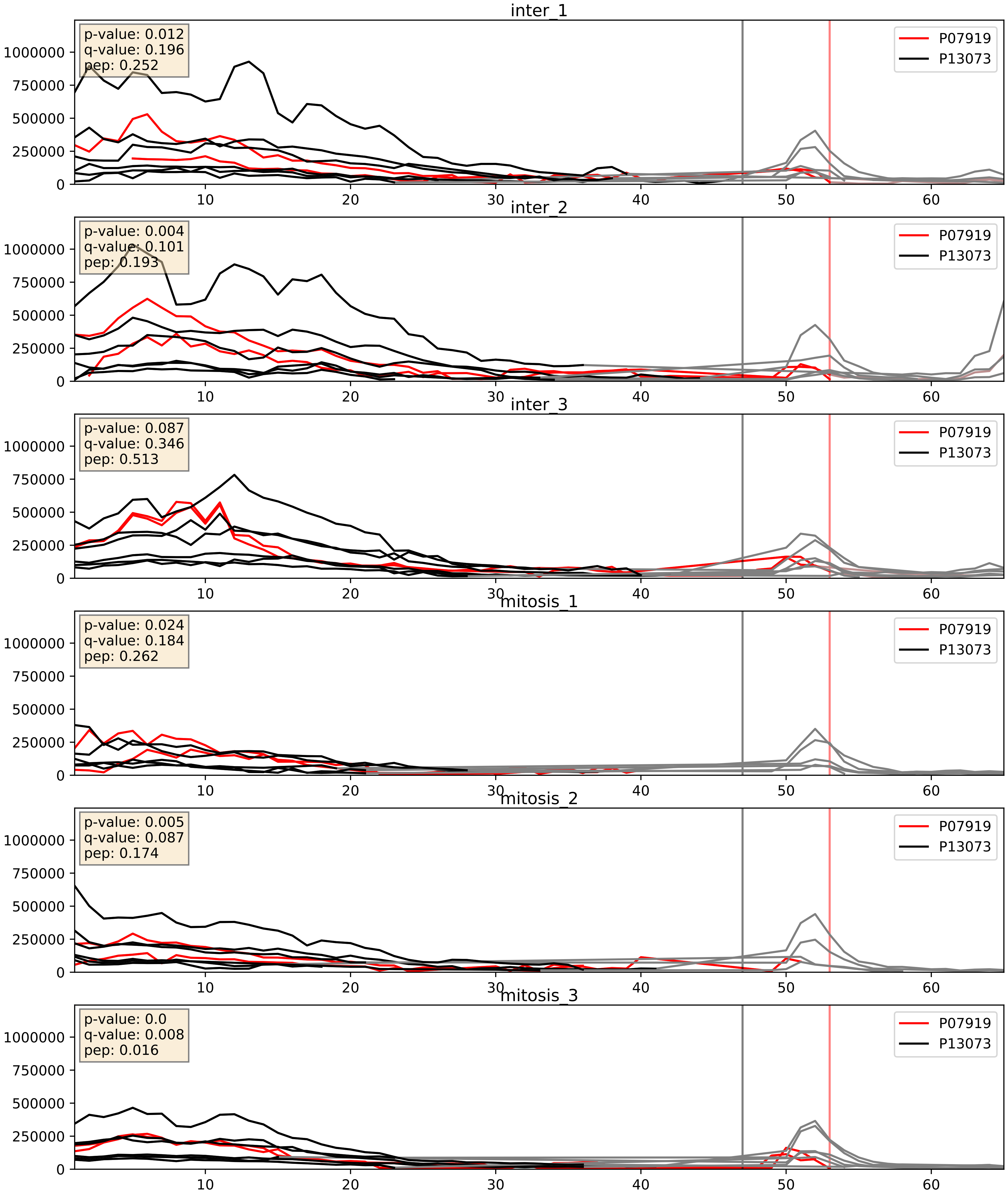


| level | condition_1 | condition_2 | pvalue | pvalue_adjusted |
|---|---|---|---|---|
| complex_abundance | mitosis | inter | 0.010248751113203294 | 0.07485435966639949 |
| interactor_abundance | mitosis | inter | 0.015834408859097590 | 0.11042162104592697 |
| interactor_ratio | mitosis | inter | 0.11281573811939599 | 0.553985317646331 |

P08574_P13073
CY1_HUMAN vs COX41_HUMAN
p-value: 0.001 q-value: 0.002

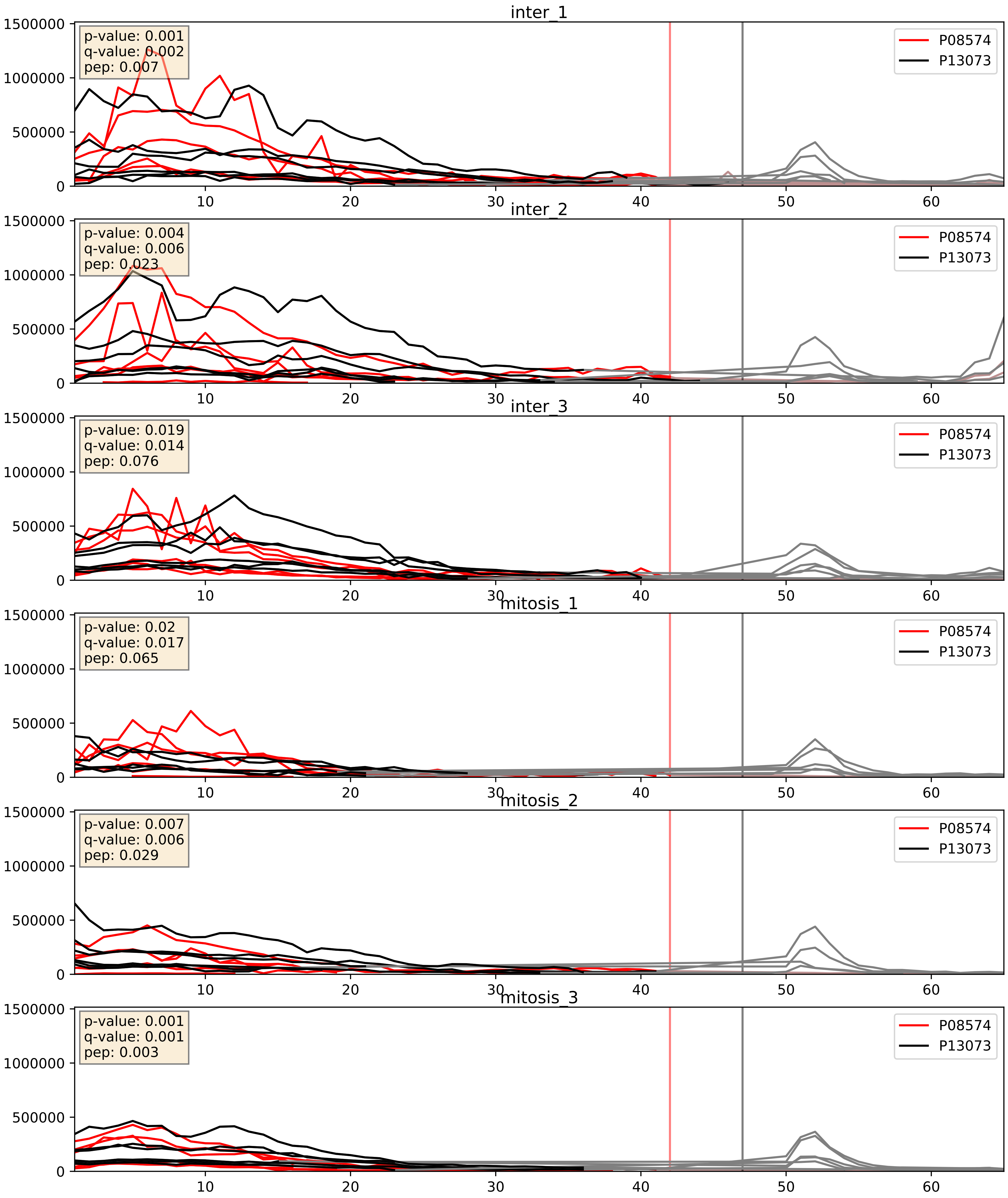


| level | condition_1 | condition_2 | pvalue | pvalue_adjusted |
|---|---|---|---|---|
| complex_abundance | mitosis | inter | 0.0024352323345376956 | 0.035755726901616935 |
| interactor_abundance | mitosis | inter | 0.006017826453889495 | 0.06595014057915963 |
| interactor_ratio | mitosis | inter | 0.4518645391568424 | 0.852724968073759 |

P10606_P13073
COX5B_HUMAN vs COX41_HUMAN
p-value: 0.004 q-value: 0.005

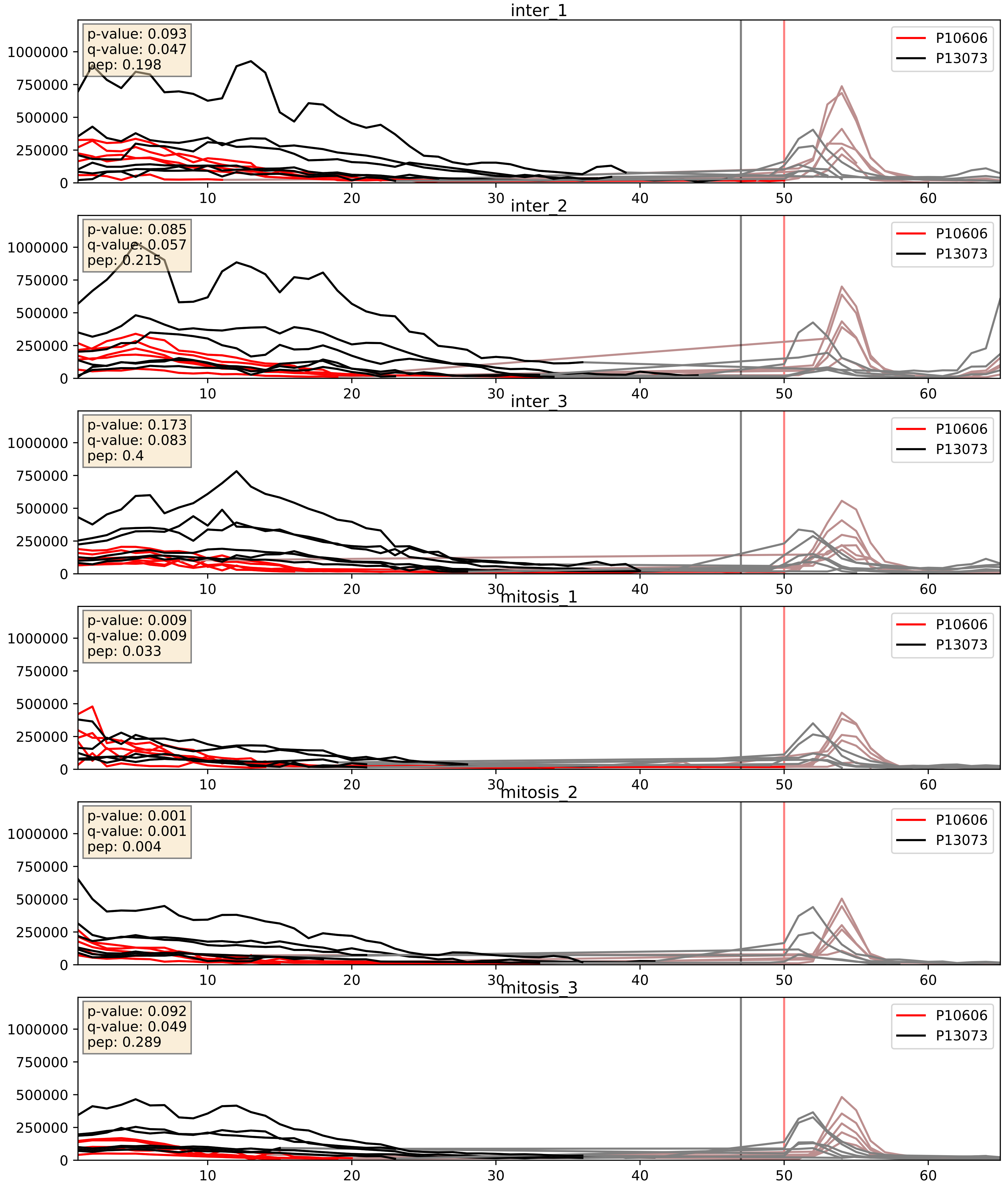


| level | condition_1 | condition_2 | pvalue | pvalue_adjusted |
|---|---|---|---|---|
| complex_abundance | mitosis | inter | 0.001172481502179381 | 0.02303709273772173 |
| interactor_abundance | mitosis | inter | 0.07779955689515294 | 0.23127772426550067 |
| interactor_ratio | mitosis | inter | 0.2788644033592743 | 0.7505446454705968 |

P13073_P18859
COX41_HUMAN vs ATP5J_HUMAN
p-value: 0.0 q-value: 0.041

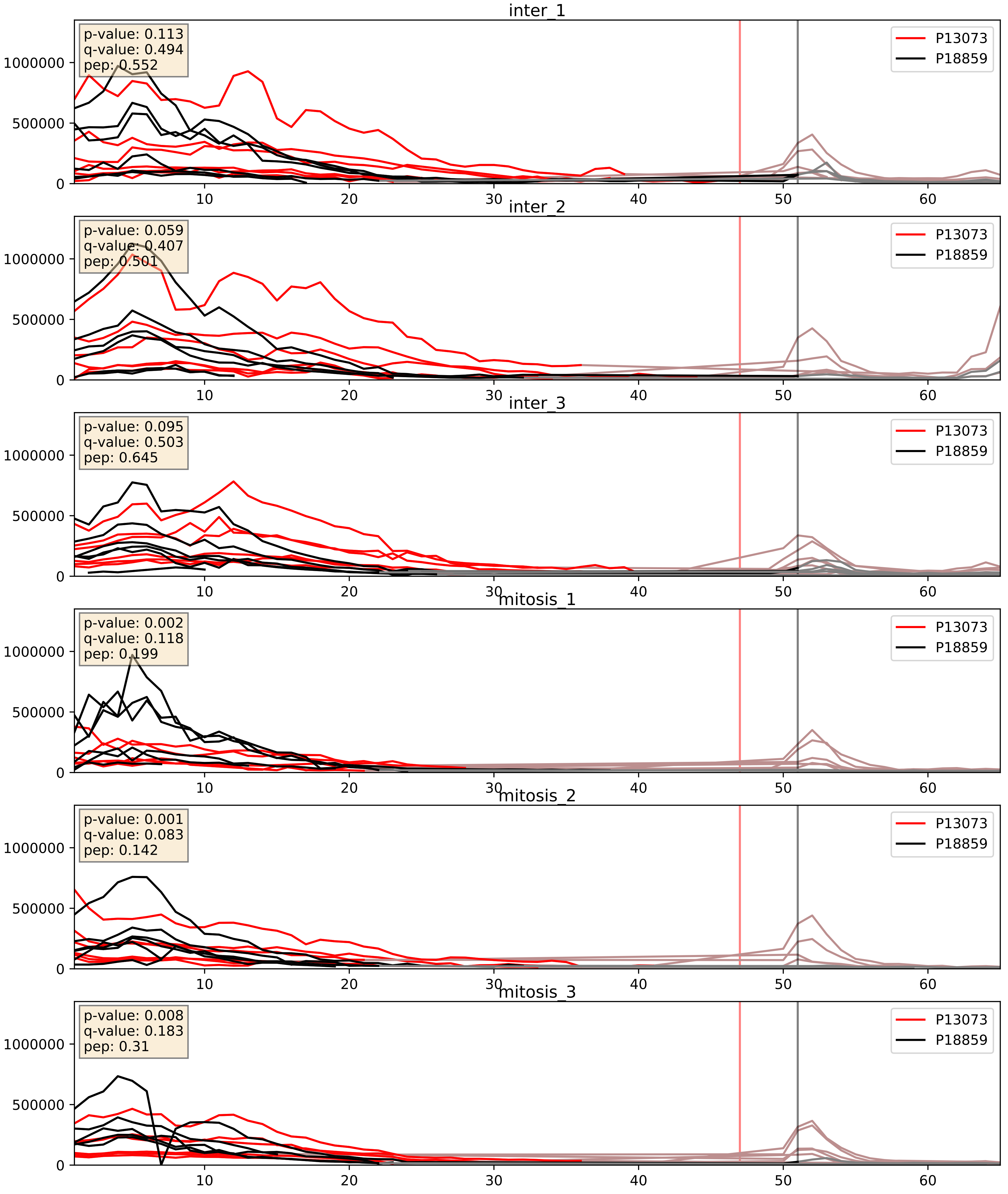


| level | condition_1 | condition_2 | pvalue | pvalue_adjusted |
|---|---|---|---|---|
| complex_abundance | mitosis | inter | 0.0025532154180830037 | 0.036547698961188146 |
| interactor_abundance | mitosis | inter | 0.02242886891826287 | 0.12329937058940194 |
| interactor_ratio | mitosis | inter | 0.19657300629598837 | 0.6709190326529746 |

P13073_P20674
COX41_HUMAN vs COX5A_HUMAN
p-value: 0.0 q-value: 0.0

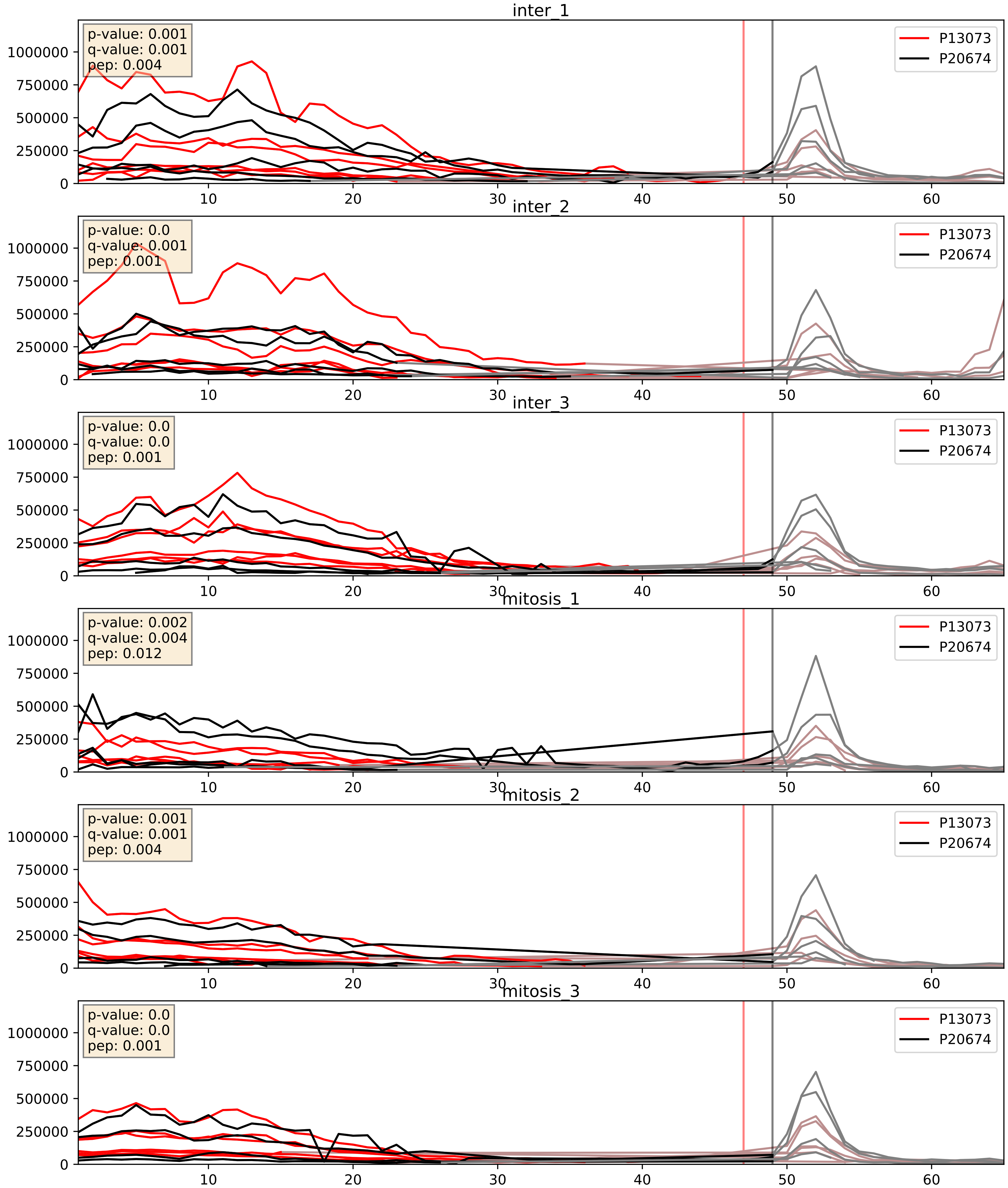


| level | condition_1 | condition_2 | pvalue | pvalue_adjusted |
|---|---|---|---|---|
| complex_abundance | mitosis | inter | 0.0025376433453244895 | 0.036409709098919725 |
| interactor_abundance | mitosis | inter | 0.024169942421986428 | 0.12811251670942622 |
| interactor_ratio | mitosis | inter | 0.24467737276607313 | 0.7204195520862359 |

P13073_P21912
COX41_HUMAN vs SDHB_HUMAN
p-value: 0.001 q-value: 0.048

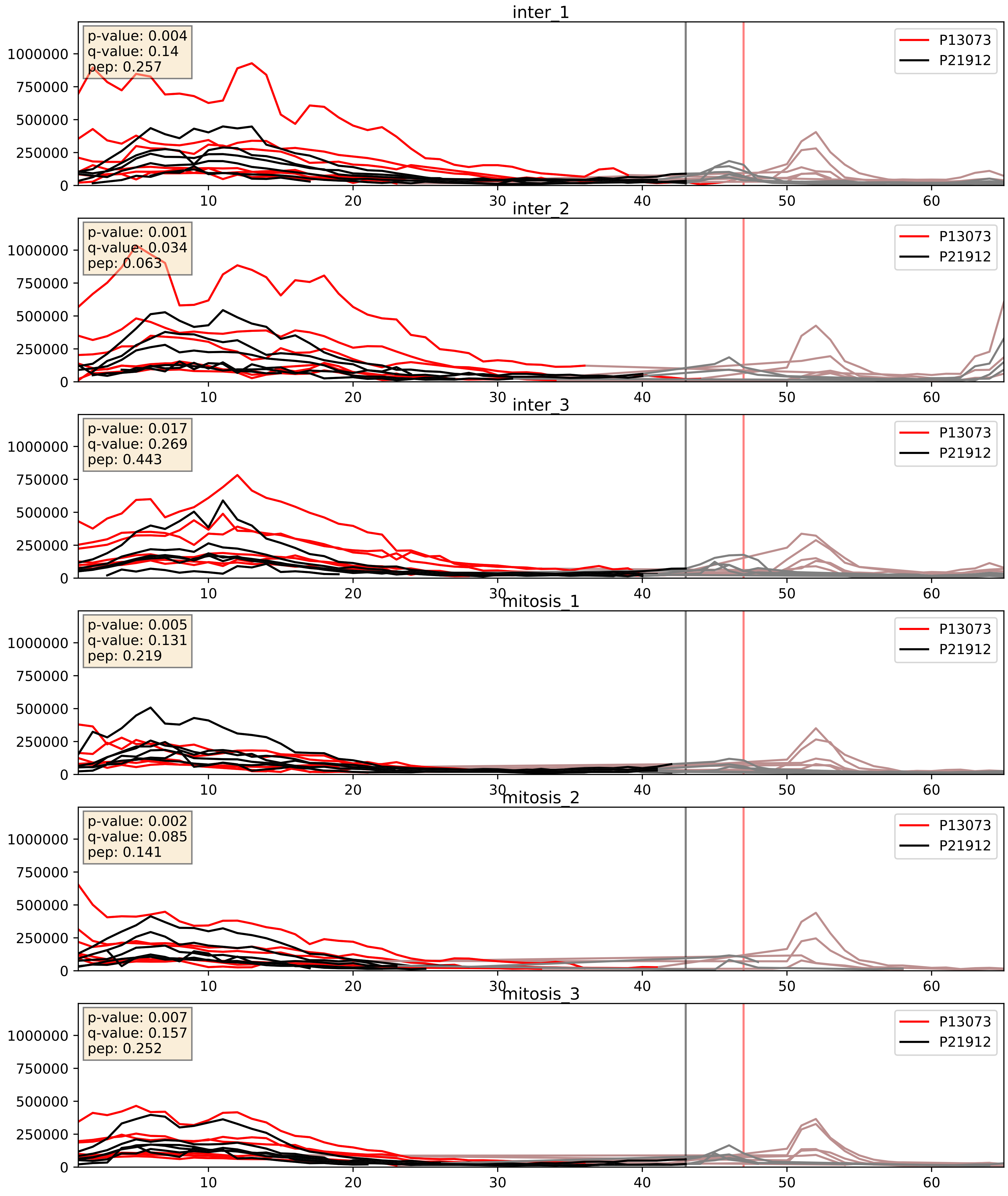


| level | condition_1 | condition_2 | pvalue | pvalue_adjusted |
|---|---|---|---|---|
| complex_abundance | mitosis | inter | 0.005389557979525 | 0.055184947732935405 |
| interactor_abundance | mitosis | inter | 0.024795500362278046 | 0.13029962525406855 |
| interactor_ratio | mitosis | inter | 0.1311008565123513 | 0.5791536040065746 |

P13073_P30049
COX41_HUMAN vs ATPD_HUMAN
p-value: 0.001 q-value: 0.015

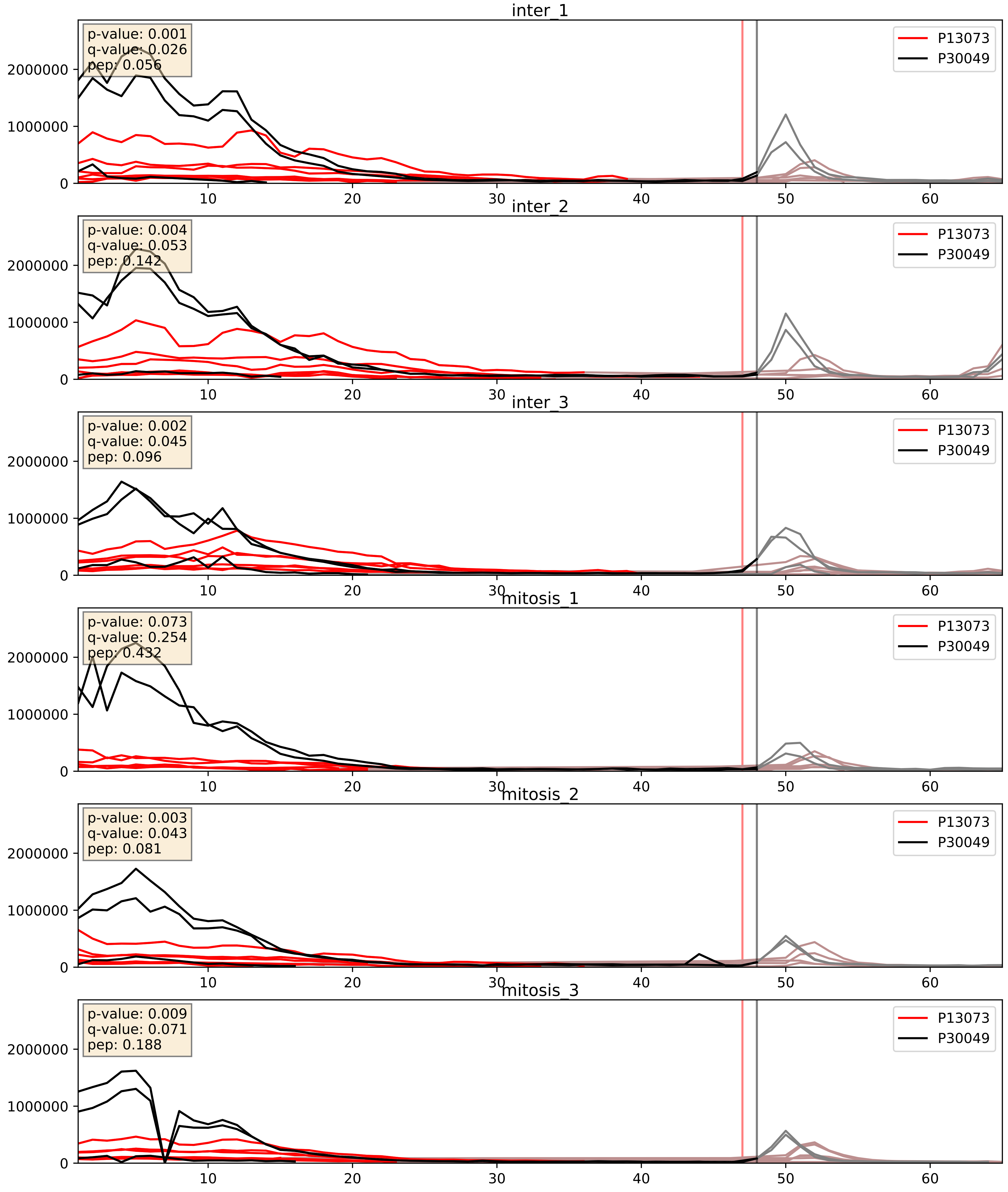


| level | condition_1 | condition_2 | pvalue | pvalue_adjusted |
|---|---|---|---|---|
| complex_abundance | mitosis | inter | 0.009389944287039965 | 0.07163807762661507 |
| interactor_abundance | mitosis | inter | 0.026369579357745133 | 0.13476035779241693 |
| interactor_ratio | mitosis | inter | 0.3361641776376856 | 0.7920475964368858 |
