## Supplementary Data 2 for "SECAT: Quantifying differential protein-protein interaction states by network-centric analysis": hela_string_000184_Q96GM5.pdf

# Q96GM5_Q96GM5

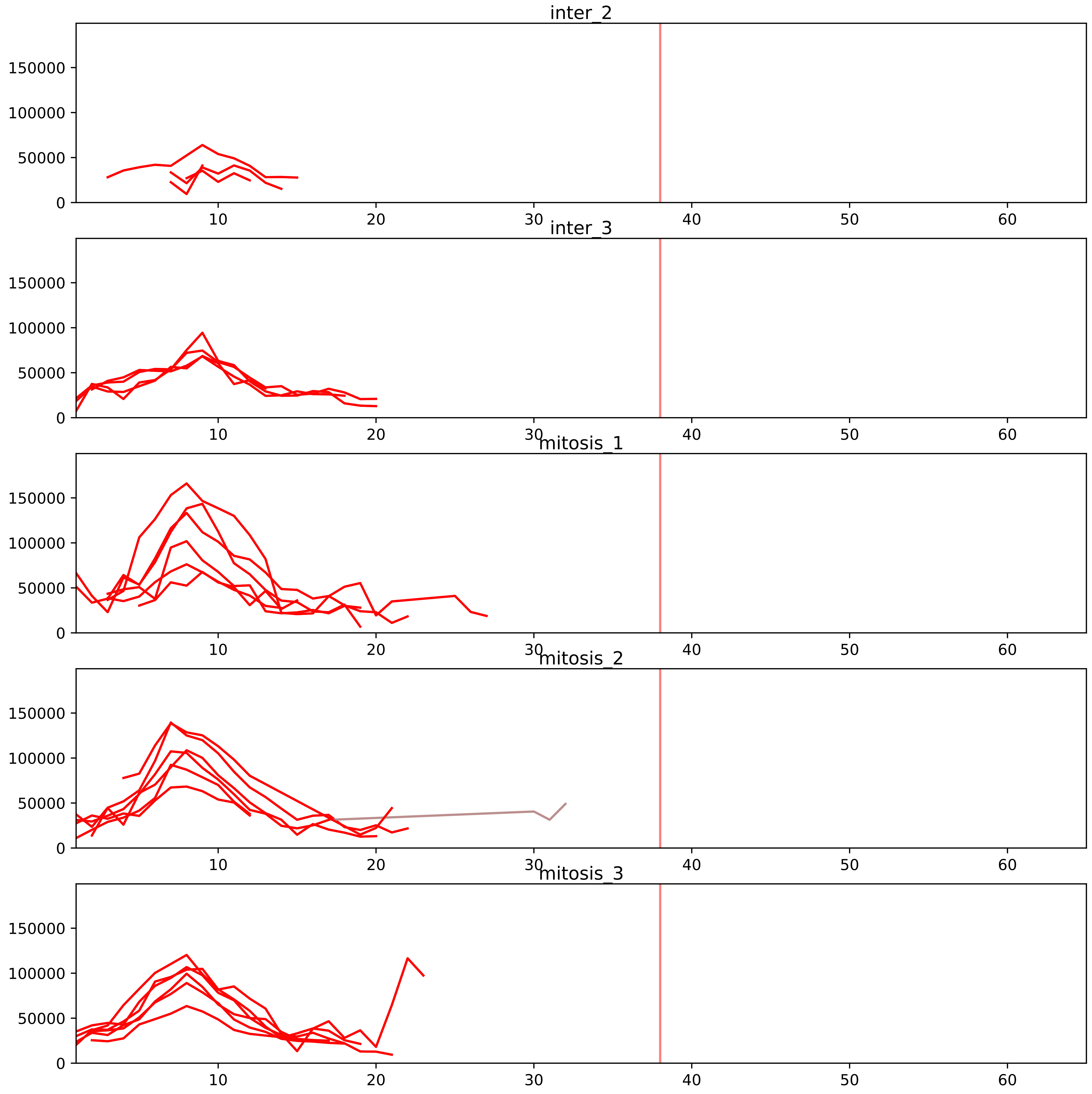


| level | condition_1 | condition_2 | pvalue | pvalue_adjusted |
|---|---|---|---|---|
| interactor_abundance | mitosis | inter | 0.00014424544131215236 | 0.002948436724638104 |
| interactor_ratio | mitosis | inter | 0.00030927694098063124 | 0.004215330158550825 |
| complex_abundance | mitosis | inter | 0.0009174272808764046 | 0.010356234336273524 |
| total_abundance | mitosis | inter | 0.011917136423098868 | 0.1355573212763089 |
| assembled_abundance | mitosis | inter | 0.01469805950415786 | 0.1387129365704898 |

O14497_Q96GM5
ARI1A_HUMAN vs SMRD1_HUMAN
p-value: 0.03 q-value: 0.022

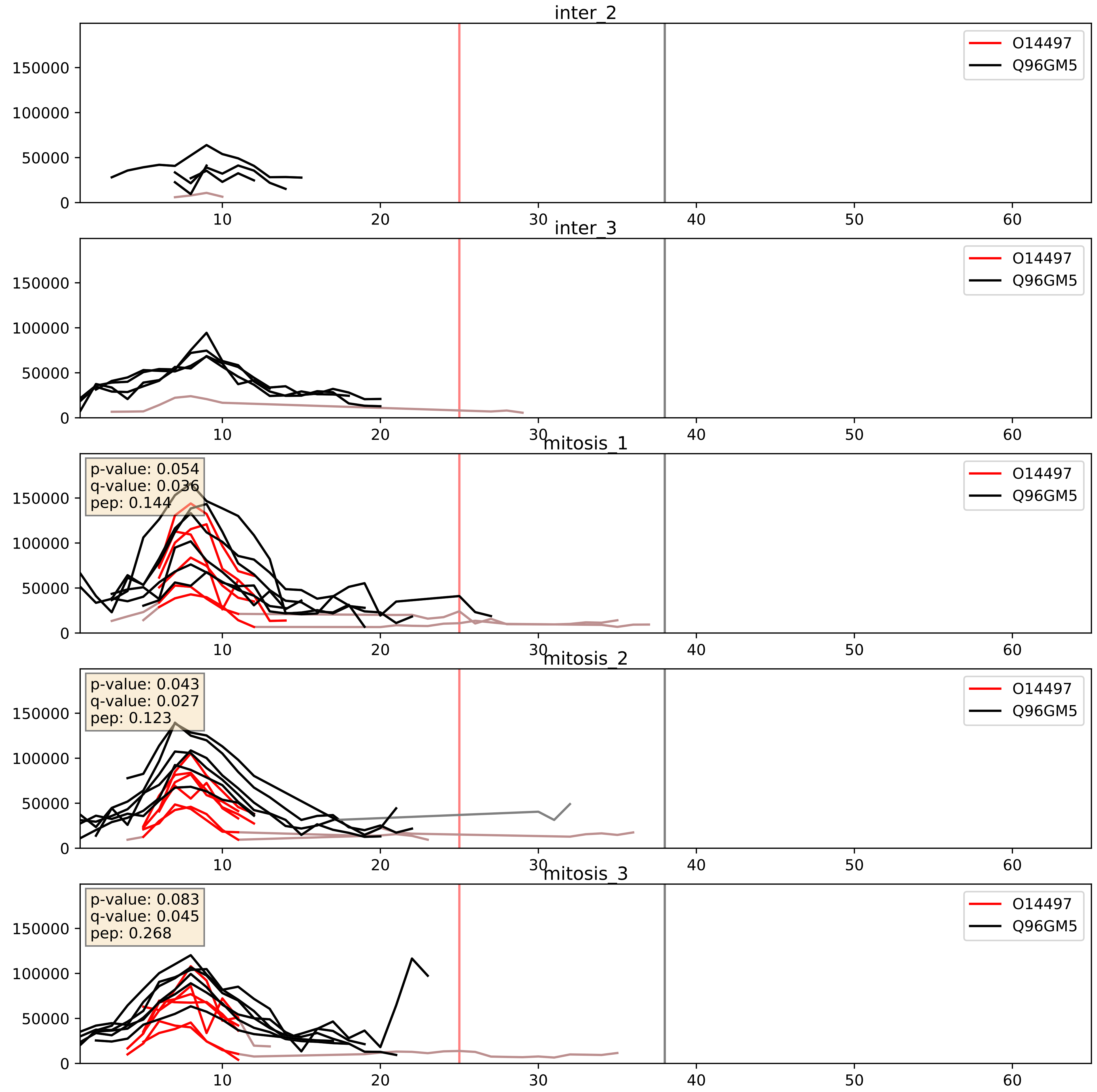


| level | condition_1 | condition_2 | pvalue | pvalue_adjusted |
|---|---|---|---|---|
| interactor_ratio | mitosis | inter | 0.0008196289026357029 | 0.03277774637652267 |
| complex_abundance | mitosis | inter | 0.00445543122107188 | 0.049788645279664145 |
| interactor_abundance | mitosis | inter | 0.012640483206095423 | 0.10027749130318384 |

O96019_Q96GM5
ACL6A_HUMAN vs SMRD1_HUMAN
p-value: 0.008 q-value: 0.007

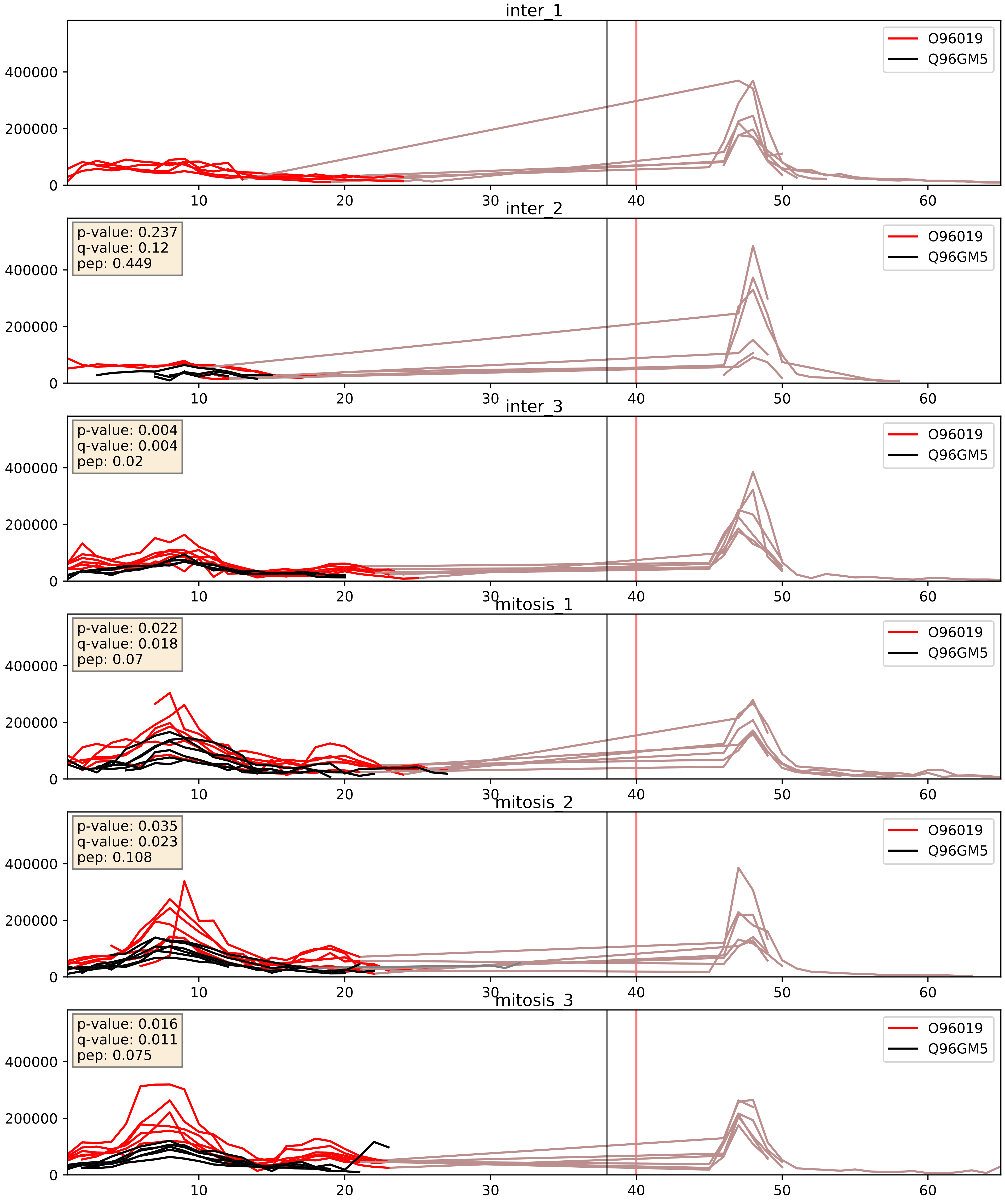


| level | condition_1 | condition_2 | pvalue | pvalue_adjusted |
|---|---|---|---|---|
| complex_abundance | mitosis | inter | 0.04039237492280713 | 0.14519139912861373 |
| interactor_abundance | mitosis | inter | 0.0854598415810031 | 0.24622559539999544 |
| interactor_ratio | mitosis | inter | 0.7353563266844123 | 0.949002778270057 |

P51531_Q96GM5
SMCA2_HUMAN vs SMRD1_HUMAN
p-value: 0.004 q-value: 0.005

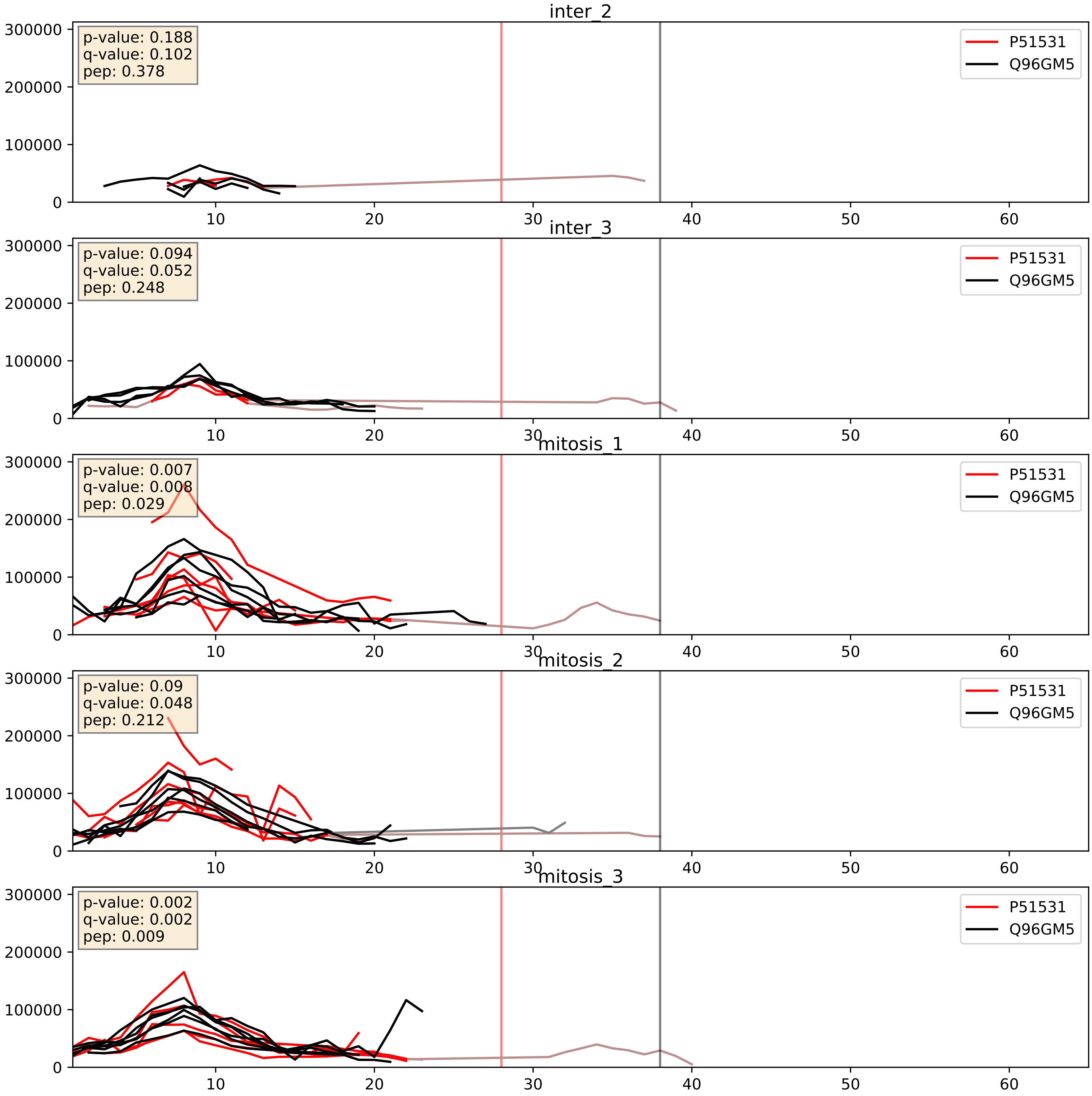


| level | condition_1 | condition_2 | pvalue | pvalue_adjusted |
|---|---|---|---|---|
| complex_abundance | mitosis | inter | 0.003988838446919328 | 0.04687907436752479 |
| interactor_abundance | mitosis | inter | 0.013657409124963755 | 0.10336642096347458 |
| interactor_ratio | mitosis | inter | 0.8054421902429661 | 0.9636037943367981 |

P51532_Q96GM5
SMCA4_HUMAN vs SMRD1_HUMAN
p-value: 0.026 q-value: 0.019

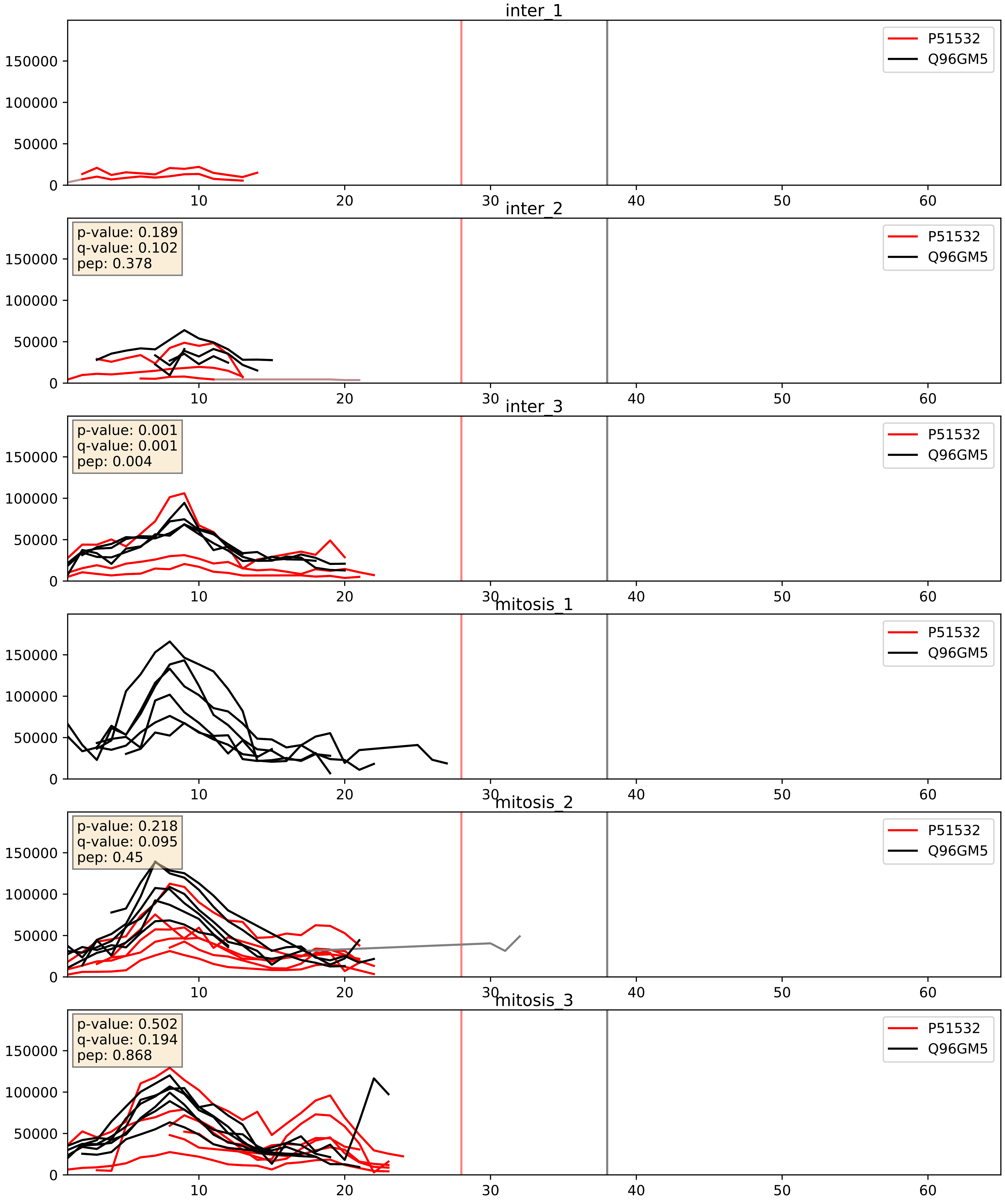


| level | condition_1 | condition_2 | pvalue | pvalue_adjusted |
|---|---|---|---|---|
| interactor_ratio | mitosis | inter | 0.25236540118785417 | 0.7251677296572153 |
| complex_abundance | mitosis | inter | 0.5989866703130391 | 0.722158577166143 |
| interactor_abundance | mitosis | inter | 0.6722259627311563 | 0.7854028855495391 |

Q8TAQ2_Q96GM5
SMRC2_HUMAN vs SMRD1_HUMAN
p-value: 0.004 q-value: 0.004

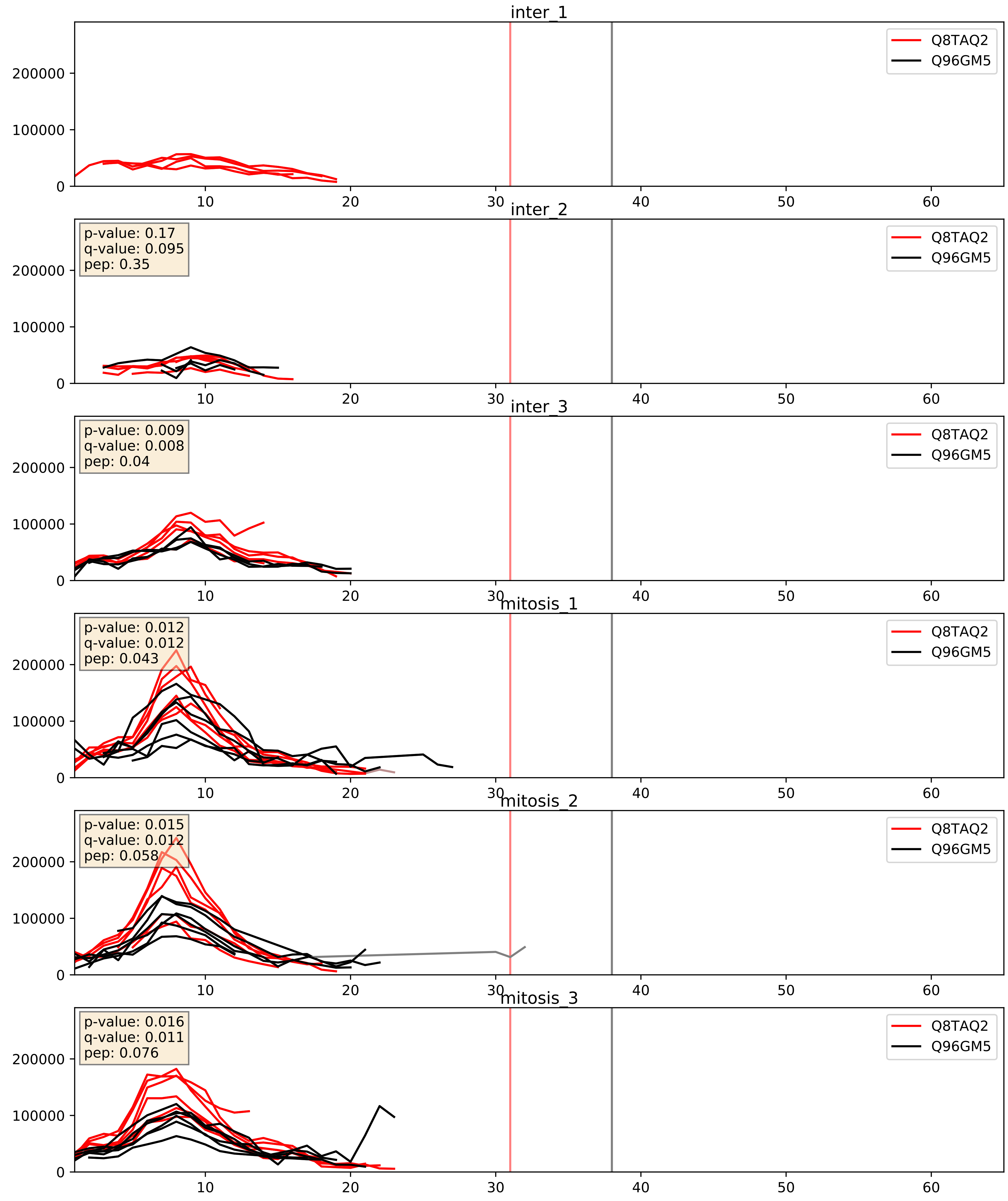


| level | condition_1 | condition_2 | pvalue | pvalue_adjusted |
|---|---|---|---|---|
| complex_abundance | mitosis | inter | 0.039208908036752275 | 0.14448905839702758 |
| interactor_abundance | mitosis | inter | 0.09491896399300535 | 0.2639407374222237 |
| interactor_ratio | mitosis | inter | 0.5537977522752682 | 0.8893358217967551 |

Q92922_Q96GM5
SMRC1_HUMAN vs SMRD1_HUMAN
p-value: 0.02 q-value: 0.016

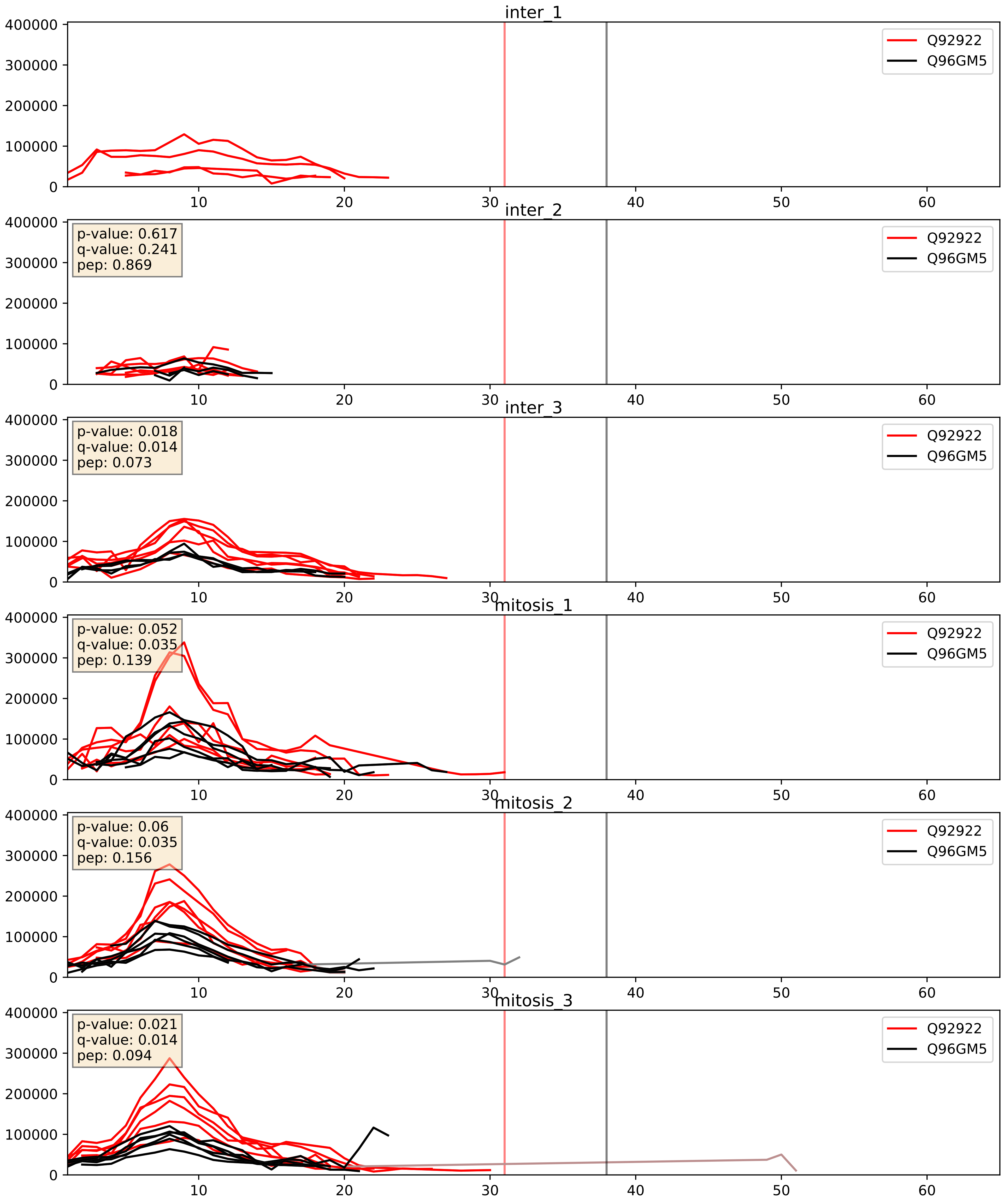


| level | condition_1 | condition_2 | pvalue | pvalue_adjusted |
|---|---|---|---|---|
| complex_abundance | mitosis | inter | 0.04777855643174045 | 0.15599228513863675 |
| interactor_abundance | mitosis | inter | 0.10152273968561054 | 0.27370850762629345 |
| interactor_ratio | mitosis | inter | 0.8269612471471082 | 0.969784012122695 |

Q92925_Q96GM5
SMRD2_HUMAN vs SMRD1_HUMAN
p-value: 0.021 q-value: 0.017

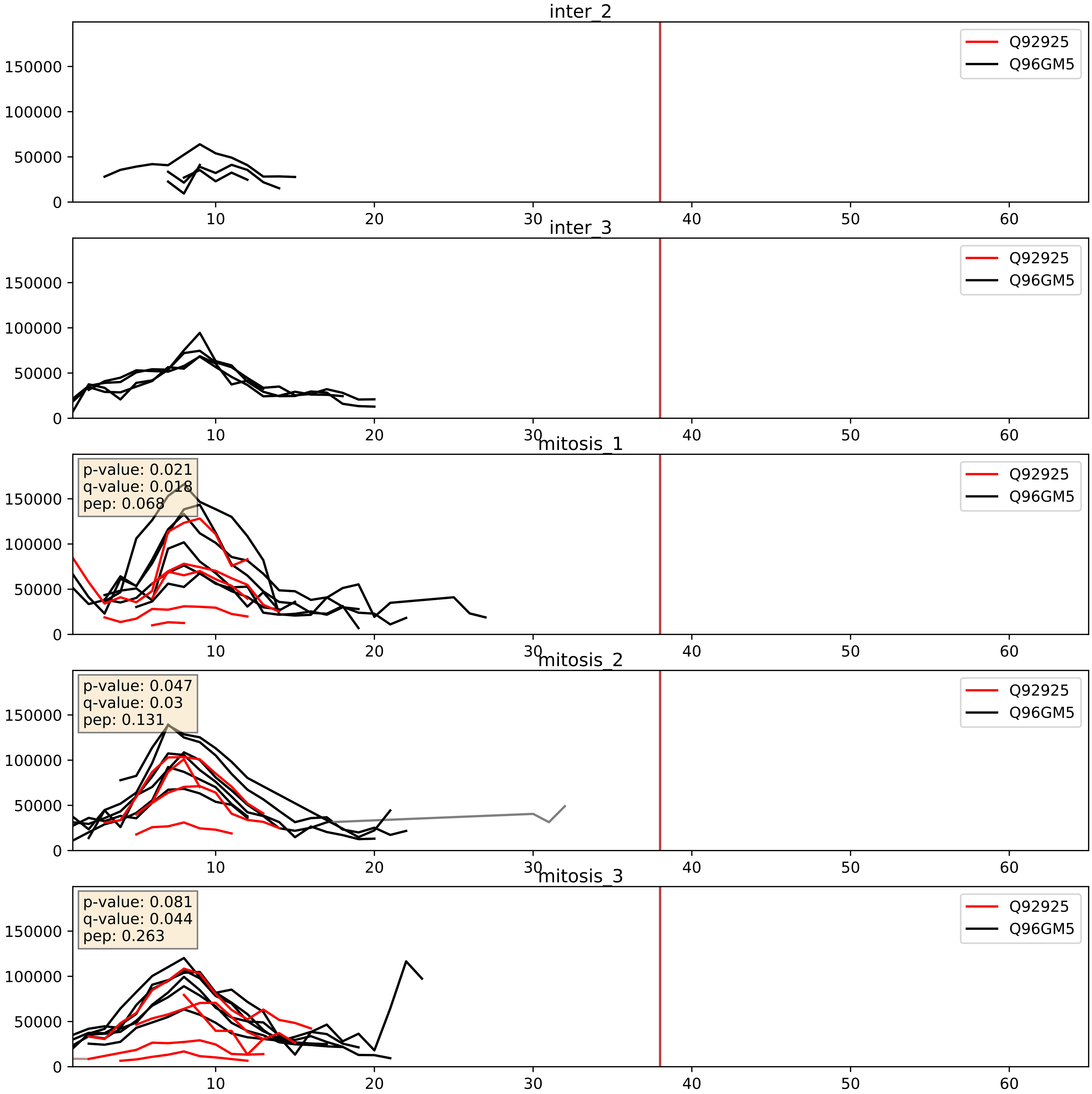


| level | condition_1 | condition_2 | pvalue | pvalue_adjusted |
|---|---|---|---|---|
| interactor_ratio | mitosis | inter | 5.661017680550811e-06 | 0.002835004461043519 |
| complex_abundance | mitosis | inter | 0.0019049156108668468 | 0.031118468757672154 |
| interactor_abundance | mitosis | inter | 0.004123135752406863 | 0.050966125690401085 |

Q969G3_Q96GM5
SMCE1_HUMAN vs SMRD1_HUMAN
p-value: 0.0 q-value: 0.0

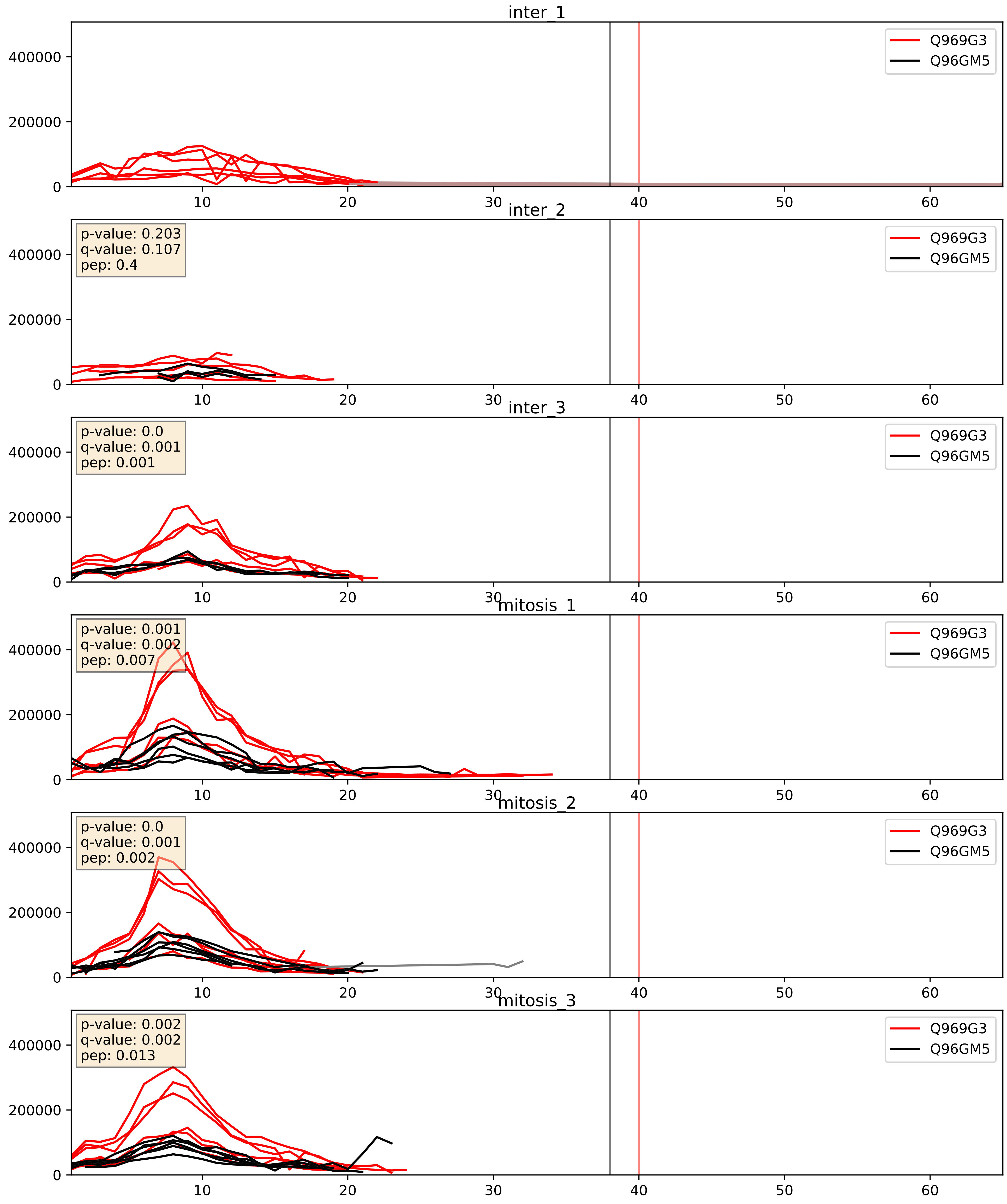


| level | condition_1 | condition_2 | pvalue | pvalue_adjusted |
|---|---|---|---|---|
| complex_abundance | mitosis | inter | 0.05012708341796544 | 0.1593936976440506 |
| interactor_abundance | mitosis | inter | 0.10625982548010601 | 0.27884246048734135 |
| interactor_ratio | mitosis | inter | 0.6739310697288592 | 0.9319765113016878 |
