## Supplementary Data 2 for "SECAT: Quantifying differential protein-protein interaction states by network-centric analysis": hela_string_000185_P51531.pdf

P51531_P51531

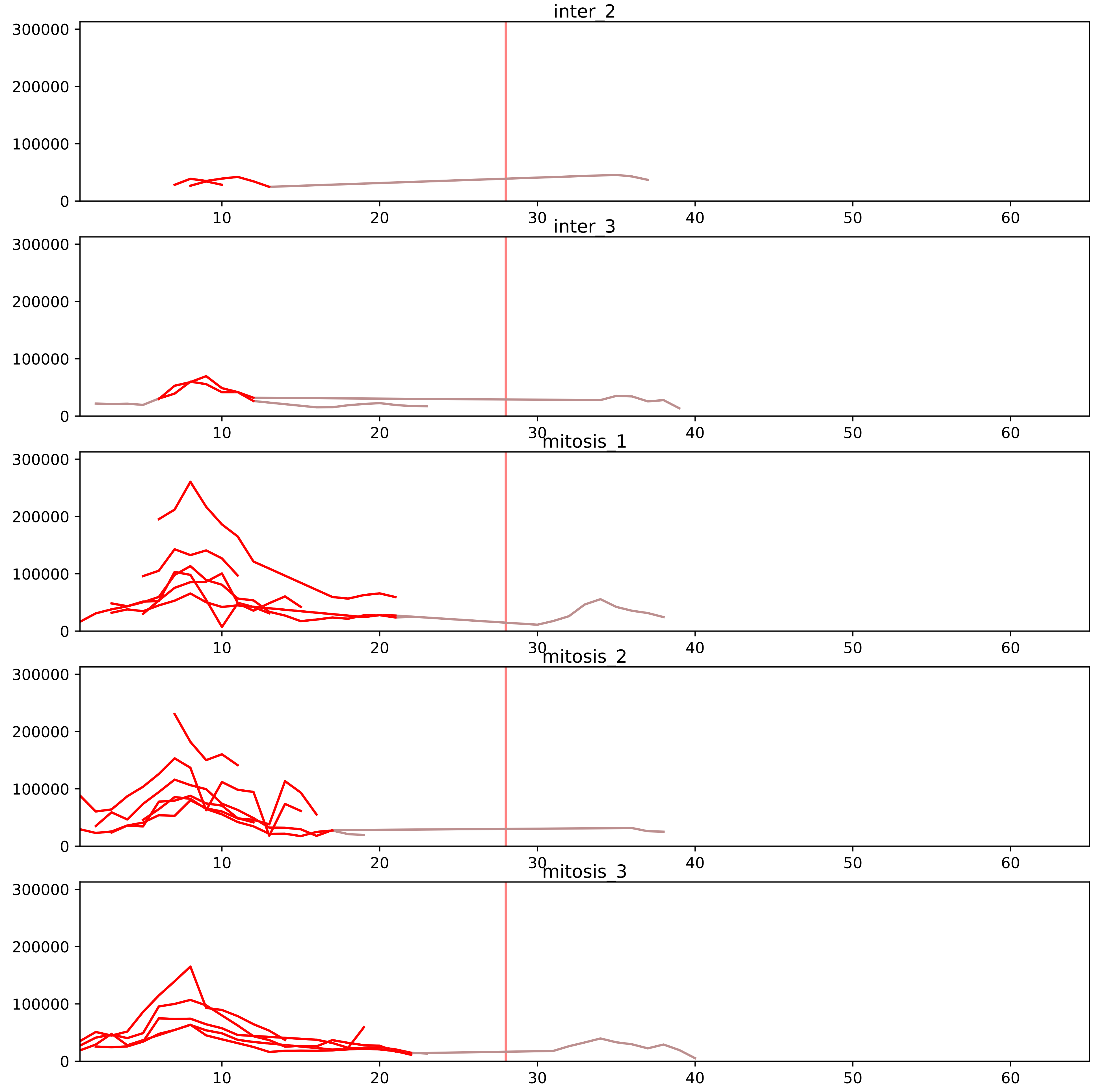


| level | condition_1 | condition_2 | pvalue | pvalue_adjusted |
|---|---|---|---|---|
| complex_abundance | mitosis | inter | 0.0001529783003852885 | 0.00293208409071803 |
| interactor_abundance | mitosis | inter | 0.0019276871190710343 | 0.018857244385255763 |
| assembled_abundance | mitosis | inter | 0.048220814615459094 | 0.26246701536623723 |
| total_abundance | mitosis | inter | 0.05407568758649025 | 0.29216412658567237 |
| monomer_abundance | mitosis | inter | 0.30705544151825115 | 0.6718670942432774 |
| interactor_ratio | mitosis | inter | 0.7948249214245131 | 0.943950409050364 |

O14497_P51531
ARI1A_HUMAN vs SMCA2_HUMAN
p-value: 0.037 q-value: 0.026

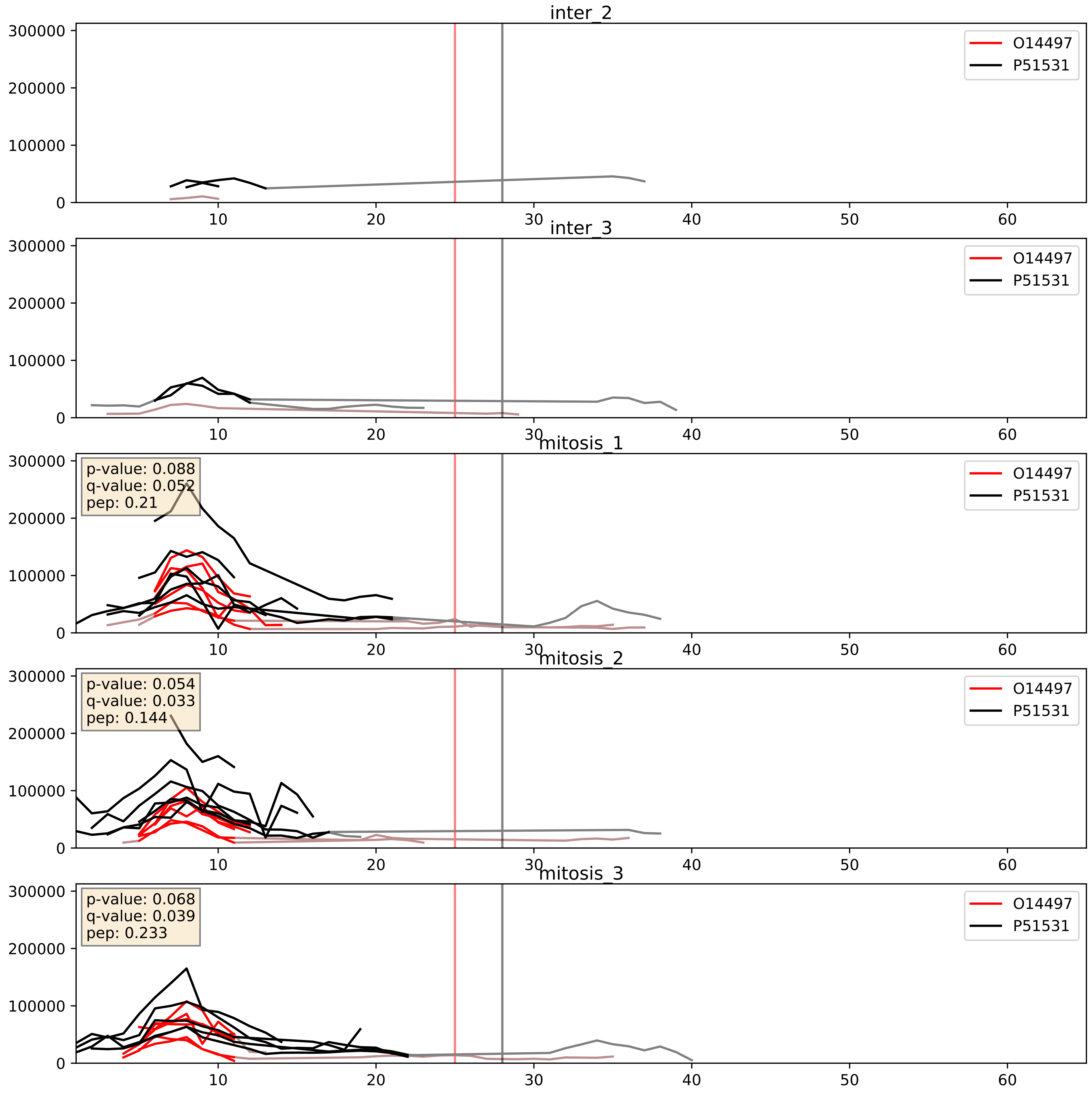


| level | condition_1 | condition_2 | pvalue | pvalue_adjusted |
|---|---|---|---|---|
| complex_abundance | mitosis | inter | 0.01207594557861336 | 0.08173689978386886 |
| interactor_abundance | mitosis | inter | 0.012640483206095423 | 0.10027749130318384 |
| interactor_ratio | mitosis | inter | 0.3232480159403212 | 0.7829618581024819 |

O96019_P51531
ACL6A_HUMAN vs SMCA2_HUMAN
p-value: 0.028 q-value: 0.02

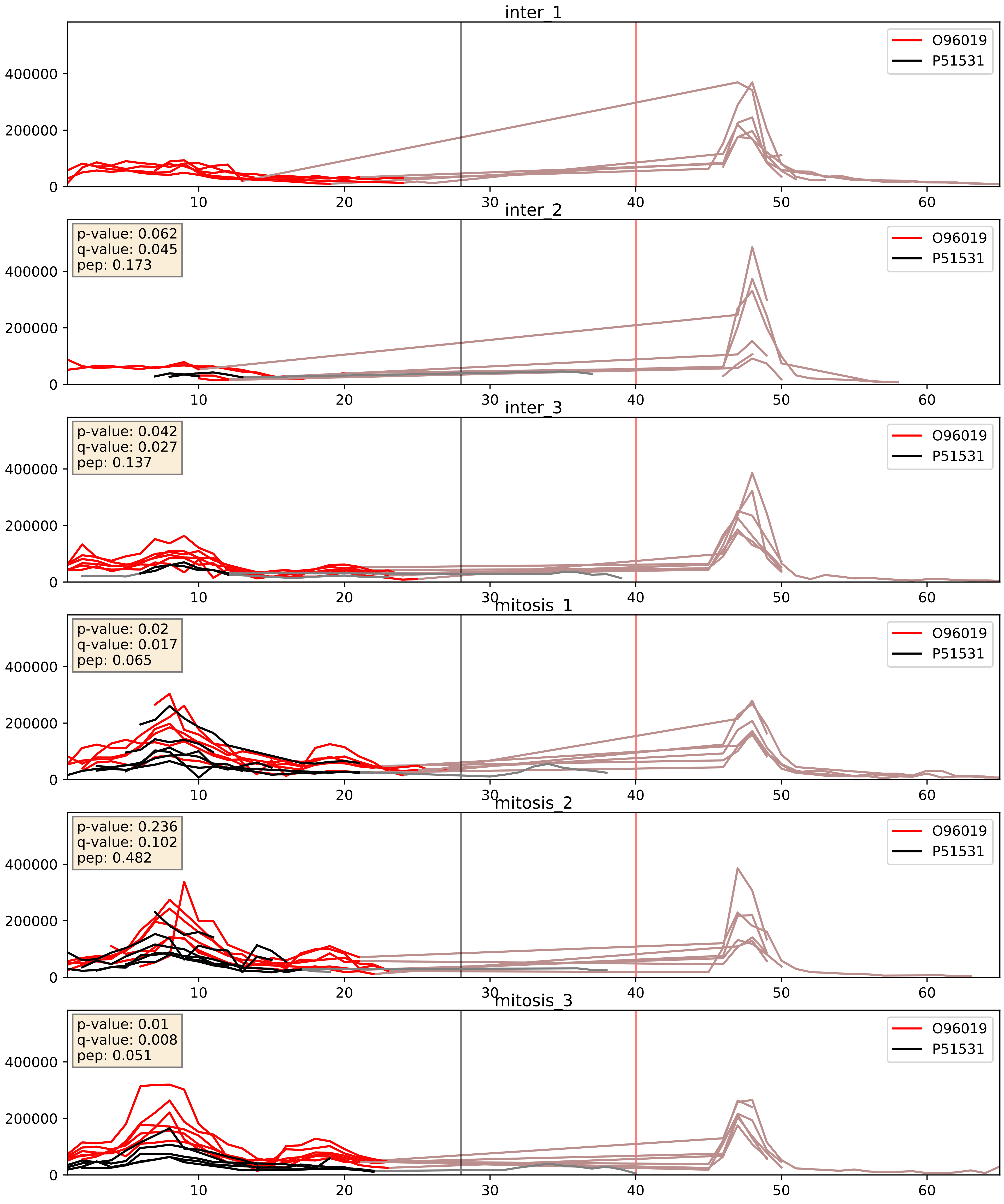


| level | condition_1 | condition_2 | pvalue | pvalue_adjusted |
|---|---|---|---|---|
| complex_abundance | mitosis | inter | 0.0050591450517678506 | 0.05313654189341448 |
| interactor_abundance | mitosis | inter | 0.02496784403535034 | 0.13071849843584032 |
| interactor_ratio | mitosis | inter | 0.6139555031434527 | 0.90762799337104 |

P51531_Q8TAQ2
SMCA2_HUMAN vs SMRC2_HUMAN
p-value: 0.001 q-value: 0.001

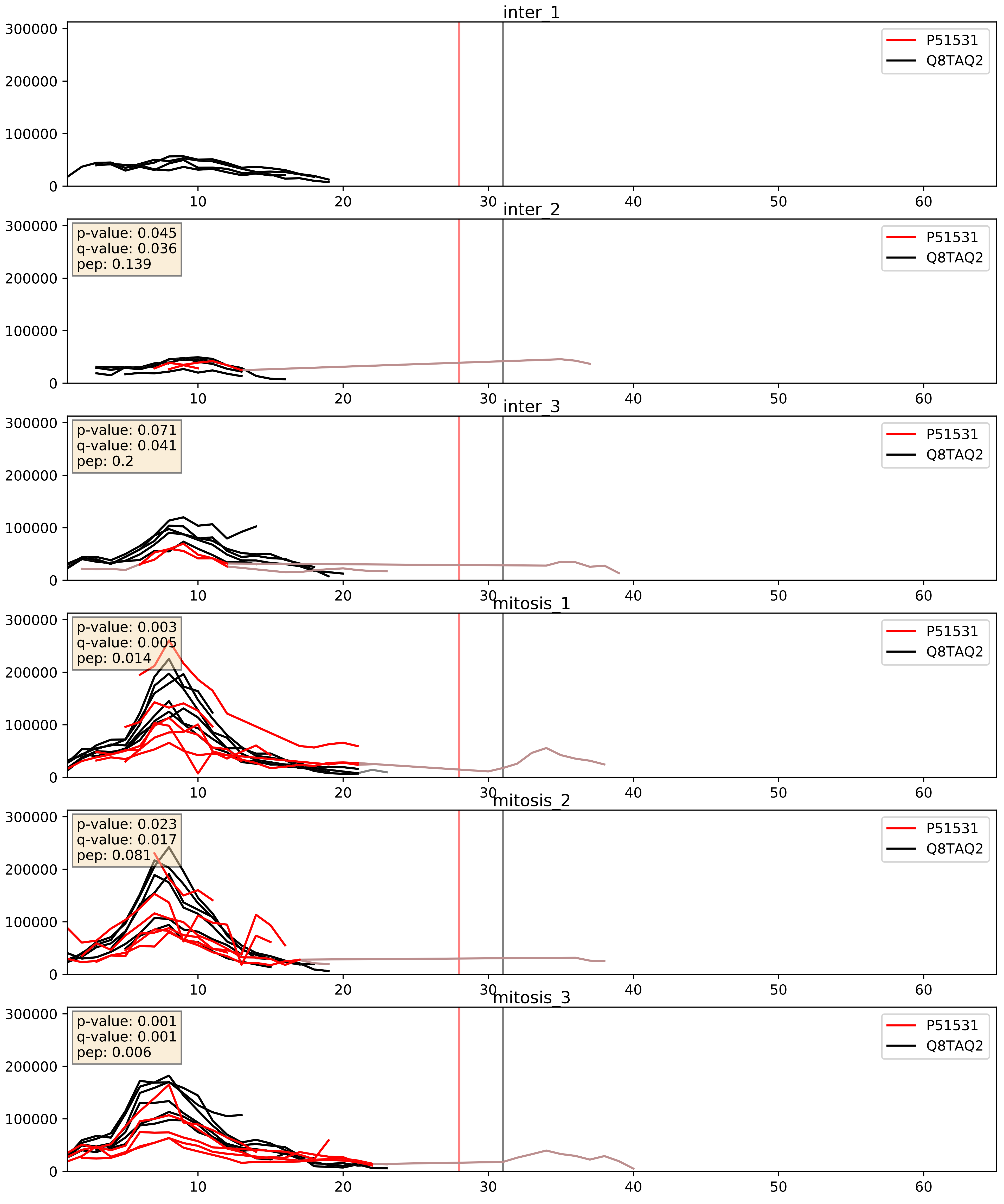


| level | condition_1 | condition_2 | pvalue | pvalue_adjusted |
|---|---|---|---|---|
| complex_abundance | mitosis | inter | 0.004971024511375018 | 0.05279244406481302 |
| interactor_abundance | mitosis | inter | 0.013657409124963755 | 0.10336642096347458 |
| interactor_ratio | mitosis | inter | 0.9846560283341652 | 0.9976654700477962 |

P51531_Q92922
SMCA2_HUMAN vs SMRC1_HUMAN
p-value: 0.024 q-value: 0.019

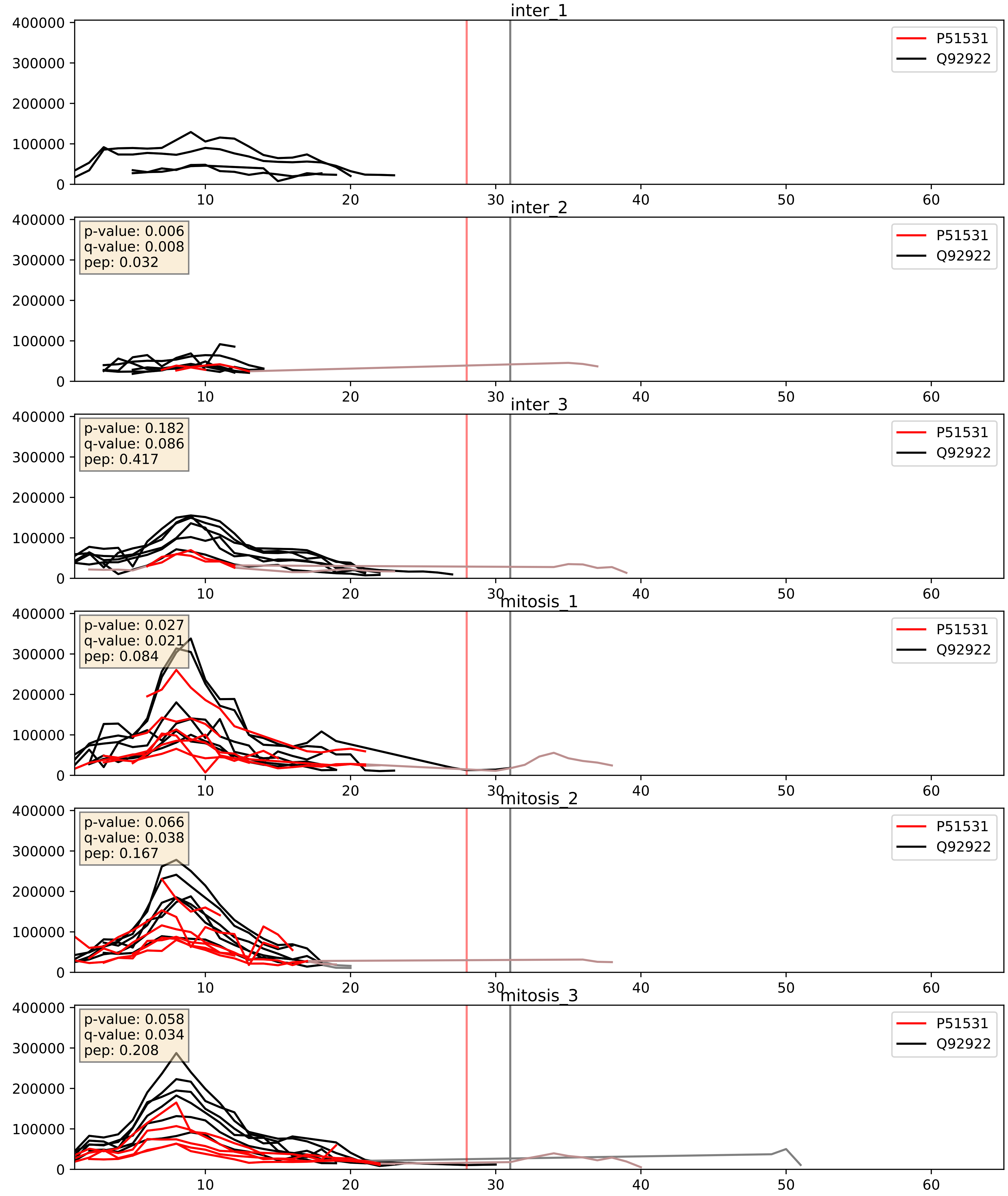


| level | condition_1 | condition_2 | pvalue | pvalue_adjusted |
|---|---|---|---|---|
| complex_abundance | mitosis | inter | 0.005693057559631294 | 0.056713450503088285 |
| interactor_abundance | mitosis | inter | 0.013657409124963755 | 0.10336642096347458 |
| interactor_ratio | mitosis | inter | 0.5531009131904417 | 0.8889886679627823 |

P51531_Q92925
SMCA2_HUMAN vs SMRD2_HUMAN
p-value: 0.085 q-value: 0.05

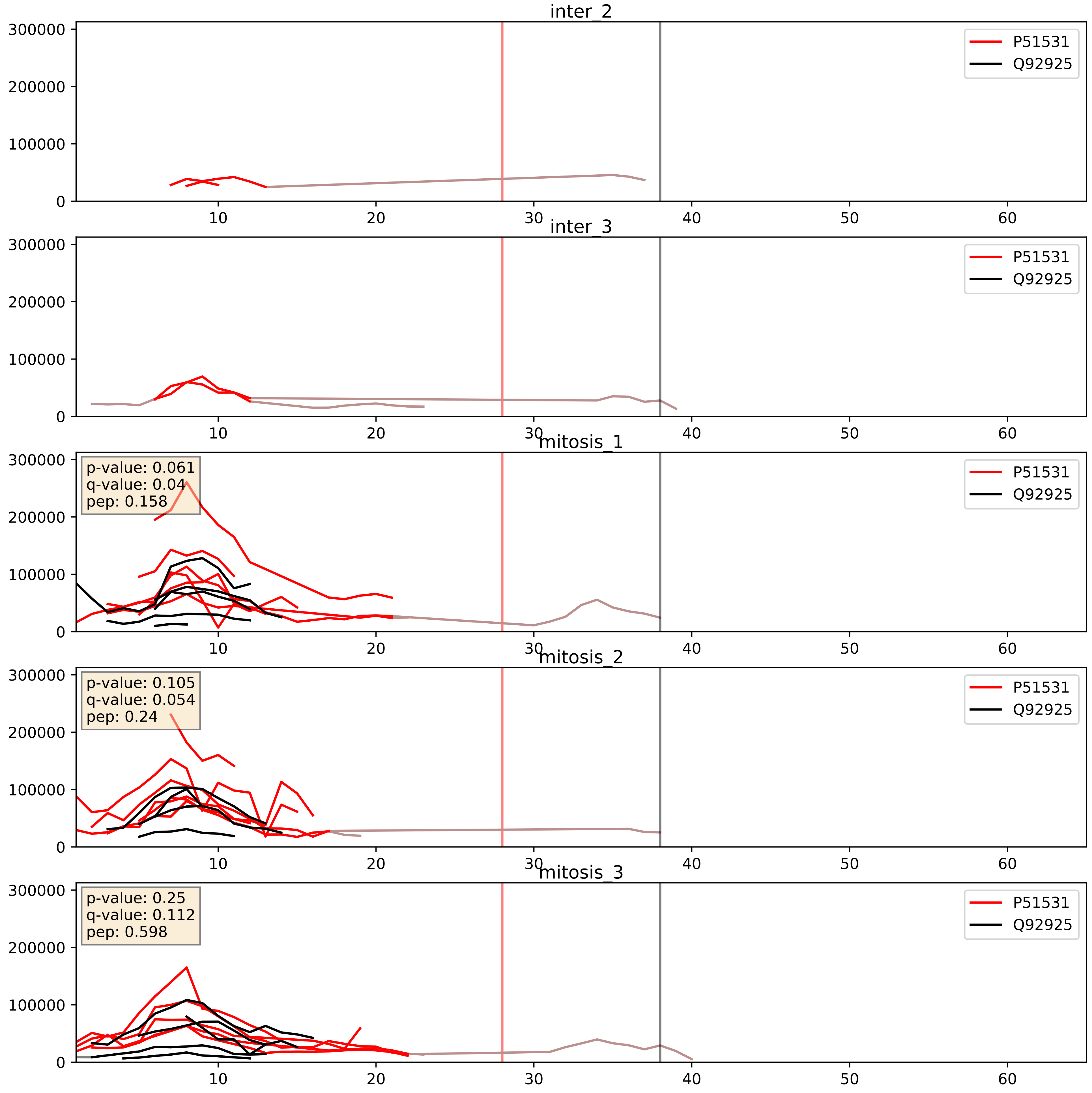


| level | condition_1 | condition_2 | pvalue | pvalue_adjusted |
|---|---|---|---|---|
| complex_abundance | mitosis | inter | 0.004621740992278264 | 0.050629452773887605 |
| interactor_abundance | mitosis | inter | 0.01019836828153345 | 0.0879625269700301 |
| interactor_ratio | mitosis | inter | 0.28817519340286885 | 0.7581846425018174 |

P51531_Q969G3
SMCA2_HUMAN vs SMCE1_HUMAN
p-value: 0.047 q-value: 0.031

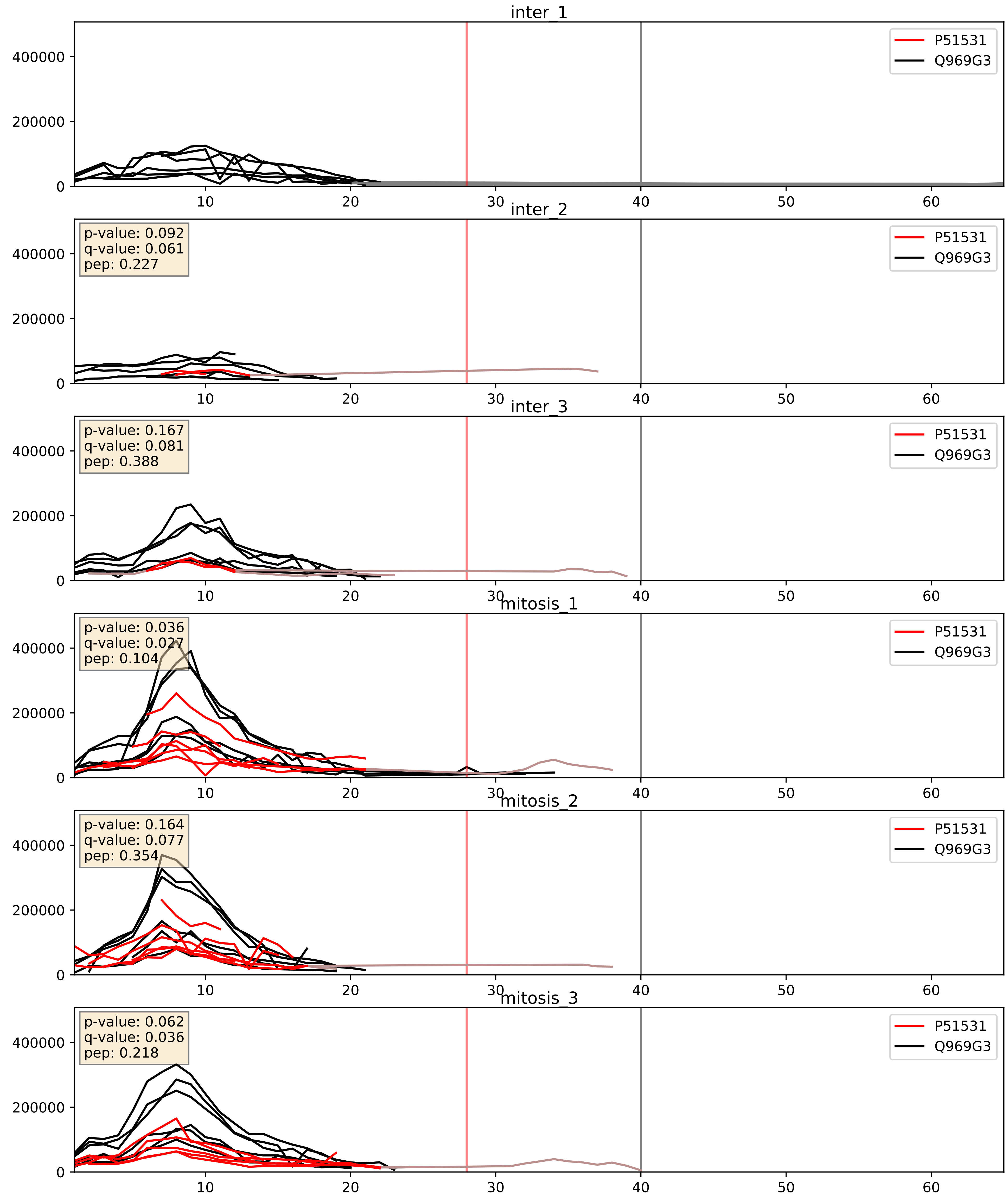


| level | condition_1 | condition_2 | pvalue | pvalue_adjusted |
|---|---|---|---|---|
| complex_abundance | mitosis | inter | 0.008183943251571598 | 0.06710589524334576 |
| interactor_abundance | mitosis | inter | 0.013657409124963755 | 0.10336642096347458 |
| interactor_ratio | mitosis | inter | 0.4403853634924024 | 0.8463612688491312 |

P51531_Q96GM5
SMCA2_HUMAN vs SMRD1_HUMAN
p-value: 0.004 q-value: 0.005

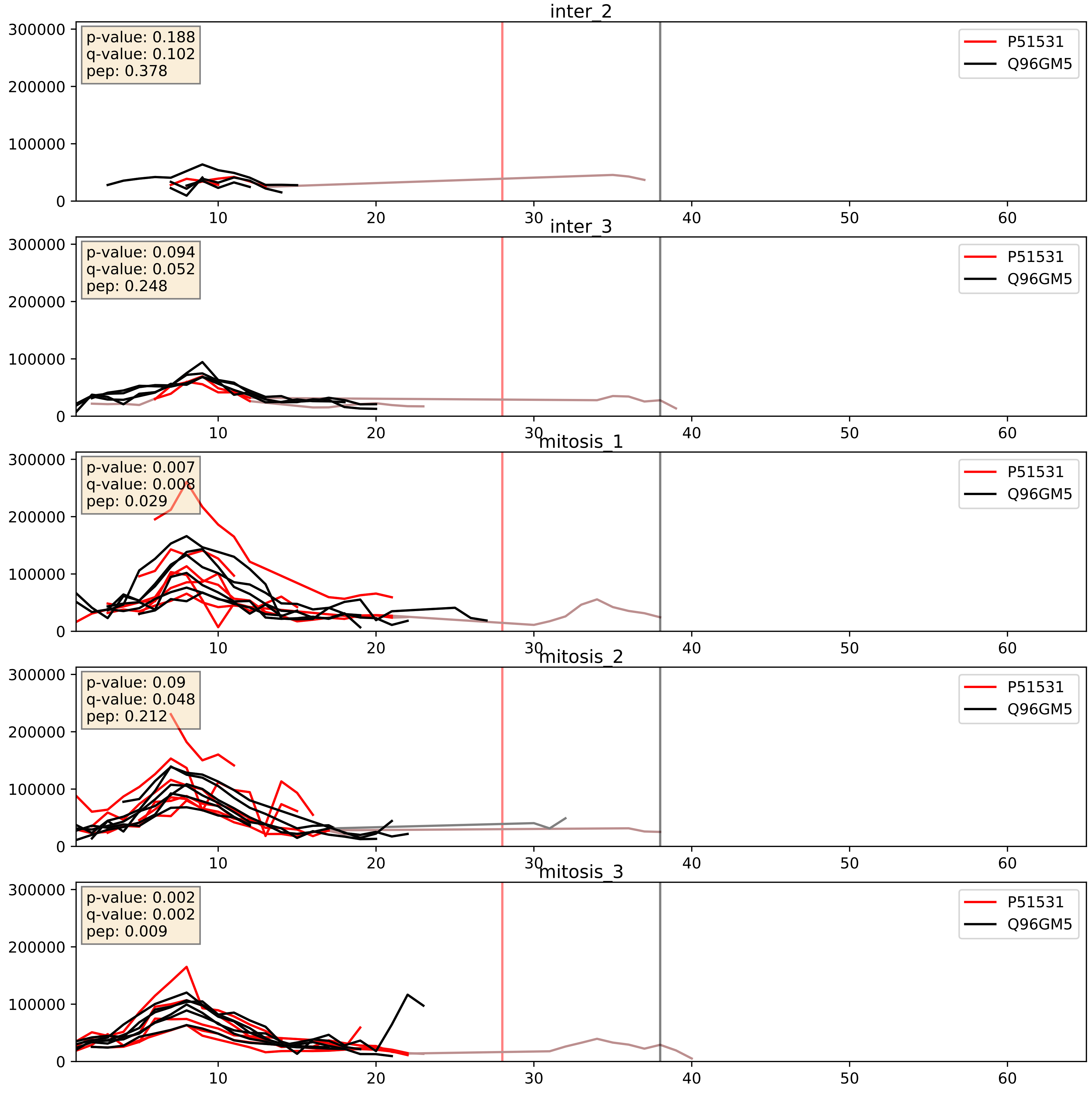


| level | condition_1 | condition_2 | pvalue | pvalue_adjusted |
|---|---|---|---|---|
| complex_abundance | mitosis | inter | 0.003988838446919328 | 0.04687907436752479 |
| interactor_abundance | mitosis | inter | 0.013657409124963755 | 0.10336642096347458 |
| interactor_ratio | mitosis | inter | 0.8054421902429661 | 0.9636037943367981 |
