## Supplementary Data 2 for "SECAT: Quantifying differential protein-protein interaction states by network-centric analysis": hela_string_000186_P38117.pdf

P38117_P38117

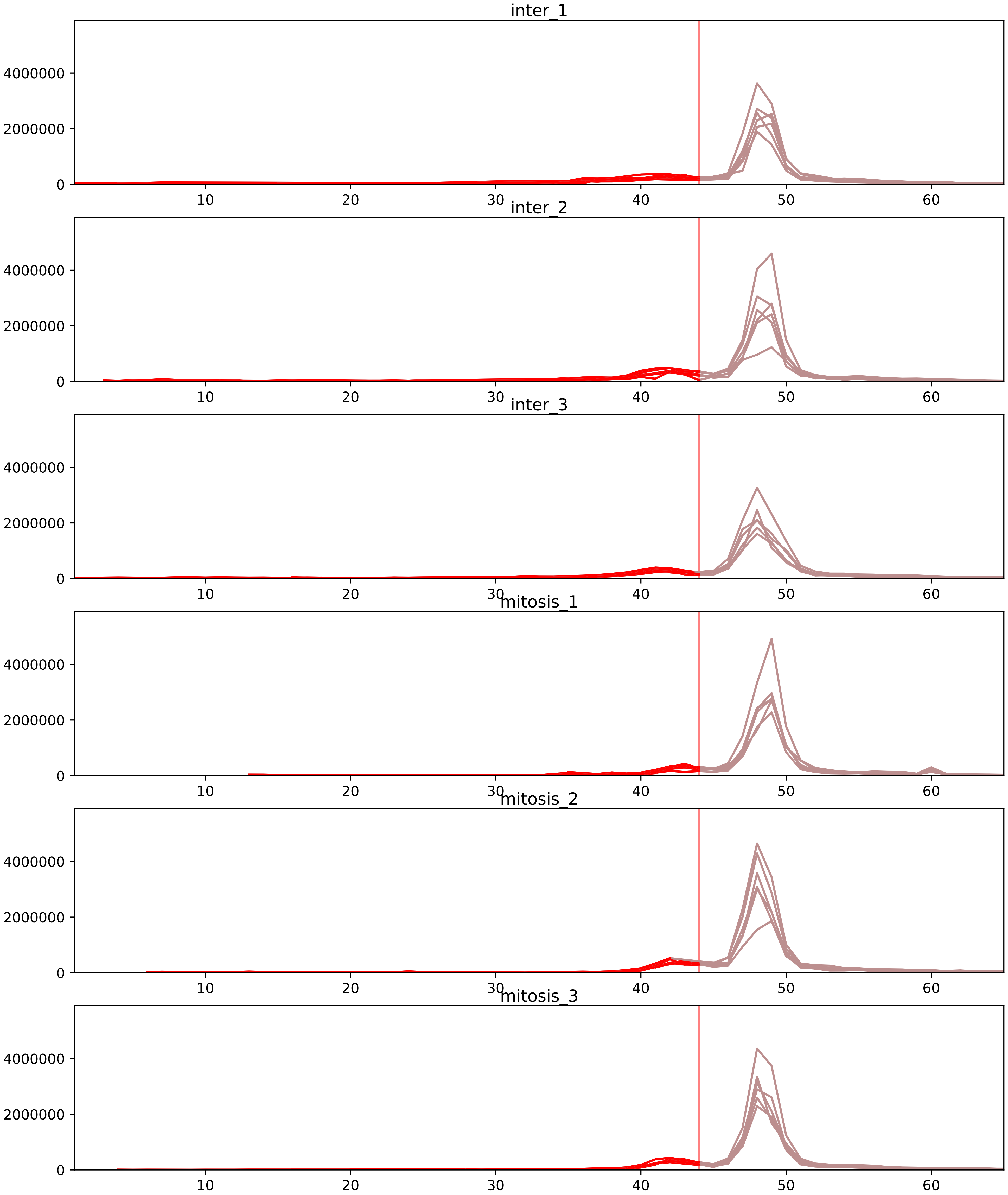


| level | condition_1 | condition_2 | pvalue | pvalue_adjusted |
|---|---|---|---|---|
| interactor_abundance | mitosis | inter | 0.00015338143620813213 | 0.0030676287241626424 |
| assembled_abundance | mitosis | inter | 0.000813826278750979 | 0.025396805405555552 |
| complex_abundance | mitosis | inter | 0.00120883895911502 | 0.011958432523639656 |
| monomer_abundance | mitosis | inter | 0.0074446564018705915 | 0.16598262670786754 |
| interactor_ratio | mitosis | inter | 0.07516951950567265 | 0.272804567831238 |
| total_abundance | mitosis | inter | 0.5520390709144615 | 0.7916035378902148 |

P13804_P38117
ETFA_HUMAN vs ETFB_HUMAN
p-value: 0.0 q-value: 0.0

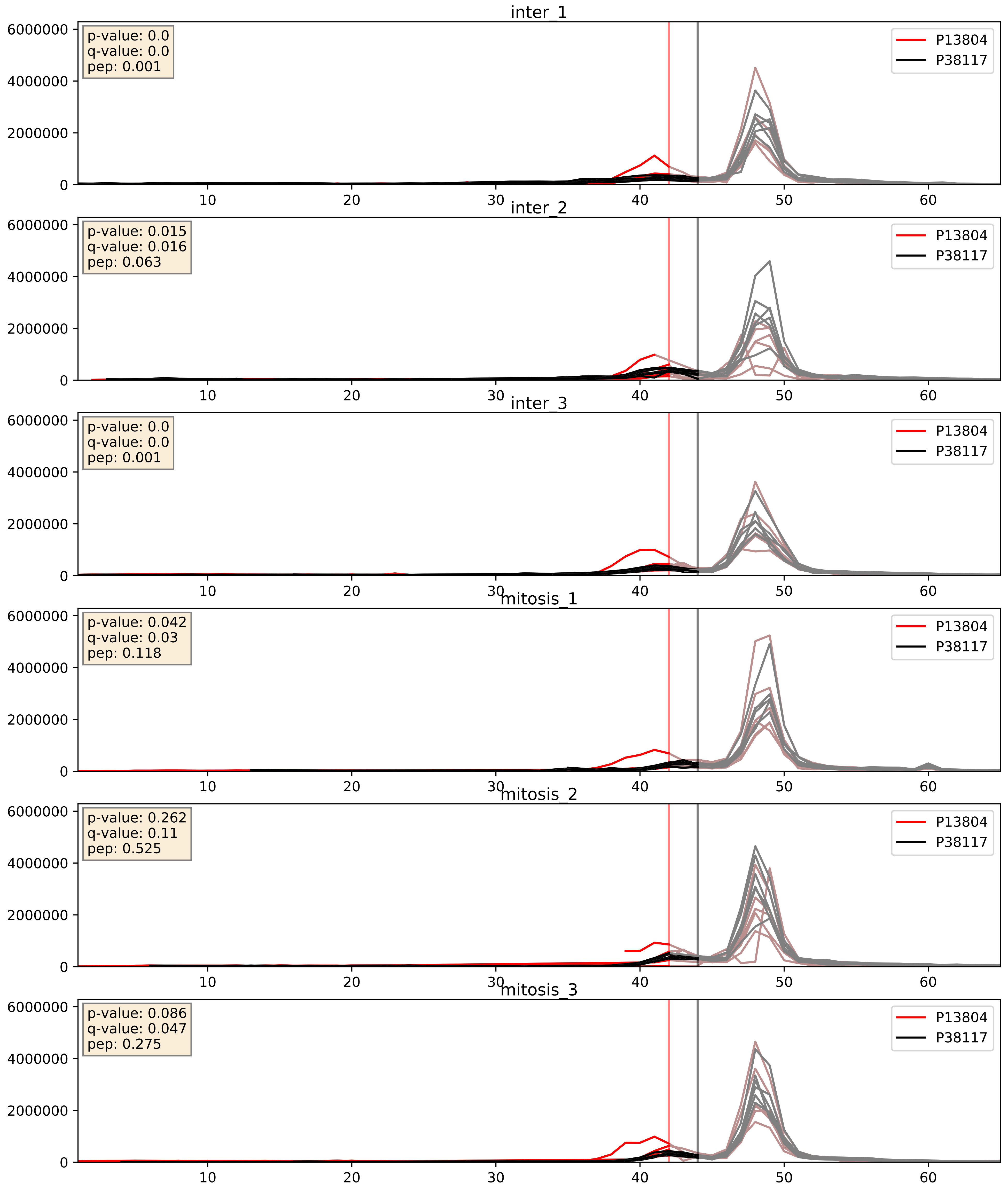


| level | condition_1 | condition_2 | pvalue | pvalue_adjusted |
|---|---|---|---|---|
| complex_abundance | mitosis | inter | 0.00120883895911502 | 0.023410998846209438 |
| interactor_abundance | mitosis | inter | 0.02890094925948924 | 0.1406836085648154 |
| interactor_ratio | mitosis | inter | 0.07516951950567265 | 0.4612552594756687 |
