## Supplementary Data 2 for "SECAT: Quantifying differential protein-protein interaction states by network-centric analysis": hela_string_000187_P11940.pdf

P11940_P11940

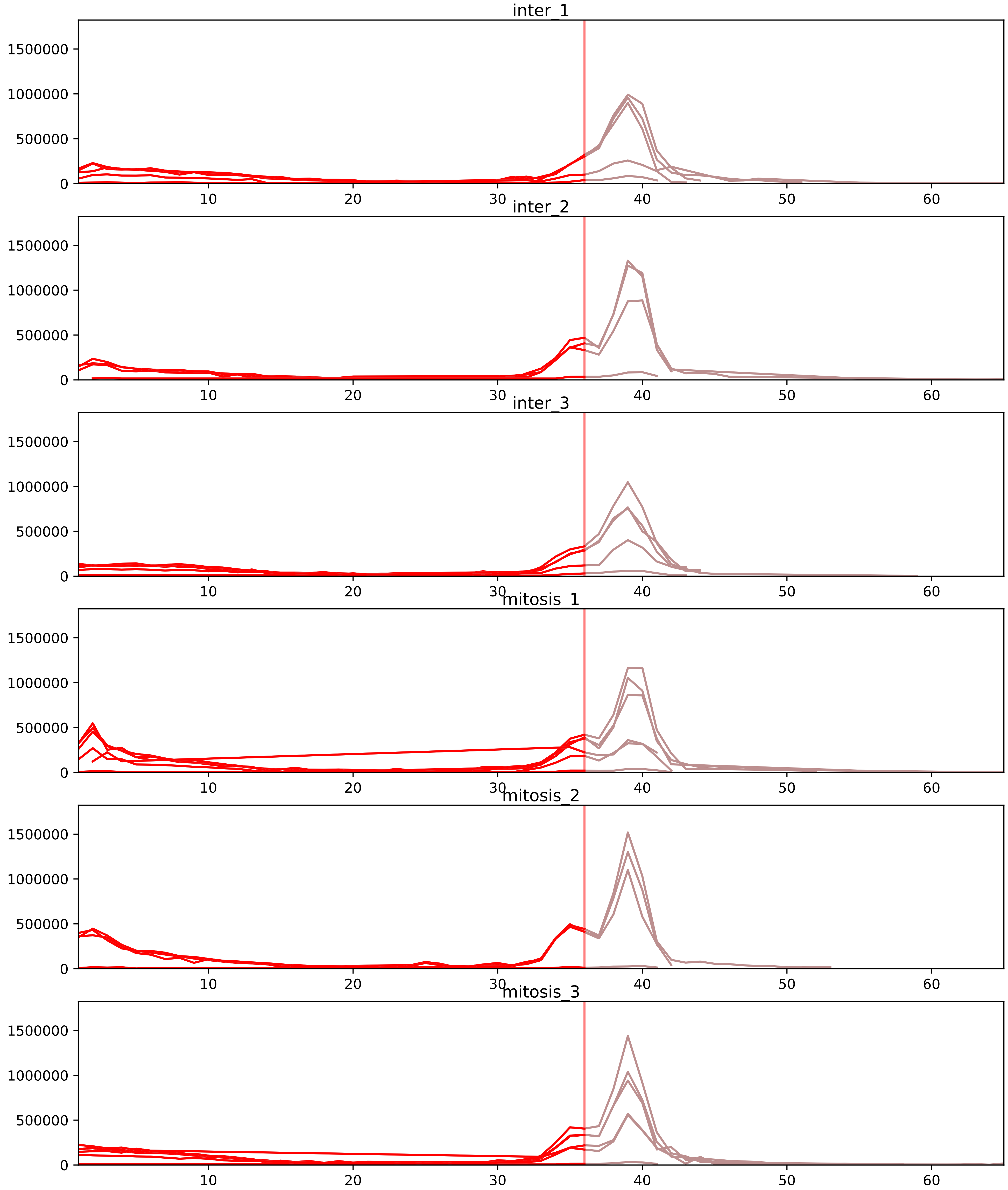


| level | condition_1 | condition_2 | pvalue | pvalue_adjusted |
|---|---|---|---|---|
| interactor_ratio | mitosis | inter | 0.00016993337253775234 | 0.0026184559792029605 |
| complex_abundance | mitosis | inter | 0.0013894223049798014 | 0.013192965785801805 |
| total_abundance | mitosis | inter | 0.031614270795032914 | 0.22522364890102928 |
| interactor_abundance | mitosis | inter | 0.0331262970797786 | 0.10923366778995093 |
| assembled_abundance | mitosis | inter | 0.04727251692982068 | 0.2606391657814966 |
| monomer_abundance | mitosis | inter | 0.22422015221783764 | 0.5988967466235329 |

P06730_P11940
IF4E_HUMAN vs PABP1_HUMAN
p-value: 0.077 q-value: 0.046

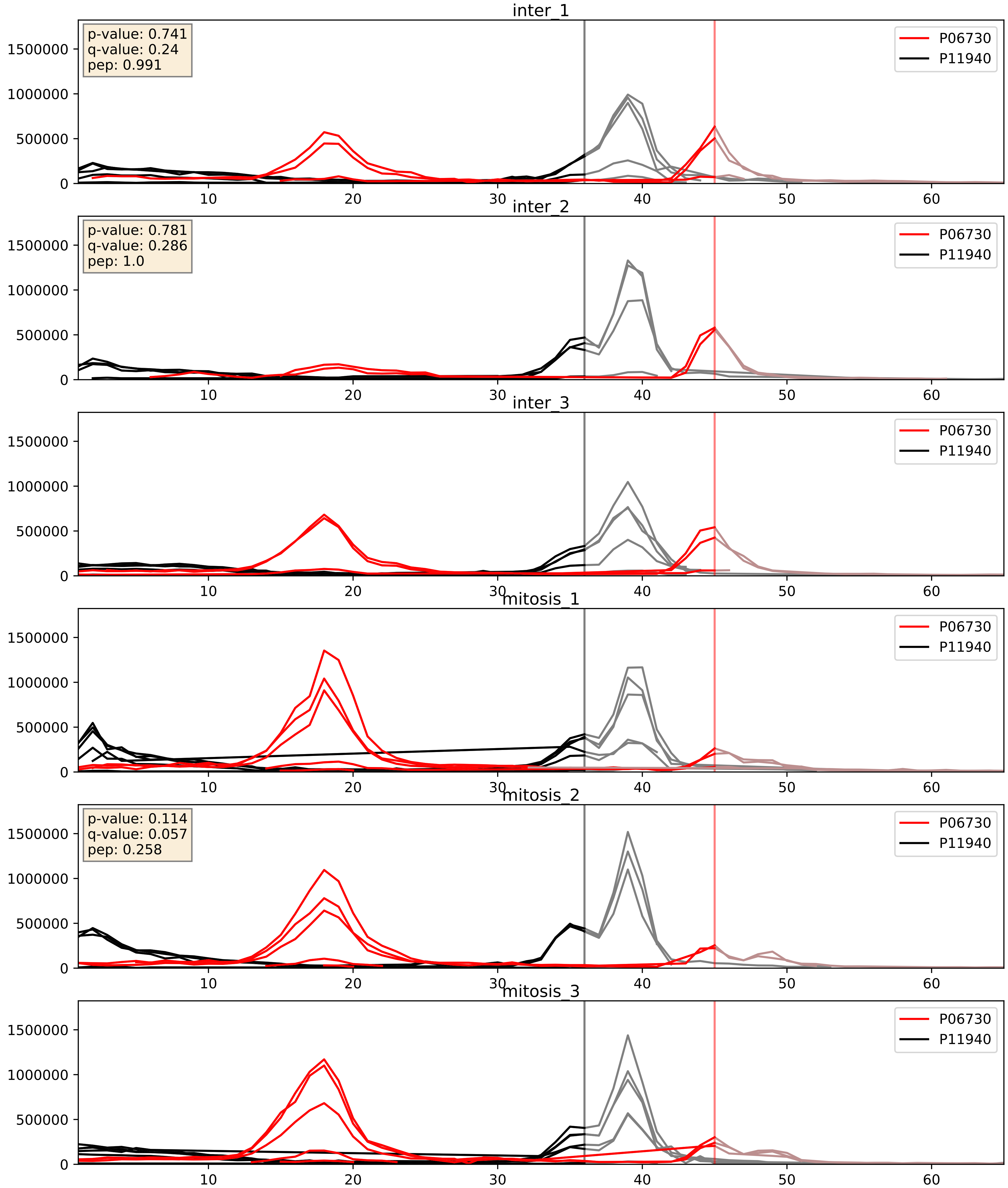


| level | condition_1 | condition_2 | pvalue | pvalue_adjusted |
|---|---|---|---|---|
| interactor_ratio | mitosis | inter | 0.005276695702534133 | 0.0974780445871482 |
| interactor_abundance | mitosis | inter | 0.008995173699102407 | 0.08274979781226932 |
| complex_abundance | mitosis | inter | 0.030746854054412083 | 0.12751204260889337 |

P11940_P15170
PABP1_HUMAN vs ERF3A_HUMAN
p-value: 0.015 q-value: 0.012

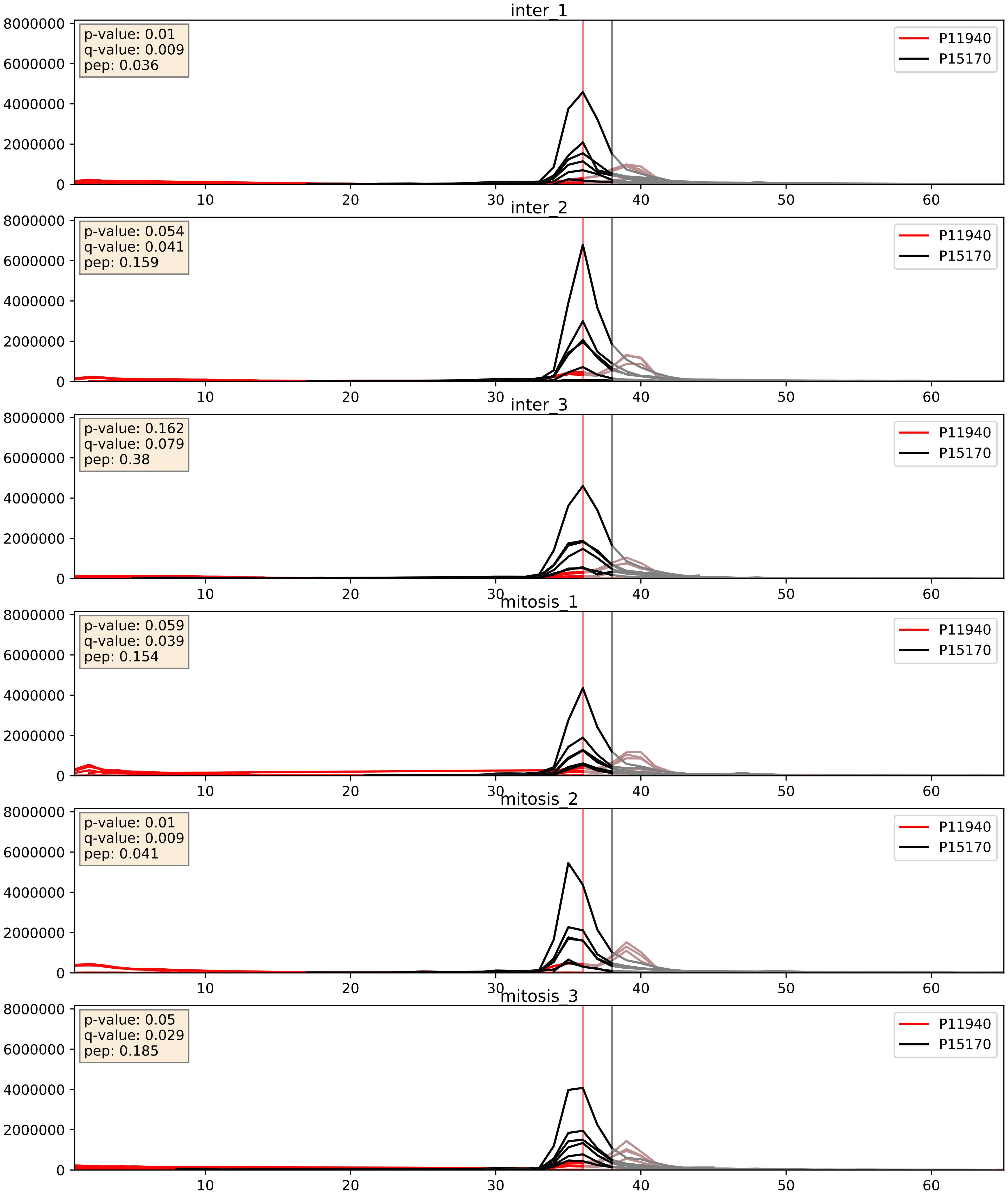


| level | condition_1 | condition_2 | pvalue | pvalue_adjusted |
|---|---|---|---|---|
| complex_abundance | mitosis | inter | 0.5037490529349358 | 0.6404410096297242 |
| interactor_abundance | mitosis | inter | 0.5653268283041282 | 0.7129047805367321 |
| interactor_ratio | mitosis | inter | 0.9651720026800956 | 0.9937294965826962 |

P11940_P17844
PABP1_HUMAN vs DDX5_HUMAN
p-value: 0.0 q-value: 0.016

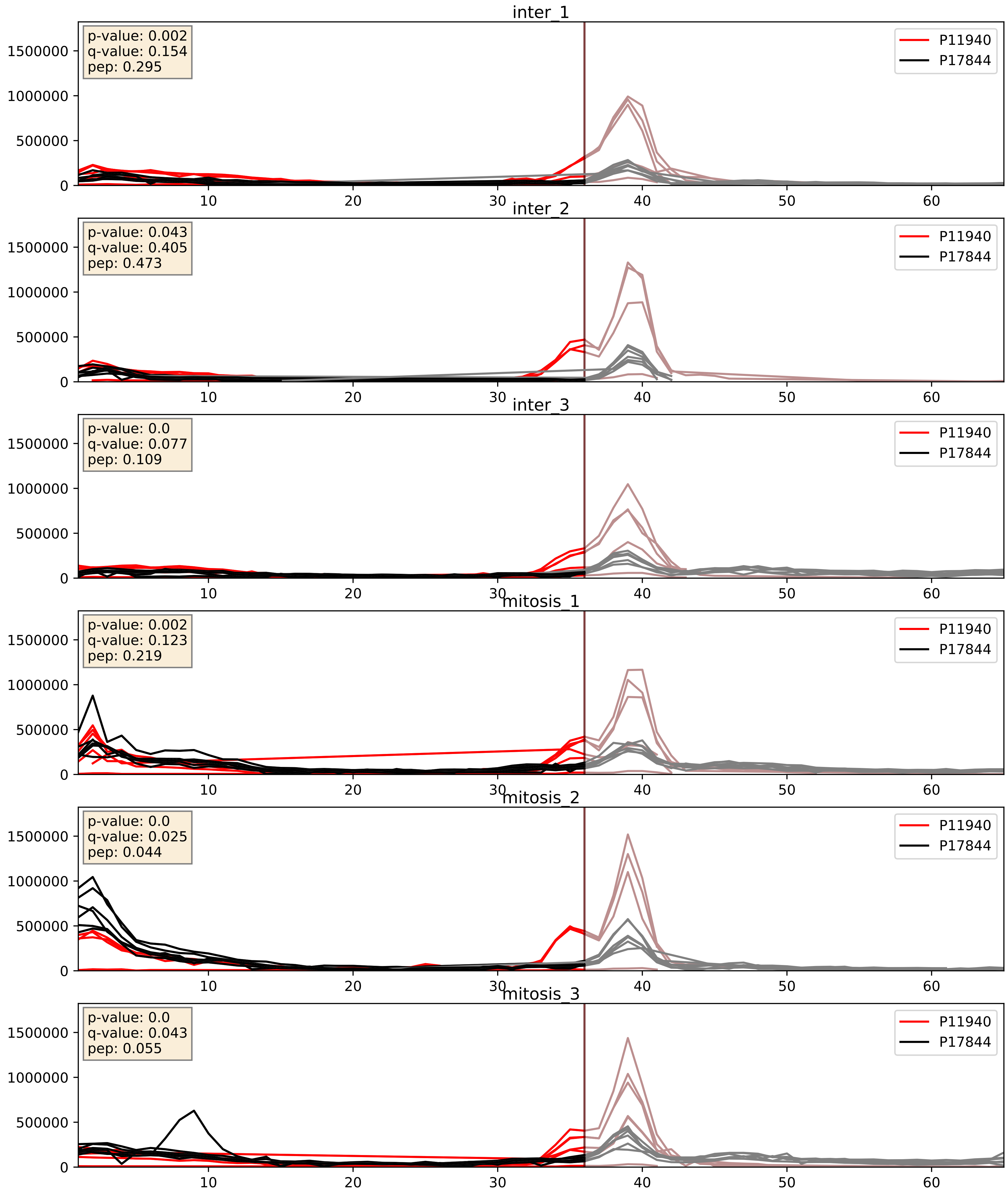


| level | condition_1 | condition_2 | pvalue | pvalue_adjusted |
|---|---|---|---|---|
| complex_abundance | mitosis | inter | 0.003636591319936893 | 0.04428054295684183 |
| interactor_ratio | mitosis | inter | 0.006113461516559377 | 0.10745632562987323 |
| interactor_abundance | mitosis | inter | 0.13490704099728298 | 0.3215381514511325 |

P11940_P18124
PABP1_HUMAN vs RL7_HUMAN
p-value: 0.017 q-value: 0.014

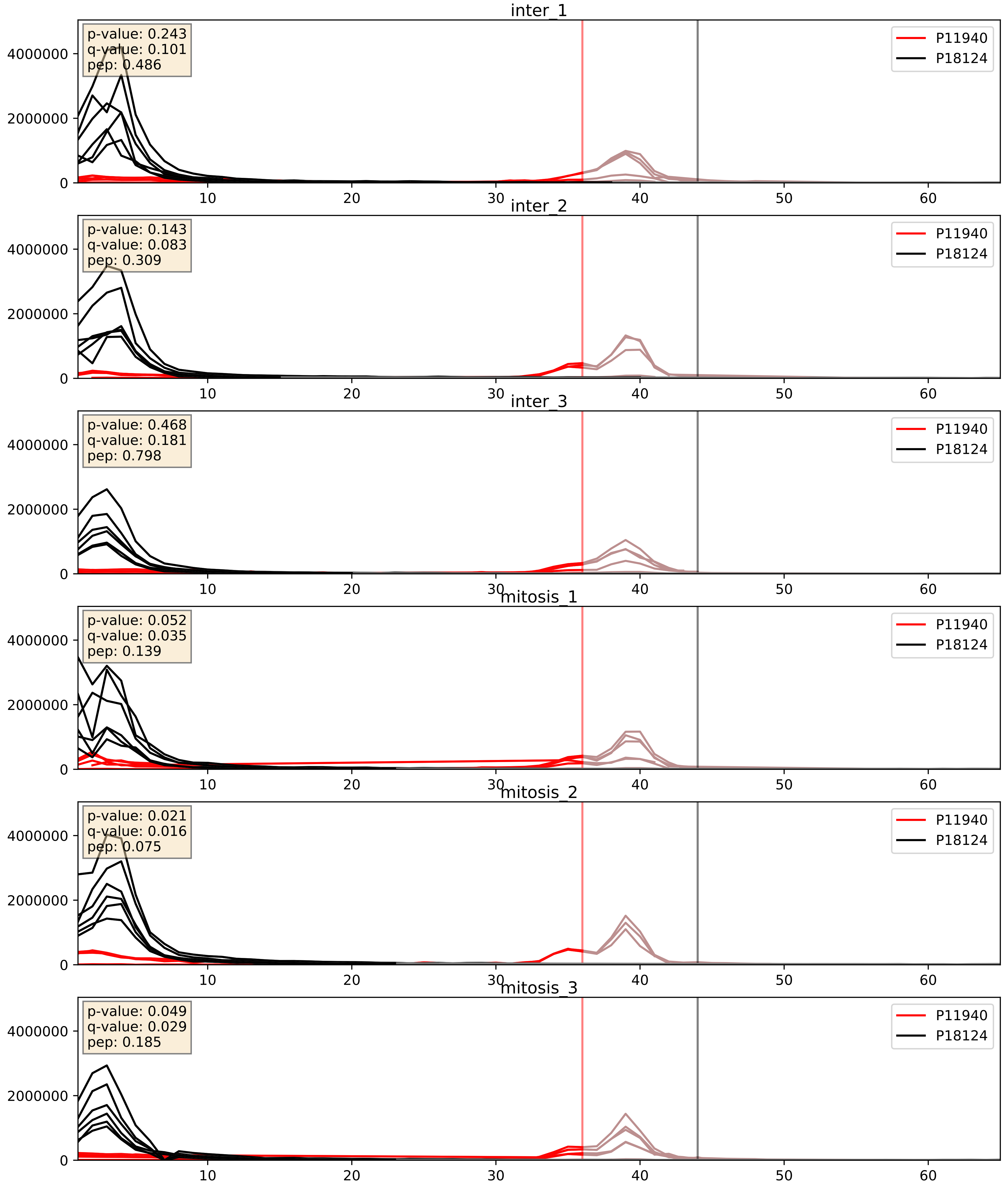


| level | condition_1 | condition_2 | pvalue | pvalue_adjusted |
|---|---|---|---|---|
| interactor_ratio | mitosis | inter | 0.0523315525000554 | 0.3780237041354213 |
| interactor_abundance | mitosis | inter | 0.15090604372175434 | 0.33732834796845007 |
| complex_abundance | mitosis | inter | 0.8756444443994904 | 0.9225251008073401 |

P11940_P26599
PABP1_HUMAN vs PTBP1_HUMAN
p-value: 0.001 q-value: 0.041

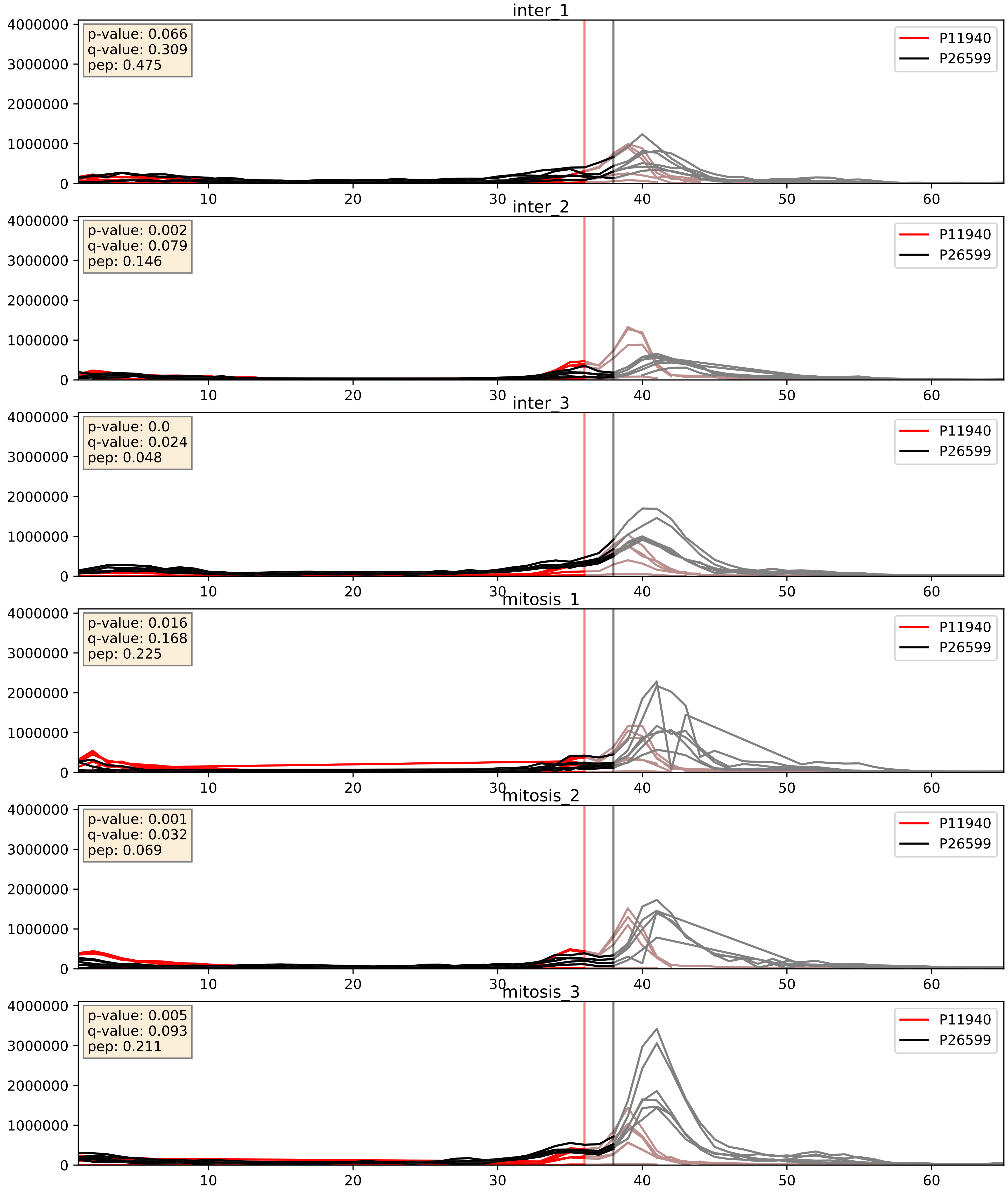


| level | condition_1 | condition_2 | pvalue | pvalue_adjusted |
|---|---|---|---|---|
| interactor_abundance | mitosis | inter | 0.14216125851351857 | 0.32880142136715784 |
| interactor_ratio | mitosis | inter | 0.29259105139089336 | 0.7623557131777331 |
| complex_abundance | mitosis | inter | 0.5262085869342336 | 0.6616253678256521 |

P11940_P36578
PABP1_HUMAN vs RL4_HUMAN
p-value: 0.001 q-value: 0.023

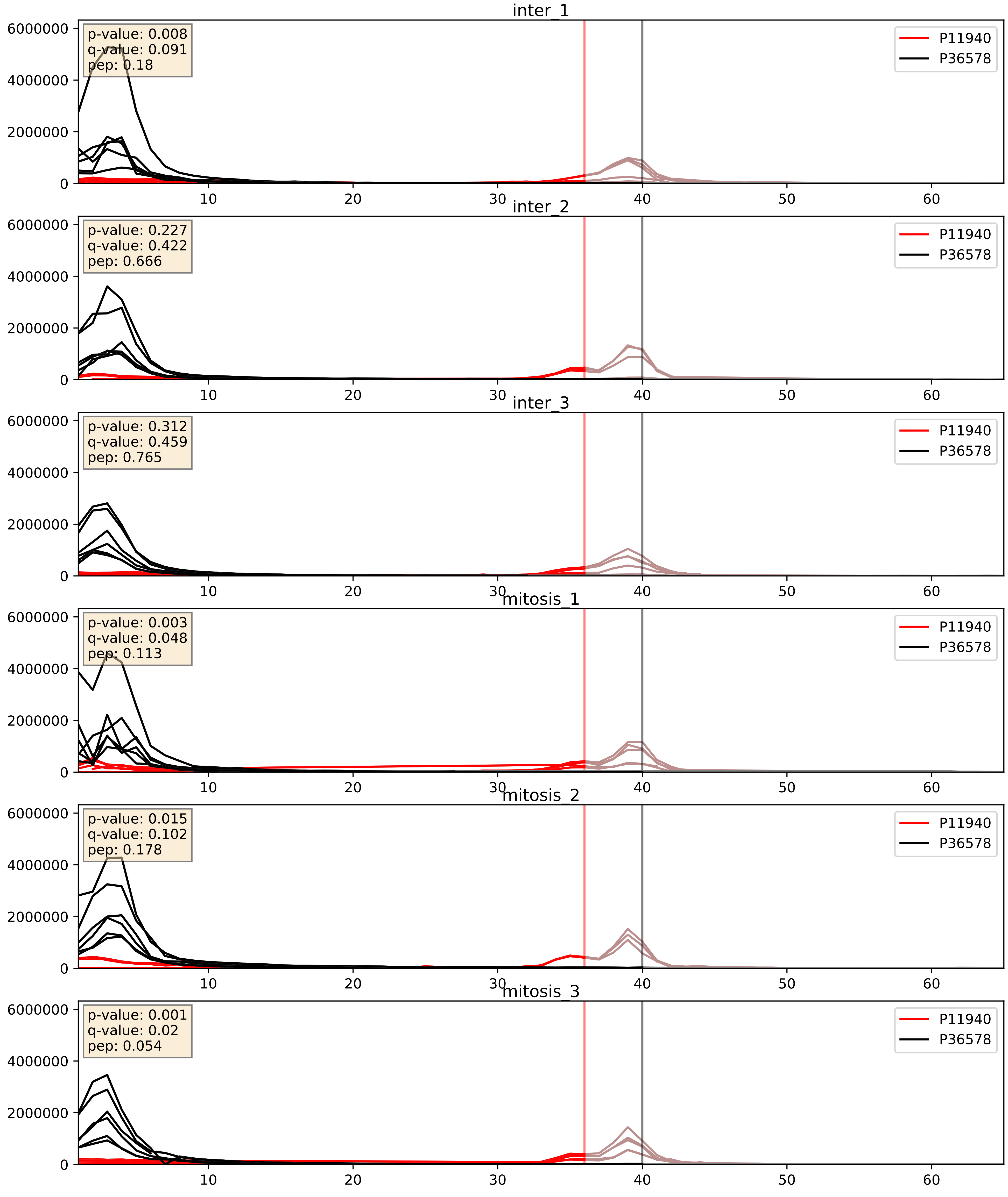


| level | condition_1 | condition_2 | pvalue | pvalue_adjusted |
|---|---|---|---|---|
| interactor_abundance | mitosis | inter | 0.15090604372175434 | 0.33732834796845007 |
| complex_abundance | mitosis | inter | 0.20994491434641527 | 0.3640122476818543 |
| interactor_ratio | mitosis | inter | 0.3785040005379135 | 0.8169425730218204 |

P11940_P39023
PABP1_HUMAN vs RL3_HUMAN
p-value: 0.002 q-value: 0.003

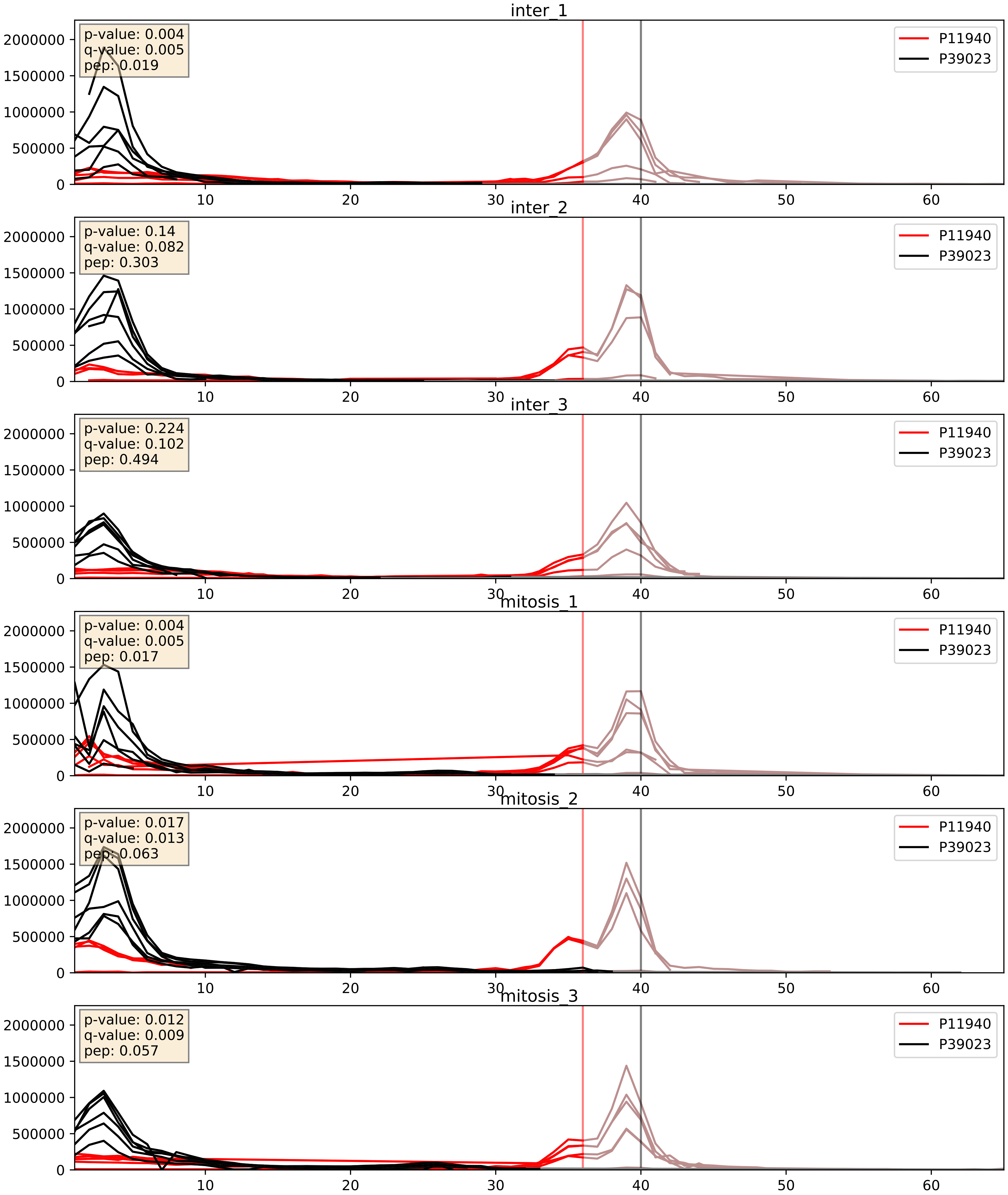


| level | condition_1 | condition_2 | pvalue | pvalue_adjusted |
|---|---|---|---|---|
| interactor_abundance | mitosis | inter | 0.10979246191660604 | 0.2847087167543616 |
| complex_abundance | mitosis | inter | 0.14603606212752657 | 0.29192772999624955 |
| interactor_ratio | mitosis | inter | 0.2750031130835264 | 0.7475191286928202 |

P11940_P46777
PABP1_HUMAN vs RL5_HUMAN
p-value: 0.006 q-value: 0.006

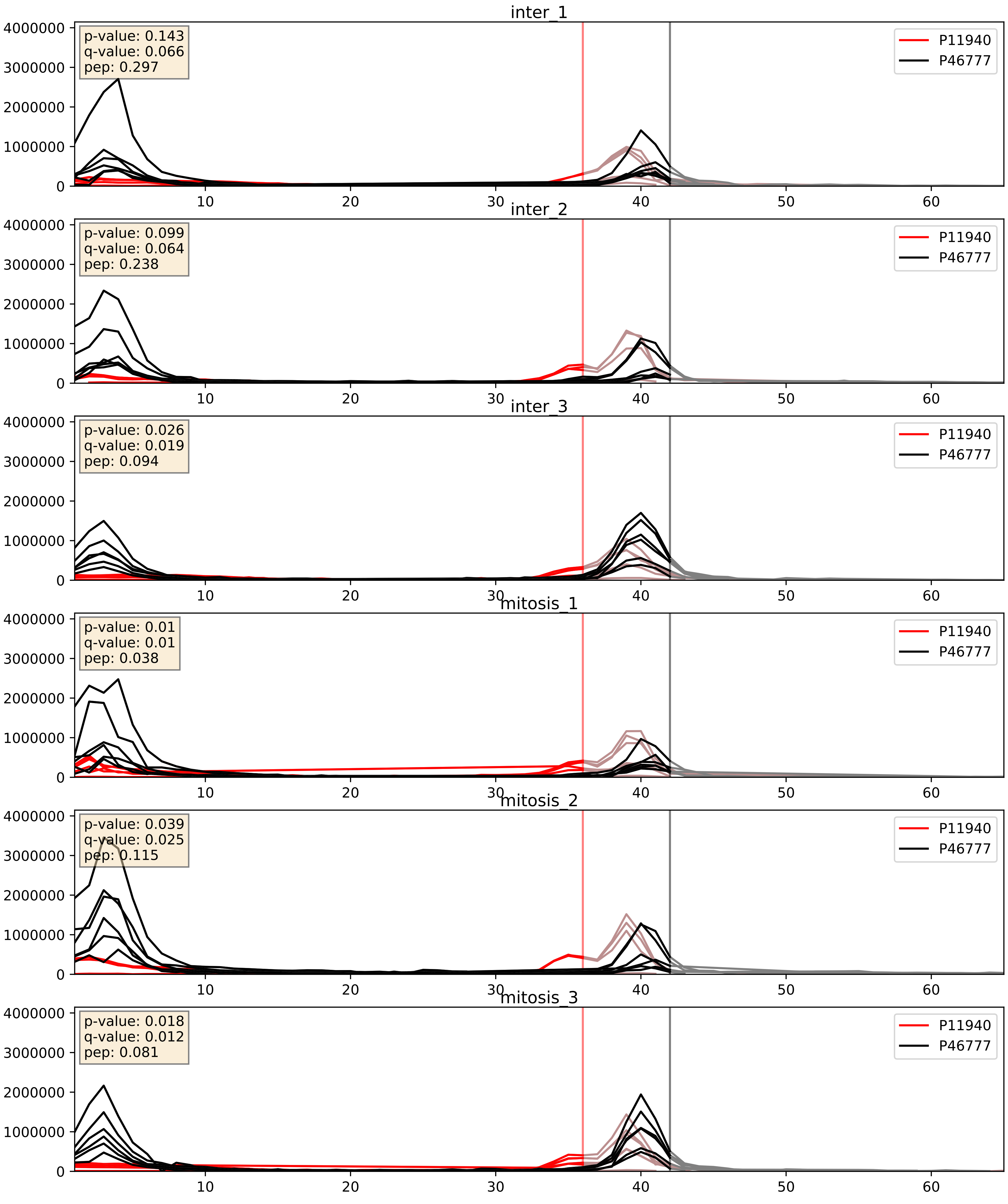


| level | condition_1 | condition_2 | pvalue | pvalue_adjusted |
|---|---|---|---|---|
| complex_abundance | mitosis | inter | 0.08744005666530409 | 0.2165134177191215 |
| interactor_abundance | mitosis | inter | 0.13289898740509534 | 0.3185259224940826 |
| interactor_ratio | mitosis | inter | 0.8236971701921706 | 0.96884652444993 |

P11940_P46779
PABP1_HUMAN vs RL28_HUMAN
p-value: 0.002 q-value: 0.03

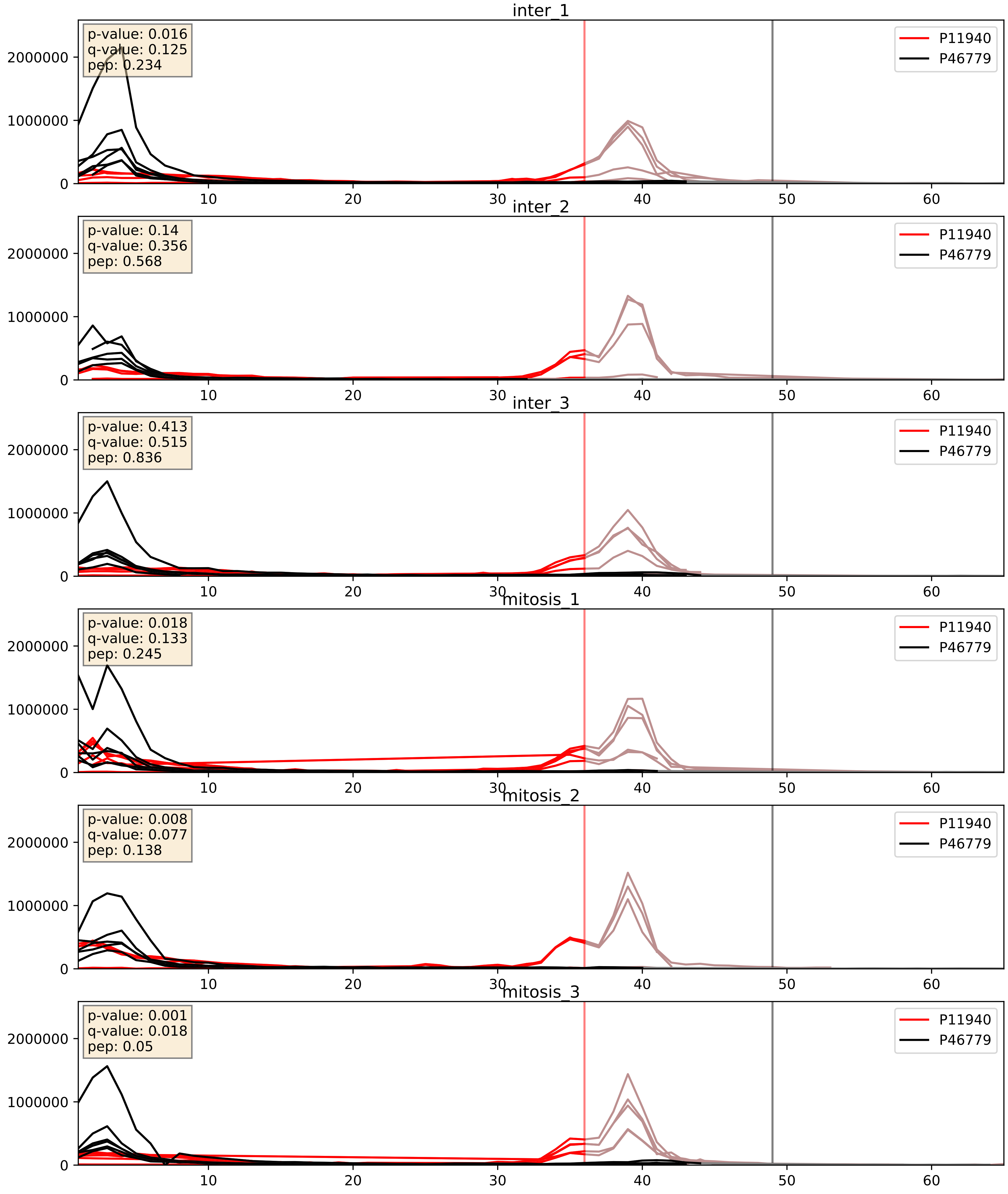


| level | condition_1 | condition_2 | pvalue | pvalue_adjusted |
|---|---|---|---|---|
| interactor_ratio | mitosis | inter | 0.0719078418302576 | 0.447008806148878 |
| interactor_abundance | mitosis | inter | 0.1703925411103742 | 0.3598133054863397 |
| complex_abundance | mitosis | inter | 0.5880292763241118 | 0.7145841290934692 |

P11940_P50914
PABP1_HUMAN vs RL14_HUMAN
p-value: 0.003 q-value: 0.047

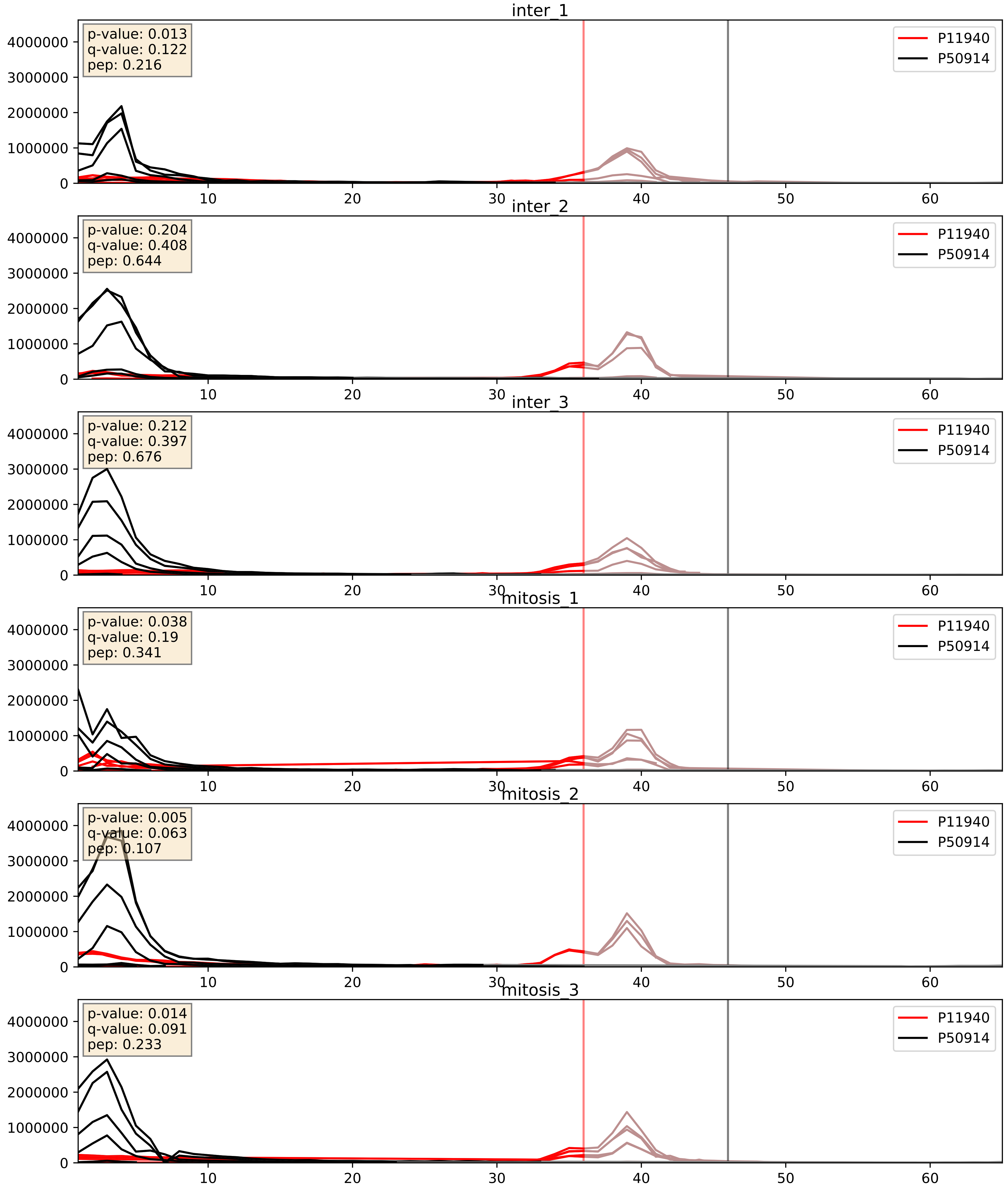


| level | condition_1 | condition_2 | pvalue | pvalue_adjusted |
|---|---|---|---|---|
| interactor_abundance | mitosis | inter | 0.38219850929418886 | 0.5748254835383039 |
| interactor_ratio | mitosis | inter | 0.5986035097059178 | 0.9019028605875125 |
| complex_abundance | mitosis | inter | 0.628373431451533 | 0.7464441539307691 |

P11940_P52597
PABP1_HUMAN vs HNRPF_HUMAN
p-value: 0.0 q-value: 0.018

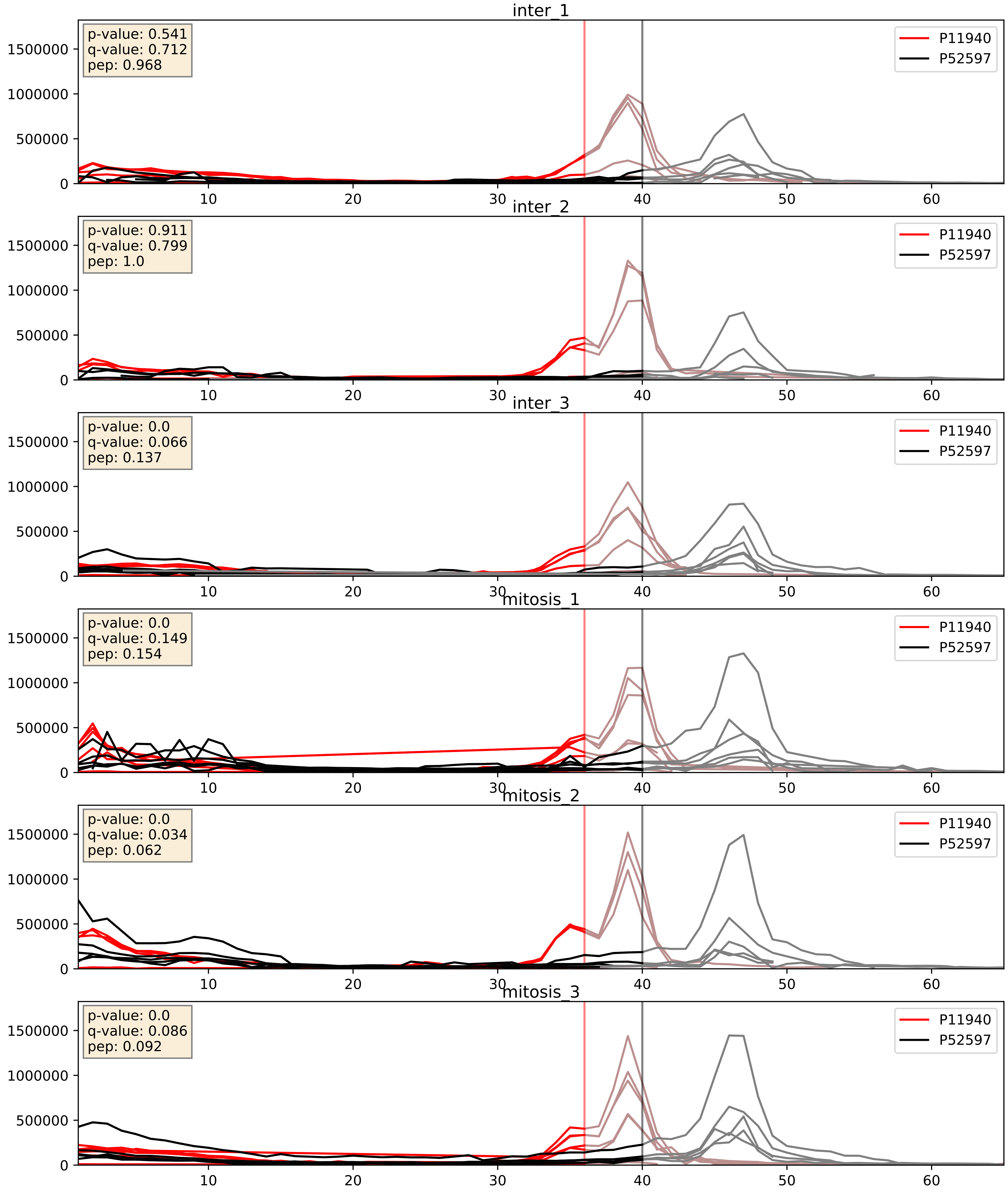


| level | condition_1 | condition_2 | pvalue | pvalue_adjusted |
|---|---|---|---|---|
| complex_abundance | mitosis | inter | 0.0061419595529173445 | 0.05828645200067901 |
| interactor_ratio | mitosis | inter | 0.016353146604940094 | 0.1886562465475569 |
| interactor_abundance | mitosis | inter | 0.15090604372175434 | 0.33732834796845007 |

P11940_P61978
PABP1_HUMAN vs HNRPK_HUMAN
p-value: 0.0 q-value: 0.023

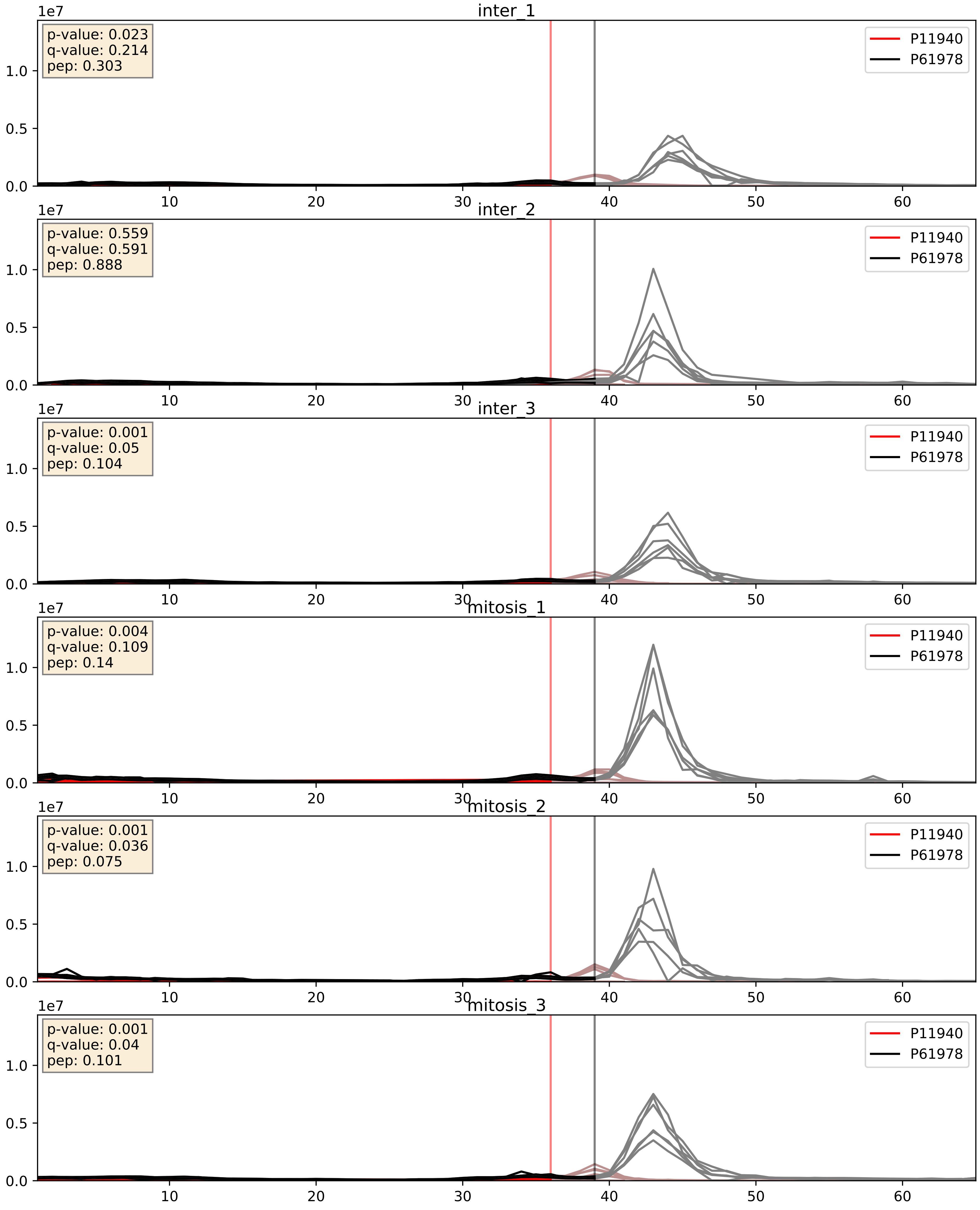


| level | condition_1 | condition_2 | pvalue | pvalue_adjusted |
|---|---|---|---|---|
| interactor_abundance | mitosis | inter | 0.15090604372175434 | 0.33732834796845007 |
| interactor_ratio | mitosis | inter | 0.6396626216060204 | 0.9203357608237295 |
| complex_abundance | mitosis | inter | 0.9759438672739589 | 0.9857075068317154 |

P11940_P62424
PABP1_HUMAN vs RL7A_HUMAN
p-value: 0.003 q-value: 0.047

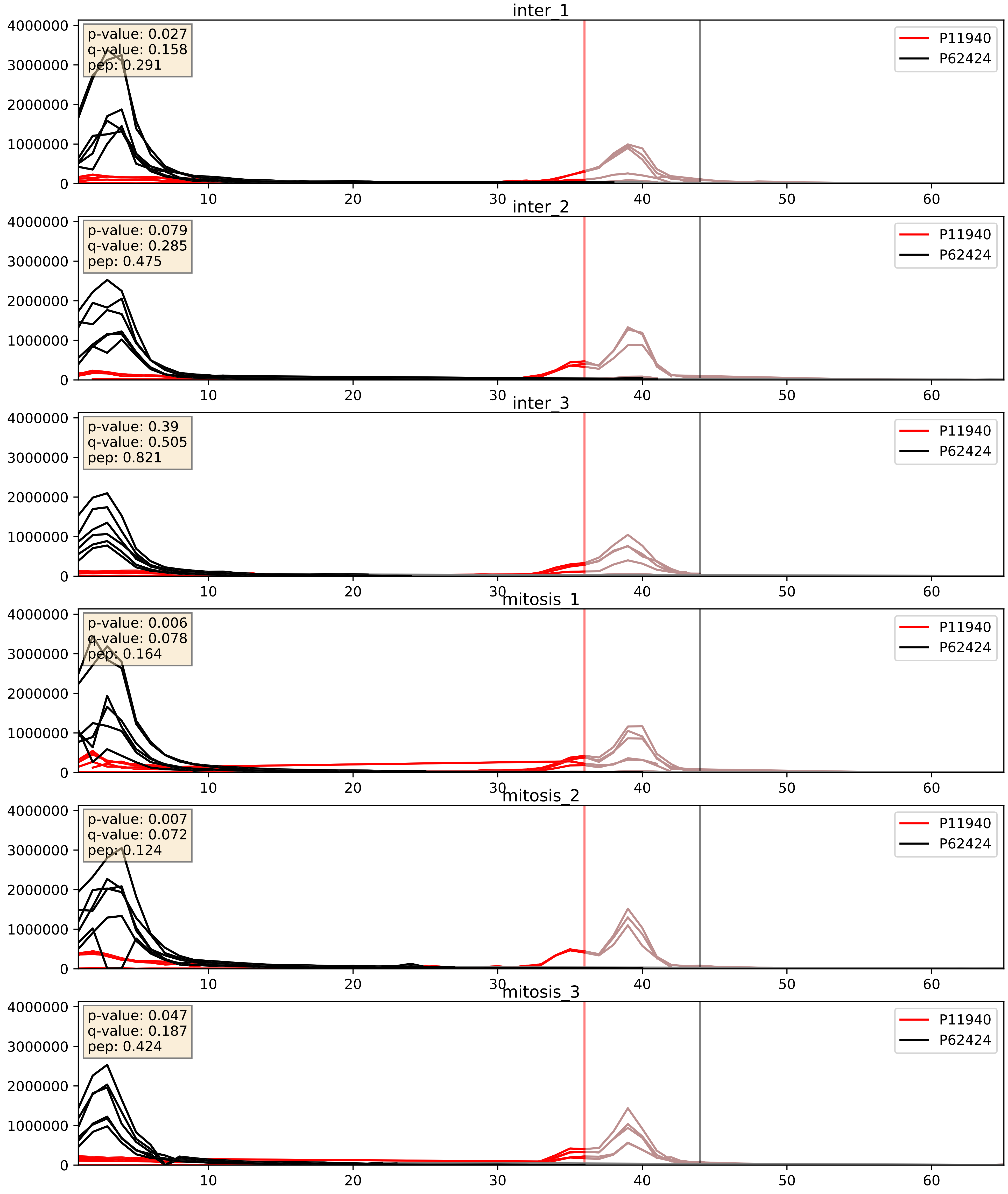


| level | condition_1 | condition_2 | pvalue | pvalue_adjusted |
|---|---|---|---|---|
| interactor_ratio | mitosis | inter | 0.1076138236161235 | 0.5447512301324761 |
| interactor_abundance | mitosis | inter | 0.21174994399655572 | 0.40715039907095185 |
| complex_abundance | mitosis | inter | 0.5033232849127958 | 0.6400902271361657 |

P11940_P62829
PABP1_HUMAN vs RL23_HUMAN
p-value: 0.003 q-value: 0.039

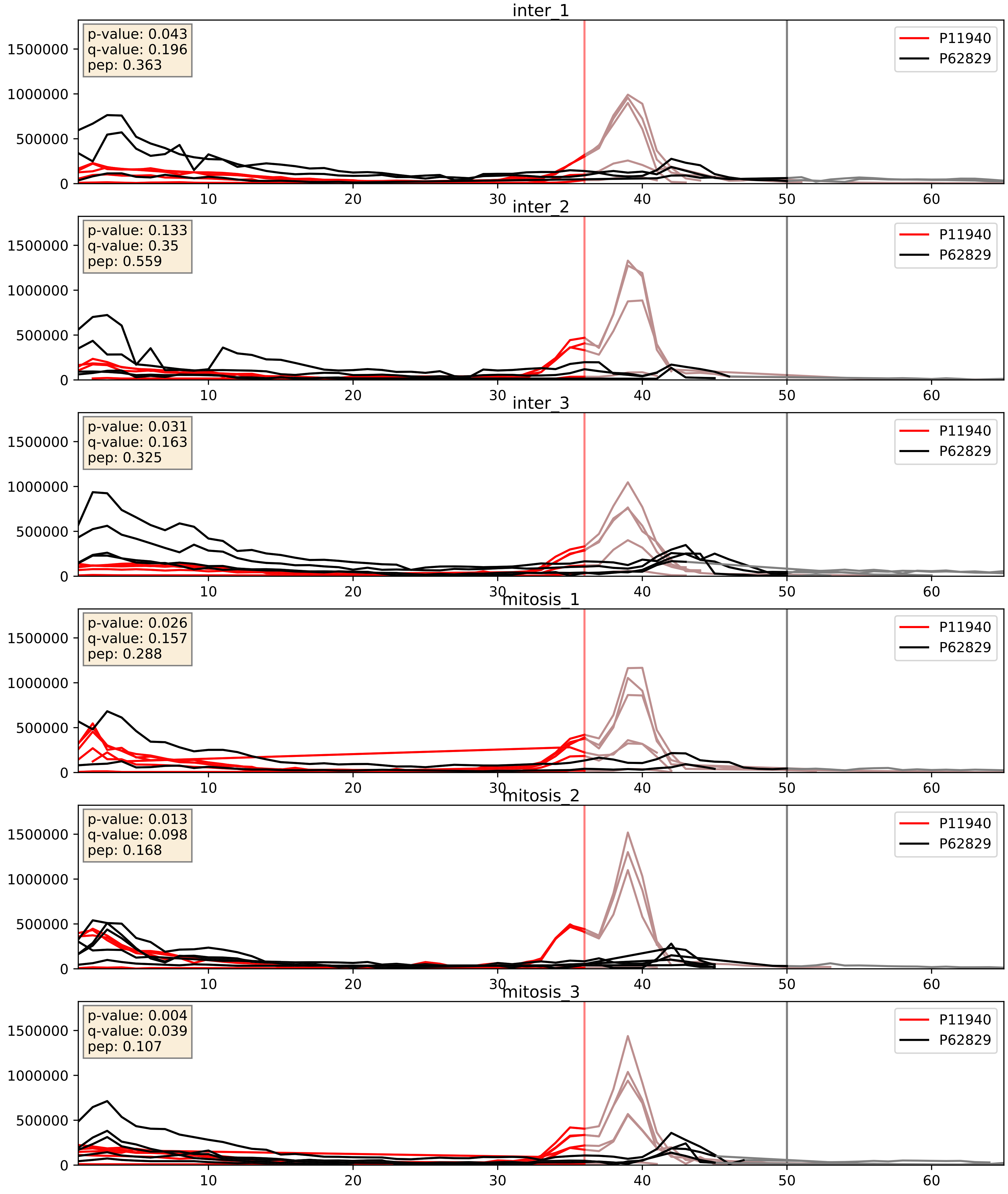


| level | condition_1 | condition_2 | pvalue | pvalue_adjusted |
|---|---|---|---|---|
| complex_abundance | mitosis | inter | 0.00659781688147954 | 0.06001839798667892 |
| interactor_ratio | mitosis | inter | 0.007455063952191203 | 0.12063392709027732 |
| interactor_abundance | mitosis | inter | 0.15090604372175434 | 0.33732834796845007 |

P11940_P62906
PABP1_HUMAN vs RL10A_HUMAN
p-value: 0.034 q-value: 0.024

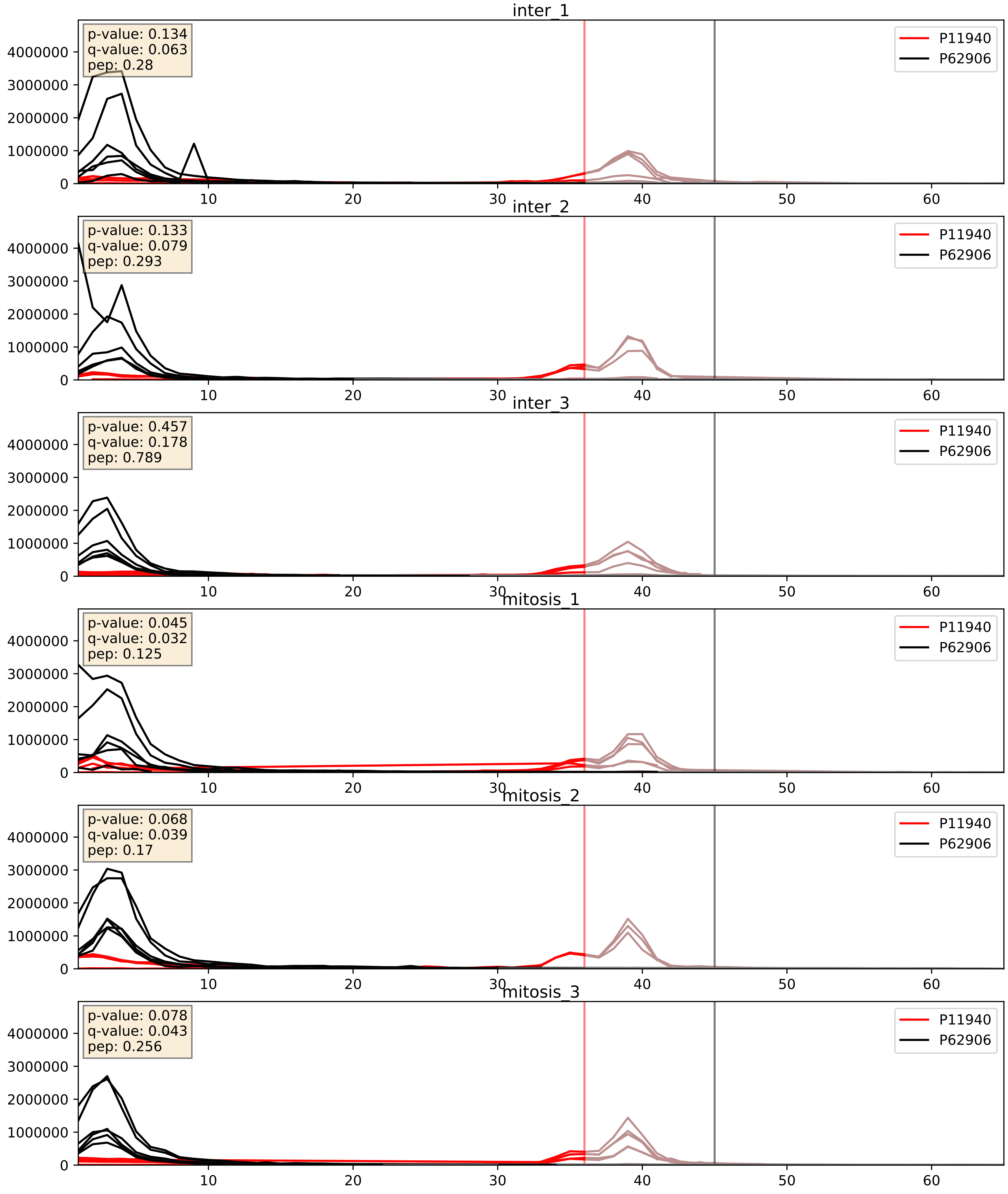


| level | condition_1 | condition_2 | pvalue | pvalue_adjusted |
|---|---|---|---|---|
| interactor_ratio | mitosis | inter | 0.13717642330363286 | 0.5873558393508659 |
| interactor_abundance | mitosis | inter | 0.22174568684515863 | 0.41786308847431103 |
| complex_abundance | mitosis | inter | 0.38602269073306805 | 0.5326732558823649 |

P11940_P62917
PABP1_HUMAN vs RL8_HUMAN
p-value: 0.004 q-value: 0.048

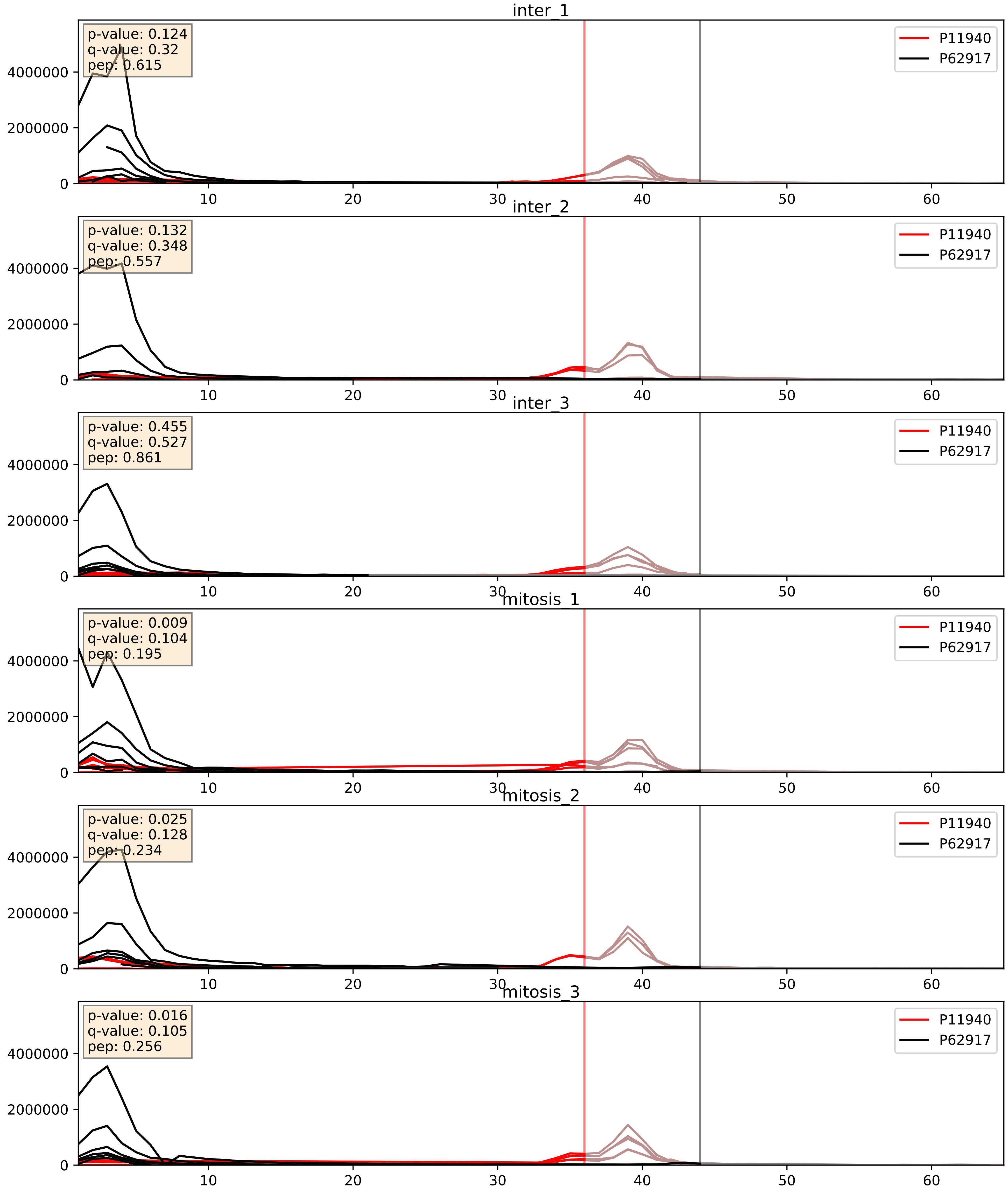


| level | condition_1 | condition_2 | pvalue | pvalue_adjusted |
|---|---|---|---|---|
| interactor_abundance | mitosis | inter | 0.06132166938843763 | 0.20248592751441813 |
| complex_abundance | mitosis | inter | 0.16032841419771246 | 0.30804508691617344 |
| interactor_ratio | mitosis | inter | 0.6692536936638933 | 0.9304647856701197 |

P11940_P62995
PABP1_HUMAN vs TRA2B_HUMAN
p-value: 0.0 q-value: 0.047

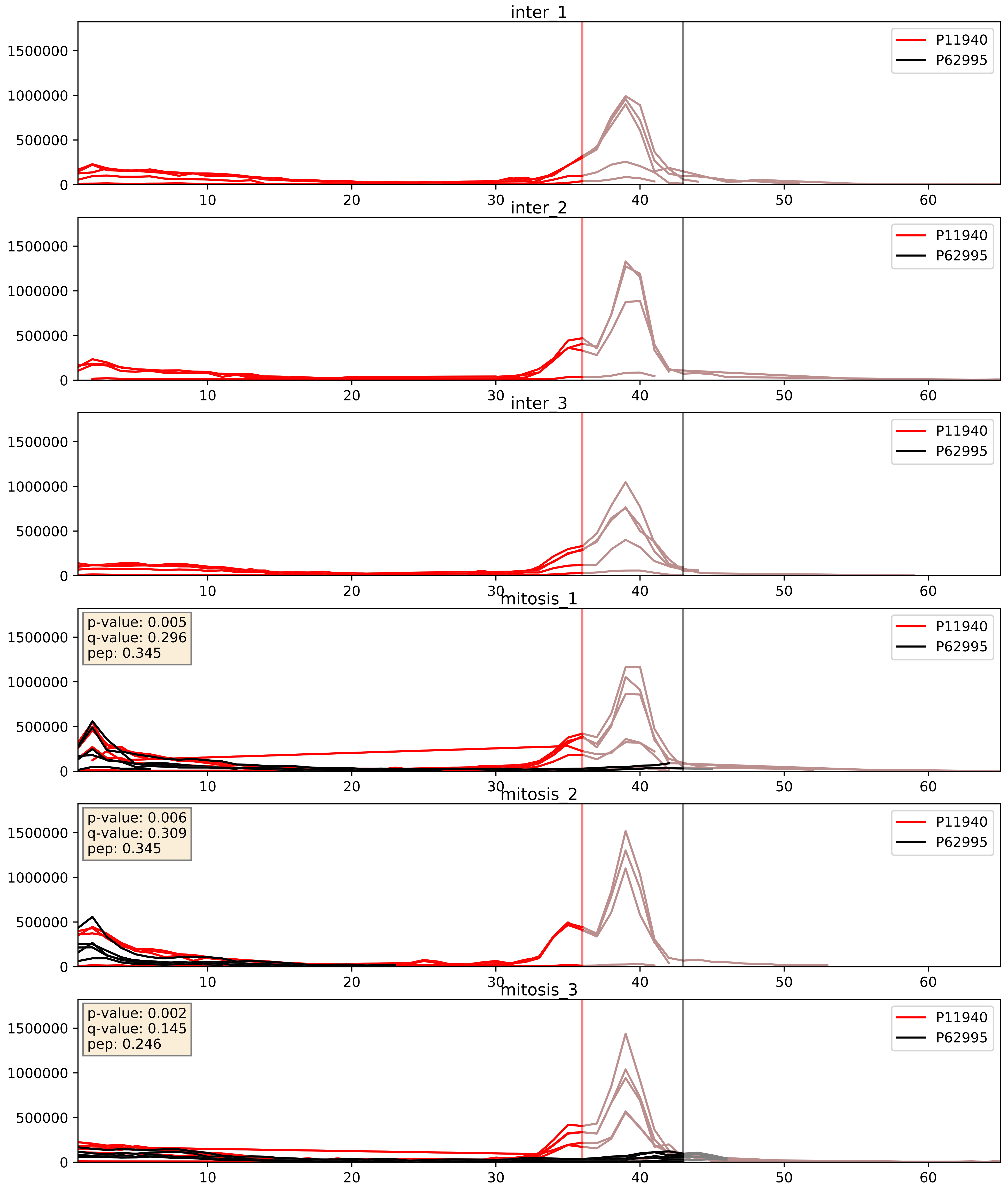


| level | condition_1 | condition_2 | pvalue | pvalue_adjusted |
|---|---|---|---|---|
| interactor_ratio | mitosis | inter | 2.219737766599098e-05 | 0.004993481478971454 |
| interactor_abundance | mitosis | inter | 7.82818796061288e-05 | 0.006980134264879818 |
| complex_abundance | mitosis | inter | 0.00015804606083051257 | 0.008563834848633118 |

P11940_Q09161
PABP1_HUMAN vs NCBP1_HUMAN
p-value: 0.0 q-value: 0.0

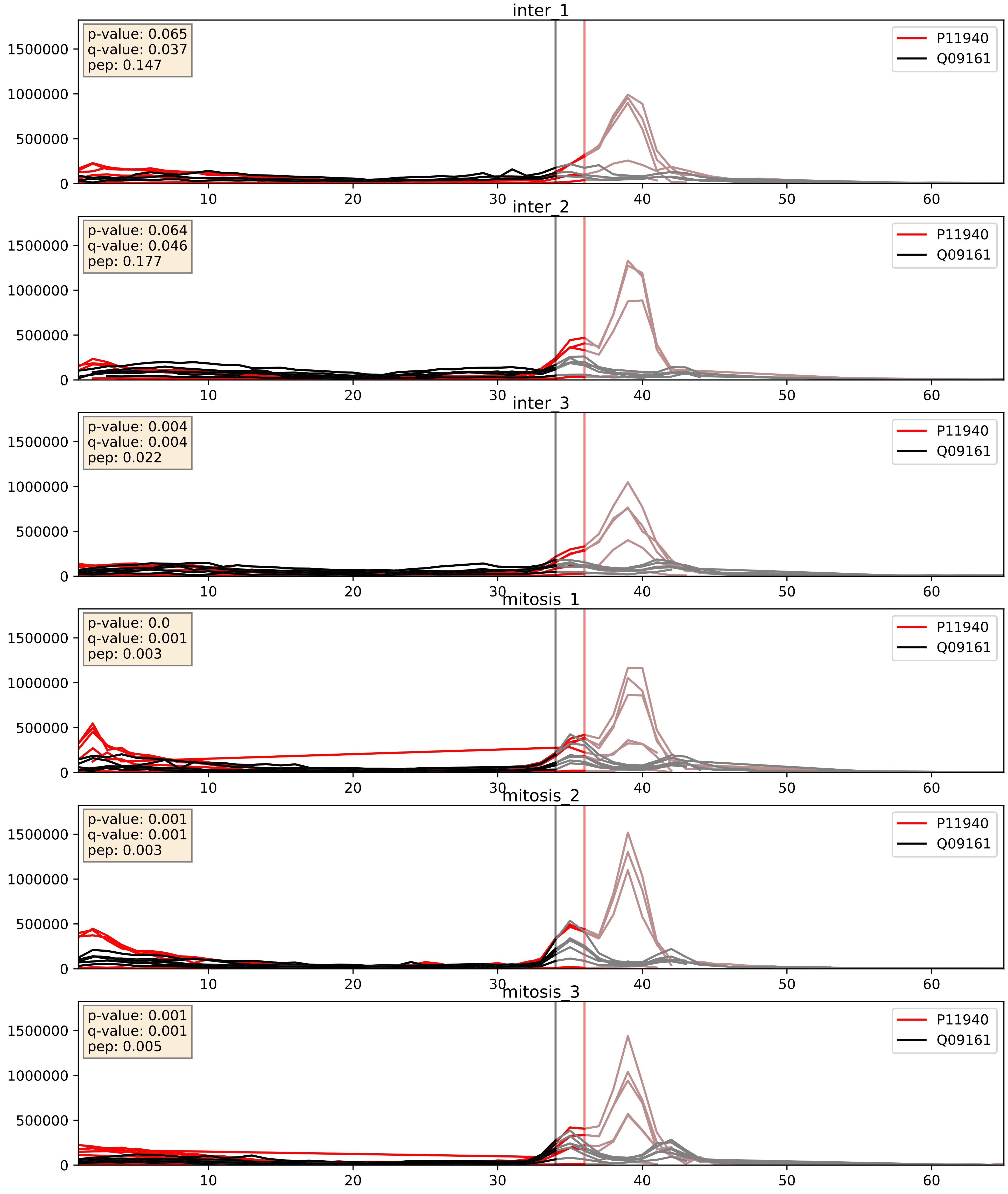


| level | condition_1 | condition_2 | pvalue | pvalue_adjusted |
|---|---|---|---|---|
| interactor_abundance | mitosis | inter | 0.24897924340370764 | 0.4426297660510358 |
| interactor_ratio | mitosis | inter | 0.3349887607442168 | 0.7914334552464135 |
| complex_abundance | mitosis | inter | 0.8602503058949373 | 0.911326116178704 |

P11940_Q10570
PABP1_HUMAN vs CPSF1_HUMAN
p-value: 0.0 q-value: 0.018

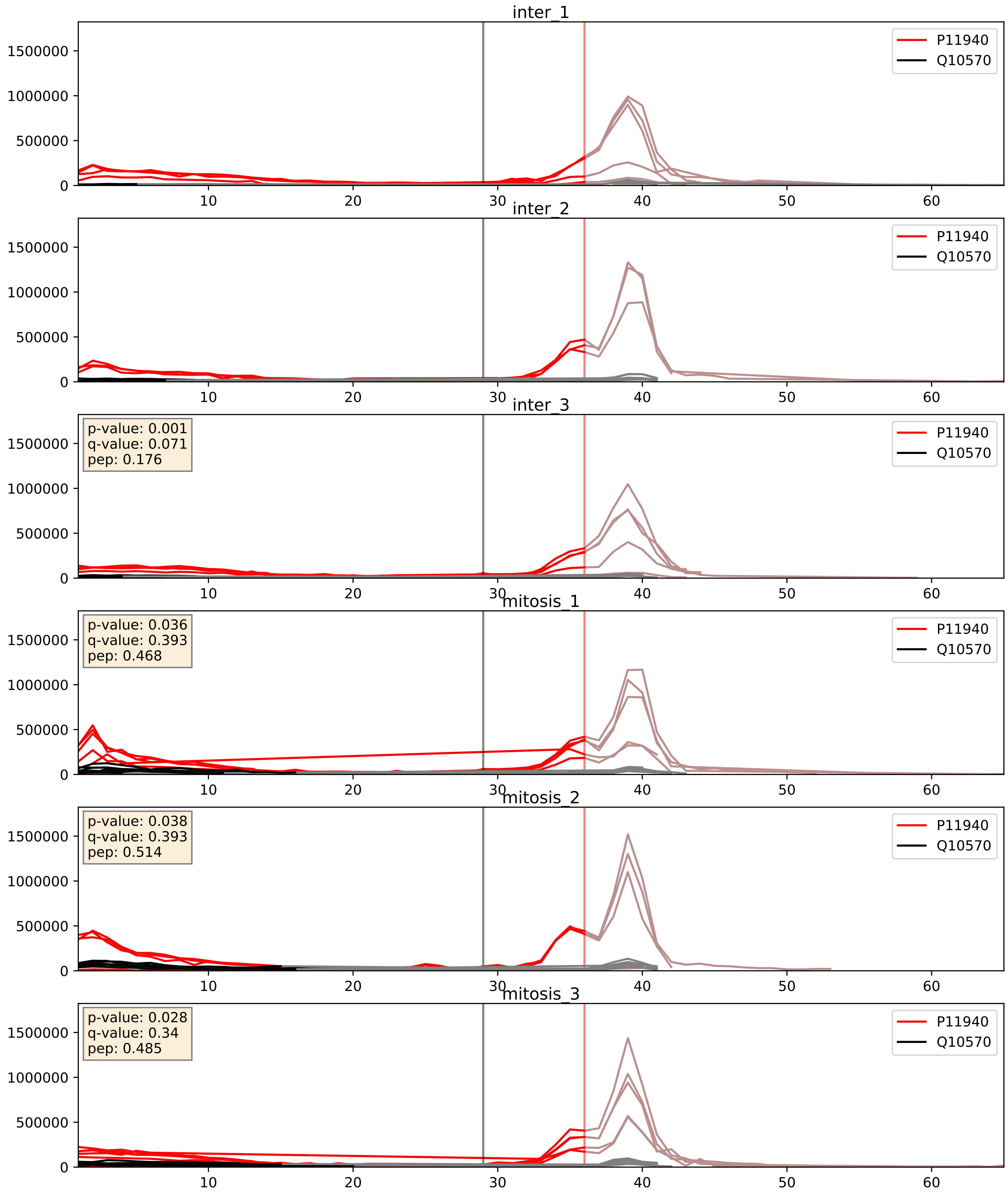


| level | condition_1 | condition_2 | pvalue | pvalue_adjusted |
|---|---|---|---|---|
| complex_abundance | mitosis | inter | 0.002346382940780768 | 0.035168510428347066 |
| interactor_abundance | mitosis | inter | 0.033644226089265476 | 0.15318860389580452 |
| interactor_ratio | mitosis | inter | 0.1927880363525544 | 0.6660896702330326 |

P11940_Q13247
PABP1_HUMAN vs SRSF6_HUMAN
p-value: 0.0 q-value: 0.047

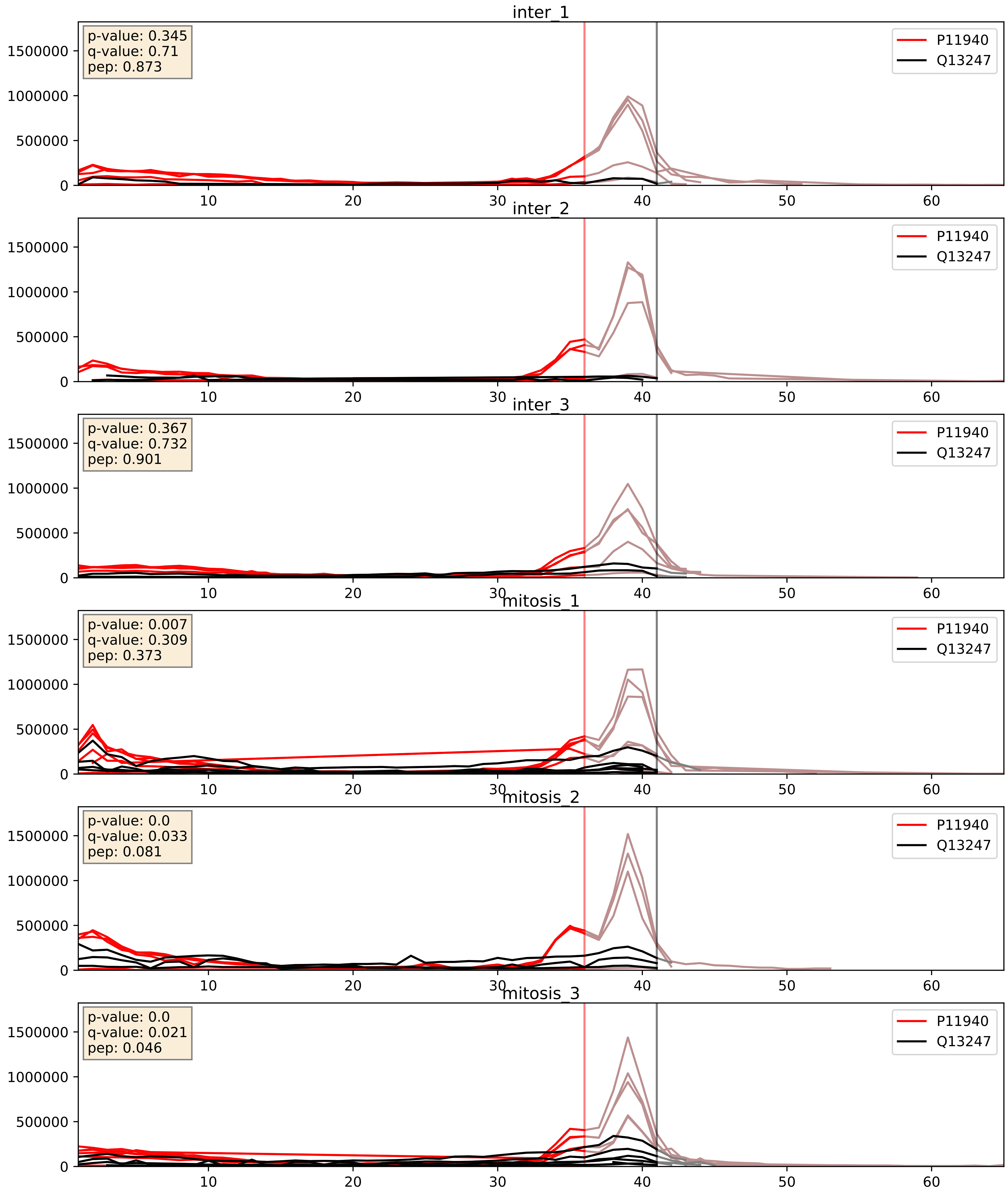


| level | condition_1 | condition_2 | pvalue | pvalue_adjusted |
|---|---|---|---|---|
| complex_abundance | mitosis | inter | 0.007836804827711203 | 0.0657361191517291 |
| interactor_abundance | mitosis | inter | 0.008345387686835815 | 0.07950642025521934 |
| interactor_ratio | mitosis | inter | 0.04176762891071437 | 0.3395355208696249 |

P11940_Q15365
PABP1_HUMAN vs PCBP1_HUMAN
p-value: 0.0 q-value: 0.023

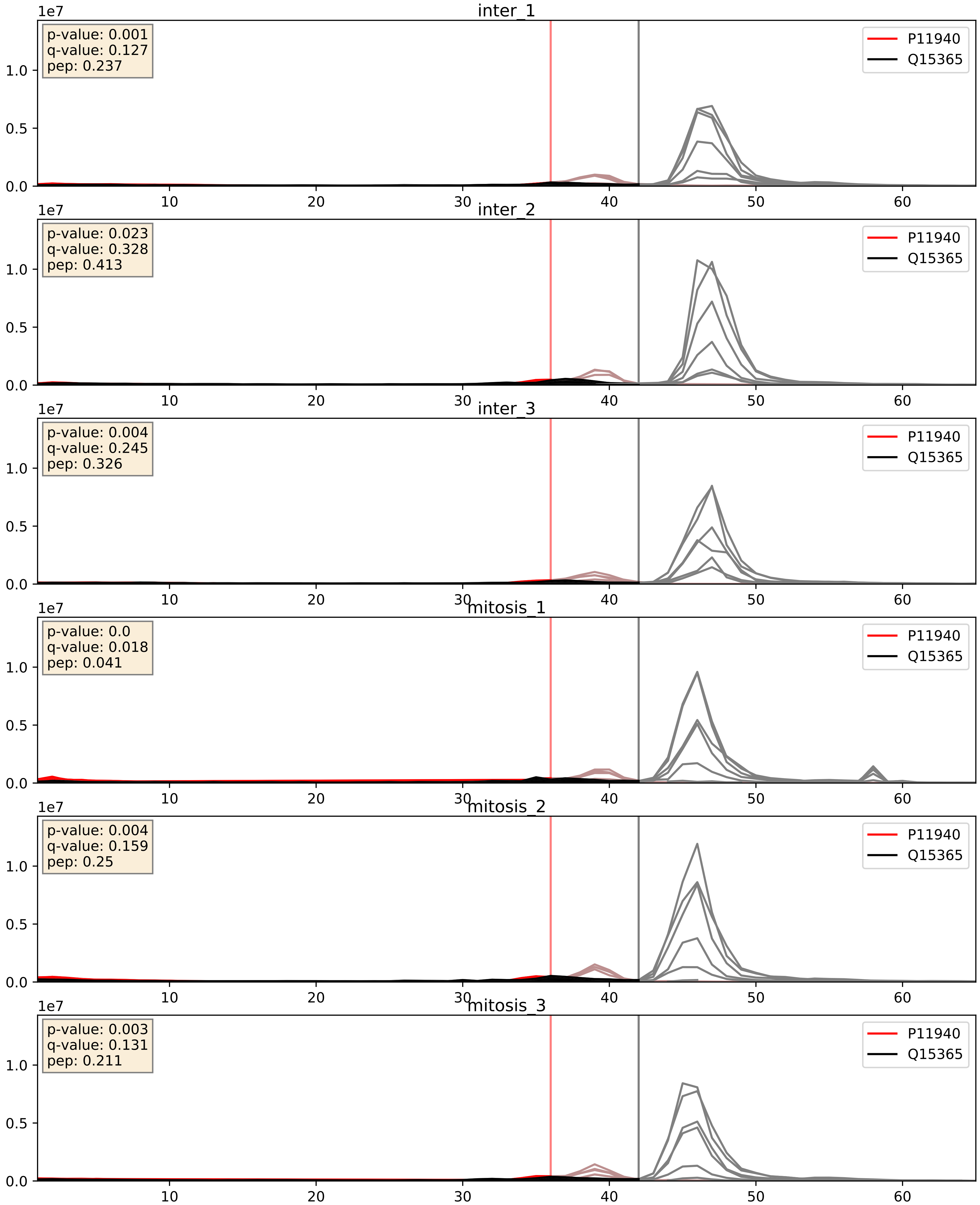


| level | condition_1 | condition_2 | pvalue | pvalue_adjusted |
|---|---|---|---|---|
| interactor_ratio | mitosis | inter | 0.06976990871901345 | 0.4375075131827156 |
| interactor_abundance | mitosis | inter | 0.15090604372175434 | 0.33732834796845007 |
| complex_abundance | mitosis | inter | 0.29680960341908563 | 0.4463615961467627 |

P11940_Q15366
PABP1_HUMAN vs PCBP2_HUMAN
p-value: 0.0 q-value: 0.026

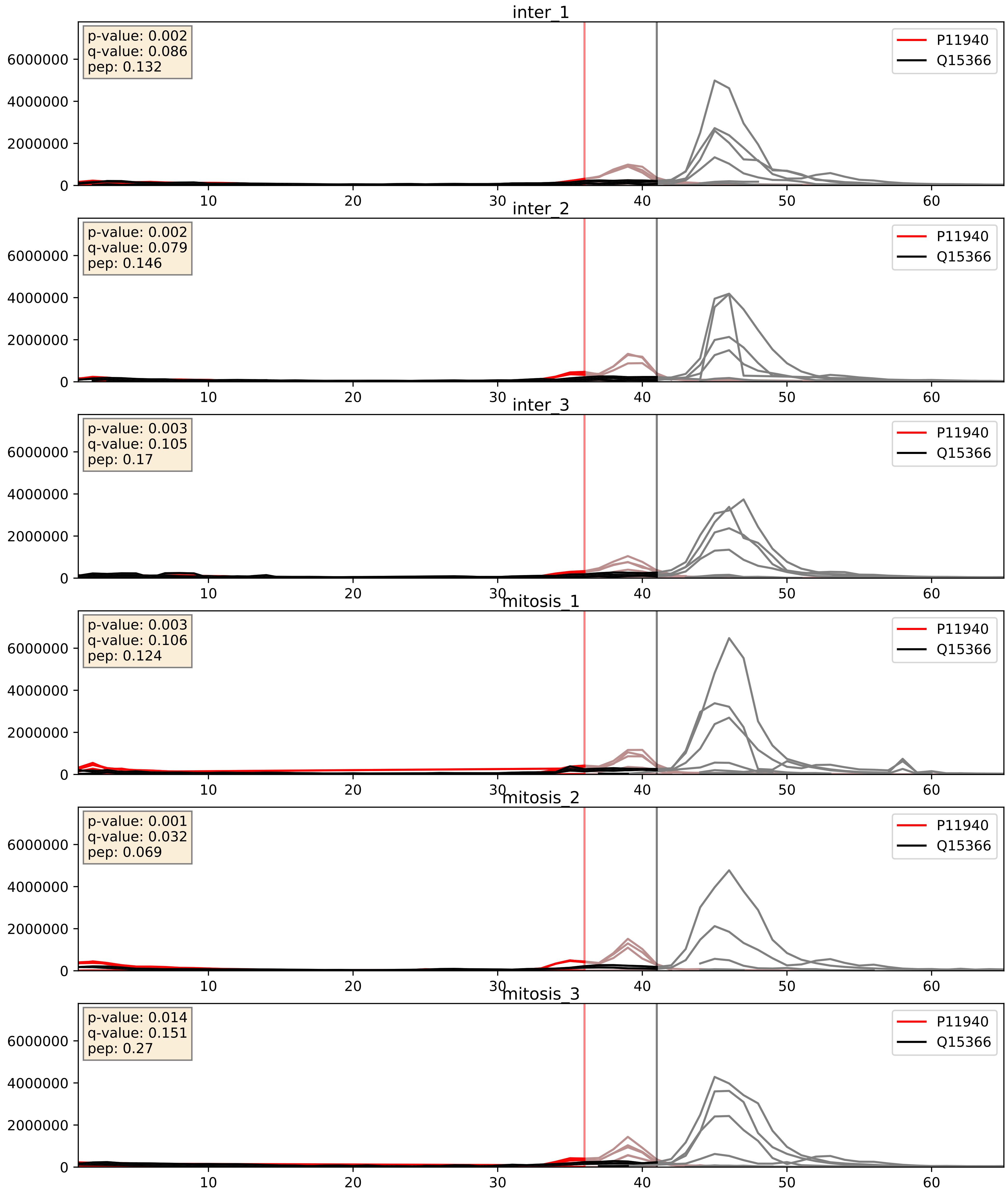


| level | condition_1 | condition_2 | pvalue | pvalue_adjusted |
|---|---|---|---|---|
| interactor_abundance | mitosis | inter | 0.15090604372175434 | 0.33732834796845007 |
| interactor_ratio | mitosis | inter | 0.3932777098083351 | 0.8261244652660977 |
| complex_abundance | mitosis | inter | 0.7295233741981375 | 0.8199474899075705 |

P11940_Q9H074
PABP1_HUMAN vs PAIP1_HUMAN
p-value: 0.03 q-value: 0.022

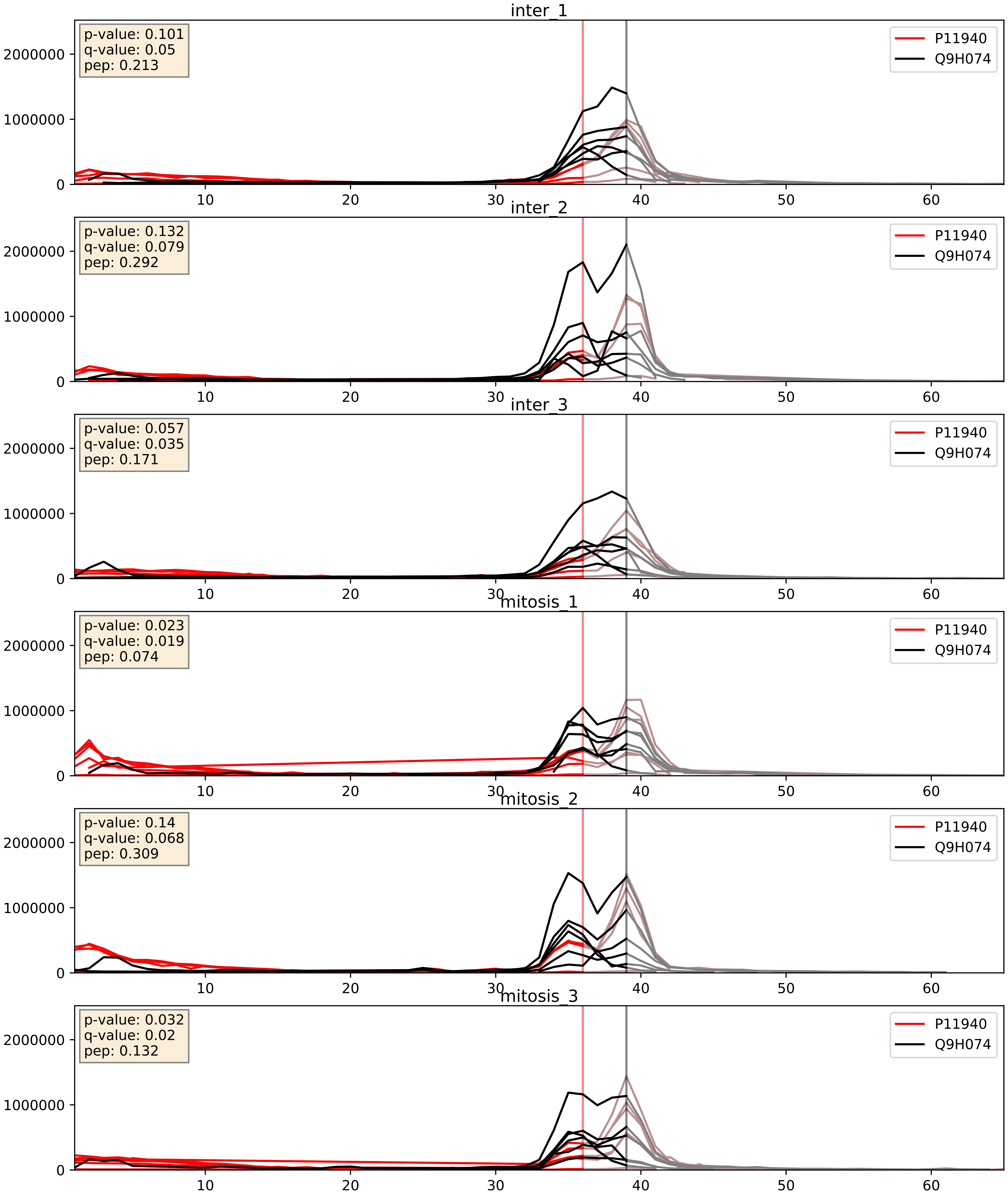


| level | condition_1 | condition_2 | pvalue | pvalue_adjusted |
|---|---|---|---|---|
| interactor_abundance | mitosis | inter | 0.45008031391935505 | 0.623412214749139 |
| interactor_ratio | mitosis | inter | 0.5553960484025842 | 0.8898061992395188 |
| complex_abundance | mitosis | inter | 0.7987628326707792 | 0.869346452340988 |
