## Supplementary Data 2 for "SECAT: Quantifying differential protein-protein interaction states by network-centric analysis": hela_string_000188_P32856.pdf

# P32856_P32856

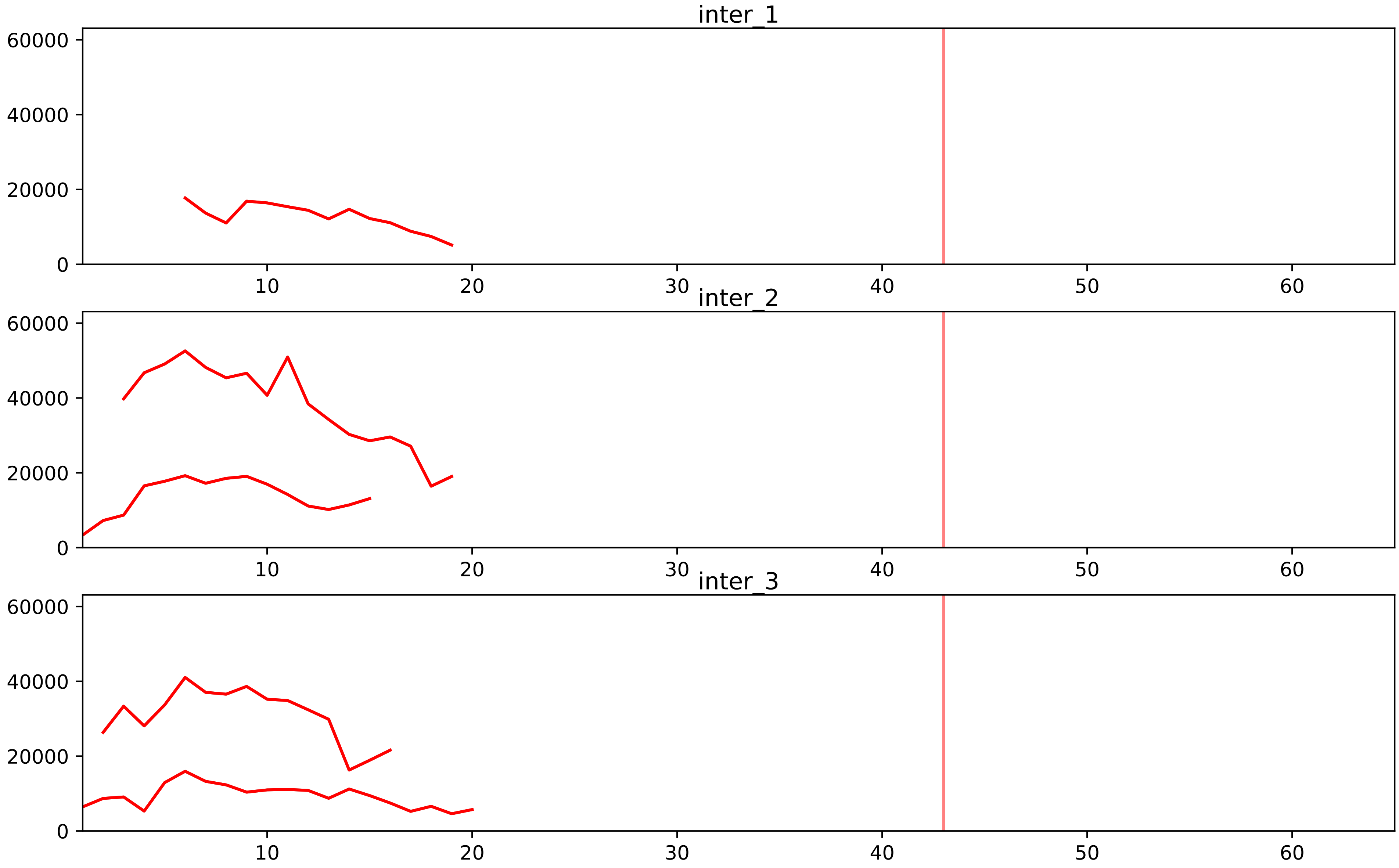


| level | condition_1 | condition_2 | pvalue | pvalue_adjusted |
|---|---|---|---|---|
| interactor_ratio | mitosis | inter | 0.00018478437099130174 | 0.002809944153917316 |
| complex_abundance | mitosis | inter | 0.001944958991355377 | 0.016491818175547898 |
| interactor_abundance | mitosis | inter | 0.004848513563914916 | 0.03328830208060987 |
| assembled_abundance | mitosis | inter | 0.00912024234292935 | 0.10420259961601182 |
| total_abundance | mitosis | inter | 0.011196062581486393 | 0.1296327302866994 |

O00161_P32856
SNP23_HUMAN vs STX2_HUMAN
p-value: 0.031 q-value: 0.022

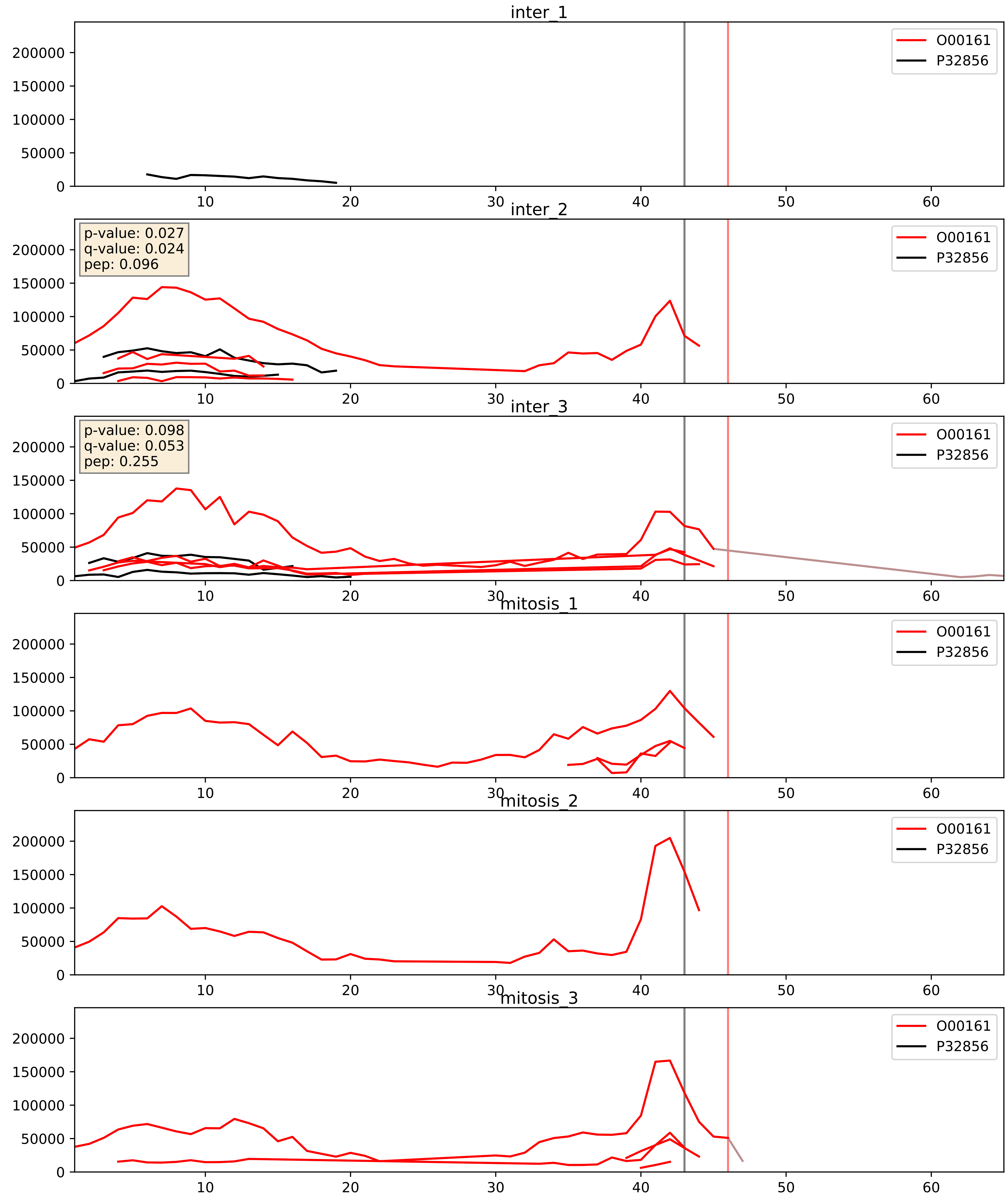


| level | condition_1 | condition_2 | pvalue | pvalue_adjusted |
|---|---|---|---|---|
| complex_abundance | mitosis | inter | 0.02589525843373415 | 0.11798182031988799 |
| interactor_abundance | mitosis | inter | 0.028138380322048524 | 0.13971260763151702 |
| interactor_ratio | mitosis | inter | 0.04420741685804907 | 0.3479294458487465 |

P32856_Q9UNK0
STX2_HUMAN vs STX8_HUMAN
p-value: 0.0 q-value: 0.026

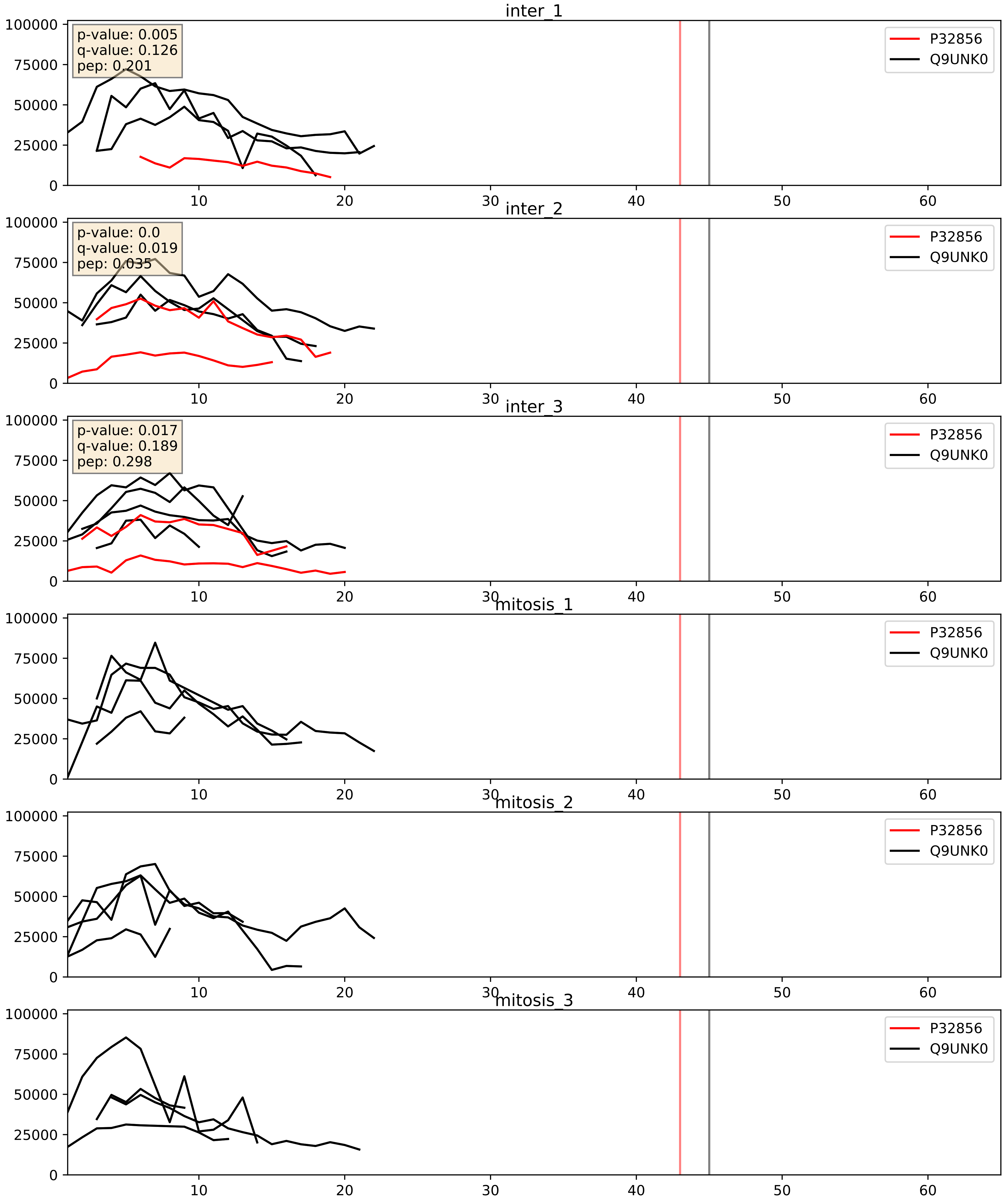


| level | condition_1 | condition_2 | pvalue | pvalue_adjusted |
|---|---|---|---|---|
| interactor_ratio | mitosis | inter | 0.0003457759019263659 | 0.019998930543849273 |
| complex_abundance | mitosis | inter | 0.0012589206953757875 | 0.0241081904975766 |
| interactor_abundance | mitosis | inter | 0.005396032441284355 | 0.06191694061312879 |
