## Supplementary Data 2 for "SECAT: Quantifying differential protein-protein interaction states by network-centric analysis": hela_string_000189_Q6UN15.pdf

# Q6UN15_Q6UN15

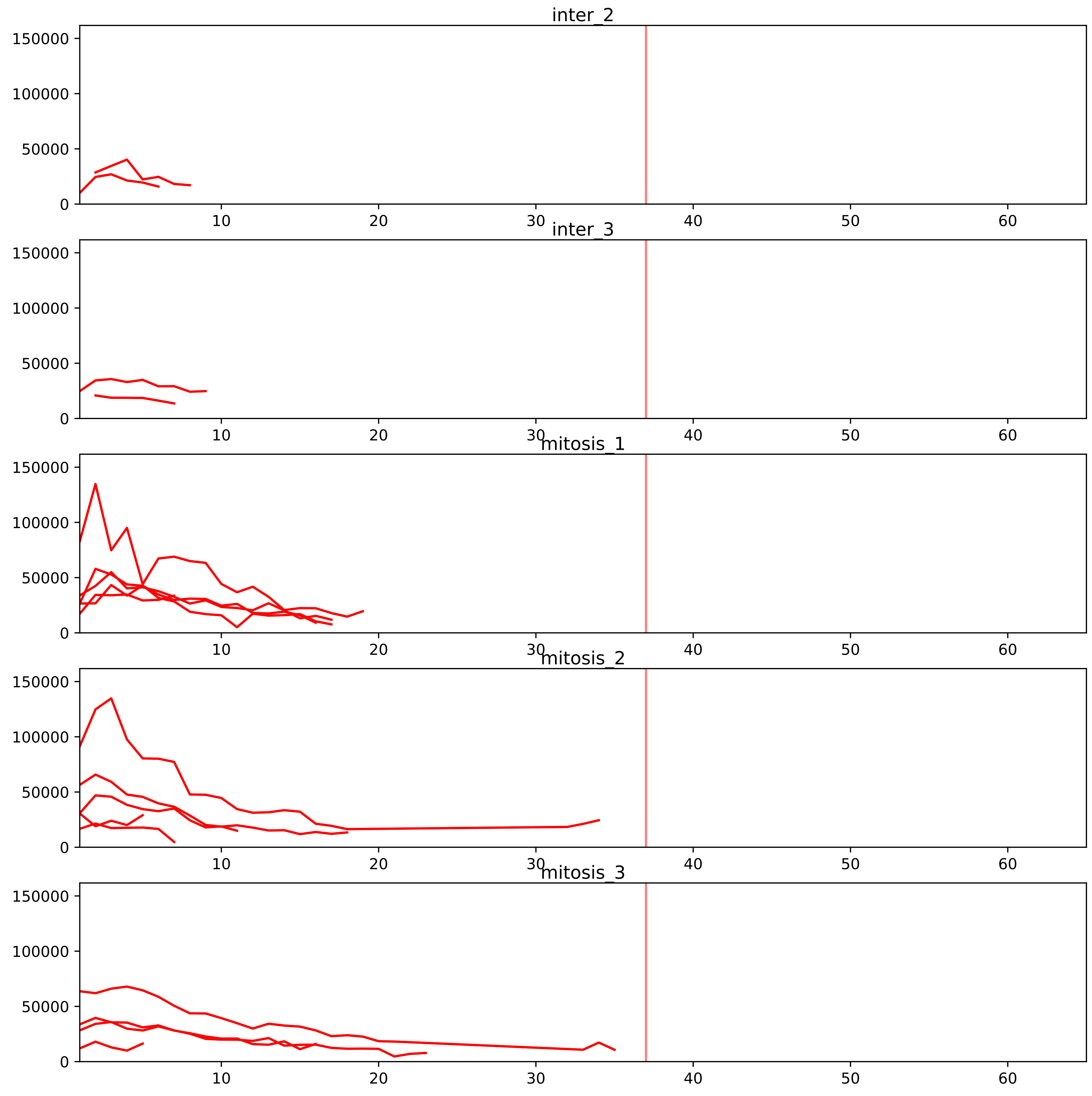


| level | condition_1 | condition_2 | pvalue | pvalue_adjusted |
|---|---|---|---|---|
| interactor_abundance | mitosis | inter | 0.00018623259479421493 | 0.00351159104032013 |
| interactor_ratio | mitosis | inter | 0.00037973060024136426 | 0.004955349676908583 |
| complex_abundance | mitosis | inter | 0.0011247098999744589 | 0.011497034533072246 |
| total_abundance | mitosis | inter | 0.005249264696182071 | 0.08323875476801054 |
| assembled_abundance | mitosis | inter | 0.0075553563468345394 | 0.09331793469809328 |

O75934_Q6UN15
SPF27_HUMAN vs FIP1_HUMAN
p-value: 0.0 q-value: 0.033

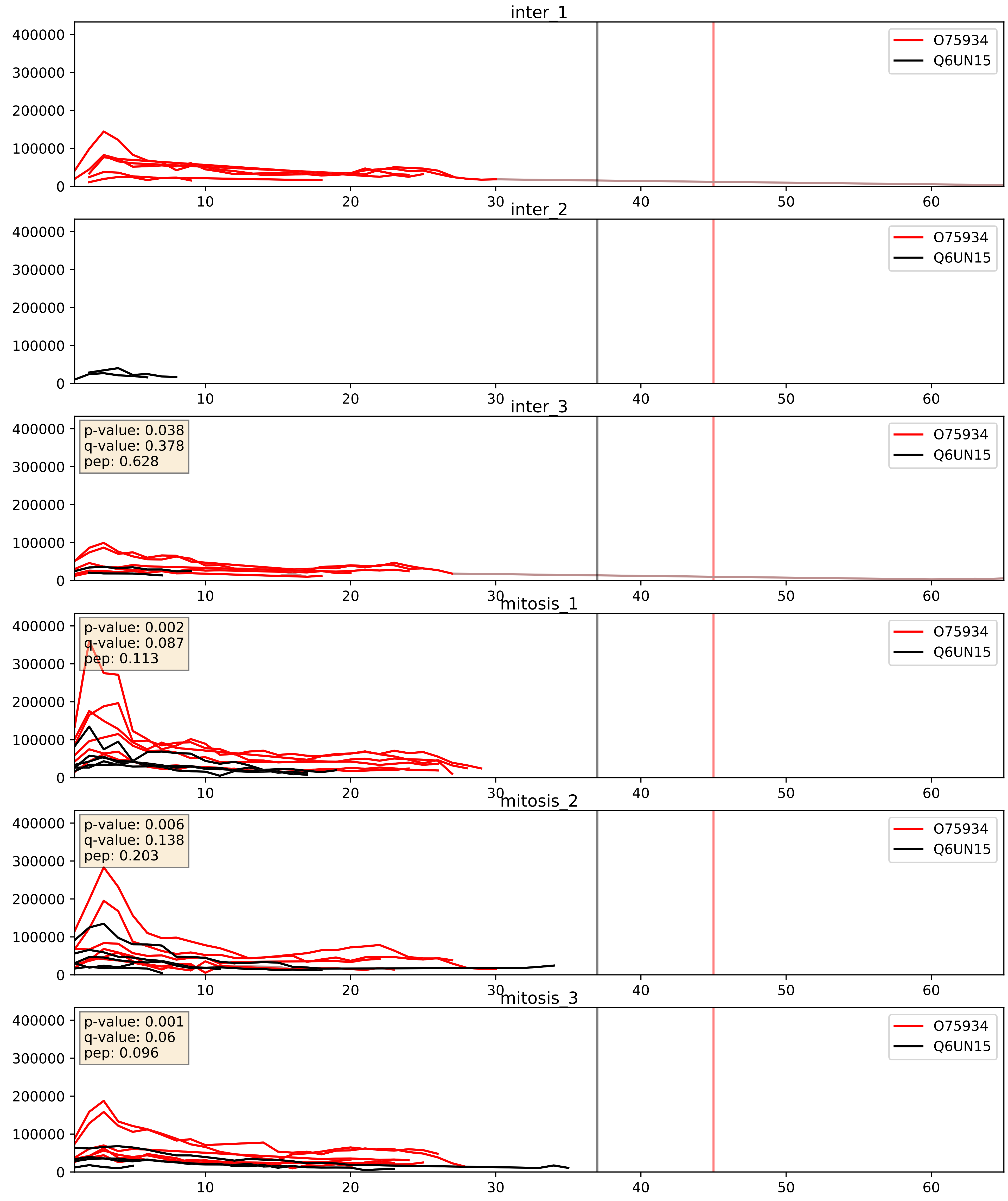


| level | condition_1 | condition_2 | pvalue | pvalue_adjusted |
|---|---|---|---|---|
| complex_abundance | mitosis | inter | 0.007783987211497894 | 0.06557020805367884 |
| interactor_abundance | mitosis | inter | 0.014772120850023046 | 0.10634933093035936 |
| interactor_ratio | mitosis | inter | 0.08024151017262679 | 0.47370160488116225 |

O95639_Q6UN15
CPSF4_HUMAN vs FIP1_HUMAN
p-value: 0.01 q-value: 0.009

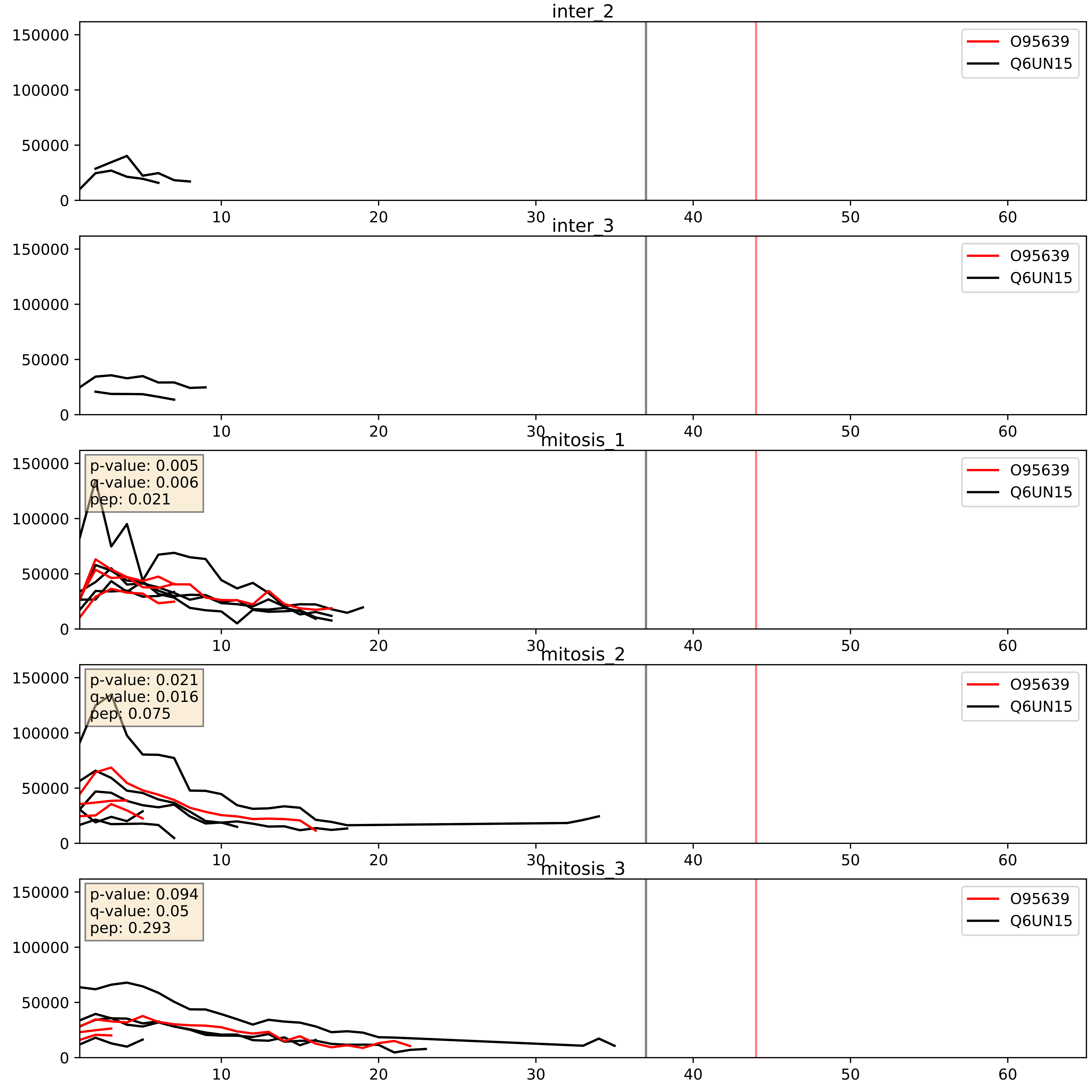


| level | condition_1 | condition_2 | pvalue | pvalue_adjusted |
|---|---|---|---|---|
| interactor_ratio | mitosis | inter | 0.00532389414285382 | 0.09758572561633554 |
| complex_abundance | mitosis | inter | 0.014845937217029401 | 0.09024336091715261 |
| interactor_abundance | mitosis | inter | 0.09608112571717416 | 0.26586534221400054 |

Q10570_Q6UN15
CPSF1_HUMAN vs FIP1_HUMAN
p-value: 0.001 q-value: 0.001

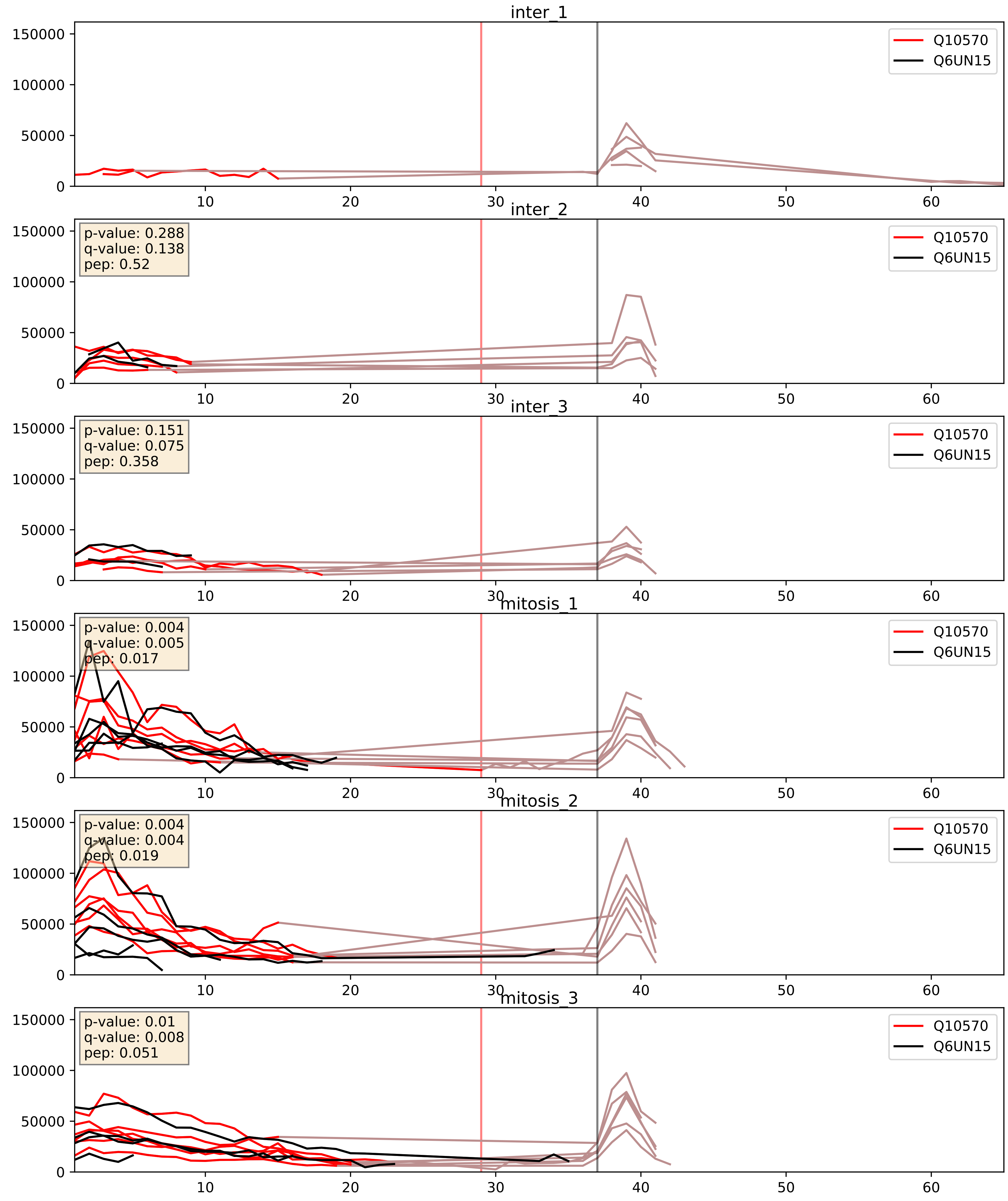


| level | condition_1 | condition_2 | pvalue | pvalue_adjusted |
|---|---|---|---|---|
| interactor_abundance | mitosis | inter | 0.00831688214137242 | 0.0793231321784378 |
| complex_abundance | mitosis | inter | 0.01014764983870757 | 0.07446889582511722 |
| interactor_ratio | mitosis | inter | 0.05863148374789707 | 0.39980872848262694 |

Q6UN15_Q92797
FIP1_HUMAN vs SYMPK_HUMAN
p-value: 0.025 q-value: 0.019

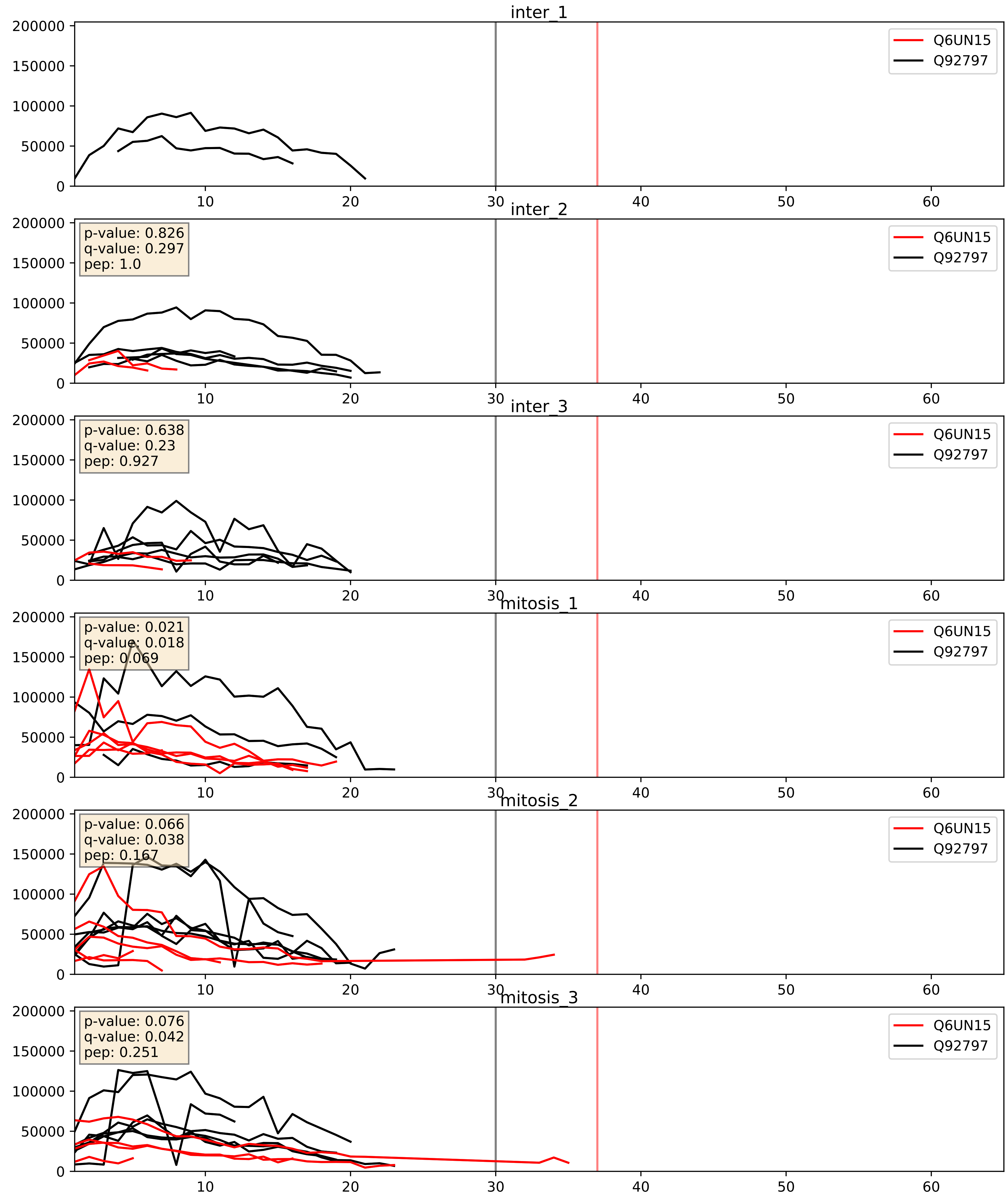


| level | condition_1 | condition_2 | pvalue | pvalue_adjusted |
|---|---|---|---|---|
| interactor_abundance | mitosis | inter | 0.0147508066463817 | 0.10628527347897926 |
| complex_abundance | mitosis | inter | 0.02263477068258683 | 0.11008729377439957 |
| interactor_ratio | mitosis | inter | 0.3550925199633546 | 0.8029332751290026 |

Q6UN15_Q9C0J8
FIP1_HUMAN vs WDR33_HUMAN
p-value: 0.065 q-value: 0.04

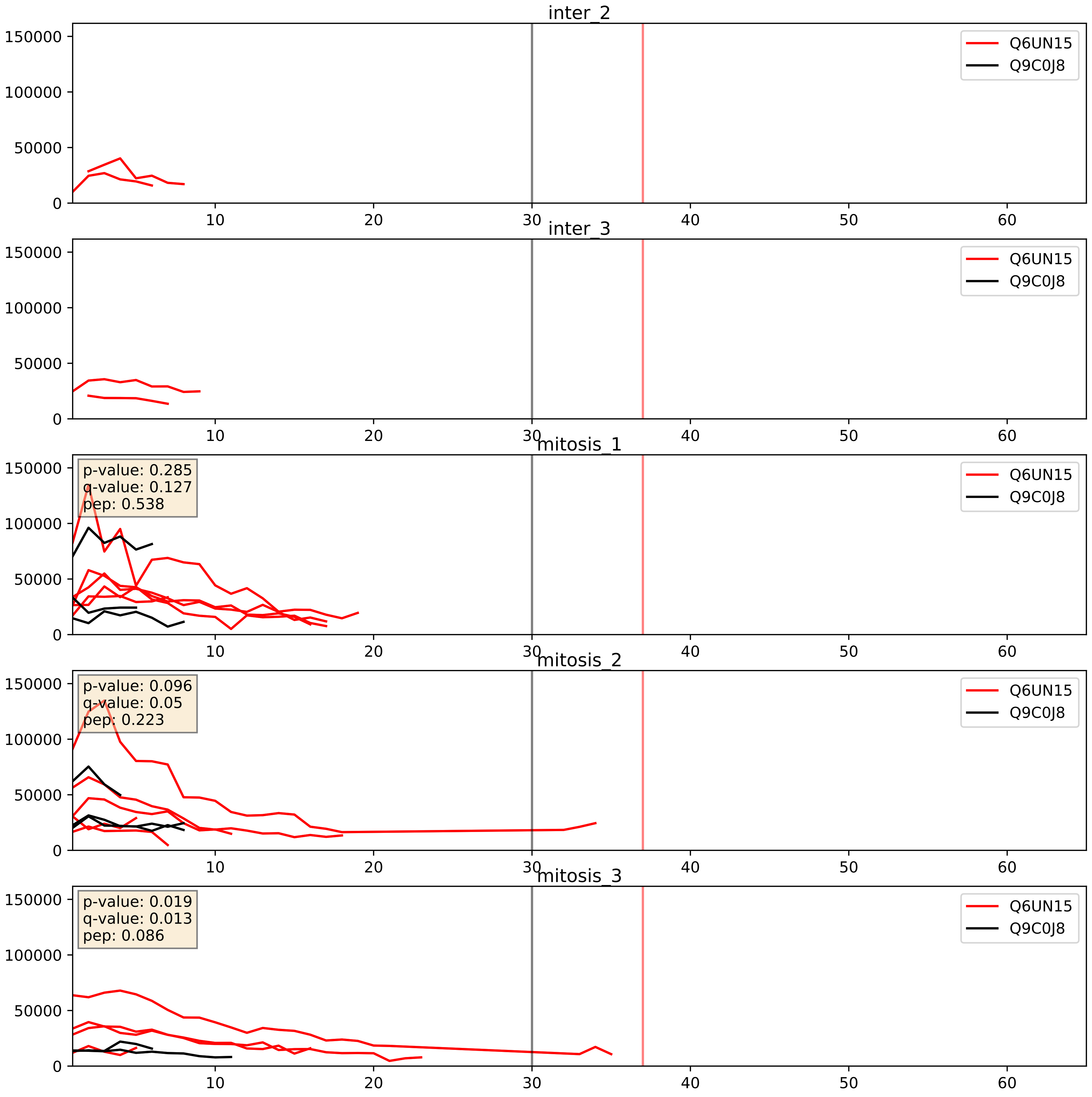


| level | condition_1 | condition_2 | pvalue | pvalue_adjusted |
|---|---|---|---|---|
| interactor_abundance | mitosis | inter | 0.005842898338158808 | 0.06478654115885933 |
| interactor_ratio | mitosis | inter | 0.03948628120844784 | 0.32783954136208876 |
| complex_abundance | mitosis | inter | 0.06924960068840534 | 0.19065640345538643 |

Q6UN15_Q9P2I0
FIP1_HUMAN vs CPSF2_HUMAN
p-value: 0.007 q-value: 0.006

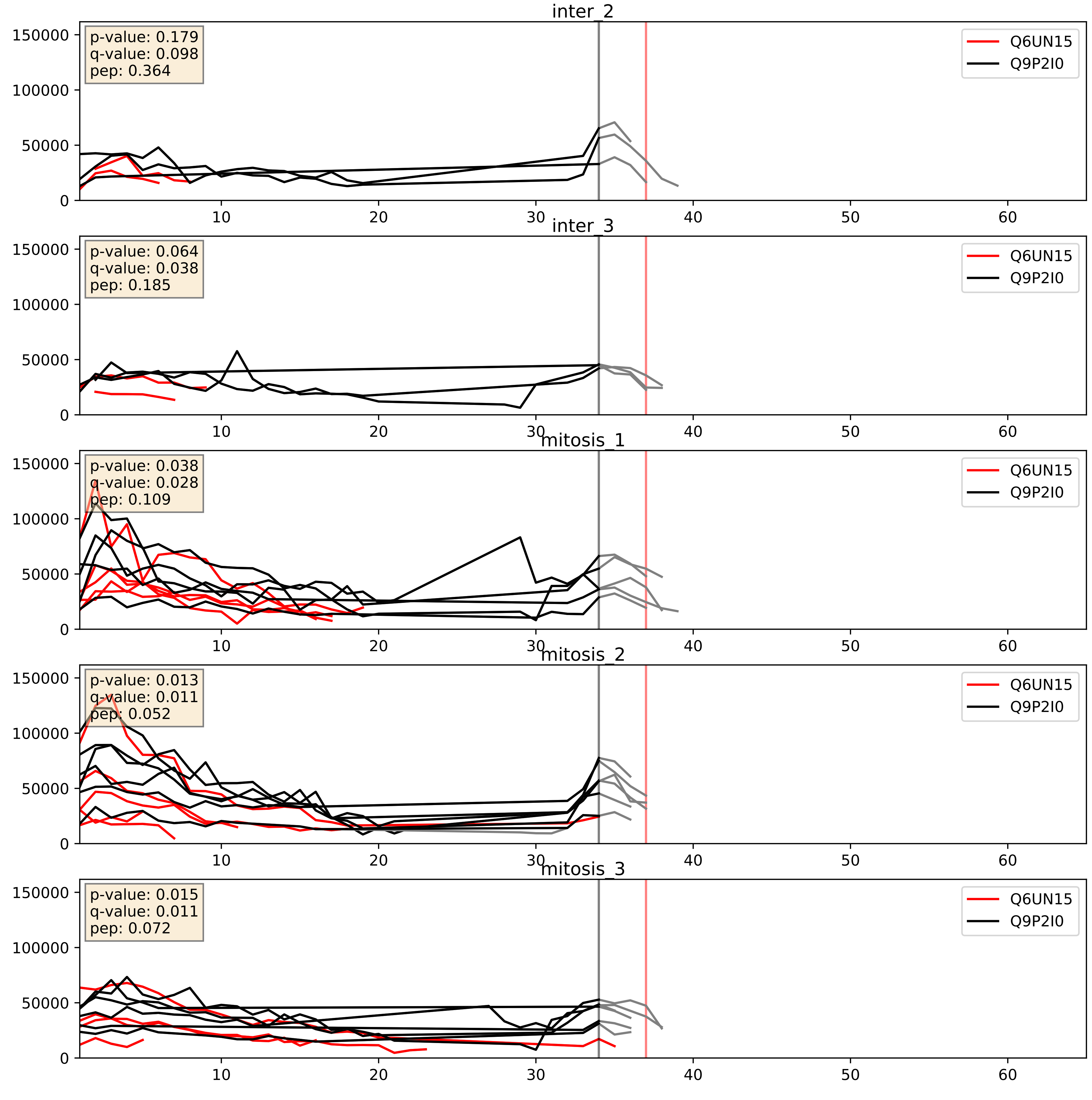


| level | condition_1 | condition_2 | pvalue | pvalue_adjusted |
|---|---|---|---|---|
| complex_abundance | mitosis | inter | 0.007845649398103807 | 0.0657361191517291 |
| interactor_abundance | mitosis | inter | 0.013852354432628004 | 0.10436859054266079 |
| interactor_ratio | mitosis | inter | 0.015100829020357177 | 0.18053505085790145 |

Q6UN15_Q9UKF6
FIP1_HUMAN vs CPSF3_HUMAN
p-value: 0.014 q-value: 0.012

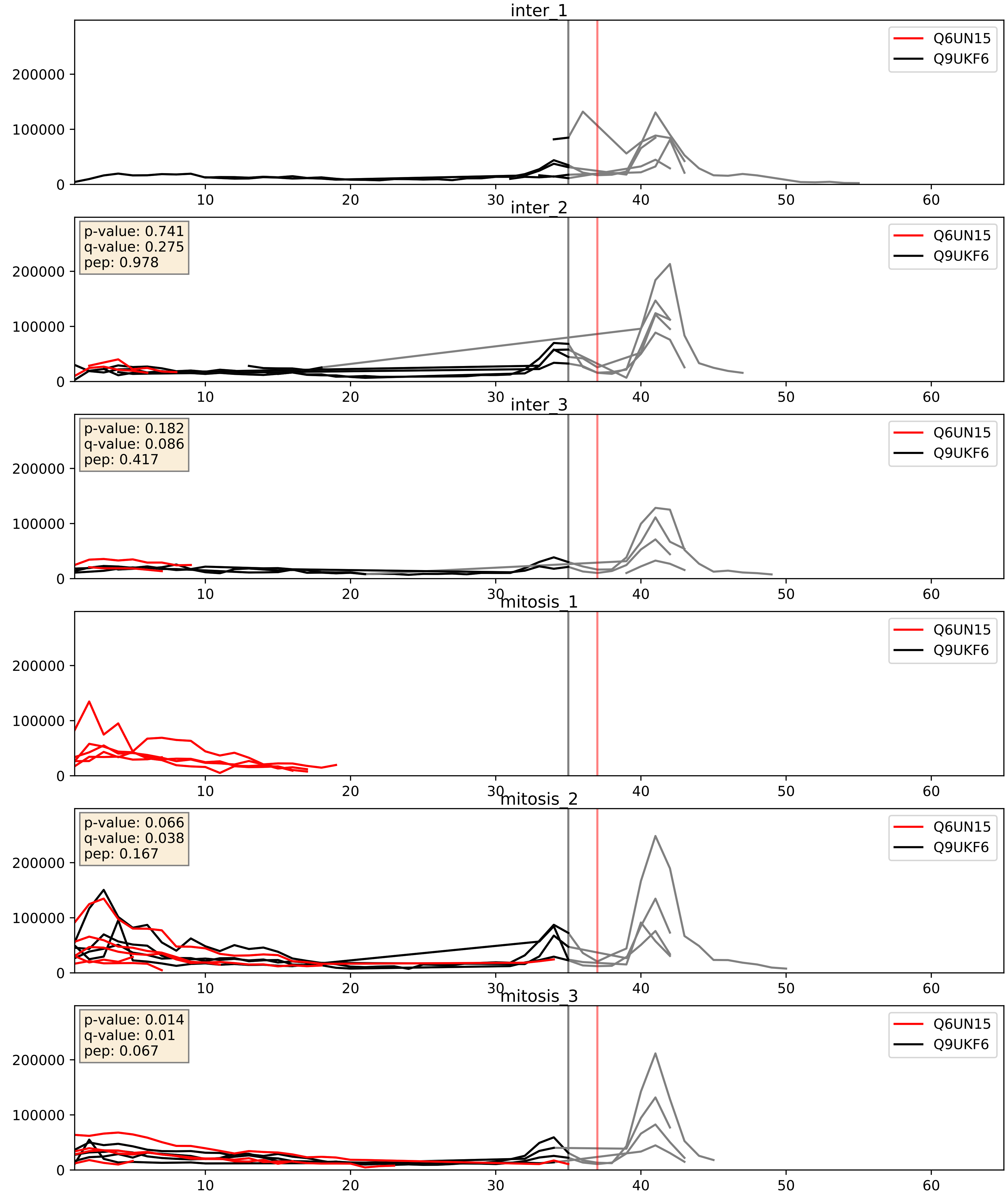


| level | condition_1 | condition_2 | pvalue | pvalue_adjusted |
|---|---|---|---|---|
| interactor_abundance | mitosis | inter | 0.6168012841297559 | 0.7473349143403655 |
| complex_abundance | mitosis | inter | 0.6779558801611931 | 0.7850787789745418 |
| interactor_ratio | mitosis | inter | 0.6781468507185926 | 0.9338400552004106 |
