## Supplementary Data 2 for "SECAT: Quantifying differential protein-protein interaction states by network-centric analysis": hela_string_000190_P56556.pdf

# P56556_P56556

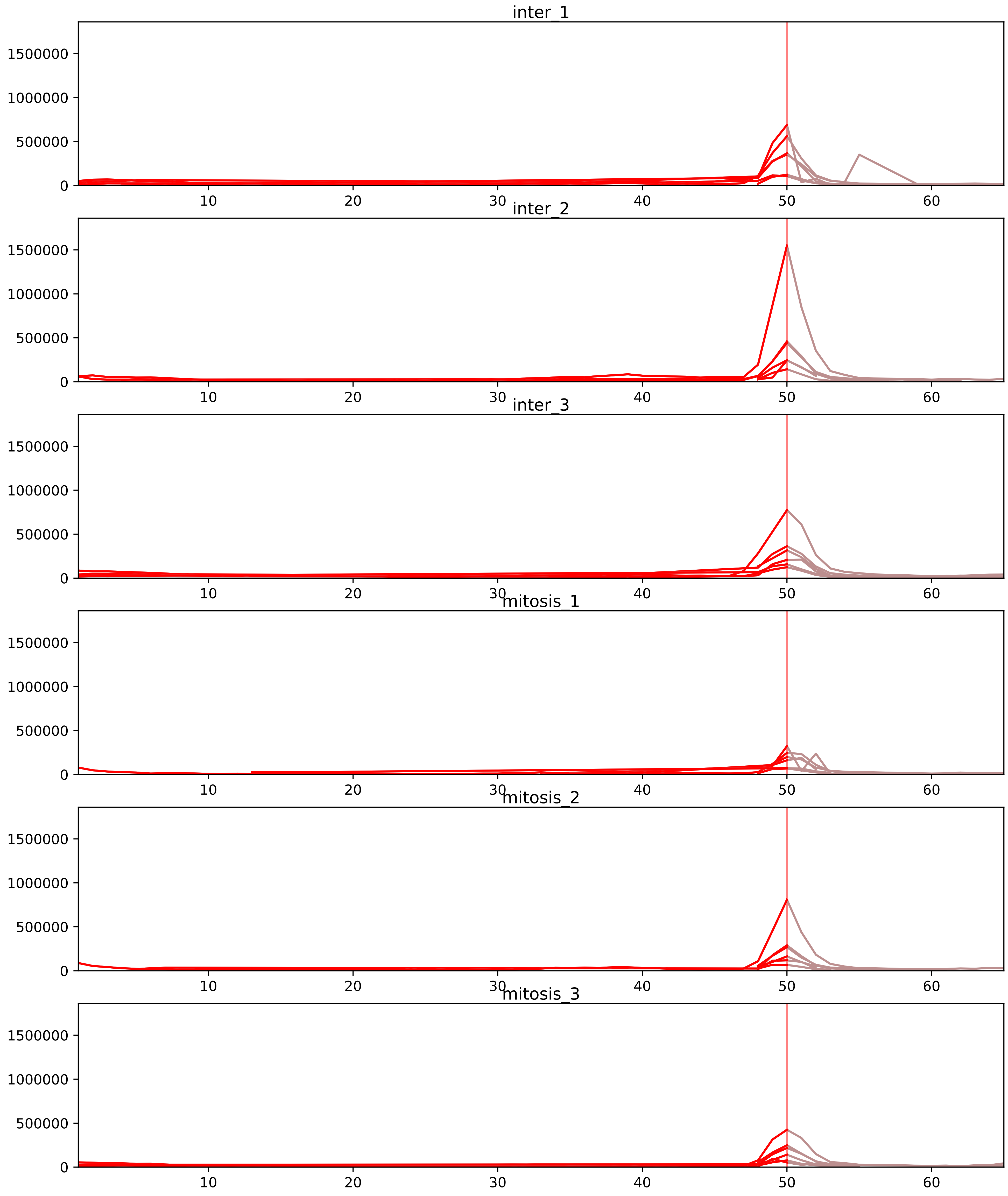


| level | condition_1 | condition_2 | pvalue | pvalue_adjusted |
|---|---|---|---|---|
| interactor_abundance | mitosis | inter | 0.0001932975922556264 | 0.0035926017146500257 |
| assembled_abundance | mitosis | inter | 0.0004479296613016665 | 0.0195958761173187 |
| complex_abundance | mitosis | inter | 0.0005081177350802482 | 0.006925456537390048 |
| total_abundance | mitosis | inter | 0.007144123284230723 | 0.0991028110082251 |
| interactor_ratio | mitosis | inter | 0.00987279082201042 | 0.062425893857385464 |
| monomer_abundance | mitosis | inter | 0.05793986928621803 | 0.3497872727142794 |

O14949_P56556
QCR8_HUMAN vs NDUA6_HUMAN
p-value: 0.077 q-value: 0.046

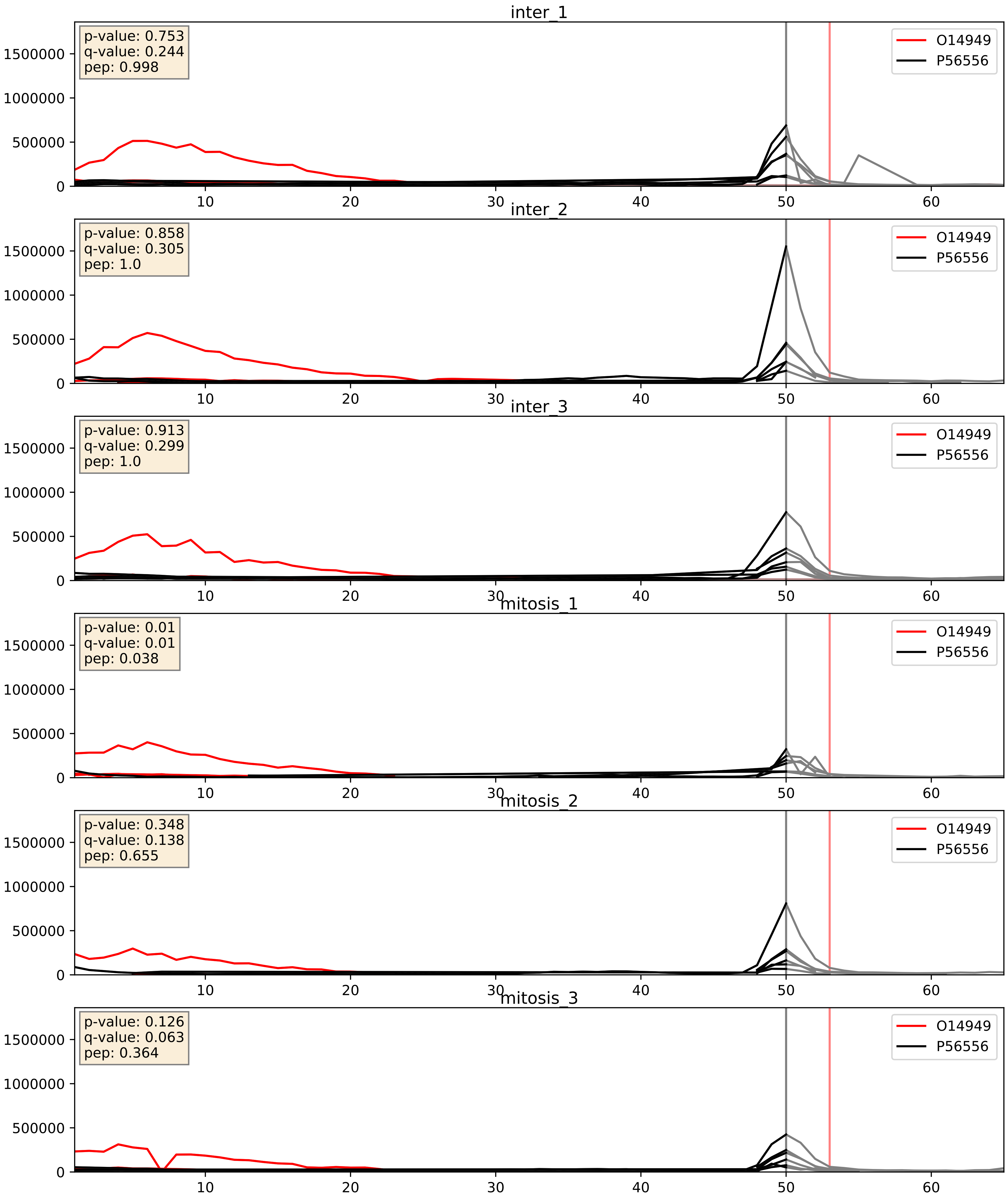


| level | condition_1 | condition_2 | pvalue | pvalue_adjusted |
|---|---|---|---|---|
| complex_abundance | mitosis | inter | 0.015264095359260698 | 0.09174685245214603 |
| interactor_abundance | mitosis | inter | 0.07386499194203666 | 0.22673988198390962 |
| interactor_ratio | mitosis | inter | 0.31467001995200305 | 0.7783421708721073 |

O43678_P56556
NDUA2_HUMAN vs NDUA6_HUMAN
p-value: 0.063 q-value: 0.039

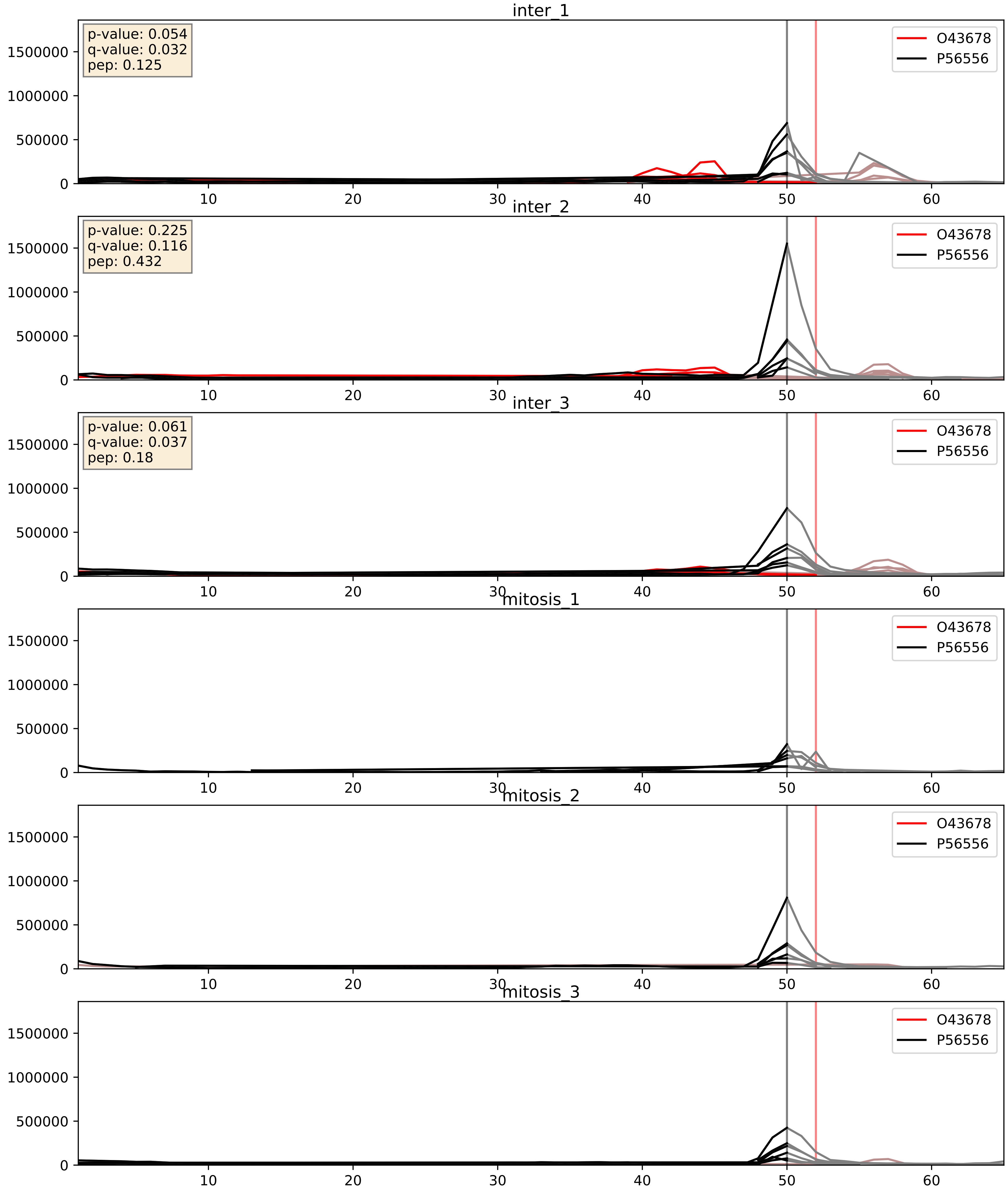


| level | condition_1 | condition_2 | pvalue | pvalue_adjusted |
|---|---|---|---|---|
| complex_abundance | mitosis | inter | 0.0010427131847453482 | 0.02161168247317235 |
| interactor_abundance | mitosis | inter | 0.005906515623201406 | 0.06511239373419707 |
| interactor_ratio | mitosis | inter | 0.15173525543561164 | 0.6067576723168497 |

O95299_P56556
NDUAA_HUMAN vs NDUA6_HUMAN
p-value: 0.064 q-value: 0.04

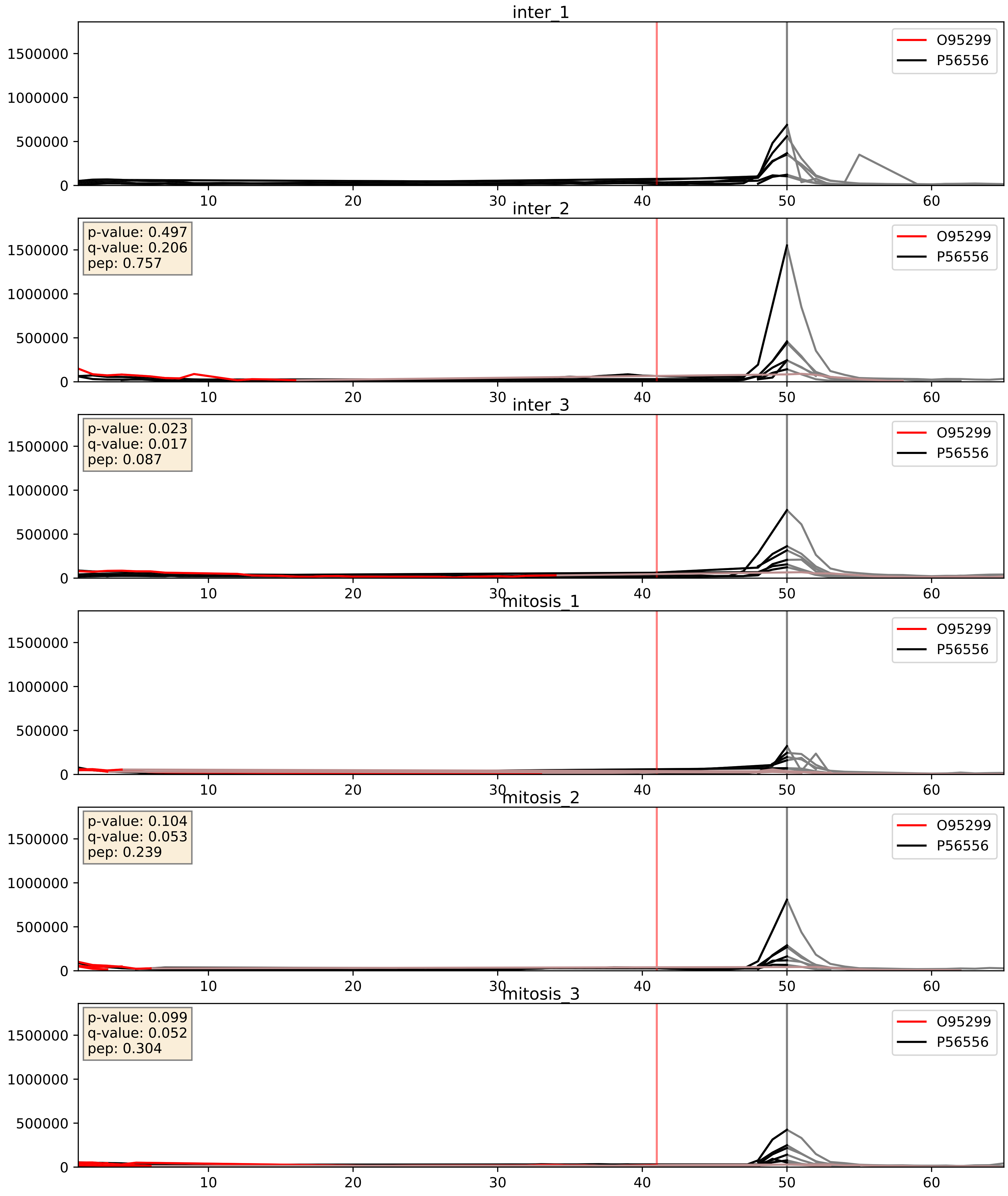


| level | condition_1 | condition_2 | pvalue | pvalue_adjusted |
|---|---|---|---|---|
| interactor_ratio | mitosis | inter | 0.03480054501247816 | 0.3021223786073155 |
| complex_abundance | mitosis | inter | 0.19508506941431797 | 0.3473950892836617 |
| interactor_abundance | mitosis | inter | 0.48614247031883095 | 0.6556967692317329 |

P20674_P56556
COX5A_HUMAN vs NDUA6_HUMAN
p-value: 0.079 q-value: 0.047

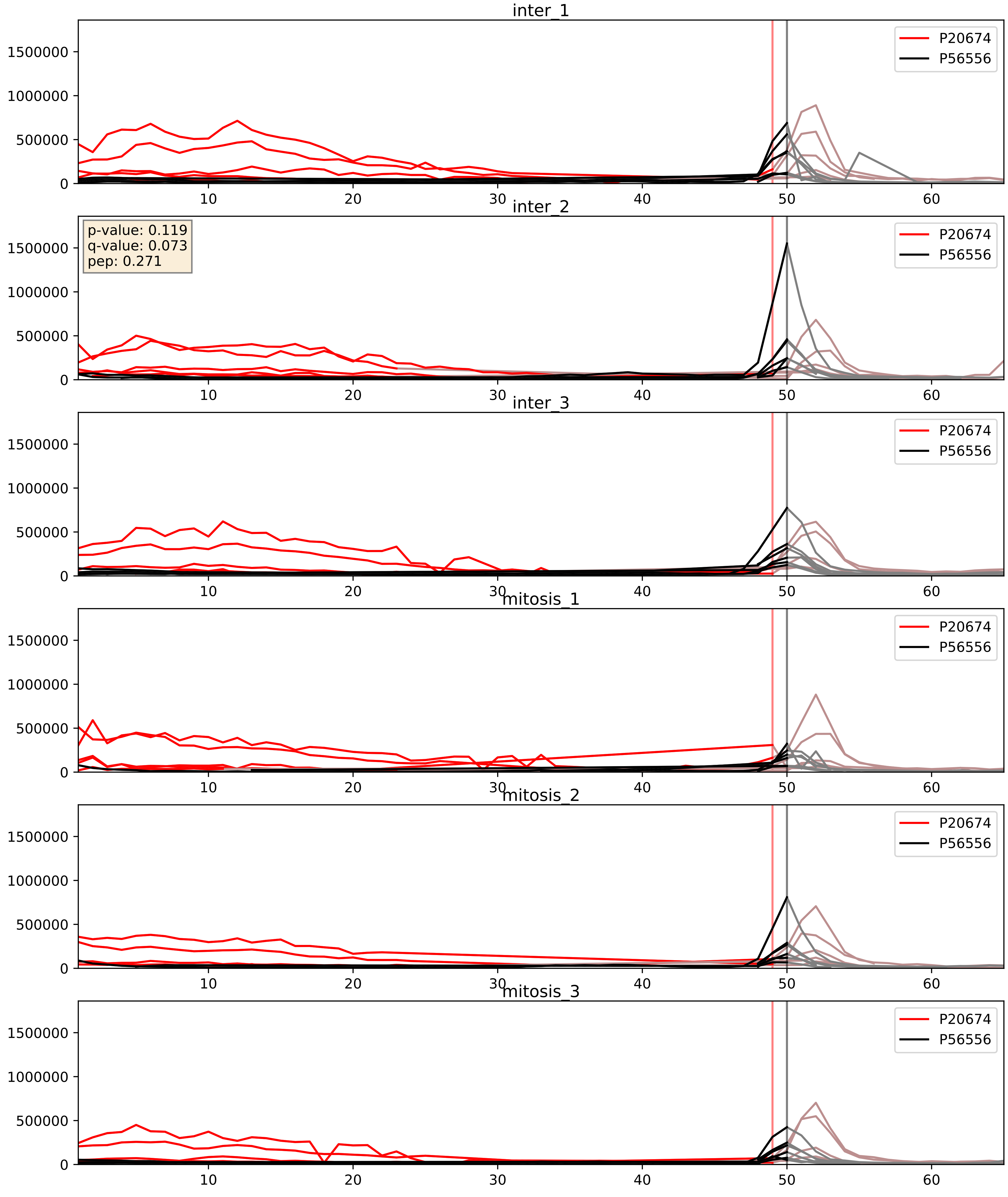


| level | condition_1 | condition_2 | pvalue | pvalue_adjusted |
|---|---|---|---|---|
| complex_abundance | mitosis | inter | 0.00808313078299871 | 0.06653038413698939 |
| interactor_abundance | mitosis | inter | 0.008547063027011064 | 0.08046463470324747 |
| interactor_ratio | mitosis | inter | 0.02440790412819327 | 0.2447639398105627 |

P28331_P56556
NDUS1_HUMAN vs NDUA6_HUMAN
p-value: 0.006 q-value: 0.006

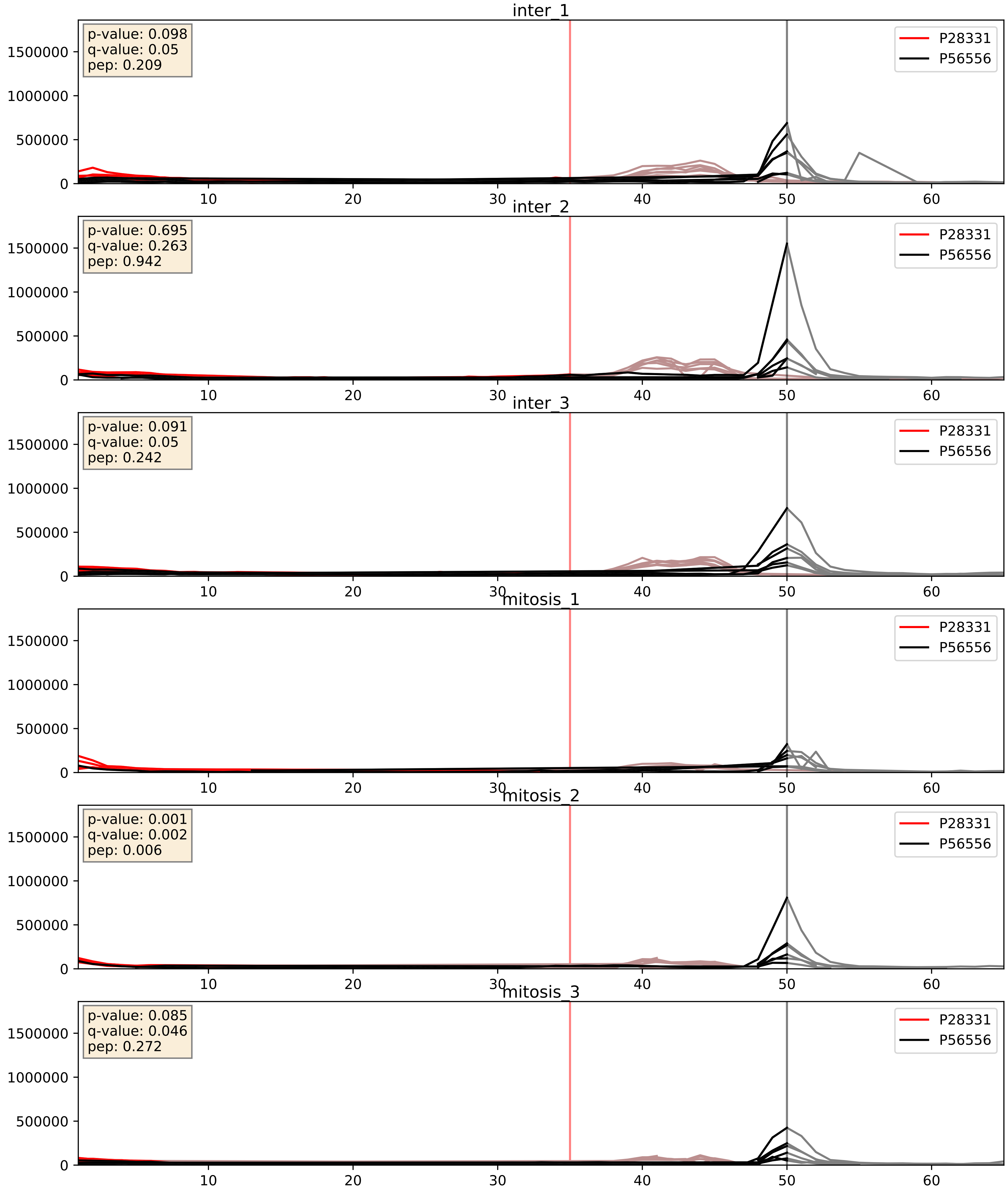


| level | condition_1 | condition_2 | pvalue | pvalue_adjusted |
|---|---|---|---|---|
| interactor_abundance | mitosis | inter | 0.002578412972883447 | 0.03806046117740333 |
| complex_abundance | mitosis | inter | 0.003507660020266425 | 0.04332694051007301 |
| interactor_ratio | mitosis | inter | 0.4030351428112066 | 0.8312490856002908 |

P56556_Q9P0J0
NDUA6_HUMAN vs NDUAD_HUMAN
p-value: 0.048 q-value: 0.031

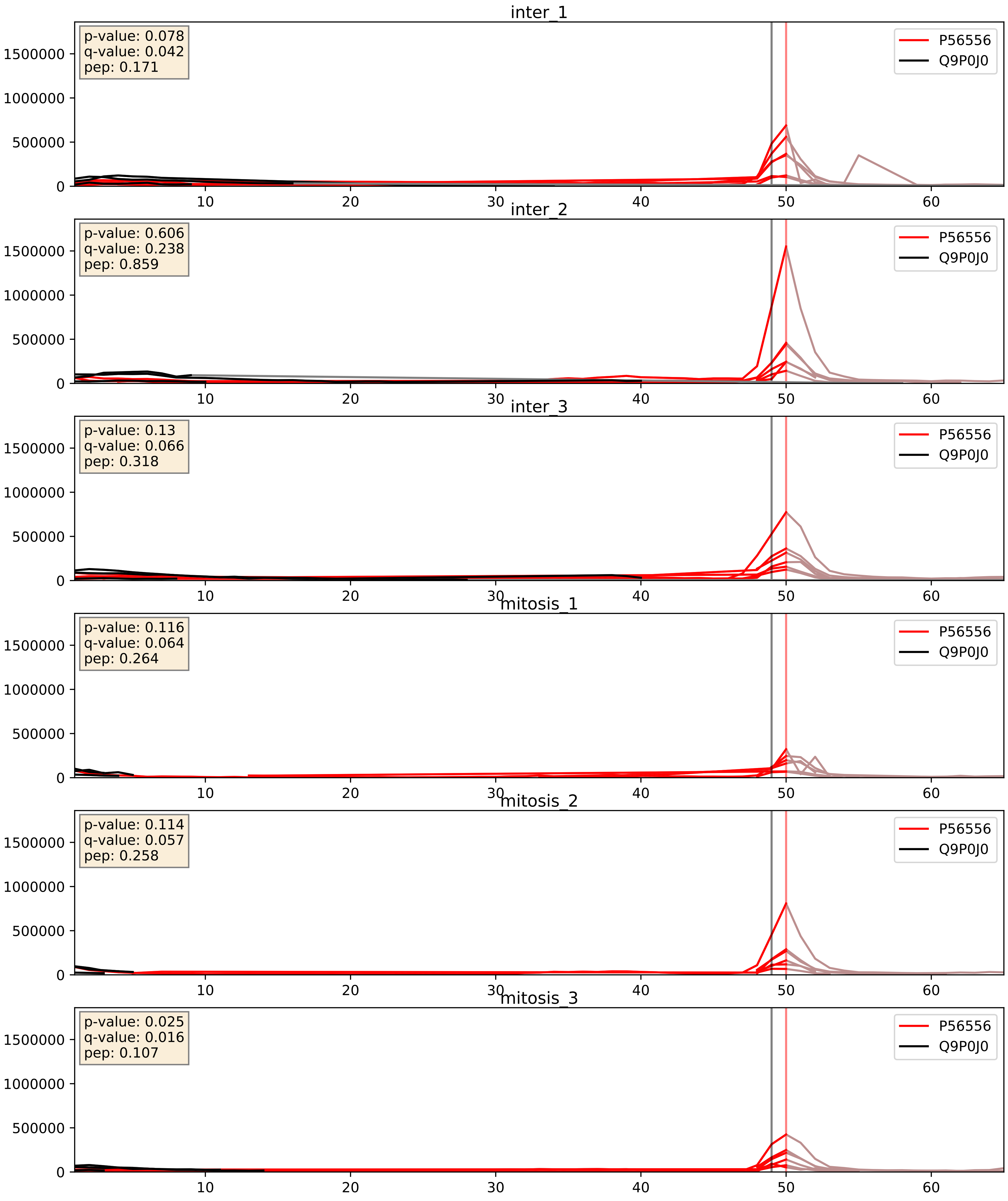


| level | condition_1 | condition_2 | pvalue | pvalue_adjusted |
|---|---|---|---|---|
| interactor_abundance | mitosis | inter | 0.0016940579623344308 | 0.030042528708242176 |
| complex_abundance | mitosis | inter | 0.0018394931343961804 | 0.030581181485790792 |
| interactor_ratio | mitosis | inter | 0.10850493173860024 | 0.5473201035252906 |
