## Supplementary Data 2 for "SECAT: Quantifying differential protein-protein interaction states by network-centric analysis": hela_string_000191_O96000.pdf

O96000_O96000

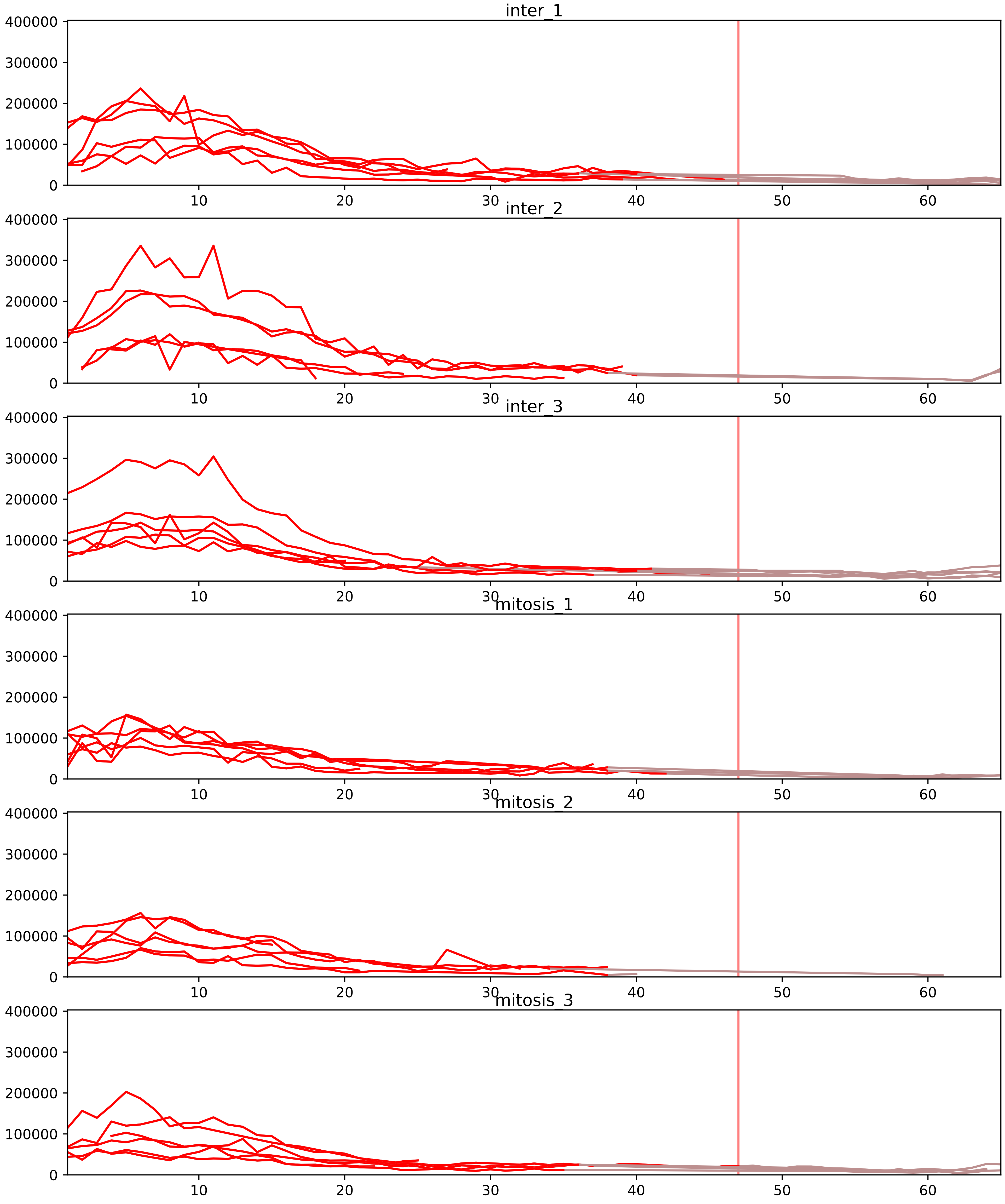


| level | condition_1 | condition_2 | pvalue | pvalue_adjusted |
|---|---|---|---|---|
| interactor_abundance | mitosis | inter | 0.00019877715730261234 | 0.003657499694368067 |
| complex_abundance | mitosis | inter | 0.0004630872323551441 | 0.006547289368558167 |
| interactor_ratio | mitosis | inter | 0.002031029127295362 | 0.0195659350482904 |
| monomer_abundance | mitosis | inter | 0.2120759311742078 | 0.5901811701101414 |
| total_abundance | mitosis | inter | 0.31383474459036037 | 0.6364019529363387 |
| assembled_abundance | mitosis | inter | 0.3286927814342444 | 0.6164231350147209 |

O00217_O96000
NDUS8_HUMAN vs NDUBA_HUMAN
p-value: 0.0 q-value: 0.0

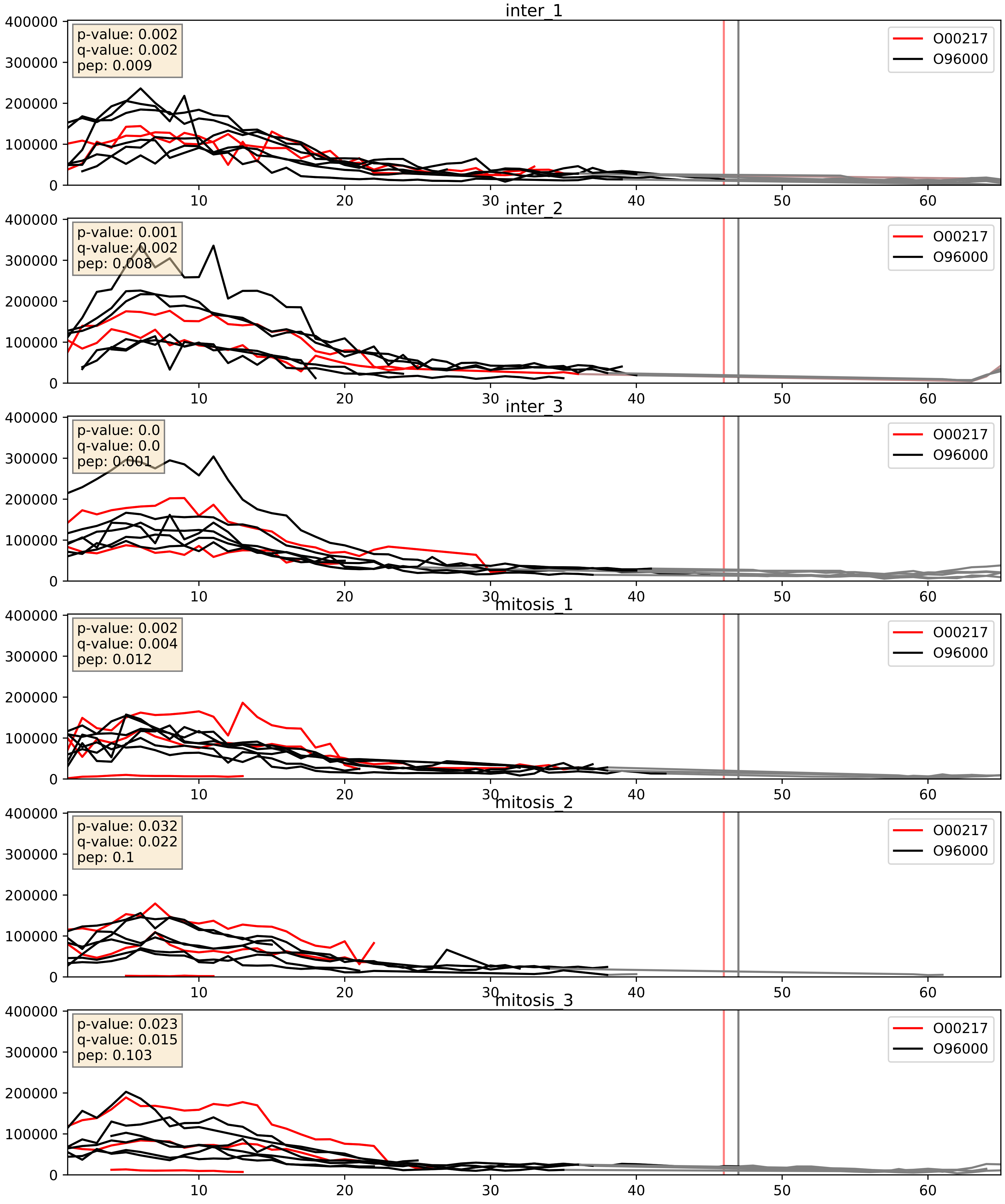


| level | condition_1 | condition_2 | pvalue | pvalue_adjusted |
|---|---|---|---|---|
| interactor_ratio | mitosis | inter | 0.0009526071034597768 | 0.035669474554159065 |
| complex_abundance | mitosis | inter | 0.03745132355060484 | 0.14122613638465967 |
| interactor_abundance | mitosis | inter | 0.4865513439277089 | 0.6558960893652855 |

O00483_O96000
NDUA4_HUMAN vs NDUBA_HUMAN
p-value: 0.002 q-value: 0.035

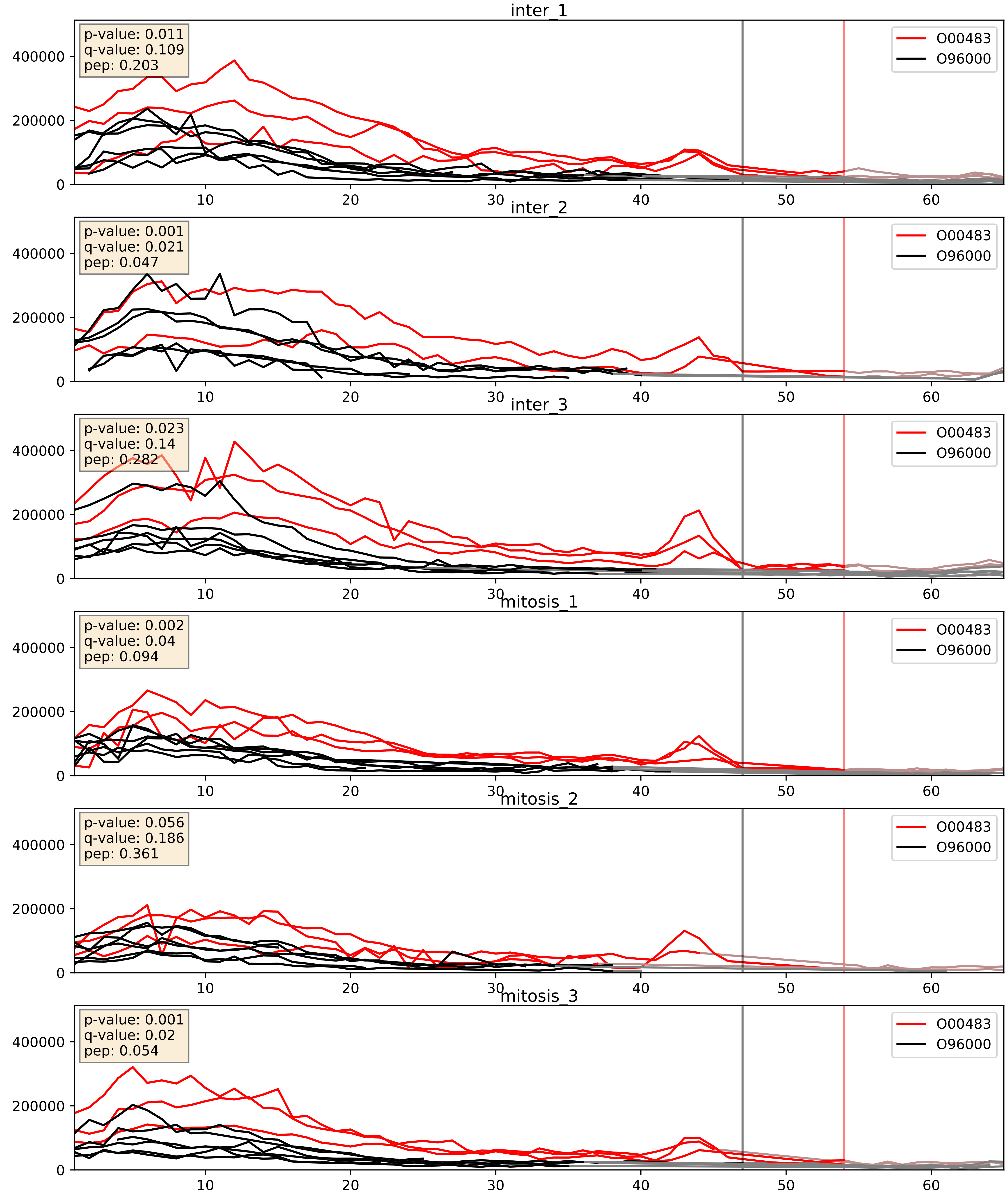


| level | condition_1 | condition_2 | pvalue | pvalue_adjusted |
|---|---|---|---|---|
| complex_abundance | mitosis | inter | 0.08962395617288287 | 0.21994869978207493 |
| interactor_ratio | mitosis | inter | 0.43547964120868093 | 0.8435878041032946 |
| interactor_abundance | mitosis | inter | 0.6231585098839934 | 0.7518304220728659 |

O14949_O96000
QCR8_HUMAN vs NDUBA_HUMAN
p-value: 0.0 q-value: 0.001

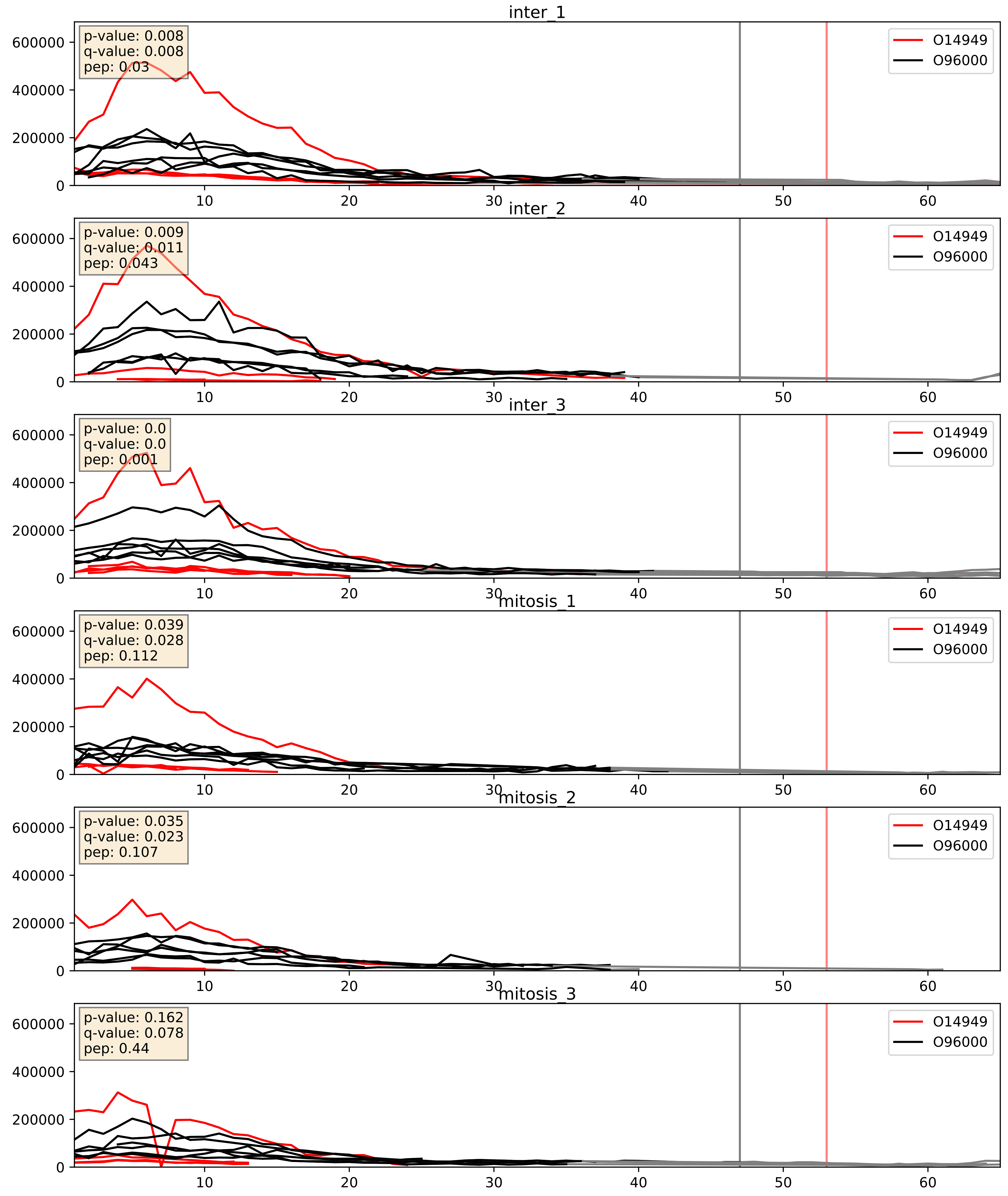


| level | condition_1 | condition_2 | pvalue | pvalue_adjusted |
|---|---|---|---|---|
| complex_abundance | mitosis | inter | 0.012811187467705593 | 0.08405635484106207 |
| interactor_abundance | mitosis | inter | 0.07386499194203666 | 0.22673988198390962 |
| interactor_ratio | mitosis | inter | 0.9566997653296503 | 0.9927224968152035 |

O43674_O96000
NDUB5_HUMAN vs NDUBA_HUMAN
p-value: 0.015 q-value: 0.013

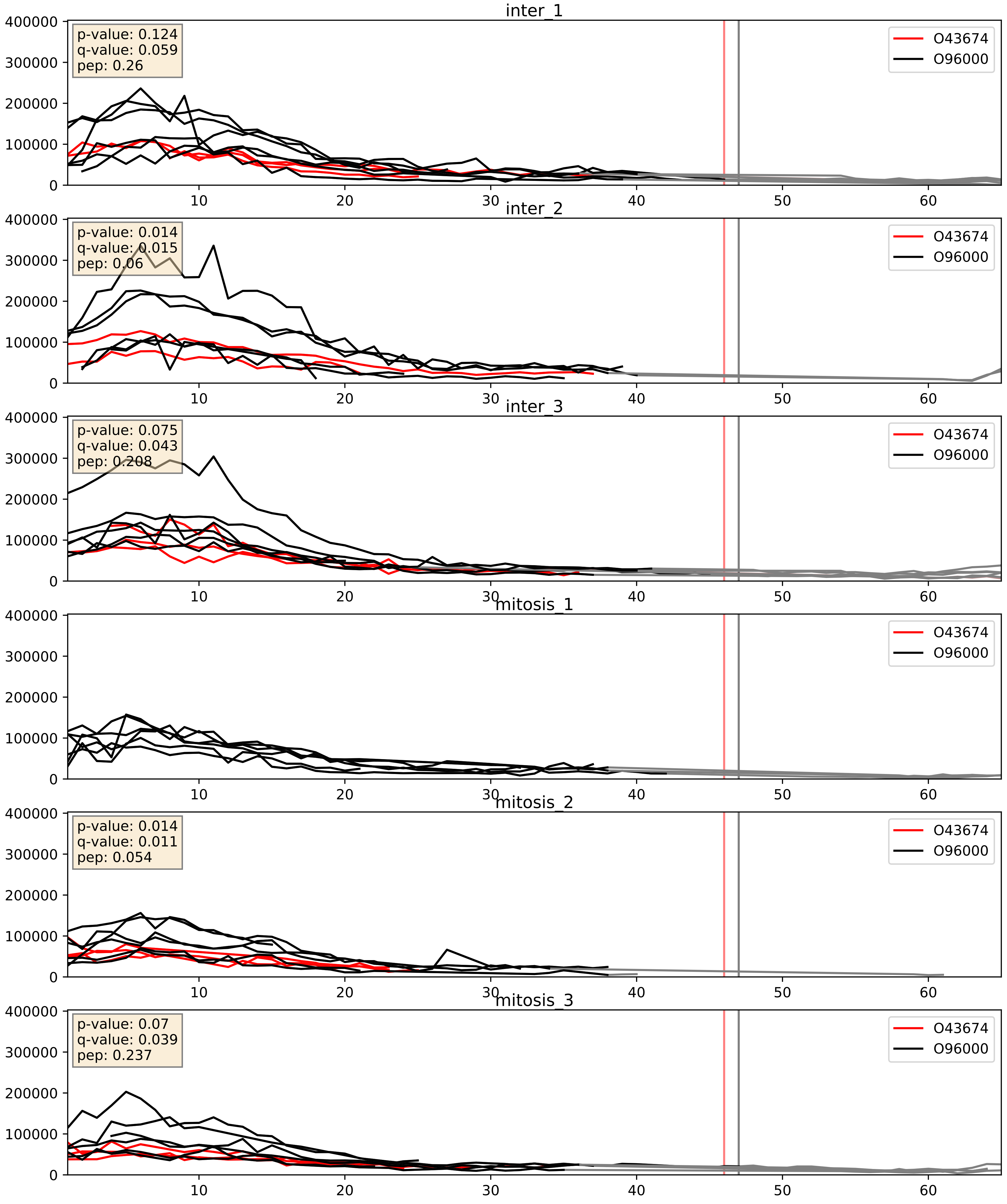


| level | condition_1 | condition_2 | pvalue | pvalue_adjusted |
|---|---|---|---|---|
| complex_abundance | mitosis | inter | 0.09253154449260821 | 0.22431889573965627 |
| interactor_abundance | mitosis | inter | 0.10848703869403525 | 0.2832110555721079 |
| interactor_ratio | mitosis | inter | 0.25377164334885527 | 0.7260311721477944 |

O75251_O96000
NDUS7_HUMAN vs NDUBA_HUMAN
p-value: 0.0 q-value: 0.0

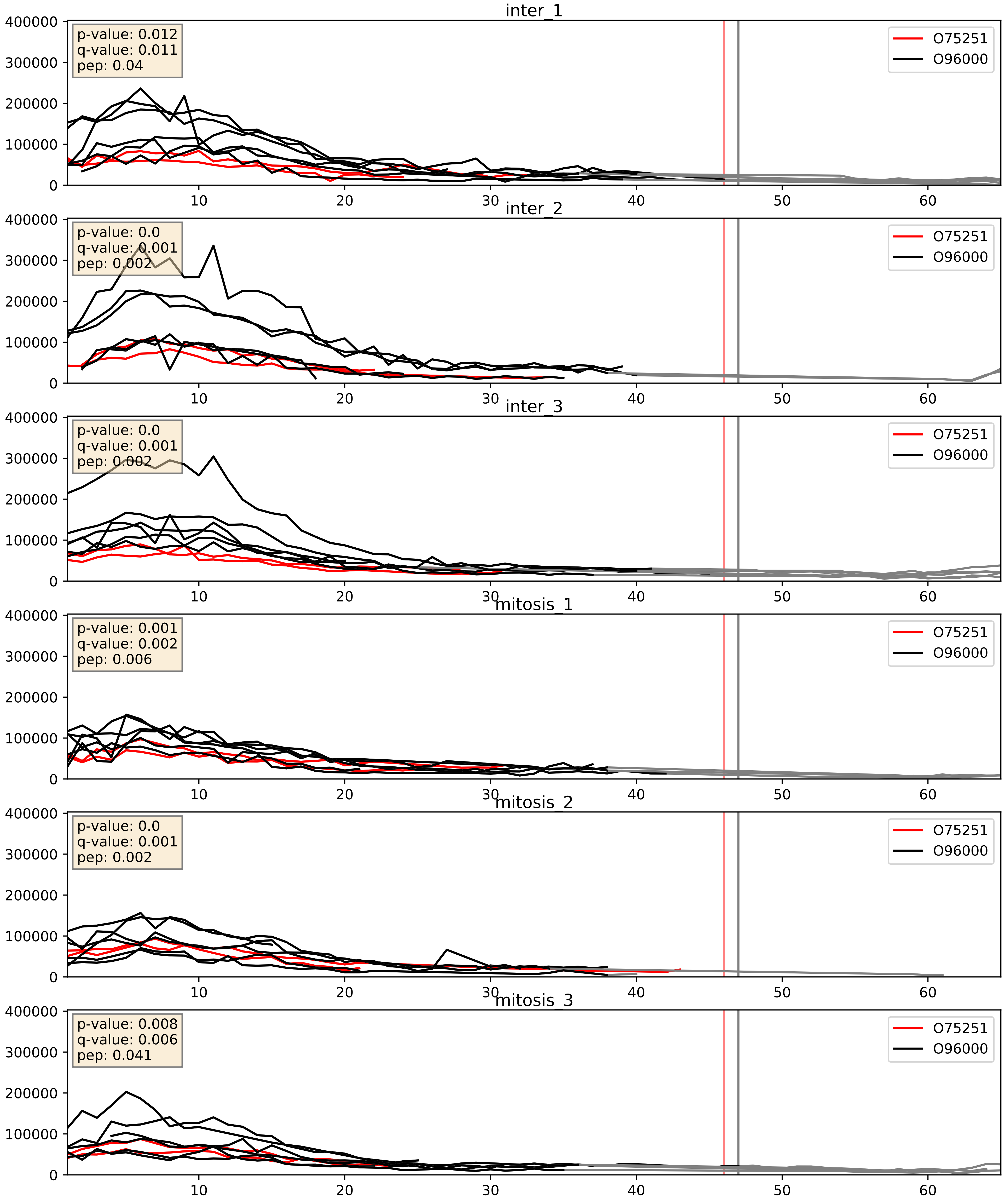


| level | condition_1 | condition_2 | pvalue | pvalue_adjusted |
|---|---|---|---|---|
| interactor_ratio | mitosis | inter | 0.0021505347076290404 | 0.05734759220344108 |
| complex_abundance | mitosis | inter | 0.004869110967109282 | 0.052397293069493756 |
| interactor_abundance | mitosis | inter | 0.016765658116067107 | 0.11277132084292767 |

O75306_O96000
NDUS2_HUMAN vs NDUBA_HUMAN
p-value: 0.007 q-value: 0.006

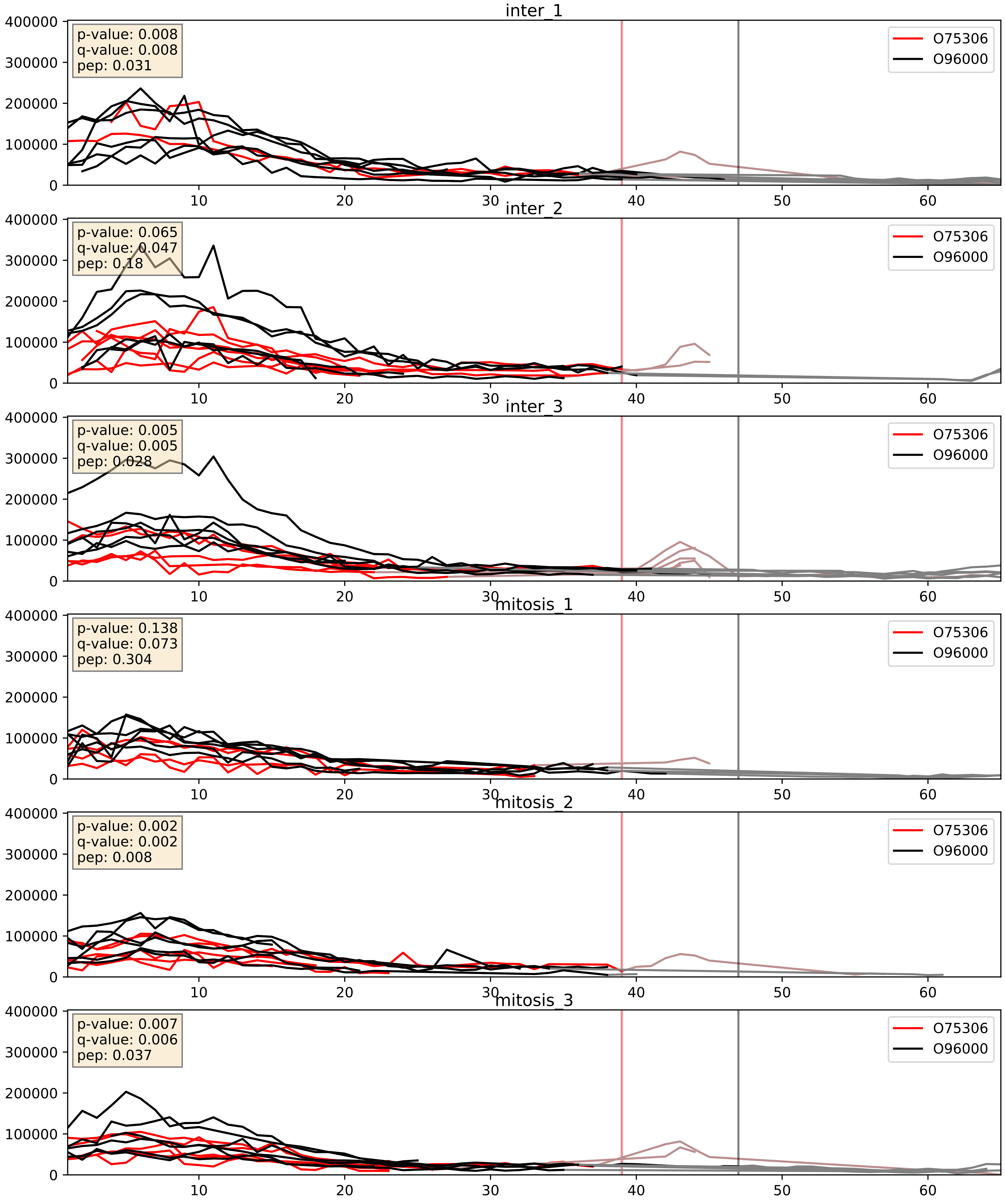


| level | condition_1 | condition_2 | pvalue | pvalue_adjusted |
|---|---|---|---|---|
| complex_abundance | mitosis | inter | 0.13840576740433103 | 0.283569499516772 |
| interactor_ratio | mitosis | inter | 0.3389910233567168 | 0.7924528012231136 |
| interactor_abundance | mitosis | inter | 0.6718484200519785 | 0.7850689339492098 |

O75489_O96000
NDUS3_HUMAN vs NDUBA_HUMAN
p-value: 0.0 q-value: 0.0

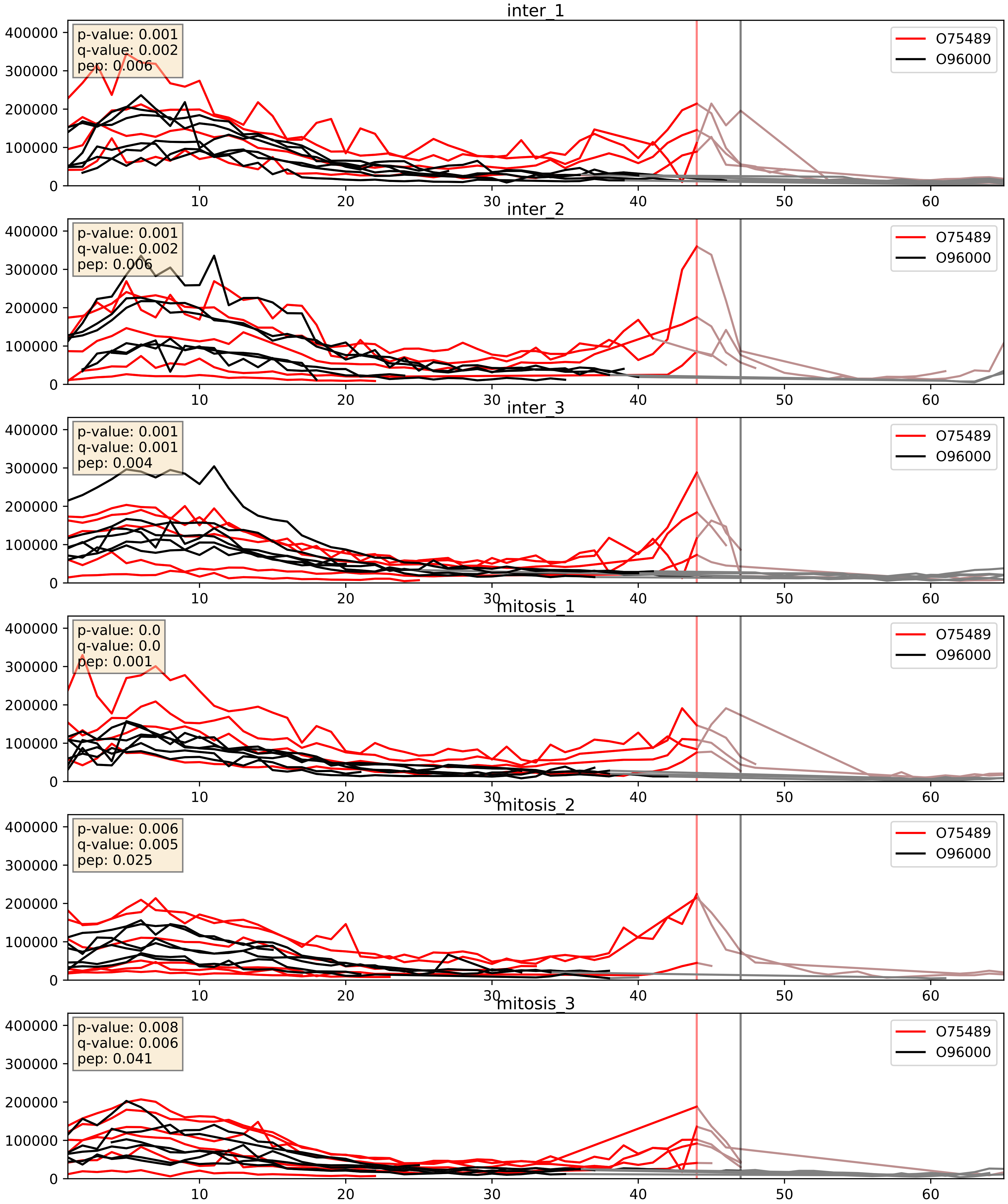


| level | condition_1 | condition_2 | pvalue | pvalue_adjusted |
|---|---|---|---|---|
| complex_abundance | mitosis | inter | 0.009718538622733542 | 0.07323124173468232 |
| interactor_ratio | mitosis | inter | 0.05739097688312565 | 0.3961608945972482 |
| interactor_abundance | mitosis | inter | 0.06442031000517522 | 0.20920098130959613 |

O75947_O96000
ATP5H_HUMAN vs NDUBA_HUMAN
p-value: 0.003 q-value: 0.039

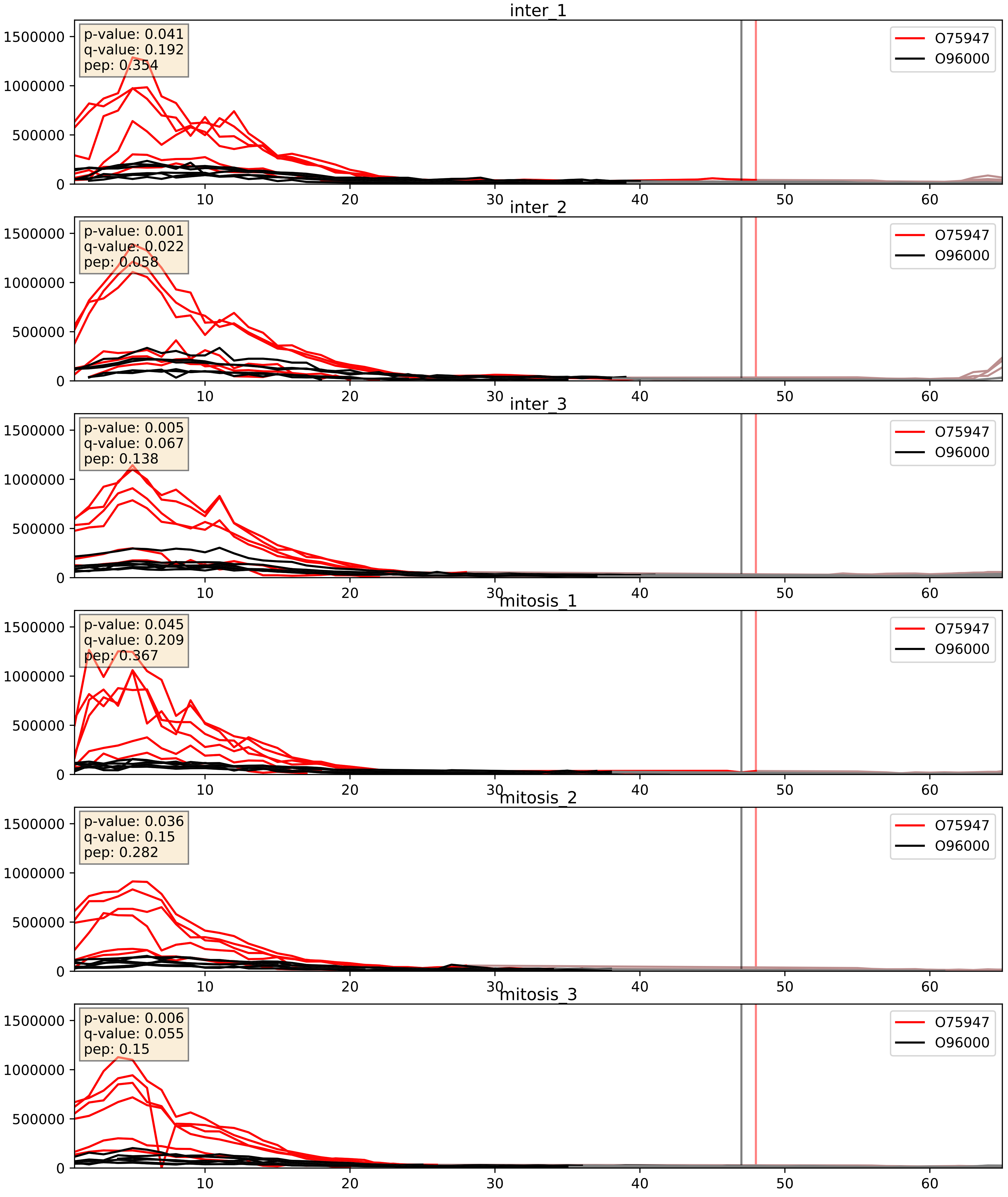


| level | condition_1 | condition_2 | pvalue | pvalue_adjusted |
|---|---|---|---|---|
| complex_abundance | mitosis | inter | 0.003939579782951728 | 0.04645014179348043 |
| interactor_abundance | mitosis | inter | 0.008426215029868284 | 0.07993154289762057 |
| interactor_ratio | mitosis | inter | 0.20046131002922904 | 0.6750388724823763 |

O75964_O96000
ATP5L_HUMAN vs NDUBA_HUMAN
p-value: 0.0 q-value: 0.023

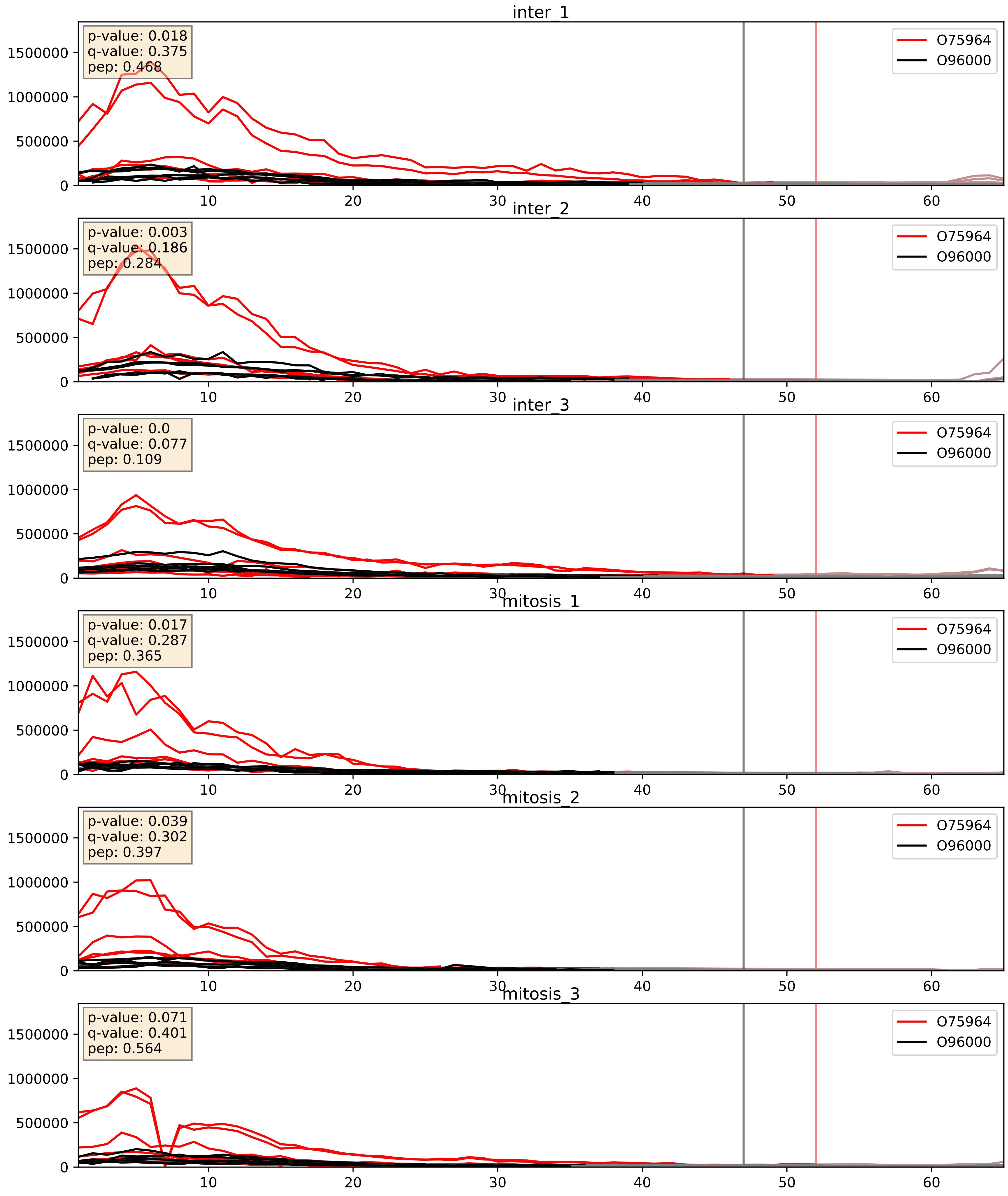


| level | condition_1 | condition_2 | pvalue | pvalue_adjusted |
|---|---|---|---|---|
| complex_abundance | mitosis | inter | 0.0029730687764829024 | 0.03988944941488032 |
| interactor_abundance | mitosis | inter | 0.007799670890406056 | 0.07671437955535183 |
| interactor_ratio | mitosis | inter | 0.6120598333070664 | 0.90762799337104 |

O95139_O96000
NDUB6_HUMAN vs NDUBA_HUMAN
p-value: 0.011 q-value: 0.01

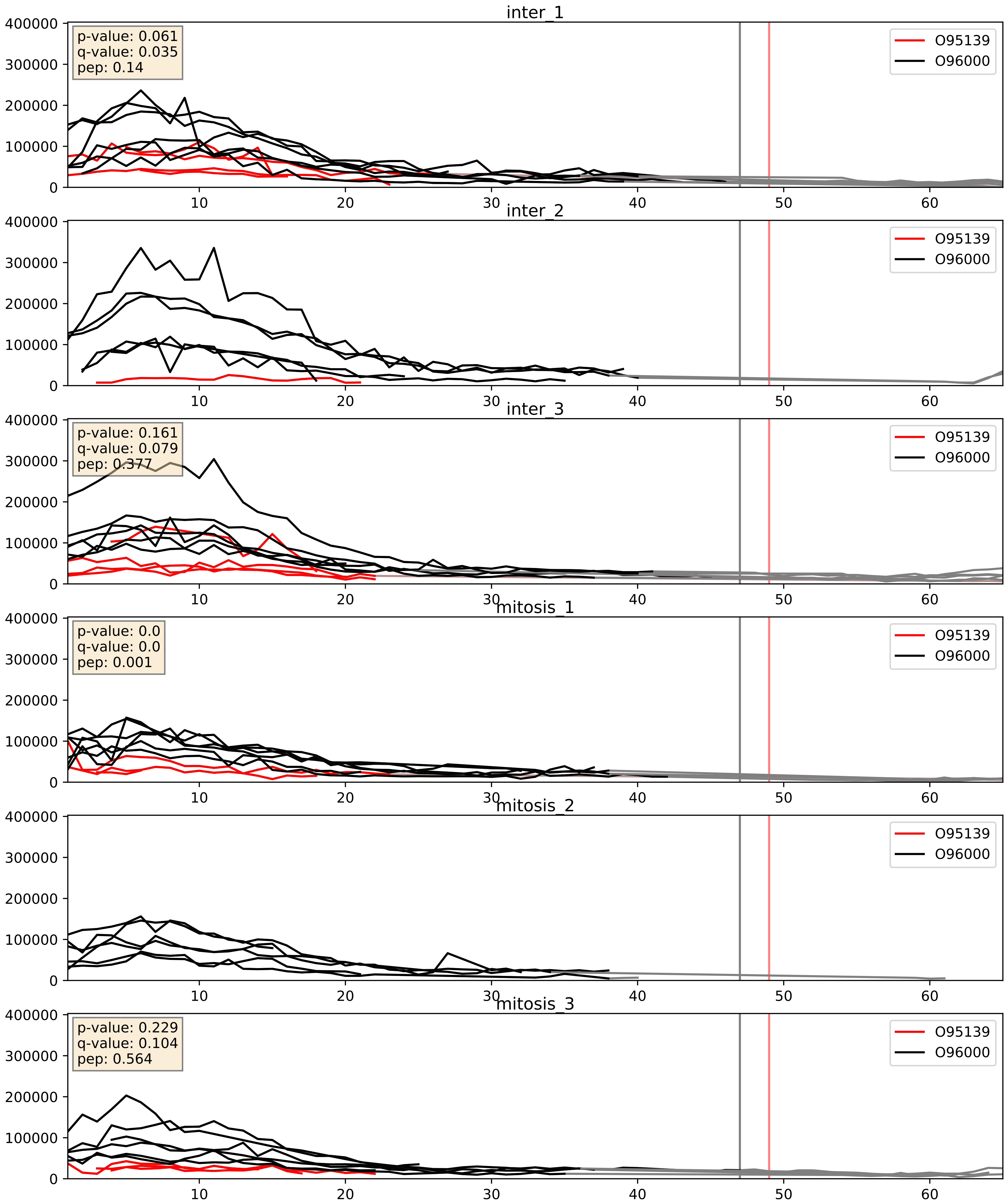


| level | condition_1 | condition_2 | pvalue | pvalue_adjusted |
|---|---|---|---|---|
| complex_abundance | mitosis | inter | 0.12672825675558308 | 0.26924643281900995 |
| interactor_abundance | mitosis | inter | 0.14414968864806146 | 0.32983729880443896 |
| interactor_ratio | mitosis | inter | 0.48080716281737923 | 0.8669092153319383 |

O95299_O96000
NDUAA_HUMAN vs NDUBA_HUMAN
p-value: 0.076 q-value: 0.046

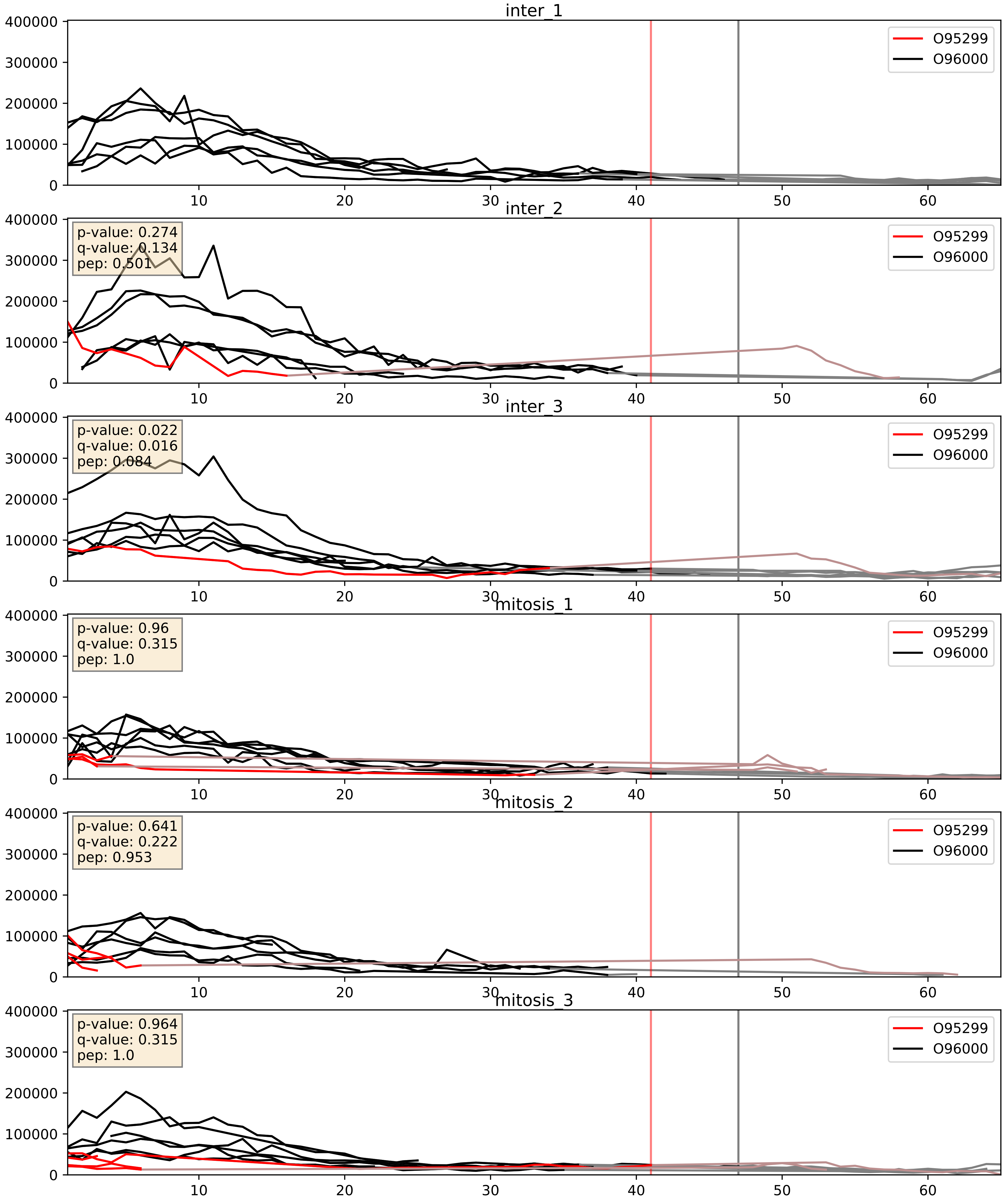


| level | condition_1 | condition_2 | pvalue | pvalue_adjusted |
|---|---|---|---|---|
| interactor_abundance | mitosis | inter | 0.48614247031883095 | 0.6556967692317329 |
| complex_abundance | mitosis | inter | 0.5120904151851724 | 0.6479910655038046 |
| interactor_ratio | mitosis | inter | 0.5793211258557531 | 0.8969051975628951 |

O96000_P07919
NDUBA_HUMAN vs QCR6_HUMAN
p-value: 0.0 q-value: 0.013

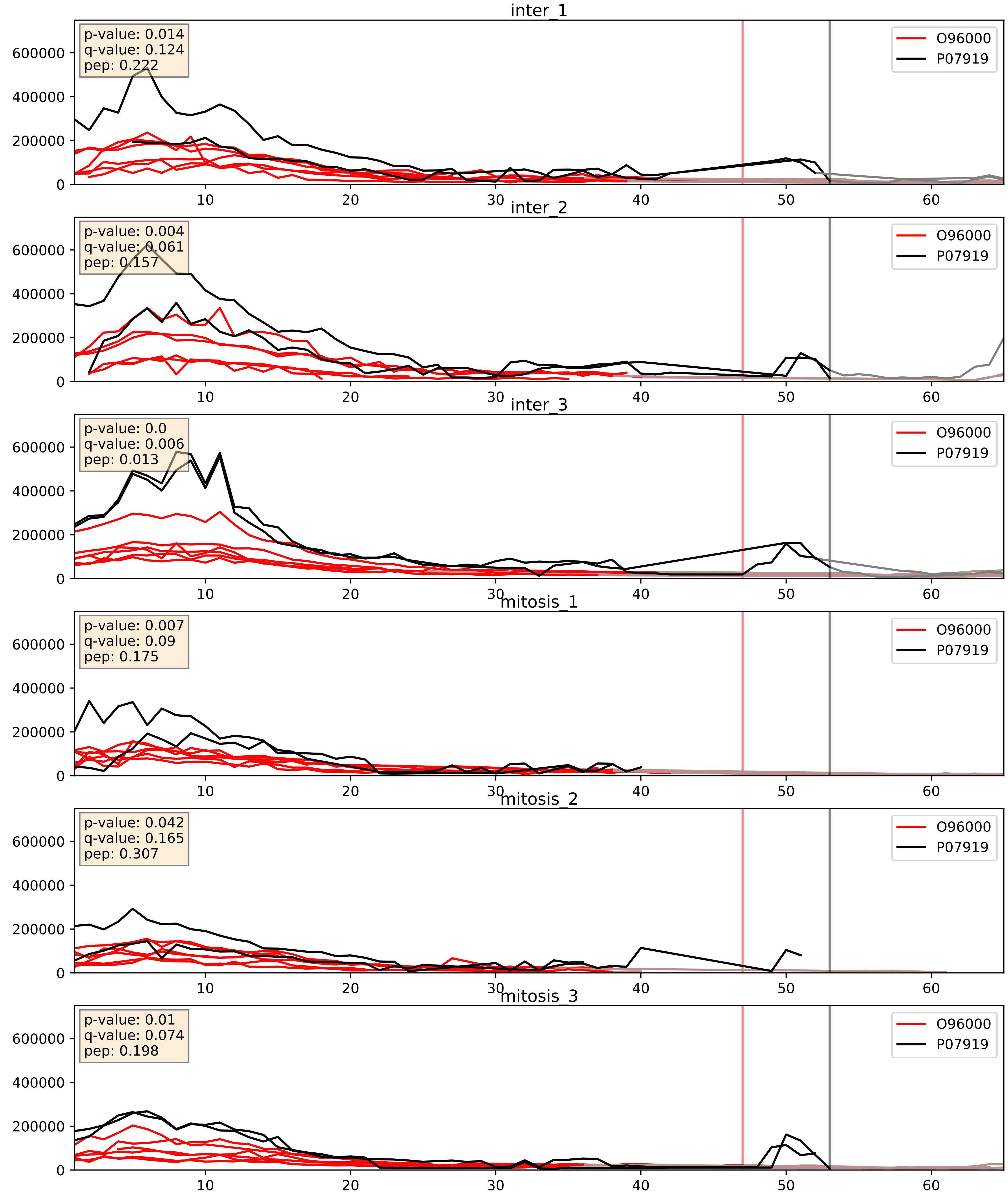


| level | condition_1 | condition_2 | pvalue | pvalue_adjusted |
|---|---|---|---|---|
| interactor_abundance | mitosis | inter | 0.0037407396194239044 | 0.047601106577632585 |
| complex_abundance | mitosis | inter | 0.0039760250442856734 | 0.04681537053519307 |
| interactor_ratio | mitosis | inter | 0.20399780328347128 | 0.6786712771498306 |

O96000_P08574
NDUBA_HUMAN vs CY1_HUMAN
p-value: 0.002 q-value: 0.03

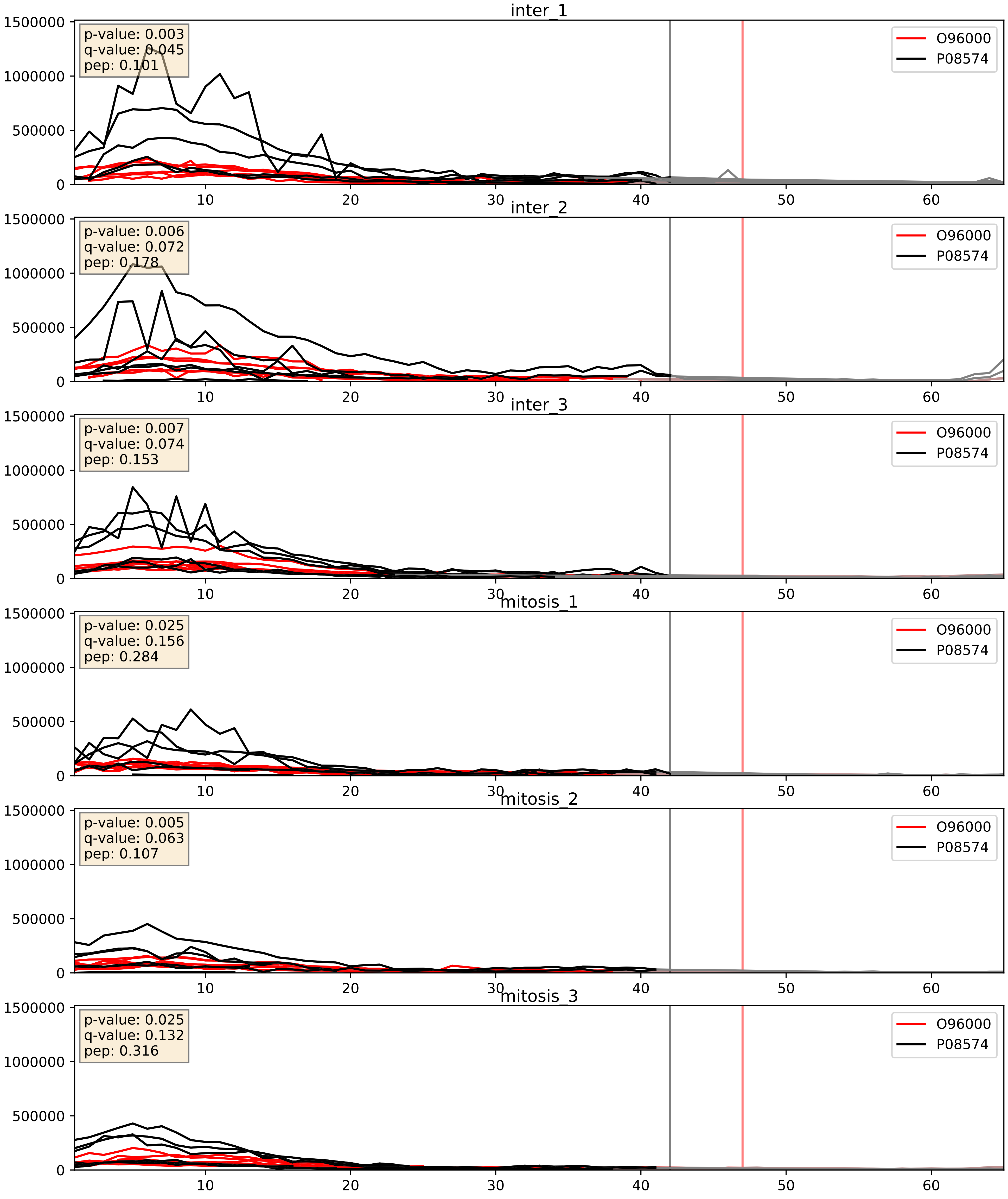


| level | condition_1 | condition_2 | pvalue | pvalue_adjusted |
|---|---|---|---|---|
| complex_abundance | mitosis | inter | 0.002334679939336067 | 0.035168510428347066 |
| interactor_abundance | mitosis | inter | 0.0030692858094850456 | 0.042465703978082026 |
| interactor_ratio | mitosis | inter | 0.3283581682329763 | 0.786600701916436 |

O96000_P13073
NDUBA_HUMAN vs COX41_HUMAN
p-value: 0.0 q-value: 0.023

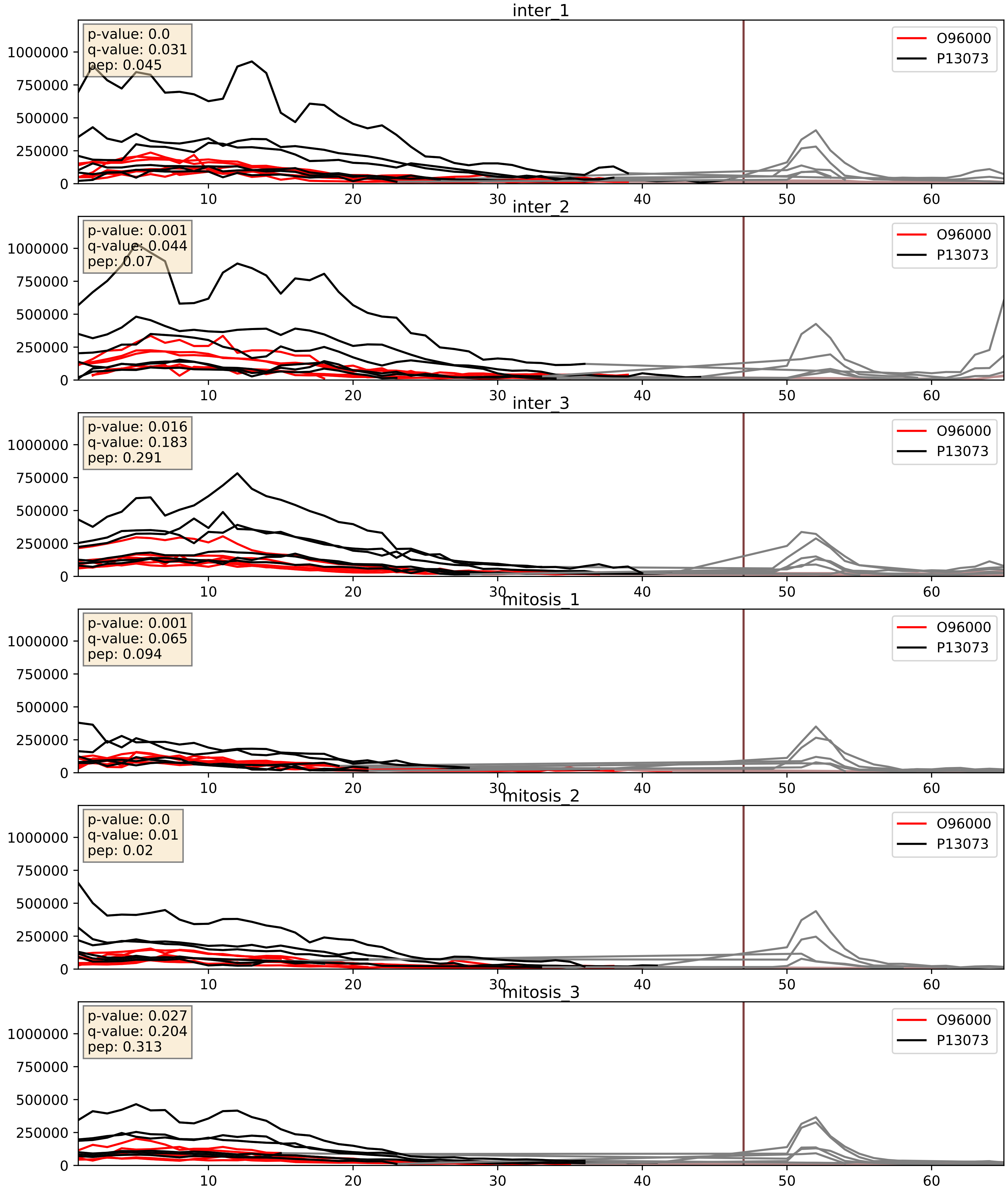


| level | condition_1 | condition_2 | pvalue | pvalue_adjusted |
|---|---|---|---|---|
| interactor_abundance | mitosis | inter | 0.0026153349281260135 | 0.038268832452578935 |
| complex_abundance | mitosis | inter | 0.00346532263349165 | 0.04302508960916016 |
| interactor_ratio | mitosis | inter | 0.25486933343791507 | 0.7264298204376385 |

O96000_P20674
NDUBA_HUMAN vs COX5A_HUMAN
p-value: 0.0 q-value: 0.013

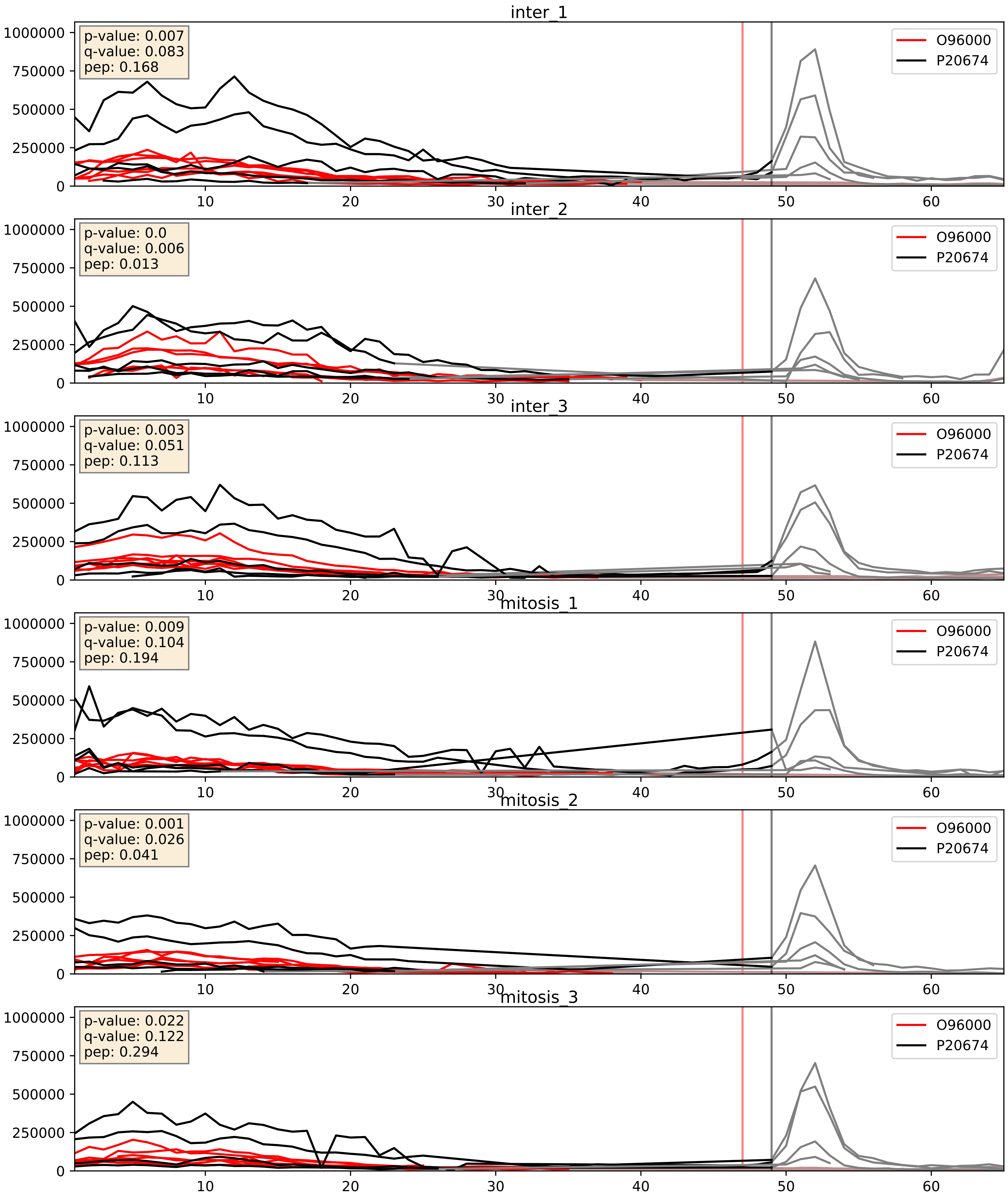


| level | condition_1 | condition_2 | pvalue | pvalue_adjusted |
|---|---|---|---|---|
| complex_abundance | mitosis | inter | 0.0035908279990228657 | 0.04393282066269434 |
| interactor_abundance | mitosis | inter | 0.004421765867454451 | 0.05334787290403116 |
| interactor_ratio | mitosis | inter | 0.6387617471795103 | 0.9197898119332029 |

O96000_P21912
NDUBA_HUMAN vs SDHB_HUMAN
p-value: 0.0 q-value: 0.023

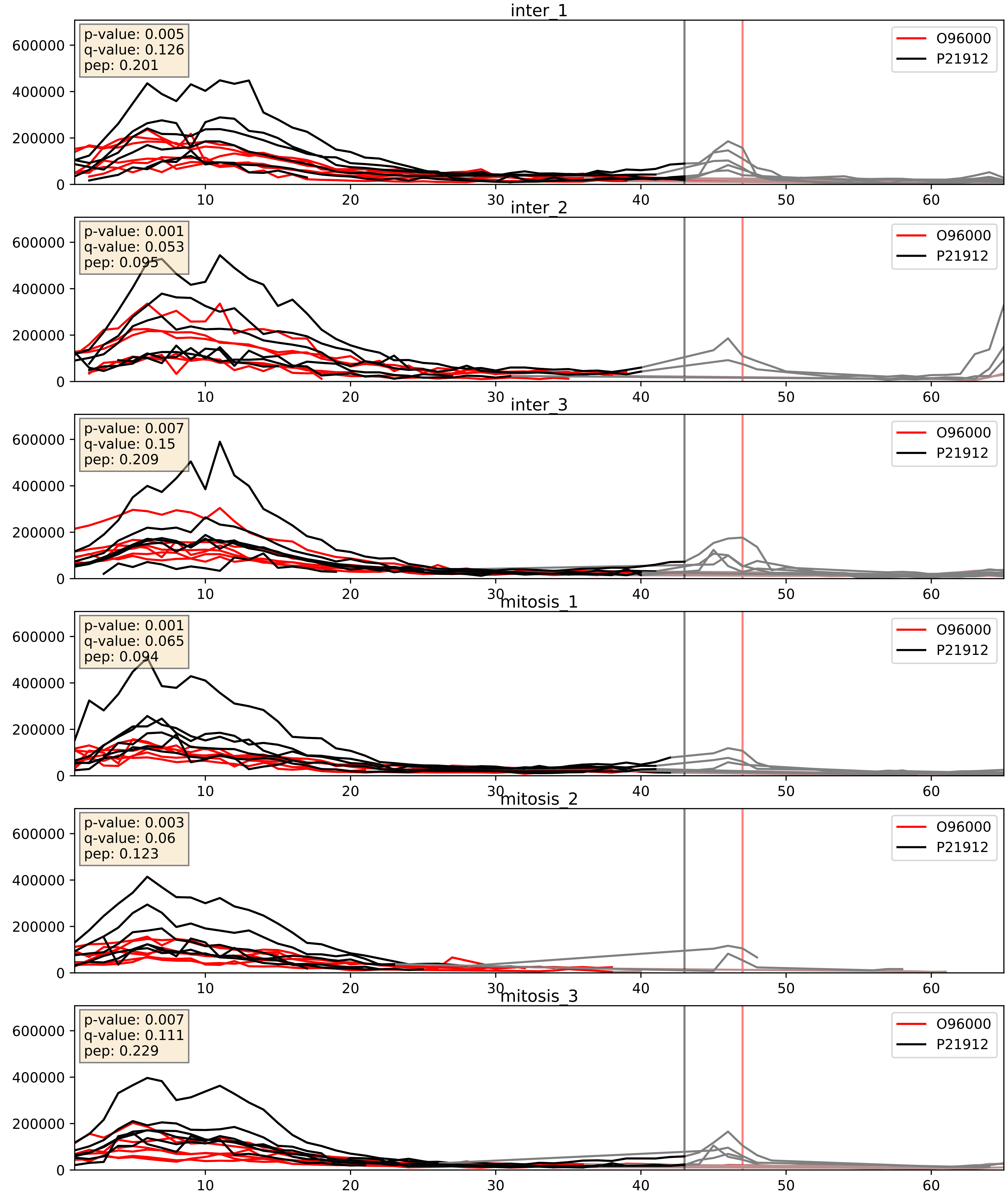


| level | condition_1 | condition_2 | pvalue | pvalue_adjusted |
|---|---|---|---|---|
| interactor_abundance | mitosis | inter | 0.004495885981421616 5 | 0.05393802943373376 |
| interactor_ratio | mitosis | inter | 0.0062007803725436465 | 0.10851698081572049 |
| complex_abundance | mitosis | inter | 0.008626823952176838 | 0.06907915157215504 |

O96000_Q16718
NDUBA_HUMAN vs NDUA5_HUMAN
p-value: 0.0 q-value: 0.001

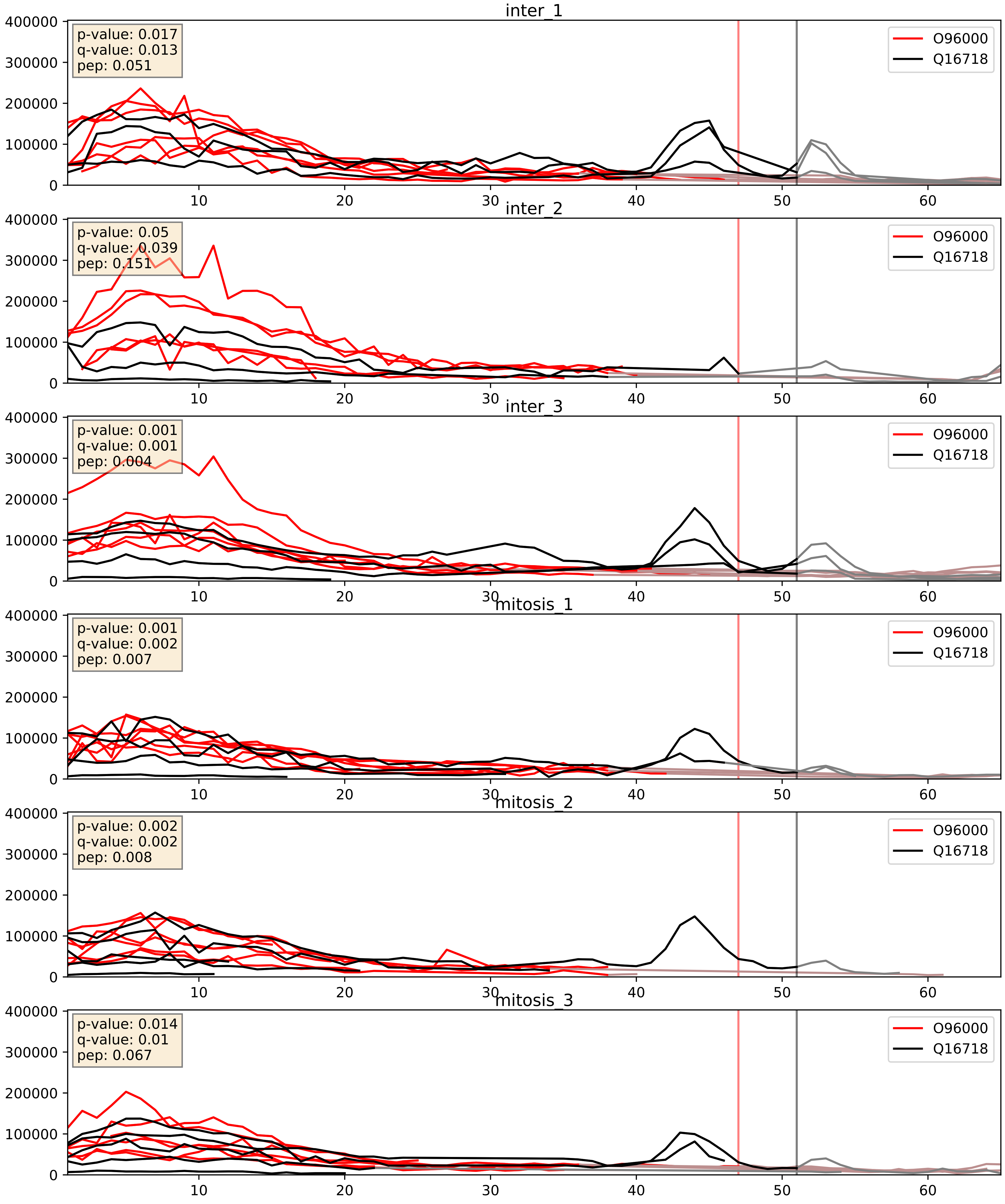


| level | condition_1 | condition_2 | pvalue | pvalue_adjusted |
|---|---|---|---|---|
| interactor_abundance | mitosis | inter | 0.0032998973550818515 | 0.04429199808372052 |
| complex_abundance | mitosis | inter | 0.04144590898197501 | 0.146541503876789 |
| interactor_ratio | mitosis | inter | 0.3393041314282266 | 0.7924528012231136 |

O96000_Q16795
NDUBA_HUMAN vs NDUA9_HUMAN
p-value: 0.016 q-value: 0.014

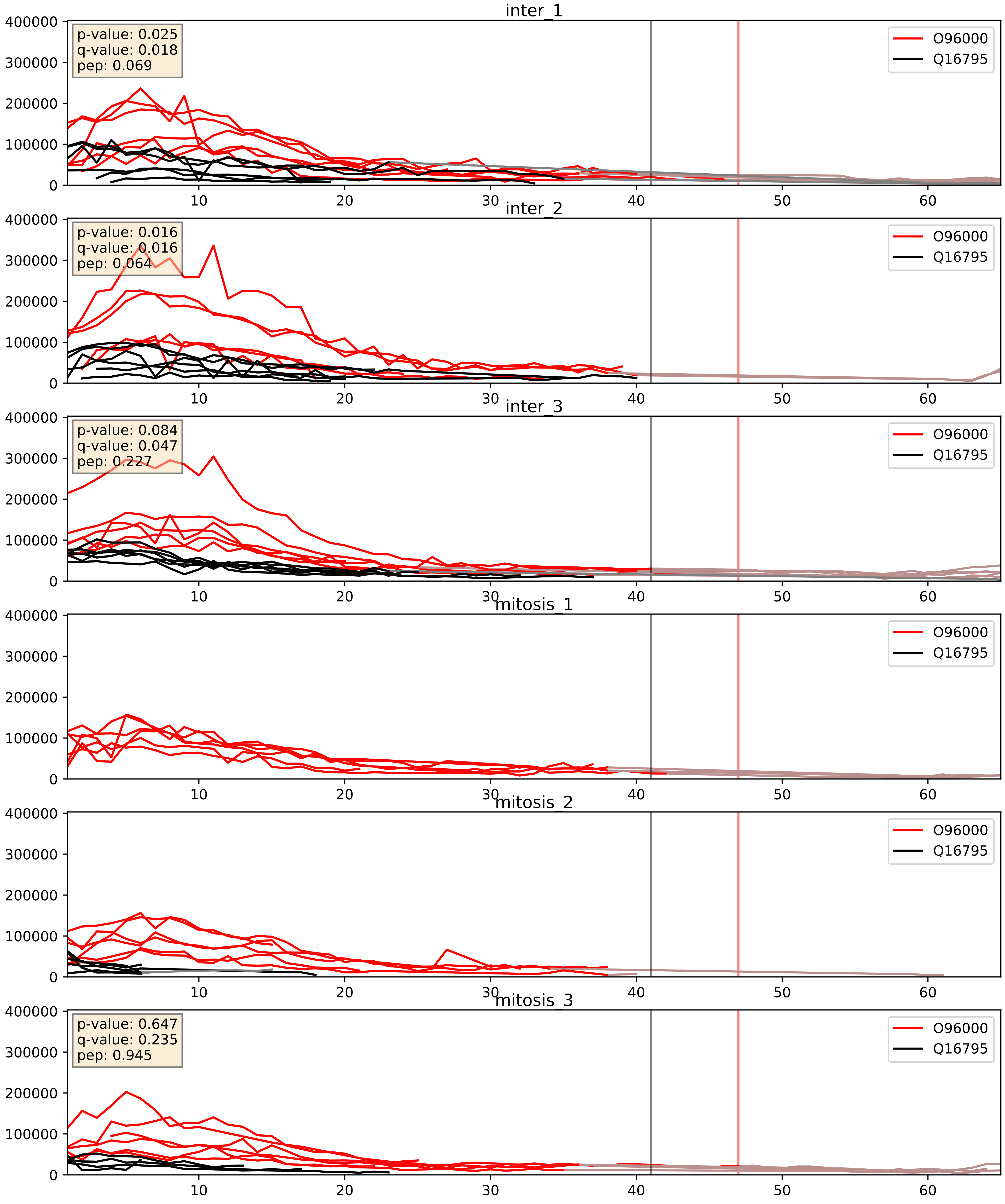


| level | condition_1 | condition_2 | pvalue | pvalue_adjusted |
|---|---|---|---|---|
| complex_abundance | mitosis | inter | 0.005866859048392097 | 0.05756820427608883 |
| interactor_abundance | mitosis | inter | 0.038734556862202 | 0.16006169767822792 |
| interactor_ratio | mitosis | inter | 0.8425742511809455 | 0.9753124530235151 |

O96000_Q9BU61
NDUBA_HUMAN vs NDUF3_HUMAN
p-value: 0.001 q-value: 0.001

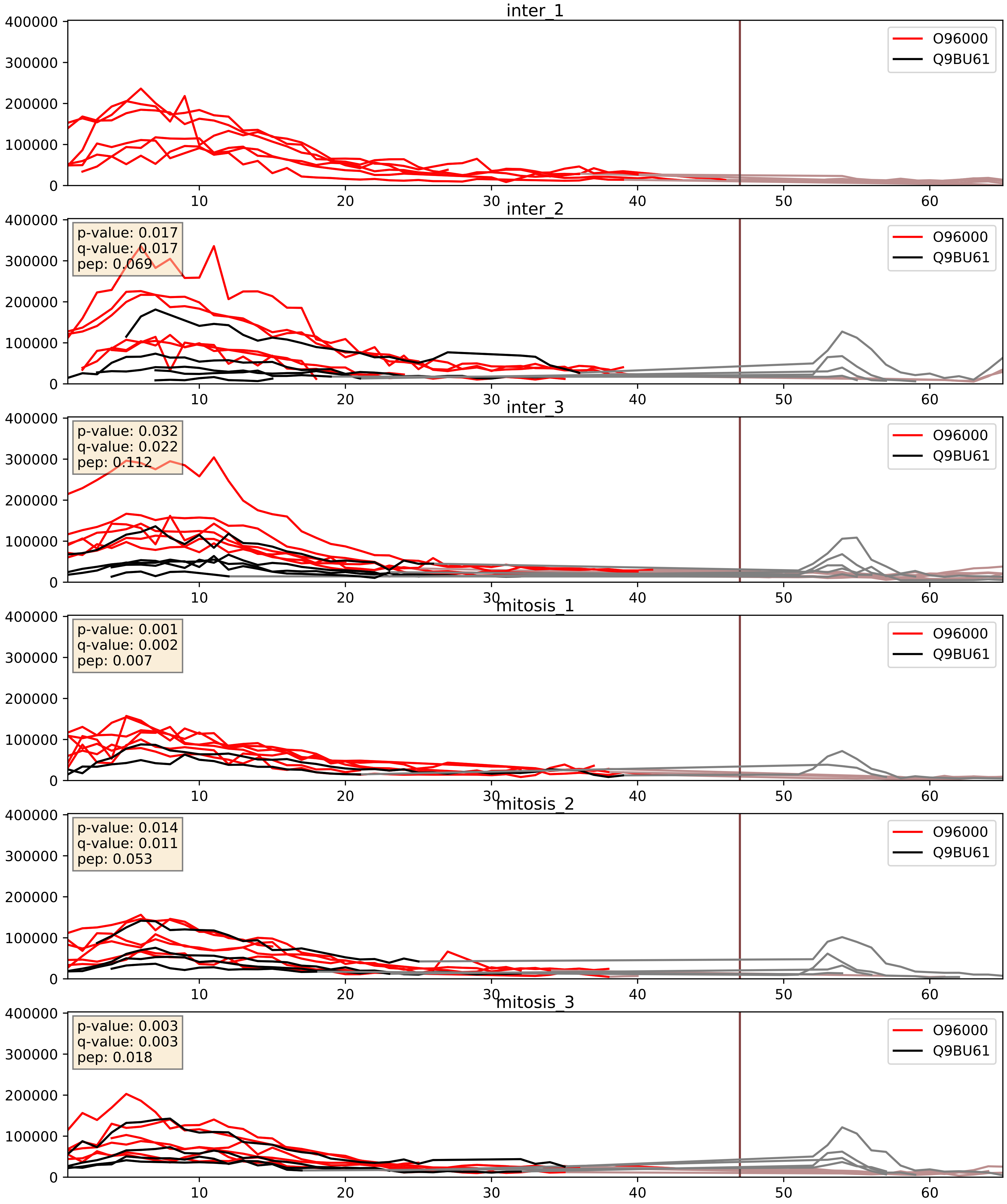


| level | condition_1 | condition_2 | pvalue | pvalue_adjusted |
|---|---|---|---|---|
| complex_abundance | mitosis | inter | 0.8220752607065606 | 0.8842629092294746 |
| interactor_abundance | mitosis | inter | 0.8238576448535243 | 0.9013575460053896 |
| interactor_ratio | mitosis | inter | 0.8665000002405782 | 0.9783867411263176 |

O96000_Q9NX14
NDUBA_HUMAN vs NDUBB_HUMAN
p-value: 0.0 q-value: 0.0

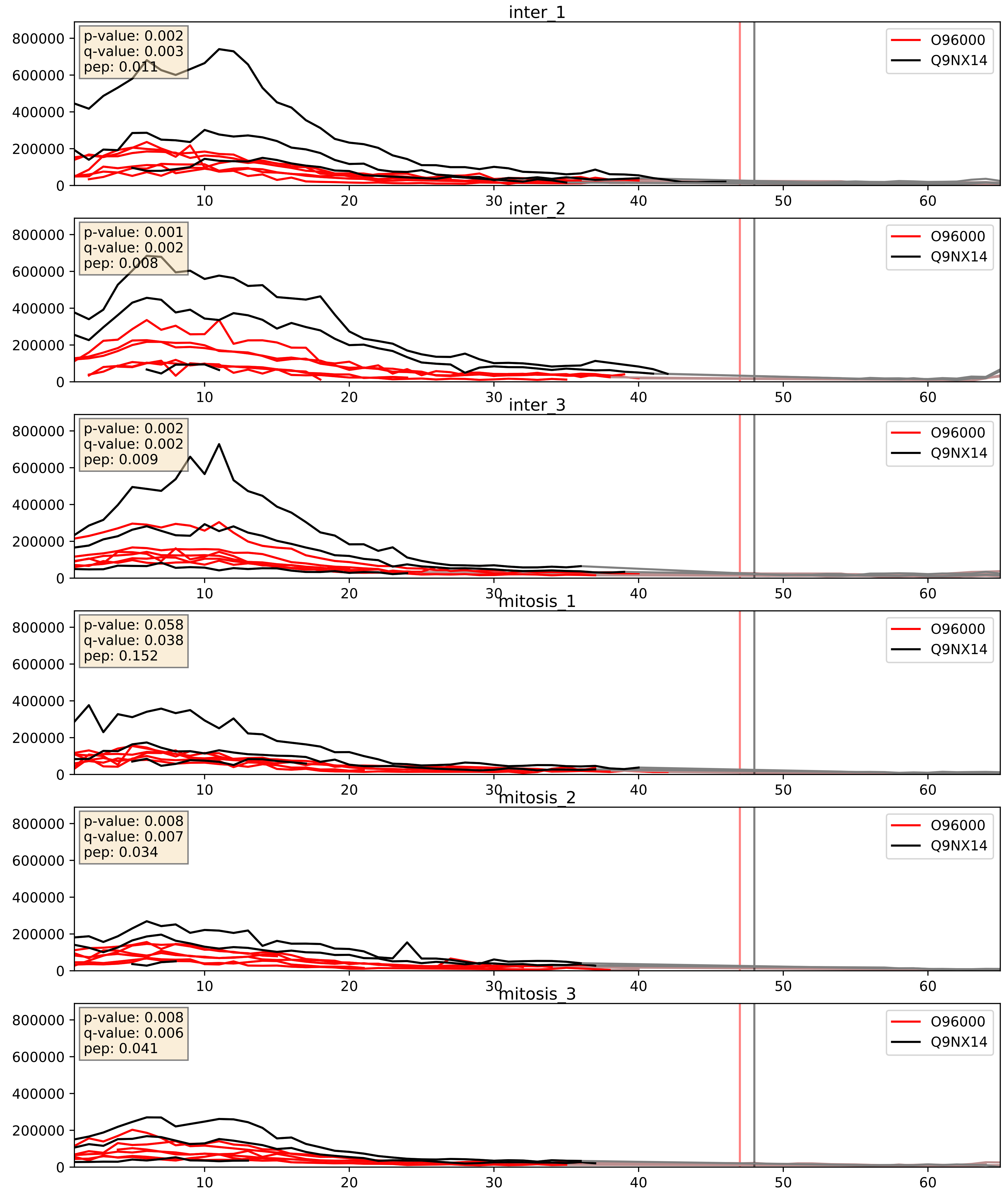


| level | condition_1 | condition_2 | pvalue | pvalue_adjusted |
|---|---|---|---|---|
| interactor_abundance | mitosis | inter | 0.003261171545258451 | 0.044100518842673526 |
| complex_abundance | mitosis | inter | 0.003872412725850539 | 0.046157376235893825 |
| interactor_ratio | mitosis | inter | 0.22262365994950983 | 0.7003665360598988 |

O96000_Q9Y375
NDUBA_HUMAN vs CIA30_HUMAN
p-value: 0.0 q-value: 0.003

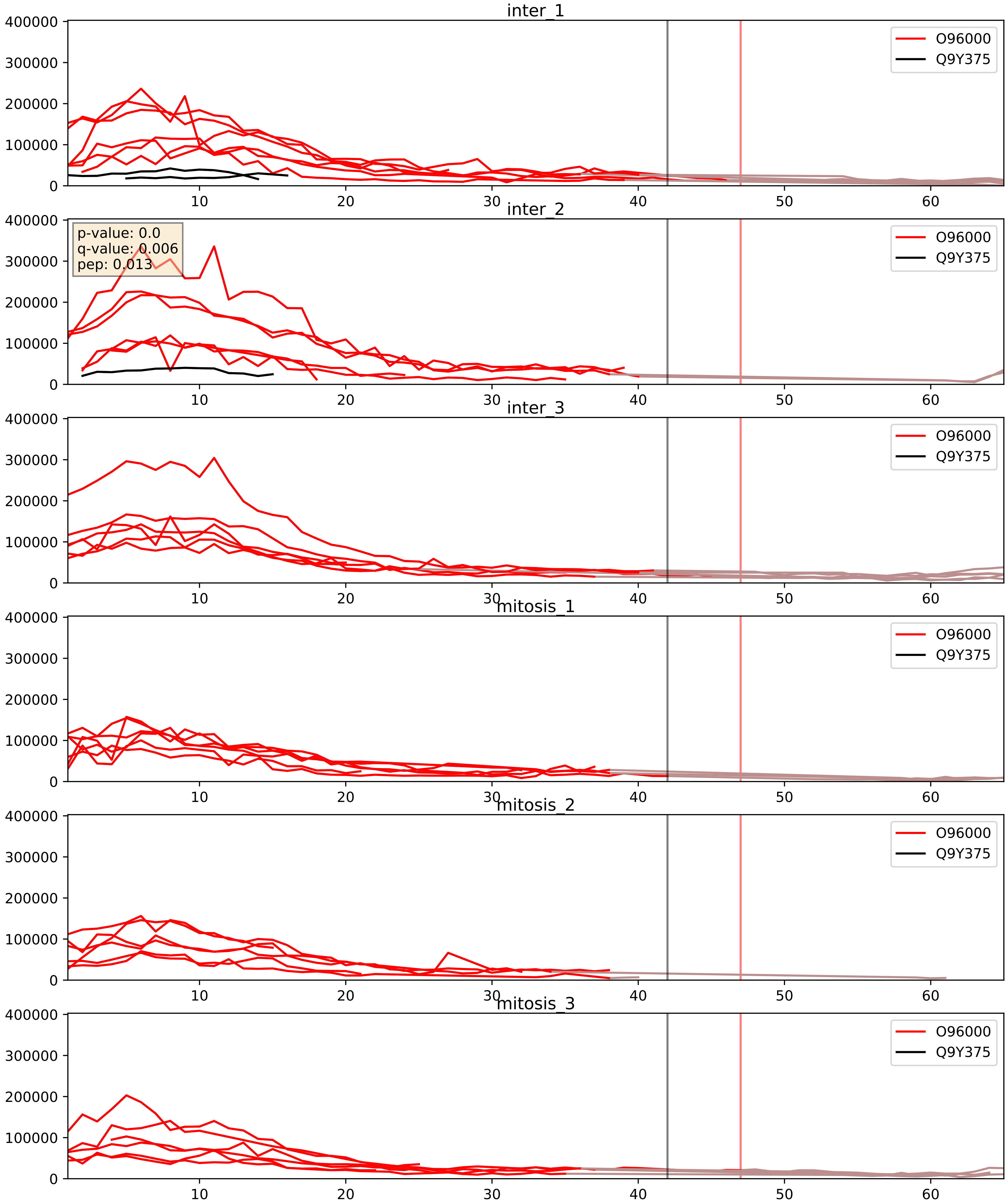


| level | condition_1 | condition_2 | pvalue | pvalue_adjusted |
|---|---|---|---|---|
| complex_abundance | mitosis | inter | 0.08025152032626533 | 0.20737287362969975 |
| interactor_abundance | mitosis | inter | 0.08672635507808389 | 0.2490831954421708 |
| interactor_ratio | mitosis | inter | 0.1043222253923626 | 0.538599667888193 |
