## Supplementary Data 2 for "SECAT: Quantifying differential protein-protein interaction states by network-centric analysis": hela_string_000192_Q92922.pdf

Q92922_Q92922

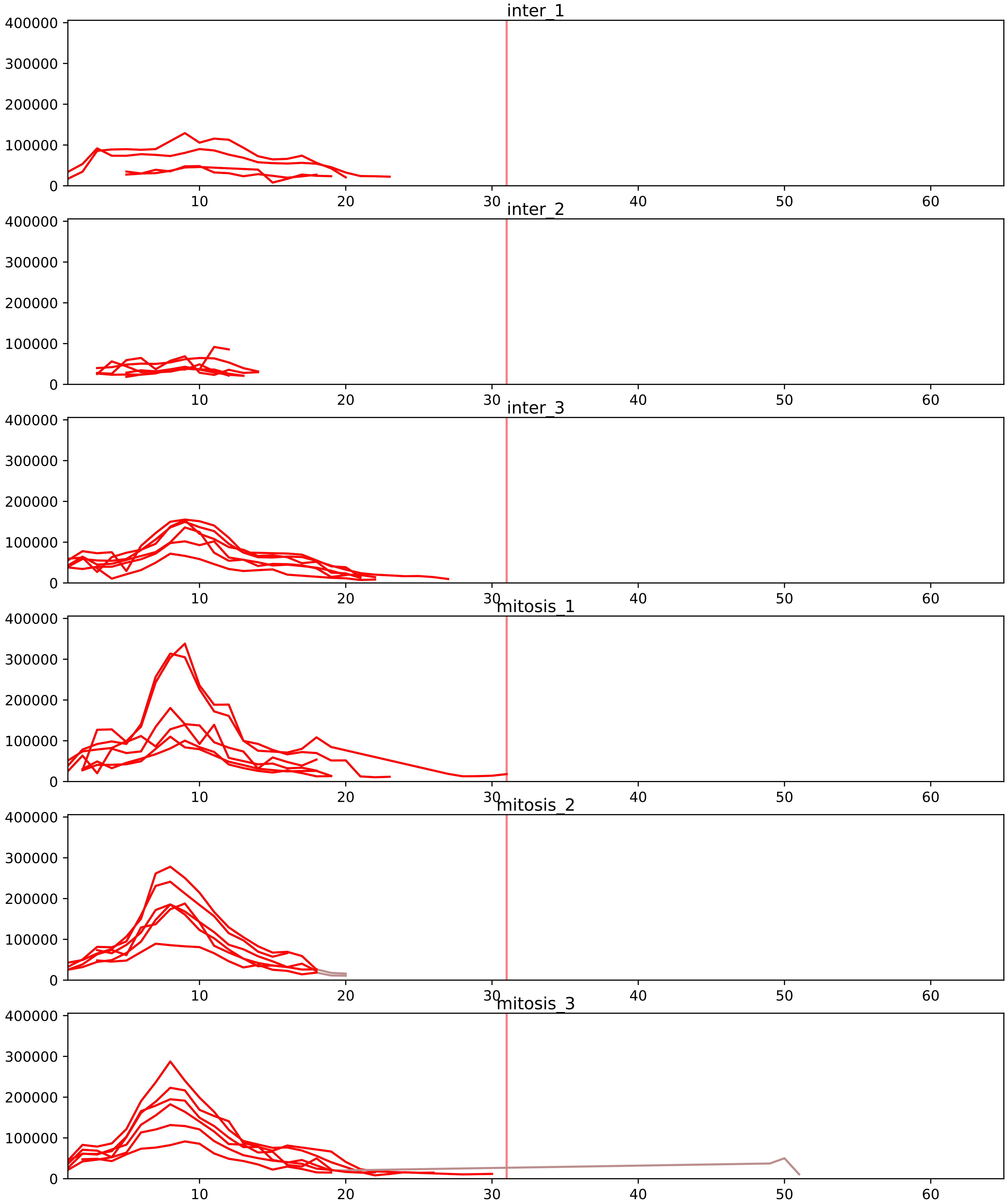


| level | condition_1 | condition_2 | pvalue | pvalue_adjusted |
|---|---|---|---|---|
| interactor_ratio | mitosis | inter | 0.00020532048253380392 | 0.0030466910311467677 |
| complex_abundance | mitosis | inter | 0.0005002792839507633 | 0.00687501553018023 |
| interactor_abundance | mitosis | inter | 0.0007035437079490738 | 0.009246574447330683 |
| monomer_abundance | mitosis | inter | 0.17163815278590833 | 0.536435242130112 |
| assembled_abundance | mitosis | inter | 0.35628373015223797 | 0.6421887334781001 |
| total_abundance | mitosis | inter | 0.4023382530993396 | 0.7023154146704781 |

O14497_Q92922
ARI1A_HUMAN vs SMRC1_HUMAN
p-value: 0.023 q-value: 0.018

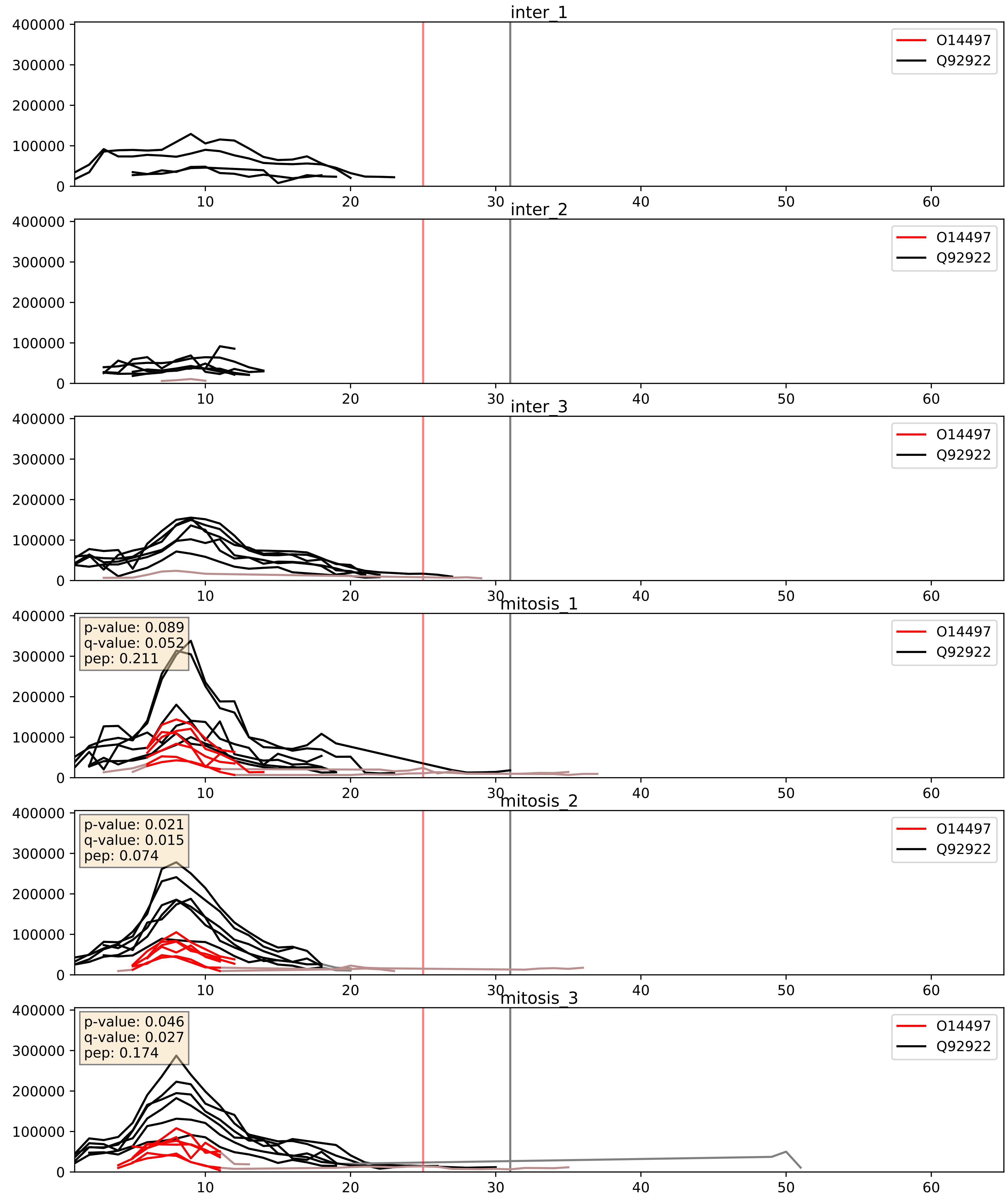


| level | condition_1 | condition_2 | pvalue | pvalue_adjusted |
|---|---|---|---|---|
| interactor_ratio | mitosis | inter | 0.0002786983396124844 | 0.017163005662466667 |
| complex_abundance | mitosis | inter | 0.0013036206631568974 | 0.024459508269436846 |
| interactor_abundance | mitosis | inter | 0.012640483206095423 | 0.10027749130318384 |

O96019_Q92922
ACL6A_HUMAN vs SMRC1_HUMAN
p-value: 0.001 q-value: 0.002

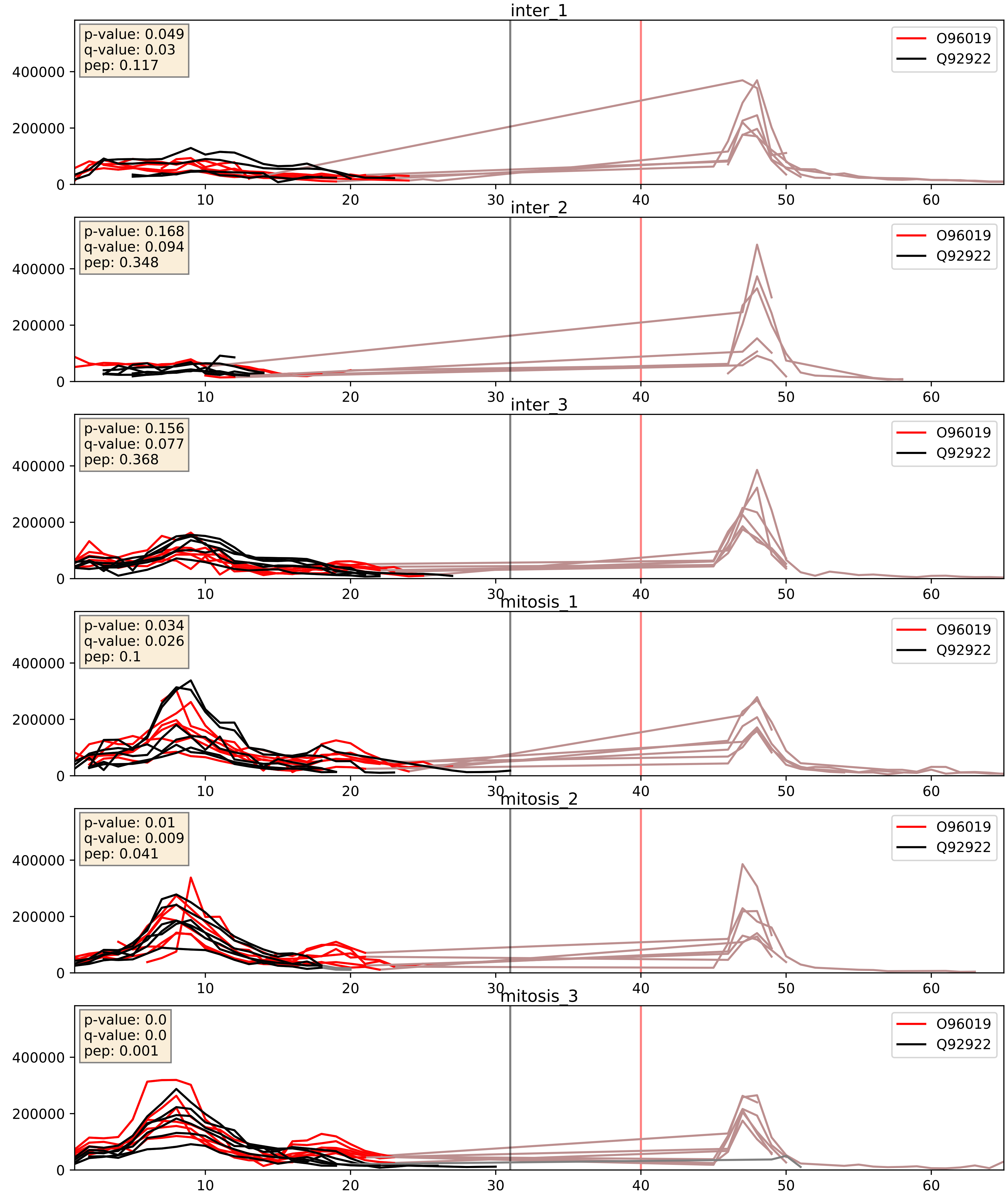


| level | condition_1 | condition_2 | pvalue | pvalue_adjusted |
|---|---|---|---|---|
| complex_abundance | mitosis | inter | 0.08346897044542251 | 0.2116393326459765 |
| interactor_abundance | mitosis | inter | 0.09277419755208206 | 0.2598223886948544 |
| interactor_ratio | mitosis | inter | 0.7994250313781059 | 0.9616467493811954 |

P51531_Q92922
SMCA2_HUMAN vs SMRC1_HUMAN
p-value: 0.024 q-value: 0.019

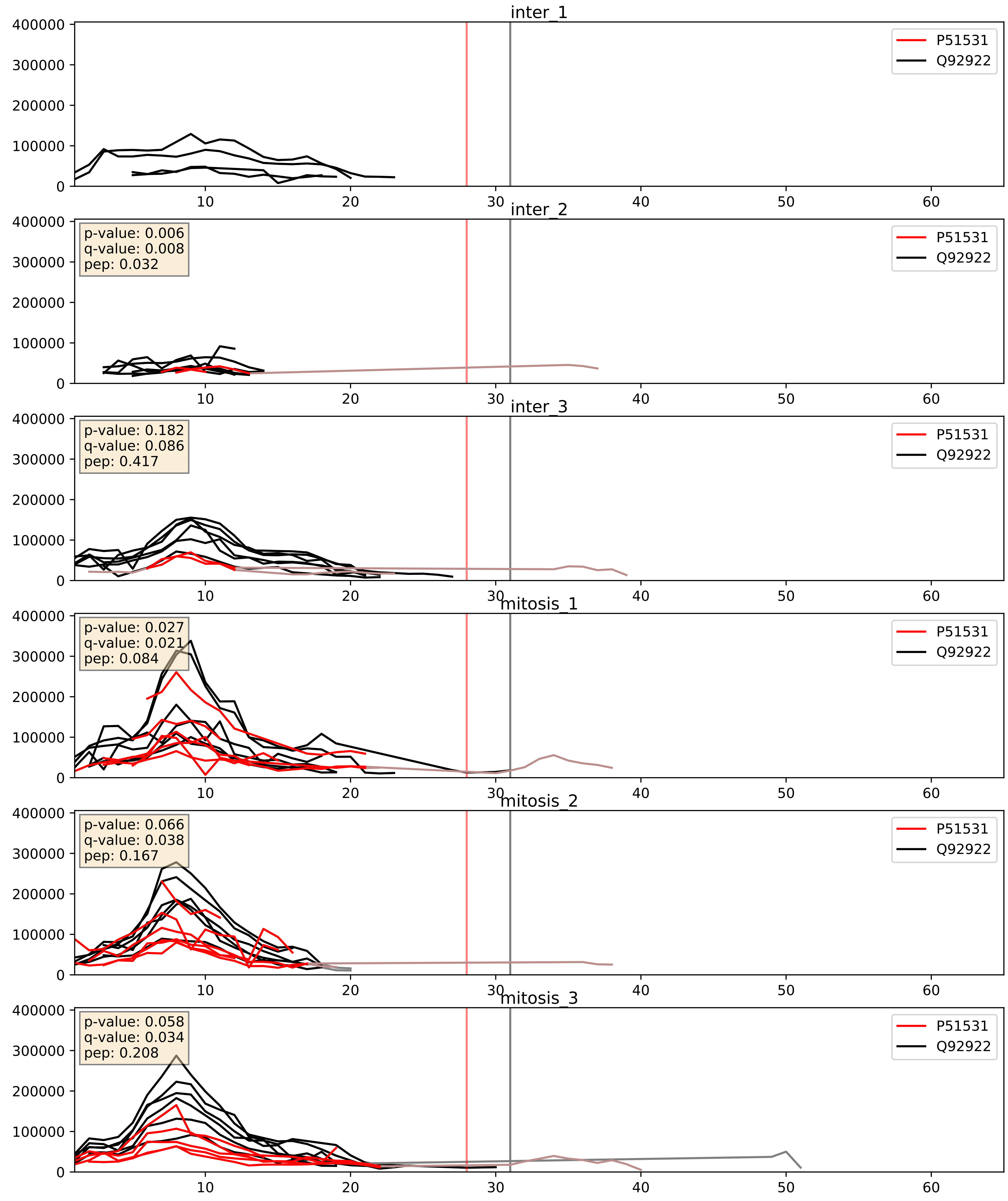


| level | condition_1 | condition_2 | pvalue | pvalue_adjusted |
|---|---|---|---|---|
| complex_abundance | mitosis | inter | 0.005693057559631294 | 0.056713450503088285 |
| interactor_abundance | mitosis | inter | 0.013657409124963755 | 0.10336642096347458 |
| interactor_ratio | mitosis | inter | 0.5531009131904417 | 0.8889886679627823 |

Q8TAQ2_Q92922
SMRC2_HUMAN vs SMRC1_HUMAN
p-value: 0.003 q-value: 0.003

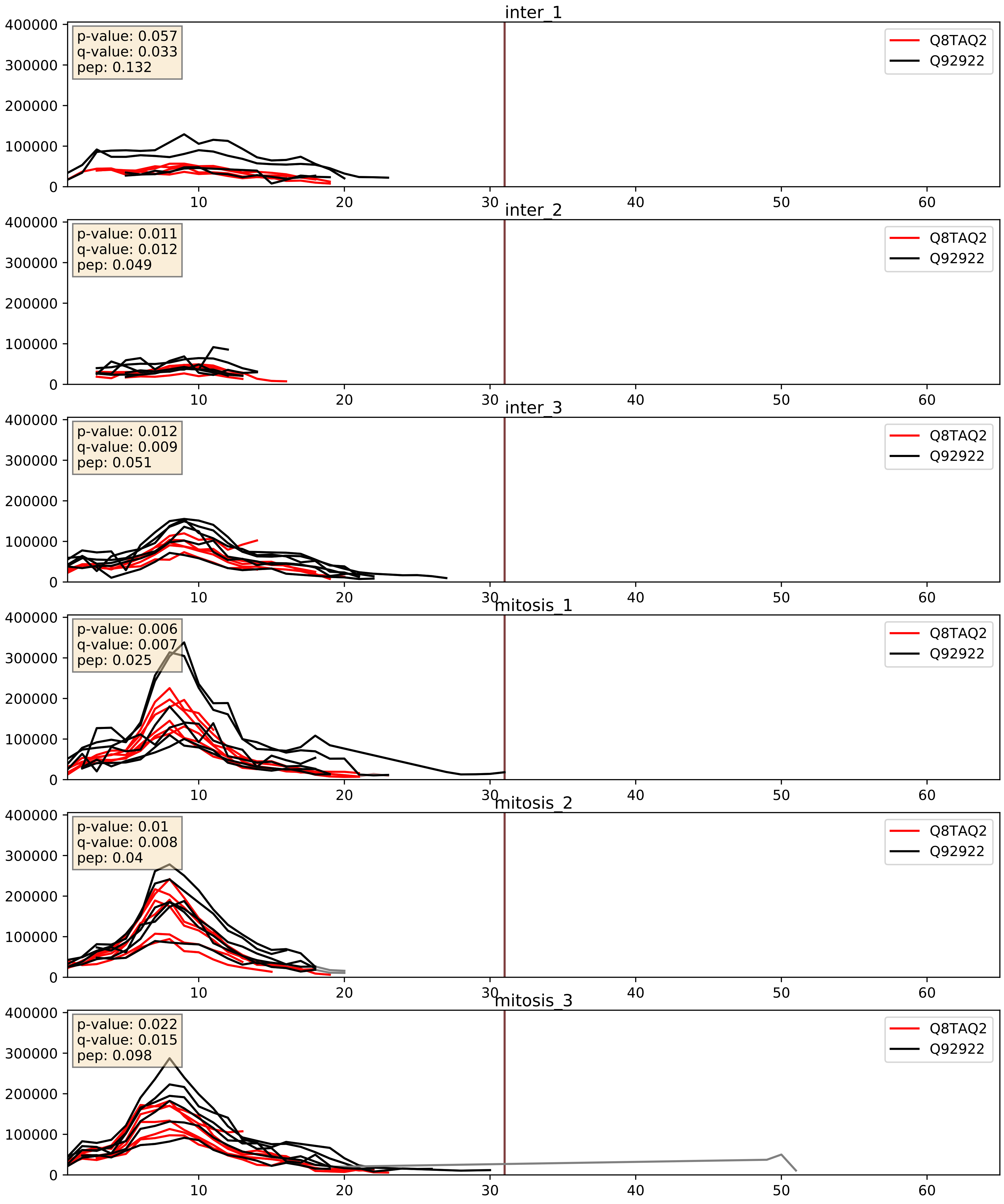


| level | condition_1 | condition_2 | pvalue | pvalue_adjusted |
|---|---|---|---|---|
| interactor_abundance | mitosis | inter | 0.01873319039954222 | 0.11790890427947162 |
| complex_abundance | mitosis | inter | 0.041723956886434606 | 0.14703872826178685 |
| interactor_ratio | mitosis | inter | 0.47647139808428074 | 0.8646587168966383 |

Q92785_Q92922
REQU_HUMAN vs SMRC1_HUMAN
p-value: 0.001 q-value: 0.023

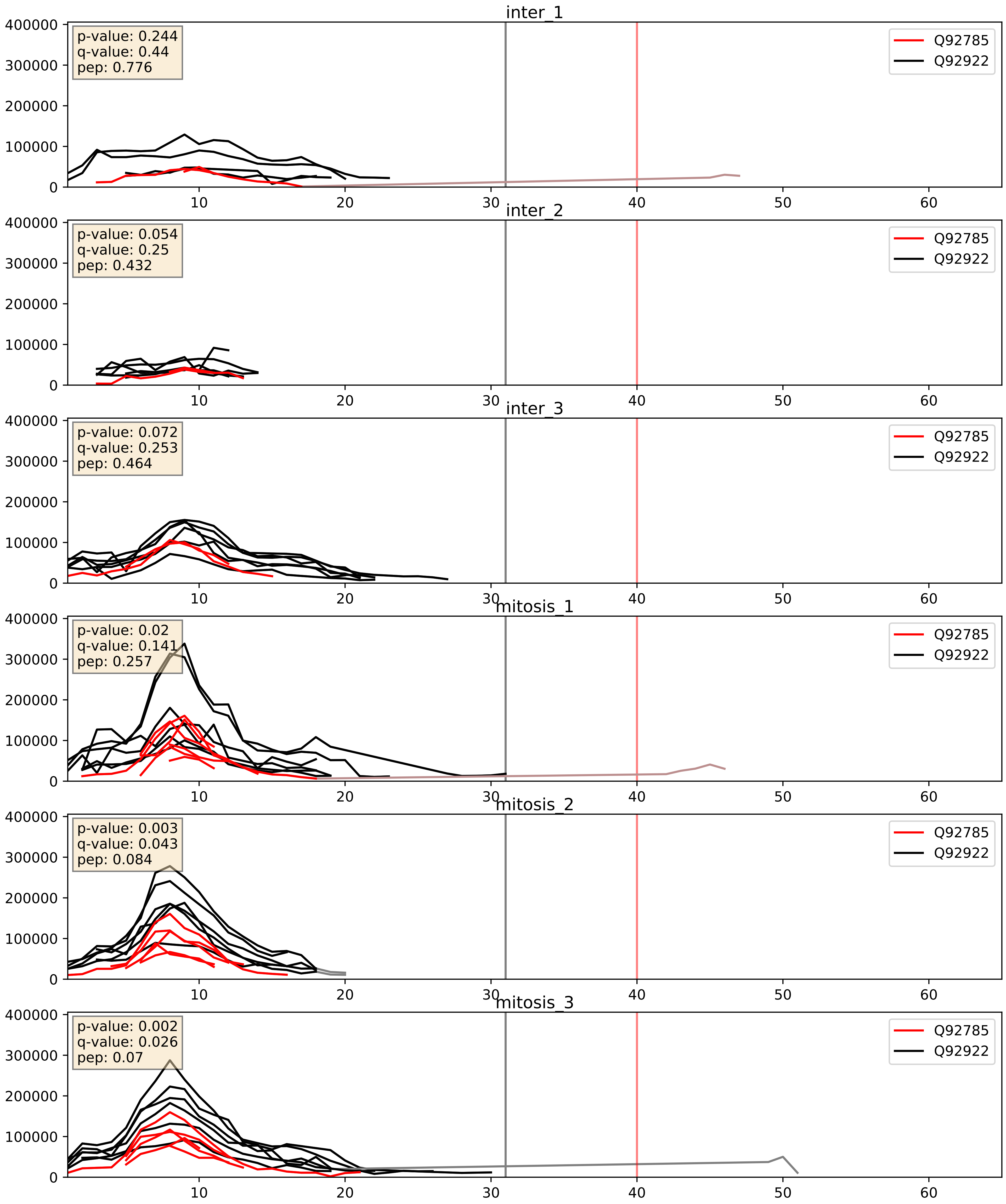


| level | condition_1 | condition_2 | pvalue | pvalue_adjusted |
|---|---|---|---|---|
| interactor_abundance | mitosis | inter | 0.0034125104988964945 | 0.04511548235131213 |
| complex_abundance | mitosis | inter | 0.013678879216987402 | 0.08692368307384146 |
| interactor_ratio | mitosis | inter | 0.2747192024977269 | 0.7473911657831552 |

Q92922_Q92925
SMRC1_HUMAN vs SMRD2_HUMAN
p-value: 0.043 q-value: 0.029

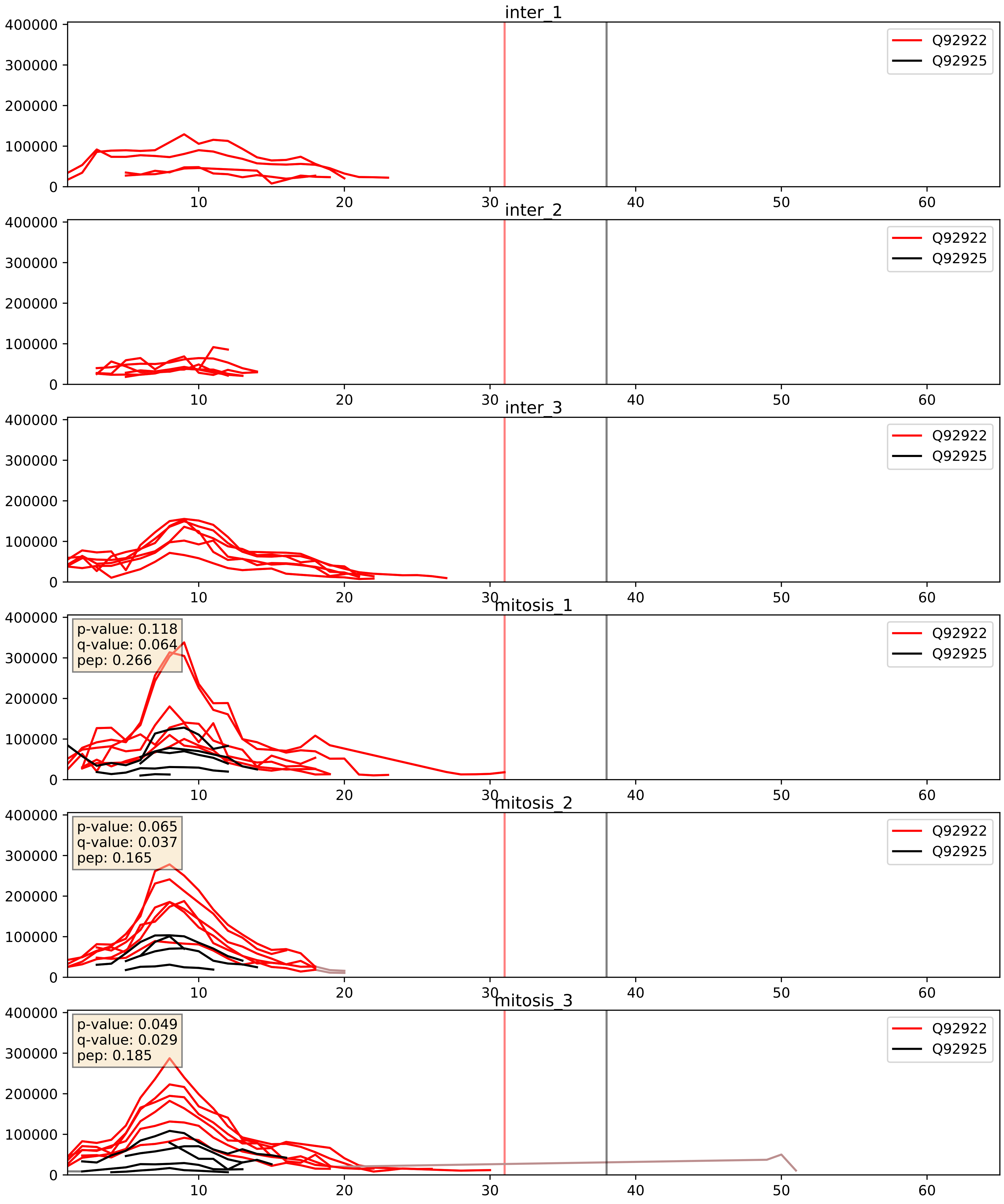


| level | condition_1 | condition_2 | pvalue | pvalue_adjusted |
|---|---|---|---|---|
| interactor_ratio | mitosis | inter | 0.00010380497123051964 | 0.010720128702277293 |
| interactor_abundance | mitosis | inter | 0.00018540588357067092 | 0.01016145604161317 |
| complex_abundance | mitosis | inter | 0.0005971387895009652 | 0.01581504091436693 |

Q92922_Q969G3
SMRC1_HUMAN vs SMCE1_HUMAN
p-value: 0.0 q-value: 0.0

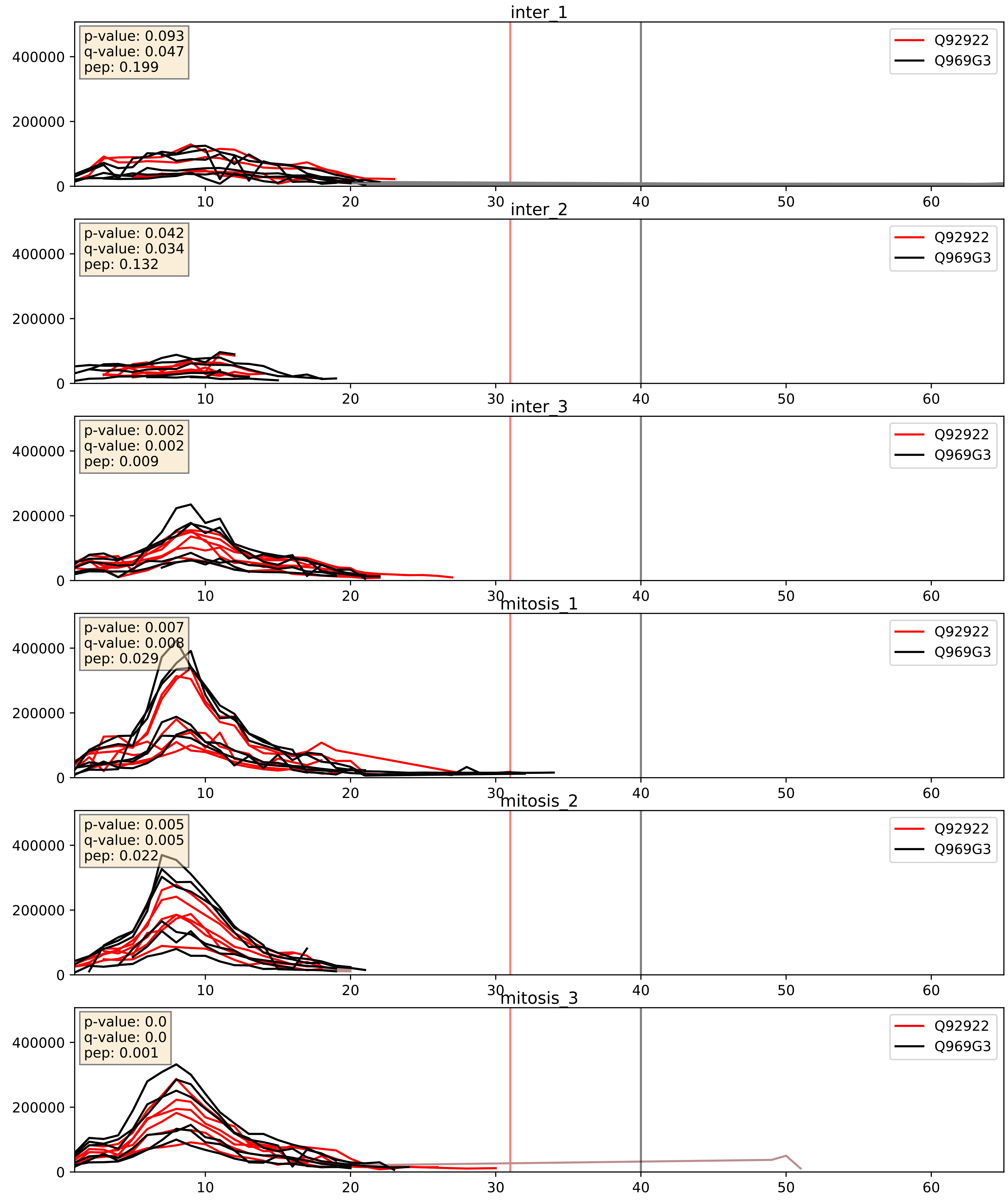


| level | condition_1 | condition_2 | pvalue | pvalue_adjusted |
|---|---|---|---|---|
| complex_abundance | mitosis | inter | 0.05577982617922177 | 0.16848105578480535 |
| interactor_abundance | mitosis | inter | 0.07736330381217478 | 0.23055134583284403 |
| interactor_ratio | mitosis | inter | 0.21333639791135564 | 0.6904192954236995 |

Q92922_Q96GM5
SMRC1_HUMAN vs SMRD1_HUMAN
p-value: 0.02 q-value: 0.016

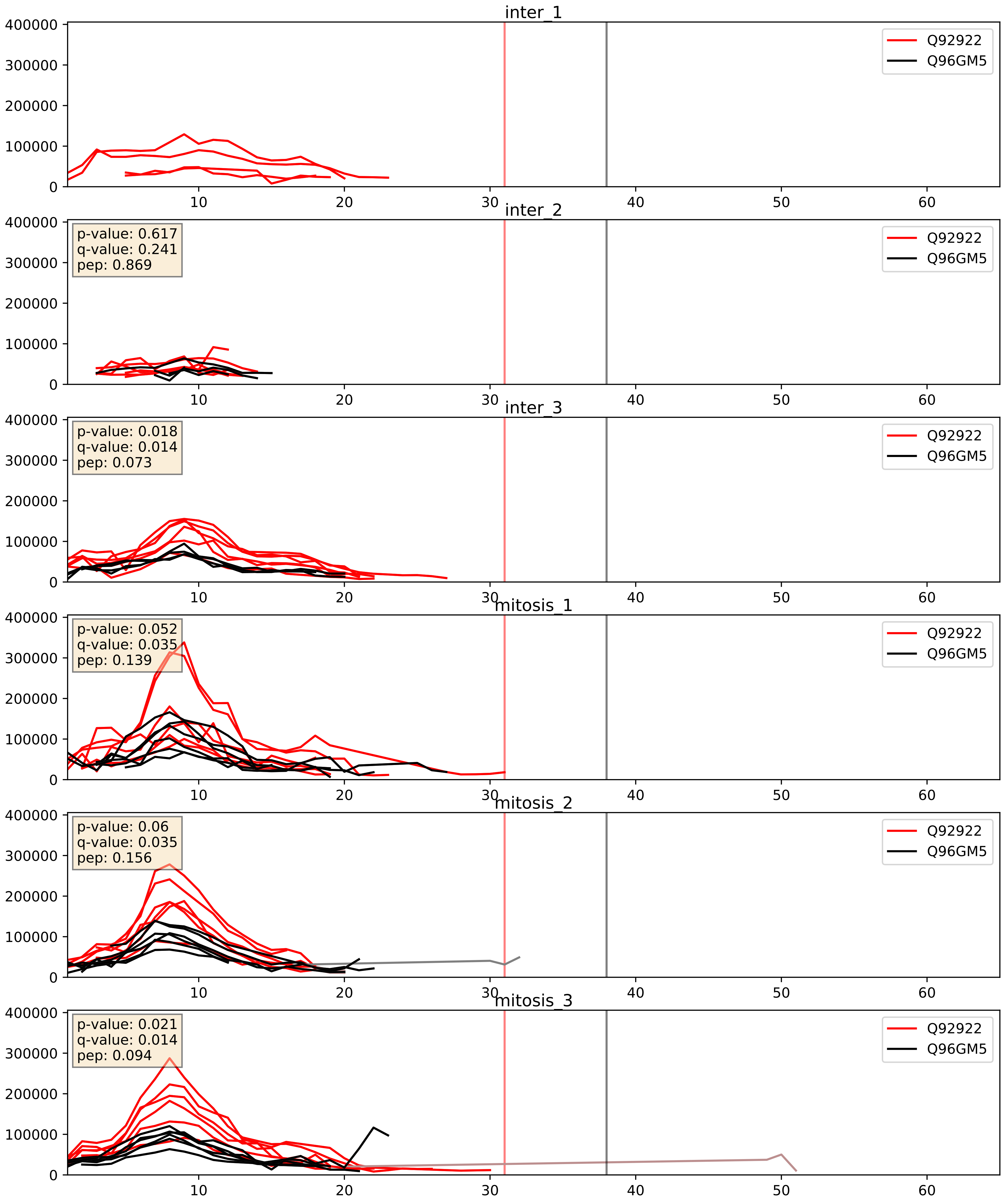


| level | condition_1 | condition_2 | pvalue | pvalue_adjusted |
|---|---|---|---|---|
| complex_abundance | mitosis | inter | 0.04777855643174045 | 0.15599228513863675 |
| interactor_abundance | mitosis | inter | 0.10152273968561054 | 0.27370850762629345 |
| interactor_ratio | mitosis | inter | 0.8269612471471082 | 0.969784012122695 |
