## Supplementary Data 2 for "SECAT: Quantifying differential protein-protein interaction states by network-centric analysis": hela_string_000193_O14548.pdf

O14548_O14548

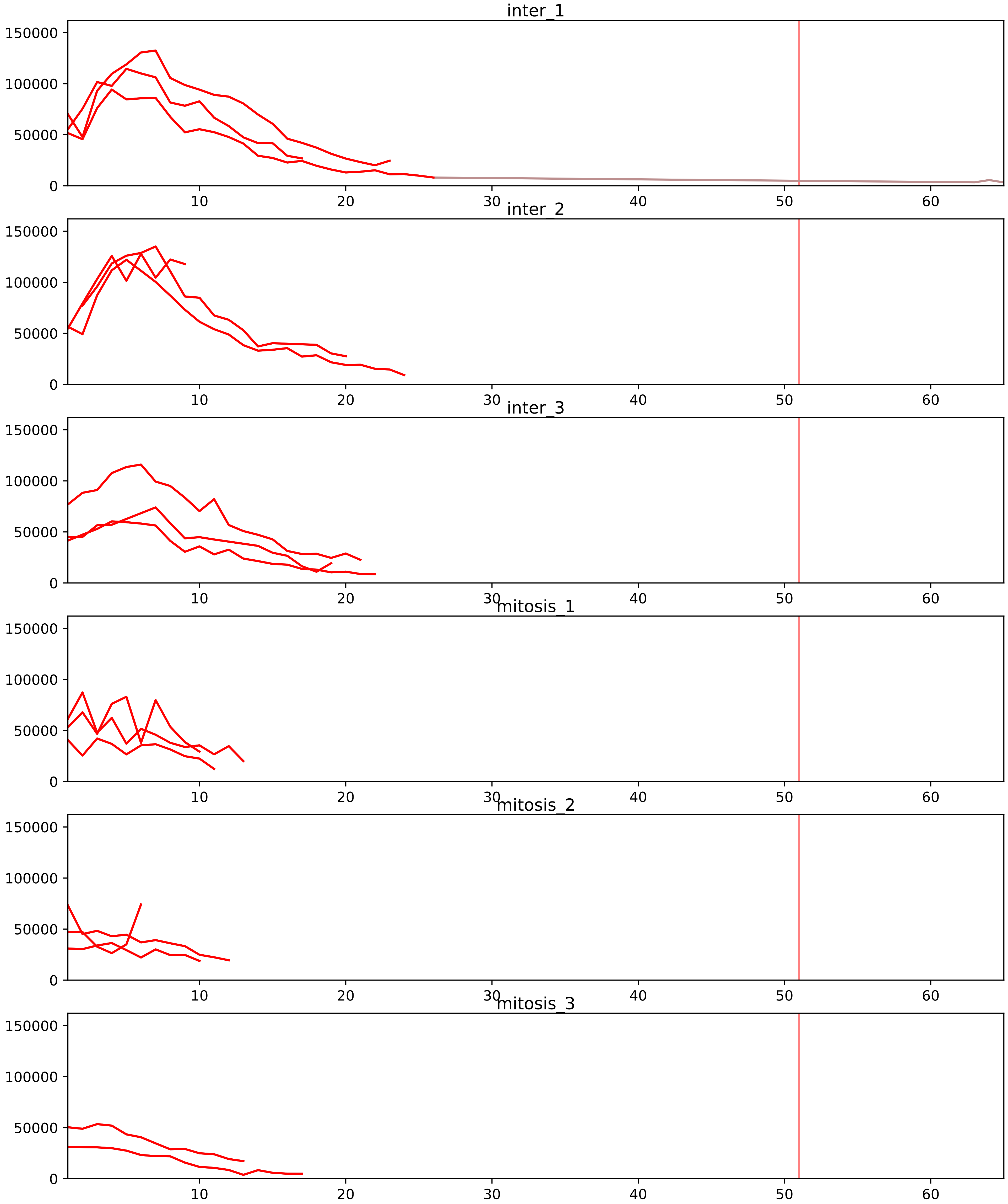


| level | condition_1 | condition_2 | pvalue | pvalue_adjusted |
|---|---|---|---|---|
| complex_abundance | mitosis | inter | 0.0002100164232376458 | 0.003794887573692578 |
| interactor_abundance | mitosis | inter | 0.0004383049086454026 | 0.006665132495103644 |
| interactor_ratio | mitosis | inter | 0.029149692138891892 | 0.13823565344216773 |
| total_abundance | mitosis | inter | 0.3122818273378753 | 0.635635760680397 |
| monomer_abundance | mitosis | inter | 0.34357785614906045 | 0.6740536609182377 |
| assembled_abundance | mitosis | inter | 0.38523839483398636 | 0.6613371783196998 |

O14548_O14949
COX7R_HUMAN vs QCR8_HUMAN
p-value: 0.002 q-value: 0.035

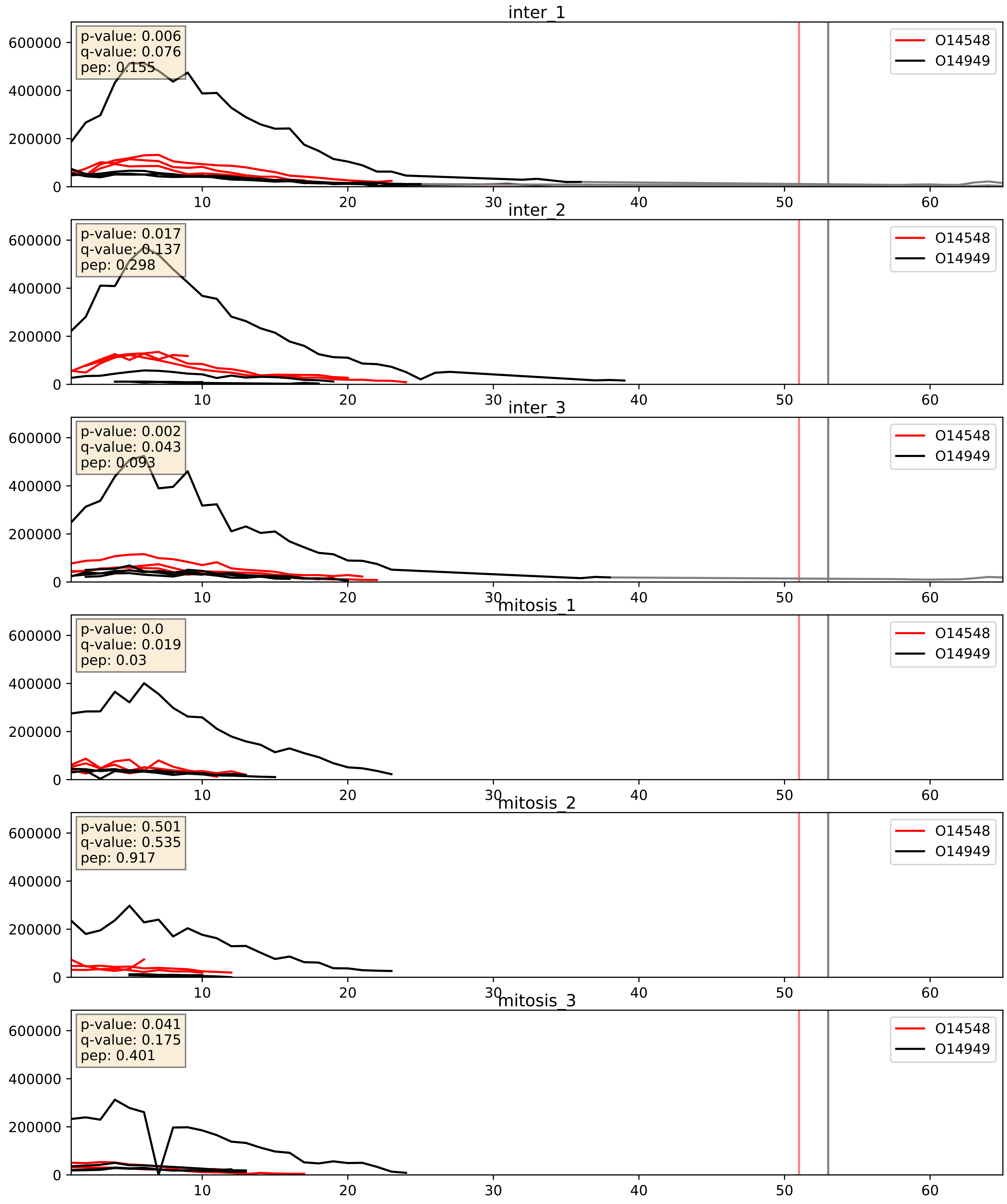


| level | condition_1 | condition_2 | pvalue | pvalue_adjusted |
|---|---|---|---|---|
| interactor_abundance | mitosis | inter | 0.012514854895198036 | 0.10000783240914232 |
| complex_abundance | mitosis | inter | 0.01658980781803441 | 0.09584596180627945 |
| interactor_ratio | mitosis | inter | 0.38235688435364557 | 0.8200889326151857 |

O14548_P09669
COX7R_HUMAN vs COX6C_HUMAN
p-value: 0.0 q-value: 0.0

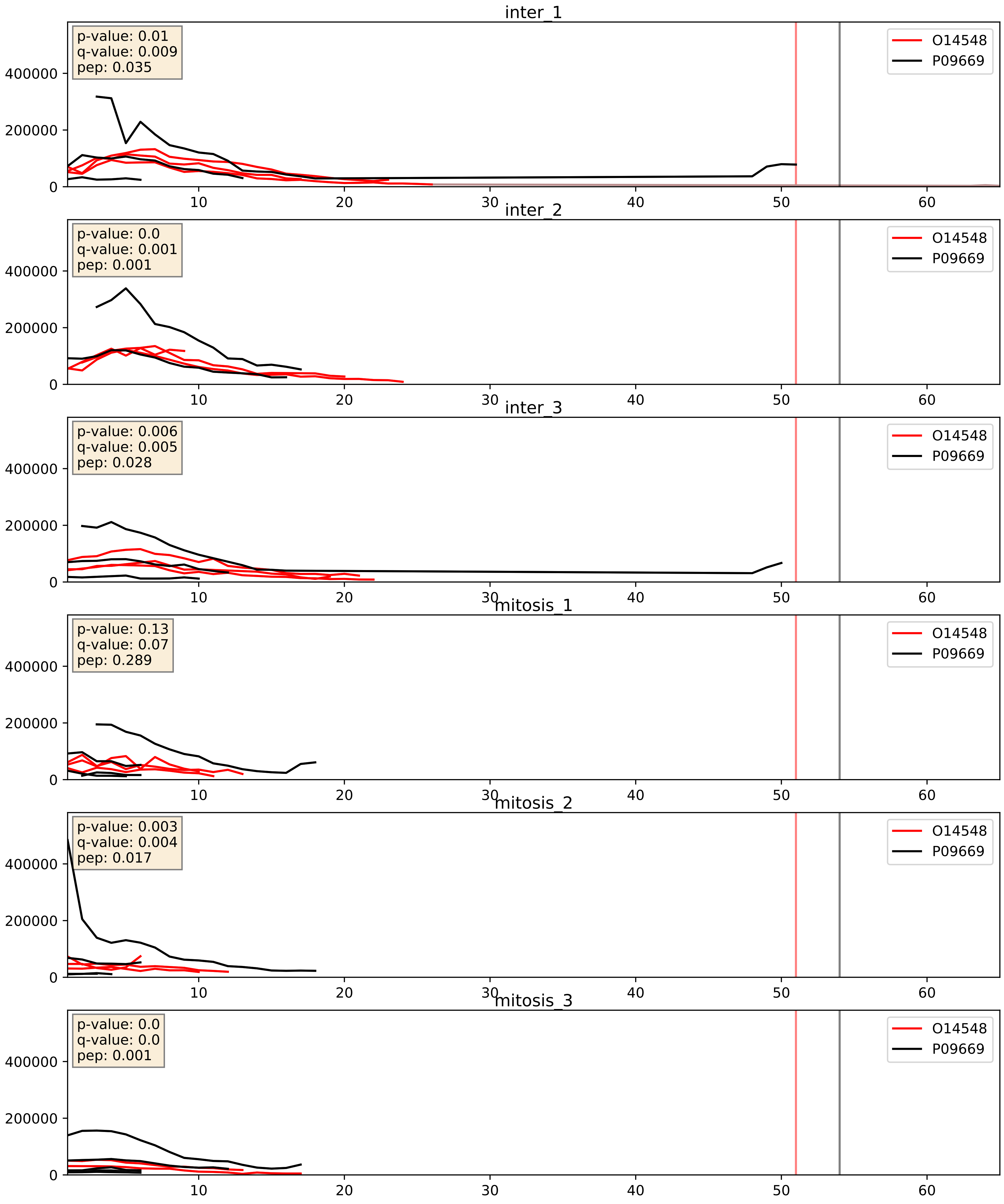


| level | condition_1 | condition_2 | pvalue | pvalue_adjusted |
|---|---|---|---|---|
| complex_abundance | mitosis | inter | 0.00828998065096952 | 0.06742444735376033 |
| interactor_abundance | mitosis | inter | 0.016084101322436272 | 0.11124909769145716 |
| interactor_ratio | mitosis | inter | 0.1824817849305413 | 0.6518307938381196 |

O14548_P10606
COX7R_HUMAN vs COX5B_HUMAN
p-value: 0.0 q-value: 0.003

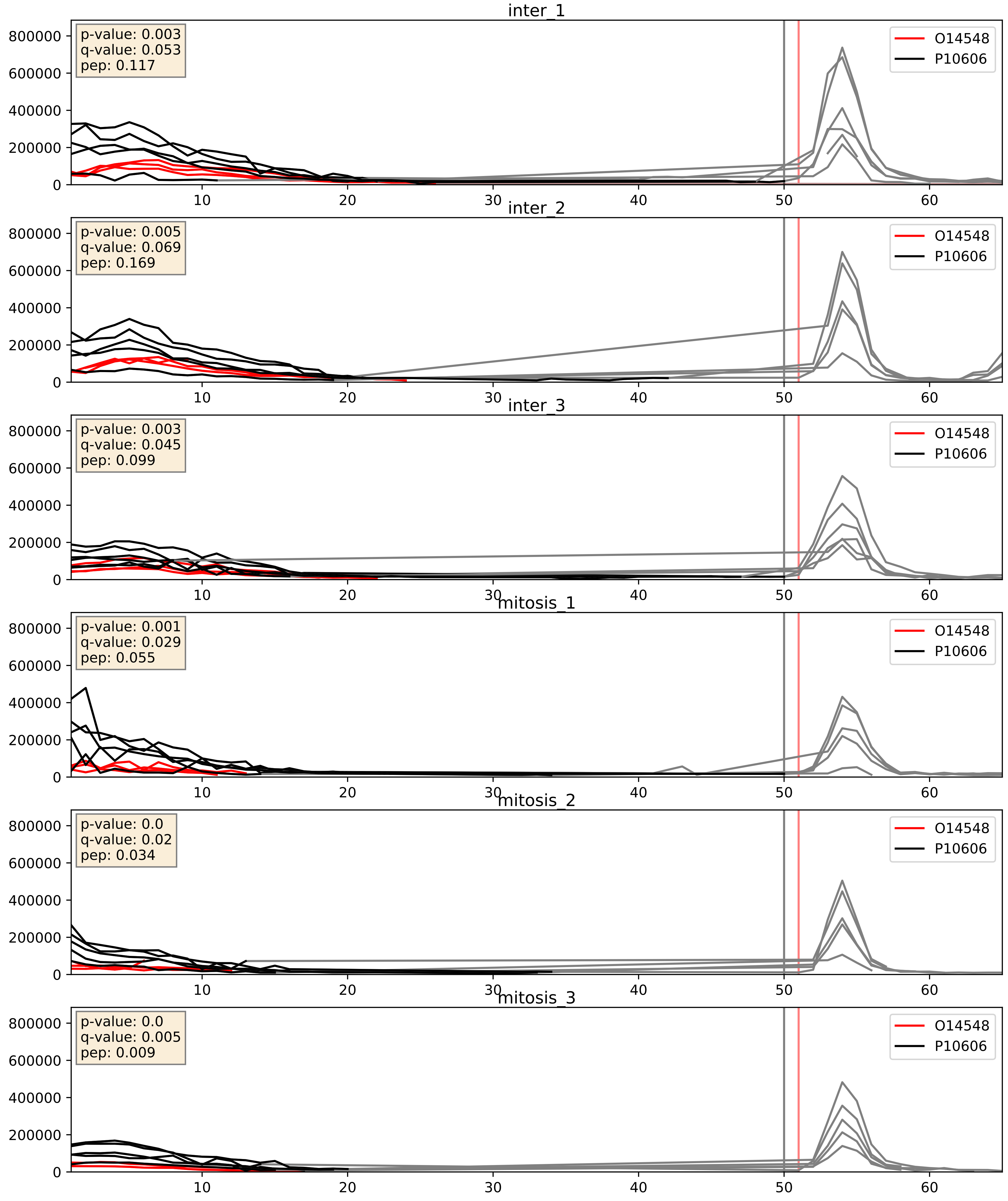


| level | condition_1 | condition_2 | pvalue | pvalue_adjusted |
|---|---|---|---|---|
| interactor_abundance | mitosis | inter | 0.012520884634712784 | 0.10000783240914232 |
| complex_abundance | mitosis | inter | 0.022456458683504935 | 0.10987189501272765 |
| interactor_ratio | mitosis | inter | 0.022930052905559373 | 0.2336681581804622 |

O14548_P14927
COX7R_HUMAN vs QCR7_HUMAN
p-value: 0.001 q-value: 0.048

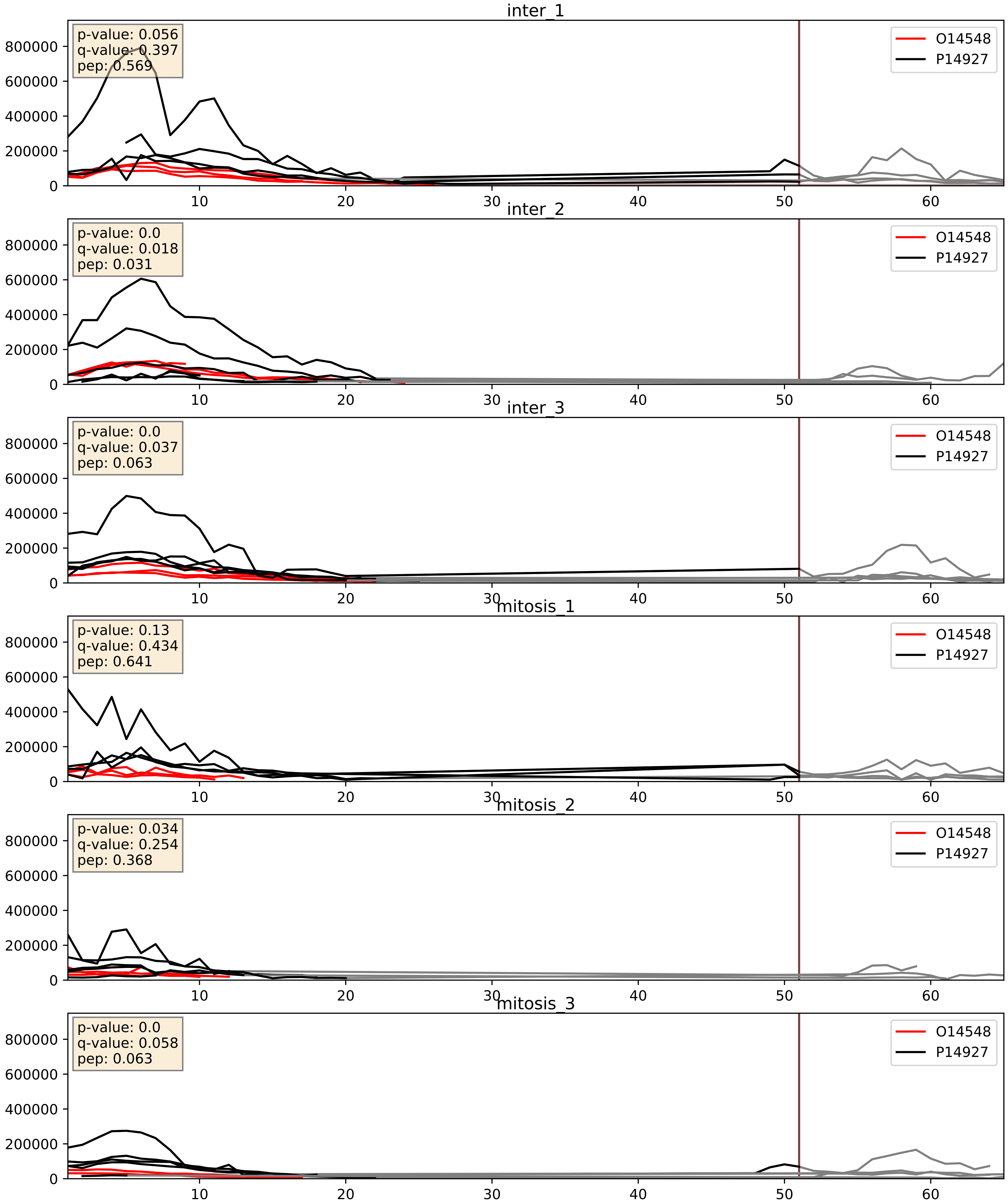


| level | condition_1 | condition_2 | pvalue | pvalue_adjusted |
|---|---|---|---|---|
| complex_abundance | mitosis | inter | 0.009937839188162696 | 0.07371585975104354 |
| interactor_abundance | mitosis | inter | 0.012524345367126235 | 0.10000783240914232 |
| interactor_ratio | mitosis | inter | 0.22098934004277596 | 0.6993230132222411 |

O14548_P15954
COX7R_HUMAN vs COX7C_HUMAN
p-value: 0.001 q-value: 0.025

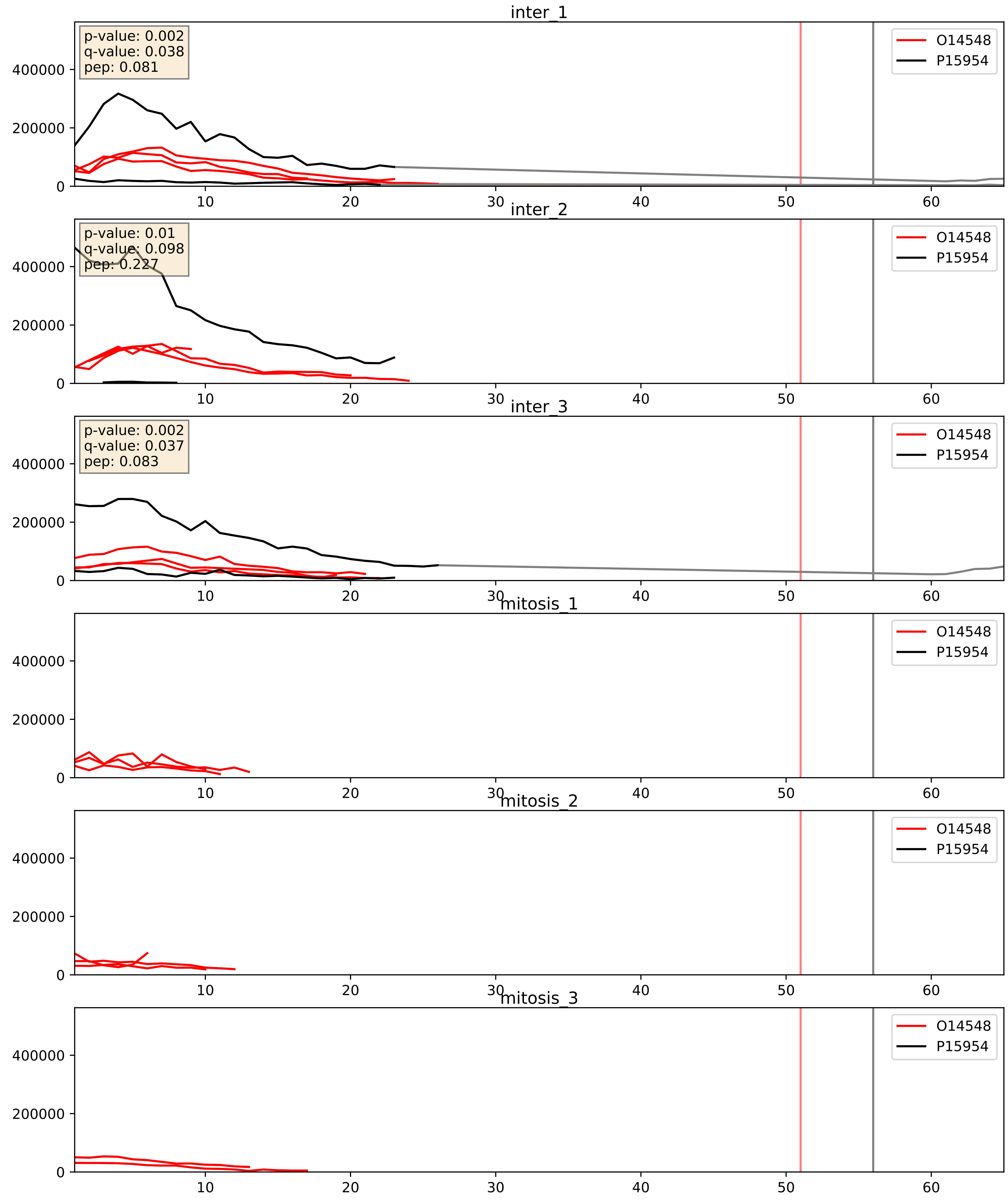


| level | condition_1 | condition_2 | pvalue | pvalue_adjusted |
|---|---|---|---|---|
| complex_abundance | mitosis | inter | 0.00021884711152757414 | 0.010087086090925927 |
| interactor_abundance | mitosis | inter | 0.0004706587247677187 | 0.015253292300429789 |
| interactor_ratio | mitosis | inter | 0.014546406987366608 | 0.17686306829592754 |

O14548_P47985
COX7R_HUMAN vs UCRI_HUMAN
p-value: 0.002 q-value: 0.03

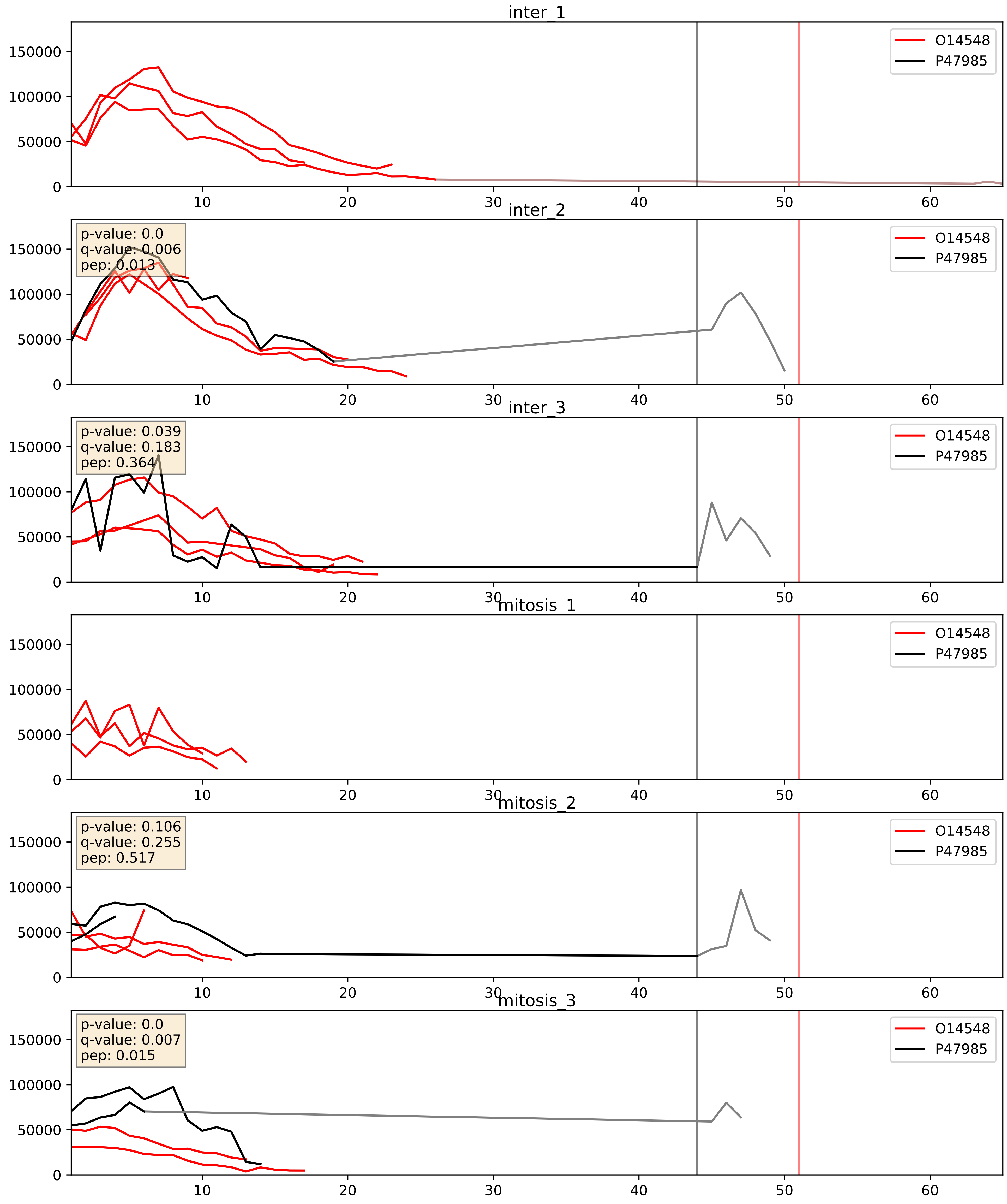


| level | condition_1 | condition_2 | pvalue | pvalue_adjusted |
|---|---|---|---|---|
| interactor_ratio | mitosis | inter | 0.07663334640763149 | 0.4648714486236808 |
| interactor_abundance | mitosis | inter | 0.15496205183180453 | 0.34179427400218604 |
| complex_abundance | mitosis | inter | 0.2642001707612497 | 0.4142039307172706 |

O14548_P56385
COX7R_HUMAN vs ATP5I_HUMAN
p-value: 0.0 q-value: 0.046

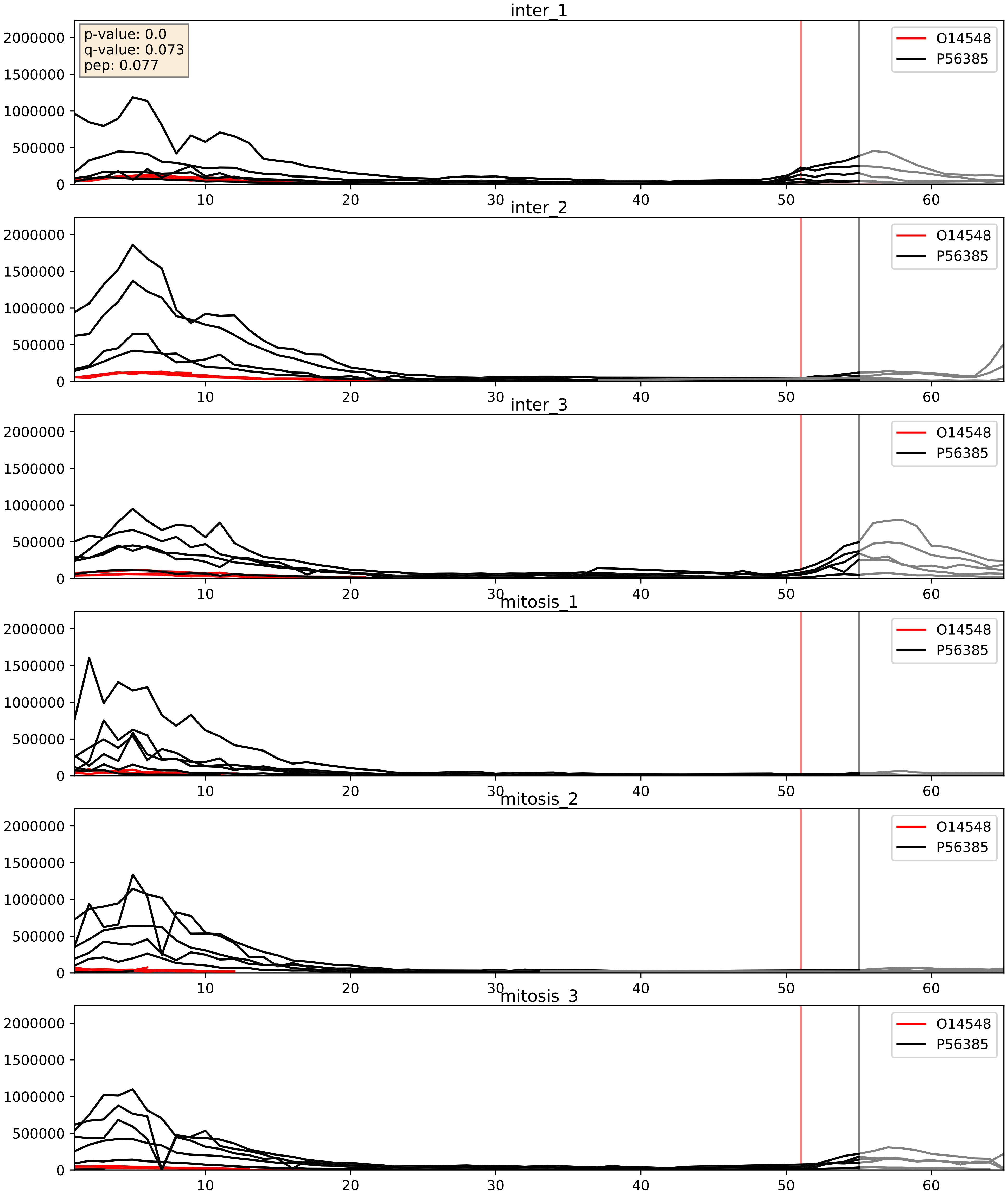


| level | condition_1 | condition_2 | pvalue | pvalue_adjusted |
|---|---|---|---|---|
| complex_abundance | mitosis | inter | 0.004430050935506871 | 0.049683977369634376 |
| interactor_abundance | mitosis | inter | 0.012520884634712784 | 0.10000783240914232 |
| interactor_ratio | mitosis | inter | 0.048677689602045764 | 0.364412066789464 |
