## Supplementary Data 2 for "SECAT: Quantifying differential protein-protein interaction states by network-centric analysis": hela_string_000194_Q5BJH7.pdf

Q5BJH7_Q5BJH7

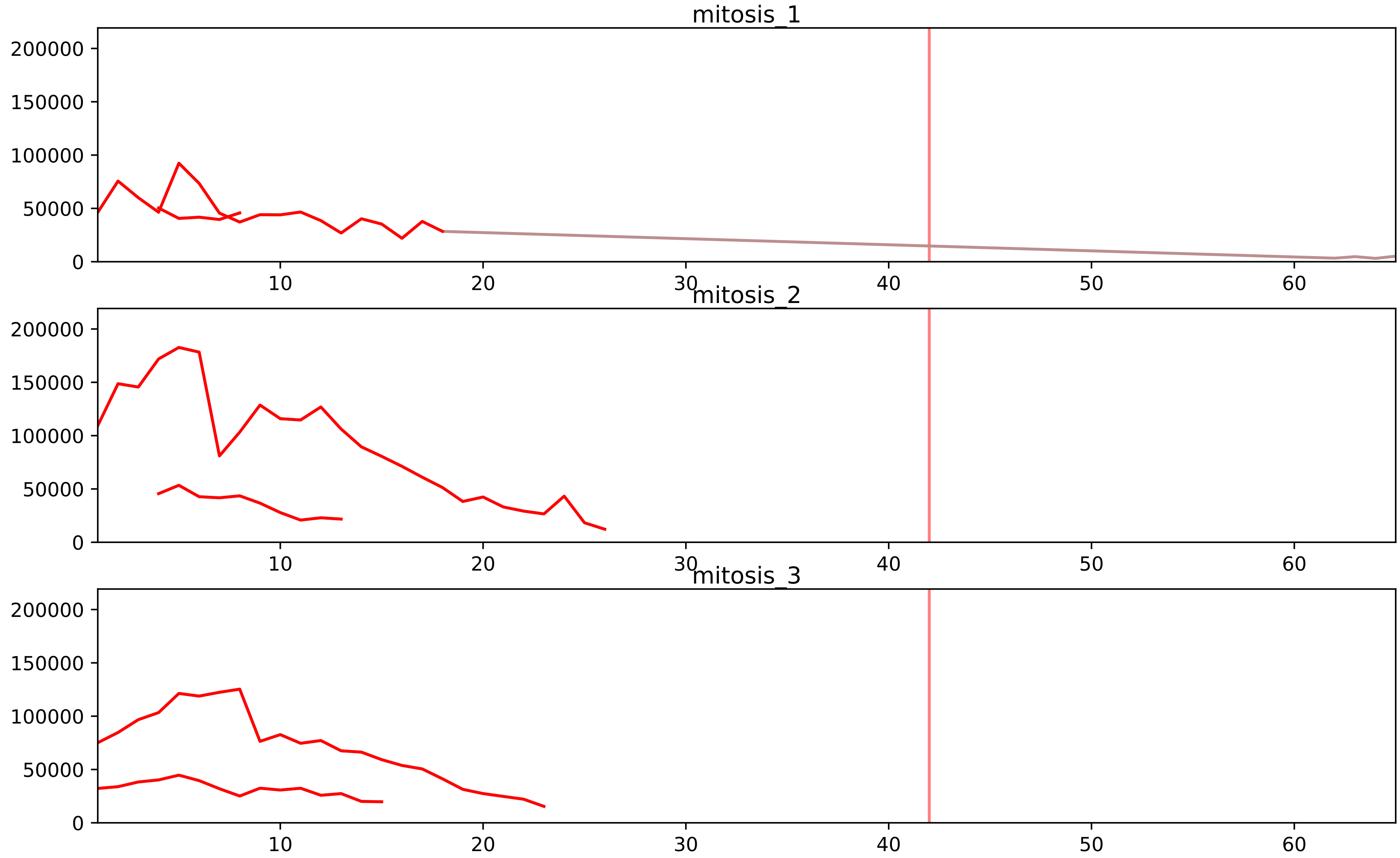


| level | condition_1 | condition_2 | pvalue | pvalue_adjusted |
|---|---|---|---|---|
| complex_abundance | mitosis | inter | 0.00022211895660948554 | 0.0038923702872519375 |
| interactor_abundance | mitosis | inter | 0.0015399695522842847 | 0.015829854615659687 |
| assembled_abundance | mitosis | inter | 0.01235848507630723 | 0.12494615257493336 |
| total_abundance | mitosis | inter | 0.012851969320205386 | 0.14285889914664981 |
| interactor_ratio | mitosis | inter | 0.35140728464106313 | 0.6594149565663059 |
| monomer_abundance | mitosis | inter | 0.44964901421802655 | 0.7255456492326616 |

Q5BJH7_Q969M3
YIF1B_HUMAN vs YIPF5_HUMAN
p-value: 0.001 q-value: 0.046

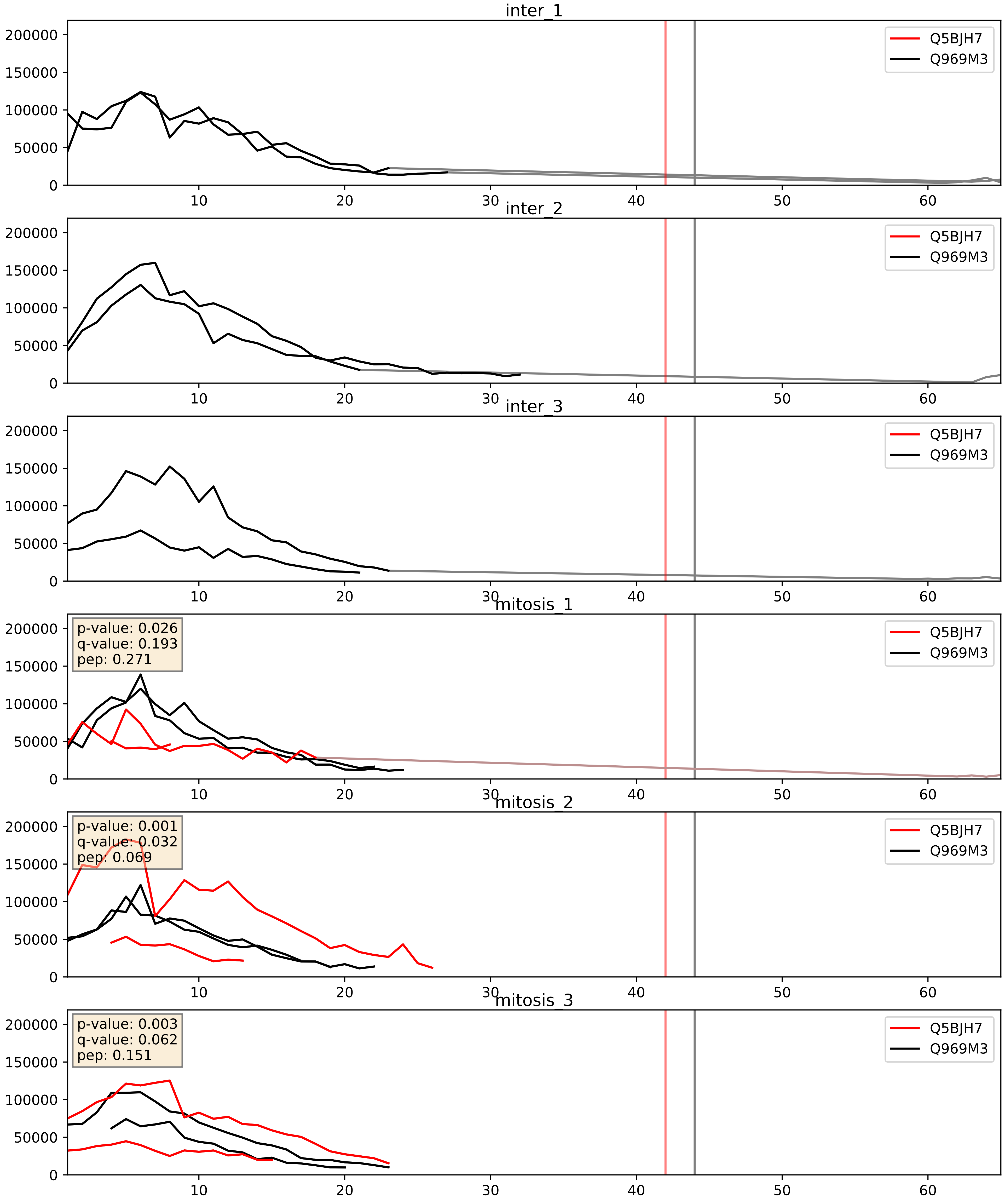


| level | condition_1 | condition_2 | pvalue | pvalue_adjusted |
|---|---|---|---|---|
| complex_abundance | mitosis | inter | 0.00022211895660948554 | 0.010113501428602107 |
| interactor_abundance | mitosis | inter | 0.0015399695522842847 | 0.028907984680718663 |
| interactor_ratio | mitosis | inter | 0.35140728464106313 | 0.8012909846903304 |
