## Supplementary Data 2 for "SECAT: Quantifying differential protein-protein interaction states by network-centric analysis": hela_string_000195_P51572.pdf

P51572_P51572

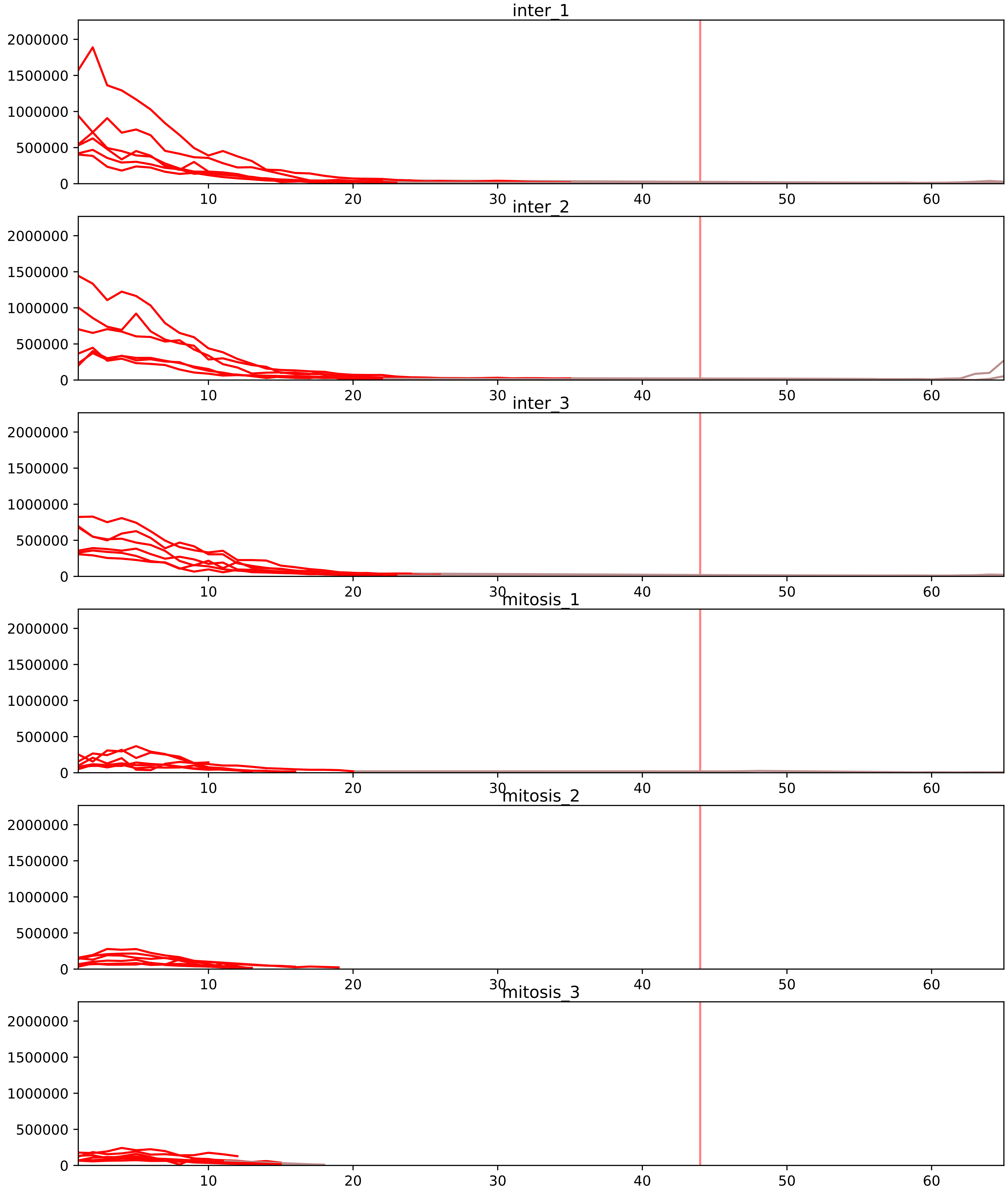


| level | condition_1 | condition_2 | pvalue | pvalue_adjusted |
|---|---|---|---|---|
| interactor_ratio | mitosis | inter | 0.00023423331011059397 | 0.0034205499254245475 |
| interactor_abundance | mitosis | inter | 0.0014127216087314219 | 0.01485375862894752 |
| complex_abundance | mitosis | inter | 0.007515969569845071 | 0.03750511614722863 |
| monomer_abundance | mitosis | inter | 0.016206195817635775 | 0.21518226668971946 |
| total_abundance | mitosis | inter | 0.31238669676337716 | 0.635635760680397 |
| assembled_abundance | mitosis | inter | 0.36979709827224405 | 0.6524709282678901 |

P51572_Q14790
BAP31_HUMAN vs CASP8_HUMAN
p-value: 0.004 q-value: 0.047

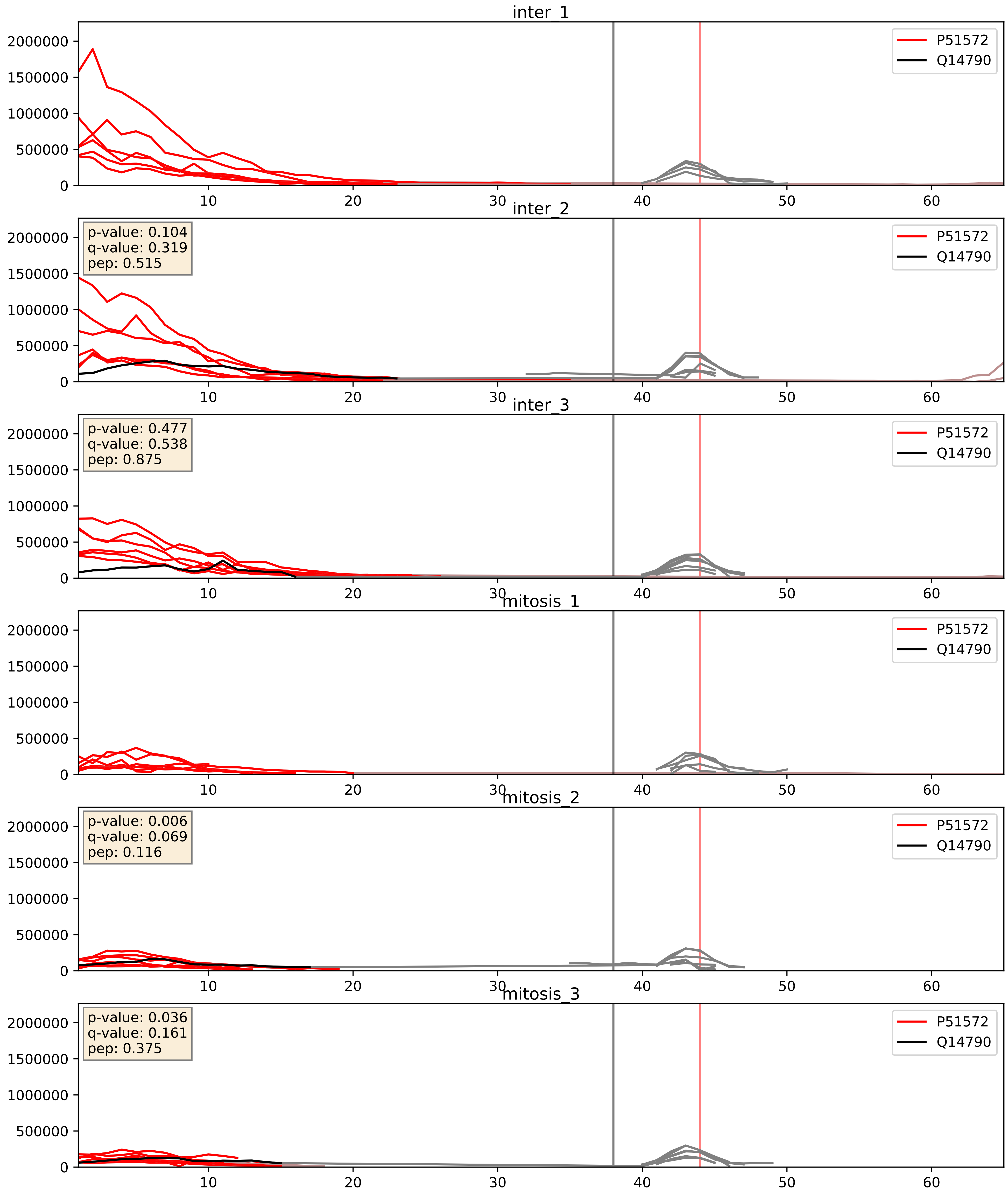


| level | condition_1 | condition_2 | pvalue | pvalue_adjusted |
|---|---|---|---|---|
| interactor_ratio | mitosis | inter | 0.4021899999618396 | 0.8311893393797354 |
| interactor_abundance | mitosis | inter | 0.44650629587583357 | 0.6220855945145077 |
| complex_abundance | mitosis | inter | 0.47500819527470256 | 0.6156980847291723 |

P51572_Q9NPA0
BAP31_HUMAN vs EMC7_HUMAN
p-value: 0.0 q-value: 0.018

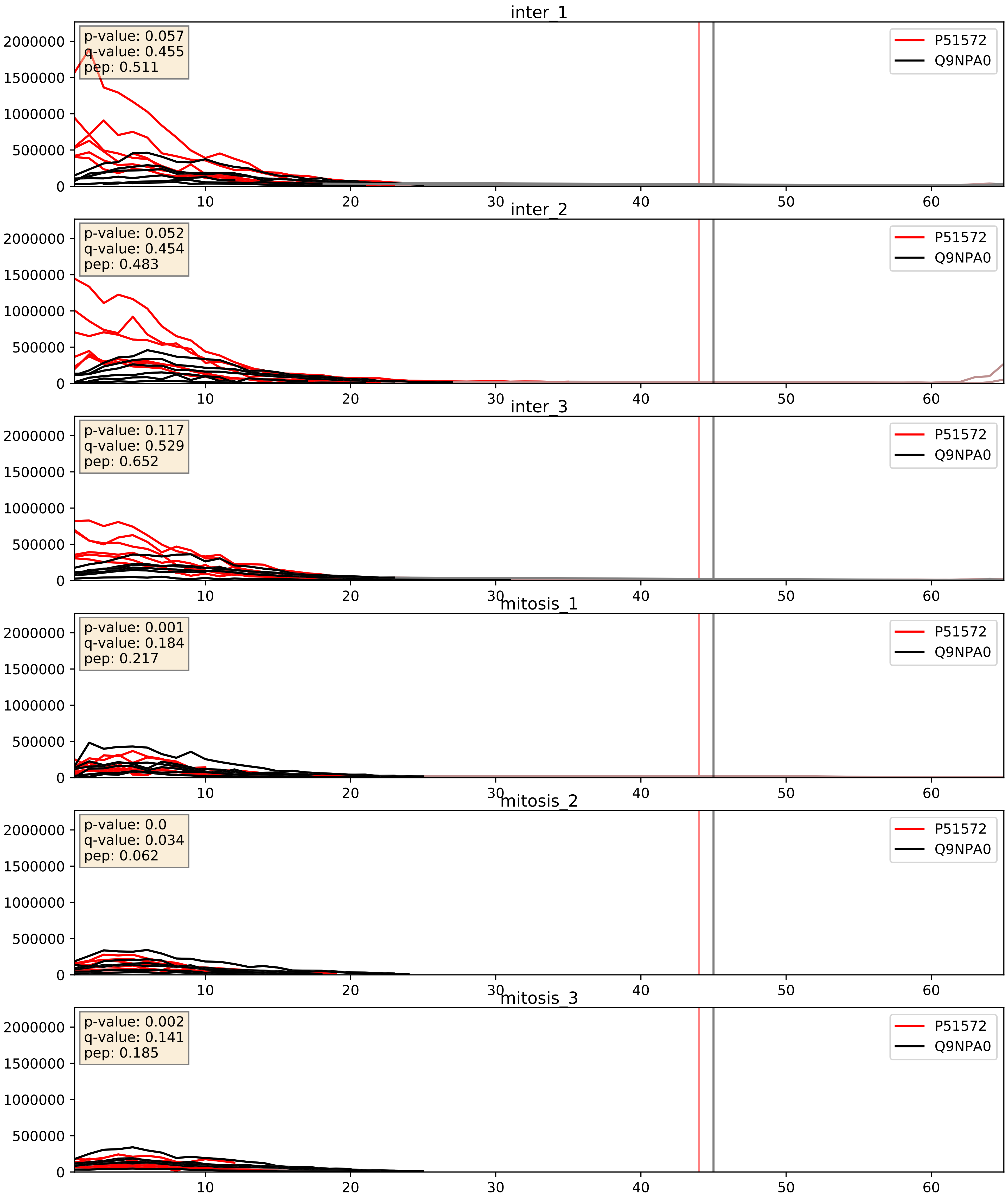


| level | condition_1 | condition_2 | pvalue | pvalue_adjusted |
|---|---|---|---|---|
| interactor_ratio | mitosis | inter | 5.961457978829829e-06 | 0.002835004461043519 |
| interactor_abundance | mitosis | inter | 0.00024214642215306928 | 0.01187905696742526 |
| complex_abundance | mitosis | inter | 0.0016414443321241888 | 0.02839616312171324 |
