## Supplementary Data 2 for "SECAT: Quantifying differential protein-protein interaction states by network-centric analysis": hela_string_000196_O95639.pdf

O95639_O95639

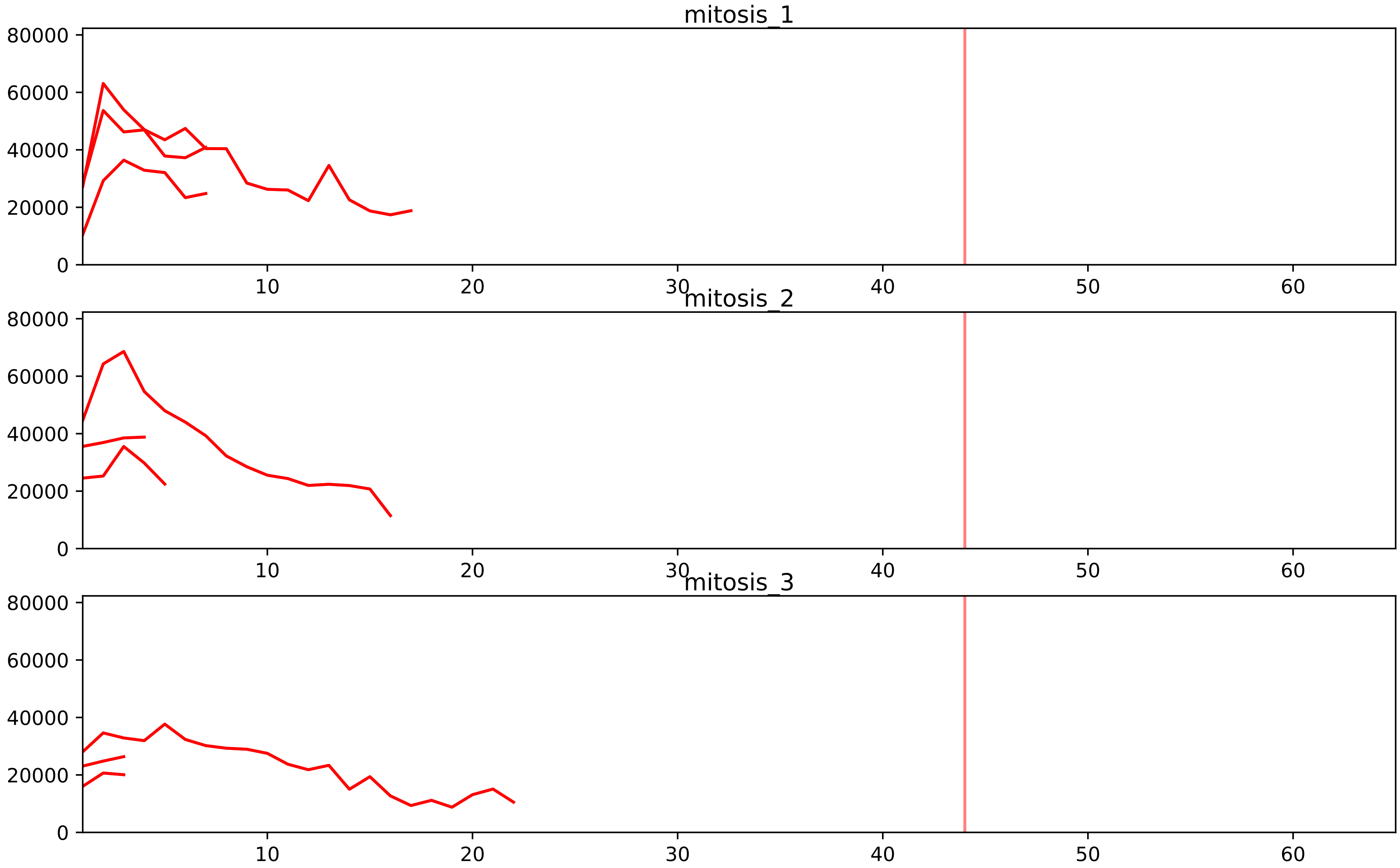


| level | condition_1 | condition_2 | pvalue | pvalue_adjusted |
|---|---|---|---|---|
| interactor_ratio | mitosis | inter | 0.00023680116704114432 | 0.003430820057918941 |
| complex_abundance | mitosis | inter | 0.0025514737338411036 | 0.019399635001105912 |
| assembled_abundance | mitosis | inter | 0.002938634703525435 | 0.053524315358764826 |
| total_abundance | mitosis | inter | 0.007710368230275846 | 0.10390002552459765 |
| interactor_abundance | mitosis | inter | 0.06798053477859406 | 0.1751879327627634 |

O95639_Q10570
CPSF4_HUMAN vs CPSF1_HUMAN
p-value: 0.04 q-value: 0.027

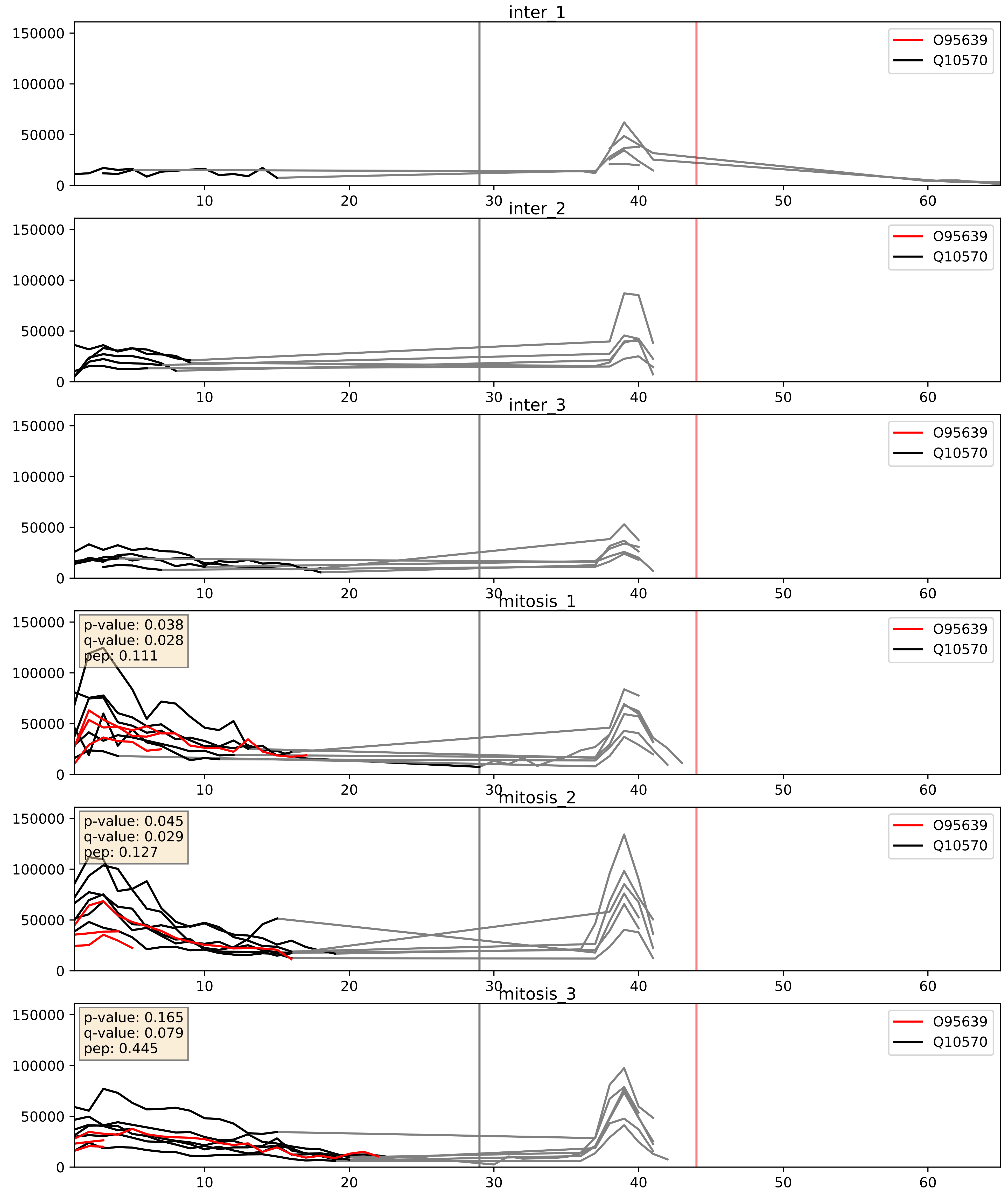


| level | condition_1 | condition_2 | pvalue | pvalue_adjusted |
|---|---|---|---|---|
| interactor_ratio | mitosis | inter | 0.002349178003548214 | 0.060752156224690976 |
| complex_abundance | mitosis | inter | 0.009086684653194026 | 0.07064670357070016 |
| interactor_abundance | mitosis | inter | 0.09608112571717416 | 0.26586534221400054 |

O95639_Q6UN15
CPSF4_HUMAN vs FIP1_HUMAN
p-value: 0.01 q-value: 0.009

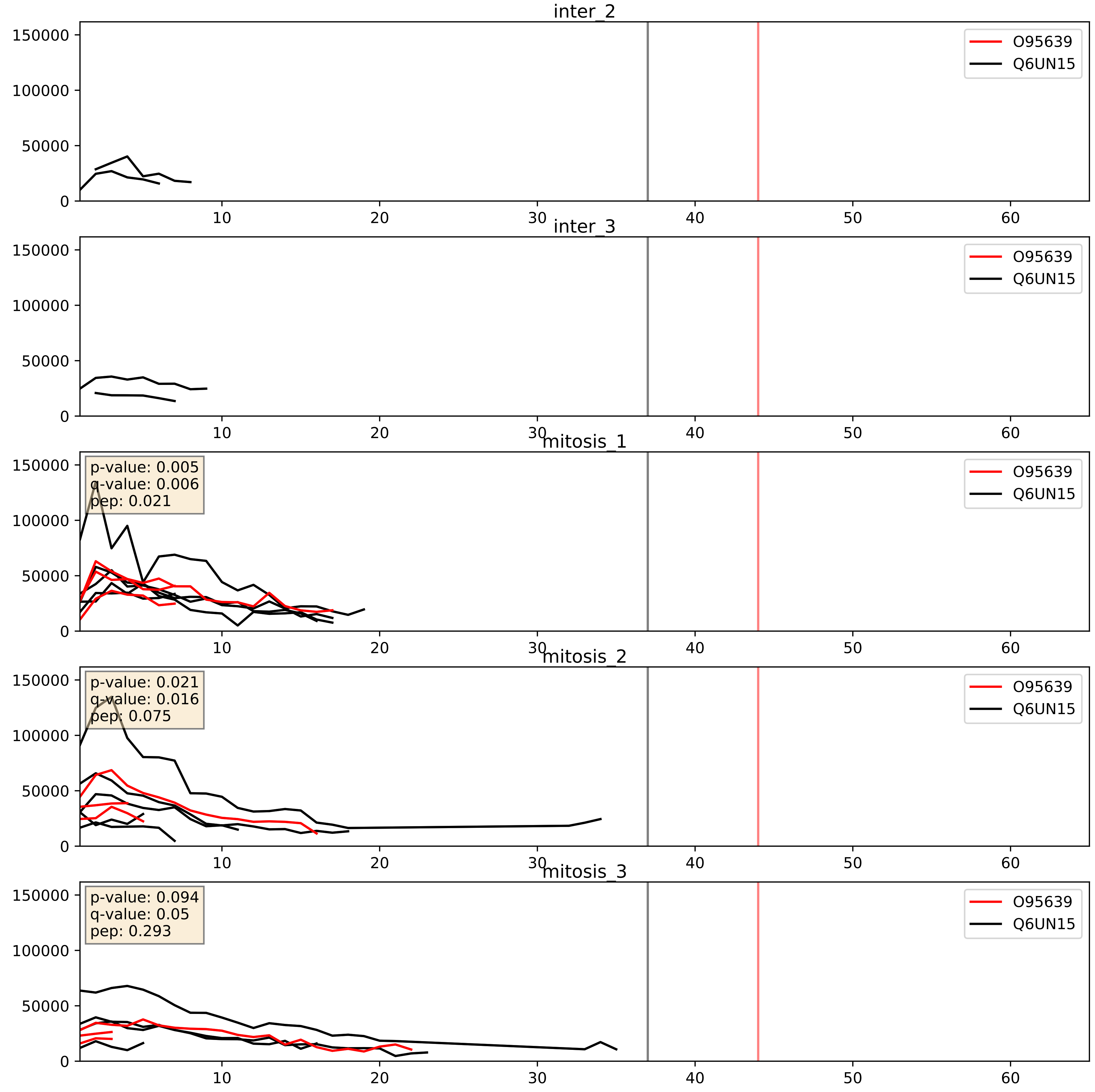


| level | condition_1 | condition_2 | pvalue | pvalue_adjusted |
|---|---|---|---|---|
| interactor_ratio | mitosis | inter | 0.00532389414285382 | 0.09758572561633554 |
| complex_abundance | mitosis | inter | 0.014845937217029401 | 0.09024336091715261 |
| interactor_abundance | mitosis | inter | 0.09608112571717416 | 0.26586534221400054 |

O95639_Q9P2I0
CPSF4_HUMAN vs CPSF2_HUMAN
p-value: 0.066 q-value: 0.041

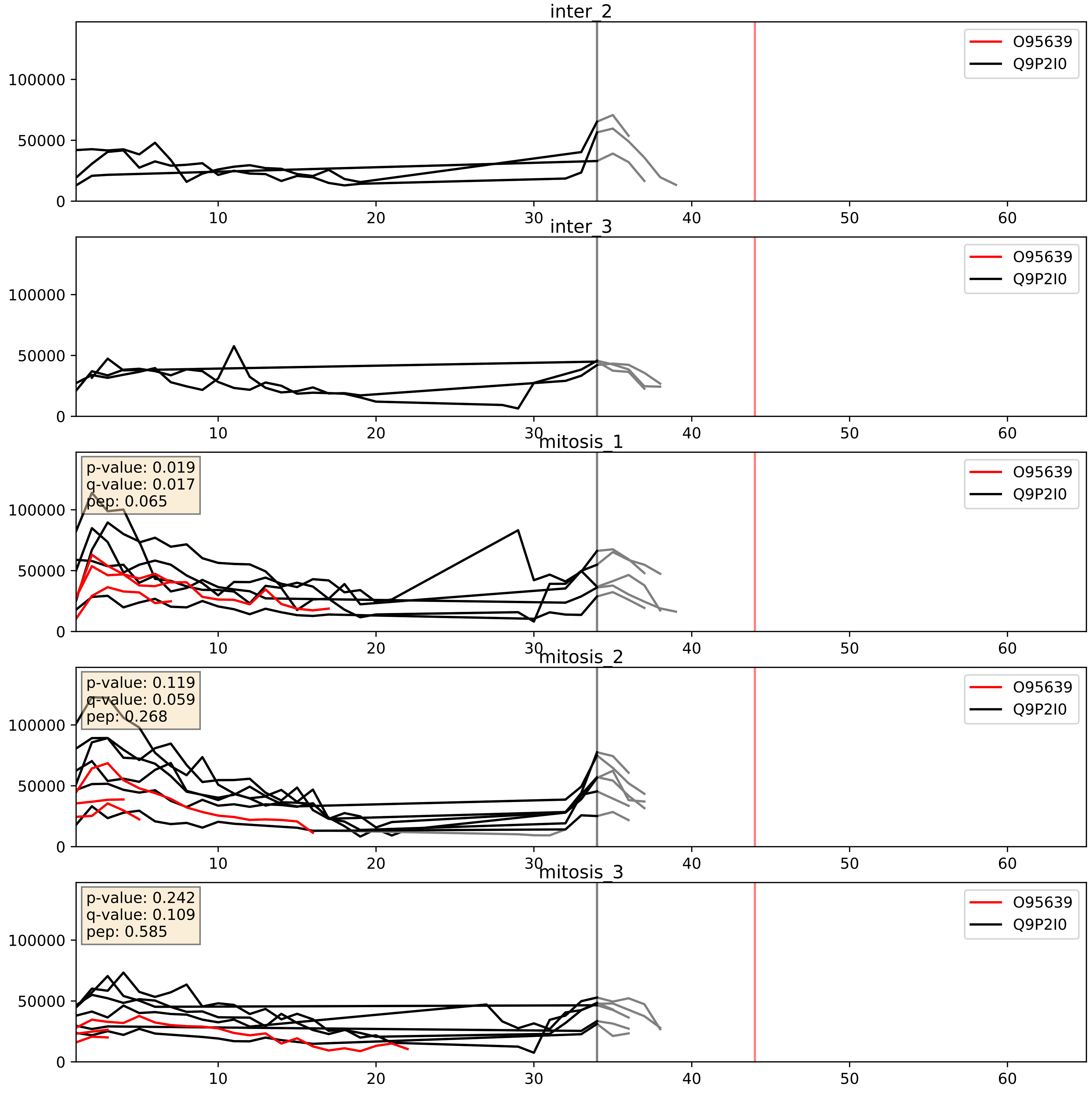


| level | condition_1 | condition_2 | pvalue | pvalue_adjusted |
|---|---|---|---|---|
| interactor_ratio | mitosis | inter | 0.0008914350338845084 | 0.034218313408302205 |
| complex_abundance | mitosis | inter | 0.007493000089970082 | 0.06444768685385507 |
| interactor_abundance | mitosis | inter | 0.09930756660188488 | 0.27089635758831565 |
