## Supplementary Data 2 for "SECAT: Quantifying differential protein-protein interaction states by network-centric analysis": hela_string_000197_O43663.pdf

O43663_O43663

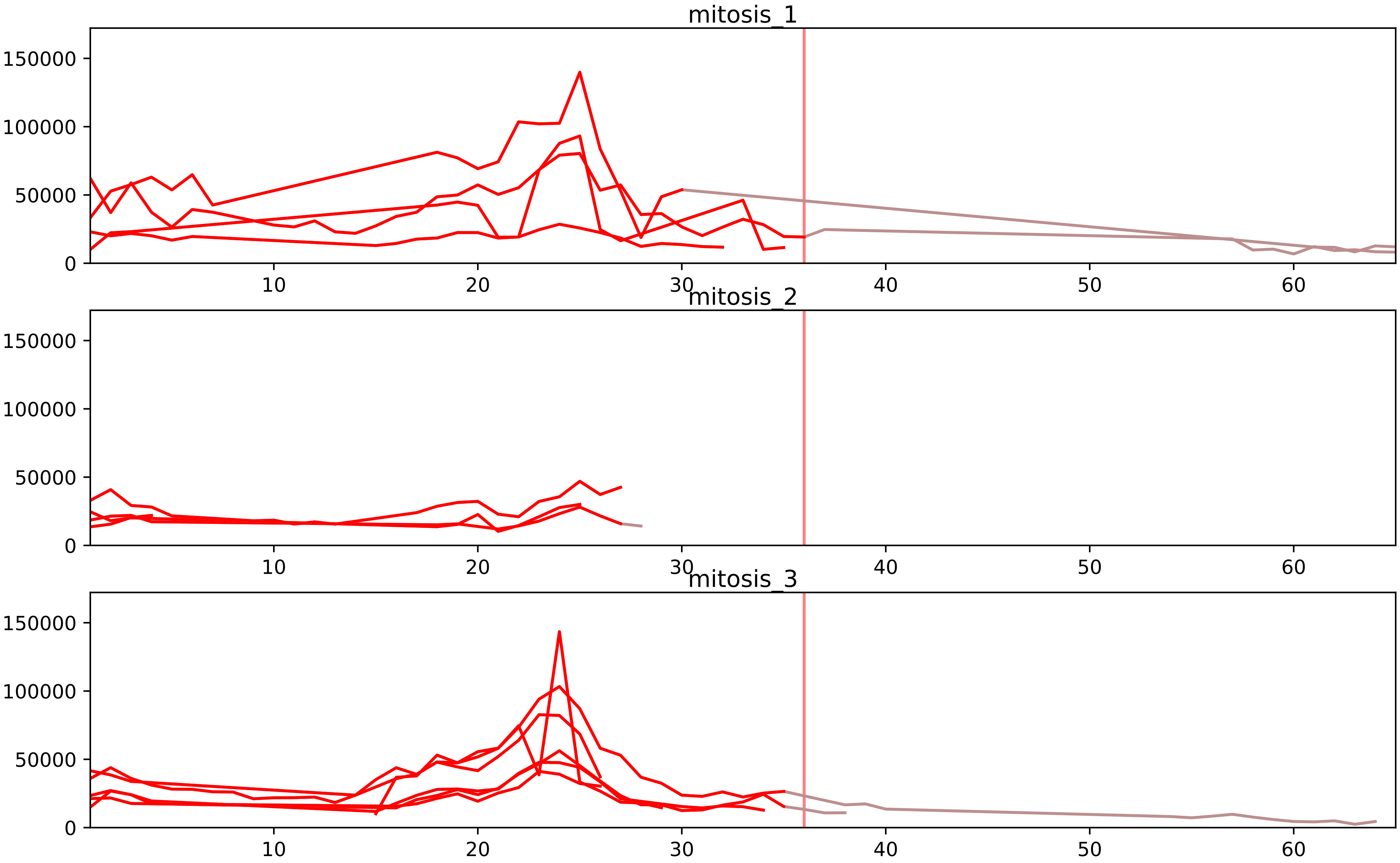


| level | condition_1 | condition_2 | pvalue | pvalue_adjusted |
|---|---|---|---|---|
| interactor_ratio | mitosis | inter | 0.00024733033683253336 | 0.003500675536706626 |
| assembled_abundance | mitosis | inter | 0.03378987078142384 | 0.22091914204238025 |
| total_abundance | mitosis | inter | 0.045471206229323616 | 0.2670056642600767 |
| complex_abundance | mitosis | inter | 0.10072858187228058 | 0.19645204292451193 |
| monomer_abundance | mitosis | inter | 0.14272340633239874 | 0.49779423054882527 |
| interactor_abundance | mitosis | inter | 0.5175837460092355 | 0.6802529233264237 |

O43663_Q15058
PRC1_HUMAN vs KIF14_HUMAN
p-value: 0.051 q-value: 0.033

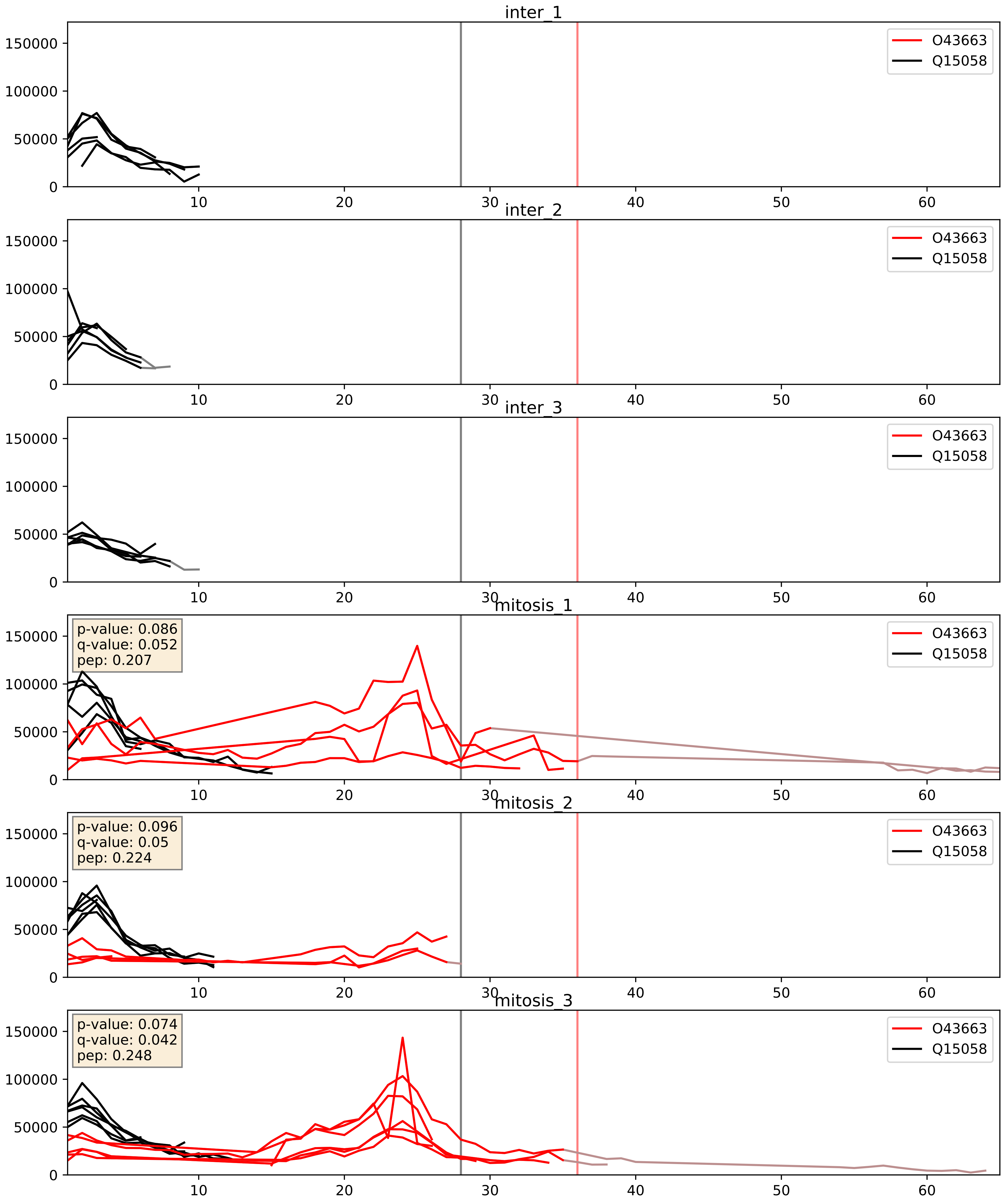


| level | condition_1 | condition_2 | pvalue | pvalue_adjusted |
|---|---|---|---|---|
| interactor_ratio | mitosis | inter | 0.00024733033683253336 | 0.016038997600655195 |
| complex_abundance | mitosis | inter | 0.10072858187228058 | 0.2354669386569353 |
| interactor_abundance | mitosis | inter | 0.5175837460092355 | 0.6804664208015752 |
