## Supplementary Data 2 for "SECAT: Quantifying differential protein-protein interaction states by network-centric analysis": hela_string_000198_Q15058.pdf

Q15058_Q15058

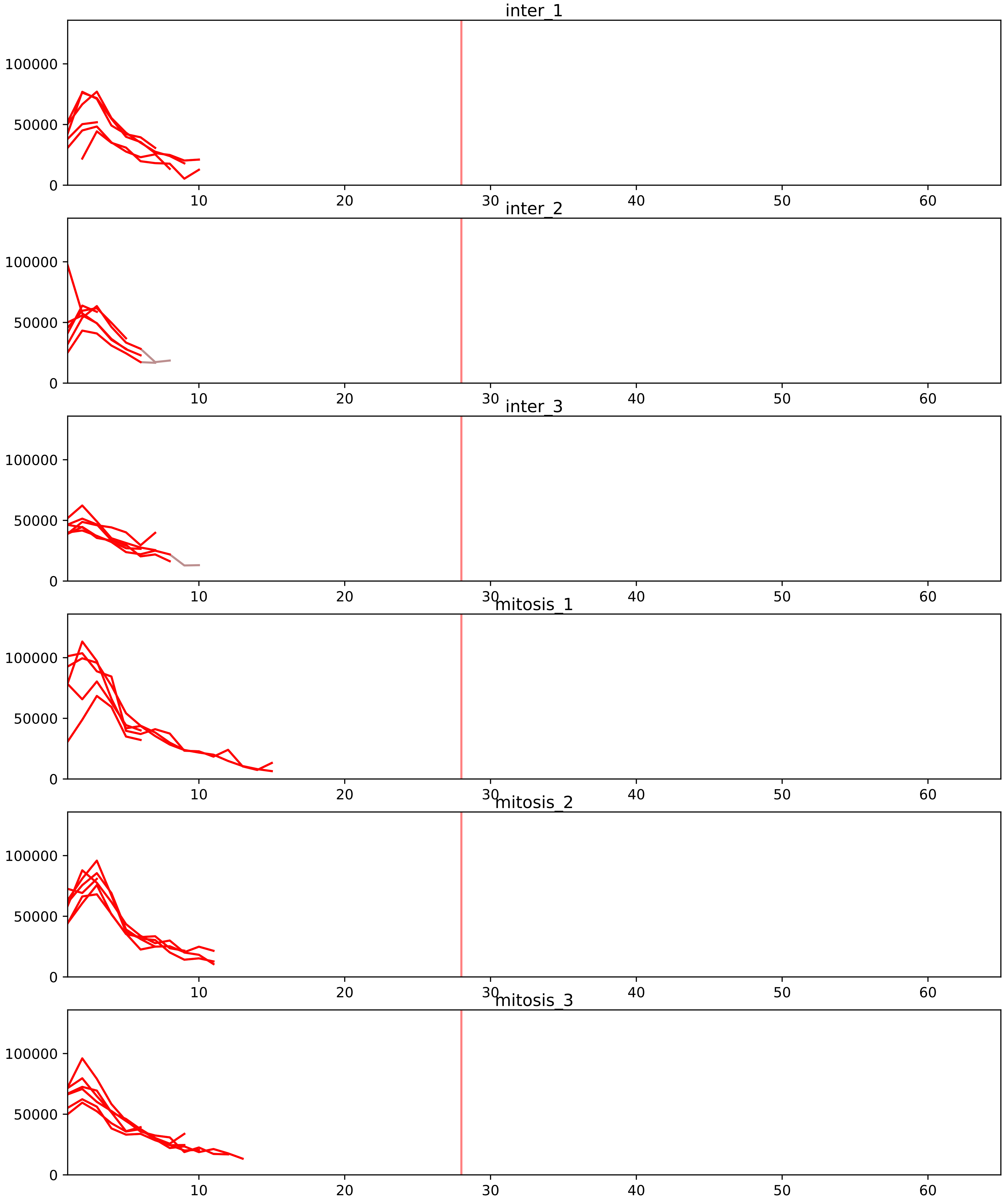


| level | condition_1 | condition_2 | pvalue | pvalue_adjusted |
|---|---|---|---|---|
| interactor_ratio | mitosis | inter | 0.0002473303368325336 | 0.003500675536706626 |
| interactor_abundance | mitosis | inter | 0.018639813633571877 | 0.07551518983979284 |
| complex_abundance | mitosis | inter | 0.10072858187228058 | 0.19645204292451193 |
| total_abundance | mitosis | inter | 0.25901924901607265 | 0.5925906412659128 |
| assembled_abundance | mitosis | inter | 0.28409765764035855 | 0.5850724968996226 |

O43663_Q15058
PRC1_HUMAN vs KIF14_HUMAN
p-value: 0.051 q-value: 0.033

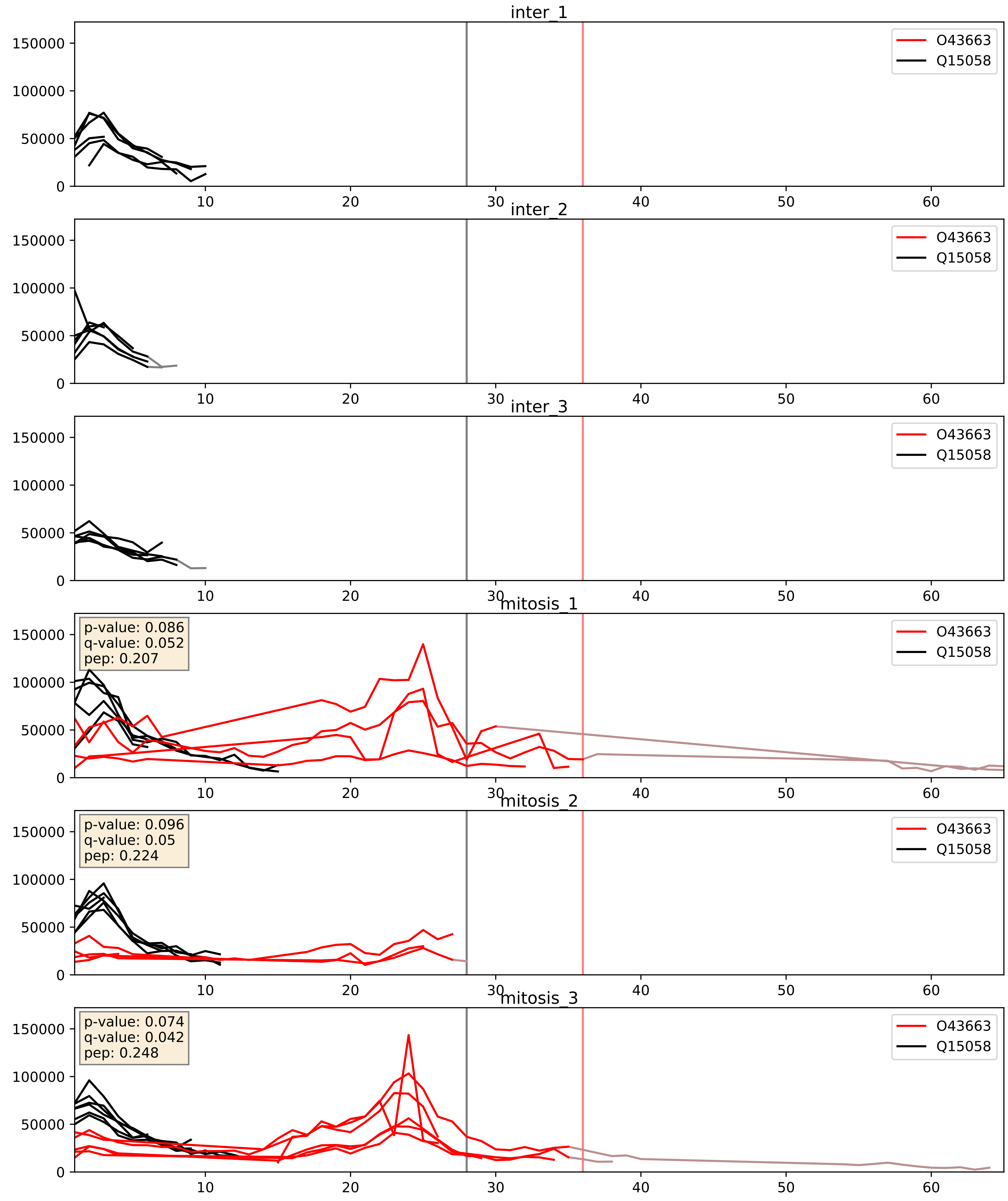


| level | condition_1 | condition_2 | pvalue | pvalue_adjusted |
|---|---|---|---|---|
| interactor_ratio | mitosis | inter | 0.00024733033683253336 | 0.016038997600655195 |
| complex_abundance | mitosis | inter | 0.10072858187228058 | 0.2354669386569353 |
| interactor_abundance | mitosis | inter | 0.5175837460092355 | 0.6804664208015752 |
