## Supplementary Data 2 for "SECAT: Quantifying differential protein-protein interaction states by network-centric analysis": hela_string_000199_O60832.pdf

O60832_O60832

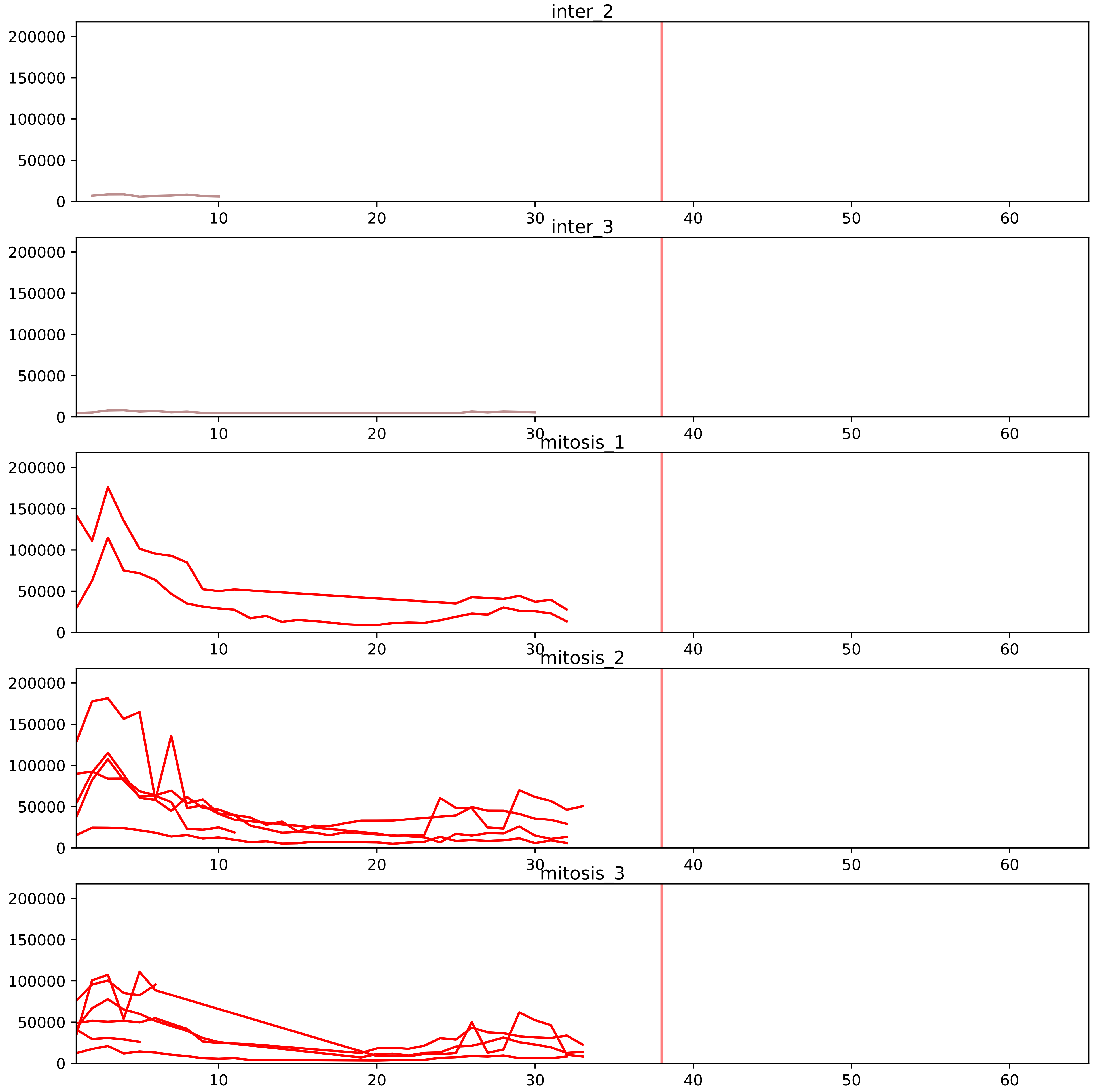


| level | condition_1 | condition_2 | pvalue | pvalue_adjusted |
|---|---|---|---|---|
| complex_abundance | mitosis | inter | 0.0002526222745112085 | 0.004330299595990477 |
| interactor_abundance | mitosis | inter | 0.02362791550279908 | 0.0878290192427279 |
| assembled_abundance | mitosis | inter | 0.039976725677557855 | 0.23808021997029047 |
| total_abundance | mitosis | inter | 0.04396672388721692 | 0.26067476584782817 |
| interactor_ratio | mitosis | inter | 0.07627116889837642 | 0.27571503098823696 |

O00567_O60832
NOP56_HUMAN vs DKC1_HUMAN
p-value: 0.008 q-value: 0.007

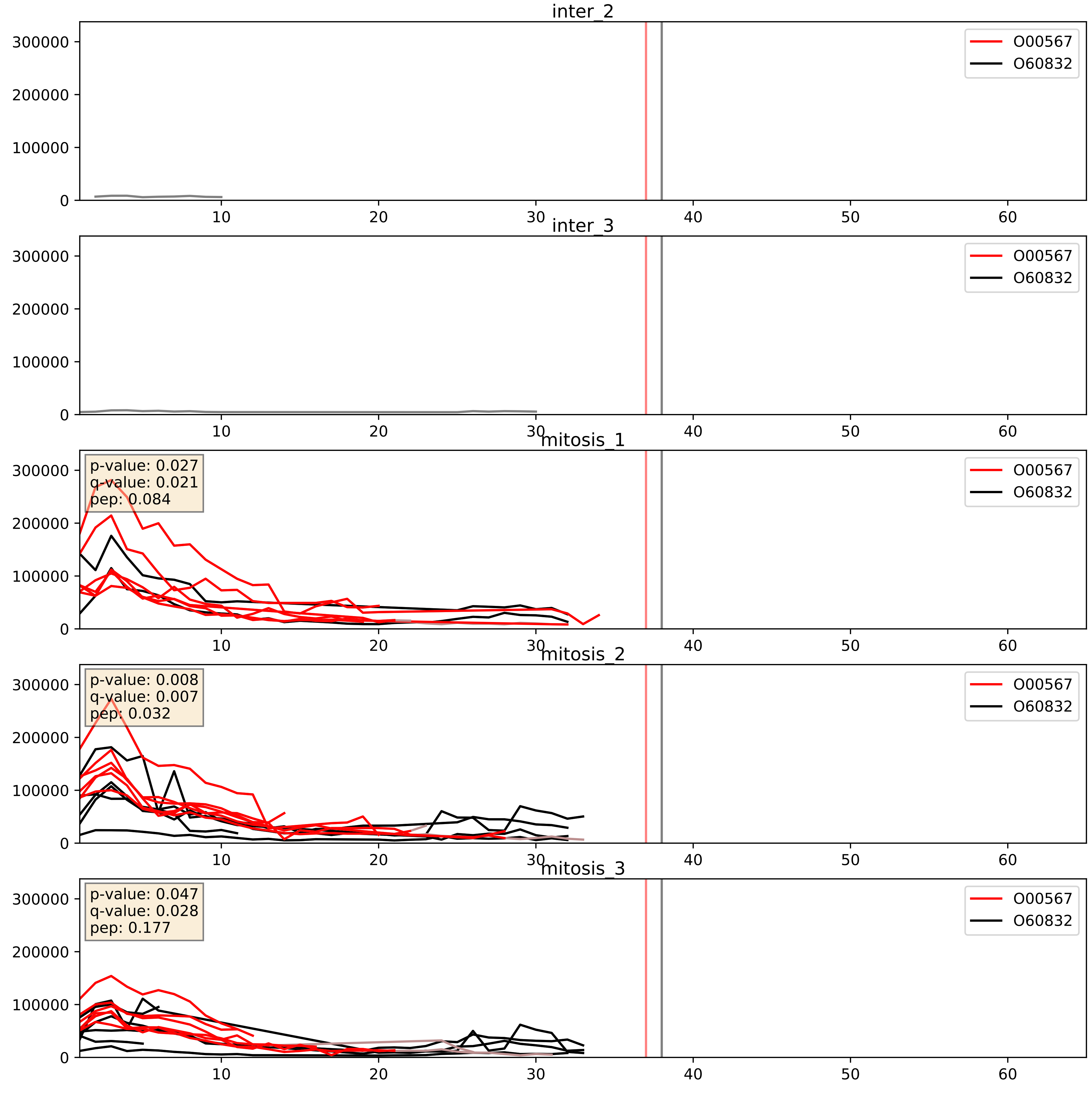


| level | condition_1 | condition_2 | pvalue | pvalue_adjusted |
|---|---|---|---|---|
| interactor_abundance | mitosis | inter | 0.0013960261808965133 | 0.02753913185123736 |
| complex_abundance | mitosis | inter | 0.0023625900336761763 | 0.035168510428347066 |
| interactor_ratio | mitosis | inter | 0.1363687124743647 | 0.5866546850118911 |

O43143_O60832
DHX15_HUMAN vs DKC1_HUMAN
p-value: 0.064 q-value: 0.04

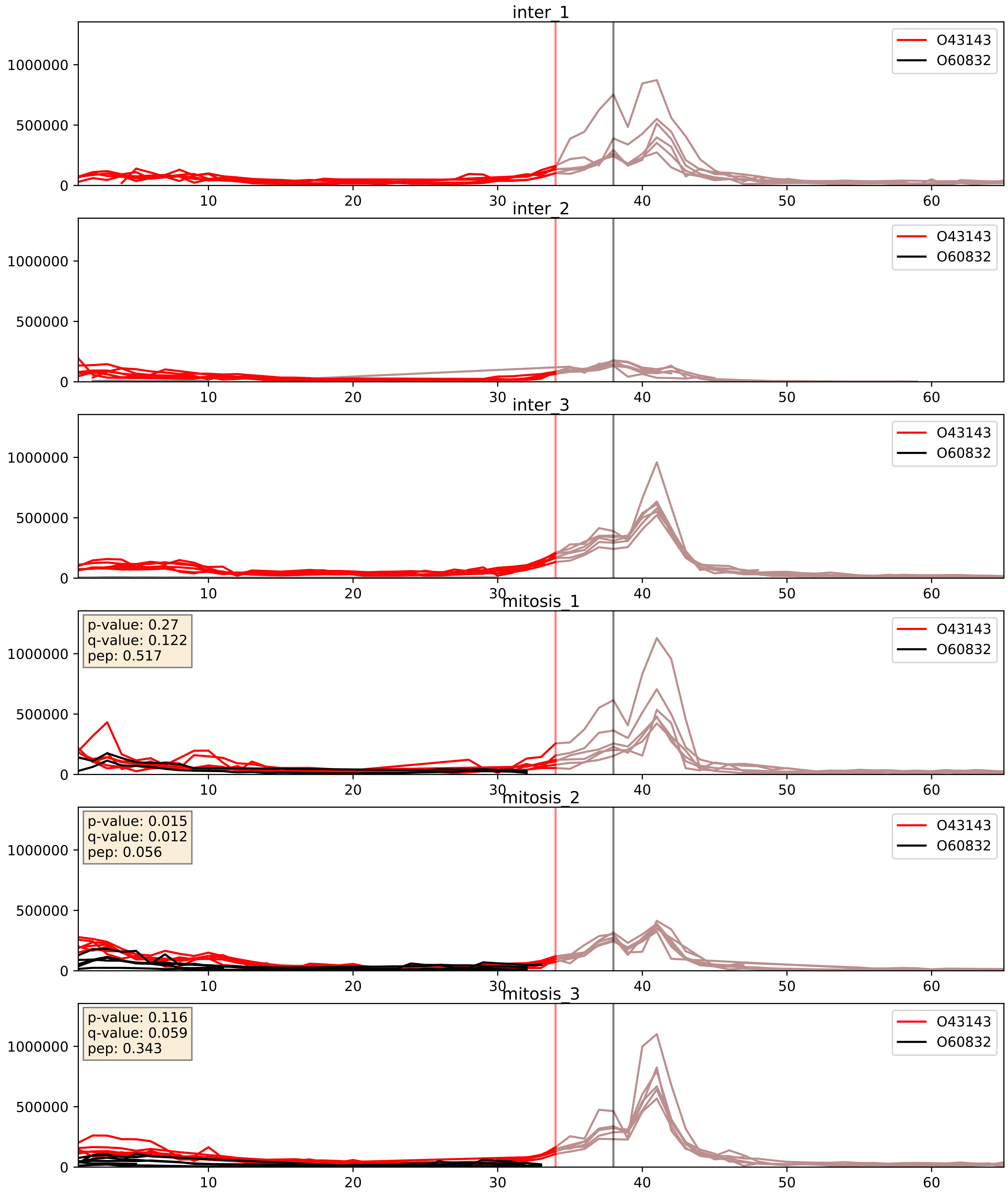


| level | condition_1 | condition_2 | pvalue | pvalue_adjusted |
|---|---|---|---|---|
| interactor_abundance | mitosis | inter | 4.356991549376841e-05 | 0.006058954526139308 |
| complex_abundance | mitosis | inter | 0.0017878025985153428 | 0.030007039692728107 |
| interactor_ratio | mitosis | inter | 0.0373689021643331 | 0.317024581295036 |

O60832_P22087
DKC1_HUMAN vs FBRL_HUMAN
p-value: 0.005 q-value: 0.005

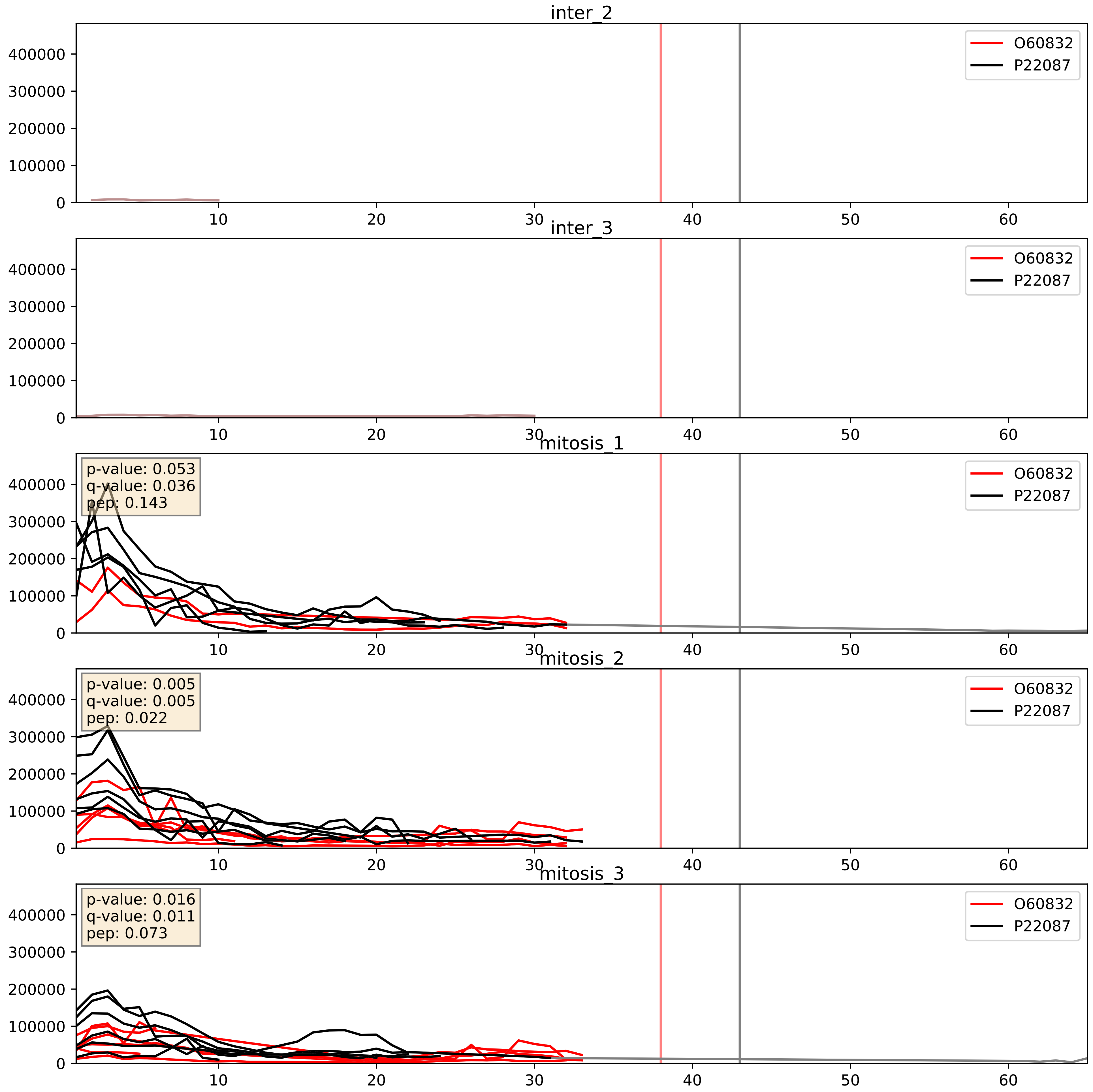


| level | condition_1 | condition_2 | pvalue | pvalue_adjusted |
|---|---|---|---|---|
| complex_abundance | mitosis | inter | 0.0015667645418376422 | 0.027691822035811595 |
| interactor_abundance | mitosis | inter | 0.050819620539780355 | 0.18065446504174412 |
| interactor_ratio | mitosis | inter | 0.07249205172524498 | 0.44970456769966505 |

O60832_P55769
DKC1_HUMAN vs NH2L1_HUMAN
p-value: 0.008 q-value: 0.007

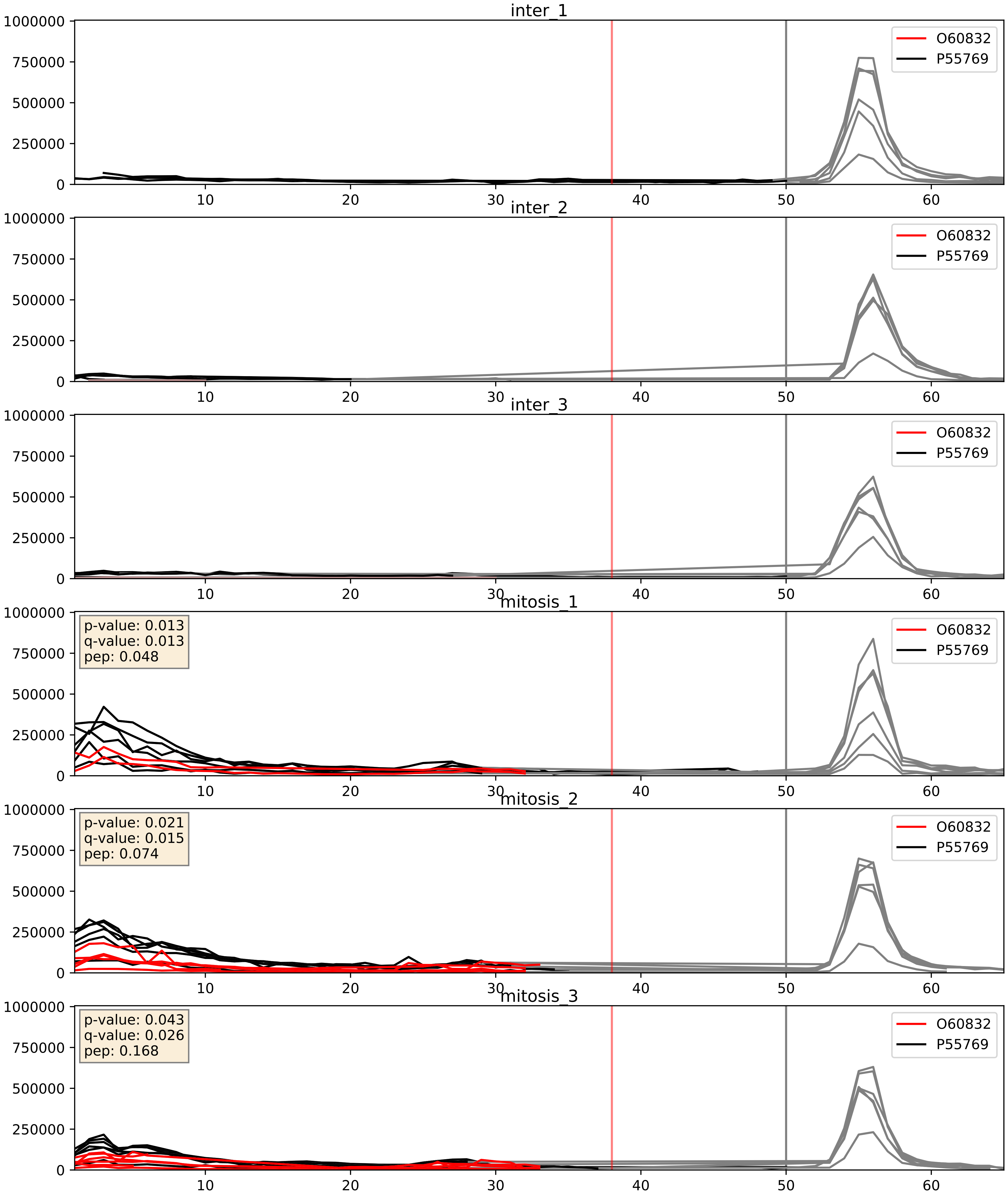


| level | condition_1 | condition_2 | pvalue | pvalue_adjusted |
|---|---|---|---|---|
| complex_abundance | mitosis | inter | 0.0010098562157940983 | 0.021187179429405588 |
| interactor_ratio | mitosis | inter | 0.043200488775377 | 0.34205635861393335 |
| interactor_abundance | mitosis | inter | 0.04824150833940766 | 0.17504902777779313 |

O60832_Q12788
DKC1_HUMAN vs TBL3_HUMAN
p-value: 0.005 q-value: 0.005

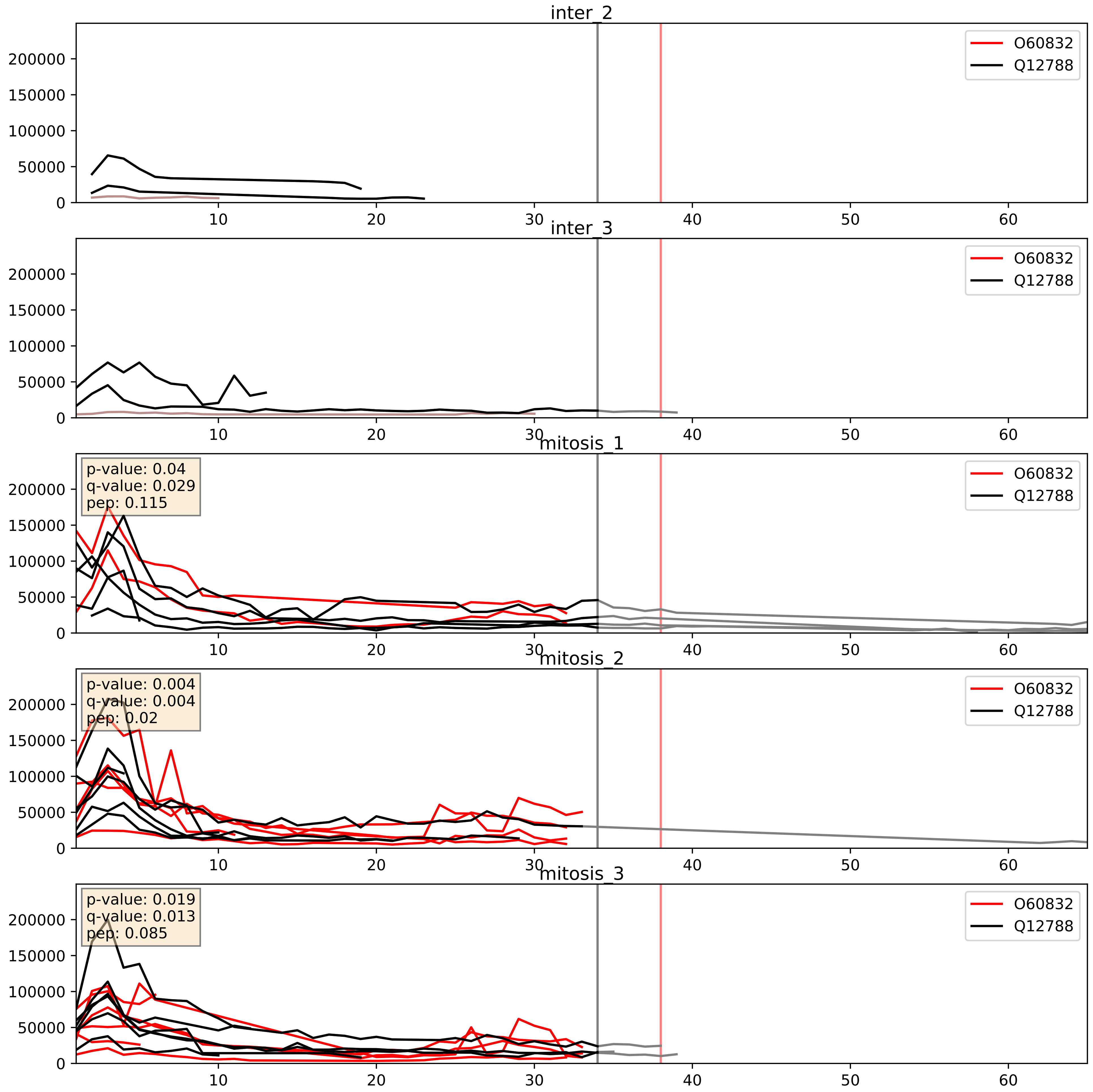


| level | condition_1 | condition_2 | pvalue | pvalue_adjusted |
|---|---|---|---|---|
| complex_abundance | mitosis | inter | 0.0037688012662694115 | 0.04531030735851989 |
| interactor_abundance | mitosis | inter | 0.04824150833940766 | 0.17504902777779313 |
| interactor_ratio | mitosis | inter | 0.4938582007919115 | 0.86984863222988 |

O60832_Q8NI36
DKC1_HUMAN vs WDR36_HUMAN
p-value: 0.004 q-value: 0.047

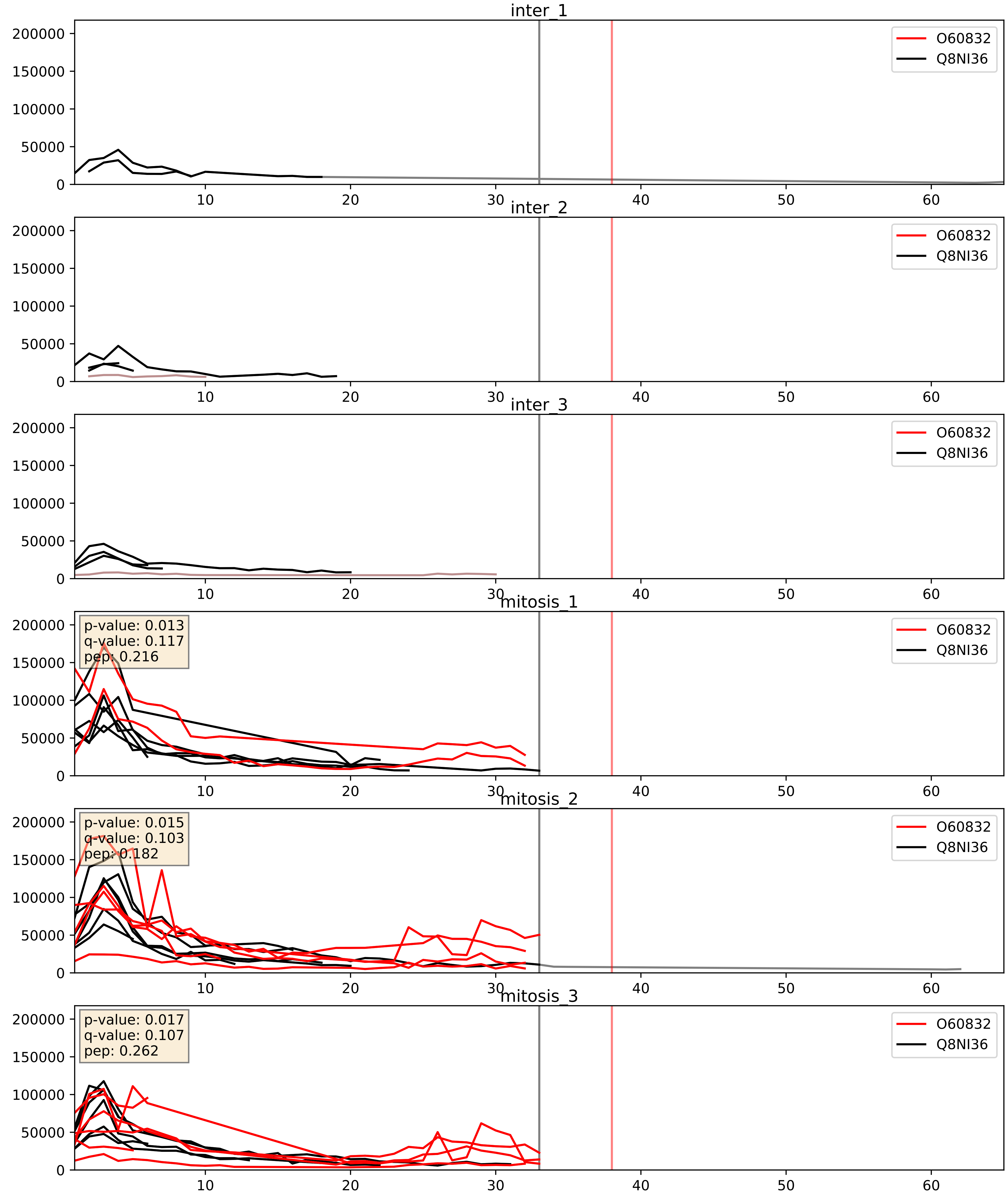


| level | condition_1 | condition_2 | pvalue | pvalue_adjusted |
|---|---|---|---|---|
| complex_abundance | mitosis | inter | 0.005657961196732316 | 0.05664578695208026 |
| interactor_abundance | mitosis | inter | 0.05267352363657362 | 0.18428385541163456 |
| interactor_ratio | mitosis | inter | 0.8878767344863951 | 0.9804177989881011 |

O60832_Q9H0A0
DKC1_HUMAN vs NAT10_HUMAN
p-value: 0.076 q-value: 0.046

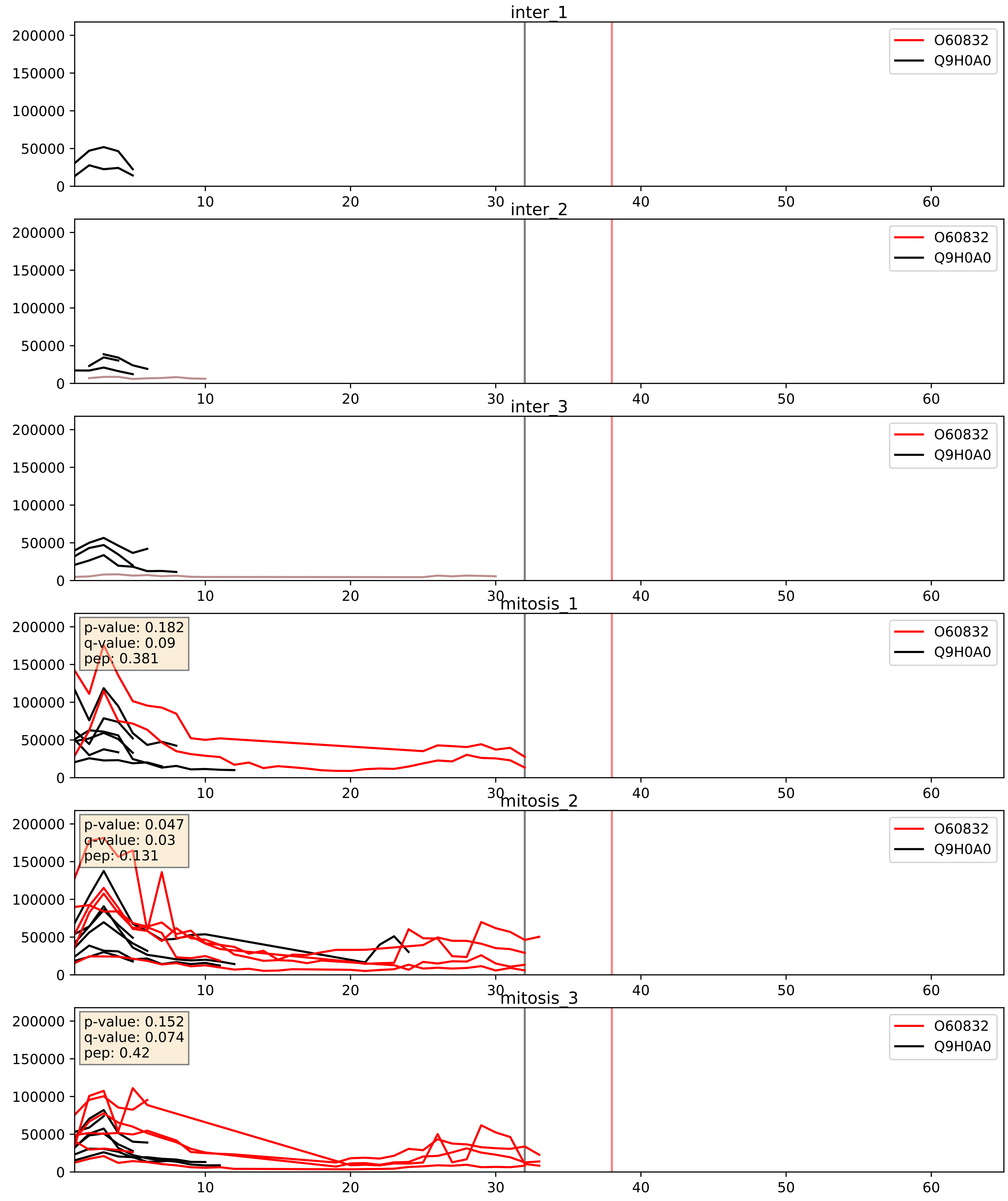


| level | condition_1 | condition_2 | pvalue | pvalue_adjusted |
|---|---|---|---|---|
| complex_abundance | mitosis | inter | 0.014138361720307944 | 0.08846811134929532 |
| interactor_abundance | mitosis | inter | 0.06548599163945651 | 0.21201213632138718 |
| interactor_ratio | mitosis | inter | 0.7825031437437657 | 0.95900240448312 |

O60832_Q9NVP1
DKC1_HUMAN vs DDX18_HUMAN
p-value: 0.076 q-value: 0.046

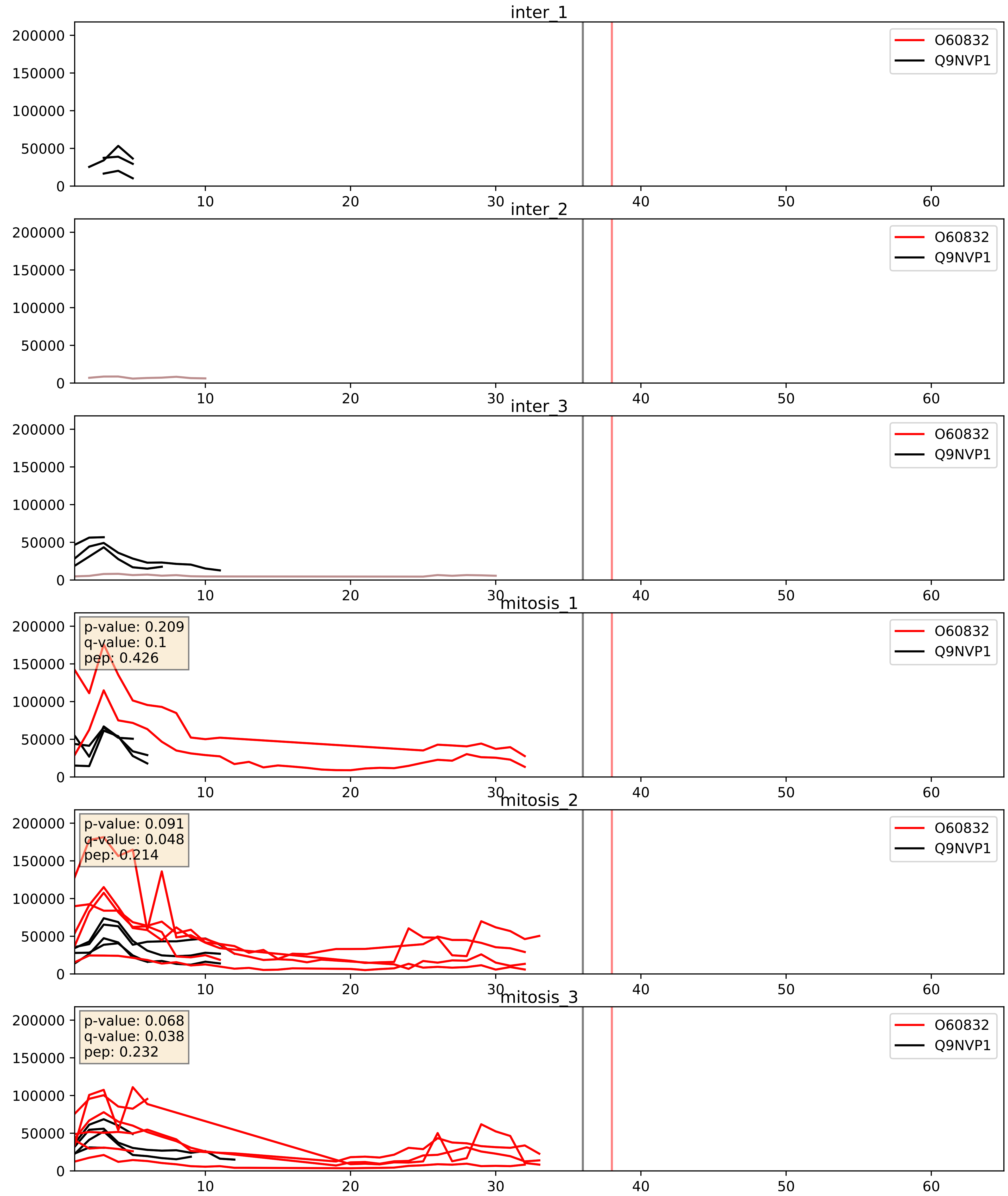


| level | condition_1 | condition_2 | pvalue | pvalue_adjusted |
|---|---|---|---|---|
| complex_abundance | mitosis | inter | 0.019061609538449552 | 0.10333589464795959 |
| interactor_abundance | mitosis | inter | 0.07954363368256376 | 0.23458863198027416 |
| interactor_ratio | mitosis | inter | 0.5961816441313526 | 0.9019028605875125 |

O60832_Q9NX24
DKC1_HUMAN vs NHP2_HUMAN
p-value: 0.003 q-value: 0.003

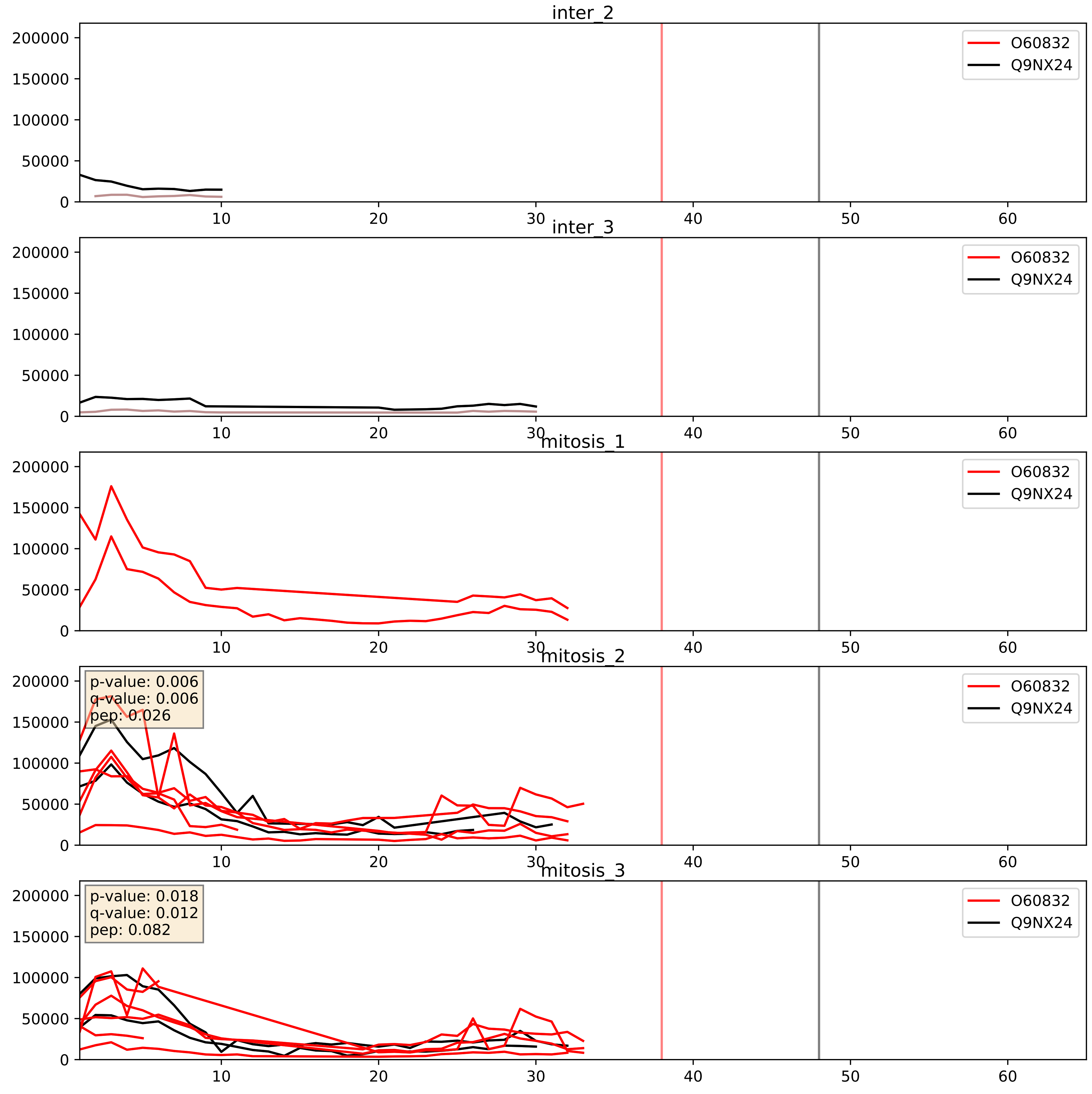


| level | condition_1 | condition_2 | pvalue | pvalue_adjusted |
|---|---|---|---|---|
| interactor_ratio | mitosis | inter | 0.17938609422294735 | 0.6462731340692043 |
| interactor_abundance | mitosis | inter | 0.18306057163109743 | 0.3757436970203683 |
| complex_abundance | mitosis | inter | 0.18394619909774976 | 0.33531940476212474 |

O60832_Q9Y2X3
DKC1_HUMAN vs NOP58_HUMAN
p-value: 0.01 q-value: 0.009

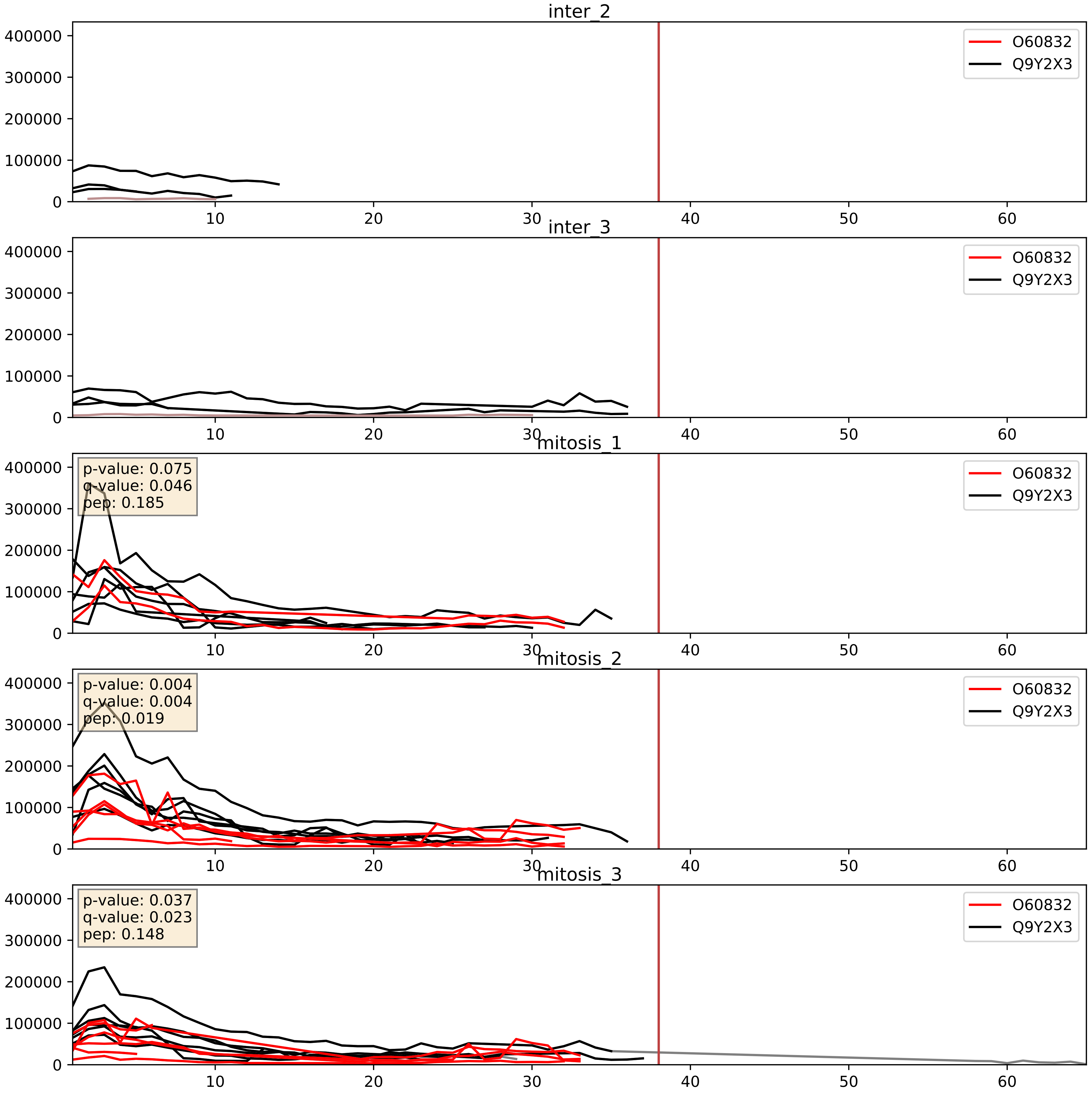


| level | condition_1 | condition_2 | pvalue | pvalue_adjusted |
|---|---|---|---|---|
| complex_abundance | mitosis | inter | 0.002344505735626372 | 0.035168510428347066 |
| interactor_abundance | mitosis | inter | 0.04900154453194635 | 0.17683525345424148 |
| interactor_ratio | mitosis | inter | 0.05935818141809091 | 0.40293896347252833 |
