## Supplementary Data 2 for "SECAT: Quantifying differential protein-protein interaction states by network-centric analysis": hela_string_000200_P19387.pdf

P19387_P19387

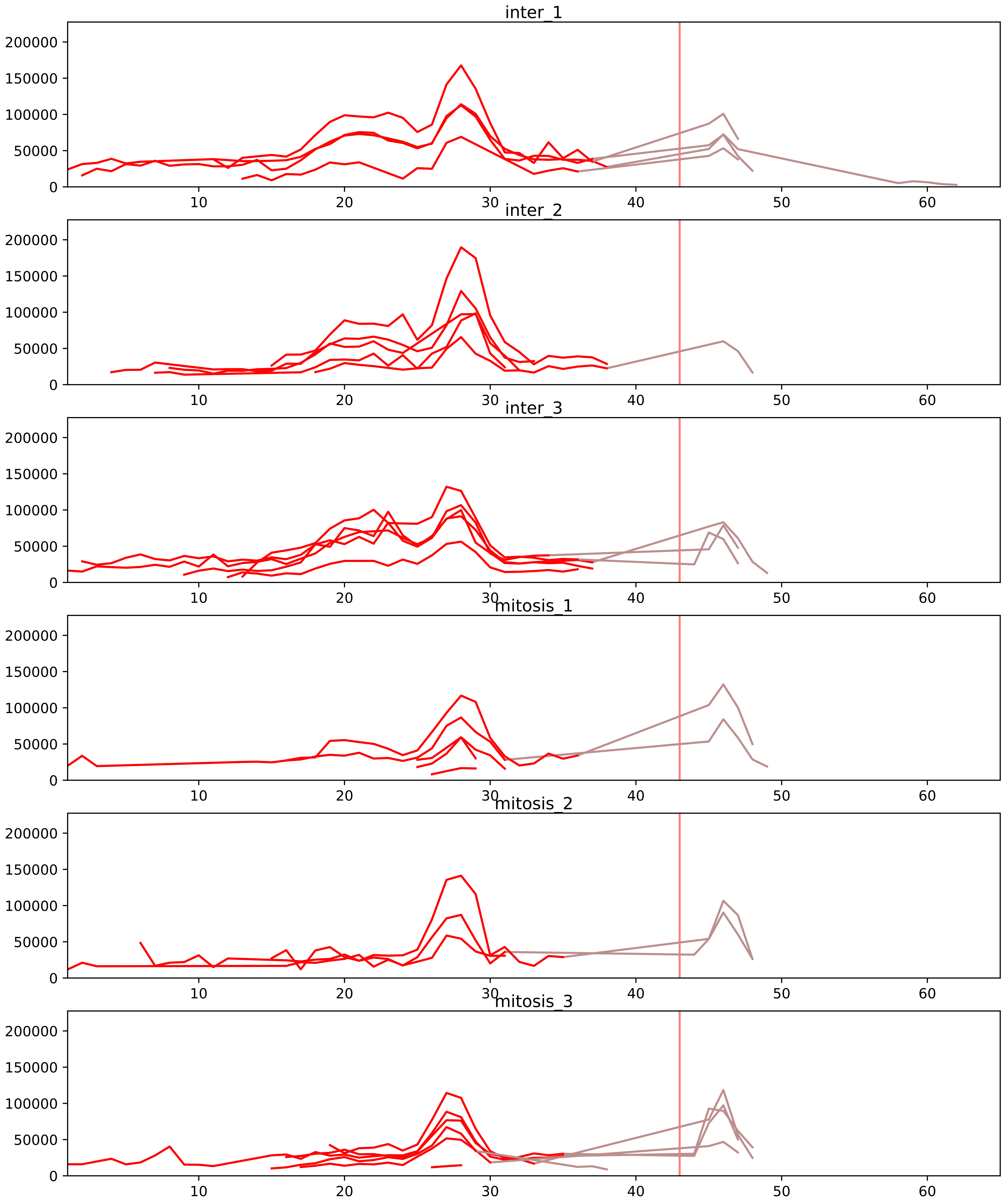


| level | condition_1 | condition_2 | pvalue | pvalue_adjusted |
|---|---|---|---|---|
| complex_abundance | mitosis | inter | 0.0002541697588950932 | 0.004330299595990477 |
| interactor_abundance | mitosis | inter | 0.0012232260669514203 | 0.013239623312885963 |
| interactor_ratio | mitosis | inter | 0.06879651741681832 | 0.26046418116655495 |
| assembled_abundance | mitosis | inter | 0.4719871026436893 | 0.7308539951952067 |
| total_abundance | mitosis | inter | 0.5704300470999222 | 0.8048945038208712 |
| monomer_abundance | mitosis | inter | 0.725238404786894 | 0.8847136478614637 |

P19387_P30876
RPB3_HUMAN vs RPB2_HUMAN
p-value: 0.006 q-value: 0.006

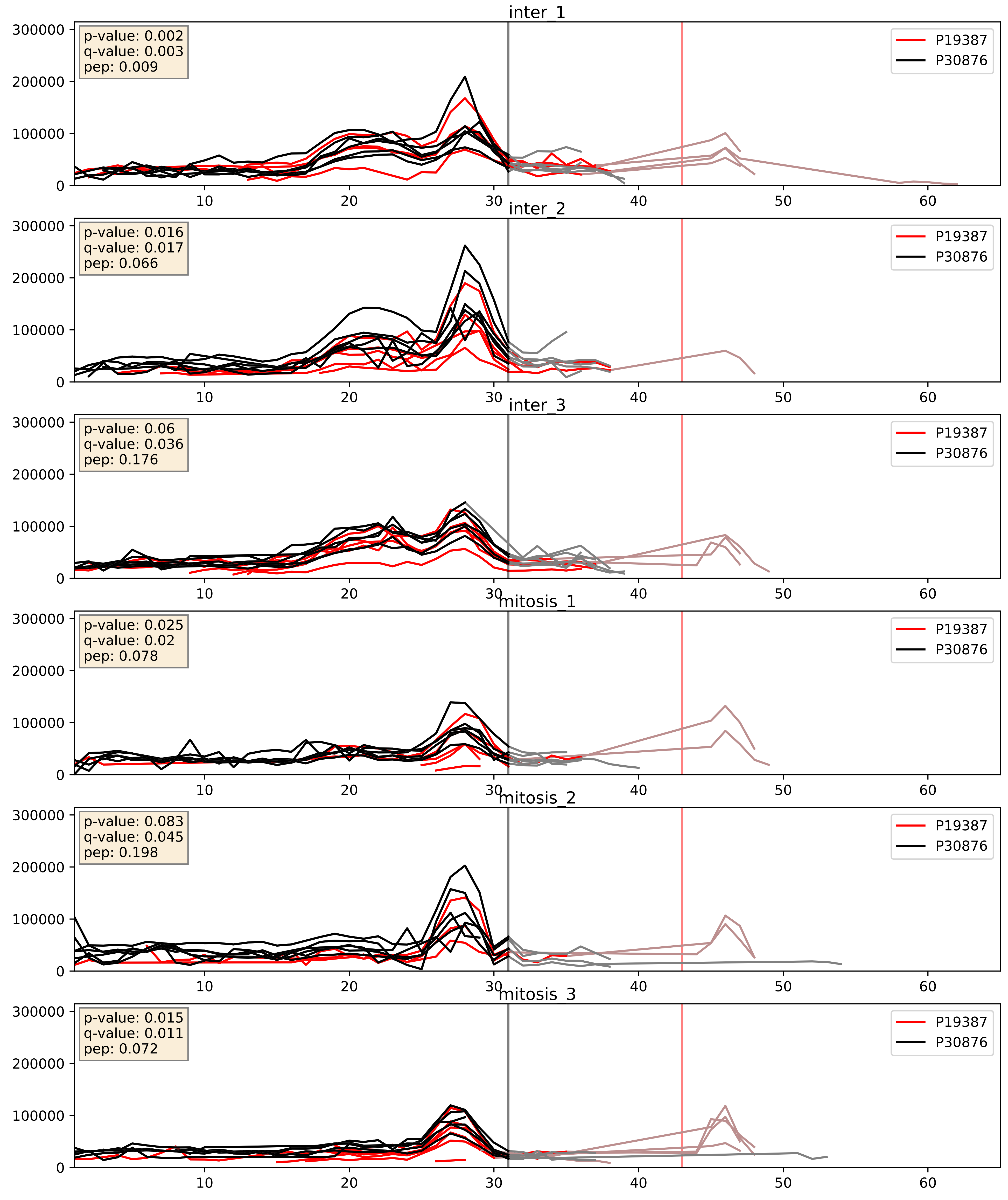


| level | condition_1 | condition_2 | pvalue | pvalue_adjusted |
|---|---|---|---|---|
| complex_abundance | mitosis | inter | 0.0002541697588950932 | 0.010665162432068616 |
| interactor_abundance | mitosis | inter | 0.0012232260669514203 | 0.02563235038703588 |
| interactor_ratio | mitosis | inter | 0.06879651741681832 | 0.4341924606131018 |
